# Variants in the *DDX6-CXCR5* autoimmune disease risk locus influence the regulatory network in immune cells and salivary gland

**DOI:** 10.1101/2023.10.05.561076

**Authors:** Mandi M. Wiley, Bhuwan Khatri, Michelle L. Joachims, Kandice L. Tessneer, Anna M. Stolarczyk, Astrid Rasmussen, Juan-Manuel Anaya, Lara A. Aqrawi, Sang-Cheol Bae, Eva Baecklund, Albin Björk, Johan G. Brun, Sara Magnusson Bucher, Nick Dand, Maija-Leena Eloranta, Fiona Engelke, Helena Forsblad-d’Elia, Cecilia Fugmann, Stuart B. Glenn, Chen Gong, Jacques-Eric Gottenberg, Daniel Hammenfors, Juliana Imgenberg-Kreuz, Janicke Liaaen Jensen, Svein Joar Auglænd Johnsen, Malin V. Jonsson, Jennifer A. Kelly, Sharmily Khanam, Kwangwoo Kim, Marika Kvarnström, Thomas Mandl, Javier Martín, David L. Morris, Gaetane Nocturne, Katrine Brække Norheim, Peter Olsson, Øyvind Palm, Jacques-Olivier Pers, Nelson L. Rhodus, Christopher Sjöwall, Kathrine Skarstein, Kimberly E. Taylor, Phil Tombleson, Gudny Ella Thorlacius, Swamy Venuturupalli, Edward M. Vital, Daniel J Wallace, Kiely M. Grundahl, Lida Radfar, Michael T. Brennan, Judith A. James, R. Hal Scofield, Patrick M. Gaffney, Lindsey A. Criswell, Roland Jonsson, Silke Appel, Per Eriksson, Simon J. Bowman, Roald Omdal, Lars Rönnblom, Blake M. Warner, Maureen Rischmueller, Torsten Witte, A. Darise Farris, Xavier Mariette, Caroline H. Shiboski, Sjögren’s International Collaborative Clinical Alliance (SICCA), Marie Wahren-Herlenius, Marta E. Alarcón-Riquelme, PRECISESADS Clinical Consortium, Wan-Fai Ng, UK Primary Sjögren’s Syndrome Registry, Kathy L. Sivils, Joel M. Guthridge, Indra Adrianto, Timothy J. Vyse, Betty P. Tsao, Gunnel Nordmark, Christopher J. Lessard

## Abstract

Fine mapping and bioinformatic analysis of the *DDX6-CXCR5* genetic risk association in Sjögren’s Disease (SjD) and Systemic Lupus Erythematosus (SLE) identified five common SNPs with functional evidence in immune cell types: rs4938573, rs57494551, rs4938572, rs4936443, rs7117261. Functional interrogation of nuclear protein binding affinity, enhancer/promoter regulatory activity, and chromatin-chromatin interactions in immune, salivary gland epithelial, and kidney epithelial cells revealed cell type-specific allelic effects for all five SNPs that expanded regulation beyond effects on *DDX6* and *CXCR5* expression. Mapping the local chromatin regulatory network revealed several additional genes of interest, including *lnc-PHLDB1-1*. Collectively, functional characterization implicated the risk alleles of these SNPs as modulators of promoter and/or enhancer activities that regulate cell type-specific expression of *DDX6*, *CXCR5*, and *lnc-PHLDB1-1*, among others. Further, these findings emphasize the importance of exploring the functional significance of SNPs in the context of complex chromatin architecture in disease-relevant cell types and tissues.

## INTRODUCTION

Sjögren’s disease (SjD) and systemic lupus erythematosus (SLE) are distinct yet related autoimmune diseases with several shared clinical features and genetic associations^1,2^. While autoimmune destruction and impaired function of the exocrine glands (e.g., lacrimal and salivary glands) are unique to SjD^3–5^, several clinical manifestations are shared, including serological features like the presence of antinuclear antibody (ANA), anti-Ro/SSA antibodies and overexpression of type I interferons^6–8^. SjD has a higher prevalence in populations of European ancestry while SLE is more frequently diagnosed in those of African descent^9–11^. SjD also has a stronger disparity in women relative to men (SjD: 9-20:1 female: male; SLE: 9-10:1 female: male)^12–17^. The complex etiologies of SjD and SLE are driven by environmental exposures in the context of genetic susceptibility^18,19^. Exploring the functional implications of shared genetic risk loci between SjD and SLE may help identify common disease mechanisms.

*DDX6-CXCR5* on chromosome 11p23.3 has been associated through candidate gene and genome-wide association studies (GWAS) with SjD and SLE of European and Asian ancestry^1,20–23^. DDX6 (DEAD-box RNA Helicase 6) is important in viral RNA recognition and modulation of type I interferon responses^23–25^. CXCR5 (C-X-C chemokine receptor type 5) is an important modulator of B and T follicular cell trafficking to peripheral lymphoid organs^26^. Loss of *CXCR5* expression has been reported in purified circulating CD19^+^ B cells isolated from SjD patients homozygous for the risk allele (T) of rs4938573^27^; a proxy in strong linkage disequilibrium (LD; *r*^2^=0.865) with the top associated SNP, rs7119038, of the *DDX6-CXCR5* SjD risk interval^21^. Loss of *CXCR5* was inversely correlated with increased homing of CXCR5^+^ cells to the salivary gland^27^, suggesting a potential SjD-specific role for this risk interval.

In this study, meta-analysis and genetic fine mapping of the *DDX6-CXCR5* risk interval of SjD and SLE were performed to refine the association signals and identify the most likely functional SNPs shared across the two diseases (**Figure 1A,B**). The *DDX6-CXCR5* locus has a complex local chromatin regulatory network including several genes, gene promoters, and regulatory elements with cell type-specific patterns of epigenetic enrichment. Deep bioinformatic analyses prioritized five SNPs for functional validation using electromobility shift assays (EMSAs), luciferase transactivation assays, and chromatin conformation capture with quantitative PCR (3C-qPCR), identifying cell type- and context-specific functional effects (**Figure 1C**). Understanding how functional SNPs located within this locus regulate the expression of *DDX6*, *CXCR5*, and/or other disease-relevant genes through interactions within the chromatin regulatory network will provide new insights into how genetic risk at the *DDX6-CXCR5* locus may modulate molecular mechanisms of autoimmunity.

**Figure 1.**
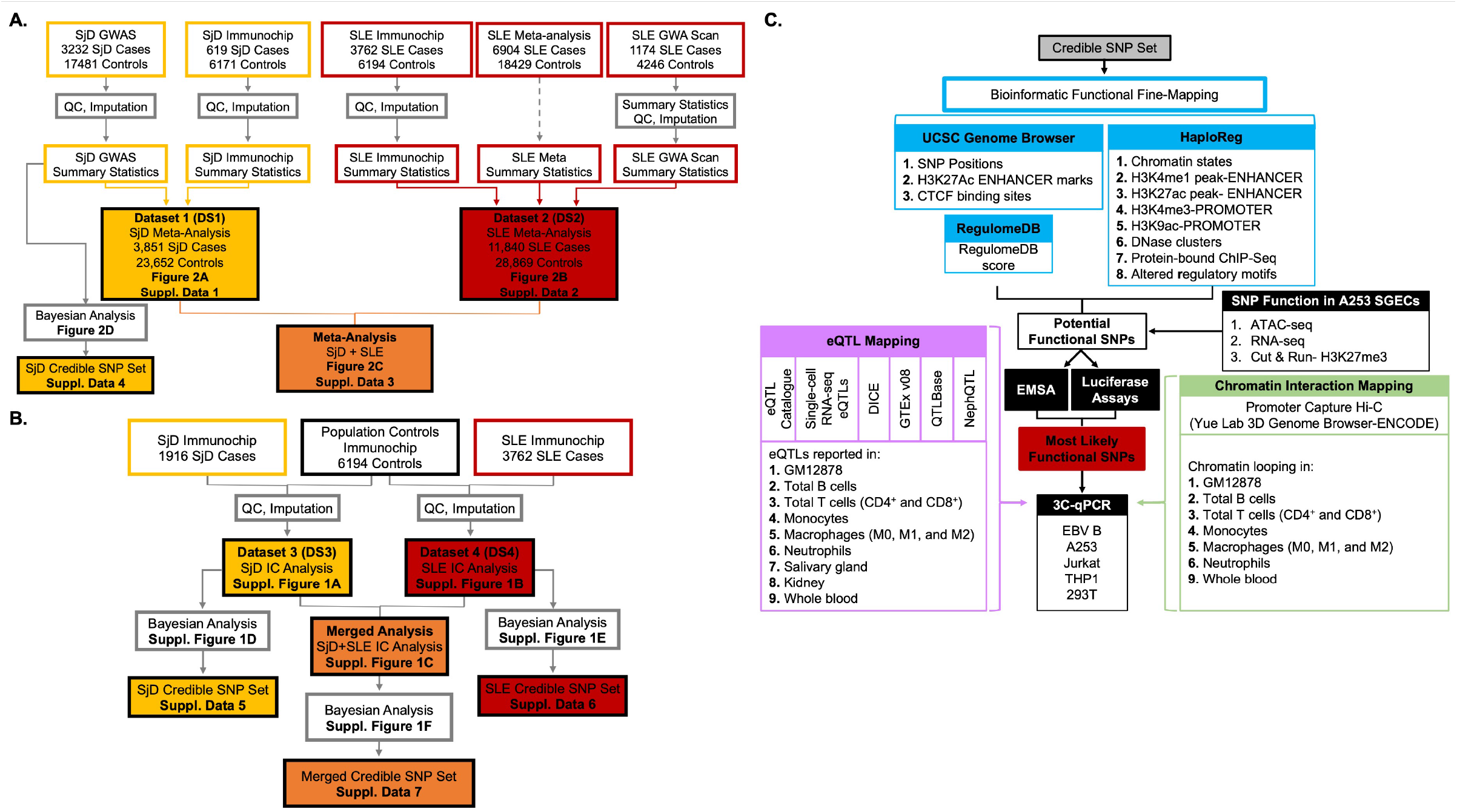
Schematic of the meta-analysis, fine mapping, and bioinformatic workflow of the shared *DDX6-CXCR5* interval associated with Sjögren’s disease (SjD) and systemic lupus erythematosus (SLE) **(A)** Composition and workflow of three datasets (DS) used in the meta-analysis of SjD (DS1; yellow), SLE (DS2; red), and SjD and SLE merged (DS1+DS2; orange). DS1 was also used to perform SNP-SjD single marker trait analysis and identify a SjD credible SNP set. **(B)** Composition and workflow of the SjD (DS3; yellow) and SLE (DS4; Red) Immunochip datasets used for SNP-single marker trait analysis and identification of credible SNP sets for SjD, SLE, and SjD and SLE merged. **(C)** Workflow of the bioinformatic fine mapping applied to the 95% credible SNP sets from the *DDX6-CXCR5* risk locus to identify, prioritize, and functionally characterize SNPs common between the SjD and SLE risk associations.

## RESULTS

### Meta-analysis and genetic fine mapping of the *DDX6-CXCR5* risk interval shows similar profiles in SjD and SLE

Meta-analysis was performed using genotype and Immunochip data from 3,851 SjD cases and 23,652 population controls of European ancestry (Dataset (DS1)) or Immunochip data and summary statistics from 11,840 SLE cases and 28,869 population controls of European or Korean ancestry (DS2), after quality control and imputation, to define the disease-specific associations on the *DDX6-CXCR5* region (**Figure 1A**)^1,20,28^. Similar association signals were observed, but different peak SNP-trait associations (i.e., index SNPs) were identified for each disease (**Figure 2A, B; Table 1; Supplemental Data 1, 2**). The index SNP, rs7481819, in SjD is positioned near the promoter region of *CXCR5* (**Figure 2A**), whereas the index SNP in SLE, rs11217045, is positioned in a shared regulatory region between *DDX6* and *CXCR5* (**Figure 2B**). The SNP, rs11217045, was also identified as the index SNP when the SjD and SLE datasets were meta-analyzed (**Figure 2C; Table 1; Supplemental Data 3**).

**Figure 2.**
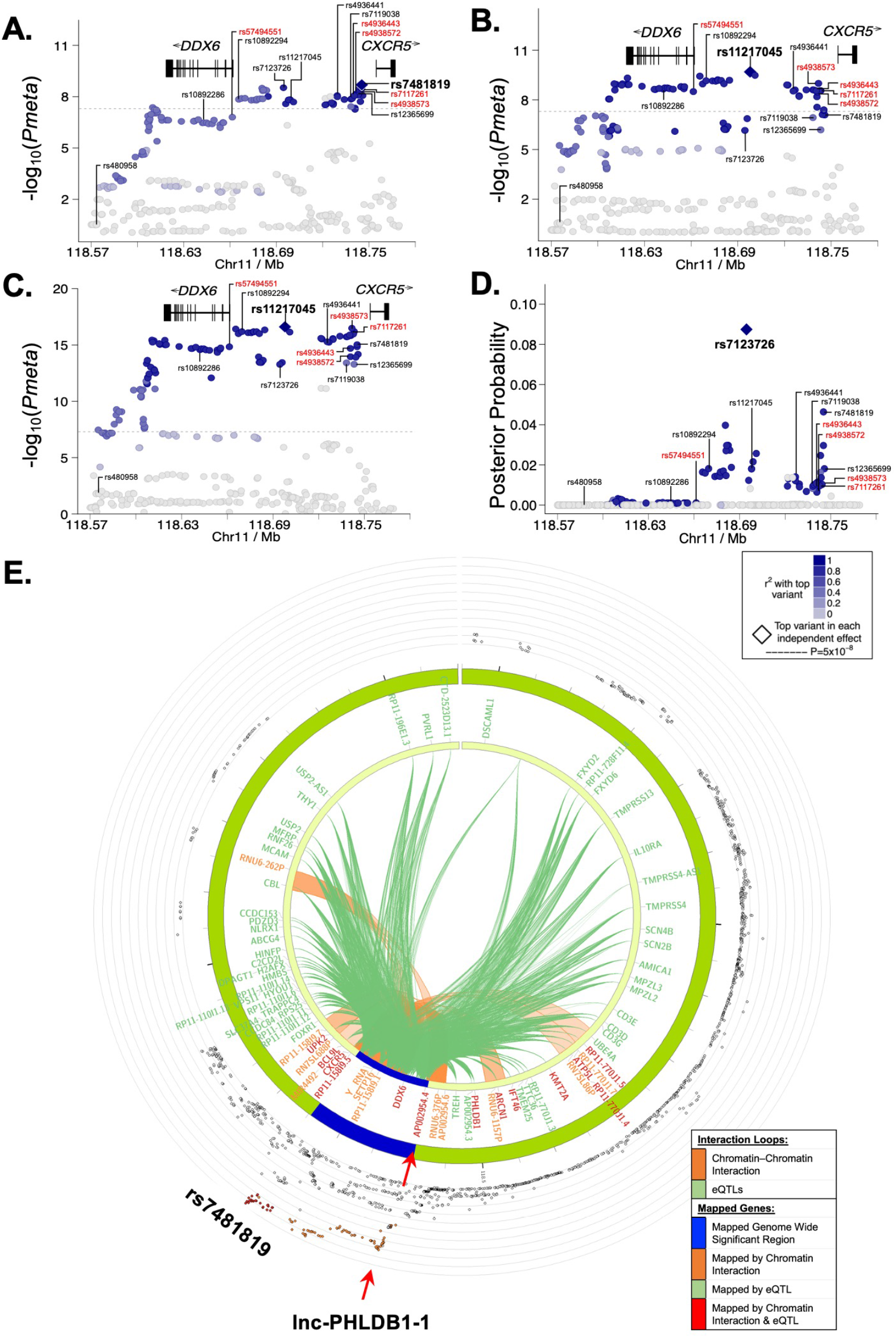
Fine mapping of the *DDX6-CXCR5* region in Sjögren’s disease (SjD) and systemic lupus erythematosus (SLE) after imputation and meta-analysis. **(A-C)** Logistic regression analysis was performed on **(A)** Dataset 1 (DS1)-SjD (3851 SjD cases; 23652 controls), **(B)** DS2-SLE (11840 SLE cases; 28869), and **(C)** DS1+DS2 (merged SjD and SLE) after quality control and imputation, identifying the top SNPs (e.g., index SNPs indicated in bold) of the *DDX6-CXCR5* region. **(D)** Posterior probability distribution of SNPs in the *DDX6-CXCR5* region of DS1-SjD, identifying rs7123726 as having the highest posterior probability. For **A-D,** SNPs prioritized for bioinformatic screening are indicated in black; five SNPs prioritized for functional characterization are labeled in red. **(E)** Circos plot showing chromatin-chromatin interactions in GM12878 EBV B cell line (orange) and reported eQTLs from different cells/tissues (green). The outer ring shows the logistic regression analysis of DS1-SjD with index SNP rs7481819 indicated.

**Table 1:** Summary Results of the Fine Mapping of the *DDX6-CXCR5* Risk Locus in SjD and SLE of European Ancestry.

|  |  |  |  |  |  |  |  |
| --- | --- | --- | --- | --- | --- | --- | --- |
| SNP rsID | Chr | Base pair | Minor | Major | SjD | SLE | SjD+SLE |
|  |  |  |  |  | Meta | Meta | Meta |
|  |  |  |  |  | P value | P value | P value |
| rs480958 | 11 | 118577990 | G | A | $8.11 \times 10^{-1}$ | $1.77 \times 10^{-1}$ | $4.19 \times 10^{-1}$ |
| rs10892286 | 11 | 118642085 | C | A | $3.39 \times 10^{-7}$ | $5.91 \times 10^{-4}$ | $2.93 \times 10^{-9}$ |
| rs57494551 | 11 | 118661398 | T | C | $1.57 \times 10^{-7}$ | $3.16 \times 10^{-9}$ | $1.45 \times 10^{-15}$ |
| rs10892294 | 11 | 118667357 | C | G | $1.26 \times 10^{-8}$ | $1.09 \times 10^{-9}$ | $9.87 \times 10^{-17}$ |
| rs7123726 | 11 | 118694297 | C | T | $2.98 \times 10^{-9}$ | $7.02 \times 10^{-7}$ | $5.31 \times 10^{-14}$ |
| rs11217045 | 11 | 118697865 | A | C | $1.54 \times 10^{-8}$ | $2.02 \times 10^{-10}$ | $2.42 \times 10^{-17}$ |
| rs4936441 | 11 | 118725660 | C | G | $2.23 \times 10^{-8}$ | $2.09 \times 10^{-1}$ | $7.42 \times 10^{-6}$ |
| rs7119038 | 11 | 118738281 | G | A | $3.77 \times 10^{-8}$ | $2.46 \times 10^{-2}$ | $8.54 \times 10^{-2}$ |
| rs4936443 | 11 | 118740864 | C | T | $4.57 \times 10^{-8}$ | $4.58 \times 10^{-2}$ | $5.79 \times 10^{-2}$ |
| rs4938572 | 11 | 118740931 | C | T | $5.48 \times 10^{-8}$ | $9.47 \times 10^{-2}$ | $3.33 \times 10^{-2}$ |
| rs7117261 | 11 | 118741157 | T | C | $8.01 \times 10^{-8}$ | $2.23 \times 10^{-1}$ | $3.75 \times 10^{-6}$ |
| rs4938573 | 11 | 118741842 | C | T | $6.69 \times 10^{-8}$ | $3.92 \times 10^{-2}$ | $3.50 \times 10^{-2}$ |
| rs12365699 | 11 | 118743286 | A | G | $7.01 \times 10^{-8}$ | $6.49 \times 10^{-7}$ | $5.17 \times 10^{-14}$ |
| rs7481819 | 11 | 118745278 | C | T | $2.02 \times 10^{-8}$ | $1.53 \times 10^{-1}$ | $7.47 \times 10^{-3}$ |

To further refine the *DDX6-CXCR5* association signals in SjD and SLE, Bayesian analyses were performed using Trinculo^29^, which leverages genotype data rather than summary statistics to calculate posterior probabilities and identify 95% credible sets of potential functional SNPs (**Figure 1A, B; Figure 2D; Supplemental Figure 1D-F; Supplemental Data 4-7**). Analysis of DS1-SjD, the largest publicly available SjD genotype dataset, identified a total of 66 SNPs in the 95% credible set with a maximum posterior probability of 0.087 (rs7123726) (**Figure 2D; Supplemental Data 4**). Use of SLE genomic summary results in the SLE meta-analysis (DS2) or merged meta-analysis (DS1+DS2) prohibited posterior probabilities calculations using Trinculo. Therefore, separate logistical regression analyses were performed using Immunochip data only (1916 SjD cases (DS3), 3,762 SLE cases (DS4), and 6,194 population controls of European ancestry), revealing similar association signals (**Figure 1B**). Analysis of DS3-SjD identified 105 SNPs in the 95% credible set with the top SNP in LD yielding a posterior probability of 0.034 (rs10790268) (**Supplemental Figure 1D; Supplemental Data 5**). Analysis of DS4-SjD identified 54 SNPs in the 95% credible set with a maximum posterior probability of 0.020 (rs76409436) (**Supplemental Figure 1E; Supplemental Data 6**). Analysis of the merged Immunochip datasets (DS3 and DS4) identified 15 SNPs in the 95% credible set and a maximum posterior probability of 0.059 (rs11217058) (**Supplemental Figure 1F; Supplemental Data 7**).

Fine mapping revealed that SjD and SLE of European or Korean ancestry exhibit similar association signals at the *DDX6-CXCR5* locus with no heterogeneity observed in the meta-analyses among the top variants (**Supplemental Data 1-3**). While there were several shared SNPs in the 95% credible set, the Bayesian analyses were unable to refine the association with sufficient posterior probability (>0.5) to predict the most likely functional SNPs. To further evaluate the co-inheritance of the index SNP and other possible functional variants within the region, the haplotype structure was examined. Co-inheritance in European ancestry (*D’* _≥_0.89) and LD (*r*^2^ =0.64-0.99) between several shared SNPs were observed (**Supplemental Figure 2A-C**) with the exception of rs480958 (*D’* ≤0.81; *r*^2^ ≤11), which was previously associated with SLE of East Asian ancestry (**Supplemental Figure 2D-F**)^22^. Strong co-inheritance of SNPs spanning the risk locus may explain why the Bayesian analyses were unable to sufficiently refine the association and predict likely functional SNPs.

### Bioinformatic interrogation identifies multiple SNPs with evidence of functionality on the *DDX6-CXCR5* interval

Bioinformatics data from multiple databases were interrogated to further refine the shared *DDX6-CXCR5* interval and identify which polymorphisms are most likely to have regulatory potential. From the top 100 SNPs in the 95% credible set from the merged Immunochip analysis (**Supplemental Data 7**), 14 SNPs exhibited bioinformatic evidence of functionally activity, and were as identified index SNPs in this study (**Table 1**) or were previously associated with SjD and/or SLE^20–22,27^ (**Supplemental Figure 3**).

Index SNPs, rs7481819, rs11217045, and rs7123726 (**Figure 2A-D**), as well as previously implicated SNPs rs10892286^20^ and rs7119038^21^, received RegulomeDB scores of 5, suggesting low functional potential (**Supplemental Figure 3; Supplemental Data 7**). HaploReg was used to assess epigenetic activity and, along with other eQTL databases (eQTL catalogue, single cell RNA-seq eQTLs, DICE, GTEx v08, QTLBase, and NephQTL), evaluate reported eQTLs in disease-relevant immune cell types, as well as salivary gland and kidney tissue. Consistently, minimal epigenetic activity or eQTLs (n<7) were reported for the index SNPs (**Supplemental Figure 3**). Previously reported SNPs, rs480958 (RegulomeDB Score: 4)^22^, rs10892294 (RegulomeDB Score: 3a)^21^, and rs4936441 (RegulomeDB Score: 3a)^22^, also had few reported eQTLs (n<8) and exhibited minimal epigenetic evidence of activity (n<56 total epigenetic promoter/enhancer peaks) (**Supplemental Figure 3; Supplemental Data 7**). Variant rs12365699 received a RegulomeDB score of 1f but only had 5 reported eQTLs and exhibited minimal evidence of epigenetic activity (n=80 epigenetic promoter/enhancer peaks) limited primarily to T cells (**Supplemental Figure 3**).

In contrast, five SNPs exhibited bioinformatic evidence of epigenetic regulation (including promoter and/or enhancer activity (n>243)), eQTL enrichment (n>36), and protein binding by ChIP-seq (n>2) across several immune cell types and tissues: rs57494551 (RegulomeDB Score: 2b) positioned in the first intron of *DDX6*, and rs4936443 (RegulomeDB Score: 4), rs4938572 (RegulomeDB Score: 1b), rs7117261 (RegulomeDB Score: 2b), and rs4938573 (RegulomeDB Score: 4) clustered in an 978 bp shared regulatory region approximately 79 kb downstream of *DDX6* and 12.5 kb upstream of *CXCR5* (**Figure 3A; Supplemental Figure 3; Supplemental Data 7**). Our comprehensive bioinformatic interrogation revealed additional eQTLs (n>36) that were not reported in RegulomeDB for four of the five SNPs (**Supplemental Figure 3**), suggesting that rs57494551, rs4936443, rs7117261, and rs4938573 should have amended RegulomeDB scores of 1c or better.

**Figure 3.**
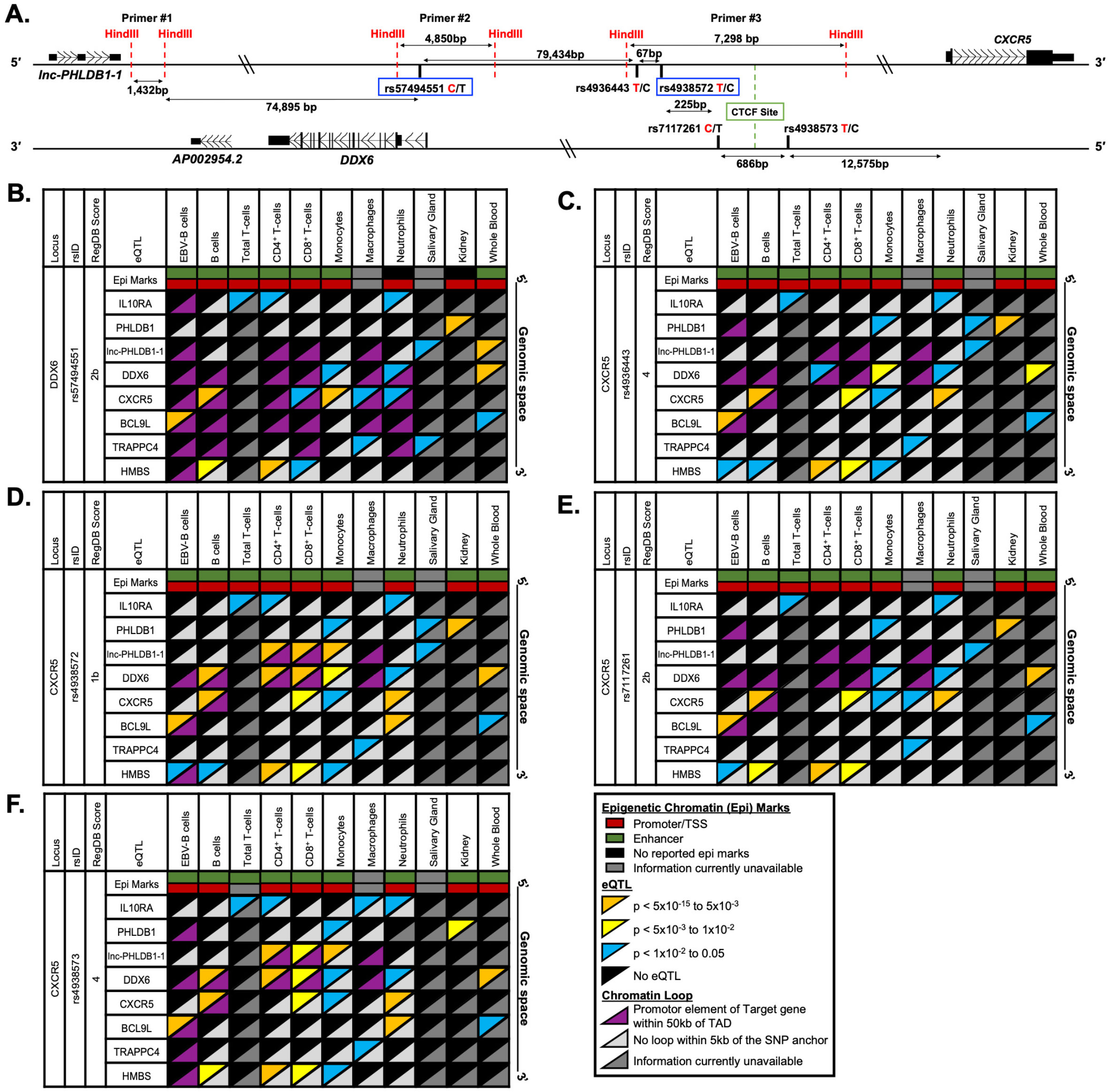
Reported expression quantitative trait loci (eQTLs) and chromatin-chromatin interactions in immune and disease-relevant tissues for the prioritized SNPs on the *DDX6-CXCR5* risk interval. **(A)** Schematic of the five prioritized SNPs arranged in genomic space. SNP rs57494551 is in the first intron of *DDX6*. The other four SNPs span 978 bp in a shared promoter/enhancer region between *DDX6* and *CXCR5*. Risk alleles are indicated by red font; non-risk alleles by black. SNPs rs57494551 and rs4938572 (blue boxes) are representative SNPs. *HindIII* restriction sites (red dotted line), CTCF site (green box), and positions of 3C-qPCR primers #1-3 are also indicated. **(B-F)** Publicly reported cell type-specific functional annotations (horizontal rectangles), select eQTLs (top triangles), and chromatin-chromatin interactions (bottom triangles) are shown for **(B)** rs57494551, **(C)** rs4936443, **(D)** rs4938572, **(E)** rs7117261, and **(F)** rs4938573 across 10 different immune cell types or disease-specific tissues (GM12878 EBV B cells and primary human B cells, CD4+ T cells, CD8+ T cells, monocytes, macrophages, neutrophils, salivary gland tissue, kidney tissue, and whole blood).

Analysis of previously reported chromatin-chromatin interactions from the GM12878 EBV B cell line and eQTLs from multiple different immune cell types revealed a complex regulatory network spanning the *DDX6-CXCR5* SjD/SLE risk locus (**Figure 2E**). Coalescence of reported eQTLs and publicly available promoter-capture Hi-C (hereafter simplified as Hi-C) data further revealed that these five SNPs are positioned in regulatory elements that may modulate gene expression beyond that of *DDX6* and *CXCR5* (**Figure 3B-F**). For example, rs57494551 is an eQTL for several transcripts in immune cells, including *DDX6* in monocytes and neutrophils, *CXCR5* in B cells, CD8^+^ T cells, monocytes, macrophages, and neutrophils, *IL10RA* in T cells and neutrophils, *BCL9L* (B cell CLL/Lymphoma 9) in B cells and whole blood, and *TRAPPC4* (transport protein particle complex subunit 4) in macrophages and minor salivary gland (**Figure 3B**). SNP rs57494551 is positioned in an intronic regulatory element that forms chromatin-chromatin interactions with the promoters of *DDX6, CXCR5,* and *BCL9L*, but not *IL10RA*, in an immune cell type and/or context specific manner, suggesting that rs57494551 may exhibit regulatory effects on *DDX6*, *CXCR5*, and *BCL9L* through modulating interactions between the intronic enhancer and the promoters of these genes. Unfortunately, publicly available Hi-C data remains unavailable for the minor salivary gland, therefore any potential coalescence between the reported *TRAPPC4* eQTL and a chromatin-chromatin interaction with the rs57494551 regulatory element could not be inferred.

Coalescence of eQTLs (*DDX6*, *CXCR5*, *IL10RA*, *TRAPPC4*) and Hi-C data were also observed for the four SNPs in the shared regulatory region between *DDX6* and *CXCR5* (**Figure 3C-F**). Interestingly, the four SNPs are also reported eQTLs for a long noncoding RNA, *lnc-PHLDB1-1*, in several immune cell types and/or the minor salivary gland (**Figure 3C-F**). Further, Hi-C data indicate cell type-specific chromatin-chromatin interactions between the regulatory region carrying rs4938572 and rs4938573 and the promoter of *lnc-PHLDB1-1* (**Figure 3D, F**). Collectively, the bioinformatic evaluation of shared polymorphisms associated with the *DDX6-CXCR5* SjD/SLE risk locus identified five SNPs with functional potential warranting further characterization.

### Prioritized SNPs demonstrate allele-specific nuclear protein complex binding

EMSAs were performed in EBV B cells, Daudi B cells, Jurkat T cells, THP1 monocytes, and/or A253 cell line originating from a submandibular gland squamous cell carcinoma to assess the cell type- and allelic effects of the five prioritized SNPs on nuclear protein binding (**Supplemental Table 1)**. Binding specificity was confirmed by competition assay with unlabeled probes. All five SNPs demonstrated varying allele- and cell type-specific nuclear protein binding (**Figure 4; Supplemental Figure 4-9**). Banding patterns varied by SNP and cell type, requiring separate analyses of multiple bands.

**Figure 4.**
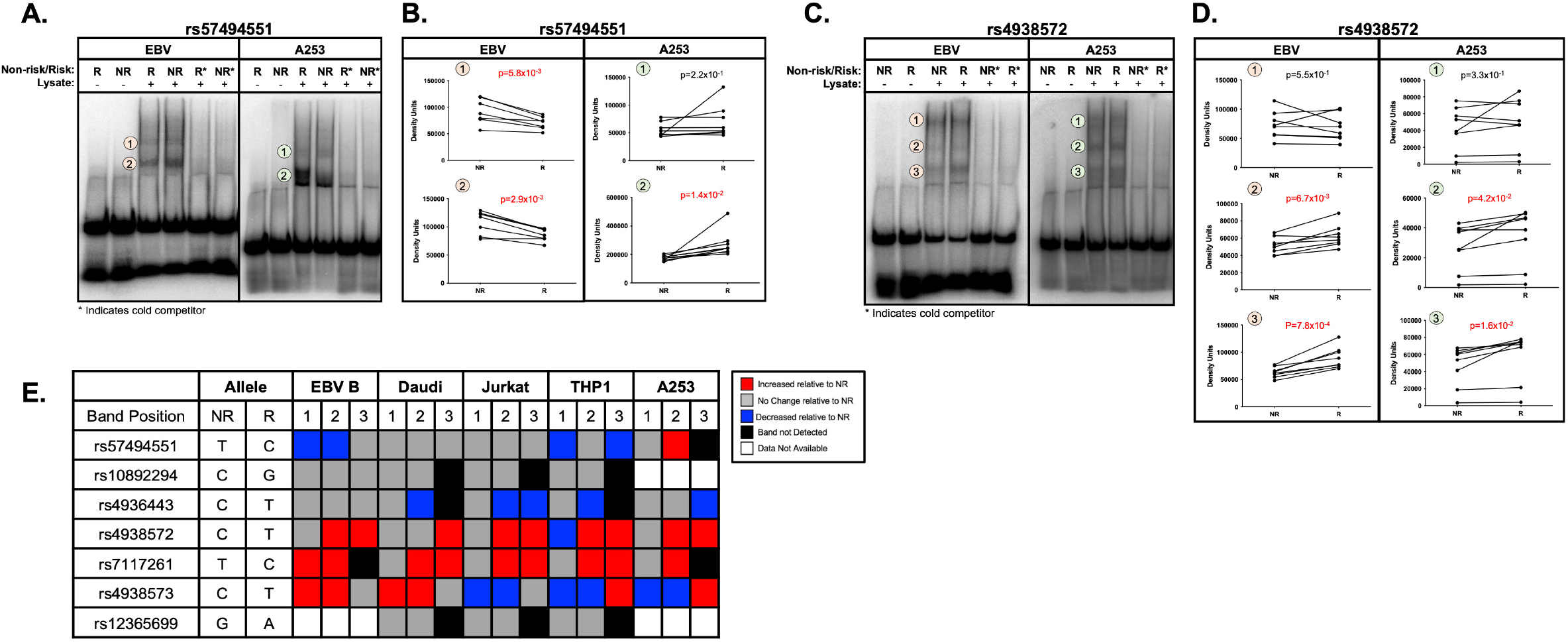
Allele- and cell-type specific differential nuclear protein affinities of the prioritized SNPs rs57494551 and rs4938572 on the shared *DDX6-CXCR5* risk region. **(A-D)** Radiolabeled electromobility shift assays (EMSA) were performed to assess the binding affinity of ribonucleoproteins isolated from EBV B or A253 cells to oligonucleotides containing the non-risk (NR) or risk (R) allele of **(A-B)** rs57494551 or **(C-D)** rs4938572. Probes incubated in the absence of nuclear lysate were used as negative control (Lanes 1, 2). Cold competitors were used to assess non-specific binding (Lanes 5, 6). Images shown in (A) and (C) are representative of n>6 biological replicates. **(B, D)** Bands indicated in (A, C) by the orange or green circles were quantified by densitometry and analyzed using paired t-test (n>6); p-values indicated. **(E)** Summary analysis of the allele-specific nuclear protein affinities of the five prioritized SNPs in EBV B, Daudi, Jurkat, THP1, and A253 cells shown in A-D and **Supplemental Figures 4-9**. Increases in binding relative to NR are shown in red; decreases relative to NR in blue; no change relative to NR in grey; no detected band in black; data not available in white.

The risk allele of rs57494551 impaired nuclear protein binding compared to the non-risk allele in EBV B cells (*P_EMSA_*<5.8×10^−3^) and THP1 monocytes (*P_EMSA_*<3.9×10^−2^), but increased binding in A253 cells (*P_EMSA_*=1.4×10^−2^) (**Figure 4A-B, 4E; Supplemental Figure 4A-B**). The risk alleles of rs4938572 (*P_EMSA_*<4.2×10^−2^) (**Figure 4C-E; Supplemental Figure 4C-D**) and rs7117261 (*P_EMSA_*<2.0×10^−3^) (**Figure 4E; Supplemental Figure 5**) increased protein binding compared to the non-risk allele in all tested immune cell lines and A253 cells. The risk allele of rs4936443 decreased nuclear protein binding compared to the non-risk allele in Daudi B cells, Jurkat T cells, THP1 monocytes, and A253 cells (*P_EMSA_*<3.9×10^−2^), but did not change binding in EBV B cells (**Figure 4E; Supplemental Figure 6**). The risk allele of rs4938573 increased protein binding in both EBV B and Daudi B cell lines (*P_EMSA_*<2.5×10^−2^) but decreased binding in the Jurkat T cells (*P_EMSA_*<3.7×10^−3^) (**Figure 4E; Supplemental Figure 7**). Interestingly, the risk allele of rs4938573 exhibited increased or decreased nuclear protein binding compared to the non-risk allele in the THP1 monocytes and A253 cells, respectively, depending on the band analyzed (**Figure 4E; Supplemental Figure 7**).

Despite cell type-specific epigenetic evidence of function in T cells, rs12365699 exhibited no change in allele-specific binding to nuclear proteins from Jurkat T cells, Daudi B, or THP1 monocytes (**Figure 4E; Supplemental Figure 8**). SNP rs10892294 also exhibited no change in allele-specific binding to nuclear proteins from Daudi B, Jurkat T, or THP1 monocytes (**Figure 4E; Supplemental Figure 9**). These results further suggest that the five prioritized SNPs are likely functional in multiple immune cell types and/or A253 cells. Further, allele-specific nuclear protein binding to rs12365699 or rs10892294 showed limited epigenetic evidence of functionality.

### Prioritized SNPs demonstrate allele-specific effects on regulatory activity

To determine if the regulatory elements carrying the prioritized SNPs exhibit allele-specific and/or cell type-specific promoter or enhancer activity, luciferase expression assays were performed using a promoter-less vector or minimal-promoter vector, respectively. The regulatory activity and allelic effects were first screened using 293T human embryonic kidney cells, which are a model for high transfection efficiency and optimal luciferase expression. Regulatory elements carrying rs57494551, rs4936443, rs4938572, or rs7117261 exhibited significantly increased enhancer and/or promoter activity compared to the empty vector control in 293T cells (**Figure 5A-D,F**). In contrast, rs10892294 and rs12365699 exhibited minimal enhancer or promoter activity above that of the empty vector in 293T cells and, therefore, were excluded from additional study (**Supplemental Figure 10**). Polymorphism rs4938573 also exhibited minimal enhancer or promoter activity in 293T cells (**Figure 5E**). However, rs4938573 exhibited strong bioinformatic evidence of function and allele-specific nuclear protein binding in EMSAs from multiple immune cell types. Further, the SjD/SLE risk allele of rs4938573 was previously reported to modulate *CXCR5* expression and CXCR5^+^ cell homing to the salivary gland^27^. For these reasons, rs4938573 was selected, along with rs57494551, rs4936443, rs4938572, and rs7117261 for further evaluation.

**Figure 5.**
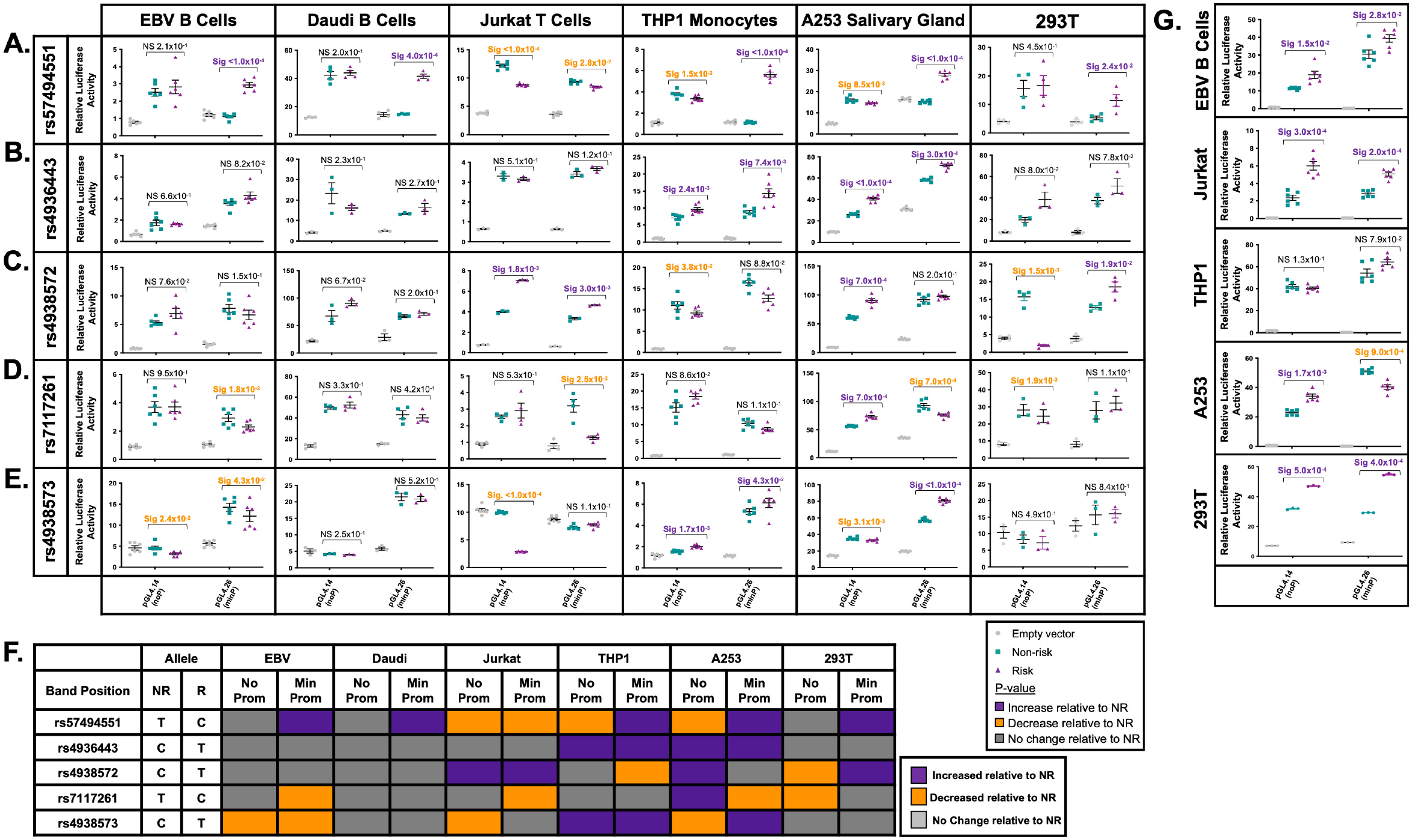
Allele- and cell-type specific promoter and enhancer activity of the prioritized SNPs on the shared *DDX6-CXCR5* risk region. **(A-E)** gBlocks carrying the non-risk or risk alleles of **(A)** rs57494551, **(B)** rs4936443, **(C)** rs4938572, **(D)** rs7117261, or **(E)** rs4938573 were cloned into a promoter-less (pGL4.14; noP) or minimal promoter (pGL4.26; minP) luciferase vector. Plasmids were transfected into EBV B, Daudi, Jurkat, THP1, A253, or 293T cells. Luciferase activity was measured after 24 hours and normalized to the Renilla transfection control and then the vector-only control; reported as Relative Luciferase Activity. Statistical comparisons were performed using a paired t-test (n>3); p-values are indicated. **(F)** Summary analysis of the allele-specific luciferase activity of the five prioritized SNPs in EBV B, Daudi, Jurkat, THP1, A253 and 293Tcells. Increases in luciferase activity relative to non-risk are shown in purple; decreases relative to non-risk in orange; no change relative to non-risk in grey. **(G)** gBlocks carrying all non-risk or all risk alleles of rs4936443, rs4938572, and rs7117261 were cloned into the promoter-less or minimal promoter above, transfected into EBV B, Jurkat, THP1, A253, or 293T cells, and luciferase activity tested as described above. Statistical comparisons were performed using a paired t-test (n>3); p-values are indicated.

Consistent with decreased nuclear protein binding in EBV B and THP1 cells, the risk allele of rs57494551 significantly decreased promoter activity compared to the non-risk allele in Jurkat (*P_luc_*<1.0×10^−4^), THP1 (*P_luc_*=1.5×10^−2^), and A253 cells (*P_luc_*=8.5×10^−3^) (**Figure 5A**). However, the risk allele of rs57494551 also significantly increased enhancer activity compared to the non-risk allele in both B cell lines (EBV B (*P_luc_*<1.0×10^−4^) and Daudi (*P_luc_*=4.0×10^−4^)), THP1 (*P_luc_*<1.0×10^−4^), A253 (*P_luc_*<1.0×10^−4^), and 293T (P_luc_=2.4×10^−2^) cells. The risk allele of rs4936443 increased promoter and enhancer activity in THP1 (*P_luc_*<2.4×10^−3^) and A253 cells (*P_luc_*<3.0×10^−4^) (**Figure 5B**). No allele specificity was observed in the EBV B, Daudi, Jurkat, or 293T cells (**Figure 5B**). The risk allele of rs4938572 increased promoter (*P_luc_*=1.8×10^−3^) and enhancer (*P_luc_*=3.0×10^−3^) activity in Jurkat cells, increased promoter activity in A253 cells (*P_luc_*=7.0×10^−4^), and increased enhancer activity in 293T cells (*P_luc_*=1.9×10^−2^), but decreased promoter activity in THP1 and 293T cells (*P_luc_*<4.0×10^−2^) (**Figure 5C**).

The risk allele of rs7117261 increased promoter activity (*P_luc_*=7.0×10^−4^) but decreased enhancer activity (*P_luc_*=7.0×10^−4^) in A253 cells, decreased promoter activity in 293T cells (*P_luc_*=1.9×10^−2^), and decreased enhancer activity in EBV B (*P_luc_*=1.8×10^−2^) and Jurkat cells (*P_luc_*=2.5×10^−2^) (**Figure 5D**). The risk allele of rs4938573 significantly decreased promoter activity in EBV B (*P_luc_*=2.4×10^−2^), Jurkat (*P_luc_*<1.0×10^−4^), and A253 (*P_luc_*=3.1×10^−3^) cells, but increased promoter activity in THP1 cells (*P_luc_*<1.7×10^−3^) (**Figure 5E**). Additionally, the rs4938573 risk allele decreased enhancer activity in EBV B cells (*P_luc_*=4.3×10^−2^), but increased enhancer activity in THP1 (*P_luc_*=4.3×10^−2^) and A253 (*P_luc_*<1.0×10^−4^) cells (**Figure 5E**). In summary, all five prioritized SNPs exhibited cell type- and allele-specific effects on promoter and/or enhancer activity with the A253 cells demonstrating the most allele-specific differences among all tested cell types; THP1 cells demonstrated the most allele-specific effects among tested immune cell lines (**Figure 5F**).

Three of the four SNPs in the regulatory region between *DDX6* and *CXCR5* (rs4936443, rs4938572, rs7117261) span a 292 bp region positioned 5’ of a CTCF binding site that separates the SNP cluster from rs4938573 (**Figure 3A**). Further, the three SNPs have similar epigenetic enrichment, reported eQTLs, and chromatin-chromatin interactions, suggesting that they likely modulate the same regulatory element. To test whether the SNPs facilitate concordant or discordant allele-specific effects on regulatory activity, the non-risk or risk alleles of the three SNPs were cloned together on the promoter-less or minimal-promoter luciferase vector. When cloned together, the risk alleles of rs4936443, rs4938572, and rs7117261 significantly increased promoter (*P_luc_*<1.5×10^−2^) and enhancer activity (*P_luc_*<2.8×10^−2^) in EBV B, Jurkat, and 293T cells relative to the non-risk alleles (**Figure 5G**). In A253 cells, the three risk alleles also significantly increased promoter activity (*P_luc_*=1.7.0×10^−3^), but impaired enhancer activity (*P_luc_*=9.0×10^−4^) (**Figure 5G**). In contrast to the allele-specific activities observed when the SNPs were separated, THP1 cells did not exhibit allele-specific differences in promoter or enhancer activity when the alleles were cloned together (**Figure 5G**).

Collectively, these findings suggest that the three clustered SNPs likely function together to modulate the haplotype- and cell type-specific regulatory activity at this locus. Further, in A253 cells, a model of salivary gland epithelial cell function, the risk alleles of rs4936443, rs4938572, and rs7117261 may suppress enhancer activity and, therefore, expression of genes that interact within the chromatin regulatory network. The opposing allelic- and cell type-specific effects observed between rs4938573 and the three SNP cluster suggest that the ∼978 bp region between *DDX6* and *CXCR5* likely has more than one regulatory element, separated by a CTCF site, that are influenced by the *DDX6-CXCR5* SjD/SLE risk haplotype.

### Chromosome architecture is similar between immune cells and salivary gland at the *DDX6-CXCR5* risk locus

Coalescence of reported Hi-C chromatin-chromatin interactions and eQTLs suggest that the five prioritized SNPs are positioned in regulatory elements that likely modulate the expression of *DDX6*, *CXCR5*, and/or other genes within the chromatin regulatory network. In further support, the SNPs are positioned in areas of open chromatin (ATAC-seq peaks), elevated promoter and/or enhancer activity (H3K4me1, H3K4me3, H3K27ac peaks), and active transcription factor binding in B cells, T cells, monocytes, macrophages, and neutrophils (**Supplemental Figures 11-16**), resulting in high IMPACT scores (**Supplemental Figure 17**). Lastly, the cell type- and allele-specific effects observed in the EMSAs and luciferase assays indicate that the risk alleles of these five variants likely influence the regulatory activity across the interval to modulate gene expression in the context of SjD and/or SLE.

To test this and address the limited availability of Hi-C data for the tested cell lines used in the functional assays, 3C-qPCR was performed in four patient-derived EBV B cell lines (two homozygous for the *DDX6-CXCR5* non-risk or risk haplotype; **Supplemental Table 2**), A253, Jurkat, THP1 and 293T cell lines using rs574994551 or rs4938572 to tag the anchor fragments determined by HindIII digestion (**Figure 3A, 6A-B**). 3C-qPCR results were also plotted in the context of cell type-specific RNA-seq, ATAC-seq, epigenetic marks, and/or Hi-C data (**Figure 6C-D; Supplemental Figures 11-16**). Publicly available epigenetic and genomic data on salivary gland tissues are limited and Hi-C or long-range chromatin interaction data are nonexistent. To provide additional insights in the salivary gland, 3C-qPCR results from the A253 cells were plotted in the context of in-house CUT & RUN (H3K27me3 pull-down primer), ATAC-seq, and RNA-seq data (**Figure 6D**).

**Figure 6.**
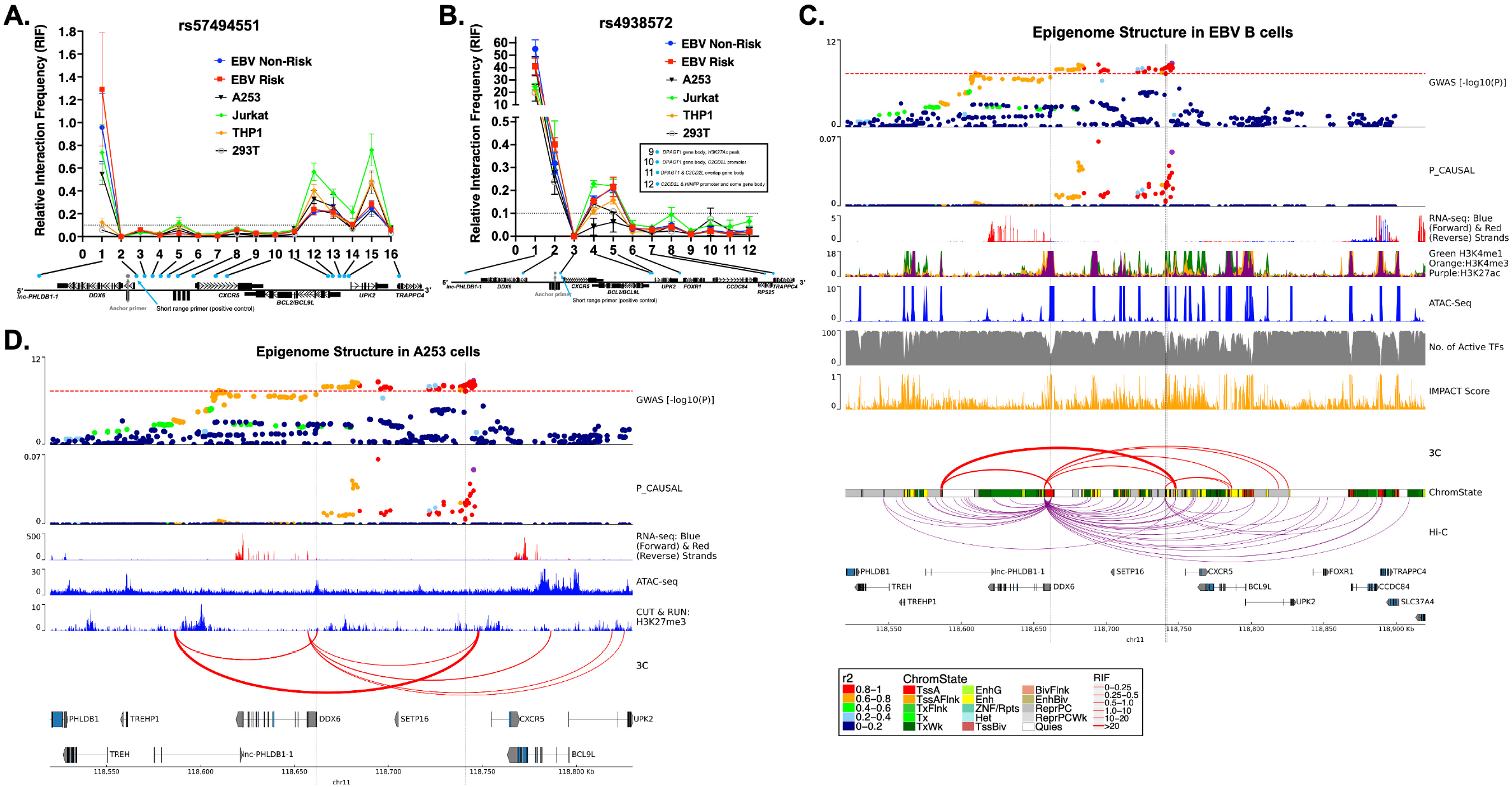
Complex chromatin architecture revealed across the *DDX6-CXCR5* region in immune cells, salivary gland, and kidney by 3C-qPCR. **(A, B)** Chromatin conformation capture with quantitative PCR (3C-qPCR) across the *DDX6-CXCR5* region where **(A)** rs57494551 or **(B)** rs4938572 is the anchor SNP (grey dot). Relative interaction frequency (RIF) is plotted relative to the primer number in 5’-3’ genomic orientation (blue dots; see **Figure 3A** for additional detail). Primers 9-12 are shown in a text box for simplicity in (B). **(C)** SjD GWAS association (top panel) and publicly available epigenomic enrichment across the *DDX6-CXCR5* region in GM12878 EBV B cells. Vertical grey lines indicate the locations of rs57494551 or rs4938572, respectively. Promoter-capture Hi-C looping (purple lines) contrasts the summary 3C-qPCR results (red lines); line thickness indicates relative interaction frequency (RIF) of the 3C data. **(D)** SjD GWAS association (top panel) across the *DDX6-CXCR5* region in A253 cells. In house ATAC-seq, CUT & RUN: H3K27me3 (Epicypher), and RNA-seq data from A253 cells are shown because of limited publicly available epigenetic data on salivary gland. Vertical grey lines indicate the locations of rs57494551 or rs4938572, respectively. Summary 3C-qPCR results are shown (red lines); line thickness indicates relative interaction frequency (RIF).

3C-qPCR designed with the anchor spanning rs57494551 revealed the highest relative interaction frequency (RIF) with the upstream promoter region of the long non-coding RNA, *lnc-PHLDB1-1*, in all tested cell types except THP1 and 293T (**Figure 6A**). Hi-C data also revealed this interaction in all examined cell types except B cells, M2 macrophage, neutrophils, and 293T cells (**Figure 6C; Supplemental Figure 11-16**). Consistent with the allele-specific increase in luciferase activity across several cell types (**Figure 5A**), the scaled effect size of reported eQTLs also indicate that the rs57494551 risk allele increases *lnc-PHLDB1-1* expression in the minor salivary gland and whole blood (**Supplemental Figure 18A**).

The rs57494551 regulatory region also interacted with the shared promoter region of *BCL9L* and *UPK2*, albeit with lower RIF, across all cell types (**Figure 6A**). Hi-C consistently showed this interaction in other immune cell types (**Figure 6C; Supplemental Figure 11-16**), as well as an additional cell type-specific (EBV B, B, and CD8^+^ T cells) interaction with the promoter of *TRAPPC4* further downstream (**Figure 6C; Supplemental Figure 11, 12B**). An interaction with *TRAPPC4* above the RIF of the short-range control was not observed in the 3C-qPCR of EBV B cells (**Figure 6A,B**). Interestingly, the scaled effect sizes of the *TRAPPC4* eQTL (minor salivary gland) and *BCL9L* eQTL (blood and EBV B cells) suggest that the rs57494551 risk allele may exhibit cell type-specific suppression of the long-range enhancer effect of this region (**Supplemental Figure 18A**). However, a discordant observation of increased luciferase enhancer activity in the context of the rs57494551 risk allele (**Figure 5A**) is indicative of a more complex regulatory mechanism that is likely allele-specific, cell type-specific, and modulated in the context of the chromatin regulatory network.

3C-qPCR designed with the anchor spanning the SNP cluster (tagged by rs4938572) also revealed the highest RIF with the promoter region of *lnc-PHLDB1-1*, in all tested cell types (**Figure 6B**) and Hi-C data from T cells (**Supplemental Figure 12**). Consistent with impaired luciferase enhancer activity in A253 cells in the context of the combined risk alleles (**Figure 5G**), the scaled effect size of *lnc-PHLDB1-1* was reduced in minor salivary gland in the context of the rs4936443, rs4938572, and rs7117261 risk alleles (**Supplemental Figure 18B-D**). The risk alleles of rs4938572 and 4938573 also exhibited decreased effect size for *lnc-PHLDB1-1* in CD4^+^ and CD8^+^ T cells and natural killer cells but increased scaled effect size in peripheral CD14^+^ monocytes (**Supplemental Figures 18C, E**). Consistently, increased luciferase enhancer activity in THP1 cells, but decreased activity in Jurkat cells, was observed in the context of rs4938573 alone (**Figures 5E**). Allele-specific enhancer activity was not observed in THP1 cells in the context of rs4938572 alone or in combination with rs4936443 and rs7117261 (**Figure 5E, G**). Given that the rs4938573 is separated from the three SNP cluster by a CTCF site, the discordant observations in observed luciferase enhancer activity and eQTL effect sizes may indicate that rs4938573 modulates the activity of a separate enhancer that forms independent interactions with the *lnc-PHLDB1-1* promoter.

Chromatin-chromatin interactions were also observed, albeit with lower RIF, between the regulatory region tagged by rs4938572 and the shared promoter region of *BCL9L* and *UPK2* in all tested cell types (**Figure 6B**) and in Hi-C data from EBV B, CD4^+^ and CD8^+^ T cells (**Figure 6C; Supplemental Figure 12**). The risk alleles of all four SNPs in this region are associated with increased expression of *BCL9L* in blood, EBV B cells, and CD16^+^ neutrophils (rs4938572 and rs4938573 only) (**Supplemental Figure 18B-E**). Interestingly, Hi-C data from B and T cells, macrophage, and neutrophils demonstrated interactions between this regulatory region and the rs57494551 regulatory region, but not the *BCL9L*-*UPK2* promoter (**Supplemental Figure 11-12, 14-15**). The risk alleles of rs4938572, rs4936443, and/or rs4938573 all exhibited reduced scaled effect size for *DDX6* in PBMCs (rs4938572, rs4936443 only), B cells (rs4938572 only), T cell, and monocyte populations (**Supplemental Figure 18B,C,E**). This finding is discordant with the observed increase in luciferase enhancer activity in EBV B and Jurkat cells in the context of three risk alleles together (rs4936443, rs4938572, and rs7117261) and rs4938573 alone (**Figure 5E, G**). Discordance between the observed allele-specific effects in the luciferase assays and the directionality of reported eQTLs may result from the potential regulatory effects of the native chromatin architecture that cannot be recapitulated in the luciferase assay, the influence of additional SNPs that are carried on *DDX6-CXCR5* risk haplotype, and additional regulation by non-coding RNAs, among others.

## DISCUSSION

This study performed fine mapping across two unique but related autoimmune diseases, SjD and SLE, to identify the most likely functional SNPs in the shared *DDX6-CXCR5* risk locus, hypothesizing that the identification and functional characterization of common SNPs may identify common disease mechanisms. Although the statistical genetic approach revealed a similar genetic association profile across the two diseases, strong co-inheritance and LD across the interval rendered it unable to identify SNPs with likely functional potential. As an alternative approach, this study leveraged publicly available datasets and databases to perform bioinformatic fine mapping of the top 100 common SNPs, identifying SNPs with cumulative evidence of transcription factor binding enrichment, epigenetic marks of regulatory activity, and reported eQTLs and their position in the chromatin regulatory network. This approach successfully prioritized five SNPs that exhibited both allele- and cell type-specific regulatory activity in subsequent functional assays and predicted the observed minimal functional activity of several additional SNPs, including the index SNPs, providing strong justification for using bioinformatic fine mapping to discern likely functional SNPs from other co-inherited non-functional SNPs carried on a disease risk haplotype. Although the statistical and bioinformatic fine mapping examined an extensive credible SNP set, as well as SNPs previously implicated in either SjD or SLE, the *DDX6-CXCR5* risk interval likely has additional functional SNPs not examined in this study.

Noncoding functional SNPs influence disease mechanisms by altering the expression of key regulatory genes^30–34^. Risk alleles can modulate transcription factor binding, activity of promoters and/or regulatory elements, and organization of the chromatin regulatory network resulting in altered long-range interactions between promoters and distant regulatory elements. Deep bioinformatic interrogation of the five selected SNPs identified three potential regulatory elements on the *DDX6-CXCR5* locus with evidence of allele- and cell type-specific promoter and/or enhancer activity that likely modulate expression of *DDX6* and *CXCR5*, as well as other genes spanning the interval through local and/or distant regulatory activity. Many of these transcripts have functional implications in established autoimmune disease mechanisms, including *DDX6*, *CXCR5*, and *BCL9L*^30,32,34^. Others, such as *lnc-PHLDB1-1*, have not been previously reported in SjD or SLE.

DDX6 is a central regulator of the viral RNA recognition pathway of the innate immune system and, as such, an important modulator of type I interferon responses^23–25,35^. Genetic ablation of *DDX6* in human cells induced global upregulation of type I interferon-stimulated genes (ISGs)^24^. Chronic upregulation of ISG expression, i.e., the interferon signature, is a hallmark of SjD and SLE^7,24,36–42^. *DDX6* is a reported eQTL of rs4936443, rs4938572, and rs4938573, where the risk alleles exhibited strong negative relative eQTL effect sizes in PBMCs and isolated B and T cell populations (**Supplemental Figure 18B-C,E**). Further, an interaction between the regulatory region carrying rs4936443, rs4938572, and rs4938573 and a region spanning part of the *DDX6* promoter was observed in the immune and A253 cell lines by 3C-qPCR (**Figure 6B**) and primary human B cells and CD4^+^ and CD8^+^ T cells by Hi-C (**Supplemental Figure 11-12**). Collectively these data suggest that the expression of *DDX6* may be regulated through a long-range interaction with the three SNP cluster (tagged by rs4938572) and/or rs4938573 enhancer regions.

The three SNP cluster enhancer and the rs4938573 enhancer are separated by a CTCF site and exhibit opposing allele-specific nuclear binding and luciferase activity across several of the cell lines tested but, due to the location of the HindIII sites used in the 3C-qPCR and Hi-C, are positioned in the same loop anchor. Consistent with the reported eQTL effect sizes, the risk allele of rs4938573 significantly impaired promoter and enhancer luciferase activity, as well as nuclear protein binding, in the EBV B and Jurkat T cells (**Figure 4,5**). Despite being a reported eQTL of *DDX6*, the risk allele of rs4938572 increased luciferase activity and nuclear protein binding across several cell types when examined separately and as part of the three SNP cluster (**Figure 4,5**). Since eQTLs are examined in the context of LD and co-inheritance, it is possible that the discordance between the functional studies and the reported eQTL effect sizes for the three SNP cluster are due to the co-inheritance between the three SNPs and rs4938573. In summary, these findings suggest that the rs4938573 risk allele may modulate a long-range interaction between the rs4938573 enhancer and the *DDX6* promoter to impair *DDX6* expression in the context of the *DDX6-CXCR5* SjD/SLE risk haplotype.

CXCR5 is expressed on the surface of B and T cell subsets. CXCR5-CXCL13 signaling activates the recruitment and infiltration of immune cells to lymphoid germinal centers and the germinal center-like structures that form in the salivary gland of SjD patients and kidneys of SLE patients^26,27,43^. SjD patients carrying the *DDX6-CXCR5* risk haplotype (tagged by the risk allele (T) of rs4938573) reportedly exhibit increased infiltration of CXCR5^+^ immune cells in the salivary gland, but lower *CXCR5* expression in circulating CXCR5^+^ cells^27^. Aqwari, et al.^27^ hypothesized that impaired *CXCR5* expression in CD19^+^ B cells may initially delay activation of the CXCR5-CXCR13 signaling axis in genetically predisposed individuals, resulting in early glandular destruction. CXCR5 was a reported eQTL for the SNPs in the three SNP cluster and rs4938573, where the risk alleles exhibited strong negative relative eQTL effect sizes in B cells, but positive effect sizes in neutrophils (**Supplemental Figure 18B-E**). CXCR5 was also a reported eQTL for rs57494551, where the risk allele exhibited increased relative expression in macrophage and neutrophils. Since rs4938573 is positioned ∼12.5 kb upstream of the CXCR5 promoter and exhibited allele-specific loss of promoter and enhancer activity in the EBV B, Jurkat, and A253 cell lines, but had increased nuclear protein binding, we hypothesize that the rs4938573 regulatory element may function as a proximal repressor to reduce *CXCR5* expression in the context of the *DDX6-CXCR5* risk haplotype. The location of HindIII restriction cut sites prohibited separation of rs4938573 from the three SNP cluster and, therefore, examination of potential long-range interactions between the three SNP cluster and the *CXCR5* promoter in this study.

Like in *DDX6*, the rs57494551 regulatory element likely modulated *CXCR5* through chromatin-chromatin interactions in immune, salivary gland, and kidney cells. Luciferase and eQTL data suggest that the rs57494551 risk allele increases enhancer activity and expression of *CXCR5* in B cells, but decreases in T cells; opposite effect of the other SNPs. Because activation and modulation of the CXCR5-CXCL13 axis is critical for adaptive immune responses, it is possible that these regulatory mechanisms work synergistically to titrate *CXCR5* expression and that genetic susceptibly from the *DDX6-CXCR5* risk haplotype dysregulates this mechanism, resulting in delayed & exacerbated signaling. SNP rs57494551 exhibited minimal evidence of *DDX6* regulation, despite being positioned within its first intron.

*Lnc-PHLDB1-1* is a previously uncharacterized lncRNA located in the promoter/enhancer of *TREH* and downstream of *DDX6.* Mined RNA-seq datasets from the Expression Atlas revealed that although not characterized, lnc-PHLDB1-1 does exhibit downregulated expression in whole blood of individuals with juvenile idiopathic arthritis (JIA), several primary and cultured immune cell types under varying inflammatory conditions, and in multiple types of cancer (**Supplemental Figure 19**)^44^. Further, rs4938572 and rs4938573 are strong eQTLs of *lnc-PHLDB1-1* in T cells and macrophages (**Figure 3E, G; Supplemental Figure 18C, E**). This lncRNA region has no CTCF binding sites or H3K27Ac peaks (data not shown) but, because the looping activity is so strong, we propose that *lnc-PHLDB1-1* may be acting as a regulatory mechanism in this risk interval. The functional significance of *lnc-PHLDB1-1* remains unclear; however, we speculate that *lnc-PHLDB1-1* may function as an enhancer lncRNA (elncRNA). In a previous lupus study, it was discovered that a functional SNP regulated allele-specific expression of IRF8 through a mechanism involving the enhancer RNA AC092723.1^45^. Similar to the findings of this study, the prioritized SNPs in this study are located in (rs57494551) or near (rs4938572, rs4936443, rs7117261, and rs4938573) an H3K27ac peak that could modulate the expression of the risk interval through *lnc-PHLDB1-1* by looping the DNA into physical proximity^45^. Identification and characterization of lncRNAs and their functions in health and human disease is of growing interest, but continues to be limited by classification standards, inclusion in publicly available bioinformatic databases, and technologies to characterize them.

*BCL9-like* (*BCL9L*) gene is a homology gene of BCL9 that functions within the Wnt and B-catenin signaling pathways to drive EMT. In breast cancer, *BCL9L* was also shown to promote T cell infiltration of the lymphoid tissue^46,47^. 3C-qPCR and Hi-C analyses suggest that the rs57494551 enhancer may form a chromatin interaction with the *BCL9L* promoter in cell lines examined. Although rs4938573 cluster also demonstrated interactions in the 3C-qPCR, the interactions were not observed by Hi-C. The eQTL analyses suggest that *BCL9L* expression is upregulated in blood and neutrophils in the context of the risk allele. Further, rs4938573 is also a reported eQTL of follicular lymphoma^48,49^. The evidence provided here suggest that rs4938573 may act as a tag SNP for rs57494551 and that risk alleles of rs57494551 may increase enhancer activity and interactions with the *BCL9L* promoter.

TRAPPC4 is a reported eQTL with elevated positive effect size in the salivary gland. Interestingly, Hi-C revealed a chromatin interaction between the rs57494551 enhancer and the promoter of *TRAPPC4* in primary B and T cell subsets. This interaction was not observed by 3C-qPCR above the RIF threshold, indicating potential important differences between cell lines and primary cells. *TRAPPC4* encodes an important regulator of autophagy^50,51^. Dysregulated autophagy and subsequent accumulation of potential autoantigens is implicated in the loss of self-tolerance in autoimmune diseases, including SjD and SLE^52^. Collectively, evidence discovered in this study suggest that the *DDX6-CXCR5* risk haplotype may play a role in the autophagic dysregulation in target tissues (salivary gland and kidney) through regulation of *TRAPPC4*.

In summary, deep interrogation of publicly available and cell type-specific bioinformatic data to prioritize and gain insights into the functionality of common SNPs spanning the *DDX6-CXCR5* SjD/SLE risk haplotypes. Functional interrogation of nuclear protein binding, enhancer/promoter activity, and interactions with the local chromatin regulatory network revealed important insights into how genetic susceptibly at this locus may influence immune cell dysregulation and loss of self-tolerance in disease target tissues through cell type- and allele-specific expression of *DDX6* and *CXCR5*, as well as other genes in the local network (*BCL9L*, *TRAPPC4*, *lnc-PHLDB1-1*) that have not been previously implicated and/or functionally characterized in autoimmune disease. Further, four of the five SNPs prioritized for functional characterization in this study have also been associated with other autoimmune diseases: rheumatoid arthritis (rs7117261, rs4938572, and rs4936443)^53^, primary biliary cholangitis (rs7117261)^54^, and follicular lymphoma (rs4938573)^48,49^. SNP rs4938573 is also a reported SLE- and SjD-associated sex-influenced eQTL^55^. The results of this study performed across multiple immune and disease-specific tissue cell types may offer important mechanistic insights into other diseases beyond autoimmunity.

## MATERIALS AND METHODS

### Study Datasets and Quality Control

Deidentified genotype, Immunochip, and genomic summary results were obtained with approval from and analyzed in accordance with the Oklahoma Medical Research Foundation Institutional Review Board. Dataset (DS) 1 included a) genotype data from 3,232 SjD case and 17,481 controls of European ancestry and b) Immunochip data from 619 SjD cases and 6,171 controls of European ancestry (**Figure 1A**)^1^. DS2 included a) Immunochip data from 3,762 SLE cases and 6,194 controls of European ancestry^20^, b) summary statistics from the meta-analysis of 6,904 SLE cases and 18,429 controls of European ancestry, and c) GWA scan summary results from 1,174 SLE cases and 4,246 controls of Korean ancestry (**Figure 1A**)^28^. DS3 included Immunochip data from an independent cohort of 1916 SjD cases analyzed with the 6,194 population controls from DS2 (**Figure 1B**)^20^. DS4 included Immunochip data from the 3,762 SLE cases and 6,194 population controls from DS2 (**Figure 1B**)^20^.

Written informed consent was obtained in accordance with the institutional review board of each respective study^1,20,28^. All SjD cases fulfilled the America-European Consensus Group (AECG) criteria for primary SjD according to clinical evaluations performed within their respective studies^56–58^. All SLE cases fulfilled at least four of the eleven American College of Rheumatology revised criteria for SLE according to clinical evaluations performed within their respective studies^59,60^.

All data were subjected to the following quality control (QC) measures as previously described^1^: i) a well-defined cluster scatter plot; ii) MAF>1%; iii) SNP call rate >95%; iv) sample call rate >95%; v) Hardy-Weinberg equilibrium test with p>0.001 and rate >95% in controls; vi) p>0.001 for differential missingness between cases and controls. Individual data with excessive heterozygosity (>5 s.d. from mean) and relatedness determined by identity-by-descent (IBD) >0.4 using PLINK (v1.9)^61^ were excluded. EIGENSTRAT^62^ was used to identify population substructure using independent genetic markers (*r*^2^<0.2 between variants).

### Imputation and Statistical Analyses

Genotype and Immunochip data were imputed for the *DDX6-CXCR5* region (chr11: 116194547-121194547; hg19). Imputation was performed using the Haplotype Reference Consortium panel version 1.1 in the Michigan Imputation Server^63^. Data were prephased using SHAPEIT and imputed using Minimac3^64,65^. Imputed variants with imputation quality score (INFO) >0.5 and that fulfilled the above QC criteria were included. GWA Scan data was imputed separately using Z-scores of the missing variants in the summary statistic. Meta-analysis summary statistics were imputed using Ssimp v.0.5.6 imputation software and the East Asian population from 1000 Genome Projects as a reference panel for LD calculation^66^. After filtering SNPs with MAF>1%, 66,397 variants were available for a meta-analysis.

Logistic regression models in PLINK (v1.9) were utilized to assess SNP-trait association in the genotype and Immunochip data, adjusting for the first 3 (or 4) principal components. Stepwise logistic regression was used to adjust for the most significant variant with a P<0.0001. Results from the logistic regressions or summary statistics were meta-analyzed using a fixed-effect model in METAL by weighing the SNP effect by sample size. Meta-analyses were performed between/among: 1) data from SjD datasets (DS1), 2) data from SLE datasets (DS2), and 3) data from merged SjD and SLE datasets (**Figure 1A**). Both Cochran’s Q test statistic and I^2^ index were used to evaluate meta-analysis heterogeneity. LocusZoom^67^ was used to plot meta-analyses results. Linkage disequilibrium (LD) and haplotype reconstruction were calculated utilizing the solid spine of the LD block with minimum *r*^2^ value of 0.8 at HAPLOVIEW (4.2)^68^.

### Fine mapping and Functional Annotations

Fine mapping was implemented using Trinculo^29^ with multinomial logistic regression in the *DDX6-CXCR5* region using genotype and/or Immunochip data (**Figure 1A – DS1; Figure 1B – DS3, DS4, Merged DS3+DS4**). For each SNP tested, population structure was accounted for by including the first four principal components as covariates. The 95% credible sets were derived from the likelihood function of the multinomial logistic model. Posterior probabilities were calculated for each SNP.

Publicly available databases were mined and bioinformatic functional fine mapping performed to further prioritize candidate functional variants in the 95% credible set. RegulomeDB^69,70^, HaploReg v.4.1^71^, and UCSC Genome Browser^72^ were used to screen the 95% credible set for likely functional SNPs (**Figure 1C**). Candidate functional SNPs in the *DDX6-CXCR5* region were further evaluated for reported cell/tissue-specific cis-eQTL (P<0.05) using FUMA^73^, the eQTL catalogue^74^, single-cell RNA sequencing eQTLs^75^, DICE (https://dice-database.org), GTEx v08 (https://gtexportal.org), QTLBase (http://www.mulinlab.org/qtlbase), and NephQTL (https://nephqtl.org) (**Figure 1C**). eQTL significance was further grouped based on p-values (p<5×10^−15^ to 5×10^−3^, p<5×10^−3^ to 1×10^−2^, and p<1×10^−2^ to 0.05), Promoter-capture Hi-C data from the 3D Genome Browser at the YUE lab, Northwestern University (http://3dgenome.fsm.northwestern.edu/chic.php), were mined for chromatin-chromatin interaction start and end coordinates within ± 5kb of each SNP (**Figure 1C**). The pyGenomeTracks tool was used to visualize chromatin interactions in genomic space^76^. Reported eQTLs and chromatin interactions were analyzed in the following cell types/tissues: GM12878 EBV B cell line and primary human total B cells, CD4^+^ and CD8^+^ T cells, monocytes, macrophages (M0, M1 and M2), neutrophils, salivary gland tissue, kidney tissue, and whole blood. Promoter-capture Hi-C data was not available for kidney and salivary gland.

### Tissue Culture and Maintenance

Patient-derived EBV B cells^77^, Daudi B, and Jurkat T cells (ATCC #CCL-213 and #TIB-151 respectively) were grown to 80% confluency using RPMI 1640 complete media (Corning #10-040-CM) supplemented with 10% fetal bovine serum (FBS) (Atlanta Biologicals, #S12450H), 1% penicillin/streptomycin, and 1% L-glutamate in 75-cm^2^ vented tissue culture flasks at 37°C under 5% CO_2_. THP1 monocytes (ATCC, #TIB-202) were grown in the above complete RPMI media supplemented with 0.05mM 2-mercaptoethanol (Sigma, #M3148). 293T human embryonic kidney epithelial cells (ATCC, #CRL-3216) were grown in complete Dulbecco’s Modified Eagle’s Medium (DMEM) (Thermo Fisher, #11995-065) supplemented with 10% FBS, 1% penicillin/streptomycin, and 1% L-glutamate at 37°C under 5% CO_2_. Adherent A253 cells (ATCC, #HTB-41) were grown in McCoy’s 5a medium (ATCC, #30-2007) supplemented with 10% FBS, 1% penicillin/streptomycin, and 1% L-glutamate at 37°C under 5% CO_2_. Cell viability was assessed using Trypan blue as a live/dead stain and visualized and counted on the Countess II Automated Cell Counter (Thermo Fisher, #AMQAX1000).

### SNP Genotyping of Tissue Cultures (TaqMan Genotyping Assay)

TaqMan assays was used to genotype Jurkat T cells, THP1 monocytes, and A253 cells. Total genomic DNA was isolated from 1×10^6^ cells using the Zymo Quick-DNA Miniprep Plus Kit (ZymoResearch, #D4068) and quantified. Taqman reactions were performed on a QuantStudio 6 qPCR machine (Thermo Fisher, #A43168) following manufacturer instructions using ∼25ng of DNA, TaqMan Genotyping Master Mix (ThermoFisher, #4371355), and TaqMan SNP Genotyping probes for rs57494551 (ThermoFisher, #4351379, Assay ID#C_90471390_10) and rs4938572 (ThermoFisher, #4351379, Assay ID#C_28012712_10) (**Supplemental Table 1**). Genotypes for each allele were compared to EBV B cell lines homozygous or heterozygous for the minor or major allele of rs57494551 and rs4938572 (**Supplemental Table 2**). A minimum of three biological replicates were analyzed in triplicate. Genotyping results were analyzed using the Design & Analysis Software package from ThermoFisher.

### Electromobility shift assay (EMSA)

Nuclear lysates from EBV B cells, Daudi B, Jurkat T, THP1 monocyte, and A253 cells were prepared using the NE-PER Nuclear and Cytoplasmic Extraction Kit (Fisher Scientific, #78833) supplemented with Halt Protease Inhibitor Cocktail 100x (Thermo Fisher, #78425), DTT (Fisher Scientific, #R0861) and PMSF (Sigma Aldrich, #P7626-5G). Complementary pairs of ∼60bp non-risk or risk oligonucleotide probes for each SNP were chemically synthesized (IDT) (**Supplemental Table 3**), annealed using 10X Oligonucleotide Annealing Buffer (100mM Tris-HCl pH 7.5, 1M NaCl, 10mM EDTA pH 7.4) in 95°C for 2 minutes, and end-labeled with (γ-P^32^) adenosine triphosphate (MP Biomedicals or Perkin Elmer, #NEG002A250U) using T4 polynucleotide kinase (NEB, #M0201S). Labeled oligos were purified using MicroSpin G-25 Columns (VWR, #95017-621), then scintillation counts were performed using Liquid Scintillation Analyzer Tri-Carb 4910 TR (Perkin Elmer). Nuclear protein extracts (5μg) were incubated for 30 minutes at room temperature with labeled probes (50,000cpm/reaction) in binding buffer (1μg poly (dI-dC), 20mM HEPES, 10% Glycerol, 1mM MgCl_2_, 0.5mM EDTA, pH 8.0, 50mM NaCl and 10mM Tris-HCl, pH 7.4). DNA-protein complexes were resolved on non-denaturing 5% acrylamide gels. Competition assays were performed by adding 100x unlabeled probe and incubating for 15 minutes on ice. Differential binding was assessed by densitometry analysis of photostimulated luminescence (PSL) using Molecular Imager PharosFX System (Bio-Rad, #170-9450) and Quantity One 1-D analysis software (Bio-Rad). Band density was normalized to the background and statistically analyzed by Student’s t-test in Prism 9 software.

### Luciferase Reporter Assay

A 500 bp gBlock (IDT) carrying the non-risk or risk allele of each SNP (**Supplemental Table 4**) were cloned into the no-promoter firefly luciferase reporter plasmid pGL4.14 [luc2/Hygro] (Promega, #E6691) or minimal promoter firefly luciferase reporter plasmid pGL4.26 [luc2/miniP/Hygro] (Promega, #E8441). Note, each gBlock had 1 SNP with risk allele in the context of the non-risk alleles for all other SNPs in close proximity. Plasmids were double digested with HindIII-HF (NEB, #R3104S) and KpnI-HF (NEB, #R3142S) restriction enzymes. A Thymidine Kinase (TK) promoter-driven Renilla luciferase plasmid pGL4.74 [hRluc/TK] (Promega, #E6921) was used as a transfection efficiency control.

Plasmids were transfected using either the 4D Nucleofector (EBV B, Daudi, THP1; Lonza, 4D-Nucleofector Core Unit #AAF-1002B and 4D-Nucleofector X Unit #AAF-1002X), the Neon Transfection System (Jurkats; ThermoFisher, #MPK5000S), or Lipofectamine 3000 Transfection Reagent (HEK 293T, A253 cells; Life Technologies, #L3000015). The SF Cell Line 4D-Nucleofector X Kit S (Lonza, #V4XC-2032) was used to transfect EBV B cells and Daudi cells; the SG Cell Line 4D-Nucleofector X Kit S (Lonza, #V4XC-3032) was used to transfect THP1 monocytes. One million cells were resuspended in 20μL of SF or SG buffer and transfected with either pGL4.14 or pGL4.26 plasmid (1μg/μL) containing the appropriate gBlock or empty vector, and pGL4.74 (125ng/μL) Renilla as a transfection control using program EH-100 (EBV B), CA-137 (Daudi), or FF-100 (THP1). After transfection, cells were rested at room temperature for 15 minutes, then 80μL of pre-warmed complete media was added and cells incubated at 37°C for 1 hour. Transfected cells were then transferred into a 96-well plate containing 100μL of pre-warmed complete media for a final volume of 200μL. Cells were collected, washed, lysed, and luciferase activity measured after 24 hours using the Dual-Luciferase Reporter Assay System (Promega, #E1980) and the Synergy H1 Hybrid Multi-Mode Microplate Reader with dual injectors (BioTek, #8041000) or the GloMax Explorer Multimode Microplate Reader (Promega, #GM3500). Relative luciferase activity (RLA) was reported as a ratio of firefly luciferase to Renilla transfection control. A minimum of three experimental replicates were plotted and analyzed by Student’s t-test.

Jurkat cells were transfected using the Neon Transfection System with the Neon Transfection System 10μL Kit (Thermo Fisher, #MPK1096) according to manufacturer instructions with modification. Briefly, 4×10^5^cells/mL were transfected with either pGL4.14 or pGL4.26 (1μg/μL) containing the appropriate gBlock or empty vector, and pGL4.74 (62.5ng/μL) Renilla transfection control in Buffer R. Cells were pulsed with a voltage of 1600v, pulse width 10, and pulse number 3. Luciferase activity was measured and analyzed after 24 hours as described above.

A253 and 293T cells were transfected with Lipofectamine 3000 Transfection Reagent following manufacturer protocols. Briefly, 8×10^4^ cells were seeded per well of a 24-well plate. After 24 hours, media was replaced with a mixture of Lipofectamine 3000 and pGL4.14 (1μg/μL) or pGL4.26 (1μg/μL) containing the appropriate gBlock or empty vector, and pGL4.74 (10ng/μL) Renilla transfection control diluted in Opti-MEM Reduced Serum Medium (Life Technologies, #31985-070). Cells were lysed and luciferase activity analyzed after 24 hours as described above.

### Chromatin Conformation Capture with Quantitative PCR (3C-qPCR)

3C was performed as described with few modifications^78^. Briefly, 1×10^6^ cells from EBV B cell lines homozygous for non-risk or risk genotypes at the SNP of interest, Jurkat T cells, or A253 cells were crosslinked with 2% formaldehyde for 10 minutes. Crosslinking was quenched with 0.125M glycine for 20 minutes then cell pellets were flash frozen in liquid nitrogen and stored at −80°C overnight. Cell pellets were thawed on ice and lysed with fresh lysis buffer (10mM TrisHCl, 0.2% NP-40, 10mM NaCl, 1x Protease Inhibitor Cocktail (Life Technologies, #78425)) for 30 minutes on ice. Nuclei were pelleted and washed with 1.25x CutSmart Buffer (New England Biolabs (NEB), #B7204S), then incubated with 0.3% SDS at 37°C for one hour and then 2% Triton X-100 at 37°C for one hour. Nuclei were incubated with 3000U of HindIII-HF (NEB, #R3104M) at 37°C overnight while shaking to digest chromatin, then ligation incubation buffer (10mg/mL BSA, 1x T4 DNA Ligase Reaction Buffer (NEB, #B0202S) and 2000U of T4 DNA Ligase (NEB, #M0202S) were added and incubated overnight on ice. 3C ligation mixtures were de-crosslinked with Proteinase-K at 65°C overnight. Samples were treated with RNaseA for 1 hour at 37°C and then purified with phenol:chloroform:isoamyl alcohol 25:24:1, twice.

Primers were designed to target 100 bp upstream of HindIII cut sites positioned in or near a Hi-C anchor; only fragments <5kb were considered (**Supplemental Table 5**). Primer efficiency was validated using three Bacterial Artificial Chromosome (BAC) clones (Thermo Fisher, #RP11-1011H15, #CTC-3245B9, #RP11-45N4) spanning the *DDX6-CXCR5* locus, mixed in equimolar concentrations. All samples were run on a QuantStudio 6 qPCR machine (Thermo Fisher, #A43168). CT values were normalized to standard curves generated from the BAC clones. Relative interaction frequency (RIF) of each 3C primer pair was calculated by comparing the interaction frequency from each 3C primer pair with the interaction frequency from the short-range interaction primer control. Graphs were plotted using Prism V9 and student’s t-test was used to calculate standard deviation.

### RNA-sequencing

A253 cells grown to 80-85% confluence were washed, trypsinized, and counted. One million cells in triplicate were collected and resuspended in 700µL of Trizol. RNA was isolated using the Direct-zol RNA Microprep kit following manufacturer instructions (Zymo Research, #R2060) and quantified by fluorometric analysis (Thermo Fisher Qubit fluorometer). mRNA was isolated and libraries made using the Swift Biosciences Rapid RNA Library kit (Swift Biosciences, #R2024) according to manufacturer instructions. Libraries were indexed using Swift Biosciences Unique Dual Indexing primers (Swift Biosciences, #X9096). Libraries were quantified, pooled, and sequenced using an Illumina NovaSeq 6000 with PE150 reads. Twenty million total sequenced reads were quality assessed using FastQC tool and cleaned for adapters using trimmomatic software (Bolger et al., 2014). QC reads were aligned to hg19 reference genome using HISAT2 (Kim et al., 2019). SAMtools was used to sort and index the aligned files into bam files (Li et al., 2009). Bam files were split by strandedness to obtain read coverage for plus(+) and minus(-) strands.

### ATAC-sequencing

In-house chromatin accessibility maps were generated using ATAC-seq data from A253 cells. A253 cells grown to 80-85% confluence were released by trypsinization, washed, and counted. Six replicates of 5×10^4^ cells were pelleted by centrifugation and resuspended in 50 µL of cold lysis buffer (10 mM Tris-HCL, pH 7.4, 10 mM NaCl, 3mM MgCL2, 0.1% IGEPAL (NP-40)) for ∼20 seconds. Nuclei were collected by centrifugation and resuspended in 50 µL transposase mix (25 µL 2x TD Buffer, 2.5 µL TDE1 transpose enzyme (Illumina Tagment DNA Enzyme and Buffer Small Kit, #20034197), and 22.5 µL nuclease-free water). After incubation at 37°C for 30 minutes, all samples were cleaned using the Qiagen MinElute Kit (Qiagen, #28006) following manufacturer instructions and eluted in 10 µL EB buffer. DNA was amplified using NEBNext Ultra II Q5 PCR Master Mix (NEB, #M0544S) with Index Primers from Illumina (IDT for Illumina DNA/RNA UD Indexes Set A (Illumina, #20027213)). PCR amplified libraries were cleaned up using AMPure XP beads (Beckman Coulter, #A63880). Libraries were quantified, pooled, and sequenced using an Illumina NovaSeq 6000 with PE150 reads. Fifty million reads from each replicate were analyzed as follows: Paired-end reads (100 bp) were sequenced, quality of the reads was determined using FASTQC tool, adapters were trimmed using trimmomatic and bowtie2 was used to align the reads to hg19 reference genome with parameters --very-sensitive -X 2000 -k 10 --threads 8 –dovetail^79^. Aligned reads were sorted by SAMtools^80^, then additional processing was performed to remove duplicate and nonuniquely mapped reads. Finally, peak calling was performed in sorted files using Genrich tool^81^.

### CUT&RUN

The CUT & RUN protocol (CUTANA ChIC/CUT & RUN Kit, EpiCypher, #14-1048) was performed according to manufacturer protocols with optimizations for A253 cells. For example, A253 cell viability increased when cells were detached with 0.05% trypsin, as opposed to manual scraping. A final concentration of 0.01% digitonin was optimal for 92-100% permeabilization of A253 cells and 2.5 µL of pAG-MNase for 2 hours was optimal sufficient cleavage of 6×10^5^ A253 cells into mononucleosome fragments of ∼300bp. Heterochromatin formation or gene inactivation were detected using antibodies against H3K4me3 (0.5 µg/sample, EpiCypher #13-0041), H2K27me3 (0.5 µg/sample, Invitrogen #MA5-11198), IgG (0.5 µg/sample, EpiCypher #13-0042). Isolated DNA pull-down fragments were quantified, and libraries prepared using NEB Ultra II DNA Library Prep Kit (NEB, #E7645S) and NEBNext Multiplex Oligos for Illumina (96 Unique Dual Index Primer Pairs Set 4) (NEB, E6446S) as index primers. Libraries were run on an Agilent TapeStation to assess quality and ensure enrichment of ∼300bp fragments (∼170bp DNA fragments + library sequencing adapters). After PCR enrichment, libraries were pooled and sequenced using Illumina NovaSeq 6000 with PE150 reads, 20 million reads per sample. Raw reads were quality controlled using FastQC and adapters were trimmed using trimmomatic tools. Clean reads were aligned to hg19 reference genome using Bowtie2. SAMtools was used to convert and sort SAM files into BAM format. Picard was used to mark and remove duplicate reads^82^. SEACR was used for peak calling^83^. Signals from sorted BAM files via bamCoverage were normalized by deeptools then computeMatrix was used to calculate overall signal distribution around the peaks center called by SEACR^84^.

### IMPACT Regulatory Annotation

To better understand binding patterns of transcription factors (TFs) around putative functional SNPs, the 707-cell type-specific IMPACT regulatory annotations resource was used^85^. IMPACT annotations for *DDX6-CXCR5* region were extracted and the total number of transcription factors with their probabilities was used to annotate SNPs in the region. pyGenomeTracks tool was then used to visualize the number of active TFs and their IMPACT scores in different cell types^76^.

### Epigenetic colocalization

Regulatory functions of putative SNPs were investigated by alignment with publicly available and in-house epigenetic data. Chromatin segmentation maps of GM12878, B, CD4^+^ T, CD8^+^ T cells, monocytes and neutrophils were obtained from the Roadmap Epigenomics Consortium [https://egg2.wustl.edu/roadmap/data/byFileType/chromhmmSegmentations/ChmmMod els/coreMarks/jointModel/final/]. RNA-seq and histone marks data of B, CD4+ T, CD8+ T cells, macrophages, monocytes, and neutrophils were obtained from BluePrint Epigenome [https://epigenomesportal.ca/tracks/Blueprint/hg19/]. Histone marks data of GM12878 and 293T, and RNA-seq data of GM12878 cell lines were obtained from ENCODE database. The RNA-seq data of 293T was from NCBI short read archive (GSM2258993).

The putative functional SNPs were colocalized in different cell types using a SNP-trait association, fine mapping, mRNA expressions, histone marks, capture Hi-C, 3C, chromatin segmentation marks, and impact annotations along with hg19 reference genome. The results were visualized using pyGenome tracks^76^.

### Data availability

SjD genome-wide association summary statistics and individual-level data used in Dataset 1 of this study were previously reported in^1^ and are available through the Databases of Genotypes and Phenotypes (dbGaP) under accession number: phs002723.v1.p1 [http://www.ncbi.nlm.nih.gov/projects/gap/cgi-bin/study.cgi?study_id=phs002723.v1.p1], phs000672.v1.p1 (n=735) [https://www.ncbi.nlm.nih.gov/projects/gap/cgi-bin/study.cgi?study_id=phs000672.v1.p1], phs000428.v2.p2 (n=8,519) [https://www.ncbi.nlm.nih.gov/projects/gap/cgi-bin/study.cgi?study_id=phs000428.v2.p2], phs000196.v3.p1 (n=995) [https://www.ncbi.nlm.nih.gov/projects/gap/cgi-bin/study.cgi?study_id=phs000196.v3.p1], phs000187.v1.p1 (n=602) [https://www.ncbi.nlm.nih.gov/projects/gap/cgi-bin/study.cgi?study_id=phs000187.v1.p1]. SLE Immunochip and genome-wide association (GWA) scan data used in Dataset 2 of this study were previously reported in^20,28^ and are available from the corresponding authors of the original manuscript in accordance with respective Institutional Review Board approval and subject consent. All remaining data were generated by coauthors in this study and will be made available upon request to the corresponding author in accordance with an established material transfer agreement. All other data presented in this study were previously published and can be accessed by URLs provided in the methods section.

## Supporting information

Compiled Supplemental Data

## ACKNOWLEDGEMENTS

We thank all the research and clinical staff, consortium investigators, study participants, and funding agencies who made this study possible.

We also thank the investigators and funding mechanisms for the following dbGaP studies:

**Phs002723.v1.p1:** Genotype data from the international Sjogren’s Genetics Network (SGENE) was reported in Khatri, et al. *Nat Commun*, 2022 and obtained through dbGAP accession number phs002723.v1.p1. The study was supported by the National Institutes of Health (NIH): R01AR073855 (Lessard); R01AR065953 (Lessard); R01AR074310 (Farris); P50AR060804 (Sivils), N01DE32636 (SICCA), HHSN26S201300057C (SICCA), U01DE028891 (SICCA), R03DE029800 (SICCA), U01HG004446 (SICCA-GWAS), P30AR070155 (SICCA-GWAS), R61AR076803 (Adrianto); NIDCR Sjögren’s Syndrome Clinic and Salivary Disorders Unit were supported by funds awarded to BMW (Z01-DE000704) by the NIDCR Division of Intramural Research at the National Institutes of Health (Warner); Birmingham NIHR Biomedical Research Centre (Bowman); Deutsche Forschungsgemeinschaft (DFG, German Research Foundation) under Germany’s Excellence Strategy – EXC 2155 – Projektnummer 390874280 (Witte); Research Council of Norway (Oslo, Norway) – Grant 240421 (Reksten), Grant 316120 (Wahren-Herlenius); Western Norway Regional Health Authority (Helse Vest) – 911807, 912043 (Omdal); Swedish Research Council for Medicine and Health (Rönnblom, Nordmark, Wahren-Herlenius); Swedish Rheumatism Association (Rönnblom, Nordmark, Wahren-Herlenius); King Gustav V’s 80-year Foundation (Nordmark); Swedish Society of Medicine (Rönnblom, Wahren-Herlenius); Swedish Cancer Society (Baecklund); The Stockholm County Council (Wahren-Herlenius); The Swedish Twin Registry is managed through the Swedish Research Council under the grant 2017-000641. The SNP&SEQ Technology Platform was supported by Science for Life Laboratory, Uppsala University, the Knut and Alice Wallenberg Foundation and the Swedish Research Council (Nordmark).

**Phs000428.v2.p2**: This study used control data from the Health and Retirement Study in dbGaP (phs000428.v2.p2) submitted by David Weir, PhD at the University of Michigan and funded by the National Institute of Aging RC2 AG036495 and RC4 AG039029.

**Phs000672.v1.p1**: Genotype data from the Sjögren’s International Collaborative Clinical Alliance (SICCA) Registry was obtained through dbGAP accession number phs000672.v1.p1. This study was supported by the National Institute of Dental and Craniofacial Research (NIDCR), the National Eye Institute, and the Office of Research on Women’s Health through contract number N01-DE-32636. Genotyping services were provided by the Center for Inherited Disease Research (CIDR). CIDR is fully funded through a federal contract from the National Institutes of Health (NIH) to the Johns Hopkins University (contract numbers HHSN268200782096C, HHSN268201100011I, HHSN268201200008I). Funds for genotyping were provided by the NIDCR through CIDR’s NIH contract. Assistance with data cleaning and imputation was provided by the University of Washington. We thank investigators from the following studies that provided DNA samples for genotyping: the Genetic Architecture of Smoking and Smoking Cessation, Collaborative Genetic Study of Nicotine Dependence (phs000404.v1.p1); Age-Related Eye Disease Study (AREDS) - Genetic Variation in Refractive Error Substudy (phs000429.v1.p1); and National Institute of Mental Health’s Human Genetics Initiative (phs000021.v3.p2, phs000167.v1.p1). We thank the many clinical collaborators and research participants who contributed to this research.

**Phs000196.v3.p1:** Investigators and Parkinson Disease patients that contributed to this Genome-wide Association Study of Parkinson Disease.

**Phs000187.v1.p1**: Research support to collect data and develop an application to support the High Density SNP Association Analysis of Melanoma project was provided by 3P50CA093459, 5P50CA097007, 5R01ES011740, and 5R01CA133996.

## Author Contributions

C.J.L.* supervised the research and serves as senior author (*); M.M.W., B.K., M.L.J., A.M.S. conceived and designed experiments under the supervision of C.J.L.*; C.J.L.*, B.K., A.R., J-M.A., L.A.A., S-C.B., E.B., A.B., J.G.B., S.M.B., N.D., M-L.E., F.E., H.F-d’E., C.F., C.G., J-E.G., D.H., J.I-K., J.L.J., S.J.A.J., M.V.J., J.A.K., S.K., K.K., M.K., T.M., J.M., D.L.M., G.Nocturne, K.B.N., P.O., Ø.P., J-O.P., N.L.R., C.S., K.S., K.E.T., P.T., G.E.T., S.V., E.M.V., D.J.W., K.M.G., L.R., M.T.B., J.A.J., R.H.S., P.M.G., L.A.C., R.J., S.A., P.E., S.J.B., R.O., L.R., B.M.W., M.R., T.W., A.D.F., X.M., C.H.S., M.W-H., M.E.A-R., W-F.N., K.L.A., J.M.G., I.A., T.J.V., B.P.T., G. Nordmark collected, characterized, and/or genotyped the Sjögren’s and/or SLE cases and population controls used in this study; S.B.G., J.A.K., K.M.G., J.M.G. manages the technology and/or data required for processing and analysis of large-scale sequencing approaches used; M.M.W., B.K., M.L.J., A.M.S. performed experiments; M.M.W., B.K., M.L.J., A.M.S. performed statistical analyses; M.M.W., B.K., M.L.J., K.L.T., A.M.S., B.P.T., G. Nordmark analyzed the data; C.J.L.*, M.M.W., B.K., K.L.T. wrote the paper; all authors critically reviewed the paper and accepted authorship responsibilities.

