## Supplementary material for "Variants in the *DDX6-CXCR5* autoimmune disease risk locus influence the regulatory network in immune cells and salivary gland": Compiled Supplemental Data

[18]Lund University, Malmö, Sweden;
[19]Instituto de Biomedicina y Parasitología López-Neyra, Granada, Spain;
[20]Université Paris-Saclay, Paris, France;
[21]Assistance Publique – Hôpitaux de Paris, Hôpital Bicêtre, Paris, France;
[22]Oslo University Hospital, Oslo, Norway;
[23]LBAI, UMR1227, University of Brest, Inserm, Brest, France;
[24]University of Minnesota Medical School, Minnesota, USA;
[25]Linköping University, Linköping, Sweden;
[26]University of California San Francisco, San Francisco, California, USA;
[27]Cedars-Sinai Medical Center, Los Angeles, California, USA;
[28]University of Leeds, Leeds, United Kingdom;
[29]University of Oklahoma College of Dentistry, Oklahoma City, Oklahoma, USA;
[30]Atrium Health Carolinas Medical Center, Charlotte, North Carolina, USA;
[31]University of Oklahoma Health Sciences Center, Oklahoma City, Oklahoma, USA;
[32]US Department of Veteran Affairs Medical Center, Oklahoma City, Oklahoma, USA;
[33]National Human Genome Research Institute, NIH, Bethesda, Maryland, USA;
[34]University Hospitals Birmingham NHS Foundation Trust, Birmingham, United Kingdom;
[35]National Institute of Dental and Craniofacial Research, Bethesda, Maryland, USA;
[36]University of Adelaide, Adelaide, South Australia;
[37]Genyo, Center for Genomics and Oncological Research, Pfizer/University of Granada/Andalusian Regional Government, Spain;
[38]NIHR Newcastle Biomedical Research Centre and NIHR Newcastle Clinical Research Facility, Newcastle upon Tyne Hospitals NHS Foundation Trust, Newcastle Upon Tyne, United Kingdom;
[39]Translational and Clinical Research Institute, Newcastle University, Newcastle Upon Tyne, United Kingdom;
[40]Center for Bioinformatics, Department of Public Health Sciences, Henry Ford Health, Detroit, Michigan, USA;
[41]Department of Medicine, College of Human Medicine, Michigan State University, East Lansing, Michigan, USA;
[42]Medical University of South Carolina, Charleston, South Carolina, USA;
[#]A list of consortia investigators and their affiliations appear as a Supplementary Note;
[†]Current Affiliation: Translational Science, The Janssen Pharmaceutical Companies of Johnson & Johnson, Spring House, Pennsylvania, USA;

**TABLE OF CONTENTS** **Page**

i. Consortium Acknowledgements and Funding* ……………………….. 5

ii. Supplemental Tables
a. Supplemental Table 1: Risk and Non-Risk Alleles of Prioritized SNPs from the *DDX6-CXCR5* SjD/SLE Risk Locus …………….. 8
b. Supplemental Table 2: Patient-Derived EBV B Cell Genotypes... 8
c. Supplemental Table 3: EMSA Probes …………………………….. 9
d. Supplemental Table 4: Luciferase Reporter Assay gBlocks ……. 10-13
e. Supplemental Table 5: 3C-qPCR Primers ………………………… 14

iii. Supplemental Figures
a. Supplemental Figure 1: Fine-mapping of the *DDX6-CXCR5* Region in SjD and SLE Immunochip After Imputation …………… 15
b. Supplemental Figure 2: Haplotype Frequency of the *DDX6-CXCR5* Risk Region in SjD ………………………………………. 16
c. Supplemental Figure 3: Functional Bioinformatic Analyses of Putative Functional SNPs in the *DDX6-CXCR5* Risk Region …… 17
d. Supplemental Figure 4: Allele- and Cell-type Specific Differential Nuclear Protein Affinities of SNPs rs57494551 and rs4938572 … 18
e. Supplemental Figure 5: Allele- and Cell-type Specific Differential Nuclear Protein Affinities of SNP rs7117261 ……………………… 19
f. Supplemental Figure 6: Allele- and Cell-type Specific Differential Nuclear Protein Affinities of SNP rs4936443 ……………………… 20
g. Supplemental Figure 7: Allele- and Cell-type Specific Differential Nuclear Protein Affinities of SNP rs4938573 ……………………… 21
h. Supplemental Figure 8: Allele- and Cell-type Specific Differential Nuclear Protein Affinities of SNP rs12365699 …………………… 22
i. Supplemental Figure 9: Allele- and Cell-type Specific Differential Nuclear Protein Affinities of SNP rs10892294 …………………… 23
j. Supplemental Figure 10: Allele-specific Promoter and Enhancer Activity of rs10892294 and rs12365699 on the *DDX6-CXCR5* Region in 293T cells ……………………………………………… 24
k. Supplemental Figure 11: Complex Chromatin Architecture Revealed Across the *DDX6-CXCR5* Region in Human Primary B Cells ……………………………………………………………… 25
l. Supplemental Figure 12: Complex Chromatin Architecture Revealed Across the *DDX6-CXCR5* Region in Primary Human T Cells ……………………………………………………………… 26
m. Supplemental Figure 13: Complex Chromatin Architecture Revealed Across the *DDX6-CXCR5* Region in Human Primary Monocytes ………………………………………………………….. 27
n. Supplemental Figure 14: Complex Chromatin Architecture Revealed Across the *DDX6-CXCR5* Region in Human Primary Macrophages ......................................................................... 28

o. Supplemental Figure 15: Complex Chromatin Architecture Revealed Across the *DDX6-CXCR5* Region in Human Primary Neutrophils …………………………………………………………… 29
p. Supplemental Figure 16: Complex Chromatin Architecture Revealed Across the *DDX6-CXCR5* Region in Human 293T Cells 30
q. Supplemental Figure 17: IMPACT Regulatory Element Probabilities and Corresponding Transcription Factor Elements in Immune Cells …………………………………………………………… 31
r. Supplemental Figure 18: eQTLs for Five Prioritized SNPs Across Blood, Salivary and Kidney Tissues, and Immune Cells ………… 32
s. Supplemental Figure 19: Expression Atlas Results for lnc-PHLDB1-1 (ENSG00000255422) …………………………………… 33

iv. Supplemental Data (see accompanying excel worksheets)
a. Supplemental Data 1: Meta-analysis of SjD GWAS and Immunochip Data in *DDX6-CXCR5* Locus (Dataset 1)
b. Supplemental Data 2: Meta-analysis of SLE Data in *DDX6-CXCR5* Locus (Dataset 2)
c. Supplemental Data 3: Meta-analysis of SjD and SLE Data in *DDX6-CXCR5* Locus (Merged Datasets 1 and 2)
d. Supplemental Data 4: Bayesian Analysis of SjD GWAS and Immunochip Data in *DDX6-CXCR5* Locus (Dataset 1)
e. Supplemental Data 5: Bayesian Analysis of SjD Immunochip Data in *DDX6-CXCR5* Locus (Dataset 3)
f. Supplemental Data 6: Bayesian Analysis of SLE Immunochip Data in *DDX6-CXCR5* Locus (Dataset 4)
g. Supplemental Data 7: Bayesian Analysis of merged SjD and SLE Immunochip Data in *DDX6-CXCR5* Locus (Merged Datasets 3 and 4) with bioinformatic analysis of the top 100 SNPs (RegulomeDB, HaploReg epimarks, and publications).

**SUPPLEMENTAL NOTE**

**Consortium Acknowledgments and Funding*:**

**A. The PRECISESADS Clinical Consortium is composed of the following members:** Lorenzo Beretta[1], Barbara Vigone[1], Jacques-Olivier Pers[2], Alain Saraux[2], Valérie Devauchelle-Pensec[2], Divi Cornec[2], Sandrine Jousse-Joulin[2], Bernard Lauwerys[3], Julie Ducreux[3], Anne-Lise Maudoux[3], Carlos Vasconcelos[4], Ana Tavares[4], Esmeralda Neves[4], Raquel Faria[4], Mariana Brandão[4], Ana Campar[4], António Marinho[4], Fátima Farinha[4], Isabel Almeida[4], Miguel Angel Gonzalez-Gay Montecón[5], Ricardo Blanco Alonso[5], Alfonso Corrales Martinez[5], Ricard Cervera[6], Ignasi Rodríguez-Pintó[6], Gerard Espinosa[6], Rik Lories[7], Ellen De Langhe[7], Nicolas Huzelmann[8], Doreen Belz[8], Torsten Witte[9], Niklas Baerlecken[9], Georg Stummvoll[10], Michael Zauner[10], Michaela Lehner[10], Eduardo Collantes[11], Rafaela Ortega-Castro[11], Mª Angeles Aguirre-Zamorano[11], Alejandro Escudero-Contreras[11], Mª Carmen Castro-Villegas[11], Norberto Ortego[12], María Concepción Fernández Roldán[12], Enrique Raya[13], Immaculada Jiménez Moleón[13], Enrique de Ramon[14], Isabel Díaz Quintero[14], Pier Luigi Meroni[15], Maria Gerosa[15], Tommaso Schioppo[15], Carolina Artusi[15], Carlo Chizzolini[16], Aleksandra Zuber[16], Donatienne Wynar[16], Laszló Kovács[17], Attila Balog[17], Magdolna Deák[17], Márta Bocskai[17], Sonja Dulic[17], Gabriella Kádár[17], Falk Hiepe[18], Velia Gerl[18], Silvia Thiel[18], Manuel Rodriguez Maresca[19], Antonio López-Berrio[19], Rocío Aguilar-Quesada[19], Héctor Navarro-Linares[19], **Marta E. Alarcon-Riquelme**[20].

[1]Referral Center for Systemic Autoimmune Diseases, Fondazione IRCCS Ca' Granda Ospedale Maggiore Policlinico di Milano, Italy; [2]Centre Hospitalier Universitaire de Brest, Hospital de la Cavale Blanche, Brest, France; [3]Pôle de pathologies rhumatismales systémiques et inflammatoires, Institut de Recherche Expérimentale et Clinique, Université catholique de Louvain, Brussels, Belgium; [4]Centro Hospitalar do Porto, Portugal; [5]Servicio Cantabro de Salud, Hospital Universitario Marqués de Valdecilla, Santander, Spain; [6]Hospital Clinic I Provicia, Institut d'Investigacions Biomèdiques August Pi i Sunyer, Barcelona, Spain; [7]Katholieke Universiteit Leuven, Belgium; [8]Klinikum der Universitaet zu Koeln, Cologne, Germany; [9]Medizinische Hochschule Hannover, Germany; [10]Medical University Vienna, Vienna, Austria; [11]Servicio Andaluz de Salud, Hospital Universitario Reina Sofía Córdoba, Spain; [12]Servicio Andaluz de Salud, Complejo hospitalario Universitario de Granada (Hospital Universitario San Cecilio), Spain; [13]Servicio Andaluz de Salud, Complejo hospitalario Universitario de Granada (Hospital Virgen de las Nieves), Spain; [14]Servicio Andaluz de Salud, Hospital Regional Universitario de Málaga, Spain; [15]Università degli studi di Milano, Milan, Italy; [16]Hospitaux Universitaires de Genève, Switzerland; [17]University of Szeged, Szeged, Hungary; [18]Charite, Berlin, Germany; [19]Andalusian Public Health System Biobank, Granada, Spain; [20]Genyo, Center for Genomics and Oncological Research, Pfizer/University of Granada/Andalusian Regional Government, Granada, Spain.

The study was approved by the following ethic committees: Comitato Etico Area 2 (Fondazione IRCCS Ca' Granda Ospedale Maggiore Policlinico di Milano and University of Milan); approval no. 425bis Nov 19, 2014, and no. 671_2018 Sep 19, 2018; Klinikum der Universitaet zu Koeln, Cologne, Germany. Geschaftsstelle Ethikkommission; Pôle de pathologies rhumatismales systémiques et inflammatoires, Institut de Recherche Expérimentale et Clinique, Université catholique de Louvain, Brussels, Belgium. Comité d`Èthique Hospitalo-Facultaire; University of Szeged, Szeged, Hungary. Csongrad Megyei Kormanyhivatal; Hospital Clinic I Provicia, Institut d'Investigacions Biomèdiques August Pi i Sunyer, Barcelona, Spain. Comité Ética de Investigación Clínica del Hospital Clínic de Barcelona. Hospital Clinic del Barcelona; Servicio Andaluz de Salud, Hospital Universitario Reina Sofía Córdoba, Spain. Comité de Ética e la Investigación de Centro de Granada (CEI – Granada); Centro Hospitalar do Porto, Portugal.

Comissao de ética para a Saude – CES do CHP; Centre Hospitalier Universitaire de Brest, Hospital de la Cavale Blanche, Avenue Tanguy Prigent 29609, Brest, France. Comite de Protection des Personnes Ouest VI; Hospitaux Universitaires de Genève, Switzerland. DEAS – Commission Cantonale d'ethique de la recherche Hopitaux universitaires de Geneve; Andalusian Public Health System Biobank, Granada, Spain; Katholieke Universiteit Leuven, Belgium. Commissie Medische Ethiek UZ KU Leuven /Onderzoek; Charite, Berlin, Germany. Ethikkommission; Medizinische Hochschule Hannover, Germany. Ethikkommission.

PRECISESADS Study was funded by the Innovative Medicines Initiative of the European Union with grant number 115565 partly supported by the EFPIA Companies (Alarcon-Riquelme).

**B. Sjögren's International Collaborative Clinical Alliance (SICCA) is composed of the following members:** Cox D[1], Jordan R[1], Lee D[1], DeSouza Y[1], Drury D[1], Do A[1], Scott L[1], Nespeco J[1], Whiteford J[1], Margaret M[1], Sack S[1], Adler I[2], Smith AC[2], Bisio AM[2], Gandolfo MS[2], Chirife AM[2], Keszler A[2], Daverio S[2], Kambo V[2], Dong Y[3], Jiang Y[3], Xu D[3], Su J[3], Du D[3], Wang H[3], Li Z[3], Xiao J[3], Wu Q[3], Zhang C[3], Meng W[3], Zhang J[3], Johansen S[4], Hamann S[4], Schiødt J[4], Holm H[4], Ibsen P[4], Manniche AM[4], Kreutzmann SP[4], and Villadsen J[4], Sugai S[5], Masaki Y[5], Sakai T[5], Shibata N[5], Honjo M[5], Kurose N[5], Nojima T[5], Kawanami T[5], Sawaki T[5], Fujimoto K[5], Odell E[6], Morgan P[6], Fernandes-Naglik L[6], Varghese-Jacob B[6], Ali S[6], Adamson M[6], Seghal S[7], Mishra R[7], Bunya V[7], Massaro-Giordano M[7], Abboud SK[7], Pinto A[7], Sia YW[7], Dow K[7], Akpek E[8], Ingrodi S[8], Henderson W[8], Gourin C[8], Keyes A[8], Srinivasan M[9], Mascarenhas J[9], Das M[9], Kumar A[9], Joshi P[9], Banushree R[9], Kim U[9], Babu B[9], Ram A[9], Saravanan R[9], Kannappan KN[9], Kalyani N[9], Criswell LA[1], Shiboski SC[1], Baer A[8], Challacombe S[6], Lanfranchi H[2], Schiødt M[4], Umehara H[5], Vivino F[7], Zhao Y[3], Dong Y[3], Greenspan D[1], Heidenreich AM[2], Helin P[4], Kirkham B[6], Kitagawa K[5], Larkin G[6], Li M[3], Lietman T[1], Lindegaard J[4], McNamara N[1], Sack K[1], Shirlaw P[6], Sugai S[5], Vollenweider C[2], Whitcher J[1], Wu A[1], Zhang S[3], Zhang W[3], Greenspan JS[1], Daniels TE[1], **Shiboski CH**[1], Criswell LA[10].

[1]University of California San Francisco, San Francisco, CA, USA; [2]University of Buenos Aires and German Hospital, Buenos Aires, Argentina; [3]Peking Union Medical College Hospital, Beijing, China; [4]Rigshospitalet, Copenhagen, Denmark; [5]Kanazawa Medical University, Ishikawa, Japan; [6]King's College London, London, UK; [7]University of Pennsylvania, Philadelphia, Pennsylvania, USA; [8]Johns Hopkins University, Baltimore, Maryland, USA; [9]Aravind Eye Hospital, Madurai, India; [10]National Human Genome Research Institute, NIH, Bethesda, Maryland, USA.

SICCA Study was funded by the National Institutes of Health (NIH): N01DE32636 (SICCA), HHSN26S201300057C (SICCA), U01DE028891 (SICCA), R03DE029800 (SICCA), U01HG004446 (SICCA-GWAS), P30AR070155 (SICCA-GWAS).

Genotype data from the Sjögren's International Collaborative Clinical Alliance (SICCA) Registry was obtained through **dbGAP accession number phs000672.v1.p1**. This study was supported by the National Institute of Dental and Craniofacial Research (NIDCR), the National Eye Institute, and the Office of Research on Women's Health through contract number N01-DE-32636. Genotyping services were provided by the Center for Inherited Disease Research (CIDR). CIDR is fully funded through a federal contract from the National Institutes of Health (NIH) to the Johns Hopkins University (contract numbers HHSN268200782096C, HHSN268201100011I, HHSN268201200008I). Funds for genotyping were provided by the NIDCR through CIDR's NIH contract. Assistance with data cleaning and imputation was provided by the University of Washington. SICCA thanks investigators from the following studies that provided DNA samples for genotyping: the Genetic Architecture of Smoking and Smoking Cessation, Collaborative

Genetic Study of Nicotine Dependence (phs000404.v1.p1); Age-Related Eye Disease Study (AREDS) - Genetic Variation in Refractive Error Substudy (phs000429.v1.p1); and National Institute of Mental Health's Human Genetics Initiative (phs000021.v3.p2, phs000167.v1.p1). SICCA thanks the many clinical collaborators and research participants who contributed to this research.

**C. The UK Primary Sjögren's Syndrome Registry is composed of the following members: Wan-Fai Ng**[1], Simon J. Bowman[2], Bridget Griffiths[3], Frances Hall[4], Elalaine C. Bacabac[5], Robert Moots[5], Kuntal Chadravarty[6], Shamin Lamabadusuriya[6], Michele Bombardieri[7], Constantino Pitzalis[7], Nurhan Sutcliffe[7], Nagui Gendi[8], Rashidat Adeniba[8], John Hamburger[9], Andrea Richards[9], Saaeha Rauz[10], Sue Brailsford[1], Joanne Logan[11], Diamuid Mulherin[11], Paul Emery[12], Alison McManus[12], Colin Pease[12], Alison Booth[13], Marian Regan[13], Theodoros Dimitroulas[14], Lucy Kadiki[14], Daljit Kaur[14], George Kitas[14], Mark Lloyd[15], Lisa Moore[15], Esther Gordon[16], Cathy Lawson[16], Monica Gupta[17], John Hunter[17], Lesley Stirton[17], Gill Ortiz[18], Elizabeth Price[18], Gavin Clunie[19], Ginny Rose[19], Sue Cuckow[19], Susan Knight[20], Deborah Symmons[20], Beverley Jones[20], Shereen Al-Ali[1], Andrew Carr[1], Katherine Collins[1], Andini Natasari[1], Philip Stocks[1], Jessica Tarn[1], Ian Corbett[3], Christine Downie[3], Suzanne Edgar[3], Marco Carrozzo[3], Francisco Figuereido[3], Heather Foggo[3], Dennis Lendrem[3], Iain Macleod[3], Philip Mawson[3], Sheryl Mitchell[3], Adrian Jones[21], Peter Lanyon[21], Alice Muir[21], Paula White[22], Steven Young-Min[22], Susan Pugmire[23],Saravanan Vadivelu[23], Annie Cooper[24], Marianne Watkins[24], Anne Field[25], Stephen Kaye[25], Devesh Mewar[25], Patricia Medcalf[25], Pamela Tomlinson[25], Debbie Whiteside[25], Neil McHugh[26], John Pauling[26], Julie James[26], Nike Olaitan[26], Mohammed Akil[27], Jayne McDermott[27], Olivia Godia[27], David Coady[28], Elizabeth Kidd[28], Lynne Palmer[28], Bhaskar Dasgupta[29], Victoria Katsande[29], Pamela Long[29], Charles Li[30], Usha Chandra[31], Kirsten MacKay[31], Stefano Fedele[32], Ada Ferenkey-Koroma[32], Ian Giles[32], David Isenberg[32], Helena Maconnell[32], Stephen Porter[32],Paul Allcoat[33], John Mclaren[33].

[1]Newcastle University, Newcastle upon Tyne, UK; [2]University Hospital Birmingham, Birmingham,UK; [3]Newcastle upon Tyne Hospitals NHS Foundation Trust, Newcastle upon Tyne, UK; [4]Addenbrooke's Hospital, Cambridge, UK; [5]Aintree University Hospitals, Liverpool, UK; [6]Barking, Havering and Redbridge NHS Trust, Barking, UK; [7]Bart and the London NHS Trust, London, UK; [8]Basildon Hospital, Basildon, UK; [9]Birmingham Dental Hospital, Brimingham, UK; [10]Birmingham & Midland Eye Centre, Birmingham, UK; [11]Cannock Chase Hospital, Cannock, UK; [12]Chapel Allerton Hospital, Leeds, Leeds, UK; [13]Derbyshire Royal Infirmary, Derby, UK; [14]Dudley Group of Hospitals NHS Foundation Trust, Dudley, UK; [15]Frimley Park Hospital, Frimley Park, UK; [16]Harrogate District Foundation Trust Hospital, Harrogate, UK; [17]Gartnavel General Hospital, Glasgow, UK; [18]Great Western Hospital, Swindon, UK; [19]Ipswich Hospital NHS Trust, Ipswich, UK; [20]Macclesfield District General Hospital & Arthritis Research UK Epidemiology Unit, Manchester, Manchester, UK; [21]Nottingham University Hospital, Nottingham, UK; [22]Portsmouth Hospitals NHS Trust, Portsmouth, UK; [23]Queen's Elizabeth Hospital, Gateshead, Gateshead, UK; [24]Royal Hampshire County Hospital, Winchester, UK; [25]Royal Liverpool University Hospital, Liverpool, UK; [26]Royal National Hospital for Rheumatic Diseases, Bath, UK; [27]Sheffield Teaching Hospitals NHS Trust, Sheffield, UK; [28]Sunderland Royal Hospital, Sunderland, UK; [29]Southend University Hospital, Southend, UK; [30]Royal Surrey Hospital, Guildford, UK; [31]Torbay Hospital, Torbay, UK; [32]University College Hospital & Eastman Dental Institute, London, UK; [33]Whyteman's Brae Hospital, Kirkaldy, Fife, UK.

The UK Primary Sjögren's Syndrome Registry was funded by the Medical Research Council (G080062; W-F.N.), and the British Sjögren's Syndrome Association (W-F.N.). This work also received infra-structure support from the NIHR Newcastle Biomedical Research Centre, Newcastle and NIHR Newcastle Clinical Research Facility.

| SNP Name | Reference Allele | Alternative Allele | Non-Risk | Risk | Notes |
|---|---|---|---|---|---|
| rs57494551 | C (major) | T (minor) | T | C | Reference is risk allele |
| rs10892294 | G (major) | C (minor) | C | G | Reference is risk allele |
| rs4936443 | C (minor) | T (major) | C | T | |
| rs4938572 | C (minor) | T (major) | C | T | |
| rs7117261 | T (minor) | C (major) | T | C | |
| rs4938573 | C (minor) | T (major) | C | T | |
| rs12365699 | G (major) | A (minor) | G | A | |

**Supplemental Table 2: Patient-Derived EBV B Cell Genotypes**

| Cell Line | Haplotype | Allele | Application Used |
|---|---|---|---|
| p1000068-2 | Heterozygous | | Luciferase & EMSA (pooled) |
| p1000069-5 | Heterozygous | | Luciferase & EMSA (pooled) |
| p1000334-2 | Heterozygous | | Luciferase & EMSA (pooled) |
| p1000181-5 | Homozygous | Minor allele (1) Non-Risk | Luciferase & EMSA (pooled), 3C |
| p1000370-0 | Homozygous | Minor allele (1) Non-Risk | 3C |
| p1000373-7 | Homozygous | Major allele (2) Risk | Luciferase & EMSA (pooled) |
| p1000333-5 | Homozygous | Major allele (2) Risk | Luciferase & EMSA (pooled), 3C |
| p1000161-0 | Homozygous | Major allele (2) Risk | 3C |

**Supplemental Table 3: EMSA Probes**

| EMSA Probe Name | | Sequence |
|---|---|---|
| rs57494551-altT-F | Non-Risk | TTCCTCCTGCCGACCCTGCTGCGCACCACATT**T**CTATATTCGCCTCCTCTCAGGCGCTCC |
| rs57494551-altT-R | Non-Risk | GGAGCGCCTGAGAGGAGGCGAATATAG**A**AATGTGGTGCGCAGCAGGGTCGGCAGGAGGAA |
| rs57494551-refC-F | Risk | TTCCTCCTGCCGACCCTGCTGCGCACCACATT**C**CTATATTCGCCTCCTCTCAGGCGCTCC |
| rs57494551-refC-R | Risk | GGAGCGCCTGAGAGGAGGCGAATATAG**G**AATGTGGTGCGCAGCAGGGTCGGCAGGAGGAA |
| rs10892294-altC-F | Non-Risk | GGTTTCCTGCCACTAGTCAATCTGCAGA**C**ACTTTTATTGATTCTTGAAAATACAACTGTG |
| rs10892294-altC-R | Non-Risk | CACAGTTGTATTTTCAAGAATCAATAAAAGT**G**TCTGCAGATTGACTAGTGGCAGGAAACC |
| rs10892294-refG-F | Risk | GGTTTCCTGCCACTAGTCAATCTGCAGA**G**ACTTTTATTGATTCTTGAAAATACAACTGTG |
| rs10892294-refG-R | Risk | CACAGTTGTATTTTCAAGAATCAATAAAAGT**C**TCTGCAGATTGACTAGTGGCAGGAAACC |
| rs4936443-refC-F | Non-Risk | AGGTTTAGTTTGCCTGGAGAGAAACAGGC**C**GGAGAGAGACTGCGGCCTCCCTAGGGTCTT |
| rs4936443-refC-R | Non-Risk | AAGACCCTAGGGAGGCCGCAGTCTCTCTCC**G**GCCTGTTTCTCTCCAGGCAAACTAAACCT |
| rs4936443-altT-F | Risk | AGGTTTAGTTTGCCTGGAGAGAAACAGGC**T**GGAGAGAGACTGCGGCCTCCCTAGGGTCTT |
| rs4936443-altT-R | Risk | AAGACCCTAGGGAGGCCGCAGTCTCTCTCC**A**GCCTGTTTCTCTCCAGGCAAACTAAACCT |
| rs4938572-refC-F | Non-Risk | CGGCAAATTCCTCCAGCTCAGTGGCTGCTGGG**C**AGCAGCACAGCCGGTTTCTCTCAAGGG |
| rs4938572-refC-R | Non-Risk | CCCTTGAGAGAAACCGGCTGTGCTGCT**G**CCCAGCAGCCACTGAGCTGGAGGAATTTGCCG |
| rs4938572-altT-F | Risk | CGGCAAATTCCTCCAGCTCAGTGGCTGCTGGG**T**AGCAGCACAGCCGGTTTCTCTCAAGGG |
| rs4938572-altT-R | Risk | CCCTTGAGAGAAACCGGCTGTGCTGCTACCCAGCAGCCACTGAGCTGGAGGAATTTGCCG |
| rs7117261-refT-F | Non-Risk | CCCTTCTTTCCTGTCCCTGGGCACTTCCC**T**GGCTGCTCTCTTTCCCCACTGCCCAGCCCA |
| rs7117261-refT-R | Non-Risk | TGGGCTGGGCAGTGGGGAAAGAGAGCAGCCAGGGAAGTGCCCAGGGACAGGAAAGAAGGG |
| rs7117261-altC-F | Risk | CCCTTCTTTCCTGTCCCTGGGCACTTCCC**C**GGCTGCTCTCTTTCCCCACTGCCCAGCCCA |
| rs7117261-altC-R | Risk | TGGGCTGGGCAGTGGGGAAAGAGAGCAGCC**G**GGGAAGTGCCCAGGGACAGGAAAGAAGGG |
| rs4938573-refC-F | Non-Risk | TCACTTGTGTAATTCATCAACAAACTTTA**C**TGAGCACCTAATAGGCACTGAGTGTTTTCG |
| rs4938573-refC-R | Non-Risk | CGAAAACACTCAGTGCCTATTAGGTGCTCA**G**TAAAGTTTGTTGATGAATTACACAAGTGA |
| rs4938573-altT-F | Risk | TCACTTGTGTAATTCATCAACAAACTTTA**T**TGAGCACCTAATAGGCACTGAGTGTTTTCG |
| rs4938573-altT-R | Risk | CGAAAACACTCAGTGCCTATTAGGTGCTCA**A**TAAAGTTTGTTGATGAATTACACAAGTGA |
| rs12365699-refG-F | Non-Risk | TCATTTGGAAACCTCTCTCGGAGGAGCTCC**G**TGATCAAGGTGCAGATGCGGCAGGTGGGC |
| rs12365699-refG-R | Non-Risk | GCCCACCTGCCGCATCTGCACCTTGATCA**C**GGAGCTCCTCCGAGAGAGGTTTCCAAATGA |
| rs12365699-altA-F | Risk | TCATTTGGAAACCTCTCTCGGAGGAGCTCC**A**TGATCAAGGTGCAGATGCGGCAGGTGGGC |
| rs12365699-altA-R | Risk | GCCCACCTGCCGCATCTGCACCTTGATCA**T**GGAGCTCCTCCGAGAGAGGTTTCCAAATGA |

**Supplemental Table 4: Luciferase Reporter Assay gBlocks**

| | |
|---|---|
| **rs57494551-refC** | ACCCTCTGGTACCTGTTATTGACCCGCAAGGCCTACTGCAATCAAGCAGCGGCCGGT<br>TTGCTTCTAAACCGAGCCCTCCAATACAGCATGTCCCTGCCGCCCCCTATAGGGCCG<br>CCTCGTACCCTATAACCTCCACCATCATCCCCCTAAGTCCTTGCCGCCCCCTTCGGCC<br>TCATATTCCCTCATCTTCGATAAAGCTACTCCGAGTACTTAGCCTGTTCCTCCTGCCGA<br>CCCTGCTGCGCACCACATT**C**CTATATTCGCCTCCTCTCAGGCGCTCCCACCCCACAC<br>AGCTGCCGACCGCCTTCCTCCCCAGGCCCGGCCAGGCCTTAGGCCTCCGCCCGAGA<br>GTCCCCCAGAGCCGGCCCGGGGGGCTCCCCACAGCCCCCAAAGCACCGCTGACCT<br>CGACCCCACCACCTCACCCCAAGCCTCGCGACTCGGGCCCGTGTCCTACCAACGAG<br>GCCACTCCCGCTCGGCACCCTCGGTCCTTTATAAGCTTTGTTGAC |
| **rs57494551-altT** | ACCCTCTGGTACCTGTTATTGACCCGCAAGGCCTACTGCAATCAAGCAGCGGCCGGT<br>TTGCTTCTAAACCGAGCCCTCCAATACAGCATGTCCCTGCCGCCCCCTATAGGGCCG<br>CCTCGTACCCTATAACCTCCACCATCATCCCCCTAAGTCCTTGCCGCCCCCTTCGGCC<br>TCATATTCCCTCATCTTCGATAAAGCTACTCCGAGTACTTAGCCTGTTCCTCCTGCCGA<br>CCCTGCTGCGCACCACATT**T**CTATATTCGCCTCCTCTCAGGCGCTCCCACCCCACACA<br>GCTGCCGACCGCCTTCCTCCCCAGGCCCGGCCAGGCCTTAGGCCTCCGCCCGAGAG<br>TCCCCCAGAGCCGGCCCGGGGGGCTCCCCACAGCCCCCAAAGCACCGCTGACCTC<br>GACCCCACCACCTCACCCCAAGCCTCGCGACTCGGGCCCGTGTCCTACCAACGAGG<br>CCACTCCCGCTCGGCACCCTCGGTCCTTTATAAGCTTTGTTGAC |
| **rs10892294-refG** | TTCTCCTTGGTACCGGGTATTTTCCAAGTTAGTTCAGGGGCAGTTGCCGAGGAATAAC<br>ACTGATGGGGGTTCACACTATGGCGATCTTGTTGAACTGCCTGATGTTGGTTTGTGTA<br>ATCTGCCCCCTTTGTGCCCAAGAACCTGTGACAAGATTCTGCTTCTGACAACCTTCTG<br>TGCAGGGGTAGCGACAGGAGTCTGAACAATCATTAAGTGTCCAGCCCTGGTTTCCTG<br>CCACTAGTCAATCTGCAGA**G**ACTTTTATTGATTCTTGAAAATACAACTGTGATTGTAGG<br>TTTGGGCCCTGGGAATGTAATTTATAGCTAAAGAATGGGCAATGTCTTCTGTCCTGAG<br>TGCCATGGTTGGATTGTGTAACATCTGTGGCTACCAGGACAAGGAAGGAACAAAGGA<br>AGACAAAAGTATTTCAGTGCAGAGATCTTCTCAGTGACCCTCCTTGCTTTTCATCTGCC<br>TGATGAATTTTAAAGACTACTTAAGCTTGTCATCAT |
| **rs10892294-altC** | TTCTCCTTGGTACCGGGTATTTTCCAAGTTAGTTCAGGGGCAGTTGCCGAGGAATAAC<br>ACTGATGGGGGTTCACACTATGGCGATCTTGTTGAACTGCCTGATGTTGGTTTGTGTA<br>ATCTGCCCCCTTTGTGCCCAAGAACCTGTGACAAGATTCTGCTTCTGACAACCTTCTG<br>TGCAGGGGTAGCGACAGGAGTCTGAACAATCATTAAGTGTCCAGCCCTGGTTTCCTG<br>CCACTAGTCAATCTGCAGA**C**ACTTTTATTGATTCTTGAAAATACAACTGTGATTGTAGG<br>TTTGGGCCCTGGGAATGTAATTTATAGCTAAAGAATGGGCAATGTCTTCTGTCCTGAG<br>TGCCATGGTTGGATTGTGTAACATCTGTGGCTACCAGGACAAGGAAGGAACAAAGGA<br>AGACAAAAGTATTTCAGTGCAGAGATCTTCTCAGTGACCCTCCTTGCTTTTCATCTGCC<br>TGATGAATTTTAAAGACTACTTAAGCTTGTCATCAT |
| **rs4936443-refC** | GCTGGAGGGTACCAAAGGCCTGGAGAGCTCCCAGCGCCCTCTGAGACATGGCTCCA<br>GGTCACACAGCCCAAAGCCTTGGCCTGTTTTGTACGTGGACGGGGCAAAGAGAACAC<br>CCTCGCCGCTTCTCTCTGCCTTTAGCAGGGCTGTAGGAAACCCCCACCAGAGACCTC<br>CAGCTTGGAGAGGAAAGAGTGGAACAGCCCTCTGGAAGCAAGTTACCCACAGGTTTA<br>GTTTGCCTGGAGAGAAACAGGC**C**GGAGAGAGACTGCGGCCTCCCTAGGGTCTTCTG<br>ACGGCAAATTCCTCCAGCTCAGTGGCTGCTGGG**C**AGCAGCACAGCCGGTTTCTCTCA<br>AGGGCACACCCCACACACCGCGTCACTGTGCACTAGCCTCAGATGACAGACAAGCCT<br>TTCACAAGACTTTTGTGGCACTGTTCATTTCTGAGACCTTCTCTATGATGAGCTCAAAC<br>TGCTTACCTCAGAGAAGAAACTGCGTGCACAAGCTTAGAAAGC<br>Green=rs4938572 |

**Supplementary Table 4 Continued**

| | |
|---|---|
| **rs4936443-altT** | GCTGGAGGGTACCAAAGGCCTGGAGAGCTCCCAGCGCCCTCTGAGACATGGCTCCA<br>GGTCACACAGCCCAAAGCCTTGGCCTGTTTTGTACGTGGACGGGGCAAAGAGAACAC<br>CCTCGCCGCTTCTCTCTGCCTTTAGCAGGGCTGTAGGAAACCCCCACCAGAGACCTC<br>CAGCTTGGAGAGGAAAGAGTGGAACAGCCCTCTGGAAGCAAGTTACCCACAGGTTTA<br>GTTTGCCTGGAGAGAAACAGGC**T**GGAGAGAGACTGCGGCCTCCCTAGGGTCTTCTGA<br>CGGCAAATTCCTCCAGCTCAGTGGCTGCTGGG**C**AGCAGCACAGCCGGTTTCTCTCAA<br>GGGCACACCCCACACACCGCGTCACTGTGCACTAGCCTCAGATGACAGACAAGCCTT<br>TCACAAGACTTTTGTGGCACTGTTCATTTCTGAGACCTTCTCTATGATGAGCTCAAACT<br>GCTTACCTCAGAGAAGAAACTGCGTGCACAAGCTTAGAAAGC<br>Green=rs4938572 |
| **rs4938572-refC** | GCCTTGGCGGTACCCTGTTTTGTACGTGGACGGGGCAAAGAGAACACCCTCGCCGCT<br>TCTCTCTGCCTTTAGCAGGGCTGTAGGAAACCCCCACCAGAGACCTCCAGCTTGGAG<br>AGGAAAGAGTGGAACAGCCCTCTGGAAGCAAGTTACCCACAGGTTTAGTTTGCCTGG<br>AGAGAAACAGGC**C**GGAGAGAGACTGCGGCCTCCCTAGGGTCTTCTGACGGCAAATT<br>CCTCCAGCTCAGTGGCTGCTGGG**C**AGCAGCACAGCCGGTTTCTCTCAAGGGCACAC<br>CCCACACACCGCGTCACTGTGCACTAGCCTCAGATGACAGACAAGCCTTTCACAAGA<br>CTTTTGTGGCACTGTTCATTTCTGAGACCTTCTCTATGATGAGCTCAAACTGCTTACCT<br>CAGAGAAGAAACTGCGTGCACAGAAAGCTGCTGAGGCCAGCTTGGGGCCCCTTCTTT<br>CCTGTCCCTGGGCACTTCCC**T**GGCTGCTCTAAGCTTCTTTCCCC<br>Red=rs4936443<br>Blue=rs7117261 |
| **rs4938572-altT** | GCCTTGGCGGTACCCTGTTTTGTACGTGGACGGGGCAAAGAGAACACCCTCGCCGCT<br>TCTCTCTGCCTTTAGCAGGGCTGTAGGAAACCCCCACCAGAGACCTCCAGCTTGGAG<br>AGGAAAGAGTGGAACAGCCCTCTGGAAGCAAGTTACCCACAGGTTTAGTTTGCCTGG<br>AGAGAAACAGGC**C**GGAGAGAGACTGCGGCCTCCCTAGGGTCTTCTGACGGCAAATT<br>CCTCCAGCTCAGTGGCTGCTGGG**T**AGCAGCACAGCCGGTTTCTCTCAAGGGCACACC<br>CCACACACCGCGTCACTGTGCACTAGCCTCAGATGACAGACAAGCCTTTCACAAGAC<br>TTTTGTGGCACTGTTCATTTCTGAGACCTTCTCTATGATGAGCTCAAACTGCTTACCTC<br>AGAGAAGAAACTGCGTGCACAGAAAGCTGCTGAGGCCAGCTTGGGGCCCCTTCTTTC<br>CTGTCCCTGGGCACTTCCC**T**GGCTGCTCTAAGCTTCTTTCCCC<br>Red=rs4936443<br>Blue=rs7117261 |
| **rs7117261-refT** | GCTCAGTGGTACCGGCTGCTGGG**C**AGCAGCACAGCCGGTTTCTCTCAAGGGCACAC<br>CCCACACACCGCGTCACTGTGCACTAGCCTCAGATGACAGACAAGCCTTTCACAAGA<br>CTTTTGTGGCACTGTTCATTTCTGAGACCTTCTCTATGATGAGCTCAAACTGCTTACCT<br>CAGAGAAGAAACTGCGTGCACAGAAAGCTGCTGAGGCCAGCTTGGGGCCCCTTCTTT<br>CCTGTCCCTGGGCACTTCCC**T**GGCTGCTCTCTTTCCCCACTGCCCAGCCCAAGGAGT<br>CCCCTCTGCAGCTGACCCGGGTTCAGCCTCCAGAACAGCGAGTTCCACAGCCCTGAA<br>GCCTGGCCATCGTCCCTTTTCTGGACACTGGACTGGTTCATAGGGCTCAGTGCCCTG<br>CGGCTTTCTCCTCCCCACCACAGGCCTGGGAGGGGCAAGAAGCACCAGTTTGTTTTC<br>TGGGTCAGCCTGCAGCAAAGAGCGACTGCAAGCTTCCTCACC<br>Green=rs4938572 |
| **rs7117261-altC** | GCTCAGTGGTACCGGCTGCTGGG**C**AGCAGCACAGCCGGTTTCTCTCAAGGGCACAC<br>CCCACACACCGCGTCACTGTGCACTAGCCTCAGATGACAGACAAGCCTTTCACAAGA<br>CTTTTGTGGCACTGTTCATTTCTGAGACCTTCTCTATGATGAGCTCAAACTGCTTACCT<br>CAGAGAAGAAACTGCGTGCACAGAAAGCTGCTGAGGCCAGCTTGGGGCCCCTTCTTT<br>CCTGTCCCTGGGCACTTCCC**C**GGCTGCTCTCTTTCCCCACTGCCCAGCCCAAGGAGT<br>CCCCTCTGCAGCTGACCCGGGTTCAGCCTCCAGAACAGCGAGTTCCACAGCCCTGAA<br>GCCTGGCCATCGTCCCTTTTCTGGACACTGGACTGGTTCATAGGGCTCAGTGCCCTG<br>CGGCTTTCTCCTCCCCACCACAGGCCTGGGAGGGGCAAGAAGCACCAGTTTGTTTTC<br>TGGGTCAGCCTGCAGCAAAGAGCGACTGCAAGCTTCCTCACC<br>Green=rs4938572 |

**Supplementary Table 4 Continued**

| | |
|---|---|
| **rs4938573-refC** | ACTACTGGTACCGATTTACATATCACACATGTGCTCATCCATCCATTCACCAAACCTCT<br>GTTGACTGCTCGCGCTGCCCTGCGTTCCTGGCACTGTGCAGGGTTATAATCTAGTAG<br>GAAAGACCTGACAAGGTCACAGATGTCAGTACTGAAAGGAAGAGTAGGGTAGATGGC<br>CACCCCACCCCCACCCCAACCTCAAAAAAGAGACTAAAAAATTACTGTCACTTGTGTA<br>ATTCATCAACAAACTTTA**C**TGAGCACCTAATAGGCACTGAGTGTTTTCGTGTATTGATC<br>ACGCATTGATCCTCACAATAACCTTTGAGATGGGTTGTGCCATTTACACAGGGCGAAG<br>AAGAGAGACTGGCCACTGTCACACAGCTACTGATATCCAAGCTGAGATCCAAAGCTC<br>CTCTGCTTCTCCTGATGAGAATGAGCACCACAGGCAGGCCACAGAAAAACACCCAGG<br>AGAGCCAAACTCAGACCCACCCCAGAAGCTTATTTCC |
| **rs4938573-altT** | ACTACTGGTACCGATTTACATATCACACATGTGCTCATCCATCCATTCACCAAACCTCT<br>GTTGACTGCTCGCGCTGCCCTGCGTTCCTGGCACTGTGCAGGGTTATAATCTAGTAG<br>GAAAGACCTGACAAGGTCACAGATGTCAGTACTGAAAGGAAGAGTAGGGTAGATGGC<br>CACCCCACCCCCACCCCAACCTCAAAAAAGAGACTAAAAAATTACTGTCACTTGTGTA<br>ATTCATCAACAAACTTTA**T**TGAGCACCTAATAGGCACTGAGTGTTTTCGTGTATTGATC<br>ACGCATTGATCCTCACAATAACCTTTGAGATGGGTTGTGCCATTTACACAGGGCGAAG<br>AAGAGAGACTGGCCACTGTCACACAGCTACTGATATCCAAGCTGAGATCCAAAGCTC<br>CTCTGCTTCTCCTGATGAGAATGAGCACCACAGGCAGGCCACAGAAAAACACCCAGG<br>AGAGCCAAACTCAGACCCACCCCAGAAGCTTATTTCC |
| **rs12365699-refG** | CACAAGAGGTACCGCCGTTGGCGGGATTTCCCATTGTCCCCCTTGGGTAGGTAAGGC<br>CAGGTGGCTGCTCCATCTCTGCCACCTCCAGCGCCGGTCCCACTGTGTCAGCAGCC<br>CTGTCCCCTACTGCTGTGTCATCAATTACTCTCAGGTGCCTGGCCCCCACCCAGTGG<br>CCCCCACCTTGCAGCCCCGAAGGCTTCCTTCCTGGGGCAGCAGGGCCGAGTCATTT<br>GGAAACCTCTCTCGGAGGAGCTCC**G**TGATCAAGGTGCAGATGCGGCAGGTGGGCCG<br>GCCTCAATCATGTCTCCAATTGCGACGGTGAATGCGGTGAGGAGTTTCGTTGGCCCA<br>TCACCTCTGCCTTGGGCCTGGCTCACTTTCACTGCTGAGTTAGTTCCACGGCCGCCTT<br>TGATGATGCCGCTTCAGCATCTTTTTTCTTCGGCGTTTCCTGCTCCTTTGTTTTCAAGG<br>TCACTCTGTCCTGCCTGCCCCACTCTGGGCCAAGCTTACCCAAG |
| **rs12365699-altA** | CACAAGAGGTACCGCCGTTGGCGGGATTTCCCATTGTCCCCCTTGGGTAGGTAAGGC<br>CAGGTGGCTGCTCCATCTCTGCCACCTCCAGCGCCGGTCCCACTGTGTCAGCAGCC<br>CTGTCCCCTACTGCTGTGTCATCAATTACTCTCAGGTGCCTGGCCCCCACCCAGTGG<br>CCCCCACCTTGCAGCCCCGAAGGCTTCCTTCCTGGGGCAGCAGGGCCGAGTCATTT<br>GGAAACCTCTCTCGGAGGAGCTCC**A**TGATCAAGGTGCAGATGCGGCAGGTGGGCCG<br>GCCTCAATCATGTCTCCAATTGCGACGGTGAATGCGGTGAGGAGTTTCGTTGGCCCA<br>TCACCTCTGCCTTGGGCCTGGCTCACTTTCACTGCTGAGTTAGTTCCACGGCCGCCTT<br>TGATGATGCCGCTTCAGCATCTTTTTTCTTCGGCGTTTCCTGCTCCTTTGTTTTCAAGG<br>TCACTCTGTCCTGCCTGCCCCACTCTGGGCCAAGCTTACCCAAG |
| **All Non-Risk** | TCTGAGAGGTACCCATGGCTCCAGGTCACACAGCCCAAAGCCTTGGCCTGTTTTGTA<br>CGTGGACGGGGCAAAGAGAACACCCTCGCCGCTTCTCTCTGCCTTTAGCAGGGCTGT<br>AGGAAACCCCCACCAGAGACCTCCAGCTTGGAGAGGAAAGAGTGGAACAGCCCTCT<br>GGAAGCAAGTTACCCACAGGTTTAGTTTGCCTGGAGAGAAACAGGC**C**GGAGAGAGAC<br>TGCGGCCTCCCTAGGGTCTTCTGACGGCAAATTCCTCCAGCTCAGTGGCTGCTGGG**C**<br>AGCAGCACAGCCGGTTTCTCTCAAGGGCACACCCCACACACCGCGTCACTGTGCACT<br>AGCCTCAGATGACAGACAAGCCTTTCACAAGACTTTTGTGGCACTGTTCATTTCTGAG<br>ACCTTCTCTATGATGAGCTCAAACTGCTTACCTCAGAGAAGAAACTGCGTGCACAGAA<br>AGCTGCTGAGGCCAGCTTGGGGCCCCTTCTTTCCTGTCCCTGGGCACTTCCC**T**GGCT<br>GCTCTCTTTCCCCACTGCCCAGCCCAAGGAGTCCCCTCTGCAGCTGACCCGGGTTCA<br>GCCTCCAGAACAGCGAGTTCCACAGCCCTGAAGCCTGGCCATCGTCCCTTTTCTGGA<br>CACTGGACTGGTTCATAGGGCTCAGTGCCCTGCGGCTTTCTCCTCCCCACCACAGGC<br>CTGGGAGGGGCAAGAAGCAAGCTTACCAGTT<br>Red=rs4936443_Non-risk (C)<br>Green=rs4938572_Non-risk (C)<br>Blue=rs7117261_Non-risk (T) |

**Supplementary Table 4 Continued**

| | |
|---|---|
| **All Risk** | TCTGAGAGGTACCCATGGCTCCAGGTCACACAGCCCAAAGCCTTGGCCTGTTTTGTA<br>CGTGGACGGGGCAAAGAGAACACCCTCGCCGCTTCTCTCTGCCTTTAGCAGGGCTGT<br>AGGAAACCCCCACCAGAGACCTCCAGCTTGGAGAGGAAAGAGTGGAACAGCCCTCT<br>GGAAGCAAGTTACCCACAGGTTTAGTTTGCCTGGAGAGAAACAGGC**T**GGAGAGAGAC<br>TGCGGCCTCCCTAGGGTCTTCTGACGGCAAATTCCTCCAGCTCAGTGGCTGCTGGG**T**<br>AGCAGCACAGCCGGTTTCTCTCAAGGGCACACCCCACACACCGCGTCACTGTGCACT<br>AGCCTCAGATGACAGACAAGCCTTTCACAAGACTTTTGTGGCACTGTTCATTTCTGAG<br>ACCTTCTCTATGATGAGCTCAAACTGCTTACCTCAGAGAAGAAACTGCGTGCACAGAA<br>AGCTGCTGAGGCCAGCTTGGGGCCCCTTCTTTCCTGTCCCTGGGCACTTCCC**C**GGCT<br>GCTCTCTTTCCCCACTGCCCAGCCCAAGGAGTCCCCTCTGCAGCTGACCCGGGTTCA<br>GCCTCCAGAACAGCGAGTTCCACAGCCCTGAAGCCTGGCCATCGTCCCTTTTCTGGA<br>CACTGGACTGGTTCATAGGGCTCAGTGCCCTGCGGCTTTCTCCTCCCCACCACAGGC<br>CTGGGAGGGGCAAGAAGCAAGCTTACCAGTT<br>Red=rs4936443_Risk (T)<br>Green=rs4938572_Risk (T)<br>Blue=rs7117261_Risk (C) |

**Supplemental Table 5: 3C-qPCR Primers**

| Primer Name | Primer Sequence |
|---|---|
| rs57494551 Anchor | CTCCACTCAAGATGGCGAAA |
| 1-rs57494551 3C Short Range Primer Control | ACTGAAGAAAGTAGGGGCGG |
| 2-rs57494551 | GACACAGGAAACCTGAGGGA |
| 3-rs57494551 | GTGCATTCATGATTGTTGCC |
| 4-rs57494551 | TTGGTGAGCTGTGATTGAGC |
| 5-rs57494551 | AGTCCCTTCCAGAGGGTTTT |
| 6-rs57494551 | TTGGTTACAGATTACACCTTGT |
| 7-rs57494551 | TCCTTCCTGATCAATGTCCC |
| 8-rs57494551 | TGAAGGTAACAGTGGCCCTT |
| 9-rs57494551 | AGGTGCATGTTGCTGTCAAG |
| 10-rs57494551 | CAAGGCTCTGGGAGAGAGG |
| 11-rs57494551 | CAAGGCTCTGGGAGAGAGG |
| 12-rs57494551 | TTAGCGTGGGATACAAAGCC |
| 13-rs57494551 | TTCCTGTATTCCAATTTCCCC |
| 14-rs57494551 | TTCTCCACTCACCCCAAACT |
| 15-rs57494551 | GAGGTGCTGGAGTATCTGGG |
| rs4938572 Anchor | TGTAATGGGGTGTTGGGTCC |
| 1-rs4938572 3C Short Range Primer Control | TCCTTCCTGATCAATGTCCC |
| 2-rs4938572 | TGACTTTGTGATCCAGCTGC |
| 3-rs4938572 | CTCCACTCAAGATGGCGAAA |
| 4-rs4938572 | CAAGGCTCTGGGAGAGAGG |
| 5-rs4938572 | TTTCCCTTCAAGAGAGCCAG |
| 6-rs4938572 | GCATAGAAAGGTGCTTTGGG |
| 7-rs4938572 | CTCTCCCCACTGAGTCCTCA |
| 8-rs4938572 | GAGGTGCTGGAGTATCTGGG |
| 9-rs4938572 | AAATCTTCCTTCCCAGCCTG |
| 10-rs4938572 | GTCTGAGGGTTCCTGAAGGA |
| 11-rs4938572 | CATATCCTGGGCCTTCACTG |
| 12-rs4938572 | AGGACAGTCAGAGAGCGTCAG |

### SUPPLEMENTAL FIGURES & LEGENDS

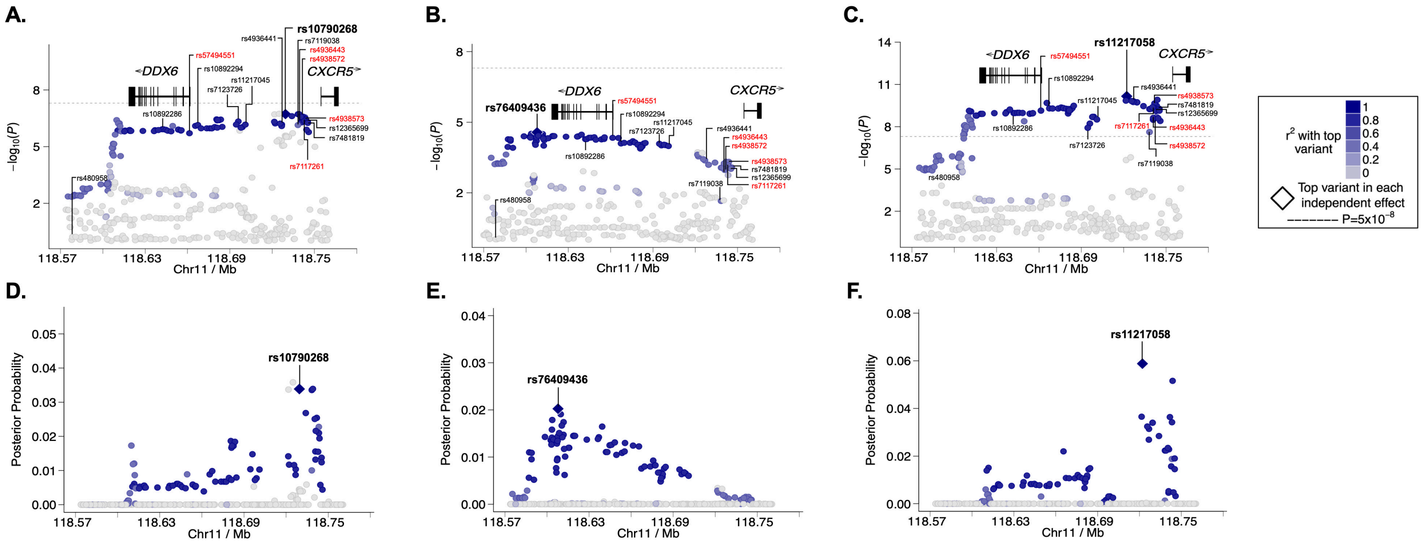


**Supplemental Figure 1. Fine mapping of the *DDX6-CXCR5* region in SjD and SLE Immunochip data after imputation.** **(A-C)** Logistic regression analysis was performed on **(A)** DS3-SjD (1916 SjD cases; 6194 controls), **(B)** DS4-SLE (3762 SLE cases; 6194 controls), and **(C)** DS3+DS4 (merged SjD and SLE) after quality control and imputation, identifying the top SNPs (e.g., index SNPs indicated in bold) of the *DDX6-CXCR5* association. SNPs prioritized for bioinformatic screening in this study are indicated in black; five SNPs prioritized for functional characterized are labeled in red. **(D-F)** Posterior probability distributions of SNPs in the *DDX6-CXCR5* region of **(D)** DS3, **(E)** DS4, and **(F)** DS3+DS4. SNPs with highest posterior probability are indicated. Pairwise ($r^2$) analysis for DS3-SjD **(A, D)** was based on the second most significant SNP because the most significant SNP was not in linkage disequilibrium (LD) with the haplotype.

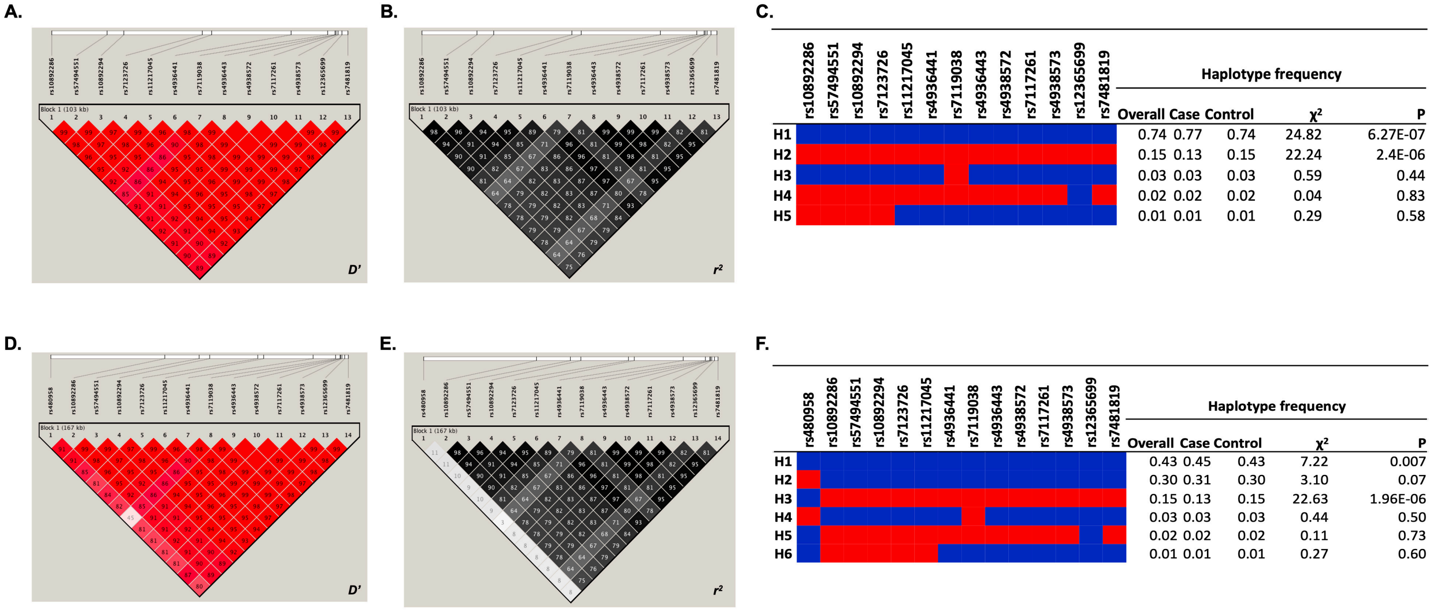


**Supplemental Figure 2. Haplotype frequency of the *DDX6-CXCR5* risk region in SjD. (A,D)** Co-inheritance (D') and **(B,E)** pairwise linkage disequilibrium ($r^2$) were assessed across the *DDX6-CXCR5* risk haplotype of SjD without **(A-C)** or with **(D-F)** rs480958. **(C,F)** Haplotype organization and frequencies of the index SNPs from the meta-analyses, SNPs previously reported as associated with SjD and/or SLE, and SNPs with strong bioinformatic functional evidence are shown.

| Variant ID | Refseq Gene | Chr | pos (hg 19) | pos (hg 38) | Ref | Alt | No. of eQTLs reported for variant | No. of protein bound by ChIP (ENCODE) | No. of regulatory motifs altered | Regulome DB score | DNase I HS clusters (% of cell types) | H3K4me1 peak - ENHANCER | H3K27ac peak - ENHANCER | H3K4me3 peak - PROMOTER | H3K9ac peak - PROMOTER |
|---|---|---|---|---|---|---|---|---|---|---|---|---|---|---|---|
| rs480958 [B] | Within *lnc-PHLDB1-1* | 11 | 118577990 | 118707281 | G | A | 2 | 0 | 8 | 4 | 0 | 24 | 16 | 6 | 6 |
| rs10892286 [A] | Within *DDX6* | 11 | 118642085 | 118771376 | A | C | 6 | 0 | 4 | 5 | 5 | 93 | 59 | 27 | 19 |
| **rs57494551** [E] | Within *DDX6* | 11 | 118661398 | 118790689 | C | T | 43 | 8 | 4 | 2b | 37 | 43 | 100 | 100 | 100 |
| rs10892294 [C] | 5.4 kb 3' of *DDX6* | 11 | 118667357 | 118796648 | G | C | 8 | 0 | 3 | 3a | 0 | 15 | 4 | 1 | 1 |
| rs7123726 [E] | 33 kb 3' of *DDX6* | 11 | 118694547 | 118823838 | T | C | 7 | 0 | 11 | 5 | 0 | 4 | 3 | 1 | 2 |
| rs11217045 [E] | 35 kb 3' of *DDX6* | 11 | 118697865 | 118827156 | C | A | 5 | 0 | 4 | 5 | 0 | 5 | 6 | 0 | 2 |
| rs4936441 [B] | 28 kb 5' of *CXCR5* | 11 | 118725660 | 118854951 | C | G | 2 | 0 | 3 | 3a | 1 | 2 | 2 | 0 | 0 |
| rs7119038 [C] | 16 kb 5' of *CXCR5* | 11 | 118738281 | 118867572 | G | A | 1 | 0 | 2 | 5 | 0 | 13 | 6 | 1 | 1 |
| **rs4936443** [E] | 14 kb 5' of *CXCR5* | 11 | 118740864 | 118870155 | C | T | 36 | 2 | 1 | 4 | 30 | 100 | 73 | 80 | 17 |
| **rs4938572** [E] | 14 kb 5' of *CXCR5* | 11 | 118740931 | 118870222 | C | T | 44 | 3 | 4 | 1b | 27 | 100 | 79 | 80 | 38 |
| **rs7117261** [E] | 14 kb 5' of *CXCR5* | 11 | 118741157 | 118870448 | T | C | 38 | 2 | 3 | 2b | 27 | 100 | 73 | 80 | 20 |
| **rs4938573** [D,E] | 13 kb 5' of *CXCR5* | 11 | 118741842 | 118871133 | C | T | 44 | 0 | 4 | 4 | 10 | 100 | 70 | 60 | 13 |
| rs12365699 [C] | 11 kb 5' of *CXCR5* | 11 | 118743286 | 118872577 | G | A | 5 | 7 | 3 | 1f | 5 | 34 | 29 | 10 | 7 |
| rs7481819 [E] | 9 kb 5' of *CXCR5* | 11 | 118745278 | 118874569 | C | T | 2 | 0 | 4 | 5 | 0 | 26 | 21 | 6 | 3 |

**Chromatin State (15-state model)**

| Variant ID | T cells (primary, cord blood) | T cells (primary, peripheral blood) | T cells (effector/memory enriched, periph., prim.) | T cells (CD8+ naïve, periph., prim.) | T cells (CD8+ memory, periph., prim.) | Th cells (CD4+, primary, peripheral blood) | Th cells (CD4+, memory, periph., prim., No.1) | Th cells (CD4+, memory, periph., prim., No.2) | Th cells (CD4+, naïve, periph., prim., NPC, CD25M, CD45RA) | Th cells (CD4+, naïve, periph., prim., NPC) | Th cells (CD4+, periph., prim., PMA/iono stim) | Th17 cells (periph., prim., PMA/iono stim.) | Treg cells (primary, peripheral blood) | B cells (primary, cord blood) | B cells (primary, peripheral blood) | NK cells (primary) | Monocytes (primary, peripheral blood) | Monocytes (primary, CD14+) | Neutrophils (primary, peripheral blood) | Mononuclear cells (primary, peripheral blood) | Hematopoietic stem cells (primary) | Hematopoietic stem cells (primary, G-CSF-mobilized Male) | Hematopoietic stem cells (primary, G-CSF-mobilized Female) | Hematopoietic stem cells (primary, short term culture) | Dnd41 T cell Leukemia Cell Line, cultured | GM12878 Lymphoblastoid Cell Line, cultured | K562 Leukemia Cell Line, cultured | Spleen | Thymus, fetal | Thymus | Kidney, fetal |
|---|---|---|---|---|---|---|---|---|---|---|---|---|---|---|---|---|---|---|---|---|---|---|---|---|---|---|---|---|---|---|---|
| rs480958 | | | | Enh | | | | | | | | | | | | | | | | | | | | | | | | | | | |
| rs10892286 | | | | | | | | | | | | | | | | | | | | | | | | | | | | | | | |
| **rs57494551** | TssA | TssA | TssA | TssA | TssA | TssA | TssA | TssA | Tx Flnk | TssA | TssA | Tx Flnk | Tx Flnk | TssA Flnk | TssA | TssA | TssA | TssA | TssA | TssA Flnk | TssA | TssA | TssA | TssA | TssA | TssA | TssA | TssA | TssA | TssA | TssA |
| rs10892294 | | | | | | | | | | | | | | | | | | | | | | | | | | | | | | | |
| rs7123726 | | | | | | | | | | | | | | Enh | Enh | | | | | | | | | | | Enh | | | | | |
| rs11217045 | | | | | | | | | | | | | | | | | | | | | | | | | | | | | | | |
| rs4936441 | | | | | | | | | | | | | | | | | | | | | | | | | | | | | | | |
| rs7119038 | | Enh | | | | | | | | | | | | | Enh | Enh | | | | | | | | | | | | | | | |
| **rs4936443** | TssA Flnk | TssA Flnk | TssA Flnk | TssA Flnk | Enh | TssA Flnk | TssA Flnk | TssA Flnk | TssA Flnk | TssA Flnk | TssA Flnk | TssA Flnk | TssA Flnk | TssA Flnk | TssA Flnk | TssA Flnk | Enh | Enh | TssA Flnk | Enh | Enh | TssA Flnk | Enh | TssA Flnk | Enh | Biv Flnk | Enh | Enh | TssA Flnk | TssA Flnk | TssA Flnk |
| **rs4938572** | TssA Flnk | TssA Flnk | TssA Flnk | TssA Flnk | Enh | TssA Flnk | TssA Flnk | TssA Flnk | TssA Flnk | TssA Flnk | TssA Flnk | TssA Flnk | TssA Flnk | TssA Flnk | TssA Flnk | TssA Flnk | Enh | Enh | TssA Flnk | Enh | Enh | TssA Flnk | Enh | TssA Flnk | Enh | Biv Flnk | Enh | Enh | TssA Flnk | TssA Flnk | TssA Flnk |
| **rs7117261** | TssA Flnk | TssA Flnk | Enh | TssA Flnk | Enh | TssA Flnk | TssA Flnk | TssA Flnk | TssA Flnk | TssA Flnk | TssA Flnk | TssA Flnk | TssA Flnk | TssA Flnk | TssA Flnk | TssA Flnk | Enh | Enh | TssA Flnk | Enh | Enh | TssA Flnk | Enh | TssA Flnk | Enh | TssA Flnk | Enh | Enh | TssA Flnk | TssA Flnk | Enh |
| **rs4938573** | Enh | Enh | Enh | TssA Flnk | TssA Flnk | Enh | Enh | TssA Flnk | Enh | TssA Flnk | Enh | Enh | Enh | TssA Flnk | TssA Flnk | Enh | Enh | | Enh | Enh | Enh | | | Enh | Enh | Enh | | Enh | Enh | Enh | Enh |
| rs12365699 | | TssA Flnk | Enh | Enh | Enh | TssA Flnk | Enh | TssA Flnk | Enh | TssA Flnk | TssA Flnk | TssA Flnk | TssA Flnk | Enh | Enh | Enh | | | | Enh | | | | | | Enh | | | Enh | Enh | |
| rs7481819 | Enh | Enh | Enh | Enh | Enh | Enh | Enh | Enh | Enh | Enh | Enh | Enh | Enh | | Enh | Enh | | | Enh | Enh | | | | | Enh | | | | Enh | | |

5' — Genomic space — 3'

[A] Langefeld *et al.*, Nat Commun., 2017 Jul17;8:16021.
[B] Yin *et al.*, Ann Rheum Dis., 2021 May; 80(5); 632-640.
[C] Lessard *et al.*, Nat Genet. 2013 Nov;45(11):1284-92.
[D] Aqrawi et al., Clin Exp Immunol. 2018 Jun;192(3):259-270.
[E] Current Study

Legend:
- Active Transcription start site (TSS) (TssA)
- Flanking Active TSS (TssA Flnk)
- Transcription at gene 5' and 3' (Tx Flnk)
- Flanking Bivalent TSS/Enh (Biv Flnk)
- Enhancer (Enh)
- Both enhancer and promoter marks present
- No epimarks
- Number of promoter epimarks >50
- Number of promoter epimarks <50
- Number of enhancer epimarks >50
- Enhancer epimarks <50
- # <10 epimarks

**Supplemental Figure 3. Functional bioinformatic analyses of putative functional SNPs in the *DDX6-CXCR5* risk region.** RegulomeDB, HaploReg 4.1, UCSC genome browser, GTEx, and ENCODE were used to assess the predicted histone marks and regulatory elements positioned at the indicated SNPs in 31 cell types and tissues. Total number of enhancer marks (H3K4me1, H3K27ac) or promoter marks (H3K4me3, H3K9ac) are also indicated. RegulomeDB scores, total number of eQTLs, proteins bound by ChIP, altered regulatory motifs, and DNase I HS clusters from HaploReg are depicted. White boxes indicate that no epimarks were found. Five SNPs (red) exhibit compounding bioinformatic evidence of function and were prioritized for functional interrogation.

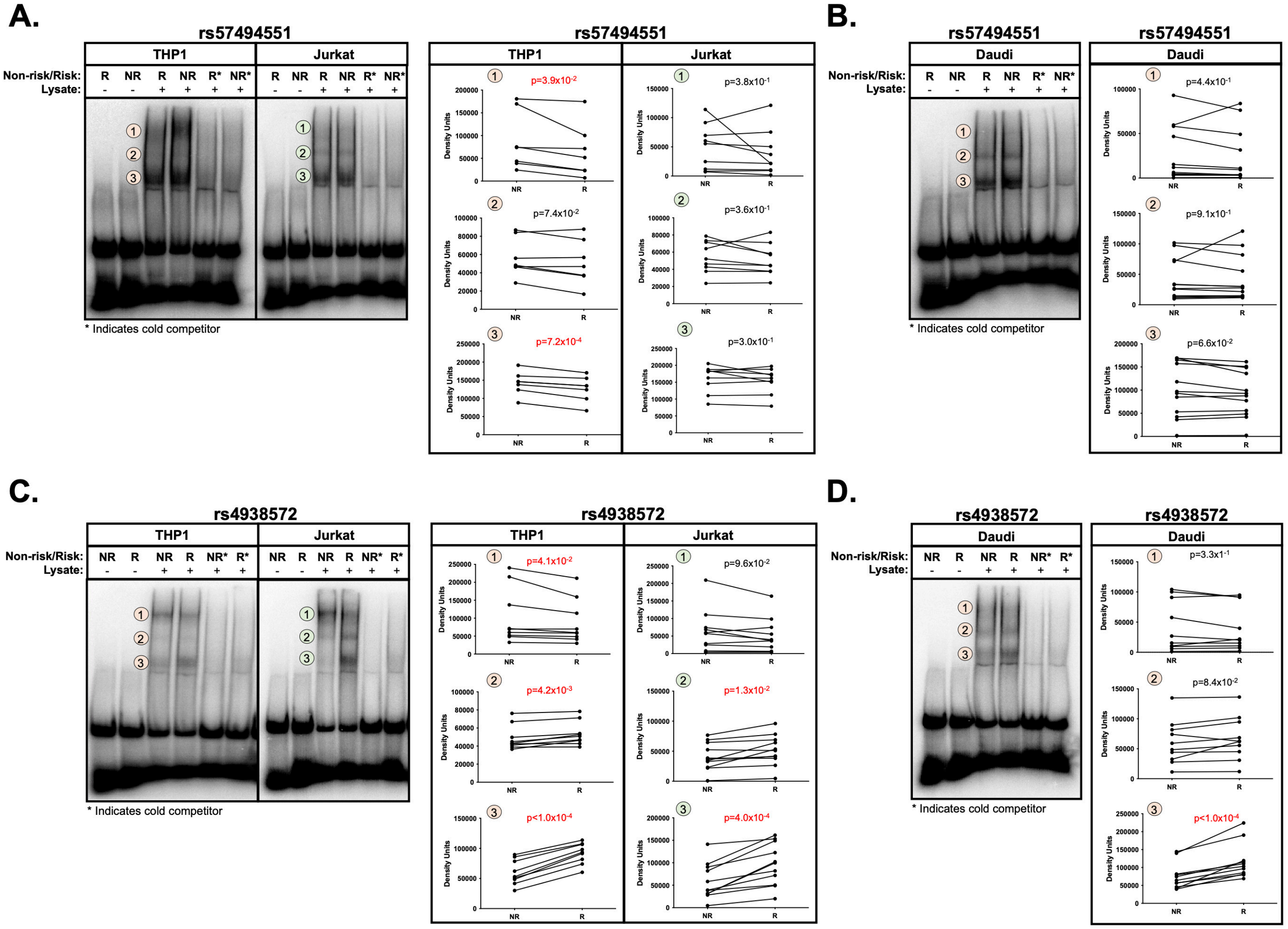


**Supplemental Figure 4. Allele- and cell-type specific differential nuclear protein affinities of SNPs rs57494551 and rs4938572. (A-B)** Radiolabeled electromobility shift assays (EMSA) were performed using oligonucleotides containing the non-risk (NR) or risk (R) allele of rs57494551 and nuclear extracts from **(A)** THP1 and Jurkat cells or **(B)** Daudi cells. **(C-D)** Radiolabeled EMSAs were performed using oligonucleotides containing the NR or R allele of rs4938572 and nuclear extracts from **(A)** THP1 and Jurkat cells or **(B)** Daudi cells. For all panels, probes incubated in the absence of nuclear lysate were used as negative control (Lanes 1, 2). Cold competitors were used to assess non-specific binding (Lanes 5, 6). Images are representative of n>6 biological replicates. Bands indicated by numbered orange or green circles were quantified by densitometry and analyzed using paired t-test; p-values are indicated.

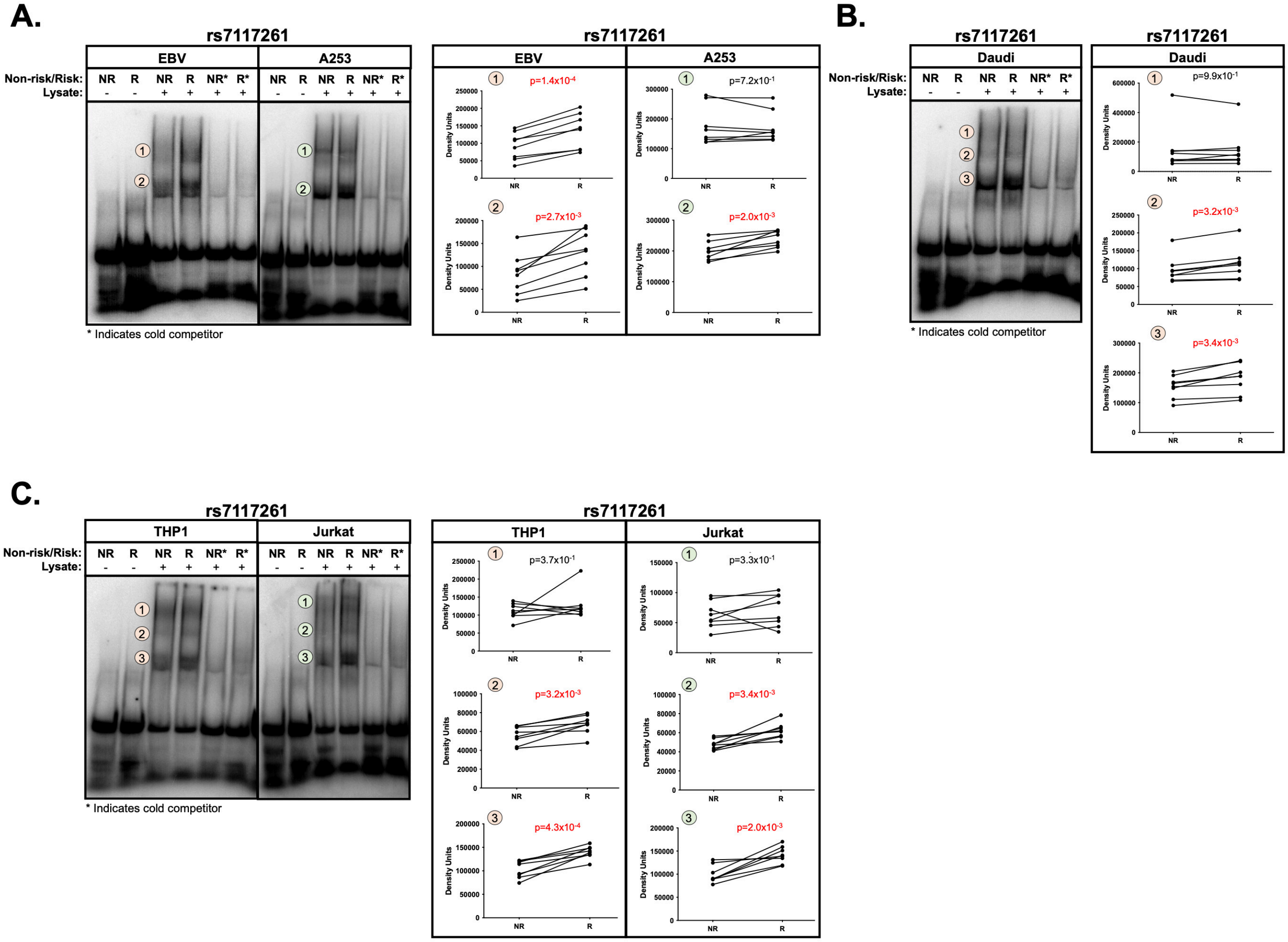


**Supplemental Figure 5. Allele- and cell-type specific differential nuclear protein affinities of SNPs rs7117261. (A-C)** Radiolabeled electromobility shift assays (EMSA) were performed using oligonucleotides containing the non-risk (NR) or risk (R) allele of rs7117261 and nuclear extracts from **(A)** EBV B and A253 cells, **(B)** Daudi cells, or **(C)** THP1 and Jurkat cells. For all panels, probes incubated in the absence of nuclear lysate were used as negative control (Lanes 1, 2). Cold competitors were used to assess non-specific binding (Lanes 5, 6). Images are representative of n>6 biological replicates. Bands indicated by numbered orange or green circles were quantified by densitometry and analyzed using paired t-test; p-values are indicated.

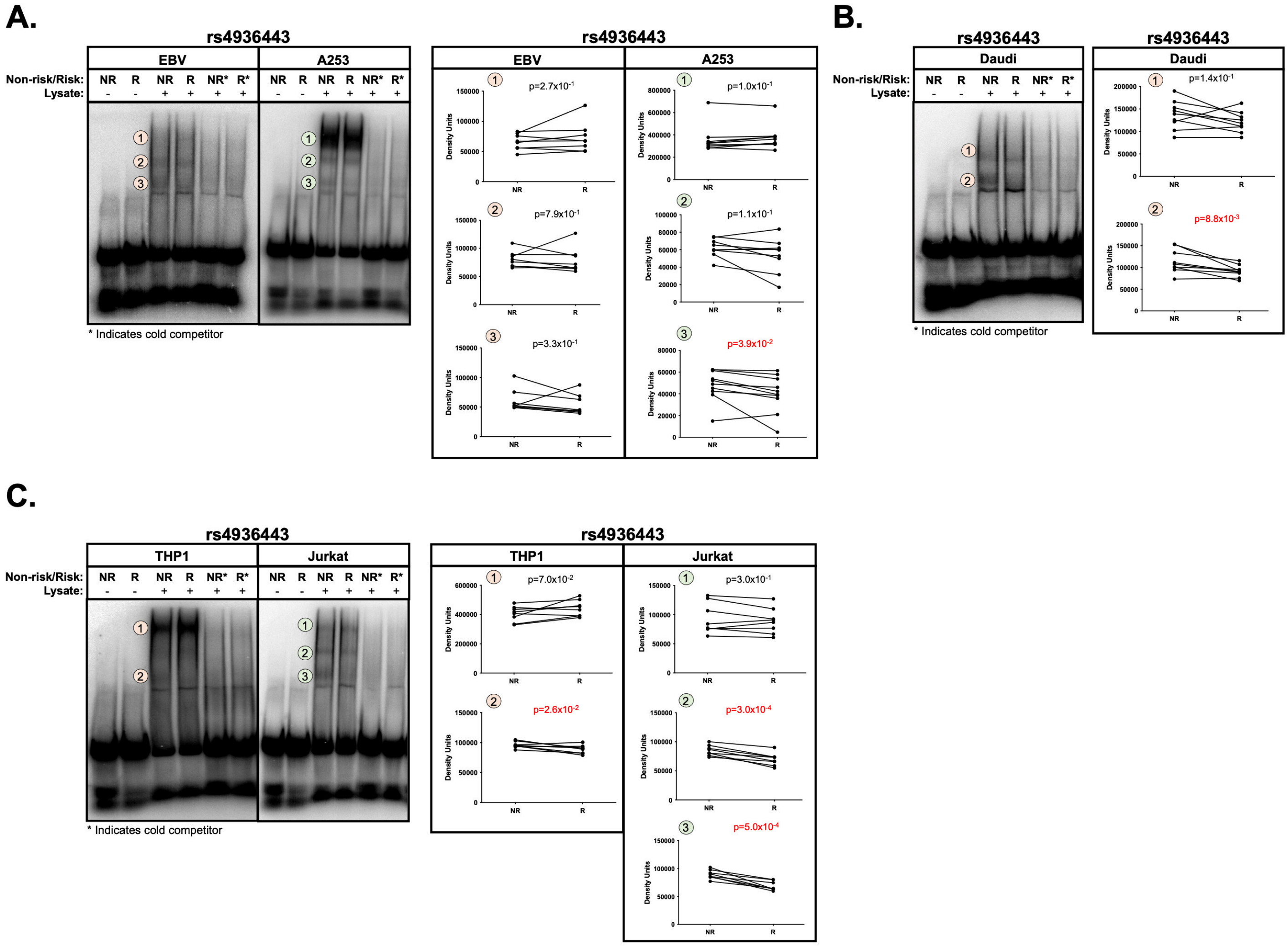


**Supplemental Figure 6. Allele- and cell-type specific differential nuclear protein affinities of SNPs rs4936443. (A-C)** Radiolabeled electromobility shift assays (EMSA) were performed using oligonucleotides containing the non-risk (NR) or risk (R) allele of rs4936443 and nuclear extracts from **(A)** EBV B and A253 cells, **(B)** Daudi cells, or **(C)** THP1 and Jurkat cells. For all panels, probes incubated in the absence of nuclear lysate were used as negative control (Lanes 1, 2). Cold competitors were used to assess non-specific binding (Lanes 5, 6). Images are representative of n>6 biological replicates. Bands indicated by numbered orange or green circles were quantified by densitometry and analyzed using paired t-test; p-values are indicated.

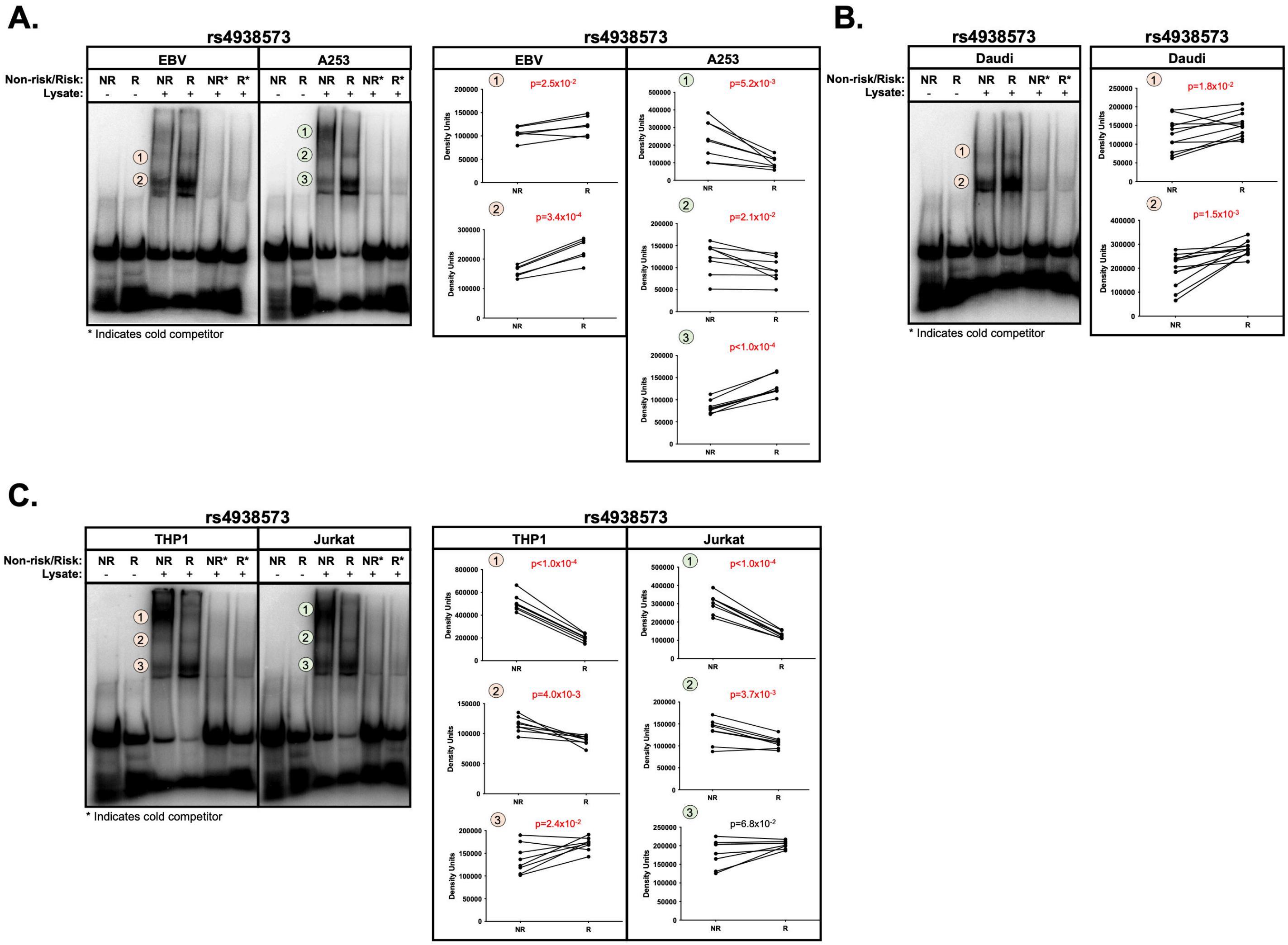


**Supplemental Figure 7. Allele- and cell-type specific differential nuclear protein affinities of SNPs rs4938573. (A-C)** Radiolabeled electromobility shift assays (EMSA) were performed using oligonucleotides containing the non-risk (NR) or risk (R) allele of rs4938573 and nuclear extracts from **(A)** EBV B and A253 cells, **(B)** Daudi cells, or **(C)** THP1 and Jurkat cells. For all panels, probes incubated in the absence of nuclear lysate were used as negative control (Lanes 1, 2). Cold competitors were used to assess non-specific binding (Lanes 5, 6). Images are representative of n>6 biological replicates. Bands indicated by numbered orange or green circles were quantified by densitometry and analyzed using paired t-test; p-values are indicated.

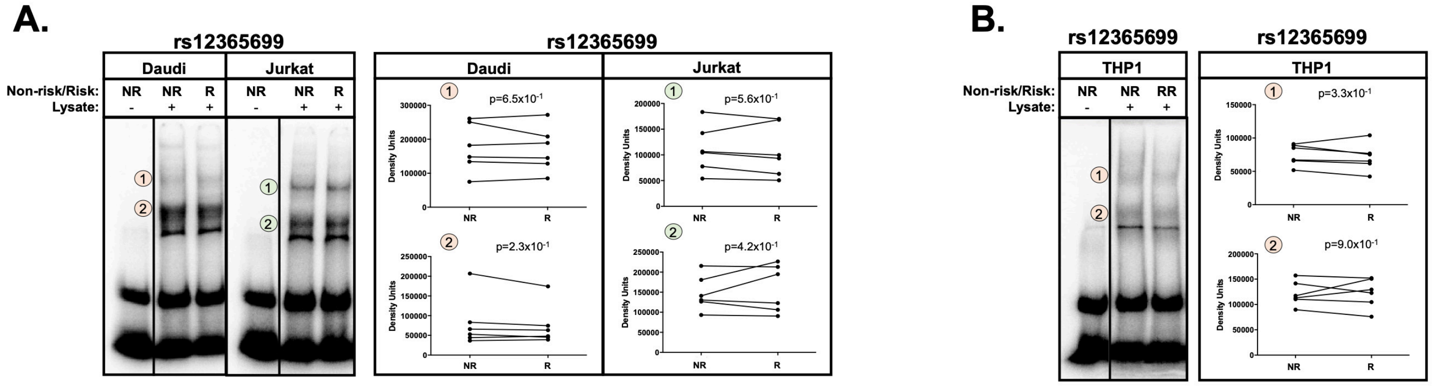


**Supplemental Figure 8. Allele- and cell-type specific differential nuclear protein affinities of SNPs rs12365699. (A-B)** Radiolabeled electromobility shift assays (EMSA) were performed using oligonucleotides containing the non-risk (NR) or risk (R) allele of rs12365699 and nuclear extracts from **(A)** Daudi and Jurkat cells or **(B)** THP 1. For all panels, probes incubated in the absence of nuclear lysate were used as negative control (Lanes 1). Images are representative of n>6 biological replicates. Bands indicated by numbered orange or green circles were quantified by densitometry and analyzed using paired t-test; p-values are indicated.

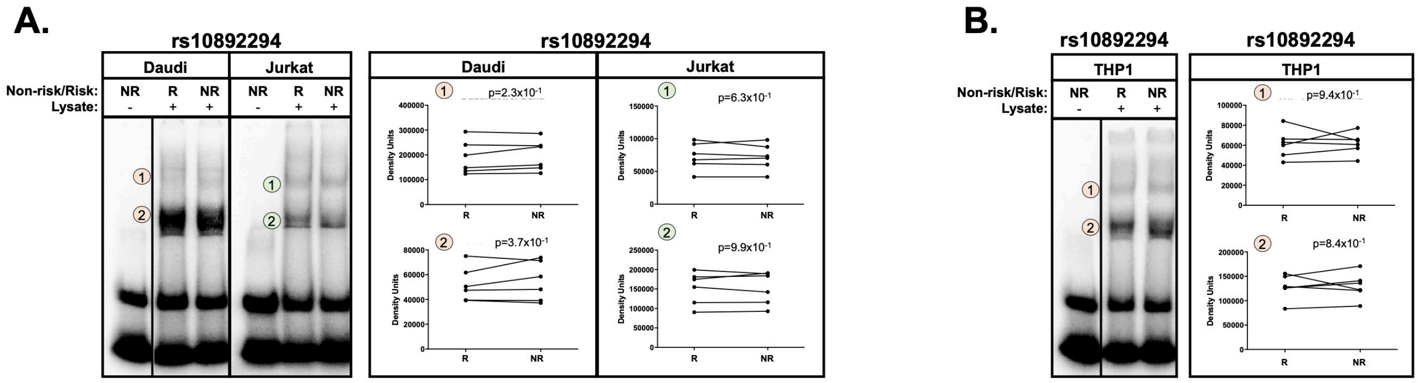


**Supplemental Figure 9. Allele- and cell-type specific differential nuclear protein affinities of SNPs rs10892294. (A-B)** Radiolabeled electromobility shift assays (EMSA) were performed using oligonucleotides containing the non-risk (NR) or risk (R) allele of rs10892294 and nuclear extracts from **(A)** Daudi and Jurkat cells or **(B)** THP1 cells. For all panels, probes incubated in the absence of nuclear lysate were used as negative control (Lanes 1). Images are representative of n>6 biological replicates. Bands indicated by numbered orange or green circles were quantified by densitometry and analyzed using paired t-test; p-values are indicated.

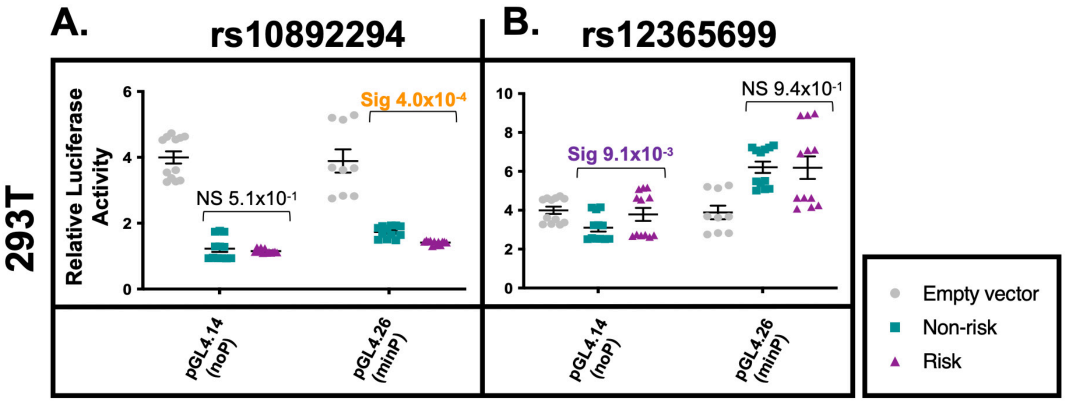


**Supplemental Figure 10. Allele-specific promoter and enhancer activity of rs10892294 and rs12365699 on the *DDX6-CXCR5* region in 293T cells.** gBlocks carrying the non-risk or risk alleles of **(A)** rs10892294 or **(B)** rs12365699 were cloned into a promoter-less (pGL4.14; noP) or minimal promoter (pGL4.26; minP) luciferase vector. Plasmids were transfected into 293T cells. Luciferase activity was measured after 24 hours and normalized to the Renilla transfection control and then the vector-only control; reported as Relative Luciferase Activity. Statistical comparisons were performed using a paired t-test; p-values are indicated.

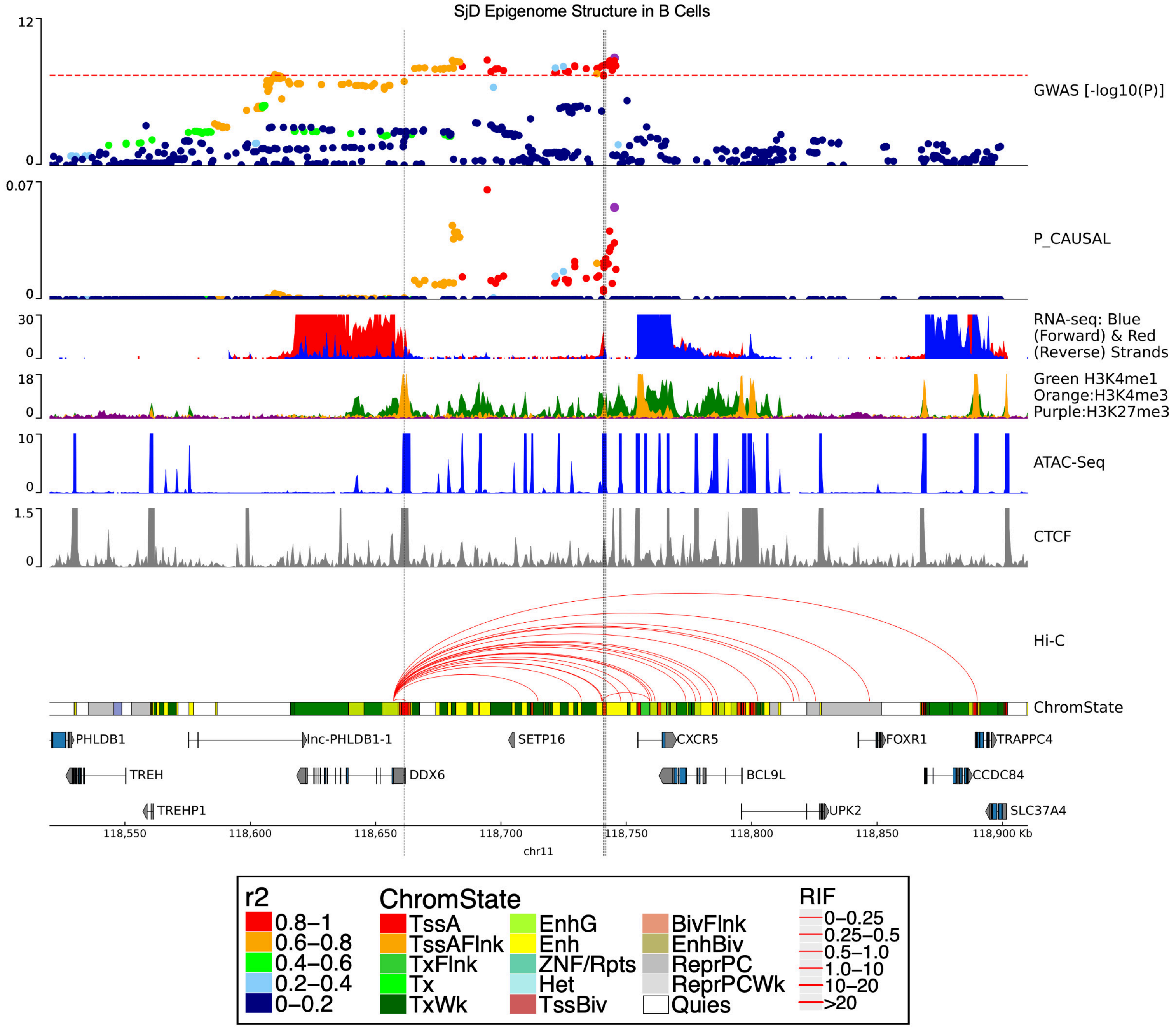


**Supplemental Figure 11. Complex chromatin architecture revealed across the *DDX6-CXCR5* region in human primary B cells.** SjD GWAS association (top panel), publicly available epigenomic enrichment, and promoter-capture Hi-C looping data (red lines) across the *DDX6-CXCR5* region in human primary B cells are shown. Vertical grey lines indicate the locations of rs57494551 or rs4938572, respectively.

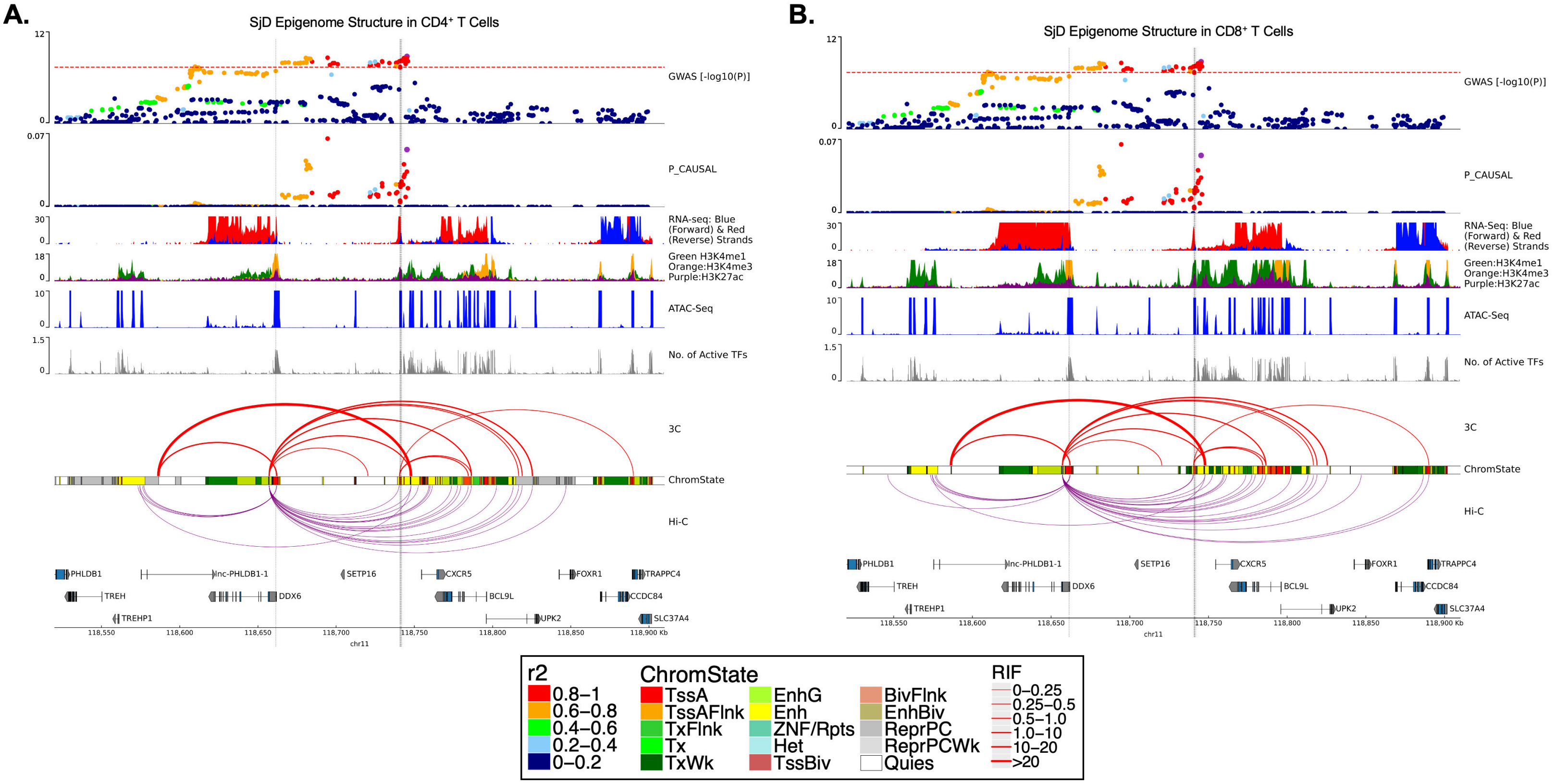


**Supplemental Figure 12. Complex chromatin architecture revealed across the *DDX6-CXCR5* region in primary human T cells.** SjD GWAS association (top panel) and publicly available epigenomic enrichment across the *DDX6-CXCR5* region in human primary **(A)** CD4+ T cells and **(B)** CD8+ T cells. Vertical grey lines indicate the locations of rs57494551 or rs4938572, respectively. Promoter-capture Hi-C looping (purple lines) contrasts the summary 3C-qPCR results reported in Figure 6 (red lines); 3C-qPCR line thickness indicates relative interaction frequency (RIF).

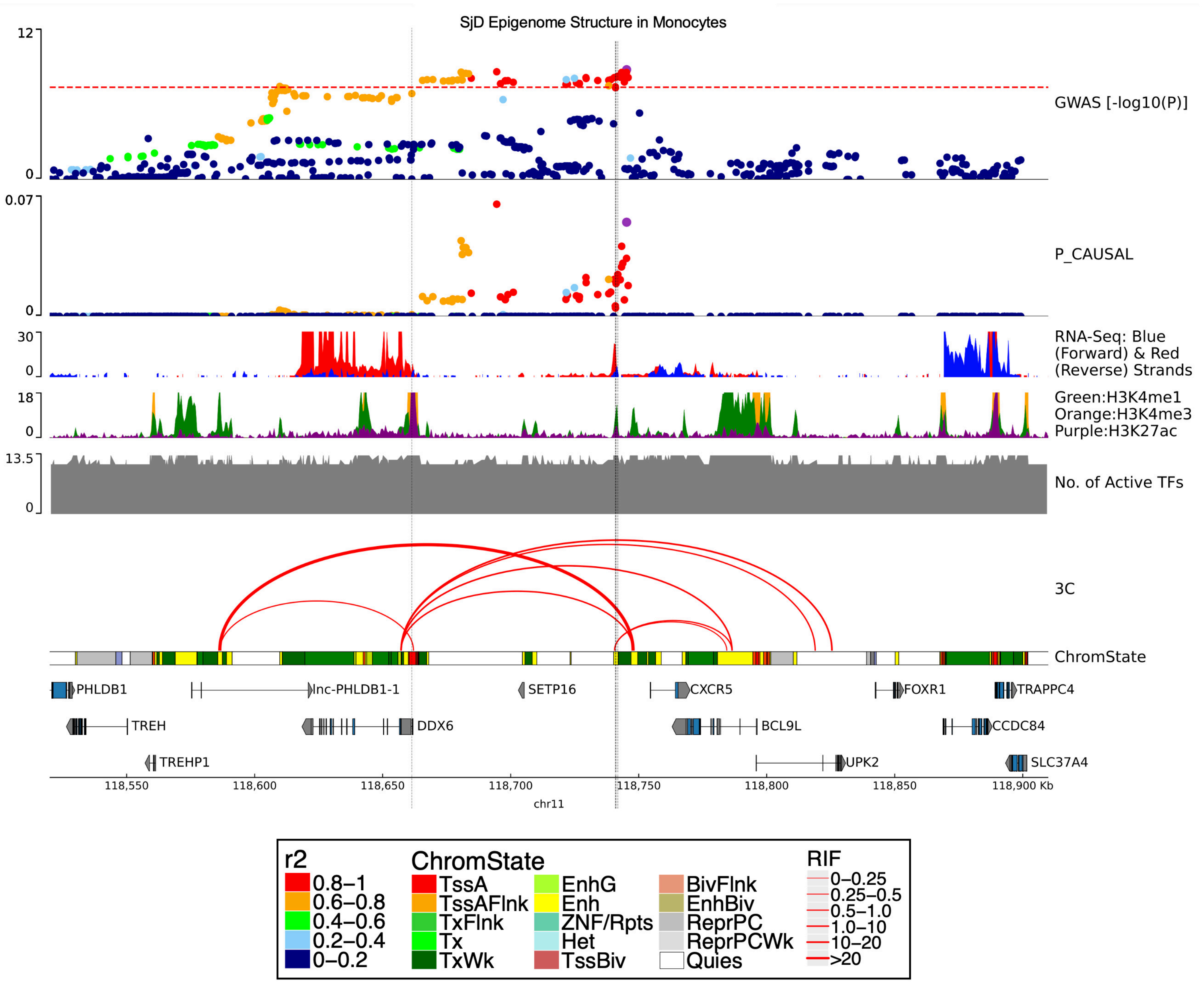


**Supplemental Figure 13. Complex chromatin architecture revealed across the *DDX6-CXCR5* region in human primary monocytes.** SjD GWAS association (top panel) and publicly available epigenomic enrichment across the *DDX6-CXCR5* region in human primary monocytes. Vertical grey lines indicate the locations of rs57494551 or rs4938572, respectively. Summary 3C-qPCR results reported in Figure 6 are also shown (red lines); line thickness indicates relative interaction frequency (RIF).

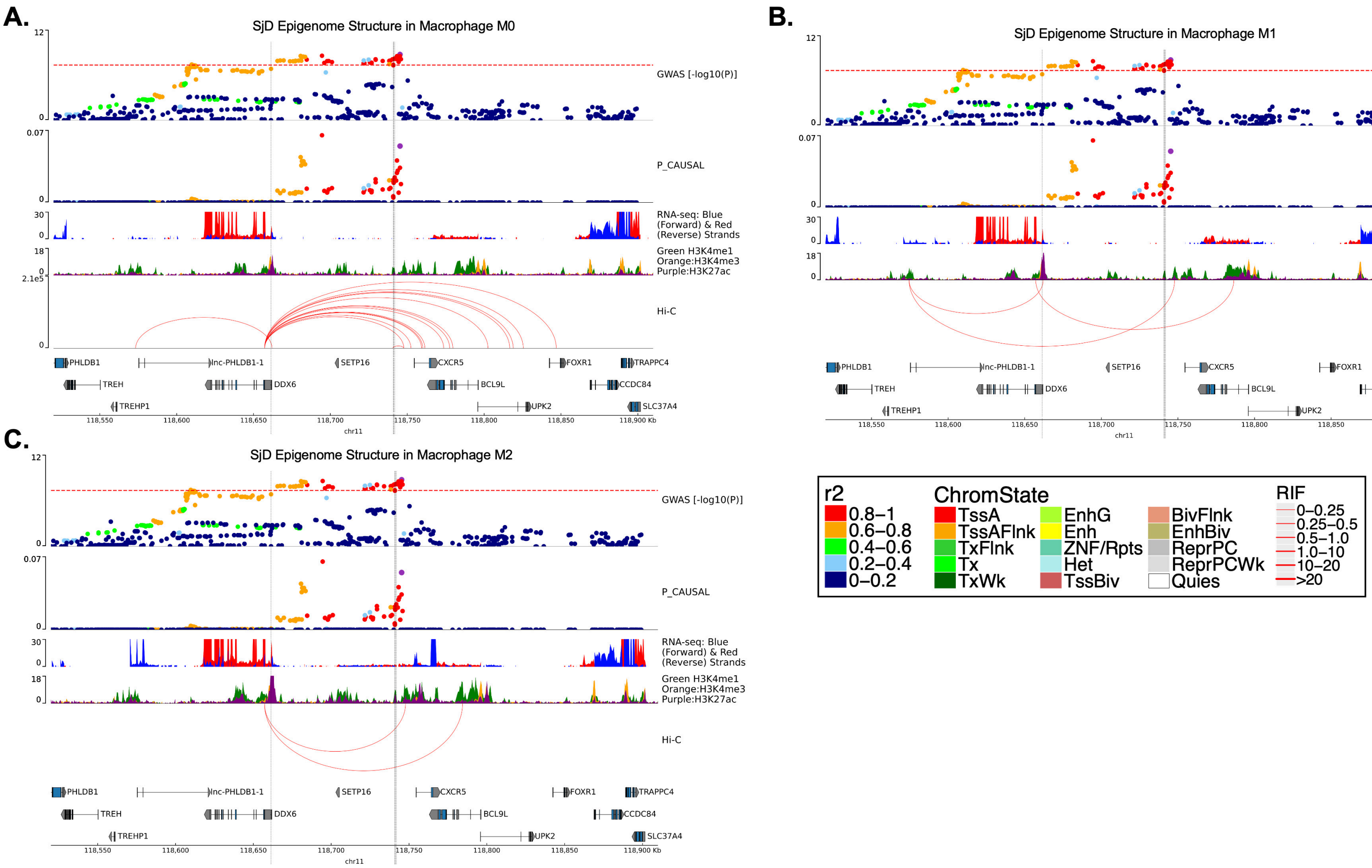


**Supplemental Figure 14. Complex chromatin architecture revealed across the *DDX6-CXCR5* region in human primary macrophages.** SjD GWAS association (top panel), publicly available epigenomic enrichment, and promoter-capture Hi-C looping data (red lines) across the *DDX6-CXCR5* region in human primary **(A)** M0, **(B)** M1, and **(C)** M2 macrophages. Vertical grey lines indicate the locations of rs57494551 or rs4938572, respectively.

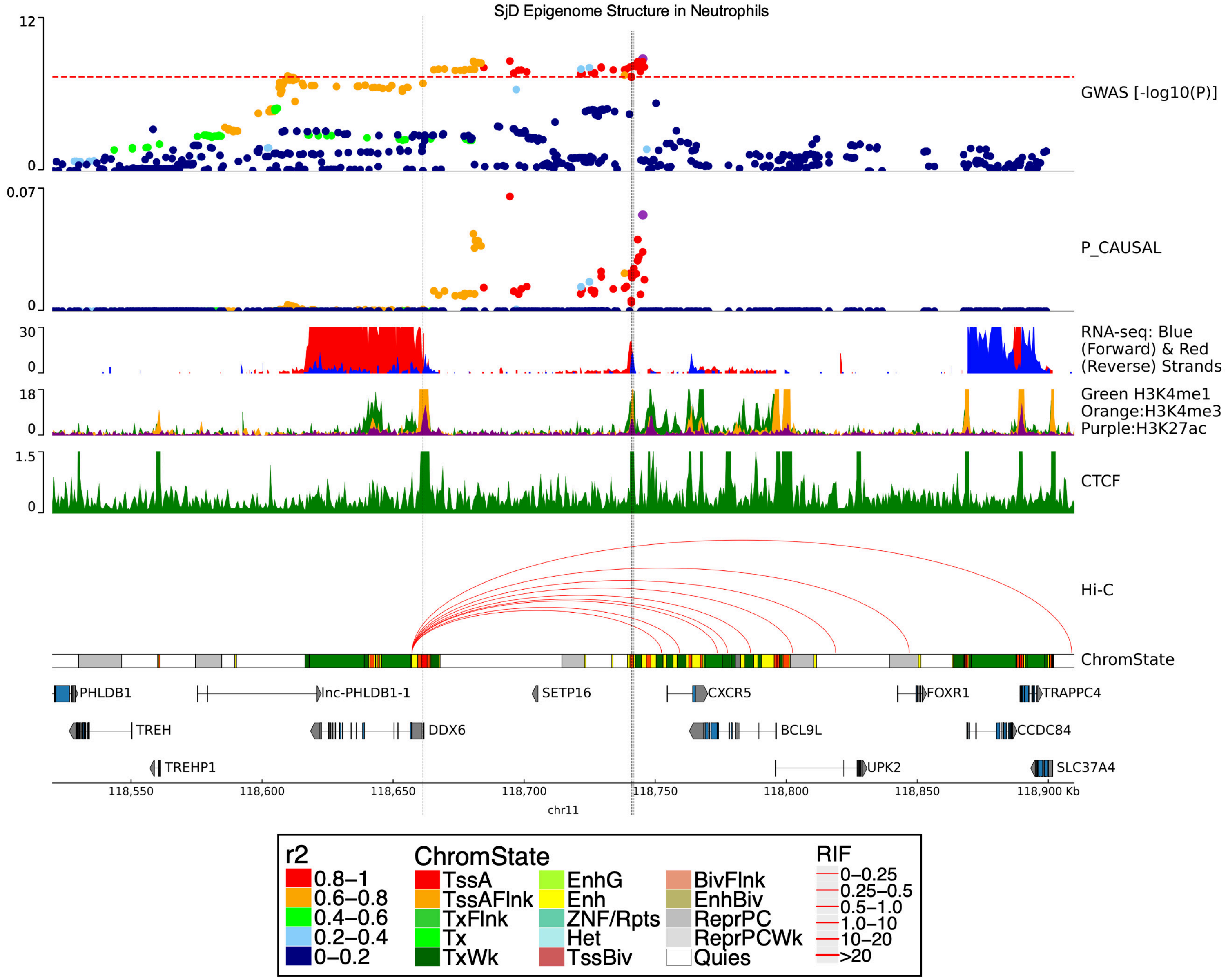


**Supplemental Figure 15. Complex chromatin architecture revealed across the *DDX6-CXCR5* region in human primary neutrophils.** SjD GWAS association (top panel), publicly available epigenomic enrichment, and promoter-capture Hi-C looping data (red lines) across the *DDX6-CXCR5* region in human neutrophils. Vertical grey lines indicate the locations of rs57494551 or rs4938572, respectively.

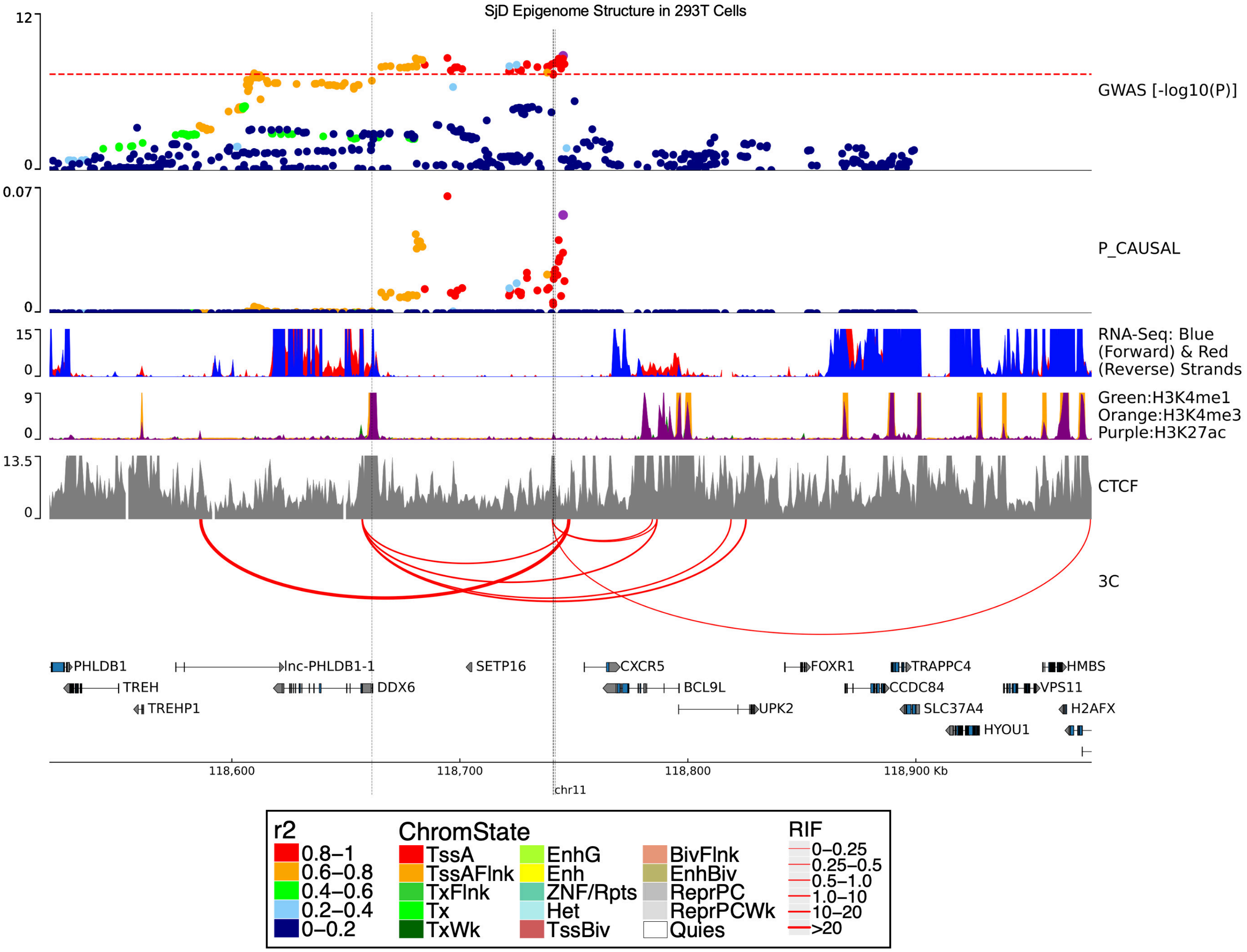


**Supplemental Figure 16. Complex chromatin architecture revealed across the *DDX6-CXCR5* region in human 293T cells.** SjD GWAS association (top panel) and publicly available epigenomic enrichment across the *DDX6-CXCR5* region in 293T cells. Vertical grey lines indicate the locations of rs57494551 or rs4938572, respectively. Summary 3C-qPCR results reported in Figure 6 are also shown (red lines); line thickness indicates relative interaction frequency (RIF).

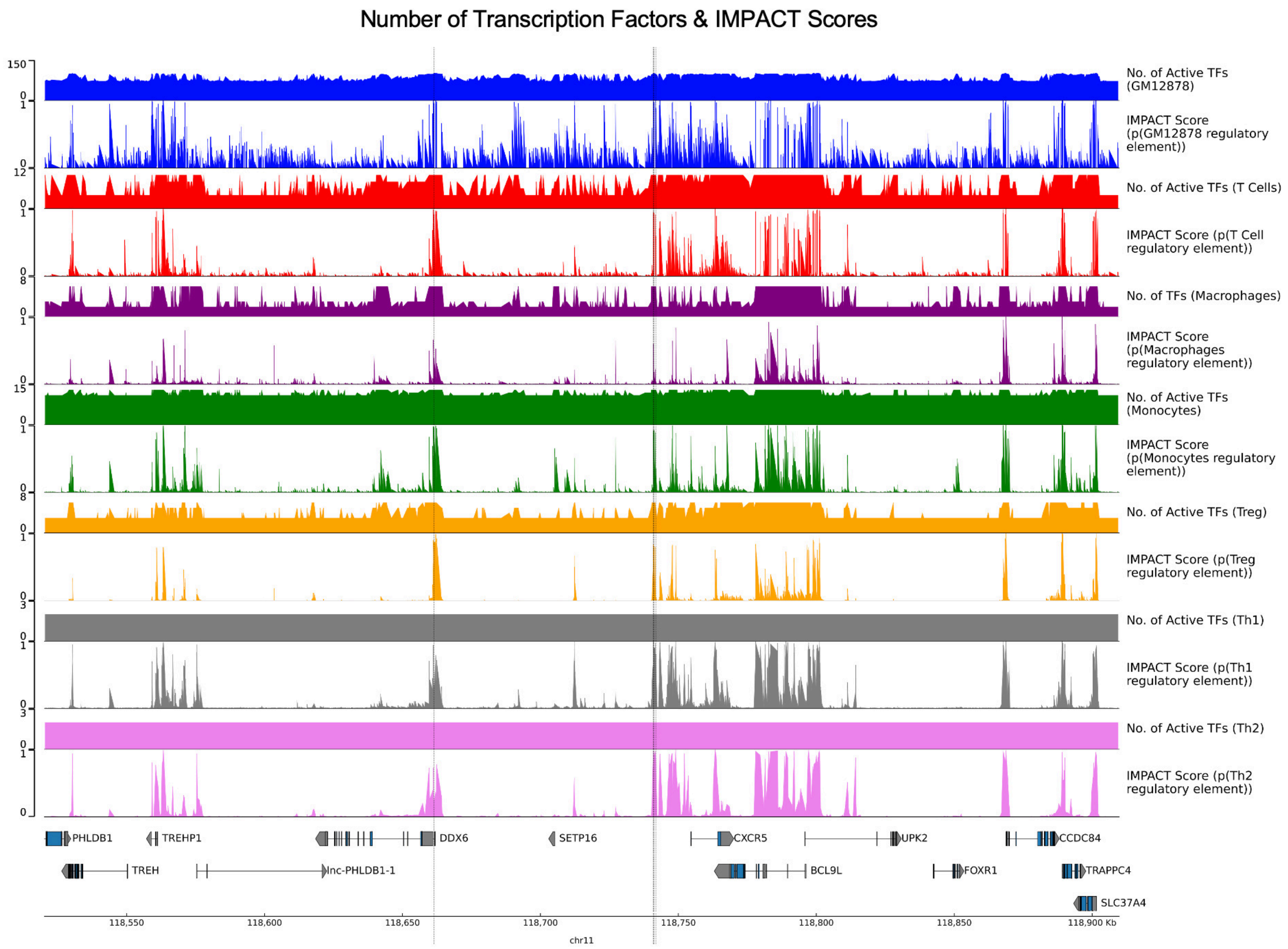


**Supplemental Figure 17. IMPACT regulatory element probabilities and corresponding transcription factor elements in immune cells.** IMPACT uses transcription factor binding sites (TF) to predict the locations of regulatory sites across the genome. Image shows predicted IMPACT regulatory elements for GM12878 B cell line and primary human T cells, macrophages, monocytes, regulatory T cells (Treg), Th1 cells, and Th2 cells. The location of the rs57494551 and rs4938572 are indicates as vertical grey lines.

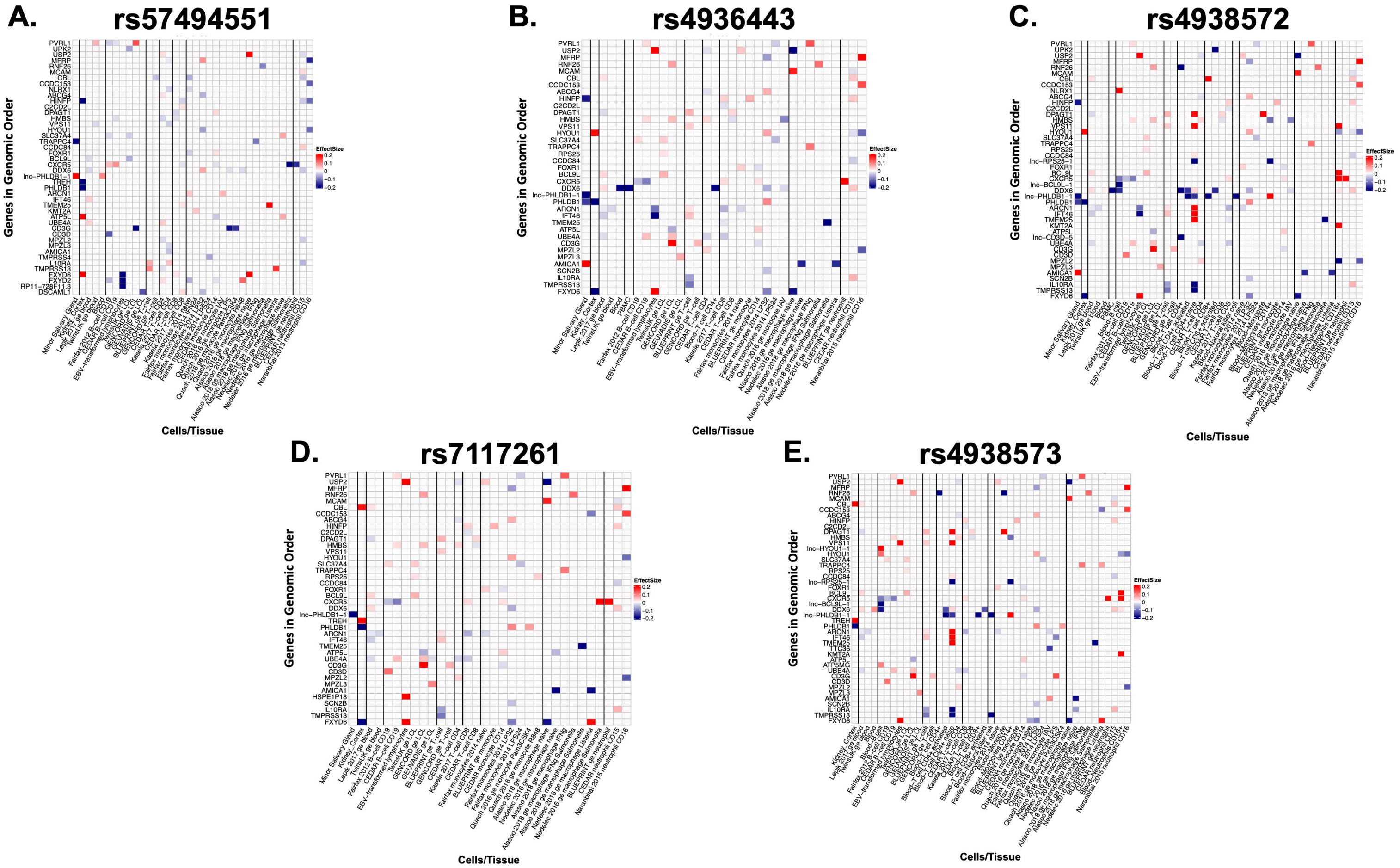


**Supplemental Figure 18. eQTLs for five prioritized SNPs across blood, salivary and kidney tissue, and immune cells.** eQTL values from GTEx in minor salivary gland, blood, kidney tissue, EBV B cells, and human primary immune cells in genes upstream and downstream of the *DDX6-CXCR5* interval for **(A)** rs57494551, **(B)** rs4936443, **(C)** rs4938572, **(D)** rs7117261, **(E)** rs4938573. Vertical lines separate different cell types. Horizontal lines highlight interesting genes such as *CXCR5*, *DDX6*, *lnc-PHLDB1-1*, *TRAPPC4*, and *IL10RA*.

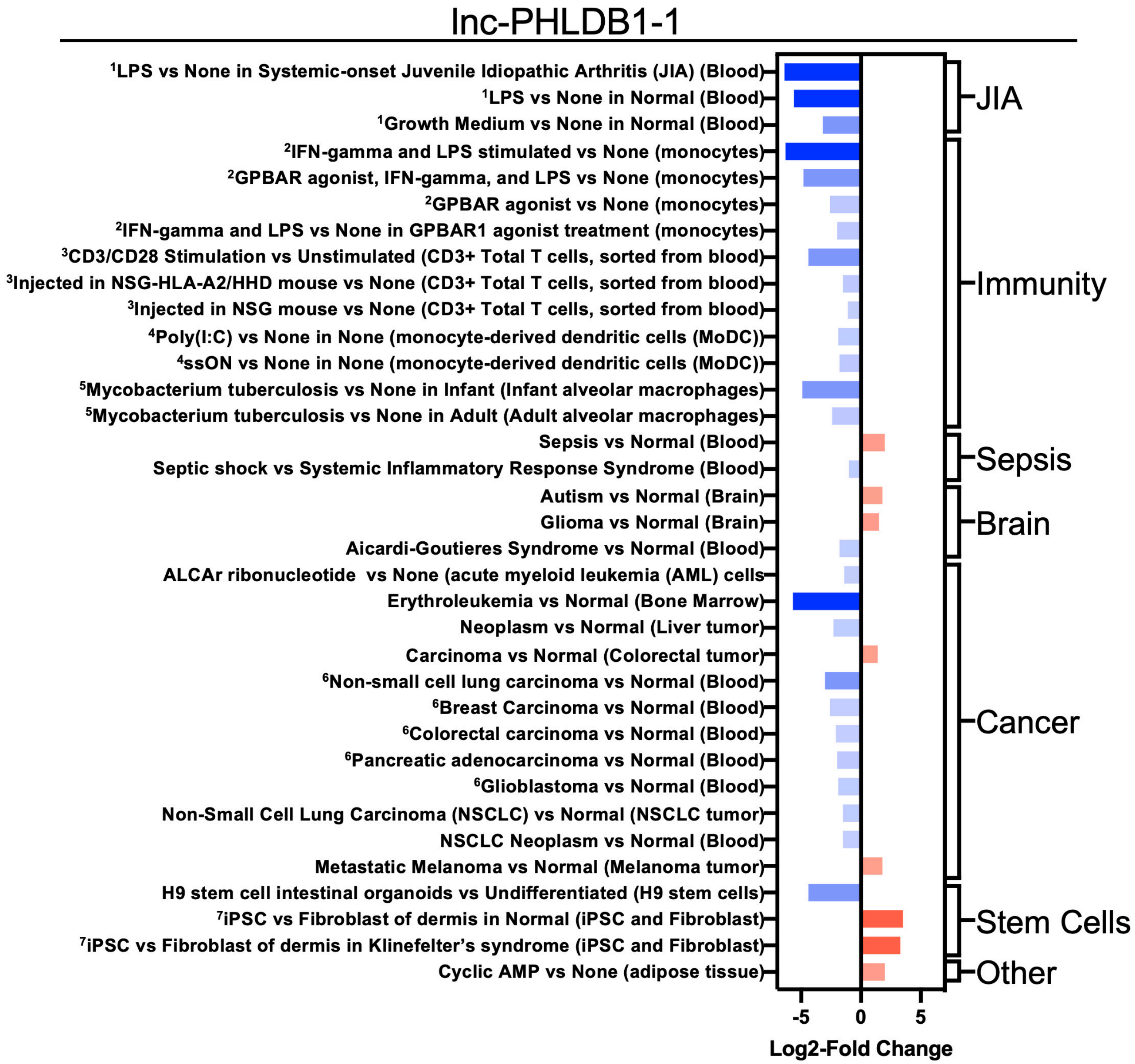


**Supplemental Figure 19. Expression Atlas results for lnc-PHLDB1-1 (ENSG00000255422).** RNA-seq results from Expression Atlas (https://www.ebi.ac.uk/gxa/home) published studies where increases in *lnc-PHLDB1-1* (ENSG00000255422) expression are in red and decreased expression are in blue. Superscripts are used to depict experiments with multiple plotted conditions.
