## Supplemental Datasets for "Variants in the *DDX6-CXCR5* autoimmune disease risk locus influence the regulatory network in immune cells and salivary gland"

**SUPPLEMENTARY DATA TABLE**

**Supplemental Data 1:** Meta-analysis of SjD GWAS and Immonochip data in DDX6-CXCR5 locus (Dataset 1).

**Supplemental Data 2:** Meta-analysis of SLE data in DDX6-CXCR5 locus (Dataset 2).

**Supplemental Data 3:** Meta-analysis of SjD and SLE data in DDX6-CXCR5 locus (Merged Datasets 1 and 2).

**Supplemental Data 4:** Bayesian analysis of SjD GWAS and Immunochip data in DDX6-CXCR5 locus (Dataset 1).

**Supplemental Data 5:** Bayesian analysis of SjD Immunochip data in DDX6-CXCR5 locus (Dataset 3).

**Supplemental Data 6:** Bayesian analysis of SLE Immunochip data in DDX6-CXCR5 locus (Dataset 4).

**Supplemental Data 7:** Bayesian analysis of merged SjD and SLE Immunochip data in DDX6-CXCR5 locus (Merged Datasets 3 and 4) with bioinformatic analysis for the top 100 SNPs (RegulomeDB Score, Haploreg epimarks, and publications)

### Supplemental Data 1: Meta-analysis of SjD GWAS and Immonochip data in DDX6-CXCR5 locus (Da

| SNP | CHR | BP(GRCh37/hg19) | A1 | A2 | Zscore | P.value | Direction | HetISq | HetChiSq | HetDf | HetPVal |
|---|---|---|---|---|---|---|---|---|---|---|---|
| rs7481819 | 11 | 118745278 | C | T | -5.996 | 2.02E-09 | -- | 0 | 0.547 | 1 | 0.4593 |
| rs7123726 | 11 | 118694547 | C | T | -5.933 | 2.98E-09 | -- | 0 | 0.133 | 1 | 0.7154 |
| rs11217040 | 11 | 118680648 | A | C | -5.923 | 3.17E-09 | -- | 0 | 0.458 | 1 | 0.4987 |
| rs4936444 | 11 | 118743338 | T | C | -5.914 | 3.35E-09 | -- | 0 | 0.752 | 1 | 0.3859 |
| rs7481797 | 11 | 118745243 | A | G | -5.914 | 3.35E-09 | -- | 0 | 0.62 | 1 | 0.4312 |
| rs11217041 | 11 | 118682350 | G | A | -5.884 | 4.01E-09 | -- | 0 | 0.537 | 1 | 0.4638 |
| rs11217042 | 11 | 118682528 | A | G | -5.884 | 4.01E-09 | -- | 0 | 0.537 | 1 | 0.4638 |
| rs7942535 | 11 | 118681464 | C | T | -5.884 | 4.01E-09 | -- | 0 | 0.537 | 1 | 0.4637 |
| rs17122453 | 11 | 118683564 | A | G | -5.874 | 4.25E-09 | -- | 0 | 0.566 | 1 | 0.452 |
| rs73005426 | 11 | 118681083 | G | A | -5.866 | 4.46E-09 | -- | 0 | 0.55 | 1 | 0.4582 |
| rs6421571 | 11 | 118743772 | T | C | -5.842 | 5.16E-09 | -- | -198.1 | 1.006 | 1 | 0.3158 |
| rs4938573 | 11 | 118741842 | C | T | -5.799 | 6.69E-09 | -- | -197.5 | 1.008 | 1 | 0.3153 |
| rs12365699 | 11 | 118743286 | A | G | -5.791 | 7.01E-09 | -- | 0 | 0.704 | 1 | 0.4015 |
| rs7117261 | 11 | 118741157 | T | C | -5.768 | 8.01E-09 | -- | 0 | 0.923 | 1 | 0.3367 |
| rs74541740 | 11 | 118742800 | A | G | -5.767 | 8.07E-09 | -- | -182.6 | 1.062 | 1 | 0.3028 |
| rs10790275 | 11 | 118745884 | G | C | -5.757 | 8.56E-09 | -- | 0 | 0.531 | 1 | 0.4662 |
| rs10790268 | 11 | 118729391 | A | G | -5.756 | 8.62E-09 | -- | 0 | 0.948 | 1 | 0.3303 |
| rs7951740 | 11 | 118740418 | G | C | -5.755 | 8.65E-09 | -- | 0 | 0.92 | 1 | 0.3374 |
| rs9736016 | 11 | 118724894 | A | T | -5.736 | 9.70E-09 | -- | 0 | 0.613 | 1 | 0.4335 |
| rs715412 | 11 | 118684610 | A | G | -5.736 | 9.72E-09 | -- | 0 | 0.454 | 1 | 0.5004 |
| rs10790269 | 11 | 118729456 | C | T | -5.728 | 1.02E-08 | -- | 0 | 0.979 | 1 | 0.3224 |
| rs11217032 | 11 | 118669605 | A | T | -5.695 | 1.23E-08 | -- | 0 | 0.381 | 1 | 0.5368 |
| rs7125066 | 11 | 118721699 | G | A | -5.692 | 1.25E-08 | -- | 0 | 0.499 | 1 | 0.4799 |
| rs10892294 | 11 | 118667357 | C | G | -5.691 | 1.26E-08 | -- | 0 | 0.448 | 1 | 0.5034 |
| rs7122669 | 11 | 118739056 | A | G | -5.691 | 1.26E-08 | -- | 0 | 0.761 | 1 | 0.3831 |
| rs28409215 | 11 | 118679304 | A | G | -5.683 | 1.32E-08 | -- | 0 | 0.393 | 1 | 0.5309 |
| rs73005423 | 11 | 118681079 | G | A | -5.681 | 1.34E-08 | -- | 0 | 0.472 | 1 | 0.4922 |
| rs11217038 | 11 | 118677355 | A | G | -5.677 | 1.37E-08 | -- | 0 | 0.397 | 1 | 0.5286 |
| rs7119044 | 11 | 118738298 | G | C | -5.675 | 1.39E-08 | -- | 0 | 0.777 | 1 | 0.3781 |
| rs10892293 | 11 | 118665575 | T | A | -5.67 | 1.42E-08 | -- | 0 | 0.384 | 1 | 0.5356 |
| rs11217074 | 11 | 118734000 | C | T | -5.666 | 1.46E-08 | -- | -191 | 1.031 | 1 | 0.3099 |
| rs11217046 | 11 | 118698907 | A | G | -5.663 | 1.49E-08 | -- | 0 | 0.236 | 1 | 0.6271 |
| rs11217045 | 11 | 118697865 | A | C | -5.657 | 1.54E-08 | -- | 0 | 0.308 | 1 | 0.5788 |
| rs73005502 | 11 | 118697906 | G | A | -5.657 | 1.54E-08 | -- | 0 | 0.308 | 1 | 0.5788 |
| rs11217036 | 11 | 118675772 | T | C | -5.654 | 1.57E-08 | -- | 0 | 0.414 | 1 | 0.52 |
| rs11217037 | 11 | 118677086 | A | G | -5.65 | 1.61E-08 | -- | 0 | 0.414 | 1 | 0.5198 |
| rs11217033 | 11 | 118673706 | C | T | -5.649 | 1.61E-08 | -- | 0 | 0.417 | 1 | 0.5182 |
| rs7117313 | 11 | 118744396 | C | G | -5.618 | 1.93E-08 | -- | 0 | 0.548 | 1 | 0.4593 |
| rs79504890 | 11 | 118701018 | A | G | -5.609 | 2.04E-08 | -- | 0 | 0.1 | 1 | 0.7522 |
| rs4936441 | 11 | 118725660 | C | G | -5.593 | 2.23E-08 | -- | 0 | 0.328 | 1 | 0.5666 |
| rs1037679725 | 11 | 118726843 | T | C | -5.574 | 2.49E-08 | -- | 0 | 0.275 | 1 | 0.6003 |
| rs11217044 | 11 | 118696022 | C | T | -5.566 | 2.60E-08 | -- | 0 | 0.321 | 1 | 0.5712 |
| rs11217058 | 11 | 118722149 | A | G | -5.551 | 2.85E-08 | -- | 0 | 0.264 | 1 | 0.6072 |
| rs11217066 | 11 | 118726753 | G | T | -5.548 | 2.89E-08 | -- | 0 | 0.294 | 1 | 0.5877 |
| rs75249660 | 11 | 118721565 | A | G | -5.539 | 3.03E-08 | -- | 0 | 0.412 | 1 | 0.5212 |
| rs7119038 | 11 | 118738281 | G | A | -5.501 | 3.77E-08 | -- | 0 | 0.395 | 1 | 0.5296 |
| rs11217006 | 11 | 118609790 | A | G | -5.48 | 4.26E-08 | -- | -174.6 | 1.093 | 1 | 0.2959 |
| rs4936443 | 11 | 118740864 | C | T | -5.467 | 4.57E-08 | -- | 0 | 0.417 | 1 | 0.5185 |
| rs4938572 | 11 | 118740931 | C | T | -5.435 | 5.48E-08 | -- | 0 | 0.405 | 1 | 0.5246 |
| rs6589684 | 11 | 118610957 | A | G | -5.431 | 5.62E-08 | -- | -196.7 | 1.011 | 1 | 0.3146 |
| rs10892279 | 11 | 118611781 | A | G | -5.426 | 5.76E-08 | -- | -162.5 | 1.143 | 1 | 0.285 |
| rs6589685 | 11 | 118612197 | C | T | -5.419 | 5.98E-08 | -- | -192.2 | 1.027 | 1 | 0.3109 |
| rs2155433 | 11 | 118612921 | A | G | -5.374 | 7.70E-08 | -- | -195.5 | 1.015 | 1 | 0.3137 |
| rs11826521 | 11 | 118610549 | T | G | -5.35 | 8.80E-08 | -- | 0 | 0.929 | 1 | 0.3351 |

| | | | | | | | | | | | |
|---|---|---|---|---|---|---|---|---|---|---|---|
| rs10892280 | 11 | 118611817 | T | C | -5.341 | 9.23E-08 | -- | 0 | 0.939 | 1 | 0.3326 |
| rs7945144 | 11 | 118608716 | G | A | -5.315 | 1.07E-07 | -- | -163.3 | 1.139 | 1 | 0.2858 |
| rs6589682 | 11 | 118606722 | T | C | -5.272 | 1.35E-07 | -- | 0 | 0.664 | 1 | 0.4151 |
| rs56758835 | 11 | 118615340 | A | G | -5.244 | 1.57E-07 | -- | 0 | 0.907 | 1 | 0.3408 |
| rs57494551 | 11 | 118661398 | T | C | -5.244 | 1.57E-07 | -- | -121.9 | 1.352 | 1 | 0.2449 |
| rs12364971 | 11 | 118607794 | G | T | -5.215 | 1.84E-07 | -- | 0 | 0.799 | 1 | 0.3715 |
| rs76409436 | 11 | 118607871 | A | G | -5.19 | 2.10E-07 | -- | 0 | 0.834 | 1 | 0.3611 |
| rs11217020 | 11 | 118639353 | A | G | -5.184 | 2.17E-07 | -- | 0 | 0.977 | 1 | 0.3229 |
| rs73003215 | 11 | 118649512 | A | G | -5.171 | 2.33E-07 | -- | -163.3 | 1.139 | 1 | 0.2858 |
| rs1048024 | 11 | 118618776 | T | A | -5.169 | 2.36E-07 | -- | -193.8 | 1.021 | 1 | 0.3122 |
| rs2004781 | 11 | 118619253 | T | C | -5.169 | 2.36E-07 | -- | -193.8 | 1.021 | 1 | 0.3122 |
| rs2077579 | 11 | 118619047 | G | T | -5.169 | 2.36E-07 | -- | -193.8 | 1.021 | 1 | 0.3122 |
| rs3889239 | 11 | 118619960 | T | C | -5.169 | 2.36E-07 | -- | -193.8 | 1.021 | 1 | 0.3122 |
| rs11217009 | 11 | 118615785 | T | C | -5.162 | 2.44E-07 | -- | 0 | 0.999 | 1 | 0.3176 |
| rs11217002 | 11 | 118607150 | T | C | -5.157 | 2.52E-07 | -- | 0 | 0.671 | 1 | 0.4127 |
| rs56267528 | 11 | 118607311 | A | C | -5.149 | 2.62E-07 | -- | 0 | 0.627 | 1 | 0.4283 |
| rs10892283 | 11 | 118636980 | G | A | -5.147 | 2.65E-07 | -- | -185.4 | 1.051 | 1 | 0.3053 |
| rs11217019 | 11 | 118636343 | A | C | -5.143 | 2.71E-07 | -- | -184.2 | 1.056 | 1 | 0.3042 |
| rs74676029 | 11 | 118628373 | T | C | -5.139 | 2.77E-07 | -- | -184 | 1.056 | 1 | 0.3041 |
| rs55894437 | 11 | 118607417 | T | C | -5.128 | 2.93E-07 | -- | 0 | 0.612 | 1 | 0.4341 |
| rs10892292 | 11 | 118655340 | C | G | -5.111 | 3.21E-07 | -- | 0 | 0.944 | 1 | 0.3313 |
| rs6589681 | 11 | 118606521 | T | C | -5.108 | 3.26E-07 | -- | 0 | 0.473 | 1 | 0.4916 |
| rs874621 | 11 | 118655645 | A | G | -5.105 | 3.31E-07 | -- | 0 | 0.946 | 1 | 0.3308 |
| rs10892287 | 11 | 118642999 | C | T | -5.105 | 3.31E-07 | -- | 0 | 0.941 | 1 | 0.332 |
| rs10892288 | 11 | 118644694 | G | A | -5.104 | 3.32E-07 | -- | 0 | 0.939 | 1 | 0.3325 |
| rs10892286 | 11 | 118642085 | C | A | -5.1 | 3.39E-07 | -- | 0 | 0.946 | 1 | 0.3307 |
| rs10892289 | 11 | 118646003 | T | C | -5.076 | 3.85E-07 | -- | 0 | 0.967 | 1 | 0.3254 |
| rs56043232 | 11 | 118648373 | T | C | -5.076 | 3.85E-07 | -- | 0 | 0.967 | 1 | 0.3254 |
| rs10892290 | 11 | 118653503 | T | G | -5.053 | 4.34E-07 | -- | 0 | 0.747 | 1 | 0.3875 |
| rs571470 | 11 | 118697012 | A | G | -5.038 | 4.70E-07 | -- | 0 | 0.043 | 1 | 0.8364 |
| rs77209083 | 11 | 118607569 | G | A | -5.006 | 5.56E-07 | -- | 0 | 0.546 | 1 | 0.4599 |
| rs10892291 | 11 | 118653517 | T | C | -4.978 | 6.44E-07 | -- | 0 | 0.588 | 1 | 0.4432 |
| rs11217001 | 11 | 118607046 | A | G | -4.896 | 9.79E-07 | -- | 0 | 0.395 | 1 | 0.5296 |
| rs6589686 | 11 | 118612565 | A | G | -4.611 | 4.01E-06 | -- | 0 | 0.644 | 1 | 0.4224 |
| rs11217078 | 11 | 118750321 | C | T | -4.543 | 5.55E-06 | -- | 0 | 0.595 | 1 | 0.4403 |
| rs10892276 | 11 | 118605628 | A | G | -4.359 | 1.31E-05 | -- | 0 | 0.007 | 1 | 0.9338 |
| rs10892277 | 11 | 118605841 | A | G | -4.354 | 1.34E-05 | -- | 0 | 0.004 | 1 | 0.9516 |
| rs11216998 | 11 | 118604837 | T | C | -4.337 | 1.44E-05 | -- | 0 | 0.012 | 1 | 0.9121 |
| rs11217072 | 11 | 118731762 | G | A | 4.328 | 1.51E-05 | ++ | 0 | 0.242 | 1 | 0.6227 |
| rs34561290 | 11 | 118731599 | T | C | 4.328 | 1.51E-05 | ++ | 0 | 0.242 | 1 | 0.6227 |
| rs10892275 | 11 | 118605617 | A | T | -4.32 | 1.56E-05 | -- | 0 | 0.004 | 1 | 0.952 |
| rs11216993 | 11 | 118603495 | A | T | -4.32 | 1.56E-05 | -- | 0 | 0.697 | 1 | 0.4038 |
| rs10892274 | 11 | 118605463 | T | C | -4.316 | 1.59E-05 | -- | 0 | 0.004 | 1 | 0.951 |
| rs73001429 | 11 | 118603165 | G | T | -4.314 | 1.60E-05 | -- | 0 | 0.529 | 1 | 0.4671 |
| rs11217070 | 11 | 118729860 | G | T | 4.31 | 1.64E-05 | ++ | 0 | 0.272 | 1 | 0.6019 |
| rs11217063 | 11 | 118725942 | C | G | 4.298 | 1.72E-05 | ++ | 0 | 0.134 | 1 | 0.7145 |
| rs10892300 | 11 | 118729210 | T | C | 4.297 | 1.73E-05 | ++ | 0 | 0.26 | 1 | 0.6103 |
| rs11217065 | 11 | 118726647 | C | T | 4.289 | 1.79E-05 | ++ | 0 | 0.137 | 1 | 0.7108 |
| rs12790489 | 11 | 118729583 | C | T | 4.286 | 1.82E-05 | ++ | 0 | 0.218 | 1 | 0.6407 |
| rs11216994 | 11 | 118603520 | T | C | -4.283 | 1.84E-05 | -- | 0 | 0.471 | 1 | 0.4926 |
| rs7929520 | 11 | 118730141 | G | A | 4.28 | 1.87E-05 | ++ | 0 | 0.221 | 1 | 0.6381 |
| rs57719838 | 11 | 118724723 | C | T | 4.28 | 1.87E-05 | ++ | 0 | 0.142 | 1 | 0.7066 |
| rs11216995 | 11 | 118603526 | A | G | -4.264 | 2.01E-05 | -- | 0 | 0.486 | 1 | 0.4856 |
| rs11216999 | 11 | 118604941 | C | T | -4.259 | 2.06E-05 | -- | 0 | 0.004 | 1 | 0.9479 |
| rs11216997 | 11 | 118604638 | T | C | -4.233 | 2.31E-05 | -- | 0 | 0.573 | 1 | 0.4491 |
| rs1174717189 | 11 | 118734521 | T | A | 4.225 | 2.38E-05 | ++ | 0 | 0.272 | 1 | 0.6021 |

| | | | | | | | | | | | |
|---|---|---|---|---|---|---|---|---|---|---|---|
| rs112561833 | 11 | 118603736 | C | A | -4.225 | 2.39E-05 | -- | 0 | 0.561 | 1 | 0.4538 |
| rs7103067 | 11 | 118723379 | A | G | 4.225 | 2.39E-05 | ++ | 0 | 0.08 | 1 | 0.7777 |
| rs10892272 | 11 | 118602707 | G | C | -4.224 | 2.40E-05 | -- | 0 | 0.511 | 1 | 0.4746 |
| rs10892273 | 11 | 118602809 | A | G | -4.22 | 2.44E-05 | -- | 0 | 0.515 | 1 | 0.473 |
| rs10892271 | 11 | 118598544 | A | G | -4.149 | 3.35E-05 | -- | 0 | 0.636 | 1 | 0.4251 |
| rs10790267 | 11 | 118727169 | T | C | 4.125 | 3.71E-05 | ++ | 0 | 0.036 | 1 | 0.8495 |
| rs149794440 | 11 | 118041740 | C | T | -4.122 | 3.76E-05 | -? | 0 | 0 | 0 | 1 |
| rs7483509 | 11 | 118740180 | A | G | 4.114 | 3.88E-05 | ++ | 0 | 0.075 | 1 | 0.7837 |
| rs11217067 | 11 | 118727332 | T | G | 4.079 | 4.52E-05 | ++ | 0 | 0.022 | 1 | 0.8827 |
| rs4936442 | 11 | 118728320 | T | C | 4.038 | 5.40E-05 | ++ | 0 | 0.029 | 1 | 0.8641 |
| rs622722 | 11 | 118711790 | A | G | 3.776 | 0.00016 | ++ | 0 | 0.006 | 1 | 0.938 |
| rs10891973 | 11 | 116287909 | A | G | -3.759 | 0.00017 | -? | 0 | 0 | 0 | 1 |
| rs10891972 | 11 | 116287783 | T | C | -3.752 | 0.00018 | -? | 0 | 0 | 0 | 1 |
| rs587816 | 11 | 118689871 | A | C | 3.583 | 0.00034 | ++ | 0 | 0.009 | 1 | 0.9236 |
| rs577121 | 11 | 118690550 | C | T | 3.568 | 0.00036 | ++ | 0 | 0.013 | 1 | 0.91 |
| rs11216972 | 11 | 118585826 | T | C | -3.536 | 0.00041 | -- | 0 | 0.166 | 1 | 0.6839 |
| rs584935 | 11 | 118696175 | A | G | 3.47 | 0.00052 | ++ | 0 | 0.02 | 1 | 0.8864 |
| rs583905 | 11 | 118695908 | G | A | 3.47 | 0.00052 | ++ | 0 | 0.024 | 1 | 0.8767 |
| rs10892263 | 11 | 118587319 | C | T | -3.467 | 0.00053 | -- | 0 | 0.077 | 1 | 0.7814 |
| rs149509653 | 11 | 118558425 | A | G | 3.446 | 0.00057 | ++ | 0 | 0.033 | 1 | 0.8562 |
| rs10892262 | 11 | 118587097 | C | G | -3.427 | 0.00061 | -- | 0 | 0.074 | 1 | 0.7859 |
| rs12421790 | 11 | 118588952 | T | C | -3.403 | 0.00067 | -- | 0 | 0.028 | 1 | 0.8673 |
| rs10790329 | 11 | 119537683 | C | T | -3.4 | 0.00067 | -? | 0 | 0 | 0 | 1 |
| rs497916 | 11 | 118758089 | T | C | -3.377 | 0.00073 | -- | 0 | 0.961 | 1 | 0.3269 |
| rs113641235 | 11 | 118700264 | T | C | 3.369 | 0.00076 | ++ | 0 | 0.001 | 1 | 0.9778 |
| rs566733 | 11 | 118608625 | G | C | 3.363 | 0.00077 | ++ | 0 | 0.133 | 1 | 0.7152 |
| rs11216980 | 11 | 118590668 | A | C | -3.363 | 0.00077 | -- | 0 | 0.005 | 1 | 0.9443 |
| rs567680 | 11 | 118608532 | G | C | 3.362 | 0.00077 | ++ | 0 | 0.154 | 1 | 0.6947 |
| rs488219 | 11 | 118620697 | C | T | 3.36 | 0.00078 | ++ | 0 | 0.031 | 1 | 0.8612 |
| rs666094 | 11 | 117489521 | C | T | 3.356 | 0.00079 | +? | 0 | 0 | 0 | 1 |
| rs546393 | 11 | 118624250 | C | T | 3.354 | 0.0008 | ++ | 0 | 0.032 | 1 | 0.8578 |
| rs632124 | 11 | 118613235 | A | T | 3.353 | 0.0008 | ++ | 0 | 0.16 | 1 | 0.689 |
| rs607125 | 11 | 118617299 | C | T | 3.336 | 0.00085 | ++ | 0 | 0.057 | 1 | 0.8115 |
| rs12421793 | 11 | 118589012 | A | C | -3.332 | 0.00086 | -- | 0 | 0.004 | 1 | 0.9474 |
| rs10128600 | 11 | 118700425 | C | T | 3.325 | 0.00089 | ++ | 0 | 0.001 | 1 | 0.9807 |
| rs11217004 | 11 | 118607622 | A | G | 3.321 | 0.0009 | ++ | 0 | 0.213 | 1 | 0.6443 |
| rs73005500 | 11 | 118697375 | C | T | 3.304 | 0.00095 | ++ | 0 | 0.004 | 1 | 0.9466 |
| rs2510901 | 11 | 118699863 | A | G | 3.288 | 0.00101 | ++ | 0 | 0 | 1 | 0.9918 |
| rs555639 | 11 | 118629031 | A | G | 3.288 | 0.00101 | ++ | 0 | 0.02 | 1 | 0.8879 |
| rs640373 | 11 | 118698160 | G | A | 3.269 | 0.00108 | ++ | 0 | 0 | 1 | 0.9992 |
| rs638893 | 11 | 118698537 | G | A | 3.268 | 0.00108 | ++ | 0 | 0 | 1 | 1 |
| rs639885 | 11 | 118698294 | A | G | 3.268 | 0.00108 | ++ | 0 | 0 | 1 | 1 |
| rs639915 | 11 | 118698274 | A | G | 3.268 | 0.00108 | ++ | 0 | 0 | 1 | 1 |
| rs2276341 | 11 | 117375461 | A | G | -3.267 | 0.00109 | -? | 0 | 0 | 0 | 1 |
| rs4938394 | 11 | 117373947 | C | T | -3.266 | 0.00109 | -? | 0 | 0 | 0 | 1 |
| rs516332 | 11 | 118698777 | G | A | 3.263 | 0.0011 | ++ | 0 | 0 | 1 | 0.9957 |
| rs2508564 | 11 | 118699744 | C | T | 3.256 | 0.00113 | ++ | 0 | 0 | 1 | 0.9979 |
| rs583213 | 11 | 118669178 | G | T | 3.221 | 0.00128 | ++ | 0 | 0.791 | 1 | 0.3739 |
| rs10892132 | 11 | 117375951 | A | G | -3.219 | 0.00129 | -? | 0 | 0 | 0 | 1 |
| rs581045 | 11 | 118632522 | T | C | 3.219 | 0.00129 | ++ | 0 | 0.01 | 1 | 0.9204 |
| rs2155394 | 11 | 117378116 | A | G | -3.218 | 0.00129 | -? | 0 | 0 | 0 | 1 |
| rs513935 | 11 | 116231840 | A | G | -3.205 | 0.00135 | -- | 30.4 | 4.307 | 1 | 0.03795 |
| rs1784543 | 11 | 118631653 | T | A | 3.202 | 0.00136 | ++ | 0 | 0.026 | 1 | 0.8727 |
| rs507890 | 11 | 118631288 | C | T | 3.202 | 0.00137 | ++ | 0 | 0.025 | 1 | 0.8732 |
| rs11216843 | 11 | 118151329 | T | A | -3.199 | 0.00138 | -? | 0 | 0 | 0 | 1 |
| rs7932097 | 11 | 117373125 | T | C | -3.178 | 0.00148 | -? | 0 | 0 | 0 | 1 |

| | | | | | | | | | | | |
|---|---|---|---|---|---|---|---|---|---|---|---|
| rs515523 | 11 | 118658075 | G | A | 3.177 | 0.00149 | ++ | 0 | 0.019 | 1 | 0.8917 |
| rs7112940 | 11 | 117373432 | G | A | -3.167 | 0.00154 | -? | 0 | 0 | 0 | 1 |
| rs582200 | 11 | 118668908 | G | A | 3.163 | 0.00156 | ++ | 0 | 0.334 | 1 | 0.5633 |
| rs555649 | 11 | 118651809 | C | T | 3.162 | 0.00157 | ++ | 0 | 0.017 | 1 | 0.8965 |
| rs472641 | 11 | 118617544 | G | A | -3.156 | 0.0016 | -- | 0 | 0.046 | 1 | 0.8303 |
| rs476246 | 11 | 118617920 | T | C | -3.154 | 0.00161 | -- | 0 | 0.042 | 1 | 0.8385 |
| rs10892256 | 11 | 118578705 | A | G | -3.151 | 0.00163 | -- | 0 | 0.447 | 1 | 0.5038 |
| rs11216960 | 11 | 118577873 | A | G | -3.151 | 0.00163 | -- | 0 | 0.447 | 1 | 0.5038 |
| rs7933859 | 11 | 118662045 | T | C | 3.15 | 0.00163 | ++ | 0 | 0.491 | 1 | 0.4835 |
| rs61901411 | 11 | 118665324 | T | C | 3.146 | 0.00165 | ++ | 0 | 0.297 | 1 | 0.5859 |
| rs4508234 | 11 | 118662203 | G | C | 3.145 | 0.00166 | ++ | 0 | 0.495 | 1 | 0.4815 |
| rs4938544 | 11 | 118665009 | A | G | 3.14 | 0.00169 | ++ | 0 | 0.273 | 1 | 0.6016 |
| rs2231449 | 11 | 118186305 | A | C | -3.138 | 0.0017 | -? | 0 | 0 | 0 | 1 |
| rs10892260 | 11 | 118583525 | C | A | -3.138 | 0.0017 | -- | 0 | 0.35 | 1 | 0.5542 |
| rs2510897 | 11 | 118644582 | C | A | 3.137 | 0.00171 | ++ | 0 | 0.017 | 1 | 0.8972 |
| rs500254 | 11 | 118678780 | G | A | 3.133 | 0.00173 | ++ | 0 | 0.259 | 1 | 0.6107 |
| rs2105782 | 11 | 118659896 | C | T | 3.133 | 0.00173 | ++ | 0 | 0.031 | 1 | 0.8608 |
| rs581062 | 11 | 118624767 | A | G | -3.13 | 0.00175 | -- | 0 | 0.053 | 1 | 0.8185 |
| rs487728 | 11 | 118621495 | G | C | -3.129 | 0.00175 | -- | 0 | 0.028 | 1 | 0.8662 |
| rs499343 | 11 | 118626511 | T | C | -3.129 | 0.00175 | -- | 0 | 0.04 | 1 | 0.8406 |
| rs10790261 | 11 | 118579747 | G | A | -3.128 | 0.00176 | -- | 0 | 0.541 | 1 | 0.4621 |
| rs10892257 | 11 | 118579327 | A | G | -3.121 | 0.0018 | -- | 0 | 0.47 | 1 | 0.4932 |
| rs6589612 | 11 | 117373055 | G | C | -3.109 | 0.00188 | -? | 0 | 0 | 0 | 1 |
| rs11216965 | 11 | 118582613 | A | C | -3.108 | 0.00188 | -- | 0 | 0.468 | 1 | 0.494 |
| rs5006102 | 11 | 118662340 | A | T | 3.108 | 0.00188 | ++ | 0 | 0.424 | 1 | 0.5151 |
| rs10892258 | 11 | 118579865 | A | G | -3.105 | 0.0019 | -- | 0 | 0.473 | 1 | 0.4917 |
| rs11216964 | 11 | 118581895 | A | G | -3.104 | 0.00191 | -- | 0 | 0.471 | 1 | 0.4927 |
| rs675003 | 11 | 118677083 | T | C | 3.103 | 0.00192 | ++ | 0 | 0.273 | 1 | 0.6013 |
| rs11216961 | 11 | 118580338 | G | C | -3.103 | 0.00192 | -- | 0 | 0.466 | 1 | 0.4947 |
| rs11216969 | 11 | 118584340 | C | T | -3.102 | 0.00192 | -- | 0 | 0.474 | 1 | 0.4912 |
| rs11216846 | 11 | 118163916 | A | G | -3.101 | 0.00193 | -? | 0 | 0 | 0 | 1 |
| rs10892259 | 11 | 118583002 | A | G | -3.088 | 0.00201 | -- | 0 | 0.482 | 1 | 0.4877 |
| rs495430 | 11 | 116235005 | A | G | -3.088 | 0.00202 | -- | 15.9 | 3.568 | 1 | 0.05889 |
| rs11216956 | 11 | 118575326 | A | G | -3.088 | 0.00202 | -- | 0 | 0.497 | 1 | 0.4809 |
| rs11216848 | 11 | 118181481 | A | C | -3.079 | 0.00208 | -? | 0 | 0 | 0 | 1 |
| rs515707 | 11 | 118685481 | T | C | 3.071 | 0.00213 | +? | 0 | 0 | 0 | 1 |
| rs10790181 | 11 | 117289347 | G | A | 3.062 | 0.0022 | +? | 0 | 0 | 0 | 1 |
| rs553906 | 11 | 117288898 | G | T | 3.062 | 0.0022 | +? | 0 | 0 | 0 | 1 |
| rs579411 | 11 | 117288387 | A | G | 3.062 | 0.0022 | +? | 0 | 0 | 0 | 1 |
| rs2124975 | 11 | 117286824 | G | A | 3.061 | 0.00221 | +? | 0 | 0 | 0 | 1 |
| rs2368164 | 11 | 117287088 | C | T | 3.061 | 0.00221 | +? | 0 | 0 | 0 | 1 |
| rs2510451 | 11 | 117287272 | G | C | 3.061 | 0.00221 | +? | 0 | 0 | 0 | 1 |
| rs2925767 | 11 | 117287230 | A | G | 3.061 | 0.00221 | +? | 0 | 0 | 0 | 1 |
| rs523159 | 11 | 117260504 | A | G | 3.057 | 0.00224 | +? | 0 | 0 | 0 | 1 |
| rs473446 | 11 | 117287942 | A | G | 3.054 | 0.00226 | +? | 0 | 0 | 0 | 1 |
| rs522383 | 11 | 117286064 | T | C | 3.054 | 0.00226 | +? | 0 | 0 | 0 | 1 |
| rs524069 | 11 | 117285907 | G | A | 3.054 | 0.00226 | +? | 0 | 0 | 0 | 1 |
| rs1309473877 | 11 | 118657311 | A | C | 3.053 | 0.00227 | ++ | 0 | 0.016 | 1 | 0.898 |
| rs512356 | 11 | 117285619 | A | G | 3.05 | 0.00229 | +? | 0 | 0 | 0 | 1 |
| rs533254 | 11 | 118701898 | A | T | 3.047 | 0.00231 | ++ | 0 | 0.06 | 1 | 0.807 |
| rs10750107 | 11 | 117379086 | C | T | -3.046 | 0.00232 | -? | 0 | 0 | 0 | 1 |
| rs11216962 | 11 | 118580813 | C | T | -3.042 | 0.00235 | -- | 0 | 0.508 | 1 | 0.476 |
| rs11216963 | 11 | 118580814 | A | G | -3.042 | 0.00235 | -- | 0 | 0.508 | 1 | 0.476 |
| rs1270801 | 11 | 117272939 | T | G | 3.039 | 0.00237 | +? | 0 | 0 | 0 | 1 |
| rs1135258 | 11 | 119210583 | T | G | 3.039 | 0.00237 | +? | 0 | 0 | 0 | 1 |
| rs524590 | 11 | 118640063 | T | A | -3.016 | 0.00256 | -- | 0 | 0.091 | 1 | 0.7633 |

| | | | | | | | | | | | |
|---|---|---|---|---|---|---|---|---|---|---|---|
| rs1170396 | 11 | 118703817 | T | A | 3.013 | 0.00259 | ++ | 0 | 0.1 | 1 | 0.7514 |
| rs2508921 | 11 | 118703484 | T | C | 3.013 | 0.00259 | ++ | 0 | 0.1 | 1 | 0.7514 |
| rs622690 | 11 | 118704265 | T | C | 3.013 | 0.00259 | ++ | 0 | 0.1 | 1 | 0.7513 |
| rs7928228 | 11 | 118703001 | T | C | 3.013 | 0.00259 | ++ | 0 | 0.1 | 1 | 0.7514 |
| rs523148 | 11 | 118682462 | T | C | 3.013 | 0.00259 | +? | 0 | 0 | 0 | 1 |
| rs493647 | 11 | 117290313 | G | A | 3.008 | 0.00263 | +? | 0 | 0 | 0 | 1 |
| rs480791 | 11 | 118704255 | G | A | 3.008 | 0.00263 | ++ | 0 | 0.099 | 1 | 0.7536 |
| rs563162 | 11 | 118944909 | T | C | -3.003 | 0.00268 | -? | 0 | 0 | 0 | 1 |
| rs607472 | 11 | 118680072 | G | C | 3 | 0.0027 | ++ | 0 | 0.001 | 1 | 0.9723 |
| rs10892147 | 11 | 117513801 | C | T | -2.999 | 0.00271 | -? | 0 | 0 | 0 | 1 |
| rs11823544 | 11 | 117285738 | A | G | 2.992 | 0.00277 | +? | 0 | 0 | 0 | 1 |
| rs511549 | 11 | 117285684 | C | A | 2.992 | 0.00277 | +? | 0 | 0 | 0 | 1 |
| rs527056 | 11 | 117285549 | G | A | 2.992 | 0.00277 | +? | 0 | 0 | 0 | 1 |
| rs649338 | 11 | 118705654 | A | G | 2.99 | 0.00279 | ++ | 0 | 0.137 | 1 | 0.7117 |
| rs4938549 | 11 | 118701597 | T | C | 2.988 | 0.00281 | ++ | 0 | 0.091 | 1 | 0.7624 |
| rs530646 | 11 | 118701668 | A | G | 2.988 | 0.00281 | ++ | 0 | 0.091 | 1 | 0.7624 |
| rs57150747 | 11 | 116291220 | A | G | -2.986 | 0.00282 | -? | 0 | 0 | 0 | 1 |
| rs534114 | 11 | 118702002 | A | G | 2.984 | 0.00285 | ++ | 0 | 0.09 | 1 | 0.764 |
| rs513327 | 11 | 117285497 | C | T | 2.978 | 0.00291 | +? | 0 | 0 | 0 | 1 |
| rs7941696 | 11 | 118702936 | G | A | 2.975 | 0.00293 | ++ | 0 | 0.047 | 1 | 0.8279 |
| rs490914 | 11 | 117475052 | T | A | 2.97 | 0.00298 | +? | 0 | 0 | 0 | 1 |
| rs4936374 | 11 | 117290898 | A | G | 2.963 | 0.00304 | +? | 0 | 0 | 0 | 1 |
| rs11606199 | 11 | 118811641 | A | G | -2.963 | 0.00305 | -- | 0 | 0.259 | 1 | 0.6109 |
| rs7941321 | 11 | 118702910 | A | G | 2.96 | 0.00307 | ++ | 0 | 0.084 | 1 | 0.7715 |
| rs598373 | 11 | 118664377 | T | C | -2.958 | 0.00309 | -- | 0 | 0.149 | 1 | 0.6994 |
| rs7928623 | 11 | 117499061 | C | T | -2.954 | 0.00313 | -? | 0 | 0 | 0 | 1 |
| rs1790189 | 11 | 118652326 | G | T | -2.952 | 0.00315 | -- | 0 | 0.036 | 1 | 0.8488 |
| rs1784293 | 11 | 118652373 | C | A | -2.949 | 0.00318 | -- | 0 | 0.037 | 1 | 0.8475 |
| rs2840350 | 11 | 117380850 | T | C | -2.942 | 0.00326 | -? | 0 | 0 | 0 | 1 |
| rs2508570 | 11 | 118678609 | A | G | -2.94 | 0.00328 | -- | 0 | 0.12 | 1 | 0.7288 |
| rs688161 | 11 | 118677710 | G | A | -2.94 | 0.00328 | -- | 0 | 0.129 | 1 | 0.7196 |
| rs680292 | 11 | 117475341 | T | G | 2.936 | 0.00333 | +? | 0 | 0 | 0 | 1 |
| rs490730 | 11 | 117289951 | T | C | 2.925 | 0.00345 | +? | 0 | 0 | 0 | 1 |
| rs1784300 | 11 | 118679629 | T | A | 2.918 | 0.00352 | ++ | 0 | 0.011 | 1 | 0.9174 |
| rs591756 | 11 | 118678881 | T | C | -2.917 | 0.00353 | -- | 0 | 0.094 | 1 | 0.7595 |
| rs2508916 | 11 | 118654651 | G | C | -2.913 | 0.00358 | -- | 0 | 0.015 | 1 | 0.9025 |
| rs4938550 | 11 | 118707171 | G | A | 2.913 | 0.00358 | ++ | 0 | 0.389 | 1 | 0.5326 |
| rs473298 | 11 | 118679387 | G | A | -2.895 | 0.00379 | -- | 0 | 0.109 | 1 | 0.7411 |
| rs516719 | 11 | 118679445 | C | T | -2.895 | 0.00379 | -- | 0 | 0.109 | 1 | 0.7411 |
| rs11823029 | 11 | 116283516 | C | T | -2.889 | 0.00387 | -? | 0 | 0 | 0 | 1 |
| rs540180 | 11 | 118653629 | T | G | -2.884 | 0.00393 | -- | 0 | 0.005 | 1 | 0.9433 |
| rs11606341 | 11 | 117431753 | T | G | -2.883 | 0.00395 | -? | 0 | 0 | 0 | 1 |
| rs528647 | 11 | 116523821 | T | C | 2.879 | 0.00399 | +? | 0 | 0 | 0 | 1 |
| rs2508569 | 11 | 118680045 | C | G | -2.876 | 0.00403 | -- | 0 | 0.083 | 1 | 0.7739 |
| rs6589610 | 11 | 117372097 | T | C | 2.875 | 0.00404 | +? | 0 | 0 | 0 | 1 |
| rs483747 | 11 | 116523167 | C | T | 2.874 | 0.00405 | +? | 0 | 0 | 0 | 1 |
| rs567576 | 11 | 116522559 | G | A | 2.874 | 0.00405 | +? | 0 | 0 | 0 | 1 |
| rs567599 | 11 | 116522565 | T | G | 2.874 | 0.00405 | +? | 0 | 0 | 0 | 1 |
| rs477014 | 11 | 118678986 | C | T | -2.872 | 0.00408 | -- | 0 | 0.084 | 1 | 0.772 |
| rs77694974 | 11 | 120492260 | T | G | 2.871 | 0.0041 | +? | 0 | 0 | 0 | 1 |
| rs3017237 | 11 | 116522625 | C | G | 2.867 | 0.00415 | +? | 0 | 0 | 0 | 1 |
| rs480028 | 11 | 116522770 | G | C | 2.867 | 0.00415 | +? | 0 | 0 | 0 | 1 |
| rs480878 | 11 | 116522854 | A | G | 2.867 | 0.00415 | +? | 0 | 0 | 0 | 1 |
| rs528732 | 11 | 116523844 | A | G | 2.866 | 0.00416 | +? | 0 | 0 | 0 | 1 |
| rs12796102 | 11 | 118662028 | T | C | 2.866 | 0.00416 | ++ | 0 | 0.039 | 1 | 0.8425 |
| rs3017778 | 11 | 116523931 | T | C | 2.858 | 0.00426 | +? | 0 | 0 | 0 | 1 |

| | | | | | | | | | | | |
|---|---|---|---|---|---|---|---|---|---|---|---|
| rs636736 | 11 | 118705145 | G | A | 2.849 | 0.00439 | ++ | 0 | 0.087 | 1 | 0.768 |
| rs3017779 | 11 | 116523988 | T | C | 2.845 | 0.00444 | +? | 0 | 0 | 0 | 1 |
| rs77416199 | 11 | 117498485 | C | T | -2.844 | 0.00445 | -? | 0 | 0 | 0 | 1 |
| rs485638 | 11 | 116523373 | C | G | 2.844 | 0.00445 | +? | 0 | 0 | 0 | 1 |
| rs10892156 | 11 | 117537864 | T | C | 2.844 | 0.00446 | +? | 0 | 0 | 0 | 1 |
| rs6589611 | 11 | 117372126 | T | A | 2.837 | 0.00456 | +? | 0 | 0 | 0 | 1 |
| rs562756 | 11 | 116521975 | C | G | 2.817 | 0.00485 | +? | 0 | 0 | 0 | 1 |
| rs951786 | 11 | 119529529 | G | A | -2.817 | 0.00485 | -? | 0 | 0 | 0 | 1 |
| rs10431082 | 11 | 117537353 | T | G | 2.814 | 0.00489 | +? | 0 | 0 | 0 | 1 |
| rs492182 | 11 | 116522164 | A | G | 2.809 | 0.00497 | +? | 0 | 0 | 0 | 1 |
| rs563917 | 11 | 116522156 | T | C | 2.809 | 0.00497 | +? | 0 | 0 | 0 | 1 |
| rs566416 | 11 | 118759610 | G | T | -2.807 | 0.005 | -- | -57.4 | 1.906 | 1 | 0.1674 |
| rs2511840 | 11 | 119208561 | C | G | -2.8 | 0.00511 | -? | 0 | 0 | 0 | 1 |
| rs563796 | 11 | 116522111 | G | A | 2.794 | 0.0052 | +? | 0 | 0 | 0 | 1 |
| rs77822543 | 11 | 117368804 | T | C | -2.794 | 0.00521 | -? | 0 | 0 | 0 | 1 |
| rs75349886 | 11 | 117496431 | G | A | -2.791 | 0.00525 | -? | 0 | 0 | 0 | 1 |
| rs624515 | 11 | 117370229 | G | T | 2.79 | 0.00526 | +? | 0 | 0 | 0 | 1 |
| rs12280210 | 11 | 116940449 | C | T | -2.788 | 0.00531 | -? | 0 | 0 | 0 | 1 |
| rs12797083 | 11 | 119205735 | C | T | 2.787 | 0.00532 | +? | 0 | 0 | 0 | 1 |
| rs4936383 | 11 | 117380108 | A | G | 2.78 | 0.00544 | +? | 0 | 0 | 0 | 1 |
| rs491022 | 11 | 116521988 | C | T | 2.776 | 0.0055 | +? | 0 | 0 | 0 | 1 |
| rs7946890 | 11 | 117494945 | C | A | -2.776 | 0.00551 | -? | 0 | 0 | 0 | 1 |
| rs565607 | 11 | 116522330 | T | C | 2.775 | 0.00552 | +? | 0 | 0 | 0 | 1 |
| rs575906 | 11 | 116521558 | G | A | 2.773 | 0.00555 | +? | 0 | 0 | 0 | 1 |
| rs4936554 | 11 | 120691285 | T | C | -2.773 | 0.00556 | -? | 0 | 0 | 0 | 1 |
| rs506222 | 11 | 116520434 | C | A | 2.772 | 0.00557 | +? | 0 | 0 | 0 | 1 |
| rs11216860 | 11 | 118240332 | C | T | -2.769 | 0.00562 | -- | 0 | 0.44 | 1 | 0.5071 |
| rs12795576 | 11 | 119205830 | G | A | 2.765 | 0.00569 | +? | 0 | 0 | 0 | 1 |
| rs1240772 | 11 | 116519129 | C | G | -2.765 | 0.0057 | -? | 0 | 0 | 0 | 1 |
| rs112525313 | 11 | 118014873 | T | C | -2.76 | 0.00578 | -- | -35.7 | 2.211 | 1 | 0.137 |
| rs948412 | 11 | 119214135 | G | A | -2.758 | 0.00582 | -? | 0 | 0 | 0 | 1 |
| rs10790191 | 11 | 117372548 | A | G | 2.751 | 0.00594 | +? | 0 | 0 | 0 | 1 |
| rs11824001 | 11 | 119203505 | T | C | 2.741 | 0.00612 | +? | 0 | 0 | 0 | 1 |
| rs533668 | 11 | 116521135 | G | T | 2.74 | 0.00614 | +? | 0 | 0 | 0 | 1 |
| rs544479 | 11 | 116520398 | C | T | 2.74 | 0.00614 | +? | 0 | 0 | 0 | 1 |
| rs549125 | 11 | 116520926 | G | C | 2.74 | 0.00614 | +? | 0 | 0 | 0 | 1 |
| rs573985 | 11 | 116521316 | A | G | 2.739 | 0.00616 | +? | 0 | 0 | 0 | 1 |
| rs577941 | 11 | 116226290 | A | G | -2.736 | 0.00622 | -- | -25.8 | 2.384 | 1 | 0.1226 |
| rs2239013 | 11 | 116693353 | T | C | -2.736 | 0.00622 | -- | 0 | 0.001 | 1 | 0.9695 |
| rs543824 | 11 | 118762218 | G | A | -2.735 | 0.00624 | -- | -69.1 | 1.774 | 1 | 0.1829 |
| rs9888246 | 11 | 119527302 | T | C | -2.734 | 0.00625 | -? | 0 | 0 | 0 | 1 |
| rs2282550 | 11 | 118220035 | C | T | -2.727 | 0.00639 | -- | 0 | 0.037 | 1 | 0.8478 |
| rs490257 | 11 | 116220570 | G | C | -2.723 | 0.00648 | -- | 51.3 | 6.155 | 1 | 0.0131 |
| rs76858861 | 11 | 117201458 | G | A | 2.721 | 0.00652 | +? | 0 | 0 | 0 | 1 |
| rs499790 | 11 | 116519739 | T | C | 2.719 | 0.00655 | +? | 0 | 0 | 0 | 1 |
| rs1561966 | 11 | 118221474 | A | G | -2.718 | 0.00657 | -- | 0 | 0.04 | 1 | 0.8413 |
| rs513533 | 11 | 116519358 | G | A | 2.715 | 0.00663 | +? | 0 | 0 | 0 | 1 |
| rs516043 | 11 | 116519584 | G | C | 2.715 | 0.00663 | +? | 0 | 0 | 0 | 1 |
| rs582179 | 11 | 117104148 | G | A | 2.714 | 0.00664 | +? | 0 | 0 | 0 | 1 |
| rs668628 | 11 | 118519213 | T | G | 2.713 | 0.00668 | ++ | 0 | 0.367 | 1 | 0.5448 |
| rs79874233 | 11 | 116636305 | C | G | 2.712 | 0.00669 | +? | 0 | 0 | 0 | 1 |
| rs476367 | 11 | 116220637 | C | T | -2.708 | 0.00677 | -- | 50.9 | 6.115 | 1 | 0.01341 |
| rs505349 | 11 | 116225004 | G | A | -2.706 | 0.00682 | -- | 54.1 | 6.536 | 1 | 0.01057 |
| rs73016221 | 11 | 118220027 | C | T | -2.705 | 0.00684 | -? | 0 | 0 | 0 | 1 |
| rs931953 | 11 | 119528905 | G | A | -2.703 | 0.00686 | -? | 0 | 0 | 0 | 1 |
| rs76457018 | 11 | 118762701 | A | G | 2.702 | 0.00689 | +- | 18.2 | 3.667 | 1 | 0.05551 |

| | | | | | | | | | | | |
|---|---|---|---|---|---|---|---|---|---|---|---|
| rs551372 | 11 | 116222588 | T | C | -2.698 | 0.00697 | -- | 55.4 | 6.721 | 1 | 0.009528 |
| rs1262892 | 11 | 116588904 | A | G | 2.694 | 0.00705 | +? | 0 | 0 | 0 | 1 |
| rs560299 | 11 | 116220063 | C | G | -2.694 | 0.00706 | -- | 51.6 | 6.204 | 1 | 0.01275 |
| rs523604 | 11 | 118755738 | G | A | -2.693 | 0.00708 | -- | 0 | 0.42 | 1 | 0.5171 |
| rs236911 | 11 | 117085270 | C | A | 2.692 | 0.00709 | +? | 0 | 0 | 0 | 1 |
| rs570448 | 11 | 116216432 | C | G | -2.692 | 0.00711 | -- | -34.6 | 2.229 | 1 | 0.1355 |
| rs11216492 | 11 | 117536758 | G | A | 2.683 | 0.0073 | +? | 0 | 0 | 0 | 1 |
| rs560476 | 11 | 116220118 | A | G | -2.681 | 0.00733 | -- | 51.2 | 6.149 | 1 | 0.01315 |
| rs4445648 | 11 | 119529524 | T | C | -2.68 | 0.00735 | -? | 0 | 0 | 0 | 1 |
| rs630923 | 11 | 118754353 | A | C | -2.68 | 0.00737 | -- | -135.9 | 1.272 | 1 | 0.2594 |
| rs558659 | 11 | 116216759 | A | G | -2.679 | 0.00738 | -- | 52.6 | 6.332 | 1 | 0.01186 |
| rs567681 | 11 | 116216713 | C | A | -2.679 | 0.00738 | -- | 52.6 | 6.332 | 1 | 0.01186 |
| rs75542613 | 11 | 116679155 | A | G | -2.679 | 0.00738 | -- | 0 | 0.023 | 1 | 0.8787 |
| rs2512160 | 11 | 116222809 | G | A | -2.679 | 0.00739 | -- | 56.9 | 6.958 | 1 | 0.008346 |
| rs4096589 | 11 | 117363891 | T | A | -2.669 | 0.0076 | -? | 0 | 0 | 0 | 1 |
| rs71480346 | 11 | 117502041 | T | C | -2.668 | 0.00764 | -? | 0 | 0 | 0 | 1 |
| rs4245201 | 11 | 119529332 | C | T | -2.666 | 0.00769 | -? | 0 | 0 | 0 | 1 |
| rs12721026 | 11 | 116706168 | G | T | -2.664 | 0.00771 | -- | 0 | 0.209 | 1 | 0.6479 |
| rs3212263 | 11 | 118216423 | G | A | -2.664 | 0.00773 | -- | 0 | 0.026 | 1 | 0.8721 |
| rs491113 | 11 | 116225111 | C | G | -2.662 | 0.00777 | -- | 54.5 | 6.587 | 1 | 0.01027 |
| rs10790182 | 11 | 117291361 | T | C | 2.661 | 0.0078 | +? | 0 | 0 | 0 | 1 |
| rs545657 | 11 | 116216809 | G | A | -2.66 | 0.00782 | -- | 53.2 | 6.406 | 1 | 0.01137 |
| rs68096541 | 11 | 117405907 | C | G | -2.657 | 0.00787 | -? | 0 | 0 | 0 | 1 |
| rs112618266 | 11 | 116726709 | T | C | -2.656 | 0.00792 | -- | 0 | 0.039 | 1 | 0.8428 |
| rs17120099 | 11 | 116735460 | C | A | -2.654 | 0.00795 | -- | 0 | 0 | 1 | 0.9826 |
| rs3212262 | 11 | 118217854 | T | G | -2.653 | 0.00798 | -? | 0 | 0 | 0 | 1 |
| rs73001406 | 11 | 118560857 | A | G | -2.651 | 0.00802 | -- | 0 | 0.408 | 1 | 0.523 |
| rs45586434 | 11 | 118828184 | A | C | -2.646 | 0.00815 | -- | 0 | 0.966 | 1 | 0.3257 |
| rs11217099 | 11 | 118809882 | A | C | -2.643 | 0.00822 | -- | 0 | 0.017 | 1 | 0.8949 |
| rs4936460 | 11 | 118912207 | T | C | -2.639 | 0.0083 | -? | 0 | 0 | 0 | 1 |
| rs948411 | 11 | 119213782 | T | A | -2.634 | 0.00843 | -? | 0 | 0 | 0 | 1 |
| rs7929496 | 11 | 117502961 | T | C | -2.629 | 0.00855 | -? | 0 | 0 | 0 | 1 |
| rs1892918 | 11 | 117531465 | C | A | -2.627 | 0.00861 | -? | 0 | 0 | 0 | 1 |
| rs472950 | 11 | 116226909 | T | C | -2.627 | 0.00861 | -- | 56.1 | 6.838 | 1 | 0.008924 |
| rs77967735 | 11 | 117453109 | G | A | -2.623 | 0.00873 | -? | 0 | 0 | 0 | 1 |
| rs4635117 | 11 | 116981001 | A | G | 2.618 | 0.00885 | +? | 0 | 0 | 0 | 1 |
| rs11216856 | 11 | 118227838 | T | C | -2.615 | 0.00892 | -- | 0 | 0.065 | 1 | 0.7994 |
| rs11217102 | 11 | 118824917 | A | G | -2.615 | 0.00892 | -- | 0 | 0.815 | 1 | 0.3668 |
| rs71480307 | 11 | 116516873 | A | G | 2.614 | 0.00895 | +? | 0 | 0 | 0 | 1 |
| rs12283648 | 11 | 120702916 | C | A | -2.601 | 0.00929 | -? | 0 | 0 | 0 | 1 |
| rs35165288 | 11 | 116517325 | C | T | 2.598 | 0.00938 | +? | 0 | 0 | 0 | 1 |
| rs2511847 | 11 | 119203359 | A | G | -2.597 | 0.0094 | -? | 0 | 0 | 0 | 1 |
| rs931954 | 11 | 119529106 | A | G | -2.597 | 0.0094 | -? | 0 | 0 | 0 | 1 |
| rs686413 | 11 | 117537710 | A | G | -2.596 | 0.00942 | -? | 0 | 0 | 0 | 1 |
| rs12275565 | 11 | 116753552 | G | A | -2.596 | 0.00945 | -- | 0 | 0.005 | 1 | 0.9417 |
| rs2276419 | 11 | 118230359 | T | C | -2.593 | 0.00952 | -? | 0 | 0 | 0 | 1 |
| rs568306 | 11 | 116223775 | A | C | -2.593 | 0.00953 | -- | -21.2 | 2.476 | 1 | 0.1156 |
| rs564046 | 11 | 116226845 | G | A | -2.592 | 0.00954 | -- | -22.3 | 2.452 | 1 | 0.1174 |
| rs12286581 | 11 | 116748744 | C | T | -2.588 | 0.00965 | -- | 0 | 0.006 | 1 | 0.9384 |
| rs12292614 | 11 | 116748515 | G | A | -2.588 | 0.00965 | -- | 0 | 0.006 | 1 | 0.9384 |
| rs12416987 | 11 | 116754265 | C | T | -2.588 | 0.00965 | -- | 0 | 0.006 | 1 | 0.9384 |
| rs75294699 | 11 | 116749072 | A | C | -2.588 | 0.00965 | -- | 0 | 0.006 | 1 | 0.9384 |
| rs2276417 | 11 | 118230223 | G | T | -2.586 | 0.00971 | -- | 0 | 0.058 | 1 | 0.8101 |
| rs10892040 | 11 | 116756986 | A | G | -2.586 | 0.00972 | -- | 0 | 0.005 | 1 | 0.9416 |
| rs11216174 | 11 | 116757248 | G | A | -2.586 | 0.00972 | -- | 0 | 0.005 | 1 | 0.9416 |
| rs11216175 | 11 | 116757398 | T | C | -2.586 | 0.00972 | -- | 0 | 0.005 | 1 | 0.9416 |

| | | | | | | | | | | | |
|---|---|---|---|---|---|---|---|---|---|---|---|
| rs12284346 | 11 | 116757793 | C | T | -2.586 | 0.00972 | -- | 0 | 0.005 | 1 | 0.9416 |
| rs2044426 | 11 | 116756183 | A | G | -2.586 | 0.00972 | -- | 0 | 0.005 | 1 | 0.9416 |
| rs12294191 | 11 | 116759824 | G | A | -2.585 | 0.00974 | -- | 0 | 0.005 | 1 | 0.9411 |
| rs4936358 | 11 | 116512477 | A | G | -2.583 | 0.00978 | -? | 0 | 0 | 0 | 1 |
| rs12278117 | 11 | 116748169 | A | G | -2.58 | 0.00988 | -- | 0 | 0.007 | 1 | 0.9347 |
| rs12279433 | 11 | 116748221 | T | G | -2.58 | 0.00988 | -- | 0 | 0.007 | 1 | 0.9347 |
| rs17120119 | 11 | 116747282 | G | C | -2.58 | 0.00988 | -- | 0 | 0.007 | 1 | 0.9347 |
| rs494459 | 11 | 118574675 | T | C | 2.578 | 0.00993 | ++ | 0 | 0.014 | 1 | 0.9046 |
| rs524409 | 11 | 118574311 | T | C | 2.578 | 0.00993 | ++ | 0 | 0.014 | 1 | 0.9046 |
| rs59607164 | 11 | 117458673 | G | A | -2.578 | 0.00993 | -? | 0 | 0 | 0 | 1 |
| rs140567534 | 11 | 120988038 | G | T | -2.576 | 0.00999 | -? | 0 | 0 | 0 | 1 |
| rs507763 | 11 | 118824683 | C | G | -2.574 | 0.01004 | -- | 0 | 0.857 | 1 | 0.3546 |
| rs594306 | 11 | 117531961 | A | T | -2.572 | 0.01012 | -? | 0 | 0 | 0 | 1 |
| rs150555490 | 11 | 116697081 | T | C | -2.571 | 0.01014 | -? | 0 | 0 | 0 | 1 |
| rs530901 | 11 | 118575011 | C | T | 2.57 | 0.01016 | ++ | 0 | 0.013 | 1 | 0.9082 |
| rs625735 | 11 | 118575419 | A | G | 2.57 | 0.01016 | ++ | 0 | 0.013 | 1 | 0.9082 |
| rs626645 | 11 | 118575180 | C | T | 2.57 | 0.01016 | ++ | 0 | 0.013 | 1 | 0.9082 |
| rs627391 | 11 | 118573916 | T | A | 2.57 | 0.01016 | ++ | 0 | 0.013 | 1 | 0.9082 |
| rs637563 | 11 | 118575479 | G | A | 2.57 | 0.01016 | ++ | 0 | 0.013 | 1 | 0.9082 |
| rs638514 | 11 | 118575693 | C | T | 2.57 | 0.01016 | ++ | 0 | 0.013 | 1 | 0.9082 |
| rs676925 | 11 | 118765688 | G | C | -2.567 | 0.01026 | -- | -193.2 | 1.023 | 1 | 0.3118 |
| rs576283 | 11 | 118573894 | G | A | 2.566 | 0.01029 | ++ | 0 | 0.014 | 1 | 0.9062 |
| rs1334757891 | 11 | 120850004 | T | C | 2.562 | 0.01041 | +? | 0 | 0 | 0 | 1 |
| rs531852 | 11 | 116228699 | C | A | -2.557 | 0.01057 | -- | 54.9 | 6.653 | 1 | 0.009898 |
| rs534437 | 11 | 116222378 | A | G | -2.553 | 0.01067 | -- | 45.9 | 5.549 | 1 | 0.01849 |
| rs78832854 | 11 | 116774475 | A | T | -2.551 | 0.01074 | -- | 0 | 0.061 | 1 | 0.8054 |
| rs7119734 | 11 | 117363066 | T | C | -2.548 | 0.01082 | -? | 0 | 0 | 0 | 1 |
| rs537080 | 11 | 118826045 | C | T | -2.548 | 0.01083 | -- | 0 | 0.974 | 1 | 0.3237 |
| rs77349713 | 11 | 116765476 | C | T | -2.548 | 0.01084 | -- | 0 | 0.062 | 1 | 0.8039 |
| rs117560784 | 11 | 117811037 | A | G | 2.547 | 0.01085 | +? | 0 | 0 | 0 | 1 |
| rs12292278 | 11 | 116758693 | C | A | -2.544 | 0.01097 | -- | 0 | 0.009 | 1 | 0.9224 |
| rs12718462 | 11 | 116705719 | C | T | -2.543 | 0.01098 | -- | 0 | 0.085 | 1 | 0.77 |
| rs117764543 | 11 | 120454129 | G | T | 2.542 | 0.01101 | +? | 0 | 0 | 0 | 1 |
| rs7130068 | 11 | 117366973 | C | T | -2.541 | 0.01104 | -? | 0 | 0 | 0 | 1 |
| rs58080742 | 11 | 117350621 | A | C | -2.541 | 0.01106 | -? | 0 | 0 | 0 | 1 |
| rs1228933145 | 11 | 117291266 | G | T | 2.54 | 0.01107 | +? | 0 | 0 | 0 | 1 |
| rs73592158 | 11 | 117291144 | A | G | 2.54 | 0.01107 | +? | 0 | 0 | 0 | 1 |
| rs61901410 | 11 | 118661197 | G | C | 2.54 | 0.01108 | ++ | -112.5 | 1.412 | 1 | 0.2347 |
| rs11820562 | 11 | 117293463 | G | A | 2.539 | 0.01111 | +? | 0 | 0 | 0 | 1 |
| rs11215924 | 11 | 116289906 | A | T | -2.536 | 0.01122 | -? | 0 | 0 | 0 | 1 |
| rs7123652 | 11 | 117377518 | C | T | -2.533 | 0.01132 | -? | 0 | 0 | 0 | 1 |
| rs139132354 | 11 | 117274521 | A | T | 2.529 | 0.01145 | +? | 0 | 0 | 0 | 1 |
| rs2076965 | 11 | 117405696 | A | T | -2.527 | 0.01152 | -? | 0 | 0 | 0 | 1 |
| rs564591 | 11 | 116223363 | G | A | -2.522 | 0.01168 | -- | 64.8 | 8.519 | 1 | 0.003514 |
| rs556437 | 11 | 118705753 | A | G | 2.521 | 0.01171 | +? | 0 | 0 | 0 | 1 |
| rs74881320 | 11 | 116765925 | C | A | -2.517 | 0.01185 | -- | 0 | 0.071 | 1 | 0.7901 |
| rs76887844 | 11 | 116281175 | T | C | -2.515 | 0.01189 | -? | 0 | 0 | 0 | 1 |
| rs10892047 | 11 | 116793693 | T | C | -2.515 | 0.0119 | -- | 0 | 0.11 | 1 | 0.7406 |
| rs79763467 | 11 | 117259782 | A | G | 2.514 | 0.01193 | +? | 0 | 0 | 0 | 1 |
| rs77280285 | 11 | 118091258 | G | C | 2.513 | 0.01197 | ++ | 45 | 5.452 | 1 | 0.01955 |
| rs7111427 | 11 | 117534083 | A | G | 2.512 | 0.01201 | +? | 0 | 0 | 0 | 1 |
| rs78049971 | 11 | 117284721 | T | C | 2.512 | 0.01202 | +? | 0 | 0 | 0 | 1 |
| rs79770727 | 11 | 117441594 | A | G | -2.511 | 0.01204 | -? | 0 | 0 | 0 | 1 |
| rs1053095947 | 11 | 116832718 | A | C | -2.508 | 0.01213 | -- | 0 | 0.112 | 1 | 0.7377 |
| rs10892050 | 11 | 116816394 | A | C | -2.508 | 0.01213 | -- | 0 | 0.112 | 1 | 0.7377 |
| rs11216193 | 11 | 116815599 | T | C | -2.508 | 0.01213 | -- | 0 | 0.112 | 1 | 0.7377 |

| | | | | | | | | | | | |
|---|---|---|---|---|---|---|---|---|---|---|---|
| rs11216194 | 11 | 116815738 | C | G | -2.508 | 0.01213 | -- | 0 | 0.112 | 1 | 0.7377 |
| rs11216197 | 11 | 116816778 | G | A | -2.508 | 0.01213 | -- | 0 | 0.112 | 1 | 0.7377 |
| rs11216199 | 11 | 116831016 | T | C | -2.508 | 0.01213 | -- | 0 | 0.112 | 1 | 0.7377 |
| rs116597701 | 11 | 116811487 | G | C | -2.508 | 0.01213 | -- | 0 | 0.112 | 1 | 0.7377 |
| rs12270837 | 11 | 116826643 | C | A | -2.508 | 0.01213 | -- | 0 | 0.112 | 1 | 0.7377 |
| rs12270974 | 11 | 116804005 | T | C | -2.508 | 0.01213 | -- | 0 | 0.112 | 1 | 0.7377 |
| rs12272491 | 11 | 116804370 | T | C | -2.508 | 0.01213 | -- | 0 | 0.112 | 1 | 0.7377 |
| rs12273769 | 11 | 116807173 | G | T | -2.508 | 0.01213 | -- | 0 | 0.112 | 1 | 0.7377 |
| rs12275355 | 11 | 116824162 | G | A | -2.508 | 0.01213 | -- | 0 | 0.112 | 1 | 0.7377 |
| rs12285956 | 11 | 116797475 | A | G | -2.508 | 0.01213 | -- | 0 | 0.112 | 1 | 0.7377 |
| rs12289974 | 11 | 116824286 | T | C | -2.508 | 0.01213 | -- | 0 | 0.112 | 1 | 0.7377 |
| rs12291040 | 11 | 116826617 | C | T | -2.508 | 0.01213 | -- | 0 | 0.112 | 1 | 0.7377 |
| rs12292371 | 11 | 116795948 | C | G | -2.508 | 0.01213 | -- | 0 | 0.112 | 1 | 0.7377 |
| rs12292434 | 11 | 116796065 | A | G | -2.508 | 0.01213 | -- | 0 | 0.112 | 1 | 0.7377 |
| rs12293755 | 11 | 116819264 | T | A | -2.508 | 0.01213 | -- | 0 | 0.112 | 1 | 0.7377 |
| rs12294657 | 11 | 116807368 | C | G | -2.508 | 0.01213 | -- | 0 | 0.112 | 1 | 0.7377 |
| rs12418735 | 11 | 116804762 | G | T | -2.508 | 0.01213 | -- | 0 | 0.112 | 1 | 0.7377 |
| rs12419047 | 11 | 116819906 | A | G | -2.508 | 0.01213 | -- | 0 | 0.112 | 1 | 0.7377 |
| rs12419437 | 11 | 116820690 | A | G | -2.508 | 0.01213 | -- | 0 | 0.112 | 1 | 0.7377 |
| rs12421834 | 11 | 116820023 | C | A | -2.508 | 0.01213 | -- | 0 | 0.112 | 1 | 0.7377 |
| rs12421902 | 11 | 116814646 | G | A | -2.508 | 0.01213 | -- | 0 | 0.112 | 1 | 0.7377 |
| rs17120157 | 11 | 116812628 | C | T | -2.508 | 0.01213 | -- | 0 | 0.112 | 1 | 0.7377 |
| rs2289890 | 11 | 116824620 | A | G | -2.508 | 0.01213 | -- | 0 | 0.112 | 1 | 0.7377 |
| rs2289892 | 11 | 116825240 | A | G | -2.508 | 0.01213 | -- | 0 | 0.112 | 1 | 0.7377 |
| rs74615346 | 11 | 116802972 | C | A | -2.508 | 0.01213 | -- | 0 | 0.112 | 1 | 0.7377 |
| rs7480278 | 11 | 116809789 | G | T | -2.508 | 0.01213 | -- | 0 | 0.112 | 1 | 0.7377 |
| rs75410648 | 11 | 116813438 | T | C | -2.508 | 0.01213 | -- | 0 | 0.112 | 1 | 0.7377 |
| rs75776964 | 11 | 116801061 | G | C | -2.508 | 0.01213 | -- | 0 | 0.112 | 1 | 0.7377 |
| rs76146269 | 11 | 116823617 | G | A | -2.508 | 0.01213 | -- | 0 | 0.112 | 1 | 0.7377 |
| rs76654964 | 11 | 116823521 | A | G | -2.508 | 0.01213 | -- | 0 | 0.112 | 1 | 0.7377 |
| rs77136879 | 11 | 116808453 | T | C | -2.508 | 0.01213 | -- | 0 | 0.112 | 1 | 0.7377 |
| rs77467134 | 11 | 116823107 | C | T | -2.508 | 0.01213 | -- | 0 | 0.112 | 1 | 0.7377 |
| rs78066444 | 11 | 116812058 | C | A | -2.508 | 0.01213 | -- | 0 | 0.112 | 1 | 0.7377 |
| rs79157715 | 11 | 116800270 | A | G | -2.508 | 0.01213 | -- | 0 | 0.112 | 1 | 0.7377 |
| rs80015203 | 11 | 116810906 | G | A | -2.508 | 0.01213 | -- | 0 | 0.112 | 1 | 0.7377 |
| rs58566212 | 11 | 116290501 | T | C | -2.508 | 0.01215 | -? | 0 | 0 | 0 | 1 |
| rs480823 | 11 | 116525730 | C | T | 2.506 | 0.0122 | +? | 0 | 0 | 0 | 1 |
| rs142174850 | 11 | 116715784 | A | G | -2.504 | 0.01226 | -- | 0 | 0.009 | 1 | 0.9243 |
| rs16867 | 11 | 117521967 | C | T | -2.503 | 0.01231 | -? | 0 | 0 | 0 | 1 |
| rs670413 | 11 | 120858914 | A | G | 2.501 | 0.0124 | +? | 0 | 0 | 0 | 1 |
| rs78692246 | 11 | 116791307 | A | G | -2.5 | 0.01242 | -- | 0 | 0.115 | 1 | 0.734 |
| rs12574158 | 11 | 117356276 | G | C | -2.498 | 0.01248 | -? | 0 | 0 | 0 | 1 |
| rs35184536 | 11 | 116778813 | G | A | -2.498 | 0.0125 | -- | 0 | 0.069 | 1 | 0.7921 |
| rs73008432 | 11 | 117378004 | T | C | 2.497 | 0.01252 | +? | 0 | 0 | 0 | 1 |
| rs12280030 | 11 | 116775196 | A | T | -2.495 | 0.01258 | -- | 0 | 0.077 | 1 | 0.7808 |
| rs17120131 | 11 | 116772486 | C | T | -2.495 | 0.01258 | -- | 0 | 0.077 | 1 | 0.7808 |
| rs17120132 | 11 | 116773318 | A | T | -2.495 | 0.01258 | -- | 0 | 0.077 | 1 | 0.7808 |
| rs17120136 | 11 | 116773761 | G | T | -2.495 | 0.01258 | -- | 0 | 0.077 | 1 | 0.7808 |
| rs151131966 | 11 | 116941555 | A | G | -2.495 | 0.0126 | -? | 0 | 0 | 0 | 1 |
| rs481843 | 11 | 116525867 | T | C | 2.495 | 0.01261 | +? | 0 | 0 | 0 | 1 |
| rs2127907 | 11 | 116981580 | A | G | 2.493 | 0.01265 | +? | 0 | 0 | 0 | 1 |
| rs6589596 | 11 | 116993181 | T | C | 2.491 | 0.01274 | +? | 0 | 0 | 0 | 1 |
| rs1942010 | 11 | 119918191 | C | T | 2.49 | 0.01278 | +? | 0 | 0 | 0 | 1 |
| rs12277181 | 11 | 116787320 | A | G | -2.489 | 0.0128 | -- | 0 | 0.079 | 1 | 0.7781 |
| rs115971514 | 11 | 116774499 | C | G | -2.488 | 0.01285 | -- | 0 | 0.08 | 1 | 0.7775 |
| rs78694306 | 11 | 116774244 | T | G | -2.488 | 0.01285 | -- | 0 | 0.08 | 1 | 0.7775 |

| | | | | | | | | | | |
|---|---|---|---|---|---|---|---|---|---|---|
| rs17120177 | 11 | 116827369 | C | T | -2.484 | 0.013 | -- | 0 | 0.033 | 1 | 0.8562 |
| rs112939258 | 11 | 118834718 | T | C | -2.483 | 0.01304 | -- | -130 | 1.304 | 1 | 0.2534 |
| rs149153426 | 11 | 116638657 | G | T | 2.482 | 0.01306 | +? | 0 | 0 | 0 | 1 |
| rs12273762 | 11 | 116777063 | A | T | -2.481 | 0.0131 | -- | 0 | 0.082 | 1 | 0.7745 |
| rs9651681 | 11 | 116778065 | C | T | -2.481 | 0.0131 | -- | 0 | 0.082 | 1 | 0.7745 |
| rs11216185 | 11 | 116782974 | G | T | -2.481 | 0.01311 | -- | 0 | 0.082 | 1 | 0.7743 |
| rs4420286 | 11 | 119922006 | A | G | 2.477 | 0.01325 | +? | 0 | 0 | 0 | 1 |
| rs1452843 | 11 | 119921117 | T | C | 2.473 | 0.01341 | +? | 0 | 0 | 0 | 1 |
| rs553449 | 11 | 116222849 | A | G | -2.472 | 0.01344 | -- | 48 | 5.771 | 1 | 0.01629 |
| rs501755 | 11 | 116224621 | T | C | -2.471 | 0.01346 | -- | -18.1 | 2.54 | 1 | 0.111 |
| rs71480306 | 11 | 116515557 | C | T | 2.471 | 0.01346 | +? | 0 | 0 | 0 | 1 |
| rs543819 | 11 | 116750073 | G | A | -2.47 | 0.01352 | -- | 0 | 0.373 | 1 | 0.5414 |
| rs1790192 | 11 | 118757191 | G | A | -2.468 | 0.01358 | -- | -74.6 | 1.718 | 1 | 0.1899 |
| rs76675432 | 11 | 118103976 | T | C | 2.468 | 0.01358 | ++ | 44.5 | 5.406 | 1 | 0.02006 |
| rs1006176 | 11 | 116788133 | C | T | -2.466 | 0.01366 | -- | 0 | 0.087 | 1 | 0.7679 |
| rs11216186 | 11 | 116784692 | C | T | -2.466 | 0.01366 | -- | 0 | 0.087 | 1 | 0.7679 |
| rs12418208 | 11 | 116787633 | T | C | -2.466 | 0.01366 | -- | 0 | 0.087 | 1 | 0.7679 |
| rs12418209 | 11 | 116787641 | T | C | -2.466 | 0.01366 | -- | 0 | 0.087 | 1 | 0.7679 |
| rs79711490 | 11 | 116785341 | T | C | -2.466 | 0.01366 | -- | 0 | 0.087 | 1 | 0.7679 |
| rs12420857 | 11 | 116829442 | T | G | -2.465 | 0.01369 | -- | 0 | 0.129 | 1 | 0.7191 |
| rs878692 | 11 | 116828939 | A | C | -2.465 | 0.01369 | -- | 0 | 0.129 | 1 | 0.7191 |
| rs2252145 | 11 | 117102648 | C | T | 2.463 | 0.01378 | +? | 0 | 0 | 0 | 1 |
| rs11216943 | 11 | 118556398 | A | G | -2.459 | 0.01392 | -- | 0 | 0.479 | 1 | 0.4891 |
| rs75216569 | 11 | 116907715 | G | A | -2.458 | 0.01395 | -- | 0 | 0.051 | 1 | 0.821 |
| rs12273545 | 11 | 116911012 | T | C | -2.455 | 0.01411 | -- | 0 | 0.052 | 1 | 0.8192 |
| rs10128570 | 11 | 116854021 | G | A | -2.453 | 0.01415 | -- | 0 | 0.057 | 1 | 0.8113 |
| rs10128571 | 11 | 116854181 | G | A | -2.453 | 0.01415 | -- | 0 | 0.057 | 1 | 0.8113 |
| rs10128690 | 11 | 116853922 | A | G | -2.453 | 0.01415 | -- | 0 | 0.057 | 1 | 0.8113 |
| rs11216209 | 11 | 116848186 | C | T | -2.453 | 0.01415 | -- | 0 | 0.057 | 1 | 0.8113 |
| rs11216212 | 11 | 116850351 | A | T | -2.453 | 0.01415 | -- | 0 | 0.057 | 1 | 0.8113 |
| rs11216215 | 11 | 116852687 | A | T | -2.453 | 0.01415 | -- | 0 | 0.057 | 1 | 0.8113 |
| rs11216217 | 11 | 116857652 | A | G | -2.453 | 0.01415 | -- | 0 | 0.057 | 1 | 0.8113 |
| rs11216218 | 11 | 116862661 | T | C | -2.453 | 0.01415 | -- | 0 | 0.057 | 1 | 0.8113 |
| rs11216219 | 11 | 116862755 | T | C | -2.453 | 0.01415 | -- | 0 | 0.057 | 1 | 0.8113 |
| rs12274771 | 11 | 116856698 | G | C | -2.453 | 0.01415 | -- | 0 | 0.057 | 1 | 0.8113 |
| rs12274773 | 11 | 116856709 | T | C | -2.453 | 0.01415 | -- | 0 | 0.057 | 1 | 0.8113 |
| rs12287768 | 11 | 116856322 | C | A | -2.453 | 0.01415 | -- | 0 | 0.057 | 1 | 0.8113 |
| rs146178826 | 11 | 116851367 | G | A | -2.453 | 0.01415 | -- | 0 | 0.057 | 1 | 0.8113 |
| rs74927399 | 11 | 116846266 | C | T | -2.453 | 0.01415 | -- | 0 | 0.057 | 1 | 0.8113 |
| rs75212922 | 11 | 116856992 | A | T | -2.453 | 0.01415 | -- | 0 | 0.057 | 1 | 0.8113 |
| rs76949321 | 11 | 116846343 | A | G | -2.453 | 0.01415 | -- | 0 | 0.057 | 1 | 0.8113 |
| rs80242245 | 11 | 116862245 | G | C | -2.453 | 0.01415 | -- | 0 | 0.057 | 1 | 0.8113 |
| rs10502220 | 11 | 116409093 | C | T | -2.453 | 0.01418 | -? | 0 | 0 | 0 | 1 |
| rs6589597 | 11 | 116997798 | C | T | 2.452 | 0.01419 | +? | 0 | 0 | 0 | 1 |
| rs149602720 | 11 | 119404234 | G | A | -2.452 | 0.01422 | -? | 0 | 0 | 0 | 1 |
| rs75890305 | 11 | 116919969 | G | A | -2.452 | 0.01422 | -- | 0 | 0.053 | 1 | 0.8179 |
| rs565578 | 11 | 116223503 | C | T | -2.451 | 0.01426 | -- | 53.8 | 6.489 | 1 | 0.01085 |
| rs478015 | 11 | 120861279 | G | T | 2.448 | 0.01436 | +? | 0 | 0 | 0 | 1 |
| rs949054 | 11 | 120795888 | C | T | 2.448 | 0.01438 | +? | 0 | 0 | 0 | 1 |
| rs10892316 | 11 | 118831755 | T | C | -2.447 | 0.01441 | -- | -158.2 | 1.162 | 1 | 0.2811 |
| rs12289032 | 11 | 116899580 | T | C | -2.447 | 0.01442 | -- | 0 | 0.054 | 1 | 0.8157 |
| rs146943088 | 11 | 116899825 | C | A | -2.447 | 0.01442 | -- | 0 | 0.054 | 1 | 0.8157 |
| rs2186844 | 11 | 116896913 | T | C | -2.447 | 0.01442 | -- | 0 | 0.054 | 1 | 0.8157 |
| rs77594377 | 11 | 116892446 | A | G | -2.447 | 0.01442 | -- | 0 | 0.054 | 1 | 0.8157 |
| rs77955134 | 11 | 116892368 | A | G | -2.447 | 0.01442 | -- | 0 | 0.054 | 1 | 0.8157 |
| rs12274590 | 11 | 116805696 | A | G | -2.445 | 0.01447 | -- | 0 | 0.138 | 1 | 0.7106 |

| | | | | | | | | | | | |
|---|---|---|---|---|---|---|---|---|---|---|---|
| rs79207844 | 11 | 116805441 | A | G | -2.445 | 0.01447 | -- | 0 | 0.138 | 1 | 0.7106 |
| rs516503 | 11 | 118580348 | T | C | 2.445 | 0.0145 | ++ | 0 | 0 | 1 | 0.9951 |
| rs596869 | 11 | 120861145 | T | A | 2.444 | 0.01453 | +? | 0 | 0 | 0 | 1 |
| rs147413777 | 11 | 117802888 | A | G | 2.444 | 0.01454 | +? | 0 | 0 | 0 | 1 |
| rs80273401 | 11 | 116908706 | C | T | -2.442 | 0.0146 | -- | 0 | 0.056 | 1 | 0.8137 |
| rs201134460 | 11 | 118550524 | T | C | -2.441 | 0.01466 | -- | 0 | 0.405 | 1 | 0.5243 |
| rs569304 | 11 | 116219447 | A | G | -2.44 | 0.01467 | -- | 51.3 | 6.159 | 1 | 0.01308 |
| rs11216250 | 11 | 116924000 | T | C | -2.44 | 0.0147 | -- | 0 | 0.056 | 1 | 0.8126 |
| rs12280172 | 11 | 116919273 | A | G | -2.44 | 0.0147 | -- | 0 | 0.056 | 1 | 0.8126 |
| rs77999164 | 11 | 116912117 | T | C | -2.44 | 0.0147 | -- | 0 | 0.056 | 1 | 0.8126 |
| rs12278943 | 11 | 116874358 | C | T | -2.439 | 0.01472 | -- | 0 | 0.061 | 1 | 0.805 |
| rs138181556 | 11 | 116879723 | A | G | -2.439 | 0.01472 | -- | 0 | 0.061 | 1 | 0.805 |
| rs10892252 | 11 | 118555692 | G | A | -2.438 | 0.01476 | -- | 0 | 0.526 | 1 | 0.4683 |
| rs598582 | 11 | 120860814 | G | A | 2.437 | 0.01481 | +? | 0 | 0 | 0 | 1 |
| rs10160799 | 11 | 116977888 | G | A | 2.437 | 0.01482 | +? | 0 | 0 | 0 | 1 |
| rs193204541 | 11 | 116550916 | G | C | -2.436 | 0.01486 | -? | 0 | 0 | 0 | 1 |
| rs536333 | 11 | 120860200 | C | T | 2.433 | 0.01497 | +? | 0 | 0 | 0 | 1 |
| rs560092 | 11 | 120860499 | G | A | 2.433 | 0.01497 | +? | 0 | 0 | 0 | 1 |
| rs611065 | 11 | 120860288 | C | A | 2.433 | 0.01497 | +? | 0 | 0 | 0 | 1 |
| rs565597 | 11 | 120861094 | C | T | 2.433 | 0.01498 | +? | 0 | 0 | 0 | 1 |
| rs552104 | 11 | 116219158 | G | T | -2.432 | 0.01503 | -- | 51.6 | 6.203 | 1 | 0.01275 |
| rs7938710 | 11 | 117292208 | T | C | 2.428 | 0.01519 | +? | 0 | 0 | 0 | 1 |
| rs34552724 | 11 | 116647152 | C | T | 2.427 | 0.01522 | +? | 0 | 0 | 0 | 1 |
| rs478081 | 11 | 118602497 | G | A | -2.427 | 0.01522 | -- | 0 | 0.004 | 1 | 0.9528 |
| rs61213686 | 11 | 117461888 | C | T | -2.424 | 0.01535 | -? | 0 | 0 | 0 | 1 |
| rs7112513 | 11 | 117037361 | A | G | 2.424 | 0.01536 | +? | 0 | 0 | 0 | 1 |
| rs10750101 | 11 | 116979689 | T | C | 2.423 | 0.0154 | +? | 0 | 0 | 0 | 1 |
| rs58949729 | 11 | 117370831 | T | G | -2.423 | 0.01541 | -? | 0 | 0 | 0 | 1 |
| rs12721025 | 11 | 116706047 | A | G | -2.422 | 0.01544 | -- | 0 | 0.063 | 1 | 0.8011 |
| rs2509656 | 11 | 119208576 | A | G | -2.422 | 0.01545 | -? | 0 | 0 | 0 | 1 |
| rs17120373 | 11 | 117003459 | G | A | -2.421 | 0.01547 | -? | 0 | 0 | 0 | 1 |
| rs12277092 | 11 | 116787327 | T | C | -2.42 | 0.01552 | -- | 0 | 0.103 | 1 | 0.7479 |
| rs656015 | 11 | 119913501 | A | G | 2.419 | 0.01556 | +? | 0 | 0 | 0 | 1 |
| rs10892121 | 11 | 117343197 | C | T | -2.419 | 0.01557 | -? | 0 | 0 | 0 | 1 |
| rs2471884 | 11 | 117010320 | T | C | 2.418 | 0.01559 | +? | 0 | 0 | 0 | 1 |
| rs2266133 | 11 | 120413124 | C | T | 2.418 | 0.01561 | +? | 0 | 0 | 0 | 1 |
| rs11218014 | 11 | 120689582 | A | G | 2.416 | 0.01568 | +? | 0 | 0 | 0 | 1 |
| rs1482242286 | 11 | 117263616 | G | T | 2.414 | 0.01576 | +? | 0 | 0 | 0 | 1 |
| rs117901629 | 11 | 116879679 | A | G | -2.413 | 0.01581 | -- | 0 | 0.008 | 1 | 0.9275 |
| rs4291686 | 11 | 118700557 | G | A | 2.411 | 0.01589 | +? | 0 | 0 | 0 | 1 |
| rs11216204 | 11 | 116839614 | A | G | -2.41 | 0.01594 | -- | 0 | 0.069 | 1 | 0.7922 |
| rs12272305 | 11 | 116839938 | A | G | -2.41 | 0.01594 | -- | 0 | 0.069 | 1 | 0.7922 |
| rs12281729 | 11 | 116838130 | C | A | -2.41 | 0.01594 | -- | 0 | 0.069 | 1 | 0.7922 |
| rs12417152 | 11 | 116839293 | C | G | -2.41 | 0.01594 | -- | 0 | 0.069 | 1 | 0.7922 |
| rs595851 | 11 | 120861385 | A | G | 2.41 | 0.01594 | +? | 0 | 0 | 0 | 1 |
| rs562059 | 11 | 118602058 | G | T | -2.41 | 0.01596 | -- | 0 | 0.004 | 1 | 0.951 |
| rs579706 | 11 | 116223405 | G | T | -2.408 | 0.01603 | -- | 54.9 | 6.652 | 1 | 0.009904 |
| rs12418324 | 11 | 116842676 | T | C | -2.407 | 0.0161 | -- | 0 | 0.07 | 1 | 0.7907 |
| rs478693 | 11 | 117263103 | C | T | 2.406 | 0.01612 | +? | 0 | 0 | 0 | 1 |
| rs77204473 | 11 | 116805064 | G | T | -2.405 | 0.01619 | -- | 0 | 0.043 | 1 | 0.8357 |
| rs4609639 | 11 | 117408890 | C | T | 2.404 | 0.0162 | +? | 0 | 0 | 0 | 1 |
| rs496058 | 11 | 118580906 | G | A | 2.402 | 0.01631 | ++ | 0 | 0.002 | 1 | 0.9672 |
| rs575636 | 11 | 116234032 | G | A | -2.396 | 0.01658 | -- | 0 | 0.422 | 1 | 0.5159 |
| rs625274 | 11 | 118699276 | T | C | 2.394 | 0.01668 | +? | 0 | 0 | 0 | 1 |
| rs7131536 | 11 | 116999446 | C | T | 2.393 | 0.01669 | +? | 0 | 0 | 0 | 1 |
| rs7131686 | 11 | 116999559 | C | T | 2.393 | 0.01669 | +? | 0 | 0 | 0 | 1 |

| | | | | | | | | | | | |
|---|---|---|---|---|---|---|---|---|---|---|---|
| rs61277802 | 11 | 117405785 | A | G | 2.392 | 0.01677 | +? | 0 | 0 | 0 | 1 |
| rs111491960 | 11 | 118833458 | G | A | 2.39 | 0.01685 | +? | 0 | 0 | 0 | 1 |
| rs2248853 | 11 | 119207446 | T | C | -2.39 | 0.01685 | -? | 0 | 0 | 0 | 1 |
| rs2248863 | 11 | 119207341 | A | G | -2.39 | 0.01685 | -? | 0 | 0 | 0 | 1 |
| rs9640 | 11 | 119209979 | A | T | -2.389 | 0.01688 | -? | 0 | 0 | 0 | 1 |
| rs607800 | 11 | 120863568 | T | C | 2.388 | 0.01696 | +? | 0 | 0 | 0 | 1 |
| rs10892319 | 11 | 118870413 | T | C | -2.387 | 0.01697 | -- | 0 | 0.771 | 1 | 0.3799 |
| rs77544215 | 11 | 116292287 | T | C | -2.384 | 0.01712 | -? | 0 | 0 | 0 | 1 |
| rs10892082 | 11 | 117039325 | T | G | 2.383 | 0.01715 | +? | 0 | 0 | 0 | 1 |
| rs474806 | 11 | 117196020 | T | C | -2.381 | 0.01725 | -? | 0 | 0 | 0 | 1 |
| rs184071181 | 11 | 118598936 | C | T | 2.38 | 0.0173 | ++ | 0 | 0.864 | 1 | 0.3525 |
| rs7940310 | 11 | 117024481 | T | C | 2.379 | 0.01734 | +? | 0 | 0 | 0 | 1 |
| rs478350 | 11 | 116233829 | T | C | -2.379 | 0.01735 | -- | 0 | 0.08 | 1 | 0.7778 |
| rs2581477 | 11 | 116218868 | A | G | -2.376 | 0.01751 | -- | 52.8 | 6.352 | 1 | 0.01172 |
| rs693448 | 11 | 116218926 | C | A | -2.376 | 0.01751 | -- | 52.8 | 6.352 | 1 | 0.01172 |
| rs1135663 | 11 | 116788837 | T | C | -2.375 | 0.01753 | -- | 0 | 0.128 | 1 | 0.7204 |
| rs7118201 | 11 | 116409112 | C | T | -2.375 | 0.01756 | -? | 0 | 0 | 0 | 1 |
| rs111358668 | 11 | 118834508 | A | G | -2.372 | 0.01771 | -- | -125.3 | 1.332 | 1 | 0.2485 |
| rs11216316 | 11 | 117081500 | C | A | 2.369 | 0.01782 | +? | 0 | 0 | 0 | 1 |
| rs78277728 | 11 | 116291283 | G | C | -2.369 | 0.01784 | -? | 0 | 0 | 0 | 1 |
| rs556194 | 11 | 116223171 | C | A | -2.368 | 0.01786 | -- | 53.6 | 6.461 | 1 | 0.01103 |
| rs79663212 | 11 | 116296917 | T | G | -2.368 | 0.01788 | -? | 0 | 0 | 0 | 1 |
| rs1320668 | 11 | 116967666 | C | T | 2.366 | 0.01798 | +? | 0 | 0 | 0 | 1 |
| rs12221530 | 11 | 120732349 | T | C | -2.365 | 0.01805 | -? | 0 | 0 | 0 | 1 |
| rs61900593 | 11 | 117937348 | G | A | 2.364 | 0.01809 | +? | 0 | 0 | 0 | 1 |
| rs7123517 | 11 | 117027755 | G | A | 2.364 | 0.01809 | +? | 0 | 0 | 0 | 1 |
| rs1395475712 | 11 | 117364120 | T | C | -2.364 | 0.0181 | -? | 0 | 0 | 0 | 1 |
| rs11215925 | 11 | 116290834 | C | T | -2.361 | 0.01824 | -? | 0 | 0 | 0 | 1 |
| rs7948838 | 11 | 116792961 | G | A | -2.36 | 0.01828 | -+ | -2.5 | 2.926 | 1 | 0.08716 |
| rs10128682 | 11 | 116863458 | T | C | -2.357 | 0.0184 | -- | 0 | 0.032 | 1 | 0.8583 |
| rs12718464 | 11 | 116707401 | A | G | -2.355 | 0.01854 | -- | 0 | 0.087 | 1 | 0.7684 |
| rs2277296 | 11 | 118550522 | C | T | -2.354 | 0.01858 | -- | 0 | 0.235 | 1 | 0.6278 |
| rs11826885 | 11 | 117401657 | T | C | 2.352 | 0.01867 | +? | 0 | 0 | 0 | 1 |
| rs11604866 | 11 | 116867514 | T | C | -2.351 | 0.01871 | -- | 0 | 0.033 | 1 | 0.8554 |
| rs74460214 | 11 | 116865207 | C | T | -2.351 | 0.01871 | -- | 0 | 0.033 | 1 | 0.8554 |
| rs75826529 | 11 | 116865100 | A | T | -2.351 | 0.01871 | -- | 0 | 0.033 | 1 | 0.8554 |
| rs79349393 | 11 | 116865367 | G | A | -2.351 | 0.01871 | -- | 0 | 0.033 | 1 | 0.8554 |
| rs7481286 | 11 | 116902372 | T | G | -2.351 | 0.01872 | -- | 0 | 0.03 | 1 | 0.8627 |
| rs541874 | 11 | 116240253 | T | C | -2.349 | 0.01884 | -- | 0 | 0.337 | 1 | 0.5615 |
| rs7930881 | 11 | 117350320 | A | G | -2.348 | 0.01885 | -? | 0 | 0 | 0 | 1 |
| rs2071381 | 11 | 118214267 | A | G | -2.347 | 0.01893 | -- | 0 | 0.399 | 1 | 0.5276 |
| rs1426560348 | 11 | 116913570 | A | C | -2.344 | 0.01907 | -- | 0 | 0.031 | 1 | 0.8596 |
| rs525493 | 11 | 117182708 | G | T | 2.344 | 0.0191 | +? | 0 | 0 | 0 | 1 |
| rs10892056 | 11 | 116870411 | C | G | -2.343 | 0.01911 | -- | 0 | 0.035 | 1 | 0.8519 |
| rs11216226 | 11 | 116878066 | G | C | -2.343 | 0.01911 | -- | 0 | 0.035 | 1 | 0.8519 |
| rs12276805 | 11 | 116878759 | C | T | -2.343 | 0.01911 | -- | 0 | 0.035 | 1 | 0.8519 |
| rs4604928 | 11 | 117044203 | C | A | 2.343 | 0.01911 | +? | 0 | 0 | 0 | 1 |
| rs4936367 | 11 | 117042377 | G | A | 2.343 | 0.01911 | +? | 0 | 0 | 0 | 1 |
| rs2155094 | 11 | 120849685 | A | G | -2.34 | 0.01928 | -? | 0 | 0 | 0 | 1 |
| rs536447 | 11 | 120860238 | T | C | 2.34 | 0.01929 | +? | 0 | 0 | 0 | 1 |
| rs12286920 | 11 | 117364821 | A | G | -2.339 | 0.01935 | -? | 0 | 0 | 0 | 1 |
| rs79870586 | 11 | 116886721 | A | G | -2.337 | 0.01944 | -- | 0 | 0.033 | 1 | 0.8564 |
| rs80054374 | 11 | 116890667 | G | A | -2.337 | 0.01944 | -- | 0 | 0.033 | 1 | 0.8564 |
| rs7122944 | 11 | 117034619 | G | T | 2.336 | 0.01947 | +? | 0 | 0 | 0 | 1 |
| rs73004983 | 11 | 119572124 | C | G | 2.336 | 0.01948 | +? | 0 | 0 | 0 | 1 |
| rs501652 | 11 | 116240625 | G | A | -2.336 | 0.01949 | -- | 0 | 0.338 | 1 | 0.561 |

| | | | | | | | | | | | |
|---|---|---|---|---|---|---|---|---|---|---|---|
| rs55965073 | 11 | 117936239 | C | T | 2.335 | 0.01956 | +? | 0 | 0 | 0 | 1 |
| rs150792202 | 11 | 118652569 | A | G | 2.334 | 0.01961 | ++ | 0 | 0.945 | 1 | 0.3311 |
| rs7950633 | 11 | 117024302 | C | T | 2.332 | 0.01968 | +? | 0 | 0 | 0 | 1 |
| rs76253444 | 11 | 118876213 | G | A | -2.331 | 0.01977 | -- | 0 | 0.901 | 1 | 0.3425 |
| rs114287696 | 11 | 117037080 | G | A | -2.33 | 0.01979 | -? | 0 | 0 | 0 | 1 |
| rs7925256 | 11 | 117045335 | T | C | 2.33 | 0.01979 | +? | 0 | 0 | 0 | 1 |
| rs55900121 | 11 | 117353812 | T | C | -2.33 | 0.01981 | -? | 0 | 0 | 0 | 1 |
| rs4938576 | 11 | 118746769 | T | G | -2.328 | 0.01989 | -- | -126.2 | 1.326 | 1 | 0.2494 |
| rs11216859 | 11 | 118238119 | C | T | -2.327 | 0.01999 | -- | 0 | 0.297 | 1 | 0.5856 |
| rs9645663 | 11 | 118873840 | C | T | -2.326 | 0.02 | -- | 0 | 0.759 | 1 | 0.3838 |
| rs117033797 | 11 | 116951943 | G | T | -2.326 | 0.02002 | -? | 0 | 0 | 0 | 1 |
| rs524968 | 11 | 117368805 | A | G | 2.325 | 0.02007 | +? | 0 | 0 | 0 | 1 |
| rs583161 | 11 | 117180921 | T | C | -2.324 | 0.02014 | -? | 0 | 0 | 0 | 1 |
| rs10892049 | 11 | 116816216 | C | G | -2.315 | 0.02059 | -+ | -6.1 | 2.827 | 1 | 0.0927 |
| rs12286405 | 11 | 116822528 | T | C | -2.315 | 0.02059 | -+ | -6.1 | 2.827 | 1 | 0.0927 |
| rs12292858 | 11 | 116813979 | C | A | -2.315 | 0.02059 | -+ | -6.1 | 2.827 | 1 | 0.0927 |
| rs12420518 | 11 | 116820088 | C | T | -2.315 | 0.02059 | -+ | -6.1 | 2.827 | 1 | 0.0927 |
| rs191655881 | 11 | 116817655 | C | A | -2.315 | 0.02059 | -+ | -6.1 | 2.827 | 1 | 0.0927 |
| rs11216190 | 11 | 116797684 | C | T | -2.314 | 0.02068 | -+ | -5.8 | 2.836 | 1 | 0.09215 |
| rs2581476 | 11 | 116218830 | A | G | -2.313 | 0.0207 | -- | 54.1 | 6.533 | 1 | 0.01059 |
| rs542821 | 11 | 117262598 | T | C | 2.313 | 0.0207 | +? | 0 | 0 | 0 | 1 |
| rs732403 | 11 | 117925812 | A | G | 2.311 | 0.02082 | +? | 0 | 0 | 0 | 1 |
| rs113611963 | 11 | 118839610 | G | T | 2.31 | 0.02089 | +? | 0 | 0 | 0 | 1 |
| rs61900594 | 11 | 117938377 | T | C | 2.31 | 0.02091 | +? | 0 | 0 | 0 | 1 |
| rs526151 | 11 | 116236775 | T | C | -2.309 | 0.02093 | -- | 0 | 0.464 | 1 | 0.4955 |
| rs140223477 | 11 | 116702630 | A | G | -2.307 | 0.02107 | -? | 0 | 0 | 0 | 1 |
| rs112746276 | 11 | 117352381 | T | A | -2.3 | 0.02145 | -? | 0 | 0 | 0 | 1 |
| rs598207 | 11 | 118765267 | C | G | -2.3 | 0.02147 | -+ | -68.1 | 1.785 | 1 | 0.1816 |
| rs2127905 | 11 | 117026374 | A | C | 2.295 | 0.02176 | +? | 0 | 0 | 0 | 1 |
| rs2234668 | 11 | 116692324 | A | G | -2.294 | 0.02178 | -? | 0 | 0 | 0 | 1 |
| rs11826012 | 11 | 117285699 | C | T | 2.294 | 0.02179 | +? | 0 | 0 | 0 | 1 |
| rs1789157 | 11 | 117285746 | G | A | 2.294 | 0.02179 | +? | 0 | 0 | 0 | 1 |
| rs75676165 | 11 | 116199019 | A | G | -2.293 | 0.02185 | -? | 0 | 0 | 0 | 1 |
| rs4331109 | 11 | 120723057 | G | A | -2.292 | 0.0219 | -? | 0 | 0 | 0 | 1 |
| rs11216479 | 11 | 117514378 | T | C | -2.291 | 0.02199 | -? | 0 | 0 | 0 | 1 |
| rs10892251 | 11 | 118543563 | T | C | -2.289 | 0.02206 | -- | 0 | 0.281 | 1 | 0.5958 |
| rs12418858 | 11 | 116328380 | A | G | 2.288 | 0.02213 | +? | 0 | 0 | 0 | 1 |
| rs2155393 | 11 | 117350336 | A | G | -2.286 | 0.02227 | -? | 0 | 0 | 0 | 1 |
| rs2370114 | 11 | 121192292 | G | A | -2.286 | 0.02228 | -? | 0 | 0 | 0 | 1 |
| rs2155392 | 11 | 117407658 | C | G | 2.285 | 0.02234 | +? | 0 | 0 | 0 | 1 |
| rs506255 | 11 | 118566069 | A | C | 2.283 | 0.02242 | +- | -27.5 | 2.352 | 1 | 0.1251 |
| rs634606 | 11 | 117381052 | T | C | 2.282 | 0.0225 | +? | 0 | 0 | 0 | 1 |
| rs7115583 | 11 | 116784376 | T | G | -2.281 | 0.02252 | -+ | 4.2 | 3.131 | 1 | 0.07682 |
| rs56198555 | 11 | 117404312 | T | C | 2.281 | 0.02254 | +? | 0 | 0 | 0 | 1 |
| rs554865 | 11 | 116219458 | T | C | -2.28 | 0.02262 | -- | 52.7 | 6.34 | 1 | 0.01181 |
| rs2270392 | 11 | 116767688 | C | T | -2.278 | 0.02273 | -+ | 6.4 | 3.205 | 1 | 0.07342 |
| rs567559 | 11 | 116239742 | C | G | -2.277 | 0.0228 | -- | 0 | 0.043 | 1 | 0.836 |
| rs7111854 | 11 | 116776826 | T | A | -2.276 | 0.02283 | -+ | 6.6 | 3.213 | 1 | 0.07306 |
| rs7128792 | 11 | 121005609 | G | A | 2.275 | 0.0229 | +? | 0 | 0 | 0 | 1 |
| rs2248540 | 11 | 120426966 | C | T | 2.274 | 0.02297 | +? | 0 | 0 | 0 | 1 |
| rs35168282 | 11 | 120715346 | A | G | -2.273 | 0.023 | -? | 0 | 0 | 0 | 1 |
| rs2252244 | 11 | 120418613 | G | A | 2.271 | 0.02317 | +? | 0 | 0 | 0 | 1 |
| rs73010348 | 11 | 117422690 | A | G | 2.269 | 0.02326 | +? | 0 | 0 | 0 | 1 |
| rs138471627 | 11 | 117296641 | T | C | 2.269 | 0.02329 | +? | 0 | 0 | 0 | 1 |
| rs11216210 | 11 | 116848329 | T | G | -2.268 | 0.02331 | -+ | 3.5 | 3.109 | 1 | 0.07784 |
| rs7358350 | 11 | 116851608 | C | T | -2.268 | 0.02331 | -+ | 3.5 | 3.109 | 1 | 0.07784 |

| | | | | | | | | | | | |
|---|---|---|---|---|---|---|---|---|---|---|---|
| rs73015719 | 11 | 117938177 | A | G | 2.267 | 0.0234 | +? | 0 | 0 | 0 | 1 |
| rs573965 | 11 | 117283471 | T | C | 2.265 | 0.02349 | +? | 0 | 0 | 0 | 1 |
| rs10892306 | 11 | 118751326 | A | T | -2.261 | 0.02373 | -- | 0 | 0.115 | 1 | 0.7343 |
| rs497902 | 11 | 117181919 | G | C | -2.26 | 0.0238 | -? | 0 | 0 | 0 | 1 |
| rs11217420 | 11 | 119583742 | A | G | -2.257 | 0.024 | -? | 0 | 0 | 0 | 1 |
| rs79459300 | 11 | 117847373 | T | C | 2.256 | 0.02406 | +? | 0 | 0 | 0 | 1 |
| rs685184 | 11 | 117489645 | T | C | 2.256 | 0.02407 | +? | 0 | 0 | 0 | 1 |
| rs61900567 | 11 | 117927488 | C | G | 2.255 | 0.02416 | +? | 0 | 0 | 0 | 1 |
| rs521099 | 11 | 117282979 | G | C | 2.253 | 0.02428 | +? | 0 | 0 | 0 | 1 |
| rs10891976 | 11 | 116308219 | T | A | 2.249 | 0.02449 | +? | 0 | 0 | 0 | 1 |
| rs61350190 | 11 | 117402325 | T | G | -2.249 | 0.02453 | -? | 0 | 0 | 0 | 1 |
| rs11216200 | 11 | 116831843 | T | C | -2.247 | 0.02463 | -+ | -2.1 | 2.939 | 1 | 0.08648 |
| rs77520149 | 11 | 116199020 | A | C | -2.245 | 0.02478 | -? | 0 | 0 | 0 | 1 |
| rs10750166 | 11 | 119678810 | C | G | 2.243 | 0.02488 | +? | 0 | 0 | 0 | 1 |
| rs10160754 | 11 | 116858994 | C | A | -2.241 | 0.025 | -+ | -1.1 | 2.966 | 1 | 0.08501 |
| rs10160755 | 11 | 116859003 | G | A | -2.241 | 0.025 | -+ | -1.1 | 2.966 | 1 | 0.08501 |
| rs512017 | 11 | 117511787 | G | A | 2.24 | 0.02511 | +? | 0 | 0 | 0 | 1 |
| rs576510 | 11 | 116234127 | G | C | -2.239 | 0.02515 | -- | 0 | 0.022 | 1 | 0.8808 |
| rs572619 | 11 | 116233643 | T | A | -2.239 | 0.02517 | -- | 0 | 0.009 | 1 | 0.9256 |
| rs2105590 | 11 | 117925668 | A | G | 2.236 | 0.02536 | +? | 0 | 0 | 0 | 1 |
| rs680465 | 11 | 118500963 | A | G | 2.235 | 0.0254 | ++ | 0 | 0.174 | 1 | 0.677 |
| rs236918 | 11 | 117091609 | C | G | 2.234 | 0.02547 | +? | 0 | 0 | 0 | 1 |
| rs11218030 | 11 | 120719994 | G | A | -2.233 | 0.02553 | -? | 0 | 0 | 0 | 1 |
| rs61900566 | 11 | 117926770 | G | A | 2.231 | 0.02567 | +? | 0 | 0 | 0 | 1 |
| rs488435 | 11 | 116237782 | T | C | -2.231 | 0.02568 | -- | 0 | 0.487 | 1 | 0.4855 |
| rs71480304 | 11 | 116485201 | A | G | 2.23 | 0.02572 | +? | 0 | 0 | 0 | 1 |
| rs613791 | 11 | 118764051 | T | C | -2.23 | 0.02574 | -+ | -17.6 | 2.552 | 1 | 0.1102 |
| rs1242229 | 11 | 117062370 | T | C | 2.23 | 0.02577 | +? | 0 | 0 | 0 | 1 |
| rs1242230 | 11 | 117062183 | T | C | 2.23 | 0.02577 | +? | 0 | 0 | 0 | 1 |
| rs4360730 | 11 | 116488748 | C | T | 2.23 | 0.02578 | +? | 0 | 0 | 0 | 1 |
| rs4565932 | 11 | 116488753 | A | G | 2.23 | 0.02578 | +? | 0 | 0 | 0 | 1 |
| rs11216452 | 11 | 117424804 | A | G | 2.227 | 0.02597 | +? | 0 | 0 | 0 | 1 |
| rs562236 | 11 | 116226662 | C | T | -2.226 | 0.02604 | -- | 51.6 | 6.204 | 1 | 0.01275 |
| rs1939128 | 11 | 117932042 | T | C | 2.225 | 0.02609 | +? | 0 | 0 | 0 | 1 |
| rs3901366 | 11 | 120408071 | A | G | 2.225 | 0.02609 | +? | 0 | 0 | 0 | 1 |
| rs78044162 | 11 | 116837089 | T | C | -2.225 | 0.0261 | -+ | -13 | 2.655 | 1 | 0.1032 |
| rs2252385 | 11 | 120417190 | C | T | 2.223 | 0.02624 | +? | 0 | 0 | 0 | 1 |
| rs45557239 | 11 | 116664654 | T | C | 2.222 | 0.02626 | +? | 0 | 0 | 0 | 1 |
| rs2096721 | 11 | 117925751 | T | C | 2.221 | 0.02635 | +? | 0 | 0 | 0 | 1 |
| rs7130708 | 11 | 117400550 | A | G | -2.218 | 0.02653 | -? | 0 | 0 | 0 | 1 |
| rs900012 | 11 | 116966435 | G | A | 2.218 | 0.02658 | +? | 0 | 0 | 0 | 1 |
| rs7103687 | 11 | 117406992 | A | C | -2.217 | 0.0266 | -? | 0 | 0 | 0 | 1 |
| rs2445936 | 11 | 117284577 | C | T | 2.216 | 0.02669 | +? | 0 | 0 | 0 | 1 |
| rs9971422 | 11 | 116837373 | C | T | -2.214 | 0.0268 | -+ | -4.9 | 2.86 | 1 | 0.0908 |
| rs113194822 | 11 | 116552860 | A | G | 2.214 | 0.02683 | +? | 0 | 0 | 0 | 1 |
| rs4938293 | 11 | 116512466 | T | C | -2.214 | 0.02685 | -? | 0 | 0 | 0 | 1 |
| rs12271319 | 11 | 116879224 | T | C | -2.213 | 0.0269 | -+ | 5.1 | 3.161 | 1 | 0.0754 |
| rs6421568 | 11 | 118655704 | C | G | 2.212 | 0.02694 | ++ | 0 | 0.958 | 1 | 0.3276 |
| rs6421569 | 11 | 118655775 | A | C | 2.212 | 0.02694 | ++ | 0 | 0.958 | 1 | 0.3276 |
| rs73592180 | 11 | 117308183 | G | A | -2.211 | 0.02704 | -? | 0 | 0 | 0 | 1 |
| rs561195 | 11 | 121011269 | A | C | 2.21 | 0.02712 | +? | 0 | 0 | 0 | 1 |
| rs34114773 | 11 | 117268683 | C | T | -2.208 | 0.02722 | -? | 0 | 0 | 0 | 1 |
| rs2508915 | 11 | 118652781 | C | G | 2.208 | 0.02728 | ++ | 0 | 0.964 | 1 | 0.3263 |
| rs78087637 | 11 | 116731852 | G | C | -2.206 | 0.02736 | -- | 0 | 0 | 1 | 0.9979 |
| rs34217740 | 11 | 121018698 | A | G | 2.203 | 0.02757 | +? | 0 | 0 | 0 | 1 |
| rs574529 | 11 | 116233857 | C | T | -2.203 | 0.02758 | -- | 0 | 0.017 | 1 | 0.8971 |

| | | | | | | | | | | | |
|---|---|---|---|---|---|---|---|---|---|---|---|
| rs12273536 | 11 | 116910971 | T | C | -2.202 | 0.02766 | -- | 0 | 0.139 | 1 | 0.709 |
| rs543692 | 11 | 118598138 | C | T | 2.2 | 0.02781 | ++ | 0 | 0.395 | 1 | 0.5299 |
| rs4936459 | 11 | 118899218 | T | C | -2.2 | 0.02782 | -- | 0 | 0.038 | 1 | 0.8448 |
| rs7112937 | 11 | 116880456 | C | T | -2.2 | 0.02782 | -+ | 4.3 | 3.135 | 1 | 0.07664 |
| rs516742 | 11 | 117192383 | G | A | 2.197 | 0.028 | +? | 0 | 0 | 0 | 1 |
| rs1784295 | 11 | 118660932 | A | G | 2.197 | 0.02801 | ++ | 0 | 0.814 | 1 | 0.3669 |
| rs1790188 | 11 | 118661055 | T | C | 2.197 | 0.02801 | ++ | 0 | 0.814 | 1 | 0.3669 |
| rs1786193 | 11 | 117997570 | A | T | -2.195 | 0.02817 | -? | 0 | 0 | 0 | 1 |
| rs117640781 | 11 | 116771226 | T | C | -2.194 | 0.02824 | -? | 0 | 0 | 0 | 1 |
| rs7942967 | 11 | 117486730 | T | C | -2.193 | 0.02833 | -? | 0 | 0 | 0 | 1 |
| rs55642294 | 11 | 117395497 | A | C | -2.191 | 0.02844 | -? | 0 | 0 | 0 | 1 |
| rs11217109 | 11 | 118853774 | C | T | -2.191 | 0.02848 | -- | 0 | 0.571 | 1 | 0.4498 |
| rs640012 | 11 | 117987573 | C | T | -2.19 | 0.02851 | -? | 0 | 0 | 0 | 1 |
| rs683842 | 11 | 117489941 | C | G | 2.19 | 0.02852 | +? | 0 | 0 | 0 | 1 |
| rs3194726 | 11 | 118279936 | C | T | -2.19 | 0.02855 | -? | 0 | 0 | 0 | 1 |
| rs113962716 | 11 | 118853021 | A | C | -2.189 | 0.02863 | -- | 0 | 0.579 | 1 | 0.4467 |
| rs2186762 | 11 | 117386067 | A | C | 2.188 | 0.02869 | +? | 0 | 0 | 0 | 1 |
| rs77543424 | 11 | 117268074 | G | C | 2.187 | 0.02871 | +? | 0 | 0 | 0 | 1 |
| rs2445939 | 11 | 117186073 | T | A | -2.184 | 0.02895 | -? | 0 | 0 | 0 | 1 |
| rs7126500 | 11 | 117399606 | T | G | -2.183 | 0.02902 | -? | 0 | 0 | 0 | 1 |
| rs11216370 | 11 | 117248832 | A | G | -2.182 | 0.02914 | -? | 0 | 0 | 0 | 1 |
| rs540904 | 11 | 117284770 | C | T | 2.181 | 0.02918 | +? | 0 | 0 | 0 | 1 |
| rs17598956 | 11 | 117903210 | C | T | 2.18 | 0.02929 | +? | 0 | 0 | 0 | 1 |
| rs115076910 | 11 | 121009383 | C | T | 2.178 | 0.02941 | +? | 0 | 0 | 0 | 1 |
| rs12279396 | 11 | 120441720 | G | A | -2.176 | 0.02953 | -? | 0 | 0 | 0 | 1 |
| rs2253023 | 11 | 120412169 | T | C | 2.176 | 0.02954 | +? | 0 | 0 | 0 | 1 |
| rs4936417 | 11 | 117903836 | C | T | 2.175 | 0.0296 | +? | 0 | 0 | 0 | 1 |
| rs1815786 | 11 | 116921390 | T | C | 2.175 | 0.02962 | +? | 0 | 0 | 0 | 1 |
| rs624428 | 11 | 121010662 | T | C | 2.173 | 0.02975 | +? | 0 | 0 | 0 | 1 |
| rs11216220 | 11 | 116863988 | C | T | -2.173 | 0.02977 | -+ | 2.6 | 3.081 | 1 | 0.07922 |
| rs538645 | 11 | 118711069 | C | T | 2.17 | 0.03004 | ++ | 0 | 0.479 | 1 | 0.4889 |
| rs7931665 | 11 | 116866240 | G | A | -2.168 | 0.03014 | -+ | 2.3 | 3.071 | 1 | 0.07971 |
| rs34002948 | 11 | 121009379 | A | G | 2.168 | 0.03019 | +? | 0 | 0 | 0 | 1 |
| rs12284696 | 11 | 116879519 | G | A | -2.167 | 0.0302 | -+ | 4.4 | 3.138 | 1 | 0.0765 |
| rs572862 | 11 | 117283634 | G | A | 2.167 | 0.03023 | +? | 0 | 0 | 0 | 1 |
| rs572884 | 11 | 117283626 | G | A | 2.167 | 0.03023 | +? | 0 | 0 | 0 | 1 |
| rs603678 | 11 | 117987391 | C | T | -2.166 | 0.0303 | -? | 0 | 0 | 0 | 1 |
| rs144466751 | 11 | 116299456 | C | T | -2.166 | 0.03033 | -? | 0 | 0 | 0 | 1 |
| rs548167 | 11 | 118648396 | A | G | 2.164 | 0.03047 | ++ | 0 | 0.84 | 1 | 0.3593 |
| rs189189723 | 11 | 117034899 | T | G | -2.164 | 0.0305 | -? | 0 | 0 | 0 | 1 |
| rs480801 | 11 | 117512831 | C | T | 2.164 | 0.0305 | +? | 0 | 0 | 0 | 1 |
| rs12282572 | 11 | 116480266 | C | T | 2.163 | 0.03057 | +? | 0 | 0 | 0 | 1 |
| rs10892057 | 11 | 116872034 | C | A | -2.162 | 0.03063 | -+ | 1.9 | 3.058 | 1 | 0.08033 |
| rs11216224 | 11 | 116875924 | C | T | -2.162 | 0.03063 | -+ | 1.9 | 3.058 | 1 | 0.08033 |
| rs11216225 | 11 | 116878014 | G | A | -2.162 | 0.03063 | -+ | 1.9 | 3.058 | 1 | 0.08033 |
| rs1940625 | 11 | 116876442 | G | A | -2.162 | 0.03063 | -+ | 1.9 | 3.058 | 1 | 0.08033 |
| rs7946390 | 11 | 116875152 | G | A | -2.162 | 0.03063 | -+ | 1.9 | 3.058 | 1 | 0.08033 |
| rs11218019 | 11 | 120703916 | C | A | -2.161 | 0.03066 | -? | 0 | 0 | 0 | 1 |
| rs477036 | 11 | 117183914 | C | T | 2.161 | 0.03067 | +? | 0 | 0 | 0 | 1 |
| rs73577182 | 11 | 119678767 | C | G | 2.161 | 0.03071 | +? | 0 | 0 | 0 | 1 |
| rs561227 | 11 | 121011258 | T | C | 2.16 | 0.03078 | +? | 0 | 0 | 0 | 1 |
| rs11823783 | 11 | 117412884 | C | T | -2.16 | 0.0308 | -? | 0 | 0 | 0 | 1 |
| rs6589577 | 11 | 116749977 | C | T | 2.16 | 0.0308 | +? | 0 | 0 | 0 | 1 |
| rs6589589 | 11 | 116932428 | T | C | 2.156 | 0.03108 | +? | 0 | 0 | 0 | 1 |
| rs7124741 | 11 | 116752219 | A | T | 2.155 | 0.03113 | +? | 0 | 0 | 0 | 1 |
| rs1351452 | 11 | 116943354 | A | C | 2.154 | 0.03121 | +? | 0 | 0 | 0 | 1 |

| | | | | | | | | | | | |
|---|---|---|---|---|---|---|---|---|---|---|---|
| rs80126726 | 11 | 118852647 | A | G | -2.154 | 0.03121 | -- | 0 | 0.591 | 1 | 0.4422 |
| rs12795310 | 11 | 120931054 | C | T | -2.153 | 0.0313 | -? | 0 | 0 | 0 | 1 |
| rs2508652 | 11 | 120405162 | T | C | 2.152 | 0.03143 | +? | 0 | 0 | 0 | 1 |
| rs4936580 | 11 | 121007777 | A | G | 2.152 | 0.03143 | +? | 0 | 0 | 0 | 1 |
| rs111698696 | 11 | 117412773 | T | C | -2.151 | 0.03148 | -? | 0 | 0 | 0 | 1 |
| rs17120139 | 11 | 116774201 | A | G | 2.15 | 0.03155 | +? | 0 | 0 | 0 | 1 |
| rs672058 | 11 | 116764021 | C | T | 2.148 | 0.03169 | +? | 0 | 0 | 0 | 1 |
| rs587263 | 11 | 118816156 | C | T | 2.148 | 0.03175 | +? | 0 | 0 | 0 | 1 |
| rs2510853 | 11 | 117410682 | A | G | -2.146 | 0.03184 | -? | 0 | 0 | 0 | 1 |
| rs530849 | 11 | 116530298 | C | T | -2.146 | 0.03188 | -? | 0 | 0 | 0 | 1 |
| rs10736487 | 11 | 116950232 | A | G | 2.144 | 0.03205 | +? | 0 | 0 | 0 | 1 |
| rs4938335 | 11 | 116947361 | A | C | 2.144 | 0.03205 | +? | 0 | 0 | 0 | 1 |
| rs535996 | 11 | 118637666 | T | C | 2.143 | 0.03213 | ++ | 0 | 0.869 | 1 | 0.3513 |
| rs375508071 | 11 | 120828334 | G | A | 2.142 | 0.03218 | +? | 0 | 0 | 0 | 1 |
| rs638463 | 11 | 121011485 | C | T | 2.142 | 0.03219 | +? | 0 | 0 | 0 | 1 |
| rs7125158 | 11 | 120521305 | C | A | -2.142 | 0.0322 | -? | 0 | 0 | 0 | 1 |
| rs529717 | 11 | 118640081 | A | G | 2.141 | 0.03224 | ++ | 0 | 0.864 | 1 | 0.3525 |
| rs11216444 | 11 | 117411410 | G | A | -2.141 | 0.03225 | -? | 0 | 0 | 0 | 1 |
| rs615697 | 11 | 121037554 | C | T | 2.14 | 0.03238 | +? | 0 | 0 | 0 | 1 |
| rs616184 | 11 | 121037670 | A | G | 2.14 | 0.03238 | +? | 0 | 0 | 0 | 1 |
| rs12797557 | 11 | 117254122 | A | G | -2.138 | 0.0325 | -? | 0 | 0 | 0 | 1 |
| rs473188 | 11 | 116812029 | C | G | 2.137 | 0.0326 | +? | 0 | 0 | 0 | 1 |
| rs59099254 | 11 | 117411717 | A | C | -2.137 | 0.03261 | -? | 0 | 0 | 0 | 1 |
| rs2508929 | 11 | 118641451 | G | T | 2.136 | 0.03266 | ++ | 0 | 0.87 | 1 | 0.351 |
| rs489985 | 11 | 121037530 | A | G | 2.136 | 0.03267 | +? | 0 | 0 | 0 | 1 |
| rs1729408 | 11 | 116674818 | G | A | -2.136 | 0.03269 | -? | 0 | 0 | 0 | 1 |
| rs75375536 | 11 | 116295218 | G | A | -2.136 | 0.0327 | -? | 0 | 0 | 0 | 1 |
| rs17599289 | 11 | 117922730 | T | C | 2.135 | 0.03275 | +? | 0 | 0 | 0 | 1 |
| rs73024367 | 11 | 117362165 | A | G | 2.133 | 0.03292 | +? | 0 | 0 | 0 | 1 |
| rs61900560 | 11 | 117908084 | C | G | 2.133 | 0.03294 | +? | 0 | 0 | 0 | 1 |
| rs553779 | 11 | 116530376 | G | A | -2.132 | 0.033 | -? | 0 | 0 | 0 | 1 |
| rs579890 | 11 | 116798963 | A | G | 2.131 | 0.03311 | +? | 0 | 0 | 0 | 1 |
| rs60851263 | 11 | 117410174 | T | C | -2.13 | 0.03317 | -? | 0 | 0 | 0 | 1 |
| rs7926330 | 11 | 117915445 | T | C | 2.13 | 0.03317 | +? | 0 | 0 | 0 | 1 |
| rs10892126 | 11 | 117355892 | C | G | -2.129 | 0.03325 | -? | 0 | 0 | 0 | 1 |
| rs741811 | 11 | 118898319 | A | G | -2.128 | 0.03335 | -- | 0 | 0.379 | 1 | 0.5384 |
| rs10502224 | 11 | 116870480 | G | A | -2.127 | 0.0334 | -+ | -0.4 | 2.989 | 1 | 0.08382 |
| rs571139 | 11 | 116237017 | A | G | -2.126 | 0.03351 | -- | 0 | 0.011 | 1 | 0.9174 |
| rs10790172 | 11 | 116927740 | G | T | 2.126 | 0.03353 | +? | 0 | 0 | 0 | 1 |
| rs580683 | 11 | 121034491 | T | C | 2.124 | 0.03368 | +? | 0 | 0 | 0 | 1 |
| rs530845 | 11 | 116530293 | T | C | -2.123 | 0.03372 | -? | 0 | 0 | 0 | 1 |
| rs17509913 | 11 | 117434487 | A | G | -2.123 | 0.03374 | -? | 0 | 0 | 0 | 1 |
| rs117158703 | 11 | 116897073 | T | C | -2.123 | 0.03375 | -? | 0 | 0 | 0 | 1 |
| rs686133 | 11 | 117373635 | G | T | 2.123 | 0.03376 | +? | 0 | 0 | 0 | 1 |
| rs57154584 | 11 | 117302152 | C | T | -2.123 | 0.03378 | -? | 0 | 0 | 0 | 1 |
| rs568688 | 11 | 118598612 | G | T | 2.122 | 0.03381 | ++ | 0 | 0.312 | 1 | 0.5764 |
| rs1008966 | 11 | 120854046 | T | C | -2.122 | 0.03382 | -? | 0 | 0 | 0 | 1 |
| rs57208438 | 11 | 117412185 | A | G | -2.122 | 0.03383 | -? | 0 | 0 | 0 | 1 |
| rs58088485 | 11 | 117411664 | A | G | -2.121 | 0.03389 | -? | 0 | 0 | 0 | 1 |
| rs10732856 | 11 | 116907113 | T | C | 2.121 | 0.03391 | +? | 0 | 0 | 0 | 1 |
| rs623052 | 11 | 117431045 | G | A | 2.12 | 0.03399 | +? | 0 | 0 | 0 | 1 |
| rs72645460 | 11 | 116771375 | G | A | 2.12 | 0.03403 | +? | 0 | 0 | 0 | 1 |
| rs78221557 | 11 | 116515528 | A | G | -2.119 | 0.03408 | -? | 0 | 0 | 0 | 1 |
| rs490296 | 11 | 116237970 | C | T | -2.118 | 0.03418 | -- | 0 | 0.017 | 1 | 0.8973 |
| rs2000615 | 11 | 116915819 | C | T | 2.116 | 0.03431 | +? | 0 | 0 | 0 | 1 |
| rs2155583 | 11 | 116915217 | A | G | 2.116 | 0.03431 | +? | 0 | 0 | 0 | 1 |

| | | | | | | | | | | | |
|---|---|---|---|---|---|---|---|---|---|---|---|
| rs2276422 | 11 | 118263388 | G | C | -2.116 | 0.03435 | -? | 0 | 0 | 0 | 1 |
| rs7115242 | 11 | 116908283 | A | G | 2.116 | 0.03435 | +? | 0 | 0 | 0 | 1 |
| rs7122110 | 11 | 121021940 | G | A | 2.115 | 0.03445 | +? | 0 | 0 | 0 | 1 |
| rs7926760 | 11 | 117222256 | C | T | -2.115 | 0.03447 | -? | 0 | 0 | 0 | 1 |
| rs1287591 | 11 | 119904686 | T | C | 2.114 | 0.03452 | +? | 0 | 0 | 0 | 1 |
| rs4356265 | 11 | 117301818 | T | C | -2.114 | 0.03452 | -? | 0 | 0 | 0 | 1 |
| rs10892237 | 11 | 118267763 | C | T | -2.113 | 0.03457 | -? | 0 | 0 | 0 | 1 |
| rs79092677 | 11 | 118265734 | G | T | -2.113 | 0.03457 | -? | 0 | 0 | 0 | 1 |
| rs11216861 | 11 | 118241516 | A | G | -2.113 | 0.03459 | -- | 0 | 0.05 | 1 | 0.8223 |
| rs4936590 | 11 | 121039234 | C | T | 2.113 | 0.03464 | +? | 0 | 0 | 0 | 1 |
| rs73008413 | 11 | 117368182 | T | A | 2.112 | 0.03466 | +? | 0 | 0 | 0 | 1 |
| rs1241658 | 11 | 116819617 | G | A | 2.112 | 0.03467 | +? | 0 | 0 | 0 | 1 |
| rs1002037 | 11 | 116516736 | C | G | -2.112 | 0.03472 | -? | 0 | 0 | 0 | 1 |
| rs6589592 | 11 | 116957907 | C | G | 2.111 | 0.03477 | +? | 0 | 0 | 0 | 1 |
| rs7130297 | 11 | 116923870 | C | T | 2.11 | 0.03484 | +? | 0 | 0 | 0 | 1 |
| rs61904786 | 11 | 116414846 | G | C | -2.11 | 0.03487 | -? | 0 | 0 | 0 | 1 |
| rs694093 | 11 | 117377135 | T | G | 2.109 | 0.03495 | +? | 0 | 0 | 0 | 1 |
| rs373191677 | 11 | 119904782 | A | G | 2.109 | 0.03496 | +? | 0 | 0 | 0 | 1 |
| rs605379 | 11 | 121011098 | T | C | 2.109 | 0.03497 | +? | 0 | 0 | 0 | 1 |
| rs10750100 | 11 | 116917244 | T | C | 2.108 | 0.035 | +? | 0 | 0 | 0 | 1 |
| rs1815787 | 11 | 116921397 | T | C | 2.108 | 0.03501 | +? | 0 | 0 | 0 | 1 |
| rs4288786 | 11 | 117033406 | A | G | -2.108 | 0.03505 | -? | 0 | 0 | 0 | 1 |
| rs641924 | 11 | 119936017 | T | C | 2.108 | 0.03505 | +? | 0 | 0 | 0 | 1 |
| rs61897703 | 11 | 119672586 | T | G | 2.108 | 0.03506 | +? | 0 | 0 | 0 | 1 |
| rs56310543 | 11 | 117905204 | A | G | 2.108 | 0.03507 | +? | 0 | 0 | 0 | 1 |
| rs570424 | 11 | 118588998 | C | T | 2.107 | 0.03511 | +? | 0 | 0 | 0 | 1 |
| rs7131196 | 11 | 118265227 | C | T | -2.106 | 0.0352 | -? | 0 | 0 | 0 | 1 |
| rs11217762 | 11 | 120071508 | A | G | -2.106 | 0.03524 | -? | 0 | 0 | 0 | 1 |
| rs12290281 | 11 | 116493633 | T | G | 2.102 | 0.03551 | +? | 0 | 0 | 0 | 1 |
| rs75871913 | 11 | 120848685 | A | G | 2.102 | 0.03551 | +? | 0 | 0 | 0 | 1 |
| rs12577702 | 11 | 117394979 | C | T | 2.102 | 0.03552 | +? | 0 | 0 | 0 | 1 |
| rs510988 | 11 | 116813804 | C | T | 2.102 | 0.03553 | +? | 0 | 0 | 0 | 1 |
| rs76948040 | 11 | 117444090 | T | C | -2.102 | 0.03554 | -? | 0 | 0 | 0 | 1 |
| rs61900561 | 11 | 117908926 | A | G | 2.102 | 0.03558 | +? | 0 | 0 | 0 | 1 |
| rs6589690 | 11 | 118632893 | A | G | 2.102 | 0.03559 | ++ | 0 | 0.968 | 1 | 0.3252 |
| rs2000614 | 11 | 116915742 | C | T | 2.101 | 0.03564 | +? | 0 | 0 | 0 | 1 |
| rs2000616 | 11 | 116916529 | C | T | 2.101 | 0.03564 | +? | 0 | 0 | 0 | 1 |
| rs4388921 | 11 | 116976778 | C | T | -2.1 | 0.03569 | -? | 0 | 0 | 0 | 1 |
| rs493134 | 11 | 116833679 | C | T | 2.1 | 0.03572 | +? | 0 | 0 | 0 | 1 |
| rs483837 | 11 | 117266466 | C | T | 2.099 | 0.0358 | +? | 0 | 0 | 0 | 1 |
| rs11215905 | 11 | 116242693 | C | T | -2.099 | 0.03581 | -- | 0 | 0.01 | 1 | 0.9205 |
| rs491111 | 11 | 116238034 | A | G | -2.099 | 0.03584 | -- | 0 | 0.014 | 1 | 0.9059 |
| rs7120982 | 11 | 118866505 | A | G | 2.099 | 0.03586 | +? | 0 | 0 | 0 | 1 |
| rs545675 | 11 | 116236469 | T | C | -2.098 | 0.03588 | -- | 0 | 0.008 | 1 | 0.93 |
| rs4938375 | 11 | 117294973 | C | T | 2.098 | 0.03589 | +? | 0 | 0 | 0 | 1 |
| rs7950364 | 11 | 116898886 | C | G | 2.097 | 0.03602 | +? | 0 | 0 | 0 | 1 |
| rs7110565 | 11 | 121062486 | C | T | 2.096 | 0.03607 | +? | 0 | 0 | 0 | 1 |
| rs596882 | 11 | 121011416 | C | G | 2.095 | 0.03619 | +? | 0 | 0 | 0 | 1 |
| rs11216864 | 11 | 118260017 | T | C | -2.094 | 0.0363 | -? | 0 | 0 | 0 | 1 |
| rs61898858 | 11 | 119304622 | G | A | -2.093 | 0.03632 | -? | 0 | 0 | 0 | 1 |
| rs640048 | 11 | 117987605 | G | T | -2.093 | 0.03633 | -? | 0 | 0 | 0 | 1 |
| rs2004927 | 11 | 117368982 | C | T | 2.093 | 0.03638 | +? | 0 | 0 | 0 | 1 |
| rs630024 | 11 | 117534353 | A | C | 2.093 | 0.03639 | +? | 0 | 0 | 0 | 1 |
| rs1176456955 | 11 | 116516966 | A | G | 2.092 | 0.03644 | +? | 0 | 0 | 0 | 1 |
| rs7940318 | 11 | 117994614 | G | A | -2.09 | 0.03664 | -? | 0 | 0 | 0 | 1 |
| rs2510876 | 11 | 120412823 | G | A | 2.089 | 0.03668 | +? | 0 | 0 | 0 | 1 |

| | | | | | | | | | | | |
|---|---|---|---|---|---|---|---|---|---|---|---|
| rs4936583 | 11 | 121015968 | C | T | 2.089 | 0.03672 | +? | 0 | 0 | 0 | 1 |
| rs75017819 | 11 | 118285789 | G | A | -2.089 | 0.03672 | -? | 0 | 0 | 0 | 1 |
| rs61897704 | 11 | 119675331 | T | C | 2.089 | 0.03675 | +? | 0 | 0 | 0 | 1 |
| rs61897705 | 11 | 119675529 | G | A | 2.089 | 0.03675 | +? | 0 | 0 | 0 | 1 |
| rs2266132 | 11 | 120413117 | T | G | 2.088 | 0.03678 | +? | 0 | 0 | 0 | 1 |
| rs10892676 | 11 | 121016098 | C | T | 2.088 | 0.03679 | +? | 0 | 0 | 0 | 1 |
| rs12275038 | 11 | 121016818 | A | G | 2.088 | 0.03679 | +? | 0 | 0 | 0 | 1 |
| rs4936584 | 11 | 121016145 | A | G | 2.088 | 0.03679 | +? | 0 | 0 | 0 | 1 |
| rs583983 | 11 | 117525125 | T | C | 2.088 | 0.0368 | +? | 0 | 0 | 0 | 1 |
| rs7113829 | 11 | 116890259 | T | C | -2.088 | 0.03682 | -- | -11 | 2.703 | 1 | 0.1002 |
| rs73008408 | 11 | 117367196 | C | A | 2.088 | 0.03683 | +? | 0 | 0 | 0 | 1 |
| rs80026097 | 11 | 116732231 | C | T | -2.086 | 0.03701 | -? | 0 | 0 | 0 | 1 |
| rs560365 | 11 | 121011342 | T | C | 2.083 | 0.03721 | +? | 0 | 0 | 0 | 1 |
| rs77110391 | 11 | 116414485 | A | G | -2.083 | 0.03729 | -? | 0 | 0 | 0 | 1 |
| rs17120152 | 11 | 116787192 | C | T | -2.081 | 0.03739 | -? | 0 | 0 | 0 | 1 |
| rs11217933 | 11 | 120503083 | A | G | 2.081 | 0.03745 | +? | 0 | 0 | 0 | 1 |
| rs61900563 | 11 | 117914450 | A | G | 2.08 | 0.03757 | +? | 0 | 0 | 0 | 1 |
| rs57300701 | 11 | 118845546 | G | C | 2.079 | 0.03759 | +? | 0 | 0 | 0 | 1 |
| rs1807116 | 11 | 117369226 | C | T | 2.079 | 0.03762 | +? | 0 | 0 | 0 | 1 |
| rs111530588 | 11 | 116922586 | A | G | -2.079 | 0.03763 | -? | 0 | 0 | 0 | 1 |
| rs4938419 | 11 | 117479686 | T | A | -2.078 | 0.03769 | -? | 0 | 0 | 0 | 1 |
| rs12794425 | 11 | 116478827 | A | G | 2.077 | 0.03782 | +? | 0 | 0 | 0 | 1 |
| rs613244 | 11 | 117513154 | A | G | 2.076 | 0.03786 | +? | 0 | 0 | 0 | 1 |
| rs12794476 | 11 | 116478904 | A | G | 2.074 | 0.03808 | +? | 0 | 0 | 0 | 1 |
| rs560401 | 11 | 121011325 | G | A | 2.073 | 0.03814 | +? | 0 | 0 | 0 | 1 |
| rs4936376 | 11 | 117294959 | T | C | 2.07 | 0.03849 | +? | 0 | 0 | 0 | 1 |
| rs55787848 | 11 | 116457294 | T | C | -2.069 | 0.03854 | -? | 0 | 0 | 0 | 1 |
| rs116582417 | 11 | 116943066 | A | G | -2.068 | 0.03867 | -? | 0 | 0 | 0 | 1 |
| rs4936421 | 11 | 117913074 | C | A | 2.068 | 0.03867 | +? | 0 | 0 | 0 | 1 |
| rs11823335 | 11 | 117350966 | T | C | -2.068 | 0.03868 | -? | 0 | 0 | 0 | 1 |
| rs34998970 | 11 | 116881126 | G | A | -2.067 | 0.0387 | -? | 0 | 0 | 0 | 1 |
| rs79862770 | 11 | 116880715 | T | C | -2.067 | 0.0387 | -? | 0 | 0 | 0 | 1 |
| rs2266134 | 11 | 120413132 | C | G | 2.067 | 0.03874 | +? | 0 | 0 | 0 | 1 |
| rs516712 | 11 | 117192376 | G | A | 2.066 | 0.03886 | +? | 0 | 0 | 0 | 1 |
| rs55730046 | 11 | 117913315 | T | C | 2.065 | 0.03897 | +? | 0 | 0 | 0 | 1 |
| rs574813 | 11 | 117258662 | G | C | 2.064 | 0.03901 | +? | 0 | 0 | 0 | 1 |
| rs574873 | 11 | 117258687 | T | G | 2.064 | 0.03901 | +? | 0 | 0 | 0 | 1 |
| rs11216183 | 11 | 116781545 | A | C | -2.064 | 0.03906 | -- | 0 | 0.093 | 1 | 0.7605 |
| rs528958 | 11 | 121012693 | T | G | 2.064 | 0.03906 | +? | 0 | 0 | 0 | 1 |
| rs581786 | 11 | 121012492 | C | T | 2.064 | 0.03906 | +? | 0 | 0 | 0 | 1 |
| rs665743 | 11 | 121012978 | A | G | 2.064 | 0.03906 | +? | 0 | 0 | 0 | 1 |
| rs688106 | 11 | 121012831 | A | T | 2.064 | 0.03906 | +? | 0 | 0 | 0 | 1 |
| rs117709132 | 11 | 116760485 | T | C | -2.063 | 0.03913 | -- | 0 | 0.078 | 1 | 0.78 |
| rs4623911 | 11 | 121018870 | G | A | 2.062 | 0.03917 | +? | 0 | 0 | 0 | 1 |
| rs4641485 | 11 | 121018999 | G | A | 2.062 | 0.03917 | +? | 0 | 0 | 0 | 1 |
| rs76882215 | 11 | 116897163 | C | T | -2.062 | 0.03921 | -? | 0 | 0 | 0 | 1 |
| rs7126416 | 11 | 117399649 | A | C | -2.061 | 0.03928 | -? | 0 | 0 | 0 | 1 |
| rs563537 | 11 | 118595207 | A | T | 2.061 | 0.03935 | +? | 0 | 0 | 0 | 1 |
| rs143793912 | 11 | 117213159 | T | C | -2.06 | 0.03941 | -? | 0 | 0 | 0 | 1 |
| rs7942406 | 11 | 117371563 | T | G | 2.059 | 0.03945 | +? | 0 | 0 | 0 | 1 |
| rs1783937 | 11 | 118609736 | A | G | 2.059 | 0.03946 | ++ | -129.2 | 1.309 | 1 | 0.2526 |
| rs12285468 | 11 | 121018488 | C | T | 2.059 | 0.0395 | +? | 0 | 0 | 0 | 1 |
| rs111519461 | 11 | 117004891 | A | T | -2.058 | 0.03961 | -? | 0 | 0 | 0 | 1 |
| rs2269397 | 11 | 117074299 | C | T | -2.058 | 0.03961 | -? | 0 | 0 | 0 | 1 |
| rs7948971 | 11 | 116202083 | G | A | -2.057 | 0.03973 | -? | 0 | 0 | 0 | 1 |
| rs45577233 | 11 | 117090225 | T | A | -2.056 | 0.03977 | -? | 0 | 0 | 0 | 1 |

| | | | | | | | | | | | |
|---|---|---|---|---|---|---|---|---|---|---|---|
| rs76567973 | 11 | 116970761 | G | C | -2.056 | 0.03978 | -? | 0 | 0 | 0 | 1 |
| rs12295025 | 11 | 116454374 | G | A | -2.056 | 0.03979 | -? | 0 | 0 | 0 | 1 |
| rs475946 | 11 | 118629143 | T | C | 2.055 | 0.03986 | ++ | 0 | 0.938 | 1 | 0.3328 |
| rs11218158 | 11 | 121014228 | G | A | 2.054 | 0.04 | +? | 0 | 0 | 0 | 1 |
| rs877538 | 11 | 121014459 | A | C | 2.054 | 0.04 | +? | 0 | 0 | 0 | 1 |
| rs4586206 | 11 | 116275617 | A | G | -2.054 | 0.04001 | -? | 0 | 0 | 0 | 1 |
| rs7114879 | 11 | 121014860 | T | C | 2.054 | 0.04002 | +? | 0 | 0 | 0 | 1 |
| rs76384483 | 11 | 116806616 | C | T | -2.052 | 0.04016 | -? | 0 | 0 | 0 | 1 |
| rs4938376 | 11 | 117295431 | G | A | 2.052 | 0.04018 | +? | 0 | 0 | 0 | 1 |
| rs4936586 | 11 | 121017863 | A | C | 2.052 | 0.04019 | +? | 0 | 0 | 0 | 1 |
| rs4936587 | 11 | 121018007 | G | A | 2.052 | 0.04019 | +? | 0 | 0 | 0 | 1 |
| rs7120647 | 11 | 121017604 | C | T | 2.052 | 0.04019 | +? | 0 | 0 | 0 | 1 |
| rs607014 | 11 | 121010788 | A | G | 2.051 | 0.04027 | +? | 0 | 0 | 0 | 1 |
| rs2510446 | 11 | 117185634 | C | T | -2.051 | 0.04028 | -? | 0 | 0 | 0 | 1 |
| rs7940750 | 11 | 117653350 | T | C | 2.051 | 0.0403 | +? | 0 | 0 | 0 | 1 |
| rs12285467 | 11 | 121018485 | C | T | 2.05 | 0.04036 | +? | 0 | 0 | 0 | 1 |
| rs144750609 | 11 | 117214617 | C | T | 2.05 | 0.04036 | +? | 0 | 0 | 0 | 1 |
| rs1474828939 | 11 | 116768491 | T | C | -2.05 | 0.04039 | -? | 0 | 0 | 0 | 1 |
| rs1020392 | 11 | 116792299 | A | G | -2.049 | 0.04048 | -? | 0 | 0 | 0 | 1 |
| rs1020393 | 11 | 116792336 | T | C | -2.049 | 0.04048 | -? | 0 | 0 | 0 | 1 |
| rs7130160 | 11 | 116789438 | C | T | -2.049 | 0.04048 | -? | 0 | 0 | 0 | 1 |
| rs78234966 | 11 | 116781699 | C | T | -2.048 | 0.04052 | -? | 0 | 0 | 0 | 1 |
| rs7114915 | 11 | 121014983 | G | A | 2.048 | 0.04054 | +? | 0 | 0 | 0 | 1 |
| rs2510877 | 11 | 120412867 | G | T | 2.048 | 0.04057 | +? | 0 | 0 | 0 | 1 |
| rs7124091 | 11 | 120520418 | G | C | -2.047 | 0.04063 | -? | 0 | 0 | 0 | 1 |
| rs6589574 | 11 | 116730638 | A | G | 2.047 | 0.04066 | +? | 0 | 0 | 0 | 1 |
| rs4936585 | 11 | 121017795 | T | G | 2.047 | 0.04069 | +? | 0 | 0 | 0 | 1 |
| rs2186781 | 11 | 118605447 | T | C | 2.046 | 0.04072 | ++ | -186.5 | 1.047 | 1 | 0.3062 |
| rs494553 | 11 | 117261853 | T | C | 2.046 | 0.04072 | +? | 0 | 0 | 0 | 1 |
| rs606674 | 11 | 121010820 | G | A | 2.046 | 0.04072 | +? | 0 | 0 | 0 | 1 |
| rs620778 | 11 | 121009800 | T | C | 2.046 | 0.04077 | +? | 0 | 0 | 0 | 1 |
| rs2187125 | 11 | 116517968 | G | T | -2.046 | 0.0408 | -? | 0 | 0 | 0 | 1 |
| rs620164 | 11 | 121010116 | A | G | 2.046 | 0.0408 | +? | 0 | 0 | 0 | 1 |
| rs4938473 | 11 | 117927550 | A | G | 2.044 | 0.04096 | +? | 0 | 0 | 0 | 1 |
| rs11606425 | 11 | 117449497 | T | A | -2.044 | 0.04098 | -- | -164.8 | 1.133 | 1 | 0.2872 |
| rs3087611 | 11 | 116645336 | T | A | -2.044 | 0.04099 | -? | 0 | 0 | 0 | 1 |
| rs7120762 | 11 | 116833399 | C | A | -2.044 | 0.041 | -? | 0 | 0 | 0 | 1 |
| rs7127416 | 11 | 116852927 | T | C | -2.044 | 0.041 | -? | 0 | 0 | 0 | 1 |
| rs2513093 | 11 | 116806687 | A | G | 2.043 | 0.04104 | +? | 0 | 0 | 0 | 1 |
| rs76636922 | 11 | 116682995 | A | G | -2.043 | 0.04106 | -- | -123.8 | 1.34 | 1 | 0.247 |
| rs11216270 | 11 | 116953644 | A | C | -2.042 | 0.04114 | -? | 0 | 0 | 0 | 1 |
| rs117043085 | 11 | 117329630 | T | C | 2.042 | 0.04116 | +? | 0 | 0 | 0 | 1 |
| rs4245168 | 11 | 116882638 | C | A | 2.042 | 0.0412 | +? | 0 | 0 | 0 | 1 |
| rs3017608 | 11 | 117189216 | A | G | 2.041 | 0.04128 | +? | 0 | 0 | 0 | 1 |
| rs115903909 | 11 | 121009384 | A | G | 2.04 | 0.04136 | +? | 0 | 0 | 0 | 1 |
| rs7108594 | 11 | 121018451 | G | A | 2.04 | 0.04138 | +? | 0 | 0 | 0 | 1 |
| rs579274 | 11 | 117288442 | C | T | 2.039 | 0.04141 | +? | 0 | 0 | 0 | 1 |
| rs7126214 | 11 | 116679711 | A | G | -2.038 | 0.04157 | -- | -97.8 | 1.517 | 1 | 0.2181 |
| rs6589578 | 11 | 116780918 | T | C | -2.038 | 0.04159 | -? | 0 | 0 | 0 | 1 |
| rs7106782 | 11 | 116782582 | G | A | -2.038 | 0.04159 | -? | 0 | 0 | 0 | 1 |
| rs1321012057 | 11 | 117711092 | A | G | -2.037 | 0.04161 | -- | 0 | 0.034 | 1 | 0.8548 |
| rs506366 | 11 | 117264256 | A | G | 2.036 | 0.04172 | +? | 0 | 0 | 0 | 1 |
| rs59283948 | 11 | 116835363 | A | G | -2.036 | 0.04174 | -? | 0 | 0 | 0 | 1 |
| rs563531 | 11 | 117414030 | A | G | -2.036 | 0.04177 | -? | 0 | 0 | 0 | 1 |
| rs10750105 | 11 | 117296160 | A | T | 2.036 | 0.04178 | +? | 0 | 0 | 0 | 1 |
| rs4935762 | 11 | 121051868 | A | G | 2.035 | 0.04181 | +? | 0 | 0 | 0 | 1 |

| | | | | | | | | | | | |
|---|---|---|---|---|---|---|---|---|---|---|---|
| rs4936591 | 11 | 121050766 | G | A | 2.035 | 0.04181 | +? | 0 | 0 | 0 | 1 |
| rs7121104 | 11 | 120520282 | T | G | -2.035 | 0.04189 | -? | 0 | 0 | 0 | 1 |
| rs559301 | 11 | 121011488 | A | G | 2.034 | 0.04199 | +? | 0 | 0 | 0 | 1 |
| rs7105356 | 11 | 121018237 | G | A | 2.033 | 0.04202 | +? | 0 | 0 | 0 | 1 |
| rs11604377 | 11 | 117449263 | C | T | -2.033 | 0.04204 | -- | -167.9 | 1.12 | 1 | 0.2899 |
| rs11216576 | 11 | 117709760 | A | G | -2.032 | 0.04211 | -- | 0 | 0.008 | 1 | 0.93 |
| rs1790187 | 11 | 118644213 | A | G | 2.032 | 0.04213 | ++ | 0 | 0.918 | 1 | 0.338 |
| rs493720 | 11 | 118604678 | T | C | 2.032 | 0.0422 | ++ | 0 | 0.924 | 1 | 0.3364 |
| rs526850 | 11 | 118605127 | G | T | 2.032 | 0.0422 | ++ | 0 | 0.924 | 1 | 0.3364 |
| rs2282565 | 11 | 121001039 | C | T | 2.03 | 0.04232 | +? | 0 | 0 | 0 | 1 |
| rs7108166 | 11 | 121001263 | G | A | 2.03 | 0.04232 | +? | 0 | 0 | 0 | 1 |
| rs11826847 | 11 | 121024461 | A | G | 2.03 | 0.04237 | +? | 0 | 0 | 0 | 1 |
| rs11216261 | 11 | 116940210 | C | T | -2.03 | 0.04239 | -? | 0 | 0 | 0 | 1 |
| rs79955870 | 11 | 116937361 | T | C | -2.03 | 0.04239 | -? | 0 | 0 | 0 | 1 |
| rs1279744828 | 11 | 117285145 | G | A | 2.03 | 0.0424 | +? | 0 | 0 | 0 | 1 |
| rs78359532 | 11 | 116653185 | C | G | -2.029 | 0.04248 | -? | 0 | 0 | 0 | 1 |
| rs56057635 | 11 | 117349401 | G | A | -2.029 | 0.0425 | -? | 0 | 0 | 0 | 1 |
| rs2282586 | 11 | 120858410 | A | G | -2.028 | 0.04252 | -? | 0 | 0 | 0 | 1 |
| rs74727906 | 11 | 120858301 | A | G | 2.028 | 0.04259 | +? | 0 | 0 | 0 | 1 |
| rs77585745 | 11 | 116872876 | C | T | -2.027 | 0.04268 | -? | 0 | 0 | 0 | 1 |
| rs7108741 | 11 | 121018507 | T | C | 2.027 | 0.0427 | +? | 0 | 0 | 0 | 1 |
| rs4420302 | 11 | 117294706 | A | G | 2.027 | 0.04271 | +? | 0 | 0 | 0 | 1 |
| rs2508653 | 11 | 120405437 | T | C | 2.026 | 0.04275 | +? | 0 | 0 | 0 | 1 |
| rs502514 | 11 | 118241995 | C | T | 2.026 | 0.04276 | +? | 0 | 0 | 0 | 1 |
| rs9919599 | 11 | 116745484 | C | G | 2.026 | 0.04278 | +? | 0 | 0 | 0 | 1 |
| rs12281358 | 11 | 116956990 | C | T | -2.025 | 0.04284 | -? | 0 | 0 | 0 | 1 |
| rs2368149 | 11 | 116955934 | A | G | -2.025 | 0.04286 | -? | 0 | 0 | 0 | 1 |
| rs541623 | 11 | 121009627 | A | G | 2.025 | 0.04288 | +? | 0 | 0 | 0 | 1 |
| rs7927736 | 11 | 120502086 | C | T | 2.025 | 0.04289 | +? | 0 | 0 | 0 | 1 |
| rs12279811 | 11 | 121059326 | T | C | 2.024 | 0.04294 | +? | 0 | 0 | 0 | 1 |
| rs620331 | 11 | 118931099 | T | C | -2.023 | 0.04304 | -? | 0 | 0 | 0 | 1 |
| rs511676 | 11 | 116824884 | T | G | 2.022 | 0.04316 | +? | 0 | 0 | 0 | 1 |
| rs4936582 | 11 | 121015763 | C | T | 2.022 | 0.04317 | +? | 0 | 0 | 0 | 1 |
| rs607999 | 11 | 121010550 | G | A | 2.021 | 0.04326 | +? | 0 | 0 | 0 | 1 |
| rs117339323 | 11 | 116548356 | C | T | 2.021 | 0.04329 | +? | 0 | 0 | 0 | 1 |
| rs2266100 | 11 | 120414586 | A | G | 2.02 | 0.04335 | +? | 0 | 0 | 0 | 1 |
| rs485570 | 11 | 117276783 | G | T | 2.02 | 0.04338 | +? | 0 | 0 | 0 | 1 |
| rs7123630 | 11 | 118995325 | T | C | 2.02 | 0.04338 | +? | 0 | 0 | 0 | 1 |
| rs79257193 | 11 | 116901175 | G | A | -2.019 | 0.04351 | -? | 0 | 0 | 0 | 1 |
| rs519000 | 11 | 116511522 | T | C | 2.017 | 0.0437 | +? | 0 | 0 | 0 | 1 |
| rs138975985 | 11 | 116890488 | T | C | -2.017 | 0.04372 | -? | 0 | 0 | 0 | 1 |
| rs7110042 | 11 | 116883716 | T | C | -2.017 | 0.04372 | -? | 0 | 0 | 0 | 1 |
| rs508487 | 11 | 117075566 | T | C | 2.016 | 0.04375 | +? | 0 | 0 | 0 | 1 |
| rs78210954 | 11 | 119008125 | C | G | 2.016 | 0.04375 | +? | 0 | 0 | 0 | 1 |
| rs4598681 | 11 | 120723872 | T | G | -2.016 | 0.04376 | -? | 0 | 0 | 0 | 1 |
| rs10082561 | 11 | 116951584 | C | A | -2.016 | 0.0438 | -? | 0 | 0 | 0 | 1 |
| rs11216265 | 11 | 116947180 | C | T | -2.016 | 0.0438 | -? | 0 | 0 | 0 | 1 |
| rs12274725 | 11 | 116946853 | T | C | -2.016 | 0.0438 | -? | 0 | 0 | 0 | 1 |
| rs12277910 | 11 | 116958533 | A | C | -2.016 | 0.0438 | -? | 0 | 0 | 0 | 1 |
| rs567988 | 11 | 117430841 | A | G | 2.016 | 0.0438 | +? | 0 | 0 | 0 | 1 |
| rs513182 | 11 | 121010427 | C | T | 2.016 | 0.04384 | +? | 0 | 0 | 0 | 1 |
| rs117978531 | 11 | 117803083 | G | T | 2.015 | 0.0439 | +? | 0 | 0 | 0 | 1 |
| rs1902624 | 11 | 117291696 | T | C | 2.015 | 0.04391 | +? | 0 | 0 | 0 | 1 |
| rs78774571 | 11 | 116280349 | T | C | -2.014 | 0.04398 | -? | 0 | 0 | 0 | 1 |
| rs112118203 | 11 | 117440076 | C | T | -2.013 | 0.0441 | -? | 0 | 0 | 0 | 1 |
| rs12271169 | 11 | 116980021 | G | C | -2.012 | 0.0442 | -? | 0 | 0 | 0 | 1 |

| | | | | | | | | | | | |
|---|---|---|---|---|---|---|---|---|---|---|---|
| rs1943656 | 11 | 121063130 | T | C | 2.012 | 0.0442 | +? | 0 | 0 | 0 | 1 |
| rs485901 | 11 | 118622245 | G | A | 2.011 | 0.04428 | ++ | -192.9 | 1.024 | 1 | 0.3115 |
| rs6589825 | 11 | 120509726 | C | T | -2.011 | 0.04432 | -? | 0 | 0 | 0 | 1 |
| rs17120293 | 11 | 116944846 | C | T | -2.011 | 0.04434 | -? | 0 | 0 | 0 | 1 |
| rs2513095 | 11 | 116806269 | T | C | 2.011 | 0.04435 | +? | 0 | 0 | 0 | 1 |
| rs80197460 | 11 | 116988790 | G | C | -2.01 | 0.04442 | -? | 0 | 0 | 0 | 1 |
| rs78135964 | 11 | 116971677 | T | G | -2.01 | 0.04445 | -? | 0 | 0 | 0 | 1 |
| rs11216863 | 11 | 118257081 | A | G | -2.01 | 0.04447 | -? | 0 | 0 | 0 | 1 |
| rs593252 | 11 | 119880519 | A | G | 2.01 | 0.04447 | +? | 0 | 0 | 0 | 1 |
| rs3802879 | 11 | 117414714 | T | G | -2.009 | 0.04458 | -? | 0 | 0 | 0 | 1 |
| rs4436552 | 11 | 121018771 | T | A | 2.009 | 0.04459 | +? | 0 | 0 | 0 | 1 |
| rs4539330 | 11 | 121018772 | C | G | 2.009 | 0.04459 | +? | 0 | 0 | 0 | 1 |
| rs10790183 | 11 | 117293516 | A | G | 2.008 | 0.04461 | +? | 0 | 0 | 0 | 1 |
| rs521910 | 11 | 118610629 | A | G | 2.008 | 0.04463 | ++ | -116.5 | 1.386 | 1 | 0.2391 |
| rs114034580 | 11 | 116811878 | G | A | -2.008 | 0.04464 | -? | 0 | 0 | 0 | 1 |
| rs76338210 | 11 | 116415864 | T | C | -2.008 | 0.04464 | -? | 0 | 0 | 0 | 1 |
| rs565083 | 11 | 117421984 | A | G | -2.008 | 0.04466 | -? | 0 | 0 | 0 | 1 |
| rs55765491 | 11 | 120404835 | G | A | 2.006 | 0.04488 | +? | 0 | 0 | 0 | 1 |
| rs1961138 | 11 | 118619423 | C | T | 2.005 | 0.04492 | ++ | -190.9 | 1.031 | 1 | 0.3098 |
| rs4499012 | 11 | 121053084 | A | G | 2.004 | 0.0451 | +? | 0 | 0 | 0 | 1 |
| rs117117171 | 11 | 118152252 | A | T | 2.003 | 0.04521 | ++ | 52.7 | 6.345 | 1 | 0.01177 |
| rs626562 | 11 | 117995697 | A | G | -2.003 | 0.04521 | -? | 0 | 0 | 0 | 1 |
| rs150492485 | 11 | 120529603 | A | G | -2.002 | 0.04532 | -? | 0 | 0 | 0 | 1 |
| rs593245 | 11 | 117183650 | T | C | 2.001 | 0.04535 | +? | 0 | 0 | 0 | 1 |
| rs2266101 | 11 | 120414838 | T | C | 2.001 | 0.04537 | +? | 0 | 0 | 0 | 1 |
| rs77138800 | 11 | 117459687 | T | C | -2.001 | 0.04537 | -? | 0 | 0 | 0 | 1 |
| rs7925881 | 11 | 117295791 | A | G | 2.001 | 0.0454 | +? | 0 | 0 | 0 | 1 |
| rs148233183 | 11 | 116972227 | A | G | -2.001 | 0.04543 | -? | 0 | 0 | 0 | 1 |
| rs684669 | 11 | 117524455 | A | C | 2 | 0.04545 | +? | 0 | 0 | 0 | 1 |
| rs640120 | 11 | 118618837 | A | C | 2 | 0.04552 | ++ | -189 | 1.038 | 1 | 0.3083 |
| rs111549567 | 11 | 119467150 | C | T | -2 | 0.04553 | -? | 0 | 0 | 0 | 1 |
| rs6589687 | 11 | 118615660 | T | A | 1.999 | 0.04562 | ++ | -188.7 | 1.039 | 1 | 0.308 |
| rs576266 | 11 | 116488133 | C | T | 1.999 | 0.04565 | +? | 0 | 0 | 0 | 1 |
| rs2298723 | 11 | 120853380 | C | A | -1.999 | 0.04566 | -? | 0 | 0 | 0 | 1 |
| rs951839 | 11 | 116409788 | A | C | -1.998 | 0.04572 | -? | 0 | 0 | 0 | 1 |
| rs74736885 | 11 | 120998225 | G | A | 1.998 | 0.04573 | +? | 0 | 0 | 0 | 1 |
| rs12418036 | 11 | 116947306 | A | G | -1.997 | 0.0458 | -? | 0 | 0 | 0 | 1 |
| rs11215942 | 11 | 116325337 | A | G | 1.996 | 0.04588 | +? | 0 | 0 | 0 | 1 |
| rs7931770 | 11 | 116866290 | C | T | 1.996 | 0.04596 | +? | 0 | 0 | 0 | 1 |
| rs7108587 | 11 | 121018391 | T | C | 1.995 | 0.04602 | +? | 0 | 0 | 0 | 1 |
| rs10892313 | 11 | 118809329 | A | G | -1.995 | 0.04603 | -- | 0 | 0.104 | 1 | 0.7472 |
| rs143138921 | 11 | 116909657 | G | T | -1.995 | 0.04606 | -? | 0 | 0 | 0 | 1 |
| rs74603899 | 11 | 116911423 | T | G | -1.995 | 0.04606 | -? | 0 | 0 | 0 | 1 |
| rs77960525 | 11 | 116912494 | A | G | -1.995 | 0.04606 | -? | 0 | 0 | 0 | 1 |
| rs73575424 | 11 | 118788517 | A | C | 1.994 | 0.04614 | +- | 27.6 | 4.142 | 1 | 0.04183 |
| rs2510878 | 11 | 120414850 | C | A | 1.994 | 0.04615 | +? | 0 | 0 | 0 | 1 |
| rs588918 | 11 | 116856642 | T | C | 1.993 | 0.04631 | +? | 0 | 0 | 0 | 1 |
| rs149576335 | 11 | 116975136 | A | G | -1.992 | 0.04639 | -? | 0 | 0 | 0 | 1 |
| rs569845 | 11 | 117268256 | T | G | 1.992 | 0.04639 | +? | 0 | 0 | 0 | 1 |
| rs76942203 | 11 | 116973247 | A | G | -1.992 | 0.04639 | -? | 0 | 0 | 0 | 1 |
| rs2105611 | 11 | 116488610 | A | G | 1.991 | 0.04646 | +? | 0 | 0 | 0 | 1 |
| rs3017773 | 11 | 116488591 | G | T | 1.991 | 0.04646 | +? | 0 | 0 | 0 | 1 |
| rs529618 | 11 | 116488978 | A | G | 1.991 | 0.04646 | +? | 0 | 0 | 0 | 1 |
| rs4323878 | 11 | 120849062 | C | G | -1.991 | 0.04649 | -? | 0 | 0 | 0 | 1 |
| rs2010611 | 11 | 116583103 | A | G | 1.991 | 0.04651 | +? | 0 | 0 | 0 | 1 |
| rs546617 | 11 | 117275815 | A | G | 1.991 | 0.04652 | +? | 0 | 0 | 0 | 1 |

| | | | | | | | | | | | |
|---|---|---|---|---|---|---|---|---|---|---|---|
| rs78641307 | 11 | 120530241 | C | A | -1.99 | 0.0466 | -? | 0 | 0 | 0 | 1 |
| rs524392 | 11 | 121066195 | A | T | 1.99 | 0.04663 | +? | 0 | 0 | 0 | 1 |
| rs640147 | 11 | 117422642 | T | C | -1.987 | 0.04691 | -? | 0 | 0 | 0 | 1 |
| rs75428021 | 11 | 116731302 | T | C | -1.986 | 0.047 | -? | 0 | 0 | 0 | 1 |
| rs61900565 | 11 | 117926624 | G | C | 1.986 | 0.04707 | +? | 0 | 0 | 0 | 1 |
| rs113213242 | 11 | 116758216 | T | C | -1.985 | 0.0471 | -? | 0 | 0 | 0 | 1 |
| rs117457463 | 11 | 116760474 | A | G | -1.985 | 0.0471 | -? | 0 | 0 | 0 | 1 |
| rs17134988 | 11 | 116758454 | A | G | -1.985 | 0.0471 | -? | 0 | 0 | 0 | 1 |
| rs568984 | 11 | 117273094 | T | G | 1.984 | 0.04727 | +? | 0 | 0 | 0 | 1 |
| rs618354 | 11 | 116700777 | C | G | -1.982 | 0.04746 | -? | 0 | 0 | 0 | 1 |
| rs535774 | 11 | 118468037 | T | C | 1.982 | 0.04748 | ++ | 0 | 0.367 | 1 | 0.5447 |
| rs73592285 | 11 | 116821415 | C | T | -1.982 | 0.04751 | -? | 0 | 0 | 0 | 1 |
| rs7111299 | 11 | 117415385 | A | C | -1.98 | 0.04765 | -? | 0 | 0 | 0 | 1 |
| rs10892631 | 11 | 120734139 | A | G | -1.98 | 0.0477 | -? | 0 | 0 | 0 | 1 |
| rs3863308 | 11 | 117293741 | G | T | 1.979 | 0.04778 | +? | 0 | 0 | 0 | 1 |
| rs925151 | 11 | 116369171 | A | G | 1.979 | 0.04782 | +? | 0 | 0 | 0 | 1 |
| rs11215928 | 11 | 116300734 | T | C | -1.979 | 0.04787 | -? | 0 | 0 | 0 | 1 |
| rs141555847 | 11 | 116767722 | A | G | -1.978 | 0.04788 | -? | 0 | 0 | 0 | 1 |
| rs76402668 | 11 | 116771204 | C | T | -1.978 | 0.04788 | -? | 0 | 0 | 0 | 1 |
| rs146523230 | 11 | 116778138 | A | G | -1.978 | 0.04792 | -? | 0 | 0 | 0 | 1 |
| rs7114963 | 11 | 116875983 | T | C | 1.978 | 0.04793 | +? | 0 | 0 | 0 | 1 |
| rs76542665 | 11 | 116587953 | A | C | 1.977 | 0.04802 | +? | 0 | 0 | 0 | 1 |
| rs74700879 | 11 | 116783142 | G | C | -1.977 | 0.04803 | -? | 0 | 0 | 0 | 1 |
| rs9971421 | 11 | 116941540 | T | C | -1.977 | 0.04808 | -? | 0 | 0 | 0 | 1 |
| rs487261 | 11 | 117205861 | T | C | 1.975 | 0.04825 | +? | 0 | 0 | 0 | 1 |
| rs2226637 | 11 | 116269990 | C | G | -1.974 | 0.04833 | -? | 0 | 0 | 0 | 1 |
| rs4516002 | 11 | 116999059 | A | G | 1.974 | 0.04833 | +? | 0 | 0 | 0 | 1 |
| rs7121347 | 11 | 116867209 | G | T | 1.974 | 0.04837 | +? | 0 | 0 | 0 | 1 |
| rs56204159 | 11 | 117925581 | T | C | 1.974 | 0.04839 | +? | 0 | 0 | 0 | 1 |
| rs78013608 | 11 | 116752045 | A | G | -1.974 | 0.04842 | -? | 0 | 0 | 0 | 1 |
| rs115255270 | 11 | 116819050 | C | T | -1.973 | 0.04847 | -? | 0 | 0 | 0 | 1 |
| rs117791268 | 11 | 116839094 | A | G | -1.973 | 0.04847 | -? | 0 | 0 | 0 | 1 |
| rs118137922 | 11 | 116842758 | A | G | -1.973 | 0.04847 | -? | 0 | 0 | 0 | 1 |
| rs12283494 | 11 | 116815717 | A | G | -1.973 | 0.04847 | -? | 0 | 0 | 0 | 1 |
| rs2289891 | 11 | 116824938 | C | A | -1.973 | 0.04847 | -? | 0 | 0 | 0 | 1 |
| rs57040912 | 11 | 116815306 | G | A | -1.973 | 0.04847 | -? | 0 | 0 | 0 | 1 |
| rs6589581 | 11 | 116828859 | T | A | -1.973 | 0.04847 | -? | 0 | 0 | 0 | 1 |
| rs7110113 | 11 | 116856152 | C | A | -1.973 | 0.04847 | -? | 0 | 0 | 0 | 1 |
| rs7112772 | 11 | 116852714 | C | T | -1.973 | 0.04847 | -? | 0 | 0 | 0 | 1 |
| rs7112773 | 11 | 116852716 | C | T | -1.973 | 0.04847 | -? | 0 | 0 | 0 | 1 |
| rs7130221 | 11 | 116828573 | A | G | -1.973 | 0.04847 | -? | 0 | 0 | 0 | 1 |
| rs74557653 | 11 | 116806367 | T | G | -1.973 | 0.04847 | -? | 0 | 0 | 0 | 1 |
| rs76223783 | 11 | 116842174 | C | T | -1.973 | 0.04847 | -? | 0 | 0 | 0 | 1 |
| rs77029349 | 11 | 116830628 | T | G | -1.973 | 0.04847 | -? | 0 | 0 | 0 | 1 |
| rs77161357 | 11 | 116826409 | C | T | -1.973 | 0.04847 | -? | 0 | 0 | 0 | 1 |
| rs77203966 | 11 | 116801299 | T | C | -1.973 | 0.04847 | -? | 0 | 0 | 0 | 1 |
| rs78241753 | 11 | 116849544 | C | G | -1.973 | 0.04847 | -? | 0 | 0 | 0 | 1 |
| rs7937585 | 11 | 116844649 | A | G | -1.973 | 0.04847 | -? | 0 | 0 | 0 | 1 |
| rs7938466 | 11 | 116829129 | C | T | -1.973 | 0.04847 | -? | 0 | 0 | 0 | 1 |
| rs7940002 | 11 | 116850485 | A | C | -1.973 | 0.04847 | -? | 0 | 0 | 0 | 1 |
| rs7940300 | 11 | 116820673 | T | A | -1.973 | 0.04847 | -? | 0 | 0 | 0 | 1 |
| rs79868224 | 11 | 116849038 | A | G | -1.973 | 0.04847 | -? | 0 | 0 | 0 | 1 |
| rs144646355 | 11 | 117052107 | A | G | -1.972 | 0.04856 | -? | 0 | 0 | 0 | 1 |
| rs61729990 | 11 | 117052532 | A | C | -1.972 | 0.04856 | -? | 0 | 0 | 0 | 1 |
| rs1786176 | 11 | 117905635 | A | G | -1.972 | 0.04863 | -? | 0 | 0 | 0 | 1 |
| rs74496689 | 11 | 116621623 | A | G | -1.972 | 0.04865 | -? | 0 | 0 | 0 | 1 |

| | | | | | | | | | | | |
|---|---|---|---|---|---|---|---|---|---|---|---|
| rs4938472 | 11 | 117926172 | A | T | 1.971 | 0.04867 | +? | 0 | 0 | 0 | 1 |
| rs73008657 | 11 | 119720735 | A | G | -1.971 | 0.04867 | -? | 0 | 0 | 0 | 1 |
| rs77535577 | 11 | 116685391 | A | G | -1.971 | 0.04868 | -- | -190.8 | 1.032 | 1 | 0.3098 |
| rs964256 | 11 | 121192650 | G | C | -1.971 | 0.04877 | -? | 0 | 0 | 0 | 1 |
| rs11217117 | 11 | 118865992 | T | C | -1.97 | 0.04879 | -? | 0 | 0 | 0 | 1 |
| rs4938589 | 11 | 118810405 | T | C | -1.97 | 0.04883 | -- | 0 | 0.212 | 1 | 0.6454 |
| rs479591 | 11 | 117429034 | G | A | 1.97 | 0.04884 | +? | 0 | 0 | 0 | 1 |
| rs3863310 | 11 | 117293929 | C | T | 1.97 | 0.04886 | +? | 0 | 0 | 0 | 1 |
| rs4935760 | 11 | 121007228 | T | A | 1.969 | 0.04891 | +? | 0 | 0 | 0 | 1 |
| rs542322 | 11 | 117421824 | A | G | -1.969 | 0.04898 | -? | 0 | 0 | 0 | 1 |
| rs4936393 | 11 | 117503394 | G | A | 1.968 | 0.04902 | +? | 0 | 0 | 0 | 1 |
| rs3889526 | 11 | 118856491 | A | C | -1.967 | 0.04914 | -? | 0 | 0 | 0 | 1 |
| rs4938610 | 11 | 118857466 | C | T | -1.967 | 0.04914 | -? | 0 | 0 | 0 | 1 |
| rs607273 | 11 | 116862818 | A | T | 1.967 | 0.04922 | +? | 0 | 0 | 0 | 1 |
| rs7112111 | 11 | 116980858 | G | T | -1.966 | 0.04927 | -? | 0 | 0 | 0 | 1 |
| rs75188049 | 11 | 116780567 | A | G | -1.966 | 0.04928 | -? | 0 | 0 | 0 | 1 |
| rs77693143 | 11 | 116780702 | G | A | -1.966 | 0.04928 | -? | 0 | 0 | 0 | 1 |
| rs10892281 | 11 | 118614397 | T | C | 1.965 | 0.04936 | ++ | -178.2 | 1.078 | 1 | 0.2991 |
| rs687100 | 11 | 117536090 | T | C | 1.964 | 0.04953 | +? | 0 | 0 | 0 | 1 |
| rs76959909 | 11 | 117444949 | A | C | -1.963 | 0.04964 | -? | 0 | 0 | 0 | 1 |
| rs116861530 | 11 | 116962661 | A | G | -1.962 | 0.04973 | -? | 0 | 0 | 0 | 1 |
| rs34899605 | 11 | 116483042 | A | G | 1.962 | 0.04975 | +? | 0 | 0 | 0 | 1 |
| rs692840 | 11 | 117509536 | A | G | 1.962 | 0.04978 | +? | 0 | 0 | 0 | 1 |
| rs10892068 | 11 | 116928546 | T | C | -1.96 | 0.05 | -? | 0 | 0 | 0 | 1 |
| rs7938234 | 11 | 116903095 | T | C | -1.959 | 0.05006 | -? | 0 | 0 | 0 | 1 |
| rs112974653 | 11 | 118807637 | T | G | -1.959 | 0.05016 | -- | 0 | 0.172 | 1 | 0.6779 |
| rs75807933 | 11 | 117444661 | A | T | -1.957 | 0.0503 | -? | 0 | 0 | 0 | 1 |
| rs624555 | 11 | 117202782 | T | C | 1.955 | 0.0506 | +? | 0 | 0 | 0 | 1 |
| rs80350987 | 11 | 119705743 | A | C | -1.954 | 0.05065 | -? | 0 | 0 | 0 | 1 |
| rs4936521 | 11 | 120459320 | G | A | -1.954 | 0.05068 | -? | 0 | 0 | 0 | 1 |
| rs557053 | 11 | 117408077 | T | C | -1.953 | 0.05077 | -? | 0 | 0 | 0 | 1 |
| rs79841374 | 11 | 116684560 | T | C | -1.953 | 0.05086 | -- | -192.7 | 1.025 | 1 | 0.3113 |
| rs6589846 | 11 | 120700147 | G | T | -1.952 | 0.05093 | -? | 0 | 0 | 0 | 1 |
| rs17491510 | 11 | 116409433 | T | C | -1.952 | 0.05098 | -? | 0 | 0 | 0 | 1 |
| rs7130952 | 11 | 121016838 | T | C | 1.951 | 0.05104 | +? | 0 | 0 | 0 | 1 |
| rs142395187 | 11 | 116976009 | G | A | -1.95 | 0.05119 | -? | 0 | 0 | 0 | 1 |
| rs17119920 | 11 | 116587889 | G | C | 1.95 | 0.05121 | +? | 0 | 0 | 0 | 1 |
| rs4938617 | 11 | 118867695 | T | C | -1.947 | 0.05156 | -- | 0 | 0.252 | 1 | 0.6158 |
| rs7114746 | 11 | 116928748 | G | T | -1.947 | 0.05158 | -? | 0 | 0 | 0 | 1 |
| rs7125464 | 11 | 116928199 | A | G | -1.947 | 0.05158 | -? | 0 | 0 | 0 | 1 |
| rs77256870 | 11 | 116891077 | G | C | -1.947 | 0.05158 | -? | 0 | 0 | 0 | 1 |
| rs7942637 | 11 | 116927101 | A | G | -1.947 | 0.05158 | -? | 0 | 0 | 0 | 1 |
| rs10892067 | 11 | 116927762 | C | G | -1.946 | 0.05162 | -? | 0 | 0 | 0 | 1 |
| rs11216577 | 11 | 117710150 | T | C | -1.946 | 0.05168 | -- | 0 | 0.009 | 1 | 0.9255 |
| rs3863311 | 11 | 117294035 | A | T | 1.945 | 0.05179 | +? | 0 | 0 | 0 | 1 |
| rs12363888 | 11 | 117713187 | T | C | -1.944 | 0.05196 | -- | 0 | 0.026 | 1 | 0.872 |
| rs74420345 | 11 | 116963277 | A | C | -1.943 | 0.05196 | -? | 0 | 0 | 0 | 1 |
| rs112119822 | 11 | 118798824 | G | A | -1.943 | 0.05199 | -- | -179.2 | 1.074 | 1 | 0.2999 |
| rs10736495 | 11 | 118607090 | C | G | 1.943 | 0.05204 | ++ | -139 | 1.255 | 1 | 0.2626 |
| rs2305827 | 11 | 117246876 | C | T | -1.943 | 0.05207 | -? | 0 | 0 | 0 | 1 |
| rs609353 | 11 | 117420356 | T | C | -1.942 | 0.05211 | -? | 0 | 0 | 0 | 1 |
| rs4938606 | 11 | 118852081 | G | A | -1.942 | 0.05218 | -? | 0 | 0 | 0 | 1 |
| rs75749368 | 11 | 120528115 | T | C | -1.941 | 0.0522 | -? | 0 | 0 | 0 | 1 |
| rs4427574 | 11 | 121006650 | A | G | 1.938 | 0.05258 | +? | 0 | 0 | 0 | 1 |
| rs4936579 | 11 | 121006387 | C | T | 1.938 | 0.05258 | +? | 0 | 0 | 0 | 1 |
| rs7947688 | 11 | 116207020 | G | A | -1.938 | 0.0526 | -? | 0 | 0 | 0 | 1 |

| | | | | | | | | | | | |
|---|---|---|---|---|---|---|---|---|---|---|---|
| rs11602493 | 11 | 120456533 | A | G | -1.938 | 0.05261 | -? | 0 | 0 | 0 | 1 |
| rs11216575 | 11 | 117709688 | G | A | -1.937 | 0.05269 | -- | 0 | 0.006 | 1 | 0.9394 |
| rs4548653 | 11 | 116903316 | A | G | -1.937 | 0.05274 | -? | 0 | 0 | 0 | 1 |
| rs112143902 | 11 | 118619948 | T | C | 1.936 | 0.05281 | ++ | 0 | 0.286 | 1 | 0.5927 |
| rs73011154 | 11 | 116448277 | C | A | -1.936 | 0.05288 | -? | 0 | 0 | 0 | 1 |
| rs10892312 | 11 | 118807740 | A | G | -1.936 | 0.05292 | -- | 0 | 0.133 | 1 | 0.7154 |
| rs79157249 | 11 | 118785982 | T | A | 1.936 | 0.05293 | +- | 21.4 | 3.819 | 1 | 0.05067 |
| rs56337383 | 11 | 120967158 | A | G | -1.934 | 0.05309 | -? | 0 | 0 | 0 | 1 |
| rs17123519 | 11 | 119853138 | T | C | 1.932 | 0.0533 | +? | 0 | 0 | 0 | 1 |
| rs7926623 | 11 | 119677955 | A | G | 1.931 | 0.05348 | +? | 0 | 0 | 0 | 1 |
| rs56751865 | 11 | 116936871 | T | C | -1.93 | 0.05359 | -? | 0 | 0 | 0 | 1 |
| rs4938614 | 11 | 118863232 | A | G | -1.93 | 0.0536 | -? | 0 | 0 | 0 | 1 |
| rs11607048 | 11 | 121152465 | G | T | -1.929 | 0.0537 | -? | 0 | 0 | 0 | 1 |
| rs3758992 | 11 | 117945171 | C | G | 1.929 | 0.0537 | +? | 0 | 0 | 0 | 1 |
| rs17492275 | 11 | 116437423 | T | C | -1.927 | 0.05394 | -? | 0 | 0 | 0 | 1 |
| rs10750171 | 11 | 120176979 | G | T | 1.927 | 0.054 | +? | 0 | 0 | 0 | 1 |
| rs78399857 | 11 | 120032251 | T | C | 1.927 | 0.05402 | +? | 0 | 0 | 0 | 1 |
| rs576390 | 11 | 117424005 | A | G | -1.926 | 0.05414 | -? | 0 | 0 | 0 | 1 |
| rs10892589 | 11 | 120509335 | T | C | -1.925 | 0.05418 | -? | 0 | 0 | 0 | 1 |
| rs539573 | 11 | 117269269 | A | T | 1.924 | 0.0544 | +? | 0 | 0 | 0 | 1 |
| rs186593252 | 11 | 116913908 | C | A | -1.923 | 0.05453 | -? | 0 | 0 | 0 | 1 |
| rs60101932 | 11 | 116916723 | A | G | -1.923 | 0.05453 | -? | 0 | 0 | 0 | 1 |
| rs74429128 | 11 | 116916984 | A | G | -1.923 | 0.05453 | -? | 0 | 0 | 0 | 1 |
| rs1902626 | 11 | 117291835 | C | A | 1.922 | 0.05455 | +? | 0 | 0 | 0 | 1 |
| rs573648 | 11 | 117424306 | A | G | -1.922 | 0.05458 | -? | 0 | 0 | 0 | 1 |
| rs11218029 | 11 | 120718086 | A | T | -1.922 | 0.05464 | -? | 0 | 0 | 0 | 1 |
| rs1939934 | 11 | 121066946 | T | C | 1.921 | 0.0547 | +? | 0 | 0 | 0 | 1 |
| rs139462136 | 11 | 116900628 | T | C | -1.921 | 0.05477 | -? | 0 | 0 | 0 | 1 |
| rs4938612 | 11 | 118863037 | C | T | -1.921 | 0.05479 | -? | 0 | 0 | 0 | 1 |
| rs4936519 | 11 | 120415158 | T | G | 1.919 | 0.05493 | +? | 0 | 0 | 0 | 1 |
| rs67120644 | 11 | 116625897 | C | T | 1.919 | 0.05504 | +? | 0 | 0 | 0 | 1 |
| rs12289097 | 11 | 116426640 | G | A | -1.918 | 0.05508 | -? | 0 | 0 | 0 | 1 |
| rs675479 | 11 | 117913430 | G | A | -1.918 | 0.05508 | -? | 0 | 0 | 0 | 1 |
| rs480120 | 11 | 117255386 | T | C | 1.916 | 0.05538 | +? | 0 | 0 | 0 | 1 |
| rs545274 | 11 | 116730833 | T | C | -1.916 | 0.05542 | -? | 0 | 0 | 0 | 1 |
| rs77892802 | 11 | 116404477 | C | T | -1.915 | 0.05548 | -? | 0 | 0 | 0 | 1 |
| rs507262 | 11 | 116491621 | G | A | 1.915 | 0.05551 | +? | 0 | 0 | 0 | 1 |
| rs534633 | 11 | 116492312 | G | A | 1.915 | 0.05554 | +? | 0 | 0 | 0 | 1 |
| rs59491594 | 11 | 116757720 | C | T | -1.913 | 0.05572 | -? | 0 | 0 | 0 | 1 |
| rs74701178 | 11 | 116763124 | C | T | -1.913 | 0.05572 | -? | 0 | 0 | 0 | 1 |
| rs759988389 | 11 | 116448341 | C | T | -1.913 | 0.05572 | -? | 0 | 0 | 0 | 1 |
| rs77718307 | 11 | 116766149 | A | G | -1.913 | 0.05572 | -? | 0 | 0 | 0 | 1 |
| rs149318234 | 11 | 116718554 | G | C | -1.913 | 0.05578 | -? | 0 | 0 | 0 | 1 |
| rs1574433 | 11 | 117347947 | T | C | -1.913 | 0.05579 | -? | 0 | 0 | 0 | 1 |
| rs562179 | 11 | 116805204 | T | C | 1.912 | 0.05584 | +? | 0 | 0 | 0 | 1 |
| rs7117408 | 11 | 118808140 | T | C | -1.912 | 0.05588 | -- | 0 | 0.123 | 1 | 0.7255 |
| rs558056 | 11 | 116530797 | T | G | 1.911 | 0.056 | +? | 0 | 0 | 0 | 1 |
| rs678327 | 11 | 117521603 | A | G | 1.91 | 0.05617 | +? | 0 | 0 | 0 | 1 |
| rs10892532 | 11 | 119951570 | G | A | 1.91 | 0.05618 | +? | 0 | 0 | 0 | 1 |
| rs11216542 | 11 | 117650725 | T | C | 1.909 | 0.0562 | +? | 0 | 0 | 0 | 1 |
| rs112422338 | 11 | 116920984 | A | G | -1.909 | 0.05622 | -? | 0 | 0 | 0 | 1 |
| rs7110034 | 11 | 118858911 | A | C | -1.909 | 0.05625 | -? | 0 | 0 | 0 | 1 |
| rs7119702 | 11 | 118607694 | G | A | 1.909 | 0.05625 | ++ | -171.6 | 1.104 | 1 | 0.2933 |
| rs4290240 | 11 | 118853488 | G | A | -1.909 | 0.05632 | -? | 0 | 0 | 0 | 1 |
| rs642215 | 11 | 117988082 | T | C | -1.909 | 0.05632 | -? | 0 | 0 | 0 | 1 |
| rs7101680 | 11 | 118799206 | A | G | -1.908 | 0.05634 | -- | -179.6 | 1.073 | 1 | 0.3003 |

| | | | | | | | | | | | |
|---|---|---|---|---|---|---|---|---|---|---|---|
| rs502816 | 11 | 116491177 | A | C | 1.907 | 0.0565 | +? | 0 | 0 | 0 | 1 |
| rs505397 | 11 | 116491428 | G | T | 1.907 | 0.0565 | +? | 0 | 0 | 0 | 1 |
| rs1452979329 | 11 | 117466540 | A | G | -1.907 | 0.05658 | -? | 0 | 0 | 0 | 1 |
| rs586763 | 11 | 118816304 | T | C | -1.906 | 0.05661 | -- | 0 | 0.348 | 1 | 0.5554 |
| rs4936534 | 11 | 120503923 | T | A | 1.904 | 0.05695 | +? | 0 | 0 | 0 | 1 |
| rs77256135 | 11 | 119813599 | A | G | -1.904 | 0.05696 | -? | 0 | 0 | 0 | 1 |
| rs112084975 | 11 | 119813015 | T | C | -1.903 | 0.057 | -? | 0 | 0 | 0 | 1 |
| rs12282154 | 11 | 118807876 | A | G | -1.903 | 0.05701 | -- | 0 | 0.12 | 1 | 0.7293 |
| rs3901285 | 11 | 120733435 | T | C | -1.903 | 0.05703 | -? | 0 | 0 | 0 | 1 |
| rs1228001593 | 11 | 116861398 | A | G | -1.902 | 0.05712 | -? | 0 | 0 | 0 | 1 |
| rs143491990 | 11 | 116859076 | T | C | -1.902 | 0.05712 | -? | 0 | 0 | 0 | 1 |
| rs59425942 | 11 | 116870159 | T | C | -1.902 | 0.05712 | -? | 0 | 0 | 0 | 1 |
| rs7122484 | 11 | 116862105 | C | G | -1.902 | 0.05712 | -? | 0 | 0 | 0 | 1 |
| rs76391131 | 11 | 116858320 | C | T | -1.902 | 0.05712 | -? | 0 | 0 | 0 | 1 |
| rs7947525 | 11 | 116862779 | T | C | -1.902 | 0.05712 | -? | 0 | 0 | 0 | 1 |
| rs11217372 | 11 | 119519217 | G | C | 1.901 | 0.05733 | +? | 0 | 0 | 0 | 1 |
| rs17311310 | 11 | 120461835 | A | G | -1.9 | 0.0574 | -? | 0 | 0 | 0 | 1 |
| rs12808437 | 11 | 116782000 | T | G | -1.899 | 0.05751 | -? | 0 | 0 | 0 | 1 |
| rs76673375 | 11 | 117794988 | C | G | 1.899 | 0.05752 | +? | 0 | 0 | 0 | 1 |
| rs12277788 | 11 | 117349482 | T | C | -1.899 | 0.05756 | -? | 0 | 0 | 0 | 1 |
| rs55741146 | 11 | 117430840 | G | C | -1.898 | 0.05772 | -? | 0 | 0 | 0 | 1 |
| rs17311317 | 11 | 120462417 | A | G | -1.898 | 0.05773 | -? | 0 | 0 | 0 | 1 |
| rs72999502 | 11 | 118556995 | C | T | -1.897 | 0.05779 | -- | 0 | 0.195 | 1 | 0.6589 |
| rs7118591 | 11 | 116908457 | A | C | -1.896 | 0.0579 | -? | 0 | 0 | 0 | 1 |
| rs112736310 | 11 | 119812958 | T | C | -1.896 | 0.05798 | -? | 0 | 0 | 0 | 1 |
| rs58354636 | 11 | 116879162 | G | A | -1.896 | 0.05799 | -? | 0 | 0 | 0 | 1 |
| rs79849088 | 11 | 116742626 | T | C | -1.896 | 0.05802 | -? | 0 | 0 | 0 | 1 |
| rs11216413 | 11 | 117344165 | A | T | -1.895 | 0.05803 | -? | 0 | 0 | 0 | 1 |
| rs2852227 | 11 | 120589179 | C | A | 1.895 | 0.05803 | +? | 0 | 0 | 0 | 1 |
| rs906541 | 11 | 116440521 | A | G | -1.894 | 0.05825 | -? | 0 | 0 | 0 | 1 |
| rs60834024 | 11 | 116985857 | A | C | -1.894 | 0.05826 | -? | 0 | 0 | 0 | 1 |
| rs78733282 | 11 | 121079940 | G | A | 1.893 | 0.05838 | +? | 0 | 0 | 0 | 1 |
| rs7928478 | 11 | 117372281 | A | G | -1.892 | 0.05843 | -? | 0 | 0 | 0 | 1 |
| rs3181 | 11 | 116907043 | A | G | -1.892 | 0.05852 | -? | 0 | 0 | 0 | 1 |
| rs4938327 | 11 | 116906728 | C | T | -1.892 | 0.05852 | -? | 0 | 0 | 0 | 1 |
| rs12280501 | 11 | 118801440 | T | C | -1.891 | 0.05857 | -- | 0 | 0.992 | 1 | 0.3193 |
| rs76666535 | 11 | 116891412 | A | G | -1.891 | 0.05857 | -? | 0 | 0 | 0 | 1 |
| rs643014 | 11 | 117988223 | G | A | -1.891 | 0.05863 | -? | 0 | 0 | 0 | 1 |
| rs7119185 | 11 | 116886096 | T | C | -1.891 | 0.05868 | -? | 0 | 0 | 0 | 1 |
| rs12269853 | 11 | 119811734 | G | C | -1.89 | 0.05875 | -? | 0 | 0 | 0 | 1 |
| rs73008653 | 11 | 119715337 | T | C | -1.89 | 0.0588 | -? | 0 | 0 | 0 | 1 |
| rs117833973 | 11 | 116730980 | T | C | -1.89 | 0.05882 | -? | 0 | 0 | 0 | 1 |
| rs117910440 | 11 | 116727547 | C | T | -1.89 | 0.05882 | -? | 0 | 0 | 0 | 1 |
| rs75385236 | 11 | 116728260 | T | C | -1.89 | 0.05882 | -? | 0 | 0 | 0 | 1 |
| rs75771311 | 11 | 116724233 | A | G | -1.89 | 0.05882 | -? | 0 | 0 | 0 | 1 |
| rs77462380 | 11 | 116723837 | T | C | -1.89 | 0.05882 | -? | 0 | 0 | 0 | 1 |
| rs78816728 | 11 | 116727871 | C | T | -1.89 | 0.05882 | -? | 0 | 0 | 0 | 1 |
| rs548638 | 11 | 116737093 | G | T | -1.889 | 0.05892 | -? | 0 | 0 | 0 | 1 |
| rs10790347 | 11 | 119679613 | T | A | 1.889 | 0.05895 | ++ | 0 | 0.414 | 1 | 0.5197 |
| rs74884652 | 11 | 118876552 | C | T | -1.888 | 0.05901 | -- | 0 | 0.331 | 1 | 0.5653 |
| rs6589677 | 11 | 118583971 | A | G | 1.888 | 0.05908 | ++ | -94.7 | 1.541 | 1 | 0.2144 |
| rs1784108 | 11 | 116505583 | T | C | 1.886 | 0.05933 | +? | 0 | 0 | 0 | 1 |
| rs621188 | 11 | 117529622 | C | T | 1.886 | 0.05936 | +? | 0 | 0 | 0 | 1 |
| rs77230637 | 11 | 117464630 | A | G | -1.885 | 0.05942 | -? | 0 | 0 | 0 | 1 |
| rs75488969 | 11 | 118712509 | T | C | 1.885 | 0.05943 | ++ | 0 | 0.167 | 1 | 0.6825 |
| rs645258 | 11 | 116801201 | A | C | 1.884 | 0.05959 | +? | 0 | 0 | 0 | 1 |

| | | | | | | | | | | | |
|---|---|---|---|---|---|---|---|---|---|---|---|
| rs4938763 | 11 | 119934934 | T | C | 1.884 | 0.0596 | +? | 0 | 0 | 0 | 1 |
| rs75212830 | 11 | 116424533 | A | G | -1.882 | 0.05986 | -? | 0 | 0 | 0 | 1 |
| rs12420361 | 11 | 116993788 | A | G | -1.881 | 0.05996 | -? | 0 | 0 | 0 | 1 |
| rs6589576 | 11 | 116735788 | C | G | 1.881 | 0.05996 | +? | 0 | 0 | 0 | 1 |
| rs149462130 | 11 | 116726181 | T | C | -1.881 | 0.06001 | -? | 0 | 0 | 0 | 1 |
| rs663166 | 11 | 117179302 | A | G | 1.879 | 0.06025 | +? | 0 | 0 | 0 | 1 |
| rs74891547 | 11 | 118811878 | C | G | -1.877 | 0.06046 | -- | 0 | 0.707 | 1 | 0.4003 |
| rs7928581 | 11 | 117271499 | C | T | 1.877 | 0.06047 | +? | 0 | 0 | 0 | 1 |
| rs2186620 | 11 | 120801185 | C | T | 1.877 | 0.06051 | +? | 0 | 0 | 0 | 1 |
| rs2509005 | 11 | 117410428 | A | G | -1.877 | 0.06054 | -? | 0 | 0 | 0 | 1 |
| rs2018774 | 11 | 119881267 | T | A | 1.876 | 0.06066 | +? | 0 | 0 | 0 | 1 |
| rs146418192 | 11 | 116761815 | C | G | -1.876 | 0.06072 | -? | 0 | 0 | 0 | 1 |
| rs10892154 | 11 | 117535724 | A | C | 1.875 | 0.06082 | +? | 0 | 0 | 0 | 1 |
| rs9943635 | 11 | 121066079 | T | C | 1.873 | 0.06104 | +? | 0 | 0 | 0 | 1 |
| rs1893176 | 11 | 119214292 | G | A | -1.873 | 0.06109 | -? | 0 | 0 | 0 | 1 |
| rs117772632 | 11 | 116813938 | A | T | -1.872 | 0.06125 | -? | 0 | 0 | 0 | 1 |
| rs147683288 | 11 | 116808658 | T | C | -1.872 | 0.06125 | -? | 0 | 0 | 0 | 1 |
| rs58123174 | 11 | 116853151 | A | C | -1.872 | 0.06125 | -? | 0 | 0 | 0 | 1 |
| rs61301656 | 11 | 116815265 | A | G | -1.872 | 0.06125 | -? | 0 | 0 | 0 | 1 |
| rs74655990 | 11 | 116843747 | C | T | -1.872 | 0.06125 | -? | 0 | 0 | 0 | 1 |
| rs74922395 | 11 | 116831357 | C | T | -1.872 | 0.06125 | -? | 0 | 0 | 0 | 1 |
| rs75176087 | 11 | 116798503 | T | C | -1.872 | 0.06125 | -? | 0 | 0 | 0 | 1 |
| rs75216082 | 11 | 116873188 | C | T | -1.872 | 0.06125 | -? | 0 | 0 | 0 | 1 |
| rs75296049 | 11 | 116838219 | C | T | -1.872 | 0.06125 | -? | 0 | 0 | 0 | 1 |
| rs76919167 | 11 | 116872657 | C | A | -1.872 | 0.06125 | -? | 0 | 0 | 0 | 1 |
| rs77322057 | 11 | 116873956 | C | T | -1.872 | 0.06125 | -? | 0 | 0 | 0 | 1 |
| rs78672214 | 11 | 116839420 | G | A | -1.872 | 0.06125 | -? | 0 | 0 | 0 | 1 |
| rs78903015 | 11 | 116876136 | T | G | -1.872 | 0.06125 | -? | 0 | 0 | 0 | 1 |
| rs79143833 | 11 | 116823800 | C | G | -1.872 | 0.06125 | -? | 0 | 0 | 0 | 1 |
| rs79523757 | 11 | 116871192 | C | T | -1.872 | 0.06125 | -? | 0 | 0 | 0 | 1 |
| rs11218160 | 11 | 121015228 | A | G | 1.871 | 0.06129 | +? | 0 | 0 | 0 | 1 |
| rs661631 | 11 | 117408179 | T | C | -1.871 | 0.06131 | -? | 0 | 0 | 0 | 1 |
| rs73008415 | 11 | 117368358 | T | C | 1.871 | 0.06132 | +? | 0 | 0 | 0 | 1 |
| rs4936528 | 11 | 120474008 | G | C | -1.871 | 0.06141 | -? | 0 | 0 | 0 | 1 |
| rs77101138 | 11 | 120528055 | G | A | -1.87 | 0.06143 | -? | 0 | 0 | 0 | 1 |
| rs495118 | 11 | 117434728 | G | A | 1.87 | 0.06147 | +? | 0 | 0 | 0 | 1 |
| rs17245685 | 11 | 120614648 | C | T | 1.869 | 0.06165 | +? | 0 | 0 | 0 | 1 |
| rs4500499 | 11 | 120445835 | A | G | -1.868 | 0.0618 | -? | 0 | 0 | 0 | 1 |
| rs11216206 | 11 | 116843425 | G | C | -1.868 | 0.06181 | -- | 0 | 0.227 | 1 | 0.6338 |
| rs147224199 | 11 | 116398570 | T | C | -1.866 | 0.06204 | -? | 0 | 0 | 0 | 1 |
| rs582285 | 11 | 117362948 | G | A | 1.865 | 0.06216 | +? | 0 | 0 | 0 | 1 |
| rs11603180 | 11 | 120457909 | A | G | -1.864 | 0.06229 | -? | 0 | 0 | 0 | 1 |
| rs1807049 | 11 | 117385531 | G | A | -1.864 | 0.06232 | -? | 0 | 0 | 0 | 1 |
| rs2509006 | 11 | 117410507 | T | C | -1.864 | 0.06234 | -? | 0 | 0 | 0 | 1 |
| rs78927965 | 11 | 119811767 | A | C | -1.863 | 0.06251 | -? | 0 | 0 | 0 | 1 |
| rs17311324 | 11 | 120462677 | T | G | -1.862 | 0.06256 | -? | 0 | 0 | 0 | 1 |
| rs75352872 | 11 | 120425668 | T | G | -1.862 | 0.06258 | -? | 0 | 0 | 0 | 1 |
| rs721783 | 11 | 116907449 | T | C | -1.862 | 0.0626 | -? | 0 | 0 | 0 | 1 |
| rs516495 | 11 | 121073219 | C | T | 1.86 | 0.06286 | +? | 0 | 0 | 0 | 1 |
| rs1241657 | 11 | 116839761 | T | C | 1.86 | 0.06291 | +? | 0 | 0 | 0 | 1 |
| rs510250 | 11 | 117430058 | G | T | 1.859 | 0.06296 | +? | 0 | 0 | 0 | 1 |
| rs530053 | 11 | 116530251 | G | A | 1.858 | 0.06314 | +? | 0 | 0 | 0 | 1 |
| rs61900590 | 11 | 117929672 | T | C | 1.858 | 0.06322 | +? | 0 | 0 | 0 | 1 |
| rs518547 | 11 | 116738046 | G | A | -1.858 | 0.06324 | -? | 0 | 0 | 0 | 1 |
| rs61900900 | 11 | 120454144 | T | C | -1.857 | 0.06325 | -? | 0 | 0 | 0 | 1 |
| rs11217091 | 11 | 118793613 | C | T | -1.856 | 0.06343 | -- | 0 | 0.482 | 1 | 0.4874 |

| | | | | | | | | | | | |
|---|---|---|---|---|---|---|---|---|---|---|---|
| rs74955808 | 11 | 116992685 | A | G | -1.855 | 0.06355 | -? | 0 | 0 | 0 | 1 |
| rs573905 | 11 | 118572267 | A | G | 1.855 | 0.06356 | ++ | 0 | 0.043 | 1 | 0.8355 |
| rs11218127 | 11 | 120925525 | G | A | -1.855 | 0.06358 | -? | 0 | 0 | 0 | 1 |
| rs201644901 | 11 | 121071633 | C | A | 1.855 | 0.06365 | +? | 0 | 0 | 0 | 1 |
| rs1573789 | 11 | 120463065 | T | C | -1.854 | 0.06372 | -? | 0 | 0 | 0 | 1 |
| rs12416941 | 11 | 116979363 | A | G | -1.854 | 0.06378 | -? | 0 | 0 | 0 | 1 |
| rs12419459 | 11 | 116996711 | T | C | -1.853 | 0.06393 | -? | 0 | 0 | 0 | 1 |
| rs10750108 | 11 | 117492181 | A | G | 1.851 | 0.06412 | +? | 0 | 0 | 0 | 1 |
| rs671553 | 11 | 117528376 | T | C | 1.851 | 0.06418 | +? | 0 | 0 | 0 | 1 |
| rs7945067 | 11 | 119678445 | T | C | 1.85 | 0.06427 | +? | 0 | 0 | 0 | 1 |
| rs10892137 | 11 | 117430479 | T | C | 1.849 | 0.06444 | +? | 0 | 0 | 0 | 1 |
| rs4512872 | 11 | 120733629 | C | A | -1.848 | 0.06463 | -? | 0 | 0 | 0 | 1 |
| rs10790169 | 11 | 116877159 | A | C | 1.847 | 0.06471 | +? | 0 | 0 | 0 | 1 |
| rs537592 | 11 | 117517424 | C | T | 1.847 | 0.06474 | +? | 0 | 0 | 0 | 1 |
| rs151215655 | 11 | 116534026 | T | C | -1.847 | 0.06476 | -? | 0 | 0 | 0 | 1 |
| rs614634 | 11 | 121076938 | T | C | 1.847 | 0.0648 | +? | 0 | 0 | 0 | 1 |
| rs74882651 | 11 | 116985191 | A | G | -1.846 | 0.06489 | -? | 0 | 0 | 0 | 1 |
| rs1784045 | 11 | 117191734 | C | T | -1.846 | 0.0649 | -? | 0 | 0 | 0 | 1 |
| rs4938294 | 11 | 116515420 | T | C | -1.843 | 0.0654 | -? | 0 | 0 | 0 | 1 |
| rs61134913 | 11 | 116298294 | G | A | -1.842 | 0.06542 | -? | 0 | 0 | 0 | 1 |
| rs12285074 | 11 | 116762028 | A | G | 1.842 | 0.06545 | +? | 0 | 0 | 0 | 1 |
| rs45478298 | 11 | 118503912 | T | C | 1.841 | 0.06563 | +- | 21.8 | 3.834 | 1 | 0.05022 |
| rs619998 | 11 | 121090162 | C | T | 1.84 | 0.06571 | +? | 0 | 0 | 0 | 1 |
| rs75967483 | 11 | 119556519 | T | C | -1.84 | 0.06572 | -? | 0 | 0 | 0 | 1 |
| rs73008405 | 11 | 117366280 | C | G | 1.84 | 0.06573 | +? | 0 | 0 | 0 | 1 |
| rs4935758 | 11 | 120862089 | T | C | -1.84 | 0.06576 | -? | 0 | 0 | 0 | 1 |
| rs4938616 | 11 | 118865465 | T | C | -1.84 | 0.06577 | -? | 0 | 0 | 0 | 1 |
| rs1944512 | 11 | 119879786 | T | G | -1.84 | 0.0658 | -? | 0 | 0 | 0 | 1 |
| rs656189 | 11 | 121073887 | A | T | 1.84 | 0.06584 | +? | 0 | 0 | 0 | 1 |
| rs55928509 | 11 | 116547924 | A | C | 1.838 | 0.06605 | +? | 0 | 0 | 0 | 1 |
| rs111677878 | 11 | 116982434 | T | C | -1.838 | 0.06607 | -? | 0 | 0 | 0 | 1 |
| rs11217941 | 11 | 120531656 | G | T | -1.838 | 0.0661 | -? | 0 | 0 | 0 | 1 |
| rs79733808 | 11 | 116237696 | A | G | -1.838 | 0.0661 | -? | 0 | 0 | 0 | 1 |
| rs4936555 | 11 | 120699899 | T | C | -1.837 | 0.06617 | -? | 0 | 0 | 0 | 1 |
| rs112791662 | 11 | 119576357 | A | G | 1.837 | 0.0662 | +? | 0 | 0 | 0 | 1 |
| rs10790214 | 11 | 117707191 | T | C | 1.836 | 0.06631 | +? | 0 | 0 | 0 | 1 |
| rs75012772 | 11 | 119807902 | A | G | -1.836 | 0.06632 | -? | 0 | 0 | 0 | 1 |
| rs1075784 | 11 | 117369450 | G | C | 1.836 | 0.06641 | +? | 0 | 0 | 0 | 1 |
| rs1940626 | 11 | 116828121 | C | T | -1.835 | 0.06644 | -? | 0 | 0 | 0 | 1 |
| rs114797458 | 11 | 117561256 | A | G | 1.835 | 0.06656 | +? | 0 | 0 | 0 | 1 |
| rs1443827410 | 11 | 117616982 | G | A | -1.832 | 0.0669 | -? | 0 | 0 | 0 | 1 |
| rs10892721 | 11 | 121189231 | T | G | -1.832 | 0.06692 | -? | 0 | 0 | 0 | 1 |
| rs1261791 | 11 | 117200638 | A | T | 1.832 | 0.06696 | +? | 0 | 0 | 0 | 1 |
| rs2508656 | 11 | 120424740 | A | G | 1.832 | 0.067 | +? | 0 | 0 | 0 | 1 |
| rs111902338 | 11 | 119814337 | A | C | -1.832 | 0.06702 | -? | 0 | 0 | 0 | 1 |
| rs75224487 | 11 | 119813733 | T | C | -1.832 | 0.06702 | -? | 0 | 0 | 0 | 1 |
| rs76661054 | 11 | 119813686 | C | T | -1.832 | 0.06702 | -? | 0 | 0 | 0 | 1 |
| rs75154936 | 11 | 118877891 | C | T | -1.831 | 0.06705 | -- | 0 | 0.261 | 1 | 0.6095 |
| rs116850963 | 11 | 120239808 | G | A | 1.83 | 0.06718 | +? | 0 | 0 | 0 | 1 |
| rs61900473 | 11 | 120011880 | A | G | 1.83 | 0.06721 | +? | 0 | 0 | 0 | 1 |
| rs75501351 | 11 | 116731744 | G | T | -1.83 | 0.06725 | -? | 0 | 0 | 0 | 1 |
| rs6589587 | 11 | 116875338 | G | T | -1.829 | 0.06733 | -? | 0 | 0 | 0 | 1 |
| rs563990 | 11 | 116285538 | G | A | 1.829 | 0.06736 | +? | 0 | 0 | 0 | 1 |
| rs114221654 | 11 | 116980679 | T | C | -1.828 | 0.0675 | -? | 0 | 0 | 0 | 1 |
| rs116770907 | 11 | 116981515 | T | C | -1.828 | 0.0675 | -? | 0 | 0 | 0 | 1 |
| rs1944495 | 11 | 119879579 | T | C | -1.828 | 0.06758 | -? | 0 | 0 | 0 | 1 |

| | | | | | | | | | | |
|---|---|---|---|---|---|---|---|---|---|---|
| rs11604129 | 11 | 120607334 | G | A | 1.828 | 0.0676 | +? | 0 | 0 | 0 | 1 |
| rs672251 | 11 | 117505239 | G | T | 1.827 | 0.06767 | +? | 0 | 0 | 0 | 1 |
| rs77763933 | 11 | 119808101 | C | G | -1.827 | 0.06769 | -? | 0 | 0 | 0 | 1 |
| rs79710878 | 11 | 119807543 | G | A | -1.827 | 0.06769 | -? | 0 | 0 | 0 | 1 |
| rs4344499 | 11 | 120734419 | G | T | -1.826 | 0.06782 | -? | 0 | 0 | 0 | 1 |
| rs533735 | 11 | 116241842 | C | A | -1.825 | 0.06793 | -- | 0 | 0.158 | 1 | 0.6906 |
| rs10892173 | 11 | 117672561 | T | C | 1.825 | 0.06797 | +? | 0 | 0 | 0 | 1 |
| rs2509007 | 11 | 117410700 | A | G | -1.825 | 0.06804 | -? | 0 | 0 | 0 | 1 |
| rs4620758 | 11 | 116945718 | A | G | -1.825 | 0.06807 | -? | 0 | 0 | 0 | 1 |
| rs7101763 | 11 | 116941985 | T | G | -1.825 | 0.06807 | -? | 0 | 0 | 0 | 1 |
| rs7119911 | 11 | 117653955 | T | G | 1.824 | 0.06809 | +? | 0 | 0 | 0 | 1 |
| rs7130213 | 11 | 116946652 | A | C | -1.824 | 0.0681 | -? | 0 | 0 | 0 | 1 |
| rs12416691 | 11 | 116996783 | T | A | -1.824 | 0.06814 | -? | 0 | 0 | 0 | 1 |
| rs12419454 | 11 | 116996631 | T | C | -1.824 | 0.06814 | -? | 0 | 0 | 0 | 1 |
| rs11216164 | 11 | 116734545 | A | G | 1.823 | 0.06833 | +? | 0 | 0 | 0 | 1 |
| rs17120344 | 11 | 116996345 | A | G | -1.822 | 0.06842 | -? | 0 | 0 | 0 | 1 |
| rs11217373 | 11 | 119519916 | G | C | 1.821 | 0.06855 | +? | 0 | 0 | 0 | 1 |
| rs2006623 | 11 | 119808535 | A | G | -1.821 | 0.06855 | -? | 0 | 0 | 0 | 1 |
| rs4938818 | 11 | 120452999 | A | G | -1.821 | 0.06855 | -? | 0 | 0 | 0 | 1 |
| rs79345551 | 11 | 119936873 | A | G | 1.821 | 0.06858 | +? | 0 | 0 | 0 | 1 |
| rs150818481 | 11 | 116749190 | A | G | -1.82 | 0.0687 | -? | 0 | 0 | 0 | 1 |
| rs12805779 | 11 | 116523063 | T | G | -1.82 | 0.06872 | -? | 0 | 0 | 0 | 1 |
| rs2004926 | 11 | 117368846 | G | A | 1.82 | 0.06874 | +? | 0 | 0 | 0 | 1 |
| rs77443589 | 11 | 119806581 | A | G | -1.82 | 0.06881 | -? | 0 | 0 | 0 | 1 |
| rs79137748 | 11 | 119806699 | C | T | -1.82 | 0.06881 | -? | 0 | 0 | 0 | 1 |
| rs7926586 | 11 | 119675931 | G | T | 1.82 | 0.06881 | +? | 0 | 0 | 0 | 1 |
| rs684973 | 11 | 117451536 | C | G | 1.819 | 0.06888 | +? | 0 | 0 | 0 | 1 |
| rs524914 | 11 | 117207122 | C | G | 1.819 | 0.06892 | +? | 0 | 0 | 0 | 1 |
| rs7929976 | 11 | 117349817 | G | A | -1.818 | 0.06901 | -? | 0 | 0 | 0 | 1 |
| rs7928007 | 11 | 117720781 | C | A | -1.817 | 0.06921 | -- | 0 | 0.059 | 1 | 0.808 |
| rs112704540 | 11 | 116818527 | G | A | -1.816 | 0.06937 | -? | 0 | 0 | 0 | 1 |
| rs2512139 | 11 | 117815564 | A | G | 1.816 | 0.06944 | +? | 0 | 0 | 0 | 1 |
| rs4938285 | 11 | 116440011 | C | T | -1.816 | 0.06944 | -? | 0 | 0 | 0 | 1 |
| rs563229 | 11 | 116288186 | G | C | 1.815 | 0.06948 | +? | 0 | 0 | 0 | 1 |
| rs1871756 | 11 | 116953645 | A | G | -1.815 | 0.06954 | -? | 0 | 0 | 0 | 1 |
| rs73008411 | 11 | 117367574 | G | A | 1.813 | 0.06976 | +? | 0 | 0 | 0 | 1 |
| rs895648 | 11 | 119677072 | C | T | 1.813 | 0.06984 | ++ | 0 | 0.254 | 1 | 0.6139 |
| rs536484 | 11 | 116280965 | C | G | 1.812 | 0.06996 | +? | 0 | 0 | 0 | 1 |
| rs6589824 | 11 | 120509621 | G | A | -1.811 | 0.07008 | -? | 0 | 0 | 0 | 1 |
| rs4938284 | 11 | 116439949 | C | T | -1.811 | 0.07013 | -? | 0 | 0 | 0 | 1 |
| rs7107702 | 11 | 116804144 | C | T | -1.811 | 0.07017 | -? | 0 | 0 | 0 | 1 |
| rs112965917 | 11 | 118725189 | A | T | 1.81 | 0.07023 | ++ | 0 | 0.024 | 1 | 0.8765 |
| rs482795 | 11 | 116227687 | G | A | -1.81 | 0.07028 | -- | 49.4 | 5.93 | 1 | 0.01489 |
| rs3819132 | 11 | 117671489 | A | G | -1.81 | 0.07029 | -? | 0 | 0 | 0 | 1 |
| rs11217090 | 11 | 118789952 | T | C | -1.81 | 0.07034 | -- | 0 | 0.295 | 1 | 0.5869 |
| rs4272793 | 11 | 119668263 | C | A | 1.806 | 0.07095 | ++ | 0 | 0.191 | 1 | 0.6624 |
| rs521187 | 11 | 117416908 | A | G | -1.805 | 0.0711 | -? | 0 | 0 | 0 | 1 |
| rs67613157 | 11 | 116514997 | T | C | -1.804 | 0.07119 | -? | 0 | 0 | 0 | 1 |
| rs1955021 | 11 | 119808052 | A | G | -1.803 | 0.07131 | -? | 0 | 0 | 0 | 1 |
| rs10790192 | 11 | 117420163 | A | G | 1.803 | 0.07134 | +? | 0 | 0 | 0 | 1 |
| rs117399581 | 11 | 117605620 | A | G | -1.802 | 0.07156 | -? | 0 | 0 | 0 | 1 |
| rs4938321 | 11 | 116852507 | T | C | -1.801 | 0.07169 | -? | 0 | 0 | 0 | 1 |
| rs76031864 | 11 | 119984598 | A | G | -1.8 | 0.0718 | -? | 0 | 0 | 0 | 1 |
| rs60171666 | 11 | 116371442 | A | G | 1.798 | 0.07213 | +? | 0 | 0 | 0 | 1 |
| rs10892054 | 11 | 116868221 | T | C | -1.797 | 0.07227 | -? | 0 | 0 | 0 | 1 |
| rs10892055 | 11 | 116868371 | A | G | -1.797 | 0.07227 | -? | 0 | 0 | 0 | 1 |

| | | | | | | | | | | | |
|---|---|---|---|---|---|---|---|---|---|---|---|
| rs7939934 | 11 | 116861268 | T | A | -1.797 | 0.07227 | -? | 0 | 0 | 0 | 1 |
| rs1815788 | 11 | 116834971 | C | T | -1.797 | 0.07239 | -? | 0 | 0 | 0 | 1 |
| rs564892 | 11 | 118602392 | A | T | 1.796 | 0.07242 | ++ | 0 | 0.708 | 1 | 0.4002 |
| rs495171 | 11 | 117338330 | C | T | -1.796 | 0.07246 | -? | 0 | 0 | 0 | 1 |
| rs509058 | 11 | 121077211 | T | C | 1.796 | 0.07248 | +? | 0 | 0 | 0 | 1 |
| rs643758 | 11 | 117531887 | T | A | 1.795 | 0.07271 | +? | 0 | 0 | 0 | 1 |
| rs4936391 | 11 | 117498605 | G | A | 1.793 | 0.07294 | +? | 0 | 0 | 0 | 1 |
| rs2510872 | 11 | 116224496 | A | G | -1.793 | 0.073 | -- | 51.1 | 6.136 | 1 | 0.01324 |
| rs143367870 | 11 | 116866944 | C | A | -1.792 | 0.0732 | -? | 0 | 0 | 0 | 1 |
| rs75292554 | 11 | 116860781 | C | T | -1.792 | 0.0732 | -? | 0 | 0 | 0 | 1 |
| rs576670 | 11 | 116285930 | G | A | -1.791 | 0.07337 | -? | 0 | 0 | 0 | 1 |
| rs557990 | 11 | 118601363 | T | C | 1.79 | 0.07345 | ++ | 0 | 0.689 | 1 | 0.4065 |
| rs12288505 | 11 | 121069225 | T | C | 1.79 | 0.07347 | +? | 0 | 0 | 0 | 1 |
| rs513548 | 11 | 120847817 | C | T | 1.79 | 0.07353 | +? | 0 | 0 | 0 | 1 |
| rs75829086 | 11 | 119820707 | G | C | 1.79 | 0.07353 | +? | 0 | 0 | 0 | 1 |
| rs553188 | 11 | 118576762 | T | C | 1.789 | 0.0736 | ++ | -141.2 | 1.244 | 1 | 0.2648 |
| rs12417881 | 11 | 117531358 | G | A | 1.789 | 0.07364 | +? | 0 | 0 | 0 | 1 |
| rs76665318 | 11 | 116864639 | C | T | -1.789 | 0.07368 | -? | 0 | 0 | 0 | 1 |
| rs497898 | 11 | 116227259 | A | G | -1.788 | 0.07378 | -- | 49.1 | 5.888 | 1 | 0.01524 |
| rs11216288 | 11 | 117004656 | T | C | 1.788 | 0.07379 | +? | 0 | 0 | 0 | 1 |
| rs74778631 | 11 | 117558945 | A | G | 1.788 | 0.07379 | +? | 0 | 0 | 0 | 1 |
| rs11218017 | 11 | 120697099 | G | A | 1.787 | 0.07389 | +? | 0 | 0 | 0 | 1 |
| rs61901099 | 11 | 119556950 | T | C | -1.787 | 0.07397 | -? | 0 | 0 | 0 | 1 |
| rs693701 | 11 | 119804461 | T | C | 1.786 | 0.07416 | +? | 0 | 0 | 0 | 1 |
| rs111345266 | 11 | 118893767 | T | C | -1.783 | 0.07456 | -- | 0 | 0.287 | 1 | 0.592 |
| rs11216545 | 11 | 117659322 | A | G | -1.783 | 0.07457 | -? | 0 | 0 | 0 | 1 |
| rs10790328 | 11 | 119506255 | G | C | -1.783 | 0.07462 | -? | 0 | 0 | 0 | 1 |
| rs11217554 | 11 | 119756941 | C | G | 1.783 | 0.07465 | +? | 0 | 0 | 0 | 1 |
| rs4938315 | 11 | 116731205 | G | A | 1.782 | 0.07478 | +? | 0 | 0 | 0 | 1 |
| rs5103 | 11 | 116692552 | G | A | -1.782 | 0.0748 | -? | 0 | 0 | 0 | 1 |
| rs638496 | 11 | 121011507 | T | C | 1.78 | 0.07506 | +? | 0 | 0 | 0 | 1 |
| rs10790356 | 11 | 119807402 | T | A | -1.78 | 0.07509 | -? | 0 | 0 | 0 | 1 |
| rs17119814 | 11 | 116424420 | G | C | -1.78 | 0.0751 | -? | 0 | 0 | 0 | 1 |
| rs45619137 | 11 | 117162284 | A | G | -1.778 | 0.07537 | -? | 0 | 0 | 0 | 1 |
| rs77018736 | 11 | 117795671 | A | G | 1.778 | 0.0754 | +? | 0 | 0 | 0 | 1 |
| rs35659242 | 11 | 120275234 | G | T | 1.778 | 0.07548 | +? | 0 | 0 | 0 | 1 |
| rs634044 | 11 | 116752558 | C | T | -1.777 | 0.07557 | -? | 0 | 0 | 0 | 1 |
| rs73008650 | 11 | 119713633 | A | G | -1.777 | 0.07558 | -? | 0 | 0 | 0 | 1 |
| rs77916839 | 11 | 118798780 | C | G | -1.777 | 0.0756 | -- | 0 | 0.792 | 1 | 0.3734 |
| rs675246 | 11 | 119880819 | T | C | 1.776 | 0.07576 | +? | 0 | 0 | 0 | 1 |
| rs4936426 | 11 | 118134597 | C | T | -1.776 | 0.07579 | -? | 0 | 0 | 0 | 1 |
| rs4938724 | 11 | 119672156 | A | G | 1.772 | 0.07643 | ++ | 0 | 0.259 | 1 | 0.6106 |
| rs663003 | 11 | 118823148 | C | T | 1.772 | 0.07643 | ++ | 0 | 0.007 | 1 | 0.9338 |
| rs498912 | 11 | 118576997 | C | T | 1.771 | 0.07649 | ++ | -143.5 | 1.232 | 1 | 0.267 |
| rs7933007 | 11 | 118730669 | C | T | 1.771 | 0.07664 | ++ | 0 | 0.399 | 1 | 0.5275 |
| rs78813250 | 11 | 120557393 | T | G | -1.77 | 0.0767 | -? | 0 | 0 | 0 | 1 |
| rs536761 | 11 | 117517366 | T | C | 1.77 | 0.07675 | +? | 0 | 0 | 0 | 1 |
| rs880004 | 11 | 119808956 | G | A | -1.77 | 0.07676 | -? | 0 | 0 | 0 | 1 |
| rs80310120 | 11 | 116357627 | T | C | 1.77 | 0.07679 | +? | 0 | 0 | 0 | 1 |
| rs1815789 | 11 | 116834953 | C | T | -1.769 | 0.07684 | -? | 0 | 0 | 0 | 1 |
| rs7941927 | 11 | 116838904 | G | A | -1.769 | 0.07684 | -? | 0 | 0 | 0 | 1 |
| rs895650 | 11 | 119806917 | T | C | -1.769 | 0.07685 | -? | 0 | 0 | 0 | 1 |
| rs7128664 | 11 | 121186914 | G | C | -1.769 | 0.07686 | -? | 0 | 0 | 0 | 1 |
| rs73578509 | 11 | 116481694 | G | C | -1.768 | 0.07712 | -? | 0 | 0 | 0 | 1 |
| rs141250514 | 11 | 116999025 | C | T | -1.767 | 0.07721 | -? | 0 | 0 | 0 | 1 |
| rs145802214 | 11 | 116998912 | C | A | -1.767 | 0.07721 | -? | 0 | 0 | 0 | 1 |

| | | | | | | | | | | | |
|---|---|---|---|---|---|---|---|---|---|---|---|
| rs4430518 | 11 | 120723433 | G | T | -1.767 | 0.07725 | -? | 0 | 0 | 0 | 1 |
| rs61900791 | 11 | 120237971 | T | C | -1.767 | 0.07731 | -? | 0 | 0 | 0 | 1 |
| rs4245037 | 11 | 120470291 | A | T | -1.766 | 0.07742 | -? | 0 | 0 | 0 | 1 |
| rs2852225 | 11 | 120588562 | T | G | 1.766 | 0.07744 | +? | 0 | 0 | 0 | 1 |
| rs1035237 | 11 | 116727850 | G | C | 1.766 | 0.07745 | +? | 0 | 0 | 0 | 1 |
| rs656083 | 11 | 117181992 | C | T | -1.766 | 0.07747 | -? | 0 | 0 | 0 | 1 |
| rs626571 | 11 | 117518038 | C | T | 1.765 | 0.07754 | +? | 0 | 0 | 0 | 1 |
| rs12575259 | 11 | 120428959 | A | G | 1.765 | 0.07756 | +? | 0 | 0 | 0 | 1 |
| rs78381558 | 11 | 116424451 | C | G | -1.765 | 0.07758 | -? | 0 | 0 | 0 | 1 |
| rs11605265 | 11 | 118805144 | A | G | -1.765 | 0.07759 | -- | -165.3 | 1.131 | 1 | 0.2876 |
| rs150308863 | 11 | 117165046 | T | C | 1.765 | 0.07759 | +? | 0 | 0 | 0 | 1 |
| rs1944502 | 11 | 119813545 | G | A | -1.764 | 0.07774 | -? | 0 | 0 | 0 | 1 |
| rs3740725 | 11 | 119859583 | A | G | 1.764 | 0.0778 | +? | 0 | 0 | 0 | 1 |
| rs11605658 | 11 | 118805220 | T | C | -1.761 | 0.0783 | -- | -179.4 | 1.074 | 1 | 0.3001 |
| rs11217617 | 11 | 119863505 | T | C | 1.761 | 0.07831 | +? | 0 | 0 | 0 | 1 |
| rs566155 | 11 | 117198638 | C | T | 1.76 | 0.07834 | +? | 0 | 0 | 0 | 1 |
| rs680908 | 11 | 117197621 | G | C | 1.76 | 0.07835 | +? | 0 | 0 | 0 | 1 |
| rs45457998 | 11 | 118492713 | G | A | 1.76 | 0.07836 | ++ | 0 | 0.109 | 1 | 0.7411 |
| rs73023056 | 11 | 116752191 | A | G | -1.76 | 0.07844 | -? | 0 | 0 | 0 | 1 |
| rs2075292 | 11 | 116732512 | G | T | 1.759 | 0.07851 | +? | 0 | 0 | 0 | 1 |
| rs10790196 | 11 | 117496475 | T | C | 1.759 | 0.07854 | +? | 0 | 0 | 0 | 1 |
| rs3863307 | 11 | 116957263 | G | C | -1.759 | 0.07855 | -? | 0 | 0 | 0 | 1 |
| rs592009 | 11 | 117987128 | G | T | -1.758 | 0.07871 | -? | 0 | 0 | 0 | 1 |
| rs74678032 | 11 | 118812559 | C | T | 1.758 | 0.07872 | ++ | 0 | 0.465 | 1 | 0.4953 |
| rs114945429 | 11 | 117004201 | T | G | -1.758 | 0.07877 | -? | 0 | 0 | 0 | 1 |
| rs12417098 | 11 | 117004853 | A | G | -1.758 | 0.07877 | -? | 0 | 0 | 0 | 1 |
| rs12421425 | 11 | 117001960 | T | C | -1.758 | 0.07877 | -? | 0 | 0 | 0 | 1 |
| rs138039720 | 11 | 116999600 | T | C | -1.758 | 0.07877 | -? | 0 | 0 | 0 | 1 |
| rs146067038 | 11 | 117000179 | G | A | -1.758 | 0.07877 | -? | 0 | 0 | 0 | 1 |
| rs17120367 | 11 | 117002828 | T | C | -1.758 | 0.07877 | -? | 0 | 0 | 0 | 1 |
| rs17120370 | 11 | 117003388 | C | T | -1.758 | 0.07877 | -? | 0 | 0 | 0 | 1 |
| rs74735277 | 11 | 117003418 | T | C | -1.758 | 0.07877 | -? | 0 | 0 | 0 | 1 |
| rs75208249 | 11 | 117003623 | C | G | -1.758 | 0.07877 | -? | 0 | 0 | 0 | 1 |
| rs668781 | 11 | 119816512 | C | T | 1.758 | 0.07878 | +? | 0 | 0 | 0 | 1 |
| rs577853 | 11 | 116226263 | T | C | -1.757 | 0.07885 | -- | 47.8 | 5.752 | 1 | 0.01647 |
| rs4936556 | 11 | 120702026 | T | C | -1.757 | 0.07893 | -? | 0 | 0 | 0 | 1 |
| rs7925794 | 11 | 116278612 | T | C | -1.757 | 0.07896 | -? | 0 | 0 | 0 | 1 |
| rs17119760 | 11 | 116358711 | A | C | 1.757 | 0.079 | +? | 0 | 0 | 0 | 1 |
| rs11599940 | 11 | 120443297 | A | G | -1.755 | 0.0792 | -? | 0 | 0 | 0 | 1 |
| rs11602332 | 11 | 117449742 | A | G | -1.755 | 0.0792 | -- | 0 | 0.865 | 1 | 0.3523 |
| rs678468 | 11 | 117413864 | C | T | -1.755 | 0.07929 | -? | 0 | 0 | 0 | 1 |
| rs7103842 | 11 | 116958028 | A | G | -1.754 | 0.0794 | -? | 0 | 0 | 0 | 1 |
| rs520333 | 11 | 117708625 | A | G | 1.753 | 0.07952 | +? | 0 | 0 | 0 | 1 |
| rs17121080 | 11 | 117554152 | A | G | 1.753 | 0.07957 | +? | 0 | 0 | 0 | 1 |
| rs7932598 | 11 | 116197556 | T | G | -1.753 | 0.07964 | -? | 0 | 0 | 0 | 1 |
| rs10892470 | 11 | 119678986 | C | T | 1.753 | 0.07967 | ++ | 0 | 0.3 | 1 | 0.5836 |
| rs80170882 | 11 | 120556274 | T | C | -1.753 | 0.07967 | -? | 0 | 0 | 0 | 1 |
| rs496547 | 11 | 118576463 | A | T | -1.752 | 0.07969 | -- | 0 | 0.001 | 1 | 0.9797 |
| rs570521 | 11 | 119733857 | G | T | -1.751 | 0.07991 | -? | 0 | 0 | 0 | 1 |
| rs555075 | 11 | 117193203 | T | C | 1.751 | 0.07999 | +? | 0 | 0 | 0 | 1 |
| rs506234 | 11 | 118540104 | G | C | 1.748 | 0.08048 | ++ | 0 | 0.18 | 1 | 0.6712 |
| rs1574249 | 11 | 120500130 | A | G | 1.748 | 0.08049 | +? | 0 | 0 | 0 | 1 |
| rs143711539 | 11 | 118804671 | A | G | -1.747 | 0.08069 | -- | -160.2 | 1.153 | 1 | 0.2829 |
| rs675522 | 11 | 117385478 | C | G | 1.746 | 0.08085 | +? | 0 | 0 | 0 | 1 |
| rs1320685 | 11 | 119932930 | C | G | -1.746 | 0.08089 | -? | 0 | 0 | 0 | 1 |
| rs4938697 | 11 | 119507468 | C | T | -1.745 | 0.0809 | -? | 0 | 0 | 0 | 1 |

| | | | | | | | | | | | |
|---|---|---|---|---|---|---|---|---|---|---|---|
| rs640888 | 11 | 117987746 | A | G | -1.744 | 0.08115 | -- | 0 | 0.016 | 1 | 0.8994 |
| rs7936516 | 11 | 120500604 | T | G | 1.743 | 0.08138 | +? | 0 | 0 | 0 | 1 |
| rs12293155 | 11 | 116423307 | G | A | -1.742 | 0.08145 | -? | 0 | 0 | 0 | 1 |
| rs652418 | 11 | 120841689 | C | T | 1.742 | 0.08152 | +? | 0 | 0 | 0 | 1 |
| rs542748 | 11 | 119797863 | C | G | 1.741 | 0.08163 | +? | 0 | 0 | 0 | 1 |
| rs11600378 | 11 | 118803842 | G | A | -1.741 | 0.08176 | -- | -158.5 | 1.16 | 1 | 0.2814 |
| rs594951 | 11 | 121011823 | C | T | 1.739 | 0.08202 | +? | 0 | 0 | 0 | 1 |
| rs4938705 | 11 | 119539324 | A | G | 1.738 | 0.08224 | +? | 0 | 0 | 0 | 1 |
| rs55897720 | 11 | 119937782 | G | A | -1.737 | 0.08231 | -? | 0 | 0 | 0 | 1 |
| rs6589616 | 11 | 117495049 | T | C | 1.737 | 0.08236 | +? | 0 | 0 | 0 | 1 |
| rs503542 | 11 | 118590743 | G | A | 1.737 | 0.08246 | +? | 0 | 0 | 0 | 1 |
| rs4938619 | 11 | 118890085 | T | C | -1.736 | 0.08253 | -- | 0 | 0.317 | 1 | 0.5736 |
| rs4447186 | 11 | 120445422 | G | A | -1.736 | 0.08257 | -? | 0 | 0 | 0 | 1 |
| rs2849167 | 11 | 116684744 | G | A | -1.735 | 0.08274 | -? | 0 | 0 | 0 | 1 |
| rs79985686 | 11 | 120425101 | C | T | 1.735 | 0.08282 | +? | 0 | 0 | 0 | 1 |
| rs2013131 | 11 | 117362198 | C | T | -1.734 | 0.08291 | -? | 0 | 0 | 0 | 1 |
| rs648389 | 11 | 118038697 | C | T | 1.733 | 0.08308 | +? | 0 | 0 | 0 | 1 |
| rs61904803 | 11 | 116429779 | C | T | -1.733 | 0.08312 | -? | 0 | 0 | 0 | 1 |
| rs527619 | 11 | 118563286 | A | G | 1.733 | 0.08318 | ++ | 0 | 0.186 | 1 | 0.6665 |
| rs11216585 | 11 | 117727584 | T | G | -1.732 | 0.08319 | -- | 0 | 0.013 | 1 | 0.9086 |
| rs12718466 | 11 | 116708663 | G | T | -1.731 | 0.0834 | -? | 0 | 0 | 0 | 1 |
| rs4938566 | 11 | 118721189 | T | A | 1.731 | 0.08343 | ++ | 0 | 0.221 | 1 | 0.6383 |
| rs4938567 | 11 | 118721249 | G | A | 1.731 | 0.08343 | ++ | 0 | 0.221 | 1 | 0.6383 |
| rs4938570 | 11 | 118722231 | G | A | 1.731 | 0.08343 | ++ | 0 | 0.221 | 1 | 0.6383 |
| rs7946961 | 11 | 118721850 | G | A | 1.731 | 0.08343 | ++ | 0 | 0.221 | 1 | 0.6383 |
| rs79365963 | 11 | 119730115 | C | T | -1.731 | 0.08351 | -? | 0 | 0 | 0 | 1 |
| rs597820 | 11 | 117911367 | C | A | -1.731 | 0.08353 | -- | 0 | 0.001 | 1 | 0.98 |
| rs61567742 | 11 | 116507206 | T | C | -1.731 | 0.08353 | -? | 0 | 0 | 0 | 1 |
| rs6589575 | 11 | 116733310 | C | T | 1.73 | 0.08358 | +? | 0 | 0 | 0 | 1 |
| rs608154 | 11 | 120838716 | G | A | 1.73 | 0.08361 | +? | 0 | 0 | 0 | 1 |
| rs11217059 | 11 | 118723087 | T | C | 1.727 | 0.08416 | ++ | 0 | 0.223 | 1 | 0.6367 |
| rs2846108 | 11 | 120588487 | T | C | 1.727 | 0.08423 | +? | 0 | 0 | 0 | 1 |
| rs3889570 | 11 | 117300040 | C | T | -1.726 | 0.08426 | -? | 0 | 0 | 0 | 1 |
| rs11217667 | 11 | 119935735 | A | C | -1.725 | 0.08446 | -? | 0 | 0 | 0 | 1 |
| rs2061293 | 11 | 117251521 | G | C | 1.725 | 0.08451 | +? | 0 | 0 | 0 | 1 |
| rs897836 | 11 | 117252489 | T | C | 1.725 | 0.08451 | +? | 0 | 0 | 0 | 1 |
| rs74651412 | 11 | 120735646 | A | G | -1.725 | 0.08455 | -? | 0 | 0 | 0 | 1 |
| rs4278519 | 11 | 120437188 | T | C | -1.725 | 0.08458 | -? | 0 | 0 | 0 | 1 |
| rs34144542 | 11 | 116774447 | G | A | -1.724 | 0.08468 | -? | 0 | 0 | 0 | 1 |
| rs718847 | 11 | 116734146 | C | T | 1.723 | 0.08488 | +? | 0 | 0 | 0 | 1 |
| rs11217918 | 11 | 120441843 | C | G | -1.722 | 0.08507 | -? | 0 | 0 | 0 | 1 |
| rs7120173 | 11 | 116829653 | G | T | -1.721 | 0.08527 | -? | 0 | 0 | 0 | 1 |
| rs80260914 | 11 | 120430224 | G | A | 1.72 | 0.08536 | +? | 0 | 0 | 0 | 1 |
| rs2509209 | 11 | 117102307 | G | A | 1.72 | 0.08537 | +? | 0 | 0 | 0 | 1 |
| rs12790243 | 11 | 117678543 | A | G | 1.72 | 0.0854 | +? | 0 | 0 | 0 | 1 |
| rs1359830167 | 11 | 117365915 | T | C | 1.72 | 0.08546 | +? | 0 | 0 | 0 | 1 |
| rs78462423 | 11 | 117551810 | A | G | 1.72 | 0.08551 | +? | 0 | 0 | 0 | 1 |
| rs79801588 | 11 | 119490788 | G | A | -1.719 | 0.08563 | -? | 0 | 0 | 0 | 1 |
| rs12575945 | 11 | 120425797 | A | C | 1.719 | 0.08564 | +? | 0 | 0 | 0 | 1 |
| rs61901398 | 11 | 120585097 | A | G | 1.718 | 0.08584 | +? | 0 | 0 | 0 | 1 |
| rs4938725 | 11 | 119674611 | G | A | 1.717 | 0.08589 | ++ | 0 | 0.244 | 1 | 0.6211 |
| rs11217118 | 11 | 118866072 | A | G | -1.717 | 0.08592 | -? | 0 | 0 | 0 | 1 |
| rs60708230 | 11 | 119935489 | A | G | -1.717 | 0.08595 | -? | 0 | 0 | 0 | 1 |
| rs4938605 | 11 | 118850707 | A | G | -1.717 | 0.08599 | -? | 0 | 0 | 0 | 1 |
| rs666191 | 11 | 118826063 | G | A | 1.717 | 0.08602 | ++ | 0 | 0 | 1 | 0.9865 |
| rs117228135 | 11 | 116225951 | A | G | -1.716 | 0.08614 | -? | 0 | 0 | 0 | 1 |

| | | | | | | | | | | | |
|---|---|---|---|---|---|---|---|---|---|---|---|
| rs72645459 | 11 | 116762157 | A | C | 1.715 | 0.08627 | +? | 0 | 0 | 0 | 1 |
| rs685308 | 11 | 117897098 | G | C | 1.714 | 0.08653 | +? | 0 | 0 | 0 | 1 |
| rs471840 | 11 | 116294634 | T | C | -1.713 | 0.08673 | -? | 0 | 0 | 0 | 1 |
| rs3781817 | 11 | 120822377 | T | C | 1.713 | 0.08677 | +? | 0 | 0 | 0 | 1 |
| rs17092713 | 11 | 117372488 | A | G | 1.712 | 0.08693 | +? | 0 | 0 | 0 | 1 |
| rs2727791 | 11 | 116684562 | C | G | -1.711 | 0.08699 | -? | 0 | 0 | 0 | 1 |
| rs7107020 | 11 | 117384412 | C | T | 1.711 | 0.08703 | +? | 0 | 0 | 0 | 1 |
| rs580965 | 11 | 116279427 | C | T | -1.711 | 0.08708 | -? | 0 | 0 | 0 | 1 |
| rs7126973 | 11 | 117503731 | T | A | 1.711 | 0.08712 | +? | 0 | 0 | 0 | 1 |
| rs1893393 | 11 | 119936507 | A | T | -1.71 | 0.08718 | -? | 0 | 0 | 0 | 1 |
| rs566848 | 11 | 118711830 | G | A | 1.71 | 0.0872 | ++ | 0 | 0.297 | 1 | 0.5859 |
| rs7103164 | 11 | 117560048 | A | G | 1.709 | 0.08742 | +? | 0 | 0 | 0 | 1 |
| rs7108237 | 11 | 116428302 | A | G | -1.709 | 0.08746 | -? | 0 | 0 | 0 | 1 |
| rs61900157 | 11 | 118824439 | A | C | 1.709 | 0.08752 | ++ | 0 | 0.027 | 1 | 0.8699 |
| rs78914704 | 11 | 119468521 | A | G | -1.708 | 0.08758 | -? | 0 | 0 | 0 | 1 |
| rs7121316 | 11 | 120431744 | A | G | 1.708 | 0.08767 | +? | 0 | 0 | 0 | 1 |
| rs477858 | 11 | 117219961 | G | A | 1.707 | 0.08779 | +? | 0 | 0 | 0 | 1 |
| rs11216586 | 11 | 117727742 | C | G | -1.707 | 0.08782 | -- | 0 | 0.017 | 1 | 0.8971 |
| rs12417682 | 11 | 117082109 | A | G | -1.707 | 0.08786 | -? | 0 | 0 | 0 | 1 |
| rs540273 | 11 | 117197018 | T | C | 1.706 | 0.08798 | +? | 0 | 0 | 0 | 1 |
| rs4938565 | 11 | 118720998 | A | G | 1.706 | 0.088 | ++ | 0 | 0.235 | 1 | 0.6281 |
| rs112201912 | 11 | 119733183 | A | G | -1.706 | 0.0881 | -? | 0 | 0 | 0 | 1 |
| rs3018285 | 11 | 120787752 | C | T | -1.704 | 0.08836 | -? | 0 | 0 | 0 | 1 |
| rs533848 | 11 | 116280688 | T | A | -1.704 | 0.08841 | -? | 0 | 0 | 0 | 1 |
| rs479455 | 11 | 119795611 | T | G | 1.703 | 0.08856 | +? | 0 | 0 | 0 | 1 |
| rs11607895 | 11 | 120606965 | T | C | 1.703 | 0.08859 | +? | 0 | 0 | 0 | 1 |
| rs12786797 | 11 | 116887751 | A | G | -1.703 | 0.08863 | -? | 0 | 0 | 0 | 1 |
| rs11218035 | 11 | 120736087 | T | C | -1.702 | 0.08876 | -? | 0 | 0 | 0 | 1 |
| rs5069 | 11 | 116708254 | A | G | -1.702 | 0.08877 | -? | 0 | 0 | 0 | 1 |
| rs625291 | 11 | 118826060 | C | T | 1.702 | 0.08877 | ++ | 0 | 0 | 1 | 0.9933 |
| rs5076 | 11 | 116707190 | A | G | -1.702 | 0.08882 | -? | 0 | 0 | 0 | 1 |
| rs4936440 | 11 | 118720684 | G | C | 1.701 | 0.08891 | ++ | 0 | 0.237 | 1 | 0.6261 |
| rs139385687 | 11 | 116687156 | C | G | -1.7 | 0.08905 | -? | 0 | 0 | 0 | 1 |
| rs10790355 | 11 | 119805958 | T | C | -1.7 | 0.0891 | -? | 0 | 0 | 0 | 1 |
| rs4938274 | 11 | 116365351 | A | G | -1.7 | 0.08912 | -? | 0 | 0 | 0 | 1 |
| rs73008407 | 11 | 117366513 | T | C | 1.7 | 0.08921 | +? | 0 | 0 | 0 | 1 |
| rs115524104 | 11 | 117727740 | C | T | -1.699 | 0.08931 | -- | 0 | 0.018 | 1 | 0.8935 |
| rs4459318 | 11 | 119541778 | T | C | 1.699 | 0.08933 | +? | 0 | 0 | 0 | 1 |
| rs4938706 | 11 | 119539364 | A | G | 1.699 | 0.0894 | +? | 0 | 0 | 0 | 1 |
| rs12283999 | 11 | 120431935 | G | C | 1.697 | 0.08961 | +? | 0 | 0 | 0 | 1 |
| rs586304 | 11 | 117902169 | T | A | -1.697 | 0.08975 | -? | 0 | 0 | 0 | 1 |
| rs7942824 | 11 | 117353294 | T | G | -1.696 | 0.08987 | -? | 0 | 0 | 0 | 1 |
| rs59626457 | 11 | 118862634 | G | A | -1.696 | 0.0899 | -? | 0 | 0 | 0 | 1 |
| rs555749 | 11 | 116530765 | T | C | -1.695 | 0.09002 | -? | 0 | 0 | 0 | 1 |
| rs7101528 | 11 | 116698665 | A | G | -1.694 | 0.09027 | -? | 0 | 0 | 0 | 1 |
| rs586112 | 11 | 119921812 | C | G | 1.694 | 0.09031 | +? | 0 | 0 | 0 | 1 |
| rs7123326 | 11 | 116696681 | A | G | -1.694 | 0.09033 | -? | 0 | 0 | 0 | 1 |
| rs632137 | 11 | 120247033 | T | C | 1.693 | 0.09043 | +? | 0 | 0 | 0 | 1 |
| rs116954303 | 11 | 118326218 | A | G | -1.693 | 0.09051 | -- | 0 | 0.648 | 1 | 0.4208 |
| rs12281009 | 11 | 117032959 | G | A | -1.692 | 0.09058 | -? | 0 | 0 | 0 | 1 |
| rs10892146 | 11 | 117500542 | T | A | 1.692 | 0.09074 | +? | 0 | 0 | 0 | 1 |
| rs4456264 | 11 | 118851138 | T | C | -1.691 | 0.09076 | -? | 0 | 0 | 0 | 1 |
| rs12718463 | 11 | 116706343 | C | T | -1.691 | 0.09079 | -? | 0 | 0 | 0 | 1 |
| rs485030 | 11 | 119029849 | T | C | -1.691 | 0.09086 | -? | 0 | 0 | 0 | 1 |
| rs79643426 | 11 | 116239462 | T | C | -1.691 | 0.09087 | -? | 0 | 0 | 0 | 1 |
| rs529373 | 11 | 119793573 | T | C | 1.691 | 0.09093 | +? | 0 | 0 | 0 | 1 |

| | | | | | | | | | | | |
|---|---|---|---|---|---|---|---|---|---|---|---|
| rs665759 | 11 | 119793489 | A | G | 1.691 | 0.09093 | +? | 0 | 0 | 0 | 1 |
| rs11217116 | 11 | 118864140 | C | T | -1.69 | 0.09099 | -? | 0 | 0 | 0 | 1 |
| rs12362865 | 11 | 118859842 | G | A | -1.69 | 0.09101 | -? | 0 | 0 | 0 | 1 |
| rs4426142 | 11 | 118824996 | G | A | 1.69 | 0.09101 | ++ | 0 | 0.023 | 1 | 0.8802 |
| rs551163 | 11 | 119795594 | A | C | 1.69 | 0.09108 | +? | 0 | 0 | 0 | 1 |
| rs10892172 | 11 | 117672474 | A | C | 1.69 | 0.09109 | +? | 0 | 0 | 0 | 1 |
| rs470679 | 11 | 118926720 | A | C | -1.689 | 0.09113 | -? | 0 | 0 | 0 | 1 |
| rs116672969 | 11 | 117549963 | A | G | 1.689 | 0.09115 | +? | 0 | 0 | 0 | 1 |
| rs500401 | 11 | 119729992 | T | C | -1.689 | 0.09118 | -? | 0 | 0 | 0 | 1 |
| rs12418705 | 11 | 117084396 | T | C | -1.689 | 0.09124 | -? | 0 | 0 | 0 | 1 |
| rs4938558 | 11 | 118717929 | T | C | 1.689 | 0.09124 | ++ | 0 | 0.227 | 1 | 0.6341 |
| rs11217112 | 11 | 118856920 | G | A | -1.689 | 0.09126 | -? | 0 | 0 | 0 | 1 |
| rs4936455 | 11 | 118861389 | T | C | -1.689 | 0.09126 | -? | 0 | 0 | 0 | 1 |
| rs7118532 | 11 | 119540470 | G | A | 1.689 | 0.09126 | +? | 0 | 0 | 0 | 1 |
| rs9794909 | 11 | 118862306 | C | G | -1.689 | 0.09126 | -? | 0 | 0 | 0 | 1 |
| rs59316586 | 11 | 118856816 | C | A | -1.688 | 0.09139 | -? | 0 | 0 | 0 | 1 |
| rs4938414 | 11 | 117464712 | A | G | -1.687 | 0.09153 | -- | 0 | 0.844 | 1 | 0.3583 |
| rs1145208 | 11 | 116637985 | C | A | 1.687 | 0.09161 | ++ | 0 | 0.003 | 1 | 0.9548 |
| rs11825496 | 11 | 118855221 | G | T | -1.687 | 0.09164 | -? | 0 | 0 | 0 | 1 |
| rs4936454 | 11 | 118859274 | T | C | -1.687 | 0.09164 | -? | 0 | 0 | 0 | 1 |
| rs45527340 | 11 | 118852526 | G | C | -1.686 | 0.0918 | -? | 0 | 0 | 0 | 1 |
| rs477113 | 11 | 119723230 | C | T | 1.686 | 0.09188 | +? | 0 | 0 | 0 | 1 |
| rs556034 | 11 | 117195136 | G | C | 1.685 | 0.09192 | +? | 0 | 0 | 0 | 1 |
| rs2156749 | 11 | 118719405 | G | C | 1.684 | 0.09215 | ++ | 0 | 0.219 | 1 | 0.6396 |
| rs2156750 | 11 | 118719254 | C | T | 1.684 | 0.09215 | ++ | 0 | 0.219 | 1 | 0.6396 |
| rs7116715 | 11 | 118719742 | C | T | 1.684 | 0.09215 | ++ | 0 | 0.219 | 1 | 0.6396 |
| rs12574810 | 11 | 116869560 | G | A | -1.683 | 0.09244 | -? | 0 | 0 | 0 | 1 |
| rs145384144 | 11 | 117017839 | C | G | -1.682 | 0.09253 | -? | 0 | 0 | 0 | 1 |
| rs67215893 | 11 | 116343033 | A | G | 1.681 | 0.09275 | +? | 0 | 0 | 0 | 1 |
| rs61901289 | 11 | 119352350 | T | G | -1.681 | 0.09278 | -? | 0 | 0 | 0 | 1 |
| rs2727788 | 11 | 116698963 | A | C | -1.681 | 0.09283 | -? | 0 | 0 | 0 | 1 |
| rs2051479 | 11 | 117953051 | G | A | -1.681 | 0.09285 | -- | 0 | 0.234 | 1 | 0.6285 |
| rs2156751 | 11 | 118718856 | G | A | 1.68 | 0.093 | ++ | 0 | 0.222 | 1 | 0.6378 |
| rs2187491 | 11 | 118718972 | C | T | 1.68 | 0.093 | ++ | 0 | 0.222 | 1 | 0.6378 |
| rs10790213 | 11 | 117706667 | T | A | -1.677 | 0.09357 | -? | 0 | 0 | 0 | 1 |
| rs471750 | 11 | 116294664 | G | A | 1.676 | 0.09368 | +? | 0 | 0 | 0 | 1 |
| rs3213459 | 11 | 117062489 | C | T | -1.676 | 0.09372 | -? | 0 | 0 | 0 | 1 |
| rs4938564 | 11 | 118720590 | T | G | 1.676 | 0.09374 | ++ | 0 | 0.252 | 1 | 0.616 |
| rs2186761 | 11 | 117467394 | A | G | -1.676 | 0.09381 | -- | 0 | 0.842 | 1 | 0.3589 |
| rs80220270 | 11 | 116680961 | C | A | -1.675 | 0.0939 | -? | 0 | 0 | 0 | 1 |
| rs544154 | 11 | 116291707 | A | G | -1.674 | 0.09414 | -? | 0 | 0 | 0 | 1 |
| rs1790191 | 11 | 118825636 | A | G | 1.674 | 0.0942 | ++ | 0 | 0.043 | 1 | 0.8366 |
| rs544096 | 11 | 118594326 | G | A | 1.674 | 0.09421 | ++ | 0 | 0.457 | 1 | 0.4991 |
| rs10892666 | 11 | 120936096 | T | G | -1.673 | 0.09424 | -? | 0 | 0 | 0 | 1 |
| rs642878 | 11 | 119907777 | A | G | 1.673 | 0.09427 | +? | 0 | 0 | 0 | 1 |
| rs2097131 | 11 | 118825069 | G | A | 1.672 | 0.09456 | ++ | 0 | 0.02 | 1 | 0.8884 |
| rs73011533 | 11 | 120890870 | A | G | -1.672 | 0.09461 | -? | 0 | 0 | 0 | 1 |
| rs2846107 | 11 | 120589674 | A | G | 1.671 | 0.09471 | +? | 0 | 0 | 0 | 1 |
| rs56316258 | 11 | 117341717 | G | C | -1.67 | 0.09486 | -? | 0 | 0 | 0 | 1 |
| rs2849174 | 11 | 116697066 | A | G | -1.67 | 0.09491 | -? | 0 | 0 | 0 | 1 |
| rs12416923 | 11 | 117340239 | A | G | 1.669 | 0.09503 | +? | 0 | 0 | 0 | 1 |
| rs661139 | 11 | 120235874 | A | T | 1.669 | 0.09508 | +? | 0 | 0 | 0 | 1 |
| rs4938670 | 11 | 119398847 | G | A | -1.668 | 0.09532 | -? | 0 | 0 | 0 | 1 |
| rs151220469 | 11 | 116613464 | A | G | -1.667 | 0.09557 | -? | 0 | 0 | 0 | 1 |
| rs874195 | 11 | 119864218 | G | A | 1.667 | 0.0956 | +? | 0 | 0 | 0 | 1 |
| rs137996620 | 11 | 118221655 | C | A | 1.666 | 0.0957 | +? | 0 | 0 | 0 | 1 |

| | | | | | | | | | | | |
|---|---|---|---|---|---|---|---|---|---|---|---|
| rs673028 | 11 | 117518200 | G | A | 1.666 | 0.09571 | +? | 0 | 0 | 0 | 1 |
| rs149421321 | 11 | 117898446 | T | C | 1.666 | 0.09575 | +? | 0 | 0 | 0 | 1 |
| rs2156755 | 11 | 118726230 | A | G | 1.665 | 0.09583 | ++ | 0 | 0.157 | 1 | 0.6918 |
| rs7943240 | 11 | 116439602 | C | T | -1.665 | 0.09584 | -? | 0 | 0 | 0 | 1 |
| rs594711 | 11 | 120992188 | T | C | -1.665 | 0.09595 | -- | 0 | 0.11 | 1 | 0.7399 |
| rs7937420 | 11 | 117953688 | A | G | 1.664 | 0.09605 | ++ | 0 | 0.413 | 1 | 0.5206 |
| rs6589617 | 11 | 117501339 | T | C | 1.664 | 0.09607 | +? | 0 | 0 | 0 | 1 |
| rs693603 | 11 | 116296243 | A | G | 1.663 | 0.09628 | +? | 0 | 0 | 0 | 1 |
| rs4936451 | 11 | 118852137 | T | C | -1.663 | 0.09631 | -? | 0 | 0 | 0 | 1 |
| rs11603573 | 11 | 119928865 | G | C | -1.663 | 0.09637 | -? | 0 | 0 | 0 | 1 |
| rs12279486 | 11 | 116428225 | T | C | -1.663 | 0.09637 | -? | 0 | 0 | 0 | 1 |
| rs34095499 | 11 | 116349950 | A | G | 1.662 | 0.09647 | +? | 0 | 0 | 0 | 1 |
| rs7103314 | 11 | 118723537 | G | A | 1.662 | 0.0966 | ++ | 0 | 0.11 | 1 | 0.7407 |
| rs11605883 | 11 | 117064776 | G | C | -1.662 | 0.09661 | -? | 0 | 0 | 0 | 1 |
| rs2000528 | 11 | 120903752 | G | A | -1.662 | 0.09661 | -? | 0 | 0 | 0 | 1 |
| rs576764 | 11 | 119795133 | G | A | 1.661 | 0.09677 | +? | 0 | 0 | 0 | 1 |
| rs12273687 | 11 | 118852641 | T | G | -1.66 | 0.09684 | -? | 0 | 0 | 0 | 1 |
| rs11217062 | 11 | 118725420 | A | G | 1.66 | 0.09688 | ++ | 0 | 0.158 | 1 | 0.6913 |
| rs17174502 | 11 | 116759655 | A | G | 1.66 | 0.09698 | +? | 0 | 0 | 0 | 1 |
| rs17475326 | 11 | 116223780 | T | A | -1.659 | 0.09711 | -? | 0 | 0 | 0 | 1 |
| rs77661993 | 11 | 117010474 | C | A | -1.658 | 0.09731 | -? | 0 | 0 | 0 | 1 |
| rs649827 | 11 | 117906759 | T | C | -1.657 | 0.0975 | -? | 0 | 0 | 0 | 1 |
| rs1942011 | 11 | 119915024 | T | C | -1.657 | 0.09753 | -? | 0 | 0 | 0 | 1 |
| rs630332 | 11 | 119885462 | A | G | 1.656 | 0.09771 | +? | 0 | 0 | 0 | 1 |
| rs7131215 | 11 | 120474370 | G | C | -1.655 | 0.09784 | -? | 0 | 0 | 0 | 1 |
| rs66991573 | 11 | 117342348 | C | A | -1.655 | 0.09792 | -? | 0 | 0 | 0 | 1 |
| rs676134 | 11 | 117179891 | T | C | 1.653 | 0.09825 | +? | 0 | 0 | 0 | 1 |
| rs67533649 | 11 | 118835272 | G | A | -1.653 | 0.09834 | -? | 0 | 0 | 0 | 1 |
| rs563828 | 11 | 117221170 | T | G | -1.651 | 0.09873 | -? | 0 | 0 | 0 | 1 |
| rs115991973 | 11 | 117010091 | A | G | -1.651 | 0.09883 | -? | 0 | 0 | 0 | 1 |
| rs116384608 | 11 | 117010123 | G | A | -1.651 | 0.09883 | -? | 0 | 0 | 0 | 1 |
| rs4936456 | 11 | 118863324 | T | C | -1.65 | 0.09904 | -? | 0 | 0 | 0 | 1 |
| rs4938613 | 11 | 118863116 | G | A | -1.65 | 0.09904 | -? | 0 | 0 | 0 | 1 |
| rs12419178 | 11 | 117050918 | T | C | -1.649 | 0.09905 | -? | 0 | 0 | 0 | 1 |
| rs12799198 | 11 | 116801422 | T | C | -1.649 | 0.09908 | -? | 0 | 0 | 0 | 1 |
| rs4938561 | 11 | 118718485 | C | T | 1.649 | 0.09912 | ++ | 0 | 0.174 | 1 | 0.6765 |
| rs496164 | 11 | 117711232 | A | G | -1.649 | 0.09914 | -? | 0 | 0 | 0 | 1 |
| rs4420279 | 11 | 118835677 | A | C | -1.648 | 0.09926 | -? | 0 | 0 | 0 | 1 |
| rs113968937 | 11 | 117329289 | G | C | -1.648 | 0.09934 | -? | 0 | 0 | 0 | 1 |
| rs6589717 | 11 | 118979026 | T | C | 1.647 | 0.09963 | +? | 0 | 0 | 0 | 1 |
| rs7940073 | 11 | 118978411 | G | C | 1.647 | 0.09963 | +? | 0 | 0 | 0 | 1 |
| rs74594162 | 11 | 117346121 | G | C | -1.646 | 0.09972 | -? | 0 | 0 | 0 | 1 |
| rs1242695 | 11 | 117200943 | T | G | -1.646 | 0.09979 | -? | 0 | 0 | 0 | 1 |
| rs2279013 | 11 | 117283091 | C | T | 1.646 | 0.09983 | +? | 0 | 0 | 0 | 1 |
| rs949055 | 11 | 120779520 | C | T | -1.645 | 0.09993 | -? | 0 | 0 | 0 | 1 |
| rs650908 | 11 | 117907041 | C | G | -1.645 | 0.09995 | -? | 0 | 0 | 0 | 1 |
| rs11827260 | 11 | 117560822 | C | G | 1.644 | 0.1002 | +? | 0 | 0 | 0 | 1 |
| rs73008672 | 11 | 119733584 | T | C | -1.644 | 0.1002 | -? | 0 | 0 | 0 | 1 |
| rs36111344 | 11 | 120437187 | G | A | -1.643 | 0.1003 | -? | 0 | 0 | 0 | 1 |
| rs56287831 | 11 | 116317789 | A | G | 1.643 | 0.1003 | +? | 0 | 0 | 0 | 1 |
| rs73000621 | 11 | 120437572 | T | C | -1.643 | 0.1003 | -? | 0 | 0 | 0 | 1 |
| rs7925845 | 11 | 116403905 | G | A | -1.643 | 0.1004 | -? | 0 | 0 | 0 | 1 |
| rs1231418 | 11 | 121079478 | C | G | 1.641 | 0.1007 | +? | 0 | 0 | 0 | 1 |
| rs7395947 | 11 | 119397680 | C | T | -1.641 | 0.1007 | -? | 0 | 0 | 0 | 1 |
| rs10502245 | 11 | 120923495 | A | G | -1.641 | 0.1009 | -? | 0 | 0 | 0 | 1 |
| rs10892665 | 11 | 120923899 | G | C | -1.641 | 0.1009 | -? | 0 | 0 | 0 | 1 |

| | | | | | | | | | | | |
|---|---|---|---|---|---|---|---|---|---|---|---|
| rs73006788 | 11 | 119663944 | T | C | -1.641 | 0.1009 | -? | 0 | 0 | 0 | 1 |
| rs540800 | 11 | 118565651 | A | G | 1.64 | 0.1011 | ++ | 0 | 0.286 | 1 | 0.5926 |
| rs11218137 | 11 | 120950752 | C | T | -1.639 | 0.1012 | -? | 0 | 0 | 0 | 1 |
| rs10790198 | 11 | 117515365 | A | G | 1.639 | 0.1013 | +? | 0 | 0 | 0 | 1 |
| rs7126193 | 11 | 118866126 | C | T | -1.639 | 0.1013 | -? | 0 | 0 | 0 | 1 |
| rs2252490 | 11 | 120417065 | C | G | 1.638 | 0.1014 | +? | 0 | 0 | 0 | 1 |
| rs2368937 | 11 | 119674095 | C | T | 1.638 | 0.1014 | ++ | 0 | 0.201 | 1 | 0.6536 |
| rs650535 | 11 | 117414385 | T | C | -1.638 | 0.1015 | -? | 0 | 0 | 0 | 1 |
| rs61905133 | 11 | 116678191 | G | A | -1.637 | 0.1016 | -? | 0 | 0 | 0 | 1 |
| rs519991 | 11 | 117711562 | C | T | -1.637 | 0.1017 | -? | 0 | 0 | 0 | 1 |
| rs2070665 | 11 | 116707684 | A | G | 1.636 | 0.1018 | +? | 0 | 0 | 0 | 1 |
| rs76883864 | 11 | 117051725 | T | C | -1.636 | 0.1018 | -? | 0 | 0 | 0 | 1 |
| rs77638281 | 11 | 118811315 | A | G | -1.634 | 0.1023 | -- | -173.7 | 1.096 | 1 | 0.2951 |
| rs12364667 | 11 | 117675286 | G | T | 1.633 | 0.1024 | +? | 0 | 0 | 0 | 1 |
| rs879412 | 11 | 116372414 | A | G | 1.633 | 0.1024 | +? | 0 | 0 | 0 | 1 |
| rs587806 | 11 | 119882908 | T | C | 1.632 | 0.1026 | +? | 0 | 0 | 0 | 1 |
| rs11217623 | 11 | 119870780 | T | C | 1.632 | 0.1027 | +? | 0 | 0 | 0 | 1 |
| rs11217924 | 11 | 120475835 | T | C | -1.632 | 0.1027 | -? | 0 | 0 | 0 | 1 |
| rs12793318 | 11 | 119703523 | A | G | -1.632 | 0.1027 | -? | 0 | 0 | 0 | 1 |
| rs2046149 | 11 | 117054709 | G | C | -1.631 | 0.1028 | -? | 0 | 0 | 0 | 1 |
| rs11216515 | 11 | 117593119 | C | T | -1.631 | 0.1029 | -? | 0 | 0 | 0 | 1 |
| rs12417961 | 11 | 117318533 | T | C | 1.631 | 0.1029 | +? | 0 | 0 | 0 | 1 |
| rs2252648 | 11 | 120415557 | C | A | 1.631 | 0.1029 | +? | 0 | 0 | 0 | 1 |
| rs11218094 | 11 | 120858815 | T | G | 1.63 | 0.103 | +? | 0 | 0 | 0 | 1 |
| rs75538174 | 11 | 119486336 | C | A | -1.63 | 0.103 | -? | 0 | 0 | 0 | 1 |
| rs10736496 | 11 | 118975113 | A | G | 1.63 | 0.1031 | +? | 0 | 0 | 0 | 1 |
| rs10790283 | 11 | 118975927 | A | G | 1.63 | 0.1031 | +? | 0 | 0 | 0 | 1 |
| rs1793178 | 11 | 117987483 | A | C | -1.63 | 0.1031 | -- | 0 | 0.016 | 1 | 0.9005 |
| rs4938628 | 11 | 118974878 | T | A | 1.63 | 0.1031 | +? | 0 | 0 | 0 | 1 |
| rs4938629 | 11 | 118975426 | C | T | 1.63 | 0.1031 | +? | 0 | 0 | 0 | 1 |
| rs10790363 | 11 | 119941807 | A | G | 1.629 | 0.1033 | +? | 0 | 0 | 0 | 1 |
| rs11216584 | 11 | 117727549 | G | A | -1.628 | 0.1035 | -- | 0 | 0.03 | 1 | 0.8615 |
| rs1312592595 | 11 | 117030633 | C | T | -1.627 | 0.1037 | -? | 0 | 0 | 0 | 1 |
| rs78064538 | 11 | 117346653 | C | G | -1.627 | 0.1037 | -? | 0 | 0 | 0 | 1 |
| rs1624780 | 11 | 118134785 | T | G | -1.625 | 0.1041 | -? | 0 | 0 | 0 | 1 |
| rs2850802 | 11 | 119881100 | A | G | 1.625 | 0.1041 | +? | 0 | 0 | 0 | 1 |
| rs3758998 | 11 | 117946941 | A | C | 1.625 | 0.1042 | +- | -1.1 | 2.967 | 1 | 0.08497 |
| rs7105963 | 11 | 120468752 | T | C | -1.625 | 0.1042 | -? | 0 | 0 | 0 | 1 |
| rs10892501 | 11 | 119818622 | G | A | 1.624 | 0.1043 | +? | 0 | 0 | 0 | 1 |
| rs7120727 | 11 | 120585020 | C | G | 1.624 | 0.1043 | +? | 0 | 0 | 0 | 1 |
| rs116941106 | 11 | 117335035 | A | G | -1.623 | 0.1046 | -? | 0 | 0 | 0 | 1 |
| rs7939112 | 11 | 120837575 | A | G | 1.623 | 0.1046 | +? | 0 | 0 | 0 | 1 |
| rs1893816 | 11 | 120798629 | T | C | 1.622 | 0.1047 | +? | 0 | 0 | 0 | 1 |
| rs2195526 | 11 | 119236751 | T | C | -1.622 | 0.1047 | -? | 0 | 0 | 0 | 1 |
| rs4936413 | 11 | 117842818 | C | T | -1.622 | 0.1048 | -- | 0 | 0.013 | 1 | 0.9081 |
| rs13377544 | 11 | 117647364 | A | T | -1.622 | 0.1049 | -? | 0 | 0 | 0 | 1 |
| rs475159 | 11 | 119732432 | T | G | -1.62 | 0.1052 | -? | 0 | 0 | 0 | 1 |
| rs4938595 | 11 | 118832582 | T | G | -1.62 | 0.1053 | -? | 0 | 0 | 0 | 1 |
| rs9787805 | 11 | 119866879 | C | G | 1.619 | 0.1054 | +? | 0 | 0 | 0 | 1 |
| rs587793 | 11 | 119882896 | C | T | 1.618 | 0.1056 | +? | 0 | 0 | 0 | 1 |
| rs117440454 | 11 | 117946849 | T | C | -1.617 | 0.1058 | -? | 0 | 0 | 0 | 1 |
| rs3016863 | 11 | 117908328 | A | G | -1.617 | 0.1058 | -- | 0 | 0.274 | 1 | 0.6009 |
| rs77719730 | 11 | 117611938 | A | G | -1.617 | 0.1058 | -? | 0 | 0 | 0 | 1 |
| rs670526 | 11 | 118938882 | C | T | -1.616 | 0.106 | -? | 0 | 0 | 0 | 1 |
| rs688099 | 11 | 121072088 | A | C | 1.616 | 0.1061 | +? | 0 | 0 | 0 | 1 |
| rs7948885 | 11 | 120834991 | A | G | 1.615 | 0.1063 | +? | 0 | 0 | 0 | 1 |

| | | | | | | | | | | | |
|---|---|---|---|---|---|---|---|---|---|---|---|
| rs895653 | 11 | 119818789 | A | T | 1.615 | 0.1063 | +? | 0 | 0 | 0 | 1 |
| rs10892023 | 11 | 116598248 | T | C | -1.614 | 0.1065 | -- | 0 | 0.071 | 1 | 0.7898 |
| rs1616199 | 11 | 118146462 | T | C | 1.614 | 0.1066 | +? | 0 | 0 | 0 | 1 |
| rs2248404 | 11 | 120428134 | A | G | 1.614 | 0.1066 | +? | 0 | 0 | 0 | 1 |
| rs471028 | 11 | 118928393 | C | G | -1.614 | 0.1066 | -? | 0 | 0 | 0 | 1 |
| rs647432 | 11 | 118932610 | T | C | -1.614 | 0.1066 | -? | 0 | 0 | 0 | 1 |
| rs117235696 | 11 | 118803193 | G | T | -1.613 | 0.1068 | -- | -126.7 | 1.323 | 1 | 0.25 |
| rs488485 | 11 | 116283172 | T | C | 1.613 | 0.1068 | +? | 0 | 0 | 0 | 1 |
| rs75524659 | 11 | 120468093 | T | C | -1.613 | 0.1068 | -? | 0 | 0 | 0 | 1 |
| rs10892298 | 11 | 118724336 | C | T | 1.612 | 0.1069 | ++ | 0 | 0.129 | 1 | 0.7195 |
| rs12273031 | 11 | 120450358 | A | G | -1.611 | 0.1071 | -? | 0 | 0 | 0 | 1 |
| rs12366015 | 11 | 116990851 | A | G | 1.611 | 0.1072 | +? | 0 | 0 | 0 | 1 |
| rs4938627 | 11 | 118974869 | A | G | 1.611 | 0.1072 | +? | 0 | 0 | 0 | 1 |
| rs2252634 | 11 | 120415823 | T | C | 1.61 | 0.1073 | +? | 0 | 0 | 0 | 1 |
| rs76388159 | 11 | 116292201 | G | A | -1.61 | 0.1074 | -? | 0 | 0 | 0 | 1 |
| rs11606480 | 11 | 117065533 | A | G | -1.61 | 0.1075 | -? | 0 | 0 | 0 | 1 |
| rs12419470 | 11 | 117061775 | T | C | -1.61 | 0.1075 | -? | 0 | 0 | 0 | 1 |
| rs1452842 | 11 | 119911542 | G | A | -1.61 | 0.1075 | -? | 0 | 0 | 0 | 1 |
| rs3781985 | 11 | 117065066 | C | G | -1.61 | 0.1075 | -? | 0 | 0 | 0 | 1 |
| rs61703672 | 11 | 117061825 | T | C | -1.61 | 0.1075 | -? | 0 | 0 | 0 | 1 |
| rs80183569 | 11 | 117065153 | A | G | -1.61 | 0.1075 | -? | 0 | 0 | 0 | 1 |
| rs11218125 | 11 | 120921109 | C | T | -1.609 | 0.1076 | -? | 0 | 0 | 0 | 1 |
| rs643295 | 11 | 117927789 | T | A | -1.609 | 0.1076 | -? | 0 | 0 | 0 | 1 |
| rs9787801 | 11 | 119866718 | A | G | 1.609 | 0.1076 | +? | 0 | 0 | 0 | 1 |
| rs4938457 | 11 | 117792611 | T | C | -1.608 | 0.1079 | -? | 0 | 0 | 0 | 1 |
| rs939274 | 11 | 118134065 | T | G | -1.608 | 0.1079 | -? | 0 | 0 | 0 | 1 |
| rs111607963 | 11 | 117502729 | T | C | -1.607 | 0.108 | -? | 0 | 0 | 0 | 1 |
| rs73580446 | 11 | 116304705 | T | G | -1.607 | 0.108 | -? | 0 | 0 | 0 | 1 |
| rs116534827 | 11 | 120568363 | G | C | 1.606 | 0.1083 | +? | 0 | 0 | 0 | 1 |
| rs61902689 | 11 | 120568477 | T | G | 1.606 | 0.1083 | +? | 0 | 0 | 0 | 1 |
| rs1944499 | 11 | 119875786 | A | G | -1.605 | 0.1084 | -? | 0 | 0 | 0 | 1 |
| rs2155465 | 11 | 121004439 | G | C | 1.605 | 0.1084 | +? | 0 | 0 | 0 | 1 |
| rs473057 | 11 | 117712887 | G | A | -1.605 | 0.1085 | -? | 0 | 0 | 0 | 1 |
| rs12419272 | 11 | 117015516 | G | C | -1.605 | 0.1086 | -? | 0 | 0 | 0 | 1 |
| rs12420127 | 11 | 117035319 | G | A | -1.605 | 0.1086 | -? | 0 | 0 | 0 | 1 |
| rs186331126 | 11 | 116823972 | G | T | -1.605 | 0.1086 | -? | 0 | 0 | 0 | 1 |
| rs646543 | 11 | 120853577 | C | T | 1.605 | 0.1086 | +? | 0 | 0 | 0 | 1 |
| rs11218135 | 11 | 120944219 | G | A | -1.604 | 0.1088 | -? | 0 | 0 | 0 | 1 |
| rs58204590 | 11 | 120946892 | G | A | -1.604 | 0.1088 | -? | 0 | 0 | 0 | 1 |
| rs687190 | 11 | 118542383 | C | T | 1.603 | 0.109 | ++ | 0 | 0.174 | 1 | 0.6763 |
| rs650104 | 11 | 117912720 | T | C | -1.602 | 0.1091 | -- | 0 | 0.234 | 1 | 0.6285 |
| rs686047 | 11 | 117908235 | A | G | -1.602 | 0.1091 | -- | 0 | 0.267 | 1 | 0.6052 |
| rs947931 | 11 | 118577614 | G | T | -1.602 | 0.1091 | -? | 0 | 0 | 0 | 1 |
| rs76406762 | 11 | 116220390 | A | C | -1.601 | 0.1093 | -? | 0 | 0 | 0 | 1 |
| rs471000 | 11 | 118936910 | T | C | -1.601 | 0.1094 | -? | 0 | 0 | 0 | 1 |
| rs1939333 | 11 | 119907974 | C | T | -1.6 | 0.1095 | -? | 0 | 0 | 0 | 1 |
| rs584620 | 11 | 119882187 | T | C | 1.6 | 0.1095 | +? | 0 | 0 | 0 | 1 |
| rs116773440 | 11 | 117039461 | C | A | -1.6 | 0.1097 | -? | 0 | 0 | 0 | 1 |
| rs142017584 | 11 | 117031363 | A | G | -1.599 | 0.1098 | -? | 0 | 0 | 0 | 1 |
| rs34051198 | 11 | 119734332 | A | G | -1.599 | 0.1098 | -? | 0 | 0 | 0 | 1 |
| rs79865588 | 11 | 117341574 | C | T | -1.599 | 0.1098 | -? | 0 | 0 | 0 | 1 |
| rs61900158 | 11 | 118831764 | A | T | -1.599 | 0.1099 | -? | 0 | 0 | 0 | 1 |
| rs3212282 | 11 | 116601945 | A | G | -1.598 | 0.11 | -- | 0 | 0.126 | 1 | 0.7226 |
| rs74330115 | 11 | 119806104 | A | G | -1.598 | 0.1101 | -? | 0 | 0 | 0 | 1 |
| rs76264435 | 11 | 119813521 | A | G | 1.597 | 0.1102 | +? | 0 | 0 | 0 | 1 |
| rs601817 | 11 | 120843834 | C | T | 1.597 | 0.1103 | +? | 0 | 0 | 0 | 1 |

| | | | | | | | | | | | |
|---|---|---|---|---|---|---|---|---|---|---|---|
| rs553062 | 11 | 117421425 | A | G | -1.596 | 0.1104 | -? | 0 | 0 | 0 | 1 |
| rs689054 | 11 | 117912731 | T | A | -1.596 | 0.1105 | -- | 0 | 0.245 | 1 | 0.6207 |
| rs1944511 | 11 | 119863584 | A | G | -1.596 | 0.1106 | -? | 0 | 0 | 0 | 1 |
| rs506842 | 11 | 119724144 | C | G | 1.596 | 0.1106 | +? | 0 | 0 | 0 | 1 |
| rs11218078 | 11 | 120824692 | G | T | 1.595 | 0.1107 | +? | 0 | 0 | 0 | 1 |
| rs527778 | 11 | 116479584 | T | G | 1.595 | 0.1107 | +? | 0 | 0 | 0 | 1 |
| rs7947524 | 11 | 117671091 | T | C | 1.595 | 0.1107 | +? | 0 | 0 | 0 | 1 |
| rs78474864 | 11 | 120847827 | A | G | -1.593 | 0.1112 | -? | 0 | 0 | 0 | 1 |
| rs11216587 | 11 | 117727965 | T | C | -1.592 | 0.1113 | -- | 0 | 0.002 | 1 | 0.962 |
| rs4938299 | 11 | 116565732 | A | G | 1.592 | 0.1114 | +? | 0 | 0 | 0 | 1 |
| rs12417984 | 11 | 117017714 | A | G | -1.591 | 0.1115 | -? | 0 | 0 | 0 | 1 |
| rs12418349 | 11 | 117025932 | G | A | -1.591 | 0.1115 | -? | 0 | 0 | 0 | 1 |
| rs12419436 | 11 | 117017455 | G | T | -1.591 | 0.1115 | -? | 0 | 0 | 0 | 1 |
| rs12420250 | 11 | 117028727 | A | G | -1.591 | 0.1115 | -? | 0 | 0 | 0 | 1 |
| rs12420725 | 11 | 117017530 | G | A | -1.591 | 0.1115 | -? | 0 | 0 | 0 | 1 |
| rs139146464 | 11 | 117019867 | G | A | -1.591 | 0.1115 | -? | 0 | 0 | 0 | 1 |
| rs139864910 | 11 | 117017762 | C | A | -1.591 | 0.1115 | -? | 0 | 0 | 0 | 1 |
| rs2306471 | 11 | 117016450 | T | C | -1.591 | 0.1115 | -? | 0 | 0 | 0 | 1 |
| rs77171490 | 11 | 117038741 | A | T | -1.591 | 0.1115 | -? | 0 | 0 | 0 | 1 |
| rs77678768 | 11 | 117040505 | C | G | -1.591 | 0.1115 | -? | 0 | 0 | 0 | 1 |
| rs78166985 | 11 | 117029785 | T | A | -1.591 | 0.1115 | -? | 0 | 0 | 0 | 1 |
| rs80353869 | 11 | 117048122 | A | G | -1.591 | 0.1115 | -? | 0 | 0 | 0 | 1 |
| rs10892123 | 11 | 117354582 | C | T | 1.591 | 0.1116 | +? | 0 | 0 | 0 | 1 |
| rs17124211 | 11 | 120574350 | G | A | 1.591 | 0.1116 | +? | 0 | 0 | 0 | 1 |
| rs35962410 | 11 | 119831163 | C | A | -1.591 | 0.1116 | -? | 0 | 0 | 0 | 1 |
| rs636283 | 11 | 118932445 | T | C | -1.591 | 0.1116 | -? | 0 | 0 | 0 | 1 |
| rs17119815 | 11 | 116429563 | G | T | -1.589 | 0.112 | -? | 0 | 0 | 0 | 1 |
| rs2155466 | 11 | 121004502 | T | C | 1.589 | 0.112 | +? | 0 | 0 | 0 | 1 |
| rs612804 | 11 | 121077373 | C | T | 1.589 | 0.112 | +? | 0 | 0 | 0 | 1 |
| rs2089647 | 11 | 119512641 | G | A | 1.587 | 0.1124 | +? | 0 | 0 | 0 | 1 |
| rs74649894 | 11 | 118136362 | G | C | 1.587 | 0.1124 | -+ | 62.8 | 8.065 | 1 | 0.004513 |
| rs651202 | 11 | 119881745 | A | G | 1.587 | 0.1125 | +? | 0 | 0 | 0 | 1 |
| rs10892148 | 11 | 117519147 | T | C | 1.587 | 0.1126 | +? | 0 | 0 | 0 | 1 |
| rs687180 | 11 | 121072293 | A | G | 1.587 | 0.1126 | +? | 0 | 0 | 0 | 1 |
| rs78341300 | 11 | 119483777 | G | A | -1.587 | 0.1126 | -? | 0 | 0 | 0 | 1 |
| rs1793170 | 11 | 118150205 | C | T | 1.586 | 0.1127 | +? | 0 | 0 | 0 | 1 |
| rs188758920 | 11 | 118858878 | A | C | -1.586 | 0.1128 | -? | 0 | 0 | 0 | 1 |
| rs2301174 | 11 | 117063709 | G | C | -1.585 | 0.1129 | -? | 0 | 0 | 0 | 1 |
| rs61901363 | 11 | 120569866 | T | C | 1.585 | 0.1129 | +? | 0 | 0 | 0 | 1 |
| rs4438048 | 11 | 119510366 | T | C | 1.585 | 0.113 | +? | 0 | 0 | 0 | 1 |
| rs616905 | 11 | 121076423 | C | T | 1.584 | 0.1131 | +? | 0 | 0 | 0 | 1 |
| rs140305819 | 11 | 118018518 | A | G | -1.584 | 0.1132 | -? | 0 | 0 | 0 | 1 |
| rs149137426 | 11 | 116682668 | A | T | 1.584 | 0.1132 | +? | 0 | 0 | 0 | 1 |
| rs1045334530 | 11 | 116301501 | C | A | 1.583 | 0.1134 | +? | 0 | 0 | 0 | 1 |
| rs10892639 | 11 | 120780402 | C | A | -1.582 | 0.1137 | -? | 0 | 0 | 0 | 1 |
| rs673768 | 11 | 118920453 | T | C | -1.582 | 0.1137 | -? | 0 | 0 | 0 | 1 |
| rs2089646 | 11 | 119512115 | G | A | 1.581 | 0.1138 | +? | 0 | 0 | 0 | 1 |
| rs61902686 | 11 | 120566375 | C | T | 1.581 | 0.1139 | +? | 0 | 0 | 0 | 1 |
| rs61902687 | 11 | 120566406 | G | A | 1.581 | 0.1139 | +? | 0 | 0 | 0 | 1 |
| rs4245199 | 11 | 119510303 | G | T | 1.58 | 0.1141 | +? | 0 | 0 | 0 | 1 |
| rs593319 | 11 | 119939806 | T | A | 1.58 | 0.1141 | +? | 0 | 0 | 0 | 1 |
| rs597253 | 11 | 119939156 | A | G | 1.58 | 0.1141 | +? | 0 | 0 | 0 | 1 |
| rs61901362 | 11 | 120569843 | C | T | 1.58 | 0.1142 | +? | 0 | 0 | 0 | 1 |
| rs11599950 | 11 | 117593368 | T | C | -1.579 | 0.1144 | -? | 0 | 0 | 0 | 1 |
| rs1316812 | 11 | 120491113 | T | C | -1.579 | 0.1144 | -? | 0 | 0 | 0 | 1 |
| rs114831648 | 11 | 117342536 | C | T | -1.578 | 0.1145 | -? | 0 | 0 | 0 | 1 |

| | | | | | | | | | | | |
|---|---|---|---|---|---|---|---|---|---|---|---|
| rs1715464 | 11 | 117987503 | A | C | -1.578 | 0.1145 | -- | 0 | 0.01 | 1 | 0.9195 |
| rs2509212 | 11 | 117214580 | G | C | -1.578 | 0.1146 | -? | 0 | 0 | 0 | 1 |
| rs12421186 | 11 | 120464049 | T | C | 1.577 | 0.1147 | +? | 0 | 0 | 0 | 1 |
| rs11216266 | 11 | 116950150 | C | T | -1.577 | 0.1148 | -? | 0 | 0 | 0 | 1 |
| rs4359220 | 11 | 120658874 | G | A | 1.577 | 0.1148 | +? | 0 | 0 | 0 | 1 |
| rs143494314 | 11 | 117023706 | T | C | -1.577 | 0.1149 | -? | 0 | 0 | 0 | 1 |
| rs150984554 | 11 | 117035016 | A | G | -1.577 | 0.1149 | -? | 0 | 0 | 0 | 1 |
| rs647975 | 11 | 118038741 | T | C | 1.577 | 0.1149 | +? | 0 | 0 | 0 | 1 |
| rs1001556897 | 11 | 117015238 | A | C | -1.576 | 0.115 | -? | 0 | 0 | 0 | 1 |
| rs34203183 | 11 | 120851138 | G | C | 1.576 | 0.115 | +? | 0 | 0 | 0 | 1 |
| rs7128009 | 11 | 120835281 | A | G | 1.576 | 0.115 | +? | 0 | 0 | 0 | 1 |
| rs11217661 | 11 | 119921007 | T | C | -1.576 | 0.1151 | -? | 0 | 0 | 0 | 1 |
| rs720313 | 11 | 119920110 | G | A | -1.576 | 0.1151 | -? | 0 | 0 | 0 | 1 |
| rs720314 | 11 | 119920056 | G | A | -1.576 | 0.1151 | -? | 0 | 0 | 0 | 1 |
| rs7933134 | 11 | 117059794 | C | G | -1.576 | 0.1151 | -? | 0 | 0 | 0 | 1 |
| rs11218060 | 11 | 120797878 | T | C | 1.574 | 0.1154 | +? | 0 | 0 | 0 | 1 |
| rs473172 | 11 | 116807970 | A | G | 1.574 | 0.1154 | +? | 0 | 0 | 0 | 1 |
| rs524209 | 11 | 120836662 | A | G | 1.574 | 0.1154 | +? | 0 | 0 | 0 | 1 |
| rs587985 | 11 | 119244095 | C | T | 1.574 | 0.1155 | +? | 0 | 0 | 0 | 1 |
| rs470324 | 11 | 118924212 | A | G | -1.574 | 0.1156 | -? | 0 | 0 | 0 | 1 |
| rs12274388 | 11 | 116946238 | T | C | -1.573 | 0.1158 | -? | 0 | 0 | 0 | 1 |
| rs2306472 | 11 | 117058062 | T | C | -1.573 | 0.1158 | -? | 0 | 0 | 0 | 1 |
| rs5072 | 11 | 116707583 | A | G | 1.573 | 0.1158 | +? | 0 | 0 | 0 | 1 |
| rs7109649 | 11 | 116943544 | T | C | -1.573 | 0.1158 | -? | 0 | 0 | 0 | 1 |
| rs7940667 | 11 | 119510644 | A | C | 1.573 | 0.1158 | +? | 0 | 0 | 0 | 1 |
| rs77998262 | 11 | 120419320 | A | G | 1.572 | 0.1159 | +? | 0 | 0 | 0 | 1 |
| rs87466 | 11 | 120981863 | G | A | -1.572 | 0.1159 | -? | 0 | 0 | 0 | 1 |
| rs640663 | 11 | 119936292 | A | G | 1.571 | 0.1161 | +? | 0 | 0 | 0 | 1 |
| rs1960536 | 11 | 119916341 | C | A | -1.571 | 0.1162 | -? | 0 | 0 | 0 | 1 |
| rs73009128 | 11 | 116333012 | T | C | 1.571 | 0.1162 | +? | 0 | 0 | 0 | 1 |
| rs12293509 | 11 | 120570595 | T | C | 1.57 | 0.1164 | +? | 0 | 0 | 0 | 1 |
| rs587308 | 11 | 119882789 | T | C | 1.57 | 0.1164 | +? | 0 | 0 | 0 | 1 |
| rs61904800 | 11 | 116424909 | A | G | -1.57 | 0.1164 | -? | 0 | 0 | 0 | 1 |
| rs2252494 | 11 | 120416923 | A | G | 1.569 | 0.1167 | +? | 0 | 0 | 0 | 1 |
| rs56221909 | 11 | 119679013 | C | G | 1.569 | 0.1167 | +? | 0 | 0 | 0 | 1 |
| rs11216240 | 11 | 116899248 | T | A | -1.568 | 0.1168 | -? | 0 | 0 | 0 | 1 |
| rs11216241 | 11 | 116899249 | C | A | -1.568 | 0.1168 | -? | 0 | 0 | 0 | 1 |
| rs10892066 | 11 | 116920927 | T | C | -1.568 | 0.1169 | -? | 0 | 0 | 0 | 1 |
| rs145723586 | 11 | 119728661 | T | C | 1.568 | 0.1169 | +? | 0 | 0 | 0 | 1 |
| rs1531706 | 11 | 116943952 | A | C | -1.568 | 0.1169 | -? | 0 | 0 | 0 | 1 |
| rs1531707 | 11 | 116944712 | A | G | -1.568 | 0.1169 | -? | 0 | 0 | 0 | 1 |
| rs7935913 | 11 | 116944348 | A | G | -1.568 | 0.1169 | -? | 0 | 0 | 0 | 1 |
| rs1876498 | 11 | 116464013 | A | G | -1.567 | 0.1171 | -? | 0 | 0 | 0 | 1 |
| rs7120429 | 11 | 116391651 | T | C | 1.567 | 0.1171 | +? | 0 | 0 | 0 | 1 |
| rs1338985819 | 11 | 117919431 | A | G | 1.567 | 0.1172 | +? | 0 | 0 | 0 | 1 |
| rs11217648 | 11 | 119904390 | G | A | -1.566 | 0.1173 | -? | 0 | 0 | 0 | 1 |
| rs582688 | 11 | 118932320 | C | T | -1.565 | 0.1175 | -? | 0 | 0 | 0 | 1 |
| rs2000527 | 11 | 120904156 | G | A | -1.564 | 0.1178 | -? | 0 | 0 | 0 | 1 |
| rs2046150 | 11 | 117053433 | A | C | -1.564 | 0.1178 | -? | 0 | 0 | 0 | 1 |
| rs12274465 | 11 | 116957189 | T | C | -1.564 | 0.1179 | -? | 0 | 0 | 0 | 1 |
| rs2298649 | 11 | 117672225 | G | A | 1.564 | 0.1179 | +? | 0 | 0 | 0 | 1 |
| rs3741279 | 11 | 117353003 | G | A | -1.564 | 0.1179 | -? | 0 | 0 | 0 | 1 |
| rs4938329 | 11 | 116928112 | T | C | -1.564 | 0.1179 | -? | 0 | 0 | 0 | 1 |
| rs7935834 | 11 | 116944437 | G | C | -1.564 | 0.1179 | -? | 0 | 0 | 0 | 1 |
| rs972534 | 11 | 119917892 | C | T | -1.564 | 0.1179 | -? | 0 | 0 | 0 | 1 |
| rs4938415 | 11 | 117467560 | A | C | -1.563 | 0.1181 | -- | 0 | 0.472 | 1 | 0.4922 |

| | | | | | | | | | | | |
|---|---|---|---|---|---|---|---|---|---|---|---|
| rs10892518 | 11 | 119905145 | T | C | -1.562 | 0.1182 | -? | 0 | 0 | 0 | 1 |
| rs11217651 | 11 | 119905245 | C | T | -1.562 | 0.1183 | -? | 0 | 0 | 0 | 1 |
| rs180339 | 11 | 116617782 | T | C | -1.562 | 0.1183 | -- | 0 | 0.721 | 1 | 0.3958 |
| rs56763747 | 11 | 120493805 | G | A | -1.562 | 0.1183 | -? | 0 | 0 | 0 | 1 |
| rs11217654 | 11 | 119914725 | G | T | -1.562 | 0.1184 | -? | 0 | 0 | 0 | 1 |
| rs11607618 | 11 | 119911205 | A | G | -1.562 | 0.1184 | -? | 0 | 0 | 0 | 1 |
| rs12790374 | 11 | 119912882 | G | A | -1.562 | 0.1184 | -? | 0 | 0 | 0 | 1 |
| rs12792065 | 11 | 119912919 | C | T | -1.562 | 0.1184 | -? | 0 | 0 | 0 | 1 |
| rs12799875 | 11 | 119910275 | T | C | -1.562 | 0.1184 | -? | 0 | 0 | 0 | 1 |
| rs12800482 | 11 | 119910556 | T | C | -1.562 | 0.1184 | -? | 0 | 0 | 0 | 1 |
| rs12800844 | 11 | 119910592 | T | G | -1.562 | 0.1184 | -? | 0 | 0 | 0 | 1 |
| rs1452841 | 11 | 119909608 | G | A | -1.562 | 0.1184 | -? | 0 | 0 | 0 | 1 |
| rs61897925 | 11 | 119910833 | A | G | -1.562 | 0.1184 | -? | 0 | 0 | 0 | 1 |
| rs2008915 | 11 | 116603134 | C | G | -1.561 | 0.1185 | -- | 0 | 0.108 | 1 | 0.7424 |
| rs61901378 | 11 | 120577241 | A | G | 1.561 | 0.1186 | +? | 0 | 0 | 0 | 1 |
| rs538359 | 11 | 116281169 | C | T | 1.56 | 0.1187 | +? | 0 | 0 | 0 | 1 |
| rs569327 | 11 | 116317745 | T | C | 1.56 | 0.1187 | +? | 0 | 0 | 0 | 1 |
| rs61903416 | 11 | 116955049 | C | T | -1.559 | 0.1189 | -? | 0 | 0 | 0 | 1 |
| rs645233 | 11 | 119938788 | G | A | 1.559 | 0.1189 | +? | 0 | 0 | 0 | 1 |
| rs11512472 | 11 | 119917309 | G | A | -1.559 | 0.119 | -? | 0 | 0 | 0 | 1 |
| rs4938630 | 11 | 118984287 | A | C | 1.559 | 0.119 | +? | 0 | 0 | 0 | 1 |
| rs11217650 | 11 | 119905180 | T | G | -1.559 | 0.1191 | -? | 0 | 0 | 0 | 1 |
| rs12279066 | 11 | 116913909 | G | C | -1.558 | 0.1192 | -? | 0 | 0 | 0 | 1 |
| rs1010260 | 11 | 119925422 | A | G | -1.557 | 0.1194 | -? | 0 | 0 | 0 | 1 |
| rs66921644 | 11 | 119835014 | A | G | -1.557 | 0.1194 | -? | 0 | 0 | 0 | 1 |
| rs10790147 | 11 | 116411836 | T | C | -1.556 | 0.1198 | -? | 0 | 0 | 0 | 1 |
| rs28737445 | 11 | 120710210 | G | A | 1.556 | 0.1198 | +? | 0 | 0 | 0 | 1 |
| rs7106662 | 11 | 116951385 | C | T | -1.556 | 0.1198 | -? | 0 | 0 | 0 | 1 |
| rs80030349 | 11 | 119804640 | G | A | -1.555 | 0.1199 | -? | 0 | 0 | 0 | 1 |
| rs61900926 | 11 | 120492000 | A | G | -1.555 | 0.12 | -? | 0 | 0 | 0 | 1 |
| rs10466590 | 11 | 116894413 | G | C | -1.554 | 0.1201 | -? | 0 | 0 | 0 | 1 |
| rs11216239 | 11 | 116895512 | G | C | -1.554 | 0.1201 | -? | 0 | 0 | 0 | 1 |
| rs7484045 | 11 | 116946890 | T | C | -1.554 | 0.1201 | -? | 0 | 0 | 0 | 1 |
| rs11493812 | 11 | 119915782 | G | A | -1.554 | 0.1202 | -? | 0 | 0 | 0 | 1 |
| rs2510639 | 11 | 120875383 | A | G | 1.554 | 0.1202 | +? | 0 | 0 | 0 | 1 |
| rs11216244 | 11 | 116912730 | T | C | -1.554 | 0.1203 | -? | 0 | 0 | 0 | 1 |
| rs11216245 | 11 | 116913202 | T | C | -1.554 | 0.1203 | -? | 0 | 0 | 0 | 1 |
| rs78312641 | 11 | 118749318 | A | T | -1.553 | 0.1204 | -+ | -126.3 | 1.326 | 1 | 0.2496 |
| rs12274778 | 11 | 120479903 | G | C | -1.553 | 0.1205 | -? | 0 | 0 | 0 | 1 |
| rs7102557 | 11 | 117745018 | A | G | 1.553 | 0.1205 | +? | 0 | 0 | 0 | 1 |
| rs7123784 | 11 | 117275887 | C | G | 1.552 | 0.1206 | +? | 0 | 0 | 0 | 1 |
| rs1379188468 | 11 | 120492853 | T | A | -1.551 | 0.1208 | -? | 0 | 0 | 0 | 1 |
| rs148945243 | 11 | 116601062 | T | C | -1.551 | 0.1208 | -? | 0 | 0 | 0 | 1 |
| rs658494 | 11 | 120984681 | T | C | -1.551 | 0.1209 | -? | 0 | 0 | 0 | 1 |
| rs73009129 | 11 | 116333791 | A | G | 1.551 | 0.1209 | +? | 0 | 0 | 0 | 1 |
| rs1567375 | 11 | 119502477 | A | C | -1.551 | 0.121 | -? | 0 | 0 | 0 | 1 |
| rs4938377 | 11 | 117296292 | G | T | 1.551 | 0.121 | +? | 0 | 0 | 0 | 1 |
| rs662146 | 11 | 119927792 | G | A | -1.551 | 0.121 | -? | 0 | 0 | 0 | 1 |
| rs7109567 | 11 | 117403017 | G | A | 1.551 | 0.121 | +? | 0 | 0 | 0 | 1 |
| rs10082609 | 11 | 116949074 | T | C | -1.55 | 0.1211 | -? | 0 | 0 | 0 | 1 |
| rs61901368 | 11 | 120571553 | G | A | 1.55 | 0.1211 | +? | 0 | 0 | 0 | 1 |
| rs591596 | 11 | 119938994 | A | G | 1.55 | 0.1212 | +? | 0 | 0 | 0 | 1 |
| rs11600552 | 11 | 116900947 | C | T | -1.549 | 0.1214 | -? | 0 | 0 | 0 | 1 |
| rs7128772 | 11 | 116889411 | A | T | -1.549 | 0.1214 | -? | 0 | 0 | 0 | 1 |
| rs10892330 | 11 | 118970875 | T | C | 1.549 | 0.1215 | +? | 0 | 0 | 0 | 1 |
| rs61901370 | 11 | 120571883 | T | C | 1.547 | 0.1218 | +? | 0 | 0 | 0 | 1 |

| | | | | | | | | | | | |
|---|---|---|---|---|---|---|---|---|---|---|---|
| rs639891 | 11 | 121094594 | T | C | 1.547 | 0.1219 | +? | 0 | 0 | 0 | 1 |
| rs7123583 | 11 | 116600021 | T | C | -1.547 | 0.1219 | -- | 0 | 0.28 | 1 | 0.5966 |
| rs10891998 | 11 | 116465292 | G | A | -1.546 | 0.122 | -? | 0 | 0 | 0 | 1 |
| rs1410186916 | 11 | 117674072 | T | A | 1.546 | 0.122 | +? | 0 | 0 | 0 | 1 |
| rs10892070 | 11 | 116951533 | A | G | 1.546 | 0.1221 | +? | 0 | 0 | 0 | 1 |
| rs7115603 | 11 | 119781153 | G | A | -1.546 | 0.1221 | -? | 0 | 0 | 0 | 1 |
| rs78915622 | 11 | 117498834 | G | A | -1.545 | 0.1224 | -? | 0 | 0 | 0 | 1 |
| rs10892310 | 11 | 118806897 | T | C | 1.544 | 0.1225 | ++ | -133.6 | 1.284 | 1 | 0.2571 |
| rs10892071 | 11 | 116954565 | T | C | -1.544 | 0.1226 | -? | 0 | 0 | 0 | 1 |
| rs10892524 | 11 | 119927321 | T | G | -1.544 | 0.1226 | -? | 0 | 0 | 0 | 1 |
| rs1939331 | 11 | 119907867 | C | G | -1.544 | 0.1226 | -? | 0 | 0 | 0 | 1 |
| rs1452840 | 11 | 119906398 | T | A | -1.543 | 0.1228 | -? | 0 | 0 | 0 | 1 |
| rs112872285 | 11 | 117668060 | T | G | 1.543 | 0.1229 | +? | 0 | 0 | 0 | 1 |
| rs1467050 | 11 | 119529365 | T | C | 1.543 | 0.1229 | +? | 0 | 0 | 0 | 1 |
| rs11216255 | 11 | 116929995 | G | A | -1.542 | 0.1231 | -? | 0 | 0 | 0 | 1 |
| rs478626 | 11 | 121088829 | A | G | 1.542 | 0.1231 | +? | 0 | 0 | 0 | 1 |
| rs637115 | 11 | 121088659 | A | G | 1.542 | 0.1231 | +? | 0 | 0 | 0 | 1 |
| rs3824976 | 11 | 120895352 | A | G | -1.541 | 0.1232 | -? | 0 | 0 | 0 | 1 |
| rs7117266 | 11 | 118838020 | A | G | -1.541 | 0.1232 | -? | 0 | 0 | 0 | 1 |
| rs74945140 | 11 | 120991571 | A | G | -1.541 | 0.1232 | -? | 0 | 0 | 0 | 1 |
| rs12273351 | 11 | 119503402 | T | G | -1.54 | 0.1236 | -? | 0 | 0 | 0 | 1 |
| rs3886960 | 11 | 116924990 | T | G | -1.54 | 0.1236 | -? | 0 | 0 | 0 | 1 |
| rs606727 | 11 | 117356858 | G | C | 1.54 | 0.1236 | +? | 0 | 0 | 0 | 1 |
| rs7108912 | 11 | 116925996 | A | G | -1.54 | 0.1236 | -? | 0 | 0 | 0 | 1 |
| rs7124872 | 11 | 116927951 | G | A | -1.54 | 0.1236 | -? | 0 | 0 | 0 | 1 |
| rs7124996 | 11 | 116928006 | T | C | -1.54 | 0.1236 | -? | 0 | 0 | 0 | 1 |
| rs723955 | 11 | 116925042 | A | G | -1.54 | 0.1236 | -? | 0 | 0 | 0 | 1 |
| rs116057214 | 11 | 119495219 | G | A | -1.539 | 0.1237 | -? | 0 | 0 | 0 | 1 |
| rs519942 | 11 | 118921306 | A | G | -1.539 | 0.1237 | -? | 0 | 0 | 0 | 1 |
| rs675783 | 11 | 119927844 | C | T | -1.539 | 0.1237 | -? | 0 | 0 | 0 | 1 |
| rs4938699 | 11 | 119510937 | C | G | 1.539 | 0.1238 | +? | 0 | 0 | 0 | 1 |
| rs61905347 | 11 | 117646968 | T | C | -1.539 | 0.1238 | -? | 0 | 0 | 0 | 1 |
| rs34100096 | 11 | 116585625 | A | G | 1.539 | 0.1239 | +? | 0 | 0 | 0 | 1 |
| rs11217655 | 11 | 119914788 | C | A | -1.538 | 0.124 | -? | 0 | 0 | 0 | 1 |
| rs74447718 | 11 | 120571314 | T | C | 1.538 | 0.124 | +? | 0 | 0 | 0 | 1 |
| rs11822344 | 11 | 118864777 | C | G | -1.538 | 0.1241 | -? | 0 | 0 | 0 | 1 |
| rs7106316 | 11 | 118441079 | G | A | 1.538 | 0.1241 | ++ | 0 | 0.163 | 1 | 0.6866 |
| rs7950093 | 11 | 116963519 | T | G | -1.538 | 0.1241 | -? | 0 | 0 | 0 | 1 |
| rs2508573 | 11 | 118555383 | T | C | 1.537 | 0.1242 | ++ | 0 | 0.405 | 1 | 0.5243 |
| rs648191 | 11 | 121088794 | A | G | 1.537 | 0.1242 | +? | 0 | 0 | 0 | 1 |
| rs12270261 | 11 | 120577413 | T | C | 1.537 | 0.1243 | +? | 0 | 0 | 0 | 1 |
| rs11216620 | 11 | 117790342 | T | C | 1.537 | 0.1244 | +? | 0 | 0 | 0 | 1 |
| rs492047 | 11 | 121058954 | A | G | 1.536 | 0.1245 | +? | 0 | 0 | 0 | 1 |
| rs670192 | 11 | 118920112 | A | G | -1.536 | 0.1245 | -? | 0 | 0 | 0 | 1 |
| rs12363889 | 11 | 119918352 | G | A | -1.536 | 0.1246 | -? | 0 | 0 | 0 | 1 |
| rs67389832 | 11 | 118839032 | A | C | -1.535 | 0.1248 | -? | 0 | 0 | 0 | 1 |
| rs10502234 | 11 | 119906313 | A | G | -1.535 | 0.1249 | -? | 0 | 0 | 0 | 1 |
| rs11605106 | 11 | 119906213 | A | G | -1.535 | 0.1249 | -? | 0 | 0 | 0 | 1 |
| rs1235340366 | 11 | 119905840 | G | C | -1.535 | 0.1249 | -? | 0 | 0 | 0 | 1 |
| rs12362929 | 11 | 119905865 | G | A | -1.535 | 0.1249 | -? | 0 | 0 | 0 | 1 |
| rs61900898 | 11 | 120421474 | T | G | 1.534 | 0.125 | +? | 0 | 0 | 0 | 1 |
| rs4938704 | 11 | 119539313 | T | C | 1.532 | 0.1255 | +? | 0 | 0 | 0 | 1 |
| rs12278795 | 11 | 120570198 | G | A | 1.531 | 0.1258 | +? | 0 | 0 | 0 | 1 |
| rs61901364 | 11 | 120569884 | T | C | 1.531 | 0.1258 | +? | 0 | 0 | 0 | 1 |
| rs10892643 | 11 | 120790963 | T | C | 1.53 | 0.1259 | +? | 0 | 0 | 0 | 1 |
| rs17311045 | 11 | 117647496 | T | C | -1.53 | 0.1259 | -? | 0 | 0 | 0 | 1 |

| | | | | | | | | | | | |
|---|---|---|---|---|---|---|---|---|---|---|---|
| rs11606672 | 11 | 117056059 | T | C | -1.53 | 0.126 | -? | 0 | 0 | 0 | 1 |
| rs675331 | 11 | 119927944 | G | A | -1.53 | 0.126 | -? | 0 | 0 | 0 | 1 |
| rs12418126 | 11 | 117336961 | G | A | 1.529 | 0.1262 | +? | 0 | 0 | 0 | 1 |
| rs11215936 | 11 | 116313516 | A | G | 1.529 | 0.1263 | +? | 0 | 0 | 0 | 1 |
| rs1203387394 | 11 | 117957631 | C | G | -1.529 | 0.1263 | -? | 0 | 0 | 0 | 1 |
| rs7951761 | 11 | 116467853 | G | A | -1.528 | 0.1264 | -? | 0 | 0 | 0 | 1 |
| rs11217366 | 11 | 119502599 | T | C | -1.527 | 0.1267 | -? | 0 | 0 | 0 | 1 |
| rs113419013 | 11 | 116870420 | T | C | -1.527 | 0.1268 | -? | 0 | 0 | 0 | 1 |
| rs514112 | 11 | 116330610 | G | A | -1.526 | 0.127 | -? | 0 | 0 | 0 | 1 |
| rs675873 | 11 | 119246708 | C | T | 1.526 | 0.127 | +? | 0 | 0 | 0 | 1 |
| rs918144 | 11 | 116633825 | T | C | -1.526 | 0.127 | -+ | 0 | 0.829 | 1 | 0.3627 |
| rs61588877 | 11 | 120792086 | G | A | -1.526 | 0.1271 | -? | 0 | 0 | 0 | 1 |
| rs118056960 | 11 | 120568248 | T | C | 1.525 | 0.1273 | +? | 0 | 0 | 0 | 1 |
| rs12288261 | 11 | 120568153 | T | C | 1.525 | 0.1273 | +? | 0 | 0 | 0 | 1 |
| rs12295200 | 11 | 120568090 | C | T | 1.525 | 0.1273 | +? | 0 | 0 | 0 | 1 |
| rs61901360 | 11 | 120568790 | T | C | 1.525 | 0.1273 | +? | 0 | 0 | 0 | 1 |
| rs78634274 | 11 | 120568318 | C | T | 1.525 | 0.1273 | +? | 0 | 0 | 0 | 1 |
| rs12291927 | 11 | 120570080 | G | C | 1.524 | 0.1274 | +? | 0 | 0 | 0 | 1 |
| rs10892468 | 11 | 119676471 | T | G | 1.523 | 0.1277 | ++ | 0 | 0.147 | 1 | 0.7017 |
| rs12293393 | 11 | 120570404 | G | C | 1.523 | 0.1277 | +? | 0 | 0 | 0 | 1 |
| rs11600946 | 11 | 120420262 | A | G | 1.522 | 0.1279 | +? | 0 | 0 | 0 | 1 |
| rs557501 | 11 | 120851221 | C | G | 1.522 | 0.1279 | +? | 0 | 0 | 0 | 1 |
| rs71480323 | 11 | 116771600 | A | G | -1.522 | 0.1279 | -? | 0 | 0 | 0 | 1 |
| rs76365645 | 11 | 119499578 | A | G | -1.522 | 0.1279 | -? | 0 | 0 | 0 | 1 |
| rs57550545 | 11 | 117560679 | C | T | 1.521 | 0.1282 | +? | 0 | 0 | 0 | 1 |
| rs627999 | 11 | 117992155 | T | C | -1.52 | 0.1285 | -? | 0 | 0 | 0 | 1 |
| rs1941410 | 11 | 120171165 | A | G | -1.52 | 0.1286 | -? | 0 | 0 | 0 | 1 |
| rs3087563 | 11 | 117707937 | C | T | -1.519 | 0.1288 | -? | 0 | 0 | 0 | 1 |
| rs568922 | 11 | 118919206 | T | C | -1.519 | 0.1288 | -? | 0 | 0 | 0 | 1 |
| rs601102 | 11 | 121057021 | C | T | 1.519 | 0.1288 | +? | 0 | 0 | 0 | 1 |
| rs61901361 | 11 | 120569355 | G | A | 1.519 | 0.1288 | +? | 0 | 0 | 0 | 1 |
| rs553637 | 11 | 117461602 | C | T | 1.518 | 0.1289 | +? | 0 | 0 | 0 | 1 |
| rs10892174 | 11 | 117672565 | C | A | 1.518 | 0.1291 | +? | 0 | 0 | 0 | 1 |
| rs588209 | 11 | 121081509 | G | C | 1.518 | 0.1291 | +? | 0 | 0 | 0 | 1 |
| rs592565 | 11 | 119939927 | A | G | 1.518 | 0.1291 | +? | 0 | 0 | 0 | 1 |
| rs7110845 | 11 | 120175749 | A | G | -1.518 | 0.1291 | -? | 0 | 0 | 0 | 1 |
| rs609272 | 11 | 117201682 | T | C | -1.517 | 0.1292 | -? | 0 | 0 | 0 | 1 |
| rs17245580 | 11 | 120480883 | G | A | -1.517 | 0.1293 | -? | 0 | 0 | 0 | 1 |
| rs947891 | 11 | 117944519 | T | C | 1.516 | 0.1295 | +- | -21.3 | 2.473 | 1 | 0.1158 |
| rs1989151 | 11 | 119926775 | A | T | -1.516 | 0.1296 | -? | 0 | 0 | 0 | 1 |
| rs1363950531 | 11 | 117461875 | G | T | 1.515 | 0.1298 | +? | 0 | 0 | 0 | 1 |
| rs481703 | 11 | 121055740 | C | T | 1.515 | 0.1298 | +? | 0 | 0 | 0 | 1 |
| rs11217556 | 11 | 119768054 | A | C | 1.514 | 0.1299 | +? | 0 | 0 | 0 | 1 |
| rs12576226 | 11 | 117464025 | G | A | 1.514 | 0.1299 | +? | 0 | 0 | 0 | 1 |
| rs2305828 | 11 | 117251143 | A | G | 1.514 | 0.1299 | +? | 0 | 0 | 0 | 1 |
| rs1893023 | 11 | 121004151 | T | C | 1.514 | 0.13 | +? | 0 | 0 | 0 | 1 |
| rs111954335 | 11 | 119797706 | T | C | -1.513 | 0.1302 | -? | 0 | 0 | 0 | 1 |
| rs112293704 | 11 | 119799826 | G | T | -1.513 | 0.1302 | -? | 0 | 0 | 0 | 1 |
| rs112297727 | 11 | 119797684 | T | C | -1.513 | 0.1302 | -? | 0 | 0 | 0 | 1 |
| rs74840179 | 11 | 119798592 | C | G | -1.513 | 0.1302 | -? | 0 | 0 | 0 | 1 |
| rs75417087 | 11 | 119801401 | T | G | -1.513 | 0.1302 | -? | 0 | 0 | 0 | 1 |
| rs754871 | 11 | 119802186 | G | A | -1.513 | 0.1302 | -? | 0 | 0 | 0 | 1 |
| rs75640250 | 11 | 119798205 | A | C | -1.513 | 0.1302 | -? | 0 | 0 | 0 | 1 |
| rs75748285 | 11 | 119798123 | C | T | -1.513 | 0.1302 | -? | 0 | 0 | 0 | 1 |
| rs76777718 | 11 | 119798178 | A | G | -1.513 | 0.1302 | -? | 0 | 0 | 0 | 1 |
| rs77109784 | 11 | 119797995 | G | A | -1.513 | 0.1302 | -? | 0 | 0 | 0 | 1 |

| | | | | | | | | | | | |
|---|---|---|---|---|---|---|---|---|---|---|---|
| rs77400160 | 11 | 119799196 | T | C | -1.513 | 0.1302 | -? | 0 | 0 | 0 | 1 |
| rs78178500 | 11 | 119800547 | A | C | -1.513 | 0.1302 | -? | 0 | 0 | 0 | 1 |
| rs78763435 | 11 | 119801962 | A | T | -1.513 | 0.1302 | -? | 0 | 0 | 0 | 1 |
| rs79048103 | 11 | 119801279 | T | C | -1.513 | 0.1302 | -? | 0 | 0 | 0 | 1 |
| rs79592175 | 11 | 119801704 | C | G | -1.513 | 0.1302 | -? | 0 | 0 | 0 | 1 |
| rs80166158 | 11 | 119799219 | A | G | -1.513 | 0.1302 | -? | 0 | 0 | 0 | 1 |
| rs11216505 | 11 | 117563295 | A | G | 1.513 | 0.1303 | +? | 0 | 0 | 0 | 1 |
| rs1447149 | 11 | 119691069 | A | G | 1.512 | 0.1305 | +? | 0 | 0 | 0 | 1 |
| rs73010304 | 11 | 117391522 | A | G | -1.512 | 0.1306 | -? | 0 | 0 | 0 | 1 |
| rs78699173 | 11 | 119922708 | A | C | -1.512 | 0.1306 | -? | 0 | 0 | 0 | 1 |
| rs113685600 | 11 | 120899875 | G | C | 1.511 | 0.1307 | +? | 0 | 0 | 0 | 1 |
| rs11216466 | 11 | 117474580 | A | C | 1.511 | 0.1308 | +? | 0 | 0 | 0 | 1 |
| rs11828031 | 11 | 119679659 | A | G | 1.511 | 0.1308 | +? | 0 | 0 | 0 | 1 |
| rs58047799 | 11 | 120489586 | G | T | -1.511 | 0.1308 | -? | 0 | 0 | 0 | 1 |
| rs10750179 | 11 | 120864593 | A | G | 1.511 | 0.1309 | +? | 0 | 0 | 0 | 1 |
| rs73012582 | 11 | 117500448 | A | G | -1.511 | 0.1309 | -? | 0 | 0 | 0 | 1 |
| rs541948 | 11 | 116470344 | T | G | -1.51 | 0.131 | -? | 0 | 0 | 0 | 1 |
| rs61901366 | 11 | 120570786 | A | G | 1.51 | 0.131 | +? | 0 | 0 | 0 | 1 |
| rs2155365 | 11 | 121013174 | C | A | 1.51 | 0.1311 | +? | 0 | 0 | 0 | 1 |
| rs60969913 | 11 | 118798154 | C | G | 1.51 | 0.1311 | +- | -37.4 | 2.183 | 1 | 0.1395 |
| rs11217360 | 11 | 119498563 | T | C | -1.509 | 0.1312 | -? | 0 | 0 | 0 | 1 |
| rs11215937 | 11 | 116313601 | G | A | 1.508 | 0.1316 | +? | 0 | 0 | 0 | 1 |
| rs7130633 | 11 | 119636704 | A | C | 1.507 | 0.1317 | +? | 0 | 0 | 0 | 1 |
| rs983567 | 11 | 116783600 | C | T | -1.507 | 0.1317 | -? | 0 | 0 | 0 | 1 |
| rs12721090 | 11 | 116700775 | T | C | -1.507 | 0.1318 | -? | 0 | 0 | 0 | 1 |
| rs7950396 | 11 | 117501271 | A | G | -1.507 | 0.1318 | -? | 0 | 0 | 0 | 1 |
| rs3889571 | 11 | 117300067 | T | G | -1.505 | 0.1322 | -? | 0 | 0 | 0 | 1 |
| rs1944518 | 11 | 120843742 | T | C | -1.505 | 0.1323 | -? | 0 | 0 | 0 | 1 |
| rs7103685 | 11 | 119523181 | C | T | -1.505 | 0.1323 | -? | 0 | 0 | 0 | 1 |
| rs7940638 | 11 | 117297415 | A | C | 1.505 | 0.1323 | +? | 0 | 0 | 0 | 1 |
| rs61901367 | 11 | 120570901 | A | G | 1.505 | 0.1324 | +? | 0 | 0 | 0 | 1 |
| rs7123336 | 11 | 116782819 | C | T | -1.505 | 0.1324 | -? | 0 | 0 | 0 | 1 |
| rs56032119 | 11 | 118797544 | G | A | 1.504 | 0.1326 | +- | -38 | 2.174 | 1 | 0.1404 |
| rs7102603 | 11 | 120853675 | T | C | -1.504 | 0.1326 | -? | 0 | 0 | 0 | 1 |
| rs6589549 | 11 | 116364066 | A | G | 1.504 | 0.1327 | +? | 0 | 0 | 0 | 1 |
| rs12285200 | 11 | 117564891 | G | A | 1.503 | 0.1328 | +? | 0 | 0 | 0 | 1 |
| rs3016346 | 11 | 116486588 | A | G | 1.503 | 0.1328 | +? | 0 | 0 | 0 | 1 |
| rs603446 | 11 | 116654435 | T | C | 1.503 | 0.1328 | ++ | 0 | 0.005 | 1 | 0.9412 |
| rs77643000 | 11 | 118749563 | C | T | -1.503 | 0.1329 | -+ | -129 | 1.31 | 1 | 0.2524 |
| rs7948548 | 11 | 116360293 | A | G | -1.503 | 0.1329 | -? | 0 | 0 | 0 | 1 |
| rs149505626 | 11 | 120571229 | A | G | 1.502 | 0.133 | +? | 0 | 0 | 0 | 1 |
| rs603455 | 11 | 117958429 | T | A | -1.502 | 0.133 | -? | 0 | 0 | 0 | 1 |
| rs11600436 | 11 | 120419868 | T | C | -1.5 | 0.1335 | -? | 0 | 0 | 0 | 1 |
| rs6589675 | 11 | 118569414 | C | G | 1.5 | 0.1335 | ++ | 0 | 0.604 | 1 | 0.4371 |
| rs111683933 | 11 | 116642049 | C | T | -1.5 | 0.1336 | -? | 0 | 0 | 0 | 1 |
| rs492257 | 11 | 116528249 | A | G | 1.5 | 0.1336 | +? | 0 | 0 | 0 | 1 |
| rs113876980 | 11 | 117667688 | A | C | 1.5 | 0.1337 | +? | 0 | 0 | 0 | 1 |
| rs11217913 | 11 | 120417553 | G | A | 1.499 | 0.1338 | +? | 0 | 0 | 0 | 1 |
| rs113271699 | 11 | 116650571 | A | C | -1.499 | 0.1339 | -? | 0 | 0 | 0 | 1 |
| rs4938440 | 11 | 117693971 | A | G | 1.498 | 0.1342 | +? | 0 | 0 | 0 | 1 |
| rs111390501 | 11 | 118797420 | G | A | 1.497 | 0.1344 | +- | -38.8 | 2.162 | 1 | 0.1415 |
| rs73575429 | 11 | 118797392 | C | T | 1.497 | 0.1344 | +- | -38.8 | 2.162 | 1 | 0.1415 |
| rs10047462 | 11 | 116722041 | G | T | 1.496 | 0.1346 | +? | 0 | 0 | 0 | 1 |
| rs10892162 | 11 | 117561265 | G | A | 1.496 | 0.1346 | +? | 0 | 0 | 0 | 1 |
| rs74584888 | 11 | 118797278 | A | G | 1.496 | 0.1347 | +- | -38.9 | 2.159 | 1 | 0.1417 |
| rs592160 | 11 | 121049136 | T | C | 1.495 | 0.1348 | +? | 0 | 0 | 0 | 1 |

| | | | | | | | | | | | |
|---|---|---|---|---|---|---|---|---|---|---|---|
| rs61901369 | 11 | 120571702 | C | T | 1.495 | 0.1348 | +? | 0 | 0 | 0 | 1 |
| rs7125270 | 11 | 119682918 | T | C | 1.495 | 0.1349 | ++ | 0 | 0.218 | 1 | 0.6407 |
| rs4938305 | 11 | 116588629 | T | C | -1.495 | 0.135 | -- | 0 | 0.154 | 1 | 0.6951 |
| rs4409845 | 11 | 119582832 | G | A | 1.494 | 0.1353 | +? | 0 | 0 | 0 | 1 |
| rs10892175 | 11 | 117674355 | A | G | 1.492 | 0.1356 | +? | 0 | 0 | 0 | 1 |
| rs11217653 | 11 | 119913541 | A | G | -1.492 | 0.1356 | -? | 0 | 0 | 0 | 1 |
| rs680730 | 11 | 117475233 | T | C | -1.492 | 0.1356 | -? | 0 | 0 | 0 | 1 |
| rs4936569 | 11 | 120825635 | C | T | 1.492 | 0.1357 | +? | 0 | 0 | 0 | 1 |
| rs75736746 | 11 | 120790814 | G | A | -1.492 | 0.1357 | -? | 0 | 0 | 0 | 1 |
| rs2513094 | 11 | 116806412 | C | T | -1.492 | 0.1358 | -? | 0 | 0 | 0 | 1 |
| rs79533157 | 11 | 119922728 | C | G | -1.491 | 0.1359 | -? | 0 | 0 | 0 | 1 |
| rs620896 | 11 | 121050913 | G | A | 1.491 | 0.136 | +? | 0 | 0 | 0 | 1 |
| rs640332 | 11 | 121050748 | C | T | 1.491 | 0.136 | +? | 0 | 0 | 0 | 1 |
| rs73009445 | 11 | 118744615 | A | G | 1.491 | 0.1361 | +- | -77.2 | 1.693 | 1 | 0.1933 |
| rs7930264 | 11 | 116870766 | A | G | -1.49 | 0.1362 | -? | 0 | 0 | 0 | 1 |
| rs7120925 | 11 | 117912465 | T | C | -1.49 | 0.1363 | -? | 0 | 0 | 0 | 1 |
| rs61697909 | 11 | 117474845 | A | G | -1.489 | 0.1364 | -? | 0 | 0 | 0 | 1 |
| rs552657 | 11 | 116530209 | T | C | 1.489 | 0.1365 | +? | 0 | 0 | 0 | 1 |
| rs1893707 | 11 | 120999631 | C | A | 1.489 | 0.1366 | +? | 0 | 0 | 0 | 1 |
| rs12288712 | 11 | 117559219 | T | C | 1.487 | 0.1369 | +? | 0 | 0 | 0 | 1 |
| rs10790255 | 11 | 118515579 | G | T | -1.487 | 0.137 | -- | 0 | 0.152 | 1 | 0.697 |
| rs12272864 | 11 | 120564637 | C | T | 1.487 | 0.1371 | +? | 0 | 0 | 0 | 1 |
| rs552580 | 11 | 121031511 | G | A | 1.486 | 0.1372 | +? | 0 | 0 | 0 | 1 |
| rs677336 | 11 | 121030381 | T | A | 1.486 | 0.1372 | +? | 0 | 0 | 0 | 1 |
| rs552787 | 11 | 116530373 | C | A | 1.486 | 0.1374 | +? | 0 | 0 | 0 | 1 |
| rs11217077 | 11 | 118746223 | C | T | 1.485 | 0.1375 | ++ | 0 | 0.131 | 1 | 0.7172 |
| rs12802202 | 11 | 116677048 | C | T | 1.485 | 0.1375 | +? | 0 | 0 | 0 | 1 |
| rs61901376 | 11 | 120573725 | G | A | 1.484 | 0.1377 | +? | 0 | 0 | 0 | 1 |
| rs681454 | 11 | 119927648 | G | C | -1.484 | 0.1377 | -? | 0 | 0 | 0 | 1 |
| rs2269398 | 11 | 117069788 | T | C | -1.484 | 0.1378 | -? | 0 | 0 | 0 | 1 |
| rs73575428 | 11 | 118797003 | G | C | 1.484 | 0.1378 | +- | -40.2 | 2.14 | 1 | 0.1435 |
| rs509171 | 11 | 116848857 | T | C | -1.483 | 0.1382 | -? | 0 | 0 | 0 | 1 |
| rs1009365 | 11 | 119504952 | T | C | -1.482 | 0.1383 | -? | 0 | 0 | 0 | 1 |
| rs117512890 | 11 | 119592965 | C | T | 1.482 | 0.1383 | +? | 0 | 0 | 0 | 1 |
| rs562281 | 11 | 121039947 | T | A | 1.482 | 0.1383 | +? | 0 | 0 | 0 | 1 |
| rs682048 | 11 | 116828320 | G | T | -1.482 | 0.1383 | -? | 0 | 0 | 0 | 1 |
| rs34637368 | 11 | 116619605 | A | G | -1.482 | 0.1384 | -? | 0 | 0 | 0 | 1 |
| rs61904804 | 11 | 116432510 | C | G | -1.482 | 0.1384 | -? | 0 | 0 | 0 | 1 |
| rs715315 | 11 | 117656774 | T | C | -1.482 | 0.1384 | -? | 0 | 0 | 0 | 1 |
| rs688641 | 11 | 121030195 | T | G | 1.481 | 0.1385 | +? | 0 | 0 | 0 | 1 |
| rs662496 | 11 | 121033105 | G | A | 1.481 | 0.1386 | +? | 0 | 0 | 0 | 1 |
| rs1317814 | 11 | 120484810 | G | A | -1.48 | 0.1388 | -? | 0 | 0 | 0 | 1 |
| rs584355 | 11 | 121035186 | A | C | 1.48 | 0.1388 | +? | 0 | 0 | 0 | 1 |
| rs4375451 | 11 | 120497208 | T | C | -1.479 | 0.1392 | -? | 0 | 0 | 0 | 1 |
| rs480761 | 11 | 120986571 | A | G | -1.479 | 0.1392 | -? | 0 | 0 | 0 | 1 |
| rs59127761 | 11 | 120493714 | G | C | -1.478 | 0.1394 | -? | 0 | 0 | 0 | 1 |
| rs6589862 | 11 | 121098681 | A | G | 1.478 | 0.1395 | +? | 0 | 0 | 0 | 1 |
| rs676426 | 11 | 120871963 | A | G | 1.478 | 0.1395 | +? | 0 | 0 | 0 | 1 |
| rs11216930 | 11 | 118488782 | C | A | -1.477 | 0.1396 | -- | 0 | 0.183 | 1 | 0.6688 |
| rs588917 | 11 | 121013909 | T | G | 1.477 | 0.1397 | +? | 0 | 0 | 0 | 1 |
| rs2846109 | 11 | 120588100 | C | T | 1.477 | 0.1398 | +? | 0 | 0 | 0 | 1 |
| rs72996698 | 11 | 119886445 | T | C | -1.476 | 0.1399 | -? | 0 | 0 | 0 | 1 |
| rs1240768 | 11 | 116516653 | C | T | 1.476 | 0.14 | +? | 0 | 0 | 0 | 1 |
| rs140657166 | 11 | 119728423 | C | A | 1.476 | 0.14 | +? | 0 | 0 | 0 | 1 |
| rs11605327 | 11 | 116823062 | A | G | 1.475 | 0.1401 | +? | 0 | 0 | 0 | 1 |
| rs12792357 | 11 | 117666595 | C | T | -1.475 | 0.1402 | -? | 0 | 0 | 0 | 1 |

| | | | | | | | | | | | |
|---|---|---|---|---|---|---|---|---|---|---|---|
| rs4252314 | 11 | 117868517 | G | A | -1.475 | 0.1402 | -- | 0 | 0.002 | 1 | 0.9638 |
| rs60905351 | 11 | 116358593 | A | G | 1.475 | 0.1402 | +? | 0 | 0 | 0 | 1 |
| rs2170317 | 11 | 117070740 | C | A | -1.475 | 0.1403 | -? | 0 | 0 | 0 | 1 |
| rs7940044 | 11 | 117435148 | A | G | 1.475 | 0.1403 | +? | 0 | 0 | 0 | 1 |
| rs10750113 | 11 | 117632609 | G | A | 1.474 | 0.1404 | +? | 0 | 0 | 0 | 1 |
| rs10790205 | 11 | 117632645 | C | T | 1.474 | 0.1404 | +? | 0 | 0 | 0 | 1 |
| rs61904810 | 11 | 116436913 | T | C | -1.474 | 0.1404 | -? | 0 | 0 | 0 | 1 |
| rs6589583 | 11 | 116863766 | T | C | -1.474 | 0.1405 | -? | 0 | 0 | 0 | 1 |
| rs587443 | 11 | 117347400 | T | C | -1.474 | 0.1406 | -? | 0 | 0 | 0 | 1 |
| rs11216386 | 11 | 117309989 | T | C | -1.473 | 0.1407 | -? | 0 | 0 | 0 | 1 |
| rs60787798 | 11 | 120482493 | T | G | -1.473 | 0.1407 | -? | 0 | 0 | 0 | 1 |
| rs689264 | 11 | 117957258 | T | C | -1.472 | 0.1409 | -? | 0 | 0 | 0 | 1 |
| rs12279718 | 11 | 119503670 | A | C | -1.472 | 0.141 | -? | 0 | 0 | 0 | 1 |
| rs61901489 | 11 | 117741933 | C | G | 1.472 | 0.141 | +? | 0 | 0 | 0 | 1 |
| rs73002650 | 11 | 120492364 | G | A | -1.472 | 0.1411 | -? | 0 | 0 | 0 | 1 |
| rs76635827 | 11 | 120529153 | C | G | 1.471 | 0.1412 | +? | 0 | 0 | 0 | 1 |
| rs61905348 | 11 | 117647145 | T | G | -1.471 | 0.1414 | -? | 0 | 0 | 0 | 1 |
| rs72999492 | 11 | 118539746 | C | A | -1.471 | 0.1414 | -- | 0 | 0.056 | 1 | 0.8131 |
| rs4344529 | 11 | 117632375 | G | A | 1.469 | 0.1417 | +? | 0 | 0 | 0 | 1 |
| rs1621211 | 11 | 120873076 | A | G | 1.469 | 0.1418 | +? | 0 | 0 | 0 | 1 |
| rs12797352 | 11 | 119518006 | G | A | -1.468 | 0.142 | -? | 0 | 0 | 0 | 1 |
| rs4442562 | 11 | 118845813 | A | G | -1.468 | 0.142 | -? | 0 | 0 | 0 | 1 |
| rs61901399 | 11 | 120586773 | T | C | 1.468 | 0.142 | +? | 0 | 0 | 0 | 1 |
| rs112444974 | 11 | 117764401 | T | G | -1.468 | 0.1421 | -? | 0 | 0 | 0 | 1 |
| rs17121096 | 11 | 117569923 | C | T | 1.468 | 0.1421 | +? | 0 | 0 | 0 | 1 |
| rs71484168 | 11 | 120309460 | T | C | 1.467 | 0.1423 | +? | 0 | 0 | 0 | 1 |
| rs141116752 | 11 | 117541718 | T | C | 1.467 | 0.1424 | +? | 0 | 0 | 0 | 1 |
| rs73580434 | 11 | 116299637 | T | A | -1.467 | 0.1424 | -? | 0 | 0 | 0 | 1 |
| rs60668211 | 11 | 117502726 | A | G | -1.467 | 0.1425 | -? | 0 | 0 | 0 | 1 |
| rs7112573 | 11 | 117069023 | C | T | -1.467 | 0.1425 | -? | 0 | 0 | 0 | 1 |
| rs10790375 | 11 | 120174671 | C | T | -1.466 | 0.1426 | -? | 0 | 0 | 0 | 1 |
| rs10892163 | 11 | 117566303 | A | G | 1.466 | 0.1427 | +? | 0 | 0 | 0 | 1 |
| rs111259707 | 11 | 119540705 | C | T | 1.466 | 0.1427 | +? | 0 | 0 | 0 | 1 |
| rs492294 | 11 | 121014403 | G | A | 1.466 | 0.1427 | +? | 0 | 0 | 0 | 1 |
| rs11827682 | 11 | 116700785 | T | C | -1.465 | 0.1428 | -? | 0 | 0 | 0 | 1 |
| rs755791 | 11 | 117433686 | C | A | 1.465 | 0.1429 | +? | 0 | 0 | 0 | 1 |
| rs10790206 | 11 | 117632764 | C | T | 1.465 | 0.143 | +? | 0 | 0 | 0 | 1 |
| rs10790207 | 11 | 117632769 | C | T | 1.465 | 0.143 | +? | 0 | 0 | 0 | 1 |
| rs520868 | 11 | 121082988 | C | T | 1.465 | 0.143 | +? | 0 | 0 | 0 | 1 |
| rs475731 | 11 | 121004224 | T | C | -1.464 | 0.1433 | -? | 0 | 0 | 0 | 1 |
| rs73012879 | 11 | 117973145 | A | G | -1.464 | 0.1433 | -? | 0 | 0 | 0 | 1 |
| rs1240778 | 11 | 116536490 | C | A | -1.463 | 0.1435 | -? | 0 | 0 | 0 | 1 |
| rs1629083 | 11 | 118126576 | T | C | 1.463 | 0.1436 | +? | 0 | 0 | 0 | 1 |
| rs565296 | 11 | 119252151 | C | T | 1.462 | 0.1437 | ++ | 0 | 0.722 | 1 | 0.3955 |
| rs78204624 | 11 | 117883397 | C | G | -1.462 | 0.1438 | -? | 0 | 0 | 0 | 1 |
| rs11600156 | 11 | 116431149 | G | A | -1.461 | 0.1439 | -? | 0 | 0 | 0 | 1 |
| rs7350 | 11 | 118965048 | A | G | -1.461 | 0.1439 | -? | 0 | 0 | 0 | 1 |
| rs582876 | 11 | 121029720 | C | T | 1.461 | 0.144 | +? | 0 | 0 | 0 | 1 |
| rs11216506 | 11 | 117564052 | G | A | 1.46 | 0.1442 | +? | 0 | 0 | 0 | 1 |
| rs2239009 | 11 | 117089722 | C | G | -1.46 | 0.1443 | -? | 0 | 0 | 0 | 1 |
| rs4558175 | 11 | 118872581 | A | G | -1.46 | 0.1444 | -- | 0 | 0.058 | 1 | 0.8101 |
| rs656439 | 11 | 119250874 | C | T | 1.459 | 0.1446 | ++ | 0 | 0.54 | 1 | 0.4625 |
| rs10502221 | 11 | 116826870 | T | C | 1.459 | 0.1447 | +? | 0 | 0 | 0 | 1 |
| rs180371 | 11 | 116592203 | G | A | -1.458 | 0.1449 | -- | 0 | 0.094 | 1 | 0.7587 |
| rs633985 | 11 | 117460030 | G | C | 1.457 | 0.1452 | +? | 0 | 0 | 0 | 1 |
| rs73002644 | 11 | 120488117 | G | C | -1.457 | 0.1452 | -? | 0 | 0 | 0 | 1 |

| | | | | | | | | | | | |
|---|---|---|---|---|---|---|---|---|---|---|---|
| rs2513097 | 11 | 116836212 | G | A | -1.456 | 0.1454 | -? | 0 | 0 | 0 | 1 |
| rs34913450 | 11 | 118020130 | C | T | 1.456 | 0.1455 | +? | 0 | 0 | 0 | 1 |
| rs76014645 | 11 | 117795717 | A | C | 1.456 | 0.1455 | +? | 0 | 0 | 0 | 1 |
| rs4936386 | 11 | 117430750 | T | C | 1.455 | 0.1456 | +? | 0 | 0 | 0 | 1 |
| rs560564 | 11 | 117195628 | C | A | 1.454 | 0.1459 | +? | 0 | 0 | 0 | 1 |
| rs75336872 | 11 | 117793169 | T | C | 1.454 | 0.1459 | +? | 0 | 0 | 0 | 1 |
| rs35314805 | 11 | 117463914 | C | T | 1.453 | 0.1461 | +? | 0 | 0 | 0 | 1 |
| rs6589834 | 11 | 120583174 | C | G | 1.452 | 0.1464 | +? | 0 | 0 | 0 | 1 |
| rs693192 | 11 | 116355678 | A | G | -1.452 | 0.1464 | -? | 0 | 0 | 0 | 1 |
| rs2239678 | 11 | 117070596 | T | C | -1.452 | 0.1465 | -? | 0 | 0 | 0 | 1 |
| rs2269399 | 11 | 117066353 | C | G | -1.452 | 0.1465 | -? | 0 | 0 | 0 | 1 |
| rs599894 | 11 | 121017741 | C | G | 1.452 | 0.1465 | +? | 0 | 0 | 0 | 1 |
| rs520328 | 11 | 117329415 | A | G | 1.451 | 0.1467 | +? | 0 | 0 | 0 | 1 |
| rs636575 | 11 | 120988600 | C | A | -1.451 | 0.1467 | -? | 0 | 0 | 0 | 1 |
| rs7108971 | 11 | 121106570 | A | T | 1.451 | 0.1468 | +? | 0 | 0 | 0 | 1 |
| rs78628783 | 11 | 117092051 | T | C | -1.451 | 0.1468 | -? | 0 | 0 | 0 | 1 |
| rs11216315 | 11 | 117080640 | G | C | -1.451 | 0.1469 | -? | 0 | 0 | 0 | 1 |
| rs11217068 | 11 | 118729283 | T | C | -1.451 | 0.1469 | -- | 0 | 0.057 | 1 | 0.811 |
| rs45499491 | 11 | 117965421 | T | C | -1.45 | 0.1471 | -? | 0 | 0 | 0 | 1 |
| rs512608 | 11 | 119253797 | G | A | 1.449 | 0.1472 | ++ | 0 | 0.705 | 1 | 0.4013 |
| rs1893394 | 11 | 119881526 | A | G | 1.449 | 0.1473 | +? | 0 | 0 | 0 | 1 |
| rs4935744 | 11 | 120496704 | G | T | -1.449 | 0.1473 | -? | 0 | 0 | 0 | 1 |
| rs61901377 | 11 | 120574079 | G | A | 1.448 | 0.1475 | +? | 0 | 0 | 0 | 1 |
| rs11217361 | 11 | 119499255 | C | G | -1.448 | 0.1476 | -? | 0 | 0 | 0 | 1 |
| rs34836631 | 11 | 117681047 | A | G | 1.448 | 0.1476 | +? | 0 | 0 | 0 | 1 |
| rs2010473 | 11 | 117433739 | C | T | 1.448 | 0.1477 | +? | 0 | 0 | 0 | 1 |
| rs117165966 | 11 | 120572772 | C | G | 1.447 | 0.1478 | +? | 0 | 0 | 0 | 1 |
| rs142600127 | 11 | 120572814 | A | G | 1.447 | 0.1478 | +? | 0 | 0 | 0 | 1 |
| rs1789403 | 11 | 116516506 | C | T | 1.447 | 0.1478 | +? | 0 | 0 | 0 | 1 |
| rs510722 | 11 | 121040370 | A | T | 1.447 | 0.1478 | +? | 0 | 0 | 0 | 1 |
| rs12280641 | 11 | 120989762 | A | G | 1.447 | 0.148 | +? | 0 | 0 | 0 | 1 |
| rs12421706 | 11 | 117403727 | T | C | 1.446 | 0.1481 | +? | 0 | 0 | 0 | 1 |
| rs626776 | 11 | 119028813 | C | G | -1.446 | 0.1481 | -? | 0 | 0 | 0 | 1 |
| rs7947185 | 11 | 118214726 | C | T | 1.446 | 0.1481 | +? | 0 | 0 | 0 | 1 |
| rs531184 | 11 | 117112526 | T | G | 1.446 | 0.1482 | +? | 0 | 0 | 0 | 1 |
| rs10892320 | 11 | 118873385 | T | C | -1.446 | 0.1483 | -- | 0 | 0.057 | 1 | 0.8109 |
| rs10892629 | 11 | 120716019 | T | C | 1.446 | 0.1483 | +? | 0 | 0 | 0 | 1 |
| rs7120237 | 11 | 120161995 | C | T | -1.444 | 0.1486 | -? | 0 | 0 | 0 | 1 |
| rs12418722 | 11 | 117320430 | T | C | 1.444 | 0.1488 | +? | 0 | 0 | 0 | 1 |
| rs11218079 | 11 | 120825043 | T | C | -1.443 | 0.149 | -? | 0 | 0 | 0 | 1 |
| rs1289103681 | 11 | 116414488 | C | T | -1.443 | 0.149 | -? | 0 | 0 | 0 | 1 |
| rs11217524 | 11 | 119692871 | C | T | 1.443 | 0.1491 | +? | 0 | 0 | 0 | 1 |
| rs11824545 | 11 | 117646692 | T | G | -1.442 | 0.1493 | -? | 0 | 0 | 0 | 1 |
| rs2855068 | 11 | 118954805 | T | C | -1.442 | 0.1493 | -? | 0 | 0 | 0 | 1 |
| rs600638 | 11 | 117957750 | T | C | -1.442 | 0.1493 | -? | 0 | 0 | 0 | 1 |
| rs80351364 | 11 | 117092394 | T | A | -1.442 | 0.1494 | -? | 0 | 0 | 0 | 1 |
| rs78566287 | 11 | 120412053 | T | C | 1.441 | 0.1495 | +? | 0 | 0 | 0 | 1 |
| rs180378 | 11 | 116588909 | A | G | -1.44 | 0.1498 | -+ | -190.6 | 1.032 | 1 | 0.3096 |
| rs61901372 | 11 | 120572387 | C | T | 1.44 | 0.1498 | +? | 0 | 0 | 0 | 1 |
| rs61901374 | 11 | 120572952 | C | T | 1.44 | 0.1498 | +? | 0 | 0 | 0 | 1 |
| rs61901375 | 11 | 120573265 | C | T | 1.44 | 0.1498 | +? | 0 | 0 | 0 | 1 |
| rs7124184 | 11 | 117655063 | T | C | 1.44 | 0.1498 | +? | 0 | 0 | 0 | 1 |
| rs78201038 | 11 | 119499316 | G | C | -1.44 | 0.1498 | -? | 0 | 0 | 0 | 1 |
| rs11216507 | 11 | 117565069 | C | T | 1.44 | 0.1499 | +? | 0 | 0 | 0 | 1 |
| rs1941635 | 11 | 117982495 | T | G | -1.44 | 0.15 | -? | 0 | 0 | 0 | 1 |
| rs527878 | 11 | 119793024 | G | A | 1.44 | 0.15 | +? | 0 | 0 | 0 | 1 |

| | | | | | | | | | | | |
|---|---|---|---|---|---|---|---|---|---|---|---|
| rs12295113 | 11 | 120577100 | T | C | 1.438 | 0.1503 | +? | 0 | 0 | 0 | 1 |
| rs79669568 | 11 | 117795611 | C | T | 1.438 | 0.1503 | +? | 0 | 0 | 0 | 1 |
| rs12277355 | 11 | 118729442 | T | C | -1.438 | 0.1504 | -- | 0 | 0.029 | 1 | 0.8638 |
| rs658654 | 11 | 119248320 | C | T | 1.438 | 0.1504 | ++ | 0 | 0.832 | 1 | 0.3617 |
| rs516226 | 11 | 116519655 | T | C | 1.438 | 0.1505 | +? | 0 | 0 | 0 | 1 |
| rs45540840 | 11 | 118486110 | A | G | -1.437 | 0.1507 | -- | 0 | 0.192 | 1 | 0.6613 |
| rs79892107 | 11 | 116306324 | T | C | -1.436 | 0.1509 | -? | 0 | 0 | 0 | 1 |
| rs113650932 | 11 | 117766665 | T | C | -1.435 | 0.1513 | -? | 0 | 0 | 0 | 1 |
| rs670906 | 11 | 119252199 | T | G | 1.435 | 0.1513 | ++ | 0 | 0.56 | 1 | 0.4543 |
| rs74385934 | 11 | 117068444 | A | G | -1.435 | 0.1514 | -? | 0 | 0 | 0 | 1 |
| rs1317815 | 11 | 120484712 | A | G | -1.434 | 0.1515 | -? | 0 | 0 | 0 | 1 |
| rs1892902 | 11 | 117669794 | T | G | -1.434 | 0.1517 | -? | 0 | 0 | 0 | 1 |
| rs628269 | 11 | 117926735 | G | A | -1.434 | 0.1517 | -? | 0 | 0 | 0 | 1 |
| rs7122817 | 11 | 117657679 | A | G | -1.434 | 0.1517 | -? | 0 | 0 | 0 | 1 |
| rs1729409 | 11 | 116673768 | A | G | -1.433 | 0.1518 | -- | 0 | 0.456 | 1 | 0.4996 |
| rs7102128 | 11 | 120505235 | A | G | 1.433 | 0.1519 | +? | 0 | 0 | 0 | 1 |
| rs61901371 | 11 | 120572361 | A | G | 1.433 | 0.152 | +? | 0 | 0 | 0 | 1 |
| rs6589805 | 11 | 120161671 | C | T | -1.433 | 0.152 | -? | 0 | 0 | 0 | 1 |
| rs2848729 | 11 | 119881097 | T | C | 1.432 | 0.1521 | +? | 0 | 0 | 0 | 1 |
| rs34163098 | 11 | 119509118 | A | G | -1.432 | 0.1521 | -? | 0 | 0 | 0 | 1 |
| rs488766 | 11 | 120990239 | G | A | -1.432 | 0.1522 | -? | 0 | 0 | 0 | 1 |
| rs55945170 | 11 | 117572395 | G | A | 1.431 | 0.1524 | +? | 0 | 0 | 0 | 1 |
| rs633770 | 11 | 121003606 | T | G | -1.431 | 0.1524 | -? | 0 | 0 | 0 | 1 |
| rs10750184 | 11 | 121160481 | A | G | -1.431 | 0.1525 | -? | 0 | 0 | 0 | 1 |
| rs11216375 | 11 | 117266094 | A | G | -1.431 | 0.1525 | -? | 0 | 0 | 0 | 1 |
| rs553370 | 11 | 117351433 | G | A | 1.43 | 0.1527 | +? | 0 | 0 | 0 | 1 |
| rs113300825 | 11 | 119544722 | T | C | -1.429 | 0.1529 | -? | 0 | 0 | 0 | 1 |
| rs633012 | 11 | 121032517 | A | G | 1.429 | 0.1531 | +? | 0 | 0 | 0 | 1 |
| rs1329232240 | 11 | 120159540 | C | T | -1.428 | 0.1532 | -? | 0 | 0 | 0 | 1 |
| rs654176 | 11 | 117961309 | G | C | -1.428 | 0.1532 | -? | 0 | 0 | 0 | 1 |
| rs7951628 | 11 | 120161117 | A | G | -1.428 | 0.1532 | -? | 0 | 0 | 0 | 1 |
| rs11601168 | 11 | 116571138 | A | T | -1.427 | 0.1536 | -? | 0 | 0 | 0 | 1 |
| rs2155854 | 11 | 118026675 | T | C | -1.427 | 0.1536 | -? | 0 | 0 | 0 | 1 |
| rs12221682 | 11 | 116578217 | G | C | 1.426 | 0.1539 | +? | 0 | 0 | 0 | 1 |
| rs11217785 | 11 | 120152591 | T | C | -1.426 | 0.154 | -? | 0 | 0 | 0 | 1 |
| rs12361671 | 11 | 117798560 | A | T | 1.426 | 0.154 | +? | 0 | 0 | 0 | 1 |
| rs2155810 | 11 | 120163869 | C | T | -1.425 | 0.1541 | -? | 0 | 0 | 0 | 1 |
| rs12278730 | 11 | 120423569 | T | C | 1.425 | 0.1542 | +? | 0 | 0 | 0 | 1 |
| rs12363277 | 11 | 118763736 | C | T | -1.425 | 0.1542 | -+ | -22.4 | 2.451 | 1 | 0.1175 |
| rs59781045 | 11 | 117095283 | T | C | -1.425 | 0.1543 | -? | 0 | 0 | 0 | 1 |
| rs527258 | 11 | 121027240 | A | T | 1.423 | 0.1546 | +? | 0 | 0 | 0 | 1 |
| rs77802172 | 11 | 120149184 | A | G | -1.423 | 0.1546 | -? | 0 | 0 | 0 | 1 |
| rs12789310 | 11 | 116197149 | G | A | 1.423 | 0.1547 | +- | -1.3 | 2.962 | 1 | 0.08525 |
| rs7128690 | 11 | 120521504 | T | G | 1.423 | 0.1547 | +? | 0 | 0 | 0 | 1 |
| rs10892687 | 11 | 121059141 | A | G | 1.423 | 0.1548 | +? | 0 | 0 | 0 | 1 |
| rs12280027 | 11 | 116399067 | G | C | -1.423 | 0.1548 | -? | 0 | 0 | 0 | 1 |
| rs11605307 | 11 | 119931576 | T | C | -1.422 | 0.1549 | -? | 0 | 0 | 0 | 1 |
| rs61740780 | 11 | 117886499 | A | G | 1.422 | 0.1549 | +? | 0 | 0 | 0 | 1 |
| rs79959350 | 11 | 120025452 | G | A | 1.422 | 0.1549 | +? | 0 | 0 | 0 | 1 |
| rs906544 | 11 | 116374399 | C | T | 1.422 | 0.1549 | +? | 0 | 0 | 0 | 1 |
| rs1944154 | 11 | 116501409 | C | T | 1.422 | 0.155 | +? | 0 | 0 | 0 | 1 |
| rs608818 | 11 | 120993071 | T | C | -1.421 | 0.1552 | -? | 0 | 0 | 0 | 1 |
| rs1787680 | 11 | 116664776 | A | T | 1.421 | 0.1553 | ++ | 0 | 0.041 | 1 | 0.8402 |
| rs1241655 | 11 | 116837808 | T | G | -1.42 | 0.1556 | -? | 0 | 0 | 0 | 1 |
| rs10892020 | 11 | 116589652 | G | C | -1.419 | 0.1558 | -- | 0 | 0.239 | 1 | 0.6249 |
| rs523549 | 11 | 116511028 | A | G | -1.419 | 0.1558 | -? | 0 | 0 | 0 | 1 |

| | | | | | | | | | | | |
|---|---|---|---|---|---|---|---|---|---|---|---|
| rs7936362 | 11 | 117635223 | A | C | -1.419 | 0.1558 | -? | 0 | 0 | 0 | 1 |
| rs3016581 | 11 | 117106116 | A | G | 1.419 | 0.1559 | +? | 0 | 0 | 0 | 1 |
| rs4582985 | 11 | 120554041 | G | C | -1.419 | 0.1559 | -? | 0 | 0 | 0 | 1 |
| rs546916 | 11 | 121035860 | T | G | 1.419 | 0.1559 | +? | 0 | 0 | 0 | 1 |
| rs7121784 | 11 | 120159052 | A | C | -1.419 | 0.1559 | -? | 0 | 0 | 0 | 1 |
| rs1263149 | 11 | 116639941 | G | A | -1.419 | 0.156 | -- | 0 | 0.364 | 1 | 0.5461 |
| rs34463064 | 11 | 120563557 | T | C | -1.419 | 0.156 | -? | 0 | 0 | 0 | 1 |
| rs1670195 | 11 | 120008468 | C | G | 1.418 | 0.1562 | +? | 0 | 0 | 0 | 1 |
| rs80183540 | 11 | 117623674 | T | C | -1.417 | 0.1565 | -? | 0 | 0 | 0 | 1 |
| rs12225548 | 11 | 118535840 | G | C | -1.416 | 0.1566 | -- | 0 | 0.278 | 1 | 0.5978 |
| rs13377448 | 11 | 116307041 | T | G | -1.416 | 0.1567 | -? | 0 | 0 | 0 | 1 |
| rs139886195 | 11 | 119804760 | A | C | -1.416 | 0.1569 | -? | 0 | 0 | 0 | 1 |
| rs11216633 | 11 | 117800269 | G | C | 1.415 | 0.1572 | +? | 0 | 0 | 0 | 1 |
| rs11216634 | 11 | 117800328 | G | A | 1.415 | 0.1572 | +? | 0 | 0 | 0 | 1 |
| rs11217076 | 11 | 118745664 | G | T | 1.413 | 0.1576 | ++ | 0 | 0.149 | 1 | 0.6992 |
| rs2542061 | 11 | 116672231 | A | G | -1.413 | 0.1576 | -- | 0 | 0.456 | 1 | 0.4996 |
| rs659042 | 11 | 120997460 | A | C | -1.413 | 0.1576 | -? | 0 | 0 | 0 | 1 |
| rs7119967 | 11 | 117646768 | A | G | -1.413 | 0.1577 | -? | 0 | 0 | 0 | 1 |
| rs663031 | 11 | 121033277 | T | C | 1.412 | 0.1579 | +? | 0 | 0 | 0 | 1 |
| rs66633375 | 11 | 119772803 | A | T | 1.412 | 0.158 | +? | 0 | 0 | 0 | 1 |
| rs1263151 | 11 | 116641028 | C | T | -1.411 | 0.1581 | -- | 0 | 0.025 | 1 | 0.8744 |
| rs576271 | 11 | 121034992 | A | T | 1.411 | 0.1581 | +? | 0 | 0 | 0 | 1 |
| rs11216937 | 11 | 118507478 | T | C | -1.411 | 0.1583 | -- | 0 | 0.355 | 1 | 0.5514 |
| rs617692 | 11 | 120986933 | T | C | -1.411 | 0.1583 | -? | 0 | 0 | 0 | 1 |
| rs4936513 | 11 | 120157168 | A | T | -1.41 | 0.1585 | -? | 0 | 0 | 0 | 1 |
| rs664059 | 11 | 116642137 | C | T | -1.41 | 0.1586 | -- | 0 | 0.284 | 1 | 0.5938 |
| rs80015904 | 11 | 118516903 | A | G | -1.409 | 0.1588 | -- | 0 | 0.368 | 1 | 0.5443 |
| rs1339633702 | 11 | 117088082 | T | C | -1.409 | 0.1589 | -? | 0 | 0 | 0 | 1 |
| rs583842 | 11 | 121001345 | C | T | -1.409 | 0.1589 | -? | 0 | 0 | 0 | 1 |
| rs114005593 | 11 | 117345114 | C | G | -1.408 | 0.159 | -? | 0 | 0 | 0 | 1 |
| rs79297003 | 11 | 117344866 | T | C | -1.408 | 0.159 | -? | 0 | 0 | 0 | 1 |
| rs11820805 | 11 | 118888831 | C | T | -1.408 | 0.1592 | -- | 0 | 0.263 | 1 | 0.6078 |
| rs10892558 | 11 | 120139487 | A | G | -1.407 | 0.1593 | -? | 0 | 0 | 0 | 1 |
| rs12806755 | 11 | 116202789 | C | T | 1.408 | 0.1593 | +- | -0.4 | 2.988 | 1 | 0.08389 |
| rs7128589 | 11 | 119775102 | C | T | 1.407 | 0.1593 | +? | 0 | 0 | 0 | 1 |
| rs71482119 | 11 | 117722478 | G | T | -1.408 | 0.1593 | -- | 0 | 0.035 | 1 | 0.8514 |
| rs566269 | 11 | 116351511 | T | C | -1.407 | 0.1594 | -? | 0 | 0 | 0 | 1 |
| rs180377 | 11 | 116589127 | T | C | -1.406 | 0.1597 | -+ | 0 | 0.993 | 1 | 0.319 |
| rs631969 | 11 | 121065886 | T | C | 1.406 | 0.1598 | +? | 0 | 0 | 0 | 1 |
| rs4936533 | 11 | 120496664 | T | C | -1.405 | 0.1599 | -? | 0 | 0 | 0 | 1 |
| rs624176 | 11 | 119893391 | T | C | 1.405 | 0.1599 | +? | 0 | 0 | 0 | 1 |
| rs2508655 | 11 | 120423595 | C | G | 1.405 | 0.16 | +? | 0 | 0 | 0 | 1 |
| rs4537790 | 11 | 117661356 | T | C | -1.405 | 0.1601 | -? | 0 | 0 | 0 | 1 |
| rs55753209 | 11 | 117392722 | A | G | -1.405 | 0.1601 | -? | 0 | 0 | 0 | 1 |
| rs61901400 | 11 | 120590901 | A | G | 1.404 | 0.1602 | +? | 0 | 0 | 0 | 1 |
| rs2174880 | 11 | 116304983 | G | A | 1.404 | 0.1604 | +? | 0 | 0 | 0 | 1 |
| rs632121 | 11 | 121003286 | T | A | -1.404 | 0.1604 | -? | 0 | 0 | 0 | 1 |
| rs511401 | 11 | 119250060 | C | G | 1.403 | 0.1605 | ++ | 0 | 0.587 | 1 | 0.4434 |
| rs12365095 | 11 | 118516774 | T | C | -1.403 | 0.1606 | -- | 0 | 0.278 | 1 | 0.5982 |
| rs12278740 | 11 | 120423601 | T | C | 1.403 | 0.1607 | +? | 0 | 0 | 0 | 1 |
| rs520805 | 11 | 121000774 | C | T | -1.403 | 0.1607 | -? | 0 | 0 | 0 | 1 |
| rs663435 | 11 | 121033374 | G | A | 1.403 | 0.1607 | +? | 0 | 0 | 0 | 1 |
| rs61906060 | 11 | 116498649 | G | A | 1.402 | 0.161 | +? | 0 | 0 | 0 | 1 |
| rs3889572 | 11 | 117300092 | T | C | -1.401 | 0.1611 | -? | 0 | 0 | 0 | 1 |
| rs10750170 | 11 | 120164102 | G | A | -1.401 | 0.1612 | -? | 0 | 0 | 0 | 1 |
| rs12280296 | 11 | 120570587 | G | A | 1.401 | 0.1613 | +? | 0 | 0 | 0 | 1 |

| | | | | | | | | | | | |
|---|---|---|---|---|---|---|---|---|---|---|---|
| rs117231130 | 11 | 116758658 | C | T | 1.4 | 0.1615 | +? | 0 | 0 | 0 | 1 |
| rs6589606 | 11 | 117092886 | T | C | -1.4 | 0.1615 | -? | 0 | 0 | 0 | 1 |
| rs675714 | 11 | 119880923 | A | G | 1.4 | 0.1616 | +? | 0 | 0 | 0 | 1 |
| rs12362152 | 11 | 118516737 | G | A | -1.399 | 0.1618 | -- | 0 | 0.259 | 1 | 0.6108 |
| rs11602764 | 11 | 118888619 | G | A | -1.398 | 0.1622 | -- | 0 | 0.262 | 1 | 0.6086 |
| rs11824502 | 11 | 117646682 | G | C | -1.398 | 0.1622 | -? | 0 | 0 | 0 | 1 |
| rs6589858 | 11 | 121021515 | C | T | 1.397 | 0.1624 | +? | 0 | 0 | 0 | 1 |
| rs12226676 | 11 | 116302939 | G | T | -1.397 | 0.1625 | -? | 0 | 0 | 0 | 1 |
| rs61906588 | 11 | 116362654 | T | C | -1.396 | 0.1628 | -? | 0 | 0 | 0 | 1 |
| rs12281528 | 11 | 116399336 | A | G | -1.395 | 0.1629 | -? | 0 | 0 | 0 | 1 |
| rs3017780 | 11 | 116352008 | G | T | -1.395 | 0.163 | -? | 0 | 0 | 0 | 1 |
| rs12273984 | 11 | 120570346 | C | T | 1.393 | 0.1636 | +? | 0 | 0 | 0 | 1 |
| rs11217514 | 11 | 119674096 | A | G | 1.392 | 0.1638 | ++ | 0 | 0.072 | 1 | 0.7883 |
| rs614192 | 11 | 119248709 | G | A | 1.392 | 0.1639 | ++ | 0 | 0.597 | 1 | 0.4395 |
| rs12271161 | 11 | 116979911 | A | G | -1.39 | 0.1645 | -? | 0 | 0 | 0 | 1 |
| rs17124632 | 11 | 120848717 | A | G | -1.39 | 0.1645 | -? | 0 | 0 | 0 | 1 |
| rs7934455 | 11 | 116359851 | C | T | -1.39 | 0.1645 | -? | 0 | 0 | 0 | 1 |
| rs11216938 | 11 | 118514625 | T | C | -1.39 | 0.1646 | -- | 0 | 0.243 | 1 | 0.6219 |
| rs684233 | 11 | 119927038 | A | G | 1.39 | 0.1646 | +? | 0 | 0 | 0 | 1 |
| rs665403 | 11 | 117307122 | T | C | -1.389 | 0.1647 | -? | 0 | 0 | 0 | 1 |
| rs495258 | 11 | 117507325 | T | C | -1.389 | 0.1648 | -? | 0 | 0 | 0 | 1 |
| rs73010609 | 11 | 119768716 | T | C | 1.389 | 0.1649 | +? | 0 | 0 | 0 | 1 |
| rs10790256 | 11 | 118534082 | T | C | -1.388 | 0.1651 | -- | 0 | 0.279 | 1 | 0.5975 |
| rs2155258 | 11 | 120657444 | G | A | 1.388 | 0.1652 | +? | 0 | 0 | 0 | 1 |
| rs61903859 | 11 | 117638305 | A | C | -1.388 | 0.1652 | -? | 0 | 0 | 0 | 1 |
| rs10790374 | 11 | 120157518 | C | T | -1.387 | 0.1653 | -? | 0 | 0 | 0 | 1 |
| rs11604214 | 11 | 119593446 | A | G | 1.387 | 0.1653 | +? | 0 | 0 | 0 | 1 |
| rs525390 | 11 | 117111214 | A | G | 1.387 | 0.1653 | +? | 0 | 0 | 0 | 1 |
| rs12273820 | 11 | 119805016 | C | T | -1.387 | 0.1654 | -? | 0 | 0 | 0 | 1 |
| rs1022080 | 11 | 119509625 | C | T | 1.386 | 0.1656 | +? | 0 | 0 | 0 | 1 |
| rs4938304 | 11 | 116588425 | T | C | -1.386 | 0.1656 | -? | 0 | 0 | 0 | 1 |
| rs17748 | 11 | 118528424 | T | C | -1.386 | 0.1659 | -- | 0 | 0.338 | 1 | 0.5612 |
| rs685425 | 11 | 117429728 | T | C | -1.385 | 0.166 | -? | 0 | 0 | 0 | 1 |
| rs12575869 | 11 | 117824206 | G | A | -1.385 | 0.1661 | -- | 0 | 0 | 1 | 0.992 |
| rs61903855 | 11 | 117605731 | C | G | -1.385 | 0.1661 | -? | 0 | 0 | 0 | 1 |
| rs7928371 | 11 | 118529127 | A | G | -1.385 | 0.1661 | -- | 0 | 0.25 | 1 | 0.6167 |
| rs651896 | 11 | 117961839 | T | C | -1.385 | 0.1662 | -? | 0 | 0 | 0 | 1 |
| rs11819854 | 11 | 117646805 | T | A | -1.384 | 0.1665 | -? | 0 | 0 | 0 | 1 |
| rs148092485 | 11 | 117397271 | A | T | -1.384 | 0.1665 | -? | 0 | 0 | 0 | 1 |
| rs2156633 | 11 | 120786892 | C | T | -1.384 | 0.1665 | -? | 0 | 0 | 0 | 1 |
| rs3825059 | 11 | 118973113 | A | C | 1.382 | 0.167 | +? | 0 | 0 | 0 | 1 |
| rs73022382 | 11 | 117352364 | A | G | -1.382 | 0.1671 | -? | 0 | 0 | 0 | 1 |
| rs112048745 | 11 | 117793222 | T | C | 1.381 | 0.1674 | +? | 0 | 0 | 0 | 1 |
| rs11217297 | 11 | 119334629 | A | G | 1.38 | 0.1676 | +? | 0 | 0 | 0 | 1 |
| rs10892657 | 11 | 120864806 | C | A | -1.38 | 0.1677 | -? | 0 | 0 | 0 | 1 |
| rs79988761 | 11 | 120126017 | A | G | 1.38 | 0.1677 | +? | 0 | 0 | 0 | 1 |
| rs2070668 | 11 | 116701153 | T | G | -1.379 | 0.1679 | -? | 0 | 0 | 0 | 1 |
| rs61907566 | 11 | 116748905 | G | C | 1.379 | 0.1679 | +? | 0 | 0 | 0 | 1 |
| rs4245211 | 11 | 120155303 | G | A | -1.379 | 0.168 | -? | 0 | 0 | 0 | 1 |
| rs73022498 | 11 | 117797120 | C | T | 1.379 | 0.168 | +? | 0 | 0 | 0 | 1 |
| rs549298 | 11 | 116354466 | G | A | -1.378 | 0.1683 | -? | 0 | 0 | 0 | 1 |
| rs11216583 | 11 | 117725057 | G | C | -1.377 | 0.1685 | -- | 0 | 0.042 | 1 | 0.8378 |
| rs4396299 | 11 | 120935369 | A | G | -1.377 | 0.1686 | -? | 0 | 0 | 0 | 1 |
| rs4936578 | 11 | 120999258 | G | A | 1.377 | 0.1686 | +? | 0 | 0 | 0 | 1 |
| rs11216658 | 11 | 117842743 | C | G | 1.376 | 0.1687 | +? | 0 | 0 | 0 | 1 |
| rs10892640 | 11 | 120789508 | G | A | 1.376 | 0.1688 | +? | 0 | 0 | 0 | 1 |

| | | | | | | | | | | | |
|---|---|---|---|---|---|---|---|---|---|---|---|
| rs77683187 | 11 | 117085210 | C | A | -1.376 | 0.1688 | -? | 0 | 0 | 0 | 1 |
| rs2298647 | 11 | 117671641 | A | T | -1.375 | 0.1691 | -? | 0 | 0 | 0 | 1 |
| rs118070092 | 11 | 119837014 | A | G | -1.375 | 0.1692 | -? | 0 | 0 | 0 | 1 |
| rs11218148 | 11 | 120992119 | A | C | 1.374 | 0.1695 | +? | 0 | 0 | 0 | 1 |
| rs4936576 | 11 | 120992859 | C | G | 1.374 | 0.1695 | +? | 0 | 0 | 0 | 1 |
| rs4936537 | 11 | 120526365 | G | A | -1.373 | 0.1699 | -? | 0 | 0 | 0 | 1 |
| rs4245041 | 11 | 120526637 | C | T | -1.372 | 0.1701 | -? | 0 | 0 | 0 | 1 |
| rs520315 | 11 | 117207626 | C | T | 1.372 | 0.1701 | +? | 0 | 0 | 0 | 1 |
| rs620848 | 11 | 117207628 | G | C | 1.372 | 0.1701 | +? | 0 | 0 | 0 | 1 |
| rs111227443 | 11 | 118590961 | T | C | 1.372 | 0.1702 | ++ | 0 | 0.18 | 1 | 0.671 |
| rs11821535 | 11 | 119518808 | A | C | -1.372 | 0.1702 | -? | 0 | 0 | 0 | 1 |
| rs10891982 | 11 | 116357955 | T | C | -1.371 | 0.1704 | -? | 0 | 0 | 0 | 1 |
| rs11601867 | 11 | 120705673 | A | G | 1.371 | 0.1704 | +? | 0 | 0 | 0 | 1 |
| rs1828121 | 11 | 116354211 | A | G | -1.369 | 0.1709 | -? | 0 | 0 | 0 | 1 |
| rs10892692 | 11 | 121103704 | G | A | 1.369 | 0.171 | +? | 0 | 0 | 0 | 1 |
| rs192734017 | 11 | 119029058 | A | G | -1.369 | 0.171 | -? | 0 | 0 | 0 | 1 |
| rs6421566 | 11 | 117671398 | G | A | -1.369 | 0.171 | -? | 0 | 0 | 0 | 1 |
| rs679607 | 11 | 121034651 | G | A | 1.369 | 0.171 | +? | 0 | 0 | 0 | 1 |
| rs7934236 | 11 | 120193511 | G | C | 1.369 | 0.171 | ++ | 0 | 0.594 | 1 | 0.4407 |
| rs543119 | 11 | 121014904 | G | A | 1.369 | 0.1711 | +? | 0 | 0 | 0 | 1 |
| rs685440 | 11 | 117429718 | T | G | -1.369 | 0.1711 | -? | 0 | 0 | 0 | 1 |
| rs7125224 | 11 | 121004851 | A | G | 1.368 | 0.1713 | +? | 0 | 0 | 0 | 1 |
| rs192113495 | 11 | 118874272 | G | T | -1.368 | 0.1714 | -- | -149 | 1.205 | 1 | 0.2723 |
| rs11217787 | 11 | 120153414 | A | T | -1.367 | 0.1715 | -? | 0 | 0 | 0 | 1 |
| rs2276065 | 11 | 118530611 | C | T | -1.367 | 0.1715 | -- | 0 | 0.114 | 1 | 0.7355 |
| rs12970 | 11 | 117074109 | A | G | -1.367 | 0.1716 | -? | 0 | 0 | 0 | 1 |
| rs10892527 | 11 | 119932137 | A | G | -1.365 | 0.1723 | -? | 0 | 0 | 0 | 1 |
| rs61898762 | 11 | 120343033 | G | T | -1.365 | 0.1723 | -? | 0 | 0 | 0 | 1 |
| rs71482116 | 11 | 117681749 | A | G | 1.365 | 0.1723 | +? | 0 | 0 | 0 | 1 |
| rs7105213 | 11 | 120505511 | T | C | 1.365 | 0.1724 | +? | 0 | 0 | 0 | 1 |
| rs611475 | 11 | 117950796 | C | T | -1.364 | 0.1725 | -? | 0 | 0 | 0 | 1 |
| rs754872 | 11 | 119802424 | T | C | 1.364 | 0.1725 | +? | 0 | 0 | 0 | 1 |
| rs893840 | 11 | 116199943 | C | T | -1.364 | 0.1725 | -? | 0 | 0 | 0 | 1 |
| rs12363946 | 11 | 116423005 | A | T | -1.364 | 0.1727 | -? | 0 | 0 | 0 | 1 |
| rs22298 | 11 | 121018954 | G | A | 1.363 | 0.1728 | +? | 0 | 0 | 0 | 1 |
| rs77776594 | 11 | 120563386 | A | G | 1.363 | 0.1728 | +? | 0 | 0 | 0 | 1 |
| rs11216551 | 11 | 117678420 | A | G | -1.363 | 0.1729 | -? | 0 | 0 | 0 | 1 |
| rs658187 | 11 | 119352570 | T | G | -1.363 | 0.1729 | -? | 0 | 0 | 0 | 1 |
| rs7129952 | 11 | 116495714 | A | T | -1.363 | 0.1729 | -? | 0 | 0 | 0 | 1 |
| rs75510322 | 11 | 120413702 | C | A | 1.363 | 0.1729 | +? | 0 | 0 | 0 | 1 |
| rs4635090 | 11 | 120844222 | A | G | -1.363 | 0.173 | -? | 0 | 0 | 0 | 1 |
| rs61906586 | 11 | 116360636 | C | T | -1.363 | 0.173 | -? | 0 | 0 | 0 | 1 |
| rs80170872 | 11 | 117909810 | A | G | 1.362 | 0.1732 | ++ | 0 | 0.005 | 1 | 0.9424 |
| rs1014034 | 11 | 117661734 | G | C | -1.361 | 0.1734 | -? | 0 | 0 | 0 | 1 |
| rs60007061 | 11 | 117937225 | A | C | 1.361 | 0.1736 | ++ | 0 | 0.007 | 1 | 0.9337 |
| rs7112429 | 11 | 117931489 | A | T | 1.36 | 0.1737 | ++ | 0 | 0.007 | 1 | 0.9335 |
| rs520332 | 11 | 120824323 | T | G | 1.36 | 0.1738 | +? | 0 | 0 | 0 | 1 |
| rs4936424 | 11 | 118009674 | T | C | -1.36 | 0.1739 | -- | 0 | 0.341 | 1 | 0.5595 |
| rs11217528 | 11 | 119701587 | C | A | 1.359 | 0.1742 | +? | 0 | 0 | 0 | 1 |
| rs1941299 | 11 | 121108907 | C | A | 1.358 | 0.1744 | +? | 0 | 0 | 0 | 1 |
| rs34189224 | 11 | 120253329 | A | G | 1.358 | 0.1745 | +? | 0 | 0 | 0 | 1 |
| rs12417676 | 11 | 116497466 | C | A | -1.357 | 0.1747 | -? | 0 | 0 | 0 | 1 |
| rs603179 | 11 | 119892194 | A | G | 1.357 | 0.1747 | +? | 0 | 0 | 0 | 1 |
| rs7940779 | 11 | 120419218 | T | C | 1.357 | 0.1747 | +? | 0 | 0 | 0 | 1 |
| rs61903185 | 11 | 120902930 | A | G | -1.357 | 0.1748 | -? | 0 | 0 | 0 | 1 |
| rs78995144 | 11 | 120867999 | A | G | -1.357 | 0.1748 | -? | 0 | 0 | 0 | 1 |

| | | | | | | | | | | | |
|---|---|---|---|---|---|---|---|---|---|---|---|
| rs76547700 | 11 | 119849683 | T | G | -1.357 | 0.1749 | -? | 0 | 0 | 0 | 1 |
| rs1787693 | 11 | 116542585 | A | G | -1.356 | 0.1751 | -? | 0 | 0 | 0 | 1 |
| rs17120753 | 11 | 117308432 | C | T | 1.356 | 0.1752 | +? | 0 | 0 | 0 | 1 |
| rs11216058 | 11 | 116501684 | A | G | -1.355 | 0.1754 | -? | 0 | 0 | 0 | 1 |
| rs649197 | 11 | 119353238 | A | G | -1.355 | 0.1754 | -? | 0 | 0 | 0 | 1 |
| rs491428 | 11 | 116353002 | T | G | -1.355 | 0.1755 | -? | 0 | 0 | 0 | 1 |
| rs490526 | 11 | 116353106 | T | A | -1.354 | 0.1756 | -? | 0 | 0 | 0 | 1 |
| rs4938270 | 11 | 116272744 | T | C | -1.354 | 0.1757 | -? | 0 | 0 | 0 | 1 |
| rs7120040 | 11 | 120349142 | G | A | -1.354 | 0.1757 | -? | 0 | 0 | 0 | 1 |
| rs7931398 | 11 | 116879676 | C | T | -1.353 | 0.1759 | -? | 0 | 0 | 0 | 1 |
| rs617345 | 11 | 117899816 | G | T | 1.353 | 0.176 | +- | -89.7 | 1.582 | 1 | 0.2085 |
| rs630842 | 11 | 119924436 | A | G | 1.353 | 0.1762 | +? | 0 | 0 | 0 | 1 |
| rs652376 | 11 | 119924214 | C | G | 1.353 | 0.1762 | +? | 0 | 0 | 0 | 1 |
| rs2276343 | 11 | 117375596 | A | G | -1.352 | 0.1763 | -? | 0 | 0 | 0 | 1 |
| rs1240773 | 11 | 116520527 | T | G | -1.351 | 0.1766 | -? | 0 | 0 | 0 | 1 |
| rs12790634 | 11 | 120176310 | A | G | 1.351 | 0.1767 | +? | 0 | 0 | 0 | 1 |
| rs12274300 | 11 | 120580038 | G | C | 1.35 | 0.1769 | +? | 0 | 0 | 0 | 1 |
| rs12285075 | 11 | 119755047 | A | G | -1.35 | 0.1769 | -? | 0 | 0 | 0 | 1 |
| rs7934267 | 11 | 117923252 | T | G | -1.35 | 0.1769 | -? | 0 | 0 | 0 | 1 |
| rs7937145 | 11 | 117923358 | A | G | -1.35 | 0.1769 | -? | 0 | 0 | 0 | 1 |
| rs947889 | 11 | 117874531 | C | T | 1.35 | 0.1769 | ++ | 0 | 0.158 | 1 | 0.691 |
| rs948467 | 11 | 117962813 | G | A | -1.35 | 0.1769 | -? | 0 | 0 | 0 | 1 |
| rs187900569 | 11 | 119337452 | C | G | 1.35 | 0.177 | +? | 0 | 0 | 0 | 1 |
| rs494560 | 11 | 118521549 | A | G | 1.35 | 0.177 | ++ | 0 | 0.153 | 1 | 0.6954 |
| rs56112767 | 11 | 117641216 | G | C | -1.349 | 0.1772 | -? | 0 | 0 | 0 | 1 |
| rs10892248 | 11 | 118501022 | A | G | -1.349 | 0.1773 | -- | 0 | 0.273 | 1 | 0.6012 |
| rs522645 | 11 | 117101574 | A | C | -1.348 | 0.1776 | -? | 0 | 0 | 0 | 1 |
| rs598650 | 11 | 120980537 | T | C | -1.348 | 0.1776 | -? | 0 | 0 | 0 | 1 |
| rs17122150 | 11 | 118271860 | G | C | -1.348 | 0.1777 | -- | 0 | 0.101 | 1 | 0.7506 |
| rs2236661 | 11 | 118499394 | G | C | -1.348 | 0.1778 | -- | 0 | 0.299 | 1 | 0.5845 |
| rs493308 | 11 | 116352789 | T | C | -1.347 | 0.1781 | -? | 0 | 0 | 0 | 1 |
| rs554912 | 11 | 117340285 | T | C | 1.347 | 0.1781 | +? | 0 | 0 | 0 | 1 |
| rs4245184 | 11 | 118510562 | A | G | 1.346 | 0.1784 | ++ | 0 | 0.123 | 1 | 0.7258 |
| rs644273 | 11 | 118511352 | T | C | 1.346 | 0.1784 | ++ | 0 | 0.123 | 1 | 0.7258 |
| rs11603533 | 11 | 120324517 | G | C | 1.345 | 0.1785 | +? | 0 | 0 | 0 | 1 |
| rs687009 | 11 | 119246567 | C | G | 1.345 | 0.1785 | ++ | 0 | 0.637 | 1 | 0.4249 |
| rs75375456 | 11 | 117430025 | T | G | 1.345 | 0.1785 | +? | 0 | 0 | 0 | 1 |
| rs10892641 | 11 | 120789800 | G | A | 1.345 | 0.1786 | +? | 0 | 0 | 0 | 1 |
| rs11218077 | 11 | 120822001 | C | T | 1.345 | 0.1787 | +? | 0 | 0 | 0 | 1 |
| rs11603023 | 11 | 118486067 | T | C | 1.345 | 0.1787 | ++ | 0 | 0.157 | 1 | 0.6924 |
| rs947990 | 11 | 116557314 | T | G | -1.345 | 0.1787 | -? | 0 | 0 | 0 | 1 |
| rs626252 | 11 | 117949861 | G | A | -1.344 | 0.1789 | -- | 0 | 0.917 | 1 | 0.3383 |
| rs576945 | 11 | 119246896 | G | A | 1.344 | 0.179 | ++ | 0 | 0.641 | 1 | 0.4234 |
| rs685916 | 11 | 117434215 | A | G | -1.344 | 0.179 | -? | 0 | 0 | 0 | 1 |
| rs74505326 | 11 | 116756690 | G | A | 1.343 | 0.1792 | +? | 0 | 0 | 0 | 1 |
| rs35021198 | 11 | 117448799 | T | C | 1.343 | 0.1794 | +? | 0 | 0 | 0 | 1 |
| rs508440 | 11 | 119745457 | G | C | -1.343 | 0.1794 | -? | 0 | 0 | 0 | 1 |
| rs78621482 | 11 | 120790033 | G | C | -1.343 | 0.1794 | -? | 0 | 0 | 0 | 1 |
| rs654423 | 11 | 118511757 | A | G | 1.342 | 0.1796 | ++ | 0 | 0.122 | 1 | 0.7274 |
| rs672499 | 11 | 119351636 | A | G | -1.342 | 0.1796 | -? | 0 | 0 | 0 | 1 |
| rs34238850 | 11 | 120462315 | T | C | 1.341 | 0.1798 | +? | 0 | 0 | 0 | 1 |
| rs556436 | 11 | 116352523 | C | T | -1.341 | 0.1799 | -? | 0 | 0 | 0 | 1 |
| rs11218103 | 11 | 120871324 | T | C | -1.341 | 0.18 | -? | 0 | 0 | 0 | 1 |
| rs12721031 | 11 | 116704505 | T | C | 1.34 | 0.1802 | +? | 0 | 0 | 0 | 1 |
| rs2370112 | 11 | 121192240 | C | T | -1.34 | 0.1802 | -? | 0 | 0 | 0 | 1 |
| rs12363335 | 11 | 119890299 | C | T | -1.34 | 0.1803 | -? | 0 | 0 | 0 | 1 |

| | | | | | | | | | | | |
|---|---|---|---|---|---|---|---|---|---|---|---|
| rs7120174 | 11 | 120349310 | C | A | -1.34 | 0.1803 | -? | 0 | 0 | 0 | 1 |
| rs613018 | 11 | 119355747 | C | T | -1.34 | 0.1804 | -? | 0 | 0 | 0 | 1 |
| rs61900897 | 11 | 120418687 | C | G | -1.339 | 0.1805 | -? | 0 | 0 | 0 | 1 |
| rs11216582 | 11 | 117724493 | A | C | -1.339 | 0.1806 | -- | 0 | 0.051 | 1 | 0.8207 |
| rs11216867 | 11 | 118262407 | G | A | -1.339 | 0.1806 | -- | 0 | 0.143 | 1 | 0.7053 |
| rs7125693 | 11 | 117670651 | C | T | -1.339 | 0.1807 | -? | 0 | 0 | 0 | 1 |
| rs2187490 | 11 | 118713180 | G | T | 1.338 | 0.1808 | ++ | 0 | 0.111 | 1 | 0.7394 |
| rs4544011 | 11 | 120656866 | G | A | 1.338 | 0.1808 | +? | 0 | 0 | 0 | 1 |
| rs7107023 | 11 | 117796722 | T | C | 1.338 | 0.1808 | +? | 0 | 0 | 0 | 1 |
| rs180322 | 11 | 116626735 | C | T | -1.338 | 0.1809 | -+ | -152.3 | 1.189 | 1 | 0.2755 |
| rs500426 | 11 | 116485319 | T | C | 1.338 | 0.181 | +? | 0 | 0 | 0 | 1 |
| rs673841 | 11 | 119351329 | G | A | -1.338 | 0.181 | -? | 0 | 0 | 0 | 1 |
| rs8301 | 11 | 118895495 | G | C | -1.337 | 0.1811 | -- | 0 | 0.027 | 1 | 0.8688 |
| rs1134734 | 11 | 118965479 | A | G | -1.337 | 0.1812 | -? | 0 | 0 | 0 | 1 |
| rs533053 | 11 | 119353577 | C | G | -1.337 | 0.1812 | -? | 0 | 0 | 0 | 1 |
| rs649557 | 11 | 119353188 | T | C | -1.337 | 0.1812 | -? | 0 | 0 | 0 | 1 |
| rs650969 | 11 | 119352890 | C | T | -1.337 | 0.1812 | -? | 0 | 0 | 0 | 1 |
| rs7936556 | 11 | 118501299 | A | G | 1.337 | 0.1812 | ++ | 0 | 0.125 | 1 | 0.7233 |
| rs12361978 | 11 | 116391272 | G | C | 1.336 | 0.1814 | +? | 0 | 0 | 0 | 1 |
| rs12362794 | 11 | 120350393 | G | T | -1.336 | 0.1816 | -? | 0 | 0 | 0 | 1 |
| rs148990923 | 11 | 118722946 | T | C | -1.335 | 0.1817 | -- | 0 | 0.274 | 1 | 0.6007 |
| rs4293153 | 11 | 116416909 | G | A | -1.336 | 0.1817 | -? | 0 | 0 | 0 | 1 |
| rs1108627 | 11 | 116369980 | T | C | 1.335 | 0.1818 | +? | 0 | 0 | 0 | 1 |
| rs11828785 | 11 | 116501918 | T | C | -1.335 | 0.1819 | -? | 0 | 0 | 0 | 1 |
| rs1185802888 | 11 | 116259493 | T | A | -1.335 | 0.1819 | -? | 0 | 0 | 0 | 1 |
| rs17124739 | 11 | 120924671 | G | A | -1.335 | 0.1819 | -? | 0 | 0 | 0 | 1 |
| rs496371 | 11 | 117722715 | T | C | 1.334 | 0.1821 | +? | 0 | 0 | 0 | 1 |
| rs7943887 | 11 | 117670282 | G | C | -1.334 | 0.1821 | -? | 0 | 0 | 0 | 1 |
| rs76347292 | 11 | 117303108 | C | T | 1.333 | 0.1824 | +? | 0 | 0 | 0 | 1 |
| rs605531 | 11 | 117120027 | G | T | 1.333 | 0.1826 | +? | 0 | 0 | 0 | 1 |
| rs547344 | 11 | 119751375 | A | G | -1.332 | 0.1827 | -? | 0 | 0 | 0 | 1 |
| rs659127 | 11 | 119913037 | A | C | 1.332 | 0.1829 | +? | 0 | 0 | 0 | 1 |
| rs548429 | 11 | 119806852 | G | A | -1.331 | 0.1832 | -? | 0 | 0 | 0 | 1 |
| rs11218106 | 11 | 120873529 | A | T | -1.331 | 0.1833 | -? | 0 | 0 | 0 | 1 |
| rs56189546 | 11 | 119435096 | G | T | -1.33 | 0.1834 | -? | 0 | 0 | 0 | 1 |
| rs11216387 | 11 | 117310022 | G | A | -1.33 | 0.1835 | -? | 0 | 0 | 0 | 1 |
| rs7125694 | 11 | 117670657 | C | T | -1.33 | 0.1835 | -? | 0 | 0 | 0 | 1 |
| rs74482798 | 11 | 120565473 | G | C | -1.329 | 0.1839 | -? | 0 | 0 | 0 | 1 |
| rs76003278 | 11 | 120565293 | C | T | -1.329 | 0.1839 | -? | 0 | 0 | 0 | 1 |
| rs545108 | 11 | 116525229 | C | G | 1.328 | 0.1842 | +? | 0 | 0 | 0 | 1 |
| rs548893 | 11 | 117193619 | G | C | 1.328 | 0.1842 | +? | 0 | 0 | 0 | 1 |
| rs142260742 | 11 | 117425169 | T | C | 1.328 | 0.1843 | +? | 0 | 0 | 0 | 1 |
| rs80350215 | 11 | 117426035 | C | T | 1.328 | 0.1843 | +? | 0 | 0 | 0 | 1 |
| rs61901380 | 11 | 120579239 | A | G | 1.327 | 0.1844 | +? | 0 | 0 | 0 | 1 |
| rs7927559 | 11 | 117845564 | G | A | -1.327 | 0.1844 | -- | 0 | 0.002 | 1 | 0.9617 |
| rs10790210 | 11 | 117662639 | G | C | 1.327 | 0.1845 | +? | 0 | 0 | 0 | 1 |
| rs10892247 | 11 | 118490076 | A | G | -1.327 | 0.1845 | -- | 0 | 0.348 | 1 | 0.555 |
| rs73008681 | 11 | 119744365 | G | C | -1.327 | 0.1846 | -? | 0 | 0 | 0 | 1 |
| rs10892619 | 11 | 120571457 | T | C | -1.326 | 0.1847 | -? | 0 | 0 | 0 | 1 |
| rs10892525 | 11 | 119930631 | T | C | -1.326 | 0.1848 | -? | 0 | 0 | 0 | 1 |
| rs4938700 | 11 | 119516805 | C | G | 1.326 | 0.1848 | +? | 0 | 0 | 0 | 1 |
| rs7106292 | 11 | 120830051 | T | C | 1.326 | 0.1848 | +? | 0 | 0 | 0 | 1 |
| rs7123228 | 11 | 120715952 | G | A | 1.326 | 0.1848 | +? | 0 | 0 | 0 | 1 |
| rs11216493 | 11 | 117542149 | T | C | -1.325 | 0.1851 | -? | 0 | 0 | 0 | 1 |
| rs11600004 | 11 | 120708132 | T | C | 1.325 | 0.1851 | +? | 0 | 0 | 0 | 1 |
| rs4936380 | 11 | 117324068 | A | G | 1.325 | 0.1851 | +? | 0 | 0 | 0 | 1 |

| | | | | | | | | | | | |
|---|---|---|---|---|---|---|---|---|---|---|---|
| rs7113753 | 11 | 118889378 | A | G | -1.325 | 0.1852 | -- | 0 | 0.342 | 1 | 0.5587 |
| rs1892903 | 11 | 117669619 | T | C | -1.325 | 0.1853 | -? | 0 | 0 | 0 | 1 |
| rs12224952 | 11 | 119736232 | T | G | -1.324 | 0.1854 | -? | 0 | 0 | 0 | 1 |
| rs61897936 | 11 | 119933153 | T | C | -1.324 | 0.1854 | -? | 0 | 0 | 0 | 1 |
| rs11216806 | 11 | 118060549 | C | T | -1.324 | 0.1856 | -+ | 17.2 | 3.625 | 1 | 0.05691 |
| rs6589547 | 11 | 116361705 | A | C | -1.323 | 0.1857 | -? | 0 | 0 | 0 | 1 |
| rs3016352 | 11 | 116521748 | G | A | 1.323 | 0.1859 | +? | 0 | 0 | 0 | 1 |
| rs600937 | 11 | 120985032 | G | A | -1.323 | 0.1859 | -- | 0 | 0.024 | 1 | 0.8776 |
| rs10892324 | 11 | 118903039 | G | T | 1.323 | 0.186 | +? | 0 | 0 | 0 | 1 |
| rs561791 | 11 | 117305418 | T | C | -1.322 | 0.1861 | -? | 0 | 0 | 0 | 1 |
| rs11217987 | 11 | 120579826 | A | T | 1.322 | 0.1863 | +? | 0 | 0 | 0 | 1 |
| rs17245326 | 11 | 119778929 | A | G | 1.322 | 0.1863 | +? | 0 | 0 | 0 | 1 |
| rs2305830 | 11 | 117266312 | G | C | -1.322 | 0.1863 | -? | 0 | 0 | 0 | 1 |
| rs6589580 | 11 | 116819873 | A | G | -1.322 | 0.1863 | -? | 0 | 0 | 0 | 1 |
| rs12363806 | 11 | 117658211 | A | G | 1.32 | 0.1867 | +? | 0 | 0 | 0 | 1 |
| rs10790280 | 11 | 118902609 | T | A | 1.32 | 0.1869 | +? | 0 | 0 | 0 | 1 |
| rs11218240 | 11 | 121151902 | T | C | -1.319 | 0.1872 | -? | 0 | 0 | 0 | 1 |
| rs3016347 | 11 | 116486664 | C | G | 1.319 | 0.1872 | +? | 0 | 0 | 0 | 1 |
| rs11216393 | 11 | 117314053 | A | G | -1.318 | 0.1875 | -? | 0 | 0 | 0 | 1 |
| rs10892322 | 11 | 118902749 | T | C | 1.317 | 0.188 | +? | 0 | 0 | 0 | 1 |
| rs12365956 | 11 | 117798431 | A | G | 1.317 | 0.188 | +? | 0 | 0 | 0 | 1 |
| rs12419866 | 11 | 119240624 | T | C | 1.317 | 0.188 | +? | 0 | 0 | 0 | 1 |
| rs73022500 | 11 | 117798073 | A | C | 1.317 | 0.188 | +? | 0 | 0 | 0 | 1 |
| rs1866683 | 11 | 116199624 | C | A | -1.316 | 0.1882 | -? | 0 | 0 | 0 | 1 |
| rs77642243 | 11 | 117795578 | C | T | 1.316 | 0.1882 | +? | 0 | 0 | 0 | 1 |
| rs7932967 | 11 | 117207665 | T | G | -1.316 | 0.1882 | -? | 0 | 0 | 0 | 1 |
| rs12292552 | 11 | 120565023 | A | G | 1.316 | 0.1883 | +? | 0 | 0 | 0 | 1 |
| rs4567478 | 11 | 120577423 | C | T | 1.315 | 0.1886 | +? | 0 | 0 | 0 | 1 |
| rs112464350 | 11 | 116539058 | A | C | 1.314 | 0.1887 | +? | 0 | 0 | 0 | 1 |
| rs4938471 | 11 | 117924986 | G | T | -1.314 | 0.1887 | -? | 0 | 0 | 0 | 1 |
| rs4499034 | 11 | 117323004 | T | C | 1.314 | 0.1889 | +? | 0 | 0 | 0 | 1 |
| rs541366 | 11 | 119744160 | C | A | -1.314 | 0.1889 | -? | 0 | 0 | 0 | 1 |
| rs56926846 | 11 | 120526291 | A | G | 1.314 | 0.1889 | +? | 0 | 0 | 0 | 1 |
| rs1940040 | 11 | 117393294 | C | G | -1.314 | 0.189 | -? | 0 | 0 | 0 | 1 |
| rs2276340 | 11 | 117391966 | A | G | -1.314 | 0.189 | -? | 0 | 0 | 0 | 1 |
| rs12800011 | 11 | 116501098 | G | A | -1.313 | 0.1892 | -? | 0 | 0 | 0 | 1 |
| rs77683570 | 11 | 118495287 | C | T | -1.313 | 0.1892 | -- | 0 | 0.347 | 1 | 0.5556 |
| rs7935609 | 11 | 116202035 | C | T | -1.313 | 0.1892 | -? | 0 | 0 | 0 | 1 |
| rs10892075 | 11 | 116982821 | T | C | -1.313 | 0.1893 | -? | 0 | 0 | 0 | 1 |
| rs138536663 | 11 | 121148641 | C | T | -1.312 | 0.1894 | -? | 0 | 0 | 0 | 1 |
| rs113708750 | 11 | 116198542 | T | C | -1.312 | 0.1895 | -? | 0 | 0 | 0 | 1 |
| rs17122154 | 11 | 118273019 | A | G | -1.312 | 0.1895 | -- | 0 | 0.109 | 1 | 0.7409 |
| rs61013776 | 11 | 120993723 | A | G | -1.312 | 0.1896 | -- | 0 | 0.286 | 1 | 0.5931 |
| rs9735317 | 11 | 118725383 | T | C | -1.312 | 0.1896 | -- | 0 | 0.027 | 1 | 0.8696 |
| rs7946325 | 11 | 120578917 | G | A | 1.311 | 0.1897 | +? | 0 | 0 | 0 | 1 |
| rs11217130 | 11 | 118902933 | T | A | 1.311 | 0.1899 | +? | 0 | 0 | 0 | 1 |
| rs3133855 | 11 | 120584851 | T | C | -1.311 | 0.19 | -? | 0 | 0 | 0 | 1 |
| rs3825060 | 11 | 118973112 | A | G | 1.311 | 0.19 | +? | 0 | 0 | 0 | 1 |
| rs75809806 | 11 | 117418077 | A | G | 1.31 | 0.1901 | +? | 0 | 0 | 0 | 1 |
| rs77342459 | 11 | 117420369 | G | C | 1.31 | 0.1901 | +? | 0 | 0 | 0 | 1 |
| rs80041541 | 11 | 117417384 | T | G | 1.31 | 0.1901 | +? | 0 | 0 | 0 | 1 |
| rs113784020 | 11 | 118559731 | T | C | 1.31 | 0.1902 | ++ | 0 | 0.168 | 1 | 0.6818 |
| rs494200 | 11 | 116352697 | A | G | -1.31 | 0.1902 | -? | 0 | 0 | 0 | 1 |
| rs60348146 | 11 | 117939413 | A | G | 1.31 | 0.1903 | ++ | 0 | 0.014 | 1 | 0.9071 |
| rs659705 | 11 | 119245349 | G | A | 1.31 | 0.1903 | ++ | 0 | 0.67 | 1 | 0.4132 |
| rs11215957 | 11 | 116355233 | C | T | -1.309 | 0.1907 | -? | 0 | 0 | 0 | 1 |

| | | | | | | | | | | | |
|---|---|---|---|---|---|---|---|---|---|---|---|
| rs61901379 | 11 | 120577940 | A | G | 1.309 | 0.1907 | +? | 0 | 0 | 0 | 1 |
| rs535831 | 11 | 119346198 | C | T | -1.308 | 0.1908 | -? | 0 | 0 | 0 | 1 |
| rs647369 | 11 | 119347074 | A | T | -1.308 | 0.1908 | -? | 0 | 0 | 0 | 1 |
| rs686882 | 11 | 121023992 | G | A | 1.307 | 0.1911 | +? | 0 | 0 | 0 | 1 |
| rs112548685 | 11 | 116548489 | T | C | -1.307 | 0.1912 | -? | 0 | 0 | 0 | 1 |
| rs483598 | 11 | 118509668 | A | G | 1.307 | 0.1912 | ++ | 0 | 0.066 | 1 | 0.7979 |
| rs57401958 | 11 | 120947699 | G | A | -1.306 | 0.1914 | -? | 0 | 0 | 0 | 1 |
| rs111494641 | 11 | 119673277 | T | C | 1.306 | 0.1915 | ++ | 0 | 0.112 | 1 | 0.7375 |
| rs4936535 | 11 | 120505771 | G | A | 1.306 | 0.1916 | +? | 0 | 0 | 0 | 1 |
| rs61903853 | 11 | 117600518 | G | T | -1.306 | 0.1917 | -? | 0 | 0 | 0 | 1 |
| rs1939004 | 11 | 120920649 | G | A | -1.304 | 0.1922 | -? | 0 | 0 | 0 | 1 |
| rs11218239 | 11 | 121149531 | G | A | -1.304 | 0.1923 | -? | 0 | 0 | 0 | 1 |
| rs4938400 | 11 | 117399604 | A | C | -1.304 | 0.1923 | -? | 0 | 0 | 0 | 1 |
| rs4938401 | 11 | 117399605 | G | T | -1.304 | 0.1923 | -? | 0 | 0 | 0 | 1 |
| rs6589713 | 11 | 118872404 | C | T | -1.304 | 0.1923 | -- | 0 | 0.019 | 1 | 0.8902 |
| rs744983 | 11 | 120166358 | G | A | -1.304 | 0.1924 | -? | 0 | 0 | 0 | 1 |
| rs7942757 | 11 | 118903651 | A | G | 1.303 | 0.1925 | +? | 0 | 0 | 0 | 1 |
| rs11216934 | 11 | 118501459 | A | G | 1.303 | 0.1926 | ++ | 0 | 0.063 | 1 | 0.8014 |
| rs12286112 | 11 | 117755580 | G | A | 1.303 | 0.1927 | +? | 0 | 0 | 0 | 1 |
| rs6589663 | 11 | 118255278 | A | G | -1.303 | 0.1927 | -- | 0 | 0.181 | 1 | 0.6707 |
| rs28781001 | 11 | 120580572 | G | A | 1.302 | 0.1929 | +? | 0 | 0 | 0 | 1 |
| rs577396 | 11 | 118499469 | C | G | 1.302 | 0.1929 | ++ | 0 | 0.073 | 1 | 0.7873 |
| rs144716047 | 11 | 120871636 | G | C | -1.302 | 0.193 | -? | 0 | 0 | 0 | 1 |
| rs667125 | 11 | 120984999 | G | A | -1.302 | 0.193 | -- | 0 | 0.028 | 1 | 0.8682 |
| rs12276408 | 11 | 120568254 | G | A | 1.301 | 0.1931 | +? | 0 | 0 | 0 | 1 |
| rs73580459 | 11 | 116309713 | A | G | -1.301 | 0.1931 | -? | 0 | 0 | 0 | 1 |
| rs10892379 | 11 | 119393309 | G | A | -1.301 | 0.1932 | -? | 0 | 0 | 0 | 1 |
| rs1045947297 | 11 | 121003604 | G | T | 1.301 | 0.1934 | +? | 0 | 0 | 0 | 1 |
| rs17120523 | 11 | 117094591 | G | A | -1.3 | 0.1937 | -? | 0 | 0 | 0 | 1 |
| rs613442 | 11 | 119355678 | T | C | -1.299 | 0.1939 | -? | 0 | 0 | 0 | 1 |
| rs619585 | 11 | 117947433 | T | C | -1.299 | 0.1939 | -- | -187.5 | 1.044 | 1 | 0.307 |
| rs671160 | 11 | 117952387 | T | C | -1.299 | 0.1939 | -? | 0 | 0 | 0 | 1 |
| rs10892323 | 11 | 118902758 | C | G | 1.298 | 0.1943 | +? | 0 | 0 | 0 | 1 |
| rs1302598824 | 11 | 117700629 | T | C | -1.298 | 0.1943 | -? | 0 | 0 | 0 | 1 |
| rs1189010881 | 11 | 117314931 | T | G | -1.297 | 0.1947 | -? | 0 | 0 | 0 | 1 |
| rs12289923 | 11 | 121112889 | G | A | -1.297 | 0.1947 | -- | 0 | 0.278 | 1 | 0.5979 |
| rs34259384 | 11 | 120860524 | T | G | 1.297 | 0.1948 | +? | 0 | 0 | 0 | 1 |
| rs620236 | 11 | 119355134 | A | T | -1.296 | 0.1949 | -? | 0 | 0 | 0 | 1 |
| rs10892204 | 11 | 117906510 | T | C | -1.296 | 0.195 | -? | 0 | 0 | 0 | 1 |
| rs883245 | 11 | 119217311 | G | A | 1.296 | 0.1951 | +? | 0 | 0 | 0 | 1 |
| rs531280 | 11 | 119353746 | C | A | -1.295 | 0.1952 | -? | 0 | 0 | 0 | 1 |
| rs2277286 | 11 | 117102706 | G | C | 1.295 | 0.1953 | +? | 0 | 0 | 0 | 1 |
| rs12276533 | 11 | 118724116 | A | G | -1.295 | 0.1954 | -- | 0 | 0.014 | 1 | 0.9051 |
| rs613406 | 11 | 119355695 | C | T | -1.294 | 0.1955 | -? | 0 | 0 | 0 | 1 |
| rs45565037 | 11 | 118496119 | A | G | -1.294 | 0.1956 | -- | 0 | 0.236 | 1 | 0.6272 |
| rs1893970 | 11 | 117432034 | T | C | 1.294 | 0.1957 | +? | 0 | 0 | 0 | 1 |
| rs587069 | 11 | 119351324 | G | A | -1.294 | 0.1957 | -? | 0 | 0 | 0 | 1 |
| rs59489957 | 11 | 118284984 | T | C | -1.294 | 0.1957 | -- | 0 | 0.203 | 1 | 0.652 |
| rs7122879 | 11 | 120897295 | T | C | -1.294 | 0.1957 | -? | 0 | 0 | 0 | 1 |
| rs61904777 | 11 | 116362669 | C | T | -1.293 | 0.1959 | -? | 0 | 0 | 0 | 1 |
| rs1508108 | 11 | 116356720 | G | A | -1.293 | 0.1961 | -? | 0 | 0 | 0 | 1 |
| rs584092 | 11 | 120823396 | T | C | 1.293 | 0.1961 | +? | 0 | 0 | 0 | 1 |
| rs61900623 | 11 | 118012623 | T | C | -1.293 | 0.1961 | -? | 0 | 0 | 0 | 1 |
| rs112550319 | 11 | 116539809 | G | C | 1.292 | 0.1962 | +? | 0 | 0 | 0 | 1 |
| rs12802672 | 11 | 116534963 | G | A | 1.292 | 0.1962 | +? | 0 | 0 | 0 | 1 |
| rs618827 | 11 | 119355438 | A | G | -1.292 | 0.1962 | -? | 0 | 0 | 0 | 1 |

| | | | | | | | | | | | |
|---|---|---|---|---|---|---|---|---|---|---|---|
| rs7121764 | 11 | 118271759 | G | A | -1.292 | 0.1962 | -- | 0 | 0.129 | 1 | 0.7195 |
| rs10891988 | 11 | 116421056 | C | T | -1.292 | 0.1964 | -? | 0 | 0 | 0 | 1 |
| rs11605788 | 11 | 116232577 | A | C | -1.292 | 0.1965 | -? | 0 | 0 | 0 | 1 |
| rs61901488 | 11 | 117731990 | T | A | 1.292 | 0.1965 | +? | 0 | 0 | 0 | 1 |
| rs11218107 | 11 | 120875634 | C | T | -1.29 | 0.1969 | -? | 0 | 0 | 0 | 1 |
| rs118003247 | 11 | 118834322 | G | A | -1.29 | 0.1969 | -? | 0 | 0 | 0 | 1 |
| rs79237143 | 11 | 120876637 | G | T | -1.29 | 0.1969 | -? | 0 | 0 | 0 | 1 |
| rs10892078 | 11 | 116993929 | T | A | 1.29 | 0.1971 | +? | 0 | 0 | 0 | 1 |
| rs525588 | 11 | 119774225 | C | G | -1.29 | 0.1971 | -? | 0 | 0 | 0 | 1 |
| rs6589689 | 11 | 118622744 | G | A | 1.289 | 0.1973 | ++ | 0 | 0.261 | 1 | 0.6094 |
| rs12805373 | 11 | 119518822 | A | T | -1.289 | 0.1974 | -? | 0 | 0 | 0 | 1 |
| rs73018291 | 11 | 118224123 | A | G | -1.289 | 0.1974 | -? | 0 | 0 | 0 | 1 |
| rs11217129 | 11 | 118902841 | T | C | 1.289 | 0.1975 | +? | 0 | 0 | 0 | 1 |
| rs1815888 | 11 | 119815915 | G | C | -1.288 | 0.1977 | -? | 0 | 0 | 0 | 1 |
| rs7936795 | 11 | 117391736 | G | A | -1.288 | 0.1977 | -? | 0 | 0 | 0 | 1 |
| rs113579428 | 11 | 118006853 | T | C | 1.288 | 0.1978 | +? | 0 | 0 | 0 | 1 |
| rs77074463 | 11 | 118748170 | C | G | -1.288 | 0.1978 | -+ | 16.1 | 3.577 | 1 | 0.05857 |
| rs11217423 | 11 | 119586134 | T | C | -1.287 | 0.1981 | -? | 0 | 0 | 0 | 1 |
| rs12791103 | 11 | 116673315 | T | G | -1.287 | 0.1982 | -+ | 0 | 0.716 | 1 | 0.3976 |
| rs7119568 | 11 | 118723753 | A | T | -1.286 | 0.1985 | -- | 0 | 0.021 | 1 | 0.8838 |
| rs56359135 | 11 | 117569931 | T | A | 1.286 | 0.1986 | +? | 0 | 0 | 0 | 1 |
| rs618523 | 11 | 121061986 | T | C | 1.283 | 0.1994 | +? | 0 | 0 | 0 | 1 |
| rs7943490 | 11 | 120845912 | A | G | -1.283 | 0.1994 | -? | 0 | 0 | 0 | 1 |
| rs11216436 | 11 | 117391374 | A | G | -1.282 | 0.1997 | -? | 0 | 0 | 0 | 1 |
| rs531273 | 11 | 119353755 | A | G | -1.282 | 0.1997 | -? | 0 | 0 | 0 | 1 |
| rs9666861 | 11 | 117829855 | G | A | -1.282 | 0.1997 | -- | 0 | 0 | 1 | 0.9984 |
| rs76429987 | 11 | 120876772 | T | C | -1.282 | 0.1998 | -? | 0 | 0 | 0 | 1 |
| rs111446534 | 11 | 117317947 | A | C | -1.282 | 0.1999 | -? | 0 | 0 | 0 | 1 |
| rs602104 | 11 | 120809383 | C | T | 1.282 | 0.1999 | +? | 0 | 0 | 0 | 1 |
| rs60385993 | 11 | 117316605 | A | G | -1.282 | 0.1999 | -? | 0 | 0 | 0 | 1 |
| rs634838 | 11 | 119354137 | T | C | -1.282 | 0.1999 | -? | 0 | 0 | 0 | 1 |
| rs4935763 | 11 | 121064513 | T | C | 1.282 | 0.2 | +? | 0 | 0 | 0 | 1 |
| rs11217977 | 11 | 120572727 | T | C | -1.28 | 0.2004 | -? | 0 | 0 | 0 | 1 |
| rs74361939 | 11 | 117547710 | A | G | -1.28 | 0.2005 | -? | 0 | 0 | 0 | 1 |
| rs73008675 | 11 | 119742642 | C | T | -1.28 | 0.2006 | -? | 0 | 0 | 0 | 1 |
| rs73008677 | 11 | 119743088 | A | G | -1.28 | 0.2006 | -? | 0 | 0 | 0 | 1 |
| rs73008679 | 11 | 119743230 | T | C | -1.28 | 0.2006 | -? | 0 | 0 | 0 | 1 |
| rs78040959 | 11 | 119324268 | G | A | 1.28 | 0.2006 | +? | 0 | 0 | 0 | 1 |
| rs35310980 | 11 | 119236410 | A | T | 1.28 | 0.2007 | +? | 0 | 0 | 0 | 1 |
| rs3802873 | 11 | 117776471 | T | C | 1.279 | 0.2008 | +? | 0 | 0 | 0 | 1 |
| rs948464 | 11 | 117966185 | G | T | -1.279 | 0.2008 | -? | 0 | 0 | 0 | 1 |
| rs948465 | 11 | 117966178 | G | A | -1.279 | 0.2008 | -? | 0 | 0 | 0 | 1 |
| rs11216093 | 11 | 116557882 | C | A | 1.279 | 0.2009 | +? | 0 | 0 | 0 | 1 |
| rs11217511 | 11 | 119671550 | C | G | 1.279 | 0.201 | +? | 0 | 0 | 0 | 1 |
| rs11605165 | 11 | 119670624 | A | G | 1.279 | 0.201 | +? | 0 | 0 | 0 | 1 |
| rs11824108 | 11 | 116501944 | G | A | -1.279 | 0.201 | -? | 0 | 0 | 0 | 1 |
| rs182928887 | 11 | 117682935 | G | A | 1.278 | 0.2011 | +? | 0 | 0 | 0 | 1 |
| rs35181812 | 11 | 120497465 | T | C | 1.278 | 0.2012 | +? | 0 | 0 | 0 | 1 |
| rs4936593 | 11 | 121064264 | A | G | 1.277 | 0.2017 | +? | 0 | 0 | 0 | 1 |
| rs118170232 | 11 | 121040934 | G | A | -1.276 | 0.202 | -? | 0 | 0 | 0 | 1 |
| rs949056 | 11 | 120779359 | G | A | -1.276 | 0.2021 | -? | 0 | 0 | 0 | 1 |
| rs750059 | 11 | 116202995 | T | G | -1.275 | 0.2022 | -? | 0 | 0 | 0 | 1 |
| rs117164485 | 11 | 116620377 | A | G | -1.275 | 0.2023 | -? | 0 | 0 | 0 | 1 |
| rs595407 | 11 | 118904015 | C | T | 1.275 | 0.2023 | +? | 0 | 0 | 0 | 1 |
| rs11216625 | 11 | 117793989 | A | C | 1.275 | 0.2024 | +? | 0 | 0 | 0 | 1 |
| rs4562840 | 11 | 120834272 | C | G | -1.275 | 0.2024 | -? | 0 | 0 | 0 | 1 |

| | | | | | | | | | | | |
|---|---|---|---|---|---|---|---|---|---|---|---|
| rs3863315 | 11 | 117313231 | C | T | -1.274 | 0.2025 | -? | 0 | 0 | 0 | 1 |
| rs1392314200 | 11 | 119929853 | T | C | 1.273 | 0.2029 | +? | 0 | 0 | 0 | 1 |
| rs10750106 | 11 | 117324964 | T | C | 1.273 | 0.203 | +? | 0 | 0 | 0 | 1 |
| rs1785899 | 11 | 119356682 | T | C | -1.272 | 0.2033 | -? | 0 | 0 | 0 | 1 |
| rs55720349 | 11 | 117796139 | C | T | 1.272 | 0.2034 | +? | 0 | 0 | 0 | 1 |
| rs10790136 | 11 | 116201751 | G | A | -1.271 | 0.2038 | -? | 0 | 0 | 0 | 1 |
| rs641620 | 11 | 117074229 | C | T | 1.271 | 0.2039 | +? | 0 | 0 | 0 | 1 |
| rs10750144 | 11 | 119337504 | A | G | -1.269 | 0.2044 | -? | 0 | 0 | 0 | 1 |
| rs504068 | 11 | 117008946 | T | C | 1.269 | 0.2044 | +? | 0 | 0 | 0 | 1 |
| rs76530395 | 11 | 117899993 | A | G | -1.269 | 0.2045 | -? | 0 | 0 | 0 | 1 |
| rs509565 | 11 | 117120473 | T | C | -1.269 | 0.2046 | -? | 0 | 0 | 0 | 1 |
| rs3016797 | 11 | 117409884 | C | G | -1.268 | 0.2049 | -? | 0 | 0 | 0 | 1 |
| rs520289 | 11 | 117350137 | A | G | -1.267 | 0.2051 | -? | 0 | 0 | 0 | 1 |
| rs6589806 | 11 | 120166817 | C | T | -1.267 | 0.2051 | -? | 0 | 0 | 0 | 1 |
| rs7121294 | 11 | 117845835 | T | C | 1.267 | 0.2051 | +? | 0 | 0 | 0 | 1 |
| rs6589568 | 11 | 116670738 | G | A | -1.267 | 0.2053 | -- | 0 | 0.378 | 1 | 0.5385 |
| rs113447691 | 11 | 119848376 | T | C | -1.266 | 0.2055 | -? | 0 | 0 | 0 | 1 |
| rs56116614 | 11 | 119673271 | G | A | 1.266 | 0.2056 | +? | 0 | 0 | 0 | 1 |
| rs56220026 | 11 | 119673235 | C | G | 1.266 | 0.2056 | +? | 0 | 0 | 0 | 1 |
| rs509728 | 11 | 116533167 | C | G | 1.265 | 0.2057 | +? | 0 | 0 | 0 | 1 |
| rs9651686 | 11 | 117967839 | G | T | -1.265 | 0.2058 | -? | 0 | 0 | 0 | 1 |
| rs115325870 | 11 | 117361165 | T | C | 1.265 | 0.2059 | +? | 0 | 0 | 0 | 1 |
| rs2508450 | 11 | 117863829 | T | C | 1.265 | 0.2059 | ++ | 0 | 0.008 | 1 | 0.927 |
| rs4526780 | 11 | 120730621 | A | G | 1.265 | 0.2059 | +? | 0 | 0 | 0 | 1 |
| rs11216647 | 11 | 117825852 | C | T | -1.264 | 0.2062 | -- | 0 | 0 | 1 | 0.9845 |
| rs2850808 | 11 | 120797604 | A | G | 1.264 | 0.2062 | +? | 0 | 0 | 0 | 1 |
| rs12281464 | 11 | 120576351 | T | A | 1.264 | 0.2063 | +? | 0 | 0 | 0 | 1 |
| rs12294858 | 11 | 120576489 | A | G | 1.264 | 0.2063 | +? | 0 | 0 | 0 | 1 |
| rs4936418 | 11 | 117904049 | T | A | -1.264 | 0.2063 | -? | 0 | 0 | 0 | 1 |
| rs549507 | 11 | 121035618 | T | C | -1.263 | 0.2066 | -? | 0 | 0 | 0 | 1 |
| rs74387077 | 11 | 119669993 | A | G | 1.263 | 0.2067 | +? | 0 | 0 | 0 | 1 |
| rs517918 | 11 | 119771968 | C | T | -1.262 | 0.2068 | -? | 0 | 0 | 0 | 1 |
| rs605805 | 11 | 117949807 | G | A | -1.262 | 0.2068 | -? | 0 | 0 | 0 | 1 |
| rs2847502 | 11 | 120118498 | T | C | -1.262 | 0.2069 | -? | 0 | 0 | 0 | 1 |
| rs1240771 | 11 | 116518889 | G | A | -1.262 | 0.207 | -? | 0 | 0 | 0 | 1 |
| rs12574613 | 11 | 117636729 | T | C | 1.262 | 0.2071 | +? | 0 | 0 | 0 | 1 |
| rs1729407 | 11 | 116677370 | C | G | -1.262 | 0.2071 | -+ | -136.5 | 1.268 | 1 | 0.2601 |
| rs2089648 | 11 | 119512770 | A | G | 1.262 | 0.2071 | +? | 0 | 0 | 0 | 1 |
| rs10892425 | 11 | 119516100 | T | C | 1.261 | 0.2074 | +? | 0 | 0 | 0 | 1 |
| rs11217131 | 11 | 118903589 | G | C | 1.261 | 0.2074 | +? | 0 | 0 | 0 | 1 |
| rs11217369 | 11 | 119515905 | T | C | 1.261 | 0.2074 | +? | 0 | 0 | 0 | 1 |
| rs12788122 | 11 | 121083183 | A | G | 1.26 | 0.2076 | +? | 0 | 0 | 0 | 1 |
| rs580948 | 11 | 119699827 | T | C | 1.26 | 0.2077 | +? | 0 | 0 | 0 | 1 |
| rs10790244 | 11 | 118059557 | C | T | -1.258 | 0.2082 | -+ | -50.4 | 1.994 | 1 | 0.1579 |
| rs6589548 | 11 | 116362136 | G | A | -1.259 | 0.2082 | -? | 0 | 0 | 0 | 1 |
| rs11217984 | 11 | 120576154 | T | C | 1.257 | 0.2086 | +? | 0 | 0 | 0 | 1 |
| rs1599810 | 11 | 118179197 | A | G | 1.257 | 0.2087 | +? | 0 | 0 | 0 | 1 |
| rs3892497 | 11 | 120411267 | A | G | -1.257 | 0.2088 | -? | 0 | 0 | 0 | 1 |
| rs1729405 | 11 | 116678760 | C | T | -1.257 | 0.2089 | -+ | -139.8 | 1.251 | 1 | 0.2634 |
| rs4936525 | 11 | 120470521 | A | C | 1.256 | 0.2091 | +? | 0 | 0 | 0 | 1 |
| rs1815889 | 11 | 119815967 | A | G | -1.256 | 0.2092 | -? | 0 | 0 | 0 | 1 |
| rs653764 | 11 | 117948080 | T | C | -1.256 | 0.2092 | -? | 0 | 0 | 0 | 1 |
| rs73009147 | 11 | 116361536 | T | C | -1.256 | 0.2092 | -? | 0 | 0 | 0 | 1 |
| rs1261672 | 11 | 116518960 | G | A | -1.255 | 0.2093 | -? | 0 | 0 | 0 | 1 |
| rs35857852 | 11 | 117448312 | C | T | 1.255 | 0.2093 | +? | 0 | 0 | 0 | 1 |
| rs10790200 | 11 | 117567949 | C | T | 1.254 | 0.2097 | +? | 0 | 0 | 0 | 1 |

| | | | | | | | | | | | |
|---|---|---|---|---|---|---|---|---|---|---|---|
| rs4488208 | 11 | 118877621 | C | G | -1.254 | 0.2097 | -? | 0 | 0 | 0 | 1 |
| rs17119839 | 11 | 116478432 | G | A | -1.254 | 0.2099 | -? | 0 | 0 | 0 | 1 |
| rs534082 | 11 | 116287337 | T | C | -1.254 | 0.21 | -? | 0 | 0 | 0 | 1 |
| rs657389 | 11 | 117670752 | A | G | -1.254 | 0.21 | -? | 0 | 0 | 0 | 1 |
| rs60798124 | 11 | 118282241 | G | A | -1.253 | 0.2101 | -- | 0 | 0.245 | 1 | 0.6209 |
| rs752796 | 11 | 117434983 | G | A | 1.253 | 0.2102 | +? | 0 | 0 | 0 | 1 |
| rs77855139 | 11 | 117455203 | G | A | 1.253 | 0.2102 | +? | 0 | 0 | 0 | 1 |
| rs78466564 | 11 | 117795494 | T | C | 1.253 | 0.2102 | +? | 0 | 0 | 0 | 1 |
| rs10892441 | 11 | 119586418 | T | C | -1.253 | 0.2103 | -? | 0 | 0 | 0 | 1 |
| rs12278852 | 11 | 117934598 | A | G | -1.253 | 0.2103 | -? | 0 | 0 | 0 | 1 |
| rs10892658 | 11 | 120868246 | T | C | 1.252 | 0.2104 | +? | 0 | 0 | 0 | 1 |
| rs626984 | 11 | 119041659 | T | C | 1.252 | 0.2105 | +? | 0 | 0 | 0 | 1 |
| rs10892426 | 11 | 119516473 | T | C | 1.252 | 0.2107 | +? | 0 | 0 | 0 | 1 |
| rs2212358 | 11 | 117914386 | A | G | -1.252 | 0.2107 | -? | 0 | 0 | 0 | 1 |
| rs1784130 | 11 | 116539412 | C | G | -1.251 | 0.2109 | -? | 0 | 0 | 0 | 1 |
| rs12806934 | 11 | 117397482 | T | C | -1.251 | 0.211 | -? | 0 | 0 | 0 | 1 |
| rs634351 | 11 | 119815248 | C | T | -1.251 | 0.211 | -? | 0 | 0 | 0 | 1 |
| rs9666004 | 11 | 121131331 | T | C | 1.25 | 0.2114 | ++ | 0 | 0.27 | 1 | 0.6032 |
| rs11605781 | 11 | 120178632 | C | T | -1.249 | 0.2117 | -? | 0 | 0 | 0 | 1 |
| rs2156120 | 11 | 116557155 | G | A | 1.248 | 0.2119 | +? | 0 | 0 | 0 | 1 |
| rs528147 | 11 | 120036845 | A | G | 1.248 | 0.2119 | +? | 0 | 0 | 0 | 1 |
| rs11217512 | 11 | 119673722 | C | G | 1.248 | 0.212 | +? | 0 | 0 | 0 | 1 |
| rs11217513 | 11 | 119673977 | C | T | 1.248 | 0.212 | +? | 0 | 0 | 0 | 1 |
| rs76070443 | 11 | 119672890 | G | A | 1.248 | 0.212 | +? | 0 | 0 | 0 | 1 |
| rs586370 | 11 | 118816353 | A | G | -1.248 | 0.2121 | -- | 0 | 0.067 | 1 | 0.7951 |
| rs4603339 | 11 | 117656916 | G | A | -1.248 | 0.2122 | -? | 0 | 0 | 0 | 1 |
| rs11216626 | 11 | 117794282 | A | T | 1.247 | 0.2124 | +? | 0 | 0 | 0 | 1 |
| rs45497100 | 11 | 117158168 | T | A | 1.247 | 0.2124 | +? | 0 | 0 | 0 | 1 |
| rs573554 | 11 | 116355860 | C | T | -1.247 | 0.2124 | -? | 0 | 0 | 0 | 1 |
| rs10750146 | 11 | 119341145 | C | T | -1.247 | 0.2125 | -? | 0 | 0 | 0 | 1 |
| rs7105816 | 11 | 117934295 | G | A | -1.247 | 0.2125 | -? | 0 | 0 | 0 | 1 |
| rs9665995 | 11 | 121131224 | T | C | 1.247 | 0.2125 | ++ | 0 | 0.272 | 1 | 0.6019 |
| rs10892620 | 11 | 120587319 | G | A | -1.246 | 0.2127 | -? | 0 | 0 | 0 | 1 |
| rs11216831 | 11 | 118117018 | A | G | 1.246 | 0.2129 | +? | 0 | 0 | 0 | 1 |
| rs2156121 | 11 | 116557142 | A | C | 1.246 | 0.2129 | +? | 0 | 0 | 0 | 1 |
| rs4938399 | 11 | 117398399 | T | C | -1.245 | 0.2133 | -? | 0 | 0 | 0 | 1 |
| rs73008683 | 11 | 119744714 | A | G | -1.245 | 0.2133 | -? | 0 | 0 | 0 | 1 |
| rs9645664 | 11 | 118873987 | C | T | -1.245 | 0.2133 | -- | 0 | 0.088 | 1 | 0.7662 |
| rs1440184 | 11 | 116203460 | A | C | -1.244 | 0.2134 | -? | 0 | 0 | 0 | 1 |
| rs1784104 | 11 | 116551538 | C | T | -1.244 | 0.2136 | -? | 0 | 0 | 0 | 1 |
| rs2846012 | 11 | 116551449 | C | T | -1.244 | 0.2136 | -? | 0 | 0 | 0 | 1 |
| rs11217980 | 11 | 120575997 | A | G | 1.243 | 0.2137 | +? | 0 | 0 | 0 | 1 |
| rs4367952 | 11 | 120475940 | T | C | 1.243 | 0.2137 | +? | 0 | 0 | 0 | 1 |
| rs6589876 | 11 | 121192382 | C | G | 1.243 | 0.2137 | +? | 0 | 0 | 0 | 1 |
| rs11826104 | 11 | 117914627 | T | C | -1.243 | 0.2139 | -? | 0 | 0 | 0 | 1 |
| rs10891987 | 11 | 116417181 | C | G | -1.243 | 0.214 | -? | 0 | 0 | 0 | 1 |
| rs11217664 | 11 | 119929632 | G | A | -1.243 | 0.214 | -? | 0 | 0 | 0 | 1 |
| rs7952568 | 11 | 119672221 | T | C | 1.243 | 0.214 | +? | 0 | 0 | 0 | 1 |
| rs10892689 | 11 | 121088158 | G | A | -1.242 | 0.2141 | -? | 0 | 0 | 0 | 1 |
| rs11216437 | 11 | 117394685 | A | G | -1.242 | 0.2143 | -? | 0 | 0 | 0 | 1 |
| rs609305 | 11 | 119392654 | G | A | -1.242 | 0.2144 | -? | 0 | 0 | 0 | 1 |
| rs7936924 | 11 | 118978319 | C | T | 1.241 | 0.2145 | +? | 0 | 0 | 0 | 1 |
| rs4553405 | 11 | 118061798 | T | C | -1.241 | 0.2146 | -+ | -35.5 | 2.215 | 1 | 0.1367 |
| rs61901382 | 11 | 120581323 | A | G | 1.241 | 0.2146 | +? | 0 | 0 | 0 | 1 |
| rs7944756 | 11 | 120581609 | A | T | 1.241 | 0.2146 | +? | 0 | 0 | 0 | 1 |
| rs56074246 | 11 | 117384450 | C | T | -1.241 | 0.2147 | -? | 0 | 0 | 0 | 1 |

| | | | | | | | | | | | |
|---|---|---|---|---|---|---|---|---|---|---|---|
| rs4423192 | 11 | 121128543 | T | C | 1.241 | 0.2148 | ++ | 0 | 0.276 | 1 | 0.5995 |
| rs7113439 | 11 | 119674742 | T | C | 1.241 | 0.2148 | ++ | 0 | 0.089 | 1 | 0.766 |
| rs73014296 | 11 | 118187216 | G | A | -1.24 | 0.2149 | -? | 0 | 0 | 0 | 1 |
| rs4936468 | 11 | 119087122 | A | G | -1.24 | 0.215 | -? | 0 | 0 | 0 | 1 |
| rs4148172 | 11 | 119030759 | G | A | -1.24 | 0.2151 | -? | 0 | 0 | 0 | 1 |
| rs551031 | 11 | 121018725 | C | T | -1.239 | 0.2152 | -? | 0 | 0 | 0 | 1 |
| rs11601887 | 11 | 120932667 | C | T | 1.239 | 0.2155 | +? | 0 | 0 | 0 | 1 |
| rs12802185 | 11 | 121005464 | A | T | -1.239 | 0.2155 | -? | 0 | 0 | 0 | 1 |
| rs1217688880 | 11 | 117845618 | C | T | 1.238 | 0.2156 | +? | 0 | 0 | 0 | 1 |
| rs597670 | 11 | 120824094 | T | C | 1.238 | 0.2156 | +? | 0 | 0 | 0 | 1 |
| rs1939981 | 11 | 117608613 | C | T | -1.238 | 0.2157 | -? | 0 | 0 | 0 | 1 |
| rs4936526 | 11 | 120470867 | A | G | 1.238 | 0.2157 | +? | 0 | 0 | 0 | 1 |
| rs56765654 | 11 | 120177530 | T | C | -1.238 | 0.2157 | -? | 0 | 0 | 0 | 1 |
| rs75432677 | 11 | 120567135 | A | G | -1.238 | 0.2157 | -? | 0 | 0 | 0 | 1 |
| rs72645479 | 11 | 120083118 | C | T | 1.238 | 0.2158 | +? | 0 | 0 | 0 | 1 |
| rs76200167 | 11 | 116307648 | A | C | 1.238 | 0.2158 | +? | 0 | 0 | 0 | 1 |
| rs7929011 | 11 | 116464194 | A | G | 1.238 | 0.2159 | +? | 0 | 0 | 0 | 1 |
| rs480009 | 11 | 116711974 | T | C | -1.236 | 0.2163 | -? | 0 | 0 | 0 | 1 |
| rs10892424 | 11 | 119515046 | T | G | 1.236 | 0.2165 | +? | 0 | 0 | 0 | 1 |
| rs1784132 | 11 | 116539921 | A | G | -1.235 | 0.2168 | -? | 0 | 0 | 0 | 1 |
| rs35412484 | 11 | 116669428 | C | T | 1.235 | 0.2168 | +? | 0 | 0 | 0 | 1 |
| rs59371082 | 11 | 116547745 | G | A | 1.235 | 0.2169 | +? | 0 | 0 | 0 | 1 |
| rs7103821 | 11 | 120435024 | C | A | -1.235 | 0.217 | -? | 0 | 0 | 0 | 1 |
| rs73009144 | 11 | 116356422 | A | G | 1.234 | 0.2171 | +? | 0 | 0 | 0 | 1 |
| rs7932330 | 11 | 116555207 | G | A | 1.234 | 0.2173 | +? | 0 | 0 | 0 | 1 |
| rs4938408 | 11 | 117434629 | A | C | 1.233 | 0.2175 | +? | 0 | 0 | 0 | 1 |
| rs593061 | 11 | 119343329 | G | A | -1.233 | 0.2175 | -? | 0 | 0 | 0 | 1 |
| rs4938469 | 11 | 117911637 | C | T | -1.233 | 0.2177 | -? | 0 | 0 | 0 | 1 |
| rs236916 | 11 | 117089628 | A | G | 1.232 | 0.2178 | +? | 0 | 0 | 0 | 1 |
| rs74954439 | 11 | 116437742 | A | G | -1.231 | 0.2183 | -? | 0 | 0 | 0 | 1 |
| rs2849169 | 11 | 116680448 | A | C | -1.231 | 0.2184 | -? | 0 | 0 | 0 | 1 |
| rs1945764 | 11 | 118182801 | G | A | 1.23 | 0.2187 | +? | 0 | 0 | 0 | 1 |
| rs61898346 | 11 | 120189261 | A | G | -1.23 | 0.2187 | -? | 0 | 0 | 0 | 1 |
| rs12273147 | 11 | 117570395 | A | G | 1.23 | 0.2188 | +? | 0 | 0 | 0 | 1 |
| rs35820582 | 11 | 116470552 | C | G | 1.229 | 0.2189 | +? | 0 | 0 | 0 | 1 |
| rs57156480 | 11 | 118896959 | A | G | -1.229 | 0.2191 | -- | 0 | 0.324 | 1 | 0.5695 |
| rs1784107 | 11 | 116549427 | T | A | -1.229 | 0.2192 | -? | 0 | 0 | 0 | 1 |
| rs7940751 | 11 | 120860425 | T | C | 1.228 | 0.2193 | +? | 0 | 0 | 0 | 1 |
| rs2253134 | 11 | 120411963 | A | C | -1.228 | 0.2195 | -? | 0 | 0 | 0 | 1 |
| rs4938474 | 11 | 117934048 | A | G | -1.228 | 0.2195 | -? | 0 | 0 | 0 | 1 |
| rs12797743 | 11 | 117775198 | G | A | 1.227 | 0.2198 | +? | 0 | 0 | 0 | 1 |
| rs655116 | 11 | 117934704 | C | G | -1.226 | 0.22 | -- | 0 | 0.562 | 1 | 0.4536 |
| rs71480349 | 11 | 117520361 | C | T | 1.227 | 0.22 | ++ | 0 | 0.568 | 1 | 0.4511 |
| rs79136224 | 11 | 117795537 | T | C | 1.227 | 0.22 | +? | 0 | 0 | 0 | 1 |
| rs4271390 | 11 | 119526136 | T | C | -1.226 | 0.2202 | -? | 0 | 0 | 0 | 1 |
| rs2156634 | 11 | 120776001 | G | A | -1.226 | 0.2203 | -? | 0 | 0 | 0 | 1 |
| rs12800036 | 11 | 120871485 | T | C | 1.225 | 0.2205 | +? | 0 | 0 | 0 | 1 |
| rs2850810 | 11 | 120785857 | A | G | -1.225 | 0.2205 | -? | 0 | 0 | 0 | 1 |
| rs750058 | 11 | 116203044 | G | C | -1.225 | 0.2205 | -? | 0 | 0 | 0 | 1 |
| rs11217982 | 11 | 120576090 | A | T | 1.225 | 0.2206 | +? | 0 | 0 | 0 | 1 |
| rs11217983 | 11 | 120576091 | C | T | 1.225 | 0.2206 | +? | 0 | 0 | 0 | 1 |
| rs1440185 | 11 | 116203721 | A | T | -1.224 | 0.2208 | -? | 0 | 0 | 0 | 1 |
| rs12281217 | 11 | 116543514 | T | C | -1.224 | 0.221 | -? | 0 | 0 | 0 | 1 |
| rs632818 | 11 | 119386473 | C | T | -1.224 | 0.221 | -? | 0 | 0 | 0 | 1 |
| rs78264252 | 11 | 117795373 | A | C | 1.224 | 0.2211 | +? | 0 | 0 | 0 | 1 |
| rs893839 | 11 | 116198680 | T | A | -1.224 | 0.2211 | -? | 0 | 0 | 0 | 1 |

| | | | | | | | | | | | |
|---|---|---|---|---|---|---|---|---|---|---|---|
| rs17530451 | 11 | 118031950 | A | C | -1.223 | 0.2215 | -? | 0 | 0 | 0 | 1 |
| rs7928550 | 11 | 120887491 | T | A | -1.223 | 0.2215 | -? | 0 | 0 | 0 | 1 |
| rs10892622 | 11 | 120624486 | G | T | -1.222 | 0.2216 | -? | 0 | 0 | 0 | 1 |
| rs11216629 | 11 | 117794650 | G | C | 1.222 | 0.2216 | +? | 0 | 0 | 0 | 1 |
| rs112732949 | 11 | 116491465 | G | A | -1.222 | 0.2216 | -? | 0 | 0 | 0 | 1 |
| rs7924647 | 11 | 118728479 | G | A | -1.222 | 0.2216 | -- | 0 | 0.016 | 1 | 0.9001 |
| rs4936422 | 11 | 117917156 | C | A | -1.222 | 0.2217 | -? | 0 | 0 | 0 | 1 |
| rs12273341 | 11 | 117931924 | T | C | -1.222 | 0.2218 | -? | 0 | 0 | 0 | 1 |
| rs7122782 | 11 | 117569523 | A | G | 1.222 | 0.2218 | +? | 0 | 0 | 0 | 1 |
| rs733762 | 11 | 119384847 | C | G | -1.221 | 0.2219 | -? | 0 | 0 | 0 | 1 |
| rs11606105 | 11 | 120706014 | T | A | 1.22 | 0.2224 | +? | 0 | 0 | 0 | 1 |
| rs6589622 | 11 | 117646330 | C | T | -1.219 | 0.2227 | -? | 0 | 0 | 0 | 1 |
| rs79517065 | 11 | 117450936 | T | C | 1.219 | 0.2227 | +? | 0 | 0 | 0 | 1 |
| rs625267 | 11 | 117932630 | T | C | -1.219 | 0.223 | -- | 0 | 0.942 | 1 | 0.3319 |
| rs6421572 | 11 | 118977926 | A | C | 1.218 | 0.2232 | +? | 0 | 0 | 0 | 1 |
| rs1787688 | 11 | 116545723 | C | T | 1.218 | 0.2234 | +? | 0 | 0 | 0 | 1 |
| rs643115 | 11 | 119788251 | T | A | 1.218 | 0.2234 | +? | 0 | 0 | 0 | 1 |
| rs6589822 | 11 | 120507564 | G | A | -1.217 | 0.2236 | -? | 0 | 0 | 0 | 1 |
| rs659092 | 11 | 117917962 | C | T | -1.217 | 0.2236 | -? | 0 | 0 | 0 | 1 |
| rs76046563 | 11 | 117795082 | G | C | 1.217 | 0.2237 | +? | 0 | 0 | 0 | 1 |
| rs7123343 | 11 | 120826623 | T | C | -1.216 | 0.2239 | -? | 0 | 0 | 0 | 1 |
| rs10891964 | 11 | 116197173 | G | T | -1.216 | 0.224 | -? | 0 | 0 | 0 | 1 |
| rs2004506 | 11 | 117700588 | A | G | 1.216 | 0.224 | +? | 0 | 0 | 0 | 1 |
| rs56168028 | 11 | 119026384 | G | A | -1.216 | 0.224 | -? | 0 | 0 | 0 | 1 |
| rs12577725 | 11 | 120500339 | T | C | 1.215 | 0.2242 | +? | 0 | 0 | 0 | 1 |
| rs5095 | 11 | 116693163 | G | A | -1.215 | 0.2242 | -? | 0 | 0 | 0 | 1 |
| rs61898761 | 11 | 120342855 | G | A | -1.215 | 0.2242 | -? | 0 | 0 | 0 | 1 |
| rs10892688 | 11 | 121082729 | C | G | -1.215 | 0.2243 | -? | 0 | 0 | 0 | 1 |
| rs1784129 | 11 | 116539233 | A | G | -1.215 | 0.2243 | -? | 0 | 0 | 0 | 1 |
| rs12801338 | 11 | 117714145 | A | G | -1.215 | 0.2245 | -- | 0 | 0.053 | 1 | 0.8174 |
| rs533203 | 11 | 117133219 | T | C | 1.215 | 0.2245 | +? | 0 | 0 | 0 | 1 |
| rs7938067 | 11 | 117924278 | C | T | -1.214 | 0.2247 | -? | 0 | 0 | 0 | 1 |
| rs17124602 | 11 | 120834199 | C | G | 1.214 | 0.2248 | +? | 0 | 0 | 0 | 1 |
| rs652469 | 11 | 117906706 | A | G | 1.214 | 0.2248 | +? | 0 | 0 | 0 | 1 |
| rs11216788 | 11 | 118034201 | A | G | 1.214 | 0.2249 | +? | 0 | 0 | 0 | 1 |
| rs139831408 | 11 | 120473450 | T | C | -1.214 | 0.2249 | -? | 0 | 0 | 0 | 1 |
| rs11216630 | 11 | 117794735 | T | A | 1.213 | 0.2253 | +? | 0 | 0 | 0 | 1 |
| rs61900795 | 11 | 120276903 | C | T | -1.212 | 0.2254 | -? | 0 | 0 | 0 | 1 |
| rs77637745 | 11 | 117795397 | A | C | 1.212 | 0.2254 | +? | 0 | 0 | 0 | 1 |
| rs7945999 | 11 | 117646291 | C | T | -1.212 | 0.2254 | -? | 0 | 0 | 0 | 1 |
| rs1786188 | 11 | 117937131 | A | G | -1.212 | 0.2255 | -- | 0 | 0.761 | 1 | 0.383 |
| rs11216807 | 11 | 118063030 | C | T | -1.212 | 0.2256 | -+ | -51.4 | 1.982 | 1 | 0.1592 |
| rs610675 | 11 | 116640094 | A | T | -1.212 | 0.2256 | -+ | -96.4 | 1.528 | 1 | 0.2164 |
| rs622604 | 11 | 116640062 | C | T | -1.212 | 0.2256 | -+ | -96.4 | 1.528 | 1 | 0.2164 |
| rs623908 | 11 | 116640368 | G | A | -1.212 | 0.2256 | -+ | -96.4 | 1.528 | 1 | 0.2164 |
| rs592828 | 11 | 117219305 | C | G | 1.212 | 0.2257 | +? | 0 | 0 | 0 | 1 |
| rs12799825 | 11 | 117448013 | A | C | 1.211 | 0.2258 | +? | 0 | 0 | 0 | 1 |
| rs617357 | 11 | 117899803 | A | G | 1.211 | 0.2258 | +- | -163.1 | 1.14 | 1 | 0.2856 |
| rs10892380 | 11 | 119400052 | A | C | -1.211 | 0.2259 | -? | 0 | 0 | 0 | 1 |
| rs623796 | 11 | 117719316 | T | C | 1.211 | 0.2259 | +? | 0 | 0 | 0 | 1 |
| rs201359394 | 11 | 118536950 | T | A | 1.21 | 0.2263 | ++ | 0 | 0.643 | 1 | 0.4225 |
| rs10892496 | 11 | 119803839 | C | A | -1.209 | 0.2265 | -? | 0 | 0 | 0 | 1 |
| rs595167 | 11 | 119853430 | T | C | 1.209 | 0.2265 | +? | 0 | 0 | 0 | 1 |
| rs61905084 | 11 | 116610294 | C | T | -1.21 | 0.2265 | -- | 0 | 0.197 | 1 | 0.657 |
| rs7125177 | 11 | 121143882 | T | C | 1.209 | 0.2265 | ++ | 0 | 0.295 | 1 | 0.5871 |
| rs11217560 | 11 | 119777403 | G | A | -1.209 | 0.2266 | -? | 0 | 0 | 0 | 1 |

| | | | | | | | | | | | |
|---|---|---|---|---|---|---|---|---|---|---|---|
| rs12281614 | 11 | 120583803 | T | C | 1.208 | 0.2269 | +? | 0 | 0 | 0 | 1 |
| rs532150 | 11 | 119353671 | C | T | -1.208 | 0.2269 | -? | 0 | 0 | 0 | 1 |
| rs11218083 | 11 | 120827003 | T | C | -1.208 | 0.227 | -? | 0 | 0 | 0 | 1 |
| rs10892188 | 11 | 117754524 | C | A | 1.208 | 0.2271 | +? | 0 | 0 | 0 | 1 |
| rs1488779268 | 11 | 118964330 | A | G | -1.208 | 0.2271 | -? | 0 | 0 | 0 | 1 |
| rs11217583 | 11 | 119801338 | T | C | -1.207 | 0.2273 | -? | 0 | 0 | 0 | 1 |
| rs117950757 | 11 | 117974397 | A | G | 1.207 | 0.2273 | +? | 0 | 0 | 0 | 1 |
| rs59868549 | 11 | 121113812 | T | A | 1.206 | 0.2277 | ++ | 0 | 0.279 | 1 | 0.5976 |
| rs1815890 | 11 | 119816115 | T | C | -1.206 | 0.2278 | -? | 0 | 0 | 0 | 1 |
| rs7104501 | 11 | 121126386 | T | A | 1.206 | 0.2278 | ++ | 0 | 0.297 | 1 | 0.5858 |
| rs17547832 | 11 | 119344568 | G | A | -1.206 | 0.228 | -? | 0 | 0 | 0 | 1 |
| rs2887176 | 11 | 117312429 | G | A | -1.206 | 0.228 | -? | 0 | 0 | 0 | 1 |
| rs10892495 | 11 | 119803564 | G | C | -1.205 | 0.2281 | -? | 0 | 0 | 0 | 1 |
| rs948379 | 11 | 121120130 | T | C | -1.205 | 0.2281 | -- | 0 | 0.015 | 1 | 0.9029 |
| rs10892010 | 11 | 116536656 | C | A | 1.205 | 0.2282 | +? | 0 | 0 | 0 | 1 |
| rs74526104 | 11 | 117446003 | G | A | 1.205 | 0.2282 | +? | 0 | 0 | 0 | 1 |
| rs10891997 | 11 | 116464675 | G | A | -1.204 | 0.2284 | -? | 0 | 0 | 0 | 1 |
| rs190143695 | 11 | 117938674 | C | A | -1.204 | 0.2284 | -? | 0 | 0 | 0 | 1 |
| rs12295930 | 11 | 121079190 | T | C | -1.204 | 0.2286 | -? | 0 | 0 | 0 | 1 |
| rs2115926 | 11 | 121145524 | A | G | 1.204 | 0.2286 | ++ | 0 | 0.298 | 1 | 0.585 |
| rs4936507 | 11 | 119934563 | A | G | -1.204 | 0.2286 | -? | 0 | 0 | 0 | 1 |
| rs80085590 | 11 | 117795232 | A | G | 1.204 | 0.2287 | +? | 0 | 0 | 0 | 1 |
| rs1787686 | 11 | 116541472 | A | G | 1.203 | 0.2288 | +? | 0 | 0 | 0 | 1 |
| rs4936523 | 11 | 120470159 | C | T | 1.203 | 0.2288 | +? | 0 | 0 | 0 | 1 |
| rs534839 | 11 | 119353388 | C | T | -1.203 | 0.2288 | -? | 0 | 0 | 0 | 1 |
| rs11216026 | 11 | 116464800 | G | A | 1.203 | 0.2289 | +? | 0 | 0 | 0 | 1 |
| rs12286892 | 11 | 119776710 | T | C | -1.203 | 0.2289 | -? | 0 | 0 | 0 | 1 |
| rs12363999 | 11 | 121144114 | T | C | 1.203 | 0.2289 | ++ | 0 | 0.299 | 1 | 0.5847 |
| rs2515798 | 11 | 117819480 | C | A | -1.203 | 0.2289 | -? | 0 | 0 | 0 | 1 |
| rs10790164 | 11 | 116699389 | A | G | 1.203 | 0.229 | +? | 0 | 0 | 0 | 1 |
| rs9667438 | 11 | 117327680 | T | C | 1.203 | 0.2291 | +? | 0 | 0 | 0 | 1 |
| rs11217584 | 11 | 119802832 | A | G | -1.202 | 0.2292 | -? | 0 | 0 | 0 | 1 |
| rs494356 | 11 | 117071970 | T | C | 1.202 | 0.2292 | +? | 0 | 0 | 0 | 1 |
| rs681569 | 11 | 119914616 | A | C | 1.202 | 0.2292 | +? | 0 | 0 | 0 | 1 |
| rs7479331 | 11 | 118829521 | T | G | -1.202 | 0.2292 | -? | 0 | 0 | 0 | 1 |
| rs750056 | 11 | 116202506 | T | C | -1.202 | 0.2292 | -? | 0 | 0 | 0 | 1 |
| rs10892485 | 11 | 119787519 | A | G | -1.202 | 0.2294 | -? | 0 | 0 | 0 | 1 |
| rs12419879 | 11 | 119240757 | T | C | 1.202 | 0.2294 | +? | 0 | 0 | 0 | 1 |
| rs71480312 | 11 | 116554088 | T | C | 1.202 | 0.2294 | +? | 0 | 0 | 0 | 1 |
| rs1941302 | 11 | 121132370 | T | A | 1.202 | 0.2295 | ++ | 0 | 0.3 | 1 | 0.5841 |
| rs7130538 | 11 | 121132553 | T | G | 1.202 | 0.2295 | ++ | 0 | 0.3 | 1 | 0.5841 |
| rs77233977 | 11 | 121057146 | T | C | 1.202 | 0.2295 | ++ | 0 | 0.717 | 1 | 0.3973 |
| rs41467444 | 11 | 119889516 | T | C | -1.201 | 0.2297 | -? | 0 | 0 | 0 | 1 |
| rs11216648 | 11 | 117830429 | G | A | -1.201 | 0.2299 | -- | 0 | 0.012 | 1 | 0.9141 |
| rs9971587 | 11 | 117131286 | A | G | 1.201 | 0.2299 | +? | 0 | 0 | 0 | 1 |
| rs644613 | 11 | 119347715 | T | C | -1.2 | 0.23 | -? | 0 | 0 | 0 | 1 |
| rs7924795 | 11 | 117924263 | T | G | -1.2 | 0.2301 | -? | 0 | 0 | 0 | 1 |
| rs11603672 | 11 | 117563212 | G | A | 1.2 | 0.2302 | +? | 0 | 0 | 0 | 1 |
| rs7105658 | 11 | 118712270 | T | C | -1.199 | 0.2306 | -- | 0 | 0.001 | 1 | 0.9775 |
| rs7114641 | 11 | 117628287 | G | A | -1.199 | 0.2306 | -? | 0 | 0 | 0 | 1 |
| rs17245664 | 11 | 120569528 | A | G | -1.199 | 0.2307 | -? | 0 | 0 | 0 | 1 |
| rs669812 | 11 | 119901422 | T | C | 1.199 | 0.2307 | +? | 0 | 0 | 0 | 1 |
| rs732480 | 11 | 120828827 | A | G | 1.199 | 0.2307 | +? | 0 | 0 | 0 | 1 |
| rs1319166 | 11 | 119942557 | C | G | -1.198 | 0.2308 | -? | 0 | 0 | 0 | 1 |
| rs633683 | 11 | 118504742 | T | C | 1.198 | 0.2308 | ++ | 0 | 0.115 | 1 | 0.734 |
| rs58856586 | 11 | 116366118 | A | G | 1.198 | 0.2309 | +? | 0 | 0 | 0 | 1 |

| | | | | | | | | | | | |
|---|---|---|---|---|---|---|---|---|---|---|---|
| rs4936600 | 11 | 121133458 | C | T | 1.198 | 0.231 | ++ | 0 | 0.302 | 1 | 0.5826 |
| rs474201 | 11 | 118958229 | G | A | 1.198 | 0.2311 | +? | 0 | 0 | 0 | 1 |
| rs4936415 | 11 | 117875313 | C | G | -1.197 | 0.2311 | -+ | 0 | 0.685 | 1 | 0.4077 |
| rs117920151 | 11 | 116759706 | C | G | 1.197 | 0.2312 | +? | 0 | 0 | 0 | 1 |
| rs4403813 | 11 | 119816331 | C | T | -1.197 | 0.2312 | -? | 0 | 0 | 0 | 1 |
| rs56895810 | 11 | 119934071 | A | C | -1.197 | 0.2314 | -? | 0 | 0 | 0 | 1 |
| rs7122510 | 11 | 118708785 | A | C | -1.197 | 0.2314 | -- | 0 | 0.046 | 1 | 0.8307 |
| rs12276440 | 11 | 120575481 | T | C | 1.197 | 0.2315 | +? | 0 | 0 | 0 | 1 |
| rs1263164 | 11 | 116675291 | C | A | -1.197 | 0.2315 | -? | 0 | 0 | 0 | 1 |
| rs6589651 | 11 | 118058156 | G | A | -1.196 | 0.2316 | -+ | -72 | 1.745 | 1 | 0.1866 |
| rs139447106 | 11 | 116737302 | T | C | -1.195 | 0.2319 | -? | 0 | 0 | 0 | 1 |
| rs4938758 | 11 | 119892030 | T | C | -1.195 | 0.2319 | -? | 0 | 0 | 0 | 1 |
| rs55644928 | 11 | 117303430 | T | C | -1.195 | 0.2319 | -? | 0 | 0 | 0 | 1 |
| rs11217579 | 11 | 119798293 | T | A | -1.195 | 0.232 | -? | 0 | 0 | 0 | 1 |
| rs34616770 | 11 | 120279132 | C | G | 1.195 | 0.2322 | ++ | 0 | 0.927 | 1 | 0.3358 |
| rs503175 | 11 | 119753926 | C | T | -1.195 | 0.2322 | -? | 0 | 0 | 0 | 1 |
| rs893837 | 11 | 116198048 | C | T | -1.194 | 0.2323 | -? | 0 | 0 | 0 | 1 |
| rs7122511 | 11 | 118708786 | A | C | -1.194 | 0.2324 | -- | 0 | 0.044 | 1 | 0.8344 |
| rs10892135 | 11 | 117392826 | A | G | -1.194 | 0.2325 | -? | 0 | 0 | 0 | 1 |
| rs540469 | 11 | 117346513 | C | T | 1.194 | 0.2325 | +? | 0 | 0 | 0 | 1 |
| rs10892645 | 11 | 120794509 | G | A | 1.193 | 0.2327 | +? | 0 | 0 | 0 | 1 |
| rs1170855838 | 11 | 119889758 | T | C | 1.193 | 0.2327 | +? | 0 | 0 | 0 | 1 |
| rs7103366 | 11 | 118057867 | G | T | -1.193 | 0.2327 | -+ | -72.4 | 1.74 | 1 | 0.1871 |
| rs554422 | 11 | 119778270 | T | C | -1.193 | 0.233 | -? | 0 | 0 | 0 | 1 |
| rs1145211 | 11 | 116557216 | A | C | 1.192 | 0.2331 | +? | 0 | 0 | 0 | 1 |
| rs752795 | 11 | 117434985 | G | C | 1.192 | 0.2331 | +? | 0 | 0 | 0 | 1 |
| rs12270409 | 11 | 120568566 | C | T | 1.192 | 0.2332 | +? | 0 | 0 | 0 | 1 |
| rs11217783 | 11 | 120151510 | T | A | -1.192 | 0.2333 | -? | 0 | 0 | 0 | 1 |
| rs1786190 | 11 | 117946720 | G | A | -1.191 | 0.2335 | -? | 0 | 0 | 0 | 1 |
| rs604411 | 11 | 117946386 | T | C | -1.191 | 0.2335 | -? | 0 | 0 | 0 | 1 |
| rs647593 | 11 | 117461576 | C | T | -1.191 | 0.2335 | -? | 0 | 0 | 0 | 1 |
| rs9630199 | 11 | 120575671 | C | T | 1.191 | 0.2338 | +? | 0 | 0 | 0 | 1 |
| rs2542062 | 11 | 116680449 | A | G | -1.19 | 0.2339 | -? | 0 | 0 | 0 | 1 |
| rs80062985 | 11 | 119817324 | T | C | -1.19 | 0.2339 | -? | 0 | 0 | 0 | 1 |
| rs10892695 | 11 | 121112943 | G | T | -1.19 | 0.234 | -- | -112.7 | 1.41 | 1 | 0.235 |
| rs580119 | 11 | 117216213 | G | T | 1.19 | 0.234 | +? | 0 | 0 | 0 | 1 |
| rs577975 | 11 | 116722910 | C | T | -1.19 | 0.2341 | -? | 0 | 0 | 0 | 1 |
| rs10219396 | 11 | 117927766 | A | G | -1.189 | 0.2345 | -? | 0 | 0 | 0 | 1 |
| rs11218115 | 11 | 120893187 | A | C | -1.189 | 0.2346 | -? | 0 | 0 | 0 | 1 |
| rs589348 | 11 | 119797699 | T | C | 1.188 | 0.2348 | +? | 0 | 0 | 0 | 1 |
| rs610775 | 11 | 117124696 | A | G | 1.188 | 0.2348 | +? | 0 | 0 | 0 | 1 |
| rs7103918 | 11 | 118061480 | T | C | -1.188 | 0.2349 | -+ | -51.2 | 1.984 | 1 | 0.159 |
| rs138897098 | 11 | 117599488 | C | T | 1.187 | 0.2351 | +? | 0 | 0 | 0 | 1 |
| rs17588728 | 11 | 117604604 | T | C | 1.187 | 0.2351 | +? | 0 | 0 | 0 | 1 |
| rs948377 | 11 | 121120917 | T | C | -1.186 | 0.2354 | -- | 0 | 0.012 | 1 | 0.9114 |
| rs10790166 | 11 | 116779499 | G | A | -1.186 | 0.2358 | -? | 0 | 0 | 0 | 1 |
| rs12275681 | 11 | 120173845 | G | A | -1.186 | 0.2358 | -? | 0 | 0 | 0 | 1 |
| rs12364606 | 11 | 117680787 | T | C | -1.185 | 0.2359 | -? | 0 | 0 | 0 | 1 |
| rs11217981 | 11 | 120576039 | A | G | 1.185 | 0.2362 | +? | 0 | 0 | 0 | 1 |
| rs36195057 | 11 | 116542247 | A | G | 1.185 | 0.2362 | +? | 0 | 0 | 0 | 1 |
| rs7101905 | 11 | 119844937 | T | C | -1.185 | 0.2362 | -? | 0 | 0 | 0 | 1 |
| rs9630200 | 11 | 120575845 | A | G | 1.185 | 0.2362 | +? | 0 | 0 | 0 | 1 |
| rs575280 | 11 | 116527728 | G | A | -1.184 | 0.2366 | -? | 0 | 0 | 0 | 1 |
| rs11821451 | 11 | 117312301 | A | G | -1.183 | 0.2368 | -? | 0 | 0 | 0 | 1 |
| rs642896 | 11 | 119353666 | G | A | -1.183 | 0.2368 | -? | 0 | 0 | 0 | 1 |
| rs7120765 | 11 | 121122021 | A | G | -1.183 | 0.2368 | -- | 0 | 0.012 | 1 | 0.9129 |

| | | | | | | | | | | | |
|---|---|---|---|---|---|---|---|---|---|---|---|
| rs75195166 | 11 | 117795189 | A | G | 1.183 | 0.2369 | +? | 0 | 0 | 0 | 1 |
| rs7928669 | 11 | 121121448 | T | C | -1.183 | 0.2369 | -- | 0 | 0.012 | 1 | 0.9131 |
| rs536141 | 11 | 117104536 | A | C | 1.183 | 0.237 | +? | 0 | 0 | 0 | 1 |
| rs74365486 | 11 | 116482449 | A | C | -1.183 | 0.237 | -? | 0 | 0 | 0 | 1 |
| rs77964702 | 11 | 117391421 | A | G | 1.182 | 0.2371 | +? | 0 | 0 | 0 | 1 |
| rs34994677 | 11 | 116472577 | G | A | 1.182 | 0.2372 | +? | 0 | 0 | 0 | 1 |
| rs487298 | 11 | 117205844 | T | G | 1.182 | 0.2372 | +? | 0 | 0 | 0 | 1 |
| rs55966937 | 11 | 120705938 | G | A | 1.182 | 0.2372 | +? | 0 | 0 | 0 | 1 |
| rs601842 | 11 | 119350396 | C | T | -1.182 | 0.2372 | -? | 0 | 0 | 0 | 1 |
| rs11217917 | 11 | 120441767 | G | C | 1.181 | 0.2375 | +? | 0 | 0 | 0 | 1 |
| rs594603 | 11 | 117923079 | T | C | -1.181 | 0.2375 | -? | 0 | 0 | 0 | 1 |
| rs4938467 | 11 | 117875171 | T | C | -1.181 | 0.2378 | -+ | 0 | 0.67 | 1 | 0.4132 |
| rs7105395 | 11 | 120152148 | T | C | 1.18 | 0.2379 | +? | 0 | 0 | 0 | 1 |
| rs611719 | 11 | 119349454 | A | G | -1.18 | 0.2381 | -? | 0 | 0 | 0 | 1 |
| rs1893817 | 11 | 120824400 | T | C | 1.179 | 0.2382 | +? | 0 | 0 | 0 | 1 |
| rs714053 | 11 | 117946138 | T | C | 1.179 | 0.2383 | ++ | 0 | 0.025 | 1 | 0.8746 |
| rs906540 | 11 | 116440439 | A | T | -1.179 | 0.2384 | -? | 0 | 0 | 0 | 1 |
| rs948102 | 11 | 117656310 | A | G | -1.179 | 0.2384 | -? | 0 | 0 | 0 | 1 |
| rs2845696 | 11 | 117947103 | G | A | -1.179 | 0.2385 | -? | 0 | 0 | 0 | 1 |
| rs1240660 | 11 | 116493738 | G | A | 1.178 | 0.2387 | +? | 0 | 0 | 0 | 1 |
| rs11216736 | 11 | 117944485 | C | T | 1.178 | 0.2389 | ++ | 0 | 0.098 | 1 | 0.7548 |
| rs576190 | 11 | 117205766 | C | T | 1.177 | 0.2391 | +? | 0 | 0 | 0 | 1 |
| rs7117594 | 11 | 118714492 | T | C | -1.177 | 0.2391 | -- | 0 | 0.005 | 1 | 0.9462 |
| rs11217854 | 11 | 120267537 | G | A | 1.177 | 0.2392 | ++ | 0 | 0.946 | 1 | 0.3307 |
| rs609530 | 11 | 119349933 | T | C | -1.177 | 0.2392 | -? | 0 | 0 | 0 | 1 |
| rs2075295 | 11 | 116628401 | C | T | -1.176 | 0.2395 | -- | 0 | 0.45 | 1 | 0.5021 |
| rs12364363 | 11 | 119500880 | C | T | -1.176 | 0.2396 | -? | 0 | 0 | 0 | 1 |
| rs11217865 | 11 | 120300327 | C | A | -1.175 | 0.2398 | -? | 0 | 0 | 0 | 1 |
| rs61898763 | 11 | 120343650 | T | C | -1.175 | 0.2398 | -? | 0 | 0 | 0 | 1 |
| rs74391474 | 11 | 117795283 | C | G | 1.175 | 0.2398 | +? | 0 | 0 | 0 | 1 |
| rs74578710 | 11 | 117795285 | A | G | 1.175 | 0.2398 | +? | 0 | 0 | 0 | 1 |
| rs1011664 | 11 | 119796458 | G | C | -1.175 | 0.24 | -? | 0 | 0 | 0 | 1 |
| rs4938738 | 11 | 119778544 | T | G | -1.175 | 0.24 | -? | 0 | 0 | 0 | 1 |
| rs145076512 | 11 | 117466661 | G | T | 1.174 | 0.2403 | +? | 0 | 0 | 0 | 1 |
| rs12361791 | 11 | 118100176 | G | T | 1.174 | 0.2404 | +? | 0 | 0 | 0 | 1 |
| rs7106859 | 11 | 121125452 | A | C | 1.174 | 0.2404 | ++ | 0 | 0.315 | 1 | 0.5745 |
| rs10219389 | 11 | 119520783 | C | A | -1.174 | 0.2405 | -? | 0 | 0 | 0 | 1 |
| rs7102869 | 11 | 120580623 | T | C | 1.174 | 0.2405 | +? | 0 | 0 | 0 | 1 |
| rs638405 | 11 | 117163824 | G | C | 1.173 | 0.2407 | +? | 0 | 0 | 0 | 1 |
| rs11216020 | 11 | 116452961 | C | G | -1.173 | 0.2408 | -? | 0 | 0 | 0 | 1 |
| rs77259443 | 11 | 117474492 | G | A | 1.173 | 0.2409 | +? | 0 | 0 | 0 | 1 |
| rs906827 | 11 | 119514129 | A | C | 1.173 | 0.2409 | +? | 0 | 0 | 0 | 1 |
| rs596770 | 11 | 119773548 | G | A | -1.172 | 0.241 | -? | 0 | 0 | 0 | 1 |
| rs662646 | 11 | 119345979 | T | C | -1.172 | 0.2411 | -? | 0 | 0 | 0 | 1 |
| rs7130866 | 11 | 119636942 | G | A | 1.172 | 0.2413 | ++ | 0 | 0.123 | 1 | 0.7256 |
| rs7121490 | 11 | 120152147 | C | T | 1.171 | 0.2414 | +? | 0 | 0 | 0 | 1 |
| rs10892628 | 11 | 120707383 | A | G | 1.171 | 0.2415 | +? | 0 | 0 | 0 | 1 |
| rs12785765 | 11 | 120890960 | C | A | -1.171 | 0.2415 | -? | 0 | 0 | 0 | 1 |
| rs7928488 | 11 | 116203828 | T | C | -1.171 | 0.2415 | -? | 0 | 0 | 0 | 1 |
| rs11217928 | 11 | 120495969 | T | C | 1.171 | 0.2416 | +? | 0 | 0 | 0 | 1 |
| rs500787 | 11 | 117725333 | T | C | 1.171 | 0.2417 | +? | 0 | 0 | 0 | 1 |
| rs11822909 | 11 | 119520995 | T | C | -1.17 | 0.2418 | -? | 0 | 0 | 0 | 1 |
| rs1320721 | 11 | 120466336 | C | G | 1.17 | 0.2419 | +? | 0 | 0 | 0 | 1 |
| rs4938815 | 11 | 120441647 | G | T | 1.17 | 0.2419 | +? | 0 | 0 | 0 | 1 |
| rs11825297 | 11 | 117642376 | T | C | -1.17 | 0.242 | -? | 0 | 0 | 0 | 1 |
| rs881517 | 11 | 116370871 | T | G | 1.17 | 0.242 | +? | 0 | 0 | 0 | 1 |

| | | | | | | | | | | | |
|---|---|---|---|---|---|---|---|---|---|---|---|
| rs12291772 | 11 | 121076227 | A | G | -1.17 | 0.2421 | -? | 0 | 0 | 0 | 1 |
| rs55973262 | 11 | 117566305 | C | G | 1.17 | 0.2421 | +? | 0 | 0 | 0 | 1 |
| rs56406822 | 11 | 120219546 | A | G | 1.17 | 0.2421 | ++ | 0 | 0.95 | 1 | 0.3296 |
| rs624882 | 11 | 119386445 | T | C | -1.17 | 0.2422 | -? | 0 | 0 | 0 | 1 |
| rs112885596 | 11 | 120833690 | C | T | -1.169 | 0.2423 | -? | 0 | 0 | 0 | 1 |
| rs2155387 | 11 | 117654176 | A | C | -1.169 | 0.2423 | -? | 0 | 0 | 0 | 1 |
| rs11217548 | 11 | 119742803 | T | C | -1.169 | 0.2425 | -? | 0 | 0 | 0 | 1 |
| rs11217562 | 11 | 119777416 | T | C | -1.168 | 0.2427 | -? | 0 | 0 | 0 | 1 |
| rs11217148 | 11 | 118957735 | T | G | 1.168 | 0.2429 | +? | 0 | 0 | 0 | 1 |
| rs11607987 | 11 | 119856928 | T | C | -1.168 | 0.2429 | -? | 0 | 0 | 0 | 1 |
| rs7947506 | 11 | 116359612 | T | C | -1.168 | 0.2429 | -? | 0 | 0 | 0 | 1 |
| rs2305826 | 11 | 117241760 | T | A | 1.167 | 0.2432 | +? | 0 | 0 | 0 | 1 |
| rs80072373 | 11 | 116485420 | G | C | -1.167 | 0.2432 | -? | 0 | 0 | 0 | 1 |
| rs147634995 | 11 | 120067199 | A | C | 1.167 | 0.2434 | +? | 0 | 0 | 0 | 1 |
| rs76785396 | 11 | 119702716 | A | T | 1.167 | 0.2434 | +? | 0 | 0 | 0 | 1 |
| rs7927356 | 11 | 120950632 | C | T | -1.167 | 0.2434 | -? | 0 | 0 | 0 | 1 |
| rs7940619 | 11 | 120950521 | A | G | -1.167 | 0.2434 | -? | 0 | 0 | 0 | 1 |
| rs77586292 | 11 | 116821525 | T | C | -1.166 | 0.2435 | -? | 0 | 0 | 0 | 1 |
| rs77963671 | 11 | 116434951 | T | C | -1.166 | 0.2435 | -? | 0 | 0 | 0 | 1 |
| rs1790469 | 11 | 119930569 | A | G | 1.166 | 0.2437 | +? | 0 | 0 | 0 | 1 |
| rs559430 | 11 | 119775597 | C | T | -1.166 | 0.2437 | -? | 0 | 0 | 0 | 1 |
| rs11218139 | 11 | 120959225 | G | A | -1.166 | 0.2438 | -? | 0 | 0 | 0 | 1 |
| rs4938397 | 11 | 117392337 | T | G | -1.166 | 0.2438 | -? | 0 | 0 | 0 | 1 |
| rs2511050 | 11 | 117338912 | A | G | 1.165 | 0.244 | +? | 0 | 0 | 0 | 1 |
| rs4936564 | 11 | 120800376 | A | G | -1.165 | 0.2442 | -+ | -182.9 | 1.06 | 1 | 0.3031 |
| rs12796185 | 11 | 120446830 | C | T | 1.164 | 0.2443 | +? | 0 | 0 | 0 | 1 |
| rs10892429 | 11 | 119548052 | A | G | 1.164 | 0.2444 | +? | 0 | 0 | 0 | 1 |
| rs487827 | 11 | 119772922 | T | G | -1.164 | 0.2445 | -? | 0 | 0 | 0 | 1 |
| rs494606 | 11 | 116714118 | G | T | -1.164 | 0.2445 | -? | 0 | 0 | 0 | 1 |
| rs115894834 | 11 | 117347073 | G | A | -1.164 | 0.2446 | -? | 0 | 0 | 0 | 1 |
| rs61902685 | 11 | 120566146 | C | T | 1.164 | 0.2446 | +? | 0 | 0 | 0 | 1 |
| rs77346757 | 11 | 117347029 | A | G | -1.164 | 0.2446 | -? | 0 | 0 | 0 | 1 |
| rs685540 | 11 | 117908079 | C | G | -1.163 | 0.2449 | -- | -188.5 | 1.04 | 1 | 0.3078 |
| rs10892635 | 11 | 120759130 | A | G | -1.162 | 0.2451 | -? | 0 | 0 | 0 | 1 |
| rs539858 | 11 | 116729013 | C | T | -1.162 | 0.2453 | -? | 0 | 0 | 0 | 1 |
| rs10892421 | 11 | 119498403 | C | T | -1.161 | 0.2455 | -? | 0 | 0 | 0 | 1 |
| rs527347 | 11 | 117203633 | G | A | 1.161 | 0.2456 | +? | 0 | 0 | 0 | 1 |
| rs61898358 | 11 | 120222480 | A | G | -1.161 | 0.2457 | -? | 0 | 0 | 0 | 1 |
| rs508479 | 11 | 119775811 | A | C | -1.16 | 0.2459 | -? | 0 | 0 | 0 | 1 |
| rs7942167 | 11 | 121121501 | G | A | -1.16 | 0.246 | -- | 0 | 0.009 | 1 | 0.9233 |
| rs4373935 | 11 | 120273301 | G | A | -1.159 | 0.2463 | -? | 0 | 0 | 0 | 1 |
| rs474867 | 11 | 120808714 | A | G | 1.159 | 0.2463 | +? | 0 | 0 | 0 | 1 |
| rs10892074 | 11 | 116980641 | T | G | 1.159 | 0.2466 | +? | 0 | 0 | 0 | 1 |
| rs11217515 | 11 | 119674099 | C | T | 1.159 | 0.2466 | +? | 0 | 0 | 0 | 1 |
| rs116904534 | 11 | 117554617 | G | A | -1.159 | 0.2466 | -? | 0 | 0 | 0 | 1 |
| rs11216785 | 11 | 118030633 | A | G | 1.158 | 0.2467 | +? | 0 | 0 | 0 | 1 |
| rs4537767 | 11 | 120273686 | G | A | -1.158 | 0.2468 | -? | 0 | 0 | 0 | 1 |
| rs67617613 | 11 | 120331916 | C | A | -1.158 | 0.247 | -? | 0 | 0 | 0 | 1 |
| rs10892422 | 11 | 119498748 | G | A | -1.157 | 0.2472 | -? | 0 | 0 | 0 | 1 |
| rs1448722646 | 11 | 120018127 | T | C | 1.157 | 0.2473 | +? | 0 | 0 | 0 | 1 |
| rs3016357 | 11 | 116555415 | T | C | 1.157 | 0.2473 | +? | 0 | 0 | 0 | 1 |
| rs10892381 | 11 | 119400817 | C | T | -1.157 | 0.2474 | -? | 0 | 0 | 0 | 1 |
| rs1893818 | 11 | 120819939 | G | A | 1.157 | 0.2474 | +? | 0 | 0 | 0 | 1 |
| rs1940039 | 11 | 117393102 | T | C | -1.157 | 0.2474 | -? | 0 | 0 | 0 | 1 |
| rs73024419 | 11 | 117801857 | T | C | 1.157 | 0.2474 | +? | 0 | 0 | 0 | 1 |
| rs76663629 | 11 | 120826260 | A | G | -1.157 | 0.2474 | -? | 0 | 0 | 0 | 1 |

| | | | | | | | | | | | |
|---|---|---|---|---|---|---|---|---|---|---|---|
| rs686624 | 11 | 118955509 | T | A | -1.156 | 0.2476 | -? | 0 | 0 | 0 | 1 |
| rs7948072 | 11 | 117312197 | G | A | -1.156 | 0.2478 | -? | 0 | 0 | 0 | 1 |
| rs12225735 | 11 | 120800039 | T | C | -1.155 | 0.2479 | -+ | -185.9 | 1.049 | 1 | 0.3056 |
| rs611279 | 11 | 119349531 | A | G | -1.155 | 0.2479 | -? | 0 | 0 | 0 | 1 |
| rs77810230 | 11 | 120170012 | C | T | -1.155 | 0.2479 | -? | 0 | 0 | 0 | 1 |
| rs112266857 | 11 | 117564845 | T | C | 1.155 | 0.248 | +? | 0 | 0 | 0 | 1 |
| rs528056 | 11 | 118317907 | T | C | 1.155 | 0.2481 | +- | 0 | 0.459 | 1 | 0.4983 |
| rs641896 | 11 | 119775435 | T | C | -1.155 | 0.2481 | -? | 0 | 0 | 0 | 1 |
| rs512443 | 11 | 117339893 | T | C | 1.155 | 0.2482 | +? | 0 | 0 | 0 | 1 |
| rs11607580 | 11 | 120713605 | G | A | 1.154 | 0.2483 | +? | 0 | 0 | 0 | 1 |
| rs6589541 | 11 | 116208396 | G | A | -1.154 | 0.2483 | -? | 0 | 0 | 0 | 1 |
| rs611616 | 11 | 119349526 | A | G | -1.154 | 0.2486 | -? | 0 | 0 | 0 | 1 |
| rs11608113 | 11 | 120714220 | G | A | 1.154 | 0.2487 | +? | 0 | 0 | 0 | 1 |
| rs512481 | 11 | 117718880 | A | G | 1.153 | 0.2488 | +? | 0 | 0 | 0 | 1 |
| rs519294 | 11 | 118573743 | A | C | 1.153 | 0.2488 | ++ | -148.4 | 1.208 | 1 | 0.2718 |
| rs4643089 | 11 | 120941583 | G | A | -1.153 | 0.2489 | -? | 0 | 0 | 0 | 1 |
| rs148838027 | 11 | 118290459 | C | G | -1.153 | 0.249 | -- | 0 | 0.354 | 1 | 0.5517 |
| rs1317586 | 11 | 119881176 | G | A | 1.153 | 0.2491 | +? | 0 | 0 | 0 | 1 |
| rs9666435 | 11 | 120310978 | C | A | -1.153 | 0.2491 | -? | 0 | 0 | 0 | 1 |
| rs116997303 | 11 | 117447871 | C | T | 1.152 | 0.2492 | +? | 0 | 0 | 0 | 1 |
| rs142639145 | 11 | 117447735 | G | A | 1.152 | 0.2492 | +? | 0 | 0 | 0 | 1 |
| rs7108596 | 11 | 120310959 | A | G | -1.152 | 0.2492 | -? | 0 | 0 | 0 | 1 |
| rs7948299 | 11 | 120316589 | C | T | -1.152 | 0.2492 | -? | 0 | 0 | 0 | 1 |
| rs1729410 | 11 | 116665661 | C | G | -1.152 | 0.2494 | -+ | -67.7 | 1.789 | 1 | 0.181 |
| rs61902688 | 11 | 120567441 | A | C | 1.152 | 0.2494 | +? | 0 | 0 | 0 | 1 |
| rs12279833 | 11 | 121119605 | C | T | -1.152 | 0.2495 | -- | 0 | 0.008 | 1 | 0.9273 |
| rs1614444 | 11 | 116547548 | G | A | -1.151 | 0.2498 | -? | 0 | 0 | 0 | 1 |
| rs112845302 | 11 | 117570168 | A | G | 1.15 | 0.2502 | +? | 0 | 0 | 0 | 1 |
| rs1482502236 | 11 | 116528047 | A | T | 1.15 | 0.2502 | +? | 0 | 0 | 0 | 1 |
| rs10892655 | 11 | 120838878 | A | G | -1.149 | 0.2505 | -? | 0 | 0 | 0 | 1 |
| rs112793334 | 11 | 117429793 | A | C | 1.149 | 0.2506 | +? | 0 | 0 | 0 | 1 |
| rs4144286 | 11 | 121126349 | A | G | 1.149 | 0.2506 | ++ | 0 | 0.334 | 1 | 0.5635 |
| rs4382920 | 11 | 121115877 | G | C | -1.148 | 0.2508 | -? | 0 | 0 | 0 | 1 |
| rs1002179 | 11 | 119344407 | C | T | -1.148 | 0.2509 | -? | 0 | 0 | 0 | 1 |
| rs10502236 | 11 | 120307742 | G | A | -1.148 | 0.2509 | -? | 0 | 0 | 0 | 1 |
| rs57912596 | 11 | 120306759 | C | T | -1.148 | 0.2509 | -? | 0 | 0 | 0 | 1 |
| rs4572163 | 11 | 117661200 | T | C | 1.148 | 0.2511 | +? | 0 | 0 | 0 | 1 |
| rs10750098 | 11 | 116705568 | G | T | 1.147 | 0.2512 | +? | 0 | 0 | 0 | 1 |
| rs145518466 | 11 | 117633482 | G | A | -1.147 | 0.2512 | -? | 0 | 0 | 0 | 1 |
| rs73587425 | 11 | 117633690 | C | G | -1.147 | 0.2512 | -? | 0 | 0 | 0 | 1 |
| rs12363974 | 11 | 120277388 | G | C | -1.147 | 0.2513 | -? | 0 | 0 | 0 | 1 |
| rs582343 | 11 | 119772693 | C | G | -1.146 | 0.2516 | -? | 0 | 0 | 0 | 1 |
| rs17245518 | 11 | 120327817 | C | T | -1.146 | 0.2517 | -? | 0 | 0 | 0 | 1 |
| rs587521 | 11 | 119351220 | G | A | -1.146 | 0.2519 | -? | 0 | 0 | 0 | 1 |
| rs635335 | 11 | 119917798 | T | C | 1.146 | 0.2519 | +? | 0 | 0 | 0 | 1 |
| rs1784042 | 11 | 117065476 | A | G | 1.146 | 0.252 | +? | 0 | 0 | 0 | 1 |
| rs505128 | 11 | 116273511 | C | T | -1.145 | 0.2521 | -? | 0 | 0 | 0 | 1 |
| rs619360 | 11 | 117168402 | T | A | 1.145 | 0.2521 | +? | 0 | 0 | 0 | 1 |
| rs4936522 | 11 | 120469504 | G | C | 1.145 | 0.2522 | +? | 0 | 0 | 0 | 1 |
| rs117839774 | 11 | 120585311 | T | C | 1.145 | 0.2523 | +? | 0 | 0 | 0 | 1 |
| rs75707401 | 11 | 117560438 | A | G | 1.144 | 0.2525 | +? | 0 | 0 | 0 | 1 |
| rs583219 | 11 | 116723171 | C | T | -1.144 | 0.2526 | -? | 0 | 0 | 0 | 1 |
| rs4938412 | 11 | 117463436 | T | G | 1.144 | 0.2528 | +? | 0 | 0 | 0 | 1 |
| rs654776 | 11 | 119346737 | A | G | -1.144 | 0.2528 | -? | 0 | 0 | 0 | 1 |
| rs76630213 | 11 | 117466663 | G | T | 1.144 | 0.2528 | +? | 0 | 0 | 0 | 1 |
| rs10892579 | 11 | 120330728 | G | A | -1.143 | 0.253 | -? | 0 | 0 | 0 | 1 |

| | | | | | | | | | | | |
|---|---|---|---|---|---|---|---|---|---|---|---|
| rs12788462 | 11 | 120151784 | C | T | -1.143 | 0.253 | -? | 0 | 0 | 0 | 1 |
| rs10790434 | 11 | 121191782 | G | C | 1.143 | 0.2531 | +? | 0 | 0 | 0 | 1 |
| rs138450618 | 11 | 117775903 | T | C | -1.143 | 0.2531 | -? | 0 | 0 | 0 | 1 |
| rs3901422 | 11 | 118837272 | A | G | -1.143 | 0.2531 | -? | 0 | 0 | 0 | 1 |
| rs529681 | 11 | 120042082 | A | G | 1.143 | 0.2531 | +? | 0 | 0 | 0 | 1 |
| rs11600723 | 11 | 120249395 | T | C | -1.142 | 0.2533 | -? | 0 | 0 | 0 | 1 |
| rs11216624 | 11 | 117792471 | A | G | -1.142 | 0.2534 | -? | 0 | 0 | 0 | 1 |
| rs4245039 | 11 | 120485547 | C | A | 1.141 | 0.2537 | +? | 0 | 0 | 0 | 1 |
| rs12361642 | 11 | 120266477 | T | C | -1.141 | 0.2538 | -? | 0 | 0 | 0 | 1 |
| rs12363073 | 11 | 117352086 | T | C | 1.141 | 0.2538 | +? | 0 | 0 | 0 | 1 |
| rs583687 | 11 | 119772958 | T | C | -1.141 | 0.2538 | -? | 0 | 0 | 0 | 1 |
| rs73020470 | 11 | 117771769 | C | T | 1.141 | 0.2539 | +? | 0 | 0 | 0 | 1 |
| rs78528013 | 11 | 120826272 | T | C | -1.141 | 0.254 | -? | 0 | 0 | 0 | 1 |
| rs7935816 | 11 | 119665842 | T | C | 1.141 | 0.254 | +? | 0 | 0 | 0 | 1 |
| rs80159023 | 11 | 119845446 | A | G | -1.141 | 0.254 | -? | 0 | 0 | 0 | 1 |
| rs2510152 | 11 | 119148573 | G | T | -1.14 | 0.2541 | -? | 0 | 0 | 0 | 1 |
| rs4245166 | 11 | 116194551 | G | A | -1.14 | 0.2541 | -? | 0 | 0 | 0 | 1 |
| rs4938603 | 11 | 118846482 | G | A | -1.14 | 0.2541 | -? | 0 | 0 | 0 | 1 |
| rs6589652 | 11 | 118058358 | G | A | -1.14 | 0.2543 | -+ | -62.5 | 1.846 | 1 | 0.1743 |
| rs947917 | 11 | 117773537 | G | A | 1.14 | 0.2543 | +? | 0 | 0 | 0 | 1 |
| rs61904847 | 11 | 116453885 | C | T | -1.14 | 0.2544 | -? | 0 | 0 | 0 | 1 |
| rs2515797 | 11 | 117818680 | G | A | -1.139 | 0.2547 | -? | 0 | 0 | 0 | 1 |
| rs34607201 | 11 | 120803734 | A | G | -1.139 | 0.2547 | -? | 0 | 0 | 0 | 1 |
| rs10892499 | 11 | 119808864 | A | G | -1.139 | 0.2549 | -? | 0 | 0 | 0 | 1 |
| rs6589819 | 11 | 120479103 | A | G | 1.138 | 0.255 | +? | 0 | 0 | 0 | 1 |
| rs73587426 | 11 | 117633791 | G | C | -1.138 | 0.255 | -? | 0 | 0 | 0 | 1 |
| rs73587428 | 11 | 117633827 | G | A | -1.138 | 0.255 | -? | 0 | 0 | 0 | 1 |
| rs73587429 | 11 | 117633869 | A | G | -1.138 | 0.255 | -? | 0 | 0 | 0 | 1 |
| rs61903857 | 11 | 117633297 | T | G | -1.138 | 0.2552 | -? | 0 | 0 | 0 | 1 |
| rs61903858 | 11 | 117633349 | A | G | -1.138 | 0.2552 | -? | 0 | 0 | 0 | 1 |
| rs74702343 | 11 | 120715471 | A | G | 1.137 | 0.2556 | +? | 0 | 0 | 0 | 1 |
| rs589925 | 11 | 118955679 | T | C | 1.137 | 0.2557 | +? | 0 | 0 | 0 | 1 |
| rs592190 | 11 | 118955314 | A | G | 1.137 | 0.2557 | +? | 0 | 0 | 0 | 1 |
| rs11216129 | 11 | 116620256 | A | C | -1.136 | 0.2559 | -+ | 0 | 0.883 | 1 | 0.3475 |
| rs2252520 | 11 | 120115570 | A | T | -1.136 | 0.2561 | -? | 0 | 0 | 0 | 1 |
| rs512393 | 11 | 119823440 | C | G | 1.136 | 0.2561 | +? | 0 | 0 | 0 | 1 |
| rs17311024 | 11 | 117644124 | A | T | -1.135 | 0.2564 | -? | 0 | 0 | 0 | 1 |
| rs602445 | 11 | 117958148 | C | T | -1.135 | 0.2565 | -? | 0 | 0 | 0 | 1 |
| rs12223674 | 11 | 117451256 | G | C | 1.134 | 0.2568 | +? | 0 | 0 | 0 | 1 |
| rs688911 | 11 | 118036532 | C | T | 1.134 | 0.2568 | +? | 0 | 0 | 0 | 1 |
| rs77806521 | 11 | 118232026 | T | C | -1.134 | 0.2568 | -? | 0 | 0 | 0 | 1 |
| rs948378 | 11 | 121120845 | A | T | -1.134 | 0.2568 | -- | 0 | 0.007 | 1 | 0.9353 |
| rs142108356 | 11 | 119749609 | G | A | -1.134 | 0.257 | -? | 0 | 0 | 0 | 1 |
| rs591252 | 11 | 120809279 | C | T | 1.134 | 0.257 | +? | 0 | 0 | 0 | 1 |
| rs11216021 | 11 | 116458660 | T | C | -1.133 | 0.2571 | -? | 0 | 0 | 0 | 1 |
| rs142724980 | 11 | 119882823 | T | C | 1.133 | 0.2573 | +? | 0 | 0 | 0 | 1 |
| rs4938502 | 11 | 118179941 | T | G | 1.133 | 0.2573 | +? | 0 | 0 | 0 | 1 |
| rs57258844 | 11 | 116538407 | C | G | 1.133 | 0.2573 | +? | 0 | 0 | 0 | 1 |
| rs77571870 | 11 | 120826325 | A | G | -1.133 | 0.2574 | -? | 0 | 0 | 0 | 1 |
| rs12796452 | 11 | 120800956 | C | G | -1.132 | 0.2575 | -+ | -193.4 | 1.023 | 1 | 0.3119 |
| rs111226844 | 11 | 117895854 | C | T | 1.131 | 0.2582 | +? | 0 | 0 | 0 | 1 |
| rs11607997 | 11 | 119941330 | T | C | -1.131 | 0.2582 | -? | 0 | 0 | 0 | 1 |
| rs4938411 | 11 | 117444884 | A | G | 1.13 | 0.2583 | +? | 0 | 0 | 0 | 1 |
| rs61903856 | 11 | 117632463 | A | G | -1.13 | 0.2583 | -? | 0 | 0 | 0 | 1 |
| rs78034012 | 11 | 116324006 | C | G | -1.13 | 0.2583 | -? | 0 | 0 | 0 | 1 |
| rs9681206 | 11 | 116539518 | T | C | 1.13 | 0.2583 | +? | 0 | 0 | 0 | 1 |

| | | | | | | | | | | | |
|---|---|---|---|---|---|---|---|---|---|---|---|
| rs12804621 | 11 | 120622385 | T | C | -1.129 | 0.2589 | -? | 0 | 0 | 0 | 1 |
| rs577166 | 11 | 117351353 | T | C | 1.129 | 0.2589 | +? | 0 | 0 | 0 | 1 |
| rs7110955 | 11 | 120323909 | G | A | -1.129 | 0.2589 | -? | 0 | 0 | 0 | 1 |
| rs11216713 | 11 | 117903942 | A | G | 1.129 | 0.2591 | ++ | 0 | 0 | 1 | 0.9836 |
| rs74236011 | 11 | 120154647 | G | T | -1.128 | 0.2593 | -? | 0 | 0 | 0 | 1 |
| rs1893460 | 11 | 116603677 | A | G | -1.128 | 0.2594 | -- | 0 | 0.049 | 1 | 0.8248 |
| rs7116690 | 11 | 118837742 | G | A | -1.128 | 0.2594 | -? | 0 | 0 | 0 | 1 |
| rs10790413 | 11 | 121113880 | C | A | -1.128 | 0.2595 | -- | 0 | 0.006 | 1 | 0.9382 |
| rs523267 | 11 | 116523279 | C | T | 1.127 | 0.2596 | +? | 0 | 0 | 0 | 1 |
| rs117083922 | 11 | 117906282 | A | G | 1.127 | 0.2597 | ++ | 0 | 0 | 1 | 1 |
| rs11217470 | 11 | 119629249 | C | T | 1.126 | 0.26 | ++ | 0 | 0.164 | 1 | 0.6857 |
| rs7924359 | 11 | 117960590 | G | C | -1.126 | 0.2601 | -- | 0 | 0.146 | 1 | 0.702 |
| rs7928767 | 11 | 117650439 | A | G | -1.126 | 0.2602 | -? | 0 | 0 | 0 | 1 |
| rs11822940 | 11 | 119520997 | C | G | -1.125 | 0.2604 | -? | 0 | 0 | 0 | 1 |
| rs7946299 | 11 | 116587849 | C | T | 1.125 | 0.2604 | +? | 0 | 0 | 0 | 1 |
| rs10892654 | 11 | 120837407 | G | A | -1.125 | 0.2605 | -? | 0 | 0 | 0 | 1 |
| rs10892497 | 11 | 119804776 | T | C | -1.125 | 0.2606 | -? | 0 | 0 | 0 | 1 |
| rs4936490 | 11 | 119506749 | A | G | 1.125 | 0.2606 | +? | 0 | 0 | 0 | 1 |
| rs555577 | 11 | 117726697 | A | G | 1.125 | 0.2606 | +? | 0 | 0 | 0 | 1 |
| rs3018521 | 11 | 117818878 | A | G | -1.125 | 0.2607 | -? | 0 | 0 | 0 | 1 |
| rs688968 | 11 | 117406739 | G | C | -1.125 | 0.2607 | -? | 0 | 0 | 0 | 1 |
| rs34028064 | 11 | 120075878 | A | C | -1.125 | 0.2608 | -? | 0 | 0 | 0 | 1 |
| rs17120893 | 11 | 117443936 | T | C | 1.124 | 0.2609 | +? | 0 | 0 | 0 | 1 |
| rs74999919 | 11 | 119829794 | G | A | -1.124 | 0.2609 | -? | 0 | 0 | 0 | 1 |
| rs1430595649 | 11 | 116435383 | T | C | -1.124 | 0.261 | -? | 0 | 0 | 0 | 1 |
| rs61904808 | 11 | 116436102 | G | A | -1.124 | 0.261 | -? | 0 | 0 | 0 | 1 |
| rs61904809 | 11 | 116436589 | T | C | -1.124 | 0.261 | -? | 0 | 0 | 0 | 1 |
| rs12794408 | 11 | 116494803 | G | A | -1.124 | 0.2612 | -? | 0 | 0 | 0 | 1 |
| rs1838947 | 11 | 119691200 | C | T | 1.124 | 0.2612 | ++ | 0 | 0.12 | 1 | 0.7295 |
| rs10160642 | 11 | 121114248 | T | C | -1.123 | 0.2615 | -- | 0 | 0.006 | 1 | 0.9403 |
| rs6589854 | 11 | 120954446 | G | T | -1.123 | 0.2615 | -? | 0 | 0 | 0 | 1 |
| rs2174876 | 11 | 116439684 | A | G | -1.123 | 0.2616 | -? | 0 | 0 | 0 | 1 |
| rs4938730 | 11 | 119689931 | C | T | 1.123 | 0.2616 | ++ | 0 | 0.049 | 1 | 0.8257 |
| rs639785 | 11 | 118062268 | T | C | -1.123 | 0.2616 | -+ | -66.4 | 1.803 | 1 | 0.1794 |
| rs73003760 | 11 | 120710519 | C | T | 1.123 | 0.2616 | +? | 0 | 0 | 0 | 1 |
| rs2276071 | 11 | 118896116 | C | A | -1.122 | 0.2617 | -- | 0 | 0.209 | 1 | 0.6472 |
| rs74442127 | 11 | 119828201 | A | G | -1.122 | 0.2617 | -? | 0 | 0 | 0 | 1 |
| rs35301288 | 11 | 119578224 | A | G | -1.122 | 0.2619 | -? | 0 | 0 | 0 | 1 |
| rs7938295 | 11 | 120644115 | A | G | 1.121 | 0.2621 | +? | 0 | 0 | 0 | 1 |
| rs59238367 | 11 | 117920100 | A | G | -1.121 | 0.2622 | -? | 0 | 0 | 0 | 1 |
| rs674576 | 11 | 117919097 | A | G | -1.121 | 0.2623 | -? | 0 | 0 | 0 | 1 |
| rs9667364 | 11 | 119785124 | G | A | -1.121 | 0.2623 | -? | 0 | 0 | 0 | 1 |
| rs180351 | 11 | 116607641 | T | C | 1.121 | 0.2625 | +- | -49.3 | 2.009 | 1 | 0.1563 |
| rs76521293 | 11 | 116253907 | C | T | -1.12 | 0.2626 | -? | 0 | 0 | 0 | 1 |
| rs4936356 | 11 | 116431521 | G | A | -1.12 | 0.2627 | -? | 0 | 0 | 0 | 1 |
| rs7929825 | 11 | 118788081 | G | C | 1.12 | 0.2628 | +- | 20.4 | 3.77 | 1 | 0.05217 |
| rs10892696 | 11 | 121114116 | C | A | -1.12 | 0.2629 | -- | 0 | 0.005 | 1 | 0.9419 |
| rs11605509 | 11 | 117396433 | C | T | -1.12 | 0.2629 | -? | 0 | 0 | 0 | 1 |
| rs12800264 | 11 | 117396269 | A | G | -1.12 | 0.2629 | -? | 0 | 0 | 0 | 1 |
| rs1941390 | 11 | 117396991 | A | G | -1.12 | 0.2629 | -? | 0 | 0 | 0 | 1 |
| rs10892364 | 11 | 119301608 | T | C | 1.119 | 0.263 | +? | 0 | 0 | 0 | 1 |
| rs2156850 | 11 | 118030186 | A | G | 1.119 | 0.263 | +? | 0 | 0 | 0 | 1 |
| rs4445646 | 11 | 120556341 | C | T | -1.119 | 0.2631 | -? | 0 | 0 | 0 | 1 |
| rs4938732 | 11 | 119690068 | T | C | 1.119 | 0.2631 | ++ | 0 | 0.048 | 1 | 0.8272 |
| rs61904807 | 11 | 116434463 | G | C | -1.118 | 0.2635 | -? | 0 | 0 | 0 | 1 |
| rs78012672 | 11 | 117443368 | A | G | 1.118 | 0.2635 | +? | 0 | 0 | 0 | 1 |

| | | | | | | | | | | | |
|---|---|---|---|---|---|---|---|---|---|---|---|
| rs11217860 | 11 | 120279620 | T | G | -1.118 | 0.2636 | -? | 0 | 0 | 0 | 1 |
| rs4938384 | 11 | 117333952 | A | G | 1.118 | 0.2636 | +? | 0 | 0 | 0 | 1 |
| rs79656669 | 11 | 117477245 | C | T | 1.118 | 0.2637 | +? | 0 | 0 | 0 | 1 |
| rs947987 | 11 | 116507107 | A | G | 1.117 | 0.2638 | +? | 0 | 0 | 0 | 1 |
| rs147189349 | 11 | 117302538 | A | G | 1.117 | 0.2642 | +? | 0 | 0 | 0 | 1 |
| rs11217608 | 11 | 119849828 | G | A | -1.116 | 0.2644 | -? | 0 | 0 | 0 | 1 |
| rs10892105 | 11 | 117312962 | G | A | -1.116 | 0.2645 | -? | 0 | 0 | 0 | 1 |
| rs643377 | 11 | 119907628 | A | G | 1.116 | 0.2645 | +? | 0 | 0 | 0 | 1 |
| rs9651694 | 11 | 119857422 | G | A | -1.116 | 0.2645 | -? | 0 | 0 | 0 | 1 |
| rs582545 | 11 | 119906741 | G | C | 1.116 | 0.2646 | +? | 0 | 0 | 0 | 1 |
| rs10892145 | 11 | 117496301 | A | G | -1.115 | 0.2648 | -? | 0 | 0 | 0 | 1 |
| rs4938288 | 11 | 116452534 | G | C | 1.115 | 0.2648 | +? | 0 | 0 | 0 | 1 |
| rs2511856 | 11 | 119144148 | G | T | -1.115 | 0.2649 | -? | 0 | 0 | 0 | 1 |
| rs518932 | 11 | 119771843 | C | T | -1.115 | 0.2649 | -? | 0 | 0 | 0 | 1 |
| rs4938737 | 11 | 119769492 | T | C | -1.114 | 0.2651 | -? | 0 | 0 | 0 | 1 |
| rs35503235 | 11 | 117392110 | G | A | 1.114 | 0.2652 | +? | 0 | 0 | 0 | 1 |
| rs4573680 | 11 | 118872741 | C | T | -1.114 | 0.2652 | -- | 0 | 0.095 | 1 | 0.7582 |
| rs7101412 | 11 | 120488476 | T | C | 1.114 | 0.2653 | +? | 0 | 0 | 0 | 1 |
| rs7117565 | 11 | 116408615 | C | T | -1.114 | 0.2653 | -? | 0 | 0 | 0 | 1 |
| rs12284644 | 11 | 120450016 | G | A | -1.113 | 0.2655 | -? | 0 | 0 | 0 | 1 |
| rs11217859 | 11 | 120278166 | C | G | -1.113 | 0.2657 | -? | 0 | 0 | 0 | 1 |
| rs2305011 | 11 | 120338040 | G | A | 1.113 | 0.2657 | ++ | 0 | 0.842 | 1 | 0.3588 |
| rs4514433 | 11 | 119492158 | C | T | -1.112 | 0.2662 | -? | 0 | 0 | 0 | 1 |
| rs11217881 | 11 | 120344551 | G | A | -1.112 | 0.2663 | -? | 0 | 0 | 0 | 1 |
| rs11217882 | 11 | 120345002 | G | T | -1.112 | 0.2663 | -? | 0 | 0 | 0 | 1 |
| rs11217883 | 11 | 120345004 | G | A | -1.112 | 0.2663 | -? | 0 | 0 | 0 | 1 |
| rs11217890 | 11 | 120357547 | A | G | -1.111 | 0.2664 | -? | 0 | 0 | 0 | 1 |
| rs11217889 | 11 | 120356317 | G | C | -1.111 | 0.2665 | -? | 0 | 0 | 0 | 1 |
| rs12420339 | 11 | 116328597 | G | T | 1.111 | 0.2666 | +? | 0 | 0 | 0 | 1 |
| rs12421652 | 11 | 116625875 | T | G | -1.111 | 0.2666 | -+ | 0 | 0.856 | 1 | 0.3549 |
| rs2276339 | 11 | 117395481 | A | G | -1.111 | 0.2666 | -? | 0 | 0 | 0 | 1 |
| rs11216539 | 11 | 117631784 | C | T | -1.11 | 0.2668 | -? | 0 | 0 | 0 | 1 |
| rs648740 | 11 | 117447373 | A | G | 1.11 | 0.2668 | +? | 0 | 0 | 0 | 1 |
| rs80294967 | 11 | 119795416 | A | G | -1.11 | 0.2668 | -? | 0 | 0 | 0 | 1 |
| rs497548 | 11 | 117343399 | T | C | 1.11 | 0.2669 | +? | 0 | 0 | 0 | 1 |
| rs503281 | 11 | 117430971 | C | G | 1.11 | 0.2671 | +? | 0 | 0 | 0 | 1 |
| rs575030 | 11 | 121092398 | C | T | 1.11 | 0.2672 | +? | 0 | 0 | 0 | 1 |
| rs579078 | 11 | 116229947 | C | G | -1.11 | 0.2672 | -? | 0 | 0 | 0 | 1 |
| rs583606 | 11 | 119860625 | C | G | 1.11 | 0.2672 | +? | 0 | 0 | 0 | 1 |
| rs7102492 | 11 | 119300712 | T | C | 1.11 | 0.2672 | +? | 0 | 0 | 0 | 1 |
| rs7105316 | 11 | 119791050 | T | C | -1.11 | 0.2672 | -? | 0 | 0 | 0 | 1 |
| rs653598 | 11 | 118036538 | C | T | 1.109 | 0.2673 | +? | 0 | 0 | 0 | 1 |
| rs4245207 | 11 | 119688442 | A | G | 1.109 | 0.2675 | ++ | 0 | 0.04 | 1 | 0.8411 |
| rs2847498 | 11 | 120113664 | C | T | -1.109 | 0.2676 | -? | 0 | 0 | 0 | 1 |
| rs2512159 | 11 | 117826191 | G | C | 1.108 | 0.2677 | ++ | 0 | 0.333 | 1 | 0.564 |
| rs5128 | 11 | 116703640 | G | C | 1.108 | 0.2677 | +? | 0 | 0 | 0 | 1 |
| rs2044545 | 11 | 119765318 | T | C | -1.108 | 0.2678 | -? | 0 | 0 | 0 | 1 |
| rs12282721 | 11 | 117151412 | A | G | -1.108 | 0.2679 | -? | 0 | 0 | 0 | 1 |
| rs61898317 | 11 | 120163184 | G | A | -1.108 | 0.268 | -? | 0 | 0 | 0 | 1 |
| rs34172399 | 11 | 120073521 | C | A | -1.107 | 0.2682 | -? | 0 | 0 | 0 | 1 |
| rs10892698 | 11 | 121118260 | C | T | -1.107 | 0.2684 | -- | 0 | 0.004 | 1 | 0.9477 |
| rs10892699 | 11 | 121119493 | C | G | -1.107 | 0.2684 | -- | 0 | 0.004 | 1 | 0.9477 |
| rs4936597 | 11 | 121118878 | G | A | -1.107 | 0.2684 | -- | 0 | 0.004 | 1 | 0.9477 |
| rs542467 | 11 | 119770931 | C | T | -1.107 | 0.2684 | -? | 0 | 0 | 0 | 1 |
| rs80138815 | 11 | 117470795 | G | A | 1.107 | 0.2684 | +? | 0 | 0 | 0 | 1 |
| rs11216714 | 11 | 117904550 | A | G | 1.106 | 0.2686 | ++ | 0 | 0 | 1 | 0.9937 |

| | | | | | | | | | | | |
|---|---|---|---|---|---|---|---|---|---|---|---|
| rs11216715 | 11 | 117905217 | T | A | 1.106 | 0.2686 | ++ | 0 | 0 | 1 | 0.9937 |
| rs11216716 | 11 | 117905394 | A | G | 1.106 | 0.2686 | ++ | 0 | 0 | 1 | 0.9937 |
| rs12223224 | 11 | 117904710 | T | C | 1.106 | 0.2686 | ++ | 0 | 0 | 1 | 0.9937 |
| rs12223683 | 11 | 117905654 | A | G | 1.106 | 0.2686 | ++ | 0 | 0 | 1 | 0.9937 |
| rs12226036 | 11 | 117904793 | G | A | 1.106 | 0.2686 | ++ | 0 | 0 | 1 | 0.9937 |
| rs77212554 | 11 | 119840963 | T | C | -1.106 | 0.2686 | -? | 0 | 0 | 0 | 1 |
| rs79075579 | 11 | 117905786 | T | C | 1.106 | 0.2686 | ++ | 0 | 0 | 1 | 0.9937 |
| rs80001712 | 11 | 117905495 | T | C | 1.106 | 0.2686 | ++ | 0 | 0 | 1 | 0.9937 |
| rs11216517 | 11 | 117595511 | T | C | -1.106 | 0.2688 | -? | 0 | 0 | 0 | 1 |
| rs603826 | 11 | 118970878 | C | A | -1.106 | 0.2688 | -? | 0 | 0 | 0 | 1 |
| rs11603340 | 11 | 117396527 | G | C | -1.106 | 0.2689 | -? | 0 | 0 | 0 | 1 |
| rs3802882 | 11 | 118889822 | G | C | 1.106 | 0.2689 | +? | 0 | 0 | 0 | 1 |
| rs1145207 | 11 | 116715688 | G | A | -1.105 | 0.269 | -? | 0 | 0 | 0 | 1 |
| rs2849179 | 11 | 116620796 | T | C | 1.105 | 0.2691 | +- | -38.7 | 2.163 | 1 | 0.1414 |
| rs61902060 | 11 | 118839828 | G | C | -1.105 | 0.2692 | -? | 0 | 0 | 0 | 1 |
| rs10790414 | 11 | 121115895 | T | G | -1.105 | 0.2693 | -- | 0 | 0.004 | 1 | 0.9486 |
| rs10790415 | 11 | 121115951 | C | G | -1.104 | 0.2694 | -- | 0 | 0.004 | 1 | 0.9487 |
| rs2276072 | 11 | 118896115 | G | A | -1.104 | 0.2696 | -- | 0 | 0.235 | 1 | 0.628 |
| rs631932 | 11 | 119768853 | C | A | -1.104 | 0.2696 | -? | 0 | 0 | 0 | 1 |
| rs599385 | 11 | 118974485 | G | A | -1.104 | 0.2697 | -? | 0 | 0 | 0 | 1 |
| rs7102696 | 11 | 119300694 | A | G | 1.104 | 0.2697 | +? | 0 | 0 | 0 | 1 |
| rs7111358 | 11 | 119089831 | A | T | -1.104 | 0.2697 | -? | 0 | 0 | 0 | 1 |
| rs10892573 | 11 | 120287803 | G | A | -1.104 | 0.2698 | -? | 0 | 0 | 0 | 1 |
| rs7947677 | 11 | 120478670 | G | T | 1.104 | 0.2698 | +? | 0 | 0 | 0 | 1 |
| rs10502226 | 11 | 117441703 | G | T | 1.103 | 0.2701 | +? | 0 | 0 | 0 | 1 |
| rs10892449 | 11 | 119628238 | G | C | 1.103 | 0.2701 | ++ | 0 | 0.153 | 1 | 0.6957 |
| rs636783 | 11 | 117539715 | C | A | -1.103 | 0.2701 | -? | 0 | 0 | 0 | 1 |
| rs7129717 | 11 | 119853069 | G | T | -1.102 | 0.2703 | -? | 0 | 0 | 0 | 1 |
| rs116006647 | 11 | 117200974 | T | C | -1.102 | 0.2704 | -? | 0 | 0 | 0 | 1 |
| rs55869319 | 11 | 120302729 | T | C | 1.102 | 0.2704 | ++ | 0 | 0.995 | 1 | 0.3185 |
| rs625404 | 11 | 117954350 | C | T | -1.102 | 0.2705 | -? | 0 | 0 | 0 | 1 |
| rs688857 | 11 | 119909507 | T | C | 1.102 | 0.2705 | +? | 0 | 0 | 0 | 1 |
| rs11217188 | 11 | 119088176 | A | G | -1.101 | 0.2707 | -? | 0 | 0 | 0 | 1 |
| rs11826230 | 11 | 119412939 | T | G | -1.101 | 0.2707 | -? | 0 | 0 | 0 | 1 |
| rs543577 | 11 | 121039069 | A | G | -1.101 | 0.2707 | -? | 0 | 0 | 0 | 1 |
| rs11217107 | 11 | 118839268 | C | G | -1.101 | 0.271 | -? | 0 | 0 | 0 | 1 |
| rs61900159 | 11 | 118839133 | C | G | -1.101 | 0.271 | -? | 0 | 0 | 0 | 1 |
| rs12575317 | 11 | 120414641 | C | T | 1.101 | 0.2711 | +? | 0 | 0 | 0 | 1 |
| rs553578 | 11 | 119778355 | G | A | -1.1 | 0.2714 | -? | 0 | 0 | 0 | 1 |
| rs7115337 | 11 | 121116773 | T | G | -1.1 | 0.2714 | -- | 0 | 0.004 | 1 | 0.9508 |
| rs11218066 | 11 | 120808928 | T | C | 1.1 | 0.2715 | +? | 0 | 0 | 0 | 1 |
| rs10160459 | 11 | 121114571 | C | T | -1.099 | 0.2716 | -- | 0 | 0.004 | 1 | 0.9511 |
| rs2849165 | 11 | 116688478 | T | C | -1.099 | 0.2716 | -? | 0 | 0 | 0 | 1 |
| rs36109901 | 11 | 119083318 | C | A | 1.099 | 0.2718 | +? | 0 | 0 | 0 | 1 |
| rs4938263 | 11 | 116210212 | G | A | -1.099 | 0.2718 | -? | 0 | 0 | 0 | 1 |
| rs7103283 | 11 | 121117803 | A | T | -1.098 | 0.272 | -- | 0 | 0.004 | 1 | 0.9515 |
| rs7103470 | 11 | 121117684 | G | T | -1.098 | 0.272 | -- | 0 | 0.004 | 1 | 0.9515 |
| rs76085613 | 11 | 119835845 | A | G | -1.098 | 0.2722 | -? | 0 | 0 | 0 | 1 |
| rs77142768 | 11 | 119835506 | G | A | -1.098 | 0.2722 | -? | 0 | 0 | 0 | 1 |
| rs9667310 | 11 | 119342678 | A | G | -1.098 | 0.2722 | -? | 0 | 0 | 0 | 1 |
| rs10160458 | 11 | 121114375 | G | A | -1.098 | 0.2723 | -- | 0 | 0.004 | 1 | 0.9518 |
| rs11218016 | 11 | 120694754 | T | C | -1.098 | 0.2723 | -? | 0 | 0 | 0 | 1 |
| rs12793150 | 11 | 118180667 | G | C | 1.098 | 0.2723 | +? | 0 | 0 | 0 | 1 |
| rs3802883 | 11 | 118889813 | G | C | 1.098 | 0.2723 | +? | 0 | 0 | 0 | 1 |
| rs522843 | 11 | 117175934 | C | T | 1.098 | 0.2724 | +? | 0 | 0 | 0 | 1 |
| rs508731 | 11 | 116244267 | A | T | -1.097 | 0.2726 | -? | 0 | 0 | 0 | 1 |

| | | | | | | | | | | | |
|---|---|---|---|---|---|---|---|---|---|---|---|
| rs604714 | 11 | 118970696 | A | C | -1.097 | 0.2727 | -? | 0 | 0 | 0 | 1 |
| rs12224853 | 11 | 117394822 | T | C | -1.097 | 0.2728 | -? | 0 | 0 | 0 | 1 |
| rs567797 | 11 | 119692211 | A | T | 1.097 | 0.2728 | ++ | 0 | 0.042 | 1 | 0.8374 |
| rs7929781 | 11 | 120441139 | T | C | -1.097 | 0.2728 | -? | 0 | 0 | 0 | 1 |
| rs1893248 | 11 | 121120755 | T | C | 1.096 | 0.2729 | ++ | 0 | 0.367 | 1 | 0.5446 |
| rs896690 | 11 | 119932578 | A | G | 1.096 | 0.2729 | +? | 0 | 0 | 0 | 1 |
| rs918143 | 11 | 116630600 | T | C | 1.096 | 0.2729 | +- | -61.1 | 1.862 | 1 | 0.1724 |
| rs73002932 | 11 | 119466427 | T | C | 1.096 | 0.2731 | +? | 0 | 0 | 0 | 1 |
| rs561830 | 11 | 119043745 | T | C | 1.096 | 0.2732 | ++ | 0 | 0.105 | 1 | 0.746 |
| rs1263176 | 11 | 116690123 | T | C | -1.095 | 0.2735 | -? | 0 | 0 | 0 | 1 |
| rs148143761 | 11 | 116950556 | G | A | -1.095 | 0.2737 | -? | 0 | 0 | 0 | 1 |
| rs1939235 | 11 | 117477867 | T | C | 1.095 | 0.2737 | +? | 0 | 0 | 0 | 1 |
| rs2298725 | 11 | 120823608 | A | G | -1.095 | 0.2737 | -? | 0 | 0 | 0 | 1 |
| rs1787687 | 11 | 116549896 | G | A | -1.094 | 0.2738 | -? | 0 | 0 | 0 | 1 |
| rs7940298 | 11 | 119635453 | T | C | 1.094 | 0.2738 | ++ | 0 | 0.149 | 1 | 0.6992 |
| rs11603366 | 11 | 117396505 | T | G | -1.094 | 0.2739 | -? | 0 | 0 | 0 | 1 |
| rs34306783 | 11 | 117841718 | T | C | 1.094 | 0.2739 | ++ | 0 | 0.651 | 1 | 0.4197 |
| rs79300298 | 11 | 119625373 | C | G | 1.094 | 0.2739 | ++ | 0 | 0.149 | 1 | 0.6993 |
| rs17540777 | 11 | 118150259 | G | T | 1.093 | 0.2744 | +? | 0 | 0 | 0 | 1 |
| rs74582224 | 11 | 117906289 | T | G | 1.093 | 0.2745 | ++ | 0 | 0 | 1 | 1 |
| rs766894 | 11 | 120328858 | C | T | -1.093 | 0.2745 | -? | 0 | 0 | 0 | 1 |
| rs78197730 | 11 | 117906408 | T | C | 1.093 | 0.2745 | ++ | 0 | 0 | 1 | 1 |
| rs79791963 | 11 | 117906406 | G | C | 1.093 | 0.2745 | ++ | 0 | 0 | 1 | 1 |
| rs61900794 | 11 | 120274893 | C | T | -1.093 | 0.2746 | -? | 0 | 0 | 0 | 1 |
| rs11217879 | 11 | 120341723 | C | T | -1.092 | 0.2747 | -? | 0 | 0 | 0 | 1 |
| rs538079 | 11 | 117342221 | G | A | -1.092 | 0.2747 | -? | 0 | 0 | 0 | 1 |
| rs11216080 | 11 | 116530998 | T | G | 1.092 | 0.2749 | +? | 0 | 0 | 0 | 1 |
| rs1440183 | 11 | 116203433 | C | T | -1.092 | 0.2749 | -? | 0 | 0 | 0 | 1 |
| rs2005529 | 11 | 120798714 | C | G | -1.092 | 0.275 | -? | 0 | 0 | 0 | 1 |
| rs77020062 | 11 | 119654894 | T | C | -1.091 | 0.2752 | -? | 0 | 0 | 0 | 1 |
| rs45539032 | 11 | 118015832 | A | G | -1.091 | 0.2753 | -? | 0 | 0 | 0 | 1 |
| rs4938800 | 11 | 120209386 | A | G | 1.091 | 0.2753 | ++ | -197.6 | 1.008 | 1 | 0.3153 |
| rs6589722 | 11 | 119085783 | C | A | -1.091 | 0.2753 | -? | 0 | 0 | 0 | 1 |
| rs7121930 | 11 | 117494246 | T | G | -1.091 | 0.2754 | -? | 0 | 0 | 0 | 1 |
| rs663191 | 11 | 117335991 | T | C | 1.09 | 0.2757 | +? | 0 | 0 | 0 | 1 |
| rs677954 | 11 | 117336947 | T | C | 1.09 | 0.2757 | +? | 0 | 0 | 0 | 1 |
| rs12294322 | 11 | 116543552 | G | A | -1.09 | 0.2758 | -? | 0 | 0 | 0 | 1 |
| rs3016355 | 11 | 116545495 | T | A | -1.09 | 0.2758 | -? | 0 | 0 | 0 | 1 |
| rs7103097 | 11 | 120728827 | A | C | 1.09 | 0.2758 | +? | 0 | 0 | 0 | 1 |
| rs7121331 | 11 | 116362253 | C | T | -1.09 | 0.2758 | -? | 0 | 0 | 0 | 1 |
| rs4938618 | 11 | 118867803 | C | G | -1.089 | 0.2762 | -- | 0 | 0.197 | 1 | 0.657 |
| rs590917 | 11 | 117902914 | T | G | -1.089 | 0.2762 | -- | 0 | 0.746 | 1 | 0.3879 |
| rs7931603 | 11 | 120965687 | A | G | -1.088 | 0.2765 | -? | 0 | 0 | 0 | 1 |
| rs616314 | 11 | 118956867 | A | T | -1.088 | 0.2766 | -? | 0 | 0 | 0 | 1 |
| rs10892581 | 11 | 120334863 | C | A | -1.088 | 0.2767 | -? | 0 | 0 | 0 | 1 |
| rs11600200 | 11 | 120334680 | G | T | -1.088 | 0.2767 | -? | 0 | 0 | 0 | 1 |
| rs113381053 | 11 | 119757989 | A | G | -1.087 | 0.277 | -? | 0 | 0 | 0 | 1 |
| rs117508216 | 11 | 120795955 | C | A | 1.087 | 0.277 | +? | 0 | 0 | 0 | 1 |
| rs12270510 | 11 | 119506018 | T | C | 1.087 | 0.277 | +? | 0 | 0 | 0 | 1 |
| rs1287394 | 11 | 120093606 | C | T | -1.087 | 0.277 | -? | 0 | 0 | 0 | 1 |
| rs10790287 | 11 | 119084914 | A | G | -1.087 | 0.2771 | -? | 0 | 0 | 0 | 1 |
| rs2887193 | 11 | 117825368 | A | G | 1.087 | 0.2771 | +? | 0 | 0 | 0 | 1 |
| rs11217871 | 11 | 120315637 | C | T | -1.087 | 0.2772 | -? | 0 | 0 | 0 | 1 |
| rs11600558 | 11 | 120310322 | C | T | -1.087 | 0.2772 | -? | 0 | 0 | 0 | 1 |
| rs61898768 | 11 | 120353237 | A | G | -1.087 | 0.2772 | -? | 0 | 0 | 0 | 1 |
| rs9625 | 11 | 120355791 | T | G | -1.087 | 0.2772 | -? | 0 | 0 | 0 | 1 |

| | | | | | | | | | | | |
|---|---|---|---|---|---|---|---|---|---|---|---|
| rs9666441 | 11 | 120311054 | G | A | -1.087 | 0.2772 | -? | 0 | 0 | 0 | 1 |
| rs12290699 | 11 | 120195702 | C | T | -1.086 | 0.2775 | -? | 0 | 0 | 0 | 1 |
| rs61898318 | 11 | 120171371 | A | G | -1.086 | 0.2775 | -? | 0 | 0 | 0 | 1 |
| rs77436344 | 11 | 119839219 | A | G | -1.086 | 0.2775 | -? | 0 | 0 | 0 | 1 |
| rs79652828 | 11 | 119838717 | T | C | -1.086 | 0.2775 | -? | 0 | 0 | 0 | 1 |
| rs7108905 | 11 | 118712427 | G | C | -1.086 | 0.2776 | -- | 0 | 0 | 1 | 0.9881 |
| rs35912781 | 11 | 120360271 | A | G | -1.086 | 0.2777 | -? | 0 | 0 | 0 | 1 |
| rs6589636 | 11 | 117824500 | C | T | 1.086 | 0.2777 | +? | 0 | 0 | 0 | 1 |
| rs6589710 | 11 | 118837460 | T | A | -1.086 | 0.2777 | -? | 0 | 0 | 0 | 1 |
| rs4936350 | 11 | 116314513 | T | C | 1.085 | 0.2778 | +? | 0 | 0 | 0 | 1 |
| rs574383 | 11 | 118573668 | C | T | 1.085 | 0.2778 | ++ | -157.8 | 1.164 | 1 | 0.2807 |
| rs3016345 | 11 | 116471264 | G | A | 1.085 | 0.2779 | +? | 0 | 0 | 0 | 1 |
| rs11217853 | 11 | 120263022 | G | A | -1.085 | 0.278 | -? | 0 | 0 | 0 | 1 |
| rs7925390 | 11 | 119455846 | T | C | 1.085 | 0.278 | +? | 0 | 0 | 0 | 1 |
| rs12574365 | 11 | 120441225 | C | T | -1.085 | 0.2781 | -? | 0 | 0 | 0 | 1 |
| rs12799401 | 11 | 117491758 | C | G | -1.085 | 0.2781 | -? | 0 | 0 | 0 | 1 |
| rs1941306 | 11 | 121115129 | C | A | -1.084 | 0.2782 | -- | 0 | 0.003 | 1 | 0.9579 |
| rs6589866 | 11 | 121115359 | A | G | -1.084 | 0.2782 | -- | 0 | 0.003 | 1 | 0.9579 |
| rs4936384 | 11 | 117398591 | T | C | -1.084 | 0.2783 | -? | 0 | 0 | 0 | 1 |
| rs150854359 | 11 | 117440220 | G | T | 1.084 | 0.2785 | +? | 0 | 0 | 0 | 1 |
| rs625464 | 11 | 118033545 | T | C | -1.084 | 0.2785 | -? | 0 | 0 | 0 | 1 |
| rs7127427 | 11 | 118805138 | A | G | -1.084 | 0.2785 | -- | 0 | 0.409 | 1 | 0.5226 |
| rs12800487 | 11 | 117396386 | A | G | -1.083 | 0.2786 | -? | 0 | 0 | 0 | 1 |
| rs11217472 | 11 | 119630763 | C | T | 1.083 | 0.2787 | ++ | 0 | 0.144 | 1 | 0.704 |
| rs3741300 | 11 | 116631690 | A | G | 1.083 | 0.2787 | +- | -53.3 | 1.956 | 1 | 0.1619 |
| rs659434 | 11 | 120984898 | G | C | 1.083 | 0.2788 | +? | 0 | 0 | 0 | 1 |
| rs2105670 | 11 | 121117142 | A | C | -1.083 | 0.2789 | -- | 0 | 0.003 | 1 | 0.9586 |
| rs118029883 | 11 | 119566235 | A | G | 1.083 | 0.279 | +? | 0 | 0 | 0 | 1 |
| rs4938604 | 11 | 118846574 | C | T | -1.082 | 0.2791 | -? | 0 | 0 | 0 | 1 |
| rs112549212 | 11 | 116495780 | C | T | -1.082 | 0.2792 | -? | 0 | 0 | 0 | 1 |
| rs180340 | 11 | 116616402 | C | T | 1.082 | 0.2792 | +- | -39.1 | 2.157 | 1 | 0.1419 |
| rs4936457 | 11 | 118875727 | G | A | -1.082 | 0.2792 | -- | 0 | 0.093 | 1 | 0.7598 |
| rs4936458 | 11 | 118875761 | G | T | -1.082 | 0.2792 | -- | 0 | 0.093 | 1 | 0.7598 |
| rs7111618 | 11 | 117848926 | G | T | -1.082 | 0.2792 | -- | 0 | 0.329 | 1 | 0.5661 |
| rs747419 | 11 | 120313539 | T | C | -1.082 | 0.2792 | -? | 0 | 0 | 0 | 1 |
| rs1784133 | 11 | 116541417 | T | C | -1.082 | 0.2793 | -? | 0 | 0 | 0 | 1 |
| rs2846016 | 11 | 116541539 | T | C | -1.082 | 0.2793 | -? | 0 | 0 | 0 | 1 |
| rs1269330 | 11 | 116701833 | A | G | -1.082 | 0.2794 | -? | 0 | 0 | 0 | 1 |
| rs34298632 | 11 | 119342143 | C | T | -1.081 | 0.2796 | -? | 0 | 0 | 0 | 1 |
| rs515871 | 11 | 117335517 | A | G | 1.081 | 0.2797 | +? | 0 | 0 | 0 | 1 |
| rs7107776 | 11 | 119080313 | T | C | -1.081 | 0.2798 | -? | 0 | 0 | 0 | 1 |
| rs139289629 | 11 | 117564084 | C | G | 1.081 | 0.2799 | +? | 0 | 0 | 0 | 1 |
| rs145253564 | 11 | 117563458 | T | C | 1.081 | 0.2799 | +? | 0 | 0 | 0 | 1 |
| rs6589865 | 11 | 121115206 | C | T | -1.08 | 0.28 | -- | 0 | 0.003 | 1 | 0.9598 |
| rs71480309 | 11 | 116529956 | T | C | 1.08 | 0.2801 | +? | 0 | 0 | 0 | 1 |
| rs77708326 | 11 | 119794526 | C | T | -1.08 | 0.2801 | -? | 0 | 0 | 0 | 1 |
| rs3758991 | 11 | 117945161 | T | C | -1.079 | 0.2806 | -- | 0 | 0.169 | 1 | 0.6809 |
| rs751370 | 11 | 120610402 | C | T | 1.079 | 0.2807 | +? | 0 | 0 | 0 | 1 |
| rs56880203 | 11 | 121190696 | G | A | 1.079 | 0.2808 | +? | 0 | 0 | 0 | 1 |
| rs12363975 | 11 | 120277392 | A | C | -1.078 | 0.2811 | -? | 0 | 0 | 0 | 1 |
| rs17120560 | 11 | 117161006 | G | C | 1.077 | 0.2813 | +? | 0 | 0 | 0 | 1 |
| rs2115645 | 11 | 119249488 | G | A | 1.077 | 0.2814 | +? | 0 | 0 | 0 | 1 |
| rs10750124 | 11 | 117964339 | C | T | -1.077 | 0.2815 | -? | 0 | 0 | 0 | 1 |
| rs2120430 | 11 | 120261934 | G | C | -1.077 | 0.2817 | -? | 0 | 0 | 0 | 1 |
| rs7129582 | 11 | 121116720 | G | A | -1.077 | 0.2817 | -- | 0 | 0.002 | 1 | 0.9615 |
| rs35401859 | 11 | 116475538 | G | T | 1.076 | 0.2819 | +? | 0 | 0 | 0 | 1 |

| | | | | | | | | | | | |
|---|---|---|---|---|---|---|---|---|---|---|---|
| rs59565018 | 11 | 120124302 | A | G | -1.076 | 0.2821 | -? | 0 | 0 | 0 | 1 |
| rs10891985 | 11 | 116392191 | T | A | 1.075 | 0.2824 | +? | 0 | 0 | 0 | 1 |
| rs7924948 | 11 | 120498024 | T | C | -1.075 | 0.2825 | -? | 0 | 0 | 0 | 1 |
| rs948468 | 11 | 117962732 | C | G | -1.074 | 0.2826 | -? | 0 | 0 | 0 | 1 |
| rs11603937 | 11 | 117367838 | A | G | 1.074 | 0.2827 | +? | 0 | 0 | 0 | 1 |
| rs12795589 | 11 | 116473704 | G | A | 1.074 | 0.2827 | +? | 0 | 0 | 0 | 1 |
| rs5141 | 11 | 116702123 | T | C | 1.074 | 0.2828 | +? | 0 | 0 | 0 | 1 |
| rs579674 | 11 | 116528224 | G | T | -1.074 | 0.2828 | -? | 0 | 0 | 0 | 1 |
| rs1268833 | 11 | 116694866 | G | A | -1.074 | 0.2829 | -? | 0 | 0 | 0 | 1 |
| rs657539 | 11 | 118572021 | T | C | 1.074 | 0.2829 | ++ | -125.5 | 1.33 | 1 | 0.2488 |
| rs3802884 | 11 | 118889810 | T | C | 1.073 | 0.2832 | +? | 0 | 0 | 0 | 1 |
| rs7128382 | 11 | 117023487 | A | C | 1.073 | 0.2832 | +? | 0 | 0 | 0 | 1 |
| rs59095933 | 11 | 118885161 | A | G | -1.073 | 0.2834 | -- | 0 | 0.327 | 1 | 0.5672 |
| rs677428 | 11 | 119849440 | T | C | -1.073 | 0.2834 | -? | 0 | 0 | 0 | 1 |
| rs7944542 | 11 | 116200726 | G | A | -1.072 | 0.2835 | -? | 0 | 0 | 0 | 1 |
| rs117315666 | 11 | 116937074 | T | G | 1.072 | 0.2837 | +? | 0 | 0 | 0 | 1 |
| rs623500 | 11 | 118828942 | G | A | 1.072 | 0.2837 | +? | 0 | 0 | 0 | 1 |
| rs3133227 | 11 | 120595653 | A | G | 1.072 | 0.2838 | +? | 0 | 0 | 0 | 1 |
| rs10892580 | 11 | 120334149 | A | G | -1.072 | 0.2839 | -? | 0 | 0 | 0 | 1 |
| rs61901404 | 11 | 120606763 | T | C | -1.072 | 0.2839 | -- | 0 | 0.271 | 1 | 0.6026 |
| rs681046 | 11 | 117452212 | C | T | 1.072 | 0.2839 | +? | 0 | 0 | 0 | 1 |
| rs871897 | 11 | 119881446 | T | C | 1.072 | 0.2839 | +? | 0 | 0 | 0 | 1 |
| rs7103570 | 11 | 118964577 | C | T | 1.071 | 0.284 | +? | 0 | 0 | 0 | 1 |
| rs78768102 | 11 | 121141095 | G | A | -1.071 | 0.284 | -? | 0 | 0 | 0 | 1 |
| rs484232 | 11 | 118708564 | T | G | 1.071 | 0.2842 | ++ | 0 | 0.472 | 1 | 0.492 |
| rs78928955 | 11 | 119823254 | C | T | -1.071 | 0.2843 | -? | 0 | 0 | 0 | 1 |
| rs11217868 | 11 | 120305787 | G | A | -1.07 | 0.2844 | -? | 0 | 0 | 0 | 1 |
| rs11217869 | 11 | 120313899 | G | T | -1.07 | 0.2844 | -? | 0 | 0 | 0 | 1 |
| rs12361340 | 11 | 120311409 | A | T | -1.07 | 0.2844 | -? | 0 | 0 | 0 | 1 |
| rs4936467 | 11 | 119086629 | T | A | -1.07 | 0.2847 | -? | 0 | 0 | 0 | 1 |
| rs7105187 | 11 | 118887581 | T | C | -1.07 | 0.2848 | -- | 0 | 0.379 | 1 | 0.5381 |
| rs7105962 | 11 | 119507477 | A | G | 1.07 | 0.2848 | +? | 0 | 0 | 0 | 1 |
| rs2253440 | 11 | 117308795 | A | G | -1.069 | 0.2849 | -? | 0 | 0 | 0 | 1 |
| rs11216045 | 11 | 116492411 | C | T | -1.069 | 0.285 | -? | 0 | 0 | 0 | 1 |
| rs11607944 | 11 | 120282566 | A | G | -1.069 | 0.285 | -? | 0 | 0 | 0 | 1 |
| rs637753 | 11 | 118041894 | C | G | -1.069 | 0.285 | -? | 0 | 0 | 0 | 1 |
| rs72999483 | 11 | 118523799 | C | T | -1.068 | 0.2853 | -- | -33.1 | 2.255 | 1 | 0.1332 |
| rs76474385 | 11 | 117581459 | C | T | 1.068 | 0.2853 | +? | 0 | 0 | 0 | 1 |
| rs10892169 | 11 | 117633054 | A | G | 1.068 | 0.2856 | +? | 0 | 0 | 0 | 1 |
| rs678914 | 11 | 119849798 | C | T | -1.068 | 0.2856 | -? | 0 | 0 | 0 | 1 |
| rs3020913 | 11 | 117831809 | G | A | 1.068 | 0.2857 | ++ | 0 | 0.582 | 1 | 0.4455 |
| rs12286723 | 11 | 118734640 | T | C | -1.067 | 0.2858 | -- | 0 | 0.011 | 1 | 0.9183 |
| rs5142 | 11 | 116701850 | T | C | 1.067 | 0.2859 | +? | 0 | 0 | 0 | 1 |
| rs12796351 | 11 | 116473789 | A | G | 1.067 | 0.2861 | +? | 0 | 0 | 0 | 1 |
| rs35777932 | 11 | 116214441 | C | T | -1.067 | 0.2862 | -- | -144.3 | 1.228 | 1 | 0.2678 |
| rs74586480 | 11 | 117906273 | G | A | 1.066 | 0.2863 | ++ | 0 | 0.001 | 1 | 0.9724 |
| rs11217656 | 11 | 119918703 | A | G | -1.066 | 0.2864 | -? | 0 | 0 | 0 | 1 |
| rs7122959 | 11 | 118804283 | G | A | -1.066 | 0.2864 | -- | 0 | 0.015 | 1 | 0.9027 |
| rs7123066 | 11 | 118804266 | T | C | -1.066 | 0.2864 | -- | 0 | 0.015 | 1 | 0.9027 |
| rs11602796 | 11 | 120268491 | A | G | -1.066 | 0.2865 | -? | 0 | 0 | 0 | 1 |
| rs11216559 | 11 | 117685169 | C | T | 1.065 | 0.2868 | ++ | 0 | 0.175 | 1 | 0.6758 |
| rs112776148 | 11 | 117683284 | C | G | 1.065 | 0.2868 | ++ | 0 | 0.185 | 1 | 0.6668 |
| rs7103418 | 11 | 120581080 | A | C | 1.065 | 0.2868 | +? | 0 | 0 | 0 | 1 |
| rs61362984 | 11 | 116450463 | G | A | -1.065 | 0.287 | -? | 0 | 0 | 0 | 1 |
| rs61898342 | 11 | 120182796 | A | G | -1.065 | 0.287 | -? | 0 | 0 | 0 | 1 |
| rs653788 | 11 | 117910111 | T | C | -1.065 | 0.287 | -? | 0 | 0 | 0 | 1 |

| | | | | | | | | | | | |
|---|---|---|---|---|---|---|---|---|---|---|---|
| rs7119619 | 11 | 120581105 | C | T | 1.065 | 0.287 | +? | 0 | 0 | 0 | 1 |
| rs60954647 | 11 | 116566933 | T | C | 1.064 | 0.2872 | +? | 0 | 0 | 0 | 1 |
| rs10790354 | 11 | 119787531 | T | C | 1.064 | 0.2873 | +? | 0 | 0 | 0 | 1 |
| rs12276890 | 11 | 121112938 | G | C | -1.064 | 0.2873 | -- | 0 | 0.546 | 1 | 0.46 |
| rs3922 | 11 | 118765600 | G | A | -1.064 | 0.2873 | -- | 0 | 0.179 | 1 | 0.6725 |
| rs9665750 | 11 | 119781695 | T | C | -1.064 | 0.2873 | -? | 0 | 0 | 0 | 1 |
| rs11608153 | 11 | 117716663 | G | A | -1.064 | 0.2874 | -? | 0 | 0 | 0 | 1 |
| rs17119975 | 11 | 116634557 | C | T | -1.064 | 0.2874 | -+ | 0 | 0.755 | 1 | 0.3848 |
| rs765814 | 11 | 119779799 | T | G | -1.063 | 0.2876 | -? | 0 | 0 | 0 | 1 |
| rs11215933 | 11 | 116312186 | A | C | 1.063 | 0.2877 | +? | 0 | 0 | 0 | 1 |
| rs78809001 | 11 | 119548402 | T | C | 1.063 | 0.2879 | +? | 0 | 0 | 0 | 1 |
| rs4936349 | 11 | 116314370 | G | A | 1.062 | 0.2881 | +? | 0 | 0 | 0 | 1 |
| rs687172 | 11 | 116731260 | A | T | -1.062 | 0.2881 | -? | 0 | 0 | 0 | 1 |
| rs528806 | 11 | 116529914 | T | C | -1.062 | 0.2882 | -? | 0 | 0 | 0 | 1 |
| rs652107 | 11 | 117909799 | G | A | -1.062 | 0.2882 | -- | -194.9 | 1.017 | 1 | 0.3132 |
| rs519356 | 11 | 117350033 | C | T | 1.062 | 0.2884 | +? | 0 | 0 | 0 | 1 |
| rs10736490 | 11 | 117964465 | T | C | -1.061 | 0.2887 | -? | 0 | 0 | 0 | 1 |
| rs10736491 | 11 | 117964490 | T | A | -1.061 | 0.2887 | -? | 0 | 0 | 0 | 1 |
| rs10790361 | 11 | 119844041 | T | C | 1.061 | 0.2887 | +? | 0 | 0 | 0 | 1 |
| rs1941629 | 11 | 117964873 | C | G | -1.061 | 0.2887 | -? | 0 | 0 | 0 | 1 |
| rs1941630 | 11 | 117964902 | C | G | -1.061 | 0.2887 | -? | 0 | 0 | 0 | 1 |
| rs7104016 | 11 | 117964598 | T | G | -1.061 | 0.2887 | -? | 0 | 0 | 0 | 1 |
| rs180329 | 11 | 116622958 | A | G | 1.06 | 0.2889 | +- | -45.2 | 2.067 | 1 | 0.1505 |
| rs12796311 | 11 | 116473730 | T | G | 1.06 | 0.2891 | +? | 0 | 0 | 0 | 1 |
| rs2512153 | 11 | 117889224 | G | A | 1.06 | 0.2891 | +- | -46.9 | 2.042 | 1 | 0.153 |
| rs486978 | 11 | 119789537 | T | G | 1.06 | 0.2891 | +? | 0 | 0 | 0 | 1 |
| rs112704487 | 11 | 119663058 | C | T | 1.06 | 0.2892 | ++ | 0 | 0.099 | 1 | 0.753 |
| rs74908422 | 11 | 120794058 | C | T | -1.059 | 0.2895 | -? | 0 | 0 | 0 | 1 |
| rs111527819 | 11 | 116481175 | A | G | -1.059 | 0.2896 | -? | 0 | 0 | 0 | 1 |
| rs7103021 | 11 | 120580795 | T | C | 1.059 | 0.2896 | +? | 0 | 0 | 0 | 1 |
| rs138159885 | 11 | 117565794 | A | G | 1.059 | 0.2897 | +? | 0 | 0 | 0 | 1 |
| rs150057425 | 11 | 117567237 | G | A | 1.059 | 0.2897 | +? | 0 | 0 | 0 | 1 |
| rs17587869 | 11 | 117565876 | T | G | 1.059 | 0.2897 | +? | 0 | 0 | 0 | 1 |
| rs17664522 | 11 | 117566372 | G | T | 1.059 | 0.2897 | +? | 0 | 0 | 0 | 1 |
| rs35664944 | 11 | 116471898 | A | G | 1.059 | 0.2897 | +? | 0 | 0 | 0 | 1 |
| rs10790291 | 11 | 119232913 | A | G | 1.059 | 0.2898 | +? | 0 | 0 | 0 | 1 |
| rs1268354 | 11 | 116690578 | C | T | -1.059 | 0.2898 | -? | 0 | 0 | 0 | 1 |
| rs540659 | 11 | 117342462 | C | T | -1.059 | 0.2898 | -? | 0 | 0 | 0 | 1 |
| rs4936549 | 11 | 120652075 | C | T | -1.058 | 0.2899 | -? | 0 | 0 | 0 | 1 |
| rs11216734 | 11 | 117921554 | A | T | -1.058 | 0.29 | -? | 0 | 0 | 0 | 1 |
| rs1263177 | 11 | 116690712 | C | T | -1.058 | 0.29 | -? | 0 | 0 | 0 | 1 |
| rs11822103 | 11 | 118869539 | A | G | -1.058 | 0.2901 | -- | 0 | 0.152 | 1 | 0.6963 |
| rs658358 | 11 | 119787190 | C | G | 1.058 | 0.2901 | +? | 0 | 0 | 0 | 1 |
| rs75195561 | 11 | 120397891 | T | C | 1.058 | 0.2901 | +? | 0 | 0 | 0 | 1 |
| rs1039094777 | 11 | 119757789 | T | C | -1.058 | 0.2902 | -? | 0 | 0 | 0 | 1 |
| rs17666283 | 11 | 117685824 | A | G | 1.058 | 0.2902 | ++ | 0 | 0.166 | 1 | 0.6835 |
| rs1209756219 | 11 | 117681924 | C | T | 1.058 | 0.2903 | ++ | 0 | 0.182 | 1 | 0.67 |
| rs2155195 | 11 | 117781618 | G | A | 1.057 | 0.2903 | +? | 0 | 0 | 0 | 1 |
| rs77045391 | 11 | 117885423 | T | C | 1.057 | 0.2903 | +? | 0 | 0 | 0 | 1 |
| rs496958 | 11 | 116528876 | C | T | -1.057 | 0.2904 | -? | 0 | 0 | 0 | 1 |
| rs7129508 | 11 | 121116786 | C | A | -1.057 | 0.2904 | -- | 0 | 0.001 | 1 | 0.9703 |
| rs116848264 | 11 | 118049817 | A | T | 1.057 | 0.2905 | +- | -133.5 | 1.285 | 1 | 0.257 |
| rs180350 | 11 | 116610048 | T | C | 1.057 | 0.2905 | +- | -34.6 | 2.23 | 1 | 0.1354 |
| rs61904846 | 11 | 116449692 | A | G | -1.057 | 0.2906 | -? | 0 | 0 | 0 | 1 |
| rs670294 | 11 | 117935840 | G | T | -1.057 | 0.2906 | -? | 0 | 0 | 0 | 1 |
| rs10892523 | 11 | 119923484 | G | A | -1.056 | 0.2908 | -? | 0 | 0 | 0 | 1 |

| | | | | | | | | | | | |
|---|---|---|---|---|---|---|---|---|---|---|---|
| rs2852238 | 11 | 120784855 | G | C | -1.056 | 0.2908 | -? | 0 | 0 | 0 | 1 |
| rs4936366 | 11 | 117026885 | G | A | 1.056 | 0.2908 | +? | 0 | 0 | 0 | 1 |
| rs12275373 | 11 | 119165129 | T | C | 1.056 | 0.2909 | +? | 0 | 0 | 0 | 1 |
| rs656287 | 11 | 118571790 | G | A | 1.055 | 0.2912 | ++ | -120.6 | 1.36 | 1 | 0.2435 |
| rs79235532 | 11 | 119382089 | A | G | 1.055 | 0.2915 | +? | 0 | 0 | 0 | 1 |
| rs550388 | 11 | 117804926 | C | T | 1.054 | 0.2917 | +? | 0 | 0 | 0 | 1 |
| rs73006585 | 11 | 116943912 | C | G | 1.054 | 0.2917 | +? | 0 | 0 | 0 | 1 |
| rs7934084 | 11 | 120478622 | T | G | 1.054 | 0.2917 | +? | 0 | 0 | 0 | 1 |
| rs4938316 | 11 | 116764582 | T | C | -1.054 | 0.2918 | -? | 0 | 0 | 0 | 1 |
| rs11217847 | 11 | 120247436 | T | C | -1.054 | 0.2919 | -? | 0 | 0 | 0 | 1 |
| rs570083 | 11 | 117343392 | A | G | 1.054 | 0.2919 | +? | 0 | 0 | 0 | 1 |
| rs12805801 | 11 | 120798482 | A | G | -1.054 | 0.292 | -+ | 0 | 0.934 | 1 | 0.3339 |
| rs611217 | 11 | 118572611 | T | C | 1.054 | 0.292 | ++ | -116 | 1.389 | 1 | 0.2386 |
| rs672113 | 11 | 119386594 | C | T | -1.054 | 0.292 | -? | 0 | 0 | 0 | 1 |
| rs7943722 | 11 | 119252448 | T | C | 1.054 | 0.292 | +? | 0 | 0 | 0 | 1 |
| rs588901 | 11 | 119882059 | C | T | 1.054 | 0.2921 | +? | 0 | 0 | 0 | 1 |
| rs7482092 | 11 | 119082706 | C | T | 1.053 | 0.2922 | +? | 0 | 0 | 0 | 1 |
| rs524467 | 11 | 116529641 | G | T | -1.053 | 0.2923 | -? | 0 | 0 | 0 | 1 |
| rs647769 | 11 | 118812512 | T | C | 1.053 | 0.2923 | ++ | -189.1 | 1.038 | 1 | 0.3083 |
| rs7129292 | 11 | 121116638 | A | G | -1.053 | 0.2923 | -- | 0 | 0.001 | 1 | 0.9722 |
| rs11217873 | 11 | 120321653 | T | C | -1.053 | 0.2925 | -? | 0 | 0 | 0 | 1 |
| rs11217861 | 11 | 120288060 | T | C | -1.052 | 0.2927 | -? | 0 | 0 | 0 | 1 |
| rs648067 | 11 | 119949927 | A | C | 1.052 | 0.2929 | +? | 0 | 0 | 0 | 1 |
| rs75210328 | 11 | 118801296 | T | G | -1.052 | 0.2929 | -- | 0 | 0.34 | 1 | 0.5597 |
| rs7926958 | 11 | 117097222 | G | T | -1.052 | 0.2929 | -? | 0 | 0 | 0 | 1 |
| rs90192 | 11 | 117059347 | C | T | 1.052 | 0.2929 | +? | 0 | 0 | 0 | 1 |
| rs12807410 | 11 | 116308355 | A | G | -1.052 | 0.293 | -? | 0 | 0 | 0 | 1 |
| rs11215985 | 11 | 116398764 | G | A | 1.051 | 0.2931 | +? | 0 | 0 | 0 | 1 |
| rs4936529 | 11 | 120476746 | T | A | 1.051 | 0.2931 | +? | 0 | 0 | 0 | 1 |
| rs664657 | 11 | 117349617 | T | C | 1.051 | 0.2931 | +? | 0 | 0 | 0 | 1 |
| rs7115562 | 11 | 117020470 | A | G | 1.051 | 0.2931 | +? | 0 | 0 | 0 | 1 |
| rs7116454 | 11 | 117021134 | A | T | 1.051 | 0.2931 | +? | 0 | 0 | 0 | 1 |
| rs1540191 | 11 | 118090432 | G | A | -1.051 | 0.2932 | -? | 0 | 0 | 0 | 1 |
| rs180344 | 11 | 116613184 | C | T | 1.051 | 0.2932 | +- | -35.6 | 2.212 | 1 | 0.1369 |
| rs190567289 | 11 | 117458098 | T | C | 1.051 | 0.2934 | +? | 0 | 0 | 0 | 1 |
| rs73578545 | 11 | 116496416 | C | T | -1.051 | 0.2934 | -? | 0 | 0 | 0 | 1 |
| rs11218075 | 11 | 120820885 | C | A | 1.05 | 0.2937 | +? | 0 | 0 | 0 | 1 |
| rs1287581 | 11 | 119948738 | T | G | 1.05 | 0.2937 | +? | 0 | 0 | 0 | 1 |
| rs514808 | 11 | 119241120 | A | G | 1.05 | 0.2938 | ++ | -188.9 | 1.039 | 1 | 0.3082 |
| rs685879 | 11 | 119341075 | G | A | -1.05 | 0.2938 | -? | 0 | 0 | 0 | 1 |
| rs7106980 | 11 | 119854322 | G | T | -1.05 | 0.2938 | -? | 0 | 0 | 0 | 1 |
| rs10892526 | 11 | 119930706 | T | C | -1.049 | 0.2941 | -? | 0 | 0 | 0 | 1 |
| rs2512152 | 11 | 117887757 | T | C | 1.049 | 0.2941 | +- | -46.9 | 2.042 | 1 | 0.153 |
| rs67551874 | 11 | 120235574 | C | T | -1.049 | 0.2941 | -? | 0 | 0 | 0 | 1 |
| rs71478235 | 11 | 116214490 | T | C | -1.049 | 0.2943 | -- | -153.9 | 1.181 | 1 | 0.2771 |
| rs56246162 | 11 | 119658388 | G | C | -1.049 | 0.2944 | -? | 0 | 0 | 0 | 1 |
| rs147019067 | 11 | 117600086 | A | C | 1.048 | 0.2945 | +? | 0 | 0 | 0 | 1 |
| rs4938343 | 11 | 117003528 | G | C | 1.048 | 0.2945 | +? | 0 | 0 | 0 | 1 |
| rs580363 | 11 | 116325156 | C | G | -1.048 | 0.2947 | -? | 0 | 0 | 0 | 1 |
| rs9610 | 11 | 117872086 | A | G | -1.047 | 0.2949 | -+ | 0 | 0.974 | 1 | 0.3237 |
| rs10892584 | 11 | 120399726 | T | G | -1.047 | 0.2951 | -? | 0 | 0 | 0 | 1 |
| rs180376 | 11 | 116590039 | C | G | 1.047 | 0.2951 | +- | -32.5 | 2.264 | 1 | 0.1324 |
| rs677027 | 11 | 119849389 | A | G | -1.047 | 0.2951 | -? | 0 | 0 | 0 | 1 |
| rs12274273 | 11 | 116471121 | C | T | 1.047 | 0.2953 | +? | 0 | 0 | 0 | 1 |
| rs12280450 | 11 | 116471149 | G | A | 1.047 | 0.2953 | +? | 0 | 0 | 0 | 1 |
| rs78991803 | 11 | 120161987 | A | G | -1.046 | 0.2955 | -? | 0 | 0 | 0 | 1 |

| | | | | | | | | | | | |
|---|---|---|---|---|---|---|---|---|---|---|---|
| rs7941202 | 11 | 117653576 | G | C | -1.046 | 0.2956 | -? | 0 | 0 | 0 | 1 |
| rs7945051 | 11 | 116201218 | C | A | -1.046 | 0.2956 | -? | 0 | 0 | 0 | 1 |
| rs115933113 | 11 | 120620840 | C | G | 1.046 | 0.2957 | +? | 0 | 0 | 0 | 1 |
| rs180346 | 11 | 116612659 | C | A | 1.045 | 0.2958 | +- | -36.2 | 2.203 | 1 | 0.1378 |
| rs142438429 | 11 | 118533042 | T | C | -1.045 | 0.296 | -- | 0 | 0.117 | 1 | 0.7321 |
| rs2510139 | 11 | 119164677 | C | T | -1.045 | 0.296 | -? | 0 | 0 | 0 | 1 |
| rs634182 | 11 | 119341038 | C | T | -1.045 | 0.296 | -? | 0 | 0 | 0 | 1 |
| rs655929 | 11 | 119893179 | C | A | 1.045 | 0.2961 | +? | 0 | 0 | 0 | 1 |
| rs2511854 | 11 | 119146659 | G | C | -1.044 | 0.2963 | -? | 0 | 0 | 0 | 1 |
| rs117414016 | 11 | 120649337 | T | G | -1.044 | 0.2964 | -? | 0 | 0 | 0 | 1 |
| rs498736 | 11 | 116485334 | C | G | -1.044 | 0.2964 | -? | 0 | 0 | 0 | 1 |
| rs544452 | 11 | 118571357 | T | G | 1.044 | 0.2965 | ++ | -122.5 | 1.348 | 1 | 0.2456 |
| rs570949 | 11 | 118571911 | T | G | 1.044 | 0.2965 | ++ | -122.5 | 1.348 | 1 | 0.2456 |
| rs658040 | 11 | 118572171 | G | A | 1.044 | 0.2965 | ++ | -122.5 | 1.348 | 1 | 0.2456 |
| rs948091 | 11 | 121091762 | G | C | 1.044 | 0.2965 | +? | 0 | 0 | 0 | 1 |
| rs12576276 | 11 | 119337109 | C | T | -1.044 | 0.2966 | -? | 0 | 0 | 0 | 1 |
| rs11216043 | 11 | 116489979 | C | T | -1.044 | 0.2967 | -? | 0 | 0 | 0 | 1 |
| rs12288877 | 11 | 117684522 | C | G | 1.044 | 0.2967 | ++ | 0 | 0.175 | 1 | 0.6758 |
| rs12793860 | 11 | 119335915 | T | G | -1.044 | 0.2967 | -? | 0 | 0 | 0 | 1 |
| rs17492366 | 11 | 116438810 | A | G | -1.043 | 0.2968 | -? | 0 | 0 | 0 | 1 |
| rs140557493 | 11 | 119474608 | G | C | 1.043 | 0.297 | +? | 0 | 0 | 0 | 1 |
| rs1939332 | 11 | 119907973 | C | T | -1.042 | 0.2973 | -? | 0 | 0 | 0 | 1 |
| rs4620730 | 11 | 119507180 | G | C | 1.042 | 0.2973 | +? | 0 | 0 | 0 | 1 |
| rs149861300 | 11 | 117362331 | C | A | 1.042 | 0.2974 | +? | 0 | 0 | 0 | 1 |
| rs58174358 | 11 | 120647015 | T | C | 1.041 | 0.2977 | +? | 0 | 0 | 0 | 1 |
| rs11604012 | 11 | 120292040 | C | T | 1.041 | 0.2978 | ++ | -171.9 | 1.104 | 1 | 0.2935 |
| rs5096 | 11 | 116693095 | G | A | -1.041 | 0.2978 | -? | 0 | 0 | 0 | 1 |
| rs5100 | 11 | 116692694 | G | A | -1.041 | 0.2978 | -? | 0 | 0 | 0 | 1 |
| rs7936503 | 11 | 119689251 | T | G | 1.041 | 0.2978 | ++ | 0 | 0.026 | 1 | 0.8716 |
| rs2509049 | 11 | 118966521 | T | C | -1.041 | 0.2979 | -? | 0 | 0 | 0 | 1 |
| rs684384 | 11 | 120977529 | A | G | 1.041 | 0.2979 | +? | 0 | 0 | 0 | 1 |
| rs11217837 | 11 | 120232766 | G | A | -1.041 | 0.298 | -? | 0 | 0 | 0 | 1 |
| rs643536 | 11 | 119788154 | T | C | 1.041 | 0.298 | +? | 0 | 0 | 0 | 1 |
| rs7118009 | 11 | 118707559 | T | C | -1.041 | 0.298 | -- | 0 | 0.003 | 1 | 0.9587 |
| rs12575295 | 11 | 119337241 | T | C | -1.04 | 0.2984 | -? | 0 | 0 | 0 | 1 |
| rs488141 | 11 | 118571168 | G | A | 1.04 | 0.2984 | ++ | -121.6 | 1.354 | 1 | 0.2446 |
| rs541631 | 11 | 118571023 | G | A | 1.04 | 0.2984 | ++ | -121.6 | 1.354 | 1 | 0.2446 |
| rs55935223 | 11 | 119337381 | C | T | -1.04 | 0.2984 | -? | 0 | 0 | 0 | 1 |
| rs57737334 | 11 | 119337436 | T | A | -1.04 | 0.2984 | -? | 0 | 0 | 0 | 1 |
| rs642224 | 11 | 118570919 | A | G | 1.04 | 0.2984 | ++ | -121.6 | 1.354 | 1 | 0.2446 |
| rs657769 | 11 | 118571316 | T | C | 1.04 | 0.2984 | ++ | -121.6 | 1.354 | 1 | 0.2446 |
| rs658676 | 11 | 118571123 | T | C | 1.04 | 0.2984 | ++ | -121.6 | 1.354 | 1 | 0.2446 |
| rs659564 | 11 | 118570914 | A | C | 1.04 | 0.2984 | ++ | -121.6 | 1.354 | 1 | 0.2446 |
| rs4611216 | 11 | 120472667 | A | G | 1.04 | 0.2985 | +? | 0 | 0 | 0 | 1 |
| rs74426327 | 11 | 119756067 | C | T | -1.04 | 0.2985 | -? | 0 | 0 | 0 | 1 |
| rs2298650 | 11 | 119155618 | T | G | 1.039 | 0.2986 | +? | 0 | 0 | 0 | 1 |
| rs57228309 | 11 | 116502706 | T | A | -1.039 | 0.2986 | -? | 0 | 0 | 0 | 1 |
| rs8551 | 11 | 118967524 | T | C | -1.039 | 0.2987 | -? | 0 | 0 | 0 | 1 |
| rs10892125 | 11 | 117355782 | C | A | -1.039 | 0.2988 | -? | 0 | 0 | 0 | 1 |
| rs2508922 | 11 | 118681088 | A | G | -1.039 | 0.2988 | -+ | 0 | 0.384 | 1 | 0.5353 |
| rs150824999 | 11 | 117564067 | T | C | 1.039 | 0.2989 | +? | 0 | 0 | 0 | 1 |
| rs4639966 | 11 | 118573519 | C | T | -1.038 | 0.2991 | -- | 0 | 0.008 | 1 | 0.9304 |
| rs2168673 | 11 | 117196901 | A | C | 1.038 | 0.2992 | +? | 0 | 0 | 0 | 1 |
| rs598696 | 11 | 121006976 | T | A | -1.038 | 0.2992 | -? | 0 | 0 | 0 | 1 |
| rs73578551 | 11 | 116498120 | T | C | -1.038 | 0.2992 | -? | 0 | 0 | 0 | 1 |
| rs1013366746 | 11 | 119501639 | T | C | 1.038 | 0.2993 | +? | 0 | 0 | 0 | 1 |

| | | | | | | | | | | | |
|---|---|---|---|---|---|---|---|---|---|---|---|
| rs1240769 | 11 | 116517591 | G | T | -1.038 | 0.2993 | -? | 0 | 0 | 0 | 1 |
| rs12419730 | 11 | 120816203 | A | G | 1.038 | 0.2995 | +? | 0 | 0 | 0 | 1 |
| rs1892975 | 11 | 120614538 | T | A | 1.038 | 0.2995 | +? | 0 | 0 | 0 | 1 |
| rs965312 | 11 | 119942972 | A | G | -1.038 | 0.2995 | -? | 0 | 0 | 0 | 1 |
| rs2368238 | 11 | 118091761 | T | G | -1.037 | 0.2996 | -? | 0 | 0 | 0 | 1 |
| rs17311233 | 11 | 120157850 | T | C | -1.037 | 0.2997 | -? | 0 | 0 | 0 | 1 |
| rs653177 | 11 | 119337199 | A | G | -1.037 | 0.2998 | -? | 0 | 0 | 0 | 1 |
| rs490460 | 11 | 117163765 | C | A | -1.037 | 0.2999 | -? | 0 | 0 | 0 | 1 |
| rs73578554 | 11 | 116501134 | G | A | -1.037 | 0.2999 | -? | 0 | 0 | 0 | 1 |
| rs7932906 | 11 | 119338245 | T | C | -1.037 | 0.2999 | -? | 0 | 0 | 0 | 1 |
| rs11217527 | 11 | 119697584 | A | G | 1.036 | 0.3 | +? | 0 | 0 | 0 | 1 |
| rs497678 | 11 | 118567881 | C | T | 1.036 | 0.3001 | ++ | -51.4 | 1.981 | 1 | 0.1592 |
| rs7123811 | 11 | 118569469 | G | T | 1.036 | 0.3001 | ++ | -120.8 | 1.359 | 1 | 0.2438 |
| rs10892199 | 11 | 117832675 | A | G | 1.036 | 0.3002 | +? | 0 | 0 | 0 | 1 |
| rs517445 | 11 | 118573535 | G | A | 1.036 | 0.3004 | ++ | -124.6 | 1.336 | 1 | 0.2478 |
| rs9326250 | 11 | 117902620 | A | C | -1.036 | 0.3004 | -? | 0 | 0 | 0 | 1 |
| rs734104 | 11 | 116700860 | C | T | 1.035 | 0.3005 | +? | 0 | 0 | 0 | 1 |
| rs80282202 | 11 | 120986075 | C | G | 1.035 | 0.3005 | +? | 0 | 0 | 0 | 1 |
| rs1546465 | 11 | 119507035 | G | T | 1.035 | 0.3007 | +? | 0 | 0 | 0 | 1 |
| rs1939667 | 11 | 120600151 | T | C | 1.035 | 0.3007 | +? | 0 | 0 | 0 | 1 |
| rs10790138 | 11 | 116209430 | C | G | -1.035 | 0.3008 | -? | 0 | 0 | 0 | 1 |
| rs7115463 | 11 | 121116824 | T | C | -1.035 | 0.3008 | -- | 0 | 0.001 | 1 | 0.9806 |
| rs1939976 | 11 | 117686829 | T | C | 1.034 | 0.3009 | ++ | 0 | 0.146 | 1 | 0.702 |
| rs55683640 | 11 | 120976311 | C | A | -1.034 | 0.301 | -? | 0 | 0 | 0 | 1 |
| rs78198334 | 11 | 116532283 | C | G | -1.034 | 0.3011 | -? | 0 | 0 | 0 | 1 |
| rs10444337 | 11 | 119929276 | A | G | -1.034 | 0.3012 | -? | 0 | 0 | 0 | 1 |
| rs56159851 | 11 | 116271858 | T | C | 1.034 | 0.3013 | +? | 0 | 0 | 0 | 1 |
| rs11216504 | 11 | 117562505 | T | C | 1.033 | 0.3014 | +? | 0 | 0 | 0 | 1 |
| rs12272981 | 11 | 118709600 | G | A | -1.033 | 0.3014 | -- | 0 | 0.002 | 1 | 0.9656 |
| rs73008610 | 11 | 119686252 | C | G | -1.033 | 0.3014 | -? | 0 | 0 | 0 | 1 |
| rs10891961 | 11 | 116196429 | C | T | -1.033 | 0.3015 | -? | 0 | 0 | 0 | 1 |
| rs1263174 | 11 | 116681243 | G | T | -1.033 | 0.3015 | -? | 0 | 0 | 0 | 1 |
| rs184243297 | 11 | 120874890 | A | G | -1.033 | 0.3015 | -? | 0 | 0 | 0 | 1 |
| rs11218235 | 11 | 121144673 | T | C | -1.033 | 0.3016 | -- | 0 | 0.294 | 1 | 0.5874 |
| rs75243489 | 11 | 116457115 | A | T | -1.033 | 0.3016 | -? | 0 | 0 | 0 | 1 |
| rs11215897 | 11 | 116196643 | G | A | -1.033 | 0.3017 | -? | 0 | 0 | 0 | 1 |
| rs58499675 | 11 | 117574923 | C | T | 1.033 | 0.3018 | +? | 0 | 0 | 0 | 1 |
| rs141169729 | 11 | 118510642 | T | C | -1.032 | 0.3019 | -- | 0 | 0.01 | 1 | 0.9204 |
| rs611733 | 11 | 118572760 | T | C | 1.032 | 0.3019 | ++ | -126.9 | 1.322 | 1 | 0.2502 |
| rs1005705 | 11 | 116549116 | C | A | -1.032 | 0.3021 | -? | 0 | 0 | 0 | 1 |
| rs108533 | 11 | 116601711 | T | A | 1.032 | 0.3021 | +- | -40.2 | 2.14 | 1 | 0.1435 |
| rs116944258 | 11 | 116546633 | A | C | -1.032 | 0.3021 | -? | 0 | 0 | 0 | 1 |
| rs117830617 | 11 | 116549971 | A | G | -1.032 | 0.3021 | -? | 0 | 0 | 0 | 1 |
| rs12793594 | 11 | 118569900 | C | T | 1.032 | 0.3021 | ++ | -119.9 | 1.364 | 1 | 0.2428 |
| rs482889 | 11 | 118570098 | T | G | 1.032 | 0.3021 | ++ | -119.9 | 1.364 | 1 | 0.2428 |
| rs523793 | 11 | 118570697 | T | C | 1.032 | 0.3021 | ++ | -119.9 | 1.364 | 1 | 0.2428 |
| rs554456 | 11 | 118570062 | C | T | 1.032 | 0.3021 | ++ | -119.9 | 1.364 | 1 | 0.2428 |
| rs7108144 | 11 | 118569694 | A | G | 1.032 | 0.3021 | ++ | -119.9 | 1.364 | 1 | 0.2428 |
| rs79493947 | 11 | 119810375 | G | C | -1.032 | 0.3021 | -? | 0 | 0 | 0 | 1 |
| rs79678269 | 11 | 116548972 | C | G | -1.032 | 0.3021 | -? | 0 | 0 | 0 | 1 |
| rs673547 | 11 | 118573339 | A | G | 1.032 | 0.3022 | ++ | -127.3 | 1.32 | 1 | 0.2506 |
| rs606683 | 11 | 117904127 | G | C | -1.032 | 0.3023 | -- | 0 | 0.189 | 1 | 0.6639 |
| rs10891999 | 11 | 116488882 | T | A | -1.031 | 0.3027 | -? | 0 | 0 | 0 | 1 |
| rs76235142 | 11 | 116480813 | A | G | -1.031 | 0.3027 | -? | 0 | 0 | 0 | 1 |
| rs573396 | 11 | 119239375 | G | A | 1.03 | 0.3028 | +? | 0 | 0 | 0 | 1 |
| rs74888816 | 11 | 118897962 | T | C | 1.029 | 0.3033 | +? | 0 | 0 | 0 | 1 |

| | | | | | | | | | | | |
|---|---|---|---|---|---|---|---|---|---|---|---|
| rs552079 | 11 | 118573022 | C | T | 1.029 | 0.3034 | ++ | -129 | 1.31 | 1 | 0.2524 |
| rs58601794 | 11 | 118884726 | C | T | -1.029 | 0.3035 | -- | 0 | 0.229 | 1 | 0.6324 |
| rs10732855 | 11 | 116209589 | C | G | -1.029 | 0.3036 | -? | 0 | 0 | 0 | 1 |
| rs10891962 | 11 | 116196666 | C | T | -1.029 | 0.3036 | -? | 0 | 0 | 0 | 1 |
| rs11215896 | 11 | 116196502 | A | C | -1.029 | 0.3036 | -? | 0 | 0 | 0 | 1 |
| rs628861 | 11 | 119925575 | A | C | -1.029 | 0.3037 | -? | 0 | 0 | 0 | 1 |
| rs1240513 | 11 | 116528202 | T | A | -1.028 | 0.3038 | -? | 0 | 0 | 0 | 1 |
| rs7932565 | 11 | 119219913 | G | A | 1.027 | 0.3043 | +? | 0 | 0 | 0 | 1 |
| rs10790231 | 11 | 117967476 | C | T | -1.027 | 0.3046 | -? | 0 | 0 | 0 | 1 |
| rs4938717 | 11 | 119576911 | T | C | -1.027 | 0.3046 | -? | 0 | 0 | 0 | 1 |
| rs612941 | 11 | 117934115 | C | A | -1.027 | 0.3046 | -? | 0 | 0 | 0 | 1 |
| rs67023719 | 11 | 117282742 | T | C | 1.027 | 0.3046 | +? | 0 | 0 | 0 | 1 |
| rs10892184 | 11 | 117707229 | T | C | -1.026 | 0.3048 | -? | 0 | 0 | 0 | 1 |
| rs7759 | 11 | 118967291 | G | A | -1.026 | 0.3049 | -? | 0 | 0 | 0 | 1 |
| rs10736483 | 11 | 116210084 | A | G | -1.026 | 0.3051 | -? | 0 | 0 | 0 | 1 |
| rs11217988 | 11 | 120581395 | C | T | 1.025 | 0.3052 | +? | 0 | 0 | 0 | 1 |
| rs1508102 | 11 | 116379889 | A | G | -1.025 | 0.3052 | -? | 0 | 0 | 0 | 1 |
| rs60651507 | 11 | 118751898 | A | G | 1.025 | 0.3052 | ++ | 0 | 0.001 | 1 | 0.9783 |
| rs61898767 | 11 | 120353153 | G | A | -1.025 | 0.3052 | -? | 0 | 0 | 0 | 1 |
| rs10891963 | 11 | 116196711 | G | T | -1.025 | 0.3053 | -? | 0 | 0 | 0 | 1 |
| rs2028960 | 11 | 119697626 | T | C | -1.025 | 0.3053 | -? | 0 | 0 | 0 | 1 |
| rs144188664 | 11 | 116541034 | T | A | -1.025 | 0.3055 | -? | 0 | 0 | 0 | 1 |
| rs144256947 | 11 | 116543025 | C | T | -1.025 | 0.3055 | -? | 0 | 0 | 0 | 1 |
| rs148462279 | 11 | 116542737 | G | A | -1.025 | 0.3055 | -? | 0 | 0 | 0 | 1 |
| rs665406 | 11 | 117347058 | C | T | -1.024 | 0.3056 | -? | 0 | 0 | 0 | 1 |
| rs73001093 | 11 | 119440387 | T | G | 1.024 | 0.3056 | +? | 0 | 0 | 0 | 1 |
| rs11828580 | 11 | 117923983 | A | G | -1.024 | 0.3057 | -? | 0 | 0 | 0 | 1 |
| rs45506400 | 11 | 117164246 | C | A | 1.024 | 0.3057 | +? | 0 | 0 | 0 | 1 |
| rs1396131 | 11 | 116395458 | C | A | -1.024 | 0.3059 | -? | 0 | 0 | 0 | 1 |
| rs11217796 | 11 | 120162250 | G | A | -1.023 | 0.3061 | -? | 0 | 0 | 0 | 1 |
| rs2051480 | 11 | 118094513 | A | G | -1.023 | 0.3063 | -? | 0 | 0 | 0 | 1 |
| rs6589721 | 11 | 119081215 | T | C | -1.023 | 0.3063 | -? | 0 | 0 | 0 | 1 |
| rs10891974 | 11 | 116302144 | T | C | -1.023 | 0.3064 | -? | 0 | 0 | 0 | 1 |
| rs11216425 | 11 | 117363426 | A | C | 1.022 | 0.3067 | +? | 0 | 0 | 0 | 1 |
| rs662318 | 11 | 119848438 | G | A | -1.022 | 0.3068 | -? | 0 | 0 | 0 | 1 |
| rs619115 | 11 | 120987253 | C | T | 1.022 | 0.307 | +? | 0 | 0 | 0 | 1 |
| rs597124 | 11 | 119949821 | T | C | 1.021 | 0.3072 | +? | 0 | 0 | 0 | 1 |
| rs600720 | 11 | 119948997 | T | C | 1.021 | 0.3072 | +? | 0 | 0 | 0 | 1 |
| rs649921 | 11 | 119949498 | T | G | 1.021 | 0.3072 | +? | 0 | 0 | 0 | 1 |
| rs662549 | 11 | 119947801 | C | T | 1.021 | 0.3072 | +? | 0 | 0 | 0 | 1 |
| rs2155811 | 11 | 120164799 | C | G | -1.021 | 0.3074 | -? | 0 | 0 | 0 | 1 |
| rs143620404 | 11 | 117941082 | T | G | 1.021 | 0.3075 | ++ | 0 | 0.01 | 1 | 0.9213 |
| rs2509851 | 11 | 118966780 | C | A | -1.02 | 0.3078 | -? | 0 | 0 | 0 | 1 |
| rs948376 | 11 | 121092505 | C | T | 1.02 | 0.3078 | +? | 0 | 0 | 0 | 1 |
| rs661621 | 11 | 119944195 | T | A | 1.02 | 0.3079 | +? | 0 | 0 | 0 | 1 |
| rs745517 | 11 | 121092052 | C | T | 1.02 | 0.3079 | +? | 0 | 0 | 0 | 1 |
| rs34068367 | 11 | 116479185 | A | C | -1.019 | 0.308 | -? | 0 | 0 | 0 | 1 |
| rs600620 | 11 | 117458539 | G | A | 1.019 | 0.308 | +? | 0 | 0 | 0 | 1 |
| rs61897924 | 11 | 119903718 | T | C | -1.019 | 0.308 | -? | 0 | 0 | 0 | 1 |
| rs17124120 | 11 | 120505628 | T | C | 1.019 | 0.3081 | +? | 0 | 0 | 0 | 1 |
| rs12276544 | 11 | 120117951 | A | T | -1.019 | 0.3082 | -? | 0 | 0 | 0 | 1 |
| rs511451 | 11 | 118568639 | T | C | 1.018 | 0.3086 | ++ | -122.1 | 1.351 | 1 | 0.2452 |
| rs61016803 | 11 | 119697261 | A | T | -1.018 | 0.3086 | -? | 0 | 0 | 0 | 1 |
| rs610226 | 11 | 117688225 | G | C | 1.018 | 0.3086 | +? | 0 | 0 | 0 | 1 |
| rs10444338 | 11 | 119929445 | C | T | -1.018 | 0.3089 | -? | 0 | 0 | 0 | 1 |
| rs638775 | 11 | 120977922 | G | C | 1.018 | 0.3089 | +? | 0 | 0 | 0 | 1 |

| | | | | | | | | | | | |
|---|---|---|---|---|---|---|---|---|---|---|---|
| rs17124305 | 11 | 120646975 | G | A | 1.017 | 0.309 | +? | 0 | 0 | 0 | 1 |
| rs12420994 | 11 | 117846083 | G | C | -1.017 | 0.3091 | -- | 0 | 0.012 | 1 | 0.9128 |
| rs3133856 | 11 | 120584612 | C | T | -1.017 | 0.3091 | -? | 0 | 0 | 0 | 1 |
| rs2929176 | 11 | 117106282 | C | T | -1.017 | 0.3092 | -? | 0 | 0 | 0 | 1 |
| rs3016582 | 11 | 117106280 | A | T | -1.017 | 0.3092 | -? | 0 | 0 | 0 | 1 |
| rs949138 | 11 | 117330669 | G | A | 1.017 | 0.3092 | +? | 0 | 0 | 0 | 1 |
| rs35508996 | 11 | 119502547 | A | G | 1.017 | 0.3093 | +? | 0 | 0 | 0 | 1 |
| rs73024427 | 11 | 117803627 | T | C | 1.017 | 0.3093 | +? | 0 | 0 | 0 | 1 |
| rs11218069 | 11 | 120816408 | A | C | 1.016 | 0.3095 | +? | 0 | 0 | 0 | 1 |
| rs56329702 | 11 | 120222475 | T | C | -1.016 | 0.3095 | -? | 0 | 0 | 0 | 1 |
| rs11217662 | 11 | 119924999 | T | A | -1.016 | 0.3096 | -? | 0 | 0 | 0 | 1 |
| rs7117145 | 11 | 116531969 | T | A | -1.016 | 0.3096 | -? | 0 | 0 | 0 | 1 |
| rs1145186 | 11 | 116528660 | A | T | -1.016 | 0.3097 | -? | 0 | 0 | 0 | 1 |
| rs35200015 | 11 | 117383215 | A | G | -1.016 | 0.3097 | -? | 0 | 0 | 0 | 1 |
| rs148108200 | 11 | 116480236 | C | T | -1.016 | 0.3098 | -? | 0 | 0 | 0 | 1 |
| rs3741301 | 11 | 116631391 | C | T | 1.016 | 0.3098 | +- | -65.9 | 1.809 | 1 | 0.1787 |
| rs1939127 | 11 | 117890681 | C | T | 1.015 | 0.3099 | +- | -49.1 | 2.012 | 1 | 0.1561 |
| rs116987336 | 11 | 117175658 | A | G | 1.015 | 0.31 | +? | 0 | 0 | 0 | 1 |
| rs4999180 | 11 | 121092522 | T | C | 1.015 | 0.31 | +? | 0 | 0 | 0 | 1 |
| rs78577252 | 11 | 116492477 | T | C | -1.015 | 0.31 | -? | 0 | 0 | 0 | 1 |
| rs10892325 | 11 | 118912702 | C | T | 1.015 | 0.3101 | +? | 0 | 0 | 0 | 1 |
| rs11607355 | 11 | 118093547 | C | T | -1.015 | 0.3101 | -? | 0 | 0 | 0 | 1 |
| rs7101613 | 11 | 117572288 | G | A | 1.015 | 0.3101 | +? | 0 | 0 | 0 | 1 |
| rs10750090 | 11 | 116209785 | C | T | -1.015 | 0.3102 | -? | 0 | 0 | 0 | 1 |
| rs948092 | 11 | 121091888 | C | T | 1.015 | 0.3102 | +? | 0 | 0 | 0 | 1 |
| rs12419497 | 11 | 119334628 | T | C | -1.015 | 0.3103 | -? | 0 | 0 | 0 | 1 |
| rs12419764 | 11 | 121134395 | G | A | -1.014 | 0.3106 | -- | 0 | 0.217 | 1 | 0.6412 |
| rs11218068 | 11 | 120816319 | C | T | 1.014 | 0.3108 | +? | 0 | 0 | 0 | 1 |
| rs12280288 | 11 | 118892940 | C | T | -1.014 | 0.3108 | -- | 0 | 0.059 | 1 | 0.8087 |
| rs12366111 | 11 | 117990553 | C | T | -1.013 | 0.3108 | -- | 0 | 0.369 | 1 | 0.5435 |
| rs10736485 | 11 | 116210182 | G | C | -1.013 | 0.3109 | -? | 0 | 0 | 0 | 1 |
| rs11217474 | 11 | 119631726 | G | C | -1.013 | 0.311 | -? | 0 | 0 | 0 | 1 |
| rs11216443 | 11 | 117409700 | A | C | 1.013 | 0.3111 | +? | 0 | 0 | 0 | 1 |
| rs675929 | 11 | 117892050 | A | G | 1.013 | 0.3111 | +- | -45.8 | 2.057 | 1 | 0.1515 |
| rs4938352 | 11 | 117027481 | G | A | 1.013 | 0.3112 | +? | 0 | 0 | 0 | 1 |
| rs1893264 | 11 | 119193721 | C | A | -1.012 | 0.3113 | -? | 0 | 0 | 0 | 1 |
| rs147507811 | 11 | 116854683 | A | C | 1.012 | 0.3115 | +? | 0 | 0 | 0 | 1 |
| rs1944496 | 11 | 119879488 | A | G | -1.012 | 0.3115 | -? | 0 | 0 | 0 | 1 |
| rs4938348 | 11 | 117021877 | G | C | 1.012 | 0.3115 | +? | 0 | 0 | 0 | 1 |
| rs12099002 | 11 | 117571475 | A | G | 1.012 | 0.3116 | +? | 0 | 0 | 0 | 1 |
| rs11827471 | 11 | 120122394 | G | A | -1.011 | 0.3118 | -? | 0 | 0 | 0 | 1 |
| rs2508449 | 11 | 117890273 | C | T | 1.011 | 0.3119 | +- | -47.6 | 2.033 | 1 | 0.1539 |
| rs56249090 | 11 | 117810532 | A | C | 1.011 | 0.3119 | +? | 0 | 0 | 0 | 1 |
| rs11217496 | 11 | 119648956 | T | G | -1.011 | 0.312 | -? | 0 | 0 | 0 | 1 |
| rs11218062 | 11 | 120804014 | C | T | -1.011 | 0.312 | -+ | 0 | 0.903 | 1 | 0.3419 |
| rs76046245 | 11 | 120193461 | A | C | -1.011 | 0.3122 | -? | 0 | 0 | 0 | 1 |
| rs2186621 | 11 | 120821263 | C | T | 1.01 | 0.3124 | +? | 0 | 0 | 0 | 1 |
| rs4936416 | 11 | 117896454 | A | G | -1.01 | 0.3124 | -? | 0 | 0 | 0 | 1 |
| rs10790151 | 11 | 116452887 | T | C | 1.01 | 0.3125 | +? | 0 | 0 | 0 | 1 |
| rs11217629 | 11 | 119880076 | A | C | -1.01 | 0.3126 | -? | 0 | 0 | 0 | 1 |
| rs11217872 | 11 | 120321201 | C | G | -1.01 | 0.3126 | -? | 0 | 0 | 0 | 1 |
| rs948466 | 11 | 117966100 | A | G | -1.01 | 0.3126 | -? | 0 | 0 | 0 | 1 |
| rs4938645 | 11 | 119163319 | G | A | 1.009 | 0.3129 | +? | 0 | 0 | 0 | 1 |
| rs61381062 | 11 | 118884708 | A | G | -1.009 | 0.313 | -- | 0 | 0.206 | 1 | 0.6499 |
| rs57877069 | 11 | 119962800 | C | G | 1.009 | 0.3132 | +? | 0 | 0 | 0 | 1 |
| rs7120309 | 11 | 117042816 | T | A | 1.009 | 0.3132 | +? | 0 | 0 | 0 | 1 |

| | | | | | | | | | | | |
|---|---|---|---|---|---|---|---|---|---|---|---|
| rs7120449 | 11 | 117042899 | T | C | 1.009 | 0.3132 | +? | 0 | 0 | 0 | 1 |
| rs7124732 | 11 | 119959273 | G | A | 1.009 | 0.3132 | +? | 0 | 0 | 0 | 1 |
| rs11215966 | 11 | 116366547 | C | T | -1.008 | 0.3133 | -? | 0 | 0 | 0 | 1 |
| rs1145192 | 11 | 116570894 | A | G | -1.008 | 0.3133 | -? | 0 | 0 | 0 | 1 |
| rs36119355 | 11 | 117845195 | T | G | 1.008 | 0.3134 | ++ | 0 | 0.522 | 1 | 0.47 |
| rs1892947 | 11 | 116530061 | A | G | -1.008 | 0.3135 | -? | 0 | 0 | 0 | 1 |
| rs2727793 | 11 | 116683374 | C | T | -1.008 | 0.3136 | -? | 0 | 0 | 0 | 1 |
| rs7930793 | 11 | 119825176 | G | T | -1.007 | 0.3137 | -? | 0 | 0 | 0 | 1 |
| rs10892471 | 11 | 119679283 | A | G | 1.007 | 0.3138 | ++ | 0 | 0.025 | 1 | 0.8744 |
| rs564989 | 11 | 117709754 | C | T | -1.007 | 0.3138 | -? | 0 | 0 | 0 | 1 |
| rs11217989 | 11 | 120583983 | C | T | 1.007 | 0.314 | +? | 0 | 0 | 0 | 1 |
| rs493721 | 11 | 116247770 | G | A | -1.007 | 0.314 | -? | 0 | 0 | 0 | 1 |
| rs536648 | 11 | 117143185 | G | C | -1.007 | 0.3141 | -? | 0 | 0 | 0 | 1 |
| rs540277 | 11 | 117362074 | T | C | 1.007 | 0.3141 | +? | 0 | 0 | 0 | 1 |
| rs4938347 | 11 | 117021759 | T | A | 1.006 | 0.3142 | +? | 0 | 0 | 0 | 1 |
| rs4938350 | 11 | 117022276 | A | G | 1.006 | 0.3142 | +? | 0 | 0 | 0 | 1 |
| rs76499613 | 11 | 119829348 | C | T | -1.006 | 0.3142 | -? | 0 | 0 | 0 | 1 |
| rs78289484 | 11 | 119870561 | T | C | 1.006 | 0.3142 | +? | 0 | 0 | 0 | 1 |
| rs2256111 | 11 | 117864047 | A | G | 1.006 | 0.3143 | ++ | 0 | 0.288 | 1 | 0.5916 |
| rs61659529 | 11 | 116533628 | G | A | -1.006 | 0.3143 | -? | 0 | 0 | 0 | 1 |
| rs75942205 | 11 | 116533595 | C | T | -1.006 | 0.3143 | -? | 0 | 0 | 0 | 1 |
| rs1263173 | 11 | 116681008 | G | A | -1.006 | 0.3145 | -? | 0 | 0 | 0 | 1 |
| rs2186882 | 11 | 119168370 | A | G | 1.006 | 0.3145 | +? | 0 | 0 | 0 | 1 |
| rs11217421 | 11 | 119583751 | C | T | -1.006 | 0.3146 | -? | 0 | 0 | 0 | 1 |
| rs554233 | 11 | 117194929 | G | A | 1.005 | 0.3147 | +? | 0 | 0 | 0 | 1 |
| rs627532 | 11 | 120826131 | C | T | -1.005 | 0.3148 | -? | 0 | 0 | 0 | 1 |
| rs7935678 | 11 | 119665733 | C | T | 1.005 | 0.3148 | +? | 0 | 0 | 0 | 1 |
| rs11218199 | 11 | 121064524 | T | C | -1.005 | 0.3149 | -? | 0 | 0 | 0 | 1 |
| rs1893969 | 11 | 117432434 | T | C | 1.005 | 0.3151 | +? | 0 | 0 | 0 | 1 |
| rs3819210 | 11 | 117389517 | G | A | -1.005 | 0.3151 | -? | 0 | 0 | 0 | 1 |
| rs11217678 | 11 | 119951347 | C | G | -1.004 | 0.3154 | -? | 0 | 0 | 0 | 1 |
| rs7126476 | 11 | 118834784 | G | A | -1.004 | 0.3154 | -? | 0 | 0 | 0 | 1 |
| rs1939666 | 11 | 120600346 | G | A | 1.004 | 0.3156 | +? | 0 | 0 | 0 | 1 |
| rs672295 | 11 | 119894514 | G | A | 1.003 | 0.3157 | +? | 0 | 0 | 0 | 1 |
| rs688587 | 11 | 118032242 | A | C | 1.003 | 0.3157 | +? | 0 | 0 | 0 | 1 |
| rs750057 | 11 | 116203045 | G | A | -1.003 | 0.3157 | -? | 0 | 0 | 0 | 1 |
| rs7926229 | 11 | 119675718 | C | T | 1.003 | 0.3157 | +? | 0 | 0 | 0 | 1 |
| rs2512156 | 11 | 117891216 | T | A | 1.003 | 0.3158 | +- | -44.7 | 2.074 | 1 | 0.1499 |
| rs489126 | 11 | 118572747 | A | G | 1.003 | 0.3159 | ++ | -148.6 | 1.207 | 1 | 0.272 |
| rs9651687 | 11 | 117967986 | A | T | -1.003 | 0.3159 | -? | 0 | 0 | 0 | 1 |
| rs2155130 | 11 | 117506565 | C | T | 1.003 | 0.3161 | +? | 0 | 0 | 0 | 1 |
| rs11215963 | 11 | 116361658 | G | C | 1.002 | 0.3162 | +? | 0 | 0 | 0 | 1 |
| rs1632755 | 11 | 116538851 | C | A | -1.002 | 0.3163 | -? | 0 | 0 | 0 | 1 |
| rs180361 | 11 | 116598649 | T | C | 1.002 | 0.3164 | +- | -37.4 | 2.183 | 1 | 0.1396 |
| rs180365 | 11 | 116596174 | T | C | 1.002 | 0.3165 | +- | -37.2 | 2.186 | 1 | 0.1392 |
| rs2071523 | 11 | 116697344 | C | T | -1.002 | 0.3165 | -? | 0 | 0 | 0 | 1 |
| rs6589826 | 11 | 120517643 | A | G | -1.002 | 0.3165 | -? | 0 | 0 | 0 | 1 |
| rs7121901 | 11 | 119824699 | T | C | -1.002 | 0.3165 | -? | 0 | 0 | 0 | 1 |
| rs10892353 | 11 | 119221449 | A | G | -1.001 | 0.3166 | -? | 0 | 0 | 0 | 1 |
| rs563372 | 11 | 119767864 | G | A | -1.001 | 0.3166 | -? | 0 | 0 | 0 | 1 |
| rs688563 | 11 | 118032228 | C | T | 1.001 | 0.3166 | +? | 0 | 0 | 0 | 1 |
| rs7927239 | 11 | 116326570 | C | G | -1.001 | 0.3167 | -? | 0 | 0 | 0 | 1 |
| rs7932145 | 11 | 119219819 | A | C | 1.001 | 0.3168 | +? | 0 | 0 | 0 | 1 |
| rs1786186 | 11 | 117945632 | T | C | -1.001 | 0.3169 | -? | 0 | 0 | 0 | 1 |
| rs17509752 | 11 | 117432779 | T | C | 1.001 | 0.317 | +? | 0 | 0 | 0 | 1 |
| rs61900957 | 11 | 118501723 | T | C | 1.001 | 0.317 | ++ | -24.3 | 2.414 | 1 | 0.1203 |

| | | | | | | | | | | | |
|---|---|---|---|---|---|---|---|---|---|---|---|
| rs12416826 | 11 | 118165955 | C | T | 1 | 0.3172 | +? | 0 | 0 | 0 | 1 |
| rs3737504 | 11 | 118869460 | G | A | -1 | 0.3172 | -- | 0 | 0.06 | 1 | 0.8072 |
| rs78347813 | 11 | 116529130 | T | C | -1 | 0.3172 | -? | 0 | 0 | 0 | 1 |
| rs2195525 | 11 | 119235404 | C | T | 1 | 0.3173 | +? | 0 | 0 | 0 | 1 |
| rs643788 | 11 | 118967758 | C | T | -1 | 0.3173 | -? | 0 | 0 | 0 | 1 |
| rs505848 | 11 | 116484741 | T | C | 0.999 | 0.3176 | +? | 0 | 0 | 0 | 1 |
| rs562422 | 11 | 117428278 | A | G | -0.999 | 0.3176 | -? | 0 | 0 | 0 | 1 |
| rs7130852 | 11 | 117571426 | G | A | 0.999 | 0.3176 | +? | 0 | 0 | 0 | 1 |
| rs1375742714 | 11 | 119146129 | C | T | 0.999 | 0.3177 | +? | 0 | 0 | 0 | 1 |
| rs1783855 | 11 | 117914504 | A | C | -0.999 | 0.3177 | -- | 0 | 0.145 | 1 | 0.7034 |
| rs2216311 | 11 | 116696043 | G | A | -0.999 | 0.3177 | -? | 0 | 0 | 0 | 1 |
| rs2887378 | 11 | 119916226 | A | G | -0.999 | 0.3177 | -? | 0 | 0 | 0 | 1 |
| rs2367794 | 11 | 116198031 | G | A | -0.999 | 0.3179 | -? | 0 | 0 | 0 | 1 |
| rs113863182 | 11 | 119220395 | T | C | 0.999 | 0.318 | +? | 0 | 0 | 0 | 1 |
| rs1508103 | 11 | 116379978 | G | A | -0.998 | 0.3183 | -? | 0 | 0 | 0 | 1 |
| rs117014628 | 11 | 116535631 | T | C | -0.998 | 0.3184 | -? | 0 | 0 | 0 | 1 |
| rs118103820 | 11 | 116537116 | G | A | -0.998 | 0.3184 | -? | 0 | 0 | 0 | 1 |
| rs140973748 | 11 | 116537798 | C | T | -0.998 | 0.3184 | -? | 0 | 0 | 0 | 1 |
| rs143380339 | 11 | 116538103 | A | C | -0.998 | 0.3184 | -? | 0 | 0 | 0 | 1 |
| rs190069353 | 11 | 116535624 | A | C | -0.998 | 0.3184 | -? | 0 | 0 | 0 | 1 |
| rs76547220 | 11 | 116534550 | G | T | -0.998 | 0.3184 | -? | 0 | 0 | 0 | 1 |
| rs77226464 | 11 | 116535435 | T | C | -0.998 | 0.3184 | -? | 0 | 0 | 0 | 1 |
| rs79063991 | 11 | 116536820 | T | C | -0.998 | 0.3184 | -? | 0 | 0 | 0 | 1 |
| rs79291519 | 11 | 116533310 | T | C | -0.997 | 0.3186 | -? | 0 | 0 | 0 | 1 |
| rs588763 | 11 | 117116781 | G | A | -0.997 | 0.3188 | -? | 0 | 0 | 0 | 1 |
| rs2515793 | 11 | 117815759 | A | G | -0.996 | 0.3191 | -? | 0 | 0 | 0 | 1 |
| rs12286525 | 11 | 117116450 | G | A | 0.996 | 0.3192 | +? | 0 | 0 | 0 | 1 |
| rs59608621 | 11 | 118885163 | G | A | -0.996 | 0.3192 | -- | 0 | 0.188 | 1 | 0.6644 |
| rs7479114 | 11 | 118568813 | T | C | 0.996 | 0.3192 | ++ | -131 | 1.299 | 1 | 0.2544 |
| rs12289816 | 11 | 116519950 | C | T | -0.996 | 0.3193 | -? | 0 | 0 | 0 | 1 |
| rs2298724 | 11 | 120823721 | G | A | -0.996 | 0.3193 | -? | 0 | 0 | 0 | 1 |
| rs56984853 | 11 | 117238478 | A | G | 0.994 | 0.3201 | +? | 0 | 0 | 0 | 1 |
| rs3133861 | 11 | 120659485 | T | G | 0.994 | 0.3202 | +? | 0 | 0 | 0 | 1 |
| rs4938736 | 11 | 119760977 | A | G | -0.994 | 0.3202 | -? | 0 | 0 | 0 | 1 |
| rs11216949 | 11 | 118568668 | T | C | 0.994 | 0.3203 | ++ | -132.7 | 1.289 | 1 | 0.2562 |
| rs12791838 | 11 | 118144209 | A | G | 0.994 | 0.3203 | +? | 0 | 0 | 0 | 1 |
| rs1630252 | 11 | 117952899 | C | T | -0.994 | 0.3204 | -? | 0 | 0 | 0 | 1 |
| rs73014974 | 11 | 118028648 | T | C | 0.994 | 0.3204 | +? | 0 | 0 | 0 | 1 |
| rs7113018 | 11 | 120804533 | G | T | 0.993 | 0.3207 | +? | 0 | 0 | 0 | 1 |
| rs117293980 | 11 | 116531475 | G | T | -0.993 | 0.3209 | -? | 0 | 0 | 0 | 1 |
| rs117889066 | 11 | 116531050 | C | G | -0.993 | 0.3209 | -? | 0 | 0 | 0 | 1 |
| rs9651691 | 11 | 119527227 | T | G | 0.993 | 0.3209 | +? | 0 | 0 | 0 | 1 |
| rs10790225 | 11 | 117805786 | T | C | 0.992 | 0.321 | ++ | -185.6 | 1.05 | 1 | 0.3054 |
| rs2100357 | 11 | 119915395 | T | C | -0.992 | 0.3212 | -? | 0 | 0 | 0 | 1 |
| rs369816532 | 11 | 118887887 | T | G | -0.992 | 0.3212 | -- | 0 | 0.371 | 1 | 0.5422 |
| rs906546 | 11 | 116389102 | T | C | 0.992 | 0.3212 | +? | 0 | 0 | 0 | 1 |
| rs79995246 | 11 | 119769724 | C | T | -0.992 | 0.3213 | -? | 0 | 0 | 0 | 1 |
| rs591221 | 11 | 121090597 | G | A | 0.992 | 0.3214 | +? | 0 | 0 | 0 | 1 |
| rs7934092 | 11 | 120643090 | A | C | 0.991 | 0.3216 | +? | 0 | 0 | 0 | 1 |
| rs523366 | 11 | 117119552 | A | G | -0.991 | 0.3218 | -? | 0 | 0 | 0 | 1 |
| rs550999 | 11 | 118811684 | G | C | 0.991 | 0.3218 | ++ | 0 | 0.895 | 1 | 0.344 |
| rs649586 | 11 | 119885674 | G | C | -0.991 | 0.3218 | -? | 0 | 0 | 0 | 1 |
| rs117170835 | 11 | 120131553 | A | G | -0.99 | 0.3222 | -? | 0 | 0 | 0 | 1 |
| rs2542063 | 11 | 116684767 | G | A | -0.99 | 0.3222 | -? | 0 | 0 | 0 | 1 |
| rs677409 | 11 | 119946763 | G | T | 0.99 | 0.3222 | +? | 0 | 0 | 0 | 1 |
| rs1396132 | 11 | 116395238 | A | T | -0.989 | 0.3225 | -? | 0 | 0 | 0 | 1 |

| | | | | | | | | | | | |
|---|---|---|---|---|---|---|---|---|---|---|---|
| rs35610956 | 11 | 118144280 | G | A | 0.989 | 0.3225 | +? | 0 | 0 | 0 | 1 |
| rs6589863 | 11 | 121098770 | C | G | 0.989 | 0.3226 | +? | 0 | 0 | 0 | 1 |
| rs678275 | 11 | 121006298 | G | C | -0.989 | 0.3226 | -? | 0 | 0 | 0 | 1 |
| rs10892505 | 11 | 119838718 | A | G | 0.989 | 0.3228 | +? | 0 | 0 | 0 | 1 |
| rs2098453 | 11 | 116696680 | C | T | -0.989 | 0.3228 | -? | 0 | 0 | 0 | 1 |
| rs4594010 | 11 | 120935558 | C | A | -0.989 | 0.3228 | -? | 0 | 0 | 0 | 1 |
| rs7127199 | 11 | 120494516 | T | C | 0.988 | 0.323 | +? | 0 | 0 | 0 | 1 |
| rs948105 | 11 | 117591654 | A | T | 0.988 | 0.3231 | +? | 0 | 0 | 0 | 1 |
| rs11217999 | 11 | 120623637 | A | C | 0.988 | 0.3232 | +? | 0 | 0 | 0 | 1 |
| rs11215908 | 11 | 116253905 | A | G | -0.988 | 0.3233 | -? | 0 | 0 | 0 | 1 |
| rs1871757 | 11 | 117030581 | T | A | 0.988 | 0.3233 | +? | 0 | 0 | 0 | 1 |
| rs2084892 | 11 | 119915651 | A | G | -0.987 | 0.3234 | -? | 0 | 0 | 0 | 1 |
| rs1784559 | 11 | 118901614 | T | A | 0.987 | 0.3235 | +? | 0 | 0 | 0 | 1 |
| rs2515800 | 11 | 117824592 | C | T | 0.987 | 0.3238 | +? | 0 | 0 | 0 | 1 |
| rs11216639 | 11 | 117805473 | T | C | 0.986 | 0.3239 | +? | 0 | 0 | 0 | 1 |
| rs180370 | 11 | 116592365 | T | C | 0.986 | 0.3242 | +- | -43.5 | 2.09 | 1 | 0.1483 |
| rs1043314 | 11 | 118915944 | G | A | 0.986 | 0.3243 | +? | 0 | 0 | 0 | 1 |
| rs4935746 | 11 | 120518653 | G | A | -0.986 | 0.3243 | -? | 0 | 0 | 0 | 1 |
| rs1278081233 | 11 | 121135004 | G | T | -0.985 | 0.3244 | -- | 0 | 0.233 | 1 | 0.6296 |
| rs633892 | 11 | 117112630 | C | A | 0.985 | 0.3246 | +? | 0 | 0 | 0 | 1 |
| rs1007369 | 11 | 119700031 | C | A | 0.984 | 0.3249 | +? | 0 | 0 | 0 | 1 |
| rs112711662 | 11 | 118882138 | T | C | 0.984 | 0.325 | +? | 0 | 0 | 0 | 1 |
| rs678837 | 11 | 117109270 | G | T | -0.984 | 0.325 | -? | 0 | 0 | 0 | 1 |
| rs12274094 | 11 | 120968462 | G | A | -0.984 | 0.3251 | -? | 0 | 0 | 0 | 1 |
| rs7952159 | 11 | 120622379 | G | A | 0.984 | 0.3252 | +? | 0 | 0 | 0 | 1 |
| rs582942 | 11 | 118515876 | T | C | -0.984 | 0.3253 | -? | 0 | 0 | 0 | 1 |
| rs3916582 | 11 | 118887888 | T | C | -0.983 | 0.3256 | -- | 0 | 0.352 | 1 | 0.5528 |
| rs75582258 | 11 | 117477761 | G | A | 0.982 | 0.3259 | +? | 0 | 0 | 0 | 1 |
| rs1786177 | 11 | 117904029 | A | G | -0.981 | 0.3264 | -- | -194.7 | 1.018 | 1 | 0.313 |
| rs7114961 | 11 | 118914388 | C | G | 0.981 | 0.3264 | +? | 0 | 0 | 0 | 1 |
| rs586407 | 11 | 119994765 | A | G | 0.981 | 0.3267 | +? | 0 | 0 | 0 | 1 |
| rs73559126 | 11 | 118880727 | A | T | 0.981 | 0.3267 | +? | 0 | 0 | 0 | 1 |
| rs2276060 | 11 | 118919132 | A | G | 0.981 | 0.3268 | +? | 0 | 0 | 0 | 1 |
| rs496391 | 11 | 116311186 | G | C | -0.98 | 0.3269 | -? | 0 | 0 | 0 | 1 |
| rs57495164 | 11 | 118095148 | T | C | -0.98 | 0.327 | -? | 0 | 0 | 0 | 1 |
| rs11607701 | 11 | 118169825 | G | A | 0.98 | 0.3271 | +? | 0 | 0 | 0 | 1 |
| rs112337443 | 11 | 118530068 | C | T | -0.979 | 0.3274 | -+ | 0 | 0.321 | 1 | 0.571 |
| rs2276083 | 11 | 119145274 | G | A | 0.979 | 0.3276 | +? | 0 | 0 | 0 | 1 |
| rs1939665 | 11 | 120600361 | T | C | 0.979 | 0.3278 | +? | 0 | 0 | 0 | 1 |
| rs948093 | 11 | 121092641 | A | T | 0.978 | 0.3279 | +? | 0 | 0 | 0 | 1 |
| rs10892001 | 11 | 116505440 | A | G | -0.978 | 0.328 | -? | 0 | 0 | 0 | 1 |
| rs9787748 | 11 | 121101075 | T | G | 0.978 | 0.328 | +? | 0 | 0 | 0 | 1 |
| rs1001973 | 11 | 119923643 | A | G | -0.978 | 0.3281 | -? | 0 | 0 | 0 | 1 |
| rs11217676 | 11 | 119951167 | G | C | -0.978 | 0.3281 | -? | 0 | 0 | 0 | 1 |
| rs1783850 | 11 | 117904561 | G | C | -0.978 | 0.3281 | -- | -192.3 | 1.026 | 1 | 0.311 |
| rs521976 | 11 | 119758831 | T | C | -0.978 | 0.3281 | -? | 0 | 0 | 0 | 1 |
| rs10437696 | 11 | 121140086 | C | T | -0.978 | 0.3282 | -- | 0 | 0.23 | 1 | 0.6317 |
| rs11217429 | 11 | 119594057 | T | G | 0.977 | 0.3285 | +? | 0 | 0 | 0 | 1 |
| rs1790475 | 11 | 119943201 | G | A | 0.977 | 0.3286 | +? | 0 | 0 | 0 | 1 |
| rs7940405 | 11 | 118142666 | G | A | 0.977 | 0.3286 | +? | 0 | 0 | 0 | 1 |
| rs948106 | 11 | 117591584 | C | T | 0.977 | 0.3287 | +? | 0 | 0 | 0 | 1 |
| rs1263171 | 11 | 116680330 | G | A | -0.977 | 0.3288 | -? | 0 | 0 | 0 | 1 |
| rs2156763 | 11 | 120624525 | G | A | 0.977 | 0.3288 | +? | 0 | 0 | 0 | 1 |
| rs56230940 | 11 | 119039028 | T | C | -0.977 | 0.3288 | -? | 0 | 0 | 0 | 1 |
| rs10790422 | 11 | 121136604 | T | C | -0.976 | 0.3291 | -- | 0 | 0.238 | 1 | 0.6257 |
| rs2849172 | 11 | 116698332 | T | C | -0.976 | 0.3291 | -? | 0 | 0 | 0 | 1 |

| | | | | | | | | | | | |
|---|---|---|---|---|---|---|---|---|---|---|---|
| rs502789 | 11 | 119784133 | A | G | 0.976 | 0.3291 | +? | 0 | 0 | 0 | 1 |
| rs12807148 | 11 | 116471492 | C | T | 0.976 | 0.3292 | +? | 0 | 0 | 0 | 1 |
| rs572063 | 11 | 117105572 | G | T | 0.976 | 0.3293 | +? | 0 | 0 | 0 | 1 |
| rs11217374 | 11 | 119520730 | G | A | 0.975 | 0.3294 | +? | 0 | 0 | 0 | 1 |
| rs55960868 | 11 | 116391666 | A | G | -0.975 | 0.3297 | -? | 0 | 0 | 0 | 1 |
| rs77973629 | 11 | 117469122 | T | G | 0.975 | 0.3297 | +? | 0 | 0 | 0 | 1 |
| rs11218071 | 11 | 120816860 | G | A | 0.975 | 0.3298 | +? | 0 | 0 | 0 | 1 |
| rs12421541 | 11 | 119922558 | T | A | -0.974 | 0.3299 | -? | 0 | 0 | 0 | 1 |
| rs61898347 | 11 | 120195689 | A | G | -0.974 | 0.3299 | -? | 0 | 0 | 0 | 1 |
| rs61150131 | 11 | 118752178 | C | T | 0.974 | 0.33 | ++ | 0 | 0.065 | 1 | 0.7985 |
| rs7950159 | 11 | 119625735 | T | C | -0.974 | 0.33 | -? | 0 | 0 | 0 | 1 |
| rs948094 | 11 | 121092679 | T | C | 0.974 | 0.3302 | +? | 0 | 0 | 0 | 1 |
| rs11216825 | 11 | 118099617 | G | A | -0.973 | 0.3304 | -? | 0 | 0 | 0 | 1 |
| rs11216826 | 11 | 118099713 | A | T | -0.973 | 0.3304 | -? | 0 | 0 | 0 | 1 |
| rs3781823 | 11 | 121057430 | A | G | -0.973 | 0.3305 | -? | 0 | 0 | 0 | 1 |
| rs11216338 | 11 | 117131719 | A | G | -0.973 | 0.3306 | -? | 0 | 0 | 0 | 1 |
| rs570371 | 11 | 118568472 | G | A | 0.973 | 0.3306 | ++ | -133.6 | 1.284 | 1 | 0.2571 |
| rs79844114 | 11 | 120334905 | G | A | 0.973 | 0.3306 | +? | 0 | 0 | 0 | 1 |
| rs11217356 | 11 | 119496258 | T | C | -0.973 | 0.3307 | -? | 0 | 0 | 0 | 1 |
| rs471814 | 11 | 119794509 | G | A | -0.973 | 0.3307 | -? | 0 | 0 | 0 | 1 |
| rs7936752 | 11 | 118978132 | G | C | -0.973 | 0.3307 | -? | 0 | 0 | 0 | 1 |
| rs585809 | 11 | 119949979 | T | C | 0.973 | 0.3308 | +? | 0 | 0 | 0 | 1 |
| rs2032397 | 11 | 118153697 | A | G | 0.972 | 0.331 | +? | 0 | 0 | 0 | 1 |
| rs2032398 | 11 | 118153708 | C | G | 0.972 | 0.331 | +? | 0 | 0 | 0 | 1 |
| rs7104819 | 11 | 118769442 | G | A | -0.972 | 0.331 | -- | 0 | 0.336 | 1 | 0.5624 |
| rs11217440 | 11 | 119611348 | G | A | 0.972 | 0.3311 | +? | 0 | 0 | 0 | 1 |
| rs11217441 | 11 | 119611360 | A | G | 0.972 | 0.3311 | +? | 0 | 0 | 0 | 1 |
| rs12801855 | 11 | 117780744 | T | C | -0.972 | 0.3311 | -? | 0 | 0 | 0 | 1 |
| rs10790180 | 11 | 117246150 | C | T | 0.972 | 0.3312 | +? | 0 | 0 | 0 | 1 |
| rs73576552 | 11 | 120623361 | T | C | 0.972 | 0.3312 | +? | 0 | 0 | 0 | 1 |
| rs117179746 | 11 | 116545251 | A | G | -0.971 | 0.3315 | -? | 0 | 0 | 0 | 1 |
| rs1893318 | 11 | 117974541 | G | A | -0.971 | 0.3315 | -? | 0 | 0 | 0 | 1 |
| rs580287 | 11 | 117758658 | G | C | -0.971 | 0.3316 | -? | 0 | 0 | 0 | 1 |
| rs61426083 | 11 | 116439118 | A | G | -0.971 | 0.3316 | -? | 0 | 0 | 0 | 1 |
| rs948463 | 11 | 117969122 | G | A | -0.971 | 0.3316 | -? | 0 | 0 | 0 | 1 |
| rs73007850 | 11 | 120800291 | C | G | 0.971 | 0.3317 | +? | 0 | 0 | 0 | 1 |
| rs2186698 | 11 | 120657934 | C | G | 0.97 | 0.3319 | +? | 0 | 0 | 0 | 1 |
| rs75137105 | 11 | 116481066 | T | C | -0.97 | 0.3319 | -? | 0 | 0 | 0 | 1 |
| rs649870 | 11 | 118971251 | G | A | -0.97 | 0.3321 | -? | 0 | 0 | 0 | 1 |
| rs664011 | 11 | 117108318 | T | A | -0.97 | 0.3321 | -? | 0 | 0 | 0 | 1 |
| rs2105609 | 11 | 116535087 | A | G | -0.97 | 0.3322 | -? | 0 | 0 | 0 | 1 |
| rs7949670 | 11 | 117530935 | A | C | 0.97 | 0.3322 | ++ | 0 | 0.288 | 1 | 0.5917 |
| rs4935747 | 11 | 120518870 | C | G | -0.969 | 0.3323 | -? | 0 | 0 | 0 | 1 |
| rs10892503 | 11 | 119826815 | G | A | -0.969 | 0.3324 | -? | 0 | 0 | 0 | 1 |
| rs11218070 | 11 | 120816711 | C | T | 0.969 | 0.3324 | +? | 0 | 0 | 0 | 1 |
| rs11218072 | 11 | 120817046 | G | A | 0.969 | 0.3324 | +? | 0 | 0 | 0 | 1 |
| rs1268353 | 11 | 116639692 | T | C | -0.969 | 0.3324 | -+ | 31.5 | 4.377 | 1 | 0.03642 |
| rs141414463 | 11 | 116598315 | T | C | 0.969 | 0.3324 | +? | 0 | 0 | 0 | 1 |
| rs10892226 | 11 | 118096618 | G | A | -0.969 | 0.3327 | -? | 0 | 0 | 0 | 1 |
| rs7943727 | 11 | 120839015 | C | T | 0.969 | 0.3327 | +? | 0 | 0 | 0 | 1 |
| rs6589777 | 11 | 119668742 | C | A | 0.968 | 0.333 | +? | 0 | 0 | 0 | 1 |
| rs1263167 | 11 | 116677723 | G | A | -0.967 | 0.3333 | -? | 0 | 0 | 0 | 1 |
| rs3132776 | 11 | 120601807 | A | G | 0.967 | 0.3333 | +? | 0 | 0 | 0 | 1 |
| rs10790358 | 11 | 119836955 | G | A | 0.967 | 0.3334 | +? | 0 | 0 | 0 | 1 |
| rs145111226 | 11 | 117570297 | T | C | 0.967 | 0.3334 | +? | 0 | 0 | 0 | 1 |
| rs147776682 | 11 | 117569803 | T | C | 0.967 | 0.3334 | +? | 0 | 0 | 0 | 1 |

| | | | | | | | | | | | |
|---|---|---|---|---|---|---|---|---|---|---|---|
| rs149084104 | 11 | 117570660 | C | G | 0.967 | 0.3334 | +? | 0 | 0 | 0 | 1 |
| rs34721671 | 11 | 116467920 | G | C | 0.967 | 0.3335 | +? | 0 | 0 | 0 | 1 |
| rs7952619 | 11 | 120914370 | C | T | -0.967 | 0.3336 | -? | 0 | 0 | 0 | 1 |
| rs61900882 | 11 | 118130199 | T | C | -0.967 | 0.3337 | -? | 0 | 0 | 0 | 1 |
| rs615144 | 11 | 118028167 | A | C | -0.966 | 0.3338 | -? | 0 | 0 | 0 | 1 |
| rs7937208 | 11 | 119229374 | T | C | 0.966 | 0.334 | +? | 0 | 0 | 0 | 1 |
| rs76741837 | 11 | 116210400 | T | C | -0.966 | 0.3341 | -? | 0 | 0 | 0 | 1 |
| rs10892002 | 11 | 116505492 | C | G | -0.966 | 0.3342 | -? | 0 | 0 | 0 | 1 |
| rs604096 | 11 | 118475098 | G | A | -0.965 | 0.3343 | -- | -187.4 | 1.044 | 1 | 0.3069 |
| rs12791282 | 11 | 120066044 | G | C | -0.965 | 0.3344 | -? | 0 | 0 | 0 | 1 |
| rs12799241 | 11 | 119980729 | A | G | -0.965 | 0.3344 | -? | 0 | 0 | 0 | 1 |
| rs17119671 | 11 | 116325220 | T | C | -0.965 | 0.3344 | -? | 0 | 0 | 0 | 1 |
| rs552176 | 11 | 117133672 | A | G | -0.965 | 0.3344 | -? | 0 | 0 | 0 | 1 |
| rs145192391 | 11 | 120334126 | T | C | -0.965 | 0.3345 | -? | 0 | 0 | 0 | 1 |
| rs3802881 | 11 | 118890022 | T | C | 0.965 | 0.3346 | +? | 0 | 0 | 0 | 1 |
| rs1597866 | 11 | 119914915 | G | A | -0.965 | 0.3347 | -? | 0 | 0 | 0 | 1 |
| rs2084893 | 11 | 119914182 | G | C | -0.965 | 0.3347 | -? | 0 | 0 | 0 | 1 |
| rs509894 | 11 | 117199870 | T | C | 0.964 | 0.3349 | +? | 0 | 0 | 0 | 1 |
| rs11216337 | 11 | 117131664 | C | T | -0.964 | 0.3351 | -? | 0 | 0 | 0 | 1 |
| rs597345 | 11 | 120988124 | G | A | 0.964 | 0.3352 | +? | 0 | 0 | 0 | 1 |
| rs7931335 | 11 | 117036805 | C | T | 0.964 | 0.3352 | +? | 0 | 0 | 0 | 1 |
| rs115791680 | 11 | 120648746 | T | C | 0.963 | 0.3353 | +? | 0 | 0 | 0 | 1 |
| rs56275759 | 11 | 120728916 | C | T | 0.963 | 0.3354 | +? | 0 | 0 | 0 | 1 |
| rs4486664 | 11 | 118569067 | C | A | 0.963 | 0.3357 | ++ | -156.7 | 1.169 | 1 | 0.2796 |
| rs11545858 | 11 | 118889073 | T | G | 0.962 | 0.3359 | +? | 0 | 0 | 0 | 1 |
| rs618166 | 11 | 119937335 | A | T | 0.962 | 0.336 | +? | 0 | 0 | 0 | 1 |
| rs6589729 | 11 | 119166651 | A | G | -0.962 | 0.336 | -? | 0 | 0 | 0 | 1 |
| rs79540430 | 11 | 120603160 | T | A | 0.962 | 0.336 | +? | 0 | 0 | 0 | 1 |
| rs11218154 | 11 | 121006301 | A | G | 0.962 | 0.3361 | +? | 0 | 0 | 0 | 1 |
| rs523518 | 11 | 116529524 | A | G | 0.962 | 0.3362 | +? | 0 | 0 | 0 | 1 |
| rs7932692 | 11 | 119220012 | G | C | 0.962 | 0.3362 | +? | 0 | 0 | 0 | 1 |
| rs4938762 | 11 | 119927043 | T | C | -0.962 | 0.3363 | -? | 0 | 0 | 0 | 1 |
| rs488931 | 11 | 117344243 | A | G | -0.961 | 0.3366 | -? | 0 | 0 | 0 | 1 |
| rs11826646 | 11 | 117890754 | C | G | 0.961 | 0.3367 | +? | 0 | 0 | 0 | 1 |
| rs77262524 | 11 | 117123218 | G | C | 0.96 | 0.3369 | +? | 0 | 0 | 0 | 1 |
| rs34224649 | 11 | 119503576 | A | G | 0.96 | 0.337 | +? | 0 | 0 | 0 | 1 |
| rs10892618 | 11 | 120568392 | C | A | -0.96 | 0.3371 | -? | 0 | 0 | 0 | 1 |
| rs12363004 | 11 | 117342333 | C | T | -0.96 | 0.3372 | -? | 0 | 0 | 0 | 1 |
| rs1945009 | 11 | 120627141 | A | G | 0.96 | 0.3372 | +? | 0 | 0 | 0 | 1 |
| rs4938416 | 11 | 117467582 | T | C | 0.959 | 0.3374 | +? | 0 | 0 | 0 | 1 |
| rs4938417 | 11 | 117467586 | C | T | 0.959 | 0.3374 | +? | 0 | 0 | 0 | 1 |
| rs56031122 | 11 | 119773520 | G | A | -0.959 | 0.3374 | -? | 0 | 0 | 0 | 1 |
| rs61899037 | 11 | 119814550 | A | G | 0.959 | 0.3374 | +? | 0 | 0 | 0 | 1 |
| rs11217660 | 11 | 119920139 | A | G | -0.959 | 0.3376 | -? | 0 | 0 | 0 | 1 |
| rs75137878 | 11 | 119945024 | A | C | -0.959 | 0.3376 | -? | 0 | 0 | 0 | 1 |
| rs76200908 | 11 | 119774817 | A | G | -0.959 | 0.3377 | -? | 0 | 0 | 0 | 1 |
| rs78755429 | 11 | 118777978 | A | C | 0.959 | 0.3377 | ++ | 0 | 0.001 | 1 | 0.9791 |
| rs650585 | 11 | 117110740 | T | C | 0.959 | 0.3378 | +? | 0 | 0 | 0 | 1 |
| rs73024501 | 11 | 117808400 | G | C | 0.958 | 0.3381 | +? | 0 | 0 | 0 | 1 |
| rs12290911 | 11 | 117773917 | T | A | 0.957 | 0.3386 | +? | 0 | 0 | 0 | 1 |
| rs637065 | 11 | 117133025 | A | G | -0.956 | 0.3391 | -? | 0 | 0 | 0 | 1 |
| rs17119659 | 11 | 116312246 | C | G | -0.956 | 0.3392 | -? | 0 | 0 | 0 | 1 |
| rs7933728 | 11 | 117923031 | A | G | -0.956 | 0.3393 | -? | 0 | 0 | 0 | 1 |
| rs12274032 | 11 | 120940461 | A | G | -0.955 | 0.3394 | -? | 0 | 0 | 0 | 1 |
| rs2071521 | 11 | 116697848 | G | A | -0.955 | 0.3395 | -? | 0 | 0 | 0 | 1 |
| rs73564406 | 11 | 119033850 | A | G | -0.955 | 0.3395 | -? | 0 | 0 | 0 | 1 |

| | | | | | | | | | | | |
|---|---|---|---|---|---|---|---|---|---|---|---|
| rs488222 | 11 | 119987576 | C | T | 0.955 | 0.3396 | +? | 0 | 0 | 0 | 1 |
| rs59711149 | 11 | 120774749 | C | T | -0.955 | 0.3396 | -? | 0 | 0 | 0 | 1 |
| rs10892668 | 11 | 120937637 | C | T | -0.955 | 0.3397 | -? | 0 | 0 | 0 | 1 |
| rs472611 | 11 | 119824102 | G | A | 0.955 | 0.3397 | +? | 0 | 0 | 0 | 1 |
| rs78830794 | 11 | 120972112 | C | T | 0.955 | 0.3397 | +? | 0 | 0 | 0 | 1 |
| rs1263166 | 11 | 116677371 | A | G | -0.954 | 0.3399 | -? | 0 | 0 | 0 | 1 |
| rs138488120 | 11 | 120144571 | A | C | 0.954 | 0.3399 | +? | 0 | 0 | 0 | 1 |
| rs35284897 | 11 | 117775895 | A | C | 0.954 | 0.3399 | +? | 0 | 0 | 0 | 1 |
| rs584231 | 11 | 117927676 | A | G | -0.954 | 0.34 | -? | 0 | 0 | 0 | 1 |
| rs7928123 | 11 | 120498337 | T | C | -0.954 | 0.3401 | -? | 0 | 0 | 0 | 1 |
| rs2509211 | 11 | 117137633 | C | T | -0.954 | 0.3402 | -? | 0 | 0 | 0 | 1 |
| rs477269 | 11 | 117138291 | G | T | -0.954 | 0.3402 | -? | 0 | 0 | 0 | 1 |
| rs507880 | 11 | 117137255 | G | T | -0.954 | 0.3402 | -? | 0 | 0 | 0 | 1 |
| rs524085 | 11 | 117137647 | G | A | -0.954 | 0.3402 | -? | 0 | 0 | 0 | 1 |
| rs540103 | 11 | 119782250 | G | A | 0.954 | 0.3403 | +? | 0 | 0 | 0 | 1 |
| rs12420795 | 11 | 118161885 | G | C | 0.953 | 0.3404 | +? | 0 | 0 | 0 | 1 |
| rs4938498 | 11 | 118162147 | G | C | 0.953 | 0.3404 | +? | 0 | 0 | 0 | 1 |
| rs4936353 | 11 | 116397649 | A | G | 0.953 | 0.3405 | +? | 0 | 0 | 0 | 1 |
| rs6589550 | 11 | 116367567 | C | A | 0.953 | 0.3407 | +? | 0 | 0 | 0 | 1 |
| rs2727789 | 11 | 116696887 | G | T | -0.952 | 0.3409 | -? | 0 | 0 | 0 | 1 |
| rs644324 | 11 | 119946171 | G | A | 0.952 | 0.341 | +? | 0 | 0 | 0 | 1 |
| rs7927785 | 11 | 116452142 | G | C | 0.952 | 0.341 | +? | 0 | 0 | 0 | 1 |
| rs4144287 | 11 | 121126285 | G | A | -0.952 | 0.3412 | -- | 0 | 0.252 | 1 | 0.616 |
| rs10892083 | 11 | 117042580 | C | G | 0.952 | 0.3413 | +? | 0 | 0 | 0 | 1 |
| rs4148173 | 11 | 119030812 | G | A | -0.952 | 0.3413 | -? | 0 | 0 | 0 | 1 |
| rs499713 | 11 | 117061961 | T | C | 0.952 | 0.3413 | +? | 0 | 0 | 0 | 1 |
| rs7118694 | 11 | 119676300 | A | G | 0.952 | 0.3413 | +? | 0 | 0 | 0 | 1 |
| rs75581341 | 11 | 117345415 | T | C | -0.952 | 0.3413 | -? | 0 | 0 | 0 | 1 |
| rs7935656 | 11 | 116364065 | C | T | 0.951 | 0.3415 | +? | 0 | 0 | 0 | 1 |
| rs79723893 | 11 | 117818515 | G | A | -0.951 | 0.3416 | -? | 0 | 0 | 0 | 1 |
| rs7939968 | 11 | 120531435 | A | T | -0.951 | 0.3418 | -? | 0 | 0 | 0 | 1 |
| rs180372 | 11 | 116591934 | A | G | 0.951 | 0.3419 | +- | -42.8 | 2.101 | 1 | 0.1472 |
| rs7928058 | 11 | 118171239 | T | G | 0.95 | 0.3419 | +? | 0 | 0 | 0 | 1 |
| rs535602 | 11 | 117107775 | A | G | -0.95 | 0.3422 | -? | 0 | 0 | 0 | 1 |
| rs7931384 | 11 | 116453047 | C | T | 0.949 | 0.3426 | +? | 0 | 0 | 0 | 1 |
| rs12280043 | 11 | 116450185 | T | A | 0.949 | 0.3428 | +? | 0 | 0 | 0 | 1 |
| rs584282 | 11 | 119945841 | G | T | 0.949 | 0.3428 | +? | 0 | 0 | 0 | 1 |
| rs11216730 | 11 | 117918702 | T | C | 0.948 | 0.3429 | ++ | 0 | 0 | 1 | 0.9888 |
| rs1145185 | 11 | 116514699 | A | G | 0.948 | 0.3429 | +? | 0 | 0 | 0 | 1 |
| rs56296109 | 11 | 118698170 | A | C | -0.948 | 0.3429 | -- | 0 | 0.352 | 1 | 0.5528 |
| rs58618446 | 11 | 117916692 | T | C | 0.948 | 0.3429 | ++ | 0 | 0 | 1 | 0.9888 |
| rs11217793 | 11 | 120161886 | T | C | -0.948 | 0.343 | -? | 0 | 0 | 0 | 1 |
| rs552879 | 11 | 117348907 | T | C | -0.948 | 0.343 | -? | 0 | 0 | 0 | 1 |
| rs11217124 | 11 | 118889132 | G | A | -0.948 | 0.3432 | -- | 0 | 0.225 | 1 | 0.635 |
| rs869638 | 11 | 118100411 | G | C | -0.948 | 0.3433 | -? | 0 | 0 | 0 | 1 |
| rs9651685 | 11 | 117967798 | C | T | -0.948 | 0.3433 | -? | 0 | 0 | 0 | 1 |
| rs55797357 | 11 | 119034374 | T | C | -0.947 | 0.3434 | -? | 0 | 0 | 0 | 1 |
| rs73001451 | 11 | 118614115 | C | T | 0.947 | 0.3437 | +- | -195.7 | 1.014 | 1 | 0.3139 |
| rs11217125 | 11 | 118889185 | C | T | -0.947 | 0.3438 | -- | 0 | 0.21 | 1 | 0.6469 |
| rs7949316 | 11 | 120495204 | T | C | 0.947 | 0.3438 | +? | 0 | 0 | 0 | 1 |
| rs1784124 | 11 | 116537618 | A | G | -0.946 | 0.3439 | -? | 0 | 0 | 0 | 1 |
| rs56703842 | 11 | 117364180 | T | C | -0.946 | 0.3439 | -? | 0 | 0 | 0 | 1 |
| rs7127949 | 11 | 119989590 | C | A | -0.946 | 0.3439 | -? | 0 | 0 | 0 | 1 |
| rs2852214 | 11 | 120613416 | A | G | 0.946 | 0.344 | +? | 0 | 0 | 0 | 1 |
| rs73008667 | 11 | 119727693 | T | C | -0.946 | 0.344 | -? | 0 | 0 | 0 | 1 |
| rs1787682 | 11 | 116539203 | G | C | -0.946 | 0.3441 | -? | 0 | 0 | 0 | 1 |

| | | | | | | | | | | | |
|---|---|---|---|---|---|---|---|---|---|---|---|
| rs4938369 | 11 | 117188120 | T | C | 0.946 | 0.3442 | +? | 0 | 0 | 0 | 1 |
| rs73557344 | 11 | 118810054 | C | G | 0.946 | 0.3442 | +- | -114.1 | 1.401 | 1 | 0.2365 |
| rs75460330 | 11 | 118809767 | A | G | 0.946 | 0.3442 | +- | -114.1 | 1.401 | 1 | 0.2365 |
| rs1263169 | 11 | 116678139 | A | T | -0.946 | 0.3444 | -? | 0 | 0 | 0 | 1 |
| rs56153022 | 11 | 117587224 | G | C | 0.945 | 0.3445 | +? | 0 | 0 | 0 | 1 |
| rs590648 | 11 | 119753355 | T | G | -0.945 | 0.3447 | -? | 0 | 0 | 0 | 1 |
| rs56784564 | 11 | 118857809 | A | G | 0.945 | 0.3449 | +? | 0 | 0 | 0 | 1 |
| rs3016242 | 11 | 120979316 | A | C | 0.944 | 0.3452 | +? | 0 | 0 | 0 | 1 |
| rs595782 | 11 | 118985151 | C | T | -0.944 | 0.3452 | -? | 0 | 0 | 0 | 1 |
| rs12807240 | 11 | 120064556 | G | A | -0.944 | 0.3453 | -? | 0 | 0 | 0 | 1 |
| rs2542051 | 11 | 116697738 | C | A | -0.944 | 0.3453 | -? | 0 | 0 | 0 | 1 |
| rs4936387 | 11 | 117435160 | T | C | -0.943 | 0.3455 | -? | 0 | 0 | 0 | 1 |
| rs60501859 | 11 | 117806383 | C | T | 0.943 | 0.3455 | +? | 0 | 0 | 0 | 1 |
| rs117341026 | 11 | 116555126 | A | G | -0.943 | 0.3456 | -? | 0 | 0 | 0 | 1 |
| rs10750165 | 11 | 119638088 | G | A | -0.943 | 0.3457 | -? | 0 | 0 | 0 | 1 |
| rs143605660 | 11 | 118707035 | A | G | 0.943 | 0.3457 | ++ | 0 | 0.475 | 1 | 0.4909 |
| rs12807541 | 11 | 120064501 | A | G | -0.943 | 0.3458 | -? | 0 | 0 | 0 | 1 |
| rs61898760 | 11 | 120330904 | G | A | -0.943 | 0.3458 | -? | 0 | 0 | 0 | 1 |
| rs4938615 | 11 | 118865319 | G | C | -0.942 | 0.346 | -? | 0 | 0 | 0 | 1 |
| rs1784131 | 11 | 116539868 | C | T | -0.942 | 0.3461 | -? | 0 | 0 | 0 | 1 |
| rs78048464 | 11 | 119761810 | T | G | -0.942 | 0.3461 | -? | 0 | 0 | 0 | 1 |
| rs11217245 | 11 | 119219311 | G | A | 0.942 | 0.3462 | +? | 0 | 0 | 0 | 1 |
| rs66879136 | 11 | 116392920 | A | C | -0.942 | 0.3464 | -? | 0 | 0 | 0 | 1 |
| rs78952625 | 11 | 120155549 | T | C | -0.942 | 0.3464 | -- | 0 | 0.151 | 1 | 0.6976 |
| rs180375 | 11 | 116591040 | T | C | 0.941 | 0.3466 | +- | -43.9 | 2.085 | 1 | 0.1487 |
| rs2136420 | 11 | 119600316 | G | A | 0.941 | 0.3468 | +? | 0 | 0 | 0 | 1 |
| rs35248712 | 11 | 119609714 | G | T | 0.94 | 0.347 | +? | 0 | 0 | 0 | 1 |
| rs4938720 | 11 | 119609846 | C | T | 0.94 | 0.347 | +? | 0 | 0 | 0 | 1 |
| rs78957222 | 11 | 119826890 | T | C | -0.94 | 0.347 | -? | 0 | 0 | 0 | 1 |
| rs6589869 | 11 | 121142453 | G | C | -0.94 | 0.3471 | -- | 0 | 0.251 | 1 | 0.6165 |
| rs10790402 | 11 | 120560994 | A | G | -0.94 | 0.3472 | -? | 0 | 0 | 0 | 1 |
| rs2071522 | 11 | 116697538 | T | C | -0.94 | 0.3472 | -? | 0 | 0 | 0 | 1 |
| rs7929799 | 11 | 119055362 | G | A | 0.94 | 0.3472 | +? | 0 | 0 | 0 | 1 |
| rs11820504 | 11 | 116529442 | C | T | 0.94 | 0.3474 | +? | 0 | 0 | 0 | 1 |
| rs61898345 | 11 | 120188964 | C | A | -0.94 | 0.3474 | -? | 0 | 0 | 0 | 1 |
| rs883247 | 11 | 119217254 | C | T | 0.94 | 0.3474 | +? | 0 | 0 | 0 | 1 |
| rs2014552 | 11 | 119504482 | A | G | 0.939 | 0.3475 | +? | 0 | 0 | 0 | 1 |
| rs587726 | 11 | 119240825 | T | C | 0.939 | 0.3476 | +? | 0 | 0 | 0 | 1 |
| rs7129284 | 11 | 117354904 | A | G | 0.939 | 0.3476 | +? | 0 | 0 | 0 | 1 |
| rs12421434 | 11 | 119603867 | C | T | 0.939 | 0.3477 | +? | 0 | 0 | 0 | 1 |
| rs12421466 | 11 | 119603983 | G | A | 0.939 | 0.3477 | +? | 0 | 0 | 0 | 1 |
| rs1837509 | 11 | 116225440 | A | G | -0.939 | 0.3477 | -- | 0 | 0.101 | 1 | 0.7501 |
| rs4938719 | 11 | 119603559 | G | A | 0.939 | 0.3477 | +? | 0 | 0 | 0 | 1 |
| rs2509210 | 11 | 117136070 | T | C | -0.939 | 0.3478 | -? | 0 | 0 | 0 | 1 |
| rs4936390 | 11 | 117498314 | A | G | 0.939 | 0.3478 | +? | 0 | 0 | 0 | 1 |
| rs4936392 | 11 | 117498656 | T | G | 0.939 | 0.3478 | +? | 0 | 0 | 0 | 1 |
| rs672177 | 11 | 117136378 | G | C | -0.939 | 0.3478 | -? | 0 | 0 | 0 | 1 |
| rs7111678 | 11 | 117135806 | T | C | -0.939 | 0.3478 | -? | 0 | 0 | 0 | 1 |
| rs78696556 | 11 | 117499830 | C | T | 0.939 | 0.3478 | +? | 0 | 0 | 0 | 1 |
| rs10750177 | 11 | 120727836 | T | A | -0.939 | 0.3479 | -? | 0 | 0 | 0 | 1 |
| rs7952478 | 11 | 120495486 | T | C | 0.939 | 0.3479 | +? | 0 | 0 | 0 | 1 |
| rs111447379 | 11 | 117116312 | T | C | 0.938 | 0.348 | +? | 0 | 0 | 0 | 1 |
| rs4938418 | 11 | 117472019 | C | T | 0.938 | 0.3481 | +? | 0 | 0 | 0 | 1 |
| rs581063 | 11 | 118767077 | C | T | -0.938 | 0.3481 | -- | 0 | 0.221 | 1 | 0.6384 |
| rs78136547 | 11 | 117470412 | G | C | 0.938 | 0.3481 | +? | 0 | 0 | 0 | 1 |
| rs78853831 | 11 | 117411223 | T | C | 0.938 | 0.3481 | +? | 0 | 0 | 0 | 1 |

| | | | | | | | | | | | |
|---|---|---|---|---|---|---|---|---|---|---|---|
| rs12787658 | 11 | 120179563 | G | A | -0.938 | 0.3483 | -? | 0 | 0 | 0 | 1 |
| rs11217818 | 11 | 120196897 | C | T | 0.937 | 0.3486 | ++ | -151.5 | 1.193 | 1 | 0.2748 |
| rs4245206 | 11 | 119609264 | A | T | 0.937 | 0.3486 | +? | 0 | 0 | 0 | 1 |
| rs8258 | 11 | 117283676 | T | C | 0.937 | 0.3486 | +? | 0 | 0 | 0 | 1 |
| rs10892446 | 11 | 119611734 | A | G | 0.937 | 0.3487 | +? | 0 | 0 | 0 | 1 |
| rs11216555 | 11 | 117684613 | A | G | 0.937 | 0.3488 | +? | 0 | 0 | 0 | 1 |
| rs2186627 | 11 | 117774146 | G | C | 0.937 | 0.349 | +? | 0 | 0 | 0 | 1 |
| rs45573242 | 11 | 118853995 | C | A | 0.937 | 0.349 | +? | 0 | 0 | 0 | 1 |
| rs7128000 | 11 | 119989380 | A | G | -0.937 | 0.349 | -? | 0 | 0 | 0 | 1 |
| rs7128133 | 11 | 119989497 | A | G | -0.937 | 0.349 | -? | 0 | 0 | 0 | 1 |
| rs560355 | 11 | 117256384 | C | G | 0.936 | 0.3492 | +? | 0 | 0 | 0 | 1 |
| rs601122 | 11 | 117757345 | T | G | -0.936 | 0.3493 | -? | 0 | 0 | 0 | 1 |
| rs675539 | 11 | 117929001 | C | G | -0.936 | 0.3493 | -? | 0 | 0 | 0 | 1 |
| rs472785 | 11 | 117134766 | C | T | -0.935 | 0.3496 | -? | 0 | 0 | 0 | 1 |
| rs540922 | 11 | 117134761 | G | A | -0.935 | 0.3496 | -? | 0 | 0 | 0 | 1 |
| rs60657140 | 11 | 119505336 | T | G | 0.935 | 0.3499 | +? | 0 | 0 | 0 | 1 |
| rs664845 | 11 | 119923652 | A | G | -0.935 | 0.3499 | -? | 0 | 0 | 0 | 1 |
| rs3889525 | 11 | 118856537 | C | G | 0.934 | 0.3501 | +? | 0 | 0 | 0 | 1 |
| rs12362636 | 11 | 117341466 | C | T | -0.934 | 0.3502 | -? | 0 | 0 | 0 | 1 |
| rs77398170 | 11 | 118792278 | G | A | -0.934 | 0.3502 | -- | 0 | 0.013 | 1 | 0.9089 |
| rs7951668 | 11 | 119383037 | T | C | -0.934 | 0.3502 | -? | 0 | 0 | 0 | 1 |
| rs11217246 | 11 | 119219360 | C | A | 0.934 | 0.3503 | +? | 0 | 0 | 0 | 1 |
| rs12291317 | 11 | 119794853 | C | T | -0.934 | 0.3505 | -? | 0 | 0 | 0 | 1 |
| rs34599893 | 11 | 120505562 | C | T | 0.933 | 0.3506 | +? | 0 | 0 | 0 | 1 |
| rs11217434 | 11 | 119602067 | C | T | 0.933 | 0.3508 | +? | 0 | 0 | 0 | 1 |
| rs1315236635 | 11 | 117357085 | G | T | -0.933 | 0.3508 | -? | 0 | 0 | 0 | 1 |
| rs2368936 | 11 | 119601779 | C | T | 0.933 | 0.3508 | +? | 0 | 0 | 0 | 1 |
| rs4938718 | 11 | 119601497 | G | C | 0.933 | 0.3508 | +? | 0 | 0 | 0 | 1 |
| rs72645469 | 11 | 119828651 | A | C | -0.933 | 0.3508 | -? | 0 | 0 | 0 | 1 |
| rs1939219 | 11 | 117686660 | C | T | 0.933 | 0.3509 | ++ | 0 | 0.112 | 1 | 0.7379 |
| rs4415772 | 11 | 121151102 | G | T | -0.933 | 0.351 | -- | 0 | 0.642 | 1 | 0.4228 |
| rs77084900 | 11 | 117906061 | C | T | 0.933 | 0.351 | ++ | 0 | 0.001 | 1 | 0.9816 |
| rs2849176 | 11 | 116696921 | T | C | -0.932 | 0.3511 | -? | 0 | 0 | 0 | 1 |
| rs61904854 | 11 | 116467100 | T | G | 0.932 | 0.3511 | +? | 0 | 0 | 0 | 1 |
| rs79679004 | 11 | 116265581 | C | T | -0.932 | 0.3511 | -? | 0 | 0 | 0 | 1 |
| rs12417736 | 11 | 119220946 | T | C | 0.932 | 0.3512 | +? | 0 | 0 | 0 | 1 |
| rs4935749 | 11 | 120518929 | A | G | -0.932 | 0.3514 | -? | 0 | 0 | 0 | 1 |
| rs657685 | 11 | 118912982 | C | G | 0.932 | 0.3514 | +? | 0 | 0 | 0 | 1 |
| rs588534 | 11 | 117068504 | A | G | 0.931 | 0.3518 | +? | 0 | 0 | 0 | 1 |
| rs10892663 | 11 | 120911203 | C | T | -0.931 | 0.352 | -? | 0 | 0 | 0 | 1 |
| rs11216726 | 11 | 117912267 | T | C | 0.931 | 0.352 | ++ | 0 | 0.001 | 1 | 0.9808 |
| rs20337 | 11 | 119502870 | A | G | 0.931 | 0.352 | +? | 0 | 0 | 0 | 1 |
| rs7480948 | 11 | 119220483 | G | C | 0.931 | 0.352 | +? | 0 | 0 | 0 | 1 |
| rs666640 | 11 | 119923259 | T | A | -0.931 | 0.3521 | -? | 0 | 0 | 0 | 1 |
| rs116958147 | 11 | 121159481 | C | T | -0.93 | 0.3522 | -? | 0 | 0 | 0 | 1 |
| rs1783846 | 11 | 117928962 | A | C | -0.93 | 0.3522 | -? | 0 | 0 | 0 | 1 |
| rs144891162 | 11 | 117821885 | T | C | -0.93 | 0.3524 | -? | 0 | 0 | 0 | 1 |
| rs7102851 | 11 | 116269470 | C | T | -0.93 | 0.3524 | -? | 0 | 0 | 0 | 1 |
| rs11216518 | 11 | 117595599 | T | G | -0.93 | 0.3525 | -? | 0 | 0 | 0 | 1 |
| rs1007530 | 11 | 119948049 | A | G | -0.929 | 0.3527 | -? | 0 | 0 | 0 | 1 |
| rs948095 | 11 | 121092828 | A | G | 0.929 | 0.3527 | +? | 0 | 0 | 0 | 1 |
| rs12421941 | 11 | 120194357 | G | A | -0.929 | 0.3528 | -? | 0 | 0 | 0 | 1 |
| rs4472950 | 11 | 120807892 | T | C | -0.929 | 0.3529 | -? | 0 | 0 | 0 | 1 |
| rs7083 | 11 | 117156552 | T | C | -0.929 | 0.3529 | -? | 0 | 0 | 0 | 1 |
| rs73024430 | 11 | 117804535 | T | C | 0.929 | 0.353 | +? | 0 | 0 | 0 | 1 |
| rs10790290 | 11 | 119218542 | T | C | 0.929 | 0.3531 | +? | 0 | 0 | 0 | 1 |

| | | | | | | | | | | | |
|---|---|---|---|---|---|---|---|---|---|---|---|
| rs10892352 | 11 | 119218553 | T | C | 0.929 | 0.3531 | +? | 0 | 0 | 0 | 1 |
| rs12575199 | 11 | 120192509 | A | G | -0.929 | 0.3531 | -? | 0 | 0 | 0 | 1 |
| rs7938191 | 11 | 117806283 | A | G | 0.928 | 0.3532 | +? | 0 | 0 | 0 | 1 |
| rs2509050 | 11 | 118977911 | G | A | -0.928 | 0.3533 | -? | 0 | 0 | 0 | 1 |
| rs4393358 | 11 | 118569097 | T | C | 0.928 | 0.3533 | ++ | -152 | 1.19 | 1 | 0.2753 |
| rs112998705 | 11 | 118890397 | G | T | 0.928 | 0.3534 | +? | 0 | 0 | 0 | 1 |
| rs2542050 | 11 | 116697117 | C | A | -0.928 | 0.3534 | -? | 0 | 0 | 0 | 1 |
| rs73010757 | 11 | 117950199 | A | G | -0.928 | 0.3534 | -- | 0 | 0.064 | 1 | 0.801 |
| rs11217437 | 11 | 119606985 | C | G | 0.928 | 0.3536 | +? | 0 | 0 | 0 | 1 |
| rs11217438 | 11 | 119607019 | C | T | 0.928 | 0.3536 | +? | 0 | 0 | 0 | 1 |
| rs7358397 | 11 | 119221078 | A | G | 0.928 | 0.3536 | +? | 0 | 0 | 0 | 1 |
| rs10466596 | 11 | 119989123 | T | G | -0.927 | 0.3537 | -? | 0 | 0 | 0 | 1 |
| rs3759001 | 11 | 118135745 | T | C | -0.927 | 0.3537 | -? | 0 | 0 | 0 | 1 |
| rs1240777 | 11 | 116536476 | C | T | -0.927 | 0.3538 | -? | 0 | 0 | 0 | 1 |
| rs35338327 | 11 | 120691871 | A | G | 0.927 | 0.3539 | +? | 0 | 0 | 0 | 1 |
| rs1876795 | 11 | 119596147 | C | G | 0.927 | 0.354 | +? | 0 | 0 | 0 | 1 |
| rs17092740 | 11 | 118117614 | G | A | -0.926 | 0.3543 | -? | 0 | 0 | 0 | 1 |
| rs12577053 | 11 | 118121421 | G | A | -0.926 | 0.3544 | -? | 0 | 0 | 0 | 1 |
| rs12577061 | 11 | 118121553 | G | C | -0.926 | 0.3544 | -? | 0 | 0 | 0 | 1 |
| rs7928348 | 11 | 119218712 | G | C | 0.926 | 0.3546 | +? | 0 | 0 | 0 | 1 |
| rs1954787 | 11 | 120663363 | T | C | 0.926 | 0.3547 | +? | 0 | 0 | 0 | 1 |
| rs474488 | 11 | 117068977 | A | G | 0.926 | 0.3547 | +? | 0 | 0 | 0 | 1 |
| rs500713 | 11 | 117320976 | C | T | -0.926 | 0.3547 | -? | 0 | 0 | 0 | 1 |
| rs11217450 | 11 | 119617639 | G | A | 0.925 | 0.3549 | ++ | 0 | 0.125 | 1 | 0.7236 |
| rs12226639 | 11 | 120963327 | A | T | -0.925 | 0.355 | -? | 0 | 0 | 0 | 1 |
| rs111804650 | 11 | 118878960 | T | C | 0.925 | 0.3552 | +? | 0 | 0 | 0 | 1 |
| rs2136421 | 11 | 119600410 | C | T | 0.925 | 0.3552 | +? | 0 | 0 | 0 | 1 |
| rs11216464 | 11 | 117456464 | T | C | 0.924 | 0.3553 | +? | 0 | 0 | 0 | 1 |
| rs12290653 | 11 | 119632975 | A | G | -0.924 | 0.3553 | -? | 0 | 0 | 0 | 1 |
| rs10790176 | 11 | 117047852 | G | T | 0.924 | 0.3554 | +? | 0 | 0 | 0 | 1 |
| rs147189138 | 11 | 121149360 | A | G | -0.924 | 0.3554 | -? | 0 | 0 | 0 | 1 |
| rs4938353 | 11 | 117046197 | G | A | 0.924 | 0.3554 | +? | 0 | 0 | 0 | 1 |
| rs473210 | 11 | 117177800 | T | C | 0.924 | 0.3555 | +? | 0 | 0 | 0 | 1 |
| rs512188 | 11 | 117126710 | A | G | -0.924 | 0.3555 | -? | 0 | 0 | 0 | 1 |
| rs10892444 | 11 | 119604854 | A | G | 0.924 | 0.3556 | +? | 0 | 0 | 0 | 1 |
| rs11217663 | 11 | 119927491 | C | T | -0.924 | 0.3556 | -? | 0 | 0 | 0 | 1 |
| rs113373988 | 11 | 119620677 | T | G | 0.924 | 0.3556 | ++ | 0 | 0.083 | 1 | 0.7728 |
| rs6589827 | 11 | 120517844 | C | T | -0.924 | 0.3556 | -? | 0 | 0 | 0 | 1 |
| rs12574203 | 11 | 119901007 | G | A | -0.923 | 0.3558 | -? | 0 | 0 | 0 | 1 |
| rs2155380 | 11 | 119080037 | G | A | 0.923 | 0.3559 | +? | 0 | 0 | 0 | 1 |
| rs630775 | 11 | 119885547 | C | A | 0.923 | 0.3559 | +? | 0 | 0 | 0 | 1 |
| rs75646928 | 11 | 120161546 | T | C | -0.923 | 0.3559 | -- | 0 | 0.141 | 1 | 0.7074 |
| rs10736484 | 11 | 116210133 | T | C | -0.923 | 0.356 | -? | 0 | 0 | 0 | 1 |
| rs1320708 | 11 | 119881259 | C | T | 0.923 | 0.356 | +? | 0 | 0 | 0 | 1 |
| rs34805238 | 11 | 121149050 | T | C | -0.923 | 0.356 | -- | 0 | 0.48 | 1 | 0.4885 |
| rs75669750 | 11 | 120130716 | C | T | -0.923 | 0.356 | -? | 0 | 0 | 0 | 1 |
| rs12269901 | 11 | 116973929 | C | G | -0.923 | 0.3561 | -? | 0 | 0 | 0 | 1 |
| rs669316 | 11 | 117931576 | T | C | -0.923 | 0.3561 | -? | 0 | 0 | 0 | 1 |
| rs670225 | 11 | 117931353 | A | G | -0.923 | 0.3561 | -? | 0 | 0 | 0 | 1 |
| rs505390 | 11 | 116316813 | A | G | -0.923 | 0.3562 | -? | 0 | 0 | 0 | 1 |
| rs75873912 | 11 | 117465314 | G | C | 0.923 | 0.3562 | +? | 0 | 0 | 0 | 1 |
| rs79944284 | 11 | 117924155 | A | G | 0.923 | 0.3562 | ++ | 0 | 0 | 1 | 0.9869 |
| rs4514426 | 11 | 120526184 | C | T | -0.922 | 0.3563 | -? | 0 | 0 | 0 | 1 |
| rs77346448 | 11 | 117473537 | C | T | 0.922 | 0.3563 | +? | 0 | 0 | 0 | 1 |
| rs2849173 | 11 | 116697321 | G | C | -0.922 | 0.3567 | -? | 0 | 0 | 0 | 1 |
| rs117041834 | 11 | 117498920 | A | T | 0.921 | 0.3568 | +? | 0 | 0 | 0 | 1 |

| | | | | | | | | | | | |
|---|---|---|---|---|---|---|---|---|---|---|---|
| rs117870205 | 11 | 117496666 | A | G | 0.921 | 0.3568 | +? | 0 | 0 | 0 | 1 |
| rs147793723 | 11 | 117496750 | C | T | 0.921 | 0.3568 | +? | 0 | 0 | 0 | 1 |
| rs584115 | 11 | 118938880 | G | A | 0.921 | 0.3568 | +? | 0 | 0 | 0 | 1 |
| rs74681797 | 11 | 117497996 | T | C | 0.921 | 0.3568 | +? | 0 | 0 | 0 | 1 |
| rs76205818 | 11 | 117499828 | A | T | 0.921 | 0.3568 | +? | 0 | 0 | 0 | 1 |
| rs7940041 | 11 | 120690039 | A | G | 0.921 | 0.3568 | +? | 0 | 0 | 0 | 1 |
| rs78221919 | 11 | 120190141 | C | T | -0.921 | 0.3569 | -? | 0 | 0 | 0 | 1 |
| rs7927959 | 11 | 119218616 | A | T | 0.921 | 0.3569 | +? | 0 | 0 | 0 | 1 |
| rs528837 | 11 | 117179176 | A | G | 0.921 | 0.357 | +? | 0 | 0 | 0 | 1 |
| rs7105874 | 11 | 120820615 | T | C | 0.921 | 0.357 | +? | 0 | 0 | 0 | 1 |
| rs35154750 | 11 | 116462431 | C | A | 0.921 | 0.3571 | +? | 0 | 0 | 0 | 1 |
| rs7130104 | 11 | 119823316 | C | T | -0.921 | 0.3571 | -? | 0 | 0 | 0 | 1 |
| rs503023 | 11 | 116270219 | A | G | -0.921 | 0.3572 | -? | 0 | 0 | 0 | 1 |
| rs567209 | 11 | 116534828 | T | C | -0.92 | 0.3574 | -? | 0 | 0 | 0 | 1 |
| rs11217122 | 11 | 118880740 | C | T | -0.92 | 0.3576 | -- | 0 | 0.011 | 1 | 0.9181 |
| rs76190886 | 11 | 117921563 | T | C | 0.92 | 0.3578 | ++ | 0 | 0.004 | 1 | 0.9517 |
| rs117028204 | 11 | 120154445 | G | C | -0.919 | 0.3579 | -- | 0 | 0.141 | 1 | 0.707 |
| rs77223730 | 11 | 120697844 | A | G | 0.919 | 0.3579 | +? | 0 | 0 | 0 | 1 |
| rs78653140 | 11 | 120697845 | T | C | 0.919 | 0.3579 | +? | 0 | 0 | 0 | 1 |
| rs11216718 | 11 | 117908922 | A | G | 0.919 | 0.358 | ++ | 0 | 0.001 | 1 | 0.9755 |
| rs11216719 | 11 | 117908985 | C | A | 0.919 | 0.358 | ++ | 0 | 0.001 | 1 | 0.9755 |
| rs11216720 | 11 | 117909101 | T | C | 0.919 | 0.358 | ++ | 0 | 0.001 | 1 | 0.9755 |
| rs11216721 | 11 | 117909531 | T | C | 0.919 | 0.358 | ++ | 0 | 0.001 | 1 | 0.9755 |
| rs11216722 | 11 | 117910244 | A | G | 0.919 | 0.358 | ++ | 0 | 0.001 | 1 | 0.9755 |
| rs11216725 | 11 | 117911450 | T | C | 0.919 | 0.358 | ++ | 0 | 0.001 | 1 | 0.9755 |
| rs12223165 | 11 | 117910017 | A | G | 0.919 | 0.358 | ++ | 0 | 0.001 | 1 | 0.9755 |
| rs12223183 | 11 | 117910116 | A | G | 0.919 | 0.358 | ++ | 0 | 0.001 | 1 | 0.9755 |
| rs12223876 | 11 | 117907713 | A | T | 0.919 | 0.358 | ++ | 0 | 0.001 | 1 | 0.9755 |
| rs513841 | 11 | 119241249 | T | A | 0.919 | 0.358 | +? | 0 | 0 | 0 | 1 |
| rs2925771 | 11 | 117118481 | T | C | 0.919 | 0.3582 | +? | 0 | 0 | 0 | 1 |
| rs11216616 | 11 | 117781192 | A | G | -0.919 | 0.3583 | -? | 0 | 0 | 0 | 1 |
| rs17119667 | 11 | 116316617 | C | T | -0.918 | 0.3586 | -? | 0 | 0 | 0 | 1 |
| rs1355191 | 11 | 119496472 | C | T | -0.918 | 0.3587 | -? | 0 | 0 | 0 | 1 |
| rs505045 | 11 | 117152540 | G | T | -0.918 | 0.3587 | -? | 0 | 0 | 0 | 1 |
| rs10892531 | 11 | 119950934 | C | T | 0.918 | 0.3588 | +? | 0 | 0 | 0 | 1 |
| rs4938452 | 11 | 117760253 | A | T | -0.918 | 0.3588 | -? | 0 | 0 | 0 | 1 |
| rs12721080 | 11 | 116699681 | A | G | -0.917 | 0.3589 | -? | 0 | 0 | 0 | 1 |
| rs12796806 | 11 | 119612572 | G | A | 0.917 | 0.3592 | +? | 0 | 0 | 0 | 1 |
| rs4310630 | 11 | 118836638 | G | A | -0.917 | 0.3594 | -? | 0 | 0 | 0 | 1 |
| rs10892423 | 11 | 119506483 | A | G | 0.916 | 0.3595 | +? | 0 | 0 | 0 | 1 |
| rs34952981 | 11 | 119620339 | T | A | -0.916 | 0.3595 | -? | 0 | 0 | 0 | 1 |
| rs4938794 | 11 | 120160262 | A | G | -0.916 | 0.3595 | -- | 0 | 0.138 | 1 | 0.7104 |
| rs77483431 | 11 | 120158933 | G | A | -0.916 | 0.3595 | -- | 0 | 0.138 | 1 | 0.7104 |
| rs11607835 | 11 | 118835258 | C | T | -0.916 | 0.3596 | -? | 0 | 0 | 0 | 1 |
| rs4525243 | 11 | 120527341 | G | A | -0.916 | 0.3596 | -? | 0 | 0 | 0 | 1 |
| rs540261 | 11 | 118947634 | T | C | 0.916 | 0.3596 | +? | 0 | 0 | 0 | 1 |
| rs2850814 | 11 | 120772254 | A | G | -0.916 | 0.3599 | -? | 0 | 0 | 0 | 1 |
| rs11218238 | 11 | 121149254 | G | A | -0.915 | 0.36 | -- | 0 | 0.541 | 1 | 0.462 |
| rs655023 | 11 | 117127620 | C | T | -0.915 | 0.36 | -? | 0 | 0 | 0 | 1 |
| rs673357 | 11 | 119940156 | G | A | 0.915 | 0.3601 | +? | 0 | 0 | 0 | 1 |
| rs3814760 | 11 | 119217556 | G | A | 0.915 | 0.3602 | +? | 0 | 0 | 0 | 1 |
| rs73024441 | 11 | 117808145 | G | A | 0.914 | 0.3605 | +? | 0 | 0 | 0 | 1 |
| rs751369 | 11 | 120610564 | G | A | 0.914 | 0.3606 | +? | 0 | 0 | 0 | 1 |
| rs1783778 | 11 | 120978982 | A | C | 0.914 | 0.3608 | +? | 0 | 0 | 0 | 1 |
| rs113137102 | 11 | 117953132 | A | C | -0.914 | 0.3609 | -? | 0 | 0 | 0 | 1 |
| rs1320679 | 11 | 119219124 | G | T | 0.914 | 0.3609 | +? | 0 | 0 | 0 | 1 |

| | | | | | | | | | | | |
|---|---|---|---|---|---|---|---|---|---|---|---|
| rs1893265 | 11 | 119218930 | C | T | 0.913 | 0.3611 | +? | 0 | 0 | 0 | 1 |
| rs538478 | 11 | 118925341 | A | G | 0.913 | 0.3611 | +? | 0 | 0 | 0 | 1 |
| rs114855326 | 11 | 117347592 | C | T | -0.913 | 0.3612 | -? | 0 | 0 | 0 | 1 |
| rs78284541 | 11 | 119847951 | T | C | -0.913 | 0.3613 | -? | 0 | 0 | 0 | 1 |
| rs1263170 | 11 | 116678413 | T | C | -0.913 | 0.3615 | -? | 0 | 0 | 0 | 1 |
| rs10444336 | 11 | 119928883 | A | C | -0.912 | 0.3616 | -? | 0 | 0 | 0 | 1 |
| rs1263168 | 11 | 116677999 | C | T | -0.912 | 0.3618 | -? | 0 | 0 | 0 | 1 |
| rs113941645 | 11 | 118545589 | G | C | 0.912 | 0.3619 | ++ | 0 | 0.705 | 1 | 0.4011 |
| rs3814754 | 11 | 119217957 | C | T | 0.912 | 0.3619 | +? | 0 | 0 | 0 | 1 |
| rs12282752 | 11 | 118793918 | C | T | -0.912 | 0.362 | -- | 0 | 0.086 | 1 | 0.7697 |
| rs12793075 | 11 | 119608316 | G | A | 0.911 | 0.3621 | +? | 0 | 0 | 0 | 1 |
| rs6589820 | 11 | 120493629 | G | A | 0.911 | 0.3623 | +? | 0 | 0 | 0 | 1 |
| rs4936496 | 11 | 119601313 | G | A | 0.911 | 0.3624 | +? | 0 | 0 | 0 | 1 |
| rs11216663 | 11 | 117848632 | T | G | 0.911 | 0.3625 | +? | 0 | 0 | 0 | 1 |
| rs113067228 | 11 | 120697913 | C | T | 0.91 | 0.3626 | +? | 0 | 0 | 0 | 1 |
| rs1893899 | 11 | 117436936 | T | G | 0.91 | 0.3626 | +? | 0 | 0 | 0 | 1 |
| rs2542052 | 11 | 116699984 | A | C | -0.91 | 0.3626 | -? | 0 | 0 | 0 | 1 |
| rs4938448 | 11 | 117757547 | C | A | -0.91 | 0.3626 | -? | 0 | 0 | 0 | 1 |
| rs1715456 | 11 | 118123517 | A | G | -0.91 | 0.3627 | -? | 0 | 0 | 0 | 1 |
| rs681311 | 11 | 120976413 | A | G | 0.91 | 0.3627 | +? | 0 | 0 | 0 | 1 |
| rs4938551 | 11 | 118707249 | C | G | 0.91 | 0.3628 | ++ | 0 | 0.227 | 1 | 0.6337 |
| rs4938451 | 11 | 117760135 | T | C | -0.91 | 0.3629 | -? | 0 | 0 | 0 | 1 |
| rs10892318 | 11 | 118848725 | A | G | -0.91 | 0.363 | -? | 0 | 0 | 0 | 1 |
| rs73564493 | 11 | 119056273 | A | G | -0.91 | 0.363 | -? | 0 | 0 | 0 | 1 |
| rs11217249 | 11 | 119220551 | G | A | 0.909 | 0.3631 | +? | 0 | 0 | 0 | 1 |
| rs2512146 | 11 | 117867008 | A | G | 0.91 | 0.3631 | ++ | 0 | 0.132 | 1 | 0.7167 |
| rs10892557 | 11 | 120138723 | T | C | -0.909 | 0.3632 | -? | 0 | 0 | 0 | 1 |
| rs34953153 | 11 | 119530129 | T | C | -0.909 | 0.3632 | -? | 0 | 0 | 0 | 1 |
| rs74235117 | 11 | 117923742 | T | C | 0.909 | 0.3632 | ++ | 0 | 0.004 | 1 | 0.947 |
| rs77731159 | 11 | 118691975 | A | T | 0.909 | 0.3632 | ++ | 0 | 0.557 | 1 | 0.4554 |
| rs10892649 | 11 | 120813148 | T | C | 0.909 | 0.3633 | +? | 0 | 0 | 0 | 1 |
| rs4938395 | 11 | 117384673 | G | A | -0.909 | 0.3633 | -? | 0 | 0 | 0 | 1 |
| rs2276122 | 11 | 117965530 | G | A | 0.909 | 0.3634 | ++ | 0 | 0.018 | 1 | 0.8945 |
| rs12364013 | 11 | 118119575 | C | T | -0.909 | 0.3635 | -? | 0 | 0 | 0 | 1 |
| rs61904853 | 11 | 116466495 | G | C | 0.909 | 0.3635 | +? | 0 | 0 | 0 | 1 |
| rs11218032 | 11 | 120724322 | A | T | 0.908 | 0.3638 | +? | 0 | 0 | 0 | 1 |
| rs10892133 | 11 | 117383694 | T | C | -0.908 | 0.364 | -? | 0 | 0 | 0 | 1 |
| rs73006796 | 11 | 119680375 | G | T | 0.908 | 0.364 | ++ | 0 | 0.019 | 1 | 0.8899 |
| rs150773762 | 11 | 119478010 | G | A | 0.907 | 0.3643 | +? | 0 | 0 | 0 | 1 |
| rs6589804 | 11 | 120161641 | A | G | -0.907 | 0.3643 | -- | 0 | 0.12 | 1 | 0.7296 |
| rs75736698 | 11 | 119823133 | A | G | -0.907 | 0.3643 | -? | 0 | 0 | 0 | 1 |
| rs10790137 | 11 | 116205687 | G | T | -0.907 | 0.3644 | -? | 0 | 0 | 0 | 1 |
| rs11217364 | 11 | 119501021 | A | G | 0.907 | 0.3644 | +? | 0 | 0 | 0 | 1 |
| rs11218074 | 11 | 120819045 | G | A | 0.907 | 0.3645 | +? | 0 | 0 | 0 | 1 |
| rs11824600 | 11 | 120406723 | A | G | -0.907 | 0.3645 | -? | 0 | 0 | 0 | 1 |
| rs4414229 | 11 | 120472569 | G | A | 0.907 | 0.3645 | +? | 0 | 0 | 0 | 1 |
| rs528508 | 11 | 117123173 | C | T | 0.907 | 0.3645 | +? | 0 | 0 | 0 | 1 |
| rs680511 | 11 | 121005824 | G | A | -0.907 | 0.3646 | -? | 0 | 0 | 0 | 1 |
| rs1783776 | 11 | 120979047 | G | A | 0.906 | 0.3647 | +? | 0 | 0 | 0 | 1 |
| rs1783777 | 11 | 120979036 | C | T | 0.906 | 0.3647 | +? | 0 | 0 | 0 | 1 |
| rs7937779 | 11 | 119957107 | G | A | -0.906 | 0.3647 | -? | 0 | 0 | 0 | 1 |
| rs12802011 | 11 | 119531135 | G | A | -0.906 | 0.3648 | -? | 0 | 0 | 0 | 1 |
| rs1939367 | 11 | 117761400 | A | C | -0.906 | 0.3648 | -? | 0 | 0 | 0 | 1 |
| rs11824351 | 11 | 120816284 | C | T | -0.906 | 0.3649 | -? | 0 | 0 | 0 | 1 |
| rs639373 | 11 | 118976719 | A | G | -0.906 | 0.3649 | -? | 0 | 0 | 0 | 1 |
| rs7478880 | 11 | 120261791 | T | C | -0.906 | 0.365 | -? | 0 | 0 | 0 | 1 |

| | | | | | | | | | | | |
|---|---|---|---|---|---|---|---|---|---|---|---|
| rs12797914 | 11 | 119613092 | G | A | 0.906 | 0.3651 | +? | 0 | 0 | 0 | 1 |
| rs548636 | 11 | 119759466 | C | T | -0.906 | 0.3651 | -? | 0 | 0 | 0 | 1 |
| rs7480302 | 11 | 116583017 | G | A | -0.906 | 0.3651 | -- | -62 | 1.852 | 1 | 0.1736 |
| rs73008513 | 11 | 117004112 | G | A | 0.906 | 0.3652 | +? | 0 | 0 | 0 | 1 |
| rs481903 | 11 | 118567733 | C | T | 0.905 | 0.3653 | ++ | -193.9 | 1.021 | 1 | 0.3123 |
| rs11218230 | 11 | 121134420 | G | T | -0.905 | 0.3654 | -- | 0 | 0.495 | 1 | 0.4818 |
| rs488538 | 11 | 117118038 | G | T | 0.905 | 0.3655 | +? | 0 | 0 | 0 | 1 |
| rs7932037 | 11 | 120259430 | A | T | -0.905 | 0.3655 | -? | 0 | 0 | 0 | 1 |
| rs7949956 | 11 | 117501072 | C | T | 0.905 | 0.3656 | +? | 0 | 0 | 0 | 1 |
| rs4938447 | 11 | 117757443 | C | A | -0.904 | 0.3658 | -? | 0 | 0 | 0 | 1 |
| rs61900655 | 11 | 118120802 | G | C | -0.904 | 0.3658 | -? | 0 | 0 | 0 | 1 |
| rs73024604 | 11 | 118120351 | C | T | -0.904 | 0.3658 | -? | 0 | 0 | 0 | 1 |
| rs73024606 | 11 | 118120366 | G | C | -0.904 | 0.3658 | -? | 0 | 0 | 0 | 1 |
| rs77492882 | 11 | 117475763 | T | C | 0.904 | 0.3662 | +? | 0 | 0 | 0 | 1 |
| rs79198917 | 11 | 117475976 | A | G | 0.904 | 0.3662 | +? | 0 | 0 | 0 | 1 |
| rs12794050 | 11 | 117244255 | A | C | -0.903 | 0.3663 | -? | 0 | 0 | 0 | 1 |
| rs12785979 | 11 | 116303839 | C | A | -0.903 | 0.3665 | -? | 0 | 0 | 0 | 1 |
| rs653330 | 11 | 117906882 | A | G | -0.903 | 0.3665 | -- | 0 | 0.851 | 1 | 0.3563 |
| rs12277255 | 11 | 119632722 | G | A | -0.903 | 0.3666 | -? | 0 | 0 | 0 | 1 |
| rs565312 | 11 | 117666712 | A | G | -0.903 | 0.3666 | -? | 0 | 0 | 0 | 1 |
| rs111932453 | 11 | 120696771 | T | C | 0.903 | 0.3667 | +? | 0 | 0 | 0 | 1 |
| rs11216740 | 11 | 117963766 | A | G | 0.903 | 0.3667 | ++ | 0 | 0.019 | 1 | 0.8916 |
| rs4936566 | 11 | 120815238 | A | G | 0.903 | 0.3667 | +? | 0 | 0 | 0 | 1 |
| rs75622430 | 11 | 120696085 | T | G | 0.903 | 0.3667 | +? | 0 | 0 | 0 | 1 |
| rs79304595 | 11 | 120697009 | G | C | 0.903 | 0.3667 | +? | 0 | 0 | 0 | 1 |
| rs9633950 | 11 | 120248249 | G | T | -0.903 | 0.3667 | -? | 0 | 0 | 0 | 1 |
| rs1784281 | 11 | 120978834 | C | A | 0.902 | 0.3668 | +? | 0 | 0 | 0 | 1 |
| rs11216136 | 11 | 116647120 | G | T | -0.902 | 0.3671 | -- | 0 | 0.196 | 1 | 0.6577 |
| rs4537792 | 11 | 117723059 | G | A | 0.902 | 0.3672 | +? | 0 | 0 | 0 | 1 |
| rs686245 | 11 | 117722587 | C | T | 0.902 | 0.3672 | +? | 0 | 0 | 0 | 1 |
| rs2226635 | 11 | 117687120 | T | C | 0.901 | 0.3676 | ++ | 0 | 0.085 | 1 | 0.7711 |
| rs4938454 | 11 | 117769598 | A | C | 0.901 | 0.3676 | +? | 0 | 0 | 0 | 1 |
| rs2851460 | 11 | 118716563 | A | G | 0.901 | 0.3678 | ++ | 0 | 0.298 | 1 | 0.585 |
| rs633983 | 11 | 117460028 | G | A | 0.901 | 0.3678 | +? | 0 | 0 | 0 | 1 |
| rs11827023 | 11 | 120406694 | C | T | -0.9 | 0.3681 | -? | 0 | 0 | 0 | 1 |
| rs56315433 | 11 | 118114576 | C | A | -0.9 | 0.3681 | -? | 0 | 0 | 0 | 1 |
| rs56356019 | 11 | 118114893 | A | G | -0.9 | 0.3681 | -? | 0 | 0 | 0 | 1 |
| rs12363923 | 11 | 120791799 | T | C | 0.9 | 0.3683 | +? | 0 | 0 | 0 | 1 |
| rs10790286 | 11 | 119054488 | T | C | 0.899 | 0.3684 | +? | 0 | 0 | 0 | 1 |
| rs3915715 | 11 | 119960260 | A | G | -0.899 | 0.3684 | -? | 0 | 0 | 0 | 1 |
| rs12800445 | 11 | 119595355 | A | C | 0.899 | 0.3687 | +? | 0 | 0 | 0 | 1 |
| rs7924314 | 11 | 117960809 | A | G | -0.899 | 0.3687 | -- | 0 | 0.024 | 1 | 0.878 |
| rs145533092 | 11 | 117472684 | A | G | 0.899 | 0.3688 | +? | 0 | 0 | 0 | 1 |
| rs666741 | 11 | 118911350 | C | A | 0.899 | 0.3689 | +? | 0 | 0 | 0 | 1 |
| rs75734044 | 11 | 117652288 | A | G | 0.899 | 0.3689 | ++ | 0 | 0.023 | 1 | 0.8792 |
| rs17124575 | 11 | 120808850 | G | A | 0.898 | 0.369 | +? | 0 | 0 | 0 | 1 |
| rs11216125 | 11 | 116612274 | A | G | -0.898 | 0.3691 | -+ | 0 | 0.69 | 1 | 0.4063 |
| rs11820112 | 11 | 116212436 | A | G | -0.898 | 0.3691 | -? | 0 | 0 | 0 | 1 |
| rs77314777 | 11 | 117495305 | A | G | 0.898 | 0.3691 | +? | 0 | 0 | 0 | 1 |
| rs113264811 | 11 | 116543111 | T | C | -0.898 | 0.3692 | -? | 0 | 0 | 0 | 1 |
| rs1551241 | 11 | 120084509 | C | T | 0.898 | 0.3692 | +? | 0 | 0 | 0 | 1 |
| rs1551242 | 11 | 120084221 | T | C | 0.898 | 0.3692 | +? | 0 | 0 | 0 | 1 |
| rs621858 | 11 | 117128971 | A | G | -0.898 | 0.3692 | -? | 0 | 0 | 0 | 1 |
| rs117604511 | 11 | 117382799 | C | A | -0.898 | 0.3694 | -? | 0 | 0 | 0 | 1 |
| rs79936530 | 11 | 119847788 | A | G | -0.898 | 0.3694 | -? | 0 | 0 | 0 | 1 |
| rs673738 | 11 | 117143581 | T | C | -0.896 | 0.37 | -? | 0 | 0 | 0 | 1 |

| | | | | | | | | | | | |
|---|---|---|---|---|---|---|---|---|---|---|---|
| rs12421814 | 11 | 119220839 | G | A | 0.896 | 0.3701 | +? | 0 | 0 | 0 | 1 |
| rs7938081 | 11 | 120701623 | C | G | 0.896 | 0.3702 | +? | 0 | 0 | 0 | 1 |
| rs531855 | 11 | 117726427 | T | C | 0.896 | 0.3704 | +? | 0 | 0 | 0 | 1 |
| rs4936565 | 11 | 120802006 | A | G | 0.896 | 0.3705 | +? | 0 | 0 | 0 | 1 |
| rs55869870 | 11 | 117446931 | A | C | 0.896 | 0.3705 | +? | 0 | 0 | 0 | 1 |
| rs11600022 | 11 | 117769430 | T | C | 0.895 | 0.3706 | +? | 0 | 0 | 0 | 1 |
| rs3016354 | 11 | 116540792 | C | T | -0.895 | 0.3707 | -? | 0 | 0 | 0 | 1 |
| rs3814759 | 11 | 119217580 | C | T | 0.894 | 0.3711 | +? | 0 | 0 | 0 | 1 |
| rs79795409 | 11 | 117229833 | T | G | 0.894 | 0.3711 | +? | 0 | 0 | 0 | 1 |
| rs11216068 | 11 | 116505239 | T | C | -0.894 | 0.3713 | -? | 0 | 0 | 0 | 1 |
| rs12789727 | 11 | 117435785 | A | C | -0.894 | 0.3714 | -- | 0 | 0.236 | 1 | 0.6275 |
| rs10892139 | 11 | 117449297 | G | A | 0.894 | 0.3715 | +? | 0 | 0 | 0 | 1 |
| rs11605293 | 11 | 116437756 | T | C | -0.894 | 0.3715 | -- | 0 | 0.544 | 1 | 0.461 |
| rs117328724 | 11 | 117378697 | T | C | 0.894 | 0.3715 | +? | 0 | 0 | 0 | 1 |
| rs10892129 | 11 | 117366111 | G | C | 0.893 | 0.3716 | +? | 0 | 0 | 0 | 1 |
| rs4462362 | 11 | 121151411 | C | G | -0.893 | 0.3717 | -- | 0 | 0.474 | 1 | 0.491 |
| rs9651695 | 11 | 120562898 | A | G | -0.893 | 0.3717 | -? | 0 | 0 | 0 | 1 |
| rs76478216 | 11 | 118605352 | C | T | -0.893 | 0.3718 | -+ | -124.6 | 1.336 | 1 | 0.2477 |
| rs686598 | 11 | 117423936 | C | G | -0.893 | 0.3719 | -? | 0 | 0 | 0 | 1 |
| rs2368242 | 11 | 118115526 | T | C | -0.893 | 0.372 | -? | 0 | 0 | 0 | 1 |
| rs10892709 | 11 | 121136915 | C | T | -0.893 | 0.3721 | -- | 0 | 0.561 | 1 | 0.4539 |
| rs619772 | 11 | 117138836 | T | A | -0.893 | 0.3721 | -? | 0 | 0 | 0 | 1 |
| rs12804550 | 11 | 118011860 | A | G | -0.892 | 0.3724 | -- | -184.8 | 1.054 | 1 | 0.3047 |
| rs4935757 | 11 | 120819657 | G | A | 0.892 | 0.3726 | +? | 0 | 0 | 0 | 1 |
| rs79357714 | 11 | 116902639 | G | A | 0.891 | 0.3727 | +? | 0 | 0 | 0 | 1 |
| rs12293490 | 11 | 121137332 | T | C | -0.891 | 0.3728 | -- | 0 | 0.557 | 1 | 0.4554 |
| rs17121997 | 11 | 118131049 | T | C | -0.891 | 0.3728 | -? | 0 | 0 | 0 | 1 |
| rs34138305 | 11 | 117376413 | A | G | 0.891 | 0.3728 | +? | 0 | 0 | 0 | 1 |
| rs7126179 | 11 | 117574062 | C | T | 0.891 | 0.3729 | +? | 0 | 0 | 0 | 1 |
| rs10892559 | 11 | 120156378 | T | C | -0.891 | 0.3731 | -? | 0 | 0 | 0 | 1 |
| rs2846015 | 11 | 116540439 | T | C | -0.891 | 0.3731 | -? | 0 | 0 | 0 | 1 |
| rs3017337 | 11 | 116540807 | T | C | -0.89 | 0.3732 | -? | 0 | 0 | 0 | 1 |
| rs10892711 | 11 | 121137543 | T | G | -0.89 | 0.3735 | -- | 0 | 0.554 | 1 | 0.4567 |
| rs10892713 | 11 | 121137629 | C | G | -0.89 | 0.3735 | -- | 0 | 0.554 | 1 | 0.4567 |
| rs10892714 | 11 | 121137690 | A | G | -0.89 | 0.3735 | -- | 0 | 0.554 | 1 | 0.4567 |
| rs2305829 | 11 | 117257876 | T | G | 0.89 | 0.3735 | +? | 0 | 0 | 0 | 1 |
| rs2508807 | 11 | 120613199 | C | T | 0.89 | 0.3735 | +? | 0 | 0 | 0 | 1 |
| rs7935472 | 11 | 121137754 | G | A | -0.89 | 0.3735 | -- | 0 | 0.554 | 1 | 0.4567 |
| rs493139 | 11 | 117145949 | T | C | -0.89 | 0.3736 | -? | 0 | 0 | 0 | 1 |
| rs2276319 | 11 | 120837821 | G | A | 0.89 | 0.3737 | +? | 0 | 0 | 0 | 1 |
| rs550319 | 11 | 117168564 | G | A | 0.89 | 0.3737 | +? | 0 | 0 | 0 | 1 |
| rs561775 | 11 | 118341212 | G | A | 0.89 | 0.3737 | +- | 0 | 0.791 | 1 | 0.3739 |
| rs644215 | 11 | 117145918 | T | A | -0.89 | 0.3737 | -? | 0 | 0 | 0 | 1 |
| rs4622278 | 11 | 120527541 | C | T | -0.889 | 0.374 | -? | 0 | 0 | 0 | 1 |
| rs1145188 | 11 | 116532314 | A | G | -0.889 | 0.3741 | -? | 0 | 0 | 0 | 1 |
| rs118110450 | 11 | 117319240 | C | A | -0.889 | 0.3741 | -? | 0 | 0 | 0 | 1 |
| rs677008 | 11 | 119240071 | A | G | 0.889 | 0.3741 | +? | 0 | 0 | 0 | 1 |
| rs116989966 | 11 | 120128055 | G | A | -0.889 | 0.3742 | -? | 0 | 0 | 0 | 1 |
| rs2252356 | 11 | 120418072 | A | G | -0.889 | 0.3742 | -? | 0 | 0 | 0 | 1 |
| rs4520 | 11 | 116701535 | T | C | 0.889 | 0.3742 | +? | 0 | 0 | 0 | 1 |
| rs10892710 | 11 | 121137123 | G | A | -0.888 | 0.3743 | -- | 0 | 0.55 | 1 | 0.4582 |
| rs2512145 | 11 | 117863032 | A | G | 0.888 | 0.3743 | ++ | 0 | 0.231 | 1 | 0.6306 |
| rs73007893 | 11 | 120823074 | A | G | 0.888 | 0.3743 | +? | 0 | 0 | 0 | 1 |
| rs495438 | 11 | 117144727 | A | C | -0.888 | 0.3744 | -? | 0 | 0 | 0 | 1 |
| rs573956 | 11 | 117143605 | G | A | -0.888 | 0.3744 | -? | 0 | 0 | 0 | 1 |
| rs7925044 | 11 | 117147987 | G | A | -0.888 | 0.3744 | -? | 0 | 0 | 0 | 1 |

| | | | | | | | | | | | |
|---|---|---|---|---|---|---|---|---|---|---|---|
| rs79297443 | 11 | 119684904 | T | C | 0.888 | 0.3744 | +? | 0 | 0 | 0 | 1 |
| rs10892507 | 11 | 119859948 | G | A | 0.888 | 0.3745 | +? | 0 | 0 | 0 | 1 |
| rs73007896 | 11 | 120823269 | T | C | 0.888 | 0.3745 | +? | 0 | 0 | 0 | 1 |
| rs17120765 | 11 | 117316546 | A | G | -0.887 | 0.375 | -? | 0 | 0 | 0 | 1 |
| rs57643685 | 11 | 116897124 | G | A | 0.887 | 0.3752 | +? | 0 | 0 | 0 | 1 |
| rs584356 | 11 | 117930428 | A | G | -0.886 | 0.3756 | -? | 0 | 0 | 0 | 1 |
| rs954319 | 11 | 117686194 | A | G | -0.886 | 0.3756 | -? | 0 | 0 | 0 | 1 |
| rs77031353 | 11 | 119839762 | A | C | -0.886 | 0.3757 | -? | 0 | 0 | 0 | 1 |
| rs7950796 | 11 | 120728304 | G | A | 0.886 | 0.3757 | +? | 0 | 0 | 0 | 1 |
| rs11600703 | 11 | 120249287 | T | C | -0.886 | 0.3758 | -? | 0 | 0 | 0 | 1 |
| rs12420378 | 11 | 120250699 | G | A | -0.886 | 0.3758 | -? | 0 | 0 | 0 | 1 |
| rs9633919 | 11 | 120249493 | A | G | -0.886 | 0.3758 | -? | 0 | 0 | 0 | 1 |
| rs1271170 | 11 | 117775345 | A | G | 0.885 | 0.3759 | +? | 0 | 0 | 0 | 1 |
| rs3814755 | 11 | 119217945 | G | A | 0.885 | 0.3759 | +? | 0 | 0 | 0 | 1 |
| rs579941 | 11 | 116279292 | T | C | 0.885 | 0.376 | +? | 0 | 0 | 0 | 1 |
| rs510435 | 11 | 118980039 | T | C | -0.884 | 0.3766 | -? | 0 | 0 | 0 | 1 |
| rs6589833 | 11 | 120561883 | G | C | -0.884 | 0.3766 | -? | 0 | 0 | 0 | 1 |
| rs12280328 | 11 | 121137458 | G | A | -0.884 | 0.3767 | -- | 0 | 0.55 | 1 | 0.4584 |
| rs12293637 | 11 | 121137477 | A | G | -0.884 | 0.3767 | -- | 0 | 0.55 | 1 | 0.4584 |
| rs12420651 | 11 | 121150839 | G | T | -0.884 | 0.3767 | -- | 0 | 0.581 | 1 | 0.4458 |
| rs112799932 | 11 | 118585146 | G | T | -0.884 | 0.3769 | -- | 0 | 0.017 | 1 | 0.8948 |
| rs4414232 | 11 | 120730638 | C | T | -0.884 | 0.3769 | -? | 0 | 0 | 0 | 1 |
| rs7950819 | 11 | 121137859 | A | T | -0.884 | 0.3769 | -- | 0 | 0.538 | 1 | 0.4634 |
| rs2155993 | 11 | 119858605 | T | C | -0.883 | 0.377 | -? | 0 | 0 | 0 | 1 |
| rs35266957 | 11 | 117411730 | T | C | 0.883 | 0.377 | +? | 0 | 0 | 0 | 1 |
| rs7118520 | 11 | 117568781 | G | A | 0.883 | 0.377 | +? | 0 | 0 | 0 | 1 |
| rs2851410 | 11 | 120399287 | T | C | -0.883 | 0.3773 | -? | 0 | 0 | 0 | 1 |
| rs3814758 | 11 | 119217818 | A | G | 0.883 | 0.3775 | +? | 0 | 0 | 0 | 1 |
| rs7935970 | 11 | 121138191 | G | A | -0.883 | 0.3775 | -- | 0 | 0.535 | 1 | 0.4645 |
| rs34150933 | 11 | 118911039 | T | C | 0.882 | 0.3776 | +? | 0 | 0 | 0 | 1 |
| rs5104 | 11 | 116692334 | C | T | 0.882 | 0.3777 | +? | 0 | 0 | 0 | 1 |
| rs73580632 | 11 | 116518901 | T | C | -0.882 | 0.3778 | -? | 0 | 0 | 0 | 1 |
| rs9943622 | 11 | 121129412 | T | G | -0.882 | 0.3778 | -- | 0 | 0.405 | 1 | 0.5244 |
| rs491292 | 11 | 117755082 | T | C | -0.882 | 0.3779 | -? | 0 | 0 | 0 | 1 |
| rs6589807 | 11 | 120166860 | A | G | -0.881 | 0.3781 | -? | 0 | 0 | 0 | 1 |
| rs7943989 | 11 | 117311169 | G | A | -0.881 | 0.3781 | -? | 0 | 0 | 0 | 1 |
| rs638415 | 11 | 117913024 | C | A | -0.881 | 0.3782 | -- | 0 | 0.697 | 1 | 0.4037 |
| rs117373043 | 11 | 117616156 | T | C | -0.881 | 0.3783 | -? | 0 | 0 | 0 | 1 |
| rs10892712 | 11 | 121137603 | C | T | -0.881 | 0.3784 | -- | 0 | 0.531 | 1 | 0.4663 |
| rs492151 | 11 | 117145815 | G | A | -0.881 | 0.3784 | -? | 0 | 0 | 0 | 1 |
| rs1374565 | 11 | 119669797 | G | A | 0.88 | 0.3789 | +? | 0 | 0 | 0 | 1 |
| rs7936408 | 11 | 121138302 | T | G | -0.88 | 0.3789 | -- | 0 | 0.529 | 1 | 0.4672 |
| rs7951282 | 11 | 121138260 | C | T | -0.88 | 0.3789 | -- | 0 | 0.529 | 1 | 0.4672 |
| rs7951286 | 11 | 121138262 | A | T | -0.88 | 0.3789 | -- | 0 | 0.529 | 1 | 0.4672 |
| rs7937927 | 11 | 119382758 | C | T | -0.88 | 0.379 | -? | 0 | 0 | 0 | 1 |
| rs11600311 | 11 | 119641843 | T | C | 0.88 | 0.3791 | ++ | 0 | 0.034 | 1 | 0.8542 |
| rs11603208 | 11 | 117290963 | T | G | -0.879 | 0.3792 | -? | 0 | 0 | 0 | 1 |
| rs3862647 | 11 | 119949832 | C | T | -0.879 | 0.3792 | -? | 0 | 0 | 0 | 1 |
| rs4938476 | 11 | 117947181 | C | A | -0.879 | 0.3794 | -- | 0 | 0.058 | 1 | 0.8105 |
| rs10790248 | 11 | 118125672 | G | A | -0.879 | 0.3795 | -? | 0 | 0 | 0 | 1 |
| rs10892701 | 11 | 121132812 | C | T | -0.879 | 0.3795 | -- | 0 | 0.483 | 1 | 0.4871 |
| rs117598476 | 11 | 120277110 | C | T | -0.879 | 0.3795 | -? | 0 | 0 | 0 | 1 |
| rs34070113 | 11 | 118911139 | G | T | 0.878 | 0.3799 | +? | 0 | 0 | 0 | 1 |
| rs7112758 | 11 | 118836821 | A | G | -0.878 | 0.3799 | -? | 0 | 0 | 0 | 1 |
| rs12802294 | 11 | 116466039 | T | C | 0.878 | 0.38 | +? | 0 | 0 | 0 | 1 |
| rs11218034 | 11 | 120726562 | A | G | 0.878 | 0.3801 | +? | 0 | 0 | 0 | 1 |

| | | | | | | | | | | | |
|---|---|---|---|---|---|---|---|---|---|---|---|
| rs472648 | 11 | 117307265 | C | T | 0.878 | 0.3801 | +? | 0 | 0 | 0 | 1 |
| rs685044 | 11 | 118942928 | A | G | 0.878 | 0.3801 | +? | 0 | 0 | 0 | 1 |
| rs7118101 | 11 | 117737382 | C | T | 0.878 | 0.3801 | +? | 0 | 0 | 0 | 1 |
| rs876798 | 11 | 117737440 | C | T | 0.878 | 0.3801 | +? | 0 | 0 | 0 | 1 |
| rs11601799 | 11 | 119659912 | A | G | 0.878 | 0.3802 | ++ | 0 | 0.021 | 1 | 0.8846 |
| rs118101513 | 11 | 119324831 | A | C | 0.878 | 0.3802 | +? | 0 | 0 | 0 | 1 |
| rs72993408 | 11 | 116225035 | T | A | -0.877 | 0.3802 | -- | 0 | 0.128 | 1 | 0.7205 |
| rs2852218 | 11 | 120618334 | G | A | 0.877 | 0.3806 | +? | 0 | 0 | 0 | 1 |
| rs4075786 | 11 | 120697162 | C | T | 0.877 | 0.3806 | +? | 0 | 0 | 0 | 1 |
| rs4076169 | 11 | 120696635 | C | T | 0.877 | 0.3806 | +? | 0 | 0 | 0 | 1 |
| rs78757430 | 11 | 119505431 | T | C | 0.877 | 0.3807 | +? | 0 | 0 | 0 | 1 |
| rs58745438 | 11 | 118848315 | C | T | -0.876 | 0.3809 | -? | 0 | 0 | 0 | 1 |
| rs12291064 | 11 | 119502703 | T | C | 0.876 | 0.3811 | +? | 0 | 0 | 0 | 1 |
| rs4245197 | 11 | 119507723 | C | T | 0.876 | 0.3813 | +? | 0 | 0 | 0 | 1 |
| rs12276124 | 11 | 121138583 | C | T | -0.875 | 0.3814 | -- | 0 | 0.604 | 1 | 0.4371 |
| rs9971517 | 11 | 121148154 | C | T | -0.875 | 0.3814 | -- | 0 | 0.535 | 1 | 0.4646 |
| rs11216022 | 11 | 116458862 | T | C | -0.875 | 0.3816 | -? | 0 | 0 | 0 | 1 |
| rs72995842 | 11 | 120630028 | T | C | 0.875 | 0.3816 | +? | 0 | 0 | 0 | 1 |
| rs1145189 | 11 | 116532856 | A | G | -0.875 | 0.3817 | -? | 0 | 0 | 0 | 1 |
| rs558590 | 11 | 117139121 | G | A | -0.875 | 0.3817 | -? | 0 | 0 | 0 | 1 |
| rs11218229 | 11 | 121134392 | G | T | -0.875 | 0.3818 | -- | 0 | 0.448 | 1 | 0.5033 |
| rs676685 | 11 | 118036942 | A | G | 0.875 | 0.3818 | +? | 0 | 0 | 0 | 1 |
| rs79164518 | 11 | 120162368 | C | T | -0.875 | 0.3818 | -- | 0 | 0.121 | 1 | 0.7283 |
| rs9667343 | 11 | 120622881 | C | G | 0.875 | 0.3818 | +? | 0 | 0 | 0 | 1 |
| rs61897706 | 11 | 119679280 | A | G | -0.874 | 0.3819 | -? | 0 | 0 | 0 | 1 |
| rs2511848 | 11 | 119201699 | A | G | -0.874 | 0.3821 | -? | 0 | 0 | 0 | 1 |
| rs35112163 | 11 | 116465552 | T | G | 0.874 | 0.3821 | +? | 0 | 0 | 0 | 1 |
| rs4938492 | 11 | 118126675 | T | C | -0.874 | 0.3821 | -? | 0 | 0 | 0 | 1 |
| rs61901381 | 11 | 120580148 | C | T | 0.874 | 0.3822 | +? | 0 | 0 | 0 | 1 |
| rs565530 | 11 | 117727735 | G | T | 0.874 | 0.3823 | +? | 0 | 0 | 0 | 1 |
| rs60790095 | 11 | 117651730 | T | C | -0.874 | 0.3823 | -- | 0 | 0.241 | 1 | 0.6237 |
| rs1104542 | 11 | 118129340 | C | T | -0.873 | 0.3824 | -? | 0 | 0 | 0 | 1 |
| rs55813570 | 11 | 117772021 | A | G | -0.873 | 0.3824 | -? | 0 | 0 | 0 | 1 |
| rs602492 | 11 | 119241743 | T | G | 0.873 | 0.3827 | +? | 0 | 0 | 0 | 1 |
| rs11602240 | 11 | 116632798 | C | T | -0.873 | 0.3828 | -+ | 0 | 0.612 | 1 | 0.4341 |
| rs7110791 | 11 | 116458278 | C | A | 0.872 | 0.3832 | +? | 0 | 0 | 0 | 1 |
| rs2512154 | 11 | 117889402 | A | G | 0.872 | 0.3834 | +- | 0 | 0.498 | 1 | 0.4802 |
| rs75693006 | 11 | 117651600 | A | G | 0.872 | 0.3835 | ++ | 0 | 0.019 | 1 | 0.8915 |
| rs4533044 | 11 | 121127518 | C | T | -0.871 | 0.3837 | -- | 0 | 0.433 | 1 | 0.5105 |
| rs4938507 | 11 | 118231584 | G | A | -0.871 | 0.3837 | -+ | 0 | 0.485 | 1 | 0.486 |
| rs56970059 | 11 | 116448578 | T | C | 0.871 | 0.3838 | +? | 0 | 0 | 0 | 1 |
| rs667584 | 11 | 118911544 | T | C | 0.871 | 0.3838 | +? | 0 | 0 | 0 | 1 |
| rs76219905 | 11 | 120699480 | T | C | 0.871 | 0.3839 | +? | 0 | 0 | 0 | 1 |
| rs2441939 | 11 | 119733445 | T | C | -0.87 | 0.3842 | -? | 0 | 0 | 0 | 1 |
| rs12804924 | 11 | 117425383 | T | C | -0.87 | 0.3845 | -? | 0 | 0 | 0 | 1 |
| rs4936425 | 11 | 118126904 | C | T | -0.869 | 0.3846 | -? | 0 | 0 | 0 | 1 |
| rs60727471 | 11 | 119624396 | C | T | 0.869 | 0.3847 | ++ | 0 | 0.066 | 1 | 0.7968 |
| rs11216827 | 11 | 118110202 | A | T | -0.869 | 0.3849 | -? | 0 | 0 | 0 | 1 |
| rs7937438 | 11 | 120731495 | G | A | -0.869 | 0.385 | -? | 0 | 0 | 0 | 1 |
| rs10892212 | 11 | 117958179 | A | T | 0.868 | 0.3852 | +? | 0 | 0 | 0 | 1 |
| rs4938302 | 11 | 116584504 | C | T | -0.868 | 0.3853 | -- | -55.3 | 1.932 | 1 | 0.1645 |
| rs10892189 | 11 | 117758431 | T | C | -0.868 | 0.3854 | -? | 0 | 0 | 0 | 1 |
| rs10892371 | 11 | 119367674 | G | A | -0.868 | 0.3854 | -? | 0 | 0 | 0 | 1 |
| rs474749 | 11 | 117455669 | A | G | -0.868 | 0.3855 | -? | 0 | 0 | 0 | 1 |
| rs11825298 | 11 | 117642382 | T | C | -0.868 | 0.3856 | -- | -45.9 | 2.056 | 1 | 0.1516 |
| rs9667661 | 11 | 121131225 | A | G | -0.868 | 0.3856 | -- | 0 | 0.507 | 1 | 0.4766 |

| | | | | | | | | | | | |
|---|---|---|---|---|---|---|---|---|---|---|---|
| rs4938703 | 11 | 119524950 | T | C | -0.867 | 0.3857 | -? | 0 | 0 | 0 | 1 |
| rs7128147 | 11 | 119989636 | T | C | -0.867 | 0.3858 | -? | 0 | 0 | 0 | 1 |
| rs1009364 | 11 | 119505025 | A | G | 0.867 | 0.386 | +? | 0 | 0 | 0 | 1 |
| rs12575964 | 11 | 117127159 | A | G | 0.867 | 0.3861 | +? | 0 | 0 | 0 | 1 |
| rs4938413 | 11 | 117464204 | G | C | -0.866 | 0.3864 | -- | 0 | 0.227 | 1 | 0.634 |
| rs12417431 | 11 | 118137383 | G | A | 0.866 | 0.3865 | +? | 0 | 0 | 0 | 1 |
| rs11216005 | 11 | 116429253 | C | A | -0.866 | 0.3866 | -? | 0 | 0 | 0 | 1 |
| rs4517527 | 11 | 121128562 | G | T | -0.866 | 0.3867 | -- | 0 | 0.435 | 1 | 0.5096 |
| rs1367711737 | 11 | 116542812 | T | G | -0.865 | 0.3869 | -? | 0 | 0 | 0 | 1 |
| rs1145187 | 11 | 116532066 | G | T | -0.865 | 0.387 | -? | 0 | 0 | 0 | 1 |
| rs117239953 | 11 | 116513188 | C | T | -0.865 | 0.387 | -? | 0 | 0 | 0 | 1 |
| rs12788483 | 11 | 117410852 | C | G | 0.865 | 0.387 | +? | 0 | 0 | 0 | 1 |
| rs1560403 | 11 | 121138891 | A | G | -0.865 | 0.3871 | -- | 0 | 0.505 | 1 | 0.4771 |
| rs4376915 | 11 | 121128752 | G | A | -0.865 | 0.3872 | -- | 0 | 0.433 | 1 | 0.5106 |
| rs498684 | 11 | 118316960 | A | G | 0.865 | 0.3872 | ++ | 0 | 0.01 | 1 | 0.9205 |
| rs2508454 | 11 | 117867397 | T | C | 0.864 | 0.3873 | ++ | 0 | 0.014 | 1 | 0.9048 |
| rs7131046 | 11 | 120238534 | A | G | -0.865 | 0.3873 | -? | 0 | 0 | 0 | 1 |
| rs112377532 | 11 | 118866550 | G | C | 0.864 | 0.3874 | +? | 0 | 0 | 0 | 1 |
| rs113032308 | 11 | 118866515 | A | G | 0.864 | 0.3874 | +? | 0 | 0 | 0 | 1 |
| rs71480357 | 11 | 117665267 | G | C | -0.864 | 0.3874 | -? | 0 | 0 | 0 | 1 |
| rs1560404 | 11 | 121139062 | C | T | -0.864 | 0.3875 | -- | 0 | 0.504 | 1 | 0.4779 |
| rs12419859 | 11 | 121127974 | G | C | -0.864 | 0.3876 | -- | 0 | 0.438 | 1 | 0.5079 |
| rs1652024 | 11 | 119242556 | C | A | 0.864 | 0.3876 | +? | 0 | 0 | 0 | 1 |
| rs4307734 | 11 | 121127491 | A | C | -0.864 | 0.3876 | -- | 0 | 0.438 | 1 | 0.5079 |
| rs4370953 | 11 | 121127870 | G | A | -0.864 | 0.3876 | -- | 0 | 0.438 | 1 | 0.5079 |
| rs4938303 | 11 | 116584987 | C | T | -0.864 | 0.3877 | -- | -54.7 | 1.939 | 1 | 0.1638 |
| rs12421355 | 11 | 121128034 | A | T | -0.863 | 0.388 | -- | 0 | 0.437 | 1 | 0.5087 |
| rs4938602 | 11 | 118846466 | A | G | -0.863 | 0.3881 | -? | 0 | 0 | 0 | 1 |
| rs78015 | 11 | 119757111 | T | G | -0.862 | 0.3885 | -? | 0 | 0 | 0 | 1 |
| rs12417414 | 11 | 121131965 | C | T | -0.862 | 0.3886 | -- | 0 | 0.509 | 1 | 0.4757 |
| rs1941304 | 11 | 121132279 | A | G | -0.862 | 0.3886 | -- | 0 | 0.509 | 1 | 0.4757 |
| rs1941305 | 11 | 121132211 | A | G | -0.862 | 0.3886 | -- | 0 | 0.509 | 1 | 0.4757 |
| rs4936599 | 11 | 121129931 | T | C | -0.862 | 0.3886 | -- | 0 | 0.435 | 1 | 0.5097 |
| rs9943676 | 11 | 121129552 | C | T | -0.862 | 0.3886 | -- | 0 | 0.435 | 1 | 0.5097 |
| rs116862564 | 11 | 120874281 | G | C | 0.862 | 0.3889 | +? | 0 | 0 | 0 | 1 |
| rs12577611 | 11 | 120897987 | C | T | -0.862 | 0.3889 | -? | 0 | 0 | 0 | 1 |
| rs1942479 | 11 | 116566117 | T | C | -0.861 | 0.389 | -? | 0 | 0 | 0 | 1 |
| rs138435357 | 11 | 120081510 | A | C | 0.861 | 0.3891 | +? | 0 | 0 | 0 | 1 |
| rs6589706 | 11 | 118747813 | A | G | 0.861 | 0.3892 | +? | 0 | 0 | 0 | 1 |
| rs7111010 | 11 | 119962890 | C | A | -0.861 | 0.3895 | -? | 0 | 0 | 0 | 1 |
| rs117445862 | 11 | 117500587 | C | T | 0.86 | 0.3896 | +? | 0 | 0 | 0 | 1 |
| rs12575215 | 11 | 117341209 | C | T | -0.86 | 0.3896 | -? | 0 | 0 | 0 | 1 |
| rs141182805 | 11 | 117500633 | G | T | 0.86 | 0.3896 | +? | 0 | 0 | 0 | 1 |
| rs17123366 | 11 | 119698359 | A | G | -0.86 | 0.3896 | -? | 0 | 0 | 0 | 1 |
| rs4938747 | 11 | 119867224 | A | G | 0.86 | 0.3896 | +? | 0 | 0 | 0 | 1 |
| rs10790373 | 11 | 120155564 | T | C | -0.86 | 0.3899 | -? | 0 | 0 | 0 | 1 |
| rs12284449 | 11 | 120148845 | G | A | -0.86 | 0.3899 | -? | 0 | 0 | 0 | 1 |
| rs4936512 | 11 | 120155336 | C | T | -0.86 | 0.3899 | -? | 0 | 0 | 0 | 1 |
| rs11218228 | 11 | 121134099 | G | A | -0.86 | 0.39 | -- | 0 | 0.45 | 1 | 0.5022 |
| rs74500359 | 11 | 117680478 | T | C | 0.859 | 0.3902 | +? | 0 | 0 | 0 | 1 |
| rs35633147 | 11 | 119633448 | T | C | -0.858 | 0.3908 | -? | 0 | 0 | 0 | 1 |
| rs495132 | 11 | 118335927 | A | G | 0.858 | 0.3908 | +- | 0 | 0.821 | 1 | 0.3649 |
| rs77101533 | 11 | 119924508 | G | A | -0.858 | 0.3908 | -? | 0 | 0 | 0 | 1 |
| rs10790197 | 11 | 117501603 | T | C | 0.858 | 0.3909 | +? | 0 | 0 | 0 | 1 |
| rs10892707 | 11 | 121136762 | T | C | -0.858 | 0.3909 | -- | 0 | 0.56 | 1 | 0.4541 |
| rs143427408 | 11 | 120910225 | T | C | 0.858 | 0.391 | +? | 0 | 0 | 0 | 1 |

| | | | | | | | | | | | |
|---|---|---|---|---|---|---|---|---|---|---|---|
| rs4938309 | 11 | 116627055 | T | C | -0.857 | 0.3912 | -+ | 0 | 0.608 | 1 | 0.4355 |
| rs76034797 | 11 | 120347586 | T | C | -0.857 | 0.3912 | -? | 0 | 0 | 0 | 1 |
| rs4340086 | 11 | 117775847 | T | A | -0.857 | 0.3915 | -? | 0 | 0 | 0 | 1 |
| rs4938439 | 11 | 117639526 | C | T | -0.857 | 0.3915 | -? | 0 | 0 | 0 | 1 |
| rs12274926 | 11 | 118295117 | C | T | 0.857 | 0.3916 | ++ | 0 | 0.014 | 1 | 0.9047 |
| rs73562872 | 11 | 119002768 | T | G | -0.857 | 0.3917 | -? | 0 | 0 | 0 | 1 |
| rs73562877 | 11 | 119010392 | A | G | -0.857 | 0.3917 | -? | 0 | 0 | 0 | 1 |
| rs498314 | 11 | 117722487 | A | G | 0.856 | 0.392 | +? | 0 | 0 | 0 | 1 |
| rs79530554 | 11 | 120627270 | T | C | 0.856 | 0.392 | +? | 0 | 0 | 0 | 1 |
| rs656199 | 11 | 117954309 | A | G | -0.856 | 0.3921 | -- | 0 | 0.132 | 1 | 0.7162 |
| rs77699630 | 11 | 117578142 | C | T | 0.855 | 0.3925 | +? | 0 | 0 | 0 | 1 |
| rs638273 | 11 | 117339926 | A | G | 0.855 | 0.3926 | +? | 0 | 0 | 0 | 1 |
| rs1940804 | 11 | 119237217 | C | T | 0.855 | 0.3927 | +? | 0 | 0 | 0 | 1 |
| rs4936601 | 11 | 121133909 | T | G | -0.855 | 0.3927 | -- | 0 | 0.52 | 1 | 0.4711 |
| rs9943623 | 11 | 121129528 | A | C | -0.855 | 0.3927 | -- | 0 | 0.503 | 1 | 0.4783 |
| rs4310650 | 11 | 116998879 | T | C | 0.855 | 0.3928 | +? | 0 | 0 | 0 | 1 |
| rs4394850 | 11 | 119504257 | C | G | 0.854 | 0.3929 | +? | 0 | 0 | 0 | 1 |
| rs895652 | 11 | 119671453 | A | G | 0.854 | 0.393 | +? | 0 | 0 | 0 | 1 |
| rs12804723 | 11 | 117425313 | T | A | -0.854 | 0.3931 | -? | 0 | 0 | 0 | 1 |
| rs57259536 | 11 | 116754753 | T | A | 0.854 | 0.3931 | +? | 0 | 0 | 0 | 1 |
| rs113684204 | 11 | 119965635 | A | T | -0.854 | 0.3932 | -? | 0 | 0 | 0 | 1 |
| rs7479050 | 11 | 120154801 | C | T | -0.854 | 0.3932 | -? | 0 | 0 | 0 | 1 |
| rs9666820 | 11 | 117327648 | C | T | 0.854 | 0.3932 | +? | 0 | 0 | 0 | 1 |
| rs10790370 | 11 | 120078393 | A | G | -0.853 | 0.3936 | -? | 0 | 0 | 0 | 1 |
| rs12272258 | 11 | 117926158 | A | G | -0.853 | 0.3936 | -? | 0 | 0 | 0 | 1 |
| rs61899015 | 11 | 119764214 | C | A | -0.853 | 0.3938 | -? | 0 | 0 | 0 | 1 |
| rs11217083 | 11 | 118770321 | G | A | -0.852 | 0.3939 | -- | 0 | 0.427 | 1 | 0.5136 |
| rs1793141 | 11 | 118108003 | G | A | -0.853 | 0.3939 | -? | 0 | 0 | 0 | 1 |
| rs2508448 | 11 | 117880688 | T | G | 0.853 | 0.3939 | +- | 0 | 0.483 | 1 | 0.487 |
| rs180327 | 11 | 116623659 | C | T | -0.852 | 0.3941 | -- | 0 | 0.027 | 1 | 0.8703 |
| rs681133 | 11 | 119798934 | T | C | 0.852 | 0.3941 | +? | 0 | 0 | 0 | 1 |
| rs536256 | 11 | 118327321 | G | A | 0.852 | 0.3943 | ++ | 0 | 0.023 | 1 | 0.8792 |
| rs4938379 | 11 | 117322782 | T | C | 0.851 | 0.3947 | +? | 0 | 0 | 0 | 1 |
| rs534845 | 11 | 116311210 | A | G | -0.851 | 0.3948 | -? | 0 | 0 | 0 | 1 |
| rs76871374 | 11 | 120699050 | G | A | 0.85 | 0.3951 | +? | 0 | 0 | 0 | 1 |
| rs12791310 | 11 | 119951884 | G | A | -0.85 | 0.3952 | -? | 0 | 0 | 0 | 1 |
| rs78097928 | 11 | 119847840 | A | G | -0.85 | 0.3952 | -? | 0 | 0 | 0 | 1 |
| rs116859831 | 11 | 116513428 | C | T | -0.85 | 0.3953 | -? | 0 | 0 | 0 | 1 |
| rs1784125 | 11 | 116537962 | C | G | -0.85 | 0.3956 | -? | 0 | 0 | 0 | 1 |
| rs7948928 | 11 | 120273224 | G | C | -0.849 | 0.3957 | -? | 0 | 0 | 0 | 1 |
| rs11217234 | 11 | 119177938 | G | A | 0.849 | 0.3959 | ++ | 0 | 0.193 | 1 | 0.6607 |
| rs7946869 | 11 | 116963312 | T | C | -0.849 | 0.3959 | -? | 0 | 0 | 0 | 1 |
| rs4259862 | 11 | 117775760 | C | T | -0.849 | 0.396 | -? | 0 | 0 | 0 | 1 |
| rs45439091 | 11 | 119054208 | T | G | -0.849 | 0.396 | -? | 0 | 0 | 0 | 1 |
| rs7128327 | 11 | 119631281 | G | A | 0.849 | 0.396 | ++ | 0 | 0.06 | 1 | 0.8059 |
| rs61899012 | 11 | 119756282 | C | G | -0.848 | 0.3962 | -? | 0 | 0 | 0 | 1 |
| rs143648302 | 11 | 117397094 | T | C | 0.848 | 0.3963 | +? | 0 | 0 | 0 | 1 |
| rs518684 | 11 | 116329917 | A | C | -0.848 | 0.3963 | -? | 0 | 0 | 0 | 1 |
| rs948415 | 11 | 119217455 | C | T | 0.848 | 0.3963 | +? | 0 | 0 | 0 | 1 |
| rs1009363 | 11 | 119505032 | G | A | 0.848 | 0.3965 | +? | 0 | 0 | 0 | 1 |
| rs116727415 | 11 | 120650530 | T | A | 0.848 | 0.3965 | +? | 0 | 0 | 0 | 1 |
| rs184841209 | 11 | 118466646 | T | G | 0.848 | 0.3965 | +- | -31.8 | 2.276 | 1 | 0.1314 |
| rs4935748 | 11 | 120518871 | A | G | -0.848 | 0.3965 | -? | 0 | 0 | 0 | 1 |
| rs7941534 | 11 | 116400382 | G | A | -0.848 | 0.3965 | -? | 0 | 0 | 0 | 1 |
| rs112186392 | 11 | 119622367 | G | A | 0.848 | 0.3966 | ++ | 0 | 0.06 | 1 | 0.8064 |
| rs948772 | 11 | 117590911 | C | G | 0.847 | 0.3969 | +? | 0 | 0 | 0 | 1 |

| | | | | | | | | | | | |
|---|---|---|---|---|---|---|---|---|---|---|---|
| rs10750111 | 11 | 117631234 | C | T | 0.847 | 0.397 | +? | 0 | 0 | 0 | 1 |
| rs10892167 | 11 | 117619027 | C | A | 0.847 | 0.397 | +? | 0 | 0 | 0 | 1 |
| rs73011741 | 11 | 117890685 | G | A | -0.847 | 0.397 | -- | 0 | 0.219 | 1 | 0.6399 |
| rs11601133 | 11 | 117424953 | G | A | -0.847 | 0.3971 | -? | 0 | 0 | 0 | 1 |
| rs1793146 | 11 | 118113045 | T | A | -0.846 | 0.3973 | -? | 0 | 0 | 0 | 1 |
| rs78764865 | 11 | 119832901 | T | C | -0.846 | 0.3973 | -? | 0 | 0 | 0 | 1 |
| rs4936399 | 11 | 117591292 | T | C | 0.846 | 0.3977 | +? | 0 | 0 | 0 | 1 |
| rs10892373 | 11 | 119368120 | T | G | -0.846 | 0.3978 | -? | 0 | 0 | 0 | 1 |
| rs490458 | 11 | 117436190 | T | C | 0.844 | 0.3986 | +? | 0 | 0 | 0 | 1 |
| rs116979369 | 11 | 117502942 | T | C | 0.844 | 0.3987 | +? | 0 | 0 | 0 | 1 |
| rs1715453 | 11 | 118125077 | T | C | -0.844 | 0.3988 | -? | 0 | 0 | 0 | 1 |
| rs1941303 | 11 | 121132347 | C | G | -0.844 | 0.3988 | -- | 0 | 0.451 | 1 | 0.5019 |
| rs73011555 | 11 | 120912139 | G | A | -0.844 | 0.3988 | -? | 0 | 0 | 0 | 1 |
| rs10892563 | 11 | 120224544 | C | T | -0.844 | 0.3989 | -? | 0 | 0 | 0 | 1 |
| rs74860176 | 11 | 117484221 | T | C | 0.843 | 0.399 | +? | 0 | 0 | 0 | 1 |
| rs74941224 | 11 | 117483626 | T | C | 0.843 | 0.399 | +? | 0 | 0 | 0 | 1 |
| rs75500945 | 11 | 117483022 | T | C | 0.843 | 0.399 | +? | 0 | 0 | 0 | 1 |
| rs114461031 | 11 | 119932617 | T | G | 0.843 | 0.3992 | +? | 0 | 0 | 0 | 1 |
| rs492471 | 11 | 118689745 | T | C | -0.843 | 0.3992 | -+ | -81.3 | 1.655 | 1 | 0.1983 |
| rs4938289 | 11 | 116458785 | T | G | -0.843 | 0.3993 | -? | 0 | 0 | 0 | 1 |
| rs2853006 | 11 | 118110977 | T | C | -0.843 | 0.3995 | -? | 0 | 0 | 0 | 1 |
| rs76517460 | 11 | 116500411 | A | G | -0.843 | 0.3995 | -? | 0 | 0 | 0 | 1 |
| rs12280323 | 11 | 121137448 | G | A | -0.842 | 0.3997 | -- | 0 | 0.528 | 1 | 0.4674 |
| rs11217679 | 11 | 119951678 | G | A | 0.842 | 0.3998 | +? | 0 | 0 | 0 | 1 |
| rs694574 | 11 | 117145053 | A | C | 0.842 | 0.3998 | +? | 0 | 0 | 0 | 1 |
| rs9971567 | 11 | 121147950 | G | A | -0.841 | 0.4002 | -- | 0 | 0.387 | 1 | 0.5336 |
| rs4938637 | 11 | 119075323 | A | G | -0.841 | 0.4003 | -? | 0 | 0 | 0 | 1 |
| rs61897958 | 11 | 119965045 | G | T | -0.841 | 0.4003 | -? | 0 | 0 | 0 | 1 |
| rs6589851 | 11 | 120835068 | T | C | 0.841 | 0.4004 | +? | 0 | 0 | 0 | 1 |
| rs7933086 | 11 | 120269308 | A | G | -0.841 | 0.4005 | -? | 0 | 0 | 0 | 1 |
| rs1056562 | 11 | 118125625 | T | C | -0.84 | 0.401 | -? | 0 | 0 | 0 | 1 |
| rs11217490 | 11 | 119645757 | A | G | 0.84 | 0.401 | ++ | 0 | 0.026 | 1 | 0.8722 |
| rs145069523 | 11 | 119844325 | G | A | -0.839 | 0.4012 | -? | 0 | 0 | 0 | 1 |
| rs1652025 | 11 | 119360083 | G | C | 0.839 | 0.4014 | +? | 0 | 0 | 0 | 1 |
| rs4604899 | 11 | 121127540 | C | T | -0.839 | 0.4014 | -- | 0 | 0.535 | 1 | 0.4645 |
| rs613808 | 11 | 116710968 | A | G | 0.839 | 0.4016 | +? | 0 | 0 | 0 | 1 |
| rs3759004 | 11 | 118136836 | T | C | 0.839 | 0.4017 | +? | 0 | 0 | 0 | 1 |
| rs75067211 | 11 | 118854372 | T | C | -0.839 | 0.4017 | -- | 0 | 0.135 | 1 | 0.7135 |
| rs17120841 | 11 | 117407435 | G | A | 0.838 | 0.4018 | +? | 0 | 0 | 0 | 1 |
| rs74735117 | 11 | 117409971 | C | T | 0.838 | 0.4018 | +? | 0 | 0 | 0 | 1 |
| rs2282537 | 11 | 120187971 | A | G | -0.838 | 0.4019 | -? | 0 | 0 | 0 | 1 |
| rs4936400 | 11 | 117592546 | A | G | 0.838 | 0.4019 | +? | 0 | 0 | 0 | 1 |
| rs1899052 | 11 | 116224964 | G | C | -0.838 | 0.402 | -? | 0 | 0 | 0 | 1 |
| rs500389 | 11 | 117100594 | G | A | 0.838 | 0.4021 | +? | 0 | 0 | 0 | 1 |
| rs79862998 | 11 | 116525319 | A | G | -0.838 | 0.4023 | -? | 0 | 0 | 0 | 1 |
| rs4935765 | 11 | 121134781 | C | T | -0.837 | 0.4025 | -- | 0 | 0.413 | 1 | 0.5206 |
| rs4936602 | 11 | 121134532 | A | G | -0.837 | 0.4025 | -- | 0 | 0.413 | 1 | 0.5206 |
| rs4938378 | 11 | 117322357 | C | T | 0.837 | 0.4025 | +? | 0 | 0 | 0 | 1 |
| rs7113731 | 11 | 118831824 | G | A | -0.837 | 0.4025 | -? | 0 | 0 | 0 | 1 |
| rs41297897 | 11 | 120838141 | A | G | -0.837 | 0.4026 | -? | 0 | 0 | 0 | 1 |
| rs11218226 | 11 | 121130758 | C | T | -0.837 | 0.4027 | -- | 0 | 0.525 | 1 | 0.4687 |
| rs59156491 | 11 | 117287082 | T | C | 0.837 | 0.4027 | +? | 0 | 0 | 0 | 1 |
| rs4936379 | 11 | 117322454 | A | C | 0.837 | 0.4028 | +? | 0 | 0 | 0 | 1 |
| rs76651078 | 11 | 118316006 | A | G | -0.836 | 0.4029 | -+ | -100.4 | 1.497 | 1 | 0.2211 |
| rs499272 | 11 | 116485496 | G | C | -0.836 | 0.403 | -? | 0 | 0 | 0 | 1 |
| rs12286677 | 11 | 117424807 | T | C | -0.836 | 0.4031 | -? | 0 | 0 | 0 | 1 |

| | | | | | | | | | | | |
|---|---|---|---|---|---|---|---|---|---|---|---|
| rs11215943 | 11 | 116331172 | T | G | 0.836 | 0.4032 | +? | 0 | 0 | 0 | 1 |
| rs12799960 | 11 | 117425206 | T | C | -0.836 | 0.4032 | -? | 0 | 0 | 0 | 1 |
| rs115767079 | 11 | 120128088 | T | C | -0.836 | 0.4034 | -? | 0 | 0 | 0 | 1 |
| rs11217462 | 11 | 119622613 | G | A | 0.835 | 0.4035 | ++ | 0 | 0.057 | 1 | 0.8119 |
| rs11607829 | 11 | 119504481 | C | G | 0.835 | 0.4036 | +? | 0 | 0 | 0 | 1 |
| rs10892700 | 11 | 121132675 | T | C | -0.835 | 0.4037 | -- | 0 | 0.458 | 1 | 0.4987 |
| rs4936604 | 11 | 121134903 | T | C | -0.835 | 0.4038 | -- | 0 | 0.409 | 1 | 0.5225 |
| rs679022 | 11 | 117125216 | C | T | 0.834 | 0.404 | +? | 0 | 0 | 0 | 1 |
| rs4635088 | 11 | 118832924 | C | G | -0.834 | 0.4042 | -? | 0 | 0 | 0 | 1 |
| rs9633947 | 11 | 119644360 | A | G | 0.834 | 0.4042 | ++ | 0 | 0.025 | 1 | 0.8747 |
| rs3781816 | 11 | 120813959 | A | G | 0.834 | 0.4043 | +? | 0 | 0 | 0 | 1 |
| rs10892708 | 11 | 121136763 | G | C | -0.834 | 0.4046 | -- | 0 | 0.582 | 1 | 0.4457 |
| rs11218231 | 11 | 121134434 | G | A | -0.834 | 0.4046 | -- | 0 | 0.406 | 1 | 0.5242 |
| rs10892651 | 11 | 120817653 | A | G | 0.833 | 0.4048 | +? | 0 | 0 | 0 | 1 |
| rs11607245 | 11 | 118843448 | T | C | -0.833 | 0.4049 | -? | 0 | 0 | 0 | 1 |
| rs574223 | 11 | 116329043 | G | T | -0.833 | 0.4051 | -? | 0 | 0 | 0 | 1 |
| rs581580 | 11 | 117538453 | G | A | -0.833 | 0.4051 | -? | 0 | 0 | 0 | 1 |
| rs113447930 | 11 | 120696251 | C | T | 0.832 | 0.4052 | +? | 0 | 0 | 0 | 1 |
| rs73003797 | 11 | 116229491 | C | T | -0.832 | 0.4052 | -- | 0 | 0.139 | 1 | 0.7091 |
| rs12279572 | 11 | 117952960 | G | A | -0.832 | 0.4053 | -- | 0 | 0.145 | 1 | 0.7034 |
| rs949050 | 11 | 119885204 | T | C | 0.832 | 0.4054 | +? | 0 | 0 | 0 | 1 |
| rs3781981 | 11 | 117256436 | C | T | -0.832 | 0.4056 | -? | 0 | 0 | 0 | 1 |
| rs4938765 | 11 | 119943485 | T | G | -0.832 | 0.4056 | -? | 0 | 0 | 0 | 1 |
| rs6589731 | 11 | 119185133 | C | T | 0.832 | 0.4056 | +? | 0 | 0 | 0 | 1 |
| rs12800734 | 11 | 120836754 | G | A | 0.831 | 0.4057 | +? | 0 | 0 | 0 | 1 |
| rs145993550 | 11 | 119499747 | A | G | 0.831 | 0.4058 | +? | 0 | 0 | 0 | 1 |
| rs1583126 | 11 | 121135578 | T | G | -0.831 | 0.4059 | -- | 0 | 0.412 | 1 | 0.5211 |
| rs7113635 | 11 | 117434146 | G | A | -0.831 | 0.4059 | -- | 0 | 0.187 | 1 | 0.6657 |
| rs4936603 | 11 | 121134854 | A | C | -0.831 | 0.406 | -- | 0 | 0.412 | 1 | 0.521 |
| rs2508948 | 11 | 118951642 | A | T | 0.831 | 0.4061 | +? | 0 | 0 | 0 | 1 |
| rs4936607 | 11 | 121135897 | G | A | -0.831 | 0.4061 | -- | 0 | 0.411 | 1 | 0.5215 |
| rs57216066 | 11 | 118861849 | A | G | 0.831 | 0.4061 | +? | 0 | 0 | 0 | 1 |
| rs10892702 | 11 | 121135196 | A | T | -0.83 | 0.4063 | -- | 0 | 0.411 | 1 | 0.5214 |
| rs10892703 | 11 | 121135199 | A | T | -0.83 | 0.4063 | -- | 0 | 0.411 | 1 | 0.5214 |
| rs10892704 | 11 | 121135240 | T | C | -0.83 | 0.4063 | -- | 0 | 0.411 | 1 | 0.5214 |
| rs10892705 | 11 | 121135286 | T | A | -0.83 | 0.4063 | -- | 0 | 0.411 | 1 | 0.5214 |
| rs12787592 | 11 | 116537974 | C | G | -0.83 | 0.4063 | -? | 0 | 0 | 0 | 1 |
| rs1583125 | 11 | 121135479 | A | C | -0.83 | 0.4063 | -- | 0 | 0.411 | 1 | 0.5214 |
| rs535327 | 11 | 118341785 | G | A | 0.831 | 0.4063 | +- | 0 | 0.727 | 1 | 0.394 |
| rs7929336 | 11 | 119961624 | C | G | -0.83 | 0.4063 | -? | 0 | 0 | 0 | 1 |
| rs113271112 | 11 | 118860598 | C | A | 0.83 | 0.4065 | +? | 0 | 0 | 0 | 1 |
| rs117470649 | 11 | 117493342 | G | A | 0.83 | 0.4065 | +? | 0 | 0 | 0 | 1 |
| rs74335840 | 11 | 117492693 | A | T | 0.83 | 0.4065 | +? | 0 | 0 | 0 | 1 |
| rs78505713 | 11 | 118860457 | T | C | 0.83 | 0.4065 | +? | 0 | 0 | 0 | 1 |
| rs117580094 | 11 | 117317855 | G | T | -0.83 | 0.4066 | -? | 0 | 0 | 0 | 1 |
| rs4936598 | 11 | 121129841 | A | G | -0.83 | 0.4066 | -- | 0 | 0.523 | 1 | 0.4695 |
| rs73020440 | 11 | 117750926 | T | C | -0.829 | 0.407 | -? | 0 | 0 | 0 | 1 |
| rs7942136 | 11 | 117760895 | C | T | -0.829 | 0.4071 | -? | 0 | 0 | 0 | 1 |
| rs499434 | 11 | 120086918 | T | C | 0.829 | 0.4073 | +? | 0 | 0 | 0 | 1 |
| rs11603274 | 11 | 120163930 | A | G | -0.828 | 0.4075 | -- | 0 | 0.09 | 1 | 0.7639 |
| rs11607289 | 11 | 120163574 | G | A | -0.828 | 0.4075 | -- | 0 | 0.09 | 1 | 0.7639 |
| rs651232 | 11 | 119764295 | T | G | -0.828 | 0.4076 | -? | 0 | 0 | 0 | 1 |
| rs11602393 | 11 | 118750967 | A | T | 0.828 | 0.4079 | +- | 27.4 | 4.134 | 1 | 0.04204 |
| rs10892570 | 11 | 120274501 | T | G | -0.827 | 0.4081 | -? | 0 | 0 | 0 | 1 |
| rs3132778 | 11 | 120611793 | C | T | -0.827 | 0.4083 | -? | 0 | 0 | 0 | 1 |
| rs76619306 | 11 | 119002868 | T | C | -0.827 | 0.4083 | -? | 0 | 0 | 0 | 1 |

| | | | | | | | | | | | |
|---|---|---|---|---|---|---|---|---|---|---|---|
| rs4936605 | 11 | 121135037 | G | A | -0.827 | 0.4084 | -- | 0 | 0.415 | 1 | 0.5194 |
| rs1573559 | 11 | 118104627 | G | A | -0.827 | 0.4085 | -? | 0 | 0 | 0 | 1 |
| rs111064597 | 11 | 116566546 | C | T | -0.826 | 0.4086 | -? | 0 | 0 | 0 | 1 |
| rs111774429 | 11 | 117335160 | T | C | 0.826 | 0.4086 | +? | 0 | 0 | 0 | 1 |
| rs4936538 | 11 | 120530316 | T | C | -0.826 | 0.4086 | -? | 0 | 0 | 0 | 1 |
| rs1583127 | 11 | 121135791 | G | A | -0.826 | 0.4087 | -- | 0 | 0.414 | 1 | 0.5199 |
| rs627291 | 11 | 118061767 | G | A | 0.826 | 0.4088 | +? | 0 | 0 | 0 | 1 |
| rs74408867 | 11 | 119558815 | C | T | -0.826 | 0.4088 | -? | 0 | 0 | 0 | 1 |
| rs114513135 | 11 | 120129129 | C | A | -0.826 | 0.4089 | -? | 0 | 0 | 0 | 1 |
| rs116667143 | 11 | 120128828 | G | C | -0.826 | 0.4089 | -? | 0 | 0 | 0 | 1 |
| rs17123747 | 11 | 120128451 | T | C | -0.826 | 0.4089 | -? | 0 | 0 | 0 | 1 |
| rs536613 | 11 | 119782588 | G | A | 0.826 | 0.4089 | +? | 0 | 0 | 0 | 1 |
| rs78314944 | 11 | 120128668 | C | G | -0.826 | 0.4089 | -? | 0 | 0 | 0 | 1 |
| rs575623 | 11 | 116278812 | C | T | 0.826 | 0.409 | +? | 0 | 0 | 0 | 1 |
| rs150628867 | 11 | 119466907 | A | G | -0.825 | 0.4091 | -? | 0 | 0 | 0 | 1 |
| rs77096386 | 11 | 117129232 | G | A | -0.825 | 0.4091 | -? | 0 | 0 | 0 | 1 |
| rs45510391 | 11 | 118936410 | A | G | -0.825 | 0.4092 | -? | 0 | 0 | 0 | 1 |
| rs61897959 | 11 | 119967825 | A | G | -0.825 | 0.4092 | -? | 0 | 0 | 0 | 1 |
| rs1941411 | 11 | 120130512 | G | A | -0.825 | 0.4094 | -? | 0 | 0 | 0 | 1 |
| rs12416965 | 11 | 121130464 | C | T | -0.825 | 0.4096 | -- | 0 | 0.51 | 1 | 0.4753 |
| rs35255208 | 11 | 121130611 | G | A | -0.825 | 0.4096 | -- | 0 | 0.51 | 1 | 0.4753 |
| rs4938497 | 11 | 118160851 | A | G | 0.824 | 0.4098 | +? | 0 | 0 | 0 | 1 |
| rs1793149 | 11 | 118127347 | A | G | -0.823 | 0.4104 | -? | 0 | 0 | 0 | 1 |
| rs4148170 | 11 | 119027545 | A | G | -0.823 | 0.4105 | -? | 0 | 0 | 0 | 1 |
| rs4938805 | 11 | 120274211 | G | A | -0.823 | 0.4106 | -? | 0 | 0 | 0 | 1 |
| rs11603307 | 11 | 120164141 | A | G | -0.822 | 0.4109 | -- | 0 | 0.088 | 1 | 0.7664 |
| rs1893317 | 11 | 117960427 | A | G | -0.822 | 0.411 | -- | 0 | 0.011 | 1 | 0.9168 |
| rs503425 | 11 | 118692727 | C | T | -0.822 | 0.411 | -+ | 0 | 0.851 | 1 | 0.3564 |
| rs947890 | 11 | 117874493 | A | G | 0.822 | 0.4112 | +- | 0 | 0.591 | 1 | 0.4422 |
| rs11217108 | 11 | 118841761 | T | C | -0.821 | 0.4114 | -? | 0 | 0 | 0 | 1 |
| rs11606461 | 11 | 120593745 | T | C | 0.821 | 0.4114 | +? | 0 | 0 | 0 | 1 |
| rs149395240 | 11 | 116480033 | T | C | -0.821 | 0.4116 | -? | 0 | 0 | 0 | 1 |
| rs10892564 | 11 | 120224650 | G | A | -0.821 | 0.4117 | -? | 0 | 0 | 0 | 1 |
| rs955917 | 11 | 118016039 | A | G | -0.821 | 0.4117 | -? | 0 | 0 | 0 | 1 |
| rs2000872 | 11 | 120520364 | G | A | -0.821 | 0.4118 | -? | 0 | 0 | 0 | 1 |
| rs514635 | 11 | 119313929 | A | G | 0.82 | 0.412 | ++ | -46.5 | 2.048 | 1 | 0.1524 |
| rs2852232 | 11 | 120606290 | G | A | -0.82 | 0.4122 | -? | 0 | 0 | 0 | 1 |
| rs7112522 | 11 | 120630452 | G | A | 0.82 | 0.4122 | +? | 0 | 0 | 0 | 1 |
| rs2845939 | 11 | 117727963 | T | C | 0.82 | 0.4124 | +? | 0 | 0 | 0 | 1 |
| rs615001 | 11 | 119242277 | T | C | 0.82 | 0.4124 | +? | 0 | 0 | 0 | 1 |
| rs880812 | 11 | 117454157 | C | G | 0.82 | 0.4125 | +? | 0 | 0 | 0 | 1 |
| rs906828 | 11 | 119456168 | T | C | 0.82 | 0.4125 | +? | 0 | 0 | 0 | 1 |
| rs4938271 | 11 | 116279191 | C | T | 0.819 | 0.4126 | +? | 0 | 0 | 0 | 1 |
| rs73023041 | 11 | 116741840 | A | G | 0.819 | 0.4126 | +? | 0 | 0 | 0 | 1 |
| rs12805320 | 11 | 117425540 | T | C | -0.819 | 0.4127 | -? | 0 | 0 | 0 | 1 |
| rs2298480 | 11 | 119002439 | G | A | -0.819 | 0.4127 | -? | 0 | 0 | 0 | 1 |
| rs77852530 | 11 | 116859675 | C | A | -0.819 | 0.4127 | -? | 0 | 0 | 0 | 1 |
| rs476130 | 11 | 117727883 | T | C | 0.819 | 0.4128 | +? | 0 | 0 | 0 | 1 |
| rs113184379 | 11 | 120695351 | G | C | 0.819 | 0.413 | +? | 0 | 0 | 0 | 1 |
| rs1793150 | 11 | 118127314 | G | A | -0.819 | 0.413 | -? | 0 | 0 | 0 | 1 |
| rs1793151 | 11 | 118126920 | C | T | -0.819 | 0.413 | -? | 0 | 0 | 0 | 1 |
| rs10892718 | 11 | 121156870 | A | C | -0.818 | 0.4133 | -? | 0 | 0 | 0 | 1 |
| rs2512144 | 11 | 117862298 | A | G | 0.818 | 0.4134 | ++ | 0 | 0.172 | 1 | 0.6783 |
| rs2077620 | 11 | 117687412 | T | G | -0.817 | 0.4138 | -? | 0 | 0 | 0 | 1 |
| rs4412764 | 11 | 119507925 | A | G | 0.817 | 0.4138 | +? | 0 | 0 | 0 | 1 |
| rs2512150 | 11 | 117883208 | C | G | 0.817 | 0.4139 | +- | 0 | 0.455 | 1 | 0.4998 |

| | | | | | | | | | | | |
|---|---|---|---|---|---|---|---|---|---|---|---|
| rs12808266 | 11 | 119649903 | T | C | -0.817 | 0.4141 | -? | 0 | 0 | 0 | 1 |
| rs1787700 | 11 | 116564408 | T | C | -0.817 | 0.4141 | -? | 0 | 0 | 0 | 1 |
| rs524936 | 11 | 118341921 | A | G | 0.817 | 0.4141 | +- | 0 | 0.744 | 1 | 0.3884 |
| rs667944 | 11 | 117128271 | G | A | 0.817 | 0.4141 | +? | 0 | 0 | 0 | 1 |
| rs11603757 | 11 | 120164658 | T | G | -0.817 | 0.4142 | -- | 0 | 0.086 | 1 | 0.769 |
| rs4148171 | 11 | 119027830 | G | A | -0.816 | 0.4143 | -? | 0 | 0 | 0 | 1 |
| rs7108508 | 11 | 119026052 | A | G | -0.816 | 0.4143 | -? | 0 | 0 | 0 | 1 |
| rs78642155 | 11 | 116514092 | C | G | -0.816 | 0.4143 | -? | 0 | 0 | 0 | 1 |
| rs17245873 | 11 | 121066137 | A | T | -0.816 | 0.4144 | -? | 0 | 0 | 0 | 1 |
| rs659471 | 11 | 119822879 | G | A | 0.816 | 0.4145 | +? | 0 | 0 | 0 | 1 |
| rs12800456 | 11 | 120063441 | G | A | -0.816 | 0.4146 | -? | 0 | 0 | 0 | 1 |
| rs1942924 | 11 | 117554338 | T | G | 0.815 | 0.4148 | +? | 0 | 0 | 0 | 1 |
| rs3802870 | 11 | 117789242 | G | C | 0.815 | 0.4149 | +? | 0 | 0 | 0 | 1 |
| rs687740 | 11 | 117177603 | C | A | -0.815 | 0.4149 | -? | 0 | 0 | 0 | 1 |
| rs11217081 | 11 | 118769032 | G | A | -0.815 | 0.415 | -- | 0 | 0.292 | 1 | 0.5887 |
| rs4252261 | 11 | 117862575 | G | T | 0.815 | 0.4151 | +- | 0 | 0.233 | 1 | 0.6293 |
| rs4604900 | 11 | 121127696 | A | T | -0.815 | 0.4151 | -- | 0 | 0.486 | 1 | 0.4859 |
| rs7104543 | 11 | 120724121 | G | A | -0.815 | 0.4151 | -? | 0 | 0 | 0 | 1 |
| rs1715449 | 11 | 118128455 | G | A | -0.814 | 0.4154 | -? | 0 | 0 | 0 | 1 |
| rs1715450 | 11 | 118128445 | C | G | -0.814 | 0.4154 | -? | 0 | 0 | 0 | 1 |
| rs12292453 | 11 | 120554278 | T | C | 0.814 | 0.4158 | +? | 0 | 0 | 0 | 1 |
| rs625447 | 11 | 117661100 | C | T | 0.813 | 0.416 | +? | 0 | 0 | 0 | 1 |
| rs147710683 | 11 | 118121356 | T | C | 0.813 | 0.4163 | +- | -185.7 | 1.05 | 1 | 0.3055 |
| rs10750181 | 11 | 121136531 | T | C | -0.813 | 0.4164 | -- | 0 | 0.446 | 1 | 0.5043 |
| rs12797625 | 11 | 120897776 | A | G | -0.813 | 0.4164 | -? | 0 | 0 | 0 | 1 |
| rs3802885 | 11 | 119032689 | C | A | -0.813 | 0.4165 | -? | 0 | 0 | 0 | 1 |
| rs117619191 | 11 | 117248283 | C | T | -0.812 | 0.4167 | -? | 0 | 0 | 0 | 1 |
| rs12362596 | 11 | 118105473 | G | C | -0.812 | 0.4167 | -? | 0 | 0 | 0 | 1 |
| rs12800503 | 11 | 117426506 | T | G | -0.812 | 0.4168 | -? | 0 | 0 | 0 | 1 |
| rs7128320 | 11 | 120835246 | G | A | 0.812 | 0.4168 | +? | 0 | 0 | 0 | 1 |
| rs10892706 | 11 | 121136756 | A | T | -0.812 | 0.417 | -- | 0 | 0.537 | 1 | 0.4637 |
| rs11600065 | 11 | 117556714 | C | A | 0.812 | 0.417 | +? | 0 | 0 | 0 | 1 |
| rs151127828 | 11 | 118206121 | A | G | 0.812 | 0.4171 | ++ | 0 | 0.294 | 1 | 0.5878 |
| rs1893038 | 11 | 117617379 | C | T | 0.811 | 0.4172 | +? | 0 | 0 | 0 | 1 |
| rs2084291 | 11 | 119697231 | G | A | 0.811 | 0.4172 | +? | 0 | 0 | 0 | 1 |
| rs57210824 | 11 | 117842344 | T | A | -0.811 | 0.4173 | -- | 0 | 0.488 | 1 | 0.4846 |
| rs574603 | 11 | 117177741 | C | T | -0.811 | 0.4173 | -? | 0 | 0 | 0 | 1 |
| rs73010962 | 11 | 116221255 | G | A | -0.811 | 0.4173 | -- | 0 | 0.125 | 1 | 0.7236 |
| rs515756 | 11 | 117201165 | G | T | 0.811 | 0.4174 | +? | 0 | 0 | 0 | 1 |
| rs10892451 | 11 | 119628589 | T | G | -0.811 | 0.4175 | -? | 0 | 0 | 0 | 1 |
| rs4936347 | 11 | 116210119 | T | C | 0.811 | 0.4175 | +? | 0 | 0 | 0 | 1 |
| rs55831600 | 11 | 119817902 | A | G | 0.811 | 0.4176 | +? | 0 | 0 | 0 | 1 |
| rs35024210 | 11 | 120817865 | A | C | 0.81 | 0.4178 | +? | 0 | 0 | 0 | 1 |
| rs17311059 | 11 | 119694933 | A | T | -0.81 | 0.418 | -? | 0 | 0 | 0 | 1 |
| rs3890562 | 11 | 118837113 | G | A | -0.81 | 0.4181 | -? | 0 | 0 | 0 | 1 |
| rs11217451 | 11 | 119618503 | A | G | 0.809 | 0.4183 | ++ | 0 | 0.05 | 1 | 0.8234 |
| rs474981 | 11 | 117008881 | A | G | 0.809 | 0.4183 | +? | 0 | 0 | 0 | 1 |
| rs45507395 | 11 | 117155696 | T | G | -0.809 | 0.4185 | -? | 0 | 0 | 0 | 1 |
| rs11607141 | 11 | 117294022 | G | T | -0.809 | 0.4186 | -? | 0 | 0 | 0 | 1 |
| rs4505073 | 11 | 119613788 | G | C | 0.809 | 0.4186 | +? | 0 | 0 | 0 | 1 |
| rs117689643 | 11 | 116514329 | T | G | -0.808 | 0.4189 | -? | 0 | 0 | 0 | 1 |
| rs652455 | 11 | 117071449 | C | T | 0.808 | 0.4189 | +? | 0 | 0 | 0 | 1 |
| rs640483 | 11 | 117770505 | G | A | -0.808 | 0.419 | -? | 0 | 0 | 0 | 1 |
| rs10750180 | 11 | 121136349 | C | T | -0.808 | 0.4191 | -- | 0 | 0.362 | 1 | 0.5476 |
| rs10790418 | 11 | 121136428 | T | C | -0.808 | 0.4191 | -- | 0 | 0.362 | 1 | 0.5476 |
| rs10790420 | 11 | 121136458 | T | C | -0.808 | 0.4191 | -- | 0 | 0.362 | 1 | 0.5476 |

| | | | | | | | | | | | |
|---|---|---|---|---|---|---|---|---|---|---|---|
| rs10790421 | 11 | 121136529 | C | T | -0.808 | 0.4191 | -- | 0 | 0.362 | 1 | 0.5476 |
| rs4936608 | 11 | 121136069 | T | C | -0.808 | 0.4191 | -- | 0 | 0.362 | 1 | 0.5476 |
| rs145811313 | 11 | 117433042 | A | C | 0.808 | 0.4193 | +? | 0 | 0 | 0 | 1 |
| rs17123976 | 11 | 120396329 | C | T | 0.807 | 0.4196 | +? | 0 | 0 | 0 | 1 |
| rs116948654 | 11 | 116514319 | C | T | -0.807 | 0.4198 | -? | 0 | 0 | 0 | 1 |
| rs73578248 | 11 | 120154200 | T | C | 0.807 | 0.4198 | +? | 0 | 0 | 0 | 1 |
| rs45445895 | 11 | 117952091 | A | G | -0.807 | 0.4199 | -? | 0 | 0 | 0 | 1 |
| rs11216396 | 11 | 117319720 | G | A | 0.806 | 0.4201 | +? | 0 | 0 | 0 | 1 |
| rs12271856 | 11 | 118791532 | T | G | -0.806 | 0.4203 | -- | 0 | 0.002 | 1 | 0.9657 |
| rs1060211 | 11 | 117039797 | G | A | 0.806 | 0.4204 | +? | 0 | 0 | 0 | 1 |
| rs6589785 | 11 | 119878605 | T | G | -0.806 | 0.4205 | -? | 0 | 0 | 0 | 1 |
| rs679556 | 11 | 119242715 | C | G | 0.806 | 0.4205 | +? | 0 | 0 | 0 | 1 |
| rs4938779 | 11 | 119966884 | C | T | -0.805 | 0.4206 | -? | 0 | 0 | 0 | 1 |
| rs7101592 | 11 | 116668867 | A | T | -0.805 | 0.4207 | -- | 0 | 0.947 | 1 | 0.3304 |
| rs476469 | 11 | 117173560 | T | C | 0.805 | 0.421 | +? | 0 | 0 | 0 | 1 |
| rs74422681 | 11 | 118312870 | G | A | -0.805 | 0.421 | -+ | -25.8 | 2.385 | 1 | 0.1225 |
| rs4935767 | 11 | 121153312 | G | A | -0.805 | 0.4211 | -? | 0 | 0 | 0 | 1 |
| rs114667713 | 11 | 120128223 | C | T | -0.804 | 0.4213 | -? | 0 | 0 | 0 | 1 |
| rs10790425 | 11 | 121139395 | T | C | -0.804 | 0.4214 | -- | 0 | 0.615 | 1 | 0.4331 |
| rs3887652 | 11 | 118856582 | C | G | -0.804 | 0.4214 | -- | 0 | 0.124 | 1 | 0.7246 |
| rs894678 | 11 | 118172520 | A | G | 0.804 | 0.4214 | +? | 0 | 0 | 0 | 1 |
| rs141859855 | 11 | 117634408 | A | G | 0.804 | 0.4216 | +? | 0 | 0 | 0 | 1 |
| rs608811 | 11 | 117538059 | A | G | -0.804 | 0.4216 | -? | 0 | 0 | 0 | 1 |
| rs58248287 | 11 | 117476375 | G | A | 0.803 | 0.4217 | +? | 0 | 0 | 0 | 1 |
| rs7939570 | 11 | 121145962 | A | C | -0.803 | 0.4217 | -- | 0 | 0.579 | 1 | 0.4467 |
| rs7103203 | 11 | 119501720 | G | A | 0.803 | 0.4218 | +? | 0 | 0 | 0 | 1 |
| rs59258276 | 11 | 117488572 | C | T | 0.803 | 0.4219 | +? | 0 | 0 | 0 | 1 |
| rs7110616 | 11 | 116458051 | T | C | 0.803 | 0.422 | +? | 0 | 0 | 0 | 1 |
| rs10790423 | 11 | 121136688 | C | A | -0.803 | 0.4221 | -- | 0 | 0.351 | 1 | 0.5537 |
| rs10892556 | 11 | 120131591 | G | A | -0.803 | 0.4221 | -? | 0 | 0 | 0 | 1 |
| rs61906081 | 11 | 116537059 | A | G | -0.803 | 0.4221 | -? | 0 | 0 | 0 | 1 |
| rs7105077 | 11 | 119760867 | G | A | -0.803 | 0.4222 | -? | 0 | 0 | 0 | 1 |
| rs12286470 | 11 | 117688517 | C | G | -0.802 | 0.4223 | -? | 0 | 0 | 0 | 1 |
| rs582973 | 11 | 119320333 | T | C | 0.802 | 0.4224 | ++ | -68.4 | 1.781 | 1 | 0.182 |
| rs11601506 | 11 | 116215109 | T | G | -0.802 | 0.4225 | -- | -67.7 | 1.789 | 1 | 0.181 |
| rs3802872 | 11 | 117776526 | G | A | -0.802 | 0.4227 | -? | 0 | 0 | 0 | 1 |
| rs1981988 | 11 | 119365380 | C | G | -0.801 | 0.4229 | -? | 0 | 0 | 0 | 1 |
| rs2041967 | 11 | 116645149 | G | A | -0.801 | 0.4229 | -- | 0 | 0.855 | 1 | 0.355 |
| rs12225131 | 11 | 120090348 | G | A | 0.801 | 0.4231 | +? | 0 | 0 | 0 | 1 |
| rs144199381 | 11 | 117384372 | T | C | 0.801 | 0.4232 | +? | 0 | 0 | 0 | 1 |
| rs17121102 | 11 | 117579111 | C | G | 0.801 | 0.4232 | +? | 0 | 0 | 0 | 1 |
| rs10892375 | 11 | 119368188 | T | C | -0.801 | 0.4234 | -? | 0 | 0 | 0 | 1 |
| rs79914174 | 11 | 116375302 | C | T | 0.801 | 0.4234 | +? | 0 | 0 | 0 | 1 |
| rs34518191 | 11 | 119680745 | G | C | -0.8 | 0.4236 | -? | 0 | 0 | 0 | 1 |
| rs10892374 | 11 | 119368153 | G | A | -0.8 | 0.4238 | -? | 0 | 0 | 0 | 1 |
| rs117843905 | 11 | 120332795 | T | C | 0.8 | 0.4239 | +? | 0 | 0 | 0 | 1 |
| rs12270574 | 11 | 119799851 | T | A | 0.8 | 0.4239 | +? | 0 | 0 | 0 | 1 |
| rs492227 | 11 | 119799829 | T | C | 0.8 | 0.4239 | +? | 0 | 0 | 0 | 1 |
| rs572287 | 11 | 119798818 | A | T | 0.8 | 0.4239 | +? | 0 | 0 | 0 | 1 |
| rs553492 | 11 | 119817670 | A | C | 0.8 | 0.424 | +? | 0 | 0 | 0 | 1 |
| rs562015 | 11 | 119992149 | T | C | 0.8 | 0.424 | +? | 0 | 0 | 0 | 1 |
| rs78238602 | 11 | 116226195 | A | G | -0.8 | 0.424 | -? | 0 | 0 | 0 | 1 |
| rs4938330 | 11 | 116928514 | G | C | -0.799 | 0.4242 | -? | 0 | 0 | 0 | 1 |
| rs607491 | 11 | 119360054 | G | C | 0.799 | 0.4244 | +? | 0 | 0 | 0 | 1 |
| rs72645461 | 11 | 117213847 | G | A | 0.798 | 0.4246 | +? | 0 | 0 | 0 | 1 |
| rs11826841 | 11 | 117790794 | G | A | -0.798 | 0.4249 | -? | 0 | 0 | 0 | 1 |

| | | | | | | | | | | | |
|---|---|---|---|---|---|---|---|---|---|---|---|
| rs35933073 | 11 | 116537846 | A | C | -0.798 | 0.4249 | -? | 0 | 0 | 0 | 1 |
| rs1000315943 | 11 | 121162486 | A | T | -0.798 | 0.425 | -? | 0 | 0 | 0 | 1 |
| rs1787714 | 11 | 116560610 | A | G | 0.798 | 0.425 | +? | 0 | 0 | 0 | 1 |
| rs20493 | 11 | 119957516 | A | G | 0.798 | 0.4251 | +? | 0 | 0 | 0 | 1 |
| rs61899038 | 11 | 119817309 | A | T | 0.797 | 0.4254 | +? | 0 | 0 | 0 | 1 |
| rs80238987 | 11 | 119634544 | A | G | 0.797 | 0.4256 | +? | 0 | 0 | 0 | 1 |
| rs117216310 | 11 | 117457472 | A | C | -0.797 | 0.4257 | -? | 0 | 0 | 0 | 1 |
| rs11216474 | 11 | 117497778 | G | A | -0.796 | 0.4258 | -? | 0 | 0 | 0 | 1 |
| rs2276342 | 11 | 117375489 | C | G | 0.796 | 0.4258 | +? | 0 | 0 | 0 | 1 |
| rs73008621 | 11 | 119693592 | G | C | -0.796 | 0.4259 | -? | 0 | 0 | 0 | 1 |
| rs584994 | 11 | 117751994 | G | T | 0.796 | 0.426 | +? | 0 | 0 | 0 | 1 |
| rs76109832 | 11 | 118755946 | T | C | 0.796 | 0.426 | ++ | 0 | 0.075 | 1 | 0.7847 |
| rs549303 | 11 | 119362888 | C | T | 0.795 | 0.4264 | +? | 0 | 0 | 0 | 1 |
| rs141047910 | 11 | 117436510 | C | G | 0.795 | 0.4268 | +? | 0 | 0 | 0 | 1 |
| rs7112232 | 11 | 119320444 | A | G | 0.795 | 0.4268 | -+ | -43.5 | 2.09 | 1 | 0.1482 |
| rs573442 | 11 | 117177920 | T | C | -0.795 | 0.4269 | -? | 0 | 0 | 0 | 1 |
| rs7116918 | 11 | 117492654 | G | A | -0.794 | 0.427 | -? | 0 | 0 | 0 | 1 |
| rs11216646 | 11 | 117824158 | G | A | -0.794 | 0.4271 | -- | 0 | 0.43 | 1 | 0.512 |
| rs3782042 | 11 | 118185919 | A | G | 0.794 | 0.4272 | +? | 0 | 0 | 0 | 1 |
| rs507622 | 11 | 119764234 | A | G | -0.794 | 0.4272 | -? | 0 | 0 | 0 | 1 |
| rs10892455 | 11 | 119647079 | C | T | 0.794 | 0.4273 | ++ | 0 | 0.018 | 1 | 0.8929 |
| rs593380 | 11 | 117538766 | C | T | -0.794 | 0.4274 | -? | 0 | 0 | 0 | 1 |
| rs7939478 | 11 | 121145976 | C | A | -0.794 | 0.4274 | -- | 0 | 0.537 | 1 | 0.4637 |
| rs12793790 | 11 | 118140372 | A | G | 0.793 | 0.4275 | +? | 0 | 0 | 0 | 1 |
| rs11217467 | 11 | 119628785 | A | G | -0.793 | 0.4276 | -? | 0 | 0 | 0 | 1 |
| rs6589695 | 11 | 118644323 | C | G | 0.793 | 0.4276 | ++ | 0 | 0.134 | 1 | 0.7143 |
| rs141270315 | 11 | 119000331 | T | C | -0.793 | 0.4277 | -? | 0 | 0 | 0 | 1 |
| rs148543195 | 11 | 117488030 | T | C | 0.793 | 0.4277 | +? | 0 | 0 | 0 | 1 |
| rs3863318 | 11 | 117811167 | A | C | 0.793 | 0.4277 | ++ | -175.3 | 1.09 | 1 | 0.2965 |
| rs79487987 | 11 | 119006788 | A | G | -0.793 | 0.4277 | -? | 0 | 0 | 0 | 1 |
| rs7123597 | 11 | 117594443 | A | C | 0.792 | 0.4282 | +? | 0 | 0 | 0 | 1 |
| rs12808105 | 11 | 117637213 | T | C | 0.792 | 0.4283 | +? | 0 | 0 | 0 | 1 |
| rs1941298 | 11 | 121108865 | T | C | 0.792 | 0.4283 | +? | 0 | 0 | 0 | 1 |
| rs34986626 | 11 | 117494151 | A | G | -0.792 | 0.4284 | -? | 0 | 0 | 0 | 1 |
| rs476636 | 11 | 120326277 | G | A | 0.792 | 0.4285 | ++ | -117.1 | 1.382 | 1 | 0.2398 |
| rs7121512 | 11 | 116455707 | G | T | 0.792 | 0.4286 | +? | 0 | 0 | 0 | 1 |
| rs74883244 | 11 | 120200571 | G | A | -0.792 | 0.4286 | -? | 0 | 0 | 0 | 1 |
| rs10790195 | 11 | 117496132 | C | T | -0.791 | 0.4287 | -? | 0 | 0 | 0 | 1 |
| rs149703780 | 11 | 118993801 | A | G | -0.791 | 0.4288 | -? | 0 | 0 | 0 | 1 |
| rs571812 | 11 | 116246514 | T | C | -0.791 | 0.4288 | -? | 0 | 0 | 0 | 1 |
| rs624588 | 11 | 118995353 | C | G | 0.791 | 0.4289 | +? | 0 | 0 | 0 | 1 |
| rs10892351 | 11 | 119218530 | T | G | 0.791 | 0.429 | +? | 0 | 0 | 0 | 1 |
| rs2257206 | 11 | 120401656 | T | C | 0.791 | 0.429 | +? | 0 | 0 | 0 | 1 |
| rs7113362 | 11 | 116454261 | A | G | 0.791 | 0.4291 | +? | 0 | 0 | 0 | 1 |
| rs7128378 | 11 | 119989630 | A | G | -0.791 | 0.4291 | -? | 0 | 0 | 0 | 1 |
| rs7948174 | 11 | 121141005 | T | G | -0.791 | 0.4291 | -- | 0 | 0.553 | 1 | 0.4572 |
| rs10790152 | 11 | 116457650 | G | A | 0.79 | 0.4293 | +? | 0 | 0 | 0 | 1 |
| rs150339153 | 11 | 118285369 | T | C | 0.79 | 0.4294 | +- | 0 | 0.343 | 1 | 0.5579 |
| rs497768 | 11 | 117750740 | C | G | 0.79 | 0.4294 | +? | 0 | 0 | 0 | 1 |
| rs6589624 | 11 | 117717297 | G | A | 0.79 | 0.4295 | +? | 0 | 0 | 0 | 1 |
| rs7115714 | 11 | 120294861 | A | G | -0.79 | 0.4295 | -? | 0 | 0 | 0 | 1 |
| rs11216655 | 11 | 117836519 | T | C | -0.79 | 0.4296 | -- | 0 | 0.526 | 1 | 0.4684 |
| rs1423902581 | 11 | 116688765 | C | A | 0.79 | 0.4298 | +? | 0 | 0 | 0 | 1 |
| rs10790241 | 11 | 118013034 | A | G | -0.789 | 0.43 | -- | 0 | 0.676 | 1 | 0.4109 |
| rs73008622 | 11 | 119693648 | T | C | -0.789 | 0.43 | -? | 0 | 0 | 0 | 1 |
| rs10892110 | 11 | 117322917 | C | A | 0.789 | 0.4301 | +? | 0 | 0 | 0 | 1 |

| | | | | | | | | | | | |
|---|---|---|---|---|---|---|---|---|---|---|---|
| rs2846097 | 11 | 120649962 | C | G | 0.789 | 0.4301 | +? | 0 | 0 | 0 | 1 |
| rs10892442 | 11 | 119592454 | G | A | 0.789 | 0.4302 | +? | 0 | 0 | 0 | 1 |
| rs12800436 | 11 | 116558802 | T | G | -0.789 | 0.4303 | -? | 0 | 0 | 0 | 1 |
| rs7130716 | 11 | 119318306 | A | G | 0.789 | 0.4303 | -+ | -42.9 | 2.1 | 1 | 0.1473 |
| rs12795905 | 11 | 118140597 | T | C | 0.789 | 0.4304 | +? | 0 | 0 | 0 | 1 |
| rs111716733 | 11 | 120784745 | T | C | -0.788 | 0.4305 | -? | 0 | 0 | 0 | 1 |
| rs11216024 | 11 | 116461333 | G | A | 0.788 | 0.4306 | +? | 0 | 0 | 0 | 1 |
| rs11216433 | 11 | 117380103 | G | A | 0.788 | 0.4306 | +? | 0 | 0 | 0 | 1 |
| rs2070669 | 11 | 116701122 | C | G | 0.788 | 0.4306 | +? | 0 | 0 | 0 | 1 |
| rs73010439 | 11 | 117132315 | G | T | 0.788 | 0.4306 | +? | 0 | 0 | 0 | 1 |
| rs76860051 | 11 | 116525306 | C | A | -0.788 | 0.4306 | -? | 0 | 0 | 0 | 1 |
| rs118176355 | 11 | 117696521 | C | G | 0.788 | 0.4307 | +? | 0 | 0 | 0 | 1 |
| rs2727784 | 11 | 116711141 | C | T | 0.788 | 0.4307 | +? | 0 | 0 | 0 | 1 |
| rs7942226 | 11 | 117617279 | C | T | 0.788 | 0.4308 | +? | 0 | 0 | 0 | 1 |
| rs12420418 | 11 | 118840857 | T | C | -0.787 | 0.4313 | -? | 0 | 0 | 0 | 1 |
| rs2017122 | 11 | 118314849 | T | C | -0.787 | 0.4314 | -+ | -110 | 1.428 | 1 | 0.232 |
| rs11606047 | 11 | 119985246 | G | A | -0.787 | 0.4315 | -? | 0 | 0 | 0 | 1 |
| rs56410112 | 11 | 118193468 | C | T | 0.787 | 0.4315 | +? | 0 | 0 | 0 | 1 |
| rs7105861 | 11 | 119070910 | G | C | -0.787 | 0.4315 | -? | 0 | 0 | 0 | 1 |
| rs621150 | 11 | 119847991 | T | C | -0.786 | 0.4318 | -? | 0 | 0 | 0 | 1 |
| rs6589737 | 11 | 119295647 | C | T | 0.786 | 0.432 | -+ | -34.8 | 2.225 | 1 | 0.1358 |
| rs641382 | 11 | 119006300 | A | C | 0.786 | 0.4321 | +? | 0 | 0 | 0 | 1 |
| rs7126377 | 11 | 119223386 | A | G | 0.785 | 0.4324 | +? | 0 | 0 | 0 | 1 |
| rs1941297 | 11 | 121108679 | T | C | 0.785 | 0.4325 | +? | 0 | 0 | 0 | 1 |
| rs652824 | 11 | 119884981 | C | T | 0.785 | 0.4325 | +? | 0 | 0 | 0 | 1 |
| rs4938307 | 11 | 116604514 | A | G | -0.785 | 0.4326 | -+ | 0 | 0.586 | 1 | 0.444 |
| rs10790331 | 11 | 119552057 | C | A | 0.785 | 0.4327 | +? | 0 | 0 | 0 | 1 |
| rs12786234 | 11 | 117638032 | A | G | 0.785 | 0.4327 | +? | 0 | 0 | 0 | 1 |
| rs3132780 | 11 | 120620159 | G | A | -0.784 | 0.4328 | -? | 0 | 0 | 0 | 1 |
| rs1366878 | 11 | 119367405 | T | C | -0.784 | 0.4329 | -? | 0 | 0 | 0 | 1 |
| rs671111 | 11 | 117988539 | A | G | 0.784 | 0.4331 | ++ | 0 | 0.088 | 1 | 0.7669 |
| rs72995856 | 11 | 120647474 | C | T | 0.784 | 0.4331 | +? | 0 | 0 | 0 | 1 |
| rs75905572 | 11 | 119005900 | G | C | -0.784 | 0.4331 | -? | 0 | 0 | 0 | 1 |
| rs2305821 | 11 | 117153420 | C | T | -0.783 | 0.4334 | -? | 0 | 0 | 0 | 1 |
| rs491812 | 11 | 117328559 | T | C | 0.783 | 0.4334 | +? | 0 | 0 | 0 | 1 |
| rs1545509 | 11 | 119927888 | A | G | -0.783 | 0.4335 | -? | 0 | 0 | 0 | 1 |
| rs3740724 | 11 | 119859475 | T | C | -0.783 | 0.4335 | -? | 0 | 0 | 0 | 1 |
| rs4938633 | 11 | 119026328 | A | G | -0.783 | 0.4335 | -? | 0 | 0 | 0 | 1 |
| rs7110192 | 11 | 116457744 | G | C | 0.783 | 0.4335 | +? | 0 | 0 | 0 | 1 |
| rs17123780 | 11 | 120157863 | A | G | -0.783 | 0.4336 | -- | 0 | 0.075 | 1 | 0.7836 |
| rs1047857 | 11 | 117158858 | G | A | -0.782 | 0.4341 | -? | 0 | 0 | 0 | 1 |
| rs187164533 | 11 | 118709275 | A | G | -0.782 | 0.4341 | -+ | 0 | 0.755 | 1 | 0.3848 |
| rs12789276 | 11 | 118140203 | G | A | 0.782 | 0.4342 | +? | 0 | 0 | 0 | 1 |
| rs6589740 | 11 | 119311477 | T | C | 0.782 | 0.4342 | -+ | -34.4 | 2.232 | 1 | 0.1352 |
| rs9645661 | 11 | 116269032 | A | G | -0.782 | 0.4342 | -? | 0 | 0 | 0 | 1 |
| rs540716 | 11 | 118685015 | T | C | -0.781 | 0.4345 | -+ | 0 | 0.719 | 1 | 0.3964 |
| rs71480344 | 11 | 117442767 | G | A | -0.781 | 0.4346 | -? | 0 | 0 | 0 | 1 |
| rs2622935 | 11 | 117010758 | A | G | 0.781 | 0.4349 | +? | 0 | 0 | 0 | 1 |
| rs2068412 | 11 | 117574296 | G | A | 0.781 | 0.435 | +? | 0 | 0 | 0 | 1 |
| rs11217852 | 11 | 120261187 | G | A | -0.78 | 0.4351 | -? | 0 | 0 | 0 | 1 |
| rs61900905 | 11 | 118185492 | C | T | -0.78 | 0.4351 | -+ | 3 | 3.094 | 1 | 0.07861 |
| rs112552940 | 11 | 118509877 | T | C | -0.78 | 0.4352 | -+ | 0 | 0.859 | 1 | 0.3539 |
| rs11602073 | 11 | 116646858 | T | C | -0.78 | 0.4352 | -+ | 0 | 0.492 | 1 | 0.4832 |
| rs11217773 | 11 | 120115450 | A | G | -0.78 | 0.4354 | -? | 0 | 0 | 0 | 1 |
| rs12274203 | 11 | 117111913 | A | C | -0.78 | 0.4354 | -? | 0 | 0 | 0 | 1 |
| rs34786601 | 11 | 118140136 | C | T | 0.78 | 0.4354 | +? | 0 | 0 | 0 | 1 |

| | | | | | | | | | | | |
|---|---|---|---|---|---|---|---|---|---|---|---|
| rs11216187 | 11 | 116787148 | T | C | 0.78 | 0.4355 | +? | 0 | 0 | 0 | 1 |
| rs11608135 | 11 | 119941819 | A | G | -0.78 | 0.4355 | -? | 0 | 0 | 0 | 1 |
| rs12286404 | 11 | 117144355 | C | T | -0.78 | 0.4356 | -? | 0 | 0 | 0 | 1 |
| rs12292027 | 11 | 117149405 | G | A | -0.78 | 0.4356 | -? | 0 | 0 | 0 | 1 |
| rs75562690 | 11 | 117655971 | T | C | 0.78 | 0.4356 | ++ | 0 | 0.007 | 1 | 0.9332 |
| rs12789293 | 11 | 117532692 | T | C | 0.779 | 0.4359 | ++ | 0 | 0.008 | 1 | 0.9266 |
| rs1944497 | 11 | 119877773 | G | A | -0.779 | 0.4359 | -? | 0 | 0 | 0 | 1 |
| rs17570978 | 11 | 116457391 | T | C | 0.779 | 0.4361 | ++ | 0 | 0.003 | 1 | 0.9565 |
| rs10892228 | 11 | 118111346 | G | A | -0.779 | 0.4362 | -? | 0 | 0 | 0 | 1 |
| rs10750140 | 11 | 119323284 | T | C | 0.778 | 0.4363 | -+ | -41.7 | 2.117 | 1 | 0.1456 |
| rs59255055 | 11 | 118296735 | A | C | 0.778 | 0.4363 | ++ | 0 | 0.018 | 1 | 0.8932 |
| rs11216678 | 11 | 117883941 | G | C | -0.778 | 0.4364 | -- | 0 | 0.168 | 1 | 0.6816 |
| rs61900902 | 11 | 118159684 | C | T | 0.778 | 0.4365 | +? | 0 | 0 | 0 | 1 |
| rs4252254 | 11 | 117861055 | T | C | 0.778 | 0.4366 | +- | 0 | 0.308 | 1 | 0.5789 |
| rs7924975 | 11 | 120233564 | C | G | -0.778 | 0.4366 | -? | 0 | 0 | 0 | 1 |
| rs10790339 | 11 | 119594525 | C | T | 0.777 | 0.4369 | +? | 0 | 0 | 0 | 1 |
| rs11606588 | 11 | 117302038 | C | G | 0.777 | 0.4369 | +? | 0 | 0 | 0 | 1 |
| rs7952138 | 11 | 118953952 | G | A | -0.777 | 0.4369 | -? | 0 | 0 | 0 | 1 |
| rs11217768 | 11 | 120086680 | A | G | 0.777 | 0.437 | +? | 0 | 0 | 0 | 1 |
| rs1942478 | 11 | 116651463 | G | T | -0.777 | 0.4371 | -+ | 0 | 0.495 | 1 | 0.4816 |
| rs12364374 | 11 | 117849268 | A | G | 0.777 | 0.4372 | ++ | 0 | 0.437 | 1 | 0.5088 |
| rs1530919 | 11 | 119905908 | A | G | -0.777 | 0.4372 | -? | 0 | 0 | 0 | 1 |
| rs3824978 | 11 | 120732508 | T | C | -0.777 | 0.4373 | -? | 0 | 0 | 0 | 1 |
| rs11607103 | 11 | 117582279 | C | T | 0.777 | 0.4374 | +? | 0 | 0 | 0 | 1 |
| rs76200019 | 11 | 120701346 | A | G | 0.776 | 0.4376 | +? | 0 | 0 | 0 | 1 |
| rs644252 | 11 | 119043500 | T | G | -0.776 | 0.4378 | -? | 0 | 0 | 0 | 1 |
| rs59005463 | 11 | 121139928 | A | G | -0.776 | 0.4379 | -? | 0 | 0 | 0 | 1 |
| rs4938754 | 11 | 119877246 | T | C | -0.776 | 0.438 | -? | 0 | 0 | 0 | 1 |
| rs616458 | 11 | 119000044 | C | G | 0.776 | 0.438 | +? | 0 | 0 | 0 | 1 |
| rs1373270 | 11 | 120324080 | A | G | -0.775 | 0.4382 | -? | 0 | 0 | 0 | 1 |
| rs549289 | 11 | 117358220 | C | A | 0.775 | 0.4383 | +? | 0 | 0 | 0 | 1 |
| rs2155169 | 11 | 119857163 | G | A | -0.775 | 0.4384 | -? | 0 | 0 | 0 | 1 |
| rs4938634 | 11 | 119026330 | C | A | -0.775 | 0.4384 | -? | 0 | 0 | 0 | 1 |
| rs7125990 | 11 | 117105726 | G | A | -0.775 | 0.4384 | -? | 0 | 0 | 0 | 1 |
| rs140967993 | 11 | 118264353 | G | A | 0.775 | 0.4385 | +- | 3.4 | 3.105 | 1 | 0.07804 |
| rs73014325 | 11 | 117196435 | A | C | -0.775 | 0.4385 | -? | 0 | 0 | 0 | 1 |
| rs652872 | 11 | 118996781 | A | G | 0.774 | 0.4387 | +? | 0 | 0 | 0 | 1 |
| rs67422215 | 11 | 117776003 | T | C | 0.774 | 0.4387 | +? | 0 | 0 | 0 | 1 |
| rs11218041 | 11 | 120749953 | G | C | -0.774 | 0.4389 | -? | 0 | 0 | 0 | 1 |
| rs117745694 | 11 | 118218491 | A | G | 0.774 | 0.4389 | ++ | 0 | 0.008 | 1 | 0.9304 |
| rs11216135 | 11 | 116646062 | G | C | -0.774 | 0.4391 | -+ | 0 | 0.486 | 1 | 0.4856 |
| rs1892904 | 11 | 117117973 | G | A | -0.774 | 0.4392 | -? | 0 | 0 | 0 | 1 |
| rs79990075 | 11 | 118267686 | T | C | 0.773 | 0.4393 | +- | 2.1 | 3.064 | 1 | 0.08005 |
| rs61897956 | 11 | 119958757 | A | G | -0.773 | 0.4394 | -? | 0 | 0 | 0 | 1 |
| rs484553 | 11 | 117201097 | T | G | 0.773 | 0.4398 | +? | 0 | 0 | 0 | 1 |
| rs4938635 | 11 | 119026460 | T | C | -0.773 | 0.4398 | -? | 0 | 0 | 0 | 1 |
| rs2096884 | 11 | 120173354 | A | G | 0.772 | 0.4399 | +? | 0 | 0 | 0 | 1 |
| rs7126048 | 11 | 121140526 | G | A | -0.772 | 0.4399 | -- | 0 | 0.578 | 1 | 0.4471 |
| rs100803 | 11 | 119005132 | T | C | 0.772 | 0.4401 | +? | 0 | 0 | 0 | 1 |
| rs662069 | 11 | 119003322 | A | C | 0.772 | 0.4401 | +? | 0 | 0 | 0 | 1 |
| rs7116825 | 11 | 116454503 | A | C | 0.772 | 0.4401 | +? | 0 | 0 | 0 | 1 |
| rs7234 | 11 | 119005485 | T | C | 0.772 | 0.4401 | +? | 0 | 0 | 0 | 1 |
| rs880487 | 11 | 120063980 | C | T | -0.772 | 0.4401 | -? | 0 | 0 | 0 | 1 |
| rs4417316 | 11 | 116652301 | T | C | -0.772 | 0.4402 | -+ | 0 | 0.491 | 1 | 0.4835 |
| rs4938636 | 11 | 119026469 | T | A | -0.772 | 0.4402 | -? | 0 | 0 | 0 | 1 |
| rs59511712 | 11 | 116945762 | T | C | -0.772 | 0.4403 | -? | 0 | 0 | 0 | 1 |

| | | | | | | | | | | | |
|---|---|---|---|---|---|---|---|---|---|---|---|
| rs1838757 | 11 | 116785083 | G | C | 0.772 | 0.4404 | +? | 0 | 0 | 0 | 1 |
| rs74886004 | 11 | 116457471 | T | C | 0.771 | 0.4404 | ++ | 0 | 0.003 | 1 | 0.9532 |
| rs12787452 | 11 | 117769938 | G | C | -0.771 | 0.4405 | -? | 0 | 0 | 0 | 1 |
| rs2715883 | 11 | 120133494 | G | A | -0.771 | 0.4405 | -? | 0 | 0 | 0 | 1 |
| rs599343 | 11 | 117994430 | A | G | 0.771 | 0.4405 | ++ | 0 | 0.089 | 1 | 0.7658 |
| rs185433085 | 11 | 117106069 | T | C | -0.771 | 0.4406 | -? | 0 | 0 | 0 | 1 |
| rs7931316 | 11 | 119585681 | C | T | -0.771 | 0.4409 | -? | 0 | 0 | 0 | 1 |
| rs10892578 | 11 | 120308290 | G | A | -0.77 | 0.4411 | -? | 0 | 0 | 0 | 1 |
| rs117236359 | 11 | 119573655 | T | C | -0.77 | 0.4412 | -? | 0 | 0 | 0 | 1 |
| rs2512148 | 11 | 117873048 | A | C | 0.77 | 0.4412 | ++ | 0 | 0.011 | 1 | 0.9148 |
| rs948028 | 11 | 120644447 | C | A | 0.77 | 0.4413 | +? | 0 | 0 | 0 | 1 |
| rs116857027 | 11 | 116525995 | T | C | -0.77 | 0.4414 | -? | 0 | 0 | 0 | 1 |
| rs1784128 | 11 | 116538689 | A | G | 0.77 | 0.4414 | +? | 0 | 0 | 0 | 1 |
| rs10790189 | 11 | 117327216 | C | A | 0.77 | 0.4415 | +? | 0 | 0 | 0 | 1 |
| rs672189 | 11 | 119004009 | T | G | 0.769 | 0.4417 | +? | 0 | 0 | 0 | 1 |
| rs7941794 | 11 | 117499026 | A | C | 0.769 | 0.4417 | +? | 0 | 0 | 0 | 1 |
| rs11218073 | 11 | 120817900 | T | C | -0.769 | 0.442 | -+ | 0 | 0.659 | 1 | 0.4171 |
| rs113460600 | 11 | 119765177 | A | G | -0.769 | 0.4421 | -? | 0 | 0 | 0 | 1 |
| rs12275546 | 11 | 118109097 | G | A | -0.769 | 0.4421 | -? | 0 | 0 | 0 | 1 |
| rs7924968 | 11 | 117619538 | A | T | 0.769 | 0.4421 | +? | 0 | 0 | 0 | 1 |
| rs7952199 | 11 | 121142126 | G | C | -0.768 | 0.4423 | -- | 0 | 0.512 | 1 | 0.4745 |
| rs10790190 | 11 | 117361447 | G | A | 0.768 | 0.4424 | +? | 0 | 0 | 0 | 1 |
| rs2512143 | 11 | 117860582 | A | G | 0.768 | 0.4424 | ++ | 0 | 0.157 | 1 | 0.6917 |
| rs12420530 | 11 | 119907605 | T | G | -0.768 | 0.4426 | -? | 0 | 0 | 0 | 1 |
| rs4531505 | 11 | 116427570 | C | G | 0.768 | 0.4426 | ++ | 0 | 0.188 | 1 | 0.6645 |
| rs10502242 | 11 | 120641004 | G | A | 0.768 | 0.4427 | +? | 0 | 0 | 0 | 1 |
| rs17124301 | 11 | 120642698 | C | T | 0.768 | 0.4427 | +? | 0 | 0 | 0 | 1 |
| rs72995844 | 11 | 120642631 | G | A | 0.768 | 0.4427 | +? | 0 | 0 | 0 | 1 |
| rs7118414 | 11 | 117811548 | C | G | -0.767 | 0.443 | -? | 0 | 0 | 0 | 1 |
| rs79876834 | 11 | 117584155 | T | C | 0.767 | 0.443 | +? | 0 | 0 | 0 | 1 |
| rs11216777 | 11 | 117997722 | G | T | -0.767 | 0.4432 | -- | 0 | 0.455 | 1 | 0.5 |
| rs35963862 | 11 | 116549897 | C | T | -0.767 | 0.4432 | -? | 0 | 0 | 0 | 1 |
| rs895649 | 11 | 119751585 | G | A | 0.767 | 0.4432 | +? | 0 | 0 | 0 | 1 |
| rs693147 | 11 | 117266754 | A | C | 0.767 | 0.4433 | +? | 0 | 0 | 0 | 1 |
| rs7951250 | 11 | 120309036 | G | A | -0.766 | 0.4437 | -? | 0 | 0 | 0 | 1 |
| rs4938657 | 11 | 119314718 | C | T | 0.766 | 0.4438 | -+ | -32.8 | 2.259 | 1 | 0.1328 |
| rs1374563 | 11 | 119762569 | A | G | -0.766 | 0.4439 | -? | 0 | 0 | 0 | 1 |
| rs11606452 | 11 | 120498485 | C | G | 0.765 | 0.4441 | +? | 0 | 0 | 0 | 1 |
| rs7128900 | 11 | 116363446 | G | T | -0.765 | 0.4441 | -? | 0 | 0 | 0 | 1 |
| rs4936515 | 11 | 120208185 | G | A | 0.765 | 0.4442 | +? | 0 | 0 | 0 | 1 |
| rs75013797 | 11 | 117200129 | T | C | 0.765 | 0.4443 | +? | 0 | 0 | 0 | 1 |
| rs7927300 | 11 | 117791132 | A | C | -0.765 | 0.4445 | -? | 0 | 0 | 0 | 1 |
| rs10790427 | 11 | 121139585 | G | A | -0.764 | 0.4448 | -- | 0 | 0.695 | 1 | 0.4044 |
| rs537153 | 11 | 119004026 | T | C | 0.764 | 0.4448 | +? | 0 | 0 | 0 | 1 |
| rs17121106 | 11 | 117581866 | C | T | 0.764 | 0.445 | +? | 0 | 0 | 0 | 1 |
| rs75160858 | 11 | 119876827 | A | G | -0.764 | 0.445 | -? | 0 | 0 | 0 | 1 |
| rs4936406 | 11 | 117618437 | C | T | 0.764 | 0.4451 | +? | 0 | 0 | 0 | 1 |
| rs56679414 | 11 | 119016547 | T | C | -0.764 | 0.4451 | -? | 0 | 0 | 0 | 1 |
| rs584057 | 11 | 117331503 | C | T | 0.763 | 0.4452 | +? | 0 | 0 | 0 | 1 |
| rs10790296 | 11 | 119310379 | A | G | 0.763 | 0.4453 | -+ | -43.3 | 2.093 | 1 | 0.148 |
| rs2070667 | 11 | 116701669 | A | G | -0.762 | 0.4459 | -? | 0 | 0 | 0 | 1 |
| rs587099 | 11 | 117509143 | G | A | -0.762 | 0.446 | -? | 0 | 0 | 0 | 1 |
| rs55927184 | 11 | 119015371 | A | G | -0.762 | 0.4463 | -? | 0 | 0 | 0 | 1 |
| rs517599 | 11 | 116329185 | C | T | -0.761 | 0.4464 | -? | 0 | 0 | 0 | 1 |
| rs10790362 | 11 | 119865290 | T | C | 0.761 | 0.4465 | +? | 0 | 0 | 0 | 1 |
| rs10892198 | 11 | 117801262 | G | C | 0.761 | 0.4466 | +? | 0 | 0 | 0 | 1 |

| | | | | | | | | | | | |
|---|---|---|---|---|---|---|---|---|---|---|---|
| rs58359402 | 11 | 120702378 | G | A | 0.761 | 0.4468 | +? | 0 | 0 | 0 | 1 |
| rs118013839 | 11 | 116561514 | A | G | -0.761 | 0.4469 | -- | 0 | 0.004 | 1 | 0.9521 |
| rs7108217 | 11 | 120183794 | T | C | -0.76 | 0.4471 | -? | 0 | 0 | 0 | 1 |
| rs7395055 | 11 | 119703507 | C | T | 0.759 | 0.4476 | +? | 0 | 0 | 0 | 1 |
| rs7930587 | 11 | 116388998 | A | T | -0.759 | 0.4478 | -? | 0 | 0 | 0 | 1 |
| rs1125618 | 11 | 119970300 | T | C | -0.759 | 0.4479 | -? | 0 | 0 | 0 | 1 |
| rs77772751 | 11 | 120872008 | A | G | -0.759 | 0.448 | -? | 0 | 0 | 0 | 1 |
| rs11606421 | 11 | 120623682 | T | C | 0.758 | 0.4482 | +? | 0 | 0 | 0 | 1 |
| rs541563 | 11 | 116329750 | A | G | -0.758 | 0.4482 | -? | 0 | 0 | 0 | 1 |
| rs55852239 | 11 | 118139755 | A | G | 0.758 | 0.4482 | +? | 0 | 0 | 0 | 1 |
| rs7117063 | 11 | 120238824 | C | T | -0.758 | 0.4482 | -? | 0 | 0 | 0 | 1 |
| rs2015914 | 11 | 120137818 | A | G | -0.758 | 0.4483 | -? | 0 | 0 | 0 | 1 |
| rs1939234 | 11 | 117557380 | T | C | 0.758 | 0.4484 | +? | 0 | 0 | 0 | 1 |
| rs876479 | 11 | 118028094 | T | C | 0.758 | 0.4485 | +? | 0 | 0 | 0 | 1 |
| rs12790585 | 11 | 120000608 | A | G | -0.758 | 0.4486 | -? | 0 | 0 | 0 | 1 |
| rs471714 | 11 | 119817579 | A | T | 0.758 | 0.4486 | +? | 0 | 0 | 0 | 1 |
| rs472665 | 11 | 119817708 | T | C | 0.758 | 0.4486 | +? | 0 | 0 | 0 | 1 |
| rs476877 | 11 | 119819087 | C | T | 0.758 | 0.4486 | +? | 0 | 0 | 0 | 1 |
| rs580318 | 11 | 119817426 | C | A | 0.758 | 0.4486 | +? | 0 | 0 | 0 | 1 |
| rs34172482 | 11 | 120278477 | G | A | 0.758 | 0.4487 | +? | 0 | 0 | 0 | 1 |
| rs7131597 | 11 | 120303939 | T | G | -0.757 | 0.4488 | -? | 0 | 0 | 0 | 1 |
| rs75050717 | 11 | 120960054 | A | G | -0.757 | 0.4488 | -? | 0 | 0 | 0 | 1 |
| rs10892498 | 11 | 119806226 | T | G | -0.757 | 0.449 | -? | 0 | 0 | 0 | 1 |
| rs755537 | 11 | 117768409 | G | A | 0.757 | 0.449 | +? | 0 | 0 | 0 | 1 |
| rs11216397 | 11 | 117319998 | C | T | 0.757 | 0.4491 | +? | 0 | 0 | 0 | 1 |
| rs688790 | 11 | 117541521 | A | T | -0.757 | 0.4492 | -? | 0 | 0 | 0 | 1 |
| rs12574466 | 11 | 119813518 | A | G | -0.757 | 0.4493 | -? | 0 | 0 | 0 | 1 |
| rs180326 | 11 | 116624703 | G | T | -0.757 | 0.4493 | -- | 0 | 0.066 | 1 | 0.797 |
| rs3802878 | 11 | 117414844 | C | A | 0.756 | 0.4497 | +? | 0 | 0 | 0 | 1 |
| rs472232 | 11 | 119701826 | A | C | 0.756 | 0.4497 | +? | 0 | 0 | 0 | 1 |
| rs73010960 | 11 | 116221203 | A | C | -0.756 | 0.4497 | -- | 0 | 0.136 | 1 | 0.7121 |
| rs12803249 | 11 | 116536418 | G | A | -0.756 | 0.4498 | -? | 0 | 0 | 0 | 1 |
| rs4298914 | 11 | 120443983 | T | C | -0.756 | 0.4498 | -? | 0 | 0 | 0 | 1 |
| rs3819250 | 11 | 118185346 | A | G | -0.755 | 0.4501 | -+ | 1.4 | 3.043 | 1 | 0.08108 |
| rs1784119 | 11 | 116562805 | T | C | 0.755 | 0.4503 | +? | 0 | 0 | 0 | 1 |
| rs476362 | 11 | 116485599 | A | G | -0.755 | 0.4503 | -? | 0 | 0 | 0 | 1 |
| rs61903699 | 11 | 117145582 | T | C | -0.755 | 0.4503 | -? | 0 | 0 | 0 | 1 |
| rs11822160 | 11 | 116577558 | A | T | -0.755 | 0.4505 | -- | 0 | 0.933 | 1 | 0.334 |
| rs79112951 | 11 | 116621972 | T | C | -0.754 | 0.4509 | -? | 0 | 0 | 0 | 1 |
| rs10790338 | 11 | 119586389 | G | T | 0.754 | 0.451 | +? | 0 | 0 | 0 | 1 |
| rs10892440 | 11 | 119584274 | T | C | 0.754 | 0.451 | +? | 0 | 0 | 0 | 1 |
| rs7931978 | 11 | 119586117 | G | A | 0.754 | 0.451 | +? | 0 | 0 | 0 | 1 |
| rs12418744 | 11 | 117084537 | A | G | -0.753 | 0.4513 | -? | 0 | 0 | 0 | 1 |
| rs17245504 | 11 | 120178753 | G | T | -0.753 | 0.4513 | -? | 0 | 0 | 0 | 1 |
| rs7107756 | 11 | 118717708 | A | G | 0.753 | 0.4514 | ++ | 0 | 0.014 | 1 | 0.9047 |
| rs12270280 | 11 | 118791150 | T | C | -0.753 | 0.4515 | -- | 0 | 0.007 | 1 | 0.9338 |
| rs11824814 | 11 | 120632841 | T | A | 0.753 | 0.4517 | +? | 0 | 0 | 0 | 1 |
| rs73562888 | 11 | 119016361 | C | T | -0.752 | 0.4521 | -? | 0 | 0 | 0 | 1 |
| rs73014900 | 11 | 116554198 | G | T | 0.752 | 0.4523 | +? | 0 | 0 | 0 | 1 |
| rs10790429 | 11 | 121147301 | A | T | -0.751 | 0.4524 | -- | 0 | 0.472 | 1 | 0.4919 |
| rs7126328 | 11 | 120244577 | T | C | -0.751 | 0.4524 | -? | 0 | 0 | 0 | 1 |
| rs4252241 | 11 | 117855295 | G | A | 0.751 | 0.4525 | +- | 0 | 0.27 | 1 | 0.6035 |
| rs191301146 | 11 | 116271393 | T | C | 0.751 | 0.4526 | +? | 0 | 0 | 0 | 1 |
| rs12278200 | 11 | 116367859 | C | G | 0.751 | 0.4527 | +? | 0 | 0 | 0 | 1 |
| rs77073823 | 11 | 117129083 | T | C | -0.751 | 0.4529 | -? | 0 | 0 | 0 | 1 |
| rs1157011088 | 11 | 120807571 | T | G | 0.75 | 0.4531 | +? | 0 | 0 | 0 | 1 |

| | | | | | | | | | | | |
|---|---|---|---|---|---|---|---|---|---|---|---|
| rs61903746 | 11 | 117255283 | T | C | -0.75 | 0.4531 | -? | 0 | 0 | 0 | 1 |
| rs17569676 | 11 | 116399249 | C | T | -0.75 | 0.4533 | -? | 0 | 0 | 0 | 1 |
| rs688415 | 11 | 117541577 | G | A | -0.749 | 0.4536 | -? | 0 | 0 | 0 | 1 |
| rs7945884 | 11 | 119062889 | C | T | -0.749 | 0.4537 | -? | 0 | 0 | 0 | 1 |
| rs11218114 | 11 | 120893054 | A | G | -0.749 | 0.4538 | -? | 0 | 0 | 0 | 1 |
| rs2298767 | 11 | 117374470 | T | C | -0.749 | 0.4539 | -? | 0 | 0 | 0 | 1 |
| rs751417 | 11 | 121164440 | G | A | -0.749 | 0.4539 | -? | 0 | 0 | 0 | 1 |
| rs9651696 | 11 | 120562930 | C | T | -0.749 | 0.4539 | -? | 0 | 0 | 0 | 1 |
| rs2298821 | 11 | 118185140 | C | T | -0.749 | 0.454 | -+ | 1 | 3.03 | 1 | 0.08174 |
| rs11826839 | 11 | 117790769 | G | A | -0.748 | 0.4542 | -? | 0 | 0 | 0 | 1 |
| rs2155808 | 11 | 120133107 | T | A | -0.748 | 0.4542 | -? | 0 | 0 | 0 | 1 |
| rs7939408 | 11 | 117845302 | T | A | 0.748 | 0.4542 | ++ | -197.4 | 1.009 | 1 | 0.3152 |
| rs7932658 | 11 | 120211212 | A | G | -0.748 | 0.4543 | -? | 0 | 0 | 0 | 1 |
| rs12804173 | 11 | 117465988 | A | T | -0.748 | 0.4546 | -- | 0 | 0.173 | 1 | 0.6775 |
| rs2508446 | 11 | 117873372 | A | G | 0.748 | 0.4546 | ++ | 0 | 0.014 | 1 | 0.9047 |
| rs748771 | 11 | 117615502 | G | A | 0.748 | 0.4546 | +? | 0 | 0 | 0 | 1 |
| rs11217841 | 11 | 120238721 | A | C | -0.748 | 0.4547 | -? | 0 | 0 | 0 | 1 |
| rs648384 | 11 | 117328700 | G | A | 0.747 | 0.455 | +? | 0 | 0 | 0 | 1 |
| rs80064625 | 11 | 119801612 | C | T | -0.747 | 0.455 | -? | 0 | 0 | 0 | 1 |
| rs12273996 | 11 | 117500940 | A | G | -0.747 | 0.4552 | -? | 0 | 0 | 0 | 1 |
| rs1145194 | 11 | 116570968 | A | C | 0.747 | 0.4553 | +? | 0 | 0 | 0 | 1 |
| rs73020495 | 11 | 117775943 | A | C | 0.747 | 0.4553 | +? | 0 | 0 | 0 | 1 |
| rs10790247 | 11 | 118113629 | T | G | -0.746 | 0.4554 | -? | 0 | 0 | 0 | 1 |
| rs12798851 | 11 | 118135973 | G | T | 0.746 | 0.4555 | -+ | 17.9 | 3.655 | 1 | 0.05589 |
| rs1596979 | 11 | 119800142 | C | A | -0.746 | 0.4555 | -? | 0 | 0 | 0 | 1 |
| rs1892894 | 11 | 120254684 | T | C | -0.746 | 0.4556 | -? | 0 | 0 | 0 | 1 |
| rs680268 | 11 | 119239689 | C | T | 0.746 | 0.4557 | +? | 0 | 0 | 0 | 1 |
| rs4938799 | 11 | 120208257 | G | T | -0.746 | 0.4558 | -? | 0 | 0 | 0 | 1 |
| rs146786560 | 11 | 117106458 | C | T | -0.746 | 0.4559 | -? | 0 | 0 | 0 | 1 |
| rs2508950 | 11 | 118993549 | T | C | 0.745 | 0.4564 | +? | 0 | 0 | 0 | 1 |
| rs7939363 | 11 | 119225134 | T | C | 0.745 | 0.4564 | +? | 0 | 0 | 0 | 1 |
| rs61010648 | 11 | 118206890 | G | C | 0.745 | 0.4565 | +? | 0 | 0 | 0 | 1 |
| rs61906102 | 11 | 116546503 | C | T | -0.744 | 0.4566 | -? | 0 | 0 | 0 | 1 |
| rs12277914 | 11 | 117733223 | T | C | 0.744 | 0.4567 | +? | 0 | 0 | 0 | 1 |
| rs2034643 | 11 | 119935789 | T | G | -0.744 | 0.4567 | -? | 0 | 0 | 0 | 1 |
| rs553080 | 11 | 116702345 | C | T | -0.744 | 0.4567 | -? | 0 | 0 | 0 | 1 |
| rs10892045 | 11 | 116784824 | A | C | 0.744 | 0.457 | +? | 0 | 0 | 0 | 1 |
| rs4938756 | 11 | 119878269 | C | T | -0.744 | 0.457 | -? | 0 | 0 | 0 | 1 |
| rs2847494 | 11 | 120133012 | T | C | -0.744 | 0.4571 | -? | 0 | 0 | 0 | 1 |
| rs12099358 | 11 | 116726048 | A | C | 0.743 | 0.4572 | +? | 0 | 0 | 0 | 1 |
| rs148189915 | 11 | 120187422 | T | A | 0.743 | 0.4572 | +? | 0 | 0 | 0 | 1 |
| rs61906104 | 11 | 116550115 | G | A | -0.743 | 0.4572 | -? | 0 | 0 | 0 | 1 |
| rs9667927 | 11 | 120622625 | C | T | 0.743 | 0.4572 | +? | 0 | 0 | 0 | 1 |
| rs61686122 | 11 | 119016539 | C | G | -0.743 | 0.4573 | -? | 0 | 0 | 0 | 1 |
| rs1793145 | 11 | 118112433 | C | T | -0.743 | 0.4574 | -? | 0 | 0 | 0 | 1 |
| rs11216631 | 11 | 117795679 | T | G | 0.743 | 0.4576 | +? | 0 | 0 | 0 | 1 |
| rs12281564 | 11 | 119923231 | A | G | 0.743 | 0.4577 | +? | 0 | 0 | 0 | 1 |
| rs1406347456 | 11 | 117709645 | G | T | 0.742 | 0.4579 | +? | 0 | 0 | 0 | 1 |
| rs67145274 | 11 | 117136277 | G | C | -0.742 | 0.4579 | -? | 0 | 0 | 0 | 1 |
| rs11607146 | 11 | 119662084 | G | A | 0.742 | 0.4581 | ++ | 0 | 0.005 | 1 | 0.9461 |
| rs1893035 | 11 | 117596086 | A | G | 0.741 | 0.4585 | +? | 0 | 0 | 0 | 1 |
| rs76589374 | 11 | 117490095 | T | G | 0.741 | 0.4585 | +? | 0 | 0 | 0 | 1 |
| rs723937 | 11 | 120312746 | A | G | -0.74 | 0.4591 | -? | 0 | 0 | 0 | 1 |
| rs11820186 | 11 | 120632745 | A | G | 0.74 | 0.4594 | +? | 0 | 0 | 0 | 1 |
| rs2850805 | 11 | 120793387 | G | A | 0.74 | 0.4594 | +? | 0 | 0 | 0 | 1 |
| rs76248354 | 11 | 118042606 | A | G | 0.74 | 0.4594 | +- | 43.8 | 5.341 | 1 | 0.02083 |

| | | | | | | | | | | | |
|---|---|---|---|---|---|---|---|---|---|---|---|
| rs10790430 | 11 | 121154837 | T | C | -0.74 | 0.4595 | -? | 0 | 0 | 0 | 1 |
| rs10790431 | 11 | 121155070 | C | T | -0.74 | 0.4595 | -? | 0 | 0 | 0 | 1 |
| rs12801136 | 11 | 116548349 | A | G | -0.74 | 0.4596 | -? | 0 | 0 | 0 | 1 |
| rs516169 | 11 | 116247262 | C | G | -0.739 | 0.4598 | -? | 0 | 0 | 0 | 1 |
| rs880745 | 11 | 119864575 | T | C | 0.739 | 0.4598 | +? | 0 | 0 | 0 | 1 |
| rs2853005 | 11 | 118120104 | C | T | -0.739 | 0.4599 | -? | 0 | 0 | 0 | 1 |
| rs56693116 | 11 | 120361394 | T | G | 0.739 | 0.4602 | +? | 0 | 0 | 0 | 1 |
| rs1551240 | 11 | 120087541 | T | C | 0.738 | 0.4605 | +? | 0 | 0 | 0 | 1 |
| rs11217840 | 11 | 120238655 | G | A | -0.738 | 0.4606 | -? | 0 | 0 | 0 | 1 |
| rs12804597 | 11 | 120391473 | A | G | 0.738 | 0.4606 | +? | 0 | 0 | 0 | 1 |
| rs7950969 | 11 | 117583898 | G | A | 0.738 | 0.4606 | +? | 0 | 0 | 0 | 1 |
| rs2846098 | 11 | 120650053 | C | T | 0.738 | 0.4607 | +? | 0 | 0 | 0 | 1 |
| rs4936388 | 11 | 117440404 | G | T | -0.738 | 0.4607 | -- | 0 | 0.157 | 1 | 0.6923 |
| rs10790239 | 11 | 118001445 | T | C | -0.737 | 0.461 | -- | 0 | 0.435 | 1 | 0.5096 |
| rs3759007 | 11 | 118137667 | T | C | 0.737 | 0.4612 | +? | 0 | 0 | 0 | 1 |
| rs4252248 | 11 | 117858983 | G | A | 0.737 | 0.4612 | ++ | 0 | 0.218 | 1 | 0.6408 |
| rs4938796 | 11 | 120184678 | A | G | 0.737 | 0.4613 | +? | 0 | 0 | 0 | 1 |
| rs1939979 | 11 | 117611162 | T | C | 0.737 | 0.4614 | +? | 0 | 0 | 0 | 1 |
| rs34180117 | 11 | 118138629 | T | C | 0.737 | 0.4614 | +? | 0 | 0 | 0 | 1 |
| rs35850218 | 11 | 118138922 | C | T | 0.737 | 0.4614 | +? | 0 | 0 | 0 | 1 |
| rs633223 | 11 | 119846524 | A | G | 0.736 | 0.4615 | +? | 0 | 0 | 0 | 1 |
| rs595049 | 11 | 116699445 | C | A | -0.736 | 0.4616 | -? | 0 | 0 | 0 | 1 |
| rs80125068 | 11 | 116996445 | G | A | 0.736 | 0.4617 | +? | 0 | 0 | 0 | 1 |
| rs145034678 | 11 | 120857142 | A | G | -0.735 | 0.4621 | -? | 0 | 0 | 0 | 1 |
| rs10892376 | 11 | 119370899 | A | C | -0.735 | 0.4622 | -? | 0 | 0 | 0 | 1 |
| rs11216832 | 11 | 118119073 | A | C | -0.735 | 0.4622 | -? | 0 | 0 | 0 | 1 |
| rs11216833 | 11 | 118119957 | T | G | -0.735 | 0.4622 | -? | 0 | 0 | 0 | 1 |
| rs76080778 | 11 | 119501013 | G | A | 0.735 | 0.4622 | +? | 0 | 0 | 0 | 1 |
| rs10892567 | 11 | 120244192 | G | A | -0.735 | 0.4623 | -? | 0 | 0 | 0 | 1 |
| rs6589829 | 11 | 120530973 | C | T | -0.735 | 0.4623 | -? | 0 | 0 | 0 | 1 |
| rs10790293 | 11 | 119304090 | A | C | 0.735 | 0.4624 | -+ | -29.7 | 2.313 | 1 | 0.1283 |
| rs4456287 | 11 | 118298252 | C | T | 0.735 | 0.4624 | ++ | 0 | 0.003 | 1 | 0.9552 |
| rs11216126 | 11 | 116617240 | C | A | -0.735 | 0.4626 | -+ | 0 | 0.524 | 1 | 0.469 |
| rs11216341 | 11 | 117134094 | T | C | -0.735 | 0.4626 | -? | 0 | 0 | 0 | 1 |
| rs117527146 | 11 | 118150882 | A | C | -0.735 | 0.4626 | -? | 0 | 0 | 0 | 1 |
| rs12295268 | 11 | 117133378 | A | G | -0.735 | 0.4626 | -? | 0 | 0 | 0 | 1 |
| rs5134 | 11 | 116702579 | C | T | -0.734 | 0.4627 | -? | 0 | 0 | 0 | 1 |
| rs35681250 | 11 | 116555386 | T | A | -0.734 | 0.4631 | -? | 0 | 0 | 0 | 1 |
| rs481473 | 11 | 116250624 | C | A | -0.734 | 0.4631 | -? | 0 | 0 | 0 | 1 |
| rs1481148807 | 11 | 116960078 | A | G | -0.733 | 0.4633 | +- | 45.2 | 5.478 | 1 | 0.01926 |
| rs10160827 | 11 | 117814506 | T | C | 0.733 | 0.4634 | ++ | -189.3 | 1.037 | 1 | 0.3085 |
| rs1573580 | 11 | 118069269 | G | C | 0.733 | 0.4634 | +? | 0 | 0 | 0 | 1 |
| rs56128937 | 11 | 118139478 | A | T | 0.733 | 0.4634 | +? | 0 | 0 | 0 | 1 |
| rs12284730 | 11 | 120560459 | T | C | 0.733 | 0.4635 | +? | 0 | 0 | 0 | 1 |
| rs1317514 | 11 | 120513914 | C | T | -0.733 | 0.4636 | -? | 0 | 0 | 0 | 1 |
| rs73008630 | 11 | 119697476 | G | A | -0.733 | 0.4636 | -? | 0 | 0 | 0 | 1 |
| rs12291550 | 11 | 116436362 | C | T | 0.733 | 0.4637 | ++ | 0 | 0.027 | 1 | 0.8694 |
| rs145749713 | 11 | 117488010 | A | G | 0.733 | 0.4637 | +? | 0 | 0 | 0 | 1 |
| rs3863317 | 11 | 117804109 | A | G | 0.733 | 0.4637 | +? | 0 | 0 | 0 | 1 |
| rs61734721 | 11 | 119692261 | C | T | -0.733 | 0.4637 | -? | 0 | 0 | 0 | 1 |
| rs3809046 | 11 | 119018766 | T | G | -0.733 | 0.4638 | -? | 0 | 0 | 0 | 1 |
| rs4936354 | 11 | 116398417 | T | G | -0.733 | 0.4638 | -? | 0 | 0 | 0 | 1 |
| rs55750089 | 11 | 119019152 | T | C | -0.733 | 0.4638 | -? | 0 | 0 | 0 | 1 |
| rs55752358 | 11 | 119019475 | T | A | -0.733 | 0.4638 | -? | 0 | 0 | 0 | 1 |
| rs56013883 | 11 | 119018774 | T | G | -0.733 | 0.4638 | -? | 0 | 0 | 0 | 1 |
| rs78951297 | 11 | 119714120 | A | G | 0.733 | 0.4638 | +? | 0 | 0 | 0 | 1 |

| | | | | | | | | | | | |
|---|---|---|---|---|---|---|---|---|---|---|---|
| rs3759006 | 11 | 118137560 | A | G | 0.732 | 0.4639 | +? | 0 | 0 | 0 | 1 |
| rs10892472 | 11 | 119683607 | A | C | 0.732 | 0.464 | +? | 0 | 0 | 0 | 1 |
| rs17092638 | 11 | 116556236 | G | C | -0.732 | 0.464 | -? | 0 | 0 | 0 | 1 |
| rs76610415 | 11 | 119900407 | A | G | -0.732 | 0.4641 | -? | 0 | 0 | 0 | 1 |
| rs7948683 | 11 | 118137911 | A | G | 0.732 | 0.4641 | +? | 0 | 0 | 0 | 1 |
| rs11601480 | 11 | 117430741 | C | A | -0.732 | 0.4643 | -? | 0 | 0 | 0 | 1 |
| rs12270661 | 11 | 117140148 | A | G | -0.731 | 0.4645 | -? | 0 | 0 | 0 | 1 |
| rs550893 | 11 | 117129617 | A | G | 0.731 | 0.4645 | +? | 0 | 0 | 0 | 1 |
| rs56966127 | 11 | 120163449 | T | G | 0.731 | 0.4646 | +? | 0 | 0 | 0 | 1 |
| rs11604424 | 11 | 116651115 | C | T | -0.731 | 0.4647 | +- | 41.3 | 5.11 | 1 | 0.02379 |
| rs7937284 | 11 | 120731363 | G | A | -0.731 | 0.465 | -? | 0 | 0 | 0 | 1 |
| rs10488782 | 11 | 117673828 | G | T | 0.73 | 0.4652 | ++ | 0 | 0.009 | 1 | 0.9224 |
| rs616889 | 11 | 119884781 | G | A | 0.73 | 0.4652 | +? | 0 | 0 | 0 | 1 |
| rs35724134 | 11 | 116306907 | A | C | -0.73 | 0.4653 | -? | 0 | 0 | 0 | 1 |
| rs4938358 | 11 | 117084091 | T | C | -0.73 | 0.4653 | -? | 0 | 0 | 0 | 1 |
| rs4938652 | 11 | 119305593 | T | C | 0.73 | 0.4653 | -+ | -29.2 | 2.321 | 1 | 0.1276 |
| rs4938653 | 11 | 119305611 | C | T | 0.73 | 0.4653 | -+ | -29.2 | 2.321 | 1 | 0.1276 |
| rs6589811 | 11 | 120235703 | T | C | -0.73 | 0.4653 | -? | 0 | 0 | 0 | 1 |
| rs57833493 | 11 | 120764256 | C | T | -0.73 | 0.4656 | -? | 0 | 0 | 0 | 1 |
| rs191917960 | 11 | 119957452 | G | C | 0.729 | 0.4657 | +? | 0 | 0 | 0 | 1 |
| rs145892119 | 11 | 119013740 | T | C | -0.729 | 0.4658 | -? | 0 | 0 | 0 | 1 |
| rs1942929 | 11 | 117605098 | C | T | 0.729 | 0.4658 | +? | 0 | 0 | 0 | 1 |
| rs1902627 | 11 | 117291975 | T | C | 0.729 | 0.4659 | +? | 0 | 0 | 0 | 1 |
| rs7104866 | 11 | 116493328 | T | C | 0.729 | 0.466 | +? | 0 | 0 | 0 | 1 |
| rs146990932 | 11 | 116392086 | T | C | 0.729 | 0.4661 | +? | 0 | 0 | 0 | 1 |
| rs12806041 | 11 | 118138998 | G | A | 0.729 | 0.4662 | +? | 0 | 0 | 0 | 1 |
| rs657317 | 11 | 119816766 | A | C | 0.729 | 0.4663 | +? | 0 | 0 | 0 | 1 |
| rs6589831 | 11 | 120558738 | T | C | -0.728 | 0.4665 | -? | 0 | 0 | 0 | 1 |
| rs7107783 | 11 | 116375253 | G | A | 0.728 | 0.4665 | +? | 0 | 0 | 0 | 1 |
| rs8192613 | 11 | 118037813 | T | G | 0.728 | 0.4667 | +? | 0 | 0 | 0 | 1 |
| rs115946185 | 11 | 117775361 | A | T | 0.728 | 0.4669 | +? | 0 | 0 | 0 | 1 |
| rs79347839 | 11 | 117775374 | A | T | 0.728 | 0.4669 | +? | 0 | 0 | 0 | 1 |
| rs482090 | 11 | 119695491 | T | C | -0.727 | 0.467 | -? | 0 | 0 | 0 | 1 |
| rs10892566 | 11 | 120241923 | C | T | -0.727 | 0.4672 | -? | 0 | 0 | 0 | 1 |
| rs74827927 | 11 | 120702968 | A | G | 0.727 | 0.4673 | +? | 0 | 0 | 0 | 1 |
| rs11216617 | 11 | 117783176 | T | G | -0.726 | 0.4676 | -? | 0 | 0 | 0 | 1 |
| rs1715437 | 11 | 118096891 | C | T | -0.726 | 0.4676 | -? | 0 | 0 | 0 | 1 |
| rs7121779 | 11 | 120213360 | T | A | -0.726 | 0.4676 | -? | 0 | 0 | 0 | 1 |
| rs11216137 | 11 | 116669828 | A | G | -0.726 | 0.4677 | -+ | 0 | 0.441 | 1 | 0.5069 |
| rs643423 | 11 | 119043656 | C | T | -0.726 | 0.4677 | -? | 0 | 0 | 0 | 1 |
| rs665597 | 11 | 117900074 | A | G | 0.726 | 0.4677 | +- | 0 | 0.684 | 1 | 0.4084 |
| rs58594771 | 11 | 120116018 | C | T | 0.726 | 0.468 | +? | 0 | 0 | 0 | 1 |
| rs7342200 | 11 | 120726066 | A | G | 0.726 | 0.4681 | +? | 0 | 0 | 0 | 1 |
| rs7952529 | 11 | 120216260 | C | A | -0.726 | 0.4681 | -? | 0 | 0 | 0 | 1 |
| rs7342199 | 11 | 120726044 | A | G | 0.725 | 0.4683 | +? | 0 | 0 | 0 | 1 |
| rs2512147 | 11 | 117872734 | T | C | 0.725 | 0.4684 | ++ | 0 | 0.015 | 1 | 0.9023 |
| rs681120 | 11 | 117173019 | C | A | -0.725 | 0.4686 | -? | 0 | 0 | 0 | 1 |
| rs7350507 | 11 | 117321606 | G | A | 0.725 | 0.4686 | +? | 0 | 0 | 0 | 1 |
| rs7924963 | 11 | 120233536 | A | G | -0.725 | 0.4687 | -? | 0 | 0 | 0 | 1 |
| rs11218131 | 11 | 120928722 | A | G | -0.724 | 0.4688 | -? | 0 | 0 | 0 | 1 |
| rs6589645 | 11 | 117971971 | G | A | -0.724 | 0.4688 | -? | 0 | 0 | 0 | 1 |
| rs117587198 | 11 | 120879138 | A | T | -0.724 | 0.469 | -? | 0 | 0 | 0 | 1 |
| rs145936741 | 11 | 116470289 | G | A | 0.724 | 0.469 | +? | 0 | 0 | 0 | 1 |
| rs11218067 | 11 | 120815567 | A | G | -0.724 | 0.4692 | -+ | 0 | 0.617 | 1 | 0.432 |
| rs11216047 | 11 | 116493715 | G | A | -0.724 | 0.4693 | -? | 0 | 0 | 0 | 1 |
| rs11216608 | 11 | 117764726 | G | A | 0.724 | 0.4693 | +? | 0 | 0 | 0 | 1 |

| | | | | | | | | | | | |
|---|---|---|---|---|---|---|---|---|---|---|---|
| rs12786734 | 11 | 120239610 | G | A | 0.723 | 0.4695 | +? | 0 | 0 | 0 | 1 |
| rs4331110 | 11 | 120730669 | A | G | -0.723 | 0.4695 | -? | 0 | 0 | 0 | 1 |
| rs7926317 | 11 | 120807747 | C | T | 0.723 | 0.4696 | +? | 0 | 0 | 0 | 1 |
| rs12799299 | 11 | 120193956 | T | C | 0.723 | 0.4697 | +? | 0 | 0 | 0 | 1 |
| rs11217250 | 11 | 119222337 | T | C | 0.723 | 0.4698 | +? | 0 | 0 | 0 | 1 |
| rs2510902 | 11 | 118700218 | A | G | -0.723 | 0.4698 | -+ | 0 | 0.566 | 1 | 0.4517 |
| rs77245011 | 11 | 117902428 | C | A | 0.723 | 0.4698 | ++ | 0 | 0.081 | 1 | 0.7758 |
| rs10790404 | 11 | 120731903 | T | C | -0.722 | 0.47 | -? | 0 | 0 | 0 | 1 |
| rs477408 | 11 | 117775056 | G | A | -0.722 | 0.4701 | -? | 0 | 0 | 0 | 1 |
| rs73578569 | 11 | 120657978 | T | C | 0.722 | 0.4702 | +? | 0 | 0 | 0 | 1 |
| rs2080586 | 11 | 116733316 | C | T | 0.722 | 0.4703 | +? | 0 | 0 | 0 | 1 |
| rs3910017 | 11 | 119295082 | A | G | 0.722 | 0.4703 | -+ | -57.4 | 1.906 | 1 | 0.1674 |
| rs7929916 | 11 | 119874301 | A | G | -0.722 | 0.4704 | -? | 0 | 0 | 0 | 1 |
| rs1480516974 | 11 | 117245988 | T | C | 0.722 | 0.4705 | +? | 0 | 0 | 0 | 1 |
| rs34955847 | 11 | 119805189 | T | C | -0.722 | 0.4705 | -? | 0 | 0 | 0 | 1 |
| rs657137 | 11 | 117441495 | T | C | -0.722 | 0.4705 | -- | 0 | 0.028 | 1 | 0.8671 |
| rs879858 | 11 | 116573120 | A | G | -0.722 | 0.4705 | -? | 0 | 0 | 0 | 1 |
| rs11217523 | 11 | 119691772 | G | A | -0.722 | 0.4706 | -? | 0 | 0 | 0 | 1 |
| rs10790380 | 11 | 120233158 | A | G | -0.721 | 0.4708 | -? | 0 | 0 | 0 | 1 |
| rs112045567 | 11 | 118804989 | G | A | 0.721 | 0.4709 | +- | -66.4 | 1.803 | 1 | 0.1794 |
| rs475178 | 11 | 119818914 | G | A | 0.721 | 0.4711 | +? | 0 | 0 | 0 | 1 |
| rs475867 | 11 | 119818937 | A | G | 0.721 | 0.4711 | +? | 0 | 0 | 0 | 1 |
| rs554338 | 11 | 119817739 | C | T | 0.721 | 0.4711 | +? | 0 | 0 | 0 | 1 |
| rs628244 | 11 | 119818657 | T | C | 0.721 | 0.4711 | +? | 0 | 0 | 0 | 1 |
| rs550764 | 11 | 119817385 | A | T | 0.721 | 0.4712 | +? | 0 | 0 | 0 | 1 |
| rs77262931 | 11 | 119968674 | A | G | -0.721 | 0.4712 | -? | 0 | 0 | 0 | 1 |
| rs688803 | 11 | 117953050 | T | C | -0.72 | 0.4713 | -? | 0 | 0 | 0 | 1 |
| rs1179920 | 11 | 118993632 | G | A | 0.72 | 0.4714 | +? | 0 | 0 | 0 | 1 |
| rs3891896 | 11 | 117291936 | G | T | -0.72 | 0.4714 | -? | 0 | 0 | 0 | 1 |
| rs1787712 | 11 | 116560738 | G | C | -0.72 | 0.4715 | -? | 0 | 0 | 0 | 1 |
| rs12294065 | 11 | 116671540 | C | G | -0.72 | 0.4716 | -+ | 0 | 0.436 | 1 | 0.5092 |
| rs4938740 | 11 | 119794036 | A | G | -0.72 | 0.4716 | -? | 0 | 0 | 0 | 1 |
| rs76861941 | 11 | 119728139 | G | A | -0.72 | 0.4716 | -? | 0 | 0 | 0 | 1 |
| rs17120652 | 11 | 117235647 | T | C | 0.72 | 0.4717 | +? | 0 | 0 | 0 | 1 |
| rs10892170 | 11 | 117633470 | G | A | -0.72 | 0.4718 | -? | 0 | 0 | 0 | 1 |
| rs11216607 | 11 | 117764641 | A | G | 0.72 | 0.4718 | +? | 0 | 0 | 0 | 1 |
| rs492517 | 11 | 116311642 | T | C | 0.72 | 0.4718 | +? | 0 | 0 | 0 | 1 |
| rs1893907 | 11 | 120524884 | G | C | -0.719 | 0.4719 | -? | 0 | 0 | 0 | 1 |
| rs4297478 | 11 | 120435854 | C | T | -0.719 | 0.4723 | -? | 0 | 0 | 0 | 1 |
| rs2515796 | 11 | 117817358 | G | A | -0.718 | 0.4725 | -? | 0 | 0 | 0 | 1 |
| rs3017229 | 11 | 116560967 | A | G | 0.718 | 0.4726 | +? | 0 | 0 | 0 | 1 |
| rs76716142 | 11 | 120695126 | T | C | 0.718 | 0.4726 | +? | 0 | 0 | 0 | 1 |
| rs10892553 | 11 | 120110775 | C | T | -0.718 | 0.4727 | -? | 0 | 0 | 0 | 1 |
| rs7107211 | 11 | 120262922 | A | G | -0.718 | 0.473 | -? | 0 | 0 | 0 | 1 |
| rs61903703 | 11 | 117169874 | A | G | -0.717 | 0.4731 | -? | 0 | 0 | 0 | 1 |
| rs117666845 | 11 | 118143062 | A | T | 0.717 | 0.4733 | ++ | 0 | 0.123 | 1 | 0.7254 |
| rs2852213 | 11 | 120789293 | T | C | 0.717 | 0.4733 | +? | 0 | 0 | 0 | 1 |
| rs35606865 | 11 | 116668843 | C | T | -0.717 | 0.4733 | -+ | 0 | 0.434 | 1 | 0.5102 |
| rs10892354 | 11 | 119238381 | T | C | 0.717 | 0.4734 | +? | 0 | 0 | 0 | 1 |
| rs6589598 | 11 | 117002403 | A | G | 0.717 | 0.4735 | +? | 0 | 0 | 0 | 1 |
| rs12803154 | 11 | 117466151 | T | C | -0.717 | 0.4736 | -- | 0 | 0.158 | 1 | 0.6906 |
| rs1941409 | 11 | 120150500 | G | A | -0.717 | 0.4736 | -? | 0 | 0 | 0 | 1 |
| rs35262372 | 11 | 118704751 | G | A | -0.717 | 0.4736 | -+ | 0 | 0.348 | 1 | 0.5552 |
| rs1793187 | 11 | 118091747 | C | G | -0.716 | 0.4737 | -? | 0 | 0 | 0 | 1 |
| rs58073046 | 11 | 120248493 | G | A | -0.716 | 0.4738 | -? | 0 | 0 | 0 | 1 |
| rs10790383 | 11 | 120344526 | T | C | -0.716 | 0.4739 | -? | 0 | 0 | 0 | 1 |

| | | | | | | | | | | | |
|---|---|---|---|---|---|---|---|---|---|---|---|
| rs1793139 | 11 | 118106875 | G | A | -0.716 | 0.474 | -? | 0 | 0 | 0 | 1 |
| rs4936410 | 11 | 117785260 | C | T | 0.716 | 0.474 | +? | 0 | 0 | 0 | 1 |
| rs7126413 | 11 | 120207405 | G | A | -0.716 | 0.4742 | -? | 0 | 0 | 0 | 1 |
| rs10892372 | 11 | 119368006 | A | G | 0.715 | 0.4747 | +? | 0 | 0 | 0 | 1 |
| rs11217821 | 11 | 120202099 | C | T | -0.715 | 0.4747 | -? | 0 | 0 | 0 | 1 |
| rs4569018 | 11 | 120733632 | C | T | -0.715 | 0.4747 | -? | 0 | 0 | 0 | 1 |
| rs58111676 | 11 | 117122831 | G | C | -0.715 | 0.4749 | -? | 0 | 0 | 0 | 1 |
| rs78972478 | 11 | 116398667 | T | C | -0.715 | 0.4749 | -? | 0 | 0 | 0 | 1 |
| rs10502233 | 11 | 119893752 | C | G | -0.714 | 0.4751 | -? | 0 | 0 | 0 | 1 |
| rs948469 | 11 | 117972721 | A | C | -0.714 | 0.4754 | -? | 0 | 0 | 0 | 1 |
| rs1944504 | 11 | 119843219 | T | C | -0.714 | 0.4755 | -? | 0 | 0 | 0 | 1 |
| rs7944850 | 11 | 119842786 | A | G | -0.713 | 0.4758 | -? | 0 | 0 | 0 | 1 |
| rs141784177 | 11 | 120916015 | G | C | 0.713 | 0.4759 | +? | 0 | 0 | 0 | 1 |
| rs1902625 | 11 | 117291793 | C | T | 0.713 | 0.476 | +? | 0 | 0 | 0 | 1 |
| rs668213 | 11 | 117577674 | G | A | 0.713 | 0.476 | +? | 0 | 0 | 0 | 1 |
| rs686992 | 11 | 117775034 | C | T | -0.713 | 0.476 | -? | 0 | 0 | 0 | 1 |
| rs1124874 | 11 | 116518008 | A | G | -0.712 | 0.4764 | -? | 0 | 0 | 0 | 1 |
| rs12293222 | 11 | 116565309 | A | G | -0.712 | 0.4765 | -? | 0 | 0 | 0 | 1 |
| rs7123209 | 11 | 117988654 | C | G | -0.712 | 0.4765 | -- | 0 | 0.24 | 1 | 0.6239 |
| rs1715438 | 11 | 118098041 | A | G | -0.712 | 0.4766 | -? | 0 | 0 | 0 | 1 |
| rs4252267 | 11 | 117863292 | A | G | 0.712 | 0.4768 | ++ | 0 | 0.141 | 1 | 0.7069 |
| rs4252270 | 11 | 117863846 | T | C | 0.712 | 0.4768 | ++ | 0 | 0.141 | 1 | 0.7069 |
| rs55713240 | 11 | 119019282 | T | C | -0.711 | 0.4769 | -? | 0 | 0 | 0 | 1 |
| rs73010950 | 11 | 116220070 | A | G | -0.711 | 0.477 | -- | 0 | 0.078 | 1 | 0.7796 |
| rs79459688 | 11 | 117582709 | G | A | 0.711 | 0.477 | +? | 0 | 0 | 0 | 1 |
| rs529411 | 11 | 116311777 | C | T | 0.711 | 0.4771 | +? | 0 | 0 | 0 | 1 |
| rs7115177 | 11 | 117239173 | C | A | -0.711 | 0.4771 | -? | 0 | 0 | 0 | 1 |
| rs4936414 | 11 | 117853613 | C | T | 0.71 | 0.4776 | ++ | 0 | 0.173 | 1 | 0.6775 |
| rs184607052 | 11 | 117482186 | G | A | 0.71 | 0.4777 | +? | 0 | 0 | 0 | 1 |
| rs7127194 | 11 | 119875060 | A | T | -0.71 | 0.4777 | -? | 0 | 0 | 0 | 1 |
| rs61906492 | 11 | 117107978 | A | G | -0.709 | 0.4781 | -? | 0 | 0 | 0 | 1 |
| rs10892582 | 11 | 120347715 | T | C | -0.709 | 0.4785 | -? | 0 | 0 | 0 | 1 |
| rs529796 | 11 | 117331864 | A | G | -0.709 | 0.4785 | -? | 0 | 0 | 0 | 1 |
| rs61905271 | 11 | 117312865 | A | G | -0.709 | 0.4785 | -? | 0 | 0 | 0 | 1 |
| rs74761525 | 11 | 117225982 | T | C | 0.709 | 0.4785 | +? | 0 | 0 | 0 | 1 |
| rs6589810 | 11 | 120233908 | A | G | -0.709 | 0.4786 | -? | 0 | 0 | 0 | 1 |
| rs715849 | 11 | 119575724 | A | G | -0.709 | 0.4786 | -? | 0 | 0 | 0 | 1 |
| rs7924772 | 11 | 120233626 | G | A | -0.709 | 0.4786 | -? | 0 | 0 | 0 | 1 |
| rs11216618 | 11 | 117784714 | C | T | -0.708 | 0.4787 | -? | 0 | 0 | 0 | 1 |
| rs486896 | 11 | 117173468 | A | G | 0.708 | 0.4788 | +? | 0 | 0 | 0 | 1 |
| rs76844341 | 11 | 116789849 | G | A | 0.708 | 0.4789 | +? | 0 | 0 | 0 | 1 |
| rs1944503 | 11 | 119813552 | A | G | -0.708 | 0.4791 | -? | 0 | 0 | 0 | 1 |
| rs7118391 | 11 | 120270709 | T | C | -0.707 | 0.4793 | -? | 0 | 0 | 0 | 1 |
| rs2277289 | 11 | 118182660 | C | T | -0.707 | 0.4795 | -+ | -6.8 | 2.81 | 1 | 0.0937 |
| rs7117321 | 11 | 120239051 | C | T | -0.707 | 0.4795 | -? | 0 | 0 | 0 | 1 |
| rs12288729 | 11 | 116463208 | C | G | 0.707 | 0.4796 | +? | 0 | 0 | 0 | 1 |
| rs117092333 | 11 | 120302456 | A | G | 0.707 | 0.4797 | +? | 0 | 0 | 0 | 1 |
| rs1793186 | 11 | 118091089 | T | C | -0.707 | 0.4798 | -? | 0 | 0 | 0 | 1 |
| rs11216399 | 11 | 117323389 | A | C | -0.706 | 0.4799 | -? | 0 | 0 | 0 | 1 |
| rs4938804 | 11 | 120273865 | A | G | -0.706 | 0.4801 | -? | 0 | 0 | 0 | 1 |
| rs559545 | 11 | 117763868 | A | C | -0.706 | 0.4801 | -? | 0 | 0 | 0 | 1 |
| rs2077276 | 11 | 117700410 | A | C | 0.706 | 0.4802 | +? | 0 | 0 | 0 | 1 |
| rs59627935 | 11 | 119307070 | C | T | -0.706 | 0.4802 | -? | 0 | 0 | 0 | 1 |
| rs117607796 | 11 | 116565603 | A | C | -0.705 | 0.4808 | -- | 0 | 0.008 | 1 | 0.9268 |
| rs7103544 | 11 | 118994516 | G | A | 0.705 | 0.4809 | +? | 0 | 0 | 0 | 1 |
| rs10892193 | 11 | 117791979 | T | C | -0.705 | 0.4811 | -? | 0 | 0 | 0 | 1 |

| | | | | | | | | | | | |
|---|---|---|---|---|---|---|---|---|---|---|---|
| rs118125468 | 11 | 120117394 | A | G | -0.705 | 0.4811 | -? | 0 | 0 | 0 | 1 |
| rs638270 | 11 | 117151378 | G | C | 0.704 | 0.4812 | +? | 0 | 0 | 0 | 1 |
| rs61898351 | 11 | 120199924 | C | T | -0.704 | 0.4813 | -? | 0 | 0 | 0 | 1 |
| rs75436216 | 11 | 119756962 | C | T | 0.704 | 0.4813 | +? | 0 | 0 | 0 | 1 |
| rs76467987 | 11 | 117390488 | C | T | 0.704 | 0.4813 | +? | 0 | 0 | 0 | 1 |
| rs1939984 | 11 | 117604158 | T | A | 0.704 | 0.4814 | +? | 0 | 0 | 0 | 1 |
| rs11216855 | 11 | 118208750 | C | T | -0.704 | 0.4815 | -- | 0 | 0.392 | 1 | 0.5313 |
| rs4354701 | 11 | 119488812 | A | G | -0.704 | 0.4816 | -? | 0 | 0 | 0 | 1 |
| rs1793162 | 11 | 118103972 | A | C | -0.703 | 0.4818 | -? | 0 | 0 | 0 | 1 |
| rs7104639 | 11 | 120239456 | G | C | -0.703 | 0.4819 | -? | 0 | 0 | 0 | 1 |
| rs1787702 | 11 | 116562288 | A | G | -0.703 | 0.4821 | -? | 0 | 0 | 0 | 1 |
| rs500943 | 11 | 116312643 | T | C | 0.702 | 0.4824 | +? | 0 | 0 | 0 | 1 |
| rs7129197 | 11 | 121107345 | T | G | 0.702 | 0.4824 | +? | 0 | 0 | 0 | 1 |
| rs4539328 | 11 | 120218243 | C | T | -0.702 | 0.4825 | -? | 0 | 0 | 0 | 1 |
| rs2370113 | 11 | 121192287 | T | G | -0.702 | 0.4826 | -? | 0 | 0 | 0 | 1 |
| rs28989504 | 11 | 117162762 | C | T | -0.701 | 0.4832 | -? | 0 | 0 | 0 | 1 |
| rs566174 | 11 | 116250106 | C | G | -0.701 | 0.4832 | -? | 0 | 0 | 0 | 1 |
| rs78058743 | 11 | 119577677 | T | C | -0.701 | 0.4832 | -? | 0 | 0 | 0 | 1 |
| rs2096882 | 11 | 120137465 | T | G | -0.701 | 0.4833 | -? | 0 | 0 | 0 | 1 |
| rs10750169 | 11 | 120136108 | C | G | -0.701 | 0.4834 | -? | 0 | 0 | 0 | 1 |
| rs56933877 | 11 | 118804643 | A | G | 0.701 | 0.4834 | +- | -62.5 | 1.846 | 1 | 0.1742 |
| rs617711 | 11 | 119366595 | T | C | 0.701 | 0.4834 | +? | 0 | 0 | 0 | 1 |
| rs10892562 | 11 | 120221618 | G | A | -0.701 | 0.4835 | -? | 0 | 0 | 0 | 1 |
| rs721487 | 11 | 117557886 | T | G | 0.701 | 0.4835 | +? | 0 | 0 | 0 | 1 |
| rs180349 | 11 | 116611827 | A | T | -0.7 | 0.4836 | -+ | 0 | 0.552 | 1 | 0.4575 |
| rs948381 | 11 | 121111458 | C | A | 0.701 | 0.4836 | +? | 0 | 0 | 0 | 1 |
| rs2515794 | 11 | 117816152 | C | A | -0.7 | 0.4837 | -? | 0 | 0 | 0 | 1 |
| rs12280992 | 11 | 119692457 | T | C | -0.7 | 0.4838 | -? | 0 | 0 | 0 | 1 |
| rs7119384 | 11 | 118994357 | T | A | 0.7 | 0.4839 | +? | 0 | 0 | 0 | 1 |
| rs80079694 | 11 | 119975192 | A | G | -0.7 | 0.4839 | -? | 0 | 0 | 0 | 1 |
| rs10892307 | 11 | 118754649 | G | C | 0.7 | 0.4842 | +- | 33.1 | 4.482 | 1 | 0.03425 |
| rs656924 | 11 | 119364461 | T | G | 0.7 | 0.4842 | +? | 0 | 0 | 0 | 1 |
| rs6589812 | 11 | 120278359 | C | T | -0.699 | 0.4843 | -? | 0 | 0 | 0 | 1 |
| rs7937612 | 11 | 120278688 | C | T | -0.699 | 0.4843 | -? | 0 | 0 | 0 | 1 |
| rs1793166 | 11 | 118116052 | A | G | 0.699 | 0.4844 | +? | 0 | 0 | 0 | 1 |
| rs2212428 | 11 | 116516587 | T | C | -0.699 | 0.4845 | -? | 0 | 0 | 0 | 1 |
| rs7940051 | 11 | 120690057 | A | G | 0.699 | 0.4845 | +? | 0 | 0 | 0 | 1 |
| rs7937697 | 11 | 120731470 | A | G | -0.699 | 0.4846 | -? | 0 | 0 | 0 | 1 |
| rs4938361 | 11 | 117099941 | C | G | 0.699 | 0.4847 | +? | 0 | 0 | 0 | 1 |
| rs79770434 | 11 | 116523165 | T | C | -0.699 | 0.4847 | -? | 0 | 0 | 0 | 1 |
| rs12366157 | 11 | 120029638 | G | A | -0.699 | 0.4848 | -? | 0 | 0 | 0 | 1 |
| rs2231445 | 11 | 118184271 | C | T | 0.699 | 0.4848 | +- | 25.6 | 4.034 | 1 | 0.04458 |
| rs2306473 | 11 | 117097952 | T | C | 0.699 | 0.4848 | +? | 0 | 0 | 0 | 1 |
| rs7928238 | 11 | 119490180 | G | A | -0.699 | 0.4848 | -? | 0 | 0 | 0 | 1 |
| rs10736497 | 11 | 119309664 | C | T | 0.698 | 0.4849 | -+ | -54.4 | 1.944 | 1 | 0.1633 |
| rs56326825 | 11 | 120690372 | C | T | 0.698 | 0.485 | +? | 0 | 0 | 0 | 1 |
| rs1558860 | 11 | 116607368 | A | C | -0.698 | 0.4851 | -- | -135.6 | 1.273 | 1 | 0.2591 |
| rs1974718 | 11 | 116606766 | G | A | -0.698 | 0.4851 | -- | -135.6 | 1.273 | 1 | 0.2591 |
| rs2229113 | 11 | 117869670 | A | G | 0.698 | 0.4851 | ++ | 0 | 0.067 | 1 | 0.7954 |
| rs521556 | 11 | 120036997 | T | C | 0.698 | 0.4851 | +? | 0 | 0 | 0 | 1 |
| rs34298380 | 11 | 120203559 | G | A | -0.698 | 0.4852 | -? | 0 | 0 | 0 | 1 |
| rs2100358 | 11 | 119939519 | A | C | -0.698 | 0.4853 | -? | 0 | 0 | 0 | 1 |
| rs608365 | 11 | 117951521 | C | G | -0.698 | 0.4853 | -? | 0 | 0 | 0 | 1 |
| rs3018382 | 11 | 118069563 | G | C | 0.697 | 0.4855 | +? | 0 | 0 | 0 | 1 |
| rs7122785 | 11 | 119222706 | G | A | 0.697 | 0.4855 | +? | 0 | 0 | 0 | 1 |
| rs906830 | 11 | 119549825 | T | C | 0.697 | 0.4855 | +? | 0 | 0 | 0 | 1 |

| | | | | | | | | | | | |
|---|---|---|---|---|---|---|---|---|---|---|---|
| rs76273814 | 11 | 120489650 | T | C | -0.697 | 0.4856 | -? | 0 | 0 | 0 | 1 |
| rs11216430 | 11 | 117376111 | C | A | 0.697 | 0.4857 | +? | 0 | 0 | 0 | 1 |
| rs45535039 | 11 | 119060963 | C | T | 0.697 | 0.4857 | +? | 0 | 0 | 0 | 1 |
| rs4938752 | 11 | 119877101 | G | C | -0.697 | 0.4857 | -? | 0 | 0 | 0 | 1 |
| rs10790378 | 11 | 120220268 | C | T | -0.697 | 0.4858 | -? | 0 | 0 | 0 | 1 |
| rs10790426 | 11 | 121139566 | C | T | -0.697 | 0.4858 | -- | 0 | 0.58 | 1 | 0.4462 |
| rs4938801 | 11 | 120223399 | T | C | -0.697 | 0.4858 | -? | 0 | 0 | 0 | 1 |
| rs7107144 | 11 | 119854514 | C | T | -0.697 | 0.486 | -? | 0 | 0 | 0 | 1 |
| rs76344204 | 11 | 121132474 | A | G | 0.697 | 0.486 | +? | 0 | 0 | 0 | 1 |
| rs10790372 | 11 | 120136078 | T | G | -0.696 | 0.4862 | -? | 0 | 0 | 0 | 1 |
| rs521053 | 11 | 119362151 | T | C | 0.696 | 0.4867 | +? | 0 | 0 | 0 | 1 |
| rs10892591 | 11 | 120521766 | T | C | -0.695 | 0.4871 | -? | 0 | 0 | 0 | 1 |
| rs78503991 | 11 | 117481112 | T | A | 0.695 | 0.4871 | +? | 0 | 0 | 0 | 1 |
| rs948026 | 11 | 120683594 | A | G | 0.695 | 0.4871 | +? | 0 | 0 | 0 | 1 |
| rs117010140 | 11 | 117773027 | A | G | -0.695 | 0.4873 | -? | 0 | 0 | 0 | 1 |
| rs3802871 | 11 | 117776648 | C | A | -0.695 | 0.4873 | -? | 0 | 0 | 0 | 1 |
| rs7940347 | 11 | 120690291 | T | G | 0.695 | 0.4873 | +? | 0 | 0 | 0 | 1 |
| rs7101415 | 11 | 120239088 | A | G | -0.694 | 0.4874 | -? | 0 | 0 | 0 | 1 |
| rs76169968 | 11 | 117082966 | A | G | -0.694 | 0.4875 | -? | 0 | 0 | 0 | 1 |
| rs17075 | 11 | 118959331 | G | A | 0.694 | 0.4876 | +? | 0 | 0 | 0 | 1 |
| rs61897696 | 11 | 119642855 | A | G | -0.694 | 0.4877 | -? | 0 | 0 | 0 | 1 |
| rs654946 | 11 | 117456034 | A | G | 0.694 | 0.4877 | +? | 0 | 0 | 0 | 1 |
| rs4482041 | 11 | 119875343 | T | C | -0.694 | 0.4878 | -? | 0 | 0 | 0 | 1 |
| rs61897867 | 11 | 119842049 | T | C | -0.694 | 0.4878 | -? | 0 | 0 | 0 | 1 |
| rs7928100 | 11 | 119377274 | A | G | -0.693 | 0.488 | -? | 0 | 0 | 0 | 1 |
| rs12806740 | 11 | 120203628 | A | G | -0.693 | 0.4882 | -? | 0 | 0 | 0 | 1 |
| rs138526953 | 11 | 118793479 | A | C | -0.693 | 0.4882 | -- | 0 | 0.107 | 1 | 0.7435 |
| rs1793163 | 11 | 118114813 | T | C | -0.693 | 0.4883 | -? | 0 | 0 | 0 | 1 |
| rs1793164 | 11 | 118114853 | T | A | -0.693 | 0.4883 | -? | 0 | 0 | 0 | 1 |
| rs2853033 | 11 | 118114344 | A | G | -0.693 | 0.4884 | -? | 0 | 0 | 0 | 1 |
| rs618627 | 11 | 119366779 | C | T | 0.693 | 0.4884 | +? | 0 | 0 | 0 | 1 |
| rs7112876 | 11 | 116363370 | C | G | -0.693 | 0.4884 | -? | 0 | 0 | 0 | 1 |
| rs1371711774 | 11 | 116368043 | T | C | 0.693 | 0.4886 | +? | 0 | 0 | 0 | 1 |
| rs668678 | 11 | 119364555 | A | G | 0.692 | 0.4888 | +? | 0 | 0 | 0 | 1 |
| rs4499013 | 11 | 119488646 | C | T | -0.692 | 0.4889 | -? | 0 | 0 | 0 | 1 |
| rs644920 | 11 | 117898231 | A | T | 0.691 | 0.4893 | +- | 0 | 0.363 | 1 | 0.5466 |
| rs73575441 | 11 | 118804919 | C | T | 0.691 | 0.4894 | +- | -63.8 | 1.832 | 1 | 0.1759 |
| rs77004360 | 11 | 120687126 | T | C | 0.691 | 0.4894 | +? | 0 | 0 | 0 | 1 |
| rs1787698 | 11 | 116564562 | T | C | 0.691 | 0.4896 | +? | 0 | 0 | 0 | 1 |
| rs1787699 | 11 | 116564553 | G | A | 0.691 | 0.4896 | +? | 0 | 0 | 0 | 1 |
| rs7939137 | 11 | 120343271 | T | C | -0.691 | 0.4896 | -? | 0 | 0 | 0 | 1 |
| rs7945424 | 11 | 119569802 | G | T | -0.691 | 0.4897 | -? | 0 | 0 | 0 | 1 |
| rs11218201 | 11 | 121086234 | C | T | -0.69 | 0.4899 | -? | 0 | 0 | 0 | 1 |
| rs12421432 | 11 | 119603847 | T | C | -0.69 | 0.49 | -? | 0 | 0 | 0 | 1 |
| rs678202 | 11 | 117899573 | G | A | 0.69 | 0.49 | +- | 0 | 0.449 | 1 | 0.503 |
| rs1047417 | 11 | 119172536 | G | A | 0.69 | 0.4901 | ++ | 0 | 0.299 | 1 | 0.5847 |
| rs4570590 | 11 | 120523139 | C | G | -0.69 | 0.4901 | -? | 0 | 0 | 0 | 1 |
| rs7950743 | 11 | 117613781 | A | G | 0.69 | 0.4902 | +? | 0 | 0 | 0 | 1 |
| rs1787703 | 11 | 116562266 | T | C | -0.69 | 0.4903 | -? | 0 | 0 | 0 | 1 |
| rs4938755 | 11 | 119878219 | G | A | -0.69 | 0.4903 | -? | 0 | 0 | 0 | 1 |
| rs556727 | 11 | 116250285 | G | A | -0.69 | 0.4903 | -? | 0 | 0 | 0 | 1 |
| rs10892511 | 11 | 119875093 | T | C | -0.69 | 0.4904 | -? | 0 | 0 | 0 | 1 |
| rs11827828 | 11 | 116755584 | A | G | 0.689 | 0.4906 | +? | 0 | 0 | 0 | 1 |
| rs2004507 | 11 | 117700725 | G | T | 0.689 | 0.4907 | +? | 0 | 0 | 0 | 1 |
| rs1533051 | 11 | 119551498 | T | C | 0.688 | 0.4912 | +? | 0 | 0 | 0 | 1 |
| rs1960036 | 11 | 116547409 | A | C | -0.688 | 0.4912 | -? | 0 | 0 | 0 | 1 |

| | | | | | | | | | | | |
|---|---|---|---|---|---|---|---|---|---|---|---|
| rs3018368 | 11 | 118113907 | G | C | -0.688 | 0.4912 | -? | 0 | 0 | 0 | 1 |
| rs11217176 | 11 | 119065044 | A | G | 0.688 | 0.4913 | +? | 0 | 0 | 0 | 1 |
| rs78390305 | 11 | 120702933 | A | G | 0.688 | 0.4913 | +? | 0 | 0 | 0 | 1 |
| rs12295634 | 11 | 118054431 | T | C | 0.688 | 0.4916 | ++ | 0 | 0.137 | 1 | 0.7117 |
| rs61898297 | 11 | 120109766 | T | C | 0.688 | 0.4916 | +? | 0 | 0 | 0 | 1 |
| rs12799449 | 11 | 116558525 | G | A | -0.688 | 0.4917 | -? | 0 | 0 | 0 | 1 |
| rs117758225 | 11 | 120165444 | T | C | -0.688 | 0.4918 | -- | 0 | 0.061 | 1 | 0.8048 |
| rs1793140 | 11 | 118107805 | A | G | -0.687 | 0.4919 | -? | 0 | 0 | 0 | 1 |
| rs4938450 | 11 | 117760117 | T | C | 0.687 | 0.492 | +? | 0 | 0 | 0 | 1 |
| rs482002 | 11 | 119330588 | T | A | 0.687 | 0.4922 | -+ | -52.9 | 1.962 | 1 | 0.1613 |
| rs59346420 | 11 | 117388656 | T | C | 0.687 | 0.4922 | +? | 0 | 0 | 0 | 1 |
| rs1790047 | 11 | 116251739 | C | T | -0.686 | 0.4924 | -? | 0 | 0 | 0 | 1 |
| rs603075 | 11 | 119365632 | C | G | 0.686 | 0.4924 | +? | 0 | 0 | 0 | 1 |
| rs689580 | 11 | 119365287 | A | G | 0.686 | 0.4924 | +? | 0 | 0 | 0 | 1 |
| rs76124005 | 11 | 116575614 | G | T | -0.687 | 0.4924 | -- | 0 | 0.003 | 1 | 0.9534 |
| rs11217803 | 11 | 120167566 | A | C | -0.686 | 0.4926 | -- | 0 | 0.013 | 1 | 0.909 |
| rs549893 | 11 | 118959732 | C | T | 0.686 | 0.4926 | +? | 0 | 0 | 0 | 1 |
| rs111551597 | 11 | 116375518 | T | C | 0.686 | 0.4927 | +? | 0 | 0 | 0 | 1 |
| rs12361680 | 11 | 119489210 | C | G | -0.686 | 0.4927 | -? | 0 | 0 | 0 | 1 |
| rs4936568 | 11 | 120825575 | A | G | 0.686 | 0.4927 | +? | 0 | 0 | 0 | 1 |
| rs116892227 | 11 | 116512747 | A | G | -0.686 | 0.493 | -? | 0 | 0 | 0 | 1 |
| rs7914 | 11 | 119180316 | A | G | -0.686 | 0.493 | -? | 0 | 0 | 0 | 1 |
| rs7931206 | 11 | 118056092 | C | T | 0.685 | 0.4932 | ++ | 0 | 0.079 | 1 | 0.7785 |
| rs7944087 | 11 | 118055733 | T | C | 0.685 | 0.4932 | ++ | 0 | 0.079 | 1 | 0.7785 |
| rs3782041 | 11 | 118184092 | T | C | 0.685 | 0.4933 | +? | 0 | 0 | 0 | 1 |
| rs12799766 | 11 | 116558427 | A | G | -0.685 | 0.4935 | -? | 0 | 0 | 0 | 1 |
| rs3759003 | 11 | 118136745 | G | T | 0.685 | 0.4936 | +? | 0 | 0 | 0 | 1 |
| rs117742492 | 11 | 116523657 | A | G | -0.684 | 0.4938 | -? | 0 | 0 | 0 | 1 |
| rs583713 | 11 | 118992177 | T | C | 0.684 | 0.4938 | +? | 0 | 0 | 0 | 1 |
| rs10892434 | 11 | 119573389 | T | C | -0.684 | 0.4941 | -? | 0 | 0 | 0 | 1 |
| rs2511841 | 11 | 119206375 | G | A | -0.684 | 0.4941 | -? | 0 | 0 | 0 | 1 |
| rs10750097 | 11 | 116664040 | G | A | -0.684 | 0.4942 | +- | -35.5 | 2.214 | 1 | 0.1368 |
| rs2512158 | 11 | 117823425 | A | G | 0.684 | 0.4942 | ++ | 0 | 0.019 | 1 | 0.8907 |
| rs6589650 | 11 | 118055217 | C | T | 0.684 | 0.4942 | ++ | 0 | 0.078 | 1 | 0.7805 |
| rs7940757 | 11 | 118055273 | T | C | 0.684 | 0.4942 | ++ | 0 | 0.078 | 1 | 0.7805 |
| rs75235704 | 11 | 117199742 | G | T | -0.683 | 0.4943 | -? | 0 | 0 | 0 | 1 |
| rs10892348 | 11 | 119194035 | T | C | -0.683 | 0.4944 | -? | 0 | 0 | 0 | 1 |
| rs35503012 | 11 | 119811841 | G | A | -0.683 | 0.4944 | -? | 0 | 0 | 0 | 1 |
| rs12222464 | 11 | 119224773 | C | T | 0.683 | 0.4945 | +? | 0 | 0 | 0 | 1 |
| rs1893392 | 11 | 119936695 | T | C | -0.683 | 0.4945 | -? | 0 | 0 | 0 | 1 |
| rs512168 | 11 | 116268668 | A | T | 0.683 | 0.4946 | +? | 0 | 0 | 0 | 1 |
| rs61906115 | 11 | 116558245 | A | T | -0.683 | 0.4947 | -? | 0 | 0 | 0 | 1 |
| rs7947338 | 11 | 117613203 | C | T | 0.683 | 0.4948 | +? | 0 | 0 | 0 | 1 |
| rs3809042 | 11 | 117749120 | G | C | -0.682 | 0.4951 | -? | 0 | 0 | 0 | 1 |
| rs582146 | 11 | 117952771 | T | G | -0.682 | 0.4951 | -? | 0 | 0 | 0 | 1 |
| rs10892108 | 11 | 117320727 | T | C | -0.682 | 0.4952 | -? | 0 | 0 | 0 | 1 |
| rs138877884 | 11 | 117135014 | A | G | -0.682 | 0.4952 | -? | 0 | 0 | 0 | 1 |
| rs12417581 | 11 | 120216906 | A | T | -0.682 | 0.4953 | -? | 0 | 0 | 0 | 1 |
| rs10892592 | 11 | 120522003 | T | C | -0.681 | 0.4956 | -? | 0 | 0 | 0 | 1 |
| rs4938362 | 11 | 117100029 | A | G | 0.681 | 0.4957 | +? | 0 | 0 | 0 | 1 |
| rs78443404 | 11 | 119935473 | C | G | 0.681 | 0.4957 | +? | 0 | 0 | 0 | 1 |
| rs11216180 | 11 | 116773378 | T | A | 0.681 | 0.4958 | +? | 0 | 0 | 0 | 1 |
| rs1715418 | 11 | 118112841 | T | A | -0.681 | 0.4958 | -? | 0 | 0 | 0 | 1 |
| rs628721 | 11 | 120987937 | G | A | -0.681 | 0.4958 | -- | 0 | 0.268 | 1 | 0.6046 |
| rs12419440 | 11 | 117318324 | C | T | 0.681 | 0.4959 | +? | 0 | 0 | 0 | 1 |
| rs2368940 | 11 | 120070770 | A | G | -0.681 | 0.4959 | -? | 0 | 0 | 0 | 1 |

| | | | | | | | | | | | |
|---|---|---|---|---|---|---|---|---|---|---|---|
| rs546734014 | 11 | 116762829 | A | G | 0.681 | 0.496 | +? | 0 | 0 | 0 | 1 |
| rs34236286 | 11 | 117642289 | C | T | 0.681 | 0.4961 | +? | 0 | 0 | 0 | 1 |
| rs518326 | 11 | 119710918 | A | T | 0.681 | 0.4961 | +? | 0 | 0 | 0 | 1 |
| rs12274740 | 11 | 118054128 | C | T | 0.68 | 0.4963 | ++ | 0 | 0.14 | 1 | 0.7085 |
| rs3759005 | 11 | 118136871 | G | A | 0.68 | 0.4963 | +? | 0 | 0 | 0 | 1 |
| rs78144220 | 11 | 117466755 | T | G | 0.68 | 0.4963 | +? | 0 | 0 | 0 | 1 |
| rs10790332 | 11 | 119553678 | T | C | 0.68 | 0.4966 | +? | 0 | 0 | 0 | 1 |
| rs3751026 | 11 | 119291688 | A | G | 0.68 | 0.4967 | -+ | -46.2 | 2.052 | 1 | 0.152 |
| rs4148167 | 11 | 119024931 | G | A | -0.68 | 0.4967 | -? | 0 | 0 | 0 | 1 |
| rs73000741 | 11 | 119935372 | A | G | -0.68 | 0.4967 | -? | 0 | 0 | 0 | 1 |
| rs73012959 | 11 | 116489644 | T | C | 0.68 | 0.4968 | +? | 0 | 0 | 0 | 1 |
| rs17123753 | 11 | 120131001 | G | C | -0.679 | 0.4969 | -? | 0 | 0 | 0 | 1 |
| rs117338034 | 11 | 116518743 | G | T | -0.679 | 0.497 | -? | 0 | 0 | 0 | 1 |
| rs74939223 | 11 | 116575880 | G | A | -0.679 | 0.497 | -- | 0 | 0.003 | 1 | 0.9568 |
| rs620566 | 11 | 119367238 | A | C | 0.679 | 0.4973 | +? | 0 | 0 | 0 | 1 |
| rs4938276 | 11 | 116393980 | T | C | -0.679 | 0.4974 | -? | 0 | 0 | 0 | 1 |
| rs77944184 | 11 | 117388657 | A | G | 0.679 | 0.4974 | +? | 0 | 0 | 0 | 1 |
| rs7944321 | 11 | 118055801 | A | G | 0.679 | 0.4974 | ++ | 0 | 0.077 | 1 | 0.7813 |
| rs7944366 | 11 | 118056047 | G | A | 0.679 | 0.4974 | ++ | 0 | 0.077 | 1 | 0.7813 |
| rs11822437 | 11 | 120637669 | A | T | 0.678 | 0.4975 | +? | 0 | 0 | 0 | 1 |
| rs4936517 | 11 | 120228286 | G | C | -0.678 | 0.4975 | -? | 0 | 0 | 0 | 1 |
| rs1145206 | 11 | 116718521 | C | G | 0.678 | 0.4978 | +? | 0 | 0 | 0 | 1 |
| rs1942925 | 11 | 117599235 | G | A | 0.678 | 0.4978 | +? | 0 | 0 | 0 | 1 |
| rs1942927 | 11 | 117599647 | G | A | 0.678 | 0.4978 | +? | 0 | 0 | 0 | 1 |
| rs948103 | 11 | 117597375 | C | A | 0.678 | 0.4978 | +? | 0 | 0 | 0 | 1 |
| rs533636 | 11 | 120036267 | A | G | 0.678 | 0.4979 | +? | 0 | 0 | 0 | 1 |
| rs11602075 | 11 | 120448992 | C | T | -0.677 | 0.4981 | -? | 0 | 0 | 0 | 1 |
| rs12293635 | 11 | 116305930 | C | G | 0.677 | 0.4981 | +? | 0 | 0 | 0 | 1 |
| rs4938503 | 11 | 118180155 | C | G | -0.677 | 0.4982 | -+ | -1.4 | 2.959 | 1 | 0.08541 |
| rs535860 | 11 | 117159878 | T | A | 0.677 | 0.4982 | +? | 0 | 0 | 0 | 1 |
| rs10892590 | 11 | 120515734 | G | A | -0.677 | 0.4983 | -? | 0 | 0 | 0 | 1 |
| rs4018665 | 11 | 119282086 | T | C | 0.677 | 0.4985 | -+ | -66.5 | 1.802 | 1 | 0.1795 |
| rs10892181 | 11 | 117699586 | G | T | 0.677 | 0.4987 | +? | 0 | 0 | 0 | 1 |
| rs12291852 | 11 | 120694915 | T | G | 0.676 | 0.4988 | +? | 0 | 0 | 0 | 1 |
| rs73575454 | 11 | 118807639 | A | G | 0.676 | 0.4988 | +- | -47.4 | 2.035 | 1 | 0.1537 |
| rs2853032 | 11 | 118114258 | G | C | -0.676 | 0.4989 | -? | 0 | 0 | 0 | 1 |
| rs58293014 | 11 | 119685035 | C | T | -0.676 | 0.4989 | -? | 0 | 0 | 0 | 1 |
| rs9888142 | 11 | 119197843 | G | A | -0.676 | 0.4989 | -? | 0 | 0 | 0 | 1 |
| rs12361632 | 11 | 120215843 | G | C | -0.676 | 0.499 | -? | 0 | 0 | 0 | 1 |
| rs1614945 | 11 | 118116522 | C | A | -0.676 | 0.499 | -? | 0 | 0 | 0 | 1 |
| rs75461585 | 11 | 118287568 | G | A | 0.676 | 0.4991 | +- | 0 | 0.932 | 1 | 0.3345 |
| rs117434119 | 11 | 119509845 | T | C | 0.676 | 0.4992 | +? | 0 | 0 | 0 | 1 |
| rs138568456 | 11 | 120915270 | G | A | -0.676 | 0.4992 | -? | 0 | 0 | 0 | 1 |
| rs73576603 | 11 | 116917885 | A | G | -0.676 | 0.4992 | +- | 46.7 | 5.634 | 1 | 0.01762 |
| rs10750147 | 11 | 119375180 | A | G | -0.676 | 0.4993 | -? | 0 | 0 | 0 | 1 |
| rs11216462 | 11 | 117456121 | C | G | 0.676 | 0.4993 | +? | 0 | 0 | 0 | 1 |
| rs2515795 | 11 | 117817276 | A | G | -0.675 | 0.4994 | -? | 0 | 0 | 0 | 1 |
| rs11217974 | 11 | 120562489 | G | A | -0.675 | 0.4995 | -? | 0 | 0 | 0 | 1 |
| rs56130750 | 11 | 119762963 | A | G | -0.675 | 0.4997 | -? | 0 | 0 | 0 | 1 |
| rs598494 | 11 | 119366884 | C | G | 0.675 | 0.4997 | +? | 0 | 0 | 0 | 1 |
| rs619180 | 11 | 119366947 | T | C | 0.675 | 0.4997 | +? | 0 | 0 | 0 | 1 |
| rs620109 | 11 | 119367127 | T | C | 0.675 | 0.4997 | +? | 0 | 0 | 0 | 1 |
| rs10790185 | 11 | 117320352 | T | C | 0.675 | 0.4998 | +? | 0 | 0 | 0 | 1 |
| rs12799480 | 11 | 120194024 | T | C | 0.675 | 0.4999 | +? | 0 | 0 | 0 | 1 |
| rs11218080 | 11 | 120825271 | T | C | 0.674 | 0.5 | +? | 0 | 0 | 0 | 1 |
| rs5093 | 11 | 116693354 | A | G | 0.674 | 0.5 | +? | 0 | 0 | 0 | 1 |

| | | | | | | | | | | | |
|---|---|---|---|---|---|---|---|---|---|---|---|
| rs4938656 | 11 | 119309436 | T | C | 0.674 | 0.5001 | -+ | -51.3 | 1.982 | 1 | 0.1591 |
| rs7113959 | 11 | 118379096 | A | T | -0.674 | 0.5002 | -+ | 0 | 0.913 | 1 | 0.3394 |
| rs11217506 | 11 | 119661809 | T | C | 0.674 | 0.5003 | ++ | 0 | 0.001 | 1 | 0.974 |
| rs146556537 | 11 | 117794456 | A | G | -0.674 | 0.5004 | -? | 0 | 0 | 0 | 1 |
| rs575021 | 11 | 119363369 | A | G | 0.674 | 0.5004 | +? | 0 | 0 | 0 | 1 |
| rs515278 | 11 | 119772239 | C | G | 0.674 | 0.5005 | +? | 0 | 0 | 0 | 1 |
| rs6589632 | 11 | 117759825 | T | C | 0.673 | 0.5007 | +? | 0 | 0 | 0 | 1 |
| rs7930658 | 11 | 117791353 | G | A | -0.673 | 0.5008 | -? | 0 | 0 | 0 | 1 |
| rs7940717 | 11 | 118055110 | T | G | 0.673 | 0.5008 | ++ | 0 | 0.077 | 1 | 0.7816 |
| rs111728839 | 11 | 120694966 | A | G | 0.673 | 0.5009 | +? | 0 | 0 | 0 | 1 |
| rs1240776 | 11 | 116536281 | T | C | 0.673 | 0.5009 | +? | 0 | 0 | 0 | 1 |
| rs1939674 | 11 | 120630546 | G | T | 0.673 | 0.501 | +? | 0 | 0 | 0 | 1 |
| rs7115526 | 11 | 117239190 | T | G | -0.673 | 0.501 | -? | 0 | 0 | 0 | 1 |
| rs1787690 | 11 | 116568516 | T | G | 0.673 | 0.5011 | +? | 0 | 0 | 0 | 1 |
| rs612979 | 11 | 117404179 | G | T | 0.672 | 0.5013 | +? | 0 | 0 | 0 | 1 |
| rs3018367 | 11 | 118119020 | T | C | -0.672 | 0.5014 | -? | 0 | 0 | 0 | 1 |
| rs73009916 | 11 | 117848134 | T | C | 0.672 | 0.5016 | +? | 0 | 0 | 0 | 1 |
| rs76477544 | 11 | 116308634 | G | A | 0.672 | 0.5017 | +? | 0 | 0 | 0 | 1 |
| rs477756 | 11 | 119702432 | C | T | 0.672 | 0.5018 | +? | 0 | 0 | 0 | 1 |
| rs1793135 | 11 | 118129705 | T | C | -0.672 | 0.5019 | -? | 0 | 0 | 0 | 1 |
| rs1418140540 | 11 | 120619909 | C | T | -0.671 | 0.502 | -? | 0 | 0 | 0 | 1 |
| rs28423005 | 11 | 118183733 | T | C | -0.671 | 0.502 | -+ | 1.8 | 3.055 | 1 | 0.08048 |
| rs139352710 | 11 | 117233378 | A | G | 0.671 | 0.5021 | +? | 0 | 0 | 0 | 1 |
| rs4938750 | 11 | 119876697 | A | G | -0.671 | 0.5021 | -? | 0 | 0 | 0 | 1 |
| rs4938753 | 11 | 119877108 | T | C | -0.671 | 0.5021 | -? | 0 | 0 | 0 | 1 |
| rs140110112 | 11 | 118629959 | A | G | 0.671 | 0.5022 | ++ | 0 | 0.247 | 1 | 0.6194 |
| rs34239965 | 11 | 119810909 | T | C | -0.671 | 0.5022 | -? | 0 | 0 | 0 | 1 |
| rs11218118 | 11 | 120896549 | G | A | -0.671 | 0.5024 | -? | 0 | 0 | 0 | 1 |
| rs10892433 | 11 | 119573259 | T | C | -0.67 | 0.5026 | -? | 0 | 0 | 0 | 1 |
| rs1715457 | 11 | 118123488 | A | G | -0.67 | 0.5026 | -? | 0 | 0 | 0 | 1 |
| rs1715458 | 11 | 118122737 | A | G | -0.67 | 0.5026 | -? | 0 | 0 | 0 | 1 |
| rs1939983 | 11 | 117604229 | G | C | 0.67 | 0.5026 | +? | 0 | 0 | 0 | 1 |
| rs7935561 | 11 | 118056722 | A | G | 0.67 | 0.5026 | ++ | 0 | 0.08 | 1 | 0.7777 |
| rs10790232 | 11 | 117976332 | A | C | -0.67 | 0.5027 | -? | 0 | 0 | 0 | 1 |
| rs10892214 | 11 | 117977388 | G | C | -0.67 | 0.5027 | -? | 0 | 0 | 0 | 1 |
| rs12361041 | 11 | 120821862 | G | A | 0.67 | 0.5028 | +? | 0 | 0 | 0 | 1 |
| rs11217815 | 11 | 120190262 | A | G | 0.67 | 0.503 | +? | 0 | 0 | 0 | 1 |
| rs4252279 | 11 | 117867187 | T | C | 0.669 | 0.5033 | +- | 0 | 0.238 | 1 | 0.6257 |
| rs947986 | 11 | 116569204 | A | G | 0.669 | 0.5035 | +? | 0 | 0 | 0 | 1 |
| rs55771168 | 11 | 119070949 | C | T | 0.669 | 0.5036 | +? | 0 | 0 | 0 | 1 |
| rs9651682 | 11 | 117240357 | A | G | -0.669 | 0.5036 | -? | 0 | 0 | 0 | 1 |
| rs11216804 | 11 | 118057320 | G | A | 0.669 | 0.5037 | ++ | 0 | 0.087 | 1 | 0.7676 |
| rs604361 | 11 | 119365917 | T | C | 0.668 | 0.5039 | +? | 0 | 0 | 0 | 1 |
| rs4936494 | 11 | 119568715 | A | G | -0.668 | 0.504 | -? | 0 | 0 | 0 | 1 |
| rs4936495 | 11 | 119568719 | C | T | -0.668 | 0.504 | -? | 0 | 0 | 0 | 1 |
| rs12796722 | 11 | 117473770 | T | C | -0.668 | 0.5041 | -+ | 0 | 0.153 | 1 | 0.6961 |
| rs579427 | 11 | 119363885 | T | C | 0.668 | 0.5041 | +? | 0 | 0 | 0 | 1 |
| rs73006770 | 11 | 119648394 | A | G | -0.668 | 0.5043 | -? | 0 | 0 | 0 | 1 |
| rs2155168 | 11 | 119876028 | T | C | -0.668 | 0.5044 | -? | 0 | 0 | 0 | 1 |
| rs885430 | 11 | 119572441 | G | A | -0.668 | 0.5044 | -? | 0 | 0 | 0 | 1 |
| rs2000874 | 11 | 120516717 | G | A | -0.667 | 0.5045 | -? | 0 | 0 | 0 | 1 |
| rs4142636 | 11 | 117604479 | T | C | 0.667 | 0.5047 | +? | 0 | 0 | 0 | 1 |
| rs1022081 | 11 | 119509746 | G | C | 0.667 | 0.5048 | +? | 0 | 0 | 0 | 1 |
| rs867899 | 11 | 116388314 | A | G | -0.667 | 0.5048 | -? | 0 | 0 | 0 | 1 |
| rs11218065 | 11 | 120808289 | T | G | -0.667 | 0.505 | -+ | 0 | 0.567 | 1 | 0.4514 |
| rs45509899 | 11 | 119217633 | T | C | 0.667 | 0.505 | +? | 0 | 0 | 0 | 1 |

| | | | | | | | | | | | |
|---|---|---|---|---|---|---|---|---|---|---|---|
| rs4936492 | 11 | 119560449 | A | G | 0.667 | 0.505 | +? | 0 | 0 | 0 | 1 |
| rs626258 | 11 | 119943918 | A | G | 0.667 | 0.505 | +? | 0 | 0 | 0 | 1 |
| rs11607356 | 11 | 116558455 | C | A | 0.666 | 0.5051 | +? | 0 | 0 | 0 | 1 |
| rs7934906 | 11 | 119488133 | T | C | -0.666 | 0.5055 | -? | 0 | 0 | 0 | 1 |
| rs17124570 | 11 | 120805056 | A | G | 0.666 | 0.5056 | +? | 0 | 0 | 0 | 1 |
| rs11217613 | 11 | 119858093 | G | A | -0.666 | 0.5057 | -? | 0 | 0 | 0 | 1 |
| rs4936561 | 11 | 120741935 | C | G | 0.666 | 0.5057 | +? | 0 | 0 | 0 | 1 |
| rs10750139 | 11 | 119065709 | G | C | -0.665 | 0.5058 | -? | 0 | 0 | 0 | 1 |
| rs673454 | 11 | 118992919 | A | C | 0.665 | 0.5058 | +? | 0 | 0 | 0 | 1 |
| rs3902979 | 11 | 120515929 | C | T | -0.665 | 0.5059 | -? | 0 | 0 | 0 | 1 |
| rs481716 | 11 | 119747312 | G | A | -0.665 | 0.5059 | -? | 0 | 0 | 0 | 1 |
| rs78740143 | 11 | 116486030 | A | G | -0.665 | 0.5059 | -? | 0 | 0 | 0 | 1 |
| rs57811256 | 11 | 117705239 | C | G | 0.665 | 0.506 | +? | 0 | 0 | 0 | 1 |
| rs629272 | 11 | 119365993 | A | G | 0.665 | 0.5061 | +? | 0 | 0 | 0 | 1 |
| rs10750172 | 11 | 120270253 | G | A | -0.665 | 0.5062 | -? | 0 | 0 | 0 | 1 |
| rs606630 | 11 | 119384079 | T | C | 0.665 | 0.5063 | +? | 0 | 0 | 0 | 1 |
| rs11216802 | 11 | 118057227 | T | C | 0.664 | 0.5065 | ++ | 0 | 0.082 | 1 | 0.775 |
| rs117275185 | 11 | 119690540 | T | C | -0.664 | 0.5065 | -? | 0 | 0 | 0 | 1 |
| rs12794946 | 11 | 120353298 | C | A | -0.664 | 0.5066 | -? | 0 | 0 | 0 | 1 |
| rs1343789 | 11 | 120516014 | C | T | -0.664 | 0.5066 | -? | 0 | 0 | 0 | 1 |
| rs7125256 | 11 | 119926054 | G | A | 0.664 | 0.5067 | +? | 0 | 0 | 0 | 1 |
| rs619054 | 11 | 116660813 | A | G | -0.664 | 0.5068 | -+ | 24 | 3.949 | 1 | 0.04691 |
| rs79446094 | 11 | 120694926 | G | A | 0.664 | 0.5068 | +? | 0 | 0 | 0 | 1 |
| rs4938638 | 11 | 119075429 | G | A | -0.664 | 0.507 | -? | 0 | 0 | 0 | 1 |
| rs7126918 | 11 | 119874855 | A | T | -0.663 | 0.5071 | -? | 0 | 0 | 0 | 1 |
| rs17245511 | 11 | 120186190 | A | G | -0.663 | 0.5073 | -? | 0 | 0 | 0 | 1 |
| rs4245210 | 11 | 119831773 | A | G | -0.663 | 0.5073 | -? | 0 | 0 | 0 | 1 |
| rs4936567 | 11 | 120819893 | T | G | -0.663 | 0.5073 | -+ | 0 | 0.564 | 1 | 0.4527 |
| rs10892508 | 11 | 119860539 | G | C | 0.663 | 0.5075 | +? | 0 | 0 | 0 | 1 |
| rs1784118 | 11 | 116565055 | T | C | 0.663 | 0.5076 | +? | 0 | 0 | 0 | 1 |
| rs1787717 | 11 | 116552766 | T | C | 0.663 | 0.5076 | +? | 0 | 0 | 0 | 1 |
| rs669226 | 11 | 117450432 | C | A | 0.662 | 0.5079 | +? | 0 | 0 | 0 | 1 |
| rs10892098 | 11 | 117238491 | T | C | -0.661 | 0.5083 | -? | 0 | 0 | 0 | 1 |
| rs12797135 | 11 | 117473944 | T | C | -0.662 | 0.5083 | -+ | 0 | 0.15 | 1 | 0.6989 |
| rs1784126 | 11 | 116538215 | T | G | 0.661 | 0.5084 | +? | 0 | 0 | 0 | 1 |
| rs1787709 | 11 | 116561328 | G | C | -0.661 | 0.5085 | -? | 0 | 0 | 0 | 1 |
| rs61582023 | 11 | 116911784 | G | A | -0.661 | 0.5085 | +- | 47.1 | 5.673 | 1 | 0.01723 |
| rs11217257 | 11 | 119238455 | G | A | 0.661 | 0.5088 | +? | 0 | 0 | 0 | 1 |
| rs35176806 | 11 | 120851210 | G | C | -0.661 | 0.5089 | -? | 0 | 0 | 0 | 1 |
| rs11216803 | 11 | 118057244 | A | C | 0.66 | 0.509 | ++ | 0 | 0.083 | 1 | 0.7733 |
| rs11216857 | 11 | 118235246 | A | G | 0.66 | 0.5093 | +- | 6.7 | 3.214 | 1 | 0.07302 |
| rs882398 | 11 | 116573838 | C | T | -0.66 | 0.5093 | -? | 0 | 0 | 0 | 1 |
| rs948775 | 11 | 117586698 | C | T | 0.66 | 0.5093 | +? | 0 | 0 | 0 | 1 |
| rs667319 | 11 | 119364258 | G | A | 0.66 | 0.5094 | +? | 0 | 0 | 0 | 1 |
| rs79481471 | 11 | 118296658 | A | G | 0.66 | 0.5096 | ++ | 0 | 0.165 | 1 | 0.6846 |
| rs1893261 | 11 | 120198093 | A | G | -0.659 | 0.5099 | -? | 0 | 0 | 0 | 1 |
| rs2508445 | 11 | 117872544 | T | G | 0.659 | 0.51 | ++ | 0 | 0.006 | 1 | 0.9364 |
| rs534761 | 11 | 117662248 | T | C | -0.659 | 0.5102 | -? | 0 | 0 | 0 | 1 |
| rs577651 | 11 | 117471825 | T | C | -0.658 | 0.5104 | -? | 0 | 0 | 0 | 1 |
| rs74661474 | 11 | 117250530 | A | G | -0.658 | 0.5106 | -? | 0 | 0 | 0 | 1 |
| rs1263160 | 11 | 116744447 | C | T | -0.657 | 0.5109 | +- | 49.8 | 5.977 | 1 | 0.0145 |
| rs3020103 | 11 | 120685533 | G | A | 0.657 | 0.5109 | +? | 0 | 0 | 0 | 1 |
| rs10892474 | 11 | 119687133 | C | T | 0.657 | 0.511 | +? | 0 | 0 | 0 | 1 |
| rs79845538 | 11 | 119686891 | C | T | 0.657 | 0.511 | +? | 0 | 0 | 0 | 1 |
| rs2305010 | 11 | 120330149 | C | G | -0.657 | 0.5112 | -? | 0 | 0 | 0 | 1 |
| rs75954141 | 11 | 120694254 | C | A | 0.657 | 0.5112 | +? | 0 | 0 | 0 | 1 |

| | | | | | | | | | | | |
|---|---|---|---|---|---|---|---|---|---|---|---|
| rs630403 | 11 | 118027074 | A | G | -0.657 | 0.5114 | -? | 0 | 0 | 0 | 1 |
| rs7126576 | 11 | 117420748 | C | T | -0.657 | 0.5114 | -? | 0 | 0 | 0 | 1 |
| rs35462788 | 11 | 119811909 | T | C | -0.657 | 0.5115 | -? | 0 | 0 | 0 | 1 |
| rs139254418 | 11 | 117380309 | G | A | 0.656 | 0.5116 | +? | 0 | 0 | 0 | 1 |
| rs45555132 | 11 | 118064030 | T | G | -0.656 | 0.5116 | -? | 0 | 0 | 0 | 1 |
| rs7118773 | 11 | 119969933 | A | G | 0.656 | 0.5116 | +? | 0 | 0 | 0 | 1 |
| rs1784127 | 11 | 116538576 | A | G | 0.656 | 0.5118 | +? | 0 | 0 | 0 | 1 |
| rs11216447 | 11 | 117416832 | A | G | 0.656 | 0.5119 | +? | 0 | 0 | 0 | 1 |
| rs7104454 | 11 | 120636167 | G | A | 0.656 | 0.512 | +? | 0 | 0 | 0 | 1 |
| rs10892647 | 11 | 120804291 | G | A | 0.656 | 0.5121 | +? | 0 | 0 | 0 | 1 |
| rs117476293 | 11 | 120898097 | C | A | 0.656 | 0.5121 | +? | 0 | 0 | 0 | 1 |
| rs4573685 | 11 | 120696297 | A | G | 0.656 | 0.5121 | +? | 0 | 0 | 0 | 1 |
| rs144553689 | 11 | 119381144 | C | T | -0.655 | 0.5123 | -? | 0 | 0 | 0 | 1 |
| rs76181827 | 11 | 120275039 | G | C | -0.655 | 0.5123 | -? | 0 | 0 | 0 | 1 |
| rs77779915 | 11 | 116232179 | T | G | -0.655 | 0.5124 | -? | 0 | 0 | 0 | 1 |
| rs11216432 | 11 | 117379572 | T | C | 0.655 | 0.5125 | +? | 0 | 0 | 0 | 1 |
| rs12271393 | 11 | 120149004 | A | G | -0.655 | 0.5125 | -? | 0 | 0 | 0 | 1 |
| rs1473177 | 11 | 116759332 | C | T | 0.655 | 0.5125 | +? | 0 | 0 | 0 | 1 |
| rs75208530 | 11 | 120694466 | G | A | 0.654 | 0.5129 | +? | 0 | 0 | 0 | 1 |
| rs117889337 | 11 | 116564493 | T | C | -0.654 | 0.5132 | -- | 0 | 0.015 | 1 | 0.9035 |
| rs61903734 | 11 | 117201056 | A | T | -0.654 | 0.5132 | -? | 0 | 0 | 0 | 1 |
| rs71482170 | 11 | 119019877 | A | G | 0.654 | 0.5132 | +? | 0 | 0 | 0 | 1 |
| rs1126924 | 11 | 118178007 | T | C | -0.654 | 0.5133 | -+ | -4.5 | 2.871 | 1 | 0.09016 |
| rs151336980 | 11 | 120470737 | T | C | -0.654 | 0.5134 | -? | 0 | 0 | 0 | 1 |
| rs74676521 | 11 | 116335869 | A | C | -0.654 | 0.5134 | -? | 0 | 0 | 0 | 1 |
| rs1942928 | 11 | 117605046 | G | A | 0.653 | 0.5136 | +? | 0 | 0 | 0 | 1 |
| rs1558861 | 11 | 116607437 | C | T | -0.653 | 0.5137 | +- | -125.2 | 1.332 | 1 | 0.2484 |
| rs35734851 | 11 | 120206746 | A | G | 0.653 | 0.5138 | +? | 0 | 0 | 0 | 1 |
| rs4938449 | 11 | 117757685 | A | T | 0.653 | 0.5138 | +? | 0 | 0 | 0 | 1 |
| rs12419495 | 11 | 117383816 | C | T | 0.652 | 0.5142 | +? | 0 | 0 | 0 | 1 |
| rs36011588 | 11 | 120196433 | G | C | -0.652 | 0.5144 | -? | 0 | 0 | 0 | 1 |
| rs60016568 | 11 | 121062777 | T | C | -0.652 | 0.5144 | -? | 0 | 0 | 0 | 1 |
| rs1784123 | 11 | 116561381 | C | T | -0.652 | 0.5145 | -? | 0 | 0 | 0 | 1 |
| rs4561221 | 11 | 120444179 | T | C | -0.651 | 0.5149 | -? | 0 | 0 | 0 | 1 |
| rs9326246 | 11 | 116611733 | C | G | -0.651 | 0.5149 | +- | -124.8 | 1.335 | 1 | 0.248 |
| rs683630 | 11 | 119365552 | T | C | 0.651 | 0.5151 | +? | 0 | 0 | 0 | 1 |
| rs74490241 | 11 | 117309494 | A | C | 0.651 | 0.5151 | +? | 0 | 0 | 0 | 1 |
| rs79423267 | 11 | 117312576 | A | G | 0.651 | 0.5151 | +? | 0 | 0 | 0 | 1 |
| rs11216379 | 11 | 117291756 | G | A | -0.651 | 0.5152 | -? | 0 | 0 | 0 | 1 |
| rs1145184 | 11 | 116513182 | A | G | 0.651 | 0.5152 | +? | 0 | 0 | 0 | 1 |
| rs888246 | 11 | 116724232 | T | C | 0.651 | 0.5153 | +? | 0 | 0 | 0 | 1 |
| rs11216622 | 11 | 117791929 | A | C | -0.65 | 0.5154 | -? | 0 | 0 | 0 | 1 |
| rs4252243 | 11 | 117855727 | T | C | 0.65 | 0.5154 | +- | 0 | 0.213 | 1 | 0.6444 |
| rs57584368 | 11 | 120783331 | C | T | -0.65 | 0.5157 | -? | 0 | 0 | 0 | 1 |
| rs111765459 | 11 | 117083373 | C | T | -0.65 | 0.5158 | -? | 0 | 0 | 0 | 1 |
| rs574387 | 11 | 118337003 | C | A | 0.65 | 0.5158 | ++ | 0 | 0.003 | 1 | 0.96 |
| rs608840 | 11 | 117150210 | A | G | 0.65 | 0.5158 | +? | 0 | 0 | 0 | 1 |
| rs10892003 | 11 | 116507479 | G | A | -0.65 | 0.5159 | -? | 0 | 0 | 0 | 1 |
| rs4537793 | 11 | 118053156 | A | C | 0.65 | 0.5159 | ++ | 0 | 0.705 | 1 | 0.4011 |
| rs11217526 | 11 | 119697028 | A | T | 0.649 | 0.5161 | +? | 0 | 0 | 0 | 1 |
| rs12273260 | 11 | 118137524 | T | C | 0.649 | 0.5161 | +- | 18.9 | 3.697 | 1 | 0.0545 |
| rs11217547 | 11 | 119742279 | A | G | 0.649 | 0.5162 | +? | 0 | 0 | 0 | 1 |
| rs4936562 | 11 | 120741937 | C | T | 0.649 | 0.5162 | +? | 0 | 0 | 0 | 1 |
| rs680210 | 11 | 117172820 | T | C | -0.649 | 0.5165 | -? | 0 | 0 | 0 | 1 |
| rs501171 | 11 | 117692394 | T | C | -0.649 | 0.5166 | -? | 0 | 0 | 0 | 1 |
| rs183832950 | 11 | 120633447 | T | G | 0.648 | 0.5168 | +? | 0 | 0 | 0 | 1 |

| | | | | | | | | | | | |
|---|---|---|---|---|---|---|---|---|---|---|---|
| rs1970750 | 11 | 118149122 | T | C | 0.648 | 0.5169 | +? | 0 | 0 | 0 | 1 |
| rs7950804 | 11 | 119683867 | G | A | 0.648 | 0.5169 | ++ | 0 | 0.003 | 1 | 0.9564 |
| rs721486 | 11 | 117557540 | G | A | 0.648 | 0.517 | +? | 0 | 0 | 0 | 1 |
| rs7950719 | 11 | 119683955 | C | T | 0.648 | 0.5172 | ++ | 0 | 0.003 | 1 | 0.9562 |
| rs891291 | 11 | 119376478 | C | G | -0.648 | 0.5172 | -? | 0 | 0 | 0 | 1 |
| rs1793165 | 11 | 118115879 | A | C | -0.648 | 0.5173 | -? | 0 | 0 | 0 | 1 |
| rs7950042 | 11 | 119683473 | A | C | 0.647 | 0.5173 | ++ | 0 | 0.003 | 1 | 0.9545 |
| rs3176905 | 11 | 118755630 | C | T | 0.647 | 0.5174 | +- | 29 | 4.223 | 1 | 0.03988 |
| rs6589796 | 11 | 119970538 | A | C | 0.647 | 0.5174 | +? | 0 | 0 | 0 | 1 |
| rs9804646 | 11 | 116665079 | T | C | -0.647 | 0.5174 | -- | 0 | 0.127 | 1 | 0.7216 |
| rs4381380 | 11 | 120273316 | A | G | -0.647 | 0.5176 | -? | 0 | 0 | 0 | 1 |
| rs11218169 | 11 | 121029733 | G | A | -0.647 | 0.5179 | -? | 0 | 0 | 0 | 1 |
| rs78053944 | 11 | 117795742 | T | C | 0.647 | 0.5179 | +? | 0 | 0 | 0 | 1 |
| rs2850804 | 11 | 120793972 | A | G | 0.646 | 0.5181 | +- | -77.7 | 1.688 | 1 | 0.1938 |
| rs11215931 | 11 | 116308090 | G | A | -0.646 | 0.5183 | -? | 0 | 0 | 0 | 1 |
| rs3935405 | 11 | 119572103 | T | C | -0.645 | 0.5189 | -? | 0 | 0 | 0 | 1 |
| rs611478 | 11 | 119784330 | A | G | 0.645 | 0.519 | +? | 0 | 0 | 0 | 1 |
| rs2512155 | 11 | 117889448 | T | C | 0.645 | 0.5192 | +- | -115 | 1.396 | 1 | 0.2375 |
| rs3018284 | 11 | 120612223 | G | T | 0.645 | 0.5192 | +? | 0 | 0 | 0 | 1 |
| rs7115491 | 11 | 117295150 | G | T | -0.645 | 0.5192 | -? | 0 | 0 | 0 | 1 |
| rs7950403 | 11 | 119683777 | A | G | 0.644 | 0.5194 | ++ | 0 | 0.003 | 1 | 0.9546 |
| rs111444487 | 11 | 119853252 | C | T | -0.644 | 0.5195 | -? | 0 | 0 | 0 | 1 |
| rs7107003 | 11 | 116532855 | C | T | 0.644 | 0.5196 | +? | 0 | 0 | 0 | 1 |
| rs891292 | 11 | 119376672 | G | A | -0.644 | 0.5196 | -? | 0 | 0 | 0 | 1 |
| rs751368 | 11 | 120610584 | T | G | 0.644 | 0.5198 | +? | 0 | 0 | 0 | 1 |
| rs689291 | 11 | 119361783 | C | G | 0.643 | 0.5199 | +? | 0 | 0 | 0 | 1 |
| rs10160375 | 11 | 117954250 | C | T | 0.643 | 0.5201 | ++ | 0 | 0.129 | 1 | 0.7194 |
| rs686593 | 11 | 119362414 | G | C | 0.643 | 0.5201 | +? | 0 | 0 | 0 | 1 |
| rs28989473 | 11 | 117179535 | A | G | 0.643 | 0.5203 | +? | 0 | 0 | 0 | 1 |
| rs12226432 | 11 | 120093920 | C | A | 0.642 | 0.5207 | +? | 0 | 0 | 0 | 1 |
| rs2155809 | 11 | 120148567 | A | T | -0.642 | 0.5207 | -? | 0 | 0 | 0 | 1 |
| rs4936368 | 11 | 117093486 | T | C | -0.642 | 0.5207 | -? | 0 | 0 | 0 | 1 |
| rs4938359 | 11 | 117093560 | G | A | -0.642 | 0.5207 | -? | 0 | 0 | 0 | 1 |
| rs4938749 | 11 | 119876516 | G | A | -0.642 | 0.5208 | -? | 0 | 0 | 0 | 1 |
| rs79219038 | 11 | 120159612 | T | C | -0.642 | 0.5209 | -? | 0 | 0 | 0 | 1 |
| rs10790224 | 11 | 117805536 | C | T | -0.642 | 0.5211 | -? | 0 | 0 | 0 | 1 |
| rs502483 | 11 | 116251791 | T | A | -0.642 | 0.5211 | -? | 0 | 0 | 0 | 1 |
| rs7105258 | 11 | 120657044 | T | C | 0.641 | 0.5212 | +? | 0 | 0 | 0 | 1 |
| rs747061 | 11 | 117330736 | G | C | 0.641 | 0.5213 | +? | 0 | 0 | 0 | 1 |
| rs4445645 | 11 | 118868750 | C | G | -0.641 | 0.5215 | -- | 0 | 0.004 | 1 | 0.9526 |
| rs11601535 | 11 | 120333801 | G | A | -0.641 | 0.5217 | -? | 0 | 0 | 0 | 1 |
| rs11600380 | 11 | 116670182 | C | T | -0.641 | 0.5218 | -+ | 0 | 0.388 | 1 | 0.5331 |
| rs61905260 | 11 | 117300380 | A | G | -0.641 | 0.5218 | -? | 0 | 0 | 0 | 1 |
| rs3133864 | 11 | 120628012 | T | C | -0.64 | 0.5219 | -? | 0 | 0 | 0 | 1 |
| rs79326407 | 11 | 116778293 | T | C | 0.64 | 0.5219 | +? | 0 | 0 | 0 | 1 |
| rs111809212 | 11 | 117099537 | C | T | 0.64 | 0.5222 | +? | 0 | 0 | 0 | 1 |
| rs17475957 | 11 | 116241411 | A | G | 0.64 | 0.5224 | +? | 0 | 0 | 0 | 1 |
| rs642548 | 11 | 120988792 | C | T | -0.64 | 0.5224 | -- | 0 | 0.293 | 1 | 0.5881 |
| rs6589620 | 11 | 117612254 | A | T | 0.64 | 0.5224 | +? | 0 | 0 | 0 | 1 |
| rs7110219 | 11 | 117420336 | A | G | -0.639 | 0.5225 | -? | 0 | 0 | 0 | 1 |
| rs74900182 | 11 | 121164050 | T | C | 0.639 | 0.5226 | +? | 0 | 0 | 0 | 1 |
| rs10892648 | 11 | 120812025 | A | C | 0.639 | 0.5228 | +? | 0 | 0 | 0 | 1 |
| rs71480317 | 11 | 116572169 | C | T | 0.639 | 0.5228 | +? | 0 | 0 | 0 | 1 |
| rs7483350 | 11 | 119040463 | G | C | -0.639 | 0.5228 | -? | 0 | 0 | 0 | 1 |
| rs470941 | 11 | 119999755 | C | T | 0.639 | 0.5229 | +? | 0 | 0 | 0 | 1 |
| rs536859 | 11 | 117362425 | A | G | 0.639 | 0.523 | +? | 0 | 0 | 0 | 1 |

| | | | | | | | | | | | |
|---|---|---|---|---|---|---|---|---|---|---|---|
| rs117754274 | 11 | 117225247 | C | G | 0.639 | 0.5231 | +? | 0 | 0 | 0 | 1 |
| rs12281635 | 11 | 120496044 | G | A | 0.638 | 0.5233 | +? | 0 | 0 | 0 | 1 |
| rs1374117 | 11 | 116709438 | T | C | 0.638 | 0.5233 | +? | 0 | 0 | 0 | 1 |
| rs113078426 | 11 | 119795996 | G | A | -0.638 | 0.5234 | -? | 0 | 0 | 0 | 1 |
| rs34458464 | 11 | 117628944 | G | A | 0.638 | 0.5235 | +? | 0 | 0 | 0 | 1 |
| rs56257632 | 11 | 116959601 | C | T | -0.638 | 0.5235 | +- | 47.7 | 5.737 | 1 | 0.01661 |
| rs71484170 | 11 | 120341938 | C | T | 0.637 | 0.524 | +? | 0 | 0 | 0 | 1 |
| rs2105610 | 11 | 116535126 | C | T | -0.637 | 0.5241 | -? | 0 | 0 | 0 | 1 |
| rs28917234 | 11 | 117186506 | A | G | 0.637 | 0.5241 | +? | 0 | 0 | 0 | 1 |
| rs7927820 | 11 | 116682156 | G | A | 0.637 | 0.5243 | +? | 0 | 0 | 0 | 1 |
| rs10892624 | 11 | 120651537 | A | G | 0.637 | 0.5244 | +? | 0 | 0 | 0 | 1 |
| rs576642 | 11 | 119987219 | C | T | -0.636 | 0.5245 | -? | 0 | 0 | 0 | 1 |
| rs949295 | 11 | 120789852 | C | T | 0.636 | 0.5245 | +? | 0 | 0 | 0 | 1 |
| rs4252246 | 11 | 117858185 | A | C | 0.636 | 0.5247 | +- | 0 | 0.231 | 1 | 0.6307 |
| rs3902978 | 11 | 120515939 | A | G | -0.636 | 0.5248 | -? | 0 | 0 | 0 | 1 |
| rs895645 | 11 | 119684321 | G | A | 0.636 | 0.525 | ++ | 0 | 0.004 | 1 | 0.9507 |
| rs895647 | 11 | 119684145 | T | G | 0.636 | 0.525 | ++ | 0 | 0.004 | 1 | 0.9507 |
| rs61122660 | 11 | 120069087 | A | G | 0.635 | 0.5253 | +? | 0 | 0 | 0 | 1 |
| rs11601826 | 11 | 120440043 | G | A | -0.635 | 0.5256 | -? | 0 | 0 | 0 | 1 |
| rs744984 | 11 | 120166410 | G | A | -0.635 | 0.5256 | -- | 0 | 0.047 | 1 | 0.8283 |
| rs1787708 | 11 | 116561393 | A | G | -0.634 | 0.5258 | -? | 0 | 0 | 0 | 1 |
| rs10750143 | 11 | 119332065 | G | A | -0.634 | 0.5259 | -? | 0 | 0 | 0 | 1 |
| rs113048801 | 11 | 118827925 | A | G | -0.634 | 0.5259 | -? | 0 | 0 | 0 | 1 |
| rs74808315 | 11 | 117107520 | G | C | -0.634 | 0.5259 | -? | 0 | 0 | 0 | 1 |
| rs1207849774 | 11 | 119989572 | A | T | 0.634 | 0.526 | +? | 0 | 0 | 0 | 1 |
| rs525028 | 11 | 116705516 | G | A | 0.634 | 0.5262 | +? | 0 | 0 | 0 | 1 |
| rs12222094 | 11 | 119163540 | C | G | 0.634 | 0.5263 | ++ | 0 | 0.339 | 1 | 0.5606 |
| rs77451506 | 11 | 117142440 | G | C | -0.634 | 0.5264 | -? | 0 | 0 | 0 | 1 |
| rs75723689 | 11 | 116468539 | A | G | 0.633 | 0.5265 | +? | 0 | 0 | 0 | 1 |
| rs592525 | 11 | 119383853 | A | G | 0.633 | 0.5266 | +? | 0 | 0 | 0 | 1 |
| rs12576431 | 11 | 116225940 | G | C | 0.633 | 0.5267 | +- | -66.4 | 1.803 | 1 | 0.1793 |
| rs3133851 | 11 | 120600950 | A | G | 0.633 | 0.5267 | +? | 0 | 0 | 0 | 1 |
| rs11217231 | 11 | 119162720 | C | G | 0.633 | 0.5268 | ++ | 0 | 0.339 | 1 | 0.5604 |
| rs4245191 | 11 | 119052826 | C | A | -0.633 | 0.5268 | -? | 0 | 0 | 0 | 1 |
| rs481400 | 11 | 119784002 | G | A | 0.633 | 0.5268 | +? | 0 | 0 | 0 | 1 |
| rs4245040 | 11 | 120520629 | T | C | -0.632 | 0.5271 | -? | 0 | 0 | 0 | 1 |
| rs531411 | 11 | 119786013 | A | G | 0.632 | 0.5271 | +? | 0 | 0 | 0 | 1 |
| rs895646 | 11 | 119684301 | T | G | 0.632 | 0.5272 | ++ | 0 | 0.004 | 1 | 0.9492 |
| rs34352231 | 11 | 117632531 | G | A | 0.632 | 0.5273 | +? | 0 | 0 | 0 | 1 |
| rs58709307 | 11 | 118877563 | T | C | -0.631 | 0.5277 | -- | 0 | 0.013 | 1 | 0.9095 |
| rs592347 | 11 | 119969030 | T | C | -0.631 | 0.5278 | -? | 0 | 0 | 0 | 1 |
| rs75808396 | 11 | 117309527 | A | G | 0.631 | 0.5279 | +? | 0 | 0 | 0 | 1 |
| rs7934690 | 11 | 117011299 | A | T | 0.631 | 0.5279 | +? | 0 | 0 | 0 | 1 |
| rs143291574 | 11 | 116365421 | T | C | -0.631 | 0.5281 | -? | 0 | 0 | 0 | 1 |
| rs523223 | 11 | 119763417 | A | G | 0.631 | 0.5283 | +? | 0 | 0 | 0 | 1 |
| rs78459638 | 11 | 116972339 | A | G | 0.631 | 0.5283 | +? | 0 | 0 | 0 | 1 |
| rs2121576 | 11 | 119683031 | G | A | 0.63 | 0.5288 | ++ | 0 | 0.005 | 1 | 0.9465 |
| rs2121577 | 11 | 119682999 | G | A | 0.63 | 0.5288 | ++ | 0 | 0.005 | 1 | 0.9465 |
| rs7936274 | 11 | 117324393 | T | C | 0.63 | 0.5289 | +? | 0 | 0 | 0 | 1 |
| rs11218081 | 11 | 120825332 | C | T | -0.629 | 0.5292 | -+ | 0 | 0.535 | 1 | 0.4644 |
| rs7937319 | 11 | 119683964 | T | C | 0.629 | 0.5293 | ++ | 0 | 0.004 | 1 | 0.9477 |
| rs78760381 | 11 | 117922310 | T | C | 0.629 | 0.5294 | +? | 0 | 0 | 0 | 1 |
| rs2239012 | 11 | 117088362 | G | A | -0.629 | 0.5295 | -? | 0 | 0 | 0 | 1 |
| rs34172923 | 11 | 120145046 | G | C | -0.629 | 0.5295 | -? | 0 | 0 | 0 | 1 |
| rs695083 | 11 | 117508837 | G | A | -0.629 | 0.5295 | -? | 0 | 0 | 0 | 1 |
| rs7396851 | 11 | 116684164 | T | C | 0.629 | 0.5296 | +? | 0 | 0 | 0 | 1 |

| | | | | | | | | | | | |
|---|---|---|---|---|---|---|---|---|---|---|---|
| rs4252249 | 11 | 117859209 | A | G | 0.628 | 0.5297 | ++ | 0 | 0.113 | 1 | 0.737 |
| rs2305008 | 11 | 120319064 | C | A | -0.628 | 0.5298 | -? | 0 | 0 | 0 | 1 |
| rs11216435 | 11 | 117388932 | T | C | -0.628 | 0.5299 | -? | 0 | 0 | 0 | 1 |
| rs117243996 | 11 | 119940917 | T | C | 0.628 | 0.53 | +? | 0 | 0 | 0 | 1 |
| rs45514102 | 11 | 119061879 | C | G | -0.628 | 0.5301 | -? | 0 | 0 | 0 | 1 |
| rs4938727 | 11 | 119684565 | C | T | 0.628 | 0.5301 | ++ | 0 | 0.004 | 1 | 0.9472 |
| rs4245213 | 11 | 120273597 | A | G | -0.628 | 0.5303 | -? | 0 | 0 | 0 | 1 |
| rs73022414 | 11 | 117787277 | C | T | 0.628 | 0.5303 | +? | 0 | 0 | 0 | 1 |
| rs2846104 | 11 | 120652371 | G | C | 0.627 | 0.5306 | +? | 0 | 0 | 0 | 1 |
| rs4938728 | 11 | 119684646 | T | C | 0.627 | 0.5306 | ++ | 0 | 0.004 | 1 | 0.9468 |
| rs10111 | 11 | 118886117 | T | C | -0.627 | 0.5307 | -- | 0 | 0.018 | 1 | 0.8926 |
| rs1790474 | 11 | 119942571 | A | G | -0.627 | 0.5307 | -? | 0 | 0 | 0 | 1 |
| rs2121575 | 11 | 119683218 | C | G | 0.627 | 0.5308 | ++ | 0 | 0.005 | 1 | 0.9451 |
| rs3825050 | 11 | 117952097 | T | C | -0.627 | 0.5308 | -? | 0 | 0 | 0 | 1 |
| rs76468794 | 11 | 116576744 | A | G | -0.627 | 0.5308 | -- | 0 | 0 | 1 | 0.9922 |
| rs6589572 | 11 | 116681563 | G | A | 0.626 | 0.5312 | +? | 0 | 0 | 0 | 1 |
| rs71486352 | 11 | 121010048 | A | T | -0.626 | 0.5312 | -? | 0 | 0 | 0 | 1 |
| rs78302438 | 11 | 120184575 | G | T | -0.626 | 0.5313 | -? | 0 | 0 | 0 | 1 |
| rs11216665 | 11 | 117851174 | G | T | 0.626 | 0.5314 | ++ | 0 | 0.245 | 1 | 0.6203 |
| rs4936511 | 11 | 120150059 | C | A | -0.626 | 0.5314 | -? | 0 | 0 | 0 | 1 |
| rs477047 | 11 | 116485583 | A | T | 0.626 | 0.5315 | +? | 0 | 0 | 0 | 1 |
| rs79053194 | 11 | 116257372 | G | T | 0.626 | 0.5315 | +? | 0 | 0 | 0 | 1 |
| rs1073635 | 11 | 119972626 | T | C | -0.625 | 0.5321 | -? | 0 | 0 | 0 | 1 |
| rs11218005 | 11 | 120652936 | T | G | 0.625 | 0.5321 | +? | 0 | 0 | 0 | 1 |
| rs189431300 | 11 | 120034758 | A | G | -0.624 | 0.5323 | -? | 0 | 0 | 0 | 1 |
| rs11218082 | 11 | 120825348 | C | G | -0.624 | 0.5324 | -+ | 0 | 0.531 | 1 | 0.4661 |
| rs17492429 | 11 | 116453270 | C | G | 0.624 | 0.5325 | ++ | 0 | 0.02 | 1 | 0.8881 |
| rs2509390 | 11 | 119196915 | C | A | -0.624 | 0.5326 | -? | 0 | 0 | 0 | 1 |
| rs75343916 | 11 | 117311365 | T | C | 0.624 | 0.5326 | +? | 0 | 0 | 0 | 1 |
| rs3882895 | 11 | 117090545 | C | T | -0.624 | 0.5327 | -? | 0 | 0 | 0 | 1 |
| rs4938434 | 11 | 117592602 | G | T | 0.624 | 0.5327 | +? | 0 | 0 | 0 | 1 |
| rs2239010 | 11 | 117089006 | C | G | -0.624 | 0.5328 | -? | 0 | 0 | 0 | 1 |
| rs2239011 | 11 | 117088994 | G | A | -0.624 | 0.5328 | -? | 0 | 0 | 0 | 1 |
| rs10750110 | 11 | 117614307 | G | T | 0.624 | 0.5329 | +? | 0 | 0 | 0 | 1 |
| rs10892481 | 11 | 119742360 | T | C | 0.623 | 0.533 | +? | 0 | 0 | 0 | 1 |
| rs504485 | 11 | 119767614 | A | C | 0.623 | 0.5332 | +? | 0 | 0 | 0 | 1 |
| rs71480343 | 11 | 117421276 | A | G | -0.623 | 0.5332 | -? | 0 | 0 | 0 | 1 |
| rs4938477 | 11 | 117956579 | G | A | 0.623 | 0.5333 | +- | 0 | 0.248 | 1 | 0.6188 |
| rs10750174 | 11 | 120292097 | T | C | -0.623 | 0.5334 | -? | 0 | 0 | 0 | 1 |
| rs10892576 | 11 | 120294423 | T | C | -0.623 | 0.5334 | -? | 0 | 0 | 0 | 1 |
| rs1787710 | 11 | 116561202 | A | G | -0.623 | 0.5334 | -? | 0 | 0 | 0 | 1 |
| rs1787711 | 11 | 116560778 | G | A | -0.623 | 0.5334 | -? | 0 | 0 | 0 | 1 |
| rs7104655 | 11 | 119577632 | A | G | 0.623 | 0.5334 | +? | 0 | 0 | 0 | 1 |
| rs671182 | 11 | 119363485 | C | T | 0.622 | 0.5337 | +? | 0 | 0 | 0 | 1 |
| rs61906112 | 11 | 116555765 | A | T | -0.622 | 0.5339 | -? | 0 | 0 | 0 | 1 |
| rs622373 | 11 | 120256845 | G | A | -0.622 | 0.5339 | -? | 0 | 0 | 0 | 1 |
| rs7113994 | 11 | 118382327 | G | A | -0.622 | 0.5339 | -+ | -174.1 | 1.095 | 1 | 0.2954 |
| rs495420 | 11 | 117456360 | C | G | 0.622 | 0.5341 | +? | 0 | 0 | 0 | 1 |
| rs4245208 | 11 | 119688712 | C | T | 0.622 | 0.5342 | ++ | 0 | 0.005 | 1 | 0.9443 |
| rs2513099 | 11 | 116811057 | C | T | -0.621 | 0.5343 | +- | 48.1 | 5.782 | 1 | 0.01619 |
| rs1979023 | 11 | 116778180 | C | T | -0.621 | 0.5344 | +- | 51.2 | 6.151 | 1 | 0.01314 |
| rs67691204 | 11 | 117775367 | A | C | 0.621 | 0.5344 | +? | 0 | 0 | 0 | 1 |
| rs12799706 | 11 | 117521490 | C | T | 0.621 | 0.5347 | ++ | -155.8 | 1.173 | 1 | 0.2788 |
| rs4529912 | 11 | 117585788 | G | A | 0.621 | 0.5347 | +? | 0 | 0 | 0 | 1 |
| rs4936398 | 11 | 117586059 | G | A | 0.621 | 0.5347 | +? | 0 | 0 | 0 | 1 |
| rs61451669 | 11 | 117853666 | C | T | 0.621 | 0.5347 | ++ | 0 | 0.085 | 1 | 0.7709 |

| | | | | | | | | | | | |
|---|---|---|---|---|---|---|---|---|---|---|---|
| rs4938806 | 11 | 120274353 | A | C | -0.621 | 0.5348 | -? | 0 | 0 | 0 | 1 |
| rs11216107 | 11 | 116579663 | C | T | -0.62 | 0.535 | -? | 0 | 0 | 0 | 1 |
| rs11600991 | 11 | 117600616 | C | T | 0.62 | 0.535 | +? | 0 | 0 | 0 | 1 |
| rs191347151 | 11 | 120702478 | T | C | 0.62 | 0.535 | +? | 0 | 0 | 0 | 1 |
| rs2368938 | 11 | 119684574 | C | T | 0.62 | 0.535 | ++ | 0 | 0.005 | 1 | 0.9437 |
| rs540780 | 11 | 117745418 | T | G | 0.62 | 0.535 | +? | 0 | 0 | 0 | 1 |
| rs11217770 | 11 | 120090735 | C | A | -0.62 | 0.5351 | -? | 0 | 0 | 0 | 1 |
| rs146525935 | 11 | 118180887 | C | T | 0.62 | 0.5352 | +- | 22.9 | 3.891 | 1 | 0.04855 |
| rs17311114 | 11 | 119872703 | C | T | -0.62 | 0.5352 | -? | 0 | 0 | 0 | 1 |
| rs676305 | 11 | 117133078 | T | C | 0.62 | 0.5352 | +? | 0 | 0 | 0 | 1 |
| rs73018464 | 11 | 117694399 | A | G | -0.62 | 0.5352 | -? | 0 | 0 | 0 | 1 |
| rs3017232 | 11 | 116560883 | T | C | -0.62 | 0.5353 | -? | 0 | 0 | 0 | 1 |
| rs623989 | 11 | 119359257 | G | T | 0.619 | 0.5357 | +? | 0 | 0 | 0 | 1 |
| rs6589571 | 11 | 116680418 | T | C | 0.619 | 0.5358 | +? | 0 | 0 | 0 | 1 |
| rs11216657 | 11 | 117839187 | C | A | 0.619 | 0.536 | ++ | 0 | 0.013 | 1 | 0.909 |
| rs1784120 | 11 | 116561859 | G | C | -0.619 | 0.536 | -? | 0 | 0 | 0 | 1 |
| rs1784122 | 11 | 116561701 | A | T | -0.619 | 0.536 | -? | 0 | 0 | 0 | 1 |
| rs1787705 | 11 | 116561938 | A | G | -0.619 | 0.536 | -? | 0 | 0 | 0 | 1 |
| rs668363 | 11 | 117144816 | C | A | 0.618 | 0.5363 | +? | 0 | 0 | 0 | 1 |
| rs2512149 | 11 | 117881601 | C | T | 0.618 | 0.5364 | +- | -143.3 | 1.233 | 1 | 0.2668 |
| rs11216440 | 11 | 117406111 | A | G | 0.618 | 0.5368 | +? | 0 | 0 | 0 | 1 |
| rs557773 | 11 | 119693206 | C | T | 0.618 | 0.5368 | ++ | 0 | 0.004 | 1 | 0.9521 |
| rs10892134 | 11 | 117386925 | T | C | 0.618 | 0.5369 | +? | 0 | 0 | 0 | 1 |
| rs6589803 | 11 | 120139700 | T | C | 0.618 | 0.5369 | +? | 0 | 0 | 0 | 1 |
| rs3802913 | 11 | 120738221 | G | C | 0.617 | 0.537 | +? | 0 | 0 | 0 | 1 |
| rs78849359 | 11 | 120565240 | T | C | 0.617 | 0.537 | +? | 0 | 0 | 0 | 1 |
| rs560320 | 11 | 116310757 | T | A | -0.617 | 0.5371 | -? | 0 | 0 | 0 | 1 |
| rs61898343 | 11 | 120182926 | C | G | -0.617 | 0.5371 | -? | 0 | 0 | 0 | 1 |
| rs10892211 | 11 | 117956409 | C | T | 0.617 | 0.5375 | +- | 0 | 0.225 | 1 | 0.6351 |
| rs2368939 | 11 | 119684789 | T | C | 0.617 | 0.5375 | ++ | 0 | 0.005 | 1 | 0.942 |
| rs11216037 | 11 | 116471044 | A | G | -0.616 | 0.5376 | -? | 0 | 0 | 0 | 1 |
| rs117498381 | 11 | 120242812 | T | A | -0.616 | 0.5378 | -? | 0 | 0 | 0 | 1 |
| rs142100881 | 11 | 119923914 | G | A | 0.616 | 0.5378 | +? | 0 | 0 | 0 | 1 |
| rs3016350 | 11 | 116474519 | T | C | -0.616 | 0.5378 | -? | 0 | 0 | 0 | 1 |
| rs11601357 | 11 | 116302248 | A | G | 0.616 | 0.5379 | +? | 0 | 0 | 0 | 1 |
| rs61905308 | 11 | 117357402 | T | C | -0.616 | 0.538 | -? | 0 | 0 | 0 | 1 |
| rs7926736 | 11 | 117549808 | A | G | -0.616 | 0.538 | -? | 0 | 0 | 0 | 1 |
| rs17121153 | 11 | 117603231 | T | C | 0.616 | 0.5382 | +? | 0 | 0 | 0 | 1 |
| rs17311414 | 11 | 120598007 | A | C | -0.615 | 0.5383 | -- | 0 | 0.729 | 1 | 0.3931 |
| rs675834 | 11 | 116737736 | A | T | -0.615 | 0.5383 | +- | 26.2 | 4.066 | 1 | 0.04376 |
| rs495981 | 11 | 119709638 | T | C | 0.615 | 0.5386 | +? | 0 | 0 | 0 | 1 |
| rs7113913 | 11 | 119941114 | T | C | -0.615 | 0.5386 | -? | 0 | 0 | 0 | 1 |
| rs79699220 | 11 | 118550841 | C | G | 0.615 | 0.5388 | ++ | 0 | 0.059 | 1 | 0.8087 |
| rs1787706 | 11 | 116561928 | A | G | -0.614 | 0.5394 | -? | 0 | 0 | 0 | 1 |
| rs4938300 | 11 | 116567619 | T | C | -0.614 | 0.5395 | -? | 0 | 0 | 0 | 1 |
| rs522848 | 11 | 116792445 | C | T | -0.614 | 0.5395 | +- | 48.3 | 5.804 | 1 | 0.01599 |
| rs6589573 | 11 | 116685766 | C | G | 0.613 | 0.5397 | +? | 0 | 0 | 0 | 1 |
| rs11218006 | 11 | 120653034 | A | G | 0.613 | 0.5401 | +? | 0 | 0 | 0 | 1 |
| rs12281008 | 11 | 120558389 | A | G | 0.613 | 0.5402 | +? | 0 | 0 | 0 | 1 |
| rs3802912 | 11 | 120738397 | A | G | 0.613 | 0.5402 | +? | 0 | 0 | 0 | 1 |
| rs11218198 | 11 | 121064444 | G | A | -0.612 | 0.5403 | -? | 0 | 0 | 0 | 1 |
| rs648374 | 11 | 119879425 | C | T | -0.612 | 0.5403 | -? | 0 | 0 | 0 | 1 |
| rs4938313 | 11 | 116671005 | A | G | -0.612 | 0.5404 | +- | -31.3 | 2.285 | 1 | 0.1306 |
| rs522742 | 11 | 117433965 | A | G | -0.612 | 0.5404 | -? | 0 | 0 | 0 | 1 |
| rs10892377 | 11 | 119381784 | A | G | -0.612 | 0.5405 | -? | 0 | 0 | 0 | 1 |
| rs12793410 | 11 | 120074907 | A | T | -0.612 | 0.5407 | -? | 0 | 0 | 0 | 1 |

| | | | | | | | | | | | |
|---|---|---|---|---|---|---|---|---|---|---|---|
| rs61903662 | 11 | 117123782 | A | G | -0.612 | 0.5407 | -? | 0 | 0 | 0 | 1 |
| rs1061332 | 11 | 121178302 | A | C | -0.611 | 0.5409 | -? | 0 | 0 | 0 | 1 |
| rs2115649 | 11 | 119267694 | A | C | 0.611 | 0.5409 | -+ | -61.2 | 1.861 | 1 | 0.1725 |
| rs113872393 | 11 | 116331421 | A | G | -0.611 | 0.541 | -? | 0 | 0 | 0 | 1 |
| rs74236028 | 11 | 120589740 | G | T | 0.611 | 0.5411 | +? | 0 | 0 | 0 | 1 |
| rs61903747 | 11 | 117281264 | A | G | -0.611 | 0.5412 | -? | 0 | 0 | 0 | 1 |
| rs11217775 | 11 | 120121323 | C | T | -0.611 | 0.5415 | -? | 0 | 0 | 0 | 1 |
| rs615166 | 11 | 118803536 | A | G | 0.61 | 0.5415 | -+ | -100.1 | 1.499 | 1 | 0.2208 |
| rs7113973 | 11 | 119940945 | A | G | -0.611 | 0.5415 | -? | 0 | 0 | 0 | 1 |
| rs35801991 | 11 | 120507687 | C | T | 0.61 | 0.5416 | +? | 0 | 0 | 0 | 1 |
| rs4581461 | 11 | 119198092 | C | T | -0.61 | 0.5416 | -? | 0 | 0 | 0 | 1 |
| rs12290383 | 11 | 120170915 | A | T | -0.61 | 0.5418 | -? | 0 | 0 | 0 | 1 |
| rs4938277 | 11 | 116397532 | T | C | -0.61 | 0.5419 | -? | 0 | 0 | 0 | 1 |
| rs10790382 | 11 | 120288372 | C | T | -0.61 | 0.542 | -? | 0 | 0 | 0 | 1 |
| rs12224796 | 11 | 117776819 | G | A | -0.61 | 0.542 | -? | 0 | 0 | 0 | 1 |
| rs4936514 | 11 | 120168201 | C | G | -0.61 | 0.5421 | -- | 0 | 0.005 | 1 | 0.9438 |
| rs57232565 | 11 | 116611457 | T | C | -0.61 | 0.5421 | +- | -47 | 2.041 | 1 | 0.1531 |
| rs6589569 | 11 | 116671476 | T | C | -0.61 | 0.5421 | +- | -36.3 | 2.2 | 1 | 0.138 |
| rs2292293 | 11 | 119548698 | G | T | 0.609 | 0.5422 | +? | 0 | 0 | 0 | 1 |
| rs73020491 | 11 | 117775043 | A | G | 0.609 | 0.5423 | +? | 0 | 0 | 0 | 1 |
| rs75618086 | 11 | 121031731 | A | T | -0.609 | 0.5423 | -? | 0 | 0 | 0 | 1 |
| rs12793354 | 11 | 120084056 | C | A | -0.609 | 0.5425 | -? | 0 | 0 | 0 | 1 |
| rs2253006 | 11 | 120111853 | C | G | -0.609 | 0.5426 | -? | 0 | 0 | 0 | 1 |
| rs12284383 | 11 | 120148717 | G | A | -0.609 | 0.5428 | -? | 0 | 0 | 0 | 1 |
| rs58848374 | 11 | 116938655 | A | G | -0.609 | 0.5428 | +- | 41.3 | 5.11 | 1 | 0.02378 |
| rs2715888 | 11 | 120124953 | A | G | -0.608 | 0.5429 | -? | 0 | 0 | 0 | 1 |
| rs11216132 | 11 | 116634585 | C | T | -0.608 | 0.543 | -+ | 10 | 3.333 | 1 | 0.0679 |
| rs891441 | 11 | 121165092 | G | T | -0.608 | 0.543 | -? | 0 | 0 | 0 | 1 |
| rs592529 | 11 | 120981195 | T | C | 0.608 | 0.5431 | +? | 0 | 0 | 0 | 1 |
| rs655824 | 11 | 120834480 | C | T | 0.608 | 0.5431 | +? | 0 | 0 | 0 | 1 |
| rs71480342 | 11 | 117421267 | T | C | -0.608 | 0.5431 | -? | 0 | 0 | 0 | 1 |
| rs644867 | 11 | 118567545 | G | A | 0.608 | 0.5432 | ++ | 0 | 0.767 | 1 | 0.3812 |
| rs4938435 | 11 | 117594364 | G | A | 0.608 | 0.5434 | +? | 0 | 0 | 0 | 1 |
| rs11217802 | 11 | 120167230 | T | C | -0.608 | 0.5435 | -- | 0 | 0.005 | 1 | 0.9448 |
| rs12225696 | 11 | 120167220 | G | A | -0.608 | 0.5435 | -- | 0 | 0.005 | 1 | 0.9448 |
| rs2156233 | 11 | 117584368 | T | C | 0.608 | 0.5435 | +? | 0 | 0 | 0 | 1 |
| rs58577143 | 11 | 118291210 | G | A | 0.608 | 0.5435 | ++ | 0 | 0.48 | 1 | 0.4882 |
| rs2044544 | 11 | 119744600 | T | C | 0.607 | 0.5436 | +? | 0 | 0 | 0 | 1 |
| rs11823300 | 11 | 120199080 | A | G | -0.607 | 0.5438 | -? | 0 | 0 | 0 | 1 |
| rs12576574 | 11 | 120898196 | T | G | -0.607 | 0.5438 | -? | 0 | 0 | 0 | 1 |
| rs12790943 | 11 | 120058623 | T | C | -0.607 | 0.5438 | -- | 0 | 0.163 | 1 | 0.6862 |
| rs12791292 | 11 | 120058640 | C | G | -0.607 | 0.5438 | -- | 0 | 0.163 | 1 | 0.6862 |
| rs947988 | 11 | 116528983 | C | G | 0.607 | 0.5438 | +? | 0 | 0 | 0 | 1 |
| rs1145210 | 11 | 116559002 | T | C | -0.607 | 0.5439 | -? | 0 | 0 | 0 | 1 |
| rs581491 | 11 | 119880792 | C | T | -0.607 | 0.5439 | -? | 0 | 0 | 0 | 1 |
| rs7928408 | 11 | 117593985 | T | C | 0.607 | 0.5439 | +? | 0 | 0 | 0 | 1 |
| rs78342015 | 11 | 118298428 | T | C | 0.607 | 0.5441 | +- | -7.7 | 2.786 | 1 | 0.09507 |
| rs10892482 | 11 | 119742989 | C | T | 0.606 | 0.5442 | +? | 0 | 0 | 0 | 1 |
| rs4142654 | 11 | 120167057 | G | A | -0.606 | 0.5443 | -- | 0 | 0.005 | 1 | 0.9453 |
| rs498889 | 11 | 117459662 | C | T | 0.606 | 0.5443 | +? | 0 | 0 | 0 | 1 |
| rs4938808 | 11 | 120336902 | T | G | -0.606 | 0.5445 | -? | 0 | 0 | 0 | 1 |
| rs629060 | 11 | 117581772 | G | T | 0.606 | 0.5447 | +? | 0 | 0 | 0 | 1 |
| rs6589619 | 11 | 117582263 | T | C | 0.606 | 0.5447 | +? | 0 | 0 | 0 | 1 |
| rs948777 | 11 | 117582054 | C | T | 0.606 | 0.5447 | +? | 0 | 0 | 0 | 1 |
| rs10892616 | 11 | 120559070 | G | A | 0.605 | 0.5449 | +? | 0 | 0 | 0 | 1 |
| rs12294046 | 11 | 120558487 | T | A | 0.605 | 0.5449 | +? | 0 | 0 | 0 | 1 |

| | | | | | | | | | | | |
|---|---|---|---|---|---|---|---|---|---|---|---|
| rs2075647 | 11 | 118037964 | C | G | 0.605 | 0.5449 | +? | 0 | 0 | 0 | 1 |
| rs12281001 | 11 | 120558364 | T | G | 0.605 | 0.545 | +? | 0 | 0 | 0 | 1 |
| rs619250 | 11 | 118064582 | A | T | 0.605 | 0.545 | +? | 0 | 0 | 0 | 1 |
| rs10892361 | 11 | 119269081 | G | A | 0.605 | 0.5453 | -+ | -58.5 | 1.892 | 1 | 0.1689 |
| rs149668068 | 11 | 118099735 | A | G | 0.605 | 0.5453 | +- | 0 | 0.868 | 1 | 0.3515 |
| rs6589856 | 11 | 120954911 | A | T | -0.605 | 0.5453 | -? | 0 | 0 | 0 | 1 |
| rs17124538 | 11 | 120783369 | G | A | -0.605 | 0.5454 | -? | 0 | 0 | 0 | 1 |
| rs72995858 | 11 | 120649949 | T | C | 0.605 | 0.5454 | +? | 0 | 0 | 0 | 1 |
| rs10892210 | 11 | 117956338 | T | G | 0.604 | 0.5455 | +- | 0 | 0.237 | 1 | 0.6264 |
| rs11217669 | 11 | 119938625 | A | G | 0.604 | 0.5456 | +? | 0 | 0 | 0 | 1 |
| rs61899022 | 11 | 119802948 | A | G | -0.604 | 0.5456 | -? | 0 | 0 | 0 | 1 |
| rs111255936 | 11 | 116710993 | T | C | 0.604 | 0.5457 | +? | 0 | 0 | 0 | 1 |
| rs4935750 | 11 | 120521066 | C | T | -0.604 | 0.5457 | -? | 0 | 0 | 0 | 1 |
| rs2509668 | 11 | 119361345 | G | A | 0.604 | 0.5458 | +? | 0 | 0 | 0 | 1 |
| rs869005 | 11 | 117614796 | T | C | 0.604 | 0.5459 | +? | 0 | 0 | 0 | 1 |
| rs609332 | 11 | 117175158 | G | C | 0.604 | 0.5461 | +? | 0 | 0 | 0 | 1 |
| rs6589832 | 11 | 120559480 | G | A | -0.604 | 0.5461 | -? | 0 | 0 | 0 | 1 |
| rs496034 | 11 | 120322079 | A | G | 0.603 | 0.5463 | +? | 0 | 0 | 0 | 1 |
| rs1263057 | 11 | 116575754 | T | C | -0.603 | 0.5464 | -? | 0 | 0 | 0 | 1 |
| rs7124930 | 11 | 120687529 | C | T | 0.603 | 0.5468 | +? | 0 | 0 | 0 | 1 |
| rs10891996 | 11 | 116457340 | A | G | 0.602 | 0.5469 | +? | 0 | 0 | 0 | 1 |
| rs2510873 | 11 | 116243868 | G | A | -0.602 | 0.5469 | -? | 0 | 0 | 0 | 1 |
| rs658274 | 11 | 117542353 | G | A | -0.602 | 0.5469 | -? | 0 | 0 | 0 | 1 |
| rs10892209 | 11 | 117956263 | T | C | 0.602 | 0.547 | +- | 0 | 0.241 | 1 | 0.6236 |
| rs11600226 | 11 | 117690423 | C | T | 0.602 | 0.547 | +? | 0 | 0 | 0 | 1 |
| rs11216140 | 11 | 116672013 | T | C | -0.602 | 0.5474 | +- | -35.5 | 2.214 | 1 | 0.1368 |
| rs35574604 | 11 | 119361057 | A | G | 0.601 | 0.5477 | +? | 0 | 0 | 0 | 1 |
| rs4245204 | 11 | 119580472 | A | G | 0.601 | 0.5477 | +? | 0 | 0 | 0 | 1 |
| rs4938478 | 11 | 117956943 | G | A | 0.601 | 0.5479 | +- | 0 | 0.303 | 1 | 0.5818 |
| rs550791 | 11 | 116322819 | T | C | 0.601 | 0.5479 | +? | 0 | 0 | 0 | 1 |
| rs10892202 | 11 | 117857338 | C | G | 0.601 | 0.548 | ++ | 0 | 0.089 | 1 | 0.7654 |
| rs519390 | 11 | 116238815 | C | T | -0.601 | 0.548 | -+ | 44.5 | 5.406 | 1 | 0.02007 |
| rs10750173 | 11 | 120288368 | C | T | -0.601 | 0.5481 | -? | 0 | 0 | 0 | 1 |
| rs1242696 | 11 | 117217391 | T | G | 0.601 | 0.5481 | +? | 0 | 0 | 0 | 1 |
| rs3017231 | 11 | 116560948 | A | G | -0.601 | 0.5481 | -? | 0 | 0 | 0 | 1 |
| rs11218140 | 11 | 120968162 | A | G | -0.6 | 0.5482 | -? | 0 | 0 | 0 | 1 |
| rs3934628 | 11 | 120707644 | T | C | 0.6 | 0.5482 | +? | 0 | 0 | 0 | 1 |
| rs58254802 | 11 | 120120523 | T | G | -0.6 | 0.5482 | -? | 0 | 0 | 0 | 1 |
| rs60758080 | 11 | 120120435 | A | G | -0.6 | 0.5482 | -? | 0 | 0 | 0 | 1 |
| rs79475413 | 11 | 119666800 | A | G | 0.6 | 0.5484 | +? | 0 | 0 | 0 | 1 |
| rs67514601 | 11 | 118189477 | G | A | -0.6 | 0.5487 | -- | 0 | 0.556 | 1 | 0.4558 |
| rs79831199 | 11 | 116236334 | A | C | -0.6 | 0.5488 | -- | 0 | 0.192 | 1 | 0.6616 |
| rs34708086 | 11 | 119331254 | C | G | -0.599 | 0.5489 | -? | 0 | 0 | 0 | 1 |
| rs73011181 | 11 | 116464824 | T | A | -0.599 | 0.5491 | -? | 0 | 0 | 0 | 1 |
| rs10892232 | 11 | 118146884 | T | A | 0.599 | 0.5492 | +- | 11.5 | 3.39 | 1 | 0.06558 |
| rs10892438 | 11 | 119579346 | T | A | -0.599 | 0.5492 | -? | 0 | 0 | 0 | 1 |
| rs11606357 | 11 | 118878222 | C | T | -0.599 | 0.5492 | -- | 0 | 0.017 | 1 | 0.8948 |
| rs34573317 | 11 | 117342672 | C | T | 0.599 | 0.5492 | +? | 0 | 0 | 0 | 1 |
| rs59814528 | 11 | 119634962 | C | T | -0.599 | 0.5492 | -? | 0 | 0 | 0 | 1 |
| rs3132781 | 11 | 120621358 | A | G | -0.599 | 0.5495 | -? | 0 | 0 | 0 | 1 |
| rs12287882 | 11 | 120558338 | C | T | 0.598 | 0.5496 | +? | 0 | 0 | 0 | 1 |
| rs2238006 | 11 | 117087792 | C | A | -0.598 | 0.5498 | -? | 0 | 0 | 0 | 1 |
| rs34198109 | 11 | 119332549 | G | C | -0.598 | 0.5501 | -? | 0 | 0 | 0 | 1 |
| rs7930786 | 11 | 116624727 | C | G | -0.597 | 0.5502 | +- | -61.4 | 1.859 | 1 | 0.1728 |
| rs480053 | 11 | 116314419 | A | G | 0.597 | 0.5505 | +? | 0 | 0 | 0 | 1 |
| rs11217552 | 11 | 119750091 | T | C | 0.597 | 0.5507 | +? | 0 | 0 | 0 | 1 |

| | | | | | | | | | | | |
|---|---|---|---|---|---|---|---|---|---|---|---|
| rs73022413 | 11 | 117787228 | C | T | 0.597 | 0.5507 | +? | 0 | 0 | 0 | 1 |
| rs17092642 | 11 | 116557248 | C | T | -0.596 | 0.5509 | -? | 0 | 0 | 0 | 1 |
| rs1079071 | 11 | 120628922 | C | T | 0.596 | 0.551 | +? | 0 | 0 | 0 | 1 |
| rs948404 | 11 | 120150533 | A | G | -0.596 | 0.551 | -? | 0 | 0 | 0 | 1 |
| rs11216403 | 11 | 117333214 | C | T | -0.596 | 0.5511 | -? | 0 | 0 | 0 | 1 |
| rs3017230 | 11 | 116560961 | C | G | -0.596 | 0.5511 | -? | 0 | 0 | 0 | 1 |
| rs4936499 | 11 | 119682612 | C | T | 0.596 | 0.5512 | ++ | 0 | 0.007 | 1 | 0.9326 |
| rs118188470 | 11 | 119692025 | C | G | 0.596 | 0.5513 | ++ | 0 | 0.047 | 1 | 0.8292 |
| rs7105363 | 11 | 120687069 | T | G | 0.596 | 0.5515 | +? | 0 | 0 | 0 | 1 |
| rs55768116 | 11 | 118108331 | A | C | 0.595 | 0.5516 | +? | 0 | 0 | 0 | 1 |
| rs4252253 | 11 | 117859925 | A | C | 0.595 | 0.5517 | +- | -191.8 | 1.028 | 1 | 0.3106 |
| rs4938436 | 11 | 117594591 | G | A | 0.595 | 0.5519 | +? | 0 | 0 | 0 | 1 |
| rs74940535 | 11 | 120182487 | C | G | -0.595 | 0.5519 | -? | 0 | 0 | 0 | 1 |
| rs518863 | 11 | 120053300 | T | C | 0.595 | 0.552 | +? | 0 | 0 | 0 | 1 |
| rs56754533 | 11 | 118188276 | A | G | -0.594 | 0.5525 | +- | -140.7 | 1.247 | 1 | 0.2642 |
| rs4252287 | 11 | 117868638 | A | G | 0.594 | 0.5526 | +- | 0 | 0.297 | 1 | 0.586 |
| rs10790365 | 11 | 119945567 | T | C | -0.594 | 0.5527 | -? | 0 | 0 | 0 | 1 |
| rs12279889 | 11 | 121119712 | T | C | 0.594 | 0.5528 | +? | 0 | 0 | 0 | 1 |
| rs11217586 | 11 | 119805109 | A | G | -0.593 | 0.553 | -? | 0 | 0 | 0 | 1 |
| rs10790203 | 11 | 117610387 | G | A | 0.593 | 0.5531 | +? | 0 | 0 | 0 | 1 |
| rs4936405 | 11 | 117611495 | G | A | 0.593 | 0.5531 | +? | 0 | 0 | 0 | 1 |
| rs56125819 | 11 | 116961617 | A | G | -0.593 | 0.5531 | +- | 48.8 | 5.86 | 1 | 0.01548 |
| rs61244196 | 11 | 116961510 | A | G | -0.593 | 0.5531 | +- | 48.8 | 5.86 | 1 | 0.01548 |
| rs2166658 | 11 | 119682874 | T | C | 0.593 | 0.5533 | ++ | 0 | 0.008 | 1 | 0.9296 |
| rs3882897 | 11 | 117091227 | A | G | -0.593 | 0.5533 | -? | 0 | 0 | 0 | 1 |
| rs11216920 | 11 | 118462554 | G | A | 0.593 | 0.5534 | +- | 0 | 0.274 | 1 | 0.6004 |
| rs10892124 | 11 | 117354818 | G | C | 0.592 | 0.5536 | +? | 0 | 0 | 0 | 1 |
| rs1309012690 | 11 | 119166047 | A | G | 0.592 | 0.5536 | ++ | 0 | 0.363 | 1 | 0.5469 |
| rs12790242 | 11 | 117842292 | A | G | 0.592 | 0.5537 | ++ | 0 | 0.012 | 1 | 0.9138 |
| rs7107887 | 11 | 117585714 | A | C | 0.592 | 0.5539 | +? | 0 | 0 | 0 | 1 |
| rs61904848 | 11 | 116454342 | C | A | 0.592 | 0.554 | +? | 0 | 0 | 0 | 1 |
| rs117273824 | 11 | 117458169 | T | C | 0.591 | 0.5542 | +? | 0 | 0 | 0 | 1 |
| rs1829 | 11 | 120981408 | T | C | 0.591 | 0.5542 | +- | -147.8 | 1.211 | 1 | 0.2712 |
| rs77080891 | 11 | 121040073 | A | G | -0.591 | 0.5543 | -? | 0 | 0 | 0 | 1 |
| rs948025 | 11 | 120679052 | T | G | 0.591 | 0.5543 | +? | 0 | 0 | 0 | 1 |
| rs10502225 | 11 | 116939724 | T | A | -0.591 | 0.5544 | +- | 48.9 | 5.866 | 1 | 0.01544 |
| rs10750091 | 11 | 116296641 | A | G | -0.591 | 0.5544 | -? | 0 | 0 | 0 | 1 |
| rs111426457 | 11 | 116931087 | G | C | -0.591 | 0.5544 | +- | 48.9 | 5.866 | 1 | 0.01544 |
| rs111641976 | 11 | 116941382 | A | G | -0.591 | 0.5544 | +- | 48.9 | 5.866 | 1 | 0.01544 |
| rs1982679 | 11 | 116936488 | A | G | -0.591 | 0.5544 | +- | 48.9 | 5.866 | 1 | 0.01544 |
| rs56188860 | 11 | 116948173 | A | T | -0.591 | 0.5544 | +- | 48.9 | 5.866 | 1 | 0.01544 |
| rs57787606 | 11 | 116942479 | A | G | -0.591 | 0.5544 | +- | 48.9 | 5.866 | 1 | 0.01544 |
| rs59752757 | 11 | 116950278 | C | A | -0.591 | 0.5544 | +- | 48.9 | 5.866 | 1 | 0.01544 |
| rs7108830 | 11 | 116948761 | A | G | -0.591 | 0.5544 | +- | 48.9 | 5.866 | 1 | 0.01544 |
| rs7121475 | 11 | 116951615 | A | G | -0.591 | 0.5544 | +- | 48.9 | 5.866 | 1 | 0.01544 |
| rs73576634 | 11 | 116944304 | T | G | -0.591 | 0.5544 | +- | 48.9 | 5.866 | 1 | 0.01544 |
| rs73576637 | 11 | 116945074 | G | A | -0.591 | 0.5544 | +- | 48.9 | 5.866 | 1 | 0.01544 |
| rs73576652 | 11 | 116950567 | A | G | -0.591 | 0.5544 | +- | 48.9 | 5.866 | 1 | 0.01544 |
| rs73576656 | 11 | 116953604 | T | C | -0.591 | 0.5544 | +- | 48.9 | 5.866 | 1 | 0.01544 |
| rs73576658 | 11 | 116953784 | A | G | -0.591 | 0.5544 | +- | 48.9 | 5.866 | 1 | 0.01544 |
| rs7946101 | 11 | 116934869 | A | G | -0.591 | 0.5544 | +- | 48.9 | 5.866 | 1 | 0.01544 |
| rs7123666 | 11 | 116667083 | A | G | -0.591 | 0.5545 | +- | -154 | 1.181 | 1 | 0.2772 |
| rs12576572 | 11 | 120922228 | T | G | -0.591 | 0.5546 | -? | 0 | 0 | 0 | 1 |
| rs3758891 | 11 | 120929197 | A | G | -0.591 | 0.5546 | -? | 0 | 0 | 0 | 1 |
| rs57826163 | 11 | 120933051 | T | G | -0.591 | 0.5546 | -? | 0 | 0 | 0 | 1 |
| rs7947565 | 11 | 120927469 | T | G | -0.591 | 0.5546 | -? | 0 | 0 | 0 | 1 |

| | | | | | | | | | | | |
|---|---|---|---|---|---|---|---|---|---|---|---|
| rs111537986 | 11 | 117763500 | T | C | 0.591 | 0.5547 | +? | 0 | 0 | 0 | 1 |
| rs17120197 | 11 | 116845104 | C | T | 0.591 | 0.5548 | +? | 0 | 0 | 0 | 1 |
| rs1787028 | 11 | 119964758 | G | C | 0.591 | 0.5548 | +? | 0 | 0 | 0 | 1 |
| rs525302 | 11 | 116762583 | C | G | -0.591 | 0.5548 | -? | 0 | 0 | 0 | 1 |
| rs11217461 | 11 | 119622391 | A | G | -0.59 | 0.555 | -? | 0 | 0 | 0 | 1 |
| rs59873384 | 11 | 116576210 | T | C | -0.59 | 0.555 | -- | 0 | 0 | 1 | 0.9974 |
| rs10892466 | 11 | 119661317 | C | G | 0.59 | 0.5551 | ++ | 0 | 0 | 1 | 0.9878 |
| rs3132782 | 11 | 120622656 | A | G | -0.59 | 0.5551 | -? | 0 | 0 | 0 | 1 |
| rs4935755 | 11 | 120735982 | T | C | 0.59 | 0.5551 | +? | 0 | 0 | 0 | 1 |
| rs470725 | 11 | 119997599 | T | C | 0.59 | 0.5552 | +? | 0 | 0 | 0 | 1 |
| rs11217520 | 11 | 119686250 | A | G | 0.59 | 0.5553 | ++ | 0 | 0.008 | 1 | 0.9299 |
| rs12575628 | 11 | 121020150 | T | C | -0.59 | 0.5553 | -? | 0 | 0 | 0 | 1 |
| rs4936501 | 11 | 119685493 | A | G | 0.59 | 0.5553 | ++ | 0 | 0.008 | 1 | 0.9299 |
| rs4936502 | 11 | 119685596 | T | C | 0.59 | 0.5553 | ++ | 0 | 0.008 | 1 | 0.9299 |
| rs501433 | 11 | 116293739 | T | C | -0.59 | 0.5554 | -? | 0 | 0 | 0 | 1 |
| rs114428118 | 11 | 120564627 | T | G | 0.589 | 0.5557 | +? | 0 | 0 | 0 | 1 |
| rs3133865 | 11 | 120623605 | A | C | -0.589 | 0.5558 | -? | 0 | 0 | 0 | 1 |
| rs4938790 | 11 | 120054202 | T | C | 0.589 | 0.5559 | +? | 0 | 0 | 0 | 1 |
| rs10892574 | 11 | 120288944 | C | G | -0.589 | 0.556 | -? | 0 | 0 | 0 | 1 |
| rs61898298 | 11 | 120119848 | T | G | -0.589 | 0.5561 | -? | 0 | 0 | 0 | 1 |
| rs11216549 | 11 | 117668104 | G | C | -0.588 | 0.5562 | -? | 0 | 0 | 0 | 1 |
| rs61593876 | 11 | 116450579 | C | T | 0.588 | 0.5562 | ++ | 0 | 0.024 | 1 | 0.8775 |
| rs141752759 | 11 | 118194647 | G | C | 0.588 | 0.5563 | ++ | 0 | 0.462 | 1 | 0.4969 |
| rs79700393 | 11 | 117551222 | A | G | 0.588 | 0.5563 | +? | 0 | 0 | 0 | 1 |
| rs1790471 | 11 | 119968219 | T | C | 0.588 | 0.5564 | +? | 0 | 0 | 0 | 1 |
| rs2854116 | 11 | 116700169 | C | T | -0.588 | 0.5564 | -? | 0 | 0 | 0 | 1 |
| rs78484152 | 11 | 119657893 | A | G | 0.588 | 0.5564 | ++ | 0 | 0 | 1 | 0.9836 |
| rs4367956 | 11 | 119490921 | A | C | 0.588 | 0.5566 | +? | 0 | 0 | 0 | 1 |
| rs12576376 | 11 | 116225239 | T | C | 0.588 | 0.5567 | +- | -73 | 1.734 | 1 | 0.1879 |
| rs2186601 | 11 | 120955028 | T | C | -0.588 | 0.5567 | -? | 0 | 0 | 0 | 1 |
| rs4936500 | 11 | 119682703 | G | C | 0.588 | 0.5567 | ++ | 0 | 0.008 | 1 | 0.9289 |
| rs7104999 | 11 | 120686999 | G | C | 0.588 | 0.5568 | +? | 0 | 0 | 0 | 1 |
| rs78062662 | 11 | 116482573 | T | C | -0.588 | 0.5568 | -? | 0 | 0 | 0 | 1 |
| rs2852224 | 11 | 120619202 | T | C | 0.587 | 0.5569 | +? | 0 | 0 | 0 | 1 |
| rs12806754 | 11 | 119784889 | A | G | -0.587 | 0.557 | -? | 0 | 0 | 0 | 1 |
| rs12575073 | 11 | 121188747 | T | C | -0.587 | 0.5571 | -? | 0 | 0 | 0 | 1 |
| rs3017228 | 11 | 116560992 | G | A | -0.587 | 0.5572 | -? | 0 | 0 | 0 | 1 |
| rs11216781 | 11 | 118012310 | A | G | -0.587 | 0.5573 | -+ | 0 | 0.119 | 1 | 0.7299 |
| rs11217521 | 11 | 119686328 | C | T | 0.587 | 0.5573 | ++ | 0 | 0.008 | 1 | 0.9285 |
| rs34570367 | 11 | 117991116 | G | A | 0.587 | 0.5573 | +- | 0 | 0.44 | 1 | 0.5073 |
| rs35676325 | 11 | 119686548 | A | G | 0.587 | 0.5573 | ++ | 0 | 0.008 | 1 | 0.9285 |
| rs653106 | 11 | 119357305 | G | A | -0.587 | 0.5573 | -? | 0 | 0 | 0 | 1 |
| rs79280888 | 11 | 116560529 | G | A | -0.587 | 0.5573 | -? | 0 | 0 | 0 | 1 |
| rs11216500 | 11 | 117555150 | C | T | 0.586 | 0.5577 | +? | 0 | 0 | 0 | 1 |
| rs75964569 | 11 | 119853613 | C | T | -0.586 | 0.5578 | -? | 0 | 0 | 0 | 1 |
| rs2277288 | 11 | 117782638 | G | A | 0.586 | 0.5579 | +? | 0 | 0 | 0 | 1 |
| rs118138924 | 11 | 116803634 | G | C | 0.586 | 0.5581 | +? | 0 | 0 | 0 | 1 |
| rs547919 | 11 | 116800154 | T | A | -0.586 | 0.5581 | +- | 49 | 5.881 | 1 | 0.0153 |
| rs7119690 | 11 | 119656902 | G | A | -0.586 | 0.5581 | -? | 0 | 0 | 0 | 1 |
| rs3133224 | 11 | 120614787 | T | C | -0.586 | 0.5582 | -? | 0 | 0 | 0 | 1 |
| rs7105286 | 11 | 120687234 | G | A | 0.586 | 0.5582 | +? | 0 | 0 | 0 | 1 |
| rs7926015 | 11 | 119261867 | T | G | 0.586 | 0.5582 | -+ | -57.7 | 1.902 | 1 | 0.1678 |
| rs11601508 | 11 | 118178757 | A | G | 0.585 | 0.5583 | +- | 21.3 | 3.813 | 1 | 0.05087 |
| rs12576759 | 11 | 120898253 | T | C | -0.585 | 0.5585 | -? | 0 | 0 | 0 | 1 |
| rs11217587 | 11 | 119809217 | C | T | -0.585 | 0.5587 | -? | 0 | 0 | 0 | 1 |
| rs11218123 | 11 | 120899492 | T | G | -0.585 | 0.5587 | -? | 0 | 0 | 0 | 1 |

| | | | | | | | | | | | |
|---|---|---|---|---|---|---|---|---|---|---|---|
| rs35004487 | 11 | 116631542 | C | G | -0.585 | 0.5587 | -? | 0 | 0 | 0 | 1 |
| rs606916 | 11 | 120982105 | G | C | 0.585 | 0.5587 | +- | -148.3 | 1.208 | 1 | 0.2717 |
| rs1202587279 | 11 | 116299004 | T | C | 0.584 | 0.559 | +? | 0 | 0 | 0 | 1 |
| rs12283240 | 11 | 120559953 | A | G | 0.584 | 0.559 | +? | 0 | 0 | 0 | 1 |
| rs948690 | 11 | 116529468 | C | T | -0.584 | 0.5591 | -? | 0 | 0 | 0 | 1 |
| rs609066 | 11 | 119879255 | G | A | -0.584 | 0.5592 | -? | 0 | 0 | 0 | 1 |
| rs475422 | 11 | 117431708 | C | T | 0.584 | 0.5593 | +? | 0 | 0 | 0 | 1 |
| rs35059164 | 11 | 119686554 | G | A | 0.584 | 0.5594 | ++ | 0 | 0.008 | 1 | 0.927 |
| rs57903286 | 11 | 116417966 | T | C | 0.584 | 0.5596 | ++ | 0 | 0.33 | 1 | 0.5659 |
| rs3181261 | 11 | 118212671 | T | C | -0.583 | 0.5597 | -? | 0 | 0 | 0 | 1 |
| rs73003745 | 11 | 120703464 | A | G | 0.583 | 0.5597 | +? | 0 | 0 | 0 | 1 |
| rs7129335 | 11 | 121107473 | A | T | -0.583 | 0.5598 | -? | 0 | 0 | 0 | 1 |
| rs180364 | 11 | 116596752 | G | A | -0.583 | 0.5599 | -+ | 0 | 0.151 | 1 | 0.6974 |
| rs34937367 | 11 | 116560524 | C | T | -0.583 | 0.5599 | -? | 0 | 0 | 0 | 1 |
| rs11218222 | 11 | 121112467 | G | C | 0.583 | 0.5601 | +? | 0 | 0 | 0 | 1 |
| rs4936403 | 11 | 117606193 | G | T | 0.582 | 0.5603 | +? | 0 | 0 | 0 | 1 |
| rs948024 | 11 | 120679042 | T | A | 0.582 | 0.5604 | +? | 0 | 0 | 0 | 1 |
| rs10892445 | 11 | 119607933 | G | A | 0.582 | 0.5605 | +? | 0 | 0 | 0 | 1 |
| rs117684324 | 11 | 116577971 | T | A | -0.582 | 0.5605 | -- | 0 | 0 | 1 | 0.9989 |
| rs61901232 | 11 | 121093389 | T | C | -0.582 | 0.5605 | -? | 0 | 0 | 0 | 1 |
| rs4936404 | 11 | 117606320 | G | A | 0.582 | 0.5606 | +? | 0 | 0 | 0 | 1 |
| rs72994672 | 11 | 119844641 | G | T | -0.582 | 0.5607 | -? | 0 | 0 | 0 | 1 |
| rs10892512 | 11 | 119878901 | A | G | -0.582 | 0.5608 | -? | 0 | 0 | 0 | 1 |
| rs6589813 | 11 | 120293490 | G | A | -0.582 | 0.5608 | -? | 0 | 0 | 0 | 1 |
| rs4938726 | 11 | 119681110 | T | C | 0.582 | 0.5609 | ++ | 0 | 0.009 | 1 | 0.9261 |
| rs634406 | 11 | 117460123 | G | C | 0.581 | 0.561 | +? | 0 | 0 | 0 | 1 |
| rs4936505 | 11 | 119740612 | T | C | 0.581 | 0.5611 | +? | 0 | 0 | 0 | 1 |
| rs3759009 | 11 | 118173850 | A | G | -0.581 | 0.5615 | -+ | -7.1 | 2.8 | 1 | 0.09423 |
| rs117388325 | 11 | 120236793 | T | C | -0.58 | 0.5617 | -? | 0 | 0 | 0 | 1 |
| rs9332852 | 11 | 118381069 | T | C | -0.58 | 0.5618 | -+ | -191.3 | 1.03 | 1 | 0.3102 |
| rs1573880 | 11 | 120465556 | T | C | -0.58 | 0.5619 | -? | 0 | 0 | 0 | 1 |
| rs2006949 | 11 | 119575104 | C | T | -0.58 | 0.5619 | -? | 0 | 0 | 0 | 1 |
| rs10892613 | 11 | 120558084 | A | G | 0.58 | 0.5622 | +? | 0 | 0 | 0 | 1 |
| rs10892614 | 11 | 120558136 | G | T | 0.58 | 0.5622 | +? | 0 | 0 | 0 | 1 |
| rs10892615 | 11 | 120558143 | T | A | 0.58 | 0.5622 | +? | 0 | 0 | 0 | 1 |
| rs11216054 | 11 | 116498951 | C | T | 0.58 | 0.5622 | +? | 0 | 0 | 0 | 1 |
| rs12286467 | 11 | 120558155 | G | T | 0.58 | 0.5622 | +? | 0 | 0 | 0 | 1 |
| rs59818386 | 11 | 116320603 | A | G | 0.58 | 0.5622 | +? | 0 | 0 | 0 | 1 |
| rs73594168 | 11 | 116899414 | T | C | -0.579 | 0.5626 | +- | 49.2 | 5.9 | 1 | 0.01514 |
| rs73594169 | 11 | 116899617 | T | C | -0.579 | 0.5626 | +- | 49.2 | 5.9 | 1 | 0.01514 |
| rs2003013 | 11 | 116372575 | A | G | -0.579 | 0.5627 | -? | 0 | 0 | 0 | 1 |
| rs1145195 | 11 | 116571250 | A | G | 0.579 | 0.5628 | +? | 0 | 0 | 0 | 1 |
| rs1941453 | 11 | 119331500 | T | C | -0.579 | 0.5628 | -? | 0 | 0 | 0 | 1 |
| rs541586 | 11 | 116729196 | C | T | -0.579 | 0.5628 | +- | 29.7 | 4.269 | 1 | 0.0388 |
| rs7115168 | 11 | 120737184 | G | A | 0.579 | 0.5629 | +? | 0 | 0 | 0 | 1 |
| rs524929 | 11 | 116260862 | G | A | 0.578 | 0.5633 | +? | 0 | 0 | 0 | 1 |
| rs17124937 | 11 | 121065198 | C | G | -0.578 | 0.5635 | -? | 0 | 0 | 0 | 1 |
| rs1787023 | 11 | 119903704 | T | C | -0.578 | 0.5635 | -? | 0 | 0 | 0 | 1 |
| rs11217365 | 11 | 119501085 | C | T | 0.577 | 0.5637 | +? | 0 | 0 | 0 | 1 |
| rs7129597 | 11 | 117555933 | T | C | 0.577 | 0.5637 | +? | 0 | 0 | 0 | 1 |
| rs61903663 | 11 | 117133242 | C | T | 0.577 | 0.5639 | +? | 0 | 0 | 0 | 1 |
| rs11216662 | 11 | 117848129 | C | A | -0.577 | 0.564 | -- | 0 | 0.534 | 1 | 0.465 |
| rs1567378 | 11 | 119480880 | G | A | 0.577 | 0.5642 | +? | 0 | 0 | 0 | 1 |
| rs11216982 | 11 | 118590864 | A | G | -0.576 | 0.5643 | -? | 0 | 0 | 0 | 1 |
| rs1245791313 | 11 | 120703432 | G | A | 0.576 | 0.5643 | +? | 0 | 0 | 0 | 1 |
| rs2212641 | 11 | 120191828 | C | T | 0.576 | 0.5644 | +? | 0 | 0 | 0 | 1 |

| | | | | | | | | | | | |
|---|---|---|---|---|---|---|---|---|---|---|---|
| rs571025 | 11 | 116292614 | T | C | -0.576 | 0.5646 | -? | 0 | 0 | 0 | 1 |
| rs602716 | 11 | 118801235 | A | G | 0.576 | 0.5647 | -+ | -100.1 | 1.499 | 1 | 0.2208 |
| rs10790141 | 11 | 116296520 | A | G | -0.576 | 0.5648 | -? | 0 | 0 | 0 | 1 |
| rs550646 | 11 | 116314641 | A | G | 0.575 | 0.565 | +? | 0 | 0 | 0 | 1 |
| rs57471964 | 11 | 117109155 | G | A | -0.575 | 0.565 | -? | 0 | 0 | 0 | 1 |
| rs964184 | 11 | 116648917 | G | C | -0.575 | 0.565 | +- | 16.1 | 3.575 | 1 | 0.05866 |
| rs139034795 | 11 | 120212377 | G | A | 0.575 | 0.5652 | +? | 0 | 0 | 0 | 1 |
| rs73020457 | 11 | 117762694 | T | C | 0.575 | 0.5652 | +? | 0 | 0 | 0 | 1 |
| rs505009 | 11 | 116293365 | A | G | -0.575 | 0.5654 | -? | 0 | 0 | 0 | 1 |
| rs78435918 | 11 | 120566473 | G | A | 0.575 | 0.5655 | +? | 0 | 0 | 0 | 1 |
| rs10892192 | 11 | 117763538 | T | C | 0.575 | 0.5656 | +? | 0 | 0 | 0 | 1 |
| rs585533 | 11 | 117927961 | A | G | -0.575 | 0.5656 | -? | 0 | 0 | 0 | 1 |
| rs118114248 | 11 | 120313999 | A | G | -0.574 | 0.5657 | -? | 0 | 0 | 0 | 1 |
| rs663319 | 11 | 121007374 | G | A | -0.574 | 0.5657 | -? | 0 | 0 | 0 | 1 |
| rs7116997 | 11 | 120060673 | C | T | -0.574 | 0.5657 | -- | 0 | 0.182 | 1 | 0.6695 |
| rs145433737 | 11 | 121117590 | A | G | 0.574 | 0.5658 | +? | 0 | 0 | 0 | 1 |
| rs4936402 | 11 | 117600876 | C | T | 0.574 | 0.5658 | +? | 0 | 0 | 0 | 1 |
| rs12798518 | 11 | 119971934 | A | G | -0.574 | 0.566 | -? | 0 | 0 | 0 | 1 |
| rs1945011 | 11 | 120621602 | A | G | 0.574 | 0.566 | +? | 0 | 0 | 0 | 1 |
| rs7928868 | 11 | 121165498 | A | G | 0.574 | 0.566 | +? | 0 | 0 | 0 | 1 |
| rs2060008 | 11 | 121179847 | A | T | -0.574 | 0.5662 | -? | 0 | 0 | 0 | 1 |
| rs3741298 | 11 | 116657561 | C | T | -0.574 | 0.5662 | +- | 51.3 | 6.157 | 1 | 0.01309 |
| rs77873962 | 11 | 117199317 | T | C | -0.574 | 0.5662 | -? | 0 | 0 | 0 | 1 |
| rs17120111 | 11 | 116744018 | T | C | 0.573 | 0.5664 | +? | 0 | 0 | 0 | 1 |
| rs4938466 | 11 | 117845068 | G | A | 0.573 | 0.5664 | ++ | 0 | 0.003 | 1 | 0.9556 |
| rs7126172 | 11 | 118567257 | G | A | 0.573 | 0.5664 | ++ | 0 | 0.825 | 1 | 0.3638 |
| rs112933729 | 11 | 117246566 | A | G | -0.573 | 0.5667 | -? | 0 | 0 | 0 | 1 |
| rs7104210 | 11 | 119073886 | G | T | 0.573 | 0.5667 | +? | 0 | 0 | 0 | 1 |
| rs4631889 | 11 | 120494034 | A | G | 0.573 | 0.5668 | +? | 0 | 0 | 0 | 1 |
| rs12278045 | 11 | 119796344 | A | G | -0.573 | 0.5669 | -? | 0 | 0 | 0 | 1 |
| rs10750117 | 11 | 117762288 | G | C | 0.572 | 0.567 | +? | 0 | 0 | 0 | 1 |
| rs11216470 | 11 | 117485422 | T | C | 0.572 | 0.567 | +? | 0 | 0 | 0 | 1 |
| rs73022412 | 11 | 117787018 | C | T | 0.572 | 0.5671 | +? | 0 | 0 | 0 | 1 |
| rs7112341 | 11 | 121110846 | A | T | -0.572 | 0.5673 | -- | 0 | 0.2 | 1 | 0.6545 |
| rs11216489 | 11 | 117535285 | A | G | 0.572 | 0.5674 | +? | 0 | 0 | 0 | 1 |
| rs4489755 | 11 | 119605480 | T | G | 0.572 | 0.5676 | +? | 0 | 0 | 0 | 1 |
| rs618923 | 11 | 116654159 | G | A | -0.572 | 0.5677 | -+ | 20.8 | 3.788 | 1 | 0.05163 |
| rs61900351 | 11 | 117801589 | A | T | 0.571 | 0.5678 | +? | 0 | 0 | 0 | 1 |
| rs11217878 | 11 | 120340383 | A | G | 0.571 | 0.5679 | +? | 0 | 0 | 0 | 1 |
| rs2276035 | 11 | 120346360 | A | G | 0.571 | 0.5679 | +? | 0 | 0 | 0 | 1 |
| rs79022222 | 11 | 121021970 | T | C | -0.571 | 0.568 | -? | 0 | 0 | 0 | 1 |
| rs11218007 | 11 | 120653294 | A | G | 0.571 | 0.5681 | +? | 0 | 0 | 0 | 1 |
| rs56311339 | 11 | 116566161 | T | C | 0.571 | 0.5681 | +? | 0 | 0 | 0 | 1 |
| rs948104 | 11 | 117596927 | G | T | 0.571 | 0.5681 | +? | 0 | 0 | 0 | 1 |
| rs111750658 | 11 | 120695729 | C | G | 0.571 | 0.5683 | +? | 0 | 0 | 0 | 1 |
| rs17120968 | 11 | 117493839 | T | C | 0.571 | 0.5683 | +? | 0 | 0 | 0 | 1 |
| rs73020492 | 11 | 117775130 | A | C | 0.57 | 0.5684 | +? | 0 | 0 | 0 | 1 |
| rs12806865 | 11 | 117404769 | G | A | 0.57 | 0.5685 | +? | 0 | 0 | 0 | 1 |
| rs1632452 | 11 | 119967615 | C | T | 0.57 | 0.5685 | +? | 0 | 0 | 0 | 1 |
| rs11823721 | 11 | 117772002 | C | T | 0.57 | 0.5686 | +? | 0 | 0 | 0 | 1 |
| rs1241550 | 11 | 116726651 | C | T | -0.57 | 0.5686 | +- | 28.7 | 4.207 | 1 | 0.04025 |
| rs61901402 | 11 | 120598054 | A | G | -0.57 | 0.5686 | -- | 0 | 0.774 | 1 | 0.3788 |
| rs633389 | 11 | 116667337 | T | C | 0.57 | 0.5686 | +- | -42.3 | 2.109 | 1 | 0.1465 |
| rs551417 | 11 | 119742372 | T | C | -0.57 | 0.5687 | -? | 0 | 0 | 0 | 1 |
| rs551662 | 11 | 117179003 | C | T | -0.57 | 0.5687 | -? | 0 | 0 | 0 | 1 |
| rs9332857 | 11 | 118389440 | T | C | -0.57 | 0.569 | -+ | -191.5 | 1.029 | 1 | 0.3104 |

| | | | | | | | | | | | |
|---|---|---|---|---|---|---|---|---|---|---|---|
| rs61904855 | 11 | 116476668 | A | C | -0.569 | 0.5691 | -? | 0 | 0 | 0 | 1 |
| rs1941408 | 11 | 120150430 | A | G | -0.569 | 0.5692 | -? | 0 | 0 | 0 | 1 |
| rs627995 | 11 | 118054324 | T | C | 0.569 | 0.5692 | +? | 0 | 0 | 0 | 1 |
| rs555179364 | 11 | 117218685 | C | T | 0.569 | 0.5696 | +? | 0 | 0 | 0 | 1 |
| rs79110667 | 11 | 121167114 | T | C | 0.568 | 0.5697 | +? | 0 | 0 | 0 | 1 |
| rs727422 | 11 | 121168178 | A | G | 0.568 | 0.5698 | +? | 0 | 0 | 0 | 1 |
| rs3863309 | 11 | 117293818 | C | T | 0.568 | 0.57 | +? | 0 | 0 | 0 | 1 |
| rs11217973 | 11 | 120553706 | T | C | 0.568 | 0.5701 | +? | 0 | 0 | 0 | 1 |
| rs7924573 | 11 | 119571342 | C | G | -0.568 | 0.5703 | -? | 0 | 0 | 0 | 1 |
| rs4372482 | 11 | 117839307 | C | T | 0.567 | 0.5709 | ++ | 0 | 0.001 | 1 | 0.9782 |
| rs12225989 | 11 | 120930947 | A | C | -0.567 | 0.571 | -? | 0 | 0 | 0 | 1 |
| rs1145193 | 11 | 116570942 | G | A | 0.566 | 0.5712 | +? | 0 | 0 | 0 | 1 |
| rs1145196 | 11 | 116572205 | T | C | 0.566 | 0.5713 | +? | 0 | 0 | 0 | 1 |
| rs7106851 | 11 | 118554259 | A | G | 0.566 | 0.5715 | ++ | 0 | 0.885 | 1 | 0.3468 |
| rs3830008 | 11 | 117093764 | C | A | -0.566 | 0.5716 | -? | 0 | 0 | 0 | 1 |
| rs11217693 | 11 | 119975273 | G | A | 0.566 | 0.5717 | +? | 0 | 0 | 0 | 1 |
| rs6589564 | 11 | 116624153 | C | G | -0.565 | 0.5718 | +- | -99.3 | 1.505 | 1 | 0.2198 |
| rs180865175 | 11 | 117250419 | G | T | 0.565 | 0.5719 | +? | 0 | 0 | 0 | 1 |
| rs478296 | 11 | 118567375 | C | T | 0.565 | 0.5719 | ++ | 0 | 0.914 | 1 | 0.3391 |
| rs12361100 | 11 | 121022231 | T | A | -0.565 | 0.572 | -? | 0 | 0 | 0 | 1 |
| rs1263499 | 11 | 117081709 | G | A | 0.565 | 0.5722 | +? | 0 | 0 | 0 | 1 |
| rs77130505 | 11 | 116448927 | T | C | 0.564 | 0.5725 | ++ | 0 | 0.028 | 1 | 0.8666 |
| rs896691 | 11 | 120048958 | C | A | 0.564 | 0.5725 | +? | 0 | 0 | 0 | 1 |
| rs11006 | 11 | 118895202 | G | A | -0.564 | 0.5726 | -- | 0 | 0.007 | 1 | 0.9321 |
| rs6589570 | 11 | 116675294 | A | T | 0.564 | 0.5726 | +? | 0 | 0 | 0 | 1 |
| rs7952366 | 11 | 119597943 | T | G | 0.564 | 0.5729 | +? | 0 | 0 | 0 | 1 |
| rs78659090 | 11 | 117558035 | G | A | 0.564 | 0.573 | +? | 0 | 0 | 0 | 1 |
| rs11217180 | 11 | 119072976 | T | C | 0.564 | 0.5731 | ++ | 0 | 0.393 | 1 | 0.5309 |
| rs578780 | 11 | 116288558 | T | C | -0.563 | 0.5731 | -? | 0 | 0 | 0 | 1 |
| rs11825655 | 11 | 117630867 | G | A | 0.563 | 0.5733 | +? | 0 | 0 | 0 | 1 |
| rs55728834 | 11 | 119682166 | C | T | 0.563 | 0.5733 | ++ | 0 | 0.008 | 1 | 0.9273 |
| rs10736502 | 11 | 119981785 | G | C | 0.563 | 0.5737 | +? | 0 | 0 | 0 | 1 |
| rs36088800 | 11 | 116223240 | C | T | 0.563 | 0.5737 | +- | -76.8 | 1.697 | 1 | 0.1927 |
| rs2156762 | 11 | 120624339 | C | T | 0.562 | 0.574 | +? | 0 | 0 | 0 | 1 |
| rs3016360 | 11 | 116559743 | C | T | -0.562 | 0.5741 | -? | 0 | 0 | 0 | 1 |
| rs2846114 | 11 | 120610144 | C | T | 0.562 | 0.5742 | +? | 0 | 0 | 0 | 1 |
| rs74341458 | 11 | 116751549 | A | G | 0.562 | 0.5742 | +? | 0 | 0 | 0 | 1 |
| rs61900350 | 11 | 117800481 | C | G | -0.562 | 0.5743 | -+ | 3.5 | 3.108 | 1 | 0.07792 |
| rs1145191 | 11 | 116570718 | G | C | 0.562 | 0.5744 | +? | 0 | 0 | 0 | 1 |
| rs1144036 | 11 | 116584421 | C | T | -0.561 | 0.5746 | -? | 0 | 0 | 0 | 1 |
| rs2155369 | 11 | 121038810 | T | C | -0.561 | 0.5746 | -? | 0 | 0 | 0 | 1 |
| rs6589802 | 11 | 119972011 | G | A | 0.561 | 0.5746 | +? | 0 | 0 | 0 | 1 |
| rs7396835 | 11 | 116684028 | T | C | 0.561 | 0.5747 | +? | 0 | 0 | 0 | 1 |
| rs3759002 | 11 | 118136160 | A | G | 0.561 | 0.5749 | -+ | -9.1 | 2.75 | 1 | 0.09725 |
| rs1939982 | 11 | 117605574 | G | A | 0.561 | 0.575 | +? | 0 | 0 | 0 | 1 |
| rs531260 | 11 | 120050515 | C | T | 0.56 | 0.5753 | +? | 0 | 0 | 0 | 1 |
| rs1240781 | 11 | 116574817 | T | C | 0.56 | 0.5756 | +? | 0 | 0 | 0 | 1 |
| rs12420005 | 11 | 119256303 | T | C | -0.56 | 0.5757 | -? | 0 | 0 | 0 | 1 |
| rs499744 | 11 | 117225033 | T | C | -0.56 | 0.5758 | -? | 0 | 0 | 0 | 1 |
| rs8192696 | 11 | 118895686 | T | C | -0.56 | 0.5758 | -- | 0 | 0.005 | 1 | 0.9423 |
| rs11216017 | 11 | 116451308 | G | C | 0.559 | 0.5759 | +? | 0 | 0 | 0 | 1 |
| rs78540725 | 11 | 119766827 | C | A | 0.559 | 0.5759 | +? | 0 | 0 | 0 | 1 |
| rs11217817 | 11 | 120191872 | G | C | 0.559 | 0.576 | +? | 0 | 0 | 0 | 1 |
| rs72994633 | 11 | 119824803 | C | T | 0.559 | 0.5763 | +? | 0 | 0 | 0 | 1 |
| rs7935369 | 11 | 119482594 | T | C | 0.559 | 0.5763 | +? | 0 | 0 | 0 | 1 |
| rs10790202 | 11 | 117597995 | C | A | 0.558 | 0.5766 | +? | 0 | 0 | 0 | 1 |

| | | | | | | | | | | | |
|---|---|---|---|---|---|---|---|---|---|---|---|
| rs116324528 | 11 | 120633977 | T | A | 0.558 | 0.5766 | +? | 0 | 0 | 0 | 1 |
| rs12292959 | 11 | 116466714 | C | T | 0.558 | 0.5766 | +? | 0 | 0 | 0 | 1 |
| rs17120244 | 11 | 116919511 | T | C | -0.558 | 0.5766 | -? | 0 | 0 | 0 | 1 |
| rs61905717 | 11 | 116919208 | A | G | -0.558 | 0.5766 | -? | 0 | 0 | 0 | 1 |
| rs11216656 | 11 | 117838400 | A | G | 0.558 | 0.5771 | ++ | 0 | 0.023 | 1 | 0.8789 |
| rs483367 | 11 | 116297397 | T | C | -0.558 | 0.5771 | -? | 0 | 0 | 0 | 1 |
| rs12577176 | 11 | 116224822 | C | T | 0.558 | 0.5772 | +- | -77.6 | 1.689 | 1 | 0.1937 |
| rs4938751 | 11 | 119876939 | C | T | -0.557 | 0.5772 | -? | 0 | 0 | 0 | 1 |
| rs79820276 | 11 | 120356832 | G | A | -0.557 | 0.5773 | -? | 0 | 0 | 0 | 1 |
| rs10437694 | 11 | 121110312 | G | T | -0.557 | 0.5776 | -- | 0 | 0.207 | 1 | 0.6488 |
| rs1787715 | 11 | 116560325 | T | C | -0.557 | 0.5776 | -? | 0 | 0 | 0 | 1 |
| rs12418662 | 11 | 119596772 | C | T | 0.557 | 0.5778 | +? | 0 | 0 | 0 | 1 |
| rs59071712 | 11 | 117557224 | A | T | 0.557 | 0.5778 | +? | 0 | 0 | 0 | 1 |
| rs79489590 | 11 | 116791391 | T | C | 0.557 | 0.5778 | +? | 0 | 0 | 0 | 1 |
| rs644498 | 11 | 118567486 | C | T | 0.556 | 0.578 | ++ | 0 | 0.708 | 1 | 0.4002 |
| rs1003081 | 11 | 118913993 | T | C | 0.556 | 0.5781 | +? | 0 | 0 | 0 | 1 |
| rs12799630 | 11 | 116393611 | T | C | 0.556 | 0.5783 | +? | 0 | 0 | 0 | 1 |
| rs11604139 | 11 | 120480291 | A | T | 0.556 | 0.5784 | +? | 0 | 0 | 0 | 1 |
| rs2252359 | 11 | 120400149 | A | G | 0.556 | 0.5784 | +? | 0 | 0 | 0 | 1 |
| rs4423191 | 11 | 120752044 | G | T | -0.556 | 0.5784 | -? | 0 | 0 | 0 | 1 |
| rs145762171 | 11 | 118748789 | A | G | 0.556 | 0.5785 | +- | 0 | 0.614 | 1 | 0.4333 |
| rs150586315 | 11 | 120636195 | A | G | 0.555 | 0.5786 | +? | 0 | 0 | 0 | 1 |
| rs10892186 | 11 | 117729274 | A | G | -0.555 | 0.5788 | -? | 0 | 0 | 0 | 1 |
| rs493602 | 11 | 116297866 | C | G | -0.555 | 0.5788 | -? | 0 | 0 | 0 | 1 |
| rs12364542 | 11 | 116468132 | A | G | -0.555 | 0.5789 | -? | 0 | 0 | 0 | 1 |
| rs180740269 | 11 | 117183277 | T | C | -0.555 | 0.5789 | -? | 0 | 0 | 0 | 1 |
| rs111313044 | 11 | 118803596 | A | G | 0.555 | 0.579 | +- | -84.5 | 1.626 | 1 | 0.2022 |
| rs73575440 | 11 | 118803160 | A | G | 0.555 | 0.579 | +- | -84.5 | 1.626 | 1 | 0.2022 |
| rs10790150 | 11 | 116450713 | T | C | 0.555 | 0.5791 | +? | 0 | 0 | 0 | 1 |
| rs113111994 | 11 | 119876350 | T | G | 0.555 | 0.5791 | +? | 0 | 0 | 0 | 1 |
| rs1240770 | 11 | 116518438 | T | G | -0.555 | 0.5791 | -? | 0 | 0 | 0 | 1 |
| rs28990986 | 11 | 118963291 | T | C | -0.554 | 0.5793 | -? | 0 | 0 | 0 | 1 |
| rs10892720 | 11 | 121169193 | T | G | -0.554 | 0.5794 | -? | 0 | 0 | 0 | 1 |
| rs1025130228 | 11 | 120480136 | T | A | 0.554 | 0.5796 | +? | 0 | 0 | 0 | 1 |
| rs11601552 | 11 | 120480191 | A | C | 0.554 | 0.5796 | +? | 0 | 0 | 0 | 1 |
| rs2231450 | 11 | 118186312 | A | C | 0.554 | 0.5796 | +- | 19.9 | 3.746 | 1 | 0.05294 |
| rs2511846 | 11 | 119203939 | C | G | -0.554 | 0.5796 | -? | 0 | 0 | 0 | 1 |
| rs7941246 | 11 | 117646354 | G | T | 0.554 | 0.5797 | +? | 0 | 0 | 0 | 1 |
| rs509712 | 11 | 116728029 | G | A | -0.554 | 0.5798 | +- | 29.4 | 4.252 | 1 | 0.0392 |
| rs11216075 | 11 | 116524832 | A | G | -0.554 | 0.5799 | -? | 0 | 0 | 0 | 1 |
| rs11215991 | 11 | 116407867 | T | C | 0.553 | 0.5801 | +? | 0 | 0 | 0 | 1 |
| rs147834459 | 11 | 116325615 | T | C | 0.553 | 0.5804 | +? | 0 | 0 | 0 | 1 |
| rs2231441 | 11 | 118175540 | G | A | -0.553 | 0.5804 | -+ | -11.2 | 2.697 | 1 | 0.1005 |
| rs1039377879 | 11 | 118068100 | T | A | 0.553 | 0.5806 | +? | 0 | 0 | 0 | 1 |
| rs1240782 | 11 | 116574584 | A | G | 0.552 | 0.5809 | +? | 0 | 0 | 0 | 1 |
| rs17124548 | 11 | 120788631 | T | A | -0.552 | 0.581 | -? | 0 | 0 | 0 | 1 |
| rs1006160 | 11 | 117739616 | A | T | 0.552 | 0.5811 | +? | 0 | 0 | 0 | 1 |
| rs7932655 | 11 | 116962563 | A | T | -0.552 | 0.5811 | -? | 0 | 0 | 0 | 1 |
| rs12421080 | 11 | 120790862 | A | T | -0.552 | 0.5812 | -? | 0 | 0 | 0 | 1 |
| rs59839888 | 11 | 116949853 | C | T | -0.551 | 0.5813 | +- | 43 | 5.259 | 1 | 0.02183 |
| rs17124764 | 11 | 120959463 | T | A | -0.551 | 0.5814 | -? | 0 | 0 | 0 | 1 |
| rs11217186 | 11 | 119083906 | T | C | 0.551 | 0.5815 | ++ | 0 | 0.383 | 1 | 0.536 |
| rs1787716 | 11 | 116560174 | A | G | -0.551 | 0.5816 | -? | 0 | 0 | 0 | 1 |
| rs1942926 | 11 | 117599333 | A | C | 0.551 | 0.5819 | +? | 0 | 0 | 0 | 1 |
| rs56016514 | 11 | 116961588 | A | G | -0.551 | 0.5819 | -? | 0 | 0 | 0 | 1 |
| rs57582454 | 11 | 116961892 | A | G | -0.551 | 0.5819 | -? | 0 | 0 | 0 | 1 |

| | | | | | | | | | | | |
|---|---|---|---|---|---|---|---|---|---|---|---|
| rs61903423 | 11 | 116962078 | A | G | -0.551 | 0.5819 | -? | 0 | 0 | 0 | 1 |
| rs79175131 | 11 | 116375719 | T | C | 0.55 | 0.5822 | +? | 0 | 0 | 0 | 1 |
| rs10750123 | 11 | 117955810 | C | T | 0.55 | 0.5824 | +- | 0 | 0.138 | 1 | 0.7107 |
| rs4938278 | 11 | 116417813 | T | G | 0.55 | 0.5826 | ++ | 0 | 0.265 | 1 | 0.6066 |
| rs73000642 | 11 | 120448042 | A | G | -0.55 | 0.5826 | -? | 0 | 0 | 0 | 1 |
| rs2155364 | 11 | 121017301 | T | C | -0.549 | 0.5828 | -? | 0 | 0 | 0 | 1 |
| rs34841529 | 11 | 116448838 | A | C | 0.549 | 0.5829 | +? | 0 | 0 | 0 | 1 |
| rs558034 | 11 | 119705210 | A | G | 0.549 | 0.5829 | +? | 0 | 0 | 0 | 1 |
| rs514649 | 11 | 119716304 | G | A | -0.549 | 0.583 | -? | 0 | 0 | 0 | 1 |
| rs633867 | 11 | 116667480 | T | C | 0.549 | 0.5833 | +- | -44.7 | 2.073 | 1 | 0.1499 |
| rs949189 | 11 | 119902167 | C | T | 0.548 | 0.5834 | +? | 0 | 0 | 0 | 1 |
| rs10892152 | 11 | 117532543 | G | A | 0.548 | 0.5835 | +? | 0 | 0 | 0 | 1 |
| rs45615131 | 11 | 117989951 | T | A | 0.548 | 0.584 | ++ | 0 | 0 | 1 | 0.9894 |
| rs470876 | 11 | 120013307 | C | T | 0.547 | 0.5842 | +? | 0 | 0 | 0 | 1 |
| rs488672 | 11 | 120047745 | C | T | 0.547 | 0.5844 | +? | 0 | 0 | 0 | 1 |
| rs79118964 | 11 | 117033808 | T | C | 0.547 | 0.5844 | +? | 0 | 0 | 0 | 1 |
| rs118086106 | 11 | 119483599 | C | T | -0.547 | 0.5846 | -? | 0 | 0 | 0 | 1 |
| rs11216023 | 11 | 116460172 | C | G | 0.546 | 0.585 | +? | 0 | 0 | 0 | 1 |
| rs3851576 | 11 | 117638172 | C | A | 0.546 | 0.5853 | +? | 0 | 0 | 0 | 1 |
| rs17124324 | 11 | 120650633 | T | A | 0.546 | 0.5854 | +? | 0 | 0 | 0 | 1 |
| rs61904806 | 11 | 116434082 | C | T | -0.546 | 0.5854 | -- | 0 | 0.004 | 1 | 0.9498 |
| rs7932398 | 11 | 119071897 | G | C | 0.545 | 0.5854 | ++ | 0 | 0.406 | 1 | 0.5241 |
| rs2298491 | 11 | 120959800 | A | G | -0.545 | 0.5855 | -? | 0 | 0 | 0 | 1 |
| rs2257198 | 11 | 120402020 | A | G | 0.545 | 0.5856 | +? | 0 | 0 | 0 | 1 |
| rs35374605 | 11 | 117630114 | C | T | 0.545 | 0.5857 | +? | 0 | 0 | 0 | 1 |
| rs11217183 | 11 | 119078886 | G | A | 0.545 | 0.5858 | ++ | 0 | 0.387 | 1 | 0.5337 |
| rs11217184 | 11 | 119078993 | G | A | 0.545 | 0.5858 | ++ | 0 | 0.387 | 1 | 0.5337 |
| rs12798188 | 11 | 119612677 | T | C | 0.545 | 0.5858 | +? | 0 | 0 | 0 | 1 |
| rs1242598 | 11 | 117129637 | T | C | 0.545 | 0.586 | +? | 0 | 0 | 0 | 1 |
| rs79293731 | 11 | 121155566 | T | C | 0.545 | 0.586 | +? | 0 | 0 | 0 | 1 |
| rs12361543 | 11 | 116467589 | G | A | -0.544 | 0.5862 | -? | 0 | 0 | 0 | 1 |
| rs12794454 | 11 | 117633286 | A | T | 0.544 | 0.5862 | +? | 0 | 0 | 0 | 1 |
| rs4938402 | 11 | 117399918 | A | G | -0.544 | 0.5862 | -? | 0 | 0 | 0 | 1 |
| rs10892000 | 11 | 116498862 | G | T | 0.544 | 0.5864 | +? | 0 | 0 | 0 | 1 |
| rs34884722 | 11 | 120479125 | G | A | 0.544 | 0.5866 | +? | 0 | 0 | 0 | 1 |
| rs113190660 | 11 | 120169537 | T | C | -0.543 | 0.5868 | -- | 0 | 0.001 | 1 | 0.974 |
| rs651474 | 11 | 117961908 | C | T | -0.543 | 0.5869 | -- | 0 | 0.003 | 1 | 0.9574 |
| rs4581455 | 11 | 119481661 | A | G | 0.542 | 0.5876 | +? | 0 | 0 | 0 | 1 |
| rs574233 | 11 | 116295758 | T | G | -0.542 | 0.5878 | -? | 0 | 0 | 0 | 1 |
| rs547761 | 11 | 116314960 | C | T | 0.542 | 0.5881 | +? | 0 | 0 | 0 | 1 |
| rs11216485 | 11 | 117523508 | G | C | -0.541 | 0.5884 | -? | 0 | 0 | 0 | 1 |
| rs542613 | 11 | 118567084 | A | G | 0.541 | 0.5884 | ++ | 0 | 0.802 | 1 | 0.3706 |
| rs10892171 | 11 | 117642030 | A | T | 0.541 | 0.5887 | +? | 0 | 0 | 0 | 1 |
| rs12272862 | 11 | 119848377 | A | G | -0.541 | 0.5888 | -? | 0 | 0 | 0 | 1 |
| rs117287138 | 11 | 116794879 | A | G | 0.54 | 0.5889 | +? | 0 | 0 | 0 | 1 |
| rs604413 | 11 | 120029497 | G | A | -0.54 | 0.5891 | -? | 0 | 0 | 0 | 1 |
| rs590005 | 11 | 120024312 | T | A | -0.54 | 0.5892 | -? | 0 | 0 | 0 | 1 |
| rs491822 | 11 | 117772848 | T | C | -0.54 | 0.5893 | -? | 0 | 0 | 0 | 1 |
| rs4938389 | 11 | 117357696 | A | G | -0.54 | 0.5893 | -? | 0 | 0 | 0 | 1 |
| rs4938661 | 11 | 119384351 | C | G | -0.54 | 0.5894 | -? | 0 | 0 | 0 | 1 |
| rs10750176 | 11 | 120557343 | G | A | -0.54 | 0.5895 | -? | 0 | 0 | 0 | 1 |
| rs602030 | 11 | 119850718 | T | C | -0.54 | 0.5895 | -? | 0 | 0 | 0 | 1 |
| rs11600568 | 11 | 120083648 | G | C | 0.539 | 0.5896 | +? | 0 | 0 | 0 | 1 |
| rs755538 | 11 | 117768454 | T | C | 0.539 | 0.5896 | +? | 0 | 0 | 0 | 1 |
| rs12576806 | 11 | 117483306 | A | G | 0.539 | 0.5898 | +? | 0 | 0 | 0 | 1 |
| rs3741280 | 11 | 117403235 | T | G | 0.539 | 0.5898 | +? | 0 | 0 | 0 | 1 |

| | | | | | | | | | | | |
|---|---|---|---|---|---|---|---|---|---|---|---|
| rs148403721 | 11 | 116757765 | A | C | -0.539 | 0.5899 | -? | 0 | 0 | 0 | 1 |
| rs11216384 | 11 | 117307002 | C | T | 0.539 | 0.59 | +? | 0 | 0 | 0 | 1 |
| rs1579885 | 11 | 119800548 | A | G | -0.539 | 0.59 | -? | 0 | 0 | 0 | 1 |
| rs28917236 | 11 | 117185892 | T | G | -0.539 | 0.59 | -? | 0 | 0 | 0 | 1 |
| rs570081 | 11 | 119740598 | G | A | -0.539 | 0.59 | -? | 0 | 0 | 0 | 1 |
| rs117785509 | 11 | 116586231 | A | G | -0.539 | 0.5901 | -- | 0 | 0.002 | 1 | 0.9664 |
| rs7940527 | 11 | 117398202 | C | G | -0.539 | 0.5901 | -? | 0 | 0 | 0 | 1 |
| rs11216416 | 11 | 117347207 | T | C | -0.538 | 0.5903 | -? | 0 | 0 | 0 | 1 |
| rs577733 | 11 | 120993066 | A | G | -0.538 | 0.5903 | -? | 0 | 0 | 0 | 1 |
| rs1240659 | 11 | 116493950 | G | T | -0.538 | 0.5904 | -? | 0 | 0 | 0 | 1 |
| rs2266788 | 11 | 116660686 | G | A | -0.538 | 0.5904 | +- | -41 | 2.128 | 1 | 0.1446 |
| rs4938793 | 11 | 120146360 | T | G | -0.538 | 0.5906 | -? | 0 | 0 | 0 | 1 |
| rs536069 | 11 | 120996292 | A | G | -0.538 | 0.5907 | -? | 0 | 0 | 0 | 1 |
| rs77196345 | 11 | 116448951 | T | C | 0.538 | 0.5907 | ++ | 0 | 0.034 | 1 | 0.8546 |
| rs113399762 | 11 | 118389698 | C | T | -0.538 | 0.5908 | -+ | -148.8 | 1.206 | 1 | 0.2722 |
| rs55987240 | 11 | 119200202 | C | A | -0.537 | 0.5911 | -? | 0 | 0 | 0 | 1 |
| rs6589674 | 11 | 118566039 | G | A | 0.537 | 0.5911 | ++ | 0 | 0.713 | 1 | 0.3984 |
| rs11821311 | 11 | 117772179 | T | C | 0.537 | 0.5914 | +? | 0 | 0 | 0 | 1 |
| rs12574794 | 11 | 119330741 | T | C | -0.537 | 0.5914 | -? | 0 | 0 | 0 | 1 |
| rs3016414 | 11 | 120670765 | G | A | 0.537 | 0.5914 | +? | 0 | 0 | 0 | 1 |
| rs7950536 | 11 | 121154118 | G | A | 0.536 | 0.5917 | +? | 0 | 0 | 0 | 1 |
| rs3133858 | 11 | 120651611 | C | T | 0.536 | 0.5918 | +? | 0 | 0 | 0 | 1 |
| rs55903632 | 11 | 121156412 | C | G | 0.536 | 0.592 | +? | 0 | 0 | 0 | 1 |
| rs56142837 | 11 | 121156037 | G | T | 0.536 | 0.592 | +? | 0 | 0 | 0 | 1 |
| rs78303474 | 11 | 121154642 | C | T | 0.536 | 0.592 | +? | 0 | 0 | 0 | 1 |
| rs11217816 | 11 | 120190997 | G | A | -0.536 | 0.5921 | -? | 0 | 0 | 0 | 1 |
| rs4589313 | 11 | 119236794 | A | G | 0.536 | 0.5921 | +? | 0 | 0 | 0 | 1 |
| rs4938317 | 11 | 116778275 | C | T | 0.536 | 0.5922 | +? | 0 | 0 | 0 | 1 |
| rs79197348 | 11 | 116238176 | G | C | -0.536 | 0.5923 | -- | 0 | 0.192 | 1 | 0.6612 |
| rs888245 | 11 | 116723737 | G | C | 0.535 | 0.5924 | +? | 0 | 0 | 0 | 1 |
| rs9667814 | 11 | 116671823 | C | G | -0.535 | 0.5925 | +- | -28.8 | 2.329 | 1 | 0.127 |
| rs2000525 | 11 | 120907482 | A | G | -0.535 | 0.5926 | -? | 0 | 0 | 0 | 1 |
| rs4245205 | 11 | 119584506 | T | C | 0.535 | 0.5926 | +? | 0 | 0 | 0 | 1 |
| rs4533047 | 11 | 119584347 | T | C | 0.535 | 0.5926 | +? | 0 | 0 | 0 | 1 |
| rs7927510 | 11 | 119584876 | A | T | 0.535 | 0.5926 | +? | 0 | 0 | 0 | 1 |
| rs7946962 | 11 | 119586216 | T | C | 0.535 | 0.5926 | +? | 0 | 0 | 0 | 1 |
| rs2852244 | 11 | 120624775 | C | G | 0.535 | 0.5927 | +? | 0 | 0 | 0 | 1 |
| rs117437529 | 11 | 116512452 | T | C | -0.535 | 0.5928 | -? | 0 | 0 | 0 | 1 |
| rs7948661 | 11 | 118341649 | C | T | -0.535 | 0.5929 | -+ | 0 | 0.806 | 1 | 0.3692 |
| rs1783861 | 11 | 117963635 | T | A | -0.535 | 0.593 | -- | 0 | 0.002 | 1 | 0.9634 |
| rs4938403 | 11 | 117409910 | A | C | -0.534 | 0.5931 | -? | 0 | 0 | 0 | 1 |
| rs11218044 | 11 | 120756908 | A | G | -0.534 | 0.5932 | -? | 0 | 0 | 0 | 1 |
| rs12419991 | 11 | 116301251 | T | C | -0.534 | 0.5932 | -? | 0 | 0 | 0 | 1 |
| rs11216034 | 11 | 116470173 | C | A | -0.534 | 0.5933 | -? | 0 | 0 | 0 | 1 |
| rs1240658 | 11 | 116496360 | C | T | -0.534 | 0.5933 | -? | 0 | 0 | 0 | 1 |
| rs635706 | 11 | 117694089 | G | A | -0.534 | 0.5936 | -? | 0 | 0 | 0 | 1 |
| rs17601549 | 11 | 117998077 | T | C | 0.534 | 0.5937 | ++ | 0 | 0.001 | 1 | 0.9749 |
| rs596911 | 11 | 120025476 | A | G | -0.533 | 0.5937 | -? | 0 | 0 | 0 | 1 |
| rs659912 | 11 | 120026241 | G | A | -0.533 | 0.5937 | -? | 0 | 0 | 0 | 1 |
| rs35288685 | 11 | 121183145 | T | C | -0.533 | 0.5938 | -? | 0 | 0 | 0 | 1 |
| rs529657 | 11 | 120042087 | C | T | 0.533 | 0.5938 | +? | 0 | 0 | 0 | 1 |
| rs11602018 | 11 | 120753572 | G | A | -0.533 | 0.594 | -? | 0 | 0 | 0 | 1 |
| rs12361746 | 11 | 119659477 | T | C | 0.533 | 0.594 | ++ | 0 | 0.002 | 1 | 0.9617 |
| rs12794726 | 11 | 118566313 | T | C | 0.533 | 0.5942 | ++ | 0 | 0.718 | 1 | 0.397 |
| rs73578608 | 11 | 120760463 | A | G | -0.533 | 0.5942 | -? | 0 | 0 | 0 | 1 |
| rs2231440 | 11 | 118175439 | A | G | -0.533 | 0.5943 | -+ | -10.5 | 2.714 | 1 | 0.09947 |

| | | | | | | | | | | | |
|---|---|---|---|---|---|---|---|---|---|---|---|
| rs2852226 | 11 | 120619586 | A | G | 0.532 | 0.5945 | +? | 0 | 0 | 0 | 1 |
| rs1567377 | 11 | 119480810 | C | T | 0.532 | 0.5946 | +? | 0 | 0 | 0 | 1 |
| rs1107149 | 11 | 119973745 | C | T | 0.532 | 0.5947 | +? | 0 | 0 | 0 | 1 |
| rs117730773 | 11 | 116547972 | T | C | -0.532 | 0.5947 | -? | 0 | 0 | 0 | 1 |
| rs894839 | 11 | 120337705 | G | A | -0.532 | 0.5947 | -? | 0 | 0 | 0 | 1 |
| rs55942271 | 11 | 118177115 | T | C | 0.532 | 0.5948 | +? | 0 | 0 | 0 | 1 |
| rs7122363 | 11 | 116449283 | T | C | 0.532 | 0.5949 | +? | 0 | 0 | 0 | 1 |
| rs11216612 | 11 | 117773893 | T | C | -0.531 | 0.5951 | -? | 0 | 0 | 0 | 1 |
| rs12146702 | 11 | 117663838 | T | C | -0.531 | 0.5953 | -? | 0 | 0 | 0 | 1 |
| rs2854117 | 11 | 116700142 | T | C | 0.531 | 0.5954 | +? | 0 | 0 | 0 | 1 |
| rs7124649 | 11 | 116948640 | C | T | -0.531 | 0.5955 | +- | 50.3 | 6.034 | 1 | 0.01403 |
| rs11216257 | 11 | 116932561 | C | T | 0.531 | 0.5956 | +? | 0 | 0 | 0 | 1 |
| rs10892208 | 11 | 117955759 | G | A | 0.531 | 0.5957 | +- | 0 | 0.131 | 1 | 0.7173 |
| rs73012993 | 11 | 116528663 | G | A | 0.531 | 0.5957 | +? | 0 | 0 | 0 | 1 |
| rs143408085 | 11 | 119580246 | A | G | 0.53 | 0.5958 | +? | 0 | 0 | 0 | 1 |
| rs6589821 | 11 | 120494089 | A | G | 0.53 | 0.5961 | +? | 0 | 0 | 0 | 1 |
| rs34250455 | 11 | 118157681 | T | C | -0.53 | 0.5963 | -+ | -3.9 | 2.887 | 1 | 0.08932 |
| rs61905132 | 11 | 116673464 | T | C | 0.53 | 0.5963 | +? | 0 | 0 | 0 | 1 |
| rs12285901 | 11 | 119235960 | A | G | 0.53 | 0.5964 | +? | 0 | 0 | 0 | 1 |
| rs3020105 | 11 | 120684447 | G | A | 0.53 | 0.5964 | +? | 0 | 0 | 0 | 1 |
| rs562780 | 11 | 118362414 | T | C | 0.53 | 0.5964 | +- | 0 | 0.862 | 1 | 0.3531 |
| rs59069289 | 11 | 118802128 | T | G | 0.53 | 0.5964 | +- | -88.7 | 1.59 | 1 | 0.2074 |
| rs59480472 | 11 | 118802386 | C | T | 0.53 | 0.5964 | +- | -88.7 | 1.59 | 1 | 0.2074 |
| rs11216157 | 11 | 116711180 | G | A | 0.529 | 0.5965 | +? | 0 | 0 | 0 | 1 |
| rs4938735 | 11 | 119747470 | T | C | 0.529 | 0.5965 | +? | 0 | 0 | 0 | 1 |
| rs76234156 | 11 | 116519181 | A | C | -0.529 | 0.5965 | -? | 0 | 0 | 0 | 1 |
| rs7947659 | 11 | 116419981 | C | A | 0.529 | 0.5965 | ++ | 0 | 0.366 | 1 | 0.5452 |
| rs11216172 | 11 | 116749878 | G | A | -0.529 | 0.5966 | -? | 0 | 0 | 0 | 1 |
| rs61900138 | 11 | 118782222 | C | T | 0.529 | 0.5966 | -+ | -22.6 | 2.447 | 1 | 0.1177 |
| rs7131629 | 11 | 119973299 | A | G | 0.529 | 0.5966 | +? | 0 | 0 | 0 | 1 |
| rs7128433 | 11 | 121156755 | T | C | 0.529 | 0.5967 | +? | 0 | 0 | 0 | 1 |
| rs3888083 | 11 | 117341163 | C | T | -0.529 | 0.5968 | -? | 0 | 0 | 0 | 1 |
| rs11216947 | 11 | 118567218 | A | G | 0.529 | 0.5969 | ++ | 0 | 0.756 | 1 | 0.3844 |
| rs10219304 | 11 | 121127685 | A | G | -0.529 | 0.597 | -? | 0 | 0 | 0 | 1 |
| rs113442800 | 11 | 119818055 | T | C | 0.529 | 0.597 | +? | 0 | 0 | 0 | 1 |
| rs111463966 | 11 | 120554375 | G | T | -0.529 | 0.5971 | -? | 0 | 0 | 0 | 1 |
| rs1786191 | 11 | 117947208 | C | T | -0.529 | 0.5971 | -? | 0 | 0 | 0 | 1 |
| rs7395060 | 11 | 119703617 | G | A | -0.529 | 0.5971 | -? | 0 | 0 | 0 | 1 |
| rs11216561 | 11 | 117685575 | C | T | 0.528 | 0.5973 | +? | 0 | 0 | 0 | 1 |
| rs3892796 | 11 | 117302910 | A | G | 0.528 | 0.5973 | +? | 0 | 0 | 0 | 1 |
| rs73020476 | 11 | 117772389 | C | T | 0.528 | 0.5973 | +? | 0 | 0 | 0 | 1 |
| rs10790218 | 11 | 117741995 | A | G | 0.528 | 0.5974 | +? | 0 | 0 | 0 | 1 |
| rs599320 | 11 | 116740659 | C | G | -0.528 | 0.5974 | +- | 35.7 | 4.667 | 1 | 0.03074 |
| rs186292996 | 11 | 117682332 | T | C | -0.528 | 0.5975 | -? | 0 | 0 | 0 | 1 |
| rs615539 | 11 | 119850725 | G | A | -0.528 | 0.5975 | -? | 0 | 0 | 0 | 1 |
| rs626225 | 11 | 120034115 | G | A | -0.528 | 0.5975 | -? | 0 | 0 | 0 | 1 |
| rs77726126 | 11 | 121153556 | G | T | 0.528 | 0.5975 | +? | 0 | 0 | 0 | 1 |
| rs7926828 | 11 | 116586423 | C | T | -0.528 | 0.5977 | -? | 0 | 0 | 0 | 1 |
| rs11602560 | 11 | 117415187 | T | C | -0.528 | 0.5978 | -? | 0 | 0 | 0 | 1 |
| rs35909519 | 11 | 117416249 | T | C | -0.528 | 0.5978 | -? | 0 | 0 | 0 | 1 |
| rs6589799 | 11 | 119971523 | C | T | 0.528 | 0.5978 | +? | 0 | 0 | 0 | 1 |
| rs1459783673 | 11 | 119480660 | A | T | 0.527 | 0.598 | +? | 0 | 0 | 0 | 1 |
| rs9332871 | 11 | 118393856 | T | G | -0.527 | 0.598 | -+ | -135.6 | 1.273 | 1 | 0.2591 |
| rs79294438 | 11 | 119341720 | T | C | 0.527 | 0.5982 | +? | 0 | 0 | 0 | 1 |
| rs111322300 | 11 | 118893878 | C | T | -0.527 | 0.5985 | -- | 0 | 0 | 1 | 0.9941 |
| rs10892177 | 11 | 117685220 | C | G | 0.526 | 0.5986 | +? | 0 | 0 | 0 | 1 |

| | | | | | | | | | | | |
|---|---|---|---|---|---|---|---|---|---|---|---|
| rs11216169 | 11 | 116744389 | C | T | 0.526 | 0.5986 | +? | 0 | 0 | 0 | 1 |
| rs11218176 | 11 | 121041724 | A | T | -0.526 | 0.5988 | -? | 0 | 0 | 0 | 1 |
| rs4938782 | 11 | 119978951 | T | C | 0.526 | 0.5988 | +? | 0 | 0 | 0 | 1 |
| rs620924 | 11 | 117947698 | A | G | -0.526 | 0.5988 | -? | 0 | 0 | 0 | 1 |
| rs75598010 | 11 | 121041584 | T | C | -0.526 | 0.5988 | -? | 0 | 0 | 0 | 1 |
| rs117507766 | 11 | 119396373 | T | C | 0.526 | 0.5989 | +? | 0 | 0 | 0 | 1 |
| rs76448372 | 11 | 118045846 | A | G | 0.526 | 0.5989 | +- | 32.1 | 4.416 | 1 | 0.03561 |
| rs76908457 | 11 | 116447795 | C | T | 0.526 | 0.5992 | ++ | 0 | 0.036 | 1 | 0.8491 |
| rs11600216 | 11 | 117690400 | C | T | 0.525 | 0.5993 | +? | 0 | 0 | 0 | 1 |
| rs1893906 | 11 | 120525032 | A | G | -0.525 | 0.5993 | -? | 0 | 0 | 0 | 1 |
| rs7930409 | 11 | 116958943 | G | A | -0.525 | 0.5994 | -? | 0 | 0 | 0 | 1 |
| rs7930783 | 11 | 116959039 | A | G | -0.525 | 0.5994 | -? | 0 | 0 | 0 | 1 |
| rs17120280 | 11 | 116942132 | A | G | 0.525 | 0.5996 | +? | 0 | 0 | 0 | 1 |
| rs11607610 | 11 | 119595348 | T | C | 0.525 | 0.5997 | +? | 0 | 0 | 0 | 1 |
| rs12798944 | 11 | 120149741 | C | T | -0.525 | 0.5997 | -? | 0 | 0 | 0 | 1 |
| rs59854309 | 11 | 116566763 | T | C | 0.525 | 0.5998 | +? | 0 | 0 | 0 | 1 |
| rs7950072 | 11 | 117525420 | G | A | -0.525 | 0.5998 | -? | 0 | 0 | 0 | 1 |
| rs117079525 | 11 | 120331909 | T | C | -0.524 | 0.6001 | -? | 0 | 0 | 0 | 1 |
| rs644287 | 11 | 118011805 | T | C | 0.524 | 0.6001 | ++ | 0 | 0.011 | 1 | 0.9159 |
| rs12294678 | 11 | 119805351 | T | C | -0.524 | 0.6002 | -? | 0 | 0 | 0 | 1 |
| rs17311212 | 11 | 120067953 | A | G | 0.524 | 0.6003 | +? | 0 | 0 | 0 | 1 |
| rs7114458 | 11 | 118566175 | A | G | 0.524 | 0.6003 | ++ | 0 | 0.726 | 1 | 0.3942 |
| rs11828621 | 11 | 120614119 | G | A | -0.524 | 0.6004 | -? | 0 | 0 | 0 | 1 |
| rs7114565 | 11 | 117963939 | A | G | -0.524 | 0.6004 | -- | 0 | 0.005 | 1 | 0.9445 |
| rs12417256 | 11 | 120337519 | C | A | -0.524 | 0.6005 | -? | 0 | 0 | 0 | 1 |
| rs4938445 | 11 | 117745003 | A | G | 0.524 | 0.6005 | +? | 0 | 0 | 0 | 1 |
| rs12295531 | 11 | 121108515 | T | C | -0.524 | 0.6006 | -- | 0 | 0.159 | 1 | 0.6905 |
| rs7114216 | 11 | 118566234 | G | A | 0.523 | 0.6006 | ++ | 0 | 0.727 | 1 | 0.394 |
| rs75051127 | 11 | 121173349 | A | G | -0.524 | 0.6006 | -? | 0 | 0 | 0 | 1 |
| rs78857232 | 11 | 116727465 | T | C | 0.524 | 0.6006 | +? | 0 | 0 | 0 | 1 |
| rs3825041 | 11 | 116631707 | T | C | -0.523 | 0.6007 | +- | -108 | 1.442 | 1 | 0.2297 |
| rs521609 | 11 | 117407301 | A | G | -0.523 | 0.6007 | -? | 0 | 0 | 0 | 1 |
| rs7928435 | 11 | 116439719 | A | C | 0.523 | 0.6007 | ++ | 0 | 0.044 | 1 | 0.8347 |
| rs7484249 | 11 | 120142118 | C | T | -0.523 | 0.6008 | -? | 0 | 0 | 0 | 1 |
| rs11215929 | 11 | 116303270 | G | A | -0.523 | 0.6011 | -? | 0 | 0 | 0 | 1 |
| rs7935488 | 11 | 119482705 | T | C | 0.523 | 0.6011 | +? | 0 | 0 | 0 | 1 |
| rs528431 | 11 | 117662909 | A | C | -0.522 | 0.6014 | -? | 0 | 0 | 0 | 1 |
| rs61903419 | 11 | 116957219 | A | G | -0.522 | 0.6015 | -? | 0 | 0 | 0 | 1 |
| rs10790236 | 11 | 117986853 | C | T | -0.522 | 0.6016 | -- | 0 | 0.008 | 1 | 0.9304 |
| rs10892102 | 11 | 117308936 | T | G | -0.522 | 0.6018 | -? | 0 | 0 | 0 | 1 |
| rs76136280 | 11 | 116788491 | T | C | 0.522 | 0.6019 | +? | 0 | 0 | 0 | 1 |
| rs1944156 | 11 | 116396613 | C | A | 0.521 | 0.6023 | +? | 0 | 0 | 0 | 1 |
| rs572126 | 11 | 118359161 | G | A | 0.521 | 0.6024 | ++ | 0 | 0.005 | 1 | 0.9428 |
| rs7130076 | 11 | 118566090 | C | T | 0.521 | 0.6024 | ++ | 0 | 0.729 | 1 | 0.3932 |
| rs1110705 | 11 | 119974597 | C | T | 0.521 | 0.6025 | +? | 0 | 0 | 0 | 1 |
| rs10892502 | 11 | 119818802 | C | T | 0.521 | 0.6026 | +? | 0 | 0 | 0 | 1 |
| rs570745 | 11 | 116329090 | A | G | -0.52 | 0.6028 | -? | 0 | 0 | 0 | 1 |
| rs7941900 | 11 | 119979007 | C | T | -0.52 | 0.6028 | -? | 0 | 0 | 0 | 1 |
| rs1787691 | 11 | 116568713 | A | G | 0.52 | 0.6032 | +? | 0 | 0 | 0 | 1 |
| rs11218031 | 11 | 120722457 | A | C | 0.52 | 0.6033 | +? | 0 | 0 | 0 | 1 |
| rs2301574 | 11 | 119063908 | T | C | 0.519 | 0.6034 | ++ | 0 | 0.71 | 1 | 0.3996 |
| rs113566564 | 11 | 118335147 | A | G | -0.519 | 0.6035 | -+ | -191.7 | 1.028 | 1 | 0.3106 |
| rs4938310 | 11 | 116629949 | T | C | -0.519 | 0.6035 | -+ | 4.8 | 3.15 | 1 | 0.07594 |
| rs7946219 | 11 | 116962759 | T | C | -0.519 | 0.6035 | -? | 0 | 0 | 0 | 1 |
| rs7946729 | 11 | 116963180 | T | C | -0.519 | 0.6035 | -? | 0 | 0 | 0 | 1 |
| rs10790211 | 11 | 117701057 | A | G | 0.519 | 0.6036 | +? | 0 | 0 | 0 | 1 |

| | | | | | | | | | | | |
|---|---|---|---|---|---|---|---|---|---|---|---|
| rs10892042 | 11 | 116760545 | A | G | 0.519 | 0.6036 | +? | 0 | 0 | 0 | 1 |
| rs3759000 | 11 | 118135449 | C | T | -0.519 | 0.6036 | -? | 0 | 0 | 0 | 1 |
| rs4938404 | 11 | 117411011 | T | C | -0.519 | 0.6036 | -? | 0 | 0 | 0 | 1 |
| rs35235858 | 11 | 118157250 | T | C | -0.519 | 0.6037 | -+ | -4.7 | 2.866 | 1 | 0.09047 |
| rs11217426 | 11 | 119590878 | T | C | 0.519 | 0.6038 | +? | 0 | 0 | 0 | 1 |
| rs17119829 | 11 | 116453571 | A | G | 0.519 | 0.6039 | ++ | 0 | 0 | 1 | 0.9866 |
| rs7946391 | 11 | 117323952 | C | T | 0.519 | 0.6039 | +? | 0 | 0 | 0 | 1 |
| rs117181393 | 11 | 118017538 | C | T | 0.519 | 0.6041 | +? | 0 | 0 | 0 | 1 |
| rs1939669 | 11 | 120592483 | A | T | 0.519 | 0.6041 | +? | 0 | 0 | 0 | 1 |
| rs79425731 | 11 | 120357341 | A | G | -0.519 | 0.6041 | -? | 0 | 0 | 0 | 1 |
| rs480793 | 11 | 116263533 | A | G | 0.518 | 0.6042 | +? | 0 | 0 | 0 | 1 |
| rs4635116 | 11 | 116417048 | C | A | 0.518 | 0.6043 | ++ | 0 | 0.284 | 1 | 0.5941 |
| rs4252251 | 11 | 117859629 | A | C | 0.518 | 0.6044 | ++ | 0 | 0.076 | 1 | 0.783 |
| rs11607569 | 11 | 118880668 | C | T | -0.518 | 0.6046 | -- | 0 | 0.018 | 1 | 0.8931 |
| rs12281156 | 11 | 117677039 | A | T | 0.518 | 0.6046 | ++ | 0 | 0.208 | 1 | 0.6486 |
| rs3133223 | 11 | 120605232 | A | G | 0.518 | 0.6046 | +? | 0 | 0 | 0 | 1 |
| rs2243935 | 11 | 117824960 | A | G | 0.518 | 0.6047 | +? | 0 | 0 | 0 | 1 |
| rs558247 | 11 | 117767498 | G | A | 0.518 | 0.6047 | +? | 0 | 0 | 0 | 1 |
| rs7129944 | 11 | 120299707 | C | T | 0.518 | 0.6047 | +? | 0 | 0 | 0 | 1 |
| rs11216158 | 11 | 116711350 | A | G | 0.518 | 0.6048 | +? | 0 | 0 | 0 | 1 |
| rs11826999 | 11 | 116553778 | A | G | -0.517 | 0.6049 | -? | 0 | 0 | 0 | 1 |
| rs76747451 | 11 | 116508833 | G | T | -0.517 | 0.6049 | -? | 0 | 0 | 0 | 1 |
| rs7949154 | 11 | 119482832 | C | T | 0.517 | 0.6049 | +? | 0 | 0 | 0 | 1 |
| rs10892101 | 11 | 117308927 | A | C | -0.517 | 0.605 | -? | 0 | 0 | 0 | 1 |
| rs10892114 | 11 | 117324662 | G | C | 0.517 | 0.6051 | +? | 0 | 0 | 0 | 1 |
| rs141023293 | 11 | 116772917 | A | G | -0.517 | 0.6051 | -? | 0 | 0 | 0 | 1 |
| rs2852242 | 11 | 120796595 | T | A | 0.517 | 0.6053 | +? | 0 | 0 | 0 | 1 |
| rs722729 | 11 | 117799804 | T | C | -0.517 | 0.6054 | -+ | 2.4 | 3.074 | 1 | 0.07954 |
| rs73004764 | 11 | 120055326 | G | C | -0.517 | 0.6054 | -? | 0 | 0 | 0 | 1 |
| rs948462 | 11 | 117969163 | C | T | -0.517 | 0.6054 | -- | 0 | 0.03 | 1 | 0.8618 |
| rs10790184 | 11 | 117319260 | A | G | -0.517 | 0.6055 | -? | 0 | 0 | 0 | 1 |
| rs12808181 | 11 | 120094400 | T | C | -0.517 | 0.6055 | -? | 0 | 0 | 0 | 1 |
| rs74660871 | 11 | 117572969 | C | T | 0.517 | 0.6055 | +? | 0 | 0 | 0 | 1 |
| rs12361715 | 11 | 119659301 | T | C | 0.516 | 0.6056 | ++ | 0 | 0 | 1 | 0.9868 |
| rs639978 | 11 | 117953403 | G | A | 0.516 | 0.6058 | +- | 0 | 0.621 | 1 | 0.4307 |
| rs11606979 | 11 | 116440214 | A | G | 0.516 | 0.6059 | ++ | 0 | 0.045 | 1 | 0.8314 |
| rs10160771 | 11 | 119809582 | A | G | -0.516 | 0.606 | -? | 0 | 0 | 0 | 1 |
| rs180357 | 11 | 116599504 | C | T | -0.516 | 0.6061 | -+ | 0 | 0.461 | 1 | 0.4973 |
| rs7941518 | 11 | 116632787 | G | A | -0.515 | 0.6062 | -+ | 4.5 | 3.142 | 1 | 0.07632 |
| rs7942054 | 11 | 116632975 | A | G | -0.515 | 0.6062 | -+ | 4.5 | 3.142 | 1 | 0.07632 |
| rs1939670 | 11 | 120679559 | C | T | 0.515 | 0.6064 | +? | 0 | 0 | 0 | 1 |
| rs2852217 | 11 | 120647226 | C | T | 0.515 | 0.6065 | +? | 0 | 0 | 0 | 1 |
| rs61905145 | 11 | 116718415 | A | G | 0.515 | 0.6065 | +? | 0 | 0 | 0 | 1 |
| rs77297809 | 11 | 116370266 | T | C | -0.515 | 0.6066 | -? | 0 | 0 | 0 | 1 |
| rs4936518 | 11 | 120311198 | T | A | 0.515 | 0.6067 | +? | 0 | 0 | 0 | 1 |
| rs10790428 | 11 | 121140993 | C | T | -0.515 | 0.6068 | -? | 0 | 0 | 0 | 1 |
| rs61179449 | 11 | 118892835 | C | G | -0.515 | 0.6068 | -- | 0 | 0 | 1 | 0.9827 |
| rs4938446 | 11 | 117745049 | A | T | 0.514 | 0.607 | +? | 0 | 0 | 0 | 1 |
| rs79099791 | 11 | 116239244 | C | T | -0.514 | 0.607 | -- | 0 | 0.264 | 1 | 0.6074 |
| rs9666263 | 11 | 119589703 | C | G | 0.514 | 0.607 | +? | 0 | 0 | 0 | 1 |
| rs111527225 | 11 | 118065830 | A | G | 0.514 | 0.6071 | -+ | -110.2 | 1.427 | 1 | 0.2322 |
| rs12792619 | 11 | 116222151 | C | T | 0.514 | 0.6071 | +- | -84.6 | 1.625 | 1 | 0.2024 |
| rs34759019 | 11 | 120479913 | A | C | 0.514 | 0.6071 | +? | 0 | 0 | 0 | 1 |
| rs10790186 | 11 | 117324868 | G | A | 0.514 | 0.6072 | +? | 0 | 0 | 0 | 1 |
| rs61901420 | 11 | 120612976 | G | A | -0.514 | 0.6072 | -? | 0 | 0 | 0 | 1 |
| rs10790388 | 11 | 120494449 | A | C | 0.514 | 0.6074 | +? | 0 | 0 | 0 | 1 |

| | | | | | | | | | | | |
|---|---|---|---|---|---|---|---|---|---|---|---|
| rs11823889 | 11 | 116960403 | A | G | -0.514 | 0.6074 | -? | 0 | 0 | 0 | 1 |
| rs1715441 | 11 | 118072181 | T | C | 0.514 | 0.6074 | +? | 0 | 0 | 0 | 1 |
| rs77657279 | 11 | 120305873 | G | A | -0.513 | 0.6079 | -? | 0 | 0 | 0 | 1 |
| rs12805061 | 11 | 116553025 | G | A | -0.513 | 0.608 | -? | 0 | 0 | 0 | 1 |
| rs12806224 | 11 | 116552815 | C | T | -0.513 | 0.608 | -? | 0 | 0 | 0 | 1 |
| rs3825051 | 11 | 118174629 | G | A | 0.513 | 0.6081 | +? | 0 | 0 | 0 | 1 |
| rs1240783 | 11 | 116572525 | C | T | 0.513 | 0.6082 | +? | 0 | 0 | 0 | 1 |
| rs5012703 | 11 | 119054989 | C | T | -0.513 | 0.6082 | -? | 0 | 0 | 0 | 1 |
| rs10790419 | 11 | 121136455 | T | C | -0.513 | 0.6083 | -? | 0 | 0 | 0 | 1 |
| rs4612814 | 11 | 120594813 | G | A | -0.513 | 0.6083 | +- | 0 | 0.834 | 1 | 0.3612 |
| rs143448490 | 11 | 119923295 | A | G | -0.512 | 0.6084 | -? | 0 | 0 | 0 | 1 |
| rs1942930 | 11 | 117607138 | C | T | 0.512 | 0.6084 | +? | 0 | 0 | 0 | 1 |
| rs7951237 | 11 | 117606937 | A | T | 0.512 | 0.6084 | +? | 0 | 0 | 0 | 1 |
| rs1715455 | 11 | 118124092 | T | C | -0.512 | 0.6086 | -? | 0 | 0 | 0 | 1 |
| rs7120665 | 11 | 118131223 | C | T | -0.512 | 0.6086 | -? | 0 | 0 | 0 | 1 |
| rs11512467 | 11 | 117835174 | T | C | 0.512 | 0.6087 | ++ | 0 | 0.033 | 1 | 0.8558 |
| rs45539536 | 11 | 116617895 | T | C | -0.512 | 0.6087 | -- | 0 | 0.041 | 1 | 0.8396 |
| rs734167 | 11 | 120829359 | A | G | -0.512 | 0.6087 | -+ | 0 | 0.441 | 1 | 0.5065 |
| rs61903661 | 11 | 117113422 | G | A | 0.511 | 0.6091 | +? | 0 | 0 | 0 | 1 |
| rs651013 | 11 | 117947454 | C | T | -0.511 | 0.6091 | -? | 0 | 0 | 0 | 1 |
| rs56083337 | 11 | 118131813 | T | A | 0.511 | 0.6092 | +? | 0 | 0 | 0 | 1 |
| rs11217866 | 11 | 120300804 | A | T | 0.511 | 0.6093 | +? | 0 | 0 | 0 | 1 |
| rs117184822 | 11 | 117989798 | T | C | -0.511 | 0.6093 | -? | 0 | 0 | 0 | 1 |
| rs4938559 | 11 | 118718061 | T | C | 0.511 | 0.6095 | ++ | 0 | 0.174 | 1 | 0.6764 |
| rs4938560 | 11 | 118718062 | G | A | 0.511 | 0.6095 | ++ | 0 | 0.174 | 1 | 0.6764 |
| rs4572168 | 11 | 116555926 | A | G | 0.511 | 0.6096 | +? | 0 | 0 | 0 | 1 |
| rs502607 | 11 | 116474899 | T | C | -0.511 | 0.6097 | -? | 0 | 0 | 0 | 1 |
| rs74984550 | 11 | 119854036 | A | G | -0.511 | 0.6097 | -? | 0 | 0 | 0 | 1 |
| rs493834 | 11 | 118567444 | G | A | 0.51 | 0.6098 | ++ | 0 | 0.757 | 1 | 0.3842 |
| rs77206548 | 11 | 116574463 | A | G | -0.51 | 0.6099 | -- | 0 | 0.002 | 1 | 0.966 |
| rs58955701 | 11 | 118892872 | T | C | -0.51 | 0.6101 | -- | 0 | 0.001 | 1 | 0.9805 |
| rs142744740 | 11 | 116688791 | A | C | 0.51 | 0.6102 | +? | 0 | 0 | 0 | 1 |
| rs75214151 | 11 | 116464362 | A | G | -0.51 | 0.6102 | -? | 0 | 0 | 0 | 1 |
| rs7937589 | 11 | 121133278 | C | G | -0.51 | 0.6102 | -? | 0 | 0 | 0 | 1 |
| rs41440247 | 11 | 117332557 | C | G | 0.51 | 0.6104 | +? | 0 | 0 | 0 | 1 |
| rs4936401 | 11 | 117595260 | C | T | 0.509 | 0.6106 | +? | 0 | 0 | 0 | 1 |
| rs531860 | 11 | 116261676 | A | G | 0.509 | 0.6106 | +? | 0 | 0 | 0 | 1 |
| rs61900894 | 11 | 120398787 | G | C | 0.509 | 0.6107 | +? | 0 | 0 | 0 | 1 |
| rs7479373 | 11 | 118171285 | C | T | -0.509 | 0.611 | -+ | -11.2 | 2.698 | 1 | 0.1005 |
| rs12798115 | 11 | 118156517 | C | G | -0.509 | 0.6111 | -+ | -4.5 | 2.87 | 1 | 0.09025 |
| rs12275106 | 11 | 121170025 | C | A | 0.508 | 0.6112 | +? | 0 | 0 | 0 | 1 |
| rs17125142 | 11 | 121170179 | G | A | 0.508 | 0.6112 | +? | 0 | 0 | 0 | 1 |
| rs7114993 | 11 | 116496310 | T | G | 0.508 | 0.6113 | +? | 0 | 0 | 0 | 1 |
| rs11217558 | 11 | 119776105 | T | G | 0.508 | 0.6114 | +? | 0 | 0 | 0 | 1 |
| rs11828763 | 11 | 116559663 | G | A | 0.508 | 0.6114 | +? | 0 | 0 | 0 | 1 |
| rs12225622 | 11 | 116553924 | T | C | 0.507 | 0.6119 | +? | 0 | 0 | 0 | 1 |
| rs1294767885 | 11 | 116780095 | T | A | 0.507 | 0.6121 | +? | 0 | 0 | 0 | 1 |
| rs1145219 | 11 | 117537799 | T | G | -0.507 | 0.6122 | -? | 0 | 0 | 0 | 1 |
| rs58290590 | 11 | 117308984 | A | G | -0.507 | 0.6122 | -? | 0 | 0 | 0 | 1 |
| rs180323 | 11 | 116626570 | G | A | -0.507 | 0.6123 | -+ | 0 | 0.258 | 1 | 0.6114 |
| rs673292 | 11 | 120019288 | A | G | -0.507 | 0.6124 | -? | 0 | 0 | 0 | 1 |
| rs4245212 | 11 | 120273353 | G | A | 0.507 | 0.6125 | +? | 0 | 0 | 0 | 1 |
| rs10466588 | 11 | 116610249 | G | A | -0.506 | 0.6127 | +- | -46.3 | 2.051 | 1 | 0.1521 |
| rs117788500 | 11 | 116613034 | C | G | -0.506 | 0.6129 | -- | 0 | 0.013 | 1 | 0.9108 |
| rs12280753 | 11 | 116613660 | T | C | -0.506 | 0.6129 | +- | -44.2 | 2.081 | 1 | 0.1492 |
| rs12577638 | 11 | 120755655 | A | G | -0.506 | 0.6131 | -? | 0 | 0 | 0 | 1 |

| | | | | | | | | | | | |
|---|---|---|---|---|---|---|---|---|---|---|---|
| rs79530380 | 11 | 116789188 | A | T | 0.506 | 0.6132 | +? | 0 | 0 | 0 | 1 |
| rs79461323 | 11 | 120065462 | G | A | 0.505 | 0.6133 | +? | 0 | 0 | 0 | 1 |
| rs17518841 | 11 | 116615244 | A | G | -0.505 | 0.6134 | -- | 0 | 0.043 | 1 | 0.8365 |
| rs4938621 | 11 | 118892277 | T | C | -0.505 | 0.6134 | -- | 0 | 0.001 | 1 | 0.9746 |
| rs11216480 | 11 | 117515473 | A | G | -0.505 | 0.6138 | -? | 0 | 0 | 0 | 1 |
| rs61905711 | 11 | 116906431 | T | C | -0.505 | 0.6139 | -? | 0 | 0 | 0 | 1 |
| rs3133846 | 11 | 120597136 | C | T | 0.504 | 0.614 | +? | 0 | 0 | 0 | 1 |
| rs1941406 | 11 | 120141165 | A | G | -0.504 | 0.6141 | -? | 0 | 0 | 0 | 1 |
| rs17120035 | 11 | 116663851 | T | C | -0.504 | 0.6142 | -+ | 0 | 0.296 | 1 | 0.5865 |
| rs3802877 | 11 | 117414948 | T | C | -0.504 | 0.6146 | -? | 0 | 0 | 0 | 1 |
| rs4500497 | 11 | 119492095 | T | C | 0.504 | 0.6146 | +? | 0 | 0 | 0 | 1 |
| rs117447786 | 11 | 119859971 | A | C | -0.503 | 0.6147 | -? | 0 | 0 | 0 | 1 |
| rs75355006 | 11 | 117332512 | C | T | 0.503 | 0.6148 | +? | 0 | 0 | 0 | 1 |
| rs4936553 | 11 | 120688599 | A | G | 0.503 | 0.615 | +? | 0 | 0 | 0 | 1 |
| rs78052530 | 11 | 116229010 | C | A | -0.503 | 0.6151 | -? | 0 | 0 | 0 | 1 |
| rs629470 | 11 | 118320553 | A | G | 0.503 | 0.6152 | ++ | 0 | 0.015 | 1 | 0.9018 |
| rs7103224 | 11 | 116663966 | A | G | -0.502 | 0.6156 | -+ | 0 | 0.3 | 1 | 0.5842 |
| rs11217530 | 11 | 119708875 | A | G | 0.502 | 0.6157 | +? | 0 | 0 | 0 | 1 |
| rs2282549 | 11 | 117539771 | C | A | 0.502 | 0.6158 | +? | 0 | 0 | 0 | 1 |
| rs561394 | 11 | 117719605 | G | A | -0.502 | 0.6158 | -? | 0 | 0 | 0 | 1 |
| rs140078194 | 11 | 116568290 | A | C | -0.501 | 0.616 | -- | 0 | 0.004 | 1 | 0.9468 |
| rs2187008 | 11 | 117954657 | G | C | 0.501 | 0.6163 | ++ | 0 | 0.004 | 1 | 0.9469 |
| rs10790353 | 11 | 119782056 | C | G | -0.501 | 0.6164 | -? | 0 | 0 | 0 | 1 |
| rs11823264 | 11 | 120294004 | A | G | 0.501 | 0.6164 | +? | 0 | 0 | 0 | 1 |
| rs906542 | 11 | 116426886 | G | A | 0.5 | 0.6169 | +? | 0 | 0 | 0 | 1 |
| rs12417661 | 11 | 121086572 | A | G | -0.5 | 0.6173 | -? | 0 | 0 | 0 | 1 |
| rs507133 | 11 | 117202437 | A | G | 0.5 | 0.6174 | +? | 0 | 0 | 0 | 1 |
| rs73018457 | 11 | 117688433 | T | C | -0.5 | 0.6174 | -? | 0 | 0 | 0 | 1 |
| rs12794618 | 11 | 120289699 | C | T | 0.499 | 0.6175 | +? | 0 | 0 | 0 | 1 |
| rs34004290 | 11 | 119684793 | C | G | -0.499 | 0.6176 | -? | 0 | 0 | 0 | 1 |
| rs34022886 | 11 | 118155804 | A | G | -0.499 | 0.6176 | -+ | -2.8 | 2.919 | 1 | 0.08756 |
| rs569 | 11 | 118894287 | G | A | -0.499 | 0.6176 | -- | 0 | 0.005 | 1 | 0.9424 |
| rs678262 | 11 | 118021740 | G | C | -0.499 | 0.6179 | -- | 0 | 0.239 | 1 | 0.6247 |
| rs538847 | 11 | 117335722 | T | C | -0.499 | 0.618 | -? | 0 | 0 | 0 | 1 |
| rs11605749 | 11 | 117784434 | G | C | 0.499 | 0.6181 | +? | 0 | 0 | 0 | 1 |
| rs1787689 | 11 | 116568382 | G | A | 0.498 | 0.6182 | +? | 0 | 0 | 0 | 1 |
| rs12786658 | 11 | 120091272 | A | G | -0.498 | 0.6183 | -? | 0 | 0 | 0 | 1 |
| rs10892670 | 11 | 120963898 | T | C | -0.498 | 0.6184 | -? | 0 | 0 | 0 | 1 |
| rs2115925 | 11 | 121181848 | G | A | -0.498 | 0.6184 | -? | 0 | 0 | 0 | 1 |
| rs149021690 | 11 | 119908975 | C | T | 0.498 | 0.6186 | +? | 0 | 0 | 0 | 1 |
| rs731512 | 11 | 120089886 | A | G | 0.498 | 0.6187 | +? | 0 | 0 | 0 | 1 |
| rs73584981 | 11 | 117658934 | C | T | 0.498 | 0.6187 | ++ | 0 | 0.005 | 1 | 0.946 |
| rs141633273 | 11 | 121181207 | T | C | 0.498 | 0.6188 | +? | 0 | 0 | 0 | 1 |
| rs7129707 | 11 | 120396485 | T | C | 0.498 | 0.6188 | +? | 0 | 0 | 0 | 1 |
| rs11605223 | 11 | 117748844 | G | C | 0.497 | 0.6189 | +? | 0 | 0 | 0 | 1 |
| rs3133845 | 11 | 120596053 | A | G | 0.497 | 0.619 | +? | 0 | 0 | 0 | 1 |
| rs57703856 | 11 | 117696211 | A | G | 0.497 | 0.619 | +? | 0 | 0 | 0 | 1 |
| rs12786505 | 11 | 119607659 | A | G | 0.497 | 0.6191 | +? | 0 | 0 | 0 | 1 |
| rs139131517 | 11 | 121176185 | T | C | 0.497 | 0.6191 | +? | 0 | 0 | 0 | 1 |
| rs623353 | 11 | 119969871 | C | A | 0.497 | 0.6191 | +? | 0 | 0 | 0 | 1 |
| rs77452664 | 11 | 117726559 | T | G | 0.497 | 0.6192 | +? | 0 | 0 | 0 | 1 |
| rs112636863 | 11 | 118333510 | C | T | -0.497 | 0.6193 | -+ | 0 | 0.801 | 1 | 0.3707 |
| rs10892143 | 11 | 117484392 | C | G | 0.497 | 0.6194 | +? | 0 | 0 | 0 | 1 |
| rs3133226 | 11 | 120592036 | C | T | 0.497 | 0.6194 | +? | 0 | 0 | 0 | 1 |
| rs7119090 | 11 | 117747110 | C | G | -0.497 | 0.6195 | -? | 0 | 0 | 0 | 1 |
| rs3758893 | 11 | 120975390 | A | G | -0.496 | 0.6196 | -? | 0 | 0 | 0 | 1 |

| | | | | | | | | | | | |
|---|---|---|---|---|---|---|---|---|---|---|---|
| rs11217052 | 11 | 118717732 | G | A | 0.496 | 0.6198 | ++ | 0 | 0.211 | 1 | 0.6463 |
| rs73003743 | 11 | 120703243 | G | A | 0.496 | 0.6199 | +? | 0 | 0 | 0 | 1 |
| rs9783385 | 11 | 120715674 | G | T | 0.496 | 0.6199 | +? | 0 | 0 | 0 | 1 |
| rs10892007 | 11 | 116527609 | T | C | 0.496 | 0.62 | +? | 0 | 0 | 0 | 1 |
| rs7106652 | 11 | 118554248 | T | C | 0.496 | 0.6201 | -+ | 0 | 0.959 | 1 | 0.3275 |
| rs10892411 | 11 | 119457828 | G | C | 0.496 | 0.6202 | +? | 0 | 0 | 0 | 1 |
| rs11218024 | 11 | 120709387 | T | C | 0.495 | 0.6204 | +? | 0 | 0 | 0 | 1 |
| rs17119701 | 11 | 116331582 | G | A | 0.495 | 0.6205 | +? | 0 | 0 | 0 | 1 |
| rs657898 | 11 | 120017974 | C | T | -0.495 | 0.6205 | -? | 0 | 0 | 0 | 1 |
| rs143811287 | 11 | 116628396 | T | C | 0.495 | 0.6208 | +? | 0 | 0 | 0 | 1 |
| rs78893389 | 11 | 116475411 | T | C | 0.494 | 0.6211 | +? | 0 | 0 | 0 | 1 |
| rs60065241 | 11 | 118883491 | A | C | -0.494 | 0.6213 | -- | 0 | 0.001 | 1 | 0.9818 |
| rs642662 | 11 | 118567020 | G | A | 0.494 | 0.6213 | -+ | 0 | 0.816 | 1 | 0.3664 |
| rs4938279 | 11 | 116419268 | A | G | 0.494 | 0.6216 | ++ | 0 | 0.6 | 1 | 0.4387 |
| rs35400250 | 11 | 116398935 | T | C | 0.493 | 0.6217 | +? | 0 | 0 | 0 | 1 |
| rs509733 | 11 | 119738623 | T | C | -0.493 | 0.6217 | -? | 0 | 0 | 0 | 1 |
| rs10750096 | 11 | 116656788 | C | A | -0.493 | 0.6218 | +- | -47.5 | 2.035 | 1 | 0.1538 |
| rs537975 | 11 | 118362869 | T | C | 0.493 | 0.6218 | ++ | 0 | 0 | 1 | 0.9844 |
| rs477908 | 11 | 119702490 | T | G | 0.493 | 0.6219 | +? | 0 | 0 | 0 | 1 |
| rs644057 | 11 | 120827609 | A | G | 0.493 | 0.6222 | +? | 0 | 0 | 0 | 1 |
| rs575535 | 11 | 117772229 | C | G | -0.492 | 0.6224 | -? | 0 | 0 | 0 | 1 |
| rs2850806 | 11 | 120792285 | T | C | 0.492 | 0.6226 | +? | 0 | 0 | 0 | 1 |
| rs7127188 | 11 | 118893023 | C | A | -0.492 | 0.6226 | -- | 0 | 0.001 | 1 | 0.9724 |
| rs7114844 | 11 | 116496187 | A | G | 0.492 | 0.6227 | +? | 0 | 0 | 0 | 1 |
| rs79751425 | 11 | 121089364 | A | G | -0.492 | 0.6227 | -? | 0 | 0 | 0 | 1 |
| rs996888 | 11 | 119428158 | A | G | 0.492 | 0.623 | +? | 0 | 0 | 0 | 1 |
| rs10892715 | 11 | 121140729 | A | G | -0.491 | 0.6231 | -? | 0 | 0 | 0 | 1 |
| rs11606521 | 11 | 120556817 | G | A | 0.491 | 0.6232 | +? | 0 | 0 | 0 | 1 |
| rs78584663 | 11 | 120556631 | G | A | 0.491 | 0.6232 | +? | 0 | 0 | 0 | 1 |
| rs7484090 | 11 | 119458695 | G | A | 0.491 | 0.6237 | +? | 0 | 0 | 0 | 1 |
| rs1508106 | 11 | 116395210 | A | G | 0.49 | 0.624 | +? | 0 | 0 | 0 | 1 |
| rs551826 | 11 | 117138403 | G | C | 0.49 | 0.624 | +? | 0 | 0 | 0 | 1 |
| rs77630867 | 11 | 119598971 | T | A | -0.49 | 0.624 | -? | 0 | 0 | 0 | 1 |
| rs142273276 | 11 | 117176075 | C | T | -0.49 | 0.6241 | -? | 0 | 0 | 0 | 1 |
| rs2000526 | 11 | 120905771 | C | T | -0.489 | 0.6245 | -? | 0 | 0 | 0 | 1 |
| rs61900474 | 11 | 120021171 | A | T | -0.489 | 0.6245 | -? | 0 | 0 | 0 | 1 |
| rs1715425 | 11 | 118077689 | C | T | 0.489 | 0.6246 | +? | 0 | 0 | 0 | 1 |
| rs598503 | 11 | 116724204 | C | T | -0.489 | 0.6246 | +- | 26.3 | 4.071 | 1 | 0.04362 |
| rs75401133 | 11 | 119824399 | T | C | -0.489 | 0.6246 | -? | 0 | 0 | 0 | 1 |
| rs57053294 | 11 | 120711869 | C | T | 0.489 | 0.6247 | +? | 0 | 0 | 0 | 1 |
| rs12362811 | 11 | 116299000 | A | G | 0.489 | 0.6248 | +? | 0 | 0 | 0 | 1 |
| rs5130 | 11 | 116703146 | C | T | 0.489 | 0.6248 | +? | 0 | 0 | 0 | 1 |
| rs4936592 | 11 | 121053985 | T | C | 0.489 | 0.6249 | +? | 0 | 0 | 0 | 1 |
| rs73005121 | 11 | 121145354 | T | A | 0.489 | 0.6249 | +? | 0 | 0 | 0 | 1 |
| rs180360 | 11 | 116598988 | G | A | -0.488 | 0.6252 | -+ | -93.9 | 1.547 | 1 | 0.2136 |
| rs12361192 | 11 | 119603642 | C | T | 0.488 | 0.6253 | +? | 0 | 0 | 0 | 1 |
| rs597583 | 11 | 117421799 | G | C | -0.488 | 0.6254 | -? | 0 | 0 | 0 | 1 |
| rs10790424 | 11 | 121137520 | T | C | -0.488 | 0.6256 | -? | 0 | 0 | 0 | 1 |
| rs17120347 | 11 | 116996539 | A | G | -0.488 | 0.6256 | -? | 0 | 0 | 0 | 1 |
| rs12788624 | 11 | 118022107 | T | C | 0.488 | 0.6258 | +? | 0 | 0 | 0 | 1 |
| rs3809059 | 11 | 120206163 | A | T | 0.487 | 0.626 | +? | 0 | 0 | 0 | 1 |
| rs4938405 | 11 | 117411254 | C | G | -0.487 | 0.626 | -? | 0 | 0 | 0 | 1 |
| rs588930 | 11 | 117581689 | C | T | 0.487 | 0.626 | +? | 0 | 0 | 0 | 1 |
| rs12272993 | 11 | 116214112 | T | G | -0.487 | 0.6263 | -? | 0 | 0 | 0 | 1 |
| rs6589766 | 11 | 119483293 | A | G | 0.487 | 0.6263 | +? | 0 | 0 | 0 | 1 |
| rs7119061 | 11 | 121126530 | G | A | -0.487 | 0.6264 | -? | 0 | 0 | 0 | 1 |

| | | | | | | | | | | | |
|---|---|---|---|---|---|---|---|---|---|---|---|
| rs73020466 | 11 | 117770990 | G | A | 0.487 | 0.6266 | +? | 0 | 0 | 0 | 1 |
| rs997956 | 11 | 117770775 | T | C | 0.487 | 0.6266 | +? | 0 | 0 | 0 | 1 |
| rs112580285 | 11 | 121049649 | G | T | 0.486 | 0.6267 | +? | 0 | 0 | 0 | 1 |
| rs4225 | 11 | 116703671 | G | T | -0.486 | 0.6267 | -? | 0 | 0 | 0 | 1 |
| rs601052 | 11 | 119850450 | C | T | -0.486 | 0.6269 | -? | 0 | 0 | 0 | 1 |
| rs12721030 | 11 | 116705278 | T | C | -0.486 | 0.627 | -? | 0 | 0 | 0 | 1 |
| rs77983701 | 11 | 116507056 | C | T | 0.486 | 0.627 | +? | 0 | 0 | 0 | 1 |
| rs610664 | 11 | 118556691 | G | A | 0.486 | 0.6272 | -+ | 0 | 0.912 | 1 | 0.3396 |
| rs11218191 | 11 | 121054619 | T | C | -0.485 | 0.6274 | -? | 0 | 0 | 0 | 1 |
| rs533646 | 11 | 118566746 | G | C | 0.485 | 0.6275 | -+ | 0 | 0.764 | 1 | 0.382 |
| rs1784106 | 11 | 116567449 | A | C | 0.485 | 0.6276 | +? | 0 | 0 | 0 | 1 |
| rs7107833 | 11 | 116396188 | C | T | -0.485 | 0.6276 | -? | 0 | 0 | 0 | 1 |
| rs11602869 | 11 | 118554448 | A | G | 0.485 | 0.6277 | -+ | 0 | 0.923 | 1 | 0.3368 |
| rs145141827 | 11 | 119919351 | T | G | -0.485 | 0.6277 | -? | 0 | 0 | 0 | 1 |
| rs1287392 | 11 | 120074732 | T | C | -0.485 | 0.6278 | -? | 0 | 0 | 0 | 1 |
| rs10892575 | 11 | 120291137 | G | C | 0.485 | 0.628 | +? | 0 | 0 | 0 | 1 |
| rs11217177 | 11 | 119065102 | C | T | 0.484 | 0.6281 | ++ | 0 | 0.597 | 1 | 0.4399 |
| rs2077334 | 11 | 116506410 | A | C | 0.484 | 0.6281 | +? | 0 | 0 | 0 | 1 |
| rs4331144 | 11 | 118554100 | G | A | 0.484 | 0.6281 | -+ | 0 | 0.971 | 1 | 0.3244 |
| rs180363 | 11 | 116597889 | C | T | -0.484 | 0.6283 | -+ | 0 | 0.106 | 1 | 0.745 |
| rs2848728 | 11 | 120021674 | T | C | -0.484 | 0.6284 | -? | 0 | 0 | 0 | 1 |
| rs4252305 | 11 | 117862979 | A | G | 0.484 | 0.6284 | +? | 0 | 0 | 0 | 1 |
| rs552035 | 11 | 116265522 | C | T | 0.484 | 0.6284 | +? | 0 | 0 | 0 | 1 |
| rs73009944 | 11 | 117863183 | A | G | 0.484 | 0.6284 | +? | 0 | 0 | 0 | 1 |
| rs73009948 | 11 | 117863307 | T | G | 0.484 | 0.6284 | +? | 0 | 0 | 0 | 1 |
| rs79879928 | 11 | 117252135 | A | G | 0.484 | 0.6284 | +? | 0 | 0 | 0 | 1 |
| rs10892109 | 11 | 117321874 | G | C | -0.484 | 0.6285 | -? | 0 | 0 | 0 | 1 |
| rs1350280779 | 11 | 119825798 | A | C | 0.484 | 0.6286 | +? | 0 | 0 | 0 | 1 |
| rs12574374 | 11 | 120119395 | A | G | -0.484 | 0.6287 | -? | 0 | 0 | 0 | 1 |
| rs11216032 | 11 | 116466894 | A | G | -0.483 | 0.6288 | -? | 0 | 0 | 0 | 1 |
| rs13377487 | 11 | 118187395 | A | T | 0.483 | 0.6288 | +- | 31.4 | 4.373 | 1 | 0.03651 |
| rs75318540 | 11 | 116683862 | T | C | 0.483 | 0.6289 | +? | 0 | 0 | 0 | 1 |
| rs1270305911 | 11 | 116915229 | T | C | -0.483 | 0.6293 | +- | 51.4 | 6.171 | 1 | 0.01299 |
| rs145339787 | 11 | 116921143 | C | T | -0.483 | 0.6293 | +- | 51.4 | 6.171 | 1 | 0.01299 |
| rs717052 | 11 | 120180479 | A | G | -0.483 | 0.6293 | -? | 0 | 0 | 0 | 1 |
| rs73594187 | 11 | 116915881 | C | T | -0.483 | 0.6293 | +- | 51.4 | 6.171 | 1 | 0.01299 |
| rs505064 | 11 | 116319221 | C | T | -0.483 | 0.6294 | -? | 0 | 0 | 0 | 1 |
| rs9332782 | 11 | 118347020 | C | T | 0.482 | 0.6295 | ++ | 0 | 0.121 | 1 | 0.7275 |
| rs78823230 | 11 | 117904328 | T | G | 0.482 | 0.6296 | ++ | 0 | 0.322 | 1 | 0.5707 |
| rs76836072 | 11 | 116653762 | T | C | -0.482 | 0.6297 | -- | 0 | 0.108 | 1 | 0.7429 |
| rs7940234 | 11 | 120531547 | C | G | -0.482 | 0.6299 | -? | 0 | 0 | 0 | 1 |
| rs6589845 | 11 | 120689035 | T | A | 0.482 | 0.6301 | +? | 0 | 0 | 0 | 1 |
| rs143660533 | 11 | 118892759 | C | A | -0.481 | 0.6304 | -- | 0 | 0 | 1 | 0.9879 |
| rs2852243 | 11 | 120796418 | C | T | 0.481 | 0.6305 | +? | 0 | 0 | 0 | 1 |
| rs11600894 | 11 | 120478676 | T | C | 0.481 | 0.6306 | +? | 0 | 0 | 0 | 1 |
| rs76820520 | 11 | 120524498 | A | G | -0.481 | 0.6306 | -? | 0 | 0 | 0 | 1 |
| rs77132149 | 11 | 120522714 | A | G | -0.481 | 0.6306 | -? | 0 | 0 | 0 | 1 |
| rs1268832 | 11 | 116575229 | T | G | 0.481 | 0.6308 | +? | 0 | 0 | 0 | 1 |
| rs73020481 | 11 | 117772955 | C | T | 0.481 | 0.6308 | +? | 0 | 0 | 0 | 1 |
| rs1793176 | 11 | 118080771 | T | A | 0.48 | 0.631 | +? | 0 | 0 | 0 | 1 |
| rs117289748 | 11 | 118892757 | G | C | -0.48 | 0.6311 | -- | 0 | 0 | 1 | 0.9893 |
| rs28989476 | 11 | 117177372 | C | T | -0.48 | 0.6311 | -? | 0 | 0 | 0 | 1 |
| rs10750138 | 11 | 119055075 | G | A | -0.48 | 0.6312 | -? | 0 | 0 | 0 | 1 |
| rs11607675 | 11 | 117333778 | A | G | 0.48 | 0.6314 | +? | 0 | 0 | 0 | 1 |
| rs2715890 | 11 | 120110219 | C | G | -0.48 | 0.6314 | -? | 0 | 0 | 0 | 1 |
| rs638903 | 11 | 120983264 | A | C | -0.48 | 0.6315 | -? | 0 | 0 | 0 | 1 |

| | | | | | | | | | | | |
|---|---|---|---|---|---|---|---|---|---|---|---|
| rs59462353 | 11 | 117690620 | C | T | 0.479 | 0.6316 | +? | 0 | 0 | 0 | 1 |
| rs113028342 | 11 | 118333537 | G | A | -0.479 | 0.6318 | -+ | 0 | 0.982 | 1 | 0.3217 |
| rs7111020 | 11 | 120494400 | T | C | 0.479 | 0.6318 | +? | 0 | 0 | 0 | 1 |
| rs111901186 | 11 | 118328052 | G | A | -0.479 | 0.632 | -+ | 0 | 0.783 | 1 | 0.3762 |
| rs138907015 | 11 | 119599641 | T | C | -0.478 | 0.6323 | -? | 0 | 0 | 0 | 1 |
| rs10790162 | 11 | 116639104 | A | G | -0.478 | 0.6324 | +- | -126.8 | 1.322 | 1 | 0.2501 |
| rs1263058 | 11 | 116575555 | A | G | 0.478 | 0.6325 | +? | 0 | 0 | 0 | 1 |
| rs73014299 | 11 | 118189965 | T | C | 0.478 | 0.6325 | +? | 0 | 0 | 0 | 1 |
| rs111982661 | 11 | 121055215 | T | C | 0.478 | 0.6328 | +? | 0 | 0 | 0 | 1 |
| rs11217330 | 11 | 119453980 | C | A | -0.478 | 0.6328 | -? | 0 | 0 | 0 | 1 |
| rs11216035 | 11 | 116470227 | C | T | -0.478 | 0.6329 | -? | 0 | 0 | 0 | 1 |
| rs57192657 | 11 | 116437360 | C | A | 0.477 | 0.633 | ++ | 0 | 0.056 | 1 | 0.8123 |
| rs57502313 | 11 | 116437347 | C | T | 0.477 | 0.633 | ++ | 0 | 0.056 | 1 | 0.8123 |
| rs58567678 | 11 | 116437333 | A | G | 0.477 | 0.633 | ++ | 0 | 0.056 | 1 | 0.8123 |
| rs526602 | 11 | 117070691 | A | G | 0.477 | 0.6331 | +? | 0 | 0 | 0 | 1 |
| rs77571570 | 11 | 116438634 | G | C | 0.477 | 0.6331 | ++ | 0 | 0.055 | 1 | 0.8142 |
| rs11825206 | 11 | 119570128 | C | T | -0.477 | 0.6333 | -? | 0 | 0 | 0 | 1 |
| rs35430963 | 11 | 119749018 | T | A | 0.477 | 0.6334 | +? | 0 | 0 | 0 | 1 |
| rs10790188 | 11 | 117325456 | A | G | -0.476 | 0.634 | -? | 0 | 0 | 0 | 1 |
| rs508994 | 11 | 116280252 | A | C | 0.476 | 0.634 | +? | 0 | 0 | 0 | 1 |
| rs594081 | 11 | 119969402 | A | C | 0.476 | 0.634 | +? | 0 | 0 | 0 | 1 |
| rs7110261 | 11 | 117810129 | A | G | 0.476 | 0.6343 | -+ | -24.1 | 2.416 | 1 | 0.1201 |
| rs602168 | 11 | 119883753 | T | C | 0.476 | 0.6344 | +? | 0 | 0 | 0 | 1 |
| rs7103239 | 11 | 117325898 | A | G | -0.476 | 0.6344 | -? | 0 | 0 | 0 | 1 |
| rs115696360 | 11 | 116304790 | C | G | 0.475 | 0.6345 | +? | 0 | 0 | 0 | 1 |
| rs1941631 | 11 | 117966597 | C | A | -0.475 | 0.6345 | -- | 0 | 0.014 | 1 | 0.9065 |
| rs2512151 | 11 | 117885694 | G | A | 0.475 | 0.6346 | +- | 0 | 0.487 | 1 | 0.4854 |
| rs3016356 | 11 | 116477772 | C | T | 0.474 | 0.6352 | +? | 0 | 0 | 0 | 1 |
| rs73010409 | 11 | 117110417 | C | T | 0.474 | 0.6352 | +? | 0 | 0 | 0 | 1 |
| rs79487385 | 11 | 116380781 | G | A | -0.474 | 0.6352 | -? | 0 | 0 | 0 | 1 |
| rs117207582 | 11 | 120200671 | T | C | 0.474 | 0.6354 | +? | 0 | 0 | 0 | 1 |
| rs73019251 | 11 | 116253169 | T | C | -0.474 | 0.6355 | -? | 0 | 0 | 0 | 1 |
| rs12286715 | 11 | 119806447 | T | C | -0.474 | 0.6356 | -? | 0 | 0 | 0 | 1 |
| rs145968026 | 11 | 118421841 | G | C | -0.474 | 0.6356 | -- | 0 | 0.017 | 1 | 0.8949 |
| rs12295249 | 11 | 119806145 | A | G | -0.474 | 0.6357 | -? | 0 | 0 | 0 | 1 |
| rs10892117 | 11 | 117325478 | C | T | -0.474 | 0.6358 | -? | 0 | 0 | 0 | 1 |
| rs1939672 | 11 | 120675122 | T | C | 0.474 | 0.6358 | +? | 0 | 0 | 0 | 1 |
| rs514143 | 11 | 118558449 | A | G | 0.474 | 0.6359 | -+ | 0 | 0.788 | 1 | 0.3747 |
| rs7927742 | 11 | 117314044 | C | T | 0.473 | 0.636 | +? | 0 | 0 | 0 | 1 |
| rs1793173 | 11 | 118074117 | A | G | 0.473 | 0.6362 | +? | 0 | 0 | 0 | 1 |
| rs1793174 | 11 | 118074337 | A | G | 0.473 | 0.6362 | +? | 0 | 0 | 0 | 1 |
| rs4938425 | 11 | 117523275 | A | G | 0.473 | 0.6362 | +? | 0 | 0 | 0 | 1 |
| rs471776 | 11 | 120050959 | A | G | 0.473 | 0.6365 | +? | 0 | 0 | 0 | 1 |
| rs546441 | 11 | 120051363 | C | A | 0.473 | 0.6365 | +? | 0 | 0 | 0 | 1 |
| rs551843 | 11 | 120052037 | A | G | 0.473 | 0.6365 | +? | 0 | 0 | 0 | 1 |
| rs659375 | 11 | 119012846 | A | T | -0.472 | 0.6369 | -? | 0 | 0 | 0 | 1 |
| rs6589565 | 11 | 116640237 | A | G | -0.472 | 0.6371 | +- | -125.4 | 1.331 | 1 | 0.2486 |
| rs2156122 | 11 | 116496623 | A | C | 0.472 | 0.6372 | +? | 0 | 0 | 0 | 1 |
| rs11218025 | 11 | 120710628 | C | T | 0.471 | 0.6373 | +? | 0 | 0 | 0 | 1 |
| rs672176 | 11 | 120829249 | A | G | 0.471 | 0.6373 | +? | 0 | 0 | 0 | 1 |
| rs7121857 | 11 | 120556389 | G | T | -0.471 | 0.6374 | -? | 0 | 0 | 0 | 1 |
| rs7940963 | 11 | 120348444 | A | G | 0.471 | 0.6375 | +? | 0 | 0 | 0 | 1 |
| rs867897 | 11 | 116388264 | G | C | 0.471 | 0.6375 | +? | 0 | 0 | 0 | 1 |
| rs571751 | 11 | 118556797 | C | A | 0.471 | 0.6376 | -+ | 0 | 0.864 | 1 | 0.3527 |
| rs621843 | 11 | 119883723 | A | G | 0.471 | 0.6376 | +? | 0 | 0 | 0 | 1 |
| rs668361 | 11 | 119883781 | A | G | 0.471 | 0.6376 | +? | 0 | 0 | 0 | 1 |

| | | | | | | | | | | | |
|---|---|---|---|---|---|---|---|---|---|---|---|
| rs948034 | 11 | 120677398 | C | T | 0.471 | 0.6378 | +? | 0 | 0 | 0 | 1 |
| rs17492380 | 11 | 116438851 | A | G | 0.47 | 0.6381 | ++ | 0 | 0.057 | 1 | 0.8111 |
| rs20494 | 11 | 119957453 | A | G | 0.47 | 0.6383 | +? | 0 | 0 | 0 | 1 |
| rs4938757 | 11 | 119878632 | T | C | -0.47 | 0.6383 | -? | 0 | 0 | 0 | 1 |
| rs78020509 | 11 | 118384811 | G | A | -0.47 | 0.6384 | -+ | -125.9 | 1.328 | 1 | 0.2491 |
| rs140479252 | 11 | 117111178 | C | T | 0.47 | 0.6385 | +? | 0 | 0 | 0 | 1 |
| rs111218525 | 11 | 116566538 | T | C | -0.47 | 0.6386 | -? | 0 | 0 | 0 | 1 |
| rs509641 | 11 | 118558952 | C | T | 0.47 | 0.6387 | -+ | 0 | 0.792 | 1 | 0.3735 |
| rs563549 | 11 | 118558953 | A | G | 0.47 | 0.6387 | -+ | 0 | 0.792 | 1 | 0.3735 |
| rs77229823 | 11 | 116380855 | G | A | -0.469 | 0.6389 | -? | 0 | 0 | 0 | 1 |
| rs10892161 | 11 | 117555644 | A | G | 0.469 | 0.639 | +? | 0 | 0 | 0 | 1 |
| rs7101554 | 11 | 118078761 | C | T | 0.469 | 0.639 | +? | 0 | 0 | 0 | 1 |
| rs581122 | 11 | 116838615 | T | C | -0.469 | 0.6391 | +- | 51.7 | 6.21 | 1 | 0.0127 |
| rs613377 | 11 | 118556084 | T | C | 0.469 | 0.6392 | -+ | 0 | 0.809 | 1 | 0.3684 |
| rs12223620 | 11 | 120709874 | G | T | 0.469 | 0.6393 | +? | 0 | 0 | 0 | 1 |
| rs11218022 | 11 | 120709008 | A | G | 0.468 | 0.6396 | +? | 0 | 0 | 0 | 1 |
| rs1475614837 | 11 | 116910320 | T | G | -0.468 | 0.6397 | +- | 51.7 | 6.212 | 1 | 0.01269 |
| rs73594175 | 11 | 116909994 | G | A | -0.468 | 0.6397 | +- | 51.7 | 6.212 | 1 | 0.01269 |
| rs73594181 | 11 | 116910427 | A | T | -0.468 | 0.6397 | +- | 51.7 | 6.212 | 1 | 0.01269 |
| rs79340485 | 11 | 116910454 | T | C | -0.468 | 0.6397 | +- | 51.7 | 6.212 | 1 | 0.01269 |
| rs563838 | 11 | 117201997 | T | C | 0.468 | 0.6398 | +? | 0 | 0 | 0 | 1 |
| rs73008690 | 11 | 119750813 | A | G | -0.468 | 0.6398 | -? | 0 | 0 | 0 | 1 |
| rs12366082 | 11 | 117380086 | G | C | 0.468 | 0.64 | +? | 0 | 0 | 0 | 1 |
| rs74743296 | 11 | 120750636 | C | A | -0.468 | 0.6401 | -? | 0 | 0 | 0 | 1 |
| rs717051 | 11 | 120180657 | T | C | -0.467 | 0.6403 | -? | 0 | 0 | 0 | 1 |
| rs4938465 | 11 | 117844785 | T | A | 0.467 | 0.6405 | ++ | 0 | 0.016 | 1 | 0.9006 |
| rs10466533 | 11 | 116612128 | G | A | -0.467 | 0.6407 | +- | -72.9 | 1.736 | 1 | 0.1877 |
| rs4938365 | 11 | 117112138 | C | A | 0.467 | 0.6407 | +? | 0 | 0 | 0 | 1 |
| rs11607656 | 11 | 117333657 | A | G | 0.467 | 0.6408 | +? | 0 | 0 | 0 | 1 |
| rs10892103 | 11 | 117309283 | A | G | -0.466 | 0.6409 | -? | 0 | 0 | 0 | 1 |
| rs4579962 | 11 | 117692334 | A | G | 0.466 | 0.6409 | +? | 0 | 0 | 0 | 1 |
| rs60524902 | 11 | 116380536 | G | T | -0.466 | 0.6409 | -? | 0 | 0 | 0 | 1 |
| rs117164546 | 11 | 119297248 | A | G | 0.466 | 0.6411 | +? | 0 | 0 | 0 | 1 |
| rs7120445 | 11 | 118879285 | C | A | -0.466 | 0.6411 | -- | 0 | 0 | 1 | 0.986 |
| rs11605069 | 11 | 118554436 | G | T | 0.466 | 0.6412 | -+ | 0 | 0.86 | 1 | 0.3537 |
| rs3133852 | 11 | 120602425 | G | T | -0.466 | 0.6412 | -? | 0 | 0 | 0 | 1 |
| rs34228131 | 11 | 119647700 | T | C | -0.466 | 0.6412 | -? | 0 | 0 | 0 | 1 |
| rs142774084 | 11 | 119652997 | A | G | 0.466 | 0.6414 | +? | 0 | 0 | 0 | 1 |
| rs7932832 | 11 | 119573524 | G | A | -0.466 | 0.6414 | -? | 0 | 0 | 0 | 1 |
| rs1320649 | 11 | 120677301 | T | C | 0.465 | 0.6418 | +? | 0 | 0 | 0 | 1 |
| rs17245340 | 11 | 119822015 | C | T | 0.465 | 0.642 | +? | 0 | 0 | 0 | 1 |
| rs2160669 | 11 | 116647607 | C | T | -0.465 | 0.6422 | +- | -75.5 | 1.71 | 1 | 0.191 |
| rs116915994 | 11 | 121142335 | T | C | 0.464 | 0.6423 | +? | 0 | 0 | 0 | 1 |
| rs11825284 | 11 | 118880012 | A | G | -0.464 | 0.6423 | -- | 0 | 0 | 1 | 0.9853 |
| rs73559119 | 11 | 118879824 | C | T | -0.464 | 0.6423 | -- | 0 | 0 | 1 | 0.9853 |
| rs56303695 | 11 | 119823216 | A | G | 0.464 | 0.6424 | +? | 0 | 0 | 0 | 1 |
| rs625513 | 11 | 118555719 | C | A | 0.464 | 0.6424 | -+ | 0 | 0.777 | 1 | 0.378 |
| rs609341 | 11 | 117341808 | T | G | -0.464 | 0.6427 | -? | 0 | 0 | 0 | 1 |
| rs12418028 | 11 | 117790674 | A | G | -0.463 | 0.6431 | -? | 0 | 0 | 0 | 1 |
| rs496288 | 11 | 116266614 | C | A | 0.463 | 0.6431 | +? | 0 | 0 | 0 | 1 |
| rs149239940 | 11 | 120826569 | A | G | -0.463 | 0.6434 | -? | 0 | 0 | 0 | 1 |
| rs117361624 | 11 | 118568427 | T | C | 0.463 | 0.6435 | ++ | 0 | 0.363 | 1 | 0.5469 |
| rs73585234 | 11 | 117618530 | T | C | 0.463 | 0.6435 | +? | 0 | 0 | 0 | 1 |
| rs525485 | 11 | 118555754 | A | G | 0.463 | 0.6436 | -+ | 0 | 0.772 | 1 | 0.3796 |
| rs558781 | 11 | 118564397 | T | G | 0.463 | 0.6436 | -+ | 0 | 0.684 | 1 | 0.4083 |
| rs56363339 | 11 | 119683659 | G | A | -0.462 | 0.6439 | -? | 0 | 0 | 0 | 1 |

| | | | | | | | | | | | |
|---|---|---|---|---|---|---|---|---|---|---|---|
| rs60657131 | 11 | 117679078 | C | T | 0.462 | 0.6439 | +? | 0 | 0 | 0 | 1 |
| rs145305155 | 11 | 117330326 | G | A | -0.462 | 0.644 | -? | 0 | 0 | 0 | 1 |
| rs638301 | 11 | 118559003 | G | A | 0.462 | 0.644 | -+ | 0 | 0.8 | 1 | 0.3712 |
| rs117547981 | 11 | 117104900 | A | G | 0.462 | 0.6441 | +? | 0 | 0 | 0 | 1 |
| rs510408 | 11 | 118558901 | C | G | 0.462 | 0.6444 | -+ | 0 | 0.8 | 1 | 0.371 |
| rs561845 | 11 | 118558827 | G | A | 0.462 | 0.6444 | -+ | 0 | 0.8 | 1 | 0.371 |
| rs638805 | 11 | 118558889 | T | G | 0.462 | 0.6444 | -+ | 0 | 0.8 | 1 | 0.371 |
| rs75960142 | 11 | 121169052 | C | T | 0.461 | 0.6445 | +? | 0 | 0 | 0 | 1 |
| rs10892568 | 11 | 120252828 | G | A | 0.461 | 0.6447 | +? | 0 | 0 | 0 | 1 |
| rs117050823 | 11 | 116782202 | G | A | 0.461 | 0.6448 | +? | 0 | 0 | 0 | 1 |
| rs626517 | 11 | 120990070 | C | G | -0.461 | 0.6448 | -? | 0 | 0 | 0 | 1 |
| rs657405 | 11 | 120989780 | T | C | -0.461 | 0.6448 | -? | 0 | 0 | 0 | 1 |
| rs201365537 | 11 | 117138456 | A | G | -0.461 | 0.6449 | -? | 0 | 0 | 0 | 1 |
| rs11216083 | 11 | 116537634 | T | C | 0.461 | 0.6451 | +? | 0 | 0 | 0 | 1 |
| rs502601 | 11 | 118565679 | A | G | 0.461 | 0.6451 | -+ | 0 | 0.789 | 1 | 0.3743 |
| rs1145183 | 11 | 116502286 | A | G | -0.46 | 0.6455 | -? | 0 | 0 | 0 | 1 |
| rs558593 | 11 | 118562520 | T | C | 0.46 | 0.6455 | -+ | 0 | 0.752 | 1 | 0.3858 |
| rs4936409 | 11 | 117694392 | G | A | -0.46 | 0.6456 | -? | 0 | 0 | 0 | 1 |
| rs4938387 | 11 | 117334519 | A | G | 0.46 | 0.6456 | +? | 0 | 0 | 0 | 1 |
| rs6589769 | 11 | 119574060 | G | C | -0.46 | 0.6456 | -? | 0 | 0 | 0 | 1 |
| rs7103799 | 11 | 117811563 | T | C | 0.46 | 0.6457 | -+ | -32.6 | 2.263 | 1 | 0.1325 |
| rs7122174 | 11 | 117842533 | A | T | 0.46 | 0.6457 | ++ | 0 | 0.004 | 1 | 0.9483 |
| rs6589774 | 11 | 119646732 | A | G | -0.46 | 0.6458 | -? | 0 | 0 | 0 | 1 |
| rs61900865 | 11 | 120392401 | T | C | 0.459 | 0.646 | +? | 0 | 0 | 0 | 1 |
| rs117268885 | 11 | 117786899 | T | C | -0.459 | 0.6461 | -? | 0 | 0 | 0 | 1 |
| rs7942396 | 11 | 121179549 | T | C | 0.459 | 0.6463 | +? | 0 | 0 | 0 | 1 |
| rs77654673 | 11 | 119364445 | C | G | 0.459 | 0.6465 | +? | 0 | 0 | 0 | 1 |
| rs558582 | 11 | 117328639 | T | C | -0.458 | 0.6467 | -? | 0 | 0 | 0 | 1 |
| rs56338557 | 11 | 116409130 | C | T | 0.458 | 0.6468 | +? | 0 | 0 | 0 | 1 |
| rs1110703 | 11 | 119974755 | G | A | 0.458 | 0.6469 | +? | 0 | 0 | 0 | 1 |
| rs11216486 | 11 | 117524000 | G | T | -0.458 | 0.6469 | -? | 0 | 0 | 0 | 1 |
| rs558907 | 11 | 118558455 | G | A | 0.458 | 0.647 | -+ | 0 | 0.804 | 1 | 0.3699 |
| rs693306 | 11 | 118558333 | G | A | 0.458 | 0.647 | -+ | 0 | 0.804 | 1 | 0.3699 |
| rs75672383 | 11 | 120790997 | T | C | -0.458 | 0.6472 | -? | 0 | 0 | 0 | 1 |
| rs59293325 | 11 | 116327447 | A | G | -0.457 | 0.6476 | -? | 0 | 0 | 0 | 1 |
| rs4936433 | 11 | 118552420 | G | A | 0.457 | 0.6477 | -+ | 0 | 0.831 | 1 | 0.362 |
| rs519848 | 11 | 118552715 | A | G | 0.457 | 0.6477 | -+ | 0 | 0.831 | 1 | 0.362 |
| rs549370 | 11 | 118552842 | A | C | 0.457 | 0.6477 | -+ | 0 | 0.831 | 1 | 0.362 |
| rs592280 | 11 | 118553218 | C | T | 0.457 | 0.6477 | -+ | 0 | 0.831 | 1 | 0.362 |
| rs654792 | 11 | 118552996 | T | G | 0.457 | 0.6477 | -+ | 0 | 0.831 | 1 | 0.362 |
| rs12291974 | 11 | 117661447 | T | C | 0.456 | 0.6481 | ++ | 0 | 0 | 1 | 0.9842 |
| rs12577361 | 11 | 117337289 | A | C | -0.456 | 0.6481 | -? | 0 | 0 | 0 | 1 |
| rs180352 | 11 | 116606630 | A | G | -0.456 | 0.6481 | -+ | -2.3 | 2.931 | 1 | 0.08688 |
| rs28989500 | 11 | 117165397 | T | C | -0.456 | 0.6481 | -? | 0 | 0 | 0 | 1 |
| rs59138930 | 11 | 118173783 | T | C | 0.456 | 0.6481 | +? | 0 | 0 | 0 | 1 |
| rs2845702 | 11 | 120113661 | G | T | -0.456 | 0.6482 | -? | 0 | 0 | 0 | 1 |
| rs7933848 | 11 | 119561362 | T | C | 0.456 | 0.6483 | +? | 0 | 0 | 0 | 1 |
| rs11218195 | 11 | 121062860 | C | A | -0.456 | 0.6485 | -? | 0 | 0 | 0 | 1 |
| rs45555231 | 11 | 116665120 | C | G | 0.456 | 0.6485 | +? | 0 | 0 | 0 | 1 |
| rs693037 | 11 | 118553326 | T | C | 0.456 | 0.6485 | -+ | 0 | 0.815 | 1 | 0.3666 |
| rs9332819 | 11 | 118365263 | T | C | -0.456 | 0.6487 | -+ | 0 | 0.892 | 1 | 0.345 |
| rs12786590 | 11 | 121161826 | G | A | -0.455 | 0.6488 | -? | 0 | 0 | 0 | 1 |
| rs2075290 | 11 | 116653296 | C | T | -0.455 | 0.6488 | +- | -75.6 | 1.709 | 1 | 0.1911 |
| rs10790233 | 11 | 117976568 | A | G | -0.455 | 0.6489 | -- | 0 | 0 | 1 | 0.9868 |
| rs1715429 | 11 | 118083664 | A | G | 0.455 | 0.6489 | +? | 0 | 0 | 0 | 1 |
| rs75128335 | 11 | 118072915 | A | G | 0.455 | 0.6492 | -+ | -143.4 | 1.232 | 1 | 0.267 |

| | | | | | | | | | | | |
|---|---|---|---|---|---|---|---|---|---|---|---|
| rs117391362 | 11 | 120785542 | A | G | 0.455 | 0.6493 | +? | 0 | 0 | 0 | 1 |
| rs118046730 | 11 | 118711376 | C | T | 0.455 | 0.6494 | -+ | -3.8 | 2.891 | 1 | 0.08906 |
| rs574217358 | 11 | 118559709 | A | G | 0.455 | 0.6494 | -+ | 0 | 0.808 | 1 | 0.3686 |
| rs753309 | 11 | 120138885 | A | G | -0.455 | 0.6494 | -? | 0 | 0 | 0 | 1 |
| rs7926959 | 11 | 118559866 | C | T | 0.455 | 0.6494 | -+ | 0 | 0.808 | 1 | 0.3686 |
| rs7926970 | 11 | 118559881 | G | T | 0.455 | 0.6494 | -+ | 0 | 0.808 | 1 | 0.3686 |
| rs1076485 | 11 | 116772441 | T | C | 0.454 | 0.6495 | +? | 0 | 0 | 0 | 1 |
| rs10790432 | 11 | 121160519 | C | T | -0.454 | 0.6496 | -? | 0 | 0 | 0 | 1 |
| rs486462 | 11 | 118559178 | C | T | 0.454 | 0.6496 | -+ | 0 | 0.808 | 1 | 0.3688 |
| rs533144 | 11 | 118559247 | A | C | 0.454 | 0.6496 | -+ | 0 | 0.808 | 1 | 0.3688 |
| rs625918 | 11 | 118559542 | T | C | 0.454 | 0.6496 | -+ | 0 | 0.808 | 1 | 0.3688 |
| rs7115581 | 11 | 121161041 | A | G | -0.454 | 0.6496 | -? | 0 | 0 | 0 | 1 |
| rs7115602 | 11 | 121161106 | A | G | -0.454 | 0.6496 | -? | 0 | 0 | 0 | 1 |
| rs518028 | 11 | 118564982 | C | T | 0.454 | 0.6497 | -+ | 0 | 0.692 | 1 | 0.4056 |
| rs56227539 | 11 | 116964469 | C | T | -0.454 | 0.6497 | -? | 0 | 0 | 0 | 1 |
| rs57950207 | 11 | 116620670 | C | T | -0.454 | 0.6497 | -- | 0 | 0.055 | 1 | 0.8138 |
| rs77112220 | 11 | 116626226 | T | C | -0.454 | 0.6497 | -- | 0 | 0.055 | 1 | 0.8138 |
| rs117792004 | 11 | 120355388 | A | T | -0.454 | 0.6498 | -? | 0 | 0 | 0 | 1 |
| rs2071702 | 11 | 118373861 | T | C | -0.454 | 0.6498 | -+ | -136.4 | 1.269 | 1 | 0.2599 |
| rs487962 | 11 | 116245350 | A | G | -0.454 | 0.6498 | -? | 0 | 0 | 0 | 1 |
| rs9332781 | 11 | 118346599 | G | A | -0.454 | 0.6499 | -+ | 0 | 0.943 | 1 | 0.3316 |
| rs1786185 | 11 | 117922143 | G | A | 0.454 | 0.65 | +? | 0 | 0 | 0 | 1 |
| rs508579 | 11 | 118559084 | C | G | 0.454 | 0.65 | -+ | 0 | 0.808 | 1 | 0.3687 |
| rs518368 | 11 | 117275561 | C | G | -0.454 | 0.65 | -? | 0 | 0 | 0 | 1 |
| rs555356 | 11 | 118558078 | T | G | 0.454 | 0.6501 | -+ | 0 | 0.808 | 1 | 0.3686 |
| rs2513091 | 11 | 116845538 | G | A | -0.453 | 0.6502 | +- | 52 | 6.254 | 1 | 0.01239 |
| rs2921655 | 11 | 116836548 | C | G | -0.453 | 0.6502 | +- | 52 | 6.254 | 1 | 0.01239 |
| rs493101 | 11 | 116833669 | A | G | -0.453 | 0.6502 | +- | 52 | 6.254 | 1 | 0.01239 |
| rs541496 | 11 | 116835362 | T | C | -0.453 | 0.6502 | +- | 52 | 6.254 | 1 | 0.01239 |
| rs622302 | 11 | 116831071 | G | A | -0.453 | 0.6502 | +- | 52 | 6.254 | 1 | 0.01239 |
| rs667119 | 11 | 116835883 | T | A | -0.453 | 0.6502 | +- | 52 | 6.254 | 1 | 0.01239 |
| rs117117915 | 11 | 117113511 | T | C | 0.453 | 0.6505 | +? | 0 | 0 | 0 | 1 |
| rs538661 | 11 | 120309851 | T | C | 0.453 | 0.6506 | +? | 0 | 0 | 0 | 1 |
| rs11216382 | 11 | 117303858 | C | T | 0.453 | 0.6508 | +? | 0 | 0 | 0 | 1 |
| rs35066934 | 11 | 117652665 | T | C | -0.453 | 0.6508 | -+ | 0 | 0.153 | 1 | 0.6954 |
| rs61903739 | 11 | 117220324 | T | C | -0.453 | 0.6508 | -? | 0 | 0 | 0 | 1 |
| rs11215970 | 11 | 116379681 | C | T | -0.453 | 0.6509 | -? | 0 | 0 | 0 | 1 |
| rs7116481 | 11 | 120238413 | T | C | 0.453 | 0.6509 | +? | 0 | 0 | 0 | 1 |
| rs67859744 | 11 | 118200657 | A | T | 0.452 | 0.651 | +? | 0 | 0 | 0 | 1 |
| rs4082031 | 11 | 116227283 | A | T | -0.452 | 0.6513 | -? | 0 | 0 | 0 | 1 |
| rs7952521 | 11 | 116385354 | A | G | 0.452 | 0.6515 | +? | 0 | 0 | 0 | 1 |
| rs12576486 | 11 | 118235490 | C | T | -0.451 | 0.6517 | -+ | 0 | 0.299 | 1 | 0.5844 |
| rs12798453 | 11 | 118561410 | G | T | 0.451 | 0.6518 | -+ | 0 | 0.695 | 1 | 0.4046 |
| rs3132786 | 11 | 118365956 | C | T | 0.451 | 0.6518 | ++ | 0 | 0 | 1 | 0.9882 |
| rs7945395 | 11 | 119569737 | G | A | -0.451 | 0.6518 | -? | 0 | 0 | 0 | 1 |
| rs12796373 | 11 | 118554939 | A | G | 0.451 | 0.6519 | -+ | 0 | 0.813 | 1 | 0.3674 |
| rs12420972 | 11 | 117337012 | A | G | 0.451 | 0.6521 | +? | 0 | 0 | 0 | 1 |
| rs590580 | 11 | 118066104 | A | G | 0.451 | 0.6521 | +? | 0 | 0 | 0 | 1 |
| rs7131534 | 11 | 118886656 | A | G | -0.451 | 0.6521 | -- | 0 | 0 | 1 | 0.9981 |
| rs2368427 | 11 | 119453770 | C | T | -0.451 | 0.6522 | -? | 0 | 0 | 0 | 1 |
| rs10750182 | 11 | 121153455 | A | T | -0.451 | 0.6523 | -? | 0 | 0 | 0 | 1 |
| rs10892044 | 11 | 116766899 | C | T | 0.45 | 0.6524 | +? | 0 | 0 | 0 | 1 |
| rs11216178 | 11 | 116766390 | T | G | 0.45 | 0.6524 | +? | 0 | 0 | 0 | 1 |
| rs2163281 | 11 | 121158682 | A | G | -0.45 | 0.6524 | -? | 0 | 0 | 0 | 1 |
| rs516153 | 11 | 118564762 | A | C | 0.45 | 0.6524 | -+ | 0 | 0.695 | 1 | 0.4044 |
| rs7128047 | 11 | 121156532 | A | G | -0.45 | 0.6524 | -? | 0 | 0 | 0 | 1 |

| | | | | | | | | | | | |
|---|---|---|---|---|---|---|---|---|---|---|---|
| rs7128342 | 11 | 121156809 | A | T | -0.45 | 0.6524 | -? | 0 | 0 | 0 | 1 |
| rs74505693 | 11 | 117668721 | A | G | 0.45 | 0.6524 | ++ | 0 | 0.005 | 1 | 0.9436 |
| rs3782040 | 11 | 118176097 | G | A | 0.45 | 0.6526 | +? | 0 | 0 | 0 | 1 |
| rs519982 | 11 | 118565231 | T | C | 0.45 | 0.6527 | -+ | 0 | 0.696 | 1 | 0.4042 |
| rs117170971 | 11 | 116605265 | A | G | -0.449 | 0.6531 | -- | 0 | 0.012 | 1 | 0.9136 |
| rs73577982 | 11 | 117337708 | A | G | 0.449 | 0.6531 | +? | 0 | 0 | 0 | 1 |
| rs10437673 | 11 | 116328809 | A | G | -0.449 | 0.6532 | -? | 0 | 0 | 0 | 1 |
| rs10790379 | 11 | 120221652 | G | A | 0.449 | 0.6532 | +? | 0 | 0 | 0 | 1 |
| rs11215953 | 11 | 116347728 | G | A | -0.449 | 0.6532 | -? | 0 | 0 | 0 | 1 |
| rs656445 | 11 | 118012164 | A | G | 0.449 | 0.6533 | ++ | 0 | 0 | 1 | 0.9834 |
| rs6589872 | 11 | 121158100 | C | T | -0.449 | 0.6533 | -? | 0 | 0 | 0 | 1 |
| rs12277299 | 11 | 119845755 | G | C | -0.449 | 0.6536 | -? | 0 | 0 | 0 | 1 |
| rs12294978 | 11 | 119805884 | A | C | -0.449 | 0.6536 | -? | 0 | 0 | 0 | 1 |
| rs11216162 | 11 | 116728277 | A | G | -0.449 | 0.6537 | -? | 0 | 0 | 0 | 1 |
| rs12417440 | 11 | 117241429 | T | C | -0.448 | 0.6538 | -? | 0 | 0 | 0 | 1 |
| rs569770 | 11 | 118584498 | T | C | 0.448 | 0.6538 | ++ | 0 | 0.46 | 1 | 0.4975 |
| rs6589770 | 11 | 119614499 | G | C | 0.448 | 0.6538 | +? | 0 | 0 | 0 | 1 |
| rs73006720 | 11 | 119609314 | C | A | 0.448 | 0.6538 | +? | 0 | 0 | 0 | 1 |
| rs34033074 | 11 | 119648831 | A | G | -0.448 | 0.654 | -? | 0 | 0 | 0 | 1 |
| rs35904674 | 11 | 116666099 | T | C | -0.448 | 0.654 | -+ | 0 | 0.205 | 1 | 0.6506 |
| rs4388901 | 11 | 119648152 | C | T | -0.448 | 0.654 | -? | 0 | 0 | 0 | 1 |
| rs1349688 | 11 | 119978608 | C | G | 0.448 | 0.6542 | +? | 0 | 0 | 0 | 1 |
| rs4938529 | 11 | 118560569 | T | C | 0.448 | 0.6542 | -+ | 0 | 0.698 | 1 | 0.4036 |
| rs79532870 | 11 | 120636299 | A | G | -0.448 | 0.6542 | -? | 0 | 0 | 0 | 1 |
| rs11216751 | 11 | 117973058 | G | T | 0.448 | 0.6545 | +? | 0 | 0 | 0 | 1 |
| rs6589567 | 11 | 116670676 | A | C | -0.447 | 0.6547 | +- | -21.3 | 2.474 | 1 | 0.1158 |
| rs471227 | 11 | 118564601 | C | T | 0.447 | 0.6548 | -+ | 0 | 0.699 | 1 | 0.4033 |
| rs508640 | 11 | 118564413 | G | A | 0.447 | 0.6548 | -+ | 0 | 0.699 | 1 | 0.4033 |
| rs582630 | 11 | 118562799 | A | C | 0.447 | 0.6548 | -+ | 0 | 0.699 | 1 | 0.4033 |
| rs585039 | 11 | 118563381 | A | C | 0.447 | 0.6548 | -+ | 0 | 0.699 | 1 | 0.4033 |
| rs12278532 | 11 | 120481773 | T | C | 0.447 | 0.6549 | +? | 0 | 0 | 0 | 1 |
| rs73022489 | 11 | 117795838 | A | G | 0.447 | 0.6549 | +- | 0 | 0.961 | 1 | 0.327 |
| rs4938708 | 11 | 119560847 | T | C | 0.447 | 0.6552 | +? | 0 | 0 | 0 | 1 |
| rs4938709 | 11 | 119561042 | A | T | 0.447 | 0.6552 | +? | 0 | 0 | 0 | 1 |
| rs10790401 | 11 | 120557543 | G | A | -0.446 | 0.6556 | -? | 0 | 0 | 0 | 1 |
| rs11217769 | 11 | 120089688 | G | A | 0.446 | 0.6556 | +? | 0 | 0 | 0 | 1 |
| rs12799314 | 11 | 118551597 | A | G | 0.446 | 0.6556 | -+ | 0 | 0.861 | 1 | 0.3535 |
| rs17440396 | 11 | 116621770 | A | G | -0.446 | 0.6556 | -- | 0 | 0.058 | 1 | 0.8102 |
| rs642497 | 11 | 118550784 | A | C | 0.446 | 0.6556 | -+ | 0 | 0.861 | 1 | 0.3535 |
| rs642530 | 11 | 118550756 | C | T | 0.446 | 0.6556 | -+ | 0 | 0.861 | 1 | 0.3535 |
| rs673770 | 11 | 118550601 | G | T | 0.446 | 0.6556 | -+ | 0 | 0.861 | 1 | 0.3535 |
| rs17124945 | 11 | 121068047 | T | C | -0.446 | 0.6559 | -? | 0 | 0 | 0 | 1 |
| rs76693355 | 11 | 120292477 | C | T | -0.446 | 0.6559 | -? | 0 | 0 | 0 | 1 |
| rs10790230 | 11 | 117954384 | A | C | 0.445 | 0.656 | ++ | 0 | 0.081 | 1 | 0.7764 |
| rs1893517 | 11 | 117522237 | A | G | 0.445 | 0.6561 | +? | 0 | 0 | 0 | 1 |
| rs6589864 | 11 | 121109489 | G | A | -0.445 | 0.6561 | -? | 0 | 0 | 0 | 1 |
| rs4614 | 11 | 118952371 | G | A | -0.445 | 0.6562 | -? | 0 | 0 | 0 | 1 |
| rs667982 | 11 | 118562767 | C | T | 0.445 | 0.6564 | -+ | 0 | 0.748 | 1 | 0.3872 |
| rs79255840 | 11 | 119933897 | T | A | 0.445 | 0.6565 | +? | 0 | 0 | 0 | 1 |
| rs7106203 | 11 | 121158551 | A | T | -0.445 | 0.6566 | -? | 0 | 0 | 0 | 1 |
| rs4938524 | 11 | 118552074 | T | G | 0.444 | 0.657 | -+ | 0 | 0.775 | 1 | 0.3788 |
| rs61903660 | 11 | 117113335 | G | A | 0.444 | 0.657 | +? | 0 | 0 | 0 | 1 |
| rs12803325 | 11 | 119643807 | T | A | -0.444 | 0.6571 | -? | 0 | 0 | 0 | 1 |
| rs4420283 | 11 | 119644249 | C | T | -0.444 | 0.6571 | -? | 0 | 0 | 0 | 1 |
| rs4938383 | 11 | 117332917 | C | G | 0.444 | 0.6571 | +? | 0 | 0 | 0 | 1 |
| rs7104562 | 11 | 119577324 | G | A | 0.444 | 0.6571 | +? | 0 | 0 | 0 | 1 |

| | | | | | | | | | | | |
|---|---|---|---|---|---|---|---|---|---|---|---|
| rs17123767 | 11 | 120146127 | A | C | -0.444 | 0.6574 | -? | 0 | 0 | 0 | 1 |
| rs557331 | 11 | 116734688 | A | G | -0.444 | 0.6574 | +- | 33.9 | 4.537 | 1 | 0.03317 |
| rs582071 | 11 | 120017875 | A | G | -0.444 | 0.6574 | -? | 0 | 0 | 0 | 1 |
| rs76598600 | 11 | 119738420 | T | C | 0.444 | 0.6574 | +? | 0 | 0 | 0 | 1 |
| rs10892384 | 11 | 119406091 | C | T | -0.443 | 0.6575 | -? | 0 | 0 | 0 | 1 |
| rs45539534 | 11 | 119236192 | T | C | 0.443 | 0.6576 | +? | 0 | 0 | 0 | 1 |
| rs11820814 | 11 | 116765700 | C | T | 0.443 | 0.6577 | +? | 0 | 0 | 0 | 1 |
| rs625790 | 11 | 119358835 | C | T | 0.443 | 0.6578 | +? | 0 | 0 | 0 | 1 |
| rs4938807 | 11 | 120331701 | A | C | 0.443 | 0.658 | +? | 0 | 0 | 0 | 1 |
| rs59097294 | 11 | 116964437 | C | T | -0.443 | 0.658 | -? | 0 | 0 | 0 | 1 |
| rs10466595 | 11 | 119989107 | G | C | -0.443 | 0.6581 | -? | 0 | 0 | 0 | 1 |
| rs41460849 | 11 | 120104859 | A | G | -0.443 | 0.6581 | -? | 0 | 0 | 0 | 1 |
| rs73578234 | 11 | 120146707 | T | C | -0.442 | 0.6582 | -? | 0 | 0 | 0 | 1 |
| rs12364375 | 11 | 117317941 | T | A | 0.442 | 0.6585 | +? | 0 | 0 | 0 | 1 |
| rs76377040 | 11 | 117666548 | G | C | 0.442 | 0.6586 | ++ | 0 | 0.009 | 1 | 0.9226 |
| rs7111290 | 11 | 119574140 | G | A | -0.442 | 0.6587 | -? | 0 | 0 | 0 | 1 |
| rs10790327 | 11 | 119500417 | T | C | -0.442 | 0.6588 | -? | 0 | 0 | 0 | 1 |
| rs73011187 | 11 | 116469461 | T | C | -0.441 | 0.6589 | -? | 0 | 0 | 0 | 1 |
| rs73014332 | 11 | 117201404 | G | A | 0.441 | 0.6589 | +? | 0 | 0 | 0 | 1 |
| rs45584835 | 11 | 118004863 | G | A | 0.441 | 0.659 | ++ | 0 | 0.006 | 1 | 0.9386 |
| rs11216475 | 11 | 117500025 | T | C | -0.441 | 0.6592 | -? | 0 | 0 | 0 | 1 |
| rs3902806 | 11 | 119572051 | G | T | -0.441 | 0.6592 | -? | 0 | 0 | 0 | 1 |
| rs10750183 | 11 | 121154721 | A | G | -0.441 | 0.6593 | -? | 0 | 0 | 0 | 1 |
| rs10892612 | 11 | 120553182 | A | G | -0.441 | 0.6593 | -? | 0 | 0 | 0 | 1 |
| rs1583124 | 11 | 121154941 | G | A | -0.441 | 0.6593 | -? | 0 | 0 | 0 | 1 |
| rs2513092 | 11 | 116844279 | C | T | -0.441 | 0.6593 | +- | 52.3 | 6.29 | 1 | 0.01214 |
| rs6589870 | 11 | 121153921 | G | A | -0.441 | 0.6593 | -? | 0 | 0 | 0 | 1 |
| rs61906076 | 11 | 116521505 | G | C | 0.441 | 0.6595 | +? | 0 | 0 | 0 | 1 |
| rs582800 | 11 | 120861967 | C | T | -0.44 | 0.6596 | -+ | 0 | 0.389 | 1 | 0.533 |
| rs486476 | 11 | 116312926 | G | C | -0.44 | 0.6597 | -? | 0 | 0 | 0 | 1 |
| rs117889729 | 11 | 120979833 | G | C | 0.44 | 0.6598 | +? | 0 | 0 | 0 | 1 |
| rs12222405 | 11 | 120090344 | T | C | 0.44 | 0.6601 | +? | 0 | 0 | 0 | 1 |
| rs12223954 | 11 | 120090552 | C | T | 0.44 | 0.6601 | +? | 0 | 0 | 0 | 1 |
| rs11217415 | 11 | 119574609 | C | T | -0.44 | 0.6602 | -? | 0 | 0 | 0 | 1 |
| rs12796257 | 11 | 119830778 | C | T | -0.439 | 0.6603 | -? | 0 | 0 | 0 | 1 |
| rs11216498 | 11 | 117552437 | G | A | 0.439 | 0.6605 | +? | 0 | 0 | 0 | 1 |
| rs1261706 | 11 | 116819991 | A | T | -0.439 | 0.6605 | +- | 52.3 | 6.295 | 1 | 0.01211 |
| rs3133352 | 11 | 116818187 | A | G | -0.439 | 0.6605 | +- | 52.3 | 6.295 | 1 | 0.01211 |
| rs587703 | 11 | 116827565 | G | A | -0.439 | 0.6605 | +- | 52.3 | 6.295 | 1 | 0.01211 |
| rs56129109 | 11 | 120594279 | C | A | -0.439 | 0.6607 | +- | 0 | 0.912 | 1 | 0.3395 |
| rs7119027 | 11 | 120142525 | G | A | -0.439 | 0.6608 | -? | 0 | 0 | 0 | 1 |
| rs682109 | 11 | 116673149 | T | C | -0.439 | 0.6609 | -+ | 8.2 | 3.267 | 1 | 0.07069 |
| rs2444240 | 11 | 120040442 | G | T | -0.439 | 0.661 | -? | 0 | 0 | 0 | 1 |
| rs10466532 | 11 | 116407823 | C | T | 0.438 | 0.6611 | +? | 0 | 0 | 0 | 1 |
| rs7103982 | 11 | 120397775 | T | C | 0.438 | 0.6611 | +? | 0 | 0 | 0 | 1 |
| rs72999491 | 11 | 118539151 | A | G | -0.438 | 0.6611 | -? | 0 | 0 | 0 | 1 |
| rs1783839 | 11 | 117905405 | C | T | -0.438 | 0.6613 | +- | 0 | 0.626 | 1 | 0.4286 |
| rs67435655 | 11 | 118196399 | C | T | 0.438 | 0.6613 | +? | 0 | 0 | 0 | 1 |
| rs10790397 | 11 | 120551569 | A | C | -0.438 | 0.6614 | -? | 0 | 0 | 0 | 1 |
| rs11217267 | 11 | 119263703 | C | T | -0.438 | 0.6614 | -? | 0 | 0 | 0 | 1 |
| rs4938268 | 11 | 116272339 | G | A | 0.438 | 0.6614 | +? | 0 | 0 | 0 | 1 |
| rs544201 | 11 | 119702015 | A | T | 0.438 | 0.6614 | +? | 0 | 0 | 0 | 1 |
| rs11217863 | 11 | 120293138 | A | G | -0.438 | 0.6617 | -? | 0 | 0 | 0 | 1 |
| rs117498846 | 11 | 117255837 | A | G | 0.437 | 0.6621 | +? | 0 | 0 | 0 | 1 |
| rs45558732 | 11 | 118402962 | T | G | 0.437 | 0.6621 | ++ | 0 | 0.033 | 1 | 0.8568 |
| rs12277700 | 11 | 119236692 | A | C | 0.437 | 0.6622 | +? | 0 | 0 | 0 | 1 |

| | | | | | | | | | | | |
|---|---|---|---|---|---|---|---|---|---|---|---|
| rs640411 | 11 | 116725303 | C | G | -0.437 | 0.6623 | +- | 43.7 | 5.333 | 1 | 0.02092 |
| rs11216591 | 11 | 117732007 | T | A | -0.437 | 0.6624 | -? | 0 | 0 | 0 | 1 |
| rs2851718 | 11 | 119995447 | C | T | 0.436 | 0.6625 | +? | 0 | 0 | 0 | 1 |
| rs2852229 | 11 | 120604533 | A | G | 0.436 | 0.6629 | +? | 0 | 0 | 0 | 1 |
| rs11217551 | 11 | 119749812 | T | G | 0.436 | 0.6631 | +? | 0 | 0 | 0 | 1 |
| rs10790335 | 11 | 119572905 | C | T | -0.435 | 0.6633 | -? | 0 | 0 | 0 | 1 |
| rs509566 | 11 | 120801460 | T | A | 0.435 | 0.6633 | +? | 0 | 0 | 0 | 1 |
| rs45460396 | 11 | 118005622 | G | T | 0.435 | 0.6635 | ++ | 0 | 0.005 | 1 | 0.9441 |
| rs11217414 | 11 | 119574558 | C | T | -0.435 | 0.6636 | -? | 0 | 0 | 0 | 1 |
| rs692750 | 11 | 118538308 | T | G | 0.435 | 0.6636 | -+ | 0 | 0.694 | 1 | 0.4049 |
| rs112900272 | 11 | 118326445 | T | C | -0.435 | 0.6638 | -+ | 0 | 0.932 | 1 | 0.3343 |
| rs78865628 | 11 | 119835382 | C | T | -0.435 | 0.6638 | -? | 0 | 0 | 0 | 1 |
| rs10892500 | 11 | 119816075 | T | C | -0.435 | 0.6639 | -? | 0 | 0 | 0 | 1 |
| rs35585361 | 11 | 117337965 | A | G | 0.434 | 0.664 | +? | 0 | 0 | 0 | 1 |
| rs6589566 | 11 | 116652423 | G | A | -0.434 | 0.6642 | +- | -117.2 | 1.381 | 1 | 0.2399 |
| rs7483863 | 11 | 116652491 | A | G | -0.434 | 0.6642 | +- | -117.2 | 1.381 | 1 | 0.2399 |
| rs12294669 | 11 | 118041972 | G | A | 0.434 | 0.6643 | +- | 0 | 0.288 | 1 | 0.5914 |
| rs12575015 | 11 | 118197645 | T | A | 0.434 | 0.6644 | +? | 0 | 0 | 0 | 1 |
| rs66962317 | 11 | 118197486 | C | T | 0.434 | 0.6644 | +? | 0 | 0 | 0 | 1 |
| rs10892588 | 11 | 120488347 | A | G | 0.434 | 0.6645 | +? | 0 | 0 | 0 | 1 |
| rs58136657 | 11 | 119610490 | T | C | 0.434 | 0.6646 | +? | 0 | 0 | 0 | 1 |
| rs12418040 | 11 | 120126782 | A | G | -0.433 | 0.6647 | -? | 0 | 0 | 0 | 1 |
| rs2155824 | 11 | 119166280 | A | T | 0.433 | 0.6647 | ++ | 0 | 0.385 | 1 | 0.535 |
| rs10892453 | 11 | 119636367 | T | C | -0.433 | 0.6648 | -? | 0 | 0 | 0 | 1 |
| rs543480 | 11 | 117771069 | T | C | -0.433 | 0.6649 | -? | 0 | 0 | 0 | 1 |
| rs1939673 | 11 | 120618079 | C | T | 0.433 | 0.6653 | +? | 0 | 0 | 0 | 1 |
| rs10160554 | 11 | 117646824 | G | A | 0.432 | 0.6654 | +? | 0 | 0 | 0 | 1 |
| rs4938741 | 11 | 119827163 | C | T | 0.432 | 0.6657 | +? | 0 | 0 | 0 | 1 |
| rs61902683 | 11 | 120555070 | G | T | 0.432 | 0.6658 | +? | 0 | 0 | 0 | 1 |
| rs885429 | 11 | 119572571 | T | G | -0.432 | 0.6658 | -? | 0 | 0 | 0 | 1 |
| rs2846122 | 11 | 120605140 | A | G | 0.432 | 0.6659 | +? | 0 | 0 | 0 | 1 |
| rs3863313 | 11 | 117312602 | G | T | -0.432 | 0.6659 | -? | 0 | 0 | 0 | 1 |
| rs17453205 | 11 | 117801635 | G | A | 0.432 | 0.666 | +? | 0 | 0 | 0 | 1 |
| rs11218175 | 11 | 121041146 | G | T | -0.432 | 0.6661 | -? | 0 | 0 | 0 | 1 |
| rs12577602 | 11 | 120755218 | T | G | -0.431 | 0.6662 | -? | 0 | 0 | 0 | 1 |
| rs686977 | 11 | 117544475 | C | T | -0.431 | 0.6662 | -? | 0 | 0 | 0 | 1 |
| rs11217875 | 11 | 120325559 | A | C | 0.431 | 0.6663 | +? | 0 | 0 | 0 | 1 |
| rs4482033 | 11 | 120713654 | G | A | 0.431 | 0.6663 | +? | 0 | 0 | 0 | 1 |
| rs7121233 | 11 | 120717153 | C | T | 0.431 | 0.6665 | +? | 0 | 0 | 0 | 1 |
| rs7952068 | 11 | 120693133 | G | C | 0.431 | 0.6665 | +? | 0 | 0 | 0 | 1 |
| rs4457755 | 11 | 121150625 | G | A | 0.431 | 0.6666 | +? | 0 | 0 | 0 | 1 |
| rs2249466 | 11 | 119181170 | T | C | -0.431 | 0.6668 | -? | 0 | 0 | 0 | 1 |
| rs11216755 | 11 | 117974937 | C | T | -0.43 | 0.6669 | -+ | 0 | 0.636 | 1 | 0.425 |
| rs582340 | 11 | 119956964 | A | G | 0.43 | 0.667 | +? | 0 | 0 | 0 | 1 |
| rs647437 | 11 | 119879217 | C | T | 0.43 | 0.667 | +? | 0 | 0 | 0 | 1 |
| rs74597362 | 11 | 121189853 | A | G | -0.43 | 0.667 | -? | 0 | 0 | 0 | 1 |
| rs11217529 | 11 | 119701941 | A | G | -0.43 | 0.6674 | -? | 0 | 0 | 0 | 1 |
| rs141734119 | 11 | 120872163 | G | A | 0.43 | 0.6675 | +? | 0 | 0 | 0 | 1 |
| rs76368626 | 11 | 119465670 | C | T | 0.43 | 0.6675 | +? | 0 | 0 | 0 | 1 |
| rs111405302 | 11 | 118385691 | C | T | -0.43 | 0.6676 | -+ | -135.2 | 1.275 | 1 | 0.2588 |
| rs111748140 | 11 | 119872269 | T | C | 0.429 | 0.6676 | +? | 0 | 0 | 0 | 1 |
| rs10892187 | 11 | 117732806 | A | G | -0.429 | 0.6682 | -? | 0 | 0 | 0 | 1 |
| rs11600071 | 11 | 119593452 | G | A | 0.428 | 0.6684 | +? | 0 | 0 | 0 | 1 |
| rs45531834 | 11 | 118096459 | T | C | 0.428 | 0.6685 | +? | 0 | 0 | 0 | 1 |
| rs523715 | 11 | 118574188 | A | G | 0.428 | 0.6685 | +? | 0 | 0 | 0 | 1 |
| rs562089 | 11 | 117710118 | C | A | -0.428 | 0.6685 | -? | 0 | 0 | 0 | 1 |

| | | | | | | | | | | | |
|---|---|---|---|---|---|---|---|---|---|---|---|
| rs687664 | 11 | 118574185 | A | C | 0.428 | 0.6685 | +? | 0 | 0 | 0 | 1 |
| rs868344 | 11 | 118005119 | C | T | 0.428 | 0.6685 | ++ | 0 | 0.005 | 1 | 0.9445 |
| rs10892201 | 11 | 117848292 | T | C | -0.428 | 0.6686 | -- | 0 | 0.468 | 1 | 0.4938 |
| rs113488707 | 11 | 118045876 | T | C | -0.428 | 0.6686 | -? | 0 | 0 | 0 | 1 |
| rs12278678 | 11 | 120481866 | A | G | 0.428 | 0.6686 | +? | 0 | 0 | 0 | 1 |
| rs79315770 | 11 | 116510542 | A | T | -0.428 | 0.6686 | -? | 0 | 0 | 0 | 1 |
| rs112473235 | 11 | 120599666 | T | C | -0.428 | 0.6687 | -? | 0 | 0 | 0 | 1 |
| rs2276045 | 11 | 118365168 | A | G | -0.428 | 0.6688 | -+ | -169 | 1.115 | 1 | 0.2909 |
| rs7101835 | 11 | 117737359 | G | A | 0.428 | 0.6688 | +? | 0 | 0 | 0 | 1 |
| rs9332809 | 11 | 118361472 | A | G | -0.428 | 0.6688 | -+ | -169 | 1.115 | 1 | 0.2909 |
| rs478371 | 11 | 118565336 | A | G | 0.428 | 0.6689 | -+ | 0 | 0.717 | 1 | 0.397 |
| rs12420386 | 11 | 116527281 | A | C | 0.427 | 0.6691 | +? | 0 | 0 | 0 | 1 |
| rs66553012 | 11 | 120067624 | G | A | 0.427 | 0.6691 | +? | 0 | 0 | 0 | 1 |
| rs4502019 | 11 | 116502834 | G | C | 0.427 | 0.6692 | +? | 0 | 0 | 0 | 1 |
| rs147710439 | 11 | 119481969 | A | G | -0.427 | 0.6693 | -? | 0 | 0 | 0 | 1 |
| rs77159892 | 11 | 116436761 | A | C | 0.427 | 0.6693 | ++ | 0 | 0.07 | 1 | 0.792 |
| rs546890 | 11 | 118557198 | C | T | 0.427 | 0.6694 | -+ | 0 | 0.727 | 1 | 0.3939 |
| rs117639653 | 11 | 117113478 | C | G | -0.427 | 0.6695 | -? | 0 | 0 | 0 | 1 |
| rs3020100 | 11 | 120666931 | G | A | 0.427 | 0.6696 | +? | 0 | 0 | 0 | 1 |
| rs113712143 | 11 | 116817785 | A | G | -0.427 | 0.6697 | +- | 52.6 | 6.331 | 1 | 0.01186 |
| rs2441218 | 11 | 116854580 | G | T | -0.427 | 0.6697 | +- | 52.6 | 6.331 | 1 | 0.01186 |
| rs4935754 | 11 | 120708690 | G | A | 0.427 | 0.6697 | +? | 0 | 0 | 0 | 1 |
| rs537431 | 11 | 116856179 | C | T | -0.427 | 0.6697 | +- | 52.6 | 6.331 | 1 | 0.01186 |
| rs548215 | 11 | 120019335 | A | G | 0.427 | 0.6697 | +? | 0 | 0 | 0 | 1 |
| rs560897 | 11 | 116862087 | T | C | -0.427 | 0.6697 | +- | 52.6 | 6.331 | 1 | 0.01186 |
| rs569568 | 11 | 116857400 | C | T | -0.427 | 0.6697 | +- | 52.6 | 6.331 | 1 | 0.01186 |
| rs607756 | 11 | 116862718 | C | T | -0.427 | 0.6697 | +- | 52.6 | 6.331 | 1 | 0.01186 |
| rs623423 | 11 | 116861519 | C | G | -0.427 | 0.6697 | +- | 52.6 | 6.331 | 1 | 0.01186 |
| rs638392 | 11 | 116860456 | A | T | -0.427 | 0.6697 | +- | 52.6 | 6.331 | 1 | 0.01186 |
| rs649084 | 11 | 116815769 | A | G | -0.427 | 0.6697 | +- | 52.6 | 6.331 | 1 | 0.01186 |
| rs73022490 | 11 | 117795859 | A | G | 0.427 | 0.6697 | +- | 0 | 0.938 | 1 | 0.3328 |
| rs6589563 | 11 | 116590787 | A | G | -0.426 | 0.6698 | +- | -92.1 | 1.561 | 1 | 0.2115 |
| rs619473 | 11 | 119342115 | C | A | 0.426 | 0.67 | -+ | -62.9 | 1.842 | 1 | 0.1747 |
| rs17491207 | 11 | 116400897 | A | G | -0.426 | 0.6702 | -? | 0 | 0 | 0 | 1 |
| rs73555296 | 11 | 118445559 | T | G | -0.426 | 0.6702 | -- | 0 | 0 | 1 | 0.9994 |
| rs495033 | 11 | 116352622 | C | T | 0.426 | 0.6703 | +? | 0 | 0 | 0 | 1 |
| rs11828220 | 11 | 118579395 | A | T | 0.426 | 0.6704 | ++ | 0 | 0.133 | 1 | 0.7156 |
| rs491575 | 11 | 116325442 | T | G | 0.425 | 0.6705 | +? | 0 | 0 | 0 | 1 |
| rs4938392 | 11 | 117359386 | T | C | -0.425 | 0.6705 | -? | 0 | 0 | 0 | 1 |
| rs74833570 | 11 | 117573319 | T | G | 0.425 | 0.6705 | +? | 0 | 0 | 0 | 1 |
| rs117432935 | 11 | 116927976 | A | G | -0.425 | 0.6708 | -+ | 69 | 9.684 | 1 | 0.001859 |
| rs75582303 | 11 | 116928302 | C | T | -0.425 | 0.6708 | -+ | 69 | 9.684 | 1 | 0.001859 |
| rs78111243 | 11 | 116925898 | C | T | -0.425 | 0.6708 | -+ | 69 | 9.684 | 1 | 0.001859 |
| rs573971 | 11 | 118393396 | A | G | 0.425 | 0.6709 | ++ | 0 | 0.034 | 1 | 0.8528 |
| rs34166664 | 11 | 116216072 | G | C | 0.424 | 0.6713 | +- | -100.5 | 1.497 | 1 | 0.2212 |
| rs61903743 | 11 | 117238526 | T | A | -0.424 | 0.6713 | -? | 0 | 0 | 0 | 1 |
| rs61903744 | 11 | 117240002 | T | C | -0.424 | 0.6713 | -? | 0 | 0 | 0 | 1 |
| rs61903745 | 11 | 117240214 | T | G | -0.424 | 0.6713 | -? | 0 | 0 | 0 | 1 |
| rs4938500 | 11 | 118166212 | A | C | 0.424 | 0.6714 | +? | 0 | 0 | 0 | 1 |
| rs150758276 | 11 | 117085874 | C | G | -0.424 | 0.6719 | -? | 0 | 0 | 0 | 1 |
| rs61905108 | 11 | 116624601 | A | C | -0.423 | 0.672 | +- | -160.9 | 1.15 | 1 | 0.2836 |
| rs75619771 | 11 | 119833476 | C | T | -0.423 | 0.672 | -? | 0 | 0 | 0 | 1 |
| rs7118517 | 11 | 120166539 | C | T | 0.423 | 0.6722 | +? | 0 | 0 | 0 | 1 |
| rs948100 | 11 | 117660275 | G | C | 0.423 | 0.6722 | ++ | 0 | 0 | 1 | 0.9994 |
| rs503473 | 11 | 120355230 | G | A | 0.423 | 0.6724 | +? | 0 | 0 | 0 | 1 |
| rs17490647 | 11 | 116351099 | C | A | 0.423 | 0.6726 | +? | 0 | 0 | 0 | 1 |

| | | | | | | | | | | | |
|---|---|---|---|---|---|---|---|---|---|---|---|
| rs12797329 | 11 | 120948832 | G | T | 0.422 | 0.6727 | +? | 0 | 0 | 0 | 1 |
| rs7944487 | 11 | 119192956 | G | A | 0.422 | 0.6727 | +? | 0 | 0 | 0 | 1 |
| rs10892234 | 11 | 118194571 | G | A | -0.422 | 0.6728 | -- | 0 | 0.17 | 1 | 0.6803 |
| rs2004901 | 11 | 120608816 | C | T | 0.422 | 0.6728 | +? | 0 | 0 | 0 | 1 |
| rs79886324 | 11 | 116585444 | A | G | -0.422 | 0.6728 | -- | 0 | 0.012 | 1 | 0.9134 |
| rs4589335 | 11 | 118539140 | T | C | 0.422 | 0.6729 | -+ | 0 | 0.706 | 1 | 0.4008 |
| rs117857960 | 11 | 116924439 | C | T | -0.422 | 0.6731 | -+ | 69.1 | 9.717 | 1 | 0.001826 |
| rs143856824 | 11 | 116922585 | A | C | -0.422 | 0.6731 | -+ | 69.1 | 9.717 | 1 | 0.001826 |
| rs3903091 | 11 | 117348145 | C | T | -0.422 | 0.6731 | -? | 0 | 0 | 0 | 1 |
| rs11217874 | 11 | 120325548 | A | G | 0.422 | 0.6733 | +? | 0 | 0 | 0 | 1 |
| rs11217256 | 11 | 119236590 | G | C | 0.421 | 0.6735 | +? | 0 | 0 | 0 | 1 |
| rs2663620 | 11 | 119427293 | A | G | 0.421 | 0.6736 | +? | 0 | 0 | 0 | 1 |
| rs143315658 | 11 | 116592248 | C | T | -0.421 | 0.6738 | -- | 0 | 0.012 | 1 | 0.9127 |
| rs2845664 | 11 | 119425258 | C | A | 0.421 | 0.6738 | +? | 0 | 0 | 0 | 1 |
| rs4935753 | 11 | 120704184 | C | G | 0.421 | 0.6738 | +? | 0 | 0 | 0 | 1 |
| rs1318166 | 11 | 118171058 | G | A | 0.421 | 0.6739 | +? | 0 | 0 | 0 | 1 |
| rs76758319 | 11 | 121192247 | A | G | -0.421 | 0.6739 | -? | 0 | 0 | 0 | 1 |
| rs61900866 | 11 | 120392508 | G | T | 0.42 | 0.6742 | +? | 0 | 0 | 0 | 1 |
| rs10502246 | 11 | 120924200 | C | T | 0.42 | 0.6745 | +? | 0 | 0 | 0 | 1 |
| rs7941217 | 11 | 118349247 | T | C | -0.42 | 0.6746 | -+ | 0 | 0.844 | 1 | 0.3583 |
| rs2186699 | 11 | 120667742 | G | T | 0.42 | 0.6747 | +? | 0 | 0 | 0 | 1 |
| rs625524 | 11 | 116674503 | A | G | 0.42 | 0.6747 | +? | 0 | 0 | 0 | 1 |
| rs139322068 | 11 | 118121180 | G | T | 0.42 | 0.6748 | -+ | -22.1 | 2.457 | 1 | 0.117 |
| rs55837479 | 11 | 116327971 | A | G | -0.419 | 0.675 | -? | 0 | 0 | 0 | 1 |
| rs76835919 | 11 | 117795852 | G | C | 0.419 | 0.675 | +- | 0 | 0.93 | 1 | 0.3349 |
| rs201373950 | 11 | 118463221 | A | T | 0.419 | 0.6755 | +- | 0 | 0.216 | 1 | 0.642 |
| rs11599994 | 11 | 116383064 | A | G | -0.418 | 0.6759 | -? | 0 | 0 | 0 | 1 |
| rs4252307 | 11 | 117863888 | G | C | 0.418 | 0.6761 | +? | 0 | 0 | 0 | 1 |
| rs9735421 | 11 | 118447597 | A | G | 0.418 | 0.6762 | +- | 0 | 0.201 | 1 | 0.6538 |
| rs9736327 | 11 | 118447580 | G | T | 0.418 | 0.6762 | +- | 0 | 0.201 | 1 | 0.6538 |
| rs471909 | 11 | 119713257 | T | A | -0.418 | 0.6763 | -? | 0 | 0 | 0 | 1 |
| rs17540708 | 11 | 118150127 | G | T | -0.417 | 0.6765 | -+ | 13.2 | 3.456 | 1 | 0.06302 |
| rs7947950 | 11 | 117312089 | G | A | -0.417 | 0.6765 | -? | 0 | 0 | 0 | 1 |
| rs562313 | 11 | 118548421 | A | G | 0.417 | 0.6767 | -+ | 0 | 0.769 | 1 | 0.3805 |
| rs600969 | 11 | 118545797 | T | C | 0.417 | 0.6767 | -+ | 0 | 0.769 | 1 | 0.3805 |
| rs7938877 | 11 | 117410237 | T | C | -0.417 | 0.6767 | -? | 0 | 0 | 0 | 1 |
| rs12797404 | 11 | 120927789 | A | G | 0.417 | 0.677 | +? | 0 | 0 | 0 | 1 |
| rs61905269 | 11 | 117309581 | A | G | 0.416 | 0.6772 | +? | 0 | 0 | 0 | 1 |
| rs35351462 | 11 | 119613090 | A | C | -0.416 | 0.6773 | -? | 0 | 0 | 0 | 1 |
| rs76523703 | 11 | 116749509 | G | T | 0.416 | 0.6773 | +? | 0 | 0 | 0 | 1 |
| rs6589755 | 11 | 119420401 | T | C | 0.416 | 0.6774 | +? | 0 | 0 | 0 | 1 |
| rs61905067 | 11 | 116578982 | G | A | 0.416 | 0.6776 | +? | 0 | 0 | 0 | 1 |
| rs10892359 | 11 | 119263794 | A | G | -0.416 | 0.6777 | -? | 0 | 0 | 0 | 1 |
| rs180348 | 11 | 116611995 | A | G | -0.415 | 0.6779 | -+ | -5.1 | 2.856 | 1 | 0.09105 |
| rs7116797 | 11 | 116707338 | A | G | 0.415 | 0.6781 | +? | 0 | 0 | 0 | 1 |
| rs12420591 | 11 | 120061195 | G | A | 0.415 | 0.6783 | +? | 0 | 0 | 0 | 1 |
| rs4077910 | 11 | 120749741 | T | C | -0.415 | 0.6783 | -? | 0 | 0 | 0 | 1 |
| rs117133249 | 11 | 116524965 | A | G | -0.415 | 0.6784 | -? | 0 | 0 | 0 | 1 |
| rs67674314 | 11 | 117757116 | C | T | -0.415 | 0.6784 | -? | 0 | 0 | 0 | 1 |
| rs2015078 | 11 | 119575185 | C | T | -0.415 | 0.6785 | -? | 0 | 0 | 0 | 1 |
| rs4938442 | 11 | 117734224 | T | C | -0.414 | 0.6786 | -? | 0 | 0 | 0 | 1 |
| rs642673 | 11 | 117118204 | A | T | 0.414 | 0.6786 | +? | 0 | 0 | 0 | 1 |
| rs11217673 | 11 | 119946041 | T | G | -0.414 | 0.6787 | -? | 0 | 0 | 0 | 1 |
| rs34211230 | 11 | 119462879 | A | G | 0.414 | 0.6787 | +? | 0 | 0 | 0 | 1 |
| rs749688 | 11 | 117984995 | C | T | -0.414 | 0.6789 | -- | 0 | 0.013 | 1 | 0.909 |
| rs10502223 | 11 | 116869827 | C | T | -0.414 | 0.679 | +- | 52.9 | 6.368 | 1 | 0.01162 |

| | | | | | | | | | | | |
|---|---|---|---|---|---|---|---|---|---|---|---|
| rs12799683 | 11 | 116232173 | A | G | -0.414 | 0.679 | +- | 0 | 0.989 | 1 | 0.3199 |
| rs73594128 | 11 | 116866609 | C | T | -0.414 | 0.679 | +- | 52.9 | 6.368 | 1 | 0.01162 |
| rs925152 | 11 | 116413688 | T | G | 0.414 | 0.679 | ++ | 0 | 0.275 | 1 | 0.5999 |
| rs12294259 | 11 | 116637146 | T | C | -0.414 | 0.6792 | +- | -75.3 | 1.711 | 1 | 0.1908 |
| rs77564847 | 11 | 116955831 | C | T | -0.414 | 0.6792 | +- | 52.9 | 6.369 | 1 | 0.01161 |
| rs12146701 | 11 | 116403238 | G | T | 0.413 | 0.6794 | +? | 0 | 0 | 0 | 1 |
| rs11216844 | 11 | 118162796 | G | A | 0.413 | 0.6796 | +? | 0 | 0 | 0 | 1 |
| rs654018 | 11 | 119343580 | A | G | 0.413 | 0.6797 | -+ | -56.6 | 1.916 | 1 | 0.1663 |
| rs7947520 | 11 | 120539020 | A | G | -0.413 | 0.6797 | -? | 0 | 0 | 0 | 1 |
| rs4252306 | 11 | 117863730 | C | G | 0.413 | 0.6798 | +? | 0 | 0 | 0 | 1 |
| rs538936 | 11 | 120980822 | A | G | -0.413 | 0.6798 | -? | 0 | 0 | 0 | 1 |
| rs551141 | 11 | 116480631 | A | G | 0.413 | 0.6798 | +? | 0 | 0 | 0 | 1 |
| rs12283803 | 11 | 119799765 | T | C | 0.413 | 0.6799 | +? | 0 | 0 | 0 | 1 |
| rs1942923 | 11 | 117554061 | C | T | 0.413 | 0.6799 | +? | 0 | 0 | 0 | 1 |
| rs1233266397 | 11 | 117334393 | C | T | 0.412 | 0.6801 | +? | 0 | 0 | 0 | 1 |
| rs7925897 | 11 | 117986152 | C | A | 0.412 | 0.6801 | ++ | 0 | 0.013 | 1 | 0.9076 |
| rs7937037 | 11 | 120706789 | C | G | 0.412 | 0.6804 | +? | 0 | 0 | 0 | 1 |
| rs10892412 | 11 | 119459317 | C | T | 0.412 | 0.6806 | +? | 0 | 0 | 0 | 1 |
| rs2187557 | 11 | 118148271 | T | C | 0.412 | 0.6807 | +? | 0 | 0 | 0 | 1 |
| rs2187126 | 11 | 116635784 | G | A | -0.411 | 0.6808 | -- | 0 | 0.068 | 1 | 0.7948 |
| rs10892301 | 11 | 118735476 | A | G | 0.411 | 0.6809 | ++ | 0 | 0.135 | 1 | 0.7129 |
| rs79436592 | 11 | 119762339 | A | G | 0.411 | 0.6811 | +? | 0 | 0 | 0 | 1 |
| rs7350537 | 11 | 116438159 | T | C | 0.411 | 0.6812 | ++ | 0 | 0.01 | 1 | 0.9217 |
| rs4554895 | 11 | 121010539 | A | C | 0.411 | 0.6813 | +? | 0 | 0 | 0 | 1 |
| rs4938301 | 11 | 116575473 | A | G | -0.411 | 0.6813 | -? | 0 | 0 | 0 | 1 |
| rs746910 | 11 | 117444044 | G | A | 0.411 | 0.6814 | +? | 0 | 0 | 0 | 1 |
| rs55843215 | 11 | 118195962 | C | T | 0.41 | 0.6815 | +? | 0 | 0 | 0 | 1 |
| rs7946423 | 11 | 116503403 | T | A | 0.41 | 0.6815 | +? | 0 | 0 | 0 | 1 |
| rs4317991 | 11 | 120696839 | A | G | 0.41 | 0.6816 | +? | 0 | 0 | 0 | 1 |
| rs2852231 | 11 | 120605882 | C | T | 0.41 | 0.6817 | +? | 0 | 0 | 0 | 1 |
| rs3935406 | 11 | 119572185 | G | A | -0.41 | 0.6817 | -? | 0 | 0 | 0 | 1 |
| rs725524 | 11 | 116510369 | A | G | -0.41 | 0.6818 | -? | 0 | 0 | 0 | 1 |
| rs10892216 | 11 | 117979744 | C | A | -0.41 | 0.6819 | -- | 0 | 0 | 1 | 0.9938 |
| rs520176 | 11 | 116471690 | T | C | -0.41 | 0.6819 | -? | 0 | 0 | 0 | 1 |
| rs79110162 | 11 | 116643011 | G | A | -0.41 | 0.6819 | -- | 0 | 0.081 | 1 | 0.7754 |
| rs11216845 | 11 | 118163216 | T | C | 0.41 | 0.6821 | +? | 0 | 0 | 0 | 1 |
| rs10892217 | 11 | 117979814 | C | T | -0.409 | 0.6822 | -- | 0 | 0 | 1 | 0.9931 |
| rs11218018 | 11 | 120703415 | G | A | 0.409 | 0.6822 | +? | 0 | 0 | 0 | 1 |
| rs60816279 | 11 | 117617127 | A | G | 0.409 | 0.6822 | +? | 0 | 0 | 0 | 1 |
| rs680671 | 11 | 117331279 | A | T | -0.409 | 0.6822 | -? | 0 | 0 | 0 | 1 |
| rs111997050 | 11 | 118387892 | T | C | -0.409 | 0.6823 | -+ | -176.3 | 1.086 | 1 | 0.2974 |
| rs7112577 | 11 | 117044603 | G | C | -0.409 | 0.6823 | -? | 0 | 0 | 0 | 1 |
| rs4936558 | 11 | 120704307 | C | T | 0.409 | 0.6825 | +? | 0 | 0 | 0 | 1 |
| rs7937921 | 11 | 117741051 | A | G | -0.409 | 0.6825 | -? | 0 | 0 | 0 | 1 |
| rs12721028 | 11 | 116705590 | G | A | 0.409 | 0.6827 | +? | 0 | 0 | 0 | 1 |
| rs34809912 | 11 | 119640427 | A | G | -0.409 | 0.6827 | -? | 0 | 0 | 0 | 1 |
| rs512588 | 11 | 116246867 | T | G | -0.409 | 0.6827 | -? | 0 | 0 | 0 | 1 |
| rs140231532 | 11 | 116697301 | T | C | 0.409 | 0.6828 | +? | 0 | 0 | 0 | 1 |
| rs78577579 | 11 | 117336708 | T | G | -0.408 | 0.683 | -? | 0 | 0 | 0 | 1 |
| rs11501117 | 11 | 117356908 | G | C | -0.408 | 0.6831 | -? | 0 | 0 | 0 | 1 |
| rs948031 | 11 | 120609495 | G | A | 0.408 | 0.6831 | +? | 0 | 0 | 0 | 1 |
| rs2846125 | 11 | 120604315 | C | T | 0.408 | 0.6833 | +? | 0 | 0 | 0 | 1 |
| rs688035 | 11 | 118802492 | G | C | 0.408 | 0.6833 | -+ | -103.7 | 1.473 | 1 | 0.2249 |
| rs1938948 | 11 | 119421419 | C | T | 0.407 | 0.6837 | +? | 0 | 0 | 0 | 1 |
| rs2845665 | 11 | 119423410 | G | A | 0.407 | 0.6837 | +? | 0 | 0 | 0 | 1 |
| rs2845666 | 11 | 119423269 | G | A | 0.407 | 0.6837 | +? | 0 | 0 | 0 | 1 |

| | | | | | | | | | | | |
|---|---|---|---|---|---|---|---|---|---|---|---|
| rs6589756 | 11 | 119423748 | C | T | 0.407 | 0.6837 | +? | 0 | 0 | 0 | 1 |
| rs7395205 | 11 | 119423822 | T | C | 0.407 | 0.6837 | +? | 0 | 0 | 0 | 1 |
| rs34420468 | 11 | 119811328 | A | G | -0.407 | 0.6838 | -? | 0 | 0 | 0 | 1 |
| rs683743 | 11 | 119962278 | C | G | 0.407 | 0.684 | +? | 0 | 0 | 0 | 1 |
| rs2850807 | 11 | 120797893 | T | A | 0.407 | 0.6841 | +? | 0 | 0 | 0 | 1 |
| rs118086079 | 11 | 117194059 | A | G | -0.407 | 0.6842 | -? | 0 | 0 | 0 | 1 |
| rs17440824 | 11 | 116634173 | T | C | -0.407 | 0.6842 | +- | -43.3 | 2.094 | 1 | 0.1479 |
| rs11216664 | 11 | 117849065 | C | G | 0.407 | 0.6843 | +? | 0 | 0 | 0 | 1 |
| rs118088243 | 11 | 121165007 | G | A | 0.406 | 0.6844 | +? | 0 | 0 | 0 | 1 |
| rs7109876 | 11 | 116848648 | A | C | 0.406 | 0.6844 | +? | 0 | 0 | 0 | 1 |
| rs7934890 | 11 | 118655356 | T | C | -0.406 | 0.6846 | -+ | 0 | 0.121 | 1 | 0.7275 |
| rs4938269 | 11 | 116272650 | C | G | -0.406 | 0.685 | -? | 0 | 0 | 0 | 1 |
| rs12805477 | 11 | 120907961 | G | C | 0.405 | 0.6852 | +? | 0 | 0 | 0 | 1 |
| rs35457276 | 11 | 120914296 | T | C | 0.405 | 0.6852 | +? | 0 | 0 | 0 | 1 |
| rs2282496 | 11 | 117864979 | T | C | 0.405 | 0.6853 | +? | 0 | 0 | 0 | 1 |
| rs11216109 | 11 | 116580815 | G | C | -0.405 | 0.6854 | -? | 0 | 0 | 0 | 1 |
| rs61902398 | 11 | 121190062 | A | G | -0.405 | 0.6854 | -? | 0 | 0 | 0 | 1 |
| rs642936 | 11 | 118019906 | C | T | -0.405 | 0.6854 | +- | 0 | 0.685 | 1 | 0.4077 |
| rs9332858 | 11 | 118390082 | C | A | -0.405 | 0.6854 | -+ | -139.6 | 1.252 | 1 | 0.2631 |
| rs1848688 | 11 | 119432440 | T | C | 0.405 | 0.6855 | +? | 0 | 0 | 0 | 1 |
| rs2155196 | 11 | 117736735 | T | C | 0.405 | 0.6855 | +? | 0 | 0 | 0 | 1 |
| rs2202884 | 11 | 119433740 | T | C | 0.405 | 0.6855 | +? | 0 | 0 | 0 | 1 |
| rs2663627 | 11 | 119433485 | G | C | 0.405 | 0.6855 | +? | 0 | 0 | 0 | 1 |
| rs71478237 | 11 | 116228118 | C | A | 0.405 | 0.6855 | +- | -104.2 | 1.47 | 1 | 0.2254 |
| rs10892569 | 11 | 120270249 | G | A | 0.405 | 0.6856 | +? | 0 | 0 | 0 | 1 |
| rs17121771 | 11 | 118000796 | T | C | 0.405 | 0.6856 | +? | 0 | 0 | 0 | 1 |
| rs74601470 | 11 | 116873382 | T | C | -0.405 | 0.6857 | -? | 0 | 0 | 0 | 1 |
| rs1787701 | 11 | 116563992 | C | G | 0.405 | 0.6858 | +? | 0 | 0 | 0 | 1 |
| rs79536557 | 11 | 116963969 | G | A | -0.404 | 0.6862 | +- | 53.1 | 6.397 | 1 | 0.01143 |
| rs17124208 | 11 | 120557814 | C | A | 0.404 | 0.6863 | +? | 0 | 0 | 0 | 1 |
| rs60972755 | 11 | 116635613 | A | C | -0.404 | 0.6863 | +- | -70.7 | 1.757 | 1 | 0.185 |
| rs1892937 | 11 | 117736770 | C | T | 0.404 | 0.6864 | +? | 0 | 0 | 0 | 1 |
| rs9332846 | 11 | 118377523 | A | G | -0.404 | 0.6864 | -+ | -121.3 | 1.355 | 1 | 0.2443 |
| rs73019249 | 11 | 116252227 | G | A | -0.403 | 0.6866 | -? | 0 | 0 | 0 | 1 |
| rs10892549 | 11 | 120075574 | T | C | -0.403 | 0.6867 | -? | 0 | 0 | 0 | 1 |
| rs3020104 | 11 | 120685242 | A | G | 0.403 | 0.6867 | +? | 0 | 0 | 0 | 1 |
| rs11605475 | 11 | 117324521 | A | G | -0.403 | 0.6868 | -? | 0 | 0 | 0 | 1 |
| rs2509004 | 11 | 117410423 | A | G | 0.403 | 0.6868 | +? | 0 | 0 | 0 | 1 |
| rs486020 | 11 | 116483126 | C | G | 0.403 | 0.687 | +? | 0 | 0 | 0 | 1 |
| rs547021 | 11 | 116329139 | C | T | 0.403 | 0.687 | +? | 0 | 0 | 0 | 1 |
| rs552040 | 11 | 119712224 | T | G | -0.403 | 0.687 | -? | 0 | 0 | 0 | 1 |
| rs1003261346 | 11 | 117359244 | A | G | -0.403 | 0.6871 | -? | 0 | 0 | 0 | 1 |
| rs10466534 | 11 | 117611292 | A | G | 0.403 | 0.6871 | +? | 0 | 0 | 0 | 1 |
| rs498689 | 11 | 117463000 | G | A | -0.403 | 0.6871 | -? | 0 | 0 | 0 | 1 |
| rs617320 | 11 | 119961694 | G | T | 0.403 | 0.6872 | +? | 0 | 0 | 0 | 1 |
| rs1942931 | 11 | 117610854 | A | G | 0.403 | 0.6873 | +? | 0 | 0 | 0 | 1 |
| rs4528371 | 11 | 117669418 | T | A | 0.402 | 0.6874 | ++ | 0 | 0.111 | 1 | 0.7391 |
| rs72999499 | 11 | 118543589 | G | C | -0.402 | 0.6877 | -? | 0 | 0 | 0 | 1 |
| rs180330 | 11 | 116622761 | A | G | -0.402 | 0.6878 | -+ | 0 | 0.198 | 1 | 0.6561 |
| rs3016611 | 11 | 116847584 | A | G | -0.402 | 0.6878 | +- | 47.2 | 5.677 | 1 | 0.01719 |
| rs12225230 | 11 | 116728630 | C | G | -0.402 | 0.6879 | -? | 0 | 0 | 0 | 1 |
| rs1893968 | 11 | 117335423 | A | G | 0.402 | 0.6879 | +? | 0 | 0 | 0 | 1 |
| rs545559 | 11 | 116481248 | A | C | 0.402 | 0.688 | +? | 0 | 0 | 0 | 1 |
| rs566254 | 11 | 116481505 | A | T | 0.402 | 0.688 | +? | 0 | 0 | 0 | 1 |
| rs11216821 | 11 | 118088362 | C | T | -0.401 | 0.6881 | -+ | 0 | 0.099 | 1 | 0.7534 |
| rs7929505 | 11 | 119647159 | C | T | -0.401 | 0.6881 | -? | 0 | 0 | 0 | 1 |

| | | | | | | | | | | | |
|---|---|---|---|---|---|---|---|---|---|---|---|
| rs2072560 | 11 | 116661826 | T | C | -0.401 | 0.6884 | +- | -77 | 1.695 | 1 | 0.1929 |
| rs76916861 | 11 | 116921802 | T | C | -0.401 | 0.6884 | -+ | 68.9 | 9.642 | 1 | 0.001901 |
| rs7950360 | 11 | 120461856 | C | G | -0.401 | 0.6885 | -? | 0 | 0 | 0 | 1 |
| rs591671 | 11 | 120878492 | A | G | -0.401 | 0.6886 | -+ | 0 | 0.361 | 1 | 0.548 |
| rs117713193 | 11 | 117263874 | T | C | 0.401 | 0.6887 | +? | 0 | 0 | 0 | 1 |
| rs1893967 | 11 | 117335148 | A | G | 0.401 | 0.6887 | +? | 0 | 0 | 0 | 1 |
| rs34505832 | 11 | 120947207 | G | A | 0.4 | 0.6888 | +? | 0 | 0 | 0 | 1 |
| rs117843763 | 11 | 116420968 | A | C | 0.4 | 0.6889 | +? | 0 | 0 | 0 | 1 |
| rs573455 | 11 | 117267884 | A | G | 0.4 | 0.6889 | +? | 0 | 0 | 0 | 1 |
| rs57643109 | 11 | 116638751 | A | G | -0.4 | 0.6889 | -- | 0 | 0.071 | 1 | 0.7899 |
| rs139872819 | 11 | 116631912 | C | T | -0.4 | 0.689 | -- | 0 | 0.071 | 1 | 0.7899 |
| rs17311407 | 11 | 120593695 | T | C | -0.4 | 0.689 | +- | 0 | 0.955 | 1 | 0.3284 |
| rs2846119 | 11 | 120605867 | C | G | 0.4 | 0.689 | +? | 0 | 0 | 0 | 1 |
| rs497653 | 11 | 116828484 | C | T | -0.4 | 0.6891 | +- | 47.2 | 5.681 | 1 | 0.01715 |
| rs4938280 | 11 | 116419442 | C | T | 0.4 | 0.6893 | ++ | 0 | 0.365 | 1 | 0.546 |
| rs80040504 | 11 | 120337712 | A | G | 0.399 | 0.6896 | +? | 0 | 0 | 0 | 1 |
| rs117579298 | 11 | 116642100 | G | A | -0.399 | 0.6897 | -- | 0 | 0.071 | 1 | 0.7895 |
| rs12226536 | 11 | 120706759 | G | C | 0.399 | 0.6897 | +? | 0 | 0 | 0 | 1 |
| rs2155197 | 11 | 117736790 | G | A | 0.399 | 0.6897 | +? | 0 | 0 | 0 | 1 |
| rs17120265 | 11 | 116932615 | G | C | -0.399 | 0.69 | +- | 53.2 | 6.412 | 1 | 0.01134 |
| rs17120271 | 11 | 116935363 | G | C | -0.399 | 0.69 | +- | 53.2 | 6.412 | 1 | 0.01134 |
| rs17120275 | 11 | 116939071 | C | T | -0.399 | 0.69 | +- | 53.2 | 6.412 | 1 | 0.01134 |
| rs73576612 | 11 | 116929666 | C | T | -0.399 | 0.69 | +- | 53.2 | 6.412 | 1 | 0.01134 |
| rs117287247 | 11 | 116470947 | T | C | -0.399 | 0.6902 | -? | 0 | 0 | 0 | 1 |
| rs7948446 | 11 | 117454506 | T | C | 0.399 | 0.6902 | +? | 0 | 0 | 0 | 1 |
| rs59810418 | 11 | 117617219 | C | G | 0.398 | 0.6903 | +? | 0 | 0 | 0 | 1 |
| rs2276434 | 11 | 117959216 | T | C | 0.398 | 0.6904 | +- | 0 | 0.15 | 1 | 0.6985 |
| rs73023350 | 11 | 118449396 | A | G | 0.398 | 0.6904 | +- | 0 | 0.233 | 1 | 0.6296 |
| rs2276435 | 11 | 117959372 | A | G | 0.398 | 0.6905 | +- | 0 | 0.15 | 1 | 0.6984 |
| rs4938803 | 11 | 120268022 | G | A | 0.398 | 0.6906 | +? | 0 | 0 | 0 | 1 |
| rs11215951 | 11 | 116344594 | T | C | 0.398 | 0.6907 | +? | 0 | 0 | 0 | 1 |
| rs7109584 | 11 | 120654400 | A | G | -0.398 | 0.6907 | -? | 0 | 0 | 0 | 1 |
| rs628418 | 11 | 116718366 | G | C | -0.397 | 0.691 | +- | 17.2 | 3.622 | 1 | 0.05702 |
| rs876797 | 11 | 117737747 | A | C | -0.397 | 0.6911 | -? | 0 | 0 | 0 | 1 |
| rs925261 | 11 | 119416327 | G | T | 0.397 | 0.6911 | +? | 0 | 0 | 0 | 1 |
| rs12223757 | 11 | 120706839 | G | A | 0.397 | 0.6912 | +? | 0 | 0 | 0 | 1 |
| rs145949765 | 11 | 116920199 | T | C | -0.397 | 0.6914 | -+ | 69 | 9.687 | 1 | 0.001856 |
| rs77775914 | 11 | 116241604 | T | C | -0.397 | 0.6915 | -- | 0 | 0.413 | 1 | 0.5203 |
| rs7122193 | 11 | 120556697 | C | T | -0.396 | 0.6918 | -? | 0 | 0 | 0 | 1 |
| rs10891966 | 11 | 116213941 | G | A | -0.396 | 0.6921 | -- | 0 | 0.441 | 1 | 0.5065 |
| rs632541 | 11 | 119960565 | G | A | 0.396 | 0.6921 | +? | 0 | 0 | 0 | 1 |
| rs1818393 | 11 | 119415314 | A | G | 0.396 | 0.6922 | +? | 0 | 0 | 0 | 1 |
| rs73010424 | 11 | 117124050 | A | G | -0.396 | 0.6922 | -? | 0 | 0 | 0 | 1 |
| rs1320720 | 11 | 120462743 | A | G | -0.396 | 0.6924 | -? | 0 | 0 | 0 | 1 |
| rs61903804 | 11 | 117720645 | G | A | -0.396 | 0.6924 | -? | 0 | 0 | 0 | 1 |
| rs76868258 | 11 | 116993292 | T | C | -0.395 | 0.6925 | -? | 0 | 0 | 0 | 1 |
| rs11217014 | 11 | 118626286 | C | T | -0.395 | 0.6926 | -+ | 0 | 0.199 | 1 | 0.6559 |
| rs138990232 | 11 | 120631792 | G | A | 0.395 | 0.6926 | +? | 0 | 0 | 0 | 1 |
| rs1784121 | 11 | 116561790 | T | G | 0.395 | 0.6926 | +? | 0 | 0 | 0 | 1 |
| rs1787704 | 11 | 116561996 | T | C | 0.395 | 0.6926 | +? | 0 | 0 | 0 | 1 |
| rs117096905 | 11 | 116467349 | T | C | -0.395 | 0.6929 | -? | 0 | 0 | 0 | 1 |
| rs35594498 | 11 | 120397162 | T | C | 0.395 | 0.693 | +? | 0 | 0 | 0 | 1 |
| rs116859176 | 11 | 116292818 | C | T | 0.394 | 0.6933 | +? | 0 | 0 | 0 | 1 |
| rs17092631 | 11 | 116476765 | C | T | 0.394 | 0.6937 | +? | 0 | 0 | 0 | 1 |
| rs11215972 | 11 | 116383138 | G | C | -0.394 | 0.6939 | -? | 0 | 0 | 0 | 1 |
| rs2070666 | 11 | 116701674 | A | T | 0.393 | 0.694 | +? | 0 | 0 | 0 | 1 |

| | | | | | | | | | | |
|---|---|---|---|---|---|---|---|---|---|---|
| rs11216404 | 11 | 117334885 | A | C | 0.393 | 0.6941 | +? | 0 | 0 | 0 | 1 |
| rs4938388 | 11 | 117334831 | C | T | 0.393 | 0.6941 | +? | 0 | 0 | 0 | 1 |
| rs45460493 | 11 | 118455465 | A | G | -0.393 | 0.6943 | -- | 0 | 0.003 | 1 | 0.9528 |
| rs2004676 | 11 | 120699625 | T | C | 0.393 | 0.6944 | +? | 0 | 0 | 0 | 1 |
| rs496248 | 11 | 120323214 | A | G | 0.393 | 0.6944 | +? | 0 | 0 | 0 | 1 |
| rs617124 | 11 | 119880367 | A | G | -0.393 | 0.6945 | -? | 0 | 0 | 0 | 1 |
| rs7112199 | 11 | 120758691 | A | G | -0.393 | 0.6945 | -? | 0 | 0 | 0 | 1 |
| rs12273529 | 11 | 119959214 | T | C | -0.393 | 0.6946 | -? | 0 | 0 | 0 | 1 |
| rs12285338 | 11 | 119959144 | G | A | -0.393 | 0.6946 | -? | 0 | 0 | 0 | 1 |
| rs7949126 | 11 | 119959362 | T | C | -0.393 | 0.6946 | -? | 0 | 0 | 0 | 1 |
| rs12276368 | 11 | 120635399 | A | G | -0.392 | 0.6947 | -? | 0 | 0 | 0 | 1 |
| rs538741 | 11 | 120980753 | T | C | -0.392 | 0.6948 | -? | 0 | 0 | 0 | 1 |
| rs77832134 | 11 | 116879636 | G | A | -0.392 | 0.6949 | -? | 0 | 0 | 0 | 1 |
| rs10502243 | 11 | 120795441 | A | G | 0.392 | 0.695 | +? | 0 | 0 | 0 | 1 |
| rs17519093 | 11 | 116629905 | A | G | -0.392 | 0.6951 | -- | 0 | 0.074 | 1 | 0.7863 |
| rs116877083 | 11 | 116634297 | C | A | -0.392 | 0.6953 | -- | 0 | 0.074 | 1 | 0.7861 |
| rs10488698 | 11 | 116633947 | A | G | -0.392 | 0.6954 | -- | 0 | 0.074 | 1 | 0.7861 |
| rs11216923 | 11 | 118465501 | G | A | 0.391 | 0.6955 | +- | 0 | 0.191 | 1 | 0.6622 |
| rs12361458 | 11 | 117317664 | T | C | 0.391 | 0.6955 | +? | 0 | 0 | 0 | 1 |
| rs150734126 | 11 | 116880018 | C | T | -0.391 | 0.6957 | +- | 47.4 | 5.706 | 1 | 0.01691 |
| rs2155260 | 11 | 120666825 | A | G | 0.391 | 0.6957 | +? | 0 | 0 | 0 | 1 |
| rs10750149 | 11 | 119417051 | T | A | 0.391 | 0.6958 | +? | 0 | 0 | 0 | 1 |
| rs11217782 | 11 | 120150936 | C | T | -0.391 | 0.6958 | -? | 0 | 0 | 0 | 1 |
| rs1605854 | 11 | 119418145 | A | G | 0.391 | 0.6958 | +? | 0 | 0 | 0 | 1 |
| rs1828280 | 11 | 119419198 | T | C | 0.391 | 0.6958 | +? | 0 | 0 | 0 | 1 |
| rs2663640 | 11 | 119418640 | C | T | 0.391 | 0.6958 | +? | 0 | 0 | 0 | 1 |
| rs608891 | 11 | 120832264 | A | G | -0.391 | 0.6958 | -+ | 0 | 0.354 | 1 | 0.5518 |
| rs9332835 | 11 | 118372923 | A | G | -0.391 | 0.6958 | -+ | -124.2 | 1.338 | 1 | 0.2473 |
| rs4936557 | 11 | 120704063 | C | T | 0.391 | 0.6959 | +? | 0 | 0 | 0 | 1 |
| rs4938691 | 11 | 119477261 | C | T | 0.39 | 0.6962 | +? | 0 | 0 | 0 | 1 |
| rs590270 | 11 | 119963787 | C | T | 0.39 | 0.6962 | +? | 0 | 0 | 0 | 1 |
| rs664579 | 11 | 119964344 | G | A | 0.39 | 0.6962 | +? | 0 | 0 | 0 | 1 |
| rs11217971 | 11 | 120551969 | A | G | -0.39 | 0.6963 | -? | 0 | 0 | 0 | 1 |
| rs11607679 | 11 | 119604844 | A | C | 0.39 | 0.6963 | +? | 0 | 0 | 0 | 1 |
| rs7126175 | 11 | 118450875 | T | C | 0.39 | 0.6964 | +- | 0 | 0.247 | 1 | 0.6195 |
| rs77325329 | 11 | 120524275 | A | G | -0.39 | 0.6964 | -? | 0 | 0 | 0 | 1 |
| rs35865949 | 11 | 118229061 | A | T | -0.39 | 0.6965 | -- | 0 | 0 | 1 | 0.983 |
| rs4435039 | 11 | 118297391 | G | A | 0.39 | 0.6966 | +- | 0 | 0.476 | 1 | 0.4904 |
| rs11217471 | 11 | 119630641 | A | G | 0.39 | 0.6967 | +? | 0 | 0 | 0 | 1 |
| rs58868870 | 11 | 118668296 | T | C | -0.39 | 0.6967 | -- | 0 | 0.039 | 1 | 0.844 |
| rs2511064 | 11 | 120611896 | A | G | 0.39 | 0.6968 | +? | 0 | 0 | 0 | 1 |
| rs570637 | 11 | 119702527 | T | C | -0.389 | 0.6971 | -? | 0 | 0 | 0 | 1 |
| rs3016351 | 11 | 116474753 | A | G | -0.389 | 0.6973 | -? | 0 | 0 | 0 | 1 |
| rs7107305 | 11 | 118368665 | G | A | -0.389 | 0.6973 | -+ | -140.1 | 1.249 | 1 | 0.2637 |
| rs7123835 | 11 | 118368822 | G | T | -0.389 | 0.6973 | -+ | -140.1 | 1.249 | 1 | 0.2637 |
| rs9735449 | 11 | 118448087 | T | C | 0.389 | 0.6976 | +- | 0 | 0.165 | 1 | 0.6849 |
| rs61903395 | 11 | 116931907 | G | A | 0.388 | 0.6979 | +? | 0 | 0 | 0 | 1 |
| rs12788928 | 11 | 119514810 | A | G | -0.388 | 0.698 | -? | 0 | 0 | 0 | 1 |
| rs2017098 | 11 | 117442388 | G | C | 0.388 | 0.698 | +? | 0 | 0 | 0 | 1 |
| rs4280006 | 11 | 121058189 | C | T | -0.388 | 0.698 | -? | 0 | 0 | 0 | 1 |
| rs10892158 | 11 | 117550767 | C | T | 0.388 | 0.6981 | +? | 0 | 0 | 0 | 1 |
| rs1320648 | 11 | 120608912 | C | T | 0.388 | 0.6981 | +? | 0 | 0 | 0 | 1 |
| rs1893262 | 11 | 120169818 | C | T | -0.388 | 0.6981 | -? | 0 | 0 | 0 | 1 |
| rs752979 | 11 | 120608964 | A | G | 0.388 | 0.6981 | +? | 0 | 0 | 0 | 1 |
| rs948032 | 11 | 120608661 | A | G | 0.388 | 0.6981 | +? | 0 | 0 | 0 | 1 |
| rs948033 | 11 | 120608222 | G | A | 0.388 | 0.6981 | +? | 0 | 0 | 0 | 1 |

| | | | | | | | | | | | |
|---|---|---|---|---|---|---|---|---|---|---|---|
| rs949298 | 11 | 120608085 | A | G | 0.388 | 0.6981 | +? | 0 | 0 | 0 | 1 |
| rs949299 | 11 | 120608142 | A | T | 0.388 | 0.6981 | +? | 0 | 0 | 0 | 1 |
| rs56314469 | 11 | 116794903 | A | G | 0.388 | 0.6983 | +? | 0 | 0 | 0 | 1 |
| rs4245203 | 11 | 119579260 | C | T | 0.387 | 0.6984 | +? | 0 | 0 | 0 | 1 |
| rs11216922 | 11 | 118465295 | G | A | 0.387 | 0.6986 | +- | 0 | 0.197 | 1 | 0.657 |
| rs3825057 | 11 | 118622985 | C | T | -0.387 | 0.6987 | -+ | 0 | 0.181 | 1 | 0.6701 |
| rs72998668 | 11 | 119904317 | T | C | 0.387 | 0.6987 | +? | 0 | 0 | 0 | 1 |
| rs10892178 | 11 | 117685343 | T | C | 0.387 | 0.6988 | +? | 0 | 0 | 0 | 1 |
| rs12416901 | 11 | 119587095 | C | T | 0.387 | 0.6989 | +? | 0 | 0 | 0 | 1 |
| rs543413 | 11 | 118550017 | C | T | 0.387 | 0.6989 | -+ | -181.5 | 1.066 | 1 | 0.302 |
| rs665008 | 11 | 119964429 | A | G | 0.387 | 0.6989 | +? | 0 | 0 | 0 | 1 |
| rs118168570 | 11 | 116322222 | A | G | 0.387 | 0.699 | +? | 0 | 0 | 0 | 1 |
| rs619744 | 11 | 117915659 | G | C | -0.387 | 0.699 | -? | 0 | 0 | 0 | 1 |
| rs11216910 | 11 | 118450046 | G | A | 0.386 | 0.6992 | +- | 0 | 0.225 | 1 | 0.6352 |
| rs3782046 | 11 | 118230739 | A | G | -0.386 | 0.6992 | -- | 0 | 0.001 | 1 | 0.9781 |
| rs59647391 | 11 | 116900956 | G | A | -0.386 | 0.6992 | +- | 53.5 | 6.448 | 1 | 0.01111 |
| rs73594141 | 11 | 116882563 | T | C | -0.386 | 0.6992 | +- | 53.5 | 6.448 | 1 | 0.01111 |
| rs73594158 | 11 | 116896441 | T | C | -0.386 | 0.6992 | +- | 53.5 | 6.448 | 1 | 0.01111 |
| rs73594162 | 11 | 116896811 | T | C | -0.386 | 0.6992 | +- | 53.5 | 6.448 | 1 | 0.01111 |
| rs73594164 | 11 | 116897613 | G | T | -0.386 | 0.6992 | +- | 53.5 | 6.448 | 1 | 0.01111 |
| rs73594172 | 11 | 116902631 | T | A | -0.386 | 0.6992 | +- | 53.5 | 6.448 | 1 | 0.01111 |
| rs11215948 | 11 | 116339688 | G | C | -0.386 | 0.6994 | -? | 0 | 0 | 0 | 1 |
| rs543327 | 11 | 118549985 | T | G | 0.386 | 0.6995 | -+ | -182.4 | 1.062 | 1 | 0.3027 |
| rs7924498 | 11 | 117960661 | A | G | 0.386 | 0.6995 | ++ | 0 | 0.015 | 1 | 0.9039 |
| rs11999 | 11 | 117690918 | C | A | 0.386 | 0.6996 | +? | 0 | 0 | 0 | 1 |
| rs2887245 | 11 | 119460505 | T | C | 0.386 | 0.6996 | +? | 0 | 0 | 0 | 1 |
| rs647099 | 11 | 118067009 | A | G | 0.386 | 0.6996 | +? | 0 | 0 | 0 | 1 |
| rs11216912 | 11 | 118452334 | C | G | 0.385 | 0.7 | +- | 0 | 0.255 | 1 | 0.6135 |
| rs11216913 | 11 | 118453681 | G | A | 0.385 | 0.7 | +- | 0 | 0.255 | 1 | 0.6135 |
| rs550164 | 11 | 120002727 | G | A | 0.385 | 0.7 | +? | 0 | 0 | 0 | 1 |
| rs12721041 | 11 | 116693871 | T | C | 0.385 | 0.7002 | +? | 0 | 0 | 0 | 1 |
| rs598746 | 11 | 118020057 | A | G | -0.385 | 0.7002 | -+ | 0 | 0.518 | 1 | 0.4717 |
| rs12365322 | 11 | 116490970 | G | T | -0.385 | 0.7003 | -? | 0 | 0 | 0 | 1 |
| rs1317723 | 11 | 119985050 | A | G | 0.385 | 0.7004 | +? | 0 | 0 | 0 | 1 |
| rs55756957 | 11 | 118747576 | G | A | 0.385 | 0.7004 | +- | 40.6 | 5.048 | 1 | 0.02466 |
| rs147864821 | 11 | 117411687 | A | G | 0.385 | 0.7005 | +? | 0 | 0 | 0 | 1 |
| rs4938716 | 11 | 119576555 | T | G | -0.385 | 0.7005 | -? | 0 | 0 | 0 | 1 |
| rs519116 | 11 | 120991298 | T | C | -0.385 | 0.7006 | -? | 0 | 0 | 0 | 1 |
| rs4936536 | 11 | 120521485 | A | G | -0.384 | 0.7007 | -? | 0 | 0 | 0 | 1 |
| rs12283211 | 11 | 120890823 | A | G | -0.384 | 0.7009 | -? | 0 | 0 | 0 | 1 |
| rs11216911 | 11 | 118451901 | C | T | 0.384 | 0.7011 | +- | 0 | 0.243 | 1 | 0.6221 |
| rs79007113 | 11 | 120039835 | A | G | 0.384 | 0.7011 | +? | 0 | 0 | 0 | 1 |
| rs10892190 | 11 | 117760079 | G | A | -0.384 | 0.7012 | -? | 0 | 0 | 0 | 1 |
| rs10892191 | 11 | 117760118 | C | G | -0.384 | 0.7012 | -? | 0 | 0 | 0 | 1 |
| rs118121169 | 11 | 116917089 | A | G | -0.384 | 0.7012 | -+ | 69.5 | 9.832 | 1 | 0.001715 |
| rs507080 | 11 | 118549582 | A | G | 0.384 | 0.7013 | -+ | -184.7 | 1.054 | 1 | 0.3046 |
| rs656575 | 11 | 118549908 | G | T | 0.384 | 0.7013 | -+ | -184.7 | 1.054 | 1 | 0.3046 |
| rs659969 | 11 | 118549803 | T | C | 0.384 | 0.7013 | -+ | -184.7 | 1.054 | 1 | 0.3046 |
| rs73580405 | 11 | 120529116 | G | A | -0.384 | 0.7013 | -? | 0 | 0 | 0 | 1 |
| rs148134535 | 11 | 116674897 | G | C | -0.383 | 0.7015 | -? | 0 | 0 | 0 | 1 |
| rs2186747 | 11 | 121037529 | G | C | -0.383 | 0.7016 | -? | 0 | 0 | 0 | 1 |
| rs573126 | 11 | 116276086 | C | A | -0.383 | 0.7016 | -? | 0 | 0 | 0 | 1 |
| rs6589579 | 11 | 116789635 | C | A | -0.383 | 0.7017 | +- | 47.6 | 5.728 | 1 | 0.01669 |
| rs17569099 | 11 | 116350128 | A | T | -0.383 | 0.7019 | -? | 0 | 0 | 0 | 1 |
| rs75542782 | 11 | 116637002 | A | G | -0.383 | 0.702 | -- | 0 | 0.076 | 1 | 0.7822 |
| rs80120780 | 11 | 117672330 | G | C | 0.383 | 0.7021 | ++ | 0 | 0.001 | 1 | 0.9739 |

| | | | | | | | | | | | |
|---|---|---|---|---|---|---|---|---|---|---|---|
| rs11217016 | 11 | 118627149 | C | A | -0.382 | 0.7022 | -+ | 0 | 0.22 | 1 | 0.639 |
| rs146877554 | 11 | 117971968 | A | G | 0.382 | 0.7022 | +? | 0 | 0 | 0 | 1 |
| rs2508806 | 11 | 120611914 | C | T | 0.382 | 0.7023 | +? | 0 | 0 | 0 | 1 |
| rs113195776 | 11 | 117853331 | A | G | 0.382 | 0.7024 | +? | 0 | 0 | 0 | 1 |
| rs144018061 | 11 | 120429035 | A | G | -0.382 | 0.7025 | -? | 0 | 0 | 0 | 1 |
| rs73003764 | 11 | 120711290 | G | A | 0.382 | 0.7025 | +? | 0 | 0 | 0 | 1 |
| rs2846121 | 11 | 120605524 | A | G | 0.382 | 0.7027 | +? | 0 | 0 | 0 | 1 |
| rs7130875 | 11 | 118613674 | C | T | -0.382 | 0.7028 | -+ | 0 | 0.18 | 1 | 0.6712 |
| rs948099 | 11 | 117660578 | A | C | 0.382 | 0.7028 | ++ | 0 | 0.003 | 1 | 0.957 |
| rs11216007 | 11 | 116432240 | C | T | 0.381 | 0.7029 | ++ | 0 | 0.005 | 1 | 0.9442 |
| rs11603831 | 11 | 120483858 | T | C | 0.381 | 0.7029 | +? | 0 | 0 | 0 | 1 |
| rs648682 | 11 | 118066582 | T | C | 0.381 | 0.7029 | +? | 0 | 0 | 0 | 1 |
| rs1628536 | 11 | 119963324 | G | A | 0.381 | 0.703 | +? | 0 | 0 | 0 | 1 |
| rs584653 | 11 | 119963001 | T | C | 0.381 | 0.703 | +? | 0 | 0 | 0 | 1 |
| rs605230 | 11 | 119962750 | C | T | 0.381 | 0.703 | +? | 0 | 0 | 0 | 1 |
| rs560919 | 11 | 120995842 | C | T | -0.381 | 0.7031 | -? | 0 | 0 | 0 | 1 |
| rs11216594 | 11 | 117736022 | A | G | 0.381 | 0.7032 | +? | 0 | 0 | 0 | 1 |
| rs2231439 | 11 | 118174885 | G | A | 0.381 | 0.7032 | +? | 0 | 0 | 0 | 1 |
| rs11217018 | 11 | 118635364 | C | T | -0.381 | 0.7033 | -+ | 0 | 0.199 | 1 | 0.6558 |
| rs11823543 | 11 | 116649135 | A | G | -0.38 | 0.7037 | +- | -62.1 | 1.851 | 1 | 0.1737 |
| rs681160 | 11 | 117304947 | C | T | 0.38 | 0.7038 | +? | 0 | 0 | 0 | 1 |
| rs73026322 | 11 | 117818505 | T | C | -0.38 | 0.7039 | -- | 0 | 0.037 | 1 | 0.847 |
| rs7925270 | 11 | 120563298 | T | C | -0.38 | 0.7039 | -? | 0 | 0 | 0 | 1 |
| rs17842763 | 11 | 118454177 | G | A | 0.38 | 0.7041 | +- | 0 | 0.252 | 1 | 0.6157 |
| rs7111646 | 11 | 120758479 | T | A | -0.38 | 0.7042 | -? | 0 | 0 | 0 | 1 |
| rs4457789 | 11 | 118539170 | T | G | 0.379 | 0.7044 | -+ | 0 | 0.862 | 1 | 0.3533 |
| rs537907 | 11 | 120078501 | C | G | 0.379 | 0.7045 | +? | 0 | 0 | 0 | 1 |
| rs540857 | 11 | 120078890 | G | A | 0.379 | 0.7045 | +? | 0 | 0 | 0 | 1 |
| rs2271249 | 11 | 119479391 | C | T | 0.379 | 0.7046 | +? | 0 | 0 | 0 | 1 |
| rs7116876 | 11 | 118371025 | C | G | -0.379 | 0.705 | -+ | -128.4 | 1.314 | 1 | 0.2517 |
| rs2155263 | 11 | 120606683 | A | G | 0.378 | 0.7053 | +? | 0 | 0 | 0 | 1 |
| rs117554512 | 11 | 120098329 | T | C | 0.378 | 0.7055 | +? | 0 | 0 | 0 | 1 |
| rs10892004 | 11 | 116514417 | T | C | -0.378 | 0.7056 | -? | 0 | 0 | 0 | 1 |
| rs595964 | 11 | 118026788 | A | G | -0.378 | 0.7058 | -- | 0 | 0 | 1 | 0.9902 |
| rs76690043 | 11 | 116801529 | G | C | 0.378 | 0.7058 | +? | 0 | 0 | 0 | 1 |
| rs7947144 | 11 | 117525255 | G | A | -0.378 | 0.7058 | -? | 0 | 0 | 0 | 1 |
| rs10892282 | 11 | 118627778 | C | G | -0.377 | 0.7061 | -+ | 0 | 0.2 | 1 | 0.6543 |
| rs605908 | 11 | 118067192 | C | A | 0.377 | 0.7062 | +? | 0 | 0 | 0 | 1 |
| rs617715 | 11 | 119961635 | A | T | 0.377 | 0.7062 | +? | 0 | 0 | 0 | 1 |
| rs4938423 | 11 | 117509797 | T | C | -0.377 | 0.7063 | -? | 0 | 0 | 0 | 1 |
| rs10892104 | 11 | 117309495 | A | G | -0.377 | 0.7064 | -? | 0 | 0 | 0 | 1 |
| rs77322791 | 11 | 116911590 | G | A | -0.377 | 0.7064 | -+ | 72 | 10.732 | 1 | 0.001053 |
| rs508023 | 11 | 116273569 | G | T | -0.377 | 0.7065 | -? | 0 | 0 | 0 | 1 |
| rs145133311 | 11 | 116886516 | T | C | -0.376 | 0.7066 | +- | 47.8 | 5.746 | 1 | 0.01652 |
| rs17120251 | 11 | 116923488 | C | T | 0.376 | 0.7066 | +? | 0 | 0 | 0 | 1 |
| rs584329 | 11 | 120994638 | T | C | -0.376 | 0.7066 | -? | 0 | 0 | 0 | 1 |
| rs659956 | 11 | 120994961 | A | G | -0.376 | 0.7066 | -? | 0 | 0 | 0 | 1 |
| rs75048292 | 11 | 116383348 | A | G | -0.376 | 0.7066 | -? | 0 | 0 | 0 | 1 |
| rs2846120 | 11 | 120605759 | T | C | 0.376 | 0.7067 | +? | 0 | 0 | 0 | 1 |
| rs2852230 | 11 | 120605736 | C | G | 0.376 | 0.7067 | +? | 0 | 0 | 0 | 1 |
| rs2845667 | 11 | 119422645 | T | C | 0.376 | 0.7068 | +? | 0 | 0 | 0 | 1 |
| rs7131676 | 11 | 119631325 | G | A | 0.376 | 0.7069 | +? | 0 | 0 | 0 | 1 |
| rs113950321 | 11 | 119609828 | C | A | 0.376 | 0.7071 | +? | 0 | 0 | 0 | 1 |
| rs77752432 | 11 | 117752769 | T | C | -0.375 | 0.7073 | -? | 0 | 0 | 0 | 1 |
| rs7105537 | 11 | 117808853 | A | G | 0.375 | 0.7074 | -+ | -35.2 | 2.219 | 1 | 0.1363 |
| rs2271250 | 11 | 119479520 | C | T | 0.375 | 0.7075 | +? | 0 | 0 | 0 | 1 |

| | | | | | | | | | | | |
|---|---|---|---|---|---|---|---|---|---|---|---|
| rs12277021 | 11 | 119452530 | C | T | -0.375 | 0.7077 | -? | 0 | 0 | 0 | 1 |
| rs2444239 | 11 | 120056416 | A | T | -0.374 | 0.7081 | -- | 0 | 0.289 | 1 | 0.5911 |
| rs73023341 | 11 | 118445585 | G | A | 0.374 | 0.7082 | +- | 0 | 0.275 | 1 | 0.6 |
| rs1939935 | 11 | 121040910 | C | T | -0.374 | 0.7083 | -? | 0 | 0 | 0 | 1 |
| rs2156637 | 11 | 120770778 | A | G | 0.374 | 0.7083 | +? | 0 | 0 | 0 | 1 |
| rs74890383 | 11 | 116428953 | A | G | 0.374 | 0.7084 | ++ | 0 | 0.093 | 1 | 0.7598 |
| rs78243160 | 11 | 116428591 | T | C | 0.374 | 0.7084 | ++ | 0 | 0.093 | 1 | 0.7598 |
| rs1786179 | 11 | 117914304 | A | T | -0.374 | 0.7085 | -? | 0 | 0 | 0 | 1 |
| rs1786180 | 11 | 117914297 | G | C | -0.374 | 0.7085 | -? | 0 | 0 | 0 | 1 |
| rs2155262 | 11 | 120610692 | A | G | 0.374 | 0.7085 | +? | 0 | 0 | 0 | 1 |
| rs10790399 | 11 | 120552067 | A | C | -0.374 | 0.7086 | -? | 0 | 0 | 0 | 1 |
| rs486394 | 11 | 116526322 | C | A | 0.374 | 0.7086 | +? | 0 | 0 | 0 | 1 |
| rs4936509 | 11 | 120041444 | C | T | -0.374 | 0.7086 | -? | 0 | 0 | 0 | 1 |
| rs17120029 | 11 | 116650118 | T | C | -0.374 | 0.7087 | +- | -61.2 | 1.862 | 1 | 0.1724 |
| rs1783853 | 11 | 117914550 | G | A | -0.374 | 0.7087 | -? | 0 | 0 | 0 | 1 |
| rs7924387 | 11 | 118954265 | A | G | -0.374 | 0.7087 | -? | 0 | 0 | 0 | 1 |
| rs34782343 | 11 | 118153511 | G | T | -0.373 | 0.7088 | -+ | 37.8 | 4.82 | 1 | 0.02814 |
| rs3016358 | 11 | 116478660 | A | G | 0.373 | 0.7091 | +? | 0 | 0 | 0 | 1 |
| rs7115634 | 11 | 118466249 | G | A | 0.373 | 0.7091 | +- | 0 | 0.15 | 1 | 0.6984 |
| rs11600715 | 11 | 116401884 | G | A | -0.373 | 0.7092 | -? | 0 | 0 | 0 | 1 |
| rs3753058 | 11 | 118221350 | T | G | -0.373 | 0.7092 | -- | 0 | 0.001 | 1 | 0.9745 |
| rs3753059 | 11 | 118221349 | C | T | -0.373 | 0.7092 | -- | 0 | 0.001 | 1 | 0.9745 |
| rs3753060 | 11 | 118221137 | T | C | -0.373 | 0.7092 | -- | 0 | 0.001 | 1 | 0.9745 |
| rs7103514 | 11 | 117820561 | G | A | -0.373 | 0.7092 | -- | 0 | 0.002 | 1 | 0.9643 |
| rs508025 | 11 | 116273570 | A | T | -0.373 | 0.7093 | -? | 0 | 0 | 0 | 1 |
| rs668400 | 11 | 119901095 | A | C | 0.372 | 0.7098 | +? | 0 | 0 | 0 | 1 |
| rs873713 | 11 | 117739916 | T | A | -0.372 | 0.7099 | -? | 0 | 0 | 0 | 1 |
| rs11216909 | 11 | 118449746 | T | C | 0.372 | 0.71 | +- | 0 | 0.217 | 1 | 0.6411 |
| rs4938482 | 11 | 117973218 | G | T | -0.372 | 0.7101 | -- | 0 | 0 | 1 | 0.9911 |
| rs61901067 | 11 | 119457720 | G | A | 0.371 | 0.7103 | +? | 0 | 0 | 0 | 1 |
| rs567610 | 11 | 116317895 | T | C | -0.371 | 0.7104 | -? | 0 | 0 | 0 | 1 |
| rs661189 | 11 | 120996969 | G | C | -0.371 | 0.7104 | -? | 0 | 0 | 0 | 1 |
| rs12362410 | 11 | 116422860 | A | G | 0.371 | 0.7105 | -+ | 0 | 0.463 | 1 | 0.496 |
| rs71484178 | 11 | 120396030 | A | G | 0.371 | 0.7109 | +? | 0 | 0 | 0 | 1 |
| rs80094129 | 11 | 121103437 | A | C | -0.371 | 0.711 | -? | 0 | 0 | 0 | 1 |
| rs76982250 | 11 | 116915242 | T | C | -0.37 | 0.7111 | -+ | 69.9 | 9.978 | 1 | 0.001584 |
| rs12365194 | 11 | 120541263 | A | G | -0.37 | 0.7114 | -? | 0 | 0 | 0 | 1 |
| rs34492753 | 11 | 119660492 | T | C | -0.37 | 0.7116 | -? | 0 | 0 | 0 | 1 |
| rs4938479 | 11 | 117956971 | A | G | 0.37 | 0.7116 | +- | 0 | 0.161 | 1 | 0.6886 |
| rs543290 | 11 | 117770998 | C | A | -0.37 | 0.7116 | -? | 0 | 0 | 0 | 1 |
| rs11218190 | 11 | 121053640 | A | G | -0.369 | 0.7118 | -? | 0 | 0 | 0 | 1 |
| rs117683585 | 11 | 117125769 | G | A | -0.369 | 0.7123 | -? | 0 | 0 | 0 | 1 |
| rs12574216 | 11 | 116239385 | G | T | -0.369 | 0.7123 | -- | 0 | 0.392 | 1 | 0.5313 |
| rs75133657 | 11 | 121184472 | C | A | 0.369 | 0.7123 | +? | 0 | 0 | 0 | 1 |
| rs2663623 | 11 | 119427624 | A | G | 0.369 | 0.7124 | +? | 0 | 0 | 0 | 1 |
| rs2845661 | 11 | 119427893 | T | G | 0.369 | 0.7124 | +? | 0 | 0 | 0 | 1 |
| rs11216906 | 11 | 118443944 | A | G | 0.368 | 0.7126 | +- | 0 | 0.272 | 1 | 0.6023 |
| rs117053908 | 11 | 117407275 | T | G | 0.368 | 0.7126 | +? | 0 | 0 | 0 | 1 |
| rs4936407 | 11 | 117638627 | T | C | 0.368 | 0.7126 | +? | 0 | 0 | 0 | 1 |
| rs56286911 | 11 | 116493201 | T | C | -0.368 | 0.7126 | -? | 0 | 0 | 0 | 1 |
| rs2846124 | 11 | 120604347 | C | A | 0.368 | 0.7127 | +? | 0 | 0 | 0 | 1 |
| rs647573 | 11 | 118067132 | C | T | 0.368 | 0.7129 | +? | 0 | 0 | 0 | 1 |
| rs11819822 | 11 | 117335165 | T | C | -0.367 | 0.7134 | -? | 0 | 0 | 0 | 1 |
| rs2276421 | 11 | 118239266 | A | G | -0.367 | 0.7134 | -- | 0 | 0.001 | 1 | 0.9721 |
| rs10892409 | 11 | 119451111 | C | A | -0.367 | 0.7135 | -? | 0 | 0 | 0 | 1 |
| rs723953 | 11 | 116924978 | C | A | 0.367 | 0.7135 | +? | 0 | 0 | 0 | 1 |

| | | | | | | | | | | | |
|---|---|---|---|---|---|---|---|---|---|---|---|
| rs624328 | 11 | 118035087 | G | A | 0.367 | 0.7136 | +? | 0 | 0 | 0 | 1 |
| rs7124498 | 11 | 118616253 | A | G | -0.367 | 0.7136 | -+ | 0 | 0.245 | 1 | 0.6204 |
| rs1605855 | 11 | 119418178 | A | C | 0.367 | 0.7139 | +? | 0 | 0 | 0 | 1 |
| rs12420596 | 11 | 116462788 | A | G | -0.366 | 0.714 | -? | 0 | 0 | 0 | 1 |
| rs11216421 | 11 | 117355493 | C | G | -0.366 | 0.7141 | -? | 0 | 0 | 0 | 1 |
| rs1629201 | 11 | 119967980 | C | T | 0.366 | 0.7141 | +? | 0 | 0 | 0 | 1 |
| rs12417015 | 11 | 116550776 | G | A | 0.366 | 0.7142 | +? | 0 | 0 | 0 | 1 |
| rs17119495 | 11 | 116224775 | C | T | -0.366 | 0.7143 | +- | -130.3 | 1.302 | 1 | 0.2538 |
| rs73022410 | 11 | 117785742 | C | G | -0.366 | 0.7143 | -? | 0 | 0 | 0 | 1 |
| rs75204992 | 11 | 117784902 | G | A | -0.366 | 0.7145 | -? | 0 | 0 | 0 | 1 |
| rs562993 | 11 | 119746467 | A | T | 0.366 | 0.7146 | +? | 0 | 0 | 0 | 1 |
| rs600484 | 11 | 119967026 | A | T | 0.366 | 0.7146 | +? | 0 | 0 | 0 | 1 |
| rs610075 | 11 | 119967383 | T | G | 0.366 | 0.7146 | +? | 0 | 0 | 0 | 1 |
| rs611656 | 11 | 119967247 | T | C | 0.366 | 0.7146 | +? | 0 | 0 | 0 | 1 |
| rs11216031 | 11 | 116466289 | C | A | 0.366 | 0.7147 | +? | 0 | 0 | 0 | 1 |
| rs112995279 | 11 | 116365711 | A | C | 0.366 | 0.7147 | +? | 0 | 0 | 0 | 1 |
| rs72645462 | 11 | 117293032 | A | T | 0.366 | 0.7147 | +? | 0 | 0 | 0 | 1 |
| rs150325557 | 11 | 117853333 | C | G | 0.365 | 0.7152 | +? | 0 | 0 | 0 | 1 |
| rs55844025 | 11 | 118446298 | T | C | 0.365 | 0.7152 | +- | 0 | 0.138 | 1 | 0.7107 |
| rs112319461 | 11 | 116493155 | A | G | -0.364 | 0.7156 | -? | 0 | 0 | 0 | 1 |
| rs648947 | 11 | 118067426 | A | G | 0.364 | 0.7156 | +? | 0 | 0 | 0 | 1 |
| rs118135828 | 11 | 119897773 | A | G | 0.364 | 0.7157 | +? | 0 | 0 | 0 | 1 |
| rs181124100 | 11 | 116464465 | T | C | 0.364 | 0.7157 | +? | 0 | 0 | 0 | 1 |
| rs592765 | 11 | 119897774 | A | C | 0.364 | 0.7157 | +? | 0 | 0 | 0 | 1 |
| rs61592631 | 11 | 116533684 | T | C | 0.364 | 0.7157 | +? | 0 | 0 | 0 | 1 |
| rs2252771 | 11 | 120414851 | G | C | -0.364 | 0.7158 | -? | 0 | 0 | 0 | 1 |
| rs4331108 | 11 | 120490713 | T | C | 0.364 | 0.7158 | +? | 0 | 0 | 0 | 1 |
| rs549406 | 11 | 119711982 | C | T | 0.364 | 0.7158 | +? | 0 | 0 | 0 | 1 |
| rs113816823 | 11 | 119612600 | G | A | 0.364 | 0.7159 | +? | 0 | 0 | 0 | 1 |
| rs607142 | 11 | 120831878 | T | C | -0.364 | 0.716 | -+ | 0 | 0.336 | 1 | 0.5622 |
| rs7106140 | 11 | 120556557 | T | G | -0.364 | 0.716 | -? | 0 | 0 | 0 | 1 |
| rs7106261 | 11 | 120556641 | A | G | -0.364 | 0.716 | -? | 0 | 0 | 0 | 1 |
| rs11217857 | 11 | 120270170 | G | A | 0.364 | 0.7161 | +? | 0 | 0 | 0 | 1 |
| rs3824943 | 11 | 117695340 | T | C | 0.364 | 0.7162 | +? | 0 | 0 | 0 | 1 |
| rs4537768 | 11 | 121192985 | T | G | -0.363 | 0.7165 | -? | 0 | 0 | 0 | 1 |
| rs2946742 | 11 | 118017093 | G | A | 0.363 | 0.7166 | ++ | 0 | 0.044 | 1 | 0.8332 |
| rs4080140 | 11 | 116676590 | A | G | 0.363 | 0.7166 | +? | 0 | 0 | 0 | 1 |
| rs56220308 | 11 | 119451380 | C | G | -0.363 | 0.7166 | -? | 0 | 0 | 0 | 1 |
| rs668021 | 11 | 120879655 | C | G | 0.363 | 0.7166 | +? | 0 | 0 | 0 | 1 |
| rs7939622 | 11 | 118087841 | A | G | -0.363 | 0.7169 | -+ | 0 | 0.085 | 1 | 0.7704 |
| rs78751065 | 11 | 117171413 | T | C | -0.362 | 0.717 | -? | 0 | 0 | 0 | 1 |
| rs10892360 | 11 | 119268984 | A | G | -0.362 | 0.7174 | -? | 0 | 0 | 0 | 1 |
| rs4565903 | 11 | 120497375 | G | C | 0.362 | 0.7174 | +? | 0 | 0 | 0 | 1 |
| rs117794242 | 11 | 120712972 | T | C | -0.362 | 0.7175 | -? | 0 | 0 | 0 | 1 |
| rs609650 | 11 | 119967484 | C | T | 0.362 | 0.7175 | +? | 0 | 0 | 0 | 1 |
| rs1629347 | 11 | 119967918 | A | C | 0.362 | 0.7177 | +? | 0 | 0 | 0 | 1 |
| rs1670196 | 11 | 119968598 | C | T | 0.362 | 0.7177 | +? | 0 | 0 | 0 | 1 |
| rs1670197 | 11 | 119968367 | C | G | 0.362 | 0.7177 | +? | 0 | 0 | 0 | 1 |
| rs1670198 | 11 | 119968311 | C | T | 0.362 | 0.7177 | +? | 0 | 0 | 0 | 1 |
| rs1787025 | 11 | 119968495 | G | T | 0.362 | 0.7177 | +? | 0 | 0 | 0 | 1 |
| rs1787026 | 11 | 119968392 | C | G | 0.362 | 0.7177 | +? | 0 | 0 | 0 | 1 |
| rs1790472 | 11 | 119967930 | G | A | 0.362 | 0.7177 | +? | 0 | 0 | 0 | 1 |
| rs639775 | 11 | 119968506 | G | C | 0.362 | 0.7177 | +? | 0 | 0 | 0 | 1 |
| rs12279393 | 11 | 117554747 | C | T | 0.361 | 0.7179 | +? | 0 | 0 | 0 | 1 |
| rs200069252 | 11 | 118354750 | A | T | -0.361 | 0.718 | -+ | -142.2 | 1.239 | 1 | 0.2657 |
| rs7118350 | 11 | 119706311 | G | C | -0.361 | 0.718 | -? | 0 | 0 | 0 | 1 |

| | | | | | | | | | | | |
|---|---|---|---|---|---|---|---|---|---|---|---|
| rs873059 | 11 | 120097498 | T | C | -0.361 | 0.718 | -? | 0 | 0 | 0 | 1 |
| rs55728331 | 11 | 117526440 | A | G | 0.361 | 0.7181 | +? | 0 | 0 | 0 | 1 |
| rs58950690 | 11 | 116543098 | C | T | 0.361 | 0.7183 | +? | 0 | 0 | 0 | 1 |
| rs11217021 | 11 | 118640828 | C | T | -0.361 | 0.7184 | -+ | 0 | 0.187 | 1 | 0.6652 |
| rs1320656 | 11 | 117739122 | C | A | 0.361 | 0.7184 | +? | 0 | 0 | 0 | 1 |
| rs7129267 | 11 | 121049690 | G | T | -0.361 | 0.7184 | -? | 0 | 0 | 0 | 1 |
| rs501763 | 11 | 116260707 | G | A | -0.36 | 0.7185 | -? | 0 | 0 | 0 | 1 |
| rs951657 | 11 | 118067621 | A | G | 0.36 | 0.7186 | +? | 0 | 0 | 0 | 1 |
| rs10790226 | 11 | 117806937 | T | C | 0.36 | 0.7187 | -+ | -32.5 | 2.264 | 1 | 0.1324 |
| rs504148 | 11 | 121003514 | T | C | -0.36 | 0.7188 | -? | 0 | 0 | 0 | 1 |
| rs616120 | 11 | 121003046 | A | G | -0.36 | 0.7188 | -? | 0 | 0 | 0 | 1 |
| rs74337439 | 11 | 118626825 | A | G | -0.36 | 0.7188 | -+ | -37.7 | 2.178 | 1 | 0.14 |
| rs2276418 | 11 | 118230264 | T | A | -0.36 | 0.7189 | -- | 0 | 0 | 1 | 0.9968 |
| rs7105943 | 11 | 120556610 | A | C | -0.36 | 0.719 | -? | 0 | 0 | 0 | 1 |
| rs11216921 | 11 | 118464815 | C | G | 0.36 | 0.7191 | +- | 0 | 0.158 | 1 | 0.691 |
| rs3181259 | 11 | 118214067 | A | G | -0.36 | 0.7191 | -- | 0 | 0.006 | 1 | 0.9378 |
| rs11217013 | 11 | 118624919 | C | G | -0.36 | 0.7192 | -+ | 0 | 0.227 | 1 | 0.6339 |
| rs180367 | 11 | 116594715 | G | A | -0.359 | 0.7193 | -+ | -7.8 | 2.783 | 1 | 0.0953 |
| rs1893900 | 11 | 117436906 | T | C | -0.359 | 0.7193 | -? | 0 | 0 | 0 | 1 |
| rs58777850 | 11 | 117317593 | C | T | 0.359 | 0.7193 | +? | 0 | 0 | 0 | 1 |
| rs11216153 | 11 | 116705100 | T | G | 0.359 | 0.7195 | +? | 0 | 0 | 0 | 1 |
| rs9332816 | 11 | 118364404 | T | G | -0.359 | 0.7195 | -+ | -155.1 | 1.176 | 1 | 0.2782 |
| rs10790217 | 11 | 117732028 | G | A | 0.359 | 0.7196 | +? | 0 | 0 | 0 | 1 |
| rs117323348 | 11 | 121193894 | G | C | -0.359 | 0.7196 | -? | 0 | 0 | 0 | 1 |
| rs2155264 | 11 | 120606577 | G | C | 0.359 | 0.7196 | +? | 0 | 0 | 0 | 1 |
| rs73588424 | 11 | 116646107 | G | A | -0.359 | 0.7197 | +- | -59.2 | 1.885 | 1 | 0.1698 |
| rs12279777 | 11 | 119451188 | T | A | -0.359 | 0.7198 | -? | 0 | 0 | 0 | 1 |
| rs10750099 | 11 | 116788699 | A | C | -0.359 | 0.7199 | +- | 45.7 | 5.526 | 1 | 0.01873 |
| rs12277899 | 11 | 119450113 | T | A | -0.359 | 0.7199 | -? | 0 | 0 | 0 | 1 |
| rs55801969 | 11 | 119841751 | G | A | -0.358 | 0.7202 | -? | 0 | 0 | 0 | 1 |
| rs80137897 | 11 | 116954760 | C | T | -0.358 | 0.7202 | +- | 48.1 | 5.779 | 1 | 0.01622 |
| rs622350 | 11 | 119883569 | A | C | 0.358 | 0.7205 | +? | 0 | 0 | 0 | 1 |
| rs71482129 | 11 | 118003694 | A | G | 0.358 | 0.7205 | ++ | 0 | 0.004 | 1 | 0.9468 |
| rs3751020 | 11 | 119479052 | C | T | 0.358 | 0.7207 | +? | 0 | 0 | 0 | 1 |
| rs651055 | 11 | 119848156 | C | G | -0.357 | 0.7208 | -? | 0 | 0 | 0 | 1 |
| rs654884 | 11 | 119967159 | T | C | 0.357 | 0.7208 | +? | 0 | 0 | 0 | 1 |
| rs12802920 | 11 | 116412160 | C | A | 0.357 | 0.7209 | ++ | 0 | 0.308 | 1 | 0.5789 |
| rs112508300 | 11 | 118385249 | G | A | 0.357 | 0.7213 | +- | 0 | 0.048 | 1 | 0.8264 |
| rs61897871 | 11 | 119875928 | G | A | -0.357 | 0.7213 | -? | 0 | 0 | 0 | 1 |
| rs629345 | 11 | 119006722 | C | T | 0.357 | 0.7213 | +? | 0 | 0 | 0 | 1 |
| rs637464 | 11 | 119968997 | G | T | 0.357 | 0.7213 | +? | 0 | 0 | 0 | 1 |
| rs10750116 | 11 | 117743822 | C | G | 0.356 | 0.7216 | +? | 0 | 0 | 0 | 1 |
| rs58270621 | 11 | 118413604 | A | G | 0.356 | 0.7216 | ++ | 0 | 0.019 | 1 | 0.8902 |
| rs12362262 | 11 | 119951702 | T | C | 0.356 | 0.7218 | +? | 0 | 0 | 0 | 1 |
| rs10790342 | 11 | 119630418 | A | G | 0.356 | 0.7219 | +? | 0 | 0 | 0 | 1 |
| rs12295555 | 11 | 119837503 | C | T | -0.356 | 0.7219 | -? | 0 | 0 | 0 | 1 |
| rs34006113 | 11 | 119838017 | T | G | -0.356 | 0.7219 | -? | 0 | 0 | 0 | 1 |
| rs56785719 | 11 | 118176623 | G | C | 0.356 | 0.722 | +? | 0 | 0 | 0 | 1 |
| rs11217505 | 11 | 119658670 | C | G | -0.356 | 0.7221 | -? | 0 | 0 | 0 | 1 |
| rs7931564 | 11 | 118469571 | G | A | 0.356 | 0.7222 | +- | 0 | 0.262 | 1 | 0.6088 |
| rs1783935 | 11 | 118539566 | T | A | 0.355 | 0.7223 | -+ | 0 | 0.931 | 1 | 0.3347 |
| rs10502219 | 11 | 116412613 | A | G | 0.355 | 0.7226 | -+ | 0 | 0.395 | 1 | 0.5295 |
| rs7110526 | 11 | 118633533 | C | T | -0.355 | 0.7226 | -+ | 0 | 0.186 | 1 | 0.6666 |
| rs1784298 | 11 | 118548665 | C | T | 0.355 | 0.7228 | -+ | 0 | 0.953 | 1 | 0.3289 |
| rs4938522 | 11 | 118548671 | T | A | 0.355 | 0.7228 | -+ | 0 | 0.953 | 1 | 0.3289 |
| rs4938523 | 11 | 118548838 | T | G | 0.355 | 0.7228 | -+ | 0 | 0.953 | 1 | 0.3289 |

| | | | | | | | | | | | |
|---|---|---|---|---|---|---|---|---|---|---|---|
| rs180366 | 11 | 116595756 | T | C | -0.355 | 0.7229 | -+ | -8.2 | 2.773 | 1 | 0.09584 |
| rs10892122 | 11 | 117353773 | A | G | -0.354 | 0.723 | -? | 0 | 0 | 0 | 1 |
| rs11216419 | 11 | 117353286 | A | G | -0.354 | 0.723 | -? | 0 | 0 | 0 | 1 |
| rs12785387 | 11 | 120221987 | T | C | 0.354 | 0.723 | +? | 0 | 0 | 0 | 1 |
| rs17092756 | 11 | 118457630 | C | G | 0.354 | 0.723 | +- | 0 | 0.312 | 1 | 0.5762 |
| rs2715882 | 11 | 120133153 | T | C | -0.354 | 0.7231 | -? | 0 | 0 | 0 | 1 |
| rs11601584 | 11 | 116571284 | C | T | 0.354 | 0.7232 | +? | 0 | 0 | 0 | 1 |
| rs12795615 | 11 | 118214562 | T | C | -0.354 | 0.7232 | -- | 0 | 0 | 1 | 0.9827 |
| rs74835404 | 11 | 117645002 | A | G | -0.354 | 0.7232 | +- | 12.3 | 3.42 | 1 | 0.0644 |
| rs2509623 | 11 | 120109967 | T | C | -0.354 | 0.7233 | -? | 0 | 0 | 0 | 1 |
| rs11217601 | 11 | 119829572 | C | A | 0.354 | 0.7234 | +? | 0 | 0 | 0 | 1 |
| rs1786183 | 11 | 118017445 | A | G | 0.354 | 0.7234 | ++ | 0 | 0.08 | 1 | 0.7778 |
| rs112910608 | 11 | 118637475 | T | C | -0.354 | 0.7237 | -+ | 0 | 0.182 | 1 | 0.6697 |
| rs34405565 | 11 | 118233424 | T | C | -0.354 | 0.7237 | -- | 0 | 0 | 1 | 0.9884 |
| rs3741311 | 11 | 117989554 | G | A | -0.354 | 0.7237 | -- | 0 | 0 | 1 | 0.9825 |
| rs573211 | 11 | 120044762 | A | G | -0.354 | 0.7237 | -? | 0 | 0 | 0 | 1 |
| rs11216917 | 11 | 118462160 | G | T | 0.353 | 0.7238 | +- | 0 | 0.253 | 1 | 0.6148 |
| rs11218028 | 11 | 120717000 | T | G | 0.353 | 0.7238 | +? | 0 | 0 | 0 | 1 |
| rs601634 | 11 | 116668511 | T | C | -0.353 | 0.7238 | -+ | 2.7 | 3.084 | 1 | 0.07907 |
| rs7928092 | 11 | 117991403 | G | C | -0.353 | 0.7239 | -- | 0 | 0 | 1 | 0.9833 |
| rs117913812 | 11 | 119773797 | C | T | 0.353 | 0.724 | +? | 0 | 0 | 0 | 1 |
| rs73615909 | 11 | 118322488 | T | C | -0.353 | 0.724 | -+ | 0 | 0.702 | 1 | 0.4021 |
| rs75819084 | 11 | 117576946 | T | C | 0.353 | 0.7242 | +? | 0 | 0 | 0 | 1 |
| rs10750167 | 11 | 119834547 | G | A | -0.353 | 0.7243 | -? | 0 | 0 | 0 | 1 |
| rs1799991 | 11 | 118957091 | A | G | -0.353 | 0.7243 | -? | 0 | 0 | 0 | 1 |
| rs4505099 | 11 | 117848645 | G | A | -0.353 | 0.7243 | +- | 0 | 0.451 | 1 | 0.5018 |
| rs9332801 | 11 | 118355642 | C | A | -0.353 | 0.7243 | -+ | -146.5 | 1.217 | 1 | 0.2699 |
| rs11217164 | 11 | 119008538 | A | G | 0.353 | 0.7244 | +? | 0 | 0 | 0 | 1 |
| rs7938794 | 11 | 117990403 | T | G | -0.352 | 0.7246 | -- | 0 | 0 | 1 | 0.9829 |
| rs948030 | 11 | 120615801 | C | G | 0.352 | 0.7246 | +? | 0 | 0 | 0 | 1 |
| rs948108 | 11 | 117738911 | A | G | 0.352 | 0.7246 | +? | 0 | 0 | 0 | 1 |
| rs17121760 | 11 | 117988797 | A | G | 0.352 | 0.7247 | +? | 0 | 0 | 0 | 1 |
| rs4938713 | 11 | 119568994 | T | C | -0.352 | 0.7247 | -? | 0 | 0 | 0 | 1 |
| rs73596628 | 11 | 117572183 | C | T | -0.352 | 0.7248 | -? | 0 | 0 | 0 | 1 |
| rs79717347 | 11 | 117721360 | G | T | -0.352 | 0.7249 | -? | 0 | 0 | 0 | 1 |
| rs11216521 | 11 | 117614122 | A | G | 0.352 | 0.725 | +? | 0 | 0 | 0 | 1 |
| rs7116903 | 11 | 117807414 | T | C | 0.352 | 0.725 | -+ | -42.9 | 2.099 | 1 | 0.1474 |
| rs117625924 | 11 | 116234916 | C | A | -0.352 | 0.7251 | -? | 0 | 0 | 0 | 1 |
| rs11824398 | 11 | 117481108 | A | G | 0.352 | 0.7252 | +? | 0 | 0 | 0 | 1 |
| rs2845659 | 11 | 119434282 | G | A | 0.352 | 0.7252 | +? | 0 | 0 | 0 | 1 |
| rs4938283 | 11 | 116432791 | T | C | 0.352 | 0.7252 | ++ | 0 | 0.277 | 1 | 0.5986 |
| rs11216025 | 11 | 116464111 | T | C | -0.351 | 0.7254 | -? | 0 | 0 | 0 | 1 |
| rs72994637 | 11 | 119828457 | A | G | 0.351 | 0.7254 | +? | 0 | 0 | 0 | 1 |
| rs694005 | 11 | 117464173 | T | C | -0.351 | 0.7258 | -? | 0 | 0 | 0 | 1 |
| rs45474998 | 11 | 118229679 | A | G | -0.351 | 0.7259 | -- | 0 | 0 | 1 | 0.9944 |
| rs526597 | 11 | 118546602 | G | A | 0.35 | 0.726 | -+ | 0 | 0.925 | 1 | 0.336 |
| rs76383990 | 11 | 116897871 | G | A | -0.35 | 0.726 | -+ | 70.6 | 10.198 | 1 | 0.001406 |
| rs2239695 | 11 | 118213632 | T | C | -0.35 | 0.7261 | -- | 0 | 0.007 | 1 | 0.9335 |
| rs10892387 | 11 | 119413149 | G | A | 0.35 | 0.7262 | +? | 0 | 0 | 0 | 1 |
| rs622860 | 11 | 117468794 | G | A | 0.35 | 0.7262 | +? | 0 | 0 | 0 | 1 |
| rs112701434 | 11 | 116797442 | C | T | -0.35 | 0.7264 | -? | 0 | 0 | 0 | 1 |
| rs1623797 | 11 | 116560447 | T | C | 0.35 | 0.7265 | +? | 0 | 0 | 0 | 1 |
| rs536031 | 11 | 117451094 | A | T | 0.35 | 0.7265 | +? | 0 | 0 | 0 | 1 |
| rs11216502 | 11 | 117557057 | C | G | 0.35 | 0.7266 | +? | 0 | 0 | 0 | 1 |
| rs688893 | 11 | 120830625 | C | T | -0.35 | 0.7266 | -+ | 0 | 0.327 | 1 | 0.5677 |
| rs73576686 | 11 | 116970006 | A | T | -0.35 | 0.7266 | +- | 54 | 6.519 | 1 | 0.01067 |

| | | | | | | | | | | | |
|---|---|---|---|---|---|---|---|---|---|---|---|
| rs144611969 | 11 | 119452937 | A | G | 0.35 | 0.7267 | +? | 0 | 0 | 0 | 1 |
| rs117235919 | 11 | 121093301 | A | G | 0.349 | 0.7268 | +? | 0 | 0 | 0 | 1 |
| rs2511857 | 11 | 119141812 | A | T | -0.349 | 0.7268 | -? | 0 | 0 | 0 | 1 |
| rs4938780 | 11 | 119976805 | T | C | 0.349 | 0.7268 | +? | 0 | 0 | 0 | 1 |
| rs7946597 | 11 | 119976523 | A | G | 0.349 | 0.7269 | +? | 0 | 0 | 0 | 1 |
| rs57438673 | 11 | 118459039 | A | G | 0.349 | 0.727 | +- | 0 | 0.268 | 1 | 0.6048 |
| rs11824032 | 11 | 120224789 | A | G | 0.349 | 0.7271 | +? | 0 | 0 | 0 | 1 |
| rs1715443 | 11 | 118017365 | G | T | 0.349 | 0.7273 | ++ | 0 | 0.032 | 1 | 0.8572 |
| rs617290 | 11 | 119006462 | G | A | 0.349 | 0.7273 | +? | 0 | 0 | 0 | 1 |
| rs11216645 | 11 | 117819863 | A | G | -0.349 | 0.7274 | -- | 0 | 0.044 | 1 | 0.8329 |
| rs4936488 | 11 | 119479922 | A | G | 0.349 | 0.7274 | +? | 0 | 0 | 0 | 1 |
| rs12787202 | 11 | 118226011 | T | C | -0.349 | 0.7275 | -- | 0 | 0.001 | 1 | 0.9809 |
| rs6589615 | 11 | 117455699 | C | G | 0.348 | 0.7275 | +? | 0 | 0 | 0 | 1 |
| rs9332793 | 11 | 118354458 | A | G | -0.348 | 0.7275 | -+ | -132.8 | 1.289 | 1 | 0.2563 |
| rs9332794 | 11 | 118354505 | T | G | -0.348 | 0.7275 | -+ | -132.8 | 1.289 | 1 | 0.2563 |
| rs10790396 | 11 | 120551561 | A | G | -0.348 | 0.7277 | -? | 0 | 0 | 0 | 1 |
| rs11216643 | 11 | 117818843 | A | T | -0.348 | 0.7277 | -- | 0 | 0.045 | 1 | 0.8327 |
| rs516655 | 11 | 117709028 | G | A | -0.348 | 0.7277 | -? | 0 | 0 | 0 | 1 |
| rs541612 | 11 | 119739796 | T | C | 0.348 | 0.7277 | +? | 0 | 0 | 0 | 1 |
| rs11216916 | 11 | 118461605 | C | T | 0.348 | 0.7278 | +- | 0 | 0.25 | 1 | 0.6169 |
| rs182784500 | 11 | 121071161 | G | A | -0.348 | 0.7278 | -? | 0 | 0 | 0 | 1 |
| rs2077623 | 11 | 120607495 | T | C | 0.348 | 0.728 | +? | 0 | 0 | 0 | 1 |
| rs505824 | 11 | 117447031 | T | C | 0.348 | 0.728 | +? | 0 | 0 | 0 | 1 |
| rs3949025 | 11 | 117954724 | A | G | 0.348 | 0.7281 | ++ | 0 | 0.262 | 1 | 0.609 |
| rs180374 | 11 | 116591418 | A | G | -0.348 | 0.7282 | -+ | -8.7 | 2.76 | 1 | 0.09665 |
| rs487177 | 11 | 118657297 | G | C | -0.348 | 0.7282 | -+ | 0 | 0.159 | 1 | 0.6898 |
| rs11601324 | 11 | 117731269 | C | G | -0.347 | 0.7283 | -? | 0 | 0 | 0 | 1 |
| rs116863015 | 11 | 120123837 | A | G | 0.347 | 0.7284 | +? | 0 | 0 | 0 | 1 |
| rs1613643 | 11 | 118068521 | T | G | 0.347 | 0.7285 | +? | 0 | 0 | 0 | 1 |
| rs61407825 | 11 | 117331806 | T | C | -0.347 | 0.7285 | -? | 0 | 0 | 0 | 1 |
| rs3863314 | 11 | 117313086 | A | G | 0.347 | 0.7286 | +? | 0 | 0 | 0 | 1 |
| rs4938795 | 11 | 120184592 | A | G | -0.347 | 0.7286 | -? | 0 | 0 | 0 | 1 |
| rs58068870 | 11 | 116948368 | C | T | -0.347 | 0.7286 | -+ | 70.7 | 10.238 | 1 | 0.001376 |
| rs948029 | 11 | 120615697 | T | A | 0.347 | 0.7286 | +? | 0 | 0 | 0 | 1 |
| rs56104144 | 11 | 118666289 | C | G | -0.347 | 0.7287 | -+ | 0 | 0.059 | 1 | 0.8086 |
| rs77320801 | 11 | 116836349 | T | C | 0.347 | 0.7287 | +? | 0 | 0 | 0 | 1 |
| rs10750157 | 11 | 119440533 | T | C | -0.347 | 0.7288 | -? | 0 | 0 | 0 | 1 |
| rs1892974 | 11 | 120614390 | T | A | 0.347 | 0.7288 | +? | 0 | 0 | 0 | 1 |
| rs11216392 | 11 | 117313174 | T | C | -0.347 | 0.7289 | -? | 0 | 0 | 0 | 1 |
| rs10892662 | 11 | 120896545 | T | G | -0.346 | 0.7292 | -? | 0 | 0 | 0 | 1 |
| rs12289253 | 11 | 118405068 | G | A | 0.346 | 0.7293 | +- | 0 | 0.041 | 1 | 0.8388 |
| rs1509464 | 11 | 119425807 | C | T | 0.346 | 0.7293 | +? | 0 | 0 | 0 | 1 |
| rs12286037 | 11 | 116652207 | T | C | -0.346 | 0.7294 | +- | -57.5 | 1.905 | 1 | 0.1675 |
| rs642280 | 11 | 120833767 | C | T | -0.346 | 0.7294 | -+ | 0 | 0.324 | 1 | 0.5691 |
| rs880615 | 11 | 117303393 | T | C | -0.346 | 0.7294 | -? | 0 | 0 | 0 | 1 |
| rs11600193 | 11 | 119607719 | C | T | 0.346 | 0.7295 | +? | 0 | 0 | 0 | 1 |
| rs4938485 | 11 | 117992620 | G | A | -0.346 | 0.7295 | -- | 0 | 0.001 | 1 | 0.9799 |
| rs17123485 | 11 | 119792173 | C | T | -0.346 | 0.7296 | -? | 0 | 0 | 0 | 1 |
| rs656382 | 11 | 119841515 | T | C | 0.346 | 0.7296 | +? | 0 | 0 | 0 | 1 |
| rs7105596 | 11 | 119656950 | C | T | 0.346 | 0.7296 | ++ | 0 | 0.012 | 1 | 0.9139 |
| rs1376996225 | 11 | 121192989 | T | G | -0.346 | 0.7297 | -? | 0 | 0 | 0 | 1 |
| rs607527 | 11 | 118538187 | T | G | 0.345 | 0.7298 | -+ | 0 | 0.862 | 1 | 0.3533 |
| rs12794546 | 11 | 121062593 | T | C | -0.345 | 0.73 | -? | 0 | 0 | 0 | 1 |
| rs2508991 | 11 | 118539522 | A | G | 0.345 | 0.73 | -+ | 0 | 0.898 | 1 | 0.3432 |
| rs495473 | 11 | 117212908 | G | A | 0.345 | 0.73 | +? | 0 | 0 | 0 | 1 |
| rs7110197 | 11 | 118228624 | C | G | -0.345 | 0.7301 | -- | 0 | 0 | 1 | 0.997 |

| | | | | | | | | | | | |
|---|---|---|---|---|---|---|---|---|---|---|---|
| rs61907602 | 11 | 116810655 | C | G | -0.345 | 0.7302 | -? | 0 | 0 | 0 | 1 |
| rs7941912 | 11 | 116815762 | A | G | -0.345 | 0.7302 | -? | 0 | 0 | 0 | 1 |
| rs4938281 | 11 | 116430875 | G | A | 0.345 | 0.7303 | +? | 0 | 0 | 0 | 1 |
| rs61747468 | 11 | 119881864 | A | G | -0.345 | 0.7303 | -? | 0 | 0 | 0 | 1 |
| rs11216914 | 11 | 118456414 | C | T | 0.345 | 0.7304 | +- | 0 | 0.271 | 1 | 0.6024 |
| rs3782045 | 11 | 118233078 | G | A | -0.345 | 0.7304 | -- | 0 | 0 | 1 | 0.9861 |
| rs1073388 | 11 | 120611792 | G | C | 0.344 | 0.7305 | +? | 0 | 0 | 0 | 1 |
| rs2187495 | 11 | 120611308 | T | C | 0.344 | 0.7305 | +? | 0 | 0 | 0 | 1 |
| rs45534632 | 11 | 118840705 | G | A | 0.344 | 0.7305 | +? | 0 | 0 | 0 | 1 |
| rs55736521 | 11 | 118840506 | T | C | 0.344 | 0.7305 | +? | 0 | 0 | 0 | 1 |
| rs620168 | 11 | 119012321 | G | C | 0.344 | 0.7306 | +? | 0 | 0 | 0 | 1 |
| rs11216006 | 11 | 116432066 | T | C | 0.344 | 0.7307 | ++ | 0 | 0.29 | 1 | 0.5902 |
| rs35305304 | 11 | 118228114 | T | C | -0.344 | 0.7307 | -- | 0 | 0 | 1 | 0.9958 |
| rs61900876 | 11 | 120395800 | C | T | 0.344 | 0.7309 | +? | 0 | 0 | 0 | 1 |
| rs504626 | 11 | 120976428 | A | G | -0.344 | 0.731 | -? | 0 | 0 | 0 | 1 |
| rs11215927 | 11 | 116300551 | C | T | -0.344 | 0.7312 | -? | 0 | 0 | 0 | 1 |
| rs117837319 | 11 | 116584447 | C | A | -0.343 | 0.7313 | -- | 0 | 0.024 | 1 | 0.8776 |
| rs1786142 | 11 | 118995481 | C | T | 0.343 | 0.7313 | +? | 0 | 0 | 0 | 1 |
| rs687329 | 11 | 121023906 | G | A | -0.343 | 0.7314 | -? | 0 | 0 | 0 | 1 |
| rs11605423 | 11 | 117618713 | A | G | 0.343 | 0.7315 | +? | 0 | 0 | 0 | 1 |
| rs12291885 | 11 | 117027942 | C | T | -0.343 | 0.7316 | -? | 0 | 0 | 0 | 1 |
| rs10790284 | 11 | 118996948 | C | A | 0.343 | 0.7318 | +? | 0 | 0 | 0 | 1 |
| rs148643334 | 11 | 118358152 | C | T | -0.343 | 0.7318 | -+ | -143.5 | 1.232 | 1 | 0.267 |
| rs3825054 | 11 | 118234191 | A | G | -0.343 | 0.732 | -- | 0 | 0 | 1 | 0.9935 |
| rs73020445 | 11 | 117752845 | T | C | -0.342 | 0.732 | -? | 0 | 0 | 0 | 1 |
| rs12275789 | 11 | 116345605 | A | G | 0.342 | 0.7321 | +? | 0 | 0 | 0 | 1 |
| rs61528732 | 11 | 120212601 | C | T | -0.342 | 0.7321 | -? | 0 | 0 | 0 | 1 |
| rs61906487 | 11 | 117088821 | T | C | -0.342 | 0.7323 | -? | 0 | 0 | 0 | 1 |
| rs61906488 | 11 | 117089505 | C | T | -0.342 | 0.7323 | -? | 0 | 0 | 0 | 1 |
| rs61906489 | 11 | 117091153 | G | C | -0.342 | 0.7323 | -? | 0 | 0 | 0 | 1 |
| rs73009920 | 11 | 117850727 | T | C | 0.342 | 0.7323 | +? | 0 | 0 | 0 | 1 |
| rs624877 | 11 | 119966457 | T | C | 0.342 | 0.7324 | +? | 0 | 0 | 0 | 1 |
| rs12286173 | 11 | 120486195 | T | C | 0.342 | 0.7326 | +? | 0 | 0 | 0 | 1 |
| rs534279 | 11 | 119358442 | G | A | -0.342 | 0.7326 | -? | 0 | 0 | 0 | 1 |
| rs10892221 | 11 | 118040642 | A | G | 0.342 | 0.7327 | +- | 0 | 0.225 | 1 | 0.635 |
| rs543312 | 11 | 117745659 | C | T | 0.342 | 0.7327 | +? | 0 | 0 | 0 | 1 |
| rs7124166 | 11 | 119630247 | C | G | 0.341 | 0.7328 | +? | 0 | 0 | 0 | 1 |
| rs17569224 | 11 | 116356327 | T | C | 0.341 | 0.7329 | +? | 0 | 0 | 0 | 1 |
| rs56160707 | 11 | 120586577 | C | T | 0.341 | 0.7331 | +? | 0 | 0 | 0 | 1 |
| rs583490 | 11 | 118022608 | A | G | -0.341 | 0.7331 | +- | 0 | 0.561 | 1 | 0.4537 |
| rs60336713 | 11 | 120054885 | T | C | -0.341 | 0.7332 | -? | 0 | 0 | 0 | 1 |
| rs657671 | 11 | 117917622 | C | T | -0.34 | 0.7336 | -? | 0 | 0 | 0 | 1 |
| rs4328225 | 11 | 119478241 | T | A | 0.34 | 0.7337 | +? | 0 | 0 | 0 | 1 |
| rs678776 | 11 | 117696744 | A | G | 0.34 | 0.7337 | +? | 0 | 0 | 0 | 1 |
| rs117052388 | 11 | 121143154 | T | C | 0.34 | 0.7338 | +? | 0 | 0 | 0 | 1 |
| rs10892222 | 11 | 118041102 | A | G | 0.34 | 0.7339 | +- | 0 | 0.228 | 1 | 0.633 |
| rs10892480 | 11 | 119737648 | A | G | 0.34 | 0.734 | +? | 0 | 0 | 0 | 1 |
| rs116981723 | 11 | 120888244 | T | C | 0.34 | 0.734 | +? | 0 | 0 | 0 | 1 |
| rs6589646 | 11 | 117977949 | T | C | -0.34 | 0.7341 | -+ | 0 | 0.455 | 1 | 0.5 |
| rs2289893 | 11 | 116720089 | T | C | 0.34 | 0.7342 | +? | 0 | 0 | 0 | 1 |
| rs3133860 | 11 | 120661178 | T | A | 0.339 | 0.7344 | +? | 0 | 0 | 0 | 1 |
| rs670 | 11 | 116708413 | T | C | 0.339 | 0.7344 | +? | 0 | 0 | 0 | 1 |
| rs118100246 | 11 | 117326419 | A | G | -0.339 | 0.7345 | -? | 0 | 0 | 0 | 1 |
| rs684613 | 11 | 120836290 | T | C | -0.339 | 0.7346 | -+ | 0 | 0.32 | 1 | 0.5718 |
| rs526573 | 11 | 119000070 | A | G | 0.339 | 0.7347 | +? | 0 | 0 | 0 | 1 |
| rs151061355 | 11 | 116586854 | A | G | -0.339 | 0.7348 | -- | 0 | 0.025 | 1 | 0.8755 |

| | | | | | | | | | | | |
|---|---|---|---|---|---|---|---|---|---|---|---|
| rs11217329 | 11 | 119453252 | A | G | -0.339 | 0.7349 | -? | 0 | 0 | 0 | 1 |
| rs1396776 | 11 | 119453579 | C | A | -0.339 | 0.7349 | -? | 0 | 0 | 0 | 1 |
| rs148406383 | 11 | 117328052 | T | C | -0.339 | 0.7349 | -? | 0 | 0 | 0 | 1 |
| rs492275 | 11 | 120076931 | T | C | 0.339 | 0.7349 | +? | 0 | 0 | 0 | 1 |
| rs57963928 | 11 | 116306360 | C | A | -0.339 | 0.7349 | -? | 0 | 0 | 0 | 1 |
| rs77822137 | 11 | 116914260 | C | T | -0.339 | 0.7349 | -+ | 71 | 10.332 | 1 | 0.001308 |
| rs520543 | 11 | 118538157 | A | G | 0.338 | 0.735 | -+ | 0 | 0.827 | 1 | 0.3631 |
| rs60112336 | 11 | 118453483 | T | C | 0.338 | 0.735 | +- | 0 | 0.186 | 1 | 0.6663 |
| rs11217602 | 11 | 119833857 | T | C | 0.338 | 0.7351 | +? | 0 | 0 | 0 | 1 |
| rs586473 | 11 | 121000784 | T | C | -0.338 | 0.7353 | -? | 0 | 0 | 0 | 1 |
| rs61905305 | 11 | 117314122 | T | C | -0.338 | 0.7354 | -? | 0 | 0 | 0 | 1 |
| rs12098917 | 11 | 117716151 | C | T | -0.338 | 0.7355 | -? | 0 | 0 | 0 | 1 |
| rs896695 | 11 | 120075939 | T | C | 0.338 | 0.7356 | +? | 0 | 0 | 0 | 1 |
| rs10790187 | 11 | 117324965 | G | A | 0.338 | 0.7357 | +? | 0 | 0 | 0 | 1 |
| rs77393043 | 11 | 116908914 | A | C | -0.338 | 0.7357 | -+ | 73.5 | 11.304 | 1 | 0.0007736 |
| rs77939786 | 11 | 116908913 | T | C | -0.338 | 0.7357 | -+ | 73.5 | 11.304 | 1 | 0.0007736 |
| rs78550859 | 11 | 116906984 | T | C | -0.338 | 0.7357 | -+ | 73.5 | 11.304 | 1 | 0.0007736 |
| rs79842631 | 11 | 116908807 | G | A | -0.338 | 0.7357 | -+ | 73.5 | 11.304 | 1 | 0.0007736 |
| rs80176870 | 11 | 116905209 | C | T | -0.338 | 0.7357 | -+ | 73.5 | 11.304 | 1 | 0.0007736 |
| rs36061634 | 11 | 118225452 | A | T | -0.337 | 0.7359 | -- | 0 | 0 | 1 | 0.9861 |
| rs149539291 | 11 | 117908503 | A | G | -0.337 | 0.7361 | -? | 0 | 0 | 0 | 1 |
| rs11600354 | 11 | 120477883 | A | G | 0.337 | 0.7362 | +? | 0 | 0 | 0 | 1 |
| rs180354 | 11 | 116599826 | T | C | -0.337 | 0.7362 | -+ | -9.9 | 2.729 | 1 | 0.09853 |
| rs180355 | 11 | 116599771 | T | C | -0.337 | 0.7362 | -+ | -9.9 | 2.729 | 1 | 0.09853 |
| rs2282495 | 11 | 117865381 | T | C | 0.337 | 0.7363 | +? | 0 | 0 | 0 | 1 |
| rs479299 | 11 | 121003848 | A | G | -0.337 | 0.7363 | -? | 0 | 0 | 0 | 1 |
| rs61900877 | 11 | 120396041 | C | G | 0.337 | 0.7364 | +? | 0 | 0 | 0 | 1 |
| rs111708971 | 11 | 116883241 | A | T | -0.336 | 0.7365 | -? | 0 | 0 | 0 | 1 |
| rs518087 | 11 | 116490583 | C | T | 0.336 | 0.7366 | +? | 0 | 0 | 0 | 1 |
| rs180324 | 11 | 116625332 | C | T | -0.336 | 0.7369 | -+ | -9.2 | 2.748 | 1 | 0.0974 |
| rs180325 | 11 | 116625294 | C | G | -0.336 | 0.7369 | -+ | -9.2 | 2.748 | 1 | 0.0974 |
| rs12225323 | 11 | 117920577 | T | C | 0.336 | 0.737 | +? | 0 | 0 | 0 | 1 |
| rs603486 | 11 | 118587347 | T | C | 0.336 | 0.7371 | +? | 0 | 0 | 0 | 1 |
| rs658922 | 11 | 119004686 | A | T | 0.335 | 0.7374 | +? | 0 | 0 | 0 | 1 |
| rs17742 | 11 | 118473385 | C | T | 0.335 | 0.7377 | +- | 0 | 0.246 | 1 | 0.62 |
| rs73576682 | 11 | 116968225 | G | A | -0.335 | 0.7377 | +- | 54.3 | 6.562 | 1 | 0.01042 |
| rs7939209 | 11 | 116969416 | C | T | -0.335 | 0.7377 | +- | 54.3 | 6.562 | 1 | 0.01042 |
| rs7939303 | 11 | 116969439 | C | T | -0.335 | 0.7377 | +- | 54.3 | 6.562 | 1 | 0.01042 |
| rs2241646 | 11 | 119229964 | G | A | 0.335 | 0.7378 | +? | 0 | 0 | 0 | 1 |
| rs11217788 | 11 | 120155118 | A | G | -0.335 | 0.7379 | -? | 0 | 0 | 0 | 1 |
| rs4938282 | 11 | 116430949 | T | C | 0.335 | 0.7379 | +? | 0 | 0 | 0 | 1 |
| rs73013776 | 11 | 117917257 | G | A | -0.335 | 0.7379 | -? | 0 | 0 | 0 | 1 |
| rs658624 | 11 | 118018767 | G | A | 0.334 | 0.738 | ++ | 0 | 0.027 | 1 | 0.8699 |
| rs80006645 | 11 | 119884588 | A | G | 0.335 | 0.738 | +? | 0 | 0 | 0 | 1 |
| rs35825681 | 11 | 119732550 | G | C | 0.334 | 0.7381 | +? | 0 | 0 | 0 | 1 |
| rs73001486 | 11 | 118629364 | A | G | -0.334 | 0.7383 | -+ | 0 | 0.249 | 1 | 0.6177 |
| rs11216925 | 11 | 118471423 | C | T | 0.334 | 0.7385 | +- | 0 | 0.302 | 1 | 0.5825 |
| rs17122113 | 11 | 118234678 | G | A | -0.334 | 0.7385 | -- | 0 | 0 | 1 | 0.9984 |
| rs73576687 | 11 | 116970285 | A | G | -0.334 | 0.7385 | +- | 54.3 | 6.565 | 1 | 0.0104 |
| rs11217689 | 11 | 119965319 | G | C | 0.334 | 0.7386 | +? | 0 | 0 | 0 | 1 |
| rs582951 | 11 | 119965482 | G | A | 0.334 | 0.7386 | +? | 0 | 0 | 0 | 1 |
| rs626302 | 11 | 119965383 | A | G | 0.334 | 0.7386 | +? | 0 | 0 | 0 | 1 |
| rs637685 | 11 | 119965639 | A | G | 0.334 | 0.7386 | +? | 0 | 0 | 0 | 1 |
| rs7936665 | 11 | 120147254 | A | G | -0.334 | 0.7386 | -? | 0 | 0 | 0 | 1 |
| rs12421460 | 11 | 119469767 | T | C | -0.333 | 0.7389 | -? | 0 | 0 | 0 | 1 |
| rs2946743 | 11 | 118017088 | G | A | 0.333 | 0.739 | ++ | 0 | 0.05 | 1 | 0.8229 |

| | | | | | | | | | | | |
|---|---|---|---|---|---|---|---|---|---|---|---|
| rs74508982 | 11 | 116301080 | T | C | -0.333 | 0.739 | -? | 0 | 0 | 0 | 1 |
| rs10750159 | 11 | 119448679 | G | A | 0.333 | 0.7391 | +? | 0 | 0 | 0 | 1 |
| rs111894427 | 11 | 116721405 | T | C | -0.333 | 0.7393 | -? | 0 | 0 | 0 | 1 |
| rs11216168 | 11 | 116741553 | A | G | 0.333 | 0.7394 | +? | 0 | 0 | 0 | 1 |
| rs670973 | 11 | 121001689 | A | C | -0.333 | 0.7394 | -? | 0 | 0 | 0 | 1 |
| rs73020446 | 11 | 117753244 | T | C | -0.332 | 0.7396 | -? | 0 | 0 | 0 | 1 |
| rs180356 | 11 | 116599534 | T | C | -0.332 | 0.7398 | -+ | -10.3 | 2.72 | 1 | 0.0991 |
| rs180358 | 11 | 116599392 | T | C | -0.332 | 0.7398 | -+ | -10.3 | 2.72 | 1 | 0.0991 |
| rs180359 | 11 | 116599281 | G | A | -0.332 | 0.7398 | -+ | -10.3 | 2.72 | 1 | 0.0991 |
| rs181268545 | 11 | 119479675 | A | G | 0.332 | 0.7398 | +? | 0 | 0 | 0 | 1 |
| rs489454 | 11 | 120077259 | A | G | 0.332 | 0.74 | +? | 0 | 0 | 0 | 1 |
| rs536743 | 11 | 120078136 | A | C | 0.332 | 0.74 | +? | 0 | 0 | 0 | 1 |
| rs1894177 | 11 | 117978264 | A | G | -0.332 | 0.7401 | -+ | 0 | 0.461 | 1 | 0.4973 |
| rs35902434 | 11 | 118222634 | C | T | -0.332 | 0.7402 | -- | 0 | 0.002 | 1 | 0.9679 |
| rs12721078 | 11 | 116699395 | A | C | -0.331 | 0.7403 | -? | 0 | 0 | 0 | 1 |
| rs2156636 | 11 | 120770782 | A | G | 0.331 | 0.7403 | +? | 0 | 0 | 0 | 1 |
| rs66899922 | 11 | 116334862 | T | G | -0.331 | 0.7403 | -? | 0 | 0 | 0 | 1 |
| rs12420903 | 11 | 120074170 | T | G | 0.331 | 0.7404 | +? | 0 | 0 | 0 | 1 |
| rs11217695 | 11 | 119977625 | T | C | -0.331 | 0.7406 | -? | 0 | 0 | 0 | 1 |
| rs73004786 | 11 | 120058833 | A | G | -0.331 | 0.7406 | -? | 0 | 0 | 0 | 1 |
| rs7125516 | 11 | 117315191 | C | T | 0.331 | 0.7409 | +? | 0 | 0 | 0 | 1 |
| rs1799993 | 11 | 118958127 | A | C | -0.33 | 0.7412 | -? | 0 | 0 | 0 | 1 |
| rs1358104428 | 11 | 119754895 | A | G | -0.33 | 0.7413 | -? | 0 | 0 | 0 | 1 |
| rs547345 | 11 | 118997869 | C | T | 0.33 | 0.7414 | +? | 0 | 0 | 0 | 1 |
| rs10892420 | 11 | 119478022 | G | A | 0.33 | 0.7415 | +? | 0 | 0 | 0 | 1 |
| rs2252222 | 11 | 120400419 | G | A | 0.33 | 0.7415 | +? | 0 | 0 | 0 | 1 |
| rs61900875 | 11 | 120395705 | G | A | 0.33 | 0.7415 | +? | 0 | 0 | 0 | 1 |
| rs1263172 | 11 | 116680918 | G | A | -0.33 | 0.7416 | -? | 0 | 0 | 0 | 1 |
| rs2845649 | 11 | 119421975 | A | G | 0.33 | 0.7416 | +? | 0 | 0 | 0 | 1 |
| rs614677 | 11 | 119884311 | T | C | 0.33 | 0.7416 | +? | 0 | 0 | 0 | 1 |
| rs61900867 | 11 | 120393284 | T | C | 0.329 | 0.7418 | +? | 0 | 0 | 0 | 1 |
| rs7924993 | 11 | 117649675 | A | G | -0.329 | 0.7419 | -? | 0 | 0 | 0 | 1 |
| rs11217906 | 11 | 120404415 | C | T | 0.329 | 0.742 | +? | 0 | 0 | 0 | 1 |
| rs12787375 | 11 | 121055191 | T | G | -0.329 | 0.7422 | -? | 0 | 0 | 0 | 1 |
| rs534427 | 11 | 119745598 | T | C | 0.329 | 0.7422 | +? | 0 | 0 | 0 | 1 |
| rs183445365 | 11 | 119263566 | A | G | -0.329 | 0.7423 | -? | 0 | 0 | 0 | 1 |
| rs504542 | 11 | 119802591 | G | T | 0.329 | 0.7423 | +? | 0 | 0 | 0 | 1 |
| rs180362 | 11 | 116598118 | G | A | -0.329 | 0.7424 | -+ | -11.7 | 2.686 | 1 | 0.1012 |
| rs1793157 | 11 | 118017434 | C | A | 0.329 | 0.7425 | ++ | 0 | 0.086 | 1 | 0.7691 |
| rs34612914 | 11 | 120553943 | G | A | 0.328 | 0.7426 | +? | 0 | 0 | 0 | 1 |
| rs61905525 | 11 | 117058854 | A | G | -0.328 | 0.7426 | -? | 0 | 0 | 0 | 1 |
| rs2852241 | 11 | 120797584 | C | T | 0.328 | 0.7429 | +? | 0 | 0 | 0 | 1 |
| rs11217326 | 11 | 119444863 | C | T | -0.327 | 0.7434 | -? | 0 | 0 | 0 | 1 |
| rs4445643 | 11 | 120497350 | T | G | 0.327 | 0.7435 | +? | 0 | 0 | 0 | 1 |
| rs61900874 | 11 | 120395521 | T | C | 0.327 | 0.7435 | +? | 0 | 0 | 0 | 1 |
| rs7103567 | 11 | 117732718 | T | C | 0.327 | 0.7437 | +? | 0 | 0 | 0 | 1 |
| rs58329329 | 11 | 117552896 | G | C | -0.327 | 0.7438 | -? | 0 | 0 | 0 | 1 |
| rs11827818 | 11 | 120198728 | G | A | 0.327 | 0.7439 | +? | 0 | 0 | 0 | 1 |
| rs17121057 | 11 | 117537132 | C | A | 0.327 | 0.7439 | +? | 0 | 0 | 0 | 1 |
| rs906822 | 11 | 119438762 | T | G | 0.327 | 0.7439 | +? | 0 | 0 | 0 | 1 |
| rs1473326 | 11 | 116823672 | C | T | 0.326 | 0.7441 | +? | 0 | 0 | 0 | 1 |
| rs10790357 | 11 | 119834038 | C | G | -0.326 | 0.7443 | -? | 0 | 0 | 0 | 1 |
| rs3133862 | 11 | 120619548 | G | T | -0.326 | 0.7444 | -? | 0 | 0 | 0 | 1 |
| rs476809 | 11 | 120080022 | G | T | 0.326 | 0.7444 | +? | 0 | 0 | 0 | 1 |
| rs477863 | 11 | 120079884 | G | T | 0.326 | 0.7444 | +? | 0 | 0 | 0 | 1 |
| rs521841 | 11 | 120080526 | C | A | 0.326 | 0.7444 | +? | 0 | 0 | 0 | 1 |

| | | | | | | | | | | | |
|---|---|---|---|---|---|---|---|---|---|---|---|
| rs74581584 | 11 | 121121331 | C | A | 0.326 | 0.7444 | +? | 0 | 0 | 0 | 1 |
| rs2228055 | 11 | 117864846 | G | A | 0.326 | 0.7445 | +? | 0 | 0 | 0 | 1 |
| rs666432 | 11 | 120003533 | T | C | 0.326 | 0.7445 | +? | 0 | 0 | 0 | 1 |
| rs180368 | 11 | 116593264 | T | C | -0.326 | 0.7446 | -+ | -10.4 | 2.719 | 1 | 0.09919 |
| rs580711 | 11 | 120809198 | G | C | -0.326 | 0.7446 | -? | 0 | 0 | 0 | 1 |
| rs78043040 | 11 | 120395934 | C | T | 0.326 | 0.7446 | +? | 0 | 0 | 0 | 1 |
| rs150342927 | 11 | 116934380 | T | C | -0.326 | 0.7447 | -+ | 71.4 | 10.478 | 1 | 0.001208 |
| rs2847499 | 11 | 120114073 | T | G | -0.326 | 0.7447 | -? | 0 | 0 | 0 | 1 |
| rs57072067 | 11 | 116939370 | G | A | -0.326 | 0.7447 | -+ | 71.4 | 10.478 | 1 | 0.001208 |
| rs603407 | 11 | 117958396 | C | T | -0.326 | 0.7448 | -- | 0 | 0 | 1 | 0.9927 |
| rs77083949 | 11 | 117490186 | T | C | 0.326 | 0.7448 | +? | 0 | 0 | 0 | 1 |
| rs7925188 | 11 | 119455748 | G | A | 0.325 | 0.7449 | +? | 0 | 0 | 0 | 1 |
| rs113054880 | 11 | 118459702 | G | C | 0.325 | 0.745 | +- | 0 | 0.254 | 1 | 0.6144 |
| rs551199 | 11 | 120999732 | A | G | -0.325 | 0.7453 | -? | 0 | 0 | 0 | 1 |
| rs643744 | 11 | 121000212 | T | G | -0.325 | 0.7453 | -? | 0 | 0 | 0 | 1 |
| rs665683 | 11 | 119884384 | C | G | 0.325 | 0.7453 | +? | 0 | 0 | 0 | 1 |
| rs11216613 | 11 | 117777599 | G | T | -0.325 | 0.7454 | -? | 0 | 0 | 0 | 1 |
| rs6589750 | 11 | 119326726 | G | A | -0.325 | 0.7455 | -? | 0 | 0 | 0 | 1 |
| rs11217350 | 11 | 119475996 | C | T | -0.324 | 0.7456 | -? | 0 | 0 | 0 | 1 |
| rs645305 | 11 | 117208747 | C | T | 0.324 | 0.7456 | +? | 0 | 0 | 0 | 1 |
| rs71484177 | 11 | 120396016 | G | A | 0.324 | 0.7459 | +? | 0 | 0 | 0 | 1 |
| rs11216100 | 11 | 116572530 | G | C | 0.324 | 0.7461 | +? | 0 | 0 | 0 | 1 |
| rs549748 | 11 | 117341617 | A | T | -0.324 | 0.7461 | -? | 0 | 0 | 0 | 1 |
| rs10790359 | 11 | 119843707 | G | A | -0.324 | 0.7462 | -? | 0 | 0 | 0 | 1 |
| rs10790360 | 11 | 119843736 | C | A | -0.324 | 0.7462 | -? | 0 | 0 | 0 | 1 |
| rs11216924 | 11 | 118470092 | G | C | 0.324 | 0.7462 | +- | 0 | 0.225 | 1 | 0.6356 |
| rs6589795 | 11 | 119970508 | T | C | 0.324 | 0.7462 | +? | 0 | 0 | 0 | 1 |
| rs74453782 | 11 | 117588461 | T | C | 0.323 | 0.7464 | +? | 0 | 0 | 0 | 1 |
| rs606970 | 11 | 119989892 | G | A | -0.323 | 0.747 | -? | 0 | 0 | 0 | 1 |
| rs493516 | 11 | 117213160 | A | G | 0.322 | 0.7471 | +? | 0 | 0 | 0 | 1 |
| rs10790398 | 11 | 120551684 | T | C | -0.322 | 0.7472 | -? | 0 | 0 | 0 | 1 |
| rs61906109 | 11 | 116554228 | A | C | 0.322 | 0.7472 | +? | 0 | 0 | 0 | 1 |
| rs76921569 | 11 | 117671216 | A | C | 0.322 | 0.7473 | +? | 0 | 0 | 0 | 1 |
| rs79046029 | 11 | 117675125 | T | C | 0.322 | 0.7473 | +? | 0 | 0 | 0 | 1 |
| rs11216566 | 11 | 117697985 | G | A | 0.322 | 0.7474 | +? | 0 | 0 | 0 | 1 |
| rs675855 | 11 | 117913374 | T | G | 0.322 | 0.7475 | +- | 0 | 0.303 | 1 | 0.582 |
| rs11216036 | 11 | 116470373 | T | A | 0.321 | 0.7479 | +? | 0 | 0 | 0 | 1 |
| rs506305 | 11 | 116520469 | T | C | -0.321 | 0.748 | -? | 0 | 0 | 0 | 1 |
| rs118179203 | 11 | 121134520 | T | C | 0.321 | 0.7481 | +? | 0 | 0 | 0 | 1 |
| rs12795183 | 11 | 119416614 | T | A | 0.321 | 0.7483 | +? | 0 | 0 | 0 | 1 |
| rs12270397 | 11 | 120248342 | T | C | 0.321 | 0.7484 | +? | 0 | 0 | 0 | 1 |
| rs2663629 | 11 | 119435180 | A | C | 0.321 | 0.7484 | +? | 0 | 0 | 0 | 1 |
| rs626466 | 11 | 119837112 | T | C | 0.32 | 0.7486 | +? | 0 | 0 | 0 | 1 |
| rs75858010 | 11 | 119790617 | T | C | 0.32 | 0.7486 | +? | 0 | 0 | 0 | 1 |
| rs11216727 | 11 | 117914652 | C | G | 0.32 | 0.7487 | +? | 0 | 0 | 0 | 1 |
| rs12807982 | 11 | 119985475 | A | C | -0.32 | 0.7487 | -? | 0 | 0 | 0 | 1 |
| rs7947434 | 11 | 117311608 | G | C | -0.32 | 0.7487 | -? | 0 | 0 | 0 | 1 |
| rs875209 | 11 | 117208270 | A | G | 0.32 | 0.7488 | +? | 0 | 0 | 0 | 1 |
| rs497960 | 11 | 116317663 | C | T | -0.32 | 0.749 | -? | 0 | 0 | 0 | 1 |
| rs536021 | 11 | 117210901 | C | T | 0.32 | 0.7491 | +? | 0 | 0 | 0 | 1 |
| rs118046012 | 11 | 116590528 | A | G | -0.32 | 0.7492 | -- | 0 | 0.028 | 1 | 0.8669 |
| rs139663973 | 11 | 116893588 | C | T | -0.32 | 0.7492 | -? | 0 | 0 | 0 | 1 |
| rs573801 | 11 | 117209268 | T | C | 0.32 | 0.7492 | +? | 0 | 0 | 0 | 1 |
| rs10736494 | 11 | 118518227 | T | C | -0.32 | 0.7493 | -+ | 0 | 0.12 | 1 | 0.7293 |
| rs1317876 | 11 | 117958909 | G | A | -0.32 | 0.7493 | -- | 0 | 0.002 | 1 | 0.9677 |
| rs3018387 | 11 | 118016981 | T | A | 0.32 | 0.7493 | ++ | 0 | 0.063 | 1 | 0.8015 |

| | | | | | | | | | | | |
|---|---|---|---|---|---|---|---|---|---|---|---|
| rs3135932 | 11 | 117864063 | G | A | 0.319 | 0.7494 | ++ | 0 | 0.193 | 1 | 0.6608 |
| rs45489394 | 11 | 118228968 | G | C | -0.319 | 0.7494 | -+ | 0 | 0.696 | 1 | 0.4042 |
| rs76814754 | 11 | 116557959 | T | C | -0.319 | 0.7494 | -? | 0 | 0 | 0 | 1 |
| rs1508099 | 11 | 116338514 | C | A | 0.319 | 0.7495 | +? | 0 | 0 | 0 | 1 |
| rs4936563 | 11 | 120793442 | T | C | -0.319 | 0.7497 | +- | 0 | 0.553 | 1 | 0.457 |
| rs11215900 | 11 | 116207032 | T | C | -0.319 | 0.7498 | -? | 0 | 0 | 0 | 1 |
| rs116083223 | 11 | 119908015 | A | G | 0.319 | 0.7498 | +? | 0 | 0 | 0 | 1 |
| rs149017065 | 11 | 116228735 | A | G | -0.319 | 0.7501 | -? | 0 | 0 | 0 | 1 |
| rs2663634 | 11 | 119438208 | T | C | -0.319 | 0.7501 | -? | 0 | 0 | 0 | 1 |
| rs475385 | 11 | 117212892 | T | C | 0.318 | 0.7503 | +? | 0 | 0 | 0 | 1 |
| rs77404358 | 11 | 120395962 | C | T | 0.318 | 0.7504 | +? | 0 | 0 | 0 | 1 |
| rs11216644 | 11 | 117819411 | T | C | -0.318 | 0.7505 | -- | 0 | 0.052 | 1 | 0.8193 |
| rs1354975 | 11 | 116352339 | T | C | -0.318 | 0.7506 | -? | 0 | 0 | 0 | 1 |
| rs56143464 | 11 | 116627056 | A | G | -0.318 | 0.7507 | +- | 1.4 | 3.044 | 1 | 0.08103 |
| rs61905112 | 11 | 116628057 | A | G | -0.318 | 0.7507 | +- | 1.4 | 3.044 | 1 | 0.08103 |
| rs12577828 | 11 | 117592582 | A | G | 0.318 | 0.7508 | +? | 0 | 0 | 0 | 1 |
| rs556550 | 11 | 116317405 | G | A | -0.317 | 0.7509 | -? | 0 | 0 | 0 | 1 |
| rs4499035 | 11 | 118610463 | T | C | -0.317 | 0.7512 | -+ | 0 | 0.326 | 1 | 0.5682 |
| rs596134 | 11 | 117984623 | C | T | -0.317 | 0.7513 | -- | 0 | 0.012 | 1 | 0.9128 |
| rs2230297 | 11 | 120745874 | T | C | -0.317 | 0.7515 | -? | 0 | 0 | 0 | 1 |
| rs140189045 | 11 | 119806408 | A | G | -0.316 | 0.7517 | -? | 0 | 0 | 0 | 1 |
| rs2715881 | 11 | 120126277 | T | C | -0.316 | 0.7517 | -? | 0 | 0 | 0 | 1 |
| rs77026903 | 11 | 117398435 | T | C | 0.316 | 0.7518 | +? | 0 | 0 | 0 | 1 |
| rs11601365 | 11 | 116214361 | G | C | -0.316 | 0.7519 | +- | 6.6 | 3.213 | 1 | 0.07306 |
| rs12278345 | 11 | 120058685 | G | T | -0.316 | 0.7519 | +- | 0 | 0.326 | 1 | 0.5682 |
| rs2851407 | 11 | 116333088 | T | C | -0.316 | 0.7519 | -? | 0 | 0 | 0 | 1 |
| rs45522239 | 11 | 118405438 | G | A | 0.316 | 0.7523 | ++ | 0 | 0 | 1 | 0.9949 |
| rs74384044 | 11 | 121138747 | A | G | 0.316 | 0.7523 | +? | 0 | 0 | 0 | 1 |
| rs1110166 | 11 | 119571975 | A | G | -0.315 | 0.7526 | -? | 0 | 0 | 0 | 1 |
| rs61906084 | 11 | 116539025 | G | C | 0.315 | 0.7526 | +? | 0 | 0 | 0 | 1 |
| rs116977890 | 11 | 121010814 | A | C | 0.315 | 0.7527 | +? | 0 | 0 | 0 | 1 |
| rs10502235 | 11 | 120212423 | G | A | 0.315 | 0.7528 | +? | 0 | 0 | 0 | 1 |
| rs533993 | 11 | 118989739 | G | A | 0.315 | 0.7528 | +? | 0 | 0 | 0 | 1 |
| rs505123 | 11 | 116319242 | T | G | -0.315 | 0.753 | -? | 0 | 0 | 0 | 1 |
| rs11216907 | 11 | 118444525 | A | G | 0.315 | 0.7531 | +- | 0 | 0.211 | 1 | 0.646 |
| rs2663632 | 11 | 119436769 | A | G | 0.314 | 0.7533 | +? | 0 | 0 | 0 | 1 |
| rs2845655 | 11 | 119437201 | T | C | 0.314 | 0.7533 | +? | 0 | 0 | 0 | 1 |
| rs7106641 | 11 | 116758101 | G | A | -0.314 | 0.7534 | -? | 0 | 0 | 0 | 1 |
| rs948181 | 11 | 117208949 | T | C | 0.314 | 0.7535 | +? | 0 | 0 | 0 | 1 |
| rs2509671 | 11 | 119175075 | A | C | -0.314 | 0.7537 | -+ | -184.2 | 1.056 | 1 | 0.3042 |
| rs474891 | 11 | 117212337 | T | G | 0.314 | 0.7537 | +? | 0 | 0 | 0 | 1 |
| rs561473 | 11 | 117212640 | C | T | 0.314 | 0.7537 | +? | 0 | 0 | 0 | 1 |
| rs150817479 | 11 | 116904279 | A | G | -0.314 | 0.7538 | -+ | 71.4 | 10.496 | 1 | 0.001196 |
| rs75467870 | 11 | 117667524 | A | G | 0.314 | 0.7538 | ++ | 0 | 0 | 1 | 0.9951 |
| rs12808008 | 11 | 119985570 | G | A | -0.313 | 0.754 | -? | 0 | 0 | 0 | 1 |
| rs112996178 | 11 | 118609598 | T | C | -0.313 | 0.7541 | -+ | 0 | 0.323 | 1 | 0.5696 |
| rs643153 | 11 | 119788216 | C | T | -0.313 | 0.7541 | -? | 0 | 0 | 0 | 1 |
| rs12576767 | 11 | 117915055 | T | C | 0.313 | 0.7542 | +? | 0 | 0 | 0 | 1 |
| rs15818 | 11 | 118952173 | G | A | -0.313 | 0.7546 | -? | 0 | 0 | 0 | 1 |
| rs4252309 | 11 | 117865859 | C | T | 0.313 | 0.7546 | +? | 0 | 0 | 0 | 1 |
| rs470370 | 11 | 120013984 | C | T | -0.313 | 0.7546 | -? | 0 | 0 | 0 | 1 |
| rs59266239 | 11 | 116911077 | T | C | 0.312 | 0.7549 | +? | 0 | 0 | 0 | 1 |
| rs6589562 | 11 | 116583439 | A | G | -0.312 | 0.7551 | -? | 0 | 0 | 0 | 1 |
| rs45548238 | 11 | 118405437 | T | C | 0.312 | 0.7552 | ++ | 0 | 0 | 1 | 0.9967 |
| rs1893029 | 11 | 118610826 | T | G | -0.312 | 0.7553 | -+ | 0 | 0.337 | 1 | 0.5618 |
| rs7103001 | 11 | 118367669 | T | A | -0.312 | 0.7554 | -+ | -160.4 | 1.152 | 1 | 0.2831 |

| | | | | | | | | | | | |
|---|---|---|---|---|---|---|---|---|---|---|---|
| rs117934631 | 11 | 120089082 | G | T | 0.311 | 0.7555 | +? | 0 | 0 | 0 | 1 |
| rs35327319 | 11 | 119262493 | G | T | -0.311 | 0.7555 | -? | 0 | 0 | 0 | 1 |
| rs10790204 | 11 | 117627333 | A | G | 0.311 | 0.7556 | +? | 0 | 0 | 0 | 1 |
| rs76116075 | 11 | 119824100 | G | A | -0.311 | 0.7556 | -? | 0 | 0 | 0 | 1 |
| rs645545 | 11 | 120853317 | A | T | -0.311 | 0.7557 | -+ | 0 | 0.302 | 1 | 0.5827 |
| rs578781 | 11 | 117210707 | A | G | 0.311 | 0.7558 | +? | 0 | 0 | 0 | 1 |
| rs17123861 | 11 | 120256337 | C | T | 0.311 | 0.7559 | +? | 0 | 0 | 0 | 1 |
| rs61905086 | 11 | 116614863 | C | T | -0.311 | 0.7561 | +- | -25.1 | 2.398 | 1 | 0.1215 |
| rs3897570 | 11 | 120395229 | C | A | 0.31 | 0.7562 | +? | 0 | 0 | 0 | 1 |
| rs61900873 | 11 | 120395252 | C | T | 0.31 | 0.7562 | +? | 0 | 0 | 0 | 1 |
| rs35661813 | 11 | 119831794 | C | A | -0.31 | 0.7563 | -? | 0 | 0 | 0 | 1 |
| rs2096834 | 11 | 119269557 | T | C | -0.31 | 0.7564 | -? | 0 | 0 | 0 | 1 |
| rs17121264 | 11 | 117690543 | C | A | 0.31 | 0.7567 | +? | 0 | 0 | 0 | 1 |
| rs17581874 | 11 | 116929384 | C | A | -0.31 | 0.7567 | -? | 0 | 0 | 0 | 1 |
| rs11600672 | 11 | 117564880 | A | G | 0.309 | 0.7572 | +? | 0 | 0 | 0 | 1 |
| rs4597100 | 11 | 117686456 | A | C | 0.309 | 0.7572 | +? | 0 | 0 | 0 | 1 |
| rs12271743 | 11 | 117026154 | C | T | -0.309 | 0.7574 | -? | 0 | 0 | 0 | 1 |
| rs4938776 | 11 | 119956631 | C | A | -0.309 | 0.7574 | -? | 0 | 0 | 0 | 1 |
| rs10790321 | 11 | 119470258 | A | G | -0.309 | 0.7575 | -? | 0 | 0 | 0 | 1 |
| rs10790322 | 11 | 119470333 | A | G | -0.309 | 0.7575 | -? | 0 | 0 | 0 | 1 |
| rs7950231 | 11 | 119470389 | G | A | -0.309 | 0.7575 | -? | 0 | 0 | 0 | 1 |
| rs56196414 | 11 | 116541268 | A | G | 0.309 | 0.7576 | +? | 0 | 0 | 0 | 1 |
| rs58208536 | 11 | 116540785 | C | G | 0.309 | 0.7576 | +? | 0 | 0 | 0 | 1 |
| rs58346469 | 11 | 116540916 | T | C | 0.309 | 0.7576 | +? | 0 | 0 | 0 | 1 |
| rs61906086 | 11 | 116539637 | G | A | 0.309 | 0.7576 | +? | 0 | 0 | 0 | 1 |
| rs61906100 | 11 | 116541656 | A | C | 0.309 | 0.7576 | +? | 0 | 0 | 0 | 1 |
| rs906820 | 11 | 119472385 | T | C | -0.309 | 0.7576 | -? | 0 | 0 | 0 | 1 |
| rs61905472 | 11 | 117017706 | T | C | -0.308 | 0.7578 | -? | 0 | 0 | 0 | 1 |
| rs4935752 | 11 | 120689119 | C | A | 0.308 | 0.7579 | +? | 0 | 0 | 0 | 1 |
| rs57879605 | 11 | 118798757 | T | C | -0.308 | 0.7579 | +- | 10.6 | 3.355 | 1 | 0.06702 |
| rs61905682 | 11 | 116837823 | C | G | -0.308 | 0.7579 | -? | 0 | 0 | 0 | 1 |
| rs649233 | 11 | 117464063 | T | C | 0.308 | 0.758 | +? | 0 | 0 | 0 | 1 |
| rs665758 | 11 | 119884319 | A | G | 0.308 | 0.758 | +? | 0 | 0 | 0 | 1 |
| rs7934734 | 11 | 120542687 | G | A | 0.308 | 0.758 | +? | 0 | 0 | 0 | 1 |
| rs10790315 | 11 | 119468951 | G | C | -0.308 | 0.7583 | -? | 0 | 0 | 0 | 1 |
| rs10790316 | 11 | 119468994 | T | C | -0.308 | 0.7583 | -? | 0 | 0 | 0 | 1 |
| rs4936378 | 11 | 117316321 | A | G | 0.308 | 0.7583 | +? | 0 | 0 | 0 | 1 |
| rs7121550 | 11 | 119469211 | A | G | -0.308 | 0.7583 | -? | 0 | 0 | 0 | 1 |
| rs10892311 | 11 | 118807735 | T | C | 0.307 | 0.7585 | ++ | 0 | 0.02 | 1 | 0.8862 |
| rs34160337 | 11 | 120393495 | G | T | 0.307 | 0.7587 | +? | 0 | 0 | 0 | 1 |
| rs4252298 | 11 | 117854977 | C | T | 0.307 | 0.7588 | +? | 0 | 0 | 0 | 1 |
| rs72999455 | 11 | 118483802 | G | A | 0.307 | 0.7588 | ++ | 0 | 0.036 | 1 | 0.8499 |
| rs649138 | 11 | 118066695 | C | T | 0.307 | 0.759 | +? | 0 | 0 | 0 | 1 |
| rs12279593 | 11 | 117533949 | A | G | 0.307 | 0.7591 | +? | 0 | 0 | 0 | 1 |
| rs17596129 | 11 | 117712849 | C | G | 0.307 | 0.7591 | +? | 0 | 0 | 0 | 1 |
| rs180347 | 11 | 116612312 | C | G | -0.307 | 0.7592 | -+ | -11.1 | 2.701 | 1 | 0.1003 |
| rs7342270 | 11 | 119843952 | A | T | -0.307 | 0.7592 | -? | 0 | 0 | 0 | 1 |
| rs11216513 | 11 | 117588733 | T | G | 0.306 | 0.7593 | +? | 0 | 0 | 0 | 1 |
| rs10790366 | 11 | 119967769 | T | G | -0.306 | 0.7594 | -? | 0 | 0 | 0 | 1 |
| rs59533604 | 11 | 117333229 | C | A | -0.306 | 0.7595 | -? | 0 | 0 | 0 | 1 |
| rs1945765 | 11 | 118182829 | A | T | 0.306 | 0.7596 | +? | 0 | 0 | 0 | 1 |
| rs676812 | 11 | 117926593 | A | G | -0.306 | 0.7597 | -? | 0 | 0 | 0 | 1 |
| rs11600274 | 11 | 120477398 | A | G | 0.305 | 0.7601 | +? | 0 | 0 | 0 | 1 |
| rs4938391 | 11 | 117358351 | A | C | -0.305 | 0.7604 | -? | 0 | 0 | 0 | 1 |
| rs73011197 | 11 | 116487300 | T | C | 0.305 | 0.7604 | +? | 0 | 0 | 0 | 1 |
| rs7478744 | 11 | 120946580 | A | G | -0.304 | 0.7609 | -? | 0 | 0 | 0 | 1 |

| | | | | | | | | | | | |
|---|---|---|---|---|---|---|---|---|---|---|---|
| rs17570295 | 11 | 116426814 | A | T | -0.304 | 0.7612 | -? | 0 | 0 | 0 | 1 |
| rs75896504 | 11 | 117784946 | A | G | -0.304 | 0.7612 | -? | 0 | 0 | 0 | 1 |
| rs598061 | 11 | 117985080 | C | A | -0.304 | 0.7613 | -- | 0 | 0.014 | 1 | 0.9068 |
| rs183901166 | 11 | 119896824 | A | G | 0.304 | 0.7614 | +? | 0 | 0 | 0 | 1 |
| rs11217712 | 11 | 120030227 | G | T | 0.303 | 0.7617 | +? | 0 | 0 | 0 | 1 |
| rs12808958 | 11 | 116229759 | A | G | -0.303 | 0.7618 | +- | -175.9 | 1.087 | 1 | 0.2971 |
| rs73003773 | 11 | 120720298 | C | T | 0.303 | 0.7618 | +? | 0 | 0 | 0 | 1 |
| rs117883320 | 11 | 119789992 | A | C | 0.303 | 0.7619 | +? | 0 | 0 | 0 | 1 |
| rs74621718 | 11 | 117590895 | G | C | 0.303 | 0.7619 | +? | 0 | 0 | 0 | 1 |
| rs78667446 | 11 | 117590919 | T | G | 0.303 | 0.7619 | +? | 0 | 0 | 0 | 1 |
| rs71478238 | 11 | 116231844 | G | T | -0.303 | 0.762 | +- | -168 | 1.12 | 1 | 0.29 |
| rs1168815 | 11 | 119018023 | C | T | 0.303 | 0.7621 | +? | 0 | 0 | 0 | 1 |
| rs7112967 | 11 | 116831912 | T | C | 0.303 | 0.7622 | +? | 0 | 0 | 0 | 1 |
| rs112571425 | 11 | 117620050 | C | T | 0.302 | 0.7624 | +? | 0 | 0 | 0 | 1 |
| rs55871712 | 11 | 117333521 | T | C | -0.302 | 0.7624 | -? | 0 | 0 | 0 | 1 |
| rs180345 | 11 | 116613017 | A | G | -0.302 | 0.7625 | -+ | -11.4 | 2.692 | 1 | 0.1008 |
| rs573827 | 11 | 117214091 | A | G | 0.302 | 0.7626 | +? | 0 | 0 | 0 | 1 |
| rs11216674 | 11 | 117880139 | T | C | 0.302 | 0.7628 | +? | 0 | 0 | 0 | 1 |
| rs59620302 | 11 | 116581159 | G | C | -0.302 | 0.7628 | -- | 0 | 0.032 | 1 | 0.8588 |
| rs188849771 | 11 | 119263574 | G | A | -0.302 | 0.7629 | -? | 0 | 0 | 0 | 1 |
| rs6589764 | 11 | 119476676 | C | T | -0.302 | 0.7629 | -? | 0 | 0 | 0 | 1 |
| rs236919 | 11 | 117095361 | T | C | 0.301 | 0.7631 | +? | 0 | 0 | 0 | 1 |
| rs7102059 | 11 | 121185726 | T | A | -0.301 | 0.7633 | -? | 0 | 0 | 0 | 1 |
| rs7121851 | 11 | 117757875 | T | C | -0.301 | 0.7634 | -? | 0 | 0 | 0 | 1 |
| rs7940175 | 11 | 117334349 | T | C | -0.301 | 0.7635 | -? | 0 | 0 | 0 | 1 |
| rs2846106 | 11 | 120601873 | G | A | 0.301 | 0.7636 | +? | 0 | 0 | 0 | 1 |
| rs140140760 | 11 | 116312961 | A | G | -0.301 | 0.7637 | -? | 0 | 0 | 0 | 1 |
| rs35778099 | 11 | 119831582 | T | C | -0.3 | 0.7639 | -? | 0 | 0 | 0 | 1 |
| rs61905474 | 11 | 117019530 | A | G | -0.3 | 0.764 | -? | 0 | 0 | 0 | 1 |
| rs61905475 | 11 | 117024435 | G | A | -0.3 | 0.764 | -? | 0 | 0 | 0 | 1 |
| rs61905477 | 11 | 117026055 | G | A | -0.3 | 0.764 | -? | 0 | 0 | 0 | 1 |
| rs6589784 | 11 | 119844270 | C | A | -0.3 | 0.764 | -? | 0 | 0 | 0 | 1 |
| rs17122278 | 11 | 118449370 | G | A | 0.3 | 0.7641 | +- | 0 | 0.228 | 1 | 0.633 |
| rs7122401 | 11 | 120183803 | A | G | -0.3 | 0.7641 | -? | 0 | 0 | 0 | 1 |
| rs80068772 | 11 | 116546161 | T | A | 0.3 | 0.7642 | +? | 0 | 0 | 0 | 1 |
| rs9163 | 11 | 117283808 | T | C | 0.3 | 0.7643 | +? | 0 | 0 | 0 | 1 |
| rs7114739 | 11 | 119455165 | C | T | 0.3 | 0.7645 | +? | 0 | 0 | 0 | 1 |
| rs1605853 | 11 | 119418040 | C | T | 0.299 | 0.7646 | +? | 0 | 0 | 0 | 1 |
| rs497043 | 11 | 117123297 | G | C | 0.299 | 0.7648 | +? | 0 | 0 | 0 | 1 |
| rs10892667 | 11 | 120936907 | C | T | -0.299 | 0.7649 | -? | 0 | 0 | 0 | 1 |
| rs2508654 | 11 | 120405520 | T | C | 0.299 | 0.7652 | +? | 0 | 0 | 0 | 1 |
| rs4252299 | 11 | 117855922 | T | G | 0.299 | 0.7652 | +? | 0 | 0 | 0 | 1 |
| rs78502545 | 11 | 118615390 | T | C | -0.299 | 0.7653 | -+ | -44.5 | 2.075 | 1 | 0.1497 |
| rs1287395 | 11 | 120085692 | C | G | 0.298 | 0.7654 | +? | 0 | 0 | 0 | 1 |
| rs59232710 | 11 | 116504984 | G | A | 0.298 | 0.7654 | +? | 0 | 0 | 0 | 1 |
| rs74533222 | 11 | 121074895 | C | T | -0.298 | 0.7654 | -? | 0 | 0 | 0 | 1 |
| rs117840927 | 11 | 121065324 | A | C | -0.298 | 0.7658 | -? | 0 | 0 | 0 | 1 |
| rs10892176 | 11 | 117685037 | C | T | 0.298 | 0.766 | +? | 0 | 0 | 0 | 1 |
| rs7110240 | 11 | 116343848 | C | T | -0.298 | 0.766 | -? | 0 | 0 | 0 | 1 |
| rs7951593 | 11 | 119840442 | T | C | -0.298 | 0.766 | -? | 0 | 0 | 0 | 1 |
| rs4938486 | 11 | 117992645 | G | A | -0.297 | 0.7663 | -- | 0 | 0.014 | 1 | 0.9061 |
| rs3894250 | 11 | 117312609 | C | A | -0.297 | 0.7667 | -? | 0 | 0 | 0 | 1 |
| rs56313145 | 11 | 119973302 | A | G | -0.297 | 0.7667 | -? | 0 | 0 | 0 | 1 |
| rs61905518 | 11 | 117040739 | G | A | -0.297 | 0.7667 | -? | 0 | 0 | 0 | 1 |
| rs665218 | 11 | 117425003 | A | G | -0.297 | 0.7667 | -? | 0 | 0 | 0 | 1 |
| rs2048246 | 11 | 119598653 | C | T | -0.297 | 0.7668 | -? | 0 | 0 | 0 | 1 |

| | | | | | | | | | | | |
|---|---|---|---|---|---|---|---|---|---|---|---|
| rs11216898 | 11 | 118432852 | C | T | 0.296 | 0.7669 | ++ | 0 | 0.007 | 1 | 0.9322 |
| rs190638475 | 11 | 121068352 | A | G | 0.296 | 0.767 | +? | 0 | 0 | 0 | 1 |
| rs113204414 | 11 | 117603352 | C | T | 0.296 | 0.7671 | +? | 0 | 0 | 0 | 1 |
| rs4938386 | 11 | 117334518 | T | C | -0.296 | 0.7671 | -? | 0 | 0 | 0 | 1 |
| rs116993355 | 11 | 119012962 | C | T | 0.296 | 0.7672 | +? | 0 | 0 | 0 | 1 |
| rs12420302 | 11 | 119299592 | G | A | -0.296 | 0.7672 | -? | 0 | 0 | 0 | 1 |
| rs144015791 | 11 | 116805874 | T | G | 0.296 | 0.7672 | +? | 0 | 0 | 0 | 1 |
| rs4936389 | 11 | 117488717 | T | C | 0.296 | 0.7672 | +? | 0 | 0 | 0 | 1 |
| rs77646493 | 11 | 116582180 | A | C | -0.296 | 0.7673 | -- | 0 | 0.033 | 1 | 0.8562 |
| rs10892355 | 11 | 119240809 | C | T | 0.296 | 0.7674 | +? | 0 | 0 | 0 | 1 |
| rs74360954 | 11 | 116582542 | T | C | -0.296 | 0.7674 | -- | 0 | 0.033 | 1 | 0.8562 |
| rs113749676 | 11 | 116712442 | T | C | -0.295 | 0.7678 | +- | 26.8 | 4.101 | 1 | 0.04287 |
| rs11601616 | 11 | 119451615 | C | T | 0.295 | 0.7678 | +? | 0 | 0 | 0 | 1 |
| rs1243255182 | 11 | 119891625 | G | A | 0.295 | 0.7678 | +? | 0 | 0 | 0 | 1 |
| rs35720648 | 11 | 119269112 | C | G | -0.295 | 0.7678 | -? | 0 | 0 | 0 | 1 |
| rs61906085 | 11 | 116539090 | G | A | 0.295 | 0.7678 | +? | 0 | 0 | 0 | 1 |
| rs671872 | 11 | 119016176 | G | A | 0.295 | 0.7678 | +? | 0 | 0 | 0 | 1 |
| rs1941636 | 11 | 117993422 | C | T | -0.295 | 0.7681 | -- | 0 | 0.003 | 1 | 0.9567 |
| rs616262 | 11 | 119851604 | G | C | 0.295 | 0.7681 | +? | 0 | 0 | 0 | 1 |
| rs10892245 | 11 | 118472354 | A | C | 0.295 | 0.7682 | +- | 0 | 0.294 | 1 | 0.5874 |
| rs45515495 | 11 | 116665025 | A | T | -0.295 | 0.7683 | +- | -192.4 | 1.026 | 1 | 0.3111 |
| rs10891965 | 11 | 116209581 | A | G | 0.294 | 0.7688 | +? | 0 | 0 | 0 | 1 |
| rs1941627 | 11 | 117989246 | G | A | -0.294 | 0.769 | -- | 0 | 0.006 | 1 | 0.9389 |
| rs77526581 | 11 | 119884520 | T | C | 0.294 | 0.769 | +? | 0 | 0 | 0 | 1 |
| rs12808254 | 11 | 118066912 | T | C | -0.294 | 0.7691 | -+ | 31.6 | 4.388 | 1 | 0.03618 |
| rs3741278 | 11 | 117352949 | A | G | -0.294 | 0.7691 | -? | 0 | 0 | 0 | 1 |
| rs10750114 | 11 | 117692170 | G | T | 0.293 | 0.7693 | +? | 0 | 0 | 0 | 1 |
| rs3138091 | 11 | 119291480 | G | A | -0.293 | 0.7693 | -? | 0 | 0 | 0 | 1 |
| rs58990403 | 11 | 116509339 | G | A | -0.293 | 0.7693 | -? | 0 | 0 | 0 | 1 |
| rs10892722 | 11 | 121194387 | A | G | -0.293 | 0.7694 | -? | 0 | 0 | 0 | 1 |
| rs12807216 | 11 | 119300681 | A | C | -0.293 | 0.7694 | -? | 0 | 0 | 0 | 1 |
| rs3138092 | 11 | 119291445 | C | T | -0.293 | 0.7694 | -? | 0 | 0 | 0 | 1 |
| rs11217325 | 11 | 119444474 | G | A | -0.293 | 0.7695 | -? | 0 | 0 | 0 | 1 |
| rs7115154 | 11 | 119840776 | A | G | -0.293 | 0.7696 | -? | 0 | 0 | 0 | 1 |
| rs10790317 | 11 | 119469009 | A | G | -0.293 | 0.7697 | -? | 0 | 0 | 0 | 1 |
| rs548381 | 11 | 117171337 | T | C | -0.293 | 0.7697 | -? | 0 | 0 | 0 | 1 |
| rs3895896 | 11 | 119961536 | A | T | -0.293 | 0.7698 | -? | 0 | 0 | 0 | 1 |
| rs56339619 | 11 | 120021546 | A | G | 0.293 | 0.7699 | +? | 0 | 0 | 0 | 1 |
| rs12365133 | 11 | 119620456 | G | A | -0.292 | 0.77 | -? | 0 | 0 | 0 | 1 |
| rs11217272 | 11 | 119279939 | G | A | 0.292 | 0.7702 | +? | 0 | 0 | 0 | 1 |
| rs662799 | 11 | 116663707 | G | A | -0.292 | 0.7702 | +- | -53.5 | 1.954 | 1 | 0.1621 |
| rs547124 | 11 | 120052554 | A | G | 0.292 | 0.7703 | +? | 0 | 0 | 0 | 1 |
| rs61898268 | 11 | 120073236 | C | A | 0.292 | 0.7703 | +? | 0 | 0 | 0 | 1 |
| rs117028190 | 11 | 117336093 | T | C | -0.292 | 0.7705 | -? | 0 | 0 | 0 | 1 |
| rs12274192 | 11 | 116652351 | G | A | -0.291 | 0.7707 | +- | -52 | 1.974 | 1 | 0.1601 |
| rs12417611 | 11 | 119299832 | A | G | -0.291 | 0.7707 | -? | 0 | 0 | 0 | 1 |
| rs78296522 | 11 | 116648776 | A | C | 0.291 | 0.7707 | +- | 0 | 0.079 | 1 | 0.7792 |
| rs12417523 | 11 | 119299408 | T | C | -0.291 | 0.7708 | -? | 0 | 0 | 0 | 1 |
| rs4385905 | 11 | 120394576 | A | T | 0.291 | 0.7709 | +? | 0 | 0 | 0 | 1 |
| rs61900871 | 11 | 120394410 | T | C | 0.291 | 0.7709 | +? | 0 | 0 | 0 | 1 |
| rs144425764 | 11 | 117710295 | C | T | -0.291 | 0.771 | -? | 0 | 0 | 0 | 1 |
| rs11217430 | 11 | 119595209 | A | G | 0.291 | 0.7712 | +? | 0 | 0 | 0 | 1 |
| rs45485302 | 11 | 118405377 | A | C | 0.29 | 0.7715 | ++ | 0 | 0 | 1 | 0.9882 |
| rs73588403 | 11 | 116628271 | T | C | -0.29 | 0.7716 | +- | 3.2 | 3.099 | 1 | 0.07833 |
| rs2444241 | 11 | 120083312 | G | A | 0.29 | 0.7717 | +? | 0 | 0 | 0 | 1 |
| rs2444243 | 11 | 120082325 | A | G | 0.29 | 0.7717 | +? | 0 | 0 | 0 | 1 |

| | | | | | | | | | | | |
|---|---|---|---|---|---|---|---|---|---|---|---|
| rs2508495 | 11 | 120082748 | C | T | 0.29 | 0.7717 | +? | 0 | 0 | 0 | 1 |
| rs2508496 | 11 | 120083078 | C | T | 0.29 | 0.7717 | +? | 0 | 0 | 0 | 1 |
| rs547990 | 11 | 120083271 | C | A | 0.29 | 0.7717 | +? | 0 | 0 | 0 | 1 |
| rs1001205 | 11 | 119292712 | T | C | -0.29 | 0.7719 | -? | 0 | 0 | 0 | 1 |
| rs76376430 | 11 | 116464209 | T | C | -0.289 | 0.7722 | -? | 0 | 0 | 0 | 1 |
| rs4544037 | 11 | 118222062 | C | T | -0.289 | 0.7724 | -+ | 0 | 0.786 | 1 | 0.3752 |
| rs45621436 | 11 | 118483912 | G | T | -0.289 | 0.7724 | -+ | 0 | 0.123 | 1 | 0.7259 |
| rs12575067 | 11 | 119065310 | A | G | -0.289 | 0.7729 | -? | 0 | 0 | 0 | 1 |
| rs3138097 | 11 | 119285008 | C | T | -0.288 | 0.773 | -? | 0 | 0 | 0 | 1 |
| rs11216668 | 11 | 117878559 | C | T | 0.288 | 0.7731 | +? | 0 | 0 | 0 | 1 |
| rs11216669 | 11 | 117878578 | T | C | 0.288 | 0.7731 | +? | 0 | 0 | 0 | 1 |
| rs11216670 | 11 | 117878659 | A | T | 0.288 | 0.7731 | +? | 0 | 0 | 0 | 1 |
| rs4938390 | 11 | 117358313 | T | C | -0.288 | 0.7734 | -? | 0 | 0 | 0 | 1 |
| rs886022 | 11 | 116583513 | A | C | 0.288 | 0.7734 | -+ | -174.9 | 1.091 | 1 | 0.2962 |
| rs11607388 | 11 | 116305597 | A | G | -0.288 | 0.7735 | -? | 0 | 0 | 0 | 1 |
| rs12417207 | 11 | 119469425 | A | C | -0.288 | 0.7735 | -? | 0 | 0 | 0 | 1 |
| rs76801336 | 11 | 116504754 | C | T | 0.288 | 0.7735 | +? | 0 | 0 | 0 | 1 |
| rs116658831 | 11 | 118970476 | A | G | 0.288 | 0.7736 | +? | 0 | 0 | 0 | 1 |
| rs2663625 | 11 | 119422334 | C | T | 0.288 | 0.7736 | +? | 0 | 0 | 0 | 1 |
| rs17121519 | 11 | 117886453 | T | C | 0.287 | 0.7738 | +? | 0 | 0 | 0 | 1 |
| rs628299 | 11 | 118018022 | G | A | -0.287 | 0.7739 | +- | 0 | 0.668 | 1 | 0.4137 |
| rs2298766 | 11 | 117332343 | T | C | -0.287 | 0.774 | -? | 0 | 0 | 0 | 1 |
| rs11216040 | 11 | 116488539 | A | G | -0.287 | 0.7741 | -? | 0 | 0 | 0 | 1 |
| rs12576975 | 11 | 117590007 | A | C | 0.287 | 0.7741 | +? | 0 | 0 | 0 | 1 |
| rs78080911 | 11 | 117590811 | G | A | 0.287 | 0.7741 | +? | 0 | 0 | 0 | 1 |
| rs7108613 | 11 | 118268613 | G | A | 0.287 | 0.7743 | +- | 0 | 0.137 | 1 | 0.7116 |
| rs61906105 | 11 | 116550528 | G | A | 0.287 | 0.7744 | +? | 0 | 0 | 0 | 1 |
| rs12293402 | 11 | 119445971 | T | C | -0.286 | 0.7746 | -? | 0 | 0 | 0 | 1 |
| rs3741324 | 11 | 118405835 | A | G | 0.286 | 0.7746 | ++ | 0 | 0 | 1 | 0.9871 |
| rs61898748 | 11 | 119445417 | T | C | -0.286 | 0.7746 | -? | 0 | 0 | 0 | 1 |
| rs77574778 | 11 | 116581800 | T | C | -0.286 | 0.7747 | -- | 0 | 0.035 | 1 | 0.8519 |
| rs7949751 | 11 | 118072373 | G | A | 0.286 | 0.7748 | +? | 0 | 0 | 0 | 1 |
| rs34003087 | 11 | 116663596 | T | C | -0.286 | 0.775 | -- | 0 | 0.106 | 1 | 0.7443 |
| rs142693807 | 11 | 116904176 | A | G | -0.286 | 0.7752 | +- | 47.4 | 5.702 | 1 | 0.01695 |
| rs1144035 | 11 | 116584317 | C | T | 0.285 | 0.7754 | -+ | -148.4 | 1.208 | 1 | 0.2718 |
| rs78470936 | 11 | 117500323 | G | T | 0.285 | 0.7754 | +? | 0 | 0 | 0 | 1 |
| rs59827403 | 11 | 117296923 | G | C | -0.285 | 0.7758 | -? | 0 | 0 | 0 | 1 |
| rs884603 | 11 | 116421659 | T | C | 0.285 | 0.7759 | -+ | 0 | 0.315 | 1 | 0.5744 |
| rs78615799 | 11 | 117400284 | T | C | 0.285 | 0.776 | +? | 0 | 0 | 0 | 1 |
| rs2847495 | 11 | 120135501 | A | C | -0.284 | 0.7761 | -? | 0 | 0 | 0 | 1 |
| rs10892017 | 11 | 116567881 | C | G | 0.284 | 0.7762 | +? | 0 | 0 | 0 | 1 |
| rs7127212 | 11 | 118953202 | T | C | -0.284 | 0.7762 | -? | 0 | 0 | 0 | 1 |
| rs75210049 | 11 | 116987964 | A | G | -0.284 | 0.7762 | -? | 0 | 0 | 0 | 1 |
| rs7938944 | 11 | 118954066 | C | T | -0.284 | 0.7762 | -? | 0 | 0 | 0 | 1 |
| rs7952700 | 11 | 118954194 | C | G | -0.284 | 0.7762 | -? | 0 | 0 | 0 | 1 |
| rs4301800 | 11 | 119019520 | A | C | 0.284 | 0.7763 | +? | 0 | 0 | 0 | 1 |
| rs646371 | 11 | 121004172 | A | G | -0.284 | 0.7763 | -? | 0 | 0 | 0 | 1 |
| rs77309894 | 11 | 119890651 | A | T | 0.284 | 0.7765 | +? | 0 | 0 | 0 | 1 |
| rs11606575 | 11 | 117237207 | G | C | -0.284 | 0.7767 | -? | 0 | 0 | 0 | 1 |
| rs143668212 | 11 | 117061197 | T | C | -0.283 | 0.7769 | -? | 0 | 0 | 0 | 1 |
| rs77086432 | 11 | 117046136 | C | T | -0.283 | 0.7769 | -? | 0 | 0 | 0 | 1 |
| rs61900461 | 11 | 119972006 | A | G | 0.283 | 0.777 | +? | 0 | 0 | 0 | 1 |
| rs7107910 | 11 | 117733559 | C | G | 0.283 | 0.7775 | +? | 0 | 0 | 0 | 1 |
| rs530291 | 11 | 119989569 | T | C | 0.282 | 0.7776 | +? | 0 | 0 | 0 | 1 |
| rs73009180 | 11 | 116417492 | T | C | -0.282 | 0.7777 | -? | 0 | 0 | 0 | 1 |
| rs1054735 | 11 | 119290034 | G | C | -0.282 | 0.7778 | -? | 0 | 0 | 0 | 1 |

| | | | | | | | | | | | |
|---|---|---|---|---|---|---|---|---|---|---|---|
| rs56133216 | 11 | 119257248 | C | T | -0.282 | 0.7779 | -? | 0 | 0 | 0 | 1 |
| rs612132 | 11 | 119014474 | T | A | 0.282 | 0.7779 | +? | 0 | 0 | 0 | 1 |
| rs10790381 | 11 | 120257495 | G | A | 0.282 | 0.778 | +? | 0 | 0 | 0 | 1 |
| rs3824977 | 11 | 120745967 | G | A | -0.282 | 0.778 | -? | 0 | 0 | 0 | 1 |
| rs61905460 | 11 | 116989524 | T | C | -0.281 | 0.7784 | -? | 0 | 0 | 0 | 1 |
| rs78825663 | 11 | 117307055 | G | C | -0.281 | 0.7785 | -? | 0 | 0 | 0 | 1 |
| rs12420850 | 11 | 120073966 | A | C | 0.281 | 0.7786 | +? | 0 | 0 | 0 | 1 |
| rs614944 | 11 | 116717663 | C | T | -0.281 | 0.7786 | +- | -15.6 | 2.596 | 1 | 0.1071 |
| rs117040938 | 11 | 119950717 | C | T | 0.281 | 0.7787 | +? | 0 | 0 | 0 | 1 |
| rs7102666 | 11 | 117589939 | C | G | 0.281 | 0.7787 | +? | 0 | 0 | 0 | 1 |
| rs74792494 | 11 | 116549408 | C | A | 0.281 | 0.7787 | +? | 0 | 0 | 0 | 1 |
| rs140393706 | 11 | 117269842 | T | C | 0.281 | 0.7788 | +? | 0 | 0 | 0 | 1 |
| rs499910 | 11 | 116763034 | G | A | -0.281 | 0.7788 | -? | 0 | 0 | 0 | 1 |
| rs12420888 | 11 | 116563822 | G | C | 0.281 | 0.7789 | +? | 0 | 0 | 0 | 1 |
| rs484988 | 11 | 117266648 | C | G | -0.28 | 0.7793 | -? | 0 | 0 | 0 | 1 |
| rs4936552 | 11 | 120688502 | T | A | 0.28 | 0.7793 | +? | 0 | 0 | 0 | 1 |
| rs11216904 | 11 | 118440863 | G | A | 0.28 | 0.7794 | +- | 0 | 0.04 | 1 | 0.8408 |
| rs470645 | 11 | 120007889 | T | G | -0.28 | 0.7794 | -? | 0 | 0 | 0 | 1 |
| rs7129848 | 11 | 119566228 | T | C | 0.28 | 0.7796 | +? | 0 | 0 | 0 | 1 |
| rs2663635 | 11 | 119438415 | C | T | 0.28 | 0.7797 | +? | 0 | 0 | 0 | 1 |
| rs7934763 | 11 | 116792357 | G | T | 0.28 | 0.7797 | +? | 0 | 0 | 0 | 1 |
| rs12224844 | 11 | 118439048 | T | A | 0.28 | 0.7799 | +- | 0 | 0.038 | 1 | 0.8455 |
| rs77216358 | 11 | 120311157 | G | A | 0.279 | 0.7799 | +? | 0 | 0 | 0 | 1 |
| rs11605568 | 11 | 116568168 | A | G | 0.279 | 0.7801 | +? | 0 | 0 | 0 | 1 |
| rs56060233 | 11 | 116540310 | T | C | 0.279 | 0.7802 | +? | 0 | 0 | 0 | 1 |
| rs67294535 | 11 | 120451223 | G | A | -0.279 | 0.7802 | -? | 0 | 0 | 0 | 1 |
| rs11217279 | 11 | 119297711 | A | G | -0.279 | 0.7803 | -? | 0 | 0 | 0 | 1 |
| rs12421915 | 11 | 119326630 | T | C | -0.279 | 0.7803 | -? | 0 | 0 | 0 | 1 |
| rs28489214 | 11 | 118993899 | G | A | 0.279 | 0.7803 | +? | 0 | 0 | 0 | 1 |
| rs629397 | 11 | 117440611 | C | G | -0.279 | 0.7803 | -? | 0 | 0 | 0 | 1 |
| rs7119828 | 11 | 119297614 | G | A | -0.279 | 0.7803 | -? | 0 | 0 | 0 | 1 |
| rs12276576 | 11 | 117672973 | G | C | 0.279 | 0.7804 | ++ | 0 | 0 | 1 | 0.9824 |
| rs140026975 | 11 | 118594669 | C | A | -0.279 | 0.7804 | -? | 0 | 0 | 0 | 1 |
| rs74724948 | 11 | 116544015 | T | A | 0.279 | 0.7804 | +? | 0 | 0 | 0 | 1 |
| rs140160504 | 11 | 119666456 | T | C | -0.279 | 0.7805 | -? | 0 | 0 | 0 | 1 |
| rs1053087454 | 11 | 117692121 | G | A | 0.279 | 0.7806 | +? | 0 | 0 | 0 | 1 |
| rs11216004 | 11 | 116426350 | T | C | 0.278 | 0.7807 | ++ | 0 | 0.124 | 1 | 0.725 |
| rs2305823 | 11 | 117306738 | C | A | 0.278 | 0.7807 | +? | 0 | 0 | 0 | 1 |
| rs10891989 | 11 | 116430203 | C | A | 0.278 | 0.7809 | ++ | 0 | 0.058 | 1 | 0.8095 |
| rs7930491 | 11 | 116427932 | A | G | 0.278 | 0.7809 | ++ | 0 | 0.007 | 1 | 0.9351 |
| rs61907605 | 11 | 116820071 | A | T | 0.278 | 0.781 | +? | 0 | 0 | 0 | 1 |
| rs35381826 | 11 | 118032438 | T | G | -0.278 | 0.7812 | -+ | 0 | 0.063 | 1 | 0.8019 |
| rs61900870 | 11 | 120394319 | G | A | 0.278 | 0.7813 | +? | 0 | 0 | 0 | 1 |
| rs61903430 | 11 | 116987342 | C | G | -0.277 | 0.7814 | -? | 0 | 0 | 0 | 1 |
| rs7106437 | 11 | 116992076 | T | C | -0.277 | 0.7814 | -? | 0 | 0 | 0 | 1 |
| rs7120515 | 11 | 116991926 | A | G | -0.277 | 0.7814 | -? | 0 | 0 | 0 | 1 |
| rs73587457 | 11 | 117647461 | G | C | -0.278 | 0.7814 | +- | 0.8 | 3.025 | 1 | 0.08201 |
| rs10790294 | 11 | 119305126 | T | C | -0.277 | 0.7815 | -? | 0 | 0 | 0 | 1 |
| rs57088249 | 11 | 119299371 | A | G | -0.277 | 0.7815 | -? | 0 | 0 | 0 | 1 |
| rs948776 | 11 | 117586623 | C | A | 0.277 | 0.7815 | +? | 0 | 0 | 0 | 1 |
| rs4450179 | 11 | 119309945 | C | T | -0.277 | 0.7818 | -? | 0 | 0 | 0 | 1 |
| rs111462461 | 11 | 117604866 | T | C | 0.277 | 0.7821 | +? | 0 | 0 | 0 | 1 |
| rs7943309 | 11 | 116773653 | A | G | -0.277 | 0.7821 | -? | 0 | 0 | 0 | 1 |
| rs508205 | 11 | 120057343 | G | A | -0.276 | 0.7822 | +- | 0 | 0.352 | 1 | 0.5529 |
| rs10160467 | 11 | 119968483 | C | G | 0.276 | 0.7824 | +? | 0 | 0 | 0 | 1 |
| rs11602570 | 11 | 117350098 | A | G | 0.276 | 0.7824 | +? | 0 | 0 | 0 | 1 |

| | | | | | | | | | | | |
|---|---|---|---|---|---|---|---|---|---|---|---|
| rs138184472 | 11 | 120486825 | C | A | -0.276 | 0.7826 | -? | 0 | 0 | 0 | 1 |
| rs45505501 | 11 | 118405305 | A | G | 0.276 | 0.7827 | ++ | 0 | 0 | 1 | 0.998 |
| rs11215990 | 11 | 116407583 | T | C | 0.276 | 0.7828 | +? | 0 | 0 | 0 | 1 |
| rs12807097 | 11 | 120242348 | A | G | 0.276 | 0.7828 | +? | 0 | 0 | 0 | 1 |
| rs7927993 | 11 | 116383211 | C | T | -0.276 | 0.7828 | -? | 0 | 0 | 0 | 1 |
| rs2000571 | 11 | 116585533 | A | G | -0.276 | 0.7829 | -? | 0 | 0 | 0 | 1 |
| rs579239 | 11 | 117425009 | C | T | -0.276 | 0.7829 | -? | 0 | 0 | 0 | 1 |
| rs7927379 | 11 | 117577925 | A | G | 0.276 | 0.7829 | +? | 0 | 0 | 0 | 1 |
| rs636518 | 11 | 119846820 | G | A | -0.275 | 0.783 | -? | 0 | 0 | 0 | 1 |
| rs11217634 | 11 | 119887151 | T | C | 0.275 | 0.7831 | +? | 0 | 0 | 0 | 1 |
| rs12272524 | 11 | 119888797 | T | C | 0.275 | 0.7831 | +? | 0 | 0 | 0 | 1 |
| rs1551238 | 11 | 119948132 | C | A | -0.275 | 0.7831 | -? | 0 | 0 | 0 | 1 |
| rs12295901 | 11 | 116347396 | A | G | 0.275 | 0.7832 | +? | 0 | 0 | 0 | 1 |
| rs180328 | 11 | 116623026 | G | A | -0.275 | 0.7832 | -+ | -13.9 | 2.633 | 1 | 0.1046 |
| rs595412 | 11 | 119015932 | T | C | 0.275 | 0.7832 | +? | 0 | 0 | 0 | 1 |
| rs10892419 | 11 | 119472574 | A | G | -0.275 | 0.7835 | -? | 0 | 0 | 0 | 1 |
| rs602594 | 11 | 118040610 | T | C | 0.275 | 0.7835 | +- | 0 | 0.282 | 1 | 0.5951 |
| rs6589775 | 11 | 119656448 | C | T | 0.275 | 0.7837 | +? | 0 | 0 | 0 | 1 |
| rs1379735938 | 11 | 116988040 | T | A | -0.274 | 0.784 | -? | 0 | 0 | 0 | 1 |
| rs142661080 | 11 | 117852972 | T | C | -0.274 | 0.784 | -? | 0 | 0 | 0 | 1 |
| rs80092204 | 11 | 116987988 | C | T | -0.274 | 0.784 | -? | 0 | 0 | 0 | 1 |
| rs1287393 | 11 | 120095872 | T | A | 0.274 | 0.7841 | +? | 0 | 0 | 0 | 1 |
| rs1429049972 | 11 | 116552062 | T | A | 0.274 | 0.7841 | +? | 0 | 0 | 0 | 1 |
| rs4936559 | 11 | 120705063 | C | T | 0.274 | 0.7841 | +? | 0 | 0 | 0 | 1 |
| rs7952043 | 11 | 116552562 | T | C | 0.274 | 0.7841 | +? | 0 | 0 | 0 | 1 |
| rs10892664 | 11 | 120911771 | A | T | -0.274 | 0.7842 | -? | 0 | 0 | 0 | 1 |
| rs61743159 | 11 | 117335686 | A | G | -0.274 | 0.7842 | -? | 0 | 0 | 0 | 1 |
| rs17122269 | 11 | 118445275 | C | T | 0.274 | 0.7843 | +- | 0 | 0.274 | 1 | 0.6006 |
| rs12417550 | 11 | 119299377 | A | G | -0.273 | 0.7845 | -? | 0 | 0 | 0 | 1 |
| rs61898266 | 11 | 120070943 | A | G | -0.273 | 0.7846 | -? | 0 | 0 | 0 | 1 |
| rs2663639 | 11 | 119418419 | G | A | 0.273 | 0.7847 | +? | 0 | 0 | 0 | 1 |
| rs3133854 | 11 | 120603801 | G | A | -0.273 | 0.7847 | -? | 0 | 0 | 0 | 1 |
| rs75949389 | 11 | 116892626 | G | A | -0.273 | 0.7847 | +- | 47.7 | 5.736 | 1 | 0.01662 |
| rs11600901 | 11 | 118031986 | A | G | -0.273 | 0.7848 | -+ | 0 | 0.084 | 1 | 0.7716 |
| rs17475804 | 11 | 116236979 | G | C | 0.273 | 0.7851 | +? | 0 | 0 | 0 | 1 |
| rs12800691 | 11 | 120145180 | C | G | -0.273 | 0.7852 | -? | 0 | 0 | 0 | 1 |
| rs1786189 | 11 | 117946691 | T | C | -0.273 | 0.7852 | -? | 0 | 0 | 0 | 1 |
| rs12785772 | 11 | 116563608 | A | G | 0.272 | 0.7853 | +? | 0 | 0 | 0 | 1 |
| rs3741323 | 11 | 118405641 | G | A | 0.272 | 0.7853 | ++ | 0 | 0 | 1 | 0.992 |
| rs1508098 | 11 | 116338899 | G | A | 0.272 | 0.7854 | +? | 0 | 0 | 0 | 1 |
| rs35238955 | 11 | 120182578 | T | G | -0.272 | 0.7854 | -? | 0 | 0 | 0 | 1 |
| rs45474901 | 11 | 118405453 | C | T | 0.272 | 0.7856 | ++ | 0 | 0 | 1 | 0.996 |
| rs4938690 | 11 | 119468661 | G | A | -0.272 | 0.7856 | -? | 0 | 0 | 0 | 1 |
| rs595483 | 11 | 119895096 | G | A | 0.272 | 0.7856 | +? | 0 | 0 | 0 | 1 |
| rs4417279 | 11 | 120543781 | T | G | -0.272 | 0.7857 | -? | 0 | 0 | 0 | 1 |
| rs1551239 | 11 | 119947953 | A | G | -0.272 | 0.7858 | -? | 0 | 0 | 0 | 1 |
| rs61905519 | 11 | 117045555 | C | T | -0.272 | 0.7858 | -? | 0 | 0 | 0 | 1 |
| rs61905521 | 11 | 117049020 | A | G | -0.272 | 0.7858 | -? | 0 | 0 | 0 | 1 |
| rs61905522 | 11 | 117049025 | A | G | -0.272 | 0.7858 | -? | 0 | 0 | 0 | 1 |
| rs11218161 | 11 | 121015711 | T | C | -0.272 | 0.7859 | -? | 0 | 0 | 0 | 1 |
| rs1145223 | 11 | 117716453 | T | C | -0.272 | 0.7859 | -? | 0 | 0 | 0 | 1 |
| rs79175883 | 11 | 116777394 | C | T | -0.272 | 0.7859 | -? | 0 | 0 | 0 | 1 |
| rs1583058 | 11 | 119269952 | C | T | -0.272 | 0.786 | -? | 0 | 0 | 0 | 1 |
| rs12801144 | 11 | 117802113 | G | A | 0.271 | 0.7861 | +? | 0 | 0 | 0 | 1 |
| rs4451741 | 11 | 120577600 | T | G | -0.271 | 0.7861 | -? | 0 | 0 | 0 | 1 |
| rs9630201 | 11 | 120577240 | A | G | -0.271 | 0.7861 | -? | 0 | 0 | 0 | 1 |

| | | | | | | | | | | | |
|---|---|---|---|---|---|---|---|---|---|---|---|
| rs4316540 | 11 | 116819241 | T | C | 0.271 | 0.7862 | +? | 0 | 0 | 0 | 1 |
| rs33929148 | 11 | 119243670 | T | C | 0.271 | 0.7864 | +? | 0 | 0 | 0 | 1 |
| rs7483081 | 11 | 118276246 | C | T | 0.271 | 0.7864 | +- | 0 | 0.14 | 1 | 0.7084 |
| rs755003 | 11 | 119439459 | A | G | 0.271 | 0.7864 | +? | 0 | 0 | 0 | 1 |
| rs9666462 | 11 | 118275997 | A | G | 0.271 | 0.7864 | +- | 0 | 0.14 | 1 | 0.7084 |
| rs7944738 | 11 | 117996119 | C | T | -0.271 | 0.7866 | -- | 0 | 0.011 | 1 | 0.9161 |
| rs1169496015 | 11 | 116960059 | G | C | -0.271 | 0.7867 | -+ | 73.6 | 11.362 | 1 | 0.0007498 |
| rs11215954 | 11 | 116348066 | T | G | 0.27 | 0.7868 | +? | 0 | 0 | 0 | 1 |
| rs1298336202 | 11 | 116640604 | T | C | 0.27 | 0.7868 | +- | -73.9 | 1.725 | 1 | 0.189 |
| rs4938341 | 11 | 117002596 | A | G | 0.27 | 0.7869 | +? | 0 | 0 | 0 | 1 |
| rs4609614 | 11 | 119448579 | A | C | -0.27 | 0.7871 | -? | 0 | 0 | 0 | 1 |
| rs59405103 | 11 | 118262305 | C | A | 0.27 | 0.7871 | +- | 0 | 0.164 | 1 | 0.6857 |
| rs141538185 | 11 | 117142625 | G | C | 0.27 | 0.7872 | +? | 0 | 0 | 0 | 1 |
| rs6589772 | 11 | 119637061 | A | C | 0.27 | 0.7872 | +? | 0 | 0 | 0 | 1 |
| rs71486349 | 11 | 120808303 | T | G | -0.27 | 0.7872 | -? | 0 | 0 | 0 | 1 |
| rs77257523 | 11 | 119884846 | C | T | 0.27 | 0.7872 | +? | 0 | 0 | 0 | 1 |
| rs77840919 | 11 | 119884816 | G | C | 0.27 | 0.7872 | +? | 0 | 0 | 0 | 1 |
| rs12280724 | 11 | 116613423 | T | G | -0.27 | 0.7873 | +- | -21.4 | 2.471 | 1 | 0.116 |
| rs118155116 | 11 | 117035228 | C | T | -0.27 | 0.7874 | -? | 0 | 0 | 0 | 1 |
| rs12576758 | 11 | 120898232 | T | C | -0.27 | 0.7875 | -? | 0 | 0 | 0 | 1 |
| rs68097814 | 11 | 120898482 | G | A | -0.27 | 0.7875 | -? | 0 | 0 | 0 | 1 |
| rs4938731 | 11 | 119689932 | A | G | -0.269 | 0.7876 | -? | 0 | 0 | 0 | 1 |
| rs637463 | 11 | 119886078 | A | T | 0.269 | 0.7876 | +? | 0 | 0 | 0 | 1 |
| rs7129877 | 11 | 119412428 | G | C | 0.269 | 0.7876 | +? | 0 | 0 | 0 | 1 |
| rs111234707 | 11 | 117912538 | A | G | -0.269 | 0.7877 | -? | 0 | 0 | 0 | 1 |
| rs17121259 | 11 | 117686567 | G | C | 0.269 | 0.7877 | +? | 0 | 0 | 0 | 1 |
| rs572608 | 11 | 118801640 | A | T | 0.269 | 0.7881 | ++ | 0 | 0 | 1 | 0.9932 |
| rs10892243 | 11 | 118441846 | C | T | 0.269 | 0.7882 | +- | 0 | 0.042 | 1 | 0.8367 |
| rs1599809 | 11 | 118179200 | A | G | 0.269 | 0.7883 | +? | 0 | 0 | 0 | 1 |
| rs12574090 | 11 | 116213060 | A | G | -0.268 | 0.7884 | -? | 0 | 0 | 0 | 1 |
| rs117259599 | 11 | 119312818 | T | C | 0.268 | 0.7885 | +? | 0 | 0 | 0 | 1 |
| rs11217495 | 11 | 119648938 | G | A | 0.268 | 0.7886 | +? | 0 | 0 | 0 | 1 |
| rs1129013 | 11 | 118989098 | C | G | 0.268 | 0.7886 | +? | 0 | 0 | 0 | 1 |
| rs61900868 | 11 | 120393597 | C | T | 0.268 | 0.7886 | +? | 0 | 0 | 0 | 1 |
| rs113337722 | 11 | 118464665 | G | A | -0.268 | 0.7888 | -- | 0 | 0.003 | 1 | 0.9549 |
| rs11604100 | 11 | 121096865 | T | C | -0.268 | 0.7888 | -? | 0 | 0 | 0 | 1 |
| rs470392 | 11 | 119994421 | T | C | -0.268 | 0.7888 | -? | 0 | 0 | 0 | 1 |
| rs584670 | 11 | 117927759 | A | G | -0.268 | 0.7889 | -? | 0 | 0 | 0 | 1 |
| rs61904859 | 11 | 116492506 | T | C | -0.268 | 0.7889 | -? | 0 | 0 | 0 | 1 |
| rs76924546 | 11 | 119885012 | A | G | 0.268 | 0.7889 | +? | 0 | 0 | 0 | 1 |
| rs11217801 | 11 | 120165942 | G | A | -0.267 | 0.7891 | -? | 0 | 0 | 0 | 1 |
| rs75310100 | 11 | 116578361 | T | G | -0.267 | 0.7892 | -? | 0 | 0 | 0 | 1 |
| rs3759012 | 11 | 118901262 | T | C | 0.267 | 0.7893 | +? | 0 | 0 | 0 | 1 |
| rs73598666 | 11 | 117591480 | T | G | 0.267 | 0.7894 | +? | 0 | 0 | 0 | 1 |
| rs78354899 | 11 | 120398577 | T | C | 0.267 | 0.7894 | +? | 0 | 0 | 0 | 1 |
| rs17567197 | 11 | 116247944 | C | A | 0.267 | 0.7895 | +? | 0 | 0 | 0 | 1 |
| rs76380380 | 11 | 117171475 | G | C | -0.267 | 0.7897 | -? | 0 | 0 | 0 | 1 |
| rs10892119 | 11 | 117331499 | A | G | -0.267 | 0.7898 | -? | 0 | 0 | 0 | 1 |
| rs111656195 | 11 | 118301375 | A | G | 0.267 | 0.7898 | -+ | 11.4 | 3.385 | 1 | 0.06579 |
| rs1244409804 | 11 | 116587146 | A | G | -0.267 | 0.7898 | -- | 0 | 0.039 | 1 | 0.843 |
| rs2510852 | 11 | 117377390 | C | G | -0.267 | 0.7898 | -? | 0 | 0 | 0 | 1 |
| rs11218155 | 11 | 121006345 | A | T | 0.266 | 0.7899 | +? | 0 | 0 | 0 | 1 |
| rs11826147 | 11 | 117613813 | C | G | 0.266 | 0.7899 | +? | 0 | 0 | 0 | 1 |
| rs12790073 | 11 | 117747277 | T | C | -0.266 | 0.7901 | -? | 0 | 0 | 0 | 1 |
| rs881122 | 11 | 117972936 | A | C | -0.266 | 0.7901 | -- | 0 | 0.123 | 1 | 0.7257 |
| rs535207 | 11 | 118408033 | G | A | 0.266 | 0.7904 | ++ | 0 | 0.003 | 1 | 0.9557 |

| | | | | | | | | | | | |
|---|---|---|---|---|---|---|---|---|---|---|---|
| rs9666463 | 11 | 118276116 | G | C | 0.266 | 0.7904 | +- | 0 | 0.138 | 1 | 0.7106 |
| rs1530604 | 11 | 119719420 | G | A | -0.266 | 0.7906 | -? | 0 | 0 | 0 | 1 |
| rs34527172 | 11 | 120394929 | C | T | 0.265 | 0.7907 | +? | 0 | 0 | 0 | 1 |
| rs35187805 | 11 | 120394910 | T | C | 0.265 | 0.7907 | +? | 0 | 0 | 0 | 1 |
| rs71484175 | 11 | 120394599 | A | G | 0.265 | 0.7907 | +? | 0 | 0 | 0 | 1 |
| rs58283940 | 11 | 116238557 | A | G | 0.265 | 0.791 | +? | 0 | 0 | 0 | 1 |
| rs2508665 | 11 | 117198789 | T | A | 0.265 | 0.7911 | +? | 0 | 0 | 0 | 1 |
| rs12805985 | 11 | 117802309 | G | A | 0.265 | 0.7912 | +? | 0 | 0 | 0 | 1 |
| rs117518542 | 11 | 116937146 | A | G | -0.264 | 0.7914 | -+ | 70.7 | 10.252 | 1 | 0.001365 |
| rs142661919 | 11 | 116937127 | G | A | -0.264 | 0.7914 | -+ | 70.7 | 10.252 | 1 | 0.001365 |
| rs11218122 | 11 | 120899458 | T | C | -0.264 | 0.7915 | -? | 0 | 0 | 0 | 1 |
| rs2845654 | 11 | 119437438 | C | A | 0.264 | 0.7915 | +? | 0 | 0 | 0 | 1 |
| rs4472969 | 11 | 117700323 | A | G | 0.264 | 0.7915 | +? | 0 | 0 | 0 | 1 |
| rs1258164206 | 11 | 117211240 | T | G | 0.264 | 0.7916 | +? | 0 | 0 | 0 | 1 |
| rs111903103 | 11 | 118308499 | T | C | 0.264 | 0.7917 | -+ | 13.9 | 3.483 | 1 | 0.062 |
| rs180369 | 11 | 116593058 | T | A | -0.264 | 0.7917 | -+ | -13.9 | 2.634 | 1 | 0.1046 |
| rs4938355 | 11 | 117053675 | G | A | 0.264 | 0.7918 | +? | 0 | 0 | 0 | 1 |
| rs111940587 | 11 | 117561070 | G | A | 0.264 | 0.7919 | +? | 0 | 0 | 0 | 1 |
| rs2097183 | 11 | 118091067 | T | C | -0.264 | 0.792 | -+ | 0 | 0.473 | 1 | 0.4918 |
| rs4936370 | 11 | 117193369 | T | G | 0.264 | 0.792 | +? | 0 | 0 | 0 | 1 |
| rs1006195 | 11 | 118958869 | T | G | -0.263 | 0.7922 | -? | 0 | 0 | 0 | 1 |
| rs74691084 | 11 | 116893790 | A | G | 0.263 | 0.7923 | +? | 0 | 0 | 0 | 1 |
| rs493678 | 11 | 120831520 | T | C | 0.263 | 0.7924 | +? | 0 | 0 | 0 | 1 |
| rs11218128 | 11 | 120926467 | A | T | -0.263 | 0.7926 | -? | 0 | 0 | 0 | 1 |
| rs511939 | 11 | 120813715 | G | C | -0.263 | 0.7926 | -? | 0 | 0 | 0 | 1 |
| rs67932968 | 11 | 120932457 | A | G | -0.263 | 0.7926 | -? | 0 | 0 | 0 | 1 |
| rs1784459 | 11 | 118938388 | A | C | 0.263 | 0.7927 | +? | 0 | 0 | 0 | 1 |
| rs1784460 | 11 | 118938371 | A | T | 0.263 | 0.7927 | +? | 0 | 0 | 0 | 1 |
| rs1786684 | 11 | 118938435 | A | G | 0.263 | 0.7927 | +? | 0 | 0 | 0 | 1 |
| rs77623050 | 11 | 116496948 | T | C | 0.263 | 0.7927 | +? | 0 | 0 | 0 | 1 |
| rs78485217 | 11 | 116496961 | A | T | 0.263 | 0.7927 | +? | 0 | 0 | 0 | 1 |
| rs12365864 | 11 | 116489036 | G | A | -0.262 | 0.7931 | -? | 0 | 0 | 0 | 1 |
| rs493693 | 11 | 116247759 | A | C | -0.262 | 0.7931 | -? | 0 | 0 | 0 | 1 |
| rs4938326 | 11 | 116905464 | C | G | 0.262 | 0.7931 | +? | 0 | 0 | 0 | 1 |
| rs2096883 | 11 | 120137544 | T | C | -0.262 | 0.7933 | -? | 0 | 0 | 0 | 1 |
| rs11216822 | 11 | 118091375 | A | G | -0.262 | 0.7935 | -+ | 0 | 0.477 | 1 | 0.4896 |
| rs11216096 | 11 | 116562580 | A | C | 0.262 | 0.7936 | +? | 0 | 0 | 0 | 1 |
| rs542987 | 11 | 116484093 | G | A | 0.262 | 0.7936 | +? | 0 | 0 | 0 | 1 |
| rs7929757 | 11 | 117612002 | C | T | 0.262 | 0.7936 | +? | 0 | 0 | 0 | 1 |
| rs10466589 | 11 | 116611985 | A | G | -0.262 | 0.7937 | +- | -20.7 | 2.486 | 1 | 0.1149 |
| rs56183153 | 11 | 119260456 | A | C | -0.262 | 0.7937 | -? | 0 | 0 | 0 | 1 |
| rs12226297 | 11 | 121185128 | A | G | -0.261 | 0.7938 | -? | 0 | 0 | 0 | 1 |
| rs2282494 | 11 | 117866868 | G | A | 0.261 | 0.7938 | +? | 0 | 0 | 0 | 1 |
| rs11825181 | 11 | 116626258 | A | G | -0.261 | 0.794 | +- | -24 | 2.42 | 1 | 0.1198 |
| rs653278 | 11 | 119899963 | T | A | 0.261 | 0.794 | +? | 0 | 0 | 0 | 1 |
| rs76260364 | 11 | 119884922 | T | C | 0.261 | 0.7942 | +? | 0 | 0 | 0 | 1 |
| rs12802321 | 11 | 119308804 | A | G | -0.261 | 0.7943 | -? | 0 | 0 | 0 | 1 |
| rs675866 | 11 | 118991019 | C | A | 0.261 | 0.7943 | +? | 0 | 0 | 0 | 1 |
| rs11216609 | 11 | 117769364 | C | T | -0.26 | 0.7946 | -? | 0 | 0 | 0 | 1 |
| rs600325 | 11 | 118041133 | C | T | 0.26 | 0.7947 | +- | 0 | 0.274 | 1 | 0.6009 |
| rs11218092 | 11 | 120857051 | A | G | -0.26 | 0.7948 | -+ | 0 | 0.271 | 1 | 0.6029 |
| rs191364497 | 11 | 120893217 | T | C | 0.26 | 0.7948 | +? | 0 | 0 | 0 | 1 |
| rs668033 | 11 | 119023107 | C | A | 0.26 | 0.7949 | +? | 0 | 0 | 0 | 1 |
| rs2853009 | 11 | 118148146 | G | A | -0.26 | 0.795 | -? | 0 | 0 | 0 | 1 |
| rs4938342 | 11 | 117003420 | G | C | 0.26 | 0.7951 | +? | 0 | 0 | 0 | 1 |
| rs57641217 | 11 | 116632956 | T | C | -0.26 | 0.7951 | +- | 5.3 | 3.168 | 1 | 0.07511 |

| | | | | | | | | | | | |
|---|---|---|---|---|---|---|---|---|---|---|---|
| rs76492199 | 11 | 117622274 | A | G | -0.26 | 0.7951 | +- | 22 | 3.845 | 1 | 0.04989 |
| rs504935 | 11 | 117446945 | C | T | 0.26 | 0.7952 | +? | 0 | 0 | 0 | 1 |
| rs4938339 | 11 | 116980594 | T | C | -0.259 | 0.7957 | -? | 0 | 0 | 0 | 1 |
| rs1078762 | 11 | 119438765 | T | C | 0.259 | 0.7959 | +? | 0 | 0 | 0 | 1 |
| rs11217281 | 11 | 119304053 | T | C | -0.258 | 0.7961 | -? | 0 | 0 | 0 | 1 |
| rs11820589 | 11 | 116633862 | A | G | -0.258 | 0.7961 | +- | -11.7 | 2.686 | 1 | 0.1012 |
| rs1901707 | 11 | 119984638 | T | C | -0.258 | 0.7961 | -? | 0 | 0 | 0 | 1 |
| rs76755469 | 11 | 121136144 | A | G | 0.258 | 0.7961 | +? | 0 | 0 | 0 | 1 |
| rs73594696 | 11 | 117561019 | C | T | 0.258 | 0.7962 | +? | 0 | 0 | 0 | 1 |
| rs78914290 | 11 | 119870442 | T | C | 0.258 | 0.7962 | +? | 0 | 0 | 0 | 1 |
| rs59590191 | 11 | 117774394 | G | A | 0.258 | 0.7963 | +? | 0 | 0 | 0 | 1 |
| rs7924856 | 11 | 120498051 | A | G | 0.258 | 0.7963 | +? | 0 | 0 | 0 | 1 |
| rs12792689 | 11 | 119259720 | G | A | -0.258 | 0.7964 | -? | 0 | 0 | 0 | 1 |
| rs676234 | 11 | 116722171 | G | A | -0.258 | 0.7964 | +- | -26.7 | 2.367 | 1 | 0.1239 |
| rs2228054 | 11 | 117864113 | A | G | 0.257 | 0.7969 | +? | 0 | 0 | 0 | 1 |
| rs73024372 | 11 | 117362820 | A | G | -0.257 | 0.7969 | -? | 0 | 0 | 0 | 1 |
| rs7930469 | 11 | 117585602 | C | T | 0.257 | 0.797 | +? | 0 | 0 | 0 | 1 |
| rs17121515 | 11 | 117882961 | A | G | 0.257 | 0.7971 | +? | 0 | 0 | 0 | 1 |
| rs7128587 | 11 | 118409302 | A | G | 0.257 | 0.7971 | ++ | 0 | 0.004 | 1 | 0.9483 |
| rs7131631 | 11 | 118409374 | A | G | 0.257 | 0.7971 | ++ | 0 | 0.004 | 1 | 0.9483 |
| rs73022041 | 11 | 118408789 | C | T | 0.257 | 0.7971 | ++ | 0 | 0.004 | 1 | 0.9483 |
| rs17123792 | 11 | 120166016 | A | G | -0.257 | 0.7973 | -? | 0 | 0 | 0 | 1 |
| rs10790343 | 11 | 119630480 | A | G | 0.257 | 0.7974 | +? | 0 | 0 | 0 | 1 |
| rs150081993 | 11 | 117337032 | T | C | -0.257 | 0.7975 | -? | 0 | 0 | 0 | 1 |
| rs583971 | 11 | 119899910 | A | G | 0.257 | 0.7975 | +? | 0 | 0 | 0 | 1 |
| rs77959607 | 11 | 119977373 | G | T | 0.257 | 0.7975 | +? | 0 | 0 | 0 | 1 |
| rs2305013 | 11 | 120340060 | T | A | 0.256 | 0.7977 | +? | 0 | 0 | 0 | 1 |
| rs10892194 | 11 | 117793239 | G | A | -0.256 | 0.7979 | -? | 0 | 0 | 0 | 1 |
| rs1305565 | 11 | 118990253 | C | T | 0.256 | 0.7979 | +? | 0 | 0 | 0 | 1 |
| rs5070 | 11 | 116708020 | A | G | 0.256 | 0.7979 | +? | 0 | 0 | 0 | 1 |
| rs71482132 | 11 | 118031211 | A | G | -0.256 | 0.798 | -+ | 0 | 0.086 | 1 | 0.7697 |
| rs11217494 | 11 | 119648830 | C | T | 0.256 | 0.7982 | +? | 0 | 0 | 0 | 1 |
| rs61903817 | 11 | 117534453 | G | C | -0.256 | 0.7982 | -? | 0 | 0 | 0 | 1 |
| rs78867156 | 11 | 120405828 | A | G | 0.256 | 0.7983 | +? | 0 | 0 | 0 | 1 |
| rs78484485 | 11 | 116598065 | A | G | -0.255 | 0.7984 | +- | 0 | 0.199 | 1 | 0.6554 |
| rs3862602 | 11 | 119296520 | G | A | -0.255 | 0.7985 | -? | 0 | 0 | 0 | 1 |
| rs73022038 | 11 | 118408471 | T | C | 0.255 | 0.7985 | ++ | 0 | 0.004 | 1 | 0.9509 |
| rs74404016 | 11 | 118408653 | T | C | 0.255 | 0.7985 | ++ | 0 | 0.004 | 1 | 0.9509 |
| rs7925835 | 11 | 116985641 | G | T | -0.255 | 0.7985 | -? | 0 | 0 | 0 | 1 |
| rs7941258 | 11 | 117392940 | T | C | 0.255 | 0.7986 | +? | 0 | 0 | 0 | 1 |
| rs7941460 | 11 | 117392987 | A | G | 0.255 | 0.7986 | +? | 0 | 0 | 0 | 1 |
| rs7944704 | 11 | 117393345 | A | G | 0.255 | 0.7986 | +? | 0 | 0 | 0 | 1 |
| rs566935 | 11 | 120055477 | C | G | 0.255 | 0.7987 | +? | 0 | 0 | 0 | 1 |
| rs7936363 | 11 | 119881024 | T | G | 0.255 | 0.7987 | +? | 0 | 0 | 0 | 1 |
| rs17122157 | 11 | 118276492 | A | G | 0.255 | 0.7988 | +- | 0 | 0.133 | 1 | 0.7152 |
| rs12289673 | 11 | 117615003 | A | G | 0.255 | 0.7989 | +? | 0 | 0 | 0 | 1 |
| rs12289727 | 11 | 117615068 | A | G | 0.255 | 0.7989 | +? | 0 | 0 | 0 | 1 |
| rs12291096 | 11 | 117615371 | T | C | 0.255 | 0.7989 | +? | 0 | 0 | 0 | 1 |
| rs2000869 | 11 | 120544871 | G | A | 0.255 | 0.7989 | +? | 0 | 0 | 0 | 1 |
| rs7107757 | 11 | 116686288 | T | C | -0.255 | 0.799 | -? | 0 | 0 | 0 | 1 |
| rs477106 | 11 | 118363424 | A | C | 0.254 | 0.7992 | +- | 0 | 0.033 | 1 | 0.8563 |
| rs7110647 | 11 | 117245133 | T | C | -0.254 | 0.7992 | -? | 0 | 0 | 0 | 1 |
| rs117195665 | 11 | 117316640 | T | G | -0.254 | 0.7997 | -? | 0 | 0 | 0 | 1 |
| rs11218213 | 11 | 121101852 | G | A | -0.253 | 0.7999 | -? | 0 | 0 | 0 | 1 |
| rs11218120 | 11 | 120898761 | G | A | -0.253 | 0.8003 | -? | 0 | 0 | 0 | 1 |
| rs11218121 | 11 | 120898784 | C | T | -0.253 | 0.8003 | -? | 0 | 0 | 0 | 1 |

| | | | | | | | | | | | |
|---|---|---|---|---|---|---|---|---|---|---|---|
| rs77953286 | 11 | 116914662 | C | G | -0.253 | 0.8004 | -+ | 72.3 | 10.835 | 1 | 0.000996 |
| rs11216886 | 11 | 118411424 | C | A | 0.253 | 0.8005 | ++ | 0 | 0.003 | 1 | 0.9545 |
| rs651348 | 11 | 117900997 | T | G | 0.253 | 0.8005 | +- | 0 | 0.355 | 1 | 0.5514 |
| rs642475 | 11 | 119899904 | C | T | 0.252 | 0.8009 | +? | 0 | 0 | 0 | 1 |
| rs61898292 | 11 | 120096056 | C | T | 0.252 | 0.801 | +? | 0 | 0 | 0 | 1 |
| rs1894006 | 11 | 119290369 | T | C | -0.252 | 0.8011 | -? | 0 | 0 | 0 | 1 |
| rs73022042 | 11 | 118408893 | A | G | 0.252 | 0.8011 | ++ | 0 | 0.005 | 1 | 0.946 |
| rs112120205 | 11 | 116899786 | A | G | 0.252 | 0.8012 | +? | 0 | 0 | 0 | 1 |
| rs609989 | 11 | 120814342 | A | G | -0.252 | 0.8012 | -? | 0 | 0 | 0 | 1 |
| rs61906103 | 11 | 116547004 | G | T | 0.252 | 0.8014 | +? | 0 | 0 | 0 | 1 |
| rs4938655 | 11 | 119306276 | A | C | -0.251 | 0.8015 | -? | 0 | 0 | 0 | 1 |
| rs61907563 | 11 | 116733565 | A | G | -0.251 | 0.8016 | -? | 0 | 0 | 0 | 1 |
| rs35407634 | 11 | 120553937 | T | G | 0.251 | 0.8018 | +? | 0 | 0 | 0 | 1 |
| rs61290697 | 11 | 118412358 | T | C | 0.251 | 0.802 | ++ | 0 | 0.002 | 1 | 0.9656 |
| rs11216487 | 11 | 117524497 | C | T | 0.25 | 0.8023 | -+ | -166.4 | 1.126 | 1 | 0.2886 |
| rs74443315 | 11 | 117583591 | G | A | 0.25 | 0.8023 | +? | 0 | 0 | 0 | 1 |
| rs59414734 | 11 | 116347944 | C | T | 0.25 | 0.8025 | +? | 0 | 0 | 0 | 1 |
| rs187236392 | 11 | 120074926 | G | A | 0.25 | 0.8027 | +? | 0 | 0 | 0 | 1 |
| rs7358294 | 11 | 119902521 | T | C | 0.25 | 0.8027 | +? | 0 | 0 | 0 | 1 |
| rs58768562 | 11 | 117896135 | T | C | 0.25 | 0.8028 | +? | 0 | 0 | 0 | 1 |
| rs73013729 | 11 | 117895163 | A | G | 0.25 | 0.8028 | +? | 0 | 0 | 0 | 1 |
| rs7942124 | 11 | 117895268 | T | A | 0.25 | 0.8028 | +? | 0 | 0 | 0 | 1 |
| rs7942146 | 11 | 117895305 | G | A | 0.25 | 0.8028 | +? | 0 | 0 | 0 | 1 |
| rs7942225 | 11 | 117895276 | T | C | 0.25 | 0.8028 | +? | 0 | 0 | 0 | 1 |
| rs7945541 | 11 | 117895712 | T | C | 0.25 | 0.8028 | +? | 0 | 0 | 0 | 1 |
| rs7945602 | 11 | 117895586 | A | G | 0.25 | 0.8028 | +? | 0 | 0 | 0 | 1 |
| rs115293580 | 11 | 120538317 | T | C | 0.249 | 0.8031 | +? | 0 | 0 | 0 | 1 |
| rs55734733 | 11 | 117894258 | A | C | 0.249 | 0.8031 | +? | 0 | 0 | 0 | 1 |
| rs55858401 | 11 | 117894371 | T | C | 0.249 | 0.8031 | +? | 0 | 0 | 0 | 1 |
| rs56029127 | 11 | 117894424 | A | G | 0.249 | 0.8031 | +? | 0 | 0 | 0 | 1 |
| rs60169792 | 11 | 117893889 | T | C | 0.249 | 0.8031 | +? | 0 | 0 | 0 | 1 |
| rs7121196 | 11 | 117729618 | A | G | -0.249 | 0.8033 | -? | 0 | 0 | 0 | 1 |
| rs4938517 | 11 | 118487673 | C | T | -0.249 | 0.8037 | -+ | 0 | 0.435 | 1 | 0.5098 |
| rs687560 | 11 | 121076285 | T | C | -0.248 | 0.8038 | -? | 0 | 0 | 0 | 1 |
| rs11216897 | 11 | 118432279 | G | A | 0.248 | 0.8039 | +- | 0 | 0.036 | 1 | 0.8495 |
| rs4245196 | 11 | 119472474 | G | A | -0.248 | 0.8039 | -? | 0 | 0 | 0 | 1 |
| rs1786141 | 11 | 118938315 | T | C | 0.248 | 0.804 | +? | 0 | 0 | 0 | 1 |
| rs59108310 | 11 | 118433729 | G | A | 0.248 | 0.804 | +- | 0 | 0.063 | 1 | 0.8017 |
| rs532815 | 11 | 116718858 | G | A | -0.248 | 0.8041 | -? | 0 | 0 | 0 | 1 |
| rs10790240 | 11 | 118009279 | G | T | -0.248 | 0.8043 | -+ | 0 | 0.024 | 1 | 0.8764 |
| rs202125027 | 11 | 119448580 | A | T | -0.248 | 0.8044 | -? | 0 | 0 | 0 | 1 |
| rs12274595 | 11 | 117692842 | C | T | 0.247 | 0.8046 | +? | 0 | 0 | 0 | 1 |
| rs11216926 | 11 | 118471996 | T | C | 0.247 | 0.8048 | +- | 0 | 0.209 | 1 | 0.6475 |
| rs4938802 | 11 | 120228989 | G | A | 0.247 | 0.8048 | +? | 0 | 0 | 0 | 1 |
| rs614825 | 11 | 118018725 | G | A | 0.247 | 0.8048 | ++ | 0 | 0.046 | 1 | 0.8294 |
| rs61355898 | 11 | 120578133 | T | C | -0.247 | 0.8049 | -? | 0 | 0 | 0 | 1 |
| rs948773 | 11 | 117586942 | A | G | 0.247 | 0.8049 | +? | 0 | 0 | 0 | 1 |
| rs948774 | 11 | 117586881 | A | T | 0.247 | 0.8049 | +? | 0 | 0 | 0 | 1 |
| rs10892452 | 11 | 119630385 | T | C | 0.247 | 0.805 | +? | 0 | 0 | 0 | 1 |
| rs4938654 | 11 | 119306183 | T | C | -0.247 | 0.805 | -? | 0 | 0 | 0 | 1 |
| rs12286604 | 11 | 118084728 | A | G | -0.247 | 0.8051 | -+ | 0 | 0.13 | 1 | 0.7181 |
| rs4625480 | 11 | 119831744 | G | A | 0.247 | 0.8051 | +? | 0 | 0 | 0 | 1 |
| rs7116106 | 11 | 120556018 | T | G | -0.247 | 0.8052 | -? | 0 | 0 | 0 | 1 |
| rs57461652 | 11 | 118146197 | A | C | -0.246 | 0.8053 | -? | 0 | 0 | 0 | 1 |
| rs580714 | 11 | 117423538 | C | A | -0.246 | 0.8053 | -? | 0 | 0 | 0 | 1 |
| rs75813208 | 11 | 117330944 | T | C | 0.246 | 0.8054 | +? | 0 | 0 | 0 | 1 |

| | | | | | | | | | | | |
|---|---|---|---|---|---|---|---|---|---|---|---|
| rs12804415 | 11 | 116232487 | G | A | -0.246 | 0.8055 | +- | -152.2 | 1.189 | 1 | 0.2755 |
| rs652449 | 11 | 119891597 | G | T | 0.246 | 0.8055 | +? | 0 | 0 | 0 | 1 |
| rs4938729 | 11 | 119687318 | C | T | -0.246 | 0.8057 | -? | 0 | 0 | 0 | 1 |
| rs12285127 | 11 | 119900754 | A | G | 0.246 | 0.8059 | +? | 0 | 0 | 0 | 1 |
| rs4576856 | 11 | 117622809 | G | T | 0.246 | 0.8059 | +- | 5.2 | 3.166 | 1 | 0.0752 |
| rs11216883 | 11 | 118410468 | G | A | 0.246 | 0.806 | ++ | 0 | 0.002 | 1 | 0.9643 |
| rs11216884 | 11 | 118410488 | C | T | 0.246 | 0.806 | ++ | 0 | 0.002 | 1 | 0.9643 |
| rs11216885 | 11 | 118410545 | G | A | 0.246 | 0.806 | ++ | 0 | 0.002 | 1 | 0.9643 |
| rs66461646 | 11 | 119319811 | C | T | -0.246 | 0.806 | -? | 0 | 0 | 0 | 1 |
| rs7943039 | 11 | 118603030 | A | G | -0.246 | 0.806 | -+ | 0 | 0.467 | 1 | 0.4946 |
| rs77341747 | 11 | 116964066 | T | C | -0.245 | 0.8061 | -+ | 76.3 | 12.678 | 1 | 0.00037 |
| rs77366224 | 11 | 116964065 | T | G | -0.245 | 0.8061 | -+ | 76.3 | 12.678 | 1 | 0.00037 |
| rs2063176 | 11 | 116428507 | G | C | 0.245 | 0.8062 | ++ | 0 | 0.009 | 1 | 0.9236 |
| rs7124100 | 11 | 118290169 | A | G | 0.245 | 0.8062 | +- | 0 | 0.253 | 1 | 0.6148 |
| rs1665777 | 11 | 119703771 | C | A | -0.245 | 0.8064 | -? | 0 | 0 | 0 | 1 |
| rs76972960 | 11 | 116916678 | C | T | 0.245 | 0.8064 | +? | 0 | 0 | 0 | 1 |
| rs1786685 | 11 | 118938957 | A | G | 0.245 | 0.8067 | +? | 0 | 0 | 0 | 1 |
| rs3212264 | 11 | 118216234 | A | C | -0.245 | 0.8067 | -+ | 0 | 0.788 | 1 | 0.3747 |
| rs45529532 | 11 | 118269674 | G | A | -0.245 | 0.8067 | -? | 0 | 0 | 0 | 1 |
| rs77252250 | 11 | 117591311 | A | C | 0.245 | 0.8067 | +? | 0 | 0 | 0 | 1 |
| rs12269936 | 11 | 119274417 | G | A | -0.244 | 0.807 | -? | 0 | 0 | 0 | 1 |
| rs4409838 | 11 | 119509911 | A | G | 0.244 | 0.807 | +? | 0 | 0 | 0 | 1 |
| rs471403 | 11 | 120816277 | A | G | -0.244 | 0.807 | -? | 0 | 0 | 0 | 1 |
| rs542903 | 11 | 119692663 | T | C | 0.244 | 0.807 | +? | 0 | 0 | 0 | 1 |
| rs57702967 | 11 | 118261736 | A | C | 0.244 | 0.8071 | +- | 0 | 0.203 | 1 | 0.6527 |
| rs11216523 | 11 | 117615875 | A | G | 0.244 | 0.8073 | +? | 0 | 0 | 0 | 1 |
| rs11216524 | 11 | 117615877 | C | G | 0.244 | 0.8073 | +? | 0 | 0 | 0 | 1 |
| rs12421359 | 11 | 119277583 | G | A | -0.244 | 0.8073 | -? | 0 | 0 | 0 | 1 |
| rs3133525 | 11 | 120671999 | A | G | 0.243 | 0.808 | +? | 0 | 0 | 0 | 1 |
| rs6589855 | 11 | 120954769 | C | T | -0.243 | 0.8081 | -? | 0 | 0 | 0 | 1 |
| rs17122021 | 11 | 118145686 | T | C | -0.243 | 0.8083 | -? | 0 | 0 | 0 | 1 |
| rs4245179 | 11 | 118144445 | G | A | -0.243 | 0.8083 | -? | 0 | 0 | 0 | 1 |
| rs4245180 | 11 | 118144512 | A | G | -0.243 | 0.8083 | -? | 0 | 0 | 0 | 1 |
| rs4938493 | 11 | 118144804 | G | T | -0.243 | 0.8083 | -? | 0 | 0 | 0 | 1 |
| rs57506367 | 11 | 118141993 | C | T | -0.243 | 0.8083 | -? | 0 | 0 | 0 | 1 |
| rs58254953 | 11 | 119472670 | A | G | -0.243 | 0.8083 | -? | 0 | 0 | 0 | 1 |
| rs607888 | 11 | 119887019 | A | G | 0.243 | 0.8083 | +? | 0 | 0 | 0 | 1 |
| rs7940047 | 11 | 118142619 | T | C | -0.243 | 0.8083 | -? | 0 | 0 | 0 | 1 |
| rs489059 | 11 | 119987517 | A | G | 0.242 | 0.8084 | +? | 0 | 0 | 0 | 1 |
| rs2048247 | 11 | 119476351 | A | G | -0.242 | 0.8086 | -? | 0 | 0 | 0 | 1 |
| rs2048248 | 11 | 119476354 | A | G | -0.242 | 0.8086 | -? | 0 | 0 | 0 | 1 |
| rs79410922 | 11 | 121101046 | G | T | -0.242 | 0.8087 | -? | 0 | 0 | 0 | 1 |
| rs35514679 | 11 | 119319701 | A | C | -0.242 | 0.8088 | -? | 0 | 0 | 0 | 1 |
| rs1789661 | 11 | 120827021 | T | C | -0.242 | 0.8089 | -+ | 0 | 0.26 | 1 | 0.6103 |
| rs1787024 | 11 | 119845535 | C | T | -0.242 | 0.809 | -? | 0 | 0 | 0 | 1 |
| rs10892458 | 11 | 119649488 | T | C | 0.242 | 0.8091 | +? | 0 | 0 | 0 | 1 |
| rs61903429 | 11 | 116986684 | A | C | -0.242 | 0.8091 | -? | 0 | 0 | 0 | 1 |
| rs7944427 | 11 | 117305461 | T | C | 0.241 | 0.8092 | +? | 0 | 0 | 0 | 1 |
| rs9804547 | 11 | 121086444 | A | T | -0.241 | 0.8092 | -? | 0 | 0 | 0 | 1 |
| rs2845653 | 11 | 119437815 | G | A | 0.241 | 0.8093 | +? | 0 | 0 | 0 | 1 |
| rs10458939 | 11 | 117898851 | A | G | 0.241 | 0.8095 | +? | 0 | 0 | 0 | 1 |
| rs11216682 | 11 | 117888564 | T | C | 0.241 | 0.8095 | +? | 0 | 0 | 0 | 1 |
| rs11216683 | 11 | 117888681 | G | A | 0.241 | 0.8095 | +? | 0 | 0 | 0 | 1 |
| rs11216685 | 11 | 117888746 | T | C | 0.241 | 0.8095 | +? | 0 | 0 | 0 | 1 |
| rs11216687 | 11 | 117889895 | A | G | 0.241 | 0.8095 | +? | 0 | 0 | 0 | 1 |
| rs11216688 | 11 | 117889922 | A | G | 0.241 | 0.8095 | +? | 0 | 0 | 0 | 1 |

| | | | | | | | | | | | |
|---|---|---|---|---|---|---|---|---|---|---|---|
| rs11216689 | 11 | 117889959 | A | C | 0.241 | 0.8095 | +? | 0 | 0 | 0 | 1 |
| rs11216690 | 11 | 117890167 | T | C | 0.241 | 0.8095 | +? | 0 | 0 | 0 | 1 |
| rs11216691 | 11 | 117890221 | G | A | 0.241 | 0.8095 | +? | 0 | 0 | 0 | 1 |
| rs11216692 | 11 | 117890295 | T | A | 0.241 | 0.8095 | +? | 0 | 0 | 0 | 1 |
| rs11216693 | 11 | 117890386 | A | C | 0.241 | 0.8095 | +? | 0 | 0 | 0 | 1 |
| rs11216694 | 11 | 117890391 | A | G | 0.241 | 0.8095 | +? | 0 | 0 | 0 | 1 |
| rs11216695 | 11 | 117890417 | A | G | 0.241 | 0.8095 | +? | 0 | 0 | 0 | 1 |
| rs11216696 | 11 | 117892322 | G | A | 0.241 | 0.8095 | +? | 0 | 0 | 0 | 1 |
| rs150204107 | 11 | 120401377 | A | G | -0.241 | 0.8095 | -? | 0 | 0 | 0 | 1 |
| rs17121524 | 11 | 117886623 | G | A | 0.241 | 0.8095 | +? | 0 | 0 | 0 | 1 |
| rs17121528 | 11 | 117889740 | T | C | 0.241 | 0.8095 | +? | 0 | 0 | 0 | 1 |
| rs4630330 | 11 | 117890857 | T | G | 0.241 | 0.8095 | +? | 0 | 0 | 0 | 1 |
| rs56118320 | 11 | 117886931 | A | C | 0.241 | 0.8095 | +? | 0 | 0 | 0 | 1 |
| rs57916935 | 11 | 117890874 | G | A | 0.241 | 0.8095 | +? | 0 | 0 | 0 | 1 |
| rs58496951 | 11 | 117890995 | C | A | 0.241 | 0.8095 | +? | 0 | 0 | 0 | 1 |
| rs58551444 | 11 | 117888418 | C | G | 0.241 | 0.8095 | +? | 0 | 0 | 0 | 1 |
| rs59198393 | 11 | 117890752 | T | C | 0.241 | 0.8095 | +? | 0 | 0 | 0 | 1 |
| rs59872579 | 11 | 117886753 | A | G | 0.241 | 0.8095 | +? | 0 | 0 | 0 | 1 |
| rs60590812 | 11 | 117892670 | C | G | 0.241 | 0.8095 | +? | 0 | 0 | 0 | 1 |
| rs61579653 | 11 | 117890989 | A | G | 0.241 | 0.8095 | +? | 0 | 0 | 0 | 1 |
| rs61598135 | 11 | 117888352 | A | T | 0.241 | 0.8095 | +? | 0 | 0 | 0 | 1 |
| rs73011704 | 11 | 117887084 | T | G | 0.241 | 0.8095 | +? | 0 | 0 | 0 | 1 |
| rs73011705 | 11 | 117887240 | A | G | 0.241 | 0.8095 | +? | 0 | 0 | 0 | 1 |
| rs73011710 | 11 | 117887667 | C | T | 0.241 | 0.8095 | +? | 0 | 0 | 0 | 1 |
| rs73011712 | 11 | 117887671 | C | T | 0.241 | 0.8095 | +? | 0 | 0 | 0 | 1 |
| rs73011721 | 11 | 117889569 | C | T | 0.241 | 0.8095 | +? | 0 | 0 | 0 | 1 |
| rs73011724 | 11 | 117889690 | A | T | 0.241 | 0.8095 | +? | 0 | 0 | 0 | 1 |
| rs73011757 | 11 | 117891787 | G | A | 0.241 | 0.8095 | +? | 0 | 0 | 0 | 1 |
| rs73013721 | 11 | 117893518 | T | G | 0.241 | 0.8095 | +? | 0 | 0 | 0 | 1 |
| rs7929175 | 11 | 117891489 | G | A | 0.241 | 0.8095 | +? | 0 | 0 | 0 | 1 |
| rs7944128 | 11 | 117891384 | C | T | 0.241 | 0.8095 | +? | 0 | 0 | 0 | 1 |
| rs7944374 | 11 | 117891582 | C | T | 0.241 | 0.8095 | +? | 0 | 0 | 0 | 1 |
| rs875025 | 11 | 116373451 | T | C | 0.241 | 0.8095 | +? | 0 | 0 | 0 | 1 |
| rs7927810 | 11 | 117991333 | A | G | -0.241 | 0.8096 | -- | 0 | 0.01 | 1 | 0.9185 |
| rs11217280 | 11 | 119304002 | G | A | -0.241 | 0.8097 | -? | 0 | 0 | 0 | 1 |
| rs12283157 | 11 | 119304238 | C | G | -0.241 | 0.8097 | -? | 0 | 0 | 0 | 1 |
| rs2511834 | 11 | 119197707 | A | G | -0.241 | 0.8097 | -? | 0 | 0 | 0 | 1 |
| rs74369487 | 11 | 117168340 | C | A | 0.241 | 0.8098 | +? | 0 | 0 | 0 | 1 |
| rs10892327 | 11 | 118939488 | G | C | 0.241 | 0.8099 | +? | 0 | 0 | 0 | 1 |
| rs949096 | 11 | 119839344 | A | G | -0.241 | 0.8099 | -? | 0 | 0 | 0 | 1 |
| rs11216676 | 11 | 117882394 | G | A | 0.24 | 0.8101 | +? | 0 | 0 | 0 | 1 |
| rs12226501 | 11 | 119831362 | G | T | 0.24 | 0.8101 | +? | 0 | 0 | 0 | 1 |
| rs57157718 | 11 | 117101445 | T | A | 0.24 | 0.8102 | +? | 0 | 0 | 0 | 1 |
| rs11216702 | 11 | 117894899 | A | T | 0.24 | 0.8103 | +? | 0 | 0 | 0 | 1 |
| rs2367970 | 11 | 116581641 | A | G | -0.24 | 0.8103 | -? | 0 | 0 | 0 | 1 |
| rs4322404 | 11 | 119420469 | C | T | 0.24 | 0.8103 | +? | 0 | 0 | 0 | 1 |
| rs61898859 | 11 | 119306752 | C | A | -0.24 | 0.8103 | -? | 0 | 0 | 0 | 1 |
| rs73013726 | 11 | 117894709 | A | G | 0.24 | 0.8103 | +? | 0 | 0 | 0 | 1 |
| rs11216902 | 11 | 118437279 | G | A | 0.24 | 0.8104 | +- | 0 | 0.063 | 1 | 0.8017 |
| rs490111 | 11 | 117426543 | G | A | 0.24 | 0.8105 | +? | 0 | 0 | 0 | 1 |
| rs480958 | 11 | 118577990 | G | A | 0.24 | 0.8106 | -+ | 0 | 0.312 | 1 | 0.5765 |
| rs9326249 | 11 | 117318148 | A | G | -0.239 | 0.8108 | -? | 0 | 0 | 0 | 1 |
| rs4938370 | 11 | 117193350 | C | A | 0.239 | 0.8109 | +? | 0 | 0 | 0 | 1 |
| rs7931873 | 11 | 119821850 | G | A | 0.239 | 0.8109 | +? | 0 | 0 | 0 | 1 |
| rs78017165 | 11 | 121100855 | A | G | -0.239 | 0.811 | -? | 0 | 0 | 0 | 1 |
| rs1939668 | 11 | 120599866 | A | G | -0.239 | 0.8111 | -? | 0 | 0 | 0 | 1 |

| | | | | | | | | | | | |
|---|---|---|---|---|---|---|---|---|---|---|---|
| rs7941150 | 11 | 117091821 | A | G | 0.239 | 0.8113 | +? | 0 | 0 | 0 | 1 |
| rs144785834 | 11 | 120095782 | C | A | 0.239 | 0.8114 | +? | 0 | 0 | 0 | 1 |
| rs73613980 | 11 | 118284650 | T | C | 0.238 | 0.8115 | +- | 0 | 0.142 | 1 | 0.706 |
| rs498872 | 11 | 118477367 | A | G | -0.238 | 0.8117 | -- | 0 | 0.109 | 1 | 0.7415 |
| rs112590920 | 11 | 118143586 | A | G | -0.238 | 0.8118 | -? | 0 | 0 | 0 | 1 |
| rs1262953 | 11 | 116579099 | A | G | 0.238 | 0.8118 | +? | 0 | 0 | 0 | 1 |
| rs7940554 | 11 | 118142930 | T | C | -0.238 | 0.8118 | -? | 0 | 0 | 0 | 1 |
| rs45600633 | 11 | 118460957 | A | G | -0.238 | 0.812 | -- | 0 | 0 | 1 | 0.9844 |
| rs1784305 | 11 | 118942048 | A | G | 0.238 | 0.8121 | +? | 0 | 0 | 0 | 1 |
| rs73005818 | 11 | 116239770 | A | G | 0.238 | 0.8121 | +? | 0 | 0 | 0 | 1 |
| rs12802944 | 11 | 116576400 | C | T | -0.238 | 0.8122 | -? | 0 | 0 | 0 | 1 |
| rs4439549 | 11 | 117780091 | A | C | 0.237 | 0.8123 | +? | 0 | 0 | 0 | 1 |
| rs526397 | 11 | 120852324 | G | T | -0.237 | 0.8124 | -+ | 0 | 0.257 | 1 | 0.6121 |
| rs6589595 | 11 | 116981335 | A | G | -0.237 | 0.8125 | -? | 0 | 0 | 0 | 1 |
| rs735007 | 11 | 119317018 | G | A | -0.237 | 0.8125 | -? | 0 | 0 | 0 | 1 |
| rs10790376 | 11 | 120182932 | A | G | -0.237 | 0.8126 | -? | 0 | 0 | 0 | 1 |
| rs10892561 | 11 | 120182869 | T | C | -0.237 | 0.8126 | -? | 0 | 0 | 0 | 1 |
| rs17500548 | 11 | 117232391 | G | A | 0.237 | 0.8126 | +? | 0 | 0 | 0 | 1 |
| rs33981325 | 11 | 121028666 | T | C | -0.237 | 0.8127 | -? | 0 | 0 | 0 | 1 |
| rs7111567 | 11 | 119316659 | T | G | -0.237 | 0.8129 | -? | 0 | 0 | 0 | 1 |
| rs4938742 | 11 | 119831351 | G | A | -0.236 | 0.8131 | -? | 0 | 0 | 0 | 1 |
| rs595359 | 11 | 118992502 | T | G | 0.236 | 0.8132 | +? | 0 | 0 | 0 | 1 |
| rs7117705 | 11 | 119492741 | C | T | -0.236 | 0.8132 | -? | 0 | 0 | 0 | 1 |
| rs7103936 | 11 | 118091636 | C | A | -0.236 | 0.8134 | -+ | 0 | 0.464 | 1 | 0.4958 |
| rs2019655 | 11 | 117699878 | T | C | -0.236 | 0.8135 | -? | 0 | 0 | 0 | 1 |
| rs10790179 | 11 | 117236319 | T | G | 0.236 | 0.8136 | +? | 0 | 0 | 0 | 1 |
| rs10892097 | 11 | 117236060 | T | C | 0.236 | 0.8136 | +? | 0 | 0 | 0 | 1 |
| rs4351828 | 11 | 119627450 | A | G | -0.236 | 0.8136 | -? | 0 | 0 | 0 | 1 |
| rs3138093 | 11 | 119291403 | C | T | -0.236 | 0.8138 | -? | 0 | 0 | 0 | 1 |
| rs56027537 | 11 | 119281397 | C | T | -0.236 | 0.8138 | -? | 0 | 0 | 0 | 1 |
| rs891293 | 11 | 119302637 | A | G | -0.236 | 0.8138 | -? | 0 | 0 | 0 | 1 |
| rs12417301 | 11 | 119284523 | A | T | -0.235 | 0.8139 | -? | 0 | 0 | 0 | 1 |
| rs11216880 | 11 | 118409536 | A | G | 0.235 | 0.8142 | ++ | 0 | 0.016 | 1 | 0.9006 |
| rs7101585 | 11 | 118409583 | A | G | 0.235 | 0.8142 | ++ | 0 | 0.016 | 1 | 0.9006 |
| rs667670 | 11 | 119885822 | C | T | 0.235 | 0.8143 | +? | 0 | 0 | 0 | 1 |
| rs11216067 | 11 | 116504377 | G | A | 0.235 | 0.8144 | +? | 0 | 0 | 0 | 1 |
| rs17476159 | 11 | 116250523 | C | T | -0.235 | 0.8145 | -? | 0 | 0 | 0 | 1 |
| rs2000873 | 11 | 120520218 | C | A | -0.234 | 0.8147 | -? | 0 | 0 | 0 | 1 |
| rs56371319 | 11 | 117042967 | A | C | 0.234 | 0.8148 | +? | 0 | 0 | 0 | 1 |
| rs58418986 | 11 | 120398613 | A | G | -0.234 | 0.8148 | -? | 0 | 0 | 0 | 1 |
| rs684913 | 11 | 120819562 | T | C | -0.234 | 0.815 | -? | 0 | 0 | 0 | 1 |
| rs1893391 | 11 | 119979732 | C | G | -0.234 | 0.8151 | -? | 0 | 0 | 0 | 1 |
| rs2187639 | 11 | 117818041 | A | C | -0.234 | 0.8151 | -+ | 0 | 0.05 | 1 | 0.8234 |
| rs58162302 | 11 | 118146199 | T | A | -0.234 | 0.8151 | -? | 0 | 0 | 0 | 1 |
| rs11608057 | 11 | 118030185 | T | C | -0.234 | 0.8152 | -+ | 0 | 0.115 | 1 | 0.735 |
| rs74996375 | 11 | 120008972 | A | C | -0.234 | 0.8152 | -? | 0 | 0 | 0 | 1 |
| rs12792929 | 11 | 118240715 | C | G | -0.234 | 0.8153 | -- | 0 | 0.011 | 1 | 0.9183 |
| rs4471466 | 11 | 116340960 | C | A | -0.234 | 0.8153 | -? | 0 | 0 | 0 | 1 |
| rs624931 | 11 | 117922486 | C | T | -0.233 | 0.8154 | -? | 0 | 0 | 0 | 1 |
| rs1614264 | 11 | 118942638 | C | A | 0.233 | 0.8155 | +? | 0 | 0 | 0 | 1 |
| rs664320 | 11 | 119947402 | A | G | -0.233 | 0.8157 | -? | 0 | 0 | 0 | 1 |
| rs150032046 | 11 | 120622042 | G | A | -0.233 | 0.8159 | -? | 0 | 0 | 0 | 1 |
| rs598804 | 11 | 120851125 | T | C | -0.233 | 0.8161 | -+ | 0 | 0.254 | 1 | 0.614 |
| rs35435609 | 11 | 118502172 | T | C | 0.232 | 0.8163 | ++ | 0 | 0.051 | 1 | 0.8212 |
| rs882856 | 11 | 120107411 | G | A | -0.232 | 0.8163 | -? | 0 | 0 | 0 | 1 |
| rs3138094 | 11 | 119288619 | G | A | -0.232 | 0.8164 | -? | 0 | 0 | 0 | 1 |

| | | | | | | | | | | | |
|---|---|---|---|---|---|---|---|---|---|---|---|
| rs60580523 | 11 | 117303242 | T | C | 0.232 | 0.8164 | +? | 0 | 0 | 0 | 1 |
| rs61899043 | 11 | 119837947 | C | A | 0.232 | 0.8164 | +? | 0 | 0 | 0 | 1 |
| rs632039 | 11 | 119885840 | C | T | 0.232 | 0.8166 | +? | 0 | 0 | 0 | 1 |
| rs74849419 | 11 | 116583685 | A | G | -0.232 | 0.8167 | +- | 0 | 0.336 | 1 | 0.562 |
| rs3138096 | 11 | 119286275 | G | C | -0.232 | 0.8168 | -? | 0 | 0 | 0 | 1 |
| rs12418678 | 11 | 117232260 | G | A | 0.232 | 0.8169 | +? | 0 | 0 | 0 | 1 |
| rs540787 | 11 | 120854479 | T | C | -0.231 | 0.817 | -+ | 0 | 0.254 | 1 | 0.6145 |
| rs11216697 | 11 | 117892362 | A | G | 0.231 | 0.8171 | +? | 0 | 0 | 0 | 1 |
| rs570373 | 11 | 120846239 | G | A | -0.231 | 0.8171 | -+ | 0 | 0.254 | 1 | 0.6145 |
| rs12793214 | 11 | 118800302 | A | C | 0.231 | 0.8173 | ++ | 0 | 0.001 | 1 | 0.9792 |
| rs76126708 | 11 | 118259379 | C | A | 0.231 | 0.8173 | +- | 0 | 0.211 | 1 | 0.6461 |
| rs10892456 | 11 | 119649342 | G | A | 0.231 | 0.8174 | +? | 0 | 0 | 0 | 1 |
| rs1429512 | 11 | 119292314 | C | T | -0.231 | 0.8174 | -? | 0 | 0 | 0 | 1 |
| rs590282 | 11 | 119881715 | A | G | 0.231 | 0.8174 | +? | 0 | 0 | 0 | 1 |
| rs59969944 | 11 | 119233110 | A | G | 0.231 | 0.8174 | +? | 0 | 0 | 0 | 1 |
| rs7111286 | 11 | 119272485 | C | G | -0.231 | 0.8174 | -? | 0 | 0 | 0 | 1 |
| rs12575027 | 11 | 119276600 | A | G | -0.231 | 0.8176 | -? | 0 | 0 | 0 | 1 |
| rs12577180 | 11 | 119276834 | C | A | -0.231 | 0.8176 | -? | 0 | 0 | 0 | 1 |
| rs4938360 | 11 | 117098436 | T | C | -0.231 | 0.8176 | -? | 0 | 0 | 0 | 1 |
| rs45480496 | 11 | 118270715 | A | G | 0.23 | 0.8177 | -+ | 0 | 0.227 | 1 | 0.6337 |
| rs77953246 | 11 | 116795987 | C | G | 0.231 | 0.8177 | +? | 0 | 0 | 0 | 1 |
| rs11216390 | 11 | 117313135 | C | G | -0.23 | 0.8178 | -? | 0 | 0 | 0 | 1 |
| rs1786690 | 11 | 118942067 | C | T | 0.23 | 0.8178 | +? | 0 | 0 | 0 | 1 |
| rs7395971 | 11 | 119693573 | C | T | 0.23 | 0.8178 | +? | 0 | 0 | 0 | 1 |
| rs10790313 | 11 | 119440066 | A | G | 0.23 | 0.8179 | +? | 0 | 0 | 0 | 1 |
| rs12364267 | 11 | 117919377 | A | G | -0.23 | 0.8184 | -? | 0 | 0 | 0 | 1 |
| rs2663643 | 11 | 119426636 | A | G | 0.23 | 0.8184 | +? | 0 | 0 | 0 | 1 |
| rs2845662 | 11 | 119427242 | C | T | 0.23 | 0.8184 | +? | 0 | 0 | 0 | 1 |
| rs11216881 | 11 | 118409539 | G | A | 0.229 | 0.8186 | ++ | 0 | 0.016 | 1 | 0.898 |
| rs180353 | 11 | 116604070 | C | T | -0.229 | 0.8186 | -+ | -18.6 | 2.529 | 1 | 0.1117 |
| rs11600793 | 11 | 119858507 | C | G | 0.229 | 0.8188 | +? | 0 | 0 | 0 | 1 |
| rs12418314 | 11 | 119301580 | T | C | -0.229 | 0.819 | -? | 0 | 0 | 0 | 1 |
| rs12418370 | 11 | 119301942 | A | G | -0.229 | 0.819 | -? | 0 | 0 | 0 | 1 |
| rs12421120 | 11 | 119301754 | C | A | -0.229 | 0.819 | -? | 0 | 0 | 0 | 1 |
| rs200041620 | 11 | 119840052 | A | G | -0.229 | 0.819 | -? | 0 | 0 | 0 | 1 |
| rs556514 | 11 | 116317399 | T | C | -0.229 | 0.819 | -? | 0 | 0 | 0 | 1 |
| rs694662 | 11 | 119895416 | T | A | 0.229 | 0.819 | +? | 0 | 0 | 0 | 1 |
| rs731392 | 11 | 119840067 | G | C | -0.229 | 0.819 | -? | 0 | 0 | 0 | 1 |
| rs10790395 | 11 | 120551419 | A | C | -0.229 | 0.8191 | -? | 0 | 0 | 0 | 1 |
| rs6589621 | 11 | 117624420 | G | C | -0.229 | 0.8192 | -? | 0 | 0 | 0 | 1 |
| rs148882737 | 11 | 117818210 | T | C | 0.228 | 0.8193 | +? | 0 | 0 | 0 | 1 |
| rs4938431 | 11 | 117587186 | G | A | 0.228 | 0.8193 | +? | 0 | 0 | 0 | 1 |
| rs4938432 | 11 | 117587345 | A | G | 0.228 | 0.8193 | +? | 0 | 0 | 0 | 1 |
| rs35095598 | 11 | 119711999 | A | G | 0.228 | 0.8196 | +? | 0 | 0 | 0 | 1 |
| rs1784461 | 11 | 118937985 | A | G | 0.228 | 0.8197 | +? | 0 | 0 | 0 | 1 |
| rs147062051 | 11 | 117014926 | A | G | -0.228 | 0.8198 | -? | 0 | 0 | 0 | 1 |
| rs181954995 | 11 | 117015290 | A | G | -0.228 | 0.8198 | -? | 0 | 0 | 0 | 1 |
| rs2846281 | 11 | 118901166 | C | G | -0.228 | 0.8198 | -? | 0 | 0 | 0 | 1 |
| rs61905473 | 11 | 117018031 | T | G | -0.228 | 0.8198 | -? | 0 | 0 | 0 | 1 |
| rs61905476 | 11 | 117024537 | G | A | -0.228 | 0.8198 | -? | 0 | 0 | 0 | 1 |
| rs1047079 | 11 | 118473231 | C | T | 0.228 | 0.82 | +- | 0 | 0.244 | 1 | 0.6214 |
| rs10892610 | 11 | 120551572 | G | A | -0.228 | 0.82 | -? | 0 | 0 | 0 | 1 |
| rs76544852 | 11 | 118487265 | T | G | -0.227 | 0.82 | -+ | 0 | 0.132 | 1 | 0.7161 |
| rs77368534 | 11 | 116362044 | G | T | 0.228 | 0.82 | +? | 0 | 0 | 0 | 1 |
| rs76841836 | 11 | 120051217 | C | G | 0.227 | 0.8202 | +? | 0 | 0 | 0 | 1 |
| rs499804 | 11 | 120852961 | A | C | -0.227 | 0.8203 | -+ | 0 | 0.251 | 1 | 0.6162 |

| | | | | | | | | | | | |
|---|---|---|---|---|---|---|---|---|---|---|---|
| rs645156 | 11 | 120853259 | T | C | -0.227 | 0.8203 | -+ | 0 | 0.251 | 1 | 0.6162 |
| rs658696 | 11 | 120853941 | A | G | -0.227 | 0.8203 | -+ | 0 | 0.251 | 1 | 0.6162 |
| rs503361 | 11 | 116321311 | C | T | -0.227 | 0.8205 | -? | 0 | 0 | 0 | 1 |
| rs147136343 | 11 | 119923305 | T | A | 0.227 | 0.8206 | +? | 0 | 0 | 0 | 1 |
| rs4608095 | 11 | 119649969 | A | G | 0.227 | 0.8207 | +? | 0 | 0 | 0 | 1 |
| rs4938357 | 11 | 117084040 | A | G | 0.227 | 0.8208 | +? | 0 | 0 | 0 | 1 |
| rs56081989 | 11 | 117085305 | A | G | 0.227 | 0.8208 | +? | 0 | 0 | 0 | 1 |
| rs67201490 | 11 | 117083004 | G | T | 0.227 | 0.8208 | +? | 0 | 0 | 0 | 1 |
| rs513083 | 11 | 116330732 | T | C | 0.226 | 0.8211 | +? | 0 | 0 | 0 | 1 |
| rs61904858 | 11 | 116492107 | G | A | -0.226 | 0.8211 | -? | 0 | 0 | 0 | 1 |
| rs80106284 | 11 | 119501909 | T | G | -0.226 | 0.8212 | -? | 0 | 0 | 0 | 1 |
| rs10892608 | 11 | 120549745 | A | G | -0.226 | 0.8213 | -? | 0 | 0 | 0 | 1 |
| rs12222723 | 11 | 119650060 | T | G | 0.226 | 0.8213 | +? | 0 | 0 | 0 | 1 |
| rs3814762 | 11 | 119216504 | T | C | -0.226 | 0.8213 | -? | 0 | 0 | 0 | 1 |
| rs7929257 | 11 | 117891475 | T | C | 0.226 | 0.8214 | +? | 0 | 0 | 0 | 1 |
| rs112152942 | 11 | 118240840 | G | C | 0.226 | 0.8215 | -+ | 1.6 | 3.05 | 1 | 0.08072 |
| rs626440 | 11 | 119898559 | T | C | 0.226 | 0.8215 | +? | 0 | 0 | 0 | 1 |
| rs11601073 | 11 | 117776747 | T | C | 0.225 | 0.8217 | +? | 0 | 0 | 0 | 1 |
| rs78925312 | 11 | 119796352 | T | C | -0.225 | 0.8217 | -? | 0 | 0 | 0 | 1 |
| rs11217476 | 11 | 119633930 | A | C | -0.225 | 0.8218 | -? | 0 | 0 | 0 | 1 |
| rs1828279 | 11 | 119419117 | C | T | 0.225 | 0.8218 | +? | 0 | 0 | 0 | 1 |
| rs11217573 | 11 | 119788395 | G | A | -0.225 | 0.8221 | -? | 0 | 0 | 0 | 1 |
| rs1060745 | 11 | 120811386 | T | G | 0.225 | 0.8222 | +? | 0 | 0 | 0 | 1 |
| rs12292921 | 11 | 116621963 | G | T | -0.225 | 0.8222 | +- | -21.6 | 2.467 | 1 | 0.1162 |
| rs1939671 | 11 | 120675437 | A | G | 0.225 | 0.8222 | +? | 0 | 0 | 0 | 1 |
| rs550974 | 11 | 118937106 | T | A | 0.225 | 0.8222 | +? | 0 | 0 | 0 | 1 |
| rs180999772 | 11 | 120074973 | A | G | 0.225 | 0.8223 | +? | 0 | 0 | 0 | 1 |
| rs61905088 | 11 | 116619867 | A | C | -0.224 | 0.8226 | +- | -15.2 | 2.604 | 1 | 0.1066 |
| rs61905526 | 11 | 117067465 | T | C | -0.224 | 0.8227 | -? | 0 | 0 | 0 | 1 |
| rs61905527 | 11 | 117069966 | T | C | -0.224 | 0.8227 | -? | 0 | 0 | 0 | 1 |
| rs2511833 | 11 | 119197379 | G | A | -0.224 | 0.8229 | -? | 0 | 0 | 0 | 1 |
| rs4454730 | 11 | 118937444 | A | G | 0.224 | 0.8229 | +? | 0 | 0 | 0 | 1 |
| rs4545564 | 11 | 118937518 | C | T | 0.224 | 0.8229 | +? | 0 | 0 | 0 | 1 |
| rs6589741 | 11 | 119311481 | A | T | -0.224 | 0.8229 | -? | 0 | 0 | 0 | 1 |
| rs6589743 | 11 | 119311933 | T | A | -0.224 | 0.8229 | -? | 0 | 0 | 0 | 1 |
| rs6589744 | 11 | 119312014 | G | A | -0.224 | 0.8229 | -? | 0 | 0 | 0 | 1 |
| rs45468196 | 11 | 118242889 | A | G | 0.224 | 0.8231 | -+ | 1.2 | 3.037 | 1 | 0.08136 |
| rs2077582 | 11 | 119880501 | T | C | 0.223 | 0.8232 | +? | 0 | 0 | 0 | 1 |
| rs638979 | 11 | 117454633 | T | C | 0.223 | 0.8232 | +? | 0 | 0 | 0 | 1 |
| rs1307145 | 11 | 118950217 | G | C | 0.223 | 0.8234 | +? | 0 | 0 | 0 | 1 |
| rs3020184 | 11 | 120670036 | A | G | 0.223 | 0.8234 | +? | 0 | 0 | 0 | 1 |
| rs7932332 | 11 | 119314765 | G | A | -0.223 | 0.8234 | -? | 0 | 0 | 0 | 1 |
| rs10892457 | 11 | 119649359 | T | C | 0.223 | 0.8236 | +? | 0 | 0 | 0 | 1 |
| rs61904849 | 11 | 116456027 | A | G | -0.223 | 0.8237 | -? | 0 | 0 | 0 | 1 |
| rs694490 | 11 | 121095669 | C | A | -0.223 | 0.8237 | -+ | 0 | 0.082 | 1 | 0.775 |
| rs11216780 | 11 | 118009385 | G | A | 0.222 | 0.824 | +? | 0 | 0 | 0 | 1 |
| rs565420 | 11 | 116481436 | A | G | 0.222 | 0.824 | +? | 0 | 0 | 0 | 1 |
| rs12280559 | 11 | 118087040 | T | C | -0.222 | 0.8241 | -+ | 0 | 0.527 | 1 | 0.4679 |
| rs1784302 | 11 | 118940957 | G | C | 0.222 | 0.8241 | +? | 0 | 0 | 0 | 1 |
| rs7119539 | 11 | 120549404 | G | A | -0.222 | 0.8243 | -? | 0 | 0 | 0 | 1 |
| rs57680824 | 11 | 117313882 | G | A | 0.222 | 0.8244 | +? | 0 | 0 | 0 | 1 |
| rs10736488 | 11 | 117285057 | G | A | 0.222 | 0.8245 | +? | 0 | 0 | 0 | 1 |
| rs7932700 | 11 | 118411713 | T | G | 0.222 | 0.8245 | ++ | 0 | 0.002 | 1 | 0.9663 |
| rs56667128 | 11 | 120097175 | A | G | 0.222 | 0.8246 | +? | 0 | 0 | 0 | 1 |
| rs7124361 | 11 | 120460445 | G | A | 0.221 | 0.8249 | +? | 0 | 0 | 0 | 1 |
| rs2075294 | 11 | 116658122 | T | G | 0.221 | 0.825 | +- | -81.6 | 1.652 | 1 | 0.1987 |

| | | | | | | | | | | | |
|---|---|---|---|---|---|---|---|---|---|---|---|
| rs645451 | 11 | 118025969 | T | C | -0.221 | 0.8251 | -? | 0 | 0 | 0 | 1 |
| rs117279127 | 11 | 117072074 | T | C | -0.221 | 0.8254 | -? | 0 | 0 | 0 | 1 |
| rs482287 | 11 | 117336108 | T | C | 0.221 | 0.8254 | +? | 0 | 0 | 0 | 1 |
| rs59455853 | 11 | 116607874 | T | A | -0.22 | 0.8257 | +- | -42.1 | 2.111 | 1 | 0.1462 |
| rs638948 | 11 | 119899143 | C | T | 0.22 | 0.8258 | +? | 0 | 0 | 0 | 1 |
| rs7951479 | 11 | 119840401 | T | C | -0.22 | 0.826 | -? | 0 | 0 | 0 | 1 |
| rs11216610 | 11 | 117769468 | A | G | -0.22 | 0.8262 | -? | 0 | 0 | 0 | 1 |
| rs3894251 | 11 | 117313577 | C | G | 0.219 | 0.8263 | +? | 0 | 0 | 0 | 1 |
| rs4938659 | 11 | 119315789 | T | C | -0.219 | 0.8263 | -? | 0 | 0 | 0 | 1 |
| rs2845708 | 11 | 120105099 | A | T | -0.219 | 0.8264 | -? | 0 | 0 | 0 | 1 |
| rs45443305 | 11 | 116645682 | A | C | 0.219 | 0.8265 | +- | -81.9 | 1.649 | 1 | 0.1991 |
| rs2015482 | 11 | 117624058 | A | G | -0.219 | 0.8266 | -? | 0 | 0 | 0 | 1 |
| rs542869 | 11 | 117225455 | A | T | 0.219 | 0.8266 | +? | 0 | 0 | 0 | 1 |
| rs4622279 | 11 | 119446118 | A | G | -0.219 | 0.8267 | -? | 0 | 0 | 0 | 1 |
| rs61906485 | 11 | 117082691 | T | G | -0.219 | 0.8267 | -? | 0 | 0 | 0 | 1 |
| rs10892609 | 11 | 120551100 | C | T | -0.219 | 0.8269 | -? | 0 | 0 | 0 | 1 |
| rs4576834 | 11 | 119542000 | T | C | 0.219 | 0.8269 | +? | 0 | 0 | 0 | 1 |
| rs57200947 | 11 | 116623674 | T | C | -0.219 | 0.8269 | +- | -20.2 | 2.496 | 1 | 0.1141 |
| rs7102643 | 11 | 119961097 | T | C | 0.219 | 0.8269 | +? | 0 | 0 | 0 | 1 |
| rs34655967 | 11 | 119792036 | A | C | 0.218 | 0.8272 | +? | 0 | 0 | 0 | 1 |
| rs10750125 | 11 | 117989989 | C | T | -0.218 | 0.8273 | -- | 0 | 0.003 | 1 | 0.9561 |
| rs12295438 | 11 | 118094921 | T | C | -0.218 | 0.8274 | -+ | 0 | 0.036 | 1 | 0.8499 |
| rs638399 | 11 | 119898995 | T | C | 0.218 | 0.8276 | +? | 0 | 0 | 0 | 1 |
| rs7102217 | 11 | 120187433 | T | C | -0.218 | 0.8277 | -? | 0 | 0 | 0 | 1 |
| rs12418475 | 11 | 119851369 | A | G | -0.217 | 0.8279 | -? | 0 | 0 | 0 | 1 |
| rs594171 | 11 | 118032974 | A | T | 0.217 | 0.828 | -+ | 0 | 0.531 | 1 | 0.4663 |
| rs12788547 | 11 | 119306878 | A | C | -0.217 | 0.8285 | -? | 0 | 0 | 0 | 1 |
| rs11825621 | 11 | 118494790 | C | T | 0.216 | 0.8286 | ++ | 0 | 0.055 | 1 | 0.8142 |
| rs11217843 | 11 | 120239937 | G | A | 0.216 | 0.8288 | +? | 0 | 0 | 0 | 1 |
| rs4938746 | 11 | 119856226 | G | A | 0.216 | 0.8288 | +? | 0 | 0 | 0 | 1 |
| rs896693 | 11 | 120067082 | A | G | -0.216 | 0.8288 | +- | 0 | 0.397 | 1 | 0.5286 |
| rs111864243 | 11 | 117291492 | C | T | 0.216 | 0.8289 | +? | 0 | 0 | 0 | 1 |
| rs112333700 | 11 | 117071505 | T | C | -0.216 | 0.829 | -? | 0 | 0 | 0 | 1 |
| rs74853011 | 11 | 117217533 | G | C | 0.216 | 0.829 | +? | 0 | 0 | 0 | 1 |
| rs76257648 | 11 | 117322099 | A | C | -0.216 | 0.829 | -? | 0 | 0 | 0 | 1 |
| rs79801428 | 11 | 116494264 | A | G | -0.216 | 0.8292 | -? | 0 | 0 | 0 | 1 |
| rs12803528 | 11 | 120905810 | A | C | 0.216 | 0.8293 | +? | 0 | 0 | 0 | 1 |
| rs180373 | 11 | 116591553 | A | C | -0.216 | 0.8293 | -+ | 0 | 0.23 | 1 | 0.6313 |
| rs2187009 | 11 | 117974484 | T | C | -0.216 | 0.8293 | -? | 0 | 0 | 0 | 1 |
| rs11216899 | 11 | 118434421 | G | A | 0.216 | 0.8294 | +- | 0 | 0.035 | 1 | 0.8519 |
| rs60753148 | 11 | 118434835 | T | C | 0.216 | 0.8294 | +- | 0 | 0.035 | 1 | 0.8519 |
| rs1232770834 | 11 | 117096652 | A | G | -0.215 | 0.8295 | -? | 0 | 0 | 0 | 1 |
| rs10892413 | 11 | 119463270 | A | G | -0.215 | 0.8296 | -? | 0 | 0 | 0 | 1 |
| rs537997 | 11 | 117451316 | T | C | 0.215 | 0.8297 | +? | 0 | 0 | 0 | 1 |
| rs1263056 | 11 | 116576415 | G | A | -0.215 | 0.8299 | +- | 0 | 0.526 | 1 | 0.4683 |
| rs59700451 | 11 | 116982928 | A | G | -0.215 | 0.8299 | -? | 0 | 0 | 0 | 1 |
| rs474490 | 11 | 117287794 | T | G | 0.215 | 0.83 | +? | 0 | 0 | 0 | 1 |
| rs11216060 | 11 | 116502267 | A | G | 0.214 | 0.8302 | +? | 0 | 0 | 0 | 1 |
| rs2845706 | 11 | 120101780 | A | G | 0.214 | 0.8302 | +? | 0 | 0 | 0 | 1 |
| rs638000 | 11 | 119885914 | T | C | 0.214 | 0.8302 | +? | 0 | 0 | 0 | 1 |
| rs11216705 | 11 | 117897020 | C | T | 0.214 | 0.8303 | +? | 0 | 0 | 0 | 1 |
| rs2276423 | 11 | 118210106 | G | C | -0.214 | 0.8303 | -+ | -91.6 | 1.566 | 1 | 0.2108 |
| rs61906106 | 11 | 116550548 | A | C | 0.214 | 0.8303 | +? | 0 | 0 | 0 | 1 |
| rs12798929 | 11 | 117776850 | T | C | 0.214 | 0.8304 | +? | 0 | 0 | 0 | 1 |
| rs2082049 | 11 | 119317208 | G | A | -0.214 | 0.8305 | -? | 0 | 0 | 0 | 1 |
| rs61903459 | 11 | 116989185 | G | A | -0.214 | 0.8305 | -? | 0 | 0 | 0 | 1 |

| | | | | | | | | | | | |
|---|---|---|---|---|---|---|---|---|---|---|---|
| rs11216366 | 11 | 117231611 | T | G | 0.214 | 0.8306 | +? | 0 | 0 | 0 | 1 |
| rs1405963027 | 11 | 119838112 | G | A | -0.214 | 0.8306 | -? | 0 | 0 | 0 | 1 |
| rs2508664 | 11 | 117229891 | T | C | 0.214 | 0.8306 | +? | 0 | 0 | 0 | 1 |
| rs9919543 | 11 | 116473707 | G | C | -0.214 | 0.8306 | -? | 0 | 0 | 0 | 1 |
| rs7483165 | 11 | 117691774 | C | T | 0.214 | 0.8308 | +? | 0 | 0 | 0 | 1 |
| rs11217316 | 11 | 119428141 | A | G | -0.213 | 0.831 | -? | 0 | 0 | 0 | 1 |
| rs11607839 | 11 | 119680383 | C | A | -0.213 | 0.831 | -? | 0 | 0 | 0 | 1 |
| rs2076765 | 11 | 120053941 | G | A | -0.213 | 0.831 | -? | 0 | 0 | 0 | 1 |
| rs10892516 | 11 | 119896830 | C | T | 0.213 | 0.8311 | +? | 0 | 0 | 0 | 1 |
| rs12362930 | 11 | 116510138 | C | T | -0.213 | 0.8311 | -? | 0 | 0 | 0 | 1 |
| rs75215496 | 11 | 121145466 | C | T | 0.213 | 0.8311 | +? | 0 | 0 | 0 | 1 |
| rs7943511 | 11 | 118096165 | T | C | -0.213 | 0.8311 | -+ | 0 | 0.043 | 1 | 0.8357 |
| rs75387178 | 11 | 117999224 | C | G | -0.213 | 0.8312 | -? | 0 | 0 | 0 | 1 |
| rs3018386 | 11 | 118016622 | G | C | -0.213 | 0.8313 | +- | 0 | 0.816 | 1 | 0.3663 |
| rs11822532 | 11 | 119690746 | A | G | 0.213 | 0.8315 | +? | 0 | 0 | 0 | 1 |
| rs2248887 | 11 | 120424000 | A | G | -0.212 | 0.8318 | -? | 0 | 0 | 0 | 1 |
| rs2845705 | 11 | 120101092 | A | T | 0.212 | 0.8318 | +? | 0 | 0 | 0 | 1 |
| rs2845650 | 11 | 119421928 | C | T | 0.212 | 0.8319 | +? | 0 | 0 | 0 | 1 |
| rs60514290 | 11 | 118246048 | C | G | 0.212 | 0.832 | +- | 0 | 0.177 | 1 | 0.6741 |
| rs7952470 | 11 | 117248268 | A | G | 0.212 | 0.832 | +? | 0 | 0 | 0 | 1 |
| rs4326835 | 11 | 118432404 | T | G | 0.212 | 0.8322 | +- | 0 | 0.015 | 1 | 0.9022 |
| rs113570704 | 11 | 117578314 | T | C | 0.212 | 0.8323 | +? | 0 | 0 | 0 | 1 |
| rs12290522 | 11 | 118486325 | T | C | -0.212 | 0.8323 | -+ | 0 | 0.485 | 1 | 0.4861 |
| rs12574878 | 11 | 117577527 | T | C | 0.212 | 0.8323 | +? | 0 | 0 | 0 | 1 |
| rs12576082 | 11 | 117577558 | A | T | 0.212 | 0.8323 | +? | 0 | 0 | 0 | 1 |
| rs12577066 | 11 | 117577909 | G | A | 0.212 | 0.8323 | +? | 0 | 0 | 0 | 1 |
| rs1310160088 | 11 | 117579516 | T | C | 0.212 | 0.8323 | +? | 0 | 0 | 0 | 1 |
| rs78696045 | 11 | 117579607 | T | A | 0.212 | 0.8323 | +? | 0 | 0 | 0 | 1 |
| rs2509121 | 11 | 118928253 | T | C | 0.212 | 0.8324 | +? | 0 | 0 | 0 | 1 |
| rs2846092 | 11 | 120641809 | A | G | 0.211 | 0.8326 | +? | 0 | 0 | 0 | 1 |
| rs10892504 | 11 | 119836087 | A | G | -0.211 | 0.8327 | -? | 0 | 0 | 0 | 1 |
| rs11606348 | 11 | 120860672 | C | T | -0.211 | 0.8327 | -? | 0 | 0 | 0 | 1 |
| rs113523070 | 11 | 117737557 | A | G | -0.211 | 0.8328 | -? | 0 | 0 | 0 | 1 |
| rs141474528 | 11 | 116952296 | G | C | -0.211 | 0.8331 | -+ | 73.2 | 11.184 | 1 | 0.000825 |
| rs59912111 | 11 | 120398707 | A | C | -0.211 | 0.8331 | -? | 0 | 0 | 0 | 1 |
| rs80068626 | 11 | 116951926 | C | T | -0.211 | 0.8331 | -+ | 73.2 | 11.184 | 1 | 0.000825 |
| rs10790389 | 11 | 120540086 | C | G | 0.21 | 0.8333 | +? | 0 | 0 | 0 | 1 |
| rs7928043 | 11 | 120498338 | A | G | 0.21 | 0.8333 | +? | 0 | 0 | 0 | 1 |
| rs7111455 | 11 | 119316614 | T | C | -0.21 | 0.8335 | -? | 0 | 0 | 0 | 1 |
| rs1848373 | 11 | 116341125 | A | T | -0.21 | 0.8336 | -? | 0 | 0 | 0 | 1 |
| rs61905715 | 11 | 116913051 | T | C | -0.21 | 0.8336 | -? | 0 | 0 | 0 | 1 |
| rs12285142 | 11 | 117557435 | T | C | 0.21 | 0.8337 | +? | 0 | 0 | 0 | 1 |
| rs12787144 | 11 | 116304152 | A | G | 0.21 | 0.8338 | +? | 0 | 0 | 0 | 1 |
| rs1793185 | 11 | 118089434 | A | C | -0.21 | 0.8338 | -+ | 0 | 0.598 | 1 | 0.4393 |
| rs11216839 | 11 | 118129583 | T | A | -0.21 | 0.8339 | -+ | 0 | 0.255 | 1 | 0.6138 |
| rs1995823 | 11 | 119446483 | T | C | 0.21 | 0.8339 | +? | 0 | 0 | 0 | 1 |
| rs4245169 | 11 | 117180905 | T | C | -0.21 | 0.834 | -? | 0 | 0 | 0 | 1 |
| rs142836613 | 11 | 116957158 | G | A | -0.209 | 0.8341 | -+ | 73 | 11.127 | 1 | 0.0008509 |
| rs1790477 | 11 | 120024827 | A | G | -0.209 | 0.8342 | -? | 0 | 0 | 0 | 1 |
| rs2155368 | 11 | 121096292 | C | A | -0.209 | 0.8342 | -+ | 0 | 0.077 | 1 | 0.7809 |
| rs11820810 | 11 | 120546088 | G | A | 0.209 | 0.8343 | +? | 0 | 0 | 0 | 1 |
| rs593598 | 11 | 119897949 | G | A | 0.209 | 0.8343 | +? | 0 | 0 | 0 | 1 |
| rs648009 | 11 | 119897964 | A | G | 0.209 | 0.8343 | +? | 0 | 0 | 0 | 1 |
| rs1168568 | 11 | 118949799 | A | G | 0.209 | 0.8344 | +? | 0 | 0 | 0 | 1 |
| rs3802866 | 11 | 118095803 | T | C | -0.209 | 0.8344 | -+ | 0 | 0.042 | 1 | 0.8377 |
| rs7943272 | 11 | 118096028 | C | A | -0.209 | 0.8344 | -+ | 0 | 0.042 | 1 | 0.8377 |

| | | | | | | | | | | | |
|---|---|---|---|---|---|---|---|---|---|---|---|
| rs1177563 | 11 | 118949083 | A | G | 0.209 | 0.8347 | +? | 0 | 0 | 0 | 1 |
| rs74498740 | 11 | 117529895 | T | C | -0.209 | 0.8348 | -? | 0 | 0 | 0 | 1 |
| rs79735246 | 11 | 117559725 | G | A | -0.208 | 0.8349 | -? | 0 | 0 | 0 | 1 |
| rs112550735 | 11 | 118303200 | C | T | 0.208 | 0.835 | -+ | 5.4 | 3.173 | 1 | 0.07488 |
| rs77728961 | 11 | 118303182 | C | T | 0.208 | 0.835 | -+ | 5.4 | 3.173 | 1 | 0.07488 |
| rs4938711 | 11 | 119566184 | T | C | 0.208 | 0.8351 | +? | 0 | 0 | 0 | 1 |
| rs61906486 | 11 | 117082865 | A | C | -0.208 | 0.8351 | -? | 0 | 0 | 0 | 1 |
| rs12419822 | 11 | 116326138 | A | C | -0.208 | 0.8353 | -? | 0 | 0 | 0 | 1 |
| rs8192614 | 11 | 118037564 | A | G | -0.208 | 0.8354 | -+ | 0 | 0.165 | 1 | 0.685 |
| rs34271655 | 11 | 121090790 | C | T | -0.208 | 0.8355 | -? | 0 | 0 | 0 | 1 |
| rs4938744 | 11 | 119836899 | A | G | -0.208 | 0.8355 | -? | 0 | 0 | 0 | 1 |
| rs4938745 | 11 | 119837342 | G | A | -0.208 | 0.8355 | -? | 0 | 0 | 0 | 1 |
| rs61899023 | 11 | 119803362 | A | G | -0.208 | 0.8355 | -? | 0 | 0 | 0 | 1 |
| rs79508602 | 11 | 119836246 | T | C | -0.208 | 0.8355 | -? | 0 | 0 | 0 | 1 |
| rs78795346 | 11 | 117297406 | A | G | 0.208 | 0.8356 | +? | 0 | 0 | 0 | 1 |
| rs61906491 | 11 | 117102200 | A | G | 0.207 | 0.8357 | +? | 0 | 0 | 0 | 1 |
| rs2509667 | 11 | 119361224 | A | C | -0.207 | 0.8358 | -? | 0 | 0 | 0 | 1 |
| rs56103633 | 11 | 119308201 | T | C | -0.207 | 0.8362 | -? | 0 | 0 | 0 | 1 |
| rs7130634 | 11 | 119309000 | A | T | -0.207 | 0.8362 | -? | 0 | 0 | 0 | 1 |
| rs74662600 | 11 | 116654959 | A | G | 0.207 | 0.8363 | +- | -84 | 1.631 | 1 | 0.2016 |
| rs7926717 | 11 | 118121399 | T | G | -0.207 | 0.8363 | -+ | 0 | 0.248 | 1 | 0.6187 |
| rs486915 | 11 | 116534288 | A | T | -0.207 | 0.8364 | -? | 0 | 0 | 0 | 1 |
| rs10892220 | 11 | 117989835 | A | T | -0.206 | 0.8365 | -- | 0 | 0.007 | 1 | 0.9352 |
| rs4258392 | 11 | 119649844 | C | T | 0.206 | 0.8368 | +? | 0 | 0 | 0 | 1 |
| rs5020633 | 11 | 119649874 | A | G | 0.206 | 0.8368 | +? | 0 | 0 | 0 | 1 |
| rs7108308 | 11 | 117586011 | G | C | 0.206 | 0.8369 | +? | 0 | 0 | 0 | 1 |
| rs1939977 | 11 | 117614720 | A | G | 0.205 | 0.8372 | +? | 0 | 0 | 0 | 1 |
| rs477389 | 11 | 120850185 | G | C | -0.206 | 0.8372 | -+ | 0 | 0.239 | 1 | 0.6249 |
| rs478978 | 11 | 120850070 | G | C | -0.205 | 0.8372 | -+ | 0 | 0.239 | 1 | 0.625 |
| rs564674 | 11 | 117227519 | T | C | 0.205 | 0.8372 | +? | 0 | 0 | 0 | 1 |
| rs61905704 | 11 | 116883665 | C | T | -0.205 | 0.8372 | -? | 0 | 0 | 0 | 1 |
| rs1287836 | 11 | 119792728 | C | A | -0.205 | 0.8373 | -? | 0 | 0 | 0 | 1 |
| rs61907561 | 11 | 116726796 | G | C | -0.205 | 0.8373 | -? | 0 | 0 | 0 | 1 |
| rs2508490 | 11 | 120099679 | G | A | 0.205 | 0.8374 | +? | 0 | 0 | 0 | 1 |
| rs670855 | 11 | 120001695 | T | C | 0.205 | 0.8374 | +? | 0 | 0 | 0 | 1 |
| rs56137014 | 11 | 117896521 | A | T | 0.205 | 0.8376 | +? | 0 | 0 | 0 | 1 |
| rs572675 | 11 | 117771888 | A | C | -0.204 | 0.838 | -? | 0 | 0 | 0 | 1 |
| rs7130266 | 11 | 119295063 | G | A | -0.204 | 0.838 | -? | 0 | 0 | 0 | 1 |
| rs10892019 | 11 | 116578223 | T | C | 0.204 | 0.8382 | -+ | -60.7 | 1.866 | 1 | 0.1719 |
| rs12797305 | 11 | 119494843 | G | A | -0.204 | 0.8382 | -? | 0 | 0 | 0 | 1 |
| rs150311062 | 11 | 117315326 | A | G | -0.204 | 0.8382 | -? | 0 | 0 | 0 | 1 |
| rs7940537 | 11 | 117656550 | A | G | -0.204 | 0.8382 | -+ | 0 | 0.072 | 1 | 0.7886 |
| rs11217771 | 11 | 120098180 | G | C | 0.204 | 0.8383 | +? | 0 | 0 | 0 | 1 |
| rs11824414 | 11 | 116348112 | G | A | 0.204 | 0.8384 | +? | 0 | 0 | 0 | 1 |
| rs67744990 | 11 | 119364296 | T | G | 0.204 | 0.8384 | +? | 0 | 0 | 0 | 1 |
| rs6589602 | 11 | 117037065 | T | C | 0.204 | 0.8385 | +? | 0 | 0 | 0 | 1 |
| rs66657611 | 11 | 117430164 | T | C | -0.204 | 0.8385 | -? | 0 | 0 | 0 | 1 |
| rs58507982 | 11 | 118146510 | T | C | -0.204 | 0.8386 | -? | 0 | 0 | 0 | 1 |
| rs5090 | 11 | 116694055 | G | C | -0.203 | 0.8388 | -? | 0 | 0 | 0 | 1 |
| rs2368426 | 11 | 119463904 | A | G | -0.203 | 0.8389 | -? | 0 | 0 | 0 | 1 |
| rs17121050 | 11 | 117533773 | A | G | -0.203 | 0.8391 | -? | 0 | 0 | 0 | 1 |
| rs524990 | 11 | 121095824 | G | T | -0.203 | 0.8391 | -+ | 0 | 0.075 | 1 | 0.7837 |
| rs59255488 | 11 | 118986118 | T | C | -0.203 | 0.8391 | -? | 0 | 0 | 0 | 1 |
| rs12792262 | 11 | 116230972 | A | G | -0.203 | 0.8393 | +- | -139.7 | 1.252 | 1 | 0.2632 |
| rs4938371 | 11 | 117193741 | T | C | 0.203 | 0.8394 | +? | 0 | 0 | 0 | 1 |
| rs11216900 | 11 | 118436079 | C | G | 0.202 | 0.8396 | +- | 0 | 0.038 | 1 | 0.8446 |

| | | | | | | | | | | | |
|---|---|---|---|---|---|---|---|---|---|---|---|
| rs10750178 | 11 | 120823528 | A | G | -0.202 | 0.8397 | -? | 0 | 0 | 0 | 1 |
| rs11216275 | 11 | 116975704 | G | T | -0.202 | 0.8398 | -? | 0 | 0 | 0 | 1 |
| rs4142639 | 11 | 117604493 | G | A | 0.202 | 0.8398 | +? | 0 | 0 | 0 | 1 |
| rs17245789 | 11 | 120904249 | G | A | 0.202 | 0.8399 | +? | 0 | 0 | 0 | 1 |
| rs2277295 | 11 | 118416253 | T | C | 0.202 | 0.84 | ++ | 0 | 0.012 | 1 | 0.9146 |
| rs17124222 | 11 | 120580969 | G | C | 0.202 | 0.8401 | +? | 0 | 0 | 0 | 1 |
| rs1185460 | 11 | 118943433 | G | A | 0.202 | 0.8402 | +? | 0 | 0 | 0 | 1 |
| rs5020632 | 11 | 119649863 | T | C | 0.202 | 0.8402 | +? | 0 | 0 | 0 | 1 |
| rs9332817 | 11 | 118365210 | C | G | 0.201 | 0.8404 | -+ | 0 | 0.197 | 1 | 0.6573 |
| rs6589542 | 11 | 116230459 | C | A | -0.201 | 0.8405 | +- | -138.2 | 1.259 | 1 | 0.2618 |
| rs7108845 | 11 | 118407523 | G | T | 0.201 | 0.8406 | ++ | 0 | 0.001 | 1 | 0.9765 |
| rs28927680 | 11 | 116619073 | G | C | -0.201 | 0.8408 | +- | -36.9 | 2.191 | 1 | 0.1388 |
| rs610831 | 11 | 119897453 | C | A | 0.201 | 0.8408 | +? | 0 | 0 | 0 | 1 |
| rs490043 | 11 | 116316173 | G | A | -0.201 | 0.8409 | -? | 0 | 0 | 0 | 1 |
| rs562966 | 11 | 120076116 | C | T | 0.201 | 0.8409 | +? | 0 | 0 | 0 | 1 |
| rs6589798 | 11 | 119971490 | T | G | 0.201 | 0.8409 | +? | 0 | 0 | 0 | 1 |
| rs12793378 | 11 | 116230993 | A | T | -0.2 | 0.8411 | +- | -139.1 | 1.255 | 1 | 0.2627 |
| rs76173441 | 11 | 119658314 | C | G | 0.2 | 0.8411 | +? | 0 | 0 | 0 | 1 |
| rs1812974 | 11 | 120221068 | T | C | 0.2 | 0.8412 | +? | 0 | 0 | 0 | 1 |
| rs4936560 | 11 | 120718009 | C | T | -0.2 | 0.8412 | -? | 0 | 0 | 0 | 1 |
| rs74948313 | 11 | 116507582 | G | A | -0.2 | 0.8413 | -? | 0 | 0 | 0 | 1 |
| rs1455831109 | 11 | 117591241 | T | C | 0.2 | 0.8414 | +? | 0 | 0 | 0 | 1 |
| rs80254203 | 11 | 120123886 | C | G | -0.2 | 0.8415 | -? | 0 | 0 | 0 | 1 |
| rs4341549 | 11 | 119637414 | G | A | 0.2 | 0.8417 | +? | 0 | 0 | 0 | 1 |
| rs75507001 | 11 | 116577694 | A | C | -0.2 | 0.8417 | -? | 0 | 0 | 0 | 1 |
| rs7950059 | 11 | 119565495 | T | C | 0.2 | 0.8417 | +? | 0 | 0 | 0 | 1 |
| rs4278525 | 11 | 119499592 | A | G | 0.2 | 0.8418 | +? | 0 | 0 | 0 | 1 |
| rs11601034 | 11 | 119789176 | T | C | -0.199 | 0.8419 | -? | 0 | 0 | 0 | 1 |
| rs35385093 | 11 | 119715710 | A | G | 0.199 | 0.8419 | +? | 0 | 0 | 0 | 1 |
| rs4938680 | 11 | 119414321 | C | G | 0.199 | 0.8419 | +? | 0 | 0 | 0 | 1 |
| rs61905718 | 11 | 116919677 | T | C | -0.199 | 0.8419 | -? | 0 | 0 | 0 | 1 |
| rs623952 | 11 | 121095863 | A | T | -0.199 | 0.8419 | -+ | 0 | 0.074 | 1 | 0.7853 |
| rs883863 | 11 | 116389017 | A | G | -0.199 | 0.842 | -? | 0 | 0 | 0 | 1 |
| rs553709 | 11 | 117726474 | C | T | -0.199 | 0.8421 | -? | 0 | 0 | 0 | 1 |
| rs671567 | 11 | 117936146 | A | G | -0.199 | 0.8421 | -? | 0 | 0 | 0 | 1 |
| rs10892206 | 11 | 117919746 | A | T | -0.199 | 0.8422 | -? | 0 | 0 | 0 | 1 |
| rs12287066 | 11 | 116662331 | T | G | -0.199 | 0.8422 | +- | -84.7 | 1.624 | 1 | 0.2025 |
| rs2242133 | 11 | 117304196 | T | C | 0.199 | 0.8422 | +? | 0 | 0 | 0 | 1 |
| rs147774547 | 11 | 119287416 | T | C | -0.199 | 0.8425 | -? | 0 | 0 | 0 | 1 |
| rs2096961 | 11 | 116503644 | A | G | 0.199 | 0.8425 | +? | 0 | 0 | 0 | 1 |
| rs7121195 | 11 | 117729616 | A | G | -0.199 | 0.8425 | -? | 0 | 0 | 0 | 1 |
| rs45555733 | 11 | 118237566 | G | C | 0.198 | 0.8427 | -+ | 3.4 | 3.105 | 1 | 0.07806 |
| rs596707 | 11 | 117371824 | C | T | -0.198 | 0.8427 | -? | 0 | 0 | 0 | 1 |
| rs10892677 | 11 | 121019288 | T | C | -0.198 | 0.8428 | -? | 0 | 0 | 0 | 1 |
| rs620602 | 11 | 120006705 | G | A | -0.198 | 0.8428 | -? | 0 | 0 | 0 | 1 |
| rs3802869 | 11 | 118034870 | A | G | 0.198 | 0.8429 | +? | 0 | 0 | 0 | 1 |
| rs748316 | 11 | 117469590 | C | T | 0.198 | 0.8429 | +? | 0 | 0 | 0 | 1 |
| rs884395 | 11 | 119485214 | G | A | -0.198 | 0.8429 | -? | 0 | 0 | 0 | 1 |
| rs588052 | 11 | 119882223 | C | T | 0.198 | 0.843 | +? | 0 | 0 | 0 | 1 |
| rs637525 | 11 | 121095137 | G | A | -0.198 | 0.843 | -+ | 0 | 0.074 | 1 | 0.7859 |
| rs2004235 | 11 | 118008212 | C | T | -0.198 | 0.8431 | -+ | 0 | 0.091 | 1 | 0.7625 |
| rs61903740 | 11 | 117226773 | A | C | -0.198 | 0.8431 | -? | 0 | 0 | 0 | 1 |
| rs61903741 | 11 | 117226941 | A | T | -0.198 | 0.8431 | -? | 0 | 0 | 0 | 1 |
| rs12785990 | 11 | 116563706 | A | G | 0.198 | 0.8432 | +? | 0 | 0 | 0 | 1 |
| rs1349689 | 11 | 119978334 | T | A | -0.198 | 0.8432 | -? | 0 | 0 | 0 | 1 |
| rs118089075 | 11 | 120398765 | T | C | 0.198 | 0.8434 | +? | 0 | 0 | 0 | 1 |

| | | | | | | | | | | | |
|---|---|---|---|---|---|---|---|---|---|---|---|
| rs1814964 | 11 | 118006957 | C | T | -0.198 | 0.8434 | -+ | 0 | 0.071 | 1 | 0.7905 |
| rs73004754 | 11 | 120053576 | A | G | -0.197 | 0.8435 | -? | 0 | 0 | 0 | 1 |
| rs11216534 | 11 | 117629468 | T | C | -0.197 | 0.8436 | +- | 14.8 | 3.519 | 1 | 0.06065 |
| rs11216535 | 11 | 117629559 | T | C | -0.197 | 0.8436 | +- | 14.8 | 3.519 | 1 | 0.06065 |
| rs17584190 | 11 | 117153815 | T | C | 0.197 | 0.8437 | +? | 0 | 0 | 0 | 1 |
| rs7944453 | 11 | 117599750 | A | G | 0.197 | 0.8437 | +? | 0 | 0 | 0 | 1 |
| rs1243337 | 11 | 117270818 | C | A | -0.197 | 0.8439 | -? | 0 | 0 | 0 | 1 |
| rs554282497 | 11 | 119495043 | C | G | 0.197 | 0.8439 | +? | 0 | 0 | 0 | 1 |
| rs117018659 | 11 | 121086259 | A | G | -0.197 | 0.844 | -? | 0 | 0 | 0 | 1 |
| rs489164 | 11 | 116316777 | C | T | -0.197 | 0.8441 | -? | 0 | 0 | 0 | 1 |
| rs9332851 | 11 | 118381018 | T | C | -0.197 | 0.8441 | -+ | 0 | 0.378 | 1 | 0.5387 |
| rs4938506 | 11 | 118205702 | A | G | -0.196 | 0.8448 | -+ | -96.8 | 1.525 | 1 | 0.2169 |
| rs67851622 | 11 | 119314499 | G | C | -0.196 | 0.8448 | -? | 0 | 0 | 0 | 1 |
| rs735006 | 11 | 119317435 | G | A | -0.196 | 0.8448 | -? | 0 | 0 | 0 | 1 |
| rs10790237 | 11 | 117989944 | G | T | -0.196 | 0.8449 | -- | 0 | 0.001 | 1 | 0.9788 |
| rs11608077 | 11 | 119462854 | G | A | -0.196 | 0.8449 | -? | 0 | 0 | 0 | 1 |
| rs1169326 | 11 | 118945185 | A | G | 0.196 | 0.8449 | +? | 0 | 0 | 0 | 1 |
| rs1184902 | 11 | 118945337 | G | A | 0.196 | 0.8449 | +? | 0 | 0 | 0 | 1 |
| rs2846095 | 11 | 120645211 | T | C | 0.196 | 0.8449 | +? | 0 | 0 | 0 | 1 |
| rs1271978583 | 11 | 119427353 | G | C | 0.196 | 0.845 | +? | 0 | 0 | 0 | 1 |
| rs2663621 | 11 | 119427309 | C | T | 0.195 | 0.8452 | +? | 0 | 0 | 0 | 1 |
| rs10892515 | 11 | 119896745 | A | T | 0.195 | 0.8454 | +? | 0 | 0 | 0 | 1 |
| rs12421142 | 11 | 120039274 | T | C | 0.195 | 0.8454 | +? | 0 | 0 | 0 | 1 |
| rs2845657 | 11 | 119429083 | T | C | 0.195 | 0.8454 | +? | 0 | 0 | 0 | 1 |
| rs56161514 | 11 | 121060783 | A | C | -0.195 | 0.8454 | -? | 0 | 0 | 0 | 1 |
| rs59565858 | 11 | 118986456 | A | G | -0.195 | 0.8454 | -? | 0 | 0 | 0 | 1 |
| rs79138835 | 11 | 117045522 | T | G | 0.195 | 0.8454 | +? | 0 | 0 | 0 | 1 |
| rs11216069 | 11 | 116507795 | C | G | -0.195 | 0.8455 | -? | 0 | 0 | 0 | 1 |
| rs11607985 | 11 | 118029798 | G | C | -0.195 | 0.8456 | -+ | 0 | 0.121 | 1 | 0.7278 |
| rs117083334 | 11 | 117252437 | G | A | 0.195 | 0.8457 | +? | 0 | 0 | 0 | 1 |
| rs470411 | 11 | 120012390 | G | C | -0.195 | 0.8457 | -? | 0 | 0 | 0 | 1 |
| rs508316 | 11 | 121085567 | C | A | -0.195 | 0.8457 | -? | 0 | 0 | 0 | 1 |
| rs12225133 | 11 | 117986722 | C | T | -0.194 | 0.8458 | -? | 0 | 0 | 0 | 1 |
| rs605742 | 11 | 119978743 | T | G | 0.194 | 0.846 | +? | 0 | 0 | 0 | 1 |
| rs625145 | 11 | 116727936 | T | A | -0.194 | 0.846 | -? | 0 | 0 | 0 | 1 |
| rs12362780 | 11 | 119762458 | A | C | 0.194 | 0.8461 | +? | 0 | 0 | 0 | 1 |
| rs595367 | 11 | 117227694 | C | T | 0.194 | 0.8463 | +? | 0 | 0 | 0 | 1 |
| rs871800 | 11 | 119485081 | C | G | -0.194 | 0.8463 | -? | 0 | 0 | 0 | 1 |
| rs11216391 | 11 | 117313136 | T | C | -0.194 | 0.8464 | -? | 0 | 0 | 0 | 1 |
| rs141076711 | 11 | 120873429 | T | G | 0.193 | 0.8467 | +? | 0 | 0 | 0 | 1 |
| rs1790476 | 11 | 120008957 | G | C | -0.193 | 0.847 | -? | 0 | 0 | 0 | 1 |
| rs189832363 | 11 | 121150291 | T | C | 0.193 | 0.847 | +? | 0 | 0 | 0 | 1 |
| rs61902684 | 11 | 120564824 | T | C | -0.193 | 0.847 | -? | 0 | 0 | 0 | 1 |
| rs11824135 | 11 | 116559553 | T | G | 0.193 | 0.8473 | +? | 0 | 0 | 0 | 1 |
| rs61903170 | 11 | 120855775 | C | T | -0.193 | 0.8473 | -+ | 0 | 0.232 | 1 | 0.6302 |
| rs80259223 | 11 | 116924318 | A | G | 0.193 | 0.8473 | +? | 0 | 0 | 0 | 1 |
| rs11216503 | 11 | 117558989 | G | A | -0.192 | 0.8474 | -? | 0 | 0 | 0 | 1 |
| rs11216677 | 11 | 117882792 | T | C | 0.192 | 0.8475 | +? | 0 | 0 | 0 | 1 |
| rs4938380 | 11 | 117323076 | A | G | 0.192 | 0.8475 | +? | 0 | 0 | 0 | 1 |
| rs7106419 | 11 | 117590322 | G | T | 0.192 | 0.8475 | +? | 0 | 0 | 0 | 1 |
| rs722058 | 11 | 119949822 | A | G | -0.192 | 0.8476 | -? | 0 | 0 | 0 | 1 |
| rs11216748 | 11 | 117970689 | A | G | 0.192 | 0.8477 | +- | 0 | 0.154 | 1 | 0.6946 |
| rs183640608 | 11 | 116877361 | G | A | 0.192 | 0.8477 | +? | 0 | 0 | 0 | 1 |
| rs7131641 | 11 | 119312750 | C | G | -0.192 | 0.8478 | -? | 0 | 0 | 0 | 1 |
| rs78036667 | 11 | 117600506 | T | C | 0.192 | 0.8478 | +? | 0 | 0 | 0 | 1 |
| rs45625638 | 11 | 117021426 | C | G | 0.192 | 0.8479 | +? | 0 | 0 | 0 | 1 |

| | | | | | | | | | | | |
|---|---|---|---|---|---|---|---|---|---|---|---|
| rs6589800 | 11 | 119971706 | C | A | 0.191 | 0.8482 | +? | 0 | 0 | 0 | 1 |
| rs113586459 | 11 | 117610811 | C | A | 0.191 | 0.8483 | +? | 0 | 0 | 0 | 1 |
| rs948027 | 11 | 120644424 | C | T | 0.191 | 0.8483 | +? | 0 | 0 | 0 | 1 |
| rs60545357 | 11 | 118281418 | C | A | 0.191 | 0.8484 | -+ | 0 | 0.155 | 1 | 0.6935 |
| rs33984246 | 11 | 116660450 | G | A | 0.191 | 0.8486 | +- | -29.5 | 2.317 | 1 | 0.128 |
| rs79212399 | 11 | 121100518 | T | G | -0.191 | 0.8487 | -? | 0 | 0 | 0 | 1 |
| rs1806 | 11 | 118076004 | C | T | 0.191 | 0.8488 | +? | 0 | 0 | 0 | 1 |
| rs17245796 | 11 | 120921962 | G | A | 0.19 | 0.849 | +? | 0 | 0 | 0 | 1 |
| rs529482 | 11 | 119786235 | G | A | -0.19 | 0.8491 | -? | 0 | 0 | 0 | 1 |
| rs72997349 | 11 | 118945502 | C | T | 0.19 | 0.8491 | +? | 0 | 0 | 0 | 1 |
| rs61409027 | 11 | 120769478 | T | C | 0.19 | 0.8492 | +? | 0 | 0 | 0 | 1 |
| rs67262752 | 11 | 118047776 | C | T | 0.19 | 0.8492 | +- | 0 | 0.474 | 1 | 0.4912 |
| rs7121573 | 11 | 117729830 | A | G | 0.19 | 0.8495 | +? | 0 | 0 | 0 | 1 |
| rs10892611 | 11 | 120551586 | T | C | -0.19 | 0.8496 | -? | 0 | 0 | 0 | 1 |
| rs2298478 | 11 | 120984087 | G | C | -0.19 | 0.8496 | -? | 0 | 0 | 0 | 1 |
| rs522098 | 11 | 116326686 | A | G | -0.189 | 0.8499 | -? | 0 | 0 | 0 | 1 |
| rs56289161 | 11 | 116535703 | T | C | 0.189 | 0.8501 | +? | 0 | 0 | 0 | 1 |
| rs56391092 | 11 | 120010550 | T | C | -0.189 | 0.8504 | -? | 0 | 0 | 0 | 1 |
| rs3825055 | 11 | 118416073 | C | A | 0.188 | 0.8505 | ++ | 0 | 0.007 | 1 | 0.9352 |
| rs624463 | 11 | 117940612 | G | A | -0.188 | 0.8505 | -? | 0 | 0 | 0 | 1 |
| rs654572 | 11 | 117940332 | C | T | -0.188 | 0.8505 | -? | 0 | 0 | 0 | 1 |
| rs61209319 | 11 | 116537880 | A | C | 0.188 | 0.8508 | +? | 0 | 0 | 0 | 1 |
| rs11216046 | 11 | 116492701 | T | G | -0.188 | 0.851 | -? | 0 | 0 | 0 | 1 |
| rs12807172 | 11 | 120497794 | T | G | 0.188 | 0.851 | +? | 0 | 0 | 0 | 1 |
| rs7120516 | 11 | 117292550 | C | G | 0.188 | 0.8511 | +? | 0 | 0 | 0 | 1 |
| rs10466594 | 11 | 119454670 | C | T | -0.187 | 0.8513 | -? | 0 | 0 | 0 | 1 |
| rs10892080 | 11 | 117035502 | A | G | 0.187 | 0.8513 | +? | 0 | 0 | 0 | 1 |
| rs146204458 | 11 | 119782437 | G | A | -0.187 | 0.8513 | -? | 0 | 0 | 0 | 1 |
| rs58164695 | 11 | 119691736 | T | C | 0.187 | 0.8513 | +? | 0 | 0 | 0 | 1 |
| rs12271703 | 11 | 116266176 | G | A | 0.187 | 0.8514 | +? | 0 | 0 | 0 | 1 |
| rs11216749 | 11 | 117970696 | C | T | 0.187 | 0.8515 | +- | 0 | 0.152 | 1 | 0.6967 |
| rs1073634 | 11 | 119972642 | T | C | 0.187 | 0.8516 | +? | 0 | 0 | 0 | 1 |
| rs1073636 | 11 | 119972448 | A | T | 0.187 | 0.8516 | +? | 0 | 0 | 0 | 1 |
| rs45501598 | 11 | 118251980 | T | A | 0.187 | 0.8516 | +- | 0 | 0.201 | 1 | 0.6537 |
| rs61906114 | 11 | 116557044 | C | G | 0.187 | 0.8518 | +? | 0 | 0 | 0 | 1 |
| rs8924 | 11 | 117988784 | A | G | -0.187 | 0.8519 | -- | 0 | 0.016 | 1 | 0.8998 |
| rs529623 | 11 | 117693255 | T | C | -0.187 | 0.852 | -? | 0 | 0 | 0 | 1 |
| rs7934404 | 11 | 120193477 | A | G | -0.187 | 0.852 | -? | 0 | 0 | 0 | 1 |
| rs7949987 | 11 | 121153421 | G | A | -0.186 | 0.8521 | -? | 0 | 0 | 0 | 1 |
| rs55920307 | 11 | 120042846 | G | A | 0.186 | 0.8523 | +? | 0 | 0 | 0 | 1 |
| rs61905081 | 11 | 116600564 | A | G | -0.186 | 0.8524 | +- | -65 | 1.818 | 1 | 0.1776 |
| rs2000870 | 11 | 120539693 | G | C | 0.186 | 0.8525 | +? | 0 | 0 | 0 | 1 |
| rs41510246 | 11 | 119836704 | A | G | -0.186 | 0.8525 | -? | 0 | 0 | 0 | 1 |
| rs649451 | 11 | 119894303 | A | G | 0.186 | 0.8525 | +? | 0 | 0 | 0 | 1 |
| rs74916845 | 11 | 119838589 | G | A | -0.186 | 0.8525 | -? | 0 | 0 | 0 | 1 |
| rs75335512 | 11 | 121106642 | C | A | -0.186 | 0.8526 | -? | 0 | 0 | 0 | 1 |
| rs80145448 | 11 | 121139014 | A | C | 0.186 | 0.8528 | +? | 0 | 0 | 0 | 1 |
| rs55986402 | 11 | 117192701 | C | A | -0.185 | 0.853 | -? | 0 | 0 | 0 | 1 |
| rs73587439 | 11 | 117644348 | A | G | -0.185 | 0.8531 | +- | 24.4 | 3.966 | 1 | 0.04643 |
| rs7939747 | 11 | 117097009 | G | A | -0.185 | 0.8531 | -? | 0 | 0 | 0 | 1 |
| rs61905307 | 11 | 117326269 | C | T | -0.185 | 0.8533 | -? | 0 | 0 | 0 | 1 |
| rs11218059 | 11 | 120787231 | T | C | -0.185 | 0.8534 | +- | 0 | 0.678 | 1 | 0.4102 |
| rs624736 | 11 | 119893208 | G | A | 0.185 | 0.8534 | +? | 0 | 0 | 0 | 1 |
| rs13377199 | 11 | 116342446 | T | C | 0.185 | 0.8536 | +? | 0 | 0 | 0 | 1 |
| rs77564141 | 11 | 119302513 | G | A | 0.184 | 0.854 | +? | 0 | 0 | 0 | 1 |
| rs117986469 | 11 | 121075781 | A | C | 0.184 | 0.8541 | +? | 0 | 0 | 0 | 1 |

| | | | | | | | | | | | |
|---|---|---|---|---|---|---|---|---|---|---|---|
| rs12291024 | 11 | 116560667 | G | T | 0.184 | 0.8541 | +? | 0 | 0 | 0 | 1 |
| rs7130303 | 11 | 117702115 | A | C | 0.184 | 0.8541 | +? | 0 | 0 | 0 | 1 |
| rs7949366 | 11 | 119959585 | T | C | 0.184 | 0.8541 | +? | 0 | 0 | 0 | 1 |
| rs61905078 | 11 | 116596309 | C | A | 0.184 | 0.8542 | +- | -43 | 2.098 | 1 | 0.1475 |
| rs11216533 | 11 | 117628729 | A | C | -0.184 | 0.8543 | +- | 15.5 | 3.549 | 1 | 0.05959 |
| rs2510146 | 11 | 119181674 | G | C | 0.184 | 0.8543 | +? | 0 | 0 | 0 | 1 |
| rs11600005 | 11 | 119781758 | A | G | -0.184 | 0.8544 | -? | 0 | 0 | 0 | 1 |
| rs55713675 | 11 | 116622722 | T | C | -0.183 | 0.8545 | +- | -17.2 | 2.56 | 1 | 0.1096 |
| rs11216527 | 11 | 117620420 | C | T | 0.183 | 0.8546 | +? | 0 | 0 | 0 | 1 |
| rs7925365 | 11 | 120637315 | G | T | -0.183 | 0.8548 | -? | 0 | 0 | 0 | 1 |
| rs7948840 | 11 | 121098423 | G | A | -0.183 | 0.8548 | -+ | 0 | 0.069 | 1 | 0.7925 |
| rs6589801 | 11 | 119971987 | T | C | 0.183 | 0.8551 | +? | 0 | 0 | 0 | 1 |
| rs596647 | 11 | 119367321 | A | C | 0.182 | 0.8552 | +? | 0 | 0 | 0 | 1 |
| rs532107 | 11 | 116311508 | G | A | -0.182 | 0.8553 | -? | 0 | 0 | 0 | 1 |
| rs56967930 | 11 | 117192528 | G | A | -0.182 | 0.8553 | -? | 0 | 0 | 0 | 1 |
| rs112357006 | 11 | 117126493 | A | G | 0.182 | 0.8556 | +? | 0 | 0 | 0 | 1 |
| rs12803171 | 11 | 117430426 | T | C | -0.182 | 0.8556 | -? | 0 | 0 | 0 | 1 |
| rs10892517 | 11 | 119902255 | T | C | 0.182 | 0.8559 | +? | 0 | 0 | 0 | 1 |
| rs3809040 | 11 | 117774829 | A | G | 0.182 | 0.8559 | +? | 0 | 0 | 0 | 1 |
| rs61415784 | 11 | 116362957 | C | A | -0.182 | 0.8559 | -? | 0 | 0 | 0 | 1 |
| rs11218105 | 11 | 120872976 | G | A | -0.181 | 0.856 | -? | 0 | 0 | 0 | 1 |
| rs56224630 | 11 | 116627097 | A | G | -0.181 | 0.8561 | +- | -6.1 | 2.826 | 1 | 0.09273 |
| rs71478236 | 11 | 116214668 | A | G | -0.181 | 0.8561 | -? | 0 | 0 | 0 | 1 |
| rs76830818 | 11 | 117083831 | T | A | -0.181 | 0.8561 | -? | 0 | 0 | 0 | 1 |
| rs10400366 | 11 | 117000955 | G | A | 0.181 | 0.8565 | +? | 0 | 0 | 0 | 1 |
| rs12574404 | 11 | 116319329 | G | C | -0.181 | 0.8566 | -? | 0 | 0 | 0 | 1 |
| rs11216675 | 11 | 117881937 | T | C | 0.181 | 0.8567 | +? | 0 | 0 | 0 | 1 |
| rs79911534 | 11 | 118232332 | A | G | -0.181 | 0.8567 | -+ | 0 | 0.027 | 1 | 0.8703 |
| rs59176377 | 11 | 118419295 | A | G | 0.181 | 0.8568 | ++ | 0 | 0 | 1 | 0.9889 |
| rs61898294 | 11 | 120104292 | A | G | -0.18 | 0.8568 | -? | 0 | 0 | 0 | 1 |
| rs12274046 | 11 | 116605070 | T | C | -0.18 | 0.8569 | +- | -41.2 | 2.125 | 1 | 0.1449 |
| rs1079118 | 11 | 118258056 | A | G | 0.18 | 0.857 | -+ | 0 | 0.148 | 1 | 0.7003 |
| rs140575350 | 11 | 116646820 | G | C | -0.18 | 0.8571 | -? | 0 | 0 | 0 | 1 |
| rs3133225 | 11 | 120624732 | A | G | 0.18 | 0.8571 | +? | 0 | 0 | 0 | 1 |
| rs11606013 | 11 | 116568225 | T | G | 0.18 | 0.8573 | +? | 0 | 0 | 0 | 1 |
| rs1177562 | 11 | 118949331 | T | C | 0.18 | 0.8573 | +? | 0 | 0 | 0 | 1 |
| rs12225399 | 11 | 118480285 | C | G | 0.18 | 0.8573 | +- | 0 | 0.13 | 1 | 0.7186 |
| rs7937973 | 11 | 117593097 | A | G | 0.18 | 0.8573 | +? | 0 | 0 | 0 | 1 |
| rs12577147 | 11 | 119218648 | A | G | 0.179 | 0.8576 | +? | 0 | 0 | 0 | 1 |
| rs66633871 | 11 | 116339980 | A | G | -0.179 | 0.8578 | -? | 0 | 0 | 0 | 1 |
| rs10750126 | 11 | 117989990 | A | G | -0.179 | 0.8579 | -- | 0 | 0.001 | 1 | 0.9727 |
| rs73003594 | 11 | 116300735 | A | G | 0.179 | 0.8579 | +? | 0 | 0 | 0 | 1 |
| rs74424292 | 11 | 116501089 | T | A | 0.179 | 0.8579 | +? | 0 | 0 | 0 | 1 |
| rs76055915 | 11 | 116501072 | T | C | 0.179 | 0.8579 | +? | 0 | 0 | 0 | 1 |
| rs61898296 | 11 | 120105300 | A | G | 0.179 | 0.858 | +? | 0 | 0 | 0 | 1 |
| rs11216105 | 11 | 116577441 | G | A | 0.178 | 0.8584 | +? | 0 | 0 | 0 | 1 |
| rs12788247 | 11 | 120954793 | G | A | -0.178 | 0.8585 | -? | 0 | 0 | 0 | 1 |
| rs60837962 | 11 | 118438448 | T | A | 0.178 | 0.8585 | +- | 0 | 0.037 | 1 | 0.8476 |
| rs476983 | 11 | 117215605 | T | C | -0.178 | 0.8586 | -? | 0 | 0 | 0 | 1 |
| rs59027088 | 11 | 119445108 | T | C | 0.178 | 0.8586 | +? | 0 | 0 | 0 | 1 |
| rs73584223 | 11 | 119444694 | A | G | 0.178 | 0.8586 | +? | 0 | 0 | 0 | 1 |
| rs603858 | 11 | 117378713 | C | G | -0.178 | 0.8587 | -? | 0 | 0 | 0 | 1 |
| rs10431083 | 11 | 117898155 | A | G | 0.178 | 0.8589 | +? | 0 | 0 | 0 | 1 |
| rs12223288 | 11 | 117898084 | G | A | 0.178 | 0.8589 | +? | 0 | 0 | 0 | 1 |
| rs192508986 | 11 | 117898042 | T | C | 0.178 | 0.8589 | +? | 0 | 0 | 0 | 1 |
| rs74430235 | 11 | 117556948 | A | G | -0.178 | 0.8589 | -? | 0 | 0 | 0 | 1 |

| | | | | | | | | | | | |
|---|---|---|---|---|---|---|---|---|---|---|---|
| rs76628593 | 11 | 117594288 | T | G | 0.178 | 0.8589 | +? | 0 | 0 | 0 | 1 |
| rs77047087 | 11 | 117594170 | A | G | 0.178 | 0.8589 | +? | 0 | 0 | 0 | 1 |
| rs78437760 | 11 | 117593858 | G | T | 0.178 | 0.8589 | +? | 0 | 0 | 0 | 1 |
| rs78908794 | 11 | 116638854 | C | T | 0.178 | 0.8589 | +- | -88.8 | 1.589 | 1 | 0.2075 |
| rs12282602 | 11 | 117628810 | G | A | -0.178 | 0.859 | +- | 15.8 | 3.562 | 1 | 0.05912 |
| rs595377 | 11 | 119898379 | T | C | 0.178 | 0.859 | +? | 0 | 0 | 0 | 1 |
| rs4142640 | 11 | 117604207 | A | G | 0.178 | 0.8591 | +? | 0 | 0 | 0 | 1 |
| rs11216059 | 11 | 116501987 | A | C | 0.177 | 0.8592 | +? | 0 | 0 | 0 | 1 |
| rs560789 | 11 | 117228892 | G | A | -0.177 | 0.8592 | -? | 0 | 0 | 0 | 1 |
| rs77746497 | 11 | 117597810 | A | G | 0.177 | 0.8592 | +? | 0 | 0 | 0 | 1 |
| rs79506234 | 11 | 117596635 | T | C | 0.177 | 0.8592 | +? | 0 | 0 | 0 | 1 |
| rs117680585 | 11 | 117048367 | T | C | 0.177 | 0.8593 | +? | 0 | 0 | 0 | 1 |
| rs74894672 | 11 | 120090711 | A | G | -0.177 | 0.8593 | -? | 0 | 0 | 0 | 1 |
| rs10892116 | 11 | 117325448 | T | C | -0.177 | 0.8594 | -? | 0 | 0 | 0 | 1 |
| rs117865451 | 11 | 116780568 | C | T | -0.177 | 0.8594 | -? | 0 | 0 | 0 | 1 |
| rs477890 | 11 | 117335074 | T | A | -0.177 | 0.8596 | -? | 0 | 0 | 0 | 1 |
| rs640421 | 11 | 117941898 | T | C | -0.177 | 0.8596 | -? | 0 | 0 | 0 | 1 |
| rs652030 | 11 | 117942125 | C | T | -0.177 | 0.8596 | -? | 0 | 0 | 0 | 1 |
| rs4936532 | 11 | 120493360 | A | G | 0.177 | 0.8598 | +? | 0 | 0 | 0 | 1 |
| rs555255 | 11 | 119701127 | G | A | 0.177 | 0.8598 | +? | 0 | 0 | 0 | 1 |
| rs79537444 | 11 | 117643958 | T | C | -0.177 | 0.8598 | +- | 24.7 | 3.985 | 1 | 0.0459 |
| rs12294242 | 11 | 116346978 | T | C | 0.176 | 0.8601 | +? | 0 | 0 | 0 | 1 |
| rs68137714 | 11 | 117025357 | G | A | 0.176 | 0.8602 | +? | 0 | 0 | 0 | 1 |
| rs7350481 | 11 | 116586283 | T | C | -0.176 | 0.8602 | +- | -18.7 | 2.528 | 1 | 0.1118 |
| rs7952087 | 11 | 119945046 | T | C | -0.176 | 0.8602 | -? | 0 | 0 | 0 | 1 |
| rs61905116 | 11 | 116649538 | G | A | -0.176 | 0.8603 | +- | -107.4 | 1.447 | 1 | 0.2291 |
| rs55820322 | 11 | 117998834 | C | A | -0.176 | 0.8604 | -? | 0 | 0 | 0 | 1 |
| rs144198120 | 11 | 117309079 | T | C | -0.176 | 0.8605 | -? | 0 | 0 | 0 | 1 |
| rs1367306 | 11 | 121142356 | T | A | 0.175 | 0.8609 | +? | 0 | 0 | 0 | 1 |
| rs1944516 | 11 | 120877786 | T | C | -0.175 | 0.8609 | -? | 0 | 0 | 0 | 1 |
| rs897837 | 11 | 117261488 | G | C | 0.175 | 0.8609 | +? | 0 | 0 | 0 | 1 |
| rs11216666 | 11 | 117876035 | C | T | 0.175 | 0.8611 | +? | 0 | 0 | 0 | 1 |
| rs10750158 | 11 | 119443669 | A | C | -0.175 | 0.8613 | -? | 0 | 0 | 0 | 1 |
| rs73005072 | 11 | 121065345 | A | C | -0.174 | 0.8615 | -? | 0 | 0 | 0 | 1 |
| rs113749645 | 11 | 120868233 | A | T | -0.174 | 0.8616 | -? | 0 | 0 | 0 | 1 |
| rs479991 | 11 | 117696958 | C | T | 0.174 | 0.8618 | +? | 0 | 0 | 0 | 1 |
| rs61898289 | 11 | 120093745 | A | G | 0.174 | 0.8618 | +? | 0 | 0 | 0 | 1 |
| rs11216385 | 11 | 117307219 | T | C | 0.174 | 0.8619 | +? | 0 | 0 | 0 | 1 |
| rs77649044 | 11 | 116501053 | C | T | 0.174 | 0.862 | +? | 0 | 0 | 0 | 1 |
| rs586053 | 11 | 118020568 | A | G | -0.174 | 0.8621 | -+ | 0 | 0.855 | 1 | 0.3551 |
| rs7124934 | 11 | 119564781 | G | A | -0.174 | 0.8622 | -? | 0 | 0 | 0 | 1 |
| rs1473324 | 11 | 116821918 | G | T | 0.173 | 0.8623 | +? | 0 | 0 | 0 | 1 |
| rs68097885 | 11 | 118596444 | A | G | -0.173 | 0.8623 | -+ | 0 | 0.32 | 1 | 0.5718 |
| rs11216029 | 11 | 116465680 | T | G | 0.173 | 0.8624 | +? | 0 | 0 | 0 | 1 |
| rs11608173 | 11 | 116568364 | A | T | 0.173 | 0.8624 | +? | 0 | 0 | 0 | 1 |
| rs2508489 | 11 | 120099658 | A | G | -0.173 | 0.8624 | -? | 0 | 0 | 0 | 1 |
| rs12417339 | 11 | 121110311 | A | G | -0.173 | 0.8625 | -? | 0 | 0 | 0 | 1 |
| rs587348 | 11 | 119895515 | T | C | 0.173 | 0.8626 | +? | 0 | 0 | 0 | 1 |
| rs11217682 | 11 | 119961707 | G | A | 0.173 | 0.8628 | +? | 0 | 0 | 0 | 1 |
| rs573052 | 11 | 116321980 | A | G | -0.173 | 0.8629 | -? | 0 | 0 | 0 | 1 |
| rs12291995 | 11 | 117587353 | C | T | 0.173 | 0.863 | +? | 0 | 0 | 0 | 1 |
| rs56152188 | 11 | 119312046 | C | T | -0.173 | 0.863 | -? | 0 | 0 | 0 | 1 |
| rs67000349 | 11 | 117842706 | G | A | 0.172 | 0.8631 | +? | 0 | 0 | 0 | 1 |
| rs11216592 | 11 | 117733364 | G | A | 0.172 | 0.8632 | +? | 0 | 0 | 0 | 1 |
| rs61901460 | 11 | 121194035 | G | C | -0.172 | 0.8632 | -? | 0 | 0 | 0 | 1 |
| rs655157 | 11 | 117942848 | T | G | -0.172 | 0.8632 | -? | 0 | 0 | 0 | 1 |

| | | | | | | | | | | | |
|---|---|---|---|---|---|---|---|---|---|---|---|
| rs2077173 | 11 | 118492591 | A | G | -0.172 | 0.8633 | -+ | 0 | 0.551 | 1 | 0.4579 |
| rs75322037 | 11 | 116479338 | G | A | -0.172 | 0.8634 | -? | 0 | 0 | 0 | 1 |
| rs4936423 | 11 | 117992718 | C | G | -0.172 | 0.8637 | -+ | 0 | 0.049 | 1 | 0.8249 |
| rs12361479 | 11 | 117555727 | T | C | -0.172 | 0.8638 | -? | 0 | 0 | 0 | 1 |
| rs7114201 | 11 | 120050113 | A | T | 0.172 | 0.8638 | +? | 0 | 0 | 0 | 1 |
| rs35287658 | 11 | 119447044 | A | G | 0.171 | 0.8639 | +? | 0 | 0 | 0 | 1 |
| rs4938336 | 11 | 116948442 | G | A | 0.171 | 0.8639 | +? | 0 | 0 | 0 | 1 |
| rs3741319 | 11 | 118415848 | C | A | 0.171 | 0.864 | ++ | 0 | 0.003 | 1 | 0.9596 |
| rs3741320 | 11 | 118415847 | C | G | 0.171 | 0.864 | ++ | 0 | 0.003 | 1 | 0.9596 |
| rs11216512 | 11 | 117584793 | G | C | 0.171 | 0.8641 | +? | 0 | 0 | 0 | 1 |
| rs2187182 | 11 | 117584761 | T | G | 0.171 | 0.8641 | +? | 0 | 0 | 0 | 1 |
| rs2187183 | 11 | 117584740 | A | G | 0.171 | 0.8641 | +? | 0 | 0 | 0 | 1 |
| rs4245171 | 11 | 117585977 | A | G | 0.171 | 0.8641 | +? | 0 | 0 | 0 | 1 |
| rs4245172 | 11 | 117585981 | C | T | 0.171 | 0.8641 | +? | 0 | 0 | 0 | 1 |
| rs4245173 | 11 | 117586129 | T | C | 0.171 | 0.8641 | +? | 0 | 0 | 0 | 1 |
| rs4255570 | 11 | 117585918 | C | T | 0.171 | 0.8641 | +? | 0 | 0 | 0 | 1 |
| rs4359247 | 11 | 117585641 | G | C | 0.171 | 0.8641 | +? | 0 | 0 | 0 | 1 |
| rs4936396 | 11 | 117585035 | A | G | 0.171 | 0.8641 | +? | 0 | 0 | 0 | 1 |
| rs4936397 | 11 | 117585370 | G | A | 0.171 | 0.8641 | +? | 0 | 0 | 0 | 1 |
| rs4938427 | 11 | 117585030 | A | G | 0.171 | 0.8641 | +? | 0 | 0 | 0 | 1 |
| rs4938429 | 11 | 117585338 | C | A | 0.171 | 0.8641 | +? | 0 | 0 | 0 | 1 |
| rs4938430 | 11 | 117586178 | C | G | 0.171 | 0.8641 | +? | 0 | 0 | 0 | 1 |
| rs7930615 | 11 | 117585523 | G | A | 0.171 | 0.8641 | +? | 0 | 0 | 0 | 1 |
| rs79078907 | 11 | 117557022 | T | C | -0.171 | 0.8644 | -? | 0 | 0 | 0 | 1 |
| rs487611 | 11 | 116330989 | C | G | 0.171 | 0.8645 | +? | 0 | 0 | 0 | 1 |
| rs680472 | 11 | 119900629 | G | C | -0.171 | 0.8646 | -? | 0 | 0 | 0 | 1 |
| rs144638086 | 11 | 117659186 | T | C | 0.17 | 0.8647 | +? | 0 | 0 | 0 | 1 |
| rs11216061 | 11 | 116502752 | C | T | 0.17 | 0.8648 | +? | 0 | 0 | 0 | 1 |
| rs10892383 | 11 | 119405750 | T | C | -0.17 | 0.865 | -? | 0 | 0 | 0 | 1 |
| rs12284089 | 11 | 118058542 | T | C | 0.17 | 0.865 | +- | 0 | 0.422 | 1 | 0.5158 |
| rs71484153 | 11 | 119792443 | C | T | 0.17 | 0.865 | +? | 0 | 0 | 0 | 1 |
| rs10892249 | 11 | 118507068 | C | T | -0.17 | 0.8651 | -+ | 0 | 0.158 | 1 | 0.6906 |
| rs73020459 | 11 | 117766606 | T | C | -0.17 | 0.8654 | -? | 0 | 0 | 0 | 1 |
| rs652346 | 11 | 117216599 | T | C | 0.169 | 0.8655 | +? | 0 | 0 | 0 | 1 |
| rs11823013 | 11 | 116839535 | C | T | 0.169 | 0.8656 | +? | 0 | 0 | 0 | 1 |
| rs73613959 | 11 | 118263054 | A | G | 0.169 | 0.8657 | -+ | 0 | 0.125 | 1 | 0.7238 |
| rs649880 | 11 | 121095030 | C | G | -0.169 | 0.8658 | -+ | 0 | 0.065 | 1 | 0.7987 |
| rs1158260 | 11 | 116356225 | T | C | 0.169 | 0.8659 | +? | 0 | 0 | 0 | 1 |
| rs17547501 | 11 | 119315417 | A | G | -0.169 | 0.866 | -? | 0 | 0 | 0 | 1 |
| rs55845693 | 11 | 119314637 | C | T | -0.169 | 0.866 | -? | 0 | 0 | 0 | 1 |
| rs61901262 | 11 | 119312851 | T | C | -0.169 | 0.866 | -? | 0 | 0 | 0 | 1 |
| rs61901263 | 11 | 119313221 | T | C | -0.169 | 0.866 | -? | 0 | 0 | 0 | 1 |
| rs61901265 | 11 | 119315316 | A | G | -0.169 | 0.866 | -? | 0 | 0 | 0 | 1 |
| rs657118 | 11 | 117954081 | G | A | 0.169 | 0.866 | +? | 0 | 0 | 0 | 1 |
| rs113770024 | 11 | 117685752 | A | G | -0.169 | 0.8661 | -? | 0 | 0 | 0 | 1 |
| rs11217646 | 11 | 119903119 | G | A | 0.168 | 0.8664 | +? | 0 | 0 | 0 | 1 |
| rs61903732 | 11 | 117193254 | G | A | 0.168 | 0.8665 | +? | 0 | 0 | 0 | 1 |
| rs11217640 | 11 | 119895939 | C | T | 0.168 | 0.8666 | +? | 0 | 0 | 0 | 1 |
| rs1473327 | 11 | 116823996 | T | A | 0.168 | 0.8666 | +? | 0 | 0 | 0 | 1 |
| rs11603616 | 11 | 118496066 | A | G | -0.168 | 0.8667 | -+ | 0 | 0.563 | 1 | 0.4529 |
| rs11217317 | 11 | 119428144 | T | C | -0.168 | 0.8668 | -? | 0 | 0 | 0 | 1 |
| rs11827322 | 11 | 117585424 | T | C | 0.168 | 0.8668 | +? | 0 | 0 | 0 | 1 |
| rs7928525 | 11 | 117118712 | T | A | -0.167 | 0.867 | -? | 0 | 0 | 0 | 1 |
| rs12803024 | 11 | 117430389 | G | C | -0.167 | 0.8671 | -? | 0 | 0 | 0 | 1 |
| rs144502429 | 11 | 119820957 | T | C | -0.167 | 0.8673 | -? | 0 | 0 | 0 | 1 |
| rs1075790 | 11 | 119983919 | A | C | -0.167 | 0.8676 | -? | 0 | 0 | 0 | 1 |

| | | | | | | | | | | | |
|---|---|---|---|---|---|---|---|---|---|---|---|
| rs61270867 | 11 | 116499500 | G | A | -0.167 | 0.8676 | -? | 0 | 0 | 0 | 1 |
| rs2735181 | 11 | 117009789 | C | T | 0.167 | 0.8677 | +? | 0 | 0 | 0 | 1 |
| rs55975161 | 11 | 117383716 | A | G | 0.167 | 0.8677 | +? | 0 | 0 | 0 | 1 |
| rs673351 | 11 | 120854938 | C | G | -0.167 | 0.8677 | -+ | 0 | 0.218 | 1 | 0.6408 |
| rs11217344 | 11 | 119468889 | A | G | 0.166 | 0.8678 | +? | 0 | 0 | 0 | 1 |
| rs12805851 | 11 | 116816884 | T | C | 0.166 | 0.8678 | +? | 0 | 0 | 0 | 1 |
| rs571001 | 11 | 118500173 | T | C | -0.166 | 0.8678 | -+ | 0 | 0.554 | 1 | 0.4569 |
| rs10790392 | 11 | 120542320 | A | G | 0.166 | 0.8679 | +? | 0 | 0 | 0 | 1 |
| rs36086368 | 11 | 117741853 | C | T | -0.166 | 0.8679 | -? | 0 | 0 | 0 | 1 |
| rs79807999 | 11 | 117554355 | G | T | -0.166 | 0.8679 | -? | 0 | 0 | 0 | 1 |
| rs11216030 | 11 | 116465734 | T | C | 0.166 | 0.868 | +? | 0 | 0 | 0 | 1 |
| rs143049161 | 11 | 117747629 | T | C | 0.166 | 0.868 | +? | 0 | 0 | 0 | 1 |
| rs7938843 | 11 | 119972861 | T | G | 0.166 | 0.868 | +? | 0 | 0 | 0 | 1 |
| rs12365185 | 11 | 119951870 | G | A | 0.166 | 0.8681 | +? | 0 | 0 | 0 | 1 |
| rs59931457 | 11 | 117528867 | A | G | 0.166 | 0.8681 | -+ | 0 | 0.994 | 1 | 0.3189 |
| rs67307131 | 11 | 118480223 | C | T | 0.166 | 0.8681 | +- | 0 | 0.159 | 1 | 0.6901 |
| rs12419204 | 11 | 118497402 | A | T | -0.166 | 0.8682 | -+ | 0 | 0.553 | 1 | 0.457 |
| rs2075548 | 11 | 117070927 | A | G | 0.166 | 0.8682 | +? | 0 | 0 | 0 | 1 |
| rs12289135 | 11 | 121085338 | C | T | -0.166 | 0.8684 | -? | 0 | 0 | 0 | 1 |
| rs80318512 | 11 | 121084717 | A | G | -0.166 | 0.8684 | -? | 0 | 0 | 0 | 1 |
| rs624717 | 11 | 119898183 | G | T | 0.165 | 0.8686 | +? | 0 | 0 | 0 | 1 |
| rs12296014 | 11 | 117628972 | A | G | -0.165 | 0.8687 | +- | 16.4 | 3.589 | 1 | 0.05818 |
| rs1876793 | 11 | 119444330 | T | C | -0.165 | 0.8687 | -? | 0 | 0 | 0 | 1 |
| rs7924895 | 11 | 117991034 | G | A | -0.165 | 0.8687 | -+ | 0 | 0.045 | 1 | 0.8315 |
| rs7946574 | 11 | 119666454 | T | C | -0.165 | 0.8687 | -? | 0 | 0 | 0 | 1 |
| rs618265 | 11 | 120028701 | A | G | -0.165 | 0.8688 | -? | 0 | 0 | 0 | 1 |
| rs7948804 | 11 | 120642807 | C | T | 0.165 | 0.8688 | +? | 0 | 0 | 0 | 1 |
| rs639484 | 11 | 119882063 | T | C | 0.165 | 0.869 | +? | 0 | 0 | 0 | 1 |
| rs494722 | 11 | 117216603 | T | A | 0.165 | 0.8691 | +? | 0 | 0 | 0 | 1 |
| rs531410 | 11 | 121094411 | A | T | -0.165 | 0.8691 | -+ | 0 | 0.064 | 1 | 0.8006 |
| rs10892636 | 11 | 120768404 | G | A | 0.165 | 0.8693 | +? | 0 | 0 | 0 | 1 |
| rs35101469 | 11 | 120550222 | G | T | -0.165 | 0.8693 | -? | 0 | 0 | 0 | 1 |
| rs10790199 | 11 | 117530473 | A | G | 0.164 | 0.8694 | +? | 0 | 0 | 0 | 1 |
| rs11216064 | 11 | 116502823 | C | G | 0.164 | 0.8694 | +? | 0 | 0 | 0 | 1 |
| rs17520254 | 11 | 116665553 | G | C | -0.164 | 0.8696 | +- | 0 | 0.157 | 1 | 0.6922 |
| rs11217507 | 11 | 119662942 | G | T | -0.164 | 0.8698 | -? | 0 | 0 | 0 | 1 |
| rs2511023 | 11 | 120009145 | A | G | 0.164 | 0.87 | +? | 0 | 0 | 0 | 1 |
| rs61903814 | 11 | 117510157 | A | G | -0.164 | 0.87 | -? | 0 | 0 | 0 | 1 |
| rs11216812 | 11 | 118071762 | A | G | 0.164 | 0.8701 | -+ | 0 | 0.506 | 1 | 0.4768 |
| rs11574653 | 11 | 119294290 | G | A | -0.163 | 0.8705 | -? | 0 | 0 | 0 | 1 |
| rs3138088 | 11 | 119293441 | G | A | -0.163 | 0.8705 | -? | 0 | 0 | 0 | 1 |
| rs3138089 | 11 | 119293397 | A | G | -0.163 | 0.8705 | -? | 0 | 0 | 0 | 1 |
| rs11216834 | 11 | 118122077 | C | T | -0.163 | 0.8707 | -+ | 0 | 0.17 | 1 | 0.68 |
| rs4936428 | 11 | 118187511 | T | C | -0.163 | 0.8707 | +- | 0 | 0.182 | 1 | 0.6696 |
| rs35705701 | 11 | 117693413 | A | G | 0.163 | 0.8708 | +? | 0 | 0 | 0 | 1 |
| rs480101 | 11 | 116314402 | T | C | 0.163 | 0.8708 | +? | 0 | 0 | 0 | 1 |
| rs117939457 | 11 | 120097897 | T | A | 0.163 | 0.8709 | +? | 0 | 0 | 0 | 1 |
| rs139060345 | 11 | 118295001 | C | T | -0.163 | 0.8709 | -? | 0 | 0 | 0 | 1 |
| rs4528369 | 11 | 116416716 | T | G | 0.163 | 0.8709 | +? | 0 | 0 | 0 | 1 |
| rs570952 | 11 | 118500193 | C | T | -0.163 | 0.8709 | -+ | 0 | 0.55 | 1 | 0.4582 |
| rs638997 | 11 | 117941554 | A | G | -0.163 | 0.8709 | -? | 0 | 0 | 0 | 1 |
| rs10892565 | 11 | 120235560 | C | T | 0.162 | 0.871 | +? | 0 | 0 | 0 | 1 |
| rs7103327 | 11 | 119443362 | T | C | 0.162 | 0.871 | +? | 0 | 0 | 0 | 1 |
| rs1304536246 | 11 | 117314320 | T | C | 0.162 | 0.8712 | +? | 0 | 0 | 0 | 1 |
| rs7103569 | 11 | 120764218 | G | A | -0.162 | 0.8713 | -? | 0 | 0 | 0 | 1 |
| rs11828506 | 11 | 117233427 | T | C | 0.162 | 0.8714 | +? | 0 | 0 | 0 | 1 |

| | | | | | | | | | | | |
|---|---|---|---|---|---|---|---|---|---|---|---|
| rs61905681 | 11 | 116836209 | C | A | 0.162 | 0.8715 | +? | 0 | 0 | 0 | 1 |
| rs7106772 | 11 | 117237186 | A | G | 0.162 | 0.8716 | +? | 0 | 0 | 0 | 1 |
| rs7109766 | 11 | 117244606 | T | A | 0.162 | 0.8716 | +? | 0 | 0 | 0 | 1 |
| rs76344095 | 11 | 117244291 | T | C | 0.162 | 0.8716 | +? | 0 | 0 | 0 | 1 |
| rs78406883 | 11 | 117253230 | T | C | 0.162 | 0.8716 | +? | 0 | 0 | 0 | 1 |
| rs673346 | 11 | 120854936 | G | A | -0.162 | 0.8717 | -+ | 0 | 0.215 | 1 | 0.6429 |
| rs11217309 | 11 | 119412000 | T | C | 0.161 | 0.8718 | +? | 0 | 0 | 0 | 1 |
| rs11824358 | 11 | 116354937 | T | C | -0.161 | 0.8718 | -? | 0 | 0 | 0 | 1 |
| rs11826857 | 11 | 116354930 | C | T | -0.161 | 0.8718 | -? | 0 | 0 | 0 | 1 |
| rs4375449 | 11 | 119406292 | A | G | -0.161 | 0.8718 | -? | 0 | 0 | 0 | 1 |
| rs675647 | 11 | 120018743 | A | G | -0.161 | 0.8718 | -? | 0 | 0 | 0 | 1 |
| rs7120565 | 11 | 117042949 | T | C | 0.161 | 0.8718 | +? | 0 | 0 | 0 | 1 |
| rs511011 | 11 | 116482895 | C | T | 0.161 | 0.8719 | +? | 0 | 0 | 0 | 1 |
| rs10790345 | 11 | 119654053 | G | A | 0.161 | 0.872 | +? | 0 | 0 | 0 | 1 |
| rs116088536 | 11 | 119884982 | G | C | 0.161 | 0.872 | +? | 0 | 0 | 0 | 1 |
| rs545613 | 11 | 116481221 | C | T | 0.161 | 0.872 | +? | 0 | 0 | 0 | 1 |
| rs117047926 | 11 | 119804674 | T | C | -0.161 | 0.8722 | -? | 0 | 0 | 0 | 1 |
| rs117933040 | 11 | 117049508 | G | A | 0.161 | 0.8722 | +? | 0 | 0 | 0 | 1 |
| rs73018318 | 11 | 117307306 | T | G | 0.161 | 0.8723 | +? | 0 | 0 | 0 | 1 |
| rs2846099 | 11 | 120650611 | T | A | 0.161 | 0.8724 | +? | 0 | 0 | 0 | 1 |
| rs60583202 | 11 | 116596464 | A | G | -0.16 | 0.8725 | +- | -77.3 | 1.692 | 1 | 0.1933 |
| rs653478 | 11 | 121087285 | A | T | -0.16 | 0.8725 | -? | 0 | 0 | 0 | 1 |
| rs473343 | 11 | 117408156 | C | G | -0.16 | 0.8726 | -? | 0 | 0 | 0 | 1 |
| rs61059752 | 11 | 118246071 | A | C | 0.16 | 0.8727 | -+ | 0 | 0.114 | 1 | 0.7359 |
| rs1027636222 | 11 | 117587085 | G | A | 0.16 | 0.8728 | +? | 0 | 0 | 0 | 1 |
| rs12361273 | 11 | 116384580 | T | C | 0.16 | 0.8728 | +? | 0 | 0 | 0 | 1 |
| rs527534 | 11 | 117689549 | T | G | -0.16 | 0.8728 | -? | 0 | 0 | 0 | 1 |
| rs1982011 | 11 | 121154437 | G | A | -0.16 | 0.8729 | -? | 0 | 0 | 0 | 1 |
| rs61906113 | 11 | 116556912 | T | C | 0.16 | 0.8732 | +? | 0 | 0 | 0 | 1 |
| rs11216706 | 11 | 117897056 | G | T | 0.159 | 0.8735 | +? | 0 | 0 | 0 | 1 |
| rs11216707 | 11 | 117897184 | G | A | 0.159 | 0.8735 | +? | 0 | 0 | 0 | 1 |
| rs1317544 | 11 | 118006930 | C | T | 0.159 | 0.8736 | +? | 0 | 0 | 0 | 1 |
| rs140763307 | 11 | 117472044 | G | A | -0.159 | 0.8736 | -? | 0 | 0 | 0 | 1 |
| rs11216814 | 11 | 118071857 | G | A | 0.159 | 0.8737 | -+ | 0 | 0.508 | 1 | 0.476 |
| rs35199514 | 11 | 120454925 | C | T | -0.159 | 0.874 | -? | 0 | 0 | 0 | 1 |
| rs3138090 | 11 | 119293216 | A | C | -0.158 | 0.8741 | -? | 0 | 0 | 0 | 1 |
| rs78559460 | 11 | 117730777 | C | A | -0.158 | 0.8743 | -? | 0 | 0 | 0 | 1 |
| rs74398561 | 11 | 118233634 | G | A | 0.158 | 0.8745 | -+ | 7.9 | 3.257 | 1 | 0.07114 |
| rs75202546 | 11 | 116961819 | A | G | -0.158 | 0.8747 | -+ | 75.7 | 12.322 | 1 | 0.0004476 |
| rs12289284 | 11 | 116338814 | A | G | -0.158 | 0.8748 | -? | 0 | 0 | 0 | 1 |
| rs595969 | 11 | 119890372 | G | C | -0.157 | 0.8751 | -? | 0 | 0 | 0 | 1 |
| rs4938788 | 11 | 120042081 | A | G | 0.157 | 0.8752 | +? | 0 | 0 | 0 | 1 |
| rs1380377167 | 11 | 120947058 | A | T | 0.157 | 0.8753 | +? | 0 | 0 | 0 | 1 |
| rs2845656 | 11 | 119429139 | C | T | 0.157 | 0.8753 | +? | 0 | 0 | 0 | 1 |
| rs674352 | 11 | 120855192 | T | C | -0.157 | 0.8753 | -+ | 0 | 0.213 | 1 | 0.6448 |
| rs12292816 | 11 | 121072114 | T | G | -0.157 | 0.8754 | -? | 0 | 0 | 0 | 1 |
| rs7928275 | 11 | 117991805 | A | G | -0.157 | 0.8754 | -+ | 0 | 0.044 | 1 | 0.8339 |
| rs7943311 | 11 | 117991817 | T | G | -0.157 | 0.8754 | -+ | 0 | 0.044 | 1 | 0.8339 |
| rs10790235 | 11 | 117977723 | G | A | -0.157 | 0.8755 | -? | 0 | 0 | 0 | 1 |
| rs12224951 | 11 | 120551964 | T | C | -0.156 | 0.8757 | -? | 0 | 0 | 0 | 1 |
| rs12792820 | 11 | 120055403 | A | T | -0.156 | 0.8758 | -? | 0 | 0 | 0 | 1 |
| rs17519079 | 11 | 116629153 | A | G | 0.156 | 0.8758 | +? | 0 | 0 | 0 | 1 |
| rs79552834 | 11 | 116953338 | C | G | -0.156 | 0.8759 | -+ | 72.5 | 10.924 | 1 | 0.0009494 |
| rs56134734 | 11 | 116833055 | A | G | 0.156 | 0.8763 | +? | 0 | 0 | 0 | 1 |
| rs61905677 | 11 | 116833655 | G | T | 0.156 | 0.8763 | +? | 0 | 0 | 0 | 1 |
| rs61905678 | 11 | 116834785 | T | C | 0.156 | 0.8763 | +? | 0 | 0 | 0 | 1 |

| | | | | | | | | | | | |
|---|---|---|---|---|---|---|---|---|---|---|---|
| rs61905680 | 11 | 116836031 | A | G | 0.156 | 0.8763 | +? | 0 | 0 | 0 | 1 |
| rs7929887 | 11 | 118428791 | G | A | 0.156 | 0.8763 | ++ | 0 | 0.007 | 1 | 0.9328 |
| rs4936363 | 11 | 116972935 | T | C | 0.155 | 0.8765 | +? | 0 | 0 | 0 | 1 |
| rs571445 | 11 | 118987971 | G | A | 0.155 | 0.8765 | +? | 0 | 0 | 0 | 1 |
| rs7110874 | 11 | 119564842 | C | T | 0.155 | 0.8765 | +? | 0 | 0 | 0 | 1 |
| rs7947704 | 11 | 119412210 | T | C | 0.155 | 0.8766 | +? | 0 | 0 | 0 | 1 |
| rs12225221 | 11 | 116438555 | T | C | 0.155 | 0.8767 | -+ | 0 | 0.18 | 1 | 0.6715 |
| rs111901638 | 11 | 116959848 | A | G | 0.155 | 0.8768 | +? | 0 | 0 | 0 | 1 |
| rs591592 | 11 | 116713566 | C | T | -0.155 | 0.8768 | -? | 0 | 0 | 0 | 1 |
| rs632153 | 11 | 116710239 | T | G | -0.155 | 0.8769 | +- | 26.3 | 4.07 | 1 | 0.04364 |
| rs1056136 | 11 | 117120998 | G | T | -0.155 | 0.8771 | -? | 0 | 0 | 0 | 1 |
| rs34089864 | 11 | 116660768 | A | G | 0.155 | 0.8771 | +- | -85.2 | 1.62 | 1 | 0.2031 |
| rs605833 | 11 | 119969700 | C | T | 0.155 | 0.8771 | +? | 0 | 0 | 0 | 1 |
| rs141740187 | 11 | 116637677 | T | G | 0.154 | 0.8772 | +- | -92.9 | 1.555 | 1 | 0.2123 |
| rs11216830 | 11 | 118115147 | A | G | -0.154 | 0.8774 | -+ | 0 | 0.16 | 1 | 0.6889 |
| rs2102405 | 11 | 116348168 | G | A | 0.154 | 0.8774 | +? | 0 | 0 | 0 | 1 |
| rs3938035 | 11 | 116348151 | G | A | 0.154 | 0.8774 | +? | 0 | 0 | 0 | 1 |
| rs10790246 | 11 | 118103542 | T | A | -0.154 | 0.8775 | -+ | 0 | 0.103 | 1 | 0.7486 |
| rs56225305 | 11 | 116625362 | A | G | -0.154 | 0.8775 | +- | -14.8 | 2.614 | 1 | 0.1059 |
| rs60067274 | 11 | 118412296 | T | A | 0.154 | 0.8775 | ++ | 0 | 0.026 | 1 | 0.872 |
| rs61898267 | 11 | 120072470 | G | C | -0.154 | 0.8775 | -? | 0 | 0 | 0 | 1 |
| rs7937334 | 11 | 118191274 | T | C | -0.154 | 0.8775 | -+ | -104.6 | 1.466 | 1 | 0.2259 |
| rs7128466 | 11 | 120491438 | A | G | 0.154 | 0.8776 | +? | 0 | 0 | 0 | 1 |
| rs1145198 | 11 | 116574336 | A | G | -0.154 | 0.8778 | -+ | -75.3 | 1.711 | 1 | 0.1909 |
| rs59253729 | 11 | 119833474 | T | C | -0.154 | 0.8778 | -? | 0 | 0 | 0 | 1 |
| rs117534497 | 11 | 119825806 | G | A | -0.154 | 0.8779 | -? | 0 | 0 | 0 | 1 |
| rs11217593 | 11 | 119822294 | A | G | -0.154 | 0.878 | -? | 0 | 0 | 0 | 1 |
| rs11217904 | 11 | 120397055 | C | T | 0.154 | 0.878 | +? | 0 | 0 | 0 | 1 |
| rs4938505 | 11 | 118205636 | A | G | 0.154 | 0.878 | +- | 0 | 0.962 | 1 | 0.3267 |
| rs12276648 | 11 | 117629056 | C | T | -0.153 | 0.8781 | +- | 17 | 3.614 | 1 | 0.05728 |
| rs10892230 | 11 | 118126682 | A | C | -0.153 | 0.8782 | -+ | 0 | 0.185 | 1 | 0.6674 |
| rs557396 | 11 | 117445258 | A | G | -0.153 | 0.8782 | -? | 0 | 0 | 0 | 1 |
| rs58089984 | 11 | 116510680 | G | T | -0.153 | 0.8782 | -? | 0 | 0 | 0 | 1 |
| rs527005 | 11 | 117486829 | G | C | -0.153 | 0.8783 | -? | 0 | 0 | 0 | 1 |
| rs582718 | 11 | 120823104 | T | C | -0.153 | 0.8783 | -? | 0 | 0 | 0 | 1 |
| rs61226783 | 11 | 120398747 | G | A | -0.153 | 0.8783 | -? | 0 | 0 | 0 | 1 |
| rs12295970 | 11 | 117628906 | T | G | -0.153 | 0.8784 | +- | 17 | 3.615 | 1 | 0.05725 |
| rs57187632 | 11 | 116353776 | A | C | 0.153 | 0.8785 | +? | 0 | 0 | 0 | 1 |
| rs73585281 | 11 | 120855386 | G | A | -0.153 | 0.8787 | -+ | 0 | 0.21 | 1 | 0.6465 |
| rs73585283 | 11 | 120855433 | A | G | -0.153 | 0.8787 | -+ | 0 | 0.21 | 1 | 0.6465 |
| rs116603457 | 11 | 119474018 | T | C | 0.152 | 0.8788 | +? | 0 | 0 | 0 | 1 |
| rs608298 | 11 | 119886924 | C | T | 0.152 | 0.8788 | +? | 0 | 0 | 0 | 1 |
| rs527813 | 11 | 121094778 | G | T | -0.152 | 0.8789 | -+ | 0 | 0.06 | 1 | 0.8061 |
| rs12274146 | 11 | 116605076 | A | G | -0.152 | 0.8792 | +- | -38.1 | 2.172 | 1 | 0.1405 |
| rs11216491 | 11 | 117536136 | T | C | -0.152 | 0.8793 | -? | 0 | 0 | 0 | 1 |
| rs542900 | 11 | 117225467 | A | T | 0.152 | 0.8793 | +? | 0 | 0 | 0 | 1 |
| rs7942259 | 11 | 119653947 | T | C | 0.152 | 0.8793 | +? | 0 | 0 | 0 | 1 |
| rs12793017 | 11 | 116858703 | G | A | -0.152 | 0.8794 | -? | 0 | 0 | 0 | 1 |
| rs61903708 | 11 | 117189201 | G | A | -0.152 | 0.8794 | -? | 0 | 0 | 0 | 1 |
| rs59049611 | 11 | 118139403 | C | T | -0.152 | 0.8795 | -? | 0 | 0 | 0 | 1 |
| rs61900932 | 11 | 118262706 | T | C | -0.152 | 0.8795 | -+ | 0 | 0.047 | 1 | 0.8281 |
| rs7949440 | 11 | 120710315 | A | G | 0.151 | 0.8796 | +? | 0 | 0 | 0 | 1 |
| rs11216708 | 11 | 117897329 | T | C | 0.151 | 0.8798 | +? | 0 | 0 | 0 | 1 |
| rs61898286 | 11 | 120092344 | C | G | 0.151 | 0.8798 | +? | 0 | 0 | 0 | 1 |
| rs557940 | 11 | 117441393 | G | A | 0.151 | 0.8799 | +? | 0 | 0 | 0 | 1 |
| rs605560 | 11 | 118067187 | C | G | 0.151 | 0.8799 | +? | 0 | 0 | 0 | 1 |

| | | | | | | | | | | | |
|---|---|---|---|---|---|---|---|---|---|---|---|
| rs552498 | 11 | 117260075 | G | A | -0.151 | 0.8801 | -? | 0 | 0 | 0 | 1 |
| rs61907594 | 11 | 116801811 | T | C | 0.151 | 0.8803 | +? | 0 | 0 | 0 | 1 |
| rs79974099 | 11 | 116802581 | A | T | 0.151 | 0.8803 | +? | 0 | 0 | 0 | 1 |
| rs112243842 | 11 | 118246021 | G | T | 0.15 | 0.8806 | -+ | -2.4 | 2.931 | 1 | 0.0869 |
| rs59294170 | 11 | 116841446 | T | A | 0.15 | 0.8807 | +? | 0 | 0 | 0 | 1 |
| rs651821 | 11 | 116662579 | C | T | -0.15 | 0.8807 | +- | -67.3 | 1.793 | 1 | 0.1806 |
| rs7109923 | 11 | 117237487 | G | A | 0.15 | 0.8808 | +? | 0 | 0 | 0 | 1 |
| rs77291360 | 11 | 120579966 | G | A | 0.15 | 0.881 | +? | 0 | 0 | 0 | 1 |
| rs609177 | 11 | 116770517 | A | G | 0.149 | 0.8813 | +? | 0 | 0 | 0 | 1 |
| rs75696653 | 11 | 117643228 | T | A | -0.149 | 0.8815 | +- | 25.9 | 4.049 | 1 | 0.04421 |
| rs10750151 | 11 | 119429736 | T | C | -0.149 | 0.8816 | -? | 0 | 0 | 0 | 1 |
| rs11216569 | 11 | 117699377 | T | C | 0.149 | 0.8817 | +? | 0 | 0 | 0 | 1 |
| rs45462696 | 11 | 117092660 | A | G | -0.149 | 0.8819 | -? | 0 | 0 | 0 | 1 |
| rs609467 | 11 | 119887725 | G | A | 0.149 | 0.8819 | +? | 0 | 0 | 0 | 1 |
| rs116966258 | 11 | 119766043 | A | G | -0.148 | 0.882 | -? | 0 | 0 | 0 | 1 |
| rs2276129 | 11 | 117973673 | A | G | 0.148 | 0.8823 | +- | 0 | 0.083 | 1 | 0.7729 |
| rs665035 | 11 | 121083812 | T | C | -0.148 | 0.8823 | -? | 0 | 0 | 0 | 1 |
| rs77984767 | 11 | 116496744 | G | A | 0.148 | 0.8823 | +? | 0 | 0 | 0 | 1 |
| rs118000999 | 11 | 116326685 | T | C | 0.148 | 0.8824 | +? | 0 | 0 | 0 | 1 |
| rs4317995 | 11 | 119622781 | A | G | 0.148 | 0.8824 | +? | 0 | 0 | 0 | 1 |
| rs4315086 | 11 | 119668265 | A | G | -0.148 | 0.8825 | -? | 0 | 0 | 0 | 1 |
| rs112646740 | 11 | 118247805 | A | T | 0.148 | 0.8827 | -+ | -2.9 | 2.915 | 1 | 0.08778 |
| rs755002 | 11 | 119439498 | T | C | 0.148 | 0.8827 | +? | 0 | 0 | 0 | 1 |
| rs5081 | 11 | 116706346 | T | A | 0.147 | 0.8828 | +- | -0.9 | 2.972 | 1 | 0.08469 |
| rs679847 | 11 | 121082861 | G | A | -0.147 | 0.8828 | -? | 0 | 0 | 0 | 1 |
| rs76867785 | 11 | 120133616 | G | A | -0.147 | 0.883 | -? | 0 | 0 | 0 | 1 |
| rs11216063 | 11 | 116502776 | A | T | 0.147 | 0.8831 | +? | 0 | 0 | 0 | 1 |
| rs507805 | 11 | 117167147 | G | A | -0.147 | 0.8831 | -? | 0 | 0 | 0 | 1 |
| rs56993455 | 11 | 119832917 | A | G | -0.147 | 0.8832 | -? | 0 | 0 | 0 | 1 |
| rs117119582 | 11 | 120181117 | C | T | -0.147 | 0.8833 | -? | 0 | 0 | 0 | 1 |
| rs61903816 | 11 | 117525308 | A | G | -0.147 | 0.8833 | -? | 0 | 0 | 0 | 1 |
| rs869789 | 11 | 117691386 | A | G | -0.147 | 0.8833 | -? | 0 | 0 | 0 | 1 |
| rs113164826 | 11 | 119664546 | C | G | -0.146 | 0.8836 | -? | 0 | 0 | 0 | 1 |
| rs7389 | 11 | 118528466 | G | T | -0.146 | 0.8836 | -+ | 0 | 0.439 | 1 | 0.5078 |
| rs77417358 | 11 | 119845184 | T | C | -0.146 | 0.8836 | -? | 0 | 0 | 0 | 1 |
| rs4938333 | 11 | 116930888 | G | A | 0.146 | 0.8837 | +? | 0 | 0 | 0 | 1 |
| rs4938338 | 11 | 116972894 | A | C | 0.146 | 0.8837 | +? | 0 | 0 | 0 | 1 |
| rs573549 | 11 | 116736679 | A | G | -0.146 | 0.8837 | -? | 0 | 0 | 0 | 1 |
| rs61903394 | 11 | 116931898 | A | G | 0.146 | 0.8837 | +? | 0 | 0 | 0 | 1 |
| rs11216055 | 11 | 116500588 | T | C | 0.146 | 0.8838 | +? | 0 | 0 | 0 | 1 |
| rs11216056 | 11 | 116500629 | A | C | 0.146 | 0.8838 | +? | 0 | 0 | 0 | 1 |
| rs11216057 | 11 | 116500768 | T | C | 0.146 | 0.8838 | +? | 0 | 0 | 0 | 1 |
| rs7108677 | 11 | 119444978 | G | C | -0.146 | 0.8838 | -? | 0 | 0 | 0 | 1 |
| rs7108679 | 11 | 119444980 | G | A | -0.146 | 0.8838 | -? | 0 | 0 | 0 | 1 |
| rs10892697 | 11 | 121115967 | C | G | 0.146 | 0.8841 | +? | 0 | 0 | 0 | 1 |
| rs2277291 | 11 | 118435756 | T | C | 0.146 | 0.8842 | +- | 0 | 0.063 | 1 | 0.802 |
| rs3016415 | 11 | 120676208 | G | T | -0.146 | 0.8842 | -? | 0 | 0 | 0 | 1 |
| rs61903169 | 11 | 120855705 | T | A | -0.146 | 0.8842 | -+ | 0 | 0.207 | 1 | 0.6494 |
| rs7119954 | 11 | 120046224 | C | T | 0.146 | 0.8842 | +? | 0 | 0 | 0 | 1 |
| rs61898281 | 11 | 120088926 | G | A | 0.146 | 0.8843 | +? | 0 | 0 | 0 | 1 |
| rs10790344 | 11 | 119639398 | G | A | 0.145 | 0.8844 | +? | 0 | 0 | 0 | 1 |
| rs61905690 | 11 | 116876168 | G | C | -0.145 | 0.8845 | -? | 0 | 0 | 0 | 1 |
| rs7949539 | 11 | 118250586 | G | A | 0.145 | 0.8845 | +- | 0 | 0.17 | 1 | 0.6802 |
| rs192958777 | 11 | 117134152 | T | C | -0.145 | 0.8846 | -? | 0 | 0 | 0 | 1 |
| rs11215978 | 11 | 116391537 | C | T | -0.145 | 0.8847 | -? | 0 | 0 | 0 | 1 |
| rs4525269 | 11 | 117722292 | C | T | -0.145 | 0.8847 | -? | 0 | 0 | 0 | 1 |

| | | | | | | | | | | | |
|---|---|---|---|---|---|---|---|---|---|---|---|
| rs4525270 | 11 | 117722501 | C | T | -0.145 | 0.8847 | -? | 0 | 0 | 0 | 1 |
| rs60044901 | 11 | 117601018 | C | G | 0.145 | 0.8849 | +? | 0 | 0 | 0 | 1 |
| rs7938470 | 11 | 116990934 | A | G | -0.145 | 0.8849 | -? | 0 | 0 | 0 | 1 |
| rs113348846 | 11 | 120123145 | A | G | -0.145 | 0.885 | -? | 0 | 0 | 0 | 1 |
| rs2276057 | 11 | 121036315 | A | G | -0.145 | 0.885 | -? | 0 | 0 | 0 | 1 |
| rs61905530 | 11 | 117077716 | A | G | -0.145 | 0.885 | -? | 0 | 0 | 0 | 1 |
| rs10750103 | 11 | 117042238 | C | T | 0.144 | 0.8852 | +? | 0 | 0 | 0 | 1 |
| rs61624438 | 11 | 116809945 | A | C | 0.144 | 0.8852 | +? | 0 | 0 | 0 | 1 |
| rs906823 | 11 | 119441311 | A | G | -0.144 | 0.8853 | -? | 0 | 0 | 0 | 1 |
| rs906824 | 11 | 119441696 | T | C | -0.144 | 0.8853 | -? | 0 | 0 | 0 | 1 |
| rs906825 | 11 | 119441804 | G | A | -0.144 | 0.8853 | -? | 0 | 0 | 0 | 1 |
| rs906826 | 11 | 119442188 | T | C | -0.144 | 0.8853 | -? | 0 | 0 | 0 | 1 |
| rs681637 | 11 | 121084190 | C | T | -0.144 | 0.8855 | -? | 0 | 0 | 0 | 1 |
| rs579163 | 11 | 116713630 | C | G | -0.144 | 0.8856 | -? | 0 | 0 | 0 | 1 |
| rs116864232 | 11 | 117830715 | C | T | -0.144 | 0.8857 | -? | 0 | 0 | 0 | 1 |
| rs10466593 | 11 | 119454588 | T | C | -0.144 | 0.8858 | -? | 0 | 0 | 0 | 1 |
| rs11216635 | 11 | 117800682 | G | A | 0.144 | 0.8858 | -+ | -160.6 | 1.151 | 1 | 0.2833 |
| rs4290266 | 11 | 118285995 | A | G | 0.144 | 0.8858 | +- | 0 | 0.166 | 1 | 0.6834 |
| rs11216122 | 11 | 116609540 | T | G | -0.143 | 0.8859 | -? | 0 | 0 | 0 | 1 |
| rs4938351 | 11 | 117022352 | A | C | 0.143 | 0.886 | +? | 0 | 0 | 0 | 1 |
| rs10790346 | 11 | 119658393 | G | C | 0.143 | 0.8865 | +? | 0 | 0 | 0 | 1 |
| rs4340085 | 11 | 116496309 | T | C | 0.143 | 0.8865 | +? | 0 | 0 | 0 | 1 |
| rs7110743 | 11 | 118825762 | A | C | -0.143 | 0.8865 | -? | 0 | 0 | 0 | 1 |
| rs73004793 | 11 | 120061255 | C | T | 0.143 | 0.8866 | +? | 0 | 0 | 0 | 1 |
| rs12275955 | 11 | 118094756 | C | T | -0.143 | 0.8867 | -+ | 0 | 0.476 | 1 | 0.4904 |
| rs56301948 | 11 | 118049880 | G | A | 0.143 | 0.8867 | +- | 0 | 0.437 | 1 | 0.5086 |
| rs57533048 | 11 | 118050014 | G | A | 0.143 | 0.8867 | +- | 0 | 0.437 | 1 | 0.5086 |
| rs59018400 | 11 | 118049811 | T | C | 0.143 | 0.8867 | +- | 0 | 0.437 | 1 | 0.5086 |
| rs61472653 | 11 | 118049759 | C | T | 0.143 | 0.8867 | +- | 0 | 0.437 | 1 | 0.5086 |
| rs7123533 | 11 | 118049257 | G | C | 0.143 | 0.8867 | +- | 0 | 0.437 | 1 | 0.5086 |
| rs7342223 | 11 | 118050348 | G | T | 0.143 | 0.8867 | +- | 0 | 0.437 | 1 | 0.5086 |
| rs7931923 | 11 | 120393776 | G | A | 0.142 | 0.8871 | +? | 0 | 0 | 0 | 1 |
| rs111430146 | 11 | 118248374 | A | C | 0.142 | 0.8873 | -+ | -4.2 | 2.88 | 1 | 0.08967 |
| rs1939664 | 11 | 120585300 | A | T | 0.142 | 0.8873 | +? | 0 | 0 | 0 | 1 |
| rs11217972 | 11 | 120552212 | T | C | -0.141 | 0.8875 | -? | 0 | 0 | 0 | 1 |
| rs61906079 | 11 | 116533660 | G | C | 0.141 | 0.8875 | +? | 0 | 0 | 0 | 1 |
| rs657706 | 11 | 119893574 | C | T | 0.141 | 0.8876 | +? | 0 | 0 | 0 | 1 |
| rs17245368 | 11 | 119879327 | T | C | -0.141 | 0.8877 | -? | 0 | 0 | 0 | 1 |
| rs61899021 | 11 | 119780975 | T | G | -0.141 | 0.8878 | -? | 0 | 0 | 0 | 1 |
| rs682795 | 11 | 118983681 | C | T | -0.141 | 0.888 | -? | 0 | 0 | 0 | 1 |
| rs587304 | 11 | 119890949 | G | C | 0.141 | 0.8881 | +? | 0 | 0 | 0 | 1 |
| rs10750152 | 11 | 119431293 | T | C | -0.14 | 0.8884 | -? | 0 | 0 | 0 | 1 |
| rs10892398 | 11 | 119431487 | C | T | -0.14 | 0.8884 | -? | 0 | 0 | 0 | 1 |
| rs11217320 | 11 | 119431655 | G | A | -0.14 | 0.8884 | -? | 0 | 0 | 0 | 1 |
| rs11217321 | 11 | 119432084 | G | A | -0.14 | 0.8884 | -? | 0 | 0 | 0 | 1 |
| rs55808044 | 11 | 119431602 | G | C | -0.14 | 0.8884 | -? | 0 | 0 | 0 | 1 |
| rs11217460 | 11 | 119622162 | G | C | 0.14 | 0.8885 | +? | 0 | 0 | 0 | 1 |
| rs513510 | 11 | 116482451 | C | T | 0.14 | 0.889 | +? | 0 | 0 | 0 | 1 |
| rs61898290 | 11 | 120094385 | A | G | 0.139 | 0.8891 | +? | 0 | 0 | 0 | 1 |
| rs73003294 | 11 | 118661376 | T | C | -0.139 | 0.8892 | +- | 0 | 0.9 | 1 | 0.3427 |
| rs10736504 | 11 | 120584018 | A | G | 0.139 | 0.8894 | +? | 0 | 0 | 0 | 1 |
| rs916493 | 11 | 119429855 | T | G | 0.139 | 0.8894 | +? | 0 | 0 | 0 | 1 |
| rs1876792 | 11 | 119444435 | G | A | -0.139 | 0.8895 | -? | 0 | 0 | 0 | 1 |
| rs11607827 | 11 | 119731345 | G | T | 0.139 | 0.8896 | +? | 0 | 0 | 0 | 1 |
| rs2663628 | 11 | 119434182 | C | A | 0.139 | 0.8897 | +? | 0 | 0 | 0 | 1 |
| rs2084898 | 11 | 120026748 | A | G | 0.139 | 0.8898 | +? | 0 | 0 | 0 | 1 |

| | | | | | | | | | | | |
|---|---|---|---|---|---|---|---|---|---|---|---|
| rs501351 | 11 | 118917814 | G | T | 0.139 | 0.8898 | +? | 0 | 0 | 0 | 1 |
| rs7130098 | 11 | 117701783 | C | G | 0.138 | 0.8899 | +? | 0 | 0 | 0 | 1 |
| rs143470629 | 11 | 119762628 | T | C | 0.138 | 0.8901 | +? | 0 | 0 | 0 | 1 |
| rs61907591 | 11 | 116796855 | A | T | 0.138 | 0.8902 | +? | 0 | 0 | 0 | 1 |
| rs113858735 | 11 | 117625225 | A | G | -0.138 | 0.8904 | +- | 27 | 4.109 | 1 | 0.04267 |
| rs3741333 | 11 | 118986972 | T | C | -0.138 | 0.8905 | -? | 0 | 0 | 0 | 1 |
| rs111354396 | 11 | 118296184 | A | G | 0.138 | 0.8906 | -+ | -15.1 | 2.607 | 1 | 0.1064 |
| rs1473325 | 11 | 116822034 | T | C | 0.138 | 0.8906 | +? | 0 | 0 | 0 | 1 |
| rs2846096 | 11 | 120645487 | G | A | 0.138 | 0.8906 | +? | 0 | 0 | 0 | 1 |
| rs3741325 | 11 | 118405989 | G | A | 0.137 | 0.8907 | ++ | 0 | 0.001 | 1 | 0.9718 |
| rs77718036 | 11 | 117964613 | T | C | 0.137 | 0.8907 | +? | 0 | 0 | 0 | 1 |
| rs10892160 | 11 | 117552640 | G | A | -0.137 | 0.8908 | -? | 0 | 0 | 0 | 1 |
| rs12574186 | 11 | 120550286 | G | A | 0.137 | 0.8909 | +? | 0 | 0 | 0 | 1 |
| rs483283 | 11 | 118506402 | C | T | -0.137 | 0.8909 | -+ | 0 | 0.509 | 1 | 0.4755 |
| rs117420503 | 11 | 119975137 | T | C | 0.137 | 0.891 | +? | 0 | 0 | 0 | 1 |
| rs4938520 | 11 | 118522338 | G | A | -0.137 | 0.891 | -+ | 0 | 0.521 | 1 | 0.4704 |
| rs558229 | 11 | 117178225 | G | A | 0.137 | 0.8911 | +? | 0 | 0 | 0 | 1 |
| rs113981653 | 11 | 120048628 | T | C | 0.137 | 0.8913 | +? | 0 | 0 | 0 | 1 |
| rs671334 | 11 | 119934058 | A | G | 0.137 | 0.8913 | +? | 0 | 0 | 0 | 1 |
| rs11217096 | 11 | 118796570 | T | C | 0.137 | 0.8914 | +- | 0 | 0.021 | 1 | 0.885 |
| rs568787 | 11 | 117218762 | A | G | -0.137 | 0.8914 | -? | 0 | 0 | 0 | 1 |
| rs1041395195 | 11 | 117225441 | T | G | 0.136 | 0.8915 | +? | 0 | 0 | 0 | 1 |
| rs12271064 | 11 | 118119211 | G | A | -0.136 | 0.8917 | -+ | 0 | 0.148 | 1 | 0.7001 |
| rs236915 | 11 | 117088450 | G | A | 0.136 | 0.8918 | +? | 0 | 0 | 0 | 1 |
| rs11218202 | 11 | 121087341 | G | A | -0.136 | 0.892 | -? | 0 | 0 | 0 | 1 |
| rs7119655 | 11 | 120549488 | G | A | -0.136 | 0.8922 | -? | 0 | 0 | 0 | 1 |
| rs1263162 | 11 | 116705021 | A | T | -0.135 | 0.8923 | +- | 4.5 | 3.142 | 1 | 0.07629 |
| rs111560312 | 11 | 116983556 | A | G | -0.135 | 0.8924 | -? | 0 | 0 | 0 | 1 |
| rs1617182 | 11 | 118068949 | C | T | 0.135 | 0.8925 | +? | 0 | 0 | 0 | 1 |
| rs568221 | 11 | 117476349 | T | C | -0.135 | 0.8926 | -? | 0 | 0 | 0 | 1 |
| rs12224987 | 11 | 120552193 | T | C | -0.135 | 0.8928 | -? | 0 | 0 | 0 | 1 |
| rs145637218 | 11 | 119805057 | A | G | -0.135 | 0.8928 | -? | 0 | 0 | 0 | 1 |
| rs1145212 | 11 | 116556865 | A | G | -0.135 | 0.8929 | -? | 0 | 0 | 0 | 1 |
| rs76604009 | 11 | 117085261 | T | C | -0.135 | 0.8929 | -? | 0 | 0 | 0 | 1 |
| rs4936493 | 11 | 119568070 | T | A | -0.135 | 0.893 | -? | 0 | 0 | 0 | 1 |
| rs4936506 | 11 | 119794035 | T | C | -0.135 | 0.893 | -? | 0 | 0 | 0 | 1 |
| rs61581699 | 11 | 118045423 | A | G | 0.135 | 0.893 | +- | 0 | 0.394 | 1 | 0.5303 |
| rs7104343 | 11 | 116529024 | T | C | -0.135 | 0.893 | -? | 0 | 0 | 0 | 1 |
| rs74332985 | 11 | 116220623 | C | T | 0.134 | 0.8931 | +? | 0 | 0 | 0 | 1 |
| rs141259801 | 11 | 117718626 | T | C | 0.134 | 0.8932 | +? | 0 | 0 | 0 | 1 |
| rs61898269 | 11 | 120073606 | C | A | -0.134 | 0.8932 | -? | 0 | 0 | 0 | 1 |
| rs87824 | 11 | 117442242 | C | G | 0.134 | 0.8932 | +? | 0 | 0 | 0 | 1 |
| rs1168569 | 11 | 118947168 | G | A | 0.134 | 0.8933 | +? | 0 | 0 | 0 | 1 |
| rs11826651 | 11 | 116920693 | A | T | 0.134 | 0.8933 | +? | 0 | 0 | 0 | 1 |
| rs12362348 | 11 | 120541508 | C | G | 0.134 | 0.8933 | +? | 0 | 0 | 0 | 1 |
| rs17566737 | 11 | 116230506 | C | T | -0.134 | 0.8935 | -? | 0 | 0 | 0 | 1 |
| rs11826742 | 11 | 116924366 | G | C | 0.134 | 0.8937 | +? | 0 | 0 | 0 | 1 |
| rs11216398 | 11 | 117323082 | T | C | -0.133 | 0.8938 | -? | 0 | 0 | 0 | 1 |
| rs11216614 | 11 | 117777929 | C | G | -0.133 | 0.8938 | -? | 0 | 0 | 0 | 1 |
| rs147893781 | 11 | 116848716 | T | C | -0.133 | 0.8938 | -? | 0 | 0 | 0 | 1 |
| rs11216790 | 11 | 118044144 | T | G | 0.133 | 0.8939 | +- | 0 | 0.381 | 1 | 0.5369 |
| rs3887268 | 11 | 119950310 | G | C | -0.133 | 0.894 | -? | 0 | 0 | 0 | 1 |
| rs570836 | 11 | 118512162 | A | G | -0.133 | 0.8942 | -+ | 0 | 0.519 | 1 | 0.4713 |
| rs10892166 | 11 | 117605839 | A | G | 0.133 | 0.8943 | +? | 0 | 0 | 0 | 1 |
| rs566880 | 11 | 119695011 | T | C | -0.133 | 0.8943 | -? | 0 | 0 | 0 | 1 |
| rs12574240 | 11 | 120569381 | C | T | -0.133 | 0.8944 | -? | 0 | 0 | 0 | 1 |

| | | | | | | | | | | | |
|---|---|---|---|---|---|---|---|---|---|---|---|
| rs7131339 | 11 | 119973082 | A | G | 0.133 | 0.8944 | +? | 0 | 0 | 0 | 1 |
| rs73588361 | 11 | 120590848 | A | G | -0.133 | 0.8945 | -? | 0 | 0 | 0 | 1 |
| rs10750154 | 11 | 119432909 | G | A | -0.132 | 0.8947 | -? | 0 | 0 | 0 | 1 |
| rs1369553921 | 11 | 117222592 | A | G | 0.132 | 0.8947 | +? | 0 | 0 | 0 | 1 |
| rs514806 | 11 | 117222942 | T | G | 0.132 | 0.8947 | +? | 0 | 0 | 0 | 1 |
| rs61900472 | 11 | 120009748 | A | G | 0.132 | 0.8947 | +? | 0 | 0 | 0 | 1 |
| rs688207 | 11 | 117222139 | T | C | 0.132 | 0.8947 | +? | 0 | 0 | 0 | 1 |
| rs35114640 | 11 | 120640009 | A | C | 0.132 | 0.8948 | +? | 0 | 0 | 0 | 1 |
| rs525914 | 11 | 119695450 | T | C | -0.132 | 0.8948 | -? | 0 | 0 | 0 | 1 |
| rs11216360 | 11 | 117213189 | A | G | 0.132 | 0.8949 | +? | 0 | 0 | 0 | 1 |
| rs11605435 | 11 | 118116496 | C | T | -0.132 | 0.8949 | -+ | 0 | 0.178 | 1 | 0.6734 |
| rs145476010 | 11 | 119943314 | A | C | 0.132 | 0.895 | +? | 0 | 0 | 0 | 1 |
| rs1876794 | 11 | 119444016 | C | G | -0.132 | 0.895 | -? | 0 | 0 | 0 | 1 |
| rs61905708 | 11 | 116886159 | T | G | -0.132 | 0.895 | -? | 0 | 0 | 0 | 1 |
| rs79973603 | 11 | 117643723 | C | T | -0.132 | 0.895 | +- | 25 | 4.002 | 1 | 0.04545 |
| rs17597688 | 11 | 117795942 | G | A | 0.132 | 0.8952 | +- | 0 | 0.785 | 1 | 0.3756 |
| rs17119549 | 11 | 116241208 | T | G | -0.132 | 0.8953 | -? | 0 | 0 | 0 | 1 |
| rs1168570 | 11 | 118947154 | C | T | 0.131 | 0.8954 | +? | 0 | 0 | 0 | 1 |
| rs12280557 | 11 | 118117118 | T | C | -0.131 | 0.8956 | -+ | 0 | 0.179 | 1 | 0.6721 |
| rs56182713 | 11 | 116558828 | A | G | 0.131 | 0.8956 | +? | 0 | 0 | 0 | 1 |
| rs7924318 | 11 | 116816630 | G | C | 0.131 | 0.8956 | +? | 0 | 0 | 0 | 1 |
| rs10502247 | 11 | 120998942 | T | C | -0.131 | 0.8958 | +- | 0 | 0.699 | 1 | 0.4031 |
| rs10790254 | 11 | 118493319 | A | G | -0.131 | 0.8959 | -+ | 0 | 0.112 | 1 | 0.7379 |
| rs80178624 | 11 | 116236271 | A | T | -0.131 | 0.8959 | +- | -176.9 | 1.084 | 1 | 0.2979 |
| rs996889 | 11 | 119428421 | G | A | 0.131 | 0.8959 | +? | 0 | 0 | 0 | 1 |
| rs11217967 | 11 | 120548795 | A | C | -0.131 | 0.896 | -? | 0 | 0 | 0 | 1 |
| rs12294117 | 11 | 117662864 | A | G | 0.131 | 0.896 | ++ | 0 | 0.01 | 1 | 0.9205 |
| rs80343155 | 11 | 117346337 | C | A | 0.131 | 0.8961 | +? | 0 | 0 | 0 | 1 |
| rs7936400 | 11 | 119655589 | C | T | 0.13 | 0.8966 | +? | 0 | 0 | 0 | 1 |
| rs17123506 | 11 | 119842708 | G | A | -0.13 | 0.8967 | -? | 0 | 0 | 0 | 1 |
| rs548877 | 11 | 118300240 | G | A | 0.13 | 0.8967 | +- | 0 | 0.062 | 1 | 0.8031 |
| rs77454413 | 11 | 120664625 | A | G | 0.13 | 0.8968 | +? | 0 | 0 | 0 | 1 |
| rs11604096 | 11 | 118012441 | C | A | 0.13 | 0.8969 | ++ | 0 | 0.001 | 1 | 0.9805 |
| rs146781782 | 11 | 117615894 | A | G | -0.13 | 0.8969 | -? | 0 | 0 | 0 | 1 |
| rs634960 | 11 | 117072963 | C | T | 0.129 | 0.897 | +? | 0 | 0 | 0 | 1 |
| rs7130531 | 11 | 116863833 | T | C | 0.129 | 0.8974 | +? | 0 | 0 | 0 | 1 |
| rs7124084 | 11 | 117235412 | T | C | 0.129 | 0.8975 | +? | 0 | 0 | 0 | 1 |
| rs11216794 | 11 | 118044740 | A | T | 0.129 | 0.8976 | +- | 0 | 0.383 | 1 | 0.5363 |
| rs35544562 | 11 | 119369816 | C | G | 0.128 | 0.8978 | +? | 0 | 0 | 0 | 1 |
| rs45448191 | 11 | 118267737 | A | T | -0.128 | 0.8981 | -+ | -21.3 | 2.474 | 1 | 0.1157 |
| rs2845710 | 11 | 117697982 | A | G | 0.128 | 0.8982 | +? | 0 | 0 | 0 | 1 |
| rs61905679 | 11 | 116834787 | T | C | 0.128 | 0.8982 | +? | 0 | 0 | 0 | 1 |
| rs75904569 | 11 | 116988313 | G | A | -0.128 | 0.8983 | -? | 0 | 0 | 0 | 1 |
| rs79281178 | 11 | 118298041 | G | C | 0.128 | 0.8983 | +- | 0 | 0.121 | 1 | 0.7282 |
| rs11608072 | 11 | 118030086 | C | G | -0.128 | 0.8984 | -+ | 0 | 0.078 | 1 | 0.7797 |
| rs679327 | 11 | 118042974 | G | A | 0.128 | 0.8984 | ++ | 0 | 0.004 | 1 | 0.948 |
| rs78054182 | 11 | 119792099 | C | T | 0.127 | 0.8988 | +? | 0 | 0 | 0 | 1 |
| rs113739713 | 11 | 118420101 | G | A | 0.127 | 0.8991 | +- | 0 | 0.007 | 1 | 0.9329 |
| rs12295676 | 11 | 118054345 | A | G | 0.127 | 0.8991 | +- | 0 | 0.398 | 1 | 0.528 |
| rs150233369 | 11 | 116629555 | A | G | 0.127 | 0.8991 | +? | 0 | 0 | 0 | 1 |
| rs61897923 | 11 | 119903434 | C | T | 0.127 | 0.8991 | +? | 0 | 0 | 0 | 1 |
| rs12419633 | 11 | 121152173 | A | G | 0.127 | 0.8993 | +? | 0 | 0 | 0 | 1 |
| rs10892219 | 11 | 117983054 | A | G | -0.126 | 0.8994 | -? | 0 | 0 | 0 | 1 |
| rs10892095 | 11 | 117213893 | G | A | 0.126 | 0.8995 | +? | 0 | 0 | 0 | 1 |
| rs7104443 | 11 | 117996885 | G | C | 0.126 | 0.8995 | +? | 0 | 0 | 0 | 1 |
| rs1715444 | 11 | 118016273 | G | T | 0.126 | 0.8997 | ++ | 0 | 0.03 | 1 | 0.8616 |

| | | | | | | | | | | | |
|---|---|---|---|---|---|---|---|---|---|---|---|
| rs12801003 | 11 | 118072716 | T | C | 0.125 | 0.9002 | -+ | 0 | 0.523 | 1 | 0.4697 |
| rs10502227 | 11 | 118147496 | C | T | 0.125 | 0.9004 | -+ | -21.4 | 2.471 | 1 | 0.116 |
| rs11603267 | 11 | 118146599 | T | G | 0.125 | 0.9004 | -+ | -21.4 | 2.471 | 1 | 0.116 |
| rs34815971 | 11 | 118301066 | G | A | 0.125 | 0.9004 | +- | 0 | 0.061 | 1 | 0.8051 |
| rs10790282 | 11 | 118973133 | G | A | -0.125 | 0.9007 | -? | 0 | 0 | 0 | 1 |
| rs11216792 | 11 | 118044450 | T | G | 0.125 | 0.9007 | +- | 0 | 0.375 | 1 | 0.5402 |
| rs510997 | 11 | 118302901 | C | T | 0.125 | 0.9007 | +- | 0 | 0.101 | 1 | 0.7505 |
| rs75403174 | 11 | 118043904 | A | C | 0.125 | 0.9008 | +- | 0 | 0.375 | 1 | 0.5402 |
| rs35650539 | 11 | 120039306 | G | C | 0.124 | 0.901 | +? | 0 | 0 | 0 | 1 |
| rs12291619 | 11 | 118062552 | A | G | -0.124 | 0.9013 | +- | 0 | 0.234 | 1 | 0.6283 |
| rs138282449 | 11 | 116331800 | C | G | -0.124 | 0.9015 | -? | 0 | 0 | 0 | 1 |
| rs7117544 | 11 | 118117938 | C | A | -0.124 | 0.9016 | -+ | 0 | 0.191 | 1 | 0.6625 |
| rs745663 | 11 | 118529595 | G | A | -0.124 | 0.9017 | -+ | 0 | 0.421 | 1 | 0.5162 |
| rs11216892 | 11 | 118424008 | T | C | 0.123 | 0.9018 | +- | 0 | 0.02 | 1 | 0.8881 |
| rs12577841 | 11 | 118194599 | T | C | 0.123 | 0.9018 | +- | 0 | 0.115 | 1 | 0.7347 |
| rs520326 | 11 | 116315660 | G | T | 0.123 | 0.9018 | +? | 0 | 0 | 0 | 1 |
| rs78160609 | 11 | 120131372 | A | G | -0.123 | 0.9018 | -? | 0 | 0 | 0 | 1 |
| rs12362311 | 11 | 120541373 | T | C | 0.123 | 0.9019 | +? | 0 | 0 | 0 | 1 |
| rs61900479 | 11 | 120040695 | T | C | 0.123 | 0.9019 | +? | 0 | 0 | 0 | 1 |
| rs1613495 | 11 | 118017645 | T | C | 0.123 | 0.902 | -+ | 0 | 0.121 | 1 | 0.728 |
| rs7934067 | 11 | 117232954 | G | A | 0.123 | 0.9021 | +? | 0 | 0 | 0 | 1 |
| rs542378 | 11 | 120855998 | T | C | -0.123 | 0.9022 | -+ | 0 | 0.195 | 1 | 0.6588 |
| rs11600129 | 11 | 117800169 | C | T | 0.123 | 0.9023 | -+ | -176.1 | 1.086 | 1 | 0.2973 |
| rs73020500 | 11 | 117777101 | T | C | 0.123 | 0.9023 | +? | 0 | 0 | 0 | 1 |
| rs591744 | 11 | 119939826 | G | A | -0.122 | 0.9026 | -? | 0 | 0 | 0 | 1 |
| rs1261744 | 11 | 117218460 | C | T | 0.122 | 0.9027 | +? | 0 | 0 | 0 | 1 |
| rs680630 | 11 | 119934081 | T | G | 0.122 | 0.9027 | +? | 0 | 0 | 0 | 1 |
| rs12225187 | 11 | 116793231 | G | A | 0.122 | 0.9029 | +? | 0 | 0 | 0 | 1 |
| rs35392431 | 11 | 117973338 | G | C | -0.122 | 0.9029 | +- | 0 | 0.052 | 1 | 0.8192 |
| rs61903398 | 11 | 116941613 | T | C | 0.122 | 0.903 | +? | 0 | 0 | 0 | 1 |
| rs61903399 | 11 | 116942475 | T | A | 0.122 | 0.903 | +? | 0 | 0 | 0 | 1 |
| rs734378 | 11 | 117597660 | T | C | 0.122 | 0.903 | +? | 0 | 0 | 0 | 1 |
| rs118155427 | 11 | 116975084 | C | G | -0.122 | 0.9031 | +- | 37.8 | 4.824 | 1 | 0.02807 |
| rs139202987 | 11 | 118474283 | G | A | -0.122 | 0.9031 | -+ | 0 | 0.034 | 1 | 0.854 |
| rs11216779 | 11 | 118002372 | T | C | -0.122 | 0.9032 | -? | 0 | 0 | 0 | 1 |
| rs12282334 | 11 | 118057783 | G | C | 0.122 | 0.9032 | +- | 0 | 0.387 | 1 | 0.5338 |
| rs57145837 | 11 | 117574499 | A | G | 0.122 | 0.9032 | +? | 0 | 0 | 0 | 1 |
| rs7934292 | 11 | 118059914 | C | T | 0.122 | 0.9032 | +- | 0 | 0.387 | 1 | 0.5338 |
| rs7935492 | 11 | 118056823 | T | C | 0.122 | 0.9032 | +- | 0 | 0.387 | 1 | 0.5338 |
| rs10488699 | 11 | 116632500 | T | C | -0.121 | 0.9035 | +- | -1.7 | 2.95 | 1 | 0.08589 |
| rs7114615 | 11 | 120704659 | T | G | 0.121 | 0.9035 | +? | 0 | 0 | 0 | 1 |
| rs11216062 | 11 | 116502768 | A | C | 0.121 | 0.9036 | +? | 0 | 0 | 0 | 1 |
| rs12365289 | 11 | 117521152 | G | C | -0.121 | 0.9037 | -? | 0 | 0 | 0 | 1 |
| rs1968541 | 11 | 116237825 | A | G | 0.121 | 0.9037 | +- | -145.3 | 1.223 | 1 | 0.2688 |
| rs10892399 | 11 | 119432144 | C | T | -0.121 | 0.9039 | -? | 0 | 0 | 0 | 1 |
| rs11217255 | 11 | 119231667 | G | A | -0.121 | 0.9039 | -? | 0 | 0 | 0 | 1 |
| rs12420446 | 11 | 116207302 | A | G | -0.121 | 0.9039 | -? | 0 | 0 | 0 | 1 |
| rs508871 | 11 | 117219770 | A | G | 0.121 | 0.9039 | +? | 0 | 0 | 0 | 1 |
| rs2115923 | 11 | 121158779 | A | G | -0.121 | 0.904 | -? | 0 | 0 | 0 | 1 |
| rs7124127 | 11 | 118006408 | C | T | -0.121 | 0.904 | -+ | 0 | 0.155 | 1 | 0.6935 |
| rs10750148 | 11 | 119412474 | A | T | 0.12 | 0.9041 | +? | 0 | 0 | 0 | 1 |
| rs11217575 | 11 | 119792847 | G | A | 0.12 | 0.9043 | +? | 0 | 0 | 0 | 1 |
| rs12286303 | 11 | 118114473 | T | C | -0.12 | 0.9045 | -+ | 0 | 0.158 | 1 | 0.6906 |
| rs11217466 | 11 | 119624993 | A | T | -0.12 | 0.9047 | -? | 0 | 0 | 0 | 1 |
| rs61898285 | 11 | 120091653 | T | C | 0.12 | 0.9048 | +? | 0 | 0 | 0 | 1 |
| rs112305884 | 11 | 116495784 | T | C | 0.119 | 0.9049 | +? | 0 | 0 | 0 | 1 |

| | | | | | | | | | | | |
|---|---|---|---|---|---|---|---|---|---|---|---|
| rs145672593 | 11 | 117010263 | T | C | 0.119 | 0.9049 | +? | 0 | 0 | 0 | 1 |
| rs75709684 | 11 | 118050958 | A | G | 0.119 | 0.905 | +- | 0 | 0.397 | 1 | 0.5287 |
| rs17092876 | 11 | 120398643 | A | G | -0.119 | 0.9053 | -? | 0 | 0 | 0 | 1 |
| rs12286567 | 11 | 118114788 | T | G | -0.119 | 0.9055 | -+ | 0 | 0.16 | 1 | 0.689 |
| rs476937 | 11 | 119697594 | A | G | -0.119 | 0.9055 | -? | 0 | 0 | 0 | 1 |
| rs73005215 | 11 | 116312418 | G | A | -0.119 | 0.9056 | -? | 0 | 0 | 0 | 1 |
| rs150762661 | 11 | 116345271 | C | A | -0.118 | 0.9058 | -? | 0 | 0 | 0 | 1 |
| rs61905687 | 11 | 116862144 | C | T | 0.118 | 0.9058 | +? | 0 | 0 | 0 | 1 |
| rs75459003 | 11 | 118053859 | C | A | 0.118 | 0.9058 | +- | 0 | 0.392 | 1 | 0.5312 |
| rs7950213 | 11 | 116868654 | A | C | 0.118 | 0.9058 | +? | 0 | 0 | 0 | 1 |
| rs35626144 | 11 | 119459523 | G | A | 0.118 | 0.9059 | +? | 0 | 0 | 0 | 1 |
| rs4344527 | 11 | 117973511 | G | A | 0.118 | 0.906 | +- | 0 | 0.089 | 1 | 0.7648 |
| rs147146093 | 11 | 117578440 | A | G | 0.118 | 0.9062 | +? | 0 | 0 | 0 | 1 |
| rs611979 | 11 | 118034223 | A | G | 0.118 | 0.9062 | -+ | 0 | 0.462 | 1 | 0.4965 |
| rs7934773 | 11 | 117579531 | A | G | 0.118 | 0.9062 | +? | 0 | 0 | 0 | 1 |
| rs2305825 | 11 | 117232492 | T | C | 0.118 | 0.9064 | +? | 0 | 0 | 0 | 1 |
| rs17591711 | 11 | 117228849 | G | A | 0.117 | 0.9066 | +? | 0 | 0 | 0 | 1 |
| rs2014537 | 11 | 117691148 | C | T | -0.117 | 0.9066 | -? | 0 | 0 | 0 | 1 |
| rs567312 | 11 | 116730738 | G | A | 0.117 | 0.9068 | +? | 0 | 0 | 0 | 1 |
| rs470606 | 11 | 120028202 | G | T | -0.117 | 0.9069 | -? | 0 | 0 | 0 | 1 |
| rs7106413 | 11 | 120769943 | T | C | 0.117 | 0.9069 | +? | 0 | 0 | 0 | 1 |
| rs501776 | 11 | 117258198 | C | T | -0.117 | 0.907 | -? | 0 | 0 | 0 | 1 |
| rs34708128 | 11 | 119691272 | T | C | 0.117 | 0.9071 | +? | 0 | 0 | 0 | 1 |
| rs601366 | 11 | 119883621 | C | T | 0.117 | 0.9072 | +? | 0 | 0 | 0 | 1 |
| rs612969 | 11 | 120989335 | G | A | -0.117 | 0.9072 | -? | 0 | 0 | 0 | 1 |
| rs552043 | 11 | 116325990 | T | C | -0.116 | 0.9074 | -? | 0 | 0 | 0 | 1 |
| rs75275400 | 11 | 117529676 | C | T | 0.116 | 0.9074 | +? | 0 | 0 | 0 | 1 |
| rs113975105 | 11 | 118836606 | G | C | -0.116 | 0.9075 | +- | -168.4 | 1.118 | 1 | 0.2904 |
| rs12787035 | 11 | 121082690 | A | G | -0.116 | 0.9077 | -? | 0 | 0 | 0 | 1 |
| rs34336740 | 11 | 117662605 | G | C | 0.116 | 0.9077 | ++ | 0 | 0.013 | 1 | 0.9107 |
| rs61372904 | 11 | 116599900 | G | A | -0.116 | 0.9078 | +- | -71.6 | 1.748 | 1 | 0.1862 |
| rs11217458 | 11 | 119620940 | A | G | 0.116 | 0.9079 | +? | 0 | 0 | 0 | 1 |
| rs12421491 | 11 | 119834296 | T | C | -0.116 | 0.908 | -? | 0 | 0 | 0 | 1 |
| rs4938340 | 11 | 116980622 | A | G | -0.116 | 0.908 | -? | 0 | 0 | 0 | 1 |
| rs61905464 | 11 | 117001384 | A | G | -0.116 | 0.908 | -? | 0 | 0 | 0 | 1 |
| rs10892483 | 11 | 119748404 | C | G | 0.115 | 0.9082 | +? | 0 | 0 | 0 | 1 |
| rs2735183 | 11 | 117015195 | C | T | 0.115 | 0.9082 | +? | 0 | 0 | 0 | 1 |
| rs11218108 | 11 | 120878916 | T | C | 0.115 | 0.9084 | +? | 0 | 0 | 0 | 1 |
| rs6589732 | 11 | 119185347 | A | G | -0.115 | 0.9084 | -? | 0 | 0 | 0 | 1 |
| rs117397221 | 11 | 120018785 | T | C | -0.115 | 0.9086 | -? | 0 | 0 | 0 | 1 |
| rs1793169 | 11 | 118070264 | G | A | 0.115 | 0.9086 | +? | 0 | 0 | 0 | 1 |
| rs11216887 | 11 | 118420736 | C | T | 0.115 | 0.9087 | +- | 0 | 0.005 | 1 | 0.9421 |
| rs4936570 | 11 | 120834573 | T | C | -0.115 | 0.9088 | -+ | 0 | 0.191 | 1 | 0.6623 |
| rs11605919 | 11 | 119902691 | C | T | 0.114 | 0.9089 | +? | 0 | 0 | 0 | 1 |
| rs543181 | 11 | 120855930 | A | G | -0.114 | 0.9091 | -+ | 0 | 0.191 | 1 | 0.6624 |
| rs488962 | 11 | 117072525 | T | C | 0.114 | 0.9092 | +? | 0 | 0 | 0 | 1 |
| rs11217133 | 11 | 118928249 | A | G | 0.114 | 0.9093 | +? | 0 | 0 | 0 | 1 |
| rs117819665 | 11 | 116393781 | T | C | -0.114 | 0.9093 | -? | 0 | 0 | 0 | 1 |
| rs660083 | 11 | 116720829 | C | G | -0.114 | 0.9095 | -? | 0 | 0 | 0 | 1 |
| rs11216357 | 11 | 117200753 | G | T | -0.113 | 0.9097 | -? | 0 | 0 | 0 | 1 |
| rs12271220 | 11 | 117664373 | A | G | 0.113 | 0.9098 | ++ | 0 | 0.012 | 1 | 0.9128 |
| rs3819108 | 11 | 117973576 | A | G | 0.113 | 0.9098 | +- | 0 | 0.085 | 1 | 0.7711 |
| rs10750155 | 11 | 119434032 | T | A | -0.113 | 0.9099 | -? | 0 | 0 | 0 | 1 |
| rs10790311 | 11 | 119433005 | T | C | -0.113 | 0.9099 | -? | 0 | 0 | 0 | 1 |
| rs10790312 | 11 | 119434275 | T | C | -0.113 | 0.9099 | -? | 0 | 0 | 0 | 1 |
| rs10892402 | 11 | 119433521 | G | A | -0.113 | 0.9099 | -? | 0 | 0 | 0 | 1 |

| | | | | | | | | | | | |
|---|---|---|---|---|---|---|---|---|---|---|---|
| rs10892403 | 11 | 119434843 | A | C | -0.113 | 0.9099 | -? | 0 | 0 | 0 | 1 |
| rs12802764 | 11 | 116474555 | A | G | -0.113 | 0.9099 | -? | 0 | 0 | 0 | 1 |
| rs2368425 | 11 | 119434468 | T | G | -0.113 | 0.9099 | -? | 0 | 0 | 0 | 1 |
| rs4018655 | 11 | 119434711 | A | G | -0.113 | 0.9099 | -? | 0 | 0 | 0 | 1 |
| rs67163042 | 11 | 118050779 | G | A | 0.113 | 0.9099 | +- | 0 | 0.415 | 1 | 0.5194 |
| rs17119713 | 11 | 116338675 | G | T | -0.113 | 0.91 | -? | 0 | 0 | 0 | 1 |
| rs142239919 | 11 | 117339203 | G | A | -0.113 | 0.9101 | -? | 0 | 0 | 0 | 1 |
| rs4938694 | 11 | 119495871 | A | G | -0.113 | 0.9101 | -? | 0 | 0 | 0 | 1 |
| rs74491080 | 11 | 120991447 | T | C | 0.113 | 0.9101 | +? | 0 | 0 | 0 | 1 |
| rs7951065 | 11 | 119495352 | C | T | -0.113 | 0.9101 | -? | 0 | 0 | 0 | 1 |
| rs7951484 | 11 | 119495527 | G | A | -0.113 | 0.9101 | -? | 0 | 0 | 0 | 1 |
| rs74830 | 11 | 117090558 | T | C | 0.113 | 0.9102 | +? | 0 | 0 | 0 | 1 |
| rs4388923 | 11 | 117973558 | A | C | 0.113 | 0.9104 | +- | 0 | 0.084 | 1 | 0.7714 |
| rs5092 | 11 | 116693464 | C | T | -0.112 | 0.9105 | -? | 0 | 0 | 0 | 1 |
| rs655499 | 11 | 118063479 | G | A | 0.112 | 0.9105 | -+ | 0 | 0.064 | 1 | 0.8001 |
| rs2845658 | 11 | 119434753 | A | G | 0.112 | 0.9106 | +? | 0 | 0 | 0 | 1 |
| rs10790223 | 11 | 117800700 | C | T | 0.112 | 0.9107 | -+ | -138.1 | 1.26 | 1 | 0.2616 |
| rs77279864 | 11 | 118017765 | T | C | 0.112 | 0.9107 | +? | 0 | 0 | 0 | 1 |
| rs12421205 | 11 | 117232156 | G | C | 0.112 | 0.9108 | +? | 0 | 0 | 0 | 1 |
| rs12421494 | 11 | 117232233 | A | G | 0.112 | 0.9108 | +? | 0 | 0 | 0 | 1 |
| rs111453105 | 11 | 116281901 | A | G | -0.112 | 0.9109 | -? | 0 | 0 | 0 | 1 |
| rs521171 | 11 | 117064693 | C | G | -0.112 | 0.9109 | -? | 0 | 0 | 0 | 1 |
| rs7104002 | 11 | 117575862 | C | T | 0.112 | 0.911 | +? | 0 | 0 | 0 | 1 |
| rs72999486 | 11 | 118526691 | T | C | 0.112 | 0.911 | ++ | 0 | 0.009 | 1 | 0.9243 |
| rs542671 | 11 | 116323001 | A | G | 0.112 | 0.9111 | +? | 0 | 0 | 0 | 1 |
| rs560801 | 11 | 116331960 | C | A | 0.112 | 0.9112 | +? | 0 | 0 | 0 | 1 |
| rs45443099 | 11 | 118254710 | G | A | 0.111 | 0.9113 | -+ | -8 | 2.778 | 1 | 0.09557 |
| rs4936463 | 11 | 118975355 | A | G | -0.111 | 0.9113 | -? | 0 | 0 | 0 | 1 |
| rs59494484 | 11 | 119445229 | T | C | 0.111 | 0.9113 | +? | 0 | 0 | 0 | 1 |
| rs3132893 | 11 | 118303822 | A | C | -0.111 | 0.9114 | +- | 0 | 0.323 | 1 | 0.5698 |
| rs4936464 | 11 | 118988745 | T | C | -0.111 | 0.912 | -? | 0 | 0 | 0 | 1 |
| rs617139 | 11 | 119880361 | T | C | 0.11 | 0.9122 | +? | 0 | 0 | 0 | 1 |
| rs7933963 | 11 | 118246906 | T | C | 0.11 | 0.9122 | -+ | 0 | 0.134 | 1 | 0.7145 |
| rs34850032 | 11 | 117797898 | G | A | 0.11 | 0.9123 | +? | 0 | 0 | 0 | 1 |
| rs74395230 | 11 | 118052662 | T | C | 0.11 | 0.9124 | +- | 0 | 0.386 | 1 | 0.5343 |
| rs4363614 | 11 | 120456605 | A | G | -0.11 | 0.9125 | -? | 0 | 0 | 0 | 1 |
| rs6589673 | 11 | 118557483 | C | A | -0.11 | 0.9125 | -+ | 0 | 0.087 | 1 | 0.7682 |
| rs11215958 | 11 | 116355310 | A | G | 0.11 | 0.9126 | +? | 0 | 0 | 0 | 1 |
| rs11216890 | 11 | 118421569 | G | C | 0.11 | 0.9126 | +- | 0 | 0.005 | 1 | 0.9443 |
| rs76105754 | 11 | 121079460 | A | G | -0.11 | 0.9126 | -? | 0 | 0 | 0 | 1 |
| rs76719908 | 11 | 121076820 | C | G | -0.11 | 0.9126 | -? | 0 | 0 | 0 | 1 |
| rs1815811 | 11 | 119059404 | G | A | -0.11 | 0.9127 | -? | 0 | 0 | 0 | 1 |
| rs7949076 | 11 | 119621691 | T | C | 0.11 | 0.9128 | +? | 0 | 0 | 0 | 1 |
| rs11216800 | 11 | 118048885 | G | A | 0.109 | 0.9129 | +- | 0 | 0.386 | 1 | 0.5346 |
| rs2077784 | 11 | 120024874 | A | G | 0.109 | 0.9129 | +? | 0 | 0 | 0 | 1 |
| rs12808524 | 11 | 116890993 | T | C | 0.109 | 0.913 | +? | 0 | 0 | 0 | 1 |
| rs1547885 | 11 | 118105222 | T | C | -0.109 | 0.913 | -+ | 0 | 0.107 | 1 | 0.7436 |
| rs664511 | 11 | 119863827 | C | G | -0.109 | 0.9131 | -? | 0 | 0 | 0 | 1 |
| rs60942267 | 11 | 117675364 | T | G | 0.109 | 0.9132 | +? | 0 | 0 | 0 | 1 |
| rs2075547 | 11 | 117068534 | A | C | 0.109 | 0.9135 | +? | 0 | 0 | 0 | 1 |
| rs10502215 | 11 | 116233398 | T | C | -0.108 | 0.9137 | -? | 0 | 0 | 0 | 1 |
| rs11217137 | 11 | 118931030 | G | A | 0.108 | 0.9137 | +? | 0 | 0 | 0 | 1 |
| rs2887244 | 11 | 119434279 | G | T | -0.108 | 0.9139 | -? | 0 | 0 | 0 | 1 |
| rs634867 | 11 | 117411041 | A | G | -0.108 | 0.9139 | -? | 0 | 0 | 0 | 1 |
| rs45508093 | 11 | 118224059 | G | A | 0.108 | 0.914 | +? | 0 | 0 | 0 | 1 |
| rs12800424 | 11 | 116558782 | A | G | 0.108 | 0.9141 | +? | 0 | 0 | 0 | 1 |

| | | | | | | | | | | | |
|---|---|---|---|---|---|---|---|---|---|---|---|
| rs10892394 | 11 | 119429614 | A | G | -0.108 | 0.9142 | -? | 0 | 0 | 0 | 1 |
| rs10892395 | 11 | 119429656 | G | A | -0.108 | 0.9142 | -? | 0 | 0 | 0 | 1 |
| rs10892396 | 11 | 119429662 | G | T | -0.108 | 0.9142 | -? | 0 | 0 | 0 | 1 |
| rs10892397 | 11 | 119430203 | G | A | -0.108 | 0.9142 | -? | 0 | 0 | 0 | 1 |
| rs11217319 | 11 | 119429923 | A | G | -0.108 | 0.9142 | -? | 0 | 0 | 0 | 1 |
| rs12274984 | 11 | 119430266 | A | G | -0.108 | 0.9142 | -? | 0 | 0 | 0 | 1 |
| rs60837248 | 11 | 117192783 | G | A | -0.107 | 0.9144 | -? | 0 | 0 | 0 | 1 |
| rs668468 | 11 | 118011776 | G | A | 0.107 | 0.9144 | ++ | 0 | 0.001 | 1 | 0.9766 |
| rs7128014 | 11 | 116384958 | A | G | 0.107 | 0.9144 | +? | 0 | 0 | 0 | 1 |
| rs4938791 | 11 | 120055170 | A | G | -0.107 | 0.9146 | -? | 0 | 0 | 0 | 1 |
| rs7113263 | 11 | 117338555 | A | G | 0.107 | 0.9146 | +? | 0 | 0 | 0 | 1 |
| rs77267645 | 11 | 117551691 | G | A | -0.107 | 0.9147 | -? | 0 | 0 | 0 | 1 |
| rs12418601 | 11 | 120052360 | T | C | 0.107 | 0.915 | +? | 0 | 0 | 0 | 1 |
| rs17496758 | 11 | 116916175 | A | G | -0.106 | 0.9152 | -? | 0 | 0 | 0 | 1 |
| rs73009178 | 11 | 116411434 | A | C | -0.106 | 0.9153 | -? | 0 | 0 | 0 | 1 |
| rs140186337 | 11 | 119564528 | G | A | -0.106 | 0.9154 | -? | 0 | 0 | 0 | 1 |
| rs79761998 | 11 | 120009308 | A | C | -0.106 | 0.9155 | -? | 0 | 0 | 0 | 1 |
| rs75830246 | 11 | 116968440 | C | T | -0.106 | 0.9157 | +- | 38.8 | 4.902 | 1 | 0.02683 |
| rs111792147 | 11 | 118259162 | A | G | 0.106 | 0.9159 | -+ | -9.3 | 2.745 | 1 | 0.09758 |
| rs10892717 | 11 | 121155632 | G | C | -0.105 | 0.916 | -? | 0 | 0 | 0 | 1 |
| rs17119743 | 11 | 116344404 | G | T | -0.105 | 0.9162 | -? | 0 | 0 | 0 | 1 |
| rs56816645 | 11 | 119768855 | G | A | 0.105 | 0.9162 | +? | 0 | 0 | 0 | 1 |
| rs6589848 | 11 | 120724018 | T | C | 0.105 | 0.9163 | +? | 0 | 0 | 0 | 1 |
| rs11216667 | 11 | 117876198 | A | C | 0.105 | 0.9164 | +? | 0 | 0 | 0 | 1 |
| rs11218200 | 11 | 121074615 | C | T | -0.105 | 0.9166 | -? | 0 | 0 | 0 | 1 |
| rs4084221 | 11 | 120405843 | A | G | -0.105 | 0.9166 | -? | 0 | 0 | 0 | 1 |
| rs56061158 | 11 | 117087118 | C | G | 0.104 | 0.9168 | +? | 0 | 0 | 0 | 1 |
| rs61897870 | 11 | 119870618 | G | C | 0.104 | 0.9169 | +? | 0 | 0 | 0 | 1 |
| rs11216117 | 11 | 116596129 | A | C | -0.104 | 0.917 | +- | -68.8 | 1.777 | 1 | 0.1825 |
| rs592979 | 11 | 119938682 | G | A | -0.104 | 0.917 | -? | 0 | 0 | 0 | 1 |
| rs671351 | 11 | 117941847 | A | C | -0.104 | 0.917 | -? | 0 | 0 | 0 | 1 |
| rs112563022 | 11 | 118262135 | T | C | 0.104 | 0.9171 | -+ | -9.6 | 2.736 | 1 | 0.09811 |
| rs73576276 | 11 | 120117335 | T | C | -0.104 | 0.9171 | -? | 0 | 0 | 0 | 1 |
| rs11216547 | 11 | 117666413 | T | C | 0.104 | 0.9172 | ++ | 0 | 0.011 | 1 | 0.9151 |
| rs56340594 | 11 | 117894618 | T | C | 0.104 | 0.9172 | +? | 0 | 0 | 0 | 1 |
| rs11216824 | 11 | 118096818 | A | G | -0.104 | 0.9175 | -+ | 0 | 0.083 | 1 | 0.7737 |
| rs596440 | 11 | 117548726 | C | T | -0.103 | 0.9177 | -? | 0 | 0 | 0 | 1 |
| rs61905463 | 11 | 116997964 | A | G | -0.103 | 0.9177 | -? | 0 | 0 | 0 | 1 |
| rs637454 | 11 | 117548858 | G | A | -0.103 | 0.9177 | -? | 0 | 0 | 0 | 1 |
| rs11603713 | 11 | 120576984 | G | A | 0.103 | 0.9179 | +? | 0 | 0 | 0 | 1 |
| rs2368961 | 11 | 119974286 | G | A | 0.103 | 0.918 | +? | 0 | 0 | 0 | 1 |
| rs76904522 | 11 | 117014863 | A | G | 0.103 | 0.918 | +? | 0 | 0 | 0 | 1 |
| rs11217962 | 11 | 120545346 | T | G | 0.103 | 0.9181 | +? | 0 | 0 | 0 | 1 |
| rs11216829 | 11 | 118112911 | A | G | -0.103 | 0.9182 | -+ | 0 | 0.192 | 1 | 0.6611 |
| rs76233351 | 11 | 119839224 | A | G | -0.103 | 0.9182 | -? | 0 | 0 | 0 | 1 |
| rs4938406 | 11 | 117427923 | A | G | -0.102 | 0.9184 | -? | 0 | 0 | 0 | 1 |
| rs631191 | 11 | 117916286 | T | C | -0.102 | 0.9185 | +- | 0 | 0.537 | 1 | 0.4635 |
| rs2084897 | 11 | 120026600 | A | G | 0.102 | 0.9189 | +? | 0 | 0 | 0 | 1 |
| rs2084899 | 11 | 120026810 | C | A | 0.102 | 0.9189 | +? | 0 | 0 | 0 | 1 |
| rs80044067 | 11 | 117880723 | A | G | -0.102 | 0.919 | -? | 0 | 0 | 0 | 1 |
| rs2845651 | 11 | 119430659 | G | A | 0.102 | 0.9191 | +? | 0 | 0 | 0 | 1 |
| rs77160958 | 11 | 119389987 | G | A | -0.102 | 0.9191 | -? | 0 | 0 | 0 | 1 |
| rs7935000 | 11 | 120542632 | A | G | 0.101 | 0.9192 | +? | 0 | 0 | 0 | 1 |
| rs74236045 | 11 | 121077362 | T | C | -0.101 | 0.9193 | -? | 0 | 0 | 0 | 1 |
| rs74942119 | 11 | 117998243 | T | C | 0.101 | 0.9193 | -+ | 0 | 0.122 | 1 | 0.727 |
| rs79660582 | 11 | 121073721 | G | T | -0.101 | 0.9193 | -? | 0 | 0 | 0 | 1 |

| | | | | | | | | | | | |
|---|---|---|---|---|---|---|---|---|---|---|---|
| rs79833470 | 11 | 121075750 | C | T | -0.101 | 0.9193 | -? | 0 | 0 | 0 | 1 |
| rs58789559 | 11 | 117568636 | A | G | -0.101 | 0.9195 | -? | 0 | 0 | 0 | 1 |
| rs111327557 | 11 | 116817784 | T | C | 0.101 | 0.9196 | +? | 0 | 0 | 0 | 1 |
| rs1317813 | 11 | 120484872 | T | C | 0.101 | 0.9196 | +? | 0 | 0 | 0 | 1 |
| rs10431079 | 11 | 116440678 | G | A | -0.101 | 0.9197 | -+ | 0 | 0.277 | 1 | 0.599 |
| rs7116650 | 11 | 120041917 | T | C | 0.101 | 0.9199 | +? | 0 | 0 | 0 | 1 |
| rs10502228 | 11 | 118113583 | T | C | -0.1 | 0.9201 | -+ | 0 | 0.196 | 1 | 0.6579 |
| rs1467051 | 11 | 119568301 | T | C | -0.1 | 0.9202 | -? | 0 | 0 | 0 | 1 |
| rs12802995 | 11 | 117527473 | A | C | -0.1 | 0.9205 | -? | 0 | 0 | 0 | 1 |
| rs2510450 | 11 | 117229902 | T | C | 0.1 | 0.9205 | +? | 0 | 0 | 0 | 1 |
| rs486366 | 11 | 117228709 | G | A | 0.1 | 0.9205 | +? | 0 | 0 | 0 | 1 |
| rs512730 | 11 | 117231057 | A | T | 0.1 | 0.9205 | +? | 0 | 0 | 0 | 1 |
| rs534296 | 11 | 117228295 | G | A | 0.1 | 0.9205 | +? | 0 | 0 | 0 | 1 |
| rs535366 | 11 | 117226616 | C | T | 0.1 | 0.9205 | +? | 0 | 0 | 0 | 1 |
| rs553303 | 11 | 117224818 | T | C | 0.1 | 0.9205 | +? | 0 | 0 | 0 | 1 |
| rs553506 | 11 | 117224887 | T | C | 0.1 | 0.9205 | +? | 0 | 0 | 0 | 1 |
| rs558854 | 11 | 117228664 | A | G | 0.1 | 0.9205 | +? | 0 | 0 | 0 | 1 |
| rs655389 | 11 | 117224878 | A | G | 0.1 | 0.9205 | +? | 0 | 0 | 0 | 1 |
| rs657666 | 11 | 117224360 | A | G | 0.1 | 0.9205 | +? | 0 | 0 | 0 | 1 |
| rs73000748 | 11 | 119937061 | T | C | 0.1 | 0.9206 | +? | 0 | 0 | 0 | 1 |
| rs10160584 | 11 | 117600136 | G | A | 0.099 | 0.9208 | +? | 0 | 0 | 0 | 1 |
| rs11828302 | 11 | 117574975 | T | A | 0.099 | 0.9208 | +? | 0 | 0 | 0 | 1 |
| rs7342287 | 11 | 118050404 | A | G | 0.099 | 0.9208 | +- | 0 | 0.405 | 1 | 0.5245 |
| rs17119878 | 11 | 116531479 | A | T | 0.099 | 0.921 | +? | 0 | 0 | 0 | 1 |
| rs10790234 | 11 | 117977678 | A | G | -0.099 | 0.9212 | -? | 0 | 0 | 0 | 1 |
| rs1080619 | 11 | 117965105 | G | A | -0.099 | 0.9212 | +- | 0 | 0.041 | 1 | 0.8402 |
| rs1080620 | 11 | 117965082 | A | G | -0.099 | 0.9212 | +- | 0 | 0.041 | 1 | 0.8402 |
| rs534421 | 11 | 116521187 | G | A | -0.099 | 0.9212 | -? | 0 | 0 | 0 | 1 |
| rs71478302 | 11 | 116451880 | G | C | -0.099 | 0.9212 | -? | 0 | 0 | 0 | 1 |
| rs1378582752 | 11 | 116789102 | C | T | 0.099 | 0.9213 | +? | 0 | 0 | 0 | 1 |
| rs4936357 | 11 | 116440163 | T | C | 0.099 | 0.9213 | -+ | 0 | 0.208 | 1 | 0.6481 |
| rs78270838 | 11 | 118072648 | G | A | 0.099 | 0.9214 | +- | 26.7 | 4.091 | 1 | 0.04311 |
| rs7950501 | 11 | 116963965 | T | C | 0.099 | 0.9214 | +? | 0 | 0 | 0 | 1 |
| rs56232713 | 11 | 120960737 | G | C | -0.099 | 0.9215 | -? | 0 | 0 | 0 | 1 |
| rs1995822 | 11 | 119446574 | A | G | -0.098 | 0.9217 | -? | 0 | 0 | 0 | 1 |
| rs10892454 | 11 | 119642827 | G | T | -0.098 | 0.9218 | -? | 0 | 0 | 0 | 1 |
| rs10892253 | 11 | 118559426 | A | C | -0.098 | 0.9221 | -+ | 0 | 0.083 | 1 | 0.7735 |
| rs74236047 | 11 | 121077823 | T | C | -0.097 | 0.9224 | -? | 0 | 0 | 0 | 1 |
| rs611960 | 11 | 118034228 | G | T | 0.097 | 0.9225 | -+ | 0 | 0.482 | 1 | 0.4875 |
| rs4936595 | 11 | 121099539 | A | C | -0.097 | 0.9226 | -+ | 0 | 0.051 | 1 | 0.8214 |
| rs3863305 | 11 | 116853124 | C | T | -0.097 | 0.9227 | -? | 0 | 0 | 0 | 1 |
| rs12419194 | 11 | 120877650 | A | G | 0.097 | 0.9229 | +? | 0 | 0 | 0 | 1 |
| rs689243 | 11 | 116722693 | G | C | -0.097 | 0.9229 | -? | 0 | 0 | 0 | 1 |
| rs11216891 | 11 | 118422234 | T | A | 0.097 | 0.923 | ++ | 0 | 0.002 | 1 | 0.9627 |
| rs11601511 | 11 | 117187551 | C | G | -0.096 | 0.9233 | -? | 0 | 0 | 0 | 1 |
| rs4996911 | 11 | 117973439 | A | G | -0.096 | 0.9233 | +- | 0 | 0.037 | 1 | 0.8469 |
| rs1261967419 | 11 | 116440036 | A | T | -0.096 | 0.9235 | -+ | 0 | 0.274 | 1 | 0.6009 |
| rs7943469 | 11 | 116439792 | G | T | -0.096 | 0.9235 | -+ | 0 | 0.274 | 1 | 0.6009 |
| rs10892596 | 11 | 120541873 | T | C | 0.096 | 0.9236 | +? | 0 | 0 | 0 | 1 |
| rs7107152 | 11 | 117056080 | A | G | -0.095 | 0.924 | -? | 0 | 0 | 0 | 1 |
| rs61898270 | 11 | 120075768 | T | A | -0.095 | 0.9241 | -? | 0 | 0 | 0 | 1 |
| rs11216815 | 11 | 118072232 | A | G | 0.095 | 0.9242 | -+ | 0 | 0.541 | 1 | 0.4621 |
| rs11217970 | 11 | 120551626 | A | G | -0.095 | 0.9243 | -? | 0 | 0 | 0 | 1 |
| rs7106296 | 11 | 116322236 | T | C | 0.095 | 0.9243 | +? | 0 | 0 | 0 | 1 |
| rs4938692 | 11 | 119494489 | T | C | -0.095 | 0.9247 | -? | 0 | 0 | 0 | 1 |
| rs74328581 | 11 | 116918046 | A | C | 0.095 | 0.9247 | +? | 0 | 0 | 0 | 1 |

| | | | | | | | | | | | |
|---|---|---|---|---|---|---|---|---|---|---|---|
| rs608093 | 11 | 117453868 | A | C | -0.094 | 0.9248 | -? | 0 | 0 | 0 | 1 |
| rs11217955 | 11 | 120540060 | T | C | 0.094 | 0.9249 | +? | 0 | 0 | 0 | 1 |
| rs12221818 | 11 | 117228519 | G | A | -0.094 | 0.9249 | -? | 0 | 0 | 0 | 1 |
| rs12419365 | 11 | 118112332 | A | C | -0.094 | 0.9249 | -- | 0 | 0.001 | 1 | 0.9793 |
| rs6589649 | 11 | 118048836 | C | T | 0.094 | 0.925 | +- | 0 | 0.401 | 1 | 0.5265 |
| rs9651688 | 11 | 118107495 | T | C | -0.094 | 0.925 | -+ | 0 | 0.114 | 1 | 0.7356 |
| rs34071565 | 11 | 119712522 | T | A | -0.094 | 0.9251 | -? | 0 | 0 | 0 | 1 |
| rs10431039 | 11 | 116440575 | T | C | 0.094 | 0.9252 | -+ | 0 | 0.211 | 1 | 0.6461 |
| rs2014536 | 11 | 117691145 | C | T | -0.094 | 0.9252 | -? | 0 | 0 | 0 | 1 |
| rs470373 | 11 | 120026271 | T | C | -0.094 | 0.9252 | -? | 0 | 0 | 0 | 1 |
| rs490592 | 11 | 116298367 | A | C | -0.094 | 0.9252 | -? | 0 | 0 | 0 | 1 |
| rs654479 | 11 | 117225134 | A | G | 0.094 | 0.9252 | +? | 0 | 0 | 0 | 1 |
| rs73019237 | 11 | 116250393 | T | C | -0.094 | 0.9254 | -? | 0 | 0 | 0 | 1 |
| rs61907596 | 11 | 116807226 | T | C | -0.093 | 0.9256 | -? | 0 | 0 | 0 | 1 |
| rs3132777 | 11 | 120603857 | A | G | -0.093 | 0.9259 | -? | 0 | 0 | 0 | 1 |
| rs57434987 | 11 | 117401292 | A | C | 0.093 | 0.926 | +? | 0 | 0 | 0 | 1 |
| rs11216028 | 11 | 116464992 | T | G | 0.093 | 0.9261 | +? | 0 | 0 | 0 | 1 |
| rs12787692 | 11 | 119885995 | A | G | -0.093 | 0.9261 | -? | 0 | 0 | 0 | 1 |
| rs642765 | 11 | 118030310 | C | G | -0.093 | 0.9261 | -+ | 0 | 0.113 | 1 | 0.7372 |
| rs2277292 | 11 | 118425419 | T | G | 0.093 | 0.9262 | +- | 0 | 0.005 | 1 | 0.9412 |
| rs2277293 | 11 | 118422868 | T | C | 0.093 | 0.9262 | +- | 0 | 0.024 | 1 | 0.8759 |
| rs11217310 | 11 | 119419568 | C | G | 0.093 | 0.9263 | +? | 0 | 0 | 0 | 1 |
| rs1295453503 | 11 | 116272249 | T | C | -0.093 | 0.9263 | -? | 0 | 0 | 0 | 1 |
| rs599411 | 11 | 117422262 | G | T | -0.093 | 0.9263 | -? | 0 | 0 | 0 | 1 |
| rs76050019 | 11 | 118051857 | C | T | 0.092 | 0.9263 | +- | 0 | 0.374 | 1 | 0.541 |
| rs77786799 | 11 | 120095847 | T | C | -0.093 | 0.9263 | -? | 0 | 0 | 0 | 1 |
| rs7927676 | 11 | 118051323 | G | C | 0.092 | 0.9263 | +- | 0 | 0.374 | 1 | 0.541 |
| rs3863304 | 11 | 116853123 | G | A | -0.092 | 0.9264 | -? | 0 | 0 | 0 | 1 |
| rs4936369 | 11 | 117141469 | C | T | 0.092 | 0.9264 | +? | 0 | 0 | 0 | 1 |
| rs1941637 | 11 | 117999443 | A | C | -0.092 | 0.9268 | -? | 0 | 0 | 0 | 1 |
| rs61905072 | 11 | 116581259 | T | C | -0.091 | 0.9271 | -? | 0 | 0 | 0 | 1 |
| rs2845712 | 11 | 117774291 | A | C | 0.091 | 0.9273 | +? | 0 | 0 | 0 | 1 |
| rs608444 | 11 | 118986630 | G | A | 0.091 | 0.9275 | +? | 0 | 0 | 0 | 1 |
| rs680424 | 11 | 117441621 | C | G | 0.091 | 0.9275 | +? | 0 | 0 | 0 | 1 |
| rs11218049 | 11 | 120770075 | G | A | 0.091 | 0.9277 | +? | 0 | 0 | 0 | 1 |
| rs11822370 | 11 | 117961734 | G | A | 0.091 | 0.9277 | +- | 0 | 0.127 | 1 | 0.7218 |
| rs1715422 | 11 | 118104577 | C | T | -0.091 | 0.9277 | -+ | 0 | 0.133 | 1 | 0.7156 |
| rs1793159 | 11 | 118104569 | A | C | -0.091 | 0.9277 | -+ | 0 | 0.133 | 1 | 0.7156 |
| rs150954345 | 11 | 119985011 | G | A | -0.091 | 0.9278 | -? | 0 | 0 | 0 | 1 |
| rs4338560 | 11 | 118143368 | T | C | 0.091 | 0.9278 | -+ | -33 | 2.255 | 1 | 0.1332 |
| rs12802402 | 11 | 120023505 | T | C | 0.09 | 0.9279 | +? | 0 | 0 | 0 | 1 |
| rs525110 | 11 | 117411392 | G | A | -0.09 | 0.928 | -? | 0 | 0 | 0 | 1 |
| rs679223 | 11 | 117412621 | A | G | -0.09 | 0.928 | -? | 0 | 0 | 0 | 1 |
| rs77346825 | 11 | 117597518 | A | G | 0.09 | 0.928 | +? | 0 | 0 | 0 | 1 |
| rs78033613 | 11 | 119371772 | A | G | -0.09 | 0.928 | -? | 0 | 0 | 0 | 1 |
| rs7125517 | 11 | 117317124 | C | A | 0.09 | 0.9282 | +? | 0 | 0 | 0 | 1 |
| rs11216828 | 11 | 118111676 | A | T | -0.09 | 0.9284 | -+ | 0 | 0.168 | 1 | 0.6821 |
| rs2186780 | 11 | 118605451 | T | C | -0.09 | 0.9284 | -- | 0 | 0 | 1 | 0.9904 |
| rs10892548 | 11 | 120074851 | A | G | 0.09 | 0.9285 | +? | 0 | 0 | 0 | 1 |
| rs11217968 | 11 | 120550229 | G | A | -0.09 | 0.9285 | -? | 0 | 0 | 0 | 1 |
| rs485778 | 11 | 116312958 | A | G | -0.09 | 0.9285 | -? | 0 | 0 | 0 | 1 |
| rs506836 | 11 | 117412453 | T | C | -0.09 | 0.9286 | -? | 0 | 0 | 0 | 1 |
| rs67877251 | 11 | 117680407 | G | A | -0.09 | 0.9286 | -? | 0 | 0 | 0 | 1 |
| rs4938693 | 11 | 119494587 | T | C | -0.089 | 0.9288 | -? | 0 | 0 | 0 | 1 |
| rs117190595 | 11 | 120887214 | T | A | -0.089 | 0.929 | -? | 0 | 0 | 0 | 1 |
| rs35251396 | 11 | 117800033 | A | G | -0.089 | 0.929 | -? | 0 | 0 | 0 | 1 |

| | | | | | | | | | | | |
|---|---|---|---|---|---|---|---|---|---|---|---|
| rs12798473 | 11 | 119494885 | T | C | -0.089 | 0.9293 | -? | 0 | 0 | 0 | 1 |
| rs79163278 | 11 | 119781157 | C | T | -0.089 | 0.9294 | -? | 0 | 0 | 0 | 1 |
| rs1171461 | 11 | 118522179 | A | G | -0.089 | 0.9295 | -+ | 0 | 0.775 | 1 | 0.3787 |
| rs636530 | 11 | 117421858 | G | A | -0.088 | 0.9295 | -? | 0 | 0 | 0 | 1 |
| rs11217950 | 11 | 120537349 | A | T | 0.088 | 0.9296 | +? | 0 | 0 | 0 | 1 |
| rs112667243 | 11 | 120122628 | T | C | -0.088 | 0.9297 | -? | 0 | 0 | 0 | 1 |
| rs504339 | 11 | 116316963 | A | G | 0.088 | 0.9298 | +? | 0 | 0 | 0 | 1 |
| rs1046727798 | 11 | 119822450 | G | A | -0.088 | 0.93 | -? | 0 | 0 | 0 | 1 |
| rs10892060 | 11 | 116881662 | C | A | 0.088 | 0.9302 | +? | 0 | 0 | 0 | 1 |
| rs73018459 | 11 | 117689383 | T | C | -0.088 | 0.9302 | -? | 0 | 0 | 0 | 1 |
| rs11217315 | 11 | 119425163 | A | G | 0.087 | 0.9304 | +? | 0 | 0 | 0 | 1 |
| rs35120633 | 11 | 116655600 | A | G | 0.087 | 0.9305 | +- | -90.1 | 1.578 | 1 | 0.2091 |
| rs4996910 | 11 | 117973454 | A | G | -0.087 | 0.9305 | +- | 0 | 0.034 | 1 | 0.853 |
| rs55857590 | 11 | 118425063 | G | A | 0.087 | 0.9307 | +- | 0 | 0.003 | 1 | 0.9549 |
| rs11216103 | 11 | 116574488 | A | G | 0.086 | 0.9311 | +? | 0 | 0 | 0 | 1 |
| rs12419025 | 11 | 120046336 | T | A | 0.086 | 0.9311 | +? | 0 | 0 | 0 | 1 |
| rs10892076 | 11 | 116988936 | G | A | -0.086 | 0.9313 | -? | 0 | 0 | 0 | 1 |
| rs17490842 | 11 | 116359615 | T | C | 0.086 | 0.9313 | +? | 0 | 0 | 0 | 1 |
| rs597686 | 11 | 117361861 | T | C | -0.086 | 0.9313 | -? | 0 | 0 | 0 | 1 |
| rs10892404 | 11 | 119435011 | C | G | -0.086 | 0.9315 | -? | 0 | 0 | 0 | 1 |
| rs10892405 | 11 | 119435047 | G | A | -0.086 | 0.9315 | -? | 0 | 0 | 0 | 1 |
| rs12785238 | 11 | 119358692 | C | T | 0.086 | 0.9315 | +? | 0 | 0 | 0 | 1 |
| rs17552226 | 11 | 118927907 | G | C | 0.086 | 0.9316 | +? | 0 | 0 | 0 | 1 |
| rs10892465 | 11 | 119659753 | C | T | -0.086 | 0.9317 | -? | 0 | 0 | 0 | 1 |
| rs11217499 | 11 | 119651089 | T | C | -0.086 | 0.9317 | -? | 0 | 0 | 0 | 1 |
| rs10750153 | 11 | 119432823 | T | C | -0.085 | 0.932 | -? | 0 | 0 | 0 | 1 |
| rs1715421 | 11 | 118105233 | C | T | -0.085 | 0.9321 | -+ | 0 | 0.13 | 1 | 0.7179 |
| rs4938687 | 11 | 119451940 | A | G | -0.085 | 0.9322 | -? | 0 | 0 | 0 | 1 |
| rs78785412 | 11 | 119844067 | T | G | -0.085 | 0.9322 | -? | 0 | 0 | 0 | 1 |
| rs494048 | 11 | 118961231 | T | C | 0.085 | 0.9323 | +? | 0 | 0 | 0 | 1 |
| rs11218096 | 11 | 120865621 | T | G | -0.085 | 0.9324 | -? | 0 | 0 | 0 | 1 |
| rs12791621 | 11 | 120025443 | G | A | 0.085 | 0.9324 | +? | 0 | 0 | 0 | 1 |
| rs544274 | 11 | 117341030 | G | A | 0.085 | 0.9324 | +? | 0 | 0 | 0 | 1 |
| rs640323 | 11 | 118010890 | C | G | 0.085 | 0.9324 | -+ | 0 | 0.069 | 1 | 0.7932 |
| rs11607627 | 11 | 116432670 | A | T | -0.085 | 0.9326 | -? | 0 | 0 | 0 | 1 |
| rs79820203 | 11 | 118051010 | C | T | 0.085 | 0.9326 | +- | 0 | 0.368 | 1 | 0.5439 |
| rs10892218 | 11 | 117982289 | T | G | 0.084 | 0.9327 | +? | 0 | 0 | 0 | 1 |
| rs1263055 | 11 | 116577043 | A | G | 0.084 | 0.9327 | +? | 0 | 0 | 0 | 1 |
| rs494965 | 11 | 120053650 | T | C | -0.084 | 0.9327 | -? | 0 | 0 | 0 | 1 |
| rs6589797 | 11 | 119971337 | C | T | -0.084 | 0.9329 | -? | 0 | 0 | 0 | 1 |
| rs11216774 | 11 | 117993836 | A | G | -0.084 | 0.933 | -? | 0 | 0 | 0 | 1 |
| rs6589557 | 11 | 116537761 | T | C | 0.084 | 0.933 | +? | 0 | 0 | 0 | 1 |
| rs6589558 | 11 | 116537942 | A | G | 0.084 | 0.933 | +? | 0 | 0 | 0 | 1 |
| rs7110327 | 11 | 117992910 | A | G | -0.084 | 0.933 | -? | 0 | 0 | 0 | 1 |
| rs11216048 | 11 | 116495059 | G | A | 0.084 | 0.9331 | +? | 0 | 0 | 0 | 1 |
| rs11216049 | 11 | 116495175 | C | T | 0.084 | 0.9331 | +? | 0 | 0 | 0 | 1 |
| rs11216050 | 11 | 116495314 | T | C | 0.084 | 0.9331 | +? | 0 | 0 | 0 | 1 |
| rs56403966 | 11 | 118425051 | A | G | 0.084 | 0.9331 | +- | 0 | 0.003 | 1 | 0.9547 |
| rs112895963 | 11 | 116937159 | T | A | 0.084 | 0.9332 | +? | 0 | 0 | 0 | 1 |
| rs35409990 | 11 | 118109196 | A | G | -0.084 | 0.9333 | -+ | 0 | 0.121 | 1 | 0.7284 |
| rs9651667 | 11 | 118108679 | G | C | -0.084 | 0.9333 | -+ | 0 | 0.121 | 1 | 0.7284 |
| rs35048090 | 11 | 117531395 | T | C | -0.084 | 0.9334 | -? | 0 | 0 | 0 | 1 |
| rs36067343 | 11 | 117531569 | G | T | -0.084 | 0.9334 | -? | 0 | 0 | 0 | 1 |
| rs11218216 | 11 | 121103527 | C | T | -0.083 | 0.9335 | -+ | 0 | 0.044 | 1 | 0.8337 |
| rs479601 | 11 | 117440171 | G | A | 0.083 | 0.9335 | +? | 0 | 0 | 0 | 1 |
| rs11217308 | 11 | 119410158 | T | C | 0.083 | 0.9336 | +? | 0 | 0 | 0 | 1 |

| | | | | | | | | | | | |
|---|---|---|---|---|---|---|---|---|---|---|---|
| rs2276123 | 11 | 117965595 | T | C | -0.082 | 0.9343 | +- | 0 | 0.044 | 1 | 0.8335 |
| rs12272004 | 11 | 116603724 | A | C | -0.082 | 0.9346 | +- | -83.7 | 1.633 | 1 | 0.2013 |
| rs56264624 | 11 | 116337814 | A | G | -0.082 | 0.9346 | -? | 0 | 0 | 0 | 1 |
| rs541407 | 11 | 116313753 | C | T | 0.082 | 0.9347 | +? | 0 | 0 | 0 | 1 |
| rs10736493 | 11 | 118518021 | G | T | -0.082 | 0.9348 | -+ | 0 | 0.172 | 1 | 0.6787 |
| rs10082646 | 11 | 120541660 | C | T | 0.082 | 0.9349 | +? | 0 | 0 | 0 | 1 |
| rs11217522 | 11 | 119690671 | T | G | -0.081 | 0.9352 | -+ | 0 | 0.148 | 1 | 0.7 |
| rs1793167 | 11 | 118102058 | T | C | -0.081 | 0.9352 | -+ | 0 | 0.483 | 1 | 0.487 |
| rs61906078 | 11 | 116529313 | A | G | 0.081 | 0.9352 | +? | 0 | 0 | 0 | 1 |
| rs73014327 | 11 | 117196583 | C | T | -0.081 | 0.9352 | -? | 0 | 0 | 0 | 1 |
| rs4936611 | 11 | 121158509 | T | C | -0.081 | 0.9353 | -? | 0 | 0 | 0 | 1 |
| rs527290 | 11 | 118515616 | T | C | -0.081 | 0.9353 | -+ | 0 | 0.524 | 1 | 0.4693 |
| rs1110704 | 11 | 119974641 | A | G | 0.081 | 0.9354 | +? | 0 | 0 | 0 | 1 |
| rs11216242 | 11 | 116901549 | G | A | -0.081 | 0.9356 | -? | 0 | 0 | 0 | 1 |
| rs10892058 | 11 | 116876621 | C | T | 0.081 | 0.9358 | +? | 0 | 0 | 0 | 1 |
| rs12365332 | 11 | 117998038 | T | C | -0.081 | 0.9358 | -? | 0 | 0 | 0 | 1 |
| rs537323 | 11 | 117226849 | T | C | 0.081 | 0.9358 | +? | 0 | 0 | 0 | 1 |
| rs626151 | 11 | 120034170 | T | G | 0.081 | 0.9358 | +? | 0 | 0 | 0 | 1 |
| rs10790257 | 11 | 118561858 | C | G | -0.08 | 0.9359 | -+ | 0 | 0.099 | 1 | 0.7528 |
| rs2727790 | 11 | 116688383 | A | G | 0.08 | 0.9359 | +- | 4.9 | 3.155 | 1 | 0.07571 |
| rs11216853 | 11 | 118200201 | T | A | 0.08 | 0.936 | -+ | 0 | 0.088 | 1 | 0.7663 |
| rs567053 | 11 | 117430931 | A | G | -0.08 | 0.9361 | -? | 0 | 0 | 0 | 1 |
| rs75893552 | 11 | 117998236 | A | G | -0.08 | 0.9362 | -? | 0 | 0 | 0 | 1 |
| rs7118777 | 11 | 119427857 | C | A | -0.08 | 0.9363 | -? | 0 | 0 | 0 | 1 |
| rs528720 | 11 | 117227686 | A | G | 0.08 | 0.9364 | +? | 0 | 0 | 0 | 1 |
| rs11217675 | 11 | 119949999 | C | T | -0.079 | 0.9367 | -? | 0 | 0 | 0 | 1 |
| rs61902671 | 11 | 120544352 | A | C | -0.079 | 0.9367 | -? | 0 | 0 | 0 | 1 |
| rs111942423 | 11 | 118429764 | G | T | 0.079 | 0.937 | +- | 0 | 0.021 | 1 | 0.8859 |
| rs75839534 | 11 | 121089547 | A | G | -0.079 | 0.937 | -? | 0 | 0 | 0 | 1 |
| rs501021 | 11 | 120000872 | T | C | -0.079 | 0.9371 | -? | 0 | 0 | 0 | 1 |
| rs77127262 | 11 | 119843716 | T | C | -0.079 | 0.9373 | -? | 0 | 0 | 0 | 1 |
| rs1190080879 | 11 | 119493873 | G | T | -0.078 | 0.9375 | -? | 0 | 0 | 0 | 1 |
| rs2622934 | 11 | 117014862 | G | T | 0.078 | 0.9377 | +? | 0 | 0 | 0 | 1 |
| rs561498 | 11 | 120002927 | C | T | 0.078 | 0.9378 | +? | 0 | 0 | 0 | 1 |
| rs79162717 | 11 | 119765750 | G | A | 0.078 | 0.9378 | +? | 0 | 0 | 0 | 1 |
| rs4938695 | 11 | 119496085 | T | C | -0.078 | 0.9379 | -? | 0 | 0 | 0 | 1 |
| rs10892593 | 11 | 120539410 | T | C | 0.078 | 0.938 | +? | 0 | 0 | 0 | 1 |
| rs10790310 | 11 | 119432935 | T | C | -0.078 | 0.9381 | -? | 0 | 0 | 0 | 1 |
| rs117114471 | 11 | 119858685 | A | G | -0.077 | 0.9383 | -? | 0 | 0 | 0 | 1 |
| rs1560409 | 11 | 121160060 | C | G | -0.077 | 0.9383 | -? | 0 | 0 | 0 | 1 |
| rs17124767 | 11 | 120963641 | C | T | 0.077 | 0.9383 | +? | 0 | 0 | 0 | 1 |
| rs4938769 | 11 | 119948566 | T | C | -0.077 | 0.9383 | -? | 0 | 0 | 0 | 1 |
| rs117080901 | 11 | 116422645 | A | G | 0.077 | 0.9384 | +? | 0 | 0 | 0 | 1 |
| rs491300 | 11 | 117442684 | G | T | 0.077 | 0.9385 | +? | 0 | 0 | 0 | 1 |
| rs568466 | 11 | 116331957 | C | T | 0.077 | 0.9385 | +? | 0 | 0 | 0 | 1 |
| rs497744 | 11 | 117368158 | T | G | -0.077 | 0.9388 | -? | 0 | 0 | 0 | 1 |
| rs1052121 | 11 | 119178050 | G | C | -0.077 | 0.9389 | -+ | 0 | 0.839 | 1 | 0.3595 |
| rs11216227 | 11 | 116878597 | C | G | 0.077 | 0.939 | +? | 0 | 0 | 0 | 1 |
| rs12281346 | 11 | 117296892 | C | G | -0.077 | 0.939 | -? | 0 | 0 | 0 | 1 |
| rs656308 | 11 | 118031206 | T | G | -0.077 | 0.939 | -+ | 0 | 0.176 | 1 | 0.6753 |
| rs12799836 | 11 | 120871452 | G | A | -0.076 | 0.9391 | -? | 0 | 0 | 0 | 1 |
| rs7127881 | 11 | 116880798 | G | A | 0.076 | 0.9391 | +? | 0 | 0 | 0 | 1 |
| rs9326253 | 11 | 119644451 | C | T | 0.076 | 0.9391 | +? | 0 | 0 | 0 | 1 |
| rs118069464 | 11 | 117313626 | C | G | -0.076 | 0.9393 | -? | 0 | 0 | 0 | 1 |
| rs55703909 | 11 | 116984866 | A | G | -0.076 | 0.9394 | -? | 0 | 0 | 0 | 1 |
| rs7121313 | 11 | 117729680 | A | G | -0.076 | 0.9395 | -? | 0 | 0 | 0 | 1 |

| | | | | | | | | | | | |
|---|---|---|---|---|---|---|---|---|---|---|---|
| rs11217954 | 11 | 120538307 | C | G | -0.076 | 0.9396 | -? | 0 | 0 | 0 | 1 |
| rs2276126 | 11 | 117965783 | G | A | -0.076 | 0.9397 | +- | 0 | 0.046 | 1 | 0.8304 |
| rs4514434 | 11 | 119492274 | C | T | -0.076 | 0.9398 | -? | 0 | 0 | 0 | 1 |
| rs56075010 | 11 | 116484610 | A | G | -0.076 | 0.9398 | -? | 0 | 0 | 0 | 1 |
| rs111993696 | 11 | 118282448 | A | G | 0.075 | 0.94 | -+ | -24.4 | 2.412 | 1 | 0.1204 |
| rs79579780 | 11 | 119405392 | G | C | -0.075 | 0.9401 | -? | 0 | 0 | 0 | 1 |
| rs2276424 | 11 | 118209960 | A | C | 0.075 | 0.9404 | -+ | 0 | 0.115 | 1 | 0.7341 |
| rs4575311 | 11 | 116904357 | G | A | -0.075 | 0.9404 | -? | 0 | 0 | 0 | 1 |
| rs1248452864 | 11 | 116872263 | G | A | 0.075 | 0.9406 | +? | 0 | 0 | 0 | 1 |
| rs17120233 | 11 | 116871223 | G | A | 0.075 | 0.9406 | +? | 0 | 0 | 0 | 1 |
| rs77250716 | 11 | 116877353 | C | T | 0.075 | 0.9406 | +? | 0 | 0 | 0 | 1 |
| rs522885 | 11 | 117282800 | G | T | 0.074 | 0.9407 | +? | 0 | 0 | 0 | 1 |
| rs11602880 | 11 | 116322769 | A | G | 0.074 | 0.9408 | +? | 0 | 0 | 0 | 1 |
| rs61601696 | 11 | 120548768 | C | T | -0.074 | 0.9408 | -? | 0 | 0 | 0 | 1 |
| rs61900463 | 11 | 119974582 | G | C | 0.074 | 0.9409 | +? | 0 | 0 | 0 | 1 |
| rs4938733 | 11 | 119732418 | C | G | -0.074 | 0.941 | -? | 0 | 0 | 0 | 1 |
| rs7131351 | 11 | 116853606 | G | A | -0.074 | 0.941 | -? | 0 | 0 | 0 | 1 |
| rs7932158 | 11 | 120550430 | C | T | -0.074 | 0.941 | -? | 0 | 0 | 0 | 1 |
| rs625573 | 11 | 119937400 | C | G | -0.074 | 0.9411 | -? | 0 | 0 | 0 | 1 |
| rs4938626 | 11 | 118974588 | T | G | -0.073 | 0.9415 | -? | 0 | 0 | 0 | 1 |
| rs7926104 | 11 | 116899782 | G | A | -0.073 | 0.9416 | -? | 0 | 0 | 0 | 1 |
| rs4936362 | 11 | 116952200 | T | C | 0.073 | 0.9417 | +? | 0 | 0 | 0 | 1 |
| rs10790238 | 11 | 117990826 | G | T | -0.073 | 0.9419 | -? | 0 | 0 | 0 | 1 |
| rs11216772 | 11 | 117992183 | A | T | -0.073 | 0.9419 | -? | 0 | 0 | 0 | 1 |
| rs12222838 | 11 | 117991381 | G | A | -0.073 | 0.9419 | -? | 0 | 0 | 0 | 1 |
| rs12361749 | 11 | 117991176 | T | A | -0.073 | 0.9419 | -? | 0 | 0 | 0 | 1 |
| rs12364660 | 11 | 117991183 | G | C | -0.073 | 0.9419 | -? | 0 | 0 | 0 | 1 |
| rs12365797 | 11 | 117993395 | A | G | -0.073 | 0.9419 | -? | 0 | 0 | 0 | 1 |
| rs12576605 | 11 | 117990003 | C | G | -0.073 | 0.9419 | -? | 0 | 0 | 0 | 1 |
| rs76043503 | 11 | 117992763 | G | A | -0.073 | 0.9419 | -? | 0 | 0 | 0 | 1 |
| rs61901429 | 11 | 120724136 | G | A | 0.073 | 0.942 | +? | 0 | 0 | 0 | 1 |
| rs7122134 | 11 | 119547005 | G | A | 0.073 | 0.942 | +? | 0 | 0 | 0 | 1 |
| rs666290 | 11 | 117476142 | T | C | -0.072 | 0.9423 | -? | 0 | 0 | 0 | 1 |
| rs7114223 | 11 | 117339276 | A | G | 0.072 | 0.9423 | +? | 0 | 0 | 0 | 1 |
| rs10891986 | 11 | 116393439 | A | T | 0.072 | 0.9424 | +? | 0 | 0 | 0 | 1 |
| rs10892594 | 11 | 120540410 | C | A | 0.072 | 0.9424 | +? | 0 | 0 | 0 | 1 |
| rs657882 | 11 | 117511951 | C | G | -0.072 | 0.9424 | -? | 0 | 0 | 0 | 1 |
| rs653008 | 11 | 118520480 | T | G | -0.072 | 0.9425 | -+ | 0 | 0.549 | 1 | 0.4587 |
| rs10892595 | 11 | 120540687 | T | C | 0.072 | 0.9427 | +? | 0 | 0 | 0 | 1 |
| rs548890 | 11 | 116312872 | A | G | 0.072 | 0.9427 | +? | 0 | 0 | 0 | 1 |
| rs11216546 | 11 | 117665021 | T | C | 0.072 | 0.9429 | -+ | 0 | 0.018 | 1 | 0.8939 |
| rs1480227044 | 11 | 116483289 | G | C | -0.072 | 0.9429 | -? | 0 | 0 | 0 | 1 |
| rs75084211 | 11 | 116482973 | A | G | -0.072 | 0.9429 | -? | 0 | 0 | 0 | 1 |
| rs79353638 | 11 | 116482683 | A | G | -0.072 | 0.9429 | -? | 0 | 0 | 0 | 1 |
| rs117013687 | 11 | 117150502 | T | C | -0.071 | 0.9434 | -? | 0 | 0 | 0 | 1 |
| rs141642186 | 11 | 119898644 | T | C | 0.071 | 0.9434 | +? | 0 | 0 | 0 | 1 |
| rs17492724 | 11 | 116480616 | C | A | -0.071 | 0.9434 | -? | 0 | 0 | 0 | 1 |
| rs56173877 | 11 | 116481208 | A | G | -0.071 | 0.9434 | -? | 0 | 0 | 0 | 1 |
| rs10892059 | 11 | 116881645 | G | A | 0.071 | 0.9435 | +? | 0 | 0 | 0 | 1 |
| rs61906117 | 11 | 116558916 | A | G | 0.071 | 0.9435 | +? | 0 | 0 | 0 | 1 |
| rs118118922 | 11 | 117935948 | T | C | 0.071 | 0.9437 | +? | 0 | 0 | 0 | 1 |
| rs9651684 | 11 | 117966994 | C | G | -0.07 | 0.9438 | +- | 0 | 0.043 | 1 | 0.835 |
| rs10892388 | 11 | 119417101 | A | G | 0.07 | 0.9439 | +? | 0 | 0 | 0 | 1 |
| rs7949789 | 11 | 120041110 | A | T | 0.07 | 0.9439 | +? | 0 | 0 | 0 | 1 |
| rs1605557 | 11 | 116342547 | G | A | 0.07 | 0.944 | +? | 0 | 0 | 0 | 1 |
| rs77610621 | 11 | 118089957 | A | G | 0.07 | 0.944 | +? | 0 | 0 | 0 | 1 |

| | | | | | | | | | | | |
|---|---|---|---|---|---|---|---|---|---|---|---|
| rs75907072 | 11 | 116233803 | A | T | 0.07 | 0.9441 | +- | -135.4 | 1.274 | 1 | 0.2589 |
| rs12273399 | 11 | 120060102 | T | C | -0.07 | 0.9442 | +- | 0 | 0.51 | 1 | 0.4752 |
| rs1353117537 | 11 | 119828505 | A | T | -0.07 | 0.9443 | -? | 0 | 0 | 0 | 1 |
| rs73613993 | 11 | 118296007 | C | T | 0.07 | 0.9443 | +- | 0 | 0.156 | 1 | 0.6931 |
| rs12276260 | 11 | 119950798 | C | G | 0.07 | 0.9444 | +? | 0 | 0 | 0 | 1 |
| rs12808972 | 11 | 120057972 | A | G | -0.07 | 0.9444 | -? | 0 | 0 | 0 | 1 |
| rs113979111 | 11 | 116983016 | A | G | -0.069 | 0.9446 | -? | 0 | 0 | 0 | 1 |
| rs11217951 | 11 | 120538183 | T | C | -0.069 | 0.9448 | -? | 0 | 0 | 0 | 1 |
| rs1158810726 | 11 | 117154091 | G | T | 0.069 | 0.9449 | +? | 0 | 0 | 0 | 1 |
| rs550767 | 11 | 116487699 | A | C | -0.069 | 0.945 | -? | 0 | 0 | 0 | 1 |
| rs74444640 | 11 | 116528544 | T | C | 0.069 | 0.945 | +? | 0 | 0 | 0 | 1 |
| rs11604327 | 11 | 120572302 | C | A | 0.069 | 0.9451 | +? | 0 | 0 | 0 | 1 |
| rs138177199 | 11 | 116371338 | G | C | 0.069 | 0.9452 | +? | 0 | 0 | 0 | 1 |
| rs35163144 | 11 | 119726313 | G | A | -0.069 | 0.9452 | -? | 0 | 0 | 0 | 1 |
| rs574778 | 11 | 116292263 | C | A | -0.069 | 0.9452 | -? | 0 | 0 | 0 | 1 |
| rs10790258 | 11 | 118561859 | T | G | -0.069 | 0.9453 | -+ | 0 | 0.095 | 1 | 0.758 |
| rs76831657 | 11 | 118051082 | G | A | 0.068 | 0.9454 | +- | 0 | 0.361 | 1 | 0.5479 |
| rs10892385 | 11 | 119412922 | A | C | -0.068 | 0.9455 | -? | 0 | 0 | 0 | 1 |
| rs3802863 | 11 | 118099186 | T | C | -0.068 | 0.9455 | -+ | 0 | 0.429 | 1 | 0.5126 |
| rs11216451 | 11 | 117424454 | G | A | -0.068 | 0.9456 | -? | 0 | 0 | 0 | 1 |
| rs11218215 | 11 | 121103497 | T | C | -0.068 | 0.9457 | -+ | 0 | 0.04 | 1 | 0.8406 |
| rs148205418 | 11 | 120071834 | G | C | -0.068 | 0.946 | -? | 0 | 0 | 0 | 1 |
| rs113784559 | 11 | 116979365 | G | C | -0.067 | 0.9465 | -? | 0 | 0 | 0 | 1 |
| rs947926 | 11 | 117539478 | C | A | 0.067 | 0.9466 | +? | 0 | 0 | 0 | 1 |
| rs112245679 | 11 | 118287622 | G | A | 0.067 | 0.9468 | -+ | -30.9 | 2.292 | 1 | 0.13 |
| rs61899017 | 11 | 119771321 | C | T | -0.067 | 0.9468 | -? | 0 | 0 | 0 | 1 |
| rs11217755 | 11 | 120053173 | A | G | 0.067 | 0.9469 | +? | 0 | 0 | 0 | 1 |
| rs76031074 | 11 | 119677419 | A | G | 0.066 | 0.9472 | +? | 0 | 0 | 0 | 1 |
| rs7946257 | 11 | 117053165 | A | G | -0.066 | 0.9473 | -? | 0 | 0 | 0 | 1 |
| rs10790167 | 11 | 116800943 | T | C | -0.066 | 0.9474 | -? | 0 | 0 | 0 | 1 |
| rs10892150 | 11 | 117524543 | C | T | -0.066 | 0.9475 | -? | 0 | 0 | 0 | 1 |
| rs504236 | 11 | 120000263 | T | C | -0.066 | 0.9475 | -? | 0 | 0 | 0 | 1 |
| rs645637 | 11 | 118505090 | A | G | -0.066 | 0.9477 | -+ | 0 | 0.474 | 1 | 0.4909 |
| rs7110837 | 11 | 118199295 | A | G | 0.066 | 0.9477 | -+ | 0 | 0.096 | 1 | 0.7568 |
| rs11602430 | 11 | 119951327 | G | C | 0.065 | 0.9478 | +? | 0 | 0 | 0 | 1 |
| rs576262 | 11 | 116764436 | C | G | 0.065 | 0.9478 | +- | 59.9 | 7.488 | 1 | 0.00621 |
| rs7129048 | 11 | 118211797 | C | T | -0.065 | 0.9478 | -+ | 11.2 | 3.376 | 1 | 0.06613 |
| rs78227947 | 11 | 117996360 | G | T | -0.065 | 0.9478 | -? | 0 | 0 | 0 | 1 |
| rs11218100 | 11 | 120867747 | T | C | -0.065 | 0.9479 | -? | 0 | 0 | 0 | 1 |
| rs6589776 | 11 | 119656449 | A | G | 0.065 | 0.9479 | +? | 0 | 0 | 0 | 1 |
| rs10736498 | 11 | 119412507 | A | G | 0.065 | 0.948 | +? | 0 | 0 | 0 | 1 |
| rs10502232 | 11 | 119790068 | G | A | 0.065 | 0.9484 | +? | 0 | 0 | 0 | 1 |
| rs12279373 | 11 | 116600400 | G | A | -0.064 | 0.9487 | +- | -80.8 | 1.659 | 1 | 0.1977 |
| rs145924639 | 11 | 119497371 | T | C | 0.064 | 0.9487 | +? | 0 | 0 | 0 | 1 |
| rs580872 | 11 | 116310858 | A | G | -0.064 | 0.9487 | -? | 0 | 0 | 0 | 1 |
| rs3741315 | 11 | 118004688 | A | G | -0.064 | 0.9488 | -+ | 0 | 0.004 | 1 | 0.9501 |
| rs10892464 | 11 | 119659720 | C | T | 0.064 | 0.9491 | +? | 0 | 0 | 0 | 1 |
| rs11216567 | 11 | 117698483 | G | A | 0.063 | 0.9495 | +? | 0 | 0 | 0 | 1 |
| rs7926282 | 11 | 118605914 | C | A | -0.063 | 0.9495 | -- | 0 | 0.001 | 1 | 0.974 |
| rs10437690 | 11 | 119422756 | G | C | 0.063 | 0.9496 | +? | 0 | 0 | 0 | 1 |
| rs61903421 | 11 | 116961802 | T | A | 0.063 | 0.9498 | +? | 0 | 0 | 0 | 1 |
| rs7950381 | 11 | 116963897 | A | C | 0.063 | 0.9498 | +? | 0 | 0 | 0 | 1 |
| rs12223789 | 11 | 117987041 | A | G | -0.063 | 0.9501 | -? | 0 | 0 | 0 | 1 |
| rs1793155 | 11 | 118098962 | A | G | -0.062 | 0.9502 | -+ | 0 | 0.432 | 1 | 0.511 |
| rs1793156 | 11 | 118098890 | C | T | -0.062 | 0.9502 | -+ | 0 | 0.432 | 1 | 0.511 |
| rs5132 | 11 | 116702778 | T | C | -0.062 | 0.9502 | +- | -3.3 | 2.904 | 1 | 0.08836 |

| | | | | | | | | | | | |
|---|---|---|---|---|---|---|---|---|---|---|---|
| rs73014407 | 11 | 117529248 | T | C | -0.062 | 0.9502 | -? | 0 | 0 | 0 | 1 |
| rs78701708 | 11 | 117315050 | C | T | -0.062 | 0.9502 | -? | 0 | 0 | 0 | 1 |
| rs12804122 | 11 | 116902616 | C | G | -0.062 | 0.9504 | -? | 0 | 0 | 0 | 1 |
| rs1508097 | 11 | 116468169 | T | A | 0.062 | 0.9506 | +? | 0 | 0 | 0 | 1 |
| rs1941626 | 11 | 117989301 | C | G | -0.062 | 0.9508 | -? | 0 | 0 | 0 | 1 |
| rs61903414 | 11 | 116946788 | T | C | -0.061 | 0.951 | -? | 0 | 0 | 0 | 1 |
| rs12795949 | 11 | 117662138 | A | G | 0.061 | 0.9513 | -+ | 0 | 0.054 | 1 | 0.8156 |
| rs71480353 | 11 | 117538907 | C | G | 0.061 | 0.9513 | +? | 0 | 0 | 0 | 1 |
| rs3825061 | 11 | 118944675 | T | C | 0.061 | 0.9514 | +? | 0 | 0 | 0 | 1 |
| rs17528237 | 11 | 117962275 | T | C | 0.061 | 0.9515 | +- | 0 | 0.117 | 1 | 0.7327 |
| rs10892328 | 11 | 118942654 | A | T | 0.061 | 0.9517 | +? | 0 | 0 | 0 | 1 |
| rs10892587 | 11 | 120450280 | A | C | -0.061 | 0.9517 | -? | 0 | 0 | 0 | 1 |
| rs688456 | 11 | 116722551 | T | G | -0.06 | 0.9518 | -? | 0 | 0 | 0 | 1 |
| rs35624176 | 11 | 116413310 | A | G | 0.06 | 0.9519 | +? | 0 | 0 | 0 | 1 |
| rs538518 | 11 | 119789044 | G | T | -0.06 | 0.9519 | -? | 0 | 0 | 0 | 1 |
| rs948460 | 11 | 117960018 | T | A | 0.06 | 0.9519 | ++ | 0 | 0.001 | 1 | 0.98 |
| rs34874964 | 11 | 117533027 | T | C | -0.06 | 0.9522 | -? | 0 | 0 | 0 | 1 |
| rs61905066 | 11 | 116571608 | G | A | -0.06 | 0.9522 | -? | 0 | 0 | 0 | 1 |
| rs11216277 | 11 | 116987046 | G | A | -0.06 | 0.9523 | -? | 0 | 0 | 0 | 1 |
| rs138697506 | 11 | 116355788 | G | A | 0.06 | 0.9523 | +? | 0 | 0 | 0 | 1 |
| rs11217953 | 11 | 120538217 | A | G | -0.06 | 0.9524 | -? | 0 | 0 | 0 | 1 |
| rs61899020 | 11 | 119778398 | T | C | -0.06 | 0.9524 | -? | 0 | 0 | 0 | 1 |
| rs10790306 | 11 | 119423690 | C | T | 0.06 | 0.9525 | +? | 0 | 0 | 0 | 1 |
| rs10790391 | 11 | 120542279 | A | G | 0.059 | 0.9526 | +? | 0 | 0 | 0 | 1 |
| rs12797009 | 11 | 118826192 | A | C | -0.059 | 0.9526 | -? | 0 | 0 | 0 | 1 |
| rs61905462 | 11 | 116994106 | A | C | -0.059 | 0.9527 | -? | 0 | 0 | 0 | 1 |
| rs17121510 | 11 | 117876155 | G | A | -0.059 | 0.9528 | -? | 0 | 0 | 0 | 1 |
| rs1715420 | 11 | 118105599 | A | T | -0.059 | 0.9531 | -+ | 0 | 0.12 | 1 | 0.7293 |
| rs7943043 | 11 | 120954436 | C | T | 0.058 | 0.9535 | +? | 0 | 0 | 0 | 1 |
| rs874069 | 11 | 120953914 | C | T | 0.058 | 0.9535 | +? | 0 | 0 | 0 | 1 |
| rs11216700 | 11 | 117894782 | T | A | 0.058 | 0.9536 | +? | 0 | 0 | 0 | 1 |
| rs11216703 | 11 | 117894927 | A | C | 0.058 | 0.9536 | +? | 0 | 0 | 0 | 1 |
| rs1892948 | 11 | 116572089 | T | C | -0.058 | 0.9536 | -? | 0 | 0 | 0 | 1 |
| rs141453720 | 11 | 119334649 | C | G | -0.058 | 0.9538 | -? | 0 | 0 | 0 | 1 |
| rs692804 | 11 | 120099368 | T | C | -0.058 | 0.9539 | -? | 0 | 0 | 0 | 1 |
| rs1260808616 | 11 | 118941596 | A | G | 0.058 | 0.954 | +? | 0 | 0 | 0 | 1 |
| rs601318 | 11 | 118519550 | G | C | -0.058 | 0.954 | -+ | 0 | 0.529 | 1 | 0.4671 |
| rs6589659 | 11 | 118191691 | A | C | 0.058 | 0.954 | -+ | 0 | 0.413 | 1 | 0.5203 |
| rs10736501 | 11 | 119437268 | G | A | 0.058 | 0.9541 | +? | 0 | 0 | 0 | 1 |
| rs11217318 | 11 | 119429848 | A | G | -0.058 | 0.9541 | -? | 0 | 0 | 0 | 1 |
| rs509809 | 11 | 119777611 | G | A | 0.058 | 0.9541 | +? | 0 | 0 | 0 | 1 |
| rs45480805 | 11 | 118267903 | A | G | 0.057 | 0.9542 | -+ | -18.1 | 2.541 | 1 | 0.1109 |
| rs664922 | 11 | 117070895 | C | A | -0.057 | 0.9543 | -? | 0 | 0 | 0 | 1 |
| rs590843 | 11 | 119991166 | T | A | -0.057 | 0.9545 | -? | 0 | 0 | 0 | 1 |
| rs17123513 | 11 | 119846442 | A | G | -0.057 | 0.9546 | -? | 0 | 0 | 0 | 1 |
| rs2508559 | 11 | 117580254 | T | C | -0.057 | 0.9546 | -? | 0 | 0 | 0 | 1 |
| rs604245 | 11 | 117580585 | A | G | -0.057 | 0.9546 | -? | 0 | 0 | 0 | 1 |
| rs655086 | 11 | 117580509 | A | G | -0.057 | 0.9546 | -? | 0 | 0 | 0 | 1 |
| rs74609299 | 11 | 119846476 | T | C | -0.057 | 0.9546 | -? | 0 | 0 | 0 | 1 |
| rs11823918 | 11 | 116960497 | C | G | 0.057 | 0.9547 | +? | 0 | 0 | 0 | 1 |
| rs6589656 | 11 | 118095996 | C | G | -0.057 | 0.9547 | -+ | 0 | 0.441 | 1 | 0.5068 |
| rs3132775 | 11 | 120585993 | G | A | -0.057 | 0.9548 | -? | 0 | 0 | 0 | 1 |
| rs112302730 | 11 | 118283383 | A | G | 0.056 | 0.9551 | -+ | -29.8 | 2.31 | 1 | 0.1285 |
| rs11217766 | 11 | 120072666 | A | C | 0.056 | 0.9552 | +? | 0 | 0 | 0 | 1 |
| rs3135506 | 11 | 116662407 | C | G | -0.056 | 0.9552 | +- | -100.8 | 1.494 | 1 | 0.2216 |
| rs595137 | 11 | 116771356 | C | T | -0.056 | 0.9553 | +- | 63 | 8.105 | 1 | 0.004413 |

| | | | | | | | | | | | |
|---|---|---|---|---|---|---|---|---|---|---|---|
| rs78509462 | 11 | 119841349 | C | T | -0.056 | 0.9553 | -? | 0 | 0 | 0 | 1 |
| rs111763202 | 11 | 118289734 | T | C | 0.056 | 0.9554 | -+ | -34.2 | 2.236 | 1 | 0.1349 |
| rs112035668 | 11 | 118290443 | G | A | 0.056 | 0.9554 | -+ | -34.2 | 2.236 | 1 | 0.1349 |
| rs3809041 | 11 | 117774560 | C | G | 0.056 | 0.9554 | +? | 0 | 0 | 0 | 1 |
| rs2000868 | 11 | 120544816 | T | C | 0.056 | 0.9555 | +? | 0 | 0 | 0 | 1 |
| rs73612281 | 11 | 118203642 | T | C | 0.056 | 0.9555 | +- | 0 | 0.912 | 1 | 0.3395 |
| rs61903418 | 11 | 116956665 | A | G | 0.056 | 0.9556 | +? | 0 | 0 | 0 | 1 |
| rs11552421 | 11 | 118430519 | T | C | 0.056 | 0.9557 | +- | 0 | 0.034 | 1 | 0.8533 |
| rs12222499 | 11 | 119814422 | T | C | -0.056 | 0.9557 | -? | 0 | 0 | 0 | 1 |
| rs544153 | 11 | 117336280 | A | G | -0.055 | 0.9558 | -? | 0 | 0 | 0 | 1 |
| rs1312372307 | 11 | 118048499 | T | C | 0.055 | 0.9561 | +- | 0 | 0.451 | 1 | 0.502 |
| rs115787357 | 11 | 117064506 | G | T | 0.055 | 0.9562 | +? | 0 | 0 | 0 | 1 |
| rs1263889 | 11 | 117473833 | A | C | 0.055 | 0.9562 | +? | 0 | 0 | 0 | 1 |
| rs11216816 | 11 | 118075152 | A | C | -0.055 | 0.9563 | -+ | 0 | 0.269 | 1 | 0.6042 |
| rs11216817 | 11 | 118075474 | T | C | -0.055 | 0.9563 | -+ | 0 | 0.269 | 1 | 0.6042 |
| rs1231238253 | 11 | 117101311 | T | C | 0.055 | 0.9563 | -+ | 73.1 | 11.152 | 1 | 0.0008395 |
| rs17092726 | 11 | 117996200 | G | A | -0.055 | 0.9565 | -? | 0 | 0 | 0 | 1 |
| rs17311121 | 11 | 119876680 | G | T | 0.054 | 0.9567 | +? | 0 | 0 | 0 | 1 |
| rs637737 | 11 | 118051646 | G | A | 0.054 | 0.9567 | +- | 0 | 0.387 | 1 | 0.5339 |
| rs34929374 | 11 | 120891614 | G | A | 0.054 | 0.9568 | +? | 0 | 0 | 0 | 1 |
| rs1064193 | 11 | 118965338 | A | T | -0.054 | 0.9569 | -? | 0 | 0 | 0 | 1 |
| rs6589644 | 11 | 117959224 | T | C | 0.054 | 0.9569 | ++ | 0 | 0 | 1 | 0.9883 |
| rs3018383 | 11 | 118093359 | A | C | -0.054 | 0.9571 | -+ | 0 | 0.438 | 1 | 0.5082 |
| rs7926574 | 11 | 119641049 | C | T | -0.054 | 0.9571 | -? | 0 | 0 | 0 | 1 |
| rs948461 | 11 | 117960138 | T | G | 0.053 | 0.9574 | ++ | 0 | 0.001 | 1 | 0.9768 |
| rs10790175 | 11 | 117034729 | T | C | -0.053 | 0.9576 | -? | 0 | 0 | 0 | 1 |
| rs17119847 | 11 | 116479752 | A | C | -0.053 | 0.9576 | -? | 0 | 0 | 0 | 1 |
| rs472948 | 11 | 120099193 | G | A | -0.053 | 0.9577 | -? | 0 | 0 | 0 | 1 |
| rs11218102 | 11 | 120868460 | C | G | -0.053 | 0.9579 | -? | 0 | 0 | 0 | 1 |
| rs514527 | 11 | 117696720 | G | A | -0.053 | 0.958 | -? | 0 | 0 | 0 | 1 |
| rs11217952 | 11 | 120538215 | T | G | -0.052 | 0.9582 | -? | 0 | 0 | 0 | 1 |
| rs2155855 | 11 | 117974600 | G | A | -0.052 | 0.9582 | -? | 0 | 0 | 0 | 1 |
| rs2155856 | 11 | 117974639 | A | C | -0.052 | 0.9582 | -? | 0 | 0 | 0 | 1 |
| rs628756 | 11 | 120987966 | C | T | -0.052 | 0.9582 | +- | 0 | 0.776 | 1 | 0.3783 |
| rs79754446 | 11 | 117997473 | G | A | -0.052 | 0.9582 | -? | 0 | 0 | 0 | 1 |
| rs11600134 | 11 | 118070788 | A | G | 0.052 | 0.9583 | -+ | 0 | 0.578 | 1 | 0.447 |
| rs11216791 | 11 | 118044360 | G | A | 0.052 | 0.9584 | +- | 0 | 0.326 | 1 | 0.568 |
| rs7130031 | 11 | 119492297 | T | C | -0.052 | 0.9585 | -? | 0 | 0 | 0 | 1 |
| rs11216027 | 11 | 116464955 | T | C | 0.052 | 0.9586 | +? | 0 | 0 | 0 | 1 |
| rs1940037 | 11 | 117436095 | T | C | 0.052 | 0.9588 | +- | 0 | 0.012 | 1 | 0.9143 |
| rs10790307 | 11 | 119424157 | C | T | 0.052 | 0.9589 | +? | 0 | 0 | 0 | 1 |
| rs78587763 | 11 | 117400684 | T | C | 0.052 | 0.9589 | +? | 0 | 0 | 0 | 1 |
| rs2120268 | 11 | 118132705 | A | G | -0.051 | 0.959 | -+ | 0 | 0.2 | 1 | 0.6548 |
| rs6589847 | 11 | 120715884 | A | G | -0.051 | 0.959 | -? | 0 | 0 | 0 | 1 |
| rs61902674 | 11 | 120548902 | A | C | -0.051 | 0.9592 | -? | 0 | 0 | 0 | 1 |
| rs11217956 | 11 | 120540795 | T | C | 0.051 | 0.9593 | +? | 0 | 0 | 0 | 1 |
| rs11602779 | 11 | 119726873 | A | G | -0.051 | 0.9593 | -? | 0 | 0 | 0 | 1 |
| rs73582578 | 11 | 116336988 | A | C | -0.051 | 0.9595 | -? | 0 | 0 | 0 | 1 |
| rs10892035 | 11 | 116694938 | A | G | -0.05 | 0.9598 | -? | 0 | 0 | 0 | 1 |
| rs60167594 | 11 | 116966878 | C | T | -0.05 | 0.9599 | -+ | 74.8 | 11.894 | 1 | 0.0005631 |
| rs144234662 | 11 | 121083487 | T | C | -0.05 | 0.96 | -? | 0 | 0 | 0 | 1 |
| rs10082698 | 11 | 120543440 | G | A | -0.05 | 0.9602 | -? | 0 | 0 | 0 | 1 |
| rs628944 | 11 | 119943275 | A | C | 0.05 | 0.9602 | +? | 0 | 0 | 0 | 1 |
| rs1022783694 | 11 | 116886114 | C | T | -0.05 | 0.9603 | -? | 0 | 0 | 0 | 1 |
| rs12808627 | 11 | 120038717 | T | C | 0.05 | 0.9605 | +? | 0 | 0 | 0 | 1 |
| rs4938354 | 11 | 117051165 | A | G | -0.05 | 0.9605 | -? | 0 | 0 | 0 | 1 |

| | | | | | | | | | | | |
|---|---|---|---|---|---|---|---|---|---|---|---|
| rs5014624 | 11 | 117731136 | A | G | 0.05 | 0.9605 | +? | 0 | 0 | 0 | 1 |
| rs11216684 | 11 | 117888685 | G | A | 0.049 | 0.9606 | +? | 0 | 0 | 0 | 1 |
| rs11216686 | 11 | 117888781 | A | T | 0.049 | 0.9606 | +? | 0 | 0 | 0 | 1 |
| rs569044 | 11 | 119999191 | A | G | -0.049 | 0.9606 | -? | 0 | 0 | 0 | 1 |
| rs73011708 | 11 | 117887528 | C | G | 0.049 | 0.9606 | +? | 0 | 0 | 0 | 1 |
| rs78640117 | 11 | 117889685 | T | C | 0.049 | 0.9606 | +? | 0 | 0 | 0 | 1 |
| rs10892315 | 11 | 118831727 | C | T | -0.049 | 0.9607 | +- | 0 | 0.814 | 1 | 0.3669 |
| rs61905337 | 11 | 117365821 | A | C | 0.049 | 0.9607 | +? | 0 | 0 | 0 | 1 |
| rs12225752 | 11 | 120546394 | G | A | 0.049 | 0.9609 | +? | 0 | 0 | 0 | 1 |
| rs72488011 | 11 | 117997360 | C | T | 0.049 | 0.9609 | +? | 0 | 0 | 0 | 1 |
| rs7928307 | 11 | 116439615 | T | C | -0.049 | 0.9609 | -+ | 0 | 0.246 | 1 | 0.6198 |
| rs10502240 | 11 | 120547912 | T | C | -0.049 | 0.961 | -? | 0 | 0 | 0 | 1 |
| rs10790394 | 11 | 120548076 | C | G | -0.049 | 0.961 | -? | 0 | 0 | 0 | 1 |
| rs10790400 | 11 | 120557091 | T | A | -0.049 | 0.961 | -? | 0 | 0 | 0 | 1 |
| rs10892605 | 11 | 120546953 | T | C | -0.049 | 0.961 | -? | 0 | 0 | 0 | 1 |
| rs11217965 | 11 | 120546914 | T | C | -0.049 | 0.961 | -? | 0 | 0 | 0 | 1 |
| rs11603586 | 11 | 120548353 | T | C | -0.049 | 0.961 | -? | 0 | 0 | 0 | 1 |
| rs11607732 | 11 | 120548318 | G | A | -0.049 | 0.961 | -? | 0 | 0 | 0 | 1 |
| rs118065597 | 11 | 116353269 | A | G | 0.049 | 0.961 | +? | 0 | 0 | 0 | 1 |
| rs4256992 | 11 | 120547174 | A | G | -0.049 | 0.961 | -? | 0 | 0 | 0 | 1 |
| rs4268524 | 11 | 120547112 | T | C | -0.049 | 0.961 | -? | 0 | 0 | 0 | 1 |
| rs4367954 | 11 | 120547109 | G | A | -0.049 | 0.961 | -? | 0 | 0 | 0 | 1 |
| rs4628679 | 11 | 120547052 | C | T | -0.049 | 0.961 | -? | 0 | 0 | 0 | 1 |
| rs7126415 | 11 | 120566535 | C | G | -0.049 | 0.961 | -? | 0 | 0 | 0 | 1 |
| rs11826038 | 11 | 116835398 | G | A | 0.049 | 0.9611 | +? | 0 | 0 | 0 | 1 |
| rs4936498 | 11 | 119680923 | G | T | -0.049 | 0.9612 | -+ | 0 | 0.164 | 1 | 0.6857 |
| rs9326248 | 11 | 116999225 | T | G | -0.049 | 0.9612 | -? | 0 | 0 | 0 | 1 |
| rs112151056 | 11 | 116919368 | A | G | 0.048 | 0.9614 | +? | 0 | 0 | 0 | 1 |
| rs11216284 | 11 | 117003060 | C | T | -0.048 | 0.9614 | -? | 0 | 0 | 0 | 1 |
| rs17120241 | 11 | 116915020 | C | T | 0.048 | 0.9614 | +? | 0 | 0 | 0 | 1 |
| rs4938328 | 11 | 116921219 | G | T | -0.048 | 0.9614 | -? | 0 | 0 | 0 | 1 |
| rs61905716 | 11 | 116919067 | T | C | 0.048 | 0.9614 | +? | 0 | 0 | 0 | 1 |
| rs61905719 | 11 | 116922714 | T | C | 0.048 | 0.9614 | +? | 0 | 0 | 0 | 1 |
| rs11216671 | 11 | 117878745 | T | C | -0.048 | 0.9615 | -? | 0 | 0 | 0 | 1 |
| rs11216672 | 11 | 117878769 | A | G | -0.048 | 0.9615 | -? | 0 | 0 | 0 | 1 |
| rs12799647 | 11 | 120032185 | G | T | 0.048 | 0.9615 | +? | 0 | 0 | 0 | 1 |
| rs7934908 | 11 | 120542603 | A | G | 0.048 | 0.9615 | +? | 0 | 0 | 0 | 1 |
| rs10892406 | 11 | 119435544 | G | A | -0.048 | 0.9616 | -? | 0 | 0 | 0 | 1 |
| rs3135507 | 11 | 116661488 | T | C | 0.048 | 0.9616 | +- | -31.9 | 2.274 | 1 | 0.1315 |
| rs10892551 | 11 | 120078286 | A | G | 0.048 | 0.9618 | +? | 0 | 0 | 0 | 1 |
| rs10892547 | 11 | 120060793 | T | G | -0.048 | 0.9619 | +- | 0 | 0.528 | 1 | 0.4674 |
| rs10790302 | 11 | 119419133 | A | G | 0.047 | 0.9622 | +? | 0 | 0 | 0 | 1 |
| rs11217311 | 11 | 119419672 | T | C | 0.047 | 0.9622 | +? | 0 | 0 | 0 | 1 |
| rs11217312 | 11 | 119419976 | C | A | 0.047 | 0.9622 | +? | 0 | 0 | 0 | 1 |
| rs59837440 | 11 | 116991842 | C | A | -0.047 | 0.9625 | -+ | 73.8 | 11.438 | 1 | 0.0007195 |
| rs117740883 | 11 | 117259904 | T | C | -0.047 | 0.9626 | -? | 0 | 0 | 0 | 1 |
| rs11216896 | 11 | 118430048 | T | C | 0.047 | 0.9628 | +- | 0 | 0.009 | 1 | 0.926 |
| rs11607159 | 11 | 120855348 | A | G | -0.047 | 0.9629 | -+ | 0 | 0.158 | 1 | 0.6908 |
| rs7944957 | 11 | 117996296 | C | T | -0.046 | 0.9629 | -+ | 0 | 0.055 | 1 | 0.8149 |
| rs10790412 | 11 | 121100193 | C | G | 0.046 | 0.963 | -+ | 0 | 0.018 | 1 | 0.8921 |
| rs7116375 | 11 | 118069979 | A | C | 0.046 | 0.963 | -+ | 0 | 0.563 | 1 | 0.4531 |
| rs76674252 | 11 | 116354727 | A | G | 0.046 | 0.9631 | +? | 0 | 0 | 0 | 1 |
| rs486876 | 11 | 119773032 | A | G | 0.046 | 0.9634 | +? | 0 | 0 | 0 | 1 |
| rs2239896 | 11 | 118983434 | T | C | -0.046 | 0.9636 | -? | 0 | 0 | 0 | 1 |
| rs34125463 | 11 | 116537393 | A | T | 0.046 | 0.9637 | +? | 0 | 0 | 0 | 1 |
| rs2852228 | 11 | 120602594 | A | T | -0.045 | 0.9639 | -? | 0 | 0 | 0 | 1 |

| | | | | | | | | | | | |
|---|---|---|---|---|---|---|---|---|---|---|---|
| rs7101652 | 11 | 118078718 | A | G | 0.045 | 0.9639 | -+ | 0 | 0.407 | 1 | 0.5233 |
| rs77751615 | 11 | 118982806 | C | A | -0.045 | 0.964 | -? | 0 | 0 | 0 | 1 |
| rs117634345 | 11 | 117065050 | A | G | -0.045 | 0.9641 | -+ | 74 | 11.537 | 1 | 0.0006823 |
| rs606151 | 11 | 119384214 | A | G | -0.045 | 0.9642 | -? | 0 | 0 | 0 | 1 |
| rs10892246 | 11 | 118481431 | A | G | 0.045 | 0.9643 | +- | 0 | 0.003 | 1 | 0.9581 |
| rs565622 | 11 | 116314866 | C | A | 0.045 | 0.9643 | +? | 0 | 0 | 0 | 1 |
| rs7123454 | 11 | 116704178 | C | A | 0.045 | 0.9644 | +? | 0 | 0 | 0 | 1 |
| rs10892358 | 11 | 119255795 | G | A | -0.045 | 0.9645 | -? | 0 | 0 | 0 | 1 |
| rs3133853 | 11 | 120603644 | C | T | 0.045 | 0.9645 | +? | 0 | 0 | 0 | 1 |
| rs608793 | 11 | 118986659 | C | T | 0.045 | 0.9645 | +? | 0 | 0 | 0 | 1 |
| rs7110179 | 11 | 120712591 | G | A | -0.044 | 0.9646 | -? | 0 | 0 | 0 | 1 |
| rs10736499 | 11 | 119429098 | C | T | -0.044 | 0.9648 | -? | 0 | 0 | 0 | 1 |
| rs10750150 | 11 | 119429080 | T | C | -0.044 | 0.9648 | -? | 0 | 0 | 0 | 1 |
| rs36108816 | 11 | 117502518 | C | G | 0.044 | 0.9648 | +? | 0 | 0 | 0 | 1 |
| rs10892065 | 11 | 116897024 | T | C | -0.044 | 0.965 | -? | 0 | 0 | 0 | 1 |
| rs11216286 | 11 | 117004087 | G | A | -0.044 | 0.9652 | -? | 0 | 0 | 0 | 1 |
| rs2298464 | 11 | 117780919 | A | G | -0.044 | 0.9652 | -? | 0 | 0 | 0 | 1 |
| rs4938675 | 11 | 119408686 | A | G | 0.044 | 0.9652 | +? | 0 | 0 | 0 | 1 |
| rs10892600 | 11 | 120545929 | G | A | -0.044 | 0.9653 | -? | 0 | 0 | 0 | 1 |
| rs10892601 | 11 | 120546027 | G | A | -0.044 | 0.9653 | -? | 0 | 0 | 0 | 1 |
| rs10892602 | 11 | 120546082 | A | G | -0.044 | 0.9653 | -? | 0 | 0 | 0 | 1 |
| rs10892603 | 11 | 120546119 | C | T | -0.044 | 0.9653 | -? | 0 | 0 | 0 | 1 |
| rs10892604 | 11 | 120546223 | T | C | -0.044 | 0.9653 | -? | 0 | 0 | 0 | 1 |
| rs11216510 | 11 | 117581589 | C | A | 0.044 | 0.9653 | +? | 0 | 0 | 0 | 1 |
| rs11217957 | 11 | 120543669 | G | A | -0.044 | 0.9653 | -? | 0 | 0 | 0 | 1 |
| rs11217959 | 11 | 120544735 | C | T | -0.044 | 0.9653 | -? | 0 | 0 | 0 | 1 |
| rs11217960 | 11 | 120545103 | T | C | -0.044 | 0.9653 | -? | 0 | 0 | 0 | 1 |
| rs11217961 | 11 | 120545219 | G | A | -0.044 | 0.9653 | -? | 0 | 0 | 0 | 1 |
| rs11217963 | 11 | 120545502 | T | C | -0.044 | 0.9653 | -? | 0 | 0 | 0 | 1 |
| rs11217964 | 11 | 120545503 | G | A | -0.044 | 0.9653 | -? | 0 | 0 | 0 | 1 |
| rs12225737 | 11 | 120546287 | C | A | -0.044 | 0.9653 | -? | 0 | 0 | 0 | 1 |
| rs61902672 | 11 | 120545098 | T | C | -0.044 | 0.9653 | -? | 0 | 0 | 0 | 1 |
| rs10790222 | 11 | 117800574 | A | G | 0.043 | 0.9654 | -+ | -146.4 | 1.217 | 1 | 0.2699 |
| rs72488012 | 11 | 117997427 | A | G | -0.043 | 0.9654 | -? | 0 | 0 | 0 | 1 |
| rs73008568 | 11 | 117065007 | G | C | 0.043 | 0.9654 | +? | 0 | 0 | 0 | 1 |
| rs79220634 | 11 | 117996402 | A | C | -0.043 | 0.9654 | -? | 0 | 0 | 0 | 1 |
| rs10431040 | 11 | 117898465 | A | G | -0.043 | 0.9655 | -? | 0 | 0 | 0 | 1 |
| rs10458984 | 11 | 117897503 | G | T | -0.043 | 0.9655 | -? | 0 | 0 | 0 | 1 |
| rs11217686 | 11 | 119963819 | A | G | 0.043 | 0.9655 | +? | 0 | 0 | 0 | 1 |
| rs474339 | 11 | 117059552 | G | A | -0.043 | 0.9656 | -? | 0 | 0 | 0 | 1 |
| rs10892478 | 11 | 119727706 | G | A | -0.043 | 0.9657 | -? | 0 | 0 | 0 | 1 |
| rs10892597 | 11 | 120542339 | T | C | 0.043 | 0.9657 | +? | 0 | 0 | 0 | 1 |
| rs61907560 | 11 | 116726087 | C | T | -0.043 | 0.9657 | -? | 0 | 0 | 0 | 1 |
| rs74236048 | 11 | 121078691 | T | C | -0.043 | 0.9657 | -? | 0 | 0 | 0 | 1 |
| rs58831762 | 11 | 119428506 | G | T | 0.043 | 0.9658 | +? | 0 | 0 | 0 | 1 |
| rs61901427 | 11 | 120715772 | G | A | -0.043 | 0.9659 | -? | 0 | 0 | 0 | 1 |
| rs7103816 | 11 | 119427346 | C | T | -0.043 | 0.9659 | -? | 0 | 0 | 0 | 1 |
| rs732481 | 11 | 120828855 | G | T | 0.043 | 0.9659 | +? | 0 | 0 | 0 | 1 |
| rs2155857 | 11 | 117974759 | A | G | -0.043 | 0.9661 | -? | 0 | 0 | 0 | 1 |
| rs579200 | 11 | 119700001 | G | T | 0.042 | 0.9663 | +? | 0 | 0 | 0 | 1 |
| rs633621 | 11 | 119866151 | A | G | -0.042 | 0.9663 | -? | 0 | 0 | 0 | 1 |
| rs117812708 | 11 | 116965555 | G | A | -0.042 | 0.9664 | -+ | 74.7 | 11.862 | 1 | 0.0005729 |
| rs482371 | 11 | 116752163 | T | C | 0.042 | 0.9664 | +? | 0 | 0 | 0 | 1 |
| rs77374709 | 11 | 116964942 | T | C | -0.042 | 0.9664 | -+ | 74.7 | 11.862 | 1 | 0.0005729 |
| rs10790390 | 11 | 120542028 | G | A | 0.042 | 0.9665 | +? | 0 | 0 | 0 | 1 |
| rs12796457 | 11 | 120855313 | T | C | -0.042 | 0.9665 | -+ | 0 | 0.156 | 1 | 0.6927 |

| | | | | | | | | | | | |
|---|---|---|---|---|---|---|---|---|---|---|---|
| rs4938481 | 11 | 117969374 | T | C | 0.042 | 0.9665 | +- | 0 | 0.15 | 1 | 0.6987 |
| rs11218212 | 11 | 121101217 | T | A | -0.042 | 0.9666 | -+ | 0 | 0.035 | 1 | 0.8523 |
| rs113839182 | 11 | 118273602 | T | C | -0.042 | 0.9668 | -+ | -39.7 | 2.147 | 1 | 0.1428 |
| rs548207 | 11 | 116489629 | G | T | -0.042 | 0.9668 | -? | 0 | 0 | 0 | 1 |
| rs10892196 | 11 | 117800083 | T | C | 0.041 | 0.9669 | -+ | -144.3 | 1.228 | 1 | 0.2678 |
| rs7934830 | 11 | 120542761 | G | A | 0.041 | 0.9669 | +? | 0 | 0 | 0 | 1 |
| rs2847699 | 11 | 117929490 | A | G | -0.041 | 0.967 | +- | 0 | 0.59 | 1 | 0.4425 |
| rs631120 | 11 | 117440267 | A | G | -0.041 | 0.967 | -? | 0 | 0 | 0 | 1 |
| rs530702 | 11 | 119983831 | A | G | -0.041 | 0.9671 | -? | 0 | 0 | 0 | 1 |
| rs1352178661 | 11 | 117024256 | A | G | -0.041 | 0.9675 | -+ | 73.5 | 11.316 | 1 | 0.0007686 |
| rs4938710 | 11 | 119566174 | C | T | 0.041 | 0.9675 | +? | 0 | 0 | 0 | 1 |
| rs73010786 | 11 | 117963955 | T | C | 0.041 | 0.9676 | +- | 0 | 0.109 | 1 | 0.7414 |
| rs519672 | 11 | 119984535 | G | C | 0.04 | 0.9677 | +? | 0 | 0 | 0 | 1 |
| rs7110879 | 11 | 116441138 | A | G | 0.04 | 0.9677 | -+ | 0 | 0.279 | 1 | 0.5973 |
| rs78080546 | 11 | 117630622 | T | A | 0.041 | 0.9677 | +- | 26.5 | 4.082 | 1 | 0.04333 |
| rs1626649 | 11 | 118071097 | T | C | 0.04 | 0.9678 | +? | 0 | 0 | 0 | 1 |
| rs10892064 | 11 | 116896553 | G | C | -0.04 | 0.9682 | -? | 0 | 0 | 0 | 1 |
| rs74428560 | 11 | 116989982 | T | C | -0.04 | 0.9683 | -? | 0 | 0 | 0 | 1 |
| rs111587825 | 11 | 118274602 | G | T | -0.039 | 0.9685 | -+ | -39 | 2.158 | 1 | 0.1418 |
| rs1866682 | 11 | 116195089 | T | C | 0.039 | 0.9685 | +? | 0 | 0 | 0 | 1 |
| rs10892182 | 11 | 117699786 | A | G | -0.039 | 0.9688 | -? | 0 | 0 | 0 | 1 |
| rs17123466 | 11 | 119771748 | T | C | 0.039 | 0.9689 | +? | 0 | 0 | 0 | 1 |
| rs55646454 | 11 | 119773103 | T | G | 0.039 | 0.9689 | +? | 0 | 0 | 0 | 1 |
| rs55895660 | 11 | 119774821 | A | G | 0.039 | 0.9689 | +? | 0 | 0 | 0 | 1 |
| rs10790212 | 11 | 117702690 | T | C | 0.039 | 0.969 | +? | 0 | 0 | 0 | 1 |
| rs76813465 | 11 | 117699173 | T | C | 0.039 | 0.969 | +? | 0 | 0 | 0 | 1 |
| rs73018483 | 11 | 117706942 | G | C | 0.039 | 0.9692 | +? | 0 | 0 | 0 | 1 |
| rs76570573 | 11 | 118543798 | T | C | 0.039 | 0.9692 | -+ | 0 | 0.155 | 1 | 0.694 |
| rs8521 | 11 | 117067699 | C | T | -0.039 | 0.9692 | -? | 0 | 0 | 0 | 1 |
| rs61905691 | 11 | 116880703 | G | A | 0.038 | 0.9693 | +? | 0 | 0 | 0 | 1 |
| rs3181258 | 11 | 118218478 | A | T | 0.038 | 0.9694 | -+ | 4.5 | 3.141 | 1 | 0.07636 |
| rs3863306 | 11 | 116853213 | C | T | -0.038 | 0.9694 | -? | 0 | 0 | 0 | 1 |
| rs499443 | 11 | 121081632 | A | G | 0.038 | 0.9694 | -+ | 0 | 0.02 | 1 | 0.8885 |
| rs60915096 | 11 | 116224575 | C | T | 0.038 | 0.9694 | +? | 0 | 0 | 0 | 1 |
| rs7110679 | 11 | 119633863 | A | G | 0.038 | 0.9694 | +? | 0 | 0 | 0 | 1 |
| rs113521822 | 11 | 118063349 | G | A | 0.038 | 0.9696 | +- | 0 | 0.223 | 1 | 0.637 |
| rs654329 | 11 | 121093676 | T | C | 0.038 | 0.9697 | -+ | 0 | 0.02 | 1 | 0.8883 |
| rs77974673 | 11 | 117491787 | T | C | 0.038 | 0.9697 | +? | 0 | 0 | 0 | 1 |
| rs12285095 | 11 | 116658031 | G | T | -0.038 | 0.9699 | +- | -72.4 | 1.74 | 1 | 0.1872 |
| rs139685157 | 11 | 117062158 | A | G | 0.038 | 0.9699 | -+ | 73.3 | 11.217 | 1 | 0.0008105 |
| rs73009131 | 11 | 116336740 | A | G | 0.038 | 0.9699 | +? | 0 | 0 | 0 | 1 |
| rs7128071 | 11 | 116968798 | A | G | 0.038 | 0.97 | +? | 0 | 0 | 0 | 1 |
| rs10892598 | 11 | 120543750 | T | G | -0.037 | 0.9701 | -? | 0 | 0 | 0 | 1 |
| rs543584 | 11 | 119984863 | C | T | 0.037 | 0.9701 | +? | 0 | 0 | 0 | 1 |
| rs11218097 | 11 | 120865697 | T | G | -0.037 | 0.9703 | -? | 0 | 0 | 0 | 1 |
| rs1944522 | 11 | 120864922 | T | C | -0.037 | 0.9703 | -? | 0 | 0 | 0 | 1 |
| rs518181 | 11 | 116772787 | C | A | 0.037 | 0.9703 | +? | 0 | 0 | 0 | 1 |
| rs10790174 | 11 | 117024264 | G | A | 0.037 | 0.9707 | +? | 0 | 0 | 0 | 1 |
| rs1243336 | 11 | 117273051 | T | C | 0.037 | 0.9707 | +? | 0 | 0 | 0 | 1 |
| rs1242127 | 11 | 117076564 | G | A | -0.036 | 0.9709 | -? | 0 | 0 | 0 | 1 |
| rs4938679 | 11 | 119411425 | T | C | 0.036 | 0.971 | +? | 0 | 0 | 0 | 1 |
| rs5014620 | 11 | 117731319 | C | A | -0.036 | 0.9711 | -? | 0 | 0 | 0 | 1 |
| rs5014621 | 11 | 117731284 | A | G | -0.036 | 0.9711 | -? | 0 | 0 | 0 | 1 |
| rs11216564 | 11 | 117695853 | A | G | -0.036 | 0.9715 | -? | 0 | 0 | 0 | 1 |
| rs1320718 | 11 | 120456829 | C | T | 0.036 | 0.9715 | +? | 0 | 0 | 0 | 1 |
| rs2136419 | 11 | 119488176 | T | C | -0.035 | 0.9717 | -? | 0 | 0 | 0 | 1 |

| | | | | | | | | | | | |
|---|---|---|---|---|---|---|---|---|---|---|---|
| rs681231 | 11 | 121094059 | A | G | 0.035 | 0.9719 | -+ | 0 | 0.02 | 1 | 0.8871 |
| rs12802652 | 11 | 120037862 | T | G | 0.035 | 0.972 | +? | 0 | 0 | 0 | 1 |
| rs7934825 | 11 | 120542709 | G | C | 0.035 | 0.9721 | +? | 0 | 0 | 0 | 1 |
| rs140158503 | 11 | 119981694 | T | C | -0.035 | 0.9722 | -? | 0 | 0 | 0 | 1 |
| rs4269933 | 11 | 119648368 | G | A | 0.035 | 0.9723 | +? | 0 | 0 | 0 | 1 |
| rs4301801 | 11 | 119648306 | G | C | 0.035 | 0.9723 | +? | 0 | 0 | 0 | 1 |
| rs4418818 | 11 | 119648005 | A | G | 0.035 | 0.9723 | +? | 0 | 0 | 0 | 1 |
| rs77487830 | 11 | 118311869 | G | A | 0.035 | 0.9723 | -+ | 16.8 | 3.606 | 1 | 0.05758 |
| rs77535829 | 11 | 118311677 | T | C | 0.035 | 0.9723 | -+ | 16.8 | 3.606 | 1 | 0.05758 |
| rs34699720 | 11 | 119712159 | A | C | -0.034 | 0.9725 | -? | 0 | 0 | 0 | 1 |
| rs79697731 | 11 | 117092808 | G | C | -0.034 | 0.9726 | -+ | 73.4 | 11.291 | 1 | 0.0007788 |
| rs10892393 | 11 | 119426651 | T | C | -0.034 | 0.9727 | -? | 0 | 0 | 0 | 1 |
| rs61905060 | 11 | 116561628 | A | G | -0.034 | 0.9727 | -? | 0 | 0 | 0 | 1 |
| rs111999954 | 11 | 119832880 | A | G | -0.034 | 0.973 | -? | 0 | 0 | 0 | 1 |
| rs12788933 | 11 | 118140184 | T | C | 0.034 | 0.9731 | -+ | -60.1 | 1.874 | 1 | 0.171 |
| rs540442 | 11 | 117688898 | G | A | -0.034 | 0.9731 | -? | 0 | 0 | 0 | 1 |
| rs79524875 | 11 | 118970682 | T | C | 0.034 | 0.9731 | +? | 0 | 0 | 0 | 1 |
| rs73022058 | 11 | 118423672 | T | C | 0.033 | 0.9733 | +- | 0 | 0.018 | 1 | 0.8942 |
| rs7117314 | 11 | 117698762 | A | G | 0.033 | 0.9734 | +? | 0 | 0 | 0 | 1 |
| rs117935983 | 11 | 116529188 | G | T | -0.033 | 0.9735 | -? | 0 | 0 | 0 | 1 |
| rs4272790 | 11 | 120445354 | T | C | 0.033 | 0.9735 | +? | 0 | 0 | 0 | 1 |
| rs75764292 | 11 | 120575606 | T | A | 0.033 | 0.9736 | +? | 0 | 0 | 0 | 1 |
| rs77445023 | 11 | 119390332 | T | G | -0.033 | 0.9736 | -? | 0 | 0 | 0 | 1 |
| rs111707199 | 11 | 118276810 | G | A | 0.033 | 0.974 | -+ | -18 | 2.543 | 1 | 0.1108 |
| rs116943892 | 11 | 116978627 | G | A | 0.033 | 0.9741 | -+ | 73.5 | 11.335 | 1 | 0.0007607 |
| rs35652094 | 11 | 120039696 | A | G | 0.032 | 0.9741 | +? | 0 | 0 | 0 | 1 |
| rs61907562 | 11 | 116727592 | A | G | -0.032 | 0.9741 | -? | 0 | 0 | 0 | 1 |
| rs1941633 | 11 | 117966677 | A | G | 0.032 | 0.9744 | +- | 0 | 0.103 | 1 | 0.7478 |
| rs2276125 | 11 | 117965646 | A | G | 0.032 | 0.9744 | +- | 0 | 0.109 | 1 | 0.741 |
| rs505142 | 11 | 120098746 | A | T | -0.032 | 0.9745 | -? | 0 | 0 | 0 | 1 |
| rs10790216 | 11 | 117731927 | C | T | -0.032 | 0.9746 | -? | 0 | 0 | 0 | 1 |
| rs12796298 | 11 | 120855341 | G | A | -0.032 | 0.9747 | -+ | 0 | 0.152 | 1 | 0.697 |
| rs11217958 | 11 | 120544575 | T | C | 0.032 | 0.9748 | +? | 0 | 0 | 0 | 1 |
| rs10892063 | 11 | 116896155 | A | C | -0.031 | 0.9749 | -? | 0 | 0 | 0 | 1 |
| rs76870367 | 11 | 119243191 | T | C | 0.031 | 0.975 | +? | 0 | 0 | 0 | 1 |
| rs74674700 | 11 | 119839599 | T | C | 0.031 | 0.9753 | +? | 0 | 0 | 0 | 1 |
| rs4408323 | 11 | 119648458 | C | A | 0.031 | 0.9754 | +? | 0 | 0 | 0 | 1 |
| rs7110823 | 11 | 116441277 | G | C | 0.031 | 0.9754 | -+ | 0 | 0.298 | 1 | 0.5853 |
| rs73005236 | 11 | 116319634 | A | G | 0.031 | 0.9754 | +? | 0 | 0 | 0 | 1 |
| rs9787799 | 11 | 121100429 | G | A | 0.031 | 0.9754 | -+ | 0 | 0.021 | 1 | 0.8851 |
| rs9787800 | 11 | 121100648 | G | A | 0.031 | 0.9754 | -+ | 0 | 0.021 | 1 | 0.8851 |
| rs484646 | 11 | 116732856 | A | G | 0.031 | 0.9755 | +? | 0 | 0 | 0 | 1 |
| rs7109251 | 11 | 119616190 | A | G | 0.031 | 0.9755 | +? | 0 | 0 | 0 | 1 |
| rs10892053 | 11 | 116857003 | C | T | -0.031 | 0.9756 | -? | 0 | 0 | 0 | 1 |
| rs11216673 | 11 | 117879926 | G | A | -0.031 | 0.9756 | -? | 0 | 0 | 0 | 1 |
| rs7125115 | 11 | 118478330 | A | G | 0.031 | 0.9756 | +- | 0 | 0.089 | 1 | 0.7657 |
| rs76900966 | 11 | 119678909 | T | G | -0.03 | 0.9757 | -? | 0 | 0 | 0 | 1 |
| rs543876 | 11 | 119750487 | C | T | -0.03 | 0.9758 | -? | 0 | 0 | 0 | 1 |
| rs78301905 | 11 | 117076336 | C | T | -0.03 | 0.976 | -+ | 73.4 | 11.275 | 1 | 0.0007856 |
| rs79278568 | 11 | 117084410 | A | G | -0.03 | 0.976 | -+ | 73.4 | 11.275 | 1 | 0.0007856 |
| rs79997323 | 11 | 117083948 | G | A | -0.03 | 0.976 | -+ | 73.4 | 11.275 | 1 | 0.0007856 |
| rs652916 | 11 | 121094021 | T | C | 0.03 | 0.9761 | -+ | 0 | 0.021 | 1 | 0.8847 |
| rs10790177 | 11 | 117048417 | A | G | -0.03 | 0.9762 | -? | 0 | 0 | 0 | 1 |
| rs651825 | 11 | 118042713 | G | T | -0.03 | 0.9763 | +- | 0 | 0.394 | 1 | 0.5304 |
| rs10750156 | 11 | 119436709 | G | A | 0.03 | 0.9764 | +? | 0 | 0 | 0 | 1 |
| rs10892408 | 11 | 119436816 | G | A | 0.03 | 0.9764 | +? | 0 | 0 | 0 | 1 |

| | | | | | | | | | | | |
|---|---|---|---|---|---|---|---|---|---|---|---|
| rs7112584 | 11 | 118069240 | A | G | 0.03 | 0.9764 | -+ | 0 | 0.457 | 1 | 0.4989 |
| rs11216071 | 11 | 116515405 | C | G | -0.029 | 0.9766 | -? | 0 | 0 | 0 | 1 |
| rs3225 | 11 | 120100520 | C | G | -0.029 | 0.9766 | -? | 0 | 0 | 0 | 1 |
| rs10892392 | 11 | 119426508 | T | C | -0.029 | 0.9767 | -? | 0 | 0 | 0 | 1 |
| rs78913295 | 11 | 119847854 | T | C | -0.029 | 0.9767 | -? | 0 | 0 | 0 | 1 |
| rs590021 | 11 | 121092625 | T | C | 0.029 | 0.9768 | -+ | 0 | 0.021 | 1 | 0.8843 |
| rs10892155 | 11 | 117537378 | T | C | -0.029 | 0.9769 | -? | 0 | 0 | 0 | 1 |
| rs11606857 | 11 | 119615828 | T | A | 0.029 | 0.9769 | +? | 0 | 0 | 0 | 1 |
| rs117734523 | 11 | 119786488 | A | G | -0.029 | 0.9769 | -? | 0 | 0 | 0 | 1 |
| rs2735182 | 11 | 117012210 | A | T | 0.029 | 0.9769 | +? | 0 | 0 | 0 | 1 |
| rs3736120 | 11 | 117032125 | G | T | -0.029 | 0.977 | -? | 0 | 0 | 0 | 1 |
| rs117649997 | 11 | 119863317 | A | G | -0.029 | 0.9771 | -? | 0 | 0 | 0 | 1 |
| rs11217290 | 11 | 119320879 | A | G | 0.028 | 0.9773 | +? | 0 | 0 | 0 | 1 |
| rs12795599 | 11 | 120036812 | A | G | 0.028 | 0.9777 | +? | 0 | 0 | 0 | 1 |
| rs10892623 | 11 | 120636628 | G | A | 0.028 | 0.9779 | +? | 0 | 0 | 0 | 1 |
| rs34980489 | 11 | 119384277 | A | T | 0.028 | 0.978 | +? | 0 | 0 | 0 | 1 |
| rs55776558 | 11 | 116527521 | C | T | 0.027 | 0.9784 | +? | 0 | 0 | 0 | 1 |
| rs12279180 | 11 | 116604032 | C | T | -0.027 | 0.9786 | +- | -75 | 1.715 | 1 | 0.1904 |
| rs79136761 | 11 | 117311507 | A | G | -0.026 | 0.9789 | -? | 0 | 0 | 0 | 1 |
| rs11216400 | 11 | 117323834 | G | A | -0.026 | 0.979 | -? | 0 | 0 | 0 | 1 |
| rs11217357 | 11 | 119497757 | T | G | -0.026 | 0.979 | -? | 0 | 0 | 0 | 1 |
| rs11217358 | 11 | 119497806 | A | G | -0.026 | 0.979 | -? | 0 | 0 | 0 | 1 |
| rs601503 | 11 | 117581240 | C | G | 0.026 | 0.9791 | +? | 0 | 0 | 0 | 1 |
| rs641569 | 11 | 117581228 | T | G | 0.026 | 0.9791 | +? | 0 | 0 | 0 | 1 |
| rs693992 | 11 | 117581785 | A | C | 0.026 | 0.9791 | +? | 0 | 0 | 0 | 1 |
| rs112468256 | 11 | 118278479 | C | T | 0.026 | 0.9792 | -+ | -19.6 | 2.508 | 1 | 0.1133 |
| rs7928668 | 11 | 117799923 | T | C | 0.026 | 0.9792 | -+ | -140.4 | 1.248 | 1 | 0.264 |
| rs112970787 | 11 | 118212235 | C | G | -0.026 | 0.9793 | -+ | 8.1 | 3.266 | 1 | 0.07074 |
| rs17123461 | 11 | 119771532 | A | C | 0.026 | 0.9794 | +? | 0 | 0 | 0 | 1 |
| rs10790308 | 11 | 119425342 | A | G | 0.026 | 0.9795 | +? | 0 | 0 | 0 | 1 |
| rs10790309 | 11 | 119426055 | A | G | 0.026 | 0.9795 | +? | 0 | 0 | 0 | 1 |
| rs10892052 | 11 | 116834313 | G | A | 0.026 | 0.9795 | +? | 0 | 0 | 0 | 1 |
| rs10892391 | 11 | 119426104 | G | T | 0.026 | 0.9795 | +? | 0 | 0 | 0 | 1 |
| rs11216394 | 11 | 117315692 | G | A | 0.026 | 0.9795 | +? | 0 | 0 | 0 | 1 |
| rs1793137 | 11 | 118016068 | C | T | 0.026 | 0.9795 | ++ | 0 | 0 | 1 | 0.9956 |
| rs61506726 | 11 | 118831335 | G | A | -0.026 | 0.9795 | +- | 0 | 0.84 | 1 | 0.3595 |
| rs10892113 | 11 | 117324522 | T | C | -0.025 | 0.9797 | -? | 0 | 0 | 0 | 1 |
| rs17092732 | 11 | 117997374 | A | G | 0.025 | 0.9797 | +? | 0 | 0 | 0 | 1 |
| rs4277146 | 11 | 117010979 | G | A | 0.025 | 0.9797 | +? | 0 | 0 | 0 | 1 |
| rs59209096 | 11 | 119427370 | A | G | 0.025 | 0.9801 | +? | 0 | 0 | 0 | 1 |
| rs10892459 | 11 | 119653883 | G | C | 0.025 | 0.9803 | +? | 0 | 0 | 0 | 1 |
| rs1466052 | 11 | 119653781 | C | T | 0.025 | 0.9803 | +? | 0 | 0 | 0 | 1 |
| rs949296 | 11 | 120649396 | A | T | -0.025 | 0.9804 | -? | 0 | 0 | 0 | 1 |
| rs11216562 | 11 | 117686732 | T | C | 0.024 | 0.9807 | +? | 0 | 0 | 0 | 1 |
| rs17122170 | 11 | 118296668 | C | G | -0.024 | 0.9807 | +- | 0 | 0.285 | 1 | 0.5931 |
| rs6589791 | 11 | 119951213 | A | C | -0.024 | 0.9808 | -? | 0 | 0 | 0 | 1 |
| rs6589781 | 11 | 119727452 | T | A | -0.024 | 0.9809 | -? | 0 | 0 | 0 | 1 |
| rs4936359 | 11 | 116893724 | A | T | 0.024 | 0.981 | +? | 0 | 0 | 0 | 1 |
| rs4936360 | 11 | 116893769 | C | G | 0.024 | 0.981 | +? | 0 | 0 | 0 | 1 |
| rs4938325 | 11 | 116898479 | A | G | 0.024 | 0.981 | +? | 0 | 0 | 0 | 1 |
| rs61905709 | 11 | 116897834 | T | C | 0.024 | 0.981 | +? | 0 | 0 | 0 | 1 |
| rs7112191 | 11 | 116902056 | A | G | 0.024 | 0.981 | +? | 0 | 0 | 0 | 1 |
| rs7125788 | 11 | 116883396 | C | T | 0.024 | 0.981 | +? | 0 | 0 | 0 | 1 |
| rs171052 | 11 | 117081763 | G | C | 0.024 | 0.9811 | +? | 0 | 0 | 0 | 1 |
| rs191030218 | 11 | 117454782 | C | T | 0.024 | 0.9811 | +? | 0 | 0 | 0 | 1 |
| rs3863316 | 11 | 117802743 | G | C | 0.024 | 0.9812 | -+ | 21.2 | 3.807 | 1 | 0.05103 |

| | | | | | | | | | | | |
|---|---|---|---|---|---|---|---|---|---|---|---|
| rs17124590 | 11 | 120828210 | T | A | 0.023 | 0.9813 | +? | 0 | 0 | 0 | 1 |
| rs55751037 | 11 | 116456659 | T | C | -0.023 | 0.9814 | +- | 0 | 0.608 | 1 | 0.4356 |
| rs3132784 | 11 | 120588584 | C | G | -0.023 | 0.9815 | -? | 0 | 0 | 0 | 1 |
| rs111849979 | 11 | 117547323 | G | A | 0.023 | 0.9816 | +? | 0 | 0 | 0 | 1 |
| rs4582984 | 11 | 119499014 | C | A | 0.023 | 0.9817 | +? | 0 | 0 | 0 | 1 |
| rs664590 | 11 | 119964351 | T | G | 0.023 | 0.9817 | +? | 0 | 0 | 0 | 1 |
| rs11216813 | 11 | 118071836 | A | G | 0.023 | 0.9818 | -+ | 0 | 0.344 | 1 | 0.5575 |
| rs6589582 | 11 | 116829200 | A | C | -0.023 | 0.9818 | -? | 0 | 0 | 0 | 1 |
| rs9787912 | 11 | 121100910 | A | G | 0.023 | 0.9818 | -+ | 0 | 0.022 | 1 | 0.8815 |
| rs489436 | 11 | 121090812 | A | C | 0.023 | 0.9819 | -+ | 0 | 0.022 | 1 | 0.8814 |
| rs543954 | 11 | 121090892 | A | G | 0.023 | 0.9819 | -+ | 0 | 0.022 | 1 | 0.8814 |
| rs588550 | 11 | 121091188 | A | G | 0.023 | 0.9819 | -+ | 0 | 0.022 | 1 | 0.8814 |
| rs618739 | 11 | 121090822 | A | G | 0.023 | 0.9819 | -+ | 0 | 0.022 | 1 | 0.8814 |
| rs10892215 | 11 | 117978103 | T | C | -0.023 | 0.982 | -? | 0 | 0 | 0 | 1 |
| rs11216765 | 11 | 117978247 | T | C | -0.023 | 0.982 | -? | 0 | 0 | 0 | 1 |
| rs4519102 | 11 | 119493539 | C | A | 0.023 | 0.982 | +? | 0 | 0 | 0 | 1 |
| rs4590884 | 11 | 120632138 | T | C | -0.023 | 0.982 | -? | 0 | 0 | 0 | 1 |
| rs10892533 | 11 | 119959410 | T | C | 0.022 | 0.9822 | +? | 0 | 0 | 0 | 1 |
| rs141512500 | 11 | 117674061 | G | A | 0.022 | 0.9822 | +? | 0 | 0 | 0 | 1 |
| rs79777938 | 11 | 116980147 | C | G | 0.022 | 0.9824 | -+ | 73.6 | 11.375 | 1 | 0.0007444 |
| rs10892317 | 11 | 118832035 | C | T | 0.022 | 0.9825 | +- | 0 | 0.885 | 1 | 0.347 |
| rs10790323 | 11 | 119498128 | A | G | -0.022 | 0.9827 | -? | 0 | 0 | 0 | 1 |
| rs117140575 | 11 | 116993291 | C | T | 0.022 | 0.9827 | -+ | 73.6 | 11.377 | 1 | 0.0007438 |
| rs12276427 | 11 | 119951111 | A | G | -0.022 | 0.9827 | -? | 0 | 0 | 0 | 1 |
| rs12417625 | 11 | 120072443 | G | A | -0.022 | 0.9827 | -? | 0 | 0 | 0 | 1 |
| rs60215110 | 11 | 116987767 | A | G | 0.022 | 0.9827 | -+ | 73.6 | 11.377 | 1 | 0.0007438 |
| rs79117087 | 11 | 116992739 | C | A | 0.022 | 0.9827 | -+ | 73.6 | 11.377 | 1 | 0.0007438 |
| rs7928437 | 11 | 119664304 | G | T | 0.022 | 0.9827 | +? | 0 | 0 | 0 | 1 |
| rs640621 | 11 | 116726769 | C | G | -0.022 | 0.9828 | -? | 0 | 0 | 0 | 1 |
| rs11216161 | 11 | 116721201 | T | C | 0.021 | 0.983 | +? | 0 | 0 | 0 | 1 |
| rs45618531 | 11 | 118087765 | T | C | 0.021 | 0.983 | -+ | -39.5 | 2.15 | 1 | 0.1426 |
| rs73005230 | 11 | 116316628 | A | G | 0.021 | 0.9831 | +? | 0 | 0 | 0 | 1 |
| rs10892509 | 11 | 119864323 | C | A | -0.021 | 0.9834 | -? | 0 | 0 | 0 | 1 |
| rs1815774 | 11 | 117731439 | C | G | -0.021 | 0.9834 | -? | 0 | 0 | 0 | 1 |
| rs57151589 | 11 | 120769382 | T | A | -0.021 | 0.9834 | -? | 0 | 0 | 0 | 1 |
| rs7396061 | 11 | 116750874 | A | C | -0.021 | 0.9835 | -? | 0 | 0 | 0 | 1 |
| rs10892461 | 11 | 119654807 | C | T | -0.021 | 0.9836 | -? | 0 | 0 | 0 | 1 |
| rs7945704 | 11 | 117235368 | A | G | -0.021 | 0.9836 | -? | 0 | 0 | 0 | 1 |
| rs527833 | 11 | 119704194 | A | G | -0.02 | 0.9837 | -? | 0 | 0 | 0 | 1 |
| rs650162 | 11 | 117574397 | T | C | 0.02 | 0.9839 | +? | 0 | 0 | 0 | 1 |
| rs140727315 | 11 | 117055893 | C | G | -0.02 | 0.984 | -+ | 73.3 | 11.236 | 1 | 0.0008021 |
| rs184936191 | 11 | 117061994 | T | A | -0.02 | 0.984 | -+ | 73.3 | 11.236 | 1 | 0.0008021 |
| rs189196849 | 11 | 117052757 | T | G | -0.02 | 0.984 | -+ | 73.3 | 11.236 | 1 | 0.0008021 |
| rs477759 | 11 | 119696165 | T | C | -0.02 | 0.984 | -+ | 0 | 0.123 | 1 | 0.7261 |
| rs59978576 | 11 | 117054029 | G | C | -0.02 | 0.984 | -+ | 73.3 | 11.236 | 1 | 0.0008021 |
| rs61268522 | 11 | 117055400 | C | A | -0.02 | 0.984 | -+ | 73.3 | 11.236 | 1 | 0.0008021 |
| rs7109733 | 11 | 119958632 | C | T | -0.02 | 0.984 | -? | 0 | 0 | 0 | 1 |
| rs74967901 | 11 | 117072970 | A | G | -0.02 | 0.984 | -+ | 73.3 | 11.236 | 1 | 0.0008021 |
| rs75525407 | 11 | 117041911 | T | C | -0.02 | 0.984 | -+ | 73.3 | 11.236 | 1 | 0.0008021 |
| rs79043792 | 11 | 117065271 | C | G | -0.02 | 0.984 | -+ | 73.3 | 11.236 | 1 | 0.0008021 |
| rs10790393 | 11 | 120548054 | T | C | 0.02 | 0.9842 | +? | 0 | 0 | 0 | 1 |
| rs11218211 | 11 | 121098347 | C | T | -0.02 | 0.9843 | -+ | 0 | 0.03 | 1 | 0.8623 |
| rs664971 | 11 | 117070857 | G | A | 0.02 | 0.9843 | +? | 0 | 0 | 0 | 1 |
| rs11218101 | 11 | 120867855 | A | G | -0.02 | 0.9844 | -? | 0 | 0 | 0 | 1 |
| rs7121898 | 11 | 116796846 | A | T | -0.019 | 0.9846 | -? | 0 | 0 | 0 | 1 |
| rs1295019 | 11 | 120098044 | A | C | -0.019 | 0.9847 | -? | 0 | 0 | 0 | 1 |

| | | | | | | | | | | | |
|---|---|---|---|---|---|---|---|---|---|---|---|
| rs1789792 | 11 | 119696100 | G | T | 0.019 | 0.9847 | +? | 0 | 0 | 0 | 1 |
| rs2282493 | 11 | 117873229 | T | C | -0.019 | 0.9847 | -? | 0 | 0 | 0 | 1 |
| rs61903426 | 11 | 116968268 | T | C | 0.019 | 0.9847 | +? | 0 | 0 | 0 | 1 |
| rs12363785 | 11 | 119965450 | A | G | -0.019 | 0.9848 | -? | 0 | 0 | 0 | 1 |
| rs947969 | 11 | 117779612 | A | G | -0.019 | 0.9848 | -? | 0 | 0 | 0 | 1 |
| rs6589792 | 11 | 119951214 | G | C | -0.019 | 0.985 | -? | 0 | 0 | 0 | 1 |
| rs7103360 | 11 | 119650105 | G | A | 0.019 | 0.985 | +? | 0 | 0 | 0 | 1 |
| rs73005128 | 11 | 121151034 | T | C | 0.019 | 0.9852 | +? | 0 | 0 | 0 | 1 |
| rs1815771 | 11 | 120656479 | A | G | 0.018 | 0.9853 | +? | 0 | 0 | 0 | 1 |
| rs2008908 | 11 | 117019010 | C | T | 0.018 | 0.9853 | +? | 0 | 0 | 0 | 1 |
| rs12787356 | 11 | 116477692 | A | G | 0.018 | 0.9854 | +? | 0 | 0 | 0 | 1 |
| rs504976 | 11 | 120000233 | C | A | 0.018 | 0.9854 | +? | 0 | 0 | 0 | 1 |
| rs80326995 | 11 | 120643050 | A | G | 0.018 | 0.9854 | +? | 0 | 0 | 0 | 1 |
| rs118000863 | 11 | 119935980 | A | G | -0.018 | 0.9855 | -? | 0 | 0 | 0 | 1 |
| rs524792 | 11 | 120830413 | C | T | -0.018 | 0.9856 | -? | 0 | 0 | 0 | 1 |
| rs11600755 | 11 | 120639087 | T | C | -0.018 | 0.9857 | -? | 0 | 0 | 0 | 1 |
| rs4310631 | 11 | 120440360 | G | C | 0.018 | 0.9858 | +? | 0 | 0 | 0 | 1 |
| rs61905712 | 11 | 116909293 | A | G | 0.018 | 0.9858 | +? | 0 | 0 | 0 | 1 |
| rs61905713 | 11 | 116909429 | T | C | 0.018 | 0.9858 | +? | 0 | 0 | 0 | 1 |
| rs10892151 | 11 | 117531731 | T | C | 0.018 | 0.9859 | -+ | -174.1 | 1.094 | 1 | 0.2955 |
| rs12289120 | 11 | 118066609 | T | C | 0.018 | 0.986 | -+ | 0 | 0.02 | 1 | 0.8869 |
| rs12574454 | 11 | 120767546 | G | A | -0.017 | 0.9861 | -? | 0 | 0 | 0 | 1 |
| rs73008656 | 11 | 119717705 | G | T | -0.017 | 0.9862 | -? | 0 | 0 | 0 | 1 |
| rs11600544 | 11 | 118076954 | C | G | 0.017 | 0.9866 | +? | 0 | 0 | 0 | 1 |
| rs1805 | 11 | 118076069 | T | G | 0.017 | 0.9866 | +? | 0 | 0 | 0 | 1 |
| rs17121871 | 11 | 118073344 | T | C | 0.017 | 0.9867 | +? | 0 | 0 | 0 | 1 |
| rs3915646 | 11 | 119956652 | C | T | -0.017 | 0.9868 | -? | 0 | 0 | 0 | 1 |
| rs634416 | 11 | 117579815 | T | C | 0.016 | 0.9869 | +? | 0 | 0 | 0 | 1 |
| rs670972 | 11 | 117579270 | A | G | 0.016 | 0.9869 | +? | 0 | 0 | 0 | 1 |
| rs879602 | 11 | 120656572 | C | T | 0.016 | 0.987 | +? | 0 | 0 | 0 | 1 |
| rs6589627 | 11 | 117730979 | C | T | 0.016 | 0.9873 | +? | 0 | 0 | 0 | 1 |
| rs6589628 | 11 | 117730983 | A | G | 0.016 | 0.9873 | +? | 0 | 0 | 0 | 1 |
| rs6589629 | 11 | 117731070 | C | G | 0.016 | 0.9873 | +? | 0 | 0 | 0 | 1 |
| rs6589630 | 11 | 117731085 | C | T | 0.016 | 0.9873 | +? | 0 | 0 | 0 | 1 |
| rs7106442 | 11 | 120770004 | T | C | -0.016 | 0.9874 | -? | 0 | 0 | 0 | 1 |
| rs61905705 | 11 | 116883776 | T | A | 0.016 | 0.9875 | +? | 0 | 0 | 0 | 1 |
| rs61905706 | 11 | 116883777 | T | G | 0.016 | 0.9875 | +? | 0 | 0 | 0 | 1 |
| rs10892407 | 11 | 119436098 | A | C | 0.016 | 0.9876 | +? | 0 | 0 | 0 | 1 |
| rs9645662 | 11 | 117974832 | A | G | 0.016 | 0.9876 | +? | 0 | 0 | 0 | 1 |
| rs10892599 | 11 | 120545811 | T | G | 0.015 | 0.9877 | +? | 0 | 0 | 0 | 1 |
| rs142742415 | 11 | 117000117 | A | G | 0.015 | 0.9877 | -+ | 73.5 | 11.303 | 1 | 0.0007738 |
| rs77983828 | 11 | 117001875 | C | T | 0.015 | 0.9877 | -+ | 73.5 | 11.303 | 1 | 0.0007738 |
| rs10892386 | 11 | 119413090 | T | C | -0.015 | 0.9878 | -? | 0 | 0 | 0 | 1 |
| rs542580 | 11 | 120827067 | T | C | 0.015 | 0.9878 | +? | 0 | 0 | 0 | 1 |
| rs11215903 | 11 | 116237754 | G | T | 0.015 | 0.988 | +? | 0 | 0 | 0 | 1 |
| rs57081718 | 11 | 120144941 | A | G | -0.015 | 0.9881 | -? | 0 | 0 | 0 | 1 |
| rs73014416 | 11 | 117535760 | A | G | -0.015 | 0.9882 | -? | 0 | 0 | 0 | 1 |
| rs533556 | 11 | 116741572 | A | C | -0.015 | 0.9884 | -? | 0 | 0 | 0 | 1 |
| rs10892538 | 11 | 119977712 | A | G | 0.014 | 0.9885 | +? | 0 | 0 | 0 | 1 |
| rs1269329 | 11 | 116755041 | T | C | 0.014 | 0.9885 | +? | 0 | 0 | 0 | 1 |
| rs73020452 | 11 | 117759895 | C | T | 0.014 | 0.9886 | +? | 0 | 0 | 0 | 1 |
| rs35968460 | 11 | 119727199 | T | C | 0.014 | 0.9888 | +? | 0 | 0 | 0 | 1 |
| rs609663 | 11 | 117201749 | G | C | -0.014 | 0.9888 | -? | 0 | 0 | 0 | 1 |
| rs11820008 | 11 | 120869570 | T | C | 0.014 | 0.9889 | +? | 0 | 0 | 0 | 1 |
| rs10502222 | 11 | 116833730 | C | T | 0.014 | 0.989 | +? | 0 | 0 | 0 | 1 |
| rs12363187 | 11 | 120544070 | T | C | -0.014 | 0.989 | -? | 0 | 0 | 0 | 1 |

| | | | | | | | | | | | |
|---|---|---|---|---|---|---|---|---|---|---|---|
| rs2306474 | 11 | 117099025 | G | A | -0.014 | 0.989 | -+ | 73.2 | 11.212 | 1 | 0.0008126 |
| rs678957 | 11 | 118050855 | T | C | -0.014 | 0.989 | +- | 0 | 0.437 | 1 | 0.5086 |
| rs61898747 | 11 | 119445010 | G | T | -0.014 | 0.9891 | -? | 0 | 0 | 0 | 1 |
| rs34178922 | 11 | 120970408 | A | G | -0.014 | 0.9892 | -? | 0 | 0 | 0 | 1 |
| rs7927088 | 11 | 117401470 | G | A | -0.014 | 0.9892 | -? | 0 | 0 | 0 | 1 |
| rs10892460 | 11 | 119654058 | C | T | 0.013 | 0.9894 | +? | 0 | 0 | 0 | 1 |
| rs236910 | 11 | 117081974 | G | A | -0.013 | 0.9896 | -? | 0 | 0 | 0 | 1 |
| rs10892079 | 11 | 117016824 | T | G | 0.013 | 0.9898 | +? | 0 | 0 | 0 | 1 |
| rs7939548 | 11 | 118212979 | A | G | -0.013 | 0.9898 | -+ | 8.2 | 3.267 | 1 | 0.0707 |
| rs11218126 | 11 | 120921866 | T | C | 0.013 | 0.9899 | +? | 0 | 0 | 0 | 1 |
| rs4391855 | 11 | 116350751 | A | G | 0.013 | 0.9899 | +? | 0 | 0 | 0 | 1 |
| rs73018334 | 11 | 117313948 | A | C | 0.013 | 0.9899 | +? | 0 | 0 | 0 | 1 |
| rs11216819 | 11 | 118084980 | G | A | -0.013 | 0.99 | -? | 0 | 0 | 0 | 1 |
| rs477337 | 11 | 120000885 | T | C | -0.012 | 0.9904 | -? | 0 | 0 | 0 | 1 |
| rs477361 | 11 | 120000873 | T | G | -0.012 | 0.9904 | -? | 0 | 0 | 0 | 1 |
| rs4936571 | 11 | 120872094 | G | A | -0.012 | 0.9904 | -? | 0 | 0 | 0 | 1 |
| rs61898283 | 11 | 120091015 | G | A | -0.012 | 0.9906 | -? | 0 | 0 | 0 | 1 |
| rs61898284 | 11 | 120091077 | A | G | -0.012 | 0.9906 | -? | 0 | 0 | 0 | 1 |
| rs61898287 | 11 | 120092774 | T | A | -0.012 | 0.9906 | -? | 0 | 0 | 0 | 1 |
| rs61903779 | 11 | 117700201 | G | A | -0.012 | 0.9906 | -? | 0 | 0 | 0 | 1 |
| rs4936477 | 11 | 119411610 | G | A | -0.012 | 0.9907 | -? | 0 | 0 | 0 | 1 |
| rs948107 | 11 | 117731643 | A | C | -0.012 | 0.9907 | -? | 0 | 0 | 0 | 1 |
| rs112583389 | 11 | 118278998 | G | T | 0.012 | 0.9908 | -+ | -20.2 | 2.495 | 1 | 0.1142 |
| rs575549 | 11 | 116315743 | A | G | -0.012 | 0.9908 | -? | 0 | 0 | 0 | 1 |
| rs1787027 | 11 | 119967661 | C | A | -0.011 | 0.9909 | -? | 0 | 0 | 0 | 1 |
| rs10502229 | 11 | 117979970 | T | G | -0.011 | 0.9911 | -? | 0 | 0 | 0 | 1 |
| rs7925677 | 11 | 119720281 | A | G | 0.011 | 0.9911 | +? | 0 | 0 | 0 | 1 |
| rs964104 | 11 | 119753972 | A | G | 0.011 | 0.9911 | +? | 0 | 0 | 0 | 1 |
| rs868066 | 11 | 116408318 | A | G | -0.011 | 0.9912 | -? | 0 | 0 | 0 | 1 |
| rs660443 | 11 | 116760693 | T | C | -0.011 | 0.9913 | -? | 0 | 0 | 0 | 1 |
| rs117303996 | 11 | 117061937 | A | G | -0.011 | 0.9916 | -+ | 73.2 | 11.2 | 1 | 0.0008181 |
| rs150881386 | 11 | 117034320 | T | C | -0.011 | 0.9916 | -+ | 73.2 | 11.2 | 1 | 0.0008181 |
| rs1944505 | 11 | 119838883 | A | G | -0.011 | 0.9916 | -? | 0 | 0 | 0 | 1 |
| rs2508560 | 11 | 117575629 | A | G | 0.01 | 0.9917 | +? | 0 | 0 | 0 | 1 |
| rs585718 | 11 | 117575841 | T | C | 0.01 | 0.9917 | +? | 0 | 0 | 0 | 1 |
| rs10047459 | 11 | 116721826 | C | T | -0.01 | 0.9918 | -? | 0 | 0 | 0 | 1 |
| rs11216818 | 11 | 118080268 | T | C | 0.01 | 0.9918 | -+ | 0 | 0.333 | 1 | 0.5639 |
| rs4306318 | 11 | 118080520 | T | C | 0.01 | 0.9918 | -+ | 0 | 0.333 | 1 | 0.5639 |
| rs35662046 | 11 | 119654326 | C | T | 0.01 | 0.9922 | +? | 0 | 0 | 0 | 1 |
| rs605374 | 11 | 120005633 | T | G | 0.01 | 0.9922 | +? | 0 | 0 | 0 | 1 |
| rs6589590 | 11 | 116936081 | A | C | -0.01 | 0.9922 | -? | 0 | 0 | 0 | 1 |
| rs664082 | 11 | 117071070 | C | G | 0.01 | 0.9922 | +? | 0 | 0 | 0 | 1 |
| rs2444242 | 11 | 120082486 | A | G | -0.01 | 0.9923 | -? | 0 | 0 | 0 | 1 |
| rs73009990 | 11 | 117882065 | A | G | 0.01 | 0.9923 | +? | 0 | 0 | 0 | 1 |
| rs7935087 | 11 | 120542998 | G | A | -0.009 | 0.9925 | -? | 0 | 0 | 0 | 1 |
| rs10892550 | 11 | 120078028 | T | G | 0.009 | 0.9926 | +? | 0 | 0 | 0 | 1 |
| rs2155194 | 11 | 117781435 | A | G | 0.009 | 0.9926 | +? | 0 | 0 | 0 | 1 |
| rs11216615 | 11 | 117777953 | G | A | -0.009 | 0.9927 | -? | 0 | 0 | 0 | 1 |
| rs17121879 | 11 | 118080133 | G | T | 0.009 | 0.9928 | +? | 0 | 0 | 0 | 1 |
| rs17121881 | 11 | 118081345 | A | T | 0.009 | 0.9928 | +? | 0 | 0 | 0 | 1 |
| rs17121882 | 11 | 118081468 | C | T | 0.009 | 0.9928 | +? | 0 | 0 | 0 | 1 |
| rs4938489 | 11 | 118079867 | G | T | 0.009 | 0.9928 | +? | 0 | 0 | 0 | 1 |
| rs4938490 | 11 | 118080977 | G | A | 0.009 | 0.9928 | +? | 0 | 0 | 0 | 1 |
| rs7945998 | 11 | 119414914 | T | C | 0.009 | 0.9928 | +? | 0 | 0 | 0 | 1 |
| rs11606569 | 11 | 119654208 | C | T | 0.009 | 0.993 | +? | 0 | 0 | 0 | 1 |
| rs12803321 | 11 | 118480115 | C | G | 0.009 | 0.993 | +- | 0 | 0.16 | 1 | 0.6892 |

| | | | | | | | | | | | |
|---|---|---|---|---|---|---|---|---|---|---|---|
| rs118143963 | 11 | 116990523 | T | A | 0.009 | 0.9931 | -+ | 73.3 | 11.224 | 1 | 0.0008077 |
| rs186927602 | 11 | 116989133 | A | G | 0.009 | 0.9931 | -+ | 73.3 | 11.224 | 1 | 0.0008077 |
| rs61899013 | 11 | 119759046 | G | T | 0.009 | 0.9931 | +? | 0 | 0 | 0 | 1 |
| rs4938817 | 11 | 120447136 | C | A | 0.009 | 0.9932 | +? | 0 | 0 | 0 | 1 |
| rs2846103 | 11 | 120652208 | C | T | 0.008 | 0.9933 | +? | 0 | 0 | 0 | 1 |
| rs657107 | 11 | 119865509 | T | C | -0.008 | 0.9933 | -? | 0 | 0 | 0 | 1 |
| rs617948 | 11 | 118981025 | A | G | 0.008 | 0.9935 | +? | 0 | 0 | 0 | 1 |
| rs674637 | 11 | 120025243 | G | A | -0.008 | 0.9935 | -? | 0 | 0 | 0 | 1 |
| rs473219 | 11 | 119697182 | C | T | 0.008 | 0.9936 | +? | 0 | 0 | 0 | 1 |
| rs530792 | 11 | 119983799 | T | C | -0.008 | 0.9936 | -? | 0 | 0 | 0 | 1 |
| rs10892072 | 11 | 116960313 | G | A | -0.008 | 0.9938 | -? | 0 | 0 | 0 | 1 |
| rs1944153 | 11 | 116532221 | C | T | 0.008 | 0.9939 | +? | 0 | 0 | 0 | 1 |
| rs2155216 | 11 | 116531739 | T | C | 0.008 | 0.9939 | +? | 0 | 0 | 0 | 1 |
| rs2186670 | 11 | 116531950 | T | C | 0.008 | 0.9939 | +? | 0 | 0 | 0 | 1 |
| rs61900956 | 11 | 120537660 | G | A | -0.008 | 0.9939 | -? | 0 | 0 | 0 | 1 |
| rs11217501 | 11 | 119652686 | C | T | 0.008 | 0.994 | +? | 0 | 0 | 0 | 1 |
| rs12295127 | 11 | 117552758 | A | G | -0.008 | 0.994 | -? | 0 | 0 | 0 | 1 |
| rs10790304 | 11 | 119420599 | T | G | 0.007 | 0.9941 | +? | 0 | 0 | 0 | 1 |
| rs10790305 | 11 | 119420615 | A | G | 0.007 | 0.9941 | +? | 0 | 0 | 0 | 1 |
| rs10892390 | 11 | 119420468 | T | C | 0.007 | 0.9941 | +? | 0 | 0 | 0 | 1 |
| rs4936365 | 11 | 117016997 | C | T | -0.007 | 0.9941 | -? | 0 | 0 | 0 | 1 |
| rs61898288 | 11 | 120093495 | A | C | 0.007 | 0.9943 | +? | 0 | 0 | 0 | 1 |
| rs10892535 | 11 | 119967718 | T | C | -0.007 | 0.9945 | -? | 0 | 0 | 0 | 1 |
| rs12802344 | 11 | 119975789 | G | T | -0.007 | 0.9945 | -? | 0 | 0 | 0 | 1 |
| rs7120706 | 11 | 116830087 | T | C | -0.007 | 0.9945 | -? | 0 | 0 | 0 | 1 |
| rs7120963 | 11 | 116830261 | T | C | -0.007 | 0.9945 | -? | 0 | 0 | 0 | 1 |
| rs881123 | 11 | 117973047 | T | C | -0.007 | 0.9945 | +- | 0 | 0.172 | 1 | 0.678 |
| rs10892048 | 11 | 116814596 | A | G | 0.006 | 0.9949 | +? | 0 | 0 | 0 | 1 |
| rs10892537 | 11 | 119977230 | T | C | -0.006 | 0.9949 | -? | 0 | 0 | 0 | 1 |
| rs75919952 | 11 | 116667545 | T | C | -0.006 | 0.9949 | +- | 0 | 0.125 | 1 | 0.7239 |
| rs5014623 | 11 | 117731191 | C | T | 0.006 | 0.995 | +? | 0 | 0 | 0 | 1 |
| rs60928822 | 11 | 117032641 | C | T | 0.006 | 0.995 | -+ | 73.1 | 11.135 | 1 | 0.000847 |
| rs7934805 | 11 | 120542648 | T | C | 0.006 | 0.9951 | +? | 0 | 0 | 0 | 1 |
| rs12286021 | 11 | 118080685 | C | T | -0.006 | 0.9952 | -+ | 0 | 0.365 | 1 | 0.5455 |
| rs75173807 | 11 | 116511766 | A | G | 0.006 | 0.9952 | +? | 0 | 0 | 0 | 1 |
| rs10790324 | 11 | 119498477 | A | G | -0.006 | 0.9954 | -? | 0 | 0 | 0 | 1 |
| rs7113932 | 11 | 116985247 | T | G | 0.006 | 0.9955 | +? | 0 | 0 | 0 | 1 |
| rs476399 | 11 | 116733728 | T | C | -0.006 | 0.9956 | -? | 0 | 0 | 0 | 1 |
| rs472446 | 11 | 119785065 | G | A | 0.005 | 0.9958 | +? | 0 | 0 | 0 | 1 |
| rs112111936 | 11 | 117045431 | G | T | -0.005 | 0.9959 | -? | 0 | 0 | 0 | 1 |
| rs475189 | 11 | 121093766 | T | C | 0.005 | 0.9959 | -+ | 0 | 0.025 | 1 | 0.8735 |
| rs4938331 | 11 | 116930682 | C | A | -0.005 | 0.9959 | -? | 0 | 0 | 0 | 1 |
| rs571429 | 11 | 121074575 | A | C | -0.005 | 0.9959 | -+ | 0 | 0.027 | 1 | 0.8688 |
| rs11217502 | 11 | 119652763 | A | G | -0.005 | 0.996 | -? | 0 | 0 | 0 | 1 |
| rs76553074 | 11 | 117084070 | C | T | 0.005 | 0.996 | -+ | 73.1 | 11.14 | 1 | 0.0008447 |
| rs117146251 | 11 | 117005207 | G | A | 0.005 | 0.9963 | -+ | 73.6 | 11.344 | 1 | 0.0007567 |
| rs150672190 | 11 | 117004023 | T | C | 0.005 | 0.9963 | -+ | 73.6 | 11.344 | 1 | 0.0007567 |
| rs12360673 | 11 | 120544237 | G | A | -0.005 | 0.9964 | -? | 0 | 0 | 0 | 1 |
| rs4331115 | 11 | 119650848 | T | C | 0.004 | 0.9966 | +? | 0 | 0 | 0 | 1 |
| rs10892674 | 11 | 121002972 | A | G | -0.004 | 0.9967 | +- | 0 | 0.826 | 1 | 0.3635 |
| rs11218153 | 11 | 121003224 | A | G | -0.004 | 0.9967 | +- | 0 | 0.826 | 1 | 0.3635 |
| rs503559 | 11 | 119986167 | T | C | -0.004 | 0.9967 | -? | 0 | 0 | 0 | 1 |
| rs7105712 | 11 | 121003358 | C | T | -0.004 | 0.9967 | +- | 0 | 0.826 | 1 | 0.3635 |
| rs76188167 | 11 | 116471494 | A | G | -0.004 | 0.9967 | -? | 0 | 0 | 0 | 1 |
| rs930430 | 11 | 116197720 | A | G | 0.004 | 0.9967 | +? | 0 | 0 | 0 | 1 |
| rs11216750 | 11 | 117972543 | T | A | -0.004 | 0.9972 | +- | 0 | 0.194 | 1 | 0.6599 |

| | | | | | | | | | | | |
|---|---|---|---|---|---|---|---|---|---|---|---|
| rs45496391 | 11 | 118477226 | C | A | 0.003 | 0.9972 | -+ | 0 | 0.079 | 1 | 0.7793 |
| rs10892415 | 11 | 119467192 | T | A | 0.003 | 0.9973 | +? | 0 | 0 | 0 | 1 |
| rs61562228 | 11 | 117053686 | A | G | 0.003 | 0.9973 | -+ | 73.6 | 11.349 | 1 | 0.0007547 |
| rs76638771 | 11 | 117552480 | C | T | -0.003 | 0.9973 | -? | 0 | 0 | 0 | 1 |
| rs61899019 | 11 | 119774394 | A | G | -0.003 | 0.9974 | -? | 0 | 0 | 0 | 1 |
| rs10790403 | 11 | 120712099 | C | T | 0.003 | 0.9976 | +? | 0 | 0 | 0 | 1 |
| rs10892448 | 11 | 119624823 | A | C | 0.003 | 0.9976 | +? | 0 | 0 | 0 | 1 |
| rs12791393 | 11 | 120058902 | T | C | 0.003 | 0.9976 | +? | 0 | 0 | 0 | 1 |
| rs1446105 | 11 | 116798544 | C | T | -0.003 | 0.9977 | -? | 0 | 0 | 0 | 1 |
| rs5014622 | 11 | 117731263 | G | A | -0.003 | 0.9977 | -? | 0 | 0 | 0 | 1 |
| rs117469840 | 11 | 117190901 | T | C | -0.003 | 0.9978 | -? | 0 | 0 | 0 | 1 |
| rs36015759 | 11 | 119216279 | A | G | 0.003 | 0.9978 | +? | 0 | 0 | 0 | 1 |
| rs79293063 | 11 | 119678083 | A | G | -0.003 | 0.9979 | -? | 0 | 0 | 0 | 1 |
| rs7951095 | 11 | 117011440 | C | A | -0.003 | 0.9979 | -? | 0 | 0 | 0 | 1 |
| rs10892534 | 11 | 119967716 | A | C | -0.003 | 0.998 | -? | 0 | 0 | 0 | 1 |
| rs117842423 | 11 | 117039208 | A | G | 0.002 | 0.998 | -+ | 73.6 | 11.353 | 1 | 0.0007534 |
| rs144762883 | 11 | 117035622 | T | C | 0.002 | 0.998 | -+ | 73.6 | 11.353 | 1 | 0.0007534 |
| rs580174 | 11 | 120100867 | G | A | -0.002 | 0.9981 | -? | 0 | 0 | 0 | 1 |
| rs59225422 | 11 | 116999303 | G | C | 0.002 | 0.9981 | -+ | 73.1 | 11.15 | 1 | 0.0008402 |
| rs59427601 | 11 | 119427371 | A | G | 0.002 | 0.9981 | +? | 0 | 0 | 0 | 1 |
| rs71484150 | 11 | 119727707 | G | T | -0.002 | 0.9981 | -? | 0 | 0 | 0 | 1 |
| rs9655 | 11 | 120100597 | G | A | -0.002 | 0.9981 | -? | 0 | 0 | 0 | 1 |
| rs7113123 | 11 | 119727494 | T | C | -0.002 | 0.9982 | -? | 0 | 0 | 0 | 1 |
| rs11822288 | 11 | 117283939 | A | G | -0.002 | 0.9983 | -? | 0 | 0 | 0 | 1 |
| rs4938344 | 11 | 117003749 | G | A | -0.002 | 0.9983 | -? | 0 | 0 | 0 | 1 |
| rs514557 | 11 | 121090430 | T | C | -0.002 | 0.9983 | -+ | 0 | 0.027 | 1 | 0.8702 |
| rs61898278 | 11 | 120085537 | G | A | 0.002 | 0.9983 | +? | 0 | 0 | 0 | 1 |
| rs61898279 | 11 | 120086365 | G | A | 0.002 | 0.9983 | +? | 0 | 0 | 0 | 1 |
| rs61898282 | 11 | 120090861 | T | C | -0.002 | 0.9983 | -? | 0 | 0 | 0 | 1 |
| rs6589790 | 11 | 119951200 | T | C | 0.002 | 0.9984 | +? | 0 | 0 | 0 | 1 |
| rs685710 | 11 | 121091284 | C | A | 0.002 | 0.9984 | -+ | 0 | 0.026 | 1 | 0.8721 |
| rs4936572 | 11 | 120872214 | A | G | -0.002 | 0.9985 | -? | 0 | 0 | 0 | 1 |
| rs585849 | 11 | 117069061 | A | G | -0.002 | 0.9985 | -? | 0 | 0 | 0 | 1 |
| rs613004 | 11 | 119967556 | T | C | -0.002 | 0.9985 | -? | 0 | 0 | 0 | 1 |
| rs1962593 | 11 | 116426097 | T | C | -0.002 | 0.9986 | -+ | 0 | 0.197 | 1 | 0.6572 |
| rs1945010 | 11 | 120617504 | T | C | 0.002 | 0.9987 | +? | 0 | 0 | 0 | 1 |
| rs10790221 | 11 | 117800326 | G | A | 0.002 | 0.9988 | -+ | -136.1 | 1.271 | 1 | 0.2596 |
| rs11604677 | 11 | 117800301 | A | G | 0.002 | 0.9988 | -+ | -136.1 | 1.271 | 1 | 0.2596 |
| rs56303423 | 11 | 119639182 | A | G | 0.002 | 0.9988 | +? | 0 | 0 | 0 | 1 |
| rs11216889 | 11 | 118421512 | A | G | 0.001 | 0.999 | +- | 0 | 0.012 | 1 | 0.9123 |
| rs4623916 | 11 | 119566825 | T | A | 0.001 | 0.999 | +? | 0 | 0 | 0 | 1 |
| rs4938332 | 11 | 116930798 | A | G | 0.001 | 0.999 | +? | 0 | 0 | 0 | 1 |
| rs4938334 | 11 | 116931042 | G | C | 0.001 | 0.999 | +? | 0 | 0 | 0 | 1 |
| rs57315752 | 11 | 117055485 | G | A | 0.001 | 0.9991 | -+ | 73.1 | 11.155 | 1 | 0.000838 |
| rs7116091 | 11 | 119497488 | C | T | 0.001 | 0.9991 | +? | 0 | 0 | 0 | 1 |
| rs112165411 | 11 | 120636109 | A | T | 0.001 | 0.9992 | +? | 0 | 0 | 0 | 1 |
| rs475724 | 11 | 119784660 | A | G | 0.001 | 0.9992 | +? | 0 | 0 | 0 | 1 |
| rs4938349 | 11 | 117021923 | C | T | 0.001 | 0.9992 | +? | 0 | 0 | 0 | 1 |
| rs5094 | 11 | 116693213 | A | G | 0.001 | 0.9992 | +- | -1.4 | 2.96 | 1 | 0.08535 |
| rs73010650 | 11 | 119796469 | A | G | -0.001 | 0.9992 | -? | 0 | 0 | 0 | 1 |
| rs506127 | 11 | 119984202 | A | G | 0.001 | 0.9993 | +? | 0 | 0 | 0 | 1 |
| rs57943482 | 11 | 120769383 | C | T | 0.001 | 0.9993 | +? | 0 | 0 | 0 | 1 |
| rs10750161 | 11 | 119498618 | A | G | 0.001 | 0.9994 | +? | 0 | 0 | 0 | 1 |
| rs56756054 | 11 | 116987421 | C | T | 0.001 | 0.9994 | -+ | 73.8 | 11.458 | 1 | 0.000712 |
| rs513944 | 11 | 120094936 | G | A | 0.001 | 0.9995 | +? | 0 | 0 | 0 | 1 |
| rs557827 | 11 | 121093832 | T | A | 0.001 | 0.9995 | -+ | 0 | 0.026 | 1 | 0.8714 |

| | | | | | | | | | | | |
|---|---|---|---|---|---|---|---|---|---|---|---|
| rs1466051 | 11 | 119653461 | C | T | 0.001 | 0.9996 | +? | 0 | 0 | 0 | 1 |
| rs7111799 | 11 | 119868171 | A | G | 0.001 | 0.9996 | +? | 0 | 0 | 0 | 1 |
| rs11216267 | 11 | 116952392 | C | T | 0 | 0.9997 | +? | 0 | 0 | 0 | 1 |
| rs2298831 | 11 | 118068752 | C | T | 0 | 0.9998 | -+ | 0 | 0.106 | 1 | 0.7447 |
| rs5026369 | 11 | 119652214 | C | G | 0 | 0.9998 | +? | 0 | 0 | 0 | 1 |
| rs59007408 | 11 | 117032934 | T | C | 0 | 0.9999 | -+ | 73.1 | 11.16 | 1 | 0.0008358 |
| rs75115775 | 11 | 117030187 | C | A | 0 | 0.9999 | -+ | 73.1 | 11.159 | 1 | 0.0008363 |
| rs116896104 | 11 | 116976598 | A | G | 0 | 1 | -+ | 74.4 | 11.696 | 1 | 0.0006263 |

**Supplemental Data 2: Meta-analysis of SLE data in DDX6-CXCR5 locus (Dataset 2).**

| SNP | CHR | BP(GRCh37/hg19) | A1 | A2 | Zscore | P.value | Direction | HetISq | HetChiSq | HetDf | HetPVal |
|---|---|---|---|---|---|---|---|---|---|---|---|
| rs11217045 | 11 | 118697865 | A | C | -6.36 | 2.02E-10 | --- | 0 | 0.873 | 2 | 0.6462 |
| rs73005502 | 11 | 118697906 | G | A | -6.358 | 2.04E-10 | --- | 0 | 0.879 | 2 | 0.6444 |
| rs11217046 | 11 | 118698907 | A | G | -6.302 | 2.93E-10 | --- | 0 | 0.999 | 2 | 0.6067 |
| rs79504890 | 11 | 118701018 | A | G | -6.284 | 3.31E-10 | --- | 0 | 0.95 | 2 | 0.6219 |
| rs10892293 | 11 | 118665575 | T | A | -6.267 | 3.69E-10 | --- | 0 | 1.347 | 2 | 0.5099 |
| rs2077579 | 11 | 118619047 | G | T | -6.226 | 4.77E-10 | --- | -66.7 | 2.399 | 2 | 0.3013 |
| rs11217037 | 11 | 118677086 | A | G | -6.189 | 6.06E-10 | --- | 0 | 1.254 | 2 | 0.5341 |
| rs715412 | 11 | 118684610 | A | G | -6.177 | 6.54E-10 | --- | 0 | 1.508 | 2 | 0.4706 |
| rs11217038 | 11 | 118677355 | A | G | -6.174 | 6.65E-10 | --- | 0 | 1.301 | 2 | 0.5217 |
| rs28409215 | 11 | 118679304 | A | G | -6.172 | 6.76E-10 | --- | 0 | 1.233 | 2 | 0.5399 |
| rs11217032 | 11 | 118669605 | A | T | -6.166 | 7.00E-10 | --- | 0 | 1.328 | 2 | 0.5149 |
| rs11217033 | 11 | 118673706 | C | T | -6.163 | 7.14E-10 | --- | 0 | 1.337 | 2 | 0.5124 |
| rs11217036 | 11 | 118675772 | T | C | -6.157 | 7.40E-10 | --- | 0 | 1.357 | 2 | 0.5075 |
| rs4938573 | 11 | 118741842 | C | T | -6.108 | 1.01E-09 | --- | 0 | 1.096 | 2 | 0.5782 |
| rs73005423 | 11 | 118681079 | G | A | -6.101 | 1.06E-09 | --- | 0 | 1.375 | 2 | 0.5027 |
| rs11826521 | 11 | 118610549 | T | G | -6.099 | 1.07E-09 | --- | -83.1 | 2.185 | 2 | 0.3353 |
| rs11217009 | 11 | 118615785 | T | C | -6.098 | 1.08E-09 | --- | 0 | 1.988 | 2 | 0.3702 |
| rs10892294 | 11 | 118667357 | C | G | -6.096 | 1.09E-09 | --- | 0 | 1.253 | 2 | 0.5345 |
| rs1048024 | 11 | 118618776 | T | A | -6.094 | 1.11E-09 | --- | -84.7 | 2.165 | 2 | 0.3387 |
| rs10892280 | 11 | 118611817 | T | C | -6.084 | 1.17E-09 | --- | -90.4 | 2.101 | 2 | 0.3498 |
| rs3889239 | 11 | 118619960 | T | C | -6.081 | 1.19E-09 | --- | -93.5 | 2.068 | 2 | 0.3557 |
| rs2004781 | 11 | 118619253 | T | C | -6.077 | 1.22E-09 | --- | -91.9 | 2.085 | 2 | 0.3526 |
| rs75249660 | 11 | 118721565 | A | G | -6.074 | 1.25E-09 | --- | 0 | 0.945 | 2 | 0.6236 |
| rs11217058 | 11 | 118722149 | A | G | -6.07 | 1.28E-09 | --- | 0 | 0.933 | 2 | 0.6272 |
| rs56758835 | 11 | 118615340 | A | G | -6.069 | 1.29E-09 | --- | 0 | 1.961 | 2 | 0.3751 |
| rs74676029 | 11 | 118628373 | T | C | -6.044 | 1.51E-09 | --- | -83.2 | 2.183 | 2 | 0.3356 |
| rs10892290 | 11 | 118653503 | T | G | -6.035 | 1.59E-09 | --- | -81.5 | 2.204 | 2 | 0.3323 |
| rs874621 | 11 | 118655645 | A | G | -6.034 | 1.60E-09 | --- | -85.4 | 2.158 | 2 | 0.3399 |
| rs10892291 | 11 | 118653517 | T | C | -6.032 | 1.62E-09 | --- | -80.6 | 2.215 | 2 | 0.3305 |
| rs10892287 | 11 | 118642999 | C | T | -6.007 | 1.89E-09 | --- | -71.6 | 2.331 | 2 | 0.3118 |
| rs10892288 | 11 | 118644694 | G | A | -6.007 | 1.89E-09 | --- | -76.1 | 2.271 | 2 | 0.3212 |
| rs11217020 | 11 | 118639353 | A | G | -5.994 | 2.04E-09 | --- | -79.8 | 2.224 | 2 | 0.3289 |
| rs10892289 | 11 | 118646003 | T | C | -5.991 | 2.08E-09 | --- | -82.6 | 2.191 | 2 | 0.3344 |
| rs10892292 | 11 | 118655340 | C | G | -5.988 | 2.12E-09 | --- | -81.6 | 2.203 | 2 | 0.3324 |
| rs10892286 | 11 | 118642085 | C | A | -5.981 | 2.22E-09 | --- | -70.9 | 2.34 | 2 | 0.3103 |
| rs56043232 | 11 | 118648373 | T | C | -5.977 | 2.27E-09 | --- | -79.4 | 2.229 | 2 | 0.328 |
| rs11217019 | 11 | 118636343 | A | C | -5.975 | 2.30E-09 | --- | -72 | 2.325 | 2 | 0.3126 |
| rs10892283 | 11 | 118636980 | G | A | -5.971 | 2.36E-09 | --- | -68 | 2.381 | 2 | 0.3041 |
| rs7119044 | 11 | 118738298 | G | C | -5.964 | 2.47E-09 | --- | 0 | 0.75 | 2 | 0.6873 |
| rs11217074 | 11 | 118734000 | C | T | -5.962 | 2.49E-09 | --- | 0 | 0.756 | 2 | 0.6851 |
| rs11217066 | 11 | 118726753 | G | T | -5.961 | 2.51E-09 | --- | 0 | 0.921 | 2 | 0.6309 |
| rs7122669 | 11 | 118739056 | A | G | -5.958 | 2.55E-09 | --- | 0 | 0.728 | 2 | 0.6948 |
| rs6589684 | 11 | 118610957 | A | G | -5.958 | 2.55E-09 | --- | 0 | 1.355 | 2 | 0.508 |
| rs74541740 | 11 | 118742800 | A | G | -5.945 | 2.77E-09 | --- | 0 | 0.814 | 2 | 0.6658 |
| rs4936444 | 11 | 118743338 | T | C | -5.944 | 2.79E-09 | --- | 0 | 0.857 | 2 | 0.6516 |
| rs7951740 | 11 | 118740418 | G | C | -5.942 | 2.82E-09 | --- | 0 | 0.841 | 2 | 0.6567 |
| rs4936441 | 11 | 118725660 | C | G | -5.935 | 2.94E-09 | --- | 0 | 0.985 | 2 | 0.6111 |
| rs57494551 | 11 | 118661398 | T | C | -5.923 | 3.16E-09 | --- | -63.2 | 2.451 | 2 | 0.2937 |
| rs7117261 | 11 | 118741157 | T | C | -5.91 | 3.42E-09 | --- | 0 | 0.84 | 2 | 0.657 |
| rs10790268 | 11 | 118729391 | A | G | -5.825 | 5.71E-09 | --- | 0 | 0.997 | 2 | 0.6074 |
| rs10790269 | 11 | 118729456 | C | T | -5.812 | 6.17E-09 | --- | 0 | 1.03 | 2 | 0.5976 |
| rs4936443 | 11 | 118740864 | C | T | -5.804 | 6.47E-09 | --- | 0 | 0.94 | 2 | 0.6249 |

| | | | | | | | | | | | |
|---|---|---|---|---|---|---|---|---|---|---|---|
| rs77209083 | 11 | 118607569 | G | A | -5.751 | 8.87E-09 | --- | 0 | 1.725 | 2 | 0.4221 |
| rs11217001 | 11 | 118607046 | A | G | -5.75 | 8.92E-09 | --- | 0 | 1.974 | 2 | 0.3726 |
| rs4938572 | 11 | 118740931 | C | T | -5.555 | 2.77E-08 | --- | 0 | 0.354 | 2 | 0.8378 |
| rs7481819 | 11 | 118745278 | C | T | -5.49 | 4.02E-08 | --- | 0 | 0.239 | 2 | 0.8874 |
| rs7481797 | 11 | 118745243 | A | G | -5.475 | 4.37E-08 | --- | 0 | 0.255 | 2 | 0.8803 |
| rs7117313 | 11 | 118744396 | C | G | -5.372 | 7.79E-08 | --- | 0 | 0.652 | 2 | 0.7216 |
| rs10790275 | 11 | 118745884 | G | C | -5.364 | 8.16E-08 | --- | 0 | 0.479 | 2 | 0.787 |
| rs10892271 | 11 | 118598544 | A | G | -5.343 | 9.14E-08 | --- | -47.9 | 2.704 | 2 | 0.2588 |
| rs10892273 | 11 | 118602809 | A | G | -5.336 | 9.51E-08 | --- | -35.7 | 2.948 | 2 | 0.229 |
| rs10892272 | 11 | 118602707 | G | C | -5.32 | 1.04E-07 | --- | -32.3 | 3.024 | 2 | 0.2205 |
| rs11216997 | 11 | 118604638 | T | C | -5.299 | 1.16E-07 | --- | -18.2 | 3.384 | 2 | 0.1841 |
| rs7119038 | 11 | 118738281 | G | A | -5.297 | 1.18E-07 | --- | 0 | 1.508 | 2 | 0.4705 |
| rs11216980 | 11 | 118590668 | A | C | -5.287 | 1.24E-07 | --- | -47 | 2.721 | 2 | 0.2565 |
| rs11217044 | 11 | 118696022 | C | T | -5.271 | 1.36E-07 | --- | 0 | 1.88 | 2 | 0.3906 |
| rs112561833 | 11 | 118603736 | C | A | -5.267 | 1.39E-07 | --- | -23.5 | 3.238 | 2 | 0.1981 |
| rs10892263 | 11 | 118587319 | C | T | -5.222 | 1.77E-07 | --- | -38 | 2.899 | 2 | 0.2347 |
| rs12421790 | 11 | 118588952 | T | C | -5.21 | 1.89E-07 | --- | -43.9 | 2.78 | 2 | 0.2491 |
| rs11217040 | 11 | 118680648 | A | C | -5.159 | 2.49E-07 | --- | -94.6 | 2.056 | 2 | 0.3578 |
| rs11217006 | 11 | 118609790 | A | G | -5.077 | 3.84E-07 | --- | -74.2 | 2.296 | 2 | 0.3173 |
| rs17122453 | 11 | 118683564 | A | G | -5.065 | 4.09E-07 | --- | 0 | 1.991 | 2 | 0.3696 |
| rs11216995 | 11 | 118603526 | A | G | -5.044 | 4.56E-07 | --- | -34 | 2.985 | 2 | 0.2248 |
| rs10892279 | 11 | 118611781 | A | G | -5.037 | 4.73E-07 | --- | 0 | 1.956 | 2 | 0.376 |
| rs6589685 | 11 | 118612197 | C | T | -5.03 | 4.90E-07 | --- | 0 | 1.878 | 2 | 0.3911 |
| rs11216994 | 11 | 118603520 | T | C | -5.025 | 5.02E-07 | --- | -36.9 | 2.921 | 2 | 0.2321 |
| rs10892262 | 11 | 118587097 | C | G | -5.024 | 5.05E-07 | --- | -59.4 | 2.51 | 2 | 0.2851 |
| rs6589686 | 11 | 118612565 | A | G | -5.024 | 5.07E-07 | --- | 0 | 1.15 | 2 | 0.5626 |
| rs73005426 | 11 | 118681083 | G | A | -5.006 | 5.57E-07 | --- | 0 | 1.707 | 2 | 0.4259 |
| rs11217042 | 11 | 118682528 | A | G | -4.997 | 5.83E-07 | --- | 0 | 1.731 | 2 | 0.4208 |
| rs73001429 | 11 | 118603165 | G | T | -4.989 | 6.08E-07 | --- | -40.1 | 2.855 | 2 | 0.2399 |
| rs7942535 | 11 | 118681464 | C | T | -4.986 | 6.17E-07 | --- | 0 | 1.744 | 2 | 0.4181 |
| rs11217041 | 11 | 118682350 | G | A | -4.984 | 6.22E-07 | --- | 0 | 1.746 | 2 | 0.4177 |
| rs12365699 | 11 | 118743286 | A | G | -4.976 | 6.49E-07 | --- | 0 | 0.647 | 2 | 0.7235 |
| rs7945144 | 11 | 118608716 | G | A | -4.965 | 6.86E-07 | --- | 0 | 1.971 | 2 | 0.3732 |
| rs2155433 | 11 | 118612921 | A | G | -4.963 | 6.95E-07 | --- | 0 | 1.92 | 2 | 0.3829 |
| rs7123726 | 11 | 118694547 | C | T | -4.961 | 7.02E-07 | --- | 0 | 1.613 | 2 | 0.4465 |
| rs12364971 | 11 | 118607794 | G | T | -4.953 | 7.32E-07 | --- | 0 | 1.921 | 2 | 0.3827 |
| rs76409436 | 11 | 118607871 | A | G | -4.945 | 7.61E-07 | --- | -94.8 | 2.053 | 2 | 0.3583 |
| rs73003215 | 11 | 118649512 | A | G | -4.935 | 8.00E-07 | --- | 0 | 1.956 | 2 | 0.376 |
| rs11216993 | 11 | 118603495 | A | T | -4.929 | 8.28E-07 | --- | -45.1 | 2.757 | 2 | 0.2519 |
| rs12421793 | 11 | 118589012 | A | C | -4.918 | 8.75E-07 | --- | -40.7 | 2.843 | 2 | 0.2413 |
| rs56267528 | 11 | 118607311 | A | C | -4.816 | 1.46E-06 | --- | 0 | 1.6 | 2 | 0.4492 |
| rs55894437 | 11 | 118607417 | T | C | -4.807 | 1.53E-06 | --- | 0 | 1.603 | 2 | 0.4486 |
| rs11217002 | 11 | 118607150 | T | C | -4.806 | 1.54E-06 | --- | 0 | 1.851 | 2 | 0.3962 |
| rs6589682 | 11 | 118606722 | T | C | -4.79 | 1.67E-06 | --- | 0 | 1.754 | 2 | 0.416 |
| rs11216956 | 11 | 118575326 | A | G | -4.536 | 5.73E-06 | --- | -86.3 | 2.147 | 2 | 0.3419 |
| rs11216972 | 11 | 118585826 | T | C | -4.508 | 6.56E-06 | --- | -38.3 | 2.892 | 2 | 0.2355 |
| rs598373 | 11 | 118664377 | T | C | -4.455 | 8.40E-06 | --- | 0 | 0.137 | 2 | 0.9336 |
| rs477014 | 11 | 118678986 | C | T | -4.424 | 9.67E-06 | --- | 0 | 0.127 | 2 | 0.9387 |
| rs476246 | 11 | 118617920 | T | C | -4.424 | 9.70E-06 | --- | 0 | 0.122 | 2 | 0.9407 |
| rs540180 | 11 | 118653629 | T | G | -4.412 | 1.03E-05 | --- | 0 | 0.111 | 2 | 0.946 |
| rs2508916 | 11 | 118654651 | G | C | -4.409 | 1.04E-05 | --- | 0 | 0.11 | 2 | 0.9467 |
| rs581062 | 11 | 118624767 | A | G | -4.406 | 1.06E-05 | --- | 0 | 0.111 | 2 | 0.9458 |
| rs472641 | 11 | 118617544 | G | A | -4.4 | 1.08E-05 | --- | 0 | 0.123 | 2 | 0.9405 |
| rs688161 | 11 | 118677710 | G | A | -4.396 | 1.10E-05 | --- | 0 | 0.109 | 2 | 0.947 |

| | | | | | | | | | | | |
|---|---|---|---|---|---|---|---|---|---|---|---|
| rs591756 | 11 | 118678881 | T | C | -4.389 | 1.14E-05 | --- | 0 | 0.106 | 2 | 0.9483 |
| rs10892259 | 11 | 118583002 | A | G | -4.386 | 1.15E-05 | --- | 0 | 1.996 | 2 | 0.3687 |
| rs1784293 | 11 | 118652373 | C | A | -4.386 | 1.16E-05 | --- | 0 | 0.099 | 2 | 0.9517 |
| rs2508570 | 11 | 118678609 | A | G | -4.382 | 1.17E-05 | --- | 0 | 0.102 | 2 | 0.9502 |
| rs473298 | 11 | 118679387 | G | A | -4.382 | 1.17E-05 | --- | 0 | 0.101 | 2 | 0.9508 |
| rs10892260 | 11 | 118583525 | C | A | -4.378 | 1.20E-05 | --- | -81.4 | 2.205 | 2 | 0.332 |
| rs11216969 | 11 | 118584340 | C | T | -4.378 | 1.20E-05 | --- | -90.1 | 2.104 | 2 | 0.3493 |
| rs516719 | 11 | 118679445 | C | T | -4.375 | 1.21E-05 | --- | 0 | 0.097 | 2 | 0.9526 |
| rs2508569 | 11 | 118680045 | C | G | -4.374 | 1.22E-05 | --- | 0 | 0.095 | 2 | 0.9538 |
| rs1790189 | 11 | 118652326 | G | T | -4.371 | 1.24E-05 | --- | 0 | 0.093 | 2 | 0.9547 |
| rs10790261 | 11 | 118579747 | G | A | -4.365 | 1.27E-05 | --- | -91 | 2.094 | 2 | 0.351 |
| rs11216960 | 11 | 118577873 | A | G | -4.361 | 1.30E-05 | --- | -77.9 | 2.249 | 2 | 0.3249 |
| rs524590 | 11 | 118640063 | T | A | -4.36 | 1.30E-05 | --- | 0 | 0.08 | 2 | 0.9607 |
| rs487728 | 11 | 118621495 | G | C | -4.353 | 1.34E-05 | --- | 0 | 0.115 | 2 | 0.944 |
| rs11216965 | 11 | 118582613 | A | C | -4.344 | 1.40E-05 | --- | -84.7 | 2.165 | 2 | 0.3387 |
| rs10892319 | 11 | 118870413 | T | C | -4.34 | 1.43E-05 | -?- | 0 | 0.032 | 1 | 0.8572 |
| rs10892256 | 11 | 118578705 | A | G | -4.336 | 1.45E-05 | --- | -84.3 | 2.17 | 2 | 0.3378 |
| rs10892257 | 11 | 118579327 | A | G | -4.335 | 1.46E-05 | --- | -83.3 | 2.182 | 2 | 0.3358 |
| rs10892258 | 11 | 118579865 | A | G | -4.33 | 1.49E-05 | --- | -85.4 | 2.157 | 2 | 0.34 |
| rs562059 | 11 | 118602058 | G | T | -4.325 | 1.52E-05 | --- | 0 | 0.416 | 2 | 0.812 |
| rs11216964 | 11 | 118581895 | A | G | -4.322 | 1.55E-05 | --- | -82 | 2.198 | 2 | 0.3332 |
| rs478081 | 11 | 118602497 | G | A | -4.321 | 1.55E-05 | --- | 0 | 0.406 | 2 | 0.8162 |
| rs4938617 | 11 | 118867695 | T | C | -4.32 | 1.56E-05 | -?- | 0 | 0.053 | 1 | 0.818 |
| rs11216961 | 11 | 118580338 | G | C | -4.319 | 1.56E-05 | --- | -83.7 | 2.177 | 2 | 0.3367 |
| rs499343 | 11 | 118626511 | T | C | -4.31 | 1.63E-05 | --- | 0 | 0.077 | 2 | 0.9624 |
| rs9645663 | 11 | 118873840 | C | T | -4.28 | 1.87E-05 | -?- | 0 | 0.014 | 1 | 0.9045 |
| rs11216962 | 11 | 118580813 | C | T | -4.278 | 1.89E-05 | --- | -94.5 | 2.056 | 2 | 0.3577 |
| rs11216963 | 11 | 118580814 | A | G | -4.278 | 1.89E-05 | --- | -94.5 | 2.056 | 2 | 0.3577 |
| rs11606199 | 11 | 118811641 | A | G | -4.272 | 1.93E-05 | -?- | 0 | 0.017 | 1 | 0.896 |
| rs11216999 | 11 | 118604941 | C | T | -4.26 | 2.05E-05 | --- | 0 | 1.349 | 2 | 0.5095 |
| rs7125066 | 11 | 118721699 | G | A | -4.216 | 2.48E-05 | --- | 31.9 | 5.877 | 2 | 0.05295 |
| rs9736016 | 11 | 118724894 | A | T | -4.155 | 3.26E-05 | --- | 30.6 | 5.768 | 2 | 0.05592 |
| rs111345266 | 11 | 118893767 | T | C | -4.144 | 3.42E-05 | -?- | 0 | 0.008 | 1 | 0.9287 |
| rs74884652 | 11 | 118876552 | C | T | -4.136 | 3.54E-05 | -?- | 0 | 0.003 | 1 | 0.9572 |
| rs4938619 | 11 | 118890085 | T | C | -4.131 | 3.62E-05 | -?- | 0 | 0.008 | 1 | 0.9282 |
| rs75154936 | 11 | 118877891 | C | T | -4.118 | 3.82E-05 | -?- | 0 | 0.001 | 1 | 0.9712 |
| rs10892247 | 11 | 118490076 | A | G | -4.113 | 3.91E-05 | --- | 0 | 1.322 | 2 | 0.5163 |
| rs76253444 | 11 | 118876213 | G | A | -4.105 | 4.05E-05 | -?- | 0 | 0 | 1 | 0.9891 |
| rs537080 | 11 | 118826045 | C | T | -4.019 | 5.84E-05 | -?- | 0 | 0.102 | 1 | 0.75 |
| rs10892316 | 11 | 118831755 | T | C | -4.016 | 5.91E-05 | -?- | 0 | 0.093 | 1 | 0.7609 |
| rs112939258 | 11 | 118834718 | T | C | -3.998 | 6.38E-05 | -?- | 0 | 0.085 | 1 | 0.77 |
| rs11216998 | 11 | 118604837 | T | C | -3.979 | 6.91E-05 | --- | 0 | 1.49 | 2 | 0.4748 |
| rs111358668 | 11 | 118834508 | A | G | -3.964 | 7.38E-05 | -?- | 0 | 0.076 | 1 | 0.7828 |
| rs45586434 | 11 | 118828184 | A | C | -3.956 | 7.63E-05 | -?- | 0 | 0.009 | 1 | 0.9228 |
| rs113962716 | 11 | 118853021 | A | C | -3.941 | 8.11E-05 | -?- | 0 | 0.085 | 1 | 0.7701 |
| rs80126726 | 11 | 118852647 | A | G | -3.938 | 8.21E-05 | -?- | 0 | 0.149 | 1 | 0.6993 |
| rs11217109 | 11 | 118853774 | C | T | -3.924 | 8.71E-05 | -?- | 0 | 0.097 | 1 | 0.756 |
| rs507763 | 11 | 118824683 | C | G | -3.918 | 8.95E-05 | -?- | 0 | 0.129 | 1 | 0.719 |
| rs12362152 | 11 | 118516737 | G | A | -3.904 | 9.47E-05 | --- | 0 | 1.191 | 2 | 0.5512 |
| rs11216937 | 11 | 118507478 | T | C | -3.89 | 0.0001004 | --- | 0 | 1.175 | 2 | 0.5558 |
| rs73001406 | 11 | 118560857 | A | G | -3.881 | 0.000104 | --- | -21.6 | 3.29 | 2 | 0.193 |
| rs80015904 | 11 | 118516903 | A | G | -3.863 | 0.0001118 | --- | 0 | 1.129 | 2 | 0.5687 |
| rs77683570 | 11 | 118495287 | C | T | -3.861 | 0.0001129 | --- | 0 | 1.168 | 2 | 0.5577 |
| rs11216938 | 11 | 118514625 | T | C | -3.858 | 0.0001141 | --- | 0 | 1.062 | 2 | 0.5881 |

| | | | | | | | | | | | |
|---|---|---|---|---|---|---|---|---|---|---|---|
| rs10892307 | 11 | 118754649 | G | C | -3.854 | 0.0001161 | --- | -91.3 | 2.09 | 2 | 0.3516 |
| rs10892277 | 11 | 118605841 | A | G | -3.852 | 0.0001171 | --- | 0 | 0.906 | 2 | 0.6358 |
| rs10892274 | 11 | 118605463 | T | C | -3.847 | 0.0001197 | --- | 0 | 0.919 | 2 | 0.6317 |
| rs45565037 | 11 | 118496119 | A | G | -3.845 | 0.0001208 | --- | 0 | 1.098 | 2 | 0.5776 |
| rs12365095 | 11 | 118516774 | T | C | -3.844 | 0.000121 | --- | 0 | 1.053 | 2 | 0.5906 |
| rs3176905 | 11 | 118755630 | C | T | -3.836 | 0.0001253 | --- | 0 | 1.651 | 2 | 0.4381 |
| rs10892251 | 11 | 118543563 | T | C | -3.826 | 0.0001303 | --- | -17.6 | 3.402 | 2 | 0.1825 |
| rs2277296 | 11 | 118550522 | C | T | -3.826 | 0.0001303 | --- | -12.1 | 3.569 | 2 | 0.1679 |
| rs7928371 | 11 | 118529127 | A | G | -3.818 | 0.0001344 | --- | 0 | 1.099 | 2 | 0.5772 |
| rs17748 | 11 | 118528424 | T | C | -3.815 | 0.0001364 | --- | 0 | 1.111 | 2 | 0.5739 |
| rs10892252 | 11 | 118555692 | G | A | -3.813 | 0.0001375 | --- | -15 | 3.479 | 2 | 0.1756 |
| rs10790256 | 11 | 118534082 | T | C | -3.81 | 0.0001391 | --- | 0 | 1.128 | 2 | 0.5691 |
| rs12225548 | 11 | 118535840 | G | C | -3.808 | 0.0001399 | --- | 0 | 1.331 | 2 | 0.5141 |
| rs2236661 | 11 | 118499394 | G | C | -3.803 | 0.0001428 | --- | 0 | 1.186 | 2 | 0.5527 |
| rs10892275 | 11 | 118605617 | A | T | -3.796 | 0.0001468 | --- | 0 | 0.844 | 2 | 0.6559 |
| rs10892276 | 11 | 118605628 | A | G | -3.794 | 0.0001485 | --- | 0 | 0.843 | 2 | 0.6561 |
| rs2276065 | 11 | 118530611 | C | T | -3.79 | 0.0001505 | --- | 0 | 1.531 | 2 | 0.465 |
| rs45540840 | 11 | 118486110 | A | G | -3.773 | 0.0001613 | --- | 0 | 1.309 | 2 | 0.5196 |
| rs496547 | 11 | 118576463 | A | T | -3.767 | 0.0001655 | --- | 0 | 0.352 | 2 | 0.8385 |
| rs11216930 | 11 | 118488782 | C | A | -3.757 | 0.0001721 | --- | 0 | 0.779 | 2 | 0.6774 |
| rs10892248 | 11 | 118501022 | A | G | -3.656 | 0.0002557 | --- | 0 | 1.325 | 2 | 0.5156 |
| rs11602393 | 11 | 118750967 | A | T | -3.562 | 0.0003683 | --- | 0 | 1.434 | 2 | 0.4882 |
| rs4938513 | 11 | 118419704 | T | C | -3.438 | 0.0005856 | --- | 0 | 0.814 | 2 | 0.6656 |
| rs11216943 | 11 | 118556398 | A | G | -3.426 | 0.0006121 | --- | 14.5 | 4.676 | 2 | 0.09651 |
| rs7943247 | 11 | 72881957 | G | A | -3.422 | 0.0006226 | --- | 45.7 | 7.363 | 2 | 0.02518 |
| rs741811 | 11 | 118898319 | A | G | -3.418 | 0.0006301 | -?- | 0 | 0.455 | 1 | 0.5001 |
| rs10790255 | 11 | 118515579 | G | T | -3.359 | 0.0007828 | --- | 0 | 1.489 | 2 | 0.475 |
| rs2276423 | 11 | 118210106 | G | C | 3.32 | 0.0008998 | +?+ | 0 | 0.007 | 1 | 0.9317 |
| rs4938506 | 11 | 118205702 | A | G | 3.281 | 0.001033 | +?+ | 0 | 0.002 | 1 | 0.9683 |
| rs7937334 | 11 | 118191274 | T | C | 3.274 | 0.001059 | +?+ | 0 | 0.004 | 1 | 0.9481 |
| rs633308 | 11 | 118459069 | G | A | -3.211 | 0.001321 | --- | 0 | 0.719 | 2 | 0.6979 |
| rs4936428 | 11 | 118187511 | T | C | 3.155 | 0.001605 | +?+ | 0 | 0.006 | 1 | 0.9384 |
| rs573905 | 11 | 118572267 | A | G | 3.152 | 0.00162 | +++ | 0 | 0.995 | 2 | 0.608 |
| rs10892306 | 11 | 118751326 | A | T | -3.147 | 0.001652 | --- | 0 | 1.626 | 2 | 0.4435 |
| rs4938576 | 11 | 118746769 | T | G | -3.138 | 0.0017 | --- | 0 | 1.074 | 2 | 0.5846 |
| rs75438046 | 11 | 118767564 | A | G | -3.132 | 0.001738 | -+- | 42.8 | 6.995 | 2 | 0.03028 |
| rs542266 | 11 | 118422240 | A | C | -3.126 | 0.001771 | --- | 0 | 0.537 | 2 | 0.7647 |
| rs2276424 | 11 | 118209960 | A | C | 3.06 | 0.002212 | +?+ | 0 | 0.077 | 1 | 0.7811 |
| rs78041268 | 11 | 118763313 | A | G | -3.05 | 0.002285 | -+- | 36.6 | 6.313 | 2 | 0.04256 |
| rs149114341 | 11 | 118783424 | A | G | -3.039 | 0.002371 | -+- | 36.7 | 6.318 | 2 | 0.04246 |
| rs61036532 | 11 | 72017321 | G | A | 3.007 | 0.002638 | +++ | 0 | 1.24 | 2 | 0.538 |
| rs77841686 | 11 | 118762073 | A | G | -2.998 | 0.002717 | -+- | 34.8 | 6.136 | 2 | 0.0465 |
| rs56076328 | 11 | 72023279 | A | G | 2.989 | 0.002795 | +++ | 0 | 1.332 | 2 | 0.5138 |
| rs11217065 | 11 | 118726647 | C | T | 2.988 | 0.002812 | +++ | -40.9 | 2.839 | 2 | 0.2418 |
| rs3750906 | 11 | 72012682 | G | A | 2.987 | 0.002815 | +++ | 0 | 1.303 | 2 | 0.5212 |
| rs586763 | 11 | 118816304 | T | C | -2.984 | 0.002841 | -?- | 0 | 0.228 | 1 | 0.6331 |
| rs11217063 | 11 | 118725942 | C | G | 2.978 | 0.002897 | +++ | -39.8 | 2.86 | 2 | 0.2393 |
| rs7110837 | 11 | 118199295 | A | G | 2.975 | 0.002932 | +?+ | 0 | 0.082 | 1 | 0.7741 |
| rs12421674 | 11 | 72056591 | T | C | 2.967 | 0.003012 | +++ | 0 | 1.632 | 2 | 0.4422 |
| rs12418637 | 11 | 72061218 | A | G | 2.966 | 0.003017 | +++ | 0 | 0.829 | 2 | 0.6608 |
| rs57719838 | 11 | 118724723 | C | T | 2.965 | 0.003024 | +++ | -36.9 | 2.922 | 2 | 0.232 |
| rs56171963 | 11 | 72009455 | G | A | 2.961 | 0.003063 | +++ | 0 | 1.361 | 2 | 0.5064 |
| rs11216853 | 11 | 118200201 | T | A | 2.959 | 0.003083 | +?+ | 0 | 0.123 | 1 | 0.7257 |
| rs56148078 | 11 | 72529358 | A | C | -2.954 | 0.003133 | +-- | 47.2 | 7.572 | 2 | 0.02268 |

| | | | | | | | | | | | |
|---|---|---|---|---|---|---|---|---|---|---|---|
| rs6589659 | 11 | 118191691 | A | C | 2.952 | 0.00316 | +?+ | 0 | 0.044 | 1 | 0.8334 |
| rs58044536 | 11 | 72011227 | G | A | 2.949 | 0.003187 | +++ | 0 | 1.247 | 2 | 0.5361 |
| rs58945323 | 11 | 72042223 | C | T | 2.948 | 0.003194 | +++ | 0 | 1.409 | 2 | 0.4943 |
| rs7103067 | 11 | 118723379 | A | G | 2.948 | 0.003194 | +++ | -32.3 | 3.023 | 2 | 0.2206 |
| rs10892246 | 11 | 118481431 | A | G | -2.946 | 0.003219 | --- | -97.2 | 2.028 | 2 | 0.3627 |
| rs633683 | 11 | 118504742 | T | C | 2.94 | 0.003281 | +?+ | -78.8 | 1.678 | 1 | 0.1952 |
| rs17162126 | 11 | 72057178 | T | A | 2.93 | 0.003395 | +++ | 0 | 1.573 | 2 | 0.4553 |
| rs11217091 | 11 | 118793613 | C | T | -2.929 | 0.003396 | -+- | 43.9 | 7.131 | 2 | 0.02828 |
| rs4936442 | 11 | 118728320 | T | C | 2.922 | 0.003473 | +++ | -44.7 | 2.765 | 2 | 0.2509 |
| rs12223292 | 11 | 72487259 | A | G | -2.917 | 0.003534 | +-- | 47.8 | 7.666 | 2 | 0.02164 |
| rs58941166 | 11 | 72083441 | A | G | 2.913 | 0.003574 | +++ | 0 | 1.001 | 2 | 0.6061 |
| rs55721826 | 11 | 72019162 | G | A | 2.913 | 0.003584 | +++ | 0 | 1.436 | 2 | 0.4877 |
| rs34782343 | 11 | 118153511 | G | T | 2.912 | 0.003589 | +?+ | -157.7 | 1.164 | 1 | 0.2806 |
| rs11217067 | 11 | 118727332 | T | G | 2.912 | 0.003592 | +++ | -45.5 | 2.749 | 2 | 0.2529 |
| rs10790267 | 11 | 118727169 | T | C | 2.912 | 0.003596 | +++ | -45.5 | 2.75 | 2 | 0.2528 |
| rs73530724 | 11 | 72049158 | G | A | 2.91 | 0.003619 | +++ | 0 | 1.439 | 2 | 0.487 |
| rs12790489 | 11 | 118729583 | C | T | 2.909 | 0.003629 | +++ | -54.4 | 2.591 | 2 | 0.2738 |
| rs11217070 | 11 | 118729860 | G | T | 2.901 | 0.003716 | +++ | -53.4 | 2.608 | 2 | 0.2715 |
| rs12416804 | 11 | 72048526 | G | T | 2.9 | 0.003735 | +++ | 0 | 1.448 | 2 | 0.4849 |
| rs3889234 | 11 | 72490986 | C | T | -2.899 | 0.003742 | +-- | 49.5 | 7.914 | 2 | 0.01912 |
| rs10892300 | 11 | 118729210 | T | C | 2.891 | 0.003843 | +++ | -56.3 | 2.56 | 2 | 0.2781 |
| rs73530718 | 11 | 72045813 | C | T | 2.891 | 0.003846 | +++ | 0 | 1.444 | 2 | 0.4857 |
| rs71482159 | 11 | 118809204 | T | C | -2.884 | 0.003922 | -?- | 0 | 0.012 | 1 | 0.9136 |
| rs11217072 | 11 | 118731762 | G | A | 2.882 | 0.003957 | +++ | -62.2 | 2.466 | 2 | 0.2915 |
| rs34561290 | 11 | 118731599 | T | C | 2.88 | 0.003974 | +++ | -62.1 | 2.468 | 2 | 0.2911 |
| rs504217 | 11 | 72006086 | T | C | 2.879 | 0.003995 | +++ | 0 | 1.217 | 2 | 0.5442 |
| rs474629 | 11 | 72006977 | C | G | 2.875 | 0.004042 | +++ | 0 | 1.069 | 2 | 0.5859 |
| rs7929520 | 11 | 118730141 | G | A | 2.872 | 0.004084 | +++ | -49.1 | 2.683 | 2 | 0.2614 |
| rs680660 | 11 | 72004714 | C | T | 2.863 | 0.004199 | +++ | 0 | 1.263 | 2 | 0.5317 |
| rs11217099 | 11 | 118809882 | A | C | -2.843 | 0.004471 | -+- | 35.5 | 6.206 | 2 | 0.04492 |
| rs17540708 | 11 | 118150127 | G | T | 2.836 | 0.004562 | +?+ | 0 | 0.798 | 1 | 0.3716 |
| rs7940062 | 11 | 72037226 | C | A | 2.834 | 0.004603 | +++ | 0 | 1.297 | 2 | 0.5229 |
| rs7483509 | 11 | 118740180 | A | G | 2.821 | 0.004784 | +++ | -73.4 | 2.307 | 2 | 0.3155 |
| rs34250455 | 11 | 118157681 | T | C | 2.821 | 0.004788 | +?+ | 0 | 0.826 | 1 | 0.3633 |
| rs35235858 | 11 | 118157250 | T | C | 2.821 | 0.004788 | +?+ | 0 | 0.826 | 1 | 0.3633 |
| rs34022886 | 11 | 118155804 | A | G | 2.818 | 0.004831 | +?+ | 0 | 0.832 | 1 | 0.3618 |
| rs12798115 | 11 | 118156517 | C | G | 2.814 | 0.004887 | +?+ | 0 | 0.838 | 1 | 0.3599 |
| rs2298821 | 11 | 118185140 | C | T | 2.81 | 0.004953 | +?+ | 0 | 0.816 | 1 | 0.3663 |
| rs622722 | 11 | 118711790 | A | G | 2.808 | 0.004981 | +++ | -13.6 | 3.523 | 2 | 0.1718 |
| rs61900905 | 11 | 118185492 | C | T | 2.805 | 0.005031 | +?+ | 0 | 0.825 | 1 | 0.3636 |
| rs11216934 | 11 | 118501459 | A | G | 2.801 | 0.005087 | +?+ | 0 | 0.136 | 1 | 0.7124 |
| rs3819250 | 11 | 118185346 | A | G | 2.788 | 0.005305 | +?+ | 0 | 0.857 | 1 | 0.3546 |
| rs7479373 | 11 | 118171285 | C | T | 2.782 | 0.005407 | +?+ | 0 | 0.974 | 1 | 0.3237 |
| rs11217090 | 11 | 118789952 | T | C | -2.773 | 0.005548 | -+- | 30.3 | 5.736 | 2 | 0.05681 |
| rs3759009 | 11 | 118173850 | A | G | 2.772 | 0.005576 | +?+ | 0 | 0.964 | 1 | 0.3263 |
| rs2231440 | 11 | 118175439 | A | G | 2.767 | 0.005661 | +?+ | 0 | 0.916 | 1 | 0.3386 |
| rs79000353 | 11 | 118420343 | A | G | -2.76 | 0.005775 | --- | -25.6 | 3.185 | 2 | 0.2034 |
| rs483598 | 11 | 118509668 | A | G | 2.759 | 0.005798 | +?+ | 0 | 0.122 | 1 | 0.7272 |
| rs2231441 | 11 | 118175540 | G | A | 2.758 | 0.005811 | +?+ | 0 | 0.983 | 1 | 0.3214 |
| rs12274090 | 11 | 72512097 | T | C | -2.754 | 0.005894 | +-- | 22.7 | 5.171 | 2 | 0.07534 |
| rs7484280 | 11 | 72167386 | A | C | 2.747 | 0.00601 | +++ | 0 | 0.145 | 2 | 0.93 |
| rs77058558 | 11 | 72065102 | A | G | 2.747 | 0.006015 | +++ | 0 | 1.09 | 2 | 0.5798 |
| rs498872 | 11 | 118477367 | A | G | -2.737 | 0.006191 | --- | 0 | 0.279 | 2 | 0.8698 |
| rs2277289 | 11 | 118182660 | C | T | 2.732 | 0.006294 | +?+ | 0 | 0.849 | 1 | 0.3567 |

| | | | | | | | | | | | |
|---|---|---|---|---|---|---|---|---|---|---|---|
| rs113573893 | 11 | 72078446 | G | T | 2.726 | 0.00642 | +++ | 0 | 0.971 | 2 | 0.6155 |
| rs60712691 | 11 | 72077961 | T | G | 2.726 | 0.00642 | +++ | 0 | 0.971 | 2 | 0.6155 |
| rs577396 | 11 | 118499469 | C | G | 2.725 | 0.006424 | +?+ | 0 | 0.124 | 1 | 0.725 |
| rs10431173 | 11 | 72512937 | G | A | -2.72 | 0.006538 | +-- | 20 | 4.999 | 2 | 0.08213 |
| rs73528886 | 11 | 72013848 | A | G | 2.706 | 0.006816 | +++ | 0 | 1.662 | 2 | 0.4356 |
| rs7944225 | 11 | 72496672 | C | G | -2.701 | 0.006909 | +-- | 24.4 | 5.292 | 2 | 0.07095 |
| rs73532667 | 11 | 72162726 | C | T | 2.699 | 0.006946 | +++ | 0 | 0.023 | 2 | 0.9884 |
| rs4938503 | 11 | 118180155 | C | G | 2.697 | 0.007003 | +?+ | 0 | 0.952 | 1 | 0.3291 |
| rs10160454 | 11 | 72496345 | C | A | -2.697 | 0.007005 | +-- | 24.3 | 5.285 | 2 | 0.0712 |
| rs576950 | 11 | 118391375 | G | A | -2.695 | 0.007041 | --- | 23.1 | 5.203 | 2 | 0.07415 |
| rs28423005 | 11 | 118183733 | T | C | 2.695 | 0.007043 | +?+ | -188.8 | 1.039 | 1 | 0.3081 |
| rs55930012 | 11 | 72079380 | C | T | 2.686 | 0.007232 | +++ | -71.4 | 2.334 | 2 | 0.3114 |
| rs10898873 | 11 | 72497713 | C | T | -2.684 | 0.007265 | +-- | 24 | 5.267 | 2 | 0.07184 |
| rs10898874 | 11 | 72497795 | G | A | -2.683 | 0.007294 | +-- | 24.4 | 5.293 | 2 | 0.07089 |
| rs7947423 | 11 | 72497001 | A | G | -2.681 | 0.007348 | +-- | 24 | 5.261 | 2 | 0.07204 |
| rs1126924 | 11 | 118178007 | T | C | 2.676 | 0.007451 | +?+ | -184.5 | 1.055 | 1 | 0.3044 |
| rs10898875 | 11 | 72497807 | C | T | -2.675 | 0.00748 | +-- | 24.3 | 5.281 | 2 | 0.07133 |
| rs7121919 | 11 | 72496087 | A | G | -2.657 | 0.007877 | +-- | 23.5 | 5.228 | 2 | 0.07323 |
| rs55756957 | 11 | 118747576 | G | A | -2.655 | 0.007938 | -?- | 0 | 0.377 | 1 | 0.5393 |
| rs77398170 | 11 | 118792278 | G | A | -2.644 | 0.008198 | -?- | 0 | 0.032 | 1 | 0.8577 |
| rs479490 | 11 | 72497321 | C | T | -2.641 | 0.008264 | +-- | 26.2 | 5.417 | 2 | 0.06663 |
| rs79769393 | 11 | 72080610 | A | G | 2.635 | 0.008404 | +++ | -77.6 | 2.252 | 2 | 0.3244 |
| rs12271856 | 11 | 118791532 | T | G | -2.628 | 0.008597 | -?- | 0 | 0.201 | 1 | 0.654 |
| rs12270280 | 11 | 118791150 | T | C | -2.617 | 0.008871 | -?- | 0 | 0.236 | 1 | 0.6272 |
| rs74766959 | 11 | 72065209 | G | A | 2.615 | 0.008927 | +++ | 0 | 1.752 | 2 | 0.4165 |
| rs9645708 | 11 | 72819318 | C | T | 2.614 | 0.008947 | -++ | 26.4 | 5.436 | 2 | 0.066 |
| rs74236665 | 11 | 72080895 | G | C | 2.613 | 0.008988 | +++ | -73.8 | 2.302 | 2 | 0.3164 |
| rs56229279 | 11 | 72013788 | G | A | 2.611 | 0.009039 | +++ | 5.2 | 4.221 | 2 | 0.1212 |
| rs75665552 | 11 | 72046487 | C | T | 2.602 | 0.009258 | +++ | -77.3 | 2.256 | 2 | 0.3238 |
| rs525549 | 11 | 118358068 | A | T | -2.597 | 0.009392 | --- | -6.6 | 3.753 | 2 | 0.1531 |
| rs8301 | 11 | 118895495 | G | C | -2.595 | 0.009449 | -?- | 0 | 0 | 1 | 0.9886 |
| rs79577700 | 11 | 72057385 | G | A | 2.595 | 0.009456 | +++ | -73.6 | 2.303 | 2 | 0.3161 |
| rs79549196 | 11 | 72057384 | T | C | 2.595 | 0.009462 | +++ | -73.6 | 2.304 | 2 | 0.3161 |
| rs79658353 | 11 | 72064041 | A | G | 2.59 | 0.009595 | +++ | -73.6 | 2.304 | 2 | 0.316 |
| rs687190 | 11 | 118542383 | C | T | 2.588 | 0.009641 | +++ | -58.2 | 2.529 | 2 | 0.2824 |
| rs12282752 | 11 | 118793918 | C | T | -2.575 | 0.01002 | -?- | 0 | 0.245 | 1 | 0.6208 |
| rs503425 | 11 | 118692727 | C | T | 2.574 | 0.01006 | +++ | 6.1 | 4.258 | 2 | 0.1189 |
| rs603486 | 11 | 118587347 | T | C | 2.57 | 0.01017 | +++ | 0 | 0.672 | 2 | 0.7147 |
| rs112974653 | 11 | 118807637 | T | G | -2.568 | 0.01022 | -?- | 0 | 0.019 | 1 | 0.8911 |
| rs73528892 | 11 | 72020144 | C | G | 2.565 | 0.01032 | +++ | -64.9 | 2.425 | 2 | 0.2974 |
| rs77638281 | 11 | 118811315 | A | G | -2.562 | 0.01041 | -?- | 0 | 0.026 | 1 | 0.8716 |
| rs480958 | 11 | 118577990 | G | A | 2.561 | 0.01044 | +++ | 0 | 0.645 | 2 | 0.7244 |
| rs140026975 | 11 | 118594669 | C | A | 2.56 | 0.01047 | +++ | 10.2 | 4.452 | 2 | 0.1079 |
| rs540716 | 11 | 118685015 | T | C | 2.558 | 0.01052 | +++ | 3.6 | 4.15 | 2 | 0.1256 |
| rs4938589 | 11 | 118810405 | T | C | -2.557 | 0.01057 | -?- | 0 | 0.032 | 1 | 0.8589 |
| rs7936556 | 11 | 118501299 | A | G | 2.556 | 0.0106 | +?+ | 0 | 0.141 | 1 | 0.7076 |
| rs68097885 | 11 | 118596444 | A | G | 2.554 | 0.01065 | +++ | 11.6 | 4.527 | 2 | 0.104 |
| rs7943039 | 11 | 118603030 | A | G | 2.55 | 0.01078 | +++ | 12.5 | 4.569 | 2 | 0.1018 |
| rs3132828 | 11 | 118431003 | A | G | -2.549 | 0.0108 | --- | 0 | 1.587 | 2 | 0.4522 |
| rs4938618 | 11 | 118867803 | C | G | -2.547 | 0.01086 | -?- | 0 | 0.013 | 1 | 0.9098 |
| rs4245184 | 11 | 118510562 | A | G | 2.541 | 0.01107 | +?+ | 0 | 0.115 | 1 | 0.7343 |
| rs4573680 | 11 | 118872741 | C | T | -2.54 | 0.01107 | -?- | 0 | 0.053 | 1 | 0.8187 |
| rs644273 | 11 | 118511352 | T | C | 2.539 | 0.01112 | +?+ | 0 | 0.116 | 1 | 0.7332 |
| rs11216982 | 11 | 118590864 | A | G | 2.538 | 0.01116 | +++ | 11.1 | 4.498 | 2 | 0.1055 |

| | | | | | | | | | | | |
|---|---|---|---|---|---|---|---|---|---|---|---|
| rs12282154 | 11 | 118807876 | A | G | -2.529 | 0.01143 | -?- | 0 | 0.01 | 1 | 0.92 |
| rs3825057 | 11 | 118622985 | C | T | 2.527 | 0.01149 | +++ | 13.3 | 4.615 | 2 | 0.09949 |
| rs11822103 | 11 | 118869539 | A | G | -2.522 | 0.01167 | -?- | 0 | 0.008 | 1 | 0.9298 |
| rs4499035 | 11 | 118610463 | T | C | 2.521 | 0.01169 | +++ | 13.3 | 4.613 | 2 | 0.0996 |
| rs11603023 | 11 | 118486067 | T | C | 2.515 | 0.01191 | +?+ | 0 | 0.177 | 1 | 0.6738 |
| rs654423 | 11 | 118511757 | A | G | 2.509 | 0.01211 | +?+ | 0 | 0.138 | 1 | 0.7106 |
| rs10892312 | 11 | 118807740 | A | G | -2.507 | 0.01216 | -?- | 0 | 0.015 | 1 | 0.9027 |
| rs4445645 | 11 | 118868750 | C | G | -2.507 | 0.01216 | -?- | 0 | 0.026 | 1 | 0.8708 |
| rs12280501 | 11 | 118801440 | T | C | -2.505 | 0.01225 | -+- | 23.2 | 5.205 | 2 | 0.07409 |
| rs4639966 | 11 | 118573519 | C | T | 2.505 | 0.01226 | +++ | -8 | 3.704 | 2 | 0.1569 |
| rs3737504 | 11 | 118869460 | G | A | -2.502 | 0.01235 | -?- | 0 | 0.017 | 1 | 0.8963 |
| rs2508922 | 11 | 118681088 | A | G | 2.5 | 0.01242 | +++ | 7.9 | 4.342 | 2 | 0.1141 |
| rs7130875 | 11 | 118613674 | C | T | 2.497 | 0.01253 | +++ | 13.8 | 4.642 | 2 | 0.09818 |
| rs10892313 | 11 | 118809329 | A | G | -2.495 | 0.01259 | -?- | 0 | 0.018 | 1 | 0.8931 |
| rs78882064 | 11 | 72084480 | A | C | 2.493 | 0.01267 | +++ | 0 | 1.814 | 2 | 0.4038 |
| rs7117408 | 11 | 118808140 | T | C | -2.488 | 0.01286 | -?- | 0 | 0.02 | 1 | 0.887 |
| rs112799932 | 11 | 118585146 | G | T | 2.486 | 0.01293 | +++ | -14.7 | 3.486 | 2 | 0.175 |
| rs7101680 | 11 | 118799206 | A | G | -2.481 | 0.01312 | -+- | 22.5 | 5.163 | 2 | 0.07566 |
| rs61563240 | 11 | 72056185 | A | G | 2.48 | 0.01313 | +++ | -42.8 | 2.801 | 2 | 0.2465 |
| rs494560 | 11 | 118521549 | A | G | 2.479 | 0.01319 | +?+ | 0 | 0.117 | 1 | 0.732 |
| rs73001486 | 11 | 118629364 | A | G | 2.478 | 0.0132 | +++ | 7.1 | 4.307 | 2 | 0.1161 |
| rs2510902 | 11 | 118700218 | A | G | 2.478 | 0.01322 | +++ | 1.5 | 4.06 | 2 | 0.1313 |
| rs112119822 | 11 | 118798824 | G | A | -2.476 | 0.01327 | -+- | 22.4 | 5.154 | 2 | 0.07602 |
| rs11235612 | 11 | 72613273 | G | A | 2.475 | 0.01334 | -++ | 37.2 | 6.373 | 2 | 0.04132 |
| rs7952074 | 11 | 72062978 | G | A | 2.472 | 0.01343 | +++ | -35.6 | 2.95 | 2 | 0.2288 |
| rs1893029 | 11 | 118610826 | T | G | 2.471 | 0.01348 | +++ | 12 | 4.545 | 2 | 0.103 |
| rs10898895 | 11 | 72741398 | A | T | 2.466 | 0.01368 | -++ | 7.4 | 4.32 | 2 | 0.1153 |
| rs11217016 | 11 | 118627149 | C | A | 2.465 | 0.01368 | +++ | 4.2 | 4.174 | 2 | 0.124 |
| rs487177 | 11 | 118657297 | G | C | 2.463 | 0.01379 | +++ | 8.9 | 4.39 | 2 | 0.1114 |
| rs75610917 | 11 | 72011276 | A | G | 2.46 | 0.01388 | +++ | 0 | 1.621 | 2 | 0.4447 |
| rs75219483 | 11 | 72008410 | T | C | 2.459 | 0.01393 | +++ | 0 | 1.62 | 2 | 0.4449 |
| rs58306819 | 11 | 72054966 | C | T | 2.451 | 0.01425 | +++ | -33.8 | 2.988 | 2 | 0.2244 |
| rs7124498 | 11 | 118616253 | A | G | 2.449 | 0.01433 | +++ | 13.7 | 4.637 | 2 | 0.09841 |
| rs6589713 | 11 | 118872404 | C | T | -2.446 | 0.01443 | -?- | 0 | 0.011 | 1 | 0.9159 |
| rs7934890 | 11 | 118655356 | T | C | 2.432 | 0.01503 | +++ | 5.7 | 4.242 | 2 | 0.1199 |
| rs57276029 | 11 | 72059859 | C | T | 2.427 | 0.0152 | +++ | -31.9 | 3.033 | 2 | 0.2195 |
| rs569770 | 11 | 118584498 | T | C | 2.426 | 0.01525 | +++ | 0 | 0.647 | 2 | 0.7238 |
| rs56104144 | 11 | 118666289 | C | G | 2.425 | 0.0153 | +++ | 17.8 | 4.865 | 2 | 0.08784 |
| rs58868870 | 11 | 118668296 | T | C | 2.412 | 0.01585 | -++ | 19 | 4.939 | 2 | 0.08463 |
| rs11217125 | 11 | 118889185 | C | T | -2.412 | 0.01586 | -?- | 0 | 0.012 | 1 | 0.9111 |
| rs74783030 | 11 | 118603615 | A | G | -2.412 | 0.01588 | --+ | 18.2 | 4.892 | 2 | 0.08665 |
| rs11217014 | 11 | 118626286 | C | T | 2.41 | 0.01597 | +++ | 7.8 | 4.34 | 2 | 0.1142 |
| rs10892282 | 11 | 118627778 | C | G | 2.405 | 0.01615 | +++ | 4.7 | 4.198 | 2 | 0.1226 |
| rs4558175 | 11 | 118872581 | A | G | -2.406 | 0.01615 | -?- | 0 | 0.099 | 1 | 0.7526 |
| rs2508204 | 11 | 72001820 | C | T | 2.401 | 0.01635 | +++ | -2.5 | 3.901 | 2 | 0.1422 |
| rs11217013 | 11 | 118624919 | C | G | 2.399 | 0.01643 | +++ | 7.9 | 4.345 | 2 | 0.1139 |
| rs12416741 | 11 | 72023828 | C | T | 2.393 | 0.0167 | +++ | -1 | 3.959 | 2 | 0.1381 |
| rs57275811 | 11 | 72127872 | G | T | 2.393 | 0.01672 | +++ | -95.9 | 2.042 | 2 | 0.3602 |
| rs4936457 | 11 | 118875727 | G | A | -2.392 | 0.01675 | -?- | 0 | 0.029 | 1 | 0.8645 |
| rs3887652 | 11 | 118856582 | C | G | -2.385 | 0.01707 | -?- | 0 | 0.009 | 1 | 0.9246 |
| rs4936458 | 11 | 118875761 | G | T | -2.381 | 0.01725 | -?- | 0 | 0.025 | 1 | 0.8732 |
| rs112996178 | 11 | 118609598 | T | C | 2.381 | 0.01729 | +++ | 14.7 | 4.69 | 2 | 0.09585 |
| rs73528897 | 11 | 72032681 | A | G | 2.378 | 0.01741 | +++ | -35.4 | 2.955 | 2 | 0.2282 |
| rs112910608 | 11 | 118637475 | T | C | 2.373 | 0.01766 | +++ | 7.5 | 4.327 | 2 | 0.1149 |

| | | | | | | | | | | | |
|---|---|---|---|---|---|---|---|---|---|---|---|
| rs57669200 | 11 | 72026863 | A | G | 2.373 | 0.01766 | +++ | 0.5 | 4.019 | 2 | 0.134 |
| rs7110526 | 11 | 118633533 | C | T | 2.368 | 0.01786 | +++ | 7.7 | 4.335 | 2 | 0.1145 |
| rs11217124 | 11 | 118889132 | G | A | -2.368 | 0.01787 | -?- | 0 | 0.013 | 1 | 0.9094 |
| rs73530754 | 11 | 72099007 | T | C | 2.363 | 0.01815 | +++ | 0 | 0.427 | 2 | 0.8077 |
| rs75184960 | 11 | 72141222 | A | C | 2.362 | 0.01818 | +++ | -96.1 | 2.039 | 2 | 0.3607 |
| rs540800 | 11 | 118565651 | A | G | 2.361 | 0.0182 | +++ | -78.8 | 2.237 | 2 | 0.3268 |
| rs34147990 | 11 | 72160112 | T | C | 2.359 | 0.01832 | +++ | 0 | 0.284 | 2 | 0.8676 |
| rs11217018 | 11 | 118635364 | C | T | 2.359 | 0.01833 | +++ | 6.6 | 4.282 | 2 | 0.1176 |
| rs7124336 | 11 | 72160271 | T | C | 2.353 | 0.01862 | +++ | 0 | 0.281 | 2 | 0.8688 |
| rs73530756 | 11 | 72101747 | T | A | 2.352 | 0.01865 | +++ | 0 | 0.428 | 2 | 0.8073 |
| rs11217021 | 11 | 118640828 | C | T | 2.352 | 0.01867 | +++ | 7.9 | 4.344 | 2 | 0.1139 |
| rs11823672 | 11 | 72160828 | G | A | 2.352 | 0.01867 | +++ | 0 | 0.182 | 2 | 0.9129 |
| rs3916582 | 11 | 118887888 | T | C | -2.35 | 0.01875 | -?- | 0 | 0.011 | 1 | 0.918 |
| rs78639187 | 11 | 72071641 | T | C | 2.347 | 0.0189 | +++ | 0 | 1.725 | 2 | 0.4222 |
| rs11600378 | 11 | 118803842 | G | A | -2.343 | 0.01912 | -?- | 0 | 0.023 | 1 | 0.8792 |
| rs117235696 | 11 | 118803193 | G | T | -2.343 | 0.01913 | -?- | 0 | 0.023 | 1 | 0.8793 |
| rs10892320 | 11 | 118873385 | T | C | -2.342 | 0.0192 | -?- | 0 | 0.063 | 1 | 0.8015 |
| rs11605265 | 11 | 118805144 | A | G | -2.342 | 0.0192 | -?- | 0 | 0.023 | 1 | 0.8803 |
| rs571470 | 11 | 118697012 | A | G | -2.338 | 0.01937 | --+ | 27 | 5.476 | 2 | 0.0647 |
| rs506234 | 11 | 118540104 | G | C | 2.329 | 0.01983 | +++ | -86.5 | 2.145 | 2 | 0.3422 |
| rs59755180 | 11 | 72073763 | C | G | 2.328 | 0.01993 | +++ | -42 | 2.817 | 2 | 0.2445 |
| rs58753751 | 11 | 72048813 | C | T | 2.328 | 0.01994 | +++ | -58.1 | 2.531 | 2 | 0.2821 |
| rs143711539 | 11 | 118804671 | A | G | -2.326 | 0.02001 | -?- | 0 | 0.018 | 1 | 0.8926 |
| rs527619 | 11 | 118563286 | A | G | 2.326 | 0.02002 | +++ | -82.9 | 2.187 | 2 | 0.3351 |
| rs11605658 | 11 | 118805220 | T | C | -2.323 | 0.02018 | -?- | 0 | 0.017 | 1 | 0.8951 |
| rs57291879 | 11 | 72041441 | A | C | 2.322 | 0.02023 | +++ | -57.5 | 2.54 | 2 | 0.2808 |
| rs2508573 | 11 | 118555383 | T | C | 2.32 | 0.02037 | +++ | -77.9 | 2.248 | 2 | 0.325 |
| rs7105187 | 11 | 118887581 | T | C | -2.312 | 0.02079 | -?- | 0 | 0.02 | 1 | 0.8878 |
| rs4291686 | 11 | 118700557 | G | A | 2.311 | 0.02083 | +++ | -15.9 | 3.45 | 2 | 0.1781 |
| rs77916839 | 11 | 118798780 | C | G | -2.308 | 0.02102 | -?- | 0 | 0.048 | 1 | 0.8263 |
| rs60849224 | 11 | 72042930 | C | T | 2.305 | 0.02114 | +++ | -55.7 | 2.57 | 2 | 0.2767 |
| rs36049191 | 11 | 72040784 | C | T | 2.305 | 0.02115 | +++ | -55.6 | 2.57 | 2 | 0.2767 |
| rs2303969 | 11 | 72536272 | A | G | 2.297 | 0.02162 | -++ | 35.7 | 6.219 | 2 | 0.04462 |
| rs625274 | 11 | 118699276 | T | C | 2.296 | 0.02166 | +++ | -20.1 | 3.33 | 2 | 0.1892 |
| rs59608621 | 11 | 118885163 | G | A | -2.29 | 0.02205 | -?- | 0 | 0.022 | 1 | 0.8826 |
| rs4936459 | 11 | 118899218 | T | C | -2.287 | 0.0222 | -+- | -1.2 | 3.955 | 2 | 0.1384 |
| rs7131534 | 11 | 118886656 | A | G | -2.286 | 0.02223 | -?- | 0 | 0.022 | 1 | 0.8829 |
| rs11825284 | 11 | 118880012 | A | G | -2.283 | 0.02244 | -?- | 0 | 0.01 | 1 | 0.9195 |
| rs492471 | 11 | 118689745 | T | C | 2.282 | 0.02251 | +++ | 0 | 0.704 | 2 | 0.7034 |
| rs77293887 | 11 | 72118937 | T | C | 2.277 | 0.0228 | +++ | 0 | 1.775 | 2 | 0.4116 |
| rs10892310 | 11 | 118806897 | T | C | 2.276 | 0.02285 | +-+ | 57.9 | 9.511 | 2 | 0.008604 |
| rs11216855 | 11 | 118208750 | C | T | -2.275 | 0.02292 | -?- | 0 | 0.455 | 1 | 0.5 |
| rs58601794 | 11 | 118884726 | C | T | -2.271 | 0.02313 | -?- | 0 | 0.025 | 1 | 0.8731 |
| rs472837 | 11 | 72007150 | T | C | 2.269 | 0.02326 | +++ | -4 | 3.846 | 2 | 0.1462 |
| rs73559119 | 11 | 118879824 | C | T | -2.268 | 0.02335 | -?- | 0 | 0.007 | 1 | 0.9316 |
| rs7120445 | 11 | 118879285 | C | A | -2.265 | 0.02348 | -?- | 0 | 0.005 | 1 | 0.9421 |
| rs590732 | 11 | 72006394 | T | C | 2.265 | 0.02352 | +++ | -3.7 | 3.859 | 2 | 0.1452 |
| rs58709307 | 11 | 118877563 | T | C | -2.264 | 0.02359 | -?- | 0 | 0.007 | 1 | 0.9319 |
| rs7113753 | 11 | 118889378 | A | G | -2.264 | 0.02359 | -?- | 0 | 0.001 | 1 | 0.9717 |
| rs55772432 | 11 | 118805385 | A | G | 2.26 | 0.02383 | +?+ | 1.9 | 3.058 | 1 | 0.08034 |
| rs61381062 | 11 | 118884708 | A | G | -2.259 | 0.02385 | -?- | 0 | 0.029 | 1 | 0.8639 |
| rs11607569 | 11 | 118880668 | C | T | -2.25 | 0.02447 | -?- | 0 | 0.009 | 1 | 0.9243 |
| rs11820805 | 11 | 118888831 | C | T | -2.249 | 0.02449 | -?- | 0 | 0.037 | 1 | 0.8483 |
| rs60081894 | 11 | 72108715 | C | A | 2.248 | 0.02458 | +++ | 0 | 0.397 | 2 | 0.82 |

| | | | | | | | | | | | |
|---|---|---|---|---|---|---|---|---|---|---|---|
| rs59095933 | 11 | 118885161 | A | G | -2.247 | 0.02466 | -?- | 0 | 0.024 | 1 | 0.8756 |
| rs10751216 | 11 | 72526123 | T | C | -2.245 | 0.02479 | --- | 0 | 0.245 | 2 | 0.8847 |
| rs7947185 | 11 | 118214726 | C | T | -2.241 | 0.02502 | --- | 0 | 1.31 | 2 | 0.5193 |
| rs78488993 | 11 | 72882539 | C | G | 2.236 | 0.02533 | +++ | 0 | 1.117 | 2 | 0.5722 |
| rs1784257 | 11 | 72007984 | C | T | 2.229 | 0.02583 | +++ | -0.7 | 3.974 | 2 | 0.1371 |
| rs35668754 | 11 | 72523110 | C | T | -2.228 | 0.02585 | --- | 0 | 0.217 | 2 | 0.8971 |
| rs6592490 | 11 | 72519756 | C | A | -2.228 | 0.02586 | --- | 0 | 0.235 | 2 | 0.889 |
| rs35343925 | 11 | 72523349 | T | A | -2.228 | 0.02587 | --- | 0 | 0.229 | 2 | 0.8917 |
| rs613791 | 11 | 118764051 | T | C | 2.227 | 0.02592 | +?+ | 0 | 0.076 | 1 | 0.7827 |
| rs35752694 | 11 | 72522881 | A | G | -2.22 | 0.02643 | --- | 0 | 0.218 | 2 | 0.8967 |
| rs11606357 | 11 | 118878222 | C | T | -2.218 | 0.02657 | -?- | 0 | 0.003 | 1 | 0.956 |
| rs1975109 | 11 | 72519217 | G | C | -2.216 | 0.02669 | --- | 0 | 0.219 | 2 | 0.8964 |
| rs10898880 | 11 | 72525111 | A | C | -2.213 | 0.02693 | --- | 0 | 0.246 | 2 | 0.8841 |
| rs10501410 | 11 | 72088806 | A | G | 2.208 | 0.02721 | +++ | 0 | 0.157 | 2 | 0.9247 |
| rs4944805 | 11 | 72522262 | G | T | -2.206 | 0.02735 | --- | 0 | 0.229 | 2 | 0.8919 |
| rs7110719 | 11 | 72521286 | C | T | -2.205 | 0.02746 | --- | 0 | 0.231 | 2 | 0.8909 |
| rs12294037 | 11 | 72807631 | T | A | -2.201 | 0.02771 | --- | -38.2 | 2.894 | 2 | 0.2353 |
| rs10751215 | 11 | 72526087 | C | T | -2.185 | 0.02892 | --- | 0 | 0.212 | 2 | 0.8994 |
| rs7952617 | 11 | 72516946 | C | T | -2.181 | 0.02918 | --- | 0 | 0.324 | 2 | 0.8506 |
| rs11602764 | 11 | 118888619 | G | A | -2.18 | 0.02929 | -?- | 0 | 0.027 | 1 | 0.8701 |
| rs11235596 | 11 | 72514163 | G | A | -2.171 | 0.02995 | --- | 0 | 0.192 | 2 | 0.9087 |
| rs111322300 | 11 | 118893878 | C | T | -2.164 | 0.03043 | --- | 0 | 1.231 | 2 | 0.5405 |
| rs3176912 | 11 | 118757788 | A | G | -2.163 | 0.03051 | -?- | 0 | 0.5 | 1 | 0.4795 |
| rs57928845 | 11 | 72149909 | T | C | 2.162 | 0.03059 | +++ | 0 | 0.732 | 2 | 0.6933 |
| rs9645664 | 11 | 118873987 | C | T | -2.162 | 0.03064 | --- | 0 | 1.311 | 2 | 0.5191 |
| rs60024067 | 11 | 72028610 | C | T | 2.158 | 0.03093 | +++ | 3.7 | 4.154 | 2 | 0.1253 |
| rs60065241 | 11 | 118883491 | A | C | -2.148 | 0.03172 | -?- | 0 | 0.03 | 1 | 0.8618 |
| rs11217122 | 11 | 118880740 | C | T | -2.147 | 0.03178 | -?- | 0 | 0.025 | 1 | 0.875 |
| rs10793057 | 11 | 72968660 | A | G | 2.147 | 0.03181 | +++ | 0 | 0.192 | 2 | 0.9086 |
| rs12280288 | 11 | 118892940 | C | T | -2.143 | 0.03213 | -?- | 0 | 0.001 | 1 | 0.9795 |
| rs10111 | 11 | 118886117 | T | C | -2.141 | 0.0323 | -?- | 0 | 0.032 | 1 | 0.8587 |
| rs6592510 | 11 | 72788823 | C | T | -2.139 | 0.03242 | +-- | -31.1 | 3.05 | 2 | 0.2176 |
| rs2276071 | 11 | 118896116 | C | A | -2.137 | 0.03258 | --- | 0 | 1.187 | 2 | 0.5524 |
| rs57879605 | 11 | 118798757 | T | C | -2.137 | 0.03261 | --- | 0 | 1.867 | 2 | 0.3931 |
| rs12294466 | 11 | 118789657 | T | C | -2.123 | 0.03372 | -+- | -3.3 | 3.873 | 2 | 0.1442 |
| rs143660533 | 11 | 118892759 | C | A | -2.117 | 0.03422 | --- | 0 | 1.178 | 2 | 0.5549 |
| rs117289748 | 11 | 118892757 | G | C | -2.116 | 0.03435 | --- | 0 | 1.175 | 2 | 0.5557 |
| rs1783584 | 11 | 72889137 | C | T | -2.112 | 0.03465 | --- | 0 | 0.197 | 2 | 0.9063 |
| rs61730467 | 11 | 118772160 | A | G | -2.105 | 0.03533 | -?- | 0 | 0.002 | 1 | 0.9613 |
| rs7926282 | 11 | 118605914 | C | A | 2.101 | 0.03563 | -++ | 52.6 | 8.441 | 2 | 0.01469 |
| rs2276072 | 11 | 118896115 | G | A | -2.1 | 0.03577 | --- | 0 | 1.131 | 2 | 0.568 |
| rs598207 | 11 | 118765267 | C | G | 2.1 | 0.03577 | +?+ | 0 | 0.221 | 1 | 0.6384 |
| rs10898877 | 11 | 72511021 | G | A | -2.099 | 0.03586 | --- | 0 | 0.142 | 2 | 0.9316 |
| rs10751213 | 11 | 72510644 | A | C | -2.094 | 0.03626 | --- | 0 | 0.137 | 2 | 0.9339 |
| rs6592489 | 11 | 72510180 | C | G | -2.093 | 0.0363 | --- | 0 | 0.136 | 2 | 0.9341 |
| rs75067211 | 11 | 118854372 | T | C | -2.093 | 0.03639 | -?- | 0 | 0.142 | 1 | 0.7064 |
| rs1784300 | 11 | 118679629 | T | A | 2.09 | 0.0366 | ++- | -3.2 | 3.877 | 2 | 0.1439 |
| rs1000620 | 11 | 72509713 | A | G | -2.089 | 0.03667 | --- | 0 | 0.132 | 2 | 0.9361 |
| rs4459332 | 11 | 72508267 | C | T | -2.087 | 0.03689 | --- | 0 | 0.132 | 2 | 0.9359 |
| rs677231 | 11 | 72504848 | A | T | -2.083 | 0.03723 | --- | 0 | 0.129 | 2 | 0.9377 |
| rs138922408 | 11 | 118356711 | T | C | -2.082 | 0.03733 | --- | 16.4 | 4.782 | 2 | 0.09154 |
| rs4944800 | 11 | 72506130 | C | T | -2.082 | 0.03737 | --- | 0 | 0.127 | 2 | 0.9385 |
| rs555649 | 11 | 118651809 | C | T | 2.08 | 0.03749 | ++- | -25 | 3.201 | 2 | 0.2018 |
| rs7925155 | 11 | 72732430 | C | G | -2.076 | 0.03793 | +-- | -33.5 | 2.997 | 2 | 0.2235 |

| | | | | | | | | | | | |
|---|---|---|---|---|---|---|---|---|---|---|---|
| rs607472 | 11 | 118680072 | G | C | 2.075 | 0.03795 | ++- | 2.1 | 4.084 | 2 | 0.1298 |
| rs947931 | 11 | 118577614 | G | T | -2.073 | 0.03817 | --- | -56.9 | 2.55 | 2 | 0.2794 |
| rs4459331 | 11 | 72507938 | C | T | -2.069 | 0.03858 | --- | 0 | 0.149 | 2 | 0.9282 |
| rs7127427 | 11 | 118805138 | A | G | -2.066 | 0.03884 | -?- | 0 | 0.22 | 1 | 0.6389 |
| rs75210328 | 11 | 118801296 | T | G | -2.063 | 0.03909 | -?- | 0 | 0.223 | 1 | 0.637 |
| rs2510897 | 11 | 118644582 | C | A | 2.062 | 0.03918 | ++- | -25.8 | 3.181 | 2 | 0.2039 |
| rs2186780 | 11 | 118605451 | T | C | 2.058 | 0.03958 | -++ | 49.8 | 7.966 | 2 | 0.01863 |
| rs12796102 | 11 | 118662028 | T | C | 2.056 | 0.03976 | ++- | -21.5 | 3.293 | 2 | 0.1927 |
| rs546393 | 11 | 118624250 | C | T | 2.054 | 0.03994 | ++- | -11.4 | 3.59 | 2 | 0.1661 |
| rs7127188 | 11 | 118893023 | C | A | -2.049 | 0.04045 | --- | 0 | 1.092 | 2 | 0.5794 |
| rs2105782 | 11 | 118659896 | C | T | 2.048 | 0.04055 | ++- | -22.4 | 3.269 | 2 | 0.195 |
| rs61179449 | 11 | 118892835 | C | G | -2.046 | 0.04076 | --- | 0 | 1.084 | 2 | 0.5817 |
| rs10431171 | 11 | 72512271 | C | G | -2.045 | 0.04085 | --- | 0 | 0.258 | 2 | 0.8789 |
| rs581045 | 11 | 118632522 | T | C | 2.045 | 0.0409 | ++- | -17.4 | 3.406 | 2 | 0.1821 |
| rs58955701 | 11 | 118892872 | T | C | -2.045 | 0.0409 | --- | 0 | 1.081 | 2 | 0.5825 |
| rs10431172 | 11 | 72512630 | T | C | -2.042 | 0.04115 | --- | 0 | 0.254 | 2 | 0.8808 |
| rs4944802 | 11 | 72510476 | G | A | -2.041 | 0.04129 | --- | 0 | 0.223 | 2 | 0.8947 |
| rs4938621 | 11 | 118892277 | T | C | -2.039 | 0.04148 | --- | 0 | 1.07 | 2 | 0.5857 |
| rs488219 | 11 | 118620697 | C | T | 2.037 | 0.04166 | ++- | -12.9 | 3.542 | 2 | 0.1702 |
| rs7929825 | 11 | 118788081 | G | C | -2.035 | 0.04189 | --- | 0 | 1.35 | 2 | 0.509 |
| rs76068633 | 11 | 118416268 | T | C | -2.032 | 0.04211 | --- | 27.9 | 5.551 | 2 | 0.06232 |
| rs555639 | 11 | 118629031 | A | G | 2.029 | 0.04244 | ++- | -18.4 | 3.378 | 2 | 0.1847 |
| rs150792202 | 11 | 118652569 | A | G | 2.029 | 0.04249 | +?+ | -20.4 | 2.492 | 1 | 0.1144 |
| rs73530785 | 11 | 72152606 | G | A | 2.021 | 0.04333 | +++ | 0 | 0.495 | 2 | 0.7807 |
| rs116976310 | 11 | 118420099 | C | A | -2.013 | 0.04414 | --- | 12.8 | 4.586 | 2 | 0.101 |
| rs6592486 | 11 | 72508609 | A | G | -2.011 | 0.04434 | --- | 0 | 0.189 | 2 | 0.9098 |
| rs515523 | 11 | 118658075 | G | A | 2.003 | 0.0452 | ++- | -12.2 | 3.564 | 2 | 0.1683 |
| rs11006 | 11 | 118895202 | G | A | -2.002 | 0.04525 | --- | 0 | 0.995 | 2 | 0.608 |
| rs74891547 | 11 | 118811878 | C | G | -2.002 | 0.04533 | -?- | 0 | 0.133 | 1 | 0.7156 |
| rs607125 | 11 | 118617299 | C | T | 1.997 | 0.04579 | ++- | -14.2 | 3.502 | 2 | 0.1736 |
| rs1864653 | 11 | 72503293 | A | G | -1.992 | 0.04641 | --- | 0 | 0.172 | 2 | 0.9176 |
| rs11235594 | 11 | 72503735 | G | A | -1.991 | 0.04649 | --- | 0 | 0.166 | 2 | 0.9203 |
| rs1784543 | 11 | 118631653 | T | A | 1.987 | 0.0469 | ++- | -16 | 3.449 | 2 | 0.1782 |
| rs11235592 | 11 | 72501277 | C | G | -1.987 | 0.04693 | --- | 0 | 0.154 | 2 | 0.9258 |
| rs507890 | 11 | 118631288 | C | T | 1.979 | 0.04784 | ++- | -18 | 3.39 | 2 | 0.1836 |
| rs485099 | 11 | 72500060 | C | T | -1.97 | 0.04881 | --- | 0 | 0.151 | 2 | 0.9271 |
| rs10892234 | 11 | 118194571 | G | A | -1.969 | 0.04893 | --- | 0 | 1.48 | 2 | 0.4771 |
| rs3862794 | 11 | 72538600 | C | T | -1.964 | 0.04954 | +-- | -1.3 | 3.948 | 2 | 0.1389 |
| rs4944797 | 11 | 72500072 | T | G | -1.963 | 0.04965 | --- | 0 | 0.173 | 2 | 0.9173 |
| rs57464210 | 11 | 72043183 | G | T | 1.961 | 0.04989 | +++ | -1.2 | 3.952 | 2 | 0.1386 |
| rs4944796 | 11 | 72499000 | C | A | -1.959 | 0.0501 | --- | 0 | 0.166 | 2 | 0.9204 |
| rs4394861 | 11 | 72800970 | C | T | -1.955 | 0.05064 | +-- | -28.1 | 3.123 | 2 | 0.2098 |
| rs79108568 | 11 | 118775524 | G | T | -1.953 | 0.05085 | -?- | 0 | 0.001 | 1 | 0.9696 |
| rs2187557 | 11 | 118148271 | T | C | -1.944 | 0.05194 | -?- | 0 | 0.023 | 1 | 0.8789 |
| rs58682117 | 11 | 72531000 | C | T | -1.931 | 0.05354 | +-- | 64.2 | 11.173 | 2 | 0.003748 |
| rs11216904 | 11 | 118440863 | G | A | 1.908 | 0.05645 | +++ | 0 | 0.488 | 2 | 0.7833 |
| rs12224844 | 11 | 118439048 | T | A | 1.896 | 0.05802 | +++ | 0 | 0.419 | 2 | 0.811 |
| rs10892243 | 11 | 118441846 | C | T | 1.889 | 0.05895 | +++ | 0 | 0.456 | 2 | 0.7962 |
| rs60837962 | 11 | 118438448 | T | A | 1.888 | 0.05899 | +++ | 0 | 0.538 | 2 | 0.7641 |
| rs10736492 | 11 | 118459269 | G | A | 1.887 | 0.05922 | +++ | 0 | 0.5 | 2 | 0.7787 |
| rs569 | 11 | 118894287 | G | A | -1.883 | 0.05972 | --- | 0 | 0.854 | 2 | 0.6525 |
| rs61506726 | 11 | 118831335 | G | A | -1.878 | 0.06038 | --- | 0 | 1.078 | 2 | 0.5832 |
| rs11216898 | 11 | 118432852 | C | T | 1.865 | 0.06218 | +++ | 0 | 0.672 | 2 | 0.7147 |
| rs60067274 | 11 | 118412296 | T | A | 1.855 | 0.06361 | +++ | 0 | 0.465 | 2 | 0.7927 |

| | | | | | | | | | | | |
|---|---|---|---|---|---|---|---|---|---|---|---|
| rs76160025 | 11 | 118782802 | T | C | -1.851 | 0.06413 | -?- | 0 | 0.271 | 1 | 0.6029 |
| rs11216897 | 11 | 118432279 | G | A | 1.851 | 0.06414 | +++ | 0 | 0.526 | 2 | 0.7687 |
| rs3741319 | 11 | 118415848 | C | A | 1.851 | 0.06424 | +++ | 0 | 0.419 | 2 | 0.8112 |
| rs11216889 | 11 | 118421512 | A | G | 1.847 | 0.06479 | +++ | 0 | 0.483 | 2 | 0.7853 |
| rs11216900 | 11 | 118436079 | C | G | 1.845 | 0.06507 | +++ | 0 | 0.632 | 2 | 0.7292 |
| rs4326835 | 11 | 118432404 | T | G | 1.844 | 0.06515 | +++ | 0 | 0.605 | 2 | 0.7391 |
| rs59108310 | 11 | 118433729 | G | A | 1.842 | 0.06542 | +++ | 0 | 0.496 | 2 | 0.7805 |
| rs566733 | 11 | 118608625 | G | C | 1.842 | 0.06551 | ++- | -29.8 | 3.082 | 2 | 0.2142 |
| rs3825055 | 11 | 118416073 | C | A | 1.841 | 0.06565 | +++ | 0 | 0.424 | 2 | 0.8091 |
| rs3741320 | 11 | 118415847 | C | G | 1.839 | 0.06593 | +++ | 0 | 0.411 | 2 | 0.8141 |
| rs11216899 | 11 | 118434421 | G | A | 1.835 | 0.06651 | +++ | 0 | 0.599 | 2 | 0.7413 |
| rs11216902 | 11 | 118437279 | G | A | 1.833 | 0.06682 | +++ | 0 | 0.483 | 2 | 0.7854 |
| rs11216890 | 11 | 118421569 | G | C | 1.83 | 0.06726 | +++ | 0 | 0.478 | 2 | 0.7876 |
| rs60753148 | 11 | 118434835 | T | C | 1.827 | 0.06769 | +++ | 0 | 0.584 | 2 | 0.7467 |
| rs11600585 | 11 | 72727840 | G | A | 1.825 | 0.06806 | -++ | 39.6 | 6.618 | 2 | 0.03655 |
| rs2277293 | 11 | 118422868 | T | C | 1.824 | 0.06816 | +++ | 0 | 0.449 | 2 | 0.7988 |
| rs2277291 | 11 | 118435756 | T | C | 1.819 | 0.06892 | +++ | 0 | 0.568 | 2 | 0.7526 |
| rs11216891 | 11 | 118422234 | T | A | 1.818 | 0.06902 | +++ | 0 | 0.468 | 2 | 0.7915 |
| rs73022058 | 11 | 118423672 | T | C | 1.818 | 0.06906 | +++ | 0 | 0.467 | 2 | 0.7918 |
| rs2277295 | 11 | 118416253 | T | C | 1.817 | 0.06929 | +++ | 0 | 0.585 | 2 | 0.7462 |
| rs11217004 | 11 | 118607622 | A | G | 1.812 | 0.06998 | ++- | -26.6 | 3.16 | 2 | 0.2059 |
| rs567680 | 11 | 118608532 | G | C | 1.81 | 0.07029 | ++- | -23.1 | 3.25 | 2 | 0.1969 |
| rs4456287 | 11 | 118298252 | C | T | 1.803 | 0.07132 | +++ | 0 | 0.627 | 2 | 0.731 |
| rs187547749 | 11 | 118439159 | T | C | -1.803 | 0.07146 | --- | 3.9 | 4.162 | 2 | 0.1248 |
| rs113739713 | 11 | 118420101 | G | A | 1.801 | 0.07167 | +++ | 0 | 0.428 | 2 | 0.8074 |
| rs12274926 | 11 | 118295117 | C | T | 1.8 | 0.07192 | +++ | 0 | 0.566 | 2 | 0.7534 |
| rs632124 | 11 | 118613235 | A | T | 1.799 | 0.07205 | ++- | -23.1 | 3.249 | 2 | 0.197 |
| rs10892245 | 11 | 118472354 | A | C | 1.798 | 0.07223 | +++ | 0 | 0.441 | 2 | 0.8022 |
| rs7929887 | 11 | 118428791 | G | A | 1.797 | 0.07232 | +++ | 0 | 0.389 | 2 | 0.8231 |
| rs2511062 | 11 | 72871055 | C | T | -1.795 | 0.0726 | +-- | -25.8 | 3.179 | 2 | 0.204 |
| rs74384601 | 11 | 118449313 | G | A | -1.793 | 0.07304 | --- | -4.7 | 3.819 | 2 | 0.1481 |
| rs12788933 | 11 | 118140184 | T | C | 1.792 | 0.07319 | +?+ | -35.7 | 2.21 | 1 | 0.1371 |
| rs7109082 | 11 | 72707441 | C | A | 1.789 | 0.07355 | -++ | 37.3 | 6.38 | 2 | 0.04118 |
| rs188843024 | 11 | 118453478 | A | G | -1.789 | 0.07356 | --- | -32.6 | 3.016 | 2 | 0.2214 |
| rs12797009 | 11 | 118826192 | A | C | 1.786 | 0.07413 | +?+ | 0 | 0.314 | 1 | 0.5751 |
| rs11216925 | 11 | 118471423 | C | T | 1.78 | 0.07502 | +++ | 0 | 0.444 | 2 | 0.8008 |
| rs187678279 | 11 | 118338067 | A | G | -1.777 | 0.07557 | --- | 22.2 | 5.138 | 2 | 0.07661 |
| rs56403966 | 11 | 118425051 | A | G | 1.774 | 0.07601 | +++ | 0 | 0.446 | 2 | 0.8 |
| rs11216926 | 11 | 118471996 | T | C | 1.772 | 0.07639 | +++ | 0 | 0.472 | 2 | 0.7898 |
| rs58270621 | 11 | 118413604 | A | G | 1.772 | 0.0764 | +++ | 0 | 0.449 | 2 | 0.7989 |
| rs141108621 | 11 | 118349211 | T | C | -1.771 | 0.07658 | --- | 13.9 | 4.648 | 2 | 0.09787 |
| rs11216909 | 11 | 118449746 | T | C | 1.769 | 0.07685 | +++ | 0 | 0.424 | 2 | 0.8088 |
| rs11216892 | 11 | 118424008 | T | C | 1.769 | 0.07694 | +++ | 0 | 0.383 | 2 | 0.8259 |
| rs55857590 | 11 | 118425063 | G | A | 1.768 | 0.07707 | +++ | 0 | 0.396 | 2 | 0.8205 |
| rs17742 | 11 | 118473385 | C | T | 1.766 | 0.07735 | +++ | 0 | 0.419 | 2 | 0.8108 |
| rs11216886 | 11 | 118411424 | C | A | 1.764 | 0.07768 | +++ | 0 | 0.505 | 2 | 0.7768 |
| rs7932700 | 11 | 118411713 | T | G | 1.764 | 0.07773 | +++ | 0 | 0.492 | 2 | 0.7817 |
| rs4338560 | 11 | 118143368 | T | C | 1.762 | 0.07805 | +?+ | -30.5 | 2.299 | 1 | 0.1294 |
| rs57156480 | 11 | 118896959 | A | G | -1.761 | 0.0782 | --- | 0 | 0.766 | 2 | 0.6819 |
| rs11216896 | 11 | 118430048 | T | C | 1.76 | 0.07833 | +++ | 0 | 0.394 | 2 | 0.8212 |
| rs3782042 | 11 | 118185919 | A | G | -1.757 | 0.07898 | -?- | -25.5 | 2.391 | 1 | 0.1221 |
| rs11552421 | 11 | 118430519 | T | C | 1.755 | 0.07925 | +++ | 0 | 0.358 | 2 | 0.8361 |
| rs1047079 | 11 | 118473231 | C | T | 1.746 | 0.08085 | +++ | 0 | 0.503 | 2 | 0.7777 |
| rs574387 | 11 | 118337003 | C | A | 1.745 | 0.08099 | +++ | 0 | 0.265 | 2 | 0.876 |

| | | | | | | | | | | | |
|---|---|---|---|---|---|---|---|---|---|---|---|
| rs3741325 | 11 | 118405989 | G | A | 1.743 | 0.08127 | +++ | 0 | 0.508 | 2 | 0.7757 |
| rs11216881 | 11 | 118409539 | G | A | 1.743 | 0.08133 | +++ | 0 | 0.455 | 2 | 0.7966 |
| rs45505501 | 11 | 118405305 | A | G | 1.742 | 0.08148 | +++ | 0 | 0.444 | 2 | 0.8008 |
| rs113054880 | 11 | 118459702 | G | C | 1.742 | 0.08158 | +++ | 0 | 0.502 | 2 | 0.7781 |
| rs11216907 | 11 | 118444525 | A | G | 1.735 | 0.08282 | +++ | 0 | 0.53 | 2 | 0.7672 |
| rs11216880 | 11 | 118409536 | A | G | 1.733 | 0.08304 | +++ | 0 | 0.452 | 2 | 0.7976 |
| rs73022038 | 11 | 118408471 | T | C | 1.732 | 0.08321 | +++ | 0 | 0.448 | 2 | 0.7993 |
| rs57438673 | 11 | 118459039 | A | G | 1.732 | 0.08329 | +++ | 0 | 0.501 | 2 | 0.7782 |
| rs11216883 | 11 | 118410468 | G | A | 1.732 | 0.08335 | +++ | 0 | 0.454 | 2 | 0.797 |
| rs11216906 | 11 | 118443944 | A | G | 1.731 | 0.08344 | +++ | 0 | 0.426 | 2 | 0.8082 |
| rs11216921 | 11 | 118464815 | C | G | 1.731 | 0.08345 | +++ | 0 | 0.466 | 2 | 0.7922 |
| rs7930148 | 11 | 118435966 | A | G | 1.73 | 0.08354 | +++ | 0 | 0.333 | 2 | 0.8466 |
| rs17122269 | 11 | 118445275 | C | T | 1.729 | 0.08382 | +++ | 0 | 0.526 | 2 | 0.7687 |
| rs2277292 | 11 | 118425419 | T | G | 1.728 | 0.08402 | +++ | 0 | 0.373 | 2 | 0.8299 |
| rs11216924 | 11 | 118470092 | G | C | 1.728 | 0.08405 | +++ | 0 | 0.526 | 2 | 0.7687 |
| rs11216912 | 11 | 118452334 | C | G | 1.728 | 0.08406 | +++ | 0 | 0.431 | 2 | 0.8061 |
| rs73022041 | 11 | 118408789 | C | T | 1.727 | 0.08412 | +++ | 0 | 0.45 | 2 | 0.7987 |
| rs73023350 | 11 | 118449396 | A | G | 1.727 | 0.08422 | +++ | 0 | 0.43 | 2 | 0.8067 |
| rs11216885 | 11 | 118410545 | G | A | 1.726 | 0.08426 | +++ | 0 | 0.444 | 2 | 0.8009 |
| rs10898881 | 11 | 72527018 | A | G | 1.726 | 0.08427 | +++ | 0 | 1.007 | 2 | 0.6043 |
| rs9736327 | 11 | 118447580 | G | T | 1.726 | 0.0843 | +++ | 0 | 0.429 | 2 | 0.807 |
| rs11216917 | 11 | 118462160 | G | T | 1.726 | 0.08431 | +++ | 0 | 0.365 | 2 | 0.8332 |
| rs7126175 | 11 | 118450875 | T | C | 1.726 | 0.08438 | +++ | 0 | 0.428 | 2 | 0.8074 |
| rs45485302 | 11 | 118405377 | A | C | 1.725 | 0.08451 | +++ | 0 | 0.415 | 2 | 0.8128 |
| rs9735421 | 11 | 118447597 | A | G | 1.724 | 0.08466 | +++ | 0 | 0.425 | 2 | 0.8085 |
| rs17092756 | 11 | 118457630 | C | G | 1.723 | 0.08497 | +++ | 0 | 0.421 | 2 | 0.8101 |
| rs11216884 | 11 | 118410488 | C | T | 1.722 | 0.08499 | +++ | 0 | 0.451 | 2 | 0.798 |
| rs45474901 | 11 | 118405453 | C | T | 1.722 | 0.08508 | +++ | 0 | 0.564 | 2 | 0.7542 |
| rs11216911 | 11 | 118451901 | C | T | 1.72 | 0.08535 | +++ | 0 | 0.419 | 2 | 0.8112 |
| rs11216923 | 11 | 118465501 | G | A | 1.72 | 0.08549 | +++ | 0 | 0.392 | 2 | 0.8221 |
| rs7131631 | 11 | 118409374 | A | G | 1.719 | 0.08552 | +++ | 0 | 0.444 | 2 | 0.8008 |
| rs45444192 | 11 | 118208083 | A | G | 1.719 | 0.0856 | +?+ | 0 | 0.144 | 1 | 0.7042 |
| rs9735449 | 11 | 118448087 | T | C | 1.719 | 0.08562 | +++ | 0 | 0.488 | 2 | 0.7835 |
| rs7128587 | 11 | 118409302 | A | G | 1.719 | 0.08566 | +++ | 0 | 0.444 | 2 | 0.8008 |
| rs61290697 | 11 | 118412358 | T | C | 1.718 | 0.08572 | +++ | 0 | 0.444 | 2 | 0.801 |
| rs73022042 | 11 | 118408893 | A | G | 1.718 | 0.08581 | +++ | 0 | 0.436 | 2 | 0.8042 |
| rs11216913 | 11 | 118453681 | G | A | 1.717 | 0.08596 | +++ | 0 | 0.469 | 2 | 0.791 |
| rs59176377 | 11 | 118419295 | A | G | 1.716 | 0.08613 | +++ | 0 | 0.366 | 2 | 0.8326 |
| rs74404016 | 11 | 118408653 | T | C | 1.716 | 0.08613 | +++ | 0 | 0.444 | 2 | 0.8011 |
| rs111942423 | 11 | 118429764 | G | T | 1.715 | 0.08628 | +++ | 0 | 0.358 | 2 | 0.836 |
| rs7101585 | 11 | 118409583 | A | G | 1.715 | 0.08635 | +++ | 0 | 0.448 | 2 | 0.7992 |
| rs17122278 | 11 | 118449370 | G | A | 1.713 | 0.08677 | +++ | 0 | 0.53 | 2 | 0.7671 |
| rs73023341 | 11 | 118445585 | G | A | 1.712 | 0.08697 | +++ | 0 | 0.373 | 2 | 0.8297 |
| rs11216922 | 11 | 118465295 | G | A | 1.711 | 0.08707 | +++ | 0 | 0.377 | 2 | 0.828 |
| rs7110743 | 11 | 118825762 | A | C | 1.711 | 0.08713 | +?+ | 0 | 0.586 | 1 | 0.4438 |
| rs4435039 | 11 | 118297391 | G | A | 1.71 | 0.08724 | +++ | 0 | 0.913 | 2 | 0.6335 |
| rs10892317 | 11 | 118832035 | C | T | -1.71 | 0.08733 | --- | 0 | 1.021 | 2 | 0.6001 |
| rs7115634 | 11 | 118466249 | G | A | 1.709 | 0.08744 | +++ | 0 | 0.49 | 2 | 0.7827 |
| rs113975105 | 11 | 118836606 | G | C | -1.708 | 0.08772 | --- | 0 | 1.082 | 2 | 0.5822 |
| rs58577143 | 11 | 118291210 | G | A | 1.706 | 0.08804 | +++ | 0 | 1.748 | 2 | 0.4174 |
| rs11216887 | 11 | 118420736 | C | T | 1.705 | 0.08826 | +++ | 0 | 0.399 | 2 | 0.819 |
| rs535207 | 11 | 118408033 | G | A | 1.703 | 0.08855 | +++ | 0 | 0.441 | 2 | 0.8023 |
| rs11216910 | 11 | 118450046 | G | A | 1.703 | 0.08856 | +++ | 0 | 0.452 | 2 | 0.7977 |
| rs55844025 | 11 | 118446298 | T | C | 1.703 | 0.0886 | +++ | 0 | 0.487 | 2 | 0.7838 |

| | | | | | | | | | | | |
|---|---|---|---|---|---|---|---|---|---|---|---|
| rs11216916 | 11 | 118461605 | C | T | 1.701 | 0.08892 | +++ | 0 | 0.341 | 2 | 0.8431 |
| rs45548238 | 11 | 118405437 | T | C | 1.7 | 0.0891 | +++ | 0 | 0.495 | 2 | 0.7809 |
| rs17842763 | 11 | 118454177 | G | A | 1.699 | 0.08923 | +++ | 0 | 0.436 | 2 | 0.804 |
| rs7931564 | 11 | 118469571 | G | A | 1.698 | 0.08948 | +++ | 0 | 0.471 | 2 | 0.7903 |
| rs59255055 | 11 | 118296735 | A | C | 1.696 | 0.0899 | +++ | 0 | 0.516 | 2 | 0.7727 |
| rs45522239 | 11 | 118405438 | G | A | 1.693 | 0.0905 | +++ | 0 | 0.491 | 2 | 0.7822 |
| rs10502227 | 11 | 118147496 | C | T | 1.692 | 0.0906 | +?+ | -49.7 | 2.003 | 1 | 0.157 |
| rs3741323 | 11 | 118405641 | G | A | 1.69 | 0.0911 | +++ | 0 | 0.424 | 2 | 0.809 |
| rs10892315 | 11 | 118831727 | C | T | -1.687 | 0.09152 | --- | 0 | 0.945 | 2 | 0.6235 |
| rs60112336 | 11 | 118453483 | T | C | 1.684 | 0.09224 | +++ | 0 | 0.509 | 2 | 0.7752 |
| rs536256 | 11 | 118327321 | G | A | 1.681 | 0.09283 | +++ | 0 | 0.174 | 2 | 0.9167 |
| rs11216919 | 11 | 118462367 | G | A | 1.678 | 0.09338 | +++ | 0 | 0.27 | 2 | 0.8739 |
| rs6592500 | 11 | 72651705 | G | A | -1.677 | 0.09361 | +-- | -2.9 | 3.889 | 2 | 0.1431 |
| rs11603267 | 11 | 118146599 | T | G | 1.669 | 0.09514 | +?+ | -56.6 | 1.916 | 1 | 0.1663 |
| rs7108845 | 11 | 118407523 | G | T | 1.661 | 0.09673 | +++ | 0 | 0.433 | 2 | 0.8053 |
| rs113971393 | 11 | 72076181 | C | T | 1.659 | 0.09703 | +++ | 0 | 1.218 | 2 | 0.544 |
| rs11216914 | 11 | 118456414 | C | T | 1.658 | 0.09723 | +++ | 0 | 0.394 | 2 | 0.8213 |
| rs6589664 | 11 | 118404804 | A | G | 1.657 | 0.09756 | +++ | 0 | 1.223 | 2 | 0.5426 |
| rs3741324 | 11 | 118405835 | A | G | 1.649 | 0.09916 | +++ | 0 | 0.47 | 2 | 0.7905 |
| rs2886599 | 11 | 72721271 | A | C | 1.643 | 0.1003 | -++ | 39.4 | 6.603 | 2 | 0.03682 |
| rs79915705 | 11 | 72016091 | C | T | 1.635 | 0.1021 | +++ | 0 | 0.52 | 2 | 0.7712 |
| rs11603313 | 11 | 72588177 | A | G | 1.634 | 0.1023 | -++ | 47 | 7.551 | 2 | 0.02292 |
| rs58571741 | 11 | 72065710 | A | C | 1.634 | 0.1024 | +++ | 0 | 1.224 | 2 | 0.5423 |
| rs4938551 | 11 | 118707249 | C | G | 1.625 | 0.1041 | -++ | 0 | 1.617 | 2 | 0.4456 |
| rs17244499 | 11 | 72629946 | G | A | 1.625 | 0.1042 | -++ | 46.2 | 7.442 | 2 | 0.02422 |
| rs8192696 | 11 | 118895686 | T | C | -1.618 | 0.1056 | --- | 0 | 0.493 | 2 | 0.7817 |
| rs548877 | 11 | 118300240 | G | A | 1.618 | 0.1057 | +++ | 0 | 0.54 | 2 | 0.7633 |
| rs666191 | 11 | 118826063 | A | G | -1.615 | 0.1064 | -?- | 0 | 0.341 | 1 | 0.5592 |
| rs11605166 | 11 | 72669777 | C | T | 1.608 | 0.1079 | -++ | 49.4 | 7.898 | 2 | 0.01927 |
| rs3814730 | 11 | 72555553 | A | G | 1.608 | 0.1079 | -++ | 48 | 7.69 | 2 | 0.02138 |
| rs566416 | 11 | 118759610 | G | T | 1.605 | 0.1086 | +-+ | -63 | 2.454 | 2 | 0.2932 |
| rs4938507 | 11 | 118231584 | G | A | 1.602 | 0.1093 | +?+ | 0 | 0.216 | 1 | 0.6421 |
| rs77074463 | 11 | 118748170 | C | G | 1.6 | 0.1095 | +++ | 0 | 0.835 | 2 | 0.6586 |
| rs498684 | 11 | 118316960 | A | G | 1.6 | 0.1096 | +++ | 0 | 0.345 | 2 | 0.8414 |
| rs625291 | 11 | 118826060 | T | C | -1.599 | 0.1097 | -?- | 0 | 0.322 | 1 | 0.5705 |
| rs79281178 | 11 | 118298041 | G | C | 1.599 | 0.1099 | +++ | 0 | 0.475 | 2 | 0.7887 |
| rs34815971 | 11 | 118301066 | G | A | 1.596 | 0.1105 | +++ | 0 | 0.552 | 2 | 0.7589 |
| rs510997 | 11 | 118302901 | C | T | 1.594 | 0.1108 | +++ | 0 | 0.586 | 2 | 0.7458 |
| rs4426142 | 11 | 118824996 | A | G | -1.594 | 0.111 | -?- | 0 | 0.152 | 1 | 0.6968 |
| rs74439784 | 11 | 72649396 | T | A | 1.593 | 0.1111 | -++ | 46.1 | 7.426 | 2 | 0.0244 |
| rs73613993 | 11 | 118296007 | C | T | 1.592 | 0.1114 | +++ | 0 | 0.533 | 2 | 0.7661 |
| rs145398037 | 11 | 72522142 | C | G | 1.59 | 0.1119 | -++ | 51.1 | 8.175 | 2 | 0.01678 |
| rs7122959 | 11 | 118804283 | G | A | -1.586 | 0.1128 | -+- | -44.6 | 2.766 | 2 | 0.2508 |
| rs45573242 | 11 | 118853995 | C | A | 1.581 | 0.1139 | +-+ | -48.4 | 2.695 | 2 | 0.2599 |
| rs7479331 | 11 | 118829521 | T | G | 1.574 | 0.1155 | +++ | -32.6 | 3.016 | 2 | 0.2213 |
| rs7123066 | 11 | 118804266 | T | C | -1.573 | 0.1158 | -+- | -47.2 | 2.718 | 2 | 0.257 |
| rs118020807 | 11 | 118759905 | A | C | -1.571 | 0.1162 | -?- | 0 | 0.569 | 1 | 0.4506 |
| rs117819392 | 11 | 72514503 | T | C | 1.57 | 0.1163 | -++ | 47.3 | 7.588 | 2 | 0.02251 |
| rs543692 | 11 | 118598138 | C | T | 1.567 | 0.1171 | ++- | 8.4 | 4.369 | 2 | 0.1126 |
| rs563537 | 11 | 118595207 | A | T | 1.563 | 0.118 | ++- | 10.1 | 4.451 | 2 | 0.108 |
| rs76793820 | 11 | 72619860 | A | G | 1.56 | 0.1187 | -++ | 47.2 | 7.582 | 2 | 0.02257 |
| rs568688 | 11 | 118598612 | G | T | 1.558 | 0.1191 | ++- | 9.1 | 4.402 | 2 | 0.1107 |
| rs2097131 | 11 | 118825069 | A | G | -1.554 | 0.1203 | -?- | 0 | 0.147 | 1 | 0.7019 |
| rs4515995 | 11 | 72687795 | G | A | -1.55 | 0.1212 | --- | 0 | 1.566 | 2 | 0.457 |

| | | | | | | | | | | | |
|---|---|---|---|---|---|---|---|---|---|---|---|
| rs566848 | 11 | 118711830 | G | A | 1.537 | 0.1243 | +++ | 0 | 0.792 | 2 | 0.6729 |
| rs75657287 | 11 | 118801518 | G | C | 1.535 | 0.1247 | +?+ | 0 | 0.129 | 1 | 0.72 |
| rs139534838 | 11 | 72506880 | C | T | 1.532 | 0.1255 | -++ | 50.1 | 8.022 | 2 | 0.01811 |
| rs61900157 | 11 | 118824439 | C | A | -1.531 | 0.1259 | -?- | 0 | 0.142 | 1 | 0.7063 |
| rs7124100 | 11 | 118290169 | A | G | 1.529 | 0.1262 | +++ | 0 | 0.677 | 2 | 0.713 |
| rs4938615 | 11 | 118865319 | G | C | 1.526 | 0.127 | +?+ | 0 | 0.943 | 1 | 0.3316 |
| rs9794909 | 11 | 118862306 | C | G | -1.521 | 0.1283 | --- | 0 | 1.112 | 2 | 0.5734 |
| rs146026283 | 11 | 118296322 | A | C | 1.52 | 0.1284 | +?+ | 0 | 0.284 | 1 | 0.5941 |
| rs113145584 | 11 | 118154798 | C | G | -1.518 | 0.129 | -?- | 0 | 0.909 | 1 | 0.3403 |
| rs11602858 | 11 | 72615069 | G | C | 1.513 | 0.1303 | -++ | 45.7 | 7.36 | 2 | 0.02523 |
| rs1059000 | 11 | 72504972 | C | G | 1.511 | 0.1308 | -++ | 49.2 | 7.871 | 2 | 0.01954 |
| rs676925 | 11 | 118765688 | G | C | 1.509 | 0.1313 | +-+ | -83.1 | 2.185 | 2 | 0.3354 |
| rs11217052 | 11 | 118717732 | G | A | 1.508 | 0.1316 | -++ | 0 | 1.774 | 2 | 0.412 |
| rs537975 | 11 | 118362869 | T | C | 1.507 | 0.1318 | +++ | 0 | 0.179 | 2 | 0.9143 |
| rs111527225 | 11 | 118065830 | A | G | 1.502 | 0.1331 | +?+ | -139.7 | 1.252 | 1 | 0.2632 |
| rs112753655 | 11 | 118154795 | A | G | -1.501 | 0.1333 | -?- | 0 | 0.942 | 1 | 0.3317 |
| rs484232 | 11 | 118708564 | T | G | 1.5 | 0.1337 | -++ | 0 | 1.75 | 2 | 0.4168 |
| rs7940405 | 11 | 118142666 | G | A | -1.493 | 0.1354 | -?- | 0 | 0.034 | 1 | 0.8536 |
| rs117392768 | 11 | 72043182 | A | T | 1.493 | 0.1355 | +?+ | 0 | 0.894 | 1 | 0.3445 |
| rs12791838 | 11 | 118144209 | A | G | -1.493 | 0.1356 | -?- | 0 | 0.032 | 1 | 0.8581 |
| rs35610956 | 11 | 118144280 | G | A | -1.491 | 0.1359 | -?- | 0 | 0.032 | 1 | 0.8571 |
| rs680465 | 11 | 118500963 | A | G | 1.491 | 0.1359 | +?- | 59.4 | 7.394 | 1 | 0.006544 |
| rs629470 | 11 | 118320553 | A | G | 1.491 | 0.136 | +++ | 0 | 0.476 | 2 | 0.7881 |
| rs56785719 | 11 | 118176623 | G | C | -1.49 | 0.1363 | -?- | 0 | 0.826 | 1 | 0.3635 |
| rs113488707 | 11 | 118045876 | T | C | 1.486 | 0.1373 | +?+ | 0 | 0.755 | 1 | 0.385 |
| rs7126193 | 11 | 118866126 | C | T | -1.486 | 0.1373 | --- | 0 | 1.322 | 2 | 0.5164 |
| rs502514 | 11 | 118241995 | C | T | -1.484 | 0.1377 | --- | 0 | 1.794 | 2 | 0.4079 |
| rs12577045 | 11 | 118151446 | A | G | -1.484 | 0.1379 | -?- | 0 | 0.79 | 1 | 0.374 |
| rs4936454 | 11 | 118859274 | T | C | -1.482 | 0.1384 | --- | 0 | 1.214 | 2 | 0.545 |
| rs117705968 | 11 | 72037907 | A | C | 1.48 | 0.1388 | +?+ | 0 | 0.87 | 1 | 0.3511 |
| rs4938498 | 11 | 118162147 | G | C | -1.48 | 0.139 | -?- | 0 | 0 | 1 | 0.987 |
| rs663003 | 11 | 118823148 | T | C | -1.479 | 0.1392 | -?- | 0 | 0.219 | 1 | 0.6397 |
| rs11217112 | 11 | 118856920 | G | A | -1.478 | 0.1394 | --- | 0 | 1.188 | 2 | 0.5522 |
| rs3759004 | 11 | 118136836 | T | C | -1.478 | 0.1394 | -?- | 0 | 0.026 | 1 | 0.8724 |
| rs12417431 | 11 | 118137383 | G | A | -1.478 | 0.1395 | -?- | 0 | 0.026 | 1 | 0.8723 |
| rs11217116 | 11 | 118864140 | C | T | -1.477 | 0.1397 | --- | 0 | 1.167 | 2 | 0.558 |
| rs4938613 | 11 | 118863116 | G | A | -1.477 | 0.1398 | --- | 0 | 1.162 | 2 | 0.5594 |
| rs111491960 | 11 | 118833458 | G | A | 1.475 | 0.1402 | +-+ | -38.6 | 2.886 | 2 | 0.2363 |
| rs117902312 | 11 | 72042303 | G | C | 1.475 | 0.1403 | +?+ | 0 | 0.86 | 1 | 0.3538 |
| rs4936456 | 11 | 118863324 | T | C | -1.473 | 0.1408 | --- | 0 | 1.167 | 2 | 0.5581 |
| rs12576119 | 11 | 118158441 | A | G | -1.47 | 0.1417 | -?- | 0 | 0.865 | 1 | 0.3523 |
| rs60545357 | 11 | 118281418 | C | A | 1.47 | 0.1417 | +++ | 0 | 0.943 | 2 | 0.624 |
| rs10892301 | 11 | 118735476 | A | G | 1.469 | 0.1419 | -++ | 0 | 1.69 | 2 | 0.4296 |
| rs11217118 | 11 | 118866072 | A | G | -1.469 | 0.1419 | --- | 0 | 1.194 | 2 | 0.5505 |
| rs12289253 | 11 | 118405068 | G | A | 1.469 | 0.1419 | +++ | 0 | 0.194 | 2 | 0.9075 |
| rs4938559 | 11 | 118718061 | T | C | 1.468 | 0.1421 | -++ | 0 | 1.716 | 2 | 0.4239 |
| rs59626457 | 11 | 118862634 | G | A | -1.466 | 0.1426 | --- | 0 | 1.143 | 2 | 0.5648 |
| rs59316586 | 11 | 118856816 | C | A | -1.464 | 0.1431 | --- | 0 | 1.237 | 2 | 0.5388 |
| rs4936455 | 11 | 118861389 | T | C | -1.464 | 0.1432 | --- | 0 | 1.237 | 2 | 0.5387 |
| rs3132893 | 11 | 118303822 | A | C | 1.463 | 0.1436 | +++ | 0 | 0.418 | 2 | 0.8116 |
| rs4938560 | 11 | 118718062 | G | A | 1.46 | 0.1444 | -++ | 0 | 1.71 | 2 | 0.4254 |
| rs12420795 | 11 | 118161885 | G | C | -1.456 | 0.1455 | -?- | 0 | 0 | 1 | 0.9963 |
| rs118115020 | 11 | 72067731 | T | C | 1.454 | 0.1458 | +?+ | -181.3 | 1.067 | 1 | 0.3017 |
| rs12362865 | 11 | 118859842 | G | A | -1.452 | 0.1464 | --- | 0 | 1.153 | 2 | 0.562 |

| | | | | | | | | | | | |
|---|---|---|---|---|---|---|---|---|---|---|---|
| rs113611963 | 11 | 118839610 | G | T | 1.45 | 0.147 | +-+ | -38.9 | 2.879 | 2 | 0.237 |
| rs570424 | 11 | 118588998 | C | T | 1.45 | 0.1472 | ++- | 11.2 | 4.506 | 2 | 0.1051 |
| rs117225442 | 11 | 72060986 | A | G | 1.448 | 0.1477 | +?+ | 0 | 0.852 | 1 | 0.3561 |
| rs12416826 | 11 | 118165955 | C | T | -1.447 | 0.148 | -?- | 0 | 0 | 1 | 0.9967 |
| rs2120269 | 11 | 118151135 | C | T | -1.445 | 0.1485 | -?- | 0 | 0.664 | 1 | 0.4153 |
| rs7928058 | 11 | 118171239 | T | G | -1.444 | 0.1487 | -?- | 0 | 0 | 1 | 0.9842 |
| rs524936 | 11 | 118341921 | A | G | 1.443 | 0.1489 | +++ | 0 | 1.417 | 2 | 0.4924 |
| rs602716 | 11 | 118801235 | A | G | 1.444 | 0.1489 | +-+ | 35.7 | 6.221 | 2 | 0.04457 |
| rs543824 | 11 | 118762218 | G | A | 1.442 | 0.1494 | +-+ | 0 | 1.961 | 2 | 0.3751 |
| rs45534632 | 11 | 118840705 | G | A | 1.44 | 0.1499 | +-+ | -45.9 | 2.741 | 2 | 0.254 |
| rs4938558 | 11 | 118717929 | T | C | 1.44 | 0.15 | +++ | 0 | 0.874 | 2 | 0.6459 |
| rs17122170 | 11 | 118296668 | C | G | 1.439 | 0.1502 | +++ | 0 | 0.275 | 2 | 0.8714 |
| rs3862791 | 11 | 72501002 | T | C | 1.439 | 0.1502 | -++ | 46.6 | 7.485 | 2 | 0.02369 |
| rs117939125 | 11 | 72062164 | G | T | 1.437 | 0.1506 | +?+ | 0 | 0.731 | 1 | 0.3924 |
| rs11607701 | 11 | 118169825 | G | A | -1.435 | 0.1513 | -?- | 0 | 0.001 | 1 | 0.9769 |
| rs11216845 | 11 | 118163216 | T | C | -1.434 | 0.1515 | -?- | 0 | 0.016 | 1 | 0.9006 |
| rs535327 | 11 | 118341785 | G | A | 1.433 | 0.1519 | +++ | 0 | 1.383 | 2 | 0.5007 |
| rs67774926 | 11 | 118157623 | G | C | -1.433 | 0.1519 | -?- | 0 | 0.828 | 1 | 0.3629 |
| rs4938614 | 11 | 118863232 | A | G | -1.433 | 0.152 | -?- | 0 | 0.628 | 1 | 0.4279 |
| rs55736521 | 11 | 118840506 | T | C | 1.432 | 0.152 | +-+ | -47.4 | 2.714 | 2 | 0.2574 |
| rs117215416 | 11 | 72046796 | C | T | 1.431 | 0.1525 | +?+ | 0 | 0.78 | 1 | 0.3771 |
| rs189530708 | 11 | 118272454 | G | C | -1.431 | 0.1526 | --+ | 13.2 | 4.611 | 2 | 0.09973 |
| rs4938494 | 11 | 118150735 | C | G | -1.429 | 0.153 | -?- | 0 | 0.69 | 1 | 0.4062 |
| rs11600727 | 11 | 72201499 | C | A | 1.425 | 0.1541 | -++ | 0 | 1.579 | 2 | 0.4541 |
| rs3782040 | 11 | 118176097 | G | A | -1.425 | 0.1541 | -?- | 0 | 0.857 | 1 | 0.3545 |
| rs2032397 | 11 | 118153697 | A | G | -1.423 | 0.1547 | -?- | 0 | 0 | 1 | 0.9974 |
| rs561775 | 11 | 118341212 | G | A | 1.422 | 0.1551 | +++ | 0 | 1.329 | 2 | 0.5144 |
| rs2032398 | 11 | 118153708 | C | G | -1.421 | 0.1553 | -?- | 0 | 0 | 1 | 0.9958 |
| rs80045923 | 11 | 72063551 | T | C | 1.42 | 0.1555 | +?+ | 0 | 0.762 | 1 | 0.3828 |
| rs11216844 | 11 | 118162796 | G | A | -1.42 | 0.1556 | -?- | 0 | 0.021 | 1 | 0.8835 |
| rs117190399 | 11 | 72056023 | T | G | 1.419 | 0.1558 | +?+ | 0 | 0.76 | 1 | 0.3833 |
| rs56140791 | 11 | 118167190 | C | T | -1.418 | 0.1562 | -?- | 0 | 0.805 | 1 | 0.3695 |
| rs4938610 | 11 | 118857466 | C | T | -1.418 | 0.1563 | -?- | 0 | 0.582 | 1 | 0.4454 |
| rs3889526 | 11 | 118856491 | A | C | -1.417 | 0.1564 | -?- | 0 | 0.558 | 1 | 0.455 |
| rs1599809 | 11 | 118179200 | A | G | -1.417 | 0.1566 | -?- | 0 | 0.676 | 1 | 0.4109 |
| rs615166 | 11 | 118803536 | A | G | 1.414 | 0.1572 | +-+ | 34.2 | 6.08 | 2 | 0.04784 |
| rs10898893 | 11 | 72676699 | C | T | -1.413 | 0.1576 | --+ | -75.6 | 2.278 | 2 | 0.3202 |
| rs587263 | 11 | 118816156 | C | T | 1.411 | 0.1581 | +++ | 0 | 1.532 | 2 | 0.4649 |
| rs7110034 | 11 | 118858911 | A | C | -1.412 | 0.1581 | -?- | 0 | 0.787 | 1 | 0.3751 |
| rs67604597 | 11 | 118154836 | G | T | -1.411 | 0.1582 | -?- | 0 | 0.799 | 1 | 0.3714 |
| rs60652614 | 11 | 118168543 | T | G | -1.408 | 0.1592 | -?- | 0 | 0.736 | 1 | 0.3909 |
| rs61706007 | 11 | 118167278 | G | T | -1.408 | 0.1592 | -?- | 0 | 0.823 | 1 | 0.3642 |
| rs688035 | 11 | 118802492 | G | C | 1.408 | 0.1592 | +-+ | 32.2 | 5.897 | 2 | 0.05243 |
| rs117613362 | 11 | 72051945 | C | A | 1.406 | 0.1598 | +?+ | 0 | 0.737 | 1 | 0.3908 |
| rs57300701 | 11 | 118845546 | G | C | 1.406 | 0.1598 | +-+ | -62.6 | 2.459 | 2 | 0.2924 |
| rs117498960 | 11 | 72065210 | C | T | 1.405 | 0.1601 | +?+ | 0 | 0.967 | 1 | 0.3255 |
| rs2851460 | 11 | 118716563 | A | G | 1.404 | 0.1602 | -++ | -92 | 2.084 | 2 | 0.3528 |
| rs55940627 | 11 | 118163920 | A | G | -1.404 | 0.1602 | -?- | 0 | 0.742 | 1 | 0.389 |
| rs56357739 | 11 | 118158507 | G | A | -1.403 | 0.1605 | -?- | 0 | 0.666 | 1 | 0.4145 |
| rs57075232 | 11 | 118159371 | T | C | -1.403 | 0.1605 | -?- | 0 | 0.688 | 1 | 0.407 |
| rs4245414 | 11 | 72188466 | A | G | 1.401 | 0.1611 | -++ | 0 | 1.808 | 2 | 0.405 |
| rs4938501 | 11 | 118166497 | C | T | -1.401 | 0.1611 | -?- | 0 | 0.8 | 1 | 0.3711 |
| rs12418805 | 11 | 72188979 | C | T | 1.401 | 0.1612 | -++ | 0 | 1.807 | 2 | 0.4051 |
| rs4938612 | 11 | 118863037 | C | T | -1.401 | 0.1613 | -?- | 0 | 0.577 | 1 | 0.4475 |

| | | | | | | | | | | | |
|---|---|---|---|---|---|---|---|---|---|---|---|
| rs4938566 | 11 | 118721189 | T | A | 1.4 | 0.1616 | +++ | 0 | 0.767 | 2 | 0.6814 |
| rs56066890 | 11 | 118169790 | C | T | -1.398 | 0.1621 | -?- | 0 | 0.729 | 1 | 0.3932 |
| rs4938565 | 11 | 118720998 | A | G | 1.398 | 0.1622 | +++ | 0 | 0.772 | 2 | 0.6797 |
| rs1970750 | 11 | 118149122 | T | C | -1.397 | 0.1624 | -?- | 0 | 0.026 | 1 | 0.8727 |
| rs59134737 | 11 | 118160959 | T | G | -1.395 | 0.1631 | -?- | 0 | 0.716 | 1 | 0.3976 |
| rs149666559 | 11 | 72042565 | A | G | 1.394 | 0.1632 | +?+ | 0 | 0.877 | 1 | 0.349 |
| rs59138930 | 11 | 118173783 | T | C | -1.394 | 0.1633 | -?- | 0 | 0.803 | 1 | 0.3703 |
| rs11217117 | 11 | 118865992 | T | C | -1.394 | 0.1635 | -?- | 0 | 0.588 | 1 | 0.4431 |
| rs12576947 | 11 | 118158055 | C | T | -1.393 | 0.1635 | -?- | 0 | 0.682 | 1 | 0.4089 |
| rs59399105 | 11 | 118168568 | A | C | -1.393 | 0.1637 | -?- | 0 | 0.762 | 1 | 0.3826 |
| rs58093803 | 11 | 118168156 | C | T | -1.391 | 0.1641 | -?- | 0 | 0.765 | 1 | 0.3819 |
| rs59450866 | 11 | 118167707 | T | C | -1.391 | 0.1641 | -?- | 0 | 0.765 | 1 | 0.3818 |
| rs7110019 | 11 | 118168617 | G | A | -1.391 | 0.1643 | -?- | 0 | 0.862 | 1 | 0.3531 |
| rs12576941 | 11 | 118158002 | C | T | -1.39 | 0.1644 | -?- | 0 | 0.687 | 1 | 0.4072 |
| rs495132 | 11 | 118335927 | A | G | 1.39 | 0.1645 | +++ | 0 | 1.344 | 2 | 0.5107 |
| rs4936440 | 11 | 118720684 | G | C | 1.39 | 0.1646 | +++ | 0 | 0.782 | 2 | 0.6763 |
| rs4938567 | 11 | 118721249 | G | A | 1.39 | 0.1647 | +++ | 0 | 0.792 | 2 | 0.6731 |
| rs12225399 | 11 | 118480285 | C | G | 1.389 | 0.1649 | +?+ | 0 | 0.228 | 1 | 0.6331 |
| rs947848 | 11 | 72895007 | T | A | -1.389 | 0.1649 | +-- | 1.6 | 4.065 | 2 | 0.131 |
| rs4943910 | 11 | 72188643 | T | C | 1.388 | 0.1651 | -++ | 0 | 1.787 | 2 | 0.4092 |
| rs7124127 | 11 | 118006408 | C | T | 1.388 | 0.1651 | +-+ | -88.7 | 2.12 | 2 | 0.3464 |
| rs3887015 | 11 | 118156410 | C | T | -1.385 | 0.166 | -?- | 0 | 0.696 | 1 | 0.4042 |
| rs56318049 | 11 | 118166608 | C | T | -1.385 | 0.166 | -?- | 0 | 0.776 | 1 | 0.3785 |
| rs66832804 | 11 | 118157049 | A | T | -1.385 | 0.166 | -?- | 0 | 0.696 | 1 | 0.4043 |
| rs7926133 | 11 | 72647296 | C | T | -1.385 | 0.166 | --+ | 0 | 1.975 | 2 | 0.3726 |
| rs573971 | 11 | 118393396 | A | G | 1.385 | 0.1661 | +++ | 0 | 0.173 | 2 | 0.9169 |
| rs4938570 | 11 | 118722231 | G | A | 1.384 | 0.1663 | +++ | 0 | 0.783 | 2 | 0.6762 |
| rs572126 | 11 | 118359161 | G | A | 1.384 | 0.1663 | +++ | 0 | 0.23 | 2 | 0.8915 |
| rs61390396 | 11 | 118156718 | T | C | -1.384 | 0.1663 | -?- | 0 | 0.697 | 1 | 0.4038 |
| rs1945765 | 11 | 118182829 | A | T | -1.384 | 0.1664 | -?- | 0 | 0.826 | 1 | 0.3634 |
| rs55910314 | 11 | 118163780 | T | C | -1.384 | 0.1664 | -?- | 0 | 0.778 | 1 | 0.3778 |
| rs4938564 | 11 | 118720590 | T | G | 1.383 | 0.1666 | +++ | 0 | 0.771 | 2 | 0.6801 |
| rs2187491 | 11 | 118718972 | C | T | 1.383 | 0.1667 | +++ | 0 | 0.721 | 2 | 0.6974 |
| rs56938099 | 11 | 118156688 | G | A | -1.383 | 0.1667 | -?- | 0 | 0.7 | 1 | 0.4029 |
| rs10892298 | 11 | 118724336 | C | T | 1.382 | 0.167 | +++ | 0 | 0.788 | 2 | 0.6744 |
| rs55899310 | 11 | 118158702 | C | T | -1.382 | 0.1671 | -?- | 0 | 0.702 | 1 | 0.4022 |
| rs7103314 | 11 | 118723537 | G | A | 1.382 | 0.1671 | +++ | 0 | 0.788 | 2 | 0.6742 |
| rs3825051 | 11 | 118174629 | G | A | -1.381 | 0.1674 | -?- | 0 | 0.835 | 1 | 0.3608 |
| rs538645 | 11 | 118711069 | C | T | 1.38 | 0.1676 | +++ | 0 | 0.891 | 2 | 0.6404 |
| rs11217059 | 11 | 118723087 | T | C | 1.38 | 0.1677 | +++ | 0 | 0.793 | 2 | 0.6726 |
| rs12576100 | 11 | 118158281 | A | G | -1.38 | 0.1677 | -?- | 0 | 0.705 | 1 | 0.4012 |
| rs12576118 | 11 | 118158440 | A | G | -1.379 | 0.1678 | -?- | 0 | 0.705 | 1 | 0.401 |
| rs4938500 | 11 | 118166212 | A | C | -1.379 | 0.1679 | -?- | 0 | 0.011 | 1 | 0.9159 |
| rs3132786 | 11 | 118365956 | C | T | 1.378 | 0.1682 | +++ | 0 | 0.213 | 2 | 0.8989 |
| rs11607671 | 11 | 72192738 | T | C | 1.377 | 0.1684 | -++ | 0 | 1.771 | 2 | 0.4125 |
| rs2156751 | 11 | 118718856 | G | A | 1.377 | 0.1686 | +++ | 0 | 0.735 | 2 | 0.6925 |
| rs4274214 | 11 | 72188003 | T | C | 1.377 | 0.1686 | -++ | 0 | 1.77 | 2 | 0.4127 |
| rs4938499 | 11 | 118162496 | C | T | -1.377 | 0.1686 | -?- | 0 | 0.791 | 1 | 0.3738 |
| rs55798741 | 11 | 118154114 | T | G | -1.376 | 0.1688 | -?- | 0 | 0.78 | 1 | 0.377 |
| rs4290240 | 11 | 118853488 | G | A | -1.376 | 0.1689 | -?- | 0 | 0.79 | 1 | 0.3741 |
| rs55806415 | 11 | 118162862 | A | G | -1.376 | 0.1689 | -?- | 0 | 0.792 | 1 | 0.3734 |
| rs58520343 | 11 | 118155000 | T | A | -1.376 | 0.1689 | -?- | 0 | 0.712 | 1 | 0.3989 |
| rs6592474 | 11 | 72184290 | A | T | 1.376 | 0.1689 | -++ | 0 | 1.769 | 2 | 0.413 |
| rs1074459 | 11 | 118155997 | C | T | -1.374 | 0.1696 | -?- | 0 | 0.715 | 1 | 0.3978 |

| | | | | | | | | | | | |
|---|---|---|---|---|---|---|---|---|---|---|---|
| rs58210741 | 11 | 118159326 | G | C | -1.373 | 0.1697 | -?- | 0 | 0.739 | 1 | 0.39 |
| rs61900902 | 11 | 118159684 | C | T | -1.373 | 0.1697 | -?- | 0 | 0 | 1 | 1 |
| rs2365385 | 11 | 72194718 | A | G | 1.373 | 0.1699 | -++ | 0 | 1.773 | 2 | 0.4122 |
| rs55815657 | 11 | 118155074 | A | G | -1.372 | 0.17 | -?- | 0 | 0.717 | 1 | 0.3971 |
| rs1318166 | 11 | 118171058 | G | A | -1.372 | 0.1701 | -?- | 0 | 0.775 | 1 | 0.3788 |
| rs67462512 | 11 | 118155336 | C | G | -1.37 | 0.1706 | -?- | 0 | 0.713 | 1 | 0.3983 |
| rs58983315 | 11 | 118167759 | A | G | -1.37 | 0.1707 | -?- | 0 | 0.686 | 1 | 0.4075 |
| rs17310153 | 11 | 72190863 | G | A | 1.37 | 0.1708 | -++ | 0 | 1.76 | 2 | 0.4148 |
| rs1074457 | 11 | 118156117 | T | A | -1.369 | 0.171 | -?- | 0 | 0.723 | 1 | 0.3951 |
| rs7946961 | 11 | 118721850 | G | A | 1.369 | 0.1711 | +++ | 0 | 0.82 | 2 | 0.6636 |
| rs11607688 | 11 | 72192830 | T | C | 1.368 | 0.1714 | -++ | 0 | 1.763 | 2 | 0.4142 |
| rs12577820 | 11 | 118164233 | T | C | -1.368 | 0.1714 | -?- | 0 | 0.807 | 1 | 0.3691 |
| rs118105218 | 11 | 72077497 | G | T | 1.366 | 0.172 | +?+ | 0 | 0.891 | 1 | 0.3451 |
| rs12576486 | 11 | 118235490 | C | T | 1.365 | 0.1722 | +++ | 0 | 0.968 | 2 | 0.6164 |
| rs2156749 | 11 | 118719405 | G | C | 1.365 | 0.1722 | +++ | 0 | 0.727 | 2 | 0.6951 |
| rs56224295 | 11 | 118161891 | A | G | -1.365 | 0.1724 | -?- | 0 | 0.756 | 1 | 0.3847 |
| rs45480496 | 11 | 118270715 | A | G | 1.364 | 0.1725 | +++ | 0 | 0.845 | 2 | 0.6554 |
| rs4938497 | 11 | 118160851 | A | G | -1.364 | 0.1727 | -?- | 0 | 0 | 1 | 0.9925 |
| rs56020103 | 11 | 118172132 | T | C | -1.363 | 0.1728 | -?- | 0 | 0.749 | 1 | 0.3869 |
| rs117652953 | 11 | 72084357 | G | A | 1.363 | 0.173 | +?+ | 0 | 0.886 | 1 | 0.3467 |
| rs11217062 | 11 | 118725420 | A | G | 1.362 | 0.1732 | +++ | 0 | 0.902 | 2 | 0.637 |
| rs56346212 | 11 | 118172342 | A | T | -1.362 | 0.1733 | -?- | 0 | 0.751 | 1 | 0.3861 |
| rs894678 | 11 | 118172520 | A | G | -1.362 | 0.1733 | -?- | 0 | 0.001 | 1 | 0.9698 |
| rs56292628 | 11 | 118163872 | A | G | -1.359 | 0.1742 | -?- | 0 | 0.823 | 1 | 0.3644 |
| rs2156750 | 11 | 118719254 | C | T | 1.357 | 0.1747 | +++ | 0 | 0.745 | 2 | 0.6889 |
| rs7480736 | 11 | 118172364 | C | T | -1.357 | 0.1747 | -?- | 0 | 0.759 | 1 | 0.3837 |
| rs73613959 | 11 | 118263054 | A | G | 1.356 | 0.1751 | +++ | 0 | 0.855 | 2 | 0.6522 |
| rs75128335 | 11 | 118072915 | A | G | 1.356 | 0.1751 | +?+ | 0 | 0.937 | 1 | 0.333 |
| rs4943907 | 11 | 72183348 | T | C | 1.354 | 0.1758 | -++ | 0 | 1.725 | 2 | 0.4221 |
| rs894677 | 11 | 118172535 | T | C | -1.353 | 0.1759 | -?- | 0 | 0.758 | 1 | 0.3839 |
| rs4936424 | 11 | 118009674 | T | C | 1.353 | 0.1762 | +-+ | -7.7 | 3.715 | 2 | 0.1561 |
| rs55789743 | 11 | 118171949 | A | C | -1.352 | 0.1763 | -?- | 0 | 0.767 | 1 | 0.381 |
| rs60618948 | 11 | 118155814 | A | G | -1.351 | 0.1767 | -?- | 0 | 0.754 | 1 | 0.3853 |
| rs3759002 | 11 | 118136160 | A | G | 1.35 | 0.177 | +?- | -19.5 | 2.51 | 1 | 0.1131 |
| rs7116715 | 11 | 118719742 | C | T | 1.347 | 0.178 | +++ | 0 | 0.729 | 2 | 0.6946 |
| rs4290266 | 11 | 118285995 | A | G | 1.338 | 0.181 | +++ | 0 | 0.436 | 2 | 0.804 |
| rs12420153 | 11 | 72195852 | A | G | 1.333 | 0.1824 | -++ | 0 | 1.71 | 2 | 0.4253 |
| rs10790239 | 11 | 118001445 | T | C | -1.332 | 0.1828 | -?- | 0 | 0.122 | 1 | 0.7274 |
| rs3759006 | 11 | 118137560 | A | G | -1.332 | 0.183 | -?- | 0 | 0.002 | 1 | 0.9613 |
| rs77643000 | 11 | 118749563 | C | T | 1.327 | 0.1845 | +++ | 0 | 0.657 | 2 | 0.72 |
| rs4938561 | 11 | 118718485 | C | T | 1.325 | 0.1851 | +++ | 0 | 0.747 | 2 | 0.6884 |
| rs34180117 | 11 | 118138629 | T | C | -1.32 | 0.187 | -?- | 0 | 0.004 | 1 | 0.9497 |
| rs3759005 | 11 | 118136871 | G | A | -1.319 | 0.1871 | -?- | 0 | 0.004 | 1 | 0.9514 |
| rs1079118 | 11 | 118258056 | A | G | 1.317 | 0.1878 | +++ | 0 | 0.736 | 2 | 0.692 |
| rs35850218 | 11 | 118138922 | C | T | -1.317 | 0.1878 | -?- | 0 | 0.004 | 1 | 0.9476 |
| rs2156755 | 11 | 118726230 | A | G | 1.317 | 0.1879 | +++ | 0 | 0.79 | 2 | 0.6736 |
| rs12795905 | 11 | 118140597 | T | C | -1.315 | 0.1886 | -?- | 0 | 0.004 | 1 | 0.9478 |
| rs61059752 | 11 | 118246071 | A | C | 1.315 | 0.1886 | +++ | 0 | 0.721 | 2 | 0.6972 |
| rs3759003 | 11 | 118136745 | G | T | -1.313 | 0.1892 | -?- | 0 | 0.002 | 1 | 0.964 |
| rs67307131 | 11 | 118480223 | C | T | 1.313 | 0.1892 | +?+ | 0 | 0.418 | 1 | 0.5179 |
| rs4938606 | 11 | 118852081 | G | A | -1.312 | 0.1894 | -?- | 0 | 0.563 | 1 | 0.453 |
| rs1790186 | 11 | 118846355 | T | C | -1.312 | 0.1895 | -?+ | -48.7 | 2.017 | 1 | 0.1556 |
| rs75896506 | 11 | 72429579 | A | G | 1.31 | 0.1903 | -++ | 12.8 | 4.586 | 2 | 0.1009 |
| rs78312641 | 11 | 118749318 | A | T | 1.31 | 0.1903 | +++ | 0 | 0.621 | 2 | 0.7332 |

| | | | | | | | | | | | |
|---|---|---|---|---|---|---|---|---|---|---|---|
| rs11607245 | 11 | 118843448 | T | C | -1.309 | 0.1905 | --- | 0 | 0.419 | 2 | 0.8111 |
| rs3759007 | 11 | 118137667 | T | C | -1.308 | 0.1908 | -?- | 0 | 0.005 | 1 | 0.9426 |
| rs543583 | 11 | 118847196 | A | G | -1.308 | 0.191 | -?+ | -49.7 | 2.004 | 1 | 0.1569 |
| rs56128937 | 11 | 118139478 | A | T | -1.307 | 0.1912 | -?- | 0 | 0.006 | 1 | 0.9397 |
| rs117088064 | 11 | 72074257 | T | C | 1.306 | 0.1915 | +?+ | 0 | 0.948 | 1 | 0.3301 |
| rs11822344 | 11 | 118864777 | C | G | -1.306 | 0.1915 | --- | 0 | 1.015 | 2 | 0.602 |
| rs117162202 | 11 | 72077463 | A | G | 1.306 | 0.1917 | +?+ | 0 | 0.947 | 1 | 0.3304 |
| rs55852239 | 11 | 118139755 | A | G | -1.306 | 0.1917 | -?- | 0 | 0.006 | 1 | 0.9405 |
| rs12789276 | 11 | 118140203 | G | A | -1.303 | 0.1927 | -?- | 0 | 0.006 | 1 | 0.9381 |
| rs34786601 | 11 | 118140136 | C | T | -1.302 | 0.1928 | -?- | 0 | 0.006 | 1 | 0.9379 |
| rs7933963 | 11 | 118246906 | T | C | 1.302 | 0.193 | +++ | 0 | 0.673 | 2 | 0.7142 |
| rs7948683 | 11 | 118137911 | A | G | -1.301 | 0.1934 | -?- | 0 | 0.006 | 1 | 0.9366 |
| rs73613980 | 11 | 118284650 | T | C | 1.299 | 0.1939 | +++ | 0 | 0.359 | 2 | 0.8356 |
| rs12793790 | 11 | 118140372 | A | G | -1.295 | 0.1952 | -?- | 0 | 0.007 | 1 | 0.9323 |
| rs1945764 | 11 | 118182801 | G | A | -1.295 | 0.1952 | --- | 0 | 0.359 | 2 | 0.8358 |
| rs4544037 | 11 | 118222062 | C | T | 1.294 | 0.1958 | +++ | 0 | 1.37 | 2 | 0.5041 |
| rs12419365 | 11 | 118112332 | A | C | 1.286 | 0.1983 | +?+ | -160.3 | 1.152 | 1 | 0.283 |
| rs3212264 | 11 | 118216234 | A | C | 1.286 | 0.1984 | +++ | 0 | 1.403 | 2 | 0.4959 |
| rs45489394 | 11 | 118228968 | G | C | 1.286 | 0.1986 | +++ | 0 | 1.339 | 2 | 0.5118 |
| rs7933007 | 11 | 118730669 | C | T | 1.284 | 0.1992 | +++ | 0 | 0.88 | 2 | 0.6442 |
| rs9666462 | 11 | 118275997 | A | G | 1.283 | 0.1995 | +++ | 0 | 0.378 | 2 | 0.8279 |
| rs11603334 | 11 | 72432985 | A | G | 1.282 | 0.1999 | -++ | 12.4 | 4.568 | 2 | 0.1019 |
| rs1599810 | 11 | 118179197 | A | G | -1.282 | 0.2 | --- | 0 | 0.375 | 2 | 0.8292 |
| rs150193966 | 11 | 72071662 | A | G | 1.271 | 0.2036 | +?+ | 0 | 0.882 | 1 | 0.3478 |
| rs117498055 | 11 | 72022810 | A | G | 1.271 | 0.2037 | +?+ | 0 | 0.56 | 1 | 0.4542 |
| rs12806041 | 11 | 118138998 | G | A | -1.271 | 0.2038 | -?- | 0 | 0.013 | 1 | 0.9109 |
| rs9666463 | 11 | 118276116 | G | C | 1.27 | 0.2042 | +++ | 0 | 0.359 | 2 | 0.8357 |
| rs117091621 | 11 | 72026473 | T | C | 1.269 | 0.2044 | +?+ | 0 | 0.558 | 1 | 0.4552 |
| rs523604 | 11 | 118755738 | G | A | -1.266 | 0.2055 | --- | 0 | 0.254 | 2 | 0.8809 |
| rs17122157 | 11 | 118276492 | A | G | 1.264 | 0.2062 | +++ | 0 | 0.358 | 2 | 0.8363 |
| rs116888156 | 11 | 72009595 | A | G | 1.261 | 0.2073 | +?+ | 0 | 0.724 | 1 | 0.3948 |
| rs7483081 | 11 | 118276246 | C | T | 1.261 | 0.2075 | +++ | 0 | 0.361 | 2 | 0.8347 |
| rs55942271 | 11 | 118177115 | T | C | -1.26 | 0.2076 | --- | 0 | 0.352 | 2 | 0.8388 |
| rs1552224 | 11 | 72433098 | C | A | 1.256 | 0.2091 | -++ | 13.8 | 4.643 | 2 | 0.09813 |
| rs4537793 | 11 | 118053156 | A | C | 1.255 | 0.2095 | +++ | 0 | 0.297 | 2 | 0.862 |
| rs4936451 | 11 | 118852137 | T | C | -1.252 | 0.2106 | --- | 0 | 0.671 | 2 | 0.715 |
| rs77342802 | 11 | 72012038 | C | G | 1.247 | 0.2125 | +?+ | 0 | 0.7 | 1 | 0.4028 |
| rs7939212 | 11 | 72021876 | T | G | 1.243 | 0.2138 | +?+ | 0 | 0.588 | 1 | 0.4433 |
| rs117524306 | 11 | 72014266 | G | A | 1.242 | 0.2143 | +?+ | 0 | 0.692 | 1 | 0.4056 |
| rs11825496 | 11 | 118855221 | G | T | -1.242 | 0.2143 | --- | 0 | 0.598 | 2 | 0.7416 |
| rs7940087 | 11 | 72021877 | T | C | 1.241 | 0.2146 | +?+ | 0 | 0.585 | 1 | 0.4445 |
| rs4456264 | 11 | 118851138 | T | C | -1.238 | 0.2157 | --- | 0 | 0.633 | 2 | 0.7288 |
| rs7129793 | 11 | 72455885 | T | C | 1.236 | 0.2165 | -++ | 31.7 | 5.855 | 2 | 0.05354 |
| rs45527340 | 11 | 118852526 | G | C | -1.235 | 0.2167 | --- | 0 | 0.633 | 2 | 0.7287 |
| rs477106 | 11 | 118363424 | A | C | 1.235 | 0.2168 | +++ | 0 | 0.283 | 2 | 0.8681 |
| rs4938616 | 11 | 118865465 | T | C | -1.233 | 0.2175 | -?- | 0 | 0.659 | 1 | 0.417 |
| rs79430446 | 11 | 72474004 | T | G | 1.229 | 0.2191 | -++ | 29.9 | 5.703 | 2 | 0.05777 |
| rs117346296 | 11 | 72025140 | A | G | 1.226 | 0.2202 | +?+ | 0 | 0.495 | 1 | 0.4817 |
| rs12273687 | 11 | 118852641 | T | G | -1.226 | 0.2203 | --- | 0 | 1 | 2 | 0.6065 |
| rs7938979 | 11 | 72021720 | C | G | 1.224 | 0.2209 | +?+ | 0 | 0.559 | 1 | 0.4546 |
| rs4936426 | 11 | 118134597 | C | T | 1.218 | 0.2232 | +?+ | 0 | 0.745 | 1 | 0.3881 |
| rs613937 | 11 | 72474839 | G | A | 1.217 | 0.2236 | -++ | 36.4 | 6.294 | 2 | 0.04299 |
| rs7125115 | 11 | 118478330 | A | G | 1.215 | 0.2242 | +++ | 0 | 0.559 | 2 | 0.7561 |
| rs528056 | 11 | 118317907 | T | C | 1.21 | 0.2264 | +++ | 0 | 0.844 | 2 | 0.6557 |

| | | | | | | | | | | | |
|---|---|---|---|---|---|---|---|---|---|---|---|
| rs74333814 | 11 | 72457487 | T | C | 1.207 | 0.2273 | -++ | 28.7 | 5.608 | 2 | 0.06056 |
| rs117237510 | 11 | 72009451 | T | C | 1.202 | 0.2295 | +?+ | 0 | 0.627 | 1 | 0.4286 |
| rs4938604 | 11 | 118846574 | C | T | -1.201 | 0.2296 | -?- | 0 | 0.448 | 1 | 0.5031 |
| rs4938502 | 11 | 118179941 | T | G | -1.2 | 0.23 | --- | 0 | 0.32 | 2 | 0.8521 |
| rs12793150 | 11 | 118180667 | G | C | -1.199 | 0.2304 | --- | 0 | 0.32 | 2 | 0.8522 |
| rs117954624 | 11 | 72028005 | G | A | 1.197 | 0.2311 | +?+ | 0 | 0.62 | 1 | 0.4311 |
| rs117999328 | 11 | 72019118 | C | T | 1.196 | 0.2318 | +?+ | 0 | 0.617 | 1 | 0.4321 |
| rs118089225 | 11 | 72018472 | T | C | 1.196 | 0.2318 | +?+ | 0 | 0.617 | 1 | 0.4321 |
| rs112337443 | 11 | 118530068 | C | T | 1.195 | 0.2321 | +-+ | 0 | 1.762 | 2 | 0.4144 |
| rs4938605 | 11 | 118850707 | A | G | -1.193 | 0.233 | --- | 0 | 0.599 | 2 | 0.741 |
| rs7108613 | 11 | 118268613 | G | A | 1.19 | 0.234 | +++ | 0 | 0.313 | 2 | 0.8552 |
| rs503542 | 11 | 118590743 | G | A | 1.19 | 0.2341 | ++- | -41.4 | 2.829 | 2 | 0.243 |
| rs2071702 | 11 | 118373861 | T | C | 1.186 | 0.2355 | +?+ | 0 | 0.407 | 1 | 0.5235 |
| rs3782041 | 11 | 118184092 | T | C | -1.182 | 0.2371 | --- | 0 | 0.272 | 2 | 0.873 |
| rs3741315 | 11 | 118004688 | A | G | 1.178 | 0.2387 | +-+ | 0 | 1.767 | 2 | 0.4133 |
| rs12420418 | 11 | 118840857 | T | C | -1.177 | 0.2393 | --- | 0 | 0.192 | 2 | 0.9086 |
| rs3890562 | 11 | 118837113 | G | A | -1.174 | 0.2405 | --- | 0 | 0.205 | 2 | 0.9027 |
| rs116951669 | 11 | 72018759 | C | T | 1.174 | 0.2406 | +?+ | 0 | 0.583 | 1 | 0.4452 |
| rs496058 | 11 | 118580906 | G | A | 1.167 | 0.2432 | ++- | -3.2 | 3.878 | 2 | 0.1439 |
| rs10892318 | 11 | 118848725 | A | G | -1.162 | 0.2452 | --- | 0 | 0.392 | 2 | 0.8219 |
| rs7103836 | 11 | 72474571 | C | G | 1.159 | 0.2466 | -++ | 27.1 | 5.49 | 2 | 0.06424 |
| rs2239695 | 11 | 118213632 | T | C | 1.158 | 0.247 | +++ | 0 | 0.67 | 2 | 0.7152 |
| rs58745438 | 11 | 118848315 | C | T | -1.154 | 0.2487 | --- | 0 | 0.397 | 2 | 0.82 |
| rs516503 | 11 | 118580348 | T | C | 1.15 | 0.25 | ++- | -2.7 | 3.896 | 2 | 0.1426 |
| rs3181259 | 11 | 118214067 | A | G | 1.15 | 0.2501 | +++ | 0 | 0.635 | 2 | 0.7278 |
| rs11217108 | 11 | 118841761 | T | C | -1.148 | 0.2509 | --- | 0 | 0.186 | 2 | 0.9114 |
| rs4938602 | 11 | 118846466 | A | G | -1.143 | 0.2529 | --- | 0 | 0.385 | 2 | 0.8251 |
| rs35865949 | 11 | 118229061 | A | T | 1.141 | 0.254 | +++ | 0 | 0.565 | 2 | 0.7538 |
| rs12795615 | 11 | 118214562 | T | C | 1.139 | 0.2548 | +++ | 0 | 0.576 | 2 | 0.7499 |
| rs7110197 | 11 | 118228624 | C | G | 1.138 | 0.255 | +++ | 0 | 0.555 | 2 | 0.7579 |
| rs1790191 | 11 | 118825636 | A | G | 1.138 | 0.2552 | +?+ | 0 | 0.001 | 1 | 0.9797 |
| rs4938603 | 11 | 118846482 | G | A | -1.138 | 0.2553 | -?- | 0 | 0.253 | 1 | 0.6152 |
| rs3782046 | 11 | 118230739 | A | G | 1.136 | 0.2558 | +++ | 0 | 0.57 | 2 | 0.752 |
| rs3753060 | 11 | 118221137 | T | C | 1.133 | 0.2573 | +++ | 0 | 0.5 | 2 | 0.7788 |
| rs11216819 | 11 | 118084980 | G | A | 1.13 | 0.2584 | +?- | -109.3 | 1.433 | 1 | 0.2312 |
| rs939274 | 11 | 118134065 | T | G | 1.129 | 0.2591 | +?- | -126.5 | 1.325 | 1 | 0.2498 |
| rs3825054 | 11 | 118234191 | A | G | 1.128 | 0.2592 | +++ | 0 | 0.565 | 2 | 0.7538 |
| rs75040111 | 11 | 118747008 | T | C | -1.127 | 0.2596 | -+- | -34.8 | 2.966 | 2 | 0.2269 |
| rs3753059 | 11 | 118221349 | C | T | 1.126 | 0.2603 | +++ | 0 | 0.507 | 2 | 0.7761 |
| rs3753058 | 11 | 118221350 | T | G | 1.125 | 0.2604 | +++ | 0 | 0.507 | 2 | 0.776 |
| rs590580 | 11 | 118066104 | A | G | 1.124 | 0.2608 | +?+ | -165.4 | 1.13 | 1 | 0.2877 |
| rs36061634 | 11 | 118225452 | A | T | 1.123 | 0.2612 | +++ | 0 | 0.57 | 2 | 0.7519 |
| rs45501598 | 11 | 118251980 | T | A | 1.121 | 0.2625 | +++ | 0 | 0.163 | 2 | 0.9217 |
| rs35305304 | 11 | 118228114 | T | C | 1.119 | 0.263 | +++ | 0 | 0.575 | 2 | 0.7502 |
| rs35902434 | 11 | 118222634 | C | T | 1.115 | 0.265 | +++ | 0 | 0.58 | 2 | 0.7482 |
| rs7949539 | 11 | 118250586 | G | A | 1.112 | 0.2664 | +++ | 0 | 0.157 | 2 | 0.9246 |
| rs494459 | 11 | 118574675 | T | C | 1.11 | 0.2671 | ++- | -4.4 | 3.83 | 2 | 0.1474 |
| rs12792929 | 11 | 118240715 | C | G | 1.105 | 0.269 | +++ | 0 | 0.469 | 2 | 0.7911 |
| rs626645 | 11 | 118575180 | C | T | 1.1 | 0.2714 | ++- | -5.6 | 3.788 | 2 | 0.1505 |
| rs60514290 | 11 | 118246048 | C | G | 1.099 | 0.2717 | +++ | 0 | 0.137 | 2 | 0.9337 |
| rs45474998 | 11 | 118229679 | A | G | 1.096 | 0.2729 | +++ | 0 | 0.459 | 2 | 0.795 |
| rs2276418 | 11 | 118230264 | T | A | 1.095 | 0.2734 | +++ | 0 | 0.46 | 2 | 0.7945 |
| rs524409 | 11 | 118574311 | T | C | 1.091 | 0.2751 | ++- | -1.8 | 3.929 | 2 | 0.1402 |
| rs7113731 | 11 | 118831824 | G | A | -1.091 | 0.2751 | --- | 0 | 0.351 | 2 | 0.8391 |

| | | | | | | | | | | | |
|---|---|---|---|---|---|---|---|---|---|---|---|
| rs186926311 | 11 | 118601426 | T | C | -1.088 | 0.2764 | --+ | -85.5 | 2.157 | 2 | 0.3401 |
| rs2276421 | 11 | 118239266 | A | G | 1.087 | 0.2769 | +++ | 0 | 0.546 | 2 | 0.7611 |
| rs76126708 | 11 | 118259379 | C | A | 1.087 | 0.2771 | +++ | 0 | 0.162 | 2 | 0.9222 |
| rs34405565 | 11 | 118233424 | T | C | 1.087 | 0.2772 | +++ | 0 | 0.469 | 2 | 0.7909 |
| rs7940757 | 11 | 118055273 | T | C | 1.086 | 0.2775 | +++ | 0 | 0.408 | 2 | 0.8155 |
| rs4635088 | 11 | 118832924 | C | G | -1.084 | 0.2782 | --- | 0 | 0.351 | 2 | 0.8391 |
| rs647769 | 11 | 118812512 | T | C | 1.082 | 0.2791 | +-+ | -49.5 | 2.675 | 2 | 0.2625 |
| rs59405103 | 11 | 118262305 | C | A | 1.081 | 0.2797 | +++ | 0 | 0.171 | 2 | 0.9182 |
| rs6589650 | 11 | 118055217 | C | T | 1.081 | 0.2798 | +++ | 0 | 0.408 | 2 | 0.8157 |
| rs11216804 | 11 | 118057320 | G | A | 1.079 | 0.2805 | +++ | 0 | 0.394 | 2 | 0.8212 |
| rs75461585 | 11 | 118287568 | G | A | -1.077 | 0.2816 | -?- | 0 | 0.093 | 1 | 0.7607 |
| rs77106637 | 11 | 72478320 | A | G | 1.076 | 0.2818 | -++ | 15.4 | 4.729 | 2 | 0.09399 |
| rs11217076 | 11 | 118745664 | G | T | 1.076 | 0.282 | ++- | 0 | 1.611 | 2 | 0.447 |
| rs3782045 | 11 | 118233078 | G | A | 1.076 | 0.282 | +++ | 0 | 0.479 | 2 | 0.7871 |
| rs625735 | 11 | 118575419 | A | G | 1.076 | 0.282 | ++- | -2.1 | 3.917 | 2 | 0.1411 |
| rs7944087 | 11 | 118055733 | T | C | 1.075 | 0.2824 | +++ | 0 | 0.395 | 2 | 0.8207 |
| rs76478216 | 11 | 118605352 | C | T | 1.075 | 0.2825 | +?- | -86.5 | 1.609 | 1 | 0.2046 |
| rs1613643 | 11 | 118068521 | T | G | 1.072 | 0.2836 | +?+ | 0 | 0.888 | 1 | 0.3461 |
| rs7944321 | 11 | 118055801 | A | G | 1.071 | 0.284 | +++ | 0 | 0.301 | 2 | 0.8603 |
| rs637563 | 11 | 118575479 | G | A | 1.071 | 0.2842 | ++- | -1.4 | 3.944 | 2 | 0.1392 |
| rs638514 | 11 | 118575693 | C | T | 1.07 | 0.2844 | ++- | -1.4 | 3.946 | 2 | 0.139 |
| rs61895726 | 11 | 72223735 | T | C | 1.068 | 0.2857 | +?+ | 0 | 0.721 | 1 | 0.3958 |
| rs530901 | 11 | 118575011 | C | T | 1.067 | 0.2859 | ++- | 1.6 | 4.064 | 2 | 0.1311 |
| rs647099 | 11 | 118067009 | A | G | 1.067 | 0.2861 | +?+ | 0 | 0.887 | 1 | 0.3463 |
| rs648947 | 11 | 118067426 | A | G | 1.066 | 0.2863 | +?+ | 0 | 0.892 | 1 | 0.3449 |
| rs76930336 | 11 | 118464764 | A | C | 1.065 | 0.2871 | +-+ | -75.4 | 2.28 | 2 | 0.3198 |
| rs6589710 | 11 | 118837460 | T | A | -1.063 | 0.2877 | --- | 0 | 0.317 | 2 | 0.8534 |
| rs7931206 | 11 | 118056092 | C | T | 1.063 | 0.2878 | +++ | 0 | 0.4 | 2 | 0.8188 |
| rs576283 | 11 | 118573894 | G | A | 1.062 | 0.2882 | ++- | -0.2 | 3.992 | 2 | 0.1359 |
| rs7126476 | 11 | 118834784 | G | A | -1.062 | 0.2882 | --- | 0 | 0.309 | 2 | 0.8569 |
| rs71482156 | 11 | 118734516 | T | A | 1.06 | 0.2891 | +?+ | 0 | 0.004 | 1 | 0.9474 |
| rs7107756 | 11 | 118717708 | A | G | 1.06 | 0.2894 | -++ | -36.5 | 2.931 | 2 | 0.2309 |
| rs11216863 | 11 | 118257081 | A | G | -1.054 | 0.2921 | -?- | 0 | 0.095 | 1 | 0.7582 |
| rs57702967 | 11 | 118261736 | A | C | 1.052 | 0.2927 | +++ | 0 | 0.146 | 2 | 0.9298 |
| rs627391 | 11 | 118573916 | T | A | 1.049 | 0.294 | ++- | -1.7 | 3.934 | 2 | 0.1398 |
| rs61895701 | 11 | 72208039 | G | T | 1.049 | 0.2941 | +?+ | 0 | 0.676 | 1 | 0.4108 |
| rs7935561 | 11 | 118056722 | A | G | 1.048 | 0.2947 | +++ | 0 | 0.304 | 2 | 0.8588 |
| rs17122113 | 11 | 118234678 | G | A | 1.047 | 0.2949 | +++ | 0 | 0.525 | 2 | 0.7693 |
| rs11216920 | 11 | 118462554 | G | A | 1.044 | 0.2964 | -++ | 0 | 1.121 | 2 | 0.571 |
| rs648682 | 11 | 118066582 | T | C | 1.044 | 0.2966 | +?+ | 0 | 0.875 | 1 | 0.3495 |
| rs7131196 | 11 | 118265227 | C | T | -1.04 | 0.2982 | -?- | 0 | 0.214 | 1 | 0.6433 |
| rs951657 | 11 | 118067621 | A | G | 1.039 | 0.2988 | +?+ | 0 | 0.947 | 1 | 0.3305 |
| rs75017819 | 11 | 118285789 | G | A | -1.034 | 0.301 | -?- | 0 | 0.192 | 1 | 0.6616 |
| rs12295634 | 11 | 118054431 | T | C | 1.034 | 0.3013 | +++ | 0 | 0.3 | 2 | 0.8606 |
| rs59489957 | 11 | 118284984 | T | C | 1.033 | 0.3014 | +++ | 0 | 0.537 | 2 | 0.7645 |
| rs7112758 | 11 | 118836821 | A | G | -1.032 | 0.3019 | --- | 0 | 0.343 | 2 | 0.8424 |
| rs7116690 | 11 | 118837742 | G | A | -1.03 | 0.3029 | --- | 0 | 0.217 | 2 | 0.897 |
| rs45496391 | 11 | 118477226 | C | A | 1.03 | 0.3031 | +?+ | 0 | 0.075 | 1 | 0.7839 |
| rs3901422 | 11 | 118837272 | A | G | -1.029 | 0.3033 | --- | 0 | 0.219 | 2 | 0.8963 |
| rs61902060 | 11 | 118839828 | G | C | -1.028 | 0.3041 | --- | 0 | 0.218 | 2 | 0.8969 |
| rs2004235 | 11 | 118008212 | C | T | 1.026 | 0.3049 | +-+ | 0 | 1.45 | 2 | 0.4844 |
| rs647573 | 11 | 118067132 | C | T | 1.025 | 0.3051 | +?+ | 0 | 0.865 | 1 | 0.3523 |
| rs11217107 | 11 | 118839268 | C | G | -1.025 | 0.3055 | --- | 0 | 0.218 | 2 | 0.8967 |
| rs4310630 | 11 | 118836638 | G | A | -1.023 | 0.3062 | --- | 0 | 0.352 | 2 | 0.8386 |

| | | | | | | | | | | | |
|---|---|---|---|---|---|---|---|---|---|---|---|
| rs3194726 | 11 | 118279936 | C | T | -1.02 | 0.3076 | -?- | 0 | 0.158 | 1 | 0.6911 |
| rs12274740 | 11 | 118054128 | C | T | 1.015 | 0.3099 | +++ | 0 | 0.308 | 2 | 0.8573 |
| rs74535623 | 11 | 118812821 | T | G | 1.014 | 0.3106 | +-+ | 0 | 1.52 | 2 | 0.4676 |
| rs60798124 | 11 | 118282241 | G | A | 1.013 | 0.3108 | +++ | 0 | 0.608 | 2 | 0.7379 |
| rs11607835 | 11 | 118835258 | C | T | -1.013 | 0.3113 | --- | 0 | 0.354 | 2 | 0.8376 |
| rs61150131 | 11 | 118752178 | C | T | 1.012 | 0.3113 | -++ | 0 | 1.399 | 2 | 0.497 |
| rs7944366 | 11 | 118056047 | G | A | 1.011 | 0.3121 | +++ | 0 | 0.326 | 2 | 0.8496 |
| rs605908 | 11 | 118067192 | C | A | 1.009 | 0.3129 | +?+ | 0 | 0.803 | 1 | 0.3701 |
| rs61900159 | 11 | 118839133 | C | G | -1.007 | 0.3137 | --- | 0 | 0.236 | 2 | 0.8887 |
| rs904469 | 11 | 72324039 | A | G | 1.008 | 0.3137 | -?+ | -73.4 | 1.73 | 1 | 0.1884 |
| rs481206 | 11 | 72497462 | T | C | 1.007 | 0.314 | ++- | -21 | 3.306 | 2 | 0.1915 |
| rs7123835 | 11 | 118368822 | G | T | 1.007 | 0.314 | +?+ | 0 | 0.363 | 1 | 0.5471 |
| rs11216803 | 11 | 118057244 | A | C | 1.006 | 0.3145 | +++ | 0 | 0.328 | 2 | 0.8486 |
| rs11216802 | 11 | 118057227 | T | C | 1.005 | 0.3149 | +++ | 0 | 0.33 | 2 | 0.848 |
| rs1814964 | 11 | 118006957 | C | T | 1.003 | 0.3156 | +-+ | 0 | 1.405 | 2 | 0.4952 |
| rs79092677 | 11 | 118265734 | G | T | -1.001 | 0.3166 | -?- | 0 | 0.173 | 1 | 0.6772 |
| rs73557345 | 11 | 118813314 | C | T | 1 | 0.3173 | +-+ | 0 | 1.531 | 2 | 0.4651 |
| rs7107305 | 11 | 118368665 | G | A | 0.999 | 0.3178 | +?+ | 0 | 0.372 | 1 | 0.5417 |
| rs11216864 | 11 | 118260017 | T | C | -0.998 | 0.3182 | -?- | 0 | 0.085 | 1 | 0.7708 |
| rs755933 | 11 | 72325482 | A | G | 0.994 | 0.3204 | -?+ | -72.3 | 1.742 | 1 | 0.1869 |
| rs7940717 | 11 | 118055110 | T | G | 0.993 | 0.3209 | +++ | 0 | 0.307 | 2 | 0.8577 |
| rs10892237 | 11 | 118267763 | C | T | -0.99 | 0.3221 | -?- | 0 | 0.183 | 1 | 0.6689 |
| rs12795307 | 11 | 72506324 | A | G | 0.987 | 0.3235 | ++- | -15.7 | 3.458 | 2 | 0.1774 |
| rs653008 | 11 | 118520480 | T | G | 0.984 | 0.3251 | ++- | 0 | 0.927 | 2 | 0.6292 |
| rs7109575 | 11 | 72463435 | A | G | 0.98 | 0.3271 | -++ | 39.2 | 6.576 | 2 | 0.03733 |
| rs519790 | 11 | 72504141 | G | C | 0.98 | 0.3273 | ++- | -7.3 | 3.726 | 2 | 0.1552 |
| rs11604096 | 11 | 118012441 | C | A | -0.978 | 0.328 | -+- | 0 | 0.62 | 2 | 0.7336 |
| rs7103001 | 11 | 118367669 | T | A | 0.977 | 0.3285 | +?+ | 0 | 0.399 | 1 | 0.5274 |
| rs2276422 | 11 | 118263388 | G | C | -0.974 | 0.3299 | -?- | 0 | 0.197 | 1 | 0.6574 |
| rs117278652 | 11 | 72093901 | A | C | 0.973 | 0.3306 | -?+ | -51.7 | 1.978 | 1 | 0.1596 |
| rs2075647 | 11 | 118037964 | C | G | -0.973 | 0.3306 | -?- | 0 | 0.471 | 1 | 0.4927 |
| rs1624780 | 11 | 118134785 | T | G | 0.972 | 0.3311 | +?- | -180.9 | 1.068 | 1 | 0.3014 |
| rs17122150 | 11 | 118271860 | G | C | 0.972 | 0.3311 | +++ | 0 | 0.597 | 2 | 0.742 |
| rs12803321 | 11 | 118480115 | C | G | 0.97 | 0.3319 | +?+ | 0 | 0.327 | 1 | 0.5674 |
| rs77280285 | 11 | 118091258 | G | C | 0.969 | 0.3326 | +?+ | 0 | 0.4 | 1 | 0.5269 |
| rs497678 | 11 | 118567881 | C | T | 0.969 | 0.3327 | ++- | 0 | 1.803 | 2 | 0.4061 |
| rs10892253 | 11 | 118559426 | A | C | 0.968 | 0.3333 | +?+ | 0 | 0.06 | 1 | 0.8068 |
| rs56296109 | 11 | 118698170 | A | C | 0.965 | 0.3344 | -?+ | 22.8 | 3.886 | 1 | 0.04868 |
| rs601318 | 11 | 118519550 | G | C | 0.964 | 0.3348 | ++- | 0 | 0.91 | 2 | 0.6344 |
| rs639785 | 11 | 118062268 | T | C | 0.964 | 0.335 | +++ | 0 | 0.116 | 2 | 0.9437 |
| rs1623316 | 11 | 118757219 | G | C | 0.963 | 0.3355 | ++- | -90.5 | 2.1 | 2 | 0.35 |
| rs7110984 | 11 | 118442234 | A | G | 0.961 | 0.3365 | +-+ | 0 | 1.975 | 2 | 0.3725 |
| rs117117171 | 11 | 118152252 | A | T | 0.959 | 0.3377 | +?+ | 0 | 0.059 | 1 | 0.8073 |
| rs7121764 | 11 | 118271759 | G | A | 0.95 | 0.3419 | +++ | 0 | 0.62 | 2 | 0.7333 |
| rs3181261 | 11 | 118212671 | T | C | 0.949 | 0.3428 | +?+ | 0 | 0.136 | 1 | 0.7126 |
| rs11603349 | 11 | 72460694 | C | T | 0.947 | 0.3437 | -++ | 36.8 | 6.327 | 2 | 0.04227 |
| rs117794131 | 11 | 72097791 | C | T | 0.946 | 0.3439 | -?+ | -57.6 | 1.904 | 1 | 0.1677 |
| rs623500 | 11 | 118828942 | G | A | 0.944 | 0.3454 | +?- | -65.5 | 1.813 | 1 | 0.1782 |
| rs11216867 | 11 | 118262407 | G | A | 0.943 | 0.3458 | +++ | 0 | 0.538 | 2 | 0.7643 |
| rs663015 | 11 | 72499035 | C | T | 0.942 | 0.346 | ++- | -16.9 | 3.42 | 2 | 0.1808 |
| rs6589673 | 11 | 118557483 | C | A | 0.942 | 0.3461 | +?+ | 0 | 0.048 | 1 | 0.8265 |
| rs7103366 | 11 | 118057867 | G | T | 0.94 | 0.3473 | +++ | 0 | 0.076 | 2 | 0.9626 |
| rs75085081 | 11 | 118801045 | A | G | 0.935 | 0.3499 | +-+ | 0 | 1.855 | 2 | 0.3955 |
| rs1791936 | 11 | 72877964 | G | A | -0.933 | 0.3508 | --- | 0 | 0.245 | 2 | 0.8849 |

| | | | | | | | | | | | |
|---|---|---|---|---|---|---|---|---|---|---|---|
| rs3781934 | 11 | 72325070 | A | G | 0.933 | 0.3509 | -?+ | -83.7 | 1.633 | 1 | 0.2013 |
| rs668468 | 11 | 118011776 | G | A | -0.932 | 0.3515 | -+- | 0 | 0.643 | 2 | 0.725 |
| rs73016221 | 11 | 118220027 | C | T | -0.926 | 0.3543 | -?- | 0 | 0.502 | 1 | 0.4786 |
| rs76675432 | 11 | 118103976 | T | C | 0.926 | 0.3546 | +?+ | 0 | 0.38 | 1 | 0.5378 |
| rs11602873 | 11 | 72460762 | T | A | 0.92 | 0.3577 | -++ | 31.7 | 5.861 | 2 | 0.05338 |
| rs73003294 | 11 | 118661376 | T | C | 0.919 | 0.3581 | -?+ | 30.1 | 4.293 | 1 | 0.03827 |
| rs77464186 | 11 | 72460398 | C | A | 0.918 | 0.3584 | -++ | 31.7 | 5.854 | 2 | 0.05356 |
| rs117822540 | 11 | 72123996 | C | T | 0.916 | 0.3597 | -?+ | -75.3 | 1.711 | 1 | 0.1909 |
| rs17122154 | 11 | 118273019 | A | G | 0.913 | 0.3611 | +++ | 0 | 0.652 | 2 | 0.7218 |
| rs9332809 | 11 | 118361472 | A | G | 0.91 | 0.363 | +?+ | 0 | 0.213 | 1 | 0.6447 |
| rs117190635 | 11 | 72104119 | A | G | 0.905 | 0.3652 | -?+ | -90.1 | 1.578 | 1 | 0.2091 |
| rs2276045 | 11 | 118365168 | A | G | 0.899 | 0.3685 | +?+ | 0 | 0.268 | 1 | 0.605 |
| rs1793170 | 11 | 118150205 | C | T | -0.899 | 0.3686 | +-- | 0 | 0.701 | 2 | 0.7045 |
| rs6589663 | 11 | 118255278 | A | G | 0.896 | 0.3703 | +++ | 0 | 0.655 | 2 | 0.7208 |
| rs619250 | 11 | 118064582 | A | T | 0.896 | 0.3704 | +?+ | 0 | 0.771 | 1 | 0.3798 |
| rs7116876 | 11 | 118371025 | C | G | 0.896 | 0.3705 | +?+ | 0 | 0.271 | 1 | 0.6024 |
| rs3781931 | 11 | 72327205 | T | C | 0.895 | 0.3707 | -?+ | -101.1 | 1.492 | 1 | 0.222 |
| rs3212262 | 11 | 118217854 | T | G | -0.893 | 0.372 | -?- | 0 | 0.551 | 1 | 0.458 |
| rs9332819 | 11 | 118365263 | T | C | 0.891 | 0.373 | +?+ | 0 | 0.248 | 1 | 0.6186 |
| rs562780 | 11 | 118362414 | T | C | 0.89 | 0.3733 | -++ | 0 | 0.856 | 2 | 0.6517 |
| rs10790240 | 11 | 118009279 | G | T | 0.889 | 0.3738 | +-+ | 0 | 1.103 | 2 | 0.5762 |
| rs9332835 | 11 | 118372923 | A | G | 0.887 | 0.3748 | +?+ | 0 | 0.28 | 1 | 0.5967 |
| rs11217077 | 11 | 118746223 | C | T | 0.885 | 0.3762 | ++- | 0 | 1.761 | 2 | 0.4145 |
| rs7933243 | 11 | 118403205 | G | T | 0.882 | 0.3778 | +-+ | 0 | 0.752 | 2 | 0.6866 |
| rs7109466 | 11 | 72114792 | T | C | 0.879 | 0.3796 | -?+ | -67.4 | 1.792 | 1 | 0.1807 |
| rs12290522 | 11 | 118486325 | T | C | 0.876 | 0.3811 | ++- | 0 | 0.92 | 2 | 0.6312 |
| rs12788624 | 11 | 118022107 | T | C | -0.876 | 0.3811 | -++ | -22.4 | 3.269 | 2 | 0.1951 |
| rs2276419 | 11 | 118230359 | T | C | -0.875 | 0.3813 | -?- | 0 | 0.589 | 1 | 0.4429 |
| rs10790254 | 11 | 118493319 | A | G | 0.874 | 0.3819 | +?+ | 0 | 0 | 1 | 0.9977 |
| rs6589651 | 11 | 118058156 | G | A | 0.874 | 0.3822 | +++ | 0 | 0.057 | 2 | 0.9721 |
| rs56754533 | 11 | 118188276 | A | G | -0.872 | 0.3833 | -?- | 0 | 0.551 | 1 | 0.4578 |
| rs9332846 | 11 | 118377523 | A | G | 0.872 | 0.3835 | +?+ | 0 | 0.197 | 1 | 0.6569 |
| rs17121871 | 11 | 118073344 | T | C | 0.871 | 0.3839 | +?+ | 0 | 0.435 | 1 | 0.5095 |
| rs113399762 | 11 | 118389698 | C | T | 0.867 | 0.3858 | +?+ | 0 | 0.35 | 1 | 0.5541 |
| rs116881498 | 11 | 72102819 | T | C | 0.866 | 0.3867 | -?+ | -73.5 | 1.729 | 1 | 0.1885 |
| rs147710683 | 11 | 118121356 | T | C | -0.865 | 0.387 | -?- | 0 | 0.486 | 1 | 0.4857 |
| rs79826955 | 11 | 72104081 | A | G | 0.865 | 0.387 | -?+ | -73.7 | 1.728 | 1 | 0.1887 |
| rs55768116 | 11 | 118108331 | A | C | -0.864 | 0.3877 | -?- | 0 | 0.329 | 1 | 0.5664 |
| rs7113994 | 11 | 118382327 | G | A | 0.861 | 0.389 | +?+ | 0 | 0.26 | 1 | 0.6103 |
| rs1616199 | 11 | 118146462 | T | C | -0.856 | 0.3918 | +-- | 0 | 0.636 | 2 | 0.7277 |
| rs1783587 | 11 | 72887612 | C | T | -0.855 | 0.3926 | ++- | 9.5 | 4.421 | 2 | 0.1097 |
| rs1632118 | 11 | 72881636 | T | C | -0.855 | 0.3928 | -+- | -44.8 | 2.762 | 2 | 0.2513 |
| rs7113959 | 11 | 118379096 | A | T | 0.853 | 0.3935 | +?+ | 0 | 0.268 | 1 | 0.6047 |
| rs45621436 | 11 | 118483912 | G | T | 0.853 | 0.3938 | +?+ | 0 | 0.005 | 1 | 0.9438 |
| rs6589652 | 11 | 118058358 | G | A | 0.853 | 0.3938 | +++ | 0 | 0.045 | 2 | 0.9779 |
| rs527290 | 11 | 118515616 | T | C | 0.851 | 0.3948 | ++- | 0 | 1.042 | 2 | 0.594 |
| rs1171461 | 11 | 118522179 | A | G | 0.85 | 0.3952 | ++- | 0 | 0.874 | 2 | 0.646 |
| rs146305387 | 11 | 118426800 | T | C | -0.849 | 0.3957 | -?+ | -177.2 | 1.082 | 1 | 0.2982 |
| rs117654328 | 11 | 72117713 | A | G | 0.849 | 0.3961 | -?+ | -75.2 | 1.712 | 1 | 0.1907 |
| rs150031150 | 11 | 72120370 | A | G | 0.849 | 0.3961 | -?+ | -75.2 | 1.712 | 1 | 0.1907 |
| rs1715444 | 11 | 118016273 | G | T | -0.846 | 0.3976 | -+- | 0 | 0.369 | 2 | 0.8316 |
| rs724673 | 11 | 72795708 | A | G | 0.838 | 0.4018 | -++ | -51.5 | 2.64 | 2 | 0.2672 |
| rs4938520 | 11 | 118522338 | G | A | 0.838 | 0.4021 | ++- | 0 | 0.787 | 2 | 0.6747 |
| rs117235147 | 11 | 72115430 | A | T | 0.838 | 0.4023 | -?+ | -78.2 | 1.684 | 1 | 0.1944 |

| | | | | | | | | | | | |
|---|---|---|---|---|---|---|---|---|---|---|---|
| rs4553405 | 11 | 118061798 | T | C | 0.837 | 0.4026 | +++ | 0 | 0.043 | 2 | 0.9785 |
| rs9332816 | 11 | 118364404 | T | G | 0.836 | 0.4029 | +?+ | 0 | 0.349 | 1 | 0.5545 |
| rs149668068 | 11 | 118099735 | A | G | -0.835 | 0.4039 | -?- | 0 | 0.358 | 1 | 0.5494 |
| rs2003368 | 11 | 72840106 | T | C | 0.832 | 0.4056 | -++ | -40.4 | 2.848 | 2 | 0.2407 |
| rs550999 | 11 | 118811684 | G | C | 0.83 | 0.4068 | +-+ | -3.7 | 3.856 | 2 | 0.1454 |
| rs148643334 | 11 | 118358152 | C | T | 0.826 | 0.4087 | +?+ | 0 | 0.42 | 1 | 0.517 |
| rs45460493 | 11 | 118455465 | A | G | 0.825 | 0.4093 | +?+ | 0 | 0.002 | 1 | 0.9689 |
| rs745663 | 11 | 118529595 | G | A | 0.823 | 0.4104 | ++- | 0 | 0.771 | 2 | 0.68 |
| rs111997050 | 11 | 118387892 | T | C | 0.815 | 0.4149 | +?+ | 0 | 0.103 | 1 | 0.748 |
| rs67514601 | 11 | 118189477 | G | A | -0.814 | 0.4155 | -?- | 0 | 0.393 | 1 | 0.531 |
| rs11235641 | 11 | 72755766 | A | G | 0.814 | 0.4159 | -++ | -39.8 | 2.861 | 2 | 0.2392 |
| rs56083337 | 11 | 118131813 | T | A | -0.812 | 0.4168 | -?+ | 0 | 0.755 | 1 | 0.385 |
| rs78020509 | 11 | 118384811 | G | A | 0.809 | 0.4186 | +?+ | 0 | 0.299 | 1 | 0.5842 |
| rs117068828 | 11 | 72139241 | A | G | 0.809 | 0.4188 | -?+ | -86.4 | 1.609 | 1 | 0.2046 |
| rs111405302 | 11 | 118385691 | C | T | 0.808 | 0.4189 | +?+ | 0 | 0.313 | 1 | 0.5759 |
| rs7389 | 11 | 118528466 | G | T | 0.806 | 0.4202 | ++- | 0 | 0.856 | 2 | 0.6519 |
| rs9332858 | 11 | 118390082 | C | A | 0.803 | 0.422 | +?+ | 0 | 0.255 | 1 | 0.6135 |
| rs4420279 | 11 | 118835677 | A | C | 0.801 | 0.4233 | -?+ | -33.1 | 2.253 | 1 | 0.1333 |
| rs117532939 | 11 | 72165516 | G | A | 0.798 | 0.4246 | -?+ | -121.3 | 1.355 | 1 | 0.2443 |
| rs117817588 | 11 | 72166041 | T | G | 0.798 | 0.4246 | -?+ | -121.3 | 1.355 | 1 | 0.2443 |
| rs570836 | 11 | 118512162 | A | G | 0.797 | 0.4255 | ++- | 0 | 0.968 | 2 | 0.6163 |
| rs1513780 | 11 | 72320500 | T | G | 0.795 | 0.4267 | -?+ | -76.9 | 1.696 | 1 | 0.1928 |
| rs630923 | 11 | 118754353 | A | C | 0.795 | 0.4269 | +-+ | -87.7 | 2.131 | 2 | 0.3445 |
| rs12419204 | 11 | 118497402 | A | T | 0.793 | 0.428 | ++- | 0 | 0.921 | 2 | 0.6309 |
| rs4944563 | 11 | 72316881 | C | A | 0.79 | 0.4294 | -?+ | -76.9 | 1.695 | 1 | 0.1929 |
| rs45460396 | 11 | 118005622 | G | T | -0.789 | 0.4299 | -+- | 0 | 0.731 | 2 | 0.6939 |
| rs10790257 | 11 | 118561858 | C | G | 0.787 | 0.4312 | +?+ | 0 | 0.004 | 1 | 0.9488 |
| rs10790258 | 11 | 118561859 | T | G | 0.787 | 0.4312 | +?+ | 0 | 0.004 | 1 | 0.9488 |
| rs4938517 | 11 | 118487673 | C | T | 0.784 | 0.4329 | ++- | 0 | 1.083 | 2 | 0.582 |
| rs4944566 | 11 | 72321872 | A | G | 0.782 | 0.4341 | -?+ | -90.3 | 1.577 | 1 | 0.2093 |
| rs649138 | 11 | 118066695 | C | T | 0.778 | 0.4363 | +?+ | 0 | 0.444 | 1 | 0.505 |
| rs45584835 | 11 | 118004863 | G | A | -0.778 | 0.4365 | -+- | 0 | 0.699 | 2 | 0.7049 |
| rs483283 | 11 | 118506402 | C | T | 0.778 | 0.4365 | ++- | 0 | 0.964 | 2 | 0.6174 |
| rs11603616 | 11 | 118496066 | A | G | 0.776 | 0.4375 | ++- | 0 | 0.987 | 2 | 0.6105 |
| rs1793185 | 11 | 118089434 | A | C | 0.776 | 0.4375 | +?- | 0 | 0.925 | 1 | 0.3363 |
| rs9332871 | 11 | 118393856 | T | G | 0.776 | 0.4377 | +?+ | 0 | 0.122 | 1 | 0.7271 |
| rs11235550 | 11 | 72331050 | C | T | 0.776 | 0.4378 | +?+ | 0 | 0 | 1 | 0.9822 |
| rs868344 | 11 | 118005119 | C | T | -0.776 | 0.4378 | -+- | 0 | 0.687 | 2 | 0.7094 |
| rs570952 | 11 | 118500193 | C | T | 0.775 | 0.4384 | ++- | 0 | 0.909 | 2 | 0.6348 |
| rs9332857 | 11 | 118389440 | T | C | 0.775 | 0.4385 | +?+ | 0 | 0.127 | 1 | 0.7215 |
| rs9332852 | 11 | 118381069 | T | C | 0.773 | 0.4395 | +?+ | 0 | 0.161 | 1 | 0.6885 |
| rs9332851 | 11 | 118381018 | T | C | 0.77 | 0.4411 | +?+ | 0 | 0.172 | 1 | 0.6783 |
| rs2231439 | 11 | 118174885 | G | A | -0.77 | 0.4412 | -?- | 0 | 0.587 | 1 | 0.4436 |
| rs67533649 | 11 | 118835272 | G | A | 0.767 | 0.4432 | -?+ | -51.4 | 1.981 | 1 | 0.1593 |
| rs458360 | 11 | 72319522 | A | C | 0.766 | 0.4435 | -?+ | -39.6 | 2.148 | 1 | 0.1427 |
| rs462491 | 11 | 72319520 | C | T | 0.766 | 0.4435 | -?+ | -39.6 | 2.148 | 1 | 0.1427 |
| rs571001 | 11 | 118500173 | T | C | 0.765 | 0.4442 | ++- | 0 | 0.961 | 2 | 0.6184 |
| rs7103936 | 11 | 118091636 | C | A | 0.765 | 0.4442 | +?- | -166.1 | 1.128 | 1 | 0.2883 |
| rs150339153 | 11 | 118285369 | T | C | -0.763 | 0.4454 | -?- | 0 | 0.141 | 1 | 0.7076 |
| rs2231449 | 11 | 118186305 | A | C | -0.76 | 0.4474 | -+- | -52.3 | 2.627 | 2 | 0.2689 |
| rs4938490 | 11 | 118080977 | G | A | 0.758 | 0.4483 | +?+ | 0 | 0.455 | 1 | 0.4998 |
| rs605560 | 11 | 118067187 | C | G | 0.755 | 0.4505 | +?+ | 0 | 0.414 | 1 | 0.5201 |
| rs2077173 | 11 | 118492591 | A | G | 0.753 | 0.4512 | ++- | 0 | 1.067 | 2 | 0.5866 |
| rs11216843 | 11 | 118151329 | T | A | -0.752 | 0.4521 | -+- | 0 | 1.722 | 2 | 0.4228 |

| | | | | | | | | | | | |
|---|---|---|---|---|---|---|---|---|---|---|---|
| rs10790244 | 11 | 118059557 | C | T | 0.747 | 0.455 | +++ | 0 | 0.021 | 2 | 0.9896 |
| rs4938595 | 11 | 118832582 | T | G | 0.746 | 0.4556 | -?+ | -37.3 | 2.184 | 1 | 0.1394 |
| rs146523897 | 11 | 72161854 | A | G | 0.745 | 0.456 | -?+ | -93.7 | 1.549 | 1 | 0.2133 |
| rs11216848 | 11 | 118181481 | A | C | -0.743 | 0.4574 | -+- | -85.2 | 2.16 | 2 | 0.3397 |
| rs2120268 | 11 | 118132705 | A | G | 0.74 | 0.4591 | +?- | 0 | 0.935 | 1 | 0.3335 |
| rs382163 | 11 | 72292796 | T | C | 0.74 | 0.4591 | -?+ | 0 | 0.907 | 1 | 0.341 |
| rs11216822 | 11 | 118091375 | A | G | 0.74 | 0.4596 | +?- | -153.7 | 1.182 | 1 | 0.2768 |
| rs2097183 | 11 | 118091067 | T | C | 0.739 | 0.4596 | +?- | -153.7 | 1.183 | 1 | 0.2768 |
| rs668628 | 11 | 118519213 | T | G | 0.736 | 0.4619 | +?- | 48.3 | 5.797 | 1 | 0.01605 |
| rs341085 | 11 | 72318882 | A | C | 0.73 | 0.4651 | -?+ | -37.2 | 2.186 | 1 | 0.1393 |
| rs17121881 | 11 | 118081345 | A | T | 0.73 | 0.4656 | +?+ | 0 | 0.49 | 1 | 0.4838 |
| rs673770 | 11 | 118550601 | G | T | 0.729 | 0.4658 | ++- | -55.4 | 2.574 | 2 | 0.276 |
| rs145968026 | 11 | 118421841 | G | C | 0.729 | 0.4661 | +?+ | 0 | 0.018 | 1 | 0.8924 |
| rs17121879 | 11 | 118080133 | G | T | 0.726 | 0.4676 | +?+ | 0 | 0.432 | 1 | 0.511 |
| rs4938489 | 11 | 118079867 | G | T | 0.726 | 0.468 | +?+ | 0 | 0.433 | 1 | 0.5106 |
| rs111227443 | 11 | 118590961 | T | C | 0.725 | 0.4684 | -?+ | -47.8 | 2.03 | 1 | 0.1543 |
| rs592280 | 11 | 118553218 | C | T | 0.725 | 0.4686 | ++- | -61.9 | 2.471 | 2 | 0.2907 |
| rs11600544 | 11 | 118076954 | C | G | 0.724 | 0.469 | +?+ | 0 | 0.417 | 1 | 0.5186 |
| rs111656195 | 11 | 118301375 | A | G | 0.724 | 0.4693 | +?- | 0 | 0.612 | 1 | 0.434 |
| rs59776769 | 11 | 72296353 | A | G | 0.724 | 0.4693 | -?+ | 0 | 0.591 | 1 | 0.4421 |
| rs1793167 | 11 | 118102058 | T | C | 0.723 | 0.4697 | +?- | -115.9 | 1.39 | 1 | 0.2385 |
| rs1805 | 11 | 118076069 | T | G | 0.722 | 0.4704 | +?+ | 0 | 0.45 | 1 | 0.5024 |
| rs151127828 | 11 | 118206121 | A | G | 0.721 | 0.4709 | +?+ | 0 | 0.002 | 1 | 0.9662 |
| rs17121882 | 11 | 118081468 | C | T | 0.721 | 0.4712 | +?+ | 0 | 0.47 | 1 | 0.4932 |
| rs76457018 | 11 | 118762701 | A | G | -0.718 | 0.4731 | +-- | 0 | 0.678 | 2 | 0.7126 |
| rs642497 | 11 | 118550784 | A | C | 0.715 | 0.4747 | ++- | -56.3 | 2.559 | 2 | 0.2782 |
| rs645637 | 11 | 118505090 | A | G | 0.713 | 0.4757 | ++- | 0 | 1.015 | 2 | 0.6019 |
| rs4331144 | 11 | 118554100 | G | A | 0.707 | 0.4795 | ++- | -46.8 | 2.725 | 2 | 0.256 |
| rs12799314 | 11 | 118551597 | A | G | 0.705 | 0.4809 | ++- | -53.6 | 2.604 | 2 | 0.272 |
| rs171021 | 11 | 72317557 | T | C | 0.703 | 0.4819 | -?+ | -157.7 | 1.164 | 1 | 0.2806 |
| rs667982 | 11 | 118562767 | C | T | 0.703 | 0.4822 | ++- | -53.3 | 2.609 | 2 | 0.2714 |
| rs586370 | 11 | 118816353 | A | G | -0.703 | 0.4823 | -?- | 0 | 0.292 | 1 | 0.5892 |
| rs112636863 | 11 | 118333510 | C | T | 0.701 | 0.4835 | +?+ | 0 | 0.422 | 1 | 0.516 |
| rs642530 | 11 | 118550756 | C | T | 0.699 | 0.4843 | ++- | -54.1 | 2.595 | 2 | 0.2732 |
| rs61900158 | 11 | 118831764 | A | T | 0.699 | 0.4849 | -?+ | -52.2 | 1.971 | 1 | 0.1603 |
| rs4936433 | 11 | 118552420 | G | A | 0.697 | 0.4857 | ++- | -54.2 | 2.594 | 2 | 0.2734 |
| rs7106652 | 11 | 118554248 | T | C | 0.697 | 0.4858 | ++- | -49.7 | 2.673 | 2 | 0.2628 |
| rs7106851 | 11 | 118554259 | A | G | 0.697 | 0.4858 | ++- | -50.2 | 2.663 | 2 | 0.2641 |
| rs600969 | 11 | 118545797 | T | C | 0.697 | 0.4859 | ++- | -51.5 | 2.64 | 2 | 0.2671 |
| rs1806 | 11 | 118076004 | C | T | 0.696 | 0.4863 | +?- | 0 | 0.747 | 1 | 0.3874 |
| rs111903103 | 11 | 118308499 | T | C | 0.694 | 0.4876 | +?+ | 0 | 0.472 | 1 | 0.4922 |
| rs11602869 | 11 | 118554448 | A | G | 0.693 | 0.4884 | ++- | -47.5 | 2.711 | 2 | 0.2578 |
| rs11605069 | 11 | 118554436 | G | T | 0.69 | 0.49 | ++- | -49.2 | 2.682 | 2 | 0.2616 |
| rs11216806 | 11 | 118060549 | C | T | 0.69 | 0.4901 | +++ | 0 | 0.062 | 2 | 0.9695 |
| rs1793155 | 11 | 118098962 | A | G | 0.689 | 0.4909 | +?- | -109.4 | 1.433 | 1 | 0.2313 |
| rs12796373 | 11 | 118554939 | A | G | 0.689 | 0.4911 | ++- | -46.5 | 2.731 | 2 | 0.2552 |
| rs12280559 | 11 | 118087040 | T | C | 0.688 | 0.4916 | +?- | -131.2 | 1.298 | 1 | 0.2546 |
| rs656575 | 11 | 118549908 | G | T | 0.688 | 0.4917 | ++- | -75.7 | 2.276 | 2 | 0.3204 |
| rs507080 | 11 | 118549582 | A | G | 0.686 | 0.4925 | ++- | -77.8 | 2.25 | 2 | 0.3246 |
| rs562313 | 11 | 118548421 | A | G | 0.686 | 0.493 | ++- | -48.6 | 2.691 | 2 | 0.2604 |
| rs549370 | 11 | 118552842 | A | C | 0.684 | 0.4939 | ++- | -50.8 | 2.653 | 2 | 0.2655 |
| rs543413 | 11 | 118550017 | C | T | 0.683 | 0.4944 | ++- | -76.8 | 2.263 | 2 | 0.3226 |
| rs3018383 | 11 | 118093359 | A | C | 0.683 | 0.4947 | +?- | -107.3 | 1.447 | 1 | 0.229 |
| rs111354396 | 11 | 118296184 | A | G | 0.682 | 0.4953 | +?- | 0 | 0.693 | 1 | 0.4052 |

| | | | | | | | | | | | |
|---|---|---|---|---|---|---|---|---|---|---|---|
| rs1793156 | 11 | 118098890 | C | T | 0.682 | 0.4954 | +?- | -107 | 1.45 | 1 | 0.2286 |
| rs11216781 | 11 | 118012310 | A | G | 0.681 | 0.496 | +++ | 0 | 0.047 | 2 | 0.977 |
| rs4938524 | 11 | 118552074 | T | G | 0.681 | 0.4961 | ++- | -48.9 | 2.686 | 2 | 0.2611 |
| rs654792 | 11 | 118552996 | T | G | 0.68 | 0.4964 | ++- | -49.8 | 2.671 | 2 | 0.2631 |
| rs111763202 | 11 | 118289734 | T | C | 0.68 | 0.4967 | +?- | 0 | 0.696 | 1 | 0.404 |
| rs112035668 | 11 | 118290443 | G | A | 0.679 | 0.4973 | +?- | 0 | 0.698 | 1 | 0.4034 |
| rs56210510 | 11 | 72333342 | C | T | 0.679 | 0.4973 | -?+ | -87 | 1.604 | 1 | 0.2054 |
| rs111901186 | 11 | 118328052 | G | A | 0.679 | 0.4974 | +?- | 0 | 0.478 | 1 | 0.4895 |
| rs7103918 | 11 | 118061480 | T | C | 0.678 | 0.498 | +++ | 0 | 0.015 | 2 | 0.9926 |
| rs519848 | 11 | 118552715 | A | G | 0.677 | 0.4985 | ++- | -48.9 | 2.686 | 2 | 0.2611 |
| rs112302730 | 11 | 118283383 | A | G | 0.676 | 0.4988 | +?- | 0 | 0.702 | 1 | 0.4021 |
| rs543327 | 11 | 118549985 | T | G | 0.675 | 0.4996 | ++- | -74.1 | 2.297 | 2 | 0.3171 |
| rs112550735 | 11 | 118303200 | C | T | 0.674 | 0.5001 | +?- | 0 | 0.738 | 1 | 0.3902 |
| rs11606719 | 11 | 118782474 | G | C | 0.674 | 0.5004 | ++- | 0 | 1.884 | 2 | 0.3898 |
| rs1784298 | 11 | 118548665 | C | T | 0.674 | 0.5004 | ++- | -73.7 | 2.303 | 2 | 0.3162 |
| rs77728961 | 11 | 118303182 | C | T | 0.674 | 0.5004 | +?- | 0 | 0.739 | 1 | 0.39 |
| rs12575015 | 11 | 118197645 | T | A | -0.674 | 0.5006 | -?- | 0 | 0.084 | 1 | 0.7716 |
| rs4938522 | 11 | 118548671 | T | A | 0.673 | 0.5012 | ++- | -73.3 | 2.308 | 2 | 0.3154 |
| rs55861597 | 11 | 72331873 | T | G | 0.672 | 0.5019 | -?+ | -127.3 | 1.32 | 1 | 0.2506 |
| rs55738331 | 11 | 72331821 | T | C | 0.67 | 0.5028 | -?+ | -127.8 | 1.317 | 1 | 0.2512 |
| rs141752759 | 11 | 118194647 | G | C | 0.67 | 0.503 | +?+ | 0 | 0 | 1 | 1 |
| rs10736494 | 11 | 118518227 | T | C | 0.669 | 0.5033 | +?+ | 0 | 0.274 | 1 | 0.6005 |
| rs67435655 | 11 | 118196399 | C | T | -0.665 | 0.5059 | -?- | 0 | 0.089 | 1 | 0.7652 |
| rs66962317 | 11 | 118197486 | C | T | -0.664 | 0.5065 | -?- | 0 | 0.09 | 1 | 0.7645 |
| rs12418930 | 11 | 72681566 | T | C | 0.663 | 0.5076 | -++ | -12.7 | 3.549 | 2 | 0.1696 |
| rs117493218 | 11 | 118027406 | C | A | 0.661 | 0.5084 | --+ | 0 | 1.311 | 2 | 0.5191 |
| rs659969 | 11 | 118549803 | T | C | 0.661 | 0.5085 | ++- | -69.8 | 2.356 | 2 | 0.308 |
| rs11216846 | 11 | 118163916 | A | G | -0.659 | 0.51 | -+- | 0 | 1.757 | 2 | 0.4154 |
| rs563549 | 11 | 118558953 | A | G | 0.653 | 0.5139 | ++- | -47.3 | 2.716 | 2 | 0.2572 |
| rs3802863 | 11 | 118099186 | T | C | 0.653 | 0.5141 | +?- | -73.2 | 1.732 | 1 | 0.1881 |
| rs4938523 | 11 | 118548838 | T | G | 0.652 | 0.5142 | ++- | -67.1 | 2.393 | 2 | 0.3022 |
| rs509641 | 11 | 118558952 | C | T | 0.652 | 0.5142 | ++- | -47.2 | 2.717 | 2 | 0.257 |
| rs112245679 | 11 | 118287622 | G | A | 0.652 | 0.5143 | +?- | 0 | 0.743 | 1 | 0.3887 |
| rs112563022 | 11 | 118262135 | T | C | 0.652 | 0.5144 | +?- | 0 | 0.86 | 1 | 0.3536 |
| rs67859744 | 11 | 118200657 | A | T | -0.652 | 0.5145 | -?- | 0 | 0.097 | 1 | 0.7549 |
| rs113028342 | 11 | 118333537 | G | A | 0.651 | 0.5153 | +?+ | 0 | 0.313 | 1 | 0.5757 |
| rs45627033 | 11 | 118766018 | T | C | 0.647 | 0.5177 | +?- | -29.7 | 2.313 | 1 | 0.1283 |
| rs510408 | 11 | 118558901 | C | G | 0.647 | 0.5179 | ++- | -48.9 | 2.686 | 2 | 0.2611 |
| rs55843215 | 11 | 118195962 | C | T | -0.645 | 0.5191 | -?- | 0 | 0.102 | 1 | 0.7496 |
| rs392565 | 11 | 72290629 | A | G | 0.645 | 0.5192 | -?+ | 0 | 0.737 | 1 | 0.3905 |
| rs143605660 | 11 | 118707035 | A | G | 0.644 | 0.5194 | -?+ | -10 | 2.728 | 1 | 0.09862 |
| rs6589656 | 11 | 118095996 | C | G | 0.644 | 0.5196 | +?- | -70.9 | 1.755 | 1 | 0.1852 |
| rs638805 | 11 | 118558889 | T | G | 0.644 | 0.5198 | ++- | -48.2 | 2.699 | 2 | 0.2593 |
| rs1617182 | 11 | 118068949 | C | T | 0.637 | 0.524 | +?- | 0 | 0.511 | 1 | 0.4747 |
| rs73612281 | 11 | 118203642 | T | C | -0.635 | 0.5255 | -?- | 0 | 0.108 | 1 | 0.7422 |
| rs111993696 | 11 | 118282448 | A | G | 0.634 | 0.5258 | +?- | 0 | 0.774 | 1 | 0.379 |
| rs4938505 | 11 | 118205636 | A | G | -0.634 | 0.526 | -?- | 0 | 0.109 | 1 | 0.7416 |
| rs12801084 | 11 | 72991793 | G | C | -0.632 | 0.5276 | --+ | -8.2 | 3.696 | 2 | 0.1576 |
| rs558781 | 11 | 118564397 | T | G | 0.63 | 0.5284 | ++- | -65.1 | 2.423 | 2 | 0.2977 |
| rs514143 | 11 | 118558449 | A | G | 0.63 | 0.5287 | ++- | -65.5 | 2.417 | 2 | 0.2986 |
| rs112583389 | 11 | 118278998 | G | T | 0.629 | 0.5295 | +?- | 0 | 0.698 | 1 | 0.4036 |
| rs11235711 | 11 | 72992343 | C | G | -0.626 | 0.5312 | --+ | -7.5 | 3.722 | 2 | 0.1555 |
| rs371862 | 11 | 72288843 | T | C | 0.626 | 0.5313 | -?+ | 0 | 0.718 | 1 | 0.3968 |
| rs625513 | 11 | 118555719 | C | A | 0.625 | 0.5318 | ++- | -65.5 | 2.418 | 2 | 0.2986 |

| | | | | | | | | | | | |
|---|---|---|---|---|---|---|---|---|---|---|---|
| rs113566564 | 11 | 118335147 | A | G | 0.625 | 0.5321 | +?+ | 0 | 0.343 | 1 | 0.5582 |
| rs45457998 | 11 | 118492713 | G | A | -0.624 | 0.5325 | +?- | 0 | 0.395 | 1 | 0.5298 |
| rs754042 | 11 | 72326275 | T | C | 0.624 | 0.5327 | --+ | 0 | 1.951 | 2 | 0.377 |
| rs146525935 | 11 | 118180887 | C | T | -0.622 | 0.5342 | -?+ | 0 | 0.449 | 1 | 0.503 |
| rs61010648 | 11 | 118206890 | G | C | -0.621 | 0.5345 | -?- | 0 | 0.123 | 1 | 0.7254 |
| rs1715421 | 11 | 118105233 | C | T | 0.621 | 0.5346 | +?- | -126.4 | 1.325 | 1 | 0.2497 |
| rs525485 | 11 | 118555754 | A | G | 0.621 | 0.5346 | ++- | -63.6 | 2.445 | 2 | 0.2945 |
| rs76651078 | 11 | 118316006 | A | G | 0.617 | 0.5371 | +?- | 0 | 0.482 | 1 | 0.4876 |
| rs1793159 | 11 | 118104569 | A | C | 0.617 | 0.5373 | +?- | -124.1 | 1.339 | 1 | 0.2472 |
| rs1715422 | 11 | 118104577 | C | T | 0.617 | 0.5375 | +?- | -123.9 | 1.34 | 1 | 0.2471 |
| rs4432047 | 11 | 72989489 | G | A | -0.617 | 0.5375 | --+ | -3.3 | 3.871 | 2 | 0.1443 |
| rs1715420 | 11 | 118105599 | A | T | 0.616 | 0.5376 | +?- | -124.6 | 1.335 | 1 | 0.2478 |
| rs1626649 | 11 | 118071097 | T | C | 0.616 | 0.5382 | +?- | 0 | 0.521 | 1 | 0.4705 |
| rs111792147 | 11 | 118259162 | A | G | 0.615 | 0.5386 | +?- | 0 | 0.931 | 1 | 0.3347 |
| rs35409990 | 11 | 118109196 | A | G | 0.614 | 0.5389 | +?- | -96.2 | 1.529 | 1 | 0.2163 |
| rs12798453 | 11 | 118561410 | G | T | 0.614 | 0.5394 | ++- | -59.6 | 2.506 | 2 | 0.2856 |
| rs111430146 | 11 | 118248374 | A | C | 0.614 | 0.5395 | +?- | 0 | 0.933 | 1 | 0.334 |
| rs555356 | 11 | 118558078 | T | G | 0.614 | 0.5395 | ++- | -61.4 | 2.478 | 2 | 0.2897 |
| rs607527 | 11 | 118538187 | T | G | 0.613 | 0.5396 | ++- | -66 | 2.409 | 2 | 0.2998 |
| rs112900272 | 11 | 118326445 | T | C | 0.612 | 0.5405 | +?- | 0 | 0.454 | 1 | 0.5004 |
| rs638301 | 11 | 118559003 | G | A | 0.612 | 0.5407 | ++- | -56.7 | 2.552 | 2 | 0.2791 |
| rs45443099 | 11 | 118254710 | G | A | 0.611 | 0.5411 | +?- | 0 | 0.906 | 1 | 0.3412 |
| rs3765638 | 11 | 72537072 | A | G | 0.61 | 0.5418 | -++ | -24.1 | 3.223 | 2 | 0.1996 |
| rs112646740 | 11 | 118247805 | A | T | 0.61 | 0.542 | +?- | 0 | 0.941 | 1 | 0.3321 |
| rs2017122 | 11 | 118314849 | T | C | 0.609 | 0.5423 | +?- | 0 | 0.437 | 1 | 0.5083 |
| rs73011440 | 11 | 118809619 | A | G | 0.609 | 0.5425 | +?+ | 0 | 0.205 | 1 | 0.6504 |
| rs11216839 | 11 | 118129583 | T | A | 0.609 | 0.5427 | +?- | -142.6 | 1.237 | 1 | 0.2661 |
| rs11216821 | 11 | 118088362 | C | T | 0.608 | 0.5434 | +?- | -189.2 | 1.037 | 1 | 0.3085 |
| rs11216859 | 11 | 118238119 | C | T | 0.608 | 0.5435 | +++ | 0 | 0.044 | 2 | 0.9782 |
| rs6589674 | 11 | 118566039 | G | A | 0.608 | 0.5435 | ++- | -67.3 | 2.391 | 2 | 0.3026 |
| rs1783935 | 11 | 118539566 | T | A | 0.607 | 0.5439 | ++- | -74.5 | 2.293 | 2 | 0.3178 |
| rs561845 | 11 | 118558827 | G | A | 0.607 | 0.544 | ++- | -50.5 | 2.658 | 2 | 0.2648 |
| rs533144 | 11 | 118559247 | A | C | 0.603 | 0.5463 | ++- | -53.8 | 2.601 | 2 | 0.2724 |
| rs486462 | 11 | 118559178 | C | T | 0.603 | 0.5464 | ++- | -55.1 | 2.579 | 2 | 0.2755 |
| rs7130076 | 11 | 118566090 | C | T | 0.603 | 0.5465 | ++- | -57.1 | 2.546 | 2 | 0.28 |
| rs876479 | 11 | 118028094 | T | C | -0.603 | 0.5465 | -?- | 0 | 0.035 | 1 | 0.8511 |
| rs508579 | 11 | 118559084 | C | G | 0.602 | 0.5471 | ++- | -58.2 | 2.529 | 2 | 0.2824 |
| rs546890 | 11 | 118557198 | C | T | 0.6 | 0.5486 | ++- | -57.5 | 2.539 | 2 | 0.281 |
| rs11216814 | 11 | 118071857 | G | A | 0.599 | 0.5492 | +?- | 0 | 0.83 | 1 | 0.3622 |
| rs12794726 | 11 | 118566313 | T | C | 0.599 | 0.5492 | ++- | -56 | 2.564 | 2 | 0.2774 |
| rs478371 | 11 | 118565336 | A | G | 0.599 | 0.5492 | ++- | -55.5 | 2.572 | 2 | 0.2764 |
| rs644498 | 11 | 118567486 | C | T | 0.599 | 0.5493 | ++- | -41.8 | 2.822 | 2 | 0.2439 |
| rs585039 | 11 | 118563381 | A | C | 0.598 | 0.5496 | ++- | -55.4 | 2.575 | 2 | 0.276 |
| rs9332801 | 11 | 118355642 | C | A | 0.598 | 0.55 | +?+ | 0 | 0.351 | 1 | 0.5537 |
| rs693306 | 11 | 118558333 | G | A | 0.597 | 0.5502 | ++- | -56.9 | 2.55 | 2 | 0.2795 |
| rs625918 | 11 | 118559542 | T | C | 0.597 | 0.5507 | ++- | -54.9 | 2.582 | 2 | 0.275 |
| rs582630 | 11 | 118562799 | A | C | 0.596 | 0.5509 | ++- | -54.8 | 2.583 | 2 | 0.2748 |
| rs558593 | 11 | 118562520 | T | C | 0.596 | 0.5511 | ++- | -55.8 | 2.568 | 2 | 0.277 |
| rs11216812 | 11 | 118071762 | A | G | 0.596 | 0.5515 | +?- | 0 | 0.836 | 1 | 0.3604 |
| rs1793137 | 11 | 118016068 | C | T | -0.595 | 0.5518 | --- | 0 | 0.238 | 2 | 0.888 |
| rs493834 | 11 | 118567444 | G | A | 0.594 | 0.5522 | ++- | -49.4 | 2.678 | 2 | 0.2622 |
| rs11235710 | 11 | 72992156 | G | C | -0.594 | 0.5523 | --+ | -6.2 | 3.768 | 2 | 0.152 |
| rs471227 | 11 | 118564601 | C | T | 0.593 | 0.5532 | ++- | -53.9 | 2.599 | 2 | 0.2727 |
| rs7926970 | 11 | 118559881 | G | T | 0.593 | 0.5534 | ++- | -55.6 | 2.57 | 2 | 0.2766 |

| | | | | | | | | | | | |
|---|---|---|---|---|---|---|---|---|---|---|---|
| rs518028 | 11 | 118564982 | C | T | 0.592 | 0.5535 | ++- | -53.8 | 2.601 | 2 | 0.2723 |
| rs1793169 | 11 | 118070264 | G | A | 0.592 | 0.5539 | +?- | 0 | 0.547 | 1 | 0.4597 |
| rs508640 | 11 | 118564413 | G | A | 0.592 | 0.5539 | ++- | -53.6 | 2.604 | 2 | 0.272 |
| rs111587825 | 11 | 118274602 | G | T | 0.592 | 0.554 | +?- | 0 | 0.653 | 1 | 0.4189 |
| rs2071381 | 11 | 118214267 | A | G | 0.592 | 0.554 | +++ | 0 | 0.047 | 2 | 0.9768 |
| rs516153 | 11 | 118564762 | A | C | 0.592 | 0.5541 | ++- | -53.5 | 2.605 | 2 | 0.2718 |
| rs4938529 | 11 | 118560569 | T | C | 0.59 | 0.5552 | ++- | -53.1 | 2.613 | 2 | 0.2708 |
| rs558907 | 11 | 118558455 | G | A | 0.589 | 0.5561 | ++- | -53.8 | 2.601 | 2 | 0.2724 |
| rs502601 | 11 | 118565679 | A | G | 0.588 | 0.5564 | ++- | -54.1 | 2.596 | 2 | 0.2731 |
| rs644867 | 11 | 118567545 | G | A | 0.587 | 0.557 | ++- | -38.2 | 2.895 | 2 | 0.2352 |
| rs571751 | 11 | 118556797 | C | A | 0.587 | 0.5572 | ++- | -54.1 | 2.596 | 2 | 0.273 |
| rs7114458 | 11 | 118566175 | A | G | 0.586 | 0.5576 | ++- | -52.6 | 2.621 | 2 | 0.2697 |
| rs35963948 | 11 | 72991186 | G | A | -0.586 | 0.5578 | --+ | -13.3 | 3.529 | 2 | 0.1713 |
| rs12801003 | 11 | 118072716 | T | C | 0.586 | 0.5581 | +?- | 0 | 0.802 | 1 | 0.3706 |
| rs610664 | 11 | 118556691 | G | A | 0.584 | 0.5589 | ++- | -53.4 | 2.608 | 2 | 0.2715 |
| rs642662 | 11 | 118567020 | G | A | 0.584 | 0.559 | ++- | -57.1 | 2.546 | 2 | 0.28 |
| rs519982 | 11 | 118565231 | T | C | 0.584 | 0.5591 | ++- | -51.6 | 2.639 | 2 | 0.2673 |
| rs140633990 | 11 | 118365883 | A | T | -0.583 | 0.5601 | --- | 0 | 0.136 | 2 | 0.9342 |
| rs7926959 | 11 | 118559866 | C | T | 0.583 | 0.5601 | ++- | -57.9 | 2.533 | 2 | 0.2819 |
| rs11216815 | 11 | 118072232 | A | G | 0.582 | 0.5603 | +?- | 0 | 0.819 | 1 | 0.3654 |
| rs533646 | 11 | 118566746 | G | C | 0.578 | 0.5631 | ++- | -53.2 | 2.611 | 2 | 0.2711 |
| rs11216807 | 11 | 118063030 | C | T | 0.578 | 0.5632 | +++ | 0 | 0.071 | 2 | 0.9651 |
| rs7117266 | 11 | 118838020 | A | G | 0.578 | 0.5634 | -?+ | -91.2 | 1.569 | 1 | 0.2104 |
| rs12275955 | 11 | 118094756 | C | T | 0.578 | 0.5635 | +?- | -78.1 | 1.684 | 1 | 0.1944 |
| rs112243842 | 11 | 118246021 | G | T | 0.577 | 0.5642 | +?- | -198.2 | 1.006 | 1 | 0.3158 |
| rs11235713 | 11 | 72997496 | G | A | -0.576 | 0.5644 | --+ | -20.5 | 3.318 | 2 | 0.1903 |
| rs112468256 | 11 | 118278479 | C | T | 0.576 | 0.5645 | +?- | 0 | 0.619 | 1 | 0.4315 |
| rs7114216 | 11 | 118566234 | G | A | 0.576 | 0.5649 | ++- | -49.8 | 2.67 | 2 | 0.2631 |
| rs111707199 | 11 | 118276810 | G | A | 0.573 | 0.5665 | +?- | 0 | 0.624 | 1 | 0.4297 |
| rs542613 | 11 | 118567084 | A | G | 0.573 | 0.5665 | ++- | -56 | 2.565 | 2 | 0.2774 |
| rs45555733 | 11 | 118237566 | G | C | 0.573 | 0.5667 | +?- | 0 | 0.98 | 1 | 0.3222 |
| rs74398561 | 11 | 118233634 | G | A | 0.573 | 0.5667 | +?- | 0 | 0.98 | 1 | 0.3222 |
| rs45448191 | 11 | 118267737 | A | T | 0.573 | 0.5669 | +?- | 0 | 0.619 | 1 | 0.4316 |
| rs12808111 | 11 | 72992714 | G | A | -0.572 | 0.567 | --+ | 1.7 | 4.071 | 2 | 0.1306 |
| rs693037 | 11 | 118553326 | T | C | 0.571 | 0.568 | ++- | -55.4 | 2.573 | 2 | 0.2762 |
| rs12804550 | 11 | 118011860 | A | G | -0.57 | 0.5689 | -?- | 0 | 0.141 | 1 | 0.7076 |
| rs73555296 | 11 | 118445559 | T | G | 0.57 | 0.569 | +?+ | 0 | 0.055 | 1 | 0.8139 |
| rs112152942 | 11 | 118240840 | G | C | 0.569 | 0.5694 | +?- | 0 | 0.988 | 1 | 0.3203 |
| rs1877652 | 11 | 72307742 | A | G | 0.569 | 0.5695 | -?+ | 0 | 0.61 | 1 | 0.4347 |
| rs67389832 | 11 | 118839032 | A | C | 0.567 | 0.5704 | -?+ | -94.4 | 1.543 | 1 | 0.2142 |
| rs2276417 | 11 | 118230223 | G | T | 0.566 | 0.5716 | +++ | 0 | 0.056 | 2 | 0.9722 |
| rs11235708 | 11 | 72985098 | A | G | -0.565 | 0.5718 | --+ | -8 | 3.705 | 2 | 0.1569 |
| rs7939548 | 11 | 118212979 | A | G | 0.564 | 0.5727 | +?- | -183 | 1.06 | 1 | 0.3032 |
| rs45468196 | 11 | 118242889 | A | G | 0.564 | 0.5729 | +?- | 0 | 0.998 | 1 | 0.3177 |
| rs11216856 | 11 | 118227838 | T | C | 0.562 | 0.5742 | +++ | 0 | 0.059 | 2 | 0.9709 |
| rs12363628 | 11 | 72987038 | C | T | -0.559 | 0.5759 | --+ | -6.8 | 3.744 | 2 | 0.1538 |
| rs12808517 | 11 | 72601276 | T | C | -0.558 | 0.5767 | ++- | 0 | 1.575 | 2 | 0.455 |
| rs12286604 | 11 | 118084728 | A | G | 0.558 | 0.577 | +?- | -162.8 | 1.141 | 1 | 0.2854 |
| rs613377 | 11 | 118556084 | T | C | 0.557 | 0.5773 | ++- | -53.2 | 2.61 | 2 | 0.2711 |
| rs7129048 | 11 | 118211797 | C | T | 0.556 | 0.5779 | +?- | -134.7 | 1.278 | 1 | 0.2582 |
| rs113839182 | 11 | 118273602 | T | C | 0.554 | 0.5796 | +?- | 0 | 0.654 | 1 | 0.4186 |
| rs12808254 | 11 | 118066912 | T | C | -0.554 | 0.5796 | +?- | 0 | 0.645 | 1 | 0.422 |
| rs2282550 | 11 | 118220035 | C | T | 0.554 | 0.5798 | +++ | 0 | 0.089 | 2 | 0.9564 |
| rs11216828 | 11 | 118111676 | A | T | 0.553 | 0.5802 | +?- | -94.6 | 1.542 | 1 | 0.2143 |

| | | | | | | | | | | | |
|---|---|---|---|---|---|---|---|---|---|---|---|
| rs7939622 | 11 | 118087841 | A | G | 0.553 | 0.5802 | +?- | -160.5 | 1.151 | 1 | 0.2832 |
| rs1980091 | 11 | 72288599 | A | G | 0.551 | 0.5816 | -?+ | 0 | 0.56 | 1 | 0.4542 |
| rs11217078 | 11 | 118750321 | C | T | -0.55 | 0.5826 | -?+ | 0 | 0.311 | 1 | 0.5768 |
| rs4442562 | 11 | 118845813 | A | G | 0.549 | 0.5833 | -?+ | -71.3 | 1.752 | 1 | 0.1857 |
| rs61751532 | 11 | 118769604 | T | C | 0.548 | 0.584 | -?+ | -149.5 | 1.202 | 1 | 0.2728 |
| rs112970787 | 11 | 118212235 | C | G | 0.547 | 0.5843 | +?- | -175.5 | 1.089 | 1 | 0.2967 |
| rs4306318 | 11 | 118080520 | T | C | 0.546 | 0.5852 | +?- | 0 | 0.776 | 1 | 0.3783 |
| rs11216947 | 11 | 118567218 | A | G | 0.545 | 0.5855 | ++- | -45.8 | 2.744 | 2 | 0.2536 |
| rs9651667 | 11 | 118108679 | G | C | 0.545 | 0.5857 | +?- | -109.9 | 1.429 | 1 | 0.2319 |
| rs3181258 | 11 | 118218478 | A | T | 0.544 | 0.5863 | +?- | -189.1 | 1.038 | 1 | 0.3084 |
| rs7126172 | 11 | 118567257 | G | A | 0.544 | 0.5865 | ++- | -49.6 | 2.673 | 2 | 0.2627 |
| rs478296 | 11 | 118567375 | C | T | 0.543 | 0.5874 | ++- | -43 | 2.797 | 2 | 0.247 |
| rs9651688 | 11 | 118107495 | T | C | 0.541 | 0.5883 | +?- | -108.6 | 1.438 | 1 | 0.2304 |
| rs10793055 | 11 | 72710761 | C | T | 0.54 | 0.5889 | --+ | 49.9 | 7.99 | 2 | 0.01841 |
| rs692750 | 11 | 118538308 | T | G | 0.54 | 0.5891 | ++- | -55.5 | 2.573 | 2 | 0.2763 |
| rs11216818 | 11 | 118080268 | T | C | 0.54 | 0.5892 | +?- | 0 | 0.786 | 1 | 0.3752 |
| rs526597 | 11 | 118546602 | G | A | 0.536 | 0.5919 | ++- | -83.6 | 2.178 | 2 | 0.3365 |
| rs4457789 | 11 | 118539170 | T | G | 0.532 | 0.5949 | ++- | -96.2 | 2.039 | 2 | 0.3608 |
| rs4589335 | 11 | 118539140 | T | C | 0.528 | 0.5976 | ++- | -52.4 | 2.625 | 2 | 0.2692 |
| rs12286303 | 11 | 118114473 | T | C | 0.527 | 0.5984 | +?- | -73.8 | 1.726 | 1 | 0.1889 |
| rs10502228 | 11 | 118113583 | T | C | 0.526 | 0.5987 | +?- | -76.9 | 1.696 | 1 | 0.1928 |
| rs11216829 | 11 | 118112911 | A | G | 0.523 | 0.6012 | +?- | -76.3 | 1.701 | 1 | 0.1921 |
| rs1547885 | 11 | 118105222 | T | C | 0.521 | 0.6025 | +?- | -97.8 | 1.517 | 1 | 0.2181 |
| rs12286021 | 11 | 118080685 | C | T | 0.519 | 0.6035 | +?- | 0 | 0.868 | 1 | 0.3514 |
| rs60969913 | 11 | 118798154 | C | G | -0.519 | 0.6036 | +-- | 0 | 0.853 | 2 | 0.6529 |
| rs7925466 | 11 | 72778113 | C | A | 0.519 | 0.6039 | --+ | 50 | 8.003 | 2 | 0.01828 |
| rs11605435 | 11 | 118116496 | C | T | 0.519 | 0.604 | +?- | -74.8 | 1.716 | 1 | 0.1902 |
| rs11216830 | 11 | 118115147 | A | G | 0.517 | 0.6049 | +?- | -74.5 | 1.72 | 1 | 0.1898 |
| rs7112584 | 11 | 118069240 | A | G | 0.517 | 0.605 | +?- | 0 | 0.73 | 1 | 0.393 |
| rs7101652 | 11 | 118078718 | A | G | 0.517 | 0.6051 | +?- | 0 | 0.789 | 1 | 0.3743 |
| rs3802866 | 11 | 118095803 | T | C | 0.517 | 0.6055 | +?- | -99.9 | 1.501 | 1 | 0.2205 |
| rs12286567 | 11 | 118114788 | T | G | 0.516 | 0.6056 | +?- | -74.2 | 1.722 | 1 | 0.1894 |
| rs7117544 | 11 | 118117938 | C | A | 0.515 | 0.6065 | +?- | -73.9 | 1.726 | 1 | 0.189 |
| rs7943511 | 11 | 118096165 | T | C | 0.515 | 0.6067 | +?- | -99.3 | 1.506 | 1 | 0.2198 |
| rs12295438 | 11 | 118094921 | T | C | 0.515 | 0.6068 | +?- | -99.2 | 1.506 | 1 | 0.2198 |
| rs426907 | 11 | 72294440 | A | G | 0.514 | 0.6072 | -?+ | 0 | 0.561 | 1 | 0.454 |
| rs1561966 | 11 | 118221474 | A | G | 0.513 | 0.6078 | +++ | 0 | 0.08 | 2 | 0.9607 |
| rs34385073 | 11 | 72987413 | C | A | -0.512 | 0.609 | --+ | -1.8 | 3.93 | 2 | 0.1402 |
| rs111390501 | 11 | 118797420 | G | A | -0.511 | 0.6091 | +-- | 0 | 0.85 | 2 | 0.6538 |
| rs10790246 | 11 | 118103542 | T | A | 0.51 | 0.6103 | +?- | -98 | 1.515 | 1 | 0.2183 |
| rs595964 | 11 | 118026788 | A | G | -0.508 | 0.6114 | -+- | 0 | 0.517 | 2 | 0.7724 |
| rs73575428 | 11 | 118797003 | G | C | -0.506 | 0.6125 | +-- | 0 | 0.857 | 2 | 0.6516 |
| rs7116594 | 11 | 72984870 | A | G | -0.506 | 0.613 | --+ | -4.7 | 3.822 | 2 | 0.1479 |
| rs12280557 | 11 | 118117118 | T | C | 0.504 | 0.6143 | +?- | -71 | 1.755 | 1 | 0.1853 |
| rs3212263 | 11 | 118216423 | G | A | 0.502 | 0.6155 | +++ | 0 | 0.087 | 2 | 0.9576 |
| rs7925928 | 11 | 72748384 | G | A | 0.502 | 0.6157 | --+ | 47.3 | 7.592 | 2 | 0.02246 |
| rs7943272 | 11 | 118096028 | C | A | 0.502 | 0.6159 | +?- | -95.1 | 1.538 | 1 | 0.215 |
| rs56032119 | 11 | 118797544 | G | A | -0.501 | 0.6162 | +-- | 0 | 0.847 | 2 | 0.6547 |
| rs45558732 | 11 | 118402962 | T | G | 0.501 | 0.6166 | +?- | 0 | 0.426 | 1 | 0.5142 |
| rs73575429 | 11 | 118797392 | C | T | -0.5 | 0.6172 | +-- | 0 | 0.847 | 2 | 0.6548 |
| rs74584888 | 11 | 118797278 | A | G | -0.499 | 0.6179 | +-- | 0 | 0.847 | 2 | 0.6549 |
| rs76448372 | 11 | 118045846 | A | G | -0.498 | 0.6186 | -?+ | 0 | 0.881 | 1 | 0.3478 |
| rs520543 | 11 | 118538157 | A | G | 0.496 | 0.6202 | ++- | -71.3 | 2.336 | 2 | 0.311 |
| rs11216834 | 11 | 118122077 | C | T | 0.494 | 0.6213 | +?- | -68.4 | 1.781 | 1 | 0.182 |

| | | | | | | | | | | | |
|---|---|---|---|---|---|---|---|---|---|---|---|
| rs7951159 | 11 | 118780950 | T | C | -0.494 | 0.6213 | -?+ | 0 | 0.788 | 1 | 0.3748 |
| rs11216861 | 11 | 118241516 | A | G | 0.493 | 0.6218 | +++ | 0 | 0.022 | 2 | 0.9891 |
| rs59049611 | 11 | 118139403 | C | T | -0.492 | 0.6226 | -?+ | -59.1 | 1.885 | 1 | 0.1697 |
| rs10892230 | 11 | 118126682 | A | C | 0.49 | 0.6243 | +?- | -67.4 | 1.792 | 1 | 0.1806 |
| rs9332793 | 11 | 118354458 | A | G | 0.488 | 0.6252 | +?- | 0 | 0.309 | 1 | 0.5784 |
| rs2365774 | 11 | 72308203 | G | A | 0.486 | 0.6269 | -?+ | 0 | 0.526 | 1 | 0.4681 |
| rs45480805 | 11 | 118267903 | A | G | 0.481 | 0.6303 | +?- | 0 | 0.637 | 1 | 0.4248 |
| rs12362596 | 11 | 118105473 | C | G | -0.48 | 0.631 | -?+ | 0 | 0.35 | 1 | 0.5543 |
| rs9332794 | 11 | 118354505 | T | G | 0.479 | 0.6319 | +?- | 0 | 0.319 | 1 | 0.572 |
| rs6589675 | 11 | 118569414 | C | G | 0.477 | 0.6333 | +?- | -100.4 | 1.497 | 1 | 0.2212 |
| rs10790241 | 11 | 118013034 | A | G | -0.472 | 0.6367 | -?- | 0 | 0.081 | 1 | 0.7756 |
| rs11604683 | 11 | 72538546 | C | T | -0.472 | 0.6369 | ++- | 0 | 1.722 | 2 | 0.4227 |
| rs79157249 | 11 | 118785982 | T | A | -0.472 | 0.6371 | +-+ | 0 | 1.032 | 2 | 0.5969 |
| rs1783937 | 11 | 118609736 | A | G | 0.47 | 0.6386 | +?- | -24.4 | 2.412 | 1 | 0.1204 |
| rs11217081 | 11 | 118769032 | G | A | -0.469 | 0.6394 | --+ | 0 | 0.333 | 2 | 0.8465 |
| rs77487830 | 11 | 118311869 | G | A | 0.467 | 0.6407 | +-- | 0 | 0.773 | 2 | 0.6794 |
| rs7116375 | 11 | 118069979 | A | C | 0.459 | 0.6462 | +?- | 0 | 0.807 | 1 | 0.3689 |
| rs581063 | 11 | 118767077 | C | T | -0.457 | 0.6475 | --+ | 0 | 0.268 | 2 | 0.8744 |
| rs644287 | 11 | 118011805 | T | C | 0.457 | 0.6478 | +-+ | 0 | 1.158 | 2 | 0.5604 |
| rs77535829 | 11 | 118311677 | T | C | 0.456 | 0.6483 | +-- | 0 | 0.782 | 2 | 0.6764 |
| rs57495164 | 11 | 118095148 | C | T | -0.456 | 0.6484 | -?+ | 0 | 0.488 | 1 | 0.485 |
| rs489126 | 11 | 118572747 | A | G | 0.452 | 0.6512 | ++- | -82.2 | 2.195 | 2 | 0.3337 |
| rs869638 | 11 | 118100411 | C | G | -0.451 | 0.6522 | -?+ | 0 | 0.5 | 1 | 0.4794 |
| rs7941321 | 11 | 118702910 | A | G | 0.449 | 0.6533 | ++- | 0 | 0.391 | 2 | 0.8224 |
| rs11216827 | 11 | 118110202 | T | A | -0.447 | 0.6548 | -?+ | 0 | 0.417 | 1 | 0.5182 |
| rs1573559 | 11 | 118104627 | A | G | -0.445 | 0.6566 | -?+ | 0 | 0.379 | 1 | 0.5381 |
| rs11217083 | 11 | 118770321 | G | A | -0.443 | 0.6577 | --+ | 0 | 0.239 | 2 | 0.8876 |
| rs11600134 | 11 | 118070788 | A | G | 0.443 | 0.6577 | +?- | 0 | 0.836 | 1 | 0.3605 |
| rs2508991 | 11 | 118539522 | A | G | 0.442 | 0.6588 | ++- | -65.4 | 2.418 | 2 | 0.2984 |
| rs2368238 | 11 | 118091761 | G | T | -0.441 | 0.6589 | -?+ | 0 | 0.411 | 1 | 0.5215 |
| rs534114 | 11 | 118702002 | A | G | 0.441 | 0.6593 | ++- | 0 | 0.447 | 2 | 0.7997 |
| rs7941696 | 11 | 118702936 | G | A | 0.441 | 0.6595 | ++- | 0 | 0.703 | 2 | 0.7036 |
| rs11216826 | 11 | 118099713 | T | A | -0.44 | 0.6596 | -?+ | 0 | 0.472 | 1 | 0.4921 |
| rs7928228 | 11 | 118703001 | T | C | 0.44 | 0.6603 | ++- | 0 | 0.436 | 2 | 0.8043 |
| rs11216825 | 11 | 118099617 | A | G | -0.439 | 0.6607 | -?+ | 0 | 0.474 | 1 | 0.4911 |
| rs12294669 | 11 | 118041972 | G | A | -0.438 | 0.6612 | +?- | -130.3 | 1.303 | 1 | 0.2537 |
| rs10892226 | 11 | 118096618 | A | G | -0.436 | 0.663 | -?+ | 0 | 0.424 | 1 | 0.515 |
| rs113521822 | 11 | 118063349 | G | A | 0.435 | 0.6638 | +?- | 0 | 0.343 | 1 | 0.5582 |
| rs1170396 | 11 | 118703817 | T | A | 0.434 | 0.6643 | ++- | 0 | 0.445 | 2 | 0.8004 |
| rs2051480 | 11 | 118094513 | G | A | -0.433 | 0.6649 | -?+ | 0 | 0.437 | 1 | 0.5086 |
| rs2508921 | 11 | 118703484 | T | C | 0.429 | 0.668 | ++- | 0 | 0.454 | 2 | 0.7968 |
| rs9332781 | 11 | 118346599 | G | A | 0.426 | 0.6704 | +?- | 0 | 0.343 | 1 | 0.558 |
| rs7936751 | 11 | 72720199 | G | T | 0.423 | 0.6722 | --+ | 46.1 | 7.415 | 2 | 0.02453 |
| rs75709684 | 11 | 118050958 | A | G | -0.42 | 0.6746 | +?- | 0 | 0.566 | 1 | 0.4519 |
| rs7948661 | 11 | 118341649 | C | T | 0.419 | 0.6749 | +?- | 0 | 0.415 | 1 | 0.5194 |
| rs112965917 | 11 | 118725189 | A | T | 0.419 | 0.6753 | +?- | 0 | 0.708 | 1 | 0.4001 |
| rs622690 | 11 | 118704265 | T | C | 0.416 | 0.6771 | ++- | 0 | 0.456 | 2 | 0.796 |
| rs77806521 | 11 | 118232026 | T | C | 0.416 | 0.6777 | +?+ | 0 | 0.031 | 1 | 0.8612 |
| rs1791049 | 11 | 72890304 | T | C | 0.415 | 0.6784 | ++- | -47.4 | 2.713 | 2 | 0.2575 |
| rs17122021 | 11 | 118145686 | T | C | -0.414 | 0.679 | -?+ | -68.7 | 1.779 | 1 | 0.1823 |
| rs544096 | 11 | 118594326 | G | A | 0.412 | 0.6803 | ++- | -53.5 | 2.605 | 2 | 0.2718 |
| rs74422681 | 11 | 118312870 | G | A | 0.411 | 0.6808 | +-+ | 0 | 1.263 | 2 | 0.5317 |
| rs480791 | 11 | 118704255 | G | A | 0.411 | 0.6813 | ++- | 0 | 0.466 | 2 | 0.792 |
| rs4938493 | 11 | 118144804 | G | T | -0.411 | 0.6813 | -?+ | -61.2 | 1.861 | 1 | 0.1725 |

| | | | | | | | | | | | |
|---|---|---|---|---|---|---|---|---|---|---|---|
| rs7127013 | 11 | 72999221 | A | G | -0.41 | 0.6818 | --+ | 2.3 | 4.094 | 2 | 0.1291 |
| rs4245180 | 11 | 118144512 | A | G | -0.409 | 0.6823 | -?+ | -67.6 | 1.79 | 1 | 0.1809 |
| rs7926717 | 11 | 118121399 | T | G | 0.409 | 0.6823 | +?- | -140.5 | 1.248 | 1 | 0.264 |
| rs10892222 | 11 | 118041102 | A | G | -0.408 | 0.6832 | +?- | -117.3 | 1.38 | 1 | 0.2401 |
| rs73615909 | 11 | 118322488 | T | C | 0.407 | 0.6839 | +?- | 0 | 0.491 | 1 | 0.4834 |
| rs11235547 | 11 | 72318831 | A | G | 0.407 | 0.6841 | -?+ | 0 | 0.798 | 1 | 0.3718 |
| rs341047 | 11 | 72301503 | A | G | 0.406 | 0.6848 | +?- | 0 | 0.459 | 1 | 0.4981 |
| rs4486664 | 11 | 118569067 | C | A | 0.405 | 0.6851 | ++- | -97.1 | 2.03 | 2 | 0.3624 |
| rs2027764 | 11 | 72998247 | G | A | -0.404 | 0.6864 | --+ | 3.9 | 4.164 | 2 | 0.1247 |
| rs4938549 | 11 | 118701597 | T | C | 0.403 | 0.687 | ++- | 0 | 0.484 | 2 | 0.7852 |
| rs6589649 | 11 | 118048836 | C | T | -0.403 | 0.6871 | +?- | 0 | 0.315 | 1 | 0.5747 |
| rs530646 | 11 | 118701668 | A | G | 0.401 | 0.6882 | ++- | 0 | 0.487 | 2 | 0.7841 |
| rs7940554 | 11 | 118142930 | T | C | -0.401 | 0.6883 | -?+ | -65.5 | 1.813 | 1 | 0.1782 |
| rs638893 | 11 | 118698537 | G | A | 0.4 | 0.6888 | ++- | 0 | 0.55 | 2 | 0.7597 |
| rs4393358 | 11 | 118569097 | T | C | 0.395 | 0.6929 | ++- | -78.3 | 2.243 | 2 | 0.3257 |
| rs1540191 | 11 | 118090432 | A | G | -0.394 | 0.6938 | -?+ | 0 | 0.492 | 1 | 0.4829 |
| rs639885 | 11 | 118698294 | A | G | 0.394 | 0.6938 | ++- | 0 | 0.563 | 2 | 0.7546 |
| rs640373 | 11 | 118698160 | G | A | 0.393 | 0.6946 | ++- | 0 | 0.565 | 2 | 0.7538 |
| rs1783583 | 11 | 72889432 | C | G | 0.392 | 0.695 | ++- | -46.6 | 2.728 | 2 | 0.2557 |
| rs7941217 | 11 | 118349247 | T | C | 0.392 | 0.695 | +?- | 0 | 0.42 | 1 | 0.5169 |
| rs341088 | 11 | 72313975 | T | C | -0.392 | 0.6954 | --+ | 0 | 1.523 | 2 | 0.467 |
| rs76248354 | 11 | 118042606 | A | G | -0.391 | 0.696 | -?+ | 0 | 0.719 | 1 | 0.3966 |
| rs11235712 | 11 | 72994199 | C | T | -0.39 | 0.6965 | --+ | 22.3 | 5.148 | 2 | 0.07622 |
| rs526850 | 11 | 118605127 | G | T | 0.389 | 0.6969 | +?- | -29.7 | 2.312 | 1 | 0.1283 |
| rs57533048 | 11 | 118050014 | G | A | -0.39 | 0.6969 | +?- | 0 | 0.397 | 1 | 0.5287 |
| rs7127705 | 11 | 72999064 | T | C | -0.389 | 0.6972 | --+ | 0 | 4.001 | 2 | 0.1353 |
| rs2368242 | 11 | 118115526 | C | T | -0.389 | 0.6974 | -?+ | 0 | 0.573 | 1 | 0.4489 |
| rs57506367 | 11 | 118141993 | C | T | -0.387 | 0.6988 | -?+ | -62.1 | 1.851 | 1 | 0.1737 |
| rs4245179 | 11 | 118144445 | G | A | -0.387 | 0.699 | -?+ | -62 | 1.852 | 1 | 0.1736 |
| rs882904 | 11 | 72997910 | G | A | -0.386 | 0.6994 | --+ | 1.8 | 4.074 | 2 | 0.1304 |
| rs639915 | 11 | 118698274 | A | G | 0.385 | 0.6999 | ++- | 0 | 0.58 | 2 | 0.7484 |
| rs11607355 | 11 | 118093547 | T | C | -0.385 | 0.7001 | -?+ | 0 | 0.436 | 1 | 0.5092 |
| rs564892 | 11 | 118602392 | A | T | 0.384 | 0.7013 | ++- | -41.3 | 2.831 | 2 | 0.2429 |
| rs7123811 | 11 | 118569469 | G | T | 0.383 | 0.7014 | ++- | -89 | 2.116 | 2 | 0.3471 |
| rs112590920 | 11 | 118143586 | A | G | -0.382 | 0.7022 | -?+ | -69 | 1.775 | 1 | 0.1828 |
| rs67262752 | 11 | 118047776 | C | T | -0.382 | 0.7025 | +?- | 0 | 0.402 | 1 | 0.5259 |
| rs7342287 | 11 | 118050404 | A | G | -0.381 | 0.703 | +?- | 0 | 0.291 | 1 | 0.5895 |
| rs7940047 | 11 | 118142619 | T | C | -0.381 | 0.7032 | -?+ | -67.6 | 1.79 | 1 | 0.181 |
| rs12577061 | 11 | 118121553 | C | G | -0.38 | 0.7043 | -?+ | 0 | 0.547 | 1 | 0.4596 |
| rs56356019 | 11 | 118114893 | G | A | -0.379 | 0.7046 | -?+ | 0 | 0.499 | 1 | 0.4799 |
| rs521910 | 11 | 118610629 | A | G | 0.377 | 0.706 | +?- | -11.6 | 2.689 | 1 | 0.1011 |
| rs497916 | 11 | 118758089 | T | C | 0.376 | 0.7072 | +++ | 0 | 0.1 | 2 | 0.9513 |
| rs517445 | 11 | 118573535 | G | A | 0.375 | 0.708 | ++- | -87.9 | 2.129 | 2 | 0.345 |
| rs57461652 | 11 | 118146197 | A | C | -0.374 | 0.7081 | -?+ | -49.5 | 2.007 | 1 | 0.1565 |
| rs11235620 | 11 | 72680931 | T | A | 0.374 | 0.7086 | ++- | 0 | 1.974 | 2 | 0.3727 |
| rs516332 | 11 | 118698777 | G | A | 0.373 | 0.7089 | ++- | 0 | 0.62 | 2 | 0.7333 |
| rs2510901 | 11 | 118699863 | A | G | 0.371 | 0.7106 | ++- | 0 | 0.609 | 2 | 0.7373 |
| rs656445 | 11 | 118012164 | A | G | 0.371 | 0.7109 | +-- | 0 | 1.073 | 2 | 0.5849 |
| rs7479114 | 11 | 118568813 | T | C | 0.37 | 0.7111 | ++- | -86 | 2.151 | 2 | 0.3412 |
| rs79699220 | 11 | 118550841 | C | G | -0.368 | 0.7125 | +?- | 0 | 0.396 | 1 | 0.5291 |
| rs12364013 | 11 | 118119575 | T | C | -0.368 | 0.7126 | -?+ | 0 | 0.563 | 1 | 0.4529 |
| rs7123533 | 11 | 118049257 | G | C | -0.367 | 0.7138 | +?- | 0 | 0.369 | 1 | 0.5437 |
| rs58162302 | 11 | 118146199 | T | A | -0.366 | 0.7141 | -?+ | -50.5 | 1.994 | 1 | 0.158 |
| rs557990 | 11 | 118601363 | T | C | 0.366 | 0.7147 | ++- | -37.6 | 2.908 | 2 | 0.2337 |

| | | | | | | | | | | | |
|---|---|---|---|---|---|---|---|---|---|---|---|
| rs611733 | 11 | 118572760 | T | C | 0.365 | 0.7153 | ++- | -89.2 | 2.114 | 2 | 0.3475 |
| rs73024604 | 11 | 118120351 | T | C | -0.364 | 0.7162 | -?+ | 0 | 0.571 | 1 | 0.45 |
| rs583213 | 11 | 118669178 | G | T | 0.363 | 0.7164 | +?- | -99.3 | 1.505 | 1 | 0.2198 |
| rs570371 | 11 | 118568472 | G | A | 0.363 | 0.7169 | ++- | -76.6 | 2.265 | 2 | 0.3222 |
| rs2508564 | 11 | 118699744 | C | T | 0.361 | 0.7178 | ++- | 0 | 0.63 | 2 | 0.7299 |
| rs56315433 | 11 | 118114576 | A | C | -0.361 | 0.718 | -?+ | 0 | 0.488 | 1 | 0.485 |
| rs611217 | 11 | 118572611 | T | C | 0.361 | 0.718 | ++- | -77.9 | 2.248 | 2 | 0.325 |
| rs657539 | 11 | 118572021 | T | C | 0.361 | 0.7184 | ++- | -91.3 | 2.091 | 2 | 0.3515 |
| rs67163042 | 11 | 118050779 | G | A | -0.361 | 0.7185 | +?- | 0 | 0.269 | 1 | 0.6039 |
| rs73024606 | 11 | 118120366 | C | G | -0.36 | 0.7187 | -?+ | 0 | 0.576 | 1 | 0.448 |
| rs61900655 | 11 | 118120802 | C | G | -0.359 | 0.7194 | -?+ | 0 | 0.577 | 1 | 0.4474 |
| rs17092740 | 11 | 118117614 | A | G | -0.359 | 0.7195 | -?+ | 0 | 0.488 | 1 | 0.4849 |
| rs59018400 | 11 | 118049811 | T | C | -0.358 | 0.7201 | +?- | 0 | 0.359 | 1 | 0.5493 |
| rs11216813 | 11 | 118071836 | A | G | 0.358 | 0.7204 | +?- | 0 | 0.737 | 1 | 0.3907 |
| rs10898872 | 11 | 72497425 | C | T | 0.357 | 0.7208 | -++ | 38.8 | 6.54 | 2 | 0.038 |
| rs604096 | 11 | 118475098 | G | A | 0.355 | 0.7224 | --+ | -13.4 | 3.528 | 2 | 0.1713 |
| rs673547 | 11 | 118573339 | A | G | 0.355 | 0.7229 | ++- | -85.7 | 2.154 | 2 | 0.3406 |
| rs481903 | 11 | 118567733 | C | T | 0.354 | 0.7232 | ++- | -82.7 | 2.189 | 2 | 0.3346 |
| rs12577053 | 11 | 118121421 | A | G | -0.353 | 0.724 | -?+ | 0 | 0.587 | 1 | 0.4438 |
| rs11216800 | 11 | 118048885 | G | A | -0.352 | 0.7248 | +?- | 0 | 0.558 | 1 | 0.455 |
| rs552079 | 11 | 118573022 | C | T | 0.352 | 0.7249 | ++- | -84.7 | 2.165 | 2 | 0.3387 |
| rs649338 | 11 | 118705654 | A | G | 0.352 | 0.7249 | ++- | 0 | 0.48 | 2 | 0.7867 |
| rs10898915 | 11 | 72998687 | A | G | -0.351 | 0.7257 | --+ | 7.2 | 4.312 | 2 | 0.1158 |
| rs498912 | 11 | 118576997 | C | T | 0.351 | 0.7258 | +?- | -55.5 | 1.93 | 1 | 0.1648 |
| rs651825 | 11 | 118042713 | G | T | 0.35 | 0.7264 | +?+ | 0 | 0.025 | 1 | 0.8744 |
| rs56301948 | 11 | 118049880 | G | A | -0.35 | 0.7267 | +?- | 0 | 0.348 | 1 | 0.5552 |
| rs7342223 | 11 | 118050348 | G | T | -0.349 | 0.7274 | +?- | 0 | 0.347 | 1 | 0.5558 |
| rs17310235 | 11 | 72607076 | G | A | 0.348 | 0.7275 | ++- | 0 | 1.311 | 2 | 0.5192 |
| rs7119702 | 11 | 118607694 | G | A | 0.348 | 0.7281 | +?- | -33.2 | 2.252 | 1 | 0.1334 |
| rs657769 | 11 | 118571316 | T | C | 0.348 | 0.7282 | ++- | -76.9 | 2.261 | 2 | 0.3229 |
| rs636736 | 11 | 118705145 | G | A | 0.346 | 0.7295 | ++- | 0 | 0.669 | 2 | 0.7156 |
| rs544452 | 11 | 118571357 | T | G | 0.346 | 0.7296 | ++- | -76.3 | 2.269 | 2 | 0.3216 |
| rs7108144 | 11 | 118569694 | A | G | 0.345 | 0.7297 | ++- | -76.2 | 2.27 | 2 | 0.3215 |
| rs523793 | 11 | 118570697 | T | C | 0.343 | 0.7313 | ++- | -75.6 | 2.278 | 2 | 0.3201 |
| rs554456 | 11 | 118570062 | C | T | 0.343 | 0.7314 | ++- | -75.5 | 2.279 | 2 | 0.32 |
| rs482889 | 11 | 118570098 | T | G | 0.342 | 0.7322 | ++- | -75.2 | 2.283 | 2 | 0.3193 |
| rs12793594 | 11 | 118569900 | C | T | 0.342 | 0.7323 | ++- | -75.1 | 2.284 | 2 | 0.3192 |
| rs488141 | 11 | 118571168 | G | A | 0.342 | 0.7323 | ++- | -75.1 | 2.284 | 2 | 0.3192 |
| rs658676 | 11 | 118571123 | T | C | 0.342 | 0.7323 | ++- | -75.1 | 2.284 | 2 | 0.3192 |
| rs541631 | 11 | 118571023 | G | A | 0.342 | 0.7324 | ++- | -75.1 | 2.284 | 2 | 0.3191 |
| rs659564 | 11 | 118570914 | A | C | 0.342 | 0.7324 | ++- | -75.1 | 2.284 | 2 | 0.3191 |
| rs642224 | 11 | 118570919 | A | G | 0.34 | 0.734 | ++- | -75.3 | 2.282 | 2 | 0.3195 |
| rs11235595 | 11 | 72512708 | C | G | 0.34 | 0.7342 | -++ | 35.5 | 6.202 | 2 | 0.04501 |
| rs628299 | 11 | 118018022 | G | A | 0.34 | 0.7342 | +-- | -89.3 | 2.113 | 2 | 0.3477 |
| rs655499 | 11 | 118063479 | G | A | 0.339 | 0.7343 | ++- | 0 | 0.192 | 2 | 0.9086 |
| rs493720 | 11 | 118604678 | T | C | 0.339 | 0.7346 | +?- | -25.3 | 2.393 | 1 | 0.1218 |
| rs656287 | 11 | 118571790 | G | A | 0.339 | 0.7348 | ++- | -88.5 | 2.122 | 2 | 0.346 |
| rs511451 | 11 | 118568639 | T | C | 0.336 | 0.7366 | ++- | -93.9 | 2.063 | 2 | 0.3565 |
| rs11216824 | 11 | 118096818 | A | G | 0.335 | 0.7376 | +?- | -105 | 1.463 | 1 | 0.2264 |
| rs113784020 | 11 | 118559731 | T | C | -0.334 | 0.7381 | +?- | 0 | 0.727 | 1 | 0.3939 |
| rs341089 | 11 | 72313462 | A | G | 0.334 | 0.7382 | +?- | 0 | 0.177 | 1 | 0.674 |
| rs7121712 | 11 | 72993695 | A | T | -0.333 | 0.739 | --+ | 13.9 | 4.645 | 2 | 0.09805 |
| rs570949 | 11 | 118571911 | T | G | 0.331 | 0.7409 | ++- | -78.1 | 2.247 | 2 | 0.3252 |
| rs58508419 | 11 | 118762102 | A | G | 0.328 | 0.7427 | +?- | 0 | 0.695 | 1 | 0.4043 |

| | | | | | | | | | | | |
|---|---|---|---|---|---|---|---|---|---|---|---|
| rs574383 | 11 | 118573668 | C | T | 0.327 | 0.7437 | ++- | -86.1 | 2.15 | 2 | 0.3413 |
| rs111313044 | 11 | 118803596 | A | G | -0.327 | 0.7439 | +-- | -58 | 2.531 | 2 | 0.2821 |
| rs10892221 | 11 | 118040642 | A | G | -0.326 | 0.7442 | +?- | -91.3 | 1.568 | 1 | 0.2104 |
| rs1715456 | 11 | 118123517 | G | A | -0.326 | 0.7443 | -?+ | 0 | 0.762 | 1 | 0.3829 |
| rs73575440 | 11 | 118803160 | A | G | -0.325 | 0.7453 | +-- | -63.8 | 2.442 | 2 | 0.2949 |
| rs4943939 | 11 | 72309540 | T | C | -0.323 | 0.7464 | --+ | 0 | 1.629 | 2 | 0.4429 |
| rs371412 | 11 | 72303491 | A | G | 0.323 | 0.747 | +?- | 0 | 0.384 | 1 | 0.5356 |
| rs12577841 | 11 | 118194599 | T | C | -0.322 | 0.7473 | -+- | 0 | 0.399 | 2 | 0.8191 |
| rs75459003 | 11 | 118053859 | C | A | -0.322 | 0.7476 | +?- | 0 | 0.514 | 1 | 0.4735 |
| rs59069289 | 11 | 118802128 | T | G | -0.321 | 0.7483 | +-- | -64.2 | 2.436 | 2 | 0.2958 |
| rs59480472 | 11 | 118802386 | C | T | -0.321 | 0.7485 | +-- | -64.2 | 2.435 | 2 | 0.2959 |
| rs642765 | 11 | 118030310 | C | G | -0.32 | 0.7487 | -?- | 0 | 0.002 | 1 | 0.9614 |
| rs11216816 | 11 | 118075152 | A | C | 0.32 | 0.7489 | +?- | 0 | 0.678 | 1 | 0.4102 |
| rs10793048 | 11 | 72580398 | T | C | 0.32 | 0.7491 | ++- | 0 | 1.454 | 2 | 0.4834 |
| rs61900882 | 11 | 118130199 | C | T | -0.317 | 0.7513 | -?+ | 0 | 0.679 | 1 | 0.41 |
| rs519294 | 11 | 118573743 | A | C | 0.315 | 0.7526 | ++- | -82.8 | 2.188 | 2 | 0.3348 |
| rs10793054 | 11 | 72662812 | A | C | 0.315 | 0.7529 | --+ | 55.7 | 9.019 | 2 | 0.011 |
| rs4488208 | 11 | 118877621 | C | G | 0.315 | 0.7529 | -?+ | 28.6 | 4.2 | 1 | 0.04042 |
| rs12295676 | 11 | 118054345 | A | G | -0.312 | 0.7548 | +?- | 0 | 0.5 | 1 | 0.4794 |
| rs78755429 | 11 | 118777978 | A | C | 0.311 | 0.7556 | +?- | -95.4 | 1.535 | 1 | 0.2153 |
| rs11216817 | 11 | 118075474 | T | C | 0.31 | 0.7568 | +?- | 0 | 0.695 | 1 | 0.4043 |
| rs74395230 | 11 | 118052662 | T | C | -0.309 | 0.7571 | +?- | 0 | 0.496 | 1 | 0.4812 |
| rs12271064 | 11 | 118119211 | G | A | 0.307 | 0.7586 | +?- | -70.1 | 1.763 | 1 | 0.1842 |
| rs500254 | 11 | 118678780 | G | A | 0.307 | 0.7589 | +?- | -101.5 | 1.489 | 1 | 0.2224 |
| rs17121997 | 11 | 118131049 | C | T | -0.305 | 0.7601 | -?+ | 0 | 0.691 | 1 | 0.406 |
| rs594171 | 11 | 118032974 | A | T | -0.303 | 0.7616 | -?- | 0 | 0.003 | 1 | 0.9531 |
| rs3018386 | 11 | 118016622 | G | C | 0.303 | 0.7619 | +-- | 0 | 1.809 | 2 | 0.4048 |
| rs1126205 | 11 | 72523794 | T | G | 0.303 | 0.762 | -++ | 37.8 | 6.433 | 2 | 0.0401 |
| rs116848264 | 11 | 118049817 | A | T | -0.302 | 0.7629 | -?+ | 0 | 0.339 | 1 | 0.5602 |
| rs75460330 | 11 | 118809767 | A | G | -0.299 | 0.7652 | +-- | -84.8 | 2.164 | 2 | 0.3389 |
| rs79820203 | 11 | 118051010 | C | T | -0.297 | 0.7662 | +?- | 0 | 0.479 | 1 | 0.4887 |
| rs7927676 | 11 | 118051323 | G | C | -0.297 | 0.7663 | +?- | 0 | 0.479 | 1 | 0.4888 |
| rs12798851 | 11 | 118135973 | G | T | 0.297 | 0.7664 | +?- | 0 | 0.798 | 1 | 0.3718 |
| rs2186781 | 11 | 118605447 | T | C | 0.296 | 0.7669 | +?- | -24.2 | 2.416 | 1 | 0.1201 |
| rs10790248 | 11 | 118125672 | A | G | -0.296 | 0.7674 | -?+ | 0 | 0.607 | 1 | 0.4359 |
| rs11235546 | 11 | 72312118 | T | G | -0.296 | 0.7674 | --+ | 0 | 1.741 | 2 | 0.4187 |
| rs687664 | 11 | 118574185 | A | C | 0.293 | 0.7692 | ++- | 0 | 1.992 | 2 | 0.3694 |
| rs523715 | 11 | 118574188 | A | G | 0.293 | 0.7695 | ++- | 0 | 1.993 | 2 | 0.3692 |
| rs73575441 | 11 | 118804919 | C | T | -0.29 | 0.7717 | +-- | -59.1 | 2.514 | 2 | 0.2845 |
| rs76050019 | 11 | 118051857 | C | T | -0.289 | 0.7729 | +?- | 0 | 0.467 | 1 | 0.4942 |
| rs56933877 | 11 | 118804643 | A | G | -0.288 | 0.7737 | +-- | -59.4 | 2.51 | 2 | 0.2851 |
| rs73557344 | 11 | 118810054 | C | G | -0.287 | 0.7744 | +-- | -86.1 | 2.149 | 2 | 0.3415 |
| rs11216949 | 11 | 118568668 | T | C | 0.286 | 0.7751 | ++- | -85.9 | 2.152 | 2 | 0.341 |
| rs4936425 | 11 | 118126904 | T | C | -0.285 | 0.7754 | -?+ | 0 | 0.675 | 1 | 0.4114 |
| rs73575454 | 11 | 118807639 | A | G | -0.285 | 0.7758 | +-- | -83.7 | 2.178 | 2 | 0.3365 |
| rs12291619 | 11 | 118062552 | A | G | -0.283 | 0.7773 | +?- | 0 | 0.528 | 1 | 0.4675 |
| rs658040 | 11 | 118572171 | G | A | 0.278 | 0.7812 | ++- | -62 | 2.469 | 2 | 0.2909 |
| rs7934292 | 11 | 118059914 | C | T | -0.277 | 0.7821 | +?- | 0 | 0.517 | 1 | 0.4721 |
| rs7933859 | 11 | 118662045 | T | C | 0.276 | 0.7828 | +?- | -124 | 1.34 | 1 | 0.2471 |
| rs4938492 | 11 | 118126675 | C | T | -0.275 | 0.7836 | -?+ | 0 | 0.644 | 1 | 0.4221 |
| rs1790192 | 11 | 118757191 | G | A | 0.274 | 0.7837 | --+ | -89.2 | 2.114 | 2 | 0.3476 |
| rs112045567 | 11 | 118804989 | G | A | -0.273 | 0.7847 | +-- | -52.2 | 2.628 | 2 | 0.2688 |
| rs553188 | 11 | 118576762 | T | C | 0.273 | 0.7847 | +?- | -32.2 | 2.269 | 1 | 0.132 |
| rs12284089 | 11 | 118058542 | T | C | -0.273 | 0.7848 | +?- | 0 | 0.512 | 1 | 0.4741 |

| | | | | | | | | | | | |
|---|---|---|---|---|---|---|---|---|---|---|---|
| rs76831657 | 11 | 118051082 | G | A | -0.272 | 0.7855 | +?- | 0 | 0.544 | 1 | 0.4608 |
| rs11608072 | 11 | 118030086 | C | G | -0.271 | 0.7865 | +?- | 0 | 0.081 | 1 | 0.7754 |
| rs637737 | 11 | 118051646 | G | A | -0.271 | 0.7867 | +?- | 0 | 0.21 | 1 | 0.6465 |
| rs1104542 | 11 | 118129340 | T | C | -0.269 | 0.788 | -?+ | 0 | 0.659 | 1 | 0.4168 |
| rs4938544 | 11 | 118665009 | A | G | 0.268 | 0.7883 | +?- | -132 | 1.293 | 1 | 0.2555 |
| rs4508234 | 11 | 118662203 | G | C | 0.267 | 0.7891 | +?- | -129.5 | 1.307 | 1 | 0.2529 |
| rs656308 | 11 | 118031206 | T | G | -0.265 | 0.7909 | -?- | 0 | 0.002 | 1 | 0.9665 |
| rs10898885 | 11 | 72571275 | T | C | 0.264 | 0.7917 | ++- | 0 | 1.901 | 2 | 0.3866 |
| rs11216860 | 11 | 118240332 | C | T | 0.263 | 0.7923 | -++ | 0 | 0.118 | 2 | 0.9425 |
| rs7935492 | 11 | 118056823 | T | C | -0.261 | 0.7938 | +?- | 0 | 0.496 | 1 | 0.4814 |
| rs11216791 | 11 | 118044360 | G | A | -0.261 | 0.7945 | +?- | 0 | 0.759 | 1 | 0.3836 |
| rs640323 | 11 | 118010890 | C | G | 0.259 | 0.7955 | +-- | 0 | 1.305 | 2 | 0.5207 |
| rs12282334 | 11 | 118057783 | G | C | -0.257 | 0.7974 | +?- | 0 | 0.489 | 1 | 0.4843 |
| rs341049 | 11 | 72298208 | T | C | -0.256 | 0.7978 | --+ | 0 | 1.893 | 2 | 0.388 |
| rs11217068 | 11 | 118729283 | T | C | -0.252 | 0.8007 | -?+ | 35.7 | 4.663 | 1 | 0.03081 |
| rs71482132 | 11 | 118031211 | A | G | -0.252 | 0.8013 | -?- | 0 | 0.003 | 1 | 0.9556 |
| rs7105658 | 11 | 118712270 | T | C | 0.249 | 0.8033 | -++ | 26.1 | 5.413 | 2 | 0.06677 |
| rs645451 | 11 | 118025969 | T | C | 0.239 | 0.8113 | -?+ | 0 | 0.134 | 1 | 0.7145 |
| rs117745694 | 11 | 118218491 | A | G | -0.238 | 0.8122 | +?- | 0 | 0.259 | 1 | 0.6108 |
| rs341086 | 11 | 72317803 | T | C | -0.236 | 0.8135 | --+ | 0 | 1.749 | 2 | 0.4171 |
| rs582200 | 11 | 118668908 | G | A | 0.233 | 0.8157 | +?- | -132.1 | 1.292 | 1 | 0.2556 |
| rs2853009 | 11 | 118148146 | G | A | -0.233 | 0.8158 | +?- | -74 | 1.724 | 1 | 0.1892 |
| rs642936 | 11 | 118019906 | C | T | 0.233 | 0.8158 | +-- | 0 | 1.647 | 2 | 0.439 |
| rs675003 | 11 | 118677083 | T | C | 0.232 | 0.8164 | +?- | -120.2 | 1.362 | 1 | 0.2431 |
| rs11235597 | 11 | 72516773 | A | G | 0.231 | 0.817 | -++ | 33 | 5.969 | 2 | 0.05055 |
| rs9735317 | 11 | 118725383 | T | C | -0.231 | 0.8173 | -?+ | 36.4 | 4.718 | 1 | 0.02985 |
| rs35381826 | 11 | 118032438 | T | G | -0.23 | 0.8182 | -?- | 0 | 0.005 | 1 | 0.9429 |
| rs73575424 | 11 | 118788517 | A | C | -0.226 | 0.8213 | +-+ | 0 | 0.92 | 2 | 0.6313 |
| rs7129921 | 11 | 72676874 | G | T | 0.224 | 0.8229 | --+ | 55.3 | 8.939 | 2 | 0.01145 |
| rs61900957 | 11 | 118501723 | T | C | -0.22 | 0.8256 | +?- | 19.3 | 3.718 | 1 | 0.05383 |
| rs11600901 | 11 | 118031986 | A | G | -0.214 | 0.8308 | -?- | 0 | 0.004 | 1 | 0.9514 |
| rs7102253 | 11 | 72792697 | C | T | -0.213 | 0.8314 | ++- | -55.8 | 2.568 | 2 | 0.277 |
| rs2276064 | 11 | 118529703 | A | G | -0.211 | 0.8328 | -+- | -53.8 | 2.601 | 2 | 0.2724 |
| rs75403174 | 11 | 118043904 | A | C | -0.211 | 0.833 | +?- | 0 | 0.92 | 1 | 0.3374 |
| rs384892 | 11 | 72309889 | A | G | 0.21 | 0.8339 | +?- | 0 | 0.297 | 1 | 0.5856 |
| rs7117594 | 11 | 118714492 | T | C | 0.206 | 0.8369 | -++ | 29.8 | 5.7 | 2 | 0.05786 |
| rs5006102 | 11 | 118662340 | A | T | 0.204 | 0.8381 | +?- | -106.1 | 1.455 | 1 | 0.2277 |
| rs11216792 | 11 | 118044450 | T | G | -0.204 | 0.8384 | +?- | 0 | 0.907 | 1 | 0.3408 |
| rs3862796 | 11 | 72705522 | T | C | -0.203 | 0.839 | -+- | -57 | 2.548 | 2 | 0.2797 |
| rs11216790 | 11 | 118044144 | T | G | -0.201 | 0.8407 | +?- | 0 | 0.912 | 1 | 0.3396 |
| rs61901410 | 11 | 118661197 | G | C | -0.2 | 0.8416 | +?- | -7.1 | 2.802 | 1 | 0.09416 |
| rs116979688 | 11 | 118032223 | T | C | 0.199 | 0.8419 | +?+ | 0 | 0.001 | 1 | 0.9778 |
| rs7924647 | 11 | 118728479 | G | A | -0.195 | 0.8453 | -?+ | 35.5 | 4.654 | 1 | 0.03098 |
| rs3759001 | 11 | 118135745 | T | C | -0.192 | 0.8475 | +?- | 0 | 0.052 | 1 | 0.819 |
| rs1790063 | 11 | 72995912 | A | G | -0.19 | 0.849 | -?- | 0 | 0.01 | 1 | 0.9199 |
| rs12289120 | 11 | 118066609 | T | C | 0.19 | 0.8494 | +?- | 0 | 0.359 | 1 | 0.5488 |
| rs10736495 | 11 | 118607090 | C | G | 0.189 | 0.8499 | +?- | -49.2 | 2.011 | 1 | 0.1562 |
| rs1629083 | 11 | 118126576 | T | C | -0.188 | 0.8512 | -?+ | 0 | 0.55 | 1 | 0.4584 |
| rs678262 | 11 | 118021740 | G | C | 0.178 | 0.8589 | +-- | -8.6 | 3.682 | 2 | 0.1587 |
| rs145762171 | 11 | 118748789 | A | G | -0.177 | 0.8598 | +?- | -76.7 | 1.697 | 1 | 0.1926 |
| rs1786183 | 11 | 118017445 | A | G | 0.172 | 0.8635 | -?+ | -119.8 | 1.365 | 1 | 0.2427 |
| rs72969876 | 11 | 72589778 | C | G | -0.166 | 0.8682 | +-- | 0 | 1.319 | 2 | 0.5171 |
| rs2187490 | 11 | 118713180 | G | T | -0.166 | 0.8684 | +?- | 0 | 0.233 | 1 | 0.6296 |
| rs12420093 | 11 | 118272963 | G | A | -0.164 | 0.8696 | --+ | 1.9 | 4.078 | 2 | 0.1302 |

| | | | | | | | | | | | |
|---|---|---|---|---|---|---|---|---|---|---|---|
| rs45555132 | 11 | 118064030 | T | G | -0.163 | 0.8704 | -?+ | 0 | 0.23 | 1 | 0.6314 |
| rs61901411 | 11 | 118665324 | T | C | 0.163 | 0.8709 | +?- | -96.7 | 1.526 | 1 | 0.2168 |
| rs61894194 | 11 | 72513028 | T | C | 0.161 | 0.8722 | -++ | 33.5 | 6.014 | 2 | 0.04944 |
| rs72979412 | 11 | 72639396 | C | T | 0.16 | 0.8726 | +?- | -157.5 | 1.165 | 1 | 0.2804 |
| rs4938550 | 11 | 118707171 | G | A | -0.158 | 0.8741 | +?- | 0 | 0.081 | 1 | 0.7754 |
| rs1628409 | 11 | 72885257 | T | C | -0.157 | 0.8749 | ++- | 0 | 1.492 | 2 | 0.4743 |
| rs12276533 | 11 | 118724116 | A | G | -0.156 | 0.8757 | -?+ | 34.9 | 4.611 | 1 | 0.03176 |
| rs658624 | 11 | 118018767 | G | A | 0.156 | 0.8759 | -?+ | -84.2 | 1.628 | 1 | 0.2019 |
| rs73005500 | 11 | 118697375 | C | T | -0.154 | 0.8778 | +?- | 0 | 0.356 | 1 | 0.5509 |
| rs1613495 | 11 | 118017645 | T | C | 0.153 | 0.8786 | -?+ | 0 | 0.779 | 1 | 0.3775 |
| rs11235617 | 11 | 72644631 | C | T | 0.151 | 0.8799 | +?- | -153.2 | 1.185 | 1 | 0.2763 |
| rs611979 | 11 | 118034223 | A | G | -0.149 | 0.8812 | -?- | 0 | 0.014 | 1 | 0.9064 |
| rs647975 | 11 | 118038741 | T | C | 0.148 | 0.882 | +?- | 0 | 0.126 | 1 | 0.7226 |
| rs1793157 | 11 | 118017434 | C | A | 0.148 | 0.8824 | -++ | 0 | 1.699 | 2 | 0.4277 |
| rs602594 | 11 | 118040610 | T | C | 0.144 | 0.8858 | +?- | 0 | 0.058 | 1 | 0.8092 |
| rs614825 | 11 | 118018725 | G | A | 0.143 | 0.886 | -?+ | -81.3 | 1.654 | 1 | 0.1984 |
| rs12272981 | 11 | 118709600 | G | A | -0.142 | 0.8873 | -?+ | 37.5 | 4.797 | 1 | 0.02851 |
| rs58507982 | 11 | 118146510 | T | C | -0.138 | 0.8904 | -?+ | -137.5 | 1.263 | 1 | 0.261 |
| rs572608 | 11 | 118801640 | A | T | 0.138 | 0.8906 | -+- | 0 | 1.256 | 2 | 0.5337 |
| rs679327 | 11 | 118042974 | G | A | 0.137 | 0.8912 | +?+ | 0 | 0.018 | 1 | 0.8939 |
| rs45508093 | 11 | 118224059 | G | A | 0.132 | 0.8947 | -?+ | 0 | 0.308 | 1 | 0.5791 |
| rs7119568 | 11 | 118723753 | A | T | -0.132 | 0.8947 | -?+ | 34.1 | 4.555 | 1 | 0.03282 |
| rs1793149 | 11 | 118127347 | A | G | 0.131 | 0.8958 | +?- | 0 | 0.513 | 1 | 0.4738 |
| rs880895 | 11 | 72884382 | A | G | -0.131 | 0.8958 | ++- | 0 | 1.561 | 2 | 0.4581 |
| rs4944014 | 11 | 72527180 | T | C | 0.13 | 0.8962 | ++- | -81.5 | 2.204 | 2 | 0.3321 |
| rs11217096 | 11 | 118796570 | T | C | -0.129 | 0.8972 | -+- | 0 | 1.591 | 2 | 0.4514 |
| rs611960 | 11 | 118034228 | G | T | -0.129 | 0.8977 | -?- | 0 | 0.008 | 1 | 0.927 |
| rs10892281 | 11 | 118614397 | T | C | 0.123 | 0.9025 | +?- | -67.8 | 1.788 | 1 | 0.1812 |
| rs7107376 | 11 | 72640991 | G | T | 0.122 | 0.9026 | +?- | -152.4 | 1.189 | 1 | 0.2756 |
| rs75488969 | 11 | 118712509 | T | C | 0.122 | 0.9027 | +?- | 0 | 0.761 | 1 | 0.3831 |
| rs11603160 | 11 | 72894273 | T | C | -0.122 | 0.9029 | -?+ | -173.5 | 1.097 | 1 | 0.2949 |
| rs1715443 | 11 | 118017365 | G | T | 0.121 | 0.9041 | -?+ | -70.6 | 1.758 | 1 | 0.1849 |
| rs12805658 | 11 | 72555300 | A | C | -0.12 | 0.9044 | ++- | -55.7 | 2.569 | 2 | 0.2768 |
| rs113641235 | 11 | 118700264 | T | C | -0.12 | 0.9049 | +-- | 0 | 0.36 | 2 | 0.8353 |
| rs583905 | 11 | 118695908 | G | A | 0.119 | 0.9049 | +?- | 0 | 0.479 | 1 | 0.4887 |
| rs3018387 | 11 | 118016981 | T | A | 0.117 | 0.9069 | -?+ | -75.5 | 1.71 | 1 | 0.191 |
| rs598746 | 11 | 118020057 | A | G | -0.112 | 0.9112 | -?+ | 0 | 0.291 | 1 | 0.5894 |
| rs6589687 | 11 | 118615660 | T | A | 0.111 | 0.912 | +?- | -64.8 | 1.82 | 1 | 0.1773 |
| rs7108905 | 11 | 118712427 | G | C | -0.11 | 0.9123 | -?+ | 37.8 | 4.821 | 1 | 0.02812 |
| rs583490 | 11 | 118022608 | A | G | 0.108 | 0.9139 | +-- | -31.6 | 3.041 | 2 | 0.2186 |
| rs1793150 | 11 | 118127314 | G | A | 0.107 | 0.9147 | +?- | 0 | 0.548 | 1 | 0.4592 |
| rs7104819 | 11 | 118769442 | G | A | 0.107 | 0.9148 | --+ | 0 | 0.556 | 2 | 0.7571 |
| rs515707 | 11 | 118685481 | T | C | 0.105 | 0.9164 | +?- | 0 | 0.429 | 1 | 0.5127 |
| rs3922 | 11 | 118765600 | G | A | 0.103 | 0.9178 | +-+ | 0 | 0.416 | 2 | 0.8123 |
| rs1783574 | 11 | 72863527 | A | G | 0.101 | 0.9197 | ++- | 0 | 0.861 | 2 | 0.6503 |
| rs8192614 | 11 | 118037564 | A | G | -0.099 | 0.9208 | -?- | 0 | 0.004 | 1 | 0.9494 |
| rs7122510 | 11 | 118708785 | A | C | -0.099 | 0.9212 | -?+ | 31.1 | 4.355 | 1 | 0.0369 |
| rs1715450 | 11 | 118128445 | C | G | 0.098 | 0.922 | +?- | 0 | 0.561 | 1 | 0.4537 |
| rs6589677 | 11 | 118583971 | A | G | 0.098 | 0.9222 | +?- | -41.2 | 2.124 | 1 | 0.145 |
| rs1715449 | 11 | 118128455 | G | A | 0.097 | 0.923 | +?- | 0 | 0.563 | 1 | 0.4529 |
| rs4944016 | 11 | 72597438 | T | C | -0.097 | 0.9231 | ++- | -60.3 | 2.495 | 2 | 0.2873 |
| rs7122511 | 11 | 118708786 | A | C | -0.096 | 0.9237 | -?+ | 31.3 | 4.368 | 1 | 0.03662 |
| rs1793151 | 11 | 118126920 | C | T | 0.094 | 0.9255 | +?- | 0 | 0.568 | 1 | 0.451 |
| rs12799445 | 11 | 72592506 | G | T | -0.093 | 0.9262 | ++- | -78.2 | 2.245 | 2 | 0.3255 |

| | | | | | | | | | | | |
|---|---|---|---|---|---|---|---|---|---|---|---|
| rs593753 | 11 | 72691440 | T | C | 0.089 | 0.9287 | ++- | -52.4 | 2.624 | 2 | 0.2693 |
| rs2298831 | 11 | 118068752 | C | T | 0.089 | 0.929 | +?- | 0 | 0.416 | 1 | 0.5189 |
| rs56027064 | 11 | 72563104 | G | T | -0.089 | 0.9293 | +-- | 0 | 1.273 | 2 | 0.5292 |
| rs341048 | 11 | 72298280 | T | C | 0.088 | 0.9295 | +?- | 0 | 0.013 | 1 | 0.9103 |
| rs10898889 | 11 | 72605382 | C | G | 0.088 | 0.9298 | ++- | -64.3 | 2.435 | 2 | 0.2959 |
| rs1056562 | 11 | 118125625 | T | C | 0.086 | 0.9312 | +?- | 0 | 0.579 | 1 | 0.4467 |
| rs577121 | 11 | 118690550 | C | T | 0.084 | 0.9333 | +?- | 0 | 0.514 | 1 | 0.4733 |
| rs180884425 | 11 | 118620195 | C | T | 0.084 | 0.9334 | +-+ | 0 | 0.065 | 2 | 0.9682 |
| rs1715453 | 11 | 118125077 | T | C | 0.081 | 0.9358 | +?- | 0 | 0.588 | 1 | 0.4433 |
| rs12286723 | 11 | 118734640 | T | C | -0.079 | 0.9369 | -?+ | 41.9 | 5.168 | 1 | 0.02301 |
| rs678957 | 11 | 118050855 | T | C | 0.078 | 0.9382 | +?+ | 0 | 0.001 | 1 | 0.9691 |
| rs533254 | 11 | 118701898 | A | T | 0.075 | 0.9403 | +?- | 0 | 0.24 | 1 | 0.6245 |
| rs11822039 | 11 | 72638704 | A | C | -0.074 | 0.9411 | ++- | -66 | 2.409 | 2 | 0.2998 |
| rs600325 | 11 | 118041133 | C | T | 0.073 | 0.9419 | +?- | 0 | 0.007 | 1 | 0.9337 |
| rs11608057 | 11 | 118030185 | T | C | -0.072 | 0.9422 | -+- | 0 | 0.103 | 2 | 0.95 |
| rs2946743 | 11 | 118017088 | G | A | 0.067 | 0.9465 | -?+ | -70.6 | 1.759 | 1 | 0.1848 |
| rs586053 | 11 | 118020568 | A | G | -0.067 | 0.9465 | -?+ | 0 | 0.233 | 1 | 0.6296 |
| rs640120 | 11 | 118618837 | A | C | 0.066 | 0.9473 | +?- | -68.5 | 1.781 | 1 | 0.1821 |
| rs4456279 | 11 | 72520003 | A | G | 0.066 | 0.9474 | -++ | 34.3 | 6.089 | 2 | 0.04763 |
| rs535996 | 11 | 118637666 | T | C | 0.065 | 0.9483 | +?- | -51 | 1.987 | 1 | 0.1587 |
| rs584935 | 11 | 118696175 | A | G | 0.065 | 0.9486 | +?- | 0 | 0.398 | 1 | 0.5284 |
| rs529717 | 11 | 118640081 | A | G | 0.063 | 0.9498 | +?- | -53 | 1.96 | 1 | 0.1615 |
| rs1961138 | 11 | 118619423 | C | T | 0.062 | 0.9502 | +?- | -67.5 | 1.791 | 1 | 0.1808 |
| rs485901 | 11 | 118622245 | G | A | 0.06 | 0.9525 | +?- | -66.8 | 1.798 | 1 | 0.1799 |
| rs577217 | 11 | 72489023 | G | A | 0.059 | 0.9527 | -++ | 38 | 6.451 | 2 | 0.03974 |
| rs1783570 | 11 | 72871654 | A | G | 0.059 | 0.9531 | ++- | -13.3 | 3.531 | 2 | 0.1711 |
| rs2508929 | 11 | 118641451 | G | T | 0.056 | 0.9551 | +?- | -51.6 | 1.979 | 1 | 0.1595 |
| rs587816 | 11 | 118689871 | A | C | 0.055 | 0.956 | +?- | 0 | 0.496 | 1 | 0.4812 |
| rs2946742 | 11 | 118017093 | G | A | 0.054 | 0.9568 | -?+ | -74.7 | 1.717 | 1 | 0.1901 |
| rs523148 | 11 | 118682462 | T | C | 0.054 | 0.957 | +?- | 0 | 0.306 | 1 | 0.58 |
| rs6421568 | 11 | 118655704 | C | G | 0.049 | 0.9613 | +?- | -49.9 | 2.001 | 1 | 0.1572 |
| rs7118009 | 11 | 118707559 | T | C | -0.047 | 0.9628 | -?+ | 25.2 | 4.011 | 1 | 0.0452 |
| rs61581699 | 11 | 118045423 | A | G | -0.042 | 0.9664 | +?- | 0 | 0.246 | 1 | 0.6202 |
| rs548167 | 11 | 118648396 | A | G | 0.039 | 0.9691 | +?- | -47.9 | 2.029 | 1 | 0.1544 |
| rs10128600 | 11 | 118700425 | C | T | 0.032 | 0.9747 | +?- | 0 | 0.362 | 1 | 0.5472 |
| rs1784295 | 11 | 118660932 | A | G | 0.032 | 0.9747 | +?- | -35.5 | 2.214 | 1 | 0.1368 |
| rs12277355 | 11 | 118729442 | T | C | -0.031 | 0.975 | -++ | 15.9 | 4.756 | 2 | 0.09275 |
| rs6589690 | 11 | 118632893 | A | G | -0.031 | 0.9755 | +?- | -55.8 | 1.926 | 1 | 0.1652 |
| rs6421569 | 11 | 118655775 | A | C | 0.03 | 0.9757 | +?- | -46.2 | 2.052 | 1 | 0.152 |
| rs2508915 | 11 | 118652781 | C | G | 0.029 | 0.9771 | +?- | -45.8 | 2.057 | 1 | 0.1515 |
| rs7946861 | 11 | 72557986 | G | A | -0.027 | 0.9786 | ++- | -62.1 | 2.468 | 2 | 0.2911 |
| rs3132744 | 11 | 72886304 | G | A | 0.027 | 0.9788 | ++- | -10.6 | 3.615 | 2 | 0.164 |
| rs11216794 | 11 | 118044740 | A | T | -0.019 | 0.9846 | +?- | 0 | 0.301 | 1 | 0.5833 |
| rs113941645 | 11 | 118545589 | G | C | 0.018 | 0.9857 | +?- | 0 | 0.212 | 1 | 0.6448 |
| rs1790188 | 11 | 118661055 | T | C | 0.015 | 0.9877 | +?- | -37 | 2.19 | 1 | 0.139 |
| rs2187494 | 11 | 72805603 | G | A | 0.015 | 0.988 | ++- | 0 | 1.833 | 2 | 0.3999 |
| rs1612659 | 11 | 72835303 | T | C | -0.009 | 0.9931 | ++- | 0 | 1.955 | 2 | 0.3762 |
| rs10892311 | 11 | 118807735 | T | C | -0.006 | 0.9949 | -+- | 0 | 1.439 | 2 | 0.487 |
| rs475946 | 11 | 118629143 | T | C | -0.004 | 0.9967 | +?- | -61.3 | 1.859 | 1 | 0.1727 |
| rs1790187 | 11 | 118644213 | A | G | 0.001 | 0.999 | +?- | -98.8 | 1.509 | 1 | 0.2192 |
| rs74678032 | 11 | 118812559 | C | T | -0.001 | 0.9994 | +?- | -50.2 | 1.997 | 1 | 0.1576 |

**Supplemental Data 3: Meta-analysis of SjD and SLE data in DDX6-CXCR5 locus (Merged Datasets**

| SNP | CHR | BP(GRCh37/hg19) | A1 | A2 | Zscore | P.value | Direction | HetISq | HetChiSq | HetDf | HetPVal |
|---|---|---|---|---|---|---|---|---|---|---|---|
| rs11217045 | 11 | 118697865 | A | C | -8.471 | 2.42E-17 | ----- | 0 | 1.872 | 4 | 0.7592 |
| rs73005502 | 11 | 118697906 | G | A | -8.47 | 2.46E-17 | ----- | 0 | 1.879 | 4 | 0.758 |
| rs4938573 | 11 | 118741842 | C | T | -8.437 | 3.26E-17 | ----- | 0 | 1.855 | 4 | 0.7623 |
| rs11217046 | 11 | 118698907 | A | G | -8.418 | 3.82E-17 | ----- | 0 | 2.15 | 4 | 0.7082 |
| rs10892293 | 11 | 118665575 | T | A | -8.418 | 3.83E-17 | ----- | 0 | 2.298 | 4 | 0.6812 |
| rs715412 | 11 | 118684610 | A | G | -8.397 | 4.58E-17 | ----- | 0 | 2.503 | 4 | 0.644 |
| rs4936444 | 11 | 118743338 | T | C | -8.357 | 6.43E-17 | ----- | 0 | 2.067 | 4 | 0.7235 |
| rs11217032 | 11 | 118669605 | A | T | -8.354 | 6.57E-17 | ----- | 0 | 2.365 | 4 | 0.669 |
| rs28409215 | 11 | 118679304 | A | G | -8.353 | 6.66E-17 | ----- | 0 | 2.239 | 4 | 0.6919 |
| rs11217038 | 11 | 118677355 | A | G | -8.352 | 6.72E-17 | ----- | 0 | 2.294 | 4 | 0.6819 |
| rs11217037 | 11 | 118677086 | A | G | -8.349 | 6.89E-17 | ----- | 0 | 2.187 | 4 | 0.7014 |
| rs79504890 | 11 | 118701018 | A | G | -8.343 | 7.25E-17 | ----- | 0 | 2.388 | 4 | 0.6648 |
| rs11217033 | 11 | 118673706 | C | T | -8.329 | 8.16E-17 | ----- | 0 | 2.281 | 4 | 0.6843 |
| rs11217036 | 11 | 118675772 | T | C | -8.327 | 8.30E-17 | ----- | 0 | 2.313 | 4 | 0.6785 |
| rs73005423 | 11 | 118681079 | G | A | -8.307 | 9.84E-17 | ----- | 0 | 2.336 | 4 | 0.6742 |
| rs10892294 | 11 | 118667357 | C | G | -8.306 | 9.87E-17 | ----- | 0 | 2.255 | 4 | 0.689 |
| rs74541740 | 11 | 118742800 | A | G | -8.296 | 1.08E-16 | ----- | 0 | 1.659 | 4 | 0.7981 |
| rs7951740 | 11 | 118740418 | G | C | -8.274 | 1.30E-16 | ----- | 0 | 1.727 | 4 | 0.7858 |
| rs7117261 | 11 | 118741157 | T | C | -8.258 | 1.48E-16 | ----- | 0 | 1.774 | 4 | 0.7773 |
| rs11217074 | 11 | 118734000 | C | T | -8.245 | 1.66E-16 | ----- | 0 | 1.462 | 4 | 0.8334 |
| rs7122669 | 11 | 118739056 | A | G | -8.233 | 1.82E-16 | ----- | 0 | 1.59 | 4 | 0.8106 |
| rs7119044 | 11 | 118738298 | G | C | -8.229 | 1.89E-16 | ----- | 0 | 1.577 | 4 | 0.8129 |
| rs75249660 | 11 | 118721565 | A | G | -8.192 | 2.56E-16 | ----- | 0 | 1.823 | 4 | 0.7683 |
| rs10790268 | 11 | 118729391 | A | G | -8.187 | 2.69E-16 | ----- | 0 | 1.987 | 4 | 0.7381 |
| rs11217058 | 11 | 118722149 | A | G | -8.175 | 2.96E-16 | ----- | 0 | 2.024 | 4 | 0.7314 |
| rs10790269 | 11 | 118729456 | C | T | -8.162 | 3.29E-16 | ----- | 0 | 1.979 | 4 | 0.7397 |
| rs11826521 | 11 | 118610549 | T | G | -8.149 | 3.66E-16 | ----- | 0 | 2.522 | 4 | 0.6407 |
| rs2077579 | 11 | 118619047 | G | T | -8.145 | 3.79E-16 | ----- | 0 | 2.56 | 4 | 0.6339 |
| rs10892280 | 11 | 118611817 | T | C | -8.133 | 4.18E-16 | ----- | 0 | 2.433 | 4 | 0.6566 |
| rs4936441 | 11 | 118725660 | C | G | -8.106 | 5.24E-16 | ----- | 0 | 2.114 | 4 | 0.7148 |
| rs6589684 | 11 | 118610957 | A | G | -8.096 | 5.66E-16 | ----- | 0 | 1.805 | 4 | 0.7715 |
| rs11217066 | 11 | 118726753 | G | T | -8.094 | 5.80E-16 | ----- | 0 | 2.026 | 4 | 0.7311 |
| rs56758835 | 11 | 118615340 | A | G | -8.059 | 7.67E-16 | ----- | 0 | 2.244 | 4 | 0.691 |
| rs1048024 | 11 | 118618776 | T | A | -8.042 | 8.82E-16 | ----- | 0 | 2.356 | 4 | 0.6707 |
| rs11217009 | 11 | 118615785 | T | C | -8.04 | 9.00E-16 | ----- | 0 | 2.181 | 4 | 0.7025 |
| rs7481819 | 11 | 118745278 | C | T | -8.035 | 9.37E-16 | ----- | 0 | 2.321 | 4 | 0.677 |
| rs3889239 | 11 | 118619960 | T | C | -8.033 | 9.53E-16 | ----- | 0 | 2.261 | 4 | 0.6878 |
| rs2004781 | 11 | 118619253 | T | C | -8.029 | 9.79E-16 | ----- | 0 | 2.28 | 4 | 0.6845 |
| rs74676029 | 11 | 118628373 | T | C | -7.988 | 1.37E-15 | ----- | 0 | 2.361 | 4 | 0.6696 |
| rs57494551 | 11 | 118661398 | T | C | -7.981 | 1.45E-15 | ----- | 0 | 2.682 | 4 | 0.6123 |
| rs7481797 | 11 | 118745243 | A | G | -7.981 | 1.45E-15 | ----- | 0 | 2.129 | 4 | 0.7121 |
| rs11217020 | 11 | 118639353 | A | G | -7.971 | 1.57E-15 | ----- | 0 | 2.469 | 4 | 0.6502 |
| rs874621 | 11 | 118655645 | A | G | -7.951 | 1.85E-15 | ----- | 0 | 2.357 | 4 | 0.6705 |
| rs4936443 | 11 | 118740864 | C | T | -7.94 | 2.03E-15 | ----- | 0 | 1.896 | 4 | 0.7549 |
| rs11217019 | 11 | 118636343 | A | C | -7.938 | 2.06E-15 | ----- | 0 | 2.527 | 4 | 0.6397 |
| rs10892283 | 11 | 118636980 | G | A | -7.936 | 2.08E-15 | ----- | 0 | 2.588 | 4 | 0.6289 |
| rs10892287 | 11 | 118642999 | C | T | -7.93 | 2.20E-15 | ----- | 0 | 2.538 | 4 | 0.6378 |
| rs10892288 | 11 | 118644694 | G | A | -7.929 | 2.21E-15 | ----- | 0 | 2.479 | 4 | 0.6484 |
| rs10892292 | 11 | 118655340 | C | G | -7.919 | 2.40E-15 | ----- | 0 | 2.418 | 4 | 0.6594 |
| rs10892286 | 11 | 118642085 | C | A | -7.907 | 2.64E-15 | ----- | 0 | 2.551 | 4 | 0.6356 |
| rs10892290 | 11 | 118653503 | T | G | -7.902 | 2.74E-15 | ----- | 0 | 2.463 | 4 | 0.6513 |
| rs10892289 | 11 | 118646003 | T | C | -7.902 | 2.75E-15 | ----- | 0 | 2.378 | 4 | 0.6665 |
| rs56043232 | 11 | 118648373 | T | C | -7.891 | 3.00E-15 | ----- | 0 | 2.421 | 4 | 0.6589 |
| rs10892291 | 11 | 118653517 | T | C | -7.838 | 4.59E-15 | ----- | 0 | 2.537 | 4 | 0.638 |

| | | | | | | | | | | | |
|---|---|---|---|---|---|---|---|---|---|---|---|
| rs10790275 | 11 | 118745884 | G | C | -7.789 | 6.74E-15 | ----- | 0 | 2.25 | 4 | 0.6899 |
| rs4938572 | 11 | 118740931 | C | T | -7.726 | 1.11E-14 | ----- | 0 | 1.472 | 4 | 0.8317 |
| rs11217040 | 11 | 118680648 | A | C | -7.723 | 1.14E-14 | ----- | -31.6 | 4.56 | 4 | 0.3355 |
| rs7117313 | 11 | 118744396 | C | G | -7.713 | 1.23E-14 | ----- | 0 | 2.135 | 4 | 0.711 |
| rs17122453 | 11 | 118683564 | A | G | -7.633 | 2.29E-14 | ----- | -34.8 | 4.45 | 4 | 0.3485 |
| rs77209083 | 11 | 118607569 | G | A | -7.633 | 2.30E-14 | ----- | 0 | 2.149 | 4 | 0.7084 |
| rs11217042 | 11 | 118682528 | A | G | -7.583 | 3.37E-14 | ----- | -37.9 | 4.352 | 4 | 0.3605 |
| rs73005426 | 11 | 118681083 | G | A | -7.581 | 3.44E-14 | ----- | -40.9 | 4.259 | 4 | 0.3721 |
| rs7942535 | 11 | 118681464 | C | T | -7.575 | 3.60E-14 | ----- | -36.9 | 4.383 | 4 | 0.3566 |
| rs11217044 | 11 | 118696022 | C | T | -7.574 | 3.62E-14 | ----- | 0 | 3.6 | 4 | 0.4628 |
| rs11217041 | 11 | 118682350 | G | A | -7.573 | 3.64E-14 | ----- | -36.7 | 4.389 | 4 | 0.3559 |
| rs7119038 | 11 | 118738281 | G | A | -7.565 | 3.88E-14 | ----- | 0 | 2.998 | 4 | 0.5582 |
| rs11217001 | 11 | 118607046 | A | G | -7.546 | 4.47E-14 | ----- | 0 | 2.455 | 4 | 0.6527 |
| rs12365699 | 11 | 118743286 | A | G | -7.528 | 5.17E-14 | ----- | 0 | 2.976 | 4 | 0.5618 |
| rs7123726 | 11 | 118694547 | C | T | -7.524 | 5.31E-14 | ----- | -21.4 | 4.942 | 4 | 0.2933 |
| rs11217006 | 11 | 118609790 | A | G | -7.45 | 9.33E-14 | ----- | 0 | 3.684 | 4 | 0.4505 |
| rs10892279 | 11 | 118611781 | A | G | -7.391 | 1.46E-13 | ----- | 0 | 3.291 | 4 | 0.5103 |
| rs6589685 | 11 | 118612197 | C | T | -7.372 | 1.68E-13 | ----- | 0 | 3.23 | 4 | 0.52 |
| rs2155433 | 11 | 118612921 | A | G | -7.291 | 3.07E-13 | ----- | 0 | 3.283 | 4 | 0.5117 |
| rs7945144 | 11 | 118608716 | G | A | -7.267 | 3.67E-13 | ----- | 0 | 3.204 | 4 | 0.5243 |
| rs12364971 | 11 | 118607794 | G | T | -7.167 | 7.65E-13 | ----- | 0 | 3.073 | 4 | 0.5457 |
| rs73003215 | 11 | 118649512 | A | G | -7.156 | 8.32E-13 | ----- | 0 | 2.985 | 4 | 0.5604 |
| rs76409436 | 11 | 118607871 | A | G | -7.15 | 8.70E-13 | ----- | 0 | 3.162 | 4 | 0.5311 |
| rs6589682 | 11 | 118606722 | T | C | -7.063 | 1.63E-12 | ----- | 0 | 3.268 | 4 | 0.5141 |
| rs11217002 | 11 | 118607150 | T | C | -7.006 | 2.45E-12 | ----- | 0 | 3.129 | 4 | 0.5365 |
| rs56267528 | 11 | 118607311 | A | C | -7.005 | 2.48E-12 | ----- | 0 | 2.872 | 4 | 0.5795 |
| rs55894437 | 11 | 118607417 | T | C | -6.983 | 2.89E-12 | ----- | 0 | 2.857 | 4 | 0.5821 |
| rs7125066 | 11 | 118721699 | G | A | -6.857 | 7.01E-12 | ----- | 37.1 | 9.534 | 4 | 0.04906 |
| rs9736016 | 11 | 118724894 | A | T | -6.849 | 7.44E-12 | ----- | 37.7 | 9.636 | 4 | 0.04703 |
| rs6589686 | 11 | 118612565 | A | G | -6.838 | 8.01E-12 | ----- | 0 | 1.526 | 4 | 0.8221 |
| rs10892273 | 11 | 118602809 | A | G | -6.828 | 8.63E-12 | ----- | 0 | 3.125 | 4 | 0.5371 |
| rs10892272 | 11 | 118602707 | G | C | -6.818 | 9.23E-12 | ----- | 0 | 3.202 | 4 | 0.5247 |
| rs11216997 | 11 | 118604638 | T | C | -6.814 | 9.50E-12 | ----- | 0 | 3.529 | 4 | 0.4734 |
| rs10892271 | 11 | 118598544 | A | G | -6.803 | 1.02E-11 | ----- | 0 | 2.818 | 4 | 0.5887 |
| rs112561833 | 11 | 118603736 | C | A | -6.783 | 1.18E-11 | ----- | 0 | 3.386 | 4 | 0.4954 |
| rs11216995 | 11 | 118603526 | A | G | -6.625 | 3.48E-11 | ----- | 0 | 3.202 | 4 | 0.5246 |
| rs11216994 | 11 | 118603520 | T | C | -6.621 | 3.58E-11 | ----- | 0 | 3.16 | 4 | 0.5315 |
| rs73001429 | 11 | 118603165 | G | T | -6.618 | 3.64E-11 | ----- | 0 | 3.086 | 4 | 0.5435 |
| rs11216993 | 11 | 118603495 | A | T | -6.592 | 4.33E-11 | ----- | 0 | 2.943 | 4 | 0.5674 |
| rs10892263 | 11 | 118587319 | C | T | -6.207 | 5.39E-10 | ----- | 0 | 3.738 | 4 | 0.4426 |
| rs12421790 | 11 | 118588952 | T | C | -6.141 | 8.21E-10 | ----- | 0 | 3.828 | 4 | 0.4298 |
| rs11216980 | 11 | 118590668 | A | C | -6.137 | 8.43E-10 | ----- | -38.4 | 4.334 | 4 | 0.3626 |
| rs10892262 | 11 | 118587097 | C | G | -6.029 | 1.65E-09 | ----- | 0 | 3.229 | 4 | 0.5203 |
| rs11216999 | 11 | 118604941 | C | T | -5.888 | 3.91E-09 | ----- | 0 | 2.966 | 4 | 0.5635 |
| rs12421793 | 11 | 118589012 | A | C | -5.854 | 4.80E-09 | ----- | 0 | 3.866 | 4 | 0.4244 |
| rs11216998 | 11 | 118604837 | T | C | -5.748 | 9.05E-09 | ----- | 0 | 3.11 | 4 | 0.5395 |
| rs11216972 | 11 | 118585826 | T | C | -5.717 | 1.08E-08 | ----- | 0 | 3.193 | 4 | 0.5261 |
| rs10892277 | 11 | 118605841 | A | G | -5.631 | 1.79E-08 | ----- | 0 | 2.995 | 4 | 0.5587 |
| rs10892274 | 11 | 118605463 | T | C | -5.604 | 2.10E-08 | ----- | 0 | 2.946 | 4 | 0.5669 |
| rs10892276 | 11 | 118605628 | A | G | -5.585 | 2.34E-08 | ----- | 0 | 3.047 | 4 | 0.55 |
| rs10892275 | 11 | 118605617 | A | T | -5.568 | 2.58E-08 | ----- | 0 | 2.928 | 4 | 0.57 |
| rs11216956 | 11 | 118575326 | A | G | -5.515 | 3.48E-08 | ----- | 0 | 2.338 | 4 | 0.6739 |
| rs11216960 | 11 | 118577873 | A | G | -5.412 | 6.24E-08 | ----- | 0 | 2.353 | 4 | 0.6712 |
| rs10790261 | 11 | 118579747 | G | A | -5.412 | 6.25E-08 | ----- | 0 | 2.181 | 4 | 0.7026 |
| rs10892260 | 11 | 118583525 | C | A | -5.404 | 6.50E-08 | ----- | 0 | 2.36 | 4 | 0.6698 |
| rs11216969 | 11 | 118584340 | C | T | -5.398 | 6.72E-08 | ----- | 0 | 2.222 | 4 | 0.695 |

| | | | | | | | | | | | |
|---|---|---|---|---|---|---|---|---|---|---|---|
| rs10892259 | 11 | 118583002 | A | G | -5.398 | 6.75E-08 | ----- | 0 | 2.121 | 4 | 0.7136 |
| rs10892256 | 11 | 118578705 | A | G | -5.393 | 6.94E-08 | ----- | 0 | 2.267 | 4 | 0.6869 |
| rs476246 | 11 | 118617920 | T | C | -5.385 | 7.23E-08 | ----- | 0 | 0.678 | 4 | 0.954 |
| rs10892257 | 11 | 118579327 | A | G | -5.376 | 7.61E-08 | ----- | 0 | 2.279 | 4 | 0.6845 |
| rs11216965 | 11 | 118582613 | A | C | -5.375 | 7.65E-08 | ----- | 0 | 2.271 | 4 | 0.6861 |
| rs472641 | 11 | 118617544 | G | A | -5.37 | 7.86E-08 | ----- | 0 | 0.652 | 4 | 0.9571 |
| rs10892258 | 11 | 118579865 | A | G | -5.364 | 8.16E-08 | ----- | 0 | 2.257 | 4 | 0.6886 |
| rs581062 | 11 | 118624767 | A | G | -5.361 | 8.29E-08 | ----- | 0 | 0.63 | 4 | 0.9596 |
| rs11216964 | 11 | 118581895 | A | G | -5.356 | 8.50E-08 | ----- | 0 | 2.296 | 4 | 0.6815 |
| rs11216961 | 11 | 118580338 | G | C | -5.353 | 8.67E-08 | ----- | 0 | 2.276 | 4 | 0.6851 |
| rs598373 | 11 | 118664377 | T | C | -5.32 | 1.04E-07 | ----- | 0 | 0.578 | 4 | 0.9655 |
| rs487728 | 11 | 118621495 | G | C | -5.31 | 1.10E-07 | ----- | 0 | 0.692 | 4 | 0.9523 |
| rs11216962 | 11 | 118580813 | C | T | -5.289 | 1.23E-07 | ----- | 0 | 2.151 | 4 | 0.7081 |
| rs11216963 | 11 | 118580814 | A | G | -5.289 | 1.23E-07 | ----- | 0 | 2.151 | 4 | 0.7081 |
| rs499343 | 11 | 118626511 | T | C | -5.282 | 1.28E-07 | ----- | 0 | 0.588 | 4 | 0.9644 |
| rs524590 | 11 | 118640063 | T | A | -5.268 | 1.38E-07 | ----- | 0 | 0.525 | 4 | 0.971 |
| rs688161 | 11 | 118677710 | G | A | -5.259 | 1.45E-07 | ----- | 0 | 0.549 | 4 | 0.9686 |
| rs2508570 | 11 | 118678609 | A | G | -5.247 | 1.55E-07 | ----- | 0 | 0.546 | 4 | 0.9689 |
| rs591756 | 11 | 118678881 | T | C | -5.231 | 1.69E-07 | ----- | 0 | 0.611 | 4 | 0.9619 |
| rs1784293 | 11 | 118652373 | C | A | -5.229 | 1.70E-07 | ----- | 0 | 0.725 | 4 | 0.9482 |
| rs477014 | 11 | 118678986 | C | T | -5.228 | 1.71E-07 | ----- | 0 | 0.702 | 4 | 0.9511 |
| rs1790189 | 11 | 118652326 | G | T | -5.219 | 1.79E-07 | ----- | 0 | 0.71 | 4 | 0.9501 |
| rs473298 | 11 | 118679387 | G | A | -5.216 | 1.83E-07 | ----- | 0 | 0.587 | 4 | 0.9645 |
| rs2508916 | 11 | 118654651 | G | C | -5.214 | 1.85E-07 | ----- | 0 | 0.869 | 4 | 0.9289 |
| rs516719 | 11 | 118679445 | C | T | -5.211 | 1.88E-07 | ----- | 0 | 0.578 | 4 | 0.9655 |
| rs2508569 | 11 | 118680045 | C | G | -5.191 | 2.09E-07 | ----- | 0 | 0.634 | 4 | 0.9593 |
| rs540180 | 11 | 118653629 | T | G | -5.189 | 2.11E-07 | ----- | 0 | 0.968 | 4 | 0.9146 |
| rs11606199 | 11 | 118811641 | A | G | -5.137 | 2.79E-07 | -?--- | 0 | 0.917 | 3 | 0.8213 |
| rs11217065 | 11 | 118726647 | C | T | 4.994 | 5.93E-07 | +++++ | -11.8 | 5.365 | 4 | 0.2519 |
| rs11217063 | 11 | 118725942 | C | G | 4.991 | 6.01E-07 | +++++ | -10.5 | 5.428 | 4 | 0.2461 |
| rs57719838 | 11 | 118724723 | C | T | 4.971 | 6.65E-07 | +++++ | -9.9 | 5.458 | 4 | 0.2435 |
| rs11217070 | 11 | 118729860 | G | T | 4.964 | 6.90E-07 | +++++ | -14.8 | 5.227 | 4 | 0.2648 |
| rs11217072 | 11 | 118731762 | G | A | 4.955 | 7.23E-07 | +++++ | -15.7 | 5.188 | 4 | 0.2686 |
| rs34561290 | 11 | 118731599 | T | C | 4.954 | 7.27E-07 | +++++ | -15.5 | 5.193 | 4 | 0.2681 |
| rs12790489 | 11 | 118729583 | C | T | 4.947 | 7.55E-07 | +++++ | -16 | 5.173 | 4 | 0.27 |
| rs10892300 | 11 | 118729210 | T | C | 4.946 | 7.56E-07 | +++++ | -16 | 5.174 | 4 | 0.2699 |
| rs7929520 | 11 | 118730141 | G | A | 4.915 | 8.89E-07 | +++++ | -12.9 | 5.316 | 4 | 0.2563 |
| rs7103067 | 11 | 118723379 | A | G | 4.909 | 9.14E-07 | +++++ | -8.2 | 5.545 | 4 | 0.2358 |
| rs10892319 | 11 | 118870413 | T | C | -4.861 | 1.17E-06 | -?--- | 0 | 1.712 | 3 | 0.6344 |
| rs571470 | 11 | 118697012 | A | G | -4.852 | 1.22E-06 | --+-- | 53.2 | 12.827 | 4 | 0.01215 |
| rs478081 | 11 | 118602497 | G | A | -4.838 | 1.31E-06 | ----- | 0 | 1.563 | 4 | 0.8154 |
| rs562059 | 11 | 118602058 | G | T | -4.832 | 1.36E-06 | ----- | 0 | 1.594 | 4 | 0.8099 |
| rs10790267 | 11 | 118727169 | T | C | 4.804 | 1.55E-06 | +++++ | -15.5 | 5.196 | 4 | 0.2678 |
| rs45586434 | 11 | 118828184 | A | C | -4.785 | 1.71E-06 | -?--- | 0 | 0.73 | 3 | 0.8662 |
| rs9645663 | 11 | 118873840 | C | T | -4.775 | 1.80E-06 | -?--- | 0 | 1.699 | 3 | 0.6371 |
| rs11217067 | 11 | 118727332 | T | G | 4.77 | 1.85E-06 | +++++ | -16.8 | 5.139 | 4 | 0.2733 |
| rs537080 | 11 | 118826045 | C | T | -4.764 | 1.90E-06 | -?--- | 0 | 1.029 | 3 | 0.7942 |
| rs4936442 | 11 | 118728320 | T | C | 4.756 | 1.97E-06 | +++++ | -19.7 | 5.014 | 4 | 0.2858 |
| rs7483509 | 11 | 118740180 | A | G | 4.742 | 2.12E-06 | +++++ | -25.4 | 4.783 | 4 | 0.3103 |
| rs112939258 | 11 | 118834718 | T | C | -4.733 | 2.21E-06 | -?--- | 0 | 1.138 | 3 | 0.7679 |
| rs73001406 | 11 | 118560857 | A | G | -4.73 | 2.24E-06 | ----- | 0 | 3.414 | 4 | 0.491 |
| rs10892316 | 11 | 118831755 | T | C | -4.709 | 2.49E-06 | -?--- | 0 | 1.193 | 3 | 0.7546 |
| rs507763 | 11 | 118824683 | C | G | -4.698 | 2.62E-06 | -?--- | 0 | 0.886 | 3 | 0.8288 |
| rs76253444 | 11 | 118876213 | G | A | -4.669 | 3.03E-06 | -?--- | 0 | 1.384 | 3 | 0.7093 |
| rs111358668 | 11 | 118834508 | A | G | -4.635 | 3.57E-06 | -?--- | 0 | 1.26 | 3 | 0.7387 |
| rs10892252 | 11 | 118555692 | G | A | -4.562 | 5.07E-06 | ----- | 0 | 3.677 | 4 | 0.4515 |

| | | | | | | | | | | | |
|---|---|---|---|---|---|---|---|---|---|---|---|
| rs2277296 | 11 | 118550522 | C | T | -4.48 | 7.45E-06 | ----- | 0 | 3.907 | 4 | 0.4187 |
| rs4938617 | 11 | 118867695 | T | C | -4.476 | 7.59E-06 | -?--- | 0 | 2.722 | 3 | 0.4366 |
| rs622722 | 11 | 118711790 | A | G | 4.467 | 7.92E-06 | +++++ | -4.9 | 5.72 | 4 | 0.2211 |
| rs10892251 | 11 | 118543563 | T | C | -4.449 | 8.63E-06 | ----- | 0 | 3.771 | 4 | 0.4379 |
| rs113962716 | 11 | 118853021 | A | C | -4.421 | 9.81E-06 | -?--- | 0 | 1.439 | 3 | 0.6964 |
| rs11217109 | 11 | 118853774 | C | T | -4.41 | 1.04E-05 | -?--- | 0 | 1.42 | 3 | 0.7009 |
| rs80126726 | 11 | 118852647 | A | G | -4.397 | 1.10E-05 | -?--- | 0 | 1.554 | 3 | 0.6698 |
| rs74884652 | 11 | 118876552 | C | T | -4.319 | 1.57E-05 | -?--- | 0 | 2.346 | 3 | 0.5039 |
| rs11216943 | 11 | 118556398 | A | G | -4.27 | 1.96E-05 | ----- | -27.3 | 4.712 | 4 | 0.3181 |
| rs75154936 | 11 | 118877891 | C | T | -4.256 | 2.08E-05 | -?--- | 0 | 2.459 | 3 | 0.4828 |
| rs111345266 | 11 | 118893767 | T | C | -4.246 | 2.18E-05 | -?--- | 0 | 2.615 | 3 | 0.4549 |
| rs4938619 | 11 | 118890085 | T | C | -4.21 | 2.56E-05 | -?--- | 0 | 2.678 | 3 | 0.4439 |
| rs10892247 | 11 | 118490076 | A | G | -4.094 | 4.25E-05 | ----- | 0 | 3.59 | 4 | 0.4643 |
| rs4938576 | 11 | 118746769 | T | G | -4.043 | 5.28E-05 | ----- | 0 | 1.325 | 4 | 0.8571 |
| rs741811 | 11 | 118898319 | A | G | -3.983 | 6.80E-05 | -?--- | 0 | 1.18 | 3 | 0.7577 |
| rs496547 | 11 | 118576463 | A | T | -3.983 | 6.81E-05 | ----- | 0 | 1.747 | 4 | 0.7821 |
| rs11216937 | 11 | 118507478 | T | C | -3.973 | 7.10E-05 | ----- | 0 | 2.867 | 4 | 0.5803 |
| rs12362152 | 11 | 118516737 | G | A | -3.962 | 7.42E-05 | ----- | 0 | 2.949 | 4 | 0.5664 |
| rs80015904 | 11 | 118516903 | A | G | -3.953 | 7.71E-05 | ----- | 0 | 2.781 | 4 | 0.5951 |
| rs12365095 | 11 | 118516774 | T | C | -3.921 | 8.81E-05 | ----- | 0 | 2.699 | 4 | 0.6093 |
| rs11216938 | 11 | 118514625 | T | C | -3.919 | 8.91E-05 | ----- | 0 | 2.768 | 4 | 0.5974 |
| rs12225548 | 11 | 118535840 | G | C | -3.902 | 9.54E-05 | ----- | 0 | 2.894 | 4 | 0.5758 |
| rs17748 | 11 | 118528424 | T | C | -3.897 | 9.75E-05 | ----- | 0 | 2.734 | 4 | 0.6032 |
| rs77683570 | 11 | 118495287 | C | T | -3.89 | 0.0001003 | ----- | 0 | 3.016 | 4 | 0.5552 |
| rs10790256 | 11 | 118534082 | T | C | -3.886 | 0.000102 | ----- | 0 | 2.748 | 4 | 0.6008 |
| rs7928371 | 11 | 118529127 | A | G | -3.886 | 0.000102 | ----- | 0 | 2.747 | 4 | 0.6011 |
| rs11216930 | 11 | 118488782 | C | A | -3.883 | 0.0001033 | ----- | 0 | 2.183 | 4 | 0.7021 |
| rs45540840 | 11 | 118486110 | A | G | -3.872 | 0.0001079 | ----- | 0 | 2.807 | 4 | 0.5905 |
| rs2236661 | 11 | 118499394 | G | C | -3.859 | 0.0001138 | ----- | 0 | 2.873 | 4 | 0.5793 |
| rs45565037 | 11 | 118496119 | A | G | -3.848 | 0.0001189 | ----- | 0 | 2.979 | 4 | 0.5613 |
| rs11217099 | 11 | 118809882 | A | C | -3.837 | 0.0001245 | -+--- | 8.6 | 6.567 | 4 | 0.1606 |
| rs2276065 | 11 | 118530611 | C | T | -3.826 | 0.00013 | ----- | 0 | 3.239 | 4 | 0.5186 |
| rs75438046 | 11 | 118767564 | A | G | -3.815 | 0.0001364 | -+--? | 29.7 | 7.113 | 3 | 0.06838 |
| rs78041268 | 11 | 118763313 | A | G | -3.806 | 0.000141 | -+--? | 23.7 | 6.556 | 3 | 0.08749 |
| rs10892306 | 11 | 118751326 | A | T | -3.761 | 0.000169 | ----- | 0 | 2.609 | 4 | 0.6252 |
| rs77841686 | 11 | 118762073 | A | G | -3.758 | 0.0001711 | -+--? | 21.9 | 6.4 | 3 | 0.09369 |
| rs10892248 | 11 | 118501022 | A | G | -3.742 | 0.0001824 | ----- | 0 | 2.783 | 4 | 0.5948 |
| rs149114341 | 11 | 118783424 | A | G | -3.714 | 0.0002041 | -+--? | 22.4 | 6.444 | 3 | 0.09192 |
| rs546393 | 11 | 118624250 | C | T | 3.609 | 0.0003075 | ++-++ | 1.2 | 6.07 | 4 | 0.194 |
| rs488219 | 11 | 118620697 | C | T | 3.6 | 0.0003185 | ++-++ | 0.9 | 6.055 | 4 | 0.1951 |
| rs10790255 | 11 | 118515579 | G | T | -3.574 | 0.0003517 | ----- | 0 | 2.361 | 4 | 0.6698 |
| rs555639 | 11 | 118629031 | A | G | 3.555 | 0.000378 | ++-++ | -5.5 | 5.686 | 4 | 0.2238 |
| rs607125 | 11 | 118617299 | C | T | 3.544 | 0.0003943 | ++-++ | 2 | 6.12 | 4 | 0.1904 |
| rs573905 | 11 | 118572267 | A | G | 3.54 | 0.0004007 | +++++ | 0 | 1.888 | 4 | 0.7564 |
| rs581045 | 11 | 118632522 | T | C | 3.532 | 0.0004126 | ++-++ | -9.4 | 5.485 | 4 | 0.2411 |
| rs586763 | 11 | 118816304 | T | C | -3.52 | 0.0004308 | -?--- | 0 | 0.723 | 3 | 0.8678 |
| rs555649 | 11 | 118651809 | C | T | 3.52 | 0.0004316 | ++-++ | -16.4 | 5.156 | 4 | 0.2717 |
| rs11217091 | 11 | 118793613 | C | T | -3.517 | 0.0004362 | -+--- | 17.5 | 7.27 | 4 | 0.1223 |
| rs2510897 | 11 | 118644582 | C | A | 3.491 | 0.000482 | ++-++ | -17.5 | 5.106 | 4 | 0.2766 |
| rs2105782 | 11 | 118659896 | C | T | 3.47 | 0.0005213 | ++-++ | -13.8 | 5.272 | 4 | 0.2605 |
| rs515523 | 11 | 118658075 | G | A | 3.468 | 0.0005252 | ++-++ | -6 | 5.661 | 4 | 0.226 |
| rs1784543 | 11 | 118631653 | T | A | 3.467 | 0.0005257 | ++-++ | -6.1 | 5.657 | 4 | 0.2263 |
| rs507890 | 11 | 118631288 | C | T | 3.46 | 0.0005396 | ++-++ | -7 | 5.607 | 4 | 0.2305 |
| rs4938513 | 11 | 118419704 | T | C | -3.449 | 0.000562 | ----? | 0 | 1.628 | 3 | 0.6532 |
| rs607472 | 11 | 118680072 | G | C | 3.433 | 0.0005972 | ++-++ | -7 | 5.608 | 4 | 0.2304 |
| rs566733 | 11 | 118608625 | G | C | 3.419 | 0.0006288 | ++-++ | 3.7 | 6.229 | 4 | 0.1827 |

| | | | | | | | | | | | |
|---|---|---|---|---|---|---|---|---|---|---|---|
| rs567680 | 11 | 118608532 | G | C | 3.389 | 0.0007017 | ++-++ | 7.7 | 6.498 | 4 | 0.1649 |
| rs1784300 | 11 | 118679629 | T | A | 3.383 | 0.0007173 | ++-++ | -12.6 | 5.328 | 4 | 0.2553 |
| rs632124 | 11 | 118613235 | A | T | 3.374 | 0.0007416 | ++-++ | 7.8 | 6.508 | 4 | 0.1643 |
| rs11217004 | 11 | 118607622 | A | G | 3.354 | 0.0007965 | ++-++ | 6.8 | 6.437 | 4 | 0.1688 |
| rs11217078 | 11 | 118750321 | C | T | -3.346 | 0.000821 | -?+-- | 53.1 | 10.654 | 3 | 0.01375 |
| rs11217090 | 11 | 118789952 | T | C | -3.343 | 0.0008293 | -+--- | -3 | 5.823 | 4 | 0.2127 |
| rs12796102 | 11 | 118662028 | T | C | 3.309 | 0.0009369 | ++-++ | -24.4 | 4.824 | 4 | 0.3058 |
| rs633308 | 11 | 118459069 | G | A | -3.277 | 0.001049 | ----? | 0 | 1.276 | 3 | 0.7347 |
| rs112119822 | 11 | 118798824 | G | A | -3.276 | 0.001052 | -+--- | -11 | 5.404 | 4 | 0.2483 |
| rs12280501 | 11 | 118801440 | T | C | -3.26 | 0.001115 | -+--- | -10.7 | 5.423 | 4 | 0.2466 |
| rs7101680 | 11 | 118799206 | A | G | -3.258 | 0.001122 | -+--- | -10.8 | 5.417 | 4 | 0.2471 |
| rs4291686 | 11 | 118700557 | G | A | 3.207 | 0.001341 | +++?+ | -15.7 | 4.321 | 3 | 0.2289 |
| rs625274 | 11 | 118699276 | T | C | 3.185 | 0.001445 | +++?+ | -19.5 | 4.186 | 3 | 0.2421 |
| rs4936459 | 11 | 118899218 | T | C | -3.146 | 0.001652 | -+--- | -44.2 | 4.161 | 4 | 0.3846 |
| rs77638281 | 11 | 118811315 | A | G | -3.12 | 0.001807 | -?--- | 0 | 0.618 | 3 | 0.8924 |
| rs112974653 | 11 | 118807637 | T | G | -3.076 | 0.0021 | -?--- | 0 | 1.164 | 3 | 0.7616 |
| rs4938589 | 11 | 118810405 | T | C | -3.067 | 0.002164 | -?--- | 0 | 1.255 | 3 | 0.7398 |
| rs10892313 | 11 | 118809329 | A | G | -3.066 | 0.002169 | -?--- | 0 | 0.928 | 3 | 0.8187 |
| rs11605265 | 11 | 118805144 | A | G | -3.057 | 0.002234 | -?--- | 0 | 0.405 | 3 | 0.9392 |
| rs542266 | 11 | 118422240 | A | C | -3.055 | 0.002247 | ----? | 0 | 1.458 | 3 | 0.6919 |
| rs11600378 | 11 | 118803842 | G | A | -3.044 | 0.002333 | -?--- | 0 | 0.436 | 3 | 0.9327 |
| rs11605658 | 11 | 118805220 | T | C | -3.036 | 0.002397 | -?--- | 0 | 0.369 | 3 | 0.9465 |
| rs143711539 | 11 | 118804671 | A | G | -3.036 | 0.0024 | -?--- | 0 | 0.417 | 3 | 0.9367 |
| rs10892312 | 11 | 118807740 | A | G | -3.026 | 0.002476 | -?--- | 0 | 1.023 | 3 | 0.7956 |
| rs12282154 | 11 | 118807876 | A | G | -3.023 | 0.002503 | -?--- | 0 | 1.01 | 3 | 0.7987 |
| rs77916839 | 11 | 118798780 | C | G | -3.009 | 0.00262 | -?--- | 0 | 0.267 | 3 | 0.966 |
| rs7117408 | 11 | 118808140 | T | C | -2.999 | 0.002712 | -?--- | 0 | 0.996 | 3 | 0.8023 |
| rs117235696 | 11 | 118803193 | G | T | -2.971 | 0.002973 | -?--- | 0 | 0.613 | 3 | 0.8934 |
| rs3176905 | 11 | 118755630 | C | T | -2.917 | 0.003539 | ----+ | 52 | 12.498 | 4 | 0.01401 |
| rs687190 | 11 | 118542383 | C | T | 2.914 | 0.003569 | +++++ | 0 | 3.481 | 4 | 0.4808 |
| rs55772432 | 11 | 118805385 | A | G | 2.91 | 0.003619 | +?++? | -29.1 | 3.098 | 2 | 0.2124 |
| rs10892307 | 11 | 118754649 | G | C | -2.909 | 0.003621 | ----+ | 55.4 | 13.452 | 4 | 0.009267 |
| rs10892310 | 11 | 118806897 | T | C | 2.893 | 0.00381 | +-+++ | 39.9 | 9.989 | 4 | 0.04062 |
| rs71482159 | 11 | 118809204 | T | C | -2.884 | 0.003922 | -?-?? | 0 | 0.012 | 1 | 0.9136 |
| rs2886599 | 11 | 72721271 | A | C | 2.849 | 0.004384 | -++++ | 24 | 7.891 | 4 | 0.09564 |
| rs150792202 | 11 | 118652569 | A | G | 2.847 | 0.004416 | +?+++ | -2.2 | 4.894 | 3 | 0.1797 |
| rs633683 | 11 | 118504742 | T | C | 2.834 | 0.004602 | +?+++ | -30.1 | 3.843 | 3 | 0.2789 |
| rs77280285 | 11 | 118091258 | G | C | 2.833 | 0.004615 | +?+++ | 1.6 | 5.082 | 3 | 0.1659 |
| rs11216934 | 11 | 118501459 | A | G | 2.823 | 0.004753 | +?+++ | 0 | 1.773 | 3 | 0.6207 |
| rs506234 | 11 | 118540104 | G | C | 2.801 | 0.005097 | +++++ | 0 | 2.962 | 4 | 0.5642 |
| rs483598 | 11 | 118509668 | A | G | 2.795 | 0.005186 | +?+++ | 0 | 1.694 | 3 | 0.6383 |
| rs527619 | 11 | 118563286 | A | G | 2.788 | 0.00531 | +++++ | 0 | 3.013 | 4 | 0.5556 |
| rs8301 | 11 | 118895495 | G | C | -2.778 | 0.005464 | -?--- | 0 | 0.833 | 3 | 0.8417 |
| rs76675432 | 11 | 118103976 | T | C | 2.77 | 0.005609 | +?+++ | 1.2 | 5.063 | 3 | 0.1672 |
| rs577396 | 11 | 118499469 | C | G | 2.766 | 0.005683 | +?+++ | 0 | 1.671 | 3 | 0.6434 |
| rs4558175 | 11 | 118872581 | A | G | -2.742 | 0.006115 | -?--- | 0 | 0.558 | 3 | 0.9059 |
| rs540800 | 11 | 118565651 | A | G | 2.741 | 0.006123 | +++++ | 0 | 3.274 | 4 | 0.5131 |
| rs523604 | 11 | 118755738 | G | A | -2.732 | 0.006299 | ----- | 0 | 2.067 | 4 | 0.7235 |
| rs11605166 | 11 | 72669777 | C | T | 2.728 | 0.006367 | -++++ | 30.2 | 8.592 | 4 | 0.07214 |
| rs3132828 | 11 | 118431003 | A | G | -2.693 | 0.007082 | ----? | 0 | 1.776 | 3 | 0.6202 |
| rs10892320 | 11 | 118873385 | T | C | -2.687 | 0.00722 | -?--- | 0 | 0.476 | 3 | 0.9242 |
| rs76793820 | 11 | 72619860 | A | G | 2.666 | 0.00767 | -++++ | 27.3 | 8.256 | 4 | 0.08264 |
| rs680465 | 11 | 118500963 | A | G | 2.657 | 0.007885 | +?-++ | 35.3 | 7.728 | 3 | 0.05197 |
| rs7936556 | 11 | 118501299 | A | G | 2.654 | 0.007957 | +?+++ | 0 | 1.542 | 3 | 0.6726 |
| rs4245184 | 11 | 118510562 | A | G | 2.65 | 0.008057 | +?+++ | 0 | 1.483 | 3 | 0.6863 |
| rs644273 | 11 | 118511352 | T | C | 2.649 | 0.008082 | +?+++ | 0 | 1.482 | 3 | 0.6865 |

| | | | | | | | | | | | |
|---|---|---|---|---|---|---|---|---|---|---|---|
| rs11820805 | 11 | 118888831 | C | T | -2.646 | 0.008138 | -?--- | 0 | 0.339 | 3 | 0.9526 |
| rs6589713 | 11 | 118872404 | C | T | -2.645 | 0.008166 | -?--- | 0 | 0.718 | 3 | 0.8689 |
| rs2508573 | 11 | 118555383 | T | C | 2.629 | 0.008552 | +++++ | 0 | 3.483 | 4 | 0.4804 |
| rs7129793 | 11 | 72455885 | T | C | 2.628 | 0.0086 | -++++ | 20.8 | 7.574 | 4 | 0.1085 |
| rs4938618 | 11 | 118867803 | C | G | -2.626 | 0.00863 | -?--- | 0 | 0.986 | 3 | 0.8048 |
| rs654423 | 11 | 118511757 | A | G | 2.625 | 0.008661 | +?+++ | 0 | 1.463 | 3 | 0.6908 |
| rs11602858 | 11 | 72615069 | G | C | 2.624 | 0.008684 | -++++ | 25.7 | 8.072 | 4 | 0.08898 |
| rs11603023 | 11 | 118486067 | T | C | 2.623 | 0.008725 | +?+++ | 0 | 1.589 | 3 | 0.662 |
| rs583213 | 11 | 118669178 | G | T | 2.619 | 0.00881 | +?-++ | 15.8 | 5.939 | 3 | 0.1146 |
| rs4573680 | 11 | 118872741 | C | T | -2.614 | 0.008955 | -?--- | 0 | 1.011 | 3 | 0.7986 |
| rs7113753 | 11 | 118889378 | A | G | -2.613 | 0.008968 | -?--- | 0 | 0.394 | 3 | 0.9414 |
| rs75896506 | 11 | 72429579 | A | G | 2.611 | 0.009033 | -++++ | -1.3 | 5.92 | 4 | 0.2052 |
| rs494560 | 11 | 118521549 | A | G | 2.601 | 0.009284 | +?+++ | 0 | 1.47 | 3 | 0.6892 |
| rs947931 | 11 | 118577614 | G | T | -2.596 | 0.009421 | ---?- | 0 | 2.673 | 3 | 0.4448 |
| rs11600585 | 11 | 72727840 | G | A | 2.595 | 0.00946 | -++++ | 12.6 | 6.867 | 4 | 0.1431 |
| rs576950 | 11 | 118391375 | G | A | -2.591 | 0.009559 | ----? | 16.9 | 6.016 | 3 | 0.1108 |
| rs11602393 | 11 | 118750967 | A | T | -2.591 | 0.009573 | ----+ | 50.9 | 12.227 | 4 | 0.01574 |
| rs11602764 | 11 | 118888619 | G | A | -2.59 | 0.009603 | -?--- | 0 | 0.286 | 3 | 0.9627 |
| rs11603334 | 11 | 72432985 | A | G | 2.584 | 0.009773 | -++++ | -1.3 | 5.923 | 4 | 0.2049 |
| rs74333814 | 11 | 72457487 | T | C | 2.582 | 0.009817 | -++++ | 17.5 | 7.277 | 4 | 0.1219 |
| rs11822103 | 11 | 118869539 | A | G | -2.578 | 0.009952 | -?--- | 0 | 0.996 | 3 | 0.8023 |
| rs10898880 | 11 | 72525111 | A | C | -2.559 | 0.01049 | ----- | 0 | 0.489 | 4 | 0.9746 |
| rs1552224 | 11 | 72433098 | C | A | 2.556 | 0.01058 | -++++ | 0.1 | 6.007 | 4 | 0.1987 |
| rs79769393 | 11 | 72080610 | A | G | 2.55 | 0.01078 | +++?+ | 0 | 2.982 | 3 | 0.3944 |
| rs35668754 | 11 | 72523110 | C | T | -2.549 | 0.0108 | ----- | 0 | 0.52 | 4 | 0.9715 |
| rs145398037 | 11 | 72522142 | C | G | 2.542 | 0.01102 | -++++ | 31.5 | 8.763 | 4 | 0.06731 |
| rs74891547 | 11 | 118811878 | C | G | -2.541 | 0.01107 | -?--- | 0 | 1.917 | 3 | 0.5898 |
| rs10751216 | 11 | 72526123 | T | C | -2.537 | 0.01117 | ----- | 0 | 0.605 | 4 | 0.9625 |
| rs35343925 | 11 | 72523349 | T | A | -2.534 | 0.01128 | ----- | 0 | 0.557 | 4 | 0.9677 |
| rs6592490 | 11 | 72519756 | C | A | -2.532 | 0.01133 | ----- | 0 | 0.567 | 4 | 0.9667 |
| rs74236665 | 11 | 72080895 | G | C | 2.53 | 0.01141 | +++?+ | -66 | 3.013 | 3 | 0.3897 |
| rs77398170 | 11 | 118792278 | G | A | -2.528 | 0.01147 | -?--- | 0 | 1.516 | 3 | 0.6786 |
| rs35752694 | 11 | 72522881 | A | G | -2.528 | 0.01149 | ----- | 0 | 0.541 | 4 | 0.9694 |
| rs525549 | 11 | 118358068 | A | T | -2.524 | 0.01161 | ----? | -12.8 | 4.431 | 3 | 0.2185 |
| rs7109082 | 11 | 72707441 | C | A | 2.524 | 0.01161 | -++++ | 8.4 | 6.55 | 4 | 0.1617 |
| rs1975109 | 11 | 72519217 | G | C | -2.523 | 0.01164 | ----- | 0 | 0.544 | 4 | 0.9691 |
| rs79577700 | 11 | 72057385 | G | A | 2.521 | 0.01169 | +++?+ | 0 | 2.981 | 3 | 0.3946 |
| rs79549196 | 11 | 72057384 | T | C | 2.521 | 0.0117 | +++?+ | 0 | 2.981 | 3 | 0.3946 |
| rs4944805 | 11 | 72522262 | G | T | -2.516 | 0.01188 | ----- | 0 | 0.549 | 4 | 0.9686 |
| rs577121 | 11 | 118690550 | C | T | 2.515 | 0.01192 | +?-++ | 27.9 | 6.938 | 3 | 0.07389 |
| rs7110719 | 11 | 72521286 | C | T | -2.514 | 0.01192 | ----- | 0 | 0.55 | 4 | 0.9684 |
| rs10751215 | 11 | 72526087 | C | T | -2.512 | 0.01199 | ----- | 0 | 0.504 | 4 | 0.9731 |
| rs75665552 | 11 | 72046487 | C | T | 2.512 | 0.012 | +++?+ | 0 | 2.984 | 3 | 0.3941 |
| rs117117171 | 11 | 118152252 | A | T | 2.511 | 0.01202 | +?+++ | 0.5 | 5.027 | 3 | 0.1698 |
| rs538645 | 11 | 118711069 | C | T | 2.508 | 0.01214 | +++++ | 0 | 1.692 | 4 | 0.7921 |
| rs79658353 | 11 | 72064041 | A | G | 2.508 | 0.01214 | +++?+ | -66.5 | 3.003 | 3 | 0.3911 |
| rs78639187 | 11 | 72071641 | T | C | 2.505 | 0.01226 | +++?+ | 0 | 1.852 | 3 | 0.6037 |
| rs117819392 | 11 | 72514503 | T | C | 2.502 | 0.01234 | -++++ | 27.4 | 8.269 | 4 | 0.08222 |
| rs139534838 | 11 | 72506880 | C | T | 2.502 | 0.01236 | -++++ | 30.9 | 8.682 | 4 | 0.06956 |
| rs587816 | 11 | 118689871 | A | C | 2.502 | 0.01237 | +?-++ | 29.4 | 7.086 | 3 | 0.0692 |
| rs9645708 | 11 | 72819318 | C | T | 2.502 | 0.01237 | -++-+ | 23.4 | 7.829 | 4 | 0.09805 |
| rs58941166 | 11 | 72083441 | A | G | 2.499 | 0.01244 | +++?- | -54.1 | 3.245 | 3 | 0.3554 |
| rs1059000 | 11 | 72504972 | C | G | 2.497 | 0.01252 | -++++ | 30 | 8.576 | 4 | 0.07263 |
| rs12282752 | 11 | 118793918 | C | T | -2.497 | 0.01252 | -?--- | 0 | 1.557 | 3 | 0.6693 |
| rs7943247 | 11 | 72881957 | G | A | -2.497 | 0.01252 | ---+- | 58.2 | 14.364 | 4 | 0.006221 |
| rs3737504 | 11 | 118869460 | G | A | -2.497 | 0.01253 | -?--- | 0 | 1.102 | 3 | 0.7765 |

| | | | | | | | | | | | |
|---|---|---|---|---|---|---|---|---|---|---|---|
| rs7103836 | 11 | 72474571 | C | G | 2.492 | 0.01272 | -++++ | 14.4 | 7.01 | 4 | 0.1353 |
| rs11235596 | 11 | 72514163 | G | A | -2.488 | 0.01286 | ----- | 0 | 0.499 | 4 | 0.9736 |
| rs4936457 | 11 | 118875727 | G | A | -2.487 | 0.0129 | -?--- | 0 | 0.833 | 3 | 0.8416 |
| rs10793057 | 11 | 72968660 | A | G | 2.484 | 0.013 | ++++? | 0 | 0.192 | 3 | 0.9789 |
| rs668628 | 11 | 118519213 | T | G | 2.484 | 0.01301 | +?-++ | 36.7 | 7.896 | 3 | 0.04822 |
| rs9645664 | 11 | 118873987 | C | T | -2.483 | 0.01301 | ----- | 0 | 1.455 | 4 | 0.8346 |
| rs583905 | 11 | 118695908 | G | A | 2.481 | 0.0131 | +?-++ | 21.9 | 6.402 | 3 | 0.09361 |
| rs4936458 | 11 | 118875761 | G | T | -2.479 | 0.01318 | -?--- | 0 | 0.815 | 3 | 0.8458 |
| rs7105187 | 11 | 118887581 | T | C | -2.479 | 0.01319 | -?--- | 0 | 0.744 | 3 | 0.8629 |
| rs7933859 | 11 | 118662045 | T | C | 2.475 | 0.01334 | +?-++ | 12.4 | 5.708 | 3 | 0.1267 |
| rs613937 | 11 | 72474839 | G | A | 2.474 | 0.01337 | -++++ | 20.3 | 7.532 | 4 | 0.1103 |
| rs111491960 | 11 | 118833458 | G | A | 2.472 | 0.01342 | +-+?+ | -7.3 | 4.661 | 3 | 0.1984 |
| rs4508234 | 11 | 118662203 | G | C | 2.466 | 0.01368 | +?-++ | 12 | 5.685 | 3 | 0.128 |
| rs494459 | 11 | 118574675 | T | C | 2.453 | 0.01416 | ++-++ | -5.2 | 5.705 | 4 | 0.2223 |
| rs543692 | 11 | 118598138 | C | T | 2.452 | 0.0142 | ++-++ | 0.7 | 6.044 | 4 | 0.1959 |
| rs500254 | 11 | 118678780 | G | A | 2.449 | 0.01431 | +?-++ | 11.7 | 5.66 | 3 | 0.1294 |
| rs3916582 | 11 | 118887888 | T | C | -2.443 | 0.01457 | -?--- | 0 | 0.887 | 3 | 0.8287 |
| rs10898877 | 11 | 72511021 | G | A | -2.442 | 0.01459 | ----- | 0 | 0.394 | 4 | 0.983 |
| rs7952617 | 11 | 72516946 | C | T | -2.441 | 0.01464 | ----- | 0 | 0.768 | 4 | 0.9426 |
| rs584935 | 11 | 118696175 | A | G | 2.44 | 0.01468 | +?-++ | 23.2 | 6.51 | 3 | 0.08926 |
| rs626645 | 11 | 118575180 | C | T | 2.44 | 0.01469 | ++-++ | -5.9 | 5.665 | 4 | 0.2256 |
| rs10751213 | 11 | 72510644 | A | C | -2.439 | 0.01473 | ----- | 0 | 0.387 | 4 | 0.9835 |
| rs524409 | 11 | 118574311 | T | C | 2.439 | 0.01473 | ++-++ | -2.8 | 5.834 | 4 | 0.2119 |
| rs6592489 | 11 | 72510180 | C | G | -2.439 | 0.01475 | ----- | 0 | 0.386 | 4 | 0.9836 |
| rs11217125 | 11 | 118889185 | C | T | -2.437 | 0.01483 | -?--- | 0 | 1 | 3 | 0.8012 |
| rs1000620 | 11 | 72509713 | A | G | -2.435 | 0.01487 | ----- | 0 | 0.38 | 4 | 0.9841 |
| rs7937334 | 11 | 118191274 | T | C | 2.435 | 0.01491 | +?++- | 20.5 | 6.288 | 3 | 0.0984 |
| rs4459332 | 11 | 72508267 | C | T | -2.434 | 0.01495 | ----- | 0 | 0.38 | 4 | 0.9841 |
| rs2276423 | 11 | 118210106 | G | C | 2.433 | 0.01496 | +?++- | 25.6 | 6.721 | 3 | 0.08134 |
| rs677231 | 11 | 72504848 | A | T | -2.433 | 0.01499 | ----- | 0 | 0.37 | 4 | 0.9848 |
| rs4944800 | 11 | 72506130 | C | T | -2.431 | 0.01504 | ----- | 0 | 0.368 | 4 | 0.985 |
| rs582200 | 11 | 118668908 | G | A | 2.43 | 0.0151 | +?-++ | 13.5 | 5.777 | 3 | 0.123 |
| rs4938544 | 11 | 118665009 | A | G | 2.429 | 0.01513 | +?-++ | 10.6 | 5.595 | 3 | 0.133 |
| rs59095933 | 11 | 118885161 | A | G | -2.427 | 0.01524 | -?--- | 0 | 0.661 | 3 | 0.8822 |
| rs78882064 | 11 | 72084480 | A | C | 2.426 | 0.01525 | +++?+ | 0 | 2.427 | 3 | 0.4886 |
| rs17244499 | 11 | 72629946 | G | A | 2.421 | 0.01546 | -++++ | 21.3 | 7.624 | 4 | 0.1064 |
| rs625735 | 11 | 118575419 | A | G | 2.421 | 0.01547 | ++-++ | -2.9 | 5.833 | 4 | 0.212 |
| rs4459331 | 11 | 72507938 | C | T | -2.419 | 0.01555 | ----- | 0 | 0.389 | 4 | 0.9834 |
| rs637563 | 11 | 118575479 | G | A | 2.417 | 0.01563 | ++-++ | -2.3 | 5.867 | 4 | 0.2093 |
| rs638514 | 11 | 118575693 | C | T | 2.417 | 0.01564 | ++-++ | -2.2 | 5.87 | 4 | 0.2091 |
| rs2276071 | 11 | 118896116 | C | A | -2.416 | 0.01567 | ----- | 0 | 1.385 | 4 | 0.8468 |
| rs12271856 | 11 | 118791532 | T | G | -2.416 | 0.01569 | -?--- | 0 | 1.92 | 3 | 0.5892 |
| rs4938506 | 11 | 118205702 | A | G | 2.415 | 0.01571 | +?++- | 23 | 6.497 | 3 | 0.0898 |
| rs530901 | 11 | 118575011 | C | T | 2.415 | 0.01575 | ++-++ | -0.1 | 5.993 | 4 | 0.1997 |
| rs113611963 | 11 | 118839610 | G | T | 2.411 | 0.01591 | +-+?+ | -11 | 4.505 | 3 | 0.2118 |
| rs568688 | 11 | 118598612 | G | T | 2.41 | 0.01596 | ++-++ | -2.7 | 5.84 | 4 | 0.2114 |
| rs11217124 | 11 | 118889132 | G | A | -2.409 | 0.01598 | -?--- | 0 | 0.94 | 3 | 0.8157 |
| rs576283 | 11 | 118573894 | G | A | 2.408 | 0.01602 | ++-++ | -1.4 | 5.918 | 4 | 0.2053 |
| rs10431171 | 11 | 72512271 | C | G | -2.406 | 0.01612 | ----- | 0 | 0.479 | 4 | 0.9755 |
| rs627391 | 11 | 118573916 | T | A | 2.401 | 0.01636 | ++-++ | -1.8 | 5.892 | 4 | 0.2073 |
| rs10431172 | 11 | 72512630 | T | C | -2.399 | 0.01645 | ----- | 0 | 0.484 | 4 | 0.9751 |
| rs4944802 | 11 | 72510476 | G | A | -2.398 | 0.0165 | ----- | 0 | 0.452 | 4 | 0.978 |
| rs58601794 | 11 | 118884726 | C | T | -2.397 | 0.01654 | -?--- | 0 | 0.727 | 3 | 0.8668 |
| rs603486 | 11 | 118587347 | T | C | 2.394 | 0.01669 | +++?+ | 0 | 1.661 | 3 | 0.6457 |
| rs3814730 | 11 | 72555553 | A | G | 2.392 | 0.01674 | -++++ | 25.4 | 8.039 | 4 | 0.09016 |
| rs75610917 | 11 | 72011276 | A | G | 2.392 | 0.01676 | +++?+ | 0 | 2.225 | 3 | 0.527 |

| | | | | | | | | | | | |
|---|---|---|---|---|---|---|---|---|---|---|---|
| rs2303969 | 11 | 72536272 | A | G | 2.391 | 0.01681 | -++-+ | 28.3 | 8.366 | 4 | 0.07904 |
| rs75219483 | 11 | 72008410 | T | C | 2.391 | 0.01681 | +++?+ | 0 | 2.224 | 3 | 0.5273 |
| rs3862791 | 11 | 72501002 | T | C | 2.389 | 0.01691 | -++++ | 25.4 | 8.048 | 4 | 0.08983 |
| rs5006102 | 11 | 118662340 | A | T | 2.386 | 0.01704 | +?-++ | 15.1 | 5.887 | 3 | 0.1172 |
| rs563537 | 11 | 118595207 | A | T | 2.384 | 0.01712 | ++-?+ | 8.4 | 5.457 | 3 | 0.1412 |
| rs11235592 | 11 | 72501277 | C | G | -2.383 | 0.01718 | ----- | 0 | 0.327 | 4 | 0.988 |
| rs516503 | 11 | 118580348 | T | C | 2.382 | 0.01722 | ++-++ | -8.7 | 5.521 | 4 | 0.2379 |
| rs6592486 | 11 | 72508609 | A | G | -2.381 | 0.01726 | ----- | 0 | 0.396 | 4 | 0.9828 |
| rs2276072 | 11 | 118896115 | G | A | -2.38 | 0.01729 | ----- | 0 | 1.326 | 4 | 0.8569 |
| rs59608621 | 11 | 118885163 | G | A | -2.379 | 0.01735 | -?--- | 0 | 0.784 | 3 | 0.8533 |
| rs675003 | 11 | 118677083 | T | C | 2.378 | 0.01738 | +?-++ | 11.7 | 5.661 | 3 | 0.1293 |
| rs79430446 | 11 | 72474004 | T | G | 2.378 | 0.01739 | -++++ | 9.8 | 6.649 | 4 | 0.1556 |
| rs11235594 | 11 | 72503735 | G | A | -2.378 | 0.0174 | ----- | 0 | 0.353 | 4 | 0.9861 |
| rs61381062 | 11 | 118884708 | A | G | -2.37 | 0.01777 | -?--- | 0 | 0.74 | 3 | 0.8637 |
| rs12294466 | 11 | 118789657 | T | C | -2.37 | 0.01779 | -+--? | -28.4 | 3.893 | 3 | 0.2732 |
| rs1864653 | 11 | 72503293 | A | G | -2.369 | 0.01781 | ----- | 0 | 0.376 | 4 | 0.9844 |
| rs496058 | 11 | 118580906 | G | A | 2.363 | 0.01812 | ++-++ | -10.6 | 5.426 | 4 | 0.2463 |
| rs61901411 | 11 | 118665324 | T | C | 2.363 | 0.01813 | +?-++ | 18.9 | 6.164 | 3 | 0.1039 |
| rs11216919 | 11 | 118462367 | G | A | 2.353 | 0.01862 | ++++? | 0 | 0.788 | 3 | 0.8524 |
| rs57275811 | 11 | 72127872 | G | T | 2.352 | 0.01869 | +++?+ | 0 | 2.55 | 3 | 0.4664 |
| rs11235612 | 11 | 72613273 | G | A | 2.35 | 0.01876 | -++-+ | 37.3 | 9.573 | 4 | 0.04827 |
| rs12270280 | 11 | 118791150 | T | C | -2.349 | 0.01883 | -?--- | 0 | 2.141 | 3 | 0.5437 |
| rs7109575 | 11 | 72463435 | A | G | 2.34 | 0.0193 | -++++ | 28 | 8.328 | 4 | 0.08026 |
| rs74439784 | 11 | 72649396 | T | A | 2.339 | 0.01931 | -++++ | 21 | 7.593 | 4 | 0.1077 |
| rs75184960 | 11 | 72141222 | A | C | 2.338 | 0.01938 | +++?+ | 0 | 2.494 | 3 | 0.4765 |
| rs11235550 | 11 | 72331050 | C | T | 2.337 | 0.01942 | +?+++ | 0 | 1.541 | 3 | 0.6729 |
| rs485099 | 11 | 72500060 | C | T | -2.334 | 0.01957 | ----- | 0 | 0.322 | 4 | 0.9884 |
| rs4944797 | 11 | 72500072 | T | G | -2.329 | 0.01987 | ----- | 0 | 0.341 | 4 | 0.987 |
| rs12274090 | 11 | 72512097 | T | C | -2.325 | 0.02006 | +--+- | 23.4 | 7.834 | 4 | 0.09783 |
| rs566848 | 11 | 118711830 | G | A | 2.325 | 0.02009 | +++++ | 0 | 0.972 | 4 | 0.914 |
| rs79000353 | 11 | 118420343 | A | G | -2.325 | 0.02009 | ---+? | 7.7 | 5.417 | 3 | 0.1437 |
| rs498872 | 11 | 118477367 | A | G | -2.321 | 0.02031 | ----- | 0 | 2.553 | 4 | 0.6351 |
| rs11603313 | 11 | 72588177 | A | G | 2.32 | 0.02035 | -++++ | 21.8 | 7.67 | 4 | 0.1044 |
| rs7127427 | 11 | 118805138 | A | G | -2.319 | 0.02041 | -?--- | 0 | 0.694 | 3 | 0.8745 |
| rs73016221 | 11 | 118220027 | C | T | -2.318 | 0.02046 | -?-?- | -21.1 | 3.302 | 2 | 0.1918 |
| rs7930148 | 11 | 118435966 | A | G | 2.316 | 0.02058 | ++++? | 0 | 0.635 | 3 | 0.8883 |
| rs113573893 | 11 | 72078446 | G | T | 2.31 | 0.02086 | +++?- | -62.8 | 3.071 | 3 | 0.3808 |
| rs60712691 | 11 | 72077961 | T | G | 2.31 | 0.02086 | +++?- | -62.8 | 3.071 | 3 | 0.3808 |
| rs4944796 | 11 | 72499000 | C | A | -2.31 | 0.0209 | ----- | 0 | 0.317 | 4 | 0.9887 |
| rs570424 | 11 | 118588998 | C | T | 2.309 | 0.02094 | ++-?+ | 12.5 | 5.716 | 3 | 0.1263 |
| rs638893 | 11 | 118698537 | G | A | 2.305 | 0.02116 | ++-++ | 1.3 | 6.08 | 4 | 0.1933 |
| rs3887652 | 11 | 118856582 | C | G | -2.301 | 0.02142 | -?--- | 0 | 1.176 | 3 | 0.7587 |
| rs639885 | 11 | 118698294 | A | G | 2.3 | 0.02146 | ++-++ | 1.8 | 6.112 | 4 | 0.191 |
| rs640373 | 11 | 118698160 | G | A | 2.3 | 0.02147 | ++-++ | 1.9 | 6.119 | 4 | 0.1904 |
| rs11603349 | 11 | 72460694 | C | T | 2.296 | 0.02166 | -++++ | 25.7 | 8.071 | 4 | 0.08901 |
| rs587263 | 11 | 118816156 | C | T | 2.296 | 0.02167 | +++?+ | 0 | 2.864 | 3 | 0.4131 |
| rs10128600 | 11 | 118700425 | C | T | 2.296 | 0.0217 | +?-++ | 18.7 | 6.147 | 3 | 0.1047 |
| rs639915 | 11 | 118698274 | A | G | 2.293 | 0.02182 | ++-++ | 2.5 | 6.151 | 4 | 0.1881 |
| rs4938610 | 11 | 118857466 | C | T | -2.293 | 0.02183 | -?-?- | 0 | 1.203 | 2 | 0.548 |
| rs3889526 | 11 | 118856491 | A | C | -2.293 | 0.02184 | -?-?- | 0 | 1.179 | 2 | 0.5545 |
| rs2510901 | 11 | 118699863 | A | G | 2.292 | 0.02188 | ++-++ | 4.8 | 6.302 | 4 | 0.1777 |
| rs75210328 | 11 | 118801296 | T | G | -2.285 | 0.02233 | -?--- | 0 | 0.706 | 3 | 0.8718 |
| rs4938614 | 11 | 118863232 | A | G | -2.284 | 0.02237 | -?-?- | 0 | 1.189 | 2 | 0.5518 |
| rs516332 | 11 | 118698777 | G | A | 2.282 | 0.02249 | ++-++ | 3.3 | 6.203 | 4 | 0.1845 |
| rs2276424 | 11 | 118209960 | A | C | 2.278 | 0.02274 | +?++- | -14.3 | 4.374 | 3 | 0.2238 |
| rs11217117 | 11 | 118865992 | T | C | -2.275 | 0.02288 | -?-?- | 0 | 1.235 | 2 | 0.5393 |

| | | | | | | | | | | | |
|---|---|---|---|---|---|---|---|---|---|---|---|
| rs77464186 | 11 | 72460398 | C | A | 2.274 | 0.02295 | -++++ | 21.5 | 7.645 | 4 | 0.1055 |
| rs7944225 | 11 | 72496672 | C | G | -2.274 | 0.02297 | +--+- | 25.6 | 8.069 | 4 | 0.08907 |
| rs2508564 | 11 | 118699744 | C | T | 2.268 | 0.02335 | ++-++ | 3.5 | 6.219 | 4 | 0.1834 |
| rs10431173 | 11 | 72512937 | G | A | -2.268 | 0.02336 | +--+- | 21.6 | 7.649 | 4 | 0.1053 |
| rs569770 | 11 | 118584498 | T | C | 2.266 | 0.02347 | +++++ | 0 | 2.062 | 4 | 0.7243 |
| rs10892246 | 11 | 118481431 | A | G | -2.263 | 0.02361 | ----+ | -7.3 | 5.589 | 4 | 0.232 |
| rs10160454 | 11 | 72496345 | C | A | -2.261 | 0.02377 | +--+- | 25.6 | 8.066 | 4 | 0.08919 |
| rs3212262 | 11 | 118217854 | T | G | -2.261 | 0.02378 | -?-?- | -22.1 | 3.276 | 2 | 0.1944 |
| rs10898873 | 11 | 72497713 | C | T | -2.259 | 0.02387 | +--+- | 25.1 | 8.015 | 4 | 0.09103 |
| rs57300701 | 11 | 118845546 | G | C | 2.257 | 0.024 | +-+?+ | -36.4 | 3.665 | 3 | 0.3 |
| rs7947423 | 11 | 72497001 | A | G | -2.256 | 0.02406 | +--+- | 25 | 8.003 | 4 | 0.09148 |
| rs10736492 | 11 | 118459269 | G | A | 2.256 | 0.02407 | ++++? | 0 | 0.518 | 3 | 0.9149 |
| rs11602873 | 11 | 72460762 | T | A | 2.256 | 0.02407 | -++++ | 21 | 7.596 | 4 | 0.1076 |
| rs7110034 | 11 | 118858911 | A | C | -2.255 | 0.02415 | -?-?- | 0 | 1.34 | 2 | 0.5117 |
| rs4938612 | 11 | 118863037 | C | T | -2.253 | 0.02429 | -?-?- | 0 | 1.154 | 2 | 0.5617 |
| rs12280288 | 11 | 118892940 | C | T | -2.252 | 0.02435 | -?--- | 0 | 0.608 | 3 | 0.8946 |
| rs11216843 | 11 | 118151329 | T | A | -2.251 | 0.02441 | -+-?- | 32.9 | 7.455 | 3 | 0.05872 |
| rs10898874 | 11 | 72497795 | G | A | -2.247 | 0.02463 | +--+- | 25.5 | 8.054 | 4 | 0.08964 |
| rs6589659 | 11 | 118191691 | A | C | 2.246 | 0.02472 | +?++- | -21.1 | 4.13 | 3 | 0.2478 |
| rs10898875 | 11 | 72497807 | C | T | -2.242 | 0.02494 | +--+- | 25.2 | 8.023 | 4 | 0.09073 |
| rs77293887 | 11 | 72118937 | T | C | 2.242 | 0.02499 | +++?+ | 0 | 2.226 | 3 | 0.5269 |
| rs7121919 | 11 | 72496087 | A | G | -2.236 | 0.02537 | +--+- | 24.3 | 7.93 | 4 | 0.09419 |
| rs2231449 | 11 | 118186305 | A | C | -2.227 | 0.02594 | -+-?- | 38.2 | 8.092 | 3 | 0.04414 |
| rs479490 | 11 | 72497321 | C | T | -2.226 | 0.02604 | +--+- | 25.8 | 8.085 | 4 | 0.0885 |
| rs4290240 | 11 | 118853488 | G | A | -2.225 | 0.02606 | -?-?- | 0 | 1.374 | 2 | 0.5032 |
| rs4938558 | 11 | 118717929 | T | C | 2.225 | 0.02609 | +++++ | 0 | 1.076 | 4 | 0.8981 |
| rs480958 | 11 | 118577990 | G | A | 2.223 | 0.02624 | ++++- | 0 | 2.633 | 4 | 0.621 |
| rs4938566 | 11 | 118721189 | T | A | 2.219 | 0.0265 | +++++ | 0 | 1.021 | 4 | 0.9065 |
| rs12418637 | 11 | 72061218 | A | G | 2.214 | 0.02682 | ++++- | -1.3 | 5.922 | 4 | 0.2051 |
| rs2276419 | 11 | 118230359 | T | C | -2.212 | 0.02698 | -?-?- | -25.6 | 3.186 | 2 | 0.2033 |
| rs34782343 | 11 | 118153511 | G | T | 2.211 | 0.02702 | +?++- | 48.5 | 9.714 | 3 | 0.02116 |
| rs4938567 | 11 | 118721249 | G | A | 2.211 | 0.02705 | +++++ | 0 | 1.052 | 4 | 0.9017 |
| rs57156480 | 11 | 118896959 | A | G | -2.209 | 0.0272 | ----- | 0 | 0.824 | 4 | 0.9353 |
| rs10898895 | 11 | 72741398 | A | T | 2.208 | 0.02727 | -++-+ | 16.9 | 7.221 | 4 | 0.1246 |
| rs4938570 | 11 | 118722231 | G | A | 2.207 | 0.02734 | +++++ | 0 | 1.047 | 4 | 0.9026 |
| rs7110837 | 11 | 118199295 | A | G | 2.206 | 0.02741 | +?++- | -20 | 4.167 | 3 | 0.244 |
| rs11216855 | 11 | 118208750 | C | T | -2.205 | 0.02746 | -?--- | 0 | 1.656 | 3 | 0.6469 |
| rs4938565 | 11 | 118720998 | A | G | 2.204 | 0.02751 | +++++ | 0 | 1.012 | 4 | 0.9079 |
| rs904469 | 11 | 72324039 | A | G | 2.203 | 0.02757 | -?+++ | 0 | 2.336 | 3 | 0.5057 |
| rs11216853 | 11 | 118200201 | T | A | 2.202 | 0.02763 | +?++- | -21.2 | 4.125 | 3 | 0.2483 |
| rs11217059 | 11 | 118723087 | T | C | 2.201 | 0.02774 | +++++ | 0 | 1.058 | 4 | 0.9009 |
| rs61036532 | 11 | 72017321 | G | A | 2.2 | 0.02783 | ++++- | 5.6 | 6.355 | 4 | 0.1741 |
| rs4445645 | 11 | 118868750 | C | G | -2.199 | 0.02784 | -?--- | 0 | 1.891 | 3 | 0.5954 |
| rs4936440 | 11 | 118720684 | G | C | 2.196 | 0.02812 | +++++ | 0 | 1.025 | 4 | 0.906 |
| rs12223292 | 11 | 72487259 | A | G | -2.195 | 0.02816 | +--+- | 49.3 | 11.836 | 4 | 0.01861 |
| rs7946961 | 11 | 118721850 | G | A | 2.195 | 0.02819 | +++++ | 0 | 1.094 | 4 | 0.8951 |
| rs4938606 | 11 | 118852081 | G | A | -2.192 | 0.02834 | -?-?- | 0 | 1.248 | 2 | 0.5358 |
| rs12421674 | 11 | 72056591 | T | C | 2.19 | 0.02853 | ++++- | 10.3 | 6.689 | 4 | 0.1532 |
| rs755933 | 11 | 72325482 | A | G | 2.188 | 0.02866 | -?+++ | 0 | 2.355 | 3 | 0.5021 |
| rs56076328 | 11 | 72023279 | A | G | 2.186 | 0.02881 | ++++- | 6.3 | 6.401 | 4 | 0.1711 |
| rs58945323 | 11 | 72042223 | C | T | 2.184 | 0.02899 | ++++- | 5.6 | 6.359 | 4 | 0.1739 |
| rs11216848 | 11 | 118181481 | A | C | -2.183 | 0.02903 | -+-?- | 32.7 | 7.426 | 3 | 0.0595 |
| rs7933007 | 11 | 118730669 | C | T | 2.18 | 0.02928 | +++++ | 0 | 1.31 | 4 | 0.8596 |
| rs4938564 | 11 | 118720590 | T | G | 2.178 | 0.02943 | +++++ | 0 | 1.003 | 4 | 0.9093 |
| rs77106637 | 11 | 72478320 | A | G | 2.175 | 0.02966 | -++++ | -5.3 | 5.701 | 4 | 0.2226 |
| rs2187491 | 11 | 118718972 | C | T | 2.174 | 0.02967 | +++++ | 0 | 0.948 | 4 | 0.9176 |

| | | | | | | | | | | | |
|---|---|---|---|---|---|---|---|---|---|---|---|
| rs2156751 | 11 | 118718856 | G | A | 2.17 | 0.03003 | +++++ | 0 | 0.966 | 4 | 0.9149 |
| rs3781934 | 11 | 72325070 | A | G | 2.166 | 0.03031 | -?+++ | 0 | 2.351 | 3 | 0.5028 |
| rs11217122 | 11 | 118880740 | C | T | -2.165 | 0.03038 | -?--- | 0 | 0.805 | 3 | 0.8484 |
| rs3750906 | 11 | 72012682 | G | A | 2.164 | 0.03048 | ++++- | 8.2 | 6.533 | 4 | 0.1627 |
| rs2156749 | 11 | 118719405 | G | C | 2.163 | 0.03055 | +++++ | 0 | 0.968 | 4 | 0.9146 |
| rs9794909 | 11 | 118862306 | C | G | -2.161 | 0.03067 | ---?- | 0 | 1.605 | 3 | 0.6582 |
| rs17162126 | 11 | 72057178 | T | A | 2.161 | 0.03068 | ++++- | 8.2 | 6.537 | 4 | 0.1625 |
| rs73005500 | 11 | 118697375 | C | T | 2.158 | 0.03091 | +?-++ | 24.7 | 6.642 | 3 | 0.08422 |
| rs2156750 | 11 | 118719254 | C | T | 2.157 | 0.03101 | +++++ | 0 | 0.991 | 4 | 0.9112 |
| rs73530724 | 11 | 72049158 | G | A | 2.15 | 0.03158 | ++++- | 5.1 | 6.323 | 4 | 0.1763 |
| rs7116715 | 11 | 118719742 | C | T | 2.149 | 0.03165 | +++++ | 0 | 0.981 | 4 | 0.9126 |
| rs56171963 | 11 | 72009455 | G | A | 2.147 | 0.03178 | ++++- | 7.6 | 6.495 | 4 | 0.1651 |
| rs57879605 | 11 | 118798757 | T | C | -2.147 | 0.0318 | ----+ | -13.8 | 5.274 | 4 | 0.2604 |
| rs12416804 | 11 | 72048526 | G | T | 2.146 | 0.03189 | ++++- | 4.4 | 6.277 | 4 | 0.1794 |
| rs7103314 | 11 | 118723537 | G | A | 2.139 | 0.03242 | +++++ | 0 | 0.991 | 4 | 0.9111 |
| rs58044536 | 11 | 72011227 | G | A | 2.138 | 0.03254 | ++++- | 5.5 | 6.349 | 4 | 0.1745 |
| rs3889234 | 11 | 72490986 | C | T | -2.137 | 0.03257 | +--+- | 51.5 | 12.383 | 4 | 0.01472 |
| rs73530718 | 11 | 72045813 | C | T | 2.135 | 0.03275 | ++++- | 4.4 | 6.278 | 4 | 0.1793 |
| rs11217062 | 11 | 118725420 | A | G | 2.134 | 0.03284 | +++++ | 0 | 1.117 | 4 | 0.8916 |
| rs504217 | 11 | 72006086 | T | C | 2.132 | 0.03299 | ++++- | -4.2 | 5.758 | 4 | 0.2179 |
| rs11217118 | 11 | 118866072 | A | G | -2.131 | 0.03311 | ---?- | 0 | 1.761 | 3 | 0.6235 |
| rs7928228 | 11 | 118703001 | T | C | 2.127 | 0.03344 | ++-++ | -13.6 | 5.283 | 4 | 0.2595 |
| rs4936454 | 11 | 118859274 | T | C | -2.127 | 0.03345 | ---?- | 0 | 1.732 | 3 | 0.6298 |
| rs11217112 | 11 | 118856920 | G | A | -2.124 | 0.03364 | ---?- | 0 | 1.711 | 3 | 0.6345 |
| rs11217116 | 11 | 118864140 | C | T | -2.124 | 0.03367 | ---?- | 0 | 1.693 | 3 | 0.6385 |
| rs1170396 | 11 | 118703817 | T | A | 2.122 | 0.0338 | ++-++ | -13.1 | 5.306 | 4 | 0.2573 |
| rs11216846 | 11 | 118163916 | A | G | -2.121 | 0.0339 | -+-?- | 31.6 | 7.31 | 3 | 0.06265 |
| rs77058558 | 11 | 72065102 | A | G | 2.121 | 0.03392 | ++++- | -19.5 | 5.023 | 4 | 0.285 |
| rs73528886 | 11 | 72013848 | A | G | 2.121 | 0.03393 | ++++- | -20.3 | 4.987 | 4 | 0.2886 |
| rs7941696 | 11 | 118702936 | G | A | 2.121 | 0.03394 | ++-++ | -13.3 | 5.295 | 4 | 0.2583 |
| rs680660 | 11 | 72004714 | C | T | 2.12 | 0.034 | ++++- | -4.1 | 5.766 | 4 | 0.2173 |
| rs75067211 | 11 | 118854372 | T | C | -2.12 | 0.03401 | -?--- | 0 | 0.864 | 3 | 0.834 |
| rs2508921 | 11 | 118703484 | T | C | 2.119 | 0.03413 | ++-++ | -12.6 | 5.327 | 4 | 0.2553 |
| rs59626457 | 11 | 118862634 | G | A | -2.118 | 0.0342 | ---?- | 0 | 1.684 | 3 | 0.6406 |
| rs10892298 | 11 | 118724336 | C | T | 2.114 | 0.03451 | +++++ | 0 | 0.957 | 4 | 0.9162 |
| rs534114 | 11 | 118702002 | A | G | 2.113 | 0.03462 | ++-++ | -16 | 5.17 | 4 | 0.2702 |
| rs4936455 | 11 | 118861389 | T | C | -2.112 | 0.03467 | ---?- | 0 | 1.771 | 3 | 0.6213 |
| rs59316586 | 11 | 118856816 | C | A | -2.112 | 0.03467 | ---?- | 0 | 1.769 | 3 | 0.6217 |
| rs622690 | 11 | 118704265 | T | C | 2.109 | 0.03495 | ++-++ | -11.9 | 5.36 | 4 | 0.2523 |
| rs474629 | 11 | 72006977 | C | G | 2.108 | 0.03501 | ++++- | -4.1 | 5.762 | 4 | 0.2176 |
| rs7941321 | 11 | 118702910 | A | G | 2.107 | 0.03515 | ++-++ | -19.9 | 5.003 | 4 | 0.287 |
| rs7126193 | 11 | 118866126 | C | T | -2.106 | 0.03519 | ---?- | 0 | 1.779 | 3 | 0.6196 |
| rs3781931 | 11 | 72327205 | T | C | 2.105 | 0.03528 | -?+++ | 0 | 2.196 | 3 | 0.5327 |
| rs4938613 | 11 | 118863116 | G | A | -2.104 | 0.03542 | ---?- | 0 | 1.638 | 3 | 0.6507 |
| rs12362865 | 11 | 118859842 | G | A | -2.103 | 0.03549 | ---?- | 0 | 1.696 | 3 | 0.6377 |
| rs2156755 | 11 | 118726230 | A | G | 2.102 | 0.03553 | +++++ | 0 | 1.036 | 4 | 0.9043 |
| rs4938561 | 11 | 118718485 | C | T | 2.102 | 0.03554 | +++++ | 0 | 0.977 | 4 | 0.9132 |
| rs480791 | 11 | 118704255 | G | A | 2.102 | 0.03556 | ++-++ | -11.9 | 5.362 | 4 | 0.2521 |
| rs4936456 | 11 | 118863324 | T | C | -2.1 | 0.03571 | ---?- | 0 | 1.646 | 3 | 0.6491 |
| rs55721826 | 11 | 72019162 | G | A | 2.1 | 0.03575 | ++++- | 8 | 6.523 | 4 | 0.1634 |
| rs3194726 | 11 | 118279936 | C | T | -2.097 | 0.03597 | -?-?- | 0 | 1.595 | 2 | 0.4505 |
| rs7940062 | 11 | 72037226 | C | A | 2.091 | 0.03652 | ++++- | -0.2 | 5.991 | 4 | 0.1998 |
| rs533254 | 11 | 118701898 | A | T | 2.087 | 0.03688 | +?-++ | 4.5 | 5.235 | 3 | 0.1554 |
| rs4938549 | 11 | 118701597 | T | C | 2.085 | 0.03703 | ++-++ | -12.9 | 5.314 | 4 | 0.2565 |
| rs530646 | 11 | 118701668 | A | G | 2.084 | 0.03714 | ++-++ | -12.8 | 5.321 | 4 | 0.2559 |
| rs140026975 | 11 | 118594669 | C | A | 2.077 | 0.03776 | +++?- | 26.1 | 6.767 | 3 | 0.07972 |

| | | | | | | | | | | | |
|---|---|---|---|---|---|---|---|---|---|---|---|
| rs4938616 | 11 | 118865465 | T | C | -2.069 | 0.03852 | -?-?- | 0 | 1.283 | 2 | 0.5264 |
| rs17540708 | 11 | 118150127 | G | T | 2.065 | 0.03888 | +?++- | 39.1 | 8.208 | 3 | 0.04191 |
| rs7131196 | 11 | 118265227 | C | T | -2.065 | 0.0389 | -?-?- | 0 | 1.467 | 2 | 0.4803 |
| rs58709307 | 11 | 118877563 | T | C | -2.053 | 0.04011 | -?--- | 0 | 1.33 | 3 | 0.722 |
| rs75017819 | 11 | 118285789 | G | A | -2.051 | 0.04031 | -?-?- | 0 | 1.42 | 2 | 0.4916 |
| rs74783030 | 11 | 118603615 | A | G | -2.05 | 0.04035 | --++? | 23.2 | 6.511 | 3 | 0.08925 |
| rs4936428 | 11 | 118187511 | T | C | 2.041 | 0.04127 | +?+-+ | 16.7 | 6.004 | 3 | 0.1114 |
| rs79092677 | 11 | 118265734 | G | T | -2.038 | 0.04156 | -?-?- | 0 | 1.49 | 2 | 0.4748 |
| rs59755180 | 11 | 72073763 | C | G | 2.037 | 0.04165 | +++?+ | -22.3 | 4.088 | 3 | 0.2522 |
| rs4944563 | 11 | 72316881 | C | A | 2.037 | 0.04166 | -?+++ | 0 | 2.542 | 3 | 0.4677 |
| rs649338 | 11 | 118705654 | A | G | 2.036 | 0.04175 | ++-++ | -8.4 | 5.533 | 4 | 0.2368 |
| rs55861597 | 11 | 72331873 | T | G | 2.036 | 0.04178 | -?+++ | 0 | 2.532 | 3 | 0.4695 |
| rs55738331 | 11 | 72331821 | T | C | 2.034 | 0.04194 | -?+++ | 0 | 2.529 | 3 | 0.47 |
| rs7484280 | 11 | 72167386 | A | C | 2.033 | 0.04204 | +++?- | -23.8 | 4.038 | 3 | 0.2574 |
| rs73532667 | 11 | 72162726 | C | T | 2.033 | 0.04205 | +++?- | -40.9 | 3.549 | 3 | 0.3145 |
| rs55756957 | 11 | 118747576 | G | A | -2.031 | 0.04225 | -?--+ | 41.1 | 8.495 | 3 | 0.03682 |
| rs10892237 | 11 | 118267763 | C | T | -2.029 | 0.0425 | -?-?- | 0 | 1.514 | 2 | 0.469 |
| rs12294037 | 11 | 72807631 | T | A | -2.029 | 0.04251 | ---+- | -24.9 | 4.805 | 4 | 0.3079 |
| rs11216864 | 11 | 118260017 | T | C | -2.024 | 0.043 | -?-?- | 0 | 1.369 | 2 | 0.5044 |
| rs11216863 | 11 | 118257081 | A | G | -2.02 | 0.04333 | -?-?- | 0 | 1.161 | 2 | 0.5596 |
| rs2276422 | 11 | 118263388 | G | C | -2.017 | 0.04368 | -?-?- | 0 | 1.554 | 2 | 0.4597 |
| rs6592510 | 11 | 72788823 | C | T | -2.013 | 0.04416 | +--+- | -26.1 | 4.76 | 4 | 0.3129 |
| rs74766959 | 11 | 72065209 | G | A | 2.013 | 0.04416 | +++?- | -4.3 | 4.793 | 3 | 0.1876 |
| rs61730467 | 11 | 118772160 | A | G | -2.008 | 0.04465 | -?--? | 0 | 0.651 | 2 | 0.7221 |
| rs59776769 | 11 | 72296353 | A | G | 2.003 | 0.04515 | -?+++ | 0 | 1.647 | 3 | 0.6489 |
| rs56148078 | 11 | 72529358 | A | C | -2.002 | 0.04529 | +--++ | 52.5 | 12.63 | 4 | 0.01323 |
| rs11606357 | 11 | 118878222 | C | T | -2.002 | 0.04534 | -?--- | 0 | 1.292 | 3 | 0.731 |
| rs111322300 | 11 | 118893878 | C | T | -1.999 | 0.04559 | ----- | 0 | 2.196 | 4 | 0.6997 |
| rs56210510 | 11 | 72333342 | C | T | 1.996 | 0.04592 | -?+++ | 0 | 2.692 | 3 | 0.4416 |
| rs1783584 | 11 | 72889137 | C | T | -1.993 | 0.04623 | ----- | 0 | 1.036 | 4 | 0.9043 |
| rs4394861 | 11 | 72800970 | C | T | -1.993 | 0.04624 | +---- | 0 | 3.885 | 4 | 0.4218 |
| rs3176912 | 11 | 118757788 | A | G | -1.988 | 0.04676 | -?--? | 0 | 1.374 | 2 | 0.503 |
| rs4944566 | 11 | 72321872 | A | G | 1.981 | 0.04755 | -?+++ | 0 | 2.343 | 3 | 0.5043 |
| rs11607569 | 11 | 118880668 | C | T | -1.969 | 0.04895 | -?--- | 0 | 1.479 | 3 | 0.6871 |
| rs34022886 | 11 | 118155804 | A | G | 1.969 | 0.049 | +?++- | 38 | 8.066 | 3 | 0.04467 |
| rs113641235 | 11 | 118700264 | T | C | 1.968 | 0.04906 | +--++ | 23.6 | 7.85 | 4 | 0.09724 |
| rs10111 | 11 | 118886117 | T | C | -1.967 | 0.04924 | -?--- | 0 | 1.158 | 3 | 0.7631 |
| rs68097885 | 11 | 118596444 | A | G | 1.966 | 0.04926 | ++++- | 20.4 | 7.534 | 4 | 0.1102 |
| rs4515995 | 11 | 72687795 | G | A | -1.963 | 0.04965 | ----+ | -13.1 | 5.304 | 4 | 0.2575 |
| rs10898881 | 11 | 72527018 | A | G | 1.958 | 0.05022 | ++++- | 0 | 2.16 | 4 | 0.7063 |
| rs636736 | 11 | 118705145 | G | A | 1.958 | 0.05027 | ++-++ | -16.3 | 5.159 | 4 | 0.2714 |
| rs12798115 | 11 | 118156517 | C | G | 1.957 | 0.05034 | +?++- | 37.9 | 8.057 | 3 | 0.04484 |
| rs35235858 | 11 | 118157250 | T | C | 1.954 | 0.05067 | +?++- | 38.3 | 8.1 | 3 | 0.04398 |
| rs1783937 | 11 | 118609736 | A | G | 1.949 | 0.05124 | +?-++ | -14.1 | 4.383 | 3 | 0.223 |
| rs34250455 | 11 | 118157681 | T | C | 1.948 | 0.0514 | +?++- | 38.7 | 8.156 | 3 | 0.0429 |
| rs1513780 | 11 | 72320500 | T | G | 1.942 | 0.0521 | -?+++ | 0 | 2.367 | 3 | 0.4999 |
| rs7943039 | 11 | 118603030 | A | G | 1.938 | 0.05259 | ++++- | 23.5 | 7.84 | 4 | 0.09763 |
| rs12274926 | 11 | 118295117 | C | T | 1.937 | 0.05278 | +++++ | 0 | 0.802 | 4 | 0.9381 |
| rs143660533 | 11 | 118892759 | C | A | -1.936 | 0.05282 | ----- | 0 | 2.144 | 4 | 0.7093 |
| rs11825284 | 11 | 118880012 | A | G | -1.935 | 0.05304 | -?--- | 0 | 1.695 | 3 | 0.638 |
| rs117289748 | 11 | 118892757 | G | C | -1.934 | 0.05308 | ----- | 0 | 2.142 | 4 | 0.7097 |
| rs484232 | 11 | 118708564 | T | G | 1.929 | 0.05371 | -++++ | 0 | 1.896 | 4 | 0.7548 |
| rs7131534 | 11 | 118886656 | A | G | -1.925 | 0.05426 | -?--- | 0 | 1.748 | 3 | 0.6264 |
| rs7479373 | 11 | 118171285 | C | T | 1.924 | 0.05431 | +?++- | 37.2 | 7.965 | 3 | 0.04673 |
| rs73559119 | 11 | 118879824 | C | T | -1.924 | 0.05438 | -?--- | 0 | 1.664 | 3 | 0.6449 |
| rs7120445 | 11 | 118879285 | C | A | -1.923 | 0.05445 | -?--- | 0 | 1.656 | 3 | 0.6467 |

| | | | | | | | | | | | |
|---|---|---|---|---|---|---|---|---|---|---|---|
| rs6589664 | 11 | 118404804 | A | G | 1.922 | 0.05457 | ++++? | 0 | 1.223 | 3 | 0.7475 |
| rs11825496 | 11 | 118855221 | G | T | -1.919 | 0.055 | ---?- | 0 | 1.303 | 3 | 0.7283 |
| rs4456264 | 11 | 118851138 | T | C | -1.918 | 0.05511 | ---?- | 0 | 1.348 | 3 | 0.7178 |
| rs4936451 | 11 | 118852137 | T | C | -1.916 | 0.05541 | ---?- | 0 | 1.334 | 3 | 0.721 |
| rs45527340 | 11 | 118852526 | G | C | -1.913 | 0.05578 | ---?- | 0 | 1.343 | 3 | 0.719 |
| rs11006 | 11 | 118895202 | G | A | -1.909 | 0.0562 | ----- | 0 | 1.684 | 4 | 0.7936 |
| rs11216982 | 11 | 118590864 | A | G | 1.909 | 0.05622 | +++?- | 34.4 | 7.623 | 3 | 0.05447 |
| rs1790191 | 11 | 118825636 | A | G | 1.908 | 0.05635 | +?+++ | 0 | 0.497 | 3 | 0.9196 |
| rs61179449 | 11 | 118892835 | C | G | -1.902 | 0.05711 | ----- | 0 | 1.915 | 4 | 0.7513 |
| rs11822344 | 11 | 118864777 | C | G | -1.9 | 0.05741 | ---?- | 0 | 1.475 | 3 | 0.688 |
| rs3862794 | 11 | 72538600 | C | T | -1.899 | 0.05752 | +--+- | -2.7 | 5.843 | 4 | 0.2112 |
| rs7122959 | 11 | 118804283 | G | A | -1.899 | 0.05752 | -+--- | 0 | 2.825 | 4 | 0.5876 |
| rs58955701 | 11 | 118892872 | T | C | -1.899 | 0.05757 | ----- | 0 | 1.915 | 4 | 0.7513 |
| rs503542 | 11 | 118590743 | G | A | 1.899 | 0.05761 | ++-?+ | -36.8 | 3.656 | 3 | 0.3011 |
| rs2231440 | 11 | 118175439 | A | G | 1.898 | 0.05765 | +?++- | 37.2 | 7.965 | 3 | 0.04675 |
| rs7127188 | 11 | 118893023 | C | A | -1.893 | 0.05831 | ----- | 0 | 1.949 | 4 | 0.7451 |
| rs4938551 | 11 | 118707249 | C | G | 1.893 | 0.05833 | -++++ | 0 | 1.729 | 4 | 0.7855 |
| rs4938621 | 11 | 118892277 | T | C | -1.893 | 0.05838 | ----- | 0 | 1.9 | 4 | 0.7542 |
| rs12273687 | 11 | 118852641 | T | G | -1.892 | 0.05852 | ---?- | 0 | 1.681 | 3 | 0.6412 |
| rs4938605 | 11 | 118850707 | A | G | -1.891 | 0.05858 | ---?- | 0 | 1.393 | 3 | 0.7073 |
| rs7123066 | 11 | 118804266 | T | C | -1.889 | 0.05888 | -+--- | 0 | 2.773 | 4 | 0.5964 |
| rs7925155 | 11 | 72732430 | C | G | -1.88 | 0.06015 | +--+- | -23.5 | 4.857 | 4 | 0.3023 |
| rs2231441 | 11 | 118175540 | G | A | 1.878 | 0.06044 | +?++- | 38 | 8.069 | 3 | 0.04461 |
| rs3759009 | 11 | 118173850 | A | G | 1.874 | 0.06094 | +?++- | 39.6 | 8.272 | 3 | 0.04071 |
| rs79108568 | 11 | 118775524 | G | T | -1.872 | 0.06115 | -?--? | 0 | 0.54 | 2 | 0.7632 |
| rs73530756 | 11 | 72101747 | T | A | 1.861 | 0.06276 | +++?- | 0 | 2.623 | 3 | 0.4534 |
| rs515707 | 11 | 118685481 | T | C | 1.859 | 0.06303 | +?-?+ | 37.7 | 6.418 | 2 | 0.0404 |
| rs73530754 | 11 | 72099007 | T | C | 1.856 | 0.06352 | +++?- | 0 | 2.71 | 3 | 0.4384 |
| rs521910 | 11 | 118610629 | A | G | 1.855 | 0.06356 | +?-++ | -4 | 4.807 | 3 | 0.1865 |
| rs4456287 | 11 | 118298252 | C | T | 1.855 | 0.06363 | +++++ | 0 | 0.982 | 4 | 0.9125 |
| rs4499035 | 11 | 118610463 | T | C | 1.854 | 0.06372 | ++++- | 24.6 | 7.959 | 4 | 0.0931 |
| rs60065241 | 11 | 118883491 | A | C | -1.852 | 0.06406 | -?--- | 0 | 1.46 | 3 | 0.6916 |
| rs10892234 | 11 | 118194571 | G | A | -1.851 | 0.06411 | ----- | 0 | 2.278 | 4 | 0.6847 |
| rs536256 | 11 | 118327321 | G | A | 1.847 | 0.06475 | +++++ | 0 | 0.336 | 4 | 0.9874 |
| rs526850 | 11 | 118605127 | G | T | 1.84 | 0.06579 | +?-++ | -21.1 | 4.13 | 3 | 0.2477 |
| rs45573242 | 11 | 118853995 | C | A | 1.837 | 0.06615 | +-+?+ | 0 | 2.695 | 3 | 0.441 |
| rs55930012 | 11 | 72079380 | C | T | 1.833 | 0.06683 | +++?- | 30.2 | 7.162 | 3 | 0.0669 |
| rs1893029 | 11 | 118610826 | T | G | 1.82 | 0.06878 | ++++- | 22.8 | 7.772 | 4 | 0.1003 |
| rs754042 | 11 | 72326275 | T | C | 1.813 | 0.06983 | --+++ | 0 | 3.474 | 4 | 0.4819 |
| rs61901410 | 11 | 118661197 | G | C | 1.812 | 0.06994 | +?-++ | 32.6 | 7.422 | 3 | 0.0596 |
| rs59255055 | 11 | 118296735 | A | C | 1.811 | 0.07014 | +++++ | 0 | 0.736 | 4 | 0.9468 |
| rs58577143 | 11 | 118291210 | G | A | 1.807 | 0.07075 | +++++ | 0 | 2.242 | 4 | 0.6914 |
| rs78488993 | 11 | 72882539 | C | G | 1.806 | 0.07088 | +++-? | 0 | 2.923 | 3 | 0.4036 |
| rs493720 | 11 | 118604678 | T | C | 1.804 | 0.0712 | +?-++ | -16.2 | 4.305 | 3 | 0.2304 |
| rs2298821 | 11 | 118185140 | C | T | 1.799 | 0.07206 | +?++- | 44.9 | 9.068 | 3 | 0.0284 |
| rs73001486 | 11 | 118629364 | A | G | 1.798 | 0.0721 | ++++- | 20.8 | 7.575 | 4 | 0.1084 |
| rs138922408 | 11 | 118356711 | T | C | -1.797 | 0.07231 | ---+? | 15.1 | 5.888 | 3 | 0.1172 |
| rs2186781 | 11 | 118605447 | T | C | 1.796 | 0.07254 | +?-++ | -10.8 | 4.514 | 3 | 0.211 |
| rs3825057 | 11 | 118622985 | C | T | 1.792 | 0.07311 | ++++- | 26.1 | 8.123 | 4 | 0.08718 |
| rs10501410 | 11 | 72088806 | A | G | 1.788 | 0.07373 | +++?- | 0 | 1.898 | 3 | 0.5939 |
| rs498684 | 11 | 118316960 | A | G | 1.784 | 0.07449 | +++++ | 0 | 0.482 | 4 | 0.9753 |
| rs523148 | 11 | 118682462 | T | C | 1.783 | 0.07454 | +?-?+ | 35.5 | 6.204 | 2 | 0.04495 |
| rs3819250 | 11 | 118185346 | A | G | 1.779 | 0.07519 | +?++- | 44.9 | 9.077 | 3 | 0.02828 |
| rs10898893 | 11 | 72676699 | C | T | -1.779 | 0.07523 | --+-+ | 9.7 | 6.646 | 4 | 0.1558 |
| rs61900905 | 11 | 118185492 | C | T | 1.777 | 0.07558 | +?++- | 45.9 | 9.239 | 3 | 0.02628 |
| rs569 | 11 | 118894287 | G | A | -1.775 | 0.07587 | ----- | 0 | 1.502 | 4 | 0.8263 |

| | | | | | | | | | | | |
|---|---|---|---|---|---|---|---|---|---|---|---|
| rs60081894 | 11 | 72108715 | C | A | 1.774 | 0.07603 | +++?- | 0 | 2.421 | 3 | 0.4897 |
| rs7130875 | 11 | 118613674 | C | T | 1.772 | 0.07646 | ++++- | 25.6 | 8.064 | 4 | 0.08928 |
| rs28423005 | 11 | 118183733 | T | C | 1.771 | 0.0765 | +?++- | 42.3 | 8.669 | 3 | 0.03404 |
| rs4938503 | 11 | 118180155 | C | G | 1.763 | 0.07782 | +?++- | 41.4 | 8.532 | 3 | 0.0362 |
| rs487177 | 11 | 118657297 | G | C | 1.762 | 0.07813 | ++++- | 21.4 | 7.631 | 4 | 0.1061 |
| rs1126924 | 11 | 118178007 | T | C | 1.76 | 0.07837 | +?++- | 40.6 | 8.416 | 3 | 0.03816 |
| rs11217076 | 11 | 118745664 | G | T | 1.76 | 0.0784 | ++-++ | 0 | 1.817 | 4 | 0.7694 |
| rs2277289 | 11 | 118182660 | C | T | 1.76 | 0.0784 | +?++- | 41.4 | 8.525 | 3 | 0.03632 |
| rs4938550 | 11 | 118707171 | G | A | 1.76 | 0.07843 | +?-++ | 15 | 5.882 | 3 | 0.1175 |
| rs574387 | 11 | 118337003 | C | A | 1.757 | 0.07899 | +++++ | 0 | 0.649 | 4 | 0.9575 |
| rs7124498 | 11 | 118616253 | A | G | 1.755 | 0.07927 | ++++- | 24.4 | 7.934 | 4 | 0.09402 |
| rs11217016 | 11 | 118627149 | C | A | 1.754 | 0.07941 | ++++- | 20.4 | 7.542 | 4 | 0.1099 |
| rs61563240 | 11 | 72056185 | A | G | 1.751 | 0.07996 | +++?- | 23.3 | 6.516 | 3 | 0.08902 |
| rs112996178 | 11 | 118609598 | T | C | 1.747 | 0.08063 | ++++- | 22.3 | 7.726 | 4 | 0.1021 |
| rs7119702 | 11 | 118607694 | G | A | 1.743 | 0.08127 | +?-++ | -22.5 | 4.083 | 3 | 0.2527 |
| rs34147990 | 11 | 72160112 | T | C | 1.741 | 0.0817 | +++?- | -57.1 | 3.183 | 3 | 0.3642 |
| rs7952074 | 11 | 72062978 | G | A | 1.739 | 0.08211 | +++?- | 25.2 | 6.687 | 3 | 0.08258 |
| rs2851460 | 11 | 118716563 | A | G | 1.728 | 0.08397 | -++++ | 0 | 2.179 | 4 | 0.7028 |
| rs6421568 | 11 | 118655704 | C | G | 1.725 | 0.08445 | +?-++ | -2.5 | 4.879 | 3 | 0.1809 |
| rs56229279 | 11 | 72013788 | G | A | 1.721 | 0.08529 | +++?- | 45.9 | 9.242 | 3 | 0.02624 |
| rs11217013 | 11 | 118624919 | C | G | 1.718 | 0.08582 | ++++- | 20.1 | 7.506 | 4 | 0.1115 |
| rs76068633 | 11 | 118416268 | T | C | -1.718 | 0.08589 | ---+? | 25.8 | 6.739 | 3 | 0.0807 |
| rs58306819 | 11 | 72054966 | C | T | 1.714 | 0.0866 | +++?- | 25.7 | 6.728 | 3 | 0.08109 |
| rs6421569 | 11 | 118655775 | A | C | 1.713 | 0.08677 | +?-++ | -0.6 | 4.973 | 3 | 0.1738 |
| rs2508915 | 11 | 118652781 | C | G | 1.709 | 0.08751 | +?-++ | -0.5 | 4.975 | 3 | 0.1736 |
| rs10892282 | 11 | 118627778 | C | G | 1.707 | 0.08776 | ++++- | 19.1 | 7.412 | 4 | 0.1157 |
| rs56104144 | 11 | 118666289 | C | G | 1.707 | 0.08782 | ++++- | 25.1 | 8.011 | 4 | 0.09118 |
| rs7124336 | 11 | 72160271 | T | C | 1.707 | 0.08788 | +++?- | -49.5 | 3.344 | 3 | 0.3416 |
| rs141108621 | 11 | 118349211 | T | C | -1.704 | 0.08833 | ----? | -0.1 | 4.996 | 3 | 0.1721 |
| rs11823672 | 11 | 72160828 | G | A | 1.702 | 0.08878 | +++?- | -53 | 3.268 | 3 | 0.3522 |
| rs7926133 | 11 | 72647296 | C | T | -1.702 | 0.08881 | --+-+ | 9.1 | 6.601 | 4 | 0.1585 |
| rs11217014 | 11 | 118626286 | C | T | 1.699 | 0.08932 | ++++- | 21.2 | 7.614 | 4 | 0.1068 |
| rs7934890 | 11 | 118655356 | T | C | 1.693 | 0.09038 | ++++- | 20.8 | 7.574 | 4 | 0.1085 |
| rs57276029 | 11 | 72059859 | C | T | 1.693 | 0.0904 | +++?- | 25.7 | 6.727 | 3 | 0.08111 |
| rs112910608 | 11 | 118637475 | T | C | 1.693 | 0.09049 | ++++- | 18.9 | 7.397 | 4 | 0.1163 |
| rs6592500 | 11 | 72651705 | G | A | -1.692 | 0.09068 | +---- | -31.1 | 4.575 | 4 | 0.3337 |
| rs7110526 | 11 | 118633533 | C | T | 1.689 | 0.09115 | ++++- | 18.9 | 7.402 | 4 | 0.1161 |
| rs1784295 | 11 | 118660932 | A | G | 1.689 | 0.09122 | +?-++ | 0.1 | 5.004 | 3 | 0.1715 |
| rs535996 | 11 | 118637666 | T | C | 1.681 | 0.09281 | +?-++ | -8.1 | 4.626 | 3 | 0.2013 |
| rs529717 | 11 | 118640081 | A | G | 1.678 | 0.09334 | +?-++ | -8.7 | 4.598 | 3 | 0.2037 |
| rs1790188 | 11 | 118661055 | T | C | 1.677 | 0.09345 | +?-++ | 0.3 | 5.017 | 3 | 0.1705 |
| rs116976310 | 11 | 118420099 | C | A | -1.677 | 0.0935 | ---+? | 14.4 | 5.841 | 3 | 0.1196 |
| rs187678279 | 11 | 118338067 | A | G | -1.676 | 0.09374 | ----? | 10.1 | 5.562 | 3 | 0.135 |
| rs57928845 | 11 | 72149909 | T | C | 1.676 | 0.09382 | +++?- | 0 | 2.756 | 3 | 0.4307 |
| rs548167 | 11 | 118648396 | A | G | 1.674 | 0.09414 | +?-++ | -5.2 | 4.751 | 3 | 0.1909 |
| rs11217021 | 11 | 118640828 | C | T | 1.674 | 0.09422 | ++++- | 18.8 | 7.393 | 4 | 0.1165 |
| rs602716 | 11 | 118801235 | A | G | 1.673 | 0.09439 | +-++- | 18.2 | 7.338 | 4 | 0.1191 |
| rs615166 | 11 | 118803536 | A | G | 1.671 | 0.09469 | +-++- | 16.2 | 7.159 | 4 | 0.1277 |
| rs2508929 | 11 | 118641451 | G | T | 1.67 | 0.09485 | +?-++ | -8.1 | 4.625 | 3 | 0.2014 |
| rs11217018 | 11 | 118635364 | C | T | 1.669 | 0.09521 | ++++- | 19 | 7.406 | 4 | 0.1159 |
| rs10736495 | 11 | 118607090 | C | G | 1.667 | 0.09547 | +?-++ | -16.4 | 4.297 | 3 | 0.2311 |
| rs58868870 | 11 | 118668296 | T | C | 1.663 | 0.09627 | -++-- | 26.7 | 8.182 | 4 | 0.08512 |
| rs498912 | 11 | 118576997 | C | T | 1.663 | 0.0964 | +?-++ | -36.7 | 3.659 | 3 | 0.3008 |
| rs11217077 | 11 | 118746223 | C | T | 1.652 | 0.09856 | ++-++ | 0 | 2.152 | 4 | 0.7077 |
| rs73528892 | 11 | 72020144 | C | G | 1.652 | 0.09861 | +++?- | 34 | 7.574 | 3 | 0.05569 |
| rs647769 | 11 | 118812512 | T | C | 1.65 | 0.09891 | +-+++ | 0 | 3.271 | 4 | 0.5136 |

| | | | | | | | | | | | |
|---|---|---|---|---|---|---|---|---|---|---|---|
| rs540716 | 11 | 118685015 | T | C | 1.645 | 0.0999 | ++++- | 35.6 | 9.318 | 4 | 0.05363 |
| rs3759002 | 11 | 118136160 | A | G | 1.645 | 0.1 | +?-+- | -6.6 | 4.692 | 3 | 0.1958 |
| rs503425 | 11 | 118692727 | C | T | 1.645 | 0.1 | ++++- | 38.2 | 9.703 | 4 | 0.04574 |
| rs76160025 | 11 | 118782802 | T | C | -1.636 | 0.1018 | -?--? | 0 | 1.067 | 2 | 0.5866 |
| rs111527225 | 11 | 118065830 | A | G | 1.634 | 0.1022 | +?++- | 0 | 2.528 | 3 | 0.4702 |
| rs6589687 | 11 | 118615660 | T | A | 1.631 | 0.1029 | +?-++ | -18.9 | 4.207 | 3 | 0.24 |
| rs61506726 | 11 | 118831335 | G | A | -1.622 | 0.1047 | ----+ | 0 | 2.814 | 4 | 0.5894 |
| rs553188 | 11 | 118576762 | T | C | 1.621 | 0.1051 | +?-++ | -20.2 | 4.161 | 3 | 0.2446 |
| rs10892281 | 11 | 118614397 | T | C | 1.62 | 0.1052 | +?-++ | -21.4 | 4.119 | 3 | 0.2489 |
| rs497678 | 11 | 118567881 | C | T | 1.616 | 0.1061 | ++-++ | 0 | 3.184 | 4 | 0.5275 |
| rs11216904 | 11 | 118440863 | G | A | 1.615 | 0.1063 | +++-+ | 0 | 1.637 | 4 | 0.8021 |
| rs58270621 | 11 | 118413604 | A | G | 1.613 | 0.1068 | +++++ | 0 | 1.133 | 4 | 0.8889 |
| rs11216898 | 11 | 118432852 | C | T | 1.611 | 0.1071 | +++++ | 0 | 1.649 | 4 | 0.8 |
| rs497916 | 11 | 118758089 | T | C | -1.608 | 0.1079 | +++-- | 40.1 | 10.022 | 4 | 0.04006 |
| rs8192696 | 11 | 118895686 | T | C | -1.607 | 0.108 | ----- | 0 | 0.847 | 4 | 0.932 |
| rs12224844 | 11 | 118439048 | T | A | 1.607 | 0.1081 | +++-+ | 0 | 1.547 | 4 | 0.8182 |
| rs4938604 | 11 | 118846574 | C | T | -1.606 | 0.1083 | -?-?- | 0 | 0.485 | 2 | 0.7848 |
| rs2510902 | 11 | 118700218 | A | G | 1.603 | 0.109 | ++++- | 31.2 | 8.719 | 4 | 0.06852 |
| rs485901 | 11 | 118622245 | G | A | 1.602 | 0.1091 | +?-++ | -16.2 | 4.305 | 3 | 0.2304 |
| rs1961138 | 11 | 118619423 | C | T | 1.601 | 0.1094 | +?-++ | -16.7 | 4.285 | 3 | 0.2323 |
| rs640120 | 11 | 118618837 | A | C | 1.6 | 0.1095 | +?-++ | -17.3 | 4.262 | 3 | 0.2346 |
| rs11216860 | 11 | 118240332 | C | T | -1.596 | 0.1104 | -++-- | -4.4 | 5.749 | 4 | 0.2187 |
| rs6589690 | 11 | 118632893 | A | G | 1.595 | 0.1108 | +?-++ | -4.9 | 4.768 | 3 | 0.1896 |
| rs10892243 | 11 | 118441846 | C | T | 1.593 | 0.1112 | +++-+ | 0 | 1.6 | 4 | 0.8087 |
| rs4938603 | 11 | 118846482 | G | A | -1.587 | 0.1125 | -?-?- | 0 | 0.328 | 2 | 0.8487 |
| rs6589677 | 11 | 118583971 | A | G | 1.587 | 0.1125 | +?-++ | -6 | 4.719 | 3 | 0.1936 |
| rs4338560 | 11 | 118143368 | T | C | 1.586 | 0.1127 | +?++- | 2.9 | 5.152 | 3 | 0.161 |
| rs7926282 | 11 | 118605914 | C | A | 1.584 | 0.1133 | -++-- | 42 | 10.353 | 4 | 0.03488 |
| rs73528897 | 11 | 72032681 | A | G | 1.581 | 0.1138 | +++?- | 28.8 | 7.022 | 3 | 0.07119 |
| rs475946 | 11 | 118629143 | T | C | 1.579 | 0.1143 | +?-++ | -10.4 | 4.528 | 3 | 0.2098 |
| rs2511062 | 11 | 72871055 | C | T | -1.579 | 0.1144 | +--+- | 14.7 | 7.036 | 4 | 0.134 |
| rs10502227 | 11 | 118147496 | C | T | 1.573 | 0.1156 | +?++- | -2.5 | 4.878 | 3 | 0.1809 |
| rs2508204 | 11 | 72001820 | C | T | 1.573 | 0.1158 | +++?- | 39.2 | 8.22 | 3 | 0.04168 |
| rs113975105 | 11 | 118836606 | G | C | -1.569 | 0.1167 | ----+ | 0 | 2.667 | 4 | 0.6149 |
| rs10790239 | 11 | 118001445 | T | C | -1.567 | 0.117 | -?--- | 0 | 0.418 | 3 | 0.9364 |
| rs1790187 | 11 | 118644213 | A | G | 1.565 | 0.1175 | +?-++ | -21.7 | 4.107 | 3 | 0.2501 |
| rs60067274 | 11 | 118412296 | T | A | 1.558 | 0.1193 | +++++ | 0 | 1.529 | 4 | 0.8215 |
| rs11603267 | 11 | 118146599 | T | G | 1.557 | 0.1195 | +?++- | -4.9 | 4.764 | 3 | 0.1899 |
| rs11216897 | 11 | 118432279 | G | A | 1.554 | 0.1202 | +++-+ | 0 | 1.636 | 4 | 0.8024 |
| rs11607245 | 11 | 118843448 | T | C | -1.55 | 0.1211 | ---?- | 0 | 0.423 | 3 | 0.9354 |
| rs57669200 | 11 | 72026863 | A | G | 1.549 | 0.1213 | +++?- | 39.5 | 8.27 | 3 | 0.04075 |
| rs2277295 | 11 | 118416253 | T | C | 1.548 | 0.1216 | +++++ | 0 | 1.541 | 4 | 0.8194 |
| rs11217052 | 11 | 118717732 | G | A | 1.547 | 0.1219 | -++++ | 0 | 2.111 | 4 | 0.7153 |
| rs58753751 | 11 | 72048813 | C | T | 1.545 | 0.1224 | +++?- | 22.5 | 6.45 | 3 | 0.09165 |
| rs12788933 | 11 | 118140184 | T | C | 1.544 | 0.1227 | +?++- | -1.8 | 4.913 | 3 | 0.1783 |
| rs688035 | 11 | 118802492 | G | C | 1.541 | 0.1234 | +-++- | 16 | 7.144 | 4 | 0.1285 |
| rs57291879 | 11 | 72041441 | A | C | 1.54 | 0.1236 | +++?- | 22.5 | 6.449 | 3 | 0.09171 |
| rs60837962 | 11 | 118438448 | T | A | 1.54 | 0.1237 | +++-+ | 0 | 1.802 | 4 | 0.7721 |
| rs11235547 | 11 | 72318831 | A | G | 1.537 | 0.1242 | -?+++ | 0 | 1.819 | 3 | 0.6107 |
| rs2186780 | 11 | 118605451 | T | C | 1.537 | 0.1242 | -++-- | 39.1 | 9.847 | 4 | 0.04309 |
| rs4326835 | 11 | 118432404 | T | G | 1.537 | 0.1242 | +++-+ | 0 | 1.702 | 4 | 0.7904 |
| rs59108310 | 11 | 118433729 | G | A | 1.537 | 0.1243 | +++-+ | 0 | 1.652 | 4 | 0.7993 |
| rs75657287 | 11 | 118801518 | G | C | 1.535 | 0.1247 | +?+?? | 0 | 0.129 | 1 | 0.72 |
| rs564892 | 11 | 118602392 | A | T | 1.533 | 0.1253 | ++-++ | -31.5 | 4.563 | 4 | 0.3352 |
| rs12416741 | 11 | 72023828 | C | T | 1.531 | 0.1257 | +++?- | 41.3 | 8.514 | 3 | 0.0365 |
| rs11216886 | 11 | 118411424 | C | A | 1.53 | 0.1259 | +++++ | 0 | 1.343 | 4 | 0.854 |

| | | | | | | | | | | | |
|---|---|---|---|---|---|---|---|---|---|---|---|
| rs3741319 | 11 | 118415848 | C | A | 1.53 | 0.1261 | +++++ | 0 | 1.535 | 4 | 0.8204 |
| rs528056 | 11 | 118317907 | T | C | 1.53 | 0.1261 | +++-+ | 0 | 1.76 | 4 | 0.7797 |
| rs3825055 | 11 | 118416073 | C | A | 1.527 | 0.1267 | +++++ | 0 | 1.521 | 4 | 0.8229 |
| rs60849224 | 11 | 72042930 | C | T | 1.525 | 0.1271 | +++?- | 22.4 | 6.445 | 3 | 0.09184 |
| rs36049191 | 11 | 72040784 | C | T | 1.525 | 0.1272 | +++?- | 22.4 | 6.445 | 3 | 0.09184 |
| rs11216902 | 11 | 118437279 | G | A | 1.525 | 0.1274 | +++-+ | 0 | 1.639 | 4 | 0.8018 |
| rs11216899 | 11 | 118434421 | G | A | 1.522 | 0.128 | +++-+ | 0 | 1.731 | 4 | 0.7851 |
| rs3741320 | 11 | 118415847 | C | G | 1.521 | 0.1284 | +++++ | 0 | 1.513 | 4 | 0.8244 |
| rs11216900 | 11 | 118436079 | C | G | 1.52 | 0.1285 | +++-+ | 0 | 1.804 | 4 | 0.7718 |
| rs11216909 | 11 | 118449746 | T | C | 1.52 | 0.1285 | +++-+ | 0 | 1.599 | 4 | 0.8089 |
| rs113145584 | 11 | 118154798 | C | G | -1.518 | 0.129 | -?-?? | 0 | 0.909 | 1 | 0.3403 |
| rs4938559 | 11 | 118718061 | T | C | 1.518 | 0.129 | -++++ | 0 | 2.002 | 4 | 0.7354 |
| rs9736327 | 11 | 118447580 | G | T | 1.518 | 0.1291 | +++-+ | 0 | 1.481 | 4 | 0.83 |
| rs45505501 | 11 | 118405305 | A | G | 1.517 | 0.1292 | +++++ | 0 | 1.253 | 4 | 0.8693 |
| rs9735421 | 11 | 118447597 | A | G | 1.516 | 0.1295 | +++-+ | 0 | 1.475 | 4 | 0.831 |
| rs60753148 | 11 | 118434835 | T | C | 1.516 | 0.1296 | +++-+ | 0 | 1.706 | 4 | 0.7896 |
| rs557990 | 11 | 118601363 | T | C | 1.513 | 0.1302 | ++-++ | -29.2 | 4.644 | 4 | 0.3258 |
| rs4938560 | 11 | 118718062 | G | A | 1.512 | 0.1306 | -++++ | 0 | 1.99 | 4 | 0.7375 |
| rs11216881 | 11 | 118409539 | G | A | 1.511 | 0.1307 | +++++ | 0 | 1.278 | 4 | 0.8651 |
| rs45485302 | 11 | 118405377 | A | C | 1.511 | 0.1308 | +++++ | 0 | 1.192 | 4 | 0.8794 |
| rs7932700 | 11 | 118411713 | T | G | 1.509 | 0.1314 | +++++ | 0 | 1.379 | 4 | 0.8479 |
| rs4537793 | 11 | 118053156 | A | C | 1.508 | 0.1316 | +++++ | 0 | 0.725 | 4 | 0.9481 |
| rs73022038 | 11 | 118408471 | T | C | 1.508 | 0.1316 | +++++ | 0 | 1.244 | 4 | 0.8707 |
| rs11216880 | 11 | 118409536 | A | G | 1.507 | 0.1319 | +++++ | 0 | 1.257 | 4 | 0.8686 |
| rs45548238 | 11 | 118405437 | T | C | 1.506 | 0.1319 | +++++ | 0 | 1.213 | 4 | 0.876 |
| rs73022041 | 11 | 118408789 | C | T | 1.506 | 0.1322 | +++++ | 0 | 1.237 | 4 | 0.872 |
| rs6589675 | 11 | 118569414 | C | G | 1.505 | 0.1323 | +?-++ | 0 | 2.314 | 3 | 0.5099 |
| rs112965917 | 11 | 118725189 | A | T | 1.503 | 0.1329 | +?-++ | 0 | 1.927 | 3 | 0.5877 |
| rs45522239 | 11 | 118405438 | G | A | 1.503 | 0.1329 | +++++ | 0 | 1.198 | 4 | 0.8784 |
| rs112753655 | 11 | 118154795 | A | G | -1.501 | 0.1333 | -?-?? | 0 | 0.942 | 1 | 0.3317 |
| rs9735449 | 11 | 118448087 | T | C | 1.501 | 0.1333 | +++-+ | 0 | 1.504 | 4 | 0.8259 |
| rs73023350 | 11 | 118449396 | A | G | 1.501 | 0.1334 | +++-+ | 0 | 1.55 | 4 | 0.8177 |
| rs7131631 | 11 | 118409374 | A | G | 1.5 | 0.1337 | +++++ | 0 | 1.223 | 4 | 0.8744 |
| rs45474901 | 11 | 118405453 | C | T | 1.499 | 0.1339 | +++++ | 0 | 1.356 | 4 | 0.8518 |
| rs7128587 | 11 | 118409302 | A | G | 1.499 | 0.1339 | +++++ | 0 | 1.222 | 4 | 0.8745 |
| rs11216883 | 11 | 118410468 | G | A | 1.499 | 0.134 | +++++ | 0 | 1.269 | 4 | 0.8667 |
| rs11216923 | 11 | 118465501 | G | A | 1.499 | 0.134 | +++-+ | 0 | 1.448 | 4 | 0.8359 |
| rs561775 | 11 | 118341212 | G | A | 1.497 | 0.1344 | +++-+ | 0 | 2.693 | 4 | 0.6105 |
| rs73022042 | 11 | 118408893 | A | G | 1.496 | 0.1347 | +++++ | 0 | 1.218 | 4 | 0.8751 |
| rs74404016 | 11 | 118408653 | T | C | 1.495 | 0.1348 | +++++ | 0 | 1.222 | 4 | 0.8745 |
| rs11216885 | 11 | 118410545 | G | A | 1.495 | 0.135 | +++++ | 0 | 1.253 | 4 | 0.8693 |
| rs11216921 | 11 | 118464815 | C | G | 1.494 | 0.1351 | +++-+ | 0 | 1.516 | 4 | 0.8237 |
| rs7126175 | 11 | 118450875 | T | C | 1.493 | 0.1355 | +++-+ | 0 | 1.577 | 4 | 0.8129 |
| rs7101585 | 11 | 118409583 | A | G | 1.493 | 0.1356 | +++++ | 0 | 1.233 | 4 | 0.8727 |
| rs11216884 | 11 | 118410488 | C | T | 1.492 | 0.1358 | +++++ | 0 | 1.256 | 4 | 0.8688 |
| rs11216925 | 11 | 118471423 | C | T | 1.492 | 0.1358 | +++-+ | 0 | 1.803 | 4 | 0.772 |
| rs61290697 | 11 | 118412358 | T | C | 1.491 | 0.1359 | +++++ | 0 | 1.238 | 4 | 0.8719 |
| rs17742 | 11 | 118473385 | C | T | 1.49 | 0.1361 | +++-+ | 0 | 1.676 | 4 | 0.7951 |
| rs535207 | 11 | 118408033 | G | A | 1.491 | 0.1361 | +++++ | 0 | 1.193 | 4 | 0.8793 |
| rs11216912 | 11 | 118452334 | C | G | 1.49 | 0.1363 | +++-+ | 0 | 1.6 | 4 | 0.8087 |
| rs11216922 | 11 | 118465295 | G | A | 1.488 | 0.1367 | +++-+ | 0 | 1.438 | 4 | 0.8376 |
| rs7115634 | 11 | 118466249 | G | A | 1.487 | 0.1369 | +++-+ | 0 | 1.488 | 4 | 0.8287 |
| rs10892315 | 11 | 118831727 | C | T | -1.487 | 0.1371 | ----+ | 0 | 2.399 | 4 | 0.6629 |
| rs11216911 | 11 | 118451901 | C | T | 1.485 | 0.1375 | +++-+ | 0 | 1.563 | 4 | 0.8154 |
| rs12577045 | 11 | 118151446 | A | G | -1.484 | 0.1379 | -?-?? | 0 | 0.79 | 1 | 0.374 |
| rs10892245 | 11 | 118472354 | A | C | 1.482 | 0.1382 | +++-+ | 0 | 1.856 | 4 | 0.7622 |

| | | | | | | | | | | | |
|---|---|---|---|---|---|---|---|---|---|---|---|
| rs11216913 | 11 | 118453681 | G | A | 1.481 | 0.1385 | +++-+ | 0 | 1.626 | 4 | 0.8041 |
| rs55844025 | 11 | 118446298 | T | C | 1.48 | 0.1388 | +++-+ | 0 | 1.467 | 4 | 0.8325 |
| rs11216906 | 11 | 118443944 | A | G | 1.479 | 0.139 | +++-+ | 0 | 1.641 | 4 | 0.8014 |
| rs11216910 | 11 | 118450046 | G | A | 1.476 | 0.1398 | +++-+ | 0 | 1.547 | 4 | 0.8183 |
| rs535327 | 11 | 118341785 | G | A | 1.475 | 0.1401 | +++-+ | 0 | 2.676 | 4 | 0.6135 |
| rs1623316 | 11 | 118757219 | G | C | 1.475 | 0.1402 | ++-+? | 0 | 2.495 | 3 | 0.4762 |
| rs75128335 | 11 | 118072915 | A | G | 1.475 | 0.1402 | +?++- | 0 | 2.04 | 3 | 0.5642 |
| rs472837 | 11 | 72007150 | T | C | 1.474 | 0.1404 | +++?- | 35.8 | 7.785 | 3 | 0.05067 |
| rs524936 | 11 | 118341921 | A | G | 1.474 | 0.1406 | +++-+ | 0 | 2.74 | 4 | 0.6022 |
| rs3741323 | 11 | 118405641 | G | A | 1.473 | 0.1407 | +++++ | 0 | 1.182 | 4 | 0.881 |
| rs7929887 | 11 | 118428791 | G | A | 1.473 | 0.1407 | +++++ | 0 | 1.48 | 4 | 0.8301 |
| rs11216890 | 11 | 118421569 | G | C | 1.473 | 0.1408 | +++-+ | 0 | 1.674 | 4 | 0.7955 |
| rs537975 | 11 | 118362869 | T | C | 1.472 | 0.1411 | +++++ | 0 | 0.528 | 4 | 0.9707 |
| rs590732 | 11 | 72006394 | T | C | 1.471 | 0.1414 | +++?- | 35.8 | 7.79 | 3 | 0.05055 |
| rs11216917 | 11 | 118462160 | G | T | 1.469 | 0.1417 | +++-+ | 0 | 1.564 | 4 | 0.8153 |
| rs12576119 | 11 | 118158441 | A | G | -1.47 | 0.1417 | -?-?? | 0 | 0.865 | 1 | 0.3523 |
| rs57438673 | 11 | 118459039 | A | G | 1.469 | 0.1419 | +++-+ | 0 | 1.733 | 4 | 0.7847 |
| rs73023341 | 11 | 118445585 | G | A | 1.467 | 0.1423 | +++-+ | 0 | 1.565 | 4 | 0.815 |
| rs10892317 | 11 | 118832035 | C | T | -1.467 | 0.1424 | ----+ | 0 | 2.678 | 4 | 0.6131 |
| rs492471 | 11 | 118689745 | T | C | 1.466 | 0.1426 | ++++- | 2.1 | 6.126 | 4 | 0.1899 |
| rs17842763 | 11 | 118454177 | G | A | 1.465 | 0.1429 | +++-+ | 0 | 1.574 | 4 | 0.8134 |
| rs113054880 | 11 | 118459702 | G | C | 1.464 | 0.1432 | +++-+ | 0 | 1.751 | 4 | 0.7814 |
| rs6589710 | 11 | 118837460 | T | A | -1.463 | 0.1433 | ---?- | 0 | 0.484 | 3 | 0.9224 |
| rs3901422 | 11 | 118837272 | A | G | -1.463 | 0.1435 | ---?- | 0 | 0.445 | 3 | 0.9309 |
| rs146026283 | 11 | 118296322 | A | C | 1.462 | 0.1437 | +?++? | 0 | 0.604 | 2 | 0.7392 |
| rs10892318 | 11 | 118848725 | A | G | -1.461 | 0.1439 | ---?- | 0 | 0.435 | 3 | 0.9329 |
| rs11216891 | 11 | 118422234 | T | A | 1.46 | 0.1444 | +++++ | 0 | 1.655 | 4 | 0.7989 |
| rs11216907 | 11 | 118444525 | A | G | 1.459 | 0.1445 | +++-+ | 0 | 1.719 | 4 | 0.7873 |
| rs113739713 | 11 | 118420101 | G | A | 1.459 | 0.1447 | +++-+ | 0 | 1.568 | 4 | 0.8146 |
| rs17092756 | 11 | 118457630 | C | G | 1.458 | 0.1449 | +++-+ | 0 | 1.701 | 4 | 0.7905 |
| rs11216924 | 11 | 118470092 | G | C | 1.457 | 0.1451 | +++-+ | 0 | 1.717 | 4 | 0.7877 |
| rs2277291 | 11 | 118435756 | T | C | 1.456 | 0.1453 | +++-+ | 0 | 1.84 | 4 | 0.7651 |
| rs7116690 | 11 | 118837742 | G | A | -1.456 | 0.1454 | ---?- | 0 | 0.43 | 3 | 0.9339 |
| rs544096 | 11 | 118594326 | G | A | 1.453 | 0.1463 | ++-++ | 0 | 3.922 | 4 | 0.4166 |
| rs495132 | 11 | 118335927 | A | G | 1.45 | 0.147 | +++-+ | 0 | 2.73 | 4 | 0.604 |
| rs10892301 | 11 | 118735476 | A | G | 1.45 | 0.1472 | -++++ | 0 | 2.05 | 4 | 0.7266 |
| rs3741324 | 11 | 118405835 | A | G | 1.449 | 0.1472 | +++++ | 0 | 1.171 | 4 | 0.8829 |
| rs4435039 | 11 | 118297391 | G | A | 1.448 | 0.1475 | +++-+ | 0 | 2.368 | 4 | 0.6685 |
| rs11216926 | 11 | 118471996 | T | C | 1.448 | 0.1477 | +++-+ | 0 | 1.786 | 4 | 0.775 |
| rs7931564 | 11 | 118469571 | G | A | 1.448 | 0.1477 | +++-+ | 0 | 1.647 | 4 | 0.8002 |
| rs11216916 | 11 | 118461605 | C | T | 1.447 | 0.1479 | +++-+ | 0 | 1.512 | 4 | 0.8245 |
| rs2120269 | 11 | 118151135 | C | T | -1.445 | 0.1485 | -?-?? | 0 | 0.664 | 1 | 0.4153 |
| rs188843024 | 11 | 118453478 | A | G | -1.443 | 0.1489 | ---+? | -19.6 | 4.179 | 3 | 0.2427 |
| rs2277293 | 11 | 118422868 | T | C | 1.443 | 0.1489 | +++-+ | 0 | 1.725 | 4 | 0.7861 |
| rs61902060 | 11 | 118839828 | G | C | -1.442 | 0.1492 | ---?- | 0 | 0.414 | 3 | 0.9374 |
| rs629470 | 11 | 118320553 | A | G | 1.441 | 0.1495 | +++++ | 0 | 0.89 | 4 | 0.926 |
| rs3741325 | 11 | 118405989 | G | A | 1.44 | 0.1498 | +++++ | 0 | 1.493 | 4 | 0.8278 |
| rs1784257 | 11 | 72007984 | C | T | 1.439 | 0.1501 | +++?- | 36.2 | 7.833 | 3 | 0.04958 |
| rs60112336 | 11 | 118453483 | T | C | 1.439 | 0.1501 | +++-+ | 0 | 1.573 | 4 | 0.8136 |
| rs11217107 | 11 | 118839268 | C | G | -1.438 | 0.1505 | ---?- | 0 | 0.412 | 3 | 0.9377 |
| rs187547749 | 11 | 118439159 | T | C | -1.437 | 0.1507 | ---+? | 7.5 | 5.408 | 3 | 0.1443 |
| rs58745438 | 11 | 118848315 | C | T | -1.437 | 0.1507 | ---?- | 0 | 0.43 | 3 | 0.934 |
| rs59176377 | 11 | 118419295 | A | G | 1.437 | 0.1507 | +++++ | 0 | 1.279 | 4 | 0.865 |
| rs67774926 | 11 | 118157623 | G | C | -1.433 | 0.1519 | -?-?? | 0 | 0.828 | 1 | 0.3629 |
| rs17122278 | 11 | 118449370 | G | A | 1.431 | 0.1525 | +++-+ | 0 | 1.735 | 4 | 0.7844 |
| rs189530708 | 11 | 118272454 | G | C | -1.431 | 0.1526 | --+?? | 13.2 | 4.611 | 2 | 0.09973 |

| | | | | | | | | | | | |
|---|---|---|---|---|---|---|---|---|---|---|---|
| rs4938494 | 11 | 118150735 | C | G | -1.429 | 0.153 | -?-?? | 0 | 0.69 | 1 | 0.4062 |
| rs74384601 | 11 | 118449313 | G | A | -1.428 | 0.1532 | ---+? | 1.1 | 5.054 | 3 | 0.1679 |
| rs12797009 | 11 | 118826192 | A | C | 1.424 | 0.1545 | +?+?- | 0 | 1.48 | 2 | 0.4772 |
| rs61900159 | 11 | 118839133 | C | G | -1.423 | 0.1548 | ---?- | 0 | 0.438 | 3 | 0.9322 |
| rs11216892 | 11 | 118424008 | T | C | 1.422 | 0.155 | +++-+ | 0 | 1.524 | 4 | 0.8224 |
| rs4938602 | 11 | 118846466 | A | G | -1.422 | 0.1551 | ---?- | 0 | 0.415 | 3 | 0.937 |
| rs7126476 | 11 | 118834784 | G | A | -1.422 | 0.1551 | ---?- | 0 | 0.423 | 3 | 0.9354 |
| rs3890562 | 11 | 118837113 | G | A | -1.421 | 0.1552 | ---?- | 0 | 0.218 | 3 | 0.9746 |
| rs17122269 | 11 | 118445275 | C | T | 1.42 | 0.1558 | +++-+ | 0 | 1.849 | 4 | 0.7634 |
| rs45534632 | 11 | 118840705 | G | A | 1.419 | 0.1558 | +-+?+ | 0 | 2.919 | 3 | 0.4043 |
| rs56140791 | 11 | 118167190 | C | T | -1.418 | 0.1562 | -?-?? | 0 | 0.805 | 1 | 0.3695 |
| rs56403966 | 11 | 118425051 | A | G | 1.416 | 0.1567 | +++-+ | 0 | 1.599 | 4 | 0.809 |
| rs7108845 | 11 | 118407523 | G | T | 1.414 | 0.1573 | +++++ | 0 | 1.233 | 4 | 0.8726 |
| rs55857590 | 11 | 118425063 | G | A | 1.413 | 0.1576 | +++-+ | 0 | 1.535 | 4 | 0.8204 |
| rs11216889 | 11 | 118421512 | A | G | 1.413 | 0.1577 | +++-+ | 0 | 1.909 | 4 | 0.7524 |
| rs55736521 | 11 | 118840506 | T | C | 1.413 | 0.1577 | +-+?+ | 0 | 2.889 | 3 | 0.4091 |
| rs12420418 | 11 | 118840857 | T | C | -1.413 | 0.1578 | ---?- | 0 | 0.2 | 3 | 0.9775 |
| rs67604597 | 11 | 118154836 | G | T | -1.411 | 0.1582 | -?-?? | 0 | 0.799 | 1 | 0.3714 |
| rs1047079 | 11 | 118473231 | C | T | 1.409 | 0.1587 | +++-+ | 0 | 1.86 | 4 | 0.7614 |
| rs60024067 | 11 | 72028610 | C | T | 1.409 | 0.1588 | +++?- | 34.8 | 7.67 | 3 | 0.05334 |
| rs11216914 | 11 | 118456414 | C | T | 1.409 | 0.159 | +++-+ | 0 | 1.55 | 4 | 0.8177 |
| rs60652614 | 11 | 118168543 | T | G | -1.408 | 0.1592 | -?-?? | 0 | 0.736 | 1 | 0.3909 |
| rs61706007 | 11 | 118167278 | G | T | -1.408 | 0.1592 | -?-?? | 0 | 0.823 | 1 | 0.3642 |
| rs73022058 | 11 | 118423672 | T | C | 1.407 | 0.1595 | +++-+ | 0 | 1.812 | 4 | 0.7702 |
| rs11217108 | 11 | 118841761 | T | C | -1.405 | 0.16 | ---?- | 0 | 0.204 | 3 | 0.9769 |
| rs2508922 | 11 | 118681088 | A | G | 1.405 | 0.16 | ++++- | 40.5 | 10.081 | 4 | 0.03908 |
| rs55940627 | 11 | 118163920 | A | G | -1.404 | 0.1602 | -?-?? | 0 | 0.742 | 1 | 0.389 |
| rs550999 | 11 | 118811684 | G | C | 1.404 | 0.1603 | +-+++ | -34.8 | 4.45 | 4 | 0.3486 |
| rs56357739 | 11 | 118158507 | G | A | -1.403 | 0.1605 | -?-?? | 0 | 0.666 | 1 | 0.4145 |
| rs57075232 | 11 | 118159371 | T | C | -1.403 | 0.1605 | -?-?? | 0 | 0.688 | 1 | 0.407 |
| rs572126 | 11 | 118359161 | G | A | 1.402 | 0.1609 | +++++ | 0 | 0.457 | 4 | 0.9776 |
| rs4938501 | 11 | 118166497 | C | T | -1.401 | 0.1611 | -?-?? | 0 | 0.8 | 1 | 0.3711 |
| rs56066890 | 11 | 118169790 | C | T | -1.398 | 0.1621 | -?-?? | 0 | 0.729 | 1 | 0.3932 |
| rs59134737 | 11 | 118160959 | T | G | -1.395 | 0.1631 | -?-?? | 0 | 0.716 | 1 | 0.3976 |
| rs12576947 | 11 | 118158055 | C | T | -1.393 | 0.1635 | -?-?? | 0 | 0.682 | 1 | 0.4089 |
| rs59399105 | 11 | 118168568 | A | C | -1.393 | 0.1637 | -?-?? | 0 | 0.762 | 1 | 0.3826 |
| rs58093803 | 11 | 118168156 | C | T | -1.391 | 0.1641 | -?-?? | 0 | 0.765 | 1 | 0.3819 |
| rs59450866 | 11 | 118167707 | T | C | -1.391 | 0.1641 | -?-?? | 0 | 0.765 | 1 | 0.3818 |
| rs7110019 | 11 | 118168617 | G | A | -1.391 | 0.1643 | -?-?? | 0 | 0.862 | 1 | 0.3531 |
| rs12576941 | 11 | 118158002 | C | T | -1.39 | 0.1644 | -?-?? | 0 | 0.687 | 1 | 0.4072 |
| rs623500 | 11 | 118828942 | G | A | 1.389 | 0.1647 | +?-?+ | 0 | 1.922 | 2 | 0.3825 |
| rs3887015 | 11 | 118156410 | C | T | -1.385 | 0.166 | -?-?? | 0 | 0.696 | 1 | 0.4042 |
| rs56318049 | 11 | 118166608 | C | T | -1.385 | 0.166 | -?-?? | 0 | 0.776 | 1 | 0.3785 |
| rs66832804 | 11 | 118157049 | A | T | -1.385 | 0.166 | -?-?? | 0 | 0.696 | 1 | 0.4043 |
| rs61390396 | 11 | 118156718 | T | C | -1.384 | 0.1663 | -?-?? | 0 | 0.697 | 1 | 0.4038 |
| rs55910314 | 11 | 118163780 | T | C | -1.384 | 0.1664 | -?-?? | 0 | 0.778 | 1 | 0.3778 |
| rs2277292 | 11 | 118425419 | T | G | 1.383 | 0.1667 | +++-+ | 0 | 1.461 | 4 | 0.8336 |
| rs56938099 | 11 | 118156688 | G | A | -1.383 | 0.1667 | -?-?? | 0 | 0.7 | 1 | 0.4029 |
| rs55899310 | 11 | 118158702 | C | T | -1.382 | 0.1671 | -?-?? | 0 | 0.702 | 1 | 0.4022 |
| rs12576100 | 11 | 118158281 | A | G | -1.38 | 0.1677 | -?-?? | 0 | 0.705 | 1 | 0.4012 |
| rs12576118 | 11 | 118158440 | A | G | -1.379 | 0.1678 | -?-?? | 0 | 0.705 | 1 | 0.401 |
| rs11216887 | 11 | 118420736 | C | T | 1.378 | 0.1681 | +++-+ | 0 | 1.424 | 4 | 0.8401 |
| rs11216896 | 11 | 118430048 | T | C | 1.377 | 0.1686 | +++-+ | 0 | 1.608 | 4 | 0.8073 |
| rs4938499 | 11 | 118162496 | C | T | -1.377 | 0.1686 | -?-?? | 0 | 0.791 | 1 | 0.3738 |
| rs79915705 | 11 | 72016091 | C | T | 1.377 | 0.1686 | +++?- | 0 | 1.302 | 3 | 0.7286 |
| rs55798741 | 11 | 118154114 | T | G | -1.376 | 0.1688 | -?-?? | 0 | 0.78 | 1 | 0.377 |

| | | | | | | | | | | | |
|---|---|---|---|---|---|---|---|---|---|---|---|
| rs55806415 | 11 | 118162862 | A | G | -1.376 | 0.1689 | -?-?? | 0 | 0.792 | 1 | 0.3734 |
| rs58520343 | 11 | 118155000 | T | A | -1.376 | 0.1689 | -?-?? | 0 | 0.712 | 1 | 0.3989 |
| rs1074459 | 11 | 118155997 | C | T | -1.374 | 0.1696 | -?-?? | 0 | 0.715 | 1 | 0.3978 |
| rs58210741 | 11 | 118159326 | G | C | -1.373 | 0.1697 | -?-?? | 0 | 0.739 | 1 | 0.39 |
| rs55815657 | 11 | 118155074 | A | G | -1.372 | 0.17 | -?-?? | 0 | 0.717 | 1 | 0.3971 |
| rs67462512 | 11 | 118155336 | C | G | -1.37 | 0.1706 | -?-?? | 0 | 0.713 | 1 | 0.3983 |
| rs111227443 | 11 | 118590961 | T | C | 1.37 | 0.1707 | -?+++ | 0 | 2.74 | 3 | 0.4334 |
| rs58983315 | 11 | 118167759 | A | G | -1.37 | 0.1707 | -?-?? | 0 | 0.686 | 1 | 0.4075 |
| rs1074457 | 11 | 118156117 | T | A | -1.369 | 0.171 | -?-?? | 0 | 0.723 | 1 | 0.3951 |
| rs12577820 | 11 | 118164233 | T | C | -1.368 | 0.1714 | -?-?? | 0 | 0.807 | 1 | 0.3691 |
| rs56224295 | 11 | 118161891 | A | G | -1.365 | 0.1724 | -?-?? | 0 | 0.756 | 1 | 0.3847 |
| rs112799932 | 11 | 118585146 | G | T | 1.364 | 0.1725 | +++-- | 30.3 | 8.602 | 4 | 0.07184 |
| rs7113731 | 11 | 118831824 | G | A | -1.364 | 0.1726 | ---?- | 0 | 0.383 | 3 | 0.9437 |
| rs56020103 | 11 | 118172132 | T | C | -1.363 | 0.1728 | -?-?? | 0 | 0.749 | 1 | 0.3869 |
| rs11552421 | 11 | 118430519 | T | C | 1.363 | 0.173 | +++-+ | 0 | 1.619 | 4 | 0.8055 |
| rs573971 | 11 | 118393396 | A | G | 1.363 | 0.173 | +++++ | 0 | 0.449 | 4 | 0.9783 |
| rs56346212 | 11 | 118172342 | A | T | -1.362 | 0.1733 | -?-?? | 0 | 0.751 | 1 | 0.3861 |
| rs56292628 | 11 | 118163872 | A | G | -1.359 | 0.1742 | -?-?? | 0 | 0.823 | 1 | 0.3644 |
| rs7480736 | 11 | 118172364 | C | T | -1.357 | 0.1747 | -?-?? | 0 | 0.759 | 1 | 0.3837 |
| rs4635088 | 11 | 118832924 | C | G | -1.356 | 0.175 | ---?- | 0 | 0.383 | 3 | 0.9437 |
| rs894677 | 11 | 118172535 | T | C | -1.353 | 0.1759 | -?-?? | 0 | 0.758 | 1 | 0.3839 |
| rs111942423 | 11 | 118429764 | G | T | 1.353 | 0.176 | +++-+ | 0 | 1.496 | 4 | 0.8273 |
| rs55789743 | 11 | 118171949 | A | C | -1.352 | 0.1763 | -?-?? | 0 | 0.767 | 1 | 0.381 |
| rs60618948 | 11 | 118155814 | A | G | -1.351 | 0.1767 | -?-?? | 0 | 0.754 | 1 | 0.3853 |
| rs2187557 | 11 | 118148271 | T | C | -1.349 | 0.1772 | -?-?+ | -86.1 | 2.149 | 2 | 0.3414 |
| rs3132786 | 11 | 118365956 | C | T | 1.345 | 0.1785 | +++++ | 0 | 0.506 | 4 | 0.9729 |
| rs4310630 | 11 | 118836638 | G | A | -1.344 | 0.1788 | ---?- | 0 | 0.432 | 3 | 0.9336 |
| rs61150131 | 11 | 118752178 | C | T | 1.336 | 0.1814 | -++++ | 0 | 1.652 | 4 | 0.7995 |
| rs11607835 | 11 | 118835258 | C | T | -1.335 | 0.1819 | ---?- | 0 | 0.437 | 3 | 0.9325 |
| rs7112758 | 11 | 118836821 | A | G | -1.333 | 0.1825 | ---?- | 0 | 0.403 | 3 | 0.9397 |
| rs60545357 | 11 | 118281418 | C | A | 1.32 | 0.1867 | ++++- | 0 | 1.551 | 4 | 0.8175 |
| rs118020807 | 11 | 118759905 | A | C | -1.317 | 0.1878 | -?--? | 0 | 1.306 | 2 | 0.5205 |
| rs12289253 | 11 | 118405068 | G | A | 1.315 | 0.1885 | +++-+ | 0 | 0.783 | 4 | 0.9407 |
| rs7110743 | 11 | 118825762 | A | C | 1.314 | 0.1887 | +?+?- | 0 | 1.806 | 2 | 0.4054 |
| rs1790186 | 11 | 118846355 | T | C | -1.312 | 0.1895 | -?+?? | -48.7 | 2.017 | 1 | 0.1556 |
| rs543583 | 11 | 118847196 | A | G | -1.308 | 0.191 | -?+?? | -49.7 | 2.004 | 1 | 0.1569 |
| rs7940757 | 11 | 118055273 | T | C | 1.305 | 0.192 | +++++ | 0 | 0.43 | 4 | 0.9799 |
| rs6589650 | 11 | 118055217 | C | T | 1.301 | 0.1934 | +++++ | 0 | 0.429 | 4 | 0.98 |
| rs7107756 | 11 | 118717708 | A | G | 1.3 | 0.1935 | -++++ | 0 | 2.945 | 4 | 0.5671 |
| rs586370 | 11 | 118816353 | A | G | -1.3 | 0.1936 | -?--- | 0 | 0.719 | 3 | 0.8687 |
| rs7944087 | 11 | 118055733 | T | C | 1.297 | 0.1945 | +++++ | 0 | 0.416 | 4 | 0.9812 |
| rs75488969 | 11 | 118712509 | T | C | 1.297 | 0.1947 | +?-++ | 0 | 2.814 | 3 | 0.4212 |
| rs11216804 | 11 | 118057320 | G | A | 1.293 | 0.196 | +++++ | 0 | 0.421 | 4 | 0.9807 |
| rs4639966 | 11 | 118573519 | C | T | 1.292 | 0.1964 | +++-- | 36.1 | 9.395 | 4 | 0.05195 |
| rs548877 | 11 | 118300240 | G | A | 1.291 | 0.1966 | +++-+ | 0 | 1.57 | 4 | 0.8142 |
| rs73530785 | 11 | 72152606 | G | A | 1.291 | 0.1968 | +++?- | -33.1 | 3.755 | 3 | 0.2891 |
| rs7944321 | 11 | 118055801 | A | G | 1.29 | 0.197 | +++++ | 0 | 0.322 | 4 | 0.9884 |
| rs58682117 | 11 | 72531000 | C | T | -1.29 | 0.1972 | +--++ | 55.2 | 13.404 | 4 | 0.009462 |
| rs7931206 | 11 | 118056092 | C | T | 1.288 | 0.1977 | +++++ | 0 | 0.419 | 4 | 0.9809 |
| rs57464210 | 11 | 72043183 | G | T | 1.285 | 0.199 | +++?- | 26.8 | 6.831 | 3 | 0.07748 |
| rs12295634 | 11 | 118054431 | T | C | 1.282 | 0.2 | +++++ | 0 | 0.335 | 4 | 0.9874 |
| rs45480496 | 11 | 118270715 | A | G | 1.276 | 0.2018 | ++++- | 0 | 1.357 | 4 | 0.8516 |
| rs34815971 | 11 | 118301066 | G | A | 1.272 | 0.2034 | +++-+ | 0 | 1.558 | 4 | 0.8163 |
| rs7935561 | 11 | 118056722 | A | G | 1.268 | 0.2049 | +++++ | 0 | 0.325 | 4 | 0.9882 |
| rs113971393 | 11 | 72076181 | C | T | 1.267 | 0.2051 | +++?- | 0 | 2.481 | 3 | 0.4787 |
| rs12274740 | 11 | 118054128 | C | T | 1.264 | 0.2063 | +++++ | 0 | 0.345 | 4 | 0.9868 |

| | | | | | | | | | | | |
|---|---|---|---|---|---|---|---|---|---|---|---|
| rs117498055 | 11 | 72022810 | A | G | 1.261 | 0.2073 | +?+?+ | 0 | 0.735 | 2 | 0.6925 |
| rs117091621 | 11 | 72026473 | T | C | 1.26 | 0.2078 | +?+?+ | 0 | 0.732 | 2 | 0.6936 |
| rs117392768 | 11 | 72043182 | A | T | 1.259 | 0.2081 | +?+?+ | 0 | 1.543 | 2 | 0.4624 |
| rs510997 | 11 | 118302901 | C | T | 1.259 | 0.2082 | +++-+ | 0 | 1.662 | 4 | 0.7977 |
| rs79281178 | 11 | 118298041 | G | C | 1.259 | 0.2082 | +++-+ | 0 | 1.583 | 4 | 0.8118 |
| rs7124100 | 11 | 118290169 | A | G | 1.251 | 0.211 | +++-+ | 0 | 1.764 | 4 | 0.779 |
| rs117705968 | 11 | 72037907 | A | C | 1.248 | 0.2119 | +?+?+ | 0 | 1.507 | 2 | 0.4707 |
| rs117902312 | 11 | 72042303 | G | C | 1.244 | 0.2135 | +?+?+ | 0 | 1.492 | 2 | 0.4743 |
| rs7944366 | 11 | 118056047 | G | A | 1.243 | 0.2138 | +++++ | 0 | 0.34 | 4 | 0.9871 |
| rs77342802 | 11 | 72012038 | C | G | 1.234 | 0.2171 | +?+?+ | 0 | 0.871 | 2 | 0.6468 |
| rs116888156 | 11 | 72009595 | A | G | 1.233 | 0.2178 | +?+?+ | 0 | 0.919 | 2 | 0.6316 |
| rs117225442 | 11 | 72060986 | A | G | 1.232 | 0.2179 | +?+?+ | 0 | 1.437 | 2 | 0.4875 |
| rs11216802 | 11 | 118057227 | T | C | 1.231 | 0.2182 | +++++ | 0 | 0.347 | 4 | 0.9866 |
| rs11216803 | 11 | 118057244 | A | C | 1.23 | 0.2187 | +++++ | 0 | 0.346 | 4 | 0.9866 |
| rs117524306 | 11 | 72014266 | G | A | 1.23 | 0.2187 | +?+?+ | 0 | 0.861 | 2 | 0.6502 |
| rs56754533 | 11 | 118188276 | A | G | -1.229 | 0.2192 | -?--+ | 0 | 1.401 | 3 | 0.7053 |
| rs1561966 | 11 | 118221474 | A | G | -1.228 | 0.2194 | +++-- | 4.2 | 6.263 | 4 | 0.1804 |
| rs143605660 | 11 | 118707035 | A | G | 1.227 | 0.2199 | -?+++ | -66.6 | 3.002 | 3 | 0.3914 |
| rs7940717 | 11 | 118055110 | T | G | 1.226 | 0.2203 | +++++ | 0 | 0.32 | 4 | 0.9885 |
| rs117346296 | 11 | 72025140 | A | G | 1.224 | 0.2208 | +?+?+ | 0 | 0.649 | 2 | 0.7229 |
| rs118115020 | 11 | 72067731 | T | C | 1.223 | 0.2213 | +?+?+ | 0 | 1.69 | 2 | 0.4296 |
| rs7947185 | 11 | 118214726 | C | T | -1.218 | 0.2233 | ---?+ | 28 | 6.942 | 3 | 0.07378 |
| rs7939212 | 11 | 72021876 | T | G | 1.217 | 0.2235 | +?+?+ | 0 | 0.774 | 2 | 0.6791 |
| rs7940087 | 11 | 72021877 | T | C | 1.216 | 0.2241 | +?+?+ | 0 | 0.77 | 2 | 0.6804 |
| rs7929825 | 11 | 118788081 | G | C | -1.214 | 0.2246 | ----+ | 33.6 | 9.04 | 4 | 0.06011 |
| rs11217068 | 11 | 118729283 | T | C | -1.213 | 0.2252 | -?+-- | 7.7 | 5.418 | 3 | 0.1436 |
| rs73613959 | 11 | 118263054 | A | G | 1.212 | 0.2255 | ++++- | 0 | 1.378 | 4 | 0.8481 |
| rs73613993 | 11 | 118296007 | C | T | 1.21 | 0.2261 | +++-+ | 0 | 1.763 | 4 | 0.7792 |
| rs3212263 | 11 | 118216423 | G | A | -1.21 | 0.2262 | +++-- | -0.1 | 5.996 | 4 | 0.1994 |
| rs117215416 | 11 | 72046796 | C | T | 1.208 | 0.2271 | +?+?+ | 0 | 1.373 | 2 | 0.5034 |
| rs117999328 | 11 | 72019118 | C | T | 1.206 | 0.2276 | +?+?+ | 0 | 0.75 | 2 | 0.6872 |
| rs118089225 | 11 | 72018472 | T | C | 1.206 | 0.2276 | +?+?+ | 0 | 0.75 | 2 | 0.6872 |
| rs2282550 | 11 | 118220035 | C | T | -1.204 | 0.2287 | +++-- | 6.5 | 6.42 | 4 | 0.1699 |
| rs12804550 | 11 | 118011860 | A | G | -1.202 | 0.2294 | -?--- | 0 | 0.87 | 3 | 0.8326 |
| rs7938979 | 11 | 72021720 | C | G | 1.202 | 0.2294 | +?+?+ | 0 | 0.736 | 2 | 0.6921 |
| rs117954624 | 11 | 72028005 | G | A | 1.201 | 0.2298 | +?+?+ | 0 | 0.761 | 2 | 0.6836 |
| rs11606719 | 11 | 118782474 | G | C | 1.197 | 0.2314 | ++-+? | 0 | 2.41 | 3 | 0.4918 |
| rs1079118 | 11 | 118258056 | A | G | 1.194 | 0.2325 | ++++- | 0 | 1.226 | 4 | 0.8738 |
| rs117237510 | 11 | 72009451 | T | C | 1.19 | 0.2339 | +?+?+ | 0 | 0.785 | 2 | 0.6754 |
| rs117939125 | 11 | 72062164 | G | T | 1.189 | 0.2346 | +?+?+ | 0 | 1.385 | 2 | 0.5002 |
| rs116951669 | 11 | 72018759 | C | T | 1.188 | 0.2347 | +?+?+ | 0 | 0.707 | 2 | 0.7023 |
| rs117190399 | 11 | 72056023 | T | G | 1.186 | 0.2356 | +?+?+ | 0 | 1.37 | 2 | 0.5041 |
| rs117613362 | 11 | 72051945 | C | A | 1.182 | 0.2373 | +?+?+ | 0 | 1.32 | 2 | 0.5169 |
| rs80045923 | 11 | 72063551 | T | C | 1.18 | 0.2378 | +?+?+ | 0 | 1.387 | 2 | 0.4998 |
| rs590580 | 11 | 118066104 | A | G | 1.178 | 0.2386 | +?+?+ | 0 | 1.209 | 2 | 0.5463 |
| rs149666559 | 11 | 72042565 | A | G | 1.178 | 0.2387 | +?+?+ | 0 | 1.438 | 2 | 0.4873 |
| rs58571741 | 11 | 72065710 | A | C | 1.173 | 0.2408 | +++?- | 0 | 2.75 | 3 | 0.4317 |
| rs61059752 | 11 | 118246071 | A | C | 1.172 | 0.2411 | ++++- | 0 | 1.216 | 4 | 0.8755 |
| rs151127828 | 11 | 118206121 | A | G | 1.164 | 0.2444 | +?+++ | 0 | 0.119 | 3 | 0.9895 |
| rs489126 | 11 | 118572747 | A | G | 1.144 | 0.2524 | ++-++ | 0 | 3.302 | 4 | 0.5086 |
| rs117498960 | 11 | 72065210 | C | T | 1.14 | 0.2544 | +?+?- | 0 | 1.641 | 2 | 0.4401 |
| rs7933963 | 11 | 118246906 | T | C | 1.136 | 0.2558 | ++++- | 0 | 1.223 | 4 | 0.8743 |
| rs519294 | 11 | 118573743 | A | C | 1.13 | 0.2584 | ++-++ | 0 | 3.548 | 4 | 0.4706 |
| rs75040111 | 11 | 118747008 | T | C | -1.127 | 0.2596 | -+-?? | -34.8 | 2.966 | 2 | 0.2269 |
| rs657539 | 11 | 118572021 | T | C | 1.126 | 0.2602 | ++-++ | 0 | 3.437 | 4 | 0.4876 |
| rs67514601 | 11 | 118189477 | G | A | -1.123 | 0.2614 | -?--- | 0 | 0.71 | 3 | 0.8709 |

| | | | | | | | | | | | |
|---|---|---|---|---|---|---|---|---|---|---|---|
| rs7123811 | 11 | 118569469 | G | T | 1.123 | 0.2616 | ++-++ | 0 | 3.435 | 4 | 0.4878 |
| rs118105218 | 11 | 72077497 | G | T | 1.119 | 0.263 | +?+?+ | 0 | 1.504 | 2 | 0.4714 |
| rs11216856 | 11 | 118227838 | T | C | -1.119 | 0.2633 | +++-- | 0.4 | 6.026 | 4 | 0.1972 |
| rs611217 | 11 | 118572611 | T | C | 1.118 | 0.2634 | ++-++ | 0 | 3.627 | 4 | 0.4588 |
| rs517445 | 11 | 118573535 | G | A | 1.114 | 0.2654 | ++-++ | 0 | 3.437 | 4 | 0.4876 |
| rs611733 | 11 | 118572760 | T | C | 1.103 | 0.2699 | ++-++ | 0 | 3.418 | 4 | 0.4905 |
| rs117652953 | 11 | 72084357 | G | A | 1.103 | 0.2701 | +?+?- | 0 | 1.527 | 2 | 0.4661 |
| rs2276417 | 11 | 118230223 | G | T | -1.1 | 0.2712 | +++-- | -1.5 | 5.911 | 4 | 0.2059 |
| rs656287 | 11 | 118571790 | G | A | 1.1 | 0.2714 | ++-++ | 0 | 3.502 | 4 | 0.4777 |
| rs544452 | 11 | 118571357 | T | G | 1.097 | 0.2725 | ++-++ | 0 | 3.622 | 4 | 0.4596 |
| rs9666462 | 11 | 118275997 | A | G | 1.097 | 0.2726 | +++-+ | 0 | 1.033 | 4 | 0.9047 |
| rs657769 | 11 | 118571316 | T | C | 1.097 | 0.2727 | ++-++ | 0 | 3.614 | 4 | 0.4608 |
| rs673547 | 11 | 118573339 | A | G | 1.095 | 0.2736 | ++-++ | 0 | 3.466 | 4 | 0.4831 |
| rs574383 | 11 | 118573668 | C | T | 1.094 | 0.2738 | ++-++ | 0 | 3.4 | 4 | 0.4932 |
| rs647099 | 11 | 118067009 | A | G | 1.094 | 0.274 | +?+?+ | 0 | 0.978 | 2 | 0.6133 |
| rs488141 | 11 | 118571168 | G | A | 1.092 | 0.2746 | ++-++ | 0 | 3.642 | 4 | 0.4566 |
| rs658676 | 11 | 118571123 | T | C | 1.092 | 0.2746 | ++-++ | 0 | 3.642 | 4 | 0.4566 |
| rs541631 | 11 | 118571023 | G | A | 1.092 | 0.2747 | ++-++ | 0 | 3.643 | 4 | 0.4565 |
| rs659564 | 11 | 118570914 | A | C | 1.092 | 0.2747 | ++-++ | 0 | 3.643 | 4 | 0.4565 |
| rs7108144 | 11 | 118569694 | A | G | 1.091 | 0.2752 | ++-++ | 0 | 3.628 | 4 | 0.4587 |
| rs642224 | 11 | 118570919 | A | G | 1.091 | 0.2754 | ++-++ | 0 | 3.643 | 4 | 0.4565 |
| rs552079 | 11 | 118573022 | C | T | 1.09 | 0.2755 | ++-++ | 0 | 3.469 | 4 | 0.4826 |
| rs523793 | 11 | 118570697 | T | C | 1.089 | 0.276 | ++-++ | 0 | 3.639 | 4 | 0.4571 |
| rs554456 | 11 | 118570062 | C | T | 1.089 | 0.276 | ++-++ | 0 | 3.639 | 4 | 0.457 |
| rs73613980 | 11 | 118284650 | T | C | 1.089 | 0.276 | +++-+ | 0 | 1.06 | 4 | 0.9006 |
| rs9735317 | 11 | 118725383 | T | C | -1.089 | 0.2762 | -?+-- | 6.2 | 5.333 | 3 | 0.149 |
| rs12793594 | 11 | 118569900 | C | T | 1.088 | 0.2764 | ++-++ | 0 | 3.645 | 4 | 0.4561 |
| rs482889 | 11 | 118570098 | T | G | 1.089 | 0.2764 | ++-++ | 0 | 3.645 | 4 | 0.4562 |
| rs570949 | 11 | 118571911 | T | G | 1.086 | 0.2776 | ++-++ | 0 | 3.615 | 4 | 0.4606 |
| rs9666463 | 11 | 118276116 | G | C | 1.084 | 0.2783 | +++-+ | 0 | 1.004 | 4 | 0.9092 |
| rs11216920 | 11 | 118462554 | G | A | 1.084 | 0.2784 | -++-+ | 0 | 1.662 | 4 | 0.7976 |
| rs7479114 | 11 | 118568813 | T | C | 1.084 | 0.2784 | ++-++ | 0 | 3.404 | 4 | 0.4926 |
| rs477106 | 11 | 118363424 | A | C | 1.082 | 0.2793 | +++-+ | 0 | 0.735 | 4 | 0.9469 |
| rs4486664 | 11 | 118569067 | C | A | 1.081 | 0.2797 | ++-++ | 0 | 3.121 | 4 | 0.5377 |
| rs648947 | 11 | 118067426 | A | G | 1.081 | 0.2797 | +?+?+ | 0 | 0.993 | 2 | 0.6086 |
| rs619250 | 11 | 118064582 | A | T | 1.081 | 0.2798 | +?+?+ | 0 | 0.772 | 2 | 0.6798 |
| rs7483081 | 11 | 118276246 | C | T | 1.08 | 0.2802 | +++-+ | 0 | 0.998 | 4 | 0.9102 |
| rs2071381 | 11 | 118214267 | A | G | -1.079 | 0.2807 | +++-- | -16.7 | 5.14 | 4 | 0.2732 |
| rs1790192 | 11 | 118757191 | G | A | -1.076 | 0.2817 | --+-- | 32.1 | 8.84 | 4 | 0.06523 |
| rs1613643 | 11 | 118068521 | T | G | 1.076 | 0.2819 | +?+?+ | 0 | 1 | 2 | 0.6064 |
| rs511451 | 11 | 118568639 | T | C | 1.075 | 0.2826 | ++-++ | 0 | 3.408 | 4 | 0.492 |
| rs17122157 | 11 | 118276492 | A | G | 1.074 | 0.2828 | +++-+ | 0 | 0.999 | 4 | 0.9099 |
| rs12798851 | 11 | 118135973 | G | T | 1.074 | 0.283 | +?-+- | -26.7 | 3.946 | 3 | 0.2674 |
| rs74678032 | 11 | 118812559 | C | T | 1.073 | 0.2833 | +?-++ | -13.6 | 4.402 | 3 | 0.2212 |
| rs648682 | 11 | 118066582 | T | C | 1.073 | 0.2835 | +?+?+ | 0 | 0.96 | 2 | 0.6187 |
| rs117088064 | 11 | 72074257 | T | C | 1.071 | 0.2843 | +?+?+ | 0 | 1.508 | 2 | 0.4704 |
| rs117162202 | 11 | 72077463 | A | G | 1.07 | 0.2846 | +?+?+ | 0 | 1.507 | 2 | 0.4708 |
| rs7124127 | 11 | 118006408 | C | T | 1.067 | 0.286 | +-++- | 0 | 3.078 | 4 | 0.5448 |
| rs570371 | 11 | 118568472 | G | A | 1.063 | 0.288 | ++-++ | 0 | 3.498 | 4 | 0.4781 |
| rs71482156 | 11 | 118734516 | T | A | 1.06 | 0.2891 | +?+?? | 0 | 0.004 | 1 | 0.9474 |
| rs1791936 | 11 | 72877964 | G | A | -1.057 | 0.2907 | ----- | 0 | 0.295 | 4 | 0.9902 |
| rs4290266 | 11 | 118285995 | A | G | 1.056 | 0.2908 | +++-+ | 0 | 1.297 | 4 | 0.8619 |
| rs951657 | 11 | 118067621 | A | G | 1.056 | 0.2908 | +?+?+ | 0 | 1.04 | 2 | 0.5944 |
| rs4393358 | 11 | 118569097 | T | C | 1.053 | 0.2922 | ++-++ | 0 | 3.342 | 4 | 0.5023 |
| rs647573 | 11 | 118067132 | C | T | 1.05 | 0.2938 | +?+?+ | 0 | 0.95 | 2 | 0.6218 |
| rs658040 | 11 | 118572171 | G | A | 1.045 | 0.2961 | ++-++ | 0 | 3.893 | 4 | 0.4207 |

| | | | | | | | | | | | |
|---|---|---|---|---|---|---|---|---|---|---|---|
| rs150193966 | 11 | 72071662 | A | G | 1.042 | 0.2974 | +?+?+ | 0 | 1.412 | 2 | 0.4937 |
| rs605908 | 11 | 118067192 | C | A | 1.042 | 0.2976 | +?+?+ | 0 | 0.879 | 2 | 0.6444 |
| rs7106851 | 11 | 118554259 | A | G | 1.041 | 0.2978 | ++-++ | 0 | 3.27 | 4 | 0.5137 |
| rs11216859 | 11 | 118238119 | C | T | -1.04 | 0.2984 | +++-- | -19 | 5.042 | 4 | 0.283 |
| rs12225399 | 11 | 118480285 | C | G | 1.038 | 0.2993 | +?+-+ | 0 | 1.241 | 3 | 0.7431 |
| rs7108613 | 11 | 118268613 | G | A | 1.036 | 0.3004 | +++-+ | 0 | 0.876 | 4 | 0.928 |
| rs111656195 | 11 | 118301375 | A | G | 1.035 | 0.3008 | +?-+- | -42 | 3.522 | 3 | 0.318 |
| rs647975 | 11 | 118038741 | T | C | 1.031 | 0.3024 | +?-?+ | 0 | 1.57 | 2 | 0.4562 |
| rs10790241 | 11 | 118013034 | A | G | -1.025 | 0.3054 | -?--- | 0 | 0.553 | 3 | 0.9071 |
| rs111903103 | 11 | 118308499 | T | C | 1.017 | 0.3092 | +?++- | -44 | 3.472 | 3 | 0.3244 |
| rs12276533 | 11 | 118724116 | A | G | -1.016 | 0.3095 | -?+-- | 5.5 | 5.293 | 3 | 0.1515 |
| rs11216949 | 11 | 118568668 | T | C | 1.016 | 0.3096 | ++-++ | 0 | 3.478 | 4 | 0.4813 |
| rs17122170 | 11 | 118296668 | C | G | 1.011 | 0.3119 | +++-+ | 0 | 1.609 | 4 | 0.8071 |
| rs56785719 | 11 | 118176623 | G | C | -1.011 | 0.312 | -?-?+ | -86.1 | 2.15 | 2 | 0.3414 |
| rs341048 | 11 | 72298280 | T | C | -1.011 | 0.3122 | +?-?- | -58.9 | 2.517 | 2 | 0.2841 |
| rs4331144 | 11 | 118554100 | G | A | 1.007 | 0.314 | ++-+- | 0 | 3.417 | 4 | 0.4906 |
| rs581063 | 11 | 118767077 | C | T | -1.005 | 0.315 | --+-- | 0 | 0.569 | 4 | 0.9664 |
| rs7106652 | 11 | 118554248 | T | C | 1.005 | 0.315 | ++-+- | 0 | 3.353 | 4 | 0.5005 |
| rs577217 | 11 | 72489023 | G | A | -1.004 | 0.3153 | -++-- | 40.3 | 10.045 | 4 | 0.03968 |
| rs1599809 | 11 | 118179200 | A | G | -1.002 | 0.3165 | -?-?+ | 0 | 1.752 | 2 | 0.4165 |
| rs141752759 | 11 | 118194647 | G | C | 1.001 | 0.3168 | +?+++ | 0 | 0.254 | 3 | 0.9684 |
| rs7119568 | 11 | 118723753 | A | T | -0.998 | 0.3182 | -?+-- | 4.8 | 5.251 | 3 | 0.1543 |
| rs7924647 | 11 | 118728479 | G | A | -0.995 | 0.3196 | -?+-- | 4.1 | 5.211 | 3 | 0.157 |
| rs481903 | 11 | 118567733 | C | T | 0.994 | 0.3201 | ++-++ | 0 | 3.167 | 4 | 0.5304 |
| rs3212264 | 11 | 118216234 | A | C | 0.994 | 0.3202 | ++++- | 0 | 2.917 | 4 | 0.5719 |
| rs11602869 | 11 | 118554448 | A | G | 0.992 | 0.3212 | ++-+- | 0 | 3.365 | 4 | 0.4987 |
| rs592280 | 11 | 118553218 | C | T | 0.991 | 0.3215 | ++-+- | 0 | 3.053 | 4 | 0.5489 |
| rs673770 | 11 | 118550601 | G | T | 0.991 | 0.3217 | ++-+- | 0 | 3.184 | 4 | 0.5275 |
| rs7110984 | 11 | 118442234 | A | G | 0.982 | 0.3262 | +-++? | 0 | 2.024 | 3 | 0.5674 |
| rs642497 | 11 | 118550784 | A | C | 0.98 | 0.3272 | ++-+- | 0 | 3.169 | 4 | 0.5299 |
| rs3782042 | 11 | 118185919 | A | G | -0.976 | 0.3291 | -?-?+ | 22.4 | 5.155 | 2 | 0.07598 |
| rs11605069 | 11 | 118554436 | G | T | 0.973 | 0.3306 | ++-+- | 0 | 3.289 | 4 | 0.5107 |
| rs4544037 | 11 | 118222062 | C | T | 0.972 | 0.3308 | ++++- | 0 | 2.967 | 4 | 0.5633 |
| rs12799314 | 11 | 118551597 | A | G | 0.972 | 0.3311 | ++-+- | 0 | 3.215 | 4 | 0.5224 |
| rs11217083 | 11 | 118770321 | G | A | -0.972 | 0.3312 | --+-- | 0 | 0.644 | 4 | 0.9581 |
| rs644867 | 11 | 118567545 | G | A | 0.971 | 0.3314 | ++-++ | 0 | 3.433 | 4 | 0.4882 |
| rs3132893 | 11 | 118303822 | A | C | 0.971 | 0.3316 | +++-+ | 0 | 1.95 | 4 | 0.745 |
| rs4936433 | 11 | 118552420 | G | A | 0.97 | 0.332 | ++-+- | 0 | 3.179 | 4 | 0.5284 |
| rs642530 | 11 | 118550756 | C | T | 0.968 | 0.3331 | ++-+- | 0 | 3.208 | 4 | 0.5237 |
| rs67307131 | 11 | 118480223 | C | T | 0.968 | 0.3332 | +?+-+ | 0 | 1.392 | 3 | 0.7075 |
| rs113488707 | 11 | 118045876 | T | C | 0.966 | 0.334 | +?+?- | -80.8 | 2.213 | 2 | 0.3308 |
| rs549370 | 11 | 118552842 | A | C | 0.96 | 0.3371 | ++-+- | 0 | 3.239 | 4 | 0.5187 |
| rs2368238 | 11 | 118091761 | G | T | -0.959 | 0.3374 | -?+?- | 0 | 0.761 | 2 | 0.6834 |
| rs667982 | 11 | 118562767 | C | T | 0.959 | 0.3375 | ++-+- | 0 | 3.128 | 4 | 0.5366 |
| rs12796373 | 11 | 118554939 | A | G | 0.958 | 0.338 | ++-+- | 0 | 3.303 | 4 | 0.5084 |
| rs654792 | 11 | 118552996 | T | G | 0.957 | 0.3386 | ++-+- | 0 | 3.257 | 4 | 0.5157 |
| rs45444192 | 11 | 118208083 | A | G | 0.956 | 0.3389 | +?+-? | -43.7 | 2.785 | 2 | 0.2485 |
| rs519848 | 11 | 118552715 | A | G | 0.954 | 0.3399 | ++-+- | 0 | 3.273 | 4 | 0.5132 |
| rs1945765 | 11 | 118182829 | A | T | -0.953 | 0.3405 | -?-?+ | 0 | 1.926 | 2 | 0.3818 |
| rs11217081 | 11 | 118769032 | G | A | -0.95 | 0.3421 | --+-- | 0 | 0.607 | 4 | 0.9623 |
| rs112550735 | 11 | 118303200 | C | T | 0.949 | 0.3425 | +?-+- | -42.5 | 3.508 | 3 | 0.3197 |
| rs77728961 | 11 | 118303182 | C | T | 0.949 | 0.3427 | +?-+- | -42.5 | 3.509 | 3 | 0.3196 |
| rs4456279 | 11 | 72520003 | A | G | -0.946 | 0.344 | -++-- | 38.1 | 9.695 | 4 | 0.04588 |
| rs76930336 | 11 | 118464764 | A | C | 0.946 | 0.3442 | +-++? | 0 | 2.521 | 3 | 0.4715 |
| rs2051480 | 11 | 118094513 | G | A | -0.944 | 0.345 | -?+?- | 0 | 0.779 | 2 | 0.6773 |
| rs4938524 | 11 | 118552074 | T | G | 0.944 | 0.3451 | ++-+- | 0 | 3.23 | 4 | 0.5202 |

| | | | | | | | | | | | |
|---|---|---|---|---|---|---|---|---|---|---|---|
| rs644498 | 11 | 118567486 | C | T | 0.943 | 0.3458 | ++-++ | 0 | 3.309 | 4 | 0.5076 |
| rs563549 | 11 | 118558953 | A | G | 0.94 | 0.3474 | ++-+- | 0 | 3.271 | 4 | 0.5135 |
| rs600969 | 11 | 118545797 | T | C | 0.94 | 0.3474 | ++-+- | 0 | 3.186 | 4 | 0.5273 |
| rs509641 | 11 | 118558952 | C | T | 0.939 | 0.3475 | ++-+- | 0 | 3.273 | 4 | 0.5133 |
| rs45489394 | 11 | 118228968 | G | C | 0.939 | 0.3476 | ++++- | 0 | 2.908 | 4 | 0.5733 |
| rs6589674 | 11 | 118566039 | G | A | 0.939 | 0.348 | ++-++ | 0 | 2.881 | 4 | 0.5779 |
| rs57495164 | 11 | 118095148 | C | T | -0.938 | 0.3481 | -?+?- | 0 | 0.776 | 2 | 0.6784 |
| rs656575 | 11 | 118549908 | G | T | 0.937 | 0.3488 | ++-+- | 0 | 3.072 | 4 | 0.5458 |
| rs543413 | 11 | 118550017 | C | T | 0.937 | 0.349 | ++-+- | 0 | 3.068 | 4 | 0.5465 |
| rs507080 | 11 | 118549582 | A | G | 0.936 | 0.3493 | ++-+- | 0 | 3.046 | 4 | 0.5501 |
| rs59405103 | 11 | 118262305 | C | A | 0.935 | 0.3497 | +++-+ | 0 | 0.702 | 4 | 0.9511 |
| rs11216845 | 11 | 118163216 | T | C | -0.935 | 0.35 | -?-?+ | 0 | 1.367 | 2 | 0.5048 |
| rs562313 | 11 | 118548421 | A | G | 0.931 | 0.3519 | ++-+- | 0 | 3.238 | 4 | 0.5189 |
| rs510408 | 11 | 118558901 | C | G | 0.931 | 0.352 | ++-+- | 0 | 3.251 | 4 | 0.5168 |
| rs12277355 | 11 | 118729442 | T | C | -0.93 | 0.3522 | -++-- | -0.2 | 5.988 | 4 | 0.2 |
| rs12794726 | 11 | 118566313 | T | C | 0.93 | 0.3526 | ++-++ | 0 | 3.06 | 4 | 0.5478 |
| rs543327 | 11 | 118549985 | T | G | 0.929 | 0.3527 | ++-+- | 0 | 3.1 | 4 | 0.5412 |
| rs638805 | 11 | 118558889 | T | G | 0.929 | 0.3531 | ++-+- | 0 | 3.265 | 4 | 0.5146 |
| rs1540191 | 11 | 118090432 | A | G | -0.928 | 0.3532 | -?+?- | 0 | 0.89 | 2 | 0.6407 |
| rs7130076 | 11 | 118566090 | C | T | 0.927 | 0.3541 | ++-++ | 0 | 3.051 | 4 | 0.5493 |
| rs7122510 | 11 | 118708785 | A | C | -0.926 | 0.3543 | -?+-- | -0.3 | 4.985 | 3 | 0.1729 |
| rs514143 | 11 | 118558449 | A | G | 0.924 | 0.3554 | ++-+- | 0 | 2.972 | 4 | 0.5625 |
| rs478296 | 11 | 118567375 | C | T | 0.924 | 0.3556 | ++-++ | 0 | 3.471 | 4 | 0.4823 |
| rs542613 | 11 | 118567084 | A | G | 0.923 | 0.3561 | ++-++ | 0 | 3.136 | 4 | 0.5353 |
| rs7126172 | 11 | 118567257 | G | A | 0.922 | 0.3566 | ++-++ | 0 | 3.273 | 4 | 0.5133 |
| rs11216826 | 11 | 118099713 | T | A | -0.922 | 0.3567 | -?+?- | 0 | 0.764 | 2 | 0.6825 |
| rs7122511 | 11 | 118708786 | A | C | -0.921 | 0.3569 | -?+-- | 0 | 4.998 | 3 | 0.1719 |
| rs11216844 | 11 | 118162796 | G | A | -0.921 | 0.3571 | -?-?+ | 0 | 1.36 | 2 | 0.5066 |
| rs11216825 | 11 | 118099617 | A | G | -0.92 | 0.3574 | -?+?- | 0 | 0.767 | 2 | 0.6814 |
| rs659969 | 11 | 118549803 | T | C | 0.916 | 0.3594 | ++-+- | 0 | 3.154 | 4 | 0.5324 |
| rs493834 | 11 | 118567444 | G | A | 0.916 | 0.3595 | ++-++ | 0 | 3.209 | 4 | 0.5235 |
| rs11216819 | 11 | 118084980 | G | A | 0.916 | 0.3599 | +?-?- | 0 | 1.873 | 2 | 0.3921 |
| rs7114458 | 11 | 118566175 | A | G | 0.915 | 0.36 | ++-++ | 0 | 3.127 | 4 | 0.5367 |
| rs869638 | 11 | 118100411 | C | G | -0.915 | 0.3601 | -?+?- | 0 | 0.764 | 2 | 0.6825 |
| rs10892226 | 11 | 118096618 | A | G | -0.915 | 0.3602 | -?+?- | 0 | 0.715 | 2 | 0.6995 |
| rs625513 | 11 | 118555719 | C | A | 0.914 | 0.3608 | ++-+- | 0 | 2.966 | 4 | 0.5635 |
| rs60514290 | 11 | 118246048 | C | G | 0.911 | 0.3623 | +++-+ | 0 | 0.737 | 4 | 0.9467 |
| rs7125115 | 11 | 118478330 | A | G | 0.911 | 0.3624 | +++-+ | 0 | 1.297 | 4 | 0.8619 |
| rs525485 | 11 | 118555754 | A | G | 0.909 | 0.3633 | ++-+- | 0 | 2.991 | 4 | 0.5594 |
| rs45501598 | 11 | 118251980 | T | A | 0.908 | 0.364 | +++-+ | 0 | 0.831 | 4 | 0.9343 |
| rs558781 | 11 | 118564397 | T | G | 0.908 | 0.364 | ++-+- | 0 | 2.894 | 4 | 0.5757 |
| rs610664 | 11 | 118556691 | G | A | 0.907 | 0.3641 | ++-+- | 0 | 3.274 | 4 | 0.5131 |
| rs76126708 | 11 | 118259379 | C | A | 0.907 | 0.3645 | +++-+ | 0 | 0.785 | 4 | 0.9404 |
| rs7114216 | 11 | 118566234 | G | A | 0.907 | 0.3646 | ++-++ | 0 | 3.18 | 4 | 0.5281 |
| rs638301 | 11 | 118559003 | G | A | 0.904 | 0.3659 | ++-+- | 0 | 3.122 | 4 | 0.5376 |
| rs3782040 | 11 | 118176097 | G | A | -0.904 | 0.3661 | -?-?+ | -75.9 | 2.274 | 2 | 0.3208 |
| rs642662 | 11 | 118567020 | G | A | 0.904 | 0.3661 | ++-+- | 0 | 3.13 | 4 | 0.5363 |
| rs555356 | 11 | 118558078 | T | G | 0.901 | 0.3674 | ++-+- | 0 | 3.056 | 4 | 0.5485 |
| rs11607355 | 11 | 118093547 | T | C | -0.901 | 0.3678 | -?+?- | 0 | 0.803 | 2 | 0.6692 |
| rs1784298 | 11 | 118548665 | C | T | 0.9 | 0.368 | ++-+- | 0 | 3.025 | 4 | 0.5537 |
| rs561845 | 11 | 118558827 | G | A | 0.9 | 0.3681 | ++-+- | 0 | 3.229 | 4 | 0.5202 |
| rs4938522 | 11 | 118548671 | T | A | 0.899 | 0.3685 | ++-+- | 0 | 3.03 | 4 | 0.5528 |
| rs571751 | 11 | 118556797 | C | A | 0.896 | 0.3701 | ++-+- | 0 | 3.223 | 4 | 0.5212 |
| rs533144 | 11 | 118559247 | A | C | 0.894 | 0.3715 | ++-+- | 0 | 3.18 | 4 | 0.5281 |
| rs486462 | 11 | 118559178 | C | T | 0.893 | 0.3716 | ++-+- | 0 | 3.158 | 4 | 0.5317 |
| rs508579 | 11 | 118559084 | C | G | 0.892 | 0.3722 | ++-+- | 0 | 3.109 | 4 | 0.5397 |

| | | | | | | | | | | | |
|---|---|---|---|---|---|---|---|---|---|---|---|
| rs693306 | 11 | 118558333 | G | A | 0.891 | 0.373 | ++-+- | 0 | 3.126 | 4 | 0.5369 |
| rs11216947 | 11 | 118567218 | A | G | 0.889 | 0.3738 | ++-++ | 0 | 3.286 | 4 | 0.5111 |
| rs57702967 | 11 | 118261736 | A | C | 0.889 | 0.3738 | +++-+ | 0 | 0.724 | 4 | 0.9484 |
| rs12798453 | 11 | 118561410 | G | T | 0.889 | 0.374 | ++-+- | 0 | 2.991 | 4 | 0.5594 |
| rs533646 | 11 | 118566746 | G | C | 0.889 | 0.374 | ++-+- | 0 | 3.155 | 4 | 0.5323 |
| rs625918 | 11 | 118559542 | T | C | 0.888 | 0.3743 | ++-+- | 0 | 3.163 | 4 | 0.5309 |
| rs558593 | 11 | 118562520 | T | C | 0.886 | 0.3755 | ++-+- | 0 | 3.101 | 4 | 0.541 |
| rs7926970 | 11 | 118559881 | G | T | 0.886 | 0.3759 | ++-+- | 0 | 3.152 | 4 | 0.5327 |
| rs502601 | 11 | 118565679 | A | G | 0.884 | 0.3767 | ++-+- | 0 | 3.162 | 4 | 0.5311 |
| rs558907 | 11 | 118558455 | G | A | 0.884 | 0.3767 | ++-+- | 0 | 3.179 | 4 | 0.5283 |
| rs3741315 | 11 | 118004688 | A | G | 0.884 | 0.3768 | +-++- | 0 | 2.382 | 4 | 0.6658 |
| rs4938523 | 11 | 118548838 | T | G | 0.884 | 0.3769 | ++-+- | 0 | 3.117 | 4 | 0.5385 |
| rs7949539 | 11 | 118250586 | G | A | 0.881 | 0.3782 | +++-+ | 0 | 0.807 | 4 | 0.9376 |
| rs4938500 | 11 | 118166212 | A | C | -0.881 | 0.3783 | -?-?+ | 0 | 1.317 | 2 | 0.5177 |
| rs112152942 | 11 | 118240840 | G | C | 0.88 | 0.3789 | +?-+- | -37.4 | 3.638 | 3 | 0.3032 |
| rs7926959 | 11 | 118559866 | C | T | 0.878 | 0.38 | ++-+- | 0 | 3.116 | 4 | 0.5385 |
| rs1318166 | 11 | 118171058 | G | A | -0.877 | 0.3804 | -?-?+ | -93.8 | 2.064 | 2 | 0.3562 |
| rs111354396 | 11 | 118296184 | A | G | 0.875 | 0.3814 | +?-+- | -65.7 | 3.018 | 3 | 0.3889 |
| rs585039 | 11 | 118563381 | A | C | 0.875 | 0.3816 | ++-+- | 0 | 3.066 | 4 | 0.5469 |
| rs59138930 | 11 | 118173783 | T | C | -0.875 | 0.3817 | -?-?+ | -82.7 | 2.189 | 2 | 0.3347 |
| rs45468196 | 11 | 118242889 | A | G | 0.874 | 0.382 | +?-+- | -37.4 | 3.639 | 3 | 0.3031 |
| rs518028 | 11 | 118564982 | C | T | 0.874 | 0.3821 | ++-+- | 0 | 3.087 | 4 | 0.5434 |
| rs582630 | 11 | 118562799 | A | C | 0.873 | 0.3824 | ++-+- | 0 | 3.075 | 4 | 0.5454 |
| rs12576486 | 11 | 118235490 | C | T | 0.873 | 0.3829 | ++++- | 0 | 2.573 | 4 | 0.6317 |
| rs516153 | 11 | 118564762 | A | C | 0.871 | 0.3835 | ++-+- | 0 | 3.094 | 4 | 0.5422 |
| rs471227 | 11 | 118564601 | C | T | 0.871 | 0.3839 | ++-+- | 0 | 3.091 | 4 | 0.5428 |
| rs11216861 | 11 | 118241516 | A | G | -0.87 | 0.3841 | +++-- | -49.1 | 4.024 | 4 | 0.4028 |
| rs693037 | 11 | 118553326 | T | C | 0.87 | 0.3842 | ++-+- | 0 | 3.165 | 4 | 0.5306 |
| rs508640 | 11 | 118564413 | G | A | 0.87 | 0.3843 | ++-+- | 0 | 3.096 | 4 | 0.542 |
| rs78755429 | 11 | 118777978 | A | C | 0.869 | 0.3846 | +?-++ | 0 | 1.796 | 3 | 0.6158 |
| rs4938529 | 11 | 118560569 | T | C | 0.869 | 0.3849 | ++-+- | 0 | 3.104 | 4 | 0.5405 |
| rs45555733 | 11 | 118237566 | G | C | 0.867 | 0.3858 | +?-+- | -35.1 | 3.701 | 3 | 0.2957 |
| rs613377 | 11 | 118556084 | T | C | 0.867 | 0.3859 | ++-+- | 0 | 3.198 | 4 | 0.5252 |
| rs11216827 | 11 | 118110202 | T | A | -0.867 | 0.3861 | -?+?- | 0 | 0.621 | 2 | 0.733 |
| rs546890 | 11 | 118557198 | C | T | 0.867 | 0.3862 | ++-+- | 0 | 3.057 | 4 | 0.5483 |
| rs478371 | 11 | 118565336 | A | G | 0.865 | 0.3868 | ++-+- | 0 | 3.082 | 4 | 0.5442 |
| rs519982 | 11 | 118565231 | T | C | 0.866 | 0.3868 | ++-+- | 0 | 3.129 | 4 | 0.5364 |
| rs61894194 | 11 | 72513028 | T | C | -0.863 | 0.3881 | -++-- | 38.8 | 9.808 | 4 | 0.04379 |
| rs12362596 | 11 | 118105473 | C | G | -0.861 | 0.3892 | -?+?- | 0 | 0.498 | 2 | 0.7794 |
| rs562780 | 11 | 118362414 | T | C | 0.859 | 0.3904 | -++-+ | 0 | 2.054 | 4 | 0.7258 |
| rs12419365 | 11 | 118112332 | A | C | 0.85 | 0.3953 | +?+-- | 0 | 2.094 | 3 | 0.5531 |
| rs146305387 | 11 | 118426800 | T | C | -0.849 | 0.3957 | -?+?? | -177.2 | 1.082 | 1 | 0.2982 |
| rs112646740 | 11 | 118247805 | A | T | 0.848 | 0.3962 | +?-+- | -41.7 | 3.529 | 3 | 0.317 |
| rs74398561 | 11 | 118233634 | G | A | 0.847 | 0.3968 | +?-+- | -29.1 | 3.872 | 3 | 0.2757 |
| rs1783935 | 11 | 118539566 | T | A | 0.847 | 0.397 | ++-+- | 0 | 3.001 | 4 | 0.5577 |
| rs947848 | 11 | 72895007 | T | A | -0.846 | 0.3976 | +--+- | 1.4 | 6.085 | 4 | 0.1929 |
| rs111430146 | 11 | 118248374 | A | C | 0.845 | 0.398 | +?-+- | -43 | 3.496 | 3 | 0.3213 |
| rs12577061 | 11 | 118121553 | C | G | -0.845 | 0.3984 | -?+?- | 0 | 0.835 | 2 | 0.6586 |
| rs653008 | 11 | 118520480 | T | G | 0.841 | 0.4004 | ++-+- | 0 | 1.742 | 4 | 0.7831 |
| rs1573559 | 11 | 118104627 | A | G | -0.84 | 0.4008 | -?+?- | 0 | 0.554 | 2 | 0.7581 |
| rs607527 | 11 | 118538187 | T | G | 0.84 | 0.401 | ++-+- | 0 | 3.061 | 4 | 0.5476 |
| rs112563022 | 11 | 118262135 | T | C | 0.839 | 0.4016 | +?-+- | -50.2 | 3.329 | 3 | 0.3436 |
| rs58508419 | 11 | 118762102 | A | G | 0.838 | 0.402 | +?-+? | 0 | 1.076 | 2 | 0.584 |
| rs630923 | 11 | 118754353 | A | C | -0.833 | 0.4047 | +-+-- | 43 | 10.522 | 4 | 0.0325 |
| rs2368242 | 11 | 118115526 | C | T | -0.833 | 0.4049 | -?+?- | 0 | 0.828 | 2 | 0.6611 |
| rs601318 | 11 | 118519550 | G | C | 0.832 | 0.4052 | ++-+- | 0 | 1.68 | 4 | 0.7944 |

| | | | | | | | | | | | |
|---|---|---|---|---|---|---|---|---|---|---|---|
| rs3825051 | 11 | 118174629 | G | A | -0.831 | 0.4058 | -?-?+ | -72.9 | 2.313 | 2 | 0.3145 |
| rs56356019 | 11 | 118114893 | G | A | -0.829 | 0.407 | -?+?- | 0 | 0.765 | 2 | 0.682 |
| rs10793054 | 11 | 72662812 | A | C | 0.828 | 0.4077 | --+-+ | 48.8 | 11.716 | 4 | 0.01959 |
| rs17092740 | 11 | 118117614 | A | G | -0.828 | 0.4077 | -?+?- | 0 | 0.789 | 2 | 0.674 |
| rs112243842 | 11 | 118246021 | G | T | 0.828 | 0.4079 | +?-+- | -38.6 | 3.607 | 3 | 0.3071 |
| rs11235597 | 11 | 72516773 | A | G | -0.828 | 0.4079 | -++-- | 39.2 | 9.867 | 4 | 0.04274 |
| rs12364013 | 11 | 118119575 | T | C | -0.825 | 0.4091 | -?+?- | 0 | 0.843 | 2 | 0.6559 |
| rs55942271 | 11 | 118177115 | T | C | -0.825 | 0.4092 | ---?+ | 0 | 1.541 | 3 | 0.6728 |
| rs12577053 | 11 | 118121421 | A | G | -0.823 | 0.4105 | -?+?- | 0 | 0.892 | 2 | 0.6403 |
| rs692750 | 11 | 118538308 | T | G | 0.822 | 0.4111 | ++-+- | 0 | 3.072 | 4 | 0.5459 |
| rs45457998 | 11 | 118492713 | G | A | 0.821 | 0.4116 | +?-++ | -50.7 | 3.318 | 3 | 0.3451 |
| rs73024604 | 11 | 118120351 | T | C | -0.819 | 0.4128 | -?+?- | 0 | 0.85 | 2 | 0.6538 |
| rs45443099 | 11 | 118254710 | G | A | 0.817 | 0.4139 | +?-+- | -47 | 3.402 | 3 | 0.3337 |
| rs61900882 | 11 | 118130199 | C | T | -0.817 | 0.414 | -?+?- | 0 | 1.046 | 2 | 0.5926 |
| rs73024606 | 11 | 118120366 | C | G | -0.816 | 0.4143 | -?+?- | 0 | 0.857 | 2 | 0.6515 |
| rs7108905 | 11 | 118712427 | G | C | -0.816 | 0.4145 | -?+-- | 6.5 | 5.346 | 3 | 0.1481 |
| rs61900655 | 11 | 118120802 | C | G | -0.815 | 0.4148 | -?+?- | 0 | 0.859 | 2 | 0.6508 |
| rs61900957 | 11 | 118501723 | T | C | 0.815 | 0.4151 | +?-++ | 23.3 | 6.517 | 3 | 0.08898 |
| rs56315433 | 11 | 118114576 | A | C | -0.815 | 0.4154 | -?+?- | 0 | 0.765 | 2 | 0.6823 |
| rs111792147 | 11 | 118259162 | A | G | 0.814 | 0.4156 | +?-+- | -47 | 3.402 | 3 | 0.3337 |
| rs649138 | 11 | 118066695 | C | T | 0.813 | 0.4164 | +?+?+ | 0 | 0.484 | 2 | 0.7851 |
| rs4589335 | 11 | 118539140 | T | C | 0.806 | 0.4203 | ++-+- | 0 | 3.138 | 4 | 0.535 |
| rs12286723 | 11 | 118734640 | T | C | -0.803 | 0.4218 | -?+-- | 11.9 | 5.678 | 3 | 0.1284 |
| rs112302730 | 11 | 118283383 | A | G | 0.798 | 0.4246 | +?-+- | 0 | 2.836 | 3 | 0.4177 |
| rs12272981 | 11 | 118709600 | G | A | -0.797 | 0.4252 | -?+-- | 4.8 | 5.251 | 3 | 0.1543 |
| rs4457789 | 11 | 118539170 | T | G | 0.797 | 0.4253 | ++-+- | 0 | 2.691 | 4 | 0.6107 |
| rs111763202 | 11 | 118289734 | T | C | 0.796 | 0.4261 | +?-+- | 0 | 2.763 | 3 | 0.4295 |
| rs112035668 | 11 | 118290443 | G | A | 0.795 | 0.4265 | +?-+- | 0 | 2.765 | 3 | 0.4293 |
| rs1715456 | 11 | 118123517 | G | A | -0.792 | 0.4284 | -?+?- | 0 | 1.069 | 2 | 0.5858 |
| rs113941645 | 11 | 118545589 | G | C | 0.791 | 0.4292 | +?-++ | 0 | 1.124 | 3 | 0.7712 |
| rs526597 | 11 | 118546602 | G | A | 0.789 | 0.4303 | ++-+- | 0 | 2.892 | 4 | 0.576 |
| rs111993696 | 11 | 118282448 | A | G | 0.788 | 0.4308 | +?-+- | 0 | 2.974 | 3 | 0.3957 |
| rs112245679 | 11 | 118287622 | G | A | 0.787 | 0.4311 | +?-+- | 0 | 2.845 | 3 | 0.4161 |
| rs45496391 | 11 | 118477226 | C | A | 0.782 | 0.4339 | +?++- | 0 | 0.602 | 3 | 0.896 |
| rs76457018 | 11 | 118762701 | A | G | 0.776 | 0.4377 | +---+ | 48.1 | 11.559 | 4 | 0.02095 |
| rs77074463 | 11 | 118748170 | C | G | 0.767 | 0.443 | ++++- | 25.4 | 8.045 | 4 | 0.08996 |
| rs1970750 | 11 | 118149122 | T | C | -0.767 | 0.4434 | -?-?+ | 0 | 1.81 | 2 | 0.4045 |
| rs17121997 | 11 | 118131049 | C | T | -0.764 | 0.445 | -?+?- | 0 | 0.995 | 2 | 0.6082 |
| rs117278652 | 11 | 72093901 | A | C | 0.762 | 0.4461 | -?+?- | -70.4 | 2.347 | 2 | 0.3093 |
| rs7479331 | 11 | 118829521 | T | G | 0.762 | 0.4461 | +++?- | 21.4 | 6.359 | 3 | 0.09539 |
| rs117822540 | 11 | 72123996 | C | T | 0.753 | 0.4514 | -?+?+ | 0 | 1.983 | 2 | 0.371 |
| rs7118009 | 11 | 118707559 | T | C | -0.753 | 0.4516 | -?+-- | -10.3 | 4.532 | 3 | 0.2094 |
| rs1171461 | 11 | 118522179 | A | G | 0.75 | 0.453 | ++-+- | 0 | 1.816 | 4 | 0.7695 |
| rs10790248 | 11 | 118125672 | A | G | -0.749 | 0.4539 | -?+?- | 0 | 0.906 | 2 | 0.6357 |
| rs112583389 | 11 | 118278998 | G | T | 0.745 | 0.4561 | +?-+- | -64.9 | 3.033 | 3 | 0.3866 |
| rs520543 | 11 | 118538157 | A | G | 0.741 | 0.4586 | ++-+- | 0 | 2.973 | 4 | 0.5623 |
| rs7925466 | 11 | 72778113 | C | A | 0.74 | 0.459 | --+-+ | 35.6 | 9.321 | 4 | 0.05356 |
| rs3181258 | 11 | 118218478 | A | T | 0.74 | 0.4596 | +?-+- | -27.3 | 3.929 | 3 | 0.2692 |
| rs2071702 | 11 | 118373861 | T | C | 0.738 | 0.4606 | +?++- | 0 | 2.745 | 3 | 0.4326 |
| rs117794131 | 11 | 72097791 | C | T | 0.735 | 0.4621 | -?+?- | -76.8 | 2.263 | 2 | 0.3226 |
| rs4936425 | 11 | 118126904 | T | C | -0.735 | 0.4624 | -?+?- | 0 | 0.972 | 2 | 0.6151 |
| rs2187490 | 11 | 118713180 | G | T | 0.734 | 0.4627 | +?-++ | 0 | 1.623 | 3 | 0.6543 |
| rs1126205 | 11 | 72523794 | T | G | -0.734 | 0.4632 | -++-- | 42.8 | 10.493 | 4 | 0.03289 |
| rs113784020 | 11 | 118559731 | T | C | 0.733 | 0.4633 | +?-++ | 0 | 2.185 | 3 | 0.535 |
| rs527290 | 11 | 118515616 | T | C | 0.73 | 0.4656 | ++-+- | 0 | 1.764 | 4 | 0.7791 |
| rs7109466 | 11 | 72114792 | T | C | 0.729 | 0.466 | -?+?+ | -96.7 | 2.033 | 2 | 0.3618 |

| | | | | | | | | | | | |
|---|---|---|---|---|---|---|---|---|---|---|---|
| rs4938492 | 11 | 118126675 | C | T | -0.729 | 0.4661 | -?+?- | 0 | 0.953 | 2 | 0.6211 |
| rs7939548 | 11 | 118212979 | A | G | 0.725 | 0.4684 | +?-+- | -21.4 | 4.119 | 3 | 0.2489 |
| rs1104542 | 11 | 118129340 | T | C | -0.724 | 0.4691 | -?+?- | 0 | 0.971 | 2 | 0.6155 |
| rs2004235 | 11 | 118008212 | C | T | 0.724 | 0.4691 | +-++- | 0 | 2.109 | 4 | 0.7158 |
| rs111707199 | 11 | 118276810 | G | A | 0.723 | 0.4696 | +?-+- | 0 | 2.974 | 3 | 0.3957 |
| rs3759004 | 11 | 118136836 | T | C | -0.723 | 0.4699 | -?-?+ | -67.3 | 2.391 | 2 | 0.3025 |
| rs17121871 | 11 | 118073344 | T | C | 0.721 | 0.4712 | +?+?+ | 0 | 0.674 | 2 | 0.7138 |
| rs112468256 | 11 | 118278479 | C | T | 0.719 | 0.4723 | +?-+- | 0 | 2.943 | 3 | 0.4005 |
| rs2508991 | 11 | 118539522 | A | G | 0.71 | 0.4778 | ++-+- | 0 | 3.127 | 4 | 0.5368 |
| rs117190635 | 11 | 72104119 | A | G | 0.707 | 0.4795 | -?+?- | 0 | 1.901 | 2 | 0.3866 |
| rs12417431 | 11 | 118137383 | G | A | -0.707 | 0.4798 | -?-?+ | -62.6 | 2.46 | 2 | 0.2922 |
| rs73575424 | 11 | 118788517 | A | C | 0.705 | 0.481 | +-+-+ | 30.2 | 8.593 | 4 | 0.07211 |
| rs117654328 | 11 | 72117713 | A | G | 0.704 | 0.4812 | -?+?+ | 0 | 1.936 | 2 | 0.3798 |
| rs150031150 | 11 | 72120370 | A | G | 0.704 | 0.4812 | -?+?+ | 0 | 1.936 | 2 | 0.3798 |
| rs112970787 | 11 | 118212235 | C | G | 0.704 | 0.4814 | +?-+- | -20.2 | 4.159 | 3 | 0.2448 |
| rs605560 | 11 | 118067187 | C | G | 0.703 | 0.4819 | +?+?+ | 0 | 0.511 | 2 | 0.7745 |
| rs4938615 | 11 | 118865319 | G | C | 0.702 | 0.4826 | +?+?- | -9.1 | 3.667 | 2 | 0.1598 |
| rs1814964 | 11 | 118006957 | C | T | 0.701 | 0.4835 | +-++- | 0 | 2.031 | 4 | 0.7301 |
| rs10793055 | 11 | 72710761 | C | T | 0.698 | 0.485 | --+-+ | 38.1 | 9.69 | 4 | 0.04599 |
| rs1790063 | 11 | 72995912 | A | G | 0.698 | 0.4851 | -?-+? | 0 | 1.745 | 2 | 0.418 |
| rs77487830 | 11 | 118311869 | G | A | 0.698 | 0.4854 | +--+- | -45.9 | 4.111 | 4 | 0.3912 |
| rs12792929 | 11 | 118240715 | C | G | 0.697 | 0.486 | +++-- | 0 | 1.27 | 4 | 0.8664 |
| rs117235147 | 11 | 72115430 | A | T | 0.695 | 0.4868 | -?+?+ | 0 | 1.902 | 2 | 0.3864 |
| rs11235595 | 11 | 72512708 | C | G | -0.695 | 0.487 | -++-- | 42.1 | 10.36 | 4 | 0.03479 |
| rs116881498 | 11 | 72102819 | T | C | 0.693 | 0.4881 | -?+?- | 0 | 1.998 | 2 | 0.3682 |
| rs79826955 | 11 | 72104081 | A | G | 0.693 | 0.4884 | -?+?- | 0 | 1.996 | 2 | 0.3686 |
| rs3759001 | 11 | 118135745 | T | C | -0.693 | 0.4886 | +?-?- | 0 | 0.47 | 2 | 0.7906 |
| rs1629083 | 11 | 118126576 | T | C | 0.691 | 0.4894 | -?+?+ | -78.1 | 2.246 | 2 | 0.3253 |
| rs7129048 | 11 | 118211797 | C | T | 0.689 | 0.4906 | +?-+- | -11.3 | 4.493 | 3 | 0.2129 |
| rs77535829 | 11 | 118311677 | T | C | 0.689 | 0.4906 | +--+- | -45.6 | 4.122 | 4 | 0.3898 |
| rs45558732 | 11 | 118402962 | T | G | 0.686 | 0.4929 | +?-++ | 0 | 0.43 | 3 | 0.9341 |
| rs4938520 | 11 | 118522338 | G | A | 0.685 | 0.4933 | ++-+- | 0 | 1.559 | 4 | 0.8161 |
| rs3759005 | 11 | 118136871 | G | A | -0.684 | 0.4937 | -?-?+ | 0 | 1.739 | 2 | 0.4193 |
| rs3782041 | 11 | 118184092 | T | C | -0.681 | 0.4957 | ---?+ | 0 | 1.674 | 3 | 0.6426 |
| rs1806 | 11 | 118076004 | C | T | 0.679 | 0.4975 | +?-?+ | 0 | 0.808 | 2 | 0.6677 |
| rs7103001 | 11 | 118367669 | T | A | 0.677 | 0.4982 | +?++- | 0 | 2.145 | 3 | 0.5429 |
| rs481206 | 11 | 72497462 | T | C | 0.677 | 0.4983 | ++--- | 0 | 3.889 | 4 | 0.4213 |
| rs3759003 | 11 | 118136745 | G | T | -0.677 | 0.4986 | -?-?+ | 0 | 1.737 | 2 | 0.4197 |
| rs45480805 | 11 | 118267903 | A | G | 0.675 | 0.4997 | +?-+- | 0 | 2.958 | 3 | 0.3982 |
| rs11604096 | 11 | 118012441 | C | A | -0.675 | 0.5 | -+-++ | 0 | 1.139 | 4 | 0.8881 |
| rs61900902 | 11 | 118159684 | C | T | -0.672 | 0.5016 | -?-?+ | -96.1 | 2.04 | 2 | 0.3607 |
| rs7110197 | 11 | 118228624 | C | G | 0.672 | 0.5017 | +++-- | 0 | 1.518 | 4 | 0.8234 |
| rs10892253 | 11 | 118559426 | A | C | 0.671 | 0.5024 | +?++- | 0 | 0.638 | 3 | 0.8876 |
| rs171021 | 11 | 72317557 | T | C | 0.671 | 0.5025 | -?+?+ | 0 | 1.237 | 2 | 0.5388 |
| rs745663 | 11 | 118529595 | G | A | 0.67 | 0.5029 | ++-+- | 0 | 1.437 | 4 | 0.8377 |
| rs12795615 | 11 | 118214562 | T | C | 0.67 | 0.5032 | +++-- | 0 | 1.55 | 4 | 0.8178 |
| rs2239695 | 11 | 118213632 | T | C | 0.669 | 0.5034 | +++-- | 0 | 1.693 | 4 | 0.7921 |
| rs35902434 | 11 | 118222634 | C | T | 0.668 | 0.5043 | +++-- | 0 | 1.488 | 4 | 0.8287 |
| rs36061634 | 11 | 118225452 | A | T | 0.667 | 0.5046 | +++-- | 0 | 1.501 | 4 | 0.8264 |
| rs4938507 | 11 | 118231584 | G | A | 0.667 | 0.5047 | +?++- | -39.7 | 3.58 | 3 | 0.3105 |
| rs3825054 | 11 | 118234191 | A | G | 0.666 | 0.5052 | +++-- | 0 | 1.512 | 4 | 0.8245 |
| rs11216812 | 11 | 118071762 | A | G | 0.665 | 0.5063 | +?-+- | 0 | 1.282 | 3 | 0.7333 |
| rs12290522 | 11 | 118486325 | T | C | 0.665 | 0.5063 | ++-+- | 0 | 1.775 | 4 | 0.777 |
| rs3759006 | 11 | 118137560 | A | G | -0.664 | 0.5064 | -?-?+ | 0 | 1.871 | 2 | 0.3924 |
| rs7925928 | 11 | 72748384 | G | A | 0.664 | 0.5064 | --+-+ | 36.2 | 9.41 | 4 | 0.05164 |
| rs11216814 | 11 | 118071857 | G | A | 0.664 | 0.5067 | +?-+- | 0 | 1.281 | 3 | 0.7336 |

| | | | | | | | | | | | |
|---|---|---|---|---|---|---|---|---|---|---|---|
| rs111587825 | 11 | 118274602 | G | T | 0.664 | 0.5068 | +?-+- | 0 | 2.723 | 3 | 0.4364 |
| rs75085081 | 11 | 118801045 | A | G | 0.664 | 0.5068 | +-+-? | 0 | 2.373 | 3 | 0.4986 |
| rs4943939 | 11 | 72309540 | T | C | -0.662 | 0.508 | --+?- | 0 | 1.879 | 3 | 0.5979 |
| rs117493218 | 11 | 118027406 | C | A | 0.661 | 0.5084 | --+?? | 0 | 1.311 | 2 | 0.5191 |
| rs3181259 | 11 | 118214067 | A | G | 0.658 | 0.5102 | +++-- | 0 | 1.66 | 4 | 0.7979 |
| rs35305304 | 11 | 118228114 | T | C | 0.658 | 0.5106 | +++-- | 0 | 1.514 | 4 | 0.8242 |
| rs4938498 | 11 | 118162147 | G | C | -0.658 | 0.5108 | -?-?+ | -50 | 2.666 | 2 | 0.2637 |
| rs644287 | 11 | 118011805 | T | C | 0.656 | 0.5116 | +-+++ | 0 | 1.222 | 4 | 0.8744 |
| rs570836 | 11 | 118512162 | A | G | 0.655 | 0.5121 | ++-+- | 0 | 1.71 | 4 | 0.7889 |
| rs7940405 | 11 | 118142666 | G | A | -0.655 | 0.5124 | -?-?+ | -43.4 | 2.789 | 2 | 0.248 |
| rs10898872 | 11 | 72497425 | C | T | -0.655 | 0.5125 | -++-+ | 44.3 | 10.767 | 4 | 0.02931 |
| rs7123835 | 11 | 118368822 | G | T | 0.654 | 0.5133 | +?++- | 0 | 2.35 | 3 | 0.503 |
| rs34180117 | 11 | 118138629 | T | C | -0.652 | 0.5143 | -?-?+ | 0 | 1.862 | 2 | 0.3941 |
| rs668468 | 11 | 118011776 | G | A | -0.651 | 0.5149 | -+-++ | 0 | 1.1 | 4 | 0.8943 |
| rs35850218 | 11 | 118138922 | C | T | -0.65 | 0.5157 | -?-?+ | 0 | 1.859 | 2 | 0.3948 |
| rs35865949 | 11 | 118229061 | A | T | 0.649 | 0.5163 | +++-- | 0 | 1.598 | 4 | 0.8092 |
| rs3782046 | 11 | 118230739 | A | G | 0.649 | 0.5164 | +++-- | 0 | 1.59 | 4 | 0.8105 |
| rs7107305 | 11 | 118368665 | G | A | 0.648 | 0.5169 | +?++- | 0 | 2.351 | 3 | 0.5028 |
| rs894678 | 11 | 118172520 | A | G | -0.648 | 0.5172 | -?-?+ | -92 | 2.083 | 2 | 0.353 |
| rs45627033 | 11 | 118766018 | T | C | 0.647 | 0.5177 | +?-?? | -29.7 | 2.313 | 1 | 0.1283 |
| rs35610956 | 11 | 118144280 | G | A | -0.646 | 0.5181 | -?-?+ | -42 | 2.817 | 2 | 0.2445 |
| rs6589673 | 11 | 118557483 | C | A | 0.646 | 0.5184 | +?++- | 0 | 0.618 | 3 | 0.8924 |
| rs12791838 | 11 | 118144209 | A | G | -0.645 | 0.519 | -?-?+ | -41.3 | 2.831 | 2 | 0.2427 |
| rs7389 | 11 | 118528466 | G | T | 0.645 | 0.519 | ++-+- | 0 | 1.55 | 4 | 0.8178 |
| rs3753060 | 11 | 118221137 | T | C | 0.645 | 0.5192 | +++-- | 0 | 1.508 | 4 | 0.8252 |
| rs56128937 | 11 | 118139478 | A | T | -0.644 | 0.5197 | -?-?+ | 0 | 1.837 | 2 | 0.3991 |
| rs3759007 | 11 | 118137667 | T | C | -0.643 | 0.5205 | -?-?+ | 0 | 1.846 | 2 | 0.3973 |
| rs3922 | 11 | 118765600 | G | A | -0.64 | 0.5224 | +-+-- | 0 | 1.328 | 4 | 0.8566 |
| rs7948683 | 11 | 118137911 | A | G | -0.639 | 0.5226 | -?-?+ | 0 | 1.825 | 2 | 0.4015 |
| rs3753059 | 11 | 118221349 | C | T | 0.639 | 0.5228 | +++-- | 0 | 1.506 | 4 | 0.8256 |
| rs3753058 | 11 | 118221350 | T | G | 0.639 | 0.523 | +++-- | 0 | 1.506 | 4 | 0.8257 |
| rs12420795 | 11 | 118161885 | G | C | -0.638 | 0.5234 | -?-?+ | -52.6 | 2.621 | 2 | 0.2697 |
| rs4938497 | 11 | 118160851 | A | G | -0.637 | 0.5238 | -?-?+ | -87.6 | 2.132 | 2 | 0.3443 |
| rs483283 | 11 | 118506402 | C | T | 0.637 | 0.5239 | ++-+- | 0 | 1.692 | 4 | 0.7922 |
| rs45474998 | 11 | 118229679 | A | G | 0.636 | 0.5245 | +++-- | 0 | 1.379 | 4 | 0.8478 |
| rs12419204 | 11 | 118497402 | A | T | 0.636 | 0.5248 | ++-+- | 0 | 1.726 | 4 | 0.786 |
| rs1791049 | 11 | 72890304 | T | C | 0.636 | 0.5251 | ++-+- | 0 | 3.742 | 4 | 0.442 |
| rs113839182 | 11 | 118273602 | T | C | 0.635 | 0.5255 | +?-+- | 0 | 2.707 | 3 | 0.439 |
| rs12801003 | 11 | 118072716 | T | C | 0.634 | 0.5263 | +?-+- | 0 | 1.281 | 3 | 0.7336 |
| rs12795307 | 11 | 72506324 | A | G | 0.634 | 0.5264 | ++--- | -47 | 4.08 | 4 | 0.3952 |
| rs7928058 | 11 | 118171239 | T | G | -0.63 | 0.5285 | -?-?+ | -54.3 | 2.592 | 2 | 0.2737 |
| rs2276418 | 11 | 118230264 | T | A | 0.629 | 0.5291 | +++-- | 0 | 1.393 | 4 | 0.8454 |
| rs1783583 | 11 | 72889432 | C | G | 0.628 | 0.5297 | ++-+- | 0 | 3.686 | 4 | 0.4502 |
| rs34405565 | 11 | 118233424 | T | C | 0.628 | 0.5299 | +++-- | 0 | 1.38 | 4 | 0.8476 |
| rs55852239 | 11 | 118139755 | A | G | -0.628 | 0.5299 | -?-?+ | 0 | 1.891 | 2 | 0.3885 |
| rs645637 | 11 | 118505090 | A | G | 0.627 | 0.5309 | ++-+- | 0 | 1.61 | 4 | 0.807 |
| rs3782045 | 11 | 118233078 | G | A | 0.626 | 0.5313 | +++-- | 0 | 1.364 | 4 | 0.8505 |
| rs117068828 | 11 | 72139241 | A | G | 0.626 | 0.5315 | -?+?- | 0 | 1.875 | 2 | 0.3916 |
| rs4938490 | 11 | 118080977 | G | A | 0.624 | 0.5324 | +?+?+ | 0 | 0.641 | 2 | 0.7259 |
| rs570952 | 11 | 118500193 | C | T | 0.624 | 0.5326 | ++-+- | 0 | 1.697 | 4 | 0.7913 |
| rs11235546 | 11 | 72312118 | T | G | -0.624 | 0.5327 | --+?- | 0 | 1.981 | 3 | 0.5765 |
| rs11603616 | 11 | 118496066 | A | G | 0.623 | 0.533 | ++-+- | 0 | 1.793 | 4 | 0.7738 |
| rs341088 | 11 | 72313975 | T | C | -0.623 | 0.533 | --+?- | 0 | 1.611 | 3 | 0.6569 |
| rs595964 | 11 | 118026788 | A | G | -0.622 | 0.534 | -+--- | 0 | 0.531 | 4 | 0.9705 |
| rs12795905 | 11 | 118140597 | T | C | -0.618 | 0.5365 | -?-?+ | 0 | 1.972 | 2 | 0.373 |
| rs12803321 | 11 | 118480115 | C | G | 0.618 | 0.5366 | +?+-+ | 0 | 1.047 | 3 | 0.7899 |

| | | | | | | | | | | | |
|---|---|---|---|---|---|---|---|---|---|---|---|
| rs12806041 | 11 | 118138998 | G | A | -0.617 | 0.5373 | -?-?+ | 0 | 1.778 | 2 | 0.4111 |
| rs571001 | 11 | 118500173 | T | C | 0.614 | 0.5389 | ++-+- | 0 | 1.75 | 4 | 0.7816 |
| rs11216815 | 11 | 118072232 | A | G | 0.613 | 0.5399 | +?-+- | 0 | 1.332 | 3 | 0.7215 |
| rs34786601 | 11 | 118140136 | C | T | -0.613 | 0.5399 | -?-?+ | 0 | 1.935 | 2 | 0.3801 |
| rs2276421 | 11 | 118239266 | A | G | 0.612 | 0.5404 | +++-- | 0 | 1.489 | 4 | 0.8285 |
| rs12789276 | 11 | 118140203 | G | A | -0.612 | 0.5405 | -?-?+ | 0 | 1.94 | 2 | 0.3791 |
| rs341049 | 11 | 72298208 | T | C | -0.61 | 0.542 | --+?- | 0 | 2.189 | 3 | 0.5341 |
| rs74535623 | 11 | 118812821 | T | G | 0.609 | 0.5423 | +-+-? | 0 | 2.466 | 3 | 0.4815 |
| rs45448191 | 11 | 118267737 | A | T | 0.609 | 0.5425 | +?-+- | -63.1 | 3.066 | 3 | 0.3816 |
| rs73011440 | 11 | 118809619 | A | G | 0.609 | 0.5425 | +?+?? | 0 | 0.205 | 1 | 0.6504 |
| rs17122113 | 11 | 118234678 | G | A | 0.608 | 0.5433 | +++-- | 0 | 1.364 | 4 | 0.8505 |
| rs7104819 | 11 | 118769442 | G | A | -0.606 | 0.5442 | --+-- | 0 | 1.481 | 4 | 0.8301 |
| rs11607701 | 11 | 118169825 | G | A | -0.606 | 0.5446 | -?-?+ | -50.8 | 2.653 | 2 | 0.2654 |
| rs12416826 | 11 | 118165955 | C | T | -0.604 | 0.546 | -?-?+ | -46.6 | 2.729 | 2 | 0.2555 |
| rs2077173 | 11 | 118492591 | A | G | 0.602 | 0.5474 | ++-+- | 0 | 1.853 | 4 | 0.7627 |
| rs11600544 | 11 | 118076954 | C | G | 0.601 | 0.5479 | +?+?+ | 0 | 0.58 | 2 | 0.7482 |
| rs17121881 | 11 | 118081345 | A | T | 0.601 | 0.5479 | +?+?+ | 0 | 0.662 | 2 | 0.7184 |
| rs2032397 | 11 | 118153697 | A | G | -0.601 | 0.5481 | -?-?+ | -53.3 | 2.609 | 2 | 0.2713 |
| rs12793790 | 11 | 118140372 | A | G | -0.599 | 0.5489 | -?-?+ | 0 | 1.955 | 2 | 0.3762 |
| rs1805 | 11 | 118076069 | T | G | 0.599 | 0.5492 | +?+?+ | 0 | 0.612 | 2 | 0.7363 |
| rs2032398 | 11 | 118153708 | C | G | -0.599 | 0.5492 | -?-?+ | -53.5 | 2.605 | 2 | 0.2718 |
| rs17121879 | 11 | 118080133 | G | T | 0.598 | 0.5496 | +?+?+ | 0 | 0.602 | 2 | 0.7402 |
| rs1617182 | 11 | 118068949 | C | T | 0.598 | 0.5497 | +?-?+ | 0 | 0.577 | 2 | 0.7493 |
| rs4938489 | 11 | 118079867 | G | T | 0.598 | 0.55 | +?+?+ | 0 | 0.602 | 2 | 0.74 |
| rs73557345 | 11 | 118813314 | C | T | 0.597 | 0.5503 | +-+-? | 0 | 2.463 | 3 | 0.482 |
| rs17121882 | 11 | 118081468 | C | T | 0.594 | 0.5528 | +?+?+ | 0 | 0.637 | 2 | 0.7274 |
| rs10790254 | 11 | 118493319 | A | G | 0.591 | 0.5545 | +?++- | 0 | 0.544 | 3 | 0.909 |
| rs73612281 | 11 | 118203642 | T | C | -0.588 | 0.5566 | -?--+ | 0 | 1.081 | 3 | 0.7817 |
| rs7116876 | 11 | 118371025 | C | G | 0.587 | 0.5569 | +?++- | 0 | 2.185 | 3 | 0.5348 |
| rs1793167 | 11 | 118102058 | T | C | 0.585 | 0.5588 | +?-+- | 0 | 2.06 | 3 | 0.56 |
| rs61751532 | 11 | 118769604 | T | C | 0.585 | 0.5588 | -?++? | 0 | 1.217 | 2 | 0.5441 |
| rs140633990 | 11 | 118365883 | A | T | -0.583 | 0.5601 | ---?? | 0 | 0.136 | 2 | 0.9342 |
| rs9332835 | 11 | 118372923 | A | G | 0.575 | 0.5652 | +?++- | 0 | 2.228 | 3 | 0.5265 |
| rs2120268 | 11 | 118132705 | A | G | 0.571 | 0.5678 | +?-+- | 0 | 1.36 | 3 | 0.715 |
| rs7117594 | 11 | 118714492 | T | C | -0.57 | 0.5686 | -++-- | 11.9 | 6.807 | 4 | 0.1464 |
| rs3018383 | 11 | 118093359 | A | C | 0.569 | 0.5695 | +?-+- | 0 | 2.03 | 3 | 0.5662 |
| rs12801084 | 11 | 72991793 | G | C | -0.567 | 0.5707 | --+-- | 0 | 3.793 | 4 | 0.4348 |
| rs10790258 | 11 | 118561859 | T | G | 0.567 | 0.5708 | +?++- | 0 | 0.402 | 3 | 0.9398 |
| rs1793155 | 11 | 118098962 | A | G | 0.566 | 0.5712 | +?-+- | 0 | 2.022 | 3 | 0.5678 |
| rs4938517 | 11 | 118487673 | C | T | 0.565 | 0.5721 | ++-+- | 0 | 1.875 | 4 | 0.7588 |
| rs10790240 | 11 | 118009279 | G | T | 0.563 | 0.5731 | +-++- | 0 | 1.661 | 4 | 0.7977 |
| rs663015 | 11 | 72499035 | C | T | 0.562 | 0.5741 | ++--- | -47.7 | 4.064 | 4 | 0.3975 |
| rs1793156 | 11 | 118098890 | C | T | 0.561 | 0.5746 | +?-+- | 0 | 2.035 | 3 | 0.5651 |
| rs11235711 | 11 | 72992343 | C | G | -0.56 | 0.5752 | --+-- | 0 | 3.817 | 4 | 0.4313 |
| rs10790257 | 11 | 118561858 | C | G | 0.56 | 0.5754 | +?++- | 0 | 0.416 | 3 | 0.937 |
| rs519790 | 11 | 72504141 | G | C | 0.56 | 0.5757 | ++--- | -34.2 | 4.472 | 4 | 0.3459 |
| rs656445 | 11 | 118012164 | A | G | 0.558 | 0.577 | +--++ | 0 | 1.101 | 4 | 0.8941 |
| rs9332846 | 11 | 118377523 | A | G | 0.557 | 0.5778 | +?++- | 0 | 2.165 | 3 | 0.5388 |
| rs148643334 | 11 | 118358152 | C | T | 0.556 | 0.5782 | +?++- | 0 | 2.143 | 3 | 0.5433 |
| rs17310235 | 11 | 72607076 | G | A | 0.552 | 0.581 | ++-+- | 0 | 1.605 | 4 | 0.8079 |
| rs11235713 | 11 | 72997496 | G | A | -0.552 | 0.5812 | --+-- | 0 | 3.376 | 4 | 0.4971 |
| rs1793169 | 11 | 118070264 | G | A | 0.55 | 0.5826 | +?-?+ | 0 | 0.608 | 2 | 0.7378 |
| rs1715444 | 11 | 118016273 | G | T | -0.549 | 0.5827 | -+-++ | 0 | 0.829 | 4 | 0.9345 |
| rs1793185 | 11 | 118089434 | A | C | 0.549 | 0.5829 | +?-+- | 0 | 1.868 | 3 | 0.6003 |
| rs9332816 | 11 | 118364404 | T | G | 0.547 | 0.5842 | +?++- | 0 | 2.054 | 3 | 0.5612 |
| rs9332809 | 11 | 118361472 | A | G | 0.547 | 0.5843 | +?++- | 0 | 2.039 | 3 | 0.5643 |

| | | | | | | | | | | | |
|---|---|---|---|---|---|---|---|---|---|---|---|
| rs7105658 | 11 | 118712270 | T | C | -0.543 | 0.5869 | -++-- | 9.3 | 6.618 | 4 | 0.1575 |
| rs2276045 | 11 | 118365168 | A | G | 0.54 | 0.5894 | +?++- | 0 | 2.083 | 3 | 0.5553 |
| rs6589656 | 11 | 118095996 | C | G | 0.54 | 0.5895 | +?-+- | 0 | 2.322 | 3 | 0.5082 |
| rs11604683 | 11 | 72538546 | C | T | -0.539 | 0.5899 | ++--- | 0 | 1.773 | 4 | 0.7774 |
| rs11235710 | 11 | 72992156 | G | C | -0.538 | 0.5905 | --+-- | 0 | 3.85 | 4 | 0.4266 |
| rs1612659 | 11 | 72835303 | T | C | -0.536 | 0.5919 | ++--- | 0 | 2.807 | 4 | 0.5906 |
| rs3802863 | 11 | 118099186 | T | C | 0.536 | 0.5919 | +?-+- | 0 | 2.304 | 3 | 0.5117 |
| rs12799445 | 11 | 72592506 | G | T | -0.531 | 0.5956 | ++-+- | 0 | 3.473 | 4 | 0.4821 |
| rs1626649 | 11 | 118071097 | T | C | 0.526 | 0.599 | +?-?+ | 0 | 0.625 | 2 | 0.7317 |
| rs4938505 | 11 | 118205636 | A | G | -0.525 | 0.5993 | -?--+ | 0 | 1.22 | 3 | 0.7481 |
| rs9332851 | 11 | 118381018 | T | C | 0.524 | 0.5999 | +?++- | 0 | 0.907 | 3 | 0.8237 |
| rs371412 | 11 | 72303491 | A | G | -0.524 | 0.6006 | +?-?- | -93.1 | 2.071 | 2 | 0.355 |
| rs12363628 | 11 | 72987038 | C | T | -0.521 | 0.6024 | --+-- | 0 | 3.813 | 4 | 0.4319 |
| rs7946861 | 11 | 72557986 | G | A | -0.519 | 0.6036 | ++-+- | 0 | 3.66 | 4 | 0.4539 |
| rs7101652 | 11 | 118078718 | A | G | 0.515 | 0.6067 | +?-+- | 0 | 1.201 | 3 | 0.7527 |
| rs12788624 | 11 | 118022107 | T | C | -0.515 | 0.6068 | -++?+ | -24.7 | 4.009 | 3 | 0.2605 |
| rs45555132 | 11 | 118064030 | T | G | -0.512 | 0.6086 | -?+?- | 0 | 0.425 | 2 | 0.8084 |
| rs7112584 | 11 | 118069240 | A | G | 0.511 | 0.6092 | +?-+- | 0 | 1.194 | 3 | 0.7545 |
| rs1945764 | 11 | 118182801 | G | A | -0.507 | 0.6123 | ---?+ | -51.9 | 3.292 | 3 | 0.3487 |
| rs7936751 | 11 | 72720199 | G | T | 0.507 | 0.6123 | --+-+ | 31.7 | 8.791 | 4 | 0.06655 |
| rs7103936 | 11 | 118091636 | C | A | 0.506 | 0.6127 | +?-+- | 0 | 1.976 | 3 | 0.5773 |
| rs384892 | 11 | 72309889 | A | G | -0.505 | 0.6135 | +?-?- | 0 | 1.458 | 2 | 0.4823 |
| rs4306318 | 11 | 118080520 | T | C | 0.5 | 0.6172 | +?-+- | 0 | 1.157 | 3 | 0.7633 |
| rs9332858 | 11 | 118390082 | C | A | 0.499 | 0.6179 | +?++- | 0 | 2.068 | 3 | 0.5585 |
| rs4432047 | 11 | 72989489 | G | A | -0.498 | 0.6183 | --+-- | -49.8 | 4.005 | 4 | 0.4054 |
| rs12808111 | 11 | 72992714 | G | A | -0.497 | 0.6194 | --+-- | -44.3 | 4.159 | 4 | 0.3849 |
| rs11216818 | 11 | 118080268 | T | C | 0.496 | 0.62 | +?-+- | 0 | 1.165 | 3 | 0.7613 |
| rs7116375 | 11 | 118069979 | A | C | 0.495 | 0.6204 | +?-+- | 0 | 1.338 | 3 | 0.7202 |
| rs7951159 | 11 | 118780950 | T | C | -0.494 | 0.6213 | -?+?? | 0 | 0.788 | 1 | 0.3748 |
| rs9332819 | 11 | 118365263 | T | C | 0.494 | 0.6213 | +?++- | 0 | 1.897 | 3 | 0.5941 |
| rs79157249 | 11 | 118785982 | T | A | 0.492 | 0.6228 | +-+-+ | 30.1 | 8.578 | 4 | 0.07255 |
| rs111997050 | 11 | 118387892 | T | C | 0.49 | 0.6239 | +?++- | 0 | 1.781 | 3 | 0.6191 |
| rs11600134 | 11 | 118070788 | A | G | 0.49 | 0.6242 | +?-+- | 0 | 1.374 | 3 | 0.7117 |
| rs12793150 | 11 | 118180667 | G | C | -0.49 | 0.6243 | ---?+ | 0 | 2.724 | 3 | 0.4362 |
| rs59049611 | 11 | 118139403 | C | T | -0.489 | 0.6246 | -?+?- | 0 | 1.911 | 2 | 0.3846 |
| rs111405302 | 11 | 118385691 | C | T | 0.488 | 0.6257 | +?++- | 0 | 2.188 | 3 | 0.5343 |
| rs11235708 | 11 | 72985098 | A | G | -0.488 | 0.6257 | --+-- | 0 | 3.794 | 4 | 0.4346 |
| rs604096 | 11 | 118475098 | G | A | -0.483 | 0.6288 | --+-- | -11.2 | 5.397 | 4 | 0.2489 |
| rs1599810 | 11 | 118179197 | A | G | -0.481 | 0.6302 | ---?+ | -48.5 | 3.366 | 3 | 0.3386 |
| rs7102253 | 11 | 72792697 | C | T | -0.48 | 0.6311 | ++-+- | 0 | 3.526 | 4 | 0.474 |
| rs34385073 | 11 | 72987413 | C | A | -0.48 | 0.6312 | --+-- | 0 | 3.982 | 4 | 0.4085 |
| rs3862796 | 11 | 72705522 | T | C | -0.48 | 0.6315 | -+-+- | 0 | 3.611 | 4 | 0.4612 |
| rs75461585 | 11 | 118287568 | G | A | -0.479 | 0.6319 | -?--+ | 0 | 2.411 | 3 | 0.4915 |
| rs7116594 | 11 | 72984870 | A | G | -0.478 | 0.6324 | --+-- | 0 | 3.872 | 4 | 0.4236 |
| rs17122021 | 11 | 118145686 | T | C | -0.478 | 0.6327 | -?+?- | 0 | 1.78 | 2 | 0.4106 |
| rs7129921 | 11 | 72676874 | G | T | 0.476 | 0.6343 | --+-+ | 43.5 | 10.613 | 4 | 0.03127 |
| rs4938493 | 11 | 118144804 | G | T | -0.475 | 0.6345 | -?+?- | 0 | 1.862 | 2 | 0.3941 |
| rs12286021 | 11 | 118080685 | C | T | 0.475 | 0.6346 | +?-+- | 0 | 1.278 | 3 | 0.7345 |
| rs35963948 | 11 | 72991186 | G | A | -0.475 | 0.6351 | --+-- | 0 | 3.649 | 4 | 0.4556 |
| rs4245180 | 11 | 118144512 | A | G | -0.474 | 0.6353 | -?+?- | 0 | 1.792 | 2 | 0.4082 |
| rs4938502 | 11 | 118179941 | T | G | -0.473 | 0.636 | ---?+ | 0 | 2.821 | 3 | 0.4201 |
| rs11216822 | 11 | 118091375 | A | G | 0.472 | 0.6367 | +?-+- | 0 | 2.052 | 3 | 0.5616 |
| rs45621436 | 11 | 118483912 | G | T | 0.471 | 0.6379 | +?++- | 0 | 0.717 | 3 | 0.8692 |
| rs2097183 | 11 | 118091067 | T | C | 0.47 | 0.6381 | +?-+- | 0 | 2.05 | 3 | 0.562 |
| rs12280559 | 11 | 118087040 | T | C | 0.469 | 0.6389 | +?-+- | 0 | 2.127 | 3 | 0.5465 |
| rs687664 | 11 | 118574185 | A | C | 0.468 | 0.6397 | ++-?+ | 0 | 2.042 | 3 | 0.5637 |

| | | | | | | | | | | | |
|---|---|---|---|---|---|---|---|---|---|---|---|
| rs523715 | 11 | 118574188 | A | G | 0.468 | 0.6399 | ++-?+ | 0 | 2.043 | 3 | 0.5635 |
| rs4488208 | 11 | 118877621 | C | G | -0.467 | 0.6404 | -?+?- | 29.3 | 5.654 | 2 | 0.05918 |
| rs78020509 | 11 | 118384811 | G | A | 0.465 | 0.642 | +?++- | 0 | 2.287 | 3 | 0.5151 |
| rs7940554 | 11 | 118142930 | T | C | -0.465 | 0.642 | -?+?- | 0 | 1.814 | 2 | 0.4037 |
| rs12805658 | 11 | 72555300 | A | C | -0.462 | 0.6442 | ++-+- | 0 | 3.378 | 4 | 0.4966 |
| rs2187494 | 11 | 72805603 | G | A | -0.46 | 0.6453 | ++--- | 0 | 2.505 | 4 | 0.6438 |
| rs1715420 | 11 | 118105599 | A | T | 0.46 | 0.6456 | +?-+- | 0 | 1.627 | 3 | 0.6532 |
| rs11235617 | 11 | 72644631 | C | T | 0.459 | 0.6464 | +?-+- | 0 | 1.604 | 3 | 0.6586 |
| rs72979412 | 11 | 72639396 | C | T | 0.458 | 0.6467 | +?-+- | 0 | 1.587 | 3 | 0.6624 |
| rs341086 | 11 | 72317803 | T | C | -0.458 | 0.6469 | --+?- | 0 | 1.852 | 3 | 0.6036 |
| rs57506367 | 11 | 118141993 | C | T | -0.456 | 0.6484 | -?+?- | 0 | 1.852 | 2 | 0.3962 |
| rs4245179 | 11 | 118144445 | G | A | -0.456 | 0.6486 | -?+?- | 0 | 1.852 | 2 | 0.396 |
| rs1632118 | 11 | 72881636 | T | C | -0.455 | 0.6492 | -+-+- | -23.2 | 4.872 | 4 | 0.3007 |
| rs7940047 | 11 | 118142619 | T | C | -0.451 | 0.6518 | -?+?- | 0 | 1.79 | 2 | 0.4086 |
| rs113399762 | 11 | 118389698 | C | T | 0.45 | 0.6528 | +?++- | 0 | 2.394 | 3 | 0.4947 |
| rs112590920 | 11 | 118143586 | A | G | -0.45 | 0.653 | -?+?- | 0 | 1.775 | 2 | 0.4116 |
| rs57461652 | 11 | 118146197 | A | C | -0.448 | 0.6541 | -?+?- | -99.3 | 2.007 | 2 | 0.3665 |
| rs1715421 | 11 | 118105233 | C | T | 0.448 | 0.6542 | +?-+- | 0 | 1.648 | 3 | 0.6486 |
| rs2075647 | 11 | 118037964 | C | G | -0.445 | 0.6565 | -?-?+ | 0 | 1.586 | 2 | 0.4525 |
| rs1793137 | 11 | 118016068 | C | T | -0.444 | 0.6568 | ---++ | 0 | 0.395 | 4 | 0.9829 |
| rs1793159 | 11 | 118104569 | A | C | 0.442 | 0.6586 | +?-+- | 0 | 1.665 | 3 | 0.6447 |
| rs35409990 | 11 | 118109196 | A | G | 0.442 | 0.6587 | +?-+- | 0 | 1.839 | 3 | 0.6065 |
| rs1715422 | 11 | 118104577 | C | T | 0.442 | 0.6588 | +?-+- | 0 | 1.666 | 3 | 0.6446 |
| rs12275955 | 11 | 118094756 | C | T | 0.439 | 0.6607 | +?-+- | 0 | 2.321 | 3 | 0.5085 |
| rs3181261 | 11 | 118212671 | T | C | 0.438 | 0.6615 | +?+?- | 0 | 1.184 | 2 | 0.5532 |
| rs382163 | 11 | 72292796 | T | C | 0.436 | 0.663 | -?+?- | 0 | 1.35 | 2 | 0.5091 |
| rs58162302 | 11 | 118146199 | T | A | -0.434 | 0.6642 | -?+?- | 0 | 1.994 | 2 | 0.369 |
| rs10793048 | 11 | 72580398 | T | C | -0.434 | 0.6643 | ++--- | 0 | 2.947 | 4 | 0.5667 |
| rs7107376 | 11 | 72640991 | G | T | 0.431 | 0.6662 | +?-+- | 0 | 1.624 | 3 | 0.654 |
| rs12808517 | 11 | 72601276 | T | C | -0.424 | 0.6713 | ++-+- | 0 | 1.805 | 4 | 0.7716 |
| rs186926311 | 11 | 118601426 | T | C | -0.424 | 0.6715 | --++? | -18 | 4.237 | 3 | 0.237 |
| rs67389832 | 11 | 118839032 | A | C | -0.423 | 0.6724 | -?+?- | 1 | 4.042 | 2 | 0.1325 |
| rs1628409 | 11 | 72885257 | T | C | 0.422 | 0.673 | ++-++ | 0 | 2.57 | 4 | 0.6322 |
| rs112337443 | 11 | 118530068 | C | T | 0.422 | 0.6731 | +-++- | -39.8 | 4.292 | 4 | 0.368 |
| rs1715453 | 11 | 118125077 | T | C | -0.421 | 0.6735 | +?-?- | 0 | 1.129 | 2 | 0.5687 |
| rs7117266 | 11 | 118838020 | A | G | -0.418 | 0.6758 | -?+?- | 2.5 | 4.104 | 2 | 0.1285 |
| rs1786183 | 11 | 118017445 | A | G | 0.415 | 0.6779 | -?+++ | 0 | 1.427 | 3 | 0.6993 |
| rs1056562 | 11 | 118125625 | T | C | -0.414 | 0.6786 | +?-?- | 0 | 1.12 | 2 | 0.5712 |
| rs2231439 | 11 | 118174885 | G | A | -0.409 | 0.6826 | -?-?+ | 0 | 1.158 | 2 | 0.5604 |
| rs11216828 | 11 | 118111676 | A | T | 0.406 | 0.685 | +?-+- | 0 | 1.859 | 3 | 0.6022 |
| rs4944016 | 11 | 72597438 | T | C | -0.404 | 0.6859 | ++-+- | 0 | 3.723 | 4 | 0.4448 |
| rs4442562 | 11 | 118845813 | A | G | -0.4 | 0.6892 | -?+?- | 1.2 | 4.049 | 2 | 0.1321 |
| rs9332871 | 11 | 118393856 | T | G | 0.398 | 0.6905 | +?++- | 0 | 2.117 | 3 | 0.5485 |
| rs1793151 | 11 | 118126920 | C | T | -0.396 | 0.6919 | +?-?- | 0 | 1.09 | 2 | 0.5798 |
| rs9651667 | 11 | 118108679 | G | C | 0.393 | 0.6946 | +?-+- | 0 | 1.7 | 3 | 0.637 |
| rs1715449 | 11 | 118128455 | G | A | -0.391 | 0.6956 | +?-?- | 0 | 1.083 | 2 | 0.5819 |
| rs1715450 | 11 | 118128445 | C | G | -0.39 | 0.6964 | +?-?- | 0 | 1.082 | 2 | 0.5821 |
| rs9332801 | 11 | 118355642 | C | A | 0.386 | 0.6991 | +?++- | 0 | 1.9 | 3 | 0.5934 |
| rs10502228 | 11 | 118113583 | T | C | 0.386 | 0.6996 | +?-+- | 0 | 2.03 | 3 | 0.5662 |
| rs1793150 | 11 | 118127314 | G | A | -0.385 | 0.7001 | +?-?- | 0 | 1.081 | 2 | 0.5824 |
| rs76448372 | 11 | 118045846 | A | G | -0.383 | 0.702 | -?+-+ | 11.9 | 5.675 | 3 | 0.1285 |
| rs146525935 | 11 | 118180887 | C | T | -0.382 | 0.7025 | -?+-+ | -0.7 | 4.964 | 3 | 0.1744 |
| rs9651688 | 11 | 118107495 | T | C | 0.381 | 0.7031 | +?-+- | 0 | 1.709 | 3 | 0.635 |
| rs11216839 | 11 | 118129583 | T | A | 0.381 | 0.7033 | +?-+- | 0 | 1.761 | 3 | 0.6235 |
| rs11216829 | 11 | 118112911 | A | G | 0.381 | 0.7034 | +?-+- | 0 | 2.032 | 3 | 0.5657 |
| rs73003294 | 11 | 118661376 | T | C | 0.379 | 0.7048 | -?+-+ | 15.4 | 5.914 | 3 | 0.1159 |

| | | | | | | | | | | | |
|---|---|---|---|---|---|---|---|---|---|---|---|
| rs7113994 | 11 | 118382327 | G | A | 0.379 | 0.705 | +?++- | 0 | 2.34 | 3 | 0.5049 |
| rs61895726 | 11 | 72223735 | T | C | 0.378 | 0.7051 | +?+-+ | 0 | 2.405 | 3 | 0.4926 |
| rs11216813 | 11 | 118071836 | A | G | 0.377 | 0.7059 | +?-+- | 0 | 1.067 | 3 | 0.7851 |
| rs117745694 | 11 | 118218491 | A | G | 0.376 | 0.7066 | +?-++ | 0 | 0.781 | 3 | 0.8541 |
| rs1980091 | 11 | 72288599 | A | G | 0.375 | 0.7075 | -?+?- | 0 | 0.74 | 2 | 0.6908 |
| rs12291619 | 11 | 118062552 | A | G | -0.374 | 0.7081 | +?--+ | 0 | 0.717 | 3 | 0.8691 |
| rs341089 | 11 | 72313462 | A | G | -0.374 | 0.7082 | +?-?- | 0 | 1.405 | 2 | 0.4953 |
| rs655499 | 11 | 118063479 | G | A | 0.373 | 0.7088 | ++-+- | 0 | 0.244 | 4 | 0.9931 |
| rs117532939 | 11 | 72165516 | G | A | 0.372 | 0.71 | -?+?- | -91.4 | 2.09 | 2 | 0.3517 |
| rs117817588 | 11 | 72166041 | T | G | 0.372 | 0.71 | -?+?- | -91.4 | 2.09 | 2 | 0.3517 |
| rs426907 | 11 | 72294440 | A | G | 0.371 | 0.7108 | -?+?- | 0 | 0.695 | 2 | 0.7066 |
| rs658624 | 11 | 118018767 | G | A | 0.369 | 0.7121 | -?+++ | 0 | 1.655 | 3 | 0.6469 |
| rs1793149 | 11 | 118127347 | A | G | -0.368 | 0.7127 | +?-?- | 0 | 1.072 | 2 | 0.585 |
| rs56083337 | 11 | 118131813 | T | A | -0.368 | 0.713 | -?+?+ | 0 | 1.54 | 2 | 0.463 |
| rs11822039 | 11 | 72638704 | A | C | -0.367 | 0.7136 | ++-+- | 0 | 3.725 | 4 | 0.4444 |
| rs76478216 | 11 | 118605352 | C | T | 0.365 | 0.7153 | +?-+- | -4.9 | 4.764 | 3 | 0.1899 |
| rs12286303 | 11 | 118114473 | T | C | 0.364 | 0.7156 | +?-+- | 0 | 2.044 | 3 | 0.5634 |
| rs1793157 | 11 | 118017434 | C | A | 0.364 | 0.716 | -++++ | 0 | 1.782 | 4 | 0.7757 |
| rs55768116 | 11 | 118108331 | A | C | -0.362 | 0.7177 | -?-?+ | 0 | 1.299 | 2 | 0.5224 |
| rs7117544 | 11 | 118117938 | C | A | 0.361 | 0.7183 | +?-+- | 0 | 2.066 | 3 | 0.5587 |
| rs12286567 | 11 | 118114788 | T | G | 0.358 | 0.7201 | +?-+- | 0 | 2.035 | 3 | 0.5653 |
| rs45460396 | 11 | 118005622 | G | T | -0.357 | 0.7207 | -+-++ | 0 | 1.42 | 4 | 0.8407 |
| rs1715443 | 11 | 118017365 | G | T | 0.357 | 0.7214 | -?+++ | 0 | 1.8 | 3 | 0.615 |
| rs11605435 | 11 | 118116496 | C | T | 0.355 | 0.7228 | +?-+- | 0 | 2.054 | 3 | 0.5612 |
| rs1547885 | 11 | 118105222 | T | C | 0.354 | 0.723 | +?-+- | 0 | 1.781 | 3 | 0.619 |
| rs371862 | 11 | 72288843 | T | C | 0.354 | 0.7231 | -?+?- | 0 | 1.058 | 2 | 0.5891 |
| rs341047 | 11 | 72301503 | A | G | -0.354 | 0.7233 | +?-?- | 0 | 1.908 | 2 | 0.3852 |
| rs61900158 | 11 | 118831764 | A | T | -0.353 | 0.7244 | -?+?- | 18.2 | 4.89 | 2 | 0.08671 |
| rs868344 | 11 | 118005119 | C | T | -0.351 | 0.7255 | -+-++ | 0 | 1.354 | 4 | 0.8522 |
| rs149668068 | 11 | 118099735 | A | G | -0.35 | 0.7262 | -?--+ | 0 | 2.166 | 3 | 0.5386 |
| rs613791 | 11 | 118764051 | T | C | 0.349 | 0.7269 | +?++- | 59.8 | 12.44 | 3 | 0.006018 |
| rs3018387 | 11 | 118016981 | T | A | 0.347 | 0.7283 | -?+++ | 0 | 1.768 | 3 | 0.622 |
| rs9332857 | 11 | 118389440 | T | C | 0.347 | 0.7284 | +?++- | 0 | 1.96 | 3 | 0.5807 |
| rs45584835 | 11 | 118004863 | G | A | -0.346 | 0.7292 | -+-++ | 0 | 1.386 | 4 | 0.8467 |
| rs12280557 | 11 | 118117118 | T | C | 0.345 | 0.7299 | +?-+- | 0 | 2.086 | 3 | 0.5548 |
| rs880895 | 11 | 72884382 | A | G | 0.345 | 0.7303 | ++-+- | 0 | 2.785 | 4 | 0.5943 |
| rs392565 | 11 | 72290629 | A | G | 0.343 | 0.7315 | -?+?- | 0 | 1.136 | 2 | 0.5667 |
| rs2853009 | 11 | 118148146 | G | A | -0.34 | 0.7337 | +?-?- | 0 | 1.73 | 2 | 0.421 |
| rs7127705 | 11 | 72999064 | T | C | -0.34 | 0.734 | --+-- | -48.4 | 4.043 | 4 | 0.4003 |
| rs9332852 | 11 | 118381069 | T | C | 0.339 | 0.7348 | +?++- | 0 | 2.01 | 3 | 0.5703 |
| rs6589649 | 11 | 118048836 | C | T | -0.338 | 0.7354 | +?--+ | 0 | 0.773 | 3 | 0.8559 |
| rs10898915 | 11 | 72998687 | A | G | -0.335 | 0.7375 | --+-- | -38.4 | 4.334 | 4 | 0.3627 |
| rs61895701 | 11 | 72208039 | G | T | 0.335 | 0.7377 | +?+-+ | 0 | 2.334 | 3 | 0.506 |
| rs11216830 | 11 | 118115147 | A | G | 0.335 | 0.7378 | +?-+- | 0 | 2.059 | 3 | 0.5602 |
| rs7127013 | 11 | 72999221 | A | G | -0.334 | 0.7382 | --+-+ | -44.3 | 4.157 | 4 | 0.3851 |
| rs3132744 | 11 | 72886304 | G | A | 0.334 | 0.7387 | ++-+- | -25.4 | 4.785 | 4 | 0.3101 |
| rs75709684 | 11 | 118050958 | A | G | -0.332 | 0.7397 | +?--+ | 0 | 1.043 | 3 | 0.7909 |
| rs543824 | 11 | 118762218 | G | A | -0.331 | 0.7403 | +-+-- | 54.5 | 13.182 | 4 | 0.01042 |
| rs67533649 | 11 | 118835272 | G | A | -0.328 | 0.7427 | -?+?- | 23 | 5.193 | 2 | 0.07452 |
| rs146523897 | 11 | 72161854 | A | G | 0.328 | 0.743 | -?+?- | -79.1 | 2.233 | 2 | 0.3274 |
| rs11235712 | 11 | 72994199 | C | T | -0.327 | 0.7434 | --+-+ | -15.4 | 5.198 | 4 | 0.2676 |
| rs45460493 | 11 | 118455465 | A | G | 0.326 | 0.7442 | +?+-- | 0 | 0.734 | 3 | 0.8652 |
| rs4938595 | 11 | 118832582 | T | G | -0.326 | 0.7445 | -?+?- | 23.9 | 5.258 | 2 | 0.07214 |
| rs2946742 | 11 | 118017093 | G | A | 0.325 | 0.7451 | -?+++ | 0 | 1.79 | 3 | 0.617 |
| rs112636863 | 11 | 118333510 | C | T | 0.322 | 0.7471 | +?++- | 0 | 1.857 | 3 | 0.6026 |
| rs10892230 | 11 | 118126682 | A | C | 0.321 | 0.7479 | +?-+- | 0 | 2.137 | 3 | 0.5444 |

| | | | | | | | | | | |
|---|---|---|---|---|---|---|---|---|---|---|
| rs1783574 | 11 | 72863527 | A | G | 0.32 | 0.7486 | ++-+- | 0 | 2.044 | 4 | 0.7276 |
| rs7113959 | 11 | 118379096 | A | T | 0.32 | 0.7487 | +?++- | 0 | 2.261 | 3 | 0.5201 |
| rs7342287 | 11 | 118050404 | A | G | -0.32 | 0.7492 | +?--+ | 0 | 0.749 | 3 | 0.8616 |
| rs10736494 | 11 | 118518227 | T | C | 0.319 | 0.7495 | +?++- | 0 | 0.842 | 3 | 0.8394 |
| rs9332793 | 11 | 118354458 | A | G | 0.318 | 0.7504 | +?-+- | 0 | 1.856 | 3 | 0.6028 |
| rs111901186 | 11 | 118328052 | G | A | 0.317 | 0.7512 | +?-+- | 0 | 1.85 | 3 | 0.6041 |
| rs113028342 | 11 | 118333537 | G | A | 0.317 | 0.7514 | +?++- | 0 | 1.848 | 3 | 0.6046 |
| rs2946743 | 11 | 118017088 | G | A | 0.316 | 0.7518 | -?+++ | 0 | 1.824 | 3 | 0.6096 |
| rs77806521 | 11 | 118232026 | T | C | -0.315 | 0.7525 | +?+?- | 0 | 1.39 | 2 | 0.4991 |
| rs112900272 | 11 | 118326445 | T | C | 0.315 | 0.7527 | +?-+- | 0 | 1.851 | 3 | 0.604 |
| rs11216834 | 11 | 118122077 | C | T | 0.315 | 0.753 | +?-+- | 0 | 2.123 | 3 | 0.5473 |
| rs10790246 | 11 | 118103542 | T | A | 0.315 | 0.7531 | +?-+- | 0 | 1.803 | 3 | 0.6144 |
| rs10898889 | 11 | 72605382 | C | G | -0.312 | 0.755 | ++-+- | 0 | 3.964 | 4 | 0.4109 |
| rs9332794 | 11 | 118354505 | T | G | 0.311 | 0.7555 | +?-+- | 0 | 1.862 | 3 | 0.6016 |
| rs7121712 | 11 | 72993695 | A | T | -0.311 | 0.7556 | --+-- | -28.6 | 4.666 | 4 | 0.3233 |
| rs614825 | 11 | 118018725 | G | A | 0.31 | 0.7566 | -?+++ | 0 | 1.686 | 3 | 0.64 |
| rs2027764 | 11 | 72998247 | G | A | -0.307 | 0.7589 | --+-+ | -41.1 | 4.252 | 4 | 0.373 |
| rs35381826 | 11 | 118032438 | T | G | -0.306 | 0.76 | -?-+- | 0 | 0.105 | 3 | 0.9913 |
| rs57533048 | 11 | 118050014 | G | A | -0.301 | 0.7636 | +?--+ | 0 | 0.915 | 3 | 0.8217 |
| rs77643000 | 11 | 118749563 | C | T | 0.3 | 0.7639 | ++++- | -1.7 | 5.897 | 4 | 0.207 |
| rs12575015 | 11 | 118197645 | T | A | -0.299 | 0.7646 | -?-?+ | 0 | 0.637 | 2 | 0.7274 |
| rs71482132 | 11 | 118031211 | A | G | -0.298 | 0.7655 | -?-+- | 0 | 0.129 | 3 | 0.9882 |
| rs4420279 | 11 | 118835677 | A | C | -0.298 | 0.7657 | -?+?- | 27.6 | 5.523 | 2 | 0.06319 |
| rs116848264 | 11 | 118049817 | A | T | 0.298 | 0.7658 | -?+-+ | 0 | 2.744 | 3 | 0.4328 |
| rs67163042 | 11 | 118050779 | G | A | -0.297 | 0.7664 | +?--+ | 0 | 0.739 | 3 | 0.864 |
| rs640323 | 11 | 118010890 | C | G | 0.296 | 0.7672 | +--+- | 0 | 1.361 | 4 | 0.851 |
| rs882904 | 11 | 72997910 | G | A | -0.293 | 0.7693 | --+-+ | -44.4 | 4.156 | 4 | 0.3854 |
| rs12286604 | 11 | 118084728 | A | G | 0.293 | 0.7696 | +?-+- | 0 | 1.558 | 3 | 0.669 |
| rs66962317 | 11 | 118197486 | C | T | -0.292 | 0.7703 | -?-?+ | 0 | 0.634 | 2 | 0.7283 |
| rs67435655 | 11 | 118196399 | C | T | -0.29 | 0.7717 | -?-?+ | 0 | 0.639 | 2 | 0.7263 |
| rs1783587 | 11 | 72887612 | C | T | -0.29 | 0.772 | ++-+- | 4.3 | 6.268 | 4 | 0.18 |
| rs11216800 | 11 | 118048885 | G | A | -0.289 | 0.7722 | +?--+ | 0 | 0.996 | 3 | 0.8022 |
| rs55843215 | 11 | 118195962 | C | T | -0.289 | 0.7722 | -?-?+ | 0 | 0.602 | 2 | 0.74 |
| rs56032119 | 11 | 118797544 | G | A | 0.285 | 0.7754 | +---+ | -10 | 5.453 | 4 | 0.2439 |
| rs11216816 | 11 | 118075152 | A | C | 0.285 | 0.7756 | +?-+- | 0 | 0.971 | 3 | 0.8082 |
| rs7123533 | 11 | 118049257 | G | C | -0.285 | 0.7759 | +?--+ | 0 | 0.88 | 3 | 0.8304 |
| rs639785 | 11 | 118062268 | T | C | 0.284 | 0.7762 | ++++- | -49 | 4.028 | 4 | 0.4023 |
| rs11600901 | 11 | 118031986 | A | G | -0.284 | 0.7767 | -?-+- | 0 | 0.128 | 3 | 0.9883 |
| rs4944014 | 11 | 72527180 | T | C | -0.283 | 0.7772 | ++-+- | 0 | 3.475 | 4 | 0.4817 |
| rs73575429 | 11 | 118797392 | C | T | 0.283 | 0.7773 | +---+ | -10.7 | 5.419 | 4 | 0.247 |
| rs74584888 | 11 | 118797278 | A | G | 0.283 | 0.7773 | +---+ | -10.9 | 5.412 | 4 | 0.2476 |
| rs59018400 | 11 | 118049811 | T | C | -0.279 | 0.7805 | +?--+ | 0 | 0.867 | 3 | 0.8335 |
| rs11216817 | 11 | 118075474 | T | C | 0.278 | 0.7812 | +?-+- | 0 | 0.986 | 3 | 0.8046 |
| rs12420093 | 11 | 118272963 | G | A | -0.278 | 0.7812 | --+-? | -21.9 | 4.101 | 3 | 0.2507 |
| rs113566564 | 11 | 118335147 | A | G | 0.275 | 0.7831 | +?++- | 0 | 1.956 | 3 | 0.5817 |
| rs60969913 | 11 | 118798154 | C | G | 0.274 | 0.7838 | +---+ | -8.9 | 5.509 | 4 | 0.2389 |
| rs111390501 | 11 | 118797420 | G | A | 0.274 | 0.7843 | +---+ | -10.3 | 5.439 | 4 | 0.2452 |
| rs502514 | 11 | 118241995 | C | T | -0.272 | 0.7853 | ---?+ | 37.7 | 8.028 | 3 | 0.04545 |
| rs56301948 | 11 | 118049880 | G | A | -0.272 | 0.7853 | +?--+ | 0 | 0.853 | 3 | 0.8367 |
| rs572608 | 11 | 118801640 | A | T | 0.272 | 0.7856 | -+-++ | 0 | 1.273 | 4 | 0.8659 |
| rs7342223 | 11 | 118050348 | G | T | -0.272 | 0.7858 | +?--+ | 0 | 0.852 | 3 | 0.837 |
| rs67859744 | 11 | 118200657 | A | T | -0.271 | 0.7864 | -?-?+ | 0 | 0.653 | 2 | 0.7213 |
| rs73575428 | 11 | 118797003 | G | C | 0.271 | 0.7864 | +---+ | -11.5 | 5.382 | 4 | 0.2503 |
| rs637737 | 11 | 118051646 | G | A | -0.27 | 0.7875 | +?--+ | 0 | 0.601 | 3 | 0.8962 |
| rs67262752 | 11 | 118047776 | C | T | -0.268 | 0.7887 | +?--+ | 0 | 0.986 | 3 | 0.8046 |
| rs1877652 | 11 | 72307742 | A | G | 0.268 | 0.789 | -?+?- | 0 | 0.978 | 2 | 0.6132 |

| | | | | | | | | | | | |
|---|---|---|---|---|---|---|---|---|---|---|---|
| rs79820203 | 11 | 118051010 | C | T | -0.265 | 0.791 | +?--+ | 0 | 0.873 | 3 | 0.8319 |
| rs75459003 | 11 | 118053859 | C | A | -0.263 | 0.7927 | +?--+ | 0 | 0.954 | 3 | 0.8123 |
| rs3802866 | 11 | 118095803 | T | C | 0.261 | 0.7944 | +?-+- | 0 | 1.786 | 3 | 0.618 |
| rs7927676 | 11 | 118051323 | G | C | -0.26 | 0.7946 | +?--+ | 0 | 0.882 | 3 | 0.8298 |
| rs74395230 | 11 | 118052662 | T | C | -0.259 | 0.7957 | +?--+ | 0 | 0.923 | 3 | 0.8199 |
| rs78312641 | 11 | 118749318 | A | T | 0.258 | 0.7968 | ++++- | 0.1 | 6.008 | 4 | 0.1986 |
| rs76831657 | 11 | 118051082 | G | A | -0.257 | 0.7971 | +?--+ | 0 | 0.918 | 3 | 0.8212 |
| rs7943511 | 11 | 118096165 | T | C | 0.257 | 0.7973 | +?-+- | 0 | 1.793 | 3 | 0.6164 |
| rs1613495 | 11 | 118017645 | T | C | 0.257 | 0.7975 | -?++- | 0 | 0.873 | 3 | 0.832 |
| rs11216791 | 11 | 118044360 | G | A | -0.254 | 0.7992 | +?--+ | 0 | 1.091 | 3 | 0.7792 |
| rs76050019 | 11 | 118051857 | C | T | -0.254 | 0.7993 | +?--+ | 0 | 0.868 | 3 | 0.8331 |
| rs10898885 | 11 | 72571275 | T | C | -0.252 | 0.8011 | ++--- | 0 | 2.662 | 4 | 0.616 |
| rs12295676 | 11 | 118054345 | A | G | -0.251 | 0.8016 | +?--+ | 0 | 0.949 | 3 | 0.8136 |
| rs12295438 | 11 | 118094921 | T | C | 0.25 | 0.8024 | +?-+- | 0 | 1.791 | 3 | 0.6168 |
| rs7943272 | 11 | 118096028 | C | A | 0.25 | 0.8025 | +?-+- | 0 | 1.813 | 3 | 0.6122 |
| rs583490 | 11 | 118022608 | A | G | -0.249 | 0.8037 | +---+ | 0 | 3.668 | 4 | 0.4528 |
| rs678262 | 11 | 118021740 | G | C | -0.248 | 0.8042 | +---- | -44.9 | 4.14 | 4 | 0.3874 |
| rs147710683 | 11 | 118121356 | T | C | -0.247 | 0.8049 | -?--+ | 0 | 2.884 | 3 | 0.4098 |
| rs113521822 | 11 | 118063349 | G | A | 0.246 | 0.8058 | +?--+ | 0 | 0.695 | 3 | 0.8743 |
| rs566416 | 11 | 118759610 | G | T | -0.241 | 0.8093 | +-+-- | 59.3 | 14.754 | 4 | 0.00524 |
| rs7926717 | 11 | 118121399 | T | G | 0.241 | 0.8098 | +?-+- | 0 | 1.648 | 3 | 0.6486 |
| rs12577841 | 11 | 118194599 | T | C | -0.23 | 0.8178 | -+--+ | 0 | 0.58 | 4 | 0.9653 |
| rs676925 | 11 | 118765688 | G | C | -0.23 | 0.8179 | +-+-- | 50.1 | 12.021 | 4 | 0.01719 |
| rs58507982 | 11 | 118146510 | T | C | -0.23 | 0.818 | -?+?- | 0 | 1.271 | 2 | 0.5297 |
| rs6592474 | 11 | 72184290 | A | T | 0.229 | 0.8191 | -++-- | -9.2 | 5.495 | 4 | 0.2402 |
| rs7934292 | 11 | 118059914 | C | T | -0.228 | 0.8197 | +?--+ | 0 | 0.944 | 3 | 0.8149 |
| rs642765 | 11 | 118030310 | C | G | -0.228 | 0.8199 | -?-+- | 0 | 0.174 | 3 | 0.9816 |
| rs11608072 | 11 | 118030086 | C | G | -0.227 | 0.8206 | +?-+- | 0 | 0.198 | 3 | 0.9779 |
| rs11216781 | 11 | 118012310 | A | G | 0.226 | 0.8209 | ++++- | 0 | 0.922 | 4 | 0.9213 |
| rs11216824 | 11 | 118096818 | A | G | 0.22 | 0.8262 | +?-+- | 0 | 1.621 | 3 | 0.6547 |
| rs7103366 | 11 | 118057867 | G | T | 0.218 | 0.8272 | ++++- | -47.2 | 4.076 | 4 | 0.3958 |
| rs7935492 | 11 | 118056823 | T | C | -0.217 | 0.828 | +?--+ | 0 | 0.919 | 3 | 0.8209 |
| rs145968026 | 11 | 118421841 | G | C | 0.217 | 0.8286 | +?+-- | 0 | 0.745 | 3 | 0.8627 |
| rs11216821 | 11 | 118088362 | C | T | 0.214 | 0.8306 | +?-+- | 0 | 1.621 | 3 | 0.6547 |
| rs12282334 | 11 | 118057783 | G | C | -0.214 | 0.8306 | +?--+ | 0 | 0.911 | 3 | 0.8227 |
| rs2365774 | 11 | 72308203 | G | A | 0.213 | 0.8314 | -?+?- | 0 | 0.819 | 2 | 0.664 |
| rs2276064 | 11 | 118529703 | A | G | -0.211 | 0.8328 | -+-?? | -53.8 | 2.601 | 2 | 0.2724 |
| rs598746 | 11 | 118020057 | A | G | -0.208 | 0.835 | -?++- | 0 | 0.927 | 3 | 0.819 |
| rs10892311 | 11 | 118807735 | T | C | 0.206 | 0.8365 | -+-++ | 0 | 1.511 | 4 | 0.8246 |
| rs642936 | 11 | 118019906 | C | T | -0.204 | 0.8383 | +---+ | 0 | 2.509 | 4 | 0.6431 |
| rs12808254 | 11 | 118066912 | T | C | -0.204 | 0.8385 | +?-+- | 7.1 | 5.385 | 3 | 0.1457 |
| rs79699220 | 11 | 118550841 | C | G | 0.204 | 0.8386 | +?-++ | 0 | 0.927 | 3 | 0.8189 |
| rs4245414 | 11 | 72188466 | A | G | 0.203 | 0.8393 | -++-- | -4.1 | 5.765 | 4 | 0.2174 |
| rs73615909 | 11 | 118322488 | T | C | 0.202 | 0.8398 | +?-+- | 0 | 1.443 | 3 | 0.6956 |
| rs12418805 | 11 | 72188979 | C | T | 0.2 | 0.8413 | -++-- | -3.9 | 5.772 | 4 | 0.2168 |
| rs76248354 | 11 | 118042606 | A | G | -0.2 | 0.8417 | -?+-+ | 25.6 | 6.72 | 3 | 0.08139 |
| rs116979688 | 11 | 118032223 | T | C | 0.199 | 0.8419 | +?+?? | 0 | 0.001 | 1 | 0.9778 |
| rs12284089 | 11 | 118058542 | T | C | -0.198 | 0.8433 | +?--+ | 0 | 0.999 | 3 | 0.8015 |
| rs7939622 | 11 | 118087841 | A | G | 0.198 | 0.8433 | +?-+- | 0 | 1.635 | 3 | 0.6515 |
| rs12271064 | 11 | 118119211 | G | A | 0.196 | 0.8448 | +?-+- | 0 | 1.986 | 3 | 0.5753 |
| rs593753 | 11 | 72691440 | T | C | -0.194 | 0.8465 | ++-+- | 0 | 3.636 | 4 | 0.4575 |
| rs602594 | 11 | 118040610 | T | C | 0.19 | 0.8489 | +?--+ | 0 | 0.401 | 3 | 0.9401 |
| rs4943910 | 11 | 72188643 | T | C | 0.19 | 0.849 | -++-- | -4.9 | 5.72 | 4 | 0.2211 |
| rs6589652 | 11 | 118058358 | G | A | 0.19 | 0.8493 | ++++- | 0 | 3.881 | 4 | 0.4223 |
| rs4274214 | 11 | 72188003 | T | C | 0.188 | 0.8513 | -++-- | -6.2 | 5.651 | 4 | 0.2268 |
| rs11607671 | 11 | 72192738 | T | C | 0.185 | 0.853 | -++-- | -5.9 | 5.663 | 4 | 0.2258 |

| | | | | | | | | | | | |
|---|---|---|---|---|---|---|---|---|---|---|---|
| rs11607688 | 11 | 72192830 | T | C | 0.178 | 0.8588 | -++-- | -6.5 | 5.631 | 4 | 0.2284 |
| rs17310153 | 11 | 72190863 | G | A | 0.178 | 0.8591 | -++-- | -6.4 | 5.641 | 4 | 0.2276 |
| rs75403174 | 11 | 118043904 | A | C | -0.178 | 0.8591 | +?--+ | 0 | 1.324 | 3 | 0.7235 |
| rs2365385 | 11 | 72194718 | A | G | 0.177 | 0.8597 | -++-- | -5.8 | 5.672 | 4 | 0.225 |
| rs11600727 | 11 | 72201499 | C | A | -0.176 | 0.8601 | -++-- | 22.8 | 7.771 | 4 | 0.1004 |
| rs12289120 | 11 | 118066609 | T | C | 0.173 | 0.8629 | +?-+- | 0 | 0.386 | 3 | 0.9431 |
| rs11216792 | 11 | 118044450 | T | G | -0.173 | 0.863 | +?--+ | 0 | 1.31 | 3 | 0.7268 |
| rs679327 | 11 | 118042974 | G | A | 0.172 | 0.8636 | +?+++ | 0 | 0.028 | 3 | 0.9988 |
| rs9332781 | 11 | 118346599 | G | A | 0.171 | 0.8641 | +?-+- | 0 | 1.644 | 3 | 0.6495 |
| rs45508093 | 11 | 118224059 | G | A | 0.17 | 0.8647 | -?+?+ | 0 | 0.308 | 2 | 0.8573 |
| rs1783570 | 11 | 72871654 | A | G | -0.168 | 0.8666 | ++-+- | 12.1 | 6.83 | 4 | 0.1452 |
| rs6589651 | 11 | 118058156 | G | A | 0.166 | 0.8684 | ++++- | 0 | 3.969 | 4 | 0.4103 |
| rs11216790 | 11 | 118044144 | T | G | -0.166 | 0.8685 | +?--+ | 0 | 1.324 | 3 | 0.7234 |
| rs12418930 | 11 | 72681566 | T | C | -0.164 | 0.8697 | -++-+ | 35.4 | 9.293 | 4 | 0.05417 |
| rs598207 | 11 | 118765267 | C | G | 0.163 | 0.8706 | +?++- | 57.2 | 11.676 | 3 | 0.008581 |
| rs656308 | 11 | 118031206 | T | G | -0.162 | 0.8713 | -?-+- | 0 | 0.227 | 3 | 0.9731 |
| rs7941217 | 11 | 118349247 | T | C | 0.161 | 0.8721 | +?-+- | 0 | 1.568 | 3 | 0.6666 |
| rs73557344 | 11 | 118810054 | C | G | 0.159 | 0.8736 | +---+ | -33.3 | 4.502 | 4 | 0.3423 |
| rs12420153 | 11 | 72195852 | A | G | 0.155 | 0.8766 | -++-- | -9.5 | 5.481 | 4 | 0.2414 |
| rs75460330 | 11 | 118809767 | A | G | 0.15 | 0.881 | +---+ | -32.5 | 4.527 | 4 | 0.3394 |
| rs663003 | 11 | 118823148 | T | C | 0.148 | 0.8824 | -?-++ | 9.6 | 5.53 | 3 | 0.1368 |
| rs10892222 | 11 | 118041102 | A | G | -0.145 | 0.8846 | +?--+ | 0 | 1.869 | 3 | 0.5999 |
| rs1624780 | 11 | 118134785 | T | G | -0.145 | 0.8849 | +?-?- | 13.7 | 4.633 | 2 | 0.09861 |
| rs11608057 | 11 | 118030185 | T | C | -0.143 | 0.8865 | -+-+- | 0 | 0.257 | 4 | 0.9924 |
| rs4553405 | 11 | 118061798 | T | C | 0.138 | 0.8905 | ++++- | -33.9 | 4.48 | 4 | 0.3449 |
| rs8192614 | 11 | 118037564 | A | G | -0.137 | 0.8911 | -?-+- | 0 | 0.203 | 3 | 0.9771 |
| rs56027064 | 11 | 72563104 | G | T | 0.136 | 0.8915 | +--+- | 0 | 1.538 | 4 | 0.8199 |
| rs59069289 | 11 | 118802128 | T | G | -0.134 | 0.8931 | +---+ | -36.6 | 4.391 | 4 | 0.3557 |
| rs59480472 | 11 | 118802386 | C | T | -0.134 | 0.8933 | +---+ | -36.7 | 4.39 | 4 | 0.3558 |
| rs600325 | 11 | 118041133 | C | T | 0.132 | 0.8948 | +?--+ | 0 | 0.336 | 3 | 0.9531 |
| rs111313044 | 11 | 118803596 | A | G | -0.126 | 0.8999 | +---+ | -31.7 | 4.556 | 4 | 0.3359 |
| rs73575440 | 11 | 118803160 | A | G | -0.124 | 0.9011 | +---+ | -34.3 | 4.466 | 4 | 0.3466 |
| rs2298831 | 11 | 118068752 | C | T | 0.123 | 0.902 | +?-+- | 0 | 0.515 | 3 | 0.9156 |
| rs724673 | 11 | 72795708 | A | G | 0.122 | 0.9029 | -++-+ | 30.6 | 8.649 | 4 | 0.07051 |
| rs4936424 | 11 | 118009674 | T | C | 0.121 | 0.9037 | +-+-- | 22.3 | 7.72 | 4 | 0.1024 |
| rs2017122 | 11 | 118314849 | T | C | 0.115 | 0.9081 | +?-+- | 0 | 2.843 | 3 | 0.4165 |
| rs12294669 | 11 | 118041972 | G | A | -0.113 | 0.9098 | +?--+ | 0 | 1.958 | 3 | 0.5811 |
| rs73555296 | 11 | 118445559 | T | G | 0.112 | 0.9108 | +?+-- | 0 | 0.549 | 3 | 0.9081 |
| rs4943907 | 11 | 72183348 | T | C | -0.112 | 0.911 | -++-- | 13.4 | 6.925 | 4 | 0.1399 |
| rs651825 | 11 | 118042713 | G | T | 0.111 | 0.9118 | +?+-+ | 0 | 0.53 | 3 | 0.9123 |
| rs2003368 | 11 | 72840106 | T | C | -0.11 | 0.9121 | -++-+ | 37.1 | 9.537 | 4 | 0.04899 |
| rs145762171 | 11 | 118748789 | A | G | 0.109 | 0.9132 | +?--+ | 0 | 2.639 | 3 | 0.4507 |
| rs150339153 | 11 | 118285369 | T | C | -0.109 | 0.9132 | -?--+ | 0 | 1.679 | 3 | 0.6416 |
| rs611960 | 11 | 118034228 | G | T | 0.104 | 0.917 | -?-+- | 0 | 0.505 | 3 | 0.9177 |
| rs60798124 | 11 | 118282241 | G | A | 0.102 | 0.9188 | +++-- | 0 | 3.44 | 4 | 0.487 |
| rs611979 | 11 | 118034223 | A | G | 0.101 | 0.9197 | -?-+- | 0 | 0.502 | 3 | 0.9184 |
| rs7948661 | 11 | 118341649 | C | T | 0.098 | 0.9218 | +?-+- | 0 | 1.673 | 3 | 0.6428 |
| rs72969876 | 11 | 72589778 | C | G | 0.096 | 0.9231 | +--+- | 0 | 1.682 | 4 | 0.7939 |
| rs3765638 | 11 | 72537072 | A | G | -0.095 | 0.9244 | -++-+ | 18.9 | 7.398 | 4 | 0.1163 |
| rs76651078 | 11 | 118316006 | A | G | 0.092 | 0.9264 | +?-+- | -63.9 | 3.051 | 3 | 0.3838 |
| rs10892221 | 11 | 118040642 | A | G | -0.086 | 0.9318 | +?--+ | 0 | 2.01 | 3 | 0.5704 |
| rs59489957 | 11 | 118284984 | T | C | 0.086 | 0.9318 | +++-- | 0 | 3.475 | 4 | 0.4816 |
| rs180884425 | 11 | 118620195 | C | T | 0.084 | 0.9334 | +-+?? | 0 | 0.065 | 2 | 0.9682 |
| rs74422681 | 11 | 118312870 | G | A | 0.083 | 0.9335 | +-++- | -34.6 | 4.458 | 4 | 0.3476 |
| rs61010648 | 11 | 118206890 | G | C | -0.077 | 0.9384 | -?-?+ | 0 | 1.058 | 2 | 0.5893 |
| rs678957 | 11 | 118050855 | T | C | -0.077 | 0.9385 | +?+-+ | 0 | 0.439 | 3 | 0.9321 |

| | | | | | | | | | | | |
|---|---|---|---|---|---|---|---|---|---|---|---|
| rs56296109 | 11 | 118698170 | A | C | -0.075 | 0.9404 | -?+-- | 17.5 | 6.064 | 3 | 0.1085 |
| rs594171 | 11 | 118032974 | A | T | 0.069 | 0.9452 | -?-+- | 0 | 0.669 | 3 | 0.8805 |
| rs645451 | 11 | 118025969 | T | C | 0.067 | 0.9463 | -?+?- | 0 | 0.235 | 2 | 0.8891 |
| rs1616199 | 11 | 118146462 | T | C | 0.065 | 0.948 | +--?+ | -26 | 3.969 | 3 | 0.2649 |
| rs7933243 | 11 | 118403205 | G | T | 0.063 | 0.9496 | +-+-? | -43.3 | 3.49 | 3 | 0.3221 |
| rs11216807 | 11 | 118063030 | C | T | -0.058 | 0.9534 | ++++- | 0 | 3.852 | 4 | 0.4264 |
| rs11235620 | 11 | 72680931 | T | A | 0.058 | 0.9534 | ++-+- | 0 | 3.41 | 4 | 0.4917 |
| rs876479 | 11 | 118028094 | T | C | -0.055 | 0.9563 | -?-?+ | 0 | 0.97 | 2 | 0.6156 |
| rs61581699 | 11 | 118045423 | A | G | -0.054 | 0.9567 | +?--+ | 0 | 0.656 | 3 | 0.8834 |
| rs61900157 | 11 | 118824439 | C | A | 0.053 | 0.9575 | -?-++ | 7.9 | 5.428 | 3 | 0.143 |
| rs11603160 | 11 | 72894273 | T | C | 0.053 | 0.9578 | -?++? | 0 | 1.179 | 2 | 0.5546 |
| rs628299 | 11 | 118018022 | G | A | -0.048 | 0.9618 | +---+ | 0 | 2.977 | 4 | 0.5617 |
| rs3018386 | 11 | 118016622 | G | C | -0.045 | 0.964 | +---+ | 0 | 2.76 | 4 | 0.5987 |
| rs10790244 | 11 | 118059557 | C | T | 0.045 | 0.9643 | ++++- | -44.4 | 4.155 | 4 | 0.3855 |
| rs73575454 | 11 | 118807639 | A | G | -0.044 | 0.9647 | +---+ | -26.3 | 4.749 | 4 | 0.314 |
| rs11216806 | 11 | 118060549 | C | T | 0.042 | 0.9662 | ++++- | -1.5 | 5.914 | 4 | 0.2057 |
| rs11217096 | 11 | 118796570 | T | C | -0.041 | 0.9675 | -+--+ | 0 | 1.645 | 4 | 0.8006 |
| rs11216794 | 11 | 118044740 | A | T | -0.041 | 0.9677 | +?--+ | 0 | 0.699 | 3 | 0.8735 |
| rs17122154 | 11 | 118273019 | A | G | -0.039 | 0.9692 | +++-- | 0 | 3.316 | 4 | 0.5065 |
| rs7103918 | 11 | 118061480 | T | C | 0.033 | 0.9734 | ++++- | 0 | 3.868 | 4 | 0.4242 |
| rs458360 | 11 | 72319522 | A | C | 0.033 | 0.9739 | -?+?- | -5.6 | 3.789 | 2 | 0.1504 |
| rs462491 | 11 | 72319520 | C | T | 0.033 | 0.9739 | -?+?- | -5.6 | 3.789 | 2 | 0.1504 |
| rs4936426 | 11 | 118134597 | C | T | -0.031 | 0.9756 | +?+?- | 25.7 | 5.381 | 2 | 0.06785 |
| rs6589663 | 11 | 118255278 | A | G | -0.031 | 0.9756 | +++-- | 0 | 3.334 | 4 | 0.5035 |
| rs625291 | 11 | 118826060 | T | C | 0.029 | 0.977 | -?-++ | 13.4 | 5.776 | 3 | 0.123 |
| rs73575441 | 11 | 118804919 | C | T | -0.027 | 0.9783 | +---+ | -22.3 | 4.907 | 4 | 0.297 |
| rs666191 | 11 | 118826063 | A | G | 0.027 | 0.9788 | -?-++ | 15.2 | 5.895 | 3 | 0.1168 |
| rs11216867 | 11 | 118262407 | G | A | -0.024 | 0.9806 | +++-- | 0 | 3.361 | 4 | 0.4993 |
| rs56933877 | 11 | 118804643 | A | G | -0.02 | 0.9838 | +---+ | -21.7 | 4.93 | 4 | 0.2945 |
| rs341085 | 11 | 72318882 | A | C | 0.02 | 0.9844 | -?+?- | -7.6 | 3.717 | 2 | 0.1559 |
| rs17122150 | 11 | 118271860 | G | C | -0.017 | 0.9863 | +++-- | 0 | 3.459 | 4 | 0.4842 |
| rs2097131 | 11 | 118825069 | A | G | 0.016 | 0.9871 | -?-++ | 7 | 5.375 | 3 | 0.1463 |
| rs1793170 | 11 | 118150205 | C | T | 0.014 | 0.9884 | +--?+ | -24.2 | 4.025 | 3 | 0.2588 |
| rs11235641 | 11 | 72755766 | A | G | -0.011 | 0.9912 | -++-+ | 26.4 | 8.151 | 4 | 0.08619 |
| rs7121764 | 11 | 118271759 | G | A | 0.007 | 0.9945 | +++-- | 0 | 3.323 | 4 | 0.5053 |
| rs939274 | 11 | 118134065 | T | G | -0.007 | 0.9946 | +?-?- | 22.8 | 5.183 | 2 | 0.07491 |
| rs112045567 | 11 | 118804989 | G | A | 0.006 | 0.9953 | +---+ | -19.4 | 5.025 | 4 | 0.2847 |
| rs586053 | 11 | 118020568 | A | G | 0.005 | 0.9957 | -?++- | 0 | 1.122 | 3 | 0.7717 |
| rs4426142 | 11 | 118824996 | A | G | -0.002 | 0.9988 | -?-++ | 10.2 | 5.571 | 3 | 0.1345 |

**Supplemental Data 4: Bayesian analysis of SjD GWAS and Immunochip data in DDX6-CXCR5 locus (Dataset 1).**

Note: CHROM=chromosome; POS=basepair position; A1=major allele; A2=minor allele; OR=odd ratio; L+P0=The log null marginal likelihood; L+P1=The log alternative marginal likelihood; BF=Bayes factor; P_CAUSAL=The probability that this variant is the causal variant at this locus;

### CREDIBLE SET

| CHROM | POS | RSID | A1 | A2 | BestModel | BestModelProb | CredibleSet |
|---|---|---|---|---|---|---|---|
| 11 | 118694547 | rs7123726 | C | T | M_Disease | 0.993406136 | M_Disease |
| 11 | 118745278 | rs7481819 | C | T | M_Disease | 0.990986701 | M_Disease |
| 11 | 118680648 | rs11217040 | A | C | M_Disease | 0.990243334 | M_Disease |
| 11 | 118682350 | rs11217041 | G | A | M_Disease | 0.988738054 | M_Disease |
| 11 | 118682528 | rs11217042 | A | G | M_Disease | 0.988738054 | M_Disease |
| 11 | 118681464 | rs7942535 | C | T | M_Disease | 0.988738054 | M_Disease |
| 11 | 118745243 | rs7481797 | A | G | M_Disease | 0.988738054 | M_Disease |
| 11 | 118683564 | rs17122453 | A | G | M_Disease | 0.98828383 | M_Disease |
| 11 | 118681083 | rs73005426 | G | A | M_Disease | 0.988225794 | M_Disease |
| 11 | 118701018 | rs79504890 | A | G | M_Disease | 0.987871565 | M_Disease |
| 11 | 118743338 | rs4936444 | T | C | M_Disease | 0.987629585 | M_Disease |
| 11 | 118698907 | rs11217046 | A | G | M_Disease | 0.986809759 | M_Disease |
| 11 | 118684610 | rs715412 | A | G | M_Disease | 0.985866875 | M_Disease |
| 11 | 118669605 | rs11217032 | A | T | M_Disease | 0.985656344 | M_Disease |
| 11 | 118745884 | rs10790275 | G | C | M_Disease | 0.985585482 | M_Disease |
| 11 | 118697865 | rs11217045 | A | C | M_Disease | 0.985585482 | M_Disease |
| 11 | 118697906 | rs73005502 | G | A | M_Disease | 0.985585482 | M_Disease |
| 11 | 118679304 | rs28409215 | A | G | M_Disease | 0.985153013 | M_Disease |
| 11 | 118677355 | rs11217038 | A | G | M_Disease | 0.98493201 | M_Disease |
| 11 | 118665575 | rs10892293 | T | A | M_Disease | 0.98493201 | M_Disease |
| 11 | 118743286 | rs12365699 | A | G | M_Disease | 0.984857625 | M_Disease |
| 11 | 118667357 | rs10892294 | C | G | M_Disease | 0.984707769 | M_Disease |
| 11 | 118681079 | rs73005423 | G | A | M_Disease | 0.984093608 | M_Disease |
| 11 | 118675772 | rs11217036 | T | C | M_Disease | 0.984015152 | M_Disease |
| 11 | 118677086 | rs11217037 | A | G | M_Disease | 0.983857095 | M_Disease |
| 11 | 118673706 | rs11217033 | C | T | M_Disease | 0.983777491 | M_Disease |
| 11 | 118726843 | rs1037679725 | T | C | M_Disease | 0.983697501 | M_Disease |
| 11 | 118721699 | rs7125066 | G | A | M_Disease | 0.983536355 | M_Disease |
| 11 | 118724894 | rs9736016 | A | T | M_Disease | 0.983455196 | M_Disease |
| 11 | 118743772 | rs6421571 | T | C | M_Disease | 0.983455196 | M_Disease |
| 11 | 118725660 | rs4936441 | C | G | M_Disease | 0.983373644 | M_Disease |
| 11 | 118722149 | rs11217058 | A | G | M_Disease | 0.983126609 | M_Disease |
| 11 | 118696022 | rs11217044 | C | T | M_Disease | 0.982621668 | M_Disease |
| 11 | 118726753 | rs11217066 | G | T | M_Disease | 0.98253608 | M_Disease |
| 11 | 118741842 | rs4938573 | C | T | M_Disease | 0.98201379 | M_Disease |
| 11 | 118741157 | rs7117261 | T | C | M_Disease | 0.981836309 | M_Disease |
| 11 | 118740418 | rs7951740 | G | C | M_Disease | 0.981385052 | M_Disease |
| 11 | 118729391 | rs10790268 | A | G | M_Disease | 0.981201486 | M_Disease |
| 11 | 118744396 | rs7117313 | C | G | M_Disease | 0.981016144 | M_Disease |
| 11 | 118739056 | rs7122669 | A | G | M_Disease | 0.980922803 | M_Disease |
| 11 | 118742800 | rs74541740 | A | G | M_Disease | 0.98035323 | M_Disease |
| 11 | 118721565 | rs75249660 | A | G | M_Disease | 0.980159694 | M_Disease |
| 11 | 118738298 | rs7119044 | G | C | M_Disease | 0.980159694 | M_Disease |
| 11 | 118729456 | rs10790269 | C | T | M_Disease | 0.979767003 | M_Disease |
| 11 | 118738281 | rs7119038 | G | A | M_Disease | 0.978958653 | M_Disease |
| 11 | 118697012 | rs571470 | A | G | M_Disease | 0.978751677 | M_Disease |
| 11 | 118740864 | rs4936443 | C | T | M_Disease | 0.977246057 | M_Disease |
| 11 | 118734000 | rs11217074 | C | T | M_Disease | 0.976683468 | M_Disease |
| 11 | 118740931 | rs4938572 | C | T | M_Disease | 0.976107311 | M_Disease |
| 11 | 118750321 | rs11217078 | C | T | M_Disease | 0.971390786 | M_Disease |
| 11 | 118609790 | rs11217006 | A | G | M_Disease | 0.966430777 | M_Disease |
| 11 | 118610957 | rs6589684 | A | G | M_Disease | 0.964939829 | M_Disease |
| 11 | 118612197 | rs6589685 | C | T | M_Disease | 0.963910622 | M_Disease |
| 11 | 118611781 | rs10892279 | A | G | M_Disease | 0.962312109 | M_Disease |
| 11 | 118606722 | rs6589682 | T | C | M_Disease | 0.961764308 | M_Disease |
| 11 | 118610549 | rs11826521 | T | G | M_Disease | 0.961208866 | M_Disease |
| 11 | 118612921 | rs2155433 | A | G | M_Disease | 0.961208866 | M_Disease |
| 11 | 118611817 | rs10892280 | T | C | M_Disease | 0.96045622 | M_Disease |
| 11 | 118606521 | rs6589681 | T | C | M_Disease | 0.95542572 | M_Disease |
| 11 | 118608716 | rs7945144 | G | A | M_Disease | 0.954566171 | M_Disease |
| 11 | 118607794 | rs12364971 | G | T | M_Disease | 0.954566171 | M_Disease |
| 11 | 118615340 | rs56758835 | A | G | M_Disease | 0.954348829 | M_Disease |
| 11 | 118607311 | rs56267528 | A | C | M_Disease | 0.953911173 | M_Disease |
| 11 | 118607150 | rs11217002 | T | C | M_Disease | 0.953469525 | M_Disease |
| 11 | 118607417 | rs55894437 | T | C | M_Disease | 0.9527995 | M_Disease |
| 11 | 118607871 | rs76409436 | A | G | M_Disease | 0.951891858 | M_Disease |
| 11 | 118041740 | rs149794440 | C | T | M_Disease | 0.948826299 | M_Disease&NULL |
| 11 | 118639353 | rs11217020 | A | G | M_Disease | 0.947846437 | M_Disease&NULL |
| 11 | 118615785 | rs11217009 | T | C | M_Disease | 0.945576564 | M_Disease&NULL |
| 11 | 118618776 | rs1048024 | T | A | M_Disease | 0.945576564 | M_Disease&NULL |
| 11 | 118619047 | rs2077579 | G | T | M_Disease | 0.945576564 | M_Disease&NULL |
| 11 | 118619253 | rs2004781 | T | C | M_Disease | 0.945576564 | M_Disease&NULL |
| 11 | 118619960 | rs3889239 | T | C | M_Disease | 0.945576564 | M_Disease&NULL |
| 11 | 118607569 | rs77209083 | G | A | M_Disease | 0.944799462 | M_Disease&NULL |
| 11 | 118661398 | rs57494551 | T | C | M_Disease | 0.944538114 | M_Disease&NULL |
| 11 | 118636980 | rs10892283 | G | A | M_Disease | 0.942945417 | M_Disease&NULL |
| 11 | 118649512 | rs73003215 | A | G | M_Disease | 0.942945417 | M_Disease&NULL |
| 11 | 118653503 | rs10892290 | T | G | M_Disease | 0.942675824 | M_Disease&NULL |
| 11 | 118655340 | rs10892292 | C | G | M_Disease | 0.942405035 | M_Disease&NULL |
| 11 | 118636343 | rs11217019 | A | C | M_Disease | 0.942405035 | M_Disease&NULL |
| 11 | 118628373 | rs74676029 | T | C | M_Disease | 0.942133045 | M_Disease&NULL |
| 11 | 118642999 | rs10892287 | C | T | M_Disease | 0.941859849 | M_Disease&NULL |
| 11 | 118644694 | rs10892288 | G | A | M_Disease | 0.941859849 | M_Disease&NULL |
| 11 | 118655645 | rs874621 | A | G | M_Disease | 0.941859849 | M_Disease&NULL |
| 11 | 118642085 | rs10892286 | C | A | M_Disease | 0.941309825 | M_Disease&NULL |
| 11 | 118653517 | rs10892291 | T | C | M_Disease | 0.941032987 | M_Disease&NULL |
| 11 | 118607046 | rs11217001 | A | G | M_Disease | 0.940754924 | M_Disease&NULL |
| 11 | 118646003 | rs10892289 | T | C | M_Disease | 0.938485818 | M_Disease&NULL |
| 11 | 118648373 | rs56043232 | T | C | M_Disease | 0.938485818 | M_Disease&NULL |
| 11 | 118605628 | rs10892276 | A | G | M_Disease | 0.926898827 | M_Disease&NULL |
| 11 | 118605841 | rs10892277 | A | G | M_Disease | 0.924839891 | M_Disease&NULL |

### FINEMAP

| CHROM | POS | RSID | A1 | A2 | OR_Disease | L+P0 | L+P1 | logBF | P_CAUSAL |
|---|---|---|---|---|---|---|---|---|---|
| 11 | 118694547 | rs7123726 | C | T | 1.19 | -7977.388 | -7967.363 | 10.026 | 0.087445691 |
| 11 | 118745278 | rs7481819 | C | T | 1.19 | -7977.388 | -7967.996 | 9.392 | 0.046386939 |
| 11 | 118680648 | rs11217040 | A | C | 1.183 | -7977.388 | -7968.151 | 9.237 | 0.039726478 |
| 11 | 118682350 | rs11217041 | G | A | 1.18 | -7977.388 | -7968.442 | 8.946 | 0.029696165 |
| 11 | 118682528 | rs11217042 | A | G | 1.18 | -7977.388 | -7968.442 | 8.946 | 0.029696165 |
| 11 | 118681464 | rs7942535 | C | T | 1.18 | -7977.388 | -7968.443 | 8.945 | 0.029666484 |
| 11 | 118745243 | rs7481797 | A | G | 1.185 | -7977.388 | -7968.443 | 8.945 | 0.029666484 |
| 11 | 118683564 | rs17122453 | A | G | 1.179 | -7977.388 | -7968.526 | 8.863 | 0.0273309 |
| 11 | 118681083 | rs73005426 | G | A | 1.179 | -7977.388 | -7968.534 | 8.854 | 0.027086025 |
| 11 | 118701018 | rs79504890 | A | G | 1.181 | -7977.388 | -7968.588 | 8.801 | 0.025687845 |
| 11 | 118743338 | rs4936444 | T | C | 1.185 | -7977.388 | -7968.628 | 8.761 | 0.02468061 |
| 11 | 118698907 | rs11217046 | A | G | 1.18 | -7977.388 | -7968.762 | 8.626 | 0.021563842 |
| 11 | 118684610 | rs715412 | A | G | 1.177 | -7977.388 | -7968.901 | 8.487 | 0.01876546 |
| 11 | 118669605 | rs11217032 | A | T | 1.176 | -7977.388 | -7968.934 | 8.454 | 0.018156306 |
| 11 | 118745884 | rs10790275 | G | C | 1.181 | -7977.388 | -7968.938 | 8.451 | 0.018101918 |
| 11 | 118697865 | rs11217045 | A | C | 1.178 | -7977.388 | -7968.94 | 8.448 | 0.018047694 |
| 11 | 118697906 | rs73005502 | G | A | 1.178 | -7977.388 | -7968.94 | 8.448 | 0.018047694 |
| 11 | 118679304 | rs28409215 | A | G | 1.175 | -7977.388 | -7969.002 | 8.386 | 0.016962719 |
| 11 | 118677355 | rs11217038 | A | G | 1.175 | -7977.388 | -7969.033 | 8.355 | 0.016444942 |
| 11 | 118665575 | rs10892293 | T | A | 1.175 | -7977.388 | -7969.035 | 8.354 | 0.016428505 |
| 11 | 118743286 | rs12365699 | A | G | 1.191 | -7977.388 | -7969.045 | 8.343 | 0.016248782 |
| 11 | 118667357 | rs10892294 | C | G | 1.175 | -7977.388 | -7969.064 | 8.324 | 0.015942969 |
| 11 | 118681079 | rs73005423 | G | A | 1.174 | -7977.388 | -7969.146 | 8.242 | 0.01468781 |
| 11 | 118675772 | rs11217036 | T | C | 1.174 | -7977.388 | -7969.153 | 8.235 | 0.014585355 |
| 11 | 118677086 | rs11217037 | A | G | 1.174 | -7977.388 | -7969.17 | 8.218 | 0.014339499 |
| 11 | 118673706 | rs11217033 | C | T | 1.174 | -7977.388 | -7969.178 | 8.21 | 0.014225241 |
| 11 | 118726843 | rs1037679725 | T | C | 1.177 | -7977.388 | -7969.193 | 8.196 | 0.014027475 |
| 11 | 118721699 | rs7125066 | G | A | 1.143 | -7977.388 | -7969.215 | 8.174 | 0.013722241 |
| 11 | 118724894 | rs9736016 | A | T | 1.143 | -7977.388 | -7969.216 | 8.172 | 0.013694824 |
| 11 | 118743772 | rs6421571 | T | C | 1.178 | -7977.388 | -7969.22 | 8.168 | 0.013640154 |
| 11 | 118725660 | rs4936441 | C | G | 1.177 | -7977.388 | -7969.229 | 8.159 | 0.013517943 |
| 11 | 118722149 | rs11217058 | A | G | 1.177 | -7977.388 | -7969.263 | 8.126 | 0.013079131 |
| 11 | 118696022 | rs11217044 | C | T | 1.172 | -7977.388 | -7969.323 | 8.065 | 0.012305151 |
| 11 | 118726753 | rs11217066 | G | T | 1.176 | -7977.388 | -7969.334 | 8.054 | 0.012170536 |
| 11 | 118741842 | rs4938573 | C | T | 1.176 | -7977.388 | -7969.39 | 7.998 | 0.011507718 |
| 11 | 118741157 | rs7117261 | T | C | 1.176 | -7977.388 | -7969.41 | 7.978 | 0.01127985 |
| 11 | 118740418 | rs7951740 | G | C | 1.176 | -7977.388 | -7969.457 | 7.931 | 0.010761962 |
| 11 | 118729391 | rs10790268 | A | G | 1.175 | -7977.388 | -7969.486 | 7.902 | 0.010454348 |
| 11 | 118744396 | rs7117313 | C | G | 1.175 | -7977.388 | -7969.503 | 7.885 | 0.010278126 |
| 11 | 118739056 | rs7122669 | A | G | 1.175 | -7977.388 | -7969.513 | 7.875 | 0.010175857 |
| 11 | 118742800 | rs74541740 | A | G | 1.174 | -7977.388 | -7969.569 | 7.819 | 0.009621671 |
| 11 | 118721565 | rs75249660 | A | G | 1.174 | -7977.388 | -7969.587 | 7.802 | 0.009459485 |
| 11 | 118738298 | rs7119044 | G | C | 1.174 | -7977.388 | -7969.595 | 7.793 | 0.009374731 |
| 11 | 118729456 | rs10790269 | C | T | 1.174 | -7977.388 | -7969.629 | 7.76 | 0.009070414 |
| 11 | 118738281 | rs7119038 | G | A | 1.163 | -7977.388 | -7969.715 | 7.673 | 0.008314641 |
| 11 | 118697012 | rs571470 | A | G | 1.138 | -7977.388 | -7969.728 | 7.66 | 0.00820725 |
| 11 | 118740864 | rs4936443 | C | T | 1.171 | -7977.388 | -7969.87 | 7.518 | 0.007120785 |
| 11 | 118734000 | rs11217074 | C | T | 1.171 | -7977.388 | -7969.918 | 7.47 | 0.00678706 |
| 11 | 118740931 | rs4938572 | C | T | 1.17 | -7977.388 | -7969.971 | 7.417 | 0.006436712 |
| 11 | 118750321 | rs11217078 | C | T | 1.139 | -7977.388 | -7970.342 | 7.046 | 0.004441614 |
| 11 | 118609790 | rs11217006 | A | G | 1.158 | -7977.388 | -7970.672 | 6.717 | 0.003196377 |
| 11 | 118610957 | rs6589684 | A | G | 1.157 | -7977.388 | -7970.766 | 6.622 | 0.002906699 |
| 11 | 118612197 | rs6589685 | C | T | 1.157 | -7977.388 | -7970.821 | 6.567 | 0.002751147 |
| 11 | 118611781 | rs10892279 | A | G | 1.156 | -7977.388 | -7970.912 | 6.476 | 0.002511846 |
| 11 | 118606722 | rs6589682 | T | C | 1.155 | -7977.388 | -7970.945 | 6.444 | 0.002432739 |
| 11 | 118610549 | rs11826521 | T | G | 1.156 | -7977.388 | -7970.967 | 6.421 | 0.002377425 |
| 11 | 118612921 | rs2155433 | A | G | 1.155 | -7977.388 | -7970.97 | 6.418 | 0.002370303 |
| 11 | 118611817 | rs10892280 | T | C | 1.156 | -7977.388 | -7971.007 | 6.381 | 0.002284205 |
| 11 | 118606521 | rs6589681 | T | C | 1.152 | -7977.388 | -7971.263 | 6.125 | 0.001768299 |
| 11 | 118608716 | rs7945144 | G | A | 1.151 | -7977.388 | -7971.297 | 6.091 | 0.001709187 |
| 11 | 118607794 | rs12364971 | G | T | 1.151 | -7977.388 | -7971.3 | 6.088 | 0.001704067 |
| 11 | 118615340 | rs56758835 | A | G | 1.153 | -7977.388 | -7971.313 | 6.076 | 0.001683741 |
| 11 | 118607311 | rs56267528 | A | C | 1.151 | -7977.388 | -7971.327 | 6.061 | 0.001658673 |
| 11 | 118607150 | rs11217002 | T | C | 1.151 | -7977.388 | -7971.354 | 6.034 | 0.001614488 |
| 11 | 118607417 | rs55894437 | T | C | 1.15 | -7977.388 | -7971.379 | 6.009 | 0.001574626 |
| 11 | 118607871 | rs76409436 | A | G | 1.15 | -7977.388 | -7971.423 | 5.966 | 0.001508352 |
| 11 | 118041740 | rs149794440 | C | T | 1.309 | -7977.388 | -7971.556 | 5.833 | 0.00132051 |
| 11 | 118639353 | rs11217020 | A | G | 1.15 | -7977.388 | -7971.588 | 5.8 | 0.001277644 |
| 11 | 118615785 | rs11217009 | T | C | 1.149 | -7977.388 | -7971.682 | 5.707 | 0.001164181 |
| 11 | 118618776 | rs1048024 | T | A | 1.149 | -7977.388 | -7971.682 | 5.707 | 0.001164181 |
| 11 | 118619047 | rs2077579 | G | T | 1.149 | -7977.388 | -7971.682 | 5.707 | 0.001164181 |
| 11 | 118619253 | rs2004781 | T | C | 1.149 | -7977.388 | -7971.682 | 5.707 | 0.001164181 |
| 11 | 118619960 | rs3889239 | T | C | 1.149 | -7977.388 | -7971.682 | 5.707 | 0.001164181 |
| 11 | 118607569 | rs77209083 | G | A | 1.147 | -7977.388 | -7971.709 | 5.679 | 0.001132036 |
| 11 | 118661398 | rs57494551 | T | C | 1.149 | -7977.388 | -7971.722 | 5.667 | 0.001118533 |
| 11 | 118636980 | rs10892283 | G | A | 1.148 | -7977.388 | -7971.784 | 5.604 | 0.001050239 |
| 11 | 118649512 | rs73003215 | A | G | 1.147 | -7977.388 | -7971.785 | 5.603 | 0.001049189 |
| 11 | 118653503 | rs10892290 | T | G | 1.148 | -7977.388 | -7971.787 | 5.601 | 0.001047093 |
| 11 | 118655340 | rs10892292 | C | G | 1.148 | -7977.388 | -7971.801 | 5.588 | 0.001033569 |
| 11 | 118636343 | rs11217019 | A | C | 1.148 | -7977.388 | -7971.801 | 5.587 | 0.001032536 |
| 11 | 118628373 | rs74676029 | T | C | 1.147 | -7977.388 | -7971.816 | 5.573 | 0.001018181 |
| 11 | 118642999 | rs10892287 | C | T | 1.147 | -7977.388 | -7971.818 | 5.57 | 0.001015131 |
| 11 | 118644694 | rs10892288 | G | A | 1.147 | -7977.388 | -7971.818 | 5.57 | 0.001015131 |
| 11 | 118655645 | rs874621 | A | G | 1.147 | -7977.388 | -7971.821 | 5.567 | 0.00101209 |
| 11 | 118642085 | rs10892286 | C | A | 1.147 | -7977.388 | -7971.838 | 5.55 | 0.00099503 |
| 11 | 118653517 | rs10892291 | T | C | 1.147 | -7977.388 | -7971.854 | 5.534 | 0.000979237 |
| 11 | 118607046 | rs11217001 | A | G | 1.146 | -7977.388 | -7971.866 | 5.523 | 0.000968524 |
| 11 | 118646003 | rs10892289 | T | C | 1.146 | -7977.388 | -7971.939 | 5.449 | 0.000899441 |
| 11 | 118648373 | rs56043232 | T | C | 1.146 | -7977.388 | -7971.939 | 5.449 | 0.000899441 |
| 11 | 118605628 | rs10892276 | A | G | 1.129 | -7977.388 | -7972.315 | 5.073 | 0.000617558 |
| 11 | 118605841 | rs10892277 | A | G | 1.128 | -7977.388 | -7972.373 | 5.016 | 0.000583342 |

| | | | | | | | |
|---|---|---|---|---|---|---|---|
| 11 | 118605617 | rs10892275 | A | T | M_Disease | 0.920926323 | M_Disease&NULL |
| 11 | 118605463 | rs10892274 | T | C | M_Disease | 0.920561451 | M_Disease&NULL |
| 11 | 116287909 | rs10891973 | A | G | M_Disease | 0.919827088 | M_Disease&NULL |
| 11 | 116287783 | rs10891972 | T | C | M_Disease | 0.919086533 | M_Disease&NULL |
| 11 | 118604941 | rs11216999 | C | T | M_Disease | 0.913725806 | M_Disease&NULL |
| 11 | 118604837 | rs11216998 | T | C | M_Disease | 0.910926472 | M_Disease&NULL |
| 11 | 118725942 | rs11217063 | C | G | M_Disease | 0.902031196 | M_Disease&NULL |
| 11 | 118726647 | rs11217065 | C | T | M_Disease | 0.900249511 | M_Disease&NULL |
| 11 | 118724723 | rs57719838 | C | T | M_Disease | 0.898439072 | M_Disease&NULL |
| 11 | 118723379 | rs7103067 | A | G | M_Disease | 0.898439072 | M_Disease&NULL |
| 11 | 118612565 | rs6589686 | A | G | M_Disease | 0.896135084 | M_Disease&NULL |
| 11 | 118731599 | rs34561290 | T | C | M_Disease | 0.896135084 | M_Disease&NULL |
| 11 | 118731762 | rs11217072 | G | A | M_Disease | 0.896135084 | M_Disease&NULL |
| 11 | 118729583 | rs12790489 | C | T | M_Disease | 0.892352574 | M_Disease&NULL |
| 11 | 118727169 | rs10790267 | T | C | M_Disease | 0.891871333 | M_Disease&NULL |
| 11 | 118729860 | rs11217070 | G | T | M_Disease | 0.890903179 | M_Disease&NULL |
| 11 | 118730141 | rs7929520 | G | A | M_Disease | 0.890903179 | M_Disease&NULL |
| 11 | 118729210 | rs10892300 | T | C | M_Disease | 0.890416255 | M_Disease&NULL |
| 11 | 118727332 | rs11217067 | T | G | M_Disease | 0.888944033 | M_Disease&NULL |
| 11 | 118740180 | rs7483509 | A | G | M_Disease | 0.883397031 | M_Disease&NULL |
| 11 | 118728320 | rs4936442 | T | C | M_Disease | 0.88027111 | M_Disease&NULL |
| 11 | 118734521 | rs1174717189 | T | A | M_Disease | 0.878681162 | M_Disease&NULL |
| 11 | 118762701 | rs76457018 | A | G | M_Disease | 0.875446642 | M_Disease&NULL |
| 11 | 117489521 | rs666094 | C | T | M_Disease | 0.868184382 | M_Disease&NULL |
| 11 | 118711790 | rs622722 | A | G | M_Disease | 0.861165006 | M_Disease&NULL |
| 11 | 118758089 | rs497916 | T | C | M_Disease | 0.861165006 | M_Disease&NULL |
| 11 | 119537683 | rs10790329 | C | T | M_Disease | 0.858148935 | M_Disease&NULL |
| 11 | 118603165 | rs73001429 | G | T | M_Disease | 0.855696866 | M_Disease&NULL |
| 11 | 118603520 | rs11216994 | T | C | M_Disease | 0.85383477 | M_Disease&NULL |
| 11 | 118603526 | rs11216995 | A | G | M_Disease | 0.84941242 | M_Disease&NULL |
| 11 | 118603495 | rs11216993 | A | T | M_Disease | 0.845534735 | M_Disease&NULL |
| 11 | 117288387 | rs579411 | A | G | M_Disease | 0.843565479 | M_Disease&NULL |
| 11 | 117288898 | rs553906 | G | T | M_Disease | 0.843565479 | M_Disease&NULL |
| 11 | 117289347 | rs10790181 | G | A | M_Disease | 0.843565479 | M_Disease&NULL |
| 11 | 117286824 | rs2124975 | G | A | M_Disease | 0.842904531 | M_Disease&NULL |
| 11 | 117287088 | rs2368164 | C | T | M_Disease | 0.842904531 | M_Disease&NULL |
| 11 | 117287230 | rs2925767 | A | G | M_Disease | 0.842904531 | M_Disease&NULL |
| 11 | 117287272 | rs2510451 | G | C | M_Disease | 0.842904531 | M_Disease&NULL |
| 11 | 117260504 | rs523159 | A | G | M_Disease | 0.842241313 | M_Disease&NULL |
| 11 | 117285907 | rs524069 | G | A | M_Disease | 0.841575821 | M_Disease&NULL |
| 11 | 117286064 | rs522383 | T | C | M_Disease | 0.841575821 | M_Disease&NULL |
| 11 | 117287942 | rs473446 | A | G | M_Disease | 0.841575821 | M_Disease&NULL |
| 11 | 117285619 | rs512356 | A | G | M_Disease | 0.840908052 | M_Disease&NULL |
| 11 | 118602707 | rs10892272 | G | C | M_Disease | 0.840238003 | M_Disease&NULL |
| 11 | 118602809 | rs10892273 | A | G | M_Disease | 0.83956567 | M_Disease&NULL |
| 11 | 117272939 | rs1270801 | T | G | M_Disease | 0.83889105 | M_Disease&NULL |
| 11 | 118604638 | rs11216997 | T | C | M_Disease | 0.837534937 | M_Disease&NULL |
| 11 | 118603736 | rs112561833 | C | A | M_Disease | 0.836853438 | M_Disease&NULL |
| 11 | 117290313 | rs493647 | G | A | M_Disease | 0.833411386 | M_Disease&NULL |
| 11 | 117475052 | rs490914 | T | A | M_Disease | 0.829911475 | M_Disease&NULL |
| 11 | 117285549 | rs527056 | G | A | M_Disease | 0.829911475 | M_Disease&NULL |
| 11 | 117285684 | rs511549 | C | A | M_Disease | 0.829911475 | M_Disease&NULL |
| 11 | 117285738 | rs11823544 | A | G | M_Disease | 0.829911475 | M_Disease&NULL |
| 11 | 118558425 | rs149509653 | A | G | M_Disease | 0.827783608 | M_Disease&NULL |
| 11 | 117285497 | rs513327 | C | T | M_Disease | 0.82706965 | M_Disease&NULL |
| 11 | 117375461 | rs2276341 | A | G | M_Disease | 0.826353353 | M_Disease&NULL |
| 11 | 118685481 | rs515707 | T | C | M_Disease | 0.826353353 | M_Disease&NULL |
| 11 | 117373947 | rs4938394 | C | T | M_Disease | 0.825634714 | M_Disease&NULL |
| 11 | 117290898 | rs4936374 | A | G | M_Disease | 0.824913732 | M_Disease&NULL |
| 11 | 118689871 | rs587816 | A | C | M_Disease | 0.824190403 | M_Disease&NULL |
| 11 | 117475341 | rs680292 | T | G | M_Disease | 0.823464725 | M_Disease&NULL |
| 11 | 118690550 | rs577121 | C | T | M_Disease | 0.819061207 | M_Disease&NULL |
| 11 | 118598544 | rs10892271 | A | G | M_Disease | 0.816827559 | M_Disease&NULL |
| 11 | 118608532 | rs567680 | G | C | M_Disease | 0.816827559 | M_Disease&NULL |
| 11 | 118607622 | rs11217004 | A | G | M_Disease | 0.816078273 | M_Disease&NULL |
| 11 | 117289951 | rs490730 | T | C | M_Disease | 0.816078273 | M_Disease&NULL |
| 11 | 118613235 | rs632124 | A | T | M_Disease | 0.815326614 | M_Disease&NULL |
| 11 | 117375951 | rs10892132 | A | G | M_Disease | 0.815326614 | M_Disease&NULL |
| 11 | 117378116 | rs2155394 | A | G | M_Disease | 0.814572581 | M_Disease&NULL |
| 11 | 120492260 | rs77694974 | T | G | M_Disease | 0.814572581 | M_Disease&NULL |
| 11 | 118608625 | rs566733 | G | C | M_Disease | 0.813816172 | M_Disease&NULL |
| 11 | 118682462 | rs523148 | T | C | M_Disease | 0.813057386 | M_Disease&NULL |
| 11 | 117373125 | rs7932097 | T | C | M_Disease | 0.805338416 | M_Disease&NULL |
| 11 | 117373432 | rs7112940 | G | A | M_Disease | 0.802976114 | M_Disease&NULL |
| 11 | 119210583 | rs1135258 | T | G | M_Disease | 0.798186778 | M_Disease&NULL |
| 11 | 116523821 | rs528647 | T | C | M_Disease | 0.797380154 | M_Disease&NULL |
| 11 | 118696175 | rs584935 | A | G | M_Disease | 0.797380154 | M_Disease&NULL |
| 11 | 118759610 | rs566416 | G | T | M_Disease | 0.796571127 | M_Disease&NULL |
| 11 | 116523167 | rs483747 | C | T | M_Disease | 0.796571127 | M_Disease&NULL |
| 11 | 116522559 | rs567576 | G | A | M_Disease | 0.796571127 | M_Disease&NULL |
| 11 | 116522565 | rs567599 | T | G | M_Disease | 0.796571127 | M_Disease&NULL |
| 11 | 118695908 | rs583905 | G | A | M_Disease | 0.795759698 | M_Disease&NULL |
| 11 | 118151329 | rs11216843 | T | A | M_Disease | 0.795759698 | M_Disease&NULL |
| 11 | 116522625 | rs3017237 | C | G | M_Disease | 0.794945865 | M_Disease&NULL |
| 11 | 116522770 | rs480028 | G | C | M_Disease | 0.794945865 | M_Disease&NULL |
| 11 | 116522854 | rs480878 | A | G | M_Disease | 0.794945865 | M_Disease&NULL |
| 11 | 116523844 | rs528732 | A | G | M_Disease | 0.794945865 | M_Disease&NULL |
| 11 | 116523931 | rs3017778 | T | C | M_Disease | 0.793310987 | M_Disease&NULL |
| 11 | 118617299 | rs607125 | C | T | M_Disease | 0.793310987 | M_Disease&NULL |
| 11 | 118620697 | rs488219 | C | T | M_Disease | 0.790012373 | M_Disease&NULL |
| 11 | 118700264 | rs113641235 | T | C | M_Disease | 0.790012373 | M_Disease&NULL |
| 11 | 116523988 | rs3017779 | T | C | M_Disease | 0.790012373 | M_Disease&NULL |
| 11 | 116523373 | rs485638 | C | G | M_Disease | 0.790012373 | M_Disease&NULL |
| 11 | 117201458 | rs76858861 | G | A | M_Disease | 0.790012373 | M_Disease&NULL |
| 11 | 118624250 | rs546393 | C | T | M_Disease | 0.789181707 | M_Disease&NULL |
| 11 | 116636305 | rs79874233 | C | G | M_Disease | 0.789181707 | M_Disease&NULL |

| | | | | | | | | | |
|---|---|---|---|---|---|---|---|---|---|
| 11 | 118605617 | rs10892275 | A | T | 1.127 | -7977.388 | -7972.481 | 4.908 | 0.000523624 |
| 11 | 118605463 | rs10892274 | T | C | 1.127 | -7977.388 | -7972.492 | 4.896 | 0.000517378 |
| 11 | 116287909 | rs10891973 | A | G | 1.153 | -7977.388 | -7972.507 | 4.882 | 0.000510185 |
| 11 | 116287783 | rs10891972 | T | C | 1.152 | -7977.388 | -7972.531 | 4.858 | 0.000498086 |
| 11 | 118604941 | rs11216999 | C | T | 1.125 | -7977.388 | -7972.667 | 4.722 | 0.000434751 |
| 11 | 118604837 | rs11216998 | T | C | 1.124 | -7977.388 | -7972.738 | 4.65 | 0.000404549 |
| 11 | 118725942 | rs11217063 | C | G | 0.9 | -7977.388 | -7972.952 | 4.436 | 0.000326612 |
| 11 | 118726647 | rs11217065 | C | T | 0.901 | -7977.388 | -7972.987 | 4.401 | 0.000315378 |
| 11 | 118724723 | rs57719838 | C | T | 0.901 | -7977.388 | -7973.026 | 4.362 | 0.000303315 |
| 11 | 118723379 | rs7103067 | A | G | 0.901 | -7977.388 | -7973.028 | 4.36 | 0.000302709 |
| 11 | 118612565 | rs6589686 | A | G | 1.136 | -7977.388 | -7973.077 | 4.312 | 0.000288523 |
| 11 | 118731599 | rs34561290 | T | C | 0.901 | -7977.388 | -7973.082 | 4.306 | 0.000286797 |
| 11 | 118731762 | rs11217072 | G | A | 0.901 | -7977.388 | -7973.082 | 4.306 | 0.000286797 |
| 11 | 118729583 | rs12790489 | C | T | 0.902 | -7977.388 | -7973.162 | 4.226 | 0.000264747 |
| 11 | 118727169 | rs10790267 | T | C | 0.902 | -7977.388 | -7973.169 | 4.219 | 0.0002629 |
| 11 | 118729860 | rs11217070 | G | T | 0.902 | -7977.388 | -7973.187 | 4.201 | 0.00025821 |
| 11 | 118730141 | rs7929520 | G | A | 0.902 | -7977.388 | -7973.187 | 4.201 | 0.00025821 |
| 11 | 118729210 | rs10892300 | T | C | 0.902 | -7977.388 | -7973.204 | 4.184 | 0.000253858 |
| 11 | 118727332 | rs11217067 | T | G | 0.902 | -7977.388 | -7973.234 | 4.154 | 0.000246355 |
| 11 | 118740180 | rs7483509 | A | G | 0.903 | -7977.388 | -7973.343 | 4.046 | 0.000221135 |
| 11 | 118728320 | rs4936442 | T | C | 0.904 | -7977.388 | -7973.396 | 3.992 | 0.00020951 |
| 11 | 118734521 | rs1174717189 | T | A | 0.904 | -7977.388 | -7973.434 | 3.954 | 0.000201698 |
| 11 | 118762701 | rs76457018 | A | G | 0.872 | -7977.388 | -7973.493 | 3.895 | 0.000190142 |
| 11 | 117489521 | rs666094 | C | T | 0.907 | -7977.388 | -7973.624 | 3.764 | 0.000166796 |
| 11 | 118711790 | rs622722 | A | G | 0.906 | -7977.388 | -7973.741 | 3.648 | 0.000148528 |
| 11 | 118758089 | rs497916 | T | C | 1.113 | -7977.388 | -7973.743 | 3.645 | 0.000148083 |
| 11 | 119537683 | rs10790329 | C | T | 1.114 | -7977.388 | -7973.794 | 3.594 | 0.00014072 |
| 11 | 118603165 | rs73001429 | G | T | 1.12 | -7977.388 | -7973.835 | 3.553 | 0.000135067 |
| 11 | 118603520 | rs11216994 | T | C | 1.12 | -7977.388 | -7973.857 | 3.532 | 0.00013226 |
| 11 | 118603526 | rs11216995 | A | G | 1.119 | -7977.388 | -7973.93 | 3.459 | 0.000122949 |
| 11 | 118603495 | rs11216993 | A | T | 1.119 | -7977.388 | -7973.99 | 3.398 | 0.000115674 |
| 11 | 117288387 | rs579411 | A | G | 0.851 | -7977.388 | -7974.024 | 3.364 | 0.000111807 |
| 11 | 117288898 | rs553906 | G | T | 0.851 | -7977.388 | -7974.024 | 3.364 | 0.000111807 |
| 11 | 117289347 | rs10790181 | G | A | 0.851 | -7977.388 | -7974.024 | 3.364 | 0.000111807 |
| 11 | 117286824 | rs2124975 | G | A | 0.851 | -7977.388 | -7974.028 | 3.361 | 0.000111472 |
| 11 | 117287088 | rs2368164 | C | T | 0.851 | -7977.388 | -7974.028 | 3.361 | 0.000111472 |
| 11 | 117287230 | rs2925767 | A | G | 0.851 | -7977.388 | -7974.028 | 3.361 | 0.000111472 |
| 11 | 117287272 | rs2510451 | G | C | 0.851 | -7977.388 | -7974.028 | 3.361 | 0.000111472 |
| 11 | 117260504 | rs523159 | A | G | 0.852 | -7977.388 | -7974.041 | 3.347 | 0.000109922 |
| 11 | 117285907 | rs524069 | G | A | 0.852 | -7977.388 | -7974.049 | 3.339 | 0.000109046 |
| 11 | 117286064 | rs522383 | T | C | 0.852 | -7977.388 | -7974.049 | 3.339 | 0.000109046 |
| 11 | 117287942 | rs473446 | A | G | 0.852 | -7977.388 | -7974.049 | 3.339 | 0.000109046 |
| 11 | 117285619 | rs512356 | A | G | 0.852 | -7977.388 | -7974.059 | 3.329 | 0.000107961 |
| 11 | 118602707 | rs10892272 | G | C | 1.117 | -7977.388 | -7974.067 | 3.321 | 0.000107101 |
| 11 | 118602809 | rs10892273 | A | G | 1.117 | -7977.388 | -7974.084 | 3.305 | 0.000105401 |
| 11 | 117272939 | rs1270801 | T | G | 0.853 | -7977.388 | -7974.09 | 3.298 | 0.000104666 |
| 11 | 118604638 | rs11216997 | T | C | 1.117 | -7977.388 | -7974.109 | 3.279 | 0.000102696 |
| 11 | 118603736 | rs112561833 | C | A | 1.117 | -7977.388 | -7974.118 | 3.271 | 0.000101878 |
| 11 | 117290313 | rs493647 | G | A | 0.854 | -7977.388 | -7974.176 | 3.212 | 9.60E-05 |
| 11 | 117475052 | rs490914 | T | A | 0.843 | -7977.388 | -7974.22 | 3.168 | 9.19E-05 |
| 11 | 117285549 | rs527056 | G | A | 0.854 | -7977.388 | -7974.223 | 3.165 | 9.16E-05 |
| 11 | 117285684 | rs511549 | C | A | 0.854 | -7977.388 | -7974.223 | 3.165 | 9.16E-05 |
| 11 | 117285738 | rs11823544 | A | G | 0.854 | -7977.388 | -7974.223 | 3.165 | 9.16E-05 |
| 11 | 118558425 | rs149509653 | A | G | 0.891 | -7977.388 | -7974.251 | 3.137 | 8.91E-05 |
| 11 | 117285497 | rs513327 | C | T | 0.855 | -7977.388 | -7974.264 | 3.124 | 8.80E-05 |
| 11 | 117375461 | rs2276341 | A | G | 1.097 | -7977.388 | -7974.272 | 3.116 | 8.72E-05 |
| 11 | 118685481 | rs515707 | T | C | 0.891 | -7977.388 | -7974.273 | 3.115 | 8.72E-05 |
| 11 | 117373947 | rs4938394 | C | T | 1.097 | -7977.388 | -7974.276 | 3.113 | 8.70E-05 |
| 11 | 117290898 | rs4936374 | A | G | 0.854 | -7977.388 | -7974.296 | 3.092 | 8.52E-05 |
| 11 | 118689871 | rs587816 | A | C | 0.891 | -7977.388 | -7974.304 | 3.085 | 8.46E-05 |
| 11 | 117475341 | rs680292 | T | G | 0.844 | -7977.388 | -7974.312 | 3.076 | 8.38E-05 |
| 11 | 118690550 | rs577121 | C | T | 0.892 | -7977.388 | -7974.368 | 3.02 | 7.93E-05 |
| 11 | 118598544 | rs10892271 | A | G | 1.113 | -7977.388 | -7974.402 | 2.986 | 7.66E-05 |
| 11 | 118608532 | rs567680 | G | C | 0.913 | -7977.388 | -7974.405 | 2.983 | 7.64E-05 |
| 11 | 118607622 | rs11217004 | A | G | 0.913 | -7977.388 | -7974.407 | 2.981 | 7.62E-05 |
| 11 | 117289951 | rs490730 | T | C | 0.857 | -7977.388 | -7974.408 | 2.98 | 7.62E-05 |
| 11 | 118613235 | rs632124 | A | T | 0.913 | -7977.388 | -7974.416 | 2.972 | 7.55E-05 |
| 11 | 117375951 | rs10892132 | A | G | 1.095 | -7977.388 | -7974.422 | 2.966 | 7.51E-05 |
| 11 | 117378116 | rs2155394 | A | G | 1.095 | -7977.388 | -7974.426 | 2.962 | 7.48E-05 |
| 11 | 120492260 | rs77694974 | T | G | 0.833 | -7977.388 | -7974.433 | 2.955 | 7.43E-05 |
| 11 | 118608625 | rs566733 | G | C | 0.913 | -7977.388 | -7974.444 | 2.945 | 7.35E-05 |
| 11 | 118682462 | rs523148 | T | C | 0.892 | -7977.388 | -7974.447 | 2.942 | 7.33E-05 |
| 11 | 117373125 | rs7932097 | T | C | 1.094 | -7977.388 | -7974.549 | 2.839 | 6.61E-05 |
| 11 | 117373432 | rs7112940 | G | A | 1.093 | -7977.388 | -7974.584 | 2.804 | 6.39E-05 |
| 11 | 119210583 | rs1135258 | T | G | 0.915 | -7977.388 | -7974.638 | 2.75 | 6.05E-05 |
| 11 | 116523821 | rs528647 | T | C | 0.876 | -7977.388 | -7974.648 | 2.74 | 5.99E-05 |
| 11 | 118696175 | rs584935 | A | G | 0.895 | -7977.388 | -7974.655 | 2.734 | 5.95E-05 |
| 11 | 118759610 | rs566416 | G | T | 1.108 | -7977.388 | -7974.657 | 2.731 | 5.94E-05 |
| 11 | 116523167 | rs483747 | C | T | 0.876 | -7977.388 | -7974.662 | 2.727 | 5.91E-05 |
| 11 | 116522559 | rs567576 | G | A | 0.876 | -7977.388 | -7974.662 | 2.726 | 5.91E-05 |
| 11 | 116522565 | rs567599 | T | G | 0.876 | -7977.388 | -7974.662 | 2.726 | 5.91E-05 |
| 11 | 118695908 | rs583905 | G | A | 0.895 | -7977.388 | -7974.673 | 2.715 | 5.84E-05 |
| 11 | 118151329 | rs11216843 | T | A | 1.248 | -7977.388 | -7974.678 | 2.711 | 5.82E-05 |
| 11 | 116522625 | rs3017237 | C | G | 0.877 | -7977.388 | -7974.682 | 2.706 | 5.79E-05 |
| 11 | 116522770 | rs480028 | G | C | 0.877 | -7977.388 | -7974.682 | 2.706 | 5.79E-05 |
| 11 | 116522854 | rs480878 | A | G | 0.877 | -7977.388 | -7974.682 | 2.706 | 5.79E-05 |
| 11 | 116523844 | rs528732 | A | G | 0.877 | -7977.388 | -7974.684 | 2.704 | 5.78E-05 |
| 11 | 116523931 | rs3017778 | T | C | 0.877 | -7977.388 | -7974.705 | 2.683 | 5.66E-05 |
| 11 | 118617299 | rs607125 | C | T | 0.916 | -7977.388 | -7974.705 | 2.683 | 5.66E-05 |
| 11 | 118620697 | rs488219 | C | T | 0.916 | -7977.388 | -7974.738 | 2.651 | 5.48E-05 |
| 11 | 118700264 | rs113641235 | T | C | 0.895 | -7977.388 | -7974.74 | 2.648 | 5.46E-05 |
| 11 | 116523988 | rs3017779 | T | C | 0.878 | -7977.388 | -7974.741 | 2.647 | 5.46E-05 |
| 11 | 116523373 | rs485638 | C | G | 0.878 | -7977.388 | -7974.744 | 2.644 | 5.44E-05 |
| 11 | 117201458 | rs76858861 | G | A | 0.779 | -7977.388 | -7974.747 | 2.641 | 5.43E-05 |
| 11 | 118624250 | rs546393 | C | T | 0.916 | -7977.388 | -7974.747 | 2.641 | 5.43E-05 |
| 11 | 116636305 | rs79874233 | C | G | 0.782 | -7977.388 | -7974.756 | 2.633 | 5.38E-05 |

| | | | | | | | |
|---|---|---|---|---|---|---|---|
| 11 | 117373055 | rs6589612 | G | C | M_Disease | 0.788348634 | M_Disease&NULL |
| 11 | 116521975 | rs562756 | C | G | M_Disease | 0.783299685 | M_Disease&NULL |
| 11 | 118186305 | rs2231449 | A | C | M_Disease | 0.782449776 | M_Disease&NULL |
| 11 | 118700425 | rs10128600 | C | T | M_Disease | 0.781597464 | M_Disease&NULL |
| 11 | 116522156 | rs563917 | T | C | M_Disease | 0.781597464 | M_Disease&NULL |
| 11 | 116522164 | rs492182 | A | G | M_Disease | 0.781597464 | M_Disease&NULL |
| 11 | 116588904 | rs1262892 | A | G | M_Disease | 0.781597464 | M_Disease&NULL |
| 11 | 116522111 | rs563796 | G | A | M_Disease | 0.778164185 | M_Disease&NULL |
| 11 | 118699863 | rs2510901 | A | G | M_Disease | 0.777299861 | M_Disease&NULL |
| 11 | 118762218 | rs543824 | G | A | M_Disease | 0.775564014 | M_Disease&NULL |
| 11 | 118944909 | rs563162 | T | C | M_Disease | 0.775564014 | M_Disease&NULL |
| 11 | 118163916 | rs11216846 | A | G | M_Disease | 0.774692493 | M_Disease&NULL |
| 11 | 116521988 | rs491022 | C | T | M_Disease | 0.773818574 | M_Disease&NULL |
| 11 | 118707171 | rs4938550 | G | A | M_Disease | 0.773818574 | M_Disease&NULL |
| 11 | 116522330 | rs565607 | T | C | M_Disease | 0.772942259 | M_Disease&NULL |
| 11 | 116521558 | rs575906 | G | A | M_Disease | 0.772942259 | M_Disease&NULL |
| 11 | 117379086 | rs10750107 | C | T | M_Disease | 0.772942259 | M_Disease&NULL |
| 11 | 116520434 | rs506222 | C | A | M_Disease | 0.772942259 | M_Disease&NULL |
| 11 | 118698160 | rs640373 | G | A | M_Disease | 0.772063549 | M_Disease&NULL |
| 11 | 118698274 | rs639915 | A | G | M_Disease | 0.772063549 | M_Disease&NULL |
| 11 | 118698294 | rs639885 | A | G | M_Disease | 0.772063549 | M_Disease&NULL |
| 11 | 118698537 | rs638893 | G | A | M_Disease | 0.772063549 | M_Disease&NULL |
| 11 | 118697375 | rs73005500 | C | T | M_Disease | 0.772063549 | M_Disease&NULL |
| 11 | 116291220 | rs57150747 | A | G | M_Disease | 0.771182446 | M_Disease&NULL |
| 11 | 118698777 | rs516332 | G | A | M_Disease | 0.770298949 | M_Disease&NULL |
| 11 | 118629031 | rs555639 | A | G | M_Disease | 0.770298949 | M_Disease&NULL |
| 11 | 117513801 | rs10892147 | C | T | M_Disease | 0.769413061 | M_Disease&NULL |
| 11 | 118699744 | rs2508564 | C | T | M_Disease | 0.769413061 | M_Disease&NULL |
| 11 | 118181481 | rs11216848 | A | C | M_Disease | 0.769413061 | M_Disease&NULL |
| 11 | 118566069 | rs506255 | A | C | M_Disease | 0.765845626 | M_Disease&NULL |
| 11 | 116520398 | rs544479 | C | T | M_Disease | 0.764947804 | M_Disease&NULL |
| 11 | 116520926 | rs549125 | G | C | M_Disease | 0.764947804 | M_Disease&NULL |
| 11 | 116521135 | rs533668 | G | T | M_Disease | 0.764947804 | M_Disease&NULL |
| 11 | 118788517 | rs73575424 | A | C | M_Disease | 0.764947804 | M_Disease&NULL |
| 11 | 116521316 | rs573985 | A | G | M_Disease | 0.764947804 | M_Disease&NULL |
| 11 | 117499061 | rs7928623 | C | T | M_Disease | 0.763145016 | M_Disease&NULL |
| 11 | 116519739 | rs499790 | T | C | M_Disease | 0.760423002 | M_Disease&NULL |
| 11 | 116519358 | rs513533 | G | A | M_Disease | 0.759510917 | M_Disease&NULL |
| 11 | 116519584 | rs516043 | G | C | M_Disease | 0.759510917 | M_Disease&NULL |
| 11 | 117811037 | rs117560784 | A | G | M_Disease | 0.759510917 | M_Disease&NULL |
| 11 | 118705654 | rs649338 | A | G | M_Disease | 0.758596462 | M_Disease&NULL |
| 11 | 117291144 | rs73592158 | A | G | M_Disease | 0.758596462 | M_Disease&NULL |
| 11 | 117291266 | rs1228933145 | G | T | M_Disease | 0.758596462 | M_Disease&NULL |
| 11 | 117293463 | rs11820562 | G | A | M_Disease | 0.757679639 | M_Disease&NULL |
| 11 | 118703001 | rs7928228 | T | C | M_Disease | 0.757679639 | M_Disease&NULL |
| 11 | 118703484 | rs2508921 | T | C | M_Disease | 0.757679639 | M_Disease&NULL |
| 11 | 118703817 | rs1170396 | T | A | M_Disease | 0.757679639 | M_Disease&NULL |
| 11 | 118704265 | rs622690 | T | C | M_Disease | 0.757679639 | M_Disease&NULL |
| 11 | 117372097 | rs6589610 | T | C | M_Disease | 0.755838899 | M_Disease&NULL |
| 11 | 118701898 | rs533254 | A | T | M_Disease | 0.755838899 | M_Disease&NULL |
| 11 | 118704255 | rs480791 | G | A | M_Disease | 0.755838899 | M_Disease&NULL |
| 11 | 117274521 | rs139132354 | A | T | M_Disease | 0.755838899 | M_Disease&NULL |
| 11 | 118590668 | rs11216980 | A | C | M_Disease | 0.754914987 | M_Disease&NULL |
| 11 | 118503912 | rs45478298 | T | C | M_Disease | 0.753988716 | M_Disease&NULL |
| 11 | 118631653 | rs1784543 | T | A | M_Disease | 0.75306009 | M_Disease&NULL |
| 11 | 118631288 | rs507890 | C | T | M_Disease | 0.75306009 | M_Disease&NULL |
| 11 | 117259782 | rs79763467 | A | G | M_Disease | 0.75306009 | M_Disease&NULL |
| 11 | 117537864 | rs10892156 | T | C | M_Disease | 0.752129111 | M_Disease&NULL |
| 11 | 117284721 | rs78049971 | T | C | M_Disease | 0.75306009 | M_Disease&NULL |
| 11 | 116792961 | rs7948838 | G | A | M_Disease | 0.750260106 | M_Disease&NULL |
| 11 | 118701597 | rs4938549 | T | C | M_Disease | 0.750260106 | M_Disease&NULL |
| 11 | 118701668 | rs530646 | A | G | M_Disease | 0.750260106 | M_Disease&NULL |
| 11 | 118632522 | rs581045 | T | C | M_Disease | 0.749322084 | M_Disease&NULL |
| 11 | 117291361 | rs10790182 | T | C | M_Disease | 0.749322084 | M_Disease&NULL |
| 11 | 117537353 | rs10431082 | T | G | M_Disease | 0.748381722 | M_Disease&NULL |
| 11 | 118702002 | rs534114 | A | G | M_Disease | 0.749322084 | M_Disease&NULL |
| 11 | 116516873 | rs71480307 | A | G | M_Disease | 0.748381722 | M_Disease&NULL |
| 11 | 117378004 | rs73008432 | T | C | M_Disease | 0.748381722 | M_Disease&NULL |
| 11 | 117085270 | rs236911 | C | A | M_Disease | 0.74743902 | M_Disease&NULL |
| 11 | 116283516 | rs11823029 | C | T | M_Disease | 0.74743902 | M_Disease&NULL |
| 11 | 116638657 | rs149153426 | G | T | M_Disease | 0.74743902 | M_Disease&NULL |
| 11 | 117372126 | rs6589611 | T | A | M_Disease | 0.745546614 | M_Disease&NULL |
| 11 | 117380850 | rs2840350 | T | C | M_Disease | 0.745546614 | M_Disease&NULL |
| 11 | 120454129 | rs117764543 | G | T | M_Disease | 0.745546614 | M_Disease&NULL |
| 11 | 116517325 | rs35165288 | C | T | M_Disease | 0.744596916 | M_Disease&NULL |
| 11 | 118658075 | rs515523 | G | A | M_Disease | 0.743644892 | M_Disease&NULL |
| 11 | 118598936 | rs184071181 | C | T | M_Disease | 0.744596916 | M_Disease&NULL |
| 11 | 118785982 | rs79157249 | T | A | M_Disease | 0.743644892 | M_Disease&NULL |
| 11 | 116981001 | rs4635117 | A | G | M_Disease | 0.742690545 | M_Disease&NULL |
| 11 | 118702910 | rs7941321 | A | G | M_Disease | 0.742690545 | M_Disease&NULL |
| 11 | 118754353 | rs630923 | A | C | M_Disease | 0.742690545 | M_Disease&NULL |
| 11 | 116767688 | rs2270392 | C | T | M_Disease | 0.74173388 | M_Disease&NULL |
| 11 | 116776826 | rs7111854 | T | A | M_Disease | 0.74173388 | M_Disease&NULL |
| 11 | 116784376 | rs7115583 | T | G | M_Disease | 0.739813607 | M_Disease&NULL |
| 11 | 118651809 | rs555649 | C | T | M_Disease | 0.739813607 | M_Disease&NULL |
| 11 | 117802888 | rs147413777 | A | G | M_Disease | 0.739813607 | M_Disease&NULL |
| 11 | 118659896 | rs2105782 | C | T | M_Disease | 0.738850006 | M_Disease&NULL |
| 11 | 116797684 | rs11216190 | C | T | M_Disease | 0.736915896 | M_Disease&NULL |
| 11 | 116813979 | rs12292858 | C | A | M_Disease | 0.736915896 | M_Disease&NULL |
| 11 | 116816216 | rs10892049 | C | G | M_Disease | 0.736915896 | M_Disease&NULL |
| 11 | 116817655 | rs191655881 | C | A | M_Disease | 0.736915896 | M_Disease&NULL |
| 11 | 116820088 | rs12420518 | C | T | M_Disease | 0.736915896 | M_Disease&NULL |
| 11 | 116822528 | rs12286405 | T | C | M_Disease | 0.736915896 | M_Disease&NULL |
| 11 | 116848329 | rs11216210 | T | G | M_Disease | 0.736915896 | M_Disease&NULL |
| 11 | 116851608 | rs7358350 | C | T | M_Disease | 0.736915896 | M_Disease&NULL |
| 11 | 117370229 | rs624515 | G | T | M_Disease | 0.735945394 | M_Disease&NULL |

| | | | | | | | | | |
|---|---|---|---|---|---|---|---|---|---|
| 11 | 117373055 | rs6589612 | G | C | 1.091 | -7977.388 | -7974.76 | 2.629 | 5.36E-05 |
| 11 | 116521975 | rs562756 | C | G | 0.879 | -7977.388 | -7974.824 | 2.564 | 5.02E-05 |
| 11 | 118186305 | rs2231449 | A | C | 1.243 | -7977.388 | -7974.839 | 2.549 | 4.95E-05 |
| 11 | 118700425 | rs10128600 | C | T | 0.897 | -7977.388 | -7974.845 | 2.543 | 4.92E-05 |
| 11 | 116522156 | rs563917 | T | C | 0.88 | -7977.388 | -7974.846 | 2.542 | 4.91E-05 |
| 11 | 116522164 | rs492182 | A | G | 0.88 | -7977.388 | -7974.846 | 2.542 | 4.91E-05 |
| 11 | 116588904 | rs1262892 | A | G | 0.834 | -7977.388 | -7974.848 | 2.541 | 4.91E-05 |
| 11 | 116522111 | rs563796 | G | A | 0.88 | -7977.388 | -7974.886 | 2.503 | 4.73E-05 |
| 11 | 118699863 | rs2510901 | A | G | 0.897 | -7977.388 | -7974.887 | 2.502 | 4.72E-05 |
| 11 | 118762218 | rs543824 | G | A | 1.104 | -7977.388 | -7974.914 | 2.474 | 4.59E-05 |
| 11 | 118944909 | rs563162 | T | C | 1.152 | -7977.388 | -7974.917 | 2.471 | 4.58E-05 |
| 11 | 118163916 | rs11216846 | A | G | 1.238 | -7977.388 | -7974.926 | 2.463 | 4.54E-05 |
| 11 | 116521988 | rs491022 | C | T | 0.881 | -7977.388 | -7974.934 | 2.454 | 4.50E-05 |
| 11 | 118707171 | rs4938550 | G | A | 0.897 | -7977.388 | -7974.936 | 2.452 | 4.49E-05 |
| 11 | 116522330 | rs565607 | T | C | 0.881 | -7977.388 | -7974.937 | 2.451 | 4.49E-05 |
| 11 | 116521558 | rs575906 | G | A | 0.881 | -7977.388 | -7974.943 | 2.446 | 4.46E-05 |
| 11 | 117379086 | rs10750107 | C | T | 1.09 | -7977.388 | -7974.943 | 2.445 | 4.46E-05 |
| 11 | 116520434 | rs506222 | C | A | 0.881 | -7977.388 | -7974.946 | 2.442 | 4.45E-05 |
| 11 | 118698160 | rs640373 | G | A | 0.898 | -7977.388 | -7974.947 | 2.441 | 4.44E-05 |
| 11 | 118698274 | rs639915 | A | G | 0.898 | -7977.388 | -7974.947 | 2.441 | 4.44E-05 |
| 11 | 118698294 | rs639885 | A | G | 0.898 | -7977.388 | -7974.947 | 2.441 | 4.44E-05 |
| 11 | 118698537 | rs638893 | G | A | 0.898 | -7977.388 | -7974.947 | 2.441 | 4.44E-05 |
| 11 | 118697375 | rs73005500 | C | T | 0.898 | -7977.388 | -7974.952 | 2.436 | 4.42E-05 |
| 11 | 116291220 | rs57150747 | A | G | 1.142 | -7977.388 | -7974.96 | 2.428 | 4.38E-05 |
| 11 | 118698777 | rs516332 | G | A | 0.898 | -7977.388 | -7974.967 | 2.421 | 4.35E-05 |
| 11 | 118629031 | rs555639 | A | G | 0.918 | -7977.388 | -7974.973 | 2.415 | 4.33E-05 |
| 11 | 117513801 | rs10892147 | C | T | 1.113 | -7977.388 | -7974.98 | 2.408 | 4.30E-05 |
| 11 | 118699744 | rs2508564 | C | T | 0.898 | -7977.388 | -7974.981 | 2.407 | 4.29E-05 |
| 11 | 118181481 | rs11216848 | A | C | 1.236 | -7977.388 | -7974.982 | 2.406 | 4.29E-05 |
| 11 | 118566069 | rs506255 | A | C | 0.882 | -7977.388 | -7975.022 | 2.366 | 4.12E-05 |
| 11 | 116520398 | rs544479 | C | T | 0.882 | -7977.388 | -7975.03 | 2.358 | 4.09E-05 |
| 11 | 116520926 | rs549125 | G | C | 0.882 | -7977.388 | -7975.03 | 2.358 | 4.09E-05 |
| 11 | 116521135 | rs533668 | G | T | 0.882 | -7977.388 | -7975.03 | 2.358 | 4.09E-05 |
| 11 | 118788517 | rs73575424 | A | C | 0.883 | -7977.388 | -7975.03 | 2.358 | 4.09E-05 |
| 11 | 116521316 | rs573985 | A | G | 0.882 | -7977.388 | -7975.032 | 2.356 | 4.08E-05 |
| 11 | 117499061 | rs7928623 | C | T | 1.142 | -7977.388 | -7975.049 | 2.339 | 4.01E-05 |
| 11 | 116519739 | rs499790 | T | C | 0.883 | -7977.388 | -7975.085 | 2.303 | 3.87E-05 |
| 11 | 116519358 | rs513533 | G | A | 0.883 | -7977.388 | -7975.095 | 2.293 | 3.83E-05 |
| 11 | 116519584 | rs516043 | G | C | 0.883 | -7977.388 | -7975.095 | 2.293 | 3.83E-05 |
| 11 | 117811037 | rs117560784 | A | G | 0.789 | -7977.388 | -7975.099 | 2.289 | 3.82E-05 |
| 11 | 118705654 | rs649338 | A | G | 0.899 | -7977.388 | -7975.1 | 2.289 | 3.82E-05 |
| 11 | 117291144 | rs73592158 | A | G | 0.796 | -7977.388 | -7975.111 | 2.277 | 3.77E-05 |
| 11 | 117291266 | rs1228933145 | G | T | 0.796 | -7977.388 | -7975.111 | 2.277 | 3.77E-05 |
| 11 | 117293463 | rs11820562 | G | A | 0.795 | -7977.388 | -7975.115 | 2.273 | 3.76E-05 |
| 11 | 118703001 | rs7928228 | T | C | 0.899 | -7977.388 | -7975.116 | 2.272 | 3.75E-05 |
| 11 | 118703484 | rs2508921 | T | C | 0.899 | -7977.388 | -7975.116 | 2.272 | 3.75E-05 |
| 11 | 118703817 | rs1170396 | T | A | 0.899 | -7977.388 | -7975.116 | 2.272 | 3.75E-05 |
| 11 | 118704265 | rs622690 | T | C | 0.899 | -7977.388 | -7975.116 | 2.272 | 3.75E-05 |
| 11 | 117372097 | rs6589610 | T | C | 0.92 | -7977.388 | -7975.127 | 2.261 | 3.71E-05 |
| 11 | 118701898 | rs533254 | A | T | 0.899 | -7977.388 | -7975.132 | 2.256 | 3.69E-05 |
| 11 | 118704255 | rs480791 | G | A | 0.9 | -7977.388 | -7975.132 | 2.256 | 3.69E-05 |
| 11 | 117274521 | rs139132354 | A | T | 0.795 | -7977.388 | -7975.137 | 2.251 | 3.67E-05 |
| 11 | 118590668 | rs11216980 | A | C | 1.104 | -7977.388 | -7975.138 | 2.25 | 3.67E-05 |
| 11 | 118503912 | rs45478298 | T | C | 0.842 | -7977.388 | -7975.153 | 2.236 | 3.62E-05 |
| 11 | 118631653 | rs1784543 | T | A | 0.92 | -7977.388 | -7975.158 | 2.23 | 3.60E-05 |
| 11 | 118631288 | rs507890 | C | T | 0.92 | -7977.388 | -7975.161 | 2.227 | 3.59E-05 |
| 11 | 117259782 | rs79763467 | A | G | 0.796 | -7977.388 | -7975.167 | 2.222 | 3.57E-05 |
| 11 | 117537864 | rs10892156 | T | C | 0.918 | -7977.388 | -7975.169 | 2.219 | 3.56E-05 |
| 11 | 117284721 | rs78049971 | T | C | 0.796 | -7977.388 | -7975.172 | 2.216 | 3.55E-05 |
| 11 | 116792961 | rs7948838 | G | A | 1.145 | -7977.388 | -7975.194 | 2.195 | 3.47E-05 |
| 11 | 118701597 | rs4938549 | T | C | 0.9 | -7977.388 | -7975.195 | 2.194 | 3.47E-05 |
| 11 | 118701668 | rs530646 | A | G | 0.9 | -7977.388 | -7975.195 | 2.194 | 3.47E-05 |
| 11 | 118632522 | rs581045 | T | C | 0.92 | -7977.388 | -7975.202 | 2.186 | 3.44E-05 |
| 11 | 117291361 | rs10790182 | T | C | 0.882 | -7977.388 | -7975.206 | 2.182 | 3.43E-05 |
| 11 | 117537353 | rs10431082 | T | G | 0.915 | -7977.388 | -7975.206 | 2.182 | 3.43E-05 |
| 11 | 118702002 | rs534114 | A | G | 0.9 | -7977.388 | -7975.206 | 2.182 | 3.43E-05 |
| 11 | 116516873 | rs71480307 | A | G | 0.869 | -7977.388 | -7975.21 | 2.178 | 3.42E-05 |
| 11 | 117378004 | rs73008432 | T | C | 0.812 | -7977.388 | -7975.214 | 2.175 | 3.40E-05 |
| 11 | 117085270 | rs236911 | C | A | 0.891 | -7977.388 | -7975.22 | 2.168 | 3.38E-05 |
| 11 | 116283516 | rs11823029 | C | T | 1.138 | -7977.388 | -7975.224 | 2.164 | 3.37E-05 |
| 11 | 116638657 | rs149153426 | G | T | 0.795 | -7977.388 | -7975.23 | 2.159 | 3.35E-05 |
| 11 | 117372126 | rs6589611 | T | A | 0.921 | -7977.388 | -7975.236 | 2.152 | 3.33E-05 |
| 11 | 117380850 | rs2840350 | T | C | 1.087 | -7977.388 | -7975.243 | 2.145 | 3.30E-05 |
| 11 | 120454129 | rs117764543 | G | T | 0.849 | -7977.388 | -7975.244 | 2.145 | 3.30E-05 |
| 11 | 116517325 | rs35165288 | C | T | 0.87 | -7977.388 | -7975.25 | 2.138 | 3.28E-05 |
| 11 | 118658075 | rs515523 | G | A | 0.92 | -7977.388 | -7975.256 | 2.133 | 3.26E-05 |
| 11 | 118598936 | rs184071181 | C | T | 0.845 | -7977.388 | -7975.256 | 2.132 | 3.26E-05 |
| 11 | 118785982 | rs79157249 | T | A | 0.886 | -7977.388 | -7975.266 | 2.123 | 3.23E-05 |
| 11 | 116981001 | rs4635117 | A | G | 0.878 | -7977.388 | -7975.27 | 2.119 | 3.22E-05 |
| 11 | 118702910 | rs7941321 | A | G | 0.901 | -7977.388 | -7975.273 | 2.115 | 3.21E-05 |
| 11 | 118754353 | rs630923 | A | C | 1.117 | -7977.388 | -7975.275 | 2.114 | 3.20E-05 |
| 11 | 116767688 | rs2270392 | C | T | 1.141 | -7977.388 | -7975.279 | 2.109 | 3.19E-05 |
| 11 | 116776826 | rs7111854 | T | A | 1.141 | -7977.388 | -7975.28 | 2.109 | 3.19E-05 |
| 11 | 116784376 | rs7115583 | T | G | 1.141 | -7977.388 | -7975.298 | 2.09 | 3.13E-05 |
| 11 | 118651809 | rs555649 | C | T | 0.921 | -7977.388 | -7975.299 | 2.089 | 3.12E-05 |
| 11 | 117802888 | rs147413777 | A | G | 0.793 | -7977.388 | -7975.305 | 2.083 | 3.11E-05 |
| 11 | 118659896 | rs2105782 | C | T | 0.921 | -7977.388 | -7975.307 | 2.081 | 3.10E-05 |
| 11 | 116797684 | rs11216190 | C | T | 1.142 | -7977.388 | -7975.334 | 2.054 | 3.02E-05 |
| 11 | 116813979 | rs12292858 | C | A | 1.142 | -7977.388 | -7975.334 | 2.054 | 3.02E-05 |
| 11 | 116816216 | rs10892049 | C | G | 1.142 | -7977.388 | -7975.334 | 2.054 | 3.02E-05 |
| 11 | 116817655 | rs191655881 | C | A | 1.142 | -7977.388 | -7975.334 | 2.054 | 3.02E-05 |
| 11 | 116820088 | rs12420518 | C | T | 1.142 | -7977.388 | -7975.334 | 2.054 | 3.02E-05 |
| 11 | 116822528 | rs12286405 | T | C | 1.142 | -7977.388 | -7975.334 | 2.054 | 3.02E-05 |
| 11 | 116848329 | rs11216210 | T | G | 1.142 | -7977.388 | -7975.334 | 2.054 | 3.02E-05 |
| 11 | 116851608 | rs7358350 | C | T | 1.142 | -7977.388 | -7975.334 | 2.054 | 3.02E-05 |
| 11 | 117370229 | rs624515 | G | T | 0.921 | -7977.388 | -7975.336 | 2.053 | 3.01E-05 |

| | | | | | | | |
|---|---|---|---|---|---|---|---|
| 11 | 117292208 | rs7938710 | T | C | M_Disease | 0.736915896 | M_Disease&NULL |
| 11 | 116647152 | rs34552724 | C | T | M_Disease | 0.736915896 | M_Disease&NULL |
| 11 | 117498485 | rs77416199 | C | T | M_Disease | 0.735945394 | M_Disease&NULL |
| 11 | 118702936 | rs7941696 | G | A | M_Disease | 0.735945394 | M_Disease&NULL |
| 11 | 118644582 | rs2510897 | C | A | M_Disease | 0.733997517 | M_Disease&NULL |
| 11 | 118587319 | rs10892263 | C | T | M_Disease | 0.733997517 | M_Disease&NULL |
| 11 | 118588952 | rs12421790 | T | C | M_Disease | 0.733020149 | M_Disease&NULL |
| 11 | 117104148 | rs582179 | G | A | M_Disease | 0.733020149 | M_Disease&NULL |
| 11 | 119205735 | rs12797083 | C | T | M_Disease | 0.733020149 | M_Disease&NULL |
| 11 | 118585826 | rs11216972 | T | C | M_Disease | 0.732040502 | M_Disease&NULL |
| 11 | 117431753 | rs11606341 | T | G | M_Disease | 0.731058579 | M_Disease&NULL |
| 11 | 117380108 | rs4936383 | A | G | M_Disease | 0.730074384 | M_Disease&NULL |
| 11 | 118589012 | rs12421793 | A | C | M_Disease | 0.730074384 | M_Disease&NULL |
| 11 | 117408890 | rs4609639 | C | T | M_Disease | 0.730074384 | M_Disease&NULL |
| 11 | 119205830 | rs12795576 | G | A | M_Disease | 0.727108216 | M_Disease&NULL |
| 11 | 116879224 | rs12271319 | T | C | M_Disease | 0.726114981 | M_Disease&NULL |
| 11 | 116831843 | rs11216200 | T | C | M_Disease | 0.726114981 | M_Disease&NULL |
| 11 | 116858994 | rs10160754 | C | A | M_Disease | 0.726114981 | M_Disease&NULL |
| 11 | 116859003 | rs10160755 | G | A | M_Disease | 0.726114981 | M_Disease&NULL |
| 11 | 118587097 | rs10892262 | C | G | M_Disease | 0.725119498 | M_Disease&NULL |
| 11 | 117368804 | rs77822543 | T | C | M_Disease | 0.723121805 | M_Disease&NULL |
| 11 | 117372548 | rs10790191 | A | G | M_Disease | 0.722119606 | M_Disease&NULL |
| 11 | 116880456 | rs7112937 | C | T | M_Disease | 0.722119606 | M_Disease&NULL |
| 11 | 116940449 | rs12280210 | C | T | M_Disease | 0.722119606 | M_Disease&NULL |
| 11 | 117496431 | rs75349886 | G | A | M_Disease | 0.722119606 | M_Disease&NULL |
| 11 | 119203505 | rs11824001 | T | C | M_Disease | 0.720108527 | M_Disease&NULL |
| 11 | 119529529 | rs951786 | G | A | M_Disease | 0.719099657 | M_Disease&NULL |
| 11 | 116525730 | rs480823 | C | T | M_Disease | 0.719099657 | M_Disease&NULL |
| 11 | 117494945 | rs7946890 | C | A | M_Disease | 0.718088575 | M_Disease&NULL |
| 11 | 118705145 | rs636736 | G | A | M_Disease | 0.717075285 | M_Disease&NULL |
| 11 | 116525867 | rs481843 | T | C | M_Disease | 0.716059794 | M_Disease&NULL |
| 11 | 116837373 | rs9971422 | C | T | M_Disease | 0.716059794 | M_Disease&NULL |
| 11 | 116879519 | rs12284696 | G | A | M_Disease | 0.715042106 | M_Disease&NULL |
| 11 | 118652569 | rs150792202 | A | G | M_Disease | 0.715042106 | M_Disease&NULL |
| 11 | 116515557 | rs71480306 | C | T | M_Disease | 0.715042106 | M_Disease&NULL |
| 11 | 116863988 | rs11216220 | C | T | M_Disease | 0.715042106 | M_Disease&NULL |
| 11 | 118657311 | rs1309473877 | A | C | M_Disease | 0.714022227 | M_Disease&NULL |
| 11 | 117536758 | rs11216492 | G | A | M_Disease | 0.714022227 | M_Disease&NULL |
| 11 | 116866240 | rs7931665 | G | A | M_Disease | 0.713000163 | M_Disease&NULL |
| 11 | 116872034 | rs10892057 | C | A | M_Disease | 0.710949503 | M_Disease&NULL |
| 11 | 116875152 | rs7946390 | G | A | M_Disease | 0.710949503 | M_Disease&NULL |
| 11 | 116875924 | rs11216224 | C | T | M_Disease | 0.710949503 | M_Disease&NULL |
| 11 | 116876442 | rs1940625 | G | A | M_Disease | 0.710949503 | M_Disease&NULL |
| 11 | 116878014 | rs11216225 | G | A | M_Disease | 0.710949503 | M_Disease&NULL |
| 11 | 119214135 | rs948412 | G | A | M_Disease | 0.710949503 | M_Disease&NULL |
| 11 | 116837089 | rs78044162 | T | C | M_Disease | 0.709920918 | M_Disease&NULL |
| 11 | 120691285 | rs4936554 | T | C | M_Disease | 0.706822221 | M_Disease&NULL |
| 11 | 119208561 | rs2511840 | C | G | M_Disease | 0.706822221 | M_Disease&NULL |
| 11 | 117296641 | rs138471627 | T | C | M_Disease | 0.706822221 | M_Disease&NULL |
| 11 | 117407658 | rs2155392 | C | G | M_Disease | 0.706822221 | M_Disease&NULL |
| 11 | 116993181 | rs6589596 | T | C | M_Disease | 0.704745698 | M_Disease&NULL |
| 11 | 118757191 | rs1790192 | G | A | M_Disease | 0.704745698 | M_Disease&NULL |
| 11 | 116981580 | rs2127907 | A | G | M_Disease | 0.704745698 | M_Disease&NULL |
| 11 | 117405785 | rs61277802 | A | G | M_Disease | 0.703704238 | M_Disease&NULL |
| 11 | 116870480 | rs10502224 | G | A | M_Disease | 0.700567142 | M_Disease&NULL |
| 11 | 120689582 | rs11218014 | A | G | M_Disease | 0.699517228 | M_Disease&NULL |
| 11 | 117847373 | rs79459300 | T | C | M_Disease | 0.699517228 | M_Disease&NULL |
| 11 | 118220027 | rs73016221 | C | T | M_Disease | 0.698465216 | M_Disease&NULL |
| 11 | 116519129 | rs1240772 | C | G | M_Disease | 0.69635493 | M_Disease&NULL |
| 11 | 116664654 | rs45557239 | T | C | M_Disease | 0.697411114 | M_Disease&NULL |
| 11 | 119527302 | rs9888246 | T | C | M_Disease | 0.69635493 | M_Disease&NULL |
| 11 | 116997798 | rs6589597 | C | T | M_Disease | 0.695296669 | M_Disease&NULL |
| 11 | 117401657 | rs11826885 | T | C | M_Disease | 0.69423634 | M_Disease&NULL |
| 11 | 118765688 | rs676925 | G | C | M_Disease | 0.693173949 | M_Disease&NULL |
| 11 | 118678780 | rs500254 | G | A | M_Disease | 0.693173949 | M_Disease&NULL |
| 11 | 118665009 | rs4938544 | A | G | M_Disease | 0.693173949 | M_Disease&NULL |
| 11 | 116552860 | rs113194822 | A | G | M_Disease | 0.693173949 | M_Disease&NULL |
| 11 | 118764051 | rs613791 | T | C | M_Disease | 0.692109504 | M_Disease&NULL |
| 11 | 118665324 | rs61901411 | T | C | M_Disease | 0.691043012 | M_Disease&NULL |
| 11 | 116977888 | rs10160799 | G | A | M_Disease | 0.691043012 | M_Disease&NULL |
| 11 | 117263616 | rs1482242286 | G | T | M_Disease | 0.691043012 | M_Disease&NULL |
| 11 | 118668908 | rs582200 | G | A | M_Disease | 0.691043012 | M_Disease&NULL |
| 11 | 117268074 | rs77543424 | G | C | M_Disease | 0.691043012 | M_Disease&NULL |
| 11 | 117263103 | rs478693 | C | T | M_Disease | 0.688903918 | M_Disease&NULL |
| 11 | 117502041 | rs71480346 | T | C | M_Disease | 0.688903918 | M_Disease&NULL |
| 11 | 117037361 | rs7112513 | A | G | M_Disease | 0.687831331 | M_Disease&NULL |
| 11 | 118680072 | rs607472 | G | C | M_Disease | 0.687831331 | M_Disease&NULL |
| 11 | 119528905 | rs931953 | G | A | M_Disease | 0.687831331 | M_Disease&NULL |
| 11 | 116979689 | rs10750101 | T | C | M_Disease | 0.687831331 | M_Disease&NULL |
| 11 | 116328380 | rs12418858 | A | G | M_Disease | 0.686756727 | M_Disease&NULL |
| 11 | 117010320 | rs2471884 | T | C | M_Disease | 0.686756727 | M_Disease&NULL |
| 11 | 117422690 | rs73010348 | A | G | M_Disease | 0.685680114 | M_Disease&NULL |
| 11 | 119678810 | rs10750166 | C | G | M_Disease | 0.685680114 | M_Disease&NULL |
| 11 | 118217854 | rs3212262 | T | G | M_Disease | 0.6846015 | M_Disease&NULL |
| 11 | 118677083 | rs675003 | T | C | M_Disease | 0.6846015 | M_Disease&NULL |
| 11 | 118912207 | rs4936460 | T | C | M_Disease | 0.683520894 | M_Disease&NULL |
| 11 | 119529524 | rs4445648 | T | C | M_Disease | 0.681353734 | M_Disease&NULL |
| 11 | 116999446 | rs7131536 | C | T | M_Disease | 0.680267197 | M_Disease&NULL |
| 11 | 116999559 | rs7131686 | C | T | M_Disease | 0.680267197 | M_Disease&NULL |
| 11 | 119572124 | rs73004983 | C | G | M_Disease | 0.680267197 | M_Disease&NULL |
| 11 | 117362165 | rs73024367 | A | G | M_Disease | 0.680267197 | M_Disease&NULL |
| 11 | 117453109 | rs77967735 | G | A | M_Disease | 0.679178699 | M_Disease&NULL |
| 11 | 117502961 | rs7929496 | T | C | M_Disease | 0.679178699 | M_Disease&NULL |
| 11 | 118662028 | rs12796102 | T | C | M_Disease | 0.67808825 | M_Disease&NULL |
| 11 | 117039325 | rs10892082 | T | G | M_Disease | 0.67808825 | M_Disease&NULL |
| 11 | 118679629 | rs1784300 | T | A | M_Disease | 0.67808825 | M_Disease&NULL |

| | | | | | | | | | |
|---|---|---|---|---|---|---|---|---|---|
| 11 | 117292208 | rs7938710 | T | C | 0.802 | -7977.388 | -7975.338 | 2.05 | 3.00E-05 |
| 11 | 116647152 | rs34552724 | C | T | 0.805 | -7977.388 | -7975.339 | 2.049 | 3.00E-05 |
| 11 | 117498485 | rs77416199 | C | T | 1.136 | -7977.388 | -7975.343 | 2.045 | 2.99E-05 |
| 11 | 118702936 | rs7941696 | G | A | 0.903 | -7977.388 | -7975.344 | 2.044 | 2.99E-05 |
| 11 | 118644582 | rs2510897 | C | A | 0.921 | -7977.388 | -7975.362 | 2.027 | 2.94E-05 |
| 11 | 118587319 | rs10892263 | C | T | 1.101 | -7977.388 | -7975.366 | 2.022 | 2.92E-05 |
| 11 | 118588952 | rs12421790 | T | C | 1.101 | -7977.388 | -7975.368 | 2.02 | 2.92E-05 |
| 11 | 117104148 | rs582179 | G | A | 0.91 | -7977.388 | -7975.371 | 2.017 | 2.91E-05 |
| 11 | 119205735 | rs12797083 | C | T | 0.922 | -7977.388 | -7975.372 | 2.017 | 2.91E-05 |
| 11 | 118585826 | rs11216972 | T | C | 1.1 | -7977.388 | -7975.383 | 2.005 | 2.87E-05 |
| 11 | 117431753 | rs11606341 | T | G | 1.09 | -7977.388 | -7975.389 | 1.999 | 2.86E-05 |
| 11 | 117380108 | rs4936383 | A | G | 0.922 | -7977.388 | -7975.396 | 1.992 | 2.84E-05 |
| 11 | 118589012 | rs12421793 | A | C | 1.1 | -7977.388 | -7975.4 | 1.989 | 2.83E-05 |
| 11 | 117408890 | rs4609639 | C | T | 0.816 | -7977.388 | -7975.403 | 1.985 | 2.82E-05 |
| 11 | 119205830 | rs12795576 | G | A | 0.922 | -7977.388 | -7975.431 | 1.957 | 2.74E-05 |
| 11 | 116879224 | rs12271319 | T | C | 1.139 | -7977.388 | -7975.44 | 1.948 | 2.71E-05 |
| 11 | 116831843 | rs11216200 | T | C | 1.139 | -7977.388 | -7975.445 | 1.943 | 2.70E-05 |
| 11 | 116858994 | rs10160754 | C | A | 1.139 | -7977.388 | -7975.447 | 1.941 | 2.69E-05 |
| 11 | 116859003 | rs10160755 | G | A | 1.139 | -7977.388 | -7975.447 | 1.941 | 2.69E-05 |
| 11 | 118587097 | rs10892262 | C | G | 1.1 | -7977.388 | -7975.453 | 1.935 | 2.68E-05 |
| 11 | 117368804 | rs77822543 | T | C | 1.146 | -7977.388 | -7975.47 | 1.918 | 2.63E-05 |
| 11 | 117372548 | rs10790191 | A | G | 0.923 | -7977.388 | -7975.475 | 1.913 | 2.62E-05 |
| 11 | 116880456 | rs7112937 | C | T | 1.138 | -7977.388 | -7975.479 | 1.91 | 2.61E-05 |
| 11 | 116940449 | rs12280210 | C | T | 1.138 | -7977.388 | -7975.482 | 1.906 | 2.60E-05 |
| 11 | 117496431 | rs75349886 | G | A | 1.132 | -7977.388 | -7975.483 | 1.906 | 2.60E-05 |
| 11 | 119203505 | rs11824001 | T | C | 0.923 | -7977.388 | -7975.499 | 1.889 | 2.56E-05 |
| 11 | 119529529 | rs951786 | G | A | 1.098 | -7977.388 | -7975.515 | 1.873 | 2.52E-05 |
| 11 | 116525730 | rs480823 | C | T | 0.879 | -7977.388 | -7975.517 | 1.872 | 2.51E-05 |
| 11 | 117494945 | rs7946890 | C | A | 1.131 | -7977.388 | -7975.522 | 1.866 | 2.50E-05 |
| 11 | 118705145 | rs636736 | G | A | 0.905 | -7977.388 | -7975.527 | 1.861 | 2.49E-05 |
| 11 | 116525867 | rs481843 | T | C | 0.879 | -7977.388 | -7975.541 | 1.847 | 2.45E-05 |
| 11 | 116837373 | rs9971422 | C | T | 1.137 | -7977.388 | -7975.547 | 1.842 | 2.44E-05 |
| 11 | 116879519 | rs12284696 | G | A | 1.137 | -7977.388 | -7975.549 | 1.839 | 2.43E-05 |
| 11 | 118652569 | rs150792202 | A | G | 0.884 | -7977.388 | -7975.551 | 1.837 | 2.43E-05 |
| 11 | 116515557 | rs71480306 | C | T | 0.875 | -7977.388 | -7975.553 | 1.835 | 2.42E-05 |
| 11 | 116863988 | rs11216220 | C | T | 1.137 | -7977.388 | -7975.557 | 1.832 | 2.42E-05 |
| 11 | 118657311 | rs1309473877 | A | C | 0.923 | -7977.388 | -7975.563 | 1.825 | 2.40E-05 |
| 11 | 117536758 | rs11216492 | G | A | 0.919 | -7977.388 | -7975.565 | 1.824 | 2.40E-05 |
| 11 | 116866240 | rs7931665 | G | A | 1.136 | -7977.388 | -7975.571 | 1.817 | 2.38E-05 |
| 11 | 116872034 | rs10892057 | C | A | 1.136 | -7977.388 | -7975.59 | 1.799 | 2.34E-05 |
| 11 | 116875152 | rs7946390 | G | A | 1.136 | -7977.388 | -7975.59 | 1.799 | 2.34E-05 |
| 11 | 116875924 | rs11216224 | C | T | 1.136 | -7977.388 | -7975.59 | 1.799 | 2.34E-05 |
| 11 | 116876442 | rs1940625 | G | A | 1.136 | -7977.388 | -7975.59 | 1.799 | 2.34E-05 |
| 11 | 116878014 | rs11216225 | G | A | 1.136 | -7977.388 | -7975.59 | 1.799 | 2.34E-05 |
| 11 | 119214135 | rs948412 | G | A | 1.116 | -7977.388 | -7975.596 | 1.793 | 2.32E-05 |
| 11 | 116837089 | rs78044162 | T | C | 1.136 | -7977.388 | -7975.601 | 1.788 | 2.31E-05 |
| 11 | 120691285 | rs4936554 | T | C | 1.097 | -7977.388 | -7975.627 | 1.761 | 2.25E-05 |
| 11 | 119208561 | rs2511840 | C | G | 1.084 | -7977.388 | -7975.631 | 1.757 | 2.24E-05 |
| 11 | 117296641 | rs138471627 | T | C | 0.805 | -7977.388 | -7975.636 | 1.753 | 2.23E-05 |
| 11 | 117407658 | rs2155392 | C | G | 0.823 | -7977.388 | -7975.641 | 1.748 | 2.22E-05 |
| 11 | 116993181 | rs6589596 | T | C | 0.891 | -7977.388 | -7975.647 | 1.741 | 2.21E-05 |
| 11 | 118757191 | rs1790192 | G | A | 1.084 | -7977.388 | -7975.648 | 1.74 | 2.20E-05 |
| 11 | 116981580 | rs2127907 | A | G | 0.892 | -7977.388 | -7975.651 | 1.737 | 2.20E-05 |
| 11 | 117405785 | rs61277802 | A | G | 0.868 | -7977.388 | -7975.662 | 1.727 | 2.18E-05 |
| 11 | 116870480 | rs10502224 | G | A | 1.133 | -7977.388 | -7975.689 | 1.699 | 2.12E-05 |
| 11 | 120689582 | rs11218014 | A | G | 0.88 | -7977.388 | -7975.703 | 1.686 | 2.09E-05 |
| 11 | 117847373 | rs79459300 | T | C | 0.829 | -7977.388 | -7975.708 | 1.68 | 2.08E-05 |
| 11 | 118220027 | rs73016221 | C | T | 1.122 | -7977.388 | -7975.713 | 1.675 | 2.07E-05 |
| 11 | 116519129 | rs1240772 | C | G | 1.082 | -7977.388 | -7975.729 | 1.659 | 2.03E-05 |
| 11 | 116664654 | rs45557239 | T | C | 0.817 | -7977.388 | -7975.731 | 1.657 | 2.03E-05 |
| 11 | 119527302 | rs9888246 | T | C | 1.095 | -7977.388 | -7975.733 | 1.656 | 2.03E-05 |
| 11 | 116997798 | rs6589597 | C | T | 0.892 | -7977.388 | -7975.74 | 1.649 | 2.01E-05 |
| 11 | 117401657 | rs11826885 | T | C | 0.87 | -7977.388 | -7975.75 | 1.639 | 1.99E-05 |
| 11 | 118765688 | rs676925 | G | C | 1.091 | -7977.388 | -7975.763 | 1.626 | 1.97E-05 |
| 11 | 118678780 | rs500254 | G | A | 0.897 | -7977.388 | -7975.765 | 1.623 | 1.96E-05 |
| 11 | 118665009 | rs4938544 | A | G | 0.897 | -7977.388 | -7975.766 | 1.622 | 1.96E-05 |
| 11 | 116552860 | rs113194822 | A | G | 0.827 | -7977.388 | -7975.772 | 1.616 | 1.95E-05 |
| 11 | 118764051 | rs613791 | T | C | 1.09 | -7977.388 | -7975.772 | 1.616 | 1.95E-05 |
| 11 | 118665324 | rs61901411 | T | C | 0.897 | -7977.388 | -7975.778 | 1.61 | 1.94E-05 |
| 11 | 116977888 | rs10160799 | G | A | 0.893 | -7977.388 | -7975.782 | 1.606 | 1.93E-05 |
| 11 | 117263616 | rs1482242286 | G | T | 0.889 | -7977.388 | -7975.783 | 1.605 | 1.93E-05 |
| 11 | 118668908 | rs582200 | G | A | 0.898 | -7977.388 | -7975.786 | 1.602 | 1.92E-05 |
| 11 | 117268074 | rs77543424 | G | C | 0.815 | -7977.388 | -7975.788 | 1.6 | 1.92E-05 |
| 11 | 117263103 | rs478693 | C | T | 0.889 | -7977.388 | -7975.802 | 1.586 | 1.89E-05 |
| 11 | 117502041 | rs71480346 | T | C | 1.12 | -7977.388 | -7975.803 | 1.585 | 1.89E-05 |
| 11 | 117037361 | rs7112513 | A | G | 0.893 | -7977.388 | -7975.807 | 1.581 | 1.88E-05 |
| 11 | 118680072 | rs607472 | G | C | 0.926 | -7977.388 | -7975.808 | 1.581 | 1.88E-05 |
| 11 | 119528905 | rs931953 | G | A | 1.094 | -7977.388 | -7975.814 | 1.575 | 1.87E-05 |
| 11 | 116979689 | rs10750101 | T | C | 0.894 | -7977.388 | -7975.815 | 1.573 | 1.86E-05 |
| 11 | 116328380 | rs12418858 | A | G | 0.863 | -7977.388 | -7975.821 | 1.567 | 1.85E-05 |
| 11 | 117010320 | rs2471884 | T | C | 0.894 | -7977.388 | -7975.823 | 1.566 | 1.85E-05 |
| 11 | 117422690 | rs73010348 | A | G | 0.86 | -7977.388 | -7975.831 | 1.558 | 1.84E-05 |
| 11 | 119678810 | rs10750166 | C | G | 0.853 | -7977.388 | -7975.834 | 1.554 | 1.83E-05 |
| 11 | 118217854 | rs3212262 | T | G | 1.12 | -7977.388 | -7975.841 | 1.548 | 1.82E-05 |
| 11 | 118677083 | rs675003 | T | C | 0.899 | -7977.388 | -7975.844 | 1.544 | 1.81E-05 |
| 11 | 118912207 | rs4936460 | T | C | 1.127 | -7977.388 | -7975.857 | 1.531 | 1.79E-05 |
| 11 | 119529524 | rs4445648 | T | C | 1.093 | -7977.388 | -7975.871 | 1.517 | 1.76E-05 |
| 11 | 116999446 | rs7131536 | C | T | 0.895 | -7977.388 | -7975.88 | 1.508 | 1.75E-05 |
| 11 | 116999559 | rs7131686 | C | T | 0.895 | -7977.388 | -7975.88 | 1.508 | 1.75E-05 |
| 11 | 119572124 | rs73004983 | C | G | 0.883 | -7977.388 | -7975.882 | 1.507 | 1.75E-05 |
| 11 | 117362165 | rs73024367 | A | G | 0.819 | -7977.388 | -7975.888 | 1.5 | 1.73E-05 |
| 11 | 117453109 | rs77967735 | G | A | 1.126 | -7977.388 | -7975.896 | 1.493 | 1.72E-05 |
| 11 | 117502961 | rs7929496 | T | C | 1.118 | -7977.388 | -7975.897 | 1.491 | 1.72E-05 |
| 11 | 118662028 | rs12796102 | T | C | 0.927 | -7977.388 | -7975.899 | 1.489 | 1.71E-05 |
| 11 | 117039325 | rs10892082 | T | G | 0.895 | -7977.388 | -7975.901 | 1.488 | 1.71E-05 |
| 11 | 118679629 | rs1784300 | T | A | 0.927 | -7977.388 | -7975.902 | 1.486 | 1.71E-05 |

| | | | | | | | |
|---|---|---|---|---|---|---|---|
| 11 | 117404312 | rs56198555 | T | C | M_Disease | 0.67808825 | M_Disease&NULL |
| 11 | 119213782 | rs948411 | T | A | M_Disease | 0.676995856 | M_Disease&NULL |
| 11 | 119529332 | rs4245201 | C | T | M_Disease | 0.676995856 | M_Disease&NULL |
| 11 | 117024481 | rs7940310 | T | C | M_Disease | 0.676995856 | M_Disease&NULL |
| 11 | 117081500 | rs11216316 | C | A | M_Disease | 0.675901528 | M_Disease&NULL |
| 11 | 120850004 | rs1334757891 | T | C | M_Disease | 0.673707099 | M_Disease&NULL |
| 11 | 117045335 | rs7925256 | T | C | M_Disease | 0.673707099 | M_Disease&NULL |
| 11 | 116967666 | rs1320668 | C | T | M_Disease | 0.673707099 | M_Disease&NULL |
| 11 | 117027755 | rs7123517 | G | A | M_Disease | 0.673707099 | M_Disease&NULL |
| 11 | 117405907 | rs68096541 | C | G | M_Disease | 0.672607017 | M_Disease&NULL |
| 11 | 118662045 | rs7933859 | T | C | M_Disease | 0.672607017 | M_Disease&NULL |
| 11 | 118765267 | rs598207 | C | G | M_Disease | 0.672607017 | M_Disease&NULL |
| 11 | 119678767 | rs73577182 | C | G | M_Disease | 0.672607017 | M_Disease&NULL |
| 11 | 118833458 | rs111491960 | G | A | M_Disease | 0.671505034 | M_Disease&NULL |
| 11 | 116516966 | rs1176456955 | A | G | M_Disease | 0.671505034 | M_Disease&NULL |
| 11 | 118662203 | rs4508234 | G | C | M_Disease | 0.67040116 | M_Disease&NULL |
| 11 | 117534083 | rs7111427 | A | G | M_Disease | 0.67040116 | M_Disease&NULL |
| 11 | 117363891 | rs4096589 | T | A | M_Disease | 0.669295403 | M_Disease&NULL |
| 11 | 118662340 | rs5006102 | A | T | M_Disease | 0.669295403 | M_Disease&NULL |
| 11 | 118230359 | rs2276419 | T | C | M_Disease | 0.669295403 | M_Disease&NULL |
| 11 | 120848685 | rs75871913 | A | G | M_Disease | 0.669295403 | M_Disease&NULL |
| 11 | 117042377 | rs4936367 | G | A | M_Disease | 0.668187772 | M_Disease&NULL |
| 11 | 117044203 | rs4604928 | C | A | M_Disease | 0.668187772 | M_Disease&NULL |
| 11 | 117458673 | rs59607164 | G | A | M_Disease | 0.667078277 | M_Disease&NULL |
| 11 | 119203359 | rs2511847 | A | G | M_Disease | 0.667078277 | M_Disease&NULL |
| 11 | 116697081 | rs150555490 | T | C | M_Disease | 0.667078277 | M_Disease&NULL |
| 11 | 117034619 | rs7122944 | G | T | M_Disease | 0.665966927 | M_Disease&NULL |
| 11 | 117262598 | rs542821 | T | C | M_Disease | 0.665966927 | M_Disease&NULL |
| 11 | 117329630 | rs117043085 | T | C | M_Disease | 0.667078277 | M_Disease&NULL |
| 11 | 117024302 | rs7950633 | C | T | M_Disease | 0.66485373 | M_Disease&NULL |
| 11 | 117102648 | rs2252145 | C | T | M_Disease | 0.66485373 | M_Disease&NULL |
| 11 | 117531465 | rs1892918 | C | A | M_Disease | 0.66485373 | M_Disease&NULL |
| 11 | 118669178 | rs583213 | G | T | M_Disease | 0.663738697 | M_Disease&NULL |
| 11 | 120702916 | rs12283648 | C | A | M_Disease | 0.663738697 | M_Disease&NULL |
| 11 | 116485201 | rs71480304 | A | G | M_Disease | 0.662621837 | M_Disease&NULL |
| 11 | 117368182 | rs73008413 | T | A | M_Disease | 0.663738697 | M_Disease&NULL |
| 11 | 120795888 | rs949054 | C | T | M_Disease | 0.662621837 | M_Disease&NULL |
| 11 | 116488748 | rs4360730 | C | T | M_Disease | 0.662621837 | M_Disease&NULL |
| 11 | 116488753 | rs4565932 | A | G | M_Disease | 0.662621837 | M_Disease&NULL |
| 11 | 118617920 | rs476246 | T | C | M_Disease | 0.662621837 | M_Disease&NULL |
| 11 | 117214617 | rs144750609 | C | T | M_Disease | 0.662621837 | M_Disease&NULL |
| 11 | 117285699 | rs11826012 | C | T | M_Disease | 0.661503159 | M_Disease&NULL |
| 11 | 117285746 | rs1789157 | G | A | M_Disease | 0.661503159 | M_Disease&NULL |
| 11 | 118617544 | rs472641 | G | A | M_Disease | 0.661503159 | M_Disease&NULL |
| 11 | 118621495 | rs487728 | G | C | M_Disease | 0.661503159 | M_Disease&NULL |
| 11 | 118705753 | rs556437 | A | G | M_Disease | 0.660382673 | M_Disease&NULL |
| 11 | 116706168 | rs12721026 | G | T | M_Disease | 0.659260388 | M_Disease&NULL |
| 11 | 117368982 | rs2004927 | C | T | M_Disease | 0.659260388 | M_Disease&NULL |
| 11 | 119529106 | rs931954 | A | G | M_Disease | 0.658136315 | M_Disease&NULL |
| 11 | 117366973 | rs7130068 | C | T | M_Disease | 0.658136315 | M_Disease&NULL |
| 11 | 120858914 | rs670413 | A | G | M_Disease | 0.658136315 | M_Disease&NULL |
| 11 | 117537710 | rs686413 | A | G | M_Disease | 0.658136315 | M_Disease&NULL |
| 11 | 117367196 | rs73008408 | C | A | M_Disease | 0.658136315 | M_Disease&NULL |
| 11 | 116548356 | rs117339323 | C | T | M_Disease | 0.658136315 | M_Disease&NULL |
| 11 | 117803083 | rs117978531 | G | T | M_Disease | 0.658136315 | M_Disease&NULL |
| 11 | 118626511 | rs499343 | T | C | M_Disease | 0.657010463 | M_Disease&NULL |
| 11 | 117369226 | rs1807116 | C | T | M_Disease | 0.655882841 | M_Disease&NULL |
| 11 | 120858301 | rs74727906 | A | G | M_Disease | 0.655882841 | M_Disease&NULL |
| 11 | 117026374 | rs2127905 | A | C | M_Disease | 0.654753461 | M_Disease&NULL |
| 11 | 119918191 | rs1942010 | C | T | M_Disease | 0.653622331 | M_Disease&NULL |
| 11 | 119675331 | rs61897704 | T | C | M_Disease | 0.654753461 | M_Disease&NULL |
| 11 | 119675529 | rs61897705 | G | A | M_Disease | 0.654753461 | M_Disease&NULL |
| 11 | 117283471 | rs573965 | T | C | M_Disease | 0.653622331 | M_Disease&NULL |
| 11 | 117405696 | rs2076965 | A | T | M_Disease | 0.653622331 | M_Disease&NULL |
| 11 | 116281175 | rs76887844 | T | C | M_Disease | 0.653622331 | M_Disease&NULL |
| 11 | 116289906 | rs11215924 | A | T | M_Disease | 0.652489462 | M_Disease&NULL |
| 11 | 118624767 | rs581062 | A | G | M_Disease | 0.652489462 | M_Disease&NULL |
| 11 | 120988038 | rs140567534 | G | T | M_Disease | 0.652489462 | M_Disease&NULL |
| 11 | 119672586 | rs61897703 | T | G | M_Disease | 0.651354865 | M_Disease&NULL |
| 11 | 118839610 | rs113611963 | G | T | M_Disease | 0.651354865 | M_Disease&NULL |
| 11 | 117282979 | rs521099 | G | C | M_Disease | 0.650218549 | M_Disease&NULL |
| 11 | 119922006 | rs4420286 | A | G | M_Disease | 0.650218549 | M_Disease&NULL |
| 11 | 117441594 | rs79770727 | A | G | M_Disease | 0.650218549 | M_Disease&NULL |
| 11 | 119936017 | rs641924 | T | C | M_Disease | 0.650218549 | M_Disease&NULL |
| 11 | 119921117 | rs1452843 | T | C | M_Disease | 0.649080524 | M_Disease&NULL |
| 11 | 117531961 | rs594306 | A | T | M_Disease | 0.649080524 | M_Disease&NULL |
| 11 | 116493633 | rs12290281 | T | G | M_Disease | 0.649080524 | M_Disease&NULL |
| 11 | 116512477 | rs4936358 | A | G | M_Disease | 0.647940802 | M_Disease&NULL |
| 11 | 116290501 | rs58566212 | T | C | M_Disease | 0.645656306 | M_Disease&NULL |
| 11 | 117363066 | rs7119734 | T | C | M_Disease | 0.644511554 | M_Disease&NULL |
| 11 | 116583103 | rs2010611 | A | G | M_Disease | 0.644511554 | M_Disease&NULL |
| 11 | 120861279 | rs478015 | G | T | M_Disease | 0.643365146 | M_Disease&NULL |
| 11 | 117350621 | rs58080742 | A | C | M_Disease | 0.643365146 | M_Disease&NULL |
| 11 | 120861145 | rs596869 | T | A | M_Disease | 0.642217093 | M_Disease&NULL |
| 11 | 117091609 | rs236918 | C | G | M_Disease | 0.642217093 | M_Disease&NULL |
| 11 | 117284577 | rs2445936 | C | T | M_Disease | 0.641067406 | M_Disease&NULL |
| 11 | 116587953 | rs76542665 | A | C | M_Disease | 0.642217093 | M_Disease&NULL |
| 11 | 117356276 | rs12574158 | G | C | M_Disease | 0.641067406 | M_Disease&NULL |
| 11 | 116480266 | rs12282572 | C | T | M_Disease | 0.641067406 | M_Disease&NULL |
| 11 | 117062183 | rs1242230 | T | C | M_Disease | 0.641067406 | M_Disease&NULL |
| 11 | 117062370 | rs1242229 | T | C | M_Disease | 0.641067406 | M_Disease&NULL |
| 11 | 120860814 | rs598582 | G | A | M_Disease | 0.639916097 | M_Disease&NULL |
| 11 | 117377518 | rs7123652 | C | T | M_Disease | 0.639916097 | M_Disease&NULL |
| 11 | 120413124 | rs2266133 | C | T | M_Disease | 0.638763175 | M_Disease&NULL |
| 11 | 120860200 | rs536333 | C | T | M_Disease | 0.638763175 | M_Disease&NULL |
| 11 | 120860288 | rs611065 | C | A | M_Disease | 0.638763175 | M_Disease&NULL |

| | | | | | | | | | |
|---|---|---|---|---|---|---|---|---|---|
| 11 | 117404312 | rs56198555 | T | C | 0.873 | -7977.388 | -7975.903 | 1.485 | 1.71E-05 |
| 11 | 119213782 | rs948411 | T | A | 1.11 | -7977.388 | -7975.908 | 1.481 | 1.70E-05 |
| 11 | 119529332 | rs4245201 | C | T | 1.093 | -7977.388 | -7975.909 | 1.479 | 1.70E-05 |
| 11 | 117024481 | rs7940310 | T | C | 0.895 | -7977.388 | -7975.911 | 1.478 | 1.70E-05 |
| 11 | 117081500 | rs11216316 | C | A | 0.894 | -7977.388 | -7975.923 | 1.465 | 1.67E-05 |
| 11 | 120850004 | rs1334757891 | T | C | 0.926 | -7977.388 | -7975.936 | 1.452 | 1.65E-05 |
| 11 | 117045335 | rs7925256 | T | C | 0.888 | -7977.388 | -7975.94 | 1.449 | 1.65E-05 |
| 11 | 116967666 | rs1320668 | C | T | 0.896 | -7977.388 | -7975.943 | 1.446 | 1.64E-05 |
| 11 | 117027755 | rs7123517 | G | A | 0.895 | -7977.388 | -7975.946 | 1.442 | 1.64E-05 |
| 11 | 117405907 | rs68096541 | C | G | 1.088 | -7977.388 | -7975.954 | 1.434 | 1.62E-05 |
| 11 | 118662045 | rs7933859 | T | C | 0.9 | -7977.388 | -7975.954 | 1.434 | 1.62E-05 |
| 11 | 118765267 | rs598207 | C | G | 1.087 | -7977.388 | -7975.955 | 1.433 | 1.62E-05 |
| 11 | 119678767 | rs73577182 | C | G | 0.849 | -7977.388 | -7975.958 | 1.43 | 1.62E-05 |
| 11 | 118833458 | rs111491960 | G | A | 0.902 | -7977.388 | -7975.963 | 1.425 | 1.61E-05 |
| 11 | 116516966 | rs1176456955 | A | G | 0.824 | -7977.388 | -7975.965 | 1.423 | 1.61E-05 |
| 11 | 118662203 | rs4508234 | G | C | 0.9 | -7977.388 | -7975.969 | 1.419 | 1.60E-05 |
| 11 | 117534083 | rs7111427 | A | G | 0.922 | -7977.388 | -7975.972 | 1.416 | 1.59E-05 |
| 11 | 117363891 | rs4096589 | T | A | 1.079 | -7977.388 | -7975.978 | 1.41 | 1.58E-05 |
| 11 | 118662340 | rs5006102 | A | T | 0.901 | -7977.388 | -7975.983 | 1.406 | 1.58E-05 |
| 11 | 118230359 | rs2276419 | T | C | 1.117 | -7977.388 | -7975.986 | 1.402 | 1.57E-05 |
| 11 | 120848685 | rs75871913 | A | G | 0.836 | -7977.388 | -7975.989 | 1.399 | 1.57E-05 |
| 11 | 117042377 | rs4936367 | G | A | 0.896 | -7977.388 | -7975.992 | 1.396 | 1.56E-05 |
| 11 | 117044203 | rs4604928 | C | A | 0.896 | -7977.388 | -7975.992 | 1.396 | 1.56E-05 |
| 11 | 117458673 | rs59607164 | G | A | 1.124 | -7977.388 | -7976.001 | 1.388 | 1.55E-05 |
| 11 | 119203359 | rs2511847 | A | G | 1.108 | -7977.388 | -7976 | 1.388 | 1.55E-05 |
| 11 | 116697081 | rs150555490 | T | C | 1.152 | -7977.388 | -7976.007 | 1.381 | 1.54E-05 |
| 11 | 117034619 | rs7122944 | G | T | 0.896 | -7977.388 | -7976.008 | 1.381 | 1.54E-05 |
| 11 | 117262598 | rs542821 | T | C | 0.892 | -7977.388 | -7976.009 | 1.379 | 1.54E-05 |
| 11 | 117329630 | rs117043085 | T | C | 0.808 | -7977.388 | -7976.013 | 1.375 | 1.53E-05 |
| 11 | 117024302 | rs7950633 | C | T | 0.897 | -7977.388 | -7976.017 | 1.371 | 1.52E-05 |
| 11 | 117102648 | rs2252145 | C | T | 0.918 | -7977.388 | -7976.017 | 1.371 | 1.52E-05 |
| 11 | 117531465 | rs1892918 | C | A | 1.088 | -7977.388 | -7976.025 | 1.363 | 1.51E-05 |
| 11 | 118669178 | rs583213 | G | T | 0.902 | -7977.388 | -7976.033 | 1.356 | 1.50E-05 |
| 11 | 120702916 | rs12283648 | C | A | 1.097 | -7977.388 | -7976.034 | 1.354 | 1.50E-05 |
| 11 | 116485201 | rs71480304 | A | G | 0.879 | -7977.388 | -7976.04 | 1.348 | 1.49E-05 |
| 11 | 117368182 | rs73008413 | T | A | 0.849 | -7977.388 | -7976.04 | 1.348 | 1.49E-05 |
| 11 | 120795888 | rs949054 | C | T | 0.917 | -7977.388 | -7976.04 | 1.348 | 1.49E-05 |
| 11 | 116488748 | rs4360730 | C | T | 0.879 | -7977.388 | -7976.041 | 1.347 | 1.49E-05 |
| 11 | 116488753 | rs4565932 | A | G | 0.879 | -7977.388 | -7976.041 | 1.347 | 1.49E-05 |
| 11 | 118617920 | rs476246 | T | C | 1.081 | -7977.388 | -7976.045 | 1.343 | 1.48E-05 |
| 11 | 117214617 | rs144750609 | C | T | 0.83 | -7977.388 | -7976.049 | 1.34 | 1.48E-05 |
| 11 | 117285699 | rs11826012 | C | T | 0.893 | -7977.388 | -7976.055 | 1.334 | 1.47E-05 |
| 11 | 117285746 | rs1789157 | G | A | 0.893 | -7977.388 | -7976.055 | 1.334 | 1.47E-05 |
| 11 | 118617544 | rs472641 | G | A | 1.081 | -7977.388 | -7976.054 | 1.334 | 1.47E-05 |
| 11 | 118621495 | rs487728 | G | C | 1.081 | -7977.388 | -7976.055 | 1.334 | 1.47E-05 |
| 11 | 118705753 | rs556437 | A | G | 0.929 | -7977.388 | -7976.064 | 1.324 | 1.45E-05 |
| 11 | 116706168 | rs12721026 | G | T | 1.141 | -7977.388 | -7976.072 | 1.316 | 1.44E-05 |
| 11 | 117368982 | rs2004927 | C | T | 0.85 | -7977.388 | -7976.076 | 1.312 | 1.44E-05 |
| 11 | 119529106 | rs931954 | A | G | 1.09 | -7977.388 | -7976.08 | 1.308 | 1.43E-05 |
| 11 | 117366973 | rs7130068 | C | T | 1.126 | -7977.388 | -7976.082 | 1.306 | 1.43E-05 |
| 11 | 120858914 | rs670413 | A | G | 0.927 | -7977.388 | -7976.082 | 1.306 | 1.43E-05 |
| 11 | 117537710 | rs686413 | A | G | 1.089 | -7977.388 | -7976.085 | 1.303 | 1.42E-05 |
| 11 | 117367196 | rs73008408 | C | A | 0.851 | -7977.388 | -7976.087 | 1.301 | 1.42E-05 |
| 11 | 116548356 | rs117339323 | C | T | 0.829 | -7977.388 | -7976.089 | 1.3 | 1.42E-05 |
| 11 | 117803083 | rs117978531 | G | T | 0.827 | -7977.388 | -7976.089 | 1.299 | 1.42E-05 |
| 11 | 118626511 | rs499343 | T | C | 1.08 | -7977.388 | -7976.096 | 1.293 | 1.41E-05 |
| 11 | 117369226 | rs1807116 | C | T | 0.851 | -7977.388 | -7976.102 | 1.286 | 1.40E-05 |
| 11 | 120858301 | rs74727906 | A | G | 0.837 | -7977.388 | -7976.11 | 1.279 | 1.39E-05 |
| 11 | 117026374 | rs2127905 | A | C | 0.899 | -7977.388 | -7976.113 | 1.275 | 1.38E-05 |
| 11 | 119918191 | rs1942010 | C | T | 0.928 | -7977.388 | -7976.117 | 1.271 | 1.38E-05 |
| 11 | 119675331 | rs61897704 | T | C | 0.857 | -7977.388 | -7976.119 | 1.269 | 1.38E-05 |
| 11 | 119675529 | rs61897705 | G | A | 0.857 | -7977.388 | -7976.119 | 1.269 | 1.38E-05 |
| 11 | 117283471 | rs573965 | T | C | 0.895 | -7977.388 | -7976.121 | 1.268 | 1.37E-05 |
| 11 | 117405696 | rs2076965 | A | T | 1.121 | -7977.388 | -7976.122 | 1.267 | 1.37E-05 |
| 11 | 116281175 | rs76887844 | T | C | 1.139 | -7977.388 | -7976.124 | 1.264 | 1.37E-05 |
| 11 | 116289906 | rs11215924 | A | T | 1.11 | -7977.388 | -7976.13 | 1.258 | 1.36E-05 |
| 11 | 118624767 | rs581062 | A | G | 1.08 | -7977.388 | -7976.13 | 1.258 | 1.36E-05 |
| 11 | 120988038 | rs140567534 | G | T | 1.193 | -7977.388 | -7976.137 | 1.251 | 1.35E-05 |
| 11 | 119672586 | rs61897703 | T | G | 0.865 | -7977.388 | -7976.141 | 1.247 | 1.35E-05 |
| 11 | 118839610 | rs113611963 | G | T | 0.905 | -7977.388 | -7976.146 | 1.243 | 1.34E-05 |
| 11 | 117282979 | rs521099 | G | C | 0.895 | -7977.388 | -7976.149 | 1.24 | 1.34E-05 |
| 11 | 119922006 | rs4420286 | A | G | 0.928 | -7977.388 | -7976.149 | 1.24 | 1.34E-05 |
| 11 | 117441594 | rs79770727 | A | G | 1.12 | -7977.388 | -7976.157 | 1.232 | 1.33E-05 |
| 11 | 119936017 | rs641924 | T | C | 0.867 | -7977.388 | -7976.156 | 1.232 | 1.33E-05 |
| 11 | 119921117 | rs1452843 | T | C | 0.929 | -7977.388 | -7976.159 | 1.229 | 1.32E-05 |
| 11 | 117531961 | rs594306 | A | T | 1.085 | -7977.388 | -7976.163 | 1.226 | 1.32E-05 |
| 11 | 116493633 | rs12290281 | T | G | 0.867 | -7977.388 | -7976.167 | 1.222 | 1.31E-05 |
| 11 | 116512477 | rs4936358 | A | G | 1.079 | -7977.388 | -7976.174 | 1.214 | 1.30E-05 |
| 11 | 116290501 | rs58566212 | T | C | 1.108 | -7977.388 | -7976.196 | 1.192 | 1.27E-05 |
| 11 | 117363066 | rs7119734 | T | C | 1.088 | -7977.388 | -7976.201 | 1.187 | 1.27E-05 |
| 11 | 116583103 | rs2010611 | A | G | 0.846 | -7977.388 | -7976.203 | 1.185 | 1.27E-05 |
| 11 | 120861279 | rs478015 | G | T | 0.929 | -7977.388 | -7976.212 | 1.176 | 1.25E-05 |
| 11 | 117350621 | rs58080742 | A | C | 1.089 | -7977.388 | -7976.213 | 1.175 | 1.25E-05 |
| 11 | 120861145 | rs596869 | T | A | 0.929 | -7977.388 | -7976.222 | 1.166 | 1.24E-05 |
| 11 | 117091609 | rs236918 | C | G | 0.899 | -7977.388 | -7976.223 | 1.165 | 1.24E-05 |
| 11 | 117284577 | rs2445936 | C | T | 0.896 | -7977.388 | -7976.228 | 1.16 | 1.23E-05 |
| 11 | 116587953 | rs76542665 | A | C | 0.847 | -7977.388 | -7976.229 | 1.159 | 1.23E-05 |
| 11 | 117356276 | rs12574158 | G | C | 1.104 | -7977.388 | -7976.231 | 1.157 | 1.23E-05 |
| 11 | 116480266 | rs12282572 | C | T | 0.888 | -7977.388 | -7976.233 | 1.155 | 1.23E-05 |
| 11 | 117062183 | rs1242230 | T | C | 0.9 | -7977.388 | -7976.236 | 1.152 | 1.22E-05 |
| 11 | 117062370 | rs1242229 | T | C | 0.9 | -7977.388 | -7976.236 | 1.152 | 1.22E-05 |
| 11 | 120860814 | rs598582 | G | A | 0.929 | -7977.388 | -7976.239 | 1.149 | 1.22E-05 |
| 11 | 117377518 | rs7123652 | C | T | 1.086 | -7977.388 | -7976.244 | 1.144 | 1.21E-05 |
| 11 | 120413124 | rs2266133 | C | T | 0.927 | -7977.388 | -7976.247 | 1.141 | 1.21E-05 |
| 11 | 120860200 | rs536333 | C | T | 0.929 | -7977.388 | -7976.249 | 1.14 | 1.21E-05 |
| 11 | 120860288 | rs611065 | C | A | 0.929 | -7977.388 | -7976.249 | 1.14 | 1.21E-05 |

| | | | | | | | |
|---|---|---|---|---|---|---|---|
| 11 | 120860499 | rs560092 | G | A | M_Disease | 0.638763175 | M_Disease&NULL |
| 11 | 120861094 | rs565597 | C | T | M_Disease | 0.638763175 | M_Disease&NULL |
| 11 | 117294973 | rs4938375 | C | T | M_Disease | 0.638763175 | M_Disease&NULL |
| 11 | 117266466 | rs483837 | C | T | M_Disease | 0.637608653 | M_Disease&NULL |
| 11 | 117394979 | rs12577702 | C | T | M_Disease | 0.63645254 | M_Disease&NULL |
| 11 | 116587889 | rs17119920 | G | C | M_Disease | 0.63645254 | M_Disease&NULL |
| 11 | 117521967 | rs16867 | C | T | M_Disease | 0.635294849 | M_Disease&NULL |
| 11 | 121079940 | rs78733282 | G | A | M_Disease | 0.63645254 | M_Disease&NULL |
| 11 | 116197149 | rs12789310 | G | A | M_Disease | 0.635294849 | M_Disease&NULL |
| 11 | 116966435 | rs900012 | G | A | M_Disease | 0.635294849 | M_Disease&NULL |
| 11 | 119913501 | rs656015 | A | G | M_Disease | 0.634135591 | M_Disease&NULL |
| 11 | 116921390 | rs1815786 | T | C | M_Disease | 0.632974777 | M_Disease&NULL |
| 11 | 117284770 | rs540904 | C | T | M_Disease | 0.632974777 | M_Disease&NULL |
| 11 | 116202789 | rs12806755 | C | T | M_Disease | 0.632974777 | M_Disease&NULL |
| 11 | 120861385 | rs595851 | A | G | M_Disease | 0.632974777 | M_Disease&NULL |
| 11 | 117294959 | rs4936376 | T | C | M_Disease | 0.631812418 | M_Disease&NULL |
| 11 | 117283626 | rs572884 | G | A | M_Disease | 0.629483112 | M_Disease&NULL |
| 11 | 117283634 | rs572862 | G | A | M_Disease | 0.629483112 | M_Disease&NULL |
| 11 | 117258662 | rs574813 | G | C | M_Disease | 0.629483112 | M_Disease&NULL |
| 11 | 117258687 | rs574873 | T | G | M_Disease | 0.629483112 | M_Disease&NULL |
| 11 | 118700557 | rs4291686 | G | A | M_Disease | 0.629483112 | M_Disease&NULL |
| 11 | 117370831 | rs58949729 | T | G | M_Disease | 0.628316188 | M_Disease&NULL |
| 11 | 117295431 | rs4938376 | G | A | M_Disease | 0.628316188 | M_Disease&NULL |
| 11 | 117003459 | rs17120373 | G | A | M_Disease | 0.627147766 | M_Disease&NULL |
| 11 | 117461888 | rs61213686 | C | T | M_Disease | 0.627147766 | M_Disease&NULL |
| 11 | 117561256 | rs114797458 | A | G | M_Disease | 0.628316188 | M_Disease&NULL |
| 11 | 120863568 | rs607800 | T | C | M_Disease | 0.625977858 | M_Disease&NULL |
| 11 | 117489645 | rs685184 | T | C | M_Disease | 0.625977858 | M_Disease&NULL |
| 11 | 117261853 | rs494553 | T | C | M_Disease | 0.624806474 | M_Disease&NULL |
| 11 | 119853138 | rs17123519 | T | C | M_Disease | 0.624806474 | M_Disease&NULL |
| 11 | 118699276 | rs625274 | T | C | M_Disease | 0.623633628 | M_Disease&NULL |
| 11 | 117296160 | rs10750105 | A | T | M_Disease | 0.624806474 | M_Disease&NULL |
| 11 | 117368358 | rs73008415 | T | C | M_Disease | 0.624806474 | M_Disease&NULL |
| 11 | 117288442 | rs579274 | C | T | M_Disease | 0.623633628 | M_Disease&NULL |
| 11 | 116409093 | rs10502220 | C | T | M_Disease | 0.623633628 | M_Disease&NULL |
| 11 | 117264256 | rs506366 | A | G | M_Disease | 0.622459331 | M_Disease&NULL |
| 11 | 116325337 | rs11215942 | A | G | M_Disease | 0.623633628 | M_Disease&NULL |
| 11 | 116749977 | rs6589577 | C | T | M_Disease | 0.622459331 | M_Disease&NULL |
| 11 | 117937348 | rs61900593 | G | A | M_Disease | 0.622459331 | M_Disease&NULL |
| 11 | 117294706 | rs4420302 | A | G | M_Disease | 0.622459331 | M_Disease&NULL |
| 11 | 116752219 | rs7124741 | A | T | M_Disease | 0.621283595 | M_Disease&NULL |
| 11 | 117285145 | rs1279744828 | G | A | M_Disease | 0.621283595 | M_Disease&NULL |
| 11 | 116478827 | rs12794425 | A | G | M_Disease | 0.621283595 | M_Disease&NULL |
| 11 | 116292287 | rs77544215 | T | C | M_Disease | 0.621283595 | M_Disease&NULL |
| 11 | 116774201 | rs17120139 | A | G | M_Disease | 0.620106432 | M_Disease&NULL |
| 11 | 116478904 | rs12794476 | A | G | M_Disease | 0.620106432 | M_Disease&NULL |
| 11 | 119880519 | rs593252 | A | G | M_Disease | 0.620106432 | M_Disease&NULL |
| 11 | 116932428 | rs6589589 | T | C | M_Disease | 0.620106432 | M_Disease&NULL |
| 11 | 119208576 | rs2509656 | A | G | M_Disease | 0.618927855 | M_Disease&NULL |
| 11 | 116943354 | rs1351452 | A | C | M_Disease | 0.618927855 | M_Disease&NULL |
| 11 | 117291696 | rs1902624 | T | C | M_Disease | 0.618927855 | M_Disease&NULL |
| 11 | 117276783 | rs485570 | G | T | M_Disease | 0.618927855 | M_Disease&NULL |
| 11 | 116941555 | rs151131966 | A | G | M_Disease | 0.620106432 | M_Disease&NULL |
| 11 | 117192383 | rs516742 | G | A | M_Disease | 0.618927855 | M_Disease&NULL |
| 11 | 116908283 | rs7115242 | A | G | M_Disease | 0.618927855 | M_Disease&NULL |
| 11 | 117366280 | rs73008405 | C | G | M_Disease | 0.618927855 | M_Disease&NULL |
| 11 | 117293516 | rs10790183 | A | G | M_Disease | 0.617747875 | M_Disease&NULL |
| 11 | 116957907 | rs6589592 | C | G | M_Disease | 0.617747875 | M_Disease&NULL |
| 11 | 121005609 | rs7128792 | G | A | M_Disease | 0.617747875 | M_Disease&NULL |
| 11 | 116291283 | rs78277728 | G | C | M_Disease | 0.617747875 | M_Disease&NULL |
| 11 | 116296917 | rs79663212 | T | G | M_Disease | 0.616566505 | M_Disease&NULL |
| 11 | 117369450 | rs1075784 | G | C | M_Disease | 0.617747875 | M_Disease&NULL |
| 11 | 116947361 | rs4938335 | A | C | M_Disease | 0.616566505 | M_Disease&NULL |
| 11 | 116950232 | rs10736487 | A | G | M_Disease | 0.616566505 | M_Disease&NULL |
| 11 | 117558945 | rs74778631 | A | G | M_Disease | 0.617747875 | M_Disease&NULL |
| 11 | 118812559 | rs74678032 | C | T | M_Disease | 0.616566505 | M_Disease&NULL |
| 11 | 117295791 | rs7925881 | A | G | M_Disease | 0.616566505 | M_Disease&NULL |
| 11 | 119008125 | rs78210954 | C | G | M_Disease | 0.615383756 | M_Disease&NULL |
| 11 | 117075566 | rs508487 | T | C | M_Disease | 0.615383756 | M_Disease&NULL |
| 11 | 117936239 | rs55965073 | C | T | M_Disease | 0.615383756 | M_Disease&NULL |
| 11 | 116290834 | rs11215925 | C | T | M_Disease | 0.615383756 | M_Disease&NULL |
| 11 | 118220035 | rs2282550 | C | T | M_Disease | 0.615383756 | M_Disease&NULL |
| 11 | 118640063 | rs524590 | T | A | M_Disease | 0.615383756 | M_Disease&NULL |
| 11 | 118652326 | rs1790189 | G | T | M_Disease | 0.614199643 | M_Disease&NULL |
| 11 | 118653629 | rs540180 | T | G | M_Disease | 0.614199643 | M_Disease&NULL |
| 11 | 118654651 | rs2508916 | G | C | M_Disease | 0.614199643 | M_Disease&NULL |
| 11 | 116798963 | rs579890 | A | G | M_Disease | 0.614199643 | M_Disease&NULL |
| 11 | 116812029 | rs473188 | C | G | M_Disease | 0.614199643 | M_Disease&NULL |
| 11 | 120032251 | rs78399857 | T | C | M_Disease | 0.615383756 | M_Disease&NULL |
| 11 | 116693353 | rs2239013 | T | C | M_Disease | 0.614199643 | M_Disease&NULL |
| 11 | 117368846 | rs2004926 | G | A | M_Disease | 0.614199643 | M_Disease&NULL |
| 11 | 118221474 | rs1561966 | A | G | M_Disease | 0.614199643 | M_Disease&NULL |
| 11 | 118652373 | rs1784293 | C | A | M_Disease | 0.614199643 | M_Disease&NULL |
| 11 | 116705719 | rs12718462 | C | T | M_Disease | 0.613014176 | M_Disease&NULL |
| 11 | 120860238 | rs536447 | T | C | M_Disease | 0.613014176 | M_Disease&NULL |
| 11 | 116771375 | rs72645460 | G | A | M_Disease | 0.613014176 | M_Disease&NULL |
| 11 | 117268256 | rs569845 | T | G | M_Disease | 0.613014176 | M_Disease&NULL |
| 11 | 117367574 | rs73008411 | G | A | M_Disease | 0.613014176 | M_Disease&NULL |
| 11 | 118038697 | rs648389 | C | T | M_Disease | 0.613014176 | M_Disease&NULL |
| 11 | 117275815 | rs546617 | A | G | M_Disease | 0.613014176 | M_Disease&NULL |
| 11 | 118995325 | rs7123630 | T | C | M_Disease | 0.613014176 | M_Disease&NULL |
| 11 | 116550916 | rs193204541 | G | C | M_Disease | 0.613014176 | M_Disease&NULL |
| 11 | 116927740 | rs10790172 | G | T | M_Disease | 0.611827369 | M_Disease&NULL |
| 11 | 117293741 | rs3863308 | G | T | M_Disease | 0.610639234 | M_Disease&NULL |
| 11 | 116907113 | rs10732856 | T | C | M_Disease | 0.610639234 | M_Disease&NULL |
| 11 | 117273094 | rs568984 | T | G | M_Disease | 0.610639234 | M_Disease&NULL |

| | | | | | | | | | |
|---|---|---|---|---|---|---|---|---|---|
| 11 | 120860499 | rs560092 | G | A | 0.929 | -7977.388 | -7976.249 | 1.14 | 1.21E-05 |
| 11 | 120861094 | rs565597 | C | T | 0.929 | -7977.388 | -7976.25 | 1.139 | 1.21E-05 |
| 11 | 117294973 | rs4938375 | C | T | 0.878 | -7977.388 | -7976.257 | 1.132 | 1.20E-05 |
| 11 | 117266466 | rs483837 | C | T | 0.879 | -7977.388 | -7976.265 | 1.123 | 1.19E-05 |
| 11 | 117394979 | rs12577702 | C | T | 0.881 | -7977.388 | -7976.276 | 1.113 | 1.18E-05 |
| 11 | 116587889 | rs17119920 | G | C | 0.849 | -7977.388 | -7976.278 | 1.111 | 1.17E-05 |
| 11 | 117521967 | rs16867 | C | T | 1.091 | -7977.388 | -7976.282 | 1.106 | 1.17E-05 |
| 11 | 121079940 | rs78733282 | G | A | 0.832 | -7977.388 | -7976.283 | 1.105 | 1.17E-05 |
| 11 | 116197149 | rs12789310 | G | A | 0.88 | -7977.388 | -7976.284 | 1.104 | 1.17E-05 |
| 11 | 116966435 | rs900012 | G | A | 0.902 | -7977.388 | -7976.285 | 1.103 | 1.17E-05 |
| 11 | 119913501 | rs656015 | A | G | 0.93 | -7977.388 | -7976.29 | 1.098 | 1.16E-05 |
| 11 | 116921390 | rs1815786 | T | C | 0.897 | -7977.388 | -7976.302 | 1.087 | 1.15E-05 |
| 11 | 117284770 | rs540904 | C | T | 0.898 | -7977.388 | -7976.302 | 1.086 | 1.15E-05 |
| 11 | 116202789 | rs12806755 | C | T | 0.881 | -7977.388 | -7976.304 | 1.084 | 1.14E-05 |
| 11 | 120861385 | rs595851 | A | G | 0.93 | -7977.388 | -7976.304 | 1.084 | 1.14E-05 |
| 11 | 117294959 | rs4936376 | T | C | 0.88 | -7977.388 | -7976.314 | 1.075 | 1.13E-05 |
| 11 | 117283626 | rs572884 | G | A | 0.898 | -7977.388 | -7976.332 | 1.057 | 1.11E-05 |
| 11 | 117283634 | rs572862 | G | A | 0.898 | -7977.388 | -7976.332 | 1.057 | 1.11E-05 |
| 11 | 117258662 | rs574813 | G | C | 0.881 | -7977.388 | -7976.334 | 1.054 | 1.11E-05 |
| 11 | 117258687 | rs574873 | T | G | 0.881 | -7977.388 | -7976.334 | 1.054 | 1.11E-05 |
| 11 | 118700557 | rs4291686 | G | A | 0.931 | -7977.388 | -7976.334 | 1.054 | 1.11E-05 |
| 11 | 117370831 | rs58949729 | T | G | 1.119 | -7977.388 | -7976.347 | 1.042 | 1.10E-05 |
| 11 | 117295431 | rs4938376 | G | A | 0.88 | -7977.388 | -7976.348 | 1.041 | 1.10E-05 |
| 11 | 117003459 | rs17120373 | G | A | 1.117 | -7977.388 | -7976.35 | 1.038 | 1.09E-05 |
| 11 | 117461888 | rs61213686 | C | T | 1.115 | -7977.388 | -7976.351 | 1.038 | 1.09E-05 |
| 11 | 117561256 | rs114797458 | A | G | 0.83 | -7977.388 | -7976.351 | 1.038 | 1.09E-05 |
| 11 | 120863568 | rs607800 | T | C | 0.93 | -7977.388 | -7976.359 | 1.03 | 1.08E-05 |
| 11 | 117489645 | rs685184 | T | C | 0.914 | -7977.388 | -7976.364 | 1.025 | 1.08E-05 |
| 11 | 117261853 | rs494553 | T | C | 0.882 | -7977.388 | -7976.369 | 1.019 | 1.07E-05 |
| 11 | 119853138 | rs17123519 | T | C | 0.86 | -7977.388 | -7976.373 | 1.015 | 1.07E-05 |
| 11 | 118699276 | rs625274 | T | C | 0.932 | -7977.388 | -7976.377 | 1.012 | 1.06E-05 |
| 11 | 117296160 | rs10750105 | A | T | 0.881 | -7977.388 | -7976.377 | 1.011 | 1.06E-05 |
| 11 | 117368358 | rs73008415 | T | C | 0.847 | -7977.388 | -7976.38 | 1.009 | 1.06E-05 |
| 11 | 117288442 | rs579274 | C | T | 0.882 | -7977.388 | -7976.383 | 1.005 | 1.06E-05 |
| 11 | 116409093 | rs10502220 | C | T | 1.092 | -7977.388 | -7976.385 | 1.003 | 1.05E-05 |
| 11 | 117264256 | rs506366 | A | G | 0.882 | -7977.388 | -7976.389 | 1 | 1.05E-05 |
| 11 | 116325337 | rs11215942 | A | G | 0.875 | -7977.388 | -7976.389 | 0.999 | 1.05E-05 |
| 11 | 116749977 | rs6589577 | C | T | 0.903 | -7977.388 | -7976.39 | 0.998 | 1.05E-05 |
| 11 | 117937348 | rs61900593 | G | A | 0.93 | -7977.388 | -7976.391 | 0.997 | 1.05E-05 |
| 11 | 117294706 | rs4420302 | A | G | 0.882 | -7977.388 | -7976.397 | 0.991 | 1.04E-05 |
| 11 | 116752219 | rs7124741 | A | T | 0.903 | -7977.388 | -7976.399 | 0.989 | 1.04E-05 |
| 11 | 117285145 | rs1279744828 | G | A | 0.883 | -7977.388 | -7976.402 | 0.986 | 1.04E-05 |
| 11 | 116478827 | rs12794425 | A | G | 0.891 | -7977.388 | -7976.404 | 0.984 | 1.03E-05 |
| 11 | 116292287 | rs77544215 | T | C | 1.13 | -7977.388 | -7976.405 | 0.983 | 1.03E-05 |
| 11 | 116774201 | rs17120139 | A | G | 0.903 | -7977.388 | -7976.408 | 0.98 | 1.03E-05 |
| 11 | 116478904 | rs12794476 | A | G | 0.891 | -7977.388 | -7976.412 | 0.977 | 1.03E-05 |
| 11 | 119880519 | rs593252 | A | G | 0.881 | -7977.388 | -7976.414 | 0.974 | 1.02E-05 |
| 11 | 116932428 | rs6589589 | T | C | 0.905 | -7977.388 | -7976.415 | 0.973 | 1.02E-05 |
| 11 | 119208576 | rs2509656 | A | G | 1.097 | -7977.388 | -7976.418 | 0.97 | 1.02E-05 |
| 11 | 116943354 | rs1351452 | A | C | 0.905 | -7977.388 | -7976.419 | 0.969 | 1.02E-05 |
| 11 | 117291696 | rs1902624 | T | C | 0.882 | -7977.388 | -7976.42 | 0.969 | 1.02E-05 |
| 11 | 117276783 | rs485570 | G | T | 0.883 | -7977.388 | -7976.42 | 0.968 | 1.02E-05 |
| 11 | 116941555 | rs151131966 | A | G | 1.198 | -7977.388 | -7976.424 | 0.964 | 1.01E-05 |
| 11 | 117192383 | rs516742 | G | A | 0.911 | -7977.388 | -7976.425 | 0.963 | 1.01E-05 |
| 11 | 116908283 | rs7115242 | A | G | 0.9 | -7977.388 | -7976.426 | 0.962 | 1.01E-05 |
| 11 | 117366280 | rs73008405 | C | G | 0.849 | -7977.388 | -7976.431 | 0.958 | 1.01E-05 |
| 11 | 117293516 | rs10790183 | A | G | 0.883 | -7977.388 | -7976.433 | 0.955 | 1.01E-05 |
| 11 | 116957907 | rs6589592 | C | G | 0.9 | -7977.388 | -7976.435 | 0.954 | 1.00E-05 |
| 11 | 121005609 | rs7128792 | G | A | 0.922 | -7977.388 | -7976.434 | 0.954 | 1.00E-05 |
| 11 | 116291283 | rs78277728 | G | C | 1.129 | -7977.388 | -7976.436 | 0.952 | 1.00E-05 |
| 11 | 116296917 | rs79663212 | T | G | 1.128 | -7977.388 | -7976.439 | 0.95 | 1.00E-05 |
| 11 | 117369450 | rs1075784 | G | C | 0.849 | -7977.388 | -7976.438 | 0.95 | 1.00E-05 |
| 11 | 116947361 | rs4938335 | A | C | 0.905 | -7977.388 | -7976.441 | 0.947 | 9.97E-06 |
| 11 | 116950232 | rs10736487 | A | G | 0.905 | -7977.388 | -7976.441 | 0.947 | 9.97E-06 |
| 11 | 117558945 | rs74778631 | A | G | 0.838 | -7977.388 | -7976.442 | 0.946 | 9.96E-06 |
| 11 | 118812559 | rs74678032 | C | T | 0.857 | -7977.388 | -7976.445 | 0.944 | 9.94E-06 |
| 11 | 117295791 | rs7925881 | A | G | 0.883 | -7977.388 | -7976.446 | 0.943 | 9.93E-06 |
| 11 | 119008125 | rs78210954 | C | G | 0.886 | -7977.388 | -7976.449 | 0.939 | 9.89E-06 |
| 11 | 117075566 | rs508487 | T | C | 0.886 | -7977.388 | -7976.451 | 0.937 | 9.87E-06 |
| 11 | 117936239 | rs55965073 | C | T | 0.93 | -7977.388 | -7976.452 | 0.937 | 9.87E-06 |
| 11 | 116290834 | rs11215925 | C | T | 1.129 | -7977.388 | -7976.453 | 0.935 | 9.85E-06 |
| 11 | 118220035 | rs2282550 | C | T | 1.077 | -7977.388 | -7976.455 | 0.934 | 9.84E-06 |
| 11 | 118640063 | rs524590 | T | A | 1.075 | -7977.388 | -7976.455 | 0.933 | 9.83E-06 |
| 11 | 118652326 | rs1790189 | G | T | 1.075 | -7977.388 | -7976.456 | 0.932 | 9.82E-06 |
| 11 | 118653629 | rs540180 | T | G | 1.075 | -7977.388 | -7976.456 | 0.932 | 9.82E-06 |
| 11 | 118654651 | rs2508916 | G | C | 1.075 | -7977.388 | -7976.456 | 0.932 | 9.82E-06 |
| 11 | 116798963 | rs579890 | A | G | 0.905 | -7977.388 | -7976.457 | 0.931 | 9.81E-06 |
| 11 | 116812029 | rs473188 | C | G | 0.905 | -7977.388 | -7976.458 | 0.931 | 9.81E-06 |
| 11 | 120032251 | rs78399857 | T | C | 0.871 | -7977.388 | -7976.457 | 0.931 | 9.81E-06 |
| 11 | 116693353 | rs2239013 | T | C | 1.131 | -7977.388 | -7976.462 | 0.926 | 9.76E-06 |
| 11 | 117368846 | rs2004926 | G | A | 0.85 | -7977.388 | -7976.464 | 0.925 | 9.75E-06 |
| 11 | 118221474 | rs1561966 | A | G | 1.077 | -7977.388 | -7976.463 | 0.925 | 9.75E-06 |
| 11 | 118652373 | rs1784293 | C | A | 1.075 | -7977.388 | -7976.464 | 0.924 | 9.75E-06 |
| 11 | 116705719 | rs12718462 | C | T | 1.13 | -7977.388 | -7976.469 | 0.92 | 9.71E-06 |
| 11 | 120860238 | rs536447 | T | C | 0.932 | -7977.388 | -7976.471 | 0.918 | 9.69E-06 |
| 11 | 116771375 | rs72645460 | G | A | 0.904 | -7977.388 | -7976.474 | 0.914 | 9.65E-06 |
| 11 | 117268256 | rs569845 | T | G | 0.884 | -7977.388 | -7976.474 | 0.914 | 9.65E-06 |
| 11 | 117367574 | rs73008411 | G | A | 0.851 | -7977.388 | -7976.474 | 0.914 | 9.65E-06 |
| 11 | 118038697 | rs648389 | C | T | 0.83 | -7977.388 | -7976.476 | 0.913 | 9.64E-06 |
| 11 | 117275815 | rs546617 | A | G | 0.885 | -7977.388 | -7976.477 | 0.912 | 9.63E-06 |
| 11 | 118995325 | rs7123630 | T | C | 0.89 | -7977.388 | -7976.478 | 0.911 | 9.62E-06 |
| 11 | 116550916 | rs193204541 | G | C | 1.189 | -7977.388 | -7976.48 | 0.908 | 9.59E-06 |
| 11 | 116927740 | rs10790172 | G | T | 0.906 | -7977.388 | -7976.48 | 0.908 | 9.59E-06 |
| 11 | 117293741 | rs3863308 | G | T | 0.884 | -7977.388 | -7976.489 | 0.9 | 9.51E-06 |
| 11 | 116907113 | rs10732856 | T | C | 0.906 | -7977.388 | -7976.489 | 0.899 | 9.50E-06 |
| 11 | 117273094 | rs568984 | T | G | 0.885 | -7977.388 | -7976.49 | 0.899 | 9.50E-06 |

| | | | | | | | |
|---|---|---|---|---|---|---|---|
| 11 | 119207341 | rs2248863 | A | G | M_Disease | 0.610639234 | M_Disease&NULL |
| 11 | 119207446 | rs2248853 | T | C | M_Disease | 0.610639234 | M_Disease&NULL |
| 11 | 120239808 | rs116850963 | G | A | M_Disease | 0.611827369 | M_Disease&NULL |
| 11 | 119209979 | rs9640 | A | T | M_Disease | 0.610639234 | M_Disease&NULL |
| 11 | 118583525 | rs10892260 | C | A | M_Disease | 0.610639234 | M_Disease&NULL |
| 11 | 117554152 | rs17121080 | A | G | M_Disease | 0.610639234 | M_Disease&NULL |
| 11 | 117368805 | rs524968 | A | G | M_Disease | 0.609449784 | M_Disease&NULL |
| 11 | 116764021 | rs672058 | C | T | M_Disease | 0.609449784 | M_Disease&NULL |
| 11 | 116915217 | rs2155583 | A | G | M_Disease | 0.609449784 | M_Disease&NULL |
| 11 | 116915819 | rs2000615 | C | T | M_Disease | 0.609449784 | M_Disease&NULL |
| 11 | 117182708 | rs525493 | G | T | M_Disease | 0.609449784 | M_Disease&NULL |
| 11 | 119820707 | rs75829086 | G | C | M_Disease | 0.609449784 | M_Disease&NULL |
| 11 | 117293929 | rs3863310 | C | T | M_Disease | 0.609449784 | M_Disease&NULL |
| 11 | 116819617 | rs1241658 | G | A | M_Disease | 0.608259031 | M_Disease&NULL |
| 11 | 117925812 | rs732403 | A | G | M_Disease | 0.608259031 | M_Disease&NULL |
| 11 | 117938377 | rs61900594 | T | C | M_Disease | 0.608259031 | M_Disease&NULL |
| 11 | 117205861 | rs487261 | T | C | M_Disease | 0.608259031 | M_Disease&NULL |
| 11 | 116923870 | rs7130297 | C | T | M_Disease | 0.608259031 | M_Disease&NULL |
| 11 | 117343197 | rs10892121 | C | T | M_Disease | 0.608259031 | M_Disease&NULL |
| 11 | 116483042 | rs34899605 | A | G | M_Disease | 0.608259031 | M_Disease&NULL |
| 11 | 116917244 | rs10750100 | T | C | M_Disease | 0.607066988 | M_Disease&NULL |
| 11 | 116921397 | rs1815787 | T | C | M_Disease | 0.607066988 | M_Disease&NULL |
| 11 | 118744615 | rs73009445 | A | G | M_Disease | 0.607066988 | M_Disease&NULL |
| 11 | 116813804 | rs510988 | C | T | M_Disease | 0.605873668 | M_Disease&NULL |
| 11 | 116833679 | rs493134 | C | T | M_Disease | 0.605873668 | M_Disease&NULL |
| 11 | 116915742 | rs2000614 | C | T | M_Disease | 0.605873668 | M_Disease&NULL |
| 11 | 116916529 | rs2000616 | C | T | M_Disease | 0.605873668 | M_Disease&NULL |
| 11 | 117179302 | rs663166 | A | G | M_Disease | 0.605873668 | M_Disease&NULL |
| 11 | 116898886 | rs7950364 | C | G | M_Disease | 0.604679085 | M_Disease&NULL |
| 11 | 117551810 | rs78462423 | A | G | M_Disease | 0.604679085 | M_Disease&NULL |
| 11 | 116730638 | rs6589574 | A | G | M_Disease | 0.60348325 | M_Disease&NULL |
| 11 | 117202782 | rs624555 | T | C | M_Disease | 0.60348325 | M_Disease&NULL |
| 11 | 119576357 | rs112791662 | A | G | M_Disease | 0.60348325 | M_Disease&NULL |
| 11 | 117294035 | rs3863311 | A | T | M_Disease | 0.60348325 | M_Disease&NULL |
| 11 | 118577873 | rs11216960 | A | G | M_Disease | 0.60348325 | M_Disease&NULL |
| 11 | 118578705 | rs10892256 | A | G | M_Disease | 0.60348325 | M_Disease&NULL |
| 11 | 117037080 | rs114287696 | G | A | M_Disease | 0.60348325 | M_Disease&NULL |
| 11 | 117489941 | rs683842 | C | G | M_Disease | 0.602286177 | M_Disease&NULL |
| 11 | 116409112 | rs7118201 | C | T | M_Disease | 0.602286177 | M_Disease&NULL |
| 11 | 117560048 | rs7103164 | A | G | M_Disease | 0.60348325 | M_Disease&NULL |
| 11 | 116951943 | rs117033797 | G | T | M_Disease | 0.602286177 | M_Disease&NULL |
| 11 | 116692324 | rs2234668 | A | G | M_Disease | 0.599888369 | M_Disease&NULL |
| 11 | 116735460 | rs17120099 | C | A | M_Disease | 0.599888369 | M_Disease&NULL |
| 11 | 120426966 | rs2248540 | C | T | M_Disease | 0.59868766 | M_Disease&NULL |
| 11 | 120732349 | rs12221530 | T | C | M_Disease | 0.59868766 | M_Disease&NULL |
| 11 | 117255386 | rs480120 | T | C | M_Disease | 0.599888369 | M_Disease&NULL |
| 11 | 117549963 | rs116672969 | A | G | M_Disease | 0.599888369 | M_Disease&NULL |
| 11 | 121018698 | rs34217740 | A | G | M_Disease | 0.59868766 | M_Disease&NULL |
| 11 | 117898446 | rs149421321 | T | C | M_Disease | 0.599888369 | M_Disease&NULL |
| 11 | 117291835 | rs1902626 | C | A | M_Disease | 0.59868766 | M_Disease&NULL |
| 11 | 118798154 | rs60969913 | C | G | M_Disease | 0.59868766 | M_Disease&NULL |
| 11 | 117381052 | rs634606 | T | C | M_Disease | 0.597485766 | M_Disease&NULL |
| 11 | 120418613 | rs2252244 | G | A | M_Disease | 0.597485766 | M_Disease&NULL |
| 11 | 117269269 | rs539573 | A | T | M_Disease | 0.597485766 | M_Disease&NULL |
| 11 | 116199019 | rs75676165 | A | G | M_Disease | 0.597485766 | M_Disease&NULL |
| 11 | 117353812 | rs55900121 | T | C | M_Disease | 0.596282699 | M_Disease&NULL |
| 11 | 117938177 | rs73015719 | A | G | M_Disease | 0.596282699 | M_Disease&NULL |
| 11 | 118227838 | rs11216856 | T | C | M_Disease | 0.596282699 | M_Disease&NULL |
| 11 | 118797544 | rs56032119 | G | A | M_Disease | 0.596282699 | M_Disease&NULL |
| 11 | 118468037 | rs535774 | T | C | M_Disease | 0.596282699 | M_Disease&NULL |
| 11 | 117365915 | rs1359830167 | T | C | M_Disease | 0.596282699 | M_Disease&NULL |
| 11 | 118216423 | rs3212263 | G | A | M_Disease | 0.595078474 | M_Disease&NULL |
| 11 | 121001039 | rs2282565 | C | T | M_Disease | 0.595078474 | M_Disease&NULL |
| 11 | 121001263 | rs7108166 | G | A | M_Disease | 0.595078474 | M_Disease&NULL |
| 11 | 118797392 | rs73575429 | C | T | M_Disease | 0.595078474 | M_Disease&NULL |
| 11 | 118797420 | rs111390501 | G | A | M_Disease | 0.595078474 | M_Disease&NULL |
| 11 | 118797278 | rs74584888 | A | G | M_Disease | 0.593873103 | M_Disease&NULL |
| 11 | 118579327 | rs10892257 | A | G | M_Disease | 0.593873103 | M_Disease&NULL |
| 11 | 117196020 | rs474806 | T | C | M_Disease | 0.593873103 | M_Disease&NULL |
| 11 | 117511787 | rs512017 | G | A | M_Disease | 0.5926666 | M_Disease&NULL |
| 11 | 119404234 | rs149602720 | G | A | M_Disease | 0.595078474 | M_Disease&NULL |
| 11 | 117946941 | rs3758998 | A | C | M_Disease | 0.5926666 | M_Disease&NULL |
| 11 | 118845546 | rs57300701 | G | C | M_Disease | 0.5926666 | M_Disease&NULL |
| 11 | 116806687 | rs2513093 | A | G | M_Disease | 0.591458978 | M_Disease&NULL |
| 11 | 121009383 | rs115076910 | C | T | M_Disease | 0.591458978 | M_Disease&NULL |
| 11 | 117366513 | rs73008407 | T | C | M_Disease | 0.5926666 | M_Disease&NULL |
| 11 | 116882638 | rs4245168 | C | A | M_Disease | 0.591458978 | M_Disease&NULL |
| 11 | 118582613 | rs11216965 | A | C | M_Disease | 0.591458978 | M_Disease&NULL |
| 11 | 118797003 | rs73575428 | G | C | M_Disease | 0.591458978 | M_Disease&NULL |
| 11 | 118678609 | rs2508570 | A | G | M_Disease | 0.590250252 | M_Disease&NULL |
| 11 | 118678881 | rs591756 | T | C | M_Disease | 0.590250252 | M_Disease&NULL |
| 11 | 121010662 | rs624428 | T | C | M_Disease | 0.590250252 | M_Disease&NULL |
| 11 | 117794988 | rs76673375 | C | G | M_Disease | 0.590250252 | M_Disease&NULL |
| 11 | 118811641 | rs11606199 | A | G | M_Disease | 0.590250252 | M_Disease&NULL |
| 11 | 117927488 | rs61900567 | C | G | M_Disease | 0.590250252 | M_Disease&NULL |
| 11 | 118579865 | rs10892258 | A | G | M_Disease | 0.590250252 | M_Disease&NULL |
| 11 | 118580338 | rs11216961 | G | C | M_Disease | 0.590250252 | M_Disease&NULL |
| 11 | 118581895 | rs11216964 | A | G | M_Disease | 0.590250252 | M_Disease&NULL |
| 11 | 117350320 | rs7930881 | A | G | M_Disease | 0.590250252 | M_Disease&NULL |
| 11 | 118664377 | rs598373 | T | C | M_Disease | 0.589040434 | M_Disease&NULL |
| 11 | 118584340 | rs11216969 | C | T | M_Disease | 0.589040434 | M_Disease&NULL |
| 11 | 121009379 | rs34002948 | A | G | M_Disease | 0.589040434 | M_Disease&NULL |
| 11 | 117352381 | rs112746276 | T | A | M_Disease | 0.589040434 | M_Disease&NULL |
| 11 | 118579747 | rs10790261 | G | A | M_Disease | 0.589040434 | M_Disease&NULL |
| 11 | 121192292 | rs2370114 | G | A | M_Disease | 0.589040434 | M_Disease&NULL |
| 11 | 116745484 | rs9919599 | C | G | M_Disease | 0.589040434 | M_Disease&NULL |

| | | | | | | | | | |
|---|---|---|---|---|---|---|---|---|---|
| 11 | 119207341 | rs2248863 | A | G | 1.096 | -7977.388 | -7976.489 | 0.899 | 9.50E-06 |
| 11 | 119207446 | rs2248853 | T | C | 1.096 | -7977.388 | -7976.489 | 0.899 | 9.50E-06 |
| 11 | 120239808 | rs116850963 | G | A | 0.857 | -7977.388 | -7976.489 | 0.899 | 9.50E-06 |
| 11 | 119209979 | rs9640 | A | T | 1.096 | -7977.388 | -7976.49 | 0.898 | 9.50E-06 |
| 11 | 118583525 | rs10892260 | C | A | 1.082 | -7977.388 | -7976.491 | 0.897 | 9.49E-06 |
| 11 | 117554152 | rs17121080 | A | G | 0.841 | -7977.388 | -7976.495 | 0.893 | 9.45E-06 |
| 11 | 117368805 | rs524968 | A | G | 0.932 | -7977.388 | -7976.496 | 0.892 | 9.44E-06 |
| 11 | 116764021 | rs672058 | C | T | 0.91 | -7977.388 | -7976.497 | 0.891 | 9.43E-06 |
| 11 | 116915217 | rs2155583 | A | G | 0.906 | -7977.388 | -7976.5 | 0.888 | 9.40E-06 |
| 11 | 116915819 | rs2000615 | C | T | 0.906 | -7977.388 | -7976.5 | 0.888 | 9.40E-06 |
| 11 | 117182708 | rs525493 | G | T | 0.934 | -7977.388 | -7976.501 | 0.887 | 9.39E-06 |
| 11 | 119820707 | rs75829086 | G | C | 0.851 | -7977.388 | -7976.504 | 0.884 | 9.36E-06 |
| 11 | 117293929 | rs3863310 | C | T | 0.885 | -7977.388 | -7976.507 | 0.882 | 9.34E-06 |
| 11 | 116819617 | rs1241658 | G | A | 0.906 | -7977.388 | -7976.507 | 0.881 | 9.34E-06 |
| 11 | 117925812 | rs732403 | A | G | 0.931 | -7977.388 | -7976.51 | 0.879 | 9.32E-06 |
| 11 | 117938377 | rs61900594 | T | C | 0.931 | -7977.388 | -7976.51 | 0.879 | 9.32E-06 |
| 11 | 117205861 | rs487261 | T | C | 0.886 | -7977.388 | -7976.512 | 0.876 | 9.29E-06 |
| 11 | 116923870 | rs7130297 | C | T | 0.907 | -7977.388 | -7976.514 | 0.875 | 9.28E-06 |
| 11 | 117343197 | rs10892121 | C | T | 1.082 | -7977.388 | -7976.514 | 0.874 | 9.27E-06 |
| 11 | 116483042 | rs34899605 | A | G | 0.884 | -7977.388 | -7976.516 | 0.872 | 9.25E-06 |
| 11 | 116917244 | rs10750100 | T | C | 0.907 | -7977.388 | -7976.518 | 0.87 | 9.23E-06 |
| 11 | 116921397 | rs1815787 | T | C | 0.907 | -7977.388 | -7976.518 | 0.87 | 9.23E-06 |
| 11 | 118744615 | rs73009445 | A | G | 0.881 | -7977.388 | -7976.519 | 0.869 | 9.22E-06 |
| 11 | 116813804 | rs510988 | C | T | 0.907 | -7977.388 | -7976.528 | 0.861 | 9.15E-06 |
| 11 | 116833679 | rs493134 | C | T | 0.907 | -7977.388 | -7976.532 | 0.856 | 9.10E-06 |
| 11 | 116915742 | rs2000614 | C | T | 0.907 | -7977.388 | -7976.533 | 0.855 | 9.10E-06 |
| 11 | 116916529 | rs2000616 | C | T | 0.907 | -7977.388 | -7976.533 | 0.855 | 9.10E-06 |
| 11 | 117179302 | rs663166 | A | G | 0.873 | -7977.388 | -7976.539 | 0.849 | 9.04E-06 |
| 11 | 116898886 | rs7950364 | C | G | 0.907 | -7977.388 | -7976.54 | 0.848 | 9.03E-06 |
| 11 | 117551810 | rs78462423 | A | G | 0.843 | -7977.388 | -7976.545 | 0.843 | 8.99E-06 |
| 11 | 116730638 | rs6589574 | A | G | 0.901 | -7977.388 | -7976.547 | 0.841 | 8.97E-06 |
| 11 | 117202782 | rs624555 | T | C | 0.887 | -7977.388 | -7976.551 | 0.838 | 8.94E-06 |
| 11 | 119576357 | rs112791662 | A | G | 0.867 | -7977.388 | -7976.55 | 0.838 | 8.94E-06 |
| 11 | 117294035 | rs3863311 | A | T | 0.886 | -7977.388 | -7976.553 | 0.836 | 8.92E-06 |
| 11 | 118577873 | rs11216960 | A | G | 1.081 | -7977.388 | -7976.553 | 0.835 | 8.92E-06 |
| 11 | 118578705 | rs10892256 | A | G | 1.081 | -7977.388 | -7976.553 | 0.835 | 8.92E-06 |
| 11 | 117037080 | rs114287696 | G | A | 1.164 | -7977.388 | -7976.561 | 0.828 | 8.85E-06 |
| 11 | 117489941 | rs683842 | C | G | 0.92 | -7977.388 | -7976.56 | 0.828 | 8.85E-06 |
| 11 | 116409112 | rs7118201 | C | T | 1.088 | -7977.388 | -7976.562 | 0.827 | 8.84E-06 |
| 11 | 117560048 | rs7103164 | A | G | 0.844 | -7977.388 | -7976.561 | 0.827 | 8.84E-06 |
| 11 | 116951943 | rs117033797 | G | T | 1.164 | -7977.388 | -7976.571 | 0.817 | 8.76E-06 |
| 11 | 116692324 | rs2234668 | A | G | 1.132 | -7977.388 | -7976.581 | 0.807 | 8.67E-06 |
| 11 | 116735460 | rs17120099 | C | A | 1.129 | -7977.388 | -7976.585 | 0.803 | 8.63E-06 |
| 11 | 120426966 | rs2248540 | C | T | 0.931 | -7977.388 | -7976.587 | 0.801 | 8.62E-06 |
| 11 | 120732349 | rs12221530 | T | C | 1.087 | -7977.388 | -7976.588 | 0.801 | 8.62E-06 |
| 11 | 117255386 | rs480120 | T | C | 0.885 | -7977.388 | -7976.588 | 0.8 | 8.61E-06 |
| 11 | 117549963 | rs116672969 | A | G | 0.845 | -7977.388 | -7976.588 | 0.8 | 8.61E-06 |
| 11 | 121018698 | rs34217740 | A | G | 0.924 | -7977.388 | -7976.588 | 0.8 | 8.61E-06 |
| 11 | 117898446 | rs149421321 | T | C | 0.84 | -7977.388 | -7976.589 | 0.799 | 8.60E-06 |
| 11 | 117291835 | rs1902626 | C | A | 0.887 | -7977.388 | -7976.594 | 0.795 | 8.57E-06 |
| 11 | 118798154 | rs60969913 | C | G | 0.904 | -7977.388 | -7976.595 | 0.794 | 8.56E-06 |
| 11 | 117381052 | rs634606 | T | C | 0.933 | -7977.388 | -7976.598 | 0.79 | 8.52E-06 |
| 11 | 120418613 | rs2252244 | G | A | 0.932 | -7977.388 | -7976.6 | 0.789 | 8.51E-06 |
| 11 | 117269269 | rs539573 | A | T | 0.888 | -7977.388 | -7976.603 | 0.785 | 8.48E-06 |
| 11 | 116199019 | rs75676165 | A | G | 1.117 | -7977.388 | -7976.605 | 0.783 | 8.46E-06 |
| 11 | 117353812 | rs55900121 | T | C | 1.096 | -7977.388 | -7976.607 | 0.781 | 8.45E-06 |
| 11 | 117938177 | rs73015719 | A | G | 0.932 | -7977.388 | -7976.607 | 0.781 | 8.45E-06 |
| 11 | 118227838 | rs11216856 | T | C | 1.075 | -7977.388 | -7976.608 | 0.781 | 8.45E-06 |
| 11 | 118797544 | rs56032119 | G | A | 0.905 | -7977.388 | -7976.608 | 0.781 | 8.45E-06 |
| 11 | 118468037 | rs535774 | T | C | 0.903 | -7977.388 | -7976.614 | 0.774 | 8.39E-06 |
| 11 | 117365915 | rs1359830167 | T | C | 0.856 | -7977.388 | -7976.617 | 0.771 | 8.36E-06 |
| 11 | 118216423 | rs3212263 | G | A | 1.075 | -7977.388 | -7976.617 | 0.771 | 8.36E-06 |
| 11 | 121001039 | rs2282565 | C | T | 0.905 | -7977.388 | -7976.62 | 0.768 | 8.34E-06 |
| 11 | 121001263 | rs7108166 | G | A | 0.905 | -7977.388 | -7976.62 | 0.768 | 8.34E-06 |
| 11 | 118797392 | rs73575429 | C | T | 0.905 | -7977.388 | -7976.624 | 0.764 | 8.30E-06 |
| 11 | 118797420 | rs111390501 | G | A | 0.905 | -7977.388 | -7976.624 | 0.764 | 8.30E-06 |
| 11 | 118797278 | rs74584888 | A | G | 0.905 | -7977.388 | -7976.627 | 0.761 | 8.28E-06 |
| 11 | 118579327 | rs10892257 | A | G | 1.08 | -7977.388 | -7976.629 | 0.759 | 8.26E-06 |
| 11 | 117196020 | rs474806 | T | C | 1.075 | -7977.388 | -7976.634 | 0.754 | 8.22E-06 |
| 11 | 117511787 | rs512017 | G | A | 0.931 | -7977.388 | -7976.637 | 0.752 | 8.21E-06 |
| 11 | 119404234 | rs149602720 | G | A | 1.199 | -7977.388 | -7976.637 | 0.752 | 8.21E-06 |
| 11 | 117946941 | rs3758998 | A | C | 0.934 | -7977.388 | -7976.638 | 0.751 | 8.20E-06 |
| 11 | 118845546 | rs57300701 | G | C | 0.913 | -7977.388 | -7976.642 | 0.747 | 8.16E-06 |
| 11 | 116806687 | rs2513093 | A | G | 0.909 | -7977.388 | -7976.647 | 0.741 | 8.12E-06 |
| 11 | 121009383 | rs115076910 | C | T | 0.925 | -7977.388 | -7976.647 | 0.741 | 8.12E-06 |
| 11 | 117366513 | rs73008407 | T | C | 0.857 | -7977.388 | -7976.65 | 0.738 | 8.09E-06 |
| 11 | 116882638 | rs4245168 | C | A | 0.909 | -7977.388 | -7976.652 | 0.736 | 8.08E-06 |
| 11 | 118582613 | rs11216965 | A | C | 1.079 | -7977.388 | -7976.653 | 0.735 | 8.07E-06 |
| 11 | 118797003 | rs73575428 | G | C | 0.905 | -7977.388 | -7976.653 | 0.735 | 8.07E-06 |
| 11 | 118678609 | rs2508570 | A | G | 1.073 | -7977.388 | -7976.659 | 0.729 | 8.02E-06 |
| 11 | 118678881 | rs591756 | T | C | 1.073 | -7977.388 | -7976.659 | 0.729 | 8.02E-06 |
| 11 | 121010662 | rs624428 | T | C | 0.925 | -7977.388 | -7976.661 | 0.727 | 8.00E-06 |
| 11 | 117794988 | rs76673375 | C | G | 0.891 | -7977.388 | -7976.663 | 0.726 | 7.99E-06 |
| 11 | 118811641 | rs11606199 | A | G | 1.09 | -7977.388 | -7976.662 | 0.726 | 7.99E-06 |
| 11 | 117927488 | rs61900567 | C | G | 0.934 | -7977.388 | -7976.663 | 0.725 | 7.99E-06 |
| 11 | 118579865 | rs10892258 | A | G | 1.079 | -7977.388 | -7976.663 | 0.725 | 7.99E-06 |
| 11 | 118580338 | rs11216961 | G | C | 1.079 | -7977.388 | -7976.663 | 0.725 | 7.99E-06 |
| 11 | 118581895 | rs11216964 | A | G | 1.079 | -7977.388 | -7976.663 | 0.725 | 7.99E-06 |
| 11 | 117350320 | rs7930881 | A | G | 1.08 | -7977.388 | -7976.666 | 0.723 | 7.97E-06 |
| 11 | 118664377 | rs598373 | T | C | 1.072 | -7977.388 | -7976.668 | 0.72 | 7.95E-06 |
| 11 | 118584340 | rs11216969 | C | T | 1.079 | -7977.388 | -7976.669 | 0.719 | 7.94E-06 |
| 11 | 121009379 | rs34002948 | A | G | 0.925 | -7977.388 | -7976.669 | 0.719 | 7.94E-06 |
| 11 | 117352381 | rs112746276 | T | A | 1.095 | -7977.388 | -7976.671 | 0.717 | 7.92E-06 |
| 11 | 118579747 | rs10790261 | G | A | 1.079 | -7977.388 | -7976.673 | 0.716 | 7.92E-06 |
| 11 | 121192292 | rs2370114 | G | A | 1.1 | -7977.388 | -7976.672 | 0.716 | 7.92E-06 |
| 11 | 116745484 | rs9919599 | C | G | 0.909 | -7977.388 | -7976.673 | 0.715 | 7.91E-06 |

| | | | | | | | |
|---|---|---|---|---|---|---|---|
| 11 | 117372488 | rs17092713 | A | G | M_Disease | 0.589040434 | M_Disease&NULL |
| 11 | 117925668 | rs2105590 | A | G | M_Disease | 0.589040434 | M_Disease&NULL |
| 11 | 118677710 | rs688161 | G | A | M_Disease | 0.589040434 | M_Disease&NULL |
| 11 | 120408071 | rs3901366 | A | G | M_Disease | 0.589040434 | M_Disease&NULL |
| 11 | 119859583 | rs3740725 | A | G | M_Disease | 0.589040434 | M_Disease&NULL |
| 11 | 116679155 | rs75542613 | A | G | M_Disease | 0.589040434 | M_Disease&NULL |
| 11 | 118230223 | rs2276417 | G | T | M_Disease | 0.587829538 | M_Disease&NULL |
| 11 | 119863505 | rs11217617 | T | C | M_Disease | 0.587829538 | M_Disease&NULL |
| 11 | 120614648 | rs17245685 | C | T | M_Disease | 0.587829538 | M_Disease&NULL |
| 11 | 120998225 | rs74736885 | G | A | M_Disease | 0.587829538 | M_Disease&NULL |
| 11 | 117926770 | rs61900566 | G | A | M_Disease | 0.587829538 | M_Disease&NULL |
| 11 | 120424740 | rs2508656 | A | G | M_Disease | 0.587829538 | M_Disease&NULL |
| 11 | 117364821 | rs12286920 | A | G | M_Disease | 0.587829538 | M_Disease&NULL |
| 11 | 116824884 | rs511676 | T | G | M_Disease | 0.586617579 | M_Disease&NULL |
| 11 | 116199020 | rs77520149 | A | C | M_Disease | 0.585404569 | M_Disease&NULL |
| 11 | 117932042 | rs1939128 | T | C | M_Disease | 0.585404569 | M_Disease&NULL |
| 11 | 118583002 | rs10892259 | A | G | M_Disease | 0.585404569 | M_Disease&NULL |
| 11 | 116682668 | rs149137426 | A | T | M_Disease | 0.586617579 | M_Disease&NULL |
| 11 | 116702630 | rs140223477 | A | G | M_Disease | 0.586617579 | M_Disease&NULL |
| 11 | 117925751 | rs2096721 | T | C | M_Disease | 0.584190523 | M_Disease&NULL |
| 11 | 120417190 | rs2252385 | C | T | M_Disease | 0.584190523 | M_Disease&NULL |
| 11 | 117364120 | rs1395475712 | T | C | M_Disease | 0.584190523 | M_Disease&NULL |
| 11 | 116806269 | rs2513095 | T | C | M_Disease | 0.584190523 | M_Disease&NULL |
| 11 | 116706047 | rs12721025 | A | G | M_Disease | 0.584190523 | M_Disease&NULL |
| 11 | 116308219 | rs10891976 | T | A | M_Disease | 0.584190523 | M_Disease&NULL |
| 11 | 118575326 | rs11216956 | A | G | M_Disease | 0.584190523 | M_Disease&NULL |
| 11 | 116488133 | rs576266 | C | T | M_Disease | 0.582975454 | M_Disease&NULL |
| 11 | 118680045 | rs2508569 | C | G | M_Disease | 0.582975454 | M_Disease&NULL |
| 11 | 118619948 | rs112143902 | T | C | M_Disease | 0.582975454 | M_Disease&NULL |
| 11 | 116715784 | rs142174850 | A | G | M_Disease | 0.582975454 | M_Disease&NULL |
| 11 | 116862818 | rs607273 | A | T | M_Disease | 0.582975454 | M_Disease&NULL |
| 11 | 117165046 | rs150308863 | T | C | M_Disease | 0.581759377 | M_Disease&NULL |
| 11 | 118678986 | rs477014 | C | T | M_Disease | 0.581759377 | M_Disease&NULL |
| 11 | 118679387 | rs473298 | G | A | M_Disease | 0.581759377 | M_Disease&NULL |
| 11 | 118679445 | rs516719 | C | T | M_Disease | 0.581759377 | M_Disease&NULL |
| 11 | 121037554 | rs615697 | C | T | M_Disease | 0.581759377 | M_Disease&NULL |
| 11 | 121037670 | rs616184 | A | G | M_Disease | 0.581759377 | M_Disease&NULL |
| 11 | 120011880 | rs61900473 | A | G | M_Disease | 0.581759377 | M_Disease&NULL |
| 11 | 116753552 | rs12275565 | G | A | M_Disease | 0.581759377 | M_Disease&NULL |
| 11 | 117795671 | rs77018736 | A | G | M_Disease | 0.581759377 | M_Disease&NULL |
| 11 | 120412169 | rs2253023 | T | C | M_Disease | 0.580542305 | M_Disease&NULL |
| 11 | 121037530 | rs489985 | A | G | M_Disease | 0.580542305 | M_Disease&NULL |
| 11 | 118042606 | rs76248354 | A | G | M_Disease | 0.580542305 | M_Disease&NULL |
| 11 | 116488591 | rs3017773 | G | T | M_Disease | 0.580542305 | M_Disease&NULL |
| 11 | 116488610 | rs2105611 | A | G | M_Disease | 0.580542305 | M_Disease&NULL |
| 11 | 116488978 | rs529618 | A | G | M_Disease | 0.580542305 | M_Disease&NULL |
| 11 | 117514378 | rs11216479 | T | C | M_Disease | 0.580542305 | M_Disease&NULL |
| 11 | 117271499 | rs7928581 | C | T | M_Disease | 0.580542305 | M_Disease&NULL |
| 11 | 116866290 | rs7931770 | C | T | M_Disease | 0.580542305 | M_Disease&NULL |
| 11 | 120723057 | rs4331109 | G | A | M_Disease | 0.579324252 | M_Disease&NULL |
| 11 | 116856642 | rs588918 | T | C | M_Disease | 0.579324252 | M_Disease&NULL |
| 11 | 116748515 | rs12292614 | G | A | M_Disease | 0.579324252 | M_Disease&NULL |
| 11 | 116748744 | rs12286581 | C | T | M_Disease | 0.579324252 | M_Disease&NULL |
| 11 | 116749072 | rs75294699 | A | C | M_Disease | 0.579324252 | M_Disease&NULL |
| 11 | 116754265 | rs12416987 | C | T | M_Disease | 0.579324252 | M_Disease&NULL |
| 11 | 116756183 | rs2044426 | A | G | M_Disease | 0.579324252 | M_Disease&NULL |
| 11 | 116756986 | rs10892040 | A | G | M_Disease | 0.579324252 | M_Disease&NULL |
| 11 | 116757248 | rs11216174 | G | A | M_Disease | 0.579324252 | M_Disease&NULL |
| 11 | 116757398 | rs11216175 | T | C | M_Disease | 0.579324252 | M_Disease&NULL |
| 11 | 116757793 | rs12284346 | C | T | M_Disease | 0.579324252 | M_Disease&NULL |
| 11 | 119881267 | rs2018774 | T | A | M_Disease | 0.579324252 | M_Disease&NULL |
| 11 | 116726709 | rs112618266 | T | C | M_Disease | 0.579324252 | M_Disease&NULL |
| 11 | 116759824 | rs12294191 | G | A | M_Disease | 0.579324252 | M_Disease&NULL |
| 11 | 120607334 | rs11604129 | G | A | M_Disease | 0.579324252 | M_Disease&NULL |
| 11 | 119583742 | rs11217420 | A | G | M_Disease | 0.578105233 | M_Disease&NULL |
| 11 | 117997570 | rs1786193 | A | T | M_Disease | 0.578105233 | M_Disease&NULL |
| 11 | 121034491 | rs580683 | T | C | M_Disease | 0.578105233 | M_Disease&NULL |
| 11 | 118598138 | rs543692 | C | T | M_Disease | 0.576885261 | M_Disease&NULL |
| 11 | 116747282 | rs17120119 | G | C | M_Disease | 0.578105233 | M_Disease&NULL |
| 11 | 116748169 | rs12278117 | A | G | M_Disease | 0.578105233 | M_Disease&NULL |
| 11 | 116748221 | rs12279433 | T | G | M_Disease | 0.578105233 | M_Disease&NULL |
| 11 | 120849685 | rs2155094 | A | G | M_Disease | 0.576885261 | M_Disease&NULL |
| 11 | 117180921 | rs583161 | T | C | M_Disease | 0.576885261 | M_Disease&NULL |
| 11 | 120428959 | rs12575259 | A | G | M_Disease | 0.576885261 | M_Disease&NULL |
| 11 | 117424804 | rs11216452 | A | G | M_Disease | 0.576885261 | M_Disease&NULL |
| 11 | 116875983 | rs7114963 | T | C | M_Disease | 0.576885261 | M_Disease&NULL |
| 11 | 121011269 | rs561195 | A | C | M_Disease | 0.575664351 | M_Disease&NULL |
| 11 | 117987573 | rs640012 | C | T | M_Disease | 0.576885261 | M_Disease&NULL |
| 11 | 119936873 | rs79345551 | A | G | M_Disease | 0.575664351 | M_Disease&NULL |
| 11 | 116707401 | rs12718464 | A | G | M_Disease | 0.575664351 | M_Disease&NULL |
| 11 | 120715346 | rs35168282 | A | G | M_Disease | 0.575664351 | M_Disease&NULL |
| 11 | 116867209 | rs7121347 | G | T | M_Disease | 0.575664351 | M_Disease&NULL |
| 11 | 121021940 | rs7122110 | G | A | M_Disease | 0.575664351 | M_Disease&NULL |
| 11 | 117200638 | rs1261791 | A | T | M_Disease | 0.575664351 | M_Disease&NULL |
| 11 | 118038741 | rs647975 | T | C | M_Disease | 0.575664351 | M_Disease&NULL |
| 11 | 121039234 | rs4936590 | C | T | M_Disease | 0.574442517 | M_Disease&NULL |
| 11 | 117903210 | rs17598956 | C | T | M_Disease | 0.573219773 | M_Disease&NULL |
| 11 | 117207122 | rs524914 | C | G | M_Disease | 0.573219773 | M_Disease&NULL |
| 11 | 117512831 | rs480801 | C | T | M_Disease | 0.573219773 | M_Disease&NULL |
| 11 | 120275234 | rs35659242 | G | T | M_Disease | 0.573219773 | M_Disease&NULL |
| 11 | 117350336 | rs2155393 | A | G | M_Disease | 0.573219773 | M_Disease&NULL |
| 11 | 120405162 | rs2508652 | T | C | M_Disease | 0.573219773 | M_Disease&NULL |
| 11 | 117903836 | rs4936417 | C | T | M_Disease | 0.571996133 | M_Disease&NULL |
| 11 | 118580813 | rs11216962 | C | T | M_Disease | 0.571996133 | M_Disease&NULL |
| 11 | 118580814 | rs11216963 | A | G | M_Disease | 0.571996133 | M_Disease&NULL |
| 11 | 117987391 | rs603678 | C | T | M_Disease | 0.571996133 | M_Disease&NULL |

| | | | | | | | | | |
|---|---|---|---|---|---|---|---|---|---|
| 11 | 117372488 | rs17092713 | A | G | 0.863 | -7977.388 | -7976.673 | 0.715 | 7.91E-06 |
| 11 | 117925668 | rs2105590 | A | G | 0.933 | -7977.388 | -7976.673 | 0.715 | 7.91E-06 |
| 11 | 118677710 | rs688161 | G | A | 1.072 | -7977.388 | -7976.674 | 0.715 | 7.91E-06 |
| 11 | 120408071 | rs3901366 | A | G | 0.932 | -7977.388 | -7976.674 | 0.714 | 7.90E-06 |
| 11 | 119859583 | rs3740725 | A | G | 0.872 | -7977.388 | -7976.676 | 0.713 | 7.89E-06 |
| 11 | 116679155 | rs75542613 | A | G | 1.12 | -7977.388 | -7976.678 | 0.711 | 7.88E-06 |
| 11 | 118230223 | rs2276417 | G | T | 1.074 | -7977.388 | -7976.68 | 0.708 | 7.85E-06 |
| 11 | 119863505 | rs11217617 | T | C | 0.872 | -7977.388 | -7976.681 | 0.708 | 7.85E-06 |
| 11 | 120614648 | rs17245685 | C | T | 0.888 | -7977.388 | -7976.681 | 0.707 | 7.84E-06 |
| 11 | 120998225 | rs74736885 | G | A | 0.906 | -7977.388 | -7976.682 | 0.707 | 7.84E-06 |
| 11 | 117926770 | rs61900566 | G | A | 0.933 | -7977.388 | -7976.684 | 0.705 | 7.83E-06 |
| 11 | 120424740 | rs2508656 | A | G | 0.883 | -7977.388 | -7976.683 | 0.705 | 7.83E-06 |
| 11 | 117364821 | rs12286920 | A | G | 1.08 | -7977.388 | -7976.685 | 0.703 | 7.81E-06 |
| 11 | 116824884 | rs511676 | T | G | 0.91 | -7977.388 | -7976.689 | 0.7 | 7.79E-06 |
| 11 | 116199020 | rs77520149 | A | C | 1.114 | -7977.388 | -7976.701 | 0.687 | 7.69E-06 |
| 11 | 117932042 | rs1939128 | T | C | 0.933 | -7977.388 | -7976.701 | 0.687 | 7.69E-06 |
| 11 | 118583002 | rs10892259 | A | G | 1.079 | -7977.388 | -7976.702 | 0.686 | 7.68E-06 |
| 11 | 116682668 | rs149137426 | A | T | 0.846 | -7977.388 | -7976.704 | 0.684 | 7.67E-06 |
| 11 | 116702630 | rs140223477 | A | G | 1.176 | -7977.388 | -7976.704 | 0.684 | 7.67E-06 |
| 11 | 117925751 | rs2096721 | T | C | 0.933 | -7977.388 | -7976.706 | 0.682 | 7.65E-06 |
| 11 | 120417190 | rs2252385 | C | T | 0.933 | -7977.388 | -7976.707 | 0.681 | 7.64E-06 |
| 11 | 117364120 | rs1395475712 | T | C | 1.07 | -7977.388 | -7976.708 | 0.68 | 7.64E-06 |
| 11 | 116806269 | rs2513095 | T | C | 0.91 | -7977.388 | -7976.712 | 0.677 | 7.61E-06 |
| 11 | 116706047 | rs12721025 | A | G | 1.123 | -7977.388 | -7976.714 | 0.674 | 7.59E-06 |
| 11 | 116308219 | rs10891976 | T | A | 0.936 | -7977.388 | -7976.715 | 0.673 | 7.58E-06 |
| 11 | 118575326 | rs11216956 | A | G | 1.078 | -7977.388 | -7976.715 | 0.673 | 7.58E-06 |
| 11 | 116488133 | rs576266 | C | T | 0.909 | -7977.388 | -7976.719 | 0.67 | 7.56E-06 |
| 11 | 118680045 | rs2508569 | C | G | 1.072 | -7977.388 | -7976.719 | 0.669 | 7.55E-06 |
| 11 | 118619948 | rs112143902 | T | C | 0.903 | -7977.388 | -7976.722 | 0.666 | 7.53E-06 |
| 11 | 116715784 | rs142174850 | A | G | 1.124 | -7977.388 | -7976.723 | 0.665 | 7.52E-06 |
| 11 | 116862818 | rs607273 | A | T | 0.906 | -7977.388 | -7976.725 | 0.664 | 7.51E-06 |
| 11 | 117165046 | rs150308863 | T | C | 0.879 | -7977.388 | -7976.731 | 0.658 | 7.47E-06 |
| 11 | 118678986 | rs477014 | C | T | 1.071 | -7977.388 | -7976.73 | 0.658 | 7.47E-06 |
| 11 | 118679387 | rs473298 | G | A | 1.071 | -7977.388 | -7976.73 | 0.658 | 7.47E-06 |
| 11 | 118679445 | rs516719 | C | T | 1.071 | -7977.388 | -7976.73 | 0.658 | 7.47E-06 |
| 11 | 121037554 | rs615697 | C | T | 0.926 | -7977.388 | -7976.731 | 0.657 | 7.46E-06 |
| 11 | 121037670 | rs616184 | A | G | 0.926 | -7977.388 | -7976.731 | 0.657 | 7.46E-06 |
| 11 | 120011880 | rs61900473 | A | G | 0.888 | -7977.388 | -7976.732 | 0.656 | 7.45E-06 |
| 11 | 116753552 | rs12275565 | G | A | 1.122 | -7977.388 | -7976.733 | 0.655 | 7.45E-06 |
| 11 | 117795671 | rs77018736 | A | G | 0.881 | -7977.388 | -7976.735 | 0.654 | 7.44E-06 |
| 11 | 120412169 | rs2253023 | T | C | 0.93 | -7977.388 | -7976.738 | 0.65 | 7.41E-06 |
| 11 | 121037530 | rs489985 | A | G | 0.926 | -7977.388 | -7976.739 | 0.649 | 7.40E-06 |
| 11 | 118042606 | rs76248354 | A | G | 0.884 | -7977.388 | -7976.74 | 0.648 | 7.39E-06 |
| 11 | 116488591 | rs3017773 | G | T | 0.91 | -7977.388 | -7976.742 | 0.646 | 7.38E-06 |
| 11 | 116488610 | rs2105611 | A | G | 0.91 | -7977.388 | -7976.742 | 0.646 | 7.38E-06 |
| 11 | 116488978 | rs529618 | A | G | 0.91 | -7977.388 | -7976.742 | 0.646 | 7.38E-06 |
| 11 | 117514378 | rs11216479 | T | C | 1.085 | -7977.388 | -7976.743 | 0.645 | 7.37E-06 |
| 11 | 117271499 | rs7928581 | C | T | 0.896 | -7977.388 | -7976.744 | 0.644 | 7.37E-06 |
| 11 | 116866290 | rs7931770 | C | T | 0.911 | -7977.388 | -7976.745 | 0.643 | 7.36E-06 |
| 11 | 120723057 | rs4331109 | G | A | 1.084 | -7977.388 | -7976.748 | 0.641 | 7.34E-06 |
| 11 | 116856642 | rs588918 | T | C | 0.911 | -7977.388 | -7976.748 | 0.64 | 7.34E-06 |
| 11 | 116748515 | rs12292614 | G | A | 1.122 | -7977.388 | -7976.749 | 0.639 | 7.33E-06 |
| 11 | 116748744 | rs12286581 | C | T | 1.122 | -7977.388 | -7976.749 | 0.639 | 7.33E-06 |
| 11 | 116749072 | rs75294699 | A | C | 1.122 | -7977.388 | -7976.749 | 0.639 | 7.33E-06 |
| 11 | 116754265 | rs12416987 | C | T | 1.122 | -7977.388 | -7976.749 | 0.639 | 7.33E-06 |
| 11 | 116756183 | rs2044426 | A | G | 1.122 | -7977.388 | -7976.749 | 0.639 | 7.33E-06 |
| 11 | 116756986 | rs10892040 | A | G | 1.122 | -7977.388 | -7976.749 | 0.639 | 7.33E-06 |
| 11 | 116757248 | rs11216174 | G | A | 1.122 | -7977.388 | -7976.749 | 0.639 | 7.33E-06 |
| 11 | 116757398 | rs11216175 | T | C | 1.122 | -7977.388 | -7976.749 | 0.639 | 7.33E-06 |
| 11 | 116757793 | rs12284346 | C | T | 1.122 | -7977.388 | -7976.749 | 0.639 | 7.33E-06 |
| 11 | 119881267 | rs2018774 | T | A | 0.896 | -7977.388 | -7976.749 | 0.639 | 7.33E-06 |
| 11 | 116726709 | rs112618266 | T | C | 1.122 | -7977.388 | -7976.751 | 0.637 | 7.31E-06 |
| 11 | 116759824 | rs12294191 | G | A | 1.122 | -7977.388 | -7976.751 | 0.637 | 7.31E-06 |
| 11 | 120607334 | rs11604129 | G | A | 0.89 | -7977.388 | -7976.756 | 0.632 | 7.28E-06 |
| 11 | 119583742 | rs11217420 | A | G | 1.094 | -7977.388 | -7976.757 | 0.631 | 7.27E-06 |
| 11 | 117997570 | rs1786193 | A | T | 1.131 | -7977.388 | -7976.762 | 0.626 | 7.23E-06 |
| 11 | 121034491 | rs580683 | T | C | 0.927 | -7977.388 | -7976.765 | 0.624 | 7.22E-06 |
| 11 | 118598138 | rs543692 | C | T | 0.936 | -7977.388 | -7976.766 | 0.623 | 7.21E-06 |
| 11 | 116747282 | rs17120119 | G | C | 1.121 | -7977.388 | -7976.766 | 0.622 | 7.20E-06 |
| 11 | 116748169 | rs12278117 | A | G | 1.121 | -7977.388 | -7976.766 | 0.622 | 7.20E-06 |
| 11 | 116748221 | rs12279433 | T | G | 1.121 | -7977.388 | -7976.766 | 0.622 | 7.20E-06 |
| 11 | 120849685 | rs2155094 | A | G | 1.069 | -7977.388 | -7976.766 | 0.622 | 7.20E-06 |
| 11 | 117180921 | rs583161 | T | C | 1.072 | -7977.388 | -7976.77 | 0.618 | 7.18E-06 |
| 11 | 120428959 | rs12575259 | A | G | 0.884 | -7977.388 | -7976.77 | 0.618 | 7.18E-06 |
| 11 | 117424804 | rs11216452 | A | G | 0.937 | -7977.388 | -7976.773 | 0.616 | 7.16E-06 |
| 11 | 116875983 | rs7114963 | T | C | 0.911 | -7977.388 | -7976.777 | 0.612 | 7.13E-06 |
| 11 | 121011269 | rs561195 | A | C | 0.936 | -7977.388 | -7976.776 | 0.612 | 7.13E-06 |
| 11 | 117987573 | rs640012 | C | T | 1.145 | -7977.388 | -7976.777 | 0.611 | 7.13E-06 |
| 11 | 119936873 | rs79345551 | A | G | 0.892 | -7977.388 | -7976.78 | 0.608 | 7.10E-06 |
| 11 | 116707401 | rs12718464 | A | G | 1.12 | -7977.388 | -7976.784 | 0.605 | 7.08E-06 |
| 11 | 120715346 | rs35168282 | A | G | 1.083 | -7977.388 | -7976.786 | 0.603 | 7.07E-06 |
| 11 | 116867209 | rs7121347 | G | T | 0.912 | -7977.388 | -7976.786 | 0.602 | 7.06E-06 |
| 11 | 121021940 | rs7122110 | G | A | 0.927 | -7977.388 | -7976.786 | 0.602 | 7.06E-06 |
| 11 | 117200638 | rs1261791 | A | T | 0.894 | -7977.388 | -7976.787 | 0.601 | 7.06E-06 |
| 11 | 118038741 | rs647975 | T | C | 0.858 | -7977.388 | -7976.787 | 0.601 | 7.06E-06 |
| 11 | 121039234 | rs4936590 | C | T | 0.927 | -7977.388 | -7976.789 | 0.6 | 7.05E-06 |
| 11 | 117903210 | rs17598956 | C | T | 0.934 | -7977.388 | -7976.798 | 0.59 | 6.98E-06 |
| 11 | 117207122 | rs524914 | C | G | 0.894 | -7977.388 | -7976.8 | 0.588 | 6.96E-06 |
| 11 | 117512831 | rs480801 | C | T | 0.933 | -7977.388 | -7976.802 | 0.587 | 6.96E-06 |
| 11 | 120275234 | rs35659242 | G | T | 0.889 | -7977.388 | -7976.801 | 0.587 | 6.96E-06 |
| 11 | 117350336 | rs2155393 | A | G | 1.077 | -7977.388 | -7976.805 | 0.584 | 6.94E-06 |
| 11 | 120405162 | rs2508652 | T | C | 0.932 | -7977.388 | -7976.806 | 0.583 | 6.93E-06 |
| 11 | 117903836 | rs4936417 | C | T | 0.934 | -7977.388 | -7976.807 | 0.581 | 6.92E-06 |
| 11 | 118580813 | rs11216962 | C | T | 1.077 | -7977.388 | -7976.809 | 0.579 | 6.90E-06 |
| 11 | 118580814 | rs11216963 | A | G | 1.077 | -7977.388 | -7976.809 | 0.579 | 6.90E-06 |
| 11 | 117987391 | rs603678 | C | T | 1.133 | -7977.388 | -7976.811 | 0.577 | 6.89E-06 |

| | | | | | | | |
|---|---|---|---|---|---|---|---|
| 11 | 118279936 | rs3194726 | C | T | M_Disease | 0.570771612 | M_Disease&NULL |
| 11 | 116299456 | rs144466751 | C | T | M_Disease | 0.570771612 | M_Disease&NULL |
| 11 | 121062486 | rs7110565 | C | T | M_Disease | 0.570771612 | M_Disease&NULL |
| 11 | 120425101 | rs79985686 | C | T | M_Disease | 0.570771612 | M_Disease&NULL |
| 11 | 118519213 | rs668628 | T | G | M_Disease | 0.569546224 | M_Disease&NULL |
| 11 | 119864218 | rs874195 | G | A | M_Disease | 0.570771612 | M_Disease&NULL |
| 11 | 119880819 | rs675246 | T | C | M_Disease | 0.569546224 | M_Disease&NULL |
| 11 | 117034899 | rs189189723 | T | G | M_Disease | 0.569546224 | M_Disease&NULL |
| 11 | 117308183 | rs73592180 | G | A | M_Disease | 0.569546224 | M_Disease&NULL |
| 11 | 117449497 | rs11606425 | T | A | M_Disease | 0.569546224 | M_Disease&NULL |
| 11 | 116758693 | rs12292278 | C | A | M_Disease | 0.568319983 | M_Disease&NULL |
| 11 | 121015968 | rs4936583 | C | T | M_Disease | 0.568319983 | M_Disease&NULL |
| 11 | 121016098 | rs10892676 | C | T | M_Disease | 0.568319983 | M_Disease&NULL |
| 11 | 121016145 | rs4936584 | A | G | M_Disease | 0.568319983 | M_Disease&NULL |
| 11 | 121016818 | rs12275038 | A | G | M_Disease | 0.568319983 | M_Disease&NULL |
| 11 | 119519217 | rs11217372 | G | C | M_Disease | 0.568319983 | M_Disease&NULL |
| 11 | 120430224 | rs80260914 | G | A | M_Disease | 0.568319983 | M_Disease&NULL |
| 11 | 120425797 | rs12575945 | A | C | M_Disease | 0.567092905 | M_Disease&NULL |
| 11 | 117449263 | rs11604377 | C | T | M_Disease | 0.565865003 | M_Disease&NULL |
| 11 | 118809882 | rs11217099 | A | C | M_Disease | 0.565865003 | M_Disease&NULL |
| 11 | 117004656 | rs11216288 | T | C | M_Disease | 0.565865003 | M_Disease&NULL |
| 11 | 117386067 | rs2186762 | A | C | M_Disease | 0.565865003 | M_Disease&NULL |
| 11 | 118574311 | rs524409 | T | C | M_Disease | 0.564636292 | M_Disease&NULL |
| 11 | 118574675 | rs494459 | T | C | M_Disease | 0.564636292 | M_Disease&NULL |
| 11 | 116771226 | rs117640781 | T | C | M_Disease | 0.565865003 | M_Disease&NULL |
| 11 | 120719994 | rs11218030 | G | A | M_Disease | 0.564636292 | M_Disease&NULL |
| 11 | 118573916 | rs627391 | T | A | M_Disease | 0.563406786 | M_Disease&NULL |
| 11 | 118575011 | rs530901 | C | T | M_Disease | 0.563406786 | M_Disease&NULL |
| 11 | 118575180 | rs626645 | C | T | M_Disease | 0.563406786 | M_Disease&NULL |
| 11 | 118575419 | rs625735 | A | G | M_Disease | 0.563406786 | M_Disease&NULL |
| 11 | 118575479 | rs637563 | G | A | M_Disease | 0.563406786 | M_Disease&NULL |
| 11 | 118575693 | rs638514 | C | T | M_Disease | 0.563406786 | M_Disease&NULL |
| 11 | 116801201 | rs645258 | A | C | M_Disease | 0.563406786 | M_Disease&NULL |
| 11 | 120521305 | rs7125158 | C | A | M_Disease | 0.564636292 | M_Disease&NULL |
| 11 | 119870780 | rs11217623 | T | C | M_Disease | 0.563406786 | M_Disease&NULL |
| 11 | 120431744 | rs7121316 | A | G | M_Disease | 0.563406786 | M_Disease&NULL |
| 11 | 121011258 | rs561227 | T | C | M_Disease | 0.563406786 | M_Disease&NULL |
| 11 | 118573894 | rs576283 | G | A | M_Disease | 0.562176501 | M_Disease&NULL |
| 11 | 118184271 | rs2231445 | C | T | M_Disease | 0.563406786 | M_Disease&NULL |
| 11 | 116491621 | rs507262 | G | A | M_Disease | 0.562176501 | M_Disease&NULL |
| 11 | 116492312 | rs534633 | G | A | M_Disease | 0.562176501 | M_Disease&NULL |
| 11 | 121012492 | rs581786 | C | T | M_Disease | 0.562176501 | M_Disease&NULL |
| 11 | 121012693 | rs528958 | T | G | M_Disease | 0.562176501 | M_Disease&NULL |
| 11 | 121012831 | rs688106 | A | T | M_Disease | 0.562176501 | M_Disease&NULL |
| 11 | 121012978 | rs665743 | A | G | M_Disease | 0.562176501 | M_Disease&NULL |
| 11 | 117922730 | rs17599289 | T | C | M_Disease | 0.562176501 | M_Disease&NULL |
| 11 | 121018870 | rs4623911 | G | A | M_Disease | 0.562176501 | M_Disease&NULL |
| 11 | 121018999 | rs4641485 | G | A | M_Disease | 0.562176501 | M_Disease&NULL |
| 11 | 119921812 | rs586112 | C | G | M_Disease | 0.562176501 | M_Disease&NULL |
| 11 | 117908084 | rs61900560 | C | G | M_Disease | 0.56094545 | M_Disease&NULL |
| 11 | 118751326 | rs10892306 | A | T | M_Disease | 0.562176501 | M_Disease&NULL |
| 11 | 116999059 | rs4516002 | A | G | M_Disease | 0.56094545 | M_Disease&NULL |
| 11 | 117197621 | rs680908 | G | C | M_Disease | 0.562176501 | M_Disease&NULL |
| 11 | 117198638 | rs566155 | C | T | M_Disease | 0.562176501 | M_Disease&NULL |
| 11 | 121018488 | rs12285468 | C | T | M_Disease | 0.56094545 | M_Disease&NULL |
| 11 | 120431935 | rs12283999 | G | C | M_Disease | 0.56094545 | M_Disease&NULL |
| 11 | 116511522 | rs519000 | T | C | M_Disease | 0.56094545 | M_Disease&NULL |
| 11 | 119866879 | rs9787805 | C | G | M_Disease | 0.56094545 | M_Disease&NULL |
| 11 | 116805204 | rs562179 | T | C | M_Disease | 0.56094545 | M_Disease&NULL |
| 11 | 117915445 | rs7926330 | T | C | M_Disease | 0.56094545 | M_Disease&NULL |
| 11 | 116295218 | rs75375536 | G | A | M_Disease | 0.56094545 | M_Disease&NULL |
| 11 | 121007777 | rs4936580 | A | G | M_Disease | 0.56094545 | M_Disease&NULL |
| 11 | 116491177 | rs502816 | A | C | M_Disease | 0.56094545 | M_Disease&NULL |
| 11 | 116491428 | rs505397 | G | T | M_Disease | 0.56094545 | M_Disease&NULL |
| 11 | 117181919 | rs497902 | G | C | M_Disease | 0.559713649 | M_Disease&NULL |
| 11 | 121014228 | rs11218158 | G | A | M_Disease | 0.559713649 | M_Disease&NULL |
| 11 | 121014459 | rs877538 | A | C | M_Disease | 0.559713649 | M_Disease&NULL |
| 11 | 121014860 | rs7114879 | T | C | M_Disease | 0.559713649 | M_Disease&NULL |
| 11 | 121010788 | rs607014 | A | G | M_Disease | 0.558481112 | M_Disease&NULL |
| 11 | 121017604 | rs7120647 | C | T | M_Disease | 0.558481112 | M_Disease&NULL |
| 11 | 121017863 | rs4936586 | A | C | M_Disease | 0.558481112 | M_Disease&NULL |
| 11 | 121018007 | rs4936587 | G | A | M_Disease | 0.558481112 | M_Disease&NULL |
| 11 | 121018485 | rs12285467 | C | T | M_Disease | 0.558481112 | M_Disease&NULL |
| 11 | 117183914 | rs477036 | C | T | M_Disease | 0.558481112 | M_Disease&NULL |
| 11 | 119866718 | rs9787801 | A | G | M_Disease | 0.559713649 | M_Disease&NULL |
| 11 | 121014983 | rs7114915 | G | A | M_Disease | 0.558481112 | M_Disease&NULL |
| 11 | 117402325 | rs61350190 | T | G | M_Disease | 0.558481112 | M_Disease&NULL |
| 11 | 121011485 | rs638463 | C | T | M_Disease | 0.558481112 | M_Disease&NULL |
| 11 | 116839761 | rs1241657 | T | C | M_Disease | 0.558481112 | M_Disease&NULL |
| 11 | 120412823 | rs2510876 | G | A | M_Disease | 0.558481112 | M_Disease&NULL |
| 11 | 117541718 | rs141116752 | T | C | M_Disease | 0.558481112 | M_Disease&NULL |
| 11 | 121010820 | rs606674 | G | A | M_Disease | 0.557247855 | M_Disease&NULL |
| 11 | 118816156 | rs587263 | C | T | M_Disease | 0.557247855 | M_Disease&NULL |
| 11 | 120413117 | rs2266132 | T | G | M_Disease | 0.557247855 | M_Disease&NULL |
| 11 | 121009800 | rs620778 | T | C | M_Disease | 0.557247855 | M_Disease&NULL |
| 11 | 121017795 | rs4936585 | T | G | M_Disease | 0.557247855 | M_Disease&NULL |
| 11 | 117430750 | rs4936386 | T | C | M_Disease | 0.558481112 | M_Disease&NULL |
| 11 | 119907777 | rs642878 | A | G | M_Disease | 0.558481112 | M_Disease&NULL |
| 11 | 121010116 | rs620164 | A | G | M_Disease | 0.557247855 | M_Disease&NULL |
| 11 | 121009384 | rs115903909 | A | G | M_Disease | 0.557247855 | M_Disease&NULL |
| 11 | 119904686 | rs1287591 | T | C | M_Disease | 0.556013891 | M_Disease&NULL |
| 11 | 116585625 | rs34100096 | A | G | M_Disease | 0.557247855 | M_Disease&NULL |
| 11 | 117987605 | rs640048 | G | T | M_Disease | 0.557247855 | M_Disease&NULL |
| 11 | 117033406 | rs4288786 | A | G | M_Disease | 0.557247855 | M_Disease&NULL |
| 11 | 121018451 | rs7108594 | G | A | M_Disease | 0.556013891 | M_Disease&NULL |
| 11 | 119304622 | rs61898858 | G | A | M_Disease | 0.556013891 | M_Disease&NULL |

| | | | | | | | | | |
|---|---|---|---|---|---|---|---|---|---|
| 11 | 118279936 | rs3194726 | C | T | 1.107 | -7977.388 | -7976.82 | 0.568 | 6.83E-06 |
| 11 | 116299456 | rs144466751 | C | T | 1.122 | -7977.388 | -7976.823 | 0.566 | 6.81E-06 |
| 11 | 121062486 | rs7110565 | C | T | 0.928 | -7977.388 | -7976.824 | 0.564 | 6.80E-06 |
| 11 | 120425101 | rs79985686 | C | T | 0.886 | -7977.388 | -7976.826 | 0.562 | 6.79E-06 |
| 11 | 118519213 | rs668628 | T | G | 0.923 | -7977.388 | -7976.827 | 0.561 | 6.78E-06 |
| 11 | 119864218 | rs874195 | G | A | 0.877 | -7977.388 | -7976.83 | 0.558 | 6.76E-06 |
| 11 | 119880819 | rs675246 | T | C | 0.891 | -7977.388 | -7976.83 | 0.558 | 6.76E-06 |
| 11 | 117034899 | rs189189723 | T | G | 1.146 | -7977.388 | -7976.831 | 0.557 | 6.75E-06 |
| 11 | 117308183 | rs73592180 | G | A | 1.095 | -7977.388 | -7976.831 | 0.557 | 6.75E-06 |
| 11 | 117449497 | rs11606425 | T | A | 1.07 | -7977.388 | -7976.831 | 0.557 | 6.75E-06 |
| 11 | 116758693 | rs12292278 | C | A | 1.119 | -7977.388 | -7976.839 | 0.549 | 6.70E-06 |
| 11 | 121015968 | rs4936583 | C | T | 0.928 | -7977.388 | -7976.84 | 0.548 | 6.69E-06 |
| 11 | 121016098 | rs10892676 | C | T | 0.928 | -7977.388 | -7976.842 | 0.546 | 6.68E-06 |
| 11 | 121016145 | rs4936584 | A | G | 0.928 | -7977.388 | -7976.842 | 0.546 | 6.68E-06 |
| 11 | 121016818 | rs12275038 | A | G | 0.928 | -7977.388 | -7976.842 | 0.546 | 6.68E-06 |
| 11 | 119519217 | rs11217372 | G | C | 0.908 | -7977.388 | -7976.845 | 0.543 | 6.66E-06 |
| 11 | 120430224 | rs80260914 | G | A | 0.886 | -7977.388 | -7976.846 | 0.543 | 6.66E-06 |
| 11 | 120425797 | rs12575945 | A | C | 0.887 | -7977.388 | -7976.853 | 0.536 | 6.61E-06 |
| 11 | 117449263 | rs11604377 | C | T | 1.07 | -7977.388 | -7976.857 | 0.531 | 6.58E-06 |
| 11 | 118809882 | rs11217099 | A | C | 1.085 | -7977.388 | -7976.858 | 0.531 | 6.58E-06 |
| 11 | 117004656 | rs11216288 | T | C | 0.896 | -7977.388 | -7976.862 | 0.527 | 6.55E-06 |
| 11 | 117386067 | rs2186762 | A | C | 0.938 | -7977.388 | -7976.865 | 0.524 | 6.53E-06 |
| 11 | 118574311 | rs524409 | T | C | 0.937 | -7977.388 | -7976.866 | 0.522 | 6.52E-06 |
| 11 | 118574675 | rs494459 | T | C | 0.937 | -7977.388 | -7976.866 | 0.522 | 6.52E-06 |
| 11 | 116771226 | rs117640781 | T | C | 1.165 | -7977.388 | -7976.87 | 0.518 | 6.49E-06 |
| 11 | 120719994 | rs11218030 | G | A | 1.082 | -7977.388 | -7976.871 | 0.517 | 6.49E-06 |
| 11 | 118573916 | rs627391 | T | A | 0.938 | -7977.388 | -7976.876 | 0.512 | 6.45E-06 |
| 11 | 118575011 | rs530901 | C | T | 0.938 | -7977.388 | -7976.876 | 0.512 | 6.45E-06 |
| 11 | 118575180 | rs626645 | C | T | 0.938 | -7977.388 | -7976.876 | 0.512 | 6.45E-06 |
| 11 | 118575419 | rs625735 | A | G | 0.938 | -7977.388 | -7976.876 | 0.512 | 6.45E-06 |
| 11 | 118575479 | rs637563 | G | A | 0.938 | -7977.388 | -7976.876 | 0.512 | 6.45E-06 |
| 11 | 118575693 | rs638514 | C | T | 0.938 | -7977.388 | -7976.876 | 0.512 | 6.45E-06 |
| 11 | 116801201 | rs645258 | A | C | 0.909 | -7977.388 | -7976.878 | 0.51 | 6.44E-06 |
| 11 | 120521305 | rs7125158 | C | A | 1.148 | -7977.388 | -7976.878 | 0.51 | 6.44E-06 |
| 11 | 119870780 | rs11217623 | T | C | 0.879 | -7977.388 | -7976.883 | 0.506 | 6.42E-06 |
| 11 | 120431744 | rs7121316 | A | G | 0.889 | -7977.388 | -7976.884 | 0.504 | 6.40E-06 |
| 11 | 121011258 | rs561227 | T | C | 0.937 | -7977.388 | -7976.885 | 0.503 | 6.40E-06 |
| 11 | 118573894 | rs576283 | G | A | 0.938 | -7977.388 | -7976.887 | 0.502 | 6.39E-06 |
| 11 | 118184271 | rs2231445 | C | T | 0.876 | -7977.388 | -7976.888 | 0.5 | 6.38E-06 |
| 11 | 116491621 | rs507262 | G | A | 0.913 | -7977.388 | -7976.891 | 0.498 | 6.36E-06 |
| 11 | 116492312 | rs534633 | G | A | 0.913 | -7977.388 | -7976.891 | 0.497 | 6.36E-06 |
| 11 | 121012492 | rs581786 | C | T | 0.929 | -7977.388 | -7976.892 | 0.497 | 6.36E-06 |
| 11 | 121012693 | rs528958 | T | G | 0.929 | -7977.388 | -7976.892 | 0.497 | 6.36E-06 |
| 11 | 121012831 | rs688106 | A | T | 0.929 | -7977.388 | -7976.892 | 0.497 | 6.36E-06 |
| 11 | 121012978 | rs665743 | A | G | 0.929 | -7977.388 | -7976.892 | 0.497 | 6.36E-06 |
| 11 | 117922730 | rs17599289 | T | C | 0.935 | -7977.388 | -7976.893 | 0.495 | 6.35E-06 |
| 11 | 121018870 | rs4623911 | G | A | 0.929 | -7977.388 | -7976.895 | 0.493 | 6.33E-06 |
| 11 | 121018999 | rs4641485 | G | A | 0.929 | -7977.388 | -7976.895 | 0.493 | 6.33E-06 |
| 11 | 119921812 | rs586112 | C | G | 0.888 | -7977.388 | -7976.896 | 0.492 | 6.33E-06 |
| 11 | 117908084 | rs61900560 | C | G | 0.935 | -7977.388 | -7976.897 | 0.491 | 6.32E-06 |
| 11 | 118751326 | rs10892306 | A | T | 1.114 | -7977.388 | -7976.897 | 0.491 | 6.32E-06 |
| 11 | 116999059 | rs4516002 | A | G | 0.92 | -7977.388 | -7976.898 | 0.49 | 6.31E-06 |
| 11 | 117197621 | rs680908 | G | C | 0.896 | -7977.388 | -7976.898 | 0.49 | 6.31E-06 |
| 11 | 117198638 | rs566155 | C | T | 0.896 | -7977.388 | -7976.898 | 0.49 | 6.31E-06 |
| 11 | 121018488 | rs12285468 | C | T | 0.929 | -7977.388 | -7976.899 | 0.489 | 6.31E-06 |
| 11 | 120431935 | rs12283999 | G | C | 0.889 | -7977.388 | -7976.901 | 0.487 | 6.30E-06 |
| 11 | 116511522 | rs519000 | T | C | 0.925 | -7977.388 | -7976.902 | 0.486 | 6.29E-06 |
| 11 | 119866879 | rs9787805 | C | G | 0.88 | -7977.388 | -7976.903 | 0.486 | 6.29E-06 |
| 11 | 116805204 | rs562179 | T | C | 0.914 | -7977.388 | -7976.903 | 0.485 | 6.28E-06 |
| 11 | 117915445 | rs7926330 | T | C | 0.935 | -7977.388 | -7976.903 | 0.485 | 6.28E-06 |
| 11 | 116295218 | rs75375536 | G | A | 1.109 | -7977.388 | -7976.904 | 0.484 | 6.28E-06 |
| 11 | 121007777 | rs4936580 | A | G | 0.937 | -7977.388 | -7976.904 | 0.484 | 6.28E-06 |
| 11 | 116491177 | rs502816 | A | C | 0.913 | -7977.388 | -7976.905 | 0.483 | 6.27E-06 |
| 11 | 116491428 | rs505397 | G | T | 0.913 | -7977.388 | -7976.905 | 0.483 | 6.27E-06 |
| 11 | 117181919 | rs497902 | G | C | 1.07 | -7977.388 | -7976.909 | 0.48 | 6.25E-06 |
| 11 | 121014228 | rs11218158 | G | A | 0.929 | -7977.388 | -7976.911 | 0.477 | 6.23E-06 |
| 11 | 121014459 | rs877538 | A | C | 0.929 | -7977.388 | -7976.911 | 0.477 | 6.23E-06 |
| 11 | 121014860 | rs7114879 | T | C | 0.929 | -7977.388 | -7976.912 | 0.477 | 6.23E-06 |
| 11 | 121010788 | rs607014 | A | G | 0.929 | -7977.388 | -7976.917 | 0.471 | 6.20E-06 |
| 11 | 121017604 | rs7120647 | C | T | 0.929 | -7977.388 | -7976.917 | 0.471 | 6.20E-06 |
| 11 | 121017863 | rs4936586 | A | C | 0.929 | -7977.388 | -7976.917 | 0.471 | 6.20E-06 |
| 11 | 121018007 | rs4936587 | G | A | 0.929 | -7977.388 | -7976.917 | 0.471 | 6.20E-06 |
| 11 | 121018485 | rs12285467 | C | T | 0.929 | -7977.388 | -7976.917 | 0.471 | 6.20E-06 |
| 11 | 117183914 | rs477036 | C | T | 0.939 | -7977.388 | -7976.918 | 0.47 | 6.19E-06 |
| 11 | 119866718 | rs9787801 | A | G | 0.88 | -7977.388 | -7976.919 | 0.47 | 6.19E-06 |
| 11 | 121014983 | rs7114915 | G | A | 0.929 | -7977.388 | -7976.923 | 0.466 | 6.16E-06 |
| 11 | 117402325 | rs61350190 | T | G | 1.071 | -7977.388 | -7976.924 | 0.464 | 6.15E-06 |
| 11 | 121011485 | rs638463 | C | T | 0.938 | -7977.388 | -7976.924 | 0.464 | 6.15E-06 |
| 11 | 116839761 | rs1241657 | T | C | 0.91 | -7977.388 | -7976.925 | 0.463 | 6.15E-06 |
| 11 | 120412823 | rs2510876 | G | A | 0.933 | -7977.388 | -7976.926 | 0.463 | 6.15E-06 |
| 11 | 117541718 | rs141116752 | T | C | 0.864 | -7977.388 | -7976.926 | 0.462 | 6.14E-06 |
| 11 | 121010820 | rs606674 | G | A | 0.929 | -7977.388 | -7976.926 | 0.462 | 6.14E-06 |
| 11 | 118816156 | rs587263 | C | T | 0.938 | -7977.388 | -7976.928 | 0.461 | 6.13E-06 |
| 11 | 120413117 | rs2266132 | T | G | 0.933 | -7977.388 | -7976.928 | 0.461 | 6.13E-06 |
| 11 | 121009800 | rs620778 | T | C | 0.929 | -7977.388 | -7976.927 | 0.461 | 6.13E-06 |
| 11 | 121017795 | rs4936585 | T | G | 0.929 | -7977.388 | -7976.927 | 0.461 | 6.13E-06 |
| 11 | 117430750 | rs4936386 | T | C | 0.863 | -7977.388 | -7976.928 | 0.46 | 6.13E-06 |
| 11 | 119907777 | rs642878 | A | G | 0.889 | -7977.388 | -7976.929 | 0.46 | 6.13E-06 |
| 11 | 121010116 | rs620164 | A | G | 0.929 | -7977.388 | -7976.928 | 0.46 | 6.13E-06 |
| 11 | 121009384 | rs115903909 | A | G | 0.929 | -7977.388 | -7976.935 | 0.453 | 6.08E-06 |
| 11 | 119904686 | rs1287591 | T | C | 0.936 | -7977.388 | -7976.937 | 0.452 | 6.08E-06 |
| 11 | 116585625 | rs34100096 | A | G | 0.874 | -7977.388 | -7976.937 | 0.451 | 6.07E-06 |
| 11 | 117987605 | rs640048 | G | T | 1.126 | -7977.388 | -7976.937 | 0.451 | 6.07E-06 |
| 11 | 117033406 | rs4288786 | A | G | 1.149 | -7977.388 | -7976.941 | 0.448 | 6.05E-06 |
| 11 | 121018451 | rs7108594 | G | A | 0.929 | -7977.388 | -7976.94 | 0.448 | 6.05E-06 |
| 11 | 119304622 | rs61898858 | G | A | 1.124 | -7977.388 | -7976.943 | 0.446 | 6.04E-06 |

| | | | | | | | |
|---|---|---|---|---|---|---|---|
| 11 | 117373635 | rs686133 | G | T | M_Disease | 0.554779235 | M_Disease&NULL |
| 11 | 118590961 | rs111227443 | T | C | M_Disease | 0.556013891 | M_Disease&NULL |
| 11 | 119904782 | rs373191677 | A | G | M_Disease | 0.554779235 | M_Disease&NULL |
| 11 | 117994614 | rs7940318 | G | A | M_Disease | 0.556013891 | M_Disease&NULL |
| 11 | 121050766 | rs4936591 | G | A | M_Disease | 0.554779235 | M_Disease&NULL |
| 11 | 121051868 | rs4935762 | A | G | M_Disease | 0.554779235 | M_Disease&NULL |
| 11 | 117905204 | rs56310543 | A | G | M_Disease | 0.554779235 | M_Disease&NULL |
| 11 | 120828334 | rs375508071 | G | A | M_Disease | 0.554779235 | M_Disease&NULL |
| 11 | 121018237 | rs7105356 | G | A | M_Disease | 0.554779235 | M_Disease&NULL |
| 11 | 120529153 | rs76635827 | C | G | M_Disease | 0.554779235 | M_Disease&NULL |
| 11 | 116787192 | rs17120152 | C | T | M_Disease | 0.554779235 | M_Disease&NULL |
| 11 | 118263388 | rs2276422 | G | C | M_Disease | 0.553543903 | M_Disease&NULL |
| 11 | 116774475 | rs78832854 | A | T | M_Disease | 0.553543903 | M_Disease&NULL |
| 11 | 120071508 | rs11217762 | A | G | M_Disease | 0.553543903 | M_Disease&NULL |
| 11 | 116732231 | rs80026097 | C | T | M_Disease | 0.554779235 | M_Disease&NULL |
| 11 | 121024461 | rs11826847 | A | G | M_Disease | 0.553543903 | M_Disease&NULL |
| 11 | 117908926 | rs61900561 | A | G | M_Disease | 0.553543903 | M_Disease&NULL |
| 11 | 118265734 | rs79092677 | G | T | M_Disease | 0.553543903 | M_Disease&NULL |
| 11 | 118267763 | rs10892237 | C | T | M_Disease | 0.553543903 | M_Disease&NULL |
| 11 | 120606965 | rs11607895 | T | C | M_Disease | 0.553543903 | M_Disease&NULL |
| 11 | 116735788 | rs6589576 | C | G | M_Disease | 0.553543903 | M_Disease&NULL |
| 11 | 121018507 | rs7108741 | T | C | M_Disease | 0.553543903 | M_Disease&NULL |
| 11 | 116515528 | rs78221557 | A | G | M_Disease | 0.553543903 | M_Disease&NULL |
| 11 | 120899875 | rs113685600 | G | C | M_Disease | 0.553543903 | M_Disease&NULL |
| 11 | 116765476 | rs77349713 | C | T | M_Disease | 0.55230791 | M_Disease&NULL |
| 11 | 121009627 | rs541623 | A | G | M_Disease | 0.55230791 | M_Disease&NULL |
| 11 | 116897073 | rs117158703 | T | C | M_Disease | 0.553543903 | M_Disease&NULL |
| 11 | 121059326 | rs12279811 | T | C | M_Disease | 0.55230791 | M_Disease&NULL |
| 11 | 120008468 | rs1670195 | C | G | M_Disease | 0.553543903 | M_Disease&NULL |
| 11 | 120413132 | rs2266134 | C | G | M_Disease | 0.55230791 | M_Disease&NULL |
| 11 | 117434487 | rs17509913 | A | G | M_Disease | 0.55230791 | M_Disease&NULL |
| 11 | 117377135 | rs694093 | T | G | M_Disease | 0.551071269 | M_Disease&NULL |
| 11 | 118265227 | rs7131196 | C | T | M_Disease | 0.551071269 | M_Disease&NULL |
| 11 | 121010550 | rs607999 | G | A | M_Disease | 0.551071269 | M_Disease&NULL |
| 11 | 121015763 | rs4936582 | C | T | M_Disease | 0.551071269 | M_Disease&NULL |
| 11 | 116922586 | rs111530588 | A | G | M_Disease | 0.55230791 | M_Disease&NULL |
| 11 | 117513154 | rs613244 | A | G | M_Disease | 0.551071269 | M_Disease&NULL |
| 11 | 117406992 | rs7103687 | A | C | M_Disease | 0.551071269 | M_Disease&NULL |
| 11 | 118580348 | rs516503 | T | C | M_Disease | 0.551071269 | M_Disease&NULL |
| 11 | 116880715 | rs79862770 | T | C | M_Disease | 0.551071269 | M_Disease&NULL |
| 11 | 116881126 | rs34998970 | G | A | M_Disease | 0.551071269 | M_Disease&NULL |
| 11 | 116565732 | rs4938299 | A | G | M_Disease | 0.551071269 | M_Disease&NULL |
| 11 | 118725189 | rs112965917 | A | T | M_Disease | 0.551071269 | M_Disease&NULL |
| 11 | 120703916 | rs11218019 | C | A | M_Disease | 0.551071269 | M_Disease&NULL |
| 11 | 118712509 | rs75488969 | T | C | M_Disease | 0.551071269 | M_Disease&NULL |
| 11 | 118598612 | rs568688 | G | T | M_Disease | 0.549833997 | M_Disease&NULL |
| 11 | 117400550 | rs7130708 | A | G | M_Disease | 0.549833997 | M_Disease&NULL |
| 11 | 121010427 | rs513182 | C | T | M_Disease | 0.549833997 | M_Disease&NULL |
| 11 | 117197018 | rs540273 | T | C | M_Disease | 0.549833997 | M_Disease&NULL |
| 11 | 116762028 | rs12285074 | A | G | M_Disease | 0.549833997 | M_Disease&NULL |
| 11 | 119519916 | rs11217373 | G | C | M_Disease | 0.549833997 | M_Disease&NULL |
| 11 | 117302152 | rs57154584 | C | T | M_Disease | 0.549833997 | M_Disease&NULL |
| 11 | 117444090 | rs76948040 | T | C | M_Disease | 0.549833997 | M_Disease&NULL |
| 11 | 121011098 | rs605379 | T | C | M_Disease | 0.549833997 | M_Disease&NULL |
| 11 | 121063130 | rs1943656 | T | C | M_Disease | 0.549833997 | M_Disease&NULL |
| 11 | 116943066 | rs116582417 | A | G | M_Disease | 0.549833997 | M_Disease&NULL |
| 11 | 116897163 | rs76882215 | C | T | M_Disease | 0.549833997 | M_Disease&NULL |
| 11 | 116827369 | rs17120177 | C | T | M_Disease | 0.548596109 | M_Disease&NULL |
| 11 | 117944519 | rs947891 | T | C | M_Disease | 0.548596109 | M_Disease&NULL |
| 11 | 118260017 | rs11216864 | T | C | M_Disease | 0.548596109 | M_Disease&NULL |
| 11 | 118264353 | rs140967993 | G | A | M_Disease | 0.549833997 | M_Disease&NULL |
| 11 | 117431045 | rs623052 | G | A | M_Disease | 0.548596109 | M_Disease&NULL |
| 11 | 121018771 | rs4436552 | T | A | M_Disease | 0.548596109 | M_Disease&NULL |
| 11 | 121018772 | rs4539330 | C | G | M_Disease | 0.548596109 | M_Disease&NULL |
| 11 | 116677048 | rs12802202 | C | T | M_Disease | 0.548596109 | M_Disease&NULL |
| 11 | 117090225 | rs45577233 | T | A | M_Disease | 0.548596109 | M_Disease&NULL |
| 11 | 118267686 | rs79990075 | T | C | M_Disease | 0.548596109 | M_Disease&NULL |
| 11 | 116768491 | rs1474828939 | T | C | M_Disease | 0.548596109 | M_Disease&NULL |
| 11 | 117914450 | rs61900563 | A | G | M_Disease | 0.547357618 | M_Disease&NULL |
| 11 | 116789438 | rs7130160 | C | T | M_Disease | 0.548596109 | M_Disease&NULL |
| 11 | 116792299 | rs1020392 | A | G | M_Disease | 0.548596109 | M_Disease&NULL |
| 11 | 116792336 | rs1020393 | T | C | M_Disease | 0.548596109 | M_Disease&NULL |
| 11 | 118285789 | rs75017819 | G | A | M_Disease | 0.547357618 | M_Disease&NULL |
| 11 | 119728661 | rs145723586 | T | C | M_Disease | 0.547357618 | M_Disease&NULL |
| 11 | 116781699 | rs78234966 | C | T | M_Disease | 0.547357618 | M_Disease&NULL |
| 11 | 117251521 | rs2061293 | G | C | M_Disease | 0.547357618 | M_Disease&NULL |
| 11 | 117252489 | rs897836 | T | C | M_Disease | 0.547357618 | M_Disease&NULL |
| 11 | 120412867 | rs2510877 | G | T | M_Disease | 0.547357618 | M_Disease&NULL |
| 11 | 121053084 | rs4499012 | A | G | M_Disease | 0.547357618 | M_Disease&NULL |
| 11 | 117004891 | rs111519461 | A | T | M_Disease | 0.547357618 | M_Disease&NULL |
| 11 | 118588998 | rs570424 | C | T | M_Disease | 0.547357618 | M_Disease&NULL |
| 11 | 118180887 | rs146525935 | C | T | M_Disease | 0.547357618 | M_Disease&NULL |
| 11 | 116970761 | rs76567973 | G | C | M_Disease | 0.547357618 | M_Disease&NULL |
| 11 | 116833399 | rs7120762 | C | A | M_Disease | 0.547357618 | M_Disease&NULL |
| 11 | 116852927 | rs7127416 | T | C | M_Disease | 0.547357618 | M_Disease&NULL |
| 11 | 117301818 | rs4356265 | T | C | M_Disease | 0.546118541 | M_Disease&NULL |
| 11 | 118137524 | rs12273260 | T | C | M_Disease | 0.547357618 | M_Disease&NULL |
| 11 | 116877159 | rs10790169 | A | C | M_Disease | 0.546118541 | M_Disease&NULL |
| 11 | 117434728 | rs495118 | G | A | M_Disease | 0.546118541 | M_Disease&NULL |
| 11 | 118580906 | rs496058 | G | A | M_Disease | 0.546118541 | M_Disease&NULL |
| 11 | 117195136 | rs556034 | G | C | M_Disease | 0.546118541 | M_Disease&NULL |
| 11 | 118049817 | rs116848264 | A | T | M_Disease | 0.546118541 | M_Disease&NULL |
| 11 | 120520418 | rs7124091 | G | C | M_Disease | 0.546118541 | M_Disease&NULL |
| 11 | 121011416 | rs596882 | C | G | M_Disease | 0.546118541 | M_Disease&NULL |
| 11 | 116233829 | rs478350 | T | C | M_Disease | 0.546118541 | M_Disease&NULL |
| 11 | 119337452 | rs187900569 | C | G | M_Disease | 0.546118541 | M_Disease&NULL |

| | | | | | | | | | |
|---|---|---|---|---|---|---|---|---|---|
| 11 | 117373635 | rs686133 | G | T | 0.937 | -7977.388 | -7976.946 | 0.442 | 6.02E-06 |
| 11 | 118590961 | rs111227443 | T | C | 0.859 | -7977.388 | -7976.948 | 0.441 | 6.01E-06 |
| 11 | 119904782 | rs373191677 | A | G | 0.936 | -7977.388 | -7976.948 | 0.441 | 6.01E-06 |
| 11 | 117994614 | rs7940318 | G | A | 1.122 | -7977.388 | -7976.948 | 0.44 | 6.01E-06 |
| 11 | 121050766 | rs4936591 | G | A | 0.93 | -7977.388 | -7976.949 | 0.439 | 6.00E-06 |
| 11 | 121051868 | rs4935762 | A | G | 0.93 | -7977.388 | -7976.949 | 0.439 | 6.00E-06 |
| 11 | 117905204 | rs56310543 | A | G | 0.936 | -7977.388 | -7976.952 | 0.437 | 5.99E-06 |
| 11 | 120828334 | rs375508071 | G | A | 0.939 | -7977.388 | -7976.953 | 0.435 | 5.98E-06 |
| 11 | 121018237 | rs7105356 | G | A | 0.93 | -7977.388 | -7976.955 | 0.434 | 5.97E-06 |
| 11 | 120529153 | rs76635827 | C | G | 0.869 | -7977.388 | -7976.958 | 0.431 | 5.95E-06 |
| 11 | 116787192 | rs17120152 | C | T | 1.136 | -7977.388 | -7976.959 | 0.429 | 5.94E-06 |
| 11 | 118263388 | rs2276422 | G | C | 1.103 | -7977.388 | -7976.959 | 0.429 | 5.94E-06 |
| 11 | 116774475 | rs78832854 | A | T | 1.115 | -7977.388 | -7976.961 | 0.428 | 5.93E-06 |
| 11 | 120071508 | rs11217762 | A | G | 1.107 | -7977.388 | -7976.96 | 0.428 | 5.93E-06 |
| 11 | 116732231 | rs80026097 | C | T | 1.142 | -7977.388 | -7976.961 | 0.427 | 5.93E-06 |
| 11 | 121024461 | rs11826847 | A | G | 0.93 | -7977.388 | -7976.962 | 0.426 | 5.92E-06 |
| 11 | 117908926 | rs61900561 | A | G | 0.936 | -7977.388 | -7976.963 | 0.425 | 5.92E-06 |
| 11 | 118265734 | rs79092677 | G | T | 1.103 | -7977.388 | -7976.964 | 0.424 | 5.91E-06 |
| 11 | 118267763 | rs10892237 | C | T | 1.103 | -7977.388 | -7976.964 | 0.424 | 5.91E-06 |
| 11 | 120606965 | rs11607895 | T | C | 0.896 | -7977.388 | -7976.965 | 0.424 | 5.91E-06 |
| 11 | 116735788 | rs6589576 | C | G | 0.915 | -7977.388 | -7976.966 | 0.423 | 5.90E-06 |
| 11 | 121018507 | rs7108741 | T | C | 0.93 | -7977.388 | -7976.965 | 0.423 | 5.90E-06 |
| 11 | 116515528 | rs78221557 | A | G | 1.157 | -7977.388 | -7976.969 | 0.42 | 5.89E-06 |
| 11 | 120899875 | rs113685600 | G | C | 0.875 | -7977.388 | -7976.969 | 0.42 | 5.89E-06 |
| 11 | 116765476 | rs77349713 | C | T | 1.114 | -7977.388 | -7976.969 | 0.419 | 5.88E-06 |
| 11 | 121009627 | rs541623 | A | G | 0.93 | -7977.388 | -7976.97 | 0.419 | 5.88E-06 |
| 11 | 116897073 | rs117158703 | T | C | 1.158 | -7977.388 | -7976.971 | 0.417 | 5.87E-06 |
| 11 | 121059326 | rs12279811 | T | C | 0.93 | -7977.388 | -7976.971 | 0.417 | 5.87E-06 |
| 11 | 120008468 | rs1670195 | C | G | 0.865 | -7977.388 | -7976.972 | 0.416 | 5.86E-06 |
| 11 | 120413132 | rs2266134 | C | G | 0.934 | -7977.388 | -7976.972 | 0.416 | 5.86E-06 |
| 11 | 117434487 | rs17509913 | A | G | 1.097 | -7977.388 | -7976.974 | 0.414 | 5.85E-06 |
| 11 | 117377135 | rs694093 | T | G | 0.937 | -7977.388 | -7976.976 | 0.412 | 5.84E-06 |
| 11 | 118265227 | rs7131196 | C | T | 1.102 | -7977.388 | -7976.978 | 0.411 | 5.83E-06 |
| 11 | 121010550 | rs607999 | G | A | 0.93 | -7977.388 | -7976.978 | 0.411 | 5.83E-06 |
| 11 | 121015763 | rs4936582 | C | T | 0.93 | -7977.388 | -7976.978 | 0.411 | 5.83E-06 |
| 11 | 116922586 | rs111530588 | A | G | 1.115 | -7977.388 | -7976.978 | 0.41 | 5.83E-06 |
| 11 | 117513154 | rs613244 | A | G | 0.935 | -7977.388 | -7976.979 | 0.409 | 5.82E-06 |
| 11 | 117406992 | rs7103687 | A | C | 1.071 | -7977.388 | -7976.981 | 0.408 | 5.82E-06 |
| 11 | 118580348 | rs516503 | T | C | 0.939 | -7977.388 | -7976.981 | 0.407 | 5.81E-06 |
| 11 | 116880715 | rs79862770 | T | C | 1.136 | -7977.388 | -7976.982 | 0.406 | 5.81E-06 |
| 11 | 116881126 | rs34998970 | G | A | 1.136 | -7977.388 | -7976.982 | 0.406 | 5.81E-06 |
| 11 | 116565732 | rs4938299 | A | G | 0.886 | -7977.388 | -7976.984 | 0.405 | 5.80E-06 |
| 11 | 118725189 | rs112965917 | A | T | 0.892 | -7977.388 | -7976.983 | 0.405 | 5.80E-06 |
| 11 | 120703916 | rs11218019 | C | A | 1.083 | -7977.388 | -7976.986 | 0.403 | 5.79E-06 |
| 11 | 118712509 | rs75488969 | T | C | 0.913 | -7977.388 | -7976.986 | 0.402 | 5.78E-06 |
| 11 | 118598612 | rs568688 | G | T | 0.939 | -7977.388 | -7976.988 | 0.401 | 5.78E-06 |
| 11 | 117400550 | rs7130708 | A | G | 1.07 | -7977.388 | -7976.988 | 0.4 | 5.77E-06 |
| 11 | 121010427 | rs513182 | C | T | 0.93 | -7977.388 | -7976.989 | 0.4 | 5.77E-06 |
| 11 | 117197018 | rs540273 | T | C | 0.899 | -7977.388 | -7976.989 | 0.399 | 5.76E-06 |
| 11 | 116762028 | rs12285074 | A | G | 0.913 | -7977.388 | -7976.991 | 0.397 | 5.75E-06 |
| 11 | 119519916 | rs11217373 | G | C | 0.911 | -7977.388 | -7976.991 | 0.397 | 5.75E-06 |
| 11 | 117302152 | rs57154584 | C | T | 1.093 | -7977.388 | -7976.993 | 0.395 | 5.74E-06 |
| 11 | 117444090 | rs76948040 | T | C | 1.1 | -7977.388 | -7976.993 | 0.395 | 5.74E-06 |
| 11 | 121011098 | rs605379 | T | C | 0.938 | -7977.388 | -7976.995 | 0.393 | 5.73E-06 |
| 11 | 121063130 | rs1943656 | T | C | 0.93 | -7977.388 | -7976.995 | 0.393 | 5.73E-06 |
| 11 | 116943066 | rs116582417 | A | G | 1.115 | -7977.388 | -7976.997 | 0.391 | 5.72E-06 |
| 11 | 116897163 | rs76882215 | C | T | 1.14 | -7977.388 | -7976.998 | 0.39 | 5.71E-06 |
| 11 | 116827369 | rs17120177 | C | T | 1.115 | -7977.388 | -7977.001 | 0.388 | 5.70E-06 |
| 11 | 117944519 | rs947891 | T | C | 0.938 | -7977.388 | -7977.001 | 0.388 | 5.70E-06 |
| 11 | 118260017 | rs11216864 | T | C | 1.102 | -7977.388 | -7977.001 | 0.388 | 5.70E-06 |
| 11 | 118264353 | rs140967993 | G | A | 0.883 | -7977.388 | -7977 | 0.388 | 5.70E-06 |
| 11 | 117431045 | rs623052 | G | A | 0.94 | -7977.388 | -7977.004 | 0.385 | 5.68E-06 |
| 11 | 121018771 | rs4436552 | T | A | 0.93 | -7977.388 | -7977.005 | 0.384 | 5.68E-06 |
| 11 | 121018772 | rs4539330 | C | G | 0.93 | -7977.388 | -7977.005 | 0.384 | 5.68E-06 |
| 11 | 116677048 | rs12802202 | C | T | 0.877 | -7977.388 | -7977.006 | 0.383 | 5.67E-06 |
| 11 | 117090225 | rs45577233 | T | A | 1.14 | -7977.388 | -7977.007 | 0.381 | 5.66E-06 |
| 11 | 118267686 | rs79990075 | T | C | 0.883 | -7977.388 | -7977.009 | 0.38 | 5.66E-06 |
| 11 | 116768491 | rs1474828939 | T | C | 1.134 | -7977.388 | -7977.009 | 0.379 | 5.65E-06 |
| 11 | 117914450 | rs61900563 | A | G | 0.937 | -7977.388 | -7977.01 | 0.379 | 5.65E-06 |
| 11 | 116789438 | rs7130160 | C | T | 1.134 | -7977.388 | -7977.011 | 0.378 | 5.65E-06 |
| 11 | 116792299 | rs1020392 | A | G | 1.134 | -7977.388 | -7977.011 | 0.378 | 5.65E-06 |
| 11 | 116792336 | rs1020393 | T | C | 1.134 | -7977.388 | -7977.011 | 0.378 | 5.65E-06 |
| 11 | 118285789 | rs75017819 | G | A | 1.101 | -7977.388 | -7977.01 | 0.378 | 5.65E-06 |
| 11 | 119728661 | rs145723586 | T | C | 0.886 | -7977.388 | -7977.01 | 0.378 | 5.65E-06 |
| 11 | 116781699 | rs78234966 | C | T | 1.134 | -7977.388 | -7977.011 | 0.377 | 5.64E-06 |
| 11 | 117251521 | rs2061293 | G | C | 0.903 | -7977.388 | -7977.011 | 0.377 | 5.64E-06 |
| 11 | 117252489 | rs897836 | T | C | 0.903 | -7977.388 | -7977.011 | 0.377 | 5.64E-06 |
| 11 | 120412867 | rs2510877 | G | T | 0.934 | -7977.388 | -7977.012 | 0.376 | 5.63E-06 |
| 11 | 121053084 | rs4499012 | A | G | 0.931 | -7977.388 | -7977.013 | 0.375 | 5.63E-06 |
| 11 | 117004891 | rs111519461 | A | T | 1.144 | -7977.388 | -7977.014 | 0.374 | 5.62E-06 |
| 11 | 118588998 | rs570424 | C | T | 0.939 | -7977.388 | -7977.015 | 0.374 | 5.62E-06 |
| 11 | 118180887 | rs146525935 | C | T | 0.881 | -7977.388 | -7977.016 | 0.372 | 5.61E-06 |
| 11 | 116970761 | rs76567973 | G | C | 1.114 | -7977.388 | -7977.018 | 0.371 | 5.61E-06 |
| 11 | 116833399 | rs7120762 | C | A | 1.134 | -7977.388 | -7977.019 | 0.369 | 5.59E-06 |
| 11 | 116852927 | rs7127416 | T | C | 1.134 | -7977.388 | -7977.019 | 0.369 | 5.59E-06 |
| 11 | 117301818 | rs4356265 | T | C | 1.09 | -7977.388 | -7977.021 | 0.368 | 5.59E-06 |
| 11 | 118137524 | rs12273260 | T | C | 0.882 | -7977.388 | -7977.02 | 0.368 | 5.59E-06 |
| 11 | 116877159 | rs10790169 | A | C | 0.916 | -7977.388 | -7977.022 | 0.367 | 5.58E-06 |
| 11 | 117434728 | rs495118 | G | A | 0.919 | -7977.388 | -7977.023 | 0.366 | 5.58E-06 |
| 11 | 118580906 | rs496058 | G | A | 0.939 | -7977.388 | -7977.023 | 0.366 | 5.58E-06 |
| 11 | 117195136 | rs556034 | G | C | 0.9 | -7977.388 | -7977.024 | 0.365 | 5.57E-06 |
| 11 | 118049817 | rs116848264 | A | T | 0.878 | -7977.388 | -7977.023 | 0.365 | 5.57E-06 |
| 11 | 120520418 | rs7124091 | G | C | 1.14 | -7977.388 | -7977.023 | 0.365 | 5.57E-06 |
| 11 | 121011416 | rs596882 | C | G | 0.939 | -7977.388 | -7977.025 | 0.364 | 5.57E-06 |
| 11 | 116233829 | rs478350 | T | C | 1.069 | -7977.388 | -7977.025 | 0.363 | 5.56E-06 |
| 11 | 119337452 | rs187900569 | C | G | 0.865 | -7977.388 | -7977.025 | 0.363 | 5.56E-06 |

| | | | | | | | |
|---|---|---|---|---|---|---|---|
| 11 | 116512466 | rs4938293 | T | C | M_Disease | 0.544878892 | M_Disease&NULL |
| 11 | 116780918 | rs6589578 | T | C | M_Disease | 0.546118541 | M_Disease&NULL |
| 11 | 116782582 | rs7106782 | G | A | M_Disease | 0.546118541 | M_Disease&NULL |
| 11 | 116879679 | rs117901629 | A | G | M_Disease | 0.546118541 | M_Disease&NULL |
| 11 | 119885462 | rs630332 | A | G | M_Disease | 0.546118541 | M_Disease&NULL |
| 11 | 117534353 | rs630024 | A | C | M_Disease | 0.544878892 | M_Disease&NULL |
| 11 | 116765925 | rs74881320 | C | A | M_Disease | 0.544878892 | M_Disease&NULL |
| 11 | 116835363 | rs59283948 | A | G | M_Disease | 0.546118541 | M_Disease&NULL |
| 11 | 121018391 | rs7108587 | T | C | M_Disease | 0.544878892 | M_Disease&NULL |
| 11 | 117486730 | rs7942967 | T | C | M_Disease | 0.544878892 | M_Disease&NULL |
| 11 | 120521504 | rs7128690 | T | G | M_Disease | 0.544878892 | M_Disease&NULL |
| 11 | 117913074 | rs4936421 | C | A | M_Disease | 0.544878892 | M_Disease&NULL |
| 11 | 120428134 | rs2248404 | A | G | M_Disease | 0.544878892 | M_Disease&NULL |
| 11 | 121066195 | rs524392 | A | T | M_Disease | 0.543638687 | M_Disease&NULL |
| 11 | 116953644 | rs11216270 | A | C | M_Disease | 0.543638687 | M_Disease&NULL |
| 11 | 117525125 | rs583983 | T | C | M_Disease | 0.543638687 | M_Disease&NULL |
| 11 | 117913315 | rs55730046 | T | C | M_Disease | 0.543638687 | M_Disease&NULL |
| 11 | 120520282 | rs7121104 | T | G | M_Disease | 0.544878892 | M_Disease&NULL |
| 11 | 117395497 | rs55642294 | A | C | M_Disease | 0.543638687 | M_Disease&NULL |
| 11 | 118748170 | rs77074463 | C | G | M_Disease | 0.543638687 | M_Disease&NULL |
| 11 | 121011342 | rs560365 | T | C | M_Disease | 0.542397941 | M_Disease&NULL |
| 11 | 117268683 | rs34114773 | C | T | M_Disease | 0.542397941 | M_Disease&NULL |
| 11 | 117361165 | rs115325870 | T | C | M_Disease | 0.543638687 | M_Disease&NULL |
| 11 | 118866505 | rs7120982 | A | G | M_Disease | 0.542397941 | M_Disease&NULL |
| 11 | 119941807 | rs10790363 | A | G | M_Disease | 0.542397941 | M_Disease&NULL |
| 11 | 116778813 | rs35184536 | G | A | M_Disease | 0.542397941 | M_Disease&NULL |
| 11 | 116901175 | rs79257193 | G | A | M_Disease | 0.542397941 | M_Disease&NULL |
| 11 | 117399606 | rs7126500 | T | G | M_Disease | 0.541156668 | M_Disease&NULL |
| 11 | 120503083 | rs11217933 | A | G | M_Disease | 0.541156668 | M_Disease&NULL |
| 11 | 116625897 | rs67120644 | C | T | M_Disease | 0.541156668 | M_Disease&NULL |
| 11 | 117790342 | rs11216620 | T | C | M_Disease | 0.542397941 | M_Disease&NULL |
| 11 | 119582832 | rs4409845 | G | A | M_Disease | 0.542397941 | M_Disease&NULL |
| 11 | 120414586 | rs2266100 | A | G | M_Disease | 0.541156668 | M_Disease&NULL |
| 11 | 116883716 | rs7110042 | T | C | M_Disease | 0.542397941 | M_Disease&NULL |
| 11 | 116890488 | rs138975985 | T | C | M_Disease | 0.542397941 | M_Disease&NULL |
| 11 | 116333012 | rs73009128 | T | C | M_Disease | 0.541156668 | M_Disease&NULL |
| 11 | 116937361 | rs79955870 | T | C | M_Disease | 0.541156668 | M_Disease&NULL |
| 11 | 116940210 | rs11216261 | C | T | M_Disease | 0.541156668 | M_Disease&NULL |
| 11 | 117248832 | rs11216370 | A | G | M_Disease | 0.541156668 | M_Disease&NULL |
| 11 | 117430025 | rs75375456 | T | G | M_Disease | 0.541156668 | M_Disease&NULL |
| 11 | 116806616 | rs76384483 | C | T | M_Disease | 0.541156668 | M_Disease&NULL |
| 11 | 118788081 | rs7929825 | G | C | M_Disease | 0.539914885 | M_Disease&NULL |
| 11 | 119593446 | rs11604214 | A | G | M_Disease | 0.541156668 | M_Disease&NULL |
| 11 | 121011325 | rs560401 | G | A | M_Disease | 0.539914885 | M_Disease&NULL |
| 11 | 116955934 | rs2368149 | A | G | M_Disease | 0.539914885 | M_Disease&NULL |
| 11 | 116956990 | rs12281358 | C | T | M_Disease | 0.539914885 | M_Disease&NULL |
| 11 | 117074299 | rs2269397 | C | T | M_Disease | 0.539914885 | M_Disease&NULL |
| 11 | 118006853 | rs113579428 | T | C | M_Disease | 0.541156668 | M_Disease&NULL |
| 11 | 118178757 | rs11601508 | A | G | M_Disease | 0.539914885 | M_Disease&NULL |
| 11 | 116772486 | rs17120131 | C | T | M_Disease | 0.539914885 | M_Disease&NULL |
| 11 | 116773318 | rs17120132 | A | T | M_Disease | 0.539914885 | M_Disease&NULL |
| 11 | 116773761 | rs17120136 | G | T | M_Disease | 0.539914885 | M_Disease&NULL |
| 11 | 116775196 | rs12280030 | A | T | M_Disease | 0.539914885 | M_Disease&NULL |
| 11 | 116811878 | rs114034580 | G | A | M_Disease | 0.539914885 | M_Disease&NULL |
| 11 | 118811878 | rs74891547 | C | G | M_Disease | 0.539914885 | M_Disease&NULL |
| 11 | 120405437 | rs2508653 | T | C | M_Disease | 0.539914885 | M_Disease&NULL |
| 11 | 119881100 | rs2850802 | A | G | M_Disease | 0.539914885 | M_Disease&NULL |
| 11 | 116907715 | rs75216569 | G | A | M_Disease | 0.539914885 | M_Disease&NULL |
| 11 | 116645336 | rs3087611 | T | A | M_Disease | 0.539914885 | M_Disease&NULL |
| 11 | 121007228 | rs4935760 | T | A | M_Disease | 0.538672605 | M_Disease&NULL |
| 11 | 116793693 | rs10892047 | T | C | M_Disease | 0.538672605 | M_Disease&NULL |
| 11 | 120419320 | rs77998262 | A | G | M_Disease | 0.538672605 | M_Disease&NULL |
| 11 | 116505583 | rs1784108 | T | C | M_Disease | 0.538672605 | M_Disease&NULL |
| 11 | 116414846 | rs61904786 | G | C | M_Disease | 0.538672605 | M_Disease&NULL |
| 11 | 116787320 | rs12277181 | A | G | M_Disease | 0.538672605 | M_Disease&NULL |
| 11 | 116911012 | rs12273545 | T | C | M_Disease | 0.538672605 | M_Disease&NULL |
| 11 | 116946853 | rs12274725 | T | C | M_Disease | 0.538672605 | M_Disease&NULL |
| 11 | 116947180 | rs11216265 | C | T | M_Disease | 0.538672605 | M_Disease&NULL |
| 11 | 116951584 | rs10082561 | C | A | M_Disease | 0.538672605 | M_Disease&NULL |
| 11 | 116958533 | rs12277910 | A | C | M_Disease | 0.538672605 | M_Disease&NULL |
| 11 | 117995697 | rs626562 | A | G | M_Disease | 0.538672605 | M_Disease&NULL |
| 11 | 116764436 | rs576262 | C | G | M_Disease | 0.538672605 | M_Disease&NULL |
| 11 | 116774244 | rs78694306 | T | G | M_Disease | 0.538672605 | M_Disease&NULL |
| 11 | 116774499 | rs115971514 | C | G | M_Disease | 0.538672605 | M_Disease&NULL |
| 11 | 117425169 | rs142260742 | T | C | M_Disease | 0.538672605 | M_Disease&NULL |
| 11 | 117426035 | rs80350215 | C | T | M_Disease | 0.538672605 | M_Disease&NULL |
| 11 | 117213159 | rs143793912 | T | C | M_Disease | 0.538672605 | M_Disease&NULL |
| 11 | 119882896 | rs587793 | C | T | M_Disease | 0.538672605 | M_Disease&NULL |
| 11 | 117403727 | rs12421706 | T | C | M_Disease | 0.538672605 | M_Disease&NULL |
| 11 | 120529603 | rs150492485 | A | G | M_Disease | 0.538672605 | M_Disease&NULL |
| 11 | 116919969 | rs75890305 | G | A | M_Disease | 0.537429845 | M_Disease&NULL |
| 11 | 117186073 | rs2445939 | T | A | M_Disease | 0.537429845 | M_Disease&NULL |
| 11 | 117340239 | rs12416923 | A | G | M_Disease | 0.537429845 | M_Disease&NULL |
| 11 | 116909657 | rs143138921 | G | T | M_Disease | 0.537429845 | M_Disease&NULL |
| 11 | 116911423 | rs74603899 | T | G | M_Disease | 0.537429845 | M_Disease&NULL |
| 11 | 116912494 | rs77960525 | A | G | M_Disease | 0.537429845 | M_Disease&NULL |
| 11 | 118763736 | rs12363277 | C | T | M_Disease | 0.537429845 | M_Disease&NULL |
| 11 | 116795948 | rs12292371 | C | G | M_Disease | 0.537429845 | M_Disease&NULL |
| 11 | 116796065 | rs12292434 | A | G | M_Disease | 0.537429845 | M_Disease&NULL |
| 11 | 116797475 | rs12285956 | A | G | M_Disease | 0.537429845 | M_Disease&NULL |
| 11 | 116800270 | rs79157715 | A | G | M_Disease | 0.537429845 | M_Disease&NULL |
| 11 | 116801061 | rs75776964 | G | C | M_Disease | 0.537429845 | M_Disease&NULL |
| 11 | 116802972 | rs74615346 | C | A | M_Disease | 0.537429845 | M_Disease&NULL |
| 11 | 116804005 | rs12270974 | T | C | M_Disease | 0.537429845 | M_Disease&NULL |
| 11 | 116804370 | rs12272491 | T | C | M_Disease | 0.537429845 | M_Disease&NULL |
| 11 | 116804762 | rs12418735 | G | T | M_Disease | 0.537429845 | M_Disease&NULL |

| | | | | | | | | | |
|---|---|---|---|---|---|---|---|---|---|
| 11 | 116512466 | rs4938293 | T | C | 1.067 | -7977.388 | -7977.026 | 0.362 | 5.56E-06 |
| 11 | 116780918 | rs6589578 | T | C | 1.133 | -7977.388 | -7977.028 | 0.36 | 5.54E-06 |
| 11 | 116782582 | rs7106782 | G | A | 1.133 | -7977.388 | -7977.028 | 0.36 | 5.54E-06 |
| 11 | 116879679 | rs117901629 | A | G | 1.114 | -7977.388 | -7977.029 | 0.36 | 5.54E-06 |
| 11 | 119885462 | rs630332 | A | G | 0.897 | -7977.388 | -7977.028 | 0.36 | 5.54E-06 |
| 11 | 117534353 | rs630024 | A | C | 0.939 | -7977.388 | -7977.03 | 0.359 | 5.54E-06 |
| 11 | 116765925 | rs74881320 | C | A | 1.112 | -7977.388 | -7977.03 | 0.358 | 5.53E-06 |
| 11 | 116835363 | rs59283948 | A | G | 1.133 | -7977.388 | -7977.03 | 0.358 | 5.53E-06 |
| 11 | 121018391 | rs7108587 | T | C | 0.931 | -7977.388 | -7977.031 | 0.357 | 5.53E-06 |
| 11 | 117486730 | rs7942967 | T | C | 1.07 | -7977.388 | -7977.034 | 0.355 | 5.52E-06 |
| 11 | 120521504 | rs7128690 | T | G | 0.874 | -7977.388 | -7977.033 | 0.355 | 5.52E-06 |
| 11 | 117913074 | rs4936421 | C | A | 0.937 | -7977.388 | -7977.034 | 0.354 | 5.51E-06 |
| 11 | 120428134 | rs2248404 | A | G | 0.894 | -7977.388 | -7977.038 | 0.351 | 5.49E-06 |
| 11 | 121066195 | rs524392 | A | T | 0.931 | -7977.388 | -7977.041 | 0.348 | 5.48E-06 |
| 11 | 116953644 | rs11216270 | A | C | 1.113 | -7977.388 | -7977.041 | 0.347 | 5.47E-06 |
| 11 | 117525125 | rs583983 | T | C | 0.939 | -7977.388 | -7977.041 | 0.347 | 5.47E-06 |
| 11 | 117913315 | rs55730046 | T | C | 0.937 | -7977.388 | -7977.041 | 0.347 | 5.47E-06 |
| 11 | 120520282 | rs7121104 | T | G | 1.139 | -7977.388 | -7977.042 | 0.346 | 5.47E-06 |
| 11 | 117395497 | rs55642294 | A | C | 1.069 | -7977.388 | -7977.043 | 0.345 | 5.46E-06 |
| 11 | 118748170 | rs77074463 | C | G | 1.105 | -7977.388 | -7977.046 | 0.342 | 5.45E-06 |
| 11 | 121011342 | rs560365 | T | C | 0.939 | -7977.388 | -7977.048 | 0.34 | 5.43E-06 |
| 11 | 117268683 | rs34114773 | C | T | 1.065 | -7977.388 | -7977.05 | 0.338 | 5.42E-06 |
| 11 | 117361165 | rs115325870 | T | C | 0.861 | -7977.388 | -7977.052 | 0.337 | 5.42E-06 |
| 11 | 118866505 | rs7120982 | A | G | 0.94 | -7977.388 | -7977.051 | 0.337 | 5.42E-06 |
| 11 | 119941807 | rs10790363 | A | G | 0.897 | -7977.388 | -7977.054 | 0.335 | 5.41E-06 |
| 11 | 116778813 | rs35184536 | G | A | 1.111 | -7977.388 | -7977.056 | 0.333 | 5.40E-06 |
| 11 | 116901175 | rs79257193 | G | A | 1.131 | -7977.388 | -7977.057 | 0.331 | 5.39E-06 |
| 11 | 117399606 | rs7126500 | T | G | 1.069 | -7977.388 | -7977.058 | 0.33 | 5.38E-06 |
| 11 | 120503083 | rs11217933 | A | G | 0.939 | -7977.388 | -7977.058 | 0.33 | 5.38E-06 |
| 11 | 116625897 | rs67120644 | C | T | 0.926 | -7977.388 | -7977.059 | 0.329 | 5.38E-06 |
| 11 | 117790342 | rs11216620 | T | C | 0.888 | -7977.388 | -7977.059 | 0.329 | 5.38E-06 |
| 11 | 119582832 | rs4409845 | G | A | 0.884 | -7977.388 | -7977.059 | 0.329 | 5.38E-06 |
| 11 | 120414586 | rs2266100 | A | G | 0.935 | -7977.388 | -7977.06 | 0.329 | 5.38E-06 |
| 11 | 116883716 | rs7110042 | T | C | 1.131 | -7977.388 | -7977.06 | 0.328 | 5.37E-06 |
| 11 | 116890488 | rs138975985 | T | C | 1.131 | -7977.388 | -7977.06 | 0.328 | 5.37E-06 |
| 11 | 116333012 | rs73009128 | T | C | 0.892 | -7977.388 | -7977.063 | 0.326 | 5.36E-06 |
| 11 | 116937361 | rs79955870 | T | C | 1.112 | -7977.388 | -7977.063 | 0.326 | 5.36E-06 |
| 11 | 116940210 | rs11216261 | C | T | 1.112 | -7977.388 | -7977.063 | 0.326 | 5.36E-06 |
| 11 | 117248832 | rs11216370 | A | G | 1.069 | -7977.388 | -7977.065 | 0.324 | 5.35E-06 |
| 11 | 117430025 | rs75375456 | T | G | 0.87 | -7977.388 | -7977.064 | 0.324 | 5.35E-06 |
| 11 | 116806616 | rs76384483 | C | T | 1.15 | -7977.388 | -7977.065 | 0.323 | 5.34E-06 |
| 11 | 118788081 | rs7929825 | G | C | 0.928 | -7977.388 | -7977.068 | 0.32 | 5.33E-06 |
| 11 | 119593446 | rs11604214 | A | G | 0.875 | -7977.388 | -7977.069 | 0.319 | 5.32E-06 |
| 11 | 121011325 | rs560401 | G | A | 0.939 | -7977.388 | -7977.07 | 0.319 | 5.32E-06 |
| 11 | 116955934 | rs2368149 | A | G | 1.112 | -7977.388 | -7977.07 | 0.318 | 5.32E-06 |
| 11 | 116956990 | rs12281358 | C | T | 1.112 | -7977.388 | -7977.07 | 0.318 | 5.32E-06 |
| 11 | 117074299 | rs2269397 | C | T | 1.098 | -7977.388 | -7977.071 | 0.317 | 5.31E-06 |
| 11 | 118006853 | rs113579428 | T | C | 0.866 | -7977.388 | -7977.071 | 0.317 | 5.31E-06 |
| 11 | 118178757 | rs11601508 | A | G | 0.884 | -7977.388 | -7977.072 | 0.317 | 5.31E-06 |
| 11 | 116772486 | rs17120131 | C | T | 1.11 | -7977.388 | -7977.072 | 0.316 | 5.31E-06 |
| 11 | 116773318 | rs17120132 | A | T | 1.11 | -7977.388 | -7977.072 | 0.316 | 5.31E-06 |
| 11 | 116773761 | rs17120136 | G | T | 1.11 | -7977.388 | -7977.072 | 0.316 | 5.31E-06 |
| 11 | 116775196 | rs12280030 | A | T | 1.11 | -7977.388 | -7977.072 | 0.316 | 5.31E-06 |
| 11 | 116811878 | rs114034580 | G | A | 1.13 | -7977.388 | -7977.073 | 0.315 | 5.30E-06 |
| 11 | 118811878 | rs74891547 | C | G | 1.101 | -7977.388 | -7977.074 | 0.315 | 5.30E-06 |
| 11 | 120405437 | rs2508653 | T | C | 0.936 | -7977.388 | -7977.073 | 0.315 | 5.30E-06 |
| 11 | 119881100 | rs2850802 | A | G | 0.899 | -7977.388 | -7977.076 | 0.313 | 5.29E-06 |
| 11 | 116907715 | rs75216569 | G | A | 1.112 | -7977.388 | -7977.078 | 0.311 | 5.28E-06 |
| 11 | 116645336 | rs3087611 | T | A | 1.15 | -7977.388 | -7977.079 | 0.309 | 5.27E-06 |
| 11 | 121007228 | rs4935760 | T | A | 0.932 | -7977.388 | -7977.08 | 0.308 | 5.26E-06 |
| 11 | 116793693 | rs10892047 | T | C | 1.112 | -7977.388 | -7977.082 | 0.307 | 5.26E-06 |
| 11 | 120419320 | rs77998262 | A | G | 0.894 | -7977.388 | -7977.082 | 0.307 | 5.26E-06 |
| 11 | 116505583 | rs1784108 | T | C | 0.924 | -7977.388 | -7977.082 | 0.306 | 5.25E-06 |
| 11 | 116414846 | rs61904786 | G | C | 1.081 | -7977.388 | -7977.085 | 0.304 | 5.24E-06 |
| 11 | 116787320 | rs12277181 | A | G | 1.11 | -7977.388 | -7977.084 | 0.304 | 5.24E-06 |
| 11 | 116911012 | rs12273545 | T | C | 1.111 | -7977.388 | -7977.085 | 0.303 | 5.24E-06 |
| 11 | 116946853 | rs12274725 | T | C | 1.111 | -7977.388 | -7977.086 | 0.303 | 5.24E-06 |
| 11 | 116947180 | rs11216265 | C | T | 1.111 | -7977.388 | -7977.086 | 0.303 | 5.24E-06 |
| 11 | 116951584 | rs10082561 | C | A | 1.111 | -7977.388 | -7977.086 | 0.303 | 5.24E-06 |
| 11 | 116958533 | rs12277910 | A | C | 1.111 | -7977.388 | -7977.086 | 0.303 | 5.24E-06 |
| 11 | 117995697 | rs626562 | A | G | 1.121 | -7977.388 | -7977.085 | 0.303 | 5.24E-06 |
| 11 | 116764436 | rs576262 | C | G | 0.879 | -7977.388 | -7977.086 | 0.302 | 5.23E-06 |
| 11 | 116774244 | rs78694306 | T | G | 1.11 | -7977.388 | -7977.087 | 0.302 | 5.23E-06 |
| 11 | 116774499 | rs115971514 | C | G | 1.11 | -7977.388 | -7977.087 | 0.302 | 5.23E-06 |
| 11 | 117425169 | rs142260742 | T | C | 0.871 | -7977.388 | -7977.087 | 0.302 | 5.23E-06 |
| 11 | 117426035 | rs80350215 | C | T | 0.871 | -7977.388 | -7977.087 | 0.302 | 5.23E-06 |
| 11 | 117213159 | rs143793912 | T | C | 1.154 | -7977.388 | -7977.089 | 0.3 | 5.22E-06 |
| 11 | 119882896 | rs587793 | C | T | 0.899 | -7977.388 | -7977.088 | 0.3 | 5.22E-06 |
| 11 | 117403727 | rs12421706 | T | C | 0.883 | -7977.388 | -7977.089 | 0.299 | 5.22E-06 |
| 11 | 120529603 | rs150492485 | A | G | 1.136 | -7977.388 | -7977.09 | 0.299 | 5.22E-06 |
| 11 | 116919969 | rs75890305 | G | A | 1.111 | -7977.388 | -7977.091 | 0.298 | 5.21E-06 |
| 11 | 117186073 | rs2445939 | T | A | 1.065 | -7977.388 | -7977.092 | 0.297 | 5.21E-06 |
| 11 | 117340239 | rs12416923 | A | G | 0.904 | -7977.388 | -7977.092 | 0.296 | 5.20E-06 |
| 11 | 116909657 | rs143138921 | G | T | 1.129 | -7977.388 | -7977.093 | 0.295 | 5.20E-06 |
| 11 | 116911423 | rs74603899 | T | G | 1.129 | -7977.388 | -7977.093 | 0.295 | 5.20E-06 |
| 11 | 116912494 | rs77960525 | A | G | 1.129 | -7977.388 | -7977.093 | 0.295 | 5.20E-06 |
| 11 | 118763736 | rs12363277 | C | T | 1.11 | -7977.388 | -7977.093 | 0.295 | 5.20E-06 |
| 11 | 116795948 | rs12292371 | C | G | 1.111 | -7977.388 | -7977.095 | 0.294 | 5.19E-06 |
| 11 | 116796065 | rs12292434 | A | G | 1.111 | -7977.388 | -7977.095 | 0.294 | 5.19E-06 |
| 11 | 116797475 | rs12285956 | A | G | 1.111 | -7977.388 | -7977.095 | 0.294 | 5.19E-06 |
| 11 | 116800270 | rs79157715 | A | G | 1.111 | -7977.388 | -7977.095 | 0.294 | 5.19E-06 |
| 11 | 116801061 | rs75776964 | G | C | 1.111 | -7977.388 | -7977.095 | 0.294 | 5.19E-06 |
| 11 | 116802972 | rs74615346 | C | A | 1.111 | -7977.388 | -7977.095 | 0.294 | 5.19E-06 |
| 11 | 116804005 | rs12270974 | T | C | 1.111 | -7977.388 | -7977.095 | 0.294 | 5.19E-06 |
| 11 | 116804370 | rs12272491 | T | C | 1.111 | -7977.388 | -7977.095 | 0.294 | 5.19E-06 |
| 11 | 116804762 | rs12418735 | G | T | 1.111 | -7977.388 | -7977.095 | 0.294 | 5.19E-06 |

| | | | | | | | |
|---|---|---|---|---|---|---|---|
| 11 | 116807173 | rs12273769 | G | T | M_Disease | 0.537429845 | M_Disease&NULL |
| 11 | 116807368 | rs12294657 | C | G | M_Disease | 0.537429845 | M_Disease&NULL |
| 11 | 116808453 | rs77136879 | T | C | M_Disease | 0.537429845 | M_Disease&NULL |
| 11 | 116809789 | rs7480278 | G | T | M_Disease | 0.537429845 | M_Disease&NULL |
| 11 | 116810906 | rs80015203 | G | A | M_Disease | 0.537429845 | M_Disease&NULL |
| 11 | 116811487 | rs116597701 | G | C | M_Disease | 0.537429845 | M_Disease&NULL |
| 11 | 116812058 | rs78066444 | C | A | M_Disease | 0.537429845 | M_Disease&NULL |
| 11 | 116812628 | rs17120157 | C | T | M_Disease | 0.537429845 | M_Disease&NULL |
| 11 | 116813438 | rs75410648 | T | C | M_Disease | 0.537429845 | M_Disease&NULL |
| 11 | 116814646 | rs12421902 | G | A | M_Disease | 0.537429845 | M_Disease&NULL |
| 11 | 116815599 | rs11216193 | T | C | M_Disease | 0.537429845 | M_Disease&NULL |
| 11 | 116815738 | rs11216194 | C | G | M_Disease | 0.537429845 | M_Disease&NULL |
| 11 | 116816394 | rs10892050 | A | C | M_Disease | 0.537429845 | M_Disease&NULL |
| 11 | 116816778 | rs11216197 | G | A | M_Disease | 0.537429845 | M_Disease&NULL |
| 11 | 116819264 | rs12293755 | T | A | M_Disease | 0.537429845 | M_Disease&NULL |
| 11 | 116819906 | rs12419047 | A | G | M_Disease | 0.537429845 | M_Disease&NULL |
| 11 | 116820023 | rs12421834 | C | A | M_Disease | 0.537429845 | M_Disease&NULL |
| 11 | 116820690 | rs12419437 | A | G | M_Disease | 0.537429845 | M_Disease&NULL |
| 11 | 116823107 | rs77467134 | C | T | M_Disease | 0.537429845 | M_Disease&NULL |
| 11 | 116823521 | rs76654964 | A | G | M_Disease | 0.537429845 | M_Disease&NULL |
| 11 | 116823617 | rs76146269 | G | A | M_Disease | 0.537429845 | M_Disease&NULL |
| 11 | 116824162 | rs12275355 | G | A | M_Disease | 0.537429845 | M_Disease&NULL |
| 11 | 116824286 | rs12289974 | T | C | M_Disease | 0.537429845 | M_Disease&NULL |
| 11 | 116824620 | rs2289890 | A | G | M_Disease | 0.537429845 | M_Disease&NULL |
| 11 | 116825240 | rs2289892 | A | G | M_Disease | 0.537429845 | M_Disease&NULL |
| 11 | 116826617 | rs12291040 | C | T | M_Disease | 0.537429845 | M_Disease&NULL |
| 11 | 116826643 | rs12270837 | C | A | M_Disease | 0.537429845 | M_Disease&NULL |
| 11 | 116831016 | rs11216199 | T | C | M_Disease | 0.537429845 | M_Disease&NULL |
| 11 | 116832718 | rs1053095947 | A | C | M_Disease | 0.537429845 | M_Disease&NULL |
| 11 | 116846266 | rs74927399 | C | T | M_Disease | 0.537429845 | M_Disease&NULL |
| 11 | 116846343 | rs76949321 | A | G | M_Disease | 0.537429845 | M_Disease&NULL |
| 11 | 116848186 | rs11216209 | C | T | M_Disease | 0.537429845 | M_Disease&NULL |
| 11 | 116850351 | rs11216212 | A | T | M_Disease | 0.537429845 | M_Disease&NULL |
| 11 | 116851367 | rs146178826 | G | A | M_Disease | 0.537429845 | M_Disease&NULL |
| 11 | 116852687 | rs11216215 | A | T | M_Disease | 0.537429845 | M_Disease&NULL |
| 11 | 116853922 | rs10128690 | A | G | M_Disease | 0.537429845 | M_Disease&NULL |
| 11 | 116854021 | rs10128570 | G | A | M_Disease | 0.537429845 | M_Disease&NULL |
| 11 | 116854181 | rs10128571 | G | A | M_Disease | 0.537429845 | M_Disease&NULL |
| 11 | 116856322 | rs12287768 | C | A | M_Disease | 0.537429845 | M_Disease&NULL |
| 11 | 116856698 | rs12274771 | G | C | M_Disease | 0.537429845 | M_Disease&NULL |
| 11 | 116856709 | rs12274773 | T | C | M_Disease | 0.537429845 | M_Disease&NULL |
| 11 | 116856992 | rs75212922 | A | T | M_Disease | 0.537429845 | M_Disease&NULL |
| 11 | 116857652 | rs11216217 | A | G | M_Disease | 0.537429845 | M_Disease&NULL |
| 11 | 116862245 | rs80242245 | G | C | M_Disease | 0.537429845 | M_Disease&NULL |
| 11 | 116862661 | rs11216218 | T | C | M_Disease | 0.537429845 | M_Disease&NULL |
| 11 | 116862755 | rs11216219 | T | C | M_Disease | 0.537429845 | M_Disease&NULL |
| 11 | 116944846 | rs17120293 | C | T | M_Disease | 0.537429845 | M_Disease&NULL |
| 11 | 116971677 | rs78135964 | T | G | M_Disease | 0.537429845 | M_Disease&NULL |
| 11 | 117927550 | rs4938473 | A | G | M_Disease | 0.53618662 | M_Disease&NULL |
| 11 | 116653185 | rs78359532 | C | G | M_Disease | 0.537429845 | M_Disease&NULL |
| 11 | 120414838 | rs2266101 | T | C | M_Disease | 0.53618662 | M_Disease&NULL |
| 11 | 117412884 | rs11823783 | C | T | M_Disease | 0.53618662 | M_Disease&NULL |
| 11 | 117886499 | rs61740780 | A | G | M_Disease | 0.537429845 | M_Disease&NULL |
| 11 | 116777063 | rs12273762 | A | T | M_Disease | 0.53618662 | M_Disease&NULL |
| 11 | 116778065 | rs9651681 | C | T | M_Disease | 0.53618662 | M_Disease&NULL |
| 11 | 116782974 | rs11216185 | G | T | M_Disease | 0.53618662 | M_Disease&NULL |
| 11 | 116872876 | rs77585745 | C | T | M_Disease | 0.537429845 | M_Disease&NULL |
| 11 | 116892368 | rs77955134 | A | G | M_Disease | 0.53618662 | M_Disease&NULL |
| 11 | 116892446 | rs77594377 | A | G | M_Disease | 0.53618662 | M_Disease&NULL |
| 11 | 116896913 | rs2186844 | T | C | M_Disease | 0.53618662 | M_Disease&NULL |
| 11 | 116899580 | rs12289032 | T | C | M_Disease | 0.53618662 | M_Disease&NULL |
| 11 | 116899825 | rs146943088 | C | A | M_Disease | 0.53618662 | M_Disease&NULL |
| 11 | 116988790 | rs80197460 | G | C | M_Disease | 0.53618662 | M_Disease&NULL |
| 11 | 118187395 | rs13377487 | A | T | M_Disease | 0.53618662 | M_Disease&NULL |
| 11 | 116280349 | rs78774571 | T | C | M_Disease | 0.53618662 | M_Disease&NULL |
| 11 | 120404835 | rs55765491 | G | A | M_Disease | 0.53618662 | M_Disease&NULL |
| 11 | 120530241 | rs78641307 | C | A | M_Disease | 0.53618662 | M_Disease&NULL |
| 11 | 116758216 | rs113213242 | T | C | M_Disease | 0.53618662 | M_Disease&NULL |
| 11 | 116758454 | rs17134988 | A | G | M_Disease | 0.53618662 | M_Disease&NULL |
| 11 | 116760474 | rs117457463 | A | G | M_Disease | 0.53618662 | M_Disease&NULL |
| 11 | 116333791 | rs73009129 | A | G | M_Disease | 0.53618662 | M_Disease&NULL |
| 11 | 117417384 | rs80041541 | T | G | M_Disease | 0.53618662 | M_Disease&NULL |
| 11 | 117418077 | rs75809806 | A | G | M_Disease | 0.53618662 | M_Disease&NULL |
| 11 | 117420369 | rs77342459 | G | C | M_Disease | 0.53618662 | M_Disease&NULL |
| 11 | 118978411 | rs7940073 | G | C | M_Disease | 0.534942945 | M_Disease&NULL |
| 11 | 118979026 | rs6589717 | T | C | M_Disease | 0.534942945 | M_Disease&NULL |
| 11 | 116731302 | rs75428021 | T | C | M_Disease | 0.53618662 | M_Disease&NULL |
| 11 | 116908706 | rs80273401 | C | T | M_Disease | 0.534942945 | M_Disease&NULL |
| 11 | 117192376 | rs516712 | G | A | M_Disease | 0.534942945 | M_Disease&NULL |
| 11 | 116791307 | rs78692246 | A | G | M_Disease | 0.534942945 | M_Disease&NULL |
| 11 | 116964065 | rs77366224 | T | G | M_Disease | 0.534942945 | M_Disease&NULL |
| 11 | 116964066 | rs77341747 | T | C | M_Disease | 0.534942945 | M_Disease&NULL |
| 11 | 116972227 | rs148233183 | A | G | M_Disease | 0.534942945 | M_Disease&NULL |
| 11 | 118595207 | rs563537 | A | T | M_Disease | 0.534942945 | M_Disease&NULL |
| 11 | 116821415 | rs73592285 | C | T | M_Disease | 0.534942945 | M_Disease&NULL |
| 11 | 120025452 | rs79959350 | G | A | M_Disease | 0.534942945 | M_Disease&NULL |
| 11 | 120526291 | rs56926846 | A | G | M_Disease | 0.53618662 | M_Disease&NULL |
| 11 | 121016838 | rs7130952 | T | C | M_Disease | 0.534942945 | M_Disease&NULL |
| 11 | 116912117 | rs77999164 | T | C | M_Disease | 0.534942945 | M_Disease&NULL |
| 11 | 116919273 | rs12280172 | A | G | M_Disease | 0.534942945 | M_Disease&NULL |
| 11 | 116924000 | rs11216250 | T | C | M_Disease | 0.534942945 | M_Disease&NULL |
| 11 | 120414850 | rs2510878 | C | A | M_Disease | 0.534942945 | M_Disease&NULL |
| 11 | 116758658 | rs117231130 | C | T | M_Disease | 0.534942945 | M_Disease&NULL |
| 11 | 117653350 | rs7940750 | T | C | M_Disease | 0.534942945 | M_Disease&NULL |
| 11 | 116947306 | rs12418036 | A | G | M_Disease | 0.534942945 | M_Disease&NULL |
| 11 | 117412773 | rs111698696 | T | C | M_Disease | 0.533698835 | M_Disease&NULL |

| | | | | | | | | | |
|---|---|---|---|---|---|---|---|---|---|
| 11 | 116807173 | rs12273769 | G | T | 1.111 | -7977.388 | -7977.095 | 0.294 | 5.19E-06 |
| 11 | 116807368 | rs12294657 | C | G | 1.111 | -7977.388 | -7977.095 | 0.294 | 5.19E-06 |
| 11 | 116808453 | rs77136879 | T | C | 1.111 | -7977.388 | -7977.095 | 0.294 | 5.19E-06 |
| 11 | 116809789 | rs7480278 | G | T | 1.111 | -7977.388 | -7977.095 | 0.294 | 5.19E-06 |
| 11 | 116810906 | rs80015203 | G | A | 1.111 | -7977.388 | -7977.095 | 0.294 | 5.19E-06 |
| 11 | 116811487 | rs116597701 | G | C | 1.111 | -7977.388 | -7977.095 | 0.294 | 5.19E-06 |
| 11 | 116812058 | rs78066444 | C | A | 1.111 | -7977.388 | -7977.095 | 0.294 | 5.19E-06 |
| 11 | 116812628 | rs17120157 | C | T | 1.111 | -7977.388 | -7977.095 | 0.294 | 5.19E-06 |
| 11 | 116813438 | rs75410648 | T | C | 1.111 | -7977.388 | -7977.095 | 0.294 | 5.19E-06 |
| 11 | 116814646 | rs12421902 | G | A | 1.111 | -7977.388 | -7977.095 | 0.294 | 5.19E-06 |
| 11 | 116815599 | rs11216193 | T | C | 1.111 | -7977.388 | -7977.095 | 0.294 | 5.19E-06 |
| 11 | 116815738 | rs11216194 | C | G | 1.111 | -7977.388 | -7977.095 | 0.294 | 5.19E-06 |
| 11 | 116816394 | rs10892050 | A | C | 1.111 | -7977.388 | -7977.095 | 0.294 | 5.19E-06 |
| 11 | 116816778 | rs11216197 | G | A | 1.111 | -7977.388 | -7977.095 | 0.294 | 5.19E-06 |
| 11 | 116819264 | rs12293755 | T | A | 1.111 | -7977.388 | -7977.095 | 0.294 | 5.19E-06 |
| 11 | 116819906 | rs12419047 | A | G | 1.111 | -7977.388 | -7977.095 | 0.294 | 5.19E-06 |
| 11 | 116820023 | rs12421834 | C | A | 1.111 | -7977.388 | -7977.095 | 0.294 | 5.19E-06 |
| 11 | 116820690 | rs12419437 | A | G | 1.111 | -7977.388 | -7977.095 | 0.294 | 5.19E-06 |
| 11 | 116823107 | rs77467134 | C | T | 1.111 | -7977.388 | -7977.095 | 0.294 | 5.19E-06 |
| 11 | 116823521 | rs76654964 | A | G | 1.111 | -7977.388 | -7977.095 | 0.294 | 5.19E-06 |
| 11 | 116823617 | rs76146269 | G | A | 1.111 | -7977.388 | -7977.095 | 0.294 | 5.19E-06 |
| 11 | 116824162 | rs12275355 | G | A | 1.111 | -7977.388 | -7977.095 | 0.294 | 5.19E-06 |
| 11 | 116824286 | rs12289974 | T | C | 1.111 | -7977.388 | -7977.095 | 0.294 | 5.19E-06 |
| 11 | 116824620 | rs2289890 | A | G | 1.111 | -7977.388 | -7977.095 | 0.294 | 5.19E-06 |
| 11 | 116825240 | rs2289892 | A | G | 1.111 | -7977.388 | -7977.095 | 0.294 | 5.19E-06 |
| 11 | 116826617 | rs12291040 | C | T | 1.111 | -7977.388 | -7977.095 | 0.294 | 5.19E-06 |
| 11 | 116826643 | rs12270837 | C | A | 1.111 | -7977.388 | -7977.095 | 0.294 | 5.19E-06 |
| 11 | 116831016 | rs11216199 | T | C | 1.111 | -7977.388 | -7977.095 | 0.294 | 5.19E-06 |
| 11 | 116832718 | rs1053095947 | A | C | 1.111 | -7977.388 | -7977.095 | 0.294 | 5.19E-06 |
| 11 | 116846266 | rs74927399 | C | T | 1.111 | -7977.388 | -7977.095 | 0.294 | 5.19E-06 |
| 11 | 116846343 | rs76949321 | A | G | 1.111 | -7977.388 | -7977.095 | 0.294 | 5.19E-06 |
| 11 | 116848186 | rs11216209 | C | T | 1.111 | -7977.388 | -7977.095 | 0.294 | 5.19E-06 |
| 11 | 116850351 | rs11216212 | A | T | 1.111 | -7977.388 | -7977.095 | 0.294 | 5.19E-06 |
| 11 | 116851367 | rs146178826 | G | A | 1.111 | -7977.388 | -7977.095 | 0.294 | 5.19E-06 |
| 11 | 116852687 | rs11216215 | A | T | 1.111 | -7977.388 | -7977.095 | 0.294 | 5.19E-06 |
| 11 | 116853922 | rs10128690 | A | G | 1.111 | -7977.388 | -7977.095 | 0.294 | 5.19E-06 |
| 11 | 116854021 | rs10128570 | G | A | 1.111 | -7977.388 | -7977.095 | 0.294 | 5.19E-06 |
| 11 | 116854181 | rs10128571 | G | A | 1.111 | -7977.388 | -7977.095 | 0.294 | 5.19E-06 |
| 11 | 116856322 | rs12287768 | C | A | 1.111 | -7977.388 | -7977.095 | 0.294 | 5.19E-06 |
| 11 | 116856698 | rs12274771 | G | C | 1.111 | -7977.388 | -7977.095 | 0.294 | 5.19E-06 |
| 11 | 116856709 | rs12274773 | T | C | 1.111 | -7977.388 | -7977.095 | 0.294 | 5.19E-06 |
| 11 | 116856992 | rs75212922 | A | T | 1.111 | -7977.388 | -7977.095 | 0.294 | 5.19E-06 |
| 11 | 116857652 | rs11216217 | A | G | 1.111 | -7977.388 | -7977.095 | 0.294 | 5.19E-06 |
| 11 | 116862245 | rs80242245 | G | C | 1.111 | -7977.388 | -7977.095 | 0.294 | 5.19E-06 |
| 11 | 116862661 | rs11216218 | T | C | 1.111 | -7977.388 | -7977.095 | 0.294 | 5.19E-06 |
| 11 | 116862755 | rs11216219 | T | C | 1.111 | -7977.388 | -7977.095 | 0.294 | 5.19E-06 |
| 11 | 116944846 | rs17120293 | C | T | 1.111 | -7977.388 | -7977.094 | 0.294 | 5.19E-06 |
| 11 | 116971677 | rs78135964 | T | G | 1.111 | -7977.388 | -7977.096 | 0.292 | 5.18E-06 |
| 11 | 117927550 | rs4938473 | A | G | 0.939 | -7977.388 | -7977.097 | 0.292 | 5.18E-06 |
| 11 | 116653185 | rs78359532 | C | G | 1.149 | -7977.388 | -7977.098 | 0.29 | 5.17E-06 |
| 11 | 120414838 | rs2266101 | T | C | 0.935 | -7977.388 | -7977.098 | 0.29 | 5.17E-06 |
| 11 | 117412884 | rs11823783 | C | T | 1.069 | -7977.388 | -7977.099 | 0.289 | 5.16E-06 |
| 11 | 117886499 | rs61740780 | A | G | 0.881 | -7977.388 | -7977.099 | 0.289 | 5.16E-06 |
| 11 | 116777063 | rs12273762 | A | T | 1.109 | -7977.388 | -7977.1 | 0.288 | 5.16E-06 |
| 11 | 116778065 | rs9651681 | C | T | 1.109 | -7977.388 | -7977.1 | 0.288 | 5.16E-06 |
| 11 | 116782974 | rs11216185 | G | T | 1.109 | -7977.388 | -7977.101 | 0.288 | 5.16E-06 |
| 11 | 116872876 | rs77585745 | C | T | 1.148 | -7977.388 | -7977.1 | 0.288 | 5.16E-06 |
| 11 | 116892368 | rs77955134 | A | G | 1.111 | -7977.388 | -7977.1 | 0.288 | 5.16E-06 |
| 11 | 116892446 | rs77594377 | A | G | 1.111 | -7977.388 | -7977.1 | 0.288 | 5.16E-06 |
| 11 | 116896913 | rs2186844 | T | C | 1.111 | -7977.388 | -7977.1 | 0.288 | 5.16E-06 |
| 11 | 116899580 | rs12289032 | T | C | 1.111 | -7977.388 | -7977.1 | 0.288 | 5.16E-06 |
| 11 | 116899825 | rs146943088 | C | A | 1.111 | -7977.388 | -7977.1 | 0.288 | 5.16E-06 |
| 11 | 116988790 | rs80197460 | G | C | 1.109 | -7977.388 | -7977.1 | 0.288 | 5.16E-06 |
| 11 | 118187395 | rs13377487 | A | T | 0.885 | -7977.388 | -7977.102 | 0.287 | 5.15E-06 |
| 11 | 116280349 | rs78774571 | T | C | 1.107 | -7977.388 | -7977.103 | 0.285 | 5.14E-06 |
| 11 | 120404835 | rs55765491 | G | A | 0.936 | -7977.388 | -7977.105 | 0.283 | 5.13E-06 |
| 11 | 120530241 | rs78641307 | C | A | 1.135 | -7977.388 | -7977.106 | 0.282 | 5.13E-06 |
| 11 | 116758216 | rs113213242 | T | C | 1.128 | -7977.388 | -7977.107 | 0.281 | 5.12E-06 |
| 11 | 116758454 | rs17134988 | A | G | 1.128 | -7977.388 | -7977.107 | 0.281 | 5.12E-06 |
| 11 | 116760474 | rs117457463 | A | G | 1.128 | -7977.388 | -7977.107 | 0.281 | 5.12E-06 |
| 11 | 116333791 | rs73009129 | A | G | 0.895 | -7977.388 | -7977.109 | 0.28 | 5.12E-06 |
| 11 | 117417384 | rs80041541 | T | G | 0.872 | -7977.388 | -7977.108 | 0.28 | 5.12E-06 |
| 11 | 117418077 | rs75809806 | A | G | 0.872 | -7977.388 | -7977.108 | 0.28 | 5.12E-06 |
| 11 | 117420369 | rs77342459 | G | C | 0.872 | -7977.388 | -7977.108 | 0.28 | 5.12E-06 |
| 11 | 118978411 | rs7940073 | G | C | 0.904 | -7977.388 | -7977.108 | 0.28 | 5.12E-06 |
| 11 | 118979026 | rs6589717 | T | C | 0.904 | -7977.388 | -7977.108 | 0.28 | 5.12E-06 |
| 11 | 116731302 | rs75428021 | T | C | 1.133 | -7977.388 | -7977.109 | 0.279 | 5.11E-06 |
| 11 | 116908706 | rs80273401 | C | T | 1.11 | -7977.388 | -7977.109 | 0.279 | 5.11E-06 |
| 11 | 117192376 | rs516712 | G | A | 0.941 | -7977.388 | -7977.111 | 0.278 | 5.11E-06 |
| 11 | 116791307 | rs78692246 | A | G | 1.11 | -7977.388 | -7977.111 | 0.277 | 5.10E-06 |
| 11 | 116964065 | rs77366224 | T | G | 1.128 | -7977.388 | -7977.111 | 0.277 | 5.10E-06 |
| 11 | 116964066 | rs77341747 | T | C | 1.128 | -7977.388 | -7977.111 | 0.277 | 5.10E-06 |
| 11 | 116972227 | rs148233183 | A | G | 1.11 | -7977.388 | -7977.112 | 0.277 | 5.10E-06 |
| 11 | 118595207 | rs563537 | A | T | 0.94 | -7977.388 | -7977.111 | 0.277 | 5.10E-06 |
| 11 | 116821415 | rs73592285 | C | T | 1.128 | -7977.388 | -7977.113 | 0.276 | 5.10E-06 |
| 11 | 120025452 | rs79959350 | G | A | 0.883 | -7977.388 | -7977.112 | 0.276 | 5.10E-06 |
| 11 | 120526291 | rs56926846 | A | G | 0.873 | -7977.388 | -7977.112 | 0.276 | 5.10E-06 |
| 11 | 121016838 | rs7130952 | T | C | 0.932 | -7977.388 | -7977.112 | 0.276 | 5.10E-06 |
| 11 | 116912117 | rs77999164 | T | C | 1.11 | -7977.388 | -7977.114 | 0.275 | 5.09E-06 |
| 11 | 116919273 | rs12280172 | A | G | 1.11 | -7977.388 | -7977.114 | 0.275 | 5.09E-06 |
| 11 | 116924000 | rs11216250 | T | C | 1.11 | -7977.388 | -7977.114 | 0.275 | 5.09E-06 |
| 11 | 120414850 | rs2510878 | C | A | 0.936 | -7977.388 | -7977.113 | 0.275 | 5.09E-06 |
| 11 | 116758658 | rs117231130 | C | T | 0.881 | -7977.388 | -7977.116 | 0.273 | 5.08E-06 |
| 11 | 117653350 | rs7940750 | T | C | 0.94 | -7977.388 | -7977.115 | 0.273 | 5.08E-06 |
| 11 | 116947306 | rs12418036 | A | G | 1.11 | -7977.388 | -7977.117 | 0.271 | 5.07E-06 |
| 11 | 117412773 | rs111698696 | T | C | 1.069 | -7977.388 | -7977.117 | 0.271 | 5.07E-06 |

| | | | | | | | |
|---|---|---|---|---|---|---|---|
| 11 | 116767722 | rs141555847 | A | G | M_Disease | 0.534942945 | M_Disease&NULL |
| 11 | 116771204 | rs76402668 | C | T | M_Disease | 0.534942945 | M_Disease&NULL |
| 11 | 116778138 | rs146523230 | A | G | M_Disease | 0.534942945 | M_Disease&NULL |
| 11 | 116783142 | rs74700879 | G | C | M_Disease | 0.534942945 | M_Disease&NULL |
| 11 | 116874358 | rs12278943 | C | T | M_Disease | 0.533698835 | M_Disease&NULL |
| 11 | 116879723 | rs138181556 | A | G | M_Disease | 0.533698835 | M_Disease&NULL |
| 11 | 118186312 | rs2231450 | A | C | M_Disease | 0.533698835 | M_Disease&NULL |
| 11 | 116285538 | rs563990 | G | A | M_Disease | 0.533698835 | M_Disease&NULL |
| 11 | 120441720 | rs12279396 | G | A | M_Disease | 0.533698835 | M_Disease&NULL |
| 11 | 116801299 | rs77203966 | T | C | M_Disease | 0.533698835 | M_Disease&NULL |
| 11 | 116806367 | rs74557653 | T | G | M_Disease | 0.533698835 | M_Disease&NULL |
| 11 | 116815306 | rs57040912 | G | A | M_Disease | 0.533698835 | M_Disease&NULL |
| 11 | 116815717 | rs12283494 | A | G | M_Disease | 0.533698835 | M_Disease&NULL |
| 11 | 116819050 | rs115255270 | C | T | M_Disease | 0.533698835 | M_Disease&NULL |
| 11 | 116820673 | rs7940300 | T | A | M_Disease | 0.533698835 | M_Disease&NULL |
| 11 | 116824938 | rs2289891 | C | A | M_Disease | 0.533698835 | M_Disease&NULL |
| 11 | 116826409 | rs77161357 | C | T | M_Disease | 0.533698835 | M_Disease&NULL |
| 11 | 116828573 | rs7130221 | A | G | M_Disease | 0.533698835 | M_Disease&NULL |
| 11 | 116828859 | rs6589581 | T | A | M_Disease | 0.533698835 | M_Disease&NULL |
| 11 | 116829129 | rs7938466 | C | T | M_Disease | 0.533698835 | M_Disease&NULL |
| 11 | 116830628 | rs77029349 | T | G | M_Disease | 0.533698835 | M_Disease&NULL |
| 11 | 116839094 | rs117791268 | A | G | M_Disease | 0.533698835 | M_Disease&NULL |
| 11 | 116842174 | rs76223783 | C | T | M_Disease | 0.533698835 | M_Disease&NULL |
| 11 | 116842758 | rs118137922 | A | G | M_Disease | 0.533698835 | M_Disease&NULL |
| 11 | 116844649 | rs7937585 | A | G | M_Disease | 0.533698835 | M_Disease&NULL |
| 11 | 116849038 | rs79868224 | A | G | M_Disease | 0.533698835 | M_Disease&NULL |
| 11 | 116849544 | rs78241753 | C | G | M_Disease | 0.533698835 | M_Disease&NULL |
| 11 | 116850485 | rs7940002 | A | C | M_Disease | 0.533698835 | M_Disease&NULL |
| 11 | 116852714 | rs7112772 | C | T | M_Disease | 0.533698835 | M_Disease&NULL |
| 11 | 116852716 | rs7112773 | C | T | M_Disease | 0.533698835 | M_Disease&NULL |
| 11 | 116856152 | rs7110113 | C | A | M_Disease | 0.533698835 | M_Disease&NULL |
| 11 | 116973247 | rs76942203 | A | G | M_Disease | 0.533698835 | M_Disease&NULL |
| 11 | 116975136 | rs149576335 | A | G | M_Disease | 0.533698835 | M_Disease&NULL |
| 11 | 116752045 | rs78013608 | A | G | M_Disease | 0.533698835 | M_Disease&NULL |
| 11 | 120589179 | rs2852227 | C | A | M_Disease | 0.532454306 | M_Disease&NULL |
| 11 | 116784692 | rs11216186 | C | T | M_Disease | 0.532454306 | M_Disease&NULL |
| 11 | 116785341 | rs79711490 | T | C | M_Disease | 0.532454306 | M_Disease&NULL |
| 11 | 116787633 | rs12418208 | T | C | M_Disease | 0.532454306 | M_Disease&NULL |
| 11 | 116787641 | rs12418209 | T | C | M_Disease | 0.532454306 | M_Disease&NULL |
| 11 | 116788133 | rs1006176 | C | T | M_Disease | 0.532454306 | M_Disease&NULL |
| 11 | 118235246 | rs11216857 | A | G | M_Disease | 0.533698835 | M_Disease&NULL |
| 11 | 119939156 | rs597253 | A | G | M_Disease | 0.532454306 | M_Disease&NULL |
| 11 | 119939806 | rs593319 | T | A | M_Disease | 0.532454306 | M_Disease&NULL |
| 11 | 118974878 | rs4938628 | T | A | M_Disease | 0.532454306 | M_Disease&NULL |
| 11 | 118975113 | rs10736496 | A | G | M_Disease | 0.532454306 | M_Disease&NULL |
| 11 | 118975426 | rs4938629 | C | T | M_Disease | 0.532454306 | M_Disease&NULL |
| 11 | 118975927 | rs10790283 | A | G | M_Disease | 0.532454306 | M_Disease&NULL |
| 11 | 116357627 | rs80310120 | T | C | M_Disease | 0.532454306 | M_Disease&NULL |
| 11 | 118241995 | rs502514 | C | T | M_Disease | 0.532454306 | M_Disease&NULL |
| 11 | 120931054 | rs12795310 | C | T | M_Disease | 0.532454306 | M_Disease&NULL |
| 11 | 116780567 | rs75188049 | A | G | M_Disease | 0.532454306 | M_Disease&NULL |
| 11 | 116780702 | rs77693143 | G | A | M_Disease | 0.532454306 | M_Disease&NULL |
| 11 | 118146884 | rs10892232 | T | A | M_Disease | 0.532454306 | M_Disease&NULL |
| 11 | 117411410 | rs11216444 | G | A | M_Disease | 0.531209373 | M_Disease&NULL |
| 11 | 119881745 | rs651202 | A | G | M_Disease | 0.532454306 | M_Disease&NULL |
| 11 | 116414485 | rs77110391 | A | G | M_Disease | 0.531209373 | M_Disease&NULL |
| 11 | 116805064 | rs77204473 | G | T | M_Disease | 0.532454306 | M_Disease&NULL |
| 11 | 117355892 | rs10892126 | C | G | M_Disease | 0.531209373 | M_Disease&NULL |
| 11 | 117371563 | rs7942406 | T | G | M_Disease | 0.531209373 | M_Disease&NULL |
| 11 | 116905209 | rs80176870 | C | T | M_Disease | 0.532454306 | M_Disease&NULL |
| 11 | 116906984 | rs78550859 | T | C | M_Disease | 0.532454306 | M_Disease&NULL |
| 11 | 116908807 | rs79842631 | G | A | M_Disease | 0.532454306 | M_Disease&NULL |
| 11 | 116908913 | rs77939786 | T | C | M_Disease | 0.532454306 | M_Disease&NULL |
| 11 | 116908914 | rs77393043 | A | C | M_Disease | 0.532454306 | M_Disease&NULL |
| 11 | 118221655 | rs137996620 | C | A | M_Disease | 0.531209373 | M_Disease&NULL |
| 11 | 121006387 | rs4936579 | C | T | M_Disease | 0.531209373 | M_Disease&NULL |
| 11 | 121006650 | rs4427574 | A | G | M_Disease | 0.531209373 | M_Disease&NULL |
| 11 | 117254122 | rs12797557 | A | G | M_Disease | 0.531209373 | M_Disease&NULL |
| 11 | 116547924 | rs55928509 | A | C | M_Disease | 0.531209373 | M_Disease&NULL |
| 11 | 119728423 | rs140657166 | C | A | M_Disease | 0.531209373 | M_Disease&NULL |
| 11 | 116903095 | rs7938234 | T | C | M_Disease | 0.531209373 | M_Disease&NULL |
| 11 | 117411717 | rs59099254 | A | C | M_Disease | 0.531209373 | M_Disease&NULL |
| 11 | 121066946 | rs1939934 | T | C | M_Disease | 0.531209373 | M_Disease&NULL |
| 11 | 116288186 | rs563229 | G | C | M_Disease | 0.529964052 | M_Disease&NULL |
| 11 | 121011488 | rs559301 | A | G | M_Disease | 0.529964052 | M_Disease&NULL |
| 11 | 118257081 | rs11216863 | A | G | M_Disease | 0.529964052 | M_Disease&NULL |
| 11 | 116911590 | rs77322791 | G | A | M_Disease | 0.529964052 | M_Disease&NULL |
| 11 | 116941540 | rs9971421 | T | C | M_Disease | 0.529964052 | M_Disease&NULL |
| 11 | 120801185 | rs2186620 | C | T | M_Disease | 0.529964052 | M_Disease&NULL |
| 11 | 116280965 | rs536484 | C | G | M_Disease | 0.529964052 | M_Disease&NULL |
| 11 | 116358711 | rs17119760 | A | C | M_Disease | 0.529964052 | M_Disease&NULL |
| 11 | 117440076 | rs112118203 | C | T | M_Disease | 0.529964052 | M_Disease&NULL |
| 11 | 116731205 | rs4938315 | G | A | M_Disease | 0.529964052 | M_Disease&NULL |
| 11 | 117795717 | rs76014645 | A | C | M_Disease | 0.529964052 | M_Disease&NULL |
| 11 | 117410174 | rs60851263 | T | C | M_Disease | 0.528718357 | M_Disease&NULL |
| 11 | 119938788 | rs645233 | G | A | M_Disease | 0.528718357 | M_Disease&NULL |
| 11 | 116891077 | rs77256870 | G | C | M_Disease | 0.528718357 | M_Disease&NULL |
| 11 | 116927101 | rs7942637 | A | G | M_Disease | 0.528718357 | M_Disease&NULL |
| 11 | 116928199 | rs7125464 | A | G | M_Disease | 0.528718357 | M_Disease&NULL |
| 11 | 116928748 | rs7114746 | G | T | M_Disease | 0.528718357 | M_Disease&NULL |
| 11 | 118974869 | rs4938627 | A | G | M_Disease | 0.528718357 | M_Disease&NULL |
| 11 | 118060549 | rs11216806 | C | T | M_Disease | 0.528718357 | M_Disease&NULL |
| 11 | 117052107 | rs144646355 | A | G | M_Disease | 0.528718357 | M_Disease&NULL |
| 11 | 117052532 | rs61729990 | A | C | M_Disease | 0.528718357 | M_Disease&NULL |
| 11 | 117318533 | rs12417961 | T | C | M_Disease | 0.528718357 | M_Disease&NULL |
| 11 | 119467150 | rs111549567 | C | T | M_Disease | 0.527472304 | M_Disease&NULL |

| | | | | | | | | | |
|---|---|---|---|---|---|---|---|---|---|
| 11 | 116767722 | rs141555847 | A | G | 1.128 | -7977.388 | -7977.118 | 0.27 | 5.07E-06 |
| 11 | 116771204 | rs76402668 | C | T | 1.128 | -7977.388 | -7977.118 | 0.27 | 5.07E-06 |
| 11 | 116778138 | rs146523230 | A | G | 1.128 | -7977.388 | -7977.118 | 0.27 | 5.07E-06 |
| 11 | 116783142 | rs74700879 | G | C | 1.128 | -7977.388 | -7977.12 | 0.268 | 5.06E-06 |
| 11 | 116874358 | rs12278943 | C | T | 1.11 | -7977.388 | -7977.122 | 0.266 | 5.05E-06 |
| 11 | 116879723 | rs138181556 | A | G | 1.11 | -7977.388 | -7977.122 | 0.266 | 5.05E-06 |
| 11 | 118186312 | rs2231450 | A | C | 0.886 | -7977.388 | -7977.122 | 0.266 | 5.05E-06 |
| 11 | 116285538 | rs563990 | G | A | 0.922 | -7977.388 | -7977.123 | 0.265 | 5.04E-06 |
| 11 | 120441720 | rs12279396 | G | A | 1.063 | -7977.388 | -7977.123 | 0.265 | 5.04E-06 |
| 11 | 116801299 | rs77203966 | T | C | 1.128 | -7977.388 | -7977.125 | 0.263 | 5.03E-06 |
| 11 | 116806367 | rs74557653 | T | G | 1.128 | -7977.388 | -7977.125 | 0.263 | 5.03E-06 |
| 11 | 116815306 | rs57040912 | G | A | 1.128 | -7977.388 | -7977.125 | 0.263 | 5.03E-06 |
| 11 | 116815717 | rs12283494 | A | G | 1.128 | -7977.388 | -7977.125 | 0.263 | 5.03E-06 |
| 11 | 116819050 | rs115255270 | C | T | 1.128 | -7977.388 | -7977.125 | 0.263 | 5.03E-06 |
| 11 | 116820673 | rs7940300 | T | A | 1.128 | -7977.388 | -7977.125 | 0.263 | 5.03E-06 |
| 11 | 116824938 | rs2289891 | C | A | 1.128 | -7977.388 | -7977.125 | 0.263 | 5.03E-06 |
| 11 | 116826409 | rs77161357 | C | T | 1.128 | -7977.388 | -7977.125 | 0.263 | 5.03E-06 |
| 11 | 116828573 | rs7130221 | A | G | 1.128 | -7977.388 | -7977.125 | 0.263 | 5.03E-06 |
| 11 | 116828859 | rs6589581 | T | A | 1.128 | -7977.388 | -7977.125 | 0.263 | 5.03E-06 |
| 11 | 116829129 | rs7938466 | C | T | 1.128 | -7977.388 | -7977.125 | 0.263 | 5.03E-06 |
| 11 | 116830628 | rs77029349 | T | G | 1.128 | -7977.388 | -7977.125 | 0.263 | 5.03E-06 |
| 11 | 116839094 | rs117791268 | A | G | 1.128 | -7977.388 | -7977.125 | 0.263 | 5.03E-06 |
| 11 | 116842174 | rs76223783 | C | T | 1.128 | -7977.388 | -7977.125 | 0.263 | 5.03E-06 |
| 11 | 116842758 | rs118137922 | A | G | 1.128 | -7977.388 | -7977.125 | 0.263 | 5.03E-06 |
| 11 | 116844649 | rs7937585 | A | G | 1.128 | -7977.388 | -7977.125 | 0.263 | 5.03E-06 |
| 11 | 116849038 | rs79868224 | A | G | 1.128 | -7977.388 | -7977.125 | 0.263 | 5.03E-06 |
| 11 | 116849544 | rs78241753 | C | G | 1.128 | -7977.388 | -7977.125 | 0.263 | 5.03E-06 |
| 11 | 116850485 | rs7940002 | A | C | 1.128 | -7977.388 | -7977.125 | 0.263 | 5.03E-06 |
| 11 | 116852714 | rs7112772 | C | T | 1.128 | -7977.388 | -7977.125 | 0.263 | 5.03E-06 |
| 11 | 116852716 | rs7112773 | C | T | 1.128 | -7977.388 | -7977.125 | 0.263 | 5.03E-06 |
| 11 | 116856152 | rs7110113 | C | A | 1.128 | -7977.388 | -7977.125 | 0.263 | 5.03E-06 |
| 11 | 116973247 | rs76942203 | A | G | 1.11 | -7977.388 | -7977.126 | 0.262 | 5.03E-06 |
| 11 | 116975136 | rs149576335 | A | G | 1.11 | -7977.388 | -7977.126 | 0.262 | 5.03E-06 |
| 11 | 116752045 | rs78013608 | A | G | 1.132 | -7977.388 | -7977.128 | 0.261 | 5.02E-06 |
| 11 | 120589179 | rs2852227 | C | A | 0.928 | -7977.388 | -7977.127 | 0.261 | 5.02E-06 |
| 11 | 116784692 | rs11216186 | C | T | 1.108 | -7977.388 | -7977.129 | 0.26 | 5.02E-06 |
| 11 | 116785341 | rs79711490 | T | C | 1.108 | -7977.388 | -7977.129 | 0.26 | 5.02E-06 |
| 11 | 116787633 | rs12418208 | T | C | 1.108 | -7977.388 | -7977.129 | 0.26 | 5.02E-06 |
| 11 | 116787641 | rs12418209 | T | C | 1.108 | -7977.388 | -7977.129 | 0.26 | 5.02E-06 |
| 11 | 116788133 | rs1006176 | C | T | 1.108 | -7977.388 | -7977.129 | 0.26 | 5.02E-06 |
| 11 | 118235246 | rs11216857 | A | G | 0.888 | -7977.388 | -7977.128 | 0.26 | 5.02E-06 |
| 11 | 119939156 | rs597253 | A | G | 0.899 | -7977.388 | -7977.13 | 0.258 | 5.01E-06 |
| 11 | 119939806 | rs593319 | T | A | 0.899 | -7977.388 | -7977.13 | 0.258 | 5.01E-06 |
| 11 | 118974878 | rs4938628 | T | A | 0.904 | -7977.388 | -7977.134 | 0.254 | 4.99E-06 |
| 11 | 118975113 | rs10736496 | A | G | 0.904 | -7977.388 | -7977.134 | 0.254 | 4.99E-06 |
| 11 | 118975426 | rs4938629 | C | T | 0.904 | -7977.388 | -7977.134 | 0.254 | 4.99E-06 |
| 11 | 118975927 | rs10790283 | A | G | 0.904 | -7977.388 | -7977.134 | 0.254 | 4.99E-06 |
| 11 | 116357627 | rs80310120 | T | C | 0.917 | -7977.388 | -7977.135 | 0.253 | 4.98E-06 |
| 11 | 118241995 | rs502514 | C | T | 0.939 | -7977.388 | -7977.135 | 0.253 | 4.98E-06 |
| 11 | 120931054 | rs12795310 | C | T | 1.066 | -7977.388 | -7977.136 | 0.253 | 4.98E-06 |
| 11 | 116780567 | rs75188049 | A | G | 1.127 | -7977.388 | -7977.136 | 0.252 | 4.98E-06 |
| 11 | 116780702 | rs77693143 | G | A | 1.127 | -7977.388 | -7977.136 | 0.252 | 4.98E-06 |
| 11 | 118146884 | rs10892232 | T | A | 0.887 | -7977.388 | -7977.137 | 0.252 | 4.98E-06 |
| 11 | 117411410 | rs11216444 | G | A | 1.069 | -7977.388 | -7977.137 | 0.251 | 4.97E-06 |
| 11 | 119881745 | rs651202 | A | G | 0.901 | -7977.388 | -7977.137 | 0.251 | 4.97E-06 |
| 11 | 116414485 | rs77110391 | A | G | 1.08 | -7977.388 | -7977.138 | 0.25 | 4.97E-06 |
| 11 | 116805064 | rs77204473 | G | T | 1.11 | -7977.388 | -7977.138 | 0.25 | 4.97E-06 |
| 11 | 117355892 | rs10892126 | C | G | 1.071 | -7977.388 | -7977.139 | 0.25 | 4.97E-06 |
| 11 | 117371563 | rs7942406 | T | G | 0.942 | -7977.388 | -7977.138 | 0.25 | 4.97E-06 |
| 11 | 116905209 | rs80176870 | C | T | 1.127 | -7977.388 | -7977.139 | 0.249 | 4.96E-06 |
| 11 | 116906984 | rs78550859 | T | C | 1.127 | -7977.388 | -7977.139 | 0.249 | 4.96E-06 |
| 11 | 116908807 | rs79842631 | G | A | 1.127 | -7977.388 | -7977.139 | 0.249 | 4.96E-06 |
| 11 | 116908913 | rs77939786 | T | C | 1.127 | -7977.388 | -7977.139 | 0.249 | 4.96E-06 |
| 11 | 116908914 | rs77393043 | A | C | 1.127 | -7977.388 | -7977.139 | 0.249 | 4.96E-06 |
| 11 | 118221655 | rs137996620 | C | A | 0.908 | -7977.388 | -7977.139 | 0.249 | 4.96E-06 |
| 11 | 121006387 | rs4936579 | C | T | 0.933 | -7977.388 | -7977.141 | 0.248 | 4.96E-06 |
| 11 | 121006650 | rs4427574 | A | G | 0.933 | -7977.388 | -7977.141 | 0.248 | 4.96E-06 |
| 11 | 117254122 | rs12797557 | A | G | 1.068 | -7977.388 | -7977.143 | 0.246 | 4.95E-06 |
| 11 | 116547924 | rs55928509 | A | C | 0.924 | -7977.388 | -7977.143 | 0.245 | 4.94E-06 |
| 11 | 119728423 | rs140657166 | C | A | 0.891 | -7977.388 | -7977.144 | 0.244 | 4.94E-06 |
| 11 | 116903095 | rs7938234 | T | C | 1.126 | -7977.388 | -7977.146 | 0.243 | 4.93E-06 |
| 11 | 117411717 | rs59099254 | A | C | 1.068 | -7977.388 | -7977.145 | 0.243 | 4.93E-06 |
| 11 | 121066946 | rs1939934 | T | C | 0.931 | -7977.388 | -7977.145 | 0.243 | 4.93E-06 |
| 11 | 116288186 | rs563229 | G | C | 0.922 | -7977.388 | -7977.149 | 0.239 | 4.91E-06 |
| 11 | 121011488 | rs559301 | A | G | 0.94 | -7977.388 | -7977.15 | 0.238 | 4.91E-06 |
| 11 | 118257081 | rs11216863 | A | G | 1.097 | -7977.388 | -7977.151 | 0.237 | 4.90E-06 |
| 11 | 116911590 | rs77322791 | G | A | 1.126 | -7977.388 | -7977.152 | 0.236 | 4.90E-06 |
| 11 | 116941540 | rs9971421 | T | C | 1.108 | -7977.388 | -7977.152 | 0.236 | 4.90E-06 |
| 11 | 120801185 | rs2186620 | C | T | 0.928 | -7977.388 | -7977.154 | 0.235 | 4.89E-06 |
| 11 | 116280965 | rs536484 | C | G | 0.923 | -7977.388 | -7977.154 | 0.234 | 4.89E-06 |
| 11 | 116358711 | rs17119760 | A | C | 0.918 | -7977.388 | -7977.155 | 0.233 | 4.88E-06 |
| 11 | 117440076 | rs112118203 | C | T | 1.095 | -7977.388 | -7977.155 | 0.233 | 4.88E-06 |
| 11 | 116731205 | rs4938315 | G | A | 0.92 | -7977.388 | -7977.156 | 0.232 | 4.88E-06 |
| 11 | 117795717 | rs76014645 | A | C | 0.891 | -7977.388 | -7977.158 | 0.231 | 4.87E-06 |
| 11 | 117410174 | rs60851263 | T | C | 1.068 | -7977.388 | -7977.159 | 0.229 | 4.86E-06 |
| 11 | 119938788 | rs645233 | G | A | 0.9 | -7977.388 | -7977.162 | 0.227 | 4.85E-06 |
| 11 | 116891077 | rs77256870 | G | C | 1.125 | -7977.388 | -7977.165 | 0.224 | 4.84E-06 |
| 11 | 116927101 | rs7942637 | A | G | 1.125 | -7977.388 | -7977.165 | 0.224 | 4.84E-06 |
| 11 | 116928199 | rs7125464 | A | G | 1.125 | -7977.388 | -7977.165 | 0.224 | 4.84E-06 |
| 11 | 116928748 | rs7114746 | G | T | 1.125 | -7977.388 | -7977.165 | 0.224 | 4.84E-06 |
| 11 | 118974869 | rs4938627 | A | G | 0.905 | -7977.388 | -7977.165 | 0.223 | 4.83E-06 |
| 11 | 118060549 | rs11216806 | C | T | 1.074 | -7977.388 | -7977.166 | 0.222 | 4.83E-06 |
| 11 | 117052107 | rs144646355 | A | G | 1.142 | -7977.388 | -7977.168 | 0.221 | 4.82E-06 |
| 11 | 117052532 | rs61729990 | A | C | 1.142 | -7977.388 | -7977.168 | 0.221 | 4.82E-06 |
| 11 | 117318533 | rs12417961 | T | C | 0.907 | -7977.388 | -7977.167 | 0.221 | 4.82E-06 |
| 11 | 119467150 | rs111549567 | C | T | 1.096 | -7977.388 | -7977.168 | 0.22 | 4.82E-06 |

| | | | | | | | |
|---|---|---|---|---|---|---|---|
| 11 | 116621623 | rs74496689 | A | G | M_Disease | 0.528718357 | M_Disease&NULL |
| 11 | 117455203 | rs77855139 | G | A | M_Disease | 0.528718357 | M_Disease&NULL |
| 11 | 117189216 | rs3017608 | A | G | M_Disease | 0.527472304 | M_Disease&NULL |
| 11 | 116457294 | rs55787848 | T | C | M_Disease | 0.527472304 | M_Disease&NULL |
| 11 | 116962661 | rs116861530 | A | G | M_Disease | 0.527472304 | M_Disease&NULL |
| 11 | 120528115 | rs75749368 | T | C | M_Disease | 0.527472304 | M_Disease&NULL |
| 11 | 116771356 | rs595137 | C | T | M_Disease | 0.527472304 | M_Disease&NULL |
| 11 | 117412185 | rs57208438 | A | G | M_Disease | 0.527472304 | M_Disease&NULL |
| 11 | 117459687 | rs77138800 | T | C | M_Disease | 0.527472304 | M_Disease&NULL |
| 11 | 117411664 | rs58088485 | A | G | M_Disease | 0.526225909 | M_Disease&NULL |
| 11 | 116828939 | rs878692 | A | C | M_Disease | 0.527472304 | M_Disease&NULL |
| 11 | 116829442 | rs12420857 | T | G | M_Disease | 0.527472304 | M_Disease&NULL |
| 11 | 116838130 | rs12281729 | C | A | M_Disease | 0.527472304 | M_Disease&NULL |
| 11 | 116839293 | rs12417152 | C | G | M_Disease | 0.527472304 | M_Disease&NULL |
| 11 | 116839614 | rs11216204 | A | G | M_Disease | 0.527472304 | M_Disease&NULL |
| 11 | 116839938 | rs12272305 | A | G | M_Disease | 0.527472304 | M_Disease&NULL |
| 11 | 120502086 | rs7927736 | C | T | M_Disease | 0.526225909 | M_Disease&NULL |
| 11 | 116928546 | rs10892068 | T | C | M_Disease | 0.526225909 | M_Disease&NULL |
| 11 | 117349401 | rs56057635 | G | A | M_Disease | 0.526225909 | M_Disease&NULL |
| 11 | 116674818 | rs1729408 | G | A | M_Disease | 0.526225909 | M_Disease&NULL |
| 11 | 116842676 | rs12418324 | T | C | M_Disease | 0.526225909 | M_Disease&NULL |
| 11 | 116863458 | rs10128682 | T | C | M_Disease | 0.526225909 | M_Disease&NULL |
| 11 | 116976778 | rs4388921 | C | T | M_Disease | 0.526225909 | M_Disease&NULL |
| 11 | 116727850 | rs1035237 | G | C | M_Disease | 0.526225909 | M_Disease&NULL |
| 11 | 116516736 | rs1002037 | C | G | M_Disease | 0.524979187 | M_Disease&NULL |
| 11 | 116530298 | rs530849 | C | T | M_Disease | 0.524979187 | M_Disease&NULL |
| 11 | 116902372 | rs7481286 | T | G | M_Disease | 0.524979187 | M_Disease&NULL |
| 11 | 117410682 | rs2510853 | A | G | M_Disease | 0.524979187 | M_Disease&NULL |
| 11 | 116756690 | rs74505326 | G | A | M_Disease | 0.526225909 | M_Disease&NULL |
| 11 | 117793169 | rs75336872 | T | C | M_Disease | 0.524979187 | M_Disease&NULL |
| 11 | 117572395 | rs55945170 | G | A | M_Disease | 0.524979187 | M_Disease&NULL |
| 11 | 116732512 | rs2075292 | G | T | M_Disease | 0.524979187 | M_Disease&NULL |
| 11 | 116865100 | rs75826529 | A | T | M_Disease | 0.524979187 | M_Disease&NULL |
| 11 | 116865207 | rs74460214 | C | T | M_Disease | 0.524979187 | M_Disease&NULL |
| 11 | 116865367 | rs79349393 | G | A | M_Disease | 0.524979187 | M_Disease&NULL |
| 11 | 116867514 | rs11604866 | T | C | M_Disease | 0.524979187 | M_Disease&NULL |
| 11 | 116976009 | rs142395187 | G | A | M_Disease | 0.524979187 | M_Disease&NULL |
| 11 | 116704505 | rs12721031 | T | C | M_Disease | 0.524979187 | M_Disease&NULL |
| 11 | 117599488 | rs138897098 | C | T | M_Disease | 0.524979187 | M_Disease&NULL |
| 11 | 117604604 | rs17588728 | T | C | M_Disease | 0.524979187 | M_Disease&NULL |
| 11 | 117434629 | rs4938408 | A | C | M_Disease | 0.524979187 | M_Disease&NULL |
| 11 | 120967158 | rs56337383 | A | G | M_Disease | 0.524979187 | M_Disease&NULL |
| 11 | 116913908 | rs186593252 | C | A | M_Disease | 0.524979187 | M_Disease&NULL |
| 11 | 116916723 | rs60101932 | A | G | M_Disease | 0.524979187 | M_Disease&NULL |
| 11 | 116916984 | rs74429128 | A | G | M_Disease | 0.524979187 | M_Disease&NULL |
| 11 | 118045846 | rs76448372 | A | G | M_Disease | 0.523732154 | M_Disease&NULL |
| 11 | 116913570 | rs1426560348 | A | C | M_Disease | 0.523732154 | M_Disease&NULL |
| 11 | 116927762 | rs10892067 | C | G | M_Disease | 0.523732154 | M_Disease&NULL |
| 11 | 119677955 | rs7926623 | A | G | M_Disease | 0.523732154 | M_Disease&NULL |
| 11 | 119705743 | rs80350987 | A | C | M_Disease | 0.523732154 | M_Disease&NULL |
| 11 | 116202083 | rs7948971 | G | A | M_Disease | 0.523732154 | M_Disease&NULL |
| 11 | 116963277 | rs74420345 | A | C | M_Disease | 0.523732154 | M_Disease&NULL |
| 11 | 116313516 | rs11215936 | A | G | M_Disease | 0.523732154 | M_Disease&NULL |
| 11 | 117926624 | rs61900565 | G | C | M_Disease | 0.522484825 | M_Disease&NULL |
| 11 | 120588487 | rs2846108 | T | C | M_Disease | 0.522484825 | M_Disease&NULL |
| 11 | 116870411 | rs10892056 | C | G | M_Disease | 0.522484825 | M_Disease&NULL |
| 11 | 116878066 | rs11216226 | G | C | M_Disease | 0.522484825 | M_Disease&NULL |
| 11 | 116878759 | rs12276805 | C | T | M_Disease | 0.522484825 | M_Disease&NULL |
| 11 | 116922585 | rs143856824 | A | C | M_Disease | 0.523732154 | M_Disease&NULL |
| 11 | 116924439 | rs117857960 | C | T | M_Disease | 0.523732154 | M_Disease&NULL |
| 11 | 116925898 | rs78111243 | C | T | M_Disease | 0.523732154 | M_Disease&NULL |
| 11 | 116927976 | rs117432935 | A | G | M_Disease | 0.523732154 | M_Disease&NULL |
| 11 | 116928302 | rs75582303 | C | T | M_Disease | 0.523732154 | M_Disease&NULL |
| 11 | 117450936 | rs79517065 | T | C | M_Disease | 0.523732154 | M_Disease&NULL |
| 11 | 116757720 | rs59491594 | C | T | M_Disease | 0.522484825 | M_Disease&NULL |
| 11 | 116763124 | rs74701178 | C | T | M_Disease | 0.522484825 | M_Disease&NULL |
| 11 | 116766149 | rs77718307 | A | G | M_Disease | 0.522484825 | M_Disease&NULL |
| 11 | 117509536 | rs692840 | A | G | M_Disease | 0.522484825 | M_Disease&NULL |
| 11 | 116718554 | rs149318234 | G | C | M_Disease | 0.522484825 | M_Disease&NULL |
| 11 | 116805441 | rs79207844 | A | G | M_Disease | 0.522484825 | M_Disease&NULL |
| 11 | 116805696 | rs12274590 | A | G | M_Disease | 0.522484825 | M_Disease&NULL |
| 11 | 120324517 | rs11603533 | G | C | M_Disease | 0.522484825 | M_Disease&NULL |
| 11 | 116762157 | rs72645459 | A | C | M_Disease | 0.522484825 | M_Disease&NULL |
| 11 | 116530376 | rs553779 | G | A | M_Disease | 0.521237215 | M_Disease&NULL |
| 11 | 116886721 | rs79870586 | A | G | M_Disease | 0.522484825 | M_Disease&NULL |
| 11 | 116890667 | rs80054374 | G | A | M_Disease | 0.522484825 | M_Disease&NULL |
| 11 | 116787327 | rs12277092 | T | C | M_Disease | 0.522484825 | M_Disease&NULL |
| 11 | 117430841 | rs567988 | A | G | M_Disease | 0.521237215 | M_Disease&NULL |
| 11 | 116960059 | rs1169496015 | G | C | M_Disease | 0.522484825 | M_Disease&NULL |
| 11 | 116920984 | rs112422338 | A | G | M_Disease | 0.522484825 | M_Disease&NULL |
| 11 | 117524455 | rs684669 | A | C | M_Disease | 0.521237215 | M_Disease&NULL |
| 11 | 118970875 | rs10892330 | T | C | M_Disease | 0.521237215 | M_Disease&NULL |
| 11 | 117446003 | rs74526104 | G | A | M_Disease | 0.521237215 | M_Disease&NULL |
| 11 | 120585097 | rs61901398 | A | G | M_Disease | 0.521237215 | M_Disease&NULL |
| 11 | 117296292 | rs4938377 | G | T | M_Disease | 0.521237215 | M_Disease&NULL |
| 11 | 116858320 | rs76391131 | C | T | M_Disease | 0.521237215 | M_Disease&NULL |
| 11 | 116859076 | rs143491990 | T | C | M_Disease | 0.521237215 | M_Disease&NULL |
| 11 | 116861398 | rs1228001593 | A | G | M_Disease | 0.521237215 | M_Disease&NULL |
| 11 | 116862105 | rs7122484 | C | G | M_Disease | 0.521237215 | M_Disease&NULL |
| 11 | 116862779 | rs7947525 | T | C | M_Disease | 0.521237215 | M_Disease&NULL |
| 11 | 116870159 | rs59425942 | T | C | M_Disease | 0.521237215 | M_Disease&NULL |
| 11 | 116936871 | rs56751865 | T | C | M_Disease | 0.521237215 | M_Disease&NULL |
| 11 | 117222256 | rs7926760 | C | T | M_Disease | 0.51998934 | M_Disease&NULL |
| 11 | 116349950 | rs34095499 | A | G | M_Disease | 0.51998934 | M_Disease&NULL |
| 11 | 116454374 | rs12295025 | G | A | M_Disease | 0.51998934 | M_Disease&NULL |
| 11 | 116731852 | rs78087637 | G | C | M_Disease | 0.521237215 | M_Disease&NULL |

| | | | | | | | | | |
|---|---|---|---|---|---|---|---|---|---|
| 11 | 116621623 | rs74496689 | A | G | 1.143 | -7977.388 | -7977.171 | 0.218 | 4.81E-06 |
| 11 | 117455203 | rs77855139 | G | A | 0.876 | -7977.388 | -7977.171 | 0.218 | 4.81E-06 |
| 11 | 117189216 | rs3017608 | A | G | 0.942 | -7977.388 | -7977.172 | 0.216 | 4.80E-06 |
| 11 | 116457294 | rs55787848 | T | C | 1.078 | -7977.388 | -7977.174 | 0.214 | 4.79E-06 |
| 11 | 116962661 | rs116861530 | A | G | 1.108 | -7977.388 | -7977.175 | 0.214 | 4.79E-06 |
| 11 | 120528115 | rs75749368 | T | C | 1.131 | -7977.388 | -7977.175 | 0.214 | 4.79E-06 |
| 11 | 116771356 | rs595137 | C | T | 0.885 | -7977.388 | -7977.175 | 0.213 | 4.79E-06 |
| 11 | 117412185 | rs57208438 | A | G | 1.068 | -7977.388 | -7977.175 | 0.213 | 4.79E-06 |
| 11 | 117459687 | rs77138800 | T | C | 1.094 | -7977.388 | -7977.175 | 0.213 | 4.79E-06 |
| 11 | 117411664 | rs58088485 | A | G | 1.068 | -7977.388 | -7977.177 | 0.212 | 4.78E-06 |
| 11 | 116828939 | rs878692 | A | C | 1.108 | -7977.388 | -7977.177 | 0.211 | 4.78E-06 |
| 11 | 116829442 | rs12420857 | T | G | 1.108 | -7977.388 | -7977.177 | 0.211 | 4.78E-06 |
| 11 | 116838130 | rs12281729 | C | A | 1.108 | -7977.388 | -7977.177 | 0.211 | 4.78E-06 |
| 11 | 116839293 | rs12417152 | C | G | 1.108 | -7977.388 | -7977.177 | 0.211 | 4.78E-06 |
| 11 | 116839614 | rs11216204 | A | G | 1.108 | -7977.388 | -7977.177 | 0.211 | 4.78E-06 |
| 11 | 116839938 | rs12272305 | A | G | 1.108 | -7977.388 | -7977.177 | 0.211 | 4.78E-06 |
| 11 | 120502086 | rs7927736 | C | T | 0.941 | -7977.388 | -7977.178 | 0.21 | 4.77E-06 |
| 11 | 116928546 | rs10892068 | T | C | 1.107 | -7977.388 | -7977.179 | 0.209 | 4.77E-06 |
| 11 | 117349401 | rs56057635 | G | A | 1.086 | -7977.388 | -7977.182 | 0.206 | 4.75E-06 |
| 11 | 116674818 | rs1729408 | G | A | 1.064 | -7977.388 | -7977.183 | 0.205 | 4.75E-06 |
| 11 | 116842676 | rs12418324 | T | C | 1.107 | -7977.388 | -7977.183 | 0.205 | 4.75E-06 |
| 11 | 116863458 | rs10128682 | T | C | 1.107 | -7977.388 | -7977.183 | 0.205 | 4.75E-06 |
| 11 | 116976778 | rs4388921 | C | T | 1.071 | -7977.388 | -7977.184 | 0.204 | 4.74E-06 |
| 11 | 116727850 | rs1035237 | G | C | 0.921 | -7977.388 | -7977.185 | 0.203 | 4.74E-06 |
| 11 | 116516736 | rs1002037 | C | G | 1.068 | -7977.388 | -7977.186 | 0.202 | 4.73E-06 |
| 11 | 116530298 | rs530849 | C | T | 1.061 | -7977.388 | -7977.187 | 0.201 | 4.73E-06 |
| 11 | 116902372 | rs7481286 | T | G | 1.107 | -7977.388 | -7977.189 | 0.199 | 4.72E-06 |
| 11 | 117410682 | rs2510853 | A | G | 1.061 | -7977.388 | -7977.189 | 0.199 | 4.72E-06 |
| 11 | 116756690 | rs74505326 | G | A | 0.885 | -7977.388 | -7977.19 | 0.198 | 4.72E-06 |
| 11 | 117793169 | rs75336872 | T | C | 0.894 | -7977.388 | -7977.191 | 0.197 | 4.71E-06 |
| 11 | 117572395 | rs55945170 | G | A | 0.892 | -7977.388 | -7977.195 | 0.194 | 4.70E-06 |
| 11 | 116732512 | rs2075292 | G | T | 0.921 | -7977.388 | -7977.195 | 0.193 | 4.69E-06 |
| 11 | 116865100 | rs75826529 | A | T | 1.107 | -7977.388 | -7977.195 | 0.193 | 4.69E-06 |
| 11 | 116865207 | rs74460214 | C | T | 1.107 | -7977.388 | -7977.195 | 0.193 | 4.69E-06 |
| 11 | 116865367 | rs79349393 | G | A | 1.107 | -7977.388 | -7977.195 | 0.193 | 4.69E-06 |
| 11 | 116867514 | rs11604866 | T | C | 1.107 | -7977.388 | -7977.195 | 0.193 | 4.69E-06 |
| 11 | 116976009 | rs142395187 | G | A | 1.107 | -7977.388 | -7977.195 | 0.193 | 4.69E-06 |
| 11 | 116704505 | rs12721031 | T | C | 0.885 | -7977.388 | -7977.197 | 0.191 | 4.68E-06 |
| 11 | 117599488 | rs138897098 | C | T | 0.874 | -7977.388 | -7977.197 | 0.191 | 4.68E-06 |
| 11 | 117604604 | rs17588728 | T | C | 0.874 | -7977.388 | -7977.197 | 0.191 | 4.68E-06 |
| 11 | 117434629 | rs4938408 | A | C | 0.878 | -7977.388 | -7977.199 | 0.19 | 4.68E-06 |
| 11 | 120967158 | rs56337383 | A | G | 1.136 | -7977.388 | -7977.198 | 0.19 | 4.68E-06 |
| 11 | 116913908 | rs186593252 | C | A | 1.123 | -7977.388 | -7977.199 | 0.189 | 4.67E-06 |
| 11 | 116916723 | rs60101932 | A | G | 1.123 | -7977.388 | -7977.199 | 0.189 | 4.67E-06 |
| 11 | 116916984 | rs74429128 | A | G | 1.123 | -7977.388 | -7977.199 | 0.189 | 4.67E-06 |
| 11 | 118045846 | rs76448372 | A | G | 0.899 | -7977.388 | -7977.201 | 0.188 | 4.67E-06 |
| 11 | 116913570 | rs1426560348 | A | C | 1.106 | -7977.388 | -7977.202 | 0.187 | 4.66E-06 |
| 11 | 116927762 | rs10892067 | C | G | 1.106 | -7977.388 | -7977.202 | 0.187 | 4.66E-06 |
| 11 | 119677955 | rs7926623 | A | G | 0.935 | -7977.388 | -7977.203 | 0.186 | 4.66E-06 |
| 11 | 119705743 | rs80350987 | A | C | 1.103 | -7977.388 | -7977.204 | 0.184 | 4.65E-06 |
| 11 | 116202083 | rs7948971 | G | A | 1.077 | -7977.388 | -7977.205 | 0.183 | 4.64E-06 |
| 11 | 116963277 | rs74420345 | A | C | 1.106 | -7977.388 | -7977.205 | 0.183 | 4.64E-06 |
| 11 | 116313516 | rs11215936 | A | G | 0.902 | -7977.388 | -7977.207 | 0.181 | 4.64E-06 |
| 11 | 117926624 | rs61900565 | G | C | 0.94 | -7977.388 | -7977.207 | 0.181 | 4.64E-06 |
| 11 | 120588487 | rs2846108 | T | C | 0.919 | -7977.388 | -7977.209 | 0.18 | 4.63E-06 |
| 11 | 116870411 | rs10892056 | C | G | 1.106 | -7977.388 | -7977.21 | 0.179 | 4.63E-06 |
| 11 | 116878066 | rs11216226 | G | C | 1.106 | -7977.388 | -7977.21 | 0.179 | 4.63E-06 |
| 11 | 116878759 | rs12276805 | C | T | 1.106 | -7977.388 | -7977.21 | 0.179 | 4.63E-06 |
| 11 | 116922585 | rs143856824 | A | C | 1.123 | -7977.388 | -7977.209 | 0.179 | 4.63E-06 |
| 11 | 116924439 | rs117857960 | C | T | 1.123 | -7977.388 | -7977.209 | 0.179 | 4.63E-06 |
| 11 | 116925898 | rs78111243 | C | T | 1.123 | -7977.388 | -7977.209 | 0.179 | 4.63E-06 |
| 11 | 116927976 | rs117432935 | A | G | 1.123 | -7977.388 | -7977.209 | 0.179 | 4.63E-06 |
| 11 | 116928302 | rs75582303 | C | T | 1.123 | -7977.388 | -7977.209 | 0.179 | 4.63E-06 |
| 11 | 117450936 | rs79517065 | T | C | 0.878 | -7977.388 | -7977.21 | 0.178 | 4.62E-06 |
| 11 | 116757720 | rs59491594 | C | T | 1.123 | -7977.388 | -7977.213 | 0.176 | 4.61E-06 |
| 11 | 116763124 | rs74701178 | C | T | 1.123 | -7977.388 | -7977.213 | 0.176 | 4.61E-06 |
| 11 | 116766149 | rs77718307 | A | G | 1.123 | -7977.388 | -7977.213 | 0.176 | 4.61E-06 |
| 11 | 117509536 | rs692840 | A | G | 0.938 | -7977.388 | -7977.213 | 0.176 | 4.61E-06 |
| 11 | 116718554 | rs149318234 | G | C | 1.122 | -7977.388 | -7977.213 | 0.175 | 4.61E-06 |
| 11 | 116805441 | rs79207844 | A | G | 1.106 | -7977.388 | -7977.213 | 0.175 | 4.61E-06 |
| 11 | 116805696 | rs12274590 | A | G | 1.106 | -7977.388 | -7977.213 | 0.175 | 4.61E-06 |
| 11 | 120324517 | rs11603533 | G | C | 0.888 | -7977.388 | -7977.213 | 0.175 | 4.61E-06 |
| 11 | 116762157 | rs72645459 | A | C | 0.918 | -7977.388 | -7977.214 | 0.174 | 4.60E-06 |
| 11 | 116530376 | rs553779 | G | A | 1.061 | -7977.388 | -7977.216 | 0.173 | 4.60E-06 |
| 11 | 116886721 | rs79870586 | A | G | 1.106 | -7977.388 | -7977.215 | 0.173 | 4.60E-06 |
| 11 | 116890667 | rs80054374 | G | A | 1.106 | -7977.388 | -7977.215 | 0.173 | 4.60E-06 |
| 11 | 116787327 | rs12277092 | T | C | 1.105 | -7977.388 | -7977.216 | 0.172 | 4.59E-06 |
| 11 | 117430841 | rs567988 | A | G | 0.942 | -7977.388 | -7977.217 | 0.172 | 4.59E-06 |
| 11 | 116960059 | rs1169496015 | G | C | 1.123 | -7977.388 | -7977.217 | 0.171 | 4.59E-06 |
| 11 | 116920984 | rs112422338 | A | G | 1.122 | -7977.388 | -7977.218 | 0.17 | 4.58E-06 |
| 11 | 117524455 | rs684669 | A | C | 0.942 | -7977.388 | -7977.221 | 0.168 | 4.58E-06 |
| 11 | 118970875 | rs10892330 | T | C | 0.905 | -7977.388 | -7977.222 | 0.166 | 4.57E-06 |
| 11 | 117446003 | rs74526104 | G | A | 0.879 | -7977.388 | -7977.224 | 0.165 | 4.56E-06 |
| 11 | 120585097 | rs61901398 | A | G | 0.919 | -7977.388 | -7977.224 | 0.164 | 4.56E-06 |
| 11 | 117296292 | rs4938377 | G | T | 0.905 | -7977.388 | -7977.228 | 0.161 | 4.54E-06 |
| 11 | 116858320 | rs76391131 | C | T | 1.122 | -7977.388 | -7977.228 | 0.16 | 4.54E-06 |
| 11 | 116859076 | rs143491990 | T | C | 1.122 | -7977.388 | -7977.228 | 0.16 | 4.54E-06 |
| 11 | 116861398 | rs1228001593 | A | G | 1.122 | -7977.388 | -7977.228 | 0.16 | 4.54E-06 |
| 11 | 116862105 | rs7122484 | C | G | 1.122 | -7977.388 | -7977.228 | 0.16 | 4.54E-06 |
| 11 | 116862779 | rs7947525 | T | C | 1.122 | -7977.388 | -7977.228 | 0.16 | 4.54E-06 |
| 11 | 116870159 | rs59425942 | T | C | 1.122 | -7977.388 | -7977.228 | 0.16 | 4.54E-06 |
| 11 | 116936871 | rs56751865 | T | C | 1.105 | -7977.388 | -7977.228 | 0.16 | 4.54E-06 |
| 11 | 117222256 | rs7926760 | C | T | 1.064 | -7977.388 | -7977.229 | 0.16 | 4.54E-06 |
| 11 | 116349950 | rs34095499 | A | G | 0.915 | -7977.388 | -7977.229 | 0.159 | 4.53E-06 |
| 11 | 116454374 | rs12295025 | G | A | 1.074 | -7977.388 | -7977.23 | 0.159 | 4.53E-06 |
| 11 | 116731852 | rs78087637 | G | C | 1.113 | -7977.388 | -7977.229 | 0.159 | 4.53E-06 |

| | | | | | | | |
|---|---|---|---|---|---|---|---|
| 11 | 120854046 | rs1008966 | T | C | M_Disease | 0.51998934 | M_Disease&NULL |
| 11 | 116900628 | rs139462136 | T | C | M_Disease | 0.521237215 | M_Disease&NULL |
| 11 | 117925581 | rs56204159 | T | C | M_Disease | 0.51998934 | M_Disease&NULL |
| 11 | 116530293 | rs530845 | T | C | M_Disease | 0.51998934 | M_Disease&NULL |
| 11 | 117926172 | rs4938472 | A | T | M_Disease | 0.518741216 | M_Disease&NULL |
| 11 | 116879162 | rs58354636 | G | A | M_Disease | 0.51998934 | M_Disease&NULL |
| 11 | 117905635 | rs1786176 | A | G | M_Disease | 0.51998934 | M_Disease&NULL |
| 11 | 116742626 | rs79849088 | T | C | M_Disease | 0.51998934 | M_Disease&NULL |
| 11 | 116313601 | rs11215937 | G | A | M_Disease | 0.518741216 | M_Disease&NULL |
| 11 | 117434985 | rs752795 | G | C | M_Disease | 0.51998934 | M_Disease&NULL |
| 11 | 117444949 | rs76959909 | A | C | M_Disease | 0.518741216 | M_Disease&NULL |
| 11 | 118931099 | rs620331 | T | C | M_Disease | 0.518741216 | M_Disease&NULL |
| 11 | 116891412 | rs76666535 | A | G | M_Disease | 0.518741216 | M_Disease&NULL |
| 11 | 116897871 | rs76383990 | G | A | M_Disease | 0.518741216 | M_Disease&NULL |
| 11 | 116914260 | rs77822137 | C | T | M_Disease | 0.518741216 | M_Disease&NULL |
| 11 | 116915242 | rs76982250 | T | C | M_Disease | 0.518741216 | M_Disease&NULL |
| 11 | 116917089 | rs118121169 | A | G | M_Disease | 0.518741216 | M_Disease&NULL |
| 11 | 116920199 | rs145949765 | T | C | M_Disease | 0.518741216 | M_Disease&NULL |
| 11 | 116921802 | rs76916861 | T | C | M_Disease | 0.518741216 | M_Disease&NULL |
| 11 | 116934380 | rs150342927 | T | C | M_Disease | 0.518741216 | M_Disease&NULL |
| 11 | 116939370 | rs57072067 | G | A | M_Disease | 0.518741216 | M_Disease&NULL |
| 11 | 116948368 | rs58068870 | C | T | M_Disease | 0.518741216 | M_Disease&NULL |
| 11 | 117350966 | rs11823335 | T | C | M_Disease | 0.518741216 | M_Disease&NULL |
| 11 | 116733310 | rs6589575 | C | T | M_Disease | 0.518741216 | M_Disease&NULL |
| 11 | 116723837 | rs77462380 | T | C | M_Disease | 0.518741216 | M_Disease&NULL |
| 11 | 116724233 | rs75771311 | A | G | M_Disease | 0.518741216 | M_Disease&NULL |
| 11 | 116727547 | rs117910440 | C | T | M_Disease | 0.518741216 | M_Disease&NULL |
| 11 | 116727871 | rs78816728 | C | T | M_Disease | 0.518741216 | M_Disease&NULL |
| 11 | 116728260 | rs75385236 | T | C | M_Disease | 0.518741216 | M_Disease&NULL |
| 11 | 116730980 | rs117833973 | T | C | M_Disease | 0.518741216 | M_Disease&NULL |
| 11 | 117795611 | rs79669568 | C | T | M_Disease | 0.518741216 | M_Disease&NULL |
| 11 | 116707684 | rs2070665 | A | G | M_Disease | 0.518741216 | M_Disease&NULL |
| 11 | 117183650 | rs593245 | T | C | M_Disease | 0.517492858 | M_Disease&NULL |
| 11 | 117479686 | rs4938419 | T | A | M_Disease | 0.517492858 | M_Disease&NULL |
| 11 | 116517968 | rs2187125 | G | T | M_Disease | 0.517492858 | M_Disease&NULL |
| 11 | 119882823 | rs142724980 | T | C | M_Disease | 0.518741216 | M_Disease&NULL |
| 11 | 117444661 | rs75807933 | A | T | M_Disease | 0.517492858 | M_Disease&NULL |
| 11 | 119679013 | rs56221909 | C | G | M_Disease | 0.517492858 | M_Disease&NULL |
| 11 | 120067199 | rs147634995 | A | C | M_Disease | 0.517492858 | M_Disease&NULL |
| 11 | 120420262 | rs11600946 | A | G | M_Disease | 0.517492858 | M_Disease&NULL |
| 11 | 116904279 | rs150817479 | A | G | M_Disease | 0.517492858 | M_Disease&NULL |
| 11 | 116734146 | rs718847 | C | T | M_Disease | 0.516244281 | M_Disease&NULL |
| 11 | 116961819 | rs75202546 | A | G | M_Disease | 0.517492858 | M_Disease&NULL |
| 11 | 116726181 | rs149462130 | T | C | M_Disease | 0.517492858 | M_Disease&NULL |
| 11 | 117466661 | rs145076512 | G | T | M_Disease | 0.517492858 | M_Disease&NULL |
| 11 | 117693971 | rs4938440 | A | G | M_Disease | 0.517492858 | M_Disease&NULL |
| 11 | 120415158 | rs4936519 | T | G | M_Disease | 0.516244281 | M_Disease&NULL |
| 11 | 121066079 | rs9943635 | T | C | M_Disease | 0.516244281 | M_Disease&NULL |
| 11 | 121073219 | rs516495 | C | T | M_Disease | 0.516244281 | M_Disease&NULL |
| 11 | 117474492 | rs77259443 | G | A | M_Disease | 0.517492858 | M_Disease&NULL |
| 11 | 119951570 | rs10892532 | G | A | M_Disease | 0.516244281 | M_Disease&NULL |
| 11 | 116275617 | rs4586206 | A | G | M_Disease | 0.516244281 | M_Disease&NULL |
| 11 | 117563458 | rs145253564 | T | C | M_Disease | 0.517492858 | M_Disease&NULL |
| 11 | 117564084 | rs139289629 | C | G | M_Disease | 0.517492858 | M_Disease&NULL |
| 11 | 116317789 | rs56287831 | A | G | M_Disease | 0.516244281 | M_Disease&NULL |
| 11 | 118984287 | rs4938630 | A | C | M_Disease | 0.516244281 | M_Disease&NULL |
| 11 | 121015228 | rs11218160 | A | G | M_Disease | 0.516244281 | M_Disease&NULL |
| 11 | 118973113 | rs3825059 | A | C | M_Disease | 0.516244281 | M_Disease&NULL |
| 11 | 119679613 | rs10790347 | T | A | M_Disease | 0.516244281 | M_Disease&NULL |
| 11 | 118240332 | rs11216860 | C | T | M_Disease | 0.514995502 | M_Disease&NULL |
| 11 | 120412053 | rs78566287 | T | C | M_Disease | 0.516244281 | M_Disease&NULL |
| 11 | 121071633 | rs201644901 | C | A | M_Disease | 0.514995502 | M_Disease&NULL |
| 11 | 116369171 | rs925151 | A | G | M_Disease | 0.514995502 | M_Disease&NULL |
| 11 | 116667337 | rs633389 | T | C | M_Disease | 0.516244281 | M_Disease&NULL |
| 11 | 116798503 | rs75176087 | T | C | M_Disease | 0.516244281 | M_Disease&NULL |
| 11 | 116808658 | rs147683288 | T | C | M_Disease | 0.516244281 | M_Disease&NULL |
| 11 | 116813938 | rs117772632 | A | T | M_Disease | 0.516244281 | M_Disease&NULL |
| 11 | 116815265 | rs61301656 | A | G | M_Disease | 0.516244281 | M_Disease&NULL |
| 11 | 116823800 | rs79143833 | C | G | M_Disease | 0.516244281 | M_Disease&NULL |
| 11 | 116831357 | rs74922395 | C | T | M_Disease | 0.516244281 | M_Disease&NULL |
| 11 | 116838219 | rs75296049 | C | T | M_Disease | 0.516244281 | M_Disease&NULL |
| 11 | 116839420 | rs78672214 | G | A | M_Disease | 0.516244281 | M_Disease&NULL |
| 11 | 116843747 | rs74655990 | C | T | M_Disease | 0.516244281 | M_Disease&NULL |
| 11 | 116853151 | rs58123174 | A | C | M_Disease | 0.516244281 | M_Disease&NULL |
| 11 | 116871192 | rs79523757 | C | T | M_Disease | 0.516244281 | M_Disease&NULL |
| 11 | 116872657 | rs76919167 | C | A | M_Disease | 0.516244281 | M_Disease&NULL |
| 11 | 116873188 | rs75216082 | C | T | M_Disease | 0.516244281 | M_Disease&NULL |
| 11 | 116873956 | rs77322057 | C | T | M_Disease | 0.516244281 | M_Disease&NULL |
| 11 | 116876136 | rs78903015 | T | G | M_Disease | 0.516244281 | M_Disease&NULL |
| 11 | 120528055 | rs77101138 | G | A | M_Disease | 0.516244281 | M_Disease&NULL |
| 11 | 117474580 | rs11216466 | A | C | M_Disease | 0.514995502 | M_Disease&NULL |
| 11 | 118466646 | rs184841209 | T | G | M_Disease | 0.514995502 | M_Disease&NULL |
| 11 | 119813015 | rs112084975 | T | C | M_Disease | 0.514995502 | M_Disease&NULL |
| 11 | 119813599 | rs77256135 | A | G | M_Disease | 0.514995502 | M_Disease&NULL |
| 11 | 120247033 | rs632137 | T | C | M_Disease | 0.514995502 | M_Disease&NULL |
| 11 | 117581459 | rs76474385 | C | T | M_Disease | 0.514995502 | M_Disease&NULL |
| 11 | 118298428 | rs78342015 | T | C | M_Disease | 0.514995502 | M_Disease&NULL |
| 11 | 120723872 | rs4598681 | T | G | M_Disease | 0.513746535 | M_Disease&NULL |
| 11 | 116398570 | rs147224199 | T | C | M_Disease | 0.514995502 | M_Disease&NULL |
| 11 | 116415864 | rs76338210 | T | C | M_Disease | 0.513746535 | M_Disease&NULL |
| 11 | 116239742 | rs567559 | C | G | M_Disease | 0.513746535 | M_Disease&NULL |
| 11 | 117503394 | rs4936393 | G | A | M_Disease | 0.513746535 | M_Disease&NULL |
| 11 | 118020130 | rs34913450 | C | T | M_Disease | 0.513746535 | M_Disease&NULL |
| 11 | 121192650 | rs964256 | G | C | M_Disease | 0.513746535 | M_Disease&NULL |
| 11 | 116300734 | rs11215928 | T | C | M_Disease | 0.513746535 | M_Disease&NULL |
| 11 | 117447735 | rs142639145 | G | A | M_Disease | 0.514995502 | M_Disease&NULL |

| | | | | | | | | | |
|---|---|---|---|---|---|---|---|---|---|
| 11 | 120854046 | rs1008966 | T | C | 1.062 | -7977.388 | -7977.23 | 0.159 | 4.53E-06 |
| 11 | 116900628 | rs139462136 | T | C | 1.137 | -7977.388 | -7977.23 | 0.158 | 4.53E-06 |
| 11 | 117925581 | rs56204159 | T | C | 0.94 | -7977.388 | -7977.23 | 0.158 | 4.53E-06 |
| 11 | 116530293 | rs530845 | T | C | 1.061 | -7977.388 | -7977.233 | 0.155 | 4.52E-06 |
| 11 | 117926172 | rs4938472 | A | T | 0.94 | -7977.388 | -7977.236 | 0.152 | 4.50E-06 |
| 11 | 116879162 | rs58354636 | G | A | 1.121 | -7977.388 | -7977.237 | 0.151 | 4.50E-06 |
| 11 | 117905635 | rs1786176 | A | G | 1.091 | -7977.388 | -7977.237 | 0.151 | 4.50E-06 |
| 11 | 116742626 | rs79849088 | T | C | 1.121 | -7977.388 | -7977.238 | 0.15 | 4.49E-06 |
| 11 | 116313601 | rs11215937 | G | A | 0.903 | -7977.388 | -7977.239 | 0.149 | 4.49E-06 |
| 11 | 117434985 | rs752795 | G | C | 0.88 | -7977.388 | -7977.243 | 0.146 | 4.48E-06 |
| 11 | 117444949 | rs76959909 | A | C | 1.092 | -7977.388 | -7977.242 | 0.146 | 4.48E-06 |
| 11 | 118931099 | rs620331 | T | C | 1.079 | -7977.388 | -7977.242 | 0.146 | 4.48E-06 |
| 11 | 116891412 | rs76666535 | A | G | 1.121 | -7977.388 | -7977.243 | 0.145 | 4.47E-06 |
| 11 | 116897871 | rs76383990 | G | A | 1.121 | -7977.388 | -7977.243 | 0.145 | 4.47E-06 |
| 11 | 116914260 | rs77822137 | C | T | 1.121 | -7977.388 | -7977.243 | 0.145 | 4.47E-06 |
| 11 | 116915242 | rs76982250 | T | C | 1.121 | -7977.388 | -7977.243 | 0.145 | 4.47E-06 |
| 11 | 116917089 | rs118121169 | A | G | 1.121 | -7977.388 | -7977.243 | 0.145 | 4.47E-06 |
| 11 | 116920199 | rs145949765 | T | C | 1.121 | -7977.388 | -7977.243 | 0.145 | 4.47E-06 |
| 11 | 116921802 | rs76916861 | T | C | 1.121 | -7977.388 | -7977.243 | 0.145 | 4.47E-06 |
| 11 | 116934380 | rs150342927 | T | C | 1.121 | -7977.388 | -7977.243 | 0.145 | 4.47E-06 |
| 11 | 116939370 | rs57072067 | G | A | 1.121 | -7977.388 | -7977.243 | 0.145 | 4.47E-06 |
| 11 | 116948368 | rs58068870 | C | T | 1.121 | -7977.388 | -7977.243 | 0.145 | 4.47E-06 |
| 11 | 117350966 | rs11823335 | T | C | 1.07 | -7977.388 | -7977.244 | 0.145 | 4.47E-06 |
| 11 | 116733310 | rs6589575 | C | T | 0.922 | -7977.388 | -7977.245 | 0.144 | 4.47E-06 |
| 11 | 116723837 | rs77462380 | T | C | 1.121 | -7977.388 | -7977.246 | 0.142 | 4.46E-06 |
| 11 | 116724233 | rs75771311 | A | G | 1.121 | -7977.388 | -7977.246 | 0.142 | 4.46E-06 |
| 11 | 116727547 | rs117910440 | C | T | 1.121 | -7977.388 | -7977.246 | 0.142 | 4.46E-06 |
| 11 | 116727871 | rs78816728 | C | T | 1.121 | -7977.388 | -7977.246 | 0.142 | 4.46E-06 |
| 11 | 116728260 | rs75385236 | T | C | 1.121 | -7977.388 | -7977.246 | 0.142 | 4.46E-06 |
| 11 | 116730980 | rs117833973 | T | C | 1.121 | -7977.388 | -7977.246 | 0.142 | 4.46E-06 |
| 11 | 117795611 | rs79669568 | C | T | 0.898 | -7977.388 | -7977.246 | 0.142 | 4.46E-06 |
| 11 | 116707684 | rs2070665 | A | G | 0.914 | -7977.388 | -7977.247 | 0.141 | 4.45E-06 |
| 11 | 117183650 | rs593245 | T | C | 0.943 | -7977.388 | -7977.249 | 0.139 | 4.44E-06 |
| 11 | 117479686 | rs4938419 | T | A | 1.068 | -7977.388 | -7977.249 | 0.139 | 4.44E-06 |
| 11 | 116517968 | rs2187125 | G | T | 1.073 | -7977.388 | -7977.251 | 0.138 | 4.44E-06 |
| 11 | 119882823 | rs142724980 | T | C | 0.878 | -7977.388 | -7977.25 | 0.138 | 4.44E-06 |
| 11 | 117444661 | rs75807933 | A | T | 1.092 | -7977.388 | -7977.252 | 0.136 | 4.43E-06 |
| 11 | 119679013 | rs56221909 | C | G | 0.909 | -7977.388 | -7977.254 | 0.134 | 4.42E-06 |
| 11 | 120067199 | rs147634995 | A | C | 0.88 | -7977.388 | -7977.254 | 0.134 | 4.42E-06 |
| 11 | 120420262 | rs11600946 | A | G | 0.905 | -7977.388 | -7977.254 | 0.134 | 4.42E-06 |
| 11 | 116904279 | rs150817479 | A | G | 1.12 | -7977.388 | -7977.256 | 0.132 | 4.41E-06 |
| 11 | 116734146 | rs718847 | C | T | 0.922 | -7977.388 | -7977.257 | 0.131 | 4.41E-06 |
| 11 | 116961819 | rs75202546 | A | G | 1.12 | -7977.388 | -7977.257 | 0.131 | 4.41E-06 |
| 11 | 116726181 | rs149462130 | T | C | 1.12 | -7977.388 | -7977.259 | 0.13 | 4.41E-06 |
| 11 | 117466661 | rs145076512 | G | T | 0.881 | -7977.388 | -7977.259 | 0.13 | 4.41E-06 |
| 11 | 117693971 | rs4938440 | A | G | 0.904 | -7977.388 | -7977.258 | 0.13 | 4.41E-06 |
| 11 | 120415158 | rs4936519 | T | G | 0.938 | -7977.388 | -7977.258 | 0.13 | 4.41E-06 |
| 11 | 121066079 | rs9943635 | T | C | 0.934 | -7977.388 | -7977.259 | 0.129 | 4.40E-06 |
| 11 | 121073219 | rs516495 | C | T | 0.933 | -7977.388 | -7977.259 | 0.129 | 4.40E-06 |
| 11 | 117474492 | rs77259443 | G | A | 0.881 | -7977.388 | -7977.26 | 0.128 | 4.40E-06 |
| 11 | 119951570 | rs10892532 | G | A | 0.937 | -7977.388 | -7977.26 | 0.128 | 4.40E-06 |
| 11 | 116275617 | rs4586206 | A | G | 1.07 | -7977.388 | -7977.263 | 0.125 | 4.38E-06 |
| 11 | 117563458 | rs145253564 | T | C | 0.877 | -7977.388 | -7977.264 | 0.125 | 4.38E-06 |
| 11 | 117564084 | rs139289629 | C | G | 0.877 | -7977.388 | -7977.264 | 0.125 | 4.38E-06 |
| 11 | 116317789 | rs56287831 | A | G | 0.916 | -7977.388 | -7977.264 | 0.124 | 4.38E-06 |
| 11 | 118984287 | rs4938630 | A | C | 0.909 | -7977.388 | -7977.265 | 0.124 | 4.38E-06 |
| 11 | 121015228 | rs11218160 | A | G | 0.934 | -7977.388 | -7977.265 | 0.124 | 4.38E-06 |
| 11 | 118973113 | rs3825059 | A | C | 0.896 | -7977.388 | -7977.265 | 0.123 | 4.37E-06 |
| 11 | 119679613 | rs10790347 | T | A | 0.941 | -7977.388 | -7977.266 | 0.123 | 4.37E-06 |
| 11 | 118240332 | rs11216860 | C | T | 1.067 | -7977.388 | -7977.267 | 0.122 | 4.37E-06 |
| 11 | 120412053 | rs78566287 | T | C | 0.9 | -7977.388 | -7977.269 | 0.12 | 4.36E-06 |
| 11 | 121071633 | rs201644901 | C | A | 0.933 | -7977.388 | -7977.27 | 0.119 | 4.36E-06 |
| 11 | 116369171 | rs925151 | A | G | 0.942 | -7977.388 | -7977.271 | 0.117 | 4.35E-06 |
| 11 | 116667337 | rs633389 | T | C | 0.885 | -7977.388 | -7977.272 | 0.117 | 4.35E-06 |
| 11 | 116798503 | rs75176087 | T | C | 1.119 | -7977.388 | -7977.271 | 0.117 | 4.35E-06 |
| 11 | 116808658 | rs147683288 | T | C | 1.119 | -7977.388 | -7977.271 | 0.117 | 4.35E-06 |
| 11 | 116813938 | rs117772632 | A | T | 1.119 | -7977.388 | -7977.271 | 0.117 | 4.35E-06 |
| 11 | 116815265 | rs61301656 | A | G | 1.119 | -7977.388 | -7977.271 | 0.117 | 4.35E-06 |
| 11 | 116823800 | rs79143833 | C | G | 1.119 | -7977.388 | -7977.271 | 0.117 | 4.35E-06 |
| 11 | 116831357 | rs74922395 | C | T | 1.119 | -7977.388 | -7977.271 | 0.117 | 4.35E-06 |
| 11 | 116838219 | rs75296049 | C | T | 1.119 | -7977.388 | -7977.271 | 0.117 | 4.35E-06 |
| 11 | 116839420 | rs78672214 | G | A | 1.119 | -7977.388 | -7977.271 | 0.117 | 4.35E-06 |
| 11 | 116843747 | rs74655990 | C | T | 1.119 | -7977.388 | -7977.271 | 0.117 | 4.35E-06 |
| 11 | 116853151 | rs58123174 | A | C | 1.119 | -7977.388 | -7977.271 | 0.117 | 4.35E-06 |
| 11 | 116871192 | rs79523757 | C | T | 1.119 | -7977.388 | -7977.271 | 0.117 | 4.35E-06 |
| 11 | 116872657 | rs76919167 | C | A | 1.119 | -7977.388 | -7977.271 | 0.117 | 4.35E-06 |
| 11 | 116873188 | rs75216082 | C | T | 1.119 | -7977.388 | -7977.271 | 0.117 | 4.35E-06 |
| 11 | 116873956 | rs77322057 | C | T | 1.119 | -7977.388 | -7977.271 | 0.117 | 4.35E-06 |
| 11 | 116876136 | rs78903015 | T | G | 1.119 | -7977.388 | -7977.271 | 0.117 | 4.35E-06 |
| 11 | 120528055 | rs77101138 | G | A | 1.124 | -7977.388 | -7977.271 | 0.117 | 4.35E-06 |
| 11 | 117474580 | rs11216466 | A | C | 0.906 | -7977.388 | -7977.273 | 0.116 | 4.34E-06 |
| 11 | 118466646 | rs184841209 | T | G | 0.904 | -7977.388 | -7977.275 | 0.114 | 4.34E-06 |
| 11 | 119813015 | rs112084975 | T | C | 1.103 | -7977.388 | -7977.274 | 0.114 | 4.34E-06 |
| 11 | 119813599 | rs77256135 | A | G | 1.103 | -7977.388 | -7977.274 | 0.114 | 4.34E-06 |
| 11 | 120247033 | rs632137 | T | C | 0.921 | -7977.388 | -7977.275 | 0.114 | 4.34E-06 |
| 11 | 117581459 | rs76474385 | C | T | 0.878 | -7977.388 | -7977.277 | 0.112 | 4.33E-06 |
| 11 | 118298428 | rs78342015 | T | C | 0.895 | -7977.388 | -7977.277 | 0.112 | 4.33E-06 |
| 11 | 120723872 | rs4598681 | T | G | 1.076 | -7977.388 | -7977.277 | 0.112 | 4.33E-06 |
| 11 | 116398570 | rs147224199 | T | C | 1.12 | -7977.388 | -7977.277 | 0.111 | 4.32E-06 |
| 11 | 116415864 | rs76338210 | T | C | 1.077 | -7977.388 | -7977.277 | 0.111 | 4.32E-06 |
| 11 | 116239742 | rs567559 | C | G | 1.065 | -7977.388 | -7977.279 | 0.109 | 4.31E-06 |
| 11 | 117503394 | rs4936393 | G | A | 0.942 | -7977.388 | -7977.279 | 0.109 | 4.31E-06 |
| 11 | 118020130 | rs34913450 | C | T | 0.903 | -7977.388 | -7977.282 | 0.106 | 4.30E-06 |
| 11 | 121192650 | rs964256 | G | C | 1.084 | -7977.388 | -7977.282 | 0.106 | 4.30E-06 |
| 11 | 116300734 | rs11215928 | T | C | 1.082 | -7977.388 | -7977.283 | 0.105 | 4.30E-06 |
| 11 | 117447735 | rs142639145 | G | A | 0.883 | -7977.388 | -7977.284 | 0.105 | 4.30E-06 |

| | | | | | | | |
|---|---|---|---|---|---|---|---|
| 11 | 117447871 | rs116997303 | C | T | M_Disease | 0.514995502 | M_Disease&NULL |
| 11 | 121076938 | rs614634 | T | C | M_Disease | 0.513746535 | M_Disease&NULL |
| 11 | 117565794 | rs138159885 | A | G | M_Disease | 0.513746535 | M_Disease&NULL |
| 11 | 117565876 | rs17587869 | T | G | M_Disease | 0.513746535 | M_Disease&NULL |
| 11 | 117566372 | rs17664522 | G | T | M_Disease | 0.513746535 | M_Disease&NULL |
| 11 | 117567237 | rs150057425 | G | A | M_Disease | 0.513746535 | M_Disease&NULL |
| 11 | 119812958 | rs112736310 | T | C | M_Disease | 0.513746535 | M_Disease&NULL |
| 11 | 120425668 | rs75352872 | T | G | M_Disease | 0.513746535 | M_Disease&NULL |
| 11 | 116985857 | rs60834024 | A | C | M_Disease | 0.512497396 | M_Disease&NULL |
| 11 | 120309460 | rs71484168 | T | C | M_Disease | 0.512497396 | M_Disease&NULL |
| 11 | 116914662 | rs77953286 | C | G | M_Disease | 0.512497396 | M_Disease&NULL |
| 11 | 119566235 | rs118029883 | A | G | M_Disease | 0.513746535 | M_Disease&NULL |
| 11 | 117463436 | rs4938412 | T | G | M_Disease | 0.513746535 | M_Disease&NULL |
| 11 | 117466663 | rs76630213 | G | T | M_Disease | 0.513746535 | M_Disease&NULL |
| 11 | 121090162 | rs619998 | C | T | M_Disease | 0.512497396 | M_Disease&NULL |
| 11 | 119720735 | rs73008657 | A | G | M_Disease | 0.512497396 | M_Disease&NULL |
| 11 | 119811734 | rs12269853 | G | C | M_Disease | 0.512497396 | M_Disease&NULL |
| 11 | 121073887 | rs656189 | A | T | M_Disease | 0.511248102 | M_Disease&NULL |
| 11 | 116761815 | rs146418192 | C | G | M_Disease | 0.512497396 | M_Disease&NULL |
| 11 | 116409788 | rs951839 | A | C | M_Disease | 0.511248102 | M_Disease&NULL |
| 11 | 116343033 | rs67215893 | A | G | M_Disease | 0.511248102 | M_Disease&NULL |
| 11 | 116667480 | rs633867 | T | C | M_Disease | 0.512497396 | M_Disease&NULL |
| 11 | 120589674 | rs2846107 | A | G | M_Disease | 0.511248102 | M_Disease&NULL |
| 11 | 117399649 | rs7126416 | A | C | M_Disease | 0.511248102 | M_Disease&NULL |
| 11 | 117429793 | rs112793334 | A | C | M_Disease | 0.511248102 | M_Disease&NULL |
| 11 | 117564067 | rs150824999 | T | C | M_Disease | 0.511248102 | M_Disease&NULL |
| 11 | 118602497 | rs478081 | G | A | M_Disease | 0.509998667 | M_Disease&NULL |
| 11 | 116951926 | rs80068626 | C | T | M_Disease | 0.511248102 | M_Disease&NULL |
| 11 | 116952296 | rs141474528 | G | C | M_Disease | 0.511248102 | M_Disease&NULL |
| 11 | 116993788 | rs12420361 | A | G | M_Disease | 0.511248102 | M_Disease&NULL |
| 11 | 117444884 | rs4938411 | A | G | M_Disease | 0.511248102 | M_Disease&NULL |
| 11 | 119548402 | rs78809001 | T | C | M_Disease | 0.511248102 | M_Disease&NULL |
| 11 | 120235874 | rs661139 | A | T | M_Disease | 0.509998667 | M_Disease&NULL |
| 11 | 117302538 | rs147189349 | A | G | M_Disease | 0.509998667 | M_Disease&NULL |
| 11 | 117429034 | rs479591 | G | A | M_Disease | 0.509998667 | M_Disease&NULL |
| 11 | 116957158 | rs142836613 | G | A | M_Disease | 0.509998667 | M_Disease&NULL |
| 11 | 117336961 | rs12418126 | G | A | M_Disease | 0.509998667 | M_Disease&NULL |
| 11 | 116759655 | rs17174502 | A | G | M_Disease | 0.509998667 | M_Disease&NULL |
| 11 | 116788837 | rs1135663 | T | C | M_Disease | 0.509998667 | M_Disease&NULL |
| 11 | 119556519 | rs75967483 | T | C | M_Disease | 0.509998667 | M_Disease&NULL |
| 11 | 120697099 | rs11218017 | G | A | M_Disease | 0.508749107 | M_Disease&NULL |
| 11 | 117536090 | rs687100 | T | C | M_Disease | 0.508749107 | M_Disease&NULL |
| 11 | 117793222 | rs112048745 | T | C | M_Disease | 0.508749107 | M_Disease&NULL |
| 11 | 117443368 | rs78012672 | A | G | M_Disease | 0.509998667 | M_Disease&NULL |
| 11 | 120795955 | rs117508216 | C | A | M_Disease | 0.509998667 | M_Disease&NULL |
| 11 | 117574923 | rs58499675 | C | T | M_Disease | 0.509998667 | M_Disease&NULL |
| 11 | 117158168 | rs45497100 | T | A | M_Disease | 0.508749107 | M_Disease&NULL |
| 11 | 117988082 | rs642215 | T | C | M_Disease | 0.508749107 | M_Disease&NULL |
| 11 | 116731744 | rs75501351 | G | T | M_Disease | 0.508749107 | M_Disease&NULL |
| 11 | 117443936 | rs17120893 | T | C | M_Disease | 0.508749107 | M_Disease&NULL |
| 11 | 117616982 | rs1443827410 | G | A | M_Disease | 0.508749107 | M_Disease&NULL |
| 11 | 117909810 | rs80170872 | A | G | M_Disease | 0.508749107 | M_Disease&NULL |
| 11 | 117477245 | rs79656669 | C | T | M_Disease | 0.508749107 | M_Disease&NULL |
| 11 | 117470795 | rs80138815 | G | A | M_Disease | 0.507499438 | M_Disease&NULL |
| 11 | 118492713 | rs45457998 | G | A | M_Disease | 0.507499438 | M_Disease&NULL |
| 11 | 118810405 | rs4938589 | T | C | M_Disease | 0.507499438 | M_Disease&NULL |
| 11 | 119811767 | rs78927965 | A | C | M_Disease | 0.507499438 | M_Disease&NULL |
| 11 | 117441703 | rs10502226 | G | T | M_Disease | 0.507499438 | M_Disease&NULL |
| 11 | 117466540 | rs1452979329 | A | G | M_Disease | 0.506249674 | M_Disease&NULL |
| 11 | 118602058 | rs562059 | G | T | M_Disease | 0.506249674 | M_Disease&NULL |
| 11 | 116937127 | rs142661919 | G | A | M_Disease | 0.507499438 | M_Disease&NULL |
| 11 | 116937146 | rs117518542 | A | G | M_Disease | 0.507499438 | M_Disease&NULL |
| 11 | 116749190 | rs150818481 | A | G | M_Disease | 0.507499438 | M_Disease&NULL |
| 11 | 117711092 | rs1321012057 | A | G | M_Disease | 0.506249674 | M_Disease&NULL |
| 11 | 120253329 | rs34189224 | A | G | M_Disease | 0.506249674 | M_Disease&NULL |
| 11 | 119529365 | rs1467050 | T | C | M_Disease | 0.506249674 | M_Disease&NULL |
| 11 | 117600086 | rs147019067 | A | C | M_Disease | 0.507499438 | M_Disease&NULL |
| 11 | 117945171 | rs3758992 | C | G | M_Disease | 0.506249674 | M_Disease&NULL |
| 11 | 116534026 | rs151215655 | T | C | M_Disease | 0.506249674 | M_Disease&NULL |
| 11 | 116992685 | rs74955808 | A | G | M_Disease | 0.506249674 | M_Disease&NULL |
| 11 | 116979363 | rs12416941 | A | G | M_Disease | 0.504999833 | M_Disease&NULL |
| 11 | 116707583 | rs5072 | A | G | M_Disease | 0.504999833 | M_Disease&NULL |
| 11 | 116996711 | rs12419459 | T | C | M_Disease | 0.504999833 | M_Disease&NULL |
| 11 | 116980021 | rs12271169 | G | C | M_Disease | 0.504999833 | M_Disease&NULL |
| 11 | 120718086 | rs11218029 | A | T | M_Disease | 0.504999833 | M_Disease&NULL |
| 11 | 117988223 | rs643014 | G | A | M_Disease | 0.504999833 | M_Disease&NULL |
| 11 | 116269990 | rs2226637 | C | G | M_Disease | 0.50374993 | M_Disease&NULL |
| 11 | 118973112 | rs3825060 | A | G | M_Disease | 0.504999833 | M_Disease&NULL |
| 11 | 117440220 | rs150854359 | G | T | M_Disease | 0.504999833 | M_Disease&NULL |
| 11 | 118121356 | rs147710683 | T | C | M_Disease | 0.504999833 | M_Disease&NULL |
| 11 | 117185634 | rs2510446 | C | T | M_Disease | 0.50374993 | M_Disease&NULL |
| 11 | 117352086 | rs12363073 | T | C | M_Disease | 0.504999833 | M_Disease&NULL |
| 11 | 120588562 | rs2852225 | T | G | M_Disease | 0.50374993 | M_Disease&NULL |
| 11 | 116985191 | rs74882651 | A | G | M_Disease | 0.50374993 | M_Disease&NULL |
| 11 | 117458098 | rs190567289 | T | C | M_Disease | 0.504999833 | M_Disease&NULL |
| 11 | 116356422 | rs73009144 | A | G | M_Disease | 0.50374993 | M_Disease&NULL |
| 11 | 117275887 | rs7123784 | C | G | M_Disease | 0.50374993 | M_Disease&NULL |
| 11 | 119382089 | rs79235532 | A | G | M_Disease | 0.50374993 | M_Disease&NULL |
| 11 | 116759706 | rs117920151 | C | G | M_Disease | 0.50374993 | M_Disease&NULL |
| 11 | 116307648 | rs76200167 | A | C | M_Disease | 0.50374993 | M_Disease&NULL |
| 11 | 117605620 | rs117399581 | A | G | M_Disease | 0.50374993 | M_Disease&NULL |
| 11 | 117569803 | rs147776682 | T | C | M_Disease | 0.50374993 | M_Disease&NULL |
| 11 | 117570297 | rs145111226 | T | C | M_Disease | 0.50374993 | M_Disease&NULL |
| 11 | 117570660 | rs149084104 | C | G | M_Disease | 0.50374993 | M_Disease&NULL |
| 11 | 116982434 | rs111677878 | T | C | M_Disease | 0.502499979 | M_Disease&NULL |
| 11 | 117464630 | rs77230637 | A | G | M_Disease | 0.502499979 | M_Disease&NULL |

| | | | | | | | | | |
|---|---|---|---|---|---|---|---|---|---|
| 11 | 117447871 | rs116997303 | C | T | 0.883 | -7977.388 | -7977.284 | 0.105 | 4.30E-06 |
| 11 | 121076938 | rs614634 | T | C | 0.934 | -7977.388 | -7977.284 | 0.104 | 4.29E-06 |
| 11 | 117565794 | rs138159885 | A | G | 0.878 | -7977.388 | -7977.286 | 0.103 | 4.29E-06 |
| 11 | 117565876 | rs17587869 | T | G | 0.878 | -7977.388 | -7977.286 | 0.103 | 4.29E-06 |
| 11 | 117566372 | rs17664522 | G | T | 0.878 | -7977.388 | -7977.286 | 0.103 | 4.29E-06 |
| 11 | 117567237 | rs150057425 | G | A | 0.878 | -7977.388 | -7977.286 | 0.103 | 4.29E-06 |
| 11 | 119812958 | rs112736310 | T | C | 1.102 | -7977.388 | -7977.286 | 0.102 | 4.28E-06 |
| 11 | 120425668 | rs75352872 | T | G | 1.129 | -7977.388 | -7977.287 | 0.102 | 4.28E-06 |
| 11 | 116985857 | rs60834024 | A | C | 1.102 | -7977.388 | -7977.289 | 0.099 | 4.27E-06 |
| 11 | 120309460 | rs71484168 | T | C | 0.904 | -7977.388 | -7977.29 | 0.098 | 4.27E-06 |
| 11 | 116914662 | rs77953286 | C | G | 1.119 | -7977.388 | -7977.292 | 0.096 | 4.26E-06 |
| 11 | 119566235 | rs118029883 | A | G | 0.88 | -7977.388 | -7977.292 | 0.096 | 4.26E-06 |
| 11 | 117463436 | rs4938412 | T | G | 0.883 | -7977.388 | -7977.293 | 0.095 | 4.25E-06 |
| 11 | 117466663 | rs76630213 | G | T | 0.883 | -7977.388 | -7977.293 | 0.095 | 4.25E-06 |
| 11 | 121090162 | rs619998 | C | T | 0.934 | -7977.388 | -7977.293 | 0.095 | 4.25E-06 |
| 11 | 119720735 | rs73008657 | A | G | 1.082 | -7977.388 | -7977.295 | 0.093 | 4.25E-06 |
| 11 | 119811734 | rs12269853 | G | C | 1.102 | -7977.388 | -7977.295 | 0.093 | 4.25E-06 |
| 11 | 121073887 | rs656189 | A | T | 0.934 | -7977.388 | -7977.298 | 0.091 | 4.24E-06 |
| 11 | 116761815 | rs146418192 | C | G | 1.134 | -7977.388 | -7977.299 | 0.089 | 4.23E-06 |
| 11 | 116409788 | rs951839 | A | C | 1.076 | -7977.388 | -7977.301 | 0.088 | 4.22E-06 |
| 11 | 116343033 | rs67215893 | A | G | 0.922 | -7977.388 | -7977.302 | 0.087 | 4.22E-06 |
| 11 | 116667480 | rs633867 | T | C | 0.887 | -7977.388 | -7977.301 | 0.087 | 4.22E-06 |
| 11 | 120589674 | rs2846107 | A | G | 0.921 | -7977.388 | -7977.301 | 0.087 | 4.22E-06 |
| 11 | 117399649 | rs7126416 | A | C | 1.065 | -7977.388 | -7977.302 | 0.086 | 4.22E-06 |
| 11 | 117429793 | rs112793334 | A | C | 0.885 | -7977.388 | -7977.304 | 0.084 | 4.21E-06 |
| 11 | 117564067 | rs150824999 | T | C | 0.88 | -7977.388 | -7977.306 | 0.083 | 4.20E-06 |
| 11 | 118602497 | rs478081 | G | A | 1.062 | -7977.388 | -7977.307 | 0.082 | 4.20E-06 |
| 11 | 116951926 | rs80068626 | C | T | 1.117 | -7977.388 | -7977.308 | 0.081 | 4.19E-06 |
| 11 | 116952296 | rs141474528 | G | C | 1.117 | -7977.388 | -7977.308 | 0.081 | 4.19E-06 |
| 11 | 116993788 | rs12420361 | A | G | 1.101 | -7977.388 | -7977.308 | 0.081 | 4.19E-06 |
| 11 | 117444884 | rs4938411 | A | G | 0.884 | -7977.388 | -7977.308 | 0.08 | 4.19E-06 |
| 11 | 119548402 | rs78809001 | T | C | 0.882 | -7977.388 | -7977.313 | 0.076 | 4.17E-06 |
| 11 | 120235874 | rs661139 | A | T | 0.922 | -7977.388 | -7977.313 | 0.075 | 4.17E-06 |
| 11 | 117302538 | rs147189349 | A | G | 0.884 | -7977.388 | -7977.314 | 0.074 | 4.17E-06 |
| 11 | 117429034 | rs479591 | G | A | 0.944 | -7977.388 | -7977.314 | 0.074 | 4.17E-06 |
| 11 | 116957158 | rs142836613 | G | A | 1.117 | -7977.388 | -7977.315 | 0.073 | 4.16E-06 |
| 11 | 117336961 | rs12418126 | G | A | 0.911 | -7977.388 | -7977.316 | 0.072 | 4.16E-06 |
| 11 | 116759655 | rs17174502 | A | G | 0.921 | -7977.388 | -7977.317 | 0.071 | 4.15E-06 |
| 11 | 116788837 | rs1135663 | T | C | 1.099 | -7977.388 | -7977.317 | 0.071 | 4.15E-06 |
| 11 | 119556519 | rs75967483 | T | C | 1.114 | -7977.388 | -7977.32 | 0.069 | 4.14E-06 |
| 11 | 120697099 | rs11218017 | G | A | 0.931 | -7977.388 | -7977.319 | 0.069 | 4.14E-06 |
| 11 | 117536090 | rs687100 | T | C | 0.944 | -7977.388 | -7977.32 | 0.068 | 4.14E-06 |
| 11 | 117793222 | rs112048745 | T | C | 0.901 | -7977.388 | -7977.32 | 0.068 | 4.14E-06 |
| 11 | 117443368 | rs78012672 | A | G | 0.885 | -7977.388 | -7977.321 | 0.067 | 4.14E-06 |
| 11 | 120795955 | rs117508216 | C | A | 0.884 | -7977.388 | -7977.324 | 0.065 | 4.13E-06 |
| 11 | 117574923 | rs58499675 | C | T | 0.882 | -7977.388 | -7977.324 | 0.064 | 4.12E-06 |
| 11 | 117158168 | rs45497100 | T | A | 0.893 | -7977.388 | -7977.328 | 0.061 | 4.11E-06 |
| 11 | 117988082 | rs642215 | T | C | 1.09 | -7977.388 | -7977.328 | 0.061 | 4.11E-06 |
| 11 | 116731744 | rs75501351 | G | T | 1.116 | -7977.388 | -7977.329 | 0.06 | 4.11E-06 |
| 11 | 117443936 | rs17120893 | T | C | 0.886 | -7977.388 | -7977.329 | 0.059 | 4.10E-06 |
| 11 | 117616982 | rs1443827410 | G | A | 1.114 | -7977.388 | -7977.329 | 0.059 | 4.10E-06 |
| 11 | 117909810 | rs80170872 | A | G | 0.891 | -7977.388 | -7977.331 | 0.057 | 4.10E-06 |
| 11 | 117477245 | rs79656669 | C | T | 0.886 | -7977.388 | -7977.332 | 0.056 | 4.09E-06 |
| 11 | 117470795 | rs80138815 | G | A | 0.886 | -7977.388 | -7977.334 | 0.054 | 4.08E-06 |
| 11 | 118492713 | rs45457998 | G | A | 0.901 | -7977.388 | -7977.336 | 0.052 | 4.07E-06 |
| 11 | 118810405 | rs4938589 | T | C | 1.083 | -7977.388 | -7977.336 | 0.052 | 4.07E-06 |
| 11 | 119811767 | rs78927965 | A | C | 1.1 | -7977.388 | -7977.337 | 0.051 | 4.07E-06 |
| 11 | 117441703 | rs10502226 | G | T | 0.886 | -7977.388 | -7977.338 | 0.05 | 4.07E-06 |
| 11 | 117466540 | rs1452979329 | A | G | 1.089 | -7977.388 | -7977.338 | 0.05 | 4.07E-06 |
| 11 | 118602058 | rs562059 | G | T | 1.061 | -7977.388 | -7977.338 | 0.05 | 4.07E-06 |
| 11 | 116937127 | rs142661919 | G | A | 1.115 | -7977.388 | -7977.34 | 0.048 | 4.06E-06 |
| 11 | 116937146 | rs117518542 | A | G | 1.115 | -7977.388 | -7977.34 | 0.048 | 4.06E-06 |
| 11 | 116749190 | rs150818481 | A | G | 1.115 | -7977.388 | -7977.341 | 0.047 | 4.05E-06 |
| 11 | 117711092 | rs1321012057 | A | G | 1.1 | -7977.388 | -7977.341 | 0.047 | 4.05E-06 |
| 11 | 120253329 | rs34189224 | A | G | 0.901 | -7977.388 | -7977.341 | 0.047 | 4.05E-06 |
| 11 | 119529365 | rs1467050 | T | C | 0.914 | -7977.388 | -7977.343 | 0.045 | 4.05E-06 |
| 11 | 117600086 | rs147019067 | A | C | 0.884 | -7977.388 | -7977.344 | 0.044 | 4.04E-06 |
| 11 | 117945171 | rs3758992 | C | G | 0.943 | -7977.388 | -7977.345 | 0.043 | 4.04E-06 |
| 11 | 116534026 | rs151215655 | T | C | 1.132 | -7977.388 | -7977.348 | 0.04 | 4.03E-06 |
| 11 | 116992685 | rs74955808 | A | G | 1.1 | -7977.388 | -7977.348 | 0.04 | 4.03E-06 |
| 11 | 116979363 | rs12416941 | A | G | 1.099 | -7977.388 | -7977.35 | 0.038 | 4.02E-06 |
| 11 | 116707583 | rs5072 | A | G | 0.917 | -7977.388 | -7977.351 | 0.037 | 4.01E-06 |
| 11 | 116996711 | rs12419459 | T | C | 1.099 | -7977.388 | -7977.352 | 0.036 | 4.01E-06 |
| 11 | 116980021 | rs12271169 | G | C | 1.068 | -7977.388 | -7977.354 | 0.034 | 4.00E-06 |
| 11 | 120718086 | rs11218029 | A | T | 1.083 | -7977.388 | -7977.355 | 0.033 | 4.00E-06 |
| 11 | 117988223 | rs643014 | G | A | 1.089 | -7977.388 | -7977.357 | 0.032 | 3.99E-06 |
| 11 | 116269990 | rs2226637 | C | G | 1.073 | -7977.388 | -7977.357 | 0.031 | 3.99E-06 |
| 11 | 118973112 | rs3825060 | A | G | 0.9 | -7977.388 | -7977.359 | 0.029 | 3.98E-06 |
| 11 | 117440220 | rs150854359 | G | T | 0.888 | -7977.388 | -7977.36 | 0.028 | 3.98E-06 |
| 11 | 118121356 | rs147710683 | T | C | 0.894 | -7977.388 | -7977.36 | 0.028 | 3.98E-06 |
| 11 | 117185634 | rs2510446 | C | T | 1.061 | -7977.388 | -7977.361 | 0.027 | 3.97E-06 |
| 11 | 117352086 | rs12363073 | T | C | 0.89 | -7977.388 | -7977.362 | 0.027 | 3.97E-06 |
| 11 | 120588562 | rs2852225 | T | G | 0.932 | -7977.388 | -7977.362 | 0.027 | 3.97E-06 |
| 11 | 116985191 | rs74882651 | A | G | 1.099 | -7977.388 | -7977.362 | 0.026 | 3.97E-06 |
| 11 | 117458098 | rs190567289 | T | C | 0.887 | -7977.388 | -7977.364 | 0.025 | 3.97E-06 |
| 11 | 116356422 | rs73009144 | A | G | 0.896 | -7977.388 | -7977.364 | 0.024 | 3.96E-06 |
| 11 | 117275887 | rs7123784 | C | G | 0.917 | -7977.388 | -7977.364 | 0.024 | 3.96E-06 |
| 11 | 119382089 | rs79235532 | A | G | 0.887 | -7977.388 | -7977.365 | 0.024 | 3.96E-06 |
| 11 | 116759706 | rs117920151 | C | G | 0.894 | -7977.388 | -7977.368 | 0.021 | 3.95E-06 |
| 11 | 116307648 | rs76200167 | A | C | 0.896 | -7977.388 | -7977.368 | 0.02 | 3.95E-06 |
| 11 | 117605620 | rs117399581 | A | G | 1.112 | -7977.388 | -7977.37 | 0.018 | 3.94E-06 |
| 11 | 117569803 | rs147776682 | T | C | 0.885 | -7977.388 | -7977.372 | 0.017 | 3.93E-06 |
| 11 | 117570297 | rs145111226 | T | C | 0.885 | -7977.388 | -7977.372 | 0.017 | 3.93E-06 |
| 11 | 117570660 | rs149084104 | C | G | 0.885 | -7977.388 | -7977.372 | 0.017 | 3.93E-06 |
| 11 | 116982434 | rs111677878 | T | C | 1.098 | -7977.388 | -7977.374 | 0.014 | 3.92E-06 |
| 11 | 117464630 | rs77230637 | A | G | 1.088 | -7977.388 | -7977.374 | 0.014 | 3.92E-06 |

| | | | | | | | |
|---|---|---|---|---|---|---|---|
| 11 | 117477867 | rs1939235 | T | C | M_Disease | 0.502499979 | M_Disease&NULL |
| 11 | 117560438 | rs75707401 | A | G | M_Disease | 0.502499979 | M_Disease&NULL |
| 11 | 121077211 | rs509058 | T | C | M_Disease | 0.502499979 | M_Disease&NULL |
| 11 | 119807902 | rs75012772 | A | G | M_Disease | 0.502499979 | M_Disease&NULL |
| 11 | 120715471 | rs74702343 | A | G | M_Disease | 0.502499979 | M_Disease&NULL |
| 11 | 116860781 | rs75292554 | C | T | M_Disease | 0.502499979 | M_Disease&NULL |
| 11 | 116866944 | rs143367870 | C | A | M_Disease | 0.502499979 | M_Disease&NULL |
| 11 | 119556950 | rs61901099 | T | C | M_Disease | 0.502499979 | M_Disease&NULL |
| 11 | 119802424 | rs754872 | T | C | M_Disease | 0.502499979 | M_Disease&NULL |
| 11 | 119510366 | rs4438048 | T | C | M_Disease | 0.501249997 | M_Disease&NULL |
| 11 | 119984598 | rs76031864 | A | G | M_Disease | 0.501249997 | M_Disease&NULL |
| 11 | 118749318 | rs78312641 | A | T | M_Disease | 0.501249997 | M_Disease&NULL |
| 11 | 116409433 | rs17491510 | T | C | M_Disease | 0.501249997 | M_Disease&NULL |
| 11 | 116864639 | rs76665318 | C | T | M_Disease | 0.501249997 | M_Disease&NULL |
| 11 | 119813686 | rs76661054 | C | T | M_Disease | 0.501249997 | M_Disease&NULL |
| 11 | 119813733 | rs75224487 | T | C | M_Disease | 0.501249997 | M_Disease&NULL |
| 11 | 119814337 | rs111902338 | A | C | M_Disease | 0.501249997 | M_Disease&NULL |
| 11 | 119512641 | rs2089647 | G | A | M_Disease | 0.501249997 | M_Disease&NULL |
| 11 | 120413702 | rs75510322 | C | A | M_Disease | 0.501249997 | M_Disease&NULL |
| 11 | 116294664 | rs471750 | G | A | M_Disease | 0.501249997 | M_Disease&NULL |
| 11 | 117338330 | rs495171 | C | T | M_Disease | 0.501249997 | M_Disease&NULL |
| 11 | 119934934 | rs4938763 | T | C | NULL | 0.5 | NULL&M_Disease |
| 11 | 116980679 | rs114221654 | T | C | NULL | 0.5 | NULL&M_Disease |
| 11 | 116981515 | rs116770907 | T | C | NULL | 0.5 | NULL&M_Disease |
| 11 | 116539058 | rs112464350 | A | C | NULL | 0.5 | NULL&M_Disease |
| 11 | 117414714 | rs3802879 | T | G | NULL | 0.5 | NULL&M_Disease |
| 11 | 117587224 | rs56153022 | G | C | M_Disease | 0.501249997 | M_Disease&NULL |
| 11 | 118807637 | rs112974653 | T | G | NULL | 0.5 | NULL&M_Disease |
| 11 | 119807543 | rs79710878 | G | A | NULL | 0.5 | NULL&M_Disease |
| 11 | 119808101 | rs77763933 | C | G | NULL | 0.5 | NULL&M_Disease |
| 11 | 116996631 | rs12419454 | T | C | NULL | 0.5 | NULL&M_Disease |
| 11 | 116996783 | rs12416691 | T | A | NULL | 0.5 | NULL&M_Disease |
| 11 | 117391421 | rs77964702 | A | G | NULL | 0.5 | NULL&M_Disease |
| 11 | 116953338 | rs79552834 | C | G | NULL | 0.5 | NULL&M_Disease |
| 11 | 119675931 | rs7926586 | G | T | NULL | 0.501249997 | NULL&M_Disease |
| 11 | 116692552 | rs5103 | G | A | NULL | 0.5 | NULL&M_Disease |
| 11 | 116996345 | rs17120344 | A | G | NULL | 0.5 | NULL&M_Disease |
| 11 | 119510644 | rs7940667 | A | C | NULL | 0.501249997 | NULL&M_Disease |
| 11 | 119512115 | rs2089646 | G | A | NULL | 0.501249997 | NULL&M_Disease |
| 11 | 116530797 | rs558056 | T | G | NULL | 0.501249997 | NULL&M_Disease |
| 11 | 119510303 | rs4245199 | G | T | NULL | 0.501249997 | NULL&M_Disease |
| 11 | 117414030 | rs563531 | A | G | NULL | 0.501249997 | NULL&M_Disease |
| 11 | 119678445 | rs7945067 | T | C | NULL | 0.501249997 | NULL&M_Disease |
| 11 | 120176979 | rs10750171 | G | T | NULL | 0.501249997 | NULL&M_Disease |
| 11 | 119806581 | rs77443589 | A | G | NULL | 0.501249997 | NULL&M_Disease |
| 11 | 119806699 | rs79137748 | C | T | NULL | 0.501249997 | NULL&M_Disease |
| 11 | 116481694 | rs73578509 | G | C | NULL | 0.5 | NULL&M_Disease |
| 11 | 116317745 | rs569327 | T | C | NULL | 0.501249997 | NULL&M_Disease |
| 11 | 116207020 | rs7947688 | G | A | NULL | 0.501249997 | NULL&M_Disease |
| 11 | 121152465 | rs11607048 | G | T | NULL | 0.501249997 | NULL&M_Disease |
| 11 | 116296243 | rs693603 | A | G | NULL | 0.502499979 | NULL&M_Disease |
| 11 | 116237696 | rs79733808 | A | G | NULL | 0.502499979 | NULL&M_Disease |
| 11 | 116750073 | rs543819 | G | A | NULL | 0.502499979 | NULL&M_Disease |
| 11 | 117795578 | rs77642243 | C | T | NULL | 0.502499979 | NULL&M_Disease |
| 11 | 119882908 | rs587806 | T | C | NULL | 0.502499979 | NULL&M_Disease |
| 11 | 117931489 | rs7112429 | A | T | NULL | 0.501249997 | NULL&M_Disease |
| 11 | 117937225 | rs60007061 | A | C | NULL | 0.501249997 | NULL&M_Disease |
| 11 | 121069225 | rs12288505 | T | C | NULL | 0.502499979 | NULL&M_Disease |
| 11 | 117899816 | rs617345 | G | T | NULL | 0.502499979 | NULL&M_Disease |
| 11 | 117521603 | rs678327 | A | G | NULL | 0.502499979 | NULL&M_Disease |
| 11 | 118560857 | rs73001406 | A | G | NULL | 0.502499979 | NULL&M_Disease |
| 11 | 120858410 | rs2282586 | A | G | NULL | 0.502499979 | NULL&M_Disease |
| 11 | 120503923 | rs4936534 | T | A | NULL | 0.50374993 | NULL&M_Disease |
| 11 | 117492181 | rs10750108 | A | G | NULL | 0.50374993 | NULL&M_Disease |
| 11 | 118809329 | rs10892313 | A | G | NULL | 0.50374993 | NULL&M_Disease |
| 11 | 116700777 | rs618354 | C | G | NULL | 0.50374993 | NULL&M_Disease |
| 11 | 116752191 | rs73023056 | A | G | NULL | 0.502499979 | NULL&M_Disease |
| 11 | 116534963 | rs12802672 | G | A | NULL | 0.50374993 | NULL&M_Disease |
| 11 | 116539809 | rs112550319 | G | C | NULL | 0.50374993 | NULL&M_Disease |
| 11 | 116448277 | rs73011154 | C | A | NULL | 0.50374993 | NULL&M_Disease |
| 11 | 116234127 | rs576510 | G | C | NULL | 0.50374993 | NULL&M_Disease |
| 11 | 116966878 | rs60167594 | C | T | NULL | 0.50374993 | NULL&M_Disease |
| 11 | 120397891 | rs75195561 | T | C | NULL | 0.50374993 | NULL&M_Disease |
| 11 | 120237971 | rs61900791 | T | C | NULL | 0.50374993 | NULL&M_Disease |
| 11 | 120557393 | rs78813250 | T | G | NULL | 0.50374993 | NULL&M_Disease |
| 11 | 117421984 | rs565083 | A | G | NULL | 0.504999833 | NULL&M_Disease |
| 11 | 120734139 | rs10892631 | A | G | NULL | 0.504999833 | NULL&M_Disease |
| 11 | 118750967 | rs11602393 | A | T | NULL | 0.504999833 | NULL&M_Disease |
| 11 | 116968440 | rs75830246 | C | T | NULL | 0.50374993 | NULL&M_Disease |
| 11 | 117650725 | rs11216542 | T | C | NULL | 0.504999833 | NULL&M_Disease |
| 11 | 120568363 | rs116534827 | G | C | NULL | 0.504999833 | NULL&M_Disease |
| 11 | 120568477 | rs61902689 | T | G | NULL | 0.504999833 | NULL&M_Disease |
| 11 | 116964942 | rs77374709 | T | C | NULL | 0.504999833 | NULL&M_Disease |
| 11 | 116965555 | rs117812708 | G | A | NULL | 0.504999833 | NULL&M_Disease |
| 11 | 119592965 | rs117512890 | C | T | NULL | 0.504999833 | NULL&M_Disease |
| 11 | 116980858 | rs7112111 | G | T | NULL | 0.506249674 | NULL&M_Disease |
| 11 | 120445422 | rs4447186 | G | A | NULL | 0.504999833 | NULL&M_Disease |
| 11 | 116437423 | rs17492275 | T | C | NULL | 0.506249674 | NULL&M_Disease |
| 11 | 117415385 | rs7111299 | A | C | NULL | 0.506249674 | NULL&M_Disease |
| 11 | 120509726 | rs6589825 | C | T | NULL | 0.506249674 | NULL&M_Disease |
| 11 | 120556274 | rs80170882 | T | C | NULL | 0.504999833 | NULL&M_Disease |
| 11 | 117517424 | rs537592 | C | T | NULL | 0.506249674 | NULL&M_Disease |
| 11 | 118755738 | rs523604 | G | A | NULL | 0.506249674 | NULL&M_Disease |
| 11 | 117775198 | rs12797743 | G | A | NULL | 0.506249674 | NULL&M_Disease |
| 11 | 117987128 | rs592009 | G | T | NULL | 0.506249674 | NULL&M_Disease |
| 11 | 116298294 | rs61134913 | G | A | NULL | 0.506249674 | NULL&M_Disease |

| | | | | | | | | | |
|---|---|---|---|---|---|---|---|---|---|
| 11 | 117477867 | rs1939235 | T | C | 0.89 | -7977.388 | -7977.374 | 0.014 | 3.92E-06 |
| 11 | 117560438 | rs75707401 | A | G | 0.892 | -7977.388 | -7977.377 | 0.012 | 3.91E-06 |
| 11 | 121077211 | rs509058 | T | C | 0.935 | -7977.388 | -7977.377 | 0.012 | 3.91E-06 |
| 11 | 119807902 | rs75012772 | A | G | 1.098 | -7977.388 | -7977.378 | 0.01 | 3.91E-06 |
| 11 | 120715471 | rs74702343 | A | G | 0.892 | -7977.388 | -7977.378 | 0.01 | 3.91E-06 |
| 11 | 116860781 | rs75292554 | C | T | 1.113 | -7977.388 | -7977.379 | 0.009 | 3.90E-06 |
| 11 | 116866944 | rs143367870 | C | A | 1.113 | -7977.388 | -7977.379 | 0.009 | 3.90E-06 |
| 11 | 119556950 | rs61901099 | T | C | 1.117 | -7977.388 | -7977.379 | 0.009 | 3.90E-06 |
| 11 | 119802424 | rs754872 | T | C | 0.905 | -7977.388 | -7977.379 | 0.009 | 3.90E-06 |
| 11 | 119510366 | rs4438048 | T | C | 0.92 | -7977.388 | -7977.381 | 0.008 | 3.90E-06 |
| 11 | 119984598 | rs76031864 | A | G | 1.108 | -7977.388 | -7977.382 | 0.007 | 3.90E-06 |
| 11 | 118749318 | rs78312641 | A | T | 1.08 | -7977.388 | -7977.382 | 0.006 | 3.89E-06 |
| 11 | 116409433 | rs17491510 | T | C | 1.074 | -7977.388 | -7977.384 | 0.005 | 3.89E-06 |
| 11 | 116864639 | rs76665318 | C | T | 1.113 | -7977.388 | -7977.383 | 0.005 | 3.89E-06 |
| 11 | 119813686 | rs76661054 | C | T | 1.098 | -7977.388 | -7977.384 | 0.005 | 3.89E-06 |
| 11 | 119813733 | rs75224487 | T | C | 1.098 | -7977.388 | -7977.384 | 0.005 | 3.89E-06 |
| 11 | 119814337 | rs111902338 | A | C | 1.098 | -7977.388 | -7977.384 | 0.005 | 3.89E-06 |
| 11 | 119512641 | rs2089647 | G | A | 0.921 | -7977.388 | -7977.384 | 0.004 | 3.88E-06 |
| 11 | 120413702 | rs75510322 | C | A | 0.905 | -7977.388 | -7977.384 | 0.004 | 3.88E-06 |
| 11 | 116294664 | rs471750 | G | A | 0.927 | -7977.388 | -7977.385 | 0.003 | 3.88E-06 |
| 11 | 117338330 | rs495171 | C | T | 1.108 | -7977.388 | -7977.387 | 0.002 | 3.88E-06 |
| 11 | 119934934 | rs4938763 | T | C | 0.942 | -7977.388 | -7977.387 | 0.001 | 3.87E-06 |
| 11 | 116980679 | rs114221654 | T | C | 1.098 | -7977.388 | -7977.389 | 0 | 3.87E-06 |
| 11 | 116981515 | rs116770907 | T | C | 1.098 | -7977.388 | -7977.389 | 0 | 3.87E-06 |
| 11 | 116539058 | rs112464350 | A | C | 0.903 | -7977.388 | -7977.39 | -0.002 | 3.86E-06 |
| 11 | 117414714 | rs3802879 | T | G | 1.065 | -7977.388 | -7977.39 | -0.002 | 3.86E-06 |
| 11 | 117587224 | rs56153022 | G | C | 0.886 | -7977.388 | -7977.392 | -0.004 | 3.85E-06 |
| 11 | 118807637 | rs112974653 | T | G | 1.081 | -7977.388 | -7977.392 | -0.004 | 3.85E-06 |
| 11 | 119807543 | rs79710878 | G | A | 1.098 | -7977.388 | -7977.392 | -0.004 | 3.85E-06 |
| 11 | 119808101 | rs77763933 | C | G | 1.098 | -7977.388 | -7977.392 | -0.004 | 3.85E-06 |
| 11 | 116996631 | rs12419454 | T | C | 1.098 | -7977.388 | -7977.395 | -0.007 | 3.84E-06 |
| 11 | 116996783 | rs12416691 | T | A | 1.098 | -7977.388 | -7977.395 | -0.007 | 3.84E-06 |
| 11 | 117391421 | rs77964702 | A | G | 0.896 | -7977.388 | -7977.396 | -0.008 | 3.84E-06 |
| 11 | 116953338 | rs79552834 | C | G | 1.112 | -7977.388 | -7977.397 | -0.009 | 3.83E-06 |
| 11 | 119675931 | rs7926586 | G | T | 0.938 | -7977.388 | -7977.397 | -0.009 | 3.83E-06 |
| 11 | 116692552 | rs5103 | G | A | 1.109 | -7977.388 | -7977.399 | -0.01 | 3.83E-06 |
| 11 | 116996345 | rs17120344 | A | G | 1.097 | -7977.388 | -7977.398 | -0.01 | 3.83E-06 |
| 11 | 119510644 | rs7940667 | A | C | 0.921 | -7977.388 | -7977.398 | -0.01 | 3.83E-06 |
| 11 | 119512115 | rs2089646 | G | A | 0.921 | -7977.388 | -7977.399 | -0.01 | 3.83E-06 |
| 11 | 116530797 | rs558056 | T | G | 0.944 | -7977.388 | -7977.4 | -0.011 | 3.83E-06 |
| 11 | 119510303 | rs4245199 | G | T | 0.921 | -7977.388 | -7977.4 | -0.011 | 3.83E-06 |
| 11 | 117414030 | rs563531 | A | G | 1.059 | -7977.388 | -7977.401 | -0.013 | 3.82E-06 |
| 11 | 119678445 | rs7945067 | T | C | 0.94 | -7977.388 | -7977.401 | -0.013 | 3.82E-06 |
| 11 | 120176979 | rs10750171 | G | T | 0.945 | -7977.388 | -7977.402 | -0.014 | 3.81E-06 |
| 11 | 119806581 | rs77443589 | A | G | 1.097 | -7977.388 | -7977.403 | -0.015 | 3.81E-06 |
| 11 | 119806699 | rs79137748 | C | T | 1.097 | -7977.388 | -7977.403 | -0.015 | 3.81E-06 |
| 11 | 116481694 | rs73578509 | G | C | 1.123 | -7977.388 | -7977.404 | -0.016 | 3.81E-06 |
| 11 | 116317745 | rs569327 | T | C | 0.92 | -7977.388 | -7977.406 | -0.017 | 3.80E-06 |
| 11 | 116207020 | rs7947688 | G | A | 1.074 | -7977.388 | -7977.406 | -0.018 | 3.80E-06 |
| 11 | 121152465 | rs11607048 | G | T | 1.075 | -7977.388 | -7977.406 | -0.018 | 3.80E-06 |
| 11 | 116296243 | rs693603 | A | G | 0.928 | -7977.388 | -7977.407 | -0.019 | 3.80E-06 |
| 11 | 116237696 | rs79733808 | A | G | 1.092 | -7977.388 | -7977.408 | -0.02 | 3.79E-06 |
| 11 | 116750073 | rs543819 | G | A | 1.091 | -7977.388 | -7977.408 | -0.02 | 3.79E-06 |
| 11 | 117795578 | rs77642243 | C | T | 0.905 | -7977.388 | -7977.41 | -0.022 | 3.78E-06 |
| 11 | 119882908 | rs587806 | T | C | 0.926 | -7977.388 | -7977.41 | -0.022 | 3.78E-06 |
| 11 | 117931489 | rs7112429 | A | T | 0.895 | -7977.388 | -7977.412 | -0.023 | 3.78E-06 |
| 11 | 117937225 | rs60007061 | A | C | 0.895 | -7977.388 | -7977.411 | -0.023 | 3.78E-06 |
| 11 | 121069225 | rs12288505 | T | C | 0.937 | -7977.388 | -7977.411 | -0.023 | 3.78E-06 |
| 11 | 117899816 | rs617345 | G | T | 0.938 | -7977.388 | -7977.413 | -0.024 | 3.78E-06 |
| 11 | 117521603 | rs678327 | A | G | 0.945 | -7977.388 | -7977.413 | -0.025 | 3.77E-06 |
| 11 | 118560857 | rs73001406 | A | G | 1.066 | -7977.388 | -7977.414 | -0.025 | 3.77E-06 |
| 11 | 120858410 | rs2282586 | A | G | 1.059 | -7977.388 | -7977.415 | -0.027 | 3.77E-06 |
| 11 | 120503923 | rs4936534 | T | A | 0.944 | -7977.388 | -7977.416 | -0.028 | 3.76E-06 |
| 11 | 117492181 | rs10750108 | A | G | 0.941 | -7977.388 | -7977.417 | -0.029 | 3.76E-06 |
| 11 | 118809329 | rs10892313 | A | G | 1.08 | -7977.388 | -7977.417 | -0.029 | 3.76E-06 |
| 11 | 116700777 | rs618354 | C | G | 1.066 | -7977.388 | -7977.419 | -0.031 | 3.75E-06 |
| 11 | 116752191 | rs73023056 | A | G | 1.11 | -7977.388 | -7977.42 | -0.032 | 3.75E-06 |
| 11 | 116534963 | rs12802672 | G | A | 0.904 | -7977.388 | -7977.421 | -0.033 | 3.74E-06 |
| 11 | 116539809 | rs112550319 | G | C | 0.904 | -7977.388 | -7977.421 | -0.033 | 3.74E-06 |
| 11 | 116448277 | rs73011154 | C | A | 1.072 | -7977.388 | -7977.422 | -0.034 | 3.74E-06 |
| 11 | 116234127 | rs576510 | G | C | 1.06 | -7977.388 | -7977.424 | -0.035 | 3.74E-06 |
| 11 | 116966878 | rs60167594 | C | T | 1.11 | -7977.388 | -7977.423 | -0.035 | 3.74E-06 |
| 11 | 120397891 | rs75195561 | T | C | 0.893 | -7977.388 | -7977.425 | -0.037 | 3.73E-06 |
| 11 | 120237971 | rs61900791 | T | C | 1.106 | -7977.388 | -7977.427 | -0.038 | 3.72E-06 |
| 11 | 120557393 | rs78813250 | T | G | 1.124 | -7977.388 | -7977.429 | -0.04 | 3.72E-06 |
| 11 | 117421984 | rs565083 | A | G | 1.061 | -7977.388 | -7977.43 | -0.041 | 3.71E-06 |
| 11 | 120734139 | rs10892631 | A | G | 1.065 | -7977.388 | -7977.429 | -0.041 | 3.71E-06 |
| 11 | 118750967 | rs11602393 | A | T | 0.934 | -7977.388 | -7977.432 | -0.043 | 3.71E-06 |
| 11 | 116968440 | rs75830246 | C | T | 0.892 | -7977.388 | -7977.432 | -0.044 | 3.70E-06 |
| 11 | 117650725 | rs11216542 | T | C | 0.945 | -7977.388 | -7977.434 | -0.045 | 3.70E-06 |
| 11 | 120568363 | rs116534827 | G | C | 0.926 | -7977.388 | -7977.434 | -0.046 | 3.69E-06 |
| 11 | 120568477 | rs61902689 | T | G | 0.926 | -7977.388 | -7977.434 | -0.046 | 3.69E-06 |
| 11 | 116964942 | rs77374709 | T | C | 1.109 | -7977.388 | -7977.435 | -0.047 | 3.69E-06 |
| 11 | 116965555 | rs117812708 | G | A | 1.109 | -7977.388 | -7977.435 | -0.047 | 3.69E-06 |
| 11 | 119592965 | rs117512890 | C | T | 0.917 | -7977.388 | -7977.435 | -0.047 | 3.69E-06 |
| 11 | 116980858 | rs7112111 | G | T | 1.066 | -7977.388 | -7977.439 | -0.051 | 3.68E-06 |
| 11 | 120445422 | rs4447186 | G | A | 1.12 | -7977.388 | -7977.442 | -0.053 | 3.67E-06 |
| 11 | 116437423 | rs17492275 | T | C | 1.071 | -7977.388 | -7977.442 | -0.054 | 3.66E-06 |
| 11 | 117415385 | rs7111299 | A | C | 1.064 | -7977.388 | -7977.442 | -0.054 | 3.66E-06 |
| 11 | 120509726 | rs6589825 | C | T | 1.059 | -7977.388 | -7977.443 | -0.055 | 3.66E-06 |
| 11 | 120556274 | rs80170882 | T | C | 1.122 | -7977.388 | -7977.445 | -0.056 | 3.66E-06 |
| 11 | 117517424 | rs537592 | C | T | 0.942 | -7977.388 | -7977.445 | -0.057 | 3.65E-06 |
| 11 | 118755738 | rs523604 | G | A | 1.058 | -7977.388 | -7977.445 | -0.057 | 3.65E-06 |
| 11 | 117775198 | rs12797743 | G | A | 0.903 | -7977.388 | -7977.446 | -0.058 | 3.65E-06 |
| 11 | 117987128 | rs592009 | G | T | 1.103 | -7977.388 | -7977.446 | -0.058 | 3.65E-06 |
| 11 | 116298294 | rs61134913 | G | A | 1.085 | -7977.388 | -7977.447 | -0.059 | 3.65E-06 |

| | | | | | | | |
|---|---|---|---|---|---|---|---|
| 11 | 116404477 | rs77892802 | C | T | NULL | 0.507499438 | NULL&M_Disease |
| 11 | 120847817 | rs513548 | C | T | NULL | 0.507499438 | NULL&M_Disease |
| 11 | 117506565 | rs2155130 | C | T | NULL | 0.506249674 | NULL&M_Disease |
| 11 | 117574062 | rs7126179 | C | T | NULL | 0.506249674 | NULL&M_Disease |
| 11 | 119510937 | rs4938699 | C | G | NULL | 0.507499438 | NULL&M_Disease |
| 11 | 118091258 | rs77280285 | G | C | NULL | 0.507499438 | NULL&M_Disease |
| 11 | 116975084 | rs118155427 | C | G | NULL | 0.507499438 | NULL&M_Disease |
| 11 | 120441843 | rs11217918 | C | G | NULL | 0.507499438 | NULL&M_Disease |
| 11 | 120585311 | rs117839774 | T | C | NULL | 0.507499438 | NULL&M_Disease |
| 11 | 116708663 | rs12718466 | G | T | NULL | 0.507499438 | NULL&M_Disease |
| 11 | 117320430 | rs12418722 | T | C | NULL | 0.507499438 | NULL&M_Disease |
| 11 | 117885423 | rs77045391 | T | C | NULL | 0.507499438 | NULL&M_Disease |
| 11 | 117929672 | rs61900590 | T | C | NULL | 0.508749107 | NULL&M_Disease |
| 11 | 118809767 | rs75460330 | A | G | NULL | 0.508749107 | NULL&M_Disease |
| 11 | 118810054 | rs73557344 | C | G | NULL | 0.508749107 | NULL&M_Disease |
| 11 | 117529622 | rs621188 | C | T | NULL | 0.508749107 | NULL&M_Disease |
| 11 | 116970285 | rs73576687 | A | G | NULL | 0.507499438 | NULL&M_Disease |
| 11 | 119882187 | rs584620 | T | C | NULL | 0.508749107 | NULL&M_Disease |
| 11 | 116448341 | rs759988389 | C | T | NULL | 0.508749107 | NULL&M_Disease |
| 11 | 116968225 | rs73576682 | G | A | NULL | 0.507499438 | NULL&M_Disease |
| 11 | 116969416 | rs7939209 | C | T | NULL | 0.507499438 | NULL&M_Disease |
| 11 | 116969439 | rs7939303 | C | T | NULL | 0.507499438 | NULL&M_Disease |
| 11 | 117065050 | rs117634345 | A | G | NULL | 0.508749107 | NULL&M_Disease |
| 11 | 118749563 | rs77643000 | C | T | NULL | 0.508749107 | NULL&M_Disease |
| 11 | 116699389 | rs10790164 | A | G | NULL | 0.508749107 | NULL&M_Disease |
| 11 | 117793989 | rs11216625 | A | C | NULL | 0.508749107 | NULL&M_Disease |
| 11 | 120569866 | rs61901363 | T | C | NULL | 0.509998667 | NULL&M_Disease |
| 11 | 116991842 | rs59837440 | C | A | NULL | 0.508749107 | NULL&M_Disease |
| 11 | 117422642 | rs640147 | T | C | NULL | 0.509998667 | NULL&M_Disease |
| 11 | 118748789 | rs145762171 | A | G | NULL | 0.508749107 | NULL&M_Disease |
| 11 | 117535724 | rs10892154 | A | C | NULL | 0.509998667 | NULL&M_Disease |
| 11 | 118807740 | rs10892312 | A | G | NULL | 0.509998667 | NULL&M_Disease |
| 11 | 116225951 | rs117228135 | A | G | NULL | 0.508749107 | NULL&M_Disease |
| 11 | 117469122 | rs77973629 | T | G | NULL | 0.508749107 | NULL&M_Disease |
| 11 | 116706346 | rs5081 | T | A | NULL | 0.508749107 | NULL&M_Disease |
| 11 | 117335160 | rs111774429 | T | C | NULL | 0.508749107 | NULL&M_Disease |
| 11 | 117449742 | rs11602332 | A | G | NULL | 0.509998667 | NULL&M_Disease |
| 11 | 120853380 | rs2298723 | C | A | NULL | 0.509998667 | NULL&M_Disease |
| 11 | 117362948 | rs582285 | G | A | NULL | 0.509998667 | NULL&M_Disease |
| 11 | 117939413 | rs60348146 | A | G | NULL | 0.509998667 | NULL&M_Disease |
| 11 | 116233643 | rs572619 | T | A | NULL | 0.511248102 | NULL&M_Disease |
| 11 | 116998912 | rs145802214 | C | A | NULL | 0.509998667 | NULL&M_Disease |
| 11 | 116999025 | rs141250514 | C | T | NULL | 0.509998667 | NULL&M_Disease |
| 11 | 117392110 | rs35503235 | G | A | NULL | 0.509998667 | NULL&M_Disease |
| 11 | 117505239 | rs672251 | G | T | NULL | 0.511248102 | NULL&M_Disease |
| 11 | 116970006 | rs73576686 | A | T | NULL | 0.509998667 | NULL&M_Disease |
| 11 | 117354582 | rs10892123 | C | T | NULL | 0.511248102 | NULL&M_Disease |
| 11 | 117411223 | rs78853831 | T | C | NULL | 0.509998667 | NULL&M_Disease |
| 11 | 119715337 | rs73008653 | T | C | NULL | 0.511248102 | NULL&M_Disease |
| 11 | 120566375 | rs61902686 | C | T | NULL | 0.511248102 | NULL&M_Disease |
| 11 | 120566406 | rs61902687 | G | A | NULL | 0.511248102 | NULL&M_Disease |
| 11 | 116507206 | rs61567742 | T | C | NULL | 0.511248102 | NULL&M_Disease |
| 11 | 116688383 | rs2727790 | A | G | NULL | 0.511248102 | NULL&M_Disease |
| 11 | 117024256 | rs1352178661 | A | G | NULL | 0.511248102 | NULL&M_Disease |
| 11 | 117467582 | rs4938416 | T | C | NULL | 0.511248102 | NULL&M_Disease |
| 11 | 117467586 | rs4938417 | C | T | NULL | 0.511248102 | NULL&M_Disease |
| 11 | 118103976 | rs76675432 | T | C | NULL | 0.511248102 | NULL&M_Disease |
| 11 | 120849062 | rs4323878 | C | G | NULL | 0.512497396 | NULL&M_Disease |
| 11 | 116903316 | rs4548653 | A | G | NULL | 0.512497396 | NULL&M_Disease |
| 11 | 117592546 | rs4936400 | A | G | NULL | 0.511248102 | NULL&M_Disease |
| 11 | 120419218 | rs7940779 | T | C | NULL | 0.512497396 | NULL&M_Disease |
| 11 | 118441079 | rs7106316 | G | A | NULL | 0.512497396 | NULL&M_Disease |
| 11 | 116426640 | rs12289097 | G | A | NULL | 0.512497396 | NULL&M_Disease |
| 11 | 116533167 | rs509728 | C | G | NULL | 0.512497396 | NULL&M_Disease |
| 11 | 116547745 | rs59371082 | G | A | NULL | 0.512497396 | NULL&M_Disease |
| 11 | 116892626 | rs75949389 | G | A | NULL | 0.511248102 | NULL&M_Disease |
| 11 | 116999600 | rs138039720 | T | C | NULL | 0.512497396 | NULL&M_Disease |
| 11 | 117000179 | rs146067038 | G | A | NULL | 0.512497396 | NULL&M_Disease |
| 11 | 117001960 | rs12421425 | T | C | NULL | 0.512497396 | NULL&M_Disease |
| 11 | 117002828 | rs17120367 | T | C | NULL | 0.512497396 | NULL&M_Disease |
| 11 | 117003388 | rs17120370 | C | T | NULL | 0.512497396 | NULL&M_Disease |
| 11 | 117003418 | rs74735277 | T | C | NULL | 0.512497396 | NULL&M_Disease |
| 11 | 117003623 | rs75208249 | C | G | NULL | 0.512497396 | NULL&M_Disease |
| 11 | 117004201 | rs114945429 | T | G | NULL | 0.512497396 | NULL&M_Disease |
| 11 | 117004853 | rs12417098 | A | G | NULL | 0.512497396 | NULL&M_Disease |
| 11 | 117477761 | rs75582258 | G | A | NULL | 0.511248102 | NULL&M_Disease |
| 11 | 117795494 | rs78466564 | T | C | NULL | 0.512497396 | NULL&M_Disease |
| 11 | 120570595 | rs12293509 | T | C | NULL | 0.512497396 | NULL&M_Disease |
| 11 | 117162284 | rs45619137 | A | G | NULL | 0.512497396 | NULL&M_Disease |
| 11 | 118661197 | rs61901410 | G | C | NULL | 0.512497396 | NULL&M_Disease |
| 11 | 116707190 | rs5076 | A | G | NULL | 0.512497396 | NULL&M_Disease |
| 11 | 116708254 | rs5069 | A | G | NULL | 0.512497396 | NULL&M_Disease |
| 11 | 116976598 | rs116896104 | A | G | NULL | 0.512497396 | NULL&M_Disease |
| 11 | 117092808 | rs79697731 | G | C | NULL | 0.512497396 | NULL&M_Disease |
| 11 | 117794282 | rs11216626 | A | T | NULL | 0.512497396 | NULL&M_Disease |
| 11 | 117498605 | rs4936391 | G | A | NULL | 0.513746535 | NULL&M_Disease |
| 11 | 119879786 | rs1944512 | T | G | NULL | 0.513746535 | NULL&M_Disease |
| 11 | 116530251 | rs530053 | G | A | NULL | 0.513746535 | NULL&M_Disease |
| 11 | 119804461 | rs693701 | T | C | NULL | 0.513746535 | NULL&M_Disease |
| 11 | 116440521 | rs906541 | A | G | NULL | 0.513746535 | NULL&M_Disease |
| 11 | 116233857 | rs574529 | C | T | NULL | 0.513746535 | NULL&M_Disease |
| 11 | 116554088 | rs71480312 | T | C | NULL | 0.513746535 | NULL&M_Disease |
| 11 | 116897124 | rs57643685 | G | A | NULL | 0.512497396 | NULL&M_Disease |
| 11 | 118614115 | rs73001451 | C | T | NULL | 0.513746535 | NULL&M_Disease |
| 11 | 119959273 | rs7124732 | G | A | NULL | 0.512497396 | NULL&M_Disease |
| 11 | 119962800 | rs57877069 | C | G | NULL | 0.512497396 | NULL&M_Disease |

| | | | | | | | | | |
|---|---|---|---|---|---|---|---|---|---|
| 11 | 116404477 | rs77892802 | C | T | 1.073 | -7977.388 | -7977.448 | -0.06 | 3.64E-06 |
| 11 | 120847817 | rs513548 | C | T | 0.939 | -7977.388 | -7977.449 | -0.06 | 3.64E-06 |
| 11 | 117506565 | rs2155130 | C | T | 0.894 | -7977.388 | -7977.449 | -0.061 | 3.64E-06 |
| 11 | 117574062 | rs7126179 | C | T | 0.892 | -7977.388 | -7977.451 | -0.063 | 3.63E-06 |
| 11 | 119510937 | rs4938699 | C | G | 0.922 | -7977.388 | -7977.451 | -0.063 | 3.63E-06 |
| 11 | 118091258 | rs77280285 | G | C | 0.895 | -7977.388 | -7977.453 | -0.065 | 3.62E-06 |
| 11 | 116975084 | rs118155427 | C | G | 0.894 | -7977.388 | -7977.455 | -0.067 | 3.62E-06 |
| 11 | 120441843 | rs11217918 | C | G | 1.119 | -7977.388 | -7977.456 | -0.067 | 3.62E-06 |
| 11 | 120585311 | rs117839774 | T | C | 0.9 | -7977.388 | -7977.457 | -0.068 | 3.61E-06 |
| 11 | 116708663 | rs12718466 | G | T | 1.108 | -7977.388 | -7977.458 | -0.069 | 3.61E-06 |
| 11 | 117320430 | rs12418722 | T | C | 0.916 | -7977.388 | -7977.457 | -0.069 | 3.61E-06 |
| 11 | 117885423 | rs77045391 | T | C | 0.897 | -7977.388 | -7977.457 | -0.069 | 3.61E-06 |
| 11 | 117929672 | rs61900590 | T | C | 0.944 | -7977.388 | -7977.458 | -0.07 | 3.61E-06 |
| 11 | 118809767 | rs75460330 | A | G | 0.914 | -7977.388 | -7977.459 | -0.07 | 3.61E-06 |
| 11 | 118810054 | rs73557344 | C | G | 0.914 | -7977.388 | -7977.459 | -0.07 | 3.61E-06 |
| 11 | 117529622 | rs621188 | C | T | 0.945 | -7977.388 | -7977.459 | -0.071 | 3.60E-06 |
| 11 | 116970285 | rs73576687 | A | G | 0.895 | -7977.388 | -7977.461 | -0.073 | 3.60E-06 |
| 11 | 119882187 | rs584620 | T | C | 0.927 | -7977.388 | -7977.461 | -0.073 | 3.60E-06 |
| 11 | 116448341 | rs759988389 | C | T | 1.071 | -7977.388 | -7977.462 | -0.074 | 3.59E-06 |
| 11 | 116968225 | rs73576682 | G | A | 0.895 | -7977.388 | -7977.462 | -0.074 | 3.59E-06 |
| 11 | 116969416 | rs7939209 | C | T | 0.895 | -7977.388 | -7977.462 | -0.074 | 3.59E-06 |
| 11 | 116969439 | rs7939303 | C | T | 0.895 | -7977.388 | -7977.462 | -0.074 | 3.59E-06 |
| 11 | 117065050 | rs117634345 | A | G | 1.107 | -7977.388 | -7977.462 | -0.074 | 3.59E-06 |
| 11 | 118749563 | rs77643000 | C | T | 1.077 | -7977.388 | -7977.463 | -0.075 | 3.59E-06 |
| 11 | 116699389 | rs10790164 | A | G | 0.904 | -7977.388 | -7977.466 | -0.078 | 3.58E-06 |
| 11 | 117793989 | rs11216625 | A | C | 0.907 | -7977.388 | -7977.466 | -0.078 | 3.58E-06 |
| 11 | 120569866 | rs61901363 | T | C | 0.926 | -7977.388 | -7977.468 | -0.079 | 3.57E-06 |
| 11 | 116991842 | rs59837440 | C | A | 1.107 | -7977.388 | -7977.469 | -0.08 | 3.57E-06 |
| 11 | 117422642 | rs640147 | T | C | 1.06 | -7977.388 | -7977.468 | -0.08 | 3.57E-06 |
| 11 | 118748789 | rs145762171 | A | G | 0.893 | -7977.388 | -7977.468 | -0.08 | 3.57E-06 |
| 11 | 117535724 | rs10892154 | A | C | 0.945 | -7977.388 | -7977.469 | -0.081 | 3.57E-06 |
| 11 | 118807740 | rs10892312 | A | G | 1.079 | -7977.388 | -7977.469 | -0.081 | 3.57E-06 |
| 11 | 116225951 | rs117228135 | A | G | 1.108 | -7977.388 | -7977.471 | -0.082 | 3.56E-06 |
| 11 | 117469122 | rs77973629 | T | G | 0.895 | -7977.388 | -7977.472 | -0.083 | 3.56E-06 |
| 11 | 116706346 | rs5081 | T | A | 0.896 | -7977.388 | -7977.472 | -0.084 | 3.56E-06 |
| 11 | 117335160 | rs111774429 | T | C | 0.893 | -7977.388 | -7977.473 | -0.085 | 3.55E-06 |
| 11 | 117449742 | rs11602332 | A | G | 1.06 | -7977.388 | -7977.474 | -0.085 | 3.55E-06 |
| 11 | 120853380 | rs2298723 | C | A | 1.058 | -7977.388 | -7977.474 | -0.085 | 3.55E-06 |
| 11 | 117362948 | rs582285 | G | A | 0.945 | -7977.388 | -7977.474 | -0.086 | 3.55E-06 |
| 11 | 117939413 | rs60348146 | A | G | 0.899 | -7977.388 | -7977.474 | -0.086 | 3.55E-06 |
| 11 | 116233643 | rs572619 | T | A | 1.059 | -7977.388 | -7977.477 | -0.089 | 3.54E-06 |
| 11 | 116998912 | rs145802214 | C | A | 1.094 | -7977.388 | -7977.478 | -0.09 | 3.54E-06 |
| 11 | 116999025 | rs141250514 | C | T | 1.094 | -7977.388 | -7977.478 | -0.09 | 3.54E-06 |
| 11 | 117392110 | rs35503235 | G | A | 0.901 | -7977.388 | -7977.478 | -0.09 | 3.54E-06 |
| 11 | 117505239 | rs672251 | G | T | 0.943 | -7977.388 | -7977.478 | -0.09 | 3.54E-06 |
| 11 | 116970006 | rs73576686 | A | T | 0.896 | -7977.388 | -7977.479 | -0.091 | 3.53E-06 |
| 11 | 117354582 | rs10892123 | C | T | 0.927 | -7977.388 | -7977.479 | -0.091 | 3.53E-06 |
| 11 | 117411223 | rs78853831 | T | C | 0.895 | -7977.388 | -7977.48 | -0.092 | 3.53E-06 |
| 11 | 119715337 | rs73008653 | T | C | 1.073 | -7977.388 | -7977.482 | -0.094 | 3.52E-06 |
| 11 | 120566375 | rs61902686 | C | T | 0.927 | -7977.388 | -7977.484 | -0.095 | 3.52E-06 |
| 11 | 120566406 | rs61902687 | G | A | 0.927 | -7977.388 | -7977.484 | -0.095 | 3.52E-06 |
| 11 | 116507206 | rs61567742 | T | C | 1.101 | -7977.388 | -7977.485 | -0.097 | 3.51E-06 |
| 11 | 116688383 | rs2727790 | A | G | 0.896 | -7977.388 | -7977.486 | -0.097 | 3.51E-06 |
| 11 | 117024256 | rs1352178661 | A | G | 1.106 | -7977.388 | -7977.487 | -0.099 | 3.50E-06 |
| 11 | 117467582 | rs4938416 | T | C | 0.897 | -7977.388 | -7977.487 | -0.099 | 3.50E-06 |
| 11 | 117467586 | rs4938417 | C | T | 0.897 | -7977.388 | -7977.487 | -0.099 | 3.50E-06 |
| 11 | 118103976 | rs76675432 | T | C | 0.897 | -7977.388 | -7977.487 | -0.099 | 3.50E-06 |
| 11 | 120849062 | rs4323878 | C | G | 1.057 | -7977.388 | -7977.488 | -0.1 | 3.50E-06 |
| 11 | 116903316 | rs4548653 | A | G | 1.065 | -7977.388 | -7977.489 | -0.101 | 3.50E-06 |
| 11 | 117592546 | rs4936400 | A | G | 0.895 | -7977.388 | -7977.49 | -0.101 | 3.50E-06 |
| 11 | 120419218 | rs7940779 | T | C | 0.914 | -7977.388 | -7977.49 | -0.101 | 3.50E-06 |
| 11 | 118441079 | rs7106316 | G | A | 0.925 | -7977.388 | -7977.49 | -0.102 | 3.49E-06 |
| 11 | 116426640 | rs12289097 | G | A | 1.068 | -7977.388 | -7977.492 | -0.103 | 3.49E-06 |
| 11 | 116533167 | rs509728 | C | G | 0.909 | -7977.388 | -7977.492 | -0.103 | 3.49E-06 |
| 11 | 116547745 | rs59371082 | G | A | 0.907 | -7977.388 | -7977.492 | -0.103 | 3.49E-06 |
| 11 | 116892626 | rs75949389 | G | A | 0.897 | -7977.388 | -7977.491 | -0.103 | 3.49E-06 |
| 11 | 116999600 | rs138039720 | T | C | 1.093 | -7977.388 | -7977.492 | -0.103 | 3.49E-06 |
| 11 | 117000179 | rs146067038 | G | A | 1.093 | -7977.388 | -7977.492 | -0.103 | 3.49E-06 |
| 11 | 117001960 | rs12421425 | T | C | 1.093 | -7977.388 | -7977.492 | -0.103 | 3.49E-06 |
| 11 | 117002828 | rs17120367 | T | C | 1.093 | -7977.388 | -7977.492 | -0.103 | 3.49E-06 |
| 11 | 117003388 | rs17120370 | C | T | 1.093 | -7977.388 | -7977.492 | -0.103 | 3.49E-06 |
| 11 | 117003418 | rs74735277 | T | C | 1.093 | -7977.388 | -7977.492 | -0.103 | 3.49E-06 |
| 11 | 117003623 | rs75208249 | C | G | 1.093 | -7977.388 | -7977.492 | -0.103 | 3.49E-06 |
| 11 | 117004201 | rs114945429 | T | G | 1.093 | -7977.388 | -7977.492 | -0.103 | 3.49E-06 |
| 11 | 117004853 | rs12417098 | A | G | 1.093 | -7977.388 | -7977.492 | -0.103 | 3.49E-06 |
| 11 | 117477761 | rs75582258 | G | A | 0.898 | -7977.388 | -7977.492 | -0.104 | 3.49E-06 |
| 11 | 117795494 | rs78466564 | T | C | 0.908 | -7977.388 | -7977.492 | -0.104 | 3.49E-06 |
| 11 | 120570595 | rs12293509 | T | C | 0.927 | -7977.388 | -7977.492 | -0.104 | 3.49E-06 |
| 11 | 117162284 | rs45619137 | A | G | 1.089 | -7977.388 | -7977.494 | -0.105 | 3.48E-06 |
| 11 | 118661197 | rs61901410 | G | C | 0.93 | -7977.388 | -7977.493 | -0.105 | 3.48E-06 |
| 11 | 116707190 | rs5076 | A | G | 1.105 | -7977.388 | -7977.496 | -0.107 | 3.48E-06 |
| 11 | 116708254 | rs5069 | A | G | 1.105 | -7977.388 | -7977.495 | -0.107 | 3.48E-06 |
| 11 | 116976598 | rs116896104 | A | G | 1.105 | -7977.388 | -7977.496 | -0.108 | 3.47E-06 |
| 11 | 117092808 | rs79697731 | G | C | 1.105 | -7977.388 | -7977.496 | -0.108 | 3.47E-06 |
| 11 | 117794282 | rs11216626 | A | T | 0.908 | -7977.388 | -7977.496 | -0.108 | 3.47E-06 |
| 11 | 117498605 | rs4936391 | G | A | 0.942 | -7977.388 | -7977.498 | -0.109 | 3.47E-06 |
| 11 | 119879786 | rs1944512 | T | G | 1.078 | -7977.388 | -7977.498 | -0.109 | 3.47E-06 |
| 11 | 116530251 | rs530053 | G | A | 0.946 | -7977.388 | -7977.499 | -0.111 | 3.46E-06 |
| 11 | 119804461 | rs693701 | T | C | 0.941 | -7977.388 | -7977.5 | -0.111 | 3.46E-06 |
| 11 | 116440521 | rs906541 | A | G | 1.07 | -7977.388 | -7977.5 | -0.112 | 3.46E-06 |
| 11 | 116233857 | rs574529 | C | T | 1.059 | -7977.388 | -7977.502 | -0.113 | 3.45E-06 |
| 11 | 116554088 | rs71480312 | T | C | 0.907 | -7977.388 | -7977.501 | -0.113 | 3.45E-06 |
| 11 | 116897124 | rs57643685 | G | A | 0.897 | -7977.388 | -7977.501 | -0.113 | 3.45E-06 |
| 11 | 118614115 | rs73001451 | C | T | 0.913 | -7977.388 | -7977.501 | -0.113 | 3.45E-06 |
| 11 | 119959273 | rs7124732 | G | A | 0.899 | -7977.388 | -7977.501 | -0.113 | 3.45E-06 |
| 11 | 119962800 | rs57877069 | C | G | 0.899 | -7977.388 | -7977.501 | -0.113 | 3.45E-06 |

| | | | | | | | |
|---|---|---|---|---|---|---|---|
| 11 | 117076336 | rs78301905 | C | T | NULL | 0.513746535 | NULL&M_Disease |
| 11 | 117083948 | rs79997323 | G | A | NULL | 0.513746535 | NULL&M_Disease |
| 11 | 117084410 | rs79278568 | A | G | NULL | 0.513746535 | NULL&M_Disease |
| 11 | 117430479 | rs10892137 | T | C | NULL | 0.513746535 | NULL&M_Disease |
| 11 | 118500963 | rs680465 | A | G | NULL | 0.513746535 | NULL&M_Disease |
| 11 | 117421824 | rs542322 | A | G | NULL | 0.513746535 | NULL&M_Disease |
| 11 | 116424533 | rs75212830 | A | G | NULL | 0.513746535 | NULL&M_Disease |
| 11 | 116687156 | rs139385687 | C | G | NULL | 0.513746535 | NULL&M_Disease |
| 11 | 116904176 | rs142693807 | A | G | NULL | 0.513746535 | NULL&M_Disease |
| 11 | 116869560 | rs12574810 | G | A | NULL | 0.513746535 | NULL&M_Disease |
| 11 | 117528376 | rs671553 | T | C | NULL | 0.514995502 | NULL&M_Disease |
| 11 | 117470412 | rs78136547 | G | C | NULL | 0.513746535 | NULL&M_Disease |
| 11 | 117472019 | rs4938418 | C | T | NULL | 0.513746535 | NULL&M_Disease |
| 11 | 120577241 | rs61901378 | A | G | NULL | 0.514995502 | NULL&M_Disease |
| 11 | 116706343 | rs12718463 | C | T | NULL | 0.513746535 | NULL&M_Disease |
| 11 | 117560822 | rs11827260 | C | G | NULL | 0.514995502 | NULL&M_Disease |
| 11 | 118622744 | rs6589689 | G | A | NULL | 0.513746535 | NULL&M_Disease |
| 11 | 119882789 | rs587308 | T | C | NULL | 0.514995502 | NULL&M_Disease |
| 11 | 116696681 | rs7123326 | A | G | NULL | 0.513746535 | NULL&M_Disease |
| 11 | 117472684 | rs145533092 | A | G | NULL | 0.513746535 | NULL&M_Disease |
| 11 | 116698665 | rs7101528 | A | G | NULL | 0.514995502 | NULL&M_Disease |
| 11 | 116613464 | rs151220469 | A | G | NULL | 0.513746535 | NULL&M_Disease |
| 11 | 118808140 | rs7117408 | T | C | NULL | 0.514995502 | NULL&M_Disease |
| 11 | 117347947 | rs1574433 | T | C | NULL | 0.514995502 | NULL&M_Disease |
| 11 | 119324831 | rs118101513 | A | C | NULL | 0.514995502 | NULL&M_Disease |
| 11 | 117041911 | rs75525407 | T | C | NULL | 0.514995502 | NULL&M_Disease |
| 11 | 117052757 | rs189196849 | T | G | NULL | 0.514995502 | NULL&M_Disease |
| 11 | 117054029 | rs59978576 | G | C | NULL | 0.514995502 | NULL&M_Disease |
| 11 | 117055400 | rs61268522 | C | A | NULL | 0.514995502 | NULL&M_Disease |
| 11 | 117055893 | rs140727315 | C | G | NULL | 0.514995502 | NULL&M_Disease |
| 11 | 117061994 | rs184936191 | T | A | NULL | 0.514995502 | NULL&M_Disease |
| 11 | 117065271 | rs79043792 | C | G | NULL | 0.514995502 | NULL&M_Disease |
| 11 | 117072970 | rs74967901 | A | G | NULL | 0.514995502 | NULL&M_Disease |
| 11 | 116987421 | rs56756054 | C | T | NULL | 0.514995502 | NULL&M_Disease |
| 11 | 119879579 | rs1944495 | T | C | NULL | 0.516244281 | NULL&M_Disease |
| 11 | 116730833 | rs545274 | T | C | NULL | 0.516244281 | NULL&M_Disease |
| 11 | 117356858 | rs606727 | G | C | NULL | 0.516244281 | NULL&M_Disease |
| 11 | 116223780 | rs17475326 | T | A | NULL | 0.514995502 | NULL&M_Disease |
| 11 | 116754753 | rs57259536 | T | A | NULL | 0.514995502 | NULL&M_Disease |
| 11 | 116882563 | rs73594141 | T | C | NULL | 0.514995502 | NULL&M_Disease |
| 11 | 116896441 | rs73594158 | T | C | NULL | 0.514995502 | NULL&M_Disease |
| 11 | 116896811 | rs73594162 | T | C | NULL | 0.514995502 | NULL&M_Disease |
| 11 | 116897613 | rs73594164 | G | T | NULL | 0.514995502 | NULL&M_Disease |
| 11 | 116900956 | rs59647391 | G | A | NULL | 0.514995502 | NULL&M_Disease |
| 11 | 116902631 | rs73594172 | T | A | NULL | 0.514995502 | NULL&M_Disease |
| 11 | 117473537 | rs77346448 | C | T | NULL | 0.514995502 | NULL&M_Disease |
| 11 | 117795537 | rs79136224 | T | C | NULL | 0.516244281 | NULL&M_Disease |
| 11 | 120841689 | rs652418 | C | T | NULL | 0.516244281 | NULL&M_Disease |
| 11 | 121189231 | rs10892721 | T | G | NULL | 0.516244281 | NULL&M_Disease |
| 11 | 120644115 | rs7938295 | A | G | NULL | 0.516244281 | NULL&M_Disease |
| 11 | 117099025 | rs2306474 | G | A | NULL | 0.516244281 | NULL&M_Disease |
| 11 | 119936292 | rs640663 | A | G | NULL | 0.516244281 | NULL&M_Disease |
| 11 | 117563295 | rs11216505 | A | G | NULL | 0.516244281 | NULL&M_Disease |
| 11 | 117579111 | rs17121102 | C | G | NULL | 0.516244281 | NULL&M_Disease |
| 11 | 118754649 | rs10892307 | G | C | NULL | 0.516244281 | NULL&M_Disease |
| 11 | 120569843 | rs61901362 | C | T | NULL | 0.516244281 | NULL&M_Disease |
| 11 | 118241516 | rs11216861 | A | G | NULL | 0.517492858 | NULL&M_Disease |
| 11 | 117430058 | rs510250 | G | T | NULL | 0.517492858 | NULL&M_Disease |
| 11 | 118865992 | rs11217117 | T | C | NULL | 0.517492858 | NULL&M_Disease |
| 11 | 120838716 | rs608154 | G | A | NULL | 0.517492858 | NULL&M_Disease |
| 11 | 117034320 | rs150881386 | T | C | NULL | 0.516244281 | NULL&M_Disease |
| 11 | 117035622 | rs144762883 | T | C | NULL | 0.516244281 | NULL&M_Disease |
| 11 | 117039208 | rs117842423 | A | G | NULL | 0.516244281 | NULL&M_Disease |
| 11 | 117061937 | rs117303996 | A | G | NULL | 0.516244281 | NULL&M_Disease |
| 11 | 117795373 | rs78264252 | A | C | NULL | 0.517492858 | NULL&M_Disease |
| 11 | 116734545 | rs11216164 | A | G | NULL | 0.517492858 | NULL&M_Disease |
| 11 | 118807876 | rs12282154 | A | G | NULL | 0.517492858 | NULL&M_Disease |
| 11 | 120571883 | rs61901370 | T | C | NULL | 0.517492858 | NULL&M_Disease |
| 11 | 117053686 | rs61562228 | A | G | NULL | 0.516244281 | NULL&M_Disease |
| 11 | 117794650 | rs11216629 | G | C | NULL | 0.517492858 | NULL&M_Disease |
| 11 | 119733857 | rs570521 | G | T | NULL | 0.517492858 | NULL&M_Disease |
| 11 | 117004023 | rs150672190 | T | C | NULL | 0.517492858 | NULL&M_Disease |
| 11 | 117005207 | rs117146251 | G | A | NULL | 0.517492858 | NULL&M_Disease |
| 11 | 119352350 | rs61901289 | T | G | NULL | 0.517492858 | NULL&M_Disease |
| 11 | 116680961 | rs80220270 | C | A | NULL | 0.517492858 | NULL&M_Disease |
| 11 | 116929666 | rs73576612 | C | T | NULL | 0.517492858 | NULL&M_Disease |
| 11 | 116932615 | rs17120265 | G | C | NULL | 0.517492858 | NULL&M_Disease |
| 11 | 116935363 | rs17120271 | G | C | NULL | 0.517492858 | NULL&M_Disease |
| 11 | 116939071 | rs17120275 | C | T | NULL | 0.517492858 | NULL&M_Disease |
| 11 | 118856491 | rs3889526 | A | C | NULL | 0.517492858 | NULL&M_Disease |
| 11 | 118857466 | rs4938610 | C | T | NULL | 0.517492858 | NULL&M_Disease |
| 11 | 119663944 | rs73006788 | T | C | NULL | 0.517492858 | NULL&M_Disease |
| 11 | 117344165 | rs11216413 | A | T | NULL | 0.518741216 | NULL&M_Disease |
| 11 | 117795082 | rs76046563 | G | C | NULL | 0.517492858 | NULL&M_Disease |
| 11 | 117346121 | rs74594162 | G | C | NULL | 0.517492858 | NULL&M_Disease |
| 11 | 117349482 | rs12277788 | T | C | NULL | 0.518741216 | NULL&M_Disease |
| 11 | 117709760 | rs11216576 | A | G | NULL | 0.518741216 | NULL&M_Disease |
| 11 | 119324268 | rs78040959 | G | A | NULL | 0.518741216 | NULL&M_Disease |
| 11 | 116963969 | rs79536557 | G | A | NULL | 0.517492858 | NULL&M_Disease |
| 11 | 118287568 | rs75461585 | G | A | NULL | 0.517492858 | NULL&M_Disease |
| 11 | 120643090 | rs7934092 | A | C | NULL | 0.517492858 | NULL&M_Disease |
| 11 | 117246876 | rs2305827 | C | T | NULL | 0.518741216 | NULL&M_Disease |
| 11 | 117475763 | rs77492882 | T | C | NULL | 0.517492858 | NULL&M_Disease |
| 11 | 117475976 | rs79198917 | A | G | NULL | 0.517492858 | NULL&M_Disease |
| 11 | 116910971 | rs12273536 | T | C | NULL | 0.518741216 | NULL&M_Disease |
| 11 | 117574296 | rs2068412 | G | A | NULL | 0.517492858 | NULL&M_Disease |

| | | | | | | | | | |
|---|---|---|---|---|---|---|---|---|---|
| 11 | 117076336 | rs78301905 | C | T | 1.105 | -7977.388 | -7977.502 | -0.114 | 3.45E-06 |
| 11 | 117083948 | rs79997323 | G | A | 1.105 | -7977.388 | -7977.502 | -0.114 | 3.45E-06 |
| 11 | 117084410 | rs79278568 | A | G | 1.105 | -7977.388 | -7977.502 | -0.114 | 3.45E-06 |
| 11 | 117430479 | rs10892137 | T | C | 0.945 | -7977.388 | -7977.502 | -0.114 | 3.45E-06 |
| 11 | 118500963 | rs680465 | A | G | 0.938 | -7977.388 | -7977.502 | -0.114 | 3.45E-06 |
| 11 | 117421824 | rs542322 | A | G | 1.06 | -7977.388 | -7977.503 | -0.115 | 3.45E-06 |
| 11 | 116424533 | rs75212830 | A | G | 1.071 | -7977.388 | -7977.504 | -0.116 | 3.44E-06 |
| 11 | 116687156 | rs139385687 | C | G | 1.103 | -7977.388 | -7977.504 | -0.116 | 3.44E-06 |
| 11 | 116904176 | rs142693807 | A | G | 0.898 | -7977.388 | -7977.505 | -0.117 | 3.44E-06 |
| 11 | 116869560 | rs12574810 | G | A | 1.107 | -7977.388 | -7977.506 | -0.118 | 3.44E-06 |
| 11 | 117528376 | rs671553 | T | C | 0.945 | -7977.388 | -7977.506 | -0.118 | 3.44E-06 |
| 11 | 117470412 | rs78136547 | G | C | 0.898 | -7977.388 | -7977.508 | -0.119 | 3.43E-06 |
| 11 | 117472019 | rs4938418 | C | T | 0.898 | -7977.388 | -7977.508 | -0.119 | 3.43E-06 |
| 11 | 120577241 | rs61901378 | A | G | 0.927 | -7977.388 | -7977.508 | -0.12 | 3.43E-06 |
| 11 | 116706343 | rs12718463 | C | T | 1.104 | -7977.388 | -7977.51 | -0.121 | 3.43E-06 |
| 11 | 117560822 | rs11827260 | C | G | 0.933 | -7977.388 | -7977.509 | -0.121 | 3.43E-06 |
| 11 | 118622744 | rs6589689 | G | A | 0.897 | -7977.388 | -7977.509 | -0.121 | 3.43E-06 |
| 11 | 119882789 | rs587308 | T | C | 0.928 | -7977.388 | -7977.51 | -0.121 | 3.43E-06 |
| 11 | 116696681 | rs7123326 | A | G | 1.103 | -7977.388 | -7977.511 | -0.122 | 3.42E-06 |
| 11 | 117472684 | rs145533092 | A | G | 0.898 | -7977.388 | -7977.51 | -0.122 | 3.42E-06 |
| 11 | 116698665 | rs7101528 | A | G | 1.102 | -7977.388 | -7977.512 | -0.124 | 3.42E-06 |
| 11 | 116613464 | rs151220469 | A | G | 1.113 | -7977.388 | -7977.513 | -0.125 | 3.41E-06 |
| 11 | 118808140 | rs7117408 | T | C | 1.077 | -7977.388 | -7977.514 | -0.125 | 3.41E-06 |
| 11 | 117347947 | rs1574433 | T | C | 1.066 | -7977.388 | -7977.515 | -0.127 | 3.41E-06 |
| 11 | 119324831 | rs118101513 | A | C | 0.898 | -7977.388 | -7977.515 | -0.127 | 3.41E-06 |
| 11 | 117041911 | rs75525407 | T | C | 1.104 | -7977.388 | -7977.516 | -0.128 | 3.40E-06 |
| 11 | 117052757 | rs189196849 | T | G | 1.104 | -7977.388 | -7977.516 | -0.128 | 3.40E-06 |
| 11 | 117054029 | rs59978576 | G | C | 1.104 | -7977.388 | -7977.516 | -0.128 | 3.40E-06 |
| 11 | 117055400 | rs61268522 | C | A | 1.104 | -7977.388 | -7977.516 | -0.128 | 3.40E-06 |
| 11 | 117055893 | rs140727315 | C | G | 1.104 | -7977.388 | -7977.516 | -0.128 | 3.40E-06 |
| 11 | 117061994 | rs184936191 | T | A | 1.104 | -7977.388 | -7977.516 | -0.128 | 3.40E-06 |
| 11 | 117065271 | rs79043792 | C | G | 1.104 | -7977.388 | -7977.516 | -0.128 | 3.40E-06 |
| 11 | 117072970 | rs74967901 | A | G | 1.104 | -7977.388 | -7977.516 | -0.128 | 3.40E-06 |
| 11 | 116987421 | rs56756054 | C | T | 1.104 | -7977.388 | -7977.518 | -0.129 | 3.40E-06 |
| 11 | 119879579 | rs1944495 | T | C | 1.078 | -7977.388 | -7977.517 | -0.129 | 3.40E-06 |
| 11 | 116730833 | rs545274 | T | C | 1.065 | -7977.388 | -7977.519 | -0.13 | 3.40E-06 |
| 11 | 117356858 | rs606727 | G | C | 0.927 | -7977.388 | -7977.52 | -0.131 | 3.39E-06 |
| 11 | 116223780 | rs17475326 | T | A | 1.109 | -7977.388 | -7977.52 | -0.132 | 3.39E-06 |
| 11 | 116754753 | rs57259536 | T | A | 0.898 | -7977.388 | -7977.52 | -0.132 | 3.39E-06 |
| 11 | 116882563 | rs73594141 | T | C | 0.899 | -7977.388 | -7977.521 | -0.133 | 3.39E-06 |
| 11 | 116896441 | rs73594158 | T | C | 0.899 | -7977.388 | -7977.521 | -0.133 | 3.39E-06 |
| 11 | 116896811 | rs73594162 | T | C | 0.899 | -7977.388 | -7977.521 | -0.133 | 3.39E-06 |
| 11 | 116897613 | rs73594164 | G | T | 0.899 | -7977.388 | -7977.521 | -0.133 | 3.39E-06 |
| 11 | 116900956 | rs59647391 | G | A | 0.899 | -7977.388 | -7977.521 | -0.133 | 3.39E-06 |
| 11 | 116902631 | rs73594172 | T | A | 0.899 | -7977.388 | -7977.521 | -0.133 | 3.39E-06 |
| 11 | 117473537 | rs77346448 | C | T | 0.899 | -7977.388 | -7977.523 | -0.135 | 3.38E-06 |
| 11 | 117795537 | rs79136224 | T | C | 0.91 | -7977.388 | -7977.523 | -0.135 | 3.38E-06 |
| 11 | 120841689 | rs652418 | C | T | 0.94 | -7977.388 | -7977.524 | -0.135 | 3.38E-06 |
| 11 | 121189231 | rs10892721 | T | G | 1.076 | -7977.388 | -7977.523 | -0.135 | 3.38E-06 |
| 11 | 120644115 | rs7938295 | A | G | 0.905 | -7977.388 | -7977.524 | -0.136 | 3.38E-06 |
| 11 | 117099025 | rs2306474 | G | A | 1.103 | -7977.388 | -7977.525 | -0.137 | 3.37E-06 |
| 11 | 119936292 | rs640663 | A | G | 0.929 | -7977.388 | -7977.525 | -0.137 | 3.37E-06 |
| 11 | 117563295 | rs11216505 | A | G | 0.926 | -7977.388 | -7977.526 | -0.138 | 3.37E-06 |
| 11 | 117579111 | rs17121102 | C | G | 0.898 | -7977.388 | -7977.526 | -0.138 | 3.37E-06 |
| 11 | 118754649 | rs10892307 | G | C | 0.935 | -7977.388 | -7977.526 | -0.138 | 3.37E-06 |
| 11 | 120569843 | rs61901362 | C | T | 0.93 | -7977.388 | -7977.526 | -0.138 | 3.37E-06 |
| 11 | 118241516 | rs11216861 | A | G | 1.061 | -7977.388 | -7977.527 | -0.139 | 3.37E-06 |
| 11 | 117430058 | rs510250 | G | T | 0.947 | -7977.388 | -7977.528 | -0.14 | 3.36E-06 |
| 11 | 118865992 | rs11217117 | T | C | 1.057 | -7977.388 | -7977.528 | -0.14 | 3.36E-06 |
| 11 | 120838716 | rs608154 | G | A | 0.939 | -7977.388 | -7977.529 | -0.14 | 3.36E-06 |
| 11 | 117034320 | rs150881386 | T | C | 1.103 | -7977.388 | -7977.53 | -0.141 | 3.36E-06 |
| 11 | 117035622 | rs144762883 | T | C | 1.103 | -7977.388 | -7977.53 | -0.141 | 3.36E-06 |
| 11 | 117039208 | rs117842423 | A | G | 1.103 | -7977.388 | -7977.53 | -0.141 | 3.36E-06 |
| 11 | 117061937 | rs117303996 | A | G | 1.103 | -7977.388 | -7977.53 | -0.141 | 3.36E-06 |
| 11 | 117795373 | rs78264252 | A | C | 0.91 | -7977.388 | -7977.53 | -0.141 | 3.36E-06 |
| 11 | 116734545 | rs11216164 | A | G | 0.945 | -7977.388 | -7977.53 | -0.142 | 3.36E-06 |
| 11 | 118807876 | rs12282154 | A | G | 1.077 | -7977.388 | -7977.53 | -0.142 | 3.36E-06 |
| 11 | 120571883 | rs61901370 | T | C | 0.928 | -7977.388 | -7977.53 | -0.142 | 3.36E-06 |
| 11 | 117053686 | rs61562228 | A | G | 1.103 | -7977.388 | -7977.531 | -0.143 | 3.35E-06 |
| 11 | 117794650 | rs11216629 | G | C | 0.91 | -7977.388 | -7977.531 | -0.143 | 3.35E-06 |
| 11 | 119733857 | rs570521 | G | T | 1.088 | -7977.388 | -7977.531 | -0.143 | 3.35E-06 |
| 11 | 117004023 | rs150672190 | T | C | 1.103 | -7977.388 | -7977.532 | -0.144 | 3.35E-06 |
| 11 | 117005207 | rs117146251 | G | A | 1.103 | -7977.388 | -7977.532 | -0.144 | 3.35E-06 |
| 11 | 119352350 | rs61901289 | T | G | 1.101 | -7977.388 | -7977.533 | -0.145 | 3.35E-06 |
| 11 | 116680961 | rs80220270 | C | A | 1.101 | -7977.388 | -7977.535 | -0.146 | 3.34E-06 |
| 11 | 116929666 | rs73576612 | C | T | 0.9 | -7977.388 | -7977.535 | -0.146 | 3.34E-06 |
| 11 | 116932615 | rs17120265 | G | C | 0.9 | -7977.388 | -7977.535 | -0.146 | 3.34E-06 |
| 11 | 116935363 | rs17120271 | G | C | 0.9 | -7977.388 | -7977.535 | -0.146 | 3.34E-06 |
| 11 | 116939071 | rs17120275 | C | T | 0.9 | -7977.388 | -7977.535 | -0.146 | 3.34E-06 |
| 11 | 118856491 | rs3889526 | A | C | 1.056 | -7977.388 | -7977.534 | -0.146 | 3.34E-06 |
| 11 | 118857466 | rs4938610 | C | T | 1.056 | -7977.388 | -7977.534 | -0.146 | 3.34E-06 |
| 11 | 119663944 | rs73006788 | T | C | 1.109 | -7977.388 | -7977.535 | -0.146 | 3.34E-06 |
| 11 | 117344165 | rs11216413 | A | T | 1.066 | -7977.388 | -7977.539 | -0.15 | 3.33E-06 |
| 11 | 117795082 | rs76046563 | G | C | 0.91 | -7977.388 | -7977.538 | -0.15 | 3.33E-06 |
| 11 | 117346121 | rs74594162 | G | C | 1.106 | -7977.388 | -7977.539 | -0.151 | 3.33E-06 |
| 11 | 117349482 | rs12277788 | T | C | 1.065 | -7977.388 | -7977.539 | -0.151 | 3.33E-06 |
| 11 | 117709760 | rs11216576 | A | G | 1.092 | -7977.388 | -7977.539 | -0.151 | 3.33E-06 |
| 11 | 119324268 | rs78040959 | G | A | 0.914 | -7977.388 | -7977.539 | -0.151 | 3.33E-06 |
| 11 | 116963969 | rs79536557 | G | A | 0.901 | -7977.388 | -7977.54 | -0.152 | 3.32E-06 |
| 11 | 118287568 | rs75461585 | G | A | 0.905 | -7977.388 | -7977.54 | -0.152 | 3.32E-06 |
| 11 | 120643090 | rs7934092 | A | C | 0.902 | -7977.388 | -7977.541 | -0.152 | 3.32E-06 |
| 11 | 117246876 | rs2305827 | C | T | 1.06 | -7977.388 | -7977.542 | -0.153 | 3.32E-06 |
| 11 | 117475763 | rs77492882 | T | C | 0.901 | -7977.388 | -7977.541 | -0.153 | 3.32E-06 |
| 11 | 117475976 | rs79198917 | A | G | 0.901 | -7977.388 | -7977.541 | -0.153 | 3.32E-06 |
| 11 | 116910971 | rs12273536 | T | C | 1.091 | -7977.388 | -7977.543 | -0.154 | 3.32E-06 |
| 11 | 117574296 | rs2068412 | G | A | 0.9 | -7977.388 | -7977.543 | -0.155 | 3.31E-06 |

| | | | | | | | |
|---|---|---|---|---|---|---|---|
| 11 | 117794735 | rs11216630 | T | A | NULL | 0.518741216 | NULL&M_Disease |
| 11 | 117795397 | rs77637745 | A | C | NULL | 0.518741216 | NULL&M_Disease |
| 11 | 120797878 | rs11218060 | T | C | NULL | 0.518741216 | NULL&M_Disease |
| 11 | 117030187 | rs75115775 | C | A | NULL | 0.518741216 | NULL&M_Disease |
| 11 | 117032934 | rs59007408 | T | C | NULL | 0.518741216 | NULL&M_Disease |
| 11 | 117329289 | rs113968937 | G | C | NULL | 0.518741216 | NULL&M_Disease |
| 11 | 117055485 | rs57315752 | G | A | NULL | 0.518741216 | NULL&M_Disease |
| 11 | 116987767 | rs60215110 | A | G | NULL | 0.518741216 | NULL&M_Disease |
| 11 | 116989133 | rs186927602 | A | G | NULL | 0.518741216 | NULL&M_Disease |
| 11 | 116990523 | rs118143963 | T | A | NULL | 0.518741216 | NULL&M_Disease |
| 11 | 116992739 | rs79117087 | C | A | NULL | 0.518741216 | NULL&M_Disease |
| 11 | 116993291 | rs117140575 | C | T | NULL | 0.518741216 | NULL&M_Disease |
| 11 | 116999303 | rs59225422 | G | C | NULL | 0.518741216 | NULL&M_Disease |
| 11 | 117000117 | rs142742415 | A | G | NULL | 0.518741216 | NULL&M_Disease |
| 11 | 117001875 | rs77983828 | C | T | NULL | 0.518741216 | NULL&M_Disease |
| 11 | 117564891 | rs12285200 | G | A | NULL | 0.518741216 | NULL&M_Disease |
| 11 | 120577413 | rs12270261 | T | C | NULL | 0.518741216 | NULL&M_Disease |
| 11 | 116980147 | rs79777938 | C | G | NULL | 0.518741216 | NULL&M_Disease |
| 11 | 116782000 | rs12808437 | T | G | NULL | 0.51998934 | NULL&M_Disease |
| 11 | 117335035 | rs116941106 | A | G | NULL | 0.518741216 | NULL&M_Disease |
| 11 | 117696521 | rs118176355 | C | G | NULL | 0.518741216 | NULL&M_Disease |
| 11 | 116854683 | rs147507811 | A | C | NULL | 0.518741216 | NULL&M_Disease |
| 11 | 116955831 | rs77564847 | C | T | NULL | 0.518741216 | NULL&M_Disease |
| 11 | 117084070 | rs76553074 | C | T | NULL | 0.51998934 | NULL&M_Disease |
| 11 | 117496475 | rs10790196 | T | C | NULL | 0.51998934 | NULL&M_Disease |
| 11 | 119816512 | rs668781 | C | T | NULL | 0.51998934 | NULL&M_Disease |
| 11 | 117420356 | rs609353 | T | C | NULL | 0.51998934 | NULL&M_Disease |
| 11 | 117498314 | rs4936390 | A | G | NULL | 0.518741216 | NULL&M_Disease |
| 11 | 117498656 | rs4936392 | T | G | NULL | 0.518741216 | NULL&M_Disease |
| 11 | 117499830 | rs78696556 | C | T | NULL | 0.518741216 | NULL&M_Disease |
| 11 | 118072648 | rs78270838 | G | A | NULL | 0.51998934 | NULL&M_Disease |
| 11 | 117032641 | rs60928822 | C | T | NULL | 0.51998934 | NULL&M_Disease |
| 11 | 117795232 | rs80085590 | A | G | NULL | 0.51998934 | NULL&M_Disease |
| 11 | 117017839 | rs145384144 | C | G | NULL | 0.51998934 | NULL&M_Disease |
| 11 | 117408077 | rs557053 | T | C | NULL | 0.521237215 | NULL&M_Disease |
| 11 | 117581866 | rs17121106 | C | T | NULL | 0.51998934 | NULL&M_Disease |
| 11 | 117903942 | rs11216713 | A | G | NULL | 0.51998934 | NULL&M_Disease |
| 11 | 120459320 | rs4936521 | G | A | NULL | 0.521237215 | NULL&M_Disease |
| 11 | 116866609 | rs73594128 | C | T | NULL | 0.51998934 | NULL&M_Disease |
| 11 | 116869827 | rs10502223 | C | T | NULL | 0.51998934 | NULL&M_Disease |
| 11 | 116954760 | rs80137897 | C | T | NULL | 0.51998934 | NULL&M_Disease |
| 11 | 120590901 | rs61901400 | A | G | NULL | 0.521237215 | NULL&M_Disease |
| 11 | 119756941 | rs11217554 | C | G | NULL | 0.521237215 | NULL&M_Disease |
| 11 | 119938994 | rs591596 | A | G | NULL | 0.521237215 | NULL&M_Disease |
| 11 | 120624525 | rs2156763 | G | A | NULL | 0.51998934 | NULL&M_Disease |
| 11 | 117346653 | rs78064538 | C | G | NULL | 0.51998934 | NULL&M_Disease |
| 11 | 120450358 | rs12273031 | A | G | NULL | 0.51998934 | NULL&M_Disease |
| 11 | 116978627 | rs116943892 | G | A | NULL | 0.521237215 | NULL&M_Disease |
| 11 | 117584155 | rs79876834 | T | C | NULL | 0.51998934 | NULL&M_Disease |
| 11 | 120700147 | rs6589846 | G | T | NULL | 0.521237215 | NULL&M_Disease |
| 11 | 116908457 | rs7118591 | A | C | NULL | 0.521237215 | NULL&M_Disease |
| 11 | 117082109 | rs12417682 | A | G | NULL | 0.521237215 | NULL&M_Disease |
| 11 | 120569884 | rs61901364 | T | C | NULL | 0.521237215 | NULL&M_Disease |
| 11 | 120570198 | rs12278795 | G | A | NULL | 0.521237215 | NULL&M_Disease |
| 11 | 116823972 | rs186331126 | G | T | NULL | 0.51998934 | NULL&M_Disease |
| 11 | 117906282 | rs117083922 | A | G | NULL | 0.521237215 | NULL&M_Disease |
| 11 | 116220390 | rs76406762 | A | C | NULL | 0.521237215 | NULL&M_Disease |
| 11 | 116283172 | rs488485 | T | C | NULL | 0.522484825 | NULL&M_Disease |
| 11 | 116528047 | rs1482502236 | A | T | NULL | 0.521237215 | NULL&M_Disease |
| 11 | 116737093 | rs548638 | G | T | NULL | 0.522484825 | NULL&M_Disease |
| 11 | 117496666 | rs117870205 | A | G | NULL | 0.521237215 | NULL&M_Disease |
| 11 | 117496750 | rs147793723 | C | T | NULL | 0.521237215 | NULL&M_Disease |
| 11 | 117497996 | rs74681797 | T | C | NULL | 0.521237215 | NULL&M_Disease |
| 11 | 117498920 | rs117041834 | A | T | NULL | 0.521237215 | NULL&M_Disease |
| 11 | 117499828 | rs76205818 | A | T | NULL | 0.521237215 | NULL&M_Disease |
| 11 | 120735646 | rs74651412 | A | G | NULL | 0.522484825 | NULL&M_Disease |
| 11 | 116906728 | rs4938327 | C | T | NULL | 0.522484825 | NULL&M_Disease |
| 11 | 116907043 | rs3181 | A | G | NULL | 0.522484825 | NULL&M_Disease |
| 11 | 117707191 | rs10790214 | T | C | NULL | 0.522484825 | NULL&M_Disease |
| 11 | 120568090 | rs12295200 | C | T | NULL | 0.522484825 | NULL&M_Disease |
| 11 | 120568153 | rs12288261 | T | C | NULL | 0.522484825 | NULL&M_Disease |
| 11 | 120568248 | rs118056960 | T | C | NULL | 0.522484825 | NULL&M_Disease |
| 11 | 120568318 | rs78634274 | C | T | NULL | 0.522484825 | NULL&M_Disease |
| 11 | 120568790 | rs61901360 | T | C | NULL | 0.522484825 | NULL&M_Disease |
| 11 | 116815769 | rs649084 | A | G | NULL | 0.521237215 | NULL&M_Disease |
| 11 | 116817785 | rs113712143 | A | G | NULL | 0.521237215 | NULL&M_Disease |
| 11 | 116854580 | rs2441218 | G | T | NULL | 0.521237215 | NULL&M_Disease |
| 11 | 116856179 | rs537431 | C | T | NULL | 0.521237215 | NULL&M_Disease |
| 11 | 116857400 | rs569568 | C | T | NULL | 0.521237215 | NULL&M_Disease |
| 11 | 116860456 | rs638392 | A | T | NULL | 0.521237215 | NULL&M_Disease |
| 11 | 116861519 | rs623423 | C | G | NULL | 0.521237215 | NULL&M_Disease |
| 11 | 116862087 | rs560897 | T | C | NULL | 0.521237215 | NULL&M_Disease |
| 11 | 116862718 | rs607756 | C | T | NULL | 0.521237215 | NULL&M_Disease |
| 11 | 116886096 | rs7119185 | T | C | NULL | 0.522484825 | NULL&M_Disease |
| 11 | 117341574 | rs79865588 | C | T | NULL | 0.521237215 | NULL&M_Disease |
| 11 | 117815564 | rs2512139 | A | G | NULL | 0.522484825 | NULL&M_Disease |
| 11 | 116693213 | rs5094 | A | G | NULL | 0.521237215 | NULL&M_Disease |
| 11 | 120570080 | rs12291927 | G | C | NULL | 0.522484825 | NULL&M_Disease |
| 11 | 120571553 | rs61901368 | G | A | NULL | 0.522484825 | NULL&M_Disease |
| 11 | 116722041 | rs10047462 | G | T | NULL | 0.522484825 | NULL&M_Disease |
| 11 | 119677072 | rs895648 | C | T | NULL | 0.522484825 | NULL&M_Disease |
| 11 | 116886516 | rs145133311 | T | C | NULL | 0.522484825 | NULL&M_Disease |
| 11 | 120570404 | rs12293393 | G | C | NULL | 0.522484825 | NULL&M_Disease |
| 11 | 120733435 | rs3901285 | T | C | NULL | 0.522484825 | NULL&M_Disease |
| 11 | 119797863 | rs542748 | C | G | NULL | 0.523732154 | NULL&M_Disease |
| 11 | 120627141 | rs1945009 | A | G | NULL | 0.522484825 | NULL&M_Disease |

| | | | | | | | | | |
|---|---|---|---|---|---|---|---|---|---|
| 11 | 117794735 | rs11216630 | T | A | 0.911 | -7977.388 | -7977.543 | -0.155 | 3.31E-06 |
| 11 | 117795397 | rs77637745 | A | C | 0.911 | -7977.388 | -7977.544 | -0.155 | 3.31E-06 |
| 11 | 120797878 | rs11218060 | T | C | 0.931 | -7977.388 | -7977.543 | -0.155 | 3.31E-06 |
| 11 | 117030187 | rs75115775 | C | A | 1.102 | -7977.388 | -7977.544 | -0.156 | 3.31E-06 |
| 11 | 117032934 | rs59007408 | T | C | 1.102 | -7977.388 | -7977.544 | -0.156 | 3.31E-06 |
| 11 | 117329289 | rs113968937 | G | C | 1.105 | -7977.388 | -7977.544 | -0.156 | 3.31E-06 |
| 11 | 117055485 | rs57315752 | G | A | 1.102 | -7977.388 | -7977.546 | -0.157 | 3.31E-06 |
| 11 | 116987767 | rs60215110 | A | G | 1.102 | -7977.388 | -7977.547 | -0.158 | 3.30E-06 |
| 11 | 116989133 | rs186927602 | A | G | 1.102 | -7977.388 | -7977.547 | -0.158 | 3.30E-06 |
| 11 | 116990523 | rs118143963 | T | A | 1.102 | -7977.388 | -7977.547 | -0.158 | 3.30E-06 |
| 11 | 116992739 | rs79117087 | C | A | 1.102 | -7977.388 | -7977.547 | -0.158 | 3.30E-06 |
| 11 | 116993291 | rs117140575 | C | T | 1.102 | -7977.388 | -7977.547 | -0.158 | 3.30E-06 |
| 11 | 116999303 | rs59225422 | G | C | 1.102 | -7977.388 | -7977.547 | -0.158 | 3.30E-06 |
| 11 | 117000117 | rs142742415 | A | G | 1.102 | -7977.388 | -7977.547 | -0.158 | 3.30E-06 |
| 11 | 117001875 | rs77983828 | C | T | 1.102 | -7977.388 | -7977.547 | -0.158 | 3.30E-06 |
| 11 | 117564891 | rs12285200 | G | A | 0.926 | -7977.388 | -7977.546 | -0.158 | 3.30E-06 |
| 11 | 120577413 | rs12270261 | T | C | 0.928 | -7977.388 | -7977.546 | -0.158 | 3.30E-06 |
| 11 | 116980147 | rs79777938 | C | G | 1.102 | -7977.388 | -7977.547 | -0.159 | 3.30E-06 |
| 11 | 116782000 | rs12808437 | T | G | 1.064 | -7977.388 | -7977.55 | -0.161 | 3.29E-06 |
| 11 | 117335035 | rs116941106 | A | G | 1.109 | -7977.388 | -7977.549 | -0.161 | 3.29E-06 |
| 11 | 117696521 | rs118176355 | C | G | 0.9 | -7977.388 | -7977.549 | -0.161 | 3.29E-06 |
| 11 | 116854683 | rs147507811 | A | C | 0.904 | -7977.388 | -7977.55 | -0.162 | 3.29E-06 |
| 11 | 116955831 | rs77564847 | C | T | 0.901 | -7977.388 | -7977.551 | -0.163 | 3.29E-06 |
| 11 | 117084070 | rs76553074 | C | T | 1.101 | -7977.388 | -7977.552 | -0.163 | 3.29E-06 |
| 11 | 117496475 | rs10790196 | T | C | 0.942 | -7977.388 | -7977.551 | -0.163 | 3.29E-06 |
| 11 | 119816512 | rs668781 | C | T | 0.942 | -7977.388 | -7977.552 | -0.163 | 3.29E-06 |
| 11 | 117420356 | rs609353 | T | C | 1.059 | -7977.388 | -7977.552 | -0.164 | 3.28E-06 |
| 11 | 117498314 | rs4936390 | A | G | 0.902 | -7977.388 | -7977.552 | -0.164 | 3.28E-06 |
| 11 | 117498656 | rs4936392 | T | G | 0.902 | -7977.388 | -7977.552 | -0.164 | 3.28E-06 |
| 11 | 117499830 | rs78696556 | C | T | 0.902 | -7977.388 | -7977.552 | -0.164 | 3.28E-06 |
| 11 | 118072648 | rs78270838 | G | A | 0.906 | -7977.388 | -7977.552 | -0.164 | 3.28E-06 |
| 11 | 117032641 | rs60928822 | C | T | 1.101 | -7977.388 | -7977.553 | -0.165 | 3.28E-06 |
| 11 | 117795232 | rs80085590 | A | G | 0.911 | -7977.388 | -7977.555 | -0.166 | 3.28E-06 |
| 11 | 117017839 | rs145384144 | C | G | 1.096 | -7977.388 | -7977.556 | -0.168 | 3.27E-06 |
| 11 | 117408077 | rs557053 | T | C | 1.057 | -7977.388 | -7977.557 | -0.169 | 3.27E-06 |
| 11 | 117581866 | rs17121106 | C | T | 0.901 | -7977.388 | -7977.557 | -0.169 | 3.27E-06 |
| 11 | 117903942 | rs11216713 | A | G | 0.904 | -7977.388 | -7977.557 | -0.169 | 3.27E-06 |
| 11 | 120459320 | rs4936521 | G | A | 1.056 | -7977.388 | -7977.557 | -0.169 | 3.27E-06 |
| 11 | 116866609 | rs73594128 | C | T | 0.902 | -7977.388 | -7977.559 | -0.17 | 3.26E-06 |
| 11 | 116869827 | rs10502223 | C | T | 0.902 | -7977.388 | -7977.559 | -0.17 | 3.26E-06 |
| 11 | 116954760 | rs80137897 | C | T | 0.902 | -7977.388 | -7977.558 | -0.17 | 3.26E-06 |
| 11 | 120590901 | rs61901400 | A | G | 0.922 | -7977.388 | -7977.558 | -0.17 | 3.26E-06 |
| 11 | 119756941 | rs11217554 | C | G | 0.944 | -7977.388 | -7977.559 | -0.171 | 3.26E-06 |
| 11 | 119938994 | rs591596 | A | G | 0.93 | -7977.388 | -7977.559 | -0.171 | 3.26E-06 |
| 11 | 120624525 | rs2156763 | G | A | 0.904 | -7977.388 | -7977.559 | -0.171 | 3.26E-06 |
| 11 | 117346653 | rs78064538 | C | G | 1.105 | -7977.388 | -7977.56 | -0.172 | 3.26E-06 |
| 11 | 120450358 | rs12273031 | A | G | 1.108 | -7977.388 | -7977.56 | -0.172 | 3.26E-06 |
| 11 | 116978627 | rs116943892 | G | A | 1.101 | -7977.388 | -7977.561 | -0.173 | 3.25E-06 |
| 11 | 117584155 | rs79876834 | T | C | 0.901 | -7977.388 | -7977.561 | -0.173 | 3.25E-06 |
| 11 | 120700147 | rs6589846 | G | T | 1.057 | -7977.388 | -7977.562 | -0.173 | 3.25E-06 |
| 11 | 116908457 | rs7118591 | A | C | 1.064 | -7977.388 | -7977.562 | -0.174 | 3.25E-06 |
| 11 | 117082109 | rs12417682 | A | G | 1.09 | -7977.388 | -7977.564 | -0.175 | 3.25E-06 |
| 11 | 120569884 | rs61901364 | T | C | 0.929 | -7977.388 | -7977.563 | -0.175 | 3.25E-06 |
| 11 | 120570198 | rs12278795 | G | A | 0.929 | -7977.388 | -7977.563 | -0.175 | 3.25E-06 |
| 11 | 116823972 | rs186331126 | G | T | 1.107 | -7977.388 | -7977.566 | -0.178 | 3.24E-06 |
| 11 | 117906282 | rs117083922 | A | G | 0.904 | -7977.388 | -7977.566 | -0.178 | 3.24E-06 |
| 11 | 116220390 | rs76406762 | A | C | 1.107 | -7977.388 | -7977.568 | -0.18 | 3.23E-06 |
| 11 | 116283172 | rs488485 | T | C | 0.934 | -7977.388 | -7977.568 | -0.18 | 3.23E-06 |
| 11 | 116528047 | rs1482502236 | A | T | 0.91 | -7977.388 | -7977.57 | -0.181 | 3.23E-06 |
| 11 | 116737093 | rs548638 | G | T | 1.064 | -7977.388 | -7977.569 | -0.181 | 3.23E-06 |
| 11 | 117496666 | rs117870205 | A | G | 0.904 | -7977.388 | -7977.569 | -0.181 | 3.23E-06 |
| 11 | 117496750 | rs147793723 | C | T | 0.904 | -7977.388 | -7977.569 | -0.181 | 3.23E-06 |
| 11 | 117497996 | rs74681797 | T | C | 0.904 | -7977.388 | -7977.569 | -0.181 | 3.23E-06 |
| 11 | 117498920 | rs117041834 | A | T | 0.904 | -7977.388 | -7977.569 | -0.181 | 3.23E-06 |
| 11 | 117499828 | rs76205818 | A | T | 0.904 | -7977.388 | -7977.569 | -0.181 | 3.23E-06 |
| 11 | 120735646 | rs74651412 | A | G | 1.087 | -7977.388 | -7977.57 | -0.181 | 3.23E-06 |
| 11 | 116906728 | rs4938327 | C | T | 1.063 | -7977.388 | -7977.57 | -0.182 | 3.22E-06 |
| 11 | 116907043 | rs3181 | A | G | 1.063 | -7977.388 | -7977.57 | -0.182 | 3.22E-06 |
| 11 | 117707191 | rs10790214 | T | C | 0.947 | -7977.388 | -7977.571 | -0.182 | 3.22E-06 |
| 11 | 120568090 | rs12295200 | C | T | 0.929 | -7977.388 | -7977.572 | -0.183 | 3.22E-06 |
| 11 | 120568153 | rs12288261 | T | C | 0.929 | -7977.388 | -7977.572 | -0.183 | 3.22E-06 |
| 11 | 120568248 | rs118056960 | T | C | 0.929 | -7977.388 | -7977.572 | -0.183 | 3.22E-06 |
| 11 | 120568318 | rs78634274 | C | T | 0.929 | -7977.388 | -7977.572 | -0.183 | 3.22E-06 |
| 11 | 120568790 | rs61901360 | T | C | 0.929 | -7977.388 | -7977.572 | -0.183 | 3.22E-06 |
| 11 | 116815769 | rs649084 | A | G | 0.903 | -7977.388 | -7977.572 | -0.184 | 3.22E-06 |
| 11 | 116817785 | rs113712143 | A | G | 0.903 | -7977.388 | -7977.572 | -0.184 | 3.22E-06 |
| 11 | 116854580 | rs2441218 | G | T | 0.903 | -7977.388 | -7977.572 | -0.184 | 3.22E-06 |
| 11 | 116856179 | rs537431 | C | T | 0.903 | -7977.388 | -7977.572 | -0.184 | 3.22E-06 |
| 11 | 116857400 | rs569568 | C | T | 0.903 | -7977.388 | -7977.572 | -0.184 | 3.22E-06 |
| 11 | 116860456 | rs638392 | A | T | 0.903 | -7977.388 | -7977.572 | -0.184 | 3.22E-06 |
| 11 | 116861519 | rs623423 | C | G | 0.903 | -7977.388 | -7977.572 | -0.184 | 3.22E-06 |
| 11 | 116862087 | rs560897 | T | C | 0.903 | -7977.388 | -7977.572 | -0.184 | 3.22E-06 |
| 11 | 116862718 | rs607756 | C | T | 0.903 | -7977.388 | -7977.572 | -0.184 | 3.22E-06 |
| 11 | 116886096 | rs7119185 | T | C | 1.063 | -7977.388 | -7977.572 | -0.184 | 3.22E-06 |
| 11 | 117341574 | rs79865588 | C | T | 1.106 | -7977.388 | -7977.572 | -0.184 | 3.22E-06 |
| 11 | 117815564 | rs2512139 | A | G | 0.946 | -7977.388 | -7977.572 | -0.184 | 3.22E-06 |
| 11 | 116693213 | rs5094 | A | G | 0.903 | -7977.388 | -7977.573 | -0.185 | 3.21E-06 |
| 11 | 120570080 | rs12291927 | G | C | 0.929 | -7977.388 | -7977.573 | -0.185 | 3.21E-06 |
| 11 | 120571553 | rs61901368 | G | A | 0.931 | -7977.388 | -7977.574 | -0.185 | 3.21E-06 |
| 11 | 116722041 | rs10047462 | G | T | 0.928 | -7977.388 | -7977.575 | -0.186 | 3.21E-06 |
| 11 | 119677072 | rs895648 | C | T | 0.947 | -7977.388 | -7977.575 | -0.186 | 3.21E-06 |
| 11 | 116886516 | rs145133311 | T | C | 0.903 | -7977.388 | -7977.576 | -0.187 | 3.21E-06 |
| 11 | 120570404 | rs12293393 | G | C | 0.929 | -7977.388 | -7977.575 | -0.187 | 3.21E-06 |
| 11 | 120733435 | rs3901285 | T | C | 1.062 | -7977.388 | -7977.575 | -0.187 | 3.21E-06 |
| 11 | 119797863 | rs542748 | C | G | 0.942 | -7977.388 | -7977.576 | -0.188 | 3.21E-06 |
| 11 | 120627141 | rs1945009 | A | G | 0.905 | -7977.388 | -7977.576 | -0.188 | 3.21E-06 |

| | | | | | | | |
|---|---|---|---|---|---|---|---|
| 11 | 117672561 | rs10892173 | T | C | NULL | 0.523732154 | NULL&M_Disease |
| 11 | 116807970 | rs473172 | A | G | NULL | 0.523732154 | NULL&M_Disease |
| 11 | 117062158 | rs139685157 | A | G | NULL | 0.522484825 | NULL&M_Disease |
| 11 | 116710239 | rs632153 | T | G | NULL | 0.522484825 | NULL&M_Disease |
| 11 | 117795189 | rs75195166 | A | G | NULL | 0.523732154 | NULL&M_Disease |
| 11 | 120569355 | rs61901361 | G | A | NULL | 0.523732154 | NULL&M_Disease |
| 11 | 116678191 | rs61905133 | G | A | NULL | 0.523732154 | NULL&M_Disease |
| 11 | 117303108 | rs76347292 | C | T | NULL | 0.523732154 | NULL&M_Disease |
| 11 | 118852081 | rs4938606 | G | A | NULL | 0.523732154 | NULL&M_Disease |
| 11 | 118858878 | rs188758920 | A | C | NULL | 0.522484825 | NULL&M_Disease |
| 11 | 116271393 | rs191301146 | T | C | NULL | 0.522484825 | NULL&M_Disease |
| 11 | 117424005 | rs576390 | A | G | NULL | 0.523732154 | NULL&M_Disease |
| 11 | 118028648 | rs73014974 | T | C | NULL | 0.523732154 | NULL&M_Disease |
| 11 | 117904550 | rs11216714 | A | G | NULL | 0.523732154 | NULL&M_Disease |
| 11 | 117904710 | rs12223224 | T | C | NULL | 0.523732154 | NULL&M_Disease |
| 11 | 117904793 | rs12226036 | G | A | NULL | 0.523732154 | NULL&M_Disease |
| 11 | 117905217 | rs11216715 | T | A | NULL | 0.523732154 | NULL&M_Disease |
| 11 | 117905394 | rs11216716 | A | G | NULL | 0.523732154 | NULL&M_Disease |
| 11 | 117905495 | rs80001712 | T | C | NULL | 0.523732154 | NULL&M_Disease |
| 11 | 117905654 | rs12223683 | A | G | NULL | 0.523732154 | NULL&M_Disease |
| 11 | 117905786 | rs79075579 | T | C | NULL | 0.523732154 | NULL&M_Disease |
| 11 | 116598315 | rs141414463 | T | C | NULL | 0.523732154 | NULL&M_Disease |
| 11 | 116774447 | rs34144542 | G | A | NULL | 0.523732154 | NULL&M_Disease |
| 11 | 117032959 | rs12281009 | G | A | NULL | 0.523732154 | NULL&M_Disease |
| 11 | 117795283 | rs74391474 | C | G | NULL | 0.523732154 | NULL&M_Disease |
| 11 | 117795285 | rs74578710 | A | G | NULL | 0.523732154 | NULL&M_Disease |
| 11 | 116818187 | rs3133352 | A | G | NULL | 0.523732154 | NULL&M_Disease |
| 11 | 116819991 | rs1261706 | A | T | NULL | 0.523732154 | NULL&M_Disease |
| 11 | 116827565 | rs587703 | G | A | NULL | 0.523732154 | NULL&M_Disease |
| 11 | 116844279 | rs2513092 | C | T | NULL | 0.523732154 | NULL&M_Disease |
| 11 | 120456533 | rs11602493 | A | G | NULL | 0.524979187 | NULL&M_Disease |
| 11 | 117424306 | rs573648 | A | G | NULL | 0.523732154 | NULL&M_Disease |
| 11 | 116788699 | rs10750099 | A | C | NULL | 0.523732154 | NULL&M_Disease |
| 11 | 116789635 | rs6589579 | C | A | NULL | 0.523732154 | NULL&M_Disease |
| 11 | 117084396 | rs12418705 | T | C | NULL | 0.524979187 | NULL&M_Disease |
| 11 | 117495049 | rs6589616 | T | C | NULL | 0.524979187 | NULL&M_Disease |
| 11 | 117617379 | rs1893038 | C | T | NULL | 0.523732154 | NULL&M_Disease |
| 11 | 117495305 | rs77314777 | A | G | NULL | 0.523732154 | NULL&M_Disease |
| 11 | 117594443 | rs7123597 | A | C | NULL | 0.523732154 | NULL&M_Disease |
| 11 | 117161006 | rs17120560 | G | C | NULL | 0.524979187 | NULL&M_Disease |
| 11 | 117464712 | rs4938414 | A | G | NULL | 0.524979187 | NULL&M_Disease |
| 11 | 118899218 | rs4936459 | T | C | NULL | 0.524979187 | NULL&M_Disease |
| 11 | 120571314 | rs74447718 | T | C | NULL | 0.524979187 | NULL&M_Disease |
| 11 | 116660450 | rs33984246 | G | A | NULL | 0.524979187 | NULL&M_Disease |
| 11 | 117219961 | rs477858 | G | A | NULL | 0.524979187 | NULL&M_Disease |
| 11 | 117974397 | rs117950757 | A | G | NULL | 0.524979187 | NULL&M_Disease |
| 11 | 116760485 | rs117709132 | T | C | NULL | 0.524979187 | NULL&M_Disease |
| 11 | 117653955 | rs7119911 | T | G | NULL | 0.524979187 | NULL&M_Disease |
| 11 | 120570786 | rs61901366 | A | G | NULL | 0.524979187 | NULL&M_Disease |
| 11 | 117342536 | rs114831648 | C | T | NULL | 0.524979187 | NULL&M_Disease |
| 11 | 117451536 | rs684973 | C | G | NULL | 0.526225909 | NULL&M_Disease |
| 11 | 117430840 | rs55741146 | G | C | NULL | 0.526225909 | NULL&M_Disease |
| 11 | 116880018 | rs150734126 | C | T | NULL | 0.524979187 | NULL&M_Disease |
| 11 | 117175658 | rs116987336 | A | G | NULL | 0.524979187 | NULL&M_Disease |
| 11 | 117362331 | rs149861300 | C | A | NULL | 0.524979187 | NULL&M_Disease |
| 11 | 117906289 | rs74582224 | T | G | NULL | 0.524979187 | NULL&M_Disease |
| 11 | 117906406 | rs79791963 | G | C | NULL | 0.524979187 | NULL&M_Disease |
| 11 | 117906408 | rs78197730 | T | C | NULL | 0.524979187 | NULL&M_Disease |
| 11 | 120925525 | rs11218127 | G | A | NULL | 0.526225909 | NULL&M_Disease |
| 11 | 117582709 | rs79459688 | G | A | NULL | 0.524979187 | NULL&M_Disease |
| 11 | 120858815 | rs11218094 | T | G | NULL | 0.526225909 | NULL&M_Disease |
| 11 | 116831071 | rs622302 | G | A | NULL | 0.524979187 | NULL&M_Disease |
| 11 | 116833669 | rs493101 | A | G | NULL | 0.524979187 | NULL&M_Disease |
| 11 | 116835362 | rs541496 | T | C | NULL | 0.524979187 | NULL&M_Disease |
| 11 | 116835883 | rs667119 | T | A | NULL | 0.524979187 | NULL&M_Disease |
| 11 | 116836548 | rs2921655 | C | G | NULL | 0.524979187 | NULL&M_Disease |
| 11 | 116845538 | rs2513091 | G | A | NULL | 0.524979187 | NULL&M_Disease |
| 11 | 117518038 | rs626571 | C | T | NULL | 0.526225909 | NULL&M_Disease |
| 11 | 116640604 | rs1298336202 | T | C | NULL | 0.524979187 | NULL&M_Disease |
| 11 | 117566303 | rs10892163 | A | G | NULL | 0.526225909 | NULL&M_Disease |
| 11 | 117569923 | rs17121096 | C | T | NULL | 0.526225909 | NULL&M_Disease |
| 11 | 117611938 | rs77719730 | A | G | NULL | 0.526225909 | NULL&M_Disease |
| 11 | 118863232 | rs4938614 | A | G | NULL | 0.526225909 | NULL&M_Disease |
| 11 | 120570901 | rs61901367 | A | G | NULL | 0.526225909 | NULL&M_Disease |
| 11 | 120585020 | rs7120727 | C | G | NULL | 0.526225909 | NULL&M_Disease |
| 11 | 117101311 | rs1231238253 | T | C | NULL | 0.526225909 | NULL&M_Disease |
| 11 | 116909994 | rs73594175 | G | A | NULL | 0.526225909 | NULL&M_Disease |
| 11 | 116910320 | rs1475614837 | T | G | NULL | 0.526225909 | NULL&M_Disease |
| 11 | 116910427 | rs73594181 | A | T | NULL | 0.526225909 | NULL&M_Disease |
| 11 | 116910454 | rs79340485 | T | C | NULL | 0.526225909 | NULL&M_Disease |
| 11 | 120571229 | rs149505626 | A | G | NULL | 0.526225909 | NULL&M_Disease |
| 11 | 118099735 | rs149668068 | A | G | NULL | 0.526225909 | NULL&M_Disease |
| 11 | 120509335 | rs10892589 | T | C | NULL | 0.527472304 | NULL&M_Disease |
| 11 | 116601062 | rs148945243 | T | C | NULL | 0.526225909 | NULL&M_Disease |
| 11 | 116828484 | rs497653 | C | T | NULL | 0.526225909 | NULL&M_Disease |
| 11 | 117062489 | rs3213459 | C | T | NULL | 0.527472304 | NULL&M_Disease |
| 11 | 116578217 | rs12221682 | G | C | NULL | 0.527472304 | NULL&M_Disease |
| 11 | 119939927 | rs592565 | A | G | NULL | 0.527472304 | NULL&M_Disease |
| 11 | 116847584 | rs3016611 | A | G | NULL | 0.526225909 | NULL&M_Disease |
| 11 | 117564052 | rs11216506 | G | A | NULL | 0.527472304 | NULL&M_Disease |
| 11 | 117467394 | rs2186761 | A | G | NULL | 0.527472304 | NULL&M_Disease |
| 11 | 121011507 | rs638496 | T | C | NULL | 0.527472304 | NULL&M_Disease |
| 11 | 116538407 | rs57258844 | C | G | NULL | 0.527472304 | NULL&M_Disease |
| 11 | 117407435 | rs17120841 | G | A | NULL | 0.526225909 | NULL&M_Disease |
| 11 | 117409971 | rs74735117 | C | T | NULL | 0.526225909 | NULL&M_Disease |
| 11 | 120647015 | rs58174358 | T | C | NULL | 0.527472304 | NULL&M_Disease |

| | | | | | | | | | |
|---|---|---|---|---|---|---|---|---|---|
| 11 | 117672561 | rs10892173 | T | C | 0.947 | -7977.388 | -7977.577 | -0.189 | 3.20E-06 |
| 11 | 116807970 | rs473172 | A | G | 0.933 | -7977.388 | -7977.578 | -0.19 | 3.20E-06 |
| 11 | 117062158 | rs139685157 | A | G | 1.1 | -7977.388 | -7977.578 | -0.19 | 3.20E-06 |
| 11 | 116710239 | rs632153 | T | G | 0.904 | -7977.388 | -7977.579 | -0.191 | 3.20E-06 |
| 11 | 117795189 | rs75195166 | A | G | 0.912 | -7977.388 | -7977.58 | -0.192 | 3.19E-06 |
| 11 | 120569355 | rs61901361 | G | A | 0.93 | -7977.388 | -7977.581 | -0.192 | 3.19E-06 |
| 11 | 116678191 | rs61905133 | G | A | 1.098 | -7977.388 | -7977.581 | -0.193 | 3.19E-06 |
| 11 | 117303108 | rs76347292 | C | T | 0.919 | -7977.388 | -7977.581 | -0.193 | 3.19E-06 |
| 11 | 118852081 | rs4938606 | G | A | 1.056 | -7977.388 | -7977.582 | -0.193 | 3.19E-06 |
| 11 | 118858878 | rs188758920 | A | C | 1.105 | -7977.388 | -7977.581 | -0.193 | 3.19E-06 |
| 11 | 116271393 | rs191301146 | T | C | 0.903 | -7977.388 | -7977.583 | -0.194 | 3.19E-06 |
| 11 | 117424005 | rs576390 | A | G | 1.058 | -7977.388 | -7977.582 | -0.194 | 3.19E-06 |
| 11 | 118028648 | rs73014974 | T | C | 0.906 | -7977.388 | -7977.582 | -0.194 | 3.19E-06 |
| 11 | 117904550 | rs11216714 | A | G | 0.906 | -7977.388 | -7977.584 | -0.195 | 3.18E-06 |
| 11 | 117904710 | rs12223224 | T | C | 0.906 | -7977.388 | -7977.584 | -0.195 | 3.18E-06 |
| 11 | 117904793 | rs12226036 | G | A | 0.906 | -7977.388 | -7977.584 | -0.195 | 3.18E-06 |
| 11 | 117905217 | rs11216715 | T | A | 0.906 | -7977.388 | -7977.584 | -0.195 | 3.18E-06 |
| 11 | 117905394 | rs11216716 | A | G | 0.906 | -7977.388 | -7977.584 | -0.195 | 3.18E-06 |
| 11 | 117905495 | rs80001712 | T | C | 0.906 | -7977.388 | -7977.584 | -0.195 | 3.18E-06 |
| 11 | 117905654 | rs12223683 | A | G | 0.906 | -7977.388 | -7977.584 | -0.195 | 3.18E-06 |
| 11 | 117905786 | rs79075579 | T | C | 0.906 | -7977.388 | -7977.584 | -0.195 | 3.18E-06 |
| 11 | 116598315 | rs141414463 | T | C | 0.906 | -7977.388 | -7977.585 | -0.197 | 3.18E-06 |
| 11 | 116774447 | rs34144542 | G | A | 1.084 | -7977.388 | -7977.585 | -0.197 | 3.18E-06 |
| 11 | 117032959 | rs12281009 | G | A | 1.089 | -7977.388 | -7977.585 | -0.197 | 3.18E-06 |
| 11 | 117795283 | rs74391474 | C | G | 0.913 | -7977.388 | -7977.586 | -0.197 | 3.18E-06 |
| 11 | 117795285 | rs74578710 | A | G | 0.913 | -7977.388 | -7977.586 | -0.197 | 3.18E-06 |
| 11 | 116818187 | rs3133352 | A | G | 0.904 | -7977.388 | -7977.586 | -0.198 | 3.17E-06 |
| 11 | 116819991 | rs1261706 | A | T | 0.904 | -7977.388 | -7977.586 | -0.198 | 3.17E-06 |
| 11 | 116827565 | rs587703 | G | A | 0.904 | -7977.388 | -7977.586 | -0.198 | 3.17E-06 |
| 11 | 116844279 | rs2513092 | C | T | 0.904 | -7977.388 | -7977.588 | -0.199 | 3.17E-06 |
| 11 | 120456533 | rs11602493 | A | G | 1.056 | -7977.388 | -7977.587 | -0.199 | 3.17E-06 |
| 11 | 117424306 | rs573648 | A | G | 1.058 | -7977.388 | -7977.589 | -0.2 | 3.17E-06 |
| 11 | 116788699 | rs10750099 | A | C | 0.905 | -7977.388 | -7977.59 | -0.202 | 3.16E-06 |
| 11 | 116789635 | rs6589579 | C | A | 0.905 | -7977.388 | -7977.59 | -0.202 | 3.16E-06 |
| 11 | 117084396 | rs12418705 | T | C | 1.089 | -7977.388 | -7977.59 | -0.202 | 3.16E-06 |
| 11 | 117495049 | rs6589616 | T | C | 0.943 | -7977.388 | -7977.59 | -0.202 | 3.16E-06 |
| 11 | 117617379 | rs1893038 | C | T | 0.904 | -7977.388 | -7977.591 | -0.202 | 3.16E-06 |
| 11 | 117495305 | rs77314777 | A | G | 0.905 | -7977.388 | -7977.592 | -0.203 | 3.16E-06 |
| 11 | 117594443 | rs7123597 | A | C | 0.904 | -7977.388 | -7977.591 | -0.203 | 3.16E-06 |
| 11 | 117161006 | rs17120560 | G | C | 0.91 | -7977.388 | -7977.592 | -0.204 | 3.15E-06 |
| 11 | 117464712 | rs4938414 | A | G | 1.058 | -7977.388 | -7977.592 | -0.204 | 3.15E-06 |
| 11 | 118899218 | rs4936459 | T | C | 1.071 | -7977.388 | -7977.592 | -0.204 | 3.15E-06 |
| 11 | 120571314 | rs74447718 | T | C | 0.931 | -7977.388 | -7977.592 | -0.204 | 3.15E-06 |
| 11 | 116660450 | rs33984246 | G | A | 0.906 | -7977.388 | -7977.594 | -0.205 | 3.15E-06 |
| 11 | 117219961 | rs477858 | G | A | 0.941 | -7977.388 | -7977.594 | -0.206 | 3.15E-06 |
| 11 | 117974397 | rs117950757 | A | G | 0.915 | -7977.388 | -7977.594 | -0.206 | 3.15E-06 |
| 11 | 116760485 | rs117709132 | T | C | 1.094 | -7977.388 | -7977.596 | -0.207 | 3.14E-06 |
| 11 | 117653955 | rs7119911 | T | G | 0.948 | -7977.388 | -7977.595 | -0.207 | 3.14E-06 |
| 11 | 120570786 | rs61901366 | A | G | 0.93 | -7977.388 | -7977.595 | -0.207 | 3.14E-06 |
| 11 | 117342536 | rs114831648 | C | T | 1.103 | -7977.388 | -7977.596 | -0.208 | 3.14E-06 |
| 11 | 117451536 | rs684973 | C | G | 0.948 | -7977.388 | -7977.596 | -0.208 | 3.14E-06 |
| 11 | 117430840 | rs55741146 | G | C | 1.06 | -7977.388 | -7977.597 | -0.209 | 3.14E-06 |
| 11 | 116880018 | rs150734126 | C | T | 0.905 | -7977.388 | -7977.599 | -0.211 | 3.13E-06 |
| 11 | 117175658 | rs116987336 | A | G | 0.908 | -7977.388 | -7977.599 | -0.211 | 3.13E-06 |
| 11 | 117362331 | rs149861300 | C | A | 0.909 | -7977.388 | -7977.599 | -0.211 | 3.13E-06 |
| 11 | 117906289 | rs74582224 | T | G | 0.907 | -7977.388 | -7977.6 | -0.211 | 3.13E-06 |
| 11 | 117906406 | rs79791963 | G | C | 0.907 | -7977.388 | -7977.6 | -0.211 | 3.13E-06 |
| 11 | 117906408 | rs78197730 | T | C | 0.907 | -7977.388 | -7977.6 | -0.211 | 3.13E-06 |
| 11 | 120925525 | rs11218127 | G | A | 1.065 | -7977.388 | -7977.599 | -0.211 | 3.13E-06 |
| 11 | 117582709 | rs79459688 | G | A | 0.905 | -7977.388 | -7977.6 | -0.212 | 3.13E-06 |
| 11 | 120858815 | rs11218094 | T | G | 0.937 | -7977.388 | -7977.6 | -0.212 | 3.13E-06 |
| 11 | 116831071 | rs622302 | G | A | 0.905 | -7977.388 | -7977.601 | -0.213 | 3.13E-06 |
| 11 | 116833669 | rs493101 | A | G | 0.905 | -7977.388 | -7977.601 | -0.213 | 3.13E-06 |
| 11 | 116835362 | rs541496 | T | C | 0.905 | -7977.388 | -7977.601 | -0.213 | 3.13E-06 |
| 11 | 116835883 | rs667119 | T | A | 0.905 | -7977.388 | -7977.601 | -0.213 | 3.13E-06 |
| 11 | 116836548 | rs2921655 | C | G | 0.905 | -7977.388 | -7977.601 | -0.213 | 3.13E-06 |
| 11 | 116845538 | rs2513091 | G | A | 0.905 | -7977.388 | -7977.601 | -0.213 | 3.13E-06 |
| 11 | 117518038 | rs626571 | C | T | 0.945 | -7977.388 | -7977.602 | -0.213 | 3.13E-06 |
| 11 | 116640604 | rs1298336202 | T | C | 0.906 | -7977.388 | -7977.603 | -0.214 | 3.12E-06 |
| 11 | 117566303 | rs10892163 | A | G | 0.928 | -7977.388 | -7977.603 | -0.214 | 3.12E-06 |
| 11 | 117569923 | rs17121096 | C | T | 0.928 | -7977.388 | -7977.603 | -0.215 | 3.12E-06 |
| 11 | 117611938 | rs77719730 | A | G | 1.097 | -7977.388 | -7977.603 | -0.215 | 3.12E-06 |
| 11 | 118863232 | rs4938614 | A | G | 1.055 | -7977.388 | -7977.603 | -0.215 | 3.12E-06 |
| 11 | 120570901 | rs61901367 | A | G | 0.93 | -7977.388 | -7977.604 | -0.215 | 3.12E-06 |
| 11 | 120585020 | rs7120727 | C | G | 0.937 | -7977.388 | -7977.603 | -0.215 | 3.12E-06 |
| 11 | 117101311 | rs1231238253 | T | C | 1.097 | -7977.388 | -7977.605 | -0.216 | 3.12E-06 |
| 11 | 116909994 | rs73594175 | G | A | 0.906 | -7977.388 | -7977.605 | -0.217 | 3.11E-06 |
| 11 | 116910320 | rs1475614837 | T | G | 0.906 | -7977.388 | -7977.605 | -0.217 | 3.11E-06 |
| 11 | 116910427 | rs73594181 | A | T | 0.906 | -7977.388 | -7977.605 | -0.217 | 3.11E-06 |
| 11 | 116910454 | rs79340485 | T | C | 0.906 | -7977.388 | -7977.605 | -0.217 | 3.11E-06 |
| 11 | 120571229 | rs149505626 | A | G | 0.93 | -7977.388 | -7977.607 | -0.218 | 3.11E-06 |
| 11 | 118099735 | rs149668068 | A | G | 0.908 | -7977.388 | -7977.607 | -0.219 | 3.11E-06 |
| 11 | 120509335 | rs10892589 | T | C | 1.056 | -7977.388 | -7977.607 | -0.219 | 3.11E-06 |
| 11 | 116601062 | rs148945243 | T | C | 1.102 | -7977.388 | -7977.608 | -0.22 | 3.10E-06 |
| 11 | 116828484 | rs497653 | C | T | 0.906 | -7977.388 | -7977.608 | -0.22 | 3.10E-06 |
| 11 | 117062489 | rs3213459 | C | T | 1.088 | -7977.388 | -7977.609 | -0.22 | 3.10E-06 |
| 11 | 116578217 | rs12221682 | G | C | 0.926 | -7977.388 | -7977.609 | -0.221 | 3.10E-06 |
| 11 | 119939927 | rs592565 | A | G | 0.931 | -7977.388 | -7977.61 | -0.221 | 3.10E-06 |
| 11 | 116847584 | rs3016611 | A | G | 0.906 | -7977.388 | -7977.61 | -0.222 | 3.10E-06 |
| 11 | 117564052 | rs11216506 | G | A | 0.928 | -7977.388 | -7977.61 | -0.222 | 3.10E-06 |
| 11 | 117467394 | rs2186761 | A | G | 1.058 | -7977.388 | -7977.611 | -0.223 | 3.09E-06 |
| 11 | 121011507 | rs638496 | T | C | 0.946 | -7977.388 | -7977.612 | -0.223 | 3.09E-06 |
| 11 | 116538407 | rs57258844 | C | G | 0.913 | -7977.388 | -7977.612 | -0.224 | 3.09E-06 |
| 11 | 117407435 | rs17120841 | G | A | 0.906 | -7977.388 | -7977.612 | -0.224 | 3.09E-06 |
| 11 | 117409971 | rs74735117 | C | T | 0.906 | -7977.388 | -7977.612 | -0.224 | 3.09E-06 |
| 11 | 120647015 | rs58174358 | T | C | 0.91 | -7977.388 | -7977.612 | -0.224 | 3.09E-06 |

| | | | | | | | |
|---|---|---|---|---|---|---|---|
| 11 | 116523063 | rs12805779 | T | G | NULL | 0.527472304 | NULL&M_Disease |
| 11 | 118807639 | rs73575454 | A | G | NULL | 0.527472304 | NULL&M_Disease |
| 11 | 116515420 | rs4938294 | T | C | NULL | 0.527472304 | NULL&M_Disease |
| 11 | 119729992 | rs500401 | T | C | NULL | 0.527472304 | NULL&M_Disease |
| 11 | 118804989 | rs112045567 | G | A | NULL | 0.527472304 | NULL&M_Disease |
| 11 | 116292201 | rs76388159 | G | A | NULL | 0.527472304 | NULL&M_Disease |
| 11 | 116371442 | rs60171666 | A | G | NULL | 0.528718357 | NULL&M_Disease |
| 11 | 116838615 | rs581122 | T | C | NULL | 0.527472304 | NULL&M_Disease |
| 11 | 118018518 | rs140305819 | A | G | NULL | 0.527472304 | NULL&M_Disease |
| 11 | 120571702 | rs61901369 | C | T | NULL | 0.528718357 | NULL&M_Disease |
| 11 | 117953688 | rs7937420 | A | G | NULL | 0.528718357 | NULL&M_Disease |
| 11 | 120334905 | rs79844114 | G | A | NULL | 0.527472304 | NULL&M_Disease |
| 11 | 116915229 | rs1270305911 | T | C | NULL | 0.527472304 | NULL&M_Disease |
| 11 | 116915881 | rs73594187 | C | T | NULL | 0.527472304 | NULL&M_Disease |
| 11 | 116921143 | rs145339787 | C | T | NULL | 0.527472304 | NULL&M_Disease |
| 11 | 118863037 | rs4938612 | C | T | NULL | 0.528718357 | NULL&M_Disease |
| 11 | 121083183 | rs12788122 | A | G | NULL | 0.528718357 | NULL&M_Disease |
| 11 | 116907449 | rs721783 | T | C | NULL | 0.528718357 | NULL&M_Disease |
| 11 | 117324068 | rs4936380 | A | G | NULL | 0.528718357 | NULL&M_Disease |
| 11 | 117913430 | rs675479 | G | A | NULL | 0.528718357 | NULL&M_Disease |
| 11 | 120630028 | rs72995842 | T | C | NULL | 0.527472304 | NULL&M_Disease |
| 11 | 116738046 | rs518547 | G | A | NULL | 0.528718357 | NULL&M_Disease |
| 11 | 119928865 | rs11603573 | G | C | NULL | 0.528718357 | NULL&M_Disease |
| 11 | 120573725 | rs61901376 | G | A | NULL | 0.528718357 | NULL&M_Disease |
| 11 | 120620840 | rs115933113 | C | G | NULL | 0.528718357 | NULL&M_Disease |
| 11 | 116358593 | rs60905351 | A | G | NULL | 0.528718357 | NULL&M_Disease |
| 11 | 117500587 | rs117445862 | C | T | NULL | 0.528718357 | NULL&M_Disease |
| 11 | 117500633 | rs141182805 | G | T | NULL | 0.528718357 | NULL&M_Disease |
| 11 | 120798629 | rs1893816 | T | C | NULL | 0.529964052 | NULL&M_Disease |
| 11 | 117064776 | rs11605883 | G | C | NULL | 0.528718357 | NULL&M_Disease |
| 11 | 117531887 | rs643758 | T | A | NULL | 0.529964052 | NULL&M_Disease |
| 11 | 118804643 | rs56933877 | A | G | NULL | 0.529964052 | NULL&M_Disease |
| 11 | 120564637 | rs12272864 | C | T | NULL | 0.529964052 | NULL&M_Disease |
| 11 | 120586773 | rs61901399 | T | C | NULL | 0.529964052 | NULL&M_Disease |
| 11 | 117010474 | rs77661993 | C | A | NULL | 0.529964052 | NULL&M_Disease |
| 11 | 117372281 | rs7928478 | A | G | NULL | 0.529964052 | NULL&M_Disease |
| 11 | 117420163 | rs10790192 | A | G | NULL | 0.529964052 | NULL&M_Disease |
| 11 | 119468521 | rs78914704 | A | G | NULL | 0.529964052 | NULL&M_Disease |
| 11 | 117476375 | rs58248287 | G | A | NULL | 0.528718357 | NULL&M_Disease |
| 11 | 117502729 | rs111607963 | T | C | NULL | 0.529964052 | NULL&M_Disease |
| 11 | 117906273 | rs74586480 | G | A | NULL | 0.529964052 | NULL&M_Disease |
| 11 | 120116018 | rs58594771 | C | T | NULL | 0.529964052 | NULL&M_Disease |
| 11 | 116237970 | rs490296 | C | T | NULL | 0.531209373 | NULL&M_Disease |
| 11 | 116530998 | rs11216080 | T | G | NULL | 0.529964052 | NULL&M_Disease |
| 11 | 119678986 | rs10892470 | C | T | NULL | 0.531209373 | NULL&M_Disease |
| 11 | 117503731 | rs7126973 | T | A | NULL | 0.531209373 | NULL&M_Disease |
| 11 | 118828184 | rs45586434 | A | C | NULL | 0.531209373 | NULL&M_Disease |
| 11 | 120646975 | rs17124305 | G | A | NULL | 0.529964052 | NULL&M_Disease |
| 11 | 117323004 | rs4499034 | T | C | NULL | 0.531209373 | NULL&M_Disease |
| 11 | 117531358 | rs12417881 | G | A | NULL | 0.531209373 | NULL&M_Disease |
| 11 | 117591654 | rs948105 | A | T | NULL | 0.529964052 | NULL&M_Disease |
| 11 | 117010091 | rs115991973 | A | G | NULL | 0.531209373 | NULL&M_Disease |
| 11 | 117010123 | rs116384608 | G | A | NULL | 0.531209373 | NULL&M_Disease |
| 11 | 117436510 | rs141047910 | C | G | NULL | 0.529964052 | NULL&M_Disease |
| 11 | 118858911 | rs7110034 | A | C | NULL | 0.531209373 | NULL&M_Disease |
| 11 | 117502942 | rs116979369 | T | C | NULL | 0.529964052 | NULL&M_Disease |
| 11 | 117745018 | rs7102557 | A | G | NULL | 0.531209373 | NULL&M_Disease |
| 11 | 118853488 | rs4290240 | G | A | NULL | 0.531209373 | NULL&M_Disease |
| 11 | 120650530 | rs116727415 | T | A | NULL | 0.529964052 | NULL&M_Disease |
| 11 | 117050918 | rs12419178 | T | C | NULL | 0.531209373 | NULL&M_Disease |
| 11 | 117385478 | rs675522 | C | G | NULL | 0.531209373 | NULL&M_Disease |
| 11 | 117483022 | rs75500945 | T | C | NULL | 0.531209373 | NULL&M_Disease |
| 11 | 117483626 | rs74941224 | T | C | NULL | 0.531209373 | NULL&M_Disease |
| 11 | 117484221 | rs74860176 | T | C | NULL | 0.531209373 | NULL&M_Disease |
| 11 | 118804919 | rs73575441 | C | T | NULL | 0.531209373 | NULL&M_Disease |
| 11 | 118824917 | rs11217102 | A | G | NULL | 0.531209373 | NULL&M_Disease |
| 11 | 117565069 | rs11216507 | C | T | NULL | 0.531209373 | NULL&M_Disease |
| 11 | 119795611 | rs479455 | T | G | NULL | 0.531209373 | NULL&M_Disease |
| 11 | 116237017 | rs571139 | A | G | NULL | 0.532454306 | NULL&M_Disease |
| 11 | 117709688 | rs11216575 | G | A | NULL | 0.531209373 | NULL&M_Disease |
| 11 | 117519147 | rs10892148 | T | C | NULL | 0.532454306 | NULL&M_Disease |
| 11 | 117710150 | rs11216577 | T | C | NULL | 0.531209373 | NULL&M_Disease |
| 11 | 117798560 | rs12361671 | A | T | NULL | 0.532454306 | NULL&M_Disease |
| 11 | 118964577 | rs7103570 | C | T | NULL | 0.531209373 | NULL&M_Disease |
| 11 | 116529956 | rs71480309 | T | C | NULL | 0.532454306 | NULL&M_Disease |
| 11 | 117591584 | rs948106 | C | T | NULL | 0.532454306 | NULL&M_Disease |
| 11 | 116642049 | rs111683933 | C | T | NULL | 0.531209373 | NULL&M_Disease |
| 11 | 117492693 | rs74335840 | A | T | NULL | 0.531209373 | NULL&M_Disease |
| 11 | 117493342 | rs117470649 | G | A | NULL | 0.531209373 | NULL&M_Disease |
| 11 | 118755630 | rs3176905 | C | T | NULL | 0.532454306 | NULL&M_Disease |
| 11 | 119672156 | rs4938724 | A | G | NULL | 0.532454306 | NULL&M_Disease |
| 11 | 119756962 | rs75436216 | C | T | NULL | 0.531209373 | NULL&M_Disease |
| 11 | 116650571 | rs113271699 | A | C | NULL | 0.531209373 | NULL&M_Disease |
| 11 | 119478010 | rs150773762 | G | A | NULL | 0.532454306 | NULL&M_Disease |
| 11 | 120890870 | rs73011533 | A | G | NULL | 0.532454306 | NULL&M_Disease |
| 11 | 120874281 | rs116862564 | G | C | NULL | 0.532454306 | NULL&M_Disease |
| 11 | 116281169 | rs538359 | C | T | NULL | 0.532454306 | NULL&M_Disease |
| 11 | 116328597 | rs12420339 | G | T | NULL | 0.532454306 | NULL&M_Disease |
| 11 | 116658122 | rs2075294 | T | G | NULL | 0.532454306 | NULL&M_Disease |
| 11 | 116870420 | rs113419013 | T | C | NULL | 0.532454306 | NULL&M_Disease |
| 11 | 117992155 | rs627999 | T | C | NULL | 0.532454306 | NULL&M_Disease |
| 11 | 120461835 | rs17311310 | A | G | NULL | 0.533698835 | NULL&M_Disease |
| 11 | 116645682 | rs45443305 | A | C | NULL | 0.532454306 | NULL&M_Disease |
| 11 | 119914616 | rs681569 | A | C | NULL | 0.532454306 | NULL&M_Disease |
| 11 | 120910225 | rs143427408 | T | C | NULL | 0.532454306 | NULL&M_Disease |
| 11 | 117051725 | rs76883864 | T | C | NULL | 0.533698835 | NULL&M_Disease |

| | | | | | | | | | |
|---|---|---|---|---|---|---|---|---|---|
| 11 | 116523063 | rs12805779 | T | G | 1.068 | -7977.388 | -7977.613 | -0.225 | 3.09E-06 |
| 11 | 118807639 | rs73575454 | A | G | 0.92 | -7977.388 | -7977.613 | -0.225 | 3.09E-06 |
| 11 | 116515420 | rs4938294 | T | C | 1.065 | -7977.388 | -7977.614 | -0.226 | 3.09E-06 |
| 11 | 119729992 | rs500401 | T | C | 1.085 | -7977.388 | -7977.614 | -0.226 | 3.09E-06 |
| 11 | 118804989 | rs112045567 | G | A | 0.92 | -7977.388 | -7977.617 | -0.228 | 3.08E-06 |
| 11 | 116292201 | rs76388159 | G | A | 1.095 | -7977.388 | -7977.617 | -0.229 | 3.08E-06 |
| 11 | 116371442 | rs60171666 | A | G | 0.947 | -7977.388 | -7977.617 | -0.229 | 3.08E-06 |
| 11 | 116838615 | rs581122 | T | C | 0.907 | -7977.388 | -7977.617 | -0.229 | 3.08E-06 |
| 11 | 118018518 | rs140305819 | A | G | 1.099 | -7977.388 | -7977.617 | -0.229 | 3.08E-06 |
| 11 | 120571702 | rs61901369 | C | T | 0.931 | -7977.388 | -7977.618 | -0.229 | 3.08E-06 |
| 11 | 117953688 | rs7937420 | A | G | 0.946 | -7977.388 | -7977.62 | -0.231 | 3.07E-06 |
| 11 | 120334905 | rs79844114 | G | A | 0.909 | -7977.388 | -7977.62 | -0.231 | 3.07E-06 |
| 11 | 116915229 | rs1270305911 | T | C | 0.907 | -7977.388 | -7977.62 | -0.232 | 3.07E-06 |
| 11 | 116915881 | rs73594187 | C | T | 0.907 | -7977.388 | -7977.62 | -0.232 | 3.07E-06 |
| 11 | 116921143 | rs145339787 | C | T | 0.907 | -7977.388 | -7977.62 | -0.232 | 3.07E-06 |
| 11 | 118863037 | rs4938612 | C | T | 1.055 | -7977.388 | -7977.621 | -0.232 | 3.07E-06 |
| 11 | 121083183 | rs12788122 | A | G | 0.919 | -7977.388 | -7977.62 | -0.232 | 3.07E-06 |
| 11 | 116907449 | rs721783 | T | C | 1.062 | -7977.388 | -7977.622 | -0.234 | 3.06E-06 |
| 11 | 117324068 | rs4936380 | A | G | 0.922 | -7977.388 | -7977.623 | -0.234 | 3.06E-06 |
| 11 | 117913430 | rs675479 | G | A | 1.055 | -7977.388 | -7977.623 | -0.234 | 3.06E-06 |
| 11 | 120630028 | rs72995842 | T | C | 0.908 | -7977.388 | -7977.622 | -0.234 | 3.06E-06 |
| 11 | 116738046 | rs518547 | G | A | 1.063 | -7977.388 | -7977.623 | -0.235 | 3.06E-06 |
| 11 | 119928865 | rs11603573 | G | C | 1.087 | -7977.388 | -7977.624 | -0.236 | 3.05E-06 |
| 11 | 120573725 | rs61901376 | G | A | 0.93 | -7977.388 | -7977.625 | -0.237 | 3.05E-06 |
| 11 | 120620840 | rs115933113 | C | G | 0.911 | -7977.388 | -7977.626 | -0.237 | 3.05E-06 |
| 11 | 116358593 | rs60905351 | A | G | 0.93 | -7977.388 | -7977.626 | -0.238 | 3.05E-06 |
| 11 | 117500587 | rs117445862 | C | T | 0.908 | -7977.388 | -7977.627 | -0.239 | 3.05E-06 |
| 11 | 117500633 | rs141182805 | G | T | 0.908 | -7977.388 | -7977.627 | -0.239 | 3.05E-06 |
| 11 | 120798629 | rs1893816 | T | C | 0.938 | -7977.388 | -7977.627 | -0.239 | 3.05E-06 |
| 11 | 117064776 | rs11605883 | G | C | 1.087 | -7977.388 | -7977.628 | -0.24 | 3.04E-06 |
| 11 | 117531887 | rs643758 | T | A | 0.948 | -7977.388 | -7977.628 | -0.24 | 3.04E-06 |
| 11 | 118804643 | rs56933877 | A | G | 0.921 | -7977.388 | -7977.629 | -0.241 | 3.04E-06 |
| 11 | 120564637 | rs12272864 | C | T | 0.931 | -7977.388 | -7977.629 | -0.241 | 3.04E-06 |
| 11 | 120586773 | rs61901399 | T | C | 0.93 | -7977.388 | -7977.63 | -0.241 | 3.04E-06 |
| 11 | 117010474 | rs77661993 | C | A | 1.087 | -7977.388 | -7977.63 | -0.242 | 3.04E-06 |
| 11 | 117372281 | rs7928478 | A | G | 1.058 | -7977.388 | -7977.63 | -0.242 | 3.04E-06 |
| 11 | 117420163 | rs10790192 | A | G | 0.948 | -7977.388 | -7977.63 | -0.242 | 3.04E-06 |
| 11 | 119468521 | rs78914704 | A | G | 1.08 | -7977.388 | -7977.631 | -0.243 | 3.03E-06 |
| 11 | 117476375 | rs58248287 | G | A | 0.908 | -7977.388 | -7977.633 | -0.244 | 3.03E-06 |
| 11 | 117502729 | rs111607963 | T | C | 1.093 | -7977.388 | -7977.633 | -0.244 | 3.03E-06 |
| 11 | 117906273 | rs74586480 | G | A | 0.909 | -7977.388 | -7977.633 | -0.245 | 3.03E-06 |
| 11 | 120116018 | rs58594771 | C | T | 0.908 | -7977.388 | -7977.635 | -0.247 | 3.02E-06 |
| 11 | 116237970 | rs490296 | C | T | 1.056 | -7977.388 | -7977.636 | -0.248 | 3.02E-06 |
| 11 | 116530998 | rs11216080 | T | G | 0.914 | -7977.388 | -7977.636 | -0.248 | 3.02E-06 |
| 11 | 119678986 | rs10892470 | C | T | 0.948 | -7977.388 | -7977.637 | -0.249 | 3.02E-06 |
| 11 | 117503731 | rs7126973 | T | A | 0.944 | -7977.388 | -7977.638 | -0.25 | 3.01E-06 |
| 11 | 118828184 | rs45586434 | A | C | 1.067 | -7977.388 | -7977.639 | -0.25 | 3.01E-06 |
| 11 | 120646975 | rs17124305 | G | A | 0.912 | -7977.388 | -7977.639 | -0.25 | 3.01E-06 |
| 11 | 117323004 | rs4499034 | T | C | 0.923 | -7977.388 | -7977.639 | -0.251 | 3.01E-06 |
| 11 | 117531358 | rs12417881 | G | A | 0.948 | -7977.388 | -7977.639 | -0.251 | 3.01E-06 |
| 11 | 117591654 | rs948105 | A | T | 0.911 | -7977.388 | -7977.639 | -0.251 | 3.01E-06 |
| 11 | 117010091 | rs115991973 | A | G | 1.086 | -7977.388 | -7977.64 | -0.252 | 3.01E-06 |
| 11 | 117010123 | rs116384608 | G | A | 1.086 | -7977.388 | -7977.64 | -0.252 | 3.01E-06 |
| 11 | 117436510 | rs141047910 | C | G | 0.909 | -7977.388 | -7977.641 | -0.253 | 3.00E-06 |
| 11 | 118858911 | rs7110034 | A | C | 1.055 | -7977.388 | -7977.641 | -0.253 | 3.00E-06 |
| 11 | 117502942 | rs116979369 | T | C | 0.909 | -7977.388 | -7977.643 | -0.254 | 3.00E-06 |
| 11 | 117745018 | rs7102557 | A | G | 0.935 | -7977.388 | -7977.642 | -0.254 | 3.00E-06 |
| 11 | 118853488 | rs4290240 | G | A | 1.055 | -7977.388 | -7977.642 | -0.254 | 3.00E-06 |
| 11 | 120650530 | rs116727415 | T | A | 0.909 | -7977.388 | -7977.643 | -0.254 | 3.00E-06 |
| 11 | 117050918 | rs12419178 | T | C | 1.086 | -7977.388 | -7977.643 | -0.255 | 3.00E-06 |
| 11 | 117385478 | rs675522 | C | G | 0.946 | -7977.388 | -7977.643 | -0.255 | 3.00E-06 |
| 11 | 117483022 | rs75500945 | T | C | 0.909 | -7977.388 | -7977.643 | -0.255 | 3.00E-06 |
| 11 | 117483626 | rs74941224 | T | C | 0.909 | -7977.388 | -7977.643 | -0.255 | 3.00E-06 |
| 11 | 117484221 | rs74860176 | T | C | 0.909 | -7977.388 | -7977.643 | -0.255 | 3.00E-06 |
| 11 | 118804919 | rs73575441 | C | T | 0.921 | -7977.388 | -7977.644 | -0.255 | 3.00E-06 |
| 11 | 118824917 | rs11217102 | A | G | 1.065 | -7977.388 | -7977.643 | -0.255 | 3.00E-06 |
| 11 | 117565069 | rs11216507 | C | T | 0.929 | -7977.388 | -7977.645 | -0.256 | 2.99E-06 |
| 11 | 119795611 | rs479455 | T | G | 0.944 | -7977.388 | -7977.644 | -0.256 | 2.99E-06 |
| 11 | 116237017 | rs571139 | A | G | 1.056 | -7977.388 | -7977.646 | -0.258 | 2.99E-06 |
| 11 | 117709688 | rs11216575 | G | A | 1.087 | -7977.388 | -7977.646 | -0.258 | 2.99E-06 |
| 11 | 117519147 | rs10892148 | T | C | 0.937 | -7977.388 | -7977.648 | -0.26 | 2.98E-06 |
| 11 | 117710150 | rs11216577 | T | C | 1.087 | -7977.388 | -7977.648 | -0.26 | 2.98E-06 |
| 11 | 117798560 | rs12361671 | A | T | 0.929 | -7977.388 | -7977.649 | -0.26 | 2.98E-06 |
| 11 | 118964577 | rs7103570 | C | T | 0.914 | -7977.388 | -7977.648 | -0.26 | 2.98E-06 |
| 11 | 116529956 | rs71480309 | T | C | 0.914 | -7977.388 | -7977.651 | -0.262 | 2.98E-06 |
| 11 | 117591584 | rs948106 | C | T | 0.912 | -7977.388 | -7977.651 | -0.263 | 2.97E-06 |
| 11 | 116642049 | rs111683933 | C | T | 1.097 | -7977.388 | -7977.653 | -0.264 | 2.97E-06 |
| 11 | 117492693 | rs74335840 | A | T | 0.91 | -7977.388 | -7977.652 | -0.264 | 2.97E-06 |
| 11 | 117493342 | rs117470649 | G | A | 0.91 | -7977.388 | -7977.652 | -0.264 | 2.97E-06 |
| 11 | 118755630 | rs3176905 | C | T | 0.938 | -7977.388 | -7977.652 | -0.264 | 2.97E-06 |
| 11 | 119672156 | rs4938724 | A | G | 0.948 | -7977.388 | -7977.652 | -0.264 | 2.97E-06 |
| 11 | 119756962 | rs75436216 | C | T | 0.91 | -7977.388 | -7977.653 | -0.264 | 2.97E-06 |
| 11 | 116650571 | rs113271699 | A | C | 1.097 | -7977.388 | -7977.653 | -0.265 | 2.97E-06 |
| 11 | 119478010 | rs150773762 | G | A | 0.911 | -7977.388 | -7977.653 | -0.265 | 2.97E-06 |
| 11 | 120890870 | rs73011533 | A | G | 1.082 | -7977.388 | -7977.654 | -0.265 | 2.97E-06 |
| 11 | 120874281 | rs116862564 | G | C | 0.91 | -7977.388 | -7977.654 | -0.266 | 2.96E-06 |
| 11 | 116281169 | rs538359 | C | T | 0.937 | -7977.388 | -7977.656 | -0.267 | 2.96E-06 |
| 11 | 116328597 | rs12420339 | G | T | 0.916 | -7977.388 | -7977.655 | -0.267 | 2.96E-06 |
| 11 | 116658122 | rs2075294 | T | G | 0.91 | -7977.388 | -7977.655 | -0.267 | 2.96E-06 |
| 11 | 116870420 | rs113419013 | T | C | 1.097 | -7977.388 | -7977.656 | -0.267 | 2.96E-06 |
| 11 | 117992155 | rs627999 | T | C | 1.097 | -7977.388 | -7977.655 | -0.267 | 2.96E-06 |
| 11 | 120461835 | rs17311310 | A | G | 1.055 | -7977.388 | -7977.656 | -0.268 | 2.96E-06 |
| 11 | 116645682 | rs45443305 | A | C | 0.91 | -7977.388 | -7977.657 | -0.269 | 2.96E-06 |
| 11 | 119914616 | rs681569 | A | C | 0.919 | -7977.388 | -7977.658 | -0.27 | 2.95E-06 |
| 11 | 120910225 | rs143427408 | T | C | 0.911 | -7977.388 | -7977.66 | -0.271 | 2.95E-06 |
| 11 | 117051725 | rs76883864 | T | C | 1.086 | -7977.388 | -7977.66 | -0.272 | 2.95E-06 |

| | | | | | | | |
|---|---|---|---|---|---|---|---|
| 11 | 120396329 | rs17123976 | C | T | NULL | 0.532454306 | NULL&M_Disease |
| 11 | 120462417 | rs17311317 | A | G | NULL | 0.533698835 | NULL&M_Disease |
| 11 | 116702778 | rs5132 | T | C | NULL | 0.532454306 | NULL&M_Disease |
| 11 | 116639692 | rs1268353 | T | C | NULL | 0.533698835 | NULL&M_Disease |
| 11 | 116948640 | rs7124649 | C | T | NULL | 0.533698835 | NULL&M_Disease |
| 11 | 117800269 | rs11216633 | G | C | NULL | 0.533698835 | NULL&M_Disease |
| 11 | 117800328 | rs11216634 | G | A | NULL | 0.533698835 | NULL&M_Disease |
| 11 | 116828121 | rs1940626 | C | T | NULL | 0.534942945 | NULL&M_Disease |
| 11 | 117517366 | rs536761 | T | C | NULL | 0.534942945 | NULL&M_Disease |
| 11 | 117708625 | rs520333 | A | G | NULL | 0.534942945 | NULL&M_Disease |
| 11 | 117922310 | rs78760381 | T | C | NULL | 0.533698835 | NULL&M_Disease |
| 11 | 117946138 | rs714053 | T | C | NULL | 0.533698835 | NULL&M_Disease |
| 11 | 119793489 | rs665759 | A | G | NULL | 0.534942945 | NULL&M_Disease |
| 11 | 119793573 | rs529373 | T | C | NULL | 0.534942945 | NULL&M_Disease |
| 11 | 116725303 | rs640411 | C | G | NULL | 0.533698835 | NULL&M_Disease |
| 11 | 119795594 | rs551163 | A | C | NULL | 0.534942945 | NULL&M_Disease |
| 11 | 120588100 | rs2846109 | C | T | NULL | 0.534942945 | NULL&M_Disease |
| 11 | 117054709 | rs2046149 | G | C | NULL | 0.534942945 | NULL&M_Disease |
| 11 | 116654959 | rs74662600 | A | G | NULL | 0.533698835 | NULL&M_Disease |
| 11 | 116712442 | rs113749676 | T | C | NULL | 0.533698835 | NULL&M_Disease |
| 11 | 119713633 | rs73008650 | A | G | NULL | 0.534942945 | NULL&M_Disease |
| 11 | 119814550 | rs61899037 | A | G | NULL | 0.533698835 | NULL&M_Disease |
| 11 | 120574350 | rs17124211 | G | A | NULL | 0.534942945 | NULL&M_Disease |
| 11 | 120710210 | rs28737445 | G | A | NULL | 0.534942945 | NULL&M_Disease |
| 11 | 116705021 | rs1263162 | A | T | NULL | 0.533698835 | NULL&M_Disease |
| 11 | 119490788 | rs79801588 | G | A | NULL | 0.534942945 | NULL&M_Disease |
| 11 | 119668263 | rs4272793 | C | A | NULL | 0.534942945 | NULL&M_Disease |
| 11 | 117030633 | rs1312592595 | C | T | NULL | 0.534942945 | NULL&M_Disease |
| 11 | 117500542 | rs10892146 | T | A | NULL | 0.534942945 | NULL&M_Disease |
| 11 | 120733629 | rs4512872 | C | A | NULL | 0.534942945 | NULL&M_Disease |
| 11 | 116424420 | rs17119814 | G | C | NULL | 0.534942945 | NULL&M_Disease |
| 11 | 116238034 | rs491111 | A | G | NULL | 0.534942945 | NULL&M_Disease |
| 11 | 120593745 | rs11606461 | T | C | NULL | 0.534942945 | NULL&M_Disease |
| 11 | 116875338 | rs6589587 | G | T | NULL | 0.53618662 | NULL&M_Disease |
| 11 | 117613781 | rs7950743 | A | G | NULL | 0.534942945 | NULL&M_Disease |
| 11 | 117897098 | rs685308 | G | C | NULL | 0.53618662 | NULL&M_Disease |
| 11 | 119922708 | rs78699173 | A | C | NULL | 0.534942945 | NULL&M_Disease |
| 11 | 116514997 | rs67613157 | T | C | NULL | 0.53618662 | NULL&M_Disease |
| 11 | 117102307 | rs2509209 | G | A | NULL | 0.53618662 | NULL&M_Disease |
| 11 | 117349817 | rs7929976 | G | A | NULL | 0.53618662 | NULL&M_Disease |
| 11 | 117308432 | rs17120753 | C | T | NULL | 0.53618662 | NULL&M_Disease |
| 11 | 119901422 | rs669812 | T | C | NULL | 0.53618662 | NULL&M_Disease |
| 11 | 120474008 | rs4936528 | G | C | NULL | 0.53618662 | NULL&M_Disease |
| 11 | 117385531 | rs1807049 | G | A | NULL | 0.53618662 | NULL&M_Disease |
| 11 | 117411730 | rs35266957 | T | C | NULL | 0.534942945 | NULL&M_Disease |
| 11 | 117414844 | rs3802878 | C | A | NULL | 0.534942945 | NULL&M_Disease |
| 11 | 119516805 | rs4938700 | C | G | NULL | 0.53618662 | NULL&M_Disease |
| 11 | 116239462 | rs79643426 | T | C | NULL | 0.53618662 | NULL&M_Disease |
| 11 | 119831163 | rs35962410 | C | A | NULL | 0.53618662 | NULL&M_Disease |
| 11 | 117944485 | rs11216736 | C | T | NULL | 0.53618662 | NULL&M_Disease |
| 11 | 116941985 | rs7101763 | T | G | NULL | 0.537429845 | NULL&M_Disease |
| 11 | 116945718 | rs4620758 | A | G | NULL | 0.537429845 | NULL&M_Disease |
| 11 | 117560679 | rs57550545 | C | T | NULL | 0.53618662 | NULL&M_Disease |
| 11 | 116946652 | rs7130213 | A | C | NULL | 0.537429845 | NULL&M_Disease |
| 11 | 117488572 | rs59258276 | C | T | NULL | 0.53618662 | NULL&M_Disease |
| 11 | 120574079 | rs61901377 | G | A | NULL | 0.537429845 | NULL&M_Disease |
| 11 | 117488030 | rs148543195 | T | C | NULL | 0.53618662 | NULL&M_Disease |
| 11 | 120572772 | rs117165966 | C | G | NULL | 0.537429845 | NULL&M_Disease |
| 11 | 120572814 | rs142600127 | A | G | NULL | 0.537429845 | NULL&M_Disease |
| 11 | 116225940 | rs12576431 | G | C | NULL | 0.537429845 | NULL&M_Disease |
| 11 | 119509625 | rs1022080 | C | T | NULL | 0.537429845 | NULL&M_Disease |
| 11 | 120126017 | rs79988761 | A | G | NULL | 0.537429845 | NULL&M_Disease |
| 11 | 119881526 | rs1893394 | A | G | NULL | 0.537429845 | NULL&M_Disease |
| 11 | 117713187 | rs12363888 | T | C | NULL | 0.537429845 | NULL&M_Disease |
| 11 | 120734419 | rs4344499 | G | T | NULL | 0.537429845 | NULL&M_Disease |
| 11 | 120837575 | rs7939112 | A | G | NULL | 0.537429845 | NULL&M_Disease |
| 11 | 120851138 | rs34203183 | G | C | NULL | 0.537429845 | NULL&M_Disease |
| 11 | 116424451 | rs78381558 | C | G | NULL | 0.538672605 | NULL&M_Disease |
| 11 | 116638854 | rs78908794 | C | T | NULL | 0.537429845 | NULL&M_Disease |
| 11 | 117061775 | rs12419470 | T | C | NULL | 0.537429845 | NULL&M_Disease |
| 11 | 117061825 | rs61703672 | T | C | NULL | 0.537429845 | NULL&M_Disease |
| 11 | 117065066 | rs3781985 | C | G | NULL | 0.537429845 | NULL&M_Disease |
| 11 | 117065153 | rs80183569 | A | G | NULL | 0.537429845 | NULL&M_Disease |
| 11 | 117065533 | rs11606480 | A | G | NULL | 0.537429845 | NULL&M_Disease |
| 11 | 117225247 | rs117754274 | C | G | NULL | 0.537429845 | NULL&M_Disease |
| 11 | 119922728 | rs79533157 | C | G | NULL | 0.537429845 | NULL&M_Disease |
| 11 | 120563557 | rs34463064 | T | C | NULL | 0.53618662 | NULL&M_Disease |
| 11 | 120834991 | rs7948885 | A | G | NULL | 0.538672605 | NULL&M_Disease |
| 11 | 119807402 | rs10790356 | T | A | NULL | 0.538672605 | NULL&M_Disease |
| 11 | 116818527 | rs112704540 | G | A | NULL | 0.538672605 | NULL&M_Disease |
| 11 | 119730115 | rs79365963 | C | T | NULL | 0.538672605 | NULL&M_Disease |
| 11 | 120577100 | rs12295113 | T | C | NULL | 0.538672605 | NULL&M_Disease |
| 11 | 120572387 | rs61901372 | C | T | NULL | 0.538672605 | NULL&M_Disease |
| 11 | 120572952 | rs61901374 | C | T | NULL | 0.538672605 | NULL&M_Disease |
| 11 | 120573265 | rs61901375 | C | T | NULL | 0.538672605 | NULL&M_Disease |
| 11 | 117015516 | rs12419272 | G | C | NULL | 0.538672605 | NULL&M_Disease |
| 11 | 117035319 | rs12420127 | G | A | NULL | 0.538672605 | NULL&M_Disease |
| 11 | 117941082 | rs143620404 | T | G | NULL | 0.537429845 | NULL&M_Disease |
| 11 | 116236469 | rs545675 | T | C | NULL | 0.538672605 | NULL&M_Disease |
| 11 | 116789849 | rs76844341 | G | A | NULL | 0.537429845 | NULL&M_Disease |
| 11 | 116953645 | rs1871756 | A | G | NULL | 0.538672605 | NULL&M_Disease |
| 11 | 118285369 | rs150339153 | T | C | NULL | 0.538672605 | NULL&M_Disease |
| 11 | 120084221 | rs1551242 | T | C | NULL | 0.538672605 | NULL&M_Disease |
| 11 | 120084509 | rs1551241 | C | T | NULL | 0.538672605 | NULL&M_Disease |
| 11 | 117384412 | rs7107020 | C | T | NULL | 0.538672605 | NULL&M_Disease |
| 11 | 120583174 | rs6589834 | C | G | NULL | 0.538672605 | NULL&M_Disease |

| | | | | | | | | | |
|---|---|---|---|---|---|---|---|---|---|
| 11 | 120396329 | rs17123976 | C | T | 0.91 | -7977.388 | -7977.661 | -0.272 | 2.95E-06 |
| 11 | 120462417 | rs17311317 | A | G | 1.055 | -7977.388 | -7977.66 | -0.272 | 2.95E-06 |
| 11 | 116702778 | rs5132 | T | C | 0.911 | -7977.388 | -7977.663 | -0.275 | 2.94E-06 |
| 11 | 116639692 | rs1268353 | T | C | 1.056 | -7977.388 | -7977.664 | -0.276 | 2.94E-06 |
| 11 | 116948640 | rs7124649 | C | T | 0.911 | -7977.388 | -7977.665 | -0.276 | 2.94E-06 |
| 11 | 117800269 | rs11216633 | G | C | 0.929 | -7977.388 | -7977.665 | -0.276 | 2.94E-06 |
| 11 | 117800328 | rs11216634 | G | A | 0.929 | -7977.388 | -7977.665 | -0.276 | 2.94E-06 |
| 11 | 116828121 | rs1940626 | C | T | 1.061 | -7977.388 | -7977.667 | -0.278 | 2.93E-06 |
| 11 | 117517366 | rs536761 | T | C | 0.948 | -7977.388 | -7977.666 | -0.278 | 2.93E-06 |
| 11 | 117708625 | rs520333 | A | G | 0.947 | -7977.388 | -7977.666 | -0.278 | 2.93E-06 |
| 11 | 117922310 | rs78760381 | T | C | 0.912 | -7977.388 | -7977.666 | -0.278 | 2.93E-06 |
| 11 | 117946138 | rs714053 | T | C | 0.912 | -7977.388 | -7977.666 | -0.278 | 2.93E-06 |
| 11 | 119793489 | rs665759 | A | G | 0.944 | -7977.388 | -7977.666 | -0.278 | 2.93E-06 |
| 11 | 119793573 | rs529373 | T | C | 0.944 | -7977.388 | -7977.666 | -0.278 | 2.93E-06 |
| 11 | 116725303 | rs640411 | C | G | 0.911 | -7977.388 | -7977.668 | -0.279 | 2.93E-06 |
| 11 | 119795594 | rs551163 | A | C | 0.944 | -7977.388 | -7977.667 | -0.279 | 2.93E-06 |
| 11 | 120588100 | rs2846109 | C | T | 0.933 | -7977.388 | -7977.668 | -0.28 | 2.92E-06 |
| 11 | 117054709 | rs2046149 | G | C | 1.085 | -7977.388 | -7977.669 | -0.281 | 2.92E-06 |
| 11 | 116654959 | rs74662600 | A | G | 0.911 | -7977.388 | -7977.67 | -0.282 | 2.92E-06 |
| 11 | 116712442 | rs113749676 | T | C | 0.911 | -7977.388 | -7977.671 | -0.282 | 2.92E-06 |
| 11 | 119713633 | rs73008650 | A | G | 1.068 | -7977.388 | -7977.671 | -0.282 | 2.92E-06 |
| 11 | 119814550 | rs61899037 | A | G | 0.913 | -7977.388 | -7977.67 | -0.282 | 2.92E-06 |
| 11 | 120574350 | rs17124211 | G | A | 0.939 | -7977.388 | -7977.67 | -0.282 | 2.92E-06 |
| 11 | 120710210 | rs28737445 | G | A | 0.937 | -7977.388 | -7977.671 | -0.282 | 2.92E-06 |
| 11 | 116705021 | rs1263162 | A | T | 0.911 | -7977.388 | -7977.671 | -0.283 | 2.91E-06 |
| 11 | 119490788 | rs79801588 | G | A | 1.074 | -7977.388 | -7977.671 | -0.283 | 2.91E-06 |
| 11 | 119668263 | rs4272793 | C | A | 0.949 | -7977.388 | -7977.671 | -0.283 | 2.91E-06 |
| 11 | 117030633 | rs1312592595 | C | T | 1.085 | -7977.388 | -7977.672 | -0.284 | 2.91E-06 |
| 11 | 117500542 | rs10892146 | T | A | 0.944 | -7977.388 | -7977.673 | -0.284 | 2.91E-06 |
| 11 | 120733629 | rs4512872 | C | A | 1.06 | -7977.388 | -7977.673 | -0.285 | 2.91E-06 |
| 11 | 116424420 | rs17119814 | G | C | 1.067 | -7977.388 | -7977.674 | -0.286 | 2.91E-06 |
| 11 | 116238034 | rs491111 | A | G | 1.056 | -7977.388 | -7977.675 | -0.287 | 2.90E-06 |
| 11 | 120593745 | rs11606461 | T | C | 0.912 | -7977.388 | -7977.675 | -0.287 | 2.90E-06 |
| 11 | 116875338 | rs6589587 | G | T | 1.061 | -7977.388 | -7977.677 | -0.288 | 2.90E-06 |
| 11 | 117613781 | rs7950743 | A | G | 0.912 | -7977.388 | -7977.678 | -0.289 | 2.90E-06 |
| 11 | 117897098 | rs685308 | G | C | 0.946 | -7977.388 | -7977.678 | -0.289 | 2.90E-06 |
| 11 | 119922708 | rs78699173 | A | C | 1.095 | -7977.388 | -7977.677 | -0.289 | 2.90E-06 |
| 11 | 116514997 | rs67613157 | T | C | 1.064 | -7977.388 | -7977.678 | -0.29 | 2.89E-06 |
| 11 | 117102307 | rs2509209 | G | A | 0.946 | -7977.388 | -7977.679 | -0.29 | 2.89E-06 |
| 11 | 117349817 | rs7929976 | G | A | 1.062 | -7977.388 | -7977.678 | -0.29 | 2.89E-06 |
| 11 | 117308432 | rs17120753 | C | T | 0.927 | -7977.388 | -7977.679 | -0.291 | 2.89E-06 |
| 11 | 119901422 | rs669812 | T | C | 0.921 | -7977.388 | -7977.679 | -0.291 | 2.89E-06 |
| 11 | 120474008 | rs4936528 | G | C | 1.056 | -7977.388 | -7977.68 | -0.291 | 2.89E-06 |
| 11 | 117385531 | rs1807049 | G | A | 1.057 | -7977.388 | -7977.681 | -0.292 | 2.89E-06 |
| 11 | 117411730 | rs35266957 | T | C | 0.913 | -7977.388 | -7977.681 | -0.293 | 2.89E-06 |
| 11 | 117414844 | rs3802878 | C | A | 0.912 | -7977.388 | -7977.681 | -0.293 | 2.89E-06 |
| 11 | 119516805 | rs4938700 | C | G | 0.926 | -7977.388 | -7977.682 | -0.294 | 2.88E-06 |
| 11 | 116239462 | rs79643426 | T | C | 1.076 | -7977.388 | -7977.684 | -0.296 | 2.88E-06 |
| 11 | 119831163 | rs35962410 | C | A | 1.087 | -7977.388 | -7977.684 | -0.296 | 2.88E-06 |
| 11 | 117944485 | rs11216736 | C | T | 0.913 | -7977.388 | -7977.686 | -0.297 | 2.87E-06 |
| 11 | 116941985 | rs7101763 | T | G | 1.061 | -7977.388 | -7977.686 | -0.298 | 2.87E-06 |
| 11 | 116945718 | rs4620758 | A | G | 1.061 | -7977.388 | -7977.686 | -0.298 | 2.87E-06 |
| 11 | 117560679 | rs57550545 | C | T | 0.936 | -7977.388 | -7977.686 | -0.298 | 2.87E-06 |
| 11 | 116946652 | rs7130213 | A | C | 1.061 | -7977.388 | -7977.687 | -0.299 | 2.87E-06 |
| 11 | 117488572 | rs59258276 | C | T | 0.913 | -7977.388 | -7977.688 | -0.299 | 2.87E-06 |
| 11 | 120574079 | rs61901377 | G | A | 0.932 | -7977.388 | -7977.687 | -0.299 | 2.87E-06 |
| 11 | 117488030 | rs148543195 | T | C | 0.913 | -7977.388 | -7977.688 | -0.3 | 2.87E-06 |
| 11 | 120572772 | rs117165966 | C | G | 0.933 | -7977.388 | -7977.69 | -0.302 | 2.86E-06 |
| 11 | 120572814 | rs142600127 | A | G | 0.933 | -7977.388 | -7977.69 | -0.302 | 2.86E-06 |
| 11 | 116225940 | rs12576431 | G | C | 0.922 | -7977.388 | -7977.691 | -0.303 | 2.86E-06 |
| 11 | 119509625 | rs1022080 | C | T | 0.93 | -7977.388 | -7977.691 | -0.303 | 2.86E-06 |
| 11 | 120126017 | rs79988761 | A | G | 0.929 | -7977.388 | -7977.692 | -0.304 | 2.85E-06 |
| 11 | 119881526 | rs1893394 | A | G | 0.933 | -7977.388 | -7977.693 | -0.305 | 2.85E-06 |
| 11 | 117713187 | rs12363888 | T | C | 1.084 | -7977.388 | -7977.695 | -0.306 | 2.85E-06 |
| 11 | 120734419 | rs4344499 | G | T | 1.06 | -7977.388 | -7977.694 | -0.306 | 2.85E-06 |
| 11 | 120837575 | rs7939112 | A | G | 0.942 | -7977.388 | -7977.694 | -0.306 | 2.85E-06 |
| 11 | 120851138 | rs34203183 | G | C | 0.94 | -7977.388 | -7977.695 | -0.307 | 2.85E-06 |
| 11 | 116424451 | rs78381558 | C | G | 1.067 | -7977.388 | -7977.697 | -0.308 | 2.84E-06 |
| 11 | 116638854 | rs78908794 | C | T | 0.913 | -7977.388 | -7977.696 | -0.308 | 2.84E-06 |
| 11 | 117061775 | rs12419470 | T | C | 1.084 | -7977.388 | -7977.697 | -0.308 | 2.84E-06 |
| 11 | 117061825 | rs61703672 | T | C | 1.084 | -7977.388 | -7977.697 | -0.308 | 2.84E-06 |
| 11 | 117065066 | rs3781985 | C | G | 1.084 | -7977.388 | -7977.697 | -0.308 | 2.84E-06 |
| 11 | 117065153 | rs80183569 | A | G | 1.084 | -7977.388 | -7977.697 | -0.308 | 2.84E-06 |
| 11 | 117065533 | rs11606480 | A | G | 1.084 | -7977.388 | -7977.697 | -0.308 | 2.84E-06 |
| 11 | 117225247 | rs117754274 | C | G | 0.914 | -7977.388 | -7977.697 | -0.308 | 2.84E-06 |
| 11 | 119922728 | rs79533157 | C | G | 1.093 | -7977.388 | -7977.696 | -0.308 | 2.84E-06 |
| 11 | 120563557 | rs34463064 | T | C | 1.087 | -7977.388 | -7977.696 | -0.308 | 2.84E-06 |
| 11 | 120834991 | rs7948885 | A | G | 0.942 | -7977.388 | -7977.698 | -0.309 | 2.84E-06 |
| 11 | 119807402 | rs10790356 | T | A | 1.065 | -7977.388 | -7977.699 | -0.31 | 2.84E-06 |
| 11 | 116818527 | rs112704540 | G | A | 1.061 | -7977.388 | -7977.7 | -0.311 | 2.83E-06 |
| 11 | 119730115 | rs79365963 | C | T | 1.07 | -7977.388 | -7977.7 | -0.311 | 2.83E-06 |
| 11 | 120577100 | rs12295113 | T | C | 0.933 | -7977.388 | -7977.699 | -0.311 | 2.83E-06 |
| 11 | 120572387 | rs61901372 | C | T | 0.933 | -7977.388 | -7977.7 | -0.312 | 2.83E-06 |
| 11 | 120572952 | rs61901374 | C | T | 0.933 | -7977.388 | -7977.7 | -0.312 | 2.83E-06 |
| 11 | 120573265 | rs61901375 | C | T | 0.933 | -7977.388 | -7977.7 | -0.312 | 2.83E-06 |
| 11 | 117015516 | rs12419272 | G | C | 1.084 | -7977.388 | -7977.701 | -0.313 | 2.83E-06 |
| 11 | 117035319 | rs12420127 | G | A | 1.084 | -7977.388 | -7977.701 | -0.313 | 2.83E-06 |
| 11 | 117941082 | rs143620404 | T | G | 0.914 | -7977.388 | -7977.702 | -0.313 | 2.83E-06 |
| 11 | 116236469 | rs545675 | T | C | 1.055 | -7977.388 | -7977.703 | -0.314 | 2.83E-06 |
| 11 | 116789849 | rs76844341 | G | A | 0.914 | -7977.388 | -7977.702 | -0.314 | 2.83E-06 |
| 11 | 116953645 | rs1871756 | A | G | 1.061 | -7977.388 | -7977.703 | -0.314 | 2.83E-06 |
| 11 | 118285369 | rs150339153 | T | C | 0.916 | -7977.388 | -7977.702 | -0.314 | 2.83E-06 |
| 11 | 120084221 | rs1551242 | T | C | 0.915 | -7977.388 | -7977.703 | -0.315 | 2.82E-06 |
| 11 | 120084509 | rs1551241 | C | T | 0.915 | -7977.388 | -7977.703 | -0.315 | 2.82E-06 |
| 11 | 117384412 | rs7107020 | C | T | 0.947 | -7977.388 | -7977.704 | -0.316 | 2.82E-06 |
| 11 | 120583174 | rs6589834 | C | G | 0.934 | -7977.388 | -7977.704 | -0.316 | 2.82E-06 |

| | | | | | | | |
|---|---|---|---|---|---|---|---|
| 11 | 120824692 | rs11218078 | G | T | NULL | 0.538672605 | NULL&M_Disease |
| 11 | 121011823 | rs594951 | C | T | NULL | 0.538672605 | NULL&M_Disease |
| 11 | 117410428 | rs2509005 | A | G | NULL | 0.538672605 | NULL&M_Disease |
| 11 | 117482186 | rs184607052 | G | A | NULL | 0.538672605 | NULL&M_Disease |
| 11 | 116899414 | rs73594168 | T | C | NULL | 0.538672605 | NULL&M_Disease |
| 11 | 116899617 | rs73594169 | T | C | NULL | 0.538672605 | NULL&M_Disease |
| 11 | 119732432 | rs475159 | T | G | NULL | 0.538672605 | NULL&M_Disease |
| 11 | 116804144 | rs7107702 | C | T | NULL | 0.539914885 | NULL&M_Disease |
| 11 | 116972339 | rs78459638 | A | G | NULL | 0.538672605 | NULL&M_Disease |
| 11 | 117031363 | rs142017584 | A | G | NULL | 0.538672605 | NULL&M_Disease |
| 11 | 117039461 | rs116773440 | C | A | NULL | 0.538672605 | NULL&M_Disease |
| 11 | 119214292 | rs1893176 | G | A | NULL | 0.539914885 | NULL&M_Disease |
| 11 | 120572361 | rs61901371 | A | G | NULL | 0.539914885 | NULL&M_Disease |
| 11 | 116660768 | rs34089864 | A | G | NULL | 0.538672605 | NULL&M_Disease |
| 11 | 118014873 | rs112525313 | T | C | NULL | 0.539914885 | NULL&M_Disease |
| 11 | 119539324 | rs4938705 | A | G | NULL | 0.539914885 | NULL&M_Disease |
| 11 | 120445835 | rs4500499 | A | G | NULL | 0.539914885 | NULL&M_Disease |
| 11 | 117408179 | rs661631 | T | C | NULL | 0.539914885 | NULL&M_Disease |
| 11 | 119881097 | rs2848729 | T | C | NULL | 0.539914885 | NULL&M_Disease |
| 11 | 117351433 | rs553370 | G | A | NULL | 0.539914885 | NULL&M_Disease |
| 11 | 117797120 | rs73022498 | C | T | NULL | 0.539914885 | NULL&M_Disease |
| 11 | 116197556 | rs7932598 | T | G | NULL | 0.541156668 | NULL&M_Disease |
| 11 | 116700775 | rs12721090 | T | C | NULL | 0.539914885 | NULL&M_Disease |
| 11 | 119795133 | rs576764 | G | A | NULL | 0.541156668 | NULL&M_Disease |
| 11 | 116931087 | rs111426457 | G | C | NULL | 0.539914885 | NULL&M_Disease |
| 11 | 116934869 | rs7946101 | A | G | NULL | 0.539914885 | NULL&M_Disease |
| 11 | 116936488 | rs1982679 | A | G | NULL | 0.539914885 | NULL&M_Disease |
| 11 | 116939724 | rs10502225 | T | A | NULL | 0.539914885 | NULL&M_Disease |
| 11 | 116941382 | rs111641976 | A | G | NULL | 0.539914885 | NULL&M_Disease |
| 11 | 116942479 | rs57787606 | A | G | NULL | 0.539914885 | NULL&M_Disease |
| 11 | 116944304 | rs73576634 | T | G | NULL | 0.539914885 | NULL&M_Disease |
| 11 | 116945074 | rs73576637 | G | A | NULL | 0.539914885 | NULL&M_Disease |
| 11 | 116948173 | rs56188860 | A | T | NULL | 0.539914885 | NULL&M_Disease |
| 11 | 116948761 | rs7108830 | A | G | NULL | 0.539914885 | NULL&M_Disease |
| 11 | 116950278 | rs59752757 | C | A | NULL | 0.539914885 | NULL&M_Disease |
| 11 | 116950567 | rs73576652 | A | G | NULL | 0.539914885 | NULL&M_Disease |
| 11 | 116951615 | rs7121475 | A | G | NULL | 0.539914885 | NULL&M_Disease |
| 11 | 116953604 | rs73576656 | T | C | NULL | 0.539914885 | NULL&M_Disease |
| 11 | 116953784 | rs73576658 | A | G | NULL | 0.539914885 | NULL&M_Disease |
| 11 | 117016450 | rs2306471 | T | C | NULL | 0.539914885 | NULL&M_Disease |
| 11 | 117017455 | rs12419436 | G | T | NULL | 0.539914885 | NULL&M_Disease |
| 11 | 117017530 | rs12420725 | G | A | NULL | 0.539914885 | NULL&M_Disease |
| 11 | 117017714 | rs12417984 | A | G | NULL | 0.539914885 | NULL&M_Disease |
| 11 | 117017762 | rs139864910 | C | A | NULL | 0.539914885 | NULL&M_Disease |
| 11 | 117019867 | rs139146464 | G | A | NULL | 0.539914885 | NULL&M_Disease |
| 11 | 117025932 | rs12418349 | G | A | NULL | 0.539914885 | NULL&M_Disease |
| 11 | 117028727 | rs12420250 | A | G | NULL | 0.539914885 | NULL&M_Disease |
| 11 | 117029785 | rs78166985 | T | A | NULL | 0.539914885 | NULL&M_Disease |
| 11 | 117038741 | rs77171490 | A | T | NULL | 0.539914885 | NULL&M_Disease |
| 11 | 117040505 | rs77678768 | C | G | NULL | 0.539914885 | NULL&M_Disease |
| 11 | 117048122 | rs80353869 | A | G | NULL | 0.539914885 | NULL&M_Disease |
| 11 | 116637677 | rs141740187 | T | G | NULL | 0.539914885 | NULL&M_Disease |
| 11 | 118061798 | rs4553405 | T | C | NULL | 0.541156668 | NULL&M_Disease |
| 11 | 120212377 | rs139034795 | G | A | NULL | 0.539914885 | NULL&M_Disease |
| 11 | 116961510 | rs61244196 | A | G | NULL | 0.539914885 | NULL&M_Disease |
| 11 | 116961617 | rs56125819 | A | G | NULL | 0.539914885 | NULL&M_Disease |
| 11 | 118824683 | rs507763 | C | G | NULL | 0.541156668 | NULL&M_Disease |
| 11 | 120457909 | rs11603180 | A | G | NULL | 0.541156668 | NULL&M_Disease |
| 11 | 120825043 | rs11218079 | T | C | NULL | 0.539914885 | NULL&M_Disease |
| 11 | 120647474 | rs72995856 | C | T | NULL | 0.539914885 | NULL&M_Disease |
| 11 | 117193203 | rs555075 | T | C | NULL | 0.541156668 | NULL&M_Disease |
| 11 | 117614307 | rs10750110 | G | T | NULL | 0.539914885 | NULL&M_Disease |
| 11 | 120462677 | rs17311324 | T | G | NULL | 0.541156668 | NULL&M_Disease |
| 11 | 116852507 | rs4938321 | T | C | NULL | 0.541156668 | NULL&M_Disease |
| 11 | 117501339 | rs6589617 | T | C | NULL | 0.541156668 | NULL&M_Disease |
| 11 | 117741933 | rs61901489 | C | G | NULL | 0.541156668 | NULL&M_Disease |
| 11 | 120036845 | rs528147 | A | G | NULL | 0.541156668 | NULL&M_Disease |
| 11 | 120414641 | rs12575317 | C | T | NULL | 0.541156668 | NULL&M_Disease |
| 11 | 116990851 | rs12366015 | A | G | NULL | 0.542397941 | NULL&M_Disease |
| 11 | 119917798 | rs635335 | T | C | NULL | 0.541156668 | NULL&M_Disease |
| 11 | 119937782 | rs55897720 | G | A | NULL | 0.541156668 | NULL&M_Disease |
| 11 | 116304705 | rs73580446 | T | G | NULL | 0.541156668 | NULL&M_Disease |
| 11 | 117063709 | rs2301174 | G | C | NULL | 0.541156668 | NULL&M_Disease |
| 11 | 117410507 | rs2509006 | T | C | NULL | 0.542397941 | NULL&M_Disease |
| 11 | 117433042 | rs145811313 | A | C | NULL | 0.541156668 | NULL&M_Disease |
| 11 | 119512770 | rs2089648 | A | G | NULL | 0.542397941 | NULL&M_Disease |
| 11 | 119682918 | rs7125270 | T | C | NULL | 0.542397941 | NULL&M_Disease |
| 11 | 119835014 | rs66921644 | A | G | NULL | 0.541156668 | NULL&M_Disease |
| 11 | 117490095 | rs76589374 | T | G | NULL | 0.541156668 | NULL&M_Disease |
| 11 | 117916692 | rs58618446 | T | C | NULL | 0.541156668 | NULL&M_Disease |
| 11 | 117918702 | rs11216730 | T | C | NULL | 0.541156668 | NULL&M_Disease |
| 11 | 121186914 | rs7128664 | G | C | NULL | 0.542397941 | NULL&M_Disease |
| 11 | 116861268 | rs7939934 | T | A | NULL | 0.542397941 | NULL&M_Disease |
| 11 | 116868221 | rs10892054 | T | C | NULL | 0.542397941 | NULL&M_Disease |
| 11 | 116868371 | rs10892055 | A | G | NULL | 0.542397941 | NULL&M_Disease |
| 11 | 117282742 | rs67023719 | T | C | NULL | 0.542397941 | NULL&M_Disease |
| 11 | 120500130 | rs1574249 | A | G | NULL | 0.542397941 | NULL&M_Disease |
| 11 | 116834971 | rs1815788 | C | T | NULL | 0.542397941 | NULL&M_Disease |
| 11 | 118660932 | rs1784295 | A | G | NULL | 0.542397941 | NULL&M_Disease |
| 11 | 118661055 | rs1790188 | T | C | NULL | 0.542397941 | NULL&M_Disease |
| 11 | 120454144 | rs61900900 | T | C | NULL | 0.542397941 | NULL&M_Disease |
| 11 | 119806104 | rs74330115 | A | G | NULL | 0.542397941 | NULL&M_Disease |
| 11 | 116423307 | rs12293155 | G | A | NULL | 0.542397941 | NULL&M_Disease |
| 11 | 118722946 | rs148990923 | T | C | NULL | 0.542397941 | NULL&M_Disease |
| 11 | 119923914 | rs142100881 | G | A | NULL | 0.542397941 | NULL&M_Disease |
| 11 | 120641004 | rs10502242 | G | A | NULL | 0.542397941 | NULL&M_Disease |

| | | | | | | | | | |
|---|---|---|---|---|---|---|---|---|---|
| 11 | 120824692 | rs11218078 | G | T | 0.941 | -7977.388 | -7977.704 | -0.316 | 2.82E-06 |
| 11 | 121011823 | rs594951 | C | T | 0.948 | -7977.388 | -7977.704 | -0.316 | 2.82E-06 |
| 11 | 117410428 | rs2509005 | A | G | 1.053 | -7977.388 | -7977.705 | -0.317 | 2.82E-06 |
| 11 | 117482186 | rs184607052 | G | A | 0.915 | -7977.388 | -7977.707 | -0.318 | 2.81E-06 |
| 11 | 116899414 | rs73594168 | T | C | 0.915 | -7977.388 | -7977.707 | -0.319 | 2.81E-06 |
| 11 | 116899617 | rs73594169 | T | C | 0.915 | -7977.388 | -7977.707 | -0.319 | 2.81E-06 |
| 11 | 119732432 | rs475159 | T | G | 1.081 | -7977.388 | -7977.708 | -0.319 | 2.81E-06 |
| 11 | 116804144 | rs7107702 | C | T | 1.061 | -7977.388 | -7977.708 | -0.32 | 2.81E-06 |
| 11 | 116972339 | rs78459638 | A | G | 0.915 | -7977.388 | -7977.708 | -0.32 | 2.81E-06 |
| 11 | 117031363 | rs142017584 | A | G | 1.083 | -7977.388 | -7977.708 | -0.32 | 2.81E-06 |
| 11 | 117039461 | rs116773440 | C | A | 1.083 | -7977.388 | -7977.708 | -0.32 | 2.81E-06 |
| 11 | 119214292 | rs1893176 | G | A | 1.054 | -7977.388 | -7977.709 | -0.32 | 2.81E-06 |
| 11 | 120572361 | rs61901371 | A | G | 0.933 | -7977.388 | -7977.711 | -0.323 | 2.80E-06 |
| 11 | 116660768 | rs34089864 | A | G | 0.915 | -7977.388 | -7977.712 | -0.324 | 2.80E-06 |
| 11 | 118014873 | rs112525313 | T | C | 1.077 | -7977.388 | -7977.714 | -0.325 | 2.79E-06 |
| 11 | 119539324 | rs4938705 | A | G | 0.949 | -7977.388 | -7977.713 | -0.325 | 2.79E-06 |
| 11 | 120445835 | rs4500499 | A | G | 1.054 | -7977.388 | -7977.714 | -0.326 | 2.79E-06 |
| 11 | 117408179 | rs661631 | T | C | 1.053 | -7977.388 | -7977.715 | -0.327 | 2.79E-06 |
| 11 | 119881097 | rs2848729 | T | C | 0.933 | -7977.388 | -7977.716 | -0.327 | 2.79E-06 |
| 11 | 117351433 | rs553370 | G | A | 0.933 | -7977.388 | -7977.716 | -0.328 | 2.79E-06 |
| 11 | 117797120 | rs73022498 | C | T | 0.931 | -7977.388 | -7977.716 | -0.328 | 2.79E-06 |
| 11 | 116197556 | rs7932598 | T | G | 1.066 | -7977.388 | -7977.718 | -0.329 | 2.78E-06 |
| 11 | 116700775 | rs12721090 | T | C | 1.09 | -7977.388 | -7977.718 | -0.329 | 2.78E-06 |
| 11 | 119795133 | rs576764 | G | A | 0.945 | -7977.388 | -7977.717 | -0.329 | 2.78E-06 |
| 11 | 116931087 | rs111426457 | G | C | 0.916 | -7977.388 | -7977.718 | -0.33 | 2.78E-06 |
| 11 | 116934869 | rs7946101 | A | G | 0.916 | -7977.388 | -7977.718 | -0.33 | 2.78E-06 |
| 11 | 116936488 | rs1982679 | A | G | 0.916 | -7977.388 | -7977.718 | -0.33 | 2.78E-06 |
| 11 | 116939724 | rs10502225 | T | A | 0.916 | -7977.388 | -7977.718 | -0.33 | 2.78E-06 |
| 11 | 116941382 | rs111641976 | A | G | 0.916 | -7977.388 | -7977.718 | -0.33 | 2.78E-06 |
| 11 | 116942479 | rs57787606 | A | G | 0.916 | -7977.388 | -7977.718 | -0.33 | 2.78E-06 |
| 11 | 116944304 | rs73576634 | T | G | 0.916 | -7977.388 | -7977.718 | -0.33 | 2.78E-06 |
| 11 | 116945074 | rs73576637 | G | A | 0.916 | -7977.388 | -7977.718 | -0.33 | 2.78E-06 |
| 11 | 116948173 | rs56188860 | A | T | 0.916 | -7977.388 | -7977.718 | -0.33 | 2.78E-06 |
| 11 | 116948761 | rs7108830 | A | G | 0.916 | -7977.388 | -7977.718 | -0.33 | 2.78E-06 |
| 11 | 116950278 | rs59752757 | C | A | 0.916 | -7977.388 | -7977.718 | -0.33 | 2.78E-06 |
| 11 | 116950567 | rs73576652 | A | G | 0.916 | -7977.388 | -7977.718 | -0.33 | 2.78E-06 |
| 11 | 116951615 | rs7121475 | A | G | 0.916 | -7977.388 | -7977.718 | -0.33 | 2.78E-06 |
| 11 | 116953604 | rs73576656 | T | C | 0.916 | -7977.388 | -7977.718 | -0.33 | 2.78E-06 |
| 11 | 116953784 | rs73576658 | A | G | 0.916 | -7977.388 | -7977.718 | -0.33 | 2.78E-06 |
| 11 | 117016450 | rs2306471 | T | C | 1.083 | -7977.388 | -7977.718 | -0.33 | 2.78E-06 |
| 11 | 117017455 | rs12419436 | G | T | 1.083 | -7977.388 | -7977.718 | -0.33 | 2.78E-06 |
| 11 | 117017530 | rs12420725 | G | A | 1.083 | -7977.388 | -7977.718 | -0.33 | 2.78E-06 |
| 11 | 117017714 | rs12417984 | A | G | 1.083 | -7977.388 | -7977.718 | -0.33 | 2.78E-06 |
| 11 | 117017762 | rs139864910 | C | A | 1.083 | -7977.388 | -7977.718 | -0.33 | 2.78E-06 |
| 11 | 117019867 | rs139146464 | G | A | 1.083 | -7977.388 | -7977.718 | -0.33 | 2.78E-06 |
| 11 | 117025932 | rs12418349 | G | A | 1.083 | -7977.388 | -7977.718 | -0.33 | 2.78E-06 |
| 11 | 117028727 | rs12420250 | A | G | 1.083 | -7977.388 | -7977.718 | -0.33 | 2.78E-06 |
| 11 | 117029785 | rs78166985 | T | A | 1.083 | -7977.388 | -7977.718 | -0.33 | 2.78E-06 |
| 11 | 117038741 | rs77171490 | A | T | 1.083 | -7977.388 | -7977.718 | -0.33 | 2.78E-06 |
| 11 | 117040505 | rs77678768 | C | G | 1.083 | -7977.388 | -7977.718 | -0.33 | 2.78E-06 |
| 11 | 117048122 | rs80353869 | A | G | 1.083 | -7977.388 | -7977.718 | -0.33 | 2.78E-06 |
| 11 | 116637677 | rs141740187 | T | G | 0.915 | -7977.388 | -7977.72 | -0.331 | 2.78E-06 |
| 11 | 118061798 | rs4553405 | T | C | 1.059 | -7977.388 | -7977.719 | -0.331 | 2.78E-06 |
| 11 | 120212377 | rs139034795 | G | A | 0.917 | -7977.388 | -7977.719 | -0.331 | 2.78E-06 |
| 11 | 116961510 | rs61244196 | A | G | 0.916 | -7977.388 | -7977.72 | -0.332 | 2.78E-06 |
| 11 | 116961617 | rs56125819 | A | G | 0.916 | -7977.388 | -7977.72 | -0.332 | 2.78E-06 |
| 11 | 118824683 | rs507763 | C | G | 1.063 | -7977.388 | -7977.72 | -0.332 | 2.78E-06 |
| 11 | 120457909 | rs11603180 | A | G | 1.054 | -7977.388 | -7977.72 | -0.332 | 2.78E-06 |
| 11 | 120825043 | rs11218079 | T | C | 1.091 | -7977.388 | -7977.72 | -0.332 | 2.78E-06 |
| 11 | 120647474 | rs72995856 | C | T | 0.916 | -7977.388 | -7977.722 | -0.333 | 2.77E-06 |
| 11 | 117193203 | rs555075 | T | C | 0.95 | -7977.388 | -7977.723 | -0.335 | 2.77E-06 |
| 11 | 117614307 | rs10750110 | G | T | 0.917 | -7977.388 | -7977.723 | -0.335 | 2.77E-06 |
| 11 | 120462677 | rs17311324 | T | G | 1.054 | -7977.388 | -7977.724 | -0.335 | 2.77E-06 |
| 11 | 116852507 | rs4938321 | T | C | 1.06 | -7977.388 | -7977.725 | -0.336 | 2.76E-06 |
| 11 | 117501339 | rs6589617 | T | C | 0.946 | -7977.388 | -7977.724 | -0.336 | 2.76E-06 |
| 11 | 117741933 | rs61901489 | C | G | 0.936 | -7977.388 | -7977.725 | -0.336 | 2.76E-06 |
| 11 | 120036845 | rs528147 | A | G | 0.926 | -7977.388 | -7977.726 | -0.337 | 2.76E-06 |
| 11 | 120414641 | rs12575317 | C | T | 0.921 | -7977.388 | -7977.726 | -0.337 | 2.76E-06 |
| 11 | 116990851 | rs12366015 | A | G | 0.943 | -7977.388 | -7977.726 | -0.338 | 2.76E-06 |
| 11 | 119917798 | rs635335 | T | C | 0.922 | -7977.388 | -7977.726 | -0.338 | 2.76E-06 |
| 11 | 119937782 | rs55897720 | G | A | 1.067 | -7977.388 | -7977.726 | -0.338 | 2.76E-06 |
| 11 | 116304705 | rs73580446 | T | G | 1.08 | -7977.388 | -7977.728 | -0.34 | 2.75E-06 |
| 11 | 117063709 | rs2301174 | G | C | 1.082 | -7977.388 | -7977.728 | -0.34 | 2.75E-06 |
| 11 | 117410507 | rs2509006 | T | C | 1.053 | -7977.388 | -7977.728 | -0.34 | 2.75E-06 |
| 11 | 117433042 | rs145811313 | A | C | 0.916 | -7977.388 | -7977.728 | -0.34 | 2.75E-06 |
| 11 | 119512770 | rs2089648 | A | G | 0.927 | -7977.388 | -7977.728 | -0.34 | 2.75E-06 |
| 11 | 119682918 | rs7125270 | T | C | 0.939 | -7977.388 | -7977.729 | -0.34 | 2.75E-06 |
| 11 | 119835014 | rs66921644 | A | G | 1.085 | -7977.388 | -7977.728 | -0.34 | 2.75E-06 |
| 11 | 117490095 | rs76589374 | T | G | 0.916 | -7977.388 | -7977.73 | -0.341 | 2.75E-06 |
| 11 | 117916692 | rs58618446 | T | C | 0.916 | -7977.388 | -7977.729 | -0.341 | 2.75E-06 |
| 11 | 117918702 | rs11216730 | T | C | 0.916 | -7977.388 | -7977.729 | -0.341 | 2.75E-06 |
| 11 | 121186914 | rs7128664 | G | C | 1.063 | -7977.388 | -7977.729 | -0.341 | 2.75E-06 |
| 11 | 116861268 | rs7939934 | T | A | 1.06 | -7977.388 | -7977.731 | -0.342 | 2.75E-06 |
| 11 | 116868221 | rs10892054 | T | C | 1.06 | -7977.388 | -7977.731 | -0.342 | 2.75E-06 |
| 11 | 116868371 | rs10892055 | A | G | 1.06 | -7977.388 | -7977.731 | -0.342 | 2.75E-06 |
| 11 | 117282742 | rs67023719 | T | C | 0.919 | -7977.388 | -7977.73 | -0.342 | 2.75E-06 |
| 11 | 120500130 | rs1574249 | A | G | 0.95 | -7977.388 | -7977.73 | -0.342 | 2.75E-06 |
| 11 | 116834971 | rs1815788 | C | T | 1.06 | -7977.388 | -7977.732 | -0.343 | 2.74E-06 |
| 11 | 118660932 | rs1784295 | A | G | 0.936 | -7977.388 | -7977.732 | -0.344 | 2.74E-06 |
| 11 | 118661055 | rs1790188 | T | C | 0.936 | -7977.388 | -7977.732 | -0.344 | 2.74E-06 |
| 11 | 120454144 | rs61900900 | T | C | 1.053 | -7977.388 | -7977.732 | -0.344 | 2.74E-06 |
| 11 | 119806104 | rs74330115 | A | G | 1.08 | -7977.388 | -7977.734 | -0.345 | 2.74E-06 |
| 11 | 116423307 | rs12293155 | G | A | 1.065 | -7977.388 | -7977.734 | -0.346 | 2.74E-06 |
| 11 | 118722946 | rs148990923 | T | C | 1.089 | -7977.388 | -7977.734 | -0.346 | 2.74E-06 |
| 11 | 119923914 | rs142100881 | G | A | 0.918 | -7977.388 | -7977.735 | -0.347 | 2.73E-06 |
| 11 | 120641004 | rs10502242 | G | A | 0.917 | -7977.388 | -7977.736 | -0.347 | 2.73E-06 |

| | | | | | | | |
|---|---|---|---|---|---|---|---|
| 11 | 120642631 | rs72995844 | G | A | NULL | 0.542397941 | NULL&M_Disease |
| 11 | 120642698 | rs17124301 | C | T | NULL | 0.542397941 | NULL&M_Disease |
| 11 | 117023706 | rs143494314 | T | C | NULL | 0.542397941 | NULL&M_Disease |
| 11 | 117035016 | rs150984554 | A | G | NULL | 0.542397941 | NULL&M_Disease |
| 11 | 118550522 | rs2277296 | C | T | NULL | 0.543638687 | NULL&M_Disease |
| 11 | 118550524 | rs201134460 | T | C | NULL | 0.543638687 | NULL&M_Disease |
| 11 | 116949853 | rs59839888 | C | T | NULL | 0.542397941 | NULL&M_Disease |
| 11 | 117015238 | rs1001556897 | A | C | NULL | 0.542397941 | NULL&M_Disease |
| 11 | 117488010 | rs145749713 | A | G | NULL | 0.542397941 | NULL&M_Disease |
| 11 | 119733183 | rs112201912 | A | G | NULL | 0.543638687 | NULL&M_Disease |
| 11 | 120463065 | rs1573789 | T | C | NULL | 0.543638687 | NULL&M_Disease |
| 11 | 120500604 | rs7936516 | T | G | NULL | 0.543638687 | NULL&M_Disease |
| 11 | 120083118 | rs72645479 | C | T | NULL | 0.543638687 | NULL&M_Disease |
| 11 | 117059794 | rs7933134 | C | G | NULL | 0.543638687 | NULL&M_Disease |
| 11 | 119544722 | rs113300825 | T | C | NULL | 0.542397941 | NULL&M_Disease |
| 11 | 116734688 | rs557331 | A | G | NULL | 0.542397941 | NULL&M_Disease |
| 11 | 117906061 | rs77084900 | C | T | NULL | 0.543638687 | NULL&M_Disease |
| 11 | 117058062 | rs2306472 | T | C | NULL | 0.543638687 | NULL&M_Disease |
| 11 | 117594591 | rs4938436 | G | A | NULL | 0.543638687 | NULL&M_Disease |
| 11 | 117924155 | rs79944284 | A | G | NULL | 0.543638687 | NULL&M_Disease |
| 11 | 117344866 | rs79297003 | T | C | NULL | 0.543638687 | NULL&M_Disease |
| 11 | 117345114 | rs114005593 | C | G | NULL | 0.543638687 | NULL&M_Disease |
| 11 | 119935735 | rs11217667 | A | C | NULL | 0.543638687 | NULL&M_Disease |
| 11 | 120632841 | rs11824814 | T | A | NULL | 0.543638687 | NULL&M_Disease |
| 11 | 120792086 | rs61588877 | G | A | NULL | 0.543638687 | NULL&M_Disease |
| 11 | 117912267 | rs11216726 | T | C | NULL | 0.543638687 | NULL&M_Disease |
| 11 | 119804760 | rs139886195 | A | C | NULL | 0.543638687 | NULL&M_Disease |
| 11 | 120565293 | rs76003278 | C | T | NULL | 0.543638687 | NULL&M_Disease |
| 11 | 120565473 | rs74482798 | G | C | NULL | 0.543638687 | NULL&M_Disease |
| 11 | 116501409 | rs1944154 | C | T | NULL | 0.544878892 | NULL&M_Disease |
| 11 | 119808535 | rs2006623 | A | G | NULL | 0.544878892 | NULL&M_Disease |
| 11 | 117191734 | rs1784045 | C | T | NULL | 0.544878892 | NULL&M_Disease |
| 11 | 117329415 | rs520328 | A | G | NULL | 0.544878892 | NULL&M_Disease |
| 11 | 119029058 | rs192734017 | A | G | NULL | 0.543638687 | NULL&M_Disease |
| 11 | 119499747 | rs145993550 | A | G | NULL | 0.544878892 | NULL&M_Disease |
| 11 | 117053433 | rs2046150 | A | C | NULL | 0.544878892 | NULL&M_Disease |
| 11 | 119674611 | rs4938725 | G | A | NULL | 0.544878892 | NULL&M_Disease |
| 11 | 116325615 | rs147834459 | T | C | NULL | 0.543638687 | NULL&M_Disease |
| 11 | 121079478 | rs1231418 | C | G | NULL | 0.544878892 | NULL&M_Disease |
| 11 | 118590743 | rs503542 | G | A | NULL | 0.544878892 | NULL&M_Disease |
| 11 | 120847827 | rs78474864 | A | G | NULL | 0.544878892 | NULL&M_Disease |
| 11 | 120853577 | rs646543 | C | T | NULL | 0.544878892 | NULL&M_Disease |
| 11 | 117907713 | rs12223876 | A | T | NULL | 0.544878892 | NULL&M_Disease |
| 11 | 117908922 | rs11216718 | A | G | NULL | 0.544878892 | NULL&M_Disease |
| 11 | 117908985 | rs11216719 | C | A | NULL | 0.544878892 | NULL&M_Disease |
| 11 | 117909101 | rs11216720 | T | C | NULL | 0.544878892 | NULL&M_Disease |
| 11 | 117909531 | rs11216721 | T | C | NULL | 0.544878892 | NULL&M_Disease |
| 11 | 117910017 | rs12223165 | A | G | NULL | 0.544878892 | NULL&M_Disease |
| 11 | 117910116 | rs12223183 | A | G | NULL | 0.544878892 | NULL&M_Disease |
| 11 | 117910244 | rs11216722 | A | G | NULL | 0.544878892 | NULL&M_Disease |
| 11 | 117911450 | rs11216725 | T | C | NULL | 0.544878892 | NULL&M_Disease |
| 11 | 118540104 | rs506234 | G | C | NULL | 0.546118541 | NULL&M_Disease |
| 11 | 118556398 | rs11216943 | A | G | NULL | 0.546118541 | NULL&M_Disease |
| 11 | 119935489 | rs60708230 | A | G | NULL | 0.544878892 | NULL&M_Disease |
| 11 | 120632745 | rs11820186 | A | G | NULL | 0.544878892 | NULL&M_Disease |
| 11 | 118713180 | rs2187490 | G | T | NULL | 0.546118541 | NULL&M_Disease |
| 11 | 119766827 | rs78540725 | C | A | NULL | 0.544878892 | NULL&M_Disease |
| 11 | 120843834 | rs601817 | C | T | NULL | 0.546118541 | NULL&M_Disease |
| 11 | 116225239 | rs12576376 | T | C | NULL | 0.544878892 | NULL&M_Disease |
| 11 | 116752558 | rs634044 | C | T | NULL | 0.546118541 | NULL&M_Disease |
| 11 | 120570587 | rs12280296 | G | A | NULL | 0.546118541 | NULL&M_Disease |
| 11 | 116781545 | rs11216183 | A | C | NULL | 0.546118541 | NULL&M_Disease |
| 11 | 117384372 | rs144199381 | T | C | NULL | 0.544878892 | NULL&M_Disease |
| 11 | 119906741 | rs582545 | G | C | NULL | 0.546118541 | NULL&M_Disease |
| 11 | 119907628 | rs643377 | A | G | NULL | 0.546118541 | NULL&M_Disease |
| 11 | 120699899 | rs4936555 | T | C | NULL | 0.546118541 | NULL&M_Disease |
| 11 | 120835281 | rs7128009 | A | G | NULL | 0.546118541 | NULL&M_Disease |
| 11 | 119880923 | rs675714 | A | G | NULL | 0.546118541 | NULL&M_Disease |
| 11 | 117164246 | rs45506400 | C | A | NULL | 0.546118541 | NULL&M_Disease |
| 11 | 117218685 | rs555179364 | C | T | NULL | 0.544878892 | NULL&M_Disease |
| 11 | 117921563 | rs76190886 | T | C | NULL | 0.546118541 | NULL&M_Disease |
| 11 | 118059557 | rs10790244 | C | T | NULL | 0.546118541 | NULL&M_Disease |
| 11 | 119236410 | rs35310980 | A | T | NULL | 0.546118541 | NULL&M_Disease |
| 11 | 120648746 | rs115791680 | T | C | NULL | 0.546118541 | NULL&M_Disease |
| 11 | 120862089 | rs4935758 | T | C | NULL | 0.546118541 | NULL&M_Disease |
| 11 | 116700785 | rs11827682 | T | C | NULL | 0.546118541 | NULL&M_Disease |
| 11 | 118865465 | rs4938616 | T | C | NULL | 0.546118541 | NULL&M_Disease |
| 11 | 116292818 | rs116859176 | C | T | NULL | 0.544878892 | NULL&M_Disease |
| 11 | 117764401 | rs112444974 | T | G | NULL | 0.546118541 | NULL&M_Disease |
| 11 | 117250419 | rs180865175 | G | T | NULL | 0.546118541 | NULL&M_Disease |
| 11 | 116429779 | rs61904803 | C | T | NULL | 0.547357618 | NULL&M_Disease |
| 11 | 116959601 | rs56257632 | C | T | NULL | 0.546118541 | NULL&M_Disease |
| 11 | 117297415 | rs7940638 | A | C | NULL | 0.547357618 | NULL&M_Disease |
| 11 | 117481112 | rs78503991 | T | A | NULL | 0.546118541 | NULL&M_Disease |
| 11 | 117647364 | rs13377544 | A | T | NULL | 0.547357618 | NULL&M_Disease |
| 11 | 119515905 | rs11217369 | T | C | NULL | 0.547357618 | NULL&M_Disease |
| 11 | 119516100 | rs10892425 | T | C | NULL | 0.547357618 | NULL&M_Disease |
| 11 | 118655704 | rs6421568 | C | G | NULL | 0.547357618 | NULL&M_Disease |
| 11 | 118655775 | rs6421569 | A | C | NULL | 0.547357618 | NULL&M_Disease |
| 11 | 118037964 | rs2075647 | C | G | NULL | 0.546118541 | NULL&M_Disease |
| 11 | 120531656 | rs11217941 | G | T | NULL | 0.547357618 | NULL&M_Disease |
| 11 | 117341717 | rs56316258 | G | C | NULL | 0.547357618 | NULL&M_Disease |
| 11 | 117518200 | rs673028 | G | A | NULL | 0.547357618 | NULL&M_Disease |
| 11 | 117600876 | rs4936402 | C | T | NULL | 0.546118541 | NULL&M_Disease |
| 11 | 116778180 | rs1979023 | C | T | NULL | 0.547357618 | NULL&M_Disease |
| 11 | 117578142 | rs77699630 | C | T | NULL | 0.547357618 | NULL&M_Disease |

| | | | | | | | | | |
|---|---|---|---|---|---|---|---|---|---|
| 11 | 120642631 | rs72995844 | G | A | 0.917 | -7977.388 | -7977.736 | -0.347 | 2.73E-06 |
| 11 | 120642698 | rs17124301 | C | T | 0.917 | -7977.388 | -7977.736 | -0.347 | 2.73E-06 |
| 11 | 117023706 | rs143494314 | T | C | 1.082 | -7977.388 | -7977.737 | -0.348 | 2.73E-06 |
| 11 | 117035016 | rs150984554 | A | G | 1.082 | -7977.388 | -7977.737 | -0.348 | 2.73E-06 |
| 11 | 118550522 | rs2277296 | C | T | 1.059 | -7977.388 | -7977.737 | -0.348 | 2.73E-06 |
| 11 | 118550524 | rs201134460 | T | C | 1.059 | -7977.388 | -7977.737 | -0.348 | 2.73E-06 |
| 11 | 116949853 | rs59839888 | C | T | 0.917 | -7977.388 | -7977.738 | -0.349 | 2.73E-06 |
| 11 | 117015238 | rs1001556897 | A | C | 1.082 | -7977.388 | -7977.738 | -0.349 | 2.73E-06 |
| 11 | 117488010 | rs145749713 | A | G | 0.917 | -7977.388 | -7977.737 | -0.349 | 2.73E-06 |
| 11 | 119733183 | rs112201912 | A | G | 1.069 | -7977.388 | -7977.738 | -0.349 | 2.73E-06 |
| 11 | 120463065 | rs1573789 | T | C | 1.053 | -7977.388 | -7977.738 | -0.35 | 2.73E-06 |
| 11 | 120500604 | rs7936516 | T | G | 0.95 | -7977.388 | -7977.738 | -0.35 | 2.73E-06 |
| 11 | 120083118 | rs72645479 | C | T | 0.926 | -7977.388 | -7977.739 | -0.351 | 2.72E-06 |
| 11 | 117059794 | rs7933134 | C | G | 1.081 | -7977.388 | -7977.741 | -0.353 | 2.72E-06 |
| 11 | 119544722 | rs113300825 | T | C | 1.089 | -7977.388 | -7977.741 | -0.353 | 2.72E-06 |
| 11 | 116734688 | rs557331 | A | G | 0.918 | -7977.388 | -7977.742 | -0.354 | 2.71E-06 |
| 11 | 117906061 | rs77084900 | C | T | 0.917 | -7977.388 | -7977.743 | -0.354 | 2.71E-06 |
| 11 | 117058062 | rs2306472 | T | C | 1.081 | -7977.388 | -7977.744 | -0.356 | 2.71E-06 |
| 11 | 117594591 | rs4938436 | G | A | 0.919 | -7977.388 | -7977.744 | -0.356 | 2.71E-06 |
| 11 | 117924155 | rs79944284 | A | G | 0.917 | -7977.388 | -7977.744 | -0.356 | 2.71E-06 |
| 11 | 117344866 | rs79297003 | T | C | 1.088 | -7977.388 | -7977.745 | -0.357 | 2.71E-06 |
| 11 | 117345114 | rs114005593 | C | G | 1.088 | -7977.388 | -7977.745 | -0.357 | 2.71E-06 |
| 11 | 119935735 | rs11217667 | A | C | 1.066 | -7977.388 | -7977.745 | -0.357 | 2.71E-06 |
| 11 | 120632841 | rs11824814 | T | A | 0.918 | -7977.388 | -7977.745 | -0.357 | 2.71E-06 |
| 11 | 120792086 | rs61588877 | G | A | 1.085 | -7977.388 | -7977.746 | -0.358 | 2.70E-06 |
| 11 | 117912267 | rs11216726 | T | C | 0.918 | -7977.388 | -7977.748 | -0.359 | 2.70E-06 |
| 11 | 119804760 | rs139886195 | A | C | 1.088 | -7977.388 | -7977.749 | -0.361 | 2.70E-06 |
| 11 | 120565293 | rs76003278 | C | T | 1.078 | -7977.388 | -7977.749 | -0.361 | 2.70E-06 |
| 11 | 120565473 | rs74482798 | G | C | 1.078 | -7977.388 | -7977.749 | -0.361 | 2.70E-06 |
| 11 | 116501409 | rs1944154 | C | T | 0.935 | -7977.388 | -7977.75 | -0.362 | 2.69E-06 |
| 11 | 119808535 | rs2006623 | A | G | 1.056 | -7977.388 | -7977.75 | -0.362 | 2.69E-06 |
| 11 | 117191734 | rs1784045 | C | T | 1.053 | -7977.388 | -7977.751 | -0.363 | 2.69E-06 |
| 11 | 117329415 | rs520328 | A | G | 0.936 | -7977.388 | -7977.752 | -0.364 | 2.69E-06 |
| 11 | 119029058 | rs192734017 | A | G | 1.084 | -7977.388 | -7977.753 | -0.364 | 2.69E-06 |
| 11 | 119499747 | rs145993550 | A | G | 0.918 | -7977.388 | -7977.752 | -0.364 | 2.69E-06 |
| 11 | 117053433 | rs2046150 | A | C | 1.081 | -7977.388 | -7977.753 | -0.365 | 2.69E-06 |
| 11 | 119674611 | rs4938725 | G | A | 0.95 | -7977.388 | -7977.753 | -0.365 | 2.69E-06 |
| 11 | 116325615 | rs147834459 | T | C | 0.921 | -7977.388 | -7977.754 | -0.366 | 2.68E-06 |
| 11 | 121079478 | rs1231418 | C | G | 0.946 | -7977.388 | -7977.754 | -0.366 | 2.68E-06 |
| 11 | 118590743 | rs503542 | G | A | 0.95 | -7977.388 | -7977.756 | -0.367 | 2.68E-06 |
| 11 | 120847827 | rs78474864 | A | G | 1.078 | -7977.388 | -7977.756 | -0.367 | 2.68E-06 |
| 11 | 120853577 | rs646543 | C | T | 0.944 | -7977.388 | -7977.755 | -0.367 | 2.68E-06 |
| 11 | 117907713 | rs12223876 | A | T | 0.918 | -7977.388 | -7977.756 | -0.368 | 2.68E-06 |
| 11 | 117908922 | rs11216718 | A | G | 0.918 | -7977.388 | -7977.756 | -0.368 | 2.68E-06 |
| 11 | 117908985 | rs11216719 | C | A | 0.918 | -7977.388 | -7977.756 | -0.368 | 2.68E-06 |
| 11 | 117909101 | rs11216720 | T | C | 0.918 | -7977.388 | -7977.756 | -0.368 | 2.68E-06 |
| 11 | 117909531 | rs11216721 | T | C | 0.918 | -7977.388 | -7977.756 | -0.368 | 2.68E-06 |
| 11 | 117910017 | rs12223165 | A | G | 0.918 | -7977.388 | -7977.756 | -0.368 | 2.68E-06 |
| 11 | 117910116 | rs12223183 | A | G | 0.918 | -7977.388 | -7977.756 | -0.368 | 2.68E-06 |
| 11 | 117910244 | rs11216722 | A | G | 0.918 | -7977.388 | -7977.756 | -0.368 | 2.68E-06 |
| 11 | 117911450 | rs11216725 | T | C | 0.918 | -7977.388 | -7977.756 | -0.368 | 2.68E-06 |
| 11 | 118540104 | rs506234 | G | C | 0.95 | -7977.388 | -7977.756 | -0.368 | 2.68E-06 |
| 11 | 118556398 | rs11216943 | A | G | 1.059 | -7977.388 | -7977.756 | -0.368 | 2.68E-06 |
| 11 | 119935489 | rs60708230 | A | G | 1.066 | -7977.388 | -7977.756 | -0.368 | 2.68E-06 |
| 11 | 120632745 | rs11820186 | A | G | 0.918 | -7977.388 | -7977.756 | -0.368 | 2.68E-06 |
| 11 | 118713180 | rs2187490 | G | T | 0.931 | -7977.388 | -7977.758 | -0.369 | 2.67E-06 |
| 11 | 119766827 | rs78540725 | C | A | 0.921 | -7977.388 | -7977.757 | -0.369 | 2.67E-06 |
| 11 | 120843834 | rs601817 | C | T | 0.944 | -7977.388 | -7977.758 | -0.369 | 2.67E-06 |
| 11 | 116225239 | rs12576376 | T | C | 0.925 | -7977.388 | -7977.758 | -0.37 | 2.67E-06 |
| 11 | 116752558 | rs634044 | C | T | 1.06 | -7977.388 | -7977.758 | -0.37 | 2.67E-06 |
| 11 | 120570587 | rs12280296 | G | A | 0.935 | -7977.388 | -7977.759 | -0.37 | 2.67E-06 |
| 11 | 116781545 | rs11216183 | A | C | 1.073 | -7977.388 | -7977.759 | -0.371 | 2.67E-06 |
| 11 | 117384372 | rs144199381 | T | C | 0.919 | -7977.388 | -7977.761 | -0.372 | 2.67E-06 |
| 11 | 119906741 | rs582545 | G | C | 0.924 | -7977.388 | -7977.761 | -0.372 | 2.67E-06 |
| 11 | 119907628 | rs643377 | A | G | 0.924 | -7977.388 | -7977.76 | -0.372 | 2.67E-06 |
| 11 | 120699899 | rs4936555 | T | C | 1.054 | -7977.388 | -7977.76 | -0.372 | 2.67E-06 |
| 11 | 120835281 | rs7128009 | A | G | 0.943 | -7977.388 | -7977.76 | -0.372 | 2.67E-06 |
| 11 | 119880923 | rs675714 | A | G | 0.935 | -7977.388 | -7977.761 | -0.373 | 2.66E-06 |
| 11 | 117164246 | rs45506400 | C | A | 0.922 | -7977.388 | -7977.762 | -0.374 | 2.66E-06 |
| 11 | 117218685 | rs555179364 | C | T | 0.921 | -7977.388 | -7977.763 | -0.374 | 2.66E-06 |
| 11 | 117921563 | rs76190886 | T | C | 0.919 | -7977.388 | -7977.763 | -0.374 | 2.66E-06 |
| 11 | 118059557 | rs10790244 | C | T | 1.058 | -7977.388 | -7977.762 | -0.374 | 2.66E-06 |
| 11 | 119236410 | rs35310980 | A | T | 0.93 | -7977.388 | -7977.762 | -0.374 | 2.66E-06 |
| 11 | 120648746 | rs115791680 | T | C | 0.92 | -7977.388 | -7977.762 | -0.374 | 2.66E-06 |
| 11 | 120862089 | rs4935758 | T | C | 1.053 | -7977.388 | -7977.763 | -0.374 | 2.66E-06 |
| 11 | 116700785 | rs11827682 | T | C | 1.086 | -7977.388 | -7977.764 | -0.375 | 2.66E-06 |
| 11 | 118865465 | rs4938616 | T | C | 1.053 | -7977.388 | -7977.763 | -0.375 | 2.66E-06 |
| 11 | 116292818 | rs116859176 | C | T | 0.926 | -7977.388 | -7977.764 | -0.376 | 2.66E-06 |
| 11 | 117764401 | rs112444974 | T | G | 1.086 | -7977.388 | -7977.764 | -0.376 | 2.66E-06 |
| 11 | 117250419 | rs180865175 | G | T | 0.921 | -7977.388 | -7977.765 | -0.377 | 2.65E-06 |
| 11 | 116429779 | rs61904803 | C | T | 1.063 | -7977.388 | -7977.768 | -0.38 | 2.65E-06 |
| 11 | 116959601 | rs56257632 | C | T | 0.92 | -7977.388 | -7977.768 | -0.38 | 2.65E-06 |
| 11 | 117297415 | rs7940638 | A | C | 0.94 | -7977.388 | -7977.768 | -0.38 | 2.65E-06 |
| 11 | 117481112 | rs78503991 | T | A | 0.92 | -7977.388 | -7977.768 | -0.38 | 2.65E-06 |
| 11 | 117647364 | rs13377544 | A | T | 1.074 | -7977.388 | -7977.768 | -0.38 | 2.65E-06 |
| 11 | 119515905 | rs11217369 | T | C | 0.929 | -7977.388 | -7977.768 | -0.38 | 2.65E-06 |
| 11 | 119516100 | rs10892425 | T | C | 0.929 | -7977.388 | -7977.768 | -0.38 | 2.65E-06 |
| 11 | 118655704 | rs6421568 | C | G | 0.937 | -7977.388 | -7977.769 | -0.381 | 2.64E-06 |
| 11 | 118655775 | rs6421569 | A | C | 0.937 | -7977.388 | -7977.769 | -0.381 | 2.64E-06 |
| 11 | 118037964 | rs2075647 | C | G | 0.921 | -7977.388 | -7977.771 | -0.382 | 2.64E-06 |
| 11 | 120531656 | rs11217941 | G | T | 1.052 | -7977.388 | -7977.77 | -0.382 | 2.64E-06 |
| 11 | 117341717 | rs56316258 | G | C | 1.069 | -7977.388 | -7977.771 | -0.383 | 2.64E-06 |
| 11 | 117518200 | rs673028 | G | A | 0.948 | -7977.388 | -7977.773 | -0.384 | 2.63E-06 |
| 11 | 117600876 | rs4936402 | C | T | 0.922 | -7977.388 | -7977.772 | -0.384 | 2.63E-06 |
| 11 | 116778180 | rs1979023 | C | T | 0.92 | -7977.388 | -7977.773 | -0.385 | 2.63E-06 |
| 11 | 117578142 | rs77699630 | C | T | 0.92 | -7977.388 | -7977.773 | -0.385 | 2.63E-06 |

| | | | | | | | |
|---|---|---|---|---|---|---|---|
| 11 | 117923742 | rs74235117 | T | C | NULL | 0.547357618 | NULL&M_Disease |
| 11 | 118563286 | rs527619 | A | G | NULL | 0.547357618 | NULL&M_Disease |
| 11 | 116800154 | rs547919 | T | A | NULL | 0.547357618 | NULL&M_Disease |
| 11 | 119909507 | rs688857 | T | C | NULL | 0.547357618 | NULL&M_Disease |
| 11 | 117796722 | rs7107023 | T | C | NULL | 0.547357618 | NULL&M_Disease |
| 11 | 116834953 | rs1815789 | C | T | NULL | 0.548596109 | NULL&M_Disease |
| 11 | 116838904 | rs7941927 | G | A | NULL | 0.548596109 | NULL&M_Disease |
| 11 | 116911784 | rs61582023 | G | A | NULL | 0.547357618 | NULL&M_Disease |
| 11 | 117316605 | rs60385993 | A | G | NULL | 0.546118541 | NULL&M_Disease |
| 11 | 117317947 | rs111446534 | A | C | NULL | 0.546118541 | NULL&M_Disease |
| 11 | 118652781 | rs2508915 | C | G | NULL | 0.548596109 | NULL&M_Disease |
| 11 | 118777978 | rs78755429 | A | C | NULL | 0.547357618 | NULL&M_Disease |
| 11 | 120836662 | rs524209 | A | G | NULL | 0.548596109 | NULL&M_Disease |
| 11 | 118117018 | rs11216831 | A | G | NULL | 0.548596109 | NULL&M_Disease |
| 11 | 116498649 | rs61906060 | G | A | NULL | 0.548596109 | NULL&M_Disease |
| 11 | 117899803 | rs617357 | A | G | NULL | 0.548596109 | NULL&M_Disease |
| 11 | 119516473 | rs10892426 | T | C | NULL | 0.548596109 | NULL&M_Disease |
| 11 | 119808052 | rs1955021 | A | G | NULL | 0.548596109 | NULL&M_Disease |
| 11 | 120916015 | rs141784177 | G | C | NULL | 0.547357618 | NULL&M_Disease |
| 11 | 119539364 | rs4938706 | A | G | NULL | 0.548596109 | NULL&M_Disease |
| 11 | 117561265 | rs10892162 | G | A | NULL | 0.548596109 | NULL&M_Disease |
| 11 | 117605574 | rs1939982 | G | A | NULL | 0.547357618 | NULL&M_Disease |
| 11 | 116548489 | rs112548685 | T | C | NULL | 0.547357618 | NULL&M_Disease |
| 11 | 119541778 | rs4459318 | T | C | NULL | 0.548596109 | NULL&M_Disease |
| 11 | 119781153 | rs7115603 | G | A | NULL | 0.548596109 | NULL&M_Disease |
| 11 | 119914788 | rs11217655 | C | A | NULL | 0.548596109 | NULL&M_Disease |
| 11 | 118648396 | rs548167 | A | G | NULL | 0.548596109 | NULL&M_Disease |
| 11 | 120790814 | rs75736746 | G | A | NULL | 0.548596109 | NULL&M_Disease |
| 11 | 118572267 | rs573905 | A | G | NULL | 0.549833997 | NULL&M_Disease |
| 11 | 118834322 | rs118003247 | G | A | NULL | 0.548596109 | NULL&M_Disease |
| 11 | 120417065 | rs2252490 | C | G | NULL | 0.549833997 | NULL&M_Disease |
| 11 | 117532692 | rs12789293 | T | C | NULL | 0.548596109 | NULL&M_Disease |
| 11 | 119936507 | rs1893393 | A | T | NULL | 0.549833997 | NULL&M_Disease |
| 11 | 117678420 | rs11216551 | A | G | NULL | 0.548596109 | NULL&M_Disease |
| 11 | 117678543 | rs12790243 | A | G | NULL | 0.549833997 | NULL&M_Disease |
| 11 | 116572169 | rs71480317 | C | T | NULL | 0.548596109 | NULL&M_Disease |
| 11 | 116917885 | rs73576603 | A | G | NULL | 0.548596109 | NULL&M_Disease |
| 11 | 117416908 | rs521187 | A | G | NULL | 0.549833997 | NULL&M_Disease |
| 11 | 117569931 | rs56359135 | T | A | NULL | 0.549833997 | NULL&M_Disease |
| 11 | 119804640 | rs80030349 | G | A | NULL | 0.549833997 | NULL&M_Disease |
| 11 | 119886445 | rs72996698 | T | C | NULL | 0.548596109 | NULL&M_Disease |
| 11 | 120563386 | rs77776594 | A | G | NULL | 0.549833997 | NULL&M_Disease |
| 11 | 120822377 | rs3781817 | T | C | NULL | 0.549833997 | NULL&M_Disease |
| 11 | 121165007 | rs118088243 | G | A | NULL | 0.548596109 | NULL&M_Disease |
| 11 | 116440011 | rs4938285 | C | T | NULL | 0.549833997 | NULL&M_Disease |
| 11 | 120332795 | rs117843905 | T | C | NULL | 0.549833997 | NULL&M_Disease |
| 11 | 116223240 | rs36088800 | C | T | NULL | 0.549833997 | NULL&M_Disease |
| 11 | 116957263 | rs3863307 | G | C | NULL | 0.549833997 | NULL&M_Disease |
| 11 | 117342348 | rs66991573 | C | A | NULL | 0.549833997 | NULL&M_Disease |
| 11 | 116996445 | rs80125068 | G | A | NULL | 0.549833997 | NULL&M_Disease |
| 11 | 117397094 | rs143648302 | T | C | NULL | 0.549833997 | NULL&M_Disease |
| 11 | 117623674 | rs80183540 | T | C | NULL | 0.549833997 | NULL&M_Disease |
| 11 | 117679078 | rs60657131 | C | T | NULL | 0.548596109 | NULL&M_Disease |
| 11 | 119515046 | rs10892424 | T | G | NULL | 0.549833997 | NULL&M_Disease |
| 11 | 120415557 | rs2252648 | C | A | NULL | 0.549833997 | NULL&M_Disease |
| 11 | 119634544 | rs80238987 | A | G | NULL | 0.549833997 | NULL&M_Disease |
| 11 | 120452999 | rs4938818 | A | G | NULL | 0.549833997 | NULL&M_Disease |
| 11 | 120871636 | rs144716047 | G | C | NULL | 0.549833997 | NULL&M_Disease |
| 11 | 117410700 | rs2509007 | A | G | NULL | 0.551071269 | NULL&M_Disease |
| 11 | 121072088 | rs688099 | A | C | NULL | 0.551071269 | NULL&M_Disease |
| 11 | 116740659 | rs599320 | C | G | NULL | 0.549833997 | NULL&M_Disease |
| 11 | 117056059 | rs11606672 | T | C | NULL | 0.549833997 | NULL&M_Disease |
| 11 | 120567135 | rs75432677 | A | G | NULL | 0.548596109 | NULL&M_Disease |
| 11 | 117766665 | rs113650932 | T | C | NULL | 0.549833997 | NULL&M_Disease |
| 11 | 116439949 | rs4938284 | C | T | NULL | 0.551071269 | NULL&M_Disease |
| 11 | 116792445 | rs522848 | C | T | NULL | 0.549833997 | NULL&M_Disease |
| 11 | 121094594 | rs639891 | T | C | NULL | 0.551071269 | NULL&M_Disease |
| 11 | 116224822 | rs12577176 | C | T | NULL | 0.551071269 | NULL&M_Disease |
| 11 | 117798073 | rs73022500 | A | C | NULL | 0.551071269 | NULL&M_Disease |
| 11 | 117798431 | rs12365956 | A | G | NULL | 0.551071269 | NULL&M_Disease |
| 11 | 117953403 | rs639978 | G | A | NULL | 0.549833997 | NULL&M_Disease |
| 11 | 119486336 | rs75538174 | C | A | NULL | 0.551071269 | NULL&M_Disease |
| 11 | 120149184 | rs77802172 | A | G | NULL | 0.549833997 | NULL&M_Disease |
| 11 | 116693354 | rs5093 | A | G | NULL | 0.549833997 | NULL&M_Disease |
| 11 | 116958028 | rs7103842 | A | G | NULL | 0.551071269 | NULL&M_Disease |
| 11 | 116744447 | rs1263160 | C | T | NULL | 0.549833997 | NULL&M_Disease |
| 11 | 119540470 | rs7118532 | G | A | NULL | 0.551071269 | NULL&M_Disease |
| 11 | 117411687 | rs147864821 | A | G | NULL | 0.549833997 | NULL&M_Disease |
| 11 | 119817902 | rs55831600 | A | G | NULL | 0.551071269 | NULL&M_Disease |
| 11 | 117771769 | rs73020470 | C | T | NULL | 0.551071269 | NULL&M_Disease |
| 11 | 117991116 | rs34570367 | G | A | NULL | 0.551071269 | NULL&M_Disease |
| 11 | 118826045 | rs537080 | C | T | NULL | 0.551071269 | NULL&M_Disease |
| 11 | 116811057 | rs2513099 | C | T | NULL | 0.551071269 | NULL&M_Disease |
| 11 | 120637669 | rs11822437 | A | T | NULL | 0.551071269 | NULL&M_Disease |
| 11 | 120972112 | rs78830794 | C | T | NULL | 0.551071269 | NULL&M_Disease |
| 11 | 116998879 | rs4310650 | T | C | NULL | 0.551071269 | NULL&M_Disease |
| 11 | 120509621 | rs6589824 | G | A | NULL | 0.55230791 | NULL&M_Disease |
| 11 | 120723433 | rs4430518 | G | T | NULL | 0.55230791 | NULL&M_Disease |
| 11 | 116285930 | rs576670 | G | A | NULL | 0.55230791 | NULL&M_Disease |
| 11 | 118926720 | rs470679 | A | C | NULL | 0.55230791 | NULL&M_Disease |
| 11 | 118555692 | rs10892252 | G | A | NULL | 0.55230791 | NULL&M_Disease |
| 11 | 120570346 | rs12273984 | C | T | NULL | 0.55230791 | NULL&M_Disease |
| 11 | 120630546 | rs1939674 | G | T | NULL | 0.55230791 | NULL&M_Disease |
| 11 | 119723230 | rs477113 | C | T | NULL | 0.55230791 | NULL&M_Disease |
| 11 | 119908975 | rs149021690 | C | T | NULL | 0.55230791 | NULL&M_Disease |
| 11 | 119592454 | rs10892442 | G | A | NULL | 0.55230791 | NULL&M_Disease |

| | | | | | | | | | |
|---|---|---|---|---|---|---|---|---|---|
| 11 | 117923742 | rs74235117 | T | C | 0.92 | -7977.388 | -7977.773 | -0.385 | 2.63E-06 |
| 11 | 118563286 | rs527619 | A | G | 0.95 | -7977.388 | -7977.774 | -0.385 | 2.63E-06 |
| 11 | 116800154 | rs547919 | T | A | 0.92 | -7977.388 | -7977.774 | -0.386 | 2.63E-06 |
| 11 | 119909507 | rs688857 | T | C | 0.924 | -7977.388 | -7977.775 | -0.387 | 2.63E-06 |
| 11 | 117796722 | rs7107023 | T | C | 0.933 | -7977.388 | -7977.776 | -0.388 | 2.62E-06 |
| 11 | 116834953 | rs1815789 | C | T | 1.059 | -7977.388 | -7977.777 | -0.389 | 2.62E-06 |
| 11 | 116838904 | rs7941927 | G | A | 1.059 | -7977.388 | -7977.777 | -0.389 | 2.62E-06 |
| 11 | 116911784 | rs61582023 | G | A | 0.921 | -7977.388 | -7977.777 | -0.389 | 2.62E-06 |
| 11 | 117316605 | rs60385993 | A | G | 1.073 | -7977.388 | -7977.777 | -0.389 | 2.62E-06 |
| 11 | 117317947 | rs111446534 | A | C | 1.073 | -7977.388 | -7977.777 | -0.389 | 2.62E-06 |
| 11 | 118652781 | rs2508915 | C | G | 0.937 | -7977.388 | -7977.777 | -0.389 | 2.62E-06 |
| 11 | 118777978 | rs78755429 | A | C | 0.92 | -7977.388 | -7977.777 | -0.389 | 2.62E-06 |
| 11 | 120836662 | rs524209 | A | G | 0.944 | -7977.388 | -7977.778 | -0.389 | 2.62E-06 |
| 11 | 118117018 | rs11216831 | A | G | 0.929 | -7977.388 | -7977.778 | -0.39 | 2.62E-06 |
| 11 | 116498649 | rs61906060 | G | A | 0.936 | -7977.388 | -7977.779 | -0.391 | 2.62E-06 |
| 11 | 117899803 | rs617357 | A | G | 0.944 | -7977.388 | -7977.781 | -0.392 | 2.61E-06 |
| 11 | 119516473 | rs10892426 | T | C | 0.93 | -7977.388 | -7977.78 | -0.392 | 2.61E-06 |
| 11 | 119808052 | rs1955021 | A | G | 1.055 | -7977.388 | -7977.78 | -0.392 | 2.61E-06 |
| 11 | 120916015 | rs141784177 | G | C | 0.92 | -7977.388 | -7977.78 | -0.392 | 2.61E-06 |
| 11 | 119539364 | rs4938706 | A | G | 0.95 | -7977.388 | -7977.782 | -0.393 | 2.61E-06 |
| 11 | 117561265 | rs10892162 | G | A | 0.94 | -7977.388 | -7977.783 | -0.394 | 2.61E-06 |
| 11 | 117605574 | rs1939982 | G | A | 0.923 | -7977.388 | -7977.782 | -0.394 | 2.61E-06 |
| 11 | 116548489 | rs112548685 | T | C | 1.078 | -7977.388 | -7977.783 | -0.395 | 2.61E-06 |
| 11 | 119541778 | rs4459318 | T | C | 0.95 | -7977.388 | -7977.784 | -0.396 | 2.60E-06 |
| 11 | 119781153 | rs7115603 | G | A | 1.079 | -7977.388 | -7977.784 | -0.396 | 2.60E-06 |
| 11 | 119914788 | rs11217655 | C | A | 1.079 | -7977.388 | -7977.785 | -0.396 | 2.60E-06 |
| 11 | 118648396 | rs548167 | A | G | 0.937 | -7977.388 | -7977.785 | -0.397 | 2.60E-06 |
| 11 | 120790814 | rs75736746 | G | A | 1.082 | -7977.388 | -7977.785 | -0.397 | 2.60E-06 |
| 11 | 118572267 | rs573905 | A | G | 0.951 | -7977.388 | -7977.787 | -0.398 | 2.60E-06 |
| 11 | 118834322 | rs118003247 | G | A | 1.076 | -7977.388 | -7977.787 | -0.398 | 2.60E-06 |
| 11 | 120417065 | rs2252490 | C | G | 0.947 | -7977.388 | -7977.786 | -0.398 | 2.60E-06 |
| 11 | 117532692 | rs12789293 | T | C | 0.921 | -7977.388 | -7977.787 | -0.399 | 2.60E-06 |
| 11 | 119936507 | rs1893393 | A | T | 1.064 | -7977.388 | -7977.788 | -0.399 | 2.60E-06 |
| 11 | 117678420 | rs11216551 | A | G | 1.083 | -7977.388 | -7977.788 | -0.4 | 2.59E-06 |
| 11 | 117678543 | rs12790243 | A | G | 0.951 | -7977.388 | -7977.788 | -0.4 | 2.59E-06 |
| 11 | 116572169 | rs71480317 | C | T | 0.922 | -7977.388 | -7977.789 | -0.401 | 2.59E-06 |
| 11 | 116917885 | rs73576603 | A | G | 0.923 | -7977.388 | -7977.789 | -0.401 | 2.59E-06 |
| 11 | 117416908 | rs521187 | A | G | 1.055 | -7977.388 | -7977.79 | -0.401 | 2.59E-06 |
| 11 | 117569931 | rs56359135 | T | A | 0.932 | -7977.388 | -7977.789 | -0.401 | 2.59E-06 |
| 11 | 119804640 | rs80030349 | G | A | 1.078 | -7977.388 | -7977.789 | -0.401 | 2.59E-06 |
| 11 | 119886445 | rs72996698 | T | C | 1.083 | -7977.388 | -7977.79 | -0.401 | 2.59E-06 |
| 11 | 120563386 | rs77776594 | A | G | 0.935 | -7977.388 | -7977.789 | -0.401 | 2.59E-06 |
| 11 | 120822377 | rs3781817 | T | C | 0.951 | -7977.388 | -7977.791 | -0.402 | 2.59E-06 |
| 11 | 121165007 | rs118088243 | G | A | 0.927 | -7977.388 | -7977.79 | -0.402 | 2.59E-06 |
| 11 | 116440011 | rs4938285 | C | T | 1.053 | -7977.388 | -7977.791 | -0.403 | 2.59E-06 |
| 11 | 120332795 | rs117843905 | T | C | 0.921 | -7977.388 | -7977.792 | -0.403 | 2.59E-06 |
| 11 | 116223240 | rs36088800 | C | T | 0.927 | -7977.388 | -7977.793 | -0.405 | 2.58E-06 |
| 11 | 116957263 | rs3863307 | G | C | 1.059 | -7977.388 | -7977.793 | -0.405 | 2.58E-06 |
| 11 | 117342348 | rs66991573 | C | A | 1.068 | -7977.388 | -7977.793 | -0.405 | 2.58E-06 |
| 11 | 116996445 | rs80125068 | G | A | 0.921 | -7977.388 | -7977.795 | -0.406 | 2.58E-06 |
| 11 | 117397094 | rs143648302 | T | C | 0.922 | -7977.388 | -7977.794 | -0.406 | 2.58E-06 |
| 11 | 117623674 | rs80183540 | T | C | 1.084 | -7977.388 | -7977.795 | -0.406 | 2.58E-06 |
| 11 | 117679078 | rs60657131 | C | T | 0.926 | -7977.388 | -7977.794 | -0.406 | 2.58E-06 |
| 11 | 119515046 | rs10892424 | T | G | 0.93 | -7977.388 | -7977.795 | -0.406 | 2.58E-06 |
| 11 | 120415557 | rs2252648 | C | A | 0.947 | -7977.388 | -7977.795 | -0.406 | 2.58E-06 |
| 11 | 119634544 | rs80238987 | A | G | 0.921 | -7977.388 | -7977.796 | -0.407 | 2.57E-06 |
| 11 | 120452999 | rs4938818 | A | G | 1.052 | -7977.388 | -7977.795 | -0.407 | 2.57E-06 |
| 11 | 120871636 | rs144716047 | G | C | 1.078 | -7977.388 | -7977.796 | -0.407 | 2.57E-06 |
| 11 | 117410700 | rs2509007 | A | G | 1.052 | -7977.388 | -7977.797 | -0.408 | 2.57E-06 |
| 11 | 121072088 | rs688099 | A | C | 0.947 | -7977.388 | -7977.796 | -0.408 | 2.57E-06 |
| 11 | 116740659 | rs599320 | C | G | 0.923 | -7977.388 | -7977.798 | -0.409 | 2.57E-06 |
| 11 | 117056059 | rs11606672 | T | C | 1.079 | -7977.388 | -7977.797 | -0.409 | 2.57E-06 |
| 11 | 120567135 | rs75432677 | A | G | 1.069 | -7977.388 | -7977.798 | -0.409 | 2.57E-06 |
| 11 | 117766665 | rs113650932 | T | C | 1.083 | -7977.388 | -7977.798 | -0.41 | 2.57E-06 |
| 11 | 116439949 | rs4938284 | C | T | 1.053 | -7977.388 | -7977.799 | -0.411 | 2.56E-06 |
| 11 | 116792445 | rs522848 | C | T | 0.922 | -7977.388 | -7977.8 | -0.411 | 2.56E-06 |
| 11 | 121094594 | rs639891 | T | C | 0.943 | -7977.388 | -7977.799 | -0.411 | 2.56E-06 |
| 11 | 116224822 | rs12577176 | C | T | 0.927 | -7977.388 | -7977.8 | -0.412 | 2.56E-06 |
| 11 | 117798073 | rs73022500 | A | C | 0.933 | -7977.388 | -7977.801 | -0.412 | 2.56E-06 |
| 11 | 117798431 | rs12365956 | A | G | 0.933 | -7977.388 | -7977.801 | -0.412 | 2.56E-06 |
| 11 | 117953403 | rs639978 | G | A | 0.922 | -7977.388 | -7977.8 | -0.412 | 2.56E-06 |
| 11 | 119486336 | rs75538174 | C | A | 1.07 | -7977.388 | -7977.8 | -0.412 | 2.56E-06 |
| 11 | 120149184 | rs77802172 | A | G | 1.083 | -7977.388 | -7977.801 | -0.412 | 2.56E-06 |
| 11 | 116693354 | rs5093 | A | G | 0.922 | -7977.388 | -7977.802 | -0.413 | 2.56E-06 |
| 11 | 116958028 | rs7103842 | A | G | 1.059 | -7977.388 | -7977.801 | -0.413 | 2.56E-06 |
| 11 | 116744447 | rs1263160 | C | T | 0.923 | -7977.388 | -7977.803 | -0.414 | 2.56E-06 |
| 11 | 119540470 | rs7118532 | G | A | 0.95 | -7977.388 | -7977.802 | -0.414 | 2.56E-06 |
| 11 | 117411687 | rs147864821 | A | G | 0.929 | -7977.388 | -7977.803 | -0.415 | 2.55E-06 |
| 11 | 119817902 | rs55831600 | A | G | 0.922 | -7977.388 | -7977.804 | -0.415 | 2.55E-06 |
| 11 | 117771769 | rs73020470 | C | T | 0.928 | -7977.388 | -7977.805 | -0.417 | 2.55E-06 |
| 11 | 117991116 | rs34570367 | G | A | 0.922 | -7977.388 | -7977.806 | -0.417 | 2.55E-06 |
| 11 | 118826045 | rs537080 | C | T | 1.061 | -7977.388 | -7977.806 | -0.417 | 2.55E-06 |
| 11 | 116811057 | rs2513099 | C | T | 0.923 | -7977.388 | -7977.807 | -0.419 | 2.54E-06 |
| 11 | 120637669 | rs11822437 | A | T | 0.923 | -7977.388 | -7977.807 | -0.419 | 2.54E-06 |
| 11 | 120972112 | rs78830794 | C | T | 0.924 | -7977.388 | -7977.807 | -0.419 | 2.54E-06 |
| 11 | 116998879 | rs4310650 | T | C | 0.923 | -7977.388 | -7977.809 | -0.42 | 2.54E-06 |
| 11 | 120509621 | rs6589824 | G | A | 1.052 | -7977.388 | -7977.809 | -0.421 | 2.54E-06 |
| 11 | 120723433 | rs4430518 | G | T | 1.057 | -7977.388 | -7977.81 | -0.422 | 2.54E-06 |
| 11 | 116285930 | rs576670 | G | A | 1.054 | -7977.388 | -7977.811 | -0.423 | 2.53E-06 |
| 11 | 118926720 | rs470679 | A | C | 1.064 | -7977.388 | -7977.812 | -0.424 | 2.53E-06 |
| 11 | 118555692 | rs10892252 | G | A | 1.057 | -7977.388 | -7977.814 | -0.425 | 2.53E-06 |
| 11 | 120570346 | rs12273984 | C | T | 0.937 | -7977.388 | -7977.813 | -0.425 | 2.53E-06 |
| 11 | 120630546 | rs1939674 | G | T | 0.923 | -7977.388 | -7977.813 | -0.425 | 2.53E-06 |
| 11 | 119723230 | rs477113 | C | T | 0.95 | -7977.388 | -7977.814 | -0.426 | 2.53E-06 |
| 11 | 119908975 | rs149021690 | C | T | 0.927 | -7977.388 | -7977.815 | -0.426 | 2.53E-06 |
| 11 | 119592454 | rs10892442 | G | A | 0.923 | -7977.388 | -7977.816 | -0.427 | 2.52E-06 |

| | | | | | | | |
|---|---|---|---|---|---|---|---|
| 11 | 120086680 | rs11217768 | A | G | NULL | 0.55230791 | NULL&M_Disease |
| 11 | 120627270 | rs79530554 | T | C | NULL | 0.55230791 | NULL&M_Disease |
| 11 | 117618530 | rs73585234 | T | C | NULL | 0.55230791 | NULL&M_Disease |
| 11 | 119817309 | rs61899038 | A | T | NULL | 0.55230791 | NULL&M_Disease |
| 11 | 119817670 | rs553492 | A | C | NULL | 0.55230791 | NULL&M_Disease |
| 11 | 116436913 | rs61904810 | T | C | NULL | 0.55230791 | NULL&M_Disease |
| 11 | 116938655 | rs58848374 | A | G | NULL | 0.55230791 | NULL&M_Disease |
| 11 | 117283091 | rs2279013 | C | T | NULL | 0.553543903 | NULL&M_Disease |
| 11 | 117889224 | rs2512153 | G | A | NULL | 0.553543903 | NULL&M_Disease |
| 11 | 119994765 | rs586407 | A | G | NULL | 0.55230791 | NULL&M_Disease |
| 11 | 117671489 | rs3819132 | A | G | NULL | 0.553543903 | NULL&M_Disease |
| 11 | 116661488 | rs3135507 | T | C | NULL | 0.553543903 | NULL&M_Disease |
| 11 | 119813521 | rs76264435 | A | G | NULL | 0.553543903 | NULL&M_Disease |
| 11 | 116331172 | rs11215943 | T | G | NULL | 0.553543903 | NULL&M_Disease |
| 11 | 117672474 | rs10892172 | A | C | NULL | 0.553543903 | NULL&M_Disease |
| 11 | 118637666 | rs535996 | T | C | NULL | 0.553543903 | NULL&M_Disease |
| 11 | 118640081 | rs529717 | A | G | NULL | 0.553543903 | NULL&M_Disease |
| 11 | 117573319 | rs74833570 | T | G | NULL | 0.55230791 | NULL&M_Disease |
| 11 | 118565651 | rs540800 | A | G | NULL | 0.553543903 | NULL&M_Disease |
| 11 | 120152148 | rs7105395 | T | C | NULL | 0.553543903 | NULL&M_Disease |
| 11 | 116937074 | rs117315666 | T | G | NULL | 0.553543903 | NULL&M_Disease |
| 11 | 117123218 | rs77262524 | G | C | NULL | 0.553543903 | NULL&M_Disease |
| 11 | 121006301 | rs11218154 | A | G | NULL | 0.553543903 | NULL&M_Disease |
| 11 | 117404179 | rs612979 | G | T | NULL | 0.553543903 | NULL&M_Disease |
| 11 | 119703523 | rs12793318 | A | G | NULL | 0.553543903 | NULL&M_Disease |
| 11 | 120415823 | rs2252634 | T | C | NULL | 0.554779235 | NULL&M_Disease |
| 11 | 120633447 | rs183832950 | T | G | NULL | 0.553543903 | NULL&M_Disease |
| 11 | 116620796 | rs2849179 | T | C | NULL | 0.554779235 | NULL&M_Disease |
| 11 | 117397271 | rs148092485 | A | T | NULL | 0.553543903 | NULL&M_Disease |
| 11 | 117675286 | rs12364667 | G | T | NULL | 0.554779235 | NULL&M_Disease |
| 11 | 119540705 | rs111259707 | C | T | NULL | 0.554779235 | NULL&M_Disease |
| 11 | 119733584 | rs73008672 | T | C | NULL | 0.554779235 | NULL&M_Disease |
| 11 | 117567949 | rs10790200 | C | T | NULL | 0.554779235 | NULL&M_Disease |
| 11 | 118063030 | rs11216807 | C | T | NULL | 0.554779235 | NULL&M_Disease |
| 11 | 118641451 | rs2508929 | G | T | NULL | 0.554779235 | NULL&M_Disease |
| 11 | 120652936 | rs11218005 | T | G | NULL | 0.553543903 | NULL&M_Disease |
| 11 | 117102706 | rs2277286 | G | C | NULL | 0.554779235 | NULL&M_Disease |
| 11 | 120152147 | rs7121490 | C | T | NULL | 0.554779235 | NULL&M_Disease |
| 11 | 116428302 | rs7108237 | A | G | NULL | 0.554779235 | NULL&M_Disease |
| 11 | 117407275 | rs117053908 | T | G | NULL | 0.553543903 | NULL&M_Disease |
| 11 | 117887757 | rs2512152 | T | C | NULL | 0.554779235 | NULL&M_Disease |
| 11 | 118146462 | rs1616199 | T | C | NULL | 0.556013891 | NULL&M_Disease |
| 11 | 118803160 | rs73575440 | A | G | NULL | 0.554779235 | NULL&M_Disease |
| 11 | 118803596 | rs111313044 | A | G | NULL | 0.554779235 | NULL&M_Disease |
| 11 | 119584274 | rs10892440 | T | C | NULL | 0.554779235 | NULL&M_Disease |
| 11 | 119586117 | rs7931978 | G | A | NULL | 0.554779235 | NULL&M_Disease |
| 11 | 119586389 | rs10790338 | G | T | NULL | 0.554779235 | NULL&M_Disease |
| 11 | 120018127 | rs1448722646 | T | C | NULL | 0.554779235 | NULL&M_Disease |
| 11 | 116392086 | rs146990932 | T | C | NULL | 0.554779235 | NULL&M_Disease |
| 11 | 120652371 | rs2846104 | G | C | NULL | 0.554779235 | NULL&M_Disease |
| 11 | 120653034 | rs11218006 | A | G | NULL | 0.554779235 | NULL&M_Disease |
| 11 | 117559219 | rs12288712 | T | C | NULL | 0.556013891 | NULL&M_Disease |
| 11 | 117957631 | rs1203387394 | C | G | NULL | 0.556013891 | NULL&M_Disease |
| 11 | 119806917 | rs895650 | T | C | NULL | 0.556013891 | NULL&M_Disease |
| 11 | 117507325 | rs495258 | T | C | NULL | 0.554779235 | NULL&M_Disease |
| 11 | 121077373 | rs612804 | C | T | NULL | 0.556013891 | NULL&M_Disease |
| 11 | 116718366 | rs628418 | G | C | NULL | 0.554779235 | NULL&M_Disease |
| 11 | 118317907 | rs528056 | T | C | NULL | 0.556013891 | NULL&M_Disease |
| 11 | 119636704 | rs7130633 | A | C | NULL | 0.556013891 | NULL&M_Disease |
| 11 | 119797684 | rs112297727 | T | C | NULL | 0.556013891 | NULL&M_Disease |
| 11 | 119797706 | rs111954335 | T | C | NULL | 0.556013891 | NULL&M_Disease |
| 11 | 119797995 | rs77109784 | G | A | NULL | 0.556013891 | NULL&M_Disease |
| 11 | 119798123 | rs75748285 | C | T | NULL | 0.556013891 | NULL&M_Disease |
| 11 | 119798178 | rs76777718 | A | G | NULL | 0.556013891 | NULL&M_Disease |
| 11 | 119798205 | rs75640250 | A | C | NULL | 0.556013891 | NULL&M_Disease |
| 11 | 119798592 | rs74840179 | C | G | NULL | 0.556013891 | NULL&M_Disease |
| 11 | 119799196 | rs77400160 | T | C | NULL | 0.556013891 | NULL&M_Disease |
| 11 | 119799219 | rs80166158 | A | G | NULL | 0.556013891 | NULL&M_Disease |
| 11 | 119799826 | rs112293704 | G | T | NULL | 0.556013891 | NULL&M_Disease |
| 11 | 119800547 | rs78178500 | A | C | NULL | 0.556013891 | NULL&M_Disease |
| 11 | 119801279 | rs79048103 | T | C | NULL | 0.556013891 | NULL&M_Disease |
| 11 | 119801401 | rs75417087 | T | G | NULL | 0.556013891 | NULL&M_Disease |
| 11 | 119801704 | rs79592175 | C | G | NULL | 0.556013891 | NULL&M_Disease |
| 11 | 119801962 | rs78763435 | A | T | NULL | 0.556013891 | NULL&M_Disease |
| 11 | 119802186 | rs754871 | G | A | NULL | 0.556013891 | NULL&M_Disease |
| 11 | 120565023 | rs12292552 | A | G | NULL | 0.556013891 | NULL&M_Disease |
| 11 | 117946849 | rs117440454 | T | C | NULL | 0.556013891 | NULL&M_Disease |
| 11 | 119818055 | rs113442800 | T | C | NULL | 0.554779235 | NULL&M_Disease |
| 11 | 121072293 | rs687180 | A | G | NULL | 0.556013891 | NULL&M_Disease |
| 11 | 118143062 | rs117666845 | A | T | NULL | 0.556013891 | NULL&M_Disease |
| 11 | 116943912 | rs73006585 | C | G | NULL | 0.556013891 | NULL&M_Disease |
| 11 | 117116312 | rs111447379 | T | C | NULL | 0.556013891 | NULL&M_Disease |
| 11 | 117582279 | rs11607103 | C | T | NULL | 0.556013891 | NULL&M_Disease |
| 11 | 118559731 | rs113784020 | T | C | NULL | 0.556013891 | NULL&M_Disease |
| 11 | 120053300 | rs518863 | T | C | NULL | 0.556013891 | NULL&M_Disease |
| 11 | 120087541 | rs1551240 | T | C | NULL | 0.556013891 | NULL&M_Disease |
| 11 | 120875383 | rs2510639 | A | G | NULL | 0.557247855 | NULL&M_Disease |
| 11 | 121040934 | rs118170232 | G | A | NULL | 0.556013891 | NULL&M_Disease |
| 11 | 119837014 | rs118070092 | A | G | NULL | 0.556013891 | NULL&M_Disease |
| 11 | 121076423 | rs616905 | C | T | NULL | 0.557247855 | NULL&M_Disease |
| 11 | 116372414 | rs879412 | A | G | NULL | 0.557247855 | NULL&M_Disease |
| 11 | 120123837 | rs116863015 | A | G | NULL | 0.556013891 | NULL&M_Disease |
| 11 | 119817426 | rs580318 | C | A | NULL | 0.556013891 | NULL&M_Disease |
| 11 | 119817579 | rs471714 | A | T | NULL | 0.556013891 | NULL&M_Disease |
| 11 | 119817708 | rs472665 | T | C | NULL | 0.556013891 | NULL&M_Disease |
| 11 | 119819087 | rs476877 | C | T | NULL | 0.556013891 | NULL&M_Disease |

| | | | | | | | | | |
|---|---|---|---|---|---|---|---|---|---|
| 11 | 120086680 | rs11217768 | A | G | 0.923 | -7977.388 | -7977.816 | -0.427 | 2.52E-06 |
| 11 | 120627270 | rs79530554 | T | C | 0.923 | -7977.388 | -7977.816 | -0.427 | 2.52E-06 |
| 11 | 117618530 | rs73585234 | T | C | 0.928 | -7977.388 | -7977.816 | -0.428 | 2.52E-06 |
| 11 | 119817309 | rs61899038 | A | T | 0.923 | -7977.388 | -7977.816 | -0.428 | 2.52E-06 |
| 11 | 119817670 | rs553492 | A | C | 0.923 | -7977.388 | -7977.816 | -0.428 | 2.52E-06 |
| 11 | 116436913 | rs61904810 | T | C | 1.08 | -7977.388 | -7977.818 | -0.43 | 2.52E-06 |
| 11 | 116938655 | rs58848374 | A | G | 0.925 | -7977.388 | -7977.819 | -0.43 | 2.52E-06 |
| 11 | 117283091 | rs2279013 | C | T | 0.949 | -7977.388 | -7977.819 | -0.431 | 2.51E-06 |
| 11 | 117889224 | rs2512153 | G | A | 0.948 | -7977.388 | -7977.82 | -0.431 | 2.51E-06 |
| 11 | 119994765 | rs586407 | A | G | 0.925 | -7977.388 | -7977.819 | -0.431 | 2.51E-06 |
| 11 | 117671489 | rs3819132 | A | G | 1.051 | -7977.388 | -7977.821 | -0.432 | 2.51E-06 |
| 11 | 116661488 | rs3135507 | T | C | 0.923 | -7977.388 | -7977.822 | -0.433 | 2.51E-06 |
| 11 | 119813521 | rs76264435 | A | G | 0.947 | -7977.388 | -7977.821 | -0.433 | 2.51E-06 |
| 11 | 116331172 | rs11215943 | T | G | 0.924 | -7977.388 | -7977.823 | -0.434 | 2.51E-06 |
| 11 | 117672474 | rs10892172 | A | C | 0.951 | -7977.388 | -7977.823 | -0.434 | 2.51E-06 |
| 11 | 118637666 | rs535996 | T | C | 0.938 | -7977.388 | -7977.822 | -0.434 | 2.51E-06 |
| 11 | 118640081 | rs529717 | A | G | 0.938 | -7977.388 | -7977.822 | -0.434 | 2.51E-06 |
| 11 | 117573319 | rs74833570 | T | G | 0.929 | -7977.388 | -7977.823 | -0.435 | 2.50E-06 |
| 11 | 118565651 | rs540800 | A | G | 0.951 | -7977.388 | -7977.823 | -0.435 | 2.50E-06 |
| 11 | 120152148 | rs7105395 | T | C | 0.93 | -7977.388 | -7977.823 | -0.435 | 2.50E-06 |
| 11 | 116937074 | rs117315666 | T | G | 0.927 | -7977.388 | -7977.824 | -0.436 | 2.50E-06 |
| 11 | 117123218 | rs77262524 | G | C | 0.925 | -7977.388 | -7977.824 | -0.436 | 2.50E-06 |
| 11 | 121006301 | rs11218154 | A | G | 0.925 | -7977.388 | -7977.824 | -0.436 | 2.50E-06 |
| 11 | 117404179 | rs612979 | G | T | 0.924 | -7977.388 | -7977.827 | -0.438 | 2.50E-06 |
| 11 | 119703523 | rs12793318 | A | G | 1.067 | -7977.388 | -7977.826 | -0.438 | 2.50E-06 |
| 11 | 120415823 | rs2252634 | T | C | 0.948 | -7977.388 | -7977.827 | -0.439 | 2.49E-06 |
| 11 | 120633447 | rs183832950 | T | G | 0.925 | -7977.388 | -7977.827 | -0.439 | 2.49E-06 |
| 11 | 116620796 | rs2849179 | T | C | 0.951 | -7977.388 | -7977.828 | -0.44 | 2.49E-06 |
| 11 | 117397271 | rs148092485 | A | T | 1.081 | -7977.388 | -7977.829 | -0.44 | 2.49E-06 |
| 11 | 117675286 | rs12364667 | G | T | 0.949 | -7977.388 | -7977.828 | -0.44 | 2.49E-06 |
| 11 | 119540705 | rs111259707 | C | T | 0.941 | -7977.388 | -7977.828 | -0.44 | 2.49E-06 |
| 11 | 119733584 | rs73008672 | T | C | 1.066 | -7977.388 | -7977.83 | -0.441 | 2.49E-06 |
| 11 | 117567949 | rs10790200 | C | T | 0.933 | -7977.388 | -7977.83 | -0.442 | 2.49E-06 |
| 11 | 118063030 | rs11216807 | C | T | 1.056 | -7977.388 | -7977.831 | -0.443 | 2.48E-06 |
| 11 | 118641451 | rs2508929 | G | T | 0.938 | -7977.388 | -7977.831 | -0.443 | 2.48E-06 |
| 11 | 120652936 | rs11218005 | T | G | 0.925 | -7977.388 | -7977.832 | -0.444 | 2.48E-06 |
| 11 | 117102706 | rs2277286 | G | C | 0.935 | -7977.388 | -7977.834 | -0.445 | 2.48E-06 |
| 11 | 120152147 | rs7121490 | C | T | 0.93 | -7977.388 | -7977.834 | -0.445 | 2.48E-06 |
| 11 | 116428302 | rs7108237 | A | G | 1.06 | -7977.388 | -7977.834 | -0.446 | 2.48E-06 |
| 11 | 117407275 | rs117053908 | T | G | 0.932 | -7977.388 | -7977.835 | -0.447 | 2.47E-06 |
| 11 | 117887757 | rs2512152 | T | C | 0.949 | -7977.388 | -7977.835 | -0.447 | 2.47E-06 |
| 11 | 118146462 | rs1616199 | T | C | 0.948 | -7977.388 | -7977.837 | -0.448 | 2.47E-06 |
| 11 | 118803160 | rs73575440 | A | G | 0.929 | -7977.388 | -7977.836 | -0.448 | 2.47E-06 |
| 11 | 118803596 | rs111313044 | A | G | 0.929 | -7977.388 | -7977.836 | -0.448 | 2.47E-06 |
| 11 | 119584274 | rs10892440 | T | C | 0.925 | -7977.388 | -7977.837 | -0.448 | 2.47E-06 |
| 11 | 119586117 | rs7931978 | G | A | 0.925 | -7977.388 | -7977.837 | -0.448 | 2.47E-06 |
| 11 | 119586389 | rs10790338 | G | T | 0.925 | -7977.388 | -7977.837 | -0.448 | 2.47E-06 |
| 11 | 120018127 | rs1448722646 | T | C | 0.93 | -7977.388 | -7977.836 | -0.448 | 2.47E-06 |
| 11 | 116392086 | rs146990932 | T | C | 0.925 | -7977.388 | -7977.837 | -0.449 | 2.47E-06 |
| 11 | 120652371 | rs2846104 | G | C | 0.926 | -7977.388 | -7977.837 | -0.449 | 2.47E-06 |
| 11 | 120653034 | rs11218006 | A | G | 0.926 | -7977.388 | -7977.838 | -0.449 | 2.47E-06 |
| 11 | 117559219 | rs12288712 | T | C | 0.943 | -7977.388 | -7977.838 | -0.45 | 2.47E-06 |
| 11 | 117957631 | rs1203387394 | C | G | 1.074 | -7977.388 | -7977.838 | -0.45 | 2.47E-06 |
| 11 | 119806917 | rs895650 | T | C | 1.054 | -7977.388 | -7977.838 | -0.45 | 2.47E-06 |
| 11 | 117507325 | rs495258 | T | C | 1.08 | -7977.388 | -7977.839 | -0.451 | 2.46E-06 |
| 11 | 121077373 | rs612804 | C | T | 0.947 | -7977.388 | -7977.839 | -0.451 | 2.46E-06 |
| 11 | 116718366 | rs628418 | G | C | 0.926 | -7977.388 | -7977.84 | -0.452 | 2.46E-06 |
| 11 | 118317907 | rs528056 | T | C | 0.937 | -7977.388 | -7977.84 | -0.452 | 2.46E-06 |
| 11 | 119636704 | rs7130633 | A | C | 0.944 | -7977.388 | -7977.84 | -0.452 | 2.46E-06 |
| 11 | 119797684 | rs112297727 | T | C | 1.075 | -7977.388 | -7977.841 | -0.453 | 2.46E-06 |
| 11 | 119797706 | rs111954335 | T | C | 1.075 | -7977.388 | -7977.841 | -0.453 | 2.46E-06 |
| 11 | 119797995 | rs77109784 | G | A | 1.075 | -7977.388 | -7977.841 | -0.453 | 2.46E-06 |
| 11 | 119798123 | rs75748285 | C | T | 1.075 | -7977.388 | -7977.841 | -0.453 | 2.46E-06 |
| 11 | 119798178 | rs76777718 | A | G | 1.075 | -7977.388 | -7977.841 | -0.453 | 2.46E-06 |
| 11 | 119798205 | rs75640250 | A | C | 1.075 | -7977.388 | -7977.841 | -0.453 | 2.46E-06 |
| 11 | 119798592 | rs74840179 | C | G | 1.075 | -7977.388 | -7977.841 | -0.453 | 2.46E-06 |
| 11 | 119799196 | rs77400160 | T | C | 1.075 | -7977.388 | -7977.841 | -0.453 | 2.46E-06 |
| 11 | 119799219 | rs80166158 | A | G | 1.075 | -7977.388 | -7977.841 | -0.453 | 2.46E-06 |
| 11 | 119799826 | rs112293704 | G | T | 1.075 | -7977.388 | -7977.841 | -0.453 | 2.46E-06 |
| 11 | 119800547 | rs78178500 | A | C | 1.075 | -7977.388 | -7977.841 | -0.453 | 2.46E-06 |
| 11 | 119801279 | rs79048103 | T | C | 1.075 | -7977.388 | -7977.841 | -0.453 | 2.46E-06 |
| 11 | 119801401 | rs75417087 | T | G | 1.075 | -7977.388 | -7977.841 | -0.453 | 2.46E-06 |
| 11 | 119801704 | rs79592175 | C | G | 1.075 | -7977.388 | -7977.841 | -0.453 | 2.46E-06 |
| 11 | 119801962 | rs78763435 | A | T | 1.075 | -7977.388 | -7977.841 | -0.453 | 2.46E-06 |
| 11 | 119802186 | rs754871 | G | A | 1.075 | -7977.388 | -7977.841 | -0.453 | 2.46E-06 |
| 11 | 120565023 | rs12292552 | A | G | 0.936 | -7977.388 | -7977.842 | -0.454 | 2.46E-06 |
| 11 | 117946849 | rs117440454 | T | C | 1.067 | -7977.388 | -7977.844 | -0.455 | 2.45E-06 |
| 11 | 119818055 | rs113442800 | T | C | 0.928 | -7977.388 | -7977.843 | -0.455 | 2.45E-06 |
| 11 | 121072293 | rs687180 | A | G | 0.947 | -7977.388 | -7977.843 | -0.455 | 2.45E-06 |
| 11 | 118143062 | rs117666845 | A | T | 0.925 | -7977.388 | -7977.845 | -0.456 | 2.45E-06 |
| 11 | 116943912 | rs73006585 | C | G | 0.928 | -7977.388 | -7977.846 | -0.457 | 2.45E-06 |
| 11 | 117116312 | rs111447379 | T | C | 0.926 | -7977.388 | -7977.845 | -0.457 | 2.45E-06 |
| 11 | 117582279 | rs11607103 | C | T | 0.925 | -7977.388 | -7977.846 | -0.457 | 2.45E-06 |
| 11 | 118559731 | rs113784020 | T | C | 0.926 | -7977.388 | -7977.845 | -0.457 | 2.45E-06 |
| 11 | 120053300 | rs518863 | T | C | 0.927 | -7977.388 | -7977.845 | -0.457 | 2.45E-06 |
| 11 | 120087541 | rs1551240 | T | C | 0.925 | -7977.388 | -7977.847 | -0.458 | 2.45E-06 |
| 11 | 120875383 | rs2510639 | A | G | 0.946 | -7977.388 | -7977.846 | -0.458 | 2.45E-06 |
| 11 | 121040934 | rs118170232 | G | A | 1.076 | -7977.388 | -7977.846 | -0.458 | 2.45E-06 |
| 11 | 119837014 | rs118070092 | A | G | 1.08 | -7977.388 | -7977.847 | -0.459 | 2.44E-06 |
| 11 | 121076423 | rs616905 | C | T | 0.948 | -7977.388 | -7977.847 | -0.459 | 2.44E-06 |
| 11 | 116372414 | rs879412 | A | G | 0.95 | -7977.388 | -7977.85 | -0.461 | 2.44E-06 |
| 11 | 120123837 | rs116863015 | A | G | 0.934 | -7977.388 | -7977.849 | -0.461 | 2.44E-06 |
| 11 | 119817426 | rs580318 | C | A | 0.926 | -7977.388 | -7977.851 | -0.462 | 2.44E-06 |
| 11 | 119817579 | rs471714 | A | T | 0.926 | -7977.388 | -7977.851 | -0.462 | 2.44E-06 |
| 11 | 119817708 | rs472665 | T | C | 0.926 | -7977.388 | -7977.851 | -0.462 | 2.44E-06 |
| 11 | 119819087 | rs476877 | C | T | 0.926 | -7977.388 | -7977.851 | -0.462 | 2.44E-06 |

| | | | | | | | |
|---|---|---|---|---|---|---|---|
| 11 | 116607641 | rs180351 | T | C | NULL | 0.557247855 | NULL&M_Disease |
| 11 | 117982495 | rs1941635 | T | G | NULL | 0.557247855 | NULL&M_Disease |
| 11 | 120991571 | rs74945140 | A | G | NULL | 0.557247855 | NULL&M_Disease |
| 11 | 117367838 | rs11603937 | A | G | NULL | 0.557247855 | NULL&M_Disease |
| 11 | 120554278 | rs12292453 | T | C | NULL | 0.557247855 | NULL&M_Disease |
| 11 | 116391651 | rs7120429 | T | C | NULL | 0.558481112 | NULL&M_Disease |
| 11 | 116829653 | rs7120173 | G | T | NULL | 0.558481112 | NULL&M_Disease |
| 11 | 120649949 | rs72995858 | T | C | NULL | 0.557247855 | NULL&M_Disease |
| 11 | 116222151 | rs12792619 | C | T | NULL | 0.558481112 | NULL&M_Disease |
| 11 | 117576946 | rs75819084 | T | C | NULL | 0.557247855 | NULL&M_Disease |
| 11 | 121169052 | rs75960142 | C | T | NULL | 0.557247855 | NULL&M_Disease |
| 11 | 116470289 | rs145936741 | G | A | NULL | 0.557247855 | NULL&M_Disease |
| 11 | 116960078 | rs1481148807 | A | G | NULL | 0.557247855 | NULL&M_Disease |
| 11 | 118061480 | rs7103918 | T | C | NULL | 0.558481112 | NULL&M_Disease |
| 11 | 116697301 | rs140231532 | T | C | NULL | 0.557247855 | NULL&M_Disease |
| 11 | 117570395 | rs12273147 | A | G | NULL | 0.558481112 | NULL&M_Disease |
| 11 | 119876350 | rs113111994 | T | G | NULL | 0.557247855 | NULL&M_Disease |
| 11 | 120470291 | rs4245037 | A | T | NULL | 0.558481112 | NULL&M_Disease |
| 11 | 117069788 | rs2269398 | T | C | NULL | 0.558481112 | NULL&M_Disease |
| 11 | 117796139 | rs55720349 | C | T | NULL | 0.558481112 | NULL&M_Disease |
| 11 | 119483777 | rs78341300 | G | A | NULL | 0.558481112 | NULL&M_Disease |
| 11 | 120042082 | rs529681 | A | G | NULL | 0.558481112 | NULL&M_Disease |
| 11 | 120653294 | rs11218007 | A | G | NULL | 0.558481112 | NULL&M_Disease |
| 11 | 116616402 | rs180340 | C | T | NULL | 0.558481112 | NULL&M_Disease |
| 11 | 116724204 | rs598503 | C | T | NULL | 0.558481112 | NULL&M_Disease |
| 11 | 118543563 | rs10892251 | T | C | NULL | 0.558481112 | NULL&M_Disease |
| 11 | 119506255 | rs10790328 | G | C | NULL | 0.558481112 | NULL&M_Disease |
| 11 | 119674096 | rs11217514 | A | G | NULL | 0.558481112 | NULL&M_Disease |
| 11 | 117659322 | rs11216545 | A | G | NULL | 0.558481112 | NULL&M_Disease |
| 11 | 120187422 | rs148189915 | T | A | NULL | 0.558481112 | NULL&M_Disease |
| 11 | 116278612 | rs7925794 | T | C | NULL | 0.559713649 | NULL&M_Disease |
| 11 | 116472577 | rs34994677 | G | A | NULL | 0.558481112 | NULL&M_Disease |
| 11 | 118100176 | rs12361791 | G | T | NULL | 0.558481112 | NULL&M_Disease |
| 11 | 120144571 | rs138488120 | A | C | NULL | 0.558481112 | NULL&M_Disease |
| 11 | 121093301 | rs117235919 | A | G | NULL | 0.558481112 | NULL&M_Disease |
| 11 | 117104900 | rs117547981 | A | G | NULL | 0.558481112 | NULL&M_Disease |
| 11 | 120302456 | rs117092333 | A | G | NULL | 0.558481112 | NULL&M_Disease |
| 11 | 116478432 | rs17119839 | G | A | NULL | 0.558481112 | NULL&M_Disease |
| 11 | 117200129 | rs75013797 | T | C | NULL | 0.558481112 | NULL&M_Disease |
| 11 | 118312870 | rs74422681 | G | A | NULL | 0.559713649 | NULL&M_Disease |
| 11 | 117958429 | rs603455 | T | A | NULL | 0.559713649 | NULL&M_Disease |
| 11 | 118802128 | rs59069289 | T | G | NULL | 0.559713649 | NULL&M_Disease |
| 11 | 118802386 | rs59480472 | C | T | NULL | 0.559713649 | NULL&M_Disease |
| 11 | 116617782 | rs180339 | T | C | NULL | 0.559713649 | NULL&M_Disease |
| 11 | 117569523 | rs7122782 | A | G | NULL | 0.559713649 | NULL&M_Disease |
| 11 | 119026384 | rs56168028 | G | A | NULL | 0.558481112 | NULL&M_Disease |
| 11 | 116633825 | rs918144 | T | C | NULL | 0.559713649 | NULL&M_Disease |
| 11 | 117179891 | rs676134 | T | C | NULL | 0.559713649 | NULL&M_Disease |
| 11 | 119514129 | rs906827 | A | C | NULL | 0.559713649 | NULL&M_Disease |
| 11 | 120825635 | rs4936569 | C | T | NULL | 0.559713649 | NULL&M_Disease |
| 11 | 117352364 | rs73022382 | A | G | NULL | 0.559713649 | NULL&M_Disease |
| 11 | 118134597 | rs4936426 | C | T | NULL | 0.559713649 | NULL&M_Disease |
| 11 | 116693871 | rs12721041 | T | C | NULL | 0.559713649 | NULL&M_Disease |
| 11 | 117378697 | rs117328724 | T | C | NULL | 0.559713649 | NULL&M_Disease |
| 11 | 117070740 | rs2170317 | C | A | NULL | 0.559713649 | NULL&M_Disease |
| 11 | 117129617 | rs550893 | A | G | NULL | 0.559713649 | NULL&M_Disease |
| 11 | 116479584 | rs527778 | T | G | NULL | 0.56094545 | NULL&M_Disease |
| 11 | 116491465 | rs112732949 | G | A | NULL | 0.559713649 | NULL&M_Disease |
| 11 | 117720781 | rs7928007 | C | A | NULL | 0.559713649 | NULL&M_Disease |
| 11 | 119816766 | rs657317 | A | C | NULL | 0.559713649 | NULL&M_Disease |
| 11 | 116610048 | rs180350 | T | C | NULL | 0.56094545 | NULL&M_Disease |
| 11 | 117646968 | rs61905347 | T | C | NULL | 0.56094545 | NULL&M_Disease |
| 11 | 117474845 | rs61697909 | A | G | NULL | 0.56094545 | NULL&M_Disease |
| 11 | 117181992 | rs656083 | C | T | NULL | 0.56094545 | NULL&M_Disease |
| 11 | 118150205 | rs1793170 | C | T | NULL | 0.56094545 | NULL&M_Disease |
| 11 | 119892194 | rs603179 | A | G | NULL | 0.56094545 | NULL&M_Disease |
| 11 | 117892050 | rs675929 | A | G | NULL | 0.56094545 | NULL&M_Disease |
| 11 | 119817739 | rs554338 | C | T | NULL | 0.56094545 | NULL&M_Disease |
| 11 | 119818657 | rs628244 | T | C | NULL | 0.56094545 | NULL&M_Disease |
| 11 | 119818914 | rs475178 | G | A | NULL | 0.56094545 | NULL&M_Disease |
| 11 | 119818937 | rs475867 | A | G | NULL | 0.56094545 | NULL&M_Disease |
| 11 | 116620377 | rs117164485 | A | G | NULL | 0.56094545 | NULL&M_Disease |
| 11 | 116887751 | rs12786797 | A | G | NULL | 0.56094545 | NULL&M_Disease |
| 11 | 117520361 | rs71480349 | C | T | NULL | 0.56094545 | NULL&M_Disease |
| 11 | 120716019 | rs10892629 | T | C | NULL | 0.56094545 | NULL&M_Disease |
| 11 | 119808956 | rs880004 | G | A | NULL | 0.56094545 | NULL&M_Disease |
| 11 | 120564627 | rs114428118 | T | G | NULL | 0.56094545 | NULL&M_Disease |
| 11 | 120851221 | rs557501 | C | G | NULL | 0.56094545 | NULL&M_Disease |
| 11 | 116242693 | rs11215905 | C | T | NULL | 0.56094545 | NULL&M_Disease |
| 11 | 116420968 | rs117843763 | A | C | NULL | 0.56094545 | NULL&M_Disease |
| 11 | 116590039 | rs180376 | C | G | NULL | 0.56094545 | NULL&M_Disease |
| 11 | 117647496 | rs17311045 | T | C | NULL | 0.56094545 | NULL&M_Disease |
| 11 | 117069023 | rs7112573 | C | T | NULL | 0.56094545 | NULL&M_Disease |
| 11 | 117537132 | rs17121057 | C | A | NULL | 0.56094545 | NULL&M_Disease |
| 11 | 118555383 | rs2508573 | T | C | NULL | 0.562176501 | NULL&M_Disease |
| 11 | 119817385 | rs550764 | A | T | NULL | 0.56094545 | NULL&M_Disease |
| 11 | 117338912 | rs2511050 | A | G | NULL | 0.562176501 | NULL&M_Disease |
| 11 | 117498834 | rs78915622 | G | A | NULL | 0.562176501 | NULL&M_Disease |
| 11 | 117515365 | rs10790198 | A | G | NULL | 0.562176501 | NULL&M_Disease |
| 11 | 116299637 | rs73580434 | T | A | NULL | 0.562176501 | NULL&M_Disease |
| 11 | 116717663 | rs614944 | C | T | NULL | 0.56094545 | NULL&M_Disease |
| 11 | 117483306 | rs12576806 | A | G | NULL | 0.56094545 | NULL&M_Disease |
| 11 | 120736087 | rs11218035 | T | C | NULL | 0.562176501 | NULL&M_Disease |
| 11 | 116728029 | rs509712 | G | A | NULL | 0.56094545 | NULL&M_Disease |
| 11 | 117588461 | rs74453782 | T | C | NULL | 0.56094545 | NULL&M_Disease |
| 11 | 117089722 | rs2239009 | C | G | NULL | 0.562176501 | NULL&M_Disease |

| | | | | | | | | | |
|---|---|---|---|---|---|---|---|---|---|
| 11 | 116607641 | rs180351 | T | C | 0.952 | -7977.388 | -7977.851 | -0.463 | 2.43E-06 |
| 11 | 117982495 | rs1941635 | T | G | 1.078 | -7977.388 | -7977.851 | -0.463 | 2.43E-06 |
| 11 | 120991571 | rs74945140 | A | G | 1.072 | -7977.388 | -7977.852 | -0.464 | 2.43E-06 |
| 11 | 117367838 | rs11603937 | A | G | 0.929 | -7977.388 | -7977.855 | -0.466 | 2.43E-06 |
| 11 | 120554278 | rs12292453 | T | C | 0.926 | -7977.388 | -7977.855 | -0.467 | 2.42E-06 |
| 11 | 116391651 | rs7120429 | T | C | 0.947 | -7977.388 | -7977.856 | -0.468 | 2.42E-06 |
| 11 | 116829653 | rs7120173 | G | T | 1.057 | -7977.388 | -7977.856 | -0.468 | 2.42E-06 |
| 11 | 120649949 | rs72995858 | T | C | 0.928 | -7977.388 | -7977.857 | -0.469 | 2.42E-06 |
| 11 | 116222151 | rs12792619 | C | T | 0.93 | -7977.388 | -7977.859 | -0.47 | 2.42E-06 |
| 11 | 117576946 | rs75819084 | T | C | 0.935 | -7977.388 | -7977.858 | -0.47 | 2.42E-06 |
| 11 | 121169052 | rs75960142 | C | T | 0.931 | -7977.388 | -7977.859 | -0.471 | 2.42E-06 |
| 11 | 116470289 | rs145936741 | G | A | 0.927 | -7977.388 | -7977.861 | -0.472 | 2.41E-06 |
| 11 | 116960078 | rs1481148807 | A | G | 0.929 | -7977.388 | -7977.86 | -0.472 | 2.41E-06 |
| 11 | 118061480 | rs7103918 | T | C | 1.056 | -7977.388 | -7977.86 | -0.472 | 2.41E-06 |
| 11 | 116697301 | rs140231532 | T | C | 0.933 | -7977.388 | -7977.861 | -0.473 | 2.41E-06 |
| 11 | 117570395 | rs12273147 | A | G | 0.934 | -7977.388 | -7977.861 | -0.473 | 2.41E-06 |
| 11 | 119876350 | rs113111994 | T | G | 0.929 | -7977.388 | -7977.861 | -0.473 | 2.41E-06 |
| 11 | 120470291 | rs4245037 | A | T | 1.053 | -7977.388 | -7977.862 | -0.473 | 2.41E-06 |
| 11 | 117069788 | rs2269398 | T | C | 1.075 | -7977.388 | -7977.863 | -0.474 | 2.41E-06 |
| 11 | 117796139 | rs55720349 | C | T | 0.936 | -7977.388 | -7977.862 | -0.474 | 2.41E-06 |
| 11 | 119483777 | rs78341300 | G | A | 1.068 | -7977.388 | -7977.862 | -0.474 | 2.41E-06 |
| 11 | 120042082 | rs529681 | A | G | 0.931 | -7977.388 | -7977.862 | -0.474 | 2.41E-06 |
| 11 | 120653294 | rs11218007 | A | G | 0.929 | -7977.388 | -7977.864 | -0.475 | 2.41E-06 |
| 11 | 116616402 | rs180340 | C | T | 0.952 | -7977.388 | -7977.864 | -0.476 | 2.40E-06 |
| 11 | 116724204 | rs598503 | C | T | 0.929 | -7977.388 | -7977.864 | -0.476 | 2.40E-06 |
| 11 | 118543563 | rs10892251 | T | C | 1.056 | -7977.388 | -7977.865 | -0.476 | 2.40E-06 |
| 11 | 119506255 | rs10790328 | G | C | 1.051 | -7977.388 | -7977.864 | -0.476 | 2.40E-06 |
| 11 | 119674096 | rs11217514 | A | G | 0.938 | -7977.388 | -7977.864 | -0.476 | 2.40E-06 |
| 11 | 117659322 | rs11216545 | A | G | 1.051 | -7977.388 | -7977.865 | -0.477 | 2.40E-06 |
| 11 | 120187422 | rs148189915 | T | A | 0.927 | -7977.388 | -7977.866 | -0.477 | 2.40E-06 |
| 11 | 116278612 | rs7925794 | T | C | 1.053 | -7977.388 | -7977.866 | -0.478 | 2.40E-06 |
| 11 | 116472577 | rs34994677 | G | A | 0.933 | -7977.388 | -7977.867 | -0.478 | 2.40E-06 |
| 11 | 118100176 | rs12361791 | G | T | 0.933 | -7977.388 | -7977.866 | -0.478 | 2.40E-06 |
| 11 | 120144571 | rs138488120 | A | C | 0.928 | -7977.388 | -7977.866 | -0.478 | 2.40E-06 |
| 11 | 121093301 | rs117235919 | A | G | 0.935 | -7977.388 | -7977.867 | -0.478 | 2.40E-06 |
| 11 | 117104900 | rs117547981 | A | G | 0.932 | -7977.388 | -7977.868 | -0.479 | 2.40E-06 |
| 11 | 120302456 | rs117092333 | A | G | 0.927 | -7977.388 | -7977.868 | -0.48 | 2.39E-06 |
| 11 | 116478432 | rs17119839 | G | A | 1.074 | -7977.388 | -7977.869 | -0.481 | 2.39E-06 |
| 11 | 117200129 | rs75013797 | T | C | 0.927 | -7977.388 | -7977.869 | -0.481 | 2.39E-06 |
| 11 | 118312870 | rs74422681 | G | A | 1.075 | -7977.388 | -7977.869 | -0.481 | 2.39E-06 |
| 11 | 117958429 | rs603455 | T | A | 1.073 | -7977.388 | -7977.87 | -0.482 | 2.39E-06 |
| 11 | 118802128 | rs59069289 | T | G | 0.93 | -7977.388 | -7977.87 | -0.482 | 2.39E-06 |
| 11 | 118802386 | rs59480472 | C | T | 0.93 | -7977.388 | -7977.87 | -0.482 | 2.39E-06 |
| 11 | 116617782 | rs180339 | T | C | 1.051 | -7977.388 | -7977.871 | -0.483 | 2.39E-06 |
| 11 | 117569523 | rs7122782 | A | G | 0.934 | -7977.388 | -7977.871 | -0.483 | 2.39E-06 |
| 11 | 119026384 | rs56168028 | G | A | 1.07 | -7977.388 | -7977.871 | -0.483 | 2.39E-06 |
| 11 | 116633825 | rs918144 | T | C | 1.051 | -7977.388 | -7977.872 | -0.484 | 2.38E-06 |
| 11 | 117179891 | rs676134 | T | C | 0.952 | -7977.388 | -7977.872 | -0.484 | 2.38E-06 |
| 11 | 119514129 | rs906827 | A | C | 0.933 | -7977.388 | -7977.872 | -0.484 | 2.38E-06 |
| 11 | 120825635 | rs4936569 | C | T | 0.945 | -7977.388 | -7977.872 | -0.484 | 2.38E-06 |
| 11 | 117352364 | rs73022382 | A | G | 1.077 | -7977.388 | -7977.873 | -0.485 | 2.38E-06 |
| 11 | 118134597 | rs4936426 | C | T | 1.051 | -7977.388 | -7977.874 | -0.486 | 2.38E-06 |
| 11 | 116693871 | rs12721041 | T | C | 0.935 | -7977.388 | -7977.876 | -0.487 | 2.38E-06 |
| 11 | 117378697 | rs117328724 | T | C | 0.928 | -7977.388 | -7977.875 | -0.487 | 2.38E-06 |
| 11 | 117070740 | rs2170317 | C | A | 1.074 | -7977.388 | -7977.877 | -0.488 | 2.37E-06 |
| 11 | 117129617 | rs550893 | A | G | 0.928 | -7977.388 | -7977.876 | -0.488 | 2.37E-06 |
| 11 | 116479584 | rs527778 | T | G | 0.949 | -7977.388 | -7977.879 | -0.49 | 2.37E-06 |
| 11 | 116491465 | rs112732949 | G | A | 1.071 | -7977.388 | -7977.878 | -0.49 | 2.37E-06 |
| 11 | 117720781 | rs7928007 | C | A | 1.075 | -7977.388 | -7977.878 | -0.49 | 2.37E-06 |
| 11 | 119816766 | rs657317 | A | C | 0.928 | -7977.388 | -7977.878 | -0.49 | 2.37E-06 |
| 11 | 116610048 | rs180350 | T | C | 0.952 | -7977.388 | -7977.88 | -0.491 | 2.37E-06 |
| 11 | 117646968 | rs61905347 | T | C | 1.07 | -7977.388 | -7977.879 | -0.491 | 2.37E-06 |
| 11 | 117474845 | rs61697909 | A | G | 1.073 | -7977.388 | -7977.88 | -0.492 | 2.36E-06 |
| 11 | 117181992 | rs656083 | C | T | 1.051 | -7977.388 | -7977.881 | -0.493 | 2.36E-06 |
| 11 | 118150205 | rs1793170 | C | T | 0.949 | -7977.388 | -7977.881 | -0.493 | 2.36E-06 |
| 11 | 119892194 | rs603179 | A | G | 0.94 | -7977.388 | -7977.882 | -0.493 | 2.36E-06 |
| 11 | 117892050 | rs675929 | A | G | 0.949 | -7977.388 | -7977.882 | -0.494 | 2.36E-06 |
| 11 | 119817739 | rs554338 | C | T | 0.928 | -7977.388 | -7977.882 | -0.494 | 2.36E-06 |
| 11 | 119818657 | rs628244 | T | C | 0.928 | -7977.388 | -7977.882 | -0.494 | 2.36E-06 |
| 11 | 119818914 | rs475178 | G | A | 0.928 | -7977.388 | -7977.882 | -0.494 | 2.36E-06 |
| 11 | 119818937 | rs475867 | A | G | 0.928 | -7977.388 | -7977.882 | -0.494 | 2.36E-06 |
| 11 | 116620377 | rs117164485 | A | G | 1.075 | -7977.388 | -7977.883 | -0.495 | 2.36E-06 |
| 11 | 116887751 | rs12786797 | A | G | 1.057 | -7977.388 | -7977.883 | -0.495 | 2.36E-06 |
| 11 | 117520361 | rs71480349 | C | T | 0.928 | -7977.388 | -7977.884 | -0.495 | 2.36E-06 |
| 11 | 120716019 | rs10892629 | T | C | 0.943 | -7977.388 | -7977.884 | -0.495 | 2.36E-06 |
| 11 | 119808956 | rs880004 | G | A | 1.051 | -7977.388 | -7977.884 | -0.496 | 2.36E-06 |
| 11 | 120564627 | rs114428118 | T | G | 0.93 | -7977.388 | -7977.885 | -0.496 | 2.36E-06 |
| 11 | 120851221 | rs557501 | C | G | 0.947 | -7977.388 | -7977.885 | -0.496 | 2.36E-06 |
| 11 | 116242693 | rs11215905 | C | T | 1.05 | -7977.388 | -7977.885 | -0.497 | 2.35E-06 |
| 11 | 116420968 | rs117843763 | A | C | 0.935 | -7977.388 | -7977.885 | -0.497 | 2.35E-06 |
| 11 | 116590039 | rs180376 | C | G | 0.952 | -7977.388 | -7977.885 | -0.497 | 2.35E-06 |
| 11 | 117647496 | rs17311045 | T | C | 1.07 | -7977.388 | -7977.886 | -0.497 | 2.35E-06 |
| 11 | 117069023 | rs7112573 | C | T | 1.073 | -7977.388 | -7977.886 | -0.498 | 2.35E-06 |
| 11 | 117537132 | rs17121057 | C | A | 0.938 | -7977.388 | -7977.886 | -0.498 | 2.35E-06 |
| 11 | 118555383 | rs2508573 | T | C | 0.952 | -7977.388 | -7977.887 | -0.499 | 2.35E-06 |
| 11 | 119817385 | rs550764 | A | T | 0.928 | -7977.388 | -7977.888 | -0.499 | 2.35E-06 |
| 11 | 117338912 | rs2511050 | A | G | 0.934 | -7977.388 | -7977.889 | -0.5 | 2.35E-06 |
| 11 | 117498834 | rs78915622 | G | A | 1.068 | -7977.388 | -7977.889 | -0.5 | 2.35E-06 |
| 11 | 117515365 | rs10790198 | A | G | 0.952 | -7977.388 | -7977.888 | -0.5 | 2.35E-06 |
| 11 | 116299637 | rs73580434 | T | A | 1.073 | -7977.388 | -7977.889 | -0.501 | 2.34E-06 |
| 11 | 116717663 | rs614944 | C | T | 0.931 | -7977.388 | -7977.889 | -0.501 | 2.34E-06 |
| 11 | 117483306 | rs12576806 | A | G | 0.931 | -7977.388 | -7977.89 | -0.501 | 2.34E-06 |
| 11 | 120736087 | rs11218035 | T | C | 1.056 | -7977.388 | -7977.889 | -0.501 | 2.34E-06 |
| 11 | 116728029 | rs509712 | G | A | 0.931 | -7977.388 | -7977.891 | -0.502 | 2.34E-06 |
| 11 | 117588461 | rs74453782 | T | C | 0.938 | -7977.388 | -7977.89 | -0.502 | 2.34E-06 |
| 11 | 117089722 | rs2239009 | C | G | 1.073 | -7977.388 | -7977.891 | -0.503 | 2.34E-06 |

| | | | | | | | |
|---|---|---|---|---|---|---|---|
| 11 | 117890273 | rs2508449 | C | T | NULL | 0.562176501 | NULL&M_Disease |
| 11 | 117890681 | rs1939127 | C | T | NULL | 0.562176501 | NULL&M_Disease |
| 11 | 117891216 | rs2512156 | T | A | NULL | 0.562176501 | NULL&M_Disease |
| 11 | 118238119 | rs11216859 | C | T | NULL | 0.562176501 | NULL&M_Disease |
| 11 | 118711069 | rs538645 | C | T | NULL | 0.562176501 | NULL&M_Disease |
| 11 | 120343033 | rs61898762 | G | T | NULL | 0.562176501 | NULL&M_Disease |
| 11 | 119734332 | rs34051198 | A | G | NULL | 0.562176501 | NULL&M_Disease |
| 11 | 116613184 | rs180344 | C | T | NULL | 0.562176501 | NULL&M_Disease |
| 11 | 117592582 | rs12577828 | A | G | NULL | 0.56094545 | NULL&M_Disease |
| 11 | 119679659 | rs11828031 | A | G | NULL | 0.562176501 | NULL&M_Disease |
| 11 | 119813545 | rs1944502 | G | A | NULL | 0.562176501 | NULL&M_Disease |
| 11 | 120890960 | rs12785765 | C | A | NULL | 0.562176501 | NULL&M_Disease |
| 11 | 117493839 | rs17120968 | T | C | NULL | 0.562176501 | NULL&M_Disease |
| 11 | 117620050 | rs112571425 | C | T | NULL | 0.562176501 | NULL&M_Disease |
| 11 | 120566473 | rs78435918 | G | A | NULL | 0.562176501 | NULL&M_Disease |
| 11 | 120702026 | rs4936556 | T | C | NULL | 0.562176501 | NULL&M_Disease |
| 11 | 119297248 | rs117164546 | A | G | NULL | 0.562176501 | NULL&M_Disease |
| 11 | 120664625 | rs77454413 | A | G | NULL | 0.562176501 | NULL&M_Disease |
| 11 | 117727584 | rs11216585 | T | G | NULL | 0.562176501 | NULL&M_Disease |
| 11 | 117938674 | rs190143695 | C | A | NULL | 0.562176501 | NULL&M_Disease |
| 11 | 120650633 | rs17124324 | T | A | NULL | 0.562176501 | NULL&M_Disease |
| 11 | 117485422 | rs11216470 | T | C | NULL | 0.562176501 | NULL&M_Disease |
| 11 | 117603352 | rs113204414 | C | T | NULL | 0.562176501 | NULL&M_Disease |
| 11 | 117605731 | rs61903855 | C | G | NULL | 0.562176501 | NULL&M_Disease |
| 11 | 120416923 | rs2252494 | A | G | NULL | 0.563406786 | NULL&M_Disease |
| 11 | 116673315 | rs12791103 | T | G | NULL | 0.563406786 | NULL&M_Disease |
| 11 | 116722171 | rs676234 | G | A | NULL | 0.562176501 | NULL&M_Disease |
| 11 | 116482449 | rs74365486 | A | C | NULL | 0.562176501 | NULL&M_Disease |
| 11 | 117092051 | rs78628783 | T | C | NULL | 0.563406786 | NULL&M_Disease |
| 11 | 117590895 | rs74621718 | G | C | NULL | 0.562176501 | NULL&M_Disease |
| 11 | 117590919 | rs78667446 | T | G | NULL | 0.562176501 | NULL&M_Disease |
| 11 | 120979833 | rs117889729 | G | C | NULL | 0.562176501 | NULL&M_Disease |
| 11 | 117070596 | rs2239678 | T | C | NULL | 0.563406786 | NULL&M_Disease |
| 11 | 117339893 | rs512443 | T | C | NULL | 0.563406786 | NULL&M_Disease |
| 11 | 119495219 | rs116057214 | G | A | NULL | 0.563406786 | NULL&M_Disease |
| 11 | 120864593 | rs10750179 | A | G | NULL | 0.563406786 | NULL&M_Disease |
| 11 | 116588909 | rs180378 | A | G | NULL | 0.563406786 | NULL&M_Disease |
| 11 | 116612659 | rs180346 | C | A | NULL | 0.563406786 | NULL&M_Disease |
| 11 | 117066353 | rs2269399 | C | G | NULL | 0.563406786 | NULL&M_Disease |
| 11 | 121010814 | rs116977890 | A | C | NULL | 0.562176501 | NULL&M_Disease |
| 11 | 116726651 | rs1241550 | C | T | NULL | 0.563406786 | NULL&M_Disease |
| 11 | 116729196 | rs541586 | C | T | NULL | 0.563406786 | NULL&M_Disease |
| 11 | 117080640 | rs11216315 | G | C | NULL | 0.563406786 | NULL&M_Disease |
| 11 | 116403905 | rs7925845 | G | A | NULL | 0.563406786 | NULL&M_Disease |
| 11 | 116428225 | rs12279486 | T | C | NULL | 0.563406786 | NULL&M_Disease |
| 11 | 118326218 | rs116954303 | A | G | NULL | 0.562176501 | NULL&M_Disease |
| 11 | 119673277 | rs111494641 | T | C | NULL | 0.563406786 | NULL&M_Disease |
| 11 | 121098681 | rs6589862 | A | G | NULL | 0.563406786 | NULL&M_Disease |
| 11 | 116485420 | rs80072373 | G | C | NULL | 0.563406786 | NULL&M_Disease |
| 11 | 117353294 | rs7942824 | T | G | NULL | 0.564636292 | NULL&M_Disease |
| 11 | 118632893 | rs6589690 | A | G | NULL | 0.564636292 | NULL&M_Disease |
| 11 | 120443297 | rs11599940 | A | G | NULL | 0.564636292 | NULL&M_Disease |
| 11 | 121088659 | rs637115 | A | G | NULL | 0.564636292 | NULL&M_Disease |
| 11 | 121088829 | rs478626 | A | G | NULL | 0.564636292 | NULL&M_Disease |
| 11 | 117604866 | rs111462461 | T | C | NULL | 0.563406786 | NULL&M_Disease |
| 11 | 119674095 | rs2368937 | C | T | NULL | 0.564636292 | NULL&M_Disease |
| 11 | 117413864 | rs678468 | C | T | NULL | 0.564636292 | NULL&M_Disease |
| 11 | 117590007 | rs12576975 | A | C | NULL | 0.563406786 | NULL&M_Disease |
| 11 | 117590811 | rs78080911 | G | A | NULL | 0.563406786 | NULL&M_Disease |
| 11 | 121117590 | rs145433737 | A | G | NULL | 0.563406786 | NULL&M_Disease |
| 11 | 117500448 | rs73012582 | A | G | NULL | 0.564636292 | NULL&M_Disease |
| 11 | 117092394 | rs80351364 | T | A | NULL | 0.564636292 | NULL&M_Disease |
| 11 | 117390488 | rs76467987 | C | T | NULL | 0.563406786 | NULL&M_Disease |
| 11 | 117789242 | rs3802870 | G | C | NULL | 0.564636292 | NULL&M_Disease |
| 11 | 117957258 | rs689264 | T | C | NULL | 0.564636292 | NULL&M_Disease |
| 11 | 118644323 | rs6589695 | C | G | NULL | 0.564636292 | NULL&M_Disease |
| 11 | 116234032 | rs575636 | G | A | NULL | 0.564636292 | NULL&M_Disease |
| 11 | 117586623 | rs948776 | C | A | NULL | 0.563406786 | NULL&M_Disease |
| 11 | 118870413 | rs10892319 | T | C | NULL | 0.564636292 | NULL&M_Disease |
| 11 | 119932930 | rs1320685 | C | G | NULL | 0.564636292 | NULL&M_Disease |
| 11 | 121088794 | rs648191 | A | G | NULL | 0.564636292 | NULL&M_Disease |
| 11 | 117351353 | rs577166 | T | C | NULL | 0.564636292 | NULL&M_Disease |
| 11 | 117501271 | rs7950396 | A | G | NULL | 0.564636292 | NULL&M_Disease |
| 11 | 116308634 | rs76477544 | G | A | NULL | 0.564636292 | NULL&M_Disease |
| 11 | 119818622 | rs10892501 | G | A | NULL | 0.565865003 | NULL&M_Disease |
| 11 | 121148641 | rs138536663 | C | T | NULL | 0.564636292 | NULL&M_Disease |
| 11 | 117583591 | rs74443315 | G | A | NULL | 0.564636292 | NULL&M_Disease |
| 11 | 118556995 | rs72999502 | C | T | NULL | 0.564636292 | NULL&M_Disease |
| 11 | 120081510 | rs138435357 | A | C | NULL | 0.564636292 | NULL&M_Disease |
| 11 | 120619586 | rs2852226 | A | G | NULL | 0.564636292 | NULL&M_Disease |
| 11 | 120936096 | rs10892666 | T | G | NULL | 0.565865003 | NULL&M_Disease |
| 11 | 121150291 | rs189832363 | T | C | NULL | 0.564636292 | NULL&M_Disease |
| 11 | 116665120 | rs45555231 | C | G | NULL | 0.564636292 | NULL&M_Disease |
| 11 | 118752178 | rs61150131 | C | T | NULL | 0.564636292 | NULL&M_Disease |
| 11 | 119762628 | rs143470629 | T | C | NULL | 0.564636292 | NULL&M_Disease |
| 11 | 116231840 | rs513935 | A | G | NULL | 0.565865003 | NULL&M_Disease |
| 11 | 116631690 | rs3741300 | A | G | NULL | 0.565865003 | NULL&M_Disease |
| 11 | 118058156 | rs6589651 | G | A | NULL | 0.565865003 | NULL&M_Disease |
| 11 | 120050959 | rs471776 | A | G | NULL | 0.564636292 | NULL&M_Disease |
| 11 | 120051363 | rs546441 | C | A | NULL | 0.564636292 | NULL&M_Disease |
| 11 | 120052037 | rs551843 | A | G | NULL | 0.564636292 | NULL&M_Disease |
| 11 | 116305930 | rs12293635 | C | G | NULL | 0.564636292 | NULL&M_Disease |
| 11 | 117591311 | rs77252250 | A | C | NULL | 0.564636292 | NULL&M_Disease |
| 11 | 118012623 | rs61900623 | T | C | NULL | 0.564636292 | NULL&M_Disease |
| 11 | 116461333 | rs11216024 | G | A | NULL | 0.565865003 | NULL&M_Disease |
| 11 | 116622958 | rs180329 | A | G | NULL | 0.565865003 | NULL&M_Disease |

| | | | | | | | | | |
|---|---|---|---|---|---|---|---|---|---|
| 11 | 117890273 | rs2508449 | C | T | 0.95 | -7977.388 | -7977.892 | -0.503 | 2.34E-06 |
| 11 | 117890681 | rs1939127 | C | T | 0.95 | -7977.388 | -7977.892 | -0.503 | 2.34E-06 |
| 11 | 117891216 | rs2512156 | T | A | 0.95 | -7977.388 | -7977.892 | -0.503 | 2.34E-06 |
| 11 | 118238119 | rs11216859 | C | T | 1.052 | -7977.388 | -7977.891 | -0.503 | 2.34E-06 |
| 11 | 118711069 | rs538645 | C | T | 0.948 | -7977.388 | -7977.891 | -0.503 | 2.34E-06 |
| 11 | 120343033 | rs61898762 | G | T | 1.076 | -7977.388 | -7977.891 | -0.503 | 2.34E-06 |
| 11 | 119734332 | rs34051198 | A | G | 1.064 | -7977.388 | -7977.892 | -0.504 | 2.34E-06 |
| 11 | 116613184 | rs180344 | C | T | 0.952 | -7977.388 | -7977.893 | -0.505 | 2.33E-06 |
| 11 | 117592582 | rs12577828 | A | G | 0.939 | -7977.388 | -7977.893 | -0.505 | 2.33E-06 |
| 11 | 119679659 | rs11828031 | A | G | 0.947 | -7977.388 | -7977.893 | -0.505 | 2.33E-06 |
| 11 | 119813545 | rs1944502 | G | A | 1.05 | -7977.388 | -7977.893 | -0.505 | 2.33E-06 |
| 11 | 120890960 | rs12785765 | C | A | 1.066 | -7977.388 | -7977.895 | -0.506 | 2.33E-06 |
| 11 | 117493839 | rs17120968 | T | C | 0.931 | -7977.388 | -7977.895 | -0.507 | 2.33E-06 |
| 11 | 117620050 | rs112571425 | C | T | 0.939 | -7977.388 | -7977.895 | -0.507 | 2.33E-06 |
| 11 | 120566473 | rs78435918 | G | A | 0.931 | -7977.388 | -7977.895 | -0.507 | 2.33E-06 |
| 11 | 120702026 | rs4936556 | T | C | 1.051 | -7977.388 | -7977.895 | -0.507 | 2.33E-06 |
| 11 | 119297248 | rs117164546 | A | G | 0.934 | -7977.388 | -7977.897 | -0.508 | 2.33E-06 |
| 11 | 120664625 | rs77454413 | A | G | 0.948 | -7977.388 | -7977.897 | -0.508 | 2.33E-06 |
| 11 | 117727584 | rs11216585 | T | G | 1.073 | -7977.388 | -7977.897 | -0.509 | 2.33E-06 |
| 11 | 117938674 | rs190143695 | C | A | 1.069 | -7977.388 | -7977.898 | -0.509 | 2.33E-06 |
| 11 | 120650633 | rs17124324 | T | A | 0.932 | -7977.388 | -7977.897 | -0.509 | 2.33E-06 |
| 11 | 117485422 | rs11216470 | T | C | 0.931 | -7977.388 | -7977.899 | -0.51 | 2.32E-06 |
| 11 | 117603352 | rs113204414 | C | T | 0.94 | -7977.388 | -7977.898 | -0.51 | 2.32E-06 |
| 11 | 117605731 | rs61903855 | C | G | 1.075 | -7977.388 | -7977.898 | -0.51 | 2.32E-06 |
| 11 | 120416923 | rs2252494 | A | G | 0.949 | -7977.388 | -7977.899 | -0.51 | 2.32E-06 |
| 11 | 116673315 | rs12791103 | T | G | 1.068 | -7977.388 | -7977.9 | -0.512 | 2.32E-06 |
| 11 | 116722171 | rs676234 | G | A | 0.932 | -7977.388 | -7977.9 | -0.512 | 2.32E-06 |
| 11 | 116482449 | rs74365486 | A | C | 1.067 | -7977.388 | -7977.902 | -0.513 | 2.32E-06 |
| 11 | 117092051 | rs78628783 | T | C | 1.073 | -7977.388 | -7977.901 | -0.513 | 2.32E-06 |
| 11 | 117590895 | rs74621718 | G | C | 0.94 | -7977.388 | -7977.901 | -0.513 | 2.32E-06 |
| 11 | 117590919 | rs78667446 | T | G | 0.94 | -7977.388 | -7977.901 | -0.513 | 2.32E-06 |
| 11 | 120979833 | rs117889729 | G | C | 0.935 | -7977.388 | -7977.901 | -0.513 | 2.32E-06 |
| 11 | 117070596 | rs2239678 | T | C | 1.072 | -7977.388 | -7977.902 | -0.514 | 2.31E-06 |
| 11 | 117339893 | rs512443 | T | C | 0.934 | -7977.388 | -7977.902 | -0.514 | 2.31E-06 |
| 11 | 119495219 | rs116057214 | G | A | 1.067 | -7977.388 | -7977.902 | -0.514 | 2.31E-06 |
| 11 | 120864593 | rs10750179 | A | G | 0.947 | -7977.388 | -7977.903 | -0.514 | 2.31E-06 |
| 11 | 116588909 | rs180378 | A | G | 1.051 | -7977.388 | -7977.903 | -0.515 | 2.31E-06 |
| 11 | 116612659 | rs180346 | C | A | 0.952 | -7977.388 | -7977.904 | -0.515 | 2.31E-06 |
| 11 | 117066353 | rs2269399 | C | G | 1.072 | -7977.388 | -7977.903 | -0.515 | 2.31E-06 |
| 11 | 121010814 | rs116977890 | A | C | 0.939 | -7977.388 | -7977.903 | -0.515 | 2.31E-06 |
| 11 | 116726651 | rs1241550 | C | T | 0.933 | -7977.388 | -7977.904 | -0.516 | 2.31E-06 |
| 11 | 116729196 | rs541586 | C | T | 0.933 | -7977.388 | -7977.904 | -0.516 | 2.31E-06 |
| 11 | 117080640 | rs11216315 | G | C | 1.072 | -7977.388 | -7977.904 | -0.516 | 2.31E-06 |
| 11 | 116403905 | rs7925845 | G | A | 1.06 | -7977.388 | -7977.905 | -0.517 | 2.31E-06 |
| 11 | 116428225 | rs12279486 | T | C | 1.058 | -7977.388 | -7977.905 | -0.517 | 2.31E-06 |
| 11 | 118326218 | rs116954303 | A | G | 1.055 | -7977.388 | -7977.905 | -0.517 | 2.31E-06 |
| 11 | 119673277 | rs111494641 | T | C | 0.939 | -7977.388 | -7977.906 | -0.517 | 2.31E-06 |
| 11 | 121098681 | rs6589862 | A | G | 0.946 | -7977.388 | -7977.906 | -0.518 | 2.30E-06 |
| 11 | 116485420 | rs80072373 | G | C | 1.066 | -7977.388 | -7977.907 | -0.519 | 2.30E-06 |
| 11 | 117353294 | rs7942824 | T | G | 1.055 | -7977.388 | -7977.907 | -0.519 | 2.30E-06 |
| 11 | 118632893 | rs6589690 | A | G | 0.94 | -7977.388 | -7977.907 | -0.519 | 2.30E-06 |
| 11 | 120443297 | rs11599940 | A | G | 1.05 | -7977.388 | -7977.907 | -0.519 | 2.30E-06 |
| 11 | 121088659 | rs637115 | A | G | 0.948 | -7977.388 | -7977.907 | -0.519 | 2.30E-06 |
| 11 | 121088829 | rs478626 | A | G | 0.948 | -7977.388 | -7977.907 | -0.519 | 2.30E-06 |
| 11 | 117604866 | rs111462461 | T | C | 0.941 | -7977.388 | -7977.908 | -0.52 | 2.30E-06 |
| 11 | 119674095 | rs2368937 | C | T | 0.953 | -7977.388 | -7977.909 | -0.52 | 2.30E-06 |
| 11 | 117413864 | rs678468 | C | T | 1.05 | -7977.388 | -7977.91 | -0.521 | 2.30E-06 |
| 11 | 117590007 | rs12576975 | A | C | 0.941 | -7977.388 | -7977.909 | -0.521 | 2.30E-06 |
| 11 | 117590811 | rs78080911 | G | A | 0.941 | -7977.388 | -7977.909 | -0.521 | 2.30E-06 |
| 11 | 121117590 | rs145433737 | A | G | 0.932 | -7977.388 | -7977.909 | -0.521 | 2.30E-06 |
| 11 | 117500448 | rs73012582 | A | G | 1.069 | -7977.388 | -7977.91 | -0.522 | 2.30E-06 |
| 11 | 117092394 | rs80351364 | T | A | 1.072 | -7977.388 | -7977.912 | -0.523 | 2.29E-06 |
| 11 | 117390488 | rs76467987 | C | T | 0.93 | -7977.388 | -7977.911 | -0.523 | 2.29E-06 |
| 11 | 117789242 | rs3802870 | G | C | 0.93 | -7977.388 | -7977.912 | -0.524 | 2.29E-06 |
| 11 | 117957258 | rs689264 | T | C | 1.07 | -7977.388 | -7977.914 | -0.525 | 2.29E-06 |
| 11 | 118644323 | rs6589695 | C | G | 0.93 | -7977.388 | -7977.914 | -0.525 | 2.29E-06 |
| 11 | 116234032 | rs575636 | G | A | 1.049 | -7977.388 | -7977.914 | -0.526 | 2.29E-06 |
| 11 | 117586623 | rs948776 | C | A | 0.942 | -7977.388 | -7977.914 | -0.526 | 2.29E-06 |
| 11 | 118870413 | rs10892319 | T | C | 1.06 | -7977.388 | -7977.915 | -0.526 | 2.29E-06 |
| 11 | 119932930 | rs1320685 | C | G | 1.051 | -7977.388 | -7977.914 | -0.526 | 2.29E-06 |
| 11 | 121088794 | rs648191 | A | G | 0.949 | -7977.388 | -7977.915 | -0.526 | 2.29E-06 |
| 11 | 117351353 | rs577166 | T | C | 0.934 | -7977.388 | -7977.915 | -0.527 | 2.28E-06 |
| 11 | 117501271 | rs7950396 | A | G | 1.068 | -7977.388 | -7977.915 | -0.527 | 2.28E-06 |
| 11 | 116308634 | rs76477544 | G | A | 0.931 | -7977.388 | -7977.916 | -0.528 | 2.28E-06 |
| 11 | 119818622 | rs10892501 | G | A | 0.952 | -7977.388 | -7977.916 | -0.528 | 2.28E-06 |
| 11 | 121148641 | rs138536663 | C | T | 1.074 | -7977.388 | -7977.916 | -0.528 | 2.28E-06 |
| 11 | 117583591 | rs74443315 | G | A | 0.943 | -7977.388 | -7977.917 | -0.529 | 2.28E-06 |
| 11 | 118556995 | rs72999502 | C | T | 1.072 | -7977.388 | -7977.917 | -0.529 | 2.28E-06 |
| 11 | 120081510 | rs138435357 | A | C | 0.93 | -7977.388 | -7977.918 | -0.529 | 2.28E-06 |
| 11 | 120619586 | rs2852226 | A | G | 0.934 | -7977.388 | -7977.918 | -0.529 | 2.28E-06 |
| 11 | 120936096 | rs10892666 | T | G | 1.056 | -7977.388 | -7977.917 | -0.529 | 2.28E-06 |
| 11 | 121150291 | rs189832363 | T | C | 0.946 | -7977.388 | -7977.917 | -0.529 | 2.28E-06 |
| 11 | 116665120 | rs45555231 | C | G | 0.936 | -7977.388 | -7977.918 | -0.53 | 2.28E-06 |
| 11 | 118752178 | rs61150131 | C | T | 0.931 | -7977.388 | -7977.918 | -0.53 | 2.28E-06 |
| 11 | 119762628 | rs143470629 | T | C | 0.949 | -7977.388 | -7977.918 | -0.53 | 2.28E-06 |
| 11 | 116231840 | rs513935 | A | G | 1.05 | -7977.388 | -7977.919 | -0.531 | 2.27E-06 |
| 11 | 116631690 | rs3741300 | A | G | 0.953 | -7977.388 | -7977.92 | -0.531 | 2.27E-06 |
| 11 | 118058156 | rs6589651 | G | A | 1.055 | -7977.388 | -7977.919 | -0.531 | 2.27E-06 |
| 11 | 120050959 | rs471776 | A | G | 0.935 | -7977.388 | -7977.919 | -0.531 | 2.27E-06 |
| 11 | 120051363 | rs546441 | C | A | 0.935 | -7977.388 | -7977.919 | -0.531 | 2.27E-06 |
| 11 | 120052037 | rs551843 | A | G | 0.935 | -7977.388 | -7977.919 | -0.531 | 2.27E-06 |
| 11 | 116305930 | rs12293635 | C | G | 0.931 | -7977.388 | -7977.92 | -0.532 | 2.27E-06 |
| 11 | 117591311 | rs77252250 | A | C | 0.944 | -7977.388 | -7977.92 | -0.532 | 2.27E-06 |
| 11 | 118012623 | rs61900623 | T | C | 1.073 | -7977.388 | -7977.921 | -0.532 | 2.27E-06 |
| 11 | 116461333 | rs11216024 | G | A | 0.931 | -7977.388 | -7977.921 | -0.533 | 2.27E-06 |
| 11 | 116622958 | rs180329 | A | G | 0.953 | -7977.388 | -7977.921 | -0.533 | 2.27E-06 |

| | | | | | | | |
|---|---|---|---|---|---|---|---|
| 11 | 117388656 | rs59346420 | T | C | NULL | 0.564636292 | NULL&M_Disease |
| 11 | 117068444 | rs74385934 | A | G | NULL | 0.565865003 | NULL&M_Disease |
| 11 | 117081709 | rs1263499 | G | A | NULL | 0.565865003 | NULL&M_Disease |
| 11 | 117958179 | rs10892212 | A | T | NULL | 0.565865003 | NULL&M_Disease |
| 11 | 118152252 | rs117117171 | A | T | NULL | 0.564636292 | NULL&M_Disease |
| 11 | 118938882 | rs670526 | C | T | NULL | 0.565865003 | NULL&M_Disease |
| 11 | 120093920 | rs12226432 | C | A | NULL | 0.565865003 | NULL&M_Disease |
| 11 | 120464049 | rs12421186 | T | C | NULL | 0.565865003 | NULL&M_Disease |
| 11 | 116240253 | rs541874 | T | C | NULL | 0.565865003 | NULL&M_Disease |
| 11 | 119652997 | rs142774084 | A | G | NULL | 0.565865003 | NULL&M_Disease |
| 11 | 116486664 | rs3016347 | C | G | NULL | 0.565865003 | NULL&M_Disease |
| 11 | 118057867 | rs7103366 | G | T | NULL | 0.565865003 | NULL&M_Disease |
| 11 | 118928393 | rs471028 | C | G | NULL | 0.565865003 | NULL&M_Disease |
| 11 | 118932610 | rs647432 | T | C | NULL | 0.565865003 | NULL&M_Disease |
| 11 | 117577527 | rs12574878 | T | C | NULL | 0.565865003 | NULL&M_Disease |
| 11 | 117577558 | rs12576082 | A | T | NULL | 0.565865003 | NULL&M_Disease |
| 11 | 117577909 | rs12577066 | G | A | NULL | 0.565865003 | NULL&M_Disease |
| 11 | 117578314 | rs113570704 | T | C | NULL | 0.565865003 | NULL&M_Disease |
| 11 | 117579516 | rs1310160088 | T | C | NULL | 0.565865003 | NULL&M_Disease |
| 11 | 117579607 | rs78696045 | T | A | NULL | 0.565865003 | NULL&M_Disease |
| 11 | 119507468 | rs4938697 | C | T | NULL | 0.567092905 | NULL&M_Disease |
| 11 | 117033808 | rs79118964 | T | C | NULL | 0.565865003 | NULL&M_Disease |
| 11 | 119577632 | rs7104655 | A | G | NULL | 0.565865003 | NULL&M_Disease |
| 11 | 116630600 | rs918143 | T | C | NULL | 0.567092905 | NULL&M_Disease |
| 11 | 116557882 | rs11216093 | C | A | NULL | 0.567092905 | NULL&M_Disease |
| 11 | 117566305 | rs55973262 | C | G | NULL | 0.567092905 | NULL&M_Disease |
| 11 | 117586881 | rs948774 | A | T | NULL | 0.565865003 | NULL&M_Disease |
| 11 | 117586942 | rs948773 | A | G | NULL | 0.565865003 | NULL&M_Disease |
| 11 | 117727742 | rs11216586 | C | G | NULL | 0.567092905 | NULL&M_Disease |
| 11 | 116226290 | rs577941 | A | G | NULL | 0.567092905 | NULL&M_Disease |
| 11 | 117113511 | rs117117915 | T | C | NULL | 0.565865003 | NULL&M_Disease |
| 11 | 118214267 | rs2071381 | A | G | NULL | 0.567092905 | NULL&M_Disease |
| 11 | 119818789 | rs895653 | A | T | NULL | 0.567092905 | NULL&M_Disease |
| 11 | 117388657 | rs77944184 | A | G | NULL | 0.567092905 | NULL&M_Disease |
| 11 | 118835272 | rs67533649 | G | A | NULL | 0.567092905 | NULL&M_Disease |
| 11 | 120473450 | rs139831408 | T | C | NULL | 0.565865003 | NULL&M_Disease |
| 11 | 116306324 | rs79892107 | T | C | NULL | 0.567092905 | NULL&M_Disease |
| 11 | 118709275 | rs187164533 | A | G | NULL | 0.565865003 | NULL&M_Disease |
| 11 | 119654894 | rs77020062 | T | C | NULL | 0.565865003 | NULL&M_Disease |
| 11 | 119768054 | rs11217556 | A | C | NULL | 0.567092905 | NULL&M_Disease |
| 11 | 116235005 | rs495430 | A | G | NULL | 0.568319983 | NULL&M_Disease |
| 11 | 117380309 | rs139254418 | G | A | NULL | 0.567092905 | NULL&M_Disease |
| 11 | 118533042 | rs142438429 | T | C | NULL | 0.567092905 | NULL&M_Disease |
| 11 | 120903752 | rs2000528 | G | A | NULL | 0.567092905 | NULL&M_Disease |
| 11 | 116683862 | rs75318540 | T | C | NULL | 0.567092905 | NULL&M_Disease |
| 11 | 116737736 | rs675834 | A | T | NULL | 0.567092905 | NULL&M_Disease |
| 11 | 117671216 | rs76921569 | A | C | NULL | 0.567092905 | NULL&M_Disease |
| 11 | 117675125 | rs79046029 | T | C | NULL | 0.567092905 | NULL&M_Disease |
| 11 | 118835677 | rs4420279 | A | C | NULL | 0.568319983 | NULL&M_Disease |
| 11 | 120790033 | rs78621482 | G | C | NULL | 0.567092905 | NULL&M_Disease |
| 11 | 117587186 | rs4938431 | G | A | NULL | 0.567092905 | NULL&M_Disease |
| 11 | 117587345 | rs4938432 | A | G | NULL | 0.567092905 | NULL&M_Disease |
| 11 | 116279427 | rs580965 | C | T | NULL | 0.568319983 | NULL&M_Disease |
| 11 | 116294634 | rs471840 | T | C | NULL | 0.568319983 | NULL&M_Disease |
| 11 | 117727740 | rs115524104 | C | T | NULL | 0.568319983 | NULL&M_Disease |
| 11 | 119691069 | rs1447149 | A | G | NULL | 0.568319983 | NULL&M_Disease |
| 11 | 116601711 | rs108533 | T | A | NULL | 0.568319983 | NULL&M_Disease |
| 11 | 117319240 | rs118110450 | C | A | NULL | 0.567092905 | NULL&M_Disease |
| 11 | 117672225 | rs2298649 | G | A | NULL | 0.568319983 | NULL&M_Disease |
| 11 | 117363426 | rs11216425 | A | C | NULL | 0.568319983 | NULL&M_Disease |
| 11 | 117957750 | rs600638 | T | C | NULL | 0.568319983 | NULL&M_Disease |
| 11 | 116240625 | rs501652 | G | A | NULL | 0.568319983 | NULL&M_Disease |
| 11 | 117481108 | rs11824398 | A | G | NULL | 0.567092905 | NULL&M_Disease |
| 11 | 117604158 | rs1939984 | T | A | NULL | 0.568319983 | NULL&M_Disease |
| 11 | 120872163 | rs141734119 | G | A | NULL | 0.567092905 | NULL&M_Disease |
| 11 | 116684744 | rs2849167 | G | A | NULL | 0.568319983 | NULL&M_Disease |
| 11 | 118936910 | rs471000 | T | C | NULL | 0.568319983 | NULL&M_Disease |
| 11 | 119509845 | rs117434119 | T | C | NULL | 0.568319983 | NULL&M_Disease |
| 11 | 119580472 | rs4245204 | A | G | NULL | 0.568319983 | NULL&M_Disease |
| 11 | 120019335 | rs548215 | A | G | NULL | 0.568319983 | NULL&M_Disease |
| 11 | 117547710 | rs74361939 | A | G | NULL | 0.568319983 | NULL&M_Disease |
| 11 | 117600506 | rs78036667 | T | C | NULL | 0.567092905 | NULL&M_Disease |
| 11 | 118751898 | rs60651507 | A | G | NULL | 0.568319983 | NULL&M_Disease |
| 11 | 121081509 | rs588209 | G | C | NULL | 0.568319983 | NULL&M_Disease |
| 11 | 117885694 | rs2512151 | G | A | NULL | 0.568319983 | NULL&M_Disease |
| 11 | 119923305 | rs147136343 | T | A | NULL | 0.568319983 | NULL&M_Disease |
| 11 | 120556631 | rs78584663 | G | A | NULL | 0.568319983 | NULL&M_Disease |
| 11 | 120556817 | rs11606521 | G | A | NULL | 0.568319983 | NULL&M_Disease |
| 11 | 117362198 | rs2013131 | C | T | NULL | 0.569546224 | NULL&M_Disease |
| 11 | 117564845 | rs112266857 | T | C | NULL | 0.568319983 | NULL&M_Disease |
| 11 | 121106570 | rs7108971 | A | T | NULL | 0.569546224 | NULL&M_Disease |
| 11 | 117586698 | rs948775 | C | T | NULL | 0.568319983 | NULL&M_Disease |
| 11 | 120421474 | rs61900898 | T | G | NULL | 0.569546224 | NULL&M_Disease |
| 11 | 121141095 | rs78768102 | G | A | NULL | 0.568319983 | NULL&M_Disease |
| 11 | 117088082 | rs1339633702 | T | C | NULL | 0.569546224 | NULL&M_Disease |
| 11 | 117610811 | rs113586459 | C | A | NULL | 0.568319983 | NULL&M_Disease |
| 11 | 118605352 | rs76478216 | C | T | NULL | 0.569546224 | NULL&M_Disease |
| 11 | 117593858 | rs78437760 | G | T | NULL | 0.568319983 | NULL&M_Disease |
| 11 | 117594170 | rs77047087 | A | G | NULL | 0.568319983 | NULL&M_Disease |
| 11 | 117594288 | rs76628593 | T | G | NULL | 0.568319983 | NULL&M_Disease |
| 11 | 117596635 | rs79506234 | T | C | NULL | 0.568319983 | NULL&M_Disease |
| 11 | 117597810 | rs77746497 | A | G | NULL | 0.568319983 | NULL&M_Disease |
| 11 | 118296658 | rs79481471 | A | G | NULL | 0.568319983 | NULL&M_Disease |
| 11 | 118629143 | rs475946 | T | C | NULL | 0.569546224 | NULL&M_Disease |
| 11 | 117233378 | rs139352710 | A | G | NULL | 0.569546224 | NULL&M_Disease |
| 11 | 116698963 | rs2727788 | A | C | NULL | 0.569546224 | NULL&M_Disease |

| | | | | | | | | | |
|---|---|---|---|---|---|---|---|---|---|
| 11 | 117388656 | rs59346420 | T | C | 0.931 | -7977.388 | -7977.921 | -0.533 | 2.27E-06 |
| 11 | 117068444 | rs74385934 | A | G | 1.071 | -7977.388 | -7977.922 | -0.534 | 2.27E-06 |
| 11 | 117081709 | rs1263499 | G | A | 0.933 | -7977.388 | -7977.922 | -0.534 | 2.27E-06 |
| 11 | 117958179 | rs10892212 | A | T | 0.931 | -7977.388 | -7977.923 | -0.534 | 2.27E-06 |
| 11 | 118152252 | rs117117171 | A | T | 0.935 | -7977.388 | -7977.922 | -0.534 | 2.27E-06 |
| 11 | 118938882 | rs670526 | C | T | 1.06 | -7977.388 | -7977.922 | -0.534 | 2.27E-06 |
| 11 | 120093920 | rs12226432 | C | A | 0.932 | -7977.388 | -7977.922 | -0.534 | 2.27E-06 |
| 11 | 120464049 | rs12421186 | T | C | 0.951 | -7977.388 | -7977.923 | -0.535 | 2.27E-06 |
| 11 | 116240253 | rs541874 | T | C | 1.049 | -7977.388 | -7977.925 | -0.536 | 2.26E-06 |
| 11 | 119652997 | rs142774084 | A | G | 0.936 | -7977.388 | -7977.924 | -0.536 | 2.26E-06 |
| 11 | 116486664 | rs3016347 | C | G | 0.941 | -7977.388 | -7977.925 | -0.537 | 2.26E-06 |
| 11 | 118057867 | rs7103366 | G | T | 1.054 | -7977.388 | -7977.925 | -0.537 | 2.26E-06 |
| 11 | 118928393 | rs471028 | C | G | 1.06 | -7977.388 | -7977.925 | -0.537 | 2.26E-06 |
| 11 | 118932610 | rs647432 | T | C | 1.06 | -7977.388 | -7977.925 | -0.537 | 2.26E-06 |
| 11 | 117577527 | rs12574878 | T | C | 0.946 | -7977.388 | -7977.926 | -0.538 | 2.26E-06 |
| 11 | 117577558 | rs12576082 | A | T | 0.946 | -7977.388 | -7977.926 | -0.538 | 2.26E-06 |
| 11 | 117577909 | rs12577066 | G | A | 0.946 | -7977.388 | -7977.926 | -0.538 | 2.26E-06 |
| 11 | 117578314 | rs113570704 | T | C | 0.946 | -7977.388 | -7977.926 | -0.538 | 2.26E-06 |
| 11 | 117579516 | rs1310160088 | T | C | 0.946 | -7977.388 | -7977.926 | -0.538 | 2.26E-06 |
| 11 | 117579607 | rs78696045 | T | A | 0.946 | -7977.388 | -7977.926 | -0.538 | 2.26E-06 |
| 11 | 119507468 | rs4938697 | C | T | 1.05 | -7977.388 | -7977.927 | -0.539 | 2.26E-06 |
| 11 | 117033808 | rs79118964 | T | C | 0.934 | -7977.388 | -7977.928 | -0.54 | 2.25E-06 |
| 11 | 119577632 | rs7104655 | A | G | 0.932 | -7977.388 | -7977.928 | -0.54 | 2.25E-06 |
| 11 | 116630600 | rs918143 | T | C | 0.953 | -7977.388 | -7977.93 | -0.541 | 2.25E-06 |
| 11 | 116557882 | rs11216093 | C | A | 0.94 | -7977.388 | -7977.93 | -0.542 | 2.25E-06 |
| 11 | 117566305 | rs55973262 | C | G | 0.936 | -7977.388 | -7977.93 | -0.542 | 2.25E-06 |
| 11 | 117586881 | rs948774 | A | T | 0.944 | -7977.388 | -7977.93 | -0.542 | 2.25E-06 |
| 11 | 117586942 | rs948773 | A | G | 0.944 | -7977.388 | -7977.93 | -0.542 | 2.25E-06 |
| 11 | 117727742 | rs11216586 | C | G | 1.071 | -7977.388 | -7977.93 | -0.542 | 2.25E-06 |
| 11 | 116226290 | rs577941 | A | G | 1.06 | -7977.388 | -7977.931 | -0.543 | 2.25E-06 |
| 11 | 117113511 | rs117117915 | T | C | 0.937 | -7977.388 | -7977.931 | -0.543 | 2.25E-06 |
| 11 | 118214267 | rs2071381 | A | G | 1.051 | -7977.388 | -7977.931 | -0.543 | 2.25E-06 |
| 11 | 119818789 | rs895653 | A | T | 0.952 | -7977.388 | -7977.932 | -0.543 | 2.25E-06 |
| 11 | 117388657 | rs77944184 | A | G | 0.932 | -7977.388 | -7977.933 | -0.544 | 2.25E-06 |
| 11 | 118835272 | rs67533649 | G | A | 1.057 | -7977.388 | -7977.932 | -0.544 | 2.25E-06 |
| 11 | 120473450 | rs139831408 | T | C | 1.069 | -7977.388 | -7977.932 | -0.544 | 2.25E-06 |
| 11 | 116306324 | rs79892107 | T | C | 1.07 | -7977.388 | -7977.934 | -0.545 | 2.24E-06 |
| 11 | 118709275 | rs187164533 | A | G | 1.061 | -7977.388 | -7977.933 | -0.545 | 2.24E-06 |
| 11 | 119654894 | rs77020062 | T | C | 1.059 | -7977.388 | -7977.934 | -0.545 | 2.24E-06 |
| 11 | 119768054 | rs11217556 | A | C | 0.949 | -7977.388 | -7977.934 | -0.546 | 2.24E-06 |
| 11 | 116235005 | rs495430 | A | G | 1.05 | -7977.388 | -7977.936 | -0.547 | 2.24E-06 |
| 11 | 117380309 | rs139254418 | G | A | 0.932 | -7977.388 | -7977.936 | -0.547 | 2.24E-06 |
| 11 | 118533042 | rs142438429 | T | C | 1.057 | -7977.388 | -7977.936 | -0.547 | 2.24E-06 |
| 11 | 120903752 | rs2000528 | G | A | 1.056 | -7977.388 | -7977.936 | -0.547 | 2.24E-06 |
| 11 | 116683862 | rs75318540 | T | C | 0.936 | -7977.388 | -7977.937 | -0.548 | 2.24E-06 |
| 11 | 116737736 | rs675834 | A | T | 0.937 | -7977.388 | -7977.939 | -0.55 | 2.23E-06 |
| 11 | 117671216 | rs76921569 | A | C | 0.942 | -7977.388 | -7977.938 | -0.55 | 2.23E-06 |
| 11 | 117675125 | rs79046029 | T | C | 0.942 | -7977.388 | -7977.938 | -0.55 | 2.23E-06 |
| 11 | 118835677 | rs4420279 | A | C | 1.057 | -7977.388 | -7977.939 | -0.55 | 2.23E-06 |
| 11 | 120790033 | rs78621482 | G | C | 1.072 | -7977.388 | -7977.938 | -0.55 | 2.23E-06 |
| 11 | 117587186 | rs4938431 | G | A | 0.946 | -7977.388 | -7977.939 | -0.551 | 2.23E-06 |
| 11 | 117587345 | rs4938432 | A | G | 0.946 | -7977.388 | -7977.939 | -0.551 | 2.23E-06 |
| 11 | 116279427 | rs580965 | C | T | 1.052 | -7977.388 | -7977.94 | -0.552 | 2.23E-06 |
| 11 | 116294634 | rs471840 | T | C | 1.052 | -7977.388 | -7977.94 | -0.552 | 2.23E-06 |
| 11 | 117727740 | rs115524104 | C | T | 1.071 | -7977.388 | -7977.94 | -0.552 | 2.23E-06 |
| 11 | 119691069 | rs1447149 | A | G | 0.949 | -7977.388 | -7977.94 | -0.552 | 2.23E-06 |
| 11 | 116601711 | rs108533 | T | A | 0.953 | -7977.388 | -7977.941 | -0.553 | 2.23E-06 |
| 11 | 117319240 | rs118110450 | C | A | 1.036 | -7977.388 | -7977.941 | -0.553 | 2.23E-06 |
| 11 | 117672225 | rs2298649 | G | A | 0.951 | -7977.388 | -7977.942 | -0.553 | 2.23E-06 |
| 11 | 117363426 | rs11216425 | A | C | 0.934 | -7977.388 | -7977.942 | -0.554 | 2.22E-06 |
| 11 | 117957750 | rs600638 | T | C | 1.069 | -7977.388 | -7977.943 | -0.554 | 2.22E-06 |
| 11 | 116240625 | rs501652 | G | A | 1.049 | -7977.388 | -7977.943 | -0.555 | 2.22E-06 |
| 11 | 117481108 | rs11824398 | A | G | 0.941 | -7977.388 | -7977.943 | -0.555 | 2.22E-06 |
| 11 | 117604158 | rs1939984 | T | A | 0.932 | -7977.388 | -7977.943 | -0.555 | 2.22E-06 |
| 11 | 120872163 | rs141734119 | G | A | 0.938 | -7977.388 | -7977.943 | -0.555 | 2.22E-06 |
| 11 | 116684744 | rs2849167 | G | A | 1.049 | -7977.388 | -7977.944 | -0.556 | 2.22E-06 |
| 11 | 118936910 | rs471000 | T | C | 1.06 | -7977.388 | -7977.944 | -0.556 | 2.22E-06 |
| 11 | 119509845 | rs117434119 | T | C | 0.933 | -7977.388 | -7977.944 | -0.556 | 2.22E-06 |
| 11 | 119580472 | rs4245204 | A | G | 0.934 | -7977.388 | -7977.945 | -0.556 | 2.22E-06 |
| 11 | 120019335 | rs548215 | A | G | 0.938 | -7977.388 | -7977.944 | -0.556 | 2.22E-06 |
| 11 | 117547710 | rs74361939 | A | G | 1.071 | -7977.388 | -7977.945 | -0.557 | 2.22E-06 |
| 11 | 117600506 | rs78036667 | T | C | 0.948 | -7977.388 | -7977.945 | -0.557 | 2.22E-06 |
| 11 | 118751898 | rs60651507 | A | G | 0.933 | -7977.388 | -7977.945 | -0.557 | 2.22E-06 |
| 11 | 121081509 | rs588209 | G | C | 0.949 | -7977.388 | -7977.945 | -0.557 | 2.22E-06 |
| 11 | 117885694 | rs2512151 | G | A | 0.932 | -7977.388 | -7977.946 | -0.558 | 2.21E-06 |
| 11 | 119923305 | rs147136343 | T | A | 0.946 | -7977.388 | -7977.946 | -0.558 | 2.21E-06 |
| 11 | 120556631 | rs78584663 | G | A | 0.937 | -7977.388 | -7977.947 | -0.558 | 2.21E-06 |
| 11 | 120556817 | rs11606521 | G | A | 0.937 | -7977.388 | -7977.947 | -0.558 | 2.21E-06 |
| 11 | 117362198 | rs2013131 | C | T | 1.049 | -7977.388 | -7977.947 | -0.559 | 2.21E-06 |
| 11 | 117564845 | rs112266857 | T | C | 0.937 | -7977.388 | -7977.947 | -0.559 | 2.21E-06 |
| 11 | 121106570 | rs7108971 | A | T | 0.947 | -7977.388 | -7977.948 | -0.559 | 2.21E-06 |
| 11 | 117586698 | rs948775 | C | T | 0.933 | -7977.388 | -7977.949 | -0.56 | 2.21E-06 |
| 11 | 120421474 | rs61900898 | T | G | 0.95 | -7977.388 | -7977.949 | -0.56 | 2.21E-06 |
| 11 | 121141095 | rs78768102 | G | A | 1.057 | -7977.388 | -7977.948 | -0.56 | 2.21E-06 |
| 11 | 117088082 | rs1339633702 | T | C | 1.07 | -7977.388 | -7977.949 | -0.561 | 2.21E-06 |
| 11 | 117610811 | rs113586459 | C | A | 0.948 | -7977.388 | -7977.95 | -0.562 | 2.21E-06 |
| 11 | 118605352 | rs76478216 | C | T | 1.071 | -7977.388 | -7977.95 | -0.562 | 2.21E-06 |
| 11 | 117593858 | rs78437760 | G | T | 0.949 | -7977.388 | -7977.951 | -0.563 | 2.20E-06 |
| 11 | 117594170 | rs77047087 | A | G | 0.949 | -7977.388 | -7977.951 | -0.563 | 2.20E-06 |
| 11 | 117594288 | rs76628593 | T | G | 0.949 | -7977.388 | -7977.951 | -0.563 | 2.20E-06 |
| 11 | 117596635 | rs79506234 | T | C | 0.949 | -7977.388 | -7977.952 | -0.563 | 2.20E-06 |
| 11 | 117597810 | rs77746497 | A | G | 0.949 | -7977.388 | -7977.952 | -0.563 | 2.20E-06 |
| 11 | 118296658 | rs79481471 | A | G | 0.941 | -7977.388 | -7977.951 | -0.563 | 2.20E-06 |
| 11 | 118629143 | rs475946 | T | C | 0.942 | -7977.388 | -7977.951 | -0.563 | 2.20E-06 |
| 11 | 117233378 | rs139352710 | A | G | 0.933 | -7977.388 | -7977.952 | -0.564 | 2.20E-06 |
| 11 | 116698963 | rs2727788 | A | C | 1.053 | -7977.388 | -7977.953 | -0.565 | 2.20E-06 |

| | | | | | | | |
|---|---|---|---|---|---|---|---|
| 11 | 117333952 | rs4938384 | A | G | NULL | 0.569546224 | NULL&M_Disease |
| 11 | 117973058 | rs11216751 | G | T | NULL | 0.569546224 | NULL&M_Disease |
| 11 | 116519655 | rs516226 | T | C | NULL | 0.569546224 | NULL&M_Disease |
| 11 | 117307306 | rs73018318 | T | G | NULL | 0.568319983 | NULL&M_Disease |
| 11 | 117383816 | rs12419495 | C | T | NULL | 0.569546224 | NULL&M_Disease |
| 11 | 119658388 | rs56246162 | G | C | NULL | 0.568319983 | NULL&M_Disease |
| 11 | 119805958 | rs10790355 | T | C | NULL | 0.569546224 | NULL&M_Disease |
| 11 | 117590322 | rs7106419 | G | T | NULL | 0.569546224 | NULL&M_Disease |
| 11 | 119724144 | rs506842 | C | G | NULL | 0.569546224 | NULL&M_Disease |
| 11 | 116216432 | rs570448 | C | G | NULL | 0.569546224 | NULL&M_Disease |
| 11 | 116578982 | rs61905067 | G | A | NULL | 0.569546224 | NULL&M_Disease |
| 11 | 117593119 | rs11216515 | C | T | NULL | 0.569546224 | NULL&M_Disease |
| 11 | 117953051 | rs2051479 | G | A | NULL | 0.570771612 | NULL&M_Disease |
| 11 | 119236751 | rs2195526 | T | C | NULL | 0.570771612 | NULL&M_Disease |
| 11 | 119334629 | rs11217297 | A | G | NULL | 0.570771612 | NULL&M_Disease |
| 11 | 117961309 | rs654176 | G | C | NULL | 0.569546224 | NULL&M_Disease |
| 11 | 120437188 | rs4278519 | T | C | NULL | 0.570771612 | NULL&M_Disease |
| 11 | 117570168 | rs112845302 | A | G | NULL | 0.570771612 | NULL&M_Disease |
| 11 | 117868517 | rs4252314 | G | A | NULL | 0.569546224 | NULL&M_Disease |
| 11 | 118232026 | rs77806521 | T | C | NULL | 0.569546224 | NULL&M_Disease |
| 11 | 119824803 | rs72994633 | C | T | NULL | 0.569546224 | NULL&M_Disease |
| 11 | 117179535 | rs28989473 | A | G | NULL | 0.569546224 | NULL&M_Disease |
| 11 | 117300040 | rs3889570 | C | T | NULL | 0.570771612 | NULL&M_Disease |
| 11 | 118932445 | rs636283 | T | C | NULL | 0.570771612 | NULL&M_Disease |
| 11 | 116198542 | rs113708750 | T | C | NULL | 0.570771612 | NULL&M_Disease |
| 11 | 116463208 | rs12288729 | C | G | NULL | 0.569546224 | NULL&M_Disease |
| 11 | 117584740 | rs2187183 | A | G | NULL | 0.569546224 | NULL&M_Disease |
| 11 | 117584761 | rs2187182 | T | G | NULL | 0.569546224 | NULL&M_Disease |
| 11 | 117584793 | rs11216512 | G | C | NULL | 0.569546224 | NULL&M_Disease |
| 11 | 117585030 | rs4938427 | A | G | NULL | 0.569546224 | NULL&M_Disease |
| 11 | 117585035 | rs4936396 | A | G | NULL | 0.569546224 | NULL&M_Disease |
| 11 | 117585338 | rs4938429 | C | A | NULL | 0.569546224 | NULL&M_Disease |
| 11 | 117585370 | rs4936397 | G | A | NULL | 0.569546224 | NULL&M_Disease |
| 11 | 117585523 | rs7930615 | G | A | NULL | 0.569546224 | NULL&M_Disease |
| 11 | 117585641 | rs4359247 | G | C | NULL | 0.569546224 | NULL&M_Disease |
| 11 | 117585918 | rs4255570 | C | T | NULL | 0.569546224 | NULL&M_Disease |
| 11 | 117585977 | rs4245171 | A | G | NULL | 0.569546224 | NULL&M_Disease |
| 11 | 117585981 | rs4245172 | C | T | NULL | 0.569546224 | NULL&M_Disease |
| 11 | 117586129 | rs4245173 | T | C | NULL | 0.569546224 | NULL&M_Disease |
| 11 | 117586178 | rs4938430 | C | G | NULL | 0.569546224 | NULL&M_Disease |
| 11 | 120171165 | rs1941410 | A | G | NULL | 0.570771612 | NULL&M_Disease |
| 11 | 120278477 | rs34172482 | G | A | NULL | 0.570771612 | NULL&M_Disease |
| 11 | 120989762 | rs12280641 | A | G | NULL | 0.570771612 | NULL&M_Disease |
| 11 | 116674503 | rs625524 | A | G | NULL | 0.569546224 | NULL&M_Disease |
| 11 | 117092886 | rs6589606 | T | C | NULL | 0.570771612 | NULL&M_Disease |
| 11 | 117881937 | rs11216675 | T | C | NULL | 0.569546224 | NULL&M_Disease |
| 11 | 120888244 | rs116981723 | T | C | NULL | 0.569546224 | NULL&M_Disease |
| 11 | 117647145 | rs61905348 | T | G | NULL | 0.570771612 | NULL&M_Disease |
| 11 | 120342855 | rs61898761 | G | A | NULL | 0.570771612 | NULL&M_Disease |
| 11 | 116778293 | rs79326407 | T | C | NULL | 0.570771612 | NULL&M_Disease |
| 11 | 120871963 | rs676426 | A | G | NULL | 0.570771612 | NULL&M_Disease |
| 11 | 116495780 | rs112549212 | C | T | NULL | 0.570771612 | NULL&M_Disease |
| 11 | 119782437 | rs146204458 | G | A | NULL | 0.569546224 | NULL&M_Disease |
| 11 | 116280688 | rs533848 | T | A | NULL | 0.570771612 | NULL&M_Disease |
| 11 | 118577614 | rs947931 | G | T | NULL | 0.570771612 | NULL&M_Disease |
| 11 | 120787752 | rs3018285 | C | T | NULL | 0.570771612 | NULL&M_Disease |
| 11 | 116801422 | rs12799198 | T | C | NULL | 0.571996133 | NULL&M_Disease |
| 11 | 117150502 | rs117013687 | T | C | NULL | 0.569546224 | NULL&M_Disease |
| 11 | 117392722 | rs55753209 | A | G | NULL | 0.570771612 | NULL&M_Disease |
| 11 | 118058358 | rs6589652 | G | A | NULL | 0.571996133 | NULL&M_Disease |
| 11 | 120630452 | rs7112522 | G | A | NULL | 0.570771612 | NULL&M_Disease |
| 11 | 116697066 | rs2849174 | A | G | NULL | 0.571996133 | NULL&M_Disease |
| 11 | 117095283 | rs59781045 | T | C | NULL | 0.570771612 | NULL&M_Disease |
| 11 | 120873076 | rs1621211 | A | G | NULL | 0.571996133 | NULL&M_Disease |
| 11 | 116557155 | rs2156120 | G | A | NULL | 0.571996133 | NULL&M_Disease |
| 11 | 116589127 | rs180377 | T | C | NULL | 0.571996133 | NULL&M_Disease |
| 11 | 117675364 | rs60942267 | T | G | NULL | 0.570771612 | NULL&M_Disease |
| 11 | 118150259 | rs17540777 | G | T | NULL | 0.571996133 | NULL&M_Disease |
| 11 | 118224123 | rs73018291 | A | G | NULL | 0.570771612 | NULL&M_Disease |
| 11 | 119658314 | rs76173441 | C | G | NULL | 0.570771612 | NULL&M_Disease |
| 11 | 120923495 | rs10502245 | A | G | NULL | 0.571996133 | NULL&M_Disease |
| 11 | 120923899 | rs10892665 | G | C | NULL | 0.571996133 | NULL&M_Disease |
| 11 | 116216072 | rs34166664 | G | C | NULL | 0.571996133 | NULL&M_Disease |
| 11 | 117597375 | rs948103 | C | A | NULL | 0.570771612 | NULL&M_Disease |
| 11 | 117599235 | rs1942925 | G | A | NULL | 0.570771612 | NULL&M_Disease |
| 11 | 117599647 | rs1942927 | G | A | NULL | 0.570771612 | NULL&M_Disease |
| 11 | 118604678 | rs493720 | T | C | NULL | 0.571996133 | NULL&M_Disease |
| 11 | 118605127 | rs526850 | G | T | NULL | 0.571996133 | NULL&M_Disease |
| 11 | 118920453 | rs673768 | T | C | NULL | 0.571996133 | NULL&M_Disease |
| 11 | 116596174 | rs180365 | T | C | NULL | 0.571996133 | NULL&M_Disease |
| 11 | 118850707 | rs4938605 | A | G | NULL | 0.571996133 | NULL&M_Disease |
| 11 | 118866072 | rs11217118 | A | G | NULL | 0.571996133 | NULL&M_Disease |
| 11 | 119452937 | rs144611969 | A | G | NULL | 0.570771612 | NULL&M_Disease |
| 11 | 120932667 | rs11601887 | C | T | NULL | 0.571996133 | NULL&M_Disease |
| 11 | 120950752 | rs11218137 | C | T | NULL | 0.571996133 | NULL&M_Disease |
| 11 | 116481175 | rs111527819 | A | G | NULL | 0.570771612 | NULL&M_Disease |
| 11 | 116598649 | rs180361 | T | C | NULL | 0.571996133 | NULL&M_Disease |
| 11 | 116365351 | rs4938274 | A | G | NULL | 0.571996133 | NULL&M_Disease |
| 11 | 116492477 | rs78577252 | T | C | NULL | 0.570771612 | NULL&M_Disease |
| 11 | 117335991 | rs663191 | T | C | NULL | 0.571996133 | NULL&M_Disease |
| 11 | 117336947 | rs677954 | T | C | NULL | 0.571996133 | NULL&M_Disease |
| 11 | 117668060 | rs112872285 | T | G | NULL | 0.571996133 | NULL&M_Disease |
| 11 | 118644213 | rs1790187 | A | G | NULL | 0.571996133 | NULL&M_Disease |
| 11 | 117458169 | rs117273824 | T | C | NULL | 0.571996133 | NULL&M_Disease |
| 11 | 119457720 | rs61901067 | G | A | NULL | 0.571996133 | NULL&M_Disease |
| 11 | 121004439 | rs2155465 | G | C | NULL | 0.571996133 | NULL&M_Disease |

| | | | | | | | | | |
|---|---|---|---|---|---|---|---|---|---|
| 11 | 117333952 | rs4938384 | A | G | 0.936 | -7977.388 | -7977.953 | -0.565 | 2.20E-06 |
| 11 | 117973058 | rs11216751 | G | T | 0.938 | -7977.388 | -7977.953 | -0.565 | 2.20E-06 |
| 11 | 116519655 | rs516226 | T | C | 0.947 | -7977.388 | -7977.954 | -0.566 | 2.20E-06 |
| 11 | 117307306 | rs73018318 | T | G | 0.95 | -7977.388 | -7977.954 | -0.566 | 2.20E-06 |
| 11 | 117383816 | rs12419495 | C | T | 0.934 | -7977.388 | -7977.955 | -0.566 | 2.20E-06 |
| 11 | 119658388 | rs56246162 | G | C | 1.055 | -7977.388 | -7977.954 | -0.566 | 2.20E-06 |
| 11 | 119805958 | rs10790355 | T | C | 1.052 | -7977.388 | -7977.954 | -0.566 | 2.20E-06 |
| 11 | 117590322 | rs7106419 | G | T | 0.949 | -7977.388 | -7977.956 | -0.567 | 2.19E-06 |
| 11 | 119724144 | rs506842 | C | G | 0.953 | -7977.388 | -7977.955 | -0.567 | 2.19E-06 |
| 11 | 116216432 | rs570448 | C | G | 1.059 | -7977.388 | -7977.956 | -0.568 | 2.19E-06 |
| 11 | 116578982 | rs61905067 | G | A | 0.94 | -7977.388 | -7977.957 | -0.568 | 2.19E-06 |
| 11 | 117593119 | rs11216515 | C | T | 1.057 | -7977.388 | -7977.956 | -0.568 | 2.19E-06 |
| 11 | 117953051 | rs2051479 | G | A | 1.052 | -7977.388 | -7977.956 | -0.568 | 2.19E-06 |
| 11 | 119236751 | rs2195526 | T | C | 1.057 | -7977.388 | -7977.957 | -0.568 | 2.19E-06 |
| 11 | 119334629 | rs11217297 | A | G | 0.945 | -7977.388 | -7977.956 | -0.568 | 2.19E-06 |
| 11 | 117961309 | rs654176 | G | C | 1.069 | -7977.388 | -7977.957 | -0.569 | 2.19E-06 |
| 11 | 120437188 | rs4278519 | T | C | 1.049 | -7977.388 | -7977.957 | -0.569 | 2.19E-06 |
| 11 | 117570168 | rs112845302 | A | G | 0.937 | -7977.388 | -7977.958 | -0.57 | 2.19E-06 |
| 11 | 117868517 | rs4252314 | G | A | 1.071 | -7977.388 | -7977.958 | -0.57 | 2.19E-06 |
| 11 | 118232026 | rs77806521 | T | C | 1.063 | -7977.388 | -7977.958 | -0.57 | 2.19E-06 |
| 11 | 119824803 | rs72994633 | C | T | 0.936 | -7977.388 | -7977.958 | -0.57 | 2.19E-06 |
| 11 | 117179535 | rs28989473 | A | G | 0.934 | -7977.388 | -7977.959 | -0.571 | 2.19E-06 |
| 11 | 117300040 | rs3889570 | C | T | 1.049 | -7977.388 | -7977.959 | -0.571 | 2.19E-06 |
| 11 | 118932445 | rs636283 | T | C | 1.059 | -7977.388 | -7977.959 | -0.571 | 2.19E-06 |
| 11 | 116198542 | rs113708750 | T | C | 1.071 | -7977.388 | -7977.96 | -0.572 | 2.18E-06 |
| 11 | 116463208 | rs12288729 | C | G | 0.934 | -7977.388 | -7977.96 | -0.572 | 2.18E-06 |
| 11 | 117584740 | rs2187183 | A | G | 0.95 | -7977.388 | -7977.961 | -0.572 | 2.18E-06 |
| 11 | 117584761 | rs2187182 | T | G | 0.95 | -7977.388 | -7977.961 | -0.572 | 2.18E-06 |
| 11 | 117584793 | rs11216512 | G | C | 0.95 | -7977.388 | -7977.961 | -0.572 | 2.18E-06 |
| 11 | 117585030 | rs4938427 | A | G | 0.95 | -7977.388 | -7977.961 | -0.572 | 2.18E-06 |
| 11 | 117585035 | rs4936396 | A | G | 0.95 | -7977.388 | -7977.961 | -0.572 | 2.18E-06 |
| 11 | 117585338 | rs4938429 | C | A | 0.95 | -7977.388 | -7977.961 | -0.572 | 2.18E-06 |
| 11 | 117585370 | rs4936397 | G | A | 0.95 | -7977.388 | -7977.961 | -0.572 | 2.18E-06 |
| 11 | 117585523 | rs7930615 | G | A | 0.95 | -7977.388 | -7977.961 | -0.572 | 2.18E-06 |
| 11 | 117585641 | rs4359247 | G | C | 0.95 | -7977.388 | -7977.961 | -0.572 | 2.18E-06 |
| 11 | 117585918 | rs4255570 | C | T | 0.95 | -7977.388 | -7977.961 | -0.572 | 2.18E-06 |
| 11 | 117585977 | rs4245171 | A | G | 0.95 | -7977.388 | -7977.961 | -0.572 | 2.18E-06 |
| 11 | 117585981 | rs4245172 | C | T | 0.95 | -7977.388 | -7977.961 | -0.572 | 2.18E-06 |
| 11 | 117586129 | rs4245173 | T | C | 0.95 | -7977.388 | -7977.961 | -0.572 | 2.18E-06 |
| 11 | 117586178 | rs4938430 | C | G | 0.95 | -7977.388 | -7977.961 | -0.572 | 2.18E-06 |
| 11 | 120171165 | rs1941410 | A | G | 1.064 | -7977.388 | -7977.96 | -0.572 | 2.18E-06 |
| 11 | 120278477 | rs34172482 | G | A | 0.933 | -7977.388 | -7977.96 | -0.572 | 2.18E-06 |
| 11 | 120989762 | rs12280641 | A | G | 0.947 | -7977.388 | -7977.96 | -0.572 | 2.18E-06 |
| 11 | 116674503 | rs625524 | A | G | 0.94 | -7977.388 | -7977.962 | -0.573 | 2.18E-06 |
| 11 | 117092886 | rs6589606 | T | C | 1.069 | -7977.388 | -7977.962 | -0.573 | 2.18E-06 |
| 11 | 117881937 | rs11216675 | T | C | 0.95 | -7977.388 | -7977.961 | -0.573 | 2.18E-06 |
| 11 | 120888244 | rs116981723 | T | C | 0.943 | -7977.388 | -7977.962 | -0.573 | 2.18E-06 |
| 11 | 117647145 | rs61905348 | T | G | 1.066 | -7977.388 | -7977.963 | -0.574 | 2.18E-06 |
| 11 | 120342855 | rs61898761 | G | A | 1.068 | -7977.388 | -7977.963 | -0.574 | 2.18E-06 |
| 11 | 116778293 | rs79326407 | T | C | 0.935 | -7977.388 | -7977.964 | -0.575 | 2.18E-06 |
| 11 | 120871963 | rs676426 | A | G | 0.949 | -7977.388 | -7977.964 | -0.575 | 2.18E-06 |
| 11 | 116495780 | rs112549212 | C | T | 1.058 | -7977.388 | -7977.965 | -0.576 | 2.17E-06 |
| 11 | 119782437 | rs146204458 | G | A | 0.972 | -7977.388 | -7977.964 | -0.576 | 2.17E-06 |
| 11 | 116280688 | rs533848 | T | A | 1.051 | -7977.388 | -7977.966 | -0.577 | 2.17E-06 |
| 11 | 118577614 | rs947931 | G | T | 1.058 | -7977.388 | -7977.966 | -0.577 | 2.17E-06 |
| 11 | 120787752 | rs3018285 | C | T | 1.051 | -7977.388 | -7977.966 | -0.577 | 2.17E-06 |
| 11 | 116801422 | rs12799198 | T | C | 1.055 | -7977.388 | -7977.967 | -0.578 | 2.17E-06 |
| 11 | 117150502 | rs117013687 | T | C | 0.965 | -7977.388 | -7977.966 | -0.578 | 2.17E-06 |
| 11 | 117392722 | rs55753209 | A | G | 1.069 | -7977.388 | -7977.967 | -0.578 | 2.17E-06 |
| 11 | 118058358 | rs6589652 | G | A | 1.054 | -7977.388 | -7977.966 | -0.578 | 2.17E-06 |
| 11 | 120630452 | rs7112522 | G | A | 0.934 | -7977.388 | -7977.966 | -0.578 | 2.17E-06 |
| 11 | 116697066 | rs2849174 | A | G | 1.053 | -7977.388 | -7977.968 | -0.579 | 2.17E-06 |
| 11 | 117095283 | rs59781045 | T | C | 1.068 | -7977.388 | -7977.967 | -0.579 | 2.17E-06 |
| 11 | 120873076 | rs1621211 | A | G | 0.948 | -7977.388 | -7977.967 | -0.579 | 2.17E-06 |
| 11 | 116557155 | rs2156120 | G | A | 0.941 | -7977.388 | -7977.968 | -0.58 | 2.17E-06 |
| 11 | 116589127 | rs180377 | T | C | 1.049 | -7977.388 | -7977.968 | -0.58 | 2.17E-06 |
| 11 | 117675364 | rs60942267 | T | G | 0.954 | -7977.388 | -7977.968 | -0.58 | 2.17E-06 |
| 11 | 118150259 | rs17540777 | G | T | 0.937 | -7977.388 | -7977.968 | -0.58 | 2.17E-06 |
| 11 | 118224123 | rs73018291 | A | G | 1.07 | -7977.388 | -7977.968 | -0.58 | 2.17E-06 |
| 11 | 119658314 | rs76173441 | C | G | 0.949 | -7977.388 | -7977.968 | -0.58 | 2.17E-06 |
| 11 | 120923495 | rs10502245 | A | G | 1.055 | -7977.388 | -7977.968 | -0.58 | 2.17E-06 |
| 11 | 120923899 | rs10892665 | G | C | 1.055 | -7977.388 | -7977.968 | -0.58 | 2.17E-06 |
| 11 | 116216072 | rs34166664 | G | C | 0.935 | -7977.388 | -7977.969 | -0.581 | 2.16E-06 |
| 11 | 117597375 | rs948103 | C | A | 0.934 | -7977.388 | -7977.969 | -0.581 | 2.16E-06 |
| 11 | 117599235 | rs1942925 | G | A | 0.934 | -7977.388 | -7977.969 | -0.581 | 2.16E-06 |
| 11 | 117599647 | rs1942927 | G | A | 0.934 | -7977.388 | -7977.969 | -0.581 | 2.16E-06 |
| 11 | 118604678 | rs493720 | T | C | 0.942 | -7977.388 | -7977.971 | -0.582 | 2.16E-06 |
| 11 | 118605127 | rs526850 | G | T | 0.942 | -7977.388 | -7977.971 | -0.582 | 2.16E-06 |
| 11 | 118920453 | rs673768 | T | C | 1.059 | -7977.388 | -7977.97 | -0.582 | 2.16E-06 |
| 11 | 116596174 | rs180365 | T | C | 0.954 | -7977.388 | -7977.971 | -0.583 | 2.16E-06 |
| 11 | 118850707 | rs4938605 | A | G | 1.049 | -7977.388 | -7977.971 | -0.583 | 2.16E-06 |
| 11 | 118866072 | rs11217118 | A | G | 1.049 | -7977.388 | -7977.971 | -0.583 | 2.16E-06 |
| 11 | 119452937 | rs144611969 | A | G | 0.943 | -7977.388 | -7977.971 | -0.583 | 2.16E-06 |
| 11 | 120932667 | rs11601887 | C | T | 0.941 | -7977.388 | -7977.971 | -0.583 | 2.16E-06 |
| 11 | 120950752 | rs11218137 | C | T | 1.055 | -7977.388 | -7977.971 | -0.583 | 2.16E-06 |
| 11 | 116481175 | rs111527819 | A | G | 1.056 | -7977.388 | -7977.972 | -0.584 | 2.16E-06 |
| 11 | 116598649 | rs180361 | T | C | 0.954 | -7977.388 | -7977.972 | -0.584 | 2.16E-06 |
| 11 | 116365351 | rs4938274 | A | G | 1.05 | -7977.388 | -7977.974 | -0.585 | 2.15E-06 |
| 11 | 116492477 | rs78577252 | T | C | 1.052 | -7977.388 | -7977.973 | -0.585 | 2.15E-06 |
| 11 | 117335991 | rs663191 | T | C | 0.937 | -7977.388 | -7977.973 | -0.585 | 2.15E-06 |
| 11 | 117336947 | rs677954 | T | C | 0.937 | -7977.388 | -7977.973 | -0.585 | 2.15E-06 |
| 11 | 117668060 | rs112872285 | T | G | 0.951 | -7977.388 | -7977.973 | -0.585 | 2.15E-06 |
| 11 | 118644213 | rs1790187 | A | G | 0.942 | -7977.388 | -7977.973 | -0.585 | 2.15E-06 |
| 11 | 117458169 | rs117273824 | T | C | 0.936 | -7977.388 | -7977.975 | -0.587 | 2.15E-06 |
| 11 | 119457720 | rs61901067 | G | A | 0.942 | -7977.388 | -7977.975 | -0.587 | 2.15E-06 |
| 11 | 121004439 | rs2155465 | G | C | 0.954 | -7977.388 | -7977.976 | -0.587 | 2.15E-06 |

| | | | | | | | |
|---|---|---|---|---|---|---|---|
| 11 | 116496416 | rs73578545 | C | T | NULL | 0.571996133 | NULL&M_Disease |
| 11 | 117600518 | rs61903853 | G | T | NULL | 0.571996133 | NULL&M_Disease |
| 11 | 121075781 | rs117986469 | A | C | NULL | 0.571996133 | NULL&M_Disease |
| 11 | 118898319 | rs741811 | A | G | NULL | 0.573219773 | NULL&M_Disease |
| 11 | 121064524 | rs11218199 | T | C | NULL | 0.571996133 | NULL&M_Disease |
| 11 | 117070691 | rs526602 | A | G | NULL | 0.571996133 | NULL&M_Disease |
| 11 | 117801857 | rs73024419 | T | C | NULL | 0.573219773 | NULL&M_Disease |
| 11 | 117842743 | rs11216658 | C | G | NULL | 0.573219773 | NULL&M_Disease |
| 11 | 119870442 | rs78914290 | T | C | NULL | 0.571996133 | NULL&M_Disease |
| 11 | 116302939 | rs12226676 | G | T | NULL | 0.573219773 | NULL&M_Disease |
| 11 | 118218491 | rs117745694 | A | G | NULL | 0.571996133 | NULL&M_Disease |
| 11 | 118542383 | rs687190 | C | T | NULL | 0.573219773 | NULL&M_Disease |
| 11 | 118831755 | rs10892316 | T | C | NULL | 0.573219773 | NULL&M_Disease |
| 11 | 119738420 | rs76598600 | T | C | NULL | 0.571996133 | NULL&M_Disease |
| 11 | 120580038 | rs12274300 | G | C | NULL | 0.573219773 | NULL&M_Disease |
| 11 | 117467560 | rs4938415 | A | C | NULL | 0.573219773 | NULL&M_Disease |
| 11 | 117727965 | rs11216587 | T | C | NULL | 0.573219773 | NULL&M_Disease |
| 11 | 118832582 | rs4938595 | T | G | NULL | 0.573219773 | NULL&M_Disease |
| 11 | 118834718 | rs112939258 | T | C | NULL | 0.573219773 | NULL&M_Disease |
| 11 | 116502706 | rs57228309 | T | A | NULL | 0.571996133 | NULL&M_Disease |
| 11 | 116684562 | rs2727791 | C | G | NULL | 0.573219773 | NULL&M_Disease |
| 11 | 116498120 | rs73578551 | T | C | NULL | 0.571996133 | NULL&M_Disease |
| 11 | 116501134 | rs73578554 | G | A | NULL | 0.571996133 | NULL&M_Disease |
| 11 | 117430971 | rs503281 | C | G | NULL | 0.573219773 | NULL&M_Disease |
| 11 | 117597518 | rs77346825 | A | G | NULL | 0.571996133 | NULL&M_Disease |
| 11 | 117910111 | rs653788 | T | C | NULL | 0.573219773 | NULL&M_Disease |
| 11 | 116771600 | rs71480323 | A | G | NULL | 0.573219773 | NULL&M_Disease |
| 11 | 117335517 | rs515871 | A | G | NULL | 0.573219773 | NULL&M_Disease |
| 11 | 118924212 | rs470324 | A | G | NULL | 0.573219773 | NULL&M_Disease |
| 11 | 119029849 | rs485030 | T | C | NULL | 0.573219773 | NULL&M_Disease |
| 11 | 117386925 | rs10892134 | T | C | NULL | 0.573219773 | NULL&M_Disease |
| 11 | 118873840 | rs9645663 | C | T | NULL | 0.574442517 | NULL&M_Disease |
| 11 | 119495043 | rs554282497 | C | G | NULL | 0.573219773 | NULL&M_Disease |
| 11 | 120208185 | rs4936515 | G | A | NULL | 0.573219773 | NULL&M_Disease |
| 11 | 116475538 | rs35401859 | G | T | NULL | 0.573219773 | NULL&M_Disease |
| 11 | 116489644 | rs73012959 | T | C | NULL | 0.573219773 | NULL&M_Disease |
| 11 | 116555207 | rs7932330 | G | A | NULL | 0.574442517 | NULL&M_Disease |
| 11 | 117085210 | rs77683187 | C | A | NULL | 0.573219773 | NULL&M_Disease |
| 11 | 118978319 | rs7936924 | C | T | NULL | 0.574442517 | NULL&M_Disease |
| 11 | 119950717 | rs117040938 | C | T | NULL | 0.573219773 | NULL&M_Disease |
| 11 | 121145466 | rs75215496 | C | T | NULL | 0.573219773 | NULL&M_Disease |
| 11 | 117309989 | rs11216386 | T | C | NULL | 0.574442517 | NULL&M_Disease |
| 11 | 117502726 | rs60668211 | A | G | NULL | 0.574442517 | NULL&M_Disease |
| 11 | 117632609 | rs10750113 | G | A | NULL | 0.574442517 | NULL&M_Disease |
| 11 | 117632645 | rs10790205 | C | T | NULL | 0.574442517 | NULL&M_Disease |
| 11 | 117731990 | rs61901488 | T | A | NULL | 0.574442517 | NULL&M_Disease |
| 11 | 119244095 | rs587985 | C | T | NULL | 0.574442517 | NULL&M_Disease |
| 11 | 119872269 | rs111748140 | T | C | NULL | 0.573219773 | NULL&M_Disease |
| 11 | 119943314 | rs145476010 | A | C | NULL | 0.573219773 | NULL&M_Disease |
| 11 | 119584347 | rs4533047 | T | C | NULL | 0.573219773 | NULL&M_Disease |
| 11 | 119584506 | rs4245205 | T | C | NULL | 0.573219773 | NULL&M_Disease |
| 11 | 119584876 | rs7927510 | A | T | NULL | 0.573219773 | NULL&M_Disease |
| 11 | 119586216 | rs7946962 | T | C | NULL | 0.573219773 | NULL&M_Disease |
| 11 | 116626735 | rs180322 | C | T | NULL | 0.574442517 | NULL&M_Disease |
| 11 | 119507035 | rs1546465 | G | T | NULL | 0.574442517 | NULL&M_Disease |
| 11 | 116473704 | rs12795589 | G | A | NULL | 0.574442517 | NULL&M_Disease |
| 11 | 120343650 | rs61898763 | T | C | NULL | 0.574442517 | NULL&M_Disease |
| 11 | 116480236 | rs148108200 | C | T | NULL | 0.573219773 | NULL&M_Disease |
| 11 | 119312818 | rs117259599 | T | C | NULL | 0.573219773 | NULL&M_Disease |
| 11 | 120912139 | rs73011555 | G | A | NULL | 0.573219773 | NULL&M_Disease |
| 11 | 116301501 | rs1045334530 | C | A | NULL | 0.574442517 | NULL&M_Disease |
| 11 | 117252135 | rs79879928 | A | G | NULL | 0.574442517 | NULL&M_Disease |
| 11 | 117714145 | rs12801338 | A | G | NULL | 0.574442517 | NULL&M_Disease |
| 11 | 116228118 | rs71478237 | C | A | NULL | 0.574442517 | NULL&M_Disease |
| 11 | 117045522 | rs79138835 | T | G | NULL | 0.574442517 | NULL&M_Disease |
| 11 | 117612254 | rs6589620 | A | T | NULL | 0.574442517 | NULL&M_Disease |
| 11 | 117632375 | rs4344529 | G | A | NULL | 0.574442517 | NULL&M_Disease |
| 11 | 118605447 | rs2186781 | T | C | NULL | 0.574442517 | NULL&M_Disease |
| 11 | 121139014 | rs80145448 | A | C | NULL | 0.574442517 | NULL&M_Disease |
| 11 | 117472044 | rs140763307 | G | A | NULL | 0.573219773 | NULL&M_Disease |
| 11 | 120601873 | rs2846106 | G | A | NULL | 0.574442517 | NULL&M_Disease |
| 11 | 117200974 | rs116006647 | T | C | NULL | 0.574442517 | NULL&M_Disease |
| 11 | 117646692 | rs11824545 | T | G | NULL | 0.574442517 | NULL&M_Disease |
| 11 | 118932320 | rs582688 | C | T | NULL | 0.575664351 | NULL&M_Disease |
| 11 | 117775903 | rs138450618 | T | C | NULL | 0.574442517 | NULL&M_Disease |
| 11 | 120557814 | rs17124208 | C | A | NULL | 0.574442517 | NULL&M_Disease |
| 11 | 116705568 | rs10750098 | G | T | NULL | 0.575664351 | NULL&M_Disease |
| 11 | 117632764 | rs10790206 | C | T | NULL | 0.575664351 | NULL&M_Disease |
| 11 | 117632769 | rs10790207 | C | T | NULL | 0.575664351 | NULL&M_Disease |
| 11 | 117671091 | rs7947524 | T | C | NULL | 0.575664351 | NULL&M_Disease |
| 11 | 118510642 | rs141169729 | T | C | NULL | 0.574442517 | NULL&M_Disease |
| 11 | 121142335 | rs116915994 | T | C | NULL | 0.574442517 | NULL&M_Disease |
| 11 | 116291707 | rs544154 | A | G | NULL | 0.575664351 | NULL&M_Disease |
| 11 | 116473789 | rs12796351 | A | G | NULL | 0.575664351 | NULL&M_Disease |
| 11 | 118062268 | rs639785 | T | C | NULL | 0.575664351 | NULL&M_Disease |
| 11 | 119466427 | rs73002932 | T | C | NULL | 0.575664351 | NULL&M_Disease |
| 11 | 121004502 | rs2155466 | T | C | NULL | 0.575664351 | NULL&M_Disease |
| 11 | 117074109 | rs12970 | A | G | NULL | 0.575664351 | NULL&M_Disease |
| 11 | 117361447 | rs10790190 | G | A | NULL | 0.575664351 | NULL&M_Disease |
| 11 | 119674742 | rs7113439 | T | C | NULL | 0.575664351 | NULL&M_Disease |
| 11 | 120893217 | rs191364497 | T | C | NULL | 0.574442517 | NULL&M_Disease |
| 11 | 117902428 | rs77245011 | C | A | NULL | 0.575664351 | NULL&M_Disease |
| 11 | 118977926 | rs6421572 | A | C | NULL | 0.575664351 | NULL&M_Disease |
| 11 | 116464465 | rs181124100 | T | C | NULL | 0.575664351 | NULL&M_Disease |
| 11 | 116676590 | rs4080140 | A | G | NULL | 0.575664351 | NULL&M_Disease |
| 11 | 117004112 | rs73008513 | G | A | NULL | 0.575664351 | NULL&M_Disease |

| | | | | | | | | | |
|---|---|---|---|---|---|---|---|---|---|
| 11 | 116496416 | rs73578545 | C | T | 1.056 | -7977.388 | -7977.977 | -0.588 | 2.15E-06 |
| 11 | 117600518 | rs61903853 | G | T | 1.07 | -7977.388 | -7977.976 | -0.588 | 2.15E-06 |
| 11 | 121075781 | rs117986469 | A | C | 0.951 | -7977.388 | -7977.977 | -0.588 | 2.15E-06 |
| 11 | 118898319 | rs741811 | A | G | 1.061 | -7977.388 | -7977.978 | -0.589 | 2.15E-06 |
| 11 | 121064524 | rs11218199 | T | C | 1.051 | -7977.388 | -7977.977 | -0.589 | 2.15E-06 |
| 11 | 117070691 | rs526602 | A | G | 0.939 | -7977.388 | -7977.978 | -0.59 | 2.14E-06 |
| 11 | 117801857 | rs73024419 | T | C | 0.939 | -7977.388 | -7977.978 | -0.59 | 2.14E-06 |
| 11 | 117842743 | rs11216658 | C | G | 0.946 | -7977.388 | -7977.979 | -0.59 | 2.14E-06 |
| 11 | 119870442 | rs78914290 | T | C | 0.947 | -7977.388 | -7977.978 | -0.59 | 2.14E-06 |
| 11 | 116302939 | rs12226676 | G | T | 1.068 | -7977.388 | -7977.979 | -0.591 | 2.14E-06 |
| 11 | 118218491 | rs117745694 | A | G | 0.936 | -7977.388 | -7977.98 | -0.591 | 2.14E-06 |
| 11 | 118542383 | rs687190 | C | T | 0.954 | -7977.388 | -7977.979 | -0.591 | 2.14E-06 |
| 11 | 118831755 | rs10892316 | T | C | 1.058 | -7977.388 | -7977.98 | -0.592 | 2.14E-06 |
| 11 | 119738420 | rs76598600 | T | C | 0.94 | -7977.388 | -7977.98 | -0.592 | 2.14E-06 |
| 11 | 120580038 | rs12274300 | G | C | 0.945 | -7977.388 | -7977.981 | -0.592 | 2.14E-06 |
| 11 | 117467560 | rs4938415 | A | C | 1.05 | -7977.388 | -7977.982 | -0.593 | 2.14E-06 |
| 11 | 117727965 | rs11216587 | T | C | 1.069 | -7977.388 | -7977.982 | -0.593 | 2.14E-06 |
| 11 | 118832582 | rs4938595 | T | G | 1.056 | -7977.388 | -7977.982 | -0.593 | 2.14E-06 |
| 11 | 118834718 | rs112939258 | T | C | 1.058 | -7977.388 | -7977.981 | -0.593 | 2.14E-06 |
| 11 | 116502706 | rs57228309 | T | A | 1.055 | -7977.388 | -7977.982 | -0.594 | 2.14E-06 |
| 11 | 116684562 | rs2727791 | C | G | 1.049 | -7977.388 | -7977.983 | -0.594 | 2.14E-06 |
| 11 | 116498120 | rs73578551 | T | C | 1.054 | -7977.388 | -7977.983 | -0.595 | 2.13E-06 |
| 11 | 116501134 | rs73578554 | G | A | 1.054 | -7977.388 | -7977.984 | -0.595 | 2.13E-06 |
| 11 | 117430971 | rs503281 | C | G | 0.938 | -7977.388 | -7977.983 | -0.595 | 2.13E-06 |
| 11 | 117597518 | rs77346825 | A | G | 0.956 | -7977.388 | -7977.983 | -0.595 | 2.13E-06 |
| 11 | 117910111 | rs653788 | T | C | 1.057 | -7977.388 | -7977.983 | -0.595 | 2.13E-06 |
| 11 | 116771600 | rs71480323 | A | G | 1.062 | -7977.388 | -7977.984 | -0.596 | 2.13E-06 |
| 11 | 117335517 | rs515871 | A | G | 0.937 | -7977.388 | -7977.984 | -0.596 | 2.13E-06 |
| 11 | 118924212 | rs470324 | A | G | 1.059 | -7977.388 | -7977.984 | -0.596 | 2.13E-06 |
| 11 | 119029849 | rs485030 | T | C | 1.05 | -7977.388 | -7977.984 | -0.596 | 2.13E-06 |
| 11 | 117386925 | rs10892134 | T | C | 0.937 | -7977.388 | -7977.987 | -0.598 | 2.13E-06 |
| 11 | 118873840 | rs9645663 | C | T | 1.058 | -7977.388 | -7977.987 | -0.598 | 2.13E-06 |
| 11 | 119495043 | rs554282497 | C | G | 0.951 | -7977.388 | -7977.987 | -0.598 | 2.13E-06 |
| 11 | 120208185 | rs4936515 | G | A | 0.935 | -7977.388 | -7977.986 | -0.598 | 2.13E-06 |
| 11 | 116475538 | rs35401859 | G | T | 0.938 | -7977.388 | -7977.987 | -0.599 | 2.13E-06 |
| 11 | 116489644 | rs73012959 | T | C | 0.936 | -7977.388 | -7977.987 | -0.599 | 2.13E-06 |
| 11 | 116555207 | rs7932330 | G | A | 0.941 | -7977.388 | -7977.987 | -0.599 | 2.13E-06 |
| 11 | 117085210 | rs77683187 | C | A | 1.068 | -7977.388 | -7977.987 | -0.599 | 2.13E-06 |
| 11 | 118978319 | rs7936924 | C | T | 0.942 | -7977.388 | -7977.987 | -0.599 | 2.13E-06 |
| 11 | 119950717 | rs117040938 | C | T | 0.947 | -7977.388 | -7977.988 | -0.599 | 2.13E-06 |
| 11 | 121145466 | rs75215496 | C | T | 0.95 | -7977.388 | -7977.987 | -0.599 | 2.13E-06 |
| 11 | 117309989 | rs11216386 | T | C | 1.064 | -7977.388 | -7977.989 | -0.6 | 2.12E-06 |
| 11 | 117502726 | rs60668211 | A | G | 1.064 | -7977.388 | -7977.988 | -0.6 | 2.12E-06 |
| 11 | 117632609 | rs10750113 | G | A | 0.95 | -7977.388 | -7977.988 | -0.6 | 2.12E-06 |
| 11 | 117632645 | rs10790205 | C | T | 0.95 | -7977.388 | -7977.988 | -0.6 | 2.12E-06 |
| 11 | 117731990 | rs61901488 | T | A | 0.943 | -7977.388 | -7977.988 | -0.6 | 2.12E-06 |
| 11 | 119244095 | rs587985 | C | T | 0.953 | -7977.388 | -7977.989 | -0.6 | 2.12E-06 |
| 11 | 119872269 | rs111748140 | T | C | 0.941 | -7977.388 | -7977.988 | -0.6 | 2.12E-06 |
| 11 | 119943314 | rs145476010 | A | C | 0.954 | -7977.388 | -7977.989 | -0.6 | 2.12E-06 |
| 11 | 119584347 | rs4533047 | T | C | 0.938 | -7977.388 | -7977.989 | -0.601 | 2.12E-06 |
| 11 | 119584506 | rs4245205 | T | C | 0.938 | -7977.388 | -7977.989 | -0.601 | 2.12E-06 |
| 11 | 119584876 | rs7927510 | A | T | 0.938 | -7977.388 | -7977.989 | -0.601 | 2.12E-06 |
| 11 | 119586216 | rs7946962 | T | C | 0.938 | -7977.388 | -7977.989 | -0.601 | 2.12E-06 |
| 11 | 116626735 | rs180322 | C | T | 1.049 | -7977.388 | -7977.99 | -0.602 | 2.12E-06 |
| 11 | 119507035 | rs1546465 | G | T | 0.937 | -7977.388 | -7977.99 | -0.602 | 2.12E-06 |
| 11 | 116473704 | rs12795589 | G | A | 0.938 | -7977.388 | -7977.992 | -0.604 | 2.11E-06 |
| 11 | 120343650 | rs61898763 | T | C | 1.065 | -7977.388 | -7977.992 | -0.604 | 2.11E-06 |
| 11 | 116480236 | rs148108200 | C | T | 1.053 | -7977.388 | -7977.994 | -0.605 | 2.11E-06 |
| 11 | 119312818 | rs117259599 | T | C | 0.948 | -7977.388 | -7977.993 | -0.605 | 2.11E-06 |
| 11 | 120912139 | rs73011555 | G | A | 1.034 | -7977.388 | -7977.994 | -0.605 | 2.11E-06 |
| 11 | 116301501 | rs1045334530 | C | A | 0.954 | -7977.388 | -7977.994 | -0.606 | 2.11E-06 |
| 11 | 117252135 | rs79879928 | A | G | 0.94 | -7977.388 | -7977.994 | -0.606 | 2.11E-06 |
| 11 | 117714145 | rs12801338 | A | G | 1.065 | -7977.388 | -7977.994 | -0.606 | 2.11E-06 |
| 11 | 116228118 | rs71478237 | C | A | 0.936 | -7977.388 | -7977.995 | -0.607 | 2.11E-06 |
| 11 | 117045522 | rs79138835 | T | G | 0.951 | -7977.388 | -7977.995 | -0.607 | 2.11E-06 |
| 11 | 117612254 | rs6589620 | A | T | 0.937 | -7977.388 | -7977.995 | -0.607 | 2.11E-06 |
| 11 | 117632375 | rs4344529 | G | A | 0.95 | -7977.388 | -7977.995 | -0.607 | 2.11E-06 |
| 11 | 118605447 | rs2186781 | T | C | 0.943 | -7977.388 | -7977.995 | -0.607 | 2.11E-06 |
| 11 | 121139014 | rs80145448 | A | C | 0.952 | -7977.388 | -7977.996 | -0.607 | 2.11E-06 |
| 11 | 117472044 | rs140763307 | G | A | 0.973 | -7977.388 | -7977.996 | -0.608 | 2.11E-06 |
| 11 | 120601873 | rs2846106 | G | A | 0.947 | -7977.388 | -7977.996 | -0.608 | 2.11E-06 |
| 11 | 117200974 | rs116006647 | T | C | 1.06 | -7977.388 | -7977.997 | -0.609 | 2.10E-06 |
| 11 | 117646692 | rs11824545 | T | G | 1.065 | -7977.388 | -7977.997 | -0.609 | 2.10E-06 |
| 11 | 118932320 | rs582688 | C | T | 1.058 | -7977.388 | -7977.997 | -0.609 | 2.10E-06 |
| 11 | 117775903 | rs138450618 | T | C | 1.063 | -7977.388 | -7977.998 | -0.61 | 2.10E-06 |
| 11 | 120557814 | rs17124208 | C | A | 0.943 | -7977.388 | -7977.998 | -0.61 | 2.10E-06 |
| 11 | 116705568 | rs10750098 | G | T | 0.94 | -7977.388 | -7977.999 | -0.611 | 2.10E-06 |
| 11 | 117632764 | rs10790206 | C | T | 0.95 | -7977.388 | -7977.999 | -0.611 | 2.10E-06 |
| 11 | 117632769 | rs10790207 | C | T | 0.95 | -7977.388 | -7977.999 | -0.611 | 2.10E-06 |
| 11 | 117671091 | rs7947524 | T | C | 0.954 | -7977.388 | -7977.999 | -0.611 | 2.10E-06 |
| 11 | 118510642 | rs141169729 | T | C | 1.046 | -7977.388 | -7978 | -0.612 | 2.10E-06 |
| 11 | 121142335 | rs116915994 | T | C | 0.941 | -7977.388 | -7978 | -0.612 | 2.10E-06 |
| 11 | 116291707 | rs544154 | A | G | 1.05 | -7977.388 | -7978.002 | -0.613 | 2.10E-06 |
| 11 | 116473789 | rs12796351 | A | G | 0.938 | -7977.388 | -7978.002 | -0.613 | 2.10E-06 |
| 11 | 118062268 | rs639785 | T | C | 1.053 | -7977.388 | -7978.001 | -0.613 | 2.10E-06 |
| 11 | 119466427 | rs73002932 | T | C | 0.939 | -7977.388 | -7978.001 | -0.613 | 2.10E-06 |
| 11 | 121004502 | rs2155466 | T | C | 0.954 | -7977.388 | -7978.002 | -0.613 | 2.10E-06 |
| 11 | 117074109 | rs12970 | A | G | 1.067 | -7977.388 | -7978.002 | -0.614 | 2.09E-06 |
| 11 | 117361447 | rs10790190 | G | A | 0.936 | -7977.388 | -7978.003 | -0.615 | 2.09E-06 |
| 11 | 119674742 | rs7113439 | T | C | 0.942 | -7977.388 | -7978.003 | -0.615 | 2.09E-06 |
| 11 | 120893217 | rs191364497 | T | C | 0.949 | -7977.388 | -7978.003 | -0.615 | 2.09E-06 |
| 11 | 117902428 | rs77245011 | C | A | 0.941 | -7977.388 | -7978.005 | -0.616 | 2.09E-06 |
| 11 | 118977926 | rs6421572 | A | C | 0.942 | -7977.388 | -7978.004 | -0.616 | 2.09E-06 |
| 11 | 116464465 | rs181124100 | T | C | 0.945 | -7977.388 | -7978.005 | -0.617 | 2.09E-06 |
| 11 | 116676590 | rs4080140 | A | G | 0.945 | -7977.388 | -7978.006 | -0.617 | 2.09E-06 |
| 11 | 117004112 | rs73008513 | G | A | 0.936 | -7977.388 | -7978.006 | -0.617 | 2.09E-06 |

| | | | | | | | |
|---|---|---|---|---|---|---|---|
| 11 | 118862634 | rs59626457 | G | A | NULL | 0.575664351 | NULL&M_Disease |
| 11 | 116471898 | rs35664944 | A | G | NULL | 0.575664351 | NULL&M_Disease |
| 11 | 117902169 | rs586304 | T | A | NULL | 0.576885261 | NULL&M_Disease |
| 11 | 117961839 | rs651896 | T | C | NULL | 0.575664351 | NULL&M_Disease |
| 11 | 119398847 | rs4938670 | G | A | NULL | 0.576885261 | NULL&M_Disease |
| 11 | 120417553 | rs11217913 | G | A | NULL | 0.576885261 | NULL&M_Disease |
| 11 | 116468539 | rs75723689 | A | G | NULL | 0.575664351 | NULL&M_Disease |
| 11 | 116473730 | rs12796311 | T | G | NULL | 0.576885261 | NULL&M_Disease |
| 11 | 117070927 | rs2075548 | A | G | NULL | 0.575664351 | NULL&M_Disease |
| 11 | 120222480 | rs61898358 | A | G | NULL | 0.575664351 | NULL&M_Disease |
| 11 | 116530765 | rs555749 | T | C | NULL | 0.576885261 | NULL&M_Disease |
| 11 | 116673464 | rs61905132 | T | C | NULL | 0.575664351 | NULL&M_Disease |
| 11 | 116902639 | rs79357714 | G | A | NULL | 0.576885261 | NULL&M_Disease |
| 11 | 117347592 | rs114855326 | C | T | NULL | 0.575664351 | NULL&M_Disease |
| 11 | 117349617 | rs664657 | T | C | NULL | 0.576885261 | NULL&M_Disease |
| 11 | 118341212 | rs561775 | G | A | NULL | 0.576885261 | NULL&M_Disease |
| 11 | 117971968 | rs146877554 | A | G | NULL | 0.575664351 | NULL&M_Disease |
| 11 | 119690540 | rs117275185 | T | C | NULL | 0.575664351 | NULL&M_Disease |
| 11 | 118316006 | rs76651078 | A | G | NULL | 0.576885261 | NULL&M_Disease |
| 11 | 118831764 | rs61900158 | A | T | NULL | 0.576885261 | NULL&M_Disease |
| 11 | 118851138 | rs4456264 | T | C | NULL | 0.576885261 | NULL&M_Disease |
| 11 | 120579239 | rs61901380 | A | G | NULL | 0.576885261 | NULL&M_Disease |
| 11 | 119341720 | rs79294438 | T | C | NULL | 0.576885261 | NULL&M_Disease |
| 11 | 120327817 | rs17245518 | C | T | NULL | 0.576885261 | NULL&M_Disease |
| 11 | 116481066 | rs75137105 | T | C | NULL | 0.576885261 | NULL&M_Disease |
| 11 | 117672565 | rs10892174 | C | A | NULL | 0.576885261 | NULL&M_Disease |
| 11 | 117818210 | rs148882737 | T | C | NULL | 0.576885261 | NULL&M_Disease |
| 11 | 118823148 | rs663003 | C | T | NULL | 0.576885261 | NULL&M_Disease |
| 11 | 118864140 | rs11217116 | C | T | NULL | 0.576885261 | NULL&M_Disease |
| 11 | 119396373 | rs117507766 | T | C | NULL | 0.576885261 | NULL&M_Disease |
| 11 | 120921109 | rs11218125 | C | T | NULL | 0.576885261 | NULL&M_Disease |
| 11 | 118206121 | rs151127828 | A | G | NULL | 0.576885261 | NULL&M_Disease |
| 11 | 118859842 | rs12362865 | G | A | NULL | 0.576885261 | NULL&M_Disease |
| 11 | 119829348 | rs76499613 | C | T | NULL | 0.576885261 | NULL&M_Disease |
| 11 | 121058954 | rs492047 | A | G | NULL | 0.576885261 | NULL&M_Disease |
| 11 | 116559663 | rs11828763 | G | A | NULL | 0.576885261 | NULL&M_Disease |
| 11 | 116843425 | rs11216206 | G | C | NULL | 0.576885261 | NULL&M_Disease |
| 11 | 117635223 | rs7936362 | A | C | NULL | 0.576885261 | NULL&M_Disease |
| 11 | 116429563 | rs17119815 | G | T | NULL | 0.578105233 | NULL&M_Disease |
| 11 | 117313626 | rs118069464 | C | G | NULL | 0.576885261 | NULL&M_Disease |
| 11 | 117594364 | rs4938435 | G | A | NULL | 0.576885261 | NULL&M_Disease |
| 11 | 118856920 | rs11217112 | G | A | NULL | 0.578105233 | NULL&M_Disease |
| 11 | 118861389 | rs4936455 | T | C | NULL | 0.578105233 | NULL&M_Disease |
| 11 | 118862306 | rs9794909 | C | G | NULL | 0.578105233 | NULL&M_Disease |
| 11 | 119679280 | rs61897706 | A | G | NULL | 0.576885261 | NULL&M_Disease |
| 11 | 119691272 | rs34708128 | T | C | NULL | 0.576885261 | NULL&M_Disease |
| 11 | 118856816 | rs59316586 | C | A | NULL | 0.578105233 | NULL&M_Disease |
| 11 | 119828457 | rs72994637 | A | G | NULL | 0.576885261 | NULL&M_Disease |
| 11 | 118187216 | rs73014296 | G | A | NULL | 0.576885261 | NULL&M_Disease |
| 11 | 119577324 | rs7104562 | G | A | NULL | 0.576885261 | NULL&M_Disease |
| 11 | 119823216 | rs56303695 | A | G | NULL | 0.576885261 | NULL&M_Disease |
| 11 | 120239610 | rs12786734 | G | A | NULL | 0.576885261 | NULL&M_Disease |
| 11 | 120579826 | rs11217987 | A | T | NULL | 0.578105233 | NULL&M_Disease |
| 11 | 116592365 | rs180370 | T | C | NULL | 0.578105233 | NULL&M_Disease |
| 11 | 118855221 | rs11825496 | G | T | NULL | 0.578105233 | NULL&M_Disease |
| 11 | 118859274 | rs4936454 | T | C | NULL | 0.578105233 | NULL&M_Disease |
| 11 | 119499578 | rs76365645 | A | G | NULL | 0.578105233 | NULL&M_Disease |
| 11 | 117883397 | rs78204624 | C | G | NULL | 0.578105233 | NULL&M_Disease |
| 11 | 118852526 | rs45527340 | G | C | NULL | 0.578105233 | NULL&M_Disease |
| 11 | 119975137 | rs117420503 | T | C | NULL | 0.576885261 | NULL&M_Disease |
| 11 | 120857142 | rs145034678 | A | G | NULL | 0.576885261 | NULL&M_Disease |
| 11 | 117706667 | rs10790213 | T | A | NULL | 0.578105233 | NULL&M_Disease |
| 11 | 117186506 | rs28917234 | A | G | NULL | 0.578105233 | NULL&M_Disease |
| 11 | 120581323 | rs61901382 | A | G | NULL | 0.578105233 | NULL&M_Disease |
| 11 | 120581609 | rs7944756 | A | T | NULL | 0.578105233 | NULL&M_Disease |
| 11 | 120658874 | rs4359220 | G | A | NULL | 0.578105233 | NULL&M_Disease |
| 11 | 120944219 | rs11218135 | G | A | NULL | 0.578105233 | NULL&M_Disease |
| 11 | 120946892 | rs58204590 | G | A | NULL | 0.578105233 | NULL&M_Disease |
| 11 | 117255837 | rs117498846 | A | G | NULL | 0.578105233 | NULL&M_Disease |
| 11 | 119692871 | rs11217524 | C | T | NULL | 0.578105233 | NULL&M_Disease |
| 11 | 119893391 | rs624176 | T | C | NULL | 0.578105233 | NULL&M_Disease |
| 11 | 121082988 | rs520868 | C | T | NULL | 0.578105233 | NULL&M_Disease |
| 11 | 116424909 | rs61904800 | A | G | NULL | 0.578105233 | NULL&M_Disease |
| 11 | 116536656 | rs10892010 | C | A | NULL | 0.578105233 | NULL&M_Disease |
| 11 | 116471121 | rs12274273 | C | T | NULL | 0.578105233 | NULL&M_Disease |
| 11 | 116471149 | rs12280450 | G | A | NULL | 0.578105233 | NULL&M_Disease |
| 11 | 117727549 | rs11216584 | G | A | NULL | 0.578105233 | NULL&M_Disease |
| 11 | 117830715 | rs116864232 | C | T | NULL | 0.578105233 | NULL&M_Disease |
| 11 | 120824323 | rs520332 | T | G | NULL | 0.579324252 | NULL&M_Disease |
| 11 | 116232577 | rs11605788 | A | C | NULL | 0.578105233 | NULL&M_Disease |
| 11 | 118889822 | rs3802882 | G | C | NULL | 0.579324252 | NULL&M_Disease |
| 11 | 120705673 | rs11601867 | A | G | NULL | 0.579324252 | NULL&M_Disease |
| 11 | 120577423 | rs4567478 | C | T | NULL | 0.579324252 | NULL&M_Disease |
| 11 | 120864806 | rs10892657 | C | A | NULL | 0.579324252 | NULL&M_Disease |
| 11 | 117448013 | rs12799825 | A | C | NULL | 0.579324252 | NULL&M_Disease |
| 11 | 117572969 | rs74660871 | C | T | NULL | 0.578105233 | NULL&M_Disease |
| 11 | 117610387 | rs10790203 | G | A | NULL | 0.578105233 | NULL&M_Disease |
| 11 | 117611495 | rs4936405 | G | A | NULL | 0.578105233 | NULL&M_Disease |
| 11 | 118622245 | rs485901 | G | A | NULL | 0.579324252 | NULL&M_Disease |
| 11 | 118689745 | rs492471 | T | C | NULL | 0.579324252 | NULL&M_Disease |
| 11 | 119915024 | rs1942011 | T | C | NULL | 0.579324252 | NULL&M_Disease |
| 11 | 117297406 | rs78795346 | A | G | NULL | 0.578105233 | NULL&M_Disease |
| 11 | 117339203 | rs142239919 | G | A | NULL | 0.578105233 | NULL&M_Disease |
| 11 | 117554617 | rs116904534 | G | A | NULL | 0.579324252 | NULL&M_Disease |
| 11 | 117593368 | rs11599950 | T | C | NULL | 0.579324252 | NULL&M_Disease |
| 11 | 116631542 | rs35004487 | C | G | NULL | 0.578105233 | NULL&M_Disease |

| | | | | | | | | | |
|---|---|---|---|---|---|---|---|---|---|
| 11 | 118862634 | rs59626457 | G | A | 1.048 | -7977.388 | -7978.005 | -0.617 | 2.09E-06 |
| 11 | 116471898 | rs35664944 | A | G | 0.938 | -7977.388 | -7978.008 | -0.619 | 2.08E-06 |
| 11 | 117902169 | rs586304 | T | A | 1.048 | -7977.388 | -7978.007 | -0.619 | 2.08E-06 |
| 11 | 117961839 | rs651896 | T | C | 1.066 | -7977.388 | -7978.008 | -0.619 | 2.08E-06 |
| 11 | 119398847 | rs4938670 | G | A | 1.05 | -7977.388 | -7978.008 | -0.619 | 2.08E-06 |
| 11 | 120417553 | rs11217913 | G | A | 0.951 | -7977.388 | -7978.007 | -0.619 | 2.08E-06 |
| 11 | 116468539 | rs75723689 | A | G | 0.938 | -7977.388 | -7978.008 | -0.62 | 2.08E-06 |
| 11 | 116473730 | rs12796311 | T | G | 0.938 | -7977.388 | -7978.009 | -0.62 | 2.08E-06 |
| 11 | 117070927 | rs2075548 | A | G | 0.954 | -7977.388 | -7978.008 | -0.62 | 2.08E-06 |
| 11 | 120222480 | rs61898358 | A | G | 1.064 | -7977.388 | -7978.008 | -0.62 | 2.08E-06 |
| 11 | 116530765 | rs555749 | T | C | 1.048 | -7977.388 | -7978.01 | -0.621 | 2.08E-06 |
| 11 | 116673464 | rs61905132 | T | C | 0.94 | -7977.388 | -7978.009 | -0.621 | 2.08E-06 |
| 11 | 116902639 | rs79357714 | G | A | 0.936 | -7977.388 | -7978.01 | -0.621 | 2.08E-06 |
| 11 | 117347592 | rs114855326 | C | T | 1.043 | -7977.388 | -7978.009 | -0.621 | 2.08E-06 |
| 11 | 117349617 | rs664657 | T | C | 0.938 | -7977.388 | -7978.009 | -0.621 | 2.08E-06 |
| 11 | 118341212 | rs561775 | G | A | 0.942 | -7977.388 | -7978.011 | -0.622 | 2.08E-06 |
| 11 | 117971968 | rs146877554 | A | G | 0.944 | -7977.388 | -7978.011 | -0.623 | 2.07E-06 |
| 11 | 119690540 | rs117275185 | T | C | 1.017 | -7977.388 | -7978.012 | -0.623 | 2.07E-06 |
| 11 | 118316006 | rs76651078 | A | G | 1.067 | -7977.388 | -7978.012 | -0.624 | 2.07E-06 |
| 11 | 118831764 | rs61900158 | A | T | 1.055 | -7977.388 | -7978.013 | -0.624 | 2.07E-06 |
| 11 | 118851138 | rs4456264 | T | C | 1.048 | -7977.388 | -7978.013 | -0.624 | 2.07E-06 |
| 11 | 120579239 | rs61901380 | A | G | 0.946 | -7977.388 | -7978.012 | -0.624 | 2.07E-06 |
| 11 | 119341720 | rs79294438 | T | C | 0.94 | -7977.388 | -7978.013 | -0.625 | 2.07E-06 |
| 11 | 120327817 | rs17245518 | C | T | 1.063 | -7977.388 | -7978.013 | -0.625 | 2.07E-06 |
| 11 | 116481066 | rs75137105 | T | C | 1.049 | -7977.388 | -7978.014 | -0.626 | 2.07E-06 |
| 11 | 117672565 | rs10892174 | C | A | 0.952 | -7977.388 | -7978.014 | -0.626 | 2.07E-06 |
| 11 | 117818210 | rs148882737 | T | C | 0.951 | -7977.388 | -7978.014 | -0.626 | 2.07E-06 |
| 11 | 118823148 | rs663003 | C | T | 0.954 | -7977.388 | -7978.014 | -0.626 | 2.07E-06 |
| 11 | 118864140 | rs11217116 | C | T | 1.048 | -7977.388 | -7978.015 | -0.626 | 2.07E-06 |
| 11 | 119396373 | rs117507766 | T | C | 0.94 | -7977.388 | -7978.014 | -0.626 | 2.07E-06 |
| 11 | 120921109 | rs11218125 | C | T | 1.054 | -7977.388 | -7978.015 | -0.626 | 2.07E-06 |
| 11 | 118206121 | rs151127828 | A | G | 0.943 | -7977.388 | -7978.016 | -0.627 | 2.07E-06 |
| 11 | 118859842 | rs12362865 | G | A | 1.048 | -7977.388 | -7978.015 | -0.627 | 2.07E-06 |
| 11 | 119829348 | rs76499613 | C | T | 1.052 | -7977.388 | -7978.015 | -0.627 | 2.07E-06 |
| 11 | 121058954 | rs492047 | A | G | 0.953 | -7977.388 | -7978.015 | -0.627 | 2.07E-06 |
| 11 | 116559663 | rs11828763 | G | A | 0.941 | -7977.388 | -7978.016 | -0.628 | 2.06E-06 |
| 11 | 116843425 | rs11216206 | G | C | 1.065 | -7977.388 | -7978.016 | -0.628 | 2.06E-06 |
| 11 | 117635223 | rs7936362 | A | C | 1.064 | -7977.388 | -7978.016 | -0.628 | 2.06E-06 |
| 11 | 116429563 | rs17119815 | G | T | 1.055 | -7977.388 | -7978.017 | -0.629 | 2.06E-06 |
| 11 | 117313626 | rs118069464 | C | G | 0.968 | -7977.388 | -7978.017 | -0.629 | 2.06E-06 |
| 11 | 117594364 | rs4938435 | G | A | 0.939 | -7977.388 | -7978.017 | -0.629 | 2.06E-06 |
| 11 | 118856920 | rs11217112 | G | A | 1.048 | -7977.388 | -7978.017 | -0.629 | 2.06E-06 |
| 11 | 118861389 | rs4936455 | T | C | 1.048 | -7977.388 | -7978.017 | -0.629 | 2.06E-06 |
| 11 | 118862306 | rs9794909 | C | G | 1.048 | -7977.388 | -7978.017 | -0.629 | 2.06E-06 |
| 11 | 119679280 | rs61897706 | A | G | 1.039 | -7977.388 | -7978.018 | -0.629 | 2.06E-06 |
| 11 | 119691272 | rs34708128 | T | C | 0.957 | -7977.388 | -7978.017 | -0.629 | 2.06E-06 |
| 11 | 118856816 | rs59316586 | C | A | 1.048 | -7977.388 | -7978.018 | -0.63 | 2.06E-06 |
| 11 | 119828457 | rs72994637 | A | G | 0.946 | -7977.388 | -7978.018 | -0.63 | 2.06E-06 |
| 11 | 118187216 | rs73014296 | G | A | 1.066 | -7977.388 | -7978.019 | -0.631 | 2.06E-06 |
| 11 | 119577324 | rs7104562 | G | A | 0.943 | -7977.388 | -7978.019 | -0.631 | 2.06E-06 |
| 11 | 119823216 | rs56303695 | A | G | 0.942 | -7977.388 | -7978.019 | -0.631 | 2.06E-06 |
| 11 | 120239610 | rs12786734 | G | A | 0.937 | -7977.388 | -7978.019 | -0.631 | 2.06E-06 |
| 11 | 120579826 | rs11217987 | A | T | 0.946 | -7977.388 | -7978.02 | -0.631 | 2.06E-06 |
| 11 | 116592365 | rs180370 | T | C | 0.954 | -7977.388 | -7978.02 | -0.632 | 2.06E-06 |
| 11 | 118855221 | rs11825496 | G | T | 1.048 | -7977.388 | -7978.02 | -0.632 | 2.06E-06 |
| 11 | 118859274 | rs4936454 | T | C | 1.048 | -7977.388 | -7978.02 | -0.632 | 2.06E-06 |
| 11 | 119499578 | rs76365645 | A | G | 1.059 | -7977.388 | -7978.02 | -0.632 | 2.06E-06 |
| 11 | 117883397 | rs78204624 | C | G | 1.062 | -7977.388 | -7978.022 | -0.633 | 2.05E-06 |
| 11 | 118852526 | rs45527340 | G | C | 1.048 | -7977.388 | -7978.021 | -0.633 | 2.05E-06 |
| 11 | 119975137 | rs117420503 | T | C | 0.956 | -7977.388 | -7978.022 | -0.633 | 2.05E-06 |
| 11 | 120857142 | rs145034678 | A | G | 1.025 | -7977.388 | -7978.022 | -0.633 | 2.05E-06 |
| 11 | 117706667 | rs10790213 | T | A | 1.049 | -7977.388 | -7978.022 | -0.634 | 2.05E-06 |
| 11 | 117186506 | rs28917234 | A | G | 0.939 | -7977.388 | -7978.023 | -0.635 | 2.05E-06 |
| 11 | 120581323 | rs61901382 | A | G | 0.943 | -7977.388 | -7978.023 | -0.635 | 2.05E-06 |
| 11 | 120581609 | rs7944756 | A | T | 0.943 | -7977.388 | -7978.023 | -0.635 | 2.05E-06 |
| 11 | 120658874 | rs4359220 | G | A | 0.955 | -7977.388 | -7978.023 | -0.635 | 2.05E-06 |
| 11 | 120944219 | rs11218135 | G | A | 1.054 | -7977.388 | -7978.023 | -0.635 | 2.05E-06 |
| 11 | 120946892 | rs58204590 | G | A | 1.054 | -7977.388 | -7978.023 | -0.635 | 2.05E-06 |
| 11 | 117255837 | rs117498846 | A | G | 0.943 | -7977.388 | -7978.024 | -0.636 | 2.05E-06 |
| 11 | 119692871 | rs11217524 | C | T | 0.95 | -7977.388 | -7978.024 | -0.636 | 2.05E-06 |
| 11 | 119893391 | rs624176 | T | C | 0.949 | -7977.388 | -7978.024 | -0.636 | 2.05E-06 |
| 11 | 121082988 | rs520868 | C | T | 0.951 | -7977.388 | -7978.025 | -0.637 | 2.05E-06 |
| 11 | 116424909 | rs61904800 | A | G | 1.056 | -7977.388 | -7978.026 | -0.638 | 2.04E-06 |
| 11 | 116536656 | rs10892010 | C | A | 0.943 | -7977.388 | -7978.027 | -0.638 | 2.04E-06 |
| 11 | 116471121 | rs12274273 | C | T | 0.939 | -7977.388 | -7978.027 | -0.639 | 2.04E-06 |
| 11 | 116471149 | rs12280450 | G | A | 0.939 | -7977.388 | -7978.027 | -0.639 | 2.04E-06 |
| 11 | 117727549 | rs11216584 | G | A | 1.066 | -7977.388 | -7978.028 | -0.639 | 2.04E-06 |
| 11 | 117830715 | rs116864232 | C | T | 0.974 | -7977.388 | -7978.027 | -0.639 | 2.04E-06 |
| 11 | 120824323 | rs520332 | T | G | 0.947 | -7977.388 | -7978.027 | -0.639 | 2.04E-06 |
| 11 | 116232577 | rs11605788 | A | C | 1.066 | -7977.388 | -7978.028 | -0.64 | 2.04E-06 |
| 11 | 118889822 | rs3802882 | G | C | 0.94 | -7977.388 | -7978.028 | -0.64 | 2.04E-06 |
| 11 | 120705673 | rs11601867 | A | G | 0.948 | -7977.388 | -7978.028 | -0.64 | 2.04E-06 |
| 11 | 120577423 | rs4567478 | C | T | 0.946 | -7977.388 | -7978.029 | -0.641 | 2.04E-06 |
| 11 | 120864806 | rs10892657 | C | A | 1.064 | -7977.388 | -7978.029 | -0.641 | 2.04E-06 |
| 11 | 117448013 | rs12799825 | A | C | 0.943 | -7977.388 | -7978.031 | -0.642 | 2.04E-06 |
| 11 | 117572969 | rs74660871 | C | T | 0.941 | -7977.388 | -7978.03 | -0.642 | 2.04E-06 |
| 11 | 117610387 | rs10790203 | G | A | 0.94 | -7977.388 | -7978.03 | -0.642 | 2.04E-06 |
| 11 | 117611495 | rs4936405 | G | A | 0.94 | -7977.388 | -7978.03 | -0.642 | 2.04E-06 |
| 11 | 118622245 | rs485901 | G | A | 0.944 | -7977.388 | -7978.03 | -0.642 | 2.04E-06 |
| 11 | 118689745 | rs492471 | T | C | 1.065 | -7977.388 | -7978.03 | -0.642 | 2.04E-06 |
| 11 | 119915024 | rs1942011 | T | C | 1.05 | -7977.388 | -7978.031 | -0.642 | 2.04E-06 |
| 11 | 117297406 | rs78795346 | A | G | 0.953 | -7977.388 | -7978.031 | -0.643 | 2.03E-06 |
| 11 | 117339203 | rs142239919 | G | A | 0.972 | -7977.388 | -7978.031 | -0.643 | 2.03E-06 |
| 11 | 117554617 | rs116904534 | G | A | 1.063 | -7977.388 | -7978.031 | -0.643 | 2.03E-06 |
| 11 | 117593368 | rs11599950 | T | C | 1.055 | -7977.388 | -7978.031 | -0.643 | 2.03E-06 |
| 11 | 116631542 | rs35004487 | C | G | 1.01 | -7977.388 | -7978.032 | -0.644 | 2.03E-06 |

| | | | | | | | |
|---|---|---|---|---|---|---|---|
| 11 | 117646768 | rs7119967 | A | G | NULL | 0.579324252 | NULL&M_Disease |
| 11 | 117848632 | rs11216663 | T | G | NULL | 0.579324252 | NULL&M_Disease |
| 11 | 118921306 | rs519942 | A | G | NULL | 0.579324252 | NULL&M_Disease |
| 11 | 117615894 | rs146781782 | A | G | NULL | 0.578105233 | NULL&M_Disease |
| 11 | 119445010 | rs61898747 | G | T | NULL | 0.578105233 | NULL&M_Disease |
| 11 | 120578917 | rs7946325 | G | A | NULL | 0.579324252 | NULL&M_Disease |
| 11 | 117398435 | rs77026903 | T | C | NULL | 0.579324252 | NULL&M_Disease |
| 11 | 118335927 | rs495132 | A | G | NULL | 0.579324252 | NULL&M_Disease |
| 11 | 119773797 | rs117913812 | C | T | NULL | 0.579324252 | NULL&M_Disease |
| 11 | 116550776 | rs12417015 | G | A | NULL | 0.579324252 | NULL&M_Disease |
| 11 | 117414385 | rs650535 | T | C | NULL | 0.580542305 | NULL&M_Disease |
| 11 | 117919431 | rs1338985819 | A | G | NULL | 0.580542305 | NULL&M_Disease |
| 11 | 118747576 | rs55756957 | G | A | NULL | 0.580542305 | NULL&M_Disease |
| 11 | 118824439 | rs61900157 | A | C | NULL | 0.580542305 | NULL&M_Disease |
| 11 | 118920112 | rs670192 | A | G | NULL | 0.580542305 | NULL&M_Disease |
| 11 | 119302513 | rs77564141 | G | A | NULL | 0.579324252 | NULL&M_Disease |
| 11 | 119927038 | rs684233 | A | G | NULL | 0.580542305 | NULL&M_Disease |
| 11 | 120474370 | rs7131215 | G | C | NULL | 0.580542305 | NULL&M_Disease |
| 11 | 116439602 | rs7943240 | C | T | NULL | 0.580542305 | NULL&M_Disease |
| 11 | 118876213 | rs76253444 | G | A | NULL | 0.580542305 | NULL&M_Disease |
| 11 | 118889813 | rs3802883 | G | C | NULL | 0.579324252 | NULL&M_Disease |
| 11 | 119957452 | rs191917960 | G | C | NULL | 0.579324252 | NULL&M_Disease |
| 11 | 120577940 | rs61901379 | A | G | NULL | 0.580542305 | NULL&M_Disease |
| 11 | 117340285 | rs554912 | T | C | NULL | 0.580542305 | NULL&M_Disease |
| 11 | 118619423 | rs1961138 | C | T | NULL | 0.580542305 | NULL&M_Disease |
| 11 | 119498563 | rs11217360 | T | C | NULL | 0.580542305 | NULL&M_Disease |
| 11 | 117342672 | rs34573317 | C | T | NULL | 0.579324252 | NULL&M_Disease |
| 11 | 117667688 | rs113876980 | A | C | NULL | 0.580542305 | NULL&M_Disease |
| 11 | 119479675 | rs181268545 | A | G | NULL | 0.579324252 | NULL&M_Disease |
| 11 | 119593452 | rs11600071 | G | A | NULL | 0.579324252 | NULL&M_Disease |
| 11 | 117315326 | rs150311062 | A | G | NULL | 0.579324252 | NULL&M_Disease |
| 11 | 116238815 | rs519390 | C | T | NULL | 0.580542305 | NULL&M_Disease |
| 11 | 116547972 | rs117730773 | T | C | NULL | 0.579324252 | NULL&M_Disease |
| 11 | 116619605 | rs34637368 | A | G | NULL | 0.580542305 | NULL&M_Disease |
| 11 | 119629249 | rs11217470 | C | T | NULL | 0.580542305 | NULL&M_Disease |
| 11 | 121057021 | rs601102 | C | T | NULL | 0.580542305 | NULL&M_Disease |
| 11 | 121184472 | rs75133657 | C | A | NULL | 0.579324252 | NULL&M_Disease |
| 11 | 117292550 | rs7120516 | C | G | NULL | 0.579324252 | NULL&M_Disease |
| 11 | 117376413 | rs34138305 | A | G | NULL | 0.580542305 | NULL&M_Disease |
| 11 | 116628396 | rs143811287 | T | C | NULL | 0.580542305 | NULL&M_Disease |
| 11 | 117347029 | rs77346757 | A | G | NULL | 0.580542305 | NULL&M_Disease |
| 11 | 117347073 | rs115894834 | G | A | NULL | 0.580542305 | NULL&M_Disease |
| 11 | 117962813 | rs948467 | G | A | NULL | 0.580542305 | NULL&M_Disease |
| 11 | 120897776 | rs12797625 | A | G | NULL | 0.580542305 | NULL&M_Disease |
| 11 | 116703640 | rs5128 | G | C | NULL | 0.580542305 | NULL&M_Disease |
| 11 | 120277110 | rs117598476 | C | T | NULL | 0.580542305 | NULL&M_Disease |
| 11 | 116366118 | rs58856586 | A | G | NULL | 0.581759377 | NULL&M_Disease |
| 11 | 118618837 | rs640120 | A | C | NULL | 0.581759377 | NULL&M_Disease |
| 11 | 120568254 | rs12276408 | G | A | NULL | 0.581759377 | NULL&M_Disease |
| 11 | 116322222 | rs118168570 | A | G | NULL | 0.580542305 | NULL&M_Disease |
| 11 | 116640062 | rs622604 | C | T | NULL | 0.581759377 | NULL&M_Disease |
| 11 | 116640094 | rs610675 | A | T | NULL | 0.581759377 | NULL&M_Disease |
| 11 | 116640368 | rs623908 | G | A | NULL | 0.581759377 | NULL&M_Disease |
| 11 | 117795742 | rs78053944 | T | C | NULL | 0.580542305 | NULL&M_Disease |
| 11 | 118615660 | rs6589687 | T | A | NULL | 0.581759377 | NULL&M_Disease |
| 11 | 119371772 | rs78033613 | A | G | NULL | 0.580542305 | NULL&M_Disease |
| 11 | 119636942 | rs7130866 | G | A | NULL | 0.581759377 | NULL&M_Disease |
| 11 | 120580572 | rs28781001 | G | A | NULL | 0.581759377 | NULL&M_Disease |
| 11 | 121055740 | rs481703 | C | T | NULL | 0.581759377 | NULL&M_Disease |
| 11 | 116951533 | rs10892070 | A | G | NULL | 0.581759377 | NULL&M_Disease |
| 11 | 117646682 | rs11824502 | G | C | NULL | 0.581759377 | NULL&M_Disease |
| 11 | 119397680 | rs7395947 | C | T | NULL | 0.581759377 | NULL&M_Disease |
| 11 | 116665661 | rs1729410 | C | G | NULL | 0.581759377 | NULL&M_Disease |
| 11 | 117693413 | rs35705701 | A | G | NULL | 0.580542305 | NULL&M_Disease |
| 11 | 118064030 | rs45555132 | T | G | NULL | 0.580542305 | NULL&M_Disease |
| 11 | 120034758 | rs189431300 | A | G | NULL | 0.580542305 | NULL&M_Disease |
| 11 | 121068352 | rs190638475 | A | G | NULL | 0.580542305 | NULL&M_Disease |
| 11 | 117952091 | rs45445895 | A | G | NULL | 0.580542305 | NULL&M_Disease |
| 11 | 117973145 | rs73012879 | A | G | NULL | 0.581759377 | NULL&M_Disease |
| 11 | 120838141 | rs41297897 | A | G | NULL | 0.580542305 | NULL&M_Disease |
| 11 | 120898097 | rs117476293 | C | A | NULL | 0.581759377 | NULL&M_Disease |
| 11 | 116304790 | rs115696360 | C | G | NULL | 0.581759377 | NULL&M_Disease |
| 11 | 116542247 | rs36195057 | A | G | NULL | 0.581759377 | NULL&M_Disease |
| 11 | 117400284 | rs78615799 | T | C | NULL | 0.580542305 | NULL&M_Disease |
| 11 | 117403017 | rs7109567 | G | A | NULL | 0.581759377 | NULL&M_Disease |
| 11 | 117674355 | rs10892175 | A | G | NULL | 0.581759377 | NULL&M_Disease |
| 11 | 118825636 | rs1790191 | A | G | NULL | 0.581759377 | NULL&M_Disease |
| 11 | 117074229 | rs641620 | C | T | NULL | 0.581759377 | NULL&M_Disease |
| 11 | 117322099 | rs76257648 | A | C | NULL | 0.580542305 | NULL&M_Disease |
| 11 | 120790963 | rs10892643 | T | C | NULL | 0.581759377 | NULL&M_Disease |
| 11 | 116791391 | rs79489590 | T | C | NULL | 0.581759377 | NULL&M_Disease |
| 11 | 117316640 | rs117195665 | T | G | NULL | 0.580542305 | NULL&M_Disease |
| 11 | 120867999 | rs78995144 | A | G | NULL | 0.581759377 | NULL&M_Disease |
| 11 | 121013174 | rs2155365 | C | A | NULL | 0.581759377 | NULL&M_Disease |
| 11 | 117324964 | rs10750106 | T | C | NULL | 0.581759377 | NULL&M_Disease |
| 11 | 117337032 | rs150081993 | T | C | NULL | 0.580542305 | NULL&M_Disease |
| 11 | 116557142 | rs2156121 | A | C | NULL | 0.582975454 | NULL&M_Disease |
| 11 | 117248283 | rs117619191 | C | T | NULL | 0.581759377 | NULL&M_Disease |
| 11 | 117953132 | rs113137102 | A | C | NULL | 0.581759377 | NULL&M_Disease |
| 11 | 118609736 | rs1783937 | A | G | NULL | 0.582975454 | NULL&M_Disease |
| 11 | 119539313 | rs4938704 | T | C | NULL | 0.582975454 | NULL&M_Disease |
| 11 | 116351099 | rs17490647 | C | A | NULL | 0.581759377 | NULL&M_Disease |
| 11 | 116460172 | rs11216023 | C | G | NULL | 0.581759377 | NULL&M_Disease |
| 11 | 116631391 | rs3741301 | C | T | NULL | 0.582975454 | NULL&M_Disease |
| 11 | 117221170 | rs563828 | T | G | NULL | 0.582975454 | NULL&M_Disease |
| 11 | 117237186 | rs7106772 | A | G | NULL | 0.581759377 | NULL&M_Disease |

| | | | | | | | | | |
|---|---|---|---|---|---|---|---|---|---|
| 11 | 117646768 | rs7119967 | A | G | 1.063 | -7977.388 | -7978.032 | -0.644 | 2.03E-06 |
| 11 | 117848632 | rs11216663 | T | G | 0.938 | -7977.388 | -7978.032 | -0.644 | 2.03E-06 |
| 11 | 118921306 | rs519942 | A | G | 1.057 | -7977.388 | -7978.033 | -0.644 | 2.03E-06 |
| 11 | 117615894 | rs146781782 | A | G | 0.973 | -7977.388 | -7978.033 | -0.645 | 2.03E-06 |
| 11 | 119445010 | rs61898747 | G | T | 0.965 | -7977.388 | -7978.033 | -0.645 | 2.03E-06 |
| 11 | 120578917 | rs7946325 | G | A | 0.946 | -7977.388 | -7978.033 | -0.645 | 2.03E-06 |
| 11 | 117398435 | rs77026903 | T | C | 0.948 | -7977.388 | -7978.035 | -0.647 | 2.03E-06 |
| 11 | 118335927 | rs495132 | A | G | 0.943 | -7977.388 | -7978.035 | -0.647 | 2.03E-06 |
| 11 | 119773797 | rs117913812 | C | T | 0.947 | -7977.388 | -7978.035 | -0.647 | 2.03E-06 |
| 11 | 116550776 | rs12417015 | G | A | 0.947 | -7977.388 | -7978.036 | -0.648 | 2.02E-06 |
| 11 | 117414385 | rs650535 | T | C | 1.051 | -7977.388 | -7978.036 | -0.648 | 2.02E-06 |
| 11 | 117919431 | rs1338985819 | A | G | 0.955 | -7977.388 | -7978.037 | -0.648 | 2.02E-06 |
| 11 | 118747576 | rs55756957 | G | A | 0.951 | -7977.388 | -7978.037 | -0.648 | 2.02E-06 |
| 11 | 118824439 | rs61900157 | A | C | 0.955 | -7977.388 | -7978.037 | -0.648 | 2.02E-06 |
| 11 | 118920112 | rs670192 | A | G | 1.057 | -7977.388 | -7978.037 | -0.648 | 2.02E-06 |
| 11 | 119302513 | rs77564141 | G | A | 0.955 | -7977.388 | -7978.037 | -0.648 | 2.02E-06 |
| 11 | 119927038 | rs684233 | A | G | 0.949 | -7977.388 | -7978.037 | -0.648 | 2.02E-06 |
| 11 | 120474370 | rs7131215 | G | C | 1.049 | -7977.388 | -7978.036 | -0.648 | 2.02E-06 |
| 11 | 116439602 | rs7943240 | C | T | 1.049 | -7977.388 | -7978.037 | -0.649 | 2.02E-06 |
| 11 | 118876213 | rs76253444 | G | A | 1.056 | -7977.388 | -7978.037 | -0.649 | 2.02E-06 |
| 11 | 118889813 | rs3802883 | G | C | 0.941 | -7977.388 | -7978.037 | -0.649 | 2.02E-06 |
| 11 | 119957452 | rs191917960 | G | C | 0.938 | -7977.388 | -7978.037 | -0.649 | 2.02E-06 |
| 11 | 120577940 | rs61901379 | A | G | 0.946 | -7977.388 | -7978.037 | -0.649 | 2.02E-06 |
| 11 | 117340285 | rs554912 | T | C | 0.947 | -7977.388 | -7978.038 | -0.65 | 2.02E-06 |
| 11 | 118619423 | rs1961138 | C | T | 0.944 | -7977.388 | -7978.039 | -0.65 | 2.02E-06 |
| 11 | 119498563 | rs11217360 | T | C | 1.058 | -7977.388 | -7978.039 | -0.651 | 2.02E-06 |
| 11 | 117342672 | rs34573317 | C | T | 0.94 | -7977.388 | -7978.04 | -0.652 | 2.02E-06 |
| 11 | 117667688 | rs113876980 | A | C | 0.953 | -7977.388 | -7978.04 | -0.652 | 2.02E-06 |
| 11 | 119479675 | rs181268545 | A | G | 0.948 | -7977.388 | -7978.04 | -0.652 | 2.02E-06 |
| 11 | 119593452 | rs11600071 | G | A | 0.945 | -7977.388 | -7978.04 | -0.652 | 2.02E-06 |
| 11 | 117315326 | rs150311062 | A | G | 0.979 | -7977.388 | -7978.042 | -0.653 | 2.01E-06 |
| 11 | 116238815 | rs519390 | C | T | 1.047 | -7977.388 | -7978.042 | -0.654 | 2.01E-06 |
| 11 | 116547972 | rs117730773 | T | C | 1.006 | -7977.388 | -7978.042 | -0.654 | 2.01E-06 |
| 11 | 116619605 | rs34637368 | A | G | 1.059 | -7977.388 | -7978.042 | -0.654 | 2.01E-06 |
| 11 | 119629249 | rs11217470 | C | T | 0.943 | -7977.388 | -7978.043 | -0.654 | 2.01E-06 |
| 11 | 121057021 | rs601102 | C | T | 0.953 | -7977.388 | -7978.042 | -0.654 | 2.01E-06 |
| 11 | 121184472 | rs75133657 | C | A | 0.947 | -7977.388 | -7978.042 | -0.654 | 2.01E-06 |
| 11 | 117292550 | rs7120516 | C | G | 0.955 | -7977.388 | -7978.043 | -0.655 | 2.01E-06 |
| 11 | 117376413 | rs34138305 | A | G | 0.939 | -7977.388 | -7978.043 | -0.655 | 2.01E-06 |
| 11 | 116628396 | rs143811287 | T | C | 0.943 | -7977.388 | -7978.045 | -0.657 | 2.01E-06 |
| 11 | 117347029 | rs77346757 | A | G | 1.062 | -7977.388 | -7978.046 | -0.657 | 2.01E-06 |
| 11 | 117347073 | rs115894834 | G | A | 1.062 | -7977.388 | -7978.046 | -0.657 | 2.01E-06 |
| 11 | 117962813 | rs948467 | G | A | 1.064 | -7977.388 | -7978.046 | -0.657 | 2.01E-06 |
| 11 | 120897776 | rs12797625 | A | G | 1.034 | -7977.388 | -7978.046 | -0.657 | 2.01E-06 |
| 11 | 116703640 | rs5128 | G | C | 0.942 | -7977.388 | -7978.047 | -0.658 | 2.00E-06 |
| 11 | 120277110 | rs117598476 | C | T | 1.04 | -7977.388 | -7978.046 | -0.658 | 2.00E-06 |
| 11 | 116366118 | rs58856586 | A | G | 0.944 | -7977.388 | -7978.048 | -0.659 | 2.00E-06 |
| 11 | 118618837 | rs640120 | A | C | 0.944 | -7977.388 | -7978.047 | -0.659 | 2.00E-06 |
| 11 | 120568254 | rs12276408 | G | A | 0.946 | -7977.388 | -7978.047 | -0.659 | 2.00E-06 |
| 11 | 116322222 | rs118168570 | A | G | 0.947 | -7977.388 | -7978.048 | -0.66 | 2.00E-06 |
| 11 | 116640062 | rs622604 | C | T | 1.048 | -7977.388 | -7978.048 | -0.66 | 2.00E-06 |
| 11 | 116640094 | rs610675 | A | T | 1.048 | -7977.388 | -7978.048 | -0.66 | 2.00E-06 |
| 11 | 116640368 | rs623908 | G | A | 1.048 | -7977.388 | -7978.048 | -0.66 | 2.00E-06 |
| 11 | 117795742 | rs78053944 | T | C | 0.94 | -7977.388 | -7978.048 | -0.66 | 2.00E-06 |
| 11 | 118615660 | rs6589687 | T | A | 0.944 | -7977.388 | -7978.048 | -0.66 | 2.00E-06 |
| 11 | 119371772 | rs78033613 | A | G | 0.971 | -7977.388 | -7978.048 | -0.66 | 2.00E-06 |
| 11 | 119636942 | rs7130866 | G | A | 0.943 | -7977.388 | -7978.048 | -0.66 | 2.00E-06 |
| 11 | 120580572 | rs28781001 | G | A | 0.947 | -7977.388 | -7978.048 | -0.66 | 2.00E-06 |
| 11 | 121055740 | rs481703 | C | T | 0.954 | -7977.388 | -7978.048 | -0.66 | 2.00E-06 |
| 11 | 116951533 | rs10892070 | A | G | 0.955 | -7977.388 | -7978.049 | -0.661 | 2.00E-06 |
| 11 | 117646682 | rs11824502 | G | C | 1.062 | -7977.388 | -7978.05 | -0.661 | 2.00E-06 |
| 11 | 119397680 | rs7395947 | C | T | 1.05 | -7977.388 | -7978.049 | -0.661 | 2.00E-06 |
| 11 | 116665661 | rs1729410 | C | G | 1.048 | -7977.388 | -7978.051 | -0.662 | 2.00E-06 |
| 11 | 117693413 | rs35705701 | A | G | 0.956 | -7977.388 | -7978.05 | -0.662 | 2.00E-06 |
| 11 | 118064030 | rs45555132 | T | G | 1.018 | -7977.388 | -7978.05 | -0.662 | 2.00E-06 |
| 11 | 120034758 | rs189431300 | A | G | 1.015 | -7977.388 | -7978.051 | -0.662 | 2.00E-06 |
| 11 | 121068352 | rs190638475 | A | G | 0.95 | -7977.388 | -7978.05 | -0.662 | 2.00E-06 |
| 11 | 117952091 | rs45445895 | A | G | 1.033 | -7977.388 | -7978.052 | -0.663 | 1.99E-06 |
| 11 | 117973145 | rs73012879 | A | G | 1.06 | -7977.388 | -7978.051 | -0.663 | 1.99E-06 |
| 11 | 120838141 | rs41297897 | A | G | 1.036 | -7977.388 | -7978.052 | -0.663 | 1.99E-06 |
| 11 | 120898097 | rs117476293 | C | A | 0.94 | -7977.388 | -7978.051 | -0.663 | 1.99E-06 |
| 11 | 116304790 | rs115696360 | C | G | 0.944 | -7977.388 | -7978.052 | -0.664 | 1.99E-06 |
| 11 | 116542247 | rs36195057 | A | G | 0.944 | -7977.388 | -7978.052 | -0.664 | 1.99E-06 |
| 11 | 117400284 | rs78615799 | T | C | 0.951 | -7977.388 | -7978.052 | -0.664 | 1.99E-06 |
| 11 | 117403017 | rs7109567 | G | A | 0.955 | -7977.388 | -7978.053 | -0.664 | 1.99E-06 |
| 11 | 117674355 | rs10892175 | A | G | 0.953 | -7977.388 | -7978.052 | -0.664 | 1.99E-06 |
| 11 | 118825636 | rs1790191 | A | G | 0.955 | -7977.388 | -7978.052 | -0.664 | 1.99E-06 |
| 11 | 117074229 | rs641620 | C | T | 0.946 | -7977.388 | -7978.053 | -0.665 | 1.99E-06 |
| 11 | 117322099 | rs76257648 | A | C | 0.981 | -7977.388 | -7978.053 | -0.665 | 1.99E-06 |
| 11 | 120790963 | rs10892643 | T | C | 0.954 | -7977.388 | -7978.053 | -0.665 | 1.99E-06 |
| 11 | 116791391 | rs79489590 | T | C | 0.942 | -7977.388 | -7978.055 | -0.666 | 1.99E-06 |
| 11 | 117316640 | rs117195665 | T | G | 0.984 | -7977.388 | -7978.055 | -0.666 | 1.99E-06 |
| 11 | 120867999 | rs78995144 | A | G | 1.063 | -7977.388 | -7978.054 | -0.666 | 1.99E-06 |
| 11 | 121013174 | rs2155365 | C | A | 0.954 | -7977.388 | -7978.055 | -0.666 | 1.99E-06 |
| 11 | 117324964 | rs10750106 | T | C | 0.946 | -7977.388 | -7978.055 | -0.667 | 1.99E-06 |
| 11 | 117337032 | rs150081993 | T | C | 0.984 | -7977.388 | -7978.055 | -0.667 | 1.99E-06 |
| 11 | 116557142 | rs2156121 | A | C | 0.945 | -7977.388 | -7978.057 | -0.668 | 1.98E-06 |
| 11 | 117248283 | rs117619191 | C | T | 1.034 | -7977.388 | -7978.056 | -0.668 | 1.98E-06 |
| 11 | 117953132 | rs113137102 | A | C | 1.044 | -7977.388 | -7978.056 | -0.668 | 1.98E-06 |
| 11 | 118609736 | rs1783937 | A | G | 0.945 | -7977.388 | -7978.057 | -0.668 | 1.98E-06 |
| 11 | 119539313 | rs4938704 | T | C | 0.954 | -7977.388 | -7978.056 | -0.668 | 1.98E-06 |
| 11 | 116351099 | rs17490647 | C | A | 0.946 | -7977.388 | -7978.057 | -0.669 | 1.98E-06 |
| 11 | 116460172 | rs11216023 | C | G | 0.942 | -7977.388 | -7978.057 | -0.669 | 1.98E-06 |
| 11 | 116631391 | rs3741301 | C | T | 0.955 | -7977.388 | -7978.057 | -0.669 | 1.98E-06 |
| 11 | 117221170 | rs563828 | T | G | 1.048 | -7977.388 | -7978.057 | -0.669 | 1.98E-06 |
| 11 | 117237186 | rs7106772 | A | G | 0.957 | -7977.388 | -7978.057 | -0.669 | 1.98E-06 |

| | | | | | | | |
|---|---|---|---|---|---|---|---|
| 11 | 117244291 | rs76344095 | T | C | NULL | 0.581759377 | NULL&M_Disease |
| 11 | 117244606 | rs7109766 | T | A | NULL | 0.581759377 | NULL&M_Disease |
| 11 | 117253230 | rs78406883 | T | C | NULL | 0.581759377 | NULL&M_Disease |
| 11 | 117335686 | rs61743159 | A | G | NULL | 0.581759377 | NULL&M_Disease |
| 11 | 118852137 | rs4936451 | T | C | NULL | 0.582975454 | NULL&M_Disease |
| 11 | 116309713 | rs73580459 | A | G | NULL | 0.581759377 | NULL&M_Disease |
| 11 | 120124302 | rs59565018 | A | G | NULL | 0.581759377 | NULL&M_Disease |
| 11 | 120779520 | rs949055 | C | T | NULL | 0.582975454 | NULL&M_Disease |
| 11 | 117908328 | rs3016863 | A | G | NULL | 0.582975454 | NULL&M_Disease |
| 11 | 120580623 | rs7102869 | T | C | NULL | 0.582975454 | NULL&M_Disease |
| 11 | 117200943 | rs1242695 | T | G | NULL | 0.582975454 | NULL&M_Disease |
| 11 | 117336093 | rs117028190 | T | C | NULL | 0.581759377 | NULL&M_Disease |
| 11 | 118919206 | rs568922 | T | C | NULL | 0.582975454 | NULL&M_Disease |
| 11 | 119246708 | rs675873 | C | T | NULL | 0.582975454 | NULL&M_Disease |
| 11 | 119604844 | rs11607679 | A | C | NULL | 0.581759377 | NULL&M_Disease |
| 11 | 120576351 | rs12281464 | T | A | NULL | 0.582975454 | NULL&M_Disease |
| 11 | 120576489 | rs12294858 | A | G | NULL | 0.582975454 | NULL&M_Disease |
| 11 | 120999258 | rs4936578 | G | A | NULL | 0.582975454 | NULL&M_Disease |
| 11 | 116236775 | rs526151 | T | C | NULL | 0.582975454 | NULL&M_Disease |
| 11 | 117530935 | rs7949670 | A | C | NULL | 0.582975454 | NULL&M_Disease |
| 11 | 117773027 | rs117010140 | A | G | NULL | 0.581759377 | NULL&M_Disease |
| 11 | 118852641 | rs12273687 | T | G | NULL | 0.582975454 | NULL&M_Disease |
| 11 | 119875786 | rs1944499 | A | G | NULL | 0.582975454 | NULL&M_Disease |
| 11 | 120603160 | rs79540430 | T | A | NULL | 0.582975454 | NULL&M_Disease |
| 11 | 120649337 | rs117414016 | T | G | NULL | 0.582975454 | NULL&M_Disease |
| 11 | 117330944 | rs75813208 | T | C | NULL | 0.582975454 | NULL&M_Disease |
| 11 | 117638305 | rs61903859 | A | C | NULL | 0.582975454 | NULL&M_Disease |
| 11 | 119944195 | rs661621 | T | A | NULL | 0.582975454 | NULL&M_Disease |
| 11 | 120992119 | rs11218148 | A | C | NULL | 0.582975454 | NULL&M_Disease |
| 11 | 120992859 | rs4936576 | C | G | NULL | 0.582975454 | NULL&M_Disease |
| 11 | 116591934 | rs180372 | A | G | NULL | 0.582975454 | NULL&M_Disease |
| 11 | 117326419 | rs118100246 | A | G | NULL | 0.581759377 | NULL&M_Disease |
| 11 | 117328052 | rs148406383 | T | C | NULL | 0.581759377 | NULL&M_Disease |
| 11 | 117965421 | rs45499491 | T | C | NULL | 0.582975454 | NULL&M_Disease |
| 11 | 118889810 | rs3802884 | T | C | NULL | 0.582975454 | NULL&M_Disease |
| 11 | 121108907 | rs1941299 | C | A | NULL | 0.582975454 | NULL&M_Disease |
| 11 | 116466714 | rs12292959 | C | T | NULL | 0.582975454 | NULL&M_Disease |
| 11 | 117330326 | rs145305155 | G | A | NULL | 0.582975454 | NULL&M_Disease |
| 11 | 117646805 | rs11819854 | T | A | NULL | 0.582975454 | NULL&M_Disease |
| 11 | 121103704 | rs10892692 | G | A | NULL | 0.582975454 | NULL&M_Disease |
| 11 | 117392940 | rs7941258 | T | C | NULL | 0.582975454 | NULL&M_Disease |
| 11 | 117392987 | rs7941460 | A | G | NULL | 0.582975454 | NULL&M_Disease |
| 11 | 117393345 | rs7944704 | A | G | NULL | 0.582975454 | NULL&M_Disease |
| 11 | 118853774 | rs11217109 | C | T | NULL | 0.584190523 | NULL&M_Disease |
| 11 | 119466907 | rs150628867 | A | G | NULL | 0.582975454 | NULL&M_Disease |
| 11 | 119684904 | rs79297443 | T | C | NULL | 0.582975454 | NULL&M_Disease |
| 11 | 119889758 | rs1170855838 | T | C | NULL | 0.584190523 | NULL&M_Disease |
| 11 | 117821885 | rs144891162 | T | C | NULL | 0.582975454 | NULL&M_Disease |
| 11 | 117906759 | rs649827 | T | C | NULL | 0.584190523 | NULL&M_Disease |
| 11 | 118314849 | rs2017122 | T | C | NULL | 0.584190523 | NULL&M_Disease |
| 11 | 120576154 | rs11217984 | T | C | NULL | 0.584190523 | NULL&M_Disease |
| 11 | 116737302 | rs139447106 | T | C | NULL | 0.584190523 | NULL&M_Disease |
| 11 | 117094591 | rs17120523 | G | A | NULL | 0.584190523 | NULL&M_Disease |
| 11 | 120860524 | rs34259384 | T | G | NULL | 0.584190523 | NULL&M_Disease |
| 11 | 116353269 | rs118065597 | A | G | NULL | 0.582975454 | NULL&M_Disease |
| 11 | 116741840 | rs73023041 | A | G | NULL | 0.584190523 | NULL&M_Disease |
| 11 | 117630622 | rs78080546 | T | A | NULL | 0.584190523 | NULL&M_Disease |
| 11 | 119609828 | rs113950321 | C | A | NULL | 0.582975454 | NULL&M_Disease |
| 11 | 120009748 | rs61900472 | A | G | NULL | 0.582975454 | NULL&M_Disease |
| 11 | 120623637 | rs11217999 | A | C | NULL | 0.584190523 | NULL&M_Disease |
| 11 | 120871324 | rs11218103 | T | C | NULL | 0.584190523 | NULL&M_Disease |
| 11 | 116677370 | rs1729407 | C | G | NULL | 0.584190523 | NULL&M_Disease |
| 11 | 121004851 | rs7125224 | A | G | NULL | 0.584190523 | NULL&M_Disease |
| 11 | 121160481 | rs10750184 | A | G | NULL | 0.584190523 | NULL&M_Disease |
| 11 | 118509877 | rs112552940 | T | C | NULL | 0.584190523 | NULL&M_Disease |
| 11 | 118824996 | rs4426142 | G | A | NULL | 0.584190523 | NULL&M_Disease |
| 11 | 118853021 | rs113962716 | A | C | NULL | 0.584190523 | NULL&M_Disease |
| 11 | 119863584 | rs1944511 | A | G | NULL | 0.584190523 | NULL&M_Disease |
| 11 | 120475835 | rs11217924 | T | C | NULL | 0.584190523 | NULL&M_Disease |
| 11 | 120715952 | rs7123228 | G | A | NULL | 0.584190523 | NULL&M_Disease |
| 11 | 117317855 | rs117580094 | G | T | NULL | 0.584190523 | NULL&M_Disease |
| 11 | 117539715 | rs636783 | C | A | NULL | 0.584190523 | NULL&M_Disease |
| 11 | 118530068 | rs112337443 | C | T | NULL | 0.584190523 | NULL&M_Disease |
| 11 | 119628238 | rs10892449 | G | C | NULL | 0.584190523 | NULL&M_Disease |
| 11 | 117464025 | rs12576226 | G | A | NULL | 0.585404569 | NULL&M_Disease |
| 11 | 117619538 | rs7924968 | A | T | NULL | 0.584190523 | NULL&M_Disease |
| 11 | 117674072 | rs1410186916 | T | A | NULL | 0.585404569 | NULL&M_Disease |
| 11 | 117683284 | rs112776148 | C | G | NULL | 0.584190523 | NULL&M_Disease |
| 11 | 120986075 | rs80282202 | C | G | NULL | 0.584190523 | NULL&M_Disease |
| 11 | 117466755 | rs78144220 | T | G | NULL | 0.584190523 | NULL&M_Disease |
| 11 | 117665267 | rs71480357 | G | C | NULL | 0.584190523 | NULL&M_Disease |
| 11 | 119612600 | rs113816823 | G | A | NULL | 0.584190523 | NULL&M_Disease |
| 11 | 119849683 | rs76547700 | T | G | NULL | 0.585404569 | NULL&M_Disease |
| 11 | 120206746 | rs35734851 | A | G | NULL | 0.584190523 | NULL&M_Disease |
| 11 | 121049136 | rs592160 | T | C | NULL | 0.585404569 | NULL&M_Disease |
| 11 | 116549408 | rs74792494 | C | A | NULL | 0.584190523 | NULL&M_Disease |
| 11 | 117366111 | rs10892129 | G | C | NULL | 0.585404569 | NULL&M_Disease |
| 11 | 117680478 | rs74500359 | T | C | NULL | 0.585404569 | NULL&M_Disease |
| 11 | 118863116 | rs4938613 | G | A | NULL | 0.585404569 | NULL&M_Disease |
| 11 | 118863324 | rs4936456 | T | C | NULL | 0.585404569 | NULL&M_Disease |
| 11 | 119497371 | rs145924639 | T | C | NULL | 0.584190523 | NULL&M_Disease |
| 11 | 119837947 | rs61899043 | C | A | NULL | 0.584190523 | NULL&M_Disease |
| 11 | 119981694 | rs140158503 | T | C | NULL | 0.584190523 | NULL&M_Disease |
| 11 | 120622042 | rs150032046 | G | A | NULL | 0.584190523 | NULL&M_Disease |
| 11 | 116470552 | rs35820582 | C | G | NULL | 0.585404569 | NULL&M_Disease |
| 11 | 116544015 | rs74724948 | T | A | NULL | 0.584190523 | NULL&M_Disease |

| | | | | | | | | | |
|---|---|---|---|---|---|---|---|---|---|
| 11 | 117244291 | rs76344095 | T | C | 0.957 | -7977.388 | -7978.057 | -0.669 | 1.98E-06 |
| 11 | 117244606 | rs7109766 | T | A | 0.957 | -7977.388 | -7978.057 | -0.669 | 1.98E-06 |
| 11 | 117253230 | rs78406883 | T | C | 0.957 | -7977.388 | -7978.057 | -0.669 | 1.98E-06 |
| 11 | 117335686 | rs61743159 | A | G | 0.985 | -7977.388 | -7978.058 | -0.669 | 1.98E-06 |
| 11 | 118852137 | rs4936451 | T | C | 1.047 | -7977.388 | -7978.058 | -0.669 | 1.98E-06 |
| 11 | 116309713 | rs73580459 | A | G | 1.064 | -7977.388 | -7978.058 | -0.67 | 1.98E-06 |
| 11 | 120124302 | rs59565018 | A | G | 1.057 | -7977.388 | -7978.058 | -0.67 | 1.98E-06 |
| 11 | 120779520 | rs949055 | C | T | 1.049 | -7977.388 | -7978.058 | -0.67 | 1.98E-06 |
| 11 | 117908328 | rs3016863 | A | G | 1.047 | -7977.388 | -7978.059 | -0.671 | 1.98E-06 |
| 11 | 120580623 | rs7102869 | T | C | 0.944 | -7977.388 | -7978.06 | -0.671 | 1.98E-06 |
| 11 | 117200943 | rs1242695 | T | G | 1.049 | -7977.388 | -7978.06 | -0.672 | 1.98E-06 |
| 11 | 117336093 | rs117028190 | T | C | 0.987 | -7977.388 | -7978.06 | -0.672 | 1.98E-06 |
| 11 | 118919206 | rs568922 | T | C | 1.056 | -7977.388 | -7978.06 | -0.672 | 1.98E-06 |
| 11 | 119246708 | rs675873 | C | T | 0.954 | -7977.388 | -7978.06 | -0.672 | 1.98E-06 |
| 11 | 119604844 | rs11607679 | A | C | 0.947 | -7977.388 | -7978.06 | -0.672 | 1.98E-06 |
| 11 | 120576351 | rs12281464 | T | A | 0.946 | -7977.388 | -7978.06 | -0.672 | 1.98E-06 |
| 11 | 120576489 | rs12294858 | A | G | 0.946 | -7977.388 | -7978.06 | -0.672 | 1.98E-06 |
| 11 | 120999258 | rs4936578 | G | A | 0.949 | -7977.388 | -7978.06 | -0.672 | 1.98E-06 |
| 11 | 116236775 | rs526151 | T | C | 1.047 | -7977.388 | -7978.061 | -0.673 | 1.97E-06 |
| 11 | 117530935 | rs7949670 | A | C | 0.942 | -7977.388 | -7978.062 | -0.673 | 1.97E-06 |
| 11 | 117773027 | rs117010140 | A | G | 1.023 | -7977.388 | -7978.061 | -0.673 | 1.97E-06 |
| 11 | 118852641 | rs12273687 | T | G | 1.047 | -7977.388 | -7978.062 | -0.673 | 1.97E-06 |
| 11 | 119875786 | rs1944499 | A | G | 1.051 | -7977.388 | -7978.061 | -0.673 | 1.97E-06 |
| 11 | 120603160 | rs79540430 | T | A | 0.94 | -7977.388 | -7978.063 | -0.675 | 1.97E-06 |
| 11 | 120649337 | rs117414016 | T | G | 1.055 | -7977.388 | -7978.063 | -0.675 | 1.97E-06 |
| 11 | 117330944 | rs75813208 | T | C | 0.953 | -7977.388 | -7978.065 | -0.676 | 1.97E-06 |
| 11 | 117638305 | rs61903859 | A | C | 1.062 | -7977.388 | -7978.065 | -0.676 | 1.97E-06 |
| 11 | 119944195 | rs661621 | T | A | 0.941 | -7977.388 | -7978.064 | -0.676 | 1.97E-06 |
| 11 | 120992119 | rs11218148 | A | C | 0.95 | -7977.388 | -7978.065 | -0.676 | 1.97E-06 |
| 11 | 120992859 | rs4936576 | C | G | 0.95 | -7977.388 | -7978.065 | -0.676 | 1.97E-06 |
| 11 | 116591934 | rs180372 | A | G | 0.955 | -7977.388 | -7978.065 | -0.677 | 1.97E-06 |
| 11 | 117326419 | rs118100246 | A | G | 0.991 | -7977.388 | -7978.065 | -0.677 | 1.97E-06 |
| 11 | 117328052 | rs148406383 | T | C | 0.991 | -7977.388 | -7978.065 | -0.677 | 1.97E-06 |
| 11 | 117965421 | rs45499491 | T | C | 1.059 | -7977.388 | -7978.066 | -0.677 | 1.97E-06 |
| 11 | 118889810 | rs3802884 | T | C | 0.942 | -7977.388 | -7978.066 | -0.677 | 1.97E-06 |
| 11 | 121108907 | rs1941299 | C | A | 0.949 | -7977.388 | -7978.065 | -0.677 | 1.97E-06 |
| 11 | 116466714 | rs12292959 | C | T | 0.943 | -7977.388 | -7978.066 | -0.678 | 1.96E-06 |
| 11 | 117330326 | rs145305155 | G | A | 1.001 | -7977.388 | -7978.066 | -0.678 | 1.96E-06 |
| 11 | 117646805 | rs11819854 | T | A | 1.062 | -7977.388 | -7978.066 | -0.678 | 1.96E-06 |
| 11 | 121103704 | rs10892692 | G | A | 0.949 | -7977.388 | -7978.066 | -0.678 | 1.96E-06 |
| 11 | 117392940 | rs7941258 | T | C | 0.953 | -7977.388 | -7978.068 | -0.679 | 1.96E-06 |
| 11 | 117392987 | rs7941460 | A | G | 0.953 | -7977.388 | -7978.068 | -0.679 | 1.96E-06 |
| 11 | 117393345 | rs7944704 | A | G | 0.953 | -7977.388 | -7978.068 | -0.679 | 1.96E-06 |
| 11 | 118853774 | rs11217109 | C | T | 1.055 | -7977.388 | -7978.067 | -0.679 | 1.96E-06 |
| 11 | 119466907 | rs150628867 | A | G | 1.036 | -7977.388 | -7978.067 | -0.679 | 1.96E-06 |
| 11 | 119684904 | rs79297443 | T | C | 0.94 | -7977.388 | -7978.067 | -0.679 | 1.96E-06 |
| 11 | 119889758 | rs1170855838 | T | C | 0.945 | -7977.388 | -7978.068 | -0.68 | 1.96E-06 |
| 11 | 117821885 | rs144891162 | T | C | 1.046 | -7977.388 | -7978.069 | -0.681 | 1.96E-06 |
| 11 | 117906759 | rs649827 | T | C | 1.047 | -7977.388 | -7978.069 | -0.681 | 1.96E-06 |
| 11 | 118314849 | rs2017122 | T | C | 1.063 | -7977.388 | -7978.069 | -0.681 | 1.96E-06 |
| 11 | 120576154 | rs11217984 | T | C | 0.946 | -7977.388 | -7978.069 | -0.681 | 1.96E-06 |
| 11 | 116737302 | rs139447106 | T | C | 1.062 | -7977.388 | -7978.07 | -0.682 | 1.96E-06 |
| 11 | 117094591 | rs17120523 | G | A | 1.063 | -7977.388 | -7978.071 | -0.682 | 1.96E-06 |
| 11 | 120860524 | rs34259384 | T | G | 0.947 | -7977.388 | -7978.071 | -0.682 | 1.96E-06 |
| 11 | 116353269 | rs118065597 | A | G | 0.964 | -7977.388 | -7978.072 | -0.683 | 1.95E-06 |
| 11 | 116741840 | rs73023041 | A | G | 0.94 | -7977.388 | -7978.071 | -0.683 | 1.95E-06 |
| 11 | 117630622 | rs78080546 | T | A | 0.942 | -7977.388 | -7978.071 | -0.683 | 1.95E-06 |
| 11 | 119609828 | rs113950321 | C | A | 0.948 | -7977.388 | -7978.072 | -0.683 | 1.95E-06 |
| 11 | 120009748 | rs61900472 | A | G | 0.959 | -7977.388 | -7978.071 | -0.683 | 1.95E-06 |
| 11 | 120623637 | rs11217999 | A | C | 0.941 | -7977.388 | -7978.072 | -0.683 | 1.95E-06 |
| 11 | 120871324 | rs11218103 | T | C | 1.062 | -7977.388 | -7978.071 | -0.683 | 1.95E-06 |
| 11 | 116677370 | rs1729407 | C | G | 1.047 | -7977.388 | -7978.073 | -0.684 | 1.95E-06 |
| 11 | 121004851 | rs7125224 | A | G | 0.95 | -7977.388 | -7978.072 | -0.684 | 1.95E-06 |
| 11 | 121160481 | rs10750184 | A | G | 1.059 | -7977.388 | -7978.072 | -0.684 | 1.95E-06 |
| 11 | 118509877 | rs112552940 | T | C | 1.06 | -7977.388 | -7978.073 | -0.685 | 1.95E-06 |
| 11 | 118824996 | rs4426142 | G | A | 0.955 | -7977.388 | -7978.073 | -0.685 | 1.95E-06 |
| 11 | 118853021 | rs113962716 | A | C | 1.055 | -7977.388 | -7978.074 | -0.685 | 1.95E-06 |
| 11 | 119863584 | rs1944511 | A | G | 1.051 | -7977.388 | -7978.073 | -0.685 | 1.95E-06 |
| 11 | 120475835 | rs11217924 | T | C | 1.049 | -7977.388 | -7978.073 | -0.685 | 1.95E-06 |
| 11 | 120715952 | rs7123228 | G | A | 0.948 | -7977.388 | -7978.074 | -0.685 | 1.95E-06 |
| 11 | 117317855 | rs117580094 | G | T | 1.036 | -7977.388 | -7978.074 | -0.686 | 1.95E-06 |
| 11 | 117539715 | rs636783 | C | A | 1.058 | -7977.388 | -7978.074 | -0.686 | 1.95E-06 |
| 11 | 118530068 | rs112337443 | C | T | 1.06 | -7977.388 | -7978.074 | -0.686 | 1.95E-06 |
| 11 | 119628238 | rs10892449 | G | C | 0.944 | -7977.388 | -7978.075 | -0.687 | 1.95E-06 |
| 11 | 117464025 | rs12576226 | G | A | 0.955 | -7977.388 | -7978.077 | -0.688 | 1.94E-06 |
| 11 | 117619538 | rs7924968 | A | T | 0.941 | -7977.388 | -7978.077 | -0.688 | 1.94E-06 |
| 11 | 117674072 | rs1410186916 | T | A | 0.956 | -7977.388 | -7978.077 | -0.688 | 1.94E-06 |
| 11 | 117683284 | rs112776148 | C | G | 0.944 | -7977.388 | -7978.077 | -0.688 | 1.94E-06 |
| 11 | 120986075 | rs80282202 | C | G | 0.942 | -7977.388 | -7978.076 | -0.688 | 1.94E-06 |
| 11 | 117466755 | rs78144220 | T | G | 0.941 | -7977.388 | -7978.078 | -0.689 | 1.94E-06 |
| 11 | 117665267 | rs71480357 | G | C | 1.04 | -7977.388 | -7978.078 | -0.69 | 1.94E-06 |
| 11 | 119612600 | rs113816823 | G | A | 0.949 | -7977.388 | -7978.078 | -0.69 | 1.94E-06 |
| 11 | 119849683 | rs76547700 | T | G | 1.061 | -7977.388 | -7978.079 | -0.69 | 1.94E-06 |
| 11 | 120206746 | rs35734851 | A | G | 0.942 | -7977.388 | -7978.078 | -0.69 | 1.94E-06 |
| 11 | 121049136 | rs592160 | T | C | 0.954 | -7977.388 | -7978.078 | -0.69 | 1.94E-06 |
| 11 | 116549408 | rs74792494 | C | A | 0.953 | -7977.388 | -7978.079 | -0.691 | 1.94E-06 |
| 11 | 117366111 | rs10892129 | G | C | 0.941 | -7977.388 | -7978.079 | -0.691 | 1.94E-06 |
| 11 | 117680478 | rs74500359 | T | C | 0.941 | -7977.388 | -7978.08 | -0.691 | 1.94E-06 |
| 11 | 118863116 | rs4938613 | G | A | 1.047 | -7977.388 | -7978.079 | -0.691 | 1.94E-06 |
| 11 | 118863324 | rs4936456 | T | C | 1.047 | -7977.388 | -7978.079 | -0.691 | 1.94E-06 |
| 11 | 119497371 | rs145924639 | T | C | 0.964 | -7977.388 | -7978.08 | -0.691 | 1.94E-06 |
| 11 | 119837947 | rs61899043 | C | A | 0.955 | -7977.388 | -7978.079 | -0.691 | 1.94E-06 |
| 11 | 119981694 | rs140158503 | T | C | 0.97 | -7977.388 | -7978.079 | -0.691 | 1.94E-06 |
| 11 | 120622042 | rs150032046 | G | A | 0.983 | -7977.388 | -7978.079 | -0.691 | 1.94E-06 |
| 11 | 116470552 | rs35820582 | C | G | 0.946 | -7977.388 | -7978.08 | -0.692 | 1.94E-06 |
| 11 | 116544015 | rs74724948 | T | A | 0.953 | -7977.388 | -7978.08 | -0.692 | 1.94E-06 |

| | | | | | | | |
|---|---|---|---|---|---|---|---|
| 11 | 119642855 | rs61897696 | A | G | NULL | 0.584190523 | NULL&M_Disease |
| 11 | 120904156 | rs2000527 | G | A | NULL | 0.585404569 | NULL&M_Disease |
| 11 | 116528249 | rs492257 | A | G | NULL | 0.585404569 | NULL&M_Disease |
| 11 | 116591040 | rs180375 | T | C | NULL | 0.585404569 | NULL&M_Disease |
| 11 | 117171413 | rs78751065 | T | C | NULL | 0.584190523 | NULL&M_Disease |
| 11 | 117685169 | rs11216559 | C | T | NULL | 0.585404569 | NULL&M_Disease |
| 11 | 117895854 | rs111226844 | C | T | NULL | 0.585404569 | NULL&M_Disease |
| 11 | 119499316 | rs78201038 | G | C | NULL | 0.585404569 | NULL&M_Disease |
| 11 | 119579260 | rs4245203 | C | T | NULL | 0.584190523 | NULL&M_Disease |
| 11 | 119587095 | rs12416901 | C | T | NULL | 0.584190523 | NULL&M_Disease |
| 11 | 120697844 | rs77223730 | A | G | NULL | 0.585404569 | NULL&M_Disease |
| 11 | 120697845 | rs78653140 | T | C | NULL | 0.585404569 | NULL&M_Disease |
| 11 | 117926735 | rs628269 | G | A | NULL | 0.585404569 | NULL&M_Disease |
| 11 | 120074926 | rs187236392 | G | A | NULL | 0.584190523 | NULL&M_Disease |
| 11 | 120873529 | rs11218106 | A | T | NULL | 0.585404569 | NULL&M_Disease |
| 11 | 116573120 | rs879858 | A | G | NULL | 0.584190523 | NULL&M_Disease |
| 11 | 117089628 | rs236916 | A | G | NULL | 0.585404569 | NULL&M_Disease |
| 11 | 118838020 | rs7117266 | A | G | NULL | 0.585404569 | NULL&M_Disease |
| 11 | 120580148 | rs61901381 | C | T | NULL | 0.585404569 | NULL&M_Disease |
| 11 | 117908235 | rs686047 | A | G | NULL | 0.585404569 | NULL&M_Disease |
| 11 | 116678760 | rs1729405 | C | T | NULL | 0.585404569 | NULL&M_Disease |
| 11 | 116782202 | rs117050823 | G | A | NULL | 0.585404569 | NULL&M_Disease |
| 11 | 117196901 | rs2168673 | A | C | NULL | 0.585404569 | NULL&M_Disease |
| 11 | 119607719 | rs11600193 | C | T | NULL | 0.585404569 | NULL&M_Disease |
| 11 | 120468752 | rs7105963 | T | C | NULL | 0.585404569 | NULL&M_Disease |
| 11 | 120822001 | rs11218077 | C | T | NULL | 0.585404569 | NULL&M_Disease |
| 11 | 121050748 | rs640332 | C | T | NULL | 0.585404569 | NULL&M_Disease |
| 11 | 121050913 | rs620896 | G | A | NULL | 0.585404569 | NULL&M_Disease |
| 11 | 116701850 | rs5142 | T | C | NULL | 0.585404569 | NULL&M_Disease |
| 11 | 116702123 | rs5141 | T | C | NULL | 0.585404569 | NULL&M_Disease |
| 11 | 117711232 | rs496164 | A | G | NULL | 0.586617579 | NULL&M_Disease |
| 11 | 120437187 | rs36111344 | G | A | NULL | 0.586617579 | NULL&M_Disease |
| 11 | 120437572 | rs73000621 | T | C | NULL | 0.586617579 | NULL&M_Disease |
| 11 | 120851210 | rs35176806 | G | C | NULL | 0.585404569 | NULL&M_Disease |
| 11 | 117907041 | rs650908 | C | G | NULL | 0.586617579 | NULL&M_Disease |
| 11 | 119625373 | rs79300298 | C | G | NULL | 0.586617579 | NULL&M_Disease |
| 11 | 120623361 | rs73576552 | T | C | NULL | 0.585404569 | NULL&M_Disease |
| 11 | 116821525 | rs77586292 | T | C | NULL | 0.585404569 | NULL&M_Disease |
| 11 | 117382799 | rs117604511 | C | A | NULL | 0.585404569 | NULL&M_Disease |
| 11 | 117681924 | rs1209756219 | C | T | NULL | 0.586617579 | NULL&M_Disease |
| 11 | 119577677 | rs78058743 | T | C | NULL | 0.585404569 | NULL&M_Disease |
| 11 | 120828210 | rs17124590 | T | A | NULL | 0.585404569 | NULL&M_Disease |
| 11 | 116465292 | rs10891998 | G | A | NULL | 0.586617579 | NULL&M_Disease |
| 11 | 116524832 | rs11216075 | A | G | NULL | 0.585404569 | NULL&M_Disease |
| 11 | 116620256 | rs11216129 | A | C | NULL | 0.585404569 | NULL&M_Disease |
| 11 | 120429035 | rs144018061 | A | G | NULL | 0.585404569 | NULL&M_Disease |
| 11 | 120708132 | rs11600004 | T | C | NULL | 0.586617579 | NULL&M_Disease |
| 11 | 116519181 | rs76234156 | A | C | NULL | 0.585404569 | NULL&M_Disease |
| 11 | 117617127 | rs60816279 | A | G | NULL | 0.585404569 | NULL&M_Disease |
| 11 | 120697913 | rs113067228 | C | T | NULL | 0.586617579 | NULL&M_Disease |
| 11 | 119635453 | rs7940298 | T | C | NULL | 0.586617579 | NULL&M_Disease |
| 11 | 120200571 | rs74883244 | G | A | NULL | 0.585404569 | NULL&M_Disease |
| 11 | 121030381 | rs677336 | T | A | NULL | 0.586617579 | NULL&M_Disease |
| 11 | 121031511 | rs552580 | G | A | NULL | 0.586617579 | NULL&M_Disease |
| 11 | 118031950 | rs17530451 | A | C | NULL | 0.586617579 | NULL&M_Disease |
| 11 | 118839032 | rs67389832 | A | C | NULL | 0.586617579 | NULL&M_Disease |
| 11 | 119717705 | rs73008656 | G | T | NULL | 0.585404569 | NULL&M_Disease |
| 11 | 119924214 | rs652376 | C | G | NULL | 0.586617579 | NULL&M_Disease |
| 11 | 119924436 | rs630842 | A | G | NULL | 0.586617579 | NULL&M_Disease |
| 11 | 119949927 | rs648067 | A | C | NULL | 0.586617579 | NULL&M_Disease |
| 11 | 120353153 | rs61898767 | G | A | NULL | 0.586617579 | NULL&M_Disease |
| 11 | 117092660 | rs45462696 | A | G | NULL | 0.585404569 | NULL&M_Disease |
| 11 | 116355788 | rs138697506 | G | A | NULL | 0.586617579 | NULL&M_Disease |
| 11 | 116950150 | rs11216266 | C | T | NULL | 0.586617579 | NULL&M_Disease |
| 11 | 117685824 | rs17666283 | A | G | NULL | 0.586617579 | NULL&M_Disease |
| 11 | 117686732 | rs11216562 | T | C | NULL | 0.586617579 | NULL&M_Disease |
| 11 | 119948738 | rs1287581 | T | G | NULL | 0.586617579 | NULL&M_Disease |
| 11 | 120159612 | rs79219038 | T | C | NULL | 0.585404569 | NULL&M_Disease |
| 11 | 117391522 | rs73010304 | A | G | NULL | 0.586617579 | NULL&M_Disease |
| 11 | 119898644 | rs141642186 | T | C | NULL | 0.586617579 | NULL&M_Disease |
| 11 | 117617219 | rs59810418 | C | G | NULL | 0.586617579 | NULL&M_Disease |
| 11 | 118614397 | rs10892281 | T | C | NULL | 0.587829538 | NULL&M_Disease |
| 11 | 118866126 | rs7126193 | C | T | NULL | 0.587829538 | NULL&M_Disease |
| 11 | 120696085 | rs75622430 | T | G | NULL | 0.586617579 | NULL&M_Disease |
| 11 | 120696771 | rs111932453 | T | C | NULL | 0.586617579 | NULL&M_Disease |
| 11 | 120697009 | rs79304595 | G | C | NULL | 0.586617579 | NULL&M_Disease |
| 11 | 118341785 | rs535327 | G | A | NULL | 0.586617579 | NULL&M_Disease |
| 11 | 116312961 | rs140140760 | A | G | NULL | 0.586617579 | NULL&M_Disease |
| 11 | 116530209 | rs552657 | T | C | NULL | 0.587829538 | NULL&M_Disease |
| 11 | 119435096 | rs56189546 | G | T | NULL | 0.587829538 | NULL&M_Disease |
| 11 | 121039947 | rs562281 | T | A | NULL | 0.587829538 | NULL&M_Disease |
| 11 | 118017538 | rs117181393 | C | T | NULL | 0.586617579 | NULL&M_Disease |
| 11 | 118136362 | rs74649894 | G | C | NULL | 0.586617579 | NULL&M_Disease |
| 11 | 121030195 | rs688641 | T | G | NULL | 0.587829538 | NULL&M_Disease |
| 11 | 116471492 | rs12807148 | C | T | NULL | 0.587829538 | NULL&M_Disease |
| 11 | 116943544 | rs7109649 | T | C | NULL | 0.587829538 | NULL&M_Disease |
| 11 | 116946238 | rs12274388 | T | C | NULL | 0.587829538 | NULL&M_Disease |
| 11 | 117628287 | rs7114641 | G | A | NULL | 0.587829538 | NULL&M_Disease |
| 11 | 121033105 | rs662496 | G | A | NULL | 0.587829538 | NULL&M_Disease |
| 11 | 117610854 | rs1942931 | A | G | NULL | 0.587829538 | NULL&M_Disease |
| 11 | 119381144 | rs144553689 | C | T | NULL | 0.586617579 | NULL&M_Disease |
| 11 | 119858685 | rs117114471 | A | G | NULL | 0.586617579 | NULL&M_Disease |
| 11 | 121035186 | rs584355 | A | C | NULL | 0.587829538 | NULL&M_Disease |
| 11 | 117562505 | rs11216504 | T | C | NULL | 0.587829538 | NULL&M_Disease |
| 11 | 118290459 | rs148838027 | C | G | NULL | 0.586617579 | NULL&M_Disease |
| 11 | 120139487 | rs10892558 | A | G | NULL | 0.587829538 | NULL&M_Disease |

| | | | | | | | | | |
|---|---|---|---|---|---|---|---|---|---|
| 11 | 119642855 | rs61897696 | A | G | 1.024 | -7977.388 | -7978.081 | -0.692 | 1.94E-06 |
| 11 | 120904156 | rs2000527 | G | A | 1.052 | -7977.388 | -7978.08 | -0.692 | 1.94E-06 |
| 11 | 116528249 | rs492257 | A | G | 0.954 | -7977.388 | -7978.082 | -0.693 | 1.93E-06 |
| 11 | 116591040 | rs180375 | T | C | 0.955 | -7977.388 | -7978.082 | -0.693 | 1.93E-06 |
| 11 | 117171413 | rs78751065 | T | C | 0.994 | -7977.388 | -7978.081 | -0.693 | 1.93E-06 |
| 11 | 117685169 | rs11216559 | C | T | 0.944 | -7977.388 | -7978.081 | -0.693 | 1.93E-06 |
| 11 | 117895854 | rs111226844 | C | T | 0.944 | -7977.388 | -7978.081 | -0.693 | 1.93E-06 |
| 11 | 119499316 | rs78201038 | G | C | 1.058 | -7977.388 | -7978.081 | -0.693 | 1.93E-06 |
| 11 | 119579260 | rs4245203 | C | T | 0.949 | -7977.388 | -7978.081 | -0.693 | 1.93E-06 |
| 11 | 119587095 | rs12416901 | C | T | 0.949 | -7977.388 | -7978.081 | -0.693 | 1.93E-06 |
| 11 | 120697844 | rs77223730 | A | G | 0.941 | -7977.388 | -7978.081 | -0.693 | 1.93E-06 |
| 11 | 120697845 | rs78653140 | T | C | 0.941 | -7977.388 | -7978.081 | -0.693 | 1.93E-06 |
| 11 | 117926735 | rs628269 | G | A | 1.059 | -7977.388 | -7978.082 | -0.694 | 1.93E-06 |
| 11 | 120074926 | rs187236392 | G | A | 0.954 | -7977.388 | -7978.082 | -0.694 | 1.93E-06 |
| 11 | 120873529 | rs11218106 | A | T | 1.062 | -7977.388 | -7978.082 | -0.694 | 1.93E-06 |
| 11 | 116573120 | rs879858 | A | G | 1.026 | -7977.388 | -7978.083 | -0.695 | 1.93E-06 |
| 11 | 117089628 | rs236916 | A | G | 0.946 | -7977.388 | -7978.083 | -0.695 | 1.93E-06 |
| 11 | 118838020 | rs7117266 | A | G | 1.053 | -7977.388 | -7978.083 | -0.695 | 1.93E-06 |
| 11 | 120580148 | rs61901381 | C | T | 0.941 | -7977.388 | -7978.083 | -0.695 | 1.93E-06 |
| 11 | 117908235 | rs686047 | A | G | 1.047 | -7977.388 | -7978.084 | -0.696 | 1.93E-06 |
| 11 | 116678760 | rs1729405 | C | T | 1.047 | -7977.388 | -7978.086 | -0.697 | 1.93E-06 |
| 11 | 116782202 | rs117050823 | G | A | 0.946 | -7977.388 | -7978.085 | -0.697 | 1.93E-06 |
| 11 | 117196901 | rs2168673 | A | C | 0.942 | -7977.388 | -7978.085 | -0.697 | 1.93E-06 |
| 11 | 119607719 | rs11600193 | C | T | 0.95 | -7977.388 | -7978.086 | -0.697 | 1.93E-06 |
| 11 | 120468752 | rs7105963 | T | C | 1.048 | -7977.388 | -7978.085 | -0.697 | 1.93E-06 |
| 11 | 120822001 | rs11218077 | C | T | 0.95 | -7977.388 | -7978.086 | -0.697 | 1.93E-06 |
| 11 | 121050748 | rs640332 | C | T | 0.954 | -7977.388 | -7978.085 | -0.697 | 1.93E-06 |
| 11 | 121050913 | rs620896 | G | A | 0.954 | -7977.388 | -7978.085 | -0.697 | 1.93E-06 |
| 11 | 116701850 | rs5142 | T | C | 0.943 | -7977.388 | -7978.087 | -0.698 | 1.92E-06 |
| 11 | 116702123 | rs5141 | T | C | 0.943 | -7977.388 | -7978.086 | -0.698 | 1.92E-06 |
| 11 | 117711232 | rs496164 | A | G | 1.046 | -7977.388 | -7978.086 | -0.698 | 1.92E-06 |
| 11 | 120437187 | rs36111344 | G | A | 1.047 | -7977.388 | -7978.086 | -0.698 | 1.92E-06 |
| 11 | 120437572 | rs73000621 | T | C | 1.047 | -7977.388 | -7978.086 | -0.698 | 1.92E-06 |
| 11 | 120851210 | rs35176806 | G | C | 1.021 | -7977.388 | -7978.086 | -0.698 | 1.92E-06 |
| 11 | 117907041 | rs650908 | C | G | 1.047 | -7977.388 | -7978.088 | -0.699 | 1.92E-06 |
| 11 | 119625373 | rs79300298 | C | G | 0.944 | -7977.388 | -7978.087 | -0.699 | 1.92E-06 |
| 11 | 120623361 | rs73576552 | T | C | 0.942 | -7977.388 | -7978.087 | -0.699 | 1.92E-06 |
| 11 | 116821525 | rs77586292 | T | C | 1.061 | -7977.388 | -7978.088 | -0.7 | 1.92E-06 |
| 11 | 117382799 | rs117604511 | C | A | 1.043 | -7977.388 | -7978.089 | -0.7 | 1.92E-06 |
| 11 | 117681924 | rs1209756219 | C | T | 0.944 | -7977.388 | -7978.088 | -0.7 | 1.92E-06 |
| 11 | 119577677 | rs78058743 | T | C | 1.025 | -7977.388 | -7978.088 | -0.7 | 1.92E-06 |
| 11 | 120828210 | rs17124590 | T | A | 0.967 | -7977.388 | -7978.088 | -0.7 | 1.92E-06 |
| 11 | 116465292 | rs10891998 | G | A | 1.053 | -7977.388 | -7978.089 | -0.701 | 1.92E-06 |
| 11 | 116524832 | rs11216075 | A | G | 1.011 | -7977.388 | -7978.089 | -0.701 | 1.92E-06 |
| 11 | 116620256 | rs11216129 | A | C | 1.057 | -7977.388 | -7978.089 | -0.701 | 1.92E-06 |
| 11 | 120429035 | rs144018061 | A | G | 0.996 | -7977.388 | -7978.09 | -0.701 | 1.92E-06 |
| 11 | 120708132 | rs11600004 | T | C | 0.949 | -7977.388 | -7978.09 | -0.701 | 1.92E-06 |
| 11 | 116519181 | rs76234156 | A | C | 1.009 | -7977.388 | -7978.091 | -0.702 | 1.92E-06 |
| 11 | 117617127 | rs60816279 | A | G | 0.948 | -7977.388 | -7978.09 | -0.702 | 1.92E-06 |
| 11 | 120697913 | rs113067228 | C | T | 0.941 | -7977.388 | -7978.09 | -0.702 | 1.92E-06 |
| 11 | 119635453 | rs7940298 | T | C | 0.945 | -7977.388 | -7978.092 | -0.703 | 1.92E-06 |
| 11 | 120200571 | rs74883244 | G | A | 1.033 | -7977.388 | -7978.091 | -0.703 | 1.92E-06 |
| 11 | 121030381 | rs677336 | T | A | 0.954 | -7977.388 | -7978.092 | -0.703 | 1.92E-06 |
| 11 | 121031511 | rs552580 | G | A | 0.954 | -7977.388 | -7978.092 | -0.703 | 1.92E-06 |
| 11 | 118031950 | rs17530451 | A | C | 1.062 | -7977.388 | -7978.092 | -0.704 | 1.91E-06 |
| 11 | 118839032 | rs67389832 | A | C | 1.053 | -7977.388 | -7978.092 | -0.704 | 1.91E-06 |
| 11 | 119717705 | rs73008656 | G | T | 0.969 | -7977.388 | -7978.092 | -0.704 | 1.91E-06 |
| 11 | 119924214 | rs652376 | C | G | 0.95 | -7977.388 | -7978.092 | -0.704 | 1.91E-06 |
| 11 | 119924436 | rs630842 | A | G | 0.95 | -7977.388 | -7978.092 | -0.704 | 1.91E-06 |
| 11 | 119949927 | rs648067 | A | C | 0.943 | -7977.388 | -7978.092 | -0.704 | 1.91E-06 |
| 11 | 120353153 | rs61898767 | G | A | 1.053 | -7977.388 | -7978.092 | -0.704 | 1.91E-06 |
| 11 | 117092660 | rs45462696 | A | G | 0.978 | -7977.388 | -7978.093 | -0.705 | 1.91E-06 |
| 11 | 116355788 | rs138697506 | G | A | 0.965 | -7977.388 | -7978.095 | -0.706 | 1.91E-06 |
| 11 | 116950150 | rs11216266 | C | T | 1.051 | -7977.388 | -7978.094 | -0.706 | 1.91E-06 |
| 11 | 117685824 | rs17666283 | A | G | 0.944 | -7977.388 | -7978.094 | -0.706 | 1.91E-06 |
| 11 | 117686732 | rs11216562 | T | C | 0.967 | -7977.388 | -7978.095 | -0.706 | 1.91E-06 |
| 11 | 119948738 | rs1287581 | T | G | 0.943 | -7977.388 | -7978.094 | -0.706 | 1.91E-06 |
| 11 | 120159612 | rs79219038 | T | C | 1.019 | -7977.388 | -7978.094 | -0.706 | 1.91E-06 |
| 11 | 117391522 | rs73010304 | A | G | 1.054 | -7977.388 | -7978.095 | -0.707 | 1.91E-06 |
| 11 | 119898644 | rs141642186 | T | C | 0.964 | -7977.388 | -7978.096 | -0.707 | 1.91E-06 |
| 11 | 117617219 | rs59810418 | C | G | 0.949 | -7977.388 | -7978.096 | -0.708 | 1.91E-06 |
| 11 | 118614397 | rs10892281 | T | C | 0.946 | -7977.388 | -7978.097 | -0.708 | 1.91E-06 |
| 11 | 118866126 | rs7126193 | C | T | 1.047 | -7977.388 | -7978.097 | -0.708 | 1.91E-06 |
| 11 | 120696085 | rs75622430 | T | G | 0.942 | -7977.388 | -7978.096 | -0.708 | 1.91E-06 |
| 11 | 120696771 | rs111932453 | T | C | 0.942 | -7977.388 | -7978.096 | -0.708 | 1.91E-06 |
| 11 | 120697009 | rs79304595 | G | C | 0.942 | -7977.388 | -7978.096 | -0.708 | 1.91E-06 |
| 11 | 118341785 | rs535327 | G | A | 0.945 | -7977.388 | -7978.097 | -0.709 | 1.90E-06 |
| 11 | 116312961 | rs140140760 | A | G | 0.99 | -7977.388 | -7978.098 | -0.71 | 1.90E-06 |
| 11 | 116530209 | rs552657 | T | C | 0.955 | -7977.388 | -7978.098 | -0.71 | 1.90E-06 |
| 11 | 119435096 | rs56189546 | G | T | 1.061 | -7977.388 | -7978.098 | -0.71 | 1.90E-06 |
| 11 | 121039947 | rs562281 | T | A | 0.955 | -7977.388 | -7978.099 | -0.71 | 1.90E-06 |
| 11 | 118017538 | rs117181393 | C | T | 0.946 | -7977.388 | -7978.099 | -0.711 | 1.90E-06 |
| 11 | 118136362 | rs74649894 | G | C | 0.971 | -7977.388 | -7978.099 | -0.711 | 1.90E-06 |
| 11 | 121030195 | rs688641 | T | G | 0.955 | -7977.388 | -7978.099 | -0.711 | 1.90E-06 |
| 11 | 116471492 | rs12807148 | C | T | 0.942 | -7977.388 | -7978.101 | -0.712 | 1.90E-06 |
| 11 | 116943544 | rs7109649 | T | C | 1.051 | -7977.388 | -7978.101 | -0.712 | 1.90E-06 |
| 11 | 116946238 | rs12274388 | T | C | 1.051 | -7977.388 | -7978.1 | -0.712 | 1.90E-06 |
| 11 | 117628287 | rs7114641 | G | A | 1.061 | -7977.388 | -7978.1 | -0.712 | 1.90E-06 |
| 11 | 121033105 | rs662496 | G | A | 0.955 | -7977.388 | -7978.1 | -0.712 | 1.90E-06 |
| 11 | 117610854 | rs1942931 | A | G | 0.949 | -7977.388 | -7978.101 | -0.713 | 1.90E-06 |
| 11 | 119381144 | rs144553689 | C | T | 1.021 | -7977.388 | -7978.102 | -0.713 | 1.90E-06 |
| 11 | 119858685 | rs117114471 | A | G | 0.974 | -7977.388 | -7978.101 | -0.713 | 1.90E-06 |
| 11 | 121035186 | rs584355 | A | C | 0.955 | -7977.388 | -7978.101 | -0.713 | 1.90E-06 |
| 11 | 117562505 | rs11216504 | T | C | 0.943 | -7977.388 | -7978.102 | -0.713 | 1.89E-06 |
| 11 | 118290459 | rs148838027 | C | G | 1.025 | -7977.388 | -7978.102 | -0.714 | 1.89E-06 |
| 11 | 120139487 | rs10892558 | A | G | 1.058 | -7977.388 | -7978.102 | -0.714 | 1.89E-06 |

| | | | | | | | |
|---|---|---|---|---|---|---|---|
| 11 | 120468093 | rs75524659 | T | C | NULL | 0.587829538 | NULL&M_Disease |
| 11 | 116530373 | rs552787 | C | A | NULL | 0.587829538 | NULL&M_Disease |
| 11 | 117461602 | rs553637 | C | T | NULL | 0.587829538 | NULL&M_Disease |
| 11 | 119630763 | rs11217472 | C | T | NULL | 0.587829538 | NULL&M_Disease |
| 11 | 119911542 | rs1452842 | G | A | NULL | 0.587829538 | NULL&M_Disease |
| 11 | 119930569 | rs1790469 | A | G | NULL | 0.587829538 | NULL&M_Disease |
| 11 | 116486588 | rs3016346 | A | G | NULL | 0.587829538 | NULL&M_Disease |
| 11 | 118041972 | rs12294669 | G | A | NULL | 0.587829538 | NULL&M_Disease |
| 11 | 121013909 | rs588917 | T | G | NULL | 0.587829538 | NULL&M_Disease |
| 11 | 117190901 | rs117469840 | T | C | NULL | 0.587829538 | NULL&M_Disease |
| 11 | 117711562 | rs519991 | C | T | NULL | 0.587829538 | NULL&M_Disease |
| 11 | 118341921 | rs524936 | A | G | NULL | 0.587829538 | NULL&M_Disease |
| 11 | 118477226 | rs45496391 | C | A | NULL | 0.587829538 | NULL&M_Disease |
| 11 | 116899248 | rs11216240 | T | A | NULL | 0.589040434 | NULL&M_Disease |
| 11 | 116899249 | rs11216241 | C | A | NULL | 0.589040434 | NULL&M_Disease |
| 11 | 117684522 | rs12288877 | C | G | NULL | 0.587829538 | NULL&M_Disease |
| 11 | 118825069 | rs2097131 | G | A | NULL | 0.589040434 | NULL&M_Disease |
| 11 | 120730621 | rs4526780 | A | G | NULL | 0.589040434 | NULL&M_Disease |
| 11 | 116920927 | rs10892066 | T | C | NULL | 0.589040434 | NULL&M_Disease |
| 11 | 116943952 | rs1531706 | A | C | NULL | 0.589040434 | NULL&M_Disease |
| 11 | 116944348 | rs7935913 | A | G | NULL | 0.589040434 | NULL&M_Disease |
| 11 | 116944712 | rs1531707 | A | G | NULL | 0.589040434 | NULL&M_Disease |
| 11 | 117641216 | rs56112767 | G | C | NULL | 0.589040434 | NULL&M_Disease |
| 11 | 117752769 | rs77752432 | T | C | NULL | 0.587829538 | NULL&M_Disease |
| 11 | 118474283 | rs139202987 | G | A | NULL | 0.587829538 | NULL&M_Disease |
| 11 | 120808303 | rs71486349 | T | G | NULL | 0.587829538 | NULL&M_Disease |
| 11 | 116660813 | rs619054 | A | G | NULL | 0.589040434 | NULL&M_Disease |
| 11 | 117611292 | rs10466534 | A | G | NULL | 0.587829538 | NULL&M_Disease |
| 11 | 117912720 | rs650104 | T | C | NULL | 0.589040434 | NULL&M_Disease |
| 11 | 117912731 | rs689054 | T | A | NULL | 0.589040434 | NULL&M_Disease |
| 11 | 118015832 | rs45539032 | A | G | NULL | 0.587829538 | NULL&M_Disease |
| 11 | 119985011 | rs150954345 | G | A | NULL | 0.587829538 | NULL&M_Disease |
| 11 | 120619202 | rs2852224 | T | C | NULL | 0.587829538 | NULL&M_Disease |
| 11 | 117804926 | rs550388 | C | T | NULL | 0.589040434 | NULL&M_Disease |
| 11 | 119745457 | rs508440 | G | C | NULL | 0.589040434 | NULL&M_Disease |
| 11 | 117590911 | rs948772 | C | G | NULL | 0.589040434 | NULL&M_Disease |
| 11 | 120576090 | rs11217982 | A | T | NULL | 0.589040434 | NULL&M_Disease |
| 11 | 120576091 | rs11217983 | C | T | NULL | 0.589040434 | NULL&M_Disease |
| 11 | 120895352 | rs3824976 | A | G | NULL | 0.589040434 | NULL&M_Disease |
| 11 | 120984681 | rs658494 | T | C | NULL | 0.589040434 | NULL&M_Disease |
| 11 | 117461875 | rs1363950531 | G | T | NULL | 0.589040434 | NULL&M_Disease |
| 11 | 117591292 | rs4936399 | T | C | NULL | 0.589040434 | NULL&M_Disease |
| 11 | 117912538 | rs111234707 | A | G | NULL | 0.587829538 | NULL&M_Disease |
| 11 | 116928112 | rs4938329 | T | C | NULL | 0.589040434 | NULL&M_Disease |
| 11 | 117526440 | rs55728331 | A | G | NULL | 0.589040434 | NULL&M_Disease |
| 11 | 117681047 | rs34836631 | A | G | NULL | 0.589040434 | NULL&M_Disease |
| 11 | 118134785 | rs1624780 | T | G | NULL | 0.589040434 | NULL&M_Disease |
| 11 | 116944437 | rs7935834 | G | C | NULL | 0.589040434 | NULL&M_Disease |
| 11 | 116957189 | rs12274465 | T | C | NULL | 0.589040434 | NULL&M_Disease |
| 11 | 117283939 | rs11822288 | A | G | NULL | 0.589040434 | NULL&M_Disease |
| 11 | 117353003 | rs3741279 | G | A | NULL | 0.589040434 | NULL&M_Disease |
| 11 | 118460957 | rs45600633 | A | G | NULL | 0.589040434 | NULL&M_Disease |
| 11 | 118518021 | rs10736493 | G | T | NULL | 0.589040434 | NULL&M_Disease |
| 11 | 118867695 | rs4938617 | T | C | NULL | 0.589040434 | NULL&M_Disease |
| 11 | 119907974 | rs1939333 | C | T | NULL | 0.590250252 | NULL&M_Disease |
| 11 | 116364066 | rs6589549 | A | G | NULL | 0.590250252 | NULL&M_Disease |
| 11 | 116539518 | rs9681206 | T | C | NULL | 0.590250252 | NULL&M_Disease |
| 11 | 117435148 | rs7940044 | A | G | NULL | 0.590250252 | NULL&M_Disease |
| 11 | 120109766 | rs61898297 | T | C | NULL | 0.589040434 | NULL&M_Disease |
| 11 | 120423569 | rs12278730 | T | C | NULL | 0.590250252 | NULL&M_Disease |
| 11 | 117251143 | rs2305828 | A | G | NULL | 0.590250252 | NULL&M_Disease |
| 11 | 117680407 | rs67877251 | G | A | NULL | 0.589040434 | NULL&M_Disease |
| 11 | 118852647 | rs80126726 | A | G | NULL | 0.590250252 | NULL&M_Disease |
| 11 | 119499255 | rs11217361 | C | G | NULL | 0.590250252 | NULL&M_Disease |
| 11 | 119768716 | rs73010609 | T | C | NULL | 0.590250252 | NULL&M_Disease |
| 11 | 119805057 | rs145637218 | A | G | NULL | 0.589040434 | NULL&M_Disease |
| 11 | 120568566 | rs12270409 | C | T | NULL | 0.590250252 | NULL&M_Disease |
| 11 | 121004151 | rs1893023 | T | C | NULL | 0.590250252 | NULL&M_Disease |
| 11 | 116955049 | rs61903416 | C | T | NULL | 0.590250252 | NULL&M_Disease |
| 11 | 117433686 | rs755791 | C | A | NULL | 0.590250252 | NULL&M_Disease |
| 11 | 118561859 | rs10790258 | T | G | NULL | 0.589040434 | NULL&M_Disease |
| 11 | 117194929 | rs554233 | G | A | NULL | 0.590250252 | NULL&M_Disease |
| 11 | 117966178 | rs948465 | G | A | NULL | 0.590250252 | NULL&M_Disease |
| 11 | 117966185 | rs948464 | G | T | NULL | 0.590250252 | NULL&M_Disease |
| 11 | 119676471 | rs10892468 | T | G | NULL | 0.590250252 | NULL&M_Disease |
| 11 | 119820957 | rs144502429 | T | C | NULL | 0.589040434 | NULL&M_Disease |
| 11 | 116913909 | rs12279066 | G | C | NULL | 0.590250252 | NULL&M_Disease |
| 11 | 118295001 | rs139060345 | C | T | NULL | 0.589040434 | NULL&M_Disease |
| 11 | 119913037 | rs659127 | A | C | NULL | 0.590250252 | NULL&M_Disease |
| 11 | 121014403 | rs492294 | G | A | NULL | 0.590250252 | NULL&M_Disease |
| 11 | 116409130 | rs56338557 | C | T | NULL | 0.590250252 | NULL&M_Disease |
| 11 | 118559426 | rs10892253 | A | C | NULL | 0.589040434 | NULL&M_Disease |
| 11 | 118561858 | rs10790257 | C | G | NULL | 0.589040434 | NULL&M_Disease |
| 11 | 118834508 | rs111358668 | A | G | NULL | 0.590250252 | NULL&M_Disease |
| 11 | 120334126 | rs145192391 | T | C | NULL | 0.590250252 | NULL&M_Disease |
| 11 | 121057146 | rs77233977 | T | C | NULL | 0.590250252 | NULL&M_Disease |
| 11 | 116226845 | rs564046 | G | A | NULL | 0.590250252 | NULL&M_Disease |
| 11 | 116625875 | rs12421652 | T | G | NULL | 0.590250252 | NULL&M_Disease |
| 11 | 118897962 | rs74888816 | T | C | NULL | 0.590250252 | NULL&M_Disease |
| 11 | 119881864 | rs61747468 | A | G | NULL | 0.589040434 | NULL&M_Disease |
| 11 | 120575997 | rs11217980 | A | G | NULL | 0.590250252 | NULL&M_Disease |
| 11 | 120633977 | rs116324528 | T | A | NULL | 0.590250252 | NULL&M_Disease |
| 11 | 121010048 | rs71486352 | A | T | NULL | 0.589040434 | NULL&M_Disease |
| 11 | 118493319 | rs10790254 | A | G | NULL | 0.590250252 | NULL&M_Disease |
| 11 | 118557483 | rs6589673 | C | A | NULL | 0.589040434 | NULL&M_Disease |
| 11 | 120636195 | rs150586315 | A | G | NULL | 0.590250252 | NULL&M_Disease |

| | | | | | | | | | |
|---|---|---|---|---|---|---|---|---|---|
| 11 | 120468093 | rs75524659 | T | C | 1.048 | -7977.388 | -7978.102 | -0.714 | 1.89E-06 |
| 11 | 116530373 | rs552787 | C | A | 0.955 | -7977.388 | -7978.103 | -0.715 | 1.89E-06 |
| 11 | 117461602 | rs553637 | C | T | 0.956 | -7977.388 | -7978.103 | -0.715 | 1.89E-06 |
| 11 | 119630763 | rs11217472 | C | T | 0.945 | -7977.388 | -7978.103 | -0.715 | 1.89E-06 |
| 11 | 119911542 | rs1452842 | G | A | 1.048 | -7977.388 | -7978.103 | -0.715 | 1.89E-06 |
| 11 | 119930569 | rs1790469 | A | G | 0.946 | -7977.388 | -7978.104 | -0.715 | 1.89E-06 |
| 11 | 116486588 | rs3016346 | A | G | 0.955 | -7977.388 | -7978.104 | -0.716 | 1.89E-06 |
| 11 | 118041972 | rs12294669 | G | A | 0.943 | -7977.388 | -7978.104 | -0.716 | 1.89E-06 |
| 11 | 121013909 | rs588917 | T | G | 0.955 | -7977.388 | -7978.105 | -0.716 | 1.89E-06 |
| 11 | 117190901 | rs117469840 | T | C | 0.969 | -7977.388 | -7978.105 | -0.717 | 1.89E-06 |
| 11 | 117711562 | rs519991 | C | T | 1.046 | -7977.388 | -7978.105 | -0.717 | 1.89E-06 |
| 11 | 118341921 | rs524936 | A | G | 0.945 | -7977.388 | -7978.105 | -0.717 | 1.89E-06 |
| 11 | 118477226 | rs45496391 | C | A | 0.978 | -7977.388 | -7978.106 | -0.717 | 1.89E-06 |
| 11 | 116899248 | rs11216240 | T | A | 1.05 | -7977.388 | -7978.106 | -0.718 | 1.89E-06 |
| 11 | 116899249 | rs11216241 | C | A | 1.05 | -7977.388 | -7978.106 | -0.718 | 1.89E-06 |
| 11 | 117684522 | rs12288877 | C | G | 0.945 | -7977.388 | -7978.106 | -0.718 | 1.89E-06 |
| 11 | 118825069 | rs2097131 | G | A | 0.956 | -7977.388 | -7978.106 | -0.718 | 1.89E-06 |
| 11 | 120730621 | rs4526780 | A | G | 0.948 | -7977.388 | -7978.107 | -0.718 | 1.89E-06 |
| 11 | 116920927 | rs10892066 | T | C | 1.05 | -7977.388 | -7978.107 | -0.719 | 1.89E-06 |
| 11 | 116943952 | rs1531706 | A | C | 1.05 | -7977.388 | -7978.108 | -0.719 | 1.88E-06 |
| 11 | 116944348 | rs7935913 | A | G | 1.05 | -7977.388 | -7978.108 | -0.719 | 1.88E-06 |
| 11 | 116944712 | rs1531707 | A | G | 1.05 | -7977.388 | -7978.108 | -0.719 | 1.88E-06 |
| 11 | 117641216 | rs56112767 | G | C | 1.059 | -7977.388 | -7978.109 | -0.72 | 1.88E-06 |
| 11 | 117752769 | rs77752432 | T | C | 0.996 | -7977.388 | -7978.108 | -0.72 | 1.88E-06 |
| 11 | 118474283 | rs139202987 | G | A | 0.982 | -7977.388 | -7978.108 | -0.72 | 1.88E-06 |
| 11 | 120808303 | rs71486349 | T | G | 0.988 | -7977.388 | -7978.108 | -0.72 | 1.88E-06 |
| 11 | 116660813 | rs619054 | A | G | 1.05 | -7977.388 | -7978.11 | -0.721 | 1.88E-06 |
| 11 | 117611292 | rs10466534 | A | G | 0.95 | -7977.388 | -7978.109 | -0.721 | 1.88E-06 |
| 11 | 117912720 | rs650104 | T | C | 1.046 | -7977.388 | -7978.109 | -0.721 | 1.88E-06 |
| 11 | 117912731 | rs689054 | T | A | 1.046 | -7977.388 | -7978.109 | -0.721 | 1.88E-06 |
| 11 | 118015832 | rs45539032 | A | G | 1.057 | -7977.388 | -7978.109 | -0.721 | 1.88E-06 |
| 11 | 119985011 | rs150954345 | G | A | 0.975 | -7977.388 | -7978.11 | -0.721 | 1.88E-06 |
| 11 | 120619202 | rs2852224 | T | C | 0.945 | -7977.388 | -7978.109 | -0.721 | 1.88E-06 |
| 11 | 117804926 | rs550388 | C | T | 0.944 | -7977.388 | -7978.111 | -0.722 | 1.88E-06 |
| 11 | 119745457 | rs508440 | G | C | 1.06 | -7977.388 | -7978.111 | -0.722 | 1.88E-06 |
| 11 | 117590911 | rs948772 | C | G | 0.942 | -7977.388 | -7978.111 | -0.723 | 1.88E-06 |
| 11 | 120576090 | rs11217982 | A | T | 0.947 | -7977.388 | -7978.111 | -0.723 | 1.88E-06 |
| 11 | 120576091 | rs11217983 | C | T | 0.947 | -7977.388 | -7978.111 | -0.723 | 1.88E-06 |
| 11 | 120895352 | rs3824976 | A | G | 1.052 | -7977.388 | -7978.112 | -0.723 | 1.88E-06 |
| 11 | 120984681 | rs658494 | T | C | 1.051 | -7977.388 | -7978.111 | -0.723 | 1.88E-06 |
| 11 | 117461875 | rs1363950531 | G | T | 0.956 | -7977.388 | -7978.112 | -0.724 | 1.88E-06 |
| 11 | 117591292 | rs4936399 | T | C | 0.942 | -7977.388 | -7978.112 | -0.724 | 1.88E-06 |
| 11 | 117912538 | rs111234707 | A | G | 0.988 | -7977.388 | -7978.112 | -0.724 | 1.88E-06 |
| 11 | 116928112 | rs4938329 | T | C | 1.05 | -7977.388 | -7978.113 | -0.725 | 1.87E-06 |
| 11 | 117526440 | rs55728331 | A | G | 0.951 | -7977.388 | -7978.114 | -0.725 | 1.87E-06 |
| 11 | 117681047 | rs34836631 | A | G | 0.954 | -7977.388 | -7978.113 | -0.725 | 1.87E-06 |
| 11 | 118134785 | rs1624780 | T | G | 1.046 | -7977.388 | -7978.114 | -0.725 | 1.87E-06 |
| 11 | 116944437 | rs7935834 | G | C | 1.05 | -7977.388 | -7978.114 | -0.726 | 1.87E-06 |
| 11 | 116957189 | rs12274465 | T | C | 1.05 | -7977.388 | -7978.114 | -0.726 | 1.87E-06 |
| 11 | 117283939 | rs11822288 | A | G | 0.97 | -7977.388 | -7978.114 | -0.726 | 1.87E-06 |
| 11 | 117353003 | rs3741279 | G | A | 1.05 | -7977.388 | -7978.115 | -0.726 | 1.87E-06 |
| 11 | 118460957 | rs45600633 | A | G | 0.984 | -7977.388 | -7978.115 | -0.727 | 1.87E-06 |
| 11 | 118518021 | rs10736493 | G | T | 0.989 | -7977.388 | -7978.117 | -0.728 | 1.87E-06 |
| 11 | 118867695 | rs4938617 | T | C | 1.056 | -7977.388 | -7978.116 | -0.728 | 1.87E-06 |
| 11 | 119907974 | rs1939333 | C | T | 1.048 | -7977.388 | -7978.117 | -0.728 | 1.87E-06 |
| 11 | 116364066 | rs6589549 | A | G | 0.956 | -7977.388 | -7978.118 | -0.729 | 1.87E-06 |
| 11 | 116539518 | rs9681206 | T | C | 0.946 | -7977.388 | -7978.117 | -0.729 | 1.87E-06 |
| 11 | 117435148 | rs7940044 | A | G | 0.955 | -7977.388 | -7978.117 | -0.729 | 1.87E-06 |
| 11 | 120109766 | rs61898297 | T | C | 0.944 | -7977.388 | -7978.117 | -0.729 | 1.87E-06 |
| 11 | 120423569 | rs12278730 | T | C | 0.953 | -7977.388 | -7978.117 | -0.729 | 1.87E-06 |
| 11 | 117251143 | rs2305828 | A | G | 0.956 | -7977.388 | -7978.119 | -0.73 | 1.86E-06 |
| 11 | 117680407 | rs67877251 | G | A | 0.975 | -7977.388 | -7978.118 | -0.73 | 1.86E-06 |
| 11 | 118852647 | rs80126726 | A | G | 1.054 | -7977.388 | -7978.119 | -0.73 | 1.86E-06 |
| 11 | 119499255 | rs11217361 | C | G | 1.056 | -7977.388 | -7978.119 | -0.73 | 1.86E-06 |
| 11 | 119768716 | rs73010609 | T | C | 0.952 | -7977.388 | -7978.118 | -0.73 | 1.86E-06 |
| 11 | 119805057 | rs145637218 | A | G | 0.979 | -7977.388 | -7978.118 | -0.73 | 1.86E-06 |
| 11 | 120568566 | rs12270409 | C | T | 0.947 | -7977.388 | -7978.118 | -0.73 | 1.86E-06 |
| 11 | 121004151 | rs1893023 | T | C | 0.956 | -7977.388 | -7978.119 | -0.73 | 1.86E-06 |
| 11 | 116955049 | rs61903416 | C | T | 1.05 | -7977.388 | -7978.12 | -0.731 | 1.86E-06 |
| 11 | 117433686 | rs755791 | C | A | 0.955 | -7977.388 | -7978.119 | -0.731 | 1.86E-06 |
| 11 | 118561859 | rs10790258 | T | G | 0.984 | -7977.388 | -7978.12 | -0.731 | 1.86E-06 |
| 11 | 117194929 | rs554233 | G | A | 0.944 | -7977.388 | -7978.12 | -0.732 | 1.86E-06 |
| 11 | 117966178 | rs948465 | G | A | 1.06 | -7977.388 | -7978.12 | -0.732 | 1.86E-06 |
| 11 | 117966185 | rs948464 | G | T | 1.06 | -7977.388 | -7978.12 | -0.732 | 1.86E-06 |
| 11 | 119676471 | rs10892468 | T | G | 0.956 | -7977.388 | -7978.12 | -0.732 | 1.86E-06 |
| 11 | 119820957 | rs144502429 | T | C | 0.981 | -7977.388 | -7978.121 | -0.732 | 1.86E-06 |
| 11 | 116913909 | rs12279066 | G | C | 1.05 | -7977.388 | -7978.121 | -0.733 | 1.86E-06 |
| 11 | 118295001 | rs139060345 | C | T | 0.981 | -7977.388 | -7978.121 | -0.733 | 1.86E-06 |
| 11 | 119913037 | rs659127 | A | C | 0.951 | -7977.388 | -7978.122 | -0.733 | 1.86E-06 |
| 11 | 121014403 | rs492294 | G | A | 0.955 | -7977.388 | -7978.121 | -0.733 | 1.86E-06 |
| 11 | 116409130 | rs56338557 | C | T | 0.949 | -7977.388 | -7978.122 | -0.734 | 1.86E-06 |
| 11 | 118559426 | rs10892253 | A | C | 0.985 | -7977.388 | -7978.122 | -0.734 | 1.86E-06 |
| 11 | 118561858 | rs10790257 | C | G | 0.985 | -7977.388 | -7978.122 | -0.734 | 1.86E-06 |
| 11 | 118834508 | rs111358668 | A | G | 1.054 | -7977.388 | -7978.122 | -0.734 | 1.86E-06 |
| 11 | 120334126 | rs145192391 | T | C | 1.049 | -7977.388 | -7978.122 | -0.734 | 1.86E-06 |
| 11 | 121057146 | rs77233977 | T | C | 0.945 | -7977.388 | -7978.122 | -0.734 | 1.86E-06 |
| 11 | 116226845 | rs564046 | G | A | 1.054 | -7977.388 | -7978.124 | -0.735 | 1.85E-06 |
| 11 | 116625875 | rs12421652 | T | G | 1.056 | -7977.388 | -7978.123 | -0.735 | 1.85E-06 |
| 11 | 118897962 | rs74888816 | T | C | 0.944 | -7977.388 | -7978.123 | -0.735 | 1.85E-06 |
| 11 | 119881864 | rs61747468 | A | G | 0.995 | -7977.388 | -7978.123 | -0.735 | 1.85E-06 |
| 11 | 120575997 | rs11217980 | A | G | 0.949 | -7977.388 | -7978.123 | -0.735 | 1.85E-06 |
| 11 | 120633977 | rs116324528 | T | A | 0.946 | -7977.388 | -7978.123 | -0.735 | 1.85E-06 |
| 11 | 121010048 | rs71486352 | A | T | 1.019 | -7977.388 | -7978.123 | -0.735 | 1.85E-06 |
| 11 | 118493319 | rs10790254 | A | G | 0.99 | -7977.388 | -7978.125 | -0.736 | 1.85E-06 |
| 11 | 118557483 | rs6589673 | C | A | 0.987 | -7977.388 | -7978.124 | -0.736 | 1.85E-06 |
| 11 | 120636195 | rs150586315 | A | G | 0.946 | -7977.388 | -7978.124 | -0.736 | 1.85E-06 |

| | | | | | | | |
|---|---|---|---|---|---|---|---|
| 11 | 120845912 | rs7943490 | A | G | NULL | 0.590250252 | NULL&M_Disease |
| 11 | 121132474 | rs76344204 | A | G | NULL | 0.590250252 | NULL&M_Disease |
| 11 | 116951385 | rs7106662 | C | T | NULL | 0.590250252 | NULL&M_Disease |
| 11 | 117964613 | rs77718036 | T | C | NULL | 0.590250252 | NULL&M_Disease |
| 11 | 120875634 | rs11218107 | C | T | NULL | 0.590250252 | NULL&M_Disease |
| 11 | 120876637 | rs79237143 | G | T | NULL | 0.590250252 | NULL&M_Disease |
| 11 | 118455465 | rs45460493 | A | G | NULL | 0.590250252 | NULL&M_Disease |
| 11 | 118507068 | rs10892249 | C | T | NULL | 0.590250252 | NULL&M_Disease |
| 11 | 118798780 | rs77916839 | C | G | NULL | 0.590250252 | NULL&M_Disease |
| 11 | 119702716 | rs76785396 | A | T | NULL | 0.590250252 | NULL&M_Disease |
| 11 | 119947801 | rs662549 | C | T | NULL | 0.590250252 | NULL&M_Disease |
| 11 | 119948997 | rs600720 | T | C | NULL | 0.590250252 | NULL&M_Disease |
| 11 | 119949498 | rs649921 | T | G | NULL | 0.590250252 | NULL&M_Disease |
| 11 | 119949821 | rs597124 | T | C | NULL | 0.590250252 | NULL&M_Disease |
| 11 | 120794058 | rs74908422 | C | T | NULL | 0.590250252 | NULL&M_Disease |
| 11 | 116571138 | rs11601168 | A | T | NULL | 0.591458978 | NULL&M_Disease |
| 11 | 116894413 | rs10466590 | G | C | NULL | 0.591458978 | NULL&M_Disease |
| 11 | 116895512 | rs11216239 | G | C | NULL | 0.591458978 | NULL&M_Disease |
| 11 | 116912730 | rs11216244 | T | C | NULL | 0.591458978 | NULL&M_Disease |
| 11 | 116913202 | rs11216245 | T | C | NULL | 0.591458978 | NULL&M_Disease |
| 11 | 116946890 | rs7484045 | T | C | NULL | 0.591458978 | NULL&M_Disease |
| 11 | 117401292 | rs57434987 | A | C | NULL | 0.590250252 | NULL&M_Disease |
| 11 | 120699480 | rs76219905 | T | C | NULL | 0.590250252 | NULL&M_Disease |
| 11 | 116223775 | rs568306 | A | C | NULL | 0.591458978 | NULL&M_Disease |
| 11 | 119481969 | rs147710439 | A | G | NULL | 0.590250252 | NULL&M_Disease |
| 11 | 120696635 | rs4076169 | C | T | NULL | 0.590250252 | NULL&M_Disease |
| 11 | 116890259 | rs7113829 | T | C | NULL | 0.590250252 | NULL&M_Disease |
| 11 | 118487265 | rs76544852 | T | G | NULL | 0.590250252 | NULL&M_Disease |
| 11 | 119863317 | rs117649997 | A | G | NULL | 0.590250252 | NULL&M_Disease |
| 11 | 120879138 | rs117587198 | A | T | NULL | 0.590250252 | NULL&M_Disease |
| 11 | 116432510 | rs61904804 | C | G | NULL | 0.591458978 | NULL&M_Disease |
| 11 | 117071970 | rs494356 | T | C | NULL | 0.591458978 | NULL&M_Disease |
| 11 | 118421841 | rs145968026 | G | C | NULL | 0.590250252 | NULL&M_Disease |
| 11 | 118483912 | rs45621436 | G | T | NULL | 0.590250252 | NULL&M_Disease |
| 11 | 120697162 | rs4075786 | C | T | NULL | 0.591458978 | NULL&M_Disease |
| 11 | 120830051 | rs7106292 | T | C | NULL | 0.591458978 | NULL&M_Disease |
| 11 | 121029720 | rs582876 | C | T | NULL | 0.591458978 | NULL&M_Disease |
| 11 | 119364445 | rs77654673 | C | G | NULL | 0.591458978 | NULL&M_Disease |
| 11 | 120981863 | rs87466 | G | A | NULL | 0.591458978 | NULL&M_Disease |
| 11 | 116949074 | rs10082609 | T | C | NULL | 0.591458978 | NULL&M_Disease |
| 11 | 117686829 | rs1939976 | T | C | NULL | 0.591458978 | NULL&M_Disease |
| 11 | 120800376 | rs4936564 | A | G | NULL | 0.591458978 | NULL&M_Disease |
| 11 | 120876772 | rs76429987 | T | C | NULL | 0.591458978 | NULL&M_Disease |
| 11 | 121055191 | rs12787375 | T | G | NULL | 0.590250252 | NULL&M_Disease |
| 11 | 116889411 | rs7128772 | A | T | NULL | 0.591458978 | NULL&M_Disease |
| 11 | 116900947 | rs11600552 | C | T | NULL | 0.591458978 | NULL&M_Disease |
| 11 | 117967839 | rs9651686 | G | T | NULL | 0.591458978 | NULL&M_Disease |
| 11 | 118087765 | rs45618531 | T | C | NULL | 0.591458978 | NULL&M_Disease |
| 11 | 118269674 | rs45529532 | G | A | NULL | 0.591458978 | NULL&M_Disease |
| 11 | 118610629 | rs521910 | A | G | NULL | 0.591458978 | NULL&M_Disease |
| 11 | 120868246 | rs10892658 | T | C | NULL | 0.591458978 | NULL&M_Disease |
| 11 | 121062593 | rs12794546 | T | C | NULL | 0.590250252 | NULL&M_Disease |
| 11 | 116234916 | rs117625924 | C | A | NULL | 0.591458978 | NULL&M_Disease |
| 11 | 117614122 | rs11216521 | A | G | NULL | 0.591458978 | NULL&M_Disease |
| 11 | 118518227 | rs10736494 | T | C | NULL | 0.591458978 | NULL&M_Disease |
| 11 | 121145354 | rs73005121 | T | A | NULL | 0.591458978 | NULL&M_Disease |
| 11 | 116629555 | rs150233369 | A | G | NULL | 0.591458978 | NULL&M_Disease |
| 11 | 117421425 | rs553062 | A | G | NULL | 0.5926666 | NULL&M_Disease |
| 11 | 117619027 | rs10892167 | C | A | NULL | 0.591458978 | NULL&M_Disease |
| 11 | 118445559 | rs73555296 | T | G | NULL | 0.591458978 | NULL&M_Disease |
| 11 | 118545589 | rs113941645 | G | C | NULL | 0.591458978 | NULL&M_Disease |
| 11 | 117380103 | rs11216433 | G | A | NULL | 0.5926666 | NULL&M_Disease |
| 11 | 116722910 | rs577975 | C | T | NULL | 0.5926666 | NULL&M_Disease |
| 11 | 118134065 | rs939274 | T | G | NULL | 0.5926666 | NULL&M_Disease |
| 11 | 118381018 | rs9332851 | T | C | NULL | 0.591458978 | NULL&M_Disease |
| 11 | 118826063 | rs666191 | G | A | NULL | 0.5926666 | NULL&M_Disease |
| 11 | 119617639 | rs11217450 | G | A | NULL | 0.5926666 | NULL&M_Disease |
| 11 | 119933153 | rs61897936 | T | C | NULL | 0.5926666 | NULL&M_Disease |
| 11 | 120538317 | rs115293580 | T | C | NULL | 0.591458978 | NULL&M_Disease |
| 11 | 117927789 | rs643295 | T | A | NULL | 0.5926666 | NULL&M_Disease |
| 11 | 116467920 | rs34721671 | G | C | NULL | 0.5926666 | NULL&M_Disease |
| 11 | 116954565 | rs10892071 | T | C | NULL | 0.5926666 | NULL&M_Disease |
| 11 | 117124050 | rs73010424 | A | G | NULL | 0.591458978 | NULL&M_Disease |
| 11 | 117125769 | rs117683585 | G | A | NULL | 0.591458978 | NULL&M_Disease |
| 11 | 117345415 | rs75581341 | T | C | NULL | 0.5926666 | NULL&M_Disease |
| 11 | 119744160 | rs541366 | C | A | NULL | 0.5926666 | NULL&M_Disease |
| 11 | 116485583 | rs477047 | A | T | NULL | 0.5926666 | NULL&M_Disease |
| 11 | 116929995 | rs11216255 | G | A | NULL | 0.5926666 | NULL&M_Disease |
| 11 | 117633482 | rs145518466 | G | A | NULL | 0.5926666 | NULL&M_Disease |
| 11 | 117633690 | rs73587425 | C | G | NULL | 0.5926666 | NULL&M_Disease |
| 11 | 117792611 | rs4938457 | T | C | NULL | 0.5926666 | NULL&M_Disease |
| 11 | 118539746 | rs72999492 | C | A | NULL | 0.5926666 | NULL&M_Disease |
| 11 | 121017741 | rs599894 | C | G | NULL | 0.5926666 | NULL&M_Disease |
| 11 | 116470344 | rs541948 | T | G | NULL | 0.5926666 | NULL&M_Disease |
| 11 | 117327680 | rs9667438 | T | C | NULL | 0.5926666 | NULL&M_Disease |
| 11 | 117433739 | rs2010473 | C | T | NULL | 0.5926666 | NULL&M_Disease |
| 11 | 118194647 | rs141752759 | G | C | NULL | 0.5926666 | NULL&M_Disease |
| 11 | 120695351 | rs113184379 | G | C | NULL | 0.5926666 | NULL&M_Disease |
| 11 | 119664546 | rs113164826 | C | G | NULL | 0.5926666 | NULL&M_Disease |
| 11 | 120575481 | rs12276440 | T | C | NULL | 0.5926666 | NULL&M_Disease |
| 11 | 120797604 | rs2850808 | A | G | NULL | 0.593873103 | NULL&M_Disease |
| 11 | 116924990 | rs3886960 | T | G | NULL | 0.593873103 | NULL&M_Disease |
| 11 | 116925042 | rs723955 | A | G | NULL | 0.593873103 | NULL&M_Disease |
| 11 | 116925996 | rs7108912 | A | G | NULL | 0.593873103 | NULL&M_Disease |
| 11 | 116927951 | rs7124872 | G | A | NULL | 0.593873103 | NULL&M_Disease |
| 11 | 116928006 | rs7124996 | T | C | NULL | 0.593873103 | NULL&M_Disease |

| | | | | | | | | | |
|---|---|---|---|---|---|---|---|---|---|
| 11 | 120845912 | rs7943490 | A | G | 1.06 | -7977.388 | -7978.124 | -0.736 | 1.85E-06 |
| 11 | 121132474 | rs76344204 | A | G | 0.944 | -7977.388 | -7978.125 | -0.736 | 1.85E-06 |
| 11 | 116951385 | rs7106662 | C | T | 1.05 | -7977.388 | -7978.126 | -0.737 | 1.85E-06 |
| 11 | 117964613 | rs77718036 | T | C | 0.962 | -7977.388 | -7978.125 | -0.737 | 1.85E-06 |
| 11 | 120875634 | rs11218107 | C | T | 1.059 | -7977.388 | -7978.125 | -0.737 | 1.85E-06 |
| 11 | 120876637 | rs79237143 | G | T | 1.059 | -7977.388 | -7978.125 | -0.737 | 1.85E-06 |
| 11 | 118455465 | rs45460493 | A | G | 0.997 | -7977.388 | -7978.127 | -0.738 | 1.85E-06 |
| 11 | 118507068 | rs10892249 | C | T | 0.995 | -7977.388 | -7978.126 | -0.738 | 1.85E-06 |
| 11 | 118798780 | rs77916839 | C | G | 1.056 | -7977.388 | -7978.126 | -0.738 | 1.85E-06 |
| 11 | 119702716 | rs76785396 | A | T | 0.947 | -7977.388 | -7978.126 | -0.738 | 1.85E-06 |
| 11 | 119947801 | rs662549 | C | T | 0.944 | -7977.388 | -7978.126 | -0.738 | 1.85E-06 |
| 11 | 119948997 | rs600720 | T | C | 0.944 | -7977.388 | -7978.126 | -0.738 | 1.85E-06 |
| 11 | 119949498 | rs649921 | T | G | 0.944 | -7977.388 | -7978.126 | -0.738 | 1.85E-06 |
| 11 | 119949821 | rs597124 | T | C | 0.944 | -7977.388 | -7978.126 | -0.738 | 1.85E-06 |
| 11 | 120794058 | rs74908422 | C | T | 1.054 | -7977.388 | -7978.127 | -0.738 | 1.85E-06 |
| 11 | 116571138 | rs11601168 | A | T | 1.056 | -7977.388 | -7978.127 | -0.739 | 1.85E-06 |
| 11 | 116894413 | rs10466590 | G | C | 1.05 | -7977.388 | -7978.127 | -0.739 | 1.85E-06 |
| 11 | 116895512 | rs11216239 | G | C | 1.05 | -7977.388 | -7978.127 | -0.739 | 1.85E-06 |
| 11 | 116912730 | rs11216244 | T | C | 1.05 | -7977.388 | -7978.128 | -0.739 | 1.85E-06 |
| 11 | 116913202 | rs11216245 | T | C | 1.05 | -7977.388 | -7978.128 | -0.739 | 1.85E-06 |
| 11 | 116946890 | rs7484045 | T | C | 1.05 | -7977.388 | -7978.127 | -0.739 | 1.85E-06 |
| 11 | 117401292 | rs57434987 | A | C | 0.965 | -7977.388 | -7978.128 | -0.739 | 1.85E-06 |
| 11 | 120699480 | rs76219905 | T | C | 0.943 | -7977.388 | -7978.127 | -0.739 | 1.85E-06 |
| 11 | 116223775 | rs568306 | A | C | 1.054 | -7977.388 | -7978.128 | -0.74 | 1.85E-06 |
| 11 | 119481969 | rs147710439 | A | G | 1.002 | -7977.388 | -7978.128 | -0.74 | 1.85E-06 |
| 11 | 120696635 | rs4076169 | C | T | 0.943 | -7977.388 | -7978.128 | -0.74 | 1.85E-06 |
| 11 | 116890259 | rs7113829 | T | C | 1.051 | -7977.388 | -7978.129 | -0.741 | 1.84E-06 |
| 11 | 118487265 | rs76544852 | T | G | 0.998 | -7977.388 | -7978.129 | -0.741 | 1.84E-06 |
| 11 | 119863317 | rs117649997 | A | G | 0.972 | -7977.388 | -7978.129 | -0.741 | 1.84E-06 |
| 11 | 120879138 | rs117587198 | A | T | 1.028 | -7977.388 | -7978.129 | -0.741 | 1.84E-06 |
| 11 | 116432510 | rs61904804 | C | G | 1.053 | -7977.388 | -7978.13 | -0.742 | 1.84E-06 |
| 11 | 117071970 | rs494356 | T | C | 0.948 | -7977.388 | -7978.131 | -0.742 | 1.84E-06 |
| 11 | 118421841 | rs145968026 | G | C | 1.006 | -7977.388 | -7978.13 | -0.742 | 1.84E-06 |
| 11 | 118483912 | rs45621436 | G | T | 1.002 | -7977.388 | -7978.13 | -0.742 | 1.84E-06 |
| 11 | 120697162 | rs4075786 | C | T | 0.943 | -7977.388 | -7978.13 | -0.742 | 1.84E-06 |
| 11 | 120830051 | rs7106292 | T | C | 0.951 | -7977.388 | -7978.13 | -0.742 | 1.84E-06 |
| 11 | 121029720 | rs582876 | C | T | 0.955 | -7977.388 | -7978.13 | -0.742 | 1.84E-06 |
| 11 | 119364445 | rs77654673 | C | G | 0.949 | -7977.388 | -7978.131 | -0.743 | 1.84E-06 |
| 11 | 120981863 | rs87466 | G | A | 1.049 | -7977.388 | -7978.131 | -0.743 | 1.84E-06 |
| 11 | 116949074 | rs10082609 | T | C | 1.05 | -7977.388 | -7978.133 | -0.745 | 1.84E-06 |
| 11 | 117686829 | rs1939976 | T | C | 0.946 | -7977.388 | -7978.133 | -0.745 | 1.84E-06 |
| 11 | 120800376 | rs4936564 | A | G | 1.051 | -7977.388 | -7978.133 | -0.745 | 1.84E-06 |
| 11 | 120876772 | rs76429987 | T | C | 1.059 | -7977.388 | -7978.133 | -0.745 | 1.84E-06 |
| 11 | 121055191 | rs12787375 | T | G | 0.994 | -7977.388 | -7978.133 | -0.745 | 1.84E-06 |
| 11 | 116889411 | rs7128772 | A | T | 1.05 | -7977.388 | -7978.134 | -0.746 | 1.83E-06 |
| 11 | 116900947 | rs11600552 | C | T | 1.05 | -7977.388 | -7978.134 | -0.746 | 1.83E-06 |
| 11 | 117967839 | rs9651686 | G | T | 1.059 | -7977.388 | -7978.134 | -0.746 | 1.83E-06 |
| 11 | 118087765 | rs45618531 | T | C | 1.027 | -7977.388 | -7978.134 | -0.746 | 1.83E-06 |
| 11 | 118269674 | rs45529532 | G | A | 0.987 | -7977.388 | -7978.134 | -0.746 | 1.83E-06 |
| 11 | 118610629 | rs521910 | A | G | 0.947 | -7977.388 | -7978.134 | -0.746 | 1.83E-06 |
| 11 | 120868246 | rs10892658 | T | C | 0.949 | -7977.388 | -7978.134 | -0.746 | 1.83E-06 |
| 11 | 121062593 | rs12794546 | T | C | 0.995 | -7977.388 | -7978.134 | -0.746 | 1.83E-06 |
| 11 | 116234916 | rs117625924 | C | A | 0.996 | -7977.388 | -7978.135 | -0.747 | 1.83E-06 |
| 11 | 117614122 | rs11216521 | A | G | 0.953 | -7977.388 | -7978.136 | -0.747 | 1.83E-06 |
| 11 | 118518227 | rs10736494 | T | C | 1.004 | -7977.388 | -7978.136 | -0.747 | 1.83E-06 |
| 11 | 121145354 | rs73005121 | T | A | 0.949 | -7977.388 | -7978.135 | -0.747 | 1.83E-06 |
| 11 | 116629555 | rs150233369 | A | G | 0.963 | -7977.388 | -7978.136 | -0.748 | 1.83E-06 |
| 11 | 117421425 | rs553062 | A | G | 1.047 | -7977.388 | -7978.136 | -0.748 | 1.83E-06 |
| 11 | 117619027 | rs10892167 | C | A | 0.944 | -7977.388 | -7978.137 | -0.749 | 1.83E-06 |
| 11 | 118445559 | rs73555296 | T | G | 0.997 | -7977.388 | -7978.138 | -0.749 | 1.83E-06 |
| 11 | 118545589 | rs113941645 | G | C | 0.952 | -7977.388 | -7978.137 | -0.749 | 1.83E-06 |
| 11 | 117380103 | rs11216433 | G | A | 0.944 | -7977.388 | -7978.139 | -0.751 | 1.83E-06 |
| 11 | 116722910 | rs577975 | C | T | 1.058 | -7977.388 | -7978.141 | -0.752 | 1.82E-06 |
| 11 | 118134065 | rs939274 | T | G | 1.046 | -7977.388 | -7978.14 | -0.752 | 1.82E-06 |
| 11 | 118381018 | rs9332851 | T | C | 1.006 | -7977.388 | -7978.14 | -0.752 | 1.82E-06 |
| 11 | 118826063 | rs666191 | G | A | 0.956 | -7977.388 | -7978.14 | -0.752 | 1.82E-06 |
| 11 | 119617639 | rs11217450 | G | A | 0.945 | -7977.388 | -7978.141 | -0.752 | 1.82E-06 |
| 11 | 119933153 | rs61897936 | T | C | 1.058 | -7977.388 | -7978.14 | -0.752 | 1.82E-06 |
| 11 | 120538317 | rs115293580 | T | C | 0.958 | -7977.388 | -7978.14 | -0.752 | 1.82E-06 |
| 11 | 117927789 | rs643295 | T | A | 1.046 | -7977.388 | -7978.141 | -0.753 | 1.82E-06 |
| 11 | 116467920 | rs34721671 | G | C | 0.945 | -7977.388 | -7978.142 | -0.754 | 1.82E-06 |
| 11 | 116954565 | rs10892071 | T | C | 1.049 | -7977.388 | -7978.142 | -0.754 | 1.82E-06 |
| 11 | 117124050 | rs73010424 | A | G | 1 | -7977.388 | -7978.143 | -0.754 | 1.82E-06 |
| 11 | 117125769 | rs117683585 | G | A | 0.998 | -7977.388 | -7978.142 | -0.754 | 1.82E-06 |
| 11 | 117345415 | rs75581341 | T | C | 1.047 | -7977.388 | -7978.142 | -0.754 | 1.82E-06 |
| 11 | 119744160 | rs541366 | C | A | 1.058 | -7977.388 | -7978.142 | -0.754 | 1.82E-06 |
| 11 | 116485583 | rs477047 | A | T | 0.946 | -7977.388 | -7978.144 | -0.756 | 1.82E-06 |
| 11 | 116929995 | rs11216255 | G | A | 1.049 | -7977.388 | -7978.144 | -0.756 | 1.82E-06 |
| 11 | 117633482 | rs145518466 | G | A | 1.057 | -7977.388 | -7978.144 | -0.756 | 1.82E-06 |
| 11 | 117633690 | rs73587425 | C | G | 1.057 | -7977.388 | -7978.144 | -0.756 | 1.82E-06 |
| 11 | 117792611 | rs4938457 | T | C | 1.046 | -7977.388 | -7978.145 | -0.756 | 1.82E-06 |
| 11 | 118539746 | rs72999492 | C | A | 1.058 | -7977.388 | -7978.144 | -0.756 | 1.82E-06 |
| 11 | 121017741 | rs599894 | C | G | 0.955 | -7977.388 | -7978.144 | -0.756 | 1.82E-06 |
| 11 | 116470344 | rs541948 | T | G | 1.051 | -7977.388 | -7978.145 | -0.757 | 1.81E-06 |
| 11 | 117327680 | rs9667438 | T | C | 0.949 | -7977.388 | -7978.145 | -0.757 | 1.81E-06 |
| 11 | 117433739 | rs2010473 | C | T | 0.955 | -7977.388 | -7978.145 | -0.757 | 1.81E-06 |
| 11 | 118194647 | rs141752759 | G | C | 0.961 | -7977.388 | -7978.145 | -0.757 | 1.81E-06 |
| 11 | 120695351 | rs113184379 | G | C | 0.944 | -7977.388 | -7978.145 | -0.757 | 1.81E-06 |
| 11 | 119664546 | rs113164826 | C | G | 0.981 | -7977.388 | -7978.146 | -0.758 | 1.81E-06 |
| 11 | 120575481 | rs12276440 | T | C | 0.949 | -7977.388 | -7978.146 | -0.758 | 1.81E-06 |
| 11 | 120797604 | rs2850808 | A | G | 0.95 | -7977.388 | -7978.146 | -0.758 | 1.81E-06 |
| 11 | 116924990 | rs3886960 | T | G | 1.049 | -7977.388 | -7978.147 | -0.759 | 1.81E-06 |
| 11 | 116925042 | rs723955 | A | G | 1.049 | -7977.388 | -7978.147 | -0.759 | 1.81E-06 |
| 11 | 116925996 | rs7108912 | A | G | 1.049 | -7977.388 | -7978.147 | -0.759 | 1.81E-06 |
| 11 | 116927951 | rs7124872 | G | A | 1.049 | -7977.388 | -7978.147 | -0.759 | 1.81E-06 |
| 11 | 116928006 | rs7124996 | T | C | 1.049 | -7977.388 | -7978.147 | -0.759 | 1.81E-06 |

| | | | | | | | |
|---|---|---|---|---|---|---|---|
| 11 | 118066912 | rs12808254 | T | C | NULL | 0.5926666 | NULL&M_Disease |
| 11 | 119870561 | rs78289484 | T | C | NULL | 0.5926666 | NULL&M_Disease |
| 11 | 120699050 | rs76871374 | G | A | NULL | 0.5926666 | NULL&M_Disease |
| 11 | 118150882 | rs117527146 | A | C | NULL | 0.5926666 | NULL&M_Disease |
| 11 | 120800039 | rs12225735 | T | C | NULL | 0.593873103 | NULL&M_Disease |
| 11 | 116437742 | rs74954439 | A | G | NULL | 0.593873103 | NULL&M_Disease |
| 11 | 117207626 | rs520315 | C | T | NULL | 0.593873103 | NULL&M_Disease |
| 11 | 117207628 | rs620848 | G | C | NULL | 0.593873103 | NULL&M_Disease |
| 11 | 119847840 | rs78097928 | A | G | NULL | 0.5926666 | NULL&M_Disease |
| 11 | 120423601 | rs12278740 | T | C | NULL | 0.593873103 | NULL&M_Disease |
| 11 | 116963519 | rs7950093 | T | G | NULL | 0.593873103 | NULL&M_Disease |
| 11 | 117803627 | rs73024427 | T | C | NULL | 0.593873103 | NULL&M_Disease |
| 11 | 121040370 | rs510722 | A | T | NULL | 0.593873103 | NULL&M_Disease |
| 11 | 117447373 | rs648740 | A | G | NULL | 0.593873103 | NULL&M_Disease |
| 11 | 117633791 | rs73587426 | G | C | NULL | 0.593873103 | NULL&M_Disease |
| 11 | 117633827 | rs73587428 | G | A | NULL | 0.593873103 | NULL&M_Disease |
| 11 | 117633869 | rs73587429 | A | G | NULL | 0.593873103 | NULL&M_Disease |
| 11 | 117848134 | rs73009916 | T | C | NULL | 0.593873103 | NULL&M_Disease |
| 11 | 120491113 | rs1316812 | T | C | NULL | 0.593873103 | NULL&M_Disease |
| 11 | 117633297 | rs61903857 | T | G | NULL | 0.593873103 | NULL&M_Disease |
| 11 | 117633349 | rs61903858 | A | G | NULL | 0.593873103 | NULL&M_Disease |
| 11 | 119920056 | rs720314 | G | A | NULL | 0.593873103 | NULL&M_Disease |
| 11 | 119920110 | rs720313 | G | A | NULL | 0.593873103 | NULL&M_Disease |
| 11 | 119921007 | rs11217661 | T | C | NULL | 0.593873103 | NULL&M_Disease |
| 11 | 120566146 | rs61902685 | C | T | NULL | 0.593873103 | NULL&M_Disease |
| 11 | 120575671 | rs9630199 | C | T | NULL | 0.593873103 | NULL&M_Disease |
| 11 | 120780402 | rs10892639 | C | A | NULL | 0.593873103 | NULL&M_Disease |
| 11 | 116330610 | rs514112 | G | A | NULL | 0.593873103 | NULL&M_Disease |
| 11 | 116516653 | rs1240768 | C | T | NULL | 0.593873103 | NULL&M_Disease |
| 11 | 117712887 | rs473057 | G | A | NULL | 0.593873103 | NULL&M_Disease |
| 11 | 117810532 | rs56249090 | A | C | NULL | 0.593873103 | NULL&M_Disease |
| 11 | 119766043 | rs116966258 | A | G | NULL | 0.593873103 | NULL&M_Disease |
| 11 | 116669428 | rs35412484 | C | T | NULL | 0.593873103 | NULL&M_Disease |
| 11 | 117112526 | rs531184 | T | G | NULL | 0.593873103 | NULL&M_Disease |
| 11 | 119030759 | rs4148172 | G | A | NULL | 0.593873103 | NULL&M_Disease |
| 11 | 120823396 | rs584092 | T | C | NULL | 0.593873103 | NULL&M_Disease |
| 11 | 120915270 | rs138568456 | G | A | NULL | 0.593873103 | NULL&M_Disease |
| 11 | 116823062 | rs11605327 | A | G | NULL | 0.595078474 | NULL&M_Disease |
| 11 | 119844325 | rs145069523 | G | A | NULL | 0.593873103 | NULL&M_Disease |
| 11 | 120200671 | rs117207582 | T | C | NULL | 0.593873103 | NULL&M_Disease |
| 11 | 120809383 | rs602104 | C | T | NULL | 0.595078474 | NULL&M_Disease |
| 11 | 118214726 | rs7947185 | C | T | NULL | 0.595078474 | NULL&M_Disease |
| 11 | 119772803 | rs66633375 | A | T | NULL | 0.595078474 | NULL&M_Disease |
| 11 | 120583803 | rs12281614 | T | C | NULL | 0.595078474 | NULL&M_Disease |
| 11 | 117008946 | rs504068 | T | C | NULL | 0.595078474 | NULL&M_Disease |
| 11 | 117401470 | rs7927088 | G | A | NULL | 0.593873103 | NULL&M_Disease |
| 11 | 117632463 | rs61903856 | A | G | NULL | 0.593873103 | NULL&M_Disease |
| 11 | 117787277 | rs73022414 | C | T | NULL | 0.593873103 | NULL&M_Disease |
| 11 | 120999631 | rs1893707 | C | A | NULL | 0.595078474 | NULL&M_Disease |
| 11 | 116431149 | rs11600156 | G | A | NULL | 0.595078474 | NULL&M_Disease |
| 11 | 117685752 | rs113770024 | A | G | NULL | 0.593873103 | NULL&M_Disease |
| 11 | 118607694 | rs7119702 | G | A | NULL | 0.595078474 | NULL&M_Disease |
| 11 | 119946763 | rs677409 | G | T | NULL | 0.595078474 | NULL&M_Disease |
| 11 | 120423595 | rs2508655 | C | G | NULL | 0.595078474 | NULL&M_Disease |
| 11 | 117613813 | rs11826147 | C | G | NULL | 0.593873103 | NULL&M_Disease |
| 11 | 118945502 | rs72997349 | C | T | NULL | 0.593873103 | NULL&M_Disease |
| 11 | 119916341 | rs1960536 | C | A | NULL | 0.595078474 | NULL&M_Disease |
| 11 | 120860425 | rs7940751 | T | C | NULL | 0.595078474 | NULL&M_Disease |
| 11 | 117952387 | rs671160 | T | C | NULL | 0.595078474 | NULL&M_Disease |
| 11 | 120575845 | rs9630200 | A | G | NULL | 0.595078474 | NULL&M_Disease |
| 11 | 120576039 | rs11217981 | A | G | NULL | 0.595078474 | NULL&M_Disease |
| 11 | 120696251 | rs113447930 | C | T | NULL | 0.595078474 | NULL&M_Disease |
| 11 | 116714118 | rs494606 | G | T | NULL | 0.595078474 | NULL&M_Disease |
| 11 | 116729013 | rs539858 | C | T | NULL | 0.595078474 | NULL&M_Disease |
| 11 | 117291492 | rs111864243 | C | T | NULL | 0.595078474 | NULL&M_Disease |
| 11 | 118601363 | rs557990 | T | C | NULL | 0.595078474 | NULL&M_Disease |
| 11 | 118602392 | rs564892 | A | T | NULL | 0.595078474 | NULL&M_Disease |
| 11 | 118954805 | rs2855068 | T | C | NULL | 0.595078474 | NULL&M_Disease |
| 11 | 119859971 | rs117447786 | A | C | NULL | 0.593873103 | NULL&M_Disease |
| 11 | 116500411 | rs76517460 | A | G | NULL | 0.595078474 | NULL&M_Disease |
| 11 | 120313999 | rs118114248 | A | G | NULL | 0.593873103 | NULL&M_Disease |
| 11 | 118607090 | rs10736495 | C | G | NULL | 0.595078474 | NULL&M_Disease |
| 11 | 119751375 | rs547344 | A | G | NULL | 0.595078474 | NULL&M_Disease |
| 11 | 119775102 | rs7128589 | C | T | NULL | 0.595078474 | NULL&M_Disease |
| 11 | 119945046 | rs7952087 | T | C | NULL | 0.595078474 | NULL&M_Disease |
| 11 | 116281901 | rs111453105 | A | G | NULL | 0.595078474 | NULL&M_Disease |
| 11 | 118826060 | rs625291 | C | T | NULL | 0.596282699 | NULL&M_Disease |
| 11 | 116588425 | rs4938304 | T | C | NULL | 0.595078474 | NULL&M_Disease |
| 11 | 117375489 | rs2276342 | C | G | NULL | 0.595078474 | NULL&M_Disease |
| 11 | 119804674 | rs117047926 | T | C | NULL | 0.595078474 | NULL&M_Disease |
| 11 | 119923295 | rs143448490 | A | G | NULL | 0.595078474 | NULL&M_Disease |
| 11 | 120712972 | rs117794242 | T | C | NULL | 0.595078474 | NULL&M_Disease |
| 11 | 116537393 | rs34125463 | A | T | NULL | 0.595078474 | NULL&M_Disease |
| 11 | 117617279 | rs7942226 | C | T | NULL | 0.595078474 | NULL&M_Disease |
| 11 | 117722587 | rs686245 | C | T | NULL | 0.595078474 | NULL&M_Disease |
| 11 | 117723059 | rs4537792 | G | A | NULL | 0.595078474 | NULL&M_Disease |
| 11 | 117747629 | rs143049161 | T | C | NULL | 0.595078474 | NULL&M_Disease |
| 11 | 117929490 | rs2847699 | A | G | NULL | 0.595078474 | NULL&M_Disease |
| 11 | 119663058 | rs112704487 | C | T | NULL | 0.595078474 | NULL&M_Disease |
| 11 | 119904390 | rs11217648 | G | A | NULL | 0.596282699 | NULL&M_Disease |
| 11 | 120567441 | rs61902688 | A | C | NULL | 0.596282699 | NULL&M_Disease |
| 11 | 117463914 | rs35314805 | C | T | NULL | 0.596282699 | NULL&M_Disease |
| 11 | 117636729 | rs12574613 | T | C | NULL | 0.596282699 | NULL&M_Disease |
| 11 | 119793024 | rs527878 | G | A | NULL | 0.596282699 | NULL&M_Disease |
| 11 | 120860672 | rs11606348 | C | T | NULL | 0.595078474 | NULL&M_Disease |
| 11 | 121003604 | rs1045947297 | G | T | NULL | 0.596282699 | NULL&M_Disease |

| | | | | | | | | | |
|---|---|---|---|---|---|---|---|---|---|
| 11 | 118066912 | rs12808254 | T | C | 1.058 | -7977.388 | -7978.147 | -0.759 | 1.81E-06 |
| 11 | 119870561 | rs78289484 | T | C | 0.945 | -7977.388 | -7978.147 | -0.759 | 1.81E-06 |
| 11 | 120699050 | rs76871374 | G | A | 0.944 | -7977.388 | -7978.147 | -0.759 | 1.81E-06 |
| 11 | 118150882 | rs117527146 | A | C | 1.03 | -7977.388 | -7978.148 | -0.76 | 1.81E-06 |
| 11 | 120800039 | rs12225735 | T | C | 1.051 | -7977.388 | -7978.149 | -0.76 | 1.81E-06 |
| 11 | 116437742 | rs74954439 | A | G | 1.058 | -7977.388 | -7978.149 | -0.761 | 1.81E-06 |
| 11 | 117207626 | rs520315 | C | T | 0.953 | -7977.388 | -7978.15 | -0.761 | 1.81E-06 |
| 11 | 117207628 | rs620848 | G | C | 0.953 | -7977.388 | -7978.15 | -0.761 | 1.81E-06 |
| 11 | 119847840 | rs78097928 | A | G | 1.04 | -7977.388 | -7978.15 | -0.761 | 1.81E-06 |
| 11 | 120423601 | rs12278740 | T | C | 0.954 | -7977.388 | -7978.149 | -0.761 | 1.81E-06 |
| 11 | 116963519 | rs7950093 | T | G | 1.049 | -7977.388 | -7978.151 | -0.762 | 1.81E-06 |
| 11 | 117803627 | rs73024427 | T | C | 0.946 | -7977.388 | -7978.15 | -0.762 | 1.81E-06 |
| 11 | 121040370 | rs510722 | A | T | 0.955 | -7977.388 | -7978.15 | -0.762 | 1.81E-06 |
| 11 | 117447373 | rs648740 | A | G | 0.947 | -7977.388 | -7978.151 | -0.763 | 1.80E-06 |
| 11 | 117633791 | rs73587426 | G | C | 1.057 | -7977.388 | -7978.152 | -0.763 | 1.80E-06 |
| 11 | 117633827 | rs73587428 | G | A | 1.057 | -7977.388 | -7978.152 | -0.763 | 1.80E-06 |
| 11 | 117633869 | rs73587429 | A | G | 1.057 | -7977.388 | -7978.152 | -0.763 | 1.80E-06 |
| 11 | 117848134 | rs73009916 | T | C | 0.946 | -7977.388 | -7978.151 | -0.763 | 1.80E-06 |
| 11 | 120491113 | rs1316812 | T | C | 1.047 | -7977.388 | -7978.152 | -0.763 | 1.80E-06 |
| 11 | 117633297 | rs61903857 | T | G | 1.057 | -7977.388 | -7978.152 | -0.764 | 1.80E-06 |
| 11 | 117633349 | rs61903858 | A | G | 1.057 | -7977.388 | -7978.152 | -0.764 | 1.80E-06 |
| 11 | 119920056 | rs720314 | G | A | 1.047 | -7977.388 | -7978.153 | -0.765 | 1.80E-06 |
| 11 | 119920110 | rs720313 | G | A | 1.047 | -7977.388 | -7978.153 | -0.765 | 1.80E-06 |
| 11 | 119921007 | rs11217661 | T | C | 1.047 | -7977.388 | -7978.153 | -0.765 | 1.80E-06 |
| 11 | 120566146 | rs61902685 | C | T | 0.948 | -7977.388 | -7978.154 | -0.765 | 1.80E-06 |
| 11 | 120575671 | rs9630199 | C | T | 0.949 | -7977.388 | -7978.154 | -0.765 | 1.80E-06 |
| 11 | 120780402 | rs10892639 | C | A | 1.047 | -7977.388 | -7978.154 | -0.765 | 1.80E-06 |
| 11 | 116330610 | rs514112 | G | A | 1.05 | -7977.388 | -7978.155 | -0.766 | 1.80E-06 |
| 11 | 116516653 | rs1240768 | C | T | 0.956 | -7977.388 | -7978.154 | -0.766 | 1.80E-06 |
| 11 | 117712887 | rs473057 | G | A | 1.045 | -7977.388 | -7978.154 | -0.766 | 1.80E-06 |
| 11 | 117810532 | rs56249090 | A | C | 0.946 | -7977.388 | -7978.154 | -0.766 | 1.80E-06 |
| 11 | 119766043 | rs116966258 | A | G | 0.981 | -7977.388 | -7978.154 | -0.766 | 1.80E-06 |
| 11 | 116669428 | rs35412484 | C | T | 0.95 | -7977.388 | -7978.156 | -0.767 | 1.80E-06 |
| 11 | 117112526 | rs531184 | T | G | 0.956 | -7977.388 | -7978.156 | -0.767 | 1.80E-06 |
| 11 | 119030759 | rs4148172 | G | A | 1.058 | -7977.388 | -7978.156 | -0.767 | 1.80E-06 |
| 11 | 120823396 | rs584092 | T | C | 0.951 | -7977.388 | -7978.155 | -0.767 | 1.80E-06 |
| 11 | 120915270 | rs138568456 | G | A | 1.025 | -7977.388 | -7978.156 | -0.767 | 1.80E-06 |
| 11 | 116823062 | rs11605327 | A | G | 0.956 | -7977.388 | -7978.156 | -0.768 | 1.79E-06 |
| 11 | 119844325 | rs145069523 | G | A | 1.039 | -7977.388 | -7978.156 | -0.768 | 1.79E-06 |
| 11 | 120200671 | rs117207582 | T | C | 0.95 | -7977.388 | -7978.157 | -0.768 | 1.79E-06 |
| 11 | 120809383 | rs602104 | C | T | 0.951 | -7977.388 | -7978.157 | -0.768 | 1.79E-06 |
| 11 | 118214726 | rs7947185 | C | T | 0.956 | -7977.388 | -7978.158 | -0.769 | 1.79E-06 |
| 11 | 119772803 | rs66633375 | A | T | 0.955 | -7977.388 | -7978.157 | -0.769 | 1.79E-06 |
| 11 | 120583803 | rs12281614 | T | C | 0.949 | -7977.388 | -7978.157 | -0.769 | 1.79E-06 |
| 11 | 117008946 | rs504068 | T | C | 0.951 | -7977.388 | -7978.158 | -0.77 | 1.79E-06 |
| 11 | 117401470 | rs7927088 | G | A | 0.973 | -7977.388 | -7978.159 | -0.77 | 1.79E-06 |
| 11 | 117632463 | rs61903856 | A | G | 1.056 | -7977.388 | -7978.158 | -0.77 | 1.79E-06 |
| 11 | 117787277 | rs73022414 | C | T | 0.947 | -7977.388 | -7978.159 | -0.77 | 1.79E-06 |
| 11 | 120999631 | rs1893707 | C | A | 0.957 | -7977.388 | -7978.158 | -0.77 | 1.79E-06 |
| 11 | 116431149 | rs11600156 | G | A | 1.052 | -7977.388 | -7978.159 | -0.771 | 1.79E-06 |
| 11 | 117685752 | rs113770024 | A | G | 0.983 | -7977.388 | -7978.159 | -0.771 | 1.79E-06 |
| 11 | 118607694 | rs7119702 | G | A | 0.948 | -7977.388 | -7978.159 | -0.771 | 1.79E-06 |
| 11 | 119946763 | rs677409 | G | T | 0.946 | -7977.388 | -7978.159 | -0.771 | 1.79E-06 |
| 11 | 120423595 | rs2508655 | C | G | 0.955 | -7977.388 | -7978.159 | -0.771 | 1.79E-06 |
| 11 | 117613813 | rs11826147 | C | G | 0.958 | -7977.388 | -7978.16 | -0.772 | 1.79E-06 |
| 11 | 118945502 | rs72997349 | C | T | 0.961 | -7977.388 | -7978.161 | -0.772 | 1.79E-06 |
| 11 | 119916341 | rs1960536 | C | A | 1.047 | -7977.388 | -7978.16 | -0.772 | 1.79E-06 |
| 11 | 120860425 | rs7940751 | T | C | 0.95 | -7977.388 | -7978.161 | -0.772 | 1.79E-06 |
| 11 | 117952387 | rs671160 | T | C | 1.057 | -7977.388 | -7978.162 | -0.773 | 1.79E-06 |
| 11 | 120575845 | rs9630200 | A | G | 0.949 | -7977.388 | -7978.161 | -0.773 | 1.79E-06 |
| 11 | 120576039 | rs11217981 | A | G | 0.949 | -7977.388 | -7978.161 | -0.773 | 1.79E-06 |
| 11 | 120696251 | rs113447930 | C | T | 0.945 | -7977.388 | -7978.161 | -0.773 | 1.79E-06 |
| 11 | 116714118 | rs494606 | G | T | 1.057 | -7977.388 | -7978.162 | -0.774 | 1.78E-06 |
| 11 | 116729013 | rs539858 | C | T | 1.057 | -7977.388 | -7978.162 | -0.774 | 1.78E-06 |
| 11 | 117291492 | rs111864243 | C | T | 0.96 | -7977.388 | -7978.162 | -0.774 | 1.78E-06 |
| 11 | 118601363 | rs557990 | T | C | 0.948 | -7977.388 | -7978.162 | -0.774 | 1.78E-06 |
| 11 | 118602392 | rs564892 | A | T | 0.948 | -7977.388 | -7978.162 | -0.774 | 1.78E-06 |
| 11 | 118954805 | rs2855068 | T | C | 1.053 | -7977.388 | -7978.162 | -0.774 | 1.78E-06 |
| 11 | 119859971 | rs117447786 | A | C | 1.01 | -7977.388 | -7978.162 | -0.774 | 1.78E-06 |
| 11 | 116500411 | rs76517460 | A | G | 1.039 | -7977.388 | -7978.163 | -0.775 | 1.78E-06 |
| 11 | 120313999 | rs118114248 | A | G | 1.016 | -7977.388 | -7978.163 | -0.775 | 1.78E-06 |
| 11 | 118607090 | rs10736495 | C | G | 0.948 | -7977.388 | -7978.164 | -0.776 | 1.78E-06 |
| 11 | 119751375 | rs547344 | A | G | 1.056 | -7977.388 | -7978.164 | -0.776 | 1.78E-06 |
| 11 | 119775102 | rs7128589 | C | T | 0.955 | -7977.388 | -7978.164 | -0.776 | 1.78E-06 |
| 11 | 119945046 | rs7952087 | T | C | 0.984 | -7977.388 | -7978.164 | -0.776 | 1.78E-06 |
| 11 | 116281901 | rs111453105 | A | G | 0.979 | -7977.388 | -7978.165 | -0.777 | 1.78E-06 |
| 11 | 118826060 | rs625291 | C | T | 0.957 | -7977.388 | -7978.166 | -0.777 | 1.78E-06 |
| 11 | 116588425 | rs4938304 | T | C | 1.055 | -7977.388 | -7978.167 | -0.778 | 1.78E-06 |
| 11 | 117375489 | rs2276342 | C | G | 0.945 | -7977.388 | -7978.166 | -0.778 | 1.78E-06 |
| 11 | 119804674 | rs117047926 | T | C | 0.983 | -7977.388 | -7978.166 | -0.778 | 1.78E-06 |
| 11 | 119923295 | rs143448490 | A | G | 1.011 | -7977.388 | -7978.166 | -0.778 | 1.78E-06 |
| 11 | 120712972 | rs117794242 | T | C | 0.998 | -7977.388 | -7978.166 | -0.778 | 1.78E-06 |
| 11 | 116537393 | rs34125463 | A | T | 0.97 | -7977.388 | -7978.167 | -0.779 | 1.77E-06 |
| 11 | 117617279 | rs7942226 | C | T | 0.946 | -7977.388 | -7978.168 | -0.779 | 1.77E-06 |
| 11 | 117722587 | rs686245 | C | T | 0.946 | -7977.388 | -7978.167 | -0.779 | 1.77E-06 |
| 11 | 117723059 | rs4537792 | G | A | 0.946 | -7977.388 | -7978.167 | -0.779 | 1.77E-06 |
| 11 | 117747629 | rs143049161 | T | C | 0.963 | -7977.388 | -7978.167 | -0.779 | 1.77E-06 |
| 11 | 117929490 | rs2847699 | A | G | 0.955 | -7977.388 | -7978.168 | -0.779 | 1.77E-06 |
| 11 | 119663058 | rs112704487 | C | T | 0.947 | -7977.388 | -7978.167 | -0.779 | 1.77E-06 |
| 11 | 119904390 | rs11217648 | G | A | 1.047 | -7977.388 | -7978.168 | -0.779 | 1.77E-06 |
| 11 | 120567441 | rs61902688 | A | C | 0.949 | -7977.388 | -7978.167 | -0.779 | 1.77E-06 |
| 11 | 117463914 | rs35314805 | C | T | 0.956 | -7977.388 | -7978.168 | -0.78 | 1.77E-06 |
| 11 | 117636729 | rs12574613 | T | C | 0.951 | -7977.388 | -7978.169 | -0.78 | 1.77E-06 |
| 11 | 119793024 | rs527878 | G | A | 0.956 | -7977.388 | -7978.168 | -0.78 | 1.77E-06 |
| 11 | 120860672 | rs11606348 | C | T | 0.987 | -7977.388 | -7978.168 | -0.78 | 1.77E-06 |
| 11 | 121003604 | rs1045947297 | G | T | 0.952 | -7977.388 | -7978.168 | -0.78 | 1.77E-06 |

| | | | | | | | |
|---|---|---|---|---|---|---|---|
| 11 | 119932617 | rs114461031 | T | G | NULL | 0.596282699 | NULL&M_Disease |
| 11 | 120871485 | rs12800036 | T | C | NULL | 0.596282699 | NULL&M_Disease |
| 11 | 121192382 | rs6589876 | C | G | NULL | 0.596282699 | NULL&M_Disease |
| 11 | 118882138 | rs112711662 | T | C | NULL | 0.596282699 | NULL&M_Disease |
| 11 | 119506749 | rs4936490 | A | G | NULL | 0.596282699 | NULL&M_Disease |
| 11 | 119917892 | rs972534 | C | T | NULL | 0.596282699 | NULL&M_Disease |
| 11 | 116307041 | rs13377448 | T | G | NULL | 0.596282699 | NULL&M_Disease |
| 11 | 116314513 | rs4936350 | T | C | NULL | 0.596282699 | NULL&M_Disease |
| 11 | 117618713 | rs11605423 | A | G | NULL | 0.596282699 | NULL&M_Disease |
| 11 | 119728139 | rs76861941 | G | A | NULL | 0.595078474 | NULL&M_Disease |
| 11 | 116331582 | rs17119701 | G | A | NULL | 0.596282699 | NULL&M_Disease |
| 11 | 116531739 | rs2155216 | T | C | NULL | 0.595078474 | NULL&M_Disease |
| 11 | 116531950 | rs2186670 | T | C | NULL | 0.595078474 | NULL&M_Disease |
| 11 | 116532221 | rs1944153 | C | T | NULL | 0.595078474 | NULL&M_Disease |
| 11 | 116950556 | rs148143761 | G | A | NULL | 0.596282699 | NULL&M_Disease |
| 11 | 119905145 | rs10892518 | T | C | NULL | 0.596282699 | NULL&M_Disease |
| 11 | 117214580 | rs2509212 | G | C | NULL | 0.596282699 | NULL&M_Disease |
| 11 | 119389987 | rs77160958 | G | A | NULL | 0.596282699 | NULL&M_Disease |
| 11 | 119905245 | rs11217651 | C | T | NULL | 0.596282699 | NULL&M_Disease |
| 11 | 120826569 | rs149239940 | A | G | NULL | 0.596282699 | NULL&M_Disease |
| 11 | 117631784 | rs11216539 | C | T | NULL | 0.596282699 | NULL&M_Disease |
| 11 | 117726427 | rs531855 | T | C | NULL | 0.596282699 | NULL&M_Disease |
| 11 | 119474608 | rs140557493 | G | C | NULL | 0.596282699 | NULL&M_Disease |
| 11 | 119909608 | rs1452841 | G | A | NULL | 0.596282699 | NULL&M_Disease |
| 11 | 119910275 | rs12799875 | T | C | NULL | 0.596282699 | NULL&M_Disease |
| 11 | 119910556 | rs12800482 | T | C | NULL | 0.596282699 | NULL&M_Disease |
| 11 | 119910592 | rs12800844 | T | G | NULL | 0.596282699 | NULL&M_Disease |
| 11 | 119910833 | rs61897925 | A | G | NULL | 0.596282699 | NULL&M_Disease |
| 11 | 119911205 | rs11607618 | A | G | NULL | 0.596282699 | NULL&M_Disease |
| 11 | 119912882 | rs12790374 | G | A | NULL | 0.596282699 | NULL&M_Disease |
| 11 | 119912919 | rs12792065 | C | T | NULL | 0.596282699 | NULL&M_Disease |
| 11 | 119914725 | rs11217654 | G | T | NULL | 0.596282699 | NULL&M_Disease |
| 11 | 117259904 | rs117740883 | T | C | NULL | 0.596282699 | NULL&M_Disease |
| 11 | 118880727 | rs73559126 | A | T | NULL | 0.596282699 | NULL&M_Disease |
| 11 | 119848376 | rs113447691 | T | C | NULL | 0.596282699 | NULL&M_Disease |
| 11 | 121032517 | rs633012 | A | G | NULL | 0.597485766 | NULL&M_Disease |
| 11 | 116464013 | rs1876498 | A | G | NULL | 0.597485766 | NULL&M_Disease |
| 11 | 117195628 | rs560564 | C | A | NULL | 0.597485766 | NULL&M_Disease |
| 11 | 117733364 | rs11216592 | G | A | NULL | 0.596282699 | NULL&M_Disease |
| 11 | 118402962 | rs45558732 | T | G | NULL | 0.596282699 | NULL&M_Disease |
| 11 | 119815248 | rs634351 | C | T | NULL | 0.596282699 | NULL&M_Disease |
| 11 | 116237782 | rs488435 | T | C | NULL | 0.597485766 | NULL&M_Disease |
| 11 | 118003694 | rs71482129 | A | G | NULL | 0.596282699 | NULL&M_Disease |
| 11 | 119917309 | rs11512472 | G | A | NULL | 0.597485766 | NULL&M_Disease |
| 11 | 119905180 | rs11217650 | T | G | NULL | 0.597485766 | NULL&M_Disease |
| 11 | 120330904 | rs61898760 | G | A | NULL | 0.596282699 | NULL&M_Disease |
| 11 | 117655063 | rs7124184 | T | C | NULL | 0.597485766 | NULL&M_Disease |
| 11 | 119949979 | rs585809 | T | C | NULL | 0.597485766 | NULL&M_Disease |
| 11 | 120493805 | rs56763747 | G | A | NULL | 0.597485766 | NULL&M_Disease |
| 11 | 116529188 | rs117935983 | G | T | NULL | 0.596282699 | NULL&M_Disease |
| 11 | 117615003 | rs12289673 | A | G | NULL | 0.596282699 | NULL&M_Disease |
| 11 | 117615068 | rs12289727 | A | G | NULL | 0.596282699 | NULL&M_Disease |
| 11 | 117615371 | rs12291096 | T | C | NULL | 0.596282699 | NULL&M_Disease |
| 11 | 119925422 | rs1010260 | A | G | NULL | 0.597485766 | NULL&M_Disease |
| 11 | 116723171 | rs583219 | C | T | NULL | 0.597485766 | NULL&M_Disease |
| 11 | 116826870 | rs10502221 | T | C | NULL | 0.597485766 | NULL&M_Disease |
| 11 | 117454782 | rs191030218 | C | T | NULL | 0.596282699 | NULL&M_Disease |
| 11 | 117805473 | rs11216639 | T | C | NULL | 0.597485766 | NULL&M_Disease |
| 11 | 119580246 | rs143408085 | A | G | NULL | 0.597485766 | NULL&M_Disease |
| 11 | 120701346 | rs76200019 | A | G | NULL | 0.597485766 | NULL&M_Disease |
| 11 | 117612002 | rs7929757 | C | T | NULL | 0.597485766 | NULL&M_Disease |
| 11 | 117787228 | rs73022413 | C | T | NULL | 0.597485766 | NULL&M_Disease |
| 11 | 116673768 | rs1729409 | A | G | NULL | 0.597485766 | NULL&M_Disease |
| 11 | 116783600 | rs983567 | C | T | NULL | 0.597485766 | NULL&M_Disease |
| 11 | 117585714 | rs7107887 | A | C | NULL | 0.597485766 | NULL&M_Disease |
| 11 | 118040642 | rs10892221 | A | G | NULL | 0.597485766 | NULL&M_Disease |
| 11 | 118041102 | rs10892222 | A | G | NULL | 0.597485766 | NULL&M_Disease |
| 11 | 119670624 | rs11605165 | A | G | NULL | 0.597485766 | NULL&M_Disease |
| 11 | 119671550 | rs11217511 | C | G | NULL | 0.597485766 | NULL&M_Disease |
| 11 | 120018785 | rs117397221 | T | C | NULL | 0.597485766 | NULL&M_Disease |
| 11 | 117310022 | rs11216387 | G | A | NULL | 0.597485766 | NULL&M_Disease |
| 11 | 118232332 | rs79911534 | A | G | NULL | 0.597485766 | NULL&M_Disease |
| 11 | 118789952 | rs11217090 | T | C | NULL | 0.597485766 | NULL&M_Disease |
| 11 | 119806408 | rs140189045 | A | G | NULL | 0.597485766 | NULL&M_Disease |
| 11 | 120800956 | rs12796452 | C | G | NULL | 0.597485766 | NULL&M_Disease |
| 11 | 118185492 | rs61900905 | C | T | NULL | 0.597485766 | NULL&M_Disease |
| 11 | 119915782 | rs11493812 | G | A | NULL | 0.597485766 | NULL&M_Disease |
| 11 | 121027240 | rs527258 | A | T | NULL | 0.597485766 | NULL&M_Disease |
| 11 | 121059141 | rs10892687 | A | G | NULL | 0.597485766 | NULL&M_Disease |
| 11 | 117615875 | rs11216523 | A | G | NULL | 0.597485766 | NULL&M_Disease |
| 11 | 117615877 | rs11216524 | C | G | NULL | 0.597485766 | NULL&M_Disease |
| 11 | 117913374 | rs675855 | T | G | NULL | 0.597485766 | NULL&M_Disease |
| 11 | 118845813 | rs4442562 | A | G | NULL | 0.59868766 | NULL&M_Disease |
| 11 | 120486825 | rs138184472 | C | A | NULL | 0.597485766 | NULL&M_Disease |
| 11 | 116782819 | rs7123336 | C | T | NULL | 0.59868766 | NULL&M_Disease |
| 11 | 119803362 | rs61899023 | A | G | NULL | 0.597485766 | NULL&M_Disease |
| 11 | 118347020 | rs9332782 | C | T | NULL | 0.597485766 | NULL&M_Disease |
| 11 | 116462431 | rs35154750 | C | A | NULL | 0.59868766 | NULL&M_Disease |
| 11 | 117618437 | rs4936406 | C | T | NULL | 0.59868766 | NULL&M_Disease |
| 11 | 119287416 | rs147774547 | T | C | NULL | 0.597485766 | NULL&M_Disease |
| 11 | 120492000 | rs61900926 | A | G | NULL | 0.59868766 | NULL&M_Disease |
| 11 | 117588733 | rs11216513 | T | G | NULL | 0.597485766 | NULL&M_Disease |
| 11 | 119927792 | rs662146 | G | A | NULL | 0.59868766 | NULL&M_Disease |
| 11 | 120177530 | rs56765654 | T | C | NULL | 0.59868766 | NULL&M_Disease |
| 11 | 121035860 | rs546916 | T | G | NULL | 0.59868766 | NULL&M_Disease |
| 11 | 117727735 | rs565530 | G | T | NULL | 0.59868766 | NULL&M_Disease |

| | | | | | | | | | |
|---|---|---|---|---|---|---|---|---|---|
| 11 | 119932617 | rs114461031 | T | G | 0.946 | -7977.388 | -7978.169 | -0.781 | 1.77E-06 |
| 11 | 120871485 | rs12800036 | T | C | 0.95 | -7977.388 | -7978.169 | -0.781 | 1.77E-06 |
| 11 | 121192382 | rs6589876 | C | G | 0.951 | -7977.388 | -7978.17 | -0.781 | 1.77E-06 |
| 11 | 118882138 | rs112711662 | T | C | 0.946 | -7977.388 | -7978.171 | -0.782 | 1.77E-06 |
| 11 | 119506749 | rs4936490 | A | G | 0.948 | -7977.388 | -7978.17 | -0.782 | 1.77E-06 |
| 11 | 119917892 | rs972534 | C | T | 1.047 | -7977.388 | -7978.171 | -0.782 | 1.77E-06 |
| 11 | 116307041 | rs13377448 | T | G | 1.053 | -7977.388 | -7978.171 | -0.783 | 1.77E-06 |
| 11 | 116314513 | rs4936350 | T | C | 0.948 | -7977.388 | -7978.171 | -0.783 | 1.77E-06 |
| 11 | 117618713 | rs11605423 | A | G | 0.955 | -7977.388 | -7978.171 | -0.783 | 1.77E-06 |
| 11 | 119728139 | rs76861941 | G | A | 1.029 | -7977.388 | -7978.171 | -0.783 | 1.77E-06 |
| 11 | 116331582 | rs17119701 | G | A | 0.95 | -7977.388 | -7978.172 | -0.784 | 1.77E-06 |
| 11 | 116531739 | rs2155216 | T | C | 0.972 | -7977.388 | -7978.172 | -0.784 | 1.77E-06 |
| 11 | 116531950 | rs2186670 | T | C | 0.972 | -7977.388 | -7978.172 | -0.784 | 1.77E-06 |
| 11 | 116532221 | rs1944153 | C | T | 0.972 | -7977.388 | -7978.172 | -0.784 | 1.77E-06 |
| 11 | 116950556 | rs148143761 | G | A | 1.054 | -7977.388 | -7978.172 | -0.784 | 1.77E-06 |
| 11 | 119905145 | rs10892518 | T | C | 1.047 | -7977.388 | -7978.173 | -0.784 | 1.77E-06 |
| 11 | 117214580 | rs2509212 | G | C | 1.046 | -7977.388 | -7978.173 | -0.785 | 1.76E-06 |
| 11 | 119389987 | rs77160958 | G | A | 0.979 | -7977.388 | -7978.173 | -0.785 | 1.76E-06 |
| 11 | 119905245 | rs11217651 | C | T | 1.047 | -7977.388 | -7978.173 | -0.785 | 1.76E-06 |
| 11 | 120826569 | rs149239940 | A | G | 1.007 | -7977.388 | -7978.174 | -0.785 | 1.76E-06 |
| 11 | 117631784 | rs11216539 | C | T | 1.055 | -7977.388 | -7978.175 | -0.786 | 1.76E-06 |
| 11 | 117726427 | rs531855 | T | C | 0.946 | -7977.388 | -7978.174 | -0.786 | 1.76E-06 |
| 11 | 119474608 | rs140557493 | G | C | 0.947 | -7977.388 | -7978.175 | -0.786 | 1.76E-06 |
| 11 | 119909608 | rs1452841 | G | A | 1.047 | -7977.388 | -7978.174 | -0.786 | 1.76E-06 |
| 11 | 119910275 | rs12799875 | T | C | 1.047 | -7977.388 | -7978.174 | -0.786 | 1.76E-06 |
| 11 | 119910556 | rs12800482 | T | C | 1.047 | -7977.388 | -7978.174 | -0.786 | 1.76E-06 |
| 11 | 119910592 | rs12800844 | T | G | 1.047 | -7977.388 | -7978.174 | -0.786 | 1.76E-06 |
| 11 | 119910833 | rs61897925 | A | G | 1.047 | -7977.388 | -7978.174 | -0.786 | 1.76E-06 |
| 11 | 119911205 | rs11607618 | A | G | 1.047 | -7977.388 | -7978.174 | -0.786 | 1.76E-06 |
| 11 | 119912882 | rs12790374 | G | A | 1.047 | -7977.388 | -7978.174 | -0.786 | 1.76E-06 |
| 11 | 119912919 | rs12792065 | C | T | 1.047 | -7977.388 | -7978.174 | -0.786 | 1.76E-06 |
| 11 | 119914725 | rs11217654 | G | T | 1.047 | -7977.388 | -7978.174 | -0.786 | 1.76E-06 |
| 11 | 117259904 | rs117740883 | T | C | 0.976 | -7977.388 | -7978.176 | -0.787 | 1.76E-06 |
| 11 | 118880727 | rs73559126 | A | T | 0.947 | -7977.388 | -7978.175 | -0.787 | 1.76E-06 |
| 11 | 119848376 | rs113447691 | T | C | 1.057 | -7977.388 | -7978.175 | -0.787 | 1.76E-06 |
| 11 | 121032517 | rs633012 | A | G | 0.956 | -7977.388 | -7978.176 | -0.787 | 1.76E-06 |
| 11 | 116464013 | rs1876498 | A | G | 1.046 | -7977.388 | -7978.177 | -0.788 | 1.76E-06 |
| 11 | 117195628 | rs560564 | C | A | 0.957 | -7977.388 | -7978.177 | -0.788 | 1.76E-06 |
| 11 | 117733364 | rs11216592 | G | A | 0.963 | -7977.388 | -7978.177 | -0.788 | 1.76E-06 |
| 11 | 118402962 | rs45558732 | T | G | 0.958 | -7977.388 | -7978.177 | -0.788 | 1.76E-06 |
| 11 | 119815248 | rs634351 | C | T | 1.057 | -7977.388 | -7978.176 | -0.788 | 1.76E-06 |
| 11 | 116237782 | rs488435 | T | C | 1.045 | -7977.388 | -7978.177 | -0.789 | 1.76E-06 |
| 11 | 118003694 | rs71482129 | A | G | 0.956 | -7977.388 | -7978.177 | -0.789 | 1.76E-06 |
| 11 | 119917309 | rs11512472 | G | A | 1.047 | -7977.388 | -7978.178 | -0.789 | 1.76E-06 |
| 11 | 119905180 | rs11217650 | T | G | 1.047 | -7977.388 | -7978.178 | -0.79 | 1.76E-06 |
| 11 | 120330904 | rs61898760 | G | A | 1.046 | -7977.388 | -7978.179 | -0.79 | 1.76E-06 |
| 11 | 117655063 | rs7124184 | T | C | 0.956 | -7977.388 | -7978.179 | -0.791 | 1.75E-06 |
| 11 | 119949979 | rs585809 | T | C | 0.947 | -7977.388 | -7978.179 | -0.791 | 1.75E-06 |
| 11 | 120493805 | rs56763747 | G | A | 1.046 | -7977.388 | -7978.179 | -0.791 | 1.75E-06 |
| 11 | 116529188 | rs117935983 | G | T | 0.975 | -7977.388 | -7978.18 | -0.792 | 1.75E-06 |
| 11 | 117615003 | rs12289673 | A | G | 0.96 | -7977.388 | -7978.181 | -0.792 | 1.75E-06 |
| 11 | 117615068 | rs12289727 | A | G | 0.96 | -7977.388 | -7978.181 | -0.792 | 1.75E-06 |
| 11 | 117615371 | rs12291096 | T | C | 0.96 | -7977.388 | -7978.181 | -0.792 | 1.75E-06 |
| 11 | 119925422 | rs1010260 | A | G | 1.047 | -7977.388 | -7978.18 | -0.792 | 1.75E-06 |
| 11 | 116723171 | rs583219 | C | T | 1.055 | -7977.388 | -7978.181 | -0.793 | 1.75E-06 |
| 11 | 116826870 | rs10502221 | T | C | 0.957 | -7977.388 | -7978.181 | -0.793 | 1.75E-06 |
| 11 | 117454782 | rs191030218 | C | T | 0.972 | -7977.388 | -7978.181 | -0.793 | 1.75E-06 |
| 11 | 117805473 | rs11216639 | T | C | 0.947 | -7977.388 | -7978.181 | -0.793 | 1.75E-06 |
| 11 | 119580246 | rs143408085 | A | G | 0.95 | -7977.388 | -7978.182 | -0.793 | 1.75E-06 |
| 11 | 120701346 | rs76200019 | A | G | 0.946 | -7977.388 | -7978.182 | -0.793 | 1.75E-06 |
| 11 | 117612002 | rs7929757 | C | T | 0.959 | -7977.388 | -7978.182 | -0.794 | 1.75E-06 |
| 11 | 117787228 | rs73022413 | C | T | 0.949 | -7977.388 | -7978.183 | -0.794 | 1.75E-06 |
| 11 | 116673768 | rs1729409 | A | G | 1.045 | -7977.388 | -7978.184 | -0.795 | 1.75E-06 |
| 11 | 116783600 | rs983567 | C | T | 1.049 | -7977.388 | -7978.183 | -0.795 | 1.75E-06 |
| 11 | 117585714 | rs7107887 | A | C | 0.949 | -7977.388 | -7978.183 | -0.795 | 1.75E-06 |
| 11 | 118040642 | rs10892221 | A | G | 0.95 | -7977.388 | -7978.183 | -0.795 | 1.75E-06 |
| 11 | 118041102 | rs10892222 | A | G | 0.95 | -7977.388 | -7978.183 | -0.795 | 1.75E-06 |
| 11 | 119670624 | rs11605165 | A | G | 0.952 | -7977.388 | -7978.184 | -0.795 | 1.75E-06 |
| 11 | 119671550 | rs11217511 | C | G | 0.952 | -7977.388 | -7978.184 | -0.795 | 1.75E-06 |
| 11 | 120018785 | rs117397221 | T | C | 0.981 | -7977.388 | -7978.183 | -0.795 | 1.75E-06 |
| 11 | 117310022 | rs11216387 | G | A | 1.055 | -7977.388 | -7978.184 | -0.796 | 1.75E-06 |
| 11 | 118232332 | rs79911534 | A | G | 0.989 | -7977.388 | -7978.184 | -0.796 | 1.75E-06 |
| 11 | 118789952 | rs11217090 | T | C | 1.056 | -7977.388 | -7978.185 | -0.796 | 1.75E-06 |
| 11 | 119806408 | rs140189045 | A | G | 0.995 | -7977.388 | -7978.184 | -0.796 | 1.75E-06 |
| 11 | 120800956 | rs12796452 | C | G | 1.05 | -7977.388 | -7978.185 | -0.796 | 1.75E-06 |
| 11 | 118185492 | rs61900905 | C | T | 1.046 | -7977.388 | -7978.185 | -0.797 | 1.74E-06 |
| 11 | 119915782 | rs11493812 | G | A | 1.046 | -7977.388 | -7978.185 | -0.797 | 1.74E-06 |
| 11 | 121027240 | rs527258 | A | T | 0.956 | -7977.388 | -7978.185 | -0.797 | 1.74E-06 |
| 11 | 121059141 | rs10892687 | A | G | 0.956 | -7977.388 | -7978.185 | -0.797 | 1.74E-06 |
| 11 | 117615875 | rs11216523 | A | G | 0.96 | -7977.388 | -7978.186 | -0.798 | 1.74E-06 |
| 11 | 117615877 | rs11216524 | C | G | 0.96 | -7977.388 | -7978.186 | -0.798 | 1.74E-06 |
| 11 | 117913374 | rs675855 | T | G | 0.95 | -7977.388 | -7978.186 | -0.798 | 1.74E-06 |
| 11 | 118845813 | rs4442562 | A | G | 1.05 | -7977.388 | -7978.186 | -0.798 | 1.74E-06 |
| 11 | 120486825 | rs138184472 | C | A | 0.992 | -7977.388 | -7978.186 | -0.798 | 1.74E-06 |
| 11 | 116782819 | rs7123336 | C | T | 1.049 | -7977.388 | -7978.187 | -0.799 | 1.74E-06 |
| 11 | 119803362 | rs61899023 | A | G | 0.987 | -7977.388 | -7978.187 | -0.799 | 1.74E-06 |
| 11 | 118347020 | rs9332782 | C | T | 0.96 | -7977.388 | -7978.188 | -0.8 | 1.74E-06 |
| 11 | 116462431 | rs35154750 | C | A | 0.947 | -7977.388 | -7978.189 | -0.801 | 1.74E-06 |
| 11 | 117618437 | rs4936406 | C | T | 0.947 | -7977.388 | -7978.189 | -0.801 | 1.74E-06 |
| 11 | 119287416 | rs147774547 | T | C | 0.987 | -7977.388 | -7978.189 | -0.801 | 1.74E-06 |
| 11 | 120492000 | rs61900926 | A | G | 1.046 | -7977.388 | -7978.189 | -0.801 | 1.74E-06 |
| 11 | 117588733 | rs11216513 | T | G | 0.958 | -7977.388 | -7978.19 | -0.802 | 1.73E-06 |
| 11 | 119927792 | rs662146 | G | A | 1.046 | -7977.388 | -7978.191 | -0.802 | 1.73E-06 |
| 11 | 120177530 | rs56765654 | T | C | 1.056 | -7977.388 | -7978.19 | -0.802 | 1.73E-06 |
| 11 | 121035860 | rs546916 | T | G | 0.956 | -7977.388 | -7978.19 | -0.802 | 1.73E-06 |
| 11 | 117727735 | rs565530 | G | T | 0.947 | -7977.388 | -7978.191 | -0.803 | 1.73E-06 |

| | | | | | | | |
|---|---|---|---|---|---|---|---|
| 11 | 118890022 | rs3802881 | T | C | NULL | 0.59868766 | NULL&M_Disease |
| 11 | 120622881 | rs9667343 | C | G | NULL | 0.59868766 | NULL&M_Disease |
| 11 | 116757765 | rs148403721 | A | C | NULL | 0.597485766 | NULL&M_Disease |
| 11 | 116772917 | rs141023293 | A | G | NULL | 0.597485766 | NULL&M_Disease |
| 11 | 118889073 | rs11545858 | T | G | NULL | 0.59868766 | NULL&M_Disease |
| 11 | 120131553 | rs117170835 | A | G | NULL | 0.59868766 | NULL&M_Disease |
| 11 | 120479903 | rs12274778 | G | C | NULL | 0.59868766 | NULL&M_Disease |
| 11 | 116241411 | rs17475957 | A | G | NULL | 0.59868766 | NULL&M_Disease |
| 11 | 116312186 | rs11215933 | A | C | NULL | 0.59868766 | NULL&M_Disease |
| 11 | 117188120 | rs4938369 | T | C | NULL | 0.59868766 | NULL&M_Disease |
| 11 | 117460030 | rs633985 | G | C | NULL | 0.59868766 | NULL&M_Disease |
| 11 | 117700588 | rs2004506 | A | G | NULL | 0.59868766 | NULL&M_Disease |
| 11 | 118805220 | rs11605658 | T | C | NULL | 0.59868766 | NULL&M_Disease |
| 11 | 120492853 | rs1379188468 | T | A | NULL | 0.59868766 | NULL&M_Disease |
| 11 | 116220623 | rs74332985 | C | T | NULL | 0.59868766 | NULL&M_Disease |
| 11 | 117106116 | rs3016581 | A | G | NULL | 0.599888369 | NULL&M_Disease |
| 11 | 118594326 | rs544096 | G | A | NULL | 0.59868766 | NULL&M_Disease |
| 11 | 116516506 | rs1789403 | C | T | NULL | 0.599888369 | NULL&M_Disease |
| 11 | 117563212 | rs11603672 | G | A | NULL | 0.59868766 | NULL&M_Disease |
| 11 | 119666456 | rs140160504 | T | C | NULL | 0.59868766 | NULL&M_Disease |
| 11 | 119742803 | rs11217548 | T | C | NULL | 0.59868766 | NULL&M_Disease |
| 11 | 116314370 | rs4936349 | G | A | NULL | 0.599888369 | NULL&M_Disease |
| 11 | 117452212 | rs681046 | C | T | NULL | 0.599888369 | NULL&M_Disease |
| 11 | 119946171 | rs644324 | G | A | NULL | 0.59868766 | NULL&M_Disease |
| 11 | 117085874 | rs150758276 | C | G | NULL | 0.59868766 | NULL&M_Disease |
| 11 | 117787018 | rs73022412 | C | T | NULL | 0.59868766 | NULL&M_Disease |
| 11 | 117795838 | rs73022489 | A | G | NULL | 0.599888369 | NULL&M_Disease |
| 11 | 117916286 | rs631191 | T | C | NULL | 0.59868766 | NULL&M_Disease |
| 11 | 120405828 | rs78867156 | A | G | NULL | 0.59868766 | NULL&M_Disease |
| 11 | 120824094 | rs597670 | T | C | NULL | 0.599888369 | NULL&M_Disease |
| 11 | 117622809 | rs4576856 | G | T | NULL | 0.599888369 | NULL&M_Disease |
| 11 | 119907867 | rs1939331 | C | G | NULL | 0.599888369 | NULL&M_Disease |
| 11 | 119929853 | rs1392314200 | T | C | NULL | 0.599888369 | NULL&M_Disease |
| 11 | 119945841 | rs584282 | G | T | NULL | 0.599888369 | NULL&M_Disease |
| 11 | 120159052 | rs7121784 | A | C | NULL | 0.599888369 | NULL&M_Disease |
| 11 | 117614720 | rs1939977 | A | G | NULL | 0.59868766 | NULL&M_Disease |
| 11 | 118805144 | rs11605265 | A | G | NULL | 0.599888369 | NULL&M_Disease |
| 11 | 119440387 | rs73001093 | T | G | NULL | 0.599888369 | NULL&M_Disease |
| 11 | 119673235 | rs56220026 | C | G | NULL | 0.599888369 | NULL&M_Disease |
| 11 | 119673271 | rs56116614 | G | A | NULL | 0.599888369 | NULL&M_Disease |
| 11 | 119906398 | rs1452840 | T | A | NULL | 0.599888369 | NULL&M_Disease |
| 11 | 119927321 | rs10892524 | T | G | NULL | 0.599888369 | NULL&M_Disease |
| 11 | 121033277 | rs663031 | T | C | NULL | 0.599888369 | NULL&M_Disease |
| 11 | 116893588 | rs139663973 | C | T | NULL | 0.59868766 | NULL&M_Disease |
| 11 | 117314053 | rs11216393 | A | G | NULL | 0.599888369 | NULL&M_Disease |
| 11 | 118126576 | rs1629083 | T | C | NULL | 0.599888369 | NULL&M_Disease |
| 11 | 118211797 | rs7129048 | C | T | NULL | 0.599888369 | NULL&M_Disease |
| 11 | 121034992 | rs576271 | A | T | NULL | 0.599888369 | NULL&M_Disease |
| 11 | 116715688 | rs1145207 | G | A | NULL | 0.599888369 | NULL&M_Disease |
| 11 | 117577925 | rs7927379 | A | G | NULL | 0.599888369 | NULL&M_Disease |
| 11 | 117603231 | rs17121153 | T | C | NULL | 0.599888369 | NULL&M_Disease |
| 11 | 119774817 | rs76200908 | A | G | NULL | 0.599888369 | NULL&M_Disease |
| 11 | 117643723 | rs79973603 | C | T | NULL | 0.599888369 | NULL&M_Disease |
| 11 | 119506018 | rs12270510 | T | C | NULL | 0.599888369 | NULL&M_Disease |
| 11 | 117312865 | rs61905271 | A | G | NULL | 0.599888369 | NULL&M_Disease |
| 11 | 118626825 | rs74337439 | A | G | NULL | 0.599888369 | NULL&M_Disease |
| 11 | 119664304 | rs7928437 | G | T | NULL | 0.599888369 | NULL&M_Disease |
| 11 | 119669993 | rs74387077 | A | G | NULL | 0.599888369 | NULL&M_Disease |
| 11 | 120448042 | rs73000642 | A | G | NULL | 0.599888369 | NULL&M_Disease |
| 11 | 119598971 | rs77630867 | T | A | NULL | 0.599888369 | NULL&M_Disease |
| 11 | 120834480 | rs655824 | C | T | NULL | 0.599888369 | NULL&M_Disease |
| 11 | 117583898 | rs7950969 | G | A | NULL | 0.599888369 | NULL&M_Disease |
| 11 | 118121180 | rs139322068 | G | T | NULL | 0.599888369 | NULL&M_Disease |
| 11 | 119599641 | rs138907015 | T | C | NULL | 0.599888369 | NULL&M_Disease |
| 11 | 119927844 | rs675783 | C | T | NULL | 0.601087879 | NULL&M_Disease |
| 11 | 116672231 | rs2542061 | A | G | NULL | 0.601087879 | NULL&M_Disease |
| 11 | 117625225 | rs113858735 | A | G | NULL | 0.601087879 | NULL&M_Disease |
| 11 | 117998243 | rs74942119 | T | C | NULL | 0.599888369 | NULL&M_Disease |
| 11 | 116365711 | rs112995279 | A | C | NULL | 0.599888369 | NULL&M_Disease |
| 11 | 116596309 | rs61905078 | C | A | NULL | 0.601087879 | NULL&M_Disease |
| 11 | 119483599 | rs118086106 | C | T | NULL | 0.599888369 | NULL&M_Disease |
| 11 | 117808400 | rs73024501 | G | C | NULL | 0.601087879 | NULL&M_Disease |
| 11 | 117908503 | rs149539291 | A | G | NULL | 0.599888369 | NULL&M_Disease |
| 11 | 117988797 | rs17121760 | A | G | NULL | 0.601087879 | NULL&M_Disease |
| 11 | 121065886 | rs631969 | T | C | NULL | 0.601087879 | NULL&M_Disease |
| 11 | 116434951 | rs77963671 | T | C | NULL | 0.601087879 | NULL&M_Disease |
| 11 | 119758831 | rs521976 | T | C | NULL | 0.601087879 | NULL&M_Disease |
| 11 | 119918352 | rs12363889 | G | A | NULL | 0.601087879 | NULL&M_Disease |
| 11 | 117643228 | rs75696653 | T | A | NULL | 0.601087879 | NULL&M_Disease |
| 11 | 117722487 | rs498314 | A | G | NULL | 0.601087879 | NULL&M_Disease |
| 11 | 120986571 | rs480761 | A | G | NULL | 0.601087879 | NULL&M_Disease |
| 11 | 119905840 | rs1235340366 | G | C | NULL | 0.601087879 | NULL&M_Disease |
| 11 | 119905865 | rs12362929 | G | A | NULL | 0.601087879 | NULL&M_Disease |
| 11 | 119906213 | rs11605106 | A | G | NULL | 0.601087879 | NULL&M_Disease |
| 11 | 119906313 | rs10502234 | A | G | NULL | 0.601087879 | NULL&M_Disease |
| 11 | 120008972 | rs74996375 | A | C | NULL | 0.601087879 | NULL&M_Disease |
| 11 | 120115450 | rs11217773 | A | G | NULL | 0.601087879 | NULL&M_Disease |
| 11 | 116806412 | rs2513094 | C | T | NULL | 0.601087879 | NULL&M_Disease |
| 11 | 117611162 | rs1939979 | T | C | NULL | 0.601087879 | NULL&M_Disease |
| 11 | 117622274 | rs76492199 | A | G | NULL | 0.601087879 | NULL&M_Disease |
| 11 | 118804671 | rs143711539 | A | G | NULL | 0.601087879 | NULL&M_Disease |
| 11 | 119523181 | rs7103685 | C | T | NULL | 0.601087879 | NULL&M_Disease |
| 11 | 121033374 | rs663435 | G | A | NULL | 0.601087879 | NULL&M_Disease |
| 11 | 118000796 | rs17121771 | T | C | NULL | 0.601087879 | NULL&M_Disease |
| 11 | 118857809 | rs56784564 | A | G | NULL | 0.601087879 | NULL&M_Disease |
| 11 | 116487300 | rs73011197 | T | C | NULL | 0.601087879 | NULL&M_Disease |

| | | | | | | | | | |
|---|---|---|---|---|---|---|---|---|---|
| 11 | 118890022 | rs3802881 | T | C | 0.947 | -7977.388 | -7978.191 | -0.803 | 1.73E-06 |
| 11 | 120622881 | rs9667343 | C | G | 0.947 | -7977.388 | -7978.192 | -0.803 | 1.73E-06 |
| 11 | 116757765 | rs148403721 | A | C | 1.014 | -7977.388 | -7978.192 | -0.804 | 1.73E-06 |
| 11 | 116772917 | rs141023293 | A | G | 1.012 | -7977.388 | -7978.192 | -0.804 | 1.73E-06 |
| 11 | 118889073 | rs11545858 | T | G | 0.947 | -7977.388 | -7978.194 | -0.805 | 1.73E-06 |
| 11 | 120131553 | rs117170835 | A | G | 1.049 | -7977.388 | -7978.193 | -0.805 | 1.73E-06 |
| 11 | 120479903 | rs12274778 | G | C | 1.046 | -7977.388 | -7978.193 | -0.805 | 1.73E-06 |
| 11 | 116241411 | rs17475957 | A | G | 0.948 | -7977.388 | -7978.194 | -0.806 | 1.73E-06 |
| 11 | 116312186 | rs11215933 | A | C | 0.948 | -7977.388 | -7978.194 | -0.806 | 1.73E-06 |
| 11 | 117188120 | rs4938369 | T | C | 0.947 | -7977.388 | -7978.195 | -0.806 | 1.73E-06 |
| 11 | 117460030 | rs633985 | G | C | 0.957 | -7977.388 | -7978.194 | -0.806 | 1.73E-06 |
| 11 | 117700588 | rs2004506 | A | G | 0.951 | -7977.388 | -7978.194 | -0.806 | 1.73E-06 |
| 11 | 118805220 | rs11605658 | T | C | 1.05 | -7977.388 | -7978.194 | -0.806 | 1.73E-06 |
| 11 | 120492853 | rs1379188468 | T | A | 1.046 | -7977.388 | -7978.194 | -0.806 | 1.73E-06 |
| 11 | 116220623 | rs74332985 | C | T | 0.966 | -7977.388 | -7978.195 | -0.807 | 1.73E-06 |
| 11 | 117106116 | rs3016581 | A | G | 0.956 | -7977.388 | -7978.196 | -0.807 | 1.73E-06 |
| 11 | 118594326 | rs544096 | G | A | 0.949 | -7977.388 | -7978.195 | -0.807 | 1.73E-06 |
| 11 | 116516506 | rs1789403 | C | T | 0.957 | -7977.388 | -7978.197 | -0.808 | 1.72E-06 |
| 11 | 117563212 | rs11603672 | G | A | 0.951 | -7977.388 | -7978.196 | -0.808 | 1.72E-06 |
| 11 | 119666456 | rs140160504 | T | C | 0.993 | -7977.388 | -7978.196 | -0.808 | 1.72E-06 |
| 11 | 119742803 | rs11217548 | T | C | 1.055 | -7977.388 | -7978.196 | -0.808 | 1.72E-06 |
| 11 | 116314370 | rs4936349 | G | A | 0.949 | -7977.388 | -7978.197 | -0.809 | 1.72E-06 |
| 11 | 117452212 | rs681046 | C | T | 0.949 | -7977.388 | -7978.197 | -0.809 | 1.72E-06 |
| 11 | 119946171 | rs644324 | G | A | 0.947 | -7977.388 | -7978.197 | -0.809 | 1.72E-06 |
| 11 | 117085874 | rs150758276 | C | G | 1.004 | -7977.388 | -7978.198 | -0.81 | 1.72E-06 |
| 11 | 117787018 | rs73022412 | C | T | 0.95 | -7977.388 | -7978.198 | -0.81 | 1.72E-06 |
| 11 | 117795838 | rs73022489 | A | G | 0.947 | -7977.388 | -7978.199 | -0.81 | 1.72E-06 |
| 11 | 117916286 | rs631191 | T | C | 0.96 | -7977.388 | -7978.199 | -0.81 | 1.72E-06 |
| 11 | 120405828 | rs78867156 | A | G | 0.96 | -7977.388 | -7978.198 | -0.81 | 1.72E-06 |
| 11 | 120824094 | rs597670 | T | C | 0.952 | -7977.388 | -7978.198 | -0.81 | 1.72E-06 |
| 11 | 117622809 | rs4576856 | G | T | 0.949 | -7977.388 | -7978.199 | -0.811 | 1.72E-06 |
| 11 | 119907867 | rs1939331 | C | G | 1.046 | -7977.388 | -7978.2 | -0.811 | 1.72E-06 |
| 11 | 119929853 | rs1392314200 | T | C | 0.953 | -7977.388 | -7978.2 | -0.811 | 1.72E-06 |
| 11 | 119945841 | rs584282 | G | T | 0.948 | -7977.388 | -7978.2 | -0.811 | 1.72E-06 |
| 11 | 120159052 | rs7121784 | A | C | 1.052 | -7977.388 | -7978.199 | -0.811 | 1.72E-06 |
| 11 | 117614720 | rs1939977 | A | G | 0.963 | -7977.388 | -7978.2 | -0.812 | 1.72E-06 |
| 11 | 118805144 | rs11605265 | A | G | 1.049 | -7977.388 | -7978.201 | -0.812 | 1.72E-06 |
| 11 | 119440387 | rs73001093 | T | G | 0.948 | -7977.388 | -7978.2 | -0.812 | 1.72E-06 |
| 11 | 119673235 | rs56220026 | C | G | 0.953 | -7977.388 | -7978.2 | -0.812 | 1.72E-06 |
| 11 | 119673271 | rs56116614 | G | A | 0.953 | -7977.388 | -7978.2 | -0.812 | 1.72E-06 |
| 11 | 119906398 | rs1452840 | T | A | 1.046 | -7977.388 | -7978.201 | -0.812 | 1.72E-06 |
| 11 | 119927321 | rs10892524 | T | G | 1.046 | -7977.388 | -7978.201 | -0.812 | 1.72E-06 |
| 11 | 121033277 | rs663031 | T | C | 0.956 | -7977.388 | -7978.2 | -0.812 | 1.72E-06 |
| 11 | 116893588 | rs139663973 | C | T | 0.996 | -7977.388 | -7978.201 | -0.813 | 1.72E-06 |
| 11 | 117314053 | rs11216393 | A | G | 1.054 | -7977.388 | -7978.202 | -0.813 | 1.72E-06 |
| 11 | 118126576 | rs1629083 | T | C | 0.958 | -7977.388 | -7978.201 | -0.813 | 1.72E-06 |
| 11 | 118211797 | rs7129048 | C | T | 1.048 | -7977.388 | -7978.202 | -0.813 | 1.72E-06 |
| 11 | 121034992 | rs576271 | A | T | 0.956 | -7977.388 | -7978.201 | -0.813 | 1.72E-06 |
| 11 | 116715688 | rs1145207 | G | A | 1.053 | -7977.388 | -7978.202 | -0.814 | 1.71E-06 |
| 11 | 117577925 | rs7927379 | A | G | 0.96 | -7977.388 | -7978.202 | -0.814 | 1.71E-06 |
| 11 | 117603231 | rs17121153 | T | C | 0.949 | -7977.388 | -7978.203 | -0.814 | 1.71E-06 |
| 11 | 119774817 | rs76200908 | A | G | 1.047 | -7977.388 | -7978.202 | -0.814 | 1.71E-06 |
| 11 | 117643723 | rs79973603 | C | T | 0.947 | -7977.388 | -7978.203 | -0.815 | 1.71E-06 |
| 11 | 119506018 | rs12270510 | T | C | 0.949 | -7977.388 | -7978.203 | -0.815 | 1.71E-06 |
| 11 | 117312865 | rs61905271 | A | G | 1.028 | -7977.388 | -7978.204 | -0.816 | 1.71E-06 |
| 11 | 118626825 | rs74337439 | A | G | 1.051 | -7977.388 | -7978.206 | -0.817 | 1.71E-06 |
| 11 | 119664304 | rs7928437 | G | T | 0.973 | -7977.388 | -7978.205 | -0.817 | 1.71E-06 |
| 11 | 119669993 | rs74387077 | A | G | 0.953 | -7977.388 | -7978.205 | -0.817 | 1.71E-06 |
| 11 | 120448042 | rs73000642 | A | G | 1.015 | -7977.388 | -7978.205 | -0.817 | 1.71E-06 |
| 11 | 119598971 | rs77630867 | T | A | 1.01 | -7977.388 | -7978.206 | -0.818 | 1.71E-06 |
| 11 | 120834480 | rs655824 | C | T | 0.95 | -7977.388 | -7978.206 | -0.818 | 1.71E-06 |
| 11 | 117583898 | rs7950969 | G | A | 0.948 | -7977.388 | -7978.207 | -0.819 | 1.71E-06 |
| 11 | 118121180 | rs139322068 | G | T | 1.004 | -7977.388 | -7978.208 | -0.819 | 1.71E-06 |
| 11 | 119599641 | rs138907015 | T | C | 1.009 | -7977.388 | -7978.207 | -0.819 | 1.71E-06 |
| 11 | 119927844 | rs675783 | C | T | 1.046 | -7977.388 | -7978.207 | -0.819 | 1.71E-06 |
| 11 | 116672231 | rs2542061 | A | G | 1.044 | -7977.388 | -7978.209 | -0.82 | 1.70E-06 |
| 11 | 117625225 | rs113858735 | A | G | 0.948 | -7977.388 | -7978.208 | -0.82 | 1.70E-06 |
| 11 | 117998243 | rs74942119 | T | C | 0.98 | -7977.388 | -7978.208 | -0.82 | 1.70E-06 |
| 11 | 116365711 | rs112995279 | A | C | 0.957 | -7977.388 | -7978.209 | -0.821 | 1.70E-06 |
| 11 | 116596309 | rs61905078 | C | A | 0.948 | -7977.388 | -7978.209 | -0.821 | 1.70E-06 |
| 11 | 119483599 | rs118086106 | C | T | 1.015 | -7977.388 | -7978.209 | -0.821 | 1.70E-06 |
| 11 | 117808400 | rs73024501 | G | C | 0.948 | -7977.388 | -7978.21 | -0.822 | 1.70E-06 |
| 11 | 117908503 | rs149539291 | A | G | 0.998 | -7977.388 | -7978.211 | -0.822 | 1.70E-06 |
| 11 | 117988797 | rs17121760 | A | G | 0.957 | -7977.388 | -7978.211 | -0.822 | 1.70E-06 |
| 11 | 121065886 | rs631969 | T | C | 0.957 | -7977.388 | -7978.211 | -0.822 | 1.70E-06 |
| 11 | 116434951 | rs77963671 | T | C | 1.054 | -7977.388 | -7978.211 | -0.823 | 1.70E-06 |
| 11 | 119758831 | rs521976 | T | C | 1.048 | -7977.388 | -7978.211 | -0.823 | 1.70E-06 |
| 11 | 119918352 | rs12363889 | G | A | 1.046 | -7977.388 | -7978.211 | -0.823 | 1.70E-06 |
| 11 | 117643228 | rs75696653 | T | A | 0.948 | -7977.388 | -7978.212 | -0.824 | 1.70E-06 |
| 11 | 117722487 | rs498314 | A | G | 0.948 | -7977.388 | -7978.213 | -0.824 | 1.70E-06 |
| 11 | 120986571 | rs480761 | A | G | 1.048 | -7977.388 | -7978.212 | -0.824 | 1.70E-06 |
| 11 | 119905840 | rs1235340366 | G | C | 1.046 | -7977.388 | -7978.213 | -0.825 | 1.70E-06 |
| 11 | 119905865 | rs12362929 | G | A | 1.046 | -7977.388 | -7978.213 | -0.825 | 1.70E-06 |
| 11 | 119906213 | rs11605106 | A | G | 1.046 | -7977.388 | -7978.213 | -0.825 | 1.70E-06 |
| 11 | 119906313 | rs10502234 | A | G | 1.046 | -7977.388 | -7978.213 | -0.825 | 1.70E-06 |
| 11 | 120008972 | rs74996375 | A | C | 0.99 | -7977.388 | -7978.213 | -0.825 | 1.70E-06 |
| 11 | 120115450 | rs11217773 | A | G | 1.034 | -7977.388 | -7978.214 | -0.825 | 1.70E-06 |
| 11 | 116806412 | rs2513094 | C | T | 1.048 | -7977.388 | -7978.214 | -0.826 | 1.69E-06 |
| 11 | 117611162 | rs1939979 | T | C | 0.948 | -7977.388 | -7978.215 | -0.826 | 1.69E-06 |
| 11 | 117622274 | rs76492199 | A | G | 0.948 | -7977.388 | -7978.214 | -0.826 | 1.69E-06 |
| 11 | 118804671 | rs143711539 | A | G | 1.048 | -7977.388 | -7978.214 | -0.826 | 1.69E-06 |
| 11 | 119523181 | rs7103685 | C | T | 1.047 | -7977.388 | -7978.214 | -0.826 | 1.69E-06 |
| 11 | 121033374 | rs663435 | G | A | 0.957 | -7977.388 | -7978.214 | -0.826 | 1.69E-06 |
| 11 | 118000796 | rs17121771 | T | C | 0.955 | -7977.388 | -7978.215 | -0.827 | 1.69E-06 |
| 11 | 118857809 | rs56784564 | A | G | 0.948 | -7977.388 | -7978.215 | -0.827 | 1.69E-06 |
| 11 | 116487300 | rs73011197 | T | C | 0.959 | -7977.388 | -7978.216 | -0.828 | 1.69E-06 |

| | | | | | | | |
|---|---|---|---|---|---|---|---|
| 11 | 116870766 | rs7930264 | A | G | NULL | 0.602286177 | NULL&M_Disease |
| 11 | 119651089 | rs11217499 | T | C | NULL | 0.601087879 | NULL&M_Disease |
| 11 | 117153815 | rs17584190 | T | C | NULL | 0.601087879 | NULL&M_Disease |
| 11 | 118803842 | rs11600378 | G | A | NULL | 0.601087879 | NULL&M_Disease |
| 11 | 119815915 | rs1815888 | G | C | NULL | 0.602286177 | NULL&M_Disease |
| 11 | 116557314 | rs947990 | T | G | NULL | 0.602286177 | NULL&M_Disease |
| 11 | 119757989 | rs113381053 | A | G | NULL | 0.602286177 | NULL&M_Disease |
| 11 | 119615828 | rs11606857 | T | A | NULL | 0.601087879 | NULL&M_Disease |
| 11 | 119927944 | rs675331 | G | A | NULL | 0.602286177 | NULL&M_Disease |
| 11 | 120828827 | rs732480 | A | G | NULL | 0.602286177 | NULL&M_Disease |
| 11 | 117199742 | rs75235704 | G | T | NULL | 0.601087879 | NULL&M_Disease |
| 11 | 117795859 | rs73022490 | A | G | NULL | 0.602286177 | NULL&M_Disease |
| 11 | 116557959 | rs76814754 | T | C | NULL | 0.602286177 | NULL&M_Disease |
| 11 | 121021515 | rs6589858 | C | T | NULL | 0.602286177 | NULL&M_Disease |
| 11 | 116411836 | rs10790147 | T | C | NULL | 0.602286177 | NULL&M_Disease |
| 11 | 117314931 | rs1189010881 | T | G | NULL | 0.602286177 | NULL&M_Disease |
| 11 | 118793613 | rs11217091 | C | T | NULL | 0.602286177 | NULL&M_Disease |
| 11 | 119672890 | rs76070443 | G | A | NULL | 0.602286177 | NULL&M_Disease |
| 11 | 119673722 | rs11217512 | C | G | NULL | 0.602286177 | NULL&M_Disease |
| 11 | 119673977 | rs11217513 | C | T | NULL | 0.602286177 | NULL&M_Disease |
| 11 | 119778929 | rs17245326 | A | G | NULL | 0.602286177 | NULL&M_Disease |
| 11 | 117615502 | rs748771 | G | A | NULL | 0.602286177 | NULL&M_Disease |
| 11 | 118853995 | rs45573242 | C | A | NULL | 0.602286177 | NULL&M_Disease |
| 11 | 118876552 | rs74884652 | C | T | NULL | 0.602286177 | NULL&M_Disease |
| 11 | 119502477 | rs1567375 | A | C | NULL | 0.602286177 | NULL&M_Disease |
| 11 | 116260707 | rs501763 | G | A | NULL | 0.602286177 | NULL&M_Disease |
| 11 | 116360293 | rs7948548 | A | G | NULL | 0.602286177 | NULL&M_Disease |
| 11 | 116422645 | rs117080901 | A | G | NULL | 0.602286177 | NULL&M_Disease |
| 11 | 117806383 | rs60501859 | C | T | NULL | 0.602286177 | NULL&M_Disease |
| 11 | 117950796 | rs611475 | C | T | NULL | 0.602286177 | NULL&M_Disease |
| 11 | 119785124 | rs9667364 | G | A | NULL | 0.602286177 | NULL&M_Disease |
| 11 | 120702968 | rs74827927 | A | G | NULL | 0.602286177 | NULL&M_Disease |
| 11 | 116701833 | rs1269330 | A | G | NULL | 0.602286177 | NULL&M_Disease |
| 11 | 116848857 | rs509171 | T | C | NULL | 0.60348325 | NULL&M_Disease |
| 11 | 118856537 | rs3889525 | C | G | NULL | 0.602286177 | NULL&M_Disease |
| 11 | 119507477 | rs7105962 | A | G | NULL | 0.602286177 | NULL&M_Disease |
| 11 | 116271858 | rs56159851 | T | C | NULL | 0.602286177 | NULL&M_Disease |
| 11 | 116637985 | rs1145208 | C | A | NULL | 0.60348325 | NULL&M_Disease |
| 11 | 116828320 | rs682048 | G | T | NULL | 0.60348325 | NULL&M_Disease |
| 11 | 117577674 | rs668213 | G | A | NULL | 0.602286177 | NULL&M_Disease |
| 11 | 118040610 | rs602594 | T | C | NULL | 0.602286177 | NULL&M_Disease |
| 11 | 118185346 | rs3819250 | A | G | NULL | 0.60348325 | NULL&M_Disease |
| 11 | 120154200 | rs73578248 | T | C | NULL | 0.602286177 | NULL&M_Disease |
| 11 | 120631792 | rs138990232 | G | A | NULL | 0.602286177 | NULL&M_Disease |
| 11 | 116674897 | rs148134535 | G | C | NULL | 0.602286177 | NULL&M_Disease |
| 11 | 118890397 | rs112998705 | G | T | NULL | 0.60348325 | NULL&M_Disease |
| 11 | 117795852 | rs76835919 | G | C | NULL | 0.60348325 | NULL&M_Disease |
| 11 | 117904328 | rs78823230 | T | G | NULL | 0.602286177 | NULL&M_Disease |
| 11 | 119768855 | rs56816645 | G | A | NULL | 0.602286177 | NULL&M_Disease |
| 11 | 116226195 | rs78238602 | A | G | NULL | 0.602286177 | NULL&M_Disease |
| 11 | 116361536 | rs73009147 | T | C | NULL | 0.60348325 | NULL&M_Disease |
| 11 | 118755946 | rs76109832 | T | C | NULL | 0.60348325 | NULL&M_Disease |
| 11 | 118793479 | rs138526953 | A | C | NULL | 0.602286177 | NULL&M_Disease |
| 11 | 119672221 | rs7952568 | T | C | NULL | 0.60348325 | NULL&M_Disease |
| 11 | 119791050 | rs7105316 | T | C | NULL | 0.60348325 | NULL&M_Disease |
| 11 | 120706014 | rs11606105 | T | A | NULL | 0.60348325 | NULL&M_Disease |
| 11 | 120833690 | rs112885596 | C | T | NULL | 0.60348325 | NULL&M_Disease |
| 11 | 116587849 | rs7946299 | C | T | NULL | 0.60348325 | NULL&M_Disease |
| 11 | 118576463 | rs496547 | A | T | NULL | 0.60348325 | NULL&M_Disease |
| 11 | 120960054 | rs75050717 | A | G | NULL | 0.60348325 | NULL&M_Disease |
| 11 | 116731260 | rs687172 | A | T | NULL | 0.60348325 | NULL&M_Disease |
| 11 | 117201404 | rs73014332 | G | A | NULL | 0.60348325 | NULL&M_Disease |
| 11 | 117521490 | rs12799706 | C | T | NULL | 0.60348325 | NULL&M_Disease |
| 11 | 118212235 | rs112970787 | C | G | NULL | 0.60348325 | NULL&M_Disease |
| 11 | 118878960 | rs111804650 | T | C | NULL | 0.60348325 | NULL&M_Disease |
| 11 | 119935372 | rs73000741 | A | G | NULL | 0.60348325 | NULL&M_Disease |
| 11 | 116375302 | rs79914174 | C | T | NULL | 0.60348325 | NULL&M_Disease |
| 11 | 116466039 | rs12802294 | T | C | NULL | 0.60348325 | NULL&M_Disease |
| 11 | 117201682 | rs609272 | T | C | NULL | 0.60348325 | NULL&M_Disease |
| 11 | 117586011 | rs7108308 | G | C | NULL | 0.60348325 | NULL&M_Disease |
| 11 | 117958148 | rs602445 | C | T | NULL | 0.60348325 | NULL&M_Disease |
| 11 | 119240624 | rs12419866 | T | C | NULL | 0.60348325 | NULL&M_Disease |
| 11 | 120097175 | rs56667128 | A | G | NULL | 0.60348325 | NULL&M_Disease |
| 11 | 120123886 | rs80254203 | C | G | NULL | 0.60348325 | NULL&M_Disease |
| 11 | 120401377 | rs150204107 | A | G | NULL | 0.60348325 | NULL&M_Disease |
| 11 | 121004224 | rs475731 | T | C | NULL | 0.60348325 | NULL&M_Disease |
| 11 | 117681749 | rs71482116 | A | G | NULL | 0.60348325 | NULL&M_Disease |
| 11 | 119924508 | rs77101533 | G | A | NULL | 0.60348325 | NULL&M_Disease |
| 11 | 120130716 | rs75669750 | C | T | NULL | 0.60348325 | NULL&M_Disease |
| 11 | 120564824 | rs61902684 | T | C | NULL | 0.60348325 | NULL&M_Disease |
| 11 | 116465552 | rs35112163 | T | G | NULL | 0.60348325 | NULL&M_Disease |
| 11 | 117596086 | rs1893035 | A | G | NULL | 0.60348325 | NULL&M_Disease |
| 11 | 120657444 | rs2155258 | G | A | NULL | 0.60348325 | NULL&M_Disease |
| 11 | 120695126 | rs76716142 | T | C | NULL | 0.60348325 | NULL&M_Disease |
| 11 | 117899993 | rs76530395 | A | G | NULL | 0.60348325 | NULL&M_Disease |
| 11 | 119765750 | rs79162717 | G | A | NULL | 0.60348325 | NULL&M_Disease |
| 11 | 120189261 | rs61898346 | A | G | NULL | 0.60348325 | NULL&M_Disease |
| 11 | 120355388 | rs117792004 | A | T | NULL | 0.60348325 | NULL&M_Disease |
| 11 | 117313948 | rs73018334 | A | C | NULL | 0.60348325 | NULL&M_Disease |
| 11 | 117542149 | rs11216493 | T | C | NULL | 0.604679085 | NULL&M_Disease |
| 11 | 117727883 | rs476130 | T | C | NULL | 0.60348325 | NULL&M_Disease |
| 11 | 117727963 | rs2845939 | T | C | NULL | 0.60348325 | NULL&M_Disease |
| 11 | 120505235 | rs7102128 | A | G | NULL | 0.604679085 | NULL&M_Disease |
| 11 | 120623682 | rs11606421 | T | C | NULL | 0.60348325 | NULL&M_Disease |
| 11 | 116453885 | rs61904847 | C | T | NULL | 0.60348325 | NULL&M_Disease |
| 11 | 116863766 | rs6589583 | T | C | NULL | 0.604679085 | NULL&M_Disease |

| | | | | | | | | | |
|---|---|---|---|---|---|---|---|---|---|
| 11 | 116870766 | rs7930264 | A | G | 1.048 | -7977.388 | -7978.216 | -0.828 | 1.69E-06 |
| 11 | 119651089 | rs11217499 | T | C | 0.98 | -7977.388 | -7978.216 | -0.828 | 1.69E-06 |
| 11 | 117153815 | rs17584190 | T | C | 0.964 | -7977.388 | -7978.218 | -0.829 | 1.69E-06 |
| 11 | 118803842 | rs11600378 | G | A | 1.047 | -7977.388 | -7978.218 | -0.83 | 1.69E-06 |
| 11 | 119815915 | rs1815888 | G | C | 1.054 | -7977.388 | -7978.218 | -0.83 | 1.69E-06 |
| 11 | 116557314 | rs947990 | T | G | 1.052 | -7977.388 | -7978.219 | -0.831 | 1.69E-06 |
| 11 | 119757989 | rs113381053 | A | G | 1.052 | -7977.388 | -7978.22 | -0.831 | 1.69E-06 |
| 11 | 119615828 | rs11606857 | T | A | 0.973 | -7977.388 | -7978.221 | -0.832 | 1.68E-06 |
| 11 | 119927944 | rs675331 | G | A | 1.046 | -7977.388 | -7978.22 | -0.832 | 1.68E-06 |
| 11 | 120828827 | rs732480 | A | G | 0.952 | -7977.388 | -7978.221 | -0.832 | 1.68E-06 |
| 11 | 117199742 | rs75235704 | G | T | 1.027 | -7977.388 | -7978.221 | -0.833 | 1.68E-06 |
| 11 | 117795859 | rs73022490 | A | G | 0.948 | -7977.388 | -7978.221 | -0.833 | 1.68E-06 |
| 11 | 116557959 | rs76814754 | T | C | 0.997 | -7977.388 | -7978.223 | -0.834 | 1.68E-06 |
| 11 | 121021515 | rs6589858 | C | T | 0.957 | -7977.388 | -7978.222 | -0.834 | 1.68E-06 |
| 11 | 116411836 | rs10790147 | T | C | 1.044 | -7977.388 | -7978.223 | -0.835 | 1.68E-06 |
| 11 | 117314931 | rs1189010881 | T | G | 1.053 | -7977.388 | -7978.223 | -0.835 | 1.68E-06 |
| 11 | 118793613 | rs11217091 | C | T | 1.054 | -7977.388 | -7978.224 | -0.835 | 1.68E-06 |
| 11 | 119672890 | rs76070443 | G | A | 0.953 | -7977.388 | -7978.223 | -0.835 | 1.68E-06 |
| 11 | 119673722 | rs11217512 | C | G | 0.953 | -7977.388 | -7978.223 | -0.835 | 1.68E-06 |
| 11 | 119673977 | rs11217513 | C | T | 0.953 | -7977.388 | -7978.223 | -0.835 | 1.68E-06 |
| 11 | 119778929 | rs17245326 | A | G | 0.955 | -7977.388 | -7978.223 | -0.836 | 1.68E-06 |
| 11 | 117615502 | rs748771 | G | A | 0.949 | -7977.388 | -7978.224 | -0.836 | 1.68E-06 |
| 11 | 118853995 | rs45573242 | C | A | 0.949 | -7977.388 | -7978.224 | -0.836 | 1.68E-06 |
| 11 | 118876552 | rs74884652 | C | T | 1.052 | -7977.388 | -7978.225 | -0.836 | 1.68E-06 |
| 11 | 119502477 | rs1567375 | A | C | 1.044 | -7977.388 | -7978.224 | -0.836 | 1.68E-06 |
| 11 | 116260707 | rs501763 | G | A | 1 | -7977.388 | -7978.226 | -0.837 | 1.67E-06 |
| 11 | 116360293 | rs7948548 | A | G | 1.046 | -7977.388 | -7978.225 | -0.837 | 1.67E-06 |
| 11 | 116422645 | rs117080901 | A | G | 0.971 | -7977.388 | -7978.225 | -0.837 | 1.67E-06 |
| 11 | 117806383 | rs60501859 | C | T | 0.949 | -7977.388 | -7978.226 | -0.837 | 1.67E-06 |
| 11 | 117950796 | rs611475 | C | T | 1.052 | -7977.388 | -7978.225 | -0.837 | 1.67E-06 |
| 11 | 119785124 | rs9667364 | G | A | 1.053 | -7977.388 | -7978.225 | -0.837 | 1.67E-06 |
| 11 | 120702968 | rs74827927 | A | G | 0.949 | -7977.388 | -7978.226 | -0.837 | 1.67E-06 |
| 11 | 116701833 | rs1269330 | A | G | 1.052 | -7977.388 | -7978.226 | -0.838 | 1.67E-06 |
| 11 | 116848857 | rs509171 | T | C | 1.047 | -7977.388 | -7978.227 | -0.838 | 1.67E-06 |
| 11 | 118856537 | rs3889525 | C | G | 0.949 | -7977.388 | -7978.227 | -0.838 | 1.67E-06 |
| 11 | 119507477 | rs7105962 | A | G | 0.95 | -7977.388 | -7978.226 | -0.838 | 1.67E-06 |
| 11 | 116271858 | rs56159851 | T | C | 0.95 | -7977.388 | -7978.227 | -0.839 | 1.67E-06 |
| 11 | 116637985 | rs1145208 | C | A | 0.958 | -7977.388 | -7978.227 | -0.839 | 1.67E-06 |
| 11 | 116828320 | rs682048 | G | T | 1.047 | -7977.388 | -7978.227 | -0.839 | 1.67E-06 |
| 11 | 117577674 | rs668213 | G | A | 0.949 | -7977.388 | -7978.228 | -0.839 | 1.67E-06 |
| 11 | 118040610 | rs602594 | T | C | 0.953 | -7977.388 | -7978.228 | -0.839 | 1.67E-06 |
| 11 | 118185346 | rs3819250 | A | G | 1.046 | -7977.388 | -7978.227 | -0.839 | 1.67E-06 |
| 11 | 120154200 | rs73578248 | T | C | 0.949 | -7977.388 | -7978.227 | -0.839 | 1.67E-06 |
| 11 | 120631792 | rs138990232 | G | A | 0.956 | -7977.388 | -7978.228 | -0.839 | 1.67E-06 |
| 11 | 116674897 | rs148134535 | G | C | 1.002 | -7977.388 | -7978.228 | -0.84 | 1.67E-06 |
| 11 | 118890397 | rs112998705 | G | T | 0.949 | -7977.388 | -7978.228 | -0.84 | 1.67E-06 |
| 11 | 117795852 | rs76835919 | G | C | 0.949 | -7977.388 | -7978.229 | -0.841 | 1.67E-06 |
| 11 | 117904328 | rs78823230 | T | G | 0.968 | -7977.388 | -7978.229 | -0.841 | 1.67E-06 |
| 11 | 119768855 | rs56816645 | G | A | 0.969 | -7977.388 | -7978.229 | -0.841 | 1.67E-06 |
| 11 | 116226195 | rs78238602 | A | G | 1.036 | -7977.388 | -7978.231 | -0.842 | 1.67E-06 |
| 11 | 116361536 | rs73009147 | T | C | 1.053 | -7977.388 | -7978.23 | -0.842 | 1.67E-06 |
| 11 | 118755946 | rs76109832 | T | C | 0.952 | -7977.388 | -7978.23 | -0.842 | 1.67E-06 |
| 11 | 118793479 | rs138526953 | A | C | 1.007 | -7977.388 | -7978.231 | -0.842 | 1.67E-06 |
| 11 | 119672221 | rs7952568 | T | C | 0.953 | -7977.388 | -7978.23 | -0.842 | 1.67E-06 |
| 11 | 119791050 | rs7105316 | T | C | 1.052 | -7977.388 | -7978.231 | -0.842 | 1.67E-06 |
| 11 | 120706014 | rs11606105 | T | A | 0.953 | -7977.388 | -7978.23 | -0.842 | 1.67E-06 |
| 11 | 120833690 | rs112885596 | C | T | 1.053 | -7977.388 | -7978.23 | -0.842 | 1.67E-06 |
| 11 | 116587849 | rs7946299 | C | T | 0.951 | -7977.388 | -7978.231 | -0.843 | 1.66E-06 |
| 11 | 118576463 | rs496547 | A | T | 1.045 | -7977.388 | -7978.231 | -0.843 | 1.66E-06 |
| 11 | 120960054 | rs75050717 | A | G | 1.033 | -7977.388 | -7978.232 | -0.843 | 1.66E-06 |
| 11 | 116731260 | rs687172 | A | T | 1.051 | -7977.388 | -7978.232 | -0.844 | 1.66E-06 |
| 11 | 117201404 | rs73014332 | G | A | 0.955 | -7977.388 | -7978.232 | -0.844 | 1.66E-06 |
| 11 | 117521490 | rs12799706 | C | T | 0.976 | -7977.388 | -7978.233 | -0.844 | 1.66E-06 |
| 11 | 118212235 | rs112970787 | C | G | 1.044 | -7977.388 | -7978.232 | -0.844 | 1.66E-06 |
| 11 | 118878960 | rs111804650 | T | C | 0.949 | -7977.388 | -7978.232 | -0.844 | 1.66E-06 |
| 11 | 119935372 | rs73000741 | A | G | 1.027 | -7977.388 | -7978.232 | -0.844 | 1.66E-06 |
| 11 | 116375302 | rs79914174 | C | T | 0.949 | -7977.388 | -7978.233 | -0.845 | 1.66E-06 |
| 11 | 116466039 | rs12802294 | T | C | 0.949 | -7977.388 | -7978.233 | -0.845 | 1.66E-06 |
| 11 | 117201682 | rs609272 | T | C | 1.045 | -7977.388 | -7978.233 | -0.845 | 1.66E-06 |
| 11 | 117586011 | rs7108308 | G | C | 0.964 | -7977.388 | -7978.234 | -0.845 | 1.66E-06 |
| 11 | 117958148 | rs602445 | C | T | 1.053 | -7977.388 | -7978.233 | -0.845 | 1.66E-06 |
| 11 | 119240624 | rs12419866 | T | C | 0.955 | -7977.388 | -7978.233 | -0.845 | 1.66E-06 |
| 11 | 120097175 | rs56667128 | A | G | 0.964 | -7977.388 | -7978.233 | -0.845 | 1.66E-06 |
| 11 | 120123886 | rs80254203 | C | G | 0.989 | -7977.388 | -7978.233 | -0.845 | 1.66E-06 |
| 11 | 120401377 | rs150204107 | A | G | 0.992 | -7977.388 | -7978.233 | -0.845 | 1.66E-06 |
| 11 | 121004224 | rs475731 | T | C | 1.048 | -7977.388 | -7978.233 | -0.845 | 1.66E-06 |
| 11 | 117681749 | rs71482116 | A | G | 0.957 | -7977.388 | -7978.234 | -0.846 | 1.66E-06 |
| 11 | 119924508 | rs77101533 | G | A | 1.04 | -7977.388 | -7978.234 | -0.846 | 1.66E-06 |
| 11 | 120130716 | rs75669750 | C | T | 1.044 | -7977.388 | -7978.234 | -0.846 | 1.66E-06 |
| 11 | 120564824 | rs61902684 | T | C | 0.988 | -7977.388 | -7978.235 | -0.846 | 1.66E-06 |
| 11 | 116465552 | rs35112163 | T | G | 0.949 | -7977.388 | -7978.235 | -0.847 | 1.66E-06 |
| 11 | 117596086 | rs1893035 | A | G | 0.949 | -7977.388 | -7978.235 | -0.847 | 1.66E-06 |
| 11 | 120657444 | rs2155258 | G | A | 0.957 | -7977.388 | -7978.236 | -0.847 | 1.66E-06 |
| 11 | 120695126 | rs76716142 | T | C | 0.95 | -7977.388 | -7978.235 | -0.847 | 1.66E-06 |
| 11 | 117899993 | rs76530395 | A | G | 1.053 | -7977.388 | -7978.236 | -0.848 | 1.66E-06 |
| 11 | 119765750 | rs79162717 | G | A | 0.971 | -7977.388 | -7978.236 | -0.848 | 1.66E-06 |
| 11 | 120189261 | rs61898346 | A | G | 1.053 | -7977.388 | -7978.237 | -0.848 | 1.66E-06 |
| 11 | 120355388 | rs117792004 | A | T | 1.008 | -7977.388 | -7978.236 | -0.848 | 1.66E-06 |
| 11 | 117313948 | rs73018334 | A | C | 0.975 | -7977.388 | -7978.237 | -0.849 | 1.65E-06 |
| 11 | 117542149 | rs11216493 | T | C | 1.052 | -7977.388 | -7978.237 | -0.849 | 1.65E-06 |
| 11 | 117727883 | rs476130 | T | C | 0.949 | -7977.388 | -7978.238 | -0.849 | 1.65E-06 |
| 11 | 117727963 | rs2845939 | T | C | 0.949 | -7977.388 | -7978.237 | -0.849 | 1.65E-06 |
| 11 | 120505235 | rs7102128 | A | G | 0.958 | -7977.388 | -7978.237 | -0.849 | 1.65E-06 |
| 11 | 120623682 | rs11606421 | T | C | 0.949 | -7977.388 | -7978.237 | -0.849 | 1.65E-06 |
| 11 | 116453885 | rs61904847 | C | T | 1.053 | -7977.388 | -7978.238 | -0.85 | 1.65E-06 |
| 11 | 116863766 | rs6589583 | T | C | 1.047 | -7977.388 | -7978.238 | -0.85 | 1.65E-06 |

| | | | | | | | |
|---|---|---|---|---|---|---|---|
| 11 | 118185140 | rs2298821 | C | T | NULL | 0.604679085 | NULL&M_Disease |
| 11 | 118212979 | rs7939548 | A | G | NULL | 0.60348325 | NULL&M_Disease |
| 11 | 118041133 | rs600325 | C | T | NULL | 0.60348325 | NULL&M_Disease |
| 11 | 118311677 | rs77535829 | T | C | NULL | 0.60348325 | NULL&M_Disease |
| 11 | 118311869 | rs77487830 | G | A | NULL | 0.60348325 | NULL&M_Disease |
| 11 | 118583971 | rs6589677 | A | G | NULL | 0.604679085 | NULL&M_Disease |
| 11 | 117111214 | rs525390 | A | G | NULL | 0.604679085 | NULL&M_Disease |
| 11 | 117806283 | rs7938191 | A | G | NULL | 0.604679085 | NULL&M_Disease |
| 11 | 119753972 | rs964104 | A | G | NULL | 0.60348325 | NULL&M_Disease |
| 11 | 120098180 | rs11217771 | G | C | NULL | 0.60348325 | NULL&M_Disease |
| 11 | 117804535 | rs73024430 | T | C | NULL | 0.604679085 | NULL&M_Disease |
| 11 | 119926775 | rs1989151 | A | T | NULL | 0.604679085 | NULL&M_Disease |
| 11 | 120834199 | rs17124602 | C | G | NULL | 0.604679085 | NULL&M_Disease |
| 11 | 116272744 | rs4938270 | T | C | NULL | 0.604679085 | NULL&M_Disease |
| 11 | 116877361 | rs183640608 | G | A | NULL | 0.604679085 | NULL&M_Disease |
| 11 | 117282800 | rs522885 | G | T | NULL | 0.604679085 | NULL&M_Disease |
| 11 | 117605098 | rs1942929 | C | T | NULL | 0.604679085 | NULL&M_Disease |
| 11 | 117643958 | rs79537444 | T | C | NULL | 0.604679085 | NULL&M_Disease |
| 11 | 117987483 | rs1793178 | A | C | NULL | 0.604679085 | NULL&M_Disease |
| 11 | 119503402 | rs12273351 | T | G | NULL | 0.604679085 | NULL&M_Disease |
| 11 | 119757789 | rs1039094777 | T | C | NULL | 0.604679085 | NULL&M_Disease |
| 11 | 116634557 | rs17119975 | C | T | NULL | 0.604679085 | NULL&M_Disease |
| 11 | 117591241 | rs1455831109 | T | C | NULL | 0.604679085 | NULL&M_Disease |
| 11 | 118864777 | rs11822344 | C | G | NULL | 0.604679085 | NULL&M_Disease |
| 11 | 117183277 | rs180740269 | T | C | NULL | 0.604679085 | NULL&M_Disease |
| 11 | 120480883 | rs17245580 | G | A | NULL | 0.604679085 | NULL&M_Disease |
| 11 | 118877891 | rs75154936 | C | T | NULL | 0.604679085 | NULL&M_Disease |
| 11 | 117199317 | rs77873962 | T | C | NULL | 0.604679085 | NULL&M_Disease |
| 11 | 117269842 | rs140393706 | T | C | NULL | 0.604679085 | NULL&M_Disease |
| 11 | 119558815 | rs74408867 | C | T | NULL | 0.604679085 | NULL&M_Disease |
| 11 | 117185892 | rs28917236 | T | G | NULL | 0.604679085 | NULL&M_Disease |
| 11 | 117321606 | rs7350507 | G | A | NULL | 0.604679085 | NULL&M_Disease |
| 11 | 117491787 | rs77974673 | T | C | NULL | 0.604679085 | NULL&M_Disease |
| 11 | 117585424 | rs11827322 | T | C | NULL | 0.604679085 | NULL&M_Disease |
| 11 | 116374399 | rs906544 | C | T | NULL | 0.605873668 | NULL&M_Disease |
| 11 | 116435383 | rs1430595649 | T | C | NULL | 0.605873668 | NULL&M_Disease |
| 11 | 116436102 | rs61904808 | G | A | NULL | 0.605873668 | NULL&M_Disease |
| 11 | 116436589 | rs61904809 | T | C | NULL | 0.605873668 | NULL&M_Disease |
| 11 | 117646330 | rs6589622 | C | T | NULL | 0.605873668 | NULL&M_Disease |
| 11 | 119771748 | rs17123466 | T | C | NULL | 0.604679085 | NULL&M_Disease |
| 11 | 119773103 | rs55646454 | T | G | NULL | 0.604679085 | NULL&M_Disease |
| 11 | 119774821 | rs55895660 | A | G | NULL | 0.604679085 | NULL&M_Disease |
| 11 | 120988600 | rs636575 | C | A | NULL | 0.605873668 | NULL&M_Disease |
| 11 | 117313231 | rs3863315 | C | T | NULL | 0.605873668 | NULL&M_Disease |
| 11 | 117599750 | rs7944453 | A | G | NULL | 0.604679085 | NULL&M_Disease |
| 11 | 119923231 | rs12281564 | A | G | NULL | 0.605873668 | NULL&M_Disease |
| 11 | 120404415 | rs11217906 | C | T | NULL | 0.604679085 | NULL&M_Disease |
| 11 | 116431521 | rs4936356 | G | A | NULL | 0.605873668 | NULL&M_Disease |
| 11 | 116476668 | rs61904855 | A | C | NULL | 0.604679085 | NULL&M_Disease |
| 11 | 116654159 | rs618923 | G | A | NULL | 0.605873668 | NULL&M_Disease |
| 11 | 117457472 | rs117216310 | A | C | NULL | 0.604679085 | NULL&M_Disease |
| 11 | 117379572 | rs11216432 | T | C | NULL | 0.605873668 | NULL&M_Disease |
| 11 | 117475233 | rs680730 | T | C | NULL | 0.605873668 | NULL&M_Disease |
| 11 | 117551222 | rs79700393 | A | G | NULL | 0.605873668 | NULL&M_Disease |
| 11 | 120489586 | rs58047799 | G | T | NULL | 0.605873668 | NULL&M_Disease |
| 11 | 116434463 | rs61904807 | G | C | NULL | 0.605873668 | NULL&M_Disease |
| 11 | 117581689 | rs588930 | C | T | NULL | 0.605873668 | NULL&M_Disease |
| 11 | 117644348 | rs73587439 | A | G | NULL | 0.605873668 | NULL&M_Disease |
| 11 | 119771532 | rs17123461 | A | C | NULL | 0.605873668 | NULL&M_Disease |
| 11 | 119786488 | rs117734523 | A | G | NULL | 0.605873668 | NULL&M_Disease |
| 11 | 120774749 | rs59711149 | C | T | NULL | 0.605873668 | NULL&M_Disease |
| 11 | 120838878 | rs10892655 | A | G | NULL | 0.605873668 | NULL&M_Disease |
| 11 | 117376111 | rs11216430 | C | A | NULL | 0.605873668 | NULL&M_Disease |
| 11 | 117889402 | rs2512154 | A | G | NULL | 0.605873668 | NULL&M_Disease |
| 11 | 119815967 | rs1815889 | A | G | NULL | 0.605873668 | NULL&M_Disease |
| 11 | 120128055 | rs116989966 | G | A | NULL | 0.605873668 | NULL&M_Disease |
| 11 | 117273051 | rs1243336 | T | C | NULL | 0.605873668 | NULL&M_Disease |
| 11 | 117604493 | rs4142639 | G | A | NULL | 0.605873668 | NULL&M_Disease |
| 11 | 117808145 | rs73024441 | G | A | NULL | 0.605873668 | NULL&M_Disease |
| 11 | 118267737 | rs45448191 | A | T | NULL | 0.605873668 | NULL&M_Disease |
| 11 | 119759046 | rs61899013 | G | T | NULL | 0.605873668 | NULL&M_Disease |
| 11 | 120580969 | rs17124222 | G | C | NULL | 0.605873668 | NULL&M_Disease |
| 11 | 117587353 | rs12291995 | C | T | NULL | 0.605873668 | NULL&M_Disease |
| 11 | 117646291 | rs7945999 | C | T | NULL | 0.605873668 | NULL&M_Disease |
| 11 | 118615390 | rs78502545 | T | C | NULL | 0.605873668 | NULL&M_Disease |
| 11 | 119774394 | rs61899019 | A | G | NULL | 0.605873668 | NULL&M_Disease |
| 11 | 116634585 | rs11216132 | C | T | NULL | 0.607066988 | NULL&M_Disease |
| 11 | 116639941 | rs1263149 | G | A | NULL | 0.607066988 | NULL&M_Disease |
| 11 | 117620420 | rs11216527 | C | T | NULL | 0.605873668 | NULL&M_Disease |
| 11 | 117674061 | rs141512500 | G | A | NULL | 0.605873668 | NULL&M_Disease |
| 11 | 119263574 | rs188849771 | G | A | NULL | 0.605873668 | NULL&M_Disease |
| 11 | 119756067 | rs74426327 | C | T | NULL | 0.605873668 | NULL&M_Disease |
| 11 | 116304983 | rs2174880 | G | A | NULL | 0.607066988 | NULL&M_Disease |
| 11 | 117686660 | rs1939219 | C | T | NULL | 0.607066988 | NULL&M_Disease |
| 11 | 117722478 | rs71482119 | G | T | NULL | 0.605873668 | NULL&M_Disease |
| 11 | 119263566 | rs183445365 | A | G | NULL | 0.605873668 | NULL&M_Disease |
| 11 | 119502599 | rs11217366 | T | C | NULL | 0.607066988 | NULL&M_Disease |
| 11 | 120554375 | rs111463966 | G | T | NULL | 0.605873668 | NULL&M_Disease |
| 11 | 120622379 | rs7952159 | G | A | NULL | 0.607066988 | NULL&M_Disease |
| 11 | 120843742 | rs1944518 | T | C | NULL | 0.607066988 | NULL&M_Disease |
| 11 | 116438810 | rs17492366 | A | G | NULL | 0.605873668 | NULL&M_Disease |
| 11 | 116479338 | rs75322037 | G | A | NULL | 0.605873668 | NULL&M_Disease |
| 11 | 117510157 | rs61903814 | A | G | NULL | 0.605873668 | NULL&M_Disease |
| 11 | 117613203 | rs7947338 | C | T | NULL | 0.605873668 | NULL&M_Disease |
| 11 | 117935948 | rs118118922 | T | C | NULL | 0.605873668 | NULL&M_Disease |
| 11 | 120827003 | rs11218083 | T | C | NULL | 0.607066988 | NULL&M_Disease |

| | | | | | | | | | |
|---|---|---|---|---|---|---|---|---|---|
| 11 | 118185140 | rs2298821 | C | T | 1.045 | -7977.388 | -7978.238 | -0.85 | 1.65E-06 |
| 11 | 118212979 | rs7939548 | A | G | 1.043 | -7977.388 | -7978.238 | -0.85 | 1.65E-06 |
| 11 | 118041133 | rs600325 | C | T | 0.954 | -7977.388 | -7978.239 | -0.851 | 1.65E-06 |
| 11 | 118311677 | rs77535829 | T | C | 1.044 | -7977.388 | -7978.239 | -0.851 | 1.65E-06 |
| 11 | 118311869 | rs77487830 | G | A | 1.044 | -7977.388 | -7978.239 | -0.851 | 1.65E-06 |
| 11 | 118583971 | rs6589677 | A | G | 0.95 | -7977.388 | -7978.24 | -0.851 | 1.65E-06 |
| 11 | 117111214 | rs525390 | A | G | 0.957 | -7977.388 | -7978.24 | -0.852 | 1.65E-06 |
| 11 | 117806283 | rs7938191 | A | G | 0.949 | -7977.388 | -7978.241 | -0.852 | 1.65E-06 |
| 11 | 119753972 | rs964104 | A | G | 0.975 | -7977.388 | -7978.24 | -0.852 | 1.65E-06 |
| 11 | 120098180 | rs11217771 | G | C | 0.965 | -7977.388 | -7978.241 | -0.852 | 1.65E-06 |
| 11 | 117804535 | rs73024430 | T | C | 0.949 | -7977.388 | -7978.241 | -0.853 | 1.65E-06 |
| 11 | 119926775 | rs1989151 | A | T | 1.045 | -7977.388 | -7978.241 | -0.853 | 1.65E-06 |
| 11 | 120834199 | rs17124602 | C | G | 0.953 | -7977.388 | -7978.241 | -0.853 | 1.65E-06 |
| 11 | 116272744 | rs4938270 | T | C | 1.051 | -7977.388 | -7978.242 | -0.854 | 1.65E-06 |
| 11 | 116877361 | rs183640608 | G | A | 0.966 | -7977.388 | -7978.242 | -0.854 | 1.65E-06 |
| 11 | 117282800 | rs522885 | G | T | 0.972 | -7977.388 | -7978.242 | -0.854 | 1.65E-06 |
| 11 | 117605098 | rs1942929 | C | T | 0.95 | -7977.388 | -7978.242 | -0.854 | 1.65E-06 |
| 11 | 117643958 | rs79537444 | T | C | 0.949 | -7977.388 | -7978.243 | -0.854 | 1.65E-06 |
| 11 | 117987483 | rs1793178 | A | C | 1.047 | -7977.388 | -7978.242 | -0.854 | 1.65E-06 |
| 11 | 119503402 | rs12273351 | T | G | 1.044 | -7977.388 | -7978.242 | -0.854 | 1.65E-06 |
| 11 | 119757789 | rs1039094777 | T | C | 1.05 | -7977.388 | -7978.242 | -0.854 | 1.65E-06 |
| 11 | 116634557 | rs17119975 | C | T | 1.051 | -7977.388 | -7978.243 | -0.855 | 1.65E-06 |
| 11 | 117591241 | rs1455831109 | T | C | 0.965 | -7977.388 | -7978.243 | -0.855 | 1.65E-06 |
| 11 | 118864777 | rs11822344 | C | G | 1.044 | -7977.388 | -7978.243 | -0.855 | 1.65E-06 |
| 11 | 117183277 | rs180740269 | T | C | 1.017 | -7977.388 | -7978.244 | -0.856 | 1.64E-06 |
| 11 | 120480883 | rs17245580 | G | A | 1.045 | -7977.388 | -7978.244 | -0.856 | 1.64E-06 |
| 11 | 118877891 | rs75154936 | C | T | 1.051 | -7977.388 | -7978.245 | -0.857 | 1.64E-06 |
| 11 | 117199317 | rs77873962 | T | C | 1.018 | -7977.388 | -7978.246 | -0.858 | 1.64E-06 |
| 11 | 117269842 | rs140393706 | T | C | 0.962 | -7977.388 | -7978.246 | -0.858 | 1.64E-06 |
| 11 | 119558815 | rs74408867 | C | T | 1.038 | -7977.388 | -7978.246 | -0.858 | 1.64E-06 |
| 11 | 117185892 | rs28917236 | T | G | 1.015 | -7977.388 | -7978.247 | -0.859 | 1.64E-06 |
| 11 | 117321606 | rs7350507 | G | A | 0.95 | -7977.388 | -7978.247 | -0.859 | 1.64E-06 |
| 11 | 117491787 | rs77974673 | T | C | 0.974 | -7977.388 | -7978.247 | -0.859 | 1.64E-06 |
| 11 | 117585424 | rs11827322 | T | C | 0.967 | -7977.388 | -7978.247 | -0.859 | 1.64E-06 |
| 11 | 116374399 | rs906544 | C | T | 0.958 | -7977.388 | -7978.248 | -0.86 | 1.64E-06 |
| 11 | 116435383 | rs1430595649 | T | C | 1.052 | -7977.388 | -7978.248 | -0.86 | 1.64E-06 |
| 11 | 116436102 | rs61904808 | G | A | 1.052 | -7977.388 | -7978.248 | -0.86 | 1.64E-06 |
| 11 | 116436589 | rs61904809 | T | C | 1.052 | -7977.388 | -7978.248 | -0.86 | 1.64E-06 |
| 11 | 117646330 | rs6589622 | C | T | 1.052 | -7977.388 | -7978.249 | -0.86 | 1.64E-06 |
| 11 | 119771748 | rs17123466 | T | C | 0.974 | -7977.388 | -7978.248 | -0.86 | 1.64E-06 |
| 11 | 119773103 | rs55646454 | T | G | 0.974 | -7977.388 | -7978.248 | -0.86 | 1.64E-06 |
| 11 | 119774821 | rs55895660 | A | G | 0.974 | -7977.388 | -7978.248 | -0.86 | 1.64E-06 |
| 11 | 120988600 | rs636575 | C | A | 1.047 | -7977.388 | -7978.249 | -0.86 | 1.64E-06 |
| 11 | 117313231 | rs3863315 | C | T | 1.052 | -7977.388 | -7978.25 | -0.861 | 1.64E-06 |
| 11 | 117599750 | rs7944453 | A | G | 0.966 | -7977.388 | -7978.25 | -0.861 | 1.64E-06 |
| 11 | 119923231 | rs12281564 | A | G | 0.95 | -7977.388 | -7978.249 | -0.861 | 1.64E-06 |
| 11 | 120404415 | rs11217906 | C | T | 0.96 | -7977.388 | -7978.249 | -0.861 | 1.64E-06 |
| 11 | 116431521 | rs4936356 | G | A | 1.052 | -7977.388 | -7978.25 | -0.862 | 1.63E-06 |
| 11 | 116476668 | rs61904855 | A | C | 1.018 | -7977.388 | -7978.25 | -0.862 | 1.63E-06 |
| 11 | 116654159 | rs618923 | G | A | 1.047 | -7977.388 | -7978.25 | -0.862 | 1.63E-06 |
| 11 | 117457472 | rs117216310 | A | C | 1.036 | -7977.388 | -7978.251 | -0.862 | 1.63E-06 |
| 11 | 117379572 | rs11216432 | T | C | 0.951 | -7977.388 | -7978.251 | -0.863 | 1.63E-06 |
| 11 | 117475233 | rs680730 | T | C | 1.046 | -7977.388 | -7978.251 | -0.863 | 1.63E-06 |
| 11 | 117551222 | rs79700393 | A | G | 0.952 | -7977.388 | -7978.252 | -0.863 | 1.63E-06 |
| 11 | 120489586 | rs58047799 | G | T | 1.045 | -7977.388 | -7978.251 | -0.863 | 1.63E-06 |
| 11 | 116434463 | rs61904807 | G | C | 1.052 | -7977.388 | -7978.252 | -0.864 | 1.63E-06 |
| 11 | 117581689 | rs588930 | C | T | 0.955 | -7977.388 | -7978.252 | -0.864 | 1.63E-06 |
| 11 | 117644348 | rs73587439 | A | G | 0.95 | -7977.388 | -7978.252 | -0.864 | 1.63E-06 |
| 11 | 119771532 | rs17123461 | A | C | 0.975 | -7977.388 | -7978.252 | -0.864 | 1.63E-06 |
| 11 | 119786488 | rs117734523 | A | G | 0.978 | -7977.388 | -7978.252 | -0.864 | 1.63E-06 |
| 11 | 120774749 | rs59711149 | C | T | 1.046 | -7977.388 | -7978.252 | -0.864 | 1.63E-06 |
| 11 | 120838878 | rs10892655 | A | G | 1.052 | -7977.388 | -7978.253 | -0.864 | 1.63E-06 |
| 11 | 117376111 | rs11216430 | C | A | 0.951 | -7977.388 | -7978.253 | -0.865 | 1.63E-06 |
| 11 | 117889402 | rs2512154 | A | G | 0.952 | -7977.388 | -7978.253 | -0.865 | 1.63E-06 |
| 11 | 119815967 | rs1815889 | A | G | 1.052 | -7977.388 | -7978.253 | -0.865 | 1.63E-06 |
| 11 | 120128055 | rs116989966 | G | A | 1.042 | -7977.388 | -7978.253 | -0.865 | 1.63E-06 |
| 11 | 117273051 | rs1243336 | T | C | 0.974 | -7977.388 | -7978.254 | -0.866 | 1.63E-06 |
| 11 | 117604493 | rs4142639 | G | A | 0.966 | -7977.388 | -7978.254 | -0.866 | 1.63E-06 |
| 11 | 117808145 | rs73024441 | G | A | 0.95 | -7977.388 | -7978.254 | -0.866 | 1.63E-06 |
| 11 | 118267737 | rs45448191 | A | T | 1.042 | -7977.388 | -7978.254 | -0.866 | 1.63E-06 |
| 11 | 119759046 | rs61899013 | G | T | 0.976 | -7977.388 | -7978.254 | -0.866 | 1.63E-06 |
| 11 | 120580969 | rs17124222 | G | C | 0.966 | -7977.388 | -7978.254 | -0.866 | 1.63E-06 |
| 11 | 117587353 | rs12291995 | C | T | 0.967 | -7977.388 | -7978.256 | -0.867 | 1.63E-06 |
| 11 | 117646291 | rs7945999 | C | T | 1.052 | -7977.388 | -7978.255 | -0.867 | 1.63E-06 |
| 11 | 118615390 | rs78502545 | T | C | 1.047 | -7977.388 | -7978.255 | -0.867 | 1.63E-06 |
| 11 | 119774394 | rs61899019 | A | G | 0.977 | -7977.388 | -7978.255 | -0.867 | 1.63E-06 |
| 11 | 116634585 | rs11216132 | C | T | 1.048 | -7977.388 | -7978.256 | -0.868 | 1.62E-06 |
| 11 | 116639941 | rs1263149 | G | A | 1.043 | -7977.388 | -7978.256 | -0.868 | 1.62E-06 |
| 11 | 117620420 | rs11216527 | C | T | 0.967 | -7977.388 | -7978.256 | -0.868 | 1.62E-06 |
| 11 | 117674061 | rs141512500 | G | A | 0.975 | -7977.388 | -7978.256 | -0.868 | 1.62E-06 |
| 11 | 119263574 | rs188849771 | G | A | 0.997 | -7977.388 | -7978.257 | -0.868 | 1.62E-06 |
| 11 | 119756067 | rs74426327 | C | T | 1.049 | -7977.388 | -7978.257 | -0.868 | 1.62E-06 |
| 11 | 116304983 | rs2174880 | G | A | 0.958 | -7977.388 | -7978.257 | -0.869 | 1.62E-06 |
| 11 | 117686660 | rs1939219 | C | T | 0.95 | -7977.388 | -7978.257 | -0.869 | 1.62E-06 |
| 11 | 117722478 | rs71482119 | G | T | 1.051 | -7977.388 | -7978.258 | -0.869 | 1.62E-06 |
| 11 | 119263566 | rs183445365 | A | G | 0.999 | -7977.388 | -7978.258 | -0.869 | 1.62E-06 |
| 11 | 119502599 | rs11217366 | T | C | 1.044 | -7977.388 | -7978.257 | -0.869 | 1.62E-06 |
| 11 | 120554375 | rs111463966 | G | T | 1.015 | -7977.388 | -7978.257 | -0.869 | 1.62E-06 |
| 11 | 120622379 | rs7952159 | G | A | 0.951 | -7977.388 | -7978.257 | -0.869 | 1.62E-06 |
| 11 | 120843742 | rs1944518 | T | C | 1.045 | -7977.388 | -7978.257 | -0.869 | 1.62E-06 |
| 11 | 116438810 | rs17492366 | A | G | 1.049 | -7977.388 | -7978.259 | -0.87 | 1.62E-06 |
| 11 | 116479338 | rs75322037 | G | A | 0.988 | -7977.388 | -7978.258 | -0.87 | 1.62E-06 |
| 11 | 117510157 | rs61903814 | A | G | 0.987 | -7977.388 | -7978.258 | -0.87 | 1.62E-06 |
| 11 | 117613203 | rs7947338 | C | T | 0.951 | -7977.388 | -7978.258 | -0.87 | 1.62E-06 |
| 11 | 117935948 | rs118118922 | T | C | 0.973 | -7977.388 | -7978.258 | -0.87 | 1.62E-06 |
| 11 | 120827003 | rs11218083 | T | C | 1.052 | -7977.388 | -7978.258 | -0.87 | 1.62E-06 |

| | | | | | | | |
|---|---|---|---|---|---|---|---|
| 11 | 117750926 | rs73020440 | T | C | NULL | 0.605873668 | NULL&M_Disease |
| 11 | 119839599 | rs74674700 | T | C | NULL | 0.605873668 | NULL&M_Disease |
| 11 | 120405520 | rs2508654 | T | C | NULL | 0.605873668 | NULL&M_Disease |
| 11 | 121066137 | rs17245873 | A | T | NULL | 0.605873668 | NULL&M_Disease |
| 11 | 117587085 | rs1027636222 | G | A | NULL | 0.605873668 | NULL&M_Disease |
| 11 | 117593097 | rs7937973 | A | G | NULL | 0.605873668 | NULL&M_Disease |
| 11 | 120462315 | rs34238850 | T | C | NULL | 0.607066988 | NULL&M_Disease |
| 11 | 117409884 | rs3016797 | C | G | NULL | 0.607066988 | NULL&M_Disease |
| 11 | 117529676 | rs75275400 | C | T | NULL | 0.605873668 | NULL&M_Disease |
| 11 | 117604207 | rs4142640 | A | G | NULL | 0.607066988 | NULL&M_Disease |
| 11 | 117912465 | rs7120925 | T | C | NULL | 0.607066988 | NULL&M_Disease |
| 11 | 119803839 | rs10892496 | C | A | NULL | 0.607066988 | NULL&M_Disease |
| 11 | 120702933 | rs78390305 | A | G | NULL | 0.607066988 | NULL&M_Disease |
| 11 | 120707383 | rs10892628 | A | G | NULL | 0.607066988 | NULL&M_Disease |
| 11 | 116836212 | rs2513097 | G | A | NULL | 0.607066988 | NULL&M_Disease |
| 11 | 117574499 | rs57145837 | A | G | NULL | 0.607066988 | NULL&M_Disease |
| 11 | 117718626 | rs141259801 | T | C | NULL | 0.607066988 | NULL&M_Disease |
| 11 | 117737557 | rs113523070 | A | G | NULL | 0.607066988 | NULL&M_Disease |
| 11 | 119781695 | rs9665750 | T | C | NULL | 0.607066988 | NULL&M_Disease |
| 11 | 119801338 | rs11217583 | T | C | NULL | 0.607066988 | NULL&M_Disease |
| 11 | 121034651 | rs679607 | G | A | NULL | 0.607066988 | NULL&M_Disease |
| 11 | 117578440 | rs147146093 | A | G | NULL | 0.607066988 | NULL&M_Disease |
| 11 | 117579531 | rs7934773 | A | G | NULL | 0.607066988 | NULL&M_Disease |
| 11 | 117709645 | rs1406347456 | G | T | NULL | 0.607066988 | NULL&M_Disease |
| 11 | 119811328 | rs34420468 | A | G | NULL | 0.607066988 | NULL&M_Disease |
| 11 | 119849828 | rs11217608 | G | A | NULL | 0.607066988 | NULL&M_Disease |
| 11 | 120687126 | rs77004360 | T | C | NULL | 0.607066988 | NULL&M_Disease |
| 11 | 119856928 | rs11607987 | T | C | NULL | 0.607066988 | NULL&M_Disease |
| 11 | 120589740 | rs74236028 | G | T | NULL | 0.607066988 | NULL&M_Disease |
| 11 | 121014904 | rs543119 | G | A | NULL | 0.607066988 | NULL&M_Disease |
| 11 | 117575862 | rs7104002 | C | T | NULL | 0.607066988 | NULL&M_Disease |
| 11 | 117954350 | rs625404 | C | T | NULL | 0.607066988 | NULL&M_Disease |
| 11 | 119803564 | rs10892495 | G | C | NULL | 0.607066988 | NULL&M_Disease |
| 11 | 120131372 | rs78160609 | A | G | NULL | 0.607066988 | NULL&M_Disease |
| 11 | 116554198 | rs73014900 | G | T | NULL | 0.607066988 | NULL&M_Disease |
| 11 | 117350033 | rs519356 | C | T | NULL | 0.607066988 | NULL&M_Disease |
| 11 | 117531731 | rs10892151 | T | C | NULL | 0.607066988 | NULL&M_Disease |
| 11 | 117874493 | rs947890 | A | G | NULL | 0.607066988 | NULL&M_Disease |
| 11 | 119778398 | rs61899020 | T | C | NULL | 0.607066988 | NULL&M_Disease |
| 11 | 120853675 | rs7102603 | T | C | NULL | 0.608259031 | NULL&M_Disease |
| 11 | 116467853 | rs7951761 | G | A | NULL | 0.608259031 | NULL&M_Disease |
| 11 | 116480631 | rs551141 | A | G | NULL | 0.607066988 | NULL&M_Disease |
| 11 | 117717297 | rs6589624 | G | A | NULL | 0.607066988 | NULL&M_Disease |
| 11 | 119780975 | rs61899021 | T | G | NULL | 0.607066988 | NULL&M_Disease |
| 11 | 117127159 | rs12575964 | A | G | NULL | 0.608259031 | NULL&M_Disease |
| 11 | 118379096 | rs7113959 | A | T | NULL | 0.607066988 | NULL&M_Disease |
| 11 | 118382327 | rs7113994 | G | A | NULL | 0.607066988 | NULL&M_Disease |
| 11 | 119711999 | rs35095598 | A | G | NULL | 0.607066988 | NULL&M_Disease |
| 11 | 119802832 | rs11217584 | A | G | NULL | 0.608259031 | NULL&M_Disease |
| 11 | 120133616 | rs76867785 | G | A | NULL | 0.607066988 | NULL&M_Disease |
| 11 | 117142625 | rs141538185 | G | C | NULL | 0.608259031 | NULL&M_Disease |
| 11 | 117574975 | rs11828302 | T | A | NULL | 0.607066988 | NULL&M_Disease |
| 11 | 117601018 | rs60044901 | C | G | NULL | 0.607066988 | NULL&M_Disease |
| 11 | 118576762 | rs553188 | T | C | NULL | 0.608259031 | NULL&M_Disease |
| 11 | 118811315 | rs77638281 | A | G | NULL | 0.608259031 | NULL&M_Disease |
| 11 | 119771321 | rs61899017 | C | T | NULL | 0.607066988 | NULL&M_Disease |
| 11 | 121018954 | rs22298 | G | A | NULL | 0.608259031 | NULL&M_Disease |
| 11 | 116483126 | rs486020 | C | G | NULL | 0.608259031 | NULL&M_Disease |
| 11 | 117722715 | rs496371 | T | C | NULL | 0.608259031 | NULL&M_Disease |
| 11 | 119573655 | rs117236359 | T | C | NULL | 0.608259031 | NULL&M_Disease |
| 11 | 121053985 | rs4936592 | T | C | NULL | 0.608259031 | NULL&M_Disease |
| 11 | 116224621 | rs501755 | T | C | NULL | 0.608259031 | NULL&M_Disease |
| 11 | 117890754 | rs11826646 | C | G | NULL | 0.608259031 | NULL&M_Disease |
| 11 | 118218478 | rs3181258 | A | T | NULL | 0.608259031 | NULL&M_Disease |
| 11 | 119913541 | rs11217653 | A | G | NULL | 0.608259031 | NULL&M_Disease |
| 11 | 120579966 | rs77291360 | G | A | NULL | 0.608259031 | NULL&M_Disease |
| 11 | 120694966 | rs111728839 | A | G | NULL | 0.608259031 | NULL&M_Disease |
| 11 | 121049649 | rs112580285 | G | T | NULL | 0.608259031 | NULL&M_Disease |
| 11 | 116481248 | rs545559 | A | C | NULL | 0.608259031 | NULL&M_Disease |
| 11 | 116481505 | rs566254 | A | T | NULL | 0.608259031 | NULL&M_Disease |
| 11 | 117524497 | rs11216487 | C | T | NULL | 0.608259031 | NULL&M_Disease |
| 11 | 117707937 | rs3087563 | C | T | NULL | 0.608259031 | NULL&M_Disease |
| 11 | 119880501 | rs2077582 | T | C | NULL | 0.608259031 | NULL&M_Disease |
| 11 | 117628906 | rs12295970 | T | G | NULL | 0.608259031 | NULL&M_Disease |
| 11 | 118072915 | rs75128335 | A | G | NULL | 0.608259031 | NULL&M_Disease |
| 11 | 119744365 | rs73008681 | G | C | NULL | 0.608259031 | NULL&M_Disease |
| 11 | 120182487 | rs74940535 | C | G | NULL | 0.608259031 | NULL&M_Disease |
| 11 | 120808714 | rs474867 | A | G | NULL | 0.608259031 | NULL&M_Disease |
| 11 | 121151902 | rs11218240 | T | C | NULL | 0.608259031 | NULL&M_Disease |
| 11 | 117629056 | rs12276648 | C | T | NULL | 0.608259031 | NULL&M_Disease |
| 11 | 120175749 | rs7110845 | A | G | NULL | 0.609449784 | NULL&M_Disease |
| 11 | 120990239 | rs488766 | G | A | NULL | 0.608259031 | NULL&M_Disease |
| 11 | 117605839 | rs10892166 | A | G | NULL | 0.608259031 | NULL&M_Disease |
| 11 | 118065830 | rs111527225 | A | G | NULL | 0.608259031 | NULL&M_Disease |
| 11 | 119798293 | rs11217579 | T | A | NULL | 0.608259031 | NULL&M_Disease |
| 11 | 120242812 | rs117498381 | T | A | NULL | 0.608259031 | NULL&M_Disease |
| 11 | 121003606 | rs633770 | T | G | NULL | 0.608259031 | NULL&M_Disease |
| 11 | 117171475 | rs76380380 | G | C | NULL | 0.608259031 | NULL&M_Disease |
| 11 | 117597660 | rs734378 | T | C | NULL | 0.608259031 | NULL&M_Disease |
| 11 | 117880688 | rs2508448 | T | G | NULL | 0.609449784 | NULL&M_Disease |
| 11 | 119699827 | rs580948 | T | C | NULL | 0.609449784 | NULL&M_Disease |
| 11 | 118723537 | rs7103314 | G | A | NULL | 0.609449784 | NULL&M_Disease |
| 11 | 121055215 | rs111982661 | T | C | NULL | 0.608259031 | NULL&M_Disease |
| 11 | 121149360 | rs147189138 | A | G | NULL | 0.608259031 | NULL&M_Disease |
| 11 | 116700860 | rs734104 | C | T | NULL | 0.609449784 | NULL&M_Disease |
| 11 | 120694926 | rs79446094 | G | A | NULL | 0.609449784 | NULL&M_Disease |

| | | | | | | | | | |
|---|---|---|---|---|---|---|---|---|---|
| 11 | 117750926 | rs73020440 | T | C | 1.038 | -7977.388 | -7978.259 | -0.871 | 1.62E-06 |
| 11 | 119839599 | rs74674700 | T | C | 0.975 | -7977.388 | -7978.26 | -0.871 | 1.62E-06 |
| 11 | 120405520 | rs2508654 | T | C | 0.962 | -7977.388 | -7978.259 | -0.871 | 1.62E-06 |
| 11 | 121066137 | rs17245873 | A | T | 1.037 | -7977.388 | -7978.259 | -0.871 | 1.62E-06 |
| 11 | 117587085 | rs1027636222 | G | A | 0.968 | -7977.388 | -7978.261 | -0.872 | 1.62E-06 |
| 11 | 117593097 | rs7937973 | A | G | 0.967 | -7977.388 | -7978.261 | -0.872 | 1.62E-06 |
| 11 | 120462315 | rs34238850 | T | C | 0.957 | -7977.388 | -7978.261 | -0.872 | 1.62E-06 |
| 11 | 117409884 | rs3016797 | C | G | 1.051 | -7977.388 | -7978.261 | -0.873 | 1.62E-06 |
| 11 | 117529676 | rs75275400 | C | T | 0.97 | -7977.388 | -7978.261 | -0.873 | 1.62E-06 |
| 11 | 117604207 | rs4142640 | A | G | 0.967 | -7977.388 | -7978.262 | -0.873 | 1.62E-06 |
| 11 | 117912465 | rs7120925 | T | C | 1.045 | -7977.388 | -7978.262 | -0.873 | 1.62E-06 |
| 11 | 119803839 | rs10892496 | C | A | 1.052 | -7977.388 | -7978.261 | -0.873 | 1.62E-06 |
| 11 | 120702933 | rs78390305 | A | G | 0.951 | -7977.388 | -7978.261 | -0.873 | 1.62E-06 |
| 11 | 120707383 | rs10892628 | A | G | 0.953 | -7977.388 | -7978.261 | -0.873 | 1.62E-06 |
| 11 | 116836212 | rs2513097 | G | A | 1.046 | -7977.388 | -7978.262 | -0.874 | 1.61E-06 |
| 11 | 117574499 | rs57145837 | A | G | 0.97 | -7977.388 | -7978.262 | -0.874 | 1.61E-06 |
| 11 | 117718626 | rs141259801 | T | C | 0.969 | -7977.388 | -7978.262 | -0.874 | 1.61E-06 |
| 11 | 117737557 | rs113523070 | A | G | 0.991 | -7977.388 | -7978.262 | -0.874 | 1.61E-06 |
| 11 | 119781695 | rs9665750 | T | C | 1.05 | -7977.388 | -7978.263 | -0.874 | 1.61E-06 |
| 11 | 119801338 | rs11217583 | T | C | 1.052 | -7977.388 | -7978.262 | -0.874 | 1.61E-06 |
| 11 | 121034651 | rs679607 | G | A | 0.958 | -7977.388 | -7978.262 | -0.874 | 1.61E-06 |
| 11 | 117578440 | rs147146093 | A | G | 0.97 | -7977.388 | -7978.263 | -0.875 | 1.61E-06 |
| 11 | 117579531 | rs7934773 | A | G | 0.97 | -7977.388 | -7978.263 | -0.875 | 1.61E-06 |
| 11 | 117709645 | rs1406347456 | G | T | 0.951 | -7977.388 | -7978.263 | -0.875 | 1.61E-06 |
| 11 | 119811328 | rs34420468 | A | G | 1.005 | -7977.388 | -7978.264 | -0.875 | 1.61E-06 |
| 11 | 119849828 | rs11217608 | G | A | 1.051 | -7977.388 | -7978.264 | -0.875 | 1.61E-06 |
| 11 | 120687126 | rs77004360 | T | C | 0.951 | -7977.388 | -7978.263 | -0.875 | 1.61E-06 |
| 11 | 119856928 | rs11607987 | T | C | 1.052 | -7977.388 | -7978.264 | -0.876 | 1.61E-06 |
| 11 | 120589740 | rs74236028 | G | T | 0.953 | -7977.388 | -7978.264 | -0.876 | 1.61E-06 |
| 11 | 121014904 | rs543119 | G | A | 0.958 | -7977.388 | -7978.264 | -0.876 | 1.61E-06 |
| 11 | 117575862 | rs7104002 | C | T | 0.971 | -7977.388 | -7978.266 | -0.877 | 1.61E-06 |
| 11 | 117954350 | rs625404 | C | T | 1.051 | -7977.388 | -7978.265 | -0.877 | 1.61E-06 |
| 11 | 119803564 | rs10892495 | G | C | 1.052 | -7977.388 | -7978.265 | -0.877 | 1.61E-06 |
| 11 | 120131372 | rs78160609 | A | G | 0.985 | -7977.388 | -7978.265 | -0.877 | 1.61E-06 |
| 11 | 116554198 | rs73014900 | G | T | 0.951 | -7977.388 | -7978.267 | -0.878 | 1.61E-06 |
| 11 | 117350033 | rs519356 | C | T | 0.952 | -7977.388 | -7978.266 | -0.878 | 1.61E-06 |
| 11 | 117531731 | rs10892151 | T | C | 1.013 | -7977.388 | -7978.266 | -0.878 | 1.61E-06 |
| 11 | 117874493 | rs947890 | A | G | 0.952 | -7977.388 | -7978.266 | -0.878 | 1.61E-06 |
| 11 | 119778398 | rs61899020 | T | C | 0.981 | -7977.388 | -7978.266 | -0.878 | 1.61E-06 |
| 11 | 120853675 | rs7102603 | T | C | 1.044 | -7977.388 | -7978.266 | -0.878 | 1.61E-06 |
| 11 | 116467853 | rs7951761 | G | A | 1.043 | -7977.388 | -7978.267 | -0.879 | 1.61E-06 |
| 11 | 116480631 | rs551141 | A | G | 0.958 | -7977.388 | -7978.267 | -0.879 | 1.61E-06 |
| 11 | 117717297 | rs6589624 | G | A | 0.951 | -7977.388 | -7978.268 | -0.879 | 1.61E-06 |
| 11 | 119780975 | rs61899021 | T | G | 0.986 | -7977.388 | -7978.267 | -0.879 | 1.61E-06 |
| 11 | 117127159 | rs12575964 | A | G | 0.951 | -7977.388 | -7978.269 | -0.88 | 1.60E-06 |
| 11 | 118379096 | rs7113959 | A | T | 1.049 | -7977.388 | -7978.268 | -0.88 | 1.60E-06 |
| 11 | 118382327 | rs7113994 | G | A | 1.049 | -7977.388 | -7978.268 | -0.88 | 1.60E-06 |
| 11 | 119711999 | rs35095598 | A | G | 0.965 | -7977.388 | -7978.269 | -0.88 | 1.60E-06 |
| 11 | 119802832 | rs11217584 | A | G | 1.052 | -7977.388 | -7978.268 | -0.88 | 1.60E-06 |
| 11 | 120133616 | rs76867785 | G | A | 0.987 | -7977.388 | -7978.269 | -0.881 | 1.60E-06 |
| 11 | 117142625 | rs141538185 | G | C | 0.963 | -7977.388 | -7978.271 | -0.882 | 1.60E-06 |
| 11 | 117574975 | rs11828302 | T | A | 0.972 | -7977.388 | -7978.271 | -0.882 | 1.60E-06 |
| 11 | 117601018 | rs60044901 | C | G | 0.969 | -7977.388 | -7978.271 | -0.882 | 1.60E-06 |
| 11 | 118576762 | rs553188 | T | C | 0.951 | -7977.388 | -7978.27 | -0.882 | 1.60E-06 |
| 11 | 118811315 | rs77638281 | A | G | 1.042 | -7977.388 | -7978.27 | -0.882 | 1.60E-06 |
| 11 | 119771321 | rs61899017 | C | T | 0.981 | -7977.388 | -7978.27 | -0.882 | 1.60E-06 |
| 11 | 121018954 | rs22298 | G | A | 0.958 | -7977.388 | -7978.27 | -0.882 | 1.60E-06 |
| 11 | 116483126 | rs486020 | C | G | 0.958 | -7977.388 | -7978.271 | -0.883 | 1.60E-06 |
| 11 | 117722715 | rs496371 | T | C | 0.957 | -7977.388 | -7978.272 | -0.883 | 1.60E-06 |
| 11 | 119573655 | rs117236359 | T | C | 1.034 | -7977.388 | -7978.271 | -0.883 | 1.60E-06 |
| 11 | 121053985 | rs4936592 | T | C | 0.956 | -7977.388 | -7978.271 | -0.883 | 1.60E-06 |
| 11 | 116224621 | rs501755 | T | C | 1.049 | -7977.388 | -7978.273 | -0.884 | 1.60E-06 |
| 11 | 117890754 | rs11826646 | C | G | 0.951 | -7977.388 | -7978.272 | -0.884 | 1.60E-06 |
| 11 | 118218478 | rs3181258 | A | T | 1.039 | -7977.388 | -7978.272 | -0.884 | 1.60E-06 |
| 11 | 119913541 | rs11217653 | A | G | 1.044 | -7977.388 | -7978.273 | -0.884 | 1.60E-06 |
| 11 | 120579966 | rs77291360 | G | A | 0.969 | -7977.388 | -7978.272 | -0.884 | 1.60E-06 |
| 11 | 120694966 | rs111728839 | A | G | 0.952 | -7977.388 | -7978.272 | -0.884 | 1.60E-06 |
| 11 | 121049649 | rs112580285 | G | T | 0.956 | -7977.388 | -7978.272 | -0.884 | 1.60E-06 |
| 11 | 116481248 | rs545559 | A | C | 0.958 | -7977.388 | -7978.274 | -0.885 | 1.60E-06 |
| 11 | 116481505 | rs566254 | A | T | 0.958 | -7977.388 | -7978.274 | -0.885 | 1.60E-06 |
| 11 | 117524497 | rs11216487 | C | T | 0.998 | -7977.388 | -7978.273 | -0.885 | 1.60E-06 |
| 11 | 117707937 | rs3087563 | C | T | 1.043 | -7977.388 | -7978.274 | -0.885 | 1.60E-06 |
| 11 | 119880501 | rs2077582 | T | C | 0.966 | -7977.388 | -7978.273 | -0.885 | 1.60E-06 |
| 11 | 117628906 | rs12295970 | T | G | 0.951 | -7977.388 | -7978.275 | -0.886 | 1.59E-06 |
| 11 | 118072915 | rs75128335 | A | G | 0.987 | -7977.388 | -7978.275 | -0.886 | 1.59E-06 |
| 11 | 119744365 | rs73008681 | G | C | 1.05 | -7977.388 | -7978.274 | -0.886 | 1.59E-06 |
| 11 | 120182487 | rs74940535 | C | G | 1.02 | -7977.388 | -7978.274 | -0.886 | 1.59E-06 |
| 11 | 120808714 | rs474867 | A | G | 0.954 | -7977.388 | -7978.275 | -0.886 | 1.59E-06 |
| 11 | 121151902 | rs11218240 | T | C | 1.05 | -7977.388 | -7978.274 | -0.886 | 1.59E-06 |
| 11 | 117629056 | rs12276648 | C | T | 0.951 | -7977.388 | -7978.276 | -0.887 | 1.59E-06 |
| 11 | 120175749 | rs7110845 | A | G | 1.043 | -7977.388 | -7978.276 | -0.887 | 1.59E-06 |
| 11 | 120990239 | rs488766 | G | A | 1.047 | -7977.388 | -7978.275 | -0.887 | 1.59E-06 |
| 11 | 117605839 | rs10892166 | A | G | 0.97 | -7977.388 | -7978.277 | -0.888 | 1.59E-06 |
| 11 | 118065830 | rs111527225 | A | G | 0.987 | -7977.388 | -7978.276 | -0.888 | 1.59E-06 |
| 11 | 119798293 | rs11217579 | T | A | 1.051 | -7977.388 | -7978.276 | -0.888 | 1.59E-06 |
| 11 | 120242812 | rs117498381 | T | A | 1.022 | -7977.388 | -7978.277 | -0.888 | 1.59E-06 |
| 11 | 121003606 | rs633770 | T | G | 1.047 | -7977.388 | -7978.276 | -0.888 | 1.59E-06 |
| 11 | 117171475 | rs76380380 | G | C | 0.995 | -7977.388 | -7978.277 | -0.889 | 1.59E-06 |
| 11 | 117597660 | rs734378 | T | C | 0.971 | -7977.388 | -7978.277 | -0.889 | 1.59E-06 |
| 11 | 117880688 | rs2508448 | T | G | 0.953 | -7977.388 | -7978.277 | -0.889 | 1.59E-06 |
| 11 | 119699827 | rs580948 | T | C | 0.956 | -7977.388 | -7978.277 | -0.889 | 1.59E-06 |
| 11 | 118723537 | rs7103314 | G | A | 0.956 | -7977.388 | -7978.278 | -0.89 | 1.59E-06 |
| 11 | 121055215 | rs111982661 | T | C | 0.956 | -7977.388 | -7978.278 | -0.89 | 1.59E-06 |
| 11 | 121149360 | rs147189138 | A | G | 1.043 | -7977.388 | -7978.278 | -0.89 | 1.59E-06 |
| 11 | 116700860 | rs734104 | C | T | 0.952 | -7977.388 | -7978.28 | -0.891 | 1.59E-06 |
| 11 | 120694926 | rs79446094 | G | A | 0.952 | -7977.388 | -7978.279 | -0.891 | 1.59E-06 |

| | | | | | | | |
|---|---|---|---|---|---|---|---|
| 11 | 116237825 | rs1968541 | A | G | NULL | 0.609449784 | NULL&M_Disease |
| 11 | 117528867 | rs59931457 | A | G | NULL | 0.608259031 | NULL&M_Disease |
| 11 | 118365210 | rs9332817 | C | G | NULL | 0.608259031 | NULL&M_Disease |
| 11 | 117263874 | rs117713193 | T | C | NULL | 0.609449784 | NULL&M_Disease |
| 11 | 117989798 | rs117184822 | T | C | NULL | 0.608259031 | NULL&M_Disease |
| 11 | 119817324 | rs80062985 | T | C | NULL | 0.609449784 | NULL&M_Disease |
| 11 | 116451880 | rs71478302 | G | C | NULL | 0.609449784 | NULL&M_Disease |
| 11 | 116803634 | rs118138924 | G | C | NULL | 0.609449784 | NULL&M_Disease |
| 11 | 117429728 | rs685425 | T | C | NULL | 0.609449784 | NULL&M_Disease |
| 11 | 117880723 | rs80044067 | A | G | NULL | 0.609449784 | NULL&M_Disease |
| 11 | 117911367 | rs597820 | C | A | NULL | 0.609449784 | NULL&M_Disease |
| 11 | 117987746 | rs640888 | A | G | NULL | 0.609449784 | NULL&M_Disease |
| 11 | 119564528 | rs140186337 | G | A | NULL | 0.609449784 | NULL&M_Disease |
| 11 | 120089082 | rs117934631 | G | T | NULL | 0.609449784 | NULL&M_Disease |
| 11 | 120683594 | rs948026 | A | G | NULL | 0.609449784 | NULL&M_Disease |
| 11 | 121159481 | rs116958147 | C | T | NULL | 0.609449784 | NULL&M_Disease |
| 11 | 117730777 | rs78559460 | C | A | NULL | 0.609449784 | NULL&M_Disease |
| 11 | 119755047 | rs12285075 | A | G | NULL | 0.609449784 | NULL&M_Disease |
| 11 | 120236793 | rs117388325 | T | C | NULL | 0.609449784 | NULL&M_Disease |
| 11 | 120837407 | rs10892654 | G | A | NULL | 0.609449784 | NULL&M_Disease |
| 11 | 118576997 | rs498912 | C | T | NULL | 0.609449784 | NULL&M_Disease |
| 11 | 119927648 | rs681454 | G | C | NULL | 0.609449784 | NULL&M_Disease |
| 11 | 120128088 | rs115767079 | T | C | NULL | 0.609449784 | NULL&M_Disease |
| 11 | 120694254 | rs75954141 | C | A | NULL | 0.609449784 | NULL&M_Disease |
| 11 | 116237754 | rs11215903 | G | T | NULL | 0.609449784 | NULL&M_Disease |
| 11 | 117320352 | rs10790185 | T | C | NULL | 0.609449784 | NULL&M_Disease |
| 11 | 120176310 | rs12790634 | A | G | NULL | 0.609449784 | NULL&M_Disease |
| 11 | 117218762 | rs568787 | A | G | NULL | 0.609449784 | NULL&M_Disease |
| 11 | 120465556 | rs1573880 | T | C | NULL | 0.609449784 | NULL&M_Disease |
| 11 | 120497208 | rs4375451 | T | C | NULL | 0.610639234 | NULL&M_Disease |
| 11 | 120576984 | rs11603713 | G | A | NULL | 0.609449784 | NULL&M_Disease |
| 11 | 120694915 | rs12291852 | T | G | NULL | 0.609449784 | NULL&M_Disease |
| 11 | 117725057 | rs11216583 | G | C | NULL | 0.609449784 | NULL&M_Disease |
| 11 | 120694466 | rs75208530 | G | A | NULL | 0.609449784 | NULL&M_Disease |
| 11 | 117303430 | rs55644928 | T | C | NULL | 0.610639234 | NULL&M_Disease |
| 11 | 117628972 | rs12296014 | A | G | NULL | 0.610639234 | NULL&M_Disease |
| 11 | 118816304 | rs586763 | T | C | NULL | 0.610639234 | NULL&M_Disease |
| 11 | 120993071 | rs608818 | T | C | NULL | 0.610639234 | NULL&M_Disease |
| 11 | 117215605 | rs476983 | T | C | NULL | 0.609449784 | NULL&M_Disease |
| 11 | 117252437 | rs117083334 | G | A | NULL | 0.609449784 | NULL&M_Disease |
| 11 | 118721189 | rs4938566 | T | A | NULL | 0.610639234 | NULL&M_Disease |
| 11 | 118721249 | rs4938567 | G | A | NULL | 0.610639234 | NULL&M_Disease |
| 11 | 118721850 | rs7946961 | G | A | NULL | 0.610639234 | NULL&M_Disease |
| 11 | 118722231 | rs4938570 | G | A | NULL | 0.610639234 | NULL&M_Disease |
| 11 | 120128451 | rs17123747 | T | C | NULL | 0.609449784 | NULL&M_Disease |
| 11 | 120128668 | rs78314944 | C | G | NULL | 0.609449784 | NULL&M_Disease |
| 11 | 120128828 | rs116667143 | G | C | NULL | 0.609449784 | NULL&M_Disease |
| 11 | 120129129 | rs114513135 | C | A | NULL | 0.609449784 | NULL&M_Disease |
| 11 | 116632500 | rs10488699 | T | C | NULL | 0.610639234 | NULL&M_Disease |
| 11 | 117604229 | rs1939983 | G | C | NULL | 0.610639234 | NULL&M_Disease |
| 11 | 120789508 | rs10892640 | G | A | NULL | 0.610639234 | NULL&M_Disease |
| 11 | 117465314 | rs75873912 | G | C | NULL | 0.610639234 | NULL&M_Disease |
| 11 | 117600136 | rs10160584 | G | A | NULL | 0.610639234 | NULL&M_Disease |
| 11 | 119844937 | rs7101905 | T | C | NULL | 0.610639234 | NULL&M_Disease |
| 11 | 120163449 | rs56966127 | T | G | NULL | 0.610639234 | NULL&M_Disease |
| 11 | 120419868 | rs11600436 | T | C | NULL | 0.610639234 | NULL&M_Disease |
| 11 | 121149531 | rs11218239 | G | A | NULL | 0.610639234 | NULL&M_Disease |
| 11 | 117228892 | rs560789 | G | A | NULL | 0.610639234 | NULL&M_Disease |
| 11 | 117581589 | rs11216510 | C | A | NULL | 0.610639234 | NULL&M_Disease |
| 11 | 117642376 | rs11825297 | T | C | NULL | 0.610639234 | NULL&M_Disease |
| 11 | 117964339 | rs10750124 | C | T | NULL | 0.610639234 | NULL&M_Disease |
| 11 | 116979911 | rs12271161 | A | G | NULL | 0.610639234 | NULL&M_Disease |
| 11 | 117581228 | rs641569 | T | G | NULL | 0.610639234 | NULL&M_Disease |
| 11 | 117581240 | rs601503 | C | G | NULL | 0.610639234 | NULL&M_Disease |
| 11 | 117581785 | rs693992 | A | C | NULL | 0.610639234 | NULL&M_Disease |
| 11 | 117604479 | rs4142636 | T | C | NULL | 0.610639234 | NULL&M_Disease |
| 11 | 120152591 | rs11217785 | T | C | NULL | 0.610639234 | NULL&M_Disease |
| 11 | 120170012 | rs77810230 | C | T | NULL | 0.610639234 | NULL&M_Disease |
| 11 | 116326685 | rs118000999 | T | C | NULL | 0.610639234 | NULL&M_Disease |
| 11 | 117574397 | rs650162 | T | C | NULL | 0.610639234 | NULL&M_Disease |
| 11 | 117962732 | rs948468 | C | G | NULL | 0.610639234 | NULL&M_Disease |
| 11 | 119890299 | rs12363335 | C | T | NULL | 0.610639234 | NULL&M_Disease |
| 11 | 116259493 | rs1185802888 | T | A | NULL | 0.610639234 | NULL&M_Disease |
| 11 | 116546633 | rs116944258 | A | C | NULL | 0.610639234 | NULL&M_Disease |
| 11 | 116548972 | rs79678269 | C | G | NULL | 0.610639234 | NULL&M_Disease |
| 11 | 116549116 | rs1005705 | C | A | NULL | 0.610639234 | NULL&M_Disease |
| 11 | 116549971 | rs117830617 | A | G | NULL | 0.610639234 | NULL&M_Disease |
| 11 | 117579270 | rs670972 | A | G | NULL | 0.610639234 | NULL&M_Disease |
| 11 | 117579815 | rs634416 | T | C | NULL | 0.610639234 | NULL&M_Disease |
| 11 | 118723087 | rs11217059 | T | C | NULL | 0.610639234 | NULL&M_Disease |
| 11 | 118866515 | rs113032308 | A | G | NULL | 0.610639234 | NULL&M_Disease |
| 11 | 118866550 | rs112377532 | G | C | NULL | 0.610639234 | NULL&M_Disease |
| 11 | 119697476 | rs73008630 | G | A | NULL | 0.610639234 | NULL&M_Disease |
| 11 | 120565240 | rs78849359 | T | C | NULL | 0.610639234 | NULL&M_Disease |
| 11 | 120705938 | rs55966937 | G | A | NULL | 0.610639234 | NULL&M_Disease |
| 11 | 116355678 | rs693192 | A | G | NULL | 0.610639234 | NULL&M_Disease |
| 11 | 116748905 | rs61907566 | G | C | NULL | 0.611827369 | NULL&M_Disease |
| 11 | 120105300 | rs61898296 | A | G | NULL | 0.610639234 | NULL&M_Disease |
| 11 | 116351511 | rs566269 | T | C | NULL | 0.611827369 | NULL&M_Disease |
| 11 | 116532283 | rs78198334 | C | G | NULL | 0.610639234 | NULL&M_Disease |
| 11 | 117575629 | rs2508560 | A | G | NULL | 0.610639234 | NULL&M_Disease |
| 11 | 117575841 | rs585718 | T | C | NULL | 0.610639234 | NULL&M_Disease |
| 11 | 119740612 | rs4936505 | T | C | NULL | 0.610639234 | NULL&M_Disease |
| 11 | 119781758 | rs11600005 | A | G | NULL | 0.610639234 | NULL&M_Disease |
| 11 | 120173845 | rs12275681 | G | A | NULL | 0.611827369 | NULL&M_Disease |
| 11 | 120493714 | rs59127761 | G | C | NULL | 0.611827369 | NULL&M_Disease |

| | | | | | | | | | |
|---|---|---|---|---|---|---|---|---|---|
| 11 | 116237825 | rs1968541 | A | G | 0.953 | -7977.388 | -7978.28 | -0.892 | 1.59E-06 |
| 11 | 117528867 | rs59931457 | A | G | 1.002 | -7977.388 | -7978.28 | -0.892 | 1.59E-06 |
| 11 | 118365210 | rs9332817 | C | G | 0.98 | -7977.388 | -7978.281 | -0.892 | 1.59E-06 |
| 11 | 117263874 | rs117713193 | T | C | 0.959 | -7977.388 | -7978.281 | -0.893 | 1.58E-06 |
| 11 | 117989798 | rs117184822 | T | C | 1.014 | -7977.388 | -7978.281 | -0.893 | 1.58E-06 |
| 11 | 119817324 | rs80062985 | T | C | 1.051 | -7977.388 | -7978.281 | -0.893 | 1.58E-06 |
| 11 | 116451880 | rs71478302 | G | C | 0.984 | -7977.388 | -7978.282 | -0.894 | 1.58E-06 |
| 11 | 116803634 | rs118138924 | G | C | 0.954 | -7977.388 | -7978.283 | -0.894 | 1.58E-06 |
| 11 | 117429728 | rs685425 | T | C | 1.048 | -7977.388 | -7978.282 | -0.894 | 1.58E-06 |
| 11 | 117880723 | rs80044067 | A | G | 0.984 | -7977.388 | -7978.282 | -0.894 | 1.58E-06 |
| 11 | 117911367 | rs597820 | C | A | 1.043 | -7977.388 | -7978.282 | -0.894 | 1.58E-06 |
| 11 | 117987746 | rs640888 | A | G | 1.046 | -7977.388 | -7978.282 | -0.894 | 1.58E-06 |
| 11 | 119564528 | rs140186337 | G | A | 0.984 | -7977.388 | -7978.282 | -0.894 | 1.58E-06 |
| 11 | 120089082 | rs117934631 | G | T | 0.962 | -7977.388 | -7978.282 | -0.894 | 1.58E-06 |
| 11 | 120683594 | rs948026 | A | G | 0.952 | -7977.388 | -7978.282 | -0.894 | 1.58E-06 |
| 11 | 121159481 | rs116958147 | C | T | 1.044 | -7977.388 | -7978.282 | -0.894 | 1.58E-06 |
| 11 | 117730777 | rs78559460 | C | A | 0.988 | -7977.388 | -7978.283 | -0.895 | 1.58E-06 |
| 11 | 119755047 | rs12285075 | A | G | 1.049 | -7977.388 | -7978.283 | -0.895 | 1.58E-06 |
| 11 | 120236793 | rs117388325 | T | C | 1.019 | -7977.388 | -7978.284 | -0.895 | 1.58E-06 |
| 11 | 120837407 | rs10892654 | G | A | 1.05 | -7977.388 | -7978.283 | -0.895 | 1.58E-06 |
| 11 | 118576997 | rs498912 | C | T | 0.952 | -7977.388 | -7978.284 | -0.896 | 1.58E-06 |
| 11 | 119927648 | rs681454 | G | C | 1.044 | -7977.388 | -7978.284 | -0.896 | 1.58E-06 |
| 11 | 120128088 | rs115767079 | T | C | 1.038 | -7977.388 | -7978.284 | -0.896 | 1.58E-06 |
| 11 | 120694254 | rs75954141 | C | A | 0.953 | -7977.388 | -7978.285 | -0.896 | 1.58E-06 |
| 11 | 116237754 | rs11215903 | G | T | 0.977 | -7977.388 | -7978.285 | -0.897 | 1.58E-06 |
| 11 | 117320352 | rs10790185 | T | C | 0.953 | -7977.388 | -7978.286 | -0.897 | 1.58E-06 |
| 11 | 120176310 | rs12790634 | A | G | 0.958 | -7977.388 | -7978.285 | -0.897 | 1.58E-06 |
| 11 | 117218762 | rs568787 | A | G | 0.987 | -7977.388 | -7978.286 | -0.898 | 1.58E-06 |
| 11 | 120465556 | rs1573880 | T | C | 1.019 | -7977.388 | -7978.286 | -0.898 | 1.58E-06 |
| 11 | 120497208 | rs4375451 | T | C | 1.044 | -7977.388 | -7978.286 | -0.898 | 1.58E-06 |
| 11 | 120576984 | rs11603713 | G | A | 0.972 | -7977.388 | -7978.286 | -0.898 | 1.58E-06 |
| 11 | 120694915 | rs12291852 | T | G | 0.953 | -7977.388 | -7978.286 | -0.898 | 1.58E-06 |
| 11 | 117725057 | rs11216583 | G | C | 1.049 | -7977.388 | -7978.287 | -0.899 | 1.57E-06 |
| 11 | 120694466 | rs75208530 | G | A | 0.953 | -7977.388 | -7978.288 | -0.899 | 1.57E-06 |
| 11 | 117303430 | rs55644928 | T | C | 1.05 | -7977.388 | -7978.288 | -0.9 | 1.57E-06 |
| 11 | 117628972 | rs12296014 | A | G | 0.952 | -7977.388 | -7978.288 | -0.9 | 1.57E-06 |
| 11 | 118816304 | rs586763 | T | C | 1.048 | -7977.388 | -7978.289 | -0.9 | 1.57E-06 |
| 11 | 120993071 | rs608818 | T | C | 1.046 | -7977.388 | -7978.288 | -0.9 | 1.57E-06 |
| 11 | 117215605 | rs476983 | T | C | 0.989 | -7977.388 | -7978.29 | -0.901 | 1.57E-06 |
| 11 | 117252437 | rs117083334 | G | A | 0.968 | -7977.388 | -7978.289 | -0.901 | 1.57E-06 |
| 11 | 118721189 | rs4938566 | T | A | 0.956 | -7977.388 | -7978.289 | -0.901 | 1.57E-06 |
| 11 | 118721249 | rs4938567 | G | A | 0.956 | -7977.388 | -7978.289 | -0.901 | 1.57E-06 |
| 11 | 118721850 | rs7946961 | G | A | 0.956 | -7977.388 | -7978.289 | -0.901 | 1.57E-06 |
| 11 | 118722231 | rs4938570 | G | A | 0.956 | -7977.388 | -7978.289 | -0.901 | 1.57E-06 |
| 11 | 120128451 | rs17123747 | T | C | 1.037 | -7977.388 | -7978.289 | -0.901 | 1.57E-06 |
| 11 | 120128668 | rs78314944 | C | G | 1.037 | -7977.388 | -7978.289 | -0.901 | 1.57E-06 |
| 11 | 120128828 | rs116667143 | G | C | 1.037 | -7977.388 | -7978.289 | -0.901 | 1.57E-06 |
| 11 | 120129129 | rs114513135 | C | A | 1.037 | -7977.388 | -7978.289 | -0.901 | 1.57E-06 |
| 11 | 116632500 | rs10488699 | T | C | 0.952 | -7977.388 | -7978.291 | -0.902 | 1.57E-06 |
| 11 | 117604229 | rs1939983 | G | C | 0.953 | -7977.388 | -7978.29 | -0.902 | 1.57E-06 |
| 11 | 120789508 | rs10892640 | G | A | 0.959 | -7977.388 | -7978.29 | -0.902 | 1.57E-06 |
| 11 | 117465314 | rs75873912 | G | C | 0.952 | -7977.388 | -7978.292 | -0.903 | 1.57E-06 |
| 11 | 117600136 | rs10160584 | G | A | 0.973 | -7977.388 | -7978.291 | -0.903 | 1.57E-06 |
| 11 | 119844937 | rs7101905 | T | C | 1.05 | -7977.388 | -7978.291 | -0.903 | 1.57E-06 |
| 11 | 120163449 | rs56966127 | T | G | 0.952 | -7977.388 | -7978.291 | -0.903 | 1.57E-06 |
| 11 | 120419868 | rs11600436 | T | C | 1.043 | -7977.388 | -7978.292 | -0.903 | 1.57E-06 |
| 11 | 121149531 | rs11218239 | G | A | 1.049 | -7977.388 | -7978.291 | -0.903 | 1.57E-06 |
| 11 | 117228892 | rs560789 | G | A | 0.989 | -7977.388 | -7978.292 | -0.904 | 1.57E-06 |
| 11 | 117581589 | rs11216510 | C | A | 0.976 | -7977.388 | -7978.293 | -0.904 | 1.57E-06 |
| 11 | 117642376 | rs11825297 | T | C | 1.05 | -7977.388 | -7978.292 | -0.904 | 1.57E-06 |
| 11 | 117964339 | rs10750124 | C | T | 1.049 | -7977.388 | -7978.292 | -0.904 | 1.57E-06 |
| 11 | 116979911 | rs12271161 | A | G | 1.047 | -7977.388 | -7978.294 | -0.905 | 1.56E-06 |
| 11 | 117581228 | rs641569 | T | G | 0.977 | -7977.388 | -7978.293 | -0.905 | 1.56E-06 |
| 11 | 117581240 | rs601503 | C | G | 0.977 | -7977.388 | -7978.293 | -0.905 | 1.56E-06 |
| 11 | 117581785 | rs693992 | A | C | 0.977 | -7977.388 | -7978.293 | -0.905 | 1.56E-06 |
| 11 | 117604479 | rs4142636 | T | C | 0.953 | -7977.388 | -7978.293 | -0.905 | 1.56E-06 |
| 11 | 120152591 | rs11217785 | T | C | 1.046 | -7977.388 | -7978.293 | -0.905 | 1.56E-06 |
| 11 | 120170012 | rs77810230 | C | T | 1.05 | -7977.388 | -7978.294 | -0.905 | 1.56E-06 |
| 11 | 116326685 | rs118000999 | T | C | 0.97 | -7977.388 | -7978.294 | -0.906 | 1.56E-06 |
| 11 | 117574397 | rs650162 | T | C | 0.977 | -7977.388 | -7978.295 | -0.906 | 1.56E-06 |
| 11 | 117962732 | rs948468 | C | G | 1.049 | -7977.388 | -7978.294 | -0.906 | 1.56E-06 |
| 11 | 119890299 | rs12363335 | C | T | 1.048 | -7977.388 | -7978.294 | -0.906 | 1.56E-06 |
| 11 | 116259493 | rs1185802888 | T | A | 1.048 | -7977.388 | -7978.295 | -0.907 | 1.56E-06 |
| 11 | 116546633 | rs116944258 | A | C | 1.047 | -7977.388 | -7978.296 | -0.907 | 1.56E-06 |
| 11 | 116548972 | rs79678269 | C | G | 1.047 | -7977.388 | -7978.296 | -0.907 | 1.56E-06 |
| 11 | 116549116 | rs1005705 | C | A | 1.047 | -7977.388 | -7978.296 | -0.907 | 1.56E-06 |
| 11 | 116549971 | rs117830617 | A | G | 1.047 | -7977.388 | -7978.296 | -0.907 | 1.56E-06 |
| 11 | 117579270 | rs670972 | A | G | 0.977 | -7977.388 | -7978.296 | -0.907 | 1.56E-06 |
| 11 | 117579815 | rs634416 | T | C | 0.977 | -7977.388 | -7978.296 | -0.907 | 1.56E-06 |
| 11 | 118723087 | rs11217059 | T | C | 0.957 | -7977.388 | -7978.296 | -0.907 | 1.56E-06 |
| 11 | 118866515 | rs113032308 | A | G | 0.952 | -7977.388 | -7978.295 | -0.907 | 1.56E-06 |
| 11 | 118866550 | rs112377532 | G | C | 0.952 | -7977.388 | -7978.295 | -0.907 | 1.56E-06 |
| 11 | 119697476 | rs73008630 | G | A | 1.031 | -7977.388 | -7978.295 | -0.907 | 1.56E-06 |
| 11 | 120565240 | rs78849359 | T | C | 0.954 | -7977.388 | -7978.295 | -0.907 | 1.56E-06 |
| 11 | 120705938 | rs55966937 | G | A | 0.955 | -7977.388 | -7978.295 | -0.907 | 1.56E-06 |
| 11 | 116355678 | rs693192 | A | G | 1.045 | -7977.388 | -7978.296 | -0.908 | 1.56E-06 |
| 11 | 116748905 | rs61907566 | G | C | 0.959 | -7977.388 | -7978.296 | -0.908 | 1.56E-06 |
| 11 | 120105300 | rs61898296 | A | G | 0.969 | -7977.388 | -7978.296 | -0.908 | 1.56E-06 |
| 11 | 116351511 | rs566269 | T | C | 1.046 | -7977.388 | -7978.297 | -0.909 | 1.56E-06 |
| 11 | 116532283 | rs78198334 | C | G | 1.047 | -7977.388 | -7978.297 | -0.909 | 1.56E-06 |
| 11 | 117575629 | rs2508560 | A | G | 0.978 | -7977.388 | -7978.297 | -0.909 | 1.56E-06 |
| 11 | 117575841 | rs585718 | T | C | 0.978 | -7977.388 | -7978.297 | -0.909 | 1.56E-06 |
| 11 | 119740612 | rs4936505 | T | C | 0.955 | -7977.388 | -7978.297 | -0.909 | 1.56E-06 |
| 11 | 119781758 | rs11600005 | A | G | 0.99 | -7977.388 | -7978.298 | -0.909 | 1.56E-06 |
| 11 | 120173845 | rs12275681 | G | A | 1.05 | -7977.388 | -7978.298 | -0.909 | 1.56E-06 |
| 11 | 120493714 | rs59127761 | G | C | 1.044 | -7977.388 | -7978.298 | -0.909 | 1.56E-06 |

| | | | | | | | |
|---|---|---|---|---|---|---|---|
| 11 | 116680448 | rs2849169 | A | C | NULL | 0.611827369 | NULL&M_Disease |
| 11 | 117662639 | rs10790210 | G | C | NULL | 0.611827369 | NULL&M_Disease |
| 11 | 118393856 | rs9332871 | T | G | NULL | 0.610639234 | NULL&M_Disease |
| 11 | 119518006 | rs12797352 | G | A | NULL | 0.611827369 | NULL&M_Disease |
| 11 | 119677419 | rs76031074 | A | G | NULL | 0.610639234 | NULL&M_Disease |
| 11 | 119769724 | rs79995246 | C | T | NULL | 0.610639234 | NULL&M_Disease |
| 11 | 119987576 | rs488222 | C | T | NULL | 0.611827369 | NULL&M_Disease |
| 11 | 117300067 | rs3889571 | T | G | NULL | 0.611827369 | NULL&M_Disease |
| 11 | 119680745 | rs34518191 | G | C | NULL | 0.610639234 | NULL&M_Disease |
| 11 | 120128223 | rs114667713 | C | T | NULL | 0.610639234 | NULL&M_Disease |
| 11 | 120997460 | rs659042 | A | C | NULL | 0.611827369 | NULL&M_Disease |
| 11 | 116449692 | rs61904846 | A | G | NULL | 0.611827369 | NULL&M_Disease |
| 11 | 116655600 | rs35120633 | A | G | NULL | 0.611827369 | NULL&M_Disease |
| 11 | 119243191 | rs76870367 | T | C | NULL | 0.610639234 | NULL&M_Disease |
| 11 | 120331909 | rs117079525 | T | C | NULL | 0.610639234 | NULL&M_Disease |
| 11 | 116541034 | rs144188664 | T | A | NULL | 0.611827369 | NULL&M_Disease |
| 11 | 116542737 | rs148462279 | G | A | NULL | 0.611827369 | NULL&M_Disease |
| 11 | 116543025 | rs144256947 | C | T | NULL | 0.611827369 | NULL&M_Disease |
| 11 | 117585788 | rs4529912 | G | A | NULL | 0.611827369 | NULL&M_Disease |
| 11 | 117586059 | rs4936398 | G | A | NULL | 0.611827369 | NULL&M_Disease |
| 11 | 117628810 | rs12282602 | G | A | NULL | 0.611827369 | NULL&M_Disease |
| 11 | 120986933 | rs617692 | T | C | NULL | 0.611827369 | NULL&M_Disease |
| 11 | 116391272 | rs12361978 | G | C | NULL | 0.611827369 | NULL&M_Disease |
| 11 | 117429718 | rs685440 | T | G | NULL | 0.611827369 | NULL&M_Disease |
| 11 | 117687120 | rs2226635 | T | C | NULL | 0.611827369 | NULL&M_Disease |
| 11 | 119620677 | rs113373988 | T | G | NULL | 0.611827369 | NULL&M_Disease |
| 11 | 120126782 | rs12418040 | A | G | NULL | 0.611827369 | NULL&M_Disease |
| 11 | 121191782 | rs10790434 | G | C | NULL | 0.611827369 | NULL&M_Disease |
| 11 | 117777101 | rs73020500 | T | C | NULL | 0.611827369 | NULL&M_Disease |
| 11 | 118389698 | rs113399762 | C | T | NULL | 0.611827369 | NULL&M_Disease |
| 11 | 118803193 | rs117235696 | G | T | NULL | 0.611827369 | NULL&M_Disease |
| 11 | 119935980 | rs118000863 | A | G | NULL | 0.611827369 | NULL&M_Disease |
| 11 | 120161995 | rs7120237 | C | T | NULL | 0.611827369 | NULL&M_Disease |
| 11 | 120484810 | rs1317814 | G | A | NULL | 0.611827369 | NULL&M_Disease |
| 11 | 120798482 | rs12805801 | A | G | NULL | 0.611827369 | NULL&M_Disease |
| 11 | 117605046 | rs1942928 | G | A | NULL | 0.611827369 | NULL&M_Disease |
| 11 | 117861055 | rs4252254 | T | C | NULL | 0.611827369 | NULL&M_Disease |
| 11 | 117964465 | rs10736490 | T | C | NULL | 0.611827369 | NULL&M_Disease |
| 11 | 117964490 | rs10736491 | T | A | NULL | 0.611827369 | NULL&M_Disease |
| 11 | 117964598 | rs7104016 | T | G | NULL | 0.611827369 | NULL&M_Disease |
| 11 | 117964873 | rs1941629 | C | G | NULL | 0.611827369 | NULL&M_Disease |
| 11 | 117964902 | rs1941630 | C | G | NULL | 0.611827369 | NULL&M_Disease |
| 11 | 119816115 | rs1815890 | T | C | NULL | 0.611827369 | NULL&M_Disease |
| 11 | 120482493 | rs60787798 | T | G | NULL | 0.611827369 | NULL&M_Disease |
| 11 | 117592602 | rs4938434 | G | T | NULL | 0.611827369 | NULL&M_Disease |
| 11 | 118381069 | rs9332852 | T | C | NULL | 0.611827369 | NULL&M_Disease |
| 11 | 118893767 | rs111345266 | T | C | NULL | 0.611827369 | NULL&M_Disease |
| 11 | 120074973 | rs180999772 | A | G | NULL | 0.611827369 | NULL&M_Disease |
| 11 | 120826260 | rs76663629 | A | G | NULL | 0.611827369 | NULL&M_Disease |
| 11 | 121001345 | rs583842 | C | T | NULL | 0.611827369 | NULL&M_Disease |
| 11 | 121193894 | rs117323348 | G | C | NULL | 0.611827369 | NULL&M_Disease |
| 11 | 116233803 | rs75907072 | A | T | NULL | 0.611827369 | NULL&M_Disease |
| 11 | 120048628 | rs113981653 | T | C | NULL | 0.611827369 | NULL&M_Disease |
| 11 | 120492364 | rs73002650 | G | A | NULL | 0.613014176 | NULL&M_Disease |
| 11 | 120575606 | rs75764292 | T | A | NULL | 0.611827369 | NULL&M_Disease |
| 11 | 120872008 | rs77772751 | A | G | NULL | 0.611827369 | NULL&M_Disease |
| 11 | 117581772 | rs629060 | G | T | NULL | 0.611827369 | NULL&M_Disease |
| 11 | 117582054 | rs948777 | C | T | NULL | 0.611827369 | NULL&M_Disease |
| 11 | 117582263 | rs6589619 | T | C | NULL | 0.611827369 | NULL&M_Disease |
| 11 | 117628729 | rs11216533 | A | C | NULL | 0.611827369 | NULL&M_Disease |
| 11 | 120069087 | rs61122660 | A | G | NULL | 0.611827369 | NULL&M_Disease |
| 11 | 120572302 | rs11604327 | C | A | NULL | 0.611827369 | NULL&M_Disease |
| 11 | 118704751 | rs35262372 | G | A | NULL | 0.611827369 | NULL&M_Disease |
| 11 | 116241208 | rs17119549 | T | G | NULL | 0.611827369 | NULL&M_Disease |
| 11 | 116837808 | rs1241655 | T | G | NULL | 0.613014176 | NULL&M_Disease |
| 11 | 117315050 | rs78701708 | C | T | NULL | 0.611827369 | NULL&M_Disease |
| 11 | 117658211 | rs12363806 | A | G | NULL | 0.613014176 | NULL&M_Disease |
| 11 | 118193468 | rs56410112 | C | T | NULL | 0.613014176 | NULL&M_Disease |
| 11 | 118278998 | rs112583389 | G | T | NULL | 0.611827369 | NULL&M_Disease |
| 11 | 119853430 | rs595167 | T | C | NULL | 0.613014176 | NULL&M_Disease |
| 11 | 120460445 | rs7124361 | G | A | NULL | 0.611827369 | NULL&M_Disease |
| 11 | 116531969 | rs7117145 | T | A | NULL | 0.613014176 | NULL&M_Disease |
| 11 | 117862575 | rs4252261 | G | T | NULL | 0.613014176 | NULL&M_Disease |
| 11 | 119810375 | rs79493947 | G | C | NULL | 0.613014176 | NULL&M_Disease |
| 11 | 120131591 | rs10892556 | G | A | NULL | 0.611827369 | NULL&M_Disease |
| 11 | 120154647 | rs74236011 | G | T | NULL | 0.613014176 | NULL&M_Disease |
| 11 | 117584368 | rs2156233 | T | C | NULL | 0.613014176 | NULL&M_Disease |
| 11 | 120874890 | rs184243297 | A | G | NULL | 0.613014176 | NULL&M_Disease |
| 11 | 121003286 | rs632121 | T | A | NULL | 0.613014176 | NULL&M_Disease |
| 11 | 117759895 | rs73020452 | C | T | NULL | 0.613014176 | NULL&M_Disease |
| 11 | 116379889 | rs1508102 | A | G | NULL | 0.613014176 | NULL&M_Disease |
| 11 | 117134152 | rs192958777 | T | C | NULL | 0.613014176 | NULL&M_Disease |
| 11 | 118233634 | rs74398561 | G | A | NULL | 0.613014176 | NULL&M_Disease |
| 11 | 118273602 | rs113839182 | T | C | NULL | 0.613014176 | NULL&M_Disease |
| 11 | 118274602 | rs111587825 | G | T | NULL | 0.613014176 | NULL&M_Disease |
| 11 | 118276810 | rs111707199 | G | A | NULL | 0.613014176 | NULL&M_Disease |
| 11 | 118278479 | rs112468256 | C | T | NULL | 0.613014176 | NULL&M_Disease |
| 11 | 118389440 | rs9332857 | T | C | NULL | 0.613014176 | NULL&M_Disease |
| 11 | 119816331 | rs4403813 | C | T | NULL | 0.613014176 | NULL&M_Disease |
| 11 | 121000774 | rs520805 | C | T | NULL | 0.613014176 | NULL&M_Disease |
| 11 | 117614796 | rs869005 | T | C | NULL | 0.613014176 | NULL&M_Disease |
| 11 | 117749120 | rs3809042 | G | C | NULL | 0.613014176 | NULL&M_Disease |
| 11 | 120702478 | rs191347151 | T | C | NULL | 0.613014176 | NULL&M_Disease |
| 11 | 120749953 | rs11218041 | G | C | NULL | 0.613014176 | NULL&M_Disease |
| 11 | 116533595 | rs75942205 | C | T | NULL | 0.613014176 | NULL&M_Disease |
| 11 | 116533628 | rs61659529 | G | A | NULL | 0.613014176 | NULL&M_Disease |

| | | | | | | | | | |
|---|---|---|---|---|---|---|---|---|---|
| 11 | 116680448 | rs2849169 | A | C | 1.05 | -7977.388 | -7978.299 | -0.91 | 1.56E-06 |
| 11 | 117662639 | rs10790210 | G | C | 0.958 | -7977.388 | -7978.299 | -0.91 | 1.56E-06 |
| 11 | 118393856 | rs9332871 | T | G | 1.047 | -7977.388 | -7978.298 | -0.91 | 1.56E-06 |
| 11 | 119518006 | rs12797352 | G | A | 1.044 | -7977.388 | -7978.298 | -0.91 | 1.56E-06 |
| 11 | 119677419 | rs76031074 | A | G | 0.975 | -7977.388 | -7978.298 | -0.91 | 1.56E-06 |
| 11 | 119769724 | rs79995246 | C | T | 1.046 | -7977.388 | -7978.299 | -0.91 | 1.56E-06 |
| 11 | 119987576 | rs488222 | C | T | 0.952 | -7977.388 | -7978.299 | -0.91 | 1.56E-06 |
| 11 | 117300067 | rs3889571 | T | G | 1.042 | -7977.388 | -7978.299 | -0.911 | 1.56E-06 |
| 11 | 119680745 | rs34518191 | G | C | 1.036 | -7977.388 | -7978.3 | -0.911 | 1.56E-06 |
| 11 | 120128223 | rs114667713 | C | T | 1.036 | -7977.388 | -7978.299 | -0.911 | 1.56E-06 |
| 11 | 120997460 | rs659042 | A | C | 1.046 | -7977.388 | -7978.299 | -0.911 | 1.56E-06 |
| 11 | 116449692 | rs61904846 | A | G | 1.048 | -7977.388 | -7978.301 | -0.912 | 1.55E-06 |
| 11 | 116655600 | rs35120633 | A | G | 0.953 | -7977.388 | -7978.3 | -0.912 | 1.55E-06 |
| 11 | 119243191 | rs76870367 | T | C | 0.977 | -7977.388 | -7978.3 | -0.912 | 1.55E-06 |
| 11 | 120331909 | rs117079525 | T | C | 1.015 | -7977.388 | -7978.3 | -0.912 | 1.55E-06 |
| 11 | 116541034 | rs144188664 | T | A | 1.047 | -7977.388 | -7978.301 | -0.913 | 1.55E-06 |
| 11 | 116542737 | rs148462279 | G | A | 1.047 | -7977.388 | -7978.301 | -0.913 | 1.55E-06 |
| 11 | 116543025 | rs144256947 | C | T | 1.047 | -7977.388 | -7978.301 | -0.913 | 1.55E-06 |
| 11 | 117585788 | rs4529912 | G | A | 0.954 | -7977.388 | -7978.301 | -0.913 | 1.55E-06 |
| 11 | 117586059 | rs4936398 | G | A | 0.954 | -7977.388 | -7978.301 | -0.913 | 1.55E-06 |
| 11 | 117628810 | rs12282602 | G | A | 0.952 | -7977.388 | -7978.301 | -0.913 | 1.55E-06 |
| 11 | 120986933 | rs617692 | T | C | 1.046 | -7977.388 | -7978.301 | -0.913 | 1.55E-06 |
| 11 | 116391272 | rs12361978 | G | C | 0.958 | -7977.388 | -7978.302 | -0.914 | 1.55E-06 |
| 11 | 117429718 | rs685440 | T | G | 1.047 | -7977.388 | -7978.302 | -0.914 | 1.55E-06 |
| 11 | 117687120 | rs2226635 | T | C | 0.952 | -7977.388 | -7978.302 | -0.914 | 1.55E-06 |
| 11 | 119620677 | rs113373988 | T | G | 0.952 | -7977.388 | -7978.302 | -0.914 | 1.55E-06 |
| 11 | 120126782 | rs12418040 | A | G | 1.008 | -7977.388 | -7978.303 | -0.914 | 1.55E-06 |
| 11 | 121191782 | rs10790434 | G | C | 0.955 | -7977.388 | -7978.303 | -0.914 | 1.55E-06 |
| 11 | 117777101 | rs73020500 | T | C | 0.972 | -7977.388 | -7978.304 | -0.915 | 1.55E-06 |
| 11 | 118389698 | rs113399762 | C | T | 1.046 | -7977.388 | -7978.304 | -0.915 | 1.55E-06 |
| 11 | 118803193 | rs117235696 | G | T | 1.037 | -7977.388 | -7978.303 | -0.915 | 1.55E-06 |
| 11 | 119935980 | rs118000863 | A | G | 0.98 | -7977.388 | -7978.304 | -0.915 | 1.55E-06 |
| 11 | 120161995 | rs7120237 | C | T | 1.045 | -7977.388 | -7978.304 | -0.915 | 1.55E-06 |
| 11 | 120484810 | rs1317814 | G | A | 1.043 | -7977.388 | -7978.303 | -0.915 | 1.55E-06 |
| 11 | 120798482 | rs12805801 | A | G | 1.046 | -7977.388 | -7978.304 | -0.915 | 1.55E-06 |
| 11 | 117605046 | rs1942928 | G | A | 0.954 | -7977.388 | -7978.304 | -0.916 | 1.55E-06 |
| 11 | 117861055 | rs4252254 | T | C | 0.953 | -7977.388 | -7978.304 | -0.916 | 1.55E-06 |
| 11 | 117964465 | rs10736490 | T | C | 1.048 | -7977.388 | -7978.305 | -0.916 | 1.55E-06 |
| 11 | 117964490 | rs10736491 | T | A | 1.048 | -7977.388 | -7978.305 | -0.916 | 1.55E-06 |
| 11 | 117964598 | rs7104016 | T | G | 1.048 | -7977.388 | -7978.305 | -0.916 | 1.55E-06 |
| 11 | 117964873 | rs1941629 | C | G | 1.048 | -7977.388 | -7978.305 | -0.916 | 1.55E-06 |
| 11 | 117964902 | rs1941630 | C | G | 1.048 | -7977.388 | -7978.305 | -0.916 | 1.55E-06 |
| 11 | 119816115 | rs1815890 | T | C | 1.05 | -7977.388 | -7978.304 | -0.916 | 1.55E-06 |
| 11 | 120482493 | rs60787798 | T | G | 1.043 | -7977.388 | -7978.305 | -0.916 | 1.55E-06 |
| 11 | 117592602 | rs4938434 | G | T | 0.954 | -7977.388 | -7978.306 | -0.917 | 1.55E-06 |
| 11 | 118381069 | rs9332852 | T | C | 1.046 | -7977.388 | -7978.306 | -0.917 | 1.55E-06 |
| 11 | 118893767 | rs111345266 | T | C | 1.049 | -7977.388 | -7978.306 | -0.917 | 1.55E-06 |
| 11 | 120074973 | rs180999772 | A | G | 0.967 | -7977.388 | -7978.305 | -0.917 | 1.55E-06 |
| 11 | 120826260 | rs76663629 | A | G | 1.049 | -7977.388 | -7978.306 | -0.917 | 1.55E-06 |
| 11 | 121001345 | rs583842 | C | T | 1.046 | -7977.388 | -7978.305 | -0.917 | 1.55E-06 |
| 11 | 121193894 | rs117323348 | G | C | 1.003 | -7977.388 | -7978.306 | -0.917 | 1.55E-06 |
| 11 | 116233803 | rs75907072 | A | T | 0.954 | -7977.388 | -7978.306 | -0.918 | 1.54E-06 |
| 11 | 120048628 | rs113981653 | T | C | 0.971 | -7977.388 | -7978.306 | -0.918 | 1.54E-06 |
| 11 | 120492364 | rs73002650 | G | A | 1.043 | -7977.388 | -7978.306 | -0.918 | 1.54E-06 |
| 11 | 120575606 | rs75764292 | T | A | 0.977 | -7977.388 | -7978.306 | -0.918 | 1.54E-06 |
| 11 | 120872008 | rs77772751 | A | G | 1.033 | -7977.388 | -7978.307 | -0.918 | 1.54E-06 |
| 11 | 117581772 | rs629060 | G | T | 0.955 | -7977.388 | -7978.307 | -0.919 | 1.54E-06 |
| 11 | 117582054 | rs948777 | C | T | 0.955 | -7977.388 | -7978.307 | -0.919 | 1.54E-06 |
| 11 | 117582263 | rs6589619 | T | C | 0.955 | -7977.388 | -7978.307 | -0.919 | 1.54E-06 |
| 11 | 117628729 | rs11216533 | A | C | 0.953 | -7977.388 | -7978.307 | -0.919 | 1.54E-06 |
| 11 | 120069087 | rs61122660 | A | G | 0.954 | -7977.388 | -7978.307 | -0.919 | 1.54E-06 |
| 11 | 120572302 | rs11604327 | C | A | 0.975 | -7977.388 | -7978.308 | -0.919 | 1.54E-06 |
| 11 | 118704751 | rs35262372 | G | A | 1.042 | -7977.388 | -7978.308 | -0.92 | 1.54E-06 |
| 11 | 116241208 | rs17119549 | T | G | 0.987 | -7977.388 | -7978.309 | -0.921 | 1.54E-06 |
| 11 | 116837808 | rs1241655 | T | G | 1.045 | -7977.388 | -7978.309 | -0.921 | 1.54E-06 |
| 11 | 117315050 | rs78701708 | C | T | 0.983 | -7977.388 | -7978.309 | -0.921 | 1.54E-06 |
| 11 | 117658211 | rs12363806 | A | G | 0.958 | -7977.388 | -7978.309 | -0.921 | 1.54E-06 |
| 11 | 118193468 | rs56410112 | C | T | 0.953 | -7977.388 | -7978.31 | -0.921 | 1.54E-06 |
| 11 | 118278998 | rs112583389 | G | T | 1.034 | -7977.388 | -7978.309 | -0.921 | 1.54E-06 |
| 11 | 119853430 | rs595167 | T | C | 0.956 | -7977.388 | -7978.309 | -0.921 | 1.54E-06 |
| 11 | 120460445 | rs7124361 | G | A | 0.967 | -7977.388 | -7978.309 | -0.921 | 1.54E-06 |
| 11 | 116531969 | rs7117145 | T | A | 1.046 | -7977.388 | -7978.311 | -0.922 | 1.54E-06 |
| 11 | 117862575 | rs4252261 | G | T | 0.953 | -7977.388 | -7978.31 | -0.922 | 1.54E-06 |
| 11 | 119810375 | rs79493947 | G | C | 1.047 | -7977.388 | -7978.31 | -0.922 | 1.54E-06 |
| 11 | 120131591 | rs10892556 | G | A | 1.036 | -7977.388 | -7978.31 | -0.922 | 1.54E-06 |
| 11 | 120154647 | rs74236011 | G | T | 1.049 | -7977.388 | -7978.311 | -0.922 | 1.54E-06 |
| 11 | 117584368 | rs2156233 | T | C | 0.955 | -7977.388 | -7978.311 | -0.923 | 1.54E-06 |
| 11 | 120874890 | rs184243297 | A | G | 1.047 | -7977.388 | -7978.312 | -0.923 | 1.54E-06 |
| 11 | 121003286 | rs632121 | T | A | 1.046 | -7977.388 | -7978.312 | -0.923 | 1.54E-06 |
| 11 | 117759895 | rs73020452 | C | T | 0.978 | -7977.388 | -7978.312 | -0.924 | 1.54E-06 |
| 11 | 116379889 | rs1508102 | A | G | 1.047 | -7977.388 | -7978.313 | -0.925 | 1.53E-06 |
| 11 | 117134152 | rs192958777 | T | C | 0.988 | -7977.388 | -7978.314 | -0.925 | 1.53E-06 |
| 11 | 118233634 | rs74398561 | G | A | 1.033 | -7977.388 | -7978.314 | -0.925 | 1.53E-06 |
| 11 | 118273602 | rs113839182 | T | C | 1.033 | -7977.388 | -7978.314 | -0.925 | 1.53E-06 |
| 11 | 118274602 | rs111587825 | G | T | 1.033 | -7977.388 | -7978.314 | -0.925 | 1.53E-06 |
| 11 | 118276810 | rs111707199 | G | A | 1.033 | -7977.388 | -7978.314 | -0.925 | 1.53E-06 |
| 11 | 118278479 | rs112468256 | C | T | 1.033 | -7977.388 | -7978.314 | -0.925 | 1.53E-06 |
| 11 | 118389440 | rs9332857 | T | C | 1.046 | -7977.388 | -7978.313 | -0.925 | 1.53E-06 |
| 11 | 119816331 | rs4403813 | C | T | 1.049 | -7977.388 | -7978.313 | -0.925 | 1.53E-06 |
| 11 | 121000774 | rs520805 | C | T | 1.046 | -7977.388 | -7978.313 | -0.925 | 1.53E-06 |
| 11 | 117614796 | rs869005 | T | C | 0.955 | -7977.388 | -7978.315 | -0.926 | 1.53E-06 |
| 11 | 117749120 | rs3809042 | G | C | 1.027 | -7977.388 | -7978.314 | -0.926 | 1.53E-06 |
| 11 | 120702478 | rs191347151 | T | C | 0.955 | -7977.388 | -7978.314 | -0.926 | 1.53E-06 |
| 11 | 120749953 | rs11218041 | G | C | 1.034 | -7977.388 | -7978.315 | -0.926 | 1.53E-06 |
| 11 | 116533595 | rs75942205 | C | T | 1.046 | -7977.388 | -7978.315 | -0.927 | 1.53E-06 |
| 11 | 116533628 | rs61659529 | G | A | 1.046 | -7977.388 | -7978.315 | -0.927 | 1.53E-06 |

| | | | | | | | |
|---|---|---|---|---|---|---|---|
| 11 | 117852972 | rs142661080 | T | C | NULL | 0.613014176 | NULL&M_Disease |
| 11 | 118726230 | rs2156755 | A | G | NULL | 0.613014176 | NULL&M_Disease |
| 11 | 120659485 | rs3133861 | T | G | NULL | 0.613014176 | NULL&M_Disease |
| 11 | 117448799 | rs35021198 | T | C | NULL | 0.614199643 | NULL&M_Disease |
| 11 | 119217633 | rs45509899 | T | C | NULL | 0.613014176 | NULL&M_Disease |
| 11 | 119935473 | rs78443404 | C | G | NULL | 0.613014176 | NULL&M_Disease |
| 11 | 116302144 | rs10891974 | T | C | NULL | 0.613014176 | NULL&M_Disease |
| 11 | 116530061 | rs1892947 | A | G | NULL | 0.613014176 | NULL&M_Disease |
| 11 | 117120027 | rs605531 | G | T | NULL | 0.614199643 | NULL&M_Disease |
| 11 | 117987503 | rs1715464 | A | C | NULL | 0.614199643 | NULL&M_Disease |
| 11 | 116621972 | rs79112951 | T | C | NULL | 0.613014176 | NULL&M_Disease |
| 11 | 117580254 | rs2508559 | T | C | NULL | 0.613014176 | NULL&M_Disease |
| 11 | 117580509 | rs655086 | A | G | NULL | 0.613014176 | NULL&M_Disease |
| 11 | 117580585 | rs604245 | A | G | NULL | 0.613014176 | NULL&M_Disease |
| 11 | 117862979 | rs4252305 | A | G | NULL | 0.613014176 | NULL&M_Disease |
| 11 | 117863183 | rs73009944 | A | G | NULL | 0.613014176 | NULL&M_Disease |
| 11 | 117863307 | rs73009948 | T | G | NULL | 0.613014176 | NULL&M_Disease |
| 11 | 119762339 | rs79436592 | A | G | NULL | 0.613014176 | NULL&M_Disease |
| 11 | 117629468 | rs11216534 | T | C | NULL | 0.614199643 | NULL&M_Disease |
| 11 | 117629559 | rs11216535 | T | C | NULL | 0.614199643 | NULL&M_Disease |
| 11 | 120161671 | rs6589805 | C | T | NULL | 0.614199643 | NULL&M_Disease |
| 11 | 120622625 | rs9667927 | C | T | NULL | 0.614199643 | NULL&M_Disease |
| 11 | 117312429 | rs2887176 | G | A | NULL | 0.614199643 | NULL&M_Disease |
| 11 | 120826272 | rs78528013 | T | C | NULL | 0.614199643 | NULL&M_Disease |
| 11 | 116529130 | rs78347813 | T | C | NULL | 0.614199643 | NULL&M_Disease |
| 11 | 116534550 | rs76547220 | G | T | NULL | 0.614199643 | NULL&M_Disease |
| 11 | 116535435 | rs77226464 | T | C | NULL | 0.614199643 | NULL&M_Disease |
| 11 | 116535624 | rs190069353 | A | C | NULL | 0.614199643 | NULL&M_Disease |
| 11 | 116535631 | rs117014628 | T | C | NULL | 0.614199643 | NULL&M_Disease |
| 11 | 116536820 | rs79063991 | T | C | NULL | 0.614199643 | NULL&M_Disease |
| 11 | 116537116 | rs118103820 | G | A | NULL | 0.614199643 | NULL&M_Disease |
| 11 | 116537798 | rs140973748 | C | T | NULL | 0.614199643 | NULL&M_Disease |
| 11 | 116538103 | rs143380339 | A | C | NULL | 0.614199643 | NULL&M_Disease |
| 11 | 116685391 | rs77535577 | A | G | NULL | 0.614199643 | NULL&M_Disease |
| 11 | 118725420 | rs11217062 | A | G | NULL | 0.614199643 | NULL&M_Disease |
| 11 | 118798824 | rs112119822 | G | A | NULL | 0.614199643 | NULL&M_Disease |
| 11 | 119504952 | rs1009365 | T | C | NULL | 0.614199643 | NULL&M_Disease |
| 11 | 119926054 | rs7125256 | G | A | NULL | 0.614199643 | NULL&M_Disease |
| 11 | 116361658 | rs11215963 | G | C | NULL | 0.614199643 | NULL&M_Disease |
| 11 | 116533310 | rs79291519 | T | C | NULL | 0.614199643 | NULL&M_Disease |
| 11 | 116642137 | rs664059 | C | T | NULL | 0.614199643 | NULL&M_Disease |
| 11 | 118267903 | rs45480805 | A | G | NULL | 0.614199643 | NULL&M_Disease |
| 11 | 120868233 | rs113749645 | A | T | NULL | 0.614199643 | NULL&M_Disease |
| 11 | 117724493 | rs11216582 | A | C | NULL | 0.614199643 | NULL&M_Disease |
| 11 | 117883208 | rs2512150 | C | G | NULL | 0.614199643 | NULL&M_Disease |
| 11 | 119805016 | rs12273820 | C | T | NULL | 0.614199643 | NULL&M_Disease |
| 11 | 120784745 | rs111716733 | T | C | NULL | 0.614199643 | NULL&M_Disease |
| 11 | 116578361 | rs75310100 | T | G | NULL | 0.614199643 | NULL&M_Disease |
| 11 | 117307122 | rs665403 | T | C | NULL | 0.614199643 | NULL&M_Disease |
| 11 | 118861849 | rs57216066 | A | G | NULL | 0.614199643 | NULL&M_Disease |
| 11 | 120159540 | rs1329232240 | C | T | NULL | 0.614199643 | NULL&M_Disease |
| 11 | 120161117 | rs7951628 | A | G | NULL | 0.614199643 | NULL&M_Disease |
| 11 | 116241842 | rs533735 | C | A | NULL | 0.614199643 | NULL&M_Disease |
| 11 | 116457115 | rs75243489 | A | T | NULL | 0.614199643 | NULL&M_Disease |
| 11 | 117059347 | rs90192 | C | T | NULL | 0.614199643 | NULL&M_Disease |
| 11 | 118183733 | rs28423005 | T | C | NULL | 0.614199643 | NULL&M_Disease |
| 11 | 118521549 | rs494560 | A | G | NULL | 0.615383756 | NULL&M_Disease |
| 11 | 118569414 | rs6589675 | C | G | NULL | 0.614199643 | NULL&M_Disease |
| 11 | 118860457 | rs78505713 | T | C | NULL | 0.614199643 | NULL&M_Disease |
| 11 | 118860598 | rs113271112 | C | A | NULL | 0.614199643 | NULL&M_Disease |
| 11 | 116789102 | rs1378582752 | C | T | NULL | 0.614199643 | NULL&M_Disease |
| 11 | 118720998 | rs4938565 | A | G | NULL | 0.615383756 | NULL&M_Disease |
| 11 | 120488117 | rs73002644 | G | C | NULL | 0.615383756 | NULL&M_Disease |
| 11 | 116531050 | rs117889066 | C | G | NULL | 0.614199643 | NULL&M_Disease |
| 11 | 116531475 | rs117293980 | G | T | NULL | 0.614199643 | NULL&M_Disease |
| 11 | 117644124 | rs17311024 | A | T | NULL | 0.614199643 | NULL&M_Disease |
| 11 | 118254710 | rs45443099 | G | A | NULL | 0.614199643 | NULL&M_Disease |
| 11 | 118259162 | rs111792147 | A | G | NULL | 0.614199643 | NULL&M_Disease |
| 11 | 118262135 | rs112563022 | T | C | NULL | 0.614199643 | NULL&M_Disease |
| 11 | 119742642 | rs73008675 | C | T | NULL | 0.615383756 | NULL&M_Disease |
| 11 | 119743088 | rs73008677 | A | G | NULL | 0.615383756 | NULL&M_Disease |
| 11 | 119743230 | rs73008679 | T | C | NULL | 0.615383756 | NULL&M_Disease |
| 11 | 120826325 | rs77571870 | A | G | NULL | 0.614199643 | NULL&M_Disease |
| 11 | 117874531 | rs947889 | C | T | NULL | 0.615383756 | NULL&M_Disease |
| 11 | 118486067 | rs11603023 | T | C | NULL | 0.615383756 | NULL&M_Disease |
| 11 | 119747470 | rs4938735 | T | C | NULL | 0.615383756 | NULL&M_Disease |
| 11 | 116375518 | rs111551597 | T | C | NULL | 0.615383756 | NULL&M_Disease |
| 11 | 117600616 | rs11600991 | C | T | NULL | 0.615383756 | NULL&M_Disease |
| 11 | 118047776 | rs67262752 | C | T | NULL | 0.615383756 | NULL&M_Disease |
| 11 | 118384811 | rs78020509 | G | A | NULL | 0.615383756 | NULL&M_Disease |
| 11 | 120163184 | rs61898317 | G | A | NULL | 0.615383756 | NULL&M_Disease |
| 11 | 120163869 | rs2155810 | C | T | NULL | 0.615383756 | NULL&M_Disease |
| 11 | 117656916 | rs4603339 | G | A | NULL | 0.615383756 | NULL&M_Disease |
| 11 | 119503670 | rs12279718 | A | C | NULL | 0.615383756 | NULL&M_Disease |
| 11 | 119761810 | rs78048464 | T | G | NULL | 0.615383756 | NULL&M_Disease |
| 11 | 120391473 | rs12804597 | A | G | NULL | 0.615383756 | NULL&M_Disease |
| 11 | 116480033 | rs149395240 | T | C | NULL | 0.615383756 | NULL&M_Disease |
| 11 | 117781435 | rs2155194 | A | G | NULL | 0.615383756 | NULL&M_Disease |
| 11 | 118182660 | rs2277289 | C | T | NULL | 0.615383756 | NULL&M_Disease |
| 11 | 118246021 | rs112243842 | G | T | NULL | 0.615383756 | NULL&M_Disease |
| 11 | 118247805 | rs112646740 | A | T | NULL | 0.615383756 | NULL&M_Disease |
| 11 | 118248374 | rs111430146 | A | C | NULL | 0.615383756 | NULL&M_Disease |
| 11 | 120789800 | rs10892641 | G | A | NULL | 0.615383756 | NULL&M_Disease |
| 11 | 116279292 | rs579941 | T | C | NULL | 0.615383756 | NULL&M_Disease |
| 11 | 116682995 | rs76636922 | A | G | NULL | 0.615383756 | NULL&M_Disease |
| 11 | 117967476 | rs10790231 | C | T | NULL | 0.615383756 | NULL&M_Disease |

| | | | | | | | | | |
|---|---|---|---|---|---|---|---|---|---|
| 11 | 117852972 | rs142661080 | T | C | 0.997 | -7977.388 | -7978.315 | -0.927 | 1.53E-06 |
| 11 | 118726230 | rs2156755 | A | G | 0.957 | -7977.388 | -7978.316 | -0.927 | 1.53E-06 |
| 11 | 120659485 | rs3133861 | T | G | 0.953 | -7977.388 | -7978.315 | -0.927 | 1.53E-06 |
| 11 | 117448799 | rs35021198 | T | C | 0.959 | -7977.388 | -7978.316 | -0.928 | 1.53E-06 |
| 11 | 119217633 | rs45509899 | T | C | 0.954 | -7977.388 | -7978.316 | -0.928 | 1.53E-06 |
| 11 | 119935473 | rs78443404 | C | G | 0.954 | -7977.388 | -7978.316 | -0.928 | 1.53E-06 |
| 11 | 116302144 | rs10891974 | T | C | 1.046 | -7977.388 | -7978.317 | -0.929 | 1.53E-06 |
| 11 | 116530061 | rs1892947 | A | G | 1.046 | -7977.388 | -7978.317 | -0.929 | 1.53E-06 |
| 11 | 117120027 | rs605531 | G | T | 0.959 | -7977.388 | -7978.317 | -0.929 | 1.53E-06 |
| 11 | 117987503 | rs1715464 | A | C | 1.045 | -7977.388 | -7978.317 | -0.929 | 1.53E-06 |
| 11 | 116621972 | rs79112951 | T | C | 1.032 | -7977.388 | -7978.318 | -0.93 | 1.53E-06 |
| 11 | 117580254 | rs2508559 | T | C | 0.983 | -7977.388 | -7978.318 | -0.93 | 1.53E-06 |
| 11 | 117580509 | rs655086 | A | G | 0.983 | -7977.388 | -7978.318 | -0.93 | 1.53E-06 |
| 11 | 117580585 | rs604245 | A | G | 0.983 | -7977.388 | -7978.318 | -0.93 | 1.53E-06 |
| 11 | 117862979 | rs4252305 | A | G | 0.958 | -7977.388 | -7978.318 | -0.93 | 1.53E-06 |
| 11 | 117863183 | rs73009944 | A | G | 0.958 | -7977.388 | -7978.318 | -0.93 | 1.53E-06 |
| 11 | 117863307 | rs73009948 | T | G | 0.958 | -7977.388 | -7978.318 | -0.93 | 1.53E-06 |
| 11 | 119762339 | rs79436592 | A | G | 0.96 | -7977.388 | -7978.318 | -0.93 | 1.53E-06 |
| 11 | 117629468 | rs11216534 | T | C | 0.953 | -7977.388 | -7978.319 | -0.931 | 1.52E-06 |
| 11 | 117629559 | rs11216535 | T | C | 0.953 | -7977.388 | -7978.319 | -0.931 | 1.52E-06 |
| 11 | 120161671 | rs6589805 | C | T | 1.044 | -7977.388 | -7978.319 | -0.931 | 1.52E-06 |
| 11 | 120622625 | rs9667927 | C | T | 0.954 | -7977.388 | -7978.32 | -0.931 | 1.52E-06 |
| 11 | 117312429 | rs2887176 | G | A | 1.049 | -7977.388 | -7978.321 | -0.932 | 1.52E-06 |
| 11 | 120826272 | rs78528013 | T | C | 1.049 | -7977.388 | -7978.32 | -0.932 | 1.52E-06 |
| 11 | 116529130 | rs78347813 | T | C | 1.045 | -7977.388 | -7978.322 | -0.933 | 1.52E-06 |
| 11 | 116534550 | rs76547220 | G | T | 1.045 | -7977.388 | -7978.321 | -0.933 | 1.52E-06 |
| 11 | 116535435 | rs77226464 | T | C | 1.045 | -7977.388 | -7978.321 | -0.933 | 1.52E-06 |
| 11 | 116535624 | rs190069353 | A | C | 1.045 | -7977.388 | -7978.321 | -0.933 | 1.52E-06 |
| 11 | 116535631 | rs117014628 | T | C | 1.045 | -7977.388 | -7978.321 | -0.933 | 1.52E-06 |
| 11 | 116536820 | rs79063991 | T | C | 1.045 | -7977.388 | -7978.321 | -0.933 | 1.52E-06 |
| 11 | 116537116 | rs118103820 | G | A | 1.045 | -7977.388 | -7978.321 | -0.933 | 1.52E-06 |
| 11 | 116537798 | rs140973748 | C | T | 1.045 | -7977.388 | -7978.321 | -0.933 | 1.52E-06 |
| 11 | 116538103 | rs143380339 | A | C | 1.045 | -7977.388 | -7978.321 | -0.933 | 1.52E-06 |
| 11 | 116685391 | rs77535577 | A | G | 1.049 | -7977.388 | -7978.321 | -0.933 | 1.52E-06 |
| 11 | 118725420 | rs11217062 | A | G | 0.957 | -7977.388 | -7978.322 | -0.933 | 1.52E-06 |
| 11 | 118798824 | rs112119822 | G | A | 1.049 | -7977.388 | -7978.321 | -0.933 | 1.52E-06 |
| 11 | 119504952 | rs1009365 | T | C | 1.042 | -7977.388 | -7978.321 | -0.933 | 1.52E-06 |
| 11 | 119926054 | rs7125256 | G | A | 0.955 | -7977.388 | -7978.321 | -0.933 | 1.52E-06 |
| 11 | 116361658 | rs11215963 | G | C | 0.954 | -7977.388 | -7978.322 | -0.934 | 1.52E-06 |
| 11 | 116533310 | rs79291519 | T | C | 1.045 | -7977.388 | -7978.322 | -0.934 | 1.52E-06 |
| 11 | 116642137 | rs664059 | C | T | 1.042 | -7977.388 | -7978.322 | -0.934 | 1.52E-06 |
| 11 | 118267903 | rs45480805 | A | G | 1.032 | -7977.388 | -7978.323 | -0.934 | 1.52E-06 |
| 11 | 120868233 | rs113749645 | A | T | 0.99 | -7977.388 | -7978.322 | -0.934 | 1.52E-06 |
| 11 | 117724493 | rs11216582 | A | C | 1.047 | -7977.388 | -7978.323 | -0.935 | 1.52E-06 |
| 11 | 117883208 | rs2512150 | C | G | 0.954 | -7977.388 | -7978.324 | -0.935 | 1.52E-06 |
| 11 | 119805016 | rs12273820 | C | T | 1.046 | -7977.388 | -7978.323 | -0.935 | 1.52E-06 |
| 11 | 120784745 | rs111716733 | T | C | 1.035 | -7977.388 | -7978.323 | -0.935 | 1.52E-06 |
| 11 | 116578361 | rs75310100 | T | G | 0.997 | -7977.388 | -7978.325 | -0.936 | 1.52E-06 |
| 11 | 117307122 | rs665403 | T | C | 1.045 | -7977.388 | -7978.324 | -0.936 | 1.52E-06 |
| 11 | 118861849 | rs57216066 | A | G | 0.953 | -7977.388 | -7978.324 | -0.936 | 1.52E-06 |
| 11 | 120159540 | rs1329232240 | C | T | 1.044 | -7977.388 | -7978.325 | -0.936 | 1.52E-06 |
| 11 | 120161117 | rs7951628 | A | G | 1.044 | -7977.388 | -7978.325 | -0.936 | 1.52E-06 |
| 11 | 116241842 | rs533735 | C | A | 1.045 | -7977.388 | -7978.325 | -0.937 | 1.52E-06 |
| 11 | 116457115 | rs75243489 | A | T | 1.046 | -7977.388 | -7978.325 | -0.937 | 1.52E-06 |
| 11 | 117059347 | rs90192 | C | T | 0.954 | -7977.388 | -7978.325 | -0.937 | 1.52E-06 |
| 11 | 118183733 | rs28423005 | T | C | 1.043 | -7977.388 | -7978.325 | -0.937 | 1.52E-06 |
| 11 | 118521549 | rs494560 | A | G | 0.96 | -7977.388 | -7978.326 | -0.937 | 1.52E-06 |
| 11 | 118569414 | rs6589675 | C | G | 0.953 | -7977.388 | -7978.326 | -0.937 | 1.52E-06 |
| 11 | 118860457 | rs78505713 | T | C | 0.953 | -7977.388 | -7978.325 | -0.937 | 1.52E-06 |
| 11 | 118860598 | rs113271112 | C | A | 0.953 | -7977.388 | -7978.325 | -0.937 | 1.52E-06 |
| 11 | 116789102 | rs1378582752 | C | T | 0.974 | -7977.388 | -7978.326 | -0.938 | 1.51E-06 |
| 11 | 118720998 | rs4938565 | A | G | 0.957 | -7977.388 | -7978.327 | -0.938 | 1.51E-06 |
| 11 | 120488117 | rs73002644 | G | C | 1.043 | -7977.388 | -7978.326 | -0.938 | 1.51E-06 |
| 11 | 116531050 | rs117889066 | C | G | 1.045 | -7977.388 | -7978.327 | -0.939 | 1.51E-06 |
| 11 | 116531475 | rs117293980 | G | T | 1.045 | -7977.388 | -7978.327 | -0.939 | 1.51E-06 |
| 11 | 117644124 | rs17311024 | A | T | 1.048 | -7977.388 | -7978.327 | -0.939 | 1.51E-06 |
| 11 | 118254710 | rs45443099 | G | A | 1.031 | -7977.388 | -7978.327 | -0.939 | 1.51E-06 |
| 11 | 118259162 | rs111792147 | A | G | 1.031 | -7977.388 | -7978.327 | -0.939 | 1.51E-06 |
| 11 | 118262135 | rs112563022 | T | C | 1.031 | -7977.388 | -7978.327 | -0.939 | 1.51E-06 |
| 11 | 119742642 | rs73008675 | C | T | 1.048 | -7977.388 | -7978.328 | -0.939 | 1.51E-06 |
| 11 | 119743088 | rs73008677 | A | G | 1.048 | -7977.388 | -7978.328 | -0.939 | 1.51E-06 |
| 11 | 119743230 | rs73008679 | T | C | 1.048 | -7977.388 | -7978.328 | -0.939 | 1.51E-06 |
| 11 | 120826325 | rs77571870 | A | G | 1.048 | -7977.388 | -7978.327 | -0.939 | 1.51E-06 |
| 11 | 117874531 | rs947889 | C | T | 0.96 | -7977.388 | -7978.329 | -0.94 | 1.51E-06 |
| 11 | 118486067 | rs11603023 | T | C | 0.96 | -7977.388 | -7978.329 | -0.94 | 1.51E-06 |
| 11 | 119747470 | rs4938735 | T | C | 0.957 | -7977.388 | -7978.329 | -0.94 | 1.51E-06 |
| 11 | 116375518 | rs111551597 | T | C | 0.955 | -7977.388 | -7978.329 | -0.941 | 1.51E-06 |
| 11 | 117600616 | rs11600991 | C | T | 0.956 | -7977.388 | -7978.329 | -0.941 | 1.51E-06 |
| 11 | 118047776 | rs67262752 | C | T | 0.958 | -7977.388 | -7978.329 | -0.941 | 1.51E-06 |
| 11 | 118384811 | rs78020509 | G | A | 1.044 | -7977.388 | -7978.329 | -0.941 | 1.51E-06 |
| 11 | 120163184 | rs61898317 | G | A | 1.048 | -7977.388 | -7978.329 | -0.941 | 1.51E-06 |
| 11 | 120163869 | rs2155810 | C | T | 1.044 | -7977.388 | -7978.329 | -0.941 | 1.51E-06 |
| 11 | 117656916 | rs4603339 | G | A | 1.048 | -7977.388 | -7978.331 | -0.942 | 1.51E-06 |
| 11 | 119503670 | rs12279718 | A | C | 1.042 | -7977.388 | -7978.33 | -0.942 | 1.51E-06 |
| 11 | 119761810 | rs78048464 | T | G | 1.043 | -7977.388 | -7978.33 | -0.942 | 1.51E-06 |
| 11 | 120391473 | rs12804597 | A | G | 0.954 | -7977.388 | -7978.33 | -0.942 | 1.51E-06 |
| 11 | 116480033 | rs149395240 | T | C | 1.037 | -7977.388 | -7978.332 | -0.943 | 1.51E-06 |
| 11 | 117781435 | rs2155194 | A | G | 0.979 | -7977.388 | -7978.332 | -0.943 | 1.51E-06 |
| 11 | 118182660 | rs2277289 | C | T | 1.043 | -7977.388 | -7978.331 | -0.943 | 1.51E-06 |
| 11 | 118246021 | rs112243842 | G | T | 1.03 | -7977.388 | -7978.331 | -0.943 | 1.51E-06 |
| 11 | 118247805 | rs112646740 | A | T | 1.03 | -7977.388 | -7978.331 | -0.943 | 1.51E-06 |
| 11 | 118248374 | rs111430146 | A | C | 1.03 | -7977.388 | -7978.331 | -0.943 | 1.51E-06 |
| 11 | 120789800 | rs10892641 | G | A | 0.96 | -7977.388 | -7978.332 | -0.943 | 1.51E-06 |
| 11 | 116279292 | rs579941 | T | C | 0.954 | -7977.388 | -7978.332 | -0.944 | 1.50E-06 |
| 11 | 116682995 | rs76636922 | A | G | 1.048 | -7977.388 | -7978.332 | -0.944 | 1.50E-06 |
| 11 | 117967476 | rs10790231 | C | T | 1.046 | -7977.388 | -7978.333 | -0.944 | 1.50E-06 |

| | | | | | | | |
|---|---|---|---|---|---|---|---|
| 11 | 119507180 | rs4620730 | G | C | NULL | 0.615383756 | NULL&M_Disease |
| 11 | 119674099 | rs11217515 | C | T | NULL | 0.615383756 | NULL&M_Disease |
| 11 | 119845446 | rs80159023 | A | G | NULL | 0.615383756 | NULL&M_Disease |
| 11 | 120470737 | rs151336980 | T | C | NULL | 0.615383756 | NULL&M_Disease |
| 11 | 116379978 | rs1508103 | G | A | NULL | 0.615383756 | NULL&M_Disease |
| 11 | 117656774 | rs715315 | T | C | NULL | 0.615383756 | NULL&M_Disease |
| 11 | 118116052 | rs1793166 | A | G | NULL | 0.615383756 | NULL&M_Disease |
| 11 | 118720684 | rs4936440 | G | C | NULL | 0.615383756 | NULL&M_Disease |
| 11 | 120505511 | rs7105213 | T | C | NULL | 0.615383756 | NULL&M_Disease |
| 11 | 120713605 | rs11607580 | G | A | NULL | 0.615383756 | NULL&M_Disease |
| 11 | 117593985 | rs7928408 | T | C | NULL | 0.615383756 | NULL&M_Disease |
| 11 | 118009385 | rs11216780 | G | A | NULL | 0.615383756 | NULL&M_Disease |
| 11 | 119937061 | rs73000748 | T | C | NULL | 0.615383756 | NULL&M_Disease |
| 11 | 120714220 | rs11608113 | G | A | NULL | 0.615383756 | NULL&M_Disease |
| 11 | 117346337 | rs80343155 | C | A | NULL | 0.615383756 | NULL&M_Disease |
| 11 | 118282448 | rs111993696 | A | G | NULL | 0.615383756 | NULL&M_Disease |
| 11 | 118283383 | rs112302730 | A | G | NULL | 0.615383756 | NULL&M_Disease |
| 11 | 118308499 | rs111903103 | T | C | NULL | 0.615383756 | NULL&M_Disease |
| 11 | 116490583 | rs518087 | C | T | NULL | 0.615383756 | NULL&M_Disease |
| 11 | 116684560 | rs79841374 | T | C | NULL | 0.615383756 | NULL&M_Disease |
| 11 | 118180155 | rs4938503 | C | G | NULL | 0.616566505 | NULL&M_Disease |
| 11 | 118303182 | rs77728961 | C | T | NULL | 0.615383756 | NULL&M_Disease |
| 11 | 118303200 | rs112550735 | C | T | NULL | 0.615383756 | NULL&M_Disease |
| 11 | 119788251 | rs643115 | T | A | NULL | 0.616566505 | NULL&M_Disease |
| 11 | 120764256 | rs57833493 | C | T | NULL | 0.615383756 | NULL&M_Disease |
| 11 | 121083487 | rs144234662 | T | C | NULL | 0.615383756 | NULL&M_Disease |
| 11 | 118237566 | rs45555733 | G | C | NULL | 0.615383756 | NULL&M_Disease |
| 11 | 120496704 | rs4935744 | G | T | NULL | 0.616566505 | NULL&M_Disease |
| 11 | 120804014 | rs11218062 | C | T | NULL | 0.616566505 | NULL&M_Disease |
| 11 | 117948080 | rs653764 | T | C | NULL | 0.616566505 | NULL&M_Disease |
| 11 | 119648394 | rs73006770 | A | G | NULL | 0.615383756 | NULL&M_Disease |
| 11 | 120935369 | rs4396299 | A | G | NULL | 0.616566505 | NULL&M_Disease |
| 11 | 116545251 | rs117179746 | A | G | NULL | 0.616566505 | NULL&M_Disease |
| 11 | 117347400 | rs587443 | T | C | NULL | 0.616566505 | NULL&M_Disease |
| 11 | 118287622 | rs112245679 | G | A | NULL | 0.616566505 | NULL&M_Disease |
| 11 | 118289734 | rs111763202 | T | C | NULL | 0.616566505 | NULL&M_Disease |
| 11 | 118290443 | rs112035668 | G | A | NULL | 0.616566505 | NULL&M_Disease |
| 11 | 118335147 | rs113566564 | A | G | NULL | 0.616566505 | NULL&M_Disease |
| 11 | 118717929 | rs4938558 | T | C | NULL | 0.616566505 | NULL&M_Disease |
| 11 | 119804776 | rs10892497 | T | C | NULL | 0.616566505 | NULL&M_Disease |
| 11 | 120796595 | rs2852242 | T | A | NULL | 0.616566505 | NULL&M_Disease |
| 11 | 121093389 | rs61901232 | T | C | NULL | 0.615383756 | NULL&M_Disease |
| 11 | 117666595 | rs12792357 | C | T | NULL | 0.616566505 | NULL&M_Disease |
| 11 | 117776471 | rs3802873 | T | C | NULL | 0.616566505 | NULL&M_Disease |
| 11 | 117777953 | rs11216615 | G | A | NULL | 0.616566505 | NULL&M_Disease |
| 11 | 118719254 | rs2156750 | C | T | NULL | 0.616566505 | NULL&M_Disease |
| 11 | 118719405 | rs2156749 | G | C | NULL | 0.616566505 | NULL&M_Disease |
| 11 | 118719742 | rs7116715 | C | T | NULL | 0.616566505 | NULL&M_Disease |
| 11 | 120809279 | rs591252 | C | T | NULL | 0.616566505 | NULL&M_Disease |
| 11 | 116535087 | rs2105609 | A | G | NULL | 0.616566505 | NULL&M_Disease |
| 11 | 117855295 | rs4252241 | G | A | NULL | 0.616566505 | NULL&M_Disease |
| 11 | 116224575 | rs60915096 | C | T | NULL | 0.616566505 | NULL&M_Disease |
| 11 | 116835398 | rs11826038 | G | A | NULL | 0.616566505 | NULL&M_Disease |
| 11 | 117547323 | rs111849979 | G | A | NULL | 0.616566505 | NULL&M_Disease |
| 11 | 117825368 | rs2887193 | A | G | NULL | 0.616566505 | NULL&M_Disease |
| 11 | 118373861 | rs2071702 | T | C | NULL | 0.616566505 | NULL&M_Disease |
| 11 | 116485319 | rs500426 | T | C | NULL | 0.616566505 | NULL&M_Disease |
| 11 | 117312301 | rs11821451 | A | G | NULL | 0.616566505 | NULL&M_Disease |
| 11 | 118296184 | rs111354396 | A | G | NULL | 0.616566505 | NULL&M_Disease |
| 11 | 118301375 | rs111656195 | A | G | NULL | 0.616566505 | NULL&M_Disease |
| 11 | 119679283 | rs10892471 | A | G | NULL | 0.616566505 | NULL&M_Disease |
| 11 | 120581080 | rs7103418 | A | C | NULL | 0.616566505 | NULL&M_Disease |
| 11 | 120581105 | rs7119619 | C | T | NULL | 0.616566505 | NULL&M_Disease |
| 11 | 116627097 | rs56224630 | A | G | NULL | 0.616566505 | NULL&M_Disease |
| 11 | 117966100 | rs948466 | A | G | NULL | 0.616566505 | NULL&M_Disease |
| 11 | 118707035 | rs143605660 | A | G | NULL | 0.616566505 | NULL&M_Disease |
| 11 | 118801440 | rs12280501 | T | C | NULL | 0.616566505 | NULL&M_Disease |
| 11 | 119835382 | rs78865628 | C | T | NULL | 0.616566505 | NULL&M_Disease |
| 11 | 120976311 | rs55683640 | C | A | NULL | 0.616566505 | NULL&M_Disease |
| 11 | 121060783 | rs56161514 | A | C | NULL | 0.616566505 | NULL&M_Disease |
| 11 | 116586283 | rs7350481 | T | C | NULL | 0.616566505 | NULL&M_Disease |
| 11 | 117391966 | rs2276340 | A | G | NULL | 0.616566505 | NULL&M_Disease |
| 11 | 117393294 | rs1940040 | C | G | NULL | 0.616566505 | NULL&M_Disease |
| 11 | 117824500 | rs6589636 | C | T | NULL | 0.616566505 | NULL&M_Disease |
| 11 | 119701587 | rs11217528 | C | A | NULL | 0.617747875 | NULL&M_Disease |
| 11 | 119832880 | rs111999954 | A | G | NULL | 0.616566505 | NULL&M_Disease |
| 11 | 121023992 | rs686882 | G | A | NULL | 0.617747875 | NULL&M_Disease |
| 11 | 118718485 | rs4938561 | C | T | NULL | 0.616566505 | NULL&M_Disease |
| 11 | 118718856 | rs2156751 | G | A | NULL | 0.617747875 | NULL&M_Disease |
| 11 | 118718972 | rs2187491 | C | T | NULL | 0.617747875 | NULL&M_Disease |
| 11 | 118827925 | rs113048801 | A | G | NULL | 0.616566505 | NULL&M_Disease |
| 11 | 118965048 | rs7350 | A | G | NULL | 0.617747875 | NULL&M_Disease |
| 11 | 119028813 | rs626776 | C | G | NULL | 0.617747875 | NULL&M_Disease |
| 11 | 119829794 | rs74999919 | G | A | NULL | 0.616566505 | NULL&M_Disease |
| 11 | 116985247 | rs7113932 | T | G | NULL | 0.616566505 | NULL&M_Disease |
| 11 | 117193619 | rs548893 | G | C | NULL | 0.617747875 | NULL&M_Disease |
| 11 | 119828201 | rs74442127 | A | G | NULL | 0.617747875 | NULL&M_Disease |
| 11 | 120157168 | rs4936513 | A | T | NULL | 0.617747875 | NULL&M_Disease |
| 11 | 120703464 | rs73003745 | A | G | NULL | 0.617747875 | NULL&M_Disease |
| 11 | 117780919 | rs2298464 | A | G | NULL | 0.616566505 | NULL&M_Disease |
| 11 | 117889448 | rs2512155 | T | C | NULL | 0.617747875 | NULL&M_Disease |
| 11 | 118058542 | rs12284089 | T | C | NULL | 0.617747875 | NULL&M_Disease |
| 11 | 118240840 | rs112152942 | G | C | NULL | 0.616566505 | NULL&M_Disease |
| 11 | 118242889 | rs45468196 | A | G | NULL | 0.616566505 | NULL&M_Disease |
| 11 | 118799206 | rs7101680 | A | G | NULL | 0.617747875 | NULL&M_Disease |
| 11 | 119789992 | rs117883320 | A | C | NULL | 0.616566505 | NULL&M_Disease |

| | | | | | | | | | |
|---|---|---|---|---|---|---|---|---|---|
| 11 | 119507180 | rs4620730 | G | C | 0.955 | -7977.388 | -7978.332 | -0.944 | 1.50E-06 |
| 11 | 119674099 | rs11217515 | C | T | 0.956 | -7977.388 | -7978.333 | -0.944 | 1.50E-06 |
| 11 | 119845446 | rs80159023 | A | G | 1.048 | -7977.388 | -7978.333 | -0.944 | 1.50E-06 |
| 11 | 120470737 | rs151336980 | T | C | 1.025 | -7977.388 | -7978.333 | -0.944 | 1.50E-06 |
| 11 | 116379978 | rs1508103 | G | A | 1.045 | -7977.388 | -7978.333 | -0.945 | 1.50E-06 |
| 11 | 117656774 | rs715315 | T | C | 1.041 | -7977.388 | -7978.333 | -0.945 | 1.50E-06 |
| 11 | 118116052 | rs1793166 | A | G | 0.955 | -7977.388 | -7978.334 | -0.945 | 1.50E-06 |
| 11 | 118720684 | rs4936440 | G | C | 0.958 | -7977.388 | -7978.334 | -0.945 | 1.50E-06 |
| 11 | 120505511 | rs7105213 | T | C | 0.96 | -7977.388 | -7978.334 | -0.945 | 1.50E-06 |
| 11 | 120713605 | rs11607580 | G | A | 0.956 | -7977.388 | -7978.333 | -0.945 | 1.50E-06 |
| 11 | 117593985 | rs7928408 | T | C | 0.956 | -7977.388 | -7978.335 | -0.946 | 1.50E-06 |
| 11 | 118009385 | rs11216780 | G | A | 0.968 | -7977.388 | -7978.335 | -0.946 | 1.50E-06 |
| 11 | 119937061 | rs73000748 | T | C | 0.974 | -7977.388 | -7978.334 | -0.946 | 1.50E-06 |
| 11 | 120714220 | rs11608113 | G | A | 0.956 | -7977.388 | -7978.334 | -0.946 | 1.50E-06 |
| 11 | 117346337 | rs80343155 | C | A | 0.973 | -7977.388 | -7978.335 | -0.947 | 1.50E-06 |
| 11 | 118282448 | rs111993696 | A | G | 1.029 | -7977.388 | -7978.336 | -0.948 | 1.50E-06 |
| 11 | 118283383 | rs112302730 | A | G | 1.029 | -7977.388 | -7978.336 | -0.948 | 1.50E-06 |
| 11 | 118308499 | rs111903103 | T | C | 1.029 | -7977.388 | -7978.336 | -0.948 | 1.50E-06 |
| 11 | 116490583 | rs518087 | C | T | 0.964 | -7977.388 | -7978.337 | -0.949 | 1.50E-06 |
| 11 | 116684560 | rs79841374 | T | C | 1.048 | -7977.388 | -7978.337 | -0.949 | 1.50E-06 |
| 11 | 118180155 | rs4938503 | C | G | 1.043 | -7977.388 | -7978.337 | -0.949 | 1.50E-06 |
| 11 | 118303182 | rs77728961 | C | T | 1.029 | -7977.388 | -7978.338 | -0.949 | 1.50E-06 |
| 11 | 118303200 | rs112550735 | C | T | 1.029 | -7977.388 | -7978.338 | -0.949 | 1.50E-06 |
| 11 | 119788251 | rs643115 | T | A | 0.957 | -7977.388 | -7978.338 | -0.949 | 1.50E-06 |
| 11 | 120764256 | rs57833493 | C | T | 1.031 | -7977.388 | -7978.338 | -0.949 | 1.50E-06 |
| 11 | 121083487 | rs144234662 | T | C | 0.983 | -7977.388 | -7978.337 | -0.949 | 1.50E-06 |
| 11 | 118237566 | rs45555733 | G | C | 1.029 | -7977.388 | -7978.338 | -0.95 | 1.50E-06 |
| 11 | 120496704 | rs4935744 | G | T | 1.043 | -7977.388 | -7978.338 | -0.95 | 1.50E-06 |
| 11 | 120804014 | rs11218062 | C | T | 1.046 | -7977.388 | -7978.338 | -0.95 | 1.50E-06 |
| 11 | 117948080 | rs653764 | T | C | 1.047 | -7977.388 | -7978.339 | -0.951 | 1.49E-06 |
| 11 | 119648394 | rs73006770 | A | G | 1.026 | -7977.388 | -7978.339 | -0.951 | 1.49E-06 |
| 11 | 120935369 | rs4396299 | A | G | 1.045 | -7977.388 | -7978.339 | -0.951 | 1.49E-06 |
| 11 | 116545251 | rs117179746 | A | G | 1.044 | -7977.388 | -7978.34 | -0.952 | 1.49E-06 |
| 11 | 117347400 | rs587443 | T | C | 1.041 | -7977.388 | -7978.341 | -0.952 | 1.49E-06 |
| 11 | 118287622 | rs112245679 | G | A | 1.028 | -7977.388 | -7978.341 | -0.952 | 1.49E-06 |
| 11 | 118289734 | rs111763202 | T | C | 1.028 | -7977.388 | -7978.341 | -0.952 | 1.49E-06 |
| 11 | 118290443 | rs112035668 | G | A | 1.028 | -7977.388 | -7978.341 | -0.952 | 1.49E-06 |
| 11 | 118335147 | rs113566564 | A | G | 1.043 | -7977.388 | -7978.34 | -0.952 | 1.49E-06 |
| 11 | 118717929 | rs4938558 | T | C | 0.958 | -7977.388 | -7978.34 | -0.952 | 1.49E-06 |
| 11 | 119804776 | rs10892497 | T | C | 1.048 | -7977.388 | -7978.341 | -0.952 | 1.49E-06 |
| 11 | 120796595 | rs2852242 | T | A | 0.958 | -7977.388 | -7978.34 | -0.952 | 1.49E-06 |
| 11 | 121093389 | rs61901232 | T | C | 1.02 | -7977.388 | -7978.34 | -0.952 | 1.49E-06 |
| 11 | 117666595 | rs12792357 | C | T | 1.041 | -7977.388 | -7978.342 | -0.953 | 1.49E-06 |
| 11 | 117776471 | rs3802873 | T | C | 0.959 | -7977.388 | -7978.341 | -0.953 | 1.49E-06 |
| 11 | 117777953 | rs11216615 | G | A | 0.981 | -7977.388 | -7978.341 | -0.953 | 1.49E-06 |
| 11 | 118719254 | rs2156750 | C | T | 0.958 | -7977.388 | -7978.341 | -0.953 | 1.49E-06 |
| 11 | 118719405 | rs2156749 | G | C | 0.958 | -7977.388 | -7978.341 | -0.953 | 1.49E-06 |
| 11 | 118719742 | rs7116715 | C | T | 0.958 | -7977.388 | -7978.341 | -0.953 | 1.49E-06 |
| 11 | 120809279 | rs591252 | C | T | 0.956 | -7977.388 | -7978.341 | -0.953 | 1.49E-06 |
| 11 | 116535087 | rs2105609 | A | G | 1.044 | -7977.388 | -7978.342 | -0.954 | 1.49E-06 |
| 11 | 117855295 | rs4252241 | G | A | 0.954 | -7977.388 | -7978.342 | -0.954 | 1.49E-06 |
| 11 | 116224575 | rs60915096 | C | T | 0.978 | -7977.388 | -7978.343 | -0.955 | 1.49E-06 |
| 11 | 116835398 | rs11826038 | G | A | 0.977 | -7977.388 | -7978.344 | -0.955 | 1.49E-06 |
| 11 | 117547323 | rs111849979 | G | A | 0.979 | -7977.388 | -7978.343 | -0.955 | 1.49E-06 |
| 11 | 117825368 | rs2887193 | A | G | 0.956 | -7977.388 | -7978.344 | -0.955 | 1.49E-06 |
| 11 | 118373861 | rs2071702 | T | C | 1.043 | -7977.388 | -7978.343 | -0.955 | 1.49E-06 |
| 11 | 116485319 | rs500426 | T | C | 0.96 | -7977.388 | -7978.344 | -0.956 | 1.49E-06 |
| 11 | 117312301 | rs11821451 | A | G | 1.048 | -7977.388 | -7978.344 | -0.956 | 1.49E-06 |
| 11 | 118296184 | rs111354396 | A | G | 1.028 | -7977.388 | -7978.345 | -0.956 | 1.49E-06 |
| 11 | 118301375 | rs111656195 | A | G | 1.028 | -7977.388 | -7978.345 | -0.956 | 1.49E-06 |
| 11 | 119679283 | rs10892471 | A | G | 0.954 | -7977.388 | -7978.344 | -0.956 | 1.49E-06 |
| 11 | 120581080 | rs7103418 | A | C | 0.955 | -7977.388 | -7978.344 | -0.956 | 1.49E-06 |
| 11 | 120581105 | rs7119619 | C | T | 0.955 | -7977.388 | -7978.344 | -0.956 | 1.49E-06 |
| 11 | 116627097 | rs56224630 | A | G | 0.955 | -7977.388 | -7978.345 | -0.957 | 1.49E-06 |
| 11 | 117966100 | rs948466 | A | G | 1.045 | -7977.388 | -7978.345 | -0.957 | 1.49E-06 |
| 11 | 118707035 | rs143605660 | A | G | 0.96 | -7977.388 | -7978.345 | -0.957 | 1.49E-06 |
| 11 | 118801440 | rs12280501 | T | C | 1.047 | -7977.388 | -7978.345 | -0.957 | 1.49E-06 |
| 11 | 119835382 | rs78865628 | C | T | 1.01 | -7977.388 | -7978.345 | -0.957 | 1.49E-06 |
| 11 | 120976311 | rs55683640 | C | A | 1.046 | -7977.388 | -7978.345 | -0.957 | 1.49E-06 |
| 11 | 121060783 | rs56161514 | A | C | 0.993 | -7977.388 | -7978.345 | -0.957 | 1.49E-06 |
| 11 | 116586283 | rs7350481 | T | C | 0.956 | -7977.388 | -7978.346 | -0.958 | 1.48E-06 |
| 11 | 117391966 | rs2276340 | A | G | 1.046 | -7977.388 | -7978.346 | -0.958 | 1.48E-06 |
| 11 | 117393294 | rs1940040 | C | G | 1.046 | -7977.388 | -7978.346 | -0.958 | 1.48E-06 |
| 11 | 117824500 | rs6589636 | C | T | 0.956 | -7977.388 | -7978.346 | -0.958 | 1.48E-06 |
| 11 | 119701587 | rs11217528 | C | A | 0.96 | -7977.388 | -7978.346 | -0.958 | 1.48E-06 |
| 11 | 119832880 | rs111999954 | A | G | 0.982 | -7977.388 | -7978.346 | -0.958 | 1.48E-06 |
| 11 | 121023992 | rs686882 | G | A | 0.959 | -7977.388 | -7978.347 | -0.958 | 1.48E-06 |
| 11 | 118718485 | rs4938561 | C | T | 0.958 | -7977.388 | -7978.347 | -0.959 | 1.48E-06 |
| 11 | 118718856 | rs2156751 | G | A | 0.958 | -7977.388 | -7978.347 | -0.959 | 1.48E-06 |
| 11 | 118718972 | rs2187491 | C | T | 0.958 | -7977.388 | -7978.347 | -0.959 | 1.48E-06 |
| 11 | 118827925 | rs113048801 | A | G | 1.024 | -7977.388 | -7978.348 | -0.959 | 1.48E-06 |
| 11 | 118965048 | rs7350 | A | G | 1.042 | -7977.388 | -7978.347 | -0.959 | 1.48E-06 |
| 11 | 119028813 | rs626776 | C | G | 1.042 | -7977.388 | -7978.347 | -0.959 | 1.48E-06 |
| 11 | 119829794 | rs74999919 | G | A | 1.047 | -7977.388 | -7978.347 | -0.959 | 1.48E-06 |
| 11 | 116985247 | rs7113932 | T | G | 0.98 | -7977.388 | -7978.348 | -0.96 | 1.48E-06 |
| 11 | 117193619 | rs548893 | G | C | 0.96 | -7977.388 | -7978.348 | -0.96 | 1.48E-06 |
| 11 | 119828201 | rs74442127 | A | G | 1.047 | -7977.388 | -7978.348 | -0.96 | 1.48E-06 |
| 11 | 120157168 | rs4936513 | A | T | 1.044 | -7977.388 | -7978.348 | -0.96 | 1.48E-06 |
| 11 | 120703464 | rs73003745 | A | G | 0.957 | -7977.388 | -7978.348 | -0.96 | 1.48E-06 |
| 11 | 117780919 | rs2298464 | A | G | 0.983 | -7977.388 | -7978.349 | -0.961 | 1.48E-06 |
| 11 | 117889448 | rs2512155 | T | C | 0.957 | -7977.388 | -7978.35 | -0.961 | 1.48E-06 |
| 11 | 118058542 | rs12284089 | T | C | 0.96 | -7977.388 | -7978.35 | -0.961 | 1.48E-06 |
| 11 | 118240840 | rs112152942 | G | C | 1.027 | -7977.388 | -7978.349 | -0.961 | 1.48E-06 |
| 11 | 118242889 | rs45468196 | A | G | 1.027 | -7977.388 | -7978.349 | -0.961 | 1.48E-06 |
| 11 | 118799206 | rs7101680 | A | G | 1.047 | -7977.388 | -7978.349 | -0.961 | 1.48E-06 |
| 11 | 119789992 | rs117883320 | A | C | 0.966 | -7977.388 | -7978.349 | -0.961 | 1.48E-06 |

| | | | | | | | |
|---|---|---|---|---|---|---|---|
| 11 | 120174671 | rs10790375 | C | T | NULL | 0.617747875 | NULL&M_Disease |
| 11 | 117967986 | rs9651687 | A | T | NULL | 0.617747875 | NULL&M_Disease |
| 11 | 118724336 | rs10892298 | C | T | NULL | 0.617747875 | NULL&M_Disease |
| 11 | 118730669 | rs7933007 | C | T | NULL | 0.617747875 | NULL&M_Disease |
| 11 | 120580795 | rs7103021 | T | C | NULL | 0.617747875 | NULL&M_Disease |
| 11 | 120657934 | rs2186698 | C | G | NULL | 0.617747875 | NULL&M_Disease |
| 11 | 120695729 | rs111750658 | C | G | NULL | 0.617747875 | NULL&M_Disease |
| 11 | 116331421 | rs113872393 | A | G | NULL | 0.617747875 | NULL&M_Disease |
| 11 | 117250530 | rs74661474 | A | G | NULL | 0.617747875 | NULL&M_Disease |
| 11 | 117266094 | rs11216375 | A | G | NULL | 0.617747875 | NULL&M_Disease |
| 11 | 117682935 | rs182928887 | G | A | NULL | 0.617747875 | NULL&M_Disease |
| 11 | 116625362 | rs56225305 | A | G | NULL | 0.617747875 | NULL&M_Disease |
| 11 | 117842818 | rs4936413 | C | T | NULL | 0.617747875 | NULL&M_Disease |
| 11 | 120497465 | rs35181812 | T | C | NULL | 0.617747875 | NULL&M_Disease |
| 11 | 116727592 | rs61907562 | A | G | NULL | 0.617747875 | NULL&M_Disease |
| 11 | 118782222 | rs61900138 | C | T | NULL | 0.617747875 | NULL&M_Disease |
| 11 | 118874272 | rs192113495 | G | T | NULL | 0.617747875 | NULL&M_Disease |
| 11 | 120801460 | rs509566 | T | A | NULL | 0.617747875 | NULL&M_Disease |
| 11 | 119795416 | rs80294967 | A | G | NULL | 0.617747875 | NULL&M_Disease |
| 11 | 116726087 | rs61907560 | C | T | NULL | 0.617747875 | NULL&M_Disease |
| 11 | 116536490 | rs1240778 | C | A | NULL | 0.618927855 | NULL&M_Disease |
| 11 | 119940917 | rs117243996 | T | C | NULL | 0.617747875 | NULL&M_Disease |
| 11 | 120484712 | rs1317815 | A | G | NULL | 0.618927855 | NULL&M_Disease |
| 11 | 117435160 | rs4936387 | T | C | NULL | 0.617747875 | NULL&M_Disease |
| 11 | 117863888 | rs4252307 | G | C | NULL | 0.617747875 | NULL&M_Disease |
| 11 | 117949807 | rs605805 | G | A | NULL | 0.618927855 | NULL&M_Disease |
| 11 | 120824400 | rs1893817 | T | C | NULL | 0.618927855 | NULL&M_Disease |
| 11 | 117129232 | rs77096386 | G | A | NULL | 0.617747875 | NULL&M_Disease |
| 11 | 117399604 | rs4938400 | A | C | NULL | 0.618927855 | NULL&M_Disease |
| 11 | 117399605 | rs4938401 | G | T | NULL | 0.618927855 | NULL&M_Disease |
| 11 | 117606193 | rs4936403 | G | T | NULL | 0.618927855 | NULL&M_Disease |
| 11 | 117606320 | rs4936404 | G | A | NULL | 0.618927855 | NULL&M_Disease |
| 11 | 117652288 | rs75734044 | A | G | NULL | 0.618927855 | NULL&M_Disease |
| 11 | 118377523 | rs9332846 | A | G | NULL | 0.617747875 | NULL&M_Disease |
| 11 | 120991447 | rs74491080 | T | C | NULL | 0.617747875 | NULL&M_Disease |
| 11 | 116555126 | rs117341026 | A | G | NULL | 0.618927855 | NULL&M_Disease |
| 11 | 117792471 | rs11216624 | A | G | NULL | 0.618927855 | NULL&M_Disease |
| 11 | 118006930 | rs1317544 | C | T | NULL | 0.617747875 | NULL&M_Disease |
| 11 | 118510562 | rs4245184 | A | G | NULL | 0.618927855 | NULL&M_Disease |
| 11 | 118511352 | rs644273 | T | C | NULL | 0.618927855 | NULL&M_Disease |
| 11 | 119796458 | rs1011664 | G | C | NULL | 0.618927855 | NULL&M_Disease |
| 11 | 120796418 | rs2852243 | C | T | NULL | 0.618927855 | NULL&M_Disease |
| 11 | 116369980 | rs1108627 | T | C | NULL | 0.618927855 | NULL&M_Disease |
| 11 | 118385249 | rs112508300 | G | A | NULL | 0.618927855 | NULL&M_Disease |
| 11 | 118711830 | rs566848 | G | A | NULL | 0.618927855 | NULL&M_Disease |
| 11 | 120164102 | rs10750170 | G | A | NULL | 0.618927855 | NULL&M_Disease |
| 11 | 120785542 | rs117391362 | A | G | NULL | 0.618927855 | NULL&M_Disease |
| 11 | 120848717 | rs17124632 | A | G | NULL | 0.618927855 | NULL&M_Disease |
| 11 | 116994106 | rs61905462 | A | C | NULL | 0.618927855 | NULL&M_Disease |
| 11 | 117196435 | rs73014325 | A | C | NULL | 0.618927855 | NULL&M_Disease |
| 11 | 117533027 | rs34874964 | T | C | NULL | 0.618927855 | NULL&M_Disease |
| 11 | 117917962 | rs659092 | C | T | NULL | 0.618927855 | NULL&M_Disease |
| 11 | 120500339 | rs12577725 | T | C | NULL | 0.618927855 | NULL&M_Disease |
| 11 | 116946788 | rs61903414 | T | C | NULL | 0.618927855 | NULL&M_Disease |
| 11 | 117596927 | rs948104 | G | T | NULL | 0.618927855 | NULL&M_Disease |
| 11 | 118385691 | rs111405302 | C | T | NULL | 0.618927855 | NULL&M_Disease |
| 11 | 119774225 | rs525588 | C | G | NULL | 0.618927855 | NULL&M_Disease |
| 11 | 117375596 | rs2276343 | A | G | NULL | 0.618927855 | NULL&M_Disease |
| 11 | 117863730 | rs4252306 | C | G | NULL | 0.618927855 | NULL&M_Disease |
| 11 | 118828942 | rs623500 | G | A | NULL | 0.618927855 | NULL&M_Disease |
| 11 | 119840963 | rs77212554 | T | C | NULL | 0.618927855 | NULL&M_Disease |
| 11 | 120902930 | rs61903185 | A | G | NULL | 0.618927855 | NULL&M_Disease |
| 11 | 117219305 | rs592828 | C | G | NULL | 0.618927855 | NULL&M_Disease |
| 11 | 117864979 | rs2282496 | T | C | NULL | 0.618927855 | NULL&M_Disease |
| 11 | 118890085 | rs4938619 | T | C | NULL | 0.618927855 | NULL&M_Disease |
| 11 | 118049257 | rs7123533 | G | C | NULL | 0.618927855 | NULL&M_Disease |
| 11 | 118049759 | rs61472653 | C | T | NULL | 0.618927855 | NULL&M_Disease |
| 11 | 118049811 | rs59018400 | T | C | NULL | 0.618927855 | NULL&M_Disease |
| 11 | 118049880 | rs56301948 | G | A | NULL | 0.618927855 | NULL&M_Disease |
| 11 | 118050014 | rs57533048 | G | A | NULL | 0.618927855 | NULL&M_Disease |
| 11 | 118050348 | rs7342223 | G | T | NULL | 0.618927855 | NULL&M_Disease |
| 11 | 118511757 | rs654423 | A | G | NULL | 0.618927855 | NULL&M_Disease |
| 11 | 116520527 | rs1240773 | T | G | NULL | 0.620106432 | NULL&M_Disease |
| 11 | 116632956 | rs57641217 | T | C | NULL | 0.618927855 | NULL&M_Disease |
| 11 | 116658031 | rs12285095 | G | T | NULL | 0.618927855 | NULL&M_Disease |
| 11 | 116807226 | rs61907596 | T | C | NULL | 0.618927855 | NULL&M_Disease |
| 11 | 117246566 | rs112933729 | A | G | NULL | 0.618927855 | NULL&M_Disease |
| 11 | 117527473 | rs12802995 | A | C | NULL | 0.618927855 | NULL&M_Disease |
| 11 | 117531395 | rs35048090 | T | C | NULL | 0.618927855 | NULL&M_Disease |
| 11 | 117531569 | rs36067343 | G | T | NULL | 0.618927855 | NULL&M_Disease |
| 11 | 119744714 | rs73008683 | A | G | NULL | 0.620106432 | NULL&M_Disease |
| 11 | 120728827 | rs7103097 | A | C | NULL | 0.618927855 | NULL&M_Disease |
| 11 | 116906431 | rs61905711 | T | C | NULL | 0.618927855 | NULL&M_Disease |
| 11 | 116997964 | rs61905463 | A | G | NULL | 0.618927855 | NULL&M_Disease |
| 11 | 118501299 | rs7936556 | A | G | NULL | 0.620106432 | NULL&M_Disease |
| 11 | 119504482 | rs2014552 | A | G | NULL | 0.620106432 | NULL&M_Disease |
| 11 | 119931576 | rs11605307 | T | C | NULL | 0.620106432 | NULL&M_Disease |
| 11 | 120577240 | rs9630201 | A | G | NULL | 0.618927855 | NULL&M_Disease |
| 11 | 120577600 | rs4451741 | T | G | NULL | 0.618927855 | NULL&M_Disease |
| 11 | 120578133 | rs61355898 | T | C | NULL | 0.618927855 | NULL&M_Disease |
| 11 | 117771888 | rs572675 | A | C | NULL | 0.618927855 | NULL&M_Disease |
| 11 | 117777929 | rs11216614 | C | G | NULL | 0.618927855 | NULL&M_Disease |
| 11 | 118372923 | rs9332835 | A | G | NULL | 0.618927855 | NULL&M_Disease |
| 11 | 117001384 | rs61905464 | A | G | NULL | 0.618927855 | NULL&M_Disease |
| 11 | 118341649 | rs7948661 | C | T | NULL | 0.620106432 | NULL&M_Disease |
| 11 | 119334649 | rs141453720 | C | G | NULL | 0.618927855 | NULL&M_Disease |

| | | | | | | | | | |
|---|---|---|---|---|---|---|---|---|---|
| 11 | 120174671 | rs10790375 | C | T | 1.041 | -7977.388 | -7978.349 | -0.961 | 1.48E-06 |
| 11 | 117967986 | rs9651687 | A | T | 1.045 | -7977.388 | -7978.35 | -0.962 | 1.48E-06 |
| 11 | 118724336 | rs10892298 | C | T | 0.958 | -7977.388 | -7978.35 | -0.962 | 1.48E-06 |
| 11 | 118730669 | rs7933007 | C | T | 0.958 | -7977.388 | -7978.35 | -0.962 | 1.48E-06 |
| 11 | 120580795 | rs7103021 | T | C | 0.956 | -7977.388 | -7978.351 | -0.962 | 1.48E-06 |
| 11 | 120657934 | rs2186698 | C | G | 0.955 | -7977.388 | -7978.351 | -0.962 | 1.48E-06 |
| 11 | 120695729 | rs111750658 | C | G | 0.958 | -7977.388 | -7978.35 | -0.962 | 1.48E-06 |
| 11 | 116331421 | rs113872393 | A | G | 1.023 | -7977.388 | -7978.351 | -0.963 | 1.48E-06 |
| 11 | 117250530 | rs74661474 | A | G | 1.026 | -7977.388 | -7978.351 | -0.963 | 1.48E-06 |
| 11 | 117266094 | rs11216375 | A | G | 1.043 | -7977.388 | -7978.351 | -0.963 | 1.48E-06 |
| 11 | 117682935 | rs182928887 | G | A | 0.959 | -7977.388 | -7978.351 | -0.963 | 1.48E-06 |
| 11 | 116625362 | rs56225305 | A | G | 0.956 | -7977.388 | -7978.352 | -0.964 | 1.48E-06 |
| 11 | 117842818 | rs4936413 | C | T | 1.041 | -7977.388 | -7978.352 | -0.964 | 1.48E-06 |
| 11 | 120497465 | rs35181812 | T | C | 0.959 | -7977.388 | -7978.353 | -0.964 | 1.48E-06 |
| 11 | 116727592 | rs61907562 | A | G | 0.983 | -7977.388 | -7978.353 | -0.965 | 1.47E-06 |
| 11 | 118782222 | rs61900138 | C | T | 1.001 | -7977.388 | -7978.353 | -0.965 | 1.47E-06 |
| 11 | 118874272 | rs192113495 | G | T | 1.025 | -7977.388 | -7978.353 | -0.965 | 1.47E-06 |
| 11 | 120801460 | rs509566 | T | A | 0.961 | -7977.388 | -7978.354 | -0.965 | 1.47E-06 |
| 11 | 119795416 | rs80294967 | A | G | 1.047 | -7977.388 | -7978.354 | -0.966 | 1.47E-06 |
| 11 | 116726087 | rs61907560 | C | T | 0.983 | -7977.388 | -7978.356 | -0.967 | 1.47E-06 |
| 11 | 116536490 | rs1240778 | C | A | 1.041 | -7977.388 | -7978.356 | -0.968 | 1.47E-06 |
| 11 | 119940917 | rs117243996 | T | C | 0.957 | -7977.388 | -7978.356 | -0.968 | 1.47E-06 |
| 11 | 120484712 | rs1317815 | A | G | 1.042 | -7977.388 | -7978.356 | -0.968 | 1.47E-06 |
| 11 | 117435160 | rs4936387 | T | C | 1.042 | -7977.388 | -7978.357 | -0.969 | 1.47E-06 |
| 11 | 117863888 | rs4252307 | G | C | 0.962 | -7977.388 | -7978.357 | -0.969 | 1.47E-06 |
| 11 | 117949807 | rs605805 | G | A | 1.046 | -7977.388 | -7978.357 | -0.969 | 1.47E-06 |
| 11 | 120824400 | rs1893817 | T | C | 0.957 | -7977.388 | -7978.357 | -0.969 | 1.47E-06 |
| 11 | 117129232 | rs77096386 | G | A | 1.036 | -7977.388 | -7978.358 | -0.97 | 1.47E-06 |
| 11 | 117399604 | rs4938400 | A | C | 1.046 | -7977.388 | -7978.358 | -0.97 | 1.47E-06 |
| 11 | 117399605 | rs4938401 | G | T | 1.046 | -7977.388 | -7978.358 | -0.97 | 1.47E-06 |
| 11 | 117606193 | rs4936403 | G | T | 0.958 | -7977.388 | -7978.359 | -0.97 | 1.47E-06 |
| 11 | 117606320 | rs4936404 | G | A | 0.958 | -7977.388 | -7978.358 | -0.97 | 1.47E-06 |
| 11 | 117652288 | rs75734044 | A | G | 0.955 | -7977.388 | -7978.358 | -0.97 | 1.47E-06 |
| 11 | 118377523 | rs9332846 | A | G | 1.042 | -7977.388 | -7978.358 | -0.97 | 1.47E-06 |
| 11 | 120991447 | rs74491080 | T | C | 0.975 | -7977.388 | -7978.359 | -0.97 | 1.47E-06 |
| 11 | 116555126 | rs117341026 | A | G | 1.042 | -7977.388 | -7978.359 | -0.971 | 1.46E-06 |
| 11 | 117792471 | rs11216624 | A | G | 1.047 | -7977.388 | -7978.359 | -0.971 | 1.46E-06 |
| 11 | 118006930 | rs1317544 | C | T | 0.972 | -7977.388 | -7978.359 | -0.971 | 1.46E-06 |
| 11 | 118510562 | rs4245184 | A | G | 0.961 | -7977.388 | -7978.359 | -0.971 | 1.46E-06 |
| 11 | 118511352 | rs644273 | T | C | 0.961 | -7977.388 | -7978.359 | -0.971 | 1.46E-06 |
| 11 | 119796458 | rs1011664 | G | C | 1.047 | -7977.388 | -7978.36 | -0.971 | 1.46E-06 |
| 11 | 120796418 | rs2852243 | C | T | 0.96 | -7977.388 | -7978.359 | -0.971 | 1.46E-06 |
| 11 | 116369980 | rs1108627 | T | C | 0.96 | -7977.388 | -7978.36 | -0.972 | 1.46E-06 |
| 11 | 118385249 | rs112508300 | G | A | 0.962 | -7977.388 | -7978.36 | -0.972 | 1.46E-06 |
| 11 | 118711830 | rs566848 | G | A | 0.958 | -7977.388 | -7978.36 | -0.972 | 1.46E-06 |
| 11 | 120164102 | rs10750170 | G | A | 1.043 | -7977.388 | -7978.36 | -0.972 | 1.46E-06 |
| 11 | 120785542 | rs117391362 | A | G | 0.961 | -7977.388 | -7978.36 | -0.972 | 1.46E-06 |
| 11 | 120848717 | rs17124632 | A | G | 1.044 | -7977.388 | -7978.36 | -0.972 | 1.46E-06 |
| 11 | 116994106 | rs61905462 | A | C | 0.984 | -7977.388 | -7978.362 | -0.973 | 1.46E-06 |
| 11 | 117196435 | rs73014325 | A | C | 1.033 | -7977.388 | -7978.361 | -0.973 | 1.46E-06 |
| 11 | 117533027 | rs34874964 | T | C | 0.984 | -7977.388 | -7978.361 | -0.973 | 1.46E-06 |
| 11 | 117917962 | rs659092 | C | T | 1.047 | -7977.388 | -7978.361 | -0.973 | 1.46E-06 |
| 11 | 120500339 | rs12577725 | T | C | 0.958 | -7977.388 | -7978.361 | -0.973 | 1.46E-06 |
| 11 | 116946788 | rs61903414 | T | C | 0.985 | -7977.388 | -7978.362 | -0.974 | 1.46E-06 |
| 11 | 117596927 | rs948104 | G | T | 0.958 | -7977.388 | -7978.363 | -0.974 | 1.46E-06 |
| 11 | 118385691 | rs111405302 | C | T | 1.042 | -7977.388 | -7978.362 | -0.974 | 1.46E-06 |
| 11 | 119774225 | rs525588 | C | G | 1.046 | -7977.388 | -7978.363 | -0.974 | 1.46E-06 |
| 11 | 117375596 | rs2276343 | A | G | 1.044 | -7977.388 | -7978.363 | -0.975 | 1.46E-06 |
| 11 | 117863730 | rs4252306 | C | G | 0.962 | -7977.388 | -7978.363 | -0.975 | 1.46E-06 |
| 11 | 118828942 | rs623500 | G | A | 0.956 | -7977.388 | -7978.364 | -0.975 | 1.46E-06 |
| 11 | 119840963 | rs77212554 | T | C | 1.046 | -7977.388 | -7978.363 | -0.975 | 1.46E-06 |
| 11 | 120902930 | rs61903185 | A | G | 1.044 | -7977.388 | -7978.364 | -0.975 | 1.46E-06 |
| 11 | 117219305 | rs592828 | C | G | 0.958 | -7977.388 | -7978.364 | -0.976 | 1.46E-06 |
| 11 | 117864979 | rs2282496 | T | C | 0.963 | -7977.388 | -7978.364 | -0.976 | 1.46E-06 |
| 11 | 118890085 | rs4938619 | T | C | 1.046 | -7977.388 | -7978.364 | -0.976 | 1.46E-06 |
| 11 | 118049257 | rs7123533 | G | C | 0.961 | -7977.388 | -7978.366 | -0.977 | 1.46E-06 |
| 11 | 118049759 | rs61472653 | C | T | 0.961 | -7977.388 | -7978.366 | -0.977 | 1.46E-06 |
| 11 | 118049811 | rs59018400 | T | C | 0.961 | -7977.388 | -7978.366 | -0.977 | 1.46E-06 |
| 11 | 118049880 | rs56301948 | G | A | 0.961 | -7977.388 | -7978.366 | -0.977 | 1.46E-06 |
| 11 | 118050014 | rs57533048 | G | A | 0.961 | -7977.388 | -7978.366 | -0.977 | 1.46E-06 |
| 11 | 118050348 | rs7342223 | G | T | 0.961 | -7977.388 | -7978.366 | -0.977 | 1.46E-06 |
| 11 | 118511757 | rs654423 | A | G | 0.961 | -7977.388 | -7978.365 | -0.977 | 1.46E-06 |
| 11 | 116520527 | rs1240773 | T | G | 1.044 | -7977.388 | -7978.367 | -0.978 | 1.45E-06 |
| 11 | 116632956 | rs57641217 | T | C | 0.957 | -7977.388 | -7978.366 | -0.978 | 1.45E-06 |
| 11 | 116658031 | rs12285095 | G | T | 0.957 | -7977.388 | -7978.366 | -0.978 | 1.45E-06 |
| 11 | 116807226 | rs61907596 | T | C | 0.987 | -7977.388 | -7978.366 | -0.978 | 1.45E-06 |
| 11 | 117246566 | rs112933729 | A | G | 1.02 | -7977.388 | -7978.366 | -0.978 | 1.45E-06 |
| 11 | 117527473 | rs12802995 | A | C | 0.987 | -7977.388 | -7978.367 | -0.978 | 1.45E-06 |
| 11 | 117531395 | rs35048090 | T | C | 0.986 | -7977.388 | -7978.366 | -0.978 | 1.45E-06 |
| 11 | 117531569 | rs36067343 | G | T | 0.986 | -7977.388 | -7978.366 | -0.978 | 1.45E-06 |
| 11 | 119744714 | rs73008683 | A | G | 1.046 | -7977.388 | -7978.367 | -0.978 | 1.45E-06 |
| 11 | 120728827 | rs7103097 | A | C | 0.956 | -7977.388 | -7978.366 | -0.978 | 1.45E-06 |
| 11 | 116906431 | rs61905711 | T | C | 1.015 | -7977.388 | -7978.367 | -0.979 | 1.45E-06 |
| 11 | 116997964 | rs61905463 | A | G | 0.987 | -7977.388 | -7978.368 | -0.979 | 1.45E-06 |
| 11 | 118501299 | rs7936556 | A | G | 0.961 | -7977.388 | -7978.367 | -0.979 | 1.45E-06 |
| 11 | 119504482 | rs2014552 | A | G | 0.955 | -7977.388 | -7978.367 | -0.979 | 1.45E-06 |
| 11 | 119931576 | rs11605307 | T | C | 1.042 | -7977.388 | -7978.367 | -0.979 | 1.45E-06 |
| 11 | 120577240 | rs9630201 | A | G | 0.998 | -7977.388 | -7978.368 | -0.979 | 1.45E-06 |
| 11 | 120577600 | rs4451741 | T | G | 0.998 | -7977.388 | -7978.368 | -0.979 | 1.45E-06 |
| 11 | 120578133 | rs61355898 | T | C | 0.997 | -7977.388 | -7978.367 | -0.979 | 1.45E-06 |
| 11 | 117771888 | rs572675 | A | C | 0.994 | -7977.388 | -7978.368 | -0.98 | 1.45E-06 |
| 11 | 117777929 | rs11216614 | C | G | 0.989 | -7977.388 | -7978.369 | -0.98 | 1.45E-06 |
| 11 | 118372923 | rs9332835 | A | G | 1.041 | -7977.388 | -7978.368 | -0.98 | 1.45E-06 |
| 11 | 117001384 | rs61905464 | A | G | 0.988 | -7977.388 | -7978.37 | -0.981 | 1.45E-06 |
| 11 | 118341649 | rs7948661 | C | T | 1.04 | -7977.388 | -7978.369 | -0.981 | 1.45E-06 |
| 11 | 119334649 | rs141453720 | C | G | 0.985 | -7977.388 | -7978.369 | -0.981 | 1.45E-06 |

| | | | | | | | |
|---|---|---|---|---|---|---|---|
| 11 | 119854322 | rs7106980 | G | T | NULL | 0.620106432 | NULL&M_Disease |
| 11 | 121142356 | rs1367306 | T | A | NULL | 0.618927855 | NULL&M_Disease |
| 11 | 116492506 | rs61904859 | T | C | NULL | 0.618927855 | NULL&M_Disease |
| 11 | 116916175 | rs17496758 | A | G | NULL | 0.618927855 | NULL&M_Disease |
| 11 | 117113478 | rs117639653 | C | G | NULL | 0.618927855 | NULL&M_Disease |
| 11 | 117432034 | rs1893970 | T | C | NULL | 0.620106432 | NULL&M_Disease |
| 11 | 117755580 | rs12286112 | G | A | NULL | 0.620106432 | NULL&M_Disease |
| 11 | 118333537 | rs113028342 | G | A | NULL | 0.620106432 | NULL&M_Disease |
| 11 | 118720590 | rs4938564 | T | G | NULL | 0.620106432 | NULL&M_Disease |
| 11 | 119624396 | rs60727471 | C | T | NULL | 0.620106432 | NULL&M_Disease |
| 11 | 119835506 | rs77142768 | G | A | NULL | 0.620106432 | NULL&M_Disease |
| 11 | 119835845 | rs76085613 | A | G | NULL | 0.620106432 | NULL&M_Disease |
| 11 | 120710519 | rs73003760 | C | T | NULL | 0.620106432 | NULL&M_Disease |
| 11 | 116679711 | rs7126214 | A | G | NULL | 0.620106432 | NULL&M_Disease |
| 11 | 117312197 | rs7948072 | G | A | NULL | 0.620106432 | NULL&M_Disease |
| 11 | 117875313 | rs4936415 | C | G | NULL | 0.620106432 | NULL&M_Disease |
| 11 | 119344568 | rs17547832 | G | A | NULL | 0.620106432 | NULL&M_Disease |
| 11 | 120092344 | rs61898286 | C | G | NULL | 0.620106432 | NULL&M_Disease |
| 11 | 120095782 | rs144785834 | C | A | NULL | 0.620106432 | NULL&M_Disease |
| 11 | 116521505 | rs61906076 | G | C | NULL | 0.620106432 | NULL&M_Disease |
| 11 | 116543098 | rs58950690 | C | T | NULL | 0.620106432 | NULL&M_Disease |
| 11 | 116699395 | rs12721078 | A | C | NULL | 0.620106432 | NULL&M_Disease |
| 11 | 116848716 | rs147893781 | T | C | NULL | 0.620106432 | NULL&M_Disease |
| 11 | 116886159 | rs61905708 | T | G | NULL | 0.620106432 | NULL&M_Disease |
| 11 | 117597995 | rs10790202 | C | A | NULL | 0.620106432 | NULL&M_Disease |
| 11 | 119505336 | rs60657140 | T | G | NULL | 0.620106432 | NULL&M_Disease |
| 11 | 119989569 | rs530291 | T | C | NULL | 0.620106432 | NULL&M_Disease |
| 11 | 120791799 | rs12363923 | T | C | NULL | 0.620106432 | NULL&M_Disease |
| 11 | 116648776 | rs78296522 | A | C | NULL | 0.620106432 | NULL&M_Disease |
| 11 | 116876168 | rs61905690 | G | C | NULL | 0.620106432 | NULL&M_Disease |
| 11 | 117077716 | rs61905530 | A | G | NULL | 0.620106432 | NULL&M_Disease |
| 11 | 117710295 | rs144425764 | C | T | NULL | 0.620106432 | NULL&M_Disease |
| 11 | 117969122 | rs948463 | G | A | NULL | 0.620106432 | NULL&M_Disease |
| 11 | 119830778 | rs12796257 | C | T | NULL | 0.620106432 | NULL&M_Disease |
| 11 | 116464194 | rs7929011 | A | G | NULL | 0.620106432 | NULL&M_Disease |
| 11 | 120470521 | rs4936525 | A | C | NULL | 0.620106432 | NULL&M_Disease |
| 11 | 120679052 | rs948025 | T | G | NULL | 0.620106432 | NULL&M_Disease |
| 11 | 116883665 | rs61905704 | C | T | NULL | 0.620106432 | NULL&M_Disease |
| 11 | 117088821 | rs61906487 | T | C | NULL | 0.620106432 | NULL&M_Disease |
| 11 | 117089505 | rs61906488 | C | T | NULL | 0.620106432 | NULL&M_Disease |
| 11 | 117091153 | rs61906489 | G | C | NULL | 0.620106432 | NULL&M_Disease |
| 11 | 117391736 | rs7936795 | G | A | NULL | 0.620106432 | NULL&M_Disease |
| 11 | 118044144 | rs11216790 | T | G | NULL | 0.620106432 | NULL&M_Disease |
| 11 | 118746769 | rs4938576 | T | G | NULL | 0.620106432 | NULL&M_Disease |
| 11 | 119853069 | rs7129717 | G | T | NULL | 0.620106432 | NULL&M_Disease |
| 11 | 121028666 | rs33981325 | T | C | NULL | 0.620106432 | NULL&M_Disease |
| 11 | 117599333 | rs1942926 | A | C | NULL | 0.620106432 | NULL&M_Disease |
| 11 | 117923079 | rs594603 | T | C | NULL | 0.620106432 | NULL&M_Disease |
| 11 | 118054345 | rs12295676 | A | G | NULL | 0.620106432 | NULL&M_Disease |
| 11 | 120656866 | rs4544011 | G | A | NULL | 0.621283595 | NULL&M_Disease |
| 11 | 117083831 | rs76830818 | T | A | NULL | 0.620106432 | NULL&M_Disease |
| 11 | 117533773 | rs17121050 | A | G | NULL | 0.620106432 | NULL&M_Disease |
| 11 | 118153511 | rs34782343 | G | T | NULL | 0.621283595 | NULL&M_Disease |
| 11 | 120157518 | rs10790374 | C | T | NULL | 0.621283595 | NULL&M_Disease |
| 11 | 121061986 | rs618523 | T | C | NULL | 0.621283595 | NULL&M_Disease |
| 11 | 121190696 | rs56880203 | G | A | NULL | 0.621283595 | NULL&M_Disease |
| 11 | 118371025 | rs7116876 | C | G | NULL | 0.620106432 | NULL&M_Disease |
| 11 | 119502870 | rs20337 | A | G | NULL | 0.621283595 | NULL&M_Disease |
| 11 | 120070943 | rs61898266 | A | G | NULL | 0.620106432 | NULL&M_Disease |
| 11 | 117014926 | rs147062051 | A | G | NULL | 0.620106432 | NULL&M_Disease |
| 11 | 117015290 | rs181954995 | A | G | NULL | 0.620106432 | NULL&M_Disease |
| 11 | 117018031 | rs61905473 | T | G | NULL | 0.620106432 | NULL&M_Disease |
| 11 | 117019530 | rs61905474 | A | G | NULL | 0.620106432 | NULL&M_Disease |
| 11 | 117024435 | rs61905475 | G | A | NULL | 0.620106432 | NULL&M_Disease |
| 11 | 117024537 | rs61905476 | G | A | NULL | 0.620106432 | NULL&M_Disease |
| 11 | 117026055 | rs61905477 | G | A | NULL | 0.620106432 | NULL&M_Disease |
| 11 | 117082691 | rs61906485 | T | G | NULL | 0.620106432 | NULL&M_Disease |
| 11 | 118361472 | rs9332809 | A | G | NULL | 0.621283595 | NULL&M_Disease |
| 11 | 118365168 | rs2276045 | A | G | NULL | 0.621283595 | NULL&M_Disease |
| 11 | 118390082 | rs9332858 | C | A | NULL | 0.621283595 | NULL&M_Disease |
| 11 | 118691975 | rs77731159 | A | T | NULL | 0.621283595 | NULL&M_Disease |
| 11 | 116558455 | rs11607356 | C | A | NULL | 0.621283595 | NULL&M_Disease |
| 11 | 116612274 | rs11216125 | A | G | NULL | 0.621283595 | NULL&M_Disease |
| 11 | 116913051 | rs61905715 | T | C | NULL | 0.620106432 | NULL&M_Disease |
| 11 | 117027942 | rs12291885 | C | T | NULL | 0.621283595 | NULL&M_Disease |
| 11 | 117082865 | rs61906486 | A | C | NULL | 0.620106432 | NULL&M_Disease |
| 11 | 118178007 | rs1126924 | T | C | NULL | 0.621283595 | NULL&M_Disease |
| 11 | 119518808 | rs11821535 | A | C | NULL | 0.621283595 | NULL&M_Disease |
| 11 | 120679042 | rs948024 | T | A | NULL | 0.621283595 | NULL&M_Disease |
| 11 | 121064513 | rs4935763 | T | C | NULL | 0.621283595 | NULL&M_Disease |
| 11 | 116780568 | rs117865451 | C | T | NULL | 0.621283595 | NULL&M_Disease |
| 11 | 117017706 | rs61905472 | T | C | NULL | 0.621283595 | NULL&M_Disease |
| 11 | 117035228 | rs118155116 | C | T | NULL | 0.621283595 | NULL&M_Disease |
| 11 | 117045555 | rs61905519 | C | T | NULL | 0.621283595 | NULL&M_Disease |
| 11 | 117046136 | rs77086432 | C | T | NULL | 0.621283595 | NULL&M_Disease |
| 11 | 117049020 | rs61905521 | A | G | NULL | 0.621283595 | NULL&M_Disease |
| 11 | 117049025 | rs61905522 | A | G | NULL | 0.621283595 | NULL&M_Disease |
| 11 | 117061197 | rs143668212 | T | C | NULL | 0.621283595 | NULL&M_Disease |
| 11 | 117067465 | rs61905526 | T | C | NULL | 0.621283595 | NULL&M_Disease |
| 11 | 117069966 | rs61905527 | T | C | NULL | 0.621283595 | NULL&M_Disease |
| 11 | 117072074 | rs117279127 | T | C | NULL | 0.621283595 | NULL&M_Disease |
| 11 | 117391374 | rs11216436 | A | G | NULL | 0.621283595 | NULL&M_Disease |
| 11 | 117752845 | rs73020445 | T | C | NULL | 0.621283595 | NULL&M_Disease |
| 11 | 118043904 | rs75403174 | A | C | NULL | 0.621283595 | NULL&M_Disease |
| 11 | 118044450 | rs11216792 | T | G | NULL | 0.621283595 | NULL&M_Disease |
| 11 | 118045423 | rs61581699 | A | G | NULL | 0.621283595 | NULL&M_Disease |

| | | | | | | | | | |
|---|---|---|---|---|---|---|---|---|---|
| 11 | 119854322 | rs7106980 | G | T | 1.045 | -7977.388 | -7978.369 | -0.981 | 1.45E-06 |
| 11 | 121142356 | rs1367306 | T | A | 0.972 | -7977.388 | -7978.369 | -0.981 | 1.45E-06 |
| 11 | 116492506 | rs61904859 | T | C | 0.998 | -7977.388 | -7978.37 | -0.982 | 1.45E-06 |
| 11 | 116916175 | rs17496758 | A | G | 0.988 | -7977.388 | -7978.37 | -0.982 | 1.45E-06 |
| 11 | 117113478 | rs117639653 | C | G | 1.009 | -7977.388 | -7978.37 | -0.982 | 1.45E-06 |
| 11 | 117432034 | rs1893970 | T | C | 0.96 | -7977.388 | -7978.37 | -0.982 | 1.45E-06 |
| 11 | 117755580 | rs12286112 | G | A | 0.96 | -7977.388 | -7978.371 | -0.982 | 1.45E-06 |
| 11 | 118333537 | rs113028342 | G | A | 1.04 | -7977.388 | -7978.37 | -0.982 | 1.45E-06 |
| 11 | 118720590 | rs4938564 | T | G | 0.958 | -7977.388 | -7978.37 | -0.982 | 1.45E-06 |
| 11 | 119624396 | rs60727471 | C | T | 0.955 | -7977.388 | -7978.371 | -0.982 | 1.45E-06 |
| 11 | 119835506 | rs77142768 | G | A | 1.046 | -7977.388 | -7978.37 | -0.982 | 1.45E-06 |
| 11 | 119835845 | rs76085613 | A | G | 1.046 | -7977.388 | -7978.37 | -0.982 | 1.45E-06 |
| 11 | 120710519 | rs73003760 | C | T | 0.957 | -7977.388 | -7978.371 | -0.982 | 1.45E-06 |
| 11 | 116679711 | rs7126214 | A | G | 1.046 | -7977.388 | -7978.371 | -0.983 | 1.45E-06 |
| 11 | 117312197 | rs7948072 | G | A | 1.046 | -7977.388 | -7978.371 | -0.983 | 1.45E-06 |
| 11 | 117875313 | rs4936415 | C | G | 1.041 | -7977.388 | -7978.372 | -0.983 | 1.45E-06 |
| 11 | 119344568 | rs17547832 | G | A | 1.046 | -7977.388 | -7978.372 | -0.983 | 1.45E-06 |
| 11 | 120092344 | rs61898286 | C | G | 0.973 | -7977.388 | -7978.371 | -0.983 | 1.45E-06 |
| 11 | 120095782 | rs144785834 | C | A | 0.969 | -7977.388 | -7978.371 | -0.983 | 1.45E-06 |
| 11 | 116521505 | rs61906076 | G | C | 0.962 | -7977.388 | -7978.372 | -0.984 | 1.45E-06 |
| 11 | 116543098 | rs58950690 | C | T | 0.964 | -7977.388 | -7978.372 | -0.984 | 1.45E-06 |
| 11 | 116699395 | rs12721078 | A | C | 1.003 | -7977.388 | -7978.372 | -0.984 | 1.45E-06 |
| 11 | 116848716 | rs147893781 | T | C | 0.989 | -7977.388 | -7978.372 | -0.984 | 1.45E-06 |
| 11 | 116886159 | rs61905708 | T | G | 0.989 | -7977.388 | -7978.372 | -0.984 | 1.45E-06 |
| 11 | 117597995 | rs10790202 | C | A | 0.959 | -7977.388 | -7978.373 | -0.984 | 1.45E-06 |
| 11 | 119505336 | rs60657140 | T | G | 0.956 | -7977.388 | -7978.372 | -0.984 | 1.45E-06 |
| 11 | 119989569 | rs530291 | T | C | 0.967 | -7977.388 | -7978.372 | -0.984 | 1.45E-06 |
| 11 | 120791799 | rs12363923 | T | C | 0.955 | -7977.388 | -7978.372 | -0.984 | 1.45E-06 |
| 11 | 116648776 | rs78296522 | A | C | 0.963 | -7977.388 | -7978.374 | -0.985 | 1.44E-06 |
| 11 | 116876168 | rs61905690 | G | C | 0.99 | -7977.388 | -7978.374 | -0.985 | 1.44E-06 |
| 11 | 117077716 | rs61905530 | A | G | 0.99 | -7977.388 | -7978.374 | -0.985 | 1.44E-06 |
| 11 | 117710295 | rs144425764 | C | T | 1 | -7977.388 | -7978.373 | -0.985 | 1.44E-06 |
| 11 | 117969122 | rs948463 | G | A | 1.043 | -7977.388 | -7978.374 | -0.985 | 1.44E-06 |
| 11 | 119830778 | rs12796257 | C | T | 1.01 | -7977.388 | -7978.374 | -0.985 | 1.44E-06 |
| 11 | 116464194 | rs7929011 | A | G | 0.959 | -7977.388 | -7978.375 | -0.986 | 1.44E-06 |
| 11 | 120470521 | rs4936525 | A | C | 0.959 | -7977.388 | -7978.375 | -0.986 | 1.44E-06 |
| 11 | 120679052 | rs948025 | T | G | 0.958 | -7977.388 | -7978.374 | -0.986 | 1.44E-06 |
| 11 | 116883665 | rs61905704 | C | T | 0.994 | -7977.388 | -7978.375 | -0.987 | 1.44E-06 |
| 11 | 117088821 | rs61906487 | T | C | 1.004 | -7977.388 | -7978.375 | -0.987 | 1.44E-06 |
| 11 | 117089505 | rs61906488 | C | T | 1.004 | -7977.388 | -7978.375 | -0.987 | 1.44E-06 |
| 11 | 117091153 | rs61906489 | G | C | 1.004 | -7977.388 | -7978.375 | -0.987 | 1.44E-06 |
| 11 | 117391736 | rs7936795 | G | A | 1.045 | -7977.388 | -7978.375 | -0.987 | 1.44E-06 |
| 11 | 118044144 | rs11216790 | T | G | 0.963 | -7977.388 | -7978.375 | -0.987 | 1.44E-06 |
| 11 | 118746769 | rs4938576 | T | G | 1.041 | -7977.388 | -7978.376 | -0.987 | 1.44E-06 |
| 11 | 119853069 | rs7129717 | G | T | 1.046 | -7977.388 | -7978.376 | -0.987 | 1.44E-06 |
| 11 | 121028666 | rs33981325 | T | C | 0.996 | -7977.388 | -7978.375 | -0.987 | 1.44E-06 |
| 11 | 117599333 | rs1942926 | A | C | 0.959 | -7977.388 | -7978.376 | -0.988 | 1.44E-06 |
| 11 | 117923079 | rs594603 | T | C | 1.046 | -7977.388 | -7978.376 | -0.988 | 1.44E-06 |
| 11 | 118054345 | rs12295676 | A | G | 0.963 | -7977.388 | -7978.376 | -0.988 | 1.44E-06 |
| 11 | 120656866 | rs4544011 | G | A | 0.961 | -7977.388 | -7978.377 | -0.988 | 1.44E-06 |
| 11 | 117083831 | rs76830818 | T | A | 0.993 | -7977.388 | -7978.377 | -0.989 | 1.44E-06 |
| 11 | 117533773 | rs17121050 | A | G | 0.994 | -7977.388 | -7978.377 | -0.989 | 1.44E-06 |
| 11 | 118153511 | rs34782343 | G | T | 1.042 | -7977.388 | -7978.377 | -0.989 | 1.44E-06 |
| 11 | 120157518 | rs10790374 | C | T | 1.043 | -7977.388 | -7978.377 | -0.989 | 1.44E-06 |
| 11 | 121061986 | rs618523 | T | C | 0.96 | -7977.388 | -7978.377 | -0.989 | 1.44E-06 |
| 11 | 121190696 | rs56880203 | G | A | 0.957 | -7977.388 | -7978.377 | -0.989 | 1.44E-06 |
| 11 | 118371025 | rs7116876 | C | G | 1.04 | -7977.388 | -7978.378 | -0.99 | 1.44E-06 |
| 11 | 119502870 | rs20337 | A | G | 0.956 | -7977.388 | -7978.379 | -0.99 | 1.44E-06 |
| 11 | 120070943 | rs61898266 | A | G | 0.999 | -7977.388 | -7978.379 | -0.99 | 1.44E-06 |
| 11 | 117014926 | rs147062051 | A | G | 0.996 | -7977.388 | -7978.379 | -0.991 | 1.44E-06 |
| 11 | 117015290 | rs181954995 | A | G | 0.996 | -7977.388 | -7978.379 | -0.991 | 1.44E-06 |
| 11 | 117018031 | rs61905473 | T | G | 0.996 | -7977.388 | -7978.379 | -0.991 | 1.44E-06 |
| 11 | 117019530 | rs61905474 | A | G | 1.001 | -7977.388 | -7978.379 | -0.991 | 1.44E-06 |
| 11 | 117024435 | rs61905475 | G | A | 1.001 | -7977.388 | -7978.379 | -0.991 | 1.44E-06 |
| 11 | 117024537 | rs61905476 | G | A | 0.996 | -7977.388 | -7978.379 | -0.991 | 1.44E-06 |
| 11 | 117026055 | rs61905477 | G | A | 1.001 | -7977.388 | -7978.379 | -0.991 | 1.44E-06 |
| 11 | 117082691 | rs61906485 | T | G | 0.995 | -7977.388 | -7978.379 | -0.991 | 1.44E-06 |
| 11 | 118361472 | rs9332809 | A | G | 1.04 | -7977.388 | -7978.379 | -0.991 | 1.44E-06 |
| 11 | 118365168 | rs2276045 | A | G | 1.04 | -7977.388 | -7978.379 | -0.991 | 1.44E-06 |
| 11 | 118390082 | rs9332858 | C | A | 1.04 | -7977.388 | -7978.379 | -0.991 | 1.44E-06 |
| 11 | 118691975 | rs77731159 | A | T | 0.963 | -7977.388 | -7978.38 | -0.991 | 1.44E-06 |
| 11 | 116558455 | rs11607356 | C | A | 0.957 | -7977.388 | -7978.381 | -0.992 | 1.43E-06 |
| 11 | 116612274 | rs11216125 | A | G | 1.046 | -7977.388 | -7978.381 | -0.992 | 1.43E-06 |
| 11 | 116913051 | rs61905715 | T | C | 0.995 | -7977.388 | -7978.38 | -0.992 | 1.43E-06 |
| 11 | 117027942 | rs12291885 | C | T | 1.004 | -7977.388 | -7978.381 | -0.992 | 1.43E-06 |
| 11 | 117082865 | rs61906486 | A | C | 0.994 | -7977.388 | -7978.38 | -0.992 | 1.43E-06 |
| 11 | 118178007 | rs1126924 | T | C | 1.042 | -7977.388 | -7978.381 | -0.992 | 1.43E-06 |
| 11 | 119518808 | rs11821535 | A | C | 1.043 | -7977.388 | -7978.38 | -0.992 | 1.43E-06 |
| 11 | 120679042 | rs948024 | T | A | 0.959 | -7977.388 | -7978.381 | -0.992 | 1.43E-06 |
| 11 | 121064513 | rs4935763 | T | C | 0.96 | -7977.388 | -7978.38 | -0.992 | 1.43E-06 |
| 11 | 116780568 | rs117865451 | C | T | 0.992 | -7977.388 | -7978.381 | -0.993 | 1.43E-06 |
| 11 | 117017706 | rs61905472 | T | C | 1.001 | -7977.388 | -7978.381 | -0.993 | 1.43E-06 |
| 11 | 117035228 | rs118155116 | C | T | 0.999 | -7977.388 | -7978.381 | -0.993 | 1.43E-06 |
| 11 | 117045555 | rs61905519 | C | T | 0.999 | -7977.388 | -7978.381 | -0.993 | 1.43E-06 |
| 11 | 117046136 | rs77086432 | C | T | 1 | -7977.388 | -7978.381 | -0.993 | 1.43E-06 |
| 11 | 117049020 | rs61905521 | A | G | 0.999 | -7977.388 | -7978.381 | -0.993 | 1.43E-06 |
| 11 | 117049025 | rs61905522 | A | G | 0.999 | -7977.388 | -7978.381 | -0.993 | 1.43E-06 |
| 11 | 117061197 | rs143668212 | T | C | 1 | -7977.388 | -7978.381 | -0.993 | 1.43E-06 |
| 11 | 117067465 | rs61905526 | T | C | 0.996 | -7977.388 | -7978.381 | -0.993 | 1.43E-06 |
| 11 | 117069966 | rs61905527 | T | C | 0.996 | -7977.388 | -7978.381 | -0.993 | 1.43E-06 |
| 11 | 117072074 | rs117279127 | T | C | 0.995 | -7977.388 | -7978.381 | -0.993 | 1.43E-06 |
| 11 | 117391374 | rs11216436 | A | G | 1.045 | -7977.388 | -7978.382 | -0.993 | 1.43E-06 |
| 11 | 117752845 | rs73020445 | T | C | 1.004 | -7977.388 | -7978.382 | -0.993 | 1.43E-06 |
| 11 | 118043904 | rs75403174 | A | C | 0.963 | -7977.388 | -7978.381 | -0.993 | 1.43E-06 |
| 11 | 118044450 | rs11216792 | T | G | 0.963 | -7977.388 | -7978.381 | -0.993 | 1.43E-06 |
| 11 | 118045423 | rs61581699 | A | G | 0.963 | -7977.388 | -7978.381 | -0.993 | 1.43E-06 |

| | | | | | | | |
|---|---|---|---|---|---|---|---|
| 11 | 118056823 | rs7935492 | T | C | NULL | 0.621283595 | NULL&M_Disease |
| 11 | 118057783 | rs12282334 | G | C | NULL | 0.621283595 | NULL&M_Disease |
| 11 | 118059914 | rs7934292 | C | T | NULL | 0.621283595 | NULL&M_Disease |
| 11 | 119794526 | rs77708326 | C | T | NULL | 0.621283595 | NULL&M_Disease |
| 11 | 119838717 | rs79652828 | T | C | NULL | 0.621283595 | NULL&M_Disease |
| 11 | 119839219 | rs77436344 | A | G | NULL | 0.621283595 | NULL&M_Disease |
| 11 | 120122628 | rs112667243 | T | C | NULL | 0.621283595 | NULL&M_Disease |
| 11 | 120144941 | rs57081718 | A | G | NULL | 0.621283595 | NULL&M_Disease |
| 11 | 120161987 | rs78991803 | A | G | NULL | 0.621283595 | NULL&M_Disease |
| 11 | 116414488 | rs1289103681 | C | T | NULL | 0.621283595 | NULL&M_Disease |
| 11 | 116521748 | rs3016352 | G | A | NULL | 0.621283595 | NULL&M_Disease |
| 11 | 116525229 | rs545108 | C | G | NULL | 0.621283595 | NULL&M_Disease |
| 11 | 116622722 | rs55713675 | T | C | NULL | 0.621283595 | NULL&M_Disease |
| 11 | 117040739 | rs61905518 | G | A | NULL | 0.621283595 | NULL&M_Disease |
| 11 | 118050958 | rs75709684 | A | G | NULL | 0.621283595 | NULL&M_Disease |
| 11 | 118053859 | rs75459003 | C | A | NULL | 0.621283595 | NULL&M_Disease |
| 11 | 118368665 | rs7107305 | G | A | NULL | 0.621283595 | NULL&M_Disease |
| 11 | 118368822 | rs7123835 | G | T | NULL | 0.621283595 | NULL&M_Disease |
| 11 | 119686252 | rs73008610 | C | G | NULL | 0.621283595 | NULL&M_Disease |
| 11 | 120349142 | rs7120040 | G | A | NULL | 0.621283595 | NULL&M_Disease |
| 11 | 116629949 | rs4938310 | T | C | NULL | 0.621283595 | NULL&M_Disease |
| 11 | 117023487 | rs7128382 | A | C | NULL | 0.621283595 | NULL&M_Disease |
| 11 | 117741853 | rs36086368 | C | T | NULL | 0.621283595 | NULL&M_Disease |
| 11 | 117849065 | rs11216664 | C | G | NULL | 0.621283595 | NULL&M_Disease |
| 11 | 119039028 | rs56230940 | T | C | NULL | 0.621283595 | NULL&M_Disease |
| 11 | 120117394 | rs118125468 | A | G | NULL | 0.621283595 | NULL&M_Disease |
| 11 | 120171371 | rs61898318 | A | G | NULL | 0.621283595 | NULL&M_Disease |
| 11 | 117058854 | rs61905525 | A | G | NULL | 0.621283595 | NULL&M_Disease |
| 11 | 117106069 | rs185433085 | T | C | NULL | 0.621283595 | NULL&M_Disease |
| 11 | 118050779 | rs67163042 | G | A | NULL | 0.621283595 | NULL&M_Disease |
| 11 | 118462554 | rs11216920 | G | A | NULL | 0.621283595 | NULL&M_Disease |
| 11 | 119240757 | rs12419879 | T | C | NULL | 0.621283595 | NULL&M_Disease |
| 11 | 120117335 | rs73576276 | T | C | NULL | 0.621283595 | NULL&M_Disease |
| 11 | 116873382 | rs74601470 | T | C | NULL | 0.621283595 | NULL&M_Disease |
| 11 | 117572288 | rs7101613 | G | A | NULL | 0.621283595 | NULL&M_Disease |
| 11 | 117651600 | rs75693006 | A | G | NULL | 0.621283595 | NULL&M_Disease |
| 11 | 118035087 | rs624328 | G | A | NULL | 0.621283595 | NULL&M_Disease |
| 11 | 118804266 | rs7123066 | T | C | NULL | 0.621283595 | NULL&M_Disease |
| 11 | 118804283 | rs7122959 | G | A | NULL | 0.621283595 | NULL&M_Disease |
| 11 | 120496664 | rs4936533 | T | C | NULL | 0.621283595 | NULL&M_Disease |
| 11 | 116879636 | rs77832134 | G | A | NULL | 0.621283595 | NULL&M_Disease |
| 11 | 116990934 | rs7938470 | A | G | NULL | 0.621283595 | NULL&M_Disease |
| 11 | 118044740 | rs11216794 | A | T | NULL | 0.621283595 | NULL&M_Disease |
| 11 | 119509118 | rs34163098 | A | G | NULL | 0.622459331 | NULL&M_Disease |
| 11 | 121064264 | rs4936593 | A | G | NULL | 0.622459331 | NULL&M_Disease |
| 11 | 116721405 | rs111894427 | T | C | NULL | 0.621283595 | NULL&M_Disease |
| 11 | 116982821 | rs10892075 | T | C | NULL | 0.622459331 | NULL&M_Disease |
| 11 | 117410852 | rs12788483 | C | G | NULL | 0.622459331 | NULL&M_Disease |
| 11 | 118052662 | rs74395230 | T | C | NULL | 0.621283595 | NULL&M_Disease |
| 11 | 118792278 | rs77398170 | G | A | NULL | 0.621283595 | NULL&M_Disease |
| 11 | 116726796 | rs61907561 | G | C | NULL | 0.621283595 | NULL&M_Disease |
| 11 | 117101574 | rs522645 | A | C | NULL | 0.622459331 | NULL&M_Disease |
| 11 | 118048885 | rs11216800 | G | A | NULL | 0.621283595 | NULL&M_Disease |
| 11 | 120155303 | rs4245211 | G | A | NULL | 0.622459331 | NULL&M_Disease |
| 11 | 116278812 | rs575623 | C | T | NULL | 0.622459331 | NULL&M_Disease |
| 11 | 116632787 | rs7941518 | G | A | NULL | 0.622459331 | NULL&M_Disease |
| 11 | 116632975 | rs7942054 | A | G | NULL | 0.622459331 | NULL&M_Disease |
| 11 | 116810655 | rs61907602 | C | G | NULL | 0.621283595 | NULL&M_Disease |
| 11 | 116815762 | rs7941912 | A | G | NULL | 0.621283595 | NULL&M_Disease |
| 11 | 118346599 | rs9332781 | G | A | NULL | 0.622459331 | NULL&M_Disease |
| 11 | 120123145 | rs113348846 | A | G | NULL | 0.621283595 | NULL&M_Disease |
| 11 | 120157850 | rs17311233 | T | C | NULL | 0.622459331 | NULL&M_Disease |
| 11 | 116646820 | rs140575350 | G | C | NULL | 0.622459331 | NULL&M_Disease |
| 11 | 116837823 | rs61905682 | C | G | NULL | 0.621283595 | NULL&M_Disease |
| 11 | 116989185 | rs61903459 | G | A | NULL | 0.621283595 | NULL&M_Disease |
| 11 | 117216213 | rs580119 | G | T | NULL | 0.622459331 | NULL&M_Disease |
| 11 | 117589939 | rs7102666 | C | G | NULL | 0.622459331 | NULL&M_Disease |
| 11 | 117753244 | rs73020446 | T | C | NULL | 0.621283595 | NULL&M_Disease |
| 11 | 117967798 | rs9651685 | C | T | NULL | 0.622459331 | NULL&M_Disease |
| 11 | 119622367 | rs112186392 | G | A | NULL | 0.622459331 | NULL&M_Disease |
| 11 | 120924671 | rs17124739 | G | A | NULL | 0.622459331 | NULL&M_Disease |
| 11 | 116354466 | rs549298 | G | A | NULL | 0.622459331 | NULL&M_Disease |
| 11 | 117996885 | rs7104443 | G | C | NULL | 0.622459331 | NULL&M_Disease |
| 11 | 119697626 | rs2028960 | T | C | NULL | 0.622459331 | NULL&M_Disease |
| 11 | 120036997 | rs521556 | T | C | NULL | 0.622459331 | NULL&M_Disease |
| 11 | 120155549 | rs78952625 | T | C | NULL | 0.622459331 | NULL&M_Disease |
| 11 | 120581395 | rs11217988 | C | T | NULL | 0.622459331 | NULL&M_Disease |
| 11 | 116475411 | rs78893389 | T | C | NULL | 0.622459331 | NULL&M_Disease |
| 11 | 116773653 | rs7943309 | A | G | NULL | 0.622459331 | NULL&M_Disease |
| 11 | 116929384 | rs17581874 | C | A | NULL | 0.622459331 | NULL&M_Disease |
| 11 | 118333510 | rs112636863 | C | T | NULL | 0.622459331 | NULL&M_Disease |
| 11 | 119412939 | rs11826230 | T | G | NULL | 0.622459331 | NULL&M_Disease |
| 11 | 116985641 | rs7925835 | G | T | NULL | 0.622459331 | NULL&M_Disease |
| 11 | 116987964 | rs75210049 | A | G | NULL | 0.622459331 | NULL&M_Disease |
| 11 | 116987988 | rs80092204 | C | T | NULL | 0.622459331 | NULL&M_Disease |
| 11 | 116988040 | rs1379735938 | T | A | NULL | 0.622459331 | NULL&M_Disease |
| 11 | 116989524 | rs61905460 | T | C | NULL | 0.622459331 | NULL&M_Disease |
| 11 | 118050404 | rs7342287 | A | G | NULL | 0.622459331 | NULL&M_Disease |
| 11 | 119771968 | rs517918 | C | T | NULL | 0.622459331 | NULL&M_Disease |
| 11 | 120583983 | rs11217989 | C | T | NULL | 0.622459331 | NULL&M_Disease |
| 11 | 120776001 | rs2156634 | G | A | NULL | 0.622459331 | NULL&M_Disease |
| 11 | 117044603 | rs7112577 | G | C | NULL | 0.622459331 | NULL&M_Disease |
| 11 | 117585602 | rs7930469 | C | T | NULL | 0.622459331 | NULL&M_Disease |
| 11 | 117591480 | rs73598666 | T | G | NULL | 0.622459331 | NULL&M_Disease |
| 11 | 117890685 | rs73011741 | G | A | NULL | 0.622459331 | NULL&M_Disease |
| 11 | 120786892 | rs2156633 | C | T | NULL | 0.622459331 | NULL&M_Disease |

| | | | | | | | | | |
|---|---|---|---|---|---|---|---|---|---|
| 11 | 118056823 | rs7935492 | T | C | 0.963 | -7977.388 | -7978.382 | -0.993 | 1.43E-06 |
| 11 | 118057783 | rs12282334 | G | C | 0.963 | -7977.388 | -7978.382 | -0.993 | 1.43E-06 |
| 11 | 118059914 | rs7934292 | C | T | 0.963 | -7977.388 | -7978.382 | -0.993 | 1.43E-06 |
| 11 | 119794526 | rs77708326 | C | T | 1.045 | -7977.388 | -7978.381 | -0.993 | 1.43E-06 |
| 11 | 119838717 | rs79652828 | T | C | 1.045 | -7977.388 | -7978.381 | -0.993 | 1.43E-06 |
| 11 | 119839219 | rs77436344 | A | G | 1.045 | -7977.388 | -7978.381 | -0.993 | 1.43E-06 |
| 11 | 120122628 | rs112667243 | T | C | 0.987 | -7977.388 | -7978.381 | -0.993 | 1.43E-06 |
| 11 | 120144941 | rs57081718 | A | G | 0.982 | -7977.388 | -7978.381 | -0.993 | 1.43E-06 |
| 11 | 120161987 | rs78991803 | A | G | 1.045 | -7977.388 | -7978.381 | -0.993 | 1.43E-06 |
| 11 | 116414488 | rs1289103681 | C | T | 1.041 | -7977.388 | -7978.382 | -0.994 | 1.43E-06 |
| 11 | 116521748 | rs3016352 | G | A | 0.961 | -7977.388 | -7978.382 | -0.994 | 1.43E-06 |
| 11 | 116525229 | rs545108 | C | G | 0.961 | -7977.388 | -7978.382 | -0.994 | 1.43E-06 |
| 11 | 116622722 | rs55713675 | T | C | 0.958 | -7977.388 | -7978.383 | -0.994 | 1.43E-06 |
| 11 | 117040739 | rs61905518 | G | A | 1.001 | -7977.388 | -7978.382 | -0.994 | 1.43E-06 |
| 11 | 118050958 | rs75709684 | A | G | 0.963 | -7977.388 | -7978.382 | -0.994 | 1.43E-06 |
| 11 | 118053859 | rs75459003 | C | A | 0.963 | -7977.388 | -7978.382 | -0.994 | 1.43E-06 |
| 11 | 118368665 | rs7107305 | G | A | 1.039 | -7977.388 | -7978.382 | -0.994 | 1.43E-06 |
| 11 | 118368822 | rs7123835 | G | T | 1.039 | -7977.388 | -7978.382 | -0.994 | 1.43E-06 |
| 11 | 119686252 | rs73008610 | C | G | 1.044 | -7977.388 | -7978.383 | -0.994 | 1.43E-06 |
| 11 | 120349142 | rs7120040 | G | A | 1.043 | -7977.388 | -7978.382 | -0.994 | 1.43E-06 |
| 11 | 116629949 | rs4938310 | T | C | 1.044 | -7977.388 | -7978.383 | -0.995 | 1.43E-06 |
| 11 | 117023487 | rs7128382 | A | C | 0.957 | -7977.388 | -7978.383 | -0.995 | 1.43E-06 |
| 11 | 117741853 | rs36086368 | C | T | 0.992 | -7977.388 | -7978.383 | -0.995 | 1.43E-06 |
| 11 | 117849065 | rs11216664 | C | G | 0.963 | -7977.388 | -7978.384 | -0.995 | 1.43E-06 |
| 11 | 119039028 | rs56230940 | T | C | 1.043 | -7977.388 | -7978.383 | -0.995 | 1.43E-06 |
| 11 | 120117394 | rs118125468 | A | G | 1.029 | -7977.388 | -7978.384 | -0.995 | 1.43E-06 |
| 11 | 120171371 | rs61898318 | A | G | 1.045 | -7977.388 | -7978.384 | -0.995 | 1.43E-06 |
| 11 | 117058854 | rs61905525 | A | G | 1.003 | -7977.388 | -7978.384 | -0.996 | 1.43E-06 |
| 11 | 117106069 | rs185433085 | T | C | 1.033 | -7977.388 | -7978.384 | -0.996 | 1.43E-06 |
| 11 | 118050779 | rs67163042 | G | A | 0.963 | -7977.388 | -7978.384 | -0.996 | 1.43E-06 |
| 11 | 118462554 | rs11216920 | G | A | 0.956 | -7977.388 | -7978.384 | -0.996 | 1.43E-06 |
| 11 | 119240757 | rs12419879 | T | C | 0.959 | -7977.388 | -7978.384 | -0.996 | 1.43E-06 |
| 11 | 120117335 | rs73576276 | T | C | 0.988 | -7977.388 | -7978.384 | -0.996 | 1.43E-06 |
| 11 | 116873382 | rs74601470 | T | C | 1.008 | -7977.388 | -7978.385 | -0.997 | 1.43E-06 |
| 11 | 117572288 | rs7101613 | G | A | 0.957 | -7977.388 | -7978.385 | -0.997 | 1.43E-06 |
| 11 | 117651600 | rs75693006 | A | G | 0.956 | -7977.388 | -7978.385 | -0.997 | 1.43E-06 |
| 11 | 118035087 | rs624328 | G | A | 0.965 | -7977.388 | -7978.385 | -0.997 | 1.43E-06 |
| 11 | 118804266 | rs7123066 | T | C | 1.038 | -7977.388 | -7978.385 | -0.997 | 1.43E-06 |
| 11 | 118804283 | rs7122959 | G | A | 1.038 | -7977.388 | -7978.385 | -0.997 | 1.43E-06 |
| 11 | 120496664 | rs4936533 | T | C | 1.042 | -7977.388 | -7978.385 | -0.997 | 1.43E-06 |
| 11 | 116879636 | rs77832134 | G | A | 1.007 | -7977.388 | -7978.386 | -0.998 | 1.43E-06 |
| 11 | 116990934 | rs7938470 | A | G | 0.991 | -7977.388 | -7978.386 | -0.998 | 1.43E-06 |
| 11 | 118044740 | rs11216794 | A | T | 0.963 | -7977.388 | -7978.387 | -0.998 | 1.43E-06 |
| 11 | 119509118 | rs34163098 | A | G | 1.041 | -7977.388 | -7978.386 | -0.998 | 1.43E-06 |
| 11 | 121064264 | rs4936593 | A | G | 0.96 | -7977.388 | -7978.386 | -0.998 | 1.43E-06 |
| 11 | 116721405 | rs111894427 | T | C | 1.003 | -7977.388 | -7978.387 | -0.999 | 1.42E-06 |
| 11 | 116982821 | rs10892075 | T | C | 1.044 | -7977.388 | -7978.387 | -0.999 | 1.42E-06 |
| 11 | 117410852 | rs12788483 | C | G | 0.956 | -7977.388 | -7978.388 | -0.999 | 1.42E-06 |
| 11 | 118052662 | rs74395230 | T | C | 0.964 | -7977.388 | -7978.387 | -0.999 | 1.42E-06 |
| 11 | 118792278 | rs77398170 | G | A | 1.032 | -7977.388 | -7978.387 | -0.999 | 1.42E-06 |
| 11 | 116726796 | rs61907561 | G | C | 0.995 | -7977.388 | -7978.388 | -1 | 1.42E-06 |
| 11 | 117101574 | rs522645 | A | C | 1.043 | -7977.388 | -7978.388 | -1 | 1.42E-06 |
| 11 | 118048885 | rs11216800 | G | A | 0.964 | -7977.388 | -7978.388 | -1 | 1.42E-06 |
| 11 | 120155303 | rs4245211 | G | A | 1.043 | -7977.388 | -7978.388 | -1 | 1.42E-06 |
| 11 | 116278812 | rs575623 | C | T | 0.956 | -7977.388 | -7978.389 | -1.001 | 1.42E-06 |
| 11 | 116632787 | rs7941518 | G | A | 1.044 | -7977.388 | -7978.389 | -1.001 | 1.42E-06 |
| 11 | 116632975 | rs7942054 | A | G | 1.044 | -7977.388 | -7978.389 | -1.001 | 1.42E-06 |
| 11 | 116810655 | rs61907602 | C | G | 1.004 | -7977.388 | -7978.389 | -1.001 | 1.42E-06 |
| 11 | 116815762 | rs7941912 | A | G | 1.004 | -7977.388 | -7978.389 | -1.001 | 1.42E-06 |
| 11 | 118346599 | rs9332781 | G | A | 1.038 | -7977.388 | -7978.389 | -1.001 | 1.42E-06 |
| 11 | 120123145 | rs113348846 | A | G | 0.991 | -7977.388 | -7978.389 | -1.001 | 1.42E-06 |
| 11 | 120157850 | rs17311233 | T | C | 1.044 | -7977.388 | -7978.389 | -1.001 | 1.42E-06 |
| 11 | 116646820 | rs140575350 | G | C | 0.993 | -7977.388 | -7978.391 | -1.002 | 1.42E-06 |
| 11 | 116837823 | rs61905682 | C | G | 1.002 | -7977.388 | -7978.39 | -1.002 | 1.42E-06 |
| 11 | 116989185 | rs61903459 | G | A | 0.995 | -7977.388 | -7978.39 | -1.002 | 1.42E-06 |
| 11 | 117216213 | rs580119 | G | T | 0.959 | -7977.388 | -7978.39 | -1.002 | 1.42E-06 |
| 11 | 117589939 | rs7102666 | C | G | 0.968 | -7977.388 | -7978.391 | -1.002 | 1.42E-06 |
| 11 | 117753244 | rs73020446 | T | C | 1.003 | -7977.388 | -7978.39 | -1.002 | 1.42E-06 |
| 11 | 117967798 | rs9651685 | C | T | 1.041 | -7977.388 | -7978.391 | -1.002 | 1.42E-06 |
| 11 | 119622367 | rs112186392 | G | A | 0.956 | -7977.388 | -7978.39 | -1.002 | 1.42E-06 |
| 11 | 120924671 | rs17124739 | G | A | 1.043 | -7977.388 | -7978.391 | -1.002 | 1.42E-06 |
| 11 | 116354466 | rs549298 | G | A | 1.042 | -7977.388 | -7978.392 | -1.003 | 1.42E-06 |
| 11 | 117996885 | rs7104443 | G | C | 0.975 | -7977.388 | -7978.392 | -1.003 | 1.42E-06 |
| 11 | 119697626 | rs2028960 | T | C | 1.044 | -7977.388 | -7978.391 | -1.003 | 1.42E-06 |
| 11 | 120036997 | rs521556 | T | C | 0.957 | -7977.388 | -7978.391 | -1.003 | 1.42E-06 |
| 11 | 120155549 | rs78952625 | T | C | 1.043 | -7977.388 | -7978.391 | -1.003 | 1.42E-06 |
| 11 | 120581395 | rs11217988 | C | T | 0.957 | -7977.388 | -7978.391 | -1.003 | 1.42E-06 |
| 11 | 116475411 | rs78893389 | T | C | 0.961 | -7977.388 | -7978.393 | -1.004 | 1.42E-06 |
| 11 | 116773653 | rs7943309 | A | G | 0.999 | -7977.388 | -7978.392 | -1.004 | 1.42E-06 |
| 11 | 116929384 | rs17581874 | C | A | 1.002 | -7977.388 | -7978.392 | -1.004 | 1.42E-06 |
| 11 | 118333510 | rs112636863 | C | T | 1.038 | -7977.388 | -7978.392 | -1.004 | 1.42E-06 |
| 11 | 119412939 | rs11826230 | T | G | 1.045 | -7977.388 | -7978.393 | -1.004 | 1.42E-06 |
| 11 | 116985641 | rs7925835 | G | T | 0.998 | -7977.388 | -7978.393 | -1.005 | 1.42E-06 |
| 11 | 116987964 | rs75210049 | A | G | 1 | -7977.388 | -7978.394 | -1.005 | 1.42E-06 |
| 11 | 116987988 | rs80092204 | C | T | 0.999 | -7977.388 | -7978.393 | -1.005 | 1.42E-06 |
| 11 | 116988040 | rs1379735938 | T | A | 0.999 | -7977.388 | -7978.393 | -1.005 | 1.42E-06 |
| 11 | 116989524 | rs61905460 | T | C | 1 | -7977.388 | -7978.393 | -1.005 | 1.42E-06 |
| 11 | 118050404 | rs7342287 | A | G | 0.964 | -7977.388 | -7978.393 | -1.005 | 1.42E-06 |
| 11 | 119771968 | rs517918 | C | T | 1.045 | -7977.388 | -7978.394 | -1.005 | 1.42E-06 |
| 11 | 120583983 | rs11217989 | C | T | 0.957 | -7977.388 | -7978.393 | -1.005 | 1.42E-06 |
| 11 | 120776001 | rs2156634 | G | A | 1.045 | -7977.388 | -7978.393 | -1.005 | 1.42E-06 |
| 11 | 117044603 | rs7112577 | G | C | 1.009 | -7977.388 | -7978.394 | -1.006 | 1.41E-06 |
| 11 | 117585602 | rs7930469 | C | T | 0.969 | -7977.388 | -7978.394 | -1.006 | 1.41E-06 |
| 11 | 117591480 | rs73598666 | T | G | 0.969 | -7977.388 | -7978.394 | -1.006 | 1.41E-06 |
| 11 | 117890685 | rs73011741 | G | A | 1.042 | -7977.388 | -7978.395 | -1.006 | 1.41E-06 |
| 11 | 120786892 | rs2156633 | C | T | 1.042 | -7977.388 | -7978.394 | -1.006 | 1.41E-06 |

| | | | | | | | |
|---|---|---|---|---|---|---|---|
| 11 | 116279191 | rs4938271 | C | T | NULL | 0.622459331 | NULL&M_Disease |
| 11 | 116312246 | rs17119659 | C | G | NULL | 0.622459331 | NULL&M_Disease |
| 11 | 116733565 | rs61907563 | A | G | NULL | 0.622459331 | NULL&M_Disease |
| 11 | 117126493 | rs112357006 | A | G | NULL | 0.622459331 | NULL&M_Disease |
| 11 | 117557886 | rs721487 | T | G | NULL | 0.622459331 | NULL&M_Disease |
| 11 | 117982289 | rs10892218 | T | G | NULL | 0.622459331 | NULL&M_Disease |
| 11 | 118354458 | rs9332793 | A | G | NULL | 0.622459331 | NULL&M_Disease |
| 11 | 118354505 | rs9332794 | T | G | NULL | 0.622459331 | NULL&M_Disease |
| 11 | 118354750 | rs200069252 | A | T | NULL | 0.622459331 | NULL&M_Disease |
| 11 | 118903039 | rs10892324 | G | T | NULL | 0.622459331 | NULL&M_Disease |
| 11 | 119631281 | rs7128327 | G | A | NULL | 0.622459331 | NULL&M_Disease |
| 11 | 119808864 | rs10892499 | A | G | NULL | 0.622459331 | NULL&M_Disease |
| 11 | 119823440 | rs512393 | C | G | NULL | 0.622459331 | NULL&M_Disease |
| 11 | 120153414 | rs11217787 | A | T | NULL | 0.622459331 | NULL&M_Disease |
| 11 | 120819939 | rs1893818 | G | A | NULL | 0.622459331 | NULL&M_Disease |
| 11 | 116539025 | rs61906084 | G | C | NULL | 0.622459331 | NULL&M_Disease |
| 11 | 116662579 | rs651821 | C | T | NULL | 0.622459331 | NULL&M_Disease |
| 11 | 118048836 | rs6589649 | C | T | NULL | 0.622459331 | NULL&M_Disease |
| 11 | 119697261 | rs61016803 | A | T | NULL | 0.622459331 | NULL&M_Disease |
| 11 | 121089364 | rs79751425 | A | G | NULL | 0.622459331 | NULL&M_Disease |
| 11 | 121164050 | rs74900182 | T | C | NULL | 0.622459331 | NULL&M_Disease |
| 11 | 117106458 | rs146786560 | C | T | NULL | 0.622459331 | NULL&M_Disease |
| 11 | 117757116 | rs67674314 | C | T | NULL | 0.622459331 | NULL&M_Disease |
| 11 | 117832675 | rs10892199 | A | G | NULL | 0.623633628 | NULL&M_Disease |
| 11 | 118387892 | rs111997050 | T | C | NULL | 0.622459331 | NULL&M_Disease |
| 11 | 120337712 | rs80040504 | A | G | NULL | 0.622459331 | NULL&M_Disease |
| 11 | 120470867 | rs4936526 | A | G | NULL | 0.623633628 | NULL&M_Disease |
| 11 | 116554228 | rs61906109 | A | C | NULL | 0.622459331 | NULL&M_Disease |
| 11 | 116628271 | rs73588403 | T | C | NULL | 0.623633628 | NULL&M_Disease |
| 11 | 117875171 | rs4938467 | T | C | NULL | 0.623633628 | NULL&M_Disease |
| 11 | 118051323 | rs7927676 | G | C | NULL | 0.622459331 | NULL&M_Disease |
| 11 | 118051857 | rs76050019 | C | T | NULL | 0.622459331 | NULL&M_Disease |
| 11 | 119684793 | rs34004290 | C | G | NULL | 0.622459331 | NULL&M_Disease |
| 11 | 116352008 | rs3017780 | G | T | NULL | 0.623633628 | NULL&M_Disease |
| 11 | 116539637 | rs61906086 | G | A | NULL | 0.622459331 | NULL&M_Disease |
| 11 | 116540785 | rs58208536 | C | G | NULL | 0.622459331 | NULL&M_Disease |
| 11 | 116540916 | rs58346469 | T | C | NULL | 0.622459331 | NULL&M_Disease |
| 11 | 116541268 | rs56196414 | A | G | NULL | 0.622459331 | NULL&M_Disease |
| 11 | 116541656 | rs61906100 | A | C | NULL | 0.622459331 | NULL&M_Disease |
| 11 | 117657679 | rs7122817 | A | G | NULL | 0.623633628 | NULL&M_Disease |
| 11 | 117865381 | rs2282495 | T | C | NULL | 0.623633628 | NULL&M_Disease |
| 11 | 117997360 | rs72488011 | C | T | NULL | 0.622459331 | NULL&M_Disease |
| 11 | 118902609 | rs10790280 | T | A | NULL | 0.623633628 | NULL&M_Disease |
| 11 | 119301608 | rs10892364 | T | C | NULL | 0.623633628 | NULL&M_Disease |
| 11 | 119776105 | rs11217558 | T | G | NULL | 0.623633628 | NULL&M_Disease |
| 11 | 120349310 | rs7120174 | C | A | NULL | 0.623633628 | NULL&M_Disease |
| 11 | 116532855 | rs7107003 | C | T | NULL | 0.623633628 | NULL&M_Disease |
| 11 | 116539090 | rs61906085 | G | A | NULL | 0.623633628 | NULL&M_Disease |
| 11 | 117026154 | rs12271743 | C | T | NULL | 0.622459331 | NULL&M_Disease |
| 11 | 117669794 | rs1892902 | T | G | NULL | 0.623633628 | NULL&M_Disease |
| 11 | 118365263 | rs9332819 | T | C | NULL | 0.623633628 | NULL&M_Disease |
| 11 | 117330669 | rs949138 | G | A | NULL | 0.623633628 | NULL&M_Disease |
| 11 | 117538907 | rs71480353 | C | G | NULL | 0.623633628 | NULL&M_Disease |
| 11 | 118063349 | rs113521822 | G | A | NULL | 0.623633628 | NULL&M_Disease |
| 11 | 118326445 | rs112900272 | T | C | NULL | 0.623633628 | NULL&M_Disease |
| 11 | 119033850 | rs73564406 | A | G | NULL | 0.623633628 | NULL&M_Disease |
| 11 | 119697584 | rs11217527 | A | G | NULL | 0.623633628 | NULL&M_Disease |
| 11 | 120797893 | rs2850807 | T | A | NULL | 0.623633628 | NULL&M_Disease |
| 11 | 117026885 | rs4936366 | G | A | NULL | 0.623633628 | NULL&M_Disease |
| 11 | 117853333 | rs150325557 | C | G | NULL | 0.623633628 | NULL&M_Disease |
| 11 | 117859925 | rs4252253 | A | C | NULL | 0.623633628 | NULL&M_Disease |
| 11 | 120475940 | rs4367952 | T | C | NULL | 0.623633628 | NULL&M_Disease |
| 11 | 116546161 | rs80068772 | T | A | NULL | 0.623633628 | NULL&M_Disease |
| 11 | 117775361 | rs115946185 | A | T | NULL | 0.623633628 | NULL&M_Disease |
| 11 | 117775374 | rs79347839 | A | T | NULL | 0.623633628 | NULL&M_Disease |
| 11 | 118051010 | rs79820203 | C | T | NULL | 0.623633628 | NULL&M_Disease |
| 11 | 118355642 | rs9332801 | C | A | NULL | 0.623633628 | NULL&M_Disease |
| 11 | 118902749 | rs10892322 | T | C | NULL | 0.623633628 | NULL&M_Disease |
| 11 | 119622613 | rs11217462 | G | A | NULL | 0.623633628 | NULL&M_Disease |
| 11 | 116662407 | rs3135506 | C | G | NULL | 0.623633628 | NULL&M_Disease |
| 11 | 117205844 | rs487298 | T | G | NULL | 0.623633628 | NULL&M_Disease |
| 11 | 117226773 | rs61903740 | A | C | NULL | 0.623633628 | NULL&M_Disease |
| 11 | 117226941 | rs61903741 | A | T | NULL | 0.623633628 | NULL&M_Disease |
| 11 | 117595260 | rs4936401 | C | T | NULL | 0.623633628 | NULL&M_Disease |
| 11 | 117645002 | rs74835404 | A | G | NULL | 0.623633628 | NULL&M_Disease |
| 11 | 118328052 | rs111901186 | G | A | NULL | 0.623633628 | NULL&M_Disease |
| 11 | 119030812 | rs4148173 | G | A | NULL | 0.623633628 | NULL&M_Disease |
| 11 | 119683659 | rs56363339 | G | A | NULL | 0.623633628 | NULL&M_Disease |
| 11 | 120350393 | rs12362794 | G | T | NULL | 0.623633628 | NULL&M_Disease |
| 11 | 117818515 | rs79723893 | G | A | NULL | 0.623633628 | NULL&M_Disease |
| 11 | 118026675 | rs2155854 | T | C | NULL | 0.623633628 | NULL&M_Disease |
| 11 | 120131001 | rs17123753 | G | C | NULL | 0.623633628 | NULL&M_Disease |
| 11 | 116478660 | rs3016358 | A | G | NULL | 0.623633628 | NULL&M_Disease |
| 11 | 117320976 | rs500713 | C | T | NULL | 0.623633628 | NULL&M_Disease |
| 11 | 117606937 | rs7951237 | A | T | NULL | 0.623633628 | NULL&M_Disease |
| 11 | 117607138 | rs1942930 | C | T | NULL | 0.623633628 | NULL&M_Disease |
| 11 | 117946386 | rs604411 | T | C | NULL | 0.623633628 | NULL&M_Disease |
| 11 | 117946720 | rs1786190 | G | A | NULL | 0.623633628 | NULL&M_Disease |
| 11 | 117976332 | rs10790232 | A | C | NULL | 0.623633628 | NULL&M_Disease |
| 11 | 117977388 | rs10892214 | G | C | NULL | 0.623633628 | NULL&M_Disease |
| 11 | 117997374 | rs17092732 | A | G | NULL | 0.623633628 | NULL&M_Disease |
| 11 | 118358152 | rs148643334 | C | T | NULL | 0.623633628 | NULL&M_Disease |
| 11 | 119034374 | rs55797357 | T | C | NULL | 0.623633628 | NULL&M_Disease |
| 11 | 120182796 | rs61898342 | A | G | NULL | 0.623633628 | NULL&M_Disease |
| 11 | 116335869 | rs74676521 | A | C | NULL | 0.623633628 | NULL&M_Disease |
| 11 | 116399067 | rs12280027 | G | C | NULL | 0.624806474 | NULL&M_Disease |

| | | | | | | | | | |
|---|---|---|---|---|---|---|---|---|---|
| 11 | 116279191 | rs4938271 | C | T | 0.957 | -7977.388 | -7978.395 | -1.007 | 1.41E-06 |
| 11 | 116312246 | rs17119659 | C | G | 1.041 | -7977.388 | -7978.395 | -1.007 | 1.41E-06 |
| 11 | 116733565 | rs61907563 | A | G | 0.998 | -7977.388 | -7978.395 | -1.007 | 1.41E-06 |
| 11 | 117126493 | rs112357006 | A | G | 0.973 | -7977.388 | -7978.395 | -1.007 | 1.41E-06 |
| 11 | 117557886 | rs721487 | T | G | 0.957 | -7977.388 | -7978.395 | -1.007 | 1.41E-06 |
| 11 | 117982289 | rs10892218 | T | G | 0.978 | -7977.388 | -7978.396 | -1.007 | 1.41E-06 |
| 11 | 118354458 | rs9332793 | A | G | 1.038 | -7977.388 | -7978.395 | -1.007 | 1.41E-06 |
| 11 | 118354505 | rs9332794 | T | G | 1.038 | -7977.388 | -7978.395 | -1.007 | 1.41E-06 |
| 11 | 118354750 | rs200069252 | A | T | 1.038 | -7977.388 | -7978.395 | -1.007 | 1.41E-06 |
| 11 | 118903039 | rs10892324 | G | T | 0.961 | -7977.388 | -7978.396 | -1.007 | 1.41E-06 |
| 11 | 119631281 | rs7128327 | G | A | 0.956 | -7977.388 | -7978.396 | -1.007 | 1.41E-06 |
| 11 | 119808864 | rs10892499 | A | G | 1.045 | -7977.388 | -7978.395 | -1.007 | 1.41E-06 |
| 11 | 119823440 | rs512393 | C | G | 0.958 | -7977.388 | -7978.395 | -1.007 | 1.41E-06 |
| 11 | 120153414 | rs11217787 | A | T | 1.042 | -7977.388 | -7978.396 | -1.007 | 1.41E-06 |
| 11 | 120819939 | rs1893818 | G | A | 0.959 | -7977.388 | -7978.395 | -1.007 | 1.41E-06 |
| 11 | 116539025 | rs61906084 | G | C | 0.967 | -7977.388 | -7978.396 | -1.008 | 1.41E-06 |
| 11 | 116662579 | rs651821 | C | T | 0.96 | -7977.388 | -7978.396 | -1.008 | 1.41E-06 |
| 11 | 118048836 | rs6589649 | C | T | 0.964 | -7977.388 | -7978.397 | -1.008 | 1.41E-06 |
| 11 | 119697261 | rs61016803 | A | T | 1.043 | -7977.388 | -7978.396 | -1.008 | 1.41E-06 |
| 11 | 121089364 | rs79751425 | A | G | 1.015 | -7977.388 | -7978.396 | -1.008 | 1.41E-06 |
| 11 | 121164050 | rs74900182 | T | C | 0.958 | -7977.388 | -7978.396 | -1.008 | 1.41E-06 |
| 11 | 117106458 | rs146786560 | C | T | 1.031 | -7977.388 | -7978.397 | -1.009 | 1.41E-06 |
| 11 | 117757116 | rs67674314 | C | T | 1.009 | -7977.388 | -7978.397 | -1.009 | 1.41E-06 |
| 11 | 117832675 | rs10892199 | A | G | 0.957 | -7977.388 | -7978.398 | -1.009 | 1.41E-06 |
| 11 | 118387892 | rs111997050 | T | C | 1.038 | -7977.388 | -7978.398 | -1.009 | 1.41E-06 |
| 11 | 120337712 | rs80040504 | A | G | 0.964 | -7977.388 | -7978.397 | -1.009 | 1.41E-06 |
| 11 | 120470867 | rs4936526 | A | G | 0.96 | -7977.388 | -7978.398 | -1.009 | 1.41E-06 |
| 11 | 116554228 | rs61906109 | A | C | 0.967 | -7977.388 | -7978.398 | -1.01 | 1.41E-06 |
| 11 | 116628271 | rs73588403 | T | C | 0.959 | -7977.388 | -7978.398 | -1.01 | 1.41E-06 |
| 11 | 117875171 | rs4938467 | T | C | 1.04 | -7977.388 | -7978.398 | -1.01 | 1.41E-06 |
| 11 | 118051323 | rs7927676 | G | C | 0.965 | -7977.388 | -7978.399 | -1.01 | 1.41E-06 |
| 11 | 118051857 | rs76050019 | C | T | 0.965 | -7977.388 | -7978.399 | -1.01 | 1.41E-06 |
| 11 | 119684793 | rs34004290 | C | G | 1.015 | -7977.388 | -7978.398 | -1.01 | 1.41E-06 |
| 11 | 116352008 | rs3017780 | G | T | 1.041 | -7977.388 | -7978.399 | -1.011 | 1.41E-06 |
| 11 | 116539637 | rs61906086 | G | A | 0.967 | -7977.388 | -7978.399 | -1.011 | 1.41E-06 |
| 11 | 116540785 | rs58208536 | C | G | 0.967 | -7977.388 | -7978.399 | -1.011 | 1.41E-06 |
| 11 | 116540916 | rs58346469 | T | C | 0.967 | -7977.388 | -7978.399 | -1.011 | 1.41E-06 |
| 11 | 116541268 | rs56196414 | A | G | 0.967 | -7977.388 | -7978.399 | -1.011 | 1.41E-06 |
| 11 | 116541656 | rs61906100 | A | C | 0.967 | -7977.388 | -7978.399 | -1.011 | 1.41E-06 |
| 11 | 117657679 | rs7122817 | A | G | 1.04 | -7977.388 | -7978.399 | -1.011 | 1.41E-06 |
| 11 | 117865381 | rs2282495 | T | C | 0.966 | -7977.388 | -7978.4 | -1.011 | 1.41E-06 |
| 11 | 117997360 | rs72488011 | C | T | 0.98 | -7977.388 | -7978.399 | -1.011 | 1.41E-06 |
| 11 | 118902609 | rs10790280 | T | A | 0.962 | -7977.388 | -7978.399 | -1.011 | 1.41E-06 |
| 11 | 119301608 | rs10892364 | T | C | 0.958 | -7977.388 | -7978.399 | -1.011 | 1.41E-06 |
| 11 | 119776105 | rs11217558 | T | G | 0.961 | -7977.388 | -7978.4 | -1.011 | 1.41E-06 |
| 11 | 120349310 | rs7120174 | C | A | 1.043 | -7977.388 | -7978.399 | -1.011 | 1.41E-06 |
| 11 | 116532855 | rs7107003 | C | T | 0.958 | -7977.388 | -7978.4 | -1.012 | 1.41E-06 |
| 11 | 116539090 | rs61906085 | G | A | 0.968 | -7977.388 | -7978.401 | -1.012 | 1.41E-06 |
| 11 | 117026154 | rs12271743 | C | T | 1.002 | -7977.388 | -7978.4 | -1.012 | 1.41E-06 |
| 11 | 117669794 | rs1892902 | T | G | 1.04 | -7977.388 | -7978.4 | -1.012 | 1.41E-06 |
| 11 | 118365263 | rs9332819 | T | C | 1.038 | -7977.388 | -7978.4 | -1.012 | 1.41E-06 |
| 11 | 117330669 | rs949138 | G | A | 0.957 | -7977.388 | -7978.401 | -1.013 | 1.40E-06 |
| 11 | 117538907 | rs71480353 | C | G | 0.979 | -7977.388 | -7978.401 | -1.013 | 1.40E-06 |
| 11 | 118063349 | rs113521822 | G | A | 0.969 | -7977.388 | -7978.402 | -1.013 | 1.40E-06 |
| 11 | 118326445 | rs112900272 | T | C | 1.037 | -7977.388 | -7978.401 | -1.013 | 1.40E-06 |
| 11 | 119033850 | rs73564406 | A | G | 1.041 | -7977.388 | -7978.401 | -1.013 | 1.40E-06 |
| 11 | 119697584 | rs11217527 | A | G | 0.957 | -7977.388 | -7978.401 | -1.013 | 1.40E-06 |
| 11 | 120797893 | rs2850807 | T | A | 0.964 | -7977.388 | -7978.401 | -1.013 | 1.40E-06 |
| 11 | 117026885 | rs4936366 | G | A | 0.958 | -7977.388 | -7978.403 | -1.014 | 1.40E-06 |
| 11 | 117853333 | rs150325557 | C | G | 0.966 | -7977.388 | -7978.402 | -1.014 | 1.40E-06 |
| 11 | 117859925 | rs4252253 | A | C | 0.957 | -7977.388 | -7978.402 | -1.014 | 1.40E-06 |
| 11 | 120475940 | rs4367952 | T | C | 0.96 | -7977.388 | -7978.402 | -1.014 | 1.40E-06 |
| 11 | 116546161 | rs80068772 | T | A | 0.968 | -7977.388 | -7978.403 | -1.015 | 1.40E-06 |
| 11 | 117775361 | rs115946185 | A | T | 0.957 | -7977.388 | -7978.403 | -1.015 | 1.40E-06 |
| 11 | 117775374 | rs79347839 | A | T | 0.957 | -7977.388 | -7978.403 | -1.015 | 1.40E-06 |
| 11 | 118051010 | rs79820203 | C | T | 0.965 | -7977.388 | -7978.404 | -1.015 | 1.40E-06 |
| 11 | 118355642 | rs9332801 | C | A | 1.037 | -7977.388 | -7978.404 | -1.015 | 1.40E-06 |
| 11 | 118902749 | rs10892322 | T | C | 0.962 | -7977.388 | -7978.404 | -1.015 | 1.40E-06 |
| 11 | 119622613 | rs11217462 | G | A | 0.957 | -7977.388 | -7978.404 | -1.015 | 1.40E-06 |
| 11 | 116662407 | rs3135506 | C | G | 0.96 | -7977.388 | -7978.404 | -1.016 | 1.40E-06 |
| 11 | 117205844 | rs487298 | T | G | 0.959 | -7977.388 | -7978.404 | -1.016 | 1.40E-06 |
| 11 | 117226773 | rs61903740 | A | C | 0.995 | -7977.388 | -7978.404 | -1.016 | 1.40E-06 |
| 11 | 117226941 | rs61903741 | A | T | 0.995 | -7977.388 | -7978.404 | -1.016 | 1.40E-06 |
| 11 | 117595260 | rs4936401 | C | T | 0.961 | -7977.388 | -7978.404 | -1.016 | 1.40E-06 |
| 11 | 117645002 | rs74835404 | A | G | 0.959 | -7977.388 | -7978.404 | -1.016 | 1.40E-06 |
| 11 | 118328052 | rs111901186 | G | A | 1.037 | -7977.388 | -7978.404 | -1.016 | 1.40E-06 |
| 11 | 119030812 | rs4148173 | G | A | 1.041 | -7977.388 | -7978.404 | -1.016 | 1.40E-06 |
| 11 | 119683659 | rs56363339 | G | A | 1.013 | -7977.388 | -7978.404 | -1.016 | 1.40E-06 |
| 11 | 120350393 | rs12362794 | G | T | 1.043 | -7977.388 | -7978.405 | -1.016 | 1.40E-06 |
| 11 | 117818515 | rs79723893 | G | A | 1.041 | -7977.388 | -7978.406 | -1.017 | 1.40E-06 |
| 11 | 118026675 | rs2155854 | T | C | 1.04 | -7977.388 | -7978.405 | -1.017 | 1.40E-06 |
| 11 | 120131001 | rs17123753 | G | C | 1.027 | -7977.388 | -7978.405 | -1.017 | 1.40E-06 |
| 11 | 116478660 | rs3016358 | A | G | 0.965 | -7977.388 | -7978.407 | -1.018 | 1.40E-06 |
| 11 | 117320976 | rs500713 | C | T | 1.04 | -7977.388 | -7978.407 | -1.018 | 1.40E-06 |
| 11 | 117606937 | rs7951237 | A | T | 0.961 | -7977.388 | -7978.406 | -1.018 | 1.40E-06 |
| 11 | 117607138 | rs1942930 | C | T | 0.961 | -7977.388 | -7978.406 | -1.018 | 1.40E-06 |
| 11 | 117946386 | rs604411 | T | C | 1.045 | -7977.388 | -7978.407 | -1.018 | 1.40E-06 |
| 11 | 117946720 | rs1786190 | G | A | 1.045 | -7977.388 | -7978.407 | -1.018 | 1.40E-06 |
| 11 | 117976332 | rs10790232 | A | C | 1.027 | -7977.388 | -7978.406 | -1.018 | 1.40E-06 |
| 11 | 117977388 | rs10892214 | G | C | 1.027 | -7977.388 | -7978.406 | -1.018 | 1.40E-06 |
| 11 | 117997374 | rs17092732 | A | G | 0.981 | -7977.388 | -7978.406 | -1.018 | 1.40E-06 |
| 11 | 118358152 | rs148643334 | C | T | 1.037 | -7977.388 | -7978.407 | -1.018 | 1.40E-06 |
| 11 | 119034374 | rs55797357 | T | C | 1.041 | -7977.388 | -7978.407 | -1.018 | 1.40E-06 |
| 11 | 120182796 | rs61898342 | A | G | 1.044 | -7977.388 | -7978.406 | -1.018 | 1.40E-06 |
| 11 | 116335869 | rs74676521 | A | C | 1.026 | -7977.388 | -7978.407 | -1.019 | 1.40E-06 |
| 11 | 116399067 | rs12280027 | G | C | 1.04 | -7977.388 | -7978.407 | -1.019 | 1.40E-06 |

| | | | | | | | |
|---|---|---|---|---|---|---|---|
| 11 | 116540310 | rs56060233 | T | C | NULL | 0.623633628 | NULL&M_Disease |
| 11 | 117020470 | rs7115562 | A | G | NULL | 0.624806474 | NULL&M_Disease |
| 11 | 117021134 | rs7116454 | A | T | NULL | 0.624806474 | NULL&M_Disease |
| 11 | 117571475 | rs12099002 | A | G | NULL | 0.624806474 | NULL&M_Disease |
| 11 | 119805189 | rs34955847 | T | C | NULL | 0.623633628 | NULL&M_Disease |
| 11 | 120036267 | rs533636 | A | G | NULL | 0.624806474 | NULL&M_Disease |
| 11 | 116604032 | rs12279180 | C | T | NULL | 0.624806474 | NULL&M_Disease |
| 11 | 116619867 | rs61905088 | A | C | NULL | 0.624806474 | NULL&M_Disease |
| 11 | 117312962 | rs10892105 | G | A | NULL | 0.624806474 | NULL&M_Disease |
| 11 | 117864846 | rs2228055 | G | A | NULL | 0.623633628 | NULL&M_Disease |
| 11 | 117986722 | rs12225133 | C | T | NULL | 0.623633628 | NULL&M_Disease |
| 11 | 119666800 | rs79475413 | A | G | NULL | 0.624806474 | NULL&M_Disease |
| 11 | 116362654 | rs61906588 | T | C | NULL | 0.624806474 | NULL&M_Disease |
| 11 | 117205766 | rs576190 | C | T | NULL | 0.624806474 | NULL&M_Disease |
| 11 | 119876680 | rs17311121 | G | T | NULL | 0.623633628 | NULL&M_Disease |
| 11 | 120161546 | rs75646928 | T | C | NULL | 0.624806474 | NULL&M_Disease |
| 11 | 116511028 | rs523549 | A | G | NULL | 0.624806474 | NULL&M_Disease |
| 11 | 117850727 | rs73009920 | T | C | NULL | 0.624806474 | NULL&M_Disease |
| 11 | 118364404 | rs9332816 | T | G | NULL | 0.624806474 | NULL&M_Disease |
| 11 | 119300712 | rs7102492 | T | C | NULL | 0.624806474 | NULL&M_Disease |
| 11 | 119496258 | rs11217356 | T | C | NULL | 0.624806474 | NULL&M_Disease |
| 11 | 120154445 | rs117028204 | G | C | NULL | 0.624806474 | NULL&M_Disease |
| 11 | 116360636 | rs61906586 | C | T | NULL | 0.624806474 | NULL&M_Disease |
| 11 | 116623674 | rs57200947 | T | C | NULL | 0.624806474 | NULL&M_Disease |
| 11 | 116633862 | rs11820589 | A | G | NULL | 0.624806474 | NULL&M_Disease |
| 11 | 117865859 | rs4252309 | C | T | NULL | 0.624806474 | NULL&M_Disease |
| 11 | 118902933 | rs11217130 | T | A | NULL | 0.624806474 | NULL&M_Disease |
| 11 | 116495714 | rs7129952 | A | T | NULL | 0.624806474 | NULL&M_Disease |
| 11 | 116511766 | rs75173807 | A | G | NULL | 0.624806474 | NULL&M_Disease |
| 11 | 116605076 | rs12274146 | A | G | NULL | 0.624806474 | NULL&M_Disease |
| 11 | 116610294 | rs61905084 | C | T | NULL | 0.624806474 | NULL&M_Disease |
| 11 | 117003528 | rs4938343 | G | C | NULL | 0.624806474 | NULL&M_Disease |
| 11 | 117122831 | rs58111676 | G | C | NULL | 0.624806474 | NULL&M_Disease |
| 11 | 117700629 | rs1302598824 | T | C | NULL | 0.624806474 | NULL&M_Disease |
| 11 | 117983054 | rs10892219 | A | G | NULL | 0.624806474 | NULL&M_Disease |
| 11 | 118051082 | rs76831657 | G | A | NULL | 0.624806474 | NULL&M_Disease |
| 11 | 120122394 | rs11827471 | G | A | NULL | 0.624806474 | NULL&M_Disease |
| 11 | 120554041 | rs4582985 | G | C | NULL | 0.624806474 | NULL&M_Disease |
| 11 | 120992188 | rs594711 | T | C | NULL | 0.624806474 | NULL&M_Disease |
| 11 | 116879676 | rs7931398 | C | T | NULL | 0.624806474 | NULL&M_Disease |
| 11 | 117085261 | rs76604009 | T | C | NULL | 0.624806474 | NULL&M_Disease |
| 11 | 117780091 | rs4439549 | A | C | NULL | 0.624806474 | NULL&M_Disease |
| 11 | 119641843 | rs11600311 | T | C | NULL | 0.624806474 | NULL&M_Disease |
| 11 | 116859675 | rs77852530 | C | A | NULL | 0.624806474 | NULL&M_Disease |
| 11 | 117434983 | rs752796 | G | A | NULL | 0.624806474 | NULL&M_Disease |
| 11 | 120505771 | rs4936535 | G | A | NULL | 0.624806474 | NULL&M_Disease |
| 11 | 116380781 | rs79487385 | G | A | NULL | 0.624806474 | NULL&M_Disease |
| 11 | 116550528 | rs61906105 | G | A | NULL | 0.624806474 | NULL&M_Disease |
| 11 | 117634408 | rs141859855 | A | G | NULL | 0.624806474 | NULL&M_Disease |
| 11 | 120158933 | rs77483431 | G | A | NULL | 0.624806474 | NULL&M_Disease |
| 11 | 120160262 | rs4938794 | A | G | NULL | 0.624806474 | NULL&M_Disease |
| 11 | 120675122 | rs1939672 | T | C | NULL | 0.624806474 | NULL&M_Disease |
| 11 | 116592203 | rs180371 | G | A | NULL | 0.625977858 | NULL&M_Disease |
| 11 | 118179197 | rs1599810 | A | G | NULL | 0.625977858 | NULL&M_Disease |
| 11 | 116359851 | rs7934455 | C | T | NULL | 0.625977858 | NULL&M_Disease |
| 11 | 116380855 | rs77229823 | G | A | NULL | 0.624806474 | NULL&M_Disease |
| 11 | 117296923 | rs59827403 | G | C | NULL | 0.624806474 | NULL&M_Disease |
| 11 | 117397482 | rs12806934 | T | C | NULL | 0.625977858 | NULL&M_Disease |
| 11 | 117881601 | rs2512149 | C | T | NULL | 0.625977858 | NULL&M_Disease |
| 11 | 117979970 | rs10502229 | T | G | NULL | 0.624806474 | NULL&M_Disease |
| 11 | 117996402 | rs79220634 | A | C | NULL | 0.624806474 | NULL&M_Disease |
| 11 | 117997427 | rs72488012 | A | G | NULL | 0.624806474 | NULL&M_Disease |
| 11 | 119300694 | rs7102696 | A | G | NULL | 0.625977858 | NULL&M_Disease |
| 11 | 116380536 | rs60524902 | G | T | NULL | 0.624806474 | NULL&M_Disease |
| 11 | 116464800 | rs11216026 | G | A | NULL | 0.625977858 | NULL&M_Disease |
| 11 | 116621963 | rs12292921 | G | T | NULL | 0.625977858 | NULL&M_Disease |
| 11 | 117155696 | rs45507395 | T | G | NULL | 0.624806474 | NULL&M_Disease |
| 11 | 118791150 | rs12270280 | T | C | NULL | 0.624806474 | NULL&M_Disease |
| 11 | 119056273 | rs73564493 | A | G | NULL | 0.624806474 | NULL&M_Disease |
| 11 | 120121323 | rs11217775 | C | T | NULL | 0.624806474 | NULL&M_Disease |
| 11 | 120586577 | rs56160707 | C | T | NULL | 0.624806474 | NULL&M_Disease |
| 11 | 116353002 | rs491428 | T | G | NULL | 0.625977858 | NULL&M_Disease |
| 11 | 116353106 | rs490526 | T | A | NULL | 0.625977858 | NULL&M_Disease |
| 11 | 116383064 | rs11599994 | A | G | NULL | 0.624806474 | NULL&M_Disease |
| 11 | 116497466 | rs12417676 | C | A | NULL | 0.625977858 | NULL&M_Disease |
| 11 | 116524965 | rs117133249 | A | G | NULL | 0.624806474 | NULL&M_Disease |
| 11 | 116701669 | rs2070667 | A | G | NULL | 0.625977858 | NULL&M_Disease |
| 11 | 117194059 | rs118086079 | A | G | NULL | 0.624806474 | NULL&M_Disease |
| 11 | 117647461 | rs73587457 | G | C | NULL | 0.625977858 | NULL&M_Disease |
| 11 | 117855922 | rs4252299 | T | G | NULL | 0.625977858 | NULL&M_Disease |
| 11 | 117989301 | rs1941626 | C | G | NULL | 0.624806474 | NULL&M_Disease |
| 11 | 117996200 | rs17092726 | G | A | NULL | 0.624806474 | NULL&M_Disease |
| 11 | 117997473 | rs79754446 | G | A | NULL | 0.624806474 | NULL&M_Disease |
| 11 | 116244267 | rs508731 | A | T | NULL | 0.625977858 | NULL&M_Disease |
| 11 | 116379681 | rs11215970 | C | T | NULL | 0.625977858 | NULL&M_Disease |
| 11 | 116552062 | rs1429049972 | T | A | NULL | 0.625977858 | NULL&M_Disease |
| 11 | 116552562 | rs7952043 | T | C | NULL | 0.625977858 | NULL&M_Disease |
| 11 | 116627056 | rs56143464 | A | G | NULL | 0.625977858 | NULL&M_Disease |
| 11 | 116628057 | rs61905112 | A | G | NULL | 0.625977858 | NULL&M_Disease |
| 11 | 117571426 | rs7130852 | G | A | NULL | 0.625977858 | NULL&M_Disease |
| 11 | 117947103 | rs2845696 | G | A | NULL | 0.625977858 | NULL&M_Disease |
| 11 | 117978103 | rs10892215 | T | C | NULL | 0.625977858 | NULL&M_Disease |
| 11 | 117978247 | rs11216765 | T | C | NULL | 0.625977858 | NULL&M_Disease |
| 11 | 118629959 | rs140110112 | A | G | NULL | 0.625977858 | NULL&M_Disease |
| 11 | 119618503 | rs11217451 | A | G | NULL | 0.625977858 | NULL&M_Disease |
| 11 | 119853252 | rs111444487 | C | T | NULL | 0.625977858 | NULL&M_Disease |

| | | | | | | | | | |
|---|---|---|---|---|---|---|---|---|---|
| 11 | 116540310 | rs56060233 | T | C | 0.969 | -7977.388 | -7978.408 | -1.019 | 1.40E-06 |
| 11 | 117020470 | rs7115562 | A | G | 0.958 | -7977.388 | -7978.408 | -1.019 | 1.40E-06 |
| 11 | 117021134 | rs7116454 | A | T | 0.958 | -7977.388 | -7978.408 | -1.019 | 1.40E-06 |
| 11 | 117571475 | rs12099002 | A | G | 0.957 | -7977.388 | -7978.407 | -1.019 | 1.40E-06 |
| 11 | 119805189 | rs34955847 | T | C | 1.03 | -7977.388 | -7978.407 | -1.019 | 1.40E-06 |
| 11 | 120036267 | rs533636 | A | G | 0.958 | -7977.388 | -7978.408 | -1.019 | 1.40E-06 |
| 11 | 116604032 | rs12279180 | C | T | 0.959 | -7977.388 | -7978.409 | -1.02 | 1.39E-06 |
| 11 | 116619867 | rs61905088 | A | C | 0.959 | -7977.388 | -7978.409 | -1.02 | 1.39E-06 |
| 11 | 117312962 | rs10892105 | G | A | 1.044 | -7977.388 | -7978.409 | -1.02 | 1.39E-06 |
| 11 | 117864846 | rs2228055 | G | A | 0.967 | -7977.388 | -7978.408 | -1.02 | 1.39E-06 |
| 11 | 117986722 | rs12225133 | C | T | 0.994 | -7977.388 | -7978.408 | -1.02 | 1.39E-06 |
| 11 | 119666800 | rs79475413 | A | G | 0.96 | -7977.388 | -7978.409 | -1.02 | 1.39E-06 |
| 11 | 116362654 | rs61906588 | T | C | 1.041 | -7977.388 | -7978.409 | -1.021 | 1.39E-06 |
| 11 | 117205766 | rs576190 | C | T | 0.959 | -7977.388 | -7978.41 | -1.021 | 1.39E-06 |
| 11 | 119876680 | rs17311121 | G | T | 0.98 | -7977.388 | -7978.409 | -1.021 | 1.39E-06 |
| 11 | 120161546 | rs75646928 | T | C | 1.042 | -7977.388 | -7978.409 | -1.021 | 1.39E-06 |
| 11 | 116511028 | rs523549 | A | G | 1.04 | -7977.388 | -7978.411 | -1.022 | 1.39E-06 |
| 11 | 117850727 | rs73009920 | T | C | 0.967 | -7977.388 | -7978.41 | -1.022 | 1.39E-06 |
| 11 | 118364404 | rs9332816 | T | G | 1.037 | -7977.388 | -7978.411 | -1.022 | 1.39E-06 |
| 11 | 119300712 | rs7102492 | T | C | 0.959 | -7977.388 | -7978.41 | -1.022 | 1.39E-06 |
| 11 | 119496258 | rs11217356 | T | C | 1.042 | -7977.388 | -7978.411 | -1.022 | 1.39E-06 |
| 11 | 120154445 | rs117028204 | G | C | 1.042 | -7977.388 | -7978.411 | -1.022 | 1.39E-06 |
| 11 | 116360636 | rs61906586 | C | T | 1.042 | -7977.388 | -7978.411 | -1.023 | 1.39E-06 |
| 11 | 116623674 | rs57200947 | T | C | 0.96 | -7977.388 | -7978.412 | -1.023 | 1.39E-06 |
| 11 | 116633862 | rs11820589 | A | G | 0.96 | -7977.388 | -7978.411 | -1.023 | 1.39E-06 |
| 11 | 117865859 | rs4252309 | C | T | 0.968 | -7977.388 | -7978.412 | -1.023 | 1.39E-06 |
| 11 | 118902933 | rs11217130 | T | A | 0.962 | -7977.388 | -7978.411 | -1.023 | 1.39E-06 |
| 11 | 116495714 | rs7129952 | A | T | 1.042 | -7977.388 | -7978.412 | -1.024 | 1.39E-06 |
| 11 | 116511766 | rs75173807 | A | G | 0.982 | -7977.388 | -7978.412 | -1.024 | 1.39E-06 |
| 11 | 116605076 | rs12274146 | A | G | 0.96 | -7977.388 | -7978.413 | -1.024 | 1.39E-06 |
| 11 | 116610294 | rs61905084 | C | T | 1.044 | -7977.388 | -7978.412 | -1.024 | 1.39E-06 |
| 11 | 117003528 | rs4938343 | G | C | 0.958 | -7977.388 | -7978.412 | -1.024 | 1.39E-06 |
| 11 | 117122831 | rs58111676 | G | C | 1.029 | -7977.388 | -7978.413 | -1.024 | 1.39E-06 |
| 11 | 117700629 | rs1302598824 | T | C | 1.043 | -7977.388 | -7978.412 | -1.024 | 1.39E-06 |
| 11 | 117983054 | rs10892219 | A | G | 0.99 | -7977.388 | -7978.412 | -1.024 | 1.39E-06 |
| 11 | 118051082 | rs76831657 | G | A | 0.966 | -7977.388 | -7978.413 | -1.024 | 1.39E-06 |
| 11 | 120122394 | rs11827471 | G | A | 1.043 | -7977.388 | -7978.412 | -1.024 | 1.39E-06 |
| 11 | 120554041 | rs4582985 | G | C | 1.04 | -7977.388 | -7978.413 | -1.024 | 1.39E-06 |
| 11 | 120992188 | rs594711 | T | C | 1.044 | -7977.388 | -7978.412 | -1.024 | 1.39E-06 |
| 11 | 116879676 | rs7931398 | C | T | 1.042 | -7977.388 | -7978.413 | -1.025 | 1.39E-06 |
| 11 | 117085261 | rs76604009 | T | C | 0.991 | -7977.388 | -7978.414 | -1.025 | 1.39E-06 |
| 11 | 117780091 | rs4439549 | A | C | 0.971 | -7977.388 | -7978.413 | -1.025 | 1.39E-06 |
| 11 | 119641843 | rs11600311 | T | C | 0.957 | -7977.388 | -7978.413 | -1.025 | 1.39E-06 |
| 11 | 116859675 | rs77852530 | C | A | 1.035 | -7977.388 | -7978.414 | -1.026 | 1.39E-06 |
| 11 | 117434983 | rs752796 | G | A | 0.961 | -7977.388 | -7978.414 | -1.026 | 1.39E-06 |
| 11 | 120505771 | rs4936535 | G | A | 0.962 | -7977.388 | -7978.414 | -1.026 | 1.39E-06 |
| 11 | 116380781 | rs79487385 | G | A | 1.014 | -7977.388 | -7978.415 | -1.027 | 1.39E-06 |
| 11 | 116550528 | rs61906105 | G | A | 0.969 | -7977.388 | -7978.416 | -1.027 | 1.39E-06 |
| 11 | 117634408 | rs141859855 | A | G | 0.957 | -7977.388 | -7978.415 | -1.027 | 1.39E-06 |
| 11 | 120158933 | rs77483431 | G | A | 1.042 | -7977.388 | -7978.415 | -1.027 | 1.39E-06 |
| 11 | 120160262 | rs4938794 | A | G | 1.042 | -7977.388 | -7978.415 | -1.027 | 1.39E-06 |
| 11 | 120675122 | rs1939672 | T | C | 0.963 | -7977.388 | -7978.416 | -1.027 | 1.39E-06 |
| 11 | 116592203 | rs180371 | G | A | 1.04 | -7977.388 | -7978.416 | -1.028 | 1.38E-06 |
| 11 | 118179197 | rs1599810 | A | G | 0.961 | -7977.388 | -7978.416 | -1.028 | 1.38E-06 |
| 11 | 116359851 | rs7934455 | C | T | 1.041 | -7977.388 | -7978.417 | -1.029 | 1.38E-06 |
| 11 | 116380855 | rs77229823 | G | A | 1.013 | -7977.388 | -7978.418 | -1.029 | 1.38E-06 |
| 11 | 117296923 | rs59827403 | G | C | 1.001 | -7977.388 | -7978.418 | -1.029 | 1.38E-06 |
| 11 | 117397482 | rs12806934 | T | C | 1.044 | -7977.388 | -7978.417 | -1.029 | 1.38E-06 |
| 11 | 117881601 | rs2512149 | C | T | 0.959 | -7977.388 | -7978.418 | -1.029 | 1.38E-06 |
| 11 | 117979970 | rs10502229 | T | G | 0.984 | -7977.388 | -7978.418 | -1.029 | 1.38E-06 |
| 11 | 117996402 | rs79220634 | A | C | 0.985 | -7977.388 | -7978.417 | -1.029 | 1.38E-06 |
| 11 | 117997427 | rs72488012 | A | G | 0.985 | -7977.388 | -7978.417 | -1.029 | 1.38E-06 |
| 11 | 119300694 | rs7102696 | A | G | 0.959 | -7977.388 | -7978.417 | -1.029 | 1.38E-06 |
| 11 | 116380536 | rs60524902 | G | T | 1.013 | -7977.388 | -7978.418 | -1.03 | 1.38E-06 |
| 11 | 116464800 | rs11216026 | G | A | 0.96 | -7977.388 | -7978.418 | -1.03 | 1.38E-06 |
| 11 | 116621963 | rs12292921 | G | T | 0.96 | -7977.388 | -7978.419 | -1.03 | 1.38E-06 |
| 11 | 117155696 | rs45507395 | T | G | 1.035 | -7977.388 | -7978.418 | -1.03 | 1.38E-06 |
| 11 | 118791150 | rs12270280 | T | C | 1.028 | -7977.388 | -7978.419 | -1.03 | 1.38E-06 |
| 11 | 119056273 | rs73564493 | A | G | 1.039 | -7977.388 | -7978.418 | -1.03 | 1.38E-06 |
| 11 | 120121323 | rs11217775 | C | T | 1.023 | -7977.388 | -7978.418 | -1.03 | 1.38E-06 |
| 11 | 120586577 | rs56160707 | C | T | 0.967 | -7977.388 | -7978.418 | -1.03 | 1.38E-06 |
| 11 | 116353002 | rs491428 | T | G | 1.042 | -7977.388 | -7978.419 | -1.031 | 1.38E-06 |
| 11 | 116353106 | rs490526 | T | A | 1.042 | -7977.388 | -7978.42 | -1.031 | 1.38E-06 |
| 11 | 116383064 | rs11599994 | A | G | 1.01 | -7977.388 | -7978.419 | -1.031 | 1.38E-06 |
| 11 | 116497466 | rs12417676 | C | A | 1.042 | -7977.388 | -7978.42 | -1.031 | 1.38E-06 |
| 11 | 116524965 | rs117133249 | A | G | 1.009 | -7977.388 | -7978.419 | -1.031 | 1.38E-06 |
| 11 | 116701669 | rs2070667 | A | G | 1.032 | -7977.388 | -7978.419 | -1.031 | 1.38E-06 |
| 11 | 117194059 | rs118086079 | A | G | 1.009 | -7977.388 | -7978.419 | -1.031 | 1.38E-06 |
| 11 | 117647461 | rs73587457 | G | C | 0.96 | -7977.388 | -7978.419 | -1.031 | 1.38E-06 |
| 11 | 117855922 | rs4252299 | T | G | 0.969 | -7977.388 | -7978.419 | -1.031 | 1.38E-06 |
| 11 | 117989301 | rs1941626 | C | G | 0.987 | -7977.388 | -7978.419 | -1.031 | 1.38E-06 |
| 11 | 117996200 | rs17092726 | G | A | 0.986 | -7977.388 | -7978.42 | -1.031 | 1.38E-06 |
| 11 | 117997473 | rs79754446 | G | A | 0.986 | -7977.388 | -7978.419 | -1.031 | 1.38E-06 |
| 11 | 116244267 | rs508731 | A | T | 1.044 | -7977.388 | -7978.42 | -1.032 | 1.38E-06 |
| 11 | 116379681 | rs11215970 | C | T | 1.012 | -7977.388 | -7978.42 | -1.032 | 1.38E-06 |
| 11 | 116552062 | rs1429049972 | T | A | 0.97 | -7977.388 | -7978.42 | -1.032 | 1.38E-06 |
| 11 | 116552562 | rs7952043 | T | C | 0.97 | -7977.388 | -7978.42 | -1.032 | 1.38E-06 |
| 11 | 116627056 | rs56143464 | A | G | 0.96 | -7977.388 | -7978.42 | -1.032 | 1.38E-06 |
| 11 | 116628057 | rs61905112 | A | G | 0.96 | -7977.388 | -7978.42 | -1.032 | 1.38E-06 |
| 11 | 117571426 | rs7130852 | G | A | 0.958 | -7977.388 | -7978.42 | -1.032 | 1.38E-06 |
| 11 | 117947103 | rs2845696 | G | A | 1.044 | -7977.388 | -7978.42 | -1.032 | 1.38E-06 |
| 11 | 117978103 | rs10892215 | T | C | 0.984 | -7977.388 | -7978.42 | -1.032 | 1.38E-06 |
| 11 | 117978247 | rs11216765 | T | C | 0.984 | -7977.388 | -7978.42 | -1.032 | 1.38E-06 |
| 11 | 118629959 | rs140110112 | A | G | 0.967 | -7977.388 | -7978.42 | -1.032 | 1.38E-06 |
| 11 | 119618503 | rs11217451 | A | G | 0.958 | -7977.388 | -7978.42 | -1.032 | 1.38E-06 |
| 11 | 119853252 | rs111444487 | C | T | 1.025 | -7977.388 | -7978.42 | -1.032 | 1.38E-06 |

| | | | | | | | |
|---|---|---|---|---|---|---|---|
| 11 | 120590848 | rs73588361 | A | G | NULL | 0.624806474 | NULL&M_Disease |
| 11 | 117867187 | rs4252279 | T | C | NULL | 0.625977858 | NULL&M_Disease |
| 11 | 117990003 | rs12576605 | C | G | NULL | 0.625977858 | NULL&M_Disease |
| 11 | 117990826 | rs10790238 | G | T | NULL | 0.625977858 | NULL&M_Disease |
| 11 | 117991176 | rs12361749 | T | A | NULL | 0.625977858 | NULL&M_Disease |
| 11 | 117991183 | rs12364660 | G | C | NULL | 0.625977858 | NULL&M_Disease |
| 11 | 117991381 | rs12222838 | G | A | NULL | 0.625977858 | NULL&M_Disease |
| 11 | 117992183 | rs11216772 | A | T | NULL | 0.625977858 | NULL&M_Disease |
| 11 | 117992763 | rs76043503 | G | A | NULL | 0.625977858 | NULL&M_Disease |
| 11 | 117993395 | rs12365797 | A | G | NULL | 0.625977858 | NULL&M_Disease |
| 11 | 117996360 | rs78227947 | G | T | NULL | 0.625977858 | NULL&M_Disease |
| 11 | 118051646 | rs637737 | G | A | NULL | 0.625977858 | NULL&M_Disease |
| 11 | 119932578 | rs896690 | A | G | NULL | 0.625977858 | NULL&M_Disease |
| 11 | 120120435 | rs60758080 | A | G | NULL | 0.625977858 | NULL&M_Disease |
| 11 | 120120523 | rs58254802 | T | G | NULL | 0.625977858 | NULL&M_Disease |
| 11 | 116316617 | rs17119667 | C | T | NULL | 0.625977858 | NULL&M_Disease |
| 11 | 116501684 | rs11216058 | A | G | NULL | 0.625977858 | NULL&M_Disease |
| 11 | 117998038 | rs12365332 | T | C | NULL | 0.625977858 | NULL&M_Disease |
| 11 | 118262407 | rs11216867 | G | A | NULL | 0.625977858 | NULL&M_Disease |
| 11 | 117359386 | rs4938392 | T | C | NULL | 0.625977858 | NULL&M_Disease |
| 11 | 117537378 | rs10892155 | T | C | NULL | 0.625977858 | NULL&M_Disease |
| 11 | 117992910 | rs7110327 | A | G | NULL | 0.625977858 | NULL&M_Disease |
| 11 | 117993836 | rs11216774 | A | G | NULL | 0.625977858 | NULL&M_Disease |
| 11 | 117998236 | rs75893552 | A | G | NULL | 0.625977858 | NULL&M_Disease |
| 11 | 118048499 | rs1312372307 | T | C | NULL | 0.625977858 | NULL&M_Disease |
| 11 | 118903651 | rs7942757 | A | G | NULL | 0.625977858 | NULL&M_Disease |
| 11 | 120515734 | rs10892590 | G | A | NULL | 0.625977858 | NULL&M_Disease |
| 11 | 116383138 | rs11215972 | G | C | NULL | 0.625977858 | NULL&M_Disease |
| 11 | 116673149 | rs682109 | T | C | NULL | 0.625977858 | NULL&M_Disease |
| 11 | 116819873 | rs6589580 | A | G | NULL | 0.625977858 | NULL&M_Disease |
| 11 | 117071505 | rs112333700 | T | C | NULL | 0.625977858 | NULL&M_Disease |
| 11 | 117398399 | rs4938399 | T | C | NULL | 0.625977858 | NULL&M_Disease |
| 11 | 118362414 | rs562780 | T | C | NULL | 0.625977858 | NULL&M_Disease |
| 11 | 118791532 | rs12271856 | T | G | NULL | 0.625977858 | NULL&M_Disease |
| 11 | 116383348 | rs75048292 | A | G | NULL | 0.625977858 | NULL&M_Disease |
| 11 | 117201056 | rs61903734 | A | T | NULL | 0.625977858 | NULL&M_Disease |
| 11 | 118284984 | rs59489957 | T | C | NULL | 0.625977858 | NULL&M_Disease |
| 11 | 118349247 | rs7941217 | T | C | NULL | 0.625977858 | NULL&M_Disease |
| 11 | 120119848 | rs61898298 | T | G | NULL | 0.625977858 | NULL&M_Disease |
| 11 | 120947699 | rs57401958 | G | A | NULL | 0.625977858 | NULL&M_Disease |
| 11 | 117135014 | rs138877884 | A | G | NULL | 0.625977858 | NULL&M_Disease |
| 11 | 117238478 | rs56984853 | A | G | NULL | 0.625977858 | NULL&M_Disease |
| 11 | 118044360 | rs11216791 | G | A | NULL | 0.625977858 | NULL&M_Disease |
| 11 | 119501909 | rs80106284 | T | G | NULL | 0.625977858 | NULL&M_Disease |
| 11 | 117394685 | rs11216437 | A | G | NULL | 0.627147766 | NULL&M_Disease |
| 11 | 117637213 | rs12808105 | T | C | NULL | 0.627147766 | NULL&M_Disease |
| 11 | 120515929 | rs3902979 | C | T | NULL | 0.625977858 | NULL&M_Disease |
| 11 | 120920649 | rs1939004 | G | A | NULL | 0.627147766 | NULL&M_Disease |
| 11 | 116375719 | rs79175131 | T | C | NULL | 0.625977858 | NULL&M_Disease |
| 11 | 118902758 | rs10892323 | C | G | NULL | 0.627147766 | NULL&M_Disease |
| 11 | 119505431 | rs78757430 | T | C | NULL | 0.627147766 | NULL&M_Disease |
| 11 | 120515939 | rs3902978 | A | G | NULL | 0.625977858 | NULL&M_Disease |
| 11 | 120670765 | rs3016414 | G | A | NULL | 0.627147766 | NULL&M_Disease |
| 11 | 116247770 | rs493721 | G | A | NULL | 0.627147766 | NULL&M_Disease |
| 11 | 116352789 | rs493308 | T | C | NULL | 0.627147766 | NULL&M_Disease |
| 11 | 116598248 | rs10892023 | T | C | NULL | 0.627147766 | NULL&M_Disease |
| 11 | 117203633 | rs527347 | G | A | NULL | 0.627147766 | NULL&M_Disease |
| 11 | 119502703 | rs12291064 | T | C | NULL | 0.627147766 | NULL&M_Disease |
| 11 | 119659912 | rs11601799 | A | G | NULL | 0.627147766 | NULL&M_Disease |
| 11 | 120684447 | rs3020105 | G | A | NULL | 0.627147766 | NULL&M_Disease |
| 11 | 116702345 | rs553080 | C | T | NULL | 0.627147766 | NULL&M_Disease |
| 11 | 117854977 | rs4252298 | C | T | NULL | 0.627147766 | NULL&M_Disease |
| 11 | 117977678 | rs10790234 | A | G | NULL | 0.627147766 | NULL&M_Disease |
| 11 | 119217311 | rs883245 | G | A | NULL | 0.627147766 | NULL&M_Disease |
| 11 | 120980537 | rs598650 | T | C | NULL | 0.627147766 | NULL&M_Disease |
| 11 | 116350751 | rs4391855 | A | G | NULL | 0.627147766 | NULL&M_Disease |
| 11 | 117158858 | rs1047857 | G | A | NULL | 0.627147766 | NULL&M_Disease |
| 11 | 117168564 | rs550319 | G | A | NULL | 0.627147766 | NULL&M_Disease |
| 11 | 117987041 | rs12223789 | A | G | NULL | 0.627147766 | NULL&M_Disease |
| 11 | 119860625 | rs583606 | C | G | NULL | 0.627147766 | NULL&M_Disease |
| 11 | 116993929 | rs10892078 | T | A | NULL | 0.627147766 | NULL&M_Disease |
| 11 | 117151412 | rs12282721 | A | G | NULL | 0.627147766 | NULL&M_Disease |
| 11 | 118255278 | rs6589663 | A | G | NULL | 0.627147766 | NULL&M_Disease |
| 11 | 116547004 | rs61906103 | G | T | NULL | 0.627147766 | NULL&M_Disease |
| 11 | 117309079 | rs144198120 | T | C | NULL | 0.627147766 | NULL&M_Disease |
| 11 | 117448312 | rs35857852 | C | T | NULL | 0.627147766 | NULL&M_Disease |
| 11 | 117899573 | rs678202 | G | A | NULL | 0.627147766 | NULL&M_Disease |
| 11 | 120893187 | rs11218115 | A | C | NULL | 0.627147766 | NULL&M_Disease |
| 11 | 116619073 | rs28927680 | G | C | NULL | 0.627147766 | NULL&M_Disease |
| 11 | 116702579 | rs5134 | C | T | NULL | 0.627147766 | NULL&M_Disease |
| 11 | 117144355 | rs12286404 | C | T | NULL | 0.627147766 | NULL&M_Disease |
| 11 | 117149405 | rs12292027 | G | A | NULL | 0.627147766 | NULL&M_Disease |
| 11 | 117638032 | rs12786234 | A | G | NULL | 0.627147766 | NULL&M_Disease |
| 11 | 116632798 | rs11602240 | C | T | NULL | 0.627147766 | NULL&M_Disease |
| 11 | 118367669 | rs7103001 | T | A | NULL | 0.627147766 | NULL&M_Disease |
| 11 | 116352523 | rs556436 | C | T | NULL | 0.628316188 | NULL&M_Disease |
| 11 | 116383211 | rs7927993 | C | T | NULL | 0.627147766 | NULL&M_Disease |
| 11 | 116605070 | rs12274046 | T | C | NULL | 0.627147766 | NULL&M_Disease |
| 11 | 116669828 | rs11216137 | A | G | NULL | 0.627147766 | NULL&M_Disease |
| 11 | 117661356 | rs4537790 | T | C | NULL | 0.628316188 | NULL&M_Disease |
| 11 | 119732550 | rs35825681 | G | C | NULL | 0.627147766 | NULL&M_Disease |
| 11 | 120054202 | rs4938790 | T | C | NULL | 0.627147766 | NULL&M_Disease |
| 11 | 120797584 | rs2852241 | C | T | NULL | 0.627147766 | NULL&M_Disease |
| 11 | 117300380 | rs61905260 | A | G | NULL | 0.627147766 | NULL&M_Disease |
| 11 | 117863292 | rs4252267 | A | G | NULL | 0.628316188 | NULL&M_Disease |
| 11 | 117863846 | rs4252270 | T | C | NULL | 0.628316188 | NULL&M_Disease |

| | | | | | | | | | |
|---|---|---|---|---|---|---|---|---|---|
| 11 | 120590848 | rs73588361 | A | G | 0.991 | -7977.388 | -7978.42 | -1.032 | 1.38E-06 |
| 11 | 117867187 | rs4252279 | T | C | 0.958 | -7977.388 | -7978.422 | -1.033 | 1.38E-06 |
| 11 | 117990003 | rs12576605 | C | G | 0.987 | -7977.388 | -7978.421 | -1.033 | 1.38E-06 |
| 11 | 117990826 | rs10790238 | G | T | 0.987 | -7977.388 | -7978.421 | -1.033 | 1.38E-06 |
| 11 | 117991176 | rs12361749 | T | A | 0.987 | -7977.388 | -7978.421 | -1.033 | 1.38E-06 |
| 11 | 117991183 | rs12364660 | G | C | 0.987 | -7977.388 | -7978.421 | -1.033 | 1.38E-06 |
| 11 | 117991381 | rs12222838 | G | A | 0.987 | -7977.388 | -7978.421 | -1.033 | 1.38E-06 |
| 11 | 117992183 | rs11216772 | A | T | 0.987 | -7977.388 | -7978.421 | -1.033 | 1.38E-06 |
| 11 | 117992763 | rs76043503 | G | A | 0.987 | -7977.388 | -7978.421 | -1.033 | 1.38E-06 |
| 11 | 117993395 | rs12365797 | A | G | 0.987 | -7977.388 | -7978.421 | -1.033 | 1.38E-06 |
| 11 | 117996360 | rs78227947 | G | T | 0.987 | -7977.388 | -7978.422 | -1.033 | 1.38E-06 |
| 11 | 118051646 | rs637737 | G | A | 0.967 | -7977.388 | -7978.422 | -1.033 | 1.38E-06 |
| 11 | 119932578 | rs896690 | A | G | 0.959 | -7977.388 | -7978.422 | -1.033 | 1.38E-06 |
| 11 | 120120435 | rs60758080 | A | G | 1.022 | -7977.388 | -7978.421 | -1.033 | 1.38E-06 |
| 11 | 120120523 | rs58254802 | T | G | 1.022 | -7977.388 | -7978.421 | -1.033 | 1.38E-06 |
| 11 | 116316617 | rs17119667 | C | T | 1.039 | -7977.388 | -7978.422 | -1.034 | 1.38E-06 |
| 11 | 116501684 | rs11216058 | A | G | 1.041 | -7977.388 | -7978.423 | -1.034 | 1.38E-06 |
| 11 | 117998038 | rs12365332 | T | C | 0.988 | -7977.388 | -7978.422 | -1.034 | 1.38E-06 |
| 11 | 118262407 | rs11216867 | G | A | 1.042 | -7977.388 | -7978.422 | -1.034 | 1.38E-06 |
| 11 | 117359386 | rs4938392 | T | C | 1.01 | -7977.388 | -7978.423 | -1.035 | 1.37E-06 |
| 11 | 117537378 | rs10892155 | T | C | 0.985 | -7977.388 | -7978.423 | -1.035 | 1.37E-06 |
| 11 | 117992910 | rs7110327 | A | G | 0.988 | -7977.388 | -7978.423 | -1.035 | 1.37E-06 |
| 11 | 117993836 | rs11216774 | A | G | 0.988 | -7977.388 | -7978.423 | -1.035 | 1.37E-06 |
| 11 | 117998236 | rs75893552 | A | G | 0.988 | -7977.388 | -7978.423 | -1.035 | 1.37E-06 |
| 11 | 118048499 | rs1312372307 | T | C | 0.966 | -7977.388 | -7978.423 | -1.035 | 1.37E-06 |
| 11 | 118903651 | rs7942757 | A | G | 0.962 | -7977.388 | -7978.423 | -1.035 | 1.37E-06 |
| 11 | 120515734 | rs10892590 | G | A | 1.027 | -7977.388 | -7978.423 | -1.035 | 1.37E-06 |
| 11 | 116383138 | rs11215972 | G | C | 1.008 | -7977.388 | -7978.425 | -1.036 | 1.37E-06 |
| 11 | 116673149 | rs682109 | T | C | 1.043 | -7977.388 | -7978.424 | -1.036 | 1.37E-06 |
| 11 | 116819873 | rs6589580 | A | G | 1.042 | -7977.388 | -7978.425 | -1.036 | 1.37E-06 |
| 11 | 117071505 | rs112333700 | T | C | 0.996 | -7977.388 | -7978.424 | -1.036 | 1.37E-06 |
| 11 | 117398399 | rs4938399 | T | C | 1.043 | -7977.388 | -7978.425 | -1.036 | 1.37E-06 |
| 11 | 118362414 | rs562780 | T | C | 0.958 | -7977.388 | -7978.424 | -1.036 | 1.37E-06 |
| 11 | 118791532 | rs12271856 | T | G | 1.027 | -7977.388 | -7978.424 | -1.036 | 1.37E-06 |
| 11 | 116383348 | rs75048292 | A | G | 1.007 | -7977.388 | -7978.426 | -1.037 | 1.37E-06 |
| 11 | 117201056 | rs61903734 | A | T | 1.026 | -7977.388 | -7978.425 | -1.037 | 1.37E-06 |
| 11 | 118284984 | rs59489957 | T | C | 1.042 | -7977.388 | -7978.426 | -1.037 | 1.37E-06 |
| 11 | 118349247 | rs7941217 | T | C | 1.035 | -7977.388 | -7978.426 | -1.037 | 1.37E-06 |
| 11 | 120119848 | rs61898298 | T | G | 1.021 | -7977.388 | -7978.425 | -1.037 | 1.37E-06 |
| 11 | 120947699 | rs57401958 | G | A | 1.042 | -7977.388 | -7978.425 | -1.037 | 1.37E-06 |
| 11 | 117135014 | rs138877884 | A | G | 1.027 | -7977.388 | -7978.426 | -1.038 | 1.37E-06 |
| 11 | 117238478 | rs56984853 | A | G | 0.958 | -7977.388 | -7978.426 | -1.038 | 1.37E-06 |
| 11 | 118044360 | rs11216791 | G | A | 0.968 | -7977.388 | -7978.426 | -1.038 | 1.37E-06 |
| 11 | 119501909 | rs80106284 | T | G | 0.997 | -7977.388 | -7978.426 | -1.038 | 1.37E-06 |
| 11 | 117394685 | rs11216437 | A | G | 1.043 | -7977.388 | -7978.427 | -1.039 | 1.37E-06 |
| 11 | 117637213 | rs12808105 | T | C | 0.958 | -7977.388 | -7978.427 | -1.039 | 1.37E-06 |
| 11 | 120515929 | rs3902979 | C | T | 1.026 | -7977.388 | -7978.428 | -1.039 | 1.37E-06 |
| 11 | 120920649 | rs1939004 | G | A | 1.042 | -7977.388 | -7978.427 | -1.039 | 1.37E-06 |
| 11 | 116375719 | rs79175131 | T | C | 0.961 | -7977.388 | -7978.428 | -1.04 | 1.37E-06 |
| 11 | 118902758 | rs10892323 | C | G | 0.962 | -7977.388 | -7978.429 | -1.04 | 1.37E-06 |
| 11 | 119505431 | rs78757430 | T | C | 0.958 | -7977.388 | -7978.428 | -1.04 | 1.37E-06 |
| 11 | 120515939 | rs3902978 | A | G | 1.024 | -7977.388 | -7978.429 | -1.04 | 1.37E-06 |
| 11 | 120670765 | rs3016414 | G | A | 0.962 | -7977.388 | -7978.428 | -1.04 | 1.37E-06 |
| 11 | 116247770 | rs493721 | G | A | 1.042 | -7977.388 | -7978.429 | -1.041 | 1.37E-06 |
| 11 | 116352789 | rs493308 | T | C | 1.041 | -7977.388 | -7978.43 | -1.041 | 1.37E-06 |
| 11 | 116598248 | rs10892023 | T | C | 1.043 | -7977.388 | -7978.43 | -1.041 | 1.37E-06 |
| 11 | 117203633 | rs527347 | G | A | 0.96 | -7977.388 | -7978.43 | -1.041 | 1.37E-06 |
| 11 | 119502703 | rs12291064 | T | C | 0.958 | -7977.388 | -7978.429 | -1.041 | 1.37E-06 |
| 11 | 119659912 | rs11601799 | A | G | 0.958 | -7977.388 | -7978.429 | -1.041 | 1.37E-06 |
| 11 | 120684447 | rs3020105 | G | A | 0.962 | -7977.388 | -7978.43 | -1.041 | 1.37E-06 |
| 11 | 116702345 | rs553080 | C | T | 1.031 | -7977.388 | -7978.43 | -1.042 | 1.36E-06 |
| 11 | 117854977 | rs4252298 | C | T | 0.969 | -7977.388 | -7978.43 | -1.042 | 1.36E-06 |
| 11 | 117977678 | rs10790234 | A | G | 0.989 | -7977.388 | -7978.43 | -1.042 | 1.36E-06 |
| 11 | 119217311 | rs883245 | G | A | 0.962 | -7977.388 | -7978.43 | -1.042 | 1.36E-06 |
| 11 | 120980537 | rs598650 | T | C | 1.041 | -7977.388 | -7978.43 | -1.042 | 1.36E-06 |
| 11 | 116350751 | rs4391855 | A | G | 0.983 | -7977.388 | -7978.431 | -1.043 | 1.36E-06 |
| 11 | 117158858 | rs1047857 | G | A | 1.033 | -7977.388 | -7978.431 | -1.043 | 1.36E-06 |
| 11 | 117168564 | rs550319 | G | A | 0.958 | -7977.388 | -7978.431 | -1.043 | 1.36E-06 |
| 11 | 117987041 | rs12223789 | A | G | 0.987 | -7977.388 | -7978.432 | -1.043 | 1.36E-06 |
| 11 | 119860625 | rs583606 | C | G | 0.959 | -7977.388 | -7978.431 | -1.043 | 1.36E-06 |
| 11 | 116993929 | rs10892078 | T | A | 0.962 | -7977.388 | -7978.433 | -1.044 | 1.36E-06 |
| 11 | 117151412 | rs12282721 | A | G | 1.043 | -7977.388 | -7978.432 | -1.044 | 1.36E-06 |
| 11 | 118255278 | rs6589663 | A | G | 1.041 | -7977.388 | -7978.432 | -1.044 | 1.36E-06 |
| 11 | 116547004 | rs61906103 | G | T | 0.971 | -7977.388 | -7978.433 | -1.045 | 1.36E-06 |
| 11 | 117309079 | rs144198120 | T | C | 0.994 | -7977.388 | -7978.433 | -1.045 | 1.36E-06 |
| 11 | 117448312 | rs35857852 | C | T | 0.962 | -7977.388 | -7978.433 | -1.045 | 1.36E-06 |
| 11 | 117899573 | rs678202 | G | A | 0.958 | -7977.388 | -7978.433 | -1.045 | 1.36E-06 |
| 11 | 120893187 | rs11218115 | A | C | 1.043 | -7977.388 | -7978.434 | -1.045 | 1.36E-06 |
| 11 | 116619073 | rs28927680 | G | C | 0.961 | -7977.388 | -7978.435 | -1.046 | 1.36E-06 |
| 11 | 116702579 | rs5134 | C | T | 1.03 | -7977.388 | -7978.434 | -1.046 | 1.36E-06 |
| 11 | 117144355 | rs12286404 | C | T | 1.033 | -7977.388 | -7978.434 | -1.046 | 1.36E-06 |
| 11 | 117149405 | rs12292027 | G | A | 1.033 | -7977.388 | -7978.434 | -1.046 | 1.36E-06 |
| 11 | 117638032 | rs12786234 | A | G | 0.958 | -7977.388 | -7978.434 | -1.046 | 1.36E-06 |
| 11 | 116632798 | rs11602240 | C | T | 1.043 | -7977.388 | -7978.435 | -1.047 | 1.36E-06 |
| 11 | 118367669 | rs7103001 | T | A | 1.034 | -7977.388 | -7978.435 | -1.047 | 1.36E-06 |
| 11 | 116352523 | rs556436 | C | T | 1.041 | -7977.388 | -7978.437 | -1.048 | 1.36E-06 |
| 11 | 116383211 | rs7927993 | C | T | 1 | -7977.388 | -7978.436 | -1.048 | 1.36E-06 |
| 11 | 116605070 | rs12274046 | T | C | 0.961 | -7977.388 | -7978.436 | -1.048 | 1.36E-06 |
| 11 | 116669828 | rs11216137 | A | G | 1.04 | -7977.388 | -7978.437 | -1.049 | 1.35E-06 |
| 11 | 117661356 | rs4537790 | T | C | 1.039 | -7977.388 | -7978.437 | -1.049 | 1.35E-06 |
| 11 | 119732550 | rs35825681 | G | C | 0.968 | -7977.388 | -7978.437 | -1.049 | 1.35E-06 |
| 11 | 120054202 | rs4938790 | T | C | 0.961 | -7977.388 | -7978.437 | -1.049 | 1.35E-06 |
| 11 | 120797584 | rs2852241 | C | T | 0.968 | -7977.388 | -7978.438 | -1.049 | 1.35E-06 |
| 11 | 117300380 | rs61905260 | A | G | 1.025 | -7977.388 | -7978.438 | -1.05 | 1.35E-06 |
| 11 | 117863292 | rs4252267 | A | G | 0.958 | -7977.388 | -7978.438 | -1.05 | 1.35E-06 |
| 11 | 117863846 | rs4252270 | T | C | 0.958 | -7977.388 | -7978.438 | -1.05 | 1.35E-06 |

| | | | | | | | |
|---|---|---|---|---|---|---|---|
| 11 | 118829521 | rs7479331 | T | G | NULL | 0.628316188 | NULL&M_Disease |
| 11 | 119240825 | rs587726 | T | C | NULL | 0.628316188 | NULL&M_Disease |
| 11 | 120161641 | rs6589804 | A | G | NULL | 0.628316188 | NULL&M_Disease |
| 11 | 120347586 | rs76034797 | T | C | NULL | 0.628316188 | NULL&M_Disease |
| 11 | 120897295 | rs7122879 | T | C | NULL | 0.628316188 | NULL&M_Disease |
| 11 | 121115877 | rs4382920 | G | C | NULL | 0.628316188 | NULL&M_Disease |
| 11 | 117153420 | rs2305821 | C | T | NULL | 0.628316188 | NULL&M_Disease |
| 11 | 117300092 | rs3889572 | T | C | NULL | 0.628316188 | NULL&M_Disease |
| 11 | 118902841 | rs11217129 | T | C | NULL | 0.628316188 | NULL&M_Disease |
| 11 | 117105726 | rs7125990 | G | A | NULL | 0.628316188 | NULL&M_Disease |
| 11 | 117557540 | rs721486 | G | A | NULL | 0.628316188 | NULL&M_Disease |
| 11 | 117682332 | rs186292996 | T | C | NULL | 0.628316188 | NULL&M_Disease |
| 11 | 118793918 | rs12282752 | C | T | NULL | 0.628316188 | NULL&M_Disease |
| 11 | 119826890 | rs78957222 | T | C | NULL | 0.628316188 | NULL&M_Disease |
| 11 | 119932137 | rs10892527 | A | G | NULL | 0.628316188 | NULL&M_Disease |
| 11 | 116357955 | rs10891982 | T | C | NULL | 0.628316188 | NULL&M_Disease |
| 11 | 116600400 | rs12279373 | G | A | NULL | 0.628316188 | NULL&M_Disease |
| 11 | 116671540 | rs12294065 | C | G | NULL | 0.628316188 | NULL&M_Disease |
| 11 | 117451256 | rs12223674 | G | C | NULL | 0.628316188 | NULL&M_Disease |
| 11 | 117866868 | rs2282494 | G | A | NULL | 0.628316188 | NULL&M_Disease |
| 11 | 117872086 | rs9610 | A | G | NULL | 0.628316188 | NULL&M_Disease |
| 11 | 118282241 | rs60798124 | G | A | NULL | 0.628316188 | NULL&M_Disease |
| 11 | 120679559 | rs1939670 | C | T | NULL | 0.628316188 | NULL&M_Disease |
| 11 | 116399336 | rs12281528 | A | G | NULL | 0.628316188 | NULL&M_Disease |
| 11 | 117107520 | rs74808315 | G | C | NULL | 0.628316188 | NULL&M_Disease |
| 11 | 117923252 | rs7934267 | T | G | NULL | 0.628316188 | NULL&M_Disease |
| 11 | 117923358 | rs7937145 | A | G | NULL | 0.628316188 | NULL&M_Disease |
| 11 | 117977723 | rs10790235 | G | A | NULL | 0.628316188 | NULL&M_Disease |
| 11 | 120181117 | rs117119582 | C | T | NULL | 0.628316188 | NULL&M_Disease |
| 11 | 120470159 | rs4936523 | C | T | NULL | 0.628316188 | NULL&M_Disease |
| 11 | 120526365 | rs4936537 | G | A | NULL | 0.628316188 | NULL&M_Disease |
| 11 | 116236271 | rs80178624 | A | T | NULL | 0.628316188 | NULL&M_Disease |
| 11 | 116550548 | rs61906106 | A | C | NULL | 0.628316188 | NULL&M_Disease |
| 11 | 116626258 | rs11825181 | A | G | NULL | 0.628316188 | NULL&M_Disease |
| 11 | 117999443 | rs1941637 | A | C | NULL | 0.628316188 | NULL&M_Disease |
| 11 | 118002372 | rs11216779 | T | C | NULL | 0.628316188 | NULL&M_Disease |
| 11 | 119787519 | rs10892485 | A | G | NULL | 0.628316188 | NULL&M_Disease |
| 11 | 120526637 | rs4245041 | C | T | NULL | 0.628316188 | NULL&M_Disease |
| 11 | 116372575 | rs2003013 | A | G | NULL | 0.628316188 | NULL&M_Disease |
| 11 | 116692334 | rs5104 | C | T | NULL | 0.628316188 | NULL&M_Disease |
| 11 | 117142440 | rs77451506 | G | C | NULL | 0.628316188 | NULL&M_Disease |
| 11 | 117145582 | rs61903699 | T | C | NULL | 0.628316188 | NULL&M_Disease |
| 11 | 117305418 | rs561791 | T | C | NULL | 0.628316188 | NULL&M_Disease |
| 11 | 121156870 | rs10892718 | A | C | NULL | 0.628316188 | NULL&M_Disease |
| 11 | 116519950 | rs12289816 | C | T | NULL | 0.628316188 | NULL&M_Disease |
| 11 | 116668843 | rs35606865 | C | T | NULL | 0.628316188 | NULL&M_Disease |
| 11 | 119777403 | rs11217560 | G | A | NULL | 0.628316188 | NULL&M_Disease |
| 11 | 116501918 | rs11828785 | T | C | NULL | 0.629483112 | NULL&M_Disease |
| 11 | 116624601 | rs61905108 | A | C | NULL | 0.628316188 | NULL&M_Disease |
| 11 | 117117973 | rs1892904 | G | A | NULL | 0.628316188 | NULL&M_Disease |
| 11 | 118805138 | rs7127427 | A | G | NULL | 0.628316188 | NULL&M_Disease |
| 11 | 119773520 | rs56031122 | G | A | NULL | 0.628316188 | NULL&M_Disease |
| 11 | 116535703 | rs56289161 | T | C | NULL | 0.628316188 | NULL&M_Disease |
| 11 | 116537880 | rs61209319 | A | C | NULL | 0.628316188 | NULL&M_Disease |
| 11 | 116611985 | rs10466589 | A | G | NULL | 0.629483112 | NULL&M_Disease |
| 11 | 116627055 | rs4938309 | T | C | NULL | 0.629483112 | NULL&M_Disease |
| 11 | 117111913 | rs12274203 | A | C | NULL | 0.629483112 | NULL&M_Disease |
| 11 | 118801296 | rs75210328 | T | G | NULL | 0.628316188 | NULL&M_Disease |
| 11 | 119727693 | rs73008667 | T | C | NULL | 0.629483112 | NULL&M_Disease |
| 11 | 117101445 | rs57157718 | T | A | NULL | 0.629483112 | NULL&M_Disease |
| 11 | 117129083 | rs77073823 | T | C | NULL | 0.629483112 | NULL&M_Disease |
| 11 | 117855727 | rs4252243 | T | C | NULL | 0.629483112 | NULL&M_Disease |
| 11 | 118034870 | rs3802869 | A | G | NULL | 0.629483112 | NULL&M_Disease |
| 11 | 118271860 | rs17122150 | G | C | NULL | 0.629483112 | NULL&M_Disease |
| 11 | 119934563 | rs4936507 | A | G | NULL | 0.629483112 | NULL&M_Disease |
| 11 | 116458785 | rs4938289 | T | G | NULL | 0.629483112 | NULL&M_Disease |
| 11 | 116596129 | rs11216117 | A | C | NULL | 0.629483112 | NULL&M_Disease |
| 11 | 117021877 | rs4938348 | G | C | NULL | 0.629483112 | NULL&M_Disease |
| 11 | 117027481 | rs4938352 | G | A | NULL | 0.629483112 | NULL&M_Disease |
| 11 | 117998834 | rs55820322 | C | A | NULL | 0.629483112 | NULL&M_Disease |
| 11 | 118182801 | rs1945764 | G | A | NULL | 0.629483112 | NULL&M_Disease |
| 11 | 120162368 | rs79164518 | C | T | NULL | 0.629483112 | NULL&M_Disease |
| 11 | 116320603 | rs59818386 | A | G | NULL | 0.629483112 | NULL&M_Disease |
| 11 | 116476765 | rs17092631 | C | T | NULL | 0.629483112 | NULL&M_Disease |
| 11 | 116646858 | rs11602073 | T | C | NULL | 0.629483112 | NULL&M_Disease |
| 11 | 116236979 | rs17475804 | G | C | NULL | 0.629483112 | NULL&M_Disease |
| 11 | 116651463 | rs1942478 | G | T | NULL | 0.629483112 | NULL&M_Disease |
| 11 | 116663707 | rs662799 | G | A | NULL | 0.629483112 | NULL&M_Disease |
| 11 | 117999224 | rs75387178 | C | G | NULL | 0.629483112 | NULL&M_Disease |
| 11 | 119776710 | rs12286892 | T | C | NULL | 0.629483112 | NULL&M_Disease |
| 11 | 120119395 | rs12574374 | A | G | NULL | 0.629483112 | NULL&M_Disease |
| 11 | 121153312 | rs4935767 | G | A | NULL | 0.629483112 | NULL&M_Disease |
| 11 | 116356327 | rs17569224 | T | C | NULL | 0.629483112 | NULL&M_Disease |
| 11 | 117136277 | rs67145274 | G | C | NULL | 0.629483112 | NULL&M_Disease |
| 11 | 117529895 | rs74498740 | T | C | NULL | 0.629483112 | NULL&M_Disease |
| 11 | 117564880 | rs11600672 | A | G | NULL | 0.629483112 | NULL&M_Disease |
| 11 | 117868638 | rs4252287 | A | G | NULL | 0.629483112 | NULL&M_Disease |
| 11 | 120844222 | rs4635090 | A | G | NULL | 0.629483112 | NULL&M_Disease |
| 11 | 116557044 | rs61906114 | C | G | NULL | 0.629483112 | NULL&M_Disease |
| 11 | 116613423 | rs12280724 | T | G | NULL | 0.629483112 | NULL&M_Disease |
| 11 | 117042816 | rs7120309 | T | A | NULL | 0.629483112 | NULL&M_Disease |
| 11 | 117042899 | rs7120449 | T | C | NULL | 0.629483112 | NULL&M_Disease |
| 11 | 117858185 | rs4252246 | A | C | NULL | 0.629483112 | NULL&M_Disease |
| 11 | 120166358 | rs744983 | G | A | NULL | 0.629483112 | NULL&M_Disease |
| 11 | 116603724 | rs12272004 | A | C | NULL | 0.629483112 | NULL&M_Disease |
| 11 | 118054324 | rs627995 | T | C | NULL | 0.629483112 | NULL&M_Disease |

| | | | | | | | | | |
|---|---|---|---|---|---|---|---|---|---|
| 11 | 118829521 | rs7479331 | T | G | 1.043 | -7977.388 | -7978.439 | -1.05 | 1.35E-06 |
| 11 | 119240825 | rs587726 | T | C | 0.958 | -7977.388 | -7978.438 | -1.05 | 1.35E-06 |
| 11 | 120161641 | rs6589804 | A | G | 1.04 | -7977.388 | -7978.438 | -1.05 | 1.35E-06 |
| 11 | 120347586 | rs76034797 | T | C | 1.036 | -7977.388 | -7978.439 | -1.051 | 1.35E-06 |
| 11 | 120897295 | rs7122879 | T | C | 1.042 | -7977.388 | -7978.439 | -1.051 | 1.35E-06 |
| 11 | 121115877 | rs4382920 | G | C | 1.043 | -7977.388 | -7978.439 | -1.051 | 1.35E-06 |
| 11 | 117153420 | rs2305821 | C | T | 1.033 | -7977.388 | -7978.441 | -1.052 | 1.35E-06 |
| 11 | 117300092 | rs3889572 | T | C | 1.039 | -7977.388 | -7978.44 | -1.052 | 1.35E-06 |
| 11 | 118902841 | rs11217129 | T | C | 0.962 | -7977.388 | -7978.44 | -1.052 | 1.35E-06 |
| 11 | 117105726 | rs7125990 | G | A | 1.032 | -7977.388 | -7978.441 | -1.053 | 1.35E-06 |
| 11 | 117557540 | rs721486 | G | A | 0.96 | -7977.388 | -7978.441 | -1.053 | 1.35E-06 |
| 11 | 117682332 | rs186292996 | T | C | 1.017 | -7977.388 | -7978.441 | -1.053 | 1.35E-06 |
| 11 | 118793918 | rs12282752 | C | T | 1.025 | -7977.388 | -7978.442 | -1.053 | 1.35E-06 |
| 11 | 119826890 | rs78957222 | T | C | 1.04 | -7977.388 | -7978.441 | -1.053 | 1.35E-06 |
| 11 | 119932137 | rs10892527 | A | G | 1.04 | -7977.388 | -7978.441 | -1.053 | 1.35E-06 |
| 11 | 116357955 | rs10891982 | T | C | 1.04 | -7977.388 | -7978.442 | -1.054 | 1.35E-06 |
| 11 | 116600400 | rs12279373 | G | A | 0.961 | -7977.388 | -7978.442 | -1.054 | 1.35E-06 |
| 11 | 116671540 | rs12294065 | C | G | 1.04 | -7977.388 | -7978.443 | -1.054 | 1.35E-06 |
| 11 | 117451256 | rs12223674 | G | C | 0.96 | -7977.388 | -7978.443 | -1.054 | 1.35E-06 |
| 11 | 117866868 | rs2282494 | G | A | 0.971 | -7977.388 | -7978.442 | -1.054 | 1.35E-06 |
| 11 | 117872086 | rs9610 | A | G | 1.039 | -7977.388 | -7978.443 | -1.054 | 1.35E-06 |
| 11 | 118282241 | rs60798124 | G | A | 1.041 | -7977.388 | -7978.442 | -1.054 | 1.35E-06 |
| 11 | 120679559 | rs1939670 | C | T | 0.963 | -7977.388 | -7978.443 | -1.054 | 1.35E-06 |
| 11 | 116399336 | rs12281528 | A | G | 1.039 | -7977.388 | -7978.443 | -1.055 | 1.35E-06 |
| 11 | 117107520 | rs74808315 | G | C | 1.024 | -7977.388 | -7978.444 | -1.055 | 1.35E-06 |
| 11 | 117923252 | rs7934267 | T | G | 1.041 | -7977.388 | -7978.443 | -1.055 | 1.35E-06 |
| 11 | 117923358 | rs7937145 | A | G | 1.041 | -7977.388 | -7978.443 | -1.055 | 1.35E-06 |
| 11 | 117977723 | rs10790235 | G | A | 0.993 | -7977.388 | -7978.443 | -1.055 | 1.35E-06 |
| 11 | 120181117 | rs117119582 | C | T | 0.992 | -7977.388 | -7978.443 | -1.055 | 1.35E-06 |
| 11 | 120470159 | rs4936523 | C | T | 0.961 | -7977.388 | -7978.443 | -1.055 | 1.35E-06 |
| 11 | 120526365 | rs4936537 | G | A | 1.04 | -7977.388 | -7978.443 | -1.055 | 1.35E-06 |
| 11 | 116236271 | rs80178624 | A | T | 0.966 | -7977.388 | -7978.444 | -1.056 | 1.35E-06 |
| 11 | 116550548 | rs61906106 | A | C | 0.973 | -7977.388 | -7978.444 | -1.056 | 1.35E-06 |
| 11 | 116626258 | rs11825181 | A | G | 0.962 | -7977.388 | -7978.445 | -1.056 | 1.35E-06 |
| 11 | 117999443 | rs1941637 | A | C | 0.989 | -7977.388 | -7978.445 | -1.056 | 1.35E-06 |
| 11 | 118002372 | rs11216779 | T | C | 0.991 | -7977.388 | -7978.445 | -1.056 | 1.35E-06 |
| 11 | 119787519 | rs10892485 | A | G | 1.043 | -7977.388 | -7978.444 | -1.056 | 1.35E-06 |
| 11 | 120526637 | rs4245041 | C | T | 1.04 | -7977.388 | -7978.444 | -1.056 | 1.35E-06 |
| 11 | 116372575 | rs2003013 | A | G | 1.021 | -7977.388 | -7978.446 | -1.057 | 1.34E-06 |
| 11 | 116692334 | rs5104 | C | T | 0.959 | -7977.388 | -7978.446 | -1.057 | 1.34E-06 |
| 11 | 117142440 | rs77451506 | G | C | 1.024 | -7977.388 | -7978.445 | -1.057 | 1.34E-06 |
| 11 | 117145582 | rs61903699 | T | C | 1.031 | -7977.388 | -7978.446 | -1.057 | 1.34E-06 |
| 11 | 117305418 | rs561791 | T | C | 1.041 | -7977.388 | -7978.445 | -1.057 | 1.34E-06 |
| 11 | 121156870 | rs10892718 | A | C | 1.035 | -7977.388 | -7978.445 | -1.057 | 1.34E-06 |
| 11 | 116519950 | rs12289816 | C | T | 1.041 | -7977.388 | -7978.446 | -1.058 | 1.34E-06 |
| 11 | 116668843 | rs35606865 | C | T | 1.04 | -7977.388 | -7978.446 | -1.058 | 1.34E-06 |
| 11 | 119777403 | rs11217560 | G | A | 1.043 | -7977.388 | -7978.446 | -1.058 | 1.34E-06 |
| 11 | 116501918 | rs11828785 | T | C | 1.041 | -7977.388 | -7978.447 | -1.059 | 1.34E-06 |
| 11 | 116624601 | rs61905108 | A | C | 0.975 | -7977.388 | -7978.447 | -1.059 | 1.34E-06 |
| 11 | 117117973 | rs1892904 | G | A | 1.032 | -7977.388 | -7978.447 | -1.059 | 1.34E-06 |
| 11 | 118805138 | rs7127427 | A | G | 1.024 | -7977.388 | -7978.447 | -1.059 | 1.34E-06 |
| 11 | 119773520 | rs56031122 | G | A | 1.04 | -7977.388 | -7978.448 | -1.059 | 1.34E-06 |
| 11 | 116535703 | rs56289161 | T | C | 0.974 | -7977.388 | -7978.448 | -1.06 | 1.34E-06 |
| 11 | 116537880 | rs61209319 | A | C | 0.974 | -7977.388 | -7978.449 | -1.06 | 1.34E-06 |
| 11 | 116611985 | rs10466589 | A | G | 0.962 | -7977.388 | -7978.448 | -1.06 | 1.34E-06 |
| 11 | 116627055 | rs4938309 | T | C | 1.043 | -7977.388 | -7978.448 | -1.06 | 1.34E-06 |
| 11 | 117111913 | rs12274203 | A | C | 1.033 | -7977.388 | -7978.449 | -1.06 | 1.34E-06 |
| 11 | 118801296 | rs75210328 | T | G | 1.024 | -7977.388 | -7978.448 | -1.06 | 1.34E-06 |
| 11 | 119727693 | rs73008667 | T | C | 1.04 | -7977.388 | -7978.448 | -1.06 | 1.34E-06 |
| 11 | 117101445 | rs57157718 | T | A | 0.972 | -7977.388 | -7978.449 | -1.061 | 1.34E-06 |
| 11 | 117129083 | rs77073823 | T | C | 1.031 | -7977.388 | -7978.45 | -1.061 | 1.34E-06 |
| 11 | 117855727 | rs4252243 | T | C | 0.959 | -7977.388 | -7978.45 | -1.061 | 1.34E-06 |
| 11 | 118034870 | rs3802869 | A | G | 0.974 | -7977.388 | -7978.449 | -1.061 | 1.34E-06 |
| 11 | 118271860 | rs17122150 | G | C | 1.041 | -7977.388 | -7978.449 | -1.061 | 1.34E-06 |
| 11 | 119934563 | rs4936507 | A | G | 1.043 | -7977.388 | -7978.45 | -1.061 | 1.34E-06 |
| 11 | 116458785 | rs4938289 | T | G | 1.036 | -7977.388 | -7978.45 | -1.062 | 1.34E-06 |
| 11 | 116596129 | rs11216117 | A | C | 0.962 | -7977.388 | -7978.45 | -1.062 | 1.34E-06 |
| 11 | 117021877 | rs4938348 | G | C | 0.959 | -7977.388 | -7978.45 | -1.062 | 1.34E-06 |
| 11 | 117027481 | rs4938352 | G | A | 0.959 | -7977.388 | -7978.45 | -1.062 | 1.34E-06 |
| 11 | 117998834 | rs55820322 | C | A | 0.994 | -7977.388 | -7978.45 | -1.062 | 1.34E-06 |
| 11 | 118182801 | rs1945764 | G | A | 0.962 | -7977.388 | -7978.451 | -1.062 | 1.34E-06 |
| 11 | 120162368 | rs79164518 | C | T | 1.039 | -7977.388 | -7978.45 | -1.062 | 1.34E-06 |
| 11 | 116320603 | rs59818386 | A | G | 0.962 | -7977.388 | -7978.451 | -1.063 | 1.34E-06 |
| 11 | 116476765 | rs17092631 | C | T | 0.967 | -7977.388 | -7978.451 | -1.063 | 1.34E-06 |
| 11 | 116646858 | rs11602073 | T | C | 1.041 | -7977.388 | -7978.451 | -1.063 | 1.34E-06 |
| 11 | 116236979 | rs17475804 | G | C | 0.971 | -7977.388 | -7978.452 | -1.064 | 1.33E-06 |
| 11 | 116651463 | rs1942478 | G | T | 1.041 | -7977.388 | -7978.452 | -1.064 | 1.33E-06 |
| 11 | 116663707 | rs662799 | G | A | 0.965 | -7977.388 | -7978.453 | -1.064 | 1.33E-06 |
| 11 | 117999224 | rs75387178 | C | G | 0.997 | -7977.388 | -7978.453 | -1.064 | 1.33E-06 |
| 11 | 119776710 | rs12286892 | T | C | 1.043 | -7977.388 | -7978.452 | -1.064 | 1.33E-06 |
| 11 | 120119395 | rs12574374 | A | G | 1.015 | -7977.388 | -7978.452 | -1.064 | 1.33E-06 |
| 11 | 121153312 | rs4935767 | G | A | 1.034 | -7977.388 | -7978.452 | -1.064 | 1.33E-06 |
| 11 | 116356327 | rs17569224 | T | C | 0.968 | -7977.388 | -7978.453 | -1.065 | 1.33E-06 |
| 11 | 117136277 | rs67145274 | G | C | 1.031 | -7977.388 | -7978.453 | -1.065 | 1.33E-06 |
| 11 | 117529895 | rs74498740 | T | C | 0.996 | -7977.388 | -7978.453 | -1.065 | 1.33E-06 |
| 11 | 117564880 | rs11600672 | A | G | 0.97 | -7977.388 | -7978.453 | -1.065 | 1.33E-06 |
| 11 | 117868638 | rs4252287 | A | G | 0.959 | -7977.388 | -7978.453 | -1.065 | 1.33E-06 |
| 11 | 120844222 | rs4635090 | A | G | 1.04 | -7977.388 | -7978.453 | -1.065 | 1.33E-06 |
| 11 | 116557044 | rs61906114 | C | G | 0.975 | -7977.388 | -7978.455 | -1.066 | 1.33E-06 |
| 11 | 116613423 | rs12280724 | T | G | 0.963 | -7977.388 | -7978.455 | -1.066 | 1.33E-06 |
| 11 | 117042816 | rs7120309 | T | A | 0.959 | -7977.388 | -7978.455 | -1.066 | 1.33E-06 |
| 11 | 117042899 | rs7120449 | T | C | 0.959 | -7977.388 | -7978.455 | -1.066 | 1.33E-06 |
| 11 | 117858185 | rs4252246 | A | C | 0.959 | -7977.388 | -7978.454 | -1.066 | 1.33E-06 |
| 11 | 120166358 | rs744983 | G | A | 1.041 | -7977.388 | -7978.455 | -1.066 | 1.33E-06 |
| 11 | 116603724 | rs12272004 | A | C | 0.962 | -7977.388 | -7978.456 | -1.067 | 1.33E-06 |
| 11 | 118054324 | rs627995 | T | C | 0.962 | -7977.388 | -7978.455 | -1.067 | 1.33E-06 |

| | | | | | | | |
|---|---|---|---|---|---|---|---|
| 11 | 119645757 | rs11217490 | A | G | NULL | 0.629483112 | NULL&M_Disease |
| 11 | 119789176 | rs11601034 | T | C | NULL | 0.629483112 | NULL&M_Disease |
| 11 | 117021759 | rs4938347 | T | A | NULL | 0.630648526 | NULL&M_Disease |
| 11 | 117022276 | rs4938350 | A | G | NULL | 0.630648526 | NULL&M_Disease |
| 11 | 117920100 | rs59238367 | A | G | NULL | 0.629483112 | NULL&M_Disease |
| 11 | 118050855 | rs678957 | T | C | NULL | 0.629483112 | NULL&M_Disease |
| 11 | 121162486 | rs1000315943 | A | T | NULL | 0.629483112 | NULL&M_Disease |
| 11 | 116646062 | rs11216135 | G | C | NULL | 0.629483112 | NULL&M_Disease |
| 11 | 116652301 | rs4417316 | T | C | NULL | 0.629483112 | NULL&M_Disease |
| 11 | 117133378 | rs12295268 | A | G | NULL | 0.629483112 | NULL&M_Disease |
| 11 | 117134094 | rs11216341 | T | C | NULL | 0.629483112 | NULL&M_Disease |
| 11 | 117442767 | rs71480344 | G | A | NULL | 0.629483112 | NULL&M_Disease |
| 11 | 117919097 | rs674576 | A | G | NULL | 0.630648526 | NULL&M_Disease |
| 11 | 119054208 | rs45439091 | T | G | NULL | 0.629483112 | NULL&M_Disease |
| 11 | 119250874 | rs656439 | C | T | NULL | 0.630648526 | NULL&M_Disease |
| 11 | 119934071 | rs56895810 | A | C | NULL | 0.630648526 | NULL&M_Disease |
| 11 | 116361705 | rs6589547 | A | C | NULL | 0.630648526 | NULL&M_Disease |
| 11 | 116654435 | rs603446 | T | C | NULL | 0.630648526 | NULL&M_Disease |
| 11 | 117140148 | rs12270661 | A | G | NULL | 0.629483112 | NULL&M_Disease |
| 11 | 118464665 | rs113337722 | G | A | NULL | 0.629483112 | NULL&M_Disease |
| 11 | 120311157 | rs77216358 | G | A | NULL | 0.629483112 | NULL&M_Disease |
| 11 | 116599900 | rs61372904 | G | A | NULL | 0.630648526 | NULL&M_Disease |
| 11 | 116607874 | rs59455853 | T | A | NULL | 0.630648526 | NULL&M_Disease |
| 11 | 117757685 | rs4938449 | A | T | NULL | 0.630648526 | NULL&M_Disease |
| 11 | 117883941 | rs11216678 | G | C | NULL | 0.630648526 | NULL&M_Disease |
| 11 | 118904015 | rs595407 | C | T | NULL | 0.630648526 | NULL&M_Disease |
| 11 | 119847951 | rs78284541 | T | C | NULL | 0.630648526 | NULL&M_Disease |
| 11 | 117169874 | rs61903703 | A | G | NULL | 0.630648526 | NULL&M_Disease |
| 11 | 119241249 | rs513841 | T | A | NULL | 0.630648526 | NULL&M_Disease |
| 11 | 120827067 | rs542580 | T | C | NULL | 0.630648526 | NULL&M_Disease |
| 11 | 116457391 | rs17570978 | T | C | NULL | 0.630648526 | NULL&M_Disease |
| 11 | 116518901 | rs73580632 | T | C | NULL | 0.630648526 | NULL&M_Disease |
| 11 | 116556912 | rs61906113 | T | C | NULL | 0.630648526 | NULL&M_Disease |
| 11 | 119788395 | rs11217573 | G | A | NULL | 0.630648526 | NULL&M_Disease |
| 11 | 116477772 | rs3016356 | C | T | NULL | 0.630648526 | NULL&M_Disease |
| 11 | 116701153 | rs2070668 | T | G | NULL | 0.630648526 | NULL&M_Disease |
| 11 | 117109155 | rs57471964 | G | A | NULL | 0.630648526 | NULL&M_Disease |
| 11 | 119075323 | rs4938637 | A | G | NULL | 0.630648526 | NULL&M_Disease |
| 11 | 119644360 | rs9633947 | A | G | NULL | 0.630648526 | NULL&M_Disease |
| 11 | 121165498 | rs7928868 | A | G | NULL | 0.630648526 | NULL&M_Disease |
| 11 | 116533660 | rs61906079 | G | C | NULL | 0.630648526 | NULL&M_Disease |
| 11 | 117343399 | rs497548 | T | C | NULL | 0.630648526 | NULL&M_Disease |
| 11 | 119352570 | rs658187 | T | G | NULL | 0.630648526 | NULL&M_Disease |
| 11 | 119985246 | rs11606047 | G | A | NULL | 0.630648526 | NULL&M_Disease |
| 11 | 120830413 | rs524792 | C | T | NULL | 0.630648526 | NULL&M_Disease |
| 11 | 117651730 | rs60790095 | T | C | NULL | 0.630648526 | NULL&M_Disease |
| 11 | 117845835 | rs7121294 | T | C | NULL | 0.630648526 | NULL&M_Disease |
| 11 | 118970476 | rs116658831 | A | G | NULL | 0.630648526 | NULL&M_Disease |
| 11 | 120522714 | rs77132149 | A | G | NULL | 0.630648526 | NULL&M_Disease |
| 11 | 120524498 | rs76820520 | A | G | NULL | 0.630648526 | NULL&M_Disease |
| 11 | 120703243 | rs73003743 | G | A | NULL | 0.630648526 | NULL&M_Disease |
| 11 | 118042713 | rs651825 | G | T | NULL | 0.630648526 | NULL&M_Disease |
| 11 | 119002768 | rs73562872 | T | G | NULL | 0.630648526 | NULL&M_Disease |
| 11 | 119010392 | rs73562877 | A | G | NULL | 0.630648526 | NULL&M_Disease |
| 11 | 117864113 | rs2228054 | A | G | NULL | 0.630648526 | NULL&M_Disease |
| 11 | 118499469 | rs577396 | C | G | NULL | 0.631812418 | NULL&M_Disease |
| 11 | 120090348 | rs12225131 | G | A | NULL | 0.630648526 | NULL&M_Disease |
| 11 | 121167114 | rs79110667 | T | C | NULL | 0.630648526 | NULL&M_Disease |
| 11 | 121168178 | rs727422 | A | G | NULL | 0.630648526 | NULL&M_Disease |
| 11 | 116239770 | rs73005818 | A | G | NULL | 0.630648526 | NULL&M_Disease |
| 11 | 116457471 | rs74886004 | T | C | NULL | 0.631812418 | NULL&M_Disease |
| 11 | 117107978 | rs61906492 | A | G | NULL | 0.630648526 | NULL&M_Disease |
| 11 | 117162762 | rs28989504 | C | T | NULL | 0.630648526 | NULL&M_Disease |
| 11 | 117880139 | rs11216674 | T | C | NULL | 0.630648526 | NULL&M_Disease |
| 11 | 119842049 | rs61897867 | T | C | NULL | 0.631812418 | NULL&M_Disease |
| 11 | 120074170 | rs12420903 | T | G | NULL | 0.630648526 | NULL&M_Disease |
| 11 | 116502286 | rs1145183 | A | G | NULL | 0.630648526 | NULL&M_Disease |
| 11 | 117824592 | rs2515800 | C | T | NULL | 0.631812418 | NULL&M_Disease |
| 11 | 119778270 | rs554422 | T | C | NULL | 0.631812418 | NULL&M_Disease |
| 11 | 119847788 | rs79936530 | A | G | NULL | 0.631812418 | NULL&M_Disease |
| 11 | 116393781 | rs117819665 | T | C | NULL | 0.631812418 | NULL&M_Disease |
| 11 | 116797442 | rs112701434 | C | T | NULL | 0.631812418 | NULL&M_Disease |
| 11 | 118509668 | rs483598 | A | G | NULL | 0.631812418 | NULL&M_Disease |
| 11 | 118550841 | rs79699220 | C | G | NULL | 0.631812418 | NULL&M_Disease |
| 11 | 120783331 | rs57584368 | C | T | NULL | 0.631812418 | NULL&M_Disease |
| 11 | 116883241 | rs111708971 | A | T | NULL | 0.631812418 | NULL&M_Disease |
| 11 | 117800481 | rs61900350 | C | G | NULL | 0.631812418 | NULL&M_Disease |
| 11 | 117898231 | rs644920 | A | T | NULL | 0.631812418 | NULL&M_Disease |
| 11 | 117971971 | rs6589645 | G | A | NULL | 0.631812418 | NULL&M_Disease |
| 11 | 120677398 | rs948034 | C | T | NULL | 0.631812418 | NULL&M_Disease |
| 11 | 116518889 | rs1240771 | G | A | NULL | 0.631812418 | NULL&M_Disease |
| 11 | 116662331 | rs12287066 | T | G | NULL | 0.631812418 | NULL&M_Disease |
| 11 | 117133219 | rs533203 | T | C | NULL | 0.631812418 | NULL&M_Disease |
| 11 | 120151510 | rs11217783 | T | A | NULL | 0.631812418 | NULL&M_Disease |
| 11 | 120441767 | rs11217917 | G | C | NULL | 0.631812418 | NULL&M_Disease |
| 11 | 116558828 | rs56182713 | A | G | NULL | 0.631812418 | NULL&M_Disease |
| 11 | 117795942 | rs17597688 | G | A | NULL | 0.631812418 | NULL&M_Disease |
| 11 | 119353238 | rs649197 | A | G | NULL | 0.631812418 | NULL&M_Disease |
| 11 | 120666931 | rs3020100 | G | A | NULL | 0.631812418 | NULL&M_Disease |
| 11 | 116352697 | rs494200 | A | G | NULL | 0.631812418 | NULL&M_Disease |
| 11 | 116363446 | rs7128900 | G | T | NULL | 0.631812418 | NULL&M_Disease |
| 11 | 116501098 | rs12800011 | G | A | NULL | 0.631812418 | NULL&M_Disease |
| 11 | 118173850 | rs3759009 | A | G | NULL | 0.631812418 | NULL&M_Disease |
| 11 | 120677301 | rs1320649 | T | C | NULL | 0.631812418 | NULL&M_Disease |
| 11 | 116542585 | rs1787693 | A | G | NULL | 0.631812418 | NULL&M_Disease |
| 11 | 116787148 | rs11216187 | T | C | NULL | 0.631812418 | NULL&M_Disease |

| | | | | | | | | | |
|---|---|---|---|---|---|---|---|---|---|
| 11 | 119645757 | rs11217490 | A | G | 0.959 | -7977.388 | -7978.455 | -1.067 | 1.33E-06 |
| 11 | 119789176 | rs11601034 | T | C | 0.996 | -7977.388 | -7978.455 | -1.067 | 1.33E-06 |
| 11 | 117021759 | rs4938347 | T | A | 0.959 | -7977.388 | -7978.457 | -1.068 | 1.33E-06 |
| 11 | 117022276 | rs4938350 | A | G | 0.959 | -7977.388 | -7978.457 | -1.068 | 1.33E-06 |
| 11 | 117920100 | rs59238367 | A | G | 1.042 | -7977.388 | -7978.457 | -1.068 | 1.33E-06 |
| 11 | 118050855 | rs678957 | T | C | 0.969 | -7977.388 | -7978.456 | -1.068 | 1.33E-06 |
| 11 | 121162486 | rs1000315943 | A | T | 1.033 | -7977.388 | -7978.456 | -1.068 | 1.33E-06 |
| 11 | 116646062 | rs11216135 | G | C | 1.041 | -7977.388 | -7978.457 | -1.069 | 1.33E-06 |
| 11 | 116652301 | rs4417316 | T | C | 1.041 | -7977.388 | -7978.457 | -1.069 | 1.33E-06 |
| 11 | 117133378 | rs12295268 | A | G | 1.03 | -7977.388 | -7978.457 | -1.069 | 1.33E-06 |
| 11 | 117134094 | rs11216341 | T | C | 1.03 | -7977.388 | -7978.457 | -1.069 | 1.33E-06 |
| 11 | 117442767 | rs71480344 | G | A | 1.033 | -7977.388 | -7978.457 | -1.069 | 1.33E-06 |
| 11 | 117919097 | rs674576 | A | G | 1.042 | -7977.388 | -7978.457 | -1.069 | 1.33E-06 |
| 11 | 119054208 | rs45439091 | T | G | 1.036 | -7977.388 | -7978.457 | -1.069 | 1.33E-06 |
| 11 | 119250874 | rs656439 | C | T | 0.959 | -7977.388 | -7978.458 | -1.069 | 1.33E-06 |
| 11 | 119934071 | rs56895810 | A | C | 1.042 | -7977.388 | -7978.457 | -1.069 | 1.33E-06 |
| 11 | 116361705 | rs6589547 | A | C | 1.04 | -7977.388 | -7978.459 | -1.07 | 1.33E-06 |
| 11 | 116654435 | rs603446 | T | C | 0.963 | -7977.388 | -7978.458 | -1.07 | 1.33E-06 |
| 11 | 117140148 | rs12270661 | A | G | 1.03 | -7977.388 | -7978.458 | -1.07 | 1.33E-06 |
| 11 | 118464665 | rs113337722 | G | A | 0.996 | -7977.388 | -7978.458 | -1.07 | 1.33E-06 |
| 11 | 120311157 | rs77216358 | G | A | 0.971 | -7977.388 | -7978.458 | -1.07 | 1.33E-06 |
| 11 | 116599900 | rs61372904 | G | A | 0.963 | -7977.388 | -7978.459 | -1.071 | 1.33E-06 |
| 11 | 116607874 | rs59455853 | T | A | 0.963 | -7977.388 | -7978.459 | -1.071 | 1.33E-06 |
| 11 | 117757685 | rs4938449 | A | T | 0.961 | -7977.388 | -7978.459 | -1.071 | 1.33E-06 |
| 11 | 117883941 | rs11216678 | G | C | 1.037 | -7977.388 | -7978.459 | -1.071 | 1.33E-06 |
| 11 | 118904015 | rs595407 | C | T | 0.963 | -7977.388 | -7978.46 | -1.071 | 1.33E-06 |
| 11 | 119847951 | rs78284541 | T | C | 1.038 | -7977.388 | -7978.459 | -1.071 | 1.33E-06 |
| 11 | 117169874 | rs61903703 | A | G | 1.029 | -7977.388 | -7978.46 | -1.072 | 1.32E-06 |
| 11 | 119241249 | rs513841 | T | A | 0.959 | -7977.388 | -7978.46 | -1.072 | 1.32E-06 |
| 11 | 120827067 | rs542580 | T | C | 0.983 | -7977.388 | -7978.461 | -1.072 | 1.32E-06 |
| 11 | 116457391 | rs17570978 | T | C | 0.961 | -7977.388 | -7978.462 | -1.073 | 1.32E-06 |
| 11 | 116518901 | rs73580632 | T | C | 1.037 | -7977.388 | -7978.461 | -1.073 | 1.32E-06 |
| 11 | 116556912 | rs61906113 | T | C | 0.976 | -7977.388 | -7978.462 | -1.073 | 1.32E-06 |
| 11 | 119788395 | rs11217573 | G | A | 0.998 | -7977.388 | -7978.462 | -1.073 | 1.32E-06 |
| 11 | 116477772 | rs3016356 | C | T | 0.965 | -7977.388 | -7978.463 | -1.074 | 1.32E-06 |
| 11 | 116701153 | rs2070668 | T | G | 1.039 | -7977.388 | -7978.462 | -1.074 | 1.32E-06 |
| 11 | 117109155 | rs57471964 | G | A | 1.021 | -7977.388 | -7978.464 | -1.075 | 1.32E-06 |
| 11 | 119075323 | rs4938637 | A | G | 1.035 | -7977.388 | -7978.463 | -1.075 | 1.32E-06 |
| 11 | 119644360 | rs9633947 | A | G | 0.959 | -7977.388 | -7978.464 | -1.075 | 1.32E-06 |
| 11 | 121165498 | rs7928868 | A | G | 0.962 | -7977.388 | -7978.463 | -1.075 | 1.32E-06 |
| 11 | 116533660 | rs61906079 | G | C | 0.977 | -7977.388 | -7978.464 | -1.076 | 1.32E-06 |
| 11 | 117343399 | rs497548 | T | C | 0.961 | -7977.388 | -7978.465 | -1.076 | 1.32E-06 |
| 11 | 119352570 | rs658187 | T | G | 1.039 | -7977.388 | -7978.464 | -1.076 | 1.32E-06 |
| 11 | 119985246 | rs11606047 | G | A | 1.033 | -7977.388 | -7978.465 | -1.076 | 1.32E-06 |
| 11 | 120830413 | rs524792 | C | T | 0.985 | -7977.388 | -7978.464 | -1.076 | 1.32E-06 |
| 11 | 117651730 | rs60790095 | T | C | 1.04 | -7977.388 | -7978.466 | -1.077 | 1.32E-06 |
| 11 | 117845835 | rs7121294 | T | C | 0.963 | -7977.388 | -7978.465 | -1.077 | 1.32E-06 |
| 11 | 118970476 | rs116658831 | A | G | 0.971 | -7977.388 | -7978.465 | -1.077 | 1.32E-06 |
| 11 | 120522714 | rs77132149 | A | G | 1.014 | -7977.388 | -7978.466 | -1.077 | 1.32E-06 |
| 11 | 120524498 | rs76820520 | A | G | 1.014 | -7977.388 | -7978.466 | -1.077 | 1.32E-06 |
| 11 | 120703243 | rs73003743 | G | A | 0.964 | -7977.388 | -7978.466 | -1.077 | 1.32E-06 |
| 11 | 118042713 | rs651825 | G | T | 0.971 | -7977.388 | -7978.466 | -1.078 | 1.32E-06 |
| 11 | 119002768 | rs73562872 | T | G | 1.036 | -7977.388 | -7978.466 | -1.078 | 1.32E-06 |
| 11 | 119010392 | rs73562877 | A | G | 1.036 | -7977.388 | -7978.466 | -1.078 | 1.32E-06 |
| 11 | 117864113 | rs2228054 | A | G | 0.972 | -7977.388 | -7978.467 | -1.079 | 1.31E-06 |
| 11 | 118499469 | rs577396 | C | G | 0.963 | -7977.388 | -7978.468 | -1.079 | 1.31E-06 |
| 11 | 120090348 | rs12225131 | G | A | 0.96 | -7977.388 | -7978.467 | -1.079 | 1.31E-06 |
| 11 | 121167114 | rs79110667 | T | C | 0.963 | -7977.388 | -7978.467 | -1.079 | 1.31E-06 |
| 11 | 121168178 | rs727422 | A | G | 0.963 | -7977.388 | -7978.467 | -1.079 | 1.31E-06 |
| 11 | 116239770 | rs73005818 | A | G | 0.973 | -7977.388 | -7978.468 | -1.08 | 1.31E-06 |
| 11 | 116457471 | rs74886004 | T | C | 0.961 | -7977.388 | -7978.468 | -1.08 | 1.31E-06 |
| 11 | 117107978 | rs61906492 | A | G | 1.029 | -7977.388 | -7978.468 | -1.08 | 1.31E-06 |
| 11 | 117162762 | rs28989504 | C | T | 1.028 | -7977.388 | -7978.468 | -1.08 | 1.31E-06 |
| 11 | 117880139 | rs11216674 | T | C | 0.97 | -7977.388 | -7978.468 | -1.08 | 1.31E-06 |
| 11 | 119842049 | rs61897867 | T | C | 1.028 | -7977.388 | -7978.469 | -1.08 | 1.31E-06 |
| 11 | 120074170 | rs12420903 | T | G | 0.969 | -7977.388 | -7978.468 | -1.08 | 1.31E-06 |
| 11 | 116502286 | rs1145183 | A | G | 1.013 | -7977.388 | -7978.469 | -1.081 | 1.31E-06 |
| 11 | 117824592 | rs2515800 | C | T | 0.96 | -7977.388 | -7978.47 | -1.081 | 1.31E-06 |
| 11 | 119778270 | rs554422 | T | C | 1.042 | -7977.388 | -7978.469 | -1.081 | 1.31E-06 |
| 11 | 119847788 | rs79936530 | A | G | 1.037 | -7977.388 | -7978.469 | -1.081 | 1.31E-06 |
| 11 | 116393781 | rs117819665 | T | C | 0.991 | -7977.388 | -7978.47 | -1.082 | 1.31E-06 |
| 11 | 116797442 | rs112701434 | C | T | 1.006 | -7977.388 | -7978.471 | -1.082 | 1.31E-06 |
| 11 | 118509668 | rs483598 | A | G | 0.963 | -7977.388 | -7978.471 | -1.082 | 1.31E-06 |
| 11 | 118550841 | rs79699220 | C | G | 0.967 | -7977.388 | -7978.47 | -1.082 | 1.31E-06 |
| 11 | 120783331 | rs57584368 | C | T | 1.025 | -7977.388 | -7978.47 | -1.082 | 1.31E-06 |
| 11 | 116883241 | rs111708971 | A | T | 1.005 | -7977.388 | -7978.471 | -1.083 | 1.31E-06 |
| 11 | 117800481 | rs61900350 | C | G | 1.039 | -7977.388 | -7978.471 | -1.083 | 1.31E-06 |
| 11 | 117898231 | rs644920 | A | T | 0.96 | -7977.388 | -7978.471 | -1.083 | 1.31E-06 |
| 11 | 117971971 | rs6589645 | G | A | 1.029 | -7977.388 | -7978.471 | -1.083 | 1.31E-06 |
| 11 | 120677398 | rs948034 | C | T | 0.965 | -7977.388 | -7978.471 | -1.083 | 1.31E-06 |
| 11 | 116518889 | rs1240771 | G | A | 1.041 | -7977.388 | -7978.473 | -1.084 | 1.31E-06 |
| 11 | 116662331 | rs12287066 | T | G | 0.965 | -7977.388 | -7978.472 | -1.084 | 1.31E-06 |
| 11 | 117133219 | rs533203 | T | C | 0.962 | -7977.388 | -7978.472 | -1.084 | 1.31E-06 |
| 11 | 120151510 | rs11217783 | T | A | 1.042 | -7977.388 | -7978.472 | -1.084 | 1.31E-06 |
| 11 | 120441767 | rs11217917 | G | C | 0.962 | -7977.388 | -7978.472 | -1.084 | 1.31E-06 |
| 11 | 116558828 | rs56182713 | A | G | 0.978 | -7977.388 | -7978.473 | -1.085 | 1.31E-06 |
| 11 | 117795942 | rs17597688 | G | A | 0.963 | -7977.388 | -7978.473 | -1.085 | 1.31E-06 |
| 11 | 119353238 | rs649197 | A | G | 1.039 | -7977.388 | -7978.474 | -1.085 | 1.31E-06 |
| 11 | 120666931 | rs3020100 | G | A | 0.966 | -7977.388 | -7978.473 | -1.085 | 1.31E-06 |
| 11 | 116352697 | rs494200 | A | G | 1.04 | -7977.388 | -7978.474 | -1.086 | 1.31E-06 |
| 11 | 116363446 | rs7128900 | G | T | 1.031 | -7977.388 | -7978.474 | -1.086 | 1.31E-06 |
| 11 | 116501098 | rs12800011 | G | A | 1.04 | -7977.388 | -7978.474 | -1.086 | 1.31E-06 |
| 11 | 118173850 | rs3759009 | A | G | 1.039 | -7977.388 | -7978.474 | -1.086 | 1.31E-06 |
| 11 | 120677301 | rs1320649 | T | C | 0.965 | -7977.388 | -7978.474 | -1.086 | 1.31E-06 |
| 11 | 116542585 | rs1787693 | A | G | 1.039 | -7977.388 | -7978.476 | -1.087 | 1.30E-06 |
| 11 | 116787148 | rs11216187 | T | C | 0.96 | -7977.388 | -7978.475 | -1.087 | 1.30E-06 |

| | | | | | | | |
|---|---|---|---|---|---|---|---|
| 11 | 117886453 | rs17121519 | T | C | NULL | 0.631812418 | NULL&M_Disease |
| 11 | 117972721 | rs948469 | A | C | NULL | 0.631812418 | NULL&M_Disease |
| 11 | 120495969 | rs11217928 | T | C | NULL | 0.631812418 | NULL&M_Disease |
| 11 | 117655971 | rs75562690 | T | C | NULL | 0.631812418 | NULL&M_Disease |
| 11 | 117878559 | rs11216668 | C | T | NULL | 0.631812418 | NULL&M_Disease |
| 11 | 117878578 | rs11216669 | T | C | NULL | 0.631812418 | NULL&M_Disease |
| 11 | 117878659 | rs11216670 | A | T | NULL | 0.631812418 | NULL&M_Disease |
| 11 | 119825806 | rs117534497 | G | A | NULL | 0.631812418 | NULL&M_Disease |
| 11 | 119839762 | rs77031353 | A | C | NULL | 0.631812418 | NULL&M_Disease |
| 11 | 120193511 | rs7934236 | G | C | NULL | 0.631812418 | NULL&M_Disease |
| 11 | 120793972 | rs2850804 | A | G | NULL | 0.632974777 | NULL&M_Disease |
| 11 | 117030581 | rs1871757 | T | A | NULL | 0.632974777 | NULL&M_Disease |
| 11 | 117168340 | rs74369487 | C | A | NULL | 0.631812418 | NULL&M_Disease |
| 11 | 117671641 | rs2298647 | A | T | NULL | 0.632974777 | NULL&M_Disease |
| 11 | 116513188 | rs117239953 | C | T | NULL | 0.632974777 | NULL&M_Disease |
| 11 | 117491758 | rs12799401 | C | G | NULL | 0.632974777 | NULL&M_Disease |
| 11 | 118322488 | rs73615909 | T | C | NULL | 0.631812418 | NULL&M_Disease |
| 11 | 118501459 | rs11216934 | A | G | NULL | 0.632974777 | NULL&M_Disease |
| 11 | 118711376 | rs118046730 | C | T | NULL | 0.631812418 | NULL&M_Disease |
| 11 | 118903589 | rs11217131 | G | C | NULL | 0.632974777 | NULL&M_Disease |
| 11 | 119792099 | rs78054182 | C | T | NULL | 0.631812418 | NULL&M_Disease |
| 11 | 116354211 | rs1828121 | A | G | NULL | 0.632974777 | NULL&M_Disease |
| 11 | 116518960 | rs1261672 | G | A | NULL | 0.632974777 | NULL&M_Disease |
| 11 | 116545723 | rs1787688 | C | T | NULL | 0.632974777 | NULL&M_Disease |
| 11 | 118273019 | rs17122154 | A | G | NULL | 0.632974777 | NULL&M_Disease |
| 11 | 119749609 | rs142108356 | G | A | NULL | 0.632974777 | NULL&M_Disease |
| 11 | 120097897 | rs117939457 | T | A | NULL | 0.631812418 | NULL&M_Disease |
| 11 | 116649538 | rs61905116 | G | A | NULL | 0.632974777 | NULL&M_Disease |
| 11 | 117795679 | rs11216631 | T | G | NULL | 0.632974777 | NULL&M_Disease |
| 11 | 116370266 | rs77297809 | T | C | NULL | 0.632974777 | NULL&M_Disease |
| 11 | 116467349 | rs117096905 | T | C | NULL | 0.632974777 | NULL&M_Disease |
| 11 | 116470947 | rs117287247 | T | C | NULL | 0.632974777 | NULL&M_Disease |
| 11 | 117307055 | rs78825663 | G | C | NULL | 0.632974777 | NULL&M_Disease |
| 11 | 117659186 | rs144638086 | T | C | NULL | 0.632974777 | NULL&M_Disease |
| 11 | 116365421 | rs143291574 | T | C | NULL | 0.632974777 | NULL&M_Disease |
| 11 | 116423005 | rs12363946 | A | T | NULL | 0.632974777 | NULL&M_Disease |
| 11 | 118271759 | rs7121764 | G | A | NULL | 0.632974777 | NULL&M_Disease |
| 11 | 119252199 | rs670906 | T | G | NULL | 0.632974777 | NULL&M_Disease |
| 11 | 119697028 | rs11217526 | A | T | NULL | 0.632974777 | NULL&M_Disease |
| 11 | 116525319 | rs79862998 | A | G | NULL | 0.632974777 | NULL&M_Disease |
| 11 | 116558782 | rs12800424 | A | G | NULL | 0.632974777 | NULL&M_Disease |
| 11 | 119002868 | rs76619306 | T | C | NULL | 0.632974777 | NULL&M_Disease |
| 11 | 116514699 | rs1145185 | A | G | NULL | 0.632974777 | NULL&M_Disease |
| 11 | 116555926 | rs4572168 | A | G | NULL | 0.632974777 | NULL&M_Disease |
| 11 | 116606630 | rs180352 | A | G | NULL | 0.632974777 | NULL&M_Disease |
| 11 | 117555150 | rs11216500 | C | T | NULL | 0.632974777 | NULL&M_Disease |
| 11 | 117900074 | rs665597 | A | G | NULL | 0.632974777 | NULL&M_Disease |
| 11 | 118746223 | rs11217077 | C | T | NULL | 0.632974777 | NULL&M_Disease |
| 11 | 119027545 | rs4148170 | A | G | NULL | 0.632974777 | NULL&M_Disease |
| 11 | 120685242 | rs3020104 | A | G | NULL | 0.632974777 | NULL&M_Disease |
| 11 | 116199943 | rs893840 | C | T | NULL | 0.632974777 | NULL&M_Disease |
| 11 | 116319634 | rs73005236 | A | G | NULL | 0.632974777 | NULL&M_Disease |
| 11 | 117671398 | rs6421566 | G | A | NULL | 0.632974777 | NULL&M_Disease |
| 11 | 119041659 | rs626984 | T | C | NULL | 0.632974777 | NULL&M_Disease |
| 11 | 120073236 | rs61898268 | C | A | NULL | 0.632974777 | NULL&M_Disease |
| 11 | 120088926 | rs61898281 | G | A | NULL | 0.632974777 | NULL&M_Disease |
| 11 | 120441647 | rs4938815 | G | T | NULL | 0.632974777 | NULL&M_Disease |
| 11 | 117176075 | rs142273276 | C | T | NULL | 0.632974777 | NULL&M_Disease |
| 11 | 120466336 | rs1320721 | C | G | NULL | 0.634135591 | NULL&M_Disease |
| 11 | 120783369 | rs17124538 | G | A | NULL | 0.632974777 | NULL&M_Disease |
| 11 | 116513428 | rs116859831 | C | T | NULL | 0.632974777 | NULL&M_Disease |
| 11 | 117350098 | rs11602570 | A | G | NULL | 0.632974777 | NULL&M_Disease |
| 11 | 117774291 | rs2845712 | A | C | NULL | 0.632974777 | NULL&M_Disease |
| 11 | 117882961 | rs17121515 | A | G | NULL | 0.632974777 | NULL&M_Disease |
| 11 | 118089957 | rs77610621 | A | G | NULL | 0.632974777 | NULL&M_Disease |
| 11 | 119778544 | rs4938738 | T | G | NULL | 0.634135591 | NULL&M_Disease |
| 11 | 121155566 | rs79293731 | T | C | NULL | 0.632974777 | NULL&M_Disease |
| 11 | 116316628 | rs73005230 | A | G | NULL | 0.632974777 | NULL&M_Disease |
| 11 | 116529313 | rs61906078 | A | G | NULL | 0.632974777 | NULL&M_Disease |
| 11 | 116553924 | rs12225622 | T | C | NULL | 0.634135591 | NULL&M_Disease |
| 11 | 117177372 | rs28989476 | C | T | NULL | 0.632974777 | NULL&M_Disease |
| 11 | 119002439 | rs2298480 | G | A | NULL | 0.632974777 | NULL&M_Disease |
| 11 | 119026052 | rs7108508 | A | G | NULL | 0.634135591 | NULL&M_Disease |
| 11 | 119027830 | rs4148171 | G | A | NULL | 0.634135591 | NULL&M_Disease |
| 11 | 119518822 | rs12805373 | A | T | NULL | 0.634135591 | NULL&M_Disease |
| 11 | 119777416 | rs11217562 | T | C | NULL | 0.634135591 | NULL&M_Disease |
| 11 | 116531479 | rs17119878 | A | T | NULL | 0.634135591 | NULL&M_Disease |
| 11 | 117131286 | rs9971587 | A | G | NULL | 0.634135591 | NULL&M_Disease |
| 11 | 119930631 | rs10892525 | T | C | NULL | 0.634135591 | NULL&M_Disease |
| 11 | 120446830 | rs12796185 | C | T | NULL | 0.634135591 | NULL&M_Disease |
| 11 | 116600564 | rs61905081 | A | G | NULL | 0.634135591 | NULL&M_Disease |
| 11 | 116614863 | rs61905086 | C | T | NULL | 0.634135591 | NULL&M_Disease |
| 11 | 116667545 | rs75919952 | T | C | NULL | 0.634135591 | NULL&M_Disease |
| 11 | 117893889 | rs60169792 | T | C | NULL | 0.634135591 | NULL&M_Disease |
| 11 | 117894258 | rs55734733 | A | C | NULL | 0.634135591 | NULL&M_Disease |
| 11 | 117894371 | rs55858401 | T | C | NULL | 0.634135591 | NULL&M_Disease |
| 11 | 117894424 | rs56029127 | A | G | NULL | 0.634135591 | NULL&M_Disease |
| 11 | 117895163 | rs73013729 | A | G | NULL | 0.634135591 | NULL&M_Disease |
| 11 | 117895268 | rs7942124 | T | A | NULL | 0.634135591 | NULL&M_Disease |
| 11 | 117895276 | rs7942225 | T | C | NULL | 0.634135591 | NULL&M_Disease |
| 11 | 117895305 | rs7942146 | G | A | NULL | 0.634135591 | NULL&M_Disease |
| 11 | 117895586 | rs7945602 | A | G | NULL | 0.634135591 | NULL&M_Disease |
| 11 | 117895712 | rs7945541 | T | C | NULL | 0.634135591 | NULL&M_Disease |
| 11 | 117896135 | rs58768562 | T | C | NULL | 0.634135591 | NULL&M_Disease |
| 11 | 117896454 | rs4936416 | A | G | NULL | 0.634135591 | NULL&M_Disease |
| 11 | 119351636 | rs672499 | A | G | NULL | 0.634135591 | NULL&M_Disease |

| | | | | | | | | | |
|---|---|---|---|---|---|---|---|---|---|
| 11 | 117886453 | rs17121519 | T | C | 0.971 | -7977.388 | -7978.475 | -1.087 | 1.30E-06 |
| 11 | 117972721 | rs948469 | A | C | 1.029 | -7977.388 | -7978.475 | -1.087 | 1.30E-06 |
| 11 | 120495969 | rs11217928 | T | C | 0.962 | -7977.388 | -7978.475 | -1.087 | 1.30E-06 |
| 11 | 117655971 | rs75562690 | T | C | 0.961 | -7977.388 | -7978.477 | -1.088 | 1.30E-06 |
| 11 | 117878559 | rs11216668 | C | T | 0.971 | -7977.388 | -7978.476 | -1.088 | 1.30E-06 |
| 11 | 117878578 | rs11216669 | T | C | 0.971 | -7977.388 | -7978.476 | -1.088 | 1.30E-06 |
| 11 | 117878659 | rs11216670 | A | T | 0.971 | -7977.388 | -7978.476 | -1.088 | 1.30E-06 |
| 11 | 119825806 | rs117534497 | G | A | 0.994 | -7977.388 | -7978.476 | -1.088 | 1.30E-06 |
| 11 | 119839762 | rs77031353 | A | C | 1.037 | -7977.388 | -7978.476 | -1.088 | 1.30E-06 |
| 11 | 120193511 | rs7934236 | G | C | 0.96 | -7977.388 | -7978.476 | -1.088 | 1.30E-06 |
| 11 | 120793972 | rs2850804 | A | G | 0.962 | -7977.388 | -7978.476 | -1.088 | 1.30E-06 |
| 11 | 117030581 | rs1871757 | T | A | 0.96 | -7977.388 | -7978.477 | -1.089 | 1.30E-06 |
| 11 | 117168340 | rs74369487 | C | A | 0.973 | -7977.388 | -7978.478 | -1.089 | 1.30E-06 |
| 11 | 117671641 | rs2298647 | A | T | 1.038 | -7977.388 | -7978.478 | -1.089 | 1.30E-06 |
| 11 | 116513188 | rs117239953 | C | T | 1.036 | -7977.388 | -7978.479 | -1.09 | 1.30E-06 |
| 11 | 117491758 | rs12799401 | C | G | 1.041 | -7977.388 | -7978.478 | -1.09 | 1.30E-06 |
| 11 | 118322488 | rs73615909 | T | C | 1.029 | -7977.388 | -7978.478 | -1.09 | 1.30E-06 |
| 11 | 118501459 | rs11216934 | A | G | 0.963 | -7977.388 | -7978.478 | -1.09 | 1.30E-06 |
| 11 | 118711376 | rs118046730 | C | T | 1.013 | -7977.388 | -7978.478 | -1.09 | 1.30E-06 |
| 11 | 118903589 | rs11217131 | G | C | 0.963 | -7977.388 | -7978.478 | -1.09 | 1.30E-06 |
| 11 | 119792099 | rs78054182 | C | T | 0.978 | -7977.388 | -7978.478 | -1.09 | 1.30E-06 |
| 11 | 116354211 | rs1828121 | A | G | 1.038 | -7977.388 | -7978.48 | -1.091 | 1.30E-06 |
| 11 | 116518960 | rs1261672 | G | A | 1.041 | -7977.388 | -7978.479 | -1.091 | 1.30E-06 |
| 11 | 116545723 | rs1787688 | C | T | 0.963 | -7977.388 | -7978.48 | -1.091 | 1.30E-06 |
| 11 | 118273019 | rs17122154 | A | G | 1.04 | -7977.388 | -7978.479 | -1.091 | 1.30E-06 |
| 11 | 119749609 | rs142108356 | G | A | 1.041 | -7977.388 | -7978.479 | -1.091 | 1.30E-06 |
| 11 | 120097897 | rs117939457 | T | A | 0.977 | -7977.388 | -7978.479 | -1.091 | 1.30E-06 |
| 11 | 116649538 | rs61905116 | G | A | 0.966 | -7977.388 | -7978.48 | -1.092 | 1.30E-06 |
| 11 | 117795679 | rs11216631 | T | G | 0.961 | -7977.388 | -7978.481 | -1.092 | 1.30E-06 |
| 11 | 116370266 | rs77297809 | T | C | 1.017 | -7977.388 | -7978.482 | -1.093 | 1.30E-06 |
| 11 | 116467349 | rs117096905 | T | C | 1.009 | -7977.388 | -7978.481 | -1.093 | 1.30E-06 |
| 11 | 116470947 | rs117287247 | T | C | 1.009 | -7977.388 | -7978.481 | -1.093 | 1.30E-06 |
| 11 | 117307055 | rs78825663 | G | C | 1.002 | -7977.388 | -7978.481 | -1.093 | 1.30E-06 |
| 11 | 117659186 | rs144638086 | T | C | 0.976 | -7977.388 | -7978.481 | -1.093 | 1.30E-06 |
| 11 | 116365421 | rs143291574 | T | C | 1.024 | -7977.388 | -7978.482 | -1.094 | 1.30E-06 |
| 11 | 116423005 | rs12363946 | A | T | 1.038 | -7977.388 | -7978.482 | -1.094 | 1.30E-06 |
| 11 | 118271759 | rs7121764 | G | A | 1.04 | -7977.388 | -7978.482 | -1.094 | 1.30E-06 |
| 11 | 119252199 | rs670906 | T | G | 0.96 | -7977.388 | -7978.482 | -1.094 | 1.30E-06 |
| 11 | 119697028 | rs11217526 | A | T | 0.962 | -7977.388 | -7978.483 | -1.094 | 1.30E-06 |
| 11 | 116525319 | rs79862998 | A | G | 1.035 | -7977.388 | -7978.484 | -1.095 | 1.29E-06 |
| 11 | 116558782 | rs12800424 | A | G | 0.979 | -7977.388 | -7978.483 | -1.095 | 1.29E-06 |
| 11 | 119002868 | rs76619306 | T | C | 1.034 | -7977.388 | -7978.483 | -1.095 | 1.29E-06 |
| 11 | 116514699 | rs1145185 | A | G | 0.96 | -7977.388 | -7978.484 | -1.096 | 1.29E-06 |
| 11 | 116555926 | rs4572168 | A | G | 0.965 | -7977.388 | -7978.484 | -1.096 | 1.29E-06 |
| 11 | 116606630 | rs180352 | A | G | 1.041 | -7977.388 | -7978.485 | -1.096 | 1.29E-06 |
| 11 | 117555150 | rs11216500 | C | T | 0.963 | -7977.388 | -7978.485 | -1.096 | 1.29E-06 |
| 11 | 117900074 | rs665597 | A | G | 0.961 | -7977.388 | -7978.484 | -1.096 | 1.29E-06 |
| 11 | 118746223 | rs11217077 | C | T | 0.961 | -7977.388 | -7978.484 | -1.096 | 1.29E-06 |
| 11 | 119027545 | rs4148170 | A | G | 1.034 | -7977.388 | -7978.484 | -1.096 | 1.29E-06 |
| 11 | 120685242 | rs3020104 | A | G | 0.968 | -7977.388 | -7978.484 | -1.096 | 1.29E-06 |
| 11 | 116199943 | rs893840 | C | T | 1.038 | -7977.388 | -7978.486 | -1.097 | 1.29E-06 |
| 11 | 116319634 | rs73005236 | A | G | 0.983 | -7977.388 | -7978.485 | -1.097 | 1.29E-06 |
| 11 | 117671398 | rs6421566 | G | A | 1.038 | -7977.388 | -7978.485 | -1.097 | 1.29E-06 |
| 11 | 119041659 | rs626984 | T | C | 0.963 | -7977.388 | -7978.485 | -1.097 | 1.29E-06 |
| 11 | 120073236 | rs61898268 | C | A | 0.971 | -7977.388 | -7978.485 | -1.097 | 1.29E-06 |
| 11 | 120088926 | rs61898281 | G | A | 0.978 | -7977.388 | -7978.485 | -1.097 | 1.29E-06 |
| 11 | 120441647 | rs4938815 | G | T | 0.962 | -7977.388 | -7978.485 | -1.097 | 1.29E-06 |
| 11 | 117176075 | rs142273276 | C | T | 1.015 | -7977.388 | -7978.486 | -1.098 | 1.29E-06 |
| 11 | 120466336 | rs1320721 | C | G | 0.962 | -7977.388 | -7978.486 | -1.098 | 1.29E-06 |
| 11 | 120783369 | rs17124538 | G | A | 1.022 | -7977.388 | -7978.486 | -1.098 | 1.29E-06 |
| 11 | 116513428 | rs116859831 | C | T | 1.035 | -7977.388 | -7978.487 | -1.099 | 1.29E-06 |
| 11 | 117350098 | rs11602570 | A | G | 0.972 | -7977.388 | -7978.487 | -1.099 | 1.29E-06 |
| 11 | 117774291 | rs2845712 | A | C | 0.98 | -7977.388 | -7978.488 | -1.099 | 1.29E-06 |
| 11 | 117882961 | rs17121515 | A | G | 0.973 | -7977.388 | -7978.487 | -1.099 | 1.29E-06 |
| 11 | 118089957 | rs77610621 | A | G | 0.981 | -7977.388 | -7978.487 | -1.099 | 1.29E-06 |
| 11 | 119778544 | rs4938738 | T | G | 1.041 | -7977.388 | -7978.487 | -1.099 | 1.29E-06 |
| 11 | 121155566 | rs79293731 | T | C | 0.964 | -7977.388 | -7978.487 | -1.099 | 1.29E-06 |
| 11 | 116316628 | rs73005230 | A | G | 0.984 | -7977.388 | -7978.488 | -1.1 | 1.29E-06 |
| 11 | 116529313 | rs61906078 | A | G | 0.981 | -7977.388 | -7978.488 | -1.1 | 1.29E-06 |
| 11 | 116553924 | rs12225622 | T | C | 0.965 | -7977.388 | -7978.488 | -1.1 | 1.29E-06 |
| 11 | 117177372 | rs28989476 | C | T | 1.015 | -7977.388 | -7978.488 | -1.1 | 1.29E-06 |
| 11 | 119002439 | rs2298480 | G | A | 1.034 | -7977.388 | -7978.488 | -1.1 | 1.29E-06 |
| 11 | 119026052 | rs7108508 | A | G | 1.034 | -7977.388 | -7978.488 | -1.1 | 1.29E-06 |
| 11 | 119027830 | rs4148171 | G | A | 1.034 | -7977.388 | -7978.488 | -1.1 | 1.29E-06 |
| 11 | 119518822 | rs12805373 | A | T | 1.04 | -7977.388 | -7978.489 | -1.1 | 1.29E-06 |
| 11 | 119777416 | rs11217562 | T | C | 1.041 | -7977.388 | -7978.488 | -1.1 | 1.29E-06 |
| 11 | 116531479 | rs17119878 | A | T | 0.98 | -7977.388 | -7978.49 | -1.101 | 1.29E-06 |
| 11 | 117131286 | rs9971587 | A | G | 0.963 | -7977.388 | -7978.489 | -1.101 | 1.29E-06 |
| 11 | 119930631 | rs10892525 | T | C | 1.039 | -7977.388 | -7978.489 | -1.101 | 1.29E-06 |
| 11 | 120446830 | rs12796185 | C | T | 0.962 | -7977.388 | -7978.489 | -1.101 | 1.29E-06 |
| 11 | 116600564 | rs61905081 | A | G | 0.965 | -7977.388 | -7978.49 | -1.102 | 1.29E-06 |
| 11 | 116614863 | rs61905086 | C | T | 0.965 | -7977.388 | -7978.49 | -1.102 | 1.29E-06 |
| 11 | 116667545 | rs75919952 | T | C | 0.977 | -7977.388 | -7978.491 | -1.102 | 1.29E-06 |
| 11 | 117893889 | rs60169792 | T | C | 0.973 | -7977.388 | -7978.491 | -1.102 | 1.29E-06 |
| 11 | 117894258 | rs55734733 | A | C | 0.973 | -7977.388 | -7978.491 | -1.102 | 1.29E-06 |
| 11 | 117894371 | rs55858401 | T | C | 0.973 | -7977.388 | -7978.491 | -1.102 | 1.29E-06 |
| 11 | 117894424 | rs56029127 | A | G | 0.973 | -7977.388 | -7978.491 | -1.102 | 1.29E-06 |
| 11 | 117895163 | rs73013729 | A | G | 0.973 | -7977.388 | -7978.49 | -1.102 | 1.29E-06 |
| 11 | 117895268 | rs7942124 | T | A | 0.973 | -7977.388 | -7978.49 | -1.102 | 1.29E-06 |
| 11 | 117895276 | rs7942225 | T | C | 0.973 | -7977.388 | -7978.49 | -1.102 | 1.29E-06 |
| 11 | 117895305 | rs7942146 | G | A | 0.973 | -7977.388 | -7978.49 | -1.102 | 1.29E-06 |
| 11 | 117895586 | rs7945602 | A | G | 0.973 | -7977.388 | -7978.49 | -1.102 | 1.29E-06 |
| 11 | 117895712 | rs7945541 | T | C | 0.973 | -7977.388 | -7978.49 | -1.102 | 1.29E-06 |
| 11 | 117896135 | rs58768562 | T | C | 0.973 | -7977.388 | -7978.49 | -1.102 | 1.29E-06 |
| 11 | 117896454 | rs4936416 | A | G | 1.04 | -7977.388 | -7978.491 | -1.102 | 1.29E-06 |
| 11 | 119351636 | rs672499 | A | G | 1.039 | -7977.388 | -7978.491 | -1.102 | 1.29E-06 |

| | | | | | | | |
|---|---|---|---|---|---|---|---|
| 11 | 116558916 | rs61906117 | A | G | NULL | 0.634135591 | NULL&M_Disease |
| 11 | 116601945 | rs3212282 | A | G | NULL | 0.634135591 | NULL&M_Disease |
| 11 | 117555933 | rs7129597 | T | C | NULL | 0.634135591 | NULL&M_Disease |
| 11 | 118579395 | rs11828220 | A | T | NULL | 0.634135591 | NULL&M_Disease |
| 11 | 119355747 | rs613018 | C | T | NULL | 0.634135591 | NULL&M_Disease |
| 11 | 120093495 | rs61898288 | A | C | NULL | 0.634135591 | NULL&M_Disease |
| 11 | 116596464 | rs60583202 | A | G | NULL | 0.634135591 | NULL&M_Disease |
| 11 | 116652351 | rs12274192 | G | A | NULL | 0.634135591 | NULL&M_Disease |
| 11 | 117220324 | rs61903739 | T | C | NULL | 0.634135591 | NULL&M_Disease |
| 11 | 117425383 | rs12804924 | T | C | NULL | 0.634135591 | NULL&M_Disease |
| 11 | 119790068 | rs10502232 | G | A | NULL | 0.634135591 | NULL&M_Disease |
| 11 | 119992149 | rs562015 | T | C | NULL | 0.634135591 | NULL&M_Disease |
| 11 | 120571457 | rs10892619 | T | C | NULL | 0.634135591 | NULL&M_Disease |
| 11 | 121154118 | rs7950536 | G | A | NULL | 0.634135591 | NULL&M_Disease |
| 11 | 117048367 | rs117680585 | T | C | NULL | 0.634135591 | NULL&M_Disease |
| 11 | 117165397 | rs28989500 | T | C | NULL | 0.634135591 | NULL&M_Disease |
| 11 | 117754524 | rs10892188 | C | A | NULL | 0.634135591 | NULL&M_Disease |
| 11 | 117898851 | rs10458939 | A | G | NULL | 0.634135591 | NULL&M_Disease |
| 11 | 117974541 | rs1893318 | G | A | NULL | 0.634135591 | NULL&M_Disease |
| 11 | 119806852 | rs548429 | G | A | NULL | 0.634135591 | NULL&M_Disease |
| 11 | 121154642 | rs78303474 | C | T | NULL | 0.634135591 | NULL&M_Disease |
| 11 | 121156037 | rs56142837 | G | T | NULL | 0.634135591 | NULL&M_Disease |
| 11 | 121156412 | rs55903632 | C | G | NULL | 0.634135591 | NULL&M_Disease |
| 11 | 116527521 | rs55776558 | C | T | NULL | 0.634135591 | NULL&M_Disease |
| 11 | 116693163 | rs5095 | G | A | NULL | 0.634135591 | NULL&M_Disease |
| 11 | 117245988 | rs1480516974 | T | C | NULL | 0.634135591 | NULL&M_Disease |
| 11 | 117661734 | rs1014034 | G | C | NULL | 0.634135591 | NULL&M_Disease |
| 11 | 117886623 | rs17121524 | G | A | NULL | 0.634135591 | NULL&M_Disease |
| 11 | 117886753 | rs59872579 | A | G | NULL | 0.634135591 | NULL&M_Disease |
| 11 | 117886931 | rs56118320 | A | C | NULL | 0.634135591 | NULL&M_Disease |
| 11 | 117887084 | rs73011704 | T | G | NULL | 0.634135591 | NULL&M_Disease |
| 11 | 117887240 | rs73011705 | A | G | NULL | 0.634135591 | NULL&M_Disease |
| 11 | 117887667 | rs73011710 | C | T | NULL | 0.634135591 | NULL&M_Disease |
| 11 | 117887671 | rs73011712 | C | T | NULL | 0.634135591 | NULL&M_Disease |
| 11 | 117888352 | rs61598135 | A | T | NULL | 0.634135591 | NULL&M_Disease |
| 11 | 117888418 | rs58551444 | C | G | NULL | 0.634135591 | NULL&M_Disease |
| 11 | 117888564 | rs11216682 | T | C | NULL | 0.634135591 | NULL&M_Disease |
| 11 | 117888681 | rs11216683 | G | A | NULL | 0.634135591 | NULL&M_Disease |
| 11 | 117888746 | rs11216685 | T | C | NULL | 0.634135591 | NULL&M_Disease |
| 11 | 117889569 | rs73011721 | C | T | NULL | 0.634135591 | NULL&M_Disease |
| 11 | 117889690 | rs73011724 | A | T | NULL | 0.634135591 | NULL&M_Disease |
| 11 | 117889740 | rs17121528 | T | C | NULL | 0.634135591 | NULL&M_Disease |
| 11 | 117889895 | rs11216687 | A | G | NULL | 0.634135591 | NULL&M_Disease |
| 11 | 117889922 | rs11216688 | A | G | NULL | 0.634135591 | NULL&M_Disease |
| 11 | 117889959 | rs11216689 | A | C | NULL | 0.634135591 | NULL&M_Disease |
| 11 | 117890167 | rs11216690 | T | C | NULL | 0.634135591 | NULL&M_Disease |
| 11 | 117890221 | rs11216691 | G | A | NULL | 0.634135591 | NULL&M_Disease |
| 11 | 117890295 | rs11216692 | T | A | NULL | 0.634135591 | NULL&M_Disease |
| 11 | 117890386 | rs11216693 | A | C | NULL | 0.634135591 | NULL&M_Disease |
| 11 | 117890391 | rs11216694 | A | G | NULL | 0.634135591 | NULL&M_Disease |
| 11 | 117890417 | rs11216695 | A | G | NULL | 0.634135591 | NULL&M_Disease |
| 11 | 117890752 | rs59198393 | T | C | NULL | 0.634135591 | NULL&M_Disease |
| 11 | 117890857 | rs4630330 | T | G | NULL | 0.634135591 | NULL&M_Disease |
| 11 | 117890874 | rs57916935 | G | A | NULL | 0.634135591 | NULL&M_Disease |
| 11 | 117890989 | rs61579653 | A | G | NULL | 0.634135591 | NULL&M_Disease |
| 11 | 117890995 | rs58496951 | C | A | NULL | 0.634135591 | NULL&M_Disease |
| 11 | 117891384 | rs7944128 | C | T | NULL | 0.634135591 | NULL&M_Disease |
| 11 | 117891489 | rs7929175 | G | A | NULL | 0.634135591 | NULL&M_Disease |
| 11 | 117891582 | rs7944374 | C | T | NULL | 0.634135591 | NULL&M_Disease |
| 11 | 117891787 | rs73011757 | G | A | NULL | 0.634135591 | NULL&M_Disease |
| 11 | 117892322 | rs11216696 | G | A | NULL | 0.634135591 | NULL&M_Disease |
| 11 | 117892670 | rs60590812 | C | G | NULL | 0.634135591 | NULL&M_Disease |
| 11 | 117893518 | rs73013721 | T | G | NULL | 0.634135591 | NULL&M_Disease |
| 11 | 117894709 | rs73013726 | A | G | NULL | 0.634135591 | NULL&M_Disease |
| 11 | 117894899 | rs11216702 | A | T | NULL | 0.634135591 | NULL&M_Disease |
| 11 | 121154837 | rs10790430 | T | C | NULL | 0.634135591 | NULL&M_Disease |
| 11 | 121155070 | rs10790431 | C | T | NULL | 0.634135591 | NULL&M_Disease |
| 11 | 118034201 | rs11216788 | A | G | NULL | 0.634135591 | NULL&M_Disease |
| 11 | 119032689 | rs3802885 | C | A | NULL | 0.634135591 | NULL&M_Disease |
| 11 | 119351329 | rs673841 | G | A | NULL | 0.634135591 | NULL&M_Disease |
| 11 | 120090861 | rs61898282 | T | C | NULL | 0.634135591 | NULL&M_Disease |
| 11 | 116356720 | rs1508108 | G | A | NULL | 0.635294849 | NULL&M_Disease |
| 11 | 116539412 | rs1784130 | C | G | NULL | 0.635294849 | NULL&M_Disease |
| 11 | 117238526 | rs61903743 | T | A | NULL | 0.634135591 | NULL&M_Disease |
| 11 | 117240002 | rs61903744 | T | C | NULL | 0.634135591 | NULL&M_Disease |
| 11 | 117240214 | rs61903745 | T | G | NULL | 0.634135591 | NULL&M_Disease |
| 11 | 117882394 | rs11216676 | G | A | NULL | 0.634135591 | NULL&M_Disease |
| 11 | 118062552 | rs12291619 | A | G | NULL | 0.634135591 | NULL&M_Disease |
| 11 | 119352890 | rs650969 | C | T | NULL | 0.635294849 | NULL&M_Disease |
| 11 | 119353188 | rs649557 | T | C | NULL | 0.635294849 | NULL&M_Disease |
| 11 | 119353577 | rs533053 | C | G | NULL | 0.635294849 | NULL&M_Disease |
| 11 | 119904317 | rs72998668 | T | C | NULL | 0.634135591 | NULL&M_Disease |
| 11 | 120089886 | rs731512 | A | G | NULL | 0.634135591 | NULL&M_Disease |
| 11 | 120667742 | rs2186699 | G | T | NULL | 0.634135591 | NULL&M_Disease |
| 11 | 116541472 | rs1787686 | A | G | NULL | 0.635294849 | NULL&M_Disease |
| 11 | 117490186 | rs77083949 | T | C | NULL | 0.634135591 | NULL&M_Disease |
| 11 | 119975192 | rs80079694 | A | G | NULL | 0.634135591 | NULL&M_Disease |
| 11 | 120071834 | rs148205418 | G | C | NULL | 0.634135591 | NULL&M_Disease |
| 11 | 120091015 | rs61898283 | G | A | NULL | 0.634135591 | NULL&M_Disease |
| 11 | 120091077 | rs61898284 | A | G | NULL | 0.634135591 | NULL&M_Disease |
| 11 | 120092774 | rs61898287 | T | A | NULL | 0.634135591 | NULL&M_Disease |
| 11 | 121156755 | rs7128433 | T | C | NULL | 0.634135591 | NULL&M_Disease |
| 11 | 116670182 | rs11600380 | C | T | NULL | 0.635294849 | NULL&M_Disease |
| 11 | 117892362 | rs11216697 | A | G | NULL | 0.634135591 | NULL&M_Disease |
| 11 | 119501720 | rs7103203 | G | A | NULL | 0.635294849 | NULL&M_Disease |
| 11 | 121153556 | rs77726126 | G | T | NULL | 0.635294849 | NULL&M_Disease |

| | | | | | | | | | |
|---|---|---|---|---|---|---|---|---|---|
| 11 | 116558916 | rs61906117 | A | G | 0.982 | -7977.388 | -7978.491 | -1.103 | 1.28E-06 |
| 11 | 116601945 | rs3212282 | A | G | 1.041 | -7977.388 | -7978.491 | -1.103 | 1.28E-06 |
| 11 | 117555933 | rs7129597 | T | C | 0.963 | -7977.388 | -7978.491 | -1.103 | 1.28E-06 |
| 11 | 118579395 | rs11828220 | A | T | 0.976 | -7977.388 | -7978.491 | -1.103 | 1.28E-06 |
| 11 | 119355747 | rs613018 | C | T | 1.039 | -7977.388 | -7978.491 | -1.103 | 1.28E-06 |
| 11 | 120093495 | rs61898288 | A | C | 0.985 | -7977.388 | -7978.492 | -1.103 | 1.28E-06 |
| 11 | 116596464 | rs60583202 | A | G | 0.965 | -7977.388 | -7978.493 | -1.104 | 1.28E-06 |
| 11 | 116652351 | rs12274192 | G | A | 0.967 | -7977.388 | -7978.493 | -1.104 | 1.28E-06 |
| 11 | 117220324 | rs61903739 | T | C | 1.013 | -7977.388 | -7978.492 | -1.104 | 1.28E-06 |
| 11 | 117425383 | rs12804924 | T | C | 1.036 | -7977.388 | -7978.492 | -1.104 | 1.28E-06 |
| 11 | 119790068 | rs10502232 | G | A | 0.982 | -7977.388 | -7978.492 | -1.104 | 1.28E-06 |
| 11 | 119992149 | rs562015 | T | C | 0.961 | -7977.388 | -7978.493 | -1.104 | 1.28E-06 |
| 11 | 120571457 | rs10892619 | T | C | 1.039 | -7977.388 | -7978.492 | -1.104 | 1.28E-06 |
| 11 | 121154118 | rs7950536 | G | A | 0.964 | -7977.388 | -7978.493 | -1.104 | 1.28E-06 |
| 11 | 117048367 | rs117680585 | T | C | 0.977 | -7977.388 | -7978.493 | -1.105 | 1.28E-06 |
| 11 | 117165397 | rs28989500 | T | C | 1.013 | -7977.388 | -7978.493 | -1.105 | 1.28E-06 |
| 11 | 117754524 | rs10892188 | C | A | 0.963 | -7977.388 | -7978.493 | -1.105 | 1.28E-06 |
| 11 | 117898851 | rs10458939 | A | G | 0.974 | -7977.388 | -7978.493 | -1.105 | 1.28E-06 |
| 11 | 117974541 | rs1893318 | G | A | 1.039 | -7977.388 | -7978.493 | -1.105 | 1.28E-06 |
| 11 | 119806852 | rs548429 | G | A | 1.039 | -7977.388 | -7978.493 | -1.105 | 1.28E-06 |
| 11 | 121154642 | rs78303474 | C | T | 0.964 | -7977.388 | -7978.493 | -1.105 | 1.28E-06 |
| 11 | 121156037 | rs56142837 | G | T | 0.964 | -7977.388 | -7978.493 | -1.105 | 1.28E-06 |
| 11 | 121156412 | rs55903632 | C | G | 0.964 | -7977.388 | -7978.493 | -1.105 | 1.28E-06 |
| 11 | 116527521 | rs55776558 | C | T | 0.984 | -7977.388 | -7978.494 | -1.106 | 1.28E-06 |
| 11 | 116693163 | rs5095 | G | A | 1.041 | -7977.388 | -7978.495 | -1.106 | 1.28E-06 |
| 11 | 117245988 | rs1480516974 | T | C | 0.961 | -7977.388 | -7978.494 | -1.106 | 1.28E-06 |
| 11 | 117661734 | rs1014034 | G | C | 1.038 | -7977.388 | -7978.495 | -1.106 | 1.28E-06 |
| 11 | 117886623 | rs17121524 | G | A | 0.974 | -7977.388 | -7978.494 | -1.106 | 1.28E-06 |
| 11 | 117886753 | rs59872579 | A | G | 0.974 | -7977.388 | -7978.494 | -1.106 | 1.28E-06 |
| 11 | 117886931 | rs56118320 | A | C | 0.974 | -7977.388 | -7978.494 | -1.106 | 1.28E-06 |
| 11 | 117887084 | rs73011704 | T | G | 0.974 | -7977.388 | -7978.494 | -1.106 | 1.28E-06 |
| 11 | 117887240 | rs73011705 | A | G | 0.974 | -7977.388 | -7978.494 | -1.106 | 1.28E-06 |
| 11 | 117887667 | rs73011710 | C | T | 0.974 | -7977.388 | -7978.494 | -1.106 | 1.28E-06 |
| 11 | 117887671 | rs73011712 | C | T | 0.974 | -7977.388 | -7978.494 | -1.106 | 1.28E-06 |
| 11 | 117888352 | rs61598135 | A | T | 0.974 | -7977.388 | -7978.494 | -1.106 | 1.28E-06 |
| 11 | 117888418 | rs58551444 | C | G | 0.974 | -7977.388 | -7978.494 | -1.106 | 1.28E-06 |
| 11 | 117888564 | rs11216682 | T | C | 0.974 | -7977.388 | -7978.494 | -1.106 | 1.28E-06 |
| 11 | 117888681 | rs11216683 | G | A | 0.974 | -7977.388 | -7978.494 | -1.106 | 1.28E-06 |
| 11 | 117888746 | rs11216685 | T | C | 0.974 | -7977.388 | -7978.494 | -1.106 | 1.28E-06 |
| 11 | 117889569 | rs73011721 | C | T | 0.974 | -7977.388 | -7978.494 | -1.106 | 1.28E-06 |
| 11 | 117889690 | rs73011724 | A | T | 0.974 | -7977.388 | -7978.494 | -1.106 | 1.28E-06 |
| 11 | 117889740 | rs17121528 | T | C | 0.974 | -7977.388 | -7978.494 | -1.106 | 1.28E-06 |
| 11 | 117889895 | rs11216687 | A | G | 0.974 | -7977.388 | -7978.494 | -1.106 | 1.28E-06 |
| 11 | 117889922 | rs11216688 | A | G | 0.974 | -7977.388 | -7978.494 | -1.106 | 1.28E-06 |
| 11 | 117889959 | rs11216689 | A | C | 0.974 | -7977.388 | -7978.494 | -1.106 | 1.28E-06 |
| 11 | 117890167 | rs11216690 | T | C | 0.974 | -7977.388 | -7978.494 | -1.106 | 1.28E-06 |
| 11 | 117890221 | rs11216691 | G | A | 0.974 | -7977.388 | -7978.494 | -1.106 | 1.28E-06 |
| 11 | 117890295 | rs11216692 | T | A | 0.974 | -7977.388 | -7978.494 | -1.106 | 1.28E-06 |
| 11 | 117890386 | rs11216693 | A | C | 0.974 | -7977.388 | -7978.494 | -1.106 | 1.28E-06 |
| 11 | 117890391 | rs11216694 | A | G | 0.974 | -7977.388 | -7978.494 | -1.106 | 1.28E-06 |
| 11 | 117890417 | rs11216695 | A | G | 0.974 | -7977.388 | -7978.494 | -1.106 | 1.28E-06 |
| 11 | 117890752 | rs59198393 | T | C | 0.974 | -7977.388 | -7978.494 | -1.106 | 1.28E-06 |
| 11 | 117890857 | rs4630330 | T | G | 0.974 | -7977.388 | -7978.494 | -1.106 | 1.28E-06 |
| 11 | 117890874 | rs57916935 | G | A | 0.974 | -7977.388 | -7978.494 | -1.106 | 1.28E-06 |
| 11 | 117890989 | rs61579653 | A | G | 0.974 | -7977.388 | -7978.494 | -1.106 | 1.28E-06 |
| 11 | 117890995 | rs58496951 | C | A | 0.974 | -7977.388 | -7978.494 | -1.106 | 1.28E-06 |
| 11 | 117891384 | rs7944128 | C | T | 0.974 | -7977.388 | -7978.494 | -1.106 | 1.28E-06 |
| 11 | 117891489 | rs7929175 | G | A | 0.974 | -7977.388 | -7978.494 | -1.106 | 1.28E-06 |
| 11 | 117891582 | rs7944374 | C | T | 0.974 | -7977.388 | -7978.494 | -1.106 | 1.28E-06 |
| 11 | 117891787 | rs73011757 | G | A | 0.974 | -7977.388 | -7978.494 | -1.106 | 1.28E-06 |
| 11 | 117892322 | rs11216696 | G | A | 0.974 | -7977.388 | -7978.494 | -1.106 | 1.28E-06 |
| 11 | 117892670 | rs60590812 | C | G | 0.974 | -7977.388 | -7978.494 | -1.106 | 1.28E-06 |
| 11 | 117893518 | rs73013721 | T | G | 0.974 | -7977.388 | -7978.494 | -1.106 | 1.28E-06 |
| 11 | 117894709 | rs73013726 | A | G | 0.974 | -7977.388 | -7978.495 | -1.106 | 1.28E-06 |
| 11 | 117894899 | rs11216702 | A | T | 0.974 | -7977.388 | -7978.495 | -1.106 | 1.28E-06 |
| 11 | 121154837 | rs10790430 | T | C | 1.03 | -7977.388 | -7978.495 | -1.106 | 1.28E-06 |
| 11 | 121155070 | rs10790431 | C | T | 1.03 | -7977.388 | -7978.495 | -1.107 | 1.28E-06 |
| 11 | 118034201 | rs11216788 | A | G | 0.963 | -7977.388 | -7978.496 | -1.107 | 1.28E-06 |
| 11 | 119032689 | rs3802885 | C | A | 1.033 | -7977.388 | -7978.496 | -1.107 | 1.28E-06 |
| 11 | 119351329 | rs673841 | G | A | 1.039 | -7977.388 | -7978.496 | -1.107 | 1.28E-06 |
| 11 | 120090861 | rs61898282 | T | C | 0.985 | -7977.388 | -7978.496 | -1.107 | 1.28E-06 |
| 11 | 116356720 | rs1508108 | G | A | 1.039 | -7977.388 | -7978.496 | -1.108 | 1.28E-06 |
| 11 | 116539412 | rs1784130 | C | G | 1.04 | -7977.388 | -7978.497 | -1.108 | 1.28E-06 |
| 11 | 117238526 | rs61903743 | T | A | 1.011 | -7977.388 | -7978.497 | -1.108 | 1.28E-06 |
| 11 | 117240002 | rs61903744 | T | C | 1.011 | -7977.388 | -7978.497 | -1.108 | 1.28E-06 |
| 11 | 117240214 | rs61903745 | T | G | 1.011 | -7977.388 | -7978.497 | -1.108 | 1.28E-06 |
| 11 | 117882394 | rs11216676 | G | A | 0.974 | -7977.388 | -7978.496 | -1.108 | 1.28E-06 |
| 11 | 118062552 | rs12291619 | A | G | 0.979 | -7977.388 | -7978.497 | -1.108 | 1.28E-06 |
| 11 | 119352890 | rs650969 | C | T | 1.038 | -7977.388 | -7978.497 | -1.108 | 1.28E-06 |
| 11 | 119353188 | rs649557 | T | C | 1.038 | -7977.388 | -7978.497 | -1.108 | 1.28E-06 |
| 11 | 119353577 | rs533053 | C | G | 1.038 | -7977.388 | -7978.497 | -1.108 | 1.28E-06 |
| 11 | 119904317 | rs72998668 | T | C | 0.969 | -7977.388 | -7978.496 | -1.108 | 1.28E-06 |
| 11 | 120089886 | rs731512 | A | G | 0.965 | -7977.388 | -7978.497 | -1.108 | 1.28E-06 |
| 11 | 120667742 | rs2186699 | G | T | 0.968 | -7977.388 | -7978.496 | -1.108 | 1.28E-06 |
| 11 | 116541472 | rs1787686 | A | G | 0.963 | -7977.388 | -7978.497 | -1.109 | 1.28E-06 |
| 11 | 117490186 | rs77083949 | T | C | 0.971 | -7977.388 | -7978.497 | -1.109 | 1.28E-06 |
| 11 | 119975192 | rs80079694 | A | G | 1.028 | -7977.388 | -7978.497 | -1.109 | 1.28E-06 |
| 11 | 120071834 | rs148205418 | G | C | 0.989 | -7977.388 | -7978.497 | -1.109 | 1.28E-06 |
| 11 | 120091015 | rs61898283 | G | A | 0.986 | -7977.388 | -7978.498 | -1.109 | 1.28E-06 |
| 11 | 120091077 | rs61898284 | A | G | 0.986 | -7977.388 | -7978.498 | -1.109 | 1.28E-06 |
| 11 | 120092774 | rs61898287 | T | A | 0.986 | -7977.388 | -7978.498 | -1.109 | 1.28E-06 |
| 11 | 121156755 | rs7128433 | T | C | 0.965 | -7977.388 | -7978.497 | -1.109 | 1.28E-06 |
| 11 | 116670182 | rs11600380 | C | T | 1.035 | -7977.388 | -7978.499 | -1.11 | 1.27E-06 |
| 11 | 117892362 | rs11216697 | A | G | 0.974 | -7977.388 | -7978.498 | -1.11 | 1.27E-06 |
| 11 | 119501720 | rs7103203 | G | A | 0.961 | -7977.388 | -7978.498 | -1.11 | 1.27E-06 |
| 11 | 121153556 | rs77726126 | G | T | 0.965 | -7977.388 | -7978.498 | -1.11 | 1.27E-06 |

| | | | | | | | |
|---|---|---|---|---|---|---|---|
| 11 | 116575614 | rs76124005 | G | T | NULL | 0.635294849 | NULL&M_Disease |
| 11 | 117049508 | rs117933040 | G | A | NULL | 0.635294849 | NULL&M_Disease |
| 11 | 117207665 | rs7932967 | T | G | NULL | 0.635294849 | NULL&M_Disease |
| 11 | 116239385 | rs12574216 | G | T | NULL | 0.635294849 | NULL&M_Disease |
| 11 | 116784824 | rs10892045 | A | C | NULL | 0.635294849 | NULL&M_Disease |
| 11 | 117266312 | rs2305830 | G | C | NULL | 0.635294849 | NULL&M_Disease |
| 11 | 117558035 | rs78659090 | G | A | NULL | 0.635294849 | NULL&M_Disease |
| 11 | 117786899 | rs117268885 | T | C | NULL | 0.635294849 | NULL&M_Disease |
| 11 | 117891475 | rs7929257 | T | C | NULL | 0.635294849 | NULL&M_Disease |
| 11 | 119759466 | rs548636 | C | T | NULL | 0.635294849 | NULL&M_Disease |
| 11 | 120073966 | rs12420850 | A | C | NULL | 0.635294849 | NULL&M_Disease |
| 11 | 120981408 | rs1829 | T | C | NULL | 0.635294849 | NULL&M_Disease |
| 11 | 116528544 | rs74444640 | T | C | NULL | 0.635294849 | NULL&M_Disease |
| 11 | 117342333 | rs12363004 | C | T | NULL | 0.635294849 | NULL&M_Disease |
| 11 | 119832901 | rs78764865 | T | C | NULL | 0.635294849 | NULL&M_Disease |
| 11 | 121071161 | rs182784500 | G | A | NULL | 0.635294849 | NULL&M_Disease |
| 11 | 116575880 | rs74939223 | G | A | NULL | 0.635294849 | NULL&M_Disease |
| 11 | 117036805 | rs7931335 | C | T | NULL | 0.635294849 | NULL&M_Disease |
| 11 | 117425313 | rs12804723 | T | A | NULL | 0.635294849 | NULL&M_Disease |
| 11 | 118231584 | rs4938507 | G | A | NULL | 0.635294849 | NULL&M_Disease |
| 11 | 118483802 | rs72999455 | G | A | NULL | 0.635294849 | NULL&M_Disease |
| 11 | 118993801 | rs149703780 | A | G | NULL | 0.635294849 | NULL&M_Disease |
| 11 | 119000331 | rs141270315 | T | C | NULL | 0.635294849 | NULL&M_Disease |
| 11 | 119006788 | rs79487987 | A | G | NULL | 0.635294849 | NULL&M_Disease |
| 11 | 120013307 | rs470876 | C | T | NULL | 0.635294849 | NULL&M_Disease |
| 11 | 120104859 | rs41460849 | A | G | NULL | 0.635294849 | NULL&M_Disease |
| 11 | 120340060 | rs2305013 | T | A | NULL | 0.635294849 | NULL&M_Disease |
| 11 | 120788631 | rs17124548 | T | A | NULL | 0.635294849 | NULL&M_Disease |
| 11 | 117897020 | rs11216705 | C | T | NULL | 0.635294849 | NULL&M_Disease |
| 11 | 118681088 | rs2508922 | A | G | NULL | 0.635294849 | NULL&M_Disease |
| 11 | 118955509 | rs686624 | T | A | NULL | 0.635294849 | NULL&M_Disease |
| 11 | 120163574 | rs11607289 | G | A | NULL | 0.635294849 | NULL&M_Disease |
| 11 | 120163930 | rs11603274 | A | G | NULL | 0.635294849 | NULL&M_Disease |
| 11 | 120790862 | rs12421080 | A | T | NULL | 0.635294849 | NULL&M_Disease |
| 11 | 116604514 | rs4938307 | A | G | NULL | 0.635294849 | NULL&M_Disease |
| 11 | 117124696 | rs610775 | A | G | NULL | 0.635294849 | NULL&M_Disease |
| 11 | 117633470 | rs10892170 | G | A | NULL | 0.635294849 | NULL&M_Disease |
| 11 | 119241743 | rs602492 | T | G | NULL | 0.635294849 | NULL&M_Disease |
| 11 | 119647079 | rs10892455 | C | T | NULL | 0.635294849 | NULL&M_Disease |
| 11 | 119675718 | rs7926229 | C | T | NULL | 0.635294849 | NULL&M_Disease |
| 11 | 116527728 | rs575280 | G | A | NULL | 0.635294849 | NULL&M_Disease |
| 11 | 117225982 | rs74761525 | T | C | NULL | 0.635294849 | NULL&M_Disease |
| 11 | 117557224 | rs59071712 | A | T | NULL | 0.635294849 | NULL&M_Disease |
| 11 | 117763500 | rs111537986 | T | C | NULL | 0.635294849 | NULL&M_Disease |
| 11 | 120166817 | rs6589806 | C | T | NULL | 0.635294849 | NULL&M_Disease |
| 11 | 120418687 | rs61900897 | C | G | NULL | 0.635294849 | NULL&M_Disease |
| 11 | 120809198 | rs580711 | G | C | NULL | 0.635294849 | NULL&M_Disease |
| 11 | 116485334 | rs498736 | C | G | NULL | 0.635294849 | NULL&M_Disease |
| 11 | 117123782 | rs61903662 | A | G | NULL | 0.635294849 | NULL&M_Disease |
| 11 | 117424953 | rs11601133 | G | A | NULL | 0.635294849 | NULL&M_Disease |
| 11 | 117974832 | rs9645662 | A | G | NULL | 0.635294849 | NULL&M_Disease |
| 11 | 119026328 | rs4938633 | A | G | NULL | 0.635294849 | NULL&M_Disease |
| 11 | 119698359 | rs17123366 | A | G | NULL | 0.635294849 | NULL&M_Disease |
| 11 | 120073606 | rs61898269 | C | A | NULL | 0.635294849 | NULL&M_Disease |
| 11 | 120157863 | rs17123780 | A | G | NULL | 0.635294849 | NULL&M_Disease |
| 11 | 121065324 | rs117840927 | A | C | NULL | 0.635294849 | NULL&M_Disease |
| 11 | 121074895 | rs74533222 | C | T | NULL | 0.635294849 | NULL&M_Disease |
| 11 | 116514092 | rs78642155 | C | G | NULL | 0.635294849 | NULL&M_Disease |
| 11 | 117896521 | rs56137014 | A | T | NULL | 0.635294849 | NULL&M_Disease |
| 11 | 119005900 | rs75905572 | G | C | NULL | 0.635294849 | NULL&M_Disease |
| 11 | 120042087 | rs529657 | C | T | NULL | 0.635294849 | NULL&M_Disease |
| 11 | 120164141 | rs11603307 | A | G | NULL | 0.635294849 | NULL&M_Disease |
| 11 | 116224964 | rs1899052 | G | C | NULL | 0.63645254 | NULL&M_Disease |
| 11 | 116312418 | rs73005215 | G | A | NULL | 0.635294849 | NULL&M_Disease |
| 11 | 116325156 | rs580363 | C | G | NULL | 0.63645254 | NULL&M_Disease |
| 11 | 117434215 | rs685916 | A | G | NULL | 0.63645254 | NULL&M_Disease |
| 11 | 117845618 | rs1217688880 | C | T | NULL | 0.63645254 | NULL&M_Disease |
| 11 | 119252151 | rs565296 | C | T | NULL | 0.63645254 | NULL&M_Disease |
| 11 | 120982105 | rs606916 | G | C | NULL | 0.63645254 | NULL&M_Disease |
| 11 | 116363370 | rs7112876 | C | G | NULL | 0.63645254 | NULL&M_Disease |
| 11 | 116603134 | rs2008915 | C | G | NULL | 0.63645254 | NULL&M_Disease |
| 11 | 117906510 | rs10892204 | T | C | NULL | 0.63645254 | NULL&M_Disease |
| 11 | 120469504 | rs4936522 | G | C | NULL | 0.63645254 | NULL&M_Disease |
| 11 | 120485547 | rs4245039 | C | A | NULL | 0.63645254 | NULL&M_Disease |
| 11 | 120666825 | rs2155260 | A | G | NULL | 0.63645254 | NULL&M_Disease |
| 11 | 120814342 | rs609989 | A | G | NULL | 0.63645254 | NULL&M_Disease |
| 11 | 120816277 | rs471403 | A | G | NULL | 0.63645254 | NULL&M_Disease |
| 11 | 120823104 | rs582718 | T | C | NULL | 0.63645254 | NULL&M_Disease |
| 11 | 116514329 | rs117689643 | T | G | NULL | 0.63645254 | NULL&M_Disease |
| 11 | 116557216 | rs1145211 | A | C | NULL | 0.63645254 | NULL&M_Disease |
| 11 | 118965479 | rs1134734 | A | G | NULL | 0.63645254 | NULL&M_Disease |
| 11 | 119649903 | rs12808266 | T | C | NULL | 0.63645254 | NULL&M_Disease |
| 11 | 120095847 | rs77786799 | T | C | NULL | 0.63645254 | NULL&M_Disease |
| 11 | 120813715 | rs511939 | G | C | NULL | 0.63645254 | NULL&M_Disease |
| 11 | 120819562 | rs684913 | T | C | NULL | 0.63645254 | NULL&M_Disease |
| 11 | 116551449 | rs2846012 | C | T | NULL | 0.63645254 | NULL&M_Disease |
| 11 | 116551538 | rs1784104 | C | T | NULL | 0.63645254 | NULL&M_Disease |
| 11 | 119026330 | rs4938634 | C | A | NULL | 0.63645254 | NULL&M_Disease |
| 11 | 119844641 | rs72994672 | G | T | NULL | 0.63645254 | NULL&M_Disease |
| 11 | 120164658 | rs11603757 | T | G | NULL | 0.63645254 | NULL&M_Disease |
| 11 | 120524275 | rs77325329 | A | G | NULL | 0.63645254 | NULL&M_Disease |
| 11 | 120823528 | rs10750178 | A | G | NULL | 0.63645254 | NULL&M_Disease |
| 11 | 121005464 | rs12802185 | A | T | NULL | 0.63645254 | NULL&M_Disease |
| 11 | 121170025 | rs12275106 | C | A | NULL | 0.63645254 | NULL&M_Disease |
| 11 | 121170179 | rs17125142 | G | A | NULL | 0.63645254 | NULL&M_Disease |
| 11 | 116525306 | rs76860051 | C | A | NULL | 0.63645254 | NULL&M_Disease |

| | | | | | | | | | |
|---|---|---|---|---|---|---|---|---|---|
| 11 | 116575614 | rs76124005 | G | T | 1.023 | -7977.388 | -7978.499 | -1.111 | 1.27E-06 |
| 11 | 117049508 | rs117933040 | G | A | 0.977 | -7977.388 | -7978.499 | -1.111 | 1.27E-06 |
| 11 | 117207665 | rs7932967 | T | G | 1.039 | -7977.388 | -7978.499 | -1.111 | 1.27E-06 |
| 11 | 116239385 | rs12574216 | G | T | 0.986 | -7977.388 | -7978.501 | -1.112 | 1.27E-06 |
| 11 | 116784824 | rs10892045 | A | C | 0.961 | -7977.388 | -7978.5 | -1.112 | 1.27E-06 |
| 11 | 117266312 | rs2305830 | G | C | 1.039 | -7977.388 | -7978.5 | -1.112 | 1.27E-06 |
| 11 | 117558035 | rs78659090 | G | A | 0.964 | -7977.388 | -7978.501 | -1.112 | 1.27E-06 |
| 11 | 117786899 | rs117268885 | T | C | 1.013 | -7977.388 | -7978.5 | -1.112 | 1.27E-06 |
| 11 | 117891475 | rs7929257 | T | C | 0.975 | -7977.388 | -7978.5 | -1.112 | 1.27E-06 |
| 11 | 119759466 | rs548636 | C | T | 1.037 | -7977.388 | -7978.5 | -1.112 | 1.27E-06 |
| 11 | 120073966 | rs12420850 | A | C | 0.972 | -7977.388 | -7978.501 | -1.112 | 1.27E-06 |
| 11 | 120981408 | rs1829 | T | C | 0.961 | -7977.388 | -7978.501 | -1.112 | 1.27E-06 |
| 11 | 116528544 | rs74444640 | T | C | 0.982 | -7977.388 | -7978.501 | -1.113 | 1.27E-06 |
| 11 | 117342333 | rs12363004 | C | T | 1.038 | -7977.388 | -7978.501 | -1.113 | 1.27E-06 |
| 11 | 119832901 | rs78764865 | T | C | 1.035 | -7977.388 | -7978.502 | -1.113 | 1.27E-06 |
| 11 | 121071161 | rs182784500 | G | A | 1.006 | -7977.388 | -7978.502 | -1.113 | 1.27E-06 |
| 11 | 116575880 | rs74939223 | G | A | 1.023 | -7977.388 | -7978.502 | -1.114 | 1.27E-06 |
| 11 | 117036805 | rs7931335 | C | T | 0.961 | -7977.388 | -7978.502 | -1.114 | 1.27E-06 |
| 11 | 117425313 | rs12804723 | T | A | 1.035 | -7977.388 | -7978.502 | -1.114 | 1.27E-06 |
| 11 | 118231584 | rs4938507 | G | A | 1.04 | -7977.388 | -7978.502 | -1.114 | 1.27E-06 |
| 11 | 118483802 | rs72999455 | G | A | 0.977 | -7977.388 | -7978.502 | -1.114 | 1.27E-06 |
| 11 | 118993801 | rs149703780 | A | G | 1.032 | -7977.388 | -7978.502 | -1.114 | 1.27E-06 |
| 11 | 119000331 | rs141270315 | T | C | 1.032 | -7977.388 | -7978.502 | -1.114 | 1.27E-06 |
| 11 | 119006788 | rs79487987 | A | G | 1.032 | -7977.388 | -7978.502 | -1.114 | 1.27E-06 |
| 11 | 120013307 | rs470876 | C | T | 0.964 | -7977.388 | -7978.503 | -1.114 | 1.27E-06 |
| 11 | 120104859 | rs41460849 | A | G | 1.012 | -7977.388 | -7978.502 | -1.114 | 1.27E-06 |
| 11 | 120340060 | rs2305013 | T | A | 0.974 | -7977.388 | -7978.502 | -1.114 | 1.27E-06 |
| 11 | 120788631 | rs17124548 | T | A | 1.019 | -7977.388 | -7978.502 | -1.114 | 1.27E-06 |
| 11 | 117897020 | rs11216705 | C | T | 0.975 | -7977.388 | -7978.504 | -1.115 | 1.27E-06 |
| 11 | 118681088 | rs2508922 | A | G | 1.04 | -7977.388 | -7978.503 | -1.115 | 1.27E-06 |
| 11 | 118955509 | rs686624 | T | A | 1.04 | -7977.388 | -7978.503 | -1.115 | 1.27E-06 |
| 11 | 120163574 | rs11607289 | G | A | 1.035 | -7977.388 | -7978.503 | -1.115 | 1.27E-06 |
| 11 | 120163930 | rs11603274 | A | G | 1.035 | -7977.388 | -7978.503 | -1.115 | 1.27E-06 |
| 11 | 120790862 | rs12421080 | A | T | 1.019 | -7977.388 | -7978.503 | -1.115 | 1.27E-06 |
| 11 | 116604514 | rs4938307 | A | G | 1.04 | -7977.388 | -7978.504 | -1.116 | 1.27E-06 |
| 11 | 117124696 | rs610775 | A | G | 0.963 | -7977.388 | -7978.505 | -1.116 | 1.27E-06 |
| 11 | 117633470 | rs10892170 | G | A | 1.029 | -7977.388 | -7978.504 | -1.116 | 1.27E-06 |
| 11 | 119241743 | rs602492 | T | G | 0.961 | -7977.388 | -7978.504 | -1.116 | 1.27E-06 |
| 11 | 119647079 | rs10892455 | C | T | 0.961 | -7977.388 | -7978.505 | -1.116 | 1.27E-06 |
| 11 | 119675718 | rs7926229 | C | T | 0.961 | -7977.388 | -7978.504 | -1.116 | 1.27E-06 |
| 11 | 116527728 | rs575280 | G | A | 1.04 | -7977.388 | -7978.505 | -1.117 | 1.27E-06 |
| 11 | 117225982 | rs74761525 | T | C | 0.962 | -7977.388 | -7978.505 | -1.117 | 1.27E-06 |
| 11 | 117557224 | rs59071712 | A | T | 0.964 | -7977.388 | -7978.505 | -1.117 | 1.27E-06 |
| 11 | 117763500 | rs111537986 | T | C | 0.964 | -7977.388 | -7978.505 | -1.117 | 1.27E-06 |
| 11 | 120166817 | rs6589806 | C | T | 1.039 | -7977.388 | -7978.505 | -1.117 | 1.27E-06 |
| 11 | 120418687 | rs61900897 | C | G | 1.038 | -7977.388 | -7978.506 | -1.117 | 1.27E-06 |
| 11 | 120809198 | rs580711 | G | C | 1.005 | -7977.388 | -7978.506 | -1.117 | 1.27E-06 |
| 11 | 116485334 | rs498736 | C | G | 1.04 | -7977.388 | -7978.506 | -1.118 | 1.26E-06 |
| 11 | 117123782 | rs61903662 | A | G | 1.023 | -7977.388 | -7978.507 | -1.118 | 1.26E-06 |
| 11 | 117424953 | rs11601133 | G | A | 1.034 | -7977.388 | -7978.506 | -1.118 | 1.26E-06 |
| 11 | 117974832 | rs9645662 | A | G | 0.985 | -7977.388 | -7978.506 | -1.118 | 1.26E-06 |
| 11 | 119026328 | rs4938633 | A | G | 1.032 | -7977.388 | -7978.507 | -1.118 | 1.26E-06 |
| 11 | 119698359 | rs17123366 | A | G | 1.035 | -7977.388 | -7978.507 | -1.118 | 1.26E-06 |
| 11 | 120073606 | rs61898269 | C | A | 0.993 | -7977.388 | -7978.506 | -1.118 | 1.26E-06 |
| 11 | 120157863 | rs17123780 | A | G | 1.033 | -7977.388 | -7978.507 | -1.118 | 1.26E-06 |
| 11 | 121065324 | rs117840927 | A | C | 1.003 | -7977.388 | -7978.506 | -1.118 | 1.26E-06 |
| 11 | 121074895 | rs74533222 | C | T | 1.003 | -7977.388 | -7978.506 | -1.118 | 1.26E-06 |
| 11 | 116514092 | rs78642155 | C | G | 1.033 | -7977.388 | -7978.507 | -1.119 | 1.26E-06 |
| 11 | 117896521 | rs56137014 | A | T | 0.976 | -7977.388 | -7978.507 | -1.119 | 1.26E-06 |
| 11 | 119005900 | rs75905572 | G | C | 1.032 | -7977.388 | -7978.507 | -1.119 | 1.26E-06 |
| 11 | 120042087 | rs529657 | C | T | 0.965 | -7977.388 | -7978.507 | -1.119 | 1.26E-06 |
| 11 | 120164141 | rs11603307 | A | G | 1.035 | -7977.388 | -7978.508 | -1.119 | 1.26E-06 |
| 11 | 116224964 | rs1899052 | G | C | 1.034 | -7977.388 | -7978.509 | -1.12 | 1.26E-06 |
| 11 | 116312418 | rs73005215 | G | A | 0.992 | -7977.388 | -7978.508 | -1.12 | 1.26E-06 |
| 11 | 116325156 | rs580363 | C | G | 1.04 | -7977.388 | -7978.508 | -1.12 | 1.26E-06 |
| 11 | 117434215 | rs685916 | A | G | 1.038 | -7977.388 | -7978.508 | -1.12 | 1.26E-06 |
| 11 | 117845618 | rs1217688880 | C | T | 0.964 | -7977.388 | -7978.508 | -1.12 | 1.26E-06 |
| 11 | 119252151 | rs565296 | C | T | 0.961 | -7977.388 | -7978.509 | -1.12 | 1.26E-06 |
| 11 | 120982105 | rs606916 | G | C | 0.962 | -7977.388 | -7978.508 | -1.12 | 1.26E-06 |
| 11 | 116363370 | rs7112876 | C | G | 1.027 | -7977.388 | -7978.509 | -1.121 | 1.26E-06 |
| 11 | 116603134 | rs2008915 | C | G | 1.04 | -7977.388 | -7978.51 | -1.121 | 1.26E-06 |
| 11 | 117906510 | rs10892204 | T | C | 1.039 | -7977.388 | -7978.509 | -1.121 | 1.26E-06 |
| 11 | 120469504 | rs4936522 | G | C | 0.963 | -7977.388 | -7978.509 | -1.121 | 1.26E-06 |
| 11 | 120485547 | rs4245039 | C | A | 0.963 | -7977.388 | -7978.51 | -1.121 | 1.26E-06 |
| 11 | 120666825 | rs2155260 | A | G | 0.969 | -7977.388 | -7978.51 | -1.121 | 1.26E-06 |
| 11 | 120814342 | rs609989 | A | G | 1 | -7977.388 | -7978.51 | -1.121 | 1.26E-06 |
| 11 | 120816277 | rs471403 | A | G | 1 | -7977.388 | -7978.51 | -1.121 | 1.26E-06 |
| 11 | 120823104 | rs582718 | T | C | 0.994 | -7977.388 | -7978.51 | -1.121 | 1.26E-06 |
| 11 | 116514329 | rs117689643 | T | G | 1.033 | -7977.388 | -7978.511 | -1.122 | 1.26E-06 |
| 11 | 116557216 | rs1145211 | A | C | 0.963 | -7977.388 | -7978.51 | -1.122 | 1.26E-06 |
| 11 | 118965479 | rs1134734 | A | G | 1.038 | -7977.388 | -7978.51 | -1.122 | 1.26E-06 |
| 11 | 119649903 | rs12808266 | T | C | 1.033 | -7977.388 | -7978.51 | -1.122 | 1.26E-06 |
| 11 | 120095847 | rs77786799 | T | C | 0.991 | -7977.388 | -7978.51 | -1.122 | 1.26E-06 |
| 11 | 120813715 | rs511939 | G | C | 1.001 | -7977.388 | -7978.51 | -1.122 | 1.26E-06 |
| 11 | 120819562 | rs684913 | T | C | 0.999 | -7977.388 | -7978.51 | -1.122 | 1.26E-06 |
| 11 | 116551449 | rs2846012 | C | T | 1.04 | -7977.388 | -7978.511 | -1.123 | 1.26E-06 |
| 11 | 116551538 | rs1784104 | C | T | 1.04 | -7977.388 | -7978.511 | -1.123 | 1.26E-06 |
| 11 | 119026330 | rs4938634 | C | A | 1.031 | -7977.388 | -7978.511 | -1.123 | 1.26E-06 |
| 11 | 119844641 | rs72994672 | G | T | 1.021 | -7977.388 | -7978.512 | -1.123 | 1.26E-06 |
| 11 | 120164658 | rs11603757 | T | G | 1.035 | -7977.388 | -7978.512 | -1.123 | 1.26E-06 |
| 11 | 120524275 | rs77325329 | A | G | 1.009 | -7977.388 | -7978.511 | -1.123 | 1.26E-06 |
| 11 | 120823528 | rs10750178 | A | G | 0.997 | -7977.388 | -7978.511 | -1.123 | 1.26E-06 |
| 11 | 121005464 | rs12802185 | A | T | 1.04 | -7977.388 | -7978.512 | -1.123 | 1.26E-06 |
| 11 | 121170025 | rs12275106 | C | A | 0.966 | -7977.388 | -7978.511 | -1.123 | 1.26E-06 |
| 11 | 121170179 | rs17125142 | G | A | 0.966 | -7977.388 | -7978.511 | -1.123 | 1.26E-06 |
| 11 | 116525306 | rs76860051 | C | A | 1.032 | -7977.388 | -7978.513 | -1.124 | 1.26E-06 |

| | | | | | | | |
|---|---|---|---|---|---|---|---|
| 11 | 117737382 | rs7118101 | C | T | NULL | 0.63645254 | NULL&M_Disease |
| 11 | 117737440 | rs876798 | C | T | NULL | 0.63645254 | NULL&M_Disease |
| 11 | 117776003 | rs67422215 | T | C | NULL | 0.63645254 | NULL&M_Disease |
| 11 | 119026460 | rs4938635 | T | C | NULL | 0.63645254 | NULL&M_Disease |
| 11 | 119026469 | rs4938636 | T | A | NULL | 0.63645254 | NULL&M_Disease |
| 11 | 119242556 | rs1652024 | C | A | NULL | 0.63645254 | NULL&M_Disease |
| 11 | 117061961 | rs499713 | T | C | NULL | 0.63645254 | NULL&M_Disease |
| 11 | 117882792 | rs11216677 | T | C | NULL | 0.63645254 | NULL&M_Disease |
| 11 | 118157681 | rs34250455 | T | C | NULL | 0.63645254 | NULL&M_Disease |
| 11 | 119731345 | rs11607827 | G | T | NULL | 0.63645254 | NULL&M_Disease |
| 11 | 120489650 | rs76273814 | T | C | NULL | 0.63645254 | NULL&M_Disease |
| 11 | 116501944 | rs11824108 | G | A | NULL | 0.63645254 | NULL&M_Disease |
| 11 | 116514319 | rs116948654 | C | T | NULL | 0.63645254 | NULL&M_Disease |
| 11 | 116527281 | rs12420386 | A | C | NULL | 0.63645254 | NULL&M_Disease |
| 11 | 116561628 | rs61905060 | A | G | NULL | 0.63645254 | NULL&M_Disease |
| 11 | 117346513 | rs540469 | C | T | NULL | 0.63645254 | NULL&M_Disease |
| 11 | 117392826 | rs10892135 | A | G | NULL | 0.63645254 | NULL&M_Disease |
| 11 | 117633054 | rs10892169 | A | G | NULL | 0.63645254 | NULL&M_Disease |
| 11 | 117859209 | rs4252249 | A | G | NULL | 0.63645254 | NULL&M_Disease |
| 11 | 118970682 | rs79524875 | T | C | NULL | 0.63645254 | NULL&M_Disease |
| 11 | 119253797 | rs512608 | G | A | NULL | 0.63645254 | NULL&M_Disease |
| 11 | 117762694 | rs73020457 | T | C | NULL | 0.63645254 | NULL&M_Disease |
| 11 | 118504742 | rs633683 | T | C | NULL | 0.63645254 | NULL&M_Disease |
| 11 | 118568427 | rs117361624 | T | C | NULL | 0.63645254 | NULL&M_Disease |
| 11 | 119897773 | rs118135828 | A | G | NULL | 0.63645254 | NULL&M_Disease |
| 11 | 119897774 | rs592765 | A | C | NULL | 0.63645254 | NULL&M_Disease |
| 11 | 120072470 | rs61898267 | G | C | NULL | 0.63645254 | NULL&M_Disease |
| 11 | 120104292 | rs61898294 | A | G | NULL | 0.63645254 | NULL&M_Disease |
| 11 | 116537634 | rs11216083 | T | C | NULL | 0.63645254 | NULL&M_Disease |
| 11 | 116577694 | rs75507001 | A | C | NULL | 0.63645254 | NULL&M_Disease |
| 11 | 117424807 | rs12286677 | T | C | NULL | 0.63645254 | NULL&M_Disease |
| 11 | 117425206 | rs12799960 | T | C | NULL | 0.63645254 | NULL&M_Disease |
| 11 | 117772002 | rs11823721 | C | T | NULL | 0.63645254 | NULL&M_Disease |
| 11 | 117898042 | rs192508986 | T | C | NULL | 0.63645254 | NULL&M_Disease |
| 11 | 117898084 | rs12223288 | G | A | NULL | 0.63645254 | NULL&M_Disease |
| 11 | 117898155 | rs10431083 | A | G | NULL | 0.63645254 | NULL&M_Disease |
| 11 | 117974759 | rs2155857 | A | G | NULL | 0.63645254 | NULL&M_Disease |
| 11 | 116680449 | rs2542062 | A | G | NULL | 0.637608653 | NULL&M_Disease |
| 11 | 121181207 | rs141633273 | T | C | NULL | 0.63645254 | NULL&M_Disease |
| 11 | 117974600 | rs2155855 | G | A | NULL | 0.63645254 | NULL&M_Disease |
| 11 | 117974639 | rs2155856 | A | C | NULL | 0.63645254 | NULL&M_Disease |
| 11 | 119016547 | rs56679414 | T | C | NULL | 0.637608653 | NULL&M_Disease |
| 11 | 121176185 | rs139131517 | T | C | NULL | 0.637608653 | NULL&M_Disease |
| 11 | 116661826 | rs2072560 | T | C | NULL | 0.637608653 | NULL&M_Disease |
| 11 | 119015371 | rs55927184 | A | G | NULL | 0.637608653 | NULL&M_Disease |
| 11 | 119250060 | rs511401 | C | G | NULL | 0.637608653 | NULL&M_Disease |
| 11 | 120075768 | rs61898270 | T | A | NULL | 0.637608653 | NULL&M_Disease |
| 11 | 117341466 | rs12362636 | C | T | NULL | 0.637608653 | NULL&M_Disease |
| 11 | 117541521 | rs688790 | A | T | NULL | 0.637608653 | NULL&M_Disease |
| 11 | 117217533 | rs74853011 | G | C | NULL | 0.637608653 | NULL&M_Disease |
| 11 | 120398765 | rs118089075 | T | C | NULL | 0.637608653 | NULL&M_Disease |
| 11 | 120811386 | rs1060745 | T | G | NULL | 0.637608653 | NULL&M_Disease |
| 11 | 121192240 | rs2370112 | C | T | NULL | 0.637608653 | NULL&M_Disease |
| 11 | 116416909 | rs4293153 | G | A | NULL | 0.637608653 | NULL&M_Disease |
| 11 | 116525995 | rs116857027 | T | C | NULL | 0.637608653 | NULL&M_Disease |
| 11 | 116539921 | rs1784132 | A | G | NULL | 0.637608653 | NULL&M_Disease |
| 11 | 117897056 | rs11216706 | G | T | NULL | 0.637608653 | NULL&M_Disease |
| 11 | 117897184 | rs11216707 | G | A | NULL | 0.637608653 | NULL&M_Disease |
| 11 | 118175540 | rs2231441 | G | A | NULL | 0.637608653 | NULL&M_Disease |
| 11 | 120085537 | rs61898278 | G | A | NULL | 0.637608653 | NULL&M_Disease |
| 11 | 120086365 | rs61898279 | G | A | NULL | 0.637608653 | NULL&M_Disease |
| 11 | 116688765 | rs1423902581 | C | A | NULL | 0.637608653 | NULL&M_Disease |
| 11 | 119756282 | rs61899012 | C | G | NULL | 0.637608653 | NULL&M_Disease |
| 11 | 120479103 | rs6589819 | A | G | NULL | 0.637608653 | NULL&M_Disease |
| 11 | 116733316 | rs2080586 | C | T | NULL | 0.637608653 | NULL&M_Disease |
| 11 | 117541577 | rs688415 | G | A | NULL | 0.637608653 | NULL&M_Disease |
| 11 | 117670651 | rs7125693 | C | T | NULL | 0.637608653 | NULL&M_Disease |
| 11 | 120151784 | rs12788462 | C | T | NULL | 0.637608653 | NULL&M_Disease |
| 11 | 120529116 | rs73580405 | G | A | NULL | 0.637608653 | NULL&M_Disease |
| 11 | 120790997 | rs75672383 | T | C | NULL | 0.637608653 | NULL&M_Disease |
| 11 | 116652207 | rs12286037 | T | C | NULL | 0.637608653 | NULL&M_Disease |
| 11 | 117799804 | rs722729 | T | C | NULL | 0.637608653 | NULL&M_Disease |
| 11 | 117897329 | rs11216708 | T | C | NULL | 0.637608653 | NULL&M_Disease |
| 11 | 117924986 | rs4938471 | G | T | NULL | 0.637608653 | NULL&M_Disease |
| 11 | 119016361 | rs73562888 | C | T | NULL | 0.637608653 | NULL&M_Disease |
| 11 | 119474018 | rs116603457 | T | C | NULL | 0.637608653 | NULL&M_Disease |
| 11 | 120584851 | rs3133855 | T | C | NULL | 0.637608653 | NULL&M_Disease |
| 11 | 116611995 | rs180348 | A | G | NULL | 0.638763175 | NULL&M_Disease |
| 11 | 117425540 | rs12805320 | T | C | NULL | 0.637608653 | NULL&M_Disease |
| 11 | 119764214 | rs61899015 | C | A | NULL | 0.637608653 | NULL&M_Disease |
| 11 | 119800142 | rs1596979 | C | A | NULL | 0.637608653 | NULL&M_Disease |
| 11 | 119801612 | rs80064625 | C | T | NULL | 0.637608653 | NULL&M_Disease |
| 11 | 119989572 | rs1207849774 | A | T | NULL | 0.637608653 | NULL&M_Disease |
| 11 | 120090344 | rs12222405 | T | C | NULL | 0.637608653 | NULL&M_Disease |
| 11 | 120090552 | rs12223954 | C | T | NULL | 0.637608653 | NULL&M_Disease |
| 11 | 116362044 | rs77368534 | G | T | NULL | 0.637608653 | NULL&M_Disease |
| 11 | 117229833 | rs79795409 | T | G | NULL | 0.638763175 | NULL&M_Disease |
| 11 | 118157250 | rs35235858 | T | C | NULL | 0.638763175 | NULL&M_Disease |
| 11 | 119665733 | rs7935678 | C | T | NULL | 0.638763175 | NULL&M_Disease |
| 11 | 116549427 | rs1784107 | T | A | NULL | 0.638763175 | NULL&M_Disease |
| 11 | 116576744 | rs76468794 | A | G | NULL | 0.638763175 | NULL&M_Disease |
| 11 | 117343392 | rs570083 | A | G | NULL | 0.638763175 | NULL&M_Disease |
| 11 | 117696211 | rs57703856 | A | G | NULL | 0.638763175 | NULL&M_Disease |
| 11 | 117974484 | rs2187009 | T | C | NULL | 0.638763175 | NULL&M_Disease |
| 11 | 119393309 | rs10892379 | G | A | NULL | 0.638763175 | NULL&M_Disease |
| 11 | 117257876 | rs2305829 | T | G | NULL | 0.638763175 | NULL&M_Disease |

| | | | | | | | | | |
|---|---|---|---|---|---|---|---|---|---|
| 11 | 117737382 | rs7118101 | C | T | 0.961 | -7977.388 | -7978.513 | -1.124 | 1.26E-06 |
| 11 | 117737440 | rs876798 | C | T | 0.961 | -7977.388 | -7978.513 | -1.124 | 1.26E-06 |
| 11 | 117776003 | rs67422215 | T | C | 0.962 | -7977.388 | -7978.512 | -1.124 | 1.26E-06 |
| 11 | 119026460 | rs4938635 | T | C | 1.031 | -7977.388 | -7978.512 | -1.124 | 1.26E-06 |
| 11 | 119026469 | rs4938636 | T | A | 1.031 | -7977.388 | -7978.513 | -1.124 | 1.26E-06 |
| 11 | 119242556 | rs1652024 | C | A | 0.961 | -7977.388 | -7978.512 | -1.124 | 1.26E-06 |
| 11 | 117061961 | rs499713 | T | C | 0.961 | -7977.388 | -7978.513 | -1.125 | 1.26E-06 |
| 11 | 117882792 | rs11216677 | T | C | 0.977 | -7977.388 | -7978.513 | -1.125 | 1.26E-06 |
| 11 | 118157681 | rs34250455 | T | C | 1.038 | -7977.388 | -7978.513 | -1.125 | 1.26E-06 |
| 11 | 119731345 | rs11607827 | G | T | 0.979 | -7977.388 | -7978.513 | -1.125 | 1.26E-06 |
| 11 | 120489650 | rs76273814 | T | C | 1.027 | -7977.388 | -7978.513 | -1.125 | 1.26E-06 |
| 11 | 116501944 | rs11824108 | G | A | 1.039 | -7977.388 | -7978.515 | -1.126 | 1.25E-06 |
| 11 | 116514319 | rs116948654 | C | T | 1.033 | -7977.388 | -7978.515 | -1.126 | 1.25E-06 |
| 11 | 116527281 | rs12420386 | A | C | 0.968 | -7977.388 | -7978.515 | -1.126 | 1.25E-06 |
| 11 | 116561628 | rs61905060 | A | G | 0.988 | -7977.388 | -7978.514 | -1.126 | 1.25E-06 |
| 11 | 117346513 | rs540469 | C | T | 0.963 | -7977.388 | -7978.514 | -1.126 | 1.25E-06 |
| 11 | 117392826 | rs10892135 | A | G | 1.04 | -7977.388 | -7978.515 | -1.126 | 1.25E-06 |
| 11 | 117633054 | rs10892169 | A | G | 0.962 | -7977.388 | -7978.514 | -1.126 | 1.25E-06 |
| 11 | 117859209 | rs4252249 | A | G | 0.962 | -7977.388 | -7978.514 | -1.126 | 1.25E-06 |
| 11 | 118970682 | rs79524875 | T | C | 0.984 | -7977.388 | -7978.514 | -1.126 | 1.25E-06 |
| 11 | 119253797 | rs512608 | G | A | 0.961 | -7977.388 | -7978.514 | -1.126 | 1.25E-06 |
| 11 | 117762694 | rs73020457 | T | C | 0.964 | -7977.388 | -7978.516 | -1.127 | 1.25E-06 |
| 11 | 118504742 | rs633683 | T | C | 0.964 | -7977.388 | -7978.515 | -1.127 | 1.25E-06 |
| 11 | 118568427 | rs117361624 | T | C | 0.981 | -7977.388 | -7978.516 | -1.127 | 1.25E-06 |
| 11 | 119897773 | rs118135828 | A | G | 0.97 | -7977.388 | -7978.515 | -1.127 | 1.25E-06 |
| 11 | 119897774 | rs592765 | A | C | 0.97 | -7977.388 | -7978.515 | -1.127 | 1.25E-06 |
| 11 | 120072470 | rs61898267 | G | C | 0.995 | -7977.388 | -7978.515 | -1.127 | 1.25E-06 |
| 11 | 120104292 | rs61898294 | A | G | 0.996 | -7977.388 | -7978.515 | -1.127 | 1.25E-06 |
| 11 | 116537634 | rs11216083 | T | C | 0.967 | -7977.388 | -7978.517 | -1.128 | 1.25E-06 |
| 11 | 116577694 | rs75507001 | A | C | 0.997 | -7977.388 | -7978.516 | -1.128 | 1.25E-06 |
| 11 | 117424807 | rs12286677 | T | C | 1.034 | -7977.388 | -7978.516 | -1.128 | 1.25E-06 |
| 11 | 117425206 | rs12799960 | T | C | 1.034 | -7977.388 | -7978.517 | -1.128 | 1.25E-06 |
| 11 | 117772002 | rs11823721 | C | T | 0.965 | -7977.388 | -7978.516 | -1.128 | 1.25E-06 |
| 11 | 117898042 | rs192508986 | T | C | 0.977 | -7977.388 | -7978.516 | -1.128 | 1.25E-06 |
| 11 | 117898084 | rs12223288 | G | A | 0.977 | -7977.388 | -7978.516 | -1.128 | 1.25E-06 |
| 11 | 117898155 | rs10431083 | A | G | 0.977 | -7977.388 | -7978.516 | -1.128 | 1.25E-06 |
| 11 | 117974759 | rs2155857 | A | G | 0.988 | -7977.388 | -7978.517 | -1.128 | 1.25E-06 |
| 11 | 116680449 | rs2542062 | A | G | 1.04 | -7977.388 | -7978.518 | -1.129 | 1.25E-06 |
| 11 | 121181207 | rs141633273 | T | C | 0.966 | -7977.388 | -7978.518 | -1.129 | 1.25E-06 |
| 11 | 117974600 | rs2155855 | G | A | 0.989 | -7977.388 | -7978.518 | -1.13 | 1.25E-06 |
| 11 | 117974639 | rs2155856 | A | C | 0.989 | -7977.388 | -7978.518 | -1.13 | 1.25E-06 |
| 11 | 119016547 | rs56679414 | T | C | 1.031 | -7977.388 | -7978.518 | -1.13 | 1.25E-06 |
| 11 | 121176185 | rs139131517 | T | C | 0.966 | -7977.388 | -7978.518 | -1.13 | 1.25E-06 |
| 11 | 116661826 | rs2072560 | T | C | 0.972 | -7977.388 | -7978.519 | -1.131 | 1.25E-06 |
| 11 | 119015371 | rs55927184 | A | G | 1.031 | -7977.388 | -7978.519 | -1.131 | 1.25E-06 |
| 11 | 119250060 | rs511401 | C | G | 0.962 | -7977.388 | -7978.519 | -1.131 | 1.25E-06 |
| 11 | 120075768 | rs61898270 | T | A | 0.991 | -7977.388 | -7978.52 | -1.131 | 1.25E-06 |
| 11 | 117341466 | rs12362636 | C | T | 1.037 | -7977.388 | -7978.52 | -1.132 | 1.25E-06 |
| 11 | 117541521 | rs688790 | A | T | 1.03 | -7977.388 | -7978.521 | -1.132 | 1.25E-06 |
| 11 | 117217533 | rs74853011 | G | C | 0.976 | -7977.388 | -7978.522 | -1.133 | 1.25E-06 |
| 11 | 120398765 | rs118089075 | T | C | 0.977 | -7977.388 | -7978.521 | -1.133 | 1.25E-06 |
| 11 | 120811386 | rs1060745 | T | G | 0.975 | -7977.388 | -7978.521 | -1.133 | 1.25E-06 |
| 11 | 121192240 | rs2370112 | C | T | 1.037 | -7977.388 | -7978.522 | -1.133 | 1.25E-06 |
| 11 | 116416909 | rs4293153 | G | A | 1.037 | -7977.388 | -7978.523 | -1.134 | 1.24E-06 |
| 11 | 116525995 | rs116857027 | T | C | 1.031 | -7977.388 | -7978.522 | -1.134 | 1.24E-06 |
| 11 | 116539921 | rs1784132 | A | G | 1.039 | -7977.388 | -7978.522 | -1.134 | 1.24E-06 |
| 11 | 117897056 | rs11216706 | G | T | 0.978 | -7977.388 | -7978.523 | -1.134 | 1.24E-06 |
| 11 | 117897184 | rs11216707 | G | A | 0.978 | -7977.388 | -7978.523 | -1.134 | 1.24E-06 |
| 11 | 118175540 | rs2231441 | G | A | 1.038 | -7977.388 | -7978.523 | -1.134 | 1.24E-06 |
| 11 | 120085537 | rs61898278 | G | A | 0.986 | -7977.388 | -7978.522 | -1.134 | 1.24E-06 |
| 11 | 120086365 | rs61898279 | G | A | 0.986 | -7977.388 | -7978.522 | -1.134 | 1.24E-06 |
| 11 | 116688765 | rs1423902581 | C | A | 0.962 | -7977.388 | -7978.523 | -1.135 | 1.24E-06 |
| 11 | 119756282 | rs61899012 | C | G | 1.034 | -7977.388 | -7978.524 | -1.135 | 1.24E-06 |
| 11 | 120479103 | rs6589819 | A | G | 0.963 | -7977.388 | -7978.523 | -1.135 | 1.24E-06 |
| 11 | 116733316 | rs2080586 | C | T | 0.963 | -7977.388 | -7978.524 | -1.136 | 1.24E-06 |
| 11 | 117541577 | rs688415 | G | A | 1.03 | -7977.388 | -7978.524 | -1.136 | 1.24E-06 |
| 11 | 117670651 | rs7125693 | C | T | 1.037 | -7977.388 | -7978.524 | -1.136 | 1.24E-06 |
| 11 | 120151784 | rs12788462 | C | T | 1.04 | -7977.388 | -7978.524 | -1.136 | 1.24E-06 |
| 11 | 120529116 | rs73580405 | G | A | 1.009 | -7977.388 | -7978.524 | -1.136 | 1.24E-06 |
| 11 | 120790997 | rs75672383 | T | C | 1.013 | -7977.388 | -7978.524 | -1.136 | 1.24E-06 |
| 11 | 116652207 | rs12286037 | T | C | 0.97 | -7977.388 | -7978.525 | -1.137 | 1.24E-06 |
| 11 | 117799804 | rs722729 | T | C | 1.038 | -7977.388 | -7978.525 | -1.137 | 1.24E-06 |
| 11 | 117897329 | rs11216708 | T | C | 0.979 | -7977.388 | -7978.525 | -1.137 | 1.24E-06 |
| 11 | 117924986 | rs4938471 | G | T | 1.038 | -7977.388 | -7978.525 | -1.137 | 1.24E-06 |
| 11 | 119016361 | rs73562888 | C | T | 1.03 | -7977.388 | -7978.525 | -1.137 | 1.24E-06 |
| 11 | 119474018 | rs116603457 | T | C | 0.979 | -7977.388 | -7978.526 | -1.137 | 1.24E-06 |
| 11 | 120584851 | rs3133855 | T | C | 1.038 | -7977.388 | -7978.526 | -1.137 | 1.24E-06 |
| 11 | 116611995 | rs180348 | A | G | 1.039 | -7977.388 | -7978.527 | -1.138 | 1.24E-06 |
| 11 | 117425540 | rs12805320 | T | C | 1.033 | -7977.388 | -7978.527 | -1.138 | 1.24E-06 |
| 11 | 119764214 | rs61899015 | C | A | 1.034 | -7977.388 | -7978.526 | -1.138 | 1.24E-06 |
| 11 | 119800142 | rs1596979 | C | A | 1.03 | -7977.388 | -7978.526 | -1.138 | 1.24E-06 |
| 11 | 119801612 | rs80064625 | C | T | 1.03 | -7977.388 | -7978.526 | -1.138 | 1.24E-06 |
| 11 | 119989572 | rs1207849774 | A | T | 0.964 | -7977.388 | -7978.527 | -1.138 | 1.24E-06 |
| 11 | 120090344 | rs12222405 | T | C | 0.968 | -7977.388 | -7978.526 | -1.138 | 1.24E-06 |
| 11 | 120090552 | rs12223954 | C | T | 0.968 | -7977.388 | -7978.526 | -1.138 | 1.24E-06 |
| 11 | 116362044 | rs77368534 | G | T | 0.976 | -7977.388 | -7978.527 | -1.139 | 1.24E-06 |
| 11 | 117229833 | rs79795409 | T | G | 0.962 | -7977.388 | -7978.527 | -1.139 | 1.24E-06 |
| 11 | 118157250 | rs35235858 | T | C | 1.038 | -7977.388 | -7978.527 | -1.139 | 1.24E-06 |
| 11 | 119665733 | rs7935678 | C | T | 0.962 | -7977.388 | -7978.528 | -1.139 | 1.24E-06 |
| 11 | 116549427 | rs1784107 | T | A | 1.039 | -7977.388 | -7978.528 | -1.14 | 1.24E-06 |
| 11 | 116576744 | rs76468794 | A | G | 1.019 | -7977.388 | -7978.528 | -1.14 | 1.24E-06 |
| 11 | 117343392 | rs570083 | A | G | 0.962 | -7977.388 | -7978.528 | -1.14 | 1.24E-06 |
| 11 | 117696211 | rs57703856 | A | G | 0.967 | -7977.388 | -7978.528 | -1.14 | 1.24E-06 |
| 11 | 117974484 | rs2187009 | T | C | 0.999 | -7977.388 | -7978.528 | -1.14 | 1.24E-06 |
| 11 | 119393309 | rs10892379 | G | A | 1.038 | -7977.388 | -7978.529 | -1.14 | 1.24E-06 |
| 11 | 117257876 | rs2305829 | T | G | 0.962 | -7977.388 | -7978.53 | -1.141 | 1.24E-06 |

| | | | | | | | |
|---|---|---|---|---|---|---|---|
| 11 | 117323389 | rs11216399 | A | C | NULL | 0.638763175 | NULL&M_Disease |
| 11 | 117670282 | rs7943887 | G | C | NULL | 0.638763175 | NULL&M_Disease |
| 11 | 117906706 | rs652469 | A | G | NULL | 0.638763175 | NULL&M_Disease |
| 11 | 119016539 | rs61686122 | C | G | NULL | 0.638763175 | NULL&M_Disease |
| 11 | 119736232 | rs12224952 | T | G | NULL | 0.638763175 | NULL&M_Disease |
| 11 | 119757111 | rs78015 | T | G | NULL | 0.638763175 | NULL&M_Disease |
| 11 | 117341209 | rs12575215 | C | T | NULL | 0.638763175 | NULL&M_Disease |
| 11 | 117426506 | rs12800503 | T | G | NULL | 0.638763175 | NULL&M_Disease |
| 11 | 118983434 | rs2239896 | T | C | NULL | 0.638763175 | NULL&M_Disease |
| 11 | 119851369 | rs12418475 | A | G | NULL | 0.638763175 | NULL&M_Disease |
| 11 | 116370871 | rs881517 | T | G | NULL | 0.638763175 | NULL&M_Disease |
| 11 | 117608613 | rs1939981 | C | T | NULL | 0.638763175 | NULL&M_Disease |
| 11 | 117688517 | rs12286470 | C | G | NULL | 0.638763175 | NULL&M_Disease |
| 11 | 118982806 | rs77751615 | C | A | NULL | 0.638763175 | NULL&M_Disease |
| 11 | 119248709 | rs614192 | G | A | NULL | 0.638763175 | NULL&M_Disease |
| 11 | 119715710 | rs35385093 | A | G | NULL | 0.638763175 | NULL&M_Disease |
| 11 | 119884588 | rs80006645 | A | G | NULL | 0.638763175 | NULL&M_Disease |
| 11 | 120711290 | rs73003764 | G | A | NULL | 0.638763175 | NULL&M_Disease |
| 11 | 116576210 | rs59873384 | T | C | NULL | 0.638763175 | NULL&M_Disease |
| 11 | 117068504 | rs588534 | A | G | NULL | 0.638763175 | NULL&M_Disease |
| 11 | 118502172 | rs35435609 | T | C | NULL | 0.638763175 | NULL&M_Disease |
| 11 | 119235960 | rs12285901 | A | G | NULL | 0.638763175 | NULL&M_Disease |
| 11 | 119346198 | rs535831 | C | T | NULL | 0.638763175 | NULL&M_Disease |
| 11 | 119347074 | rs647369 | A | T | NULL | 0.638763175 | NULL&M_Disease |
| 11 | 119797699 | rs589348 | T | C | NULL | 0.638763175 | NULL&M_Disease |
| 11 | 120794509 | rs10892645 | G | A | NULL | 0.638763175 | NULL&M_Disease |
| 11 | 121188747 | rs12575073 | T | C | NULL | 0.638763175 | NULL&M_Disease |
| 11 | 116493155 | rs112319461 | A | G | NULL | 0.638763175 | NULL&M_Disease |
| 11 | 116543514 | rs12281217 | T | C | NULL | 0.638763175 | NULL&M_Disease |
| 11 | 116646107 | rs73588424 | G | A | NULL | 0.638763175 | NULL&M_Disease |
| 11 | 116668511 | rs601634 | T | C | NULL | 0.638763175 | NULL&M_Disease |
| 11 | 119018766 | rs3809046 | T | G | NULL | 0.638763175 | NULL&M_Disease |
| 11 | 119018774 | rs56013883 | T | G | NULL | 0.638763175 | NULL&M_Disease |
| 11 | 119019152 | rs55750089 | T | C | NULL | 0.638763175 | NULL&M_Disease |
| 11 | 119019475 | rs55752358 | T | A | NULL | 0.638763175 | NULL&M_Disease |
| 11 | 120275039 | rs76181827 | G | C | NULL | 0.638763175 | NULL&M_Disease |
| 11 | 116577971 | rs117684324 | T | A | NULL | 0.638763175 | NULL&M_Disease |
| 11 | 117132315 | rs73010439 | G | T | NULL | 0.638763175 | NULL&M_Disease |
| 11 | 117670657 | rs7125694 | C | T | NULL | 0.638763175 | NULL&M_Disease |
| 11 | 118150127 | rs17540708 | G | T | NULL | 0.638763175 | NULL&M_Disease |
| 11 | 118974588 | rs4938626 | T | G | NULL | 0.638763175 | NULL&M_Disease |
| 11 | 119249488 | rs2115645 | G | A | NULL | 0.638763175 | NULL&M_Disease |
| 11 | 119665842 | rs7935816 | T | C | NULL | 0.638763175 | NULL&M_Disease |
| 11 | 119773548 | rs596770 | G | A | NULL | 0.638763175 | NULL&M_Disease |
| 11 | 116528983 | rs947988 | C | G | NULL | 0.638763175 | NULL&M_Disease |
| 11 | 117853666 | rs61451669 | C | T | NULL | 0.638763175 | NULL&M_Disease |
| 11 | 119908015 | rs116083223 | A | G | NULL | 0.638763175 | NULL&M_Disease |
| 11 | 120779359 | rs949056 | G | A | NULL | 0.638763175 | NULL&M_Disease |
| 11 | 116362136 | rs6589548 | G | A | NULL | 0.638763175 | NULL&M_Disease |
| 11 | 117775943 | rs73020495 | A | C | NULL | 0.638763175 | NULL&M_Disease |
| 11 | 116253907 | rs76521293 | C | T | NULL | 0.639916097 | NULL&M_Disease |
| 11 | 116268668 | rs512168 | A | T | NULL | 0.639916097 | NULL&M_Disease |
| 11 | 116362669 | rs61904777 | C | T | NULL | 0.639916097 | NULL&M_Disease |
| 11 | 117624058 | rs2015482 | A | G | NULL | 0.638763175 | NULL&M_Disease |
| 11 | 117772179 | rs11821311 | T | C | NULL | 0.639916097 | NULL&M_Disease |
| 11 | 118156517 | rs12798115 | C | G | NULL | 0.639916097 | NULL&M_Disease |
| 11 | 118494790 | rs11825621 | C | T | NULL | 0.638763175 | NULL&M_Disease |
| 11 | 119013740 | rs145892119 | T | C | NULL | 0.638763175 | NULL&M_Disease |
| 11 | 120488476 | rs7101412 | T | C | NULL | 0.639916097 | NULL&M_Disease |
| 11 | 117068977 | rs474488 | A | G | NULL | 0.639916097 | NULL&M_Disease |
| 11 | 118155804 | rs34022886 | A | G | NULL | 0.639916097 | NULL&M_Disease |
| 11 | 118526691 | rs72999486 | T | C | NULL | 0.638763175 | NULL&M_Disease |
| 11 | 118975355 | rs4936463 | A | G | NULL | 0.638763175 | NULL&M_Disease |
| 11 | 119530129 | rs34953153 | T | C | NULL | 0.639916097 | NULL&M_Disease |
| 11 | 116230506 | rs17566737 | C | T | NULL | 0.639916097 | NULL&M_Disease |
| 11 | 117768454 | rs755538 | T | C | NULL | 0.639916097 | NULL&M_Disease |
| 11 | 117894782 | rs11216700 | T | A | NULL | 0.639916097 | NULL&M_Disease |
| 11 | 117894927 | rs11216703 | A | C | NULL | 0.639916097 | NULL&M_Disease |
| 11 | 119355678 | rs613442 | T | C | NULL | 0.639916097 | NULL&M_Disease |
| 11 | 119794036 | rs4938740 | A | G | NULL | 0.639916097 | NULL&M_Disease |
| 11 | 116650118 | rs17120029 | T | C | NULL | 0.639916097 | NULL&M_Disease |
| 11 | 118175439 | rs2231440 | A | G | NULL | 0.639916097 | NULL&M_Disease |
| 11 | 118988745 | rs4936464 | T | C | NULL | 0.639916097 | NULL&M_Disease |
| 11 | 121179549 | rs7942396 | T | C | NULL | 0.639916097 | NULL&M_Disease |
| 11 | 116611457 | rs57232565 | T | C | NULL | 0.639916097 | NULL&M_Disease |
| 11 | 117568781 | rs7118520 | G | A | NULL | 0.639916097 | NULL&M_Disease |
| 11 | 117935840 | rs670294 | G | T | NULL | 0.639916097 | NULL&M_Disease |
| 11 | 118729442 | rs12277355 | T | C | NULL | 0.639916097 | NULL&M_Disease |
| 11 | 118986972 | rs3741333 | T | C | NULL | 0.639916097 | NULL&M_Disease |
| 11 | 119769492 | rs4938737 | T | C | NULL | 0.639916097 | NULL&M_Disease |
| 11 | 119775597 | rs559430 | C | T | NULL | 0.639916097 | NULL&M_Disease |
| 11 | 116471264 | rs3016345 | G | A | NULL | 0.639916097 | NULL&M_Disease |
| 11 | 116529442 | rs11820504 | C | T | NULL | 0.639916097 | NULL&M_Disease |
| 11 | 116539233 | rs1784129 | A | G | NULL | 0.639916097 | NULL&M_Disease |
| 11 | 117046197 | rs4938353 | G | A | NULL | 0.639916097 | NULL&M_Disease |
| 11 | 117047852 | rs10790176 | G | T | NULL | 0.639916097 | NULL&M_Disease |
| 11 | 117669619 | rs1892903 | T | C | NULL | 0.639916097 | NULL&M_Disease |
| 11 | 117716663 | rs11608153 | G | A | NULL | 0.639916097 | NULL&M_Disease |
| 11 | 117887528 | rs73011708 | C | G | NULL | 0.639916097 | NULL&M_Disease |
| 11 | 117888685 | rs11216684 | G | A | NULL | 0.639916097 | NULL&M_Disease |
| 11 | 117888781 | rs11216686 | A | T | NULL | 0.639916097 | NULL&M_Disease |
| 11 | 117889685 | rs78640117 | T | C | NULL | 0.639916097 | NULL&M_Disease |
| 11 | 119727707 | rs71484150 | G | T | NULL | 0.639916097 | NULL&M_Disease |
| 11 | 117392337 | rs4938397 | T | G | NULL | 0.639916097 | NULL&M_Disease |
| 11 | 117712849 | rs17596129 | C | G | NULL | 0.639916097 | NULL&M_Disease |
| 11 | 118692727 | rs503425 | C | T | NULL | 0.639916097 | NULL&M_Disease |

| | | | | | | | | | |
|---|---|---|---|---|---|---|---|---|---|
| 11 | 117323389 | rs11216399 | A | C | 1.028 | -7977.388 | -7978.529 | -1.141 | 1.24E-06 |
| 11 | 117670282 | rs7943887 | G | C | 1.037 | -7977.388 | -7978.529 | -1.141 | 1.24E-06 |
| 11 | 117906706 | rs652469 | A | G | 0.964 | -7977.388 | -7978.529 | -1.141 | 1.24E-06 |
| 11 | 119016539 | rs61686122 | C | G | 1.029 | -7977.388 | -7978.529 | -1.141 | 1.24E-06 |
| 11 | 119736232 | rs12224952 | T | G | 1.037 | -7977.388 | -7978.53 | -1.141 | 1.24E-06 |
| 11 | 119757111 | rs78015 | T | G | 1.034 | -7977.388 | -7978.53 | -1.141 | 1.24E-06 |
| 11 | 117341209 | rs12575215 | C | T | 1.034 | -7977.388 | -7978.53 | -1.142 | 1.23E-06 |
| 11 | 117426506 | rs12800503 | T | G | 1.033 | -7977.388 | -7978.531 | -1.142 | 1.23E-06 |
| 11 | 118983434 | rs2239896 | T | C | 0.989 | -7977.388 | -7978.53 | -1.142 | 1.23E-06 |
| 11 | 119851369 | rs12418475 | A | G | 0.999 | -7977.388 | -7978.531 | -1.142 | 1.23E-06 |
| 11 | 116370871 | rs881517 | T | G | 0.964 | -7977.388 | -7978.531 | -1.143 | 1.23E-06 |
| 11 | 117608613 | rs1939981 | C | T | 1.039 | -7977.388 | -7978.532 | -1.143 | 1.23E-06 |
| 11 | 117688517 | rs12286470 | C | G | 1.032 | -7977.388 | -7978.531 | -1.143 | 1.23E-06 |
| 11 | 118982806 | rs77751615 | C | A | 0.989 | -7977.388 | -7978.532 | -1.143 | 1.23E-06 |
| 11 | 119248709 | rs614192 | G | A | 0.962 | -7977.388 | -7978.531 | -1.143 | 1.23E-06 |
| 11 | 119715710 | rs35385093 | A | G | 0.977 | -7977.388 | -7978.531 | -1.143 | 1.23E-06 |
| 11 | 119884588 | rs80006645 | A | G | 0.972 | -7977.388 | -7978.532 | -1.143 | 1.23E-06 |
| 11 | 120711290 | rs73003764 | G | A | 0.97 | -7977.388 | -7978.531 | -1.143 | 1.23E-06 |
| 11 | 116576210 | rs59873384 | T | C | 1.017 | -7977.388 | -7978.532 | -1.144 | 1.23E-06 |
| 11 | 117068504 | rs588534 | A | G | 0.962 | -7977.388 | -7978.532 | -1.144 | 1.23E-06 |
| 11 | 118502172 | rs35435609 | T | C | 0.982 | -7977.388 | -7978.532 | -1.144 | 1.23E-06 |
| 11 | 119235960 | rs12285901 | A | G | 0.966 | -7977.388 | -7978.532 | -1.144 | 1.23E-06 |
| 11 | 119346198 | rs535831 | C | T | 1.038 | -7977.388 | -7978.533 | -1.144 | 1.23E-06 |
| 11 | 119347074 | rs647369 | A | T | 1.038 | -7977.388 | -7978.533 | -1.144 | 1.23E-06 |
| 11 | 119797699 | rs589348 | T | C | 0.964 | -7977.388 | -7978.532 | -1.144 | 1.23E-06 |
| 11 | 120794509 | rs10892645 | G | A | 0.964 | -7977.388 | -7978.532 | -1.144 | 1.23E-06 |
| 11 | 121188747 | rs12575073 | T | C | 1.021 | -7977.388 | -7978.532 | -1.144 | 1.23E-06 |
| 11 | 116493155 | rs112319461 | A | G | 1.008 | -7977.388 | -7978.533 | -1.145 | 1.23E-06 |
| 11 | 116543514 | rs12281217 | T | C | 1.039 | -7977.388 | -7978.533 | -1.145 | 1.23E-06 |
| 11 | 116646107 | rs73588424 | G | A | 0.97 | -7977.388 | -7978.533 | -1.145 | 1.23E-06 |
| 11 | 116668511 | rs601634 | T | C | 1.039 | -7977.388 | -7978.533 | -1.145 | 1.23E-06 |
| 11 | 119018766 | rs3809046 | T | G | 1.029 | -7977.388 | -7978.534 | -1.145 | 1.23E-06 |
| 11 | 119018774 | rs56013883 | T | G | 1.029 | -7977.388 | -7978.534 | -1.145 | 1.23E-06 |
| 11 | 119019152 | rs55750089 | T | C | 1.029 | -7977.388 | -7978.534 | -1.145 | 1.23E-06 |
| 11 | 119019475 | rs55752358 | T | A | 1.029 | -7977.388 | -7978.534 | -1.145 | 1.23E-06 |
| 11 | 120275039 | rs76181827 | G | C | 1.025 | -7977.388 | -7978.534 | -1.145 | 1.23E-06 |
| 11 | 116577971 | rs117684324 | T | A | 1.016 | -7977.388 | -7978.534 | -1.146 | 1.23E-06 |
| 11 | 117132315 | rs73010439 | G | T | 0.962 | -7977.388 | -7978.535 | -1.146 | 1.23E-06 |
| 11 | 117670657 | rs7125694 | C | T | 1.037 | -7977.388 | -7978.535 | -1.146 | 1.23E-06 |
| 11 | 118150127 | rs17540708 | G | T | 1.038 | -7977.388 | -7978.534 | -1.146 | 1.23E-06 |
| 11 | 118974588 | rs4938626 | T | G | 0.99 | -7977.388 | -7978.534 | -1.146 | 1.23E-06 |
| 11 | 119249488 | rs2115645 | G | A | 0.963 | -7977.388 | -7978.534 | -1.146 | 1.23E-06 |
| 11 | 119665842 | rs7935816 | T | C | 0.964 | -7977.388 | -7978.535 | -1.146 | 1.23E-06 |
| 11 | 119773548 | rs596770 | G | A | 1.039 | -7977.388 | -7978.534 | -1.146 | 1.23E-06 |
| 11 | 116528983 | rs947988 | C | G | 0.965 | -7977.388 | -7978.535 | -1.147 | 1.23E-06 |
| 11 | 117853666 | rs61451669 | C | T | 0.963 | -7977.388 | -7978.535 | -1.147 | 1.23E-06 |
| 11 | 119908015 | rs116083223 | A | G | 0.972 | -7977.388 | -7978.535 | -1.147 | 1.23E-06 |
| 11 | 120779359 | rs949056 | G | A | 1.038 | -7977.388 | -7978.536 | -1.147 | 1.23E-06 |
| 11 | 116362136 | rs6589548 | G | A | 1.038 | -7977.388 | -7978.536 | -1.148 | 1.23E-06 |
| 11 | 117775943 | rs73020495 | A | C | 0.963 | -7977.388 | -7978.536 | -1.148 | 1.23E-06 |
| 11 | 116253907 | rs76521293 | C | T | 1.039 | -7977.388 | -7978.538 | -1.149 | 1.23E-06 |
| 11 | 116268668 | rs512168 | A | T | 0.963 | -7977.388 | -7978.538 | -1.149 | 1.23E-06 |
| 11 | 116362669 | rs61904777 | C | T | 1.038 | -7977.388 | -7978.537 | -1.149 | 1.23E-06 |
| 11 | 117624058 | rs2015482 | A | G | 0.999 | -7977.388 | -7978.537 | -1.149 | 1.23E-06 |
| 11 | 117772179 | rs11821311 | T | C | 0.966 | -7977.388 | -7978.538 | -1.149 | 1.23E-06 |
| 11 | 118156517 | rs12798115 | C | G | 1.038 | -7977.388 | -7978.538 | -1.149 | 1.23E-06 |
| 11 | 118494790 | rs11825621 | C | T | 0.983 | -7977.388 | -7978.537 | -1.149 | 1.23E-06 |
| 11 | 119013740 | rs145892119 | T | C | 1.029 | -7977.388 | -7978.537 | -1.149 | 1.23E-06 |
| 11 | 120488476 | rs7101412 | T | C | 0.963 | -7977.388 | -7978.537 | -1.149 | 1.23E-06 |
| 11 | 117068977 | rs474488 | A | G | 0.962 | -7977.388 | -7978.538 | -1.15 | 1.22E-06 |
| 11 | 118155804 | rs34022886 | A | G | 1.038 | -7977.388 | -7978.539 | -1.15 | 1.22E-06 |
| 11 | 118526691 | rs72999486 | T | C | 0.984 | -7977.388 | -7978.538 | -1.15 | 1.22E-06 |
| 11 | 118975355 | rs4936463 | A | G | 0.993 | -7977.388 | -7978.538 | -1.15 | 1.22E-06 |
| 11 | 119530129 | rs34953153 | T | C | 1.036 | -7977.388 | -7978.538 | -1.15 | 1.22E-06 |
| 11 | 116230506 | rs17566737 | C | T | 0.994 | -7977.388 | -7978.54 | -1.151 | 1.22E-06 |
| 11 | 117768454 | rs755538 | T | C | 0.966 | -7977.388 | -7978.54 | -1.151 | 1.22E-06 |
| 11 | 117894782 | rs11216700 | T | A | 0.984 | -7977.388 | -7978.54 | -1.151 | 1.22E-06 |
| 11 | 117894927 | rs11216703 | A | C | 0.984 | -7977.388 | -7978.54 | -1.151 | 1.22E-06 |
| 11 | 119355678 | rs613442 | T | C | 1.037 | -7977.388 | -7978.54 | -1.151 | 1.22E-06 |
| 11 | 119794036 | rs4938740 | A | G | 1.028 | -7977.388 | -7978.539 | -1.151 | 1.22E-06 |
| 11 | 116650118 | rs17120029 | T | C | 0.971 | -7977.388 | -7978.54 | -1.152 | 1.22E-06 |
| 11 | 118175439 | rs2231440 | A | G | 1.038 | -7977.388 | -7978.54 | -1.152 | 1.22E-06 |
| 11 | 118988745 | rs4936464 | T | C | 0.993 | -7977.388 | -7978.54 | -1.152 | 1.22E-06 |
| 11 | 121179549 | rs7942396 | T | C | 0.968 | -7977.388 | -7978.54 | -1.152 | 1.22E-06 |
| 11 | 116611457 | rs57232565 | T | C | 0.978 | -7977.388 | -7978.541 | -1.153 | 1.22E-06 |
| 11 | 117568781 | rs7118520 | G | A | 0.962 | -7977.388 | -7978.542 | -1.153 | 1.22E-06 |
| 11 | 117935840 | rs670294 | G | T | 1.039 | -7977.388 | -7978.542 | -1.153 | 1.22E-06 |
| 11 | 118729442 | rs12277355 | T | C | 1.039 | -7977.388 | -7978.542 | -1.153 | 1.22E-06 |
| 11 | 118986972 | rs3741333 | T | C | 0.994 | -7977.388 | -7978.541 | -1.153 | 1.22E-06 |
| 11 | 119769492 | rs4938737 | T | C | 1.039 | -7977.388 | -7978.541 | -1.153 | 1.22E-06 |
| 11 | 119775597 | rs559430 | C | T | 1.039 | -7977.388 | -7978.541 | -1.153 | 1.22E-06 |
| 11 | 116471264 | rs3016345 | G | A | 0.963 | -7977.388 | -7978.542 | -1.154 | 1.22E-06 |
| 11 | 116529442 | rs11820504 | C | T | 0.962 | -7977.388 | -7978.542 | -1.154 | 1.22E-06 |
| 11 | 116539233 | rs1784129 | A | G | 1.039 | -7977.388 | -7978.543 | -1.154 | 1.22E-06 |
| 11 | 117046197 | rs4938353 | G | A | 0.962 | -7977.388 | -7978.542 | -1.154 | 1.22E-06 |
| 11 | 117047852 | rs10790176 | G | T | 0.962 | -7977.388 | -7978.542 | -1.154 | 1.22E-06 |
| 11 | 117669619 | rs1892903 | T | C | 1.037 | -7977.388 | -7978.542 | -1.154 | 1.22E-06 |
| 11 | 117716663 | rs11608153 | G | A | 1.039 | -7977.388 | -7978.542 | -1.154 | 1.22E-06 |
| 11 | 117887528 | rs73011708 | C | G | 0.984 | -7977.388 | -7978.542 | -1.154 | 1.22E-06 |
| 11 | 117888685 | rs11216684 | G | A | 0.984 | -7977.388 | -7978.542 | -1.154 | 1.22E-06 |
| 11 | 117888781 | rs11216686 | A | T | 0.984 | -7977.388 | -7978.542 | -1.154 | 1.22E-06 |
| 11 | 117889685 | rs78640117 | T | C | 0.984 | -7977.388 | -7978.542 | -1.154 | 1.22E-06 |
| 11 | 119727707 | rs71484150 | G | T | 0.987 | -7977.388 | -7978.542 | -1.154 | 1.22E-06 |
| 11 | 117392337 | rs4938397 | T | G | 1.039 | -7977.388 | -7978.543 | -1.155 | 1.22E-06 |
| 11 | 117712849 | rs17596129 | C | G | 0.973 | -7977.388 | -7978.543 | -1.155 | 1.22E-06 |
| 11 | 118692727 | rs503425 | C | T | 1.039 | -7977.388 | -7978.543 | -1.155 | 1.22E-06 |

| | | | | | | | |
|---|---|---|---|---|---|---|---|
| 11 | 118986118 | rs59255488 | T | C | NULL | 0.639916097 | NULL&M_Disease |
| 11 | 118986456 | rs59565858 | A | G | NULL | 0.639916097 | NULL&M_Disease |
| 11 | 119355134 | rs620236 | A | T | NULL | 0.639916097 | NULL&M_Disease |
| 11 | 119772922 | rs487827 | T | G | NULL | 0.639916097 | NULL&M_Disease |
| 11 | 119896824 | rs183901166 | A | G | NULL | 0.639916097 | NULL&M_Disease |
| 11 | 116649135 | rs11823543 | A | G | NULL | 0.639916097 | NULL&M_Disease |
| 11 | 116711974 | rs480009 | T | C | NULL | 0.639916097 | NULL&M_Disease |
| 11 | 116762583 | rs525302 | C | G | NULL | 0.639916097 | NULL&M_Disease |
| 11 | 117772389 | rs73020476 | C | T | NULL | 0.639916097 | NULL&M_Disease |
| 11 | 116436362 | rs12291550 | C | T | NULL | 0.639916097 | NULL&M_Disease |
| 11 | 117857338 | rs10892202 | C | G | NULL | 0.639916097 | NULL&M_Disease |
| 11 | 119019282 | rs55713240 | T | C | NULL | 0.639916097 | NULL&M_Disease |
| 11 | 119355695 | rs613406 | C | T | NULL | 0.639916097 | NULL&M_Disease |
| 11 | 120276903 | rs61900795 | C | T | NULL | 0.639916097 | NULL&M_Disease |
| 11 | 116287337 | rs534082 | T | C | NULL | 0.639916097 | NULL&M_Disease |
| 11 | 116762829 | rs546734014 | A | G | NULL | 0.639916097 | NULL&M_Disease |
| 11 | 117719316 | rs623796 | T | C | NULL | 0.641067406 | NULL&M_Disease |
| 11 | 119775811 | rs508479 | A | C | NULL | 0.641067406 | NULL&M_Disease |
| 11 | 120165444 | rs117758225 | T | C | NULL | 0.639916097 | NULL&M_Disease |
| 11 | 120670036 | rs3020184 | A | G | NULL | 0.639916097 | NULL&M_Disease |
| 11 | 120675437 | rs1939671 | A | G | NULL | 0.639916097 | NULL&M_Disease |
| 11 | 120808928 | rs11218066 | T | C | NULL | 0.639916097 | NULL&M_Disease |
| 11 | 116199624 | rs1866683 | C | A | NULL | 0.641067406 | NULL&M_Disease |
| 11 | 116228735 | rs149017065 | A | G | NULL | 0.639916097 | NULL&M_Disease |
| 11 | 116233398 | rs10502215 | T | C | NULL | 0.639916097 | NULL&M_Disease |
| 11 | 116858703 | rs12793017 | G | A | NULL | 0.639916097 | NULL&M_Disease |
| 11 | 119353746 | rs531280 | C | A | NULL | 0.641067406 | NULL&M_Disease |
| 11 | 119355438 | rs618827 | A | G | NULL | 0.641067406 | NULL&M_Disease |
| 11 | 119712159 | rs34699720 | A | C | NULL | 0.639916097 | NULL&M_Disease |
| 11 | 119765318 | rs2044545 | T | C | NULL | 0.641067406 | NULL&M_Disease |
| 11 | 119796352 | rs78925312 | T | C | NULL | 0.639916097 | NULL&M_Disease |
| 11 | 120572727 | rs11217977 | T | C | NULL | 0.641067406 | NULL&M_Disease |
| 11 | 117689383 | rs73018459 | T | C | NULL | 0.639916097 | NULL&M_Disease |
| 11 | 117904049 | rs4936418 | T | A | NULL | 0.641067406 | NULL&M_Disease |
| 11 | 118958229 | rs474201 | G | A | NULL | 0.641067406 | NULL&M_Disease |
| 11 | 117642289 | rs34236286 | C | T | NULL | 0.641067406 | NULL&M_Disease |
| 11 | 117882065 | rs73009990 | A | G | NULL | 0.641067406 | NULL&M_Disease |
| 11 | 119351324 | rs587069 | G | A | NULL | 0.641067406 | NULL&M_Disease |
| 11 | 119884520 | rs77526581 | T | C | NULL | 0.641067406 | NULL&M_Disease |
| 11 | 116202035 | rs7935609 | C | T | NULL | 0.641067406 | NULL&M_Disease |
| 11 | 116353776 | rs57187632 | A | C | NULL | 0.641067406 | NULL&M_Disease |
| 11 | 117104536 | rs536141 | A | C | NULL | 0.641067406 | NULL&M_Disease |
| 11 | 120118498 | rs2847502 | T | C | NULL | 0.641067406 | NULL&M_Disease |
| 11 | 116561514 | rs118013839 | A | G | NULL | 0.641067406 | NULL&M_Disease |
| 11 | 116609540 | rs11216122 | T | G | NULL | 0.641067406 | NULL&M_Disease |
| 11 | 117873229 | rs2282493 | T | C | NULL | 0.641067406 | NULL&M_Disease |
| 11 | 119252448 | rs7943722 | T | C | NULL | 0.641067406 | NULL&M_Disease |
| 11 | 117393102 | rs1940039 | T | C | NULL | 0.641067406 | NULL&M_Disease |
| 11 | 119501013 | rs76080778 | G | A | NULL | 0.641067406 | NULL&M_Disease |
| 11 | 119768853 | rs631932 | C | A | NULL | 0.641067406 | NULL&M_Disease |
| 11 | 119775435 | rs641896 | T | C | NULL | 0.641067406 | NULL&M_Disease |
| 11 | 119890651 | rs77309894 | A | T | NULL | 0.641067406 | NULL&M_Disease |
| 11 | 116555415 | rs3016357 | T | C | NULL | 0.641067406 | NULL&M_Disease |
| 11 | 117336708 | rs78577579 | T | G | NULL | 0.641067406 | NULL&M_Disease |
| 11 | 119242277 | rs615001 | T | C | NULL | 0.641067406 | NULL&M_Disease |
| 11 | 119726313 | rs35163144 | G | A | NULL | 0.641067406 | NULL&M_Disease |
| 11 | 119857422 | rs9651694 | G | A | NULL | 0.641067406 | NULL&M_Disease |
| 11 | 116574463 | rs77206548 | A | G | NULL | 0.641067406 | NULL&M_Disease |
| 11 | 116773378 | rs11216180 | T | A | NULL | 0.641067406 | NULL&M_Disease |
| 11 | 117770998 | rs543290 | C | A | NULL | 0.641067406 | NULL&M_Disease |
| 11 | 121107473 | rs7129335 | A | T | NULL | 0.641067406 | NULL&M_Disease |
| 11 | 117163824 | rs638405 | G | C | NULL | 0.641067406 | NULL&M_Disease |
| 11 | 117406739 | rs688968 | G | C | NULL | 0.641067406 | NULL&M_Disease |
| 11 | 119548052 | rs10892429 | A | G | NULL | 0.641067406 | NULL&M_Disease |
| 11 | 119764295 | rs651232 | T | G | NULL | 0.641067406 | NULL&M_Disease |
| 11 | 116355233 | rs11215957 | C | T | NULL | 0.642217093 | NULL&M_Disease |
| 11 | 116635613 | rs60972755 | A | C | NULL | 0.641067406 | NULL&M_Disease |
| 11 | 117879926 | rs11216673 | G | A | NULL | 0.641067406 | NULL&M_Disease |
| 11 | 119232913 | rs10790291 | A | G | NULL | 0.641067406 | NULL&M_Disease |
| 11 | 119248320 | rs658654 | C | T | NULL | 0.642217093 | NULL&M_Disease |
| 11 | 119531135 | rs12802011 | G | A | NULL | 0.641067406 | NULL&M_Disease |
| 11 | 119887151 | rs11217634 | T | C | NULL | 0.641067406 | NULL&M_Disease |
| 11 | 119888797 | rs12272524 | T | C | NULL | 0.641067406 | NULL&M_Disease |
| 11 | 116246514 | rs571812 | T | C | NULL | 0.641067406 | NULL&M_Disease |
| 11 | 116518008 | rs1124874 | A | G | NULL | 0.641067406 | NULL&M_Disease |
| 11 | 117914386 | rs2212358 | A | G | NULL | 0.642217093 | NULL&M_Disease |
| 11 | 119024931 | rs4148167 | G | A | NULL | 0.641067406 | NULL&M_Disease |
| 11 | 119458695 | rs7484090 | G | A | NULL | 0.641067406 | NULL&M_Disease |
| 11 | 119727199 | rs35968460 | T | C | NULL | 0.641067406 | NULL&M_Disease |
| 11 | 120671999 | rs3133525 | A | G | NULL | 0.641067406 | NULL&M_Disease |
| 11 | 120715772 | rs61901427 | G | A | NULL | 0.641067406 | NULL&M_Disease |
| 11 | 117316546 | rs17120765 | A | G | NULL | 0.642217093 | NULL&M_Disease |
| 11 | 117897503 | rs10458984 | G | T | NULL | 0.641067406 | NULL&M_Disease |
| 11 | 117898465 | rs10431040 | A | G | NULL | 0.641067406 | NULL&M_Disease |
| 11 | 119720281 | rs7925677 | A | G | NULL | 0.641067406 | NULL&M_Disease |
| 11 | 119884816 | rs77840919 | G | C | NULL | 0.642217093 | NULL&M_Disease |
| 11 | 119884846 | rs77257523 | C | T | NULL | 0.642217093 | NULL&M_Disease |
| 11 | 119885012 | rs76924546 | A | G | NULL | 0.641067406 | NULL&M_Disease |
| 11 | 120398787 | rs61900894 | G | C | NULL | 0.642217093 | NULL&M_Disease |
| 11 | 116359612 | rs7947506 | T | C | NULL | 0.642217093 | NULL&M_Disease |
| 11 | 117624420 | rs6589621 | G | C | NULL | 0.642217093 | NULL&M_Disease |
| 11 | 117878745 | rs11216671 | T | C | NULL | 0.642217093 | NULL&M_Disease |
| 11 | 117878769 | rs11216672 | A | G | NULL | 0.642217093 | NULL&M_Disease |
| 11 | 119676300 | rs7118694 | A | G | NULL | 0.642217093 | NULL&M_Disease |
| 11 | 116347728 | rs11215953 | G | A | NULL | 0.642217093 | NULL&M_Disease |
| 11 | 116523165 | rs79770434 | T | C | NULL | 0.642217093 | NULL&M_Disease |

| | | | | | | | | | |
|---|---|---|---|---|---|---|---|---|---|
| 11 | 118986118 | rs59255488 | T | C | 0.998 | -7977.388 | -7978.544 | -1.155 | 1.22E-06 |
| 11 | 118986456 | rs59565858 | A | G | 0.998 | -7977.388 | -7978.543 | -1.155 | 1.22E-06 |
| 11 | 119355134 | rs620236 | A | T | 1.037 | -7977.388 | -7978.543 | -1.155 | 1.22E-06 |
| 11 | 119772922 | rs487827 | T | G | 1.039 | -7977.388 | -7978.543 | -1.155 | 1.22E-06 |
| 11 | 119896824 | rs183901166 | A | G | 0.973 | -7977.388 | -7978.544 | -1.155 | 1.22E-06 |
| 11 | 116649135 | rs11823543 | A | G | 0.972 | -7977.388 | -7978.544 | -1.156 | 1.22E-06 |
| 11 | 116711974 | rs480009 | T | C | 1.038 | -7977.388 | -7978.544 | -1.156 | 1.22E-06 |
| 11 | 116762583 | rs525302 | C | G | 1.021 | -7977.388 | -7978.545 | -1.156 | 1.22E-06 |
| 11 | 117772389 | rs73020476 | C | T | 0.966 | -7977.388 | -7978.544 | -1.156 | 1.22E-06 |
| 11 | 116436362 | rs12291550 | C | T | 0.963 | -7977.388 | -7978.546 | -1.157 | 1.22E-06 |
| 11 | 117857338 | rs10892202 | C | G | 0.964 | -7977.388 | -7978.545 | -1.157 | 1.22E-06 |
| 11 | 119019282 | rs55713240 | T | C | 1.028 | -7977.388 | -7978.545 | -1.157 | 1.22E-06 |
| 11 | 119355695 | rs613406 | C | T | 1.037 | -7977.388 | -7978.545 | -1.157 | 1.22E-06 |
| 11 | 120276903 | rs61900795 | C | T | 1.038 | -7977.388 | -7978.545 | -1.157 | 1.22E-06 |
| 11 | 116287337 | rs534082 | T | C | 1.038 | -7977.388 | -7978.546 | -1.158 | 1.22E-06 |
| 11 | 116762829 | rs546734014 | A | G | 0.964 | -7977.388 | -7978.546 | -1.158 | 1.22E-06 |
| 11 | 117719316 | rs623796 | T | C | 0.965 | -7977.388 | -7978.546 | -1.158 | 1.22E-06 |
| 11 | 119775811 | rs508479 | A | C | 1.039 | -7977.388 | -7978.547 | -1.158 | 1.22E-06 |
| 11 | 120165444 | rs117758225 | T | C | 1.028 | -7977.388 | -7978.547 | -1.158 | 1.22E-06 |
| 11 | 120670036 | rs3020184 | A | G | 0.976 | -7977.388 | -7978.546 | -1.158 | 1.22E-06 |
| 11 | 120675437 | rs1939671 | A | G | 0.976 | -7977.388 | -7978.547 | -1.158 | 1.22E-06 |
| 11 | 120808928 | rs11218066 | T | C | 0.963 | -7977.388 | -7978.546 | -1.158 | 1.22E-06 |
| 11 | 116199624 | rs1866683 | C | A | 1.037 | -7977.388 | -7978.547 | -1.159 | 1.21E-06 |
| 11 | 116228735 | rs149017065 | A | G | 1.005 | -7977.388 | -7978.548 | -1.159 | 1.21E-06 |
| 11 | 116233398 | rs10502215 | T | C | 0.993 | -7977.388 | -7978.548 | -1.159 | 1.21E-06 |
| 11 | 116858703 | rs12793017 | G | A | 0.995 | -7977.388 | -7978.547 | -1.159 | 1.21E-06 |
| 11 | 119353746 | rs531280 | C | A | 1.037 | -7977.388 | -7978.548 | -1.159 | 1.21E-06 |
| 11 | 119355438 | rs618827 | A | G | 1.037 | -7977.388 | -7978.547 | -1.159 | 1.21E-06 |
| 11 | 119712159 | rs34699720 | A | C | 0.989 | -7977.388 | -7978.547 | -1.159 | 1.21E-06 |
| 11 | 119765318 | rs2044545 | T | C | 1.039 | -7977.388 | -7978.548 | -1.159 | 1.21E-06 |
| 11 | 119796352 | rs78925312 | T | C | 0.999 | -7977.388 | -7978.547 | -1.159 | 1.21E-06 |
| 11 | 120572727 | rs11217977 | T | C | 1.037 | -7977.388 | -7978.547 | -1.159 | 1.21E-06 |
| 11 | 117689383 | rs73018459 | T | C | 0.992 | -7977.388 | -7978.549 | -1.16 | 1.21E-06 |
| 11 | 117904049 | rs4936418 | T | A | 1.038 | -7977.388 | -7978.548 | -1.16 | 1.21E-06 |
| 11 | 118958229 | rs474201 | G | A | 0.965 | -7977.388 | -7978.548 | -1.16 | 1.21E-06 |
| 11 | 117642289 | rs34236286 | C | T | 0.964 | -7977.388 | -7978.549 | -1.161 | 1.21E-06 |
| 11 | 117882065 | rs73009990 | A | G | 0.986 | -7977.388 | -7978.55 | -1.161 | 1.21E-06 |
| 11 | 119351324 | rs587069 | G | A | 1.037 | -7977.388 | -7978.549 | -1.161 | 1.21E-06 |
| 11 | 119884520 | rs77526581 | T | C | 0.974 | -7977.388 | -7978.549 | -1.161 | 1.21E-06 |
| 11 | 116202035 | rs7935609 | C | T | 1.037 | -7977.388 | -7978.55 | -1.162 | 1.21E-06 |
| 11 | 116353776 | rs57187632 | A | C | 0.979 | -7977.388 | -7978.55 | -1.162 | 1.21E-06 |
| 11 | 117104536 | rs536141 | A | C | 0.965 | -7977.388 | -7978.551 | -1.162 | 1.21E-06 |
| 11 | 120118498 | rs2847502 | T | C | 1.038 | -7977.388 | -7978.551 | -1.162 | 1.21E-06 |
| 11 | 116561514 | rs118013839 | A | G | 1.023 | -7977.388 | -7978.551 | -1.163 | 1.21E-06 |
| 11 | 116609540 | rs11216122 | T | G | 0.995 | -7977.388 | -7978.551 | -1.163 | 1.21E-06 |
| 11 | 117873229 | rs2282493 | T | C | 0.988 | -7977.388 | -7978.552 | -1.163 | 1.21E-06 |
| 11 | 119252448 | rs7943722 | T | C | 0.963 | -7977.388 | -7978.552 | -1.163 | 1.21E-06 |
| 11 | 117393102 | rs1940039 | T | C | 1.038 | -7977.388 | -7978.553 | -1.163 | 1.21E-06 |
| 11 | 119501013 | rs76080778 | G | A | 0.963 | -7977.388 | -7978.553 | -1.164 | 1.21E-06 |
| 11 | 119768853 | rs631932 | C | A | 1.039 | -7977.388 | -7978.552 | -1.164 | 1.21E-06 |
| 11 | 119775435 | rs641896 | T | C | 1.039 | -7977.388 | -7978.552 | -1.164 | 1.21E-06 |
| 11 | 119890651 | rs77309894 | A | T | 0.974 | -7977.388 | -7978.552 | -1.164 | 1.21E-06 |
| 11 | 116555415 | rs3016357 | T | C | 0.964 | -7977.388 | -7978.554 | -1.165 | 1.21E-06 |
| 11 | 117336708 | rs78577579 | T | G | 1.011 | -7977.388 | -7978.553 | -1.165 | 1.21E-06 |
| 11 | 119242277 | rs615001 | T | C | 0.963 | -7977.388 | -7978.553 | -1.165 | 1.21E-06 |
| 11 | 119726313 | rs35163144 | G | A | 0.991 | -7977.388 | -7978.553 | -1.165 | 1.21E-06 |
| 11 | 119857422 | rs9651694 | G | A | 1.038 | -7977.388 | -7978.553 | -1.165 | 1.21E-06 |
| 11 | 116574463 | rs77206548 | A | G | 1.011 | -7977.388 | -7978.554 | -1.166 | 1.21E-06 |
| 11 | 116773378 | rs11216180 | T | A | 0.964 | -7977.388 | -7978.554 | -1.166 | 1.21E-06 |
| 11 | 117770998 | rs543290 | C | A | 1.008 | -7977.388 | -7978.554 | -1.166 | 1.21E-06 |
| 11 | 121107473 | rs7129335 | A | T | 1.021 | -7977.388 | -7978.554 | -1.166 | 1.21E-06 |
| 11 | 117163824 | rs638405 | G | C | 0.965 | -7977.388 | -7978.555 | -1.167 | 1.20E-06 |
| 11 | 117406739 | rs688968 | G | C | 1.038 | -7977.388 | -7978.556 | -1.167 | 1.20E-06 |
| 11 | 119548052 | rs10892429 | A | G | 0.964 | -7977.388 | -7978.555 | -1.167 | 1.20E-06 |
| 11 | 119764295 | rs651232 | T | G | 1.033 | -7977.388 | -7978.555 | -1.167 | 1.20E-06 |
| 11 | 116355233 | rs11215957 | C | T | 1.036 | -7977.388 | -7978.556 | -1.168 | 1.20E-06 |
| 11 | 116635613 | rs60972755 | A | C | 0.973 | -7977.388 | -7978.556 | -1.168 | 1.20E-06 |
| 11 | 117879926 | rs11216673 | G | A | 0.989 | -7977.388 | -7978.556 | -1.168 | 1.20E-06 |
| 11 | 119232913 | rs10790291 | A | G | 0.963 | -7977.388 | -7978.556 | -1.168 | 1.20E-06 |
| 11 | 119248320 | rs658654 | C | T | 0.963 | -7977.388 | -7978.557 | -1.168 | 1.20E-06 |
| 11 | 119531135 | rs12802011 | G | A | 1.035 | -7977.388 | -7978.557 | -1.168 | 1.20E-06 |
| 11 | 119887151 | rs11217634 | T | C | 0.975 | -7977.388 | -7978.556 | -1.168 | 1.20E-06 |
| 11 | 119888797 | rs12272524 | T | C | 0.975 | -7977.388 | -7978.556 | -1.168 | 1.20E-06 |
| 11 | 116246514 | rs571812 | T | C | 1.031 | -7977.388 | -7978.557 | -1.169 | 1.20E-06 |
| 11 | 116518008 | rs1124874 | A | G | 1.028 | -7977.388 | -7978.557 | -1.169 | 1.20E-06 |
| 11 | 117914386 | rs2212358 | A | G | 1.037 | -7977.388 | -7978.558 | -1.169 | 1.20E-06 |
| 11 | 119024931 | rs4148167 | G | A | 1.026 | -7977.388 | -7978.557 | -1.169 | 1.20E-06 |
| 11 | 119458695 | rs7484090 | G | A | 0.968 | -7977.388 | -7978.557 | -1.169 | 1.20E-06 |
| 11 | 119727199 | rs35968460 | T | C | 0.986 | -7977.388 | -7978.557 | -1.169 | 1.20E-06 |
| 11 | 120671999 | rs3133525 | A | G | 0.976 | -7977.388 | -7978.557 | -1.169 | 1.20E-06 |
| 11 | 120715772 | rs61901427 | G | A | 0.989 | -7977.388 | -7978.557 | -1.169 | 1.20E-06 |
| 11 | 117316546 | rs17120765 | A | G | 1.035 | -7977.388 | -7978.559 | -1.17 | 1.20E-06 |
| 11 | 117897503 | rs10458984 | G | T | 0.989 | -7977.388 | -7978.559 | -1.17 | 1.20E-06 |
| 11 | 117898465 | rs10431040 | A | G | 0.989 | -7977.388 | -7978.559 | -1.17 | 1.20E-06 |
| 11 | 119720281 | rs7925677 | A | G | 0.987 | -7977.388 | -7978.558 | -1.17 | 1.20E-06 |
| 11 | 119884816 | rs77840919 | G | C | 0.975 | -7977.388 | -7978.558 | -1.17 | 1.20E-06 |
| 11 | 119884846 | rs77257523 | C | T | 0.975 | -7977.388 | -7978.558 | -1.17 | 1.20E-06 |
| 11 | 119885012 | rs76924546 | A | G | 0.975 | -7977.388 | -7978.558 | -1.17 | 1.20E-06 |
| 11 | 120398787 | rs61900894 | G | C | 0.967 | -7977.388 | -7978.558 | -1.17 | 1.20E-06 |
| 11 | 116359612 | rs7947506 | T | C | 1.038 | -7977.388 | -7978.56 | -1.171 | 1.20E-06 |
| 11 | 117624420 | rs6589621 | G | C | 1 | -7977.388 | -7978.559 | -1.171 | 1.20E-06 |
| 11 | 117878745 | rs11216671 | T | C | 0.99 | -7977.388 | -7978.559 | -1.171 | 1.20E-06 |
| 11 | 117878769 | rs11216672 | A | G | 0.99 | -7977.388 | -7978.559 | -1.171 | 1.20E-06 |
| 11 | 119676300 | rs7118694 | A | G | 0.963 | -7977.388 | -7978.56 | -1.171 | 1.20E-06 |
| 11 | 116347728 | rs11215953 | G | A | 1.013 | -7977.388 | -7978.561 | -1.172 | 1.20E-06 |
| 11 | 116523165 | rs79770434 | T | C | 1.027 | -7977.388 | -7978.56 | -1.172 | 1.20E-06 |

| | | | | | | | |
|---|---|---|---|---|---|---|---|
| 11 | 117934115 | rs612941 | C | A | NULL | 0.642217093 | NULL&M_Disease |
| 11 | 119772693 | rs582343 | C | G | NULL | 0.642217093 | NULL&M_Disease |
| 11 | 120834272 | rs4562840 | C | G | NULL | 0.642217093 | NULL&M_Disease |
| 11 | 116516587 | rs2212428 | T | C | NULL | 0.642217093 | NULL&M_Disease |
| 11 | 119662084 | rs11607146 | G | A | NULL | 0.642217093 | NULL&M_Disease |
| 11 | 120873429 | rs141076711 | T | G | NULL | 0.642217093 | NULL&M_Disease |
| 11 | 118936410 | rs45510391 | A | G | NULL | 0.642217093 | NULL&M_Disease |
| 11 | 119884922 | rs76260364 | T | C | NULL | 0.642217093 | NULL&M_Disease |
| 11 | 116352622 | rs495033 | C | T | NULL | 0.642217093 | NULL&M_Disease |
| 11 | 116641028 | rs1263151 | C | T | NULL | 0.642217093 | NULL&M_Disease |
| 11 | 117952899 | rs1630252 | C | T | NULL | 0.642217093 | NULL&M_Disease |
| 11 | 118030633 | rs11216785 | A | G | NULL | 0.642217093 | NULL&M_Disease |
| 11 | 119083318 | rs36109901 | C | A | NULL | 0.642217093 | NULL&M_Disease |
| 11 | 119353755 | rs531273 | A | G | NULL | 0.642217093 | NULL&M_Disease |
| 11 | 119354137 | rs634838 | T | C | NULL | 0.642217093 | NULL&M_Disease |
| 11 | 119526136 | rs4271390 | T | C | NULL | 0.642217093 | NULL&M_Disease |
| 11 | 119937335 | rs618166 | A | T | NULL | 0.642217093 | NULL&M_Disease |
| 11 | 116637146 | rs12294259 | T | C | NULL | 0.642217093 | NULL&M_Disease |
| 11 | 117894618 | rs56340594 | T | C | NULL | 0.642217093 | NULL&M_Disease |
| 11 | 118543798 | rs76570573 | T | C | NULL | 0.642217093 | NULL&M_Disease |
| 11 | 118729283 | rs11217068 | T | C | NULL | 0.642217093 | NULL&M_Disease |
| 11 | 120086918 | rs499434 | T | C | NULL | 0.642217093 | NULL&M_Disease |
| 11 | 121173349 | rs75051127 | A | G | NULL | 0.642217093 | NULL&M_Disease |
| 11 | 116507107 | rs947987 | A | G | NULL | 0.642217093 | NULL&M_Disease |
| 11 | 116583685 | rs74849419 | A | G | NULL | 0.642217093 | NULL&M_Disease |
| 11 | 117934598 | rs12278852 | A | G | NULL | 0.642217093 | NULL&M_Disease |
| 11 | 119668742 | rs6589777 | C | A | NULL | 0.642217093 | NULL&M_Disease |
| 11 | 119772958 | rs583687 | T | C | NULL | 0.642217093 | NULL&M_Disease |
| 11 | 119902521 | rs7358294 | T | C | NULL | 0.642217093 | NULL&M_Disease |
| 11 | 116359615 | rs17490842 | T | C | NULL | 0.642217093 | NULL&M_Disease |
| 11 | 116523657 | rs117742492 | A | G | NULL | 0.642217093 | NULL&M_Disease |
| 11 | 117471825 | rs577651 | T | C | NULL | 0.642217093 | NULL&M_Disease |
| 11 | 117914627 | rs11826104 | T | C | NULL | 0.643365146 | NULL&M_Disease |
| 11 | 118171285 | rs7479373 | C | T | NULL | 0.643365146 | NULL&M_Disease |
| 11 | 118745664 | rs11217076 | G | T | NULL | 0.643365146 | NULL&M_Disease |
| 11 | 120478670 | rs7947677 | G | T | NULL | 0.643365146 | NULL&M_Disease |
| 11 | 117690543 | rs17121264 | C | A | NULL | 0.643365146 | NULL&M_Disease |
| 11 | 116371338 | rs138177199 | G | C | NULL | 0.643365146 | NULL&M_Disease |
| 11 | 116512747 | rs116892227 | A | G | NULL | 0.643365146 | NULL&M_Disease |
| 11 | 117770775 | rs997956 | T | C | NULL | 0.643365146 | NULL&M_Disease |
| 11 | 117770990 | rs73020466 | G | A | NULL | 0.643365146 | NULL&M_Disease |
| 11 | 119061879 | rs45514102 | C | G | NULL | 0.643365146 | NULL&M_Disease |
| 11 | 119428158 | rs996888 | A | G | NULL | 0.643365146 | NULL&M_Disease |
| 11 | 119712522 | rs34071565 | T | A | NULL | 0.643365146 | NULL&M_Disease |
| 11 | 119876827 | rs75160858 | A | G | NULL | 0.643365146 | NULL&M_Disease |
| 11 | 119900754 | rs12285127 | A | G | NULL | 0.643365146 | NULL&M_Disease |
| 11 | 120166539 | rs7118517 | C | T | NULL | 0.643365146 | NULL&M_Disease |
| 11 | 120356832 | rs79820276 | G | A | NULL | 0.643365146 | NULL&M_Disease |
| 11 | 120720298 | rs73003773 | C | T | NULL | 0.643365146 | NULL&M_Disease |
| 11 | 116257372 | rs79053194 | G | T | NULL | 0.643365146 | NULL&M_Disease |
| 11 | 116512452 | rs117437529 | T | C | NULL | 0.643365146 | NULL&M_Disease |
| 11 | 116617240 | rs11216126 | C | A | NULL | 0.643365146 | NULL&M_Disease |
| 11 | 117325448 | rs10892116 | T | C | NULL | 0.643365146 | NULL&M_Disease |
| 11 | 119586134 | rs11217423 | T | C | NULL | 0.643365146 | NULL&M_Disease |
| 11 | 119859948 | rs10892507 | G | A | NULL | 0.643365146 | NULL&M_Disease |
| 11 | 116565603 | rs117607796 | A | C | NULL | 0.643365146 | NULL&M_Disease |
| 11 | 118956867 | rs616314 | A | T | NULL | 0.643365146 | NULL&M_Disease |
| 11 | 116421056 | rs10891988 | C | T | NULL | 0.643365146 | NULL&M_Disease |
| 11 | 116518743 | rs117338034 | G | T | NULL | 0.643365146 | NULL&M_Disease |
| 11 | 117934295 | rs7105816 | G | A | NULL | 0.643365146 | NULL&M_Disease |
| 11 | 119356682 | rs1785899 | T | C | NULL | 0.643365146 | NULL&M_Disease |
| 11 | 120090711 | rs74894672 | A | G | NULL | 0.643365146 | NULL&M_Disease |
| 11 | 116515405 | rs11216071 | C | G | NULL | 0.643365146 | NULL&M_Disease |
| 11 | 117430741 | rs11601480 | C | A | NULL | 0.643365146 | NULL&M_Disease |
| 11 | 117772955 | rs73020481 | C | T | NULL | 0.643365146 | NULL&M_Disease |
| 11 | 118045876 | rs113488707 | T | C | NULL | 0.643365146 | NULL&M_Disease |
| 11 | 116508833 | rs76747451 | G | T | NULL | 0.643365146 | NULL&M_Disease |
| 11 | 116510138 | rs12362930 | C | T | NULL | 0.643365146 | NULL&M_Disease |
| 11 | 116656788 | rs10750096 | C | A | NULL | 0.643365146 | NULL&M_Disease |
| 11 | 116724232 | rs888246 | T | C | NULL | 0.643365146 | NULL&M_Disease |
| 11 | 117430931 | rs567053 | A | G | NULL | 0.643365146 | NULL&M_Disease |
| 11 | 117568636 | rs58789559 | A | G | NULL | 0.643365146 | NULL&M_Disease |
| 11 | 117914652 | rs11216727 | C | G | NULL | 0.643365146 | NULL&M_Disease |
| 11 | 116354727 | rs76674252 | A | G | NULL | 0.643365146 | NULL&M_Disease |
| 11 | 116533684 | rs61592631 | T | C | NULL | 0.643365146 | NULL&M_Disease |
| 11 | 117287082 | rs59156491 | T | C | NULL | 0.644511554 | NULL&M_Disease |
| 11 | 117458539 | rs600620 | G | A | NULL | 0.644511554 | NULL&M_Disease |
| 11 | 119242715 | rs679556 | C | G | NULL | 0.644511554 | NULL&M_Disease |
| 11 | 119337504 | rs10750144 | A | G | NULL | 0.644511554 | NULL&M_Disease |
| 11 | 119465670 | rs76368626 | C | T | NULL | 0.643365146 | NULL&M_Disease |
| 11 | 119764234 | rs507622 | A | G | NULL | 0.643365146 | NULL&M_Disease |
| 11 | 119787531 | rs10790354 | T | C | NULL | 0.644511554 | NULL&M_Disease |
| 11 | 119795996 | rs113078426 | G | A | NULL | 0.643365146 | NULL&M_Disease |
| 11 | 121086234 | rs11218201 | C | T | NULL | 0.643365146 | NULL&M_Disease |
| 11 | 117362074 | rs540277 | T | C | NULL | 0.644511554 | NULL&M_Disease |
| 11 | 118179941 | rs4938502 | T | G | NULL | 0.644511554 | NULL&M_Disease |
| 11 | 119765177 | rs113460600 | A | G | NULL | 0.644511554 | NULL&M_Disease |
| 11 | 116453270 | rs17492429 | C | G | NULL | 0.644511554 | NULL&M_Disease |
| 11 | 117595511 | rs11216517 | T | C | NULL | 0.644511554 | NULL&M_Disease |
| 11 | 117915055 | rs12576767 | T | C | NULL | 0.644511554 | NULL&M_Disease |
| 11 | 119246896 | rs576945 | G | A | NULL | 0.644511554 | NULL&M_Disease |
| 11 | 120392401 | rs61900865 | T | C | NULL | 0.644511554 | NULL&M_Disease |
| 11 | 116660686 | rs2266788 | G | A | NULL | 0.644511554 | NULL&M_Disease |
| 11 | 117228849 | rs17591711 | G | A | NULL | 0.644511554 | NULL&M_Disease |
| 11 | 119243670 | rs33929148 | T | C | NULL | 0.644511554 | NULL&M_Disease |
| 11 | 120093606 | rs1287394 | C | T | NULL | 0.644511554 | NULL&M_Disease |

| | | | | | | | | | |
|---|---|---|---|---|---|---|---|---|---|
| 11 | 117934115 | rs612941 | C | A | 1.038 | -7977.388 | -7978.56 | -1.172 | 1.20E-06 |
| 11 | 119772693 | rs582343 | C | G | 1.038 | -7977.388 | -7978.561 | -1.172 | 1.20E-06 |
| 11 | 120834272 | rs4562840 | C | G | 1.037 | -7977.388 | -7978.561 | -1.172 | 1.20E-06 |
| 11 | 116516587 | rs2212428 | T | C | 1.027 | -7977.388 | -7978.562 | -1.173 | 1.20E-06 |
| 11 | 119662084 | rs11607146 | G | A | 0.964 | -7977.388 | -7978.561 | -1.173 | 1.20E-06 |
| 11 | 120873429 | rs141076711 | T | G | 0.978 | -7977.388 | -7978.561 | -1.173 | 1.20E-06 |
| 11 | 118936410 | rs45510391 | A | G | 1.032 | -7977.388 | -7978.563 | -1.174 | 1.20E-06 |
| 11 | 119884922 | rs76260364 | T | C | 0.975 | -7977.388 | -7978.562 | -1.174 | 1.20E-06 |
| 11 | 116352622 | rs495033 | C | T | 0.97 | -7977.388 | -7978.564 | -1.175 | 1.19E-06 |
| 11 | 116641028 | rs1263151 | C | T | 1.036 | -7977.388 | -7978.563 | -1.175 | 1.19E-06 |
| 11 | 117952899 | rs1630252 | C | T | 1.037 | -7977.388 | -7978.564 | -1.175 | 1.19E-06 |
| 11 | 118030633 | rs11216785 | A | G | 0.965 | -7977.388 | -7978.563 | -1.175 | 1.19E-06 |
| 11 | 119083318 | rs36109901 | C | A | 0.964 | -7977.388 | -7978.563 | -1.175 | 1.19E-06 |
| 11 | 119353755 | rs531273 | A | G | 1.037 | -7977.388 | -7978.563 | -1.175 | 1.19E-06 |
| 11 | 119354137 | rs634838 | T | C | 1.037 | -7977.388 | -7978.563 | -1.175 | 1.19E-06 |
| 11 | 119526136 | rs4271390 | T | C | 1.038 | -7977.388 | -7978.563 | -1.175 | 1.19E-06 |
| 11 | 119937335 | rs618166 | A | T | 0.963 | -7977.388 | -7978.563 | -1.175 | 1.19E-06 |
| 11 | 116637146 | rs12294259 | T | C | 0.974 | -7977.388 | -7978.564 | -1.176 | 1.19E-06 |
| 11 | 117894618 | rs56340594 | T | C | 0.982 | -7977.388 | -7978.564 | -1.176 | 1.19E-06 |
| 11 | 118543798 | rs76570573 | T | C | 0.996 | -7977.388 | -7978.564 | -1.176 | 1.19E-06 |
| 11 | 118729283 | rs11217068 | T | C | 1.038 | -7977.388 | -7978.564 | -1.176 | 1.19E-06 |
| 11 | 120086918 | rs499434 | T | C | 0.963 | -7977.388 | -7978.565 | -1.176 | 1.19E-06 |
| 11 | 121173349 | rs75051127 | A | G | 1.017 | -7977.388 | -7978.564 | -1.176 | 1.19E-06 |
| 11 | 116507107 | rs947987 | A | G | 0.964 | -7977.388 | -7978.566 | -1.177 | 1.19E-06 |
| 11 | 116583685 | rs74849419 | A | G | 0.983 | -7977.388 | -7978.565 | -1.177 | 1.19E-06 |
| 11 | 117934598 | rs12278852 | A | G | 1.037 | -7977.388 | -7978.565 | -1.177 | 1.19E-06 |
| 11 | 119668742 | rs6589777 | C | A | 0.963 | -7977.388 | -7978.565 | -1.177 | 1.19E-06 |
| 11 | 119772958 | rs583687 | T | C | 1.038 | -7977.388 | -7978.566 | -1.178 | 1.19E-06 |
| 11 | 119902521 | rs7358294 | T | C | 0.976 | -7977.388 | -7978.566 | -1.178 | 1.19E-06 |
| 11 | 116359615 | rs17490842 | T | C | 0.983 | -7977.388 | -7978.567 | -1.179 | 1.19E-06 |
| 11 | 116523657 | rs117742492 | A | G | 1.026 | -7977.388 | -7978.567 | -1.179 | 1.19E-06 |
| 11 | 117471825 | rs577651 | T | C | 1.025 | -7977.388 | -7978.567 | -1.179 | 1.19E-06 |
| 11 | 117914627 | rs11826104 | T | C | 1.037 | -7977.388 | -7978.568 | -1.179 | 1.19E-06 |
| 11 | 118171285 | rs7479373 | C | T | 1.037 | -7977.388 | -7978.567 | -1.179 | 1.19E-06 |
| 11 | 118745664 | rs11217076 | G | T | 0.964 | -7977.388 | -7978.567 | -1.179 | 1.19E-06 |
| 11 | 120478670 | rs7947677 | G | T | 0.964 | -7977.388 | -7978.567 | -1.179 | 1.19E-06 |
| 11 | 117690543 | rs17121264 | C | A | 0.974 | -7977.388 | -7978.568 | -1.18 | 1.19E-06 |
| 11 | 116371338 | rs138177199 | G | C | 0.984 | -7977.388 | -7978.569 | -1.181 | 1.19E-06 |
| 11 | 116512747 | rs116892227 | A | G | 1.026 | -7977.388 | -7978.569 | -1.181 | 1.19E-06 |
| 11 | 117770775 | rs997956 | T | C | 0.968 | -7977.388 | -7978.569 | -1.181 | 1.19E-06 |
| 11 | 117770990 | rs73020466 | G | A | 0.968 | -7977.388 | -7978.569 | -1.181 | 1.19E-06 |
| 11 | 119061879 | rs45514102 | C | G | 1.023 | -7977.388 | -7978.57 | -1.181 | 1.19E-06 |
| 11 | 119428158 | rs996888 | A | G | 0.968 | -7977.388 | -7978.569 | -1.181 | 1.19E-06 |
| 11 | 119712522 | rs34071565 | T | A | 0.992 | -7977.388 | -7978.569 | -1.181 | 1.19E-06 |
| 11 | 119876827 | rs75160858 | A | G | 1.03 | -7977.388 | -7978.569 | -1.181 | 1.19E-06 |
| 11 | 119900754 | rs12285127 | A | G | 0.976 | -7977.388 | -7978.57 | -1.181 | 1.19E-06 |
| 11 | 120166539 | rs7118517 | C | T | 0.97 | -7977.388 | -7978.569 | -1.181 | 1.19E-06 |
| 11 | 120356832 | rs79820276 | G | A | 1.019 | -7977.388 | -7978.569 | -1.181 | 1.19E-06 |
| 11 | 120720298 | rs73003773 | C | T | 0.974 | -7977.388 | -7978.569 | -1.181 | 1.19E-06 |
| 11 | 116257372 | rs79053194 | G | T | 0.966 | -7977.388 | -7978.57 | -1.182 | 1.19E-06 |
| 11 | 116512452 | rs117437529 | T | C | 1.018 | -7977.388 | -7978.57 | -1.182 | 1.19E-06 |
| 11 | 116617240 | rs11216126 | C | A | 1.037 | -7977.388 | -7978.57 | -1.182 | 1.19E-06 |
| 11 | 117325448 | rs10892116 | T | C | 0.997 | -7977.388 | -7978.57 | -1.182 | 1.19E-06 |
| 11 | 119586134 | rs11217423 | T | C | 1.036 | -7977.388 | -7978.57 | -1.182 | 1.19E-06 |
| 11 | 119859948 | rs10892507 | G | A | 0.963 | -7977.388 | -7978.57 | -1.182 | 1.19E-06 |
| 11 | 116565603 | rs117607796 | A | C | 1.02 | -7977.388 | -7978.572 | -1.183 | 1.19E-06 |
| 11 | 118956867 | rs616314 | A | T | 1.038 | -7977.388 | -7978.572 | -1.183 | 1.19E-06 |
| 11 | 116421056 | rs10891988 | C | T | 1.036 | -7977.388 | -7978.572 | -1.184 | 1.18E-06 |
| 11 | 116518743 | rs117338034 | G | T | 1.026 | -7977.388 | -7978.572 | -1.184 | 1.18E-06 |
| 11 | 117934295 | rs7105816 | G | A | 1.037 | -7977.388 | -7978.572 | -1.184 | 1.18E-06 |
| 11 | 119356682 | rs1785899 | T | C | 1.037 | -7977.388 | -7978.572 | -1.184 | 1.18E-06 |
| 11 | 120090711 | rs74894672 | A | G | 0.997 | -7977.388 | -7978.572 | -1.184 | 1.18E-06 |
| 11 | 116515405 | rs11216071 | C | G | 0.989 | -7977.388 | -7978.574 | -1.185 | 1.18E-06 |
| 11 | 117430741 | rs11601480 | C | A | 1.028 | -7977.388 | -7978.573 | -1.185 | 1.18E-06 |
| 11 | 117772955 | rs73020481 | C | T | 0.969 | -7977.388 | -7978.573 | -1.185 | 1.18E-06 |
| 11 | 118045876 | rs113488707 | T | C | 1.012 | -7977.388 | -7978.575 | -1.186 | 1.18E-06 |
| 11 | 116508833 | rs76747451 | G | T | 1.017 | -7977.388 | -7978.576 | -1.188 | 1.18E-06 |
| 11 | 116510138 | rs12362930 | C | T | 0.999 | -7977.388 | -7978.576 | -1.188 | 1.18E-06 |
| 11 | 116656788 | rs10750096 | C | A | 0.975 | -7977.388 | -7978.576 | -1.188 | 1.18E-06 |
| 11 | 116724232 | rs888246 | T | C | 0.965 | -7977.388 | -7978.576 | -1.188 | 1.18E-06 |
| 11 | 117430931 | rs567053 | A | G | 0.992 | -7977.388 | -7978.576 | -1.188 | 1.18E-06 |
| 11 | 117568636 | rs58789559 | A | G | 0.993 | -7977.388 | -7978.576 | -1.188 | 1.18E-06 |
| 11 | 117914652 | rs11216727 | C | G | 0.974 | -7977.388 | -7978.576 | -1.188 | 1.18E-06 |
| 11 | 116354727 | rs76674252 | A | G | 0.985 | -7977.388 | -7978.577 | -1.189 | 1.18E-06 |
| 11 | 116533684 | rs61592631 | T | C | 0.972 | -7977.388 | -7978.577 | -1.189 | 1.18E-06 |
| 11 | 117287082 | rs59156491 | T | C | 0.964 | -7977.388 | -7978.577 | -1.189 | 1.18E-06 |
| 11 | 117458539 | rs600620 | G | A | 0.964 | -7977.388 | -7978.577 | -1.189 | 1.18E-06 |
| 11 | 119242715 | rs679556 | C | G | 0.964 | -7977.388 | -7978.577 | -1.189 | 1.18E-06 |
| 11 | 119337504 | rs10750144 | A | G | 1.036 | -7977.388 | -7978.578 | -1.189 | 1.18E-06 |
| 11 | 119465670 | rs76368626 | C | T | 0.97 | -7977.388 | -7978.577 | -1.189 | 1.18E-06 |
| 11 | 119764234 | rs507622 | A | G | 1.031 | -7977.388 | -7978.577 | -1.189 | 1.18E-06 |
| 11 | 119787531 | rs10790354 | T | C | 0.964 | -7977.388 | -7978.578 | -1.189 | 1.18E-06 |
| 11 | 119795996 | rs113078426 | G | A | 1.024 | -7977.388 | -7978.577 | -1.189 | 1.18E-06 |
| 11 | 121086234 | rs11218201 | C | T | 1.026 | -7977.388 | -7978.578 | -1.189 | 1.18E-06 |
| 11 | 117362074 | rs540277 | T | C | 0.964 | -7977.388 | -7978.578 | -1.19 | 1.18E-06 |
| 11 | 118179941 | rs4938502 | T | G | 0.965 | -7977.388 | -7978.579 | -1.19 | 1.18E-06 |
| 11 | 119765177 | rs113460600 | A | G | 1.03 | -7977.388 | -7978.578 | -1.19 | 1.18E-06 |
| 11 | 116453270 | rs17492429 | C | G | 0.969 | -7977.388 | -7978.579 | -1.191 | 1.18E-06 |
| 11 | 117595511 | rs11216517 | T | C | 1.037 | -7977.388 | -7978.58 | -1.191 | 1.18E-06 |
| 11 | 117915055 | rs12576767 | T | C | 0.974 | -7977.388 | -7978.579 | -1.191 | 1.18E-06 |
| 11 | 119246896 | rs576945 | G | A | 0.964 | -7977.388 | -7978.58 | -1.191 | 1.18E-06 |
| 11 | 120392401 | rs61900865 | T | C | 0.969 | -7977.388 | -7978.58 | -1.192 | 1.17E-06 |
| 11 | 116660686 | rs2266788 | G | A | 0.976 | -7977.388 | -7978.581 | -1.193 | 1.17E-06 |
| 11 | 117228849 | rs17591711 | G | A | 0.982 | -7977.388 | -7978.581 | -1.193 | 1.17E-06 |
| 11 | 119243670 | rs33929148 | T | C | 0.976 | -7977.388 | -7978.581 | -1.193 | 1.17E-06 |
| 11 | 120093606 | rs1287394 | C | T | 1.037 | -7977.388 | -7978.581 | -1.193 | 1.17E-06 |

| | | | | | | | |
|---|---|---|---|---|---|---|---|
| 11 | 116759332 | rs1473177 | C | T | NULL | 0.644511554 | NULL&M_Disease |
| 11 | 119789537 | rs486978 | T | G | NULL | 0.644511554 | NULL&M_Disease |
| 11 | 117911637 | rs4938469 | C | T | NULL | 0.644511554 | NULL&M_Disease |
| 11 | 120166410 | rs744984 | G | A | NULL | 0.644511554 | NULL&M_Disease |
| 11 | 116586231 | rs117785509 | A | G | NULL | 0.644511554 | NULL&M_Disease |
| 11 | 117071449 | rs652455 | C | T | NULL | 0.644511554 | NULL&M_Disease |
| 11 | 118957735 | rs11217148 | T | G | NULL | 0.644511554 | NULL&M_Disease |
| 11 | 119787190 | rs658358 | C | G | NULL | 0.644511554 | NULL&M_Disease |
| 11 | 119884982 | rs116088536 | G | C | NULL | 0.644511554 | NULL&M_Disease |
| 11 | 120167566 | rs11217803 | A | C | NULL | 0.644511554 | NULL&M_Disease |
| 11 | 120300327 | rs11217865 | C | A | NULL | 0.644511554 | NULL&M_Disease |
| 11 | 116493738 | rs1240660 | G | A | NULL | 0.644511554 | NULL&M_Disease |
| 11 | 117670752 | rs657389 | A | G | NULL | 0.644511554 | NULL&M_Disease |
| 11 | 116528663 | rs73012993 | G | A | NULL | 0.644511554 | NULL&M_Disease |
| 11 | 117241760 | rs2305826 | T | A | NULL | 0.645656306 | NULL&M_Disease |
| 11 | 117920577 | rs12225323 | T | C | NULL | 0.644511554 | NULL&M_Disease |
| 11 | 120357341 | rs79425731 | A | G | NULL | 0.644511554 | NULL&M_Disease |
| 11 | 120411963 | rs2253134 | A | C | NULL | 0.645656306 | NULL&M_Disease |
| 11 | 116247262 | rs516169 | C | G | NULL | 0.644511554 | NULL&M_Disease |
| 11 | 116345605 | rs12275789 | A | G | NULL | 0.644511554 | NULL&M_Disease |
| 11 | 116663851 | rs17120035 | T | C | NULL | 0.644511554 | NULL&M_Disease |
| 11 | 116663966 | rs7103224 | A | G | NULL | 0.644511554 | NULL&M_Disease |
| 11 | 117168402 | rs619360 | T | A | NULL | 0.645656306 | NULL&M_Disease |
| 11 | 117354904 | rs7129284 | A | G | NULL | 0.645656306 | NULL&M_Disease |
| 11 | 119236692 | rs12277700 | A | C | NULL | 0.644511554 | NULL&M_Disease |
| 11 | 116653296 | rs2075290 | C | T | NULL | 0.644511554 | NULL&M_Disease |
| 11 | 119246567 | rs687009 | C | G | NULL | 0.645656306 | NULL&M_Disease |
| 11 | 120072666 | rs11217766 | A | C | NULL | 0.644511554 | NULL&M_Disease |
| 11 | 120305873 | rs77657279 | G | A | NULL | 0.645656306 | NULL&M_Disease |
| 11 | 121190062 | rs61902398 | A | G | NULL | 0.645656306 | NULL&M_Disease |
| 11 | 117151378 | rs638270 | G | C | NULL | 0.645656306 | NULL&M_Disease |
| 11 | 119867224 | rs4938747 | A | G | NULL | 0.645656306 | NULL&M_Disease |
| 11 | 119929632 | rs11217664 | G | A | NULL | 0.645656306 | NULL&M_Disease |
| 11 | 116564493 | rs117889337 | T | C | NULL | 0.645656306 | NULL&M_Disease |
| 11 | 116665079 | rs9804646 | T | C | NULL | 0.645656306 | NULL&M_Disease |
| 11 | 117396269 | rs12800264 | A | G | NULL | 0.645656306 | NULL&M_Disease |
| 11 | 117396433 | rs11605509 | C | T | NULL | 0.645656306 | NULL&M_Disease |
| 11 | 117396991 | rs1941390 | A | G | NULL | 0.645656306 | NULL&M_Disease |
| 11 | 119779799 | rs765814 | T | G | NULL | 0.645656306 | NULL&M_Disease |
| 11 | 119957516 | rs20493 | A | G | NULL | 0.645656306 | NULL&M_Disease |
| 11 | 116647607 | rs2160669 | C | T | NULL | 0.645656306 | NULL&M_Disease |
| 11 | 117639526 | rs4938439 | C | T | NULL | 0.645656306 | NULL&M_Disease |
| 11 | 120816203 | rs12419730 | A | G | NULL | 0.645656306 | NULL&M_Disease |
| 11 | 119771843 | rs518932 | C | T | NULL | 0.645656306 | NULL&M_Disease |
| 11 | 121109489 | rs6589864 | G | A | NULL | 0.645656306 | NULL&M_Disease |
| 11 | 117934048 | rs4938474 | A | G | NULL | 0.645656306 | NULL&M_Disease |
| 11 | 119694933 | rs17311059 | A | T | NULL | 0.645656306 | NULL&M_Disease |
| 11 | 117173468 | rs486896 | A | G | NULL | 0.645656306 | NULL&M_Disease |
| 11 | 117725333 | rs500787 | T | C | NULL | 0.645656306 | NULL&M_Disease |
| 11 | 119823254 | rs78928955 | C | T | NULL | 0.645656306 | NULL&M_Disease |
| 11 | 120212601 | rs61528732 | C | T | NULL | 0.645656306 | NULL&M_Disease |
| 11 | 121143154 | rs117052388 | T | C | NULL | 0.645656306 | NULL&M_Disease |
| 11 | 116202995 | rs750059 | T | G | NULL | 0.646799393 | NULL&M_Disease |
| 11 | 116453571 | rs17119829 | A | G | NULL | 0.645656306 | NULL&M_Disease |
| 11 | 116585444 | rs79886324 | A | G | NULL | 0.645656306 | NULL&M_Disease |
| 11 | 116594715 | rs180367 | G | A | NULL | 0.645656306 | NULL&M_Disease |
| 11 | 117917156 | rs4936422 | C | A | NULL | 0.645656306 | NULL&M_Disease |
| 11 | 120397775 | rs7103982 | T | C | NULL | 0.645656306 | NULL&M_Disease |
| 11 | 120887491 | rs7928550 | T | A | NULL | 0.645656306 | NULL&M_Disease |
| 11 | 116232179 | rs77779915 | T | G | NULL | 0.645656306 | NULL&M_Disease |
| 11 | 116980641 | rs10892074 | T | G | NULL | 0.646799393 | NULL&M_Disease |
| 11 | 117695340 | rs3824943 | T | C | NULL | 0.646799393 | NULL&M_Disease |
| 11 | 118953952 | rs7952138 | G | A | NULL | 0.645656306 | NULL&M_Disease |
| 11 | 119236590 | rs11217256 | G | C | NULL | 0.645656306 | NULL&M_Disease |
| 11 | 119392654 | rs609305 | G | A | NULL | 0.646799393 | NULL&M_Disease |
| 11 | 120505628 | rs17124120 | T | C | NULL | 0.646799393 | NULL&M_Disease |
| 11 | 116779499 | rs10790166 | G | A | NULL | 0.646799393 | NULL&M_Disease |
| 11 | 119923484 | rs10892523 | G | A | NULL | 0.646799393 | NULL&M_Disease |
| 11 | 116510542 | rs79315770 | A | T | NULL | 0.646799393 | NULL&M_Disease |
| 11 | 116581259 | rs61905072 | T | C | NULL | 0.645656306 | NULL&M_Disease |
| 11 | 117255283 | rs61903746 | T | C | NULL | 0.646799393 | NULL&M_Disease |
| 11 | 117395481 | rs2276339 | A | G | NULL | 0.646799393 | NULL&M_Disease |
| 11 | 117726559 | rs77452664 | T | G | NULL | 0.646799393 | NULL&M_Disease |
| 11 | 119462879 | rs34211230 | A | G | NULL | 0.646799393 | NULL&M_Disease |
| 11 | 116362957 | rs61415784 | C | A | NULL | 0.646799393 | NULL&M_Disease |
| 11 | 116474519 | rs3016350 | T | C | NULL | 0.646799393 | NULL&M_Disease |
| 11 | 116581800 | rs77574778 | T | C | NULL | 0.646799393 | NULL&M_Disease |
| 11 | 116582180 | rs77646493 | A | C | NULL | 0.646799393 | NULL&M_Disease |
| 11 | 116582542 | rs74360954 | T | C | NULL | 0.646799393 | NULL&M_Disease |
| 11 | 116584447 | rs117837319 | C | A | NULL | 0.646799393 | NULL&M_Disease |
| 11 | 117781618 | rs2155195 | G | A | NULL | 0.646799393 | NULL&M_Disease |
| 11 | 119425258 | rs2845664 | C | A | NULL | 0.646799393 | NULL&M_Disease |
| 11 | 119427293 | rs2663620 | A | G | NULL | 0.646799393 | NULL&M_Disease |
| 11 | 120610402 | rs751370 | C | T | NULL | 0.646799393 | NULL&M_Disease |
| 11 | 116450579 | rs61593876 | C | T | NULL | 0.646799393 | NULL&M_Disease |
| 11 | 116479752 | rs17119847 | A | C | NULL | 0.646799393 | NULL&M_Disease |
| 11 | 116581159 | rs59620302 | G | C | NULL | 0.646799393 | NULL&M_Disease |
| 11 | 116652423 | rs6589566 | G | A | NULL | 0.646799393 | NULL&M_Disease |
| 11 | 116652491 | rs7483863 | A | G | NULL | 0.646799393 | NULL&M_Disease |
| 11 | 117931924 | rs12273341 | T | C | NULL | 0.646799393 | NULL&M_Disease |
| 11 | 119400052 | rs10892380 | A | C | NULL | 0.646799393 | NULL&M_Disease |
| 11 | 119770931 | rs542467 | C | T | NULL | 0.646799393 | NULL&M_Disease |
| 11 | 116201751 | rs10790136 | G | A | NULL | 0.646799393 | NULL&M_Disease |
| 11 | 116510369 | rs725524 | A | G | NULL | 0.646799393 | NULL&M_Disease |
| 11 | 116568290 | rs140078194 | A | C | NULL | 0.646799393 | NULL&M_Disease |
| 11 | 116595756 | rs180366 | T | C | NULL | 0.646799393 | NULL&M_Disease |

| | | | | | | | | | |
|---|---|---|---|---|---|---|---|---|---|
| 11 | 116759332 | rs1473177 | C | T | 0.966 | -7977.388 | -7978.582 | -1.194 | 1.17E-06 |
| 11 | 119789537 | rs486978 | T | G | 0.964 | -7977.388 | -7978.582 | -1.194 | 1.17E-06 |
| 11 | 117911637 | rs4938469 | C | T | 1.037 | -7977.388 | -7978.584 | -1.195 | 1.17E-06 |
| 11 | 120166410 | rs744984 | G | A | 1.025 | -7977.388 | -7978.583 | -1.195 | 1.17E-06 |
| 11 | 116586231 | rs117785509 | A | G | 1.013 | -7977.388 | -7978.584 | -1.196 | 1.17E-06 |
| 11 | 117071449 | rs652455 | C | T | 0.964 | -7977.388 | -7978.584 | -1.196 | 1.17E-06 |
| 11 | 118957735 | rs11217148 | T | G | 0.965 | -7977.388 | -7978.584 | -1.196 | 1.17E-06 |
| 11 | 119787190 | rs658358 | C | G | 0.964 | -7977.388 | -7978.584 | -1.196 | 1.17E-06 |
| 11 | 119884982 | rs116088536 | G | C | 0.98 | -7977.388 | -7978.584 | -1.196 | 1.17E-06 |
| 11 | 120167566 | rs11217803 | A | C | 1.024 | -7977.388 | -7978.585 | -1.196 | 1.17E-06 |
| 11 | 120300327 | rs11217865 | C | A | 1.037 | -7977.388 | -7978.584 | -1.196 | 1.17E-06 |
| 11 | 116493738 | rs1240660 | G | A | 0.966 | -7977.388 | -7978.585 | -1.197 | 1.17E-06 |
| 11 | 117670752 | rs657389 | A | G | 1.036 | -7977.388 | -7978.585 | -1.197 | 1.17E-06 |
| 11 | 116528663 | rs73012993 | G | A | 0.968 | -7977.388 | -7978.587 | -1.198 | 1.17E-06 |
| 11 | 117241760 | rs2305826 | T | A | 0.965 | -7977.388 | -7978.586 | -1.198 | 1.17E-06 |
| 11 | 117920577 | rs12225323 | T | C | 0.973 | -7977.388 | -7978.586 | -1.198 | 1.17E-06 |
| 11 | 120357341 | rs79425731 | A | G | 1.017 | -7977.388 | -7978.586 | -1.198 | 1.17E-06 |
| 11 | 120411963 | rs2253134 | A | C | 1.037 | -7977.388 | -7978.587 | -1.198 | 1.17E-06 |
| 11 | 116247262 | rs516169 | C | G | 1.028 | -7977.388 | -7978.587 | -1.199 | 1.17E-06 |
| 11 | 116345605 | rs12275789 | A | G | 0.973 | -7977.388 | -7978.587 | -1.199 | 1.17E-06 |
| 11 | 116663851 | rs17120035 | T | C | 1.027 | -7977.388 | -7978.588 | -1.199 | 1.17E-06 |
| 11 | 116663966 | rs7103224 | A | G | 1.027 | -7977.388 | -7978.588 | -1.199 | 1.17E-06 |
| 11 | 117168402 | rs619360 | T | A | 0.965 | -7977.388 | -7978.587 | -1.199 | 1.17E-06 |
| 11 | 117354904 | rs7129284 | A | G | 0.964 | -7977.388 | -7978.588 | -1.199 | 1.17E-06 |
| 11 | 119236692 | rs12277700 | A | C | 0.97 | -7977.388 | -7978.587 | -1.199 | 1.17E-06 |
| 11 | 116653296 | rs2075290 | C | T | 0.976 | -7977.388 | -7978.588 | -1.2 | 1.17E-06 |
| 11 | 119246567 | rs687009 | C | G | 0.964 | -7977.388 | -7978.589 | -1.2 | 1.17E-06 |
| 11 | 120072666 | rs11217766 | A | C | 0.985 | -7977.388 | -7978.588 | -1.2 | 1.17E-06 |
| 11 | 120305873 | rs77657279 | G | A | 1.017 | -7977.388 | -7978.589 | -1.2 | 1.17E-06 |
| 11 | 121190062 | rs61902398 | A | G | 1.011 | -7977.388 | -7978.589 | -1.2 | 1.17E-06 |
| 11 | 117151378 | rs638270 | G | C | 0.965 | -7977.388 | -7978.59 | -1.201 | 1.16E-06 |
| 11 | 119867224 | rs4938747 | A | G | 0.964 | -7977.388 | -7978.589 | -1.201 | 1.16E-06 |
| 11 | 119929632 | rs11217664 | G | A | 1.036 | -7977.388 | -7978.589 | -1.201 | 1.16E-06 |
| 11 | 116564493 | rs117889337 | T | C | 1.017 | -7977.388 | -7978.591 | -1.202 | 1.16E-06 |
| 11 | 116665079 | rs9804646 | T | C | 1.028 | -7977.388 | -7978.591 | -1.202 | 1.16E-06 |
| 11 | 117396269 | rs12800264 | A | G | 1.037 | -7977.388 | -7978.591 | -1.202 | 1.16E-06 |
| 11 | 117396433 | rs11605509 | C | T | 1.037 | -7977.388 | -7978.591 | -1.202 | 1.16E-06 |
| 11 | 117396991 | rs1941390 | A | G | 1.037 | -7977.388 | -7978.591 | -1.202 | 1.16E-06 |
| 11 | 119779799 | rs765814 | T | G | 1.037 | -7977.388 | -7978.59 | -1.202 | 1.16E-06 |
| 11 | 119957516 | rs20493 | A | G | 0.964 | -7977.388 | -7978.59 | -1.202 | 1.16E-06 |
| 11 | 116647607 | rs2160669 | C | T | 0.977 | -7977.388 | -7978.591 | -1.203 | 1.16E-06 |
| 11 | 117639526 | rs4938439 | C | T | 1.033 | -7977.388 | -7978.591 | -1.203 | 1.16E-06 |
| 11 | 120816203 | rs12419730 | A | G | 0.965 | -7977.388 | -7978.591 | -1.203 | 1.16E-06 |
| 11 | 119771843 | rs518932 | C | T | 1.037 | -7977.388 | -7978.592 | -1.204 | 1.16E-06 |
| 11 | 121109489 | rs6589864 | G | A | 1.013 | -7977.388 | -7978.592 | -1.204 | 1.16E-06 |
| 11 | 117934048 | rs4938474 | A | G | 1.036 | -7977.388 | -7978.593 | -1.205 | 1.16E-06 |
| 11 | 119694933 | rs17311059 | A | T | 1.031 | -7977.388 | -7978.593 | -1.205 | 1.16E-06 |
| 11 | 117173468 | rs486896 | A | G | 0.965 | -7977.388 | -7978.594 | -1.206 | 1.16E-06 |
| 11 | 117725333 | rs500787 | T | C | 0.966 | -7977.388 | -7978.594 | -1.206 | 1.16E-06 |
| 11 | 119823254 | rs78928955 | C | T | 1.037 | -7977.388 | -7978.594 | -1.206 | 1.16E-06 |
| 11 | 120212601 | rs61528732 | C | T | 1.007 | -7977.388 | -7978.594 | -1.206 | 1.16E-06 |
| 11 | 121143154 | rs117052388 | T | C | 0.974 | -7977.388 | -7978.594 | -1.206 | 1.16E-06 |
| 11 | 116202995 | rs750059 | T | G | 1.036 | -7977.388 | -7978.596 | -1.207 | 1.16E-06 |
| 11 | 116453571 | rs17119829 | A | G | 0.971 | -7977.388 | -7978.596 | -1.207 | 1.16E-06 |
| 11 | 116585444 | rs79886324 | A | G | 1.005 | -7977.388 | -7978.595 | -1.207 | 1.16E-06 |
| 11 | 116594715 | rs180367 | G | A | 1.037 | -7977.388 | -7978.595 | -1.207 | 1.16E-06 |
| 11 | 117917156 | rs4936422 | C | A | 1.036 | -7977.388 | -7978.596 | -1.207 | 1.16E-06 |
| 11 | 120397775 | rs7103982 | T | C | 0.971 | -7977.388 | -7978.595 | -1.207 | 1.16E-06 |
| 11 | 120887491 | rs7928550 | T | A | 1.036 | -7977.388 | -7978.595 | -1.207 | 1.16E-06 |
| 11 | 116232179 | rs77779915 | T | G | 1.024 | -7977.388 | -7978.597 | -1.208 | 1.16E-06 |
| 11 | 116980641 | rs10892074 | T | G | 0.966 | -7977.388 | -7978.596 | -1.208 | 1.16E-06 |
| 11 | 117695340 | rs3824943 | T | C | 0.973 | -7977.388 | -7978.596 | -1.208 | 1.16E-06 |
| 11 | 118953952 | rs7952138 | G | A | 1.03 | -7977.388 | -7978.596 | -1.208 | 1.16E-06 |
| 11 | 119236590 | rs11217256 | G | C | 0.971 | -7977.388 | -7978.596 | -1.208 | 1.16E-06 |
| 11 | 119392654 | rs609305 | G | A | 1.036 | -7977.388 | -7978.597 | -1.208 | 1.16E-06 |
| 11 | 120505628 | rs17124120 | T | C | 0.965 | -7977.388 | -7978.596 | -1.208 | 1.16E-06 |
| 11 | 116779499 | rs10790166 | G | A | 1.037 | -7977.388 | -7978.598 | -1.209 | 1.15E-06 |
| 11 | 119923484 | rs10892523 | G | A | 1.037 | -7977.388 | -7978.597 | -1.209 | 1.15E-06 |
| 11 | 116510542 | rs79315770 | A | T | 1.012 | -7977.388 | -7978.598 | -1.21 | 1.15E-06 |
| 11 | 116581259 | rs61905072 | T | C | 0.993 | -7977.388 | -7978.598 | -1.21 | 1.15E-06 |
| 11 | 117255283 | rs61903746 | T | C | 1.029 | -7977.388 | -7978.598 | -1.21 | 1.15E-06 |
| 11 | 117395481 | rs2276339 | A | G | 1.037 | -7977.388 | -7978.598 | -1.21 | 1.15E-06 |
| 11 | 117726559 | rs77452664 | T | G | 0.969 | -7977.388 | -7978.598 | -1.21 | 1.15E-06 |
| 11 | 119462879 | rs34211230 | A | G | 0.971 | -7977.388 | -7978.598 | -1.21 | 1.15E-06 |
| 11 | 116362957 | rs61415784 | C | A | 0.998 | -7977.388 | -7978.599 | -1.211 | 1.15E-06 |
| 11 | 116474519 | rs3016350 | T | C | 1.022 | -7977.388 | -7978.599 | -1.211 | 1.15E-06 |
| 11 | 116581800 | rs77574778 | T | C | 0.996 | -7977.388 | -7978.599 | -1.211 | 1.15E-06 |
| 11 | 116582180 | rs77646493 | A | C | 0.997 | -7977.388 | -7978.6 | -1.211 | 1.15E-06 |
| 11 | 116582542 | rs74360954 | T | C | 0.997 | -7977.388 | -7978.6 | -1.211 | 1.15E-06 |
| 11 | 116584447 | rs117837319 | C | A | 1 | -7977.388 | -7978.599 | -1.211 | 1.15E-06 |
| 11 | 117781618 | rs2155195 | G | A | 0.965 | -7977.388 | -7978.599 | -1.211 | 1.15E-06 |
| 11 | 119425258 | rs2845664 | C | A | 0.971 | -7977.388 | -7978.599 | -1.211 | 1.15E-06 |
| 11 | 119427293 | rs2663620 | A | G | 0.971 | -7977.388 | -7978.599 | -1.211 | 1.15E-06 |
| 11 | 120610402 | rs751370 | C | T | 0.965 | -7977.388 | -7978.599 | -1.211 | 1.15E-06 |
| 11 | 116450579 | rs61593876 | C | T | 0.971 | -7977.388 | -7978.6 | -1.212 | 1.15E-06 |
| 11 | 116479752 | rs17119847 | A | C | 0.991 | -7977.388 | -7978.6 | -1.212 | 1.15E-06 |
| 11 | 116581159 | rs59620302 | G | C | 0.997 | -7977.388 | -7978.6 | -1.212 | 1.15E-06 |
| 11 | 116652423 | rs6589566 | G | A | 0.979 | -7977.388 | -7978.6 | -1.212 | 1.15E-06 |
| 11 | 116652491 | rs7483863 | A | G | 0.979 | -7977.388 | -7978.6 | -1.212 | 1.15E-06 |
| 11 | 117931924 | rs12273341 | T | C | 1.036 | -7977.388 | -7978.6 | -1.212 | 1.15E-06 |
| 11 | 119400052 | rs10892380 | A | C | 1.036 | -7977.388 | -7978.6 | -1.212 | 1.15E-06 |
| 11 | 119770931 | rs542467 | C | T | 1.037 | -7977.388 | -7978.6 | -1.212 | 1.15E-06 |
| 11 | 116201751 | rs10790136 | G | A | 1.035 | -7977.388 | -7978.601 | -1.213 | 1.15E-06 |
| 11 | 116510369 | rs725524 | A | G | 1.011 | -7977.388 | -7978.602 | -1.213 | 1.15E-06 |
| 11 | 116568290 | rs140078194 | A | C | 1.011 | -7977.388 | -7978.602 | -1.213 | 1.15E-06 |
| 11 | 116595756 | rs180366 | T | C | 1.037 | -7977.388 | -7978.601 | -1.213 | 1.15E-06 |

| | | | | | | | |
|---|---|---|---|---|---|---|---|
| 11 | 117065476 | rs1784042 | A | G | NULL | 0.646799393 | NULL&M_Disease |
| 11 | 117175934 | rs522843 | C | T | NULL | 0.646799393 | NULL&M_Disease |
| 11 | 119693592 | rs73008621 | G | C | NULL | 0.646799393 | NULL&M_Disease |
| 11 | 121092398 | rs575030 | C | T | NULL | 0.646799393 | NULL&M_Disease |
| 11 | 121121331 | rs74581584 | C | A | NULL | 0.646799393 | NULL&M_Disease |
| 11 | 116598065 | rs78484485 | A | G | NULL | 0.646799393 | NULL&M_Disease |
| 11 | 116605265 | rs117170971 | A | G | NULL | 0.646799393 | NULL&M_Disease |
| 11 | 116613034 | rs117788500 | C | G | NULL | 0.646799393 | NULL&M_Disease |
| 11 | 117859629 | rs4252251 | A | C | NULL | 0.646799393 | NULL&M_Disease |
| 11 | 119420401 | rs6589755 | T | C | NULL | 0.646799393 | NULL&M_Disease |
| 11 | 119823316 | rs7130104 | C | T | NULL | 0.646799393 | NULL&M_Disease |
| 11 | 120072443 | rs12417625 | G | A | NULL | 0.646799393 | NULL&M_Disease |
| 11 | 120273301 | rs4373935 | G | A | NULL | 0.646799393 | NULL&M_Disease |
| 11 | 116480616 | rs17492724 | C | A | NULL | 0.646799393 | NULL&M_Disease |
| 11 | 116481208 | rs56173877 | A | G | NULL | 0.646799393 | NULL&M_Disease |
| 11 | 116482683 | rs79353638 | A | G | NULL | 0.646799393 | NULL&M_Disease |
| 11 | 116482973 | rs75084211 | A | G | NULL | 0.646799393 | NULL&M_Disease |
| 11 | 116483289 | rs1480227044 | G | C | NULL | 0.646799393 | NULL&M_Disease |
| 11 | 116484610 | rs56075010 | A | G | NULL | 0.646799393 | NULL&M_Disease |
| 11 | 116592248 | rs143315658 | C | T | NULL | 0.646799393 | NULL&M_Disease |
| 11 | 116603677 | rs1893460 | A | G | NULL | 0.646799393 | NULL&M_Disease |
| 11 | 117120473 | rs509565 | T | C | NULL | 0.646799393 | NULL&M_Disease |
| 11 | 117332557 | rs41440247 | C | G | NULL | 0.646799393 | NULL&M_Disease |
| 11 | 117661200 | rs4572163 | T | C | NULL | 0.646799393 | NULL&M_Disease |
| 11 | 119221449 | rs10892353 | A | G | NULL | 0.646799393 | NULL&M_Disease |
| 11 | 119788154 | rs643536 | T | C | NULL | 0.646799393 | NULL&M_Disease |
| 11 | 119930706 | rs10892526 | T | C | NULL | 0.646799393 | NULL&M_Disease |
| 11 | 120273686 | rs4537767 | G | A | NULL | 0.646799393 | NULL&M_Disease |
| 11 | 120392508 | rs61900866 | G | T | NULL | 0.646799393 | NULL&M_Disease |
| 11 | 121035618 | rs549507 | T | C | NULL | 0.646799393 | NULL&M_Disease |
| 11 | 121134520 | rs118179203 | T | C | NULL | 0.646799393 | NULL&M_Disease |
| 11 | 116336740 | rs73009131 | A | G | NULL | 0.646799393 | NULL&M_Disease |
| 11 | 116527609 | rs10892007 | T | C | NULL | 0.646799393 | NULL&M_Disease |
| 11 | 116587146 | rs1244409804 | A | G | NULL | 0.646799393 | NULL&M_Disease |
| 11 | 117396527 | rs11603340 | G | C | NULL | 0.646799393 | NULL&M_Disease |
| 11 | 117945632 | rs1786186 | T | C | NULL | 0.646799393 | NULL&M_Disease |
| 11 | 119341145 | rs10750146 | C | T | NULL | 0.646799393 | NULL&M_Disease |
| 11 | 119432440 | rs1848688 | T | C | NULL | 0.646799393 | NULL&M_Disease |
| 11 | 119433485 | rs2663627 | G | C | NULL | 0.646799393 | NULL&M_Disease |
| 11 | 119433740 | rs2202884 | T | C | NULL | 0.646799393 | NULL&M_Disease |
| 11 | 119753926 | rs503175 | C | T | NULL | 0.646799393 | NULL&M_Disease |
| 11 | 119957107 | rs7937779 | G | A | NULL | 0.646799393 | NULL&M_Disease |
| 11 | 120331916 | rs67617613 | C | A | NULL | 0.646799393 | NULL&M_Disease |
| 11 | 119693648 | rs73008622 | T | C | NULL | 0.646799393 | NULL&M_Disease |
| 11 | 120193461 | rs76046245 | A | C | NULL | 0.646799393 | NULL&M_Disease |
| 11 | 120411267 | rs3892497 | A | G | NULL | 0.646799393 | NULL&M_Disease |
| 11 | 116210400 | rs76741837 | T | C | NULL | 0.646799393 | NULL&M_Disease |
| 11 | 117246150 | rs10790180 | C | T | NULL | 0.647940802 | NULL&M_Disease |
| 11 | 117384450 | rs56074246 | C | T | NULL | 0.647940802 | NULL&M_Disease |
| 11 | 118017765 | rs77279864 | T | C | NULL | 0.646799393 | NULL&M_Disease |
| 11 | 119421419 | rs1938948 | C | T | NULL | 0.646799393 | NULL&M_Disease |
| 11 | 119423269 | rs2845666 | G | A | NULL | 0.646799393 | NULL&M_Disease |
| 11 | 119423410 | rs2845665 | G | A | NULL | 0.646799393 | NULL&M_Disease |
| 11 | 119423748 | rs6589756 | C | T | NULL | 0.646799393 | NULL&M_Disease |
| 11 | 119423822 | rs7395205 | T | C | NULL | 0.646799393 | NULL&M_Disease |
| 11 | 121088158 | rs10892689 | G | A | NULL | 0.647940802 | NULL&M_Disease |
| 11 | 121138747 | rs74384044 | A | G | NULL | 0.646799393 | NULL&M_Disease |
| 11 | 116586854 | rs151061355 | A | G | NULL | 0.646799393 | NULL&M_Disease |
| 11 | 116588629 | rs4938305 | T | C | NULL | 0.647940802 | NULL&M_Disease |
| 11 | 116590528 | rs118046012 | A | G | NULL | 0.646799393 | NULL&M_Disease |
| 11 | 116617895 | rs45539536 | T | C | NULL | 0.646799393 | NULL&M_Disease |
| 11 | 117332512 | rs75355006 | C | T | NULL | 0.647940802 | NULL&M_Disease |
| 11 | 117350137 | rs520289 | A | G | NULL | 0.647940802 | NULL&M_Disease |
| 11 | 119416327 | rs925261 | G | T | NULL | 0.646799393 | NULL&M_Disease |
| 11 | 119778355 | rs553578 | G | A | NULL | 0.647940802 | NULL&M_Disease |
| 11 | 120587319 | rs10892620 | G | A | NULL | 0.647940802 | NULL&M_Disease |
| 11 | 116229947 | rs579078 | C | G | NULL | 0.647940802 | NULL&M_Disease |
| 11 | 116615244 | rs17518841 | A | G | NULL | 0.646799393 | NULL&M_Disease |
| 11 | 118030186 | rs2156850 | A | G | NULL | 0.647940802 | NULL&M_Disease |
| 11 | 119907973 | rs1939332 | C | T | NULL | 0.647940802 | NULL&M_Disease |
| 11 | 120341938 | rs71484170 | C | T | NULL | 0.647940802 | NULL&M_Disease |
| 11 | 116210119 | rs4936347 | T | C | NULL | 0.647940802 | NULL&M_Disease |
| 11 | 116486030 | rs78740143 | A | G | NULL | 0.647940802 | NULL&M_Disease |
| 11 | 116591418 | rs180374 | A | G | NULL | 0.647940802 | NULL&M_Disease |
| 11 | 116675291 | rs1263164 | C | A | NULL | 0.647940802 | NULL&M_Disease |
| 11 | 119165129 | rs12275373 | T | C | NULL | 0.647940802 | NULL&M_Disease |
| 11 | 119881176 | rs1317586 | G | A | NULL | 0.647940802 | NULL&M_Disease |
| 11 | 119940156 | rs673357 | G | A | NULL | 0.647940802 | NULL&M_Disease |
| 11 | 119960260 | rs3915715 | A | G | NULL | 0.647940802 | NULL&M_Disease |
| 11 | 120310978 | rs9666435 | C | A | NULL | 0.647940802 | NULL&M_Disease |
| 11 | 116764582 | rs4938316 | T | C | NULL | 0.647940802 | NULL&M_Disease |
| 11 | 117159878 | rs535860 | T | A | NULL | 0.647940802 | NULL&M_Disease |
| 11 | 119586418 | rs10892441 | T | C | NULL | 0.647940802 | NULL&M_Disease |
| 11 | 120310959 | rs7108596 | A | G | NULL | 0.647940802 | NULL&M_Disease |
| 11 | 120316589 | rs7948299 | C | T | NULL | 0.647940802 | NULL&M_Disease |
| 11 | 120560459 | rs12284730 | T | C | NULL | 0.647940802 | NULL&M_Disease |
| 11 | 116559553 | rs11824135 | T | G | NULL | 0.647940802 | NULL&M_Disease |
| 11 | 116688478 | rs2849165 | T | C | NULL | 0.647940802 | NULL&M_Disease |
| 11 | 117400684 | rs78587763 | T | C | NULL | 0.647940802 | NULL&M_Disease |
| 11 | 119415314 | rs1818393 | A | G | NULL | 0.647940802 | NULL&M_Disease |
| 11 | 120173354 | rs2096884 | A | G | NULL | 0.647940802 | NULL&M_Disease |
| 11 | 116694055 | rs5090 | G | C | NULL | 0.647940802 | NULL&M_Disease |
| 11 | 117394822 | rs12224853 | T | C | NULL | 0.647940802 | NULL&M_Disease |
| 11 | 119417051 | rs10750149 | T | A | NULL | 0.647940802 | NULL&M_Disease |
| 11 | 119418145 | rs1605854 | A | G | NULL | 0.647940802 | NULL&M_Disease |
| 11 | 119418640 | rs2663640 | C | T | NULL | 0.647940802 | NULL&M_Disease |

| | | | | | | | | | |
|---|---|---|---|---|---|---|---|---|---|
| 11 | 117065476 | rs1784042 | A | G | 0.966 | -7977.388 | -7978.601 | -1.213 | 1.15E-06 |
| 11 | 117175934 | rs522843 | C | T | 0.965 | -7977.388 | -7978.602 | -1.213 | 1.15E-06 |
| 11 | 119693592 | rs73008621 | G | C | 1.03 | -7977.388 | -7978.601 | -1.213 | 1.15E-06 |
| 11 | 121092398 | rs575030 | C | T | 0.965 | -7977.388 | -7978.601 | -1.213 | 1.15E-06 |
| 11 | 121121331 | rs74581584 | C | A | 0.974 | -7977.388 | -7978.602 | -1.213 | 1.15E-06 |
| 11 | 116598065 | rs78484485 | A | G | 0.988 | -7977.388 | -7978.603 | -1.214 | 1.15E-06 |
| 11 | 116605265 | rs117170971 | A | G | 1.007 | -7977.388 | -7978.602 | -1.214 | 1.15E-06 |
| 11 | 116613034 | rs117788500 | C | G | 1.009 | -7977.388 | -7978.602 | -1.214 | 1.15E-06 |
| 11 | 117859629 | rs4252251 | A | C | 0.967 | -7977.388 | -7978.602 | -1.214 | 1.15E-06 |
| 11 | 119420401 | rs6589755 | T | C | 0.971 | -7977.388 | -7978.602 | -1.214 | 1.15E-06 |
| 11 | 119823316 | rs7130104 | C | T | 1.034 | -7977.388 | -7978.602 | -1.214 | 1.15E-06 |
| 11 | 120072443 | rs12417625 | G | A | 0.989 | -7977.388 | -7978.602 | -1.214 | 1.15E-06 |
| 11 | 120273301 | rs4373935 | G | A | 1.037 | -7977.388 | -7978.602 | -1.214 | 1.15E-06 |
| 11 | 116480616 | rs17492724 | C | A | 0.992 | -7977.388 | -7978.603 | -1.215 | 1.15E-06 |
| 11 | 116481208 | rs56173877 | A | G | 0.992 | -7977.388 | -7978.603 | -1.215 | 1.15E-06 |
| 11 | 116482683 | rs79353638 | A | G | 0.992 | -7977.388 | -7978.603 | -1.215 | 1.15E-06 |
| 11 | 116482973 | rs75084211 | A | G | 0.992 | -7977.388 | -7978.603 | -1.215 | 1.15E-06 |
| 11 | 116483289 | rs1480227044 | G | C | 0.992 | -7977.388 | -7978.603 | -1.215 | 1.15E-06 |
| 11 | 116484610 | rs56075010 | A | G | 0.992 | -7977.388 | -7978.603 | -1.215 | 1.15E-06 |
| 11 | 116592248 | rs143315658 | C | T | 1.005 | -7977.388 | -7978.603 | -1.215 | 1.15E-06 |
| 11 | 116603677 | rs1893460 | A | G | 1.037 | -7977.388 | -7978.603 | -1.215 | 1.15E-06 |
| 11 | 117120473 | rs509565 | T | C | 1.035 | -7977.388 | -7978.604 | -1.215 | 1.15E-06 |
| 11 | 117332557 | rs41440247 | C | G | 0.969 | -7977.388 | -7978.603 | -1.215 | 1.15E-06 |
| 11 | 117661200 | rs4572163 | T | C | 0.966 | -7977.388 | -7978.603 | -1.215 | 1.15E-06 |
| 11 | 119221449 | rs10892353 | A | G | 1.036 | -7977.388 | -7978.603 | -1.215 | 1.15E-06 |
| 11 | 119788154 | rs643536 | T | C | 0.965 | -7977.388 | -7978.603 | -1.215 | 1.15E-06 |
| 11 | 119930706 | rs10892526 | T | C | 1.036 | -7977.388 | -7978.603 | -1.215 | 1.15E-06 |
| 11 | 120273686 | rs4537767 | G | A | 1.037 | -7977.388 | -7978.603 | -1.215 | 1.15E-06 |
| 11 | 120392508 | rs61900866 | G | T | 0.971 | -7977.388 | -7978.604 | -1.215 | 1.15E-06 |
| 11 | 121035618 | rs549507 | T | C | 1.035 | -7977.388 | -7978.603 | -1.215 | 1.15E-06 |
| 11 | 121134520 | rs118179203 | T | C | 0.975 | -7977.388 | -7978.604 | -1.215 | 1.15E-06 |
| 11 | 116336740 | rs73009131 | A | G | 0.986 | -7977.388 | -7978.604 | -1.216 | 1.15E-06 |
| 11 | 116527609 | rs10892007 | T | C | 0.969 | -7977.388 | -7978.604 | -1.216 | 1.15E-06 |
| 11 | 116587146 | rs1244409804 | A | G | 0.995 | -7977.388 | -7978.604 | -1.216 | 1.15E-06 |
| 11 | 117396527 | rs11603340 | G | C | 1.037 | -7977.388 | -7978.604 | -1.216 | 1.15E-06 |
| 11 | 117945632 | rs1786186 | T | C | 1.036 | -7977.388 | -7978.604 | -1.216 | 1.15E-06 |
| 11 | 119341145 | rs10750146 | C | T | 1.036 | -7977.388 | -7978.604 | -1.216 | 1.15E-06 |
| 11 | 119432440 | rs1848688 | T | C | 0.972 | -7977.388 | -7978.605 | -1.216 | 1.15E-06 |
| 11 | 119433485 | rs2663627 | G | C | 0.972 | -7977.388 | -7978.605 | -1.216 | 1.15E-06 |
| 11 | 119433740 | rs2202884 | T | C | 0.972 | -7977.388 | -7978.605 | -1.216 | 1.15E-06 |
| 11 | 119753926 | rs503175 | C | T | 1.036 | -7977.388 | -7978.604 | -1.216 | 1.15E-06 |
| 11 | 119957107 | rs7937779 | G | A | 1.034 | -7977.388 | -7978.605 | -1.216 | 1.15E-06 |
| 11 | 120331916 | rs67617613 | C | A | 1.036 | -7977.388 | -7978.604 | -1.216 | 1.15E-06 |
| 11 | 119693648 | rs73008622 | T | C | 1.03 | -7977.388 | -7978.605 | -1.217 | 1.15E-06 |
| 11 | 120193461 | rs76046245 | A | C | 1.036 | -7977.388 | -7978.605 | -1.217 | 1.15E-06 |
| 11 | 120411267 | rs3892497 | A | G | 1.035 | -7977.388 | -7978.605 | -1.217 | 1.15E-06 |
| 11 | 116210400 | rs76741837 | T | C | 1.035 | -7977.388 | -7978.606 | -1.218 | 1.14E-06 |
| 11 | 117246150 | rs10790180 | C | T | 0.965 | -7977.388 | -7978.607 | -1.218 | 1.14E-06 |
| 11 | 117384450 | rs56074246 | C | T | 1.036 | -7977.388 | -7978.606 | -1.218 | 1.14E-06 |
| 11 | 118017765 | rs77279864 | T | C | 0.983 | -7977.388 | -7978.607 | -1.218 | 1.14E-06 |
| 11 | 119421419 | rs1938948 | C | T | 0.972 | -7977.388 | -7978.607 | -1.218 | 1.14E-06 |
| 11 | 119423269 | rs2845666 | G | A | 0.972 | -7977.388 | -7978.607 | -1.218 | 1.14E-06 |
| 11 | 119423410 | rs2845665 | G | A | 0.972 | -7977.388 | -7978.607 | -1.218 | 1.14E-06 |
| 11 | 119423748 | rs6589756 | C | T | 0.972 | -7977.388 | -7978.607 | -1.218 | 1.14E-06 |
| 11 | 119423822 | rs7395205 | T | C | 0.972 | -7977.388 | -7978.607 | -1.218 | 1.14E-06 |
| 11 | 121088158 | rs10892689 | G | A | 1.036 | -7977.388 | -7978.606 | -1.218 | 1.14E-06 |
| 11 | 121138747 | rs74384044 | A | G | 0.975 | -7977.388 | -7978.606 | -1.218 | 1.14E-06 |
| 11 | 116586854 | rs151061355 | A | G | 1 | -7977.388 | -7978.607 | -1.219 | 1.14E-06 |
| 11 | 116588629 | rs4938305 | T | C | 1.036 | -7977.388 | -7978.607 | -1.219 | 1.14E-06 |
| 11 | 116590528 | rs118046012 | A | G | 0.999 | -7977.388 | -7978.608 | -1.219 | 1.14E-06 |
| 11 | 116617895 | rs45539536 | T | C | 1.007 | -7977.388 | -7978.607 | -1.219 | 1.14E-06 |
| 11 | 117332512 | rs75355006 | C | T | 0.969 | -7977.388 | -7978.607 | -1.219 | 1.14E-06 |
| 11 | 117350137 | rs520289 | A | G | 1.035 | -7977.388 | -7978.607 | -1.219 | 1.14E-06 |
| 11 | 119416327 | rs925261 | G | T | 0.972 | -7977.388 | -7978.607 | -1.219 | 1.14E-06 |
| 11 | 119778355 | rs553578 | G | A | 1.036 | -7977.388 | -7978.608 | -1.219 | 1.14E-06 |
| 11 | 120587319 | rs10892620 | G | A | 1.036 | -7977.388 | -7978.607 | -1.219 | 1.14E-06 |
| 11 | 116229947 | rs579078 | C | G | 1.036 | -7977.388 | -7978.608 | -1.22 | 1.14E-06 |
| 11 | 116615244 | rs17518841 | A | G | 1.007 | -7977.388 | -7978.608 | -1.22 | 1.14E-06 |
| 11 | 118030186 | rs2156850 | A | G | 0.966 | -7977.388 | -7978.608 | -1.22 | 1.14E-06 |
| 11 | 119907973 | rs1939332 | C | T | 1.036 | -7977.388 | -7978.608 | -1.22 | 1.14E-06 |
| 11 | 120341938 | rs71484170 | C | T | 0.967 | -7977.388 | -7978.609 | -1.22 | 1.14E-06 |
| 11 | 116210119 | rs4936347 | T | C | 0.965 | -7977.388 | -7978.609 | -1.221 | 1.14E-06 |
| 11 | 116486030 | rs78740143 | A | G | 1.025 | -7977.388 | -7978.61 | -1.221 | 1.14E-06 |
| 11 | 116591418 | rs180374 | A | G | 1.036 | -7977.388 | -7978.609 | -1.221 | 1.14E-06 |
| 11 | 116675291 | rs1263164 | C | A | 1.036 | -7977.388 | -7978.609 | -1.221 | 1.14E-06 |
| 11 | 119165129 | rs12275373 | T | C | 0.965 | -7977.388 | -7978.61 | -1.221 | 1.14E-06 |
| 11 | 119881176 | rs1317586 | G | A | 0.966 | -7977.388 | -7978.61 | -1.221 | 1.14E-06 |
| 11 | 119940156 | rs673357 | G | A | 0.965 | -7977.388 | -7978.61 | -1.221 | 1.14E-06 |
| 11 | 119960260 | rs3915715 | A | G | 1.033 | -7977.388 | -7978.61 | -1.221 | 1.14E-06 |
| 11 | 120310978 | rs9666435 | C | A | 1.036 | -7977.388 | -7978.61 | -1.221 | 1.14E-06 |
| 11 | 116764582 | rs4938316 | T | C | 1.036 | -7977.388 | -7978.611 | -1.222 | 1.14E-06 |
| 11 | 117159878 | rs535860 | T | A | 0.966 | -7977.388 | -7978.61 | -1.222 | 1.14E-06 |
| 11 | 119586418 | rs10892441 | T | C | 1.035 | -7977.388 | -7978.611 | -1.222 | 1.14E-06 |
| 11 | 120310959 | rs7108596 | A | G | 1.036 | -7977.388 | -7978.61 | -1.222 | 1.14E-06 |
| 11 | 120316589 | rs7948299 | C | T | 1.036 | -7977.388 | -7978.61 | -1.222 | 1.14E-06 |
| 11 | 120560459 | rs12284730 | T | C | 0.966 | -7977.388 | -7978.61 | -1.222 | 1.14E-06 |
| 11 | 116559553 | rs11824135 | T | G | 0.98 | -7977.388 | -7978.611 | -1.223 | 1.14E-06 |
| 11 | 116688478 | rs2849165 | T | C | 1.036 | -7977.388 | -7978.611 | -1.223 | 1.14E-06 |
| 11 | 117400684 | rs78587763 | T | C | 0.986 | -7977.388 | -7978.612 | -1.223 | 1.14E-06 |
| 11 | 119415314 | rs1818393 | A | G | 0.972 | -7977.388 | -7978.611 | -1.223 | 1.14E-06 |
| 11 | 120173354 | rs2096884 | A | G | 0.965 | -7977.388 | -7978.611 | -1.223 | 1.14E-06 |
| 11 | 116694055 | rs5090 | G | C | 0.999 | -7977.388 | -7978.612 | -1.224 | 1.14E-06 |
| 11 | 117394822 | rs12224853 | T | C | 1.036 | -7977.388 | -7978.612 | -1.224 | 1.14E-06 |
| 11 | 119417051 | rs10750149 | T | A | 0.973 | -7977.388 | -7978.612 | -1.224 | 1.14E-06 |
| 11 | 119418145 | rs1605854 | A | G | 0.973 | -7977.388 | -7978.612 | -1.224 | 1.14E-06 |
| 11 | 119418640 | rs2663640 | C | T | 0.973 | -7977.388 | -7978.612 | -1.224 | 1.14E-06 |

| | | | | | | | |
|---|---|---|---|---|---|---|---|
| 11 | 119419198 | rs1828280 | T | C | NULL | 0.647940802 | NULL&M_Disease |
| 11 | 119823133 | rs75736698 | A | G | NULL | 0.647940802 | NULL&M_Disease |
| 11 | 116260862 | rs524929 | G | A | NULL | 0.647940802 | NULL&M_Disease |
| 11 | 116389102 | rs906546 | T | C | NULL | 0.647940802 | NULL&M_Disease |
| 11 | 116663596 | rs34003087 | T | C | NULL | 0.647940802 | NULL&M_Disease |
| 11 | 119343329 | rs593061 | G | A | NULL | 0.647940802 | NULL&M_Disease |
| 11 | 120168201 | rs4936514 | C | G | NULL | 0.647940802 | NULL&M_Disease |
| 11 | 120277388 | rs12363974 | G | C | NULL | 0.647940802 | NULL&M_Disease |
| 11 | 120816408 | rs11218069 | A | C | NULL | 0.647940802 | NULL&M_Disease |
| 11 | 116448927 | rs77130505 | T | C | NULL | 0.647940802 | NULL&M_Disease |
| 11 | 119082706 | rs7482092 | C | T | NULL | 0.647940802 | NULL&M_Disease |
| 11 | 120306759 | rs57912596 | C | T | NULL | 0.647940802 | NULL&M_Disease |
| 11 | 120307742 | rs10502236 | G | A | NULL | 0.647940802 | NULL&M_Disease |
| 11 | 120785857 | rs2850810 | A | G | NULL | 0.647940802 | NULL&M_Disease |
| 11 | 116247759 | rs493693 | A | C | NULL | 0.647940802 | NULL&M_Disease |
| 11 | 116620670 | rs57950207 | C | T | NULL | 0.647940802 | NULL&M_Disease |
| 11 | 116621770 | rs17440396 | A | G | NULL | 0.647940802 | NULL&M_Disease |
| 11 | 116626226 | rs77112220 | T | C | NULL | 0.647940802 | NULL&M_Disease |
| 11 | 116640237 | rs6589565 | A | G | NULL | 0.647940802 | NULL&M_Disease |
| 11 | 117396505 | rs11603366 | T | G | NULL | 0.647940802 | NULL&M_Disease |
| 11 | 117420748 | rs7126576 | C | T | NULL | 0.647940802 | NULL&M_Disease |
| 11 | 117773537 | rs947917 | G | A | NULL | 0.647940802 | NULL&M_Disease |
| 11 | 117804109 | rs3863317 | A | G | NULL | 0.647940802 | NULL&M_Disease |
| 11 | 116665553 | rs17520254 | G | C | NULL | 0.647940802 | NULL&M_Disease |
| 11 | 117718880 | rs512481 | A | G | NULL | 0.649080524 | NULL&M_Disease |
| 11 | 118685015 | rs540716 | T | C | NULL | 0.647940802 | NULL&M_Disease |
| 11 | 120816319 | rs11218068 | C | T | NULL | 0.649080524 | NULL&M_Disease |
| 11 | 119509911 | rs4409838 | A | G | NULL | 0.647940802 | NULL&M_Disease |
| 11 | 119880076 | rs11217629 | A | C | NULL | 0.649080524 | NULL&M_Disease |
| 11 | 116629905 | rs17519093 | A | G | NULL | 0.647940802 | NULL&M_Disease |
| 11 | 116631912 | rs139872819 | C | T | NULL | 0.647940802 | NULL&M_Disease |
| 11 | 116633947 | rs10488698 | A | G | NULL | 0.647940802 | NULL&M_Disease |
| 11 | 116634297 | rs116877083 | C | A | NULL | 0.647940802 | NULL&M_Disease |
| 11 | 116635784 | rs2187126 | G | A | NULL | 0.647940802 | NULL&M_Disease |
| 11 | 116637002 | rs75542782 | A | G | NULL | 0.647940802 | NULL&M_Disease |
| 11 | 116638751 | rs57643109 | A | G | NULL | 0.647940802 | NULL&M_Disease |
| 11 | 116639104 | rs10790162 | A | G | NULL | 0.649080524 | NULL&M_Disease |
| 11 | 116642100 | rs117579298 | G | A | NULL | 0.647940802 | NULL&M_Disease |
| 11 | 116643011 | rs79110162 | G | A | NULL | 0.647940802 | NULL&M_Disease |
| 11 | 116653762 | rs76836072 | T | C | NULL | 0.649080524 | NULL&M_Disease |
| 11 | 118877621 | rs4488208 | C | G | NULL | 0.649080524 | NULL&M_Disease |
| 11 | 119386473 | rs632818 | C | T | NULL | 0.649080524 | NULL&M_Disease |
| 11 | 119520783 | rs10219389 | C | A | NULL | 0.649080524 | NULL&M_Disease |
| 11 | 120397162 | rs35594498 | T | C | NULL | 0.649080524 | NULL&M_Disease |
| 11 | 121110311 | rs12417339 | A | G | NULL | 0.649080524 | NULL&M_Disease |
| 11 | 116613660 | rs12280753 | T | C | NULL | 0.649080524 | NULL&M_Disease |
| 11 | 116624727 | rs7930786 | C | G | NULL | 0.649080524 | NULL&M_Disease |
| 11 | 116631707 | rs3825041 | T | C | NULL | 0.649080524 | NULL&M_Disease |
| 11 | 117420336 | rs7110219 | A | G | NULL | 0.649080524 | NULL&M_Disease |
| 11 | 117673828 | rs10488782 | G | T | NULL | 0.649080524 | NULL&M_Disease |
| 11 | 118180667 | rs12793150 | G | C | NULL | 0.649080524 | NULL&M_Disease |
| 11 | 120330728 | rs10892579 | G | A | NULL | 0.649080524 | NULL&M_Disease |
| 11 | 120451223 | rs67294535 | G | A | NULL | 0.649080524 | NULL&M_Disease |
| 11 | 120478622 | rs7934084 | T | G | NULL | 0.649080524 | NULL&M_Disease |
| 11 | 120817900 | rs11218073 | T | C | NULL | 0.649080524 | NULL&M_Disease |
| 11 | 120826623 | rs7123343 | T | C | NULL | 0.649080524 | NULL&M_Disease |
| 11 | 116214112 | rs12272993 | T | G | NULL | 0.649080524 | NULL&M_Disease |
| 11 | 116666099 | rs35904674 | T | C | NULL | 0.649080524 | NULL&M_Disease |
| 11 | 117561070 | rs111940587 | G | A | NULL | 0.649080524 | NULL&M_Disease |
| 11 | 118955314 | rs592190 | A | G | NULL | 0.649080524 | NULL&M_Disease |
| 11 | 118955679 | rs589925 | T | C | NULL | 0.649080524 | NULL&M_Disease |
| 11 | 119422645 | rs2845667 | T | C | NULL | 0.649080524 | NULL&M_Disease |
| 11 | 120249395 | rs11600723 | T | C | NULL | 0.649080524 | NULL&M_Disease |
| 11 | 120649962 | rs2846097 | C | G | NULL | 0.649080524 | NULL&M_Disease |
| 11 | 116464675 | rs10891997 | G | A | NULL | 0.649080524 | NULL&M_Disease |
| 11 | 117333778 | rs11607675 | A | G | NULL | 0.649080524 | NULL&M_Disease |
| 11 | 119520995 | rs11822909 | T | C | NULL | 0.649080524 | NULL&M_Disease |
| 11 | 120167220 | rs12225696 | G | A | NULL | 0.649080524 | NULL&M_Disease |
| 11 | 120167230 | rs11217802 | T | C | NULL | 0.649080524 | NULL&M_Disease |
| 11 | 120266477 | rs12361642 | T | C | NULL | 0.649080524 | NULL&M_Disease |
| 11 | 121103437 | rs80094129 | A | C | NULL | 0.649080524 | NULL&M_Disease |
| 11 | 116574488 | rs11216103 | A | G | NULL | 0.649080524 | NULL&M_Disease |
| 11 | 117561019 | rs73594696 | C | T | NULL | 0.649080524 | NULL&M_Disease |
| 11 | 120167057 | rs4142654 | G | A | NULL | 0.649080524 | NULL&M_Disease |
| 11 | 120820885 | rs11218075 | C | A | NULL | 0.649080524 | NULL&M_Disease |
| 11 | 116599771 | rs180355 | T | C | NULL | 0.649080524 | NULL&M_Disease |
| 11 | 116599826 | rs180354 | T | C | NULL | 0.649080524 | NULL&M_Disease |
| 11 | 116625294 | rs180325 | C | G | NULL | 0.649080524 | NULL&M_Disease |
| 11 | 116625332 | rs180324 | C | T | NULL | 0.649080524 | NULL&M_Disease |
| 11 | 119245349 | rs659705 | G | A | NULL | 0.649080524 | NULL&M_Disease |
| 11 | 120454925 | rs35199514 | C | T | NULL | 0.649080524 | NULL&M_Disease |
| 11 | 120558389 | rs12281008 | A | G | NULL | 0.649080524 | NULL&M_Disease |
| 11 | 116528224 | rs579674 | G | T | NULL | 0.649080524 | NULL&M_Disease |
| 11 | 116664776 | rs1787680 | A | T | NULL | 0.649080524 | NULL&M_Disease |
| 11 | 117302910 | rs3892796 | A | G | NULL | 0.649080524 | NULL&M_Disease |
| 11 | 117398591 | rs4936384 | T | C | NULL | 0.649080524 | NULL&M_Disease |
| 11 | 117572183 | rs73596628 | C | T | NULL | 0.649080524 | NULL&M_Disease |
| 11 | 119418178 | rs1605855 | A | C | NULL | 0.649080524 | NULL&M_Disease |
| 11 | 120078501 | rs537907 | C | G | NULL | 0.649080524 | NULL&M_Disease |
| 11 | 120078890 | rs540857 | G | A | NULL | 0.649080524 | NULL&M_Disease |
| 11 | 116624153 | rs6589564 | C | G | NULL | 0.649080524 | NULL&M_Disease |
| 11 | 117337289 | rs12577361 | A | C | NULL | 0.649080524 | NULL&M_Disease |
| 11 | 117396386 | rs12800487 | A | G | NULL | 0.649080524 | NULL&M_Disease |
| 11 | 119950934 | rs10892531 | C | T | NULL | 0.649080524 | NULL&M_Disease |
| 11 | 116523279 | rs523267 | C | T | NULL | 0.650218549 | NULL&M_Disease |
| 11 | 117927676 | rs584231 | A | G | NULL | 0.649080524 | NULL&M_Disease |

| | | | | | | | | | |
|---|---|---|---|---|---|---|---|---|---|
| 11 | 119419198 | rs1828280 | T | C | 0.973 | -7977.388 | -7978.612 | -1.224 | 1.14E-06 |
| 11 | 119823133 | rs75736698 | A | G | 1.034 | -7977.388 | -7978.612 | -1.224 | 1.14E-06 |
| 11 | 116260862 | rs524929 | G | A | 0.968 | -7977.388 | -7978.613 | -1.225 | 1.14E-06 |
| 11 | 116389102 | rs906546 | T | C | 0.965 | -7977.388 | -7978.613 | -1.225 | 1.14E-06 |
| 11 | 116663596 | rs34003087 | T | C | 0.993 | -7977.388 | -7978.613 | -1.225 | 1.14E-06 |
| 11 | 119343329 | rs593061 | G | A | 1.036 | -7977.388 | -7978.613 | -1.225 | 1.14E-06 |
| 11 | 120168201 | rs4936514 | C | G | 1.02 | -7977.388 | -7978.613 | -1.225 | 1.14E-06 |
| 11 | 120277388 | rs12363974 | G | C | 1.036 | -7977.388 | -7978.614 | -1.225 | 1.14E-06 |
| 11 | 120816408 | rs11218069 | A | C | 0.965 | -7977.388 | -7978.614 | -1.225 | 1.14E-06 |
| 11 | 116448927 | rs77130505 | T | C | 0.972 | -7977.388 | -7978.614 | -1.226 | 1.14E-06 |
| 11 | 119082706 | rs7482092 | C | T | 0.965 | -7977.388 | -7978.614 | -1.226 | 1.14E-06 |
| 11 | 120306759 | rs57912596 | C | T | 1.036 | -7977.388 | -7978.614 | -1.226 | 1.14E-06 |
| 11 | 120307742 | rs10502236 | G | A | 1.036 | -7977.388 | -7978.614 | -1.226 | 1.14E-06 |
| 11 | 120785857 | rs2850810 | A | G | 1.036 | -7977.388 | -7978.615 | -1.226 | 1.14E-06 |
| 11 | 116247759 | rs493693 | A | C | 1.003 | -7977.388 | -7978.615 | -1.227 | 1.13E-06 |
| 11 | 116620670 | rs57950207 | C | T | 1.004 | -7977.388 | -7978.615 | -1.227 | 1.13E-06 |
| 11 | 116621770 | rs17440396 | A | G | 1.003 | -7977.388 | -7978.616 | -1.227 | 1.13E-06 |
| 11 | 116626226 | rs77112220 | T | C | 1.004 | -7977.388 | -7978.615 | -1.227 | 1.13E-06 |
| 11 | 116640237 | rs6589565 | A | G | 0.981 | -7977.388 | -7978.616 | -1.227 | 1.13E-06 |
| 11 | 117396505 | rs11603366 | T | G | 1.036 | -7977.388 | -7978.615 | -1.227 | 1.13E-06 |
| 11 | 117420748 | rs7126576 | C | T | 1.024 | -7977.388 | -7978.615 | -1.227 | 1.13E-06 |
| 11 | 117773537 | rs947917 | G | A | 0.966 | -7977.388 | -7978.615 | -1.227 | 1.13E-06 |
| 11 | 117804109 | rs3863317 | A | G | 0.966 | -7977.388 | -7978.616 | -1.227 | 1.13E-06 |
| 11 | 116665553 | rs17520254 | G | C | 0.986 | -7977.388 | -7978.616 | -1.228 | 1.13E-06 |
| 11 | 117718880 | rs512481 | A | G | 0.966 | -7977.388 | -7978.616 | -1.228 | 1.13E-06 |
| 11 | 118685015 | rs540716 | T | C | 1.036 | -7977.388 | -7978.616 | -1.228 | 1.13E-06 |
| 11 | 120816319 | rs11218068 | C | T | 0.965 | -7977.388 | -7978.617 | -1.228 | 1.13E-06 |
| 11 | 119509911 | rs4409838 | A | G | 0.978 | -7977.388 | -7978.617 | -1.229 | 1.13E-06 |
| 11 | 119880076 | rs11217629 | A | C | 1.035 | -7977.388 | -7978.618 | -1.229 | 1.13E-06 |
| 11 | 116629905 | rs17519093 | A | G | 1 | -7977.388 | -7978.618 | -1.23 | 1.13E-06 |
| 11 | 116631912 | rs139872819 | C | T | 1 | -7977.388 | -7978.618 | -1.23 | 1.13E-06 |
| 11 | 116633947 | rs10488698 | A | G | 0.999 | -7977.388 | -7978.618 | -1.23 | 1.13E-06 |
| 11 | 116634297 | rs116877083 | C | A | 0.999 | -7977.388 | -7978.618 | -1.23 | 1.13E-06 |
| 11 | 116635784 | rs2187126 | G | A | 1.001 | -7977.388 | -7978.618 | -1.23 | 1.13E-06 |
| 11 | 116637002 | rs75542782 | A | G | 0.999 | -7977.388 | -7978.618 | -1.23 | 1.13E-06 |
| 11 | 116638751 | rs57643109 | A | G | 1 | -7977.388 | -7978.618 | -1.23 | 1.13E-06 |
| 11 | 116639104 | rs10790162 | A | G | 0.981 | -7977.388 | -7978.618 | -1.23 | 1.13E-06 |
| 11 | 116642100 | rs117579298 | G | A | 1 | -7977.388 | -7978.618 | -1.23 | 1.13E-06 |
| 11 | 116643011 | rs79110162 | G | A | 1 | -7977.388 | -7978.618 | -1.23 | 1.13E-06 |
| 11 | 116653762 | rs76836072 | T | C | 1.002 | -7977.388 | -7978.619 | -1.23 | 1.13E-06 |
| 11 | 118877621 | rs4488208 | C | G | 1.035 | -7977.388 | -7978.618 | -1.23 | 1.13E-06 |
| 11 | 119386473 | rs632818 | C | T | 1.035 | -7977.388 | -7978.618 | -1.23 | 1.13E-06 |
| 11 | 119520783 | rs10219389 | C | A | 1.036 | -7977.388 | -7978.619 | -1.23 | 1.13E-06 |
| 11 | 120397162 | rs35594498 | T | C | 0.973 | -7977.388 | -7978.618 | -1.23 | 1.13E-06 |
| 11 | 121110311 | rs12417339 | A | G | 0.998 | -7977.388 | -7978.619 | -1.23 | 1.13E-06 |
| 11 | 116613660 | rs12280753 | T | C | 0.977 | -7977.388 | -7978.619 | -1.231 | 1.13E-06 |
| 11 | 116624727 | rs7930786 | C | G | 0.981 | -7977.388 | -7978.619 | -1.231 | 1.13E-06 |
| 11 | 116631707 | rs3825041 | T | C | 0.982 | -7977.388 | -7978.619 | -1.231 | 1.13E-06 |
| 11 | 117420336 | rs7110219 | A | G | 1.023 | -7977.388 | -7978.619 | -1.231 | 1.13E-06 |
| 11 | 117673828 | rs10488782 | G | T | 0.968 | -7977.388 | -7978.619 | -1.231 | 1.13E-06 |
| 11 | 118180667 | rs12793150 | G | C | 0.966 | -7977.388 | -7978.619 | -1.231 | 1.13E-06 |
| 11 | 120330728 | rs10892579 | G | A | 1.036 | -7977.388 | -7978.619 | -1.231 | 1.13E-06 |
| 11 | 120451223 | rs67294535 | G | A | 1.004 | -7977.388 | -7978.619 | -1.231 | 1.13E-06 |
| 11 | 120478622 | rs7934084 | T | G | 0.966 | -7977.388 | -7978.62 | -1.231 | 1.13E-06 |
| 11 | 120817900 | rs11218073 | T | C | 1.036 | -7977.388 | -7978.619 | -1.231 | 1.13E-06 |
| 11 | 120826623 | rs7123343 | T | C | 1.035 | -7977.388 | -7978.619 | -1.231 | 1.13E-06 |
| 11 | 116214112 | rs12272993 | T | G | 1.015 | -7977.388 | -7978.62 | -1.232 | 1.13E-06 |
| 11 | 116666099 | rs35904674 | T | C | 1.022 | -7977.388 | -7978.62 | -1.232 | 1.13E-06 |
| 11 | 117561070 | rs111940587 | G | A | 0.977 | -7977.388 | -7978.62 | -1.232 | 1.13E-06 |
| 11 | 118955314 | rs592190 | A | G | 0.966 | -7977.388 | -7978.62 | -1.232 | 1.13E-06 |
| 11 | 118955679 | rs589925 | T | C | 0.966 | -7977.388 | -7978.62 | -1.232 | 1.13E-06 |
| 11 | 119422645 | rs2845667 | T | C | 0.973 | -7977.388 | -7978.62 | -1.232 | 1.13E-06 |
| 11 | 120249395 | rs11600723 | T | C | 1.036 | -7977.388 | -7978.62 | -1.232 | 1.13E-06 |
| 11 | 120649962 | rs2846097 | C | G | 0.966 | -7977.388 | -7978.62 | -1.232 | 1.13E-06 |
| 11 | 116464675 | rs10891997 | G | A | 1.036 | -7977.388 | -7978.621 | -1.233 | 1.13E-06 |
| 11 | 117333778 | rs11607675 | A | G | 0.97 | -7977.388 | -7978.622 | -1.233 | 1.13E-06 |
| 11 | 119520995 | rs11822909 | T | C | 1.036 | -7977.388 | -7978.621 | -1.233 | 1.13E-06 |
| 11 | 120167220 | rs12225696 | G | A | 1.019 | -7977.388 | -7978.622 | -1.233 | 1.13E-06 |
| 11 | 120167230 | rs11217802 | T | C | 1.019 | -7977.388 | -7978.622 | -1.233 | 1.13E-06 |
| 11 | 120266477 | rs12361642 | T | C | 1.036 | -7977.388 | -7978.621 | -1.233 | 1.13E-06 |
| 11 | 121103437 | rs80094129 | A | C | 1.009 | -7977.388 | -7978.621 | -1.233 | 1.13E-06 |
| 11 | 116574488 | rs11216103 | A | G | 0.985 | -7977.388 | -7978.622 | -1.234 | 1.13E-06 |
| 11 | 117561019 | rs73594696 | C | T | 0.977 | -7977.388 | -7978.622 | -1.234 | 1.13E-06 |
| 11 | 120167057 | rs4142654 | G | A | 1.019 | -7977.388 | -7978.622 | -1.234 | 1.13E-06 |
| 11 | 120820885 | rs11218075 | C | A | 0.966 | -7977.388 | -7978.623 | -1.234 | 1.13E-06 |
| 11 | 116599771 | rs180355 | T | C | 1.036 | -7977.388 | -7978.623 | -1.235 | 1.12E-06 |
| 11 | 116599826 | rs180354 | T | C | 1.036 | -7977.388 | -7978.623 | -1.235 | 1.12E-06 |
| 11 | 116625294 | rs180325 | C | G | 1.036 | -7977.388 | -7978.623 | -1.235 | 1.12E-06 |
| 11 | 116625332 | rs180324 | C | T | 1.036 | -7977.388 | -7978.623 | -1.235 | 1.12E-06 |
| 11 | 119245349 | rs659705 | G | A | 0.966 | -7977.388 | -7978.623 | -1.235 | 1.12E-06 |
| 11 | 120454925 | rs35199514 | C | T | 0.997 | -7977.388 | -7978.623 | -1.235 | 1.12E-06 |
| 11 | 120558389 | rs12281008 | A | G | 0.968 | -7977.388 | -7978.623 | -1.235 | 1.12E-06 |
| 11 | 116528224 | rs579674 | G | T | 1.036 | -7977.388 | -7978.624 | -1.236 | 1.12E-06 |
| 11 | 116664776 | rs1787680 | A | T | 0.966 | -7977.388 | -7978.624 | -1.236 | 1.12E-06 |
| 11 | 117302910 | rs3892796 | A | G | 0.969 | -7977.388 | -7978.625 | -1.236 | 1.12E-06 |
| 11 | 117398591 | rs4936384 | T | C | 1.036 | -7977.388 | -7978.624 | -1.236 | 1.12E-06 |
| 11 | 117572183 | rs73596628 | C | T | 1.008 | -7977.388 | -7978.624 | -1.236 | 1.12E-06 |
| 11 | 119418178 | rs1605855 | A | C | 0.974 | -7977.388 | -7978.624 | -1.236 | 1.12E-06 |
| 11 | 120078501 | rs537907 | C | G | 0.973 | -7977.388 | -7978.625 | -1.236 | 1.12E-06 |
| 11 | 120078890 | rs540857 | G | A | 0.973 | -7977.388 | -7978.625 | -1.236 | 1.12E-06 |
| 11 | 116624153 | rs6589564 | C | G | 0.983 | -7977.388 | -7978.625 | -1.237 | 1.12E-06 |
| 11 | 117337289 | rs12577361 | A | C | 1.014 | -7977.388 | -7978.626 | -1.237 | 1.12E-06 |
| 11 | 117396386 | rs12800487 | A | G | 1.036 | -7977.388 | -7978.626 | -1.237 | 1.12E-06 |
| 11 | 119950934 | rs10892531 | C | T | 0.965 | -7977.388 | -7978.626 | -1.237 | 1.12E-06 |
| 11 | 116523279 | rs523267 | C | T | 0.966 | -7977.388 | -7978.626 | -1.238 | 1.12E-06 |
| 11 | 117927676 | rs584231 | A | G | 1.034 | -7977.388 | -7978.626 | -1.238 | 1.12E-06 |

| | | | | | | | |
|---|---|---|---|---|---|---|---|
| 11 | 119427624 | rs2663623 | A | G | NULL | 0.649080524 | NULL&M_Disease |
| 11 | 119427893 | rs2845661 | T | G | NULL | 0.649080524 | NULL&M_Disease |
| 11 | 120476746 | rs4936529 | T | A | NULL | 0.650218549 | NULL&M_Disease |
| 11 | 116407867 | rs11215991 | T | C | NULL | 0.650218549 | NULL&M_Disease |
| 11 | 117800033 | rs35251396 | A | G | NULL | 0.649080524 | NULL&M_Disease |
| 11 | 119903718 | rs61897924 | T | C | NULL | 0.650218549 | NULL&M_Disease |
| 11 | 116448951 | rs77196345 | T | C | NULL | 0.650218549 | NULL&M_Disease |
| 11 | 116599281 | rs180359 | G | A | NULL | 0.650218549 | NULL&M_Disease |
| 11 | 116599392 | rs180358 | T | C | NULL | 0.650218549 | NULL&M_Disease |
| 11 | 116599534 | rs180356 | T | C | NULL | 0.650218549 | NULL&M_Disease |
| 11 | 117864047 | rs2256111 | A | G | NULL | 0.650218549 | NULL&M_Disease |
| 11 | 119155618 | rs2298650 | T | G | NULL | 0.650218549 | NULL&M_Disease |
| 11 | 120558364 | rs12281001 | T | G | NULL | 0.650218549 | NULL&M_Disease |
| 11 | 120558487 | rs12294046 | T | A | NULL | 0.650218549 | NULL&M_Disease |
| 11 | 120559070 | rs10892616 | G | A | NULL | 0.650218549 | NULL&M_Disease |
| 11 | 120599666 | rs112473235 | T | C | NULL | 0.650218549 | NULL&M_Disease |
| 11 | 117082966 | rs76169968 | A | G | NULL | 0.650218549 | NULL&M_Disease |
| 11 | 117213847 | rs72645461 | G | A | NULL | 0.650218549 | NULL&M_Disease |
| 11 | 117333657 | rs11607656 | A | G | NULL | 0.650218549 | NULL&M_Disease |
| 11 | 120396030 | rs71484178 | A | G | NULL | 0.650218549 | NULL&M_Disease |
| 11 | 116355860 | rs573554 | C | T | NULL | 0.650218549 | NULL&M_Disease |
| 11 | 117421276 | rs71480343 | A | G | NULL | 0.650218549 | NULL&M_Disease |
| 11 | 120178632 | rs11605781 | C | T | NULL | 0.650218549 | NULL&M_Disease |
| 11 | 121018725 | rs551031 | C | T | NULL | 0.650218549 | NULL&M_Disease |
| 11 | 116263533 | rs480793 | A | G | NULL | 0.650218549 | NULL&M_Disease |
| 11 | 116392191 | rs10891985 | T | A | NULL | 0.650218549 | NULL&M_Disease |
| 11 | 117150210 | rs608840 | A | G | NULL | 0.650218549 | NULL&M_Disease |
| 11 | 119434282 | rs2845659 | G | A | NULL | 0.650218549 | NULL&M_Disease |
| 11 | 119792173 | rs17123485 | C | T | NULL | 0.650218549 | NULL&M_Disease |
| 11 | 119841349 | rs78509462 | C | T | NULL | 0.650218549 | NULL&M_Disease |
| 11 | 119890372 | rs595969 | G | C | NULL | 0.650218549 | NULL&M_Disease |
| 11 | 120820615 | rs7105874 | T | C | NULL | 0.650218549 | NULL&M_Disease |
| 11 | 121078691 | rs74236048 | T | C | NULL | 0.650218549 | NULL&M_Disease |
| 11 | 116203460 | rs1440184 | A | C | NULL | 0.650218549 | NULL&M_Disease |
| 11 | 117116450 | rs12286525 | G | A | NULL | 0.650218549 | NULL&M_Disease |
| 11 | 119413149 | rs10892387 | G | A | NULL | 0.650218549 | NULL&M_Disease |
| 11 | 119924999 | rs11217662 | T | A | NULL | 0.650218549 | NULL&M_Disease |
| 11 | 119942557 | rs1319166 | C | G | NULL | 0.650218549 | NULL&M_Disease |
| 11 | 121089547 | rs75839534 | A | G | NULL | 0.650218549 | NULL&M_Disease |
| 11 | 121136144 | rs76755469 | A | G | NULL | 0.650218549 | NULL&M_Disease |
| 11 | 117334519 | rs4938387 | A | G | NULL | 0.650218549 | NULL&M_Disease |
| 11 | 117733223 | rs12277914 | T | C | NULL | 0.650218549 | NULL&M_Disease |
| 11 | 117775367 | rs67691204 | A | C | NULL | 0.650218549 | NULL&M_Disease |
| 11 | 117785742 | rs73022410 | C | G | NULL | 0.650218549 | NULL&M_Disease |
| 11 | 118724116 | rs12276533 | A | G | NULL | 0.650218549 | NULL&M_Disease |
| 11 | 120323909 | rs7110955 | G | A | NULL | 0.650218549 | NULL&M_Disease |
| 11 | 120558338 | rs12287882 | C | T | NULL | 0.650218549 | NULL&M_Disease |
| 11 | 121100855 | rs78017165 | A | G | NULL | 0.650218549 | NULL&M_Disease |
| 11 | 121101046 | rs79410922 | G | T | NULL | 0.650218549 | NULL&M_Disease |
| 11 | 116417181 | rs10891987 | C | G | NULL | 0.650218549 | NULL&M_Disease |
| 11 | 116593264 | rs180368 | T | C | NULL | 0.650218549 | NULL&M_Disease |
| 11 | 117784902 | rs75204992 | G | A | NULL | 0.650218549 | NULL&M_Disease |
| 11 | 119933897 | rs79255840 | T | A | NULL | 0.650218549 | NULL&M_Disease |
| 11 | 120472667 | rs4611216 | A | G | NULL | 0.650218549 | NULL&M_Disease |
| 11 | 120624486 | rs10892622 | G | T | NULL | 0.650218549 | NULL&M_Disease |
| 11 | 116324006 | rs78034012 | C | G | NULL | 0.650218549 | NULL&M_Disease |
| 11 | 116474899 | rs502607 | T | C | NULL | 0.650218549 | NULL&M_Disease |
| 11 | 116529914 | rs528806 | T | C | NULL | 0.650218549 | NULL&M_Disease |
| 11 | 117421267 | rs71480342 | T | C | NULL | 0.650218549 | NULL&M_Disease |
| 11 | 117544475 | rs686977 | C | T | NULL | 0.650218549 | NULL&M_Disease |
| 11 | 119692261 | rs61734721 | C | T | NULL | 0.650218549 | NULL&M_Disease |
| 11 | 121100518 | rs79212399 | T | G | NULL | 0.650218549 | NULL&M_Disease |
| 11 | 121106642 | rs75335512 | C | A | NULL | 0.650218549 | NULL&M_Disease |
| 11 | 116447795 | rs76908457 | C | T | NULL | 0.650218549 | NULL&M_Disease |
| 11 | 116547548 | rs1614444 | G | A | NULL | 0.651354865 | NULL&M_Disease |
| 11 | 116598118 | rs180362 | G | A | NULL | 0.651354865 | NULL&M_Disease |
| 11 | 116726048 | rs12099358 | A | C | NULL | 0.650218549 | NULL&M_Disease |
| 11 | 117927766 | rs10219396 | A | G | NULL | 0.651354865 | NULL&M_Disease |
| 11 | 118036532 | rs688911 | C | T | NULL | 0.651354865 | NULL&M_Disease |
| 11 | 119690746 | rs11822532 | A | G | NULL | 0.650218549 | NULL&M_Disease |
| 11 | 120169537 | rs113190660 | T | C | NULL | 0.650218549 | NULL&M_Disease |
| 11 | 121073721 | rs79660582 | G | T | NULL | 0.650218549 | NULL&M_Disease |
| 11 | 121075750 | rs79833470 | C | T | NULL | 0.650218549 | NULL&M_Disease |
| 11 | 121077362 | rs74236045 | T | C | NULL | 0.650218549 | NULL&M_Disease |
| 11 | 121096865 | rs11604100 | T | C | NULL | 0.650218549 | NULL&M_Disease |
| 11 | 116629153 | rs17519079 | A | G | NULL | 0.650218549 | NULL&M_Disease |
| 11 | 119962890 | rs7111010 | C | A | NULL | 0.651354865 | NULL&M_Disease |
| 11 | 121076820 | rs76719908 | C | G | NULL | 0.650218549 | NULL&M_Disease |
| 11 | 121077823 | rs74236047 | T | C | NULL | 0.650218549 | NULL&M_Disease |
| 11 | 121079460 | rs76105754 | A | G | NULL | 0.650218549 | NULL&M_Disease |
| 11 | 121082729 | rs10892688 | C | G | NULL | 0.651354865 | NULL&M_Disease |
| 11 | 119425807 | rs1509464 | C | T | NULL | 0.651354865 | NULL&M_Disease |
| 11 | 119972006 | rs61900461 | A | G | NULL | 0.651354865 | NULL&M_Disease |
| 11 | 120941583 | rs4643089 | G | A | NULL | 0.651354865 | NULL&M_Disease |
| 11 | 116439719 | rs7928435 | A | C | NULL | 0.651354865 | NULL&M_Disease |
| 11 | 116528876 | rs496958 | C | T | NULL | 0.651354865 | NULL&M_Disease |
| 11 | 116751549 | rs74341458 | A | G | NULL | 0.651354865 | NULL&M_Disease |
| 11 | 121074615 | rs11218200 | C | T | NULL | 0.651354865 | NULL&M_Disease |
| 11 | 117337012 | rs12420972 | A | G | NULL | 0.651354865 | NULL&M_Disease |
| 11 | 117337708 | rs73577982 | A | G | NULL | 0.651354865 | NULL&M_Disease |
| 11 | 117929001 | rs675539 | C | G | NULL | 0.651354865 | NULL&M_Disease |
| 11 | 118725383 | rs9735317 | T | C | NULL | 0.651354865 | NULL&M_Disease |
| 11 | 119384847 | rs733762 | C | G | NULL | 0.651354865 | NULL&M_Disease |
| 11 | 119421975 | rs2845649 | A | G | NULL | 0.651354865 | NULL&M_Disease |
| 11 | 120098329 | rs117554512 | T | C | NULL | 0.651354865 | NULL&M_Disease |
| 11 | 121087341 | rs11218202 | G | A | NULL | 0.651354865 | NULL&M_Disease |

| | | | | | | | | | |
|---|---|---|---|---|---|---|---|---|---|
| 11 | 119427624 | rs2663623 | A | G | 0.974 | -7977.388 | -7978.626 | -1.238 | 1.12E-06 |
| 11 | 119427893 | rs2845661 | T | G | 0.974 | -7977.388 | -7978.626 | -1.238 | 1.12E-06 |
| 11 | 120476746 | rs4936529 | T | A | 0.966 | -7977.388 | -7978.626 | -1.238 | 1.12E-06 |
| 11 | 116407867 | rs11215991 | T | C | 0.969 | -7977.388 | -7978.628 | -1.239 | 1.12E-06 |
| 11 | 117800033 | rs35251396 | A | G | 0.993 | -7977.388 | -7978.627 | -1.239 | 1.12E-06 |
| 11 | 119903718 | rs61897924 | T | C | 1.035 | -7977.388 | -7978.628 | -1.239 | 1.12E-06 |
| 11 | 116448951 | rs77196345 | T | C | 0.974 | -7977.388 | -7978.629 | -1.24 | 1.12E-06 |
| 11 | 116599281 | rs180359 | G | A | 1.036 | -7977.388 | -7978.629 | -1.24 | 1.12E-06 |
| 11 | 116599392 | rs180358 | T | C | 1.036 | -7977.388 | -7978.629 | -1.24 | 1.12E-06 |
| 11 | 116599534 | rs180356 | T | C | 1.036 | -7977.388 | -7978.629 | -1.24 | 1.12E-06 |
| 11 | 117864047 | rs2256111 | A | G | 0.967 | -7977.388 | -7978.628 | -1.24 | 1.12E-06 |
| 11 | 119155618 | rs2298650 | T | G | 0.966 | -7977.388 | -7978.628 | -1.24 | 1.12E-06 |
| 11 | 120558364 | rs12281001 | T | G | 0.968 | -7977.388 | -7978.629 | -1.24 | 1.12E-06 |
| 11 | 120558487 | rs12294046 | T | A | 0.968 | -7977.388 | -7978.628 | -1.24 | 1.12E-06 |
| 11 | 120559070 | rs10892616 | G | A | 0.968 | -7977.388 | -7978.628 | -1.24 | 1.12E-06 |
| 11 | 120599666 | rs112473235 | T | C | 1.012 | -7977.388 | -7978.628 | -1.24 | 1.12E-06 |
| 11 | 117082966 | rs76169968 | A | G | 1.026 | -7977.388 | -7978.629 | -1.241 | 1.12E-06 |
| 11 | 117213847 | rs72645461 | G | A | 0.966 | -7977.388 | -7978.63 | -1.241 | 1.12E-06 |
| 11 | 117333657 | rs11607656 | A | G | 0.971 | -7977.388 | -7978.629 | -1.241 | 1.12E-06 |
| 11 | 120396030 | rs71484178 | A | G | 0.974 | -7977.388 | -7978.629 | -1.241 | 1.12E-06 |
| 11 | 116355860 | rs573554 | C | T | 1.035 | -7977.388 | -7978.63 | -1.242 | 1.12E-06 |
| 11 | 117421276 | rs71480343 | A | G | 1.022 | -7977.388 | -7978.63 | -1.242 | 1.12E-06 |
| 11 | 120178632 | rs11605781 | C | T | 1.035 | -7977.388 | -7978.63 | -1.242 | 1.12E-06 |
| 11 | 121018725 | rs551031 | C | T | 1.035 | -7977.388 | -7978.63 | -1.242 | 1.12E-06 |
| 11 | 116263533 | rs480793 | A | G | 0.97 | -7977.388 | -7978.631 | -1.243 | 1.12E-06 |
| 11 | 116392191 | rs10891985 | T | A | 0.966 | -7977.388 | -7978.631 | -1.243 | 1.12E-06 |
| 11 | 117150210 | rs608840 | A | G | 0.967 | -7977.388 | -7978.631 | -1.243 | 1.12E-06 |
| 11 | 119434282 | rs2845659 | G | A | 0.974 | -7977.388 | -7978.631 | -1.243 | 1.12E-06 |
| 11 | 119792173 | rs17123485 | C | T | 1.008 | -7977.388 | -7978.631 | -1.243 | 1.12E-06 |
| 11 | 119841349 | rs78509462 | C | T | 0.992 | -7977.388 | -7978.632 | -1.243 | 1.12E-06 |
| 11 | 119890372 | rs595969 | G | C | 0.997 | -7977.388 | -7978.631 | -1.243 | 1.12E-06 |
| 11 | 120820615 | rs7105874 | T | C | 0.966 | -7977.388 | -7978.632 | -1.243 | 1.12E-06 |
| 11 | 121078691 | rs74236048 | T | C | 0.991 | -7977.388 | -7978.631 | -1.243 | 1.12E-06 |
| 11 | 116203460 | rs1440184 | A | C | 1.035 | -7977.388 | -7978.632 | -1.244 | 1.11E-06 |
| 11 | 117116450 | rs12286525 | G | A | 0.966 | -7977.388 | -7978.633 | -1.244 | 1.11E-06 |
| 11 | 119413149 | rs10892387 | G | A | 0.974 | -7977.388 | -7978.633 | -1.244 | 1.11E-06 |
| 11 | 119924999 | rs11217662 | T | A | 1.035 | -7977.388 | -7978.633 | -1.244 | 1.11E-06 |
| 11 | 119942557 | rs1319166 | C | G | 1.035 | -7977.388 | -7978.632 | -1.244 | 1.11E-06 |
| 11 | 121089547 | rs75839534 | A | G | 0.993 | -7977.388 | -7978.633 | -1.244 | 1.11E-06 |
| 11 | 121136144 | rs76755469 | A | G | 0.978 | -7977.388 | -7978.632 | -1.244 | 1.11E-06 |
| 11 | 117334519 | rs4938387 | A | G | 0.971 | -7977.388 | -7978.634 | -1.245 | 1.11E-06 |
| 11 | 117733223 | rs12277914 | T | C | 0.966 | -7977.388 | -7978.633 | -1.245 | 1.11E-06 |
| 11 | 117775367 | rs67691204 | A | C | 0.968 | -7977.388 | -7978.633 | -1.245 | 1.11E-06 |
| 11 | 117785742 | rs73022410 | C | G | 1.009 | -7977.388 | -7978.633 | -1.245 | 1.11E-06 |
| 11 | 118724116 | rs12276533 | A | G | 1.035 | -7977.388 | -7978.634 | -1.245 | 1.11E-06 |
| 11 | 120323909 | rs7110955 | G | A | 1.035 | -7977.388 | -7978.634 | -1.245 | 1.11E-06 |
| 11 | 120558338 | rs12287882 | C | T | 0.968 | -7977.388 | -7978.633 | -1.245 | 1.11E-06 |
| 11 | 121100855 | rs78017165 | A | G | 1.002 | -7977.388 | -7978.634 | -1.245 | 1.11E-06 |
| 11 | 121101046 | rs79410922 | G | T | 1.002 | -7977.388 | -7978.634 | -1.245 | 1.11E-06 |
| 11 | 116417181 | rs10891987 | C | G | 1.035 | -7977.388 | -7978.634 | -1.246 | 1.11E-06 |
| 11 | 116593264 | rs180368 | T | C | 1.035 | -7977.388 | -7978.634 | -1.246 | 1.11E-06 |
| 11 | 117784902 | rs75204992 | G | A | 1.009 | -7977.388 | -7978.635 | -1.246 | 1.11E-06 |
| 11 | 119933897 | rs79255840 | T | A | 0.972 | -7977.388 | -7978.634 | -1.246 | 1.11E-06 |
| 11 | 120472667 | rs4611216 | A | G | 0.966 | -7977.388 | -7978.635 | -1.246 | 1.11E-06 |
| 11 | 120624486 | rs10892622 | G | T | 1.035 | -7977.388 | -7978.634 | -1.246 | 1.11E-06 |
| 11 | 116324006 | rs78034012 | C | G | 1.035 | -7977.388 | -7978.635 | -1.247 | 1.11E-06 |
| 11 | 116474899 | rs502607 | T | C | 1.017 | -7977.388 | -7978.635 | -1.247 | 1.11E-06 |
| 11 | 116529914 | rs528806 | T | C | 1.035 | -7977.388 | -7978.635 | -1.247 | 1.11E-06 |
| 11 | 117421267 | rs71480342 | T | C | 1.022 | -7977.388 | -7978.635 | -1.247 | 1.11E-06 |
| 11 | 117544475 | rs686977 | C | T | 1.012 | -7977.388 | -7978.636 | -1.247 | 1.11E-06 |
| 11 | 119692261 | rs61734721 | C | T | 1.027 | -7977.388 | -7978.635 | -1.247 | 1.11E-06 |
| 11 | 121100518 | rs79212399 | T | G | 0.999 | -7977.388 | -7978.635 | -1.247 | 1.11E-06 |
| 11 | 121106642 | rs75335512 | C | A | 0.999 | -7977.388 | -7978.635 | -1.247 | 1.11E-06 |
| 11 | 116447795 | rs76908457 | C | T | 0.974 | -7977.388 | -7978.636 | -1.248 | 1.11E-06 |
| 11 | 116547548 | rs1614444 | G | A | 1.035 | -7977.388 | -7978.636 | -1.248 | 1.11E-06 |
| 11 | 116598118 | rs180362 | G | A | 1.035 | -7977.388 | -7978.637 | -1.248 | 1.11E-06 |
| 11 | 116726048 | rs12099358 | A | C | 0.966 | -7977.388 | -7978.636 | -1.248 | 1.11E-06 |
| 11 | 117927766 | rs10219396 | A | G | 1.035 | -7977.388 | -7978.636 | -1.248 | 1.11E-06 |
| 11 | 118036532 | rs688911 | C | T | 0.967 | -7977.388 | -7978.637 | -1.248 | 1.11E-06 |
| 11 | 119690746 | rs11822532 | A | G | 0.98 | -7977.388 | -7978.636 | -1.248 | 1.11E-06 |
| 11 | 120169537 | rs113190660 | T | C | 1.015 | -7977.388 | -7978.636 | -1.248 | 1.11E-06 |
| 11 | 121073721 | rs79660582 | G | T | 0.994 | -7977.388 | -7978.636 | -1.248 | 1.11E-06 |
| 11 | 121075750 | rs79833470 | C | T | 0.994 | -7977.388 | -7978.636 | -1.248 | 1.11E-06 |
| 11 | 121077362 | rs74236045 | T | C | 0.994 | -7977.388 | -7978.636 | -1.248 | 1.11E-06 |
| 11 | 121096865 | rs11604100 | T | C | 1.003 | -7977.388 | -7978.636 | -1.248 | 1.11E-06 |
| 11 | 116629153 | rs17519079 | A | G | 0.982 | -7977.388 | -7978.637 | -1.249 | 1.11E-06 |
| 11 | 119962890 | rs7111010 | C | A | 1.032 | -7977.388 | -7978.637 | -1.249 | 1.11E-06 |
| 11 | 121076820 | rs76719908 | C | G | 0.995 | -7977.388 | -7978.637 | -1.249 | 1.11E-06 |
| 11 | 121077823 | rs74236047 | T | C | 0.994 | -7977.388 | -7978.637 | -1.249 | 1.11E-06 |
| 11 | 121079460 | rs76105754 | A | G | 0.995 | -7977.388 | -7978.637 | -1.249 | 1.11E-06 |
| 11 | 121082729 | rs10892688 | C | G | 1.035 | -7977.388 | -7978.637 | -1.249 | 1.11E-06 |
| 11 | 119425807 | rs1509464 | C | T | 0.975 | -7977.388 | -7978.638 | -1.25 | 1.11E-06 |
| 11 | 119972006 | rs61900461 | A | G | 0.977 | -7977.388 | -7978.638 | -1.25 | 1.11E-06 |
| 11 | 120941583 | rs4643089 | G | A | 1.035 | -7977.388 | -7978.638 | -1.25 | 1.11E-06 |
| 11 | 116439719 | rs7928435 | A | C | 0.975 | -7977.388 | -7978.639 | -1.251 | 1.11E-06 |
| 11 | 116528876 | rs496958 | C | T | 1.035 | -7977.388 | -7978.64 | -1.251 | 1.11E-06 |
| 11 | 116751549 | rs74341458 | A | G | 0.969 | -7977.388 | -7978.639 | -1.251 | 1.11E-06 |
| 11 | 121074615 | rs11218200 | C | T | 0.994 | -7977.388 | -7978.64 | -1.251 | 1.11E-06 |
| 11 | 117337012 | rs12420972 | A | G | 0.972 | -7977.388 | -7978.64 | -1.252 | 1.11E-06 |
| 11 | 117337708 | rs73577982 | A | G | 0.972 | -7977.388 | -7978.64 | -1.252 | 1.11E-06 |
| 11 | 117929001 | rs675539 | C | G | 1.033 | -7977.388 | -7978.64 | -1.252 | 1.11E-06 |
| 11 | 118725383 | rs9735317 | T | C | 1.035 | -7977.388 | -7978.641 | -1.252 | 1.11E-06 |
| 11 | 119384847 | rs733762 | C | G | 1.035 | -7977.388 | -7978.64 | -1.252 | 1.11E-06 |
| 11 | 119421975 | rs2845649 | A | G | 0.975 | -7977.388 | -7978.64 | -1.252 | 1.11E-06 |
| 11 | 120098329 | rs117554512 | T | C | 0.974 | -7977.388 | -7978.641 | -1.252 | 1.11E-06 |
| 11 | 121087341 | rs11218202 | G | A | 0.996 | -7977.388 | -7978.64 | -1.252 | 1.11E-06 |

| | | | | | | | |
|---|---|---|---|---|---|---|---|
| 11 | 117332917 | rs4938383 | C | G | NULL | 0.651354865 | NULL&M_Disease |
| 11 | 117924278 | rs7938067 | C | T | NULL | 0.651354865 | NULL&M_Disease |
| 11 | 119438762 | rs906822 | T | G | NULL | 0.651354865 | NULL&M_Disease |
| 11 | 119844041 | rs10790361 | T | C | NULL | 0.651354865 | NULL&M_Disease |
| 11 | 119956964 | rs582340 | A | G | NULL | 0.651354865 | NULL&M_Disease |
| 11 | 119965635 | rs113684204 | A | T | NULL | 0.651354865 | NULL&M_Disease |
| 11 | 117551691 | rs77267645 | G | A | NULL | 0.651354865 | NULL&M_Disease |
| 11 | 117784946 | rs75896504 | A | G | NULL | 0.651354865 | NULL&M_Disease |
| 11 | 117931353 | rs670225 | A | G | NULL | 0.651354865 | NULL&M_Disease |
| 11 | 117931576 | rs669316 | T | C | NULL | 0.651354865 | NULL&M_Disease |
| 11 | 119813518 | rs12574466 | A | G | NULL | 0.651354865 | NULL&M_Disease |
| 11 | 120138723 | rs10892557 | T | C | NULL | 0.651354865 | NULL&M_Disease |
| 11 | 121072114 | rs12292816 | T | G | NULL | 0.651354865 | NULL&M_Disease |
| 11 | 121084717 | rs80318512 | A | G | NULL | 0.651354865 | NULL&M_Disease |
| 11 | 121085338 | rs12289135 | C | T | NULL | 0.651354865 | NULL&M_Disease |
| 11 | 121086444 | rs9804547 | A | T | NULL | 0.651354865 | NULL&M_Disease |
| 11 | 116440214 | rs11606979 | A | G | NULL | 0.651354865 | NULL&M_Disease |
| 11 | 117335165 | rs11819822 | T | C | NULL | 0.651354865 | NULL&M_Disease |
| 11 | 117559725 | rs79735246 | G | A | NULL | 0.651354865 | NULL&M_Disease |
| 11 | 119087122 | rs4936468 | A | G | NULL | 0.651354865 | NULL&M_Disease |
| 11 | 119691772 | rs11217523 | G | A | NULL | 0.651354865 | NULL&M_Disease |
| 11 | 120075939 | rs896695 | T | C | NULL | 0.651354865 | NULL&M_Disease |
| 11 | 120076931 | rs492275 | T | C | NULL | 0.651354865 | NULL&M_Disease |
| 11 | 120279620 | rs11217860 | T | G | NULL | 0.651354865 | NULL&M_Disease |
| 11 | 120435024 | rs7103821 | C | A | NULL | 0.651354865 | NULL&M_Disease |
| 11 | 120559953 | rs12283240 | A | G | NULL | 0.651354865 | NULL&M_Disease |
| 11 | 120813148 | rs10892649 | T | C | NULL | 0.651354865 | NULL&M_Disease |
| 11 | 120893054 | rs11218114 | A | G | NULL | 0.651354865 | NULL&M_Disease |
| 11 | 116529641 | rs524467 | G | T | NULL | 0.651354865 | NULL&M_Disease |
| 11 | 116794879 | rs117287138 | A | G | NULL | 0.651354865 | NULL&M_Disease |
| 11 | 117552896 | rs58329329 | G | C | NULL | 0.651354865 | NULL&M_Disease |
| 11 | 117928962 | rs1783846 | A | C | NULL | 0.651354865 | NULL&M_Disease |
| 11 | 120395800 | rs61900876 | C | T | NULL | 0.651354865 | NULL&M_Disease |
| 11 | 121086259 | rs117018659 | A | G | NULL | 0.651354865 | NULL&M_Disease |
| 11 | 117500323 | rs78470936 | G | T | NULL | 0.651354865 | NULL&M_Disease |
| 11 | 117554355 | rs79807999 | G | T | NULL | 0.651354865 | NULL&M_Disease |
| 11 | 117555727 | rs12361479 | T | C | NULL | 0.651354865 | NULL&M_Disease |
| 11 | 117556948 | rs74430235 | A | G | NULL | 0.651354865 | NULL&M_Disease |
| 11 | 117557022 | rs79078907 | T | C | NULL | 0.651354865 | NULL&M_Disease |
| 11 | 119435180 | rs2663629 | A | C | NULL | 0.651354865 | NULL&M_Disease |
| 11 | 121194035 | rs61901460 | G | C | NULL | 0.651354865 | NULL&M_Disease |
| 11 | 116214361 | rs11601365 | G | C | NULL | 0.651354865 | NULL&M_Disease |
| 11 | 116480813 | rs76235142 | A | G | NULL | 0.652489462 | NULL&M_Disease |
| 11 | 117494246 | rs7121930 | T | G | NULL | 0.652489462 | NULL&M_Disease |
| 11 | 117666712 | rs565312 | A | G | NULL | 0.651354865 | NULL&M_Disease |
| 11 | 119691736 | rs58164695 | T | C | NULL | 0.651354865 | NULL&M_Disease |
| 11 | 119760977 | rs4938736 | A | G | NULL | 0.652489462 | NULL&M_Disease |
| 11 | 116265581 | rs79679004 | C | T | NULL | 0.652489462 | NULL&M_Disease |
| 11 | 119448679 | rs10750159 | G | A | NULL | 0.651354865 | NULL&M_Disease |
| 11 | 119853613 | rs75964569 | C | T | NULL | 0.651354865 | NULL&M_Disease |
| 11 | 120077259 | rs489454 | A | G | NULL | 0.651354865 | NULL&M_Disease |
| 11 | 120078136 | rs536743 | A | C | NULL | 0.651354865 | NULL&M_Disease |
| 11 | 120396041 | rs61900877 | C | G | NULL | 0.652489462 | NULL&M_Disease |
| 11 | 120569528 | rs17245664 | A | G | NULL | 0.652489462 | NULL&M_Disease |
| 11 | 116610249 | rs10466588 | G | A | NULL | 0.652489462 | NULL&M_Disease |
| 11 | 117039797 | rs1060211 | G | A | NULL | 0.652489462 | NULL&M_Disease |
| 11 | 117331806 | rs61407825 | T | C | NULL | 0.652489462 | NULL&M_Disease |
| 11 | 117337965 | rs35585361 | A | G | NULL | 0.652489462 | NULL&M_Disease |
| 11 | 117461576 | rs647593 | C | T | NULL | 0.652489462 | NULL&M_Disease |
| 11 | 119436769 | rs2663632 | A | G | NULL | 0.652489462 | NULL&M_Disease |
| 11 | 119437201 | rs2845655 | T | C | NULL | 0.652489462 | NULL&M_Disease |
| 11 | 119692025 | rs118188470 | C | G | NULL | 0.652489462 | NULL&M_Disease |
| 11 | 120278166 | rs11217859 | C | G | NULL | 0.652489462 | NULL&M_Disease |
| 11 | 120357547 | rs11217890 | A | G | NULL | 0.652489462 | NULL&M_Disease |
| 11 | 121090597 | rs591221 | G | A | NULL | 0.652489462 | NULL&M_Disease |
| 11 | 116452534 | rs4938288 | G | C | NULL | 0.652489462 | NULL&M_Disease |
| 11 | 117331503 | rs584057 | C | T | NULL | 0.652489462 | NULL&M_Disease |
| 11 | 117726697 | rs555577 | A | G | NULL | 0.652489462 | NULL&M_Disease |
| 11 | 117960590 | rs7924359 | G | C | NULL | 0.652489462 | NULL&M_Disease |
| 11 | 119669797 | rs1374565 | G | A | NULL | 0.652489462 | NULL&M_Disease |
| 11 | 120344551 | rs11217881 | G | A | NULL | 0.652489462 | NULL&M_Disease |
| 11 | 120345002 | rs11217882 | G | T | NULL | 0.652489462 | NULL&M_Disease |
| 11 | 120345004 | rs11217883 | G | A | NULL | 0.652489462 | NULL&M_Disease |
| 11 | 120356317 | rs11217889 | G | C | NULL | 0.652489462 | NULL&M_Disease |
| 11 | 120393284 | rs61900867 | T | C | NULL | 0.652489462 | NULL&M_Disease |
| 11 | 121079190 | rs12295930 | T | C | NULL | 0.652489462 | NULL&M_Disease |
| 11 | 117775043 | rs73020491 | A | G | NULL | 0.652489462 | NULL&M_Disease |
| 11 | 120079884 | rs477863 | G | T | NULL | 0.652489462 | NULL&M_Disease |
| 11 | 120080022 | rs476809 | G | T | NULL | 0.652489462 | NULL&M_Disease |
| 11 | 120080526 | rs521841 | C | A | NULL | 0.652489462 | NULL&M_Disease |
| 11 | 116393611 | rs12799630 | T | C | NULL | 0.652489462 | NULL&M_Disease |
| 11 | 119825176 | rs7930793 | G | T | NULL | 0.652489462 | NULL&M_Disease |
| 11 | 119885995 | rs12787692 | A | G | NULL | 0.652489462 | NULL&M_Disease |
| 11 | 119965045 | rs61897958 | G | T | NULL | 0.652489462 | NULL&M_Disease |
| 11 | 120395521 | rs61900874 | T | C | NULL | 0.652489462 | NULL&M_Disease |
| 11 | 120395705 | rs61900875 | G | A | NULL | 0.652489462 | NULL&M_Disease |
| 11 | 116238557 | rs58283940 | A | G | NULL | 0.652489462 | NULL&M_Disease |
| 11 | 116326570 | rs7927239 | C | G | NULL | 0.652489462 | NULL&M_Disease |
| 11 | 116566161 | rs56311339 | T | C | NULL | 0.652489462 | NULL&M_Disease |
| 11 | 117133078 | rs676305 | T | C | NULL | 0.652489462 | NULL&M_Disease |
| 11 | 117333229 | rs59533604 | C | A | NULL | 0.652489462 | NULL&M_Disease |
| 11 | 119692457 | rs12280992 | T | C | NULL | 0.652489462 | NULL&M_Disease |
| 11 | 119700031 | rs1007369 | C | A | NULL | 0.652489462 | NULL&M_Disease |
| 11 | 120117951 | rs12276544 | A | T | NULL | 0.652489462 | NULL&M_Disease |
| 11 | 116572089 | rs1892948 | T | C | NULL | 0.652489462 | NULL&M_Disease |
| 11 | 116612128 | rs10466533 | G | A | NULL | 0.652489462 | NULL&M_Disease |

| | | | | | | | | | |
|---|---|---|---|---|---|---|---|---|---|
| 11 | 117332917 | rs4938383 | C | G | 0.972 | -7977.388 | -7978.641 | -1.253 | 1.10E-06 |
| 11 | 117924278 | rs7938067 | C | T | 1.035 | -7977.388 | -7978.642 | -1.253 | 1.10E-06 |
| 11 | 119438762 | rs906822 | T | G | 0.976 | -7977.388 | -7978.641 | -1.253 | 1.10E-06 |
| 11 | 119844041 | rs10790361 | T | C | 0.966 | -7977.388 | -7978.642 | -1.253 | 1.10E-06 |
| 11 | 119956964 | rs582340 | A | G | 0.972 | -7977.388 | -7978.641 | -1.253 | 1.10E-06 |
| 11 | 119965635 | rs113684204 | A | T | 1.031 | -7977.388 | -7978.642 | -1.253 | 1.10E-06 |
| 11 | 117551691 | rs77267645 | G | A | 0.995 | -7977.388 | -7978.642 | -1.254 | 1.10E-06 |
| 11 | 117784946 | rs75896504 | A | G | 1.005 | -7977.388 | -7978.643 | -1.254 | 1.10E-06 |
| 11 | 117931353 | rs670225 | A | G | 1.033 | -7977.388 | -7978.642 | -1.254 | 1.10E-06 |
| 11 | 117931576 | rs669316 | T | C | 1.033 | -7977.388 | -7978.642 | -1.254 | 1.10E-06 |
| 11 | 119813518 | rs12574466 | A | G | 1.028 | -7977.388 | -7978.642 | -1.254 | 1.10E-06 |
| 11 | 120138723 | rs10892557 | T | C | 1.033 | -7977.388 | -7978.643 | -1.254 | 1.10E-06 |
| 11 | 121072114 | rs12292816 | T | G | 0.997 | -7977.388 | -7978.642 | -1.254 | 1.10E-06 |
| 11 | 121084717 | rs80318512 | A | G | 0.998 | -7977.388 | -7978.643 | -1.254 | 1.10E-06 |
| 11 | 121085338 | rs12289135 | C | T | 0.998 | -7977.388 | -7978.643 | -1.254 | 1.10E-06 |
| 11 | 121086444 | rs9804547 | A | T | 1.002 | -7977.388 | -7978.643 | -1.254 | 1.10E-06 |
| 11 | 116440214 | rs11606979 | A | G | 0.975 | -7977.388 | -7978.643 | -1.255 | 1.10E-06 |
| 11 | 117335165 | rs11819822 | T | C | 1.009 | -7977.388 | -7978.644 | -1.255 | 1.10E-06 |
| 11 | 117559725 | rs79735246 | G | A | 1 | -7977.388 | -7978.643 | -1.255 | 1.10E-06 |
| 11 | 119087122 | rs4936468 | A | G | 1.034 | -7977.388 | -7978.643 | -1.255 | 1.10E-06 |
| 11 | 119691772 | rs11217523 | G | A | 1.027 | -7977.388 | -7978.643 | -1.255 | 1.10E-06 |
| 11 | 120075939 | rs896695 | T | C | 0.975 | -7977.388 | -7978.643 | -1.255 | 1.10E-06 |
| 11 | 120076931 | rs492275 | T | C | 0.975 | -7977.388 | -7978.644 | -1.255 | 1.10E-06 |
| 11 | 120279620 | rs11217860 | T | G | 1.035 | -7977.388 | -7978.643 | -1.255 | 1.10E-06 |
| 11 | 120435024 | rs7103821 | C | A | 1.034 | -7977.388 | -7978.644 | -1.255 | 1.10E-06 |
| 11 | 120559953 | rs12283240 | A | G | 0.969 | -7977.388 | -7978.643 | -1.255 | 1.10E-06 |
| 11 | 120813148 | rs10892649 | T | C | 0.966 | -7977.388 | -7978.643 | -1.255 | 1.10E-06 |
| 11 | 120893054 | rs11218114 | A | G | 1.028 | -7977.388 | -7978.643 | -1.255 | 1.10E-06 |
| 11 | 116529641 | rs524467 | G | T | 1.035 | -7977.388 | -7978.644 | -1.256 | 1.10E-06 |
| 11 | 116794879 | rs117287138 | A | G | 0.97 | -7977.388 | -7978.645 | -1.256 | 1.10E-06 |
| 11 | 117552896 | rs58329329 | G | C | 1.007 | -7977.388 | -7978.644 | -1.256 | 1.10E-06 |
| 11 | 117928962 | rs1783846 | A | C | 1.033 | -7977.388 | -7978.645 | -1.256 | 1.10E-06 |
| 11 | 120395800 | rs61900876 | C | T | 0.975 | -7977.388 | -7978.644 | -1.256 | 1.10E-06 |
| 11 | 121086259 | rs117018659 | A | G | 0.999 | -7977.388 | -7978.644 | -1.256 | 1.10E-06 |
| 11 | 117500323 | rs78470936 | G | T | 0.977 | -7977.388 | -7978.645 | -1.257 | 1.10E-06 |
| 11 | 117554355 | rs79807999 | G | T | 0.998 | -7977.388 | -7978.645 | -1.257 | 1.10E-06 |
| 11 | 117555727 | rs12361479 | T | C | 0.998 | -7977.388 | -7978.645 | -1.257 | 1.10E-06 |
| 11 | 117556948 | rs74430235 | A | G | 0.998 | -7977.388 | -7978.646 | -1.257 | 1.10E-06 |
| 11 | 117557022 | rs79078907 | T | C | 0.998 | -7977.388 | -7978.646 | -1.257 | 1.10E-06 |
| 11 | 119435180 | rs2663629 | A | C | 0.976 | -7977.388 | -7978.645 | -1.257 | 1.10E-06 |
| 11 | 121194035 | rs61901460 | G | C | 0.998 | -7977.388 | -7978.646 | -1.257 | 1.10E-06 |
| 11 | 116214361 | rs11601365 | G | C | 0.968 | -7977.388 | -7978.646 | -1.258 | 1.10E-06 |
| 11 | 116480813 | rs76235142 | A | G | 1.035 | -7977.388 | -7978.646 | -1.258 | 1.10E-06 |
| 11 | 117494246 | rs7121930 | T | G | 1.035 | -7977.388 | -7978.646 | -1.258 | 1.10E-06 |
| 11 | 117666712 | rs565312 | A | G | 1.033 | -7977.388 | -7978.646 | -1.258 | 1.10E-06 |
| 11 | 119691736 | rs58164695 | T | C | 0.981 | -7977.388 | -7978.646 | -1.258 | 1.10E-06 |
| 11 | 119760977 | rs4938736 | A | G | 1.034 | -7977.388 | -7978.647 | -1.258 | 1.10E-06 |
| 11 | 116265581 | rs79679004 | C | T | 1.033 | -7977.388 | -7978.647 | -1.259 | 1.10E-06 |
| 11 | 119448679 | rs10750159 | G | A | 0.975 | -7977.388 | -7978.647 | -1.259 | 1.10E-06 |
| 11 | 119853613 | rs75964569 | C | T | 1.02 | -7977.388 | -7978.647 | -1.259 | 1.10E-06 |
| 11 | 120077259 | rs489454 | A | G | 0.976 | -7977.388 | -7978.647 | -1.259 | 1.10E-06 |
| 11 | 120078136 | rs536743 | A | C | 0.976 | -7977.388 | -7978.647 | -1.259 | 1.10E-06 |
| 11 | 120396041 | rs61900877 | C | G | 0.975 | -7977.388 | -7978.648 | -1.259 | 1.10E-06 |
| 11 | 120569528 | rs17245664 | A | G | 1.035 | -7977.388 | -7978.647 | -1.259 | 1.10E-06 |
| 11 | 116610249 | rs10466588 | G | A | 0.978 | -7977.388 | -7978.649 | -1.26 | 1.10E-06 |
| 11 | 117039797 | rs1060211 | G | A | 0.966 | -7977.388 | -7978.649 | -1.26 | 1.10E-06 |
| 11 | 117331806 | rs61407825 | T | C | 1.008 | -7977.388 | -7978.648 | -1.26 | 1.10E-06 |
| 11 | 117337965 | rs35585361 | A | G | 0.972 | -7977.388 | -7978.649 | -1.26 | 1.10E-06 |
| 11 | 117461576 | rs647593 | C | T | 1.035 | -7977.388 | -7978.648 | -1.26 | 1.10E-06 |
| 11 | 119436769 | rs2663632 | A | G | 0.976 | -7977.388 | -7978.648 | -1.26 | 1.10E-06 |
| 11 | 119437201 | rs2845655 | T | C | 0.976 | -7977.388 | -7978.648 | -1.26 | 1.10E-06 |
| 11 | 119692025 | rs118188470 | C | G | 0.973 | -7977.388 | -7978.648 | -1.26 | 1.10E-06 |
| 11 | 120278166 | rs11217859 | C | G | 1.035 | -7977.388 | -7978.648 | -1.26 | 1.10E-06 |
| 11 | 120357547 | rs11217890 | A | G | 1.035 | -7977.388 | -7978.649 | -1.26 | 1.10E-06 |
| 11 | 121090597 | rs591221 | G | A | 0.966 | -7977.388 | -7978.649 | -1.26 | 1.10E-06 |
| 11 | 116452534 | rs4938288 | G | C | 0.967 | -7977.388 | -7978.649 | -1.261 | 1.10E-06 |
| 11 | 117331503 | rs584057 | C | T | 0.967 | -7977.388 | -7978.649 | -1.261 | 1.10E-06 |
| 11 | 117726697 | rs555577 | A | G | 0.967 | -7977.388 | -7978.649 | -1.261 | 1.10E-06 |
| 11 | 117960590 | rs7924359 | G | C | 1.035 | -7977.388 | -7978.65 | -1.261 | 1.10E-06 |
| 11 | 119669797 | rs1374565 | G | A | 0.966 | -7977.388 | -7978.649 | -1.261 | 1.10E-06 |
| 11 | 120344551 | rs11217881 | G | A | 1.035 | -7977.388 | -7978.649 | -1.261 | 1.10E-06 |
| 11 | 120345002 | rs11217882 | G | T | 1.035 | -7977.388 | -7978.649 | -1.261 | 1.10E-06 |
| 11 | 120345004 | rs11217883 | G | A | 1.035 | -7977.388 | -7978.649 | -1.261 | 1.10E-06 |
| 11 | 120356317 | rs11217889 | G | C | 1.035 | -7977.388 | -7978.649 | -1.261 | 1.10E-06 |
| 11 | 120393284 | rs61900867 | T | C | 0.976 | -7977.388 | -7978.65 | -1.261 | 1.10E-06 |
| 11 | 121079190 | rs12295930 | T | C | 1.034 | -7977.388 | -7978.65 | -1.261 | 1.10E-06 |
| 11 | 117775043 | rs73020491 | A | G | 0.969 | -7977.388 | -7978.65 | -1.262 | 1.10E-06 |
| 11 | 120079884 | rs477863 | G | T | 0.976 | -7977.388 | -7978.65 | -1.262 | 1.10E-06 |
| 11 | 120080022 | rs476809 | G | T | 0.976 | -7977.388 | -7978.65 | -1.262 | 1.10E-06 |
| 11 | 120080526 | rs521841 | C | A | 0.976 | -7977.388 | -7978.65 | -1.262 | 1.10E-06 |
| 11 | 116393611 | rs12799630 | T | C | 0.97 | -7977.388 | -7978.651 | -1.263 | 1.09E-06 |
| 11 | 119825176 | rs7930793 | G | T | 1.034 | -7977.388 | -7978.651 | -1.263 | 1.09E-06 |
| 11 | 119885995 | rs12787692 | A | G | 0.994 | -7977.388 | -7978.652 | -1.263 | 1.09E-06 |
| 11 | 119965045 | rs61897958 | G | T | 1.031 | -7977.388 | -7978.651 | -1.263 | 1.09E-06 |
| 11 | 120395521 | rs61900874 | T | C | 0.976 | -7977.388 | -7978.651 | -1.263 | 1.09E-06 |
| 11 | 120395705 | rs61900875 | G | A | 0.976 | -7977.388 | -7978.651 | -1.263 | 1.09E-06 |
| 11 | 116238557 | rs58283940 | A | G | 0.978 | -7977.388 | -7978.653 | -1.264 | 1.09E-06 |
| 11 | 116326570 | rs7927239 | C | G | 1.034 | -7977.388 | -7978.652 | -1.264 | 1.09E-06 |
| 11 | 116566161 | rs56311339 | T | C | 0.969 | -7977.388 | -7978.653 | -1.264 | 1.09E-06 |
| 11 | 117133078 | rs676305 | T | C | 0.969 | -7977.388 | -7978.653 | -1.264 | 1.09E-06 |
| 11 | 117333229 | rs59533604 | C | A | 1.006 | -7977.388 | -7978.652 | -1.264 | 1.09E-06 |
| 11 | 119692457 | rs12280992 | T | C | 1.026 | -7977.388 | -7978.652 | -1.264 | 1.09E-06 |
| 11 | 119700031 | rs1007369 | C | A | 0.966 | -7977.388 | -7978.652 | -1.264 | 1.09E-06 |
| 11 | 120117951 | rs12276544 | A | T | 1.034 | -7977.388 | -7978.652 | -1.264 | 1.09E-06 |
| 11 | 116572089 | rs1892948 | T | C | 0.992 | -7977.388 | -7978.654 | -1.265 | 1.09E-06 |
| 11 | 116612128 | rs10466533 | G | A | 0.979 | -7977.388 | -7978.654 | -1.265 | 1.09E-06 |

| | | | | | | | |
|---|---|---|---|---|---|---|---|
| 11 | 117144816 | rs668363 | C | A | NULL | 0.652489462 | NULL&M_Disease |
| 11 | 117334349 | rs7940175 | T | C | NULL | 0.652489462 | NULL&M_Disease |
| 11 | 117945161 | rs3758991 | T | C | NULL | 0.652489462 | NULL&M_Disease |
| 11 | 118723753 | rs7119568 | A | T | NULL | 0.652489462 | NULL&M_Disease |
| 11 | 120395934 | rs78043040 | C | T | NULL | 0.652489462 | NULL&M_Disease |
| 11 | 120396016 | rs71484177 | G | A | NULL | 0.652489462 | NULL&M_Disease |
| 11 | 120558084 | rs10892613 | A | G | NULL | 0.652489462 | NULL&M_Disease |
| 11 | 120558136 | rs10892614 | G | T | NULL | 0.652489462 | NULL&M_Disease |
| 11 | 120558143 | rs10892615 | T | A | NULL | 0.652489462 | NULL&M_Disease |
| 11 | 120558155 | rs12286467 | G | T | NULL | 0.652489462 | NULL&M_Disease |
| 11 | 116203044 | rs750058 | G | C | NULL | 0.652489462 | NULL&M_Disease |
| 11 | 116479185 | rs34068367 | A | C | NULL | 0.652489462 | NULL&M_Disease |
| 11 | 116571608 | rs61905066 | G | A | NULL | 0.652489462 | NULL&M_Disease |
| 11 | 116611733 | rs9326246 | C | G | NULL | 0.652489462 | NULL&M_Disease |
| 11 | 117848926 | rs7111618 | G | T | NULL | 0.652489462 | NULL&M_Disease |
| 11 | 118872581 | rs4558175 | A | G | NULL | 0.652489462 | NULL&M_Disease |
| 11 | 120759130 | rs10892635 | A | G | NULL | 0.652489462 | NULL&M_Disease |
| 11 | 116203721 | rs1440185 | A | T | NULL | 0.652489462 | NULL&M_Disease |
| 11 | 116607437 | rs1558861 | C | T | NULL | 0.652489462 | NULL&M_Disease |
| 11 | 116612312 | rs180347 | C | G | NULL | 0.652489462 | NULL&M_Disease |
| 11 | 117105572 | rs572063 | G | T | NULL | 0.652489462 | NULL&M_Disease |
| 11 | 117321874 | rs10892109 | G | C | NULL | 0.652489462 | NULL&M_Disease |
| 11 | 116198680 | rs893839 | T | A | NULL | 0.653622331 | NULL&M_Disease |
| 11 | 116261676 | rs531860 | A | G | NULL | 0.652489462 | NULL&M_Disease |
| 11 | 118964330 | rs1488779268 | A | G | NULL | 0.653622331 | NULL&M_Disease |
| 11 | 119762458 | rs12362780 | A | C | NULL | 0.652489462 | NULL&M_Disease |
| 11 | 119824699 | rs7121901 | T | C | NULL | 0.652489462 | NULL&M_Disease |
| 11 | 120395962 | rs77404358 | C | T | NULL | 0.652489462 | NULL&M_Disease |
| 11 | 117924263 | rs7924795 | T | G | NULL | 0.653622331 | NULL&M_Disease |
| 11 | 119347715 | rs644613 | T | C | NULL | 0.653622331 | NULL&M_Disease |
| 11 | 119961624 | rs7929336 | C | G | NULL | 0.653622331 | NULL&M_Disease |
| 11 | 120287803 | rs10892573 | G | A | NULL | 0.653622331 | NULL&M_Disease |
| 11 | 116243868 | rs2510873 | G | A | NULL | 0.653622331 | NULL&M_Disease |
| 11 | 116789188 | rs79530380 | A | T | NULL | 0.653622331 | NULL&M_Disease |
| 11 | 119960565 | rs632541 | G | A | NULL | 0.653622331 | NULL&M_Disease |
| 11 | 116613017 | rs180345 | A | G | NULL | 0.653622331 | NULL&M_Disease |
| 11 | 116709438 | rs1374117 | T | C | NULL | 0.653622331 | NULL&M_Disease |
| 11 | 116785083 | rs1838757 | G | C | NULL | 0.653622331 | NULL&M_Disease |
| 11 | 118698170 | rs56296109 | A | C | NULL | 0.653622331 | NULL&M_Disease |
| 11 | 120393495 | rs34160337 | G | T | NULL | 0.653622331 | NULL&M_Disease |
| 11 | 120816860 | rs11218071 | G | A | NULL | 0.653622331 | NULL&M_Disease |
| 11 | 120985032 | rs600937 | G | A | NULL | 0.653622331 | NULL&M_Disease |
| 11 | 116590787 | rs6589563 | A | G | NULL | 0.653622331 | NULL&M_Disease |
| 11 | 116816884 | rs12805851 | T | C | NULL | 0.653622331 | NULL&M_Disease |
| 11 | 117334393 | rs1233266397 | C | T | NULL | 0.653622331 | NULL&M_Disease |
| 11 | 117663838 | rs12146702 | T | C | NULL | 0.653622331 | NULL&M_Disease |
| 11 | 119163319 | rs4938645 | G | A | NULL | 0.653622331 | NULL&M_Disease |
| 11 | 119451615 | rs11601616 | C | T | NULL | 0.653622331 | NULL&M_Disease |
| 11 | 119967825 | rs61897959 | A | G | NULL | 0.653622331 | NULL&M_Disease |
| 11 | 120395229 | rs3897570 | C | A | NULL | 0.653622331 | NULL&M_Disease |
| 11 | 120395252 | rs61900873 | C | T | NULL | 0.653622331 | NULL&M_Disease |
| 11 | 120644447 | rs948028 | C | A | NULL | 0.653622331 | NULL&M_Disease |
| 11 | 116682156 | rs7927820 | G | A | NULL | 0.653622331 | NULL&M_Disease |
| 11 | 116788491 | rs76136280 | T | C | NULL | 0.653622331 | NULL&M_Disease |
| 11 | 120085692 | rs1287395 | C | G | NULL | 0.653622331 | NULL&M_Disease |
| 11 | 116437333 | rs58567678 | A | G | NULL | 0.653622331 | NULL&M_Disease |
| 11 | 116437347 | rs57502313 | C | T | NULL | 0.653622331 | NULL&M_Disease |
| 11 | 116437360 | rs57192657 | C | A | NULL | 0.653622331 | NULL&M_Disease |
| 11 | 116438634 | rs77571570 | G | C | NULL | 0.653622331 | NULL&M_Disease |
| 11 | 116606766 | rs1974718 | G | A | NULL | 0.653622331 | NULL&M_Disease |
| 11 | 116607368 | rs1558860 | A | C | NULL | 0.653622331 | NULL&M_Disease |
| 11 | 117409910 | rs4938403 | A | C | NULL | 0.653622331 | NULL&M_Disease |
| 11 | 119342678 | rs9667310 | A | G | NULL | 0.653622331 | NULL&M_Disease |
| 11 | 117281264 | rs61903747 | A | G | NULL | 0.653622331 | NULL&M_Disease |
| 11 | 117415187 | rs11602560 | T | C | NULL | 0.653622331 | NULL&M_Disease |
| 11 | 117416249 | rs35909519 | T | C | NULL | 0.653622331 | NULL&M_Disease |
| 11 | 119168370 | rs2186882 | A | G | NULL | 0.653622331 | NULL&M_Disease |
| 11 | 119455846 | rs7925390 | T | C | NULL | 0.653622331 | NULL&M_Disease |
| 11 | 120188964 | rs61898345 | C | A | NULL | 0.653622331 | NULL&M_Disease |
| 11 | 120507564 | rs6589822 | G | A | NULL | 0.653622331 | NULL&M_Disease |
| 11 | 116432240 | rs11216007 | C | T | NULL | 0.653622331 | NULL&M_Disease |
| 11 | 117325478 | rs10892117 | C | T | NULL | 0.653622331 | NULL&M_Disease |
| 11 | 118036538 | rs653598 | C | T | NULL | 0.653622331 | NULL&M_Disease |
| 11 | 118816353 | rs586370 | A | G | NULL | 0.653622331 | NULL&M_Disease |
| 11 | 120816711 | rs11218070 | C | T | NULL | 0.653622331 | NULL&M_Disease |
| 11 | 120817046 | rs11218072 | G | A | NULL | 0.653622331 | NULL&M_Disease |
| 11 | 120821263 | rs2186621 | C | T | NULL | 0.653622331 | NULL&M_Disease |
| 11 | 120924200 | rs10502246 | C | T | NULL | 0.653622331 | NULL&M_Disease |
| 11 | 120948832 | rs12797329 | G | T | NULL | 0.653622331 | NULL&M_Disease |
| 11 | 116197173 | rs10891964 | G | T | NULL | 0.653622331 | NULL&M_Disease |
| 11 | 116684164 | rs7396851 | T | C | NULL | 0.653622331 | NULL&M_Disease |
| 11 | 117325456 | rs10790188 | A | G | NULL | 0.653622331 | NULL&M_Disease |
| 11 | 119353671 | rs532150 | C | T | NULL | 0.653622331 | NULL&M_Disease |
| 11 | 120082325 | rs2444243 | A | G | NULL | 0.653622331 | NULL&M_Disease |
| 11 | 120082748 | rs2508495 | C | T | NULL | 0.653622331 | NULL&M_Disease |
| 11 | 120083078 | rs2508496 | C | T | NULL | 0.653622331 | NULL&M_Disease |
| 11 | 120083271 | rs547990 | C | A | NULL | 0.653622331 | NULL&M_Disease |
| 11 | 120083312 | rs2444241 | G | A | NULL | 0.653622331 | NULL&M_Disease |
| 11 | 116438851 | rs17492380 | A | G | NULL | 0.653622331 | NULL&M_Disease |
| 11 | 116528202 | rs1240513 | T | A | NULL | 0.653622331 | NULL&M_Disease |
| 11 | 117335148 | rs1893967 | A | G | NULL | 0.653622331 | NULL&M_Disease |
| 11 | 117335423 | rs1893968 | A | G | NULL | 0.653622331 | NULL&M_Disease |
| 11 | 117411011 | rs4938404 | T | C | NULL | 0.653622331 | NULL&M_Disease |
| 11 | 119520997 | rs11822940 | C | G | NULL | 0.654753461 | NULL&M_Disease |
| 11 | 120394410 | rs61900871 | T | C | NULL | 0.653622331 | NULL&M_Disease |
| 11 | 120394576 | rs4385905 | A | T | NULL | 0.653622331 | NULL&M_Disease |

| | | | | | | | | | |
|---|---|---|---|---|---|---|---|---|---|
| 11 | 117144816 | rs668363 | C | A | 0.969 | -7977.388 | -7978.653 | -1.265 | 1.09E-06 |
| 11 | 117334349 | rs7940175 | T | C | 1.005 | -7977.388 | -7978.654 | -1.265 | 1.09E-06 |
| 11 | 117945161 | rs3758991 | T | C | 1.035 | -7977.388 | -7978.653 | -1.265 | 1.09E-06 |
| 11 | 118723753 | rs7119568 | A | T | 1.035 | -7977.388 | -7978.653 | -1.265 | 1.09E-06 |
| 11 | 120395934 | rs78043040 | C | T | 0.976 | -7977.388 | -7978.653 | -1.265 | 1.09E-06 |
| 11 | 120396016 | rs71484177 | G | A | 0.976 | -7977.388 | -7978.654 | -1.265 | 1.09E-06 |
| 11 | 120558084 | rs10892613 | A | G | 0.969 | -7977.388 | -7978.653 | -1.265 | 1.09E-06 |
| 11 | 120558136 | rs10892614 | G | T | 0.969 | -7977.388 | -7978.653 | -1.265 | 1.09E-06 |
| 11 | 120558143 | rs10892615 | T | A | 0.969 | -7977.388 | -7978.653 | -1.265 | 1.09E-06 |
| 11 | 120558155 | rs12286467 | G | T | 0.969 | -7977.388 | -7978.653 | -1.265 | 1.09E-06 |
| 11 | 116203044 | rs750058 | G | C | 1.034 | -7977.388 | -7978.654 | -1.266 | 1.09E-06 |
| 11 | 116479185 | rs34068367 | A | C | 1.034 | -7977.388 | -7978.654 | -1.266 | 1.09E-06 |
| 11 | 116571608 | rs61905066 | G | A | 0.992 | -7977.388 | -7978.654 | -1.266 | 1.09E-06 |
| 11 | 116611733 | rs9326246 | C | G | 0.989 | -7977.388 | -7978.655 | -1.266 | 1.09E-06 |
| 11 | 117848926 | rs7111618 | G | T | 1.034 | -7977.388 | -7978.655 | -1.266 | 1.09E-06 |
| 11 | 118872581 | rs4558175 | A | G | 1.035 | -7977.388 | -7978.654 | -1.266 | 1.09E-06 |
| 11 | 120759130 | rs10892635 | A | G | 1.035 | -7977.388 | -7978.654 | -1.266 | 1.09E-06 |
| 11 | 116203721 | rs1440185 | A | T | 1.034 | -7977.388 | -7978.655 | -1.267 | 1.09E-06 |
| 11 | 116607437 | rs1558861 | C | T | 0.989 | -7977.388 | -7978.655 | -1.267 | 1.09E-06 |
| 11 | 116612312 | rs180347 | C | G | 1.035 | -7977.388 | -7978.655 | -1.267 | 1.09E-06 |
| 11 | 117105572 | rs572063 | G | T | 0.966 | -7977.388 | -7978.656 | -1.267 | 1.09E-06 |
| 11 | 117321874 | rs10892109 | G | C | 1.015 | -7977.388 | -7978.655 | -1.267 | 1.09E-06 |
| 11 | 116198680 | rs893839 | T | A | 1.034 | -7977.388 | -7978.657 | -1.268 | 1.09E-06 |
| 11 | 116261676 | rs531860 | A | G | 0.971 | -7977.388 | -7978.657 | -1.268 | 1.09E-06 |
| 11 | 118964330 | rs1488779268 | A | G | 1.034 | -7977.388 | -7978.656 | -1.268 | 1.09E-06 |
| 11 | 119762458 | rs12362780 | A | C | 0.981 | -7977.388 | -7978.656 | -1.268 | 1.09E-06 |
| 11 | 119824699 | rs7121901 | T | C | 1.034 | -7977.388 | -7978.656 | -1.268 | 1.09E-06 |
| 11 | 120395962 | rs77404358 | C | T | 0.976 | -7977.388 | -7978.657 | -1.268 | 1.09E-06 |
| 11 | 117924263 | rs7924795 | T | G | 1.034 | -7977.388 | -7978.658 | -1.269 | 1.09E-06 |
| 11 | 119347715 | rs644613 | T | C | 1.034 | -7977.388 | -7978.658 | -1.269 | 1.09E-06 |
| 11 | 119961624 | rs7929336 | C | G | 1.03 | -7977.388 | -7978.657 | -1.269 | 1.09E-06 |
| 11 | 120287803 | rs10892573 | G | A | 1.035 | -7977.388 | -7978.657 | -1.269 | 1.09E-06 |
| 11 | 116243868 | rs2510873 | G | A | 1.021 | -7977.388 | -7978.658 | -1.27 | 1.09E-06 |
| 11 | 116789188 | rs79530380 | A | T | 0.971 | -7977.388 | -7978.659 | -1.27 | 1.09E-06 |
| 11 | 119960565 | rs632541 | G | A | 0.974 | -7977.388 | -7978.658 | -1.27 | 1.09E-06 |
| 11 | 116613017 | rs180345 | A | G | 1.034 | -7977.388 | -7978.66 | -1.271 | 1.09E-06 |
| 11 | 116709438 | rs1374117 | T | C | 0.968 | -7977.388 | -7978.659 | -1.271 | 1.09E-06 |
| 11 | 116785083 | rs1838757 | G | C | 0.967 | -7977.388 | -7978.659 | -1.271 | 1.09E-06 |
| 11 | 118698170 | rs56296109 | A | C | 1.018 | -7977.388 | -7978.66 | -1.271 | 1.09E-06 |
| 11 | 120393495 | rs34160337 | G | T | 0.977 | -7977.388 | -7978.66 | -1.271 | 1.09E-06 |
| 11 | 120816860 | rs11218071 | G | A | 0.967 | -7977.388 | -7978.659 | -1.271 | 1.09E-06 |
| 11 | 120985032 | rs600937 | G | A | 1.035 | -7977.388 | -7978.659 | -1.271 | 1.09E-06 |
| 11 | 116590787 | rs6589563 | A | G | 0.979 | -7977.388 | -7978.661 | -1.272 | 1.08E-06 |
| 11 | 116816884 | rs12805851 | T | C | 0.982 | -7977.388 | -7978.66 | -1.272 | 1.08E-06 |
| 11 | 117334393 | rs1233266397 | C | T | 0.973 | -7977.388 | -7978.661 | -1.272 | 1.08E-06 |
| 11 | 117663838 | rs12146702 | T | C | 1.018 | -7977.388 | -7978.66 | -1.272 | 1.08E-06 |
| 11 | 119163319 | rs4938645 | G | A | 0.967 | -7977.388 | -7978.661 | -1.272 | 1.08E-06 |
| 11 | 119451615 | rs11601616 | C | T | 0.977 | -7977.388 | -7978.66 | -1.272 | 1.08E-06 |
| 11 | 119967825 | rs61897959 | A | G | 1.03 | -7977.388 | -7978.661 | -1.272 | 1.08E-06 |
| 11 | 120395229 | rs3897570 | C | A | 0.977 | -7977.388 | -7978.66 | -1.272 | 1.08E-06 |
| 11 | 120395252 | rs61900873 | C | T | 0.977 | -7977.388 | -7978.66 | -1.272 | 1.08E-06 |
| 11 | 120644447 | rs948028 | C | A | 0.967 | -7977.388 | -7978.661 | -1.272 | 1.08E-06 |
| 11 | 116682156 | rs7927820 | G | A | 0.969 | -7977.388 | -7978.661 | -1.273 | 1.08E-06 |
| 11 | 116788491 | rs76136280 | T | C | 0.971 | -7977.388 | -7978.662 | -1.273 | 1.08E-06 |
| 11 | 120085692 | rs1287395 | C | G | 0.977 | -7977.388 | -7978.662 | -1.273 | 1.08E-06 |
| 11 | 116437333 | rs58567678 | A | G | 0.977 | -7977.388 | -7978.662 | -1.274 | 1.08E-06 |
| 11 | 116437347 | rs57502313 | C | T | 0.977 | -7977.388 | -7978.662 | -1.274 | 1.08E-06 |
| 11 | 116437360 | rs57192657 | C | A | 0.977 | -7977.388 | -7978.662 | -1.274 | 1.08E-06 |
| 11 | 116438634 | rs77571570 | G | C | 0.977 | -7977.388 | -7978.662 | -1.274 | 1.08E-06 |
| 11 | 116606766 | rs1974718 | G | A | 0.992 | -7977.388 | -7978.663 | -1.274 | 1.08E-06 |
| 11 | 116607368 | rs1558860 | A | C | 0.992 | -7977.388 | -7978.663 | -1.274 | 1.08E-06 |
| 11 | 117409910 | rs4938403 | A | C | 1.018 | -7977.388 | -7978.663 | -1.274 | 1.08E-06 |
| 11 | 119342678 | rs9667310 | A | G | 1.034 | -7977.388 | -7978.663 | -1.274 | 1.08E-06 |
| 11 | 117281264 | rs61903747 | A | G | 1.022 | -7977.388 | -7978.663 | -1.275 | 1.08E-06 |
| 11 | 117415187 | rs11602560 | T | C | 1.017 | -7977.388 | -7978.664 | -1.275 | 1.08E-06 |
| 11 | 117416249 | rs35909519 | T | C | 1.017 | -7977.388 | -7978.664 | -1.275 | 1.08E-06 |
| 11 | 119168370 | rs2186882 | A | G | 0.967 | -7977.388 | -7978.664 | -1.275 | 1.08E-06 |
| 11 | 119455846 | rs7925390 | T | C | 0.967 | -7977.388 | -7978.664 | -1.275 | 1.08E-06 |
| 11 | 120188964 | rs61898345 | C | A | 1.033 | -7977.388 | -7978.663 | -1.275 | 1.08E-06 |
| 11 | 120507564 | rs6589822 | G | A | 1.034 | -7977.388 | -7978.663 | -1.275 | 1.08E-06 |
| 11 | 116432240 | rs11216007 | C | T | 0.977 | -7977.388 | -7978.665 | -1.276 | 1.08E-06 |
| 11 | 117325478 | rs10892117 | C | T | 1.015 | -7977.388 | -7978.664 | -1.276 | 1.08E-06 |
| 11 | 118036538 | rs653598 | C | T | 0.967 | -7977.388 | -7978.665 | -1.276 | 1.08E-06 |
| 11 | 118816353 | rs586370 | A | G | 1.034 | -7977.388 | -7978.664 | -1.276 | 1.08E-06 |
| 11 | 120816711 | rs11218070 | C | T | 0.967 | -7977.388 | -7978.664 | -1.276 | 1.08E-06 |
| 11 | 120817046 | rs11218072 | G | A | 0.967 | -7977.388 | -7978.664 | -1.276 | 1.08E-06 |
| 11 | 120821263 | rs2186621 | C | T | 0.967 | -7977.388 | -7978.665 | -1.276 | 1.08E-06 |
| 11 | 120924200 | rs10502246 | C | T | 0.973 | -7977.388 | -7978.665 | -1.276 | 1.08E-06 |
| 11 | 120948832 | rs12797329 | G | T | 0.973 | -7977.388 | -7978.664 | -1.276 | 1.08E-06 |
| 11 | 116197173 | rs10891964 | G | T | 1.034 | -7977.388 | -7978.666 | -1.277 | 1.08E-06 |
| 11 | 116684164 | rs7396851 | T | C | 0.969 | -7977.388 | -7978.665 | -1.277 | 1.08E-06 |
| 11 | 117325456 | rs10790188 | A | G | 1.015 | -7977.388 | -7978.666 | -1.277 | 1.08E-06 |
| 11 | 119353671 | rs532150 | C | T | 1.034 | -7977.388 | -7978.665 | -1.277 | 1.08E-06 |
| 11 | 120082325 | rs2444243 | A | G | 0.977 | -7977.388 | -7978.665 | -1.277 | 1.08E-06 |
| 11 | 120082748 | rs2508495 | C | T | 0.977 | -7977.388 | -7978.665 | -1.277 | 1.08E-06 |
| 11 | 120083078 | rs2508496 | C | T | 0.977 | -7977.388 | -7978.665 | -1.277 | 1.08E-06 |
| 11 | 120083271 | rs547990 | C | A | 0.977 | -7977.388 | -7978.665 | -1.277 | 1.08E-06 |
| 11 | 120083312 | rs2444241 | G | A | 0.977 | -7977.388 | -7978.665 | -1.277 | 1.08E-06 |
| 11 | 116438851 | rs17492380 | A | G | 0.978 | -7977.388 | -7978.666 | -1.278 | 1.08E-06 |
| 11 | 116528202 | rs1240513 | T | A | 1.034 | -7977.388 | -7978.666 | -1.278 | 1.08E-06 |
| 11 | 117335148 | rs1893967 | A | G | 0.974 | -7977.388 | -7978.667 | -1.278 | 1.08E-06 |
| 11 | 117335423 | rs1893968 | A | G | 0.974 | -7977.388 | -7978.666 | -1.278 | 1.08E-06 |
| 11 | 117411011 | rs4938404 | T | C | 1.017 | -7977.388 | -7978.666 | -1.278 | 1.08E-06 |
| 11 | 119520997 | rs11822940 | C | G | 1.034 | -7977.388 | -7978.666 | -1.278 | 1.08E-06 |
| 11 | 120394410 | rs61900871 | T | C | 0.977 | -7977.388 | -7978.666 | -1.278 | 1.08E-06 |
| 11 | 120394576 | rs4385905 | A | T | 0.977 | -7977.388 | -7978.666 | -1.278 | 1.08E-06 |

| | | | | | | | |
|---|---|---|---|---|---|---|---|
| 11 | 120815567 | rs11218067 | A | G | NULL | 0.654753461 | NULL&M_Disease |
| 11 | 118873385 | rs10892320 | T | C | NULL | 0.654753461 | NULL&M_Disease |
| 11 | 120170915 | rs12290383 | A | T | NULL | 0.654753461 | NULL&M_Disease |
| 11 | 120328858 | rs766894 | C | T | NULL | 0.654753461 | NULL&M_Disease |
| 11 | 120927789 | rs12797404 | A | G | NULL | 0.654753461 | NULL&M_Disease |
| 11 | 116681563 | rs6589572 | G | A | NULL | 0.654753461 | NULL&M_Disease |
| 11 | 117380086 | rs12366082 | G | C | NULL | 0.654753461 | NULL&M_Disease |
| 11 | 118963291 | rs28990986 | T | C | NULL | 0.654753461 | NULL&M_Disease |
| 11 | 119400817 | rs10892381 | C | T | NULL | 0.654753461 | NULL&M_Disease |
| 11 | 119657893 | rs78484152 | A | G | NULL | 0.654753461 | NULL&M_Disease |
| 11 | 120341723 | rs11217879 | C | T | NULL | 0.654753461 | NULL&M_Disease |
| 11 | 116212436 | rs11820112 | A | G | NULL | 0.654753461 | NULL&M_Disease |
| 11 | 116493201 | rs56286911 | T | C | NULL | 0.654753461 | NULL&M_Disease |
| 11 | 116755584 | rs11827828 | A | G | NULL | 0.654753461 | NULL&M_Disease |
| 11 | 117775130 | rs73020492 | A | C | NULL | 0.654753461 | NULL&M_Disease |
| 11 | 117930428 | rs584356 | A | G | NULL | 0.654753461 | NULL&M_Disease |
| 11 | 118212671 | rs3181261 | T | C | NULL | 0.654753461 | NULL&M_Disease |
| 11 | 118712270 | rs7105658 | T | C | NULL | 0.654753461 | NULL&M_Disease |
| 11 | 119945024 | rs75137878 | A | C | NULL | 0.654753461 | NULL&M_Disease |
| 11 | 120274893 | rs61900794 | C | T | NULL | 0.654753461 | NULL&M_Disease |
| 11 | 120975390 | rs3758893 | A | G | NULL | 0.654753461 | NULL&M_Disease |
| 11 | 116485496 | rs499272 | G | C | NULL | 0.654753461 | NULL&M_Disease |
| 11 | 116744389 | rs11216169 | C | T | NULL | 0.654753461 | NULL&M_Disease |
| 11 | 117334831 | rs4938388 | C | T | NULL | 0.654753461 | NULL&M_Disease |
| 11 | 117334885 | rs11216404 | A | C | NULL | 0.654753461 | NULL&M_Disease |
| 11 | 117414948 | rs3802877 | T | C | NULL | 0.654753461 | NULL&M_Disease |
| 11 | 117441393 | rs557940 | G | A | NULL | 0.654753461 | NULL&M_Disease |
| 11 | 117453868 | rs608093 | A | C | NULL | 0.654753461 | NULL&M_Disease |
| 11 | 119353388 | rs534839 | C | T | NULL | 0.654753461 | NULL&M_Disease |
| 11 | 119685035 | rs58293014 | C | T | NULL | 0.654753461 | NULL&M_Disease |
| 11 | 119943201 | rs1790475 | G | A | NULL | 0.654753461 | NULL&M_Disease |
| 11 | 116396613 | rs1944156 | C | A | NULL | 0.654753461 | NULL&M_Disease |
| 11 | 116680418 | rs6589571 | T | C | NULL | 0.654753461 | NULL&M_Disease |
| 11 | 117133242 | rs61903663 | C | T | NULL | 0.654753461 | NULL&M_Disease |
| 11 | 118025969 | rs645451 | T | C | NULL | 0.654753461 | NULL&M_Disease |
| 11 | 119350396 | rs601842 | C | T | NULL | 0.654753461 | NULL&M_Disease |
| 11 | 119784133 | rs502789 | A | G | NULL | 0.654753461 | NULL&M_Disease |
| 11 | 120190141 | rs78221919 | C | T | NULL | 0.654753461 | NULL&M_Disease |
| 11 | 120194357 | rs12421941 | G | A | NULL | 0.654753461 | NULL&M_Disease |
| 11 | 116265522 | rs552035 | C | T | NULL | 0.654753461 | NULL&M_Disease |
| 11 | 116432670 | rs11607627 | A | T | NULL | 0.654753461 | NULL&M_Disease |
| 11 | 117175158 | rs609332 | G | C | NULL | 0.654753461 | NULL&M_Disease |
| 11 | 117445258 | rs557396 | A | G | NULL | 0.654753461 | NULL&M_Disease |
| 11 | 117542353 | rs658274 | G | A | NULL | 0.654753461 | NULL&M_Disease |
| 11 | 119146129 | rs1375742714 | C | T | NULL | 0.654753461 | NULL&M_Disease |
| 11 | 119854036 | rs74984550 | A | G | NULL | 0.654753461 | NULL&M_Disease |
| 11 | 120192509 | rs12575199 | A | G | NULL | 0.654753461 | NULL&M_Disease |
| 11 | 120334680 | rs11600200 | G | T | NULL | 0.654753461 | NULL&M_Disease |
| 11 | 120334863 | rs10892581 | C | A | NULL | 0.654753461 | NULL&M_Disease |
| 11 | 120394319 | rs61900870 | G | A | NULL | 0.654753461 | NULL&M_Disease |
| 11 | 120401656 | rs2257206 | T | C | NULL | 0.654753461 | NULL&M_Disease |
| 11 | 117440611 | rs629397 | C | G | NULL | 0.654753461 | NULL&M_Disease |
| 11 | 117658934 | rs73584981 | C | T | NULL | 0.654753461 | NULL&M_Disease |
| 11 | 119349454 | rs611719 | A | G | NULL | 0.654753461 | NULL&M_Disease |
| 11 | 119966884 | rs4938779 | C | T | NULL | 0.654753461 | NULL&M_Disease |
| 11 | 120310322 | rs11600558 | C | T | NULL | 0.654753461 | NULL&M_Disease |
| 11 | 120311054 | rs9666441 | G | A | NULL | 0.654753461 | NULL&M_Disease |
| 11 | 120315637 | rs11217871 | C | T | NULL | 0.654753461 | NULL&M_Disease |
| 11 | 120353237 | rs61898768 | A | G | NULL | 0.654753461 | NULL&M_Disease |
| 11 | 120355791 | rs9625 | T | G | NULL | 0.654753461 | NULL&M_Disease |
| 11 | 120907961 | rs12805477 | G | C | NULL | 0.654753461 | NULL&M_Disease |
| 11 | 120914296 | rs35457276 | T | C | NULL | 0.654753461 | NULL&M_Disease |
| 11 | 116685766 | rs6589573 | C | G | NULL | 0.654753461 | NULL&M_Disease |
| 11 | 117442242 | rs87824 | C | G | NULL | 0.654753461 | NULL&M_Disease |
| 11 | 119386445 | rs624882 | T | C | NULL | 0.654753461 | NULL&M_Disease |
| 11 | 120162250 | rs11217796 | G | A | NULL | 0.654753461 | NULL&M_Disease |
| 11 | 120360271 | rs35912781 | A | G | NULL | 0.654753461 | NULL&M_Disease |
| 11 | 116413310 | rs35624176 | A | G | NULL | 0.654753461 | NULL&M_Disease |
| 11 | 116474753 | rs3016351 | A | G | NULL | 0.654753461 | NULL&M_Disease |
| 11 | 116777394 | rs79175883 | C | T | NULL | 0.654753461 | NULL&M_Disease |
| 11 | 120095872 | rs1287393 | T | A | NULL | 0.654753461 | NULL&M_Disease |
| 11 | 120950521 | rs7940619 | A | G | NULL | 0.654753461 | NULL&M_Disease |
| 11 | 120950632 | rs7927356 | C | T | NULL | 0.654753461 | NULL&M_Disease |
| 11 | 121098770 | rs6589863 | C | G | NULL | 0.654753461 | NULL&M_Disease |
| 11 | 117411254 | rs4938405 | C | G | NULL | 0.654753461 | NULL&M_Disease |
| 11 | 118865319 | rs4938615 | G | C | NULL | 0.654753461 | NULL&M_Disease |
| 11 | 119349933 | rs609530 | T | C | NULL | 0.655882841 | NULL&M_Disease |
| 11 | 119501085 | rs11217365 | C | T | NULL | 0.654753461 | NULL&M_Disease |
| 11 | 120263022 | rs11217853 | G | A | NULL | 0.655882841 | NULL&M_Disease |
| 11 | 120947207 | rs34505832 | G | A | NULL | 0.654753461 | NULL&M_Disease |
| 11 | 117853331 | rs113195776 | A | G | NULL | 0.655882841 | NULL&M_Disease |
| 11 | 119762963 | rs56130750 | A | G | NULL | 0.654753461 | NULL&M_Disease |
| 11 | 120164799 | rs2155811 | C | G | NULL | 0.655882841 | NULL&M_Disease |
| 11 | 120195702 | rs12290699 | C | T | NULL | 0.655882841 | NULL&M_Disease |
| 11 | 120139700 | rs6589803 | T | C | NULL | 0.655882841 | NULL&M_Disease |
| 11 | 120313539 | rs747419 | T | C | NULL | 0.655882841 | NULL&M_Disease |
| 11 | 120393597 | rs61900868 | C | T | NULL | 0.655882841 | NULL&M_Disease |
| 11 | 120394599 | rs71484175 | A | G | NULL | 0.655882841 | NULL&M_Disease |
| 11 | 120394910 | rs35187805 | T | C | NULL | 0.655882841 | NULL&M_Disease |
| 11 | 120394929 | rs34527172 | C | T | NULL | 0.655882841 | NULL&M_Disease |
| 11 | 120804533 | rs7113018 | G | T | NULL | 0.655882841 | NULL&M_Disease |
| 11 | 120959225 | rs11218139 | G | A | NULL | 0.655882841 | NULL&M_Disease |
| 11 | 116202506 | rs750056 | T | C | NULL | 0.655882841 | NULL&M_Disease |
| 11 | 119342143 | rs34298632 | C | T | NULL | 0.655882841 | NULL&M_Disease |
| 11 | 120494516 | rs7127199 | T | C | NULL | 0.655882841 | NULL&M_Disease |
| 11 | 120928722 | rs11218131 | A | G | NULL | 0.655882841 | NULL&M_Disease |

| | | | | | | | | | |
|---|---|---|---|---|---|---|---|---|---|
| 11 | 120815567 | rs11218067 | A | G | 1.034 | -7977.388 | -7978.666 | -1.278 | 1.08E-06 |
| 11 | 118873385 | rs10892320 | T | C | 1.034 | -7977.388 | -7978.667 | -1.279 | 1.08E-06 |
| 11 | 120170915 | rs12290383 | A | T | 1.021 | -7977.388 | -7978.668 | -1.279 | 1.08E-06 |
| 11 | 120328858 | rs766894 | C | T | 1.034 | -7977.388 | -7978.668 | -1.279 | 1.08E-06 |
| 11 | 120927789 | rs12797404 | A | G | 0.974 | -7977.388 | -7978.668 | -1.279 | 1.08E-06 |
| 11 | 116681563 | rs6589572 | G | A | 0.969 | -7977.388 | -7978.668 | -1.28 | 1.08E-06 |
| 11 | 117380086 | rs12366082 | G | C | 0.972 | -7977.388 | -7978.668 | -1.28 | 1.08E-06 |
| 11 | 118963291 | rs28990986 | T | C | 1.019 | -7977.388 | -7978.669 | -1.28 | 1.08E-06 |
| 11 | 119400817 | rs10892381 | C | T | 1.034 | -7977.388 | -7978.668 | -1.28 | 1.08E-06 |
| 11 | 119657893 | rs78484152 | A | G | 0.971 | -7977.388 | -7978.669 | -1.28 | 1.08E-06 |
| 11 | 120341723 | rs11217879 | C | T | 1.034 | -7977.388 | -7978.668 | -1.28 | 1.08E-06 |
| 11 | 116212436 | rs11820112 | A | G | 1.032 | -7977.388 | -7978.669 | -1.281 | 1.07E-06 |
| 11 | 116493201 | rs56286911 | T | C | 1.009 | -7977.388 | -7978.67 | -1.281 | 1.07E-06 |
| 11 | 116755584 | rs11827828 | A | G | 0.968 | -7977.388 | -7978.669 | -1.281 | 1.07E-06 |
| 11 | 117775130 | rs73020492 | A | C | 0.97 | -7977.388 | -7978.669 | -1.281 | 1.07E-06 |
| 11 | 117930428 | rs584356 | A | G | 1.032 | -7977.388 | -7978.67 | -1.281 | 1.07E-06 |
| 11 | 118212671 | rs3181261 | T | C | 1.02 | -7977.388 | -7978.67 | -1.281 | 1.07E-06 |
| 11 | 118712270 | rs7105658 | T | C | 1.034 | -7977.388 | -7978.67 | -1.281 | 1.07E-06 |
| 11 | 119945024 | rs75137878 | A | C | 1.033 | -7977.388 | -7978.67 | -1.281 | 1.07E-06 |
| 11 | 120274893 | rs61900794 | C | T | 1.034 | -7977.388 | -7978.669 | -1.281 | 1.07E-06 |
| 11 | 120975390 | rs3758893 | A | G | 1.016 | -7977.388 | -7978.669 | -1.281 | 1.07E-06 |
| 11 | 116485496 | rs499272 | G | C | 1.03 | -7977.388 | -7978.67 | -1.282 | 1.07E-06 |
| 11 | 116744389 | rs11216169 | C | T | 0.971 | -7977.388 | -7978.671 | -1.282 | 1.07E-06 |
| 11 | 117334831 | rs4938388 | C | T | 0.974 | -7977.388 | -7978.671 | -1.282 | 1.07E-06 |
| 11 | 117334885 | rs11216404 | A | C | 0.974 | -7977.388 | -7978.671 | -1.282 | 1.07E-06 |
| 11 | 117414948 | rs3802877 | T | C | 1.016 | -7977.388 | -7978.671 | -1.282 | 1.07E-06 |
| 11 | 117441393 | rs557940 | G | A | 0.983 | -7977.388 | -7978.67 | -1.282 | 1.07E-06 |
| 11 | 117453868 | rs608093 | A | C | 0.994 | -7977.388 | -7978.67 | -1.282 | 1.07E-06 |
| 11 | 119353388 | rs534839 | C | T | 1.034 | -7977.388 | -7978.671 | -1.282 | 1.07E-06 |
| 11 | 119685035 | rs58293014 | C | T | 1.024 | -7977.388 | -7978.67 | -1.282 | 1.07E-06 |
| 11 | 119943201 | rs1790475 | G | A | 0.967 | -7977.388 | -7978.67 | -1.282 | 1.07E-06 |
| 11 | 116396613 | rs1944156 | C | A | 0.971 | -7977.388 | -7978.671 | -1.283 | 1.07E-06 |
| 11 | 116680418 | rs6589571 | T | C | 0.969 | -7977.388 | -7978.672 | -1.283 | 1.07E-06 |
| 11 | 117133242 | rs61903663 | C | T | 0.97 | -7977.388 | -7978.671 | -1.283 | 1.07E-06 |
| 11 | 118025969 | rs645451 | T | C | 1.001 | -7977.388 | -7978.671 | -1.283 | 1.07E-06 |
| 11 | 119350396 | rs601842 | C | T | 1.034 | -7977.388 | -7978.671 | -1.283 | 1.07E-06 |
| 11 | 119784133 | rs502789 | A | G | 0.967 | -7977.388 | -7978.671 | -1.283 | 1.07E-06 |
| 11 | 120190141 | rs78221919 | C | T | 1.032 | -7977.388 | -7978.672 | -1.283 | 1.07E-06 |
| 11 | 120194357 | rs12421941 | G | A | 1.032 | -7977.388 | -7978.671 | -1.283 | 1.07E-06 |
| 11 | 116265522 | rs552035 | C | T | 0.972 | -7977.388 | -7978.673 | -1.284 | 1.07E-06 |
| 11 | 116432670 | rs11607627 | A | T | 0.994 | -7977.388 | -7978.672 | -1.284 | 1.07E-06 |
| 11 | 117175158 | rs609332 | G | C | 0.969 | -7977.388 | -7978.672 | -1.284 | 1.07E-06 |
| 11 | 117445258 | rs557396 | A | G | 0.998 | -7977.388 | -7978.673 | -1.284 | 1.07E-06 |
| 11 | 117542353 | rs658274 | G | A | 1.021 | -7977.388 | -7978.673 | -1.284 | 1.07E-06 |
| 11 | 119146129 | rs1375742714 | C | T | 0.967 | -7977.388 | -7978.672 | -1.284 | 1.07E-06 |
| 11 | 119854036 | rs74984550 | A | G | 1.017 | -7977.388 | -7978.672 | -1.284 | 1.07E-06 |
| 11 | 120192509 | rs12575199 | A | G | 1.032 | -7977.388 | -7978.672 | -1.284 | 1.07E-06 |
| 11 | 120334680 | rs11600200 | G | T | 1.034 | -7977.388 | -7978.672 | -1.284 | 1.07E-06 |
| 11 | 120334863 | rs10892581 | C | A | 1.034 | -7977.388 | -7978.672 | -1.284 | 1.07E-06 |
| 11 | 120394319 | rs61900870 | G | A | 0.978 | -7977.388 | -7978.672 | -1.284 | 1.07E-06 |
| 11 | 120401656 | rs2257206 | T | C | 0.967 | -7977.388 | -7978.672 | -1.284 | 1.07E-06 |
| 11 | 117440611 | rs629397 | C | G | 1.004 | -7977.388 | -7978.674 | -1.285 | 1.07E-06 |
| 11 | 117658934 | rs73584981 | C | T | 0.972 | -7977.388 | -7978.673 | -1.285 | 1.07E-06 |
| 11 | 119349454 | rs611719 | A | G | 1.034 | -7977.388 | -7978.674 | -1.285 | 1.07E-06 |
| 11 | 119966884 | rs4938779 | C | T | 1.029 | -7977.388 | -7978.674 | -1.285 | 1.07E-06 |
| 11 | 120310322 | rs11600558 | C | T | 1.034 | -7977.388 | -7978.674 | -1.285 | 1.07E-06 |
| 11 | 120311054 | rs9666441 | G | A | 1.034 | -7977.388 | -7978.674 | -1.285 | 1.07E-06 |
| 11 | 120315637 | rs11217871 | C | T | 1.034 | -7977.388 | -7978.674 | -1.285 | 1.07E-06 |
| 11 | 120353237 | rs61898768 | A | G | 1.034 | -7977.388 | -7978.673 | -1.285 | 1.07E-06 |
| 11 | 120355791 | rs9625 | T | G | 1.034 | -7977.388 | -7978.673 | -1.285 | 1.07E-06 |
| 11 | 120907961 | rs12805477 | G | C | 0.974 | -7977.388 | -7978.673 | -1.285 | 1.07E-06 |
| 11 | 120914296 | rs35457276 | T | C | 0.974 | -7977.388 | -7978.673 | -1.285 | 1.07E-06 |
| 11 | 116685766 | rs6589573 | C | G | 0.969 | -7977.388 | -7978.674 | -1.286 | 1.07E-06 |
| 11 | 117442242 | rs87824 | C | G | 0.984 | -7977.388 | -7978.674 | -1.286 | 1.07E-06 |
| 11 | 119386445 | rs624882 | T | C | 1.034 | -7977.388 | -7978.674 | -1.286 | 1.07E-06 |
| 11 | 120162250 | rs11217796 | G | A | 1.034 | -7977.388 | -7978.675 | -1.286 | 1.07E-06 |
| 11 | 120360271 | rs35912781 | A | G | 1.034 | -7977.388 | -7978.674 | -1.286 | 1.07E-06 |
| 11 | 116413310 | rs35624176 | A | G | 0.987 | -7977.388 | -7978.675 | -1.287 | 1.07E-06 |
| 11 | 116474753 | rs3016351 | A | G | 1.01 | -7977.388 | -7978.675 | -1.287 | 1.07E-06 |
| 11 | 116777394 | rs79175883 | C | T | 1.004 | -7977.388 | -7978.675 | -1.287 | 1.07E-06 |
| 11 | 120095872 | rs1287393 | T | A | 0.978 | -7977.388 | -7978.675 | -1.287 | 1.07E-06 |
| 11 | 120950521 | rs7940619 | A | G | 1.034 | -7977.388 | -7978.675 | -1.287 | 1.07E-06 |
| 11 | 120950632 | rs7927356 | C | T | 1.034 | -7977.388 | -7978.675 | -1.287 | 1.07E-06 |
| 11 | 121098770 | rs6589863 | C | G | 0.967 | -7977.388 | -7978.676 | -1.287 | 1.07E-06 |
| 11 | 117411254 | rs4938405 | C | G | 1.015 | -7977.388 | -7978.676 | -1.288 | 1.07E-06 |
| 11 | 118865319 | rs4938615 | G | C | 1.033 | -7977.388 | -7978.676 | -1.288 | 1.07E-06 |
| 11 | 119349933 | rs609530 | T | C | 1.034 | -7977.388 | -7978.676 | -1.288 | 1.07E-06 |
| 11 | 119501085 | rs11217365 | C | T | 0.97 | -7977.388 | -7978.676 | -1.288 | 1.07E-06 |
| 11 | 120263022 | rs11217853 | G | A | 1.034 | -7977.388 | -7978.676 | -1.288 | 1.07E-06 |
| 11 | 120947207 | rs34505832 | G | A | 0.974 | -7977.388 | -7978.676 | -1.288 | 1.07E-06 |
| 11 | 117853331 | rs113195776 | A | G | 0.975 | -7977.388 | -7978.678 | -1.289 | 1.07E-06 |
| 11 | 119762963 | rs56130750 | A | G | 1.024 | -7977.388 | -7978.677 | -1.289 | 1.07E-06 |
| 11 | 120164799 | rs2155811 | C | G | 1.034 | -7977.388 | -7978.678 | -1.289 | 1.07E-06 |
| 11 | 120195702 | rs12290699 | C | T | 1.034 | -7977.388 | -7978.677 | -1.289 | 1.07E-06 |
| 11 | 120139700 | rs6589803 | T | C | 0.969 | -7977.388 | -7978.678 | -1.29 | 1.06E-06 |
| 11 | 120313539 | rs747419 | T | C | 1.034 | -7977.388 | -7978.678 | -1.29 | 1.06E-06 |
| 11 | 120393597 | rs61900868 | C | T | 0.979 | -7977.388 | -7978.678 | -1.29 | 1.06E-06 |
| 11 | 120394599 | rs71484175 | A | G | 0.979 | -7977.388 | -7978.679 | -1.29 | 1.06E-06 |
| 11 | 120394910 | rs35187805 | T | C | 0.979 | -7977.388 | -7978.679 | -1.29 | 1.06E-06 |
| 11 | 120394929 | rs34527172 | C | T | 0.979 | -7977.388 | -7978.679 | -1.29 | 1.06E-06 |
| 11 | 120804533 | rs7113018 | G | T | 0.967 | -7977.388 | -7978.678 | -1.29 | 1.06E-06 |
| 11 | 120959225 | rs11218139 | G | A | 1.034 | -7977.388 | -7978.678 | -1.29 | 1.06E-06 |
| 11 | 116202506 | rs750056 | T | C | 1.033 | -7977.388 | -7978.68 | -1.291 | 1.06E-06 |
| 11 | 119342143 | rs34298632 | C | T | 1.034 | -7977.388 | -7978.679 | -1.291 | 1.06E-06 |
| 11 | 120494516 | rs7127199 | T | C | 0.967 | -7977.388 | -7978.679 | -1.291 | 1.06E-06 |
| 11 | 120928722 | rs11218131 | A | G | 1.026 | -7977.388 | -7978.679 | -1.291 | 1.06E-06 |

| | | | | | | | |
|---|---|---|---|---|---|---|---|
| 11 | 117654176 | rs2155387 | A | C | NULL | 0.655882841 | NULL&M_Disease |
| 11 | 117819480 | rs2515798 | C | A | NULL | 0.655882841 | NULL&M_Disease |
| 11 | 119012962 | rs116993355 | C | T | NULL | 0.655882841 | NULL&M_Disease |
| 11 | 120277392 | rs12363975 | A | C | NULL | 0.655882841 | NULL&M_Disease |
| 11 | 120398577 | rs78354899 | T | C | NULL | 0.655882841 | NULL&M_Disease |
| 11 | 120984999 | rs667125 | G | A | NULL | 0.655882841 | NULL&M_Disease |
| 11 | 116325220 | rs17119671 | T | C | NULL | 0.655882841 | NULL&M_Disease |
| 11 | 117308795 | rs2253440 | A | G | NULL | 0.655882841 | NULL&M_Disease |
| 11 | 119345979 | rs662646 | T | C | NULL | 0.655882841 | NULL&M_Disease |
| 11 | 121181848 | rs2115925 | G | A | NULL | 0.655882841 | NULL&M_Disease |
| 11 | 116395458 | rs1396131 | C | A | NULL | 0.655882841 | NULL&M_Disease |
| 11 | 119889516 | rs41467444 | T | C | NULL | 0.655882841 | NULL&M_Disease |
| 11 | 116426814 | rs17570295 | A | T | NULL | 0.655882841 | NULL&M_Disease |
| 11 | 119881446 | rs871897 | T | C | NULL | 0.655882841 | NULL&M_Disease |
| 11 | 119970300 | rs1125618 | T | C | NULL | 0.655882841 | NULL&M_Disease |
| 11 | 116266614 | rs496288 | C | A | NULL | 0.655882841 | NULL&M_Disease |
| 11 | 117383215 | rs35200015 | A | G | NULL | 0.655882841 | NULL&M_Disease |
| 11 | 117824206 | rs12575869 | G | A | NULL | 0.655882841 | NULL&M_Disease |
| 11 | 119229374 | rs7937208 | T | C | NULL | 0.655882841 | NULL&M_Disease |
| 11 | 120261934 | rs2120430 | G | C | NULL | 0.655882841 | NULL&M_Disease |
| 11 | 116436761 | rs77159892 | A | C | NULL | 0.655882841 | NULL&M_Disease |
| 11 | 116494803 | rs12794408 | G | A | NULL | 0.655882841 | NULL&M_Disease |
| 11 | 117818505 | rs73026322 | T | C | NULL | 0.655882841 | NULL&M_Disease |
| 11 | 118515579 | rs10790255 | G | T | NULL | 0.655882841 | NULL&M_Disease |
| 11 | 120505562 | rs34599893 | C | T | NULL | 0.655882841 | NULL&M_Disease |
| 11 | 120650053 | rs2846098 | C | T | NULL | 0.655882841 | NULL&M_Disease |
| 11 | 116225440 | rs1837509 | A | G | NULL | 0.655882841 | NULL&M_Disease |
| 11 | 116710993 | rs111255936 | T | C | NULL | 0.655882841 | NULL&M_Disease |
| 11 | 117652665 | rs35066934 | T | C | NULL | 0.655882841 | NULL&M_Disease |
| 11 | 119826815 | rs10892503 | G | A | NULL | 0.655882841 | NULL&M_Disease |
| 11 | 120111853 | rs2253006 | C | G | NULL | 0.655882841 | NULL&M_Disease |
| 11 | 121076227 | rs12291772 | A | G | NULL | 0.657010463 | NULL&M_Disease |
| 11 | 117389517 | rs3819210 | G | A | NULL | 0.657010463 | NULL&M_Disease |
| 11 | 117818843 | rs11216643 | A | T | NULL | 0.655882841 | NULL&M_Disease |
| 11 | 117819411 | rs11216644 | T | C | NULL | 0.655882841 | NULL&M_Disease |
| 11 | 117819863 | rs11216645 | A | G | NULL | 0.655882841 | NULL&M_Disease |
| 11 | 120334149 | rs10892580 | A | G | NULL | 0.657010463 | NULL&M_Disease |
| 11 | 121101075 | rs9787748 | T | G | NULL | 0.657010463 | NULL&M_Disease |
| 11 | 116395210 | rs1508106 | A | G | NULL | 0.657010463 | NULL&M_Disease |
| 11 | 116398935 | rs35400250 | T | C | NULL | 0.657010463 | NULL&M_Disease |
| 11 | 117441621 | rs680424 | C | G | NULL | 0.657010463 | NULL&M_Disease |
| 11 | 119892030 | rs4938758 | T | C | NULL | 0.657010463 | NULL&M_Disease |
| 11 | 120311409 | rs12361340 | A | T | NULL | 0.657010463 | NULL&M_Disease |
| 11 | 120313899 | rs11217869 | G | T | NULL | 0.657010463 | NULL&M_Disease |
| 11 | 116198048 | rs893837 | C | T | NULL | 0.657010463 | NULL&M_Disease |
| 11 | 116331800 | rs138282449 | C | G | NULL | 0.657010463 | NULL&M_Disease |
| 11 | 117554338 | rs1942924 | T | G | NULL | 0.657010463 | NULL&M_Disease |
| 11 | 120146127 | rs17123767 | A | C | NULL | 0.657010463 | NULL&M_Disease |
| 11 | 120146707 | rs73578234 | T | C | NULL | 0.657010463 | NULL&M_Disease |
| 11 | 120305787 | rs11217868 | G | A | NULL | 0.657010463 | NULL&M_Disease |
| 11 | 116452961 | rs11216020 | C | G | NULL | 0.657010463 | NULL&M_Disease |
| 11 | 116623026 | rs180328 | G | A | NULL | 0.657010463 | NULL&M_Disease |
| 11 | 116665025 | rs45515495 | A | T | NULL | 0.657010463 | NULL&M_Disease |
| 11 | 119865290 | rs10790362 | T | C | NULL | 0.657010463 | NULL&M_Disease |
| 11 | 120282566 | rs11607944 | A | G | NULL | 0.657010463 | NULL&M_Disease |
| 11 | 117668721 | rs74505693 | A | G | NULL | 0.657010463 | NULL&M_Disease |
| 11 | 119145274 | rs2276083 | G | A | NULL | 0.657010463 | NULL&M_Disease |
| 11 | 119689931 | rs4938730 | C | T | NULL | 0.657010463 | NULL&M_Disease |
| 11 | 119958757 | rs61897956 | A | G | NULL | 0.657010463 | NULL&M_Disease |
| 11 | 116300735 | rs73003594 | A | G | NULL | 0.657010463 | NULL&M_Disease |
| 11 | 116347396 | rs12295901 | A | G | NULL | 0.657010463 | NULL&M_Disease |
| 11 | 116549896 | rs1787687 | G | A | NULL | 0.657010463 | NULL&M_Disease |
| 11 | 116593058 | rs180369 | T | A | NULL | 0.657010463 | NULL&M_Disease |
| 11 | 118728479 | rs7924647 | G | A | NULL | 0.657010463 | NULL&M_Disease |
| 11 | 119446483 | rs1995823 | T | C | NULL | 0.657010463 | NULL&M_Disease |
| 11 | 116342446 | rs13377199 | T | C | NULL | 0.657010463 | NULL&M_Disease |
| 11 | 117556714 | rs11600065 | C | A | NULL | 0.657010463 | NULL&M_Disease |
| 11 | 119353666 | rs642896 | G | A | NULL | 0.657010463 | NULL&M_Disease |
| 11 | 116385354 | rs7952521 | A | G | NULL | 0.657010463 | NULL&M_Disease |
| 11 | 117235647 | rs17120652 | T | C | NULL | 0.657010463 | NULL&M_Disease |
| 11 | 118488782 | rs11216930 | C | A | NULL | 0.657010463 | NULL&M_Disease |
| 11 | 119879488 | rs1944496 | A | G | NULL | 0.657010463 | NULL&M_Disease |
| 11 | 120268491 | rs11602796 | A | G | NULL | 0.657010463 | NULL&M_Disease |
| 11 | 117687412 | rs2077620 | T | G | NULL | 0.657010463 | NULL&M_Disease |
| 11 | 119690068 | rs4938732 | T | C | NULL | 0.657010463 | NULL&M_Disease |
| 11 | 119872703 | rs17311114 | C | T | NULL | 0.657010463 | NULL&M_Disease |
| 11 | 120096056 | rs61898292 | C | T | NULL | 0.657010463 | NULL&M_Disease |
| 11 | 120963327 | rs12226639 | A | T | NULL | 0.657010463 | NULL&M_Disease |
| 11 | 117410237 | rs7938877 | T | C | NULL | 0.657010463 | NULL&M_Disease |
| 11 | 120073521 | rs34172399 | C | A | NULL | 0.658136315 | NULL&M_Disease |
| 11 | 120656479 | rs1815771 | A | G | NULL | 0.657010463 | NULL&M_Disease |
| 11 | 116543552 | rs12294322 | G | A | NULL | 0.658136315 | NULL&M_Disease |
| 11 | 116545495 | rs3016355 | T | A | NULL | 0.658136315 | NULL&M_Disease |
| 11 | 117179003 | rs551662 | C | T | NULL | 0.657010463 | NULL&M_Disease |
| 11 | 119509746 | rs1022081 | G | C | NULL | 0.658136315 | NULL&M_Disease |
| 11 | 119767864 | rs563372 | G | A | NULL | 0.658136315 | NULL&M_Disease |
| 11 | 120150936 | rs11217782 | C | T | NULL | 0.658136315 | NULL&M_Disease |
| 11 | 120361394 | rs56693116 | T | G | NULL | 0.658136315 | NULL&M_Disease |
| 11 | 121189853 | rs74597362 | A | G | NULL | 0.657010463 | NULL&M_Disease |
| 11 | 117364180 | rs56703842 | T | C | NULL | 0.658136315 | NULL&M_Disease |
| 11 | 120115570 | rs2252520 | A | T | NULL | 0.658136315 | NULL&M_Disease |
| 11 | 120800291 | rs73007850 | C | G | NULL | 0.658136315 | NULL&M_Disease |
| 11 | 120984898 | rs659434 | G | C | NULL | 0.658136315 | NULL&M_Disease |
| 11 | 121192247 | rs76758319 | A | G | NULL | 0.658136315 | NULL&M_Disease |
| 11 | 116600021 | rs7123583 | T | C | NULL | 0.658136315 | NULL&M_Disease |
| 11 | 117432434 | rs1893969 | T | C | NULL | 0.658136315 | NULL&M_Disease |

| | | | | | | | | | |
|---|---|---|---|---|---|---|---|---|---|
| 11 | 117654176 | rs2155387 | A | C | 1.034 | -7977.388 | -7978.68 | -1.292 | 1.06E-06 |
| 11 | 117819480 | rs2515798 | C | A | 1.033 | -7977.388 | -7978.68 | -1.292 | 1.06E-06 |
| 11 | 119012962 | rs116993355 | C | T | 0.978 | -7977.388 | -7978.68 | -1.292 | 1.06E-06 |
| 11 | 120277392 | rs12363975 | A | C | 1.034 | -7977.388 | -7978.681 | -1.292 | 1.06E-06 |
| 11 | 120398577 | rs78354899 | T | C | 0.979 | -7977.388 | -7978.68 | -1.292 | 1.06E-06 |
| 11 | 120984999 | rs667125 | G | A | 1.034 | -7977.388 | -7978.681 | -1.292 | 1.06E-06 |
| 11 | 116325220 | rs17119671 | T | C | 1.033 | -7977.388 | -7978.682 | -1.293 | 1.06E-06 |
| 11 | 117308795 | rs2253440 | A | G | 1.034 | -7977.388 | -7978.681 | -1.293 | 1.06E-06 |
| 11 | 119345979 | rs662646 | T | C | 1.034 | -7977.388 | -7978.682 | -1.293 | 1.06E-06 |
| 11 | 121181848 | rs2115925 | G | A | 1.016 | -7977.388 | -7978.681 | -1.293 | 1.06E-06 |
| 11 | 116395458 | rs1396131 | C | A | 1.033 | -7977.388 | -7978.682 | -1.294 | 1.06E-06 |
| 11 | 119889516 | rs41467444 | T | C | 1.033 | -7977.388 | -7978.682 | -1.294 | 1.06E-06 |
| 11 | 116426814 | rs17570295 | A | T | 1.006 | -7977.388 | -7978.683 | -1.295 | 1.06E-06 |
| 11 | 119881446 | rs871897 | T | C | 0.968 | -7977.388 | -7978.683 | -1.295 | 1.06E-06 |
| 11 | 119970300 | rs1125618 | T | C | 1.027 | -7977.388 | -7978.684 | -1.295 | 1.06E-06 |
| 11 | 116266614 | rs496288 | C | A | 0.973 | -7977.388 | -7978.684 | -1.296 | 1.06E-06 |
| 11 | 117383215 | rs35200015 | A | G | 1.033 | -7977.388 | -7978.685 | -1.296 | 1.06E-06 |
| 11 | 117824206 | rs12575869 | G | A | 1.033 | -7977.388 | -7978.685 | -1.296 | 1.06E-06 |
| 11 | 119229374 | rs7937208 | T | C | 0.967 | -7977.388 | -7978.684 | -1.296 | 1.06E-06 |
| 11 | 120261934 | rs2120430 | G | C | 1.034 | -7977.388 | -7978.685 | -1.296 | 1.06E-06 |
| 11 | 116436761 | rs77159892 | A | C | 0.98 | -7977.388 | -7978.686 | -1.297 | 1.06E-06 |
| 11 | 116494803 | rs12794408 | G | A | 1.034 | -7977.388 | -7978.685 | -1.297 | 1.06E-06 |
| 11 | 117818505 | rs73026322 | T | C | 1.002 | -7977.388 | -7978.685 | -1.297 | 1.06E-06 |
| 11 | 118515579 | rs10790255 | G | T | 1.034 | -7977.388 | -7978.686 | -1.297 | 1.06E-06 |
| 11 | 120505562 | rs34599893 | C | T | 0.967 | -7977.388 | -7978.685 | -1.297 | 1.06E-06 |
| 11 | 120650053 | rs2846098 | C | T | 0.968 | -7977.388 | -7978.686 | -1.297 | 1.06E-06 |
| 11 | 116225440 | rs1837509 | A | G | 1.023 | -7977.388 | -7978.686 | -1.298 | 1.06E-06 |
| 11 | 116710993 | rs111255936 | T | C | 0.97 | -7977.388 | -7978.686 | -1.298 | 1.06E-06 |
| 11 | 117652665 | rs35066934 | T | C | 1.02 | -7977.388 | -7978.686 | -1.298 | 1.06E-06 |
| 11 | 119826815 | rs10892503 | G | A | 1.033 | -7977.388 | -7978.686 | -1.298 | 1.06E-06 |
| 11 | 120111853 | rs2253006 | C | G | 1.021 | -7977.388 | -7978.687 | -1.298 | 1.06E-06 |
| 11 | 121076227 | rs12291772 | A | G | 1.033 | -7977.388 | -7978.687 | -1.298 | 1.06E-06 |
| 11 | 117389517 | rs3819210 | G | A | 1.033 | -7977.388 | -7978.687 | -1.299 | 1.06E-06 |
| 11 | 117818843 | rs11216643 | A | T | 1 | -7977.388 | -7978.687 | -1.299 | 1.06E-06 |
| 11 | 117819411 | rs11216644 | T | C | 0.998 | -7977.388 | -7978.688 | -1.299 | 1.06E-06 |
| 11 | 117819863 | rs11216645 | A | G | 1 | -7977.388 | -7978.687 | -1.299 | 1.06E-06 |
| 11 | 120334149 | rs10892580 | A | G | 1.034 | -7977.388 | -7978.688 | -1.299 | 1.06E-06 |
| 11 | 121101075 | rs9787748 | T | G | 0.967 | -7977.388 | -7978.688 | -1.299 | 1.06E-06 |
| 11 | 116395210 | rs1508106 | A | G | 0.972 | -7977.388 | -7978.688 | -1.3 | 1.05E-06 |
| 11 | 116398935 | rs35400250 | T | C | 0.972 | -7977.388 | -7978.688 | -1.3 | 1.05E-06 |
| 11 | 117441621 | rs680424 | C | G | 0.986 | -7977.388 | -7978.688 | -1.3 | 1.05E-06 |
| 11 | 119892030 | rs4938758 | T | C | 1.033 | -7977.388 | -7978.688 | -1.3 | 1.05E-06 |
| 11 | 120311409 | rs12361340 | A | T | 1.033 | -7977.388 | -7978.689 | -1.3 | 1.05E-06 |
| 11 | 120313899 | rs11217869 | G | T | 1.033 | -7977.388 | -7978.689 | -1.3 | 1.05E-06 |
| 11 | 116198048 | rs893837 | C | T | 1.033 | -7977.388 | -7978.689 | -1.301 | 1.05E-06 |
| 11 | 116331800 | rs138282449 | C | G | 0.996 | -7977.388 | -7978.689 | -1.301 | 1.05E-06 |
| 11 | 117554338 | rs1942924 | T | G | 0.968 | -7977.388 | -7978.689 | -1.301 | 1.05E-06 |
| 11 | 120146127 | rs17123767 | A | C | 1.013 | -7977.388 | -7978.689 | -1.301 | 1.05E-06 |
| 11 | 120146707 | rs73578234 | T | C | 1.013 | -7977.388 | -7978.689 | -1.301 | 1.05E-06 |
| 11 | 120305787 | rs11217868 | G | A | 1.033 | -7977.388 | -7978.689 | -1.301 | 1.05E-06 |
| 11 | 116452961 | rs11216020 | C | G | 1.033 | -7977.388 | -7978.691 | -1.302 | 1.05E-06 |
| 11 | 116623026 | rs180328 | G | A | 1.033 | -7977.388 | -7978.69 | -1.302 | 1.05E-06 |
| 11 | 116665025 | rs45515495 | A | T | 0.98 | -7977.388 | -7978.691 | -1.302 | 1.05E-06 |
| 11 | 119865290 | rs10790362 | T | C | 0.968 | -7977.388 | -7978.691 | -1.302 | 1.05E-06 |
| 11 | 120282566 | rs11607944 | A | G | 1.033 | -7977.388 | -7978.69 | -1.302 | 1.05E-06 |
| 11 | 117668721 | rs74505693 | A | G | 0.974 | -7977.388 | -7978.691 | -1.303 | 1.05E-06 |
| 11 | 119145274 | rs2276083 | G | A | 0.968 | -7977.388 | -7978.692 | -1.303 | 1.05E-06 |
| 11 | 119689931 | rs4938730 | C | T | 0.968 | -7977.388 | -7978.691 | -1.303 | 1.05E-06 |
| 11 | 119958757 | rs61897956 | A | G | 1.028 | -7977.388 | -7978.692 | -1.303 | 1.05E-06 |
| 11 | 116300735 | rs73003594 | A | G | 0.982 | -7977.388 | -7978.692 | -1.304 | 1.05E-06 |
| 11 | 116347396 | rs12295901 | A | G | 0.979 | -7977.388 | -7978.692 | -1.304 | 1.05E-06 |
| 11 | 116549896 | rs1787687 | G | A | 1.033 | -7977.388 | -7978.693 | -1.304 | 1.05E-06 |
| 11 | 116593058 | rs180369 | T | A | 1.033 | -7977.388 | -7978.692 | -1.304 | 1.05E-06 |
| 11 | 118728479 | rs7924647 | G | A | 1.033 | -7977.388 | -7978.693 | -1.304 | 1.05E-06 |
| 11 | 119446483 | rs1995823 | T | C | 0.981 | -7977.388 | -7978.692 | -1.304 | 1.05E-06 |
| 11 | 116342446 | rs13377199 | T | C | 0.982 | -7977.388 | -7978.693 | -1.305 | 1.05E-06 |
| 11 | 117556714 | rs11600065 | C | A | 0.968 | -7977.388 | -7978.693 | -1.305 | 1.05E-06 |
| 11 | 119353666 | rs642896 | G | A | 1.033 | -7977.388 | -7978.693 | -1.305 | 1.05E-06 |
| 11 | 116385354 | rs7952521 | A | G | 0.973 | -7977.388 | -7978.694 | -1.306 | 1.05E-06 |
| 11 | 117235647 | rs17120652 | T | C | 0.969 | -7977.388 | -7978.694 | -1.306 | 1.05E-06 |
| 11 | 118488782 | rs11216930 | C | A | 1.033 | -7977.388 | -7978.694 | -1.306 | 1.05E-06 |
| 11 | 119879488 | rs1944496 | A | G | 1.033 | -7977.388 | -7978.694 | -1.306 | 1.05E-06 |
| 11 | 120268491 | rs11602796 | A | G | 1.033 | -7977.388 | -7978.695 | -1.306 | 1.05E-06 |
| 11 | 117687412 | rs2077620 | T | G | 1.029 | -7977.388 | -7978.695 | -1.307 | 1.05E-06 |
| 11 | 119690068 | rs4938732 | T | C | 0.968 | -7977.388 | -7978.696 | -1.307 | 1.05E-06 |
| 11 | 119872703 | rs17311114 | C | T | 1.022 | -7977.388 | -7978.695 | -1.307 | 1.05E-06 |
| 11 | 120096056 | rs61898292 | C | T | 0.98 | -7977.388 | -7978.696 | -1.307 | 1.05E-06 |
| 11 | 120963327 | rs12226639 | A | T | 1.032 | -7977.388 | -7978.695 | -1.307 | 1.05E-06 |
| 11 | 117410237 | rs7938877 | T | C | 1.012 | -7977.388 | -7978.696 | -1.308 | 1.05E-06 |
| 11 | 120073521 | rs34172399 | C | A | 1.033 | -7977.388 | -7978.696 | -1.308 | 1.05E-06 |
| 11 | 120656479 | rs1815771 | A | G | 0.989 | -7977.388 | -7978.696 | -1.308 | 1.05E-06 |
| 11 | 116543552 | rs12294322 | G | A | 1.033 | -7977.388 | -7978.697 | -1.309 | 1.04E-06 |
| 11 | 116545495 | rs3016355 | T | A | 1.033 | -7977.388 | -7978.697 | -1.309 | 1.04E-06 |
| 11 | 117179003 | rs551662 | C | T | 1.019 | -7977.388 | -7978.697 | -1.309 | 1.04E-06 |
| 11 | 119509746 | rs1022081 | G | C | 0.969 | -7977.388 | -7978.697 | -1.309 | 1.04E-06 |
| 11 | 119767864 | rs563372 | G | A | 1.033 | -7977.388 | -7978.698 | -1.309 | 1.04E-06 |
| 11 | 120150936 | rs11217782 | C | T | 1.01 | -7977.388 | -7978.698 | -1.309 | 1.04E-06 |
| 11 | 120361394 | rs56693116 | T | G | 0.969 | -7977.388 | -7978.697 | -1.309 | 1.04E-06 |
| 11 | 121189853 | rs74597362 | A | G | 1.012 | -7977.388 | -7978.698 | -1.309 | 1.04E-06 |
| 11 | 117364180 | rs56703842 | T | C | 1.032 | -7977.388 | -7978.699 | -1.31 | 1.04E-06 |
| 11 | 120115570 | rs2252520 | A | T | 1.033 | -7977.388 | -7978.698 | -1.31 | 1.04E-06 |
| 11 | 120800291 | rs73007850 | C | G | 0.968 | -7977.388 | -7978.698 | -1.31 | 1.04E-06 |
| 11 | 120984898 | rs659434 | G | C | 0.968 | -7977.388 | -7978.698 | -1.31 | 1.04E-06 |
| 11 | 121192247 | rs76758319 | A | G | 1.012 | -7977.388 | -7978.699 | -1.31 | 1.04E-06 |
| 11 | 116600021 | rs7123583 | T | C | 1.033 | -7977.388 | -7978.699 | -1.311 | 1.04E-06 |
| 11 | 117432434 | rs1893969 | T | C | 0.968 | -7977.388 | -7978.699 | -1.311 | 1.04E-06 |

| | | | | | | | |
|---|---|---|---|---|---|---|---|
| 11 | 119349531 | rs611279 | A | G | NULL | 0.658136315 | NULL&M_Disease |
| 11 | 117460028 | rs633983 | G | A | NULL | 0.658136315 | NULL&M_Disease |
| 11 | 118297391 | rs4435039 | G | A | NULL | 0.658136315 | NULL&M_Disease |
| 11 | 120595653 | rs3133227 | A | G | NULL | 0.658136315 | NULL&M_Disease |
| 11 | 116407823 | rs10466532 | C | T | NULL | 0.658136315 | NULL&M_Disease |
| 11 | 118189965 | rs73014299 | T | C | NULL | 0.658136315 | NULL&M_Disease |
| 11 | 119349526 | rs611616 | A | G | NULL | 0.658136315 | NULL&M_Disease |
| 11 | 116347944 | rs59414734 | C | T | NULL | 0.658136315 | NULL&M_Disease |
| 11 | 116848648 | rs7109876 | A | C | NULL | 0.658136315 | NULL&M_Disease |
| 11 | 117314044 | rs7927742 | C | T | NULL | 0.658136315 | NULL&M_Disease |
| 11 | 119444694 | rs73584223 | A | G | NULL | 0.658136315 | NULL&M_Disease |
| 11 | 119445108 | rs59027088 | T | C | NULL | 0.658136315 | NULL&M_Disease |
| 11 | 119878632 | rs4938757 | T | C | NULL | 0.658136315 | NULL&M_Disease |
| 11 | 119951347 | rs11217678 | C | G | NULL | 0.658136315 | NULL&M_Disease |
| 11 | 117347207 | rs11216416 | T | C | NULL | 0.658136315 | NULL&M_Disease |
| 11 | 117432779 | rs17509752 | T | C | NULL | 0.658136315 | NULL&M_Disease |
| 11 | 118714492 | rs7117594 | T | C | NULL | 0.658136315 | NULL&M_Disease |
| 11 | 116566933 | rs60954647 | T | C | NULL | 0.658136315 | NULL&M_Disease |
| 11 | 117008881 | rs474981 | A | G | NULL | 0.658136315 | NULL&M_Disease |
| 11 | 117129637 | rs1242598 | T | C | NULL | 0.658136315 | NULL&M_Disease |
| 11 | 117632531 | rs34352231 | G | A | NULL | 0.658136315 | NULL&M_Disease |
| 11 | 119447044 | rs35287658 | A | G | NULL | 0.658136315 | NULL&M_Disease |
| 11 | 119806226 | rs10892498 | T | G | NULL | 0.658136315 | NULL&M_Disease |
| 11 | 116467100 | rs61904854 | T | G | NULL | 0.658136315 | NULL&M_Disease |
| 11 | 116541417 | rs1784133 | T | C | NULL | 0.658136315 | NULL&M_Disease |
| 11 | 116541539 | rs2846016 | T | C | NULL | 0.658136315 | NULL&M_Disease |
| 11 | 117680787 | rs12364606 | T | C | NULL | 0.658136315 | NULL&M_Disease |
| 11 | 119412210 | rs7947704 | T | C | NULL | 0.658136315 | NULL&M_Disease |
| 11 | 119901007 | rs12574203 | G | A | NULL | 0.658136315 | NULL&M_Disease |
| 11 | 120247436 | rs11217847 | T | C | NULL | 0.658136315 | NULL&M_Disease |
| 11 | 120321653 | rs11217873 | T | C | NULL | 0.658136315 | NULL&M_Disease |
| 11 | 116280252 | rs508994 | A | C | NULL | 0.658136315 | NULL&M_Disease |
| 11 | 116398764 | rs11215985 | G | A | NULL | 0.659260388 | NULL&M_Disease |
| 11 | 116482573 | rs78062662 | T | C | NULL | 0.658136315 | NULL&M_Disease |
| 11 | 117357402 | rs61905308 | T | C | NULL | 0.658136315 | NULL&M_Disease |
| 11 | 117656310 | rs948102 | A | G | NULL | 0.659260388 | NULL&M_Disease |
| 11 | 120093745 | rs61898289 | A | G | NULL | 0.658136315 | NULL&M_Disease |
| 11 | 120288060 | rs11217861 | T | C | NULL | 0.658136315 | NULL&M_Disease |
| 11 | 116350128 | rs17569099 | A | T | NULL | 0.658136315 | NULL&M_Disease |
| 11 | 116428591 | rs78243160 | T | C | NULL | 0.658136315 | NULL&M_Disease |
| 11 | 116428953 | rs74890383 | A | G | NULL | 0.658136315 | NULL&M_Disease |
| 11 | 116440439 | rs906540 | A | T | NULL | 0.659260388 | NULL&M_Disease |
| 11 | 116675294 | rs6589570 | A | T | NULL | 0.659260388 | NULL&M_Disease |
| 11 | 117917257 | rs73013776 | G | A | NULL | 0.658136315 | NULL&M_Disease |
| 11 | 119344407 | rs1002179 | C | T | NULL | 0.659260388 | NULL&M_Disease |
| 11 | 119661809 | rs11217506 | T | C | NULL | 0.659260388 | NULL&M_Disease |
| 11 | 117099941 | rs4938361 | C | G | NULL | 0.659260388 | NULL&M_Disease |
| 11 | 117118481 | rs2925771 | T | C | NULL | 0.659260388 | NULL&M_Disease |
| 11 | 117661447 | rs12291974 | T | C | NULL | 0.659260388 | NULL&M_Disease |
| 11 | 118700218 | rs2510902 | A | G | NULL | 0.659260388 | NULL&M_Disease |
| 11 | 119864575 | rs880745 | T | C | NULL | 0.659260388 | NULL&M_Disease |
| 11 | 120147254 | rs7936665 | A | G | NULL | 0.659260388 | NULL&M_Disease |
| 11 | 120897987 | rs12577611 | C | T | NULL | 0.659260388 | NULL&M_Disease |
| 11 | 117688225 | rs610226 | G | C | NULL | 0.659260388 | NULL&M_Disease |
| 11 | 119351220 | rs587521 | G | A | NULL | 0.659260388 | NULL&M_Disease |
| 11 | 119443362 | rs7103327 | T | C | NULL | 0.659260388 | NULL&M_Disease |
| 11 | 119941330 | rs11607997 | T | C | NULL | 0.659260388 | NULL&M_Disease |
| 11 | 117097952 | rs2306473 | T | C | NULL | 0.659260388 | NULL&M_Disease |
| 11 | 118445585 | rs73023341 | G | A | NULL | 0.659260388 | NULL&M_Disease |
| 11 | 118447580 | rs9736327 | G | T | NULL | 0.659260388 | NULL&M_Disease |
| 11 | 118447597 | rs9735421 | A | G | NULL | 0.659260388 | NULL&M_Disease |
| 11 | 118449396 | rs73023350 | A | G | NULL | 0.659260388 | NULL&M_Disease |
| 11 | 118450875 | rs7126175 | T | C | NULL | 0.659260388 | NULL&M_Disease |
| 11 | 118452334 | rs11216912 | C | G | NULL | 0.659260388 | NULL&M_Disease |
| 11 | 118453681 | rs11216913 | G | A | NULL | 0.659260388 | NULL&M_Disease |
| 11 | 118457630 | rs17092756 | C | G | NULL | 0.659260388 | NULL&M_Disease |
| 11 | 119341075 | rs685879 | G | A | NULL | 0.659260388 | NULL&M_Disease |
| 11 | 120235574 | rs67551874 | C | T | NULL | 0.659260388 | NULL&M_Disease |
| 11 | 116684028 | rs7396835 | T | C | NULL | 0.659260388 | NULL&M_Disease |
| 11 | 119688442 | rs4245207 | A | G | NULL | 0.659260388 | NULL&M_Disease |
| 11 | 116395238 | rs1396132 | A | T | NULL | 0.659260388 | NULL&M_Disease |
| 11 | 119346737 | rs654776 | A | G | NULL | 0.659260388 | NULL&M_Disease |
| 11 | 117820561 | rs7103514 | G | A | NULL | 0.659260388 | NULL&M_Disease |
| 11 | 119498403 | rs10892421 | C | T | NULL | 0.659260388 | NULL&M_Disease |
| 11 | 119948049 | rs1007530 | A | G | NULL | 0.659260388 | NULL&M_Disease |
| 11 | 116203828 | rs7928488 | T | C | NULL | 0.659260388 | NULL&M_Disease |
| 11 | 116589652 | rs10892020 | G | C | NULL | 0.659260388 | NULL&M_Disease |
| 11 | 118443944 | rs11216906 | A | G | NULL | 0.659260388 | NULL&M_Disease |
| 11 | 118454177 | rs17842763 | G | A | NULL | 0.659260388 | NULL&M_Disease |
| 11 | 119335915 | rs12793860 | T | G | NULL | 0.659260388 | NULL&M_Disease |
| 11 | 119337109 | rs12576276 | C | T | NULL | 0.659260388 | NULL&M_Disease |
| 11 | 119341038 | rs634182 | C | T | NULL | 0.659260388 | NULL&M_Disease |
| 11 | 119500880 | rs12364363 | C | T | NULL | 0.659260388 | NULL&M_Disease |
| 11 | 116456027 | rs61904849 | A | G | NULL | 0.659260388 | NULL&M_Disease |
| 11 | 117863829 | rs2508450 | T | C | NULL | 0.660382673 | NULL&M_Disease |
| 11 | 118165955 | rs12416826 | C | T | NULL | 0.659260388 | NULL&M_Disease |
| 11 | 118451901 | rs11216911 | C | T | NULL | 0.659260388 | NULL&M_Disease |
| 11 | 119784889 | rs12806754 | A | G | NULL | 0.659260388 | NULL&M_Disease |
| 11 | 119828651 | rs72645469 | A | C | NULL | 0.659260388 | NULL&M_Disease |
| 11 | 120009308 | rs79761998 | A | C | NULL | 0.659260388 | NULL&M_Disease |
| 11 | 116344594 | rs11215951 | T | C | NULL | 0.659260388 | NULL&M_Disease |
| 11 | 117650439 | rs7928767 | A | G | NULL | 0.660382673 | NULL&M_Disease |
| 11 | 119240071 | rs677008 | A | G | NULL | 0.660382673 | NULL&M_Disease |
| 11 | 116801529 | rs76690043 | G | C | NULL | 0.660382673 | NULL&M_Disease |
| 11 | 117428278 | rs562422 | A | G | NULL | 0.660382673 | NULL&M_Disease |
| 11 | 117919377 | rs12364267 | A | G | NULL | 0.660382673 | NULL&M_Disease |

| | | | | | | | | | |
|---|---|---|---|---|---|---|---|---|---|
| 11 | 119349531 | rs611279 | A | G | 1.033 | -7977.388 | -7978.7 | -1.311 | 1.04E-06 |
| 11 | 117460028 | rs633983 | G | A | 0.968 | -7977.388 | -7978.701 | -1.312 | 1.04E-06 |
| 11 | 118297391 | rs4435039 | G | A | 0.969 | -7977.388 | -7978.701 | -1.312 | 1.04E-06 |
| 11 | 120595653 | rs3133227 | A | G | 0.968 | -7977.388 | -7978.7 | -1.312 | 1.04E-06 |
| 11 | 116407823 | rs10466532 | C | T | 0.974 | -7977.388 | -7978.701 | -1.313 | 1.04E-06 |
| 11 | 118189965 | rs73014299 | T | C | 0.973 | -7977.388 | -7978.701 | -1.313 | 1.04E-06 |
| 11 | 119349526 | rs611616 | A | G | 1.033 | -7977.388 | -7978.701 | -1.313 | 1.04E-06 |
| 11 | 116347944 | rs59414734 | C | T | 0.98 | -7977.388 | -7978.703 | -1.314 | 1.04E-06 |
| 11 | 116848648 | rs7109876 | A | C | 0.975 | -7977.388 | -7978.702 | -1.314 | 1.04E-06 |
| 11 | 117314044 | rs7927742 | C | T | 0.973 | -7977.388 | -7978.702 | -1.314 | 1.04E-06 |
| 11 | 119444694 | rs73584223 | A | G | 0.983 | -7977.388 | -7978.702 | -1.314 | 1.04E-06 |
| 11 | 119445108 | rs59027088 | T | C | 0.983 | -7977.388 | -7978.702 | -1.314 | 1.04E-06 |
| 11 | 119878632 | rs4938757 | T | C | 1.014 | -7977.388 | -7978.703 | -1.314 | 1.04E-06 |
| 11 | 119951347 | rs11217678 | C | G | 1.033 | -7977.388 | -7978.703 | -1.314 | 1.04E-06 |
| 11 | 117347207 | rs11216416 | T | C | 1.018 | -7977.388 | -7978.703 | -1.315 | 1.04E-06 |
| 11 | 117432779 | rs17509752 | T | C | 0.968 | -7977.388 | -7978.703 | -1.315 | 1.04E-06 |
| 11 | 118714492 | rs7117594 | T | C | 1.032 | -7977.388 | -7978.703 | -1.315 | 1.04E-06 |
| 11 | 116566933 | rs60954647 | T | C | 0.968 | -7977.388 | -7978.704 | -1.316 | 1.04E-06 |
| 11 | 117008881 | rs474981 | A | G | 0.968 | -7977.388 | -7978.704 | -1.316 | 1.04E-06 |
| 11 | 117129637 | rs1242598 | T | C | 0.972 | -7977.388 | -7978.704 | -1.316 | 1.04E-06 |
| 11 | 117632531 | rs34352231 | G | A | 0.97 | -7977.388 | -7978.705 | -1.316 | 1.04E-06 |
| 11 | 119447044 | rs35287658 | A | G | 0.983 | -7977.388 | -7978.705 | -1.316 | 1.04E-06 |
| 11 | 119806226 | rs10892498 | T | G | 1.027 | -7977.388 | -7978.705 | -1.316 | 1.04E-06 |
| 11 | 116467100 | rs61904854 | T | G | 0.968 | -7977.388 | -7978.705 | -1.317 | 1.04E-06 |
| 11 | 116541417 | rs1784133 | T | C | 1.033 | -7977.388 | -7978.705 | -1.317 | 1.04E-06 |
| 11 | 116541539 | rs2846016 | T | C | 1.033 | -7977.388 | -7978.705 | -1.317 | 1.04E-06 |
| 11 | 117680787 | rs12364606 | T | C | 1.033 | -7977.388 | -7978.706 | -1.317 | 1.04E-06 |
| 11 | 119412210 | rs7947704 | T | C | 0.984 | -7977.388 | -7978.705 | -1.317 | 1.04E-06 |
| 11 | 119901007 | rs12574203 | G | A | 1.031 | -7977.388 | -7978.705 | -1.317 | 1.04E-06 |
| 11 | 120247436 | rs11217847 | T | C | 1.033 | -7977.388 | -7978.705 | -1.317 | 1.04E-06 |
| 11 | 120321653 | rs11217873 | T | C | 1.033 | -7977.388 | -7978.705 | -1.317 | 1.04E-06 |
| 11 | 116280252 | rs508994 | A | C | 0.973 | -7977.388 | -7978.706 | -1.318 | 1.04E-06 |
| 11 | 116398764 | rs11215985 | G | A | 0.968 | -7977.388 | -7978.706 | -1.318 | 1.04E-06 |
| 11 | 116482573 | rs78062662 | T | C | 1.02 | -7977.388 | -7978.706 | -1.318 | 1.04E-06 |
| 11 | 117357402 | rs61905308 | T | C | 1.021 | -7977.388 | -7978.706 | -1.318 | 1.04E-06 |
| 11 | 117656310 | rs948102 | A | G | 1.033 | -7977.388 | -7978.707 | -1.318 | 1.04E-06 |
| 11 | 120093745 | rs61898289 | A | G | 0.983 | -7977.388 | -7978.706 | -1.318 | 1.04E-06 |
| 11 | 120288060 | rs11217861 | T | C | 1.033 | -7977.388 | -7978.706 | -1.318 | 1.04E-06 |
| 11 | 116350128 | rs17569099 | A | T | 1.01 | -7977.388 | -7978.707 | -1.319 | 1.03E-06 |
| 11 | 116428591 | rs78243160 | T | C | 0.983 | -7977.388 | -7978.707 | -1.319 | 1.03E-06 |
| 11 | 116428953 | rs74890383 | A | G | 0.983 | -7977.388 | -7978.707 | -1.319 | 1.03E-06 |
| 11 | 116440439 | rs906540 | A | T | 1.033 | -7977.388 | -7978.707 | -1.319 | 1.03E-06 |
| 11 | 116675294 | rs6589570 | A | T | 0.971 | -7977.388 | -7978.707 | -1.319 | 1.03E-06 |
| 11 | 117917257 | rs73013776 | G | A | 1.007 | -7977.388 | -7978.707 | -1.319 | 1.03E-06 |
| 11 | 119344407 | rs1002179 | C | T | 1.033 | -7977.388 | -7978.707 | -1.319 | 1.03E-06 |
| 11 | 119661809 | rs11217506 | T | C | 0.971 | -7977.388 | -7978.708 | -1.319 | 1.03E-06 |
| 11 | 117099941 | rs4938361 | C | G | 0.969 | -7977.388 | -7978.708 | -1.32 | 1.03E-06 |
| 11 | 117118481 | rs2925771 | T | C | 0.968 | -7977.388 | -7978.708 | -1.32 | 1.03E-06 |
| 11 | 117661447 | rs12291974 | T | C | 0.975 | -7977.388 | -7978.709 | -1.32 | 1.03E-06 |
| 11 | 118700218 | rs2510902 | A | G | 1.032 | -7977.388 | -7978.708 | -1.32 | 1.03E-06 |
| 11 | 119864575 | rs880745 | T | C | 0.969 | -7977.388 | -7978.709 | -1.32 | 1.03E-06 |
| 11 | 120147254 | rs7936665 | A | G | 1.007 | -7977.388 | -7978.708 | -1.32 | 1.03E-06 |
| 11 | 120897987 | rs12577611 | C | T | 1.03 | -7977.388 | -7978.708 | -1.32 | 1.03E-06 |
| 11 | 117688225 | rs610226 | G | C | 0.968 | -7977.388 | -7978.71 | -1.321 | 1.03E-06 |
| 11 | 119351220 | rs587521 | G | A | 1.033 | -7977.388 | -7978.709 | -1.321 | 1.03E-06 |
| 11 | 119443362 | rs7103327 | T | C | 0.983 | -7977.388 | -7978.709 | -1.321 | 1.03E-06 |
| 11 | 119941330 | rs11607997 | T | C | 1.033 | -7977.388 | -7978.709 | -1.321 | 1.03E-06 |
| 11 | 117097952 | rs2306473 | T | C | 0.969 | -7977.388 | -7978.71 | -1.322 | 1.03E-06 |
| 11 | 118445585 | rs73023341 | G | A | 0.971 | -7977.388 | -7978.71 | -1.322 | 1.03E-06 |
| 11 | 118447580 | rs9736327 | G | T | 0.971 | -7977.388 | -7978.71 | -1.322 | 1.03E-06 |
| 11 | 118447597 | rs9735421 | A | G | 0.971 | -7977.388 | -7978.71 | -1.322 | 1.03E-06 |
| 11 | 118449396 | rs73023350 | A | G | 0.971 | -7977.388 | -7978.71 | -1.322 | 1.03E-06 |
| 11 | 118450875 | rs7126175 | T | C | 0.971 | -7977.388 | -7978.71 | -1.322 | 1.03E-06 |
| 11 | 118452334 | rs11216912 | C | G | 0.971 | -7977.388 | -7978.71 | -1.322 | 1.03E-06 |
| 11 | 118453681 | rs11216913 | G | A | 0.971 | -7977.388 | -7978.71 | -1.322 | 1.03E-06 |
| 11 | 118457630 | rs17092756 | C | G | 0.971 | -7977.388 | -7978.71 | -1.322 | 1.03E-06 |
| 11 | 119341075 | rs685879 | G | A | 1.033 | -7977.388 | -7978.71 | -1.322 | 1.03E-06 |
| 11 | 120235574 | rs67551874 | C | T | 1.033 | -7977.388 | -7978.71 | -1.322 | 1.03E-06 |
| 11 | 116684028 | rs7396835 | T | C | 0.972 | -7977.388 | -7978.711 | -1.323 | 1.03E-06 |
| 11 | 119688442 | rs4245207 | A | G | 0.969 | -7977.388 | -7978.712 | -1.323 | 1.03E-06 |
| 11 | 116395238 | rs1396132 | A | T | 1.032 | -7977.388 | -7978.712 | -1.324 | 1.03E-06 |
| 11 | 119346737 | rs654776 | A | G | 1.033 | -7977.388 | -7978.712 | -1.324 | 1.03E-06 |
| 11 | 117820561 | rs7103514 | G | A | 1.006 | -7977.388 | -7978.713 | -1.325 | 1.03E-06 |
| 11 | 119498403 | rs10892421 | C | T | 1.033 | -7977.388 | -7978.714 | -1.325 | 1.03E-06 |
| 11 | 119948049 | rs1007530 | A | G | 1.031 | -7977.388 | -7978.713 | -1.325 | 1.03E-06 |
| 11 | 116203828 | rs7928488 | T | C | 1.032 | -7977.388 | -7978.714 | -1.326 | 1.03E-06 |
| 11 | 116589652 | rs10892020 | G | C | 1.032 | -7977.388 | -7978.714 | -1.326 | 1.03E-06 |
| 11 | 118443944 | rs11216906 | A | G | 0.971 | -7977.388 | -7978.714 | -1.326 | 1.03E-06 |
| 11 | 118454177 | rs17842763 | G | A | 0.971 | -7977.388 | -7978.714 | -1.326 | 1.03E-06 |
| 11 | 119335915 | rs12793860 | T | G | 1.033 | -7977.388 | -7978.714 | -1.326 | 1.03E-06 |
| 11 | 119337109 | rs12576276 | C | T | 1.033 | -7977.388 | -7978.714 | -1.326 | 1.03E-06 |
| 11 | 119341038 | rs634182 | C | T | 1.033 | -7977.388 | -7978.714 | -1.326 | 1.03E-06 |
| 11 | 119500880 | rs12364363 | C | T | 1.032 | -7977.388 | -7978.714 | -1.326 | 1.03E-06 |
| 11 | 116456027 | rs61904849 | A | G | 1.002 | -7977.388 | -7978.715 | -1.327 | 1.03E-06 |
| 11 | 117863829 | rs2508450 | T | C | 0.969 | -7977.388 | -7978.716 | -1.327 | 1.03E-06 |
| 11 | 118165955 | rs12416826 | C | T | 0.968 | -7977.388 | -7978.716 | -1.327 | 1.03E-06 |
| 11 | 118451901 | rs11216911 | C | T | 0.971 | -7977.388 | -7978.715 | -1.327 | 1.03E-06 |
| 11 | 119784889 | rs12806754 | A | G | 1.02 | -7977.388 | -7978.715 | -1.327 | 1.03E-06 |
| 11 | 119828651 | rs72645469 | A | C | 1.031 | -7977.388 | -7978.715 | -1.327 | 1.03E-06 |
| 11 | 120009308 | rs79761998 | A | C | 0.996 | -7977.388 | -7978.715 | -1.327 | 1.03E-06 |
| 11 | 116344594 | rs11215951 | T | C | 0.976 | -7977.388 | -7978.716 | -1.328 | 1.03E-06 |
| 11 | 117650439 | rs7928767 | A | G | 1.033 | -7977.388 | -7978.716 | -1.328 | 1.03E-06 |
| 11 | 119240071 | rs677008 | A | G | 0.968 | -7977.388 | -7978.717 | -1.328 | 1.03E-06 |
| 11 | 116801529 | rs76690043 | G | C | 0.976 | -7977.388 | -7978.717 | -1.329 | 1.02E-06 |
| 11 | 117428278 | rs562422 | A | G | 1.032 | -7977.388 | -7978.718 | -1.329 | 1.02E-06 |
| 11 | 117919377 | rs12364267 | A | G | 1.002 | -7977.388 | -7978.718 | -1.329 | 1.02E-06 |

| | | | | | | | |
|---|---|---|---|---|---|---|---|
| 11 | 119337241 | rs12575295 | T | C | NULL | 0.660382673 | NULL&M_Disease |
| 11 | 119337381 | rs55935223 | C | T | NULL | 0.660382673 | NULL&M_Disease |
| 11 | 119337436 | rs57737334 | T | A | NULL | 0.660382673 | NULL&M_Disease |
| 11 | 119973302 | rs56313145 | A | G | NULL | 0.659260388 | NULL&M_Disease |
| 11 | 120094385 | rs61898290 | A | G | NULL | 0.659260388 | NULL&M_Disease |
| 11 | 120232766 | rs11217837 | G | A | NULL | 0.660382673 | NULL&M_Disease |
| 11 | 116225035 | rs72993408 | T | A | NULL | 0.660382673 | NULL&M_Disease |
| 11 | 116348112 | rs11824414 | G | A | NULL | 0.660382673 | NULL&M_Disease |
| 11 | 116893790 | rs74691084 | A | G | NULL | 0.660382673 | NULL&M_Disease |
| 11 | 117919746 | rs10892206 | A | T | NULL | 0.660382673 | NULL&M_Disease |
| 11 | 119498748 | rs10892422 | G | A | NULL | 0.660382673 | NULL&M_Disease |
| 11 | 117113422 | rs61903661 | G | A | NULL | 0.660382673 | NULL&M_Disease |
| 11 | 117747277 | rs12790073 | T | C | NULL | 0.660382673 | NULL&M_Disease |
| 11 | 118450046 | rs11216910 | G | A | NULL | 0.660382673 | NULL&M_Disease |
| 11 | 120495204 | rs7949316 | T | C | NULL | 0.660382673 | NULL&M_Disease |
| 11 | 120803734 | rs34607201 | A | G | NULL | 0.660382673 | NULL&M_Disease |
| 11 | 120977529 | rs684384 | A | G | NULL | 0.660382673 | NULL&M_Disease |
| 11 | 118895495 | rs8301 | G | C | NULL | 0.660382673 | NULL&M_Disease |
| 11 | 119338245 | rs7932906 | T | C | NULL | 0.660382673 | NULL&M_Disease |
| 11 | 119838718 | rs10892505 | A | G | NULL | 0.660382673 | NULL&M_Disease |
| 11 | 120067953 | rs17311212 | A | G | NULL | 0.660382673 | NULL&M_Disease |
| 11 | 116485599 | rs476362 | A | G | NULL | 0.660382673 | NULL&M_Disease |
| 11 | 117672330 | rs80120780 | G | C | NULL | 0.660382673 | NULL&M_Disease |
| 11 | 119337199 | rs653177 | A | G | NULL | 0.660382673 | NULL&M_Disease |
| 11 | 119733445 | rs2441939 | T | C | NULL | 0.660382673 | NULL&M_Disease |
| 11 | 120802006 | rs4936565 | A | G | NULL | 0.660382673 | NULL&M_Disease |
| 11 | 117100029 | rs4938362 | A | G | NULL | 0.660382673 | NULL&M_Disease |
| 11 | 117118038 | rs488538 | G | T | NULL | 0.660382673 | NULL&M_Disease |
| 11 | 117525308 | rs61903816 | A | G | NULL | 0.660382673 | NULL&M_Disease |
| 11 | 118144209 | rs12791838 | A | G | NULL | 0.660382673 | NULL&M_Disease |
| 11 | 119824100 | rs76116075 | G | A | NULL | 0.660382673 | NULL&M_Disease |
| 11 | 119882059 | rs588901 | C | T | NULL | 0.660382673 | NULL&M_Disease |
| 11 | 119968674 | rs77262931 | A | G | NULL | 0.660382673 | NULL&M_Disease |
| 11 | 120091653 | rs61898285 | T | C | NULL | 0.660382673 | NULL&M_Disease |
| 11 | 120808289 | rs11218065 | T | G | NULL | 0.660382673 | NULL&M_Disease |
| 11 | 116273511 | rs505128 | C | T | NULL | 0.660382673 | NULL&M_Disease |
| 11 | 116430203 | rs10891989 | C | A | NULL | 0.660382673 | NULL&M_Disease |
| 11 | 116698332 | rs2849172 | T | C | NULL | 0.660382673 | NULL&M_Disease |
| 11 | 118096459 | rs45531834 | T | C | NULL | 0.660382673 | NULL&M_Disease |
| 11 | 119445229 | rs59494484 | T | C | NULL | 0.660382673 | NULL&M_Disease |
| 11 | 119678083 | rs79293063 | A | G | NULL | 0.660382673 | NULL&M_Disease |
| 11 | 120954446 | rs6589854 | G | T | NULL | 0.660382673 | NULL&M_Disease |
| 11 | 116391666 | rs55960868 | A | G | NULL | 0.660382673 | NULL&M_Disease |
| 11 | 119692211 | rs567797 | A | T | NULL | 0.660382673 | NULL&M_Disease |
| 11 | 119714120 | rs78951297 | A | G | NULL | 0.660382673 | NULL&M_Disease |
| 11 | 117010758 | rs2622935 | A | G | NULL | 0.660382673 | NULL&M_Disease |
| 11 | 117339926 | rs638273 | A | G | NULL | 0.660382673 | NULL&M_Disease |
| 11 | 117780744 | rs12801855 | T | C | NULL | 0.660382673 | NULL&M_Disease |
| 11 | 118459039 | rs57438673 | A | G | NULL | 0.660382673 | NULL&M_Disease |
| 11 | 118469571 | rs7931564 | G | A | NULL | 0.660382673 | NULL&M_Disease |
| 11 | 118471423 | rs11216925 | C | T | NULL | 0.660382673 | NULL&M_Disease |
| 11 | 119661317 | rs10892466 | C | G | NULL | 0.660382673 | NULL&M_Disease |
| 11 | 120279132 | rs34616770 | C | G | NULL | 0.660382673 | NULL&M_Disease |
| 11 | 116346978 | rs12294242 | T | C | NULL | 0.660382673 | NULL&M_Disease |
| 11 | 117660275 | rs948100 | G | C | NULL | 0.660382673 | NULL&M_Disease |
| 11 | 119977625 | rs11217695 | T | C | NULL | 0.660382673 | NULL&M_Disease |
| 11 | 120815238 | rs4936566 | A | G | NULL | 0.661503159 | NULL&M_Disease |
| 11 | 120819045 | rs11218074 | G | A | NULL | 0.661503159 | NULL&M_Disease |
| 11 | 120819893 | rs4936567 | T | G | NULL | 0.660382673 | NULL&M_Disease |
| 11 | 118144280 | rs35610956 | G | A | NULL | 0.661503159 | NULL&M_Disease |
| 11 | 119843716 | rs77127262 | T | C | NULL | 0.660382673 | NULL&M_Disease |
| 11 | 120495486 | rs7952478 | T | C | NULL | 0.661503159 | NULL&M_Disease |
| 11 | 116466495 | rs61904853 | G | C | NULL | 0.661503159 | NULL&M_Disease |
| 11 | 116677723 | rs1263167 | G | A | NULL | 0.661503159 | NULL&M_Disease |
| 11 | 118456414 | rs11216914 | C | T | NULL | 0.661503159 | NULL&M_Disease |
| 11 | 119678909 | rs76900966 | T | G | NULL | 0.661503159 | NULL&M_Disease |
| 11 | 119844067 | rs78785412 | T | G | NULL | 0.661503159 | NULL&M_Disease |
| 11 | 117660578 | rs948099 | A | C | NULL | 0.661503159 | NULL&M_Disease |
| 11 | 118462160 | rs11216917 | G | T | NULL | 0.661503159 | NULL&M_Disease |
| 11 | 118465295 | rs11216922 | G | A | NULL | 0.661503159 | NULL&M_Disease |
| 11 | 118465501 | rs11216923 | G | A | NULL | 0.661503159 | NULL&M_Disease |
| 11 | 118486110 | rs45540840 | A | G | NULL | 0.661503159 | NULL&M_Disease |
| 11 | 119836704 | rs41510246 | A | G | NULL | 0.661503159 | NULL&M_Disease |
| 11 | 119838589 | rs74916845 | G | A | NULL | 0.661503159 | NULL&M_Disease |
| 11 | 116836349 | rs77320801 | T | C | NULL | 0.661503159 | NULL&M_Disease |
| 11 | 118449746 | rs11216909 | T | C | NULL | 0.661503159 | NULL&M_Disease |
| 11 | 118712427 | rs7108905 | G | C | NULL | 0.661503159 | NULL&M_Disease |
| 11 | 119836246 | rs79508602 | T | C | NULL | 0.661503159 | NULL&M_Disease |
| 11 | 119836899 | rs4938744 | A | G | NULL | 0.661503159 | NULL&M_Disease |
| 11 | 119837342 | rs4938745 | G | A | NULL | 0.661503159 | NULL&M_Disease |
| 11 | 119838112 | rs1405963027 | G | A | NULL | 0.661503159 | NULL&M_Disease |
| 11 | 116345271 | rs150762661 | C | A | NULL | 0.661503159 | NULL&M_Disease |
| 11 | 117773917 | rs12290911 | T | A | NULL | 0.661503159 | NULL&M_Disease |
| 11 | 119219913 | rs7932565 | G | A | NULL | 0.661503159 | NULL&M_Disease |
| 11 | 119831351 | rs4938742 | G | A | NULL | 0.661503159 | NULL&M_Disease |
| 11 | 119845184 | rs77417358 | T | C | NULL | 0.661503159 | NULL&M_Disease |
| 11 | 120222475 | rs56329702 | T | C | NULL | 0.661503159 | NULL&M_Disease |
| 11 | 116208396 | rs6589541 | G | A | NULL | 0.661503159 | NULL&M_Disease |
| 11 | 118461605 | rs11216916 | C | T | NULL | 0.661503159 | NULL&M_Disease |
| 11 | 119405392 | rs79579780 | G | C | NULL | 0.661503159 | NULL&M_Disease |
| 11 | 119875928 | rs61897871 | G | A | NULL | 0.661503159 | NULL&M_Disease |
| 11 | 120065462 | rs79461323 | G | A | NULL | 0.661503159 | NULL&M_Disease |
| 11 | 116348151 | rs3938035 | G | A | NULL | 0.661503159 | NULL&M_Disease |
| 11 | 116348168 | rs2102405 | G | A | NULL | 0.661503159 | NULL&M_Disease |
| 11 | 116701535 | rs4520 | T | C | NULL | 0.661503159 | NULL&M_Disease |
| 11 | 117374470 | rs2298767 | T | C | NULL | 0.661503159 | NULL&M_Disease |

| | | | | | | | | | |
|---|---|---|---|---|---|---|---|---|---|
| 11 | 119337241 | rs12575295 | T | C | 1.032 | -7977.388 | -7978.718 | -1.329 | 1.02E-06 |
| 11 | 119337381 | rs55935223 | C | T | 1.032 | -7977.388 | -7978.718 | -1.329 | 1.02E-06 |
| 11 | 119337436 | rs57737334 | T | A | 1.032 | -7977.388 | -7978.718 | -1.329 | 1.02E-06 |
| 11 | 119973302 | rs56313145 | A | G | 1.006 | -7977.388 | -7978.717 | -1.329 | 1.02E-06 |
| 11 | 120094385 | rs61898290 | A | G | 0.985 | -7977.388 | -7978.717 | -1.329 | 1.02E-06 |
| 11 | 120232766 | rs11217837 | G | A | 1.032 | -7977.388 | -7978.718 | -1.329 | 1.02E-06 |
| 11 | 116225035 | rs72993408 | T | A | 1.02 | -7977.388 | -7978.718 | -1.33 | 1.02E-06 |
| 11 | 116348112 | rs11824414 | G | A | 0.982 | -7977.388 | -7978.718 | -1.33 | 1.02E-06 |
| 11 | 116893790 | rs74691084 | A | G | 0.98 | -7977.388 | -7978.719 | -1.33 | 1.02E-06 |
| 11 | 117919746 | rs10892206 | A | T | 1.001 | -7977.388 | -7978.718 | -1.33 | 1.02E-06 |
| 11 | 119498748 | rs10892422 | G | A | 1.032 | -7977.388 | -7978.719 | -1.33 | 1.02E-06 |
| 11 | 117113422 | rs61903661 | G | A | 0.973 | -7977.388 | -7978.719 | -1.331 | 1.02E-06 |
| 11 | 117747277 | rs12790073 | T | C | 1.004 | -7977.388 | -7978.719 | -1.331 | 1.02E-06 |
| 11 | 118450046 | rs11216910 | G | A | 0.972 | -7977.388 | -7978.719 | -1.331 | 1.02E-06 |
| 11 | 120495204 | rs7949316 | T | C | 0.968 | -7977.388 | -7978.719 | -1.331 | 1.02E-06 |
| 11 | 120803734 | rs34607201 | A | G | 1.032 | -7977.388 | -7978.719 | -1.331 | 1.02E-06 |
| 11 | 120977529 | rs684384 | A | G | 0.969 | -7977.388 | -7978.719 | -1.331 | 1.02E-06 |
| 11 | 118895495 | rs8301 | G | C | 1.032 | -7977.388 | -7978.721 | -1.332 | 1.02E-06 |
| 11 | 119338245 | rs7932906 | T | C | 1.032 | -7977.388 | -7978.721 | -1.332 | 1.02E-06 |
| 11 | 119838718 | rs10892505 | A | G | 0.968 | -7977.388 | -7978.72 | -1.332 | 1.02E-06 |
| 11 | 120067953 | rs17311212 | A | G | 0.973 | -7977.388 | -7978.72 | -1.332 | 1.02E-06 |
| 11 | 116485599 | rs476362 | A | G | 1.027 | -7977.388 | -7978.721 | -1.333 | 1.02E-06 |
| 11 | 117672330 | rs80120780 | G | C | 0.977 | -7977.388 | -7978.722 | -1.333 | 1.02E-06 |
| 11 | 119337199 | rs653177 | A | G | 1.032 | -7977.388 | -7978.721 | -1.333 | 1.02E-06 |
| 11 | 119733445 | rs2441939 | T | C | 1.03 | -7977.388 | -7978.721 | -1.333 | 1.02E-06 |
| 11 | 120802006 | rs4936565 | A | G | 0.968 | -7977.388 | -7978.722 | -1.333 | 1.02E-06 |
| 11 | 117100029 | rs4938362 | A | G | 0.97 | -7977.388 | -7978.722 | -1.334 | 1.02E-06 |
| 11 | 117118038 | rs488538 | G | T | 0.968 | -7977.388 | -7978.722 | -1.334 | 1.02E-06 |
| 11 | 117525308 | rs61903816 | A | G | 0.998 | -7977.388 | -7978.722 | -1.334 | 1.02E-06 |
| 11 | 118144209 | rs12791838 | A | G | 0.969 | -7977.388 | -7978.722 | -1.334 | 1.02E-06 |
| 11 | 119824100 | rs76116075 | G | A | 1.006 | -7977.388 | -7978.722 | -1.334 | 1.02E-06 |
| 11 | 119882059 | rs588901 | C | T | 0.969 | -7977.388 | -7978.722 | -1.334 | 1.02E-06 |
| 11 | 119968674 | rs77262931 | A | G | 1.025 | -7977.388 | -7978.722 | -1.334 | 1.02E-06 |
| 11 | 120091653 | rs61898285 | T | C | 0.986 | -7977.388 | -7978.722 | -1.334 | 1.02E-06 |
| 11 | 120808289 | rs11218065 | T | G | 1.031 | -7977.388 | -7978.723 | -1.334 | 1.02E-06 |
| 11 | 116273511 | rs505128 | C | T | 1.032 | -7977.388 | -7978.723 | -1.335 | 1.02E-06 |
| 11 | 116430203 | rs10891989 | C | A | 0.986 | -7977.388 | -7978.724 | -1.335 | 1.02E-06 |
| 11 | 116698332 | rs2849172 | T | C | 1.032 | -7977.388 | -7978.723 | -1.335 | 1.02E-06 |
| 11 | 118096459 | rs45531834 | T | C | 0.975 | -7977.388 | -7978.723 | -1.335 | 1.02E-06 |
| 11 | 119445229 | rs59494484 | T | C | 0.986 | -7977.388 | -7978.723 | -1.335 | 1.02E-06 |
| 11 | 119678083 | rs79293063 | A | G | 0.991 | -7977.388 | -7978.724 | -1.335 | 1.02E-06 |
| 11 | 120954446 | rs6589854 | G | T | 1.032 | -7977.388 | -7978.723 | -1.335 | 1.02E-06 |
| 11 | 116391666 | rs55960868 | A | G | 1.032 | -7977.388 | -7978.724 | -1.336 | 1.02E-06 |
| 11 | 119692211 | rs567797 | A | T | 0.969 | -7977.388 | -7978.724 | -1.336 | 1.02E-06 |
| 11 | 119714120 | rs78951297 | A | G | 0.969 | -7977.388 | -7978.724 | -1.336 | 1.02E-06 |
| 11 | 117010758 | rs2622935 | A | G | 0.969 | -7977.388 | -7978.725 | -1.337 | 1.02E-06 |
| 11 | 117339926 | rs638273 | A | G | 0.969 | -7977.388 | -7978.725 | -1.337 | 1.02E-06 |
| 11 | 117780744 | rs12801855 | T | C | 1.032 | -7977.388 | -7978.725 | -1.337 | 1.02E-06 |
| 11 | 118459039 | rs57438673 | A | G | 0.972 | -7977.388 | -7978.725 | -1.337 | 1.02E-06 |
| 11 | 118469571 | rs7931564 | G | A | 0.972 | -7977.388 | -7978.725 | -1.337 | 1.02E-06 |
| 11 | 118471423 | rs11216925 | C | T | 0.972 | -7977.388 | -7978.725 | -1.337 | 1.02E-06 |
| 11 | 119661317 | rs10892466 | C | G | 0.973 | -7977.388 | -7978.725 | -1.337 | 1.02E-06 |
| 11 | 120279132 | rs34616770 | C | G | 0.972 | -7977.388 | -7978.726 | -1.337 | 1.02E-06 |
| 11 | 116346978 | rs12294242 | T | C | 0.983 | -7977.388 | -7978.727 | -1.338 | 1.01E-06 |
| 11 | 117660275 | rs948100 | G | C | 0.977 | -7977.388 | -7978.726 | -1.338 | 1.01E-06 |
| 11 | 119977625 | rs11217695 | T | C | 1.007 | -7977.388 | -7978.726 | -1.338 | 1.01E-06 |
| 11 | 120815238 | rs4936566 | A | G | 0.969 | -7977.388 | -7978.727 | -1.338 | 1.01E-06 |
| 11 | 120819045 | rs11218074 | G | A | 0.969 | -7977.388 | -7978.727 | -1.338 | 1.01E-06 |
| 11 | 120819893 | rs4936567 | T | G | 1.031 | -7977.388 | -7978.726 | -1.338 | 1.01E-06 |
| 11 | 118144280 | rs35610956 | G | A | 0.969 | -7977.388 | -7978.727 | -1.339 | 1.01E-06 |
| 11 | 119843716 | rs77127262 | T | C | 0.995 | -7977.388 | -7978.727 | -1.339 | 1.01E-06 |
| 11 | 120495486 | rs7952478 | T | C | 0.969 | -7977.388 | -7978.727 | -1.339 | 1.01E-06 |
| 11 | 116466495 | rs61904853 | G | C | 0.969 | -7977.388 | -7978.729 | -1.34 | 1.01E-06 |
| 11 | 116677723 | rs1263167 | G | A | 1.031 | -7977.388 | -7978.728 | -1.34 | 1.01E-06 |
| 11 | 118456414 | rs11216914 | C | T | 0.972 | -7977.388 | -7978.728 | -1.34 | 1.01E-06 |
| 11 | 119678909 | rs76900966 | T | G | 0.992 | -7977.388 | -7978.728 | -1.34 | 1.01E-06 |
| 11 | 119844067 | rs78785412 | T | G | 0.995 | -7977.388 | -7978.728 | -1.34 | 1.01E-06 |
| 11 | 117660578 | rs948099 | A | C | 0.977 | -7977.388 | -7978.729 | -1.341 | 1.01E-06 |
| 11 | 118462160 | rs11216917 | G | T | 0.972 | -7977.388 | -7978.729 | -1.341 | 1.01E-06 |
| 11 | 118465295 | rs11216922 | G | A | 0.972 | -7977.388 | -7978.729 | -1.341 | 1.01E-06 |
| 11 | 118465501 | rs11216923 | G | A | 0.972 | -7977.388 | -7978.729 | -1.341 | 1.01E-06 |
| 11 | 118486110 | rs45540840 | A | G | 1.032 | -7977.388 | -7978.729 | -1.341 | 1.01E-06 |
| 11 | 119836704 | rs41510246 | A | G | 1 | -7977.388 | -7978.73 | -1.341 | 1.01E-06 |
| 11 | 119838589 | rs74916845 | G | A | 1 | -7977.388 | -7978.73 | -1.341 | 1.01E-06 |
| 11 | 116836349 | rs77320801 | T | C | 0.977 | -7977.388 | -7978.73 | -1.342 | 1.01E-06 |
| 11 | 118449746 | rs11216909 | T | C | 0.972 | -7977.388 | -7978.731 | -1.342 | 1.01E-06 |
| 11 | 118712427 | rs7108905 | G | C | 1.031 | -7977.388 | -7978.731 | -1.342 | 1.01E-06 |
| 11 | 119836246 | rs79508602 | T | C | 1.001 | -7977.388 | -7978.73 | -1.342 | 1.01E-06 |
| 11 | 119836899 | rs4938744 | A | G | 1.001 | -7977.388 | -7978.73 | -1.342 | 1.01E-06 |
| 11 | 119837342 | rs4938745 | G | A | 1.001 | -7977.388 | -7978.73 | -1.342 | 1.01E-06 |
| 11 | 119838112 | rs1405963027 | G | A | 1.001 | -7977.388 | -7978.73 | -1.342 | 1.01E-06 |
| 11 | 116345271 | rs150762661 | C | A | 0.997 | -7977.388 | -7978.732 | -1.343 | 1.01E-06 |
| 11 | 117773917 | rs12290911 | T | A | 0.969 | -7977.388 | -7978.731 | -1.343 | 1.01E-06 |
| 11 | 119219913 | rs7932565 | G | A | 0.969 | -7977.388 | -7978.731 | -1.343 | 1.01E-06 |
| 11 | 119831351 | rs4938742 | G | A | 1.003 | -7977.388 | -7978.731 | -1.343 | 1.01E-06 |
| 11 | 119845184 | rs77417358 | T | C | 0.998 | -7977.388 | -7978.731 | -1.343 | 1.01E-06 |
| 11 | 120222475 | rs56329702 | T | C | 1.032 | -7977.388 | -7978.732 | -1.343 | 1.01E-06 |
| 11 | 116208396 | rs6589541 | G | A | 1.032 | -7977.388 | -7978.733 | -1.344 | 1.01E-06 |
| 11 | 118461605 | rs11216916 | C | T | 0.972 | -7977.388 | -7978.733 | -1.345 | 1.01E-06 |
| 11 | 119405392 | rs79579780 | G | C | 0.995 | -7977.388 | -7978.733 | -1.345 | 1.01E-06 |
| 11 | 119875928 | rs61897871 | G | A | 1.009 | -7977.388 | -7978.733 | -1.345 | 1.01E-06 |
| 11 | 120065462 | rs79461323 | G | A | 0.973 | -7977.388 | -7978.733 | -1.345 | 1.01E-06 |
| 11 | 116348151 | rs3938035 | G | A | 0.984 | -7977.388 | -7978.734 | -1.346 | 1.01E-06 |
| 11 | 116348168 | rs2102405 | G | A | 0.984 | -7977.388 | -7978.734 | -1.346 | 1.01E-06 |
| 11 | 116701535 | rs4520 | T | C | 0.969 | -7977.388 | -7978.734 | -1.346 | 1.01E-06 |
| 11 | 117374470 | rs2298767 | T | C | 1.026 | -7977.388 | -7978.734 | -1.346 | 1.01E-06 |

| | | | | | | | |
|---|---|---|---|---|---|---|---|
| 11 | 118515876 | rs582942 | T | C | NULL | 0.661503159 | NULL&M_Disease |
| 11 | 119893179 | rs655929 | C | A | NULL | 0.661503159 | NULL&M_Disease |
| 11 | 120622385 | rs12804621 | T | C | NULL | 0.661503159 | NULL&M_Disease |
| 11 | 121091762 | rs948091 | G | C | NULL | 0.661503159 | NULL&M_Disease |
| 11 | 116229491 | rs73003797 | C | T | NULL | 0.661503159 | NULL&M_Disease |
| 11 | 116707338 | rs7116797 | A | G | NULL | 0.661503159 | NULL&M_Disease |
| 11 | 117688433 | rs73018457 | T | C | NULL | 0.661503159 | NULL&M_Disease |
| 11 | 117775895 | rs35284897 | A | C | NULL | 0.662621837 | NULL&M_Disease |
| 11 | 118169825 | rs11607701 | G | A | NULL | 0.662621837 | NULL&M_Disease |
| 11 | 120651537 | rs10892624 | A | G | NULL | 0.662621837 | NULL&M_Disease |
| 11 | 116426350 | rs11216004 | T | C | NULL | 0.662621837 | NULL&M_Disease |
| 11 | 117953050 | rs688803 | T | C | NULL | 0.662621837 | NULL&M_Disease |
| 11 | 118448087 | rs9735449 | T | C | NULL | 0.662621837 | NULL&M_Disease |
| 11 | 120161886 | rs11217793 | T | C | NULL | 0.662621837 | NULL&M_Disease |
| 11 | 120187971 | rs2282537 | A | G | NULL | 0.662621837 | NULL&M_Disease |
| 11 | 120614538 | rs1892975 | T | A | NULL | 0.662621837 | NULL&M_Disease |
| 11 | 117557380 | rs1939234 | T | C | NULL | 0.662621837 | NULL&M_Disease |
| 11 | 117707229 | rs10892184 | T | C | NULL | 0.662621837 | NULL&M_Disease |
| 11 | 120267537 | rs11217854 | G | A | NULL | 0.662621837 | NULL&M_Disease |
| 11 | 120600151 | rs1939667 | T | C | NULL | 0.662621837 | NULL&M_Disease |
| 11 | 116367859 | rs12278200 | C | G | NULL | 0.662621837 | NULL&M_Disease |
| 11 | 116677371 | rs1263166 | A | G | NULL | 0.662621837 | NULL&M_Disease |
| 11 | 116718521 | rs1145206 | C | G | NULL | 0.662621837 | NULL&M_Disease |
| 11 | 116823672 | rs1473326 | C | T | NULL | 0.662621837 | NULL&M_Disease |
| 11 | 118142666 | rs7940405 | G | A | NULL | 0.662621837 | NULL&M_Disease |
| 11 | 118872404 | rs6589713 | C | T | NULL | 0.662621837 | NULL&M_Disease |
| 11 | 119514810 | rs12788928 | A | G | NULL | 0.662621837 | NULL&M_Disease |
| 11 | 119880361 | rs617139 | T | C | NULL | 0.662621837 | NULL&M_Disease |
| 11 | 116221255 | rs73010962 | G | A | NULL | 0.662621837 | NULL&M_Disease |
| 11 | 116342547 | rs1605557 | G | A | NULL | 0.662621837 | NULL&M_Disease |
| 11 | 119239375 | rs573396 | G | A | NULL | 0.662621837 | NULL&M_Disease |
| 11 | 119334628 | rs12419497 | T | C | NULL | 0.662621837 | NULL&M_Disease |
| 11 | 120636167 | rs7104454 | G | A | NULL | 0.662621837 | NULL&M_Disease |
| 11 | 120798714 | rs2005529 | C | G | NULL | 0.662621837 | NULL&M_Disease |
| 11 | 117110417 | rs73010409 | C | T | NULL | 0.662621837 | NULL&M_Disease |
| 11 | 117700201 | rs61903779 | G | A | NULL | 0.662621837 | NULL&M_Disease |
| 11 | 120082486 | rs2444242 | A | G | NULL | 0.662621837 | NULL&M_Disease |
| 11 | 120169818 | rs1893262 | C | T | NULL | 0.662621837 | NULL&M_Disease |
| 11 | 120556341 | rs4445646 | C | T | NULL | 0.662621837 | NULL&M_Disease |
| 11 | 120823074 | rs73007893 | A | G | NULL | 0.662621837 | NULL&M_Disease |
| 11 | 120823269 | rs73007896 | T | C | NULL | 0.662621837 | NULL&M_Disease |
| 11 | 120905810 | rs12803528 | A | C | NULL | 0.662621837 | NULL&M_Disease |
| 11 | 116403238 | rs12146701 | G | T | NULL | 0.662621837 | NULL&M_Disease |
| 11 | 116711350 | rs11216158 | A | G | NULL | 0.662621837 | NULL&M_Disease |
| 11 | 118459702 | rs113054880 | G | C | NULL | 0.662621837 | NULL&M_Disease |
| 11 | 118473385 | rs17742 | C | T | NULL | 0.662621837 | NULL&M_Disease |
| 11 | 118837272 | rs3901422 | A | G | NULL | 0.662621837 | NULL&M_Disease |
| 11 | 119836955 | rs10790358 | G | A | NULL | 0.662621837 | NULL&M_Disease |
| 11 | 120248493 | rs58073046 | G | A | NULL | 0.662621837 | NULL&M_Disease |
| 11 | 116328809 | rs10437673 | A | G | NULL | 0.662621837 | NULL&M_Disease |
| 11 | 117534453 | rs61903817 | G | C | NULL | 0.662621837 | NULL&M_Disease |
| 11 | 118530611 | rs2276065 | C | T | NULL | 0.662621837 | NULL&M_Disease |
| 11 | 118846482 | rs4938603 | G | A | NULL | 0.662621837 | NULL&M_Disease |
| 11 | 119951167 | rs11217676 | G | C | NULL | 0.662621837 | NULL&M_Disease |
| 11 | 120113661 | rs2845702 | G | T | NULL | 0.662621837 | NULL&M_Disease |
| 11 | 120113664 | rs2847498 | C | T | NULL | 0.662621837 | NULL&M_Disease |
| 11 | 120977922 | rs638775 | G | C | NULL | 0.662621837 | NULL&M_Disease |
| 11 | 116711180 | rs11216157 | G | A | NULL | 0.662621837 | NULL&M_Disease |
| 11 | 117111178 | rs140479252 | C | T | NULL | 0.662621837 | NULL&M_Disease |
| 11 | 117358220 | rs549289 | C | A | NULL | 0.662621837 | NULL&M_Disease |
| 11 | 118153697 | rs2032397 | A | G | NULL | 0.662621837 | NULL&M_Disease |
| 11 | 118153708 | rs2032398 | C | G | NULL | 0.662621837 | NULL&M_Disease |
| 11 | 118708785 | rs7122510 | A | C | NULL | 0.662621837 | NULL&M_Disease |
| 11 | 118708786 | rs7122511 | A | C | NULL | 0.662621837 | NULL&M_Disease |
| 11 | 119501639 | rs1013366746 | T | C | NULL | 0.662621837 | NULL&M_Disease |
| 11 | 120321201 | rs11217872 | C | G | NULL | 0.662621837 | NULL&M_Disease |
| 11 | 117112138 | rs4938365 | C | A | NULL | 0.662621837 | NULL&M_Disease |
| 11 | 117595599 | rs11216518 | T | G | NULL | 0.662621837 | NULL&M_Disease |
| 11 | 116535126 | rs2105610 | C | T | NULL | 0.662621837 | NULL&M_Disease |
| 11 | 116604070 | rs180353 | C | T | NULL | 0.663738697 | NULL&M_Disease |
| 11 | 117698483 | rs11216567 | G | A | NULL | 0.662621837 | NULL&M_Disease |
| 11 | 118316960 | rs498684 | A | G | NULL | 0.663738697 | NULL&M_Disease |
| 11 | 119824399 | rs75401133 | T | C | NULL | 0.662621837 | NULL&M_Disease |
| 11 | 120904249 | rs17245789 | G | A | NULL | 0.662621837 | NULL&M_Disease |
| 11 | 116513182 | rs1145184 | A | G | NULL | 0.663738697 | NULL&M_Disease |
| 11 | 119527227 | rs9651691 | T | G | NULL | 0.663738697 | NULL&M_Disease |
| 11 | 120823608 | rs2298725 | A | G | NULL | 0.663738697 | NULL&M_Disease |
| 11 | 116194551 | rs4245166 | G | A | NULL | 0.663738697 | NULL&M_Disease |
| 11 | 116484093 | rs542987 | G | A | NULL | 0.663738697 | NULL&M_Disease |
| 11 | 117199870 | rs509894 | T | C | NULL | 0.663738697 | NULL&M_Disease |
| 11 | 117409700 | rs11216443 | A | C | NULL | 0.663738697 | NULL&M_Disease |
| 11 | 119080037 | rs2155380 | G | A | NULL | 0.663738697 | NULL&M_Disease |
| 11 | 120694754 | rs11218016 | T | C | NULL | 0.663738697 | NULL&M_Disease |
| 11 | 117705239 | rs57811256 | C | G | NULL | 0.663738697 | NULL&M_Disease |
| 11 | 118472354 | rs10892245 | A | C | NULL | 0.663738697 | NULL&M_Disease |
| 11 | 116227283 | rs4082031 | A | T | NULL | 0.663738697 | NULL&M_Disease |
| 11 | 116247944 | rs17567197 | C | A | NULL | 0.663738697 | NULL&M_Disease |
| 11 | 116306360 | rs57963928 | C | A | NULL | 0.663738697 | NULL&M_Disease |
| 11 | 117818680 | rs2515797 | G | A | NULL | 0.663738697 | NULL&M_Disease |
| 11 | 117951521 | rs608365 | C | G | NULL | 0.663738697 | NULL&M_Disease |
| 11 | 120921962 | rs17245796 | G | A | NULL | 0.663738697 | NULL&M_Disease |
| 11 | 116392920 | rs66879136 | A | C | NULL | 0.663738697 | NULL&M_Disease |
| 11 | 116458660 | rs11216021 | T | C | NULL | 0.663738697 | NULL&M_Disease |
| 11 | 117099537 | rs111809212 | C | T | NULL | 0.663738697 | NULL&M_Disease |
| 11 | 117666548 | rs76377040 | G | C | NULL | 0.663738697 | NULL&M_Disease |
| 11 | 117630867 | rs11825655 | G | A | NULL | 0.663738697 | NULL&M_Disease |

| | | | | | | | | | |
|---|---|---|---|---|---|---|---|---|---|
| 11 | 118515876 | rs582942 | T | C | 1.031 | -7977.388 | -7978.734 | -1.346 | 1.01E-06 |
| 11 | 119893179 | rs655929 | C | A | 0.969 | -7977.388 | -7978.735 | -1.346 | 1.01E-06 |
| 11 | 120622385 | rs12804621 | T | C | 1.032 | -7977.388 | -7978.735 | -1.346 | 1.01E-06 |
| 11 | 121091762 | rs948091 | G | C | 0.969 | -7977.388 | -7978.734 | -1.346 | 1.01E-06 |
| 11 | 116229491 | rs73003797 | C | T | 1.018 | -7977.388 | -7978.735 | -1.347 | 1.01E-06 |
| 11 | 116707338 | rs7116797 | A | G | 0.976 | -7977.388 | -7978.736 | -1.347 | 1.01E-06 |
| 11 | 117688433 | rs73018457 | T | C | 1.016 | -7977.388 | -7978.736 | -1.348 | 1.00E-06 |
| 11 | 117775895 | rs35284897 | A | C | 0.969 | -7977.388 | -7978.736 | -1.348 | 1.00E-06 |
| 11 | 118169825 | rs11607701 | G | A | 0.969 | -7977.388 | -7978.737 | -1.348 | 1.00E-06 |
| 11 | 120651537 | rs10892624 | A | G | 0.971 | -7977.388 | -7978.736 | -1.348 | 1.00E-06 |
| 11 | 116426350 | rs11216004 | T | C | 0.988 | -7977.388 | -7978.738 | -1.349 | 1.00E-06 |
| 11 | 117953050 | rs688803 | T | C | 1.025 | -7977.388 | -7978.737 | -1.349 | 1.00E-06 |
| 11 | 118448087 | rs9735449 | T | C | 0.973 | -7977.388 | -7978.738 | -1.349 | 1.00E-06 |
| 11 | 120161886 | rs11217793 | T | C | 1.031 | -7977.388 | -7978.738 | -1.349 | 1.00E-06 |
| 11 | 120187971 | rs2282537 | A | G | 1.029 | -7977.388 | -7978.737 | -1.349 | 1.00E-06 |
| 11 | 120614538 | rs1892975 | T | A | 0.969 | -7977.388 | -7978.737 | -1.349 | 1.00E-06 |
| 11 | 117557380 | rs1939234 | T | C | 0.97 | -7977.388 | -7978.738 | -1.35 | 1.00E-06 |
| 11 | 117707229 | rs10892184 | T | C | 1.032 | -7977.388 | -7978.738 | -1.35 | 1.00E-06 |
| 11 | 120267537 | rs11217854 | G | A | 0.973 | -7977.388 | -7978.738 | -1.35 | 1.00E-06 |
| 11 | 120600151 | rs1939667 | T | C | 0.969 | -7977.388 | -7978.738 | -1.35 | 1.00E-06 |
| 11 | 116367859 | rs12278200 | C | G | 0.97 | -7977.388 | -7978.739 | -1.351 | 1.00E-06 |
| 11 | 116677371 | rs1263166 | A | G | 1.031 | -7977.388 | -7978.739 | -1.351 | 1.00E-06 |
| 11 | 116718521 | rs1145206 | C | G | 0.971 | -7977.388 | -7978.739 | -1.351 | 1.00E-06 |
| 11 | 116823672 | rs1473326 | C | T | 0.978 | -7977.388 | -7978.739 | -1.351 | 1.00E-06 |
| 11 | 118142666 | rs7940405 | G | A | 0.969 | -7977.388 | -7978.74 | -1.351 | 1.00E-06 |
| 11 | 118872404 | rs6589713 | C | T | 1.032 | -7977.388 | -7978.739 | -1.351 | 1.00E-06 |
| 11 | 119514810 | rs12788928 | A | G | 1.01 | -7977.388 | -7978.74 | -1.351 | 1.00E-06 |
| 11 | 119880361 | rs617139 | T | C | 0.986 | -7977.388 | -7978.739 | -1.351 | 1.00E-06 |
| 11 | 116221255 | rs73010962 | G | A | 1.017 | -7977.388 | -7978.74 | -1.352 | 1.00E-06 |
| 11 | 116342547 | rs1605557 | G | A | 0.988 | -7977.388 | -7978.74 | -1.352 | 1.00E-06 |
| 11 | 119239375 | rs573396 | G | A | 0.969 | -7977.388 | -7978.74 | -1.352 | 1.00E-06 |
| 11 | 119334628 | rs12419497 | T | C | 1.032 | -7977.388 | -7978.74 | -1.352 | 1.00E-06 |
| 11 | 120636167 | rs7104454 | G | A | 0.971 | -7977.388 | -7978.74 | -1.352 | 1.00E-06 |
| 11 | 120798714 | rs2005529 | C | G | 1.032 | -7977.388 | -7978.74 | -1.352 | 1.00E-06 |
| 11 | 117110417 | rs73010409 | C | T | 0.974 | -7977.388 | -7978.741 | -1.353 | 1.00E-06 |
| 11 | 117700201 | rs61903779 | G | A | 0.992 | -7977.388 | -7978.742 | -1.353 | 1.00E-06 |
| 11 | 120082486 | rs2444242 | A | G | 0.992 | -7977.388 | -7978.741 | -1.353 | 1.00E-06 |
| 11 | 120169818 | rs1893262 | C | T | 1.01 | -7977.388 | -7978.742 | -1.353 | 1.00E-06 |
| 11 | 120556341 | rs4445646 | C | T | 1.032 | -7977.388 | -7978.741 | -1.353 | 1.00E-06 |
| 11 | 120823074 | rs73007893 | A | G | 0.969 | -7977.388 | -7978.742 | -1.353 | 1.00E-06 |
| 11 | 120823269 | rs73007896 | T | C | 0.969 | -7977.388 | -7978.742 | -1.353 | 1.00E-06 |
| 11 | 120905810 | rs12803528 | A | C | 0.982 | -7977.388 | -7978.742 | -1.353 | 1.00E-06 |
| 11 | 116403238 | rs12146701 | G | T | 0.976 | -7977.388 | -7978.743 | -1.354 | 9.99E-07 |
| 11 | 116711350 | rs11216158 | A | G | 0.973 | -7977.388 | -7978.742 | -1.354 | 9.99E-07 |
| 11 | 118459702 | rs113054880 | G | C | 0.973 | -7977.388 | -7978.743 | -1.354 | 9.99E-07 |
| 11 | 118473385 | rs17742 | C | T | 0.973 | -7977.388 | -7978.742 | -1.354 | 9.99E-07 |
| 11 | 118837272 | rs3901422 | A | G | 1.032 | -7977.388 | -7978.742 | -1.354 | 9.99E-07 |
| 11 | 119836955 | rs10790358 | G | A | 0.969 | -7977.388 | -7978.743 | -1.354 | 9.99E-07 |
| 11 | 120248493 | rs58073046 | G | A | 1.025 | -7977.388 | -7978.743 | -1.354 | 9.99E-07 |
| 11 | 116328809 | rs10437673 | A | G | 1.013 | -7977.388 | -7978.743 | -1.355 | 9.98E-07 |
| 11 | 117534453 | rs61903817 | G | C | 1.004 | -7977.388 | -7978.744 | -1.355 | 9.98E-07 |
| 11 | 118530611 | rs2276065 | C | T | 1.031 | -7977.388 | -7978.744 | -1.355 | 9.98E-07 |
| 11 | 118846482 | rs4938603 | G | A | 1.032 | -7977.388 | -7978.744 | -1.355 | 9.98E-07 |
| 11 | 119951167 | rs11217676 | G | C | 1.031 | -7977.388 | -7978.743 | -1.355 | 9.98E-07 |
| 11 | 120113661 | rs2845702 | G | T | 1.014 | -7977.388 | -7978.743 | -1.355 | 9.98E-07 |
| 11 | 120113664 | rs2847498 | C | T | 1.032 | -7977.388 | -7978.743 | -1.355 | 9.98E-07 |
| 11 | 120977922 | rs638775 | G | C | 0.969 | -7977.388 | -7978.744 | -1.355 | 9.98E-07 |
| 11 | 116711180 | rs11216157 | G | A | 0.973 | -7977.388 | -7978.744 | -1.356 | 9.97E-07 |
| 11 | 117111178 | rs140479252 | C | T | 0.974 | -7977.388 | -7978.744 | -1.356 | 9.97E-07 |
| 11 | 117358220 | rs549289 | C | A | 0.97 | -7977.388 | -7978.744 | -1.356 | 9.97E-07 |
| 11 | 118153697 | rs2032397 | A | G | 0.969 | -7977.388 | -7978.745 | -1.356 | 9.97E-07 |
| 11 | 118153708 | rs2032398 | C | G | 0.969 | -7977.388 | -7978.745 | -1.356 | 9.97E-07 |
| 11 | 118708785 | rs7122510 | A | C | 1.03 | -7977.388 | -7978.744 | -1.356 | 9.97E-07 |
| 11 | 118708786 | rs7122511 | A | C | 1.03 | -7977.388 | -7978.744 | -1.356 | 9.97E-07 |
| 11 | 119501639 | rs1013366746 | T | C | 0.969 | -7977.388 | -7978.744 | -1.356 | 9.97E-07 |
| 11 | 120321201 | rs11217872 | C | G | 1.031 | -7977.388 | -7978.744 | -1.356 | 9.97E-07 |
| 11 | 117112138 | rs4938365 | C | A | 0.975 | -7977.388 | -7978.745 | -1.357 | 9.96E-07 |
| 11 | 117595599 | rs11216518 | T | G | 1.03 | -7977.388 | -7978.745 | -1.357 | 9.96E-07 |
| 11 | 116535126 | rs2105610 | C | T | 1.022 | -7977.388 | -7978.746 | -1.358 | 9.95E-07 |
| 11 | 116604070 | rs180353 | C | T | 1.031 | -7977.388 | -7978.747 | -1.358 | 9.95E-07 |
| 11 | 117698483 | rs11216567 | G | A | 0.988 | -7977.388 | -7978.746 | -1.358 | 9.95E-07 |
| 11 | 118316960 | rs498684 | A | G | 0.97 | -7977.388 | -7978.746 | -1.358 | 9.95E-07 |
| 11 | 119824399 | rs75401133 | T | C | 1.015 | -7977.388 | -7978.746 | -1.358 | 9.95E-07 |
| 11 | 120904249 | rs17245789 | G | A | 0.983 | -7977.388 | -7978.746 | -1.358 | 9.95E-07 |
| 11 | 116513182 | rs1145184 | A | G | 0.971 | -7977.388 | -7978.748 | -1.359 | 9.94E-07 |
| 11 | 119527227 | rs9651691 | T | G | 0.969 | -7977.388 | -7978.748 | -1.359 | 9.94E-07 |
| 11 | 120823608 | rs2298725 | A | G | 1.032 | -7977.388 | -7978.747 | -1.359 | 9.94E-07 |
| 11 | 116194551 | rs4245166 | G | A | 1.031 | -7977.388 | -7978.748 | -1.36 | 9.93E-07 |
| 11 | 116484093 | rs542987 | G | A | 0.981 | -7977.388 | -7978.749 | -1.36 | 9.93E-07 |
| 11 | 117199870 | rs509894 | T | C | 0.969 | -7977.388 | -7978.749 | -1.36 | 9.93E-07 |
| 11 | 117409700 | rs11216443 | A | C | 0.969 | -7977.388 | -7978.748 | -1.36 | 9.93E-07 |
| 11 | 119080037 | rs2155380 | G | A | 0.969 | -7977.388 | -7978.748 | -1.36 | 9.93E-07 |
| 11 | 120694754 | rs11218016 | T | C | 1.032 | -7977.388 | -7978.748 | -1.36 | 9.93E-07 |
| 11 | 117705239 | rs57811256 | C | G | 0.971 | -7977.388 | -7978.749 | -1.361 | 9.92E-07 |
| 11 | 118472354 | rs10892245 | A | C | 0.973 | -7977.388 | -7978.749 | -1.361 | 9.92E-07 |
| 11 | 116227283 | rs4082031 | A | T | 1.013 | -7977.388 | -7978.75 | -1.362 | 9.91E-07 |
| 11 | 116247944 | rs17567197 | C | A | 0.981 | -7977.388 | -7978.751 | -1.362 | 9.91E-07 |
| 11 | 116306360 | rs57963928 | C | A | 1.008 | -7977.388 | -7978.751 | -1.362 | 9.91E-07 |
| 11 | 117818680 | rs2515797 | G | A | 1.031 | -7977.388 | -7978.75 | -1.362 | 9.91E-07 |
| 11 | 117951521 | rs608365 | C | G | 1.024 | -7977.388 | -7978.75 | -1.362 | 9.91E-07 |
| 11 | 120921962 | rs17245796 | G | A | 0.983 | -7977.388 | -7978.751 | -1.362 | 9.91E-07 |
| 11 | 116392920 | rs66879136 | A | C | 1.03 | -7977.388 | -7978.752 | -1.363 | 9.90E-07 |
| 11 | 116458660 | rs11216021 | T | C | 1.031 | -7977.388 | -7978.751 | -1.363 | 9.90E-07 |
| 11 | 117099537 | rs111809212 | C | T | 0.971 | -7977.388 | -7978.751 | -1.363 | 9.90E-07 |
| 11 | 117666548 | rs76377040 | G | C | 0.976 | -7977.388 | -7978.752 | -1.363 | 9.90E-07 |
| 11 | 117630867 | rs11825655 | G | A | 0.973 | -7977.388 | -7978.752 | -1.364 | 9.89E-07 |

| | | | | | | | |
|---|---|---|---|---|---|---|---|
| 11 | 118466249 | rs7115634 | G | A | NULL | 0.663738697 | NULL&M_Disease |
| 11 | 118470092 | rs11216924 | G | C | NULL | 0.663738697 | NULL&M_Disease |
| 11 | 120784855 | rs2852238 | G | C | NULL | 0.663738697 | NULL&M_Disease |
| 11 | 121065345 | rs73005072 | A | C | NULL | 0.663738697 | NULL&M_Disease |
| 11 | 116805874 | rs144015791 | T | G | NULL | 0.663738697 | NULL&M_Disease |
| 11 | 117774146 | rs2186627 | G | C | NULL | 0.663738697 | NULL&M_Disease |
| 11 | 119949832 | rs3862647 | C | T | NULL | 0.663738697 | NULL&M_Disease |
| 11 | 119995447 | rs2851718 | C | T | NULL | 0.663738697 | NULL&M_Disease |
| 11 | 120498485 | rs11606452 | C | G | NULL | 0.663738697 | NULL&M_Disease |
| 11 | 120772254 | rs2850814 | A | G | NULL | 0.663738697 | NULL&M_Disease |
| 11 | 120819657 | rs4935757 | G | A | NULL | 0.663738697 | NULL&M_Disease |
| 11 | 116534828 | rs567209 | T | C | NULL | 0.663738697 | NULL&M_Disease |
| 11 | 116744018 | rs17120111 | T | C | NULL | 0.663738697 | NULL&M_Disease |
| 11 | 119148573 | rs2510152 | G | T | NULL | 0.663738697 | NULL&M_Disease |
| 11 | 119943918 | rs626258 | A | G | NULL | 0.663738697 | NULL&M_Disease |
| 11 | 120555070 | rs61902683 | G | T | NULL | 0.663738697 | NULL&M_Disease |
| 11 | 121086572 | rs12417661 | A | G | NULL | 0.663738697 | NULL&M_Disease |
| 11 | 116220070 | rs73010950 | A | G | NULL | 0.663738697 | NULL&M_Disease |
| 11 | 117656550 | rs7940537 | A | G | NULL | 0.663738697 | NULL&M_Disease |
| 11 | 118709600 | rs12272981 | G | A | NULL | 0.663738697 | NULL&M_Disease |
| 11 | 120493629 | rs6589820 | G | A | NULL | 0.663738697 | NULL&M_Disease |
| 11 | 120965687 | rs7931603 | A | G | NULL | 0.663738697 | NULL&M_Disease |
| 11 | 116492411 | rs11216045 | C | T | NULL | 0.66485373 | NULL&M_Disease |
| 11 | 116570894 | rs1145192 | A | G | NULL | 0.663738697 | NULL&M_Disease |
| 11 | 117167147 | rs507805 | G | A | NULL | 0.663738697 | NULL&M_Disease |
| 11 | 117348145 | rs3903091 | C | T | NULL | 0.663738697 | NULL&M_Disease |
| 11 | 117525420 | rs7950072 | G | A | NULL | 0.663738697 | NULL&M_Disease |
| 11 | 117709754 | rs564989 | C | T | NULL | 0.66485373 | NULL&M_Disease |
| 11 | 118041894 | rs637753 | C | G | NULL | 0.66485373 | NULL&M_Disease |
| 11 | 118453483 | rs60112336 | T | C | NULL | 0.663738697 | NULL&M_Disease |
| 11 | 118464815 | rs11216921 | C | G | NULL | 0.663738697 | NULL&M_Disease |
| 11 | 119578224 | rs35301288 | A | G | NULL | 0.66485373 | NULL&M_Disease |
| 11 | 120611793 | rs3132778 | C | T | NULL | 0.663738697 | NULL&M_Disease |
| 11 | 120825332 | rs11218081 | C | T | NULL | 0.66485373 | NULL&M_Disease |
| 11 | 116305597 | rs11607388 | A | G | NULL | 0.663738697 | NULL&M_Disease |
| 11 | 116329139 | rs547021 | C | T | NULL | 0.66485373 | NULL&M_Disease |
| 11 | 120094936 | rs513944 | G | A | NULL | 0.663738697 | NULL&M_Disease |
| 11 | 116221203 | rs73010960 | A | C | NULL | 0.66485373 | NULL&M_Disease |
| 11 | 117383694 | rs10892133 | T | C | NULL | 0.66485373 | NULL&M_Disease |
| 11 | 117384673 | rs4938395 | G | A | NULL | 0.66485373 | NULL&M_Disease |
| 11 | 117952771 | rs582146 | T | G | NULL | 0.66485373 | NULL&M_Disease |
| 11 | 118837742 | rs7116690 | G | A | NULL | 0.66485373 | NULL&M_Disease |
| 11 | 119219819 | rs7932145 | A | C | NULL | 0.66485373 | NULL&M_Disease |
| 11 | 119241120 | rs514808 | A | G | NULL | 0.66485373 | NULL&M_Disease |
| 11 | 120606763 | rs61901404 | T | C | NULL | 0.66485373 | NULL&M_Disease |
| 11 | 116362253 | rs7121331 | C | T | NULL | 0.66485373 | NULL&M_Disease |
| 11 | 117113335 | rs61903660 | G | A | NULL | 0.66485373 | NULL&M_Disease |
| 11 | 120039835 | rs79007113 | A | G | NULL | 0.66485373 | NULL&M_Disease |
| 11 | 116220570 | rs490257 | G | C | NULL | 0.66485373 | NULL&M_Disease |
| 11 | 116408615 | rs7117565 | C | T | NULL | 0.66485373 | NULL&M_Disease |
| 11 | 117356908 | rs11501117 | G | C | NULL | 0.66485373 | NULL&M_Disease |
| 11 | 117845564 | rs7927559 | G | A | NULL | 0.66485373 | NULL&M_Disease |
| 11 | 118444525 | rs11216907 | A | G | NULL | 0.66485373 | NULL&M_Disease |
| 11 | 119220395 | rs113863182 | T | C | NULL | 0.66485373 | NULL&M_Disease |
| 11 | 121092052 | rs745517 | C | T | NULL | 0.66485373 | NULL&M_Disease |
| 11 | 121092505 | rs948376 | C | T | NULL | 0.66485373 | NULL&M_Disease |
| 11 | 116484741 | rs505848 | T | C | NULL | 0.66485373 | NULL&M_Disease |
| 11 | 116598988 | rs180360 | G | A | NULL | 0.66485373 | NULL&M_Disease |
| 11 | 118445275 | rs17122269 | C | T | NULL | 0.66485373 | NULL&M_Disease |
| 11 | 118446298 | rs55844025 | T | C | NULL | 0.66485373 | NULL&M_Disease |
| 11 | 118449370 | rs17122278 | G | A | NULL | 0.66485373 | NULL&M_Disease |
| 11 | 120075878 | rs34028064 | A | C | NULL | 0.66485373 | NULL&M_Disease |
| 11 | 120696297 | rs4573685 | A | G | NULL | 0.66485373 | NULL&M_Disease |
| 11 | 120825348 | rs11218082 | C | G | NULL | 0.66485373 | NULL&M_Disease |
| 11 | 120947058 | rs1380377167 | A | T | NULL | 0.66485373 | NULL&M_Disease |
| 11 | 117435785 | rs12789727 | A | C | NULL | 0.66485373 | NULL&M_Disease |
| 11 | 118912702 | rs10892325 | C | T | NULL | 0.66485373 | NULL&M_Disease |
| 11 | 119659301 | rs12361715 | T | C | NULL | 0.66485373 | NULL&M_Disease |
| 11 | 119659477 | rs12361746 | T | C | NULL | 0.66485373 | NULL&M_Disease |
| 11 | 116452887 | rs10790151 | T | C | NULL | 0.66485373 | NULL&M_Disease |
| 11 | 116481436 | rs565420 | A | G | NULL | 0.66485373 | NULL&M_Disease |
| 11 | 117630114 | rs35374605 | C | T | NULL | 0.66485373 | NULL&M_Disease |
| 11 | 117633286 | rs12794454 | A | T | NULL | 0.66485373 | NULL&M_Disease |
| 11 | 117690423 | rs11600226 | C | T | NULL | 0.66485373 | NULL&M_Disease |
| 11 | 118161885 | rs12420795 | G | C | NULL | 0.66485373 | NULL&M_Disease |
| 11 | 118162147 | rs4938498 | G | C | NULL | 0.66485373 | NULL&M_Disease |
| 11 | 119701826 | rs472232 | A | C | NULL | 0.66485373 | NULL&M_Disease |
| 11 | 120338040 | rs2305011 | G | A | NULL | 0.66485373 | NULL&M_Disease |
| 11 | 117312576 | rs79423267 | A | G | NULL | 0.66485373 | NULL&M_Disease |
| 11 | 118022107 | rs12788624 | T | C | NULL | 0.66485373 | NULL&M_Disease |
| 11 | 119824102 | rs472611 | G | A | NULL | 0.66485373 | NULL&M_Disease |
| 11 | 120484872 | rs1317813 | T | C | NULL | 0.66485373 | NULL&M_Disease |
| 11 | 120987253 | rs619115 | C | T | NULL | 0.66485373 | NULL&M_Disease |
| 11 | 117515473 | rs11216480 | A | G | NULL | 0.66485373 | NULL&M_Disease |
| 11 | 117818878 | rs3018521 | A | G | NULL | 0.66485373 | NULL&M_Disease |
| 11 | 120114073 | rs2847499 | T | G | NULL | 0.66485373 | NULL&M_Disease |
| 11 | 121091888 | rs948092 | C | T | NULL | 0.665966927 | NULL&M_Disease |
| 11 | 121092522 | rs4999180 | T | C | NULL | 0.66485373 | NULL&M_Disease |
| 11 | 116229010 | rs78052530 | C | A | NULL | 0.66485373 | NULL&M_Disease |
| 11 | 116690123 | rs1263176 | T | C | NULL | 0.665966927 | NULL&M_Disease |
| 11 | 118171239 | rs7928058 | T | G | NULL | 0.665966927 | NULL&M_Disease |
| 11 | 120109967 | rs2509623 | T | C | NULL | 0.66485373 | NULL&M_Disease |
| 11 | 120610564 | rs751369 | G | A | NULL | 0.665966927 | NULL&M_Disease |
| 11 | 116439684 | rs2174876 | A | G | NULL | 0.665966927 | NULL&M_Disease |
| 11 | 116517591 | rs1240769 | G | T | NULL | 0.665966927 | NULL&M_Disease |
| 11 | 117112630 | rs633892 | C | A | NULL | 0.665966927 | NULL&M_Disease |

| | | | | | | | | | |
|---|---|---|---|---|---|---|---|---|---|
| 11 | 118466249 | rs7115634 | G | A | 0.974 | -7977.388 | -7978.752 | -1.364 | 9.89E-07 |
| 11 | 118470092 | rs11216924 | G | C | 0.974 | -7977.388 | -7978.752 | -1.364 | 9.89E-07 |
| 11 | 120784855 | rs2852238 | G | C | 1.031 | -7977.388 | -7978.752 | -1.364 | 9.89E-07 |
| 11 | 121065345 | rs73005072 | A | C | 1 | -7977.388 | -7978.752 | -1.364 | 9.89E-07 |
| 11 | 116805874 | rs144015791 | T | G | 0.98 | -7977.388 | -7978.753 | -1.365 | 9.88E-07 |
| 11 | 117774146 | rs2186627 | G | C | 0.969 | -7977.388 | -7978.753 | -1.365 | 9.88E-07 |
| 11 | 119949832 | rs3862647 | C | T | 1.029 | -7977.388 | -7978.753 | -1.365 | 9.88E-07 |
| 11 | 119995447 | rs2851718 | C | T | 0.976 | -7977.388 | -7978.753 | -1.365 | 9.88E-07 |
| 11 | 120498485 | rs11606452 | C | G | 0.97 | -7977.388 | -7978.753 | -1.365 | 9.88E-07 |
| 11 | 120772254 | rs2850814 | A | G | 1.03 | -7977.388 | -7978.754 | -1.365 | 9.88E-07 |
| 11 | 120819657 | rs4935757 | G | A | 0.969 | -7977.388 | -7978.753 | -1.365 | 9.88E-07 |
| 11 | 116534828 | rs567209 | T | C | 1.03 | -7977.388 | -7978.754 | -1.366 | 9.87E-07 |
| 11 | 116744018 | rs17120111 | T | C | 0.973 | -7977.388 | -7978.754 | -1.366 | 9.87E-07 |
| 11 | 119148573 | rs2510152 | G | T | 1.031 | -7977.388 | -7978.754 | -1.366 | 9.87E-07 |
| 11 | 119943918 | rs626258 | A | G | 0.971 | -7977.388 | -7978.754 | -1.366 | 9.87E-07 |
| 11 | 120555070 | rs61902683 | G | T | 0.976 | -7977.388 | -7978.754 | -1.366 | 9.87E-07 |
| 11 | 121086572 | rs12417661 | A | G | 1.016 | -7977.388 | -7978.754 | -1.366 | 9.87E-07 |
| 11 | 116220070 | rs73010950 | A | G | 1.015 | -7977.388 | -7978.755 | -1.367 | 9.86E-07 |
| 11 | 117656550 | rs7940537 | A | G | 1.007 | -7977.388 | -7978.756 | -1.367 | 9.86E-07 |
| 11 | 118709600 | rs12272981 | G | A | 1.03 | -7977.388 | -7978.755 | -1.367 | 9.86E-07 |
| 11 | 120493629 | rs6589820 | G | A | 0.969 | -7977.388 | -7978.755 | -1.367 | 9.86E-07 |
| 11 | 120965687 | rs7931603 | A | G | 1.031 | -7977.388 | -7978.756 | -1.367 | 9.86E-07 |
| 11 | 116492411 | rs11216045 | C | T | 1.031 | -7977.388 | -7978.756 | -1.368 | 9.85E-07 |
| 11 | 116570894 | rs1145192 | A | G | 1.031 | -7977.388 | -7978.756 | -1.368 | 9.85E-07 |
| 11 | 117167147 | rs507805 | G | A | 0.998 | -7977.388 | -7978.756 | -1.368 | 9.85E-07 |
| 11 | 117348145 | rs3903091 | C | T | 1.012 | -7977.388 | -7978.757 | -1.368 | 9.85E-07 |
| 11 | 117525420 | rs7950072 | G | A | 1.017 | -7977.388 | -7978.756 | -1.368 | 9.85E-07 |
| 11 | 117709754 | rs564989 | C | T | 1.031 | -7977.388 | -7978.757 | -1.368 | 9.85E-07 |
| 11 | 118041894 | rs637753 | C | G | 1.031 | -7977.388 | -7978.756 | -1.368 | 9.85E-07 |
| 11 | 118453483 | rs60112336 | T | C | 0.974 | -7977.388 | -7978.756 | -1.368 | 9.85E-07 |
| 11 | 118464815 | rs11216921 | C | G | 0.974 | -7977.388 | -7978.756 | -1.368 | 9.85E-07 |
| 11 | 119578224 | rs35301288 | A | G | 1.031 | -7977.388 | -7978.756 | -1.368 | 9.85E-07 |
| 11 | 120611793 | rs3132778 | C | T | 1.028 | -7977.388 | -7978.756 | -1.368 | 9.85E-07 |
| 11 | 120825332 | rs11218081 | C | T | 1.03 | -7977.388 | -7978.757 | -1.368 | 9.85E-07 |
| 11 | 116305597 | rs11607388 | A | G | 1.005 | -7977.388 | -7978.757 | -1.369 | 9.84E-07 |
| 11 | 116329139 | rs547021 | C | T | 0.977 | -7977.388 | -7978.757 | -1.369 | 9.84E-07 |
| 11 | 120094936 | rs513944 | G | A | 0.992 | -7977.388 | -7978.757 | -1.369 | 9.84E-07 |
| 11 | 116221203 | rs73010960 | A | C | 1.014 | -7977.388 | -7978.759 | -1.37 | 9.83E-07 |
| 11 | 117383694 | rs10892133 | T | C | 1.03 | -7977.388 | -7978.758 | -1.37 | 9.83E-07 |
| 11 | 117384673 | rs4938395 | G | A | 1.03 | -7977.388 | -7978.758 | -1.37 | 9.83E-07 |
| 11 | 117952771 | rs582146 | T | G | 1.023 | -7977.388 | -7978.758 | -1.37 | 9.83E-07 |
| 11 | 118837742 | rs7116690 | G | A | 1.031 | -7977.388 | -7978.758 | -1.37 | 9.83E-07 |
| 11 | 119219819 | rs7932145 | A | C | 0.97 | -7977.388 | -7978.758 | -1.37 | 9.83E-07 |
| 11 | 119241120 | rs514808 | A | G | 0.977 | -7977.388 | -7978.758 | -1.37 | 9.83E-07 |
| 11 | 120606763 | rs61901404 | T | C | 1.023 | -7977.388 | -7978.758 | -1.37 | 9.83E-07 |
| 11 | 116362253 | rs7121331 | C | T | 1.031 | -7977.388 | -7978.76 | -1.371 | 9.82E-07 |
| 11 | 117113335 | rs61903660 | G | A | 0.976 | -7977.388 | -7978.759 | -1.371 | 9.82E-07 |
| 11 | 120039835 | rs79007113 | A | G | 0.977 | -7977.388 | -7978.759 | -1.371 | 9.82E-07 |
| 11 | 116220570 | rs490257 | G | C | 1.031 | -7977.388 | -7978.761 | -1.372 | 9.81E-07 |
| 11 | 116408615 | rs7117565 | C | T | 1.031 | -7977.388 | -7978.76 | -1.372 | 9.81E-07 |
| 11 | 117356908 | rs11501117 | G | C | 1.011 | -7977.388 | -7978.76 | -1.372 | 9.81E-07 |
| 11 | 117845564 | rs7927559 | G | A | 1.031 | -7977.388 | -7978.76 | -1.372 | 9.81E-07 |
| 11 | 118444525 | rs11216907 | A | G | 0.974 | -7977.388 | -7978.761 | -1.372 | 9.81E-07 |
| 11 | 119220395 | rs113863182 | T | C | 0.97 | -7977.388 | -7978.76 | -1.372 | 9.81E-07 |
| 11 | 121092052 | rs745517 | C | T | 0.97 | -7977.388 | -7978.761 | -1.372 | 9.81E-07 |
| 11 | 121092505 | rs948376 | C | T | 0.97 | -7977.388 | -7978.76 | -1.372 | 9.81E-07 |
| 11 | 116484741 | rs505848 | T | C | 0.97 | -7977.388 | -7978.762 | -1.373 | 9.80E-07 |
| 11 | 116598988 | rs180360 | G | A | 1.031 | -7977.388 | -7978.761 | -1.373 | 9.80E-07 |
| 11 | 118445275 | rs17122269 | C | T | 0.974 | -7977.388 | -7978.762 | -1.373 | 9.80E-07 |
| 11 | 118446298 | rs55844025 | T | C | 0.974 | -7977.388 | -7978.761 | -1.373 | 9.80E-07 |
| 11 | 118449370 | rs17122278 | G | A | 0.974 | -7977.388 | -7978.762 | -1.373 | 9.80E-07 |
| 11 | 120075878 | rs34028064 | A | C | 1.031 | -7977.388 | -7978.761 | -1.373 | 9.80E-07 |
| 11 | 120696297 | rs4573685 | A | G | 0.971 | -7977.388 | -7978.761 | -1.373 | 9.80E-07 |
| 11 | 120825348 | rs11218082 | C | G | 1.029 | -7977.388 | -7978.761 | -1.373 | 9.80E-07 |
| 11 | 120947058 | rs1380377167 | A | T | 0.985 | -7977.388 | -7978.761 | -1.373 | 9.80E-07 |
| 11 | 117435785 | rs12789727 | A | C | 1.031 | -7977.388 | -7978.762 | -1.374 | 9.79E-07 |
| 11 | 118912702 | rs10892325 | C | T | 0.97 | -7977.388 | -7978.762 | -1.374 | 9.79E-07 |
| 11 | 119659301 | rs12361715 | T | C | 0.976 | -7977.388 | -7978.762 | -1.374 | 9.79E-07 |
| 11 | 119659477 | rs12361746 | T | C | 0.976 | -7977.388 | -7978.763 | -1.374 | 9.79E-07 |
| 11 | 116452887 | rs10790151 | T | C | 0.97 | -7977.388 | -7978.763 | -1.375 | 9.78E-07 |
| 11 | 116481436 | rs565420 | A | G | 0.983 | -7977.388 | -7978.763 | -1.375 | 9.78E-07 |
| 11 | 117630114 | rs35374605 | C | T | 0.973 | -7977.388 | -7978.763 | -1.375 | 9.78E-07 |
| 11 | 117633286 | rs12794454 | A | T | 0.973 | -7977.388 | -7978.763 | -1.375 | 9.78E-07 |
| 11 | 117690423 | rs11600226 | C | T | 0.972 | -7977.388 | -7978.763 | -1.375 | 9.78E-07 |
| 11 | 118161885 | rs12420795 | G | C | 0.97 | -7977.388 | -7978.764 | -1.375 | 9.78E-07 |
| 11 | 118162147 | rs4938498 | G | C | 0.97 | -7977.388 | -7978.764 | -1.375 | 9.78E-07 |
| 11 | 119701826 | rs472232 | A | C | 0.97 | -7977.388 | -7978.764 | -1.375 | 9.78E-07 |
| 11 | 120338040 | rs2305011 | G | A | 0.974 | -7977.388 | -7978.763 | -1.375 | 9.78E-07 |
| 11 | 117312576 | rs79423267 | A | G | 0.972 | -7977.388 | -7978.764 | -1.376 | 9.77E-07 |
| 11 | 118022107 | rs12788624 | T | C | 0.975 | -7977.388 | -7978.764 | -1.376 | 9.77E-07 |
| 11 | 119824102 | rs472611 | G | A | 0.97 | -7977.388 | -7978.764 | -1.376 | 9.77E-07 |
| 11 | 120484872 | rs1317813 | T | C | 0.987 | -7977.388 | -7978.764 | -1.376 | 9.77E-07 |
| 11 | 120987253 | rs619115 | C | T | 0.97 | -7977.388 | -7978.764 | -1.376 | 9.77E-07 |
| 11 | 117515473 | rs11216480 | A | G | 1.016 | -7977.388 | -7978.766 | -1.377 | 9.76E-07 |
| 11 | 117818878 | rs3018521 | A | G | 1.031 | -7977.388 | -7978.765 | -1.377 | 9.76E-07 |
| 11 | 120114073 | rs2847499 | T | G | 1.007 | -7977.388 | -7978.765 | -1.377 | 9.76E-07 |
| 11 | 121091888 | rs948092 | C | T | 0.97 | -7977.388 | -7978.766 | -1.377 | 9.76E-07 |
| 11 | 121092522 | rs4999180 | T | C | 0.97 | -7977.388 | -7978.765 | -1.377 | 9.76E-07 |
| 11 | 116229010 | rs78052530 | C | A | 1.016 | -7977.388 | -7978.766 | -1.378 | 9.75E-07 |
| 11 | 116690123 | rs1263176 | T | C | 1.031 | -7977.388 | -7978.766 | -1.378 | 9.75E-07 |
| 11 | 118171239 | rs7928058 | T | G | 0.97 | -7977.388 | -7978.766 | -1.378 | 9.75E-07 |
| 11 | 120109967 | rs2509623 | T | C | 1.009 | -7977.388 | -7978.766 | -1.378 | 9.75E-07 |
| 11 | 120610564 | rs751369 | G | A | 0.97 | -7977.388 | -7978.766 | -1.378 | 9.75E-07 |
| 11 | 116439684 | rs2174876 | A | G | 1.031 | -7977.388 | -7978.767 | -1.379 | 9.74E-07 |
| 11 | 116517591 | rs1240769 | G | T | 1.031 | -7977.388 | -7978.767 | -1.379 | 9.74E-07 |
| 11 | 117112630 | rs633892 | C | A | 0.97 | -7977.388 | -7978.768 | -1.379 | 9.74E-07 |

| | | | | | | | |
|---|---|---|---|---|---|---|---|
| 11 | 119502547 | rs35508996 | A | G | NULL | 0.665966927 | NULL&M_Disease |
| 11 | 119794853 | rs12291317 | C | T | NULL | 0.665966927 | NULL&M_Disease |
| 11 | 119833476 | rs75619771 | C | T | NULL | 0.665966927 | NULL&M_Disease |
| 11 | 116253905 | rs11215908 | A | G | NULL | 0.665966927 | NULL&M_Disease |
| 11 | 116905464 | rs4938326 | C | G | NULL | 0.665966927 | NULL&M_Disease |
| 11 | 117309494 | rs74490241 | A | C | NULL | 0.665966927 | NULL&M_Disease |
| 11 | 117667524 | rs75467870 | A | G | NULL | 0.665966927 | NULL&M_Disease |
| 11 | 117781192 | rs11216616 | A | G | NULL | 0.665966927 | NULL&M_Disease |
| 11 | 119492158 | rs4514433 | C | T | NULL | 0.665966927 | NULL&M_Disease |
| 11 | 121120130 | rs948379 | T | C | NULL | 0.665966927 | NULL&M_Disease |
| 11 | 116220637 | rs476367 | C | T | NULL | 0.665966927 | NULL&M_Disease |
| 11 | 117290963 | rs11603208 | T | G | NULL | 0.665966927 | NULL&M_Disease |
| 11 | 117355493 | rs11216421 | C | G | NULL | 0.665966927 | NULL&M_Disease |
| 11 | 116670676 | rs6589567 | A | C | NULL | 0.665966927 | NULL&M_Disease |
| 11 | 116699681 | rs12721080 | A | G | NULL | 0.665966927 | NULL&M_Disease |
| 11 | 117672973 | rs12276576 | G | C | NULL | 0.665966927 | NULL&M_Disease |
| 11 | 119918703 | rs11217656 | A | G | NULL | 0.665966927 | NULL&M_Disease |
| 11 | 119979007 | rs7941900 | C | T | NULL | 0.665966927 | NULL&M_Disease |
| 11 | 120219546 | rs56406822 | A | G | NULL | 0.665966927 | NULL&M_Disease |
| 11 | 120654400 | rs7109584 | A | G | NULL | 0.665966927 | NULL&M_Disease |
| 11 | 116681243 | rs1263174 | G | T | NULL | 0.665966927 | NULL&M_Disease |
| 11 | 116899786 | rs112120205 | A | G | NULL | 0.665966927 | NULL&M_Disease |
| 11 | 117353286 | rs11216419 | A | G | NULL | 0.665966927 | NULL&M_Disease |
| 11 | 117353773 | rs10892122 | A | G | NULL | 0.665966927 | NULL&M_Disease |
| 11 | 120105099 | rs2845708 | A | T | NULL | 0.665966927 | NULL&M_Disease |
| 11 | 120600346 | rs1939666 | G | A | NULL | 0.665966927 | NULL&M_Disease |
| 11 | 116657561 | rs3741298 | C | T | NULL | 0.665966927 | NULL&M_Disease |
| 11 | 116795987 | rs77953246 | C | G | NULL | 0.665966927 | NULL&M_Disease |
| 11 | 117829855 | rs9666861 | G | A | NULL | 0.665966927 | NULL&M_Disease |
| 11 | 118061767 | rs627291 | G | A | NULL | 0.665966927 | NULL&M_Disease |
| 11 | 119235404 | rs2195525 | C | T | NULL | 0.665966927 | NULL&M_Disease |
| 11 | 120003533 | rs666432 | T | C | NULL | 0.665966927 | NULL&M_Disease |
| 11 | 120472569 | rs4414229 | G | A | NULL | 0.665966927 | NULL&M_Disease |
| 11 | 121119712 | rs12279889 | T | C | NULL | 0.665966927 | NULL&M_Disease |
| 11 | 116916678 | rs76972960 | C | T | NULL | 0.665966927 | NULL&M_Disease |
| 11 | 118327321 | rs536256 | G | A | NULL | 0.665966927 | NULL&M_Disease |
| 11 | 120078393 | rs10790370 | A | G | NULL | 0.665966927 | NULL&M_Disease |
| 11 | 120195689 | rs61898347 | A | G | NULL | 0.665966927 | NULL&M_Disease |
| 11 | 120636299 | rs79532870 | A | G | NULL | 0.665966927 | NULL&M_Disease |
| 11 | 116241604 | rs77775914 | T | C | NULL | 0.665966927 | NULL&M_Disease |
| 11 | 117362425 | rs536859 | A | G | NULL | 0.665966927 | NULL&M_Disease |
| 11 | 117496301 | rs10892145 | A | G | NULL | 0.665966927 | NULL&M_Disease |
| 11 | 118970878 | rs603826 | C | A | NULL | 0.665966927 | NULL&M_Disease |
| 11 | 116778275 | rs4938317 | C | T | NULL | 0.667078277 | NULL&M_Disease |
| 11 | 116942132 | rs17120280 | A | G | NULL | 0.665966927 | NULL&M_Disease |
| 11 | 117002403 | rs6589598 | A | G | NULL | 0.667078277 | NULL&M_Disease |
| 11 | 118974485 | rs599385 | G | A | NULL | 0.667078277 | NULL&M_Disease |
| 11 | 120067624 | rs66553012 | G | A | NULL | 0.665966927 | NULL&M_Disease |
| 11 | 120450016 | rs12284644 | G | A | NULL | 0.667078277 | NULL&M_Disease |
| 11 | 119980729 | rs12799241 | A | G | NULL | 0.667078277 | NULL&M_Disease |
| 11 | 120808850 | rs17124575 | G | A | NULL | 0.667078277 | NULL&M_Disease |
| 11 | 117118204 | rs642673 | A | T | NULL | 0.667078277 | NULL&M_Disease |
| 11 | 117772021 | rs55813570 | A | G | NULL | 0.667078277 | NULL&M_Disease |
| 11 | 119894514 | rs672295 | G | A | NULL | 0.667078277 | NULL&M_Disease |
| 11 | 116489979 | rs11216043 | C | T | NULL | 0.667078277 | NULL&M_Disease |
| 11 | 116760545 | rs10892042 | A | G | NULL | 0.667078277 | NULL&M_Disease |
| 11 | 117045431 | rs112111936 | G | T | NULL | 0.667078277 | NULL&M_Disease |
| 11 | 117358351 | rs4938391 | A | C | NULL | 0.667078277 | NULL&M_Disease |
| 11 | 117524000 | rs11216486 | G | T | NULL | 0.667078277 | NULL&M_Disease |
| 11 | 119193721 | rs1893264 | C | A | NULL | 0.667078277 | NULL&M_Disease |
| 11 | 119520730 | rs11217374 | G | A | NULL | 0.667078277 | NULL&M_Disease |
| 11 | 121039069 | rs543577 | A | G | NULL | 0.667078277 | NULL&M_Disease |
| 11 | 116772441 | rs1076485 | T | C | NULL | 0.667078277 | NULL&M_Disease |
| 11 | 116963312 | rs7946869 | T | C | NULL | 0.667078277 | NULL&M_Disease |
| 11 | 117097222 | rs7926958 | G | T | NULL | 0.667078277 | NULL&M_Disease |
| 11 | 117358313 | rs4938390 | T | C | NULL | 0.667078277 | NULL&M_Disease |
| 11 | 117825852 | rs11216647 | C | T | NULL | 0.667078277 | NULL&M_Disease |
| 11 | 118032242 | rs688587 | A | C | NULL | 0.667078277 | NULL&M_Disease |
| 11 | 120178753 | rs17245504 | G | T | NULL | 0.667078277 | NULL&M_Disease |
| 11 | 116253169 | rs73019251 | T | C | NULL | 0.667078277 | NULL&M_Disease |
| 11 | 116407583 | rs11215990 | T | C | NULL | 0.667078277 | NULL&M_Disease |
| 11 | 117145053 | rs694574 | A | C | NULL | 0.667078277 | NULL&M_Disease |
| 11 | 117352949 | rs3741278 | A | G | NULL | 0.667078277 | NULL&M_Disease |
| 11 | 117464204 | rs4938413 | G | C | NULL | 0.667078277 | NULL&M_Disease |
| 11 | 117616156 | rs117373043 | T | C | NULL | 0.667078277 | NULL&M_Disease |
| 11 | 117921554 | rs11216734 | A | T | NULL | 0.667078277 | NULL&M_Disease |
| 11 | 119144148 | rs2511856 | G | T | NULL | 0.667078277 | NULL&M_Disease |
| 11 | 117309527 | rs75808396 | A | G | NULL | 0.667078277 | NULL&M_Disease |
| 11 | 117311365 | rs75343916 | T | C | NULL | 0.667078277 | NULL&M_Disease |
| 11 | 118032228 | rs688563 | C | T | NULL | 0.667078277 | NULL&M_Disease |
| 11 | 118839828 | rs61902060 | G | C | NULL | 0.667078277 | NULL&M_Disease |
| 11 | 116481221 | rs545613 | C | T | NULL | 0.667078277 | NULL&M_Disease |
| 11 | 116482895 | rs511011 | C | T | NULL | 0.667078277 | NULL&M_Disease |
| 11 | 116727465 | rs78857232 | T | C | NULL | 0.667078277 | NULL&M_Disease |
| 11 | 118970696 | rs604714 | A | C | NULL | 0.667078277 | NULL&M_Disease |
| 11 | 116408318 | rs868066 | A | G | NULL | 0.667078277 | NULL&M_Disease |
| 11 | 116437756 | rs11605293 | T | C | NULL | 0.667078277 | NULL&M_Disease |
| 11 | 118535840 | rs12225548 | G | C | NULL | 0.667078277 | NULL&M_Disease |
| 11 | 119671453 | rs895652 | A | G | NULL | 0.667078277 | NULL&M_Disease |
| 11 | 119900407 | rs76610415 | A | G | NULL | 0.667078277 | NULL&M_Disease |
| 11 | 120302729 | rs55869319 | T | C | NULL | 0.667078277 | NULL&M_Disease |
| 11 | 118501723 | rs61900957 | T | C | NULL | 0.667078277 | NULL&M_Disease |
| 11 | 116322819 | rs550791 | T | C | NULL | 0.667078277 | NULL&M_Disease |
| 11 | 118839133 | rs61900159 | C | G | NULL | 0.668187772 | NULL&M_Disease |
| 11 | 118839268 | rs11217107 | C | G | NULL | 0.668187772 | NULL&M_Disease |
| 11 | 116694866 | rs1268833 | G | A | NULL | 0.668187772 | NULL&M_Disease |

| | | | | | | | | | |
|---|---|---|---|---|---|---|---|---|---|
| 11 | 119502547 | rs35508996 | A | G | 0.97 | -7977.388 | -7978.767 | -1.379 | 9.74E-07 |
| 11 | 119794853 | rs12291317 | C | T | 1.03 | -7977.388 | -7978.768 | -1.379 | 9.74E-07 |
| 11 | 119833476 | rs75619771 | C | T | 1.012 | -7977.388 | -7978.768 | -1.379 | 9.74E-07 |
| 11 | 116253905 | rs11215908 | A | G | 1.03 | -7977.388 | -7978.769 | -1.38 | 9.73E-07 |
| 11 | 116905464 | rs4938326 | C | G | 0.981 | -7977.388 | -7978.768 | -1.38 | 9.73E-07 |
| 11 | 117309494 | rs74490241 | A | C | 0.972 | -7977.388 | -7978.768 | -1.38 | 9.73E-07 |
| 11 | 117667524 | rs75467870 | A | G | 0.981 | -7977.388 | -7978.768 | -1.38 | 9.73E-07 |
| 11 | 117781192 | rs11216616 | A | G | 1.03 | -7977.388 | -7978.768 | -1.38 | 9.73E-07 |
| 11 | 119492158 | rs4514433 | C | T | 1.031 | -7977.388 | -7978.768 | -1.38 | 9.73E-07 |
| 11 | 121120130 | rs948379 | T | C | 1.031 | -7977.388 | -7978.769 | -1.38 | 9.73E-07 |
| 11 | 116220637 | rs476367 | C | T | 1.031 | -7977.388 | -7978.77 | -1.381 | 9.72E-07 |
| 11 | 117290963 | rs11603208 | T | G | 1.029 | -7977.388 | -7978.77 | -1.381 | 9.72E-07 |
| 11 | 117355493 | rs11216421 | C | G | 1.009 | -7977.388 | -7978.769 | -1.381 | 9.72E-07 |
| 11 | 116670676 | rs6589567 | A | C | 0.977 | -7977.388 | -7978.77 | -1.382 | 9.71E-07 |
| 11 | 116699681 | rs12721080 | A | G | 1.029 | -7977.388 | -7978.77 | -1.382 | 9.71E-07 |
| 11 | 117672973 | rs12276576 | G | C | 0.982 | -7977.388 | -7978.77 | -1.382 | 9.71E-07 |
| 11 | 119918703 | rs11217656 | A | G | 1.031 | -7977.388 | -7978.77 | -1.382 | 9.71E-07 |
| 11 | 119979007 | rs7941900 | C | T | 1.017 | -7977.388 | -7978.77 | -1.382 | 9.71E-07 |
| 11 | 120219546 | rs56406822 | A | G | 0.974 | -7977.388 | -7978.771 | -1.382 | 9.71E-07 |
| 11 | 120654400 | rs7109584 | A | G | 1.011 | -7977.388 | -7978.77 | -1.382 | 9.71E-07 |
| 11 | 116681243 | rs1263174 | G | T | 1.031 | -7977.388 | -7978.771 | -1.383 | 9.70E-07 |
| 11 | 116899786 | rs112120205 | A | G | 0.982 | -7977.388 | -7978.771 | -1.383 | 9.70E-07 |
| 11 | 117353286 | rs11216419 | A | G | 1.009 | -7977.388 | -7978.771 | -1.383 | 9.70E-07 |
| 11 | 117353773 | rs10892122 | A | G | 1.009 | -7977.388 | -7978.771 | -1.383 | 9.70E-07 |
| 11 | 120105099 | rs2845708 | A | T | 1.002 | -7977.388 | -7978.772 | -1.383 | 9.70E-07 |
| 11 | 120600346 | rs1939666 | G | A | 0.97 | -7977.388 | -7978.771 | -1.383 | 9.70E-07 |
| 11 | 116657561 | rs3741298 | C | T | 0.971 | -7977.388 | -7978.772 | -1.384 | 9.69E-07 |
| 11 | 116795987 | rs77953246 | C | G | 0.982 | -7977.388 | -7978.772 | -1.384 | 9.69E-07 |
| 11 | 117829855 | rs9666861 | G | A | 1.031 | -7977.388 | -7978.773 | -1.384 | 9.69E-07 |
| 11 | 118061767 | rs627291 | G | A | 0.97 | -7977.388 | -7978.772 | -1.384 | 9.69E-07 |
| 11 | 119235404 | rs2195525 | C | T | 0.97 | -7977.388 | -7978.773 | -1.384 | 9.69E-07 |
| 11 | 120003533 | rs666432 | T | C | 0.979 | -7977.388 | -7978.773 | -1.384 | 9.69E-07 |
| 11 | 120472569 | rs4414229 | G | A | 0.97 | -7977.388 | -7978.772 | -1.384 | 9.69E-07 |
| 11 | 121119712 | rs12279889 | T | C | 0.973 | -7977.388 | -7978.773 | -1.384 | 9.69E-07 |
| 11 | 116916678 | rs76972960 | C | T | 0.982 | -7977.388 | -7978.774 | -1.386 | 9.67E-07 |
| 11 | 118327321 | rs536256 | G | A | 0.972 | -7977.388 | -7978.775 | -1.386 | 9.67E-07 |
| 11 | 120078393 | rs10790370 | A | G | 1.028 | -7977.388 | -7978.774 | -1.386 | 9.67E-07 |
| 11 | 120195689 | rs61898347 | A | G | 1.03 | -7977.388 | -7978.775 | -1.386 | 9.67E-07 |
| 11 | 120636299 | rs79532870 | A | G | 1.013 | -7977.388 | -7978.774 | -1.386 | 9.67E-07 |
| 11 | 116241604 | rs77775914 | T | C | 0.993 | -7977.388 | -7978.776 | -1.387 | 9.66E-07 |
| 11 | 117362425 | rs536859 | A | G | 0.972 | -7977.388 | -7978.776 | -1.387 | 9.66E-07 |
| 11 | 117496301 | rs10892145 | A | G | 1.031 | -7977.388 | -7978.775 | -1.387 | 9.66E-07 |
| 11 | 118970878 | rs603826 | C | A | 1.031 | -7977.388 | -7978.775 | -1.387 | 9.66E-07 |
| 11 | 116778275 | rs4938317 | C | T | 0.974 | -7977.388 | -7978.777 | -1.388 | 9.65E-07 |
| 11 | 116942132 | rs17120280 | A | G | 0.974 | -7977.388 | -7978.777 | -1.388 | 9.65E-07 |
| 11 | 117002403 | rs6589598 | A | G | 0.971 | -7977.388 | -7978.776 | -1.388 | 9.65E-07 |
| 11 | 118974485 | rs599385 | G | A | 1.031 | -7977.388 | -7978.777 | -1.388 | 9.65E-07 |
| 11 | 120067624 | rs66553012 | G | A | 0.976 | -7977.388 | -7978.776 | -1.388 | 9.65E-07 |
| 11 | 120450016 | rs12284644 | G | A | 1.031 | -7977.388 | -7978.776 | -1.388 | 9.65E-07 |
| 11 | 119980729 | rs12799241 | A | G | 1.03 | -7977.388 | -7978.777 | -1.389 | 9.64E-07 |
| 11 | 120808850 | rs17124575 | G | A | 0.97 | -7977.388 | -7978.777 | -1.389 | 9.64E-07 |
| 11 | 117118204 | rs642673 | A | T | 0.977 | -7977.388 | -7978.779 | -1.39 | 9.63E-07 |
| 11 | 117772021 | rs55813570 | A | G | 1.028 | -7977.388 | -7978.778 | -1.39 | 9.63E-07 |
| 11 | 119894514 | rs672295 | G | A | 0.97 | -7977.388 | -7978.778 | -1.39 | 9.63E-07 |
| 11 | 116489979 | rs11216043 | C | T | 1.031 | -7977.388 | -7978.78 | -1.391 | 9.62E-07 |
| 11 | 116760545 | rs10892042 | A | G | 0.974 | -7977.388 | -7978.779 | -1.391 | 9.62E-07 |
| 11 | 117045431 | rs112111936 | G | T | 0.992 | -7977.388 | -7978.779 | -1.391 | 9.62E-07 |
| 11 | 117358351 | rs4938391 | A | C | 1.006 | -7977.388 | -7978.78 | -1.391 | 9.62E-07 |
| 11 | 117524000 | rs11216486 | G | T | 1.014 | -7977.388 | -7978.779 | -1.391 | 9.62E-07 |
| 11 | 119193721 | rs1893264 | C | A | 1.03 | -7977.388 | -7978.78 | -1.391 | 9.62E-07 |
| 11 | 119520730 | rs11217374 | G | A | 0.97 | -7977.388 | -7978.779 | -1.391 | 9.62E-07 |
| 11 | 121039069 | rs543577 | A | G | 1.031 | -7977.388 | -7978.78 | -1.391 | 9.62E-07 |
| 11 | 116772441 | rs1076485 | T | C | 0.976 | -7977.388 | -7978.78 | -1.392 | 9.62E-07 |
| 11 | 116963312 | rs7946869 | T | C | 1.028 | -7977.388 | -7978.781 | -1.392 | 9.62E-07 |
| 11 | 117097222 | rs7926958 | G | T | 1.031 | -7977.388 | -7978.78 | -1.392 | 9.62E-07 |
| 11 | 117358313 | rs4938390 | T | C | 1.006 | -7977.388 | -7978.78 | -1.392 | 9.62E-07 |
| 11 | 117825852 | rs11216647 | C | T | 1.031 | -7977.388 | -7978.781 | -1.392 | 9.62E-07 |
| 11 | 118032242 | rs688587 | A | C | 0.97 | -7977.388 | -7978.78 | -1.392 | 9.62E-07 |
| 11 | 120178753 | rs17245504 | G | T | 1.025 | -7977.388 | -7978.78 | -1.392 | 9.62E-07 |
| 11 | 116253169 | rs73019251 | T | C | 1.014 | -7977.388 | -7978.782 | -1.393 | 9.61E-07 |
| 11 | 116407583 | rs11215990 | T | C | 0.981 | -7977.388 | -7978.781 | -1.393 | 9.61E-07 |
| 11 | 117145053 | rs694574 | A | C | 0.97 | -7977.388 | -7978.781 | -1.393 | 9.61E-07 |
| 11 | 117352949 | rs3741278 | A | G | 1.006 | -7977.388 | -7978.782 | -1.393 | 9.61E-07 |
| 11 | 117464204 | rs4938413 | G | C | 1.03 | -7977.388 | -7978.781 | -1.393 | 9.61E-07 |
| 11 | 117616156 | rs117373043 | T | C | 1.029 | -7977.388 | -7978.781 | -1.393 | 9.61E-07 |
| 11 | 117921554 | rs11216734 | A | T | 1.031 | -7977.388 | -7978.781 | -1.393 | 9.61E-07 |
| 11 | 119144148 | rs2511856 | G | T | 1.031 | -7977.388 | -7978.781 | -1.393 | 9.61E-07 |
| 11 | 117309527 | rs75808396 | A | G | 0.972 | -7977.388 | -7978.782 | -1.394 | 9.60E-07 |
| 11 | 117311365 | rs75343916 | T | C | 0.973 | -7977.388 | -7978.783 | -1.394 | 9.60E-07 |
| 11 | 118032228 | rs688563 | C | T | 0.97 | -7977.388 | -7978.782 | -1.394 | 9.60E-07 |
| 11 | 118839828 | rs61902060 | G | C | 1.031 | -7977.388 | -7978.782 | -1.394 | 9.60E-07 |
| 11 | 116481221 | rs545613 | C | T | 0.985 | -7977.388 | -7978.784 | -1.395 | 9.59E-07 |
| 11 | 116482895 | rs511011 | C | T | 0.985 | -7977.388 | -7978.783 | -1.395 | 9.59E-07 |
| 11 | 116727465 | rs78857232 | T | C | 0.974 | -7977.388 | -7978.784 | -1.395 | 9.59E-07 |
| 11 | 118970696 | rs604714 | A | C | 1.03 | -7977.388 | -7978.784 | -1.395 | 9.59E-07 |
| 11 | 116408318 | rs868066 | A | G | 0.992 | -7977.388 | -7978.784 | -1.396 | 9.58E-07 |
| 11 | 116437756 | rs11605293 | T | C | 1.011 | -7977.388 | -7978.784 | -1.396 | 9.58E-07 |
| 11 | 118535840 | rs12225548 | G | C | 1.03 | -7977.388 | -7978.785 | -1.396 | 9.58E-07 |
| 11 | 119671453 | rs895652 | A | G | 0.971 | -7977.388 | -7978.785 | -1.396 | 9.58E-07 |
| 11 | 119900407 | rs76610415 | A | G | 1.025 | -7977.388 | -7978.784 | -1.396 | 9.58E-07 |
| 11 | 120302729 | rs55869319 | T | C | 0.976 | -7977.388 | -7978.785 | -1.396 | 9.58E-07 |
| 11 | 118501723 | rs61900957 | T | C | 0.988 | -7977.388 | -7978.786 | -1.397 | 9.57E-07 |
| 11 | 116322819 | rs550791 | T | C | 0.973 | -7977.388 | -7978.786 | -1.398 | 9.56E-07 |
| 11 | 118839133 | rs61900159 | C | G | 1.03 | -7977.388 | -7978.786 | -1.398 | 9.56E-07 |
| 11 | 118839268 | rs11217107 | C | G | 1.03 | -7977.388 | -7978.786 | -1.398 | 9.56E-07 |
| 11 | 116694866 | rs1268833 | G | A | 1.03 | -7977.388 | -7978.788 | -1.399 | 9.55E-07 |

| | | | | | | | |
|---|---|---|---|---|---|---|---|
| 11 | 116723737 | rs888245 | G | C | NULL | 0.668187772 | NULL&M_Disease |
| 11 | 118707559 | rs7118009 | T | C | NULL | 0.668187772 | NULL&M_Disease |
| 11 | 119813552 | rs1944503 | A | G | NULL | 0.668187772 | NULL&M_Disease |
| 11 | 120635399 | rs12276368 | A | G | NULL | 0.668187772 | NULL&M_Disease |
| 11 | 117125216 | rs679022 | C | T | NULL | 0.668187772 | NULL&M_Disease |
| 11 | 117769598 | rs4938454 | A | C | NULL | 0.668187772 | NULL&M_Disease |
| 11 | 119978743 | rs605742 | T | G | NULL | 0.668187772 | NULL&M_Disease |
| 11 | 120029638 | rs12366157 | G | A | NULL | 0.668187772 | NULL&M_Disease |
| 11 | 116210212 | rs4938263 | G | A | NULL | 0.668187772 | NULL&M_Disease |
| 11 | 116252227 | rs73019249 | G | A | NULL | 0.668187772 | NULL&M_Disease |
| 11 | 116482451 | rs513510 | C | T | NULL | 0.668187772 | NULL&M_Disease |
| 11 | 116507056 | rs77983701 | C | T | NULL | 0.668187772 | NULL&M_Disease |
| 11 | 118514625 | rs11216938 | T | C | NULL | 0.668187772 | NULL&M_Disease |
| 11 | 118516737 | rs12362152 | G | A | NULL | 0.668187772 | NULL&M_Disease |
| 11 | 118901614 | rs1784559 | T | A | NULL | 0.668187772 | NULL&M_Disease |
| 11 | 119796469 | rs73010650 | A | G | NULL | 0.668187772 | NULL&M_Disease |
| 11 | 119975273 | rs11217693 | G | A | NULL | 0.668187772 | NULL&M_Disease |
| 11 | 120813959 | rs3781816 | A | G | NULL | 0.668187772 | NULL&M_Disease |
| 11 | 120839015 | rs7943727 | C | T | NULL | 0.668187772 | NULL&M_Disease |
| 11 | 120887214 | rs117190595 | T | A | NULL | 0.668187772 | NULL&M_Disease |
| 11 | 121120917 | rs948377 | T | C | NULL | 0.668187772 | NULL&M_Disease |
| 11 | 116220063 | rs560299 | C | G | NULL | 0.668187772 | NULL&M_Disease |
| 11 | 117293818 | rs3863309 | C | T | NULL | 0.668187772 | NULL&M_Disease |
| 11 | 118915944 | rs1043314 | G | A | NULL | 0.668187772 | NULL&M_Disease |
| 11 | 119043745 | rs561830 | T | C | NULL | 0.668187772 | NULL&M_Disease |
| 11 | 120817653 | rs10892651 | A | G | NULL | 0.668187772 | NULL&M_Disease |
| 11 | 116488882 | rs10891999 | T | A | NULL | 0.668187772 | NULL&M_Disease |
| 11 | 118016039 | rs955917 | A | G | NULL | 0.668187772 | NULL&M_Disease |
| 11 | 119089831 | rs7111358 | A | T | NULL | 0.668187772 | NULL&M_Disease |
| 11 | 119689251 | rs7936503 | T | G | NULL | 0.668187772 | NULL&M_Disease |
| 11 | 119942972 | rs965312 | A | G | NULL | 0.668187772 | NULL&M_Disease |
| 11 | 118473231 | rs1047079 | C | T | NULL | 0.668187772 | NULL&M_Disease |
| 11 | 119307070 | rs59627935 | C | T | NULL | 0.668187772 | NULL&M_Disease |
| 11 | 119489210 | rs12361680 | C | G | NULL | 0.668187772 | NULL&M_Disease |
| 11 | 120399726 | rs10892584 | T | G | NULL | 0.668187772 | NULL&M_Disease |
| 11 | 120441139 | rs7929781 | T | C | NULL | 0.668187772 | NULL&M_Disease |
| 11 | 116236334 | rs79831199 | A | C | NULL | 0.668187772 | NULL&M_Disease |
| 11 | 116683374 | rs2727793 | C | T | NULL | 0.668187772 | NULL&M_Disease |
| 11 | 116780095 | rs1294767885 | T | A | NULL | 0.668187772 | NULL&M_Disease |
| 11 | 117110740 | rs650585 | T | C | NULL | 0.668187772 | NULL&M_Disease |
| 11 | 117341030 | rs544274 | G | A | NULL | 0.668187772 | NULL&M_Disease |
| 11 | 118471996 | rs11216926 | T | C | NULL | 0.668187772 | NULL&M_Disease |
| 11 | 118516774 | rs12365095 | T | C | NULL | 0.668187772 | NULL&M_Disease |
| 11 | 119088176 | rs11217188 | A | G | NULL | 0.668187772 | NULL&M_Disease |
| 11 | 121122021 | rs7120765 | A | G | NULL | 0.668187772 | NULL&M_Disease |
| 11 | 116681008 | rs1263173 | G | A | NULL | 0.668187772 | NULL&M_Disease |
| 11 | 117769430 | rs11600022 | T | C | NULL | 0.668187772 | NULL&M_Disease |
| 11 | 119893752 | rs10502233 | C | G | NULL | 0.668187772 | NULL&M_Disease |
| 11 | 121121448 | rs7928669 | T | C | NULL | 0.668187772 | NULL&M_Disease |
| 11 | 116996539 | rs17120347 | A | G | NULL | 0.668187772 | NULL&M_Disease |
| 11 | 118529127 | rs7928371 | A | G | NULL | 0.668187772 | NULL&M_Disease |
| 11 | 118914388 | rs7114961 | C | G | NULL | 0.669295403 | NULL&M_Disease |
| 11 | 118919132 | rs2276060 | A | G | NULL | 0.669295403 | NULL&M_Disease |
| 11 | 120600361 | rs1939665 | T | C | NULL | 0.669295403 | NULL&M_Disease |
| 11 | 120728916 | rs56275759 | C | T | NULL | 0.669295403 | NULL&M_Disease |
| 11 | 116220118 | rs560476 | A | G | NULL | 0.669295403 | NULL&M_Disease |
| 11 | 117684613 | rs11216555 | A | G | NULL | 0.669295403 | NULL&M_Disease |
| 11 | 119386594 | rs672113 | C | T | NULL | 0.669295403 | NULL&M_Disease |
| 11 | 120021546 | rs56339619 | A | G | NULL | 0.669295403 | NULL&M_Disease |
| 11 | 116203433 | rs1440183 | C | T | NULL | 0.669295403 | NULL&M_Disease |
| 11 | 116718415 | rs61905145 | A | G | NULL | 0.669295403 | NULL&M_Disease |
| 11 | 118734640 | rs12286723 | T | C | NULL | 0.669295403 | NULL&M_Disease |
| 11 | 120657978 | rs73578569 | T | C | NULL | 0.669295403 | NULL&M_Disease |
| 11 | 116250624 | rs481473 | C | A | NULL | 0.669295403 | NULL&M_Disease |
| 11 | 116529524 | rs523518 | A | G | NULL | 0.669295403 | NULL&M_Disease |
| 11 | 116537618 | rs1784124 | A | G | NULL | 0.669295403 | NULL&M_Disease |
| 11 | 116924318 | rs80259223 | A | G | NULL | 0.669295403 | NULL&M_Disease |
| 11 | 117313174 | rs11216392 | T | C | NULL | 0.669295403 | NULL&M_Disease |
| 11 | 117342221 | rs538079 | G | A | NULL | 0.669295403 | NULL&M_Disease |
| 11 | 119220012 | rs7932692 | G | C | NULL | 0.669295403 | NULL&M_Disease |
| 11 | 119594057 | rs11217429 | T | G | NULL | 0.669295403 | NULL&M_Disease |
| 11 | 116703146 | rs5130 | C | T | NULL | 0.669295403 | NULL&M_Disease |
| 11 | 117430389 | rs12803024 | G | C | NULL | 0.669295403 | NULL&M_Disease |
| 11 | 117436936 | rs1893899 | T | G | NULL | 0.669295403 | NULL&M_Disease |
| 11 | 118463221 | rs201373950 | A | T | NULL | 0.669295403 | NULL&M_Disease |
| 11 | 119811841 | rs35503012 | G | A | NULL | 0.669295403 | NULL&M_Disease |
| 11 | 119929276 | rs10444337 | A | G | NULL | 0.669295403 | NULL&M_Disease |
| 11 | 116238176 | rs79197348 | G | C | NULL | 0.669295403 | NULL&M_Disease |
| 11 | 117430164 | rs66657611 | T | C | NULL | 0.669295403 | NULL&M_Disease |
| 11 | 117430426 | rs12803171 | T | C | NULL | 0.669295403 | NULL&M_Disease |
| 11 | 118837460 | rs6589710 | T | A | NULL | 0.669295403 | NULL&M_Disease |
| 11 | 116450463 | rs61362984 | G | A | NULL | 0.669295403 | NULL&M_Disease |
| 11 | 116506410 | rs2077334 | A | C | NULL | 0.669295403 | NULL&M_Disease |
| 11 | 116690578 | rs1268354 | C | T | NULL | 0.669295403 | NULL&M_Disease |
| 11 | 116690712 | rs1263177 | C | T | NULL | 0.669295403 | NULL&M_Disease |
| 11 | 117283676 | rs8258 | T | C | NULL | 0.669295403 | NULL&M_Disease |
| 11 | 117424454 | rs11216451 | G | A | NULL | 0.669295403 | NULL&M_Disease |
| 11 | 117427923 | rs4938406 | A | G | NULL | 0.669295403 | NULL&M_Disease |
| 11 | 117446931 | rs55869870 | A | C | NULL | 0.669295403 | NULL&M_Disease |
| 11 | 118846574 | rs4938604 | C | T | NULL | 0.669295403 | NULL&M_Disease |
| 11 | 121092641 | rs948093 | A | T | NULL | 0.669295403 | NULL&M_Disease |
| 11 | 117256384 | rs560355 | C | G | NULL | 0.669295403 | NULL&M_Disease |
| 11 | 118534082 | rs10790256 | T | C | NULL | 0.669295403 | NULL&M_Disease |
| 11 | 119085783 | rs6589722 | C | A | NULL | 0.669295403 | NULL&M_Disease |
| 11 | 119279939 | rs11217272 | G | A | NULL | 0.669295403 | NULL&M_Disease |
| 11 | 119611348 | rs11217440 | G | A | NULL | 0.669295403 | NULL&M_Disease |

| | | | | | | | | | |
|---|---|---|---|---|---|---|---|---|---|
| 11 | 116723737 | rs888245 | G | C | 0.974 | -7977.388 | -7978.787 | -1.399 | 9.55E-07 |
| 11 | 118707559 | rs7118009 | T | C | 1.028 | -7977.388 | -7978.788 | -1.399 | 9.55E-07 |
| 11 | 119813552 | rs1944503 | A | G | 1.024 | -7977.388 | -7978.787 | -1.399 | 9.55E-07 |
| 11 | 120635399 | rs12276368 | A | G | 1.011 | -7977.388 | -7978.788 | -1.4 | 9.54E-07 |
| 11 | 117125216 | rs679022 | C | T | 0.971 | -7977.388 | -7978.789 | -1.401 | 9.53E-07 |
| 11 | 117769598 | rs4938454 | A | C | 0.971 | -7977.388 | -7978.789 | -1.401 | 9.53E-07 |
| 11 | 119978743 | rs605742 | T | G | 0.984 | -7977.388 | -7978.789 | -1.401 | 9.53E-07 |
| 11 | 120029638 | rs12366157 | G | A | 1.023 | -7977.388 | -7978.789 | -1.401 | 9.53E-07 |
| 11 | 116210212 | rs4938263 | G | A | 1.03 | -7977.388 | -7978.79 | -1.402 | 9.52E-07 |
| 11 | 116252227 | rs73019249 | G | A | 1.011 | -7977.388 | -7978.791 | -1.402 | 9.52E-07 |
| 11 | 116482451 | rs513510 | C | T | 0.986 | -7977.388 | -7978.79 | -1.402 | 9.52E-07 |
| 11 | 116507056 | rs77983701 | C | T | 0.975 | -7977.388 | -7978.79 | -1.402 | 9.52E-07 |
| 11 | 118514625 | rs11216938 | T | C | 1.03 | -7977.388 | -7978.79 | -1.402 | 9.52E-07 |
| 11 | 118516737 | rs12362152 | G | A | 1.03 | -7977.388 | -7978.79 | -1.402 | 9.52E-07 |
| 11 | 118901614 | rs1784559 | T | A | 0.971 | -7977.388 | -7978.79 | -1.402 | 9.52E-07 |
| 11 | 119796469 | rs73010650 | A | G | 0.992 | -7977.388 | -7978.79 | -1.402 | 9.52E-07 |
| 11 | 119975273 | rs11217693 | G | A | 0.974 | -7977.388 | -7978.79 | -1.402 | 9.52E-07 |
| 11 | 120813959 | rs3781816 | A | G | 0.971 | -7977.388 | -7978.79 | -1.402 | 9.52E-07 |
| 11 | 120839015 | rs7943727 | C | T | 0.971 | -7977.388 | -7978.79 | -1.402 | 9.52E-07 |
| 11 | 120887214 | rs117190595 | T | A | 0.996 | -7977.388 | -7978.79 | -1.402 | 9.52E-07 |
| 11 | 121120917 | rs948377 | T | C | 1.03 | -7977.388 | -7978.79 | -1.402 | 9.52E-07 |
| 11 | 116220063 | rs560299 | C | G | 1.03 | -7977.388 | -7978.792 | -1.403 | 9.51E-07 |
| 11 | 117293818 | rs3863309 | C | T | 0.974 | -7977.388 | -7978.791 | -1.403 | 9.51E-07 |
| 11 | 118915944 | rs1043314 | G | A | 0.971 | -7977.388 | -7978.792 | -1.403 | 9.51E-07 |
| 11 | 119043745 | rs561830 | T | C | 0.971 | -7977.388 | -7978.792 | -1.403 | 9.51E-07 |
| 11 | 120817653 | rs10892651 | A | G | 0.971 | -7977.388 | -7978.791 | -1.403 | 9.51E-07 |
| 11 | 116488882 | rs10891999 | T | A | 1.03 | -7977.388 | -7978.792 | -1.404 | 9.50E-07 |
| 11 | 118016039 | rs955917 | A | G | 1.027 | -7977.388 | -7978.793 | -1.404 | 9.50E-07 |
| 11 | 119089831 | rs7111358 | A | T | 1.03 | -7977.388 | -7978.792 | -1.404 | 9.50E-07 |
| 11 | 119689251 | rs7936503 | T | G | 0.971 | -7977.388 | -7978.792 | -1.404 | 9.50E-07 |
| 11 | 119942972 | rs965312 | A | G | 1.03 | -7977.388 | -7978.792 | -1.404 | 9.50E-07 |
| 11 | 118473231 | rs1047079 | C | T | 0.977 | -7977.388 | -7978.793 | -1.405 | 9.49E-07 |
| 11 | 119307070 | rs59627935 | C | T | 1.024 | -7977.388 | -7978.793 | -1.405 | 9.49E-07 |
| 11 | 119489210 | rs12361680 | C | G | 1.023 | -7977.388 | -7978.794 | -1.405 | 9.49E-07 |
| 11 | 120399726 | rs10892584 | T | G | 1.03 | -7977.388 | -7978.793 | -1.405 | 9.49E-07 |
| 11 | 120441139 | rs7929781 | T | C | 1.03 | -7977.388 | -7978.793 | -1.405 | 9.49E-07 |
| 11 | 116236334 | rs79831199 | A | C | 1.006 | -7977.388 | -7978.795 | -1.406 | 9.48E-07 |
| 11 | 116683374 | rs2727793 | C | T | 1.03 | -7977.388 | -7978.794 | -1.406 | 9.48E-07 |
| 11 | 116780095 | rs1294767885 | T | A | 0.975 | -7977.388 | -7978.794 | -1.406 | 9.48E-07 |
| 11 | 117110740 | rs650585 | T | C | 0.971 | -7977.388 | -7978.794 | -1.406 | 9.48E-07 |
| 11 | 117341030 | rs544274 | G | A | 0.989 | -7977.388 | -7978.794 | -1.406 | 9.48E-07 |
| 11 | 118471996 | rs11216926 | T | C | 0.977 | -7977.388 | -7978.795 | -1.406 | 9.48E-07 |
| 11 | 118516774 | rs12365095 | T | C | 1.029 | -7977.388 | -7978.795 | -1.406 | 9.48E-07 |
| 11 | 119088176 | rs11217188 | A | G | 1.03 | -7977.388 | -7978.795 | -1.406 | 9.48E-07 |
| 11 | 121122021 | rs7120765 | A | G | 1.03 | -7977.388 | -7978.794 | -1.406 | 9.48E-07 |
| 11 | 116681008 | rs1263173 | G | A | 1.03 | -7977.388 | -7978.795 | -1.407 | 9.47E-07 |
| 11 | 117769430 | rs11600022 | T | C | 0.971 | -7977.388 | -7978.795 | -1.407 | 9.47E-07 |
| 11 | 119893752 | rs10502233 | C | G | 1.024 | -7977.388 | -7978.795 | -1.407 | 9.47E-07 |
| 11 | 121121448 | rs7928669 | T | C | 1.03 | -7977.388 | -7978.795 | -1.407 | 9.47E-07 |
| 11 | 116996539 | rs17120347 | A | G | 1.015 | -7977.388 | -7978.796 | -1.408 | 9.46E-07 |
| 11 | 118529127 | rs7928371 | A | G | 1.029 | -7977.388 | -7978.796 | -1.408 | 9.46E-07 |
| 11 | 118914388 | rs7114961 | C | G | 0.971 | -7977.388 | -7978.796 | -1.408 | 9.46E-07 |
| 11 | 118919132 | rs2276060 | A | G | 0.971 | -7977.388 | -7978.796 | -1.408 | 9.46E-07 |
| 11 | 120600361 | rs1939665 | T | C | 0.971 | -7977.388 | -7978.797 | -1.408 | 9.46E-07 |
| 11 | 120728916 | rs56275759 | C | T | 0.971 | -7977.388 | -7978.796 | -1.408 | 9.46E-07 |
| 11 | 116220118 | rs560476 | A | G | 1.03 | -7977.388 | -7978.797 | -1.409 | 9.45E-07 |
| 11 | 117684613 | rs11216555 | A | G | 0.971 | -7977.388 | -7978.798 | -1.409 | 9.45E-07 |
| 11 | 119386594 | rs672113 | C | T | 1.03 | -7977.388 | -7978.797 | -1.409 | 9.45E-07 |
| 11 | 120021546 | rs56339619 | A | G | 0.981 | -7977.388 | -7978.798 | -1.409 | 9.45E-07 |
| 11 | 116203433 | rs1440183 | C | T | 1.03 | -7977.388 | -7978.798 | -1.41 | 9.44E-07 |
| 11 | 116718415 | rs61905145 | A | G | 0.975 | -7977.388 | -7978.798 | -1.41 | 9.44E-07 |
| 11 | 118734640 | rs12286723 | T | C | 1.028 | -7977.388 | -7978.799 | -1.41 | 9.44E-07 |
| 11 | 120657978 | rs73578569 | T | C | 0.972 | -7977.388 | -7978.799 | -1.41 | 9.44E-07 |
| 11 | 116250624 | rs481473 | C | A | 1.024 | -7977.388 | -7978.799 | -1.411 | 9.43E-07 |
| 11 | 116529524 | rs523518 | A | G | 0.971 | -7977.388 | -7978.8 | -1.411 | 9.43E-07 |
| 11 | 116537618 | rs1784124 | A | G | 1.029 | -7977.388 | -7978.799 | -1.411 | 9.43E-07 |
| 11 | 116924318 | rs80259223 | A | G | 0.984 | -7977.388 | -7978.799 | -1.411 | 9.43E-07 |
| 11 | 117313174 | rs11216392 | T | C | 1.008 | -7977.388 | -7978.799 | -1.411 | 9.43E-07 |
| 11 | 117342221 | rs538079 | G | A | 1.03 | -7977.388 | -7978.8 | -1.411 | 9.43E-07 |
| 11 | 119220012 | rs7932692 | G | C | 0.971 | -7977.388 | -7978.799 | -1.411 | 9.43E-07 |
| 11 | 119594057 | rs11217429 | T | G | 0.971 | -7977.388 | -7978.8 | -1.411 | 9.43E-07 |
| 11 | 116703146 | rs5130 | C | T | 0.976 | -7977.388 | -7978.8 | -1.412 | 9.42E-07 |
| 11 | 117430389 | rs12803024 | G | C | 1 | -7977.388 | -7978.801 | -1.412 | 9.42E-07 |
| 11 | 117436936 | rs1893899 | T | G | 0.971 | -7977.388 | -7978.8 | -1.412 | 9.42E-07 |
| 11 | 118463221 | rs201373950 | A | T | 0.974 | -7977.388 | -7978.8 | -1.412 | 9.42E-07 |
| 11 | 119811841 | rs35503012 | G | A | 1.023 | -7977.388 | -7978.8 | -1.412 | 9.42E-07 |
| 11 | 119929276 | rs10444337 | A | G | 1.03 | -7977.388 | -7978.8 | -1.412 | 9.42E-07 |
| 11 | 116238176 | rs79197348 | G | C | 1.004 | -7977.388 | -7978.801 | -1.413 | 9.42E-07 |
| 11 | 117430164 | rs66657611 | T | C | 1.002 | -7977.388 | -7978.801 | -1.413 | 9.42E-07 |
| 11 | 117430426 | rs12803171 | T | C | 1.001 | -7977.388 | -7978.801 | -1.413 | 9.42E-07 |
| 11 | 118837460 | rs6589710 | T | A | 1.03 | -7977.388 | -7978.801 | -1.413 | 9.42E-07 |
| 11 | 116450463 | rs61362984 | G | A | 1.03 | -7977.388 | -7978.803 | -1.414 | 9.41E-07 |
| 11 | 116506410 | rs2077334 | A | C | 0.976 | -7977.388 | -7978.802 | -1.414 | 9.41E-07 |
| 11 | 116690578 | rs1268354 | C | T | 1.03 | -7977.388 | -7978.802 | -1.414 | 9.41E-07 |
| 11 | 116690712 | rs1263177 | C | T | 1.03 | -7977.388 | -7978.803 | -1.414 | 9.41E-07 |
| 11 | 117283676 | rs8258 | T | C | 0.971 | -7977.388 | -7978.803 | -1.415 | 9.40E-07 |
| 11 | 117424454 | rs11216451 | G | A | 0.995 | -7977.388 | -7978.803 | -1.415 | 9.40E-07 |
| 11 | 117427923 | rs4938406 | A | G | 0.997 | -7977.388 | -7978.804 | -1.415 | 9.40E-07 |
| 11 | 117446931 | rs55869870 | A | C | 0.971 | -7977.388 | -7978.803 | -1.415 | 9.40E-07 |
| 11 | 118846574 | rs4938604 | C | T | 1.03 | -7977.388 | -7978.803 | -1.415 | 9.40E-07 |
| 11 | 121092641 | rs948093 | A | T | 0.971 | -7977.388 | -7978.803 | -1.415 | 9.40E-07 |
| 11 | 117256384 | rs560355 | C | G | 0.971 | -7977.388 | -7978.804 | -1.416 | 9.39E-07 |
| 11 | 118534082 | rs10790256 | T | C | 1.029 | -7977.388 | -7978.805 | -1.417 | 9.38E-07 |
| 11 | 119085783 | rs6589722 | C | A | 1.03 | -7977.388 | -7978.805 | -1.417 | 9.38E-07 |
| 11 | 119279939 | rs11217272 | G | A | 0.981 | -7977.388 | -7978.806 | -1.417 | 9.38E-07 |
| 11 | 119611348 | rs11217440 | G | A | 0.971 | -7977.388 | -7978.805 | -1.417 | 9.38E-07 |

| | | | | | | | |
|---|---|---|---|---|---|---|---|
| 11 | 119611360 | rs11217441 | A | G | NULL | 0.669295403 | NULL&M_Disease |
| 11 | 119925575 | rs628861 | A | C | NULL | 0.669295403 | NULL&M_Disease |
| 11 | 120441225 | rs12574365 | C | T | NULL | 0.669295403 | NULL&M_Disease |
| 11 | 120993723 | rs61013776 | A | G | NULL | 0.669295403 | NULL&M_Disease |
| 11 | 117128271 | rs667944 | G | A | NULL | 0.67040116 | NULL&M_Disease |
| 11 | 117525255 | rs7947144 | G | A | NULL | 0.669295403 | NULL&M_Disease |
| 11 | 117631234 | rs10750111 | C | T | NULL | 0.67040116 | NULL&M_Disease |
| 11 | 117653576 | rs7941202 | G | C | NULL | 0.67040116 | NULL&M_Disease |
| 11 | 119810909 | rs34239965 | T | C | NULL | 0.669295403 | NULL&M_Disease |
| 11 | 118033545 | rs625464 | T | C | NULL | 0.67040116 | NULL&M_Disease |
| 11 | 118290169 | rs7124100 | A | G | NULL | 0.67040116 | NULL&M_Disease |
| 11 | 116222378 | rs534437 | A | G | NULL | 0.67040116 | NULL&M_Disease |
| 11 | 117042580 | rs10892083 | C | G | NULL | 0.67040116 | NULL&M_Disease |
| 11 | 120601807 | rs3132776 | A | G | NULL | 0.67040116 | NULL&M_Disease |
| 11 | 121092679 | rs948094 | T | C | NULL | 0.67040116 | NULL&M_Disease |
| 11 | 116213941 | rs10891966 | G | A | NULL | 0.67040116 | NULL&M_Disease |
| 11 | 116648917 | rs964184 | G | C | NULL | 0.67040116 | NULL&M_Disease |
| 11 | 117294022 | rs11607141 | G | T | NULL | 0.67040116 | NULL&M_Disease |
| 11 | 116529024 | rs7104343 | T | C | NULL | 0.67040116 | NULL&M_Disease |
| 11 | 116684767 | rs2542063 | G | A | NULL | 0.67040116 | NULL&M_Disease |
| 11 | 116918046 | rs74328581 | A | C | NULL | 0.67040116 | NULL&M_Disease |
| 11 | 119084914 | rs10790287 | A | G | NULL | 0.67040116 | NULL&M_Disease |
| 11 | 120061195 | rs12420591 | G | A | NULL | 0.67040116 | NULL&M_Disease |
| 11 | 120155336 | rs4936512 | C | T | NULL | 0.67040116 | NULL&M_Disease |
| 11 | 120155564 | rs10790373 | T | C | NULL | 0.67040116 | NULL&M_Disease |
| 11 | 117449297 | rs10892139 | G | A | NULL | 0.67040116 | NULL&M_Disease |
| 11 | 116250106 | rs566174 | C | G | NULL | 0.67040116 | NULL&M_Disease |
| 11 | 116439118 | rs61426083 | A | G | NULL | 0.67040116 | NULL&M_Disease |
| 11 | 118661376 | rs73003294 | T | C | NULL | 0.67040116 | NULL&M_Disease |
| 11 | 119697231 | rs2084291 | G | A | NULL | 0.67040116 | NULL&M_Disease |
| 11 | 120643050 | rs80326995 | A | G | NULL | 0.67040116 | NULL&M_Disease |
| 11 | 120817865 | rs35024210 | A | C | NULL | 0.67040116 | NULL&M_Disease |
| 11 | 121029733 | rs11218169 | G | A | NULL | 0.67040116 | NULL&M_Disease |
| 11 | 121062777 | rs60016568 | T | C | NULL | 0.67040116 | NULL&M_Disease |
| 11 | 116334862 | rs66899922 | T | G | NULL | 0.67040116 | NULL&M_Disease |
| 11 | 116336988 | rs73582578 | A | C | NULL | 0.67040116 | NULL&M_Disease |
| 11 | 117923983 | rs11828580 | A | G | NULL | 0.67040116 | NULL&M_Disease |
| 11 | 119811909 | rs35462788 | T | C | NULL | 0.67040116 | NULL&M_Disease |
| 11 | 116225004 | rs505349 | G | A | NULL | 0.67040116 | NULL&M_Disease |
| 11 | 116373451 | rs875025 | T | C | NULL | 0.67040116 | NULL&M_Disease |
| 11 | 117690400 | rs11600216 | C | T | NULL | 0.67040116 | NULL&M_Disease |
| 11 | 120154801 | rs7479050 | C | T | NULL | 0.67040116 | NULL&M_Disease |
| 11 | 116337814 | rs56264624 | A | G | NULL | 0.67040116 | NULL&M_Disease |
| 11 | 117539771 | rs2282549 | C | A | NULL | 0.67040116 | NULL&M_Disease |
| 11 | 119760867 | rs7105077 | G | A | NULL | 0.67040116 | NULL&M_Disease |
| 11 | 116972894 | rs4938338 | A | C | NULL | 0.67040116 | NULL&M_Disease |
| 11 | 119080313 | rs7107776 | T | C | NULL | 0.671505034 | NULL&M_Disease |
| 11 | 119929445 | rs10444338 | C | T | NULL | 0.671505034 | NULL&M_Disease |
| 11 | 116216713 | rs567681 | C | A | NULL | 0.671505034 | NULL&M_Disease |
| 11 | 116216759 | rs558659 | A | G | NULL | 0.671505034 | NULL&M_Disease |
| 11 | 116338675 | rs17119713 | G | T | NULL | 0.671505034 | NULL&M_Disease |
| 11 | 116667083 | rs7123666 | A | G | NULL | 0.671505034 | NULL&M_Disease |
| 11 | 118507478 | rs11216937 | T | C | NULL | 0.671505034 | NULL&M_Disease |
| 11 | 116200726 | rs7944542 | G | A | NULL | 0.671505034 | NULL&M_Disease |
| 11 | 116250285 | rs556727 | G | A | NULL | 0.671505034 | NULL&M_Disease |
| 11 | 117700410 | rs2077276 | A | C | NULL | 0.671505034 | NULL&M_Disease |
| 11 | 119219311 | rs11217245 | G | A | NULL | 0.671505034 | NULL&M_Disease |
| 11 | 119782250 | rs540103 | G | A | NULL | 0.671505034 | NULL&M_Disease |
| 11 | 116367567 | rs6589550 | C | A | NULL | 0.671505034 | NULL&M_Disease |
| 11 | 116634173 | rs17440824 | T | C | NULL | 0.671505034 | NULL&M_Disease |
| 11 | 116671823 | rs9667814 | C | G | NULL | 0.671505034 | NULL&M_Disease |
| 11 | 116692694 | rs5100 | G | A | NULL | 0.671505034 | NULL&M_Disease |
| 11 | 116693095 | rs5096 | G | A | NULL | 0.671505034 | NULL&M_Disease |
| 11 | 116766390 | rs11216178 | T | G | NULL | 0.671505034 | NULL&M_Disease |
| 11 | 116766899 | rs10892044 | C | T | NULL | 0.671505034 | NULL&M_Disease |
| 11 | 117775345 | rs1271170 | A | G | NULL | 0.671505034 | NULL&M_Disease |
| 11 | 118798757 | rs57879605 | T | C | NULL | 0.671505034 | NULL&M_Disease |
| 11 | 120661178 | rs3133860 | T | A | NULL | 0.671505034 | NULL&M_Disease |
| 11 | 116308355 | rs12807410 | A | G | NULL | 0.671505034 | NULL&M_Disease |
| 11 | 120209386 | rs4938800 | A | G | NULL | 0.671505034 | NULL&M_Disease |
| 11 | 120340383 | rs11217878 | A | G | NULL | 0.671505034 | NULL&M_Disease |
| 11 | 120346360 | rs2276035 | A | G | NULL | 0.671505034 | NULL&M_Disease |
| 11 | 120979316 | rs3016242 | A | C | NULL | 0.671505034 | NULL&M_Disease |
| 11 | 121121501 | rs7942167 | G | A | NULL | 0.671505034 | NULL&M_Disease |
| 11 | 116341125 | rs1848373 | A | T | NULL | 0.671505034 | NULL&M_Disease |
| 11 | 116538851 | rs1632755 | C | A | NULL | 0.671505034 | NULL&M_Disease |
| 11 | 116680330 | rs1263171 | G | A | NULL | 0.671505034 | NULL&M_Disease |
| 11 | 116993292 | rs76868258 | T | C | NULL | 0.671505034 | NULL&M_Disease |
| 11 | 117416832 | rs11216447 | A | G | NULL | 0.671505034 | NULL&M_Disease |
| 11 | 119847854 | rs78913295 | T | C | NULL | 0.671505034 | NULL&M_Disease |
| 11 | 120498024 | rs7924948 | T | C | NULL | 0.671505034 | NULL&M_Disease |
| 11 | 120795441 | rs10502243 | A | G | NULL | 0.671505034 | NULL&M_Disease |
| 11 | 116452142 | rs7927785 | G | C | NULL | 0.671505034 | NULL&M_Disease |
| 11 | 117770505 | rs640483 | G | A | NULL | 0.671505034 | NULL&M_Disease |
| 11 | 119849440 | rs677428 | T | C | NULL | 0.671505034 | NULL&M_Disease |
| 11 | 116765700 | rs11820814 | C | T | NULL | 0.671505034 | NULL&M_Disease |
| 11 | 118516903 | rs80015904 | A | G | NULL | 0.671505034 | NULL&M_Disease |
| 11 | 119217254 | rs883247 | C | T | NULL | 0.671505034 | NULL&M_Disease |
| 11 | 120024312 | rs590005 | T | A | NULL | 0.671505034 | NULL&M_Disease |
| 11 | 120029497 | rs604413 | G | A | NULL | 0.671505034 | NULL&M_Disease |
| 11 | 120988124 | rs597345 | G | A | NULL | 0.671505034 | NULL&M_Disease |
| 11 | 116250523 | rs17476159 | C | T | NULL | 0.671505034 | NULL&M_Disease |
| 11 | 116364065 | rs7935656 | C | T | NULL | 0.671505034 | NULL&M_Disease |
| 11 | 116537974 | rs12787592 | C | G | NULL | 0.671505034 | NULL&M_Disease |
| 11 | 118112332 | rs12419365 | A | C | NULL | 0.671505034 | NULL&M_Disease |
| 11 | 118261736 | rs57702967 | A | C | NULL | 0.671505034 | NULL&M_Disease |

| | | | | | | | | | |
|---|---|---|---|---|---|---|---|---|---|
| 11 | 119611360 | rs11217441 | A | G | 0.971 | -7977.388 | -7978.805 | -1.417 | 9.38E-07 |
| 11 | 119925575 | rs628861 | A | C | 1.03 | -7977.388 | -7978.806 | -1.417 | 9.38E-07 |
| 11 | 120441225 | rs12574365 | C | T | 1.03 | -7977.388 | -7978.805 | -1.417 | 9.38E-07 |
| 11 | 120993723 | rs61013776 | A | G | 1.028 | -7977.388 | -7978.805 | -1.417 | 9.38E-07 |
| 11 | 117128271 | rs667944 | G | A | 0.971 | -7977.388 | -7978.806 | -1.418 | 9.37E-07 |
| 11 | 117525255 | rs7947144 | G | A | 1.01 | -7977.388 | -7978.807 | -1.418 | 9.37E-07 |
| 11 | 117631234 | rs10750111 | C | T | 0.971 | -7977.388 | -7978.806 | -1.418 | 9.37E-07 |
| 11 | 117653576 | rs7941202 | G | C | 1.03 | -7977.388 | -7978.806 | -1.418 | 9.37E-07 |
| 11 | 119810909 | rs34239965 | T | C | 1.022 | -7977.388 | -7978.807 | -1.418 | 9.37E-07 |
| 11 | 118033545 | rs625464 | T | C | 1.03 | -7977.388 | -7978.807 | -1.419 | 9.36E-07 |
| 11 | 118290169 | rs7124100 | A | G | 0.976 | -7977.388 | -7978.807 | -1.419 | 9.36E-07 |
| 11 | 116222378 | rs534437 | A | G | 1.03 | -7977.388 | -7978.809 | -1.42 | 9.35E-07 |
| 11 | 117042580 | rs10892083 | C | G | 0.971 | -7977.388 | -7978.808 | -1.42 | 9.35E-07 |
| 11 | 120601807 | rs3132776 | A | G | 0.971 | -7977.388 | -7978.809 | -1.42 | 9.35E-07 |
| 11 | 121092679 | rs948094 | T | C | 0.971 | -7977.388 | -7978.808 | -1.42 | 9.35E-07 |
| 11 | 116213941 | rs10891966 | G | A | 0.993 | -7977.388 | -7978.809 | -1.421 | 9.34E-07 |
| 11 | 116648917 | rs964184 | G | C | 0.977 | -7977.388 | -7978.809 | -1.421 | 9.34E-07 |
| 11 | 117294022 | rs11607141 | G | T | 1.026 | -7977.388 | -7978.809 | -1.421 | 9.34E-07 |
| 11 | 116529024 | rs7104343 | T | C | 0.998 | -7977.388 | -7978.811 | -1.422 | 9.33E-07 |
| 11 | 116684767 | rs2542063 | G | A | 1.029 | -7977.388 | -7978.81 | -1.422 | 9.33E-07 |
| 11 | 116918046 | rs74328581 | A | C | 0.988 | -7977.388 | -7978.81 | -1.422 | 9.33E-07 |
| 11 | 119084914 | rs10790287 | A | G | 1.03 | -7977.388 | -7978.81 | -1.422 | 9.33E-07 |
| 11 | 120061195 | rs12420591 | G | A | 0.978 | -7977.388 | -7978.81 | -1.422 | 9.33E-07 |
| 11 | 120155336 | rs4936512 | C | T | 1.027 | -7977.388 | -7978.81 | -1.422 | 9.33E-07 |
| 11 | 120155564 | rs10790373 | T | C | 1.027 | -7977.388 | -7978.81 | -1.422 | 9.33E-07 |
| 11 | 117449297 | rs10892139 | G | A | 0.971 | -7977.388 | -7978.811 | -1.423 | 9.32E-07 |
| 11 | 116250106 | rs566174 | C | G | 1.023 | -7977.388 | -7978.812 | -1.424 | 9.31E-07 |
| 11 | 116439118 | rs61426083 | A | G | 1.029 | -7977.388 | -7978.812 | -1.424 | 9.31E-07 |
| 11 | 118661376 | rs73003294 | T | C | 0.98 | -7977.388 | -7978.813 | -1.424 | 9.31E-07 |
| 11 | 119697231 | rs2084291 | G | A | 0.972 | -7977.388 | -7978.812 | -1.424 | 9.31E-07 |
| 11 | 120643050 | rs80326995 | A | G | 0.992 | -7977.388 | -7978.812 | -1.424 | 9.31E-07 |
| 11 | 120817865 | rs35024210 | A | C | 0.972 | -7977.388 | -7978.813 | -1.424 | 9.31E-07 |
| 11 | 121029733 | rs11218169 | G | A | 1.021 | -7977.388 | -7978.813 | -1.424 | 9.31E-07 |
| 11 | 121062777 | rs60016568 | T | C | 1.021 | -7977.388 | -7978.812 | -1.424 | 9.31E-07 |
| 11 | 116334862 | rs66899922 | T | G | 1.008 | -7977.388 | -7978.813 | -1.425 | 9.30E-07 |
| 11 | 116336988 | rs73582578 | A | C | 0.995 | -7977.388 | -7978.814 | -1.425 | 9.30E-07 |
| 11 | 117923983 | rs11828580 | A | G | 1.029 | -7977.388 | -7978.814 | -1.425 | 9.30E-07 |
| 11 | 119811909 | rs35462788 | T | C | 1.022 | -7977.388 | -7978.814 | -1.425 | 9.30E-07 |
| 11 | 116225004 | rs505349 | G | A | 1.03 | -7977.388 | -7978.815 | -1.426 | 9.29E-07 |
| 11 | 116373451 | rs875025 | T | C | 0.983 | -7977.388 | -7978.814 | -1.426 | 9.29E-07 |
| 11 | 117690400 | rs11600216 | C | T | 0.975 | -7977.388 | -7978.814 | -1.426 | 9.29E-07 |
| 11 | 120154801 | rs7479050 | C | T | 1.027 | -7977.388 | -7978.814 | -1.426 | 9.29E-07 |
| 11 | 116337814 | rs56264624 | A | G | 0.996 | -7977.388 | -7978.816 | -1.427 | 9.28E-07 |
| 11 | 117539771 | rs2282549 | C | A | 0.976 | -7977.388 | -7978.816 | -1.427 | 9.28E-07 |
| 11 | 119760867 | rs7105077 | G | A | 1.026 | -7977.388 | -7978.816 | -1.427 | 9.28E-07 |
| 11 | 116972894 | rs4938338 | A | C | 0.987 | -7977.388 | -7978.817 | -1.428 | 9.28E-07 |
| 11 | 119080313 | rs7107776 | T | C | 1.03 | -7977.388 | -7978.816 | -1.428 | 9.28E-07 |
| 11 | 119929445 | rs10444338 | C | T | 1.029 | -7977.388 | -7978.816 | -1.428 | 9.28E-07 |
| 11 | 116216713 | rs567681 | C | A | 1.029 | -7977.388 | -7978.817 | -1.429 | 9.27E-07 |
| 11 | 116216759 | rs558659 | A | G | 1.029 | -7977.388 | -7978.817 | -1.429 | 9.27E-07 |
| 11 | 116338675 | rs17119713 | G | T | 0.998 | -7977.388 | -7978.818 | -1.429 | 9.27E-07 |
| 11 | 116667083 | rs7123666 | A | G | 0.991 | -7977.388 | -7978.818 | -1.429 | 9.27E-07 |
| 11 | 118507478 | rs11216937 | T | C | 1.028 | -7977.388 | -7978.817 | -1.429 | 9.27E-07 |
| 11 | 116200726 | rs7944542 | G | A | 1.029 | -7977.388 | -7978.818 | -1.43 | 9.26E-07 |
| 11 | 116250285 | rs556727 | G | A | 1.023 | -7977.388 | -7978.818 | -1.43 | 9.26E-07 |
| 11 | 117700410 | rs2077276 | A | C | 0.973 | -7977.388 | -7978.818 | -1.43 | 9.26E-07 |
| 11 | 119219311 | rs11217245 | G | A | 0.971 | -7977.388 | -7978.818 | -1.43 | 9.26E-07 |
| 11 | 119782250 | rs540103 | G | A | 0.971 | -7977.388 | -7978.818 | -1.43 | 9.26E-07 |
| 11 | 116367567 | rs6589550 | C | A | 0.971 | -7977.388 | -7978.819 | -1.431 | 9.25E-07 |
| 11 | 116634173 | rs17440824 | T | C | 0.979 | -7977.388 | -7978.82 | -1.431 | 9.25E-07 |
| 11 | 116671823 | rs9667814 | C | G | 0.981 | -7977.388 | -7978.82 | -1.431 | 9.25E-07 |
| 11 | 116692694 | rs5100 | G | A | 1.029 | -7977.388 | -7978.819 | -1.431 | 9.25E-07 |
| 11 | 116693095 | rs5096 | G | A | 1.029 | -7977.388 | -7978.819 | -1.431 | 9.25E-07 |
| 11 | 116766390 | rs11216178 | T | G | 0.977 | -7977.388 | -7978.82 | -1.431 | 9.25E-07 |
| 11 | 116766899 | rs10892044 | C | T | 0.977 | -7977.388 | -7978.82 | -1.431 | 9.25E-07 |
| 11 | 117775345 | rs1271170 | A | G | 0.971 | -7977.388 | -7978.819 | -1.431 | 9.25E-07 |
| 11 | 118798757 | rs57879605 | T | C | 0.973 | -7977.388 | -7978.819 | -1.431 | 9.25E-07 |
| 11 | 120661178 | rs3133860 | T | A | 0.98 | -7977.388 | -7978.82 | -1.431 | 9.25E-07 |
| 11 | 116308355 | rs12807410 | A | G | 1.029 | -7977.388 | -7978.82 | -1.432 | 9.24E-07 |
| 11 | 120209386 | rs4938800 | A | G | 0.977 | -7977.388 | -7978.82 | -1.432 | 9.24E-07 |
| 11 | 120340383 | rs11217878 | A | G | 0.975 | -7977.388 | -7978.821 | -1.432 | 9.24E-07 |
| 11 | 120346360 | rs2276035 | A | G | 0.975 | -7977.388 | -7978.821 | -1.432 | 9.24E-07 |
| 11 | 120979316 | rs3016242 | A | C | 0.971 | -7977.388 | -7978.82 | -1.432 | 9.24E-07 |
| 11 | 121121501 | rs7942167 | G | A | 1.029 | -7977.388 | -7978.82 | -1.432 | 9.24E-07 |
| 11 | 116341125 | rs1848373 | A | T | 1.002 | -7977.388 | -7978.821 | -1.433 | 9.23E-07 |
| 11 | 116538851 | rs1632755 | C | A | 1.029 | -7977.388 | -7978.822 | -1.433 | 9.23E-07 |
| 11 | 116680330 | rs1263171 | G | A | 1.029 | -7977.388 | -7978.821 | -1.433 | 9.23E-07 |
| 11 | 116993292 | rs76868258 | T | C | 1.011 | -7977.388 | -7978.822 | -1.433 | 9.23E-07 |
| 11 | 117416832 | rs11216447 | A | G | 0.973 | -7977.388 | -7978.821 | -1.433 | 9.23E-07 |
| 11 | 119847854 | rs78913295 | T | C | 0.994 | -7977.388 | -7978.821 | -1.433 | 9.23E-07 |
| 11 | 120498024 | rs7924948 | T | C | 1.029 | -7977.388 | -7978.821 | -1.433 | 9.23E-07 |
| 11 | 120795441 | rs10502243 | A | G | 0.979 | -7977.388 | -7978.821 | -1.433 | 9.23E-07 |
| 11 | 116452142 | rs7927785 | G | C | 0.971 | -7977.388 | -7978.822 | -1.434 | 9.22E-07 |
| 11 | 117770505 | rs640483 | G | A | 1.026 | -7977.388 | -7978.822 | -1.434 | 9.22E-07 |
| 11 | 119849440 | rs677428 | T | C | 1.029 | -7977.388 | -7978.822 | -1.434 | 9.22E-07 |
| 11 | 116765700 | rs11820814 | C | T | 0.977 | -7977.388 | -7978.824 | -1.435 | 9.21E-07 |
| 11 | 118516903 | rs80015904 | A | G | 1.028 | -7977.388 | -7978.823 | -1.435 | 9.21E-07 |
| 11 | 119217254 | rs883247 | C | T | 0.971 | -7977.388 | -7978.823 | -1.435 | 9.21E-07 |
| 11 | 120024312 | rs590005 | T | A | 1.017 | -7977.388 | -7978.823 | -1.435 | 9.21E-07 |
| 11 | 120029497 | rs604413 | G | A | 1.017 | -7977.388 | -7978.823 | -1.435 | 9.21E-07 |
| 11 | 120988124 | rs597345 | G | A | 0.972 | -7977.388 | -7978.824 | -1.435 | 9.21E-07 |
| 11 | 116250523 | rs17476159 | C | T | 1.003 | -7977.388 | -7978.824 | -1.436 | 9.20E-07 |
| 11 | 116364065 | rs7935656 | C | T | 0.972 | -7977.388 | -7978.824 | -1.436 | 9.20E-07 |
| 11 | 116537974 | rs12787592 | C | G | 1.026 | -7977.388 | -7978.824 | -1.436 | 9.20E-07 |
| 11 | 118112332 | rs12419365 | A | C | 0.997 | -7977.388 | -7978.824 | -1.436 | 9.20E-07 |
| 11 | 118261736 | rs57702967 | A | C | 0.978 | -7977.388 | -7978.824 | -1.436 | 9.20E-07 |

| | | | | | | | |
|---|---|---|---|---|---|---|---|
| 11 | 118262305 | rs59405103 | C | A | NULL | 0.671505034 | NULL&M_Disease |
| 11 | 119834296 | rs12421491 | T | C | NULL | 0.671505034 | NULL&M_Disease |
| 11 | 119846442 | rs17123513 | A | G | NULL | 0.671505034 | NULL&M_Disease |
| 11 | 119846476 | rs74609299 | T | C | NULL | 0.671505034 | NULL&M_Disease |
| 11 | 120652075 | rs4936549 | C | T | NULL | 0.671505034 | NULL&M_Disease |
| 11 | 120960737 | rs56232713 | G | C | NULL | 0.671505034 | NULL&M_Disease |
| 11 | 116450185 | rs12280043 | T | A | NULL | 0.671505034 | NULL&M_Disease |
| 11 | 116453047 | rs7931384 | C | T | NULL | 0.671505034 | NULL&M_Disease |
| 11 | 117123173 | rs528508 | C | T | NULL | 0.671505034 | NULL&M_Disease |
| 11 | 118888831 | rs11820805 | C | T | NULL | 0.671505034 | NULL&M_Disease |
| 11 | 120025476 | rs596911 | A | G | NULL | 0.671505034 | NULL&M_Disease |
| 11 | 120026241 | rs659912 | G | A | NULL | 0.671505034 | NULL&M_Disease |
| 11 | 120055326 | rs73004764 | G | C | NULL | 0.671505034 | NULL&M_Disease |
| 11 | 116250393 | rs73019237 | T | C | NULL | 0.671505034 | NULL&M_Disease |
| 11 | 116611827 | rs180349 | A | T | NULL | 0.672607017 | NULL&M_Disease |
| 11 | 116710968 | rs613808 | A | G | NULL | 0.672607017 | NULL&M_Disease |
| 11 | 117902620 | rs9326250 | A | C | NULL | 0.672607017 | NULL&M_Disease |
| 11 | 118268613 | rs7108613 | G | A | NULL | 0.671505034 | NULL&M_Disease |
| 11 | 119086629 | rs4936467 | T | A | NULL | 0.672607017 | NULL&M_Disease |
| 11 | 119219360 | rs11217246 | C | A | NULL | 0.672607017 | NULL&M_Disease |
| 11 | 119828505 | rs1353117537 | A | T | NULL | 0.671505034 | NULL&M_Disease |
| 11 | 116397649 | rs4936353 | A | G | NULL | 0.672607017 | NULL&M_Disease |
| 11 | 119220946 | rs12417736 | T | C | NULL | 0.672607017 | NULL&M_Disease |
| 11 | 119503576 | rs34224649 | A | G | NULL | 0.672607017 | NULL&M_Disease |
| 11 | 119747312 | rs481716 | G | A | NULL | 0.672607017 | NULL&M_Disease |
| 11 | 119849798 | rs678914 | C | T | NULL | 0.672607017 | NULL&M_Disease |
| 11 | 120292040 | rs11604012 | C | T | NULL | 0.672607017 | NULL&M_Disease |
| 11 | 120651611 | rs3133858 | C | T | NULL | 0.672607017 | NULL&M_Disease |
| 11 | 116558802 | rs12800436 | T | G | NULL | 0.672607017 | NULL&M_Disease |
| 11 | 117201165 | rs515756 | G | T | NULL | 0.672607017 | NULL&M_Disease |
| 11 | 118259379 | rs76126708 | C | A | NULL | 0.672607017 | NULL&M_Disease |
| 11 | 118528424 | rs17748 | T | C | NULL | 0.672607017 | NULL&M_Disease |
| 11 | 119839224 | rs76233351 | A | G | NULL | 0.672607017 | NULL&M_Disease |
| 11 | 116688791 | rs142744740 | A | C | NULL | 0.672607017 | NULL&M_Disease |
| 11 | 117123297 | rs497043 | G | C | NULL | 0.672607017 | NULL&M_Disease |
| 11 | 117365821 | rs61905337 | A | C | NULL | 0.672607017 | NULL&M_Disease |
| 11 | 119220483 | rs7480948 | G | C | NULL | 0.672607017 | NULL&M_Disease |
| 11 | 119832917 | rs56993455 | A | G | NULL | 0.672607017 | NULL&M_Disease |
| 11 | 119833474 | rs59253729 | T | C | NULL | 0.672607017 | NULL&M_Disease |
| 11 | 120724136 | rs61901429 | G | A | NULL | 0.672607017 | NULL&M_Disease |
| 11 | 116338814 | rs12289284 | A | G | NULL | 0.672607017 | NULL&M_Disease |
| 11 | 116340960 | rs4471466 | C | A | NULL | 0.672607017 | NULL&M_Disease |
| 11 | 116507795 | rs11216069 | C | G | NULL | 0.672607017 | NULL&M_Disease |
| 11 | 117177800 | rs473210 | T | C | NULL | 0.672607017 | NULL&M_Disease |
| 11 | 119848377 | rs12272862 | A | G | NULL | 0.672607017 | NULL&M_Disease |
| 11 | 121031731 | rs75618086 | A | T | NULL | 0.672607017 | NULL&M_Disease |
| 11 | 121064444 | rs11218198 | G | A | NULL | 0.672607017 | NULL&M_Disease |
| 11 | 121119605 | rs12279833 | C | T | NULL | 0.672607017 | NULL&M_Disease |
| 11 | 116339980 | rs66633871 | A | G | NULL | 0.672607017 | NULL&M_Disease |
| 11 | 117434146 | rs7113635 | G | A | NULL | 0.672607017 | NULL&M_Disease |
| 11 | 117557435 | rs12285142 | T | C | NULL | 0.672607017 | NULL&M_Disease |
| 11 | 118275997 | rs9666462 | A | G | NULL | 0.672607017 | NULL&M_Disease |
| 11 | 118276246 | rs7483081 | C | T | NULL | 0.672607017 | NULL&M_Disease |
| 11 | 118501022 | rs10892248 | A | G | NULL | 0.672607017 | NULL&M_Disease |
| 11 | 119218542 | rs10790290 | T | C | NULL | 0.672607017 | NULL&M_Disease |
| 11 | 119218553 | rs10892352 | T | C | NULL | 0.672607017 | NULL&M_Disease |
| 11 | 119221078 | rs7358397 | A | G | NULL | 0.672607017 | NULL&M_Disease |
| 11 | 116932561 | rs11216257 | C | T | NULL | 0.672607017 | NULL&M_Disease |
| 11 | 118888619 | rs11602764 | G | A | NULL | 0.672607017 | NULL&M_Disease |
| 11 | 119916226 | rs2887378 | A | G | NULL | 0.672607017 | NULL&M_Disease |
| 11 | 120613416 | rs2852214 | A | G | NULL | 0.672607017 | NULL&M_Disease |
| 11 | 116326686 | rs522098 | A | G | NULL | 0.672607017 | NULL&M_Disease |
| 11 | 116468169 | rs1508097 | T | A | NULL | 0.672607017 | NULL&M_Disease |
| 11 | 116945762 | rs59511712 | T | C | NULL | 0.672607017 | NULL&M_Disease |
| 11 | 117179176 | rs528837 | A | G | NULL | 0.672607017 | NULL&M_Disease |
| 11 | 117244255 | rs12794050 | A | C | NULL | 0.672607017 | NULL&M_Disease |
| 11 | 117342462 | rs540659 | C | T | NULL | 0.672607017 | NULL&M_Disease |
| 11 | 120963641 | rs17124767 | C | T | NULL | 0.672607017 | NULL&M_Disease |
| 11 | 117362820 | rs73024372 | A | G | NULL | 0.672607017 | NULL&M_Disease |
| 11 | 117769938 | rs12787452 | G | C | NULL | 0.672607017 | NULL&M_Disease |
| 11 | 118276116 | rs9666463 | G | C | NULL | 0.672607017 | NULL&M_Disease |
| 11 | 119218712 | rs7928348 | G | C | NULL | 0.672607017 | NULL&M_Disease |
| 11 | 119802948 | rs61899022 | A | G | NULL | 0.672607017 | NULL&M_Disease |
| 11 | 120690039 | rs7940041 | A | G | NULL | 0.672607017 | NULL&M_Disease |
| 11 | 120970408 | rs34178922 | A | G | NULL | 0.672607017 | NULL&M_Disease |
| 11 | 118031986 | rs11600901 | A | G | NULL | 0.672607017 | NULL&M_Disease |
| 11 | 119600316 | rs2136420 | G | A | NULL | 0.672607017 | NULL&M_Disease |
| 11 | 120019288 | rs673292 | A | G | NULL | 0.672607017 | NULL&M_Disease |
| 11 | 118037564 | rs8192614 | A | G | NULL | 0.672607017 | NULL&M_Disease |
| 11 | 119609714 | rs35248712 | G | T | NULL | 0.673707099 | NULL&M_Disease |
| 11 | 119609846 | rs4938720 | C | T | NULL | 0.673707099 | NULL&M_Disease |
| 11 | 120399287 | rs2851410 | T | C | NULL | 0.672607017 | NULL&M_Disease |
| 11 | 120620159 | rs3132780 | G | A | NULL | 0.672607017 | NULL&M_Disease |
| 11 | 116384958 | rs7128014 | A | G | NULL | 0.673707099 | NULL&M_Disease |
| 11 | 117900997 | rs651348 | T | G | NULL | 0.673707099 | NULL&M_Disease |
| 11 | 119603559 | rs4938719 | G | A | NULL | 0.673707099 | NULL&M_Disease |
| 11 | 119603867 | rs12421434 | C | T | NULL | 0.673707099 | NULL&M_Disease |
| 11 | 119603983 | rs12421466 | G | A | NULL | 0.673707099 | NULL&M_Disease |
| 11 | 120953914 | rs874069 | C | T | NULL | 0.673707099 | NULL&M_Disease |
| 11 | 120954436 | rs7943043 | C | T | NULL | 0.673707099 | NULL&M_Disease |
| 11 | 121020150 | rs12575628 | T | C | NULL | 0.673707099 | NULL&M_Disease |
| 11 | 116222588 | rs551372 | T | C | NULL | 0.673707099 | NULL&M_Disease |
| 11 | 118030185 | rs11608057 | T | C | NULL | 0.673707099 | NULL&M_Disease |
| 11 | 118405068 | rs12289253 | G | A | NULL | 0.673707099 | NULL&M_Disease |
| 11 | 118440863 | rs11216904 | G | A | NULL | 0.673707099 | NULL&M_Disease |
| 11 | 118873987 | rs9645664 | C | T | NULL | 0.673707099 | NULL&M_Disease |

| | | | | | | | | | |
|---|---|---|---|---|---|---|---|---|---|
| 11 | 118262305 | rs59405103 | C | A | 0.978 | -7977.388 | -7978.824 | -1.436 | 9.20E-07 |
| 11 | 119834296 | rs12421491 | T | C | 0.998 | -7977.388 | -7978.825 | -1.436 | 9.20E-07 |
| 11 | 119846442 | rs17123513 | A | G | 0.995 | -7977.388 | -7978.824 | -1.436 | 9.20E-07 |
| 11 | 119846476 | rs74609299 | T | C | 0.995 | -7977.388 | -7978.824 | -1.436 | 9.20E-07 |
| 11 | 120652075 | rs4936549 | C | T | 1.029 | -7977.388 | -7978.824 | -1.436 | 9.20E-07 |
| 11 | 120960737 | rs56232713 | G | C | 0.997 | -7977.388 | -7978.825 | -1.436 | 9.20E-07 |
| 11 | 116450185 | rs12280043 | T | A | 0.972 | -7977.388 | -7978.825 | -1.437 | 9.19E-07 |
| 11 | 116453047 | rs7931384 | C | T | 0.972 | -7977.388 | -7978.825 | -1.437 | 9.19E-07 |
| 11 | 117123173 | rs528508 | C | T | 0.972 | -7977.388 | -7978.825 | -1.437 | 9.19E-07 |
| 11 | 118888831 | rs11820805 | C | T | 1.029 | -7977.388 | -7978.825 | -1.437 | 9.19E-07 |
| 11 | 120025476 | rs596911 | A | G | 1.017 | -7977.388 | -7978.825 | -1.437 | 9.19E-07 |
| 11 | 120026241 | rs659912 | G | A | 1.017 | -7977.388 | -7978.825 | -1.437 | 9.19E-07 |
| 11 | 120055326 | rs73004764 | G | C | 1.016 | -7977.388 | -7978.825 | -1.437 | 9.19E-07 |
| 11 | 116250393 | rs73019237 | T | C | 0.997 | -7977.388 | -7978.826 | -1.438 | 9.18E-07 |
| 11 | 116611827 | rs180349 | A | T | 1.029 | -7977.388 | -7978.826 | -1.438 | 9.18E-07 |
| 11 | 116710968 | rs613808 | A | G | 0.972 | -7977.388 | -7978.827 | -1.438 | 9.18E-07 |
| 11 | 117902620 | rs9326250 | A | C | 1.029 | -7977.388 | -7978.826 | -1.438 | 9.18E-07 |
| 11 | 118268613 | rs7108613 | G | A | 0.978 | -7977.388 | -7978.826 | -1.438 | 9.18E-07 |
| 11 | 119086629 | rs4936467 | T | A | 1.029 | -7977.388 | -7978.827 | -1.438 | 9.18E-07 |
| 11 | 119219360 | rs11217246 | C | A | 0.972 | -7977.388 | -7978.826 | -1.438 | 9.18E-07 |
| 11 | 119828505 | rs1353117537 | A | T | 0.996 | -7977.388 | -7978.827 | -1.438 | 9.18E-07 |
| 11 | 116397649 | rs4936353 | A | G | 0.972 | -7977.388 | -7978.827 | -1.439 | 9.17E-07 |
| 11 | 119220946 | rs12417736 | T | C | 0.972 | -7977.388 | -7978.827 | -1.439 | 9.17E-07 |
| 11 | 119503576 | rs34224649 | A | G | 0.972 | -7977.388 | -7978.827 | -1.439 | 9.17E-07 |
| 11 | 119747312 | rs481716 | G | A | 1.022 | -7977.388 | -7978.827 | -1.439 | 9.17E-07 |
| 11 | 119849798 | rs678914 | C | T | 1.029 | -7977.388 | -7978.827 | -1.439 | 9.17E-07 |
| 11 | 120292040 | rs11604012 | C | T | 0.979 | -7977.388 | -7978.827 | -1.439 | 9.17E-07 |
| 11 | 120651611 | rs3133858 | C | T | 0.975 | -7977.388 | -7978.827 | -1.439 | 9.17E-07 |
| 11 | 116558802 | rs12800436 | T | G | 1.025 | -7977.388 | -7978.829 | -1.44 | 9.16E-07 |
| 11 | 117201165 | rs515756 | G | T | 0.972 | -7977.388 | -7978.829 | -1.44 | 9.16E-07 |
| 11 | 118259379 | rs76126708 | C | A | 0.978 | -7977.388 | -7978.828 | -1.44 | 9.16E-07 |
| 11 | 118528424 | rs17748 | T | C | 1.028 | -7977.388 | -7978.828 | -1.44 | 9.16E-07 |
| 11 | 119839224 | rs76233351 | A | G | 0.997 | -7977.388 | -7978.828 | -1.44 | 9.16E-07 |
| 11 | 116688791 | rs142744740 | A | C | 0.976 | -7977.388 | -7978.829 | -1.441 | 9.16E-07 |
| 11 | 117123297 | rs497043 | G | C | 0.982 | -7977.388 | -7978.829 | -1.441 | 9.16E-07 |
| 11 | 117365821 | rs61905337 | A | C | 0.991 | -7977.388 | -7978.83 | -1.441 | 9.16E-07 |
| 11 | 119220483 | rs7480948 | G | C | 0.972 | -7977.388 | -7978.829 | -1.441 | 9.16E-07 |
| 11 | 119832917 | rs56993455 | A | G | 0.999 | -7977.388 | -7978.829 | -1.441 | 9.16E-07 |
| 11 | 119833474 | rs59253729 | T | C | 1 | -7977.388 | -7978.829 | -1.441 | 9.16E-07 |
| 11 | 120724136 | rs61901429 | G | A | 0.99 | -7977.388 | -7978.829 | -1.441 | 9.16E-07 |
| 11 | 116338814 | rs12289284 | A | G | 1 | -7977.388 | -7978.831 | -1.442 | 9.15E-07 |
| 11 | 116340960 | rs4471466 | C | A | 1.003 | -7977.388 | -7978.83 | -1.442 | 9.15E-07 |
| 11 | 116507795 | rs11216069 | C | G | 1.001 | -7977.388 | -7978.83 | -1.442 | 9.15E-07 |
| 11 | 117177800 | rs473210 | T | C | 0.972 | -7977.388 | -7978.83 | -1.442 | 9.15E-07 |
| 11 | 119848377 | rs12272862 | A | G | 1.017 | -7977.388 | -7978.83 | -1.442 | 9.15E-07 |
| 11 | 121031731 | rs75618086 | A | T | 1.02 | -7977.388 | -7978.831 | -1.442 | 9.15E-07 |
| 11 | 121064444 | rs11218198 | G | A | 1.02 | -7977.388 | -7978.831 | -1.442 | 9.15E-07 |
| 11 | 121119605 | rs12279833 | C | T | 1.029 | -7977.388 | -7978.83 | -1.442 | 9.15E-07 |
| 11 | 116339980 | rs66633871 | A | G | 1.001 | -7977.388 | -7978.831 | -1.443 | 9.14E-07 |
| 11 | 117434146 | rs7113635 | G | A | 1.028 | -7977.388 | -7978.832 | -1.443 | 9.14E-07 |
| 11 | 117557435 | rs12285142 | T | C | 0.985 | -7977.388 | -7978.831 | -1.443 | 9.14E-07 |
| 11 | 118275997 | rs9666462 | A | G | 0.978 | -7977.388 | -7978.831 | -1.443 | 9.14E-07 |
| 11 | 118276246 | rs7483081 | C | T | 0.978 | -7977.388 | -7978.831 | -1.443 | 9.14E-07 |
| 11 | 118501022 | rs10892248 | A | G | 1.028 | -7977.388 | -7978.831 | -1.443 | 9.14E-07 |
| 11 | 119218542 | rs10790290 | T | C | 0.972 | -7977.388 | -7978.831 | -1.443 | 9.14E-07 |
| 11 | 119218553 | rs10892352 | T | C | 0.972 | -7977.388 | -7978.831 | -1.443 | 9.14E-07 |
| 11 | 119221078 | rs7358397 | A | G | 0.972 | -7977.388 | -7978.832 | -1.443 | 9.14E-07 |
| 11 | 116932561 | rs11216257 | C | T | 0.976 | -7977.388 | -7978.833 | -1.444 | 9.13E-07 |
| 11 | 118888619 | rs11602764 | G | A | 1.028 | -7977.388 | -7978.832 | -1.444 | 9.13E-07 |
| 11 | 119916226 | rs2887378 | A | G | 1.029 | -7977.388 | -7978.833 | -1.444 | 9.13E-07 |
| 11 | 120613416 | rs2852214 | A | G | 0.972 | -7977.388 | -7978.832 | -1.444 | 9.13E-07 |
| 11 | 116326686 | rs522098 | A | G | 1.001 | -7977.388 | -7978.833 | -1.445 | 9.12E-07 |
| 11 | 116468169 | rs1508097 | T | A | 0.99 | -7977.388 | -7978.833 | -1.445 | 9.12E-07 |
| 11 | 116945762 | rs59511712 | T | C | 1.025 | -7977.388 | -7978.833 | -1.445 | 9.12E-07 |
| 11 | 117179176 | rs528837 | A | G | 0.972 | -7977.388 | -7978.833 | -1.445 | 9.12E-07 |
| 11 | 117244255 | rs12794050 | A | C | 1.028 | -7977.388 | -7978.834 | -1.445 | 9.12E-07 |
| 11 | 117342462 | rs540659 | C | T | 1.029 | -7977.388 | -7978.833 | -1.445 | 9.12E-07 |
| 11 | 120963641 | rs17124767 | C | T | 0.99 | -7977.388 | -7978.833 | -1.445 | 9.12E-07 |
| 11 | 117362820 | rs73024372 | A | G | 1.004 | -7977.388 | -7978.834 | -1.446 | 9.11E-07 |
| 11 | 117769938 | rs12787452 | G | C | 1.025 | -7977.388 | -7978.834 | -1.446 | 9.11E-07 |
| 11 | 118276116 | rs9666463 | G | C | 0.978 | -7977.388 | -7978.835 | -1.446 | 9.11E-07 |
| 11 | 119218712 | rs7928348 | G | C | 0.972 | -7977.388 | -7978.834 | -1.446 | 9.11E-07 |
| 11 | 119802948 | rs61899022 | A | G | 1.019 | -7977.388 | -7978.834 | -1.446 | 9.11E-07 |
| 11 | 120690039 | rs7940041 | A | G | 0.972 | -7977.388 | -7978.834 | -1.446 | 9.11E-07 |
| 11 | 120970408 | rs34178922 | A | G | 0.993 | -7977.388 | -7978.834 | -1.446 | 9.11E-07 |
| 11 | 118031986 | rs11600901 | A | G | 1.01 | -7977.388 | -7978.836 | -1.447 | 9.10E-07 |
| 11 | 119600316 | rs2136420 | G | A | 0.972 | -7977.388 | -7978.835 | -1.447 | 9.10E-07 |
| 11 | 120019288 | rs673292 | A | G | 1.016 | -7977.388 | -7978.835 | -1.447 | 9.10E-07 |
| 11 | 118037564 | rs8192614 | A | G | 1.01 | -7977.388 | -7978.836 | -1.448 | 9.09E-07 |
| 11 | 119609714 | rs35248712 | G | T | 0.972 | -7977.388 | -7978.836 | -1.448 | 9.09E-07 |
| 11 | 119609846 | rs4938720 | C | T | 0.972 | -7977.388 | -7978.836 | -1.448 | 9.09E-07 |
| 11 | 120399287 | rs2851410 | T | C | 1.027 | -7977.388 | -7978.836 | -1.448 | 9.09E-07 |
| 11 | 120620159 | rs3132780 | G | A | 1.025 | -7977.388 | -7978.836 | -1.448 | 9.09E-07 |
| 11 | 116384958 | rs7128014 | A | G | 0.988 | -7977.388 | -7978.837 | -1.449 | 9.08E-07 |
| 11 | 117900997 | rs651348 | T | G | 0.976 | -7977.388 | -7978.837 | -1.449 | 9.08E-07 |
| 11 | 119603559 | rs4938719 | G | A | 0.972 | -7977.388 | -7978.837 | -1.449 | 9.08E-07 |
| 11 | 119603867 | rs12421434 | C | T | 0.972 | -7977.388 | -7978.837 | -1.449 | 9.08E-07 |
| 11 | 119603983 | rs12421466 | G | A | 0.972 | -7977.388 | -7978.837 | -1.449 | 9.08E-07 |
| 11 | 120953914 | rs874069 | C | T | 0.99 | -7977.388 | -7978.838 | -1.449 | 9.08E-07 |
| 11 | 120954436 | rs7943043 | C | T | 0.99 | -7977.388 | -7978.838 | -1.449 | 9.08E-07 |
| 11 | 121020150 | rs12575628 | T | C | 1.019 | -7977.388 | -7978.838 | -1.449 | 9.08E-07 |
| 11 | 116222588 | rs551372 | T | C | 1.029 | -7977.388 | -7978.838 | -1.45 | 9.07E-07 |
| 11 | 118030185 | rs11608057 | T | C | 1.01 | -7977.388 | -7978.839 | -1.45 | 9.07E-07 |
| 11 | 118405068 | rs12289253 | G | A | 0.979 | -7977.388 | -7978.838 | -1.45 | 9.07E-07 |
| 11 | 118440863 | rs11216904 | G | A | 0.98 | -7977.388 | -7978.838 | -1.45 | 9.07E-07 |
| 11 | 118873987 | rs9645664 | C | T | 1.028 | -7977.388 | -7978.839 | -1.45 | 9.07E-07 |

| | | | | | | | |
|---|---|---|---|---|---|---|---|
| 11 | 119055362 | rs7929799 | G | A | NULL | 0.673707099 | NULL&M_Disease |
| 11 | 119218616 | rs7927959 | A | T | NULL | 0.673707099 | NULL&M_Disease |
| 11 | 119762569 | rs1374563 | A | G | NULL | 0.673707099 | NULL&M_Disease |
| 11 | 120017974 | rs657898 | C | T | NULL | 0.673707099 | NULL&M_Disease |
| 11 | 120691871 | rs35338327 | A | G | NULL | 0.673707099 | NULL&M_Disease |
| 11 | 121112889 | rs12289923 | G | A | NULL | 0.673707099 | NULL&M_Disease |
| 11 | 117163765 | rs490460 | C | A | NULL | 0.673707099 | NULL&M_Disease |
| 11 | 117755082 | rs491292 | T | C | NULL | 0.673707099 | NULL&M_Disease |
| 11 | 118031211 | rs71482132 | A | G | NULL | 0.673707099 | NULL&M_Disease |
| 11 | 118032438 | rs35381826 | T | G | NULL | 0.673707099 | NULL&M_Disease |
| 11 | 118433729 | rs59108310 | G | A | NULL | 0.673707099 | NULL&M_Disease |
| 11 | 118439048 | rs12224844 | T | A | NULL | 0.673707099 | NULL&M_Disease |
| 11 | 119609264 | rs4245206 | A | T | NULL | 0.673707099 | NULL&M_Disease |
| 11 | 119611734 | rs10892446 | A | G | NULL | 0.673707099 | NULL&M_Disease |
| 11 | 119842708 | rs17123506 | G | A | NULL | 0.673707099 | NULL&M_Disease |
| 11 | 119881259 | rs1320708 | C | T | NULL | 0.673707099 | NULL&M_Disease |
| 11 | 119915395 | rs2100357 | T | C | NULL | 0.673707099 | NULL&M_Disease |
| 11 | 121040073 | rs77080891 | A | G | NULL | 0.673707099 | NULL&M_Disease |
| 11 | 116216809 | rs545657 | G | A | NULL | 0.673707099 | NULL&M_Disease |
| 11 | 117456034 | rs654946 | A | G | NULL | 0.673707099 | NULL&M_Disease |
| 11 | 117523508 | rs11216485 | G | C | NULL | 0.673707099 | NULL&M_Disease |
| 11 | 120021171 | rs61900474 | A | T | NULL | 0.673707099 | NULL&M_Disease |
| 11 | 120400149 | rs2252359 | A | G | NULL | 0.673707099 | NULL&M_Disease |
| 11 | 116536476 | rs1240777 | C | T | NULL | 0.673707099 | NULL&M_Disease |
| 11 | 118276492 | rs17122157 | A | G | NULL | 0.673707099 | NULL&M_Disease |
| 11 | 118441846 | rs10892243 | C | T | NULL | 0.673707099 | NULL&M_Disease |
| 11 | 118499394 | rs2236661 | G | C | NULL | 0.673707099 | NULL&M_Disease |
| 11 | 119343580 | rs654018 | A | G | NULL | 0.673707099 | NULL&M_Disease |
| 11 | 119798934 | rs681133 | T | C | NULL | 0.673707099 | NULL&M_Disease |
| 11 | 121006976 | rs598696 | T | A | NULL | 0.673707099 | NULL&M_Disease |
| 11 | 116400382 | rs7941534 | G | A | NULL | 0.673707099 | NULL&M_Disease |
| 11 | 117690620 | rs59462353 | C | T | NULL | 0.673707099 | NULL&M_Disease |
| 11 | 118026788 | rs595964 | A | G | NULL | 0.673707099 | NULL&M_Disease |
| 11 | 118437279 | rs11216902 | G | A | NULL | 0.673707099 | NULL&M_Disease |
| 11 | 120021674 | rs2848728 | T | C | NULL | 0.673707099 | NULL&M_Disease |
| 11 | 116464992 | rs11216028 | T | G | NULL | 0.673707099 | NULL&M_Disease |
| 11 | 116537059 | rs61906081 | A | G | NULL | 0.673707099 | NULL&M_Disease |
| 11 | 117802309 | rs12805985 | G | A | NULL | 0.673707099 | NULL&M_Disease |
| 11 | 118090432 | rs1540191 | G | A | NULL | 0.673707099 | NULL&M_Disease |
| 11 | 119601497 | rs4938718 | G | C | NULL | 0.673707099 | NULL&M_Disease |
| 11 | 119601779 | rs2368936 | C | T | NULL | 0.673707099 | NULL&M_Disease |
| 11 | 119602067 | rs11217434 | C | T | NULL | 0.673707099 | NULL&M_Disease |
| 11 | 119915651 | rs2084892 | A | G | NULL | 0.673707099 | NULL&M_Disease |
| 11 | 120935558 | rs4594010 | C | A | NULL | 0.673707099 | NULL&M_Disease |
| 11 | 116201218 | rs7945051 | C | A | NULL | 0.673707099 | NULL&M_Disease |
| 11 | 116749509 | rs76523703 | G | T | NULL | 0.673707099 | NULL&M_Disease |
| 11 | 117802113 | rs12801144 | G | A | NULL | 0.673707099 | NULL&M_Disease |
| 11 | 118029798 | rs11607985 | G | C | NULL | 0.673707099 | NULL&M_Disease |
| 11 | 118284650 | rs73613980 | T | C | NULL | 0.673707099 | NULL&M_Disease |
| 11 | 119342115 | rs619473 | C | A | NULL | 0.673707099 | NULL&M_Disease |
| 11 | 120155118 | rs11217788 | A | G | NULL | 0.673707099 | NULL&M_Disease |
| 11 | 121021970 | rs79022222 | T | C | NULL | 0.673707099 | NULL&M_Disease |
| 11 | 118295117 | rs12274926 | C | T | NULL | 0.673707099 | NULL&M_Disease |
| 11 | 118912982 | rs657685 | C | G | NULL | 0.674805273 | NULL&M_Disease |
| 11 | 119217556 | rs3814760 | G | A | NULL | 0.673707099 | NULL&M_Disease |
| 11 | 119219124 | rs1320679 | G | T | NULL | 0.673707099 | NULL&M_Disease |
| 11 | 119223386 | rs7126377 | A | G | NULL | 0.673707099 | NULL&M_Disease |
| 11 | 120584612 | rs3133856 | C | T | NULL | 0.673707099 | NULL&M_Disease |
| 11 | 120954793 | rs12788247 | G | A | NULL | 0.673707099 | NULL&M_Disease |
| 11 | 116537846 | rs35933073 | A | C | NULL | 0.674805273 | NULL&M_Disease |
| 11 | 116989982 | rs74428560 | T | C | NULL | 0.673707099 | NULL&M_Disease |
| 11 | 118246048 | rs60514290 | C | G | NULL | 0.673707099 | NULL&M_Disease |
| 11 | 119218930 | rs1893265 | C | T | NULL | 0.674805273 | NULL&M_Disease |
| 11 | 120402020 | rs2257198 | A | G | NULL | 0.674805273 | NULL&M_Disease |
| 11 | 121065198 | rs17124937 | C | G | NULL | 0.673707099 | NULL&M_Disease |
| 11 | 116671005 | rs4938313 | A | G | NULL | 0.674805273 | NULL&M_Disease |
| 11 | 116672013 | rs11216140 | T | C | NULL | 0.674805273 | NULL&M_Disease |
| 11 | 116678139 | rs1263169 | A | T | NULL | 0.674805273 | NULL&M_Disease |
| 11 | 118137383 | rs12417431 | G | A | NULL | 0.674805273 | NULL&M_Disease |
| 11 | 119809217 | rs11217587 | C | T | NULL | 0.674805273 | NULL&M_Disease |
| 11 | 119849389 | rs677027 | A | G | NULL | 0.674805273 | NULL&M_Disease |
| 11 | 121022231 | rs12361100 | T | A | NULL | 0.674805273 | NULL&M_Disease |
| 11 | 116356225 | rs1158260 | T | C | NULL | 0.674805273 | NULL&M_Disease |
| 11 | 116577558 | rs11822160 | A | T | NULL | 0.674805273 | NULL&M_Disease |
| 11 | 119217957 | rs3814754 | C | T | NULL | 0.674805273 | NULL&M_Disease |
| 11 | 119796344 | rs12278045 | A | G | NULL | 0.674805273 | NULL&M_Disease |
| 11 | 119851604 | rs616262 | G | C | NULL | 0.674805273 | NULL&M_Disease |
| 11 | 117549808 | rs7926736 | A | G | NULL | 0.674805273 | NULL&M_Disease |
| 11 | 118251980 | rs45501598 | T | A | NULL | 0.674805273 | NULL&M_Disease |
| 11 | 119220551 | rs11217249 | G | A | NULL | 0.674805273 | NULL&M_Disease |
| 11 | 119596147 | rs1876795 | C | G | NULL | 0.674805273 | NULL&M_Disease |
| 11 | 119606985 | rs11217437 | C | G | NULL | 0.674805273 | NULL&M_Disease |
| 11 | 119607019 | rs11217438 | C | T | NULL | 0.674805273 | NULL&M_Disease |
| 11 | 120724322 | rs11218032 | A | T | NULL | 0.674805273 | NULL&M_Disease |
| 11 | 120978982 | rs1783778 | A | C | NULL | 0.674805273 | NULL&M_Disease |
| 11 | 121120845 | rs948378 | A | T | NULL | 0.674805273 | NULL&M_Disease |
| 11 | 116983016 | rs113979111 | A | G | NULL | 0.674805273 | NULL&M_Disease |
| 11 | 117456464 | rs11216464 | T | C | NULL | 0.674805273 | NULL&M_Disease |
| 11 | 118432279 | rs11216897 | G | A | NULL | 0.674805273 | NULL&M_Disease |
| 11 | 119239689 | rs680268 | C | T | NULL | 0.674805273 | NULL&M_Disease |
| 11 | 119790617 | rs75858010 | T | C | NULL | 0.674805273 | NULL&M_Disease |
| 11 | 120823721 | rs2298724 | G | A | NULL | 0.674805273 | NULL&M_Disease |
| 11 | 116427570 | rs4531505 | C | G | NULL | 0.674805273 | NULL&M_Disease |
| 11 | 116458278 | rs7110791 | C | A | NULL | 0.674805273 | NULL&M_Disease |
| 11 | 116532314 | rs1145188 | A | G | NULL | 0.674805273 | NULL&M_Disease |
| 11 | 116671476 | rs6589569 | T | C | NULL | 0.674805273 | NULL&M_Disease |

| | | | | | | | | | |
|---|---|---|---|---|---|---|---|---|---|
| 11 | 119055362 | rs7929799 | G | A | 0.972 | -7977.388 | -7978.838 | -1.45 | 9.07E-07 |
| 11 | 119218616 | rs7927959 | A | T | 0.972 | -7977.388 | -7978.838 | -1.45 | 9.07E-07 |
| 11 | 119762569 | rs1374563 | A | G | 1.025 | -7977.388 | -7978.839 | -1.45 | 9.07E-07 |
| 11 | 120017974 | rs657898 | C | T | 1.015 | -7977.388 | -7978.839 | -1.45 | 9.07E-07 |
| 11 | 120691871 | rs35338327 | A | G | 0.972 | -7977.388 | -7978.838 | -1.45 | 9.07E-07 |
| 11 | 121112889 | rs12289923 | G | A | 1.027 | -7977.388 | -7978.839 | -1.45 | 9.07E-07 |
| 11 | 117163765 | rs490460 | C | A | 1.029 | -7977.388 | -7978.839 | -1.451 | 9.06E-07 |
| 11 | 117755082 | rs491292 | T | C | 1.027 | -7977.388 | -7978.839 | -1.451 | 9.06E-07 |
| 11 | 118031211 | rs71482132 | A | G | 1.01 | -7977.388 | -7978.839 | -1.451 | 9.06E-07 |
| 11 | 118032438 | rs35381826 | T | G | 1.009 | -7977.388 | -7978.84 | -1.451 | 9.06E-07 |
| 11 | 118433729 | rs59108310 | G | A | 0.981 | -7977.388 | -7978.839 | -1.451 | 9.06E-07 |
| 11 | 118439048 | rs12224844 | T | A | 0.981 | -7977.388 | -7978.84 | -1.451 | 9.06E-07 |
| 11 | 119609264 | rs4245206 | A | T | 0.972 | -7977.388 | -7978.839 | -1.451 | 9.06E-07 |
| 11 | 119611734 | rs10892446 | A | G | 0.972 | -7977.388 | -7978.84 | -1.451 | 9.06E-07 |
| 11 | 119842708 | rs17123506 | G | A | 0.999 | -7977.388 | -7978.839 | -1.451 | 9.06E-07 |
| 11 | 119881259 | rs1320708 | C | T | 0.972 | -7977.388 | -7978.839 | -1.451 | 9.06E-07 |
| 11 | 119915395 | rs2100357 | T | C | 1.029 | -7977.388 | -7978.839 | -1.451 | 9.06E-07 |
| 11 | 121040073 | rs77080891 | A | G | 1.019 | -7977.388 | -7978.839 | -1.451 | 9.06E-07 |
| 11 | 116216809 | rs545657 | G | A | 1.029 | -7977.388 | -7978.841 | -1.452 | 9.06E-07 |
| 11 | 117456034 | rs654946 | A | G | 0.973 | -7977.388 | -7978.84 | -1.452 | 9.06E-07 |
| 11 | 117523508 | rs11216485 | G | C | 1.017 | -7977.388 | -7978.84 | -1.452 | 9.06E-07 |
| 11 | 120021171 | rs61900474 | A | T | 1.015 | -7977.388 | -7978.841 | -1.452 | 9.06E-07 |
| 11 | 120400149 | rs2252359 | A | G | 0.975 | -7977.388 | -7978.84 | -1.452 | 9.06E-07 |
| 11 | 116536476 | rs1240777 | C | T | 1.028 | -7977.388 | -7978.841 | -1.453 | 9.05E-07 |
| 11 | 118276492 | rs17122157 | A | G | 0.979 | -7977.388 | -7978.841 | -1.453 | 9.05E-07 |
| 11 | 118441846 | rs10892243 | C | T | 0.981 | -7977.388 | -7978.841 | -1.453 | 9.05E-07 |
| 11 | 118499394 | rs2236661 | G | C | 1.027 | -7977.388 | -7978.841 | -1.453 | 9.05E-07 |
| 11 | 119343580 | rs654018 | A | G | 1.008 | -7977.388 | -7978.841 | -1.453 | 9.05E-07 |
| 11 | 119798934 | rs681133 | T | C | 0.972 | -7977.388 | -7978.841 | -1.453 | 9.05E-07 |
| 11 | 121006976 | rs598696 | T | A | 1.029 | -7977.388 | -7978.841 | -1.453 | 9.05E-07 |
| 11 | 116400382 | rs7941534 | G | A | 1.026 | -7977.388 | -7978.843 | -1.454 | 9.04E-07 |
| 11 | 117690620 | rs59462353 | C | T | 0.977 | -7977.388 | -7978.842 | -1.454 | 9.04E-07 |
| 11 | 118026788 | rs595964 | A | G | 1.008 | -7977.388 | -7978.842 | -1.454 | 9.04E-07 |
| 11 | 118437279 | rs11216902 | G | A | 0.981 | -7977.388 | -7978.842 | -1.454 | 9.04E-07 |
| 11 | 120021674 | rs2848728 | T | C | 1.015 | -7977.388 | -7978.842 | -1.454 | 9.04E-07 |
| 11 | 116464992 | rs11216028 | T | G | 0.989 | -7977.388 | -7978.843 | -1.455 | 9.03E-07 |
| 11 | 116537059 | rs61906081 | A | G | 1.025 | -7977.388 | -7978.843 | -1.455 | 9.03E-07 |
| 11 | 117802309 | rs12805985 | G | A | 0.983 | -7977.388 | -7978.843 | -1.455 | 9.03E-07 |
| 11 | 118090432 | rs1540191 | G | A | 1.029 | -7977.388 | -7978.843 | -1.455 | 9.03E-07 |
| 11 | 119601497 | rs4938718 | G | C | 0.972 | -7977.388 | -7978.843 | -1.455 | 9.03E-07 |
| 11 | 119601779 | rs2368936 | C | T | 0.972 | -7977.388 | -7978.843 | -1.455 | 9.03E-07 |
| 11 | 119602067 | rs11217434 | C | T | 0.972 | -7977.388 | -7978.843 | -1.455 | 9.03E-07 |
| 11 | 119915651 | rs2084892 | A | G | 1.028 | -7977.388 | -7978.843 | -1.455 | 9.03E-07 |
| 11 | 120935558 | rs4594010 | C | A | 1.028 | -7977.388 | -7978.843 | -1.455 | 9.03E-07 |
| 11 | 116201218 | rs7945051 | C | A | 1.029 | -7977.388 | -7978.844 | -1.456 | 9.02E-07 |
| 11 | 116749509 | rs76523703 | G | T | 0.979 | -7977.388 | -7978.845 | -1.456 | 9.02E-07 |
| 11 | 117802113 | rs12801144 | G | A | 0.983 | -7977.388 | -7978.844 | -1.456 | 9.02E-07 |
| 11 | 118029798 | rs11607985 | G | C | 1.008 | -7977.388 | -7978.844 | -1.456 | 9.02E-07 |
| 11 | 118284650 | rs73613980 | T | C | 0.979 | -7977.388 | -7978.844 | -1.456 | 9.02E-07 |
| 11 | 119342115 | rs619473 | C | A | 1.007 | -7977.388 | -7978.845 | -1.456 | 9.02E-07 |
| 11 | 120155118 | rs11217788 | A | G | 1.008 | -7977.388 | -7978.844 | -1.456 | 9.02E-07 |
| 11 | 121021970 | rs79022222 | T | C | 1.018 | -7977.388 | -7978.845 | -1.456 | 9.02E-07 |
| 11 | 118295117 | rs12274926 | C | T | 0.974 | -7977.388 | -7978.845 | -1.457 | 9.01E-07 |
| 11 | 118912982 | rs657685 | C | G | 0.972 | -7977.388 | -7978.846 | -1.457 | 9.01E-07 |
| 11 | 119217556 | rs3814760 | G | A | 0.972 | -7977.388 | -7978.846 | -1.457 | 9.01E-07 |
| 11 | 119219124 | rs1320679 | G | T | 0.972 | -7977.388 | -7978.846 | -1.457 | 9.01E-07 |
| 11 | 119223386 | rs7126377 | A | G | 0.973 | -7977.388 | -7978.845 | -1.457 | 9.01E-07 |
| 11 | 120584612 | rs3133856 | C | T | 1.029 | -7977.388 | -7978.845 | -1.457 | 9.01E-07 |
| 11 | 120954793 | rs12788247 | G | A | 1.001 | -7977.388 | -7978.846 | -1.457 | 9.01E-07 |
| 11 | 116537846 | rs35933073 | A | C | 1.025 | -7977.388 | -7978.846 | -1.458 | 9.00E-07 |
| 11 | 116989982 | rs74428560 | T | C | 0.995 | -7977.388 | -7978.846 | -1.458 | 9.00E-07 |
| 11 | 118246048 | rs60514290 | C | G | 0.979 | -7977.388 | -7978.846 | -1.458 | 9.00E-07 |
| 11 | 119218930 | rs1893265 | C | T | 0.972 | -7977.388 | -7978.846 | -1.458 | 9.00E-07 |
| 11 | 120402020 | rs2257198 | A | G | 0.976 | -7977.388 | -7978.847 | -1.458 | 9.00E-07 |
| 11 | 121065198 | rs17124937 | C | G | 1.018 | -7977.388 | -7978.846 | -1.458 | 9.00E-07 |
| 11 | 116671005 | rs4938313 | A | G | 0.985 | -7977.388 | -7978.848 | -1.459 | 8.99E-07 |
| 11 | 116672013 | rs11216140 | T | C | 0.985 | -7977.388 | -7978.848 | -1.459 | 8.99E-07 |
| 11 | 116678139 | rs1263169 | A | T | 1.028 | -7977.388 | -7978.848 | -1.459 | 8.99E-07 |
| 11 | 118137383 | rs12417431 | G | A | 0.972 | -7977.388 | -7978.847 | -1.459 | 8.99E-07 |
| 11 | 119809217 | rs11217587 | C | T | 1.019 | -7977.388 | -7978.847 | -1.459 | 8.99E-07 |
| 11 | 119849389 | rs677027 | A | G | 1.029 | -7977.388 | -7978.847 | -1.459 | 8.99E-07 |
| 11 | 121022231 | rs12361100 | T | A | 1.018 | -7977.388 | -7978.847 | -1.459 | 8.99E-07 |
| 11 | 116356225 | rs1158260 | T | C | 0.986 | -7977.388 | -7978.848 | -1.46 | 8.98E-07 |
| 11 | 116577558 | rs11822160 | A | T | 1.001 | -7977.388 | -7978.848 | -1.46 | 8.98E-07 |
| 11 | 119217957 | rs3814754 | C | T | 0.972 | -7977.388 | -7978.848 | -1.46 | 8.98E-07 |
| 11 | 119796344 | rs12278045 | A | G | 1.018 | -7977.388 | -7978.848 | -1.46 | 8.98E-07 |
| 11 | 119851604 | rs616262 | G | C | 0.982 | -7977.388 | -7978.848 | -1.46 | 8.98E-07 |
| 11 | 117549808 | rs7926736 | A | G | 1.02 | -7977.388 | -7978.849 | -1.461 | 8.97E-07 |
| 11 | 118251980 | rs45501598 | T | A | 0.98 | -7977.388 | -7978.85 | -1.461 | 8.97E-07 |
| 11 | 119220551 | rs11217249 | G | A | 0.972 | -7977.388 | -7978.85 | -1.461 | 8.97E-07 |
| 11 | 119596147 | rs1876795 | C | G | 0.972 | -7977.388 | -7978.849 | -1.461 | 8.97E-07 |
| 11 | 119606985 | rs11217437 | C | G | 0.972 | -7977.388 | -7978.849 | -1.461 | 8.97E-07 |
| 11 | 119607019 | rs11217438 | C | T | 0.972 | -7977.388 | -7978.849 | -1.461 | 8.97E-07 |
| 11 | 120724322 | rs11218032 | A | T | 0.972 | -7977.388 | -7978.85 | -1.461 | 8.97E-07 |
| 11 | 120978982 | rs1783778 | A | C | 0.972 | -7977.388 | -7978.849 | -1.461 | 8.97E-07 |
| 11 | 121120845 | rs948378 | A | T | 1.028 | -7977.388 | -7978.849 | -1.461 | 8.97E-07 |
| 11 | 116983016 | rs113979111 | A | G | 0.996 | -7977.388 | -7978.851 | -1.462 | 8.97E-07 |
| 11 | 117456464 | rs11216464 | T | C | 0.972 | -7977.388 | -7978.85 | -1.462 | 8.97E-07 |
| 11 | 118432279 | rs11216897 | G | A | 0.982 | -7977.388 | -7978.851 | -1.462 | 8.97E-07 |
| 11 | 119239689 | rs680268 | C | T | 0.973 | -7977.388 | -7978.851 | -1.462 | 8.97E-07 |
| 11 | 119790617 | rs75858010 | T | C | 0.981 | -7977.388 | -7978.85 | -1.462 | 8.97E-07 |
| 11 | 120823721 | rs2298724 | G | A | 1.028 | -7977.388 | -7978.851 | -1.462 | 8.97E-07 |
| 11 | 116427570 | rs4531505 | C | G | 0.972 | -7977.388 | -7978.851 | -1.463 | 8.96E-07 |
| 11 | 116458278 | rs7110791 | C | A | 0.972 | -7977.388 | -7978.851 | -1.463 | 8.96E-07 |
| 11 | 116532314 | rs1145188 | A | G | 1.027 | -7977.388 | -7978.852 | -1.463 | 8.96E-07 |
| 11 | 116671476 | rs6589569 | T | C | 0.985 | -7977.388 | -7978.851 | -1.463 | 8.96E-07 |

| | | | | | | | |
|---|---|---|---|---|---|---|---|
| 11 | 116845104 | rs17120197 | C | T | NULL | 0.674805273 | NULL&M_Disease |
| 11 | 116979365 | rs113784559 | G | C | NULL | 0.674805273 | NULL&M_Disease |
| 11 | 116983556 | rs111560312 | A | G | NULL | 0.674805273 | NULL&M_Disease |
| 11 | 116984866 | rs55703909 | A | G | NULL | 0.674805273 | NULL&M_Disease |
| 11 | 117355782 | rs10892125 | C | A | NULL | 0.674805273 | NULL&M_Disease |
| 11 | 119146659 | rs2511854 | G | C | NULL | 0.674805273 | NULL&M_Disease |
| 11 | 119164677 | rs2510139 | C | T | NULL | 0.674805273 | NULL&M_Disease |
| 11 | 119576911 | rs4938717 | T | C | NULL | 0.674805273 | NULL&M_Disease |
| 11 | 119600410 | rs2136421 | C | T | NULL | 0.674805273 | NULL&M_Disease |
| 11 | 119923643 | rs1001973 | A | G | NULL | 0.674805273 | NULL&M_Disease |
| 11 | 119978608 | rs1349688 | C | G | NULL | 0.674805273 | NULL&M_Disease |
| 11 | 120976413 | rs681311 | A | G | NULL | 0.674805273 | NULL&M_Disease |
| 11 | 121038810 | rs2155369 | T | C | NULL | 0.674805273 | NULL&M_Disease |
| 11 | 121092828 | rs948095 | A | G | NULL | 0.674805273 | NULL&M_Disease |
| 11 | 116794903 | rs56314469 | A | G | NULL | 0.674805273 | NULL&M_Disease |
| 11 | 118938880 | rs584115 | G | A | NULL | 0.674805273 | NULL&M_Disease |
| 11 | 119604854 | rs10892444 | A | G | NULL | 0.674805273 | NULL&M_Disease |
| 11 | 117774829 | rs3809040 | A | G | NULL | 0.674805273 | NULL&M_Disease |
| 11 | 118432852 | rs11216898 | C | T | NULL | 0.674805273 | NULL&M_Disease |
| 11 | 118967291 | rs7759 | G | A | NULL | 0.674805273 | NULL&M_Disease |
| 11 | 119818802 | rs10892502 | C | T | NULL | 0.674805273 | NULL&M_Disease |
| 11 | 119885547 | rs630775 | C | A | NULL | 0.674805273 | NULL&M_Disease |
| 11 | 120165942 | rs11217801 | G | A | NULL | 0.674805273 | NULL&M_Disease |
| 11 | 120299707 | rs7129944 | C | T | NULL | 0.674805273 | NULL&M_Disease |
| 11 | 120657044 | rs7105258 | T | C | NULL | 0.674805273 | NULL&M_Disease |
| 11 | 116209430 | rs10790138 | C | G | NULL | 0.674805273 | NULL&M_Disease |
| 11 | 116701122 | rs2070669 | C | G | NULL | 0.674805273 | NULL&M_Disease |
| 11 | 118069269 | rs1573580 | G | C | NULL | 0.674805273 | NULL&M_Disease |
| 11 | 118496119 | rs45565037 | A | G | NULL | 0.674805273 | NULL&M_Disease |
| 11 | 118966521 | rs2509049 | T | C | NULL | 0.674805273 | NULL&M_Disease |
| 11 | 119922558 | rs12421541 | T | A | NULL | 0.674805273 | NULL&M_Disease |
| 11 | 120663363 | rs1954787 | T | C | NULL | 0.674805273 | NULL&M_Disease |
| 11 | 121017301 | rs2155364 | T | C | NULL | 0.674805273 | NULL&M_Disease |
| 11 | 120017875 | rs582071 | A | G | NULL | 0.674805273 | NULL&M_Disease |
| 11 | 120166016 | rs17123792 | A | G | NULL | 0.674805273 | NULL&M_Disease |
| 11 | 120186190 | rs17245511 | A | G | NULL | 0.674805273 | NULL&M_Disease |
| 11 | 116225111 | rs491113 | C | G | NULL | 0.675901528 | NULL&M_Disease |
| 11 | 116697344 | rs2071523 | C | T | NULL | 0.675901528 | NULL&M_Disease |
| 11 | 118091761 | rs2368238 | T | G | NULL | 0.675901528 | NULL&M_Disease |
| 11 | 118967524 | rs8551 | T | C | NULL | 0.675901528 | NULL&M_Disease |
| 11 | 120968462 | rs12274094 | G | A | NULL | 0.675901528 | NULL&M_Disease |
| 11 | 120979036 | rs1783777 | C | T | NULL | 0.675901528 | NULL&M_Disease |
| 11 | 120979047 | rs1783776 | G | A | NULL | 0.675901528 | NULL&M_Disease |
| 11 | 121113880 | rs10790413 | C | A | NULL | 0.675901528 | NULL&M_Disease |
| 11 | 116196429 | rs10891961 | C | T | NULL | 0.675901528 | NULL&M_Disease |
| 11 | 116196643 | rs11215897 | G | A | NULL | 0.675901528 | NULL&M_Disease |
| 11 | 116696043 | rs2216311 | G | A | NULL | 0.675901528 | NULL&M_Disease |
| 11 | 118947634 | rs540261 | T | C | NULL | 0.675901528 | NULL&M_Disease |
| 11 | 120051217 | rs76841836 | C | G | NULL | 0.675901528 | NULL&M_Disease |
| 11 | 120292477 | rs76693355 | C | T | NULL | 0.675901528 | NULL&M_Disease |
| 11 | 120300804 | rs11217866 | A | T | NULL | 0.675901528 | NULL&M_Disease |
| 11 | 120517643 | rs6589826 | A | G | NULL | 0.675901528 | NULL&M_Disease |
| 11 | 118966780 | rs2509851 | C | A | NULL | 0.675901528 | NULL&M_Disease |
| 11 | 117950199 | rs73010757 | A | G | NULL | 0.675901528 | NULL&M_Disease |
| 11 | 118320553 | rs629470 | A | G | NULL | 0.675901528 | NULL&M_Disease |
| 11 | 118337003 | rs574387 | C | A | NULL | 0.675901528 | NULL&M_Disease |
| 11 | 119612572 | rs12796806 | G | A | NULL | 0.675901528 | NULL&M_Disease |
| 11 | 120293138 | rs11217863 | A | G | NULL | 0.675901528 | NULL&M_Disease |
| 11 | 116209589 | rs10732855 | C | G | NULL | 0.675901528 | NULL&M_Disease |
| 11 | 118925341 | rs538478 | A | G | NULL | 0.675901528 | NULL&M_Disease |
| 11 | 120978834 | rs1784281 | C | A | NULL | 0.675901528 | NULL&M_Disease |
| 11 | 116196502 | rs11215896 | A | C | NULL | 0.675901528 | NULL&M_Disease |
| 11 | 116196666 | rs10891962 | C | T | NULL | 0.675901528 | NULL&M_Disease |
| 11 | 116355310 | rs11215958 | A | G | NULL | 0.675901528 | NULL&M_Disease |
| 11 | 118034228 | rs611960 | G | T | NULL | 0.675901528 | NULL&M_Disease |
| 11 | 118434421 | rs11216899 | G | A | NULL | 0.675901528 | NULL&M_Disease |
| 11 | 118434835 | rs60753148 | T | C | NULL | 0.675901528 | NULL&M_Disease |
| 11 | 120066044 | rs12791282 | G | C | NULL | 0.675901528 | NULL&M_Disease |
| 11 | 121114248 | rs10160642 | T | C | NULL | 0.675901528 | NULL&M_Disease |
| 11 | 116466289 | rs11216031 | C | A | NULL | 0.675901528 | NULL&M_Disease |
| 11 | 116532856 | rs1145189 | A | G | NULL | 0.675901528 | NULL&M_Disease |
| 11 | 117410423 | rs2509004 | A | G | NULL | 0.675901528 | NULL&M_Disease |
| 11 | 119220839 | rs12421814 | G | A | NULL | 0.675901528 | NULL&M_Disease |
| 11 | 119800548 | rs1579885 | A | G | NULL | 0.675901528 | NULL&M_Disease |
| 11 | 119951678 | rs11217679 | G | A | NULL | 0.675901528 | NULL&M_Disease |
| 11 | 120166860 | rs6589807 | A | G | NULL | 0.675901528 | NULL&M_Disease |
| 11 | 120914370 | rs7952619 | C | T | NULL | 0.675901528 | NULL&M_Disease |
| 11 | 116210084 | rs10736483 | A | G | NULL | 0.675901528 | NULL&M_Disease |
| 11 | 117347058 | rs665406 | C | T | NULL | 0.675901528 | NULL&M_Disease |
| 11 | 118034223 | rs611979 | A | G | NULL | 0.675901528 | NULL&M_Disease |
| 11 | 119683607 | rs10892472 | A | C | NULL | 0.675901528 | NULL&M_Disease |
| 11 | 119845755 | rs12277299 | G | C | NULL | 0.675901528 | NULL&M_Disease |
| 11 | 119914182 | rs2084893 | G | C | NULL | 0.675901528 | NULL&M_Disease |
| 11 | 119914915 | rs1597866 | G | A | NULL | 0.675901528 | NULL&M_Disease |
| 11 | 120294004 | rs11823264 | A | G | NULL | 0.675901528 | NULL&M_Disease |
| 11 | 120311198 | rs4936518 | T | A | NULL | 0.675901528 | NULL&M_Disease |
| 11 | 116196711 | rs10891963 | G | T | NULL | 0.675901528 | NULL&M_Disease |
| 11 | 118436079 | rs11216900 | C | G | NULL | 0.675901528 | NULL&M_Disease |
| 11 | 119217580 | rs3814759 | C | T | NULL | 0.675901528 | NULL&M_Disease |
| 11 | 119601313 | rs4936496 | G | A | NULL | 0.675901528 | NULL&M_Disease |
| 11 | 119608316 | rs12793075 | G | A | NULL | 0.675901528 | NULL&M_Disease |
| 11 | 120289699 | rs12794618 | C | T | NULL | 0.675901528 | NULL&M_Disease |
| 11 | 121114116 | rs10892696 | C | A | NULL | 0.675901528 | NULL&M_Disease |
| 11 | 119631726 | rs11217474 | G | C | NULL | 0.675901528 | NULL&M_Disease |
| 11 | 119927043 | rs4938762 | T | C | NULL | 0.675901528 | NULL&M_Disease |
| 11 | 120728304 | rs7950796 | G | A | NULL | 0.676995856 | NULL&M_Disease |

| | | | | | | | | | |
|---|---|---|---|---|---|---|---|---|---|
| 11 | 116845104 | rs17120197 | C | T | 0.975 | -7977.388 | -7978.851 | -1.463 | 8.96E-07 |
| 11 | 116979365 | rs113784559 | G | C | 0.996 | -7977.388 | -7978.851 | -1.463 | 8.96E-07 |
| 11 | 116983556 | rs111560312 | A | G | 0.999 | -7977.388 | -7978.852 | -1.463 | 8.96E-07 |
| 11 | 116984866 | rs55703909 | A | G | 0.996 | -7977.388 | -7978.851 | -1.463 | 8.96E-07 |
| 11 | 117355782 | rs10892125 | C | A | 1.028 | -7977.388 | -7978.851 | -1.463 | 8.96E-07 |
| 11 | 119146659 | rs2511854 | G | C | 1.028 | -7977.388 | -7978.852 | -1.463 | 8.96E-07 |
| 11 | 119164677 | rs2510139 | C | T | 1.028 | -7977.388 | -7978.851 | -1.463 | 8.96E-07 |
| 11 | 119576911 | rs4938717 | T | C | 1.028 | -7977.388 | -7978.852 | -1.463 | 8.96E-07 |
| 11 | 119600410 | rs2136421 | C | T | 0.972 | -7977.388 | -7978.851 | -1.463 | 8.96E-07 |
| 11 | 119923643 | rs1001973 | A | G | 1.028 | -7977.388 | -7978.851 | -1.463 | 8.96E-07 |
| 11 | 119978608 | rs1349688 | C | G | 0.978 | -7977.388 | -7978.851 | -1.463 | 8.96E-07 |
| 11 | 120976413 | rs681311 | A | G | 0.972 | -7977.388 | -7978.851 | -1.463 | 8.96E-07 |
| 11 | 121038810 | rs2155369 | T | C | 1.018 | -7977.388 | -7978.851 | -1.463 | 8.96E-07 |
| 11 | 121092828 | rs948095 | A | G | 0.972 | -7977.388 | -7978.851 | -1.463 | 8.96E-07 |
| 11 | 116794903 | rs56314469 | A | G | 0.98 | -7977.388 | -7978.852 | -1.464 | 8.95E-07 |
| 11 | 118938880 | rs584115 | G | A | 0.972 | -7977.388 | -7978.852 | -1.464 | 8.95E-07 |
| 11 | 119604854 | rs10892444 | A | G | 0.972 | -7977.388 | -7978.852 | -1.464 | 8.95E-07 |
| 11 | 117774829 | rs3809040 | A | G | 0.986 | -7977.388 | -7978.854 | -1.465 | 8.94E-07 |
| 11 | 118432852 | rs11216898 | C | T | 0.982 | -7977.388 | -7978.853 | -1.465 | 8.94E-07 |
| 11 | 118967291 | rs7759 | G | A | 1.028 | -7977.388 | -7978.853 | -1.465 | 8.94E-07 |
| 11 | 119818802 | rs10892502 | C | T | 0.976 | -7977.388 | -7978.853 | -1.465 | 8.94E-07 |
| 11 | 119885547 | rs630775 | C | A | 0.972 | -7977.388 | -7978.853 | -1.465 | 8.94E-07 |
| 11 | 120165942 | rs11217801 | G | A | 1.005 | -7977.388 | -7978.854 | -1.465 | 8.94E-07 |
| 11 | 120299707 | rs7129944 | C | T | 0.976 | -7977.388 | -7978.853 | -1.465 | 8.94E-07 |
| 11 | 120657044 | rs7105258 | T | C | 0.974 | -7977.388 | -7978.853 | -1.465 | 8.94E-07 |
| 11 | 116209430 | rs10790138 | C | G | 1.028 | -7977.388 | -7978.854 | -1.466 | 8.93E-07 |
| 11 | 116701122 | rs2070669 | C | G | 0.973 | -7977.388 | -7978.854 | -1.466 | 8.93E-07 |
| 11 | 118069269 | rs1573580 | G | C | 0.973 | -7977.388 | -7978.854 | -1.466 | 8.93E-07 |
| 11 | 118496119 | rs45565037 | A | G | 1.027 | -7977.388 | -7978.854 | -1.466 | 8.93E-07 |
| 11 | 118966521 | rs2509049 | T | C | 1.028 | -7977.388 | -7978.854 | -1.466 | 8.93E-07 |
| 11 | 119922558 | rs12421541 | T | A | 1.028 | -7977.388 | -7978.855 | -1.466 | 8.93E-07 |
| 11 | 120663363 | rs1954787 | T | C | 0.972 | -7977.388 | -7978.854 | -1.466 | 8.93E-07 |
| 11 | 121017301 | rs2155364 | T | C | 1.017 | -7977.388 | -7978.854 | -1.466 | 8.93E-07 |
| 11 | 120017875 | rs582071 | A | G | 1.013 | -7977.388 | -7978.855 | -1.467 | 8.92E-07 |
| 11 | 120166016 | rs17123792 | A | G | 1.005 | -7977.388 | -7978.855 | -1.467 | 8.92E-07 |
| 11 | 120186190 | rs17245511 | A | G | 1.021 | -7977.388 | -7978.855 | -1.467 | 8.92E-07 |
| 11 | 116225111 | rs491113 | C | G | 1.028 | -7977.388 | -7978.856 | -1.468 | 8.91E-07 |
| 11 | 116697344 | rs2071523 | C | T | 1.028 | -7977.388 | -7978.856 | -1.468 | 8.91E-07 |
| 11 | 118091761 | rs2368238 | T | G | 1.028 | -7977.388 | -7978.856 | -1.468 | 8.91E-07 |
| 11 | 118967524 | rs8551 | T | C | 1.028 | -7977.388 | -7978.856 | -1.468 | 8.91E-07 |
| 11 | 120968462 | rs12274094 | G | A | 1.028 | -7977.388 | -7978.856 | -1.468 | 8.91E-07 |
| 11 | 120979036 | rs1783777 | C | T | 0.972 | -7977.388 | -7978.856 | -1.468 | 8.91E-07 |
| 11 | 120979047 | rs1783776 | G | A | 0.972 | -7977.388 | -7978.856 | -1.468 | 8.91E-07 |
| 11 | 121113880 | rs10790413 | C | A | 1.028 | -7977.388 | -7978.856 | -1.468 | 8.91E-07 |
| 11 | 116196429 | rs10891961 | C | T | 1.028 | -7977.388 | -7978.857 | -1.469 | 8.90E-07 |
| 11 | 116196643 | rs11215897 | G | A | 1.028 | -7977.388 | -7978.857 | -1.469 | 8.90E-07 |
| 11 | 116696043 | rs2216311 | G | A | 1.028 | -7977.388 | -7978.857 | -1.469 | 8.90E-07 |
| 11 | 118947634 | rs540261 | T | C | 0.972 | -7977.388 | -7978.857 | -1.469 | 8.90E-07 |
| 11 | 120051217 | rs76841836 | C | G | 0.985 | -7977.388 | -7978.858 | -1.469 | 8.90E-07 |
| 11 | 120292477 | rs76693355 | C | T | 1.013 | -7977.388 | -7978.858 | -1.469 | 8.90E-07 |
| 11 | 120300804 | rs11217866 | A | T | 0.977 | -7977.388 | -7978.857 | -1.469 | 8.90E-07 |
| 11 | 120517643 | rs6589826 | A | G | 1.028 | -7977.388 | -7978.857 | -1.469 | 8.90E-07 |
| 11 | 118966780 | rs2509851 | C | A | 1.028 | -7977.388 | -7978.858 | -1.47 | 8.89E-07 |
| 11 | 117950199 | rs73010757 | A | G | 1.027 | -7977.388 | -7978.859 | -1.471 | 8.88E-07 |
| 11 | 118320553 | rs629470 | A | G | 0.977 | -7977.388 | -7978.859 | -1.471 | 8.88E-07 |
| 11 | 118337003 | rs574387 | C | A | 0.976 | -7977.388 | -7978.86 | -1.471 | 8.88E-07 |
| 11 | 119612572 | rs12796806 | G | A | 0.973 | -7977.388 | -7978.859 | -1.471 | 8.88E-07 |
| 11 | 120293138 | rs11217863 | A | G | 1.013 | -7977.388 | -7978.86 | -1.471 | 8.88E-07 |
| 11 | 116209589 | rs10732855 | C | G | 1.028 | -7977.388 | -7978.86 | -1.472 | 8.88E-07 |
| 11 | 118925341 | rs538478 | A | G | 0.973 | -7977.388 | -7978.86 | -1.472 | 8.88E-07 |
| 11 | 120978834 | rs1784281 | C | A | 0.973 | -7977.388 | -7978.86 | -1.472 | 8.88E-07 |
| 11 | 116196502 | rs11215896 | A | C | 1.028 | -7977.388 | -7978.861 | -1.473 | 8.87E-07 |
| 11 | 116196666 | rs10891962 | C | T | 1.028 | -7977.388 | -7978.861 | -1.473 | 8.87E-07 |
| 11 | 116355310 | rs11215958 | A | G | 0.989 | -7977.388 | -7978.861 | -1.473 | 8.87E-07 |
| 11 | 118034228 | rs611960 | G | T | 1.005 | -7977.388 | -7978.861 | -1.473 | 8.87E-07 |
| 11 | 118434421 | rs11216899 | G | A | 0.983 | -7977.388 | -7978.862 | -1.473 | 8.87E-07 |
| 11 | 118434835 | rs60753148 | T | C | 0.983 | -7977.388 | -7978.862 | -1.473 | 8.87E-07 |
| 11 | 120066044 | rs12791282 | G | C | 1.028 | -7977.388 | -7978.861 | -1.473 | 8.87E-07 |
| 11 | 121114248 | rs10160642 | T | C | 1.028 | -7977.388 | -7978.861 | -1.473 | 8.87E-07 |
| 11 | 116466289 | rs11216031 | C | A | 0.98 | -7977.388 | -7978.862 | -1.474 | 8.86E-07 |
| 11 | 116532856 | rs1145189 | A | G | 1.026 | -7977.388 | -7978.862 | -1.474 | 8.86E-07 |
| 11 | 117410423 | rs2509004 | A | G | 0.979 | -7977.388 | -7978.862 | -1.474 | 8.86E-07 |
| 11 | 119220839 | rs12421814 | G | A | 0.973 | -7977.388 | -7978.862 | -1.474 | 8.86E-07 |
| 11 | 119800548 | rs1579885 | A | G | 1.017 | -7977.388 | -7978.862 | -1.474 | 8.86E-07 |
| 11 | 119951678 | rs11217679 | G | A | 0.973 | -7977.388 | -7978.862 | -1.474 | 8.86E-07 |
| 11 | 120166860 | rs6589807 | A | G | 1.027 | -7977.388 | -7978.862 | -1.474 | 8.86E-07 |
| 11 | 120914370 | rs7952619 | C | T | 1.028 | -7977.388 | -7978.862 | -1.474 | 8.86E-07 |
| 11 | 116210084 | rs10736483 | A | G | 1.028 | -7977.388 | -7978.864 | -1.475 | 8.85E-07 |
| 11 | 117347058 | rs665406 | C | T | 1.028 | -7977.388 | -7978.863 | -1.475 | 8.85E-07 |
| 11 | 118034223 | rs611979 | A | G | 1.004 | -7977.388 | -7978.863 | -1.475 | 8.85E-07 |
| 11 | 119683607 | rs10892472 | A | C | 0.974 | -7977.388 | -7978.863 | -1.475 | 8.85E-07 |
| 11 | 119845755 | rs12277299 | G | C | 1.013 | -7977.388 | -7978.863 | -1.475 | 8.85E-07 |
| 11 | 119914182 | rs2084893 | G | C | 1.028 | -7977.388 | -7978.863 | -1.475 | 8.85E-07 |
| 11 | 119914915 | rs1597866 | G | A | 1.028 | -7977.388 | -7978.863 | -1.475 | 8.85E-07 |
| 11 | 120294004 | rs11823264 | A | G | 0.977 | -7977.388 | -7978.863 | -1.475 | 8.85E-07 |
| 11 | 120311198 | rs4936518 | T | A | 0.977 | -7977.388 | -7978.863 | -1.475 | 8.85E-07 |
| 11 | 116196711 | rs10891963 | G | T | 1.028 | -7977.388 | -7978.864 | -1.476 | 8.84E-07 |
| 11 | 118436079 | rs11216900 | C | G | 0.983 | -7977.388 | -7978.865 | -1.476 | 8.84E-07 |
| 11 | 119217580 | rs3814759 | C | T | 0.973 | -7977.388 | -7978.865 | -1.476 | 8.84E-07 |
| 11 | 119601313 | rs4936496 | G | A | 0.973 | -7977.388 | -7978.865 | -1.476 | 8.84E-07 |
| 11 | 119608316 | rs12793075 | G | A | 0.973 | -7977.388 | -7978.864 | -1.476 | 8.84E-07 |
| 11 | 120289699 | rs12794618 | C | T | 0.977 | -7977.388 | -7978.865 | -1.476 | 8.84E-07 |
| 11 | 121114116 | rs10892696 | C | A | 1.028 | -7977.388 | -7978.865 | -1.476 | 8.84E-07 |
| 11 | 119631726 | rs11217474 | G | C | 1.028 | -7977.388 | -7978.865 | -1.477 | 8.83E-07 |
| 11 | 119927043 | rs4938762 | T | C | 1.028 | -7977.388 | -7978.865 | -1.477 | 8.83E-07 |
| 11 | 120728304 | rs7950796 | G | A | 0.973 | -7977.388 | -7978.866 | -1.477 | 8.83E-07 |

| | | | | | | | |
|---|---|---|---|---|---|---|---|
| 11 | 120837821 | rs2276319 | G | A | NULL | 0.675901528 | NULL&M_Disease |
| 11 | 121041584 | rs75598010 | T | C | NULL | 0.675901528 | NULL&M_Disease |
| 11 | 121041724 | rs11218176 | A | T | NULL | 0.675901528 | NULL&M_Disease |
| 11 | 116471494 | rs76188167 | A | G | NULL | 0.675901528 | NULL&M_Disease |
| 11 | 116477692 | rs12787356 | A | G | NULL | 0.675901528 | NULL&M_Disease |
| 11 | 119390332 | rs77445023 | T | G | NULL | 0.675901528 | NULL&M_Disease |
| 11 | 120640009 | rs35114640 | A | C | NULL | 0.675901528 | NULL&M_Disease |
| 11 | 120826131 | rs627532 | C | T | NULL | 0.676995856 | NULL&M_Disease |
| 11 | 116549897 | rs35963862 | C | T | NULL | 0.676995856 | NULL&M_Disease |
| 11 | 116696680 | rs2098453 | C | T | NULL | 0.676995856 | NULL&M_Disease |
| 11 | 119999191 | rs569044 | A | G | NULL | 0.676995856 | NULL&M_Disease |
| 11 | 117311169 | rs7943989 | G | A | NULL | 0.676995856 | NULL&M_Disease |
| 11 | 117876155 | rs17121510 | G | A | NULL | 0.676995856 | NULL&M_Disease |
| 11 | 119920139 | rs11217660 | A | G | NULL | 0.676995856 | NULL&M_Disease |
| 11 | 116266176 | rs12271703 | G | A | NULL | 0.676995856 | NULL&M_Disease |
| 11 | 116333088 | rs2851407 | T | C | NULL | 0.676995856 | NULL&M_Disease |
| 11 | 117106280 | rs3016582 | A | T | NULL | 0.676995856 | NULL&M_Disease |
| 11 | 117106282 | rs2929176 | C | T | NULL | 0.676995856 | NULL&M_Disease |
| 11 | 119613092 | rs12797914 | G | A | NULL | 0.676995856 | NULL&M_Disease |
| 11 | 119831744 | rs4625480 | G | A | NULL | 0.676995856 | NULL&M_Disease |
| 11 | 120054885 | rs60336713 | T | C | NULL | 0.676995856 | NULL&M_Disease |
| 11 | 120614787 | rs3133224 | T | C | NULL | 0.676995856 | NULL&M_Disease |
| 11 | 116269470 | rs7102851 | C | T | NULL | 0.676995856 | NULL&M_Disease |
| 11 | 116393439 | rs10891986 | A | T | NULL | 0.676995856 | NULL&M_Disease |
| 11 | 116532066 | rs1145187 | G | T | NULL | 0.676995856 | NULL&M_Disease |
| 11 | 116539203 | rs1787682 | G | C | NULL | 0.676995856 | NULL&M_Disease |
| 11 | 118094513 | rs2051480 | A | G | NULL | 0.676995856 | NULL&M_Disease |
| 11 | 119805351 | rs12294678 | T | C | NULL | 0.676995856 | NULL&M_Disease |
| 11 | 119848438 | rs662318 | G | A | NULL | 0.676995856 | NULL&M_Disease |
| 11 | 120042846 | rs55920307 | G | A | NULL | 0.676995856 | NULL&M_Disease |
| 11 | 116311186 | rs496391 | G | C | NULL | 0.676995856 | NULL&M_Disease |
| 11 | 118136836 | rs3759004 | T | C | NULL | 0.676995856 | NULL&M_Disease |
| 11 | 118435756 | rs2277291 | T | C | NULL | 0.676995856 | NULL&M_Disease |
| 11 | 118438448 | rs60837962 | T | A | NULL | 0.676995856 | NULL&M_Disease |
| 11 | 118490076 | rs10892247 | A | G | NULL | 0.676995856 | NULL&M_Disease |
| 11 | 119506483 | rs10892423 | A | G | NULL | 0.676995856 | NULL&M_Disease |
| 11 | 119831362 | rs12226501 | G | T | NULL | 0.676995856 | NULL&M_Disease |
| 11 | 120024827 | rs1790477 | A | G | NULL | 0.676995856 | NULL&M_Disease |
| 11 | 116457340 | rs10891996 | A | G | NULL | 0.676995856 | NULL&M_Disease |
| 11 | 116831912 | rs7112967 | T | C | NULL | 0.676995856 | NULL&M_Disease |
| 11 | 119081215 | rs6589721 | T | C | NULL | 0.676995856 | NULL&M_Disease |
| 11 | 119320333 | rs582973 | T | C | NULL | 0.676995856 | NULL&M_Disease |
| 11 | 119648956 | rs11217496 | T | G | NULL | 0.676995856 | NULL&M_Disease |
| 11 | 120726562 | rs11218034 | A | G | NULL | 0.676995856 | NULL&M_Disease |
| 11 | 120940461 | rs12274032 | A | G | NULL | 0.676995856 | NULL&M_Disease |
| 11 | 116209785 | rs10750090 | C | T | NULL | 0.676995856 | NULL&M_Disease |
| 11 | 116539868 | rs1784131 | C | T | NULL | 0.676995856 | NULL&M_Disease |
| 11 | 117934704 | rs655116 | C | G | NULL | 0.676995856 | NULL&M_Disease |
| 11 | 118432404 | rs4326835 | T | G | NULL | 0.676995856 | NULL&M_Disease |
| 11 | 119217945 | rs3814755 | G | A | NULL | 0.676995856 | NULL&M_Disease |
| 11 | 119742372 | rs551417 | T | C | NULL | 0.676995856 | NULL&M_Disease |
| 11 | 120291137 | rs10892575 | G | C | NULL | 0.676995856 | NULL&M_Disease |
| 11 | 120937637 | rs10892668 | C | T | NULL | 0.676995856 | NULL&M_Disease |
| 11 | 116210182 | rs10736485 | G | C | NULL | 0.676995856 | NULL&M_Disease |
| 11 | 116528660 | rs1145186 | A | T | NULL | 0.676995856 | NULL&M_Disease |
| 11 | 120518653 | rs4935746 | G | A | NULL | 0.676995856 | NULL&M_Disease |
| 11 | 120896549 | rs11218118 | G | A | NULL | 0.676995856 | NULL&M_Disease |
| 11 | 116222809 | rs2512160 | G | A | NULL | 0.67808825 | NULL&M_Disease |
| 11 | 116384580 | rs12361273 | T | C | NULL | 0.676995856 | NULL&M_Disease |
| 11 | 116474555 | rs12802764 | A | G | NULL | 0.676995856 | NULL&M_Disease |
| 11 | 116677999 | rs1263168 | C | T | NULL | 0.676995856 | NULL&M_Disease |
| 11 | 116678413 | rs1263170 | T | C | NULL | 0.676995856 | NULL&M_Disease |
| 11 | 117949861 | rs626252 | G | A | NULL | 0.676995856 | NULL&M_Disease |
| 11 | 117952960 | rs12279572 | G | A | NULL | 0.676995856 | NULL&M_Disease |
| 11 | 118250586 | rs7949539 | G | A | NULL | 0.676995856 | NULL&M_Disease |
| 11 | 119217818 | rs3814758 | A | G | NULL | 0.676995856 | NULL&M_Disease |
| 11 | 119501021 | rs11217364 | A | G | NULL | 0.676995856 | NULL&M_Disease |
| 11 | 119583751 | rs11217421 | C | T | NULL | 0.676995856 | NULL&M_Disease |
| 11 | 119595355 | rs12800445 | A | C | NULL | 0.676995856 | NULL&M_Disease |
| 11 | 119809582 | rs10160771 | A | G | NULL | 0.676995856 | NULL&M_Disease |
| 11 | 120050113 | rs7114201 | A | T | NULL | 0.676995856 | NULL&M_Disease |
| 11 | 120184575 | rs78302438 | G | T | NULL | 0.676995856 | NULL&M_Disease |
| 11 | 117501072 | rs7949956 | C | T | NULL | 0.67808825 | NULL&M_Disease |
| 11 | 118285995 | rs4290266 | A | G | NULL | 0.676995856 | NULL&M_Disease |
| 11 | 118405438 | rs45522239 | G | A | NULL | 0.67808825 | NULL&M_Disease |
| 11 | 119054488 | rs10790286 | T | C | NULL | 0.67808825 | NULL&M_Disease |
| 11 | 120614119 | rs11828621 | G | A | NULL | 0.676995856 | NULL&M_Disease |
| 11 | 120642807 | rs7948804 | C | T | NULL | 0.676995856 | NULL&M_Disease |
| 11 | 116230993 | rs12793378 | A | T | NULL | 0.67808825 | NULL&M_Disease |
| 11 | 116566117 | rs1942479 | T | C | NULL | 0.67808825 | NULL&M_Disease |
| 11 | 116701674 | rs2070666 | A | T | NULL | 0.67808825 | NULL&M_Disease |
| 11 | 117774394 | rs59590191 | G | A | NULL | 0.67808825 | NULL&M_Disease |
| 11 | 117824158 | rs11216646 | G | A | NULL | 0.67808825 | NULL&M_Disease |
| 11 | 118093547 | rs11607355 | C | T | NULL | 0.67808825 | NULL&M_Disease |
| 11 | 118911350 | rs666741 | C | A | NULL | 0.67808825 | NULL&M_Disease |
| 11 | 119225134 | rs7939363 | T | C | NULL | 0.67808825 | NULL&M_Disease |
| 11 | 116230972 | rs12792262 | A | G | NULL | 0.67808825 | NULL&M_Disease |
| 11 | 116270219 | rs503023 | A | G | NULL | 0.67808825 | NULL&M_Disease |
| 11 | 118405437 | rs45548238 | T | C | NULL | 0.67808825 | NULL&M_Disease |
| 11 | 119313929 | rs514635 | A | G | NULL | 0.67808825 | NULL&M_Disease |
| 11 | 119710918 | rs518326 | A | T | NULL | 0.67808825 | NULL&M_Disease |
| 11 | 120182926 | rs61898343 | C | G | NULL | 0.67808825 | NULL&M_Disease |
| 11 | 120701623 | rs7938081 | C | G | NULL | 0.67808825 | NULL&M_Disease |
| 11 | 120807892 | rs4472950 | T | C | NULL | 0.67808825 | NULL&M_Disease |
| 11 | 121118260 | rs10892698 | C | T | NULL | 0.67808825 | NULL&M_Disease |
| 11 | 121118878 | rs4936597 | G | A | NULL | 0.67808825 | NULL&M_Disease |

| | | | | | | | | | |
|---|---|---|---|---|---|---|---|---|---|
| 11 | 120837821 | rs2276319 | G | A | 0.973 | -7977.388 | -7978.865 | -1.477 | 8.83E-07 |
| 11 | 121041584 | rs75598010 | T | C | 1.016 | -7977.388 | -7978.865 | -1.477 | 8.83E-07 |
| 11 | 121041724 | rs11218176 | A | T | 1.016 | -7977.388 | -7978.865 | -1.477 | 8.83E-07 |
| 11 | 116471494 | rs76188167 | A | G | 0.993 | -7977.388 | -7978.866 | -1.478 | 8.82E-07 |
| 11 | 116477692 | rs12787356 | A | G | 0.993 | -7977.388 | -7978.866 | -1.478 | 8.82E-07 |
| 11 | 119390332 | rs77445023 | T | G | 0.995 | -7977.388 | -7978.866 | -1.478 | 8.82E-07 |
| 11 | 120640009 | rs35114640 | A | C | 0.988 | -7977.388 | -7978.866 | -1.478 | 8.82E-07 |
| 11 | 120826131 | rs627532 | C | T | 1.028 | -7977.388 | -7978.866 | -1.478 | 8.82E-07 |
| 11 | 116549897 | rs35963862 | C | T | 1.024 | -7977.388 | -7978.867 | -1.479 | 8.81E-07 |
| 11 | 116696680 | rs2098453 | C | T | 1.028 | -7977.388 | -7978.867 | -1.479 | 8.81E-07 |
| 11 | 119999191 | rs569044 | A | G | 0.995 | -7977.388 | -7978.867 | -1.479 | 8.81E-07 |
| 11 | 117311169 | rs7943989 | G | A | 1.026 | -7977.388 | -7978.869 | -1.48 | 8.81E-07 |
| 11 | 117876155 | rs17121510 | G | A | 0.996 | -7977.388 | -7978.868 | -1.48 | 8.81E-07 |
| 11 | 119920139 | rs11217660 | A | G | 1.027 | -7977.388 | -7978.868 | -1.48 | 8.81E-07 |
| 11 | 116266176 | rs12271703 | G | A | 0.986 | -7977.388 | -7978.869 | -1.481 | 8.80E-07 |
| 11 | 116333088 | rs2851407 | T | C | 1.007 | -7977.388 | -7978.869 | -1.481 | 8.80E-07 |
| 11 | 117106280 | rs3016582 | A | T | 1.028 | -7977.388 | -7978.869 | -1.481 | 8.80E-07 |
| 11 | 117106282 | rs2929176 | C | T | 1.028 | -7977.388 | -7978.869 | -1.481 | 8.80E-07 |
| 11 | 119613092 | rs12797914 | G | A | 0.973 | -7977.388 | -7978.87 | -1.481 | 8.80E-07 |
| 11 | 119831744 | rs4625480 | G | A | 0.984 | -7977.388 | -7978.869 | -1.481 | 8.80E-07 |
| 11 | 120054885 | rs60336713 | T | C | 1.008 | -7977.388 | -7978.869 | -1.481 | 8.80E-07 |
| 11 | 120614787 | rs3133224 | T | C | 1.018 | -7977.388 | -7978.869 | -1.481 | 8.80E-07 |
| 11 | 116269470 | rs7102851 | C | T | 1.027 | -7977.388 | -7978.871 | -1.482 | 8.79E-07 |
| 11 | 116393439 | rs10891986 | A | T | 0.99 | -7977.388 | -7978.87 | -1.482 | 8.79E-07 |
| 11 | 116532066 | rs1145187 | G | T | 1.026 | -7977.388 | -7978.87 | -1.482 | 8.79E-07 |
| 11 | 116539203 | rs1787682 | G | C | 1.027 | -7977.388 | -7978.87 | -1.482 | 8.79E-07 |
| 11 | 118094513 | rs2051480 | A | G | 1.028 | -7977.388 | -7978.87 | -1.482 | 8.79E-07 |
| 11 | 119805351 | rs12294678 | T | C | 1.016 | -7977.388 | -7978.87 | -1.482 | 8.79E-07 |
| 11 | 119848438 | rs662318 | G | A | 1.028 | -7977.388 | -7978.87 | -1.482 | 8.79E-07 |
| 11 | 120042846 | rs55920307 | G | A | 0.986 | -7977.388 | -7978.87 | -1.482 | 8.79E-07 |
| 11 | 116311186 | rs496391 | G | C | 1.028 | -7977.388 | -7978.871 | -1.483 | 8.78E-07 |
| 11 | 118136836 | rs3759004 | T | C | 0.973 | -7977.388 | -7978.872 | -1.483 | 8.78E-07 |
| 11 | 118435756 | rs2277291 | T | C | 0.984 | -7977.388 | -7978.871 | -1.483 | 8.78E-07 |
| 11 | 118438448 | rs60837962 | T | A | 0.984 | -7977.388 | -7978.872 | -1.483 | 8.78E-07 |
| 11 | 118490076 | rs10892247 | A | G | 1.026 | -7977.388 | -7978.871 | -1.483 | 8.78E-07 |
| 11 | 119506483 | rs10892423 | A | G | 0.973 | -7977.388 | -7978.871 | -1.483 | 8.78E-07 |
| 11 | 119831362 | rs12226501 | G | T | 0.984 | -7977.388 | -7978.872 | -1.483 | 8.78E-07 |
| 11 | 120024827 | rs1790477 | A | G | 1.002 | -7977.388 | -7978.871 | -1.483 | 8.78E-07 |
| 11 | 116457340 | rs10891996 | A | G | 0.975 | -7977.388 | -7978.872 | -1.484 | 8.77E-07 |
| 11 | 116831912 | rs7112967 | T | C | 0.982 | -7977.388 | -7978.872 | -1.484 | 8.77E-07 |
| 11 | 119081215 | rs6589721 | T | C | 1.028 | -7977.388 | -7978.873 | -1.484 | 8.77E-07 |
| 11 | 119320333 | rs582973 | T | C | 0.992 | -7977.388 | -7978.873 | -1.484 | 8.77E-07 |
| 11 | 119648956 | rs11217496 | T | G | 1.028 | -7977.388 | -7978.873 | -1.484 | 8.77E-07 |
| 11 | 120726562 | rs11218034 | A | G | 0.973 | -7977.388 | -7978.872 | -1.484 | 8.77E-07 |
| 11 | 120940461 | rs12274032 | A | G | 1.027 | -7977.388 | -7978.873 | -1.484 | 8.77E-07 |
| 11 | 116209785 | rs10750090 | C | T | 1.028 | -7977.388 | -7978.874 | -1.485 | 8.76E-07 |
| 11 | 116539868 | rs1784131 | C | T | 1.027 | -7977.388 | -7978.874 | -1.485 | 8.76E-07 |
| 11 | 117934704 | rs655116 | C | G | 1.022 | -7977.388 | -7978.873 | -1.485 | 8.76E-07 |
| 11 | 118432404 | rs4326835 | T | G | 0.984 | -7977.388 | -7978.873 | -1.485 | 8.76E-07 |
| 11 | 119217945 | rs3814755 | G | A | 0.973 | -7977.388 | -7978.873 | -1.485 | 8.76E-07 |
| 11 | 119742372 | rs551417 | T | C | 1.018 | -7977.388 | -7978.873 | -1.485 | 8.76E-07 |
| 11 | 120291137 | rs10892575 | G | C | 0.978 | -7977.388 | -7978.873 | -1.485 | 8.76E-07 |
| 11 | 120937637 | rs10892668 | C | T | 1.027 | -7977.388 | -7978.873 | -1.485 | 8.76E-07 |
| 11 | 116210182 | rs10736485 | G | C | 1.028 | -7977.388 | -7978.875 | -1.486 | 8.75E-07 |
| 11 | 116528660 | rs1145186 | A | T | 1.028 | -7977.388 | -7978.874 | -1.486 | 8.75E-07 |
| 11 | 120518653 | rs4935746 | G | A | 1.028 | -7977.388 | -7978.874 | -1.486 | 8.75E-07 |
| 11 | 120896549 | rs11218118 | G | A | 1.021 | -7977.388 | -7978.875 | -1.486 | 8.75E-07 |
| 11 | 116222809 | rs2512160 | G | A | 1.028 | -7977.388 | -7978.876 | -1.487 | 8.74E-07 |
| 11 | 116384580 | rs12361273 | T | C | 0.987 | -7977.388 | -7978.875 | -1.487 | 8.74E-07 |
| 11 | 116474555 | rs12802764 | A | G | 0.998 | -7977.388 | -7978.876 | -1.487 | 8.74E-07 |
| 11 | 116677999 | rs1263168 | C | T | 1.027 | -7977.388 | -7978.875 | -1.487 | 8.74E-07 |
| 11 | 116678413 | rs1263170 | T | C | 1.027 | -7977.388 | -7978.876 | -1.487 | 8.74E-07 |
| 11 | 117949861 | rs626252 | G | A | 1.022 | -7977.388 | -7978.875 | -1.487 | 8.74E-07 |
| 11 | 117952960 | rs12279572 | G | A | 1.027 | -7977.388 | -7978.875 | -1.487 | 8.74E-07 |
| 11 | 118250586 | rs7949539 | G | A | 0.982 | -7977.388 | -7978.875 | -1.487 | 8.74E-07 |
| 11 | 119217818 | rs3814758 | A | G | 0.973 | -7977.388 | -7978.876 | -1.487 | 8.74E-07 |
| 11 | 119501021 | rs11217364 | A | G | 0.973 | -7977.388 | -7978.875 | -1.487 | 8.74E-07 |
| 11 | 119583751 | rs11217421 | C | T | 1.028 | -7977.388 | -7978.875 | -1.487 | 8.74E-07 |
| 11 | 119595355 | rs12800445 | A | C | 0.973 | -7977.388 | -7978.875 | -1.487 | 8.74E-07 |
| 11 | 119809582 | rs10160771 | A | G | 1.016 | -7977.388 | -7978.875 | -1.487 | 8.74E-07 |
| 11 | 120050113 | rs7114201 | A | T | 0.987 | -7977.388 | -7978.876 | -1.487 | 8.74E-07 |
| 11 | 120184575 | rs78302438 | G | T | 1.02 | -7977.388 | -7978.875 | -1.487 | 8.74E-07 |
| 11 | 117501072 | rs7949956 | C | T | 0.973 | -7977.388 | -7978.876 | -1.488 | 8.74E-07 |
| 11 | 118285995 | rs4290266 | A | G | 0.982 | -7977.388 | -7978.877 | -1.488 | 8.74E-07 |
| 11 | 118405438 | rs45522239 | G | A | 0.983 | -7977.388 | -7978.877 | -1.488 | 8.74E-07 |
| 11 | 119054488 | rs10790286 | T | C | 0.973 | -7977.388 | -7978.876 | -1.488 | 8.74E-07 |
| 11 | 120614119 | rs11828621 | G | A | 1.016 | -7977.388 | -7978.876 | -1.488 | 8.74E-07 |
| 11 | 120642807 | rs7948804 | C | T | 0.987 | -7977.388 | -7978.876 | -1.488 | 8.74E-07 |
| 11 | 116230993 | rs12793378 | A | T | 0.98 | -7977.388 | -7978.877 | -1.489 | 8.73E-07 |
| 11 | 116566117 | rs1942479 | T | C | 1.026 | -7977.388 | -7978.877 | -1.489 | 8.73E-07 |
| 11 | 116701674 | rs2070666 | A | T | 0.98 | -7977.388 | -7978.877 | -1.489 | 8.73E-07 |
| 11 | 117774394 | rs59590191 | G | A | 0.984 | -7977.388 | -7978.877 | -1.489 | 8.73E-07 |
| 11 | 117824158 | rs11216646 | G | A | 1.009 | -7977.388 | -7978.877 | -1.489 | 8.73E-07 |
| 11 | 118093547 | rs11607355 | C | T | 1.028 | -7977.388 | -7978.877 | -1.489 | 8.73E-07 |
| 11 | 118911350 | rs666741 | C | A | 0.973 | -7977.388 | -7978.877 | -1.489 | 8.73E-07 |
| 11 | 119225134 | rs7939363 | T | C | 0.974 | -7977.388 | -7978.877 | -1.489 | 8.73E-07 |
| 11 | 116230972 | rs12792262 | A | G | 0.98 | -7977.388 | -7978.878 | -1.49 | 8.72E-07 |
| 11 | 116270219 | rs503023 | A | G | 1.027 | -7977.388 | -7978.878 | -1.49 | 8.72E-07 |
| 11 | 118405437 | rs45548238 | T | C | 0.983 | -7977.388 | -7978.878 | -1.49 | 8.72E-07 |
| 11 | 119313929 | rs514635 | A | G | 0.993 | -7977.388 | -7978.878 | -1.49 | 8.72E-07 |
| 11 | 119710918 | rs518326 | A | T | 0.975 | -7977.388 | -7978.879 | -1.49 | 8.72E-07 |
| 11 | 120182926 | rs61898343 | C | G | 1.019 | -7977.388 | -7978.878 | -1.49 | 8.72E-07 |
| 11 | 120701623 | rs7938081 | C | G | 0.973 | -7977.388 | -7978.878 | -1.49 | 8.72E-07 |
| 11 | 120807892 | rs4472950 | T | C | 1.027 | -7977.388 | -7978.878 | -1.49 | 8.72E-07 |
| 11 | 121118260 | rs10892698 | C | T | 1.028 | -7977.388 | -7978.878 | -1.49 | 8.72E-07 |
| 11 | 121118878 | rs4936597 | G | A | 1.028 | -7977.388 | -7978.878 | -1.49 | 8.72E-07 |

| | | | | | | | |
|---|---|---|---|---|---|---|---|
| 11 | 121119493 | rs10892699 | C | G | NULL | 0.67808825 | NULL&M_Disease |
| 11 | 116230459 | rs6589542 | C | A | NULL | 0.67808825 | NULL&M_Disease |
| 11 | 119691200 | rs1838947 | C | T | NULL | 0.67808825 | NULL&M_Disease |
| 11 | 120064556 | rs12807240 | G | A | NULL | 0.67808825 | NULL&M_Disease |
| 11 | 116388998 | rs7930587 | A | T | NULL | 0.67808825 | NULL&M_Disease |
| 11 | 118495287 | rs77683570 | C | T | NULL | 0.67808825 | NULL&M_Disease |
| 11 | 120048958 | rs896691 | C | A | NULL | 0.67808825 | NULL&M_Disease |
| 11 | 120064501 | rs12807541 | A | G | NULL | 0.67808825 | NULL&M_Disease |
| 11 | 120612976 | rs61901420 | G | A | NULL | 0.67808825 | NULL&M_Disease |
| 11 | 120770778 | rs2156637 | A | G | NULL | 0.67808825 | NULL&M_Disease |
| 11 | 121054619 | rs11218191 | T | C | NULL | 0.67808825 | NULL&M_Disease |
| 11 | 121115895 | rs10790414 | T | G | NULL | 0.67808825 | NULL&M_Disease |
| 11 | 121115951 | rs10790415 | C | G | NULL | 0.67808825 | NULL&M_Disease |
| 11 | 116546503 | rs61906102 | C | T | NULL | 0.67808825 | NULL&M_Disease |
| 11 | 116550115 | rs61906104 | G | A | NULL | 0.67808825 | NULL&M_Disease |
| 11 | 117143185 | rs536648 | G | C | NULL | 0.67808825 | NULL&M_Disease |
| 11 | 117354818 | rs10892124 | G | C | NULL | 0.67808825 | NULL&M_Disease |
| 11 | 118031206 | rs656308 | T | G | NULL | 0.67808825 | NULL&M_Disease |
| 11 | 118206890 | rs61010648 | G | C | NULL | 0.67808825 | NULL&M_Disease |
| 11 | 118834784 | rs7126476 | G | A | NULL | 0.67808825 | NULL&M_Disease |
| 11 | 116366547 | rs11215966 | C | T | NULL | 0.67808825 | NULL&M_Disease |
| 11 | 116514417 | rs10892004 | T | C | NULL | 0.67808825 | NULL&M_Disease |
| 11 | 116820071 | rs61907605 | A | T | NULL | 0.67808825 | NULL&M_Disease |
| 11 | 118069563 | rs3018382 | G | C | NULL | 0.67808825 | NULL&M_Disease |
| 11 | 119295082 | rs3910017 | A | G | NULL | 0.67808825 | NULL&M_Disease |
| 11 | 119295647 | rs6589737 | C | T | NULL | 0.67808825 | NULL&M_Disease |
| 11 | 120039274 | rs12421142 | T | C | NULL | 0.67808825 | NULL&M_Disease |
| 11 | 117450432 | rs669226 | C | A | NULL | 0.67808825 | NULL&M_Disease |
| 11 | 119282086 | rs4018665 | T | C | NULL | 0.67808825 | NULL&M_Disease |
| 11 | 119320444 | rs7112232 | A | G | NULL | 0.67808825 | NULL&M_Disease |
| 11 | 119961536 | rs3895896 | A | T | NULL | 0.67808825 | NULL&M_Disease |
| 11 | 120829359 | rs734167 | A | G | NULL | 0.67808825 | NULL&M_Disease |
| 11 | 116203045 | rs750057 | G | A | NULL | 0.67808825 | NULL&M_Disease |
| 11 | 116548349 | rs12801136 | A | G | NULL | 0.67808825 | NULL&M_Disease |
| 11 | 116819241 | rs4316540 | T | C | NULL | 0.67808825 | NULL&M_Disease |
| 11 | 118030310 | rs642765 | C | G | NULL | 0.67808825 | NULL&M_Disease |
| 11 | 118573519 | rs4639966 | C | T | NULL | 0.67808825 | NULL&M_Disease |
| 11 | 119318306 | rs7130716 | A | G | NULL | 0.67808825 | NULL&M_Disease |
| 11 | 119330588 | rs482002 | T | A | NULL | 0.67808825 | NULL&M_Disease |
| 11 | 120026748 | rs2084898 | A | G | NULL | 0.67808825 | NULL&M_Disease |
| 11 | 120752044 | rs4423191 | G | T | NULL | 0.67808825 | NULL&M_Disease |
| 11 | 116928514 | rs4938330 | G | C | NULL | 0.67808825 | NULL&M_Disease |
| 11 | 117842344 | rs57210824 | T | A | NULL | 0.67808825 | NULL&M_Disease |
| 11 | 118030086 | rs11608072 | C | G | NULL | 0.67808825 | NULL&M_Disease |
| 11 | 119261867 | rs7926015 | T | G | NULL | 0.67808825 | NULL&M_Disease |
| 11 | 119267694 | rs2115649 | A | C | NULL | 0.67808825 | NULL&M_Disease |
| 11 | 119269081 | rs10892361 | G | A | NULL | 0.67808825 | NULL&M_Disease |
| 11 | 119309664 | rs10736497 | C | T | NULL | 0.67808825 | NULL&M_Disease |
| 11 | 119323284 | rs10750140 | T | C | NULL | 0.67808825 | NULL&M_Disease |
| 11 | 119361224 | rs2509667 | A | C | NULL | 0.67808825 | NULL&M_Disease |
| 11 | 119841751 | rs55801969 | G | A | NULL | 0.67808825 | NULL&M_Disease |
| 11 | 120053576 | rs73004754 | A | G | NULL | 0.67808825 | NULL&M_Disease |
| 11 | 121116773 | rs7115337 | T | G | NULL | 0.67808825 | NULL&M_Disease |
| 11 | 117760079 | rs10892190 | G | A | NULL | 0.67808825 | NULL&M_Disease |
| 11 | 117760118 | rs10892191 | C | G | NULL | 0.67808825 | NULL&M_Disease |
| 11 | 117774560 | rs3809041 | C | G | NULL | 0.67808825 | NULL&M_Disease |
| 11 | 118405377 | rs45485302 | A | C | NULL | 0.67808825 | NULL&M_Disease |
| 11 | 119237217 | rs1940804 | C | T | NULL | 0.679178699 | NULL&M_Disease |
| 11 | 119310379 | rs10790296 | A | G | NULL | 0.67808825 | NULL&M_Disease |
| 11 | 119951327 | rs11602430 | G | C | NULL | 0.67808825 | NULL&M_Disease |
| 11 | 120042081 | rs4938788 | A | G | NULL | 0.679178699 | NULL&M_Disease |
| 11 | 120613199 | rs2508807 | C | T | NULL | 0.679178699 | NULL&M_Disease |
| 11 | 120636628 | rs10892623 | G | A | NULL | 0.67808825 | NULL&M_Disease |
| 11 | 121114571 | rs10160459 | C | T | NULL | 0.679178699 | NULL&M_Disease |
| 11 | 121117684 | rs7103470 | G | T | NULL | 0.679178699 | NULL&M_Disease |
| 11 | 121117803 | rs7103283 | A | T | NULL | 0.679178699 | NULL&M_Disease |
| 11 | 116555386 | rs35681250 | T | A | NULL | 0.679178699 | NULL&M_Disease |
| 11 | 116973929 | rs12269901 | C | G | NULL | 0.679178699 | NULL&M_Disease |
| 11 | 117465988 | rs12804173 | A | T | NULL | 0.679178699 | NULL&M_Disease |
| 11 | 118405835 | rs3741324 | A | G | NULL | 0.679178699 | NULL&M_Disease |
| 11 | 118413604 | rs58270621 | A | G | NULL | 0.679178699 | NULL&M_Disease |
| 11 | 119309436 | rs4938656 | T | C | NULL | 0.679178699 | NULL&M_Disease |
| 11 | 119311477 | rs6589740 | T | C | NULL | 0.679178699 | NULL&M_Disease |
| 11 | 119740598 | rs570081 | G | A | NULL | 0.679178699 | NULL&M_Disease |
| 11 | 119798818 | rs572287 | A | T | NULL | 0.679178699 | NULL&M_Disease |
| 11 | 119799829 | rs492227 | T | C | NULL | 0.679178699 | NULL&M_Disease |
| 11 | 119799851 | rs12270574 | T | A | NULL | 0.679178699 | NULL&M_Disease |
| 11 | 119837503 | rs12295555 | C | T | NULL | 0.679178699 | NULL&M_Disease |
| 11 | 119838017 | rs34006113 | T | G | NULL | 0.679178699 | NULL&M_Disease |
| 11 | 119965450 | rs12363785 | A | G | NULL | 0.679178699 | NULL&M_Disease |
| 11 | 120058833 | rs73004786 | A | G | NULL | 0.679178699 | NULL&M_Disease |
| 11 | 120518870 | rs4935747 | C | G | NULL | 0.679178699 | NULL&M_Disease |
| 11 | 121006298 | rs678275 | G | C | NULL | 0.679178699 | NULL&M_Disease |
| 11 | 121114375 | rs10160458 | G | A | NULL | 0.679178699 | NULL&M_Disease |
| 11 | 116198031 | rs2367794 | G | A | NULL | 0.679178699 | NULL&M_Disease |
| 11 | 117116781 | rs588763 | G | A | NULL | 0.679178699 | NULL&M_Disease |
| 11 | 117836519 | rs11216655 | T | C | NULL | 0.679178699 | NULL&M_Disease |
| 11 | 119291688 | rs3751026 | A | G | NULL | 0.679178699 | NULL&M_Disease |
| 11 | 119792036 | rs34655967 | A | C | NULL | 0.679178699 | NULL&M_Disease |
| 11 | 119806447 | rs12286715 | T | C | NULL | 0.679178699 | NULL&M_Disease |
| 11 | 120568392 | rs10892618 | C | A | NULL | 0.679178699 | NULL&M_Disease |
| 11 | 120632138 | rs4590884 | T | C | NULL | 0.679178699 | NULL&M_Disease |
| 11 | 116556236 | rs17092638 | G | C | NULL | 0.679178699 | NULL&M_Disease |
| 11 | 117815759 | rs2515793 | A | G | NULL | 0.679178699 | NULL&M_Disease |
| 11 | 118298041 | rs79281178 | G | C | NULL | 0.679178699 | NULL&M_Disease |
| 11 | 119304090 | rs10790293 | A | C | NULL | 0.679178699 | NULL&M_Disease |

| | | | | | | | | | |
|---|---|---|---|---|---|---|---|---|---|
| 11 | 121119493 | rs10892699 | C | G | 1.028 | -7977.388 | -7978.878 | -1.49 | 8.72E-07 |
| 11 | 116230459 | rs6589542 | C | A | 0.98 | -7977.388 | -7978.879 | -1.491 | 8.71E-07 |
| 11 | 119691200 | rs1838947 | C | T | 0.973 | -7977.388 | -7978.879 | -1.491 | 8.71E-07 |
| 11 | 120064556 | rs12807240 | G | A | 1.027 | -7977.388 | -7978.879 | -1.491 | 8.71E-07 |
| 11 | 116388998 | rs7930587 | A | T | 1.024 | -7977.388 | -7978.88 | -1.492 | 8.70E-07 |
| 11 | 118495287 | rs77683570 | C | T | 1.026 | -7977.388 | -7978.88 | -1.492 | 8.70E-07 |
| 11 | 120048958 | rs896691 | C | A | 0.976 | -7977.388 | -7978.881 | -1.492 | 8.70E-07 |
| 11 | 120064501 | rs12807541 | A | G | 1.027 | -7977.388 | -7978.88 | -1.492 | 8.70E-07 |
| 11 | 120612976 | rs61901420 | G | A | 1.016 | -7977.388 | -7978.88 | -1.492 | 8.70E-07 |
| 11 | 120770778 | rs2156637 | A | G | 0.981 | -7977.388 | -7978.88 | -1.492 | 8.70E-07 |
| 11 | 121054619 | rs11218191 | T | C | 1.014 | -7977.388 | -7978.88 | -1.492 | 8.70E-07 |
| 11 | 121115895 | rs10790414 | T | G | 1.027 | -7977.388 | -7978.88 | -1.492 | 8.70E-07 |
| 11 | 121115951 | rs10790415 | C | G | 1.027 | -7977.388 | -7978.88 | -1.492 | 8.70E-07 |
| 11 | 116546503 | rs61906102 | C | T | 1.023 | -7977.388 | -7978.881 | -1.493 | 8.69E-07 |
| 11 | 116550115 | rs61906104 | G | A | 1.023 | -7977.388 | -7978.882 | -1.493 | 8.69E-07 |
| 11 | 117143185 | rs536648 | G | C | 1.028 | -7977.388 | -7978.881 | -1.493 | 8.69E-07 |
| 11 | 117354818 | rs10892124 | G | C | 0.976 | -7977.388 | -7978.881 | -1.493 | 8.69E-07 |
| 11 | 118031206 | rs656308 | T | G | 1.005 | -7977.388 | -7978.881 | -1.493 | 8.69E-07 |
| 11 | 118206890 | rs61010648 | G | C | 0.974 | -7977.388 | -7978.881 | -1.493 | 8.69E-07 |
| 11 | 118834784 | rs7126476 | G | A | 1.028 | -7977.388 | -7978.881 | -1.493 | 8.69E-07 |
| 11 | 116366547 | rs11215966 | C | T | 1.028 | -7977.388 | -7978.883 | -1.494 | 8.68E-07 |
| 11 | 116514417 | rs10892004 | T | C | 1.01 | -7977.388 | -7978.882 | -1.494 | 8.68E-07 |
| 11 | 116820071 | rs61907605 | A | T | 0.983 | -7977.388 | -7978.882 | -1.494 | 8.68E-07 |
| 11 | 118069563 | rs3018382 | G | C | 0.974 | -7977.388 | -7978.883 | -1.494 | 8.68E-07 |
| 11 | 119295082 | rs3910017 | A | G | 0.996 | -7977.388 | -7978.883 | -1.494 | 8.68E-07 |
| 11 | 119295647 | rs6589737 | C | T | 0.996 | -7977.388 | -7978.883 | -1.494 | 8.68E-07 |
| 11 | 120039274 | rs12421142 | T | C | 0.986 | -7977.388 | -7978.883 | -1.494 | 8.68E-07 |
| 11 | 117450432 | rs669226 | C | A | 0.975 | -7977.388 | -7978.884 | -1.495 | 8.67E-07 |
| 11 | 119282086 | rs4018665 | T | C | 0.997 | -7977.388 | -7978.884 | -1.495 | 8.67E-07 |
| 11 | 119320444 | rs7112232 | A | G | 0.995 | -7977.388 | -7978.884 | -1.495 | 8.67E-07 |
| 11 | 119961536 | rs3895896 | A | T | 1.006 | -7977.388 | -7978.883 | -1.495 | 8.67E-07 |
| 11 | 120829359 | rs734167 | A | G | 1.024 | -7977.388 | -7978.883 | -1.495 | 8.67E-07 |
| 11 | 116203045 | rs750057 | G | A | 1.027 | -7977.388 | -7978.885 | -1.496 | 8.67E-07 |
| 11 | 116548349 | rs12801136 | A | G | 1.023 | -7977.388 | -7978.884 | -1.496 | 8.67E-07 |
| 11 | 116819241 | rs4316540 | T | C | 0.984 | -7977.388 | -7978.885 | -1.496 | 8.67E-07 |
| 11 | 118030310 | rs642765 | C | G | 1.004 | -7977.388 | -7978.884 | -1.496 | 8.67E-07 |
| 11 | 118573519 | rs4639966 | C | T | 1.026 | -7977.388 | -7978.884 | -1.496 | 8.67E-07 |
| 11 | 119318306 | rs7130716 | A | G | 0.995 | -7977.388 | -7978.884 | -1.496 | 8.67E-07 |
| 11 | 119330588 | rs482002 | T | A | 0.998 | -7977.388 | -7978.884 | -1.496 | 8.67E-07 |
| 11 | 120026748 | rs2084898 | A | G | 0.988 | -7977.388 | -7978.884 | -1.496 | 8.67E-07 |
| 11 | 120752044 | rs4423191 | G | T | 1.017 | -7977.388 | -7978.884 | -1.496 | 8.67E-07 |
| 11 | 116928514 | rs4938330 | G | C | 1.025 | -7977.388 | -7978.885 | -1.497 | 8.66E-07 |
| 11 | 117842344 | rs57210824 | T | A | 1.009 | -7977.388 | -7978.885 | -1.497 | 8.66E-07 |
| 11 | 118030086 | rs11608072 | C | G | 1.004 | -7977.388 | -7978.886 | -1.497 | 8.66E-07 |
| 11 | 119261867 | rs7926015 | T | G | 1.001 | -7977.388 | -7978.885 | -1.497 | 8.66E-07 |
| 11 | 119267694 | rs2115649 | A | C | 1 | -7977.388 | -7978.885 | -1.497 | 8.66E-07 |
| 11 | 119269081 | rs10892361 | G | A | 1 | -7977.388 | -7978.885 | -1.497 | 8.66E-07 |
| 11 | 119309664 | rs10736497 | C | T | 0.997 | -7977.388 | -7978.886 | -1.497 | 8.66E-07 |
| 11 | 119323284 | rs10750140 | T | C | 0.996 | -7977.388 | -7978.886 | -1.497 | 8.66E-07 |
| 11 | 119361224 | rs2509667 | A | C | 1.003 | -7977.388 | -7978.886 | -1.497 | 8.66E-07 |
| 11 | 119841751 | rs55801969 | G | A | 1.009 | -7977.388 | -7978.886 | -1.497 | 8.66E-07 |
| 11 | 120053576 | rs73004754 | A | G | 1.002 | -7977.388 | -7978.885 | -1.497 | 8.66E-07 |
| 11 | 121116773 | rs7115337 | T | G | 1.027 | -7977.388 | -7978.885 | -1.497 | 8.66E-07 |
| 11 | 117760079 | rs10892190 | G | A | 1.01 | -7977.388 | -7978.886 | -1.498 | 8.65E-07 |
| 11 | 117760118 | rs10892191 | C | G | 1.01 | -7977.388 | -7978.886 | -1.498 | 8.65E-07 |
| 11 | 117774560 | rs3809041 | C | G | 0.991 | -7977.388 | -7978.887 | -1.498 | 8.65E-07 |
| 11 | 118405377 | rs45485302 | A | C | 0.984 | -7977.388 | -7978.886 | -1.498 | 8.65E-07 |
| 11 | 119237217 | rs1940804 | C | T | 0.973 | -7977.388 | -7978.886 | -1.498 | 8.65E-07 |
| 11 | 119310379 | rs10790296 | A | G | 0.996 | -7977.388 | -7978.887 | -1.498 | 8.65E-07 |
| 11 | 119951327 | rs11602430 | G | C | 0.991 | -7977.388 | -7978.887 | -1.498 | 8.65E-07 |
| 11 | 120042081 | rs4938788 | A | G | 0.988 | -7977.388 | -7978.887 | -1.498 | 8.65E-07 |
| 11 | 120613199 | rs2508807 | C | T | 0.973 | -7977.388 | -7978.886 | -1.498 | 8.65E-07 |
| 11 | 120636628 | rs10892623 | G | A | 0.992 | -7977.388 | -7978.886 | -1.498 | 8.65E-07 |
| 11 | 121114571 | rs10160459 | C | T | 1.027 | -7977.388 | -7978.886 | -1.498 | 8.65E-07 |
| 11 | 121117684 | rs7103470 | G | T | 1.027 | -7977.388 | -7978.886 | -1.498 | 8.65E-07 |
| 11 | 121117803 | rs7103283 | A | T | 1.027 | -7977.388 | -7978.886 | -1.498 | 8.65E-07 |
| 11 | 116555386 | rs35681250 | T | A | 1.023 | -7977.388 | -7978.888 | -1.499 | 8.64E-07 |
| 11 | 116973929 | rs12269901 | C | G | 1.027 | -7977.388 | -7978.887 | -1.499 | 8.64E-07 |
| 11 | 117465988 | rs12804173 | A | T | 1.026 | -7977.388 | -7978.888 | -1.499 | 8.64E-07 |
| 11 | 118405835 | rs3741324 | A | G | 0.984 | -7977.388 | -7978.888 | -1.499 | 8.64E-07 |
| 11 | 118413604 | rs58270621 | A | G | 0.985 | -7977.388 | -7978.887 | -1.499 | 8.64E-07 |
| 11 | 119309436 | rs4938656 | T | C | 0.998 | -7977.388 | -7978.888 | -1.499 | 8.64E-07 |
| 11 | 119311477 | rs6589740 | T | C | 0.996 | -7977.388 | -7978.888 | -1.499 | 8.64E-07 |
| 11 | 119740598 | rs570081 | G | A | 1.016 | -7977.388 | -7978.887 | -1.499 | 8.64E-07 |
| 11 | 119798818 | rs572287 | A | T | 0.974 | -7977.388 | -7978.887 | -1.499 | 8.64E-07 |
| 11 | 119799829 | rs492227 | T | C | 0.974 | -7977.388 | -7978.887 | -1.499 | 8.64E-07 |
| 11 | 119799851 | rs12270574 | T | A | 0.974 | -7977.388 | -7978.887 | -1.499 | 8.64E-07 |
| 11 | 119837503 | rs12295555 | C | T | 1.009 | -7977.388 | -7978.887 | -1.499 | 8.64E-07 |
| 11 | 119838017 | rs34006113 | T | G | 1.009 | -7977.388 | -7978.887 | -1.499 | 8.64E-07 |
| 11 | 119965450 | rs12363785 | A | G | 0.994 | -7977.388 | -7978.887 | -1.499 | 8.64E-07 |
| 11 | 120058833 | rs73004786 | A | G | 1.008 | -7977.388 | -7978.887 | -1.499 | 8.64E-07 |
| 11 | 120518870 | rs4935747 | C | G | 1.027 | -7977.388 | -7978.887 | -1.499 | 8.64E-07 |
| 11 | 121006298 | rs678275 | G | C | 1.027 | -7977.388 | -7978.887 | -1.499 | 8.64E-07 |
| 11 | 121114375 | rs10160458 | G | A | 1.027 | -7977.388 | -7978.887 | -1.499 | 8.64E-07 |
| 11 | 116198031 | rs2367794 | G | A | 1.027 | -7977.388 | -7978.889 | -1.5 | 8.63E-07 |
| 11 | 117116781 | rs588763 | G | A | 1.027 | -7977.388 | -7978.888 | -1.5 | 8.63E-07 |
| 11 | 117836519 | rs11216655 | T | C | 1.008 | -7977.388 | -7978.888 | -1.5 | 8.63E-07 |
| 11 | 119291688 | rs3751026 | A | G | 0.999 | -7977.388 | -7978.889 | -1.5 | 8.63E-07 |
| 11 | 119792036 | rs34655967 | A | C | 0.985 | -7977.388 | -7978.889 | -1.5 | 8.63E-07 |
| 11 | 119806447 | rs12286715 | T | C | 1.014 | -7977.388 | -7978.889 | -1.5 | 8.63E-07 |
| 11 | 120568392 | rs10892618 | C | A | 1.027 | -7977.388 | -7978.888 | -1.5 | 8.63E-07 |
| 11 | 120632138 | rs4590884 | T | C | 0.995 | -7977.388 | -7978.888 | -1.5 | 8.63E-07 |
| 11 | 116556236 | rs17092638 | G | C | 1.023 | -7977.388 | -7978.889 | -1.501 | 8.62E-07 |
| 11 | 117815759 | rs2515793 | A | G | 1.027 | -7977.388 | -7978.89 | -1.501 | 8.62E-07 |
| 11 | 118298041 | rs79281178 | G | C | 0.983 | -7977.388 | -7978.89 | -1.501 | 8.62E-07 |
| 11 | 119304090 | rs10790293 | A | C | 0.999 | -7977.388 | -7978.889 | -1.501 | 8.62E-07 |

| | | | | | | | |
|---|---|---|---|---|---|---|---|
| 11 | 119305593 | rs4938652 | T | C | NULL | 0.679178699 | NULL&M_Disease |
| 11 | 119305611 | rs4938653 | C | T | NULL | 0.679178699 | NULL&M_Disease |
| 11 | 119314718 | rs4938657 | C | T | NULL | 0.679178699 | NULL&M_Disease |
| 11 | 119806145 | rs12295249 | A | G | NULL | 0.679178699 | NULL&M_Disease |
| 11 | 119923652 | rs664845 | A | G | NULL | 0.679178699 | NULL&M_Disease |
| 11 | 119956631 | rs4938776 | C | A | NULL | 0.679178699 | NULL&M_Disease |
| 11 | 120046224 | rs7119954 | C | T | NULL | 0.679178699 | NULL&M_Disease |
| 11 | 120619909 | rs1418140540 | C | T | NULL | 0.679178699 | NULL&M_Disease |
| 11 | 120636109 | rs112165411 | A | T | NULL | 0.679178699 | NULL&M_Disease |
| 11 | 116741553 | rs11216168 | A | G | NULL | 0.679178699 | NULL&M_Disease |
| 11 | 116931907 | rs61903395 | G | A | NULL | 0.679178699 | NULL&M_Disease |
| 11 | 117501603 | rs10790197 | T | C | NULL | 0.679178699 | NULL&M_Disease |
| 11 | 117842706 | rs67000349 | G | A | NULL | 0.679178699 | NULL&M_Disease |
| 11 | 119753355 | rs590648 | T | G | NULL | 0.679178699 | NULL&M_Disease |
| 11 | 119963819 | rs11217686 | A | G | NULL | 0.679178699 | NULL&M_Disease |
| 11 | 116504984 | rs59232710 | G | A | NULL | 0.679178699 | NULL&M_Disease |
| 11 | 117307265 | rs472648 | C | T | NULL | 0.679178699 | NULL&M_Disease |
| 11 | 117904127 | rs606683 | G | C | NULL | 0.679178699 | NULL&M_Disease |
| 11 | 117914504 | rs1783855 | A | C | NULL | 0.679178699 | NULL&M_Disease |
| 11 | 118032974 | rs594171 | A | T | NULL | 0.679178699 | NULL&M_Disease |
| 11 | 118405453 | rs45474901 | C | T | NULL | 0.679178699 | NULL&M_Disease |
| 11 | 119858507 | rs11600793 | C | G | NULL | 0.679178699 | NULL&M_Disease |
| 11 | 120024874 | rs2077784 | A | G | NULL | 0.679178699 | NULL&M_Disease |
| 11 | 117303858 | rs11216382 | C | T | NULL | 0.679178699 | NULL&M_Disease |
| 11 | 118405305 | rs45505501 | A | G | NULL | 0.679178699 | NULL&M_Disease |
| 11 | 118405641 | rs3741323 | G | A | NULL | 0.679178699 | NULL&M_Disease |
| 11 | 118911039 | rs34150933 | T | C | NULL | 0.679178699 | NULL&M_Disease |
| 11 | 119782056 | rs10790353 | C | G | NULL | 0.679178699 | NULL&M_Disease |
| 11 | 119794509 | rs471814 | G | A | NULL | 0.679178699 | NULL&M_Disease |
| 11 | 119805109 | rs11217586 | A | G | NULL | 0.679178699 | NULL&M_Disease |
| 11 | 119923259 | rs666640 | T | A | NULL | 0.679178699 | NULL&M_Disease |
| 11 | 119959410 | rs10892533 | T | C | NULL | 0.679178699 | NULL&M_Disease |
| 11 | 120100597 | rs9655 | G | A | NULL | 0.679178699 | NULL&M_Disease |
| 11 | 120100867 | rs580174 | G | A | NULL | 0.679178699 | NULL&M_Disease |
| 11 | 117119552 | rs523366 | A | G | NULL | 0.679178699 | NULL&M_Disease |
| 11 | 117309581 | rs61905269 | A | G | NULL | 0.679178699 | NULL&M_Disease |
| 11 | 118160851 | rs4938497 | A | G | NULL | 0.679178699 | NULL&M_Disease |
| 11 | 118942928 | rs685044 | A | G | NULL | 0.679178699 | NULL&M_Disease |
| 11 | 118967758 | rs643788 | C | T | NULL | 0.679178699 | NULL&M_Disease |
| 11 | 119360083 | rs1652025 | G | C | NULL | 0.679178699 | NULL&M_Disease |
| 11 | 119831794 | rs35661813 | C | A | NULL | 0.679178699 | NULL&M_Disease |
| 11 | 120026600 | rs2084897 | A | G | NULL | 0.679178699 | NULL&M_Disease |
| 11 | 120026810 | rs2084899 | C | A | NULL | 0.679178699 | NULL&M_Disease |
| 11 | 120756908 | rs11218044 | A | G | NULL | 0.679178699 | NULL&M_Disease |
| 11 | 120760463 | rs73578608 | A | G | NULL | 0.679178699 | NULL&M_Disease |
| 11 | 120911203 | rs10892663 | C | T | NULL | 0.679178699 | NULL&M_Disease |
| 11 | 121068047 | rs17124945 | T | C | NULL | 0.679178699 | NULL&M_Disease |
| 11 | 116222849 | rs553449 | A | G | NULL | 0.679178699 | NULL&M_Disease |
| 11 | 117830429 | rs11216648 | G | A | NULL | 0.679178699 | NULL&M_Disease |
| 11 | 117988539 | rs671111 | A | G | NULL | 0.679178699 | NULL&M_Disease |
| 11 | 120098044 | rs1295019 | A | C | NULL | 0.679178699 | NULL&M_Disease |
| 11 | 116417492 | rs73009180 | T | C | NULL | 0.679178699 | NULL&M_Disease |
| 11 | 116504754 | rs76801336 | C | T | NULL | 0.679178699 | NULL&M_Disease |
| 11 | 116670738 | rs6589568 | G | A | NULL | 0.679178699 | NULL&M_Disease |
| 11 | 116923488 | rs17120251 | C | T | NULL | 0.679178699 | NULL&M_Disease |
| 11 | 119831582 | rs35778099 | T | C | NULL | 0.679178699 | NULL&M_Disease |
| 11 | 119885674 | rs649586 | G | C | NULL | 0.679178699 | NULL&M_Disease |
| 11 | 120040695 | rs61900479 | T | C | NULL | 0.679178699 | NULL&M_Disease |
| 11 | 120098746 | rs505142 | A | T | NULL | 0.679178699 | NULL&M_Disease |
| 11 | 120100520 | rs3225 | C | G | NULL | 0.679178699 | NULL&M_Disease |
| 11 | 120753572 | rs11602018 | G | A | NULL | 0.679178699 | NULL&M_Disease |
| 11 | 116411434 | rs73009178 | A | C | NULL | 0.679178699 | NULL&M_Disease |
| 11 | 116705590 | rs12721028 | G | A | NULL | 0.680267197 | NULL&M_Disease |
| 11 | 118302901 | rs510997 | C | T | NULL | 0.679178699 | NULL&M_Disease |
| 11 | 118523799 | rs72999483 | C | T | NULL | 0.679178699 | NULL&M_Disease |
| 11 | 118911139 | rs34070113 | G | T | NULL | 0.680267197 | NULL&M_Disease |
| 11 | 120331701 | rs4938807 | A | C | NULL | 0.679178699 | NULL&M_Disease |
| 11 | 120639087 | rs11600755 | T | C | NULL | 0.679178699 | NULL&M_Disease |
| 11 | 116697848 | rs2071521 | G | A | NULL | 0.680267197 | NULL&M_Disease |
| 11 | 117694399 | rs73018464 | A | G | NULL | 0.680267197 | NULL&M_Disease |
| 11 | 117923031 | rs7933728 | A | G | NULL | 0.680267197 | NULL&M_Disease |
| 11 | 119222337 | rs11217250 | T | C | NULL | 0.680267197 | NULL&M_Disease |
| 11 | 119805884 | rs12294978 | A | C | NULL | 0.680267197 | NULL&M_Disease |
| 11 | 119840776 | rs7115154 | A | G | NULL | 0.680267197 | NULL&M_Disease |
| 11 | 119927491 | rs11217663 | C | T | NULL | 0.680267197 | NULL&M_Disease |
| 11 | 120061255 | rs73004793 | C | T | NULL | 0.680267197 | NULL&M_Disease |
| 11 | 121057430 | rs3781823 | A | G | NULL | 0.680267197 | NULL&M_Disease |
| 11 | 116226909 | rs472950 | T | C | NULL | 0.680267197 | NULL&M_Disease |
| 11 | 116696887 | rs2727789 | G | T | NULL | 0.680267197 | NULL&M_Disease |
| 11 | 117456360 | rs495420 | C | G | NULL | 0.680267197 | NULL&M_Disease |
| 11 | 119943485 | rs4938765 | T | G | NULL | 0.680267197 | NULL&M_Disease |
| 11 | 120099193 | rs472948 | G | A | NULL | 0.680267197 | NULL&M_Disease |
| 11 | 120099368 | rs692804 | T | C | NULL | 0.680267197 | NULL&M_Disease |
| 11 | 120618334 | rs2852218 | G | A | NULL | 0.680267197 | NULL&M_Disease |
| 11 | 116448578 | rs56970059 | T | C | NULL | 0.680267197 | NULL&M_Disease |
| 11 | 117109270 | rs678837 | G | T | NULL | 0.680267197 | NULL&M_Disease |
| 11 | 117293032 | rs72645462 | A | T | NULL | 0.680267197 | NULL&M_Disease |
| 11 | 118296735 | rs59255055 | A | C | NULL | 0.680267197 | NULL&M_Disease |
| 11 | 119201699 | rs2511848 | A | G | NULL | 0.680267197 | NULL&M_Disease |
| 11 | 119625735 | rs7950159 | T | C | NULL | 0.680267197 | NULL&M_Disease |
| 11 | 120770782 | rs2156636 | A | G | NULL | 0.680267197 | NULL&M_Disease |
| 11 | 116792357 | rs7934763 | G | T | NULL | 0.680267197 | NULL&M_Disease |
| 11 | 116924978 | rs723953 | C | A | NULL | 0.680267197 | NULL&M_Disease |
| 11 | 117436190 | rs490458 | T | C | NULL | 0.680267197 | NULL&M_Disease |
| 11 | 118298252 | rs4456287 | C | T | NULL | 0.680267197 | NULL&M_Disease |
| 11 | 120041917 | rs7116650 | T | C | NULL | 0.680267197 | NULL&M_Disease |

| | | | | | | | | | |
|---|---|---|---|---|---|---|---|---|---|
| 11 | 119305593 | rs4938652 | T | C | 0.999 | -7977.388 | -7978.889 | -1.501 | 8.62E-07 |
| 11 | 119305611 | rs4938653 | C | T | 0.999 | -7977.388 | -7978.889 | -1.501 | 8.62E-07 |
| 11 | 119314718 | rs4938657 | C | T | 0.997 | -7977.388 | -7978.889 | -1.501 | 8.62E-07 |
| 11 | 119806145 | rs12295249 | A | G | 1.014 | -7977.388 | -7978.889 | -1.501 | 8.62E-07 |
| 11 | 119923652 | rs664845 | A | G | 1.027 | -7977.388 | -7978.889 | -1.501 | 8.62E-07 |
| 11 | 119956631 | rs4938776 | C | A | 1.007 | -7977.388 | -7978.89 | -1.501 | 8.62E-07 |
| 11 | 120046224 | rs7119954 | C | T | 0.988 | -7977.388 | -7978.89 | -1.501 | 8.62E-07 |
| 11 | 120619909 | rs1418140540 | C | T | 1.021 | -7977.388 | -7978.89 | -1.501 | 8.62E-07 |
| 11 | 120636109 | rs112165411 | A | T | 0.994 | -7977.388 | -7978.89 | -1.501 | 8.62E-07 |
| 11 | 116741553 | rs11216168 | A | G | 0.982 | -7977.388 | -7978.89 | -1.502 | 8.61E-07 |
| 11 | 116931907 | rs61903395 | G | A | 0.98 | -7977.388 | -7978.89 | -1.502 | 8.61E-07 |
| 11 | 117501603 | rs10790197 | T | C | 0.974 | -7977.388 | -7978.891 | -1.502 | 8.61E-07 |
| 11 | 117842706 | rs67000349 | G | A | 0.987 | -7977.388 | -7978.89 | -1.502 | 8.61E-07 |
| 11 | 119753355 | rs590648 | T | G | 1.027 | -7977.388 | -7978.891 | -1.502 | 8.61E-07 |
| 11 | 119963819 | rs11217686 | A | G | 0.992 | -7977.388 | -7978.891 | -1.502 | 8.61E-07 |
| 11 | 116504984 | rs59232710 | G | A | 0.983 | -7977.388 | -7978.891 | -1.503 | 8.61E-07 |
| 11 | 117307265 | rs472648 | C | T | 0.974 | -7977.388 | -7978.892 | -1.503 | 8.61E-07 |
| 11 | 117904127 | rs606683 | G | C | 1.021 | -7977.388 | -7978.891 | -1.503 | 8.61E-07 |
| 11 | 117914504 | rs1783855 | A | C | 1.021 | -7977.388 | -7978.891 | -1.503 | 8.61E-07 |
| 11 | 118032974 | rs594171 | A | T | 1.001 | -7977.388 | -7978.891 | -1.503 | 8.61E-07 |
| 11 | 118405453 | rs45474901 | C | T | 0.985 | -7977.388 | -7978.892 | -1.503 | 8.61E-07 |
| 11 | 119858507 | rs11600793 | C | G | 0.985 | -7977.388 | -7978.891 | -1.503 | 8.61E-07 |
| 11 | 120024874 | rs2077784 | A | G | 0.989 | -7977.388 | -7978.891 | -1.503 | 8.61E-07 |
| 11 | 117303858 | rs11216382 | C | T | 0.979 | -7977.388 | -7978.892 | -1.504 | 8.60E-07 |
| 11 | 118405305 | rs45505501 | A | G | 0.985 | -7977.388 | -7978.892 | -1.504 | 8.60E-07 |
| 11 | 118405641 | rs3741323 | G | A | 0.985 | -7977.388 | -7978.892 | -1.504 | 8.60E-07 |
| 11 | 118911039 | rs34150933 | T | C | 0.974 | -7977.388 | -7978.892 | -1.504 | 8.60E-07 |
| 11 | 119782056 | rs10790353 | C | G | 1.015 | -7977.388 | -7978.893 | -1.504 | 8.60E-07 |
| 11 | 119794509 | rs471814 | G | A | 1.027 | -7977.388 | -7978.893 | -1.504 | 8.60E-07 |
| 11 | 119805109 | rs11217586 | A | G | 1.018 | -7977.388 | -7978.892 | -1.504 | 8.60E-07 |
| 11 | 119923259 | rs666640 | T | A | 1.027 | -7977.388 | -7978.892 | -1.504 | 8.60E-07 |
| 11 | 119959410 | rs10892533 | T | C | 0.993 | -7977.388 | -7978.893 | -1.504 | 8.60E-07 |
| 11 | 120100597 | rs9655 | G | A | 0.994 | -7977.388 | -7978.892 | -1.504 | 8.60E-07 |
| 11 | 120100867 | rs580174 | G | A | 0.994 | -7977.388 | -7978.892 | -1.504 | 8.60E-07 |
| 11 | 117119552 | rs523366 | A | G | 1.027 | -7977.388 | -7978.893 | -1.505 | 8.59E-07 |
| 11 | 117309581 | rs61905269 | A | G | 0.98 | -7977.388 | -7978.894 | -1.505 | 8.59E-07 |
| 11 | 118160851 | rs4938497 | A | G | 0.974 | -7977.388 | -7978.894 | -1.505 | 8.59E-07 |
| 11 | 118942928 | rs685044 | A | G | 0.974 | -7977.388 | -7978.893 | -1.505 | 8.59E-07 |
| 11 | 118967758 | rs643788 | C | T | 1.027 | -7977.388 | -7978.893 | -1.505 | 8.59E-07 |
| 11 | 119360083 | rs1652025 | G | C | 0.974 | -7977.388 | -7978.893 | -1.505 | 8.59E-07 |
| 11 | 119831794 | rs35661813 | C | A | 1.007 | -7977.388 | -7978.893 | -1.505 | 8.59E-07 |
| 11 | 120026600 | rs2084897 | A | G | 0.99 | -7977.388 | -7978.893 | -1.505 | 8.59E-07 |
| 11 | 120026810 | rs2084899 | C | A | 0.99 | -7977.388 | -7978.893 | -1.505 | 8.59E-07 |
| 11 | 120756908 | rs11218044 | A | G | 1.016 | -7977.388 | -7978.893 | -1.505 | 8.59E-07 |
| 11 | 120760463 | rs73578608 | A | G | 1.016 | -7977.388 | -7978.893 | -1.505 | 8.59E-07 |
| 11 | 120911203 | rs10892663 | C | T | 1.027 | -7977.388 | -7978.893 | -1.505 | 8.59E-07 |
| 11 | 121068047 | rs17124945 | T | C | 1.013 | -7977.388 | -7978.894 | -1.505 | 8.59E-07 |
| 11 | 116222849 | rs553449 | A | G | 1.027 | -7977.388 | -7978.894 | -1.506 | 8.58E-07 |
| 11 | 117830429 | rs11216648 | G | A | 1.027 | -7977.388 | -7978.894 | -1.506 | 8.58E-07 |
| 11 | 117988539 | rs671111 | A | G | 0.974 | -7977.388 | -7978.894 | -1.506 | 8.58E-07 |
| 11 | 120098044 | rs1295019 | A | C | 0.995 | -7977.388 | -7978.894 | -1.506 | 8.58E-07 |
| 11 | 116417492 | rs73009180 | T | C | 1.006 | -7977.388 | -7978.896 | -1.507 | 8.57E-07 |
| 11 | 116504754 | rs76801336 | C | T | 0.983 | -7977.388 | -7978.895 | -1.507 | 8.57E-07 |
| 11 | 116670738 | rs6589568 | G | A | 1.024 | -7977.388 | -7978.896 | -1.507 | 8.57E-07 |
| 11 | 116923488 | rs17120251 | C | T | 0.981 | -7977.388 | -7978.896 | -1.507 | 8.57E-07 |
| 11 | 119831582 | rs35778099 | T | C | 1.007 | -7977.388 | -7978.895 | -1.507 | 8.57E-07 |
| 11 | 119885674 | rs649586 | G | C | 1.027 | -7977.388 | -7978.895 | -1.507 | 8.57E-07 |
| 11 | 120040695 | rs61900479 | T | C | 0.989 | -7977.388 | -7978.896 | -1.507 | 8.57E-07 |
| 11 | 120098746 | rs505142 | A | T | 0.995 | -7977.388 | -7978.895 | -1.507 | 8.57E-07 |
| 11 | 120100520 | rs3225 | C | G | 0.995 | -7977.388 | -7978.895 | -1.507 | 8.57E-07 |
| 11 | 120753572 | rs11602018 | G | A | 1.016 | -7977.388 | -7978.896 | -1.507 | 8.57E-07 |
| 11 | 116411434 | rs73009178 | A | C | 0.998 | -7977.388 | -7978.896 | -1.508 | 8.56E-07 |
| 11 | 116705590 | rs12721028 | G | A | 0.98 | -7977.388 | -7978.897 | -1.508 | 8.56E-07 |
| 11 | 118302901 | rs510997 | C | T | 0.984 | -7977.388 | -7978.896 | -1.508 | 8.56E-07 |
| 11 | 118523799 | rs72999483 | C | T | 1.001 | -7977.388 | -7978.896 | -1.508 | 8.56E-07 |
| 11 | 118911139 | rs34070113 | G | T | 0.974 | -7977.388 | -7978.896 | -1.508 | 8.56E-07 |
| 11 | 120331701 | rs4938807 | A | C | 0.979 | -7977.388 | -7978.896 | -1.508 | 8.56E-07 |
| 11 | 120639087 | rs11600755 | T | C | 0.994 | -7977.388 | -7978.897 | -1.508 | 8.56E-07 |
| 11 | 116697848 | rs2071521 | G | A | 1.027 | -7977.388 | -7978.897 | -1.509 | 8.55E-07 |
| 11 | 117694399 | rs73018464 | A | G | 1.019 | -7977.388 | -7978.898 | -1.509 | 8.55E-07 |
| 11 | 117923031 | rs7933728 | A | G | 1.027 | -7977.388 | -7978.897 | -1.509 | 8.55E-07 |
| 11 | 119222337 | rs11217250 | T | C | 0.975 | -7977.388 | -7978.897 | -1.509 | 8.55E-07 |
| 11 | 119805884 | rs12294978 | A | C | 1.013 | -7977.388 | -7978.897 | -1.509 | 8.55E-07 |
| 11 | 119840776 | rs7115154 | A | G | 1.006 | -7977.388 | -7978.898 | -1.509 | 8.55E-07 |
| 11 | 119927491 | rs11217663 | C | T | 1.026 | -7977.388 | -7978.897 | -1.509 | 8.55E-07 |
| 11 | 120061255 | rs73004793 | C | T | 0.988 | -7977.388 | -7978.898 | -1.509 | 8.55E-07 |
| 11 | 121057430 | rs3781823 | A | G | 1.027 | -7977.388 | -7978.897 | -1.509 | 8.55E-07 |
| 11 | 116226909 | rs472950 | T | C | 1.027 | -7977.388 | -7978.898 | -1.51 | 8.55E-07 |
| 11 | 116696887 | rs2727789 | G | T | 1.027 | -7977.388 | -7978.899 | -1.51 | 8.55E-07 |
| 11 | 117456360 | rs495420 | C | G | 0.976 | -7977.388 | -7978.898 | -1.51 | 8.55E-07 |
| 11 | 119943485 | rs4938765 | T | G | 1.025 | -7977.388 | -7978.898 | -1.51 | 8.55E-07 |
| 11 | 120099193 | rs472948 | G | A | 0.996 | -7977.388 | -7978.898 | -1.51 | 8.55E-07 |
| 11 | 120099368 | rs692804 | T | C | 0.996 | -7977.388 | -7978.898 | -1.51 | 8.55E-07 |
| 11 | 120618334 | rs2852218 | G | A | 0.974 | -7977.388 | -7978.898 | -1.51 | 8.55E-07 |
| 11 | 116448578 | rs56970059 | T | C | 0.974 | -7977.388 | -7978.899 | -1.511 | 8.54E-07 |
| 11 | 117109270 | rs678837 | G | T | 1.027 | -7977.388 | -7978.899 | -1.511 | 8.54E-07 |
| 11 | 117293032 | rs72645462 | A | T | 0.981 | -7977.388 | -7978.9 | -1.511 | 8.54E-07 |
| 11 | 118296735 | rs59255055 | A | C | 0.976 | -7977.388 | -7978.899 | -1.511 | 8.54E-07 |
| 11 | 119201699 | rs2511848 | A | G | 1.026 | -7977.388 | -7978.899 | -1.511 | 8.54E-07 |
| 11 | 119625735 | rs7950159 | T | C | 1.027 | -7977.388 | -7978.9 | -1.511 | 8.54E-07 |
| 11 | 120770782 | rs2156636 | A | G | 0.982 | -7977.388 | -7978.899 | -1.511 | 8.54E-07 |
| 11 | 116792357 | rs7934763 | G | T | 0.984 | -7977.388 | -7978.901 | -1.512 | 8.53E-07 |
| 11 | 116924978 | rs723953 | C | A | 0.981 | -7977.388 | -7978.9 | -1.512 | 8.53E-07 |
| 11 | 117436190 | rs490458 | T | C | 0.974 | -7977.388 | -7978.9 | -1.512 | 8.53E-07 |
| 11 | 118298252 | rs4456287 | C | T | 0.976 | -7977.388 | -7978.9 | -1.512 | 8.53E-07 |
| 11 | 120041917 | rs7116650 | T | C | 0.99 | -7977.388 | -7978.9 | -1.512 | 8.53E-07 |

| | | | | | | | |
|---|---|---|---|---|---|---|---|
| 11 | 120052360 | rs12418601 | T | C | NULL | 0.680267197 | NULL&M_Disease |
| 11 | 120418072 | rs2252356 | A | G | NULL | 0.680267197 | NULL&M_Disease |
| 11 | 116451308 | rs11216017 | G | C | NULL | 0.680267197 | NULL&M_Disease |
| 11 | 117494151 | rs34986626 | A | G | NULL | 0.680267197 | NULL&M_Disease |
| 11 | 119185133 | rs6589731 | C | T | NULL | 0.680267197 | NULL&M_Disease |
| 11 | 119738623 | rs509733 | T | C | NULL | 0.680267197 | NULL&M_Disease |
| 11 | 120052554 | rs547124 | A | G | NULL | 0.680267197 | NULL&M_Disease |
| 11 | 120196897 | rs11217818 | C | T | NULL | 0.680267197 | NULL&M_Disease |
| 11 | 120807571 | rs1157011088 | T | G | NULL | 0.680267197 | NULL&M_Disease |
| 11 | 121115129 | rs1941306 | C | A | NULL | 0.680267197 | NULL&M_Disease |
| 11 | 121115359 | rs6589866 | A | G | NULL | 0.680267197 | NULL&M_Disease |
| 11 | 116566546 | rs111064597 | C | T | NULL | 0.680267197 | NULL&M_Disease |
| 11 | 117994430 | rs599343 | A | G | NULL | 0.680267197 | NULL&M_Disease |
| 11 | 118036942 | rs676685 | A | G | NULL | 0.680267197 | NULL&M_Disease |
| 11 | 118296668 | rs17122170 | C | G | NULL | 0.680267197 | NULL&M_Disease |
| 11 | 118911544 | rs667584 | T | C | NULL | 0.680267197 | NULL&M_Disease |
| 11 | 118978132 | rs7936752 | G | C | NULL | 0.680267197 | NULL&M_Disease |
| 11 | 120325559 | rs11217875 | A | C | NULL | 0.680267197 | NULL&M_Disease |
| 11 | 120755655 | rs12577638 | A | G | NULL | 0.680267197 | NULL&M_Disease |
| 11 | 121117142 | rs2105670 | A | C | NULL | 0.680267197 | NULL&M_Disease |
| 11 | 117295150 | rs7115491 | G | T | NULL | 0.680267197 | NULL&M_Disease |
| 11 | 117848292 | rs10892201 | T | C | NULL | 0.680267197 | NULL&M_Disease |
| 11 | 118296007 | rs73613993 | C | T | NULL | 0.680267197 | NULL&M_Disease |
| 11 | 118806897 | rs10892310 | T | C | NULL | 0.680267197 | NULL&M_Disease |
| 11 | 116232487 | rs12804415 | G | A | NULL | 0.680267197 | NULL&M_Disease |
| 11 | 116450713 | rs10790150 | T | C | NULL | 0.680267197 | NULL&M_Disease |
| 11 | 118416073 | rs3825055 | C | A | NULL | 0.680267197 | NULL&M_Disease |
| 11 | 119197707 | rs2511834 | A | G | NULL | 0.680267197 | NULL&M_Disease |
| 11 | 120039306 | rs35650539 | G | C | NULL | 0.680267197 | NULL&M_Disease |
| 11 | 116505440 | rs10892001 | A | G | NULL | 0.680267197 | NULL&M_Disease |
| 11 | 117102200 | rs61906491 | A | G | NULL | 0.680267197 | NULL&M_Disease |
| 11 | 117291756 | rs11216379 | G | A | NULL | 0.680267197 | NULL&M_Disease |
| 11 | 117496132 | rs10790195 | C | T | NULL | 0.680267197 | NULL&M_Disease |
| 11 | 118408033 | rs535207 | G | A | NULL | 0.680267197 | NULL&M_Disease |
| 11 | 119197379 | rs2511833 | G | A | NULL | 0.680267197 | NULL&M_Disease |
| 11 | 119507723 | rs4245197 | C | T | NULL | 0.681353734 | NULL&M_Disease |
| 11 | 119524950 | rs4938703 | T | C | NULL | 0.680267197 | NULL&M_Disease |
| 11 | 120099658 | rs2508489 | A | G | NULL | 0.680267197 | NULL&M_Disease |
| 11 | 120628012 | rs3133864 | T | C | NULL | 0.680267197 | NULL&M_Disease |
| 11 | 120637315 | rs7925365 | G | T | NULL | 0.680267197 | NULL&M_Disease |
| 11 | 121041146 | rs11218175 | G | T | NULL | 0.680267197 | NULL&M_Disease |
| 11 | 121115206 | rs6589865 | C | T | NULL | 0.680267197 | NULL&M_Disease |
| 11 | 117947181 | rs4938476 | C | A | NULL | 0.681353734 | NULL&M_Disease |
| 11 | 119217455 | rs948415 | C | T | NULL | 0.681353734 | NULL&M_Disease |
| 11 | 119928883 | rs10444336 | A | C | NULL | 0.681353734 | NULL&M_Disease |
| 11 | 116399249 | rs17569676 | C | T | NULL | 0.681353734 | NULL&M_Disease |
| 11 | 116496948 | rs77623050 | T | C | NULL | 0.681353734 | NULL&M_Disease |
| 11 | 116496961 | rs78485217 | A | T | NULL | 0.681353734 | NULL&M_Disease |
| 11 | 116697738 | rs2542051 | C | A | NULL | 0.681353734 | NULL&M_Disease |
| 11 | 116704178 | rs7123454 | C | A | NULL | 0.681353734 | NULL&M_Disease |
| 11 | 118300240 | rs548877 | G | A | NULL | 0.681353734 | NULL&M_Disease |
| 11 | 118412358 | rs61290697 | T | C | NULL | 0.681353734 | NULL&M_Disease |
| 11 | 119989590 | rs7127949 | C | A | NULL | 0.681353734 | NULL&M_Disease |
| 11 | 120325548 | rs11217874 | A | G | NULL | 0.681353734 | NULL&M_Disease |
| 11 | 120898196 | rs12576574 | T | G | NULL | 0.681353734 | NULL&M_Disease |
| 11 | 117769364 | rs11216609 | C | T | NULL | 0.681353734 | NULL&M_Disease |
| 11 | 117769468 | rs11216610 | A | G | NULL | 0.681353734 | NULL&M_Disease |
| 11 | 119750813 | rs73008690 | A | G | NULL | 0.681353734 | NULL&M_Disease |
| 11 | 120053173 | rs11217755 | A | G | NULL | 0.681353734 | NULL&M_Disease |
| 11 | 120076116 | rs562966 | C | T | NULL | 0.681353734 | NULL&M_Disease |
| 11 | 120644424 | rs948027 | C | T | NULL | 0.681353734 | NULL&M_Disease |
| 11 | 116448838 | rs34841529 | A | C | NULL | 0.681353734 | NULL&M_Disease |
| 11 | 116697538 | rs2071522 | T | C | NULL | 0.681353734 | NULL&M_Disease |
| 11 | 117131719 | rs11216338 | A | G | NULL | 0.681353734 | NULL&M_Disease |
| 11 | 117319720 | rs11216396 | G | A | NULL | 0.681353734 | NULL&M_Disease |
| 11 | 117348907 | rs552879 | T | C | NULL | 0.681353734 | NULL&M_Disease |
| 11 | 118095148 | rs57495164 | T | C | NULL | 0.681353734 | NULL&M_Disease |
| 11 | 118301066 | rs34815971 | G | A | NULL | 0.681353734 | NULL&M_Disease |
| 11 | 118408471 | rs73022038 | T | C | NULL | 0.681353734 | NULL&M_Disease |
| 11 | 118408653 | rs74404016 | T | C | NULL | 0.681353734 | NULL&M_Disease |
| 11 | 118408789 | rs73022041 | C | T | NULL | 0.681353734 | NULL&M_Disease |
| 11 | 118409302 | rs7128587 | A | G | NULL | 0.681353734 | NULL&M_Disease |
| 11 | 118409374 | rs7131631 | A | G | NULL | 0.681353734 | NULL&M_Disease |
| 11 | 118410468 | rs11216883 | G | A | NULL | 0.681353734 | NULL&M_Disease |
| 11 | 118410488 | rs11216884 | C | T | NULL | 0.681353734 | NULL&M_Disease |
| 11 | 118410545 | rs11216885 | G | A | NULL | 0.681353734 | NULL&M_Disease |
| 11 | 118411424 | rs11216886 | C | A | NULL | 0.681353734 | NULL&M_Disease |
| 11 | 119709638 | rs495981 | T | C | NULL | 0.681353734 | NULL&M_Disease |
| 11 | 120034170 | rs626151 | T | G | NULL | 0.681353734 | NULL&M_Disease |
| 11 | 120954911 | rs6589856 | A | T | NULL | 0.681353734 | NULL&M_Disease |
| 11 | 121116720 | rs7129582 | G | A | NULL | 0.681353734 | NULL&M_Disease |
| 11 | 116543111 | rs113264811 | T | C | NULL | 0.681353734 | NULL&M_Disease |
| 11 | 116564408 | rs1787700 | T | C | NULL | 0.681353734 | NULL&M_Disease |
| 11 | 117783176 | rs11216617 | T | G | NULL | 0.681353734 | NULL&M_Disease |
| 11 | 117848645 | rs4505099 | G | A | NULL | 0.681353734 | NULL&M_Disease |
| 11 | 118172520 | rs894678 | A | G | NULL | 0.681353734 | NULL&M_Disease |
| 11 | 119362888 | rs549303 | C | T | NULL | 0.681353734 | NULL&M_Disease |
| 11 | 120835068 | rs6589851 | T | C | NULL | 0.681353734 | NULL&M_Disease |
| 11 | 117848129 | rs11216662 | C | A | NULL | 0.681353734 | NULL&M_Disease |
| 11 | 117947433 | rs619585 | T | C | NULL | 0.681353734 | NULL&M_Disease |
| 11 | 118408893 | rs73022042 | A | G | NULL | 0.681353734 | NULL&M_Disease |
| 11 | 120006705 | rs620602 | G | A | NULL | 0.681353734 | NULL&M_Disease |
| 11 | 120018743 | rs675647 | A | G | NULL | 0.681353734 | NULL&M_Disease |
| 11 | 116458051 | rs7110616 | T | C | NULL | 0.681353734 | NULL&M_Disease |
| 11 | 116540792 | rs3016354 | C | T | NULL | 0.681353734 | NULL&M_Disease |
| 11 | 117100594 | rs500389 | G | A | NULL | 0.681353734 | NULL&M_Disease |

| | | | | | | | | | |
|---|---|---|---|---|---|---|---|---|---|
| 11 | 120052360 | rs12418601 | T | C | 0.99 | -7977.388 | -7978.9 | -1.512 | 8.53E-07 |
| 11 | 120418072 | rs2252356 | A | G | 1.026 | -7977.388 | -7978.9 | -1.512 | 8.53E-07 |
| 11 | 116451308 | rs11216017 | G | C | 0.977 | -7977.388 | -7978.901 | -1.513 | 8.52E-07 |
| 11 | 117494151 | rs34986626 | A | G | 1.024 | -7977.388 | -7978.902 | -1.513 | 8.52E-07 |
| 11 | 119185133 | rs6589731 | C | T | 0.974 | -7977.388 | -7978.902 | -1.513 | 8.52E-07 |
| 11 | 119738623 | rs509733 | T | C | 1.015 | -7977.388 | -7978.902 | -1.513 | 8.52E-07 |
| 11 | 120052554 | rs547124 | A | G | 0.983 | -7977.388 | -7978.902 | -1.513 | 8.52E-07 |
| 11 | 120196897 | rs11217818 | C | T | 0.984 | -7977.388 | -7978.902 | -1.513 | 8.52E-07 |
| 11 | 120807571 | rs1157011088 | T | G | 0.974 | -7977.388 | -7978.902 | -1.513 | 8.52E-07 |
| 11 | 121115129 | rs1941306 | C | A | 1.027 | -7977.388 | -7978.901 | -1.513 | 8.52E-07 |
| 11 | 121115359 | rs6589866 | A | G | 1.027 | -7977.388 | -7978.901 | -1.513 | 8.52E-07 |
| 11 | 116566546 | rs111064597 | C | T | 1.025 | -7977.388 | -7978.903 | -1.514 | 8.51E-07 |
| 11 | 117994430 | rs599343 | A | G | 0.974 | -7977.388 | -7978.902 | -1.514 | 8.51E-07 |
| 11 | 118036942 | rs676685 | A | G | 0.974 | -7977.388 | -7978.903 | -1.514 | 8.51E-07 |
| 11 | 118296668 | rs17122170 | C | G | 0.985 | -7977.388 | -7978.902 | -1.514 | 8.51E-07 |
| 11 | 118911544 | rs667584 | T | C | 0.974 | -7977.388 | -7978.902 | -1.514 | 8.51E-07 |
| 11 | 118978132 | rs7936752 | G | C | 1.027 | -7977.388 | -7978.903 | -1.514 | 8.51E-07 |
| 11 | 120325559 | rs11217875 | A | C | 0.98 | -7977.388 | -7978.902 | -1.514 | 8.51E-07 |
| 11 | 120755655 | rs12577638 | A | G | 1.015 | -7977.388 | -7978.902 | -1.514 | 8.51E-07 |
| 11 | 121117142 | rs2105670 | A | C | 1.027 | -7977.388 | -7978.902 | -1.514 | 8.51E-07 |
| 11 | 117295150 | rs7115491 | G | T | 1.02 | -7977.388 | -7978.904 | -1.515 | 8.50E-07 |
| 11 | 117848292 | rs10892201 | T | C | 0.995 | -7977.388 | -7978.904 | -1.515 | 8.50E-07 |
| 11 | 118296007 | rs73613993 | C | T | 0.985 | -7977.388 | -7978.903 | -1.515 | 8.50E-07 |
| 11 | 118806897 | rs10892310 | T | C | 0.974 | -7977.388 | -7978.904 | -1.515 | 8.50E-07 |
| 11 | 116232487 | rs12804415 | G | A | 0.982 | -7977.388 | -7978.904 | -1.516 | 8.49E-07 |
| 11 | 116450713 | rs10790150 | T | C | 0.977 | -7977.388 | -7978.904 | -1.516 | 8.49E-07 |
| 11 | 118416073 | rs3825055 | C | A | 0.986 | -7977.388 | -7978.904 | -1.516 | 8.49E-07 |
| 11 | 119197707 | rs2511834 | A | G | 1.004 | -7977.388 | -7978.904 | -1.516 | 8.49E-07 |
| 11 | 120039306 | rs35650539 | G | C | 0.989 | -7977.388 | -7978.905 | -1.516 | 8.49E-07 |
| 11 | 116505440 | rs10892001 | A | G | 1.027 | -7977.388 | -7978.905 | -1.517 | 8.49E-07 |
| 11 | 117102200 | rs61906491 | A | G | 0.986 | -7977.388 | -7978.905 | -1.517 | 8.49E-07 |
| 11 | 117291756 | rs11216379 | G | A | 1.02 | -7977.388 | -7978.905 | -1.517 | 8.49E-07 |
| 11 | 117496132 | rs10790195 | C | T | 1.024 | -7977.388 | -7978.905 | -1.517 | 8.49E-07 |
| 11 | 118408033 | rs535207 | G | A | 0.986 | -7977.388 | -7978.905 | -1.517 | 8.49E-07 |
| 11 | 119197379 | rs2511833 | G | A | 1.003 | -7977.388 | -7978.906 | -1.517 | 8.49E-07 |
| 11 | 119507723 | rs4245197 | C | T | 0.974 | -7977.388 | -7978.906 | -1.517 | 8.49E-07 |
| 11 | 119524950 | rs4938703 | T | C | 1.025 | -7977.388 | -7978.906 | -1.517 | 8.49E-07 |
| 11 | 120099658 | rs2508489 | A | G | 1.001 | -7977.388 | -7978.905 | -1.517 | 8.49E-07 |
| 11 | 120628012 | rs3133864 | T | C | 1.02 | -7977.388 | -7978.906 | -1.517 | 8.49E-07 |
| 11 | 120637315 | rs7925365 | G | T | 1.002 | -7977.388 | -7978.905 | -1.517 | 8.49E-07 |
| 11 | 121041146 | rs11218175 | G | T | 1.012 | -7977.388 | -7978.906 | -1.517 | 8.49E-07 |
| 11 | 121115206 | rs6589865 | C | T | 1.027 | -7977.388 | -7978.905 | -1.517 | 8.49E-07 |
| 11 | 117947181 | rs4938476 | C | A | 1.026 | -7977.388 | -7978.907 | -1.518 | 8.48E-07 |
| 11 | 119217455 | rs948415 | C | T | 0.974 | -7977.388 | -7978.907 | -1.518 | 8.48E-07 |
| 11 | 119928883 | rs10444336 | A | C | 1.026 | -7977.388 | -7978.907 | -1.518 | 8.48E-07 |
| 11 | 116399249 | rs17569676 | C | T | 1.023 | -7977.388 | -7978.907 | -1.519 | 8.47E-07 |
| 11 | 116496948 | rs77623050 | T | C | 0.984 | -7977.388 | -7978.907 | -1.519 | 8.47E-07 |
| 11 | 116496961 | rs78485217 | A | T | 0.984 | -7977.388 | -7978.907 | -1.519 | 8.47E-07 |
| 11 | 116697738 | rs2542051 | C | A | 1.026 | -7977.388 | -7978.907 | -1.519 | 8.47E-07 |
| 11 | 116704178 | rs7123454 | C | A | 0.992 | -7977.388 | -7978.908 | -1.519 | 8.47E-07 |
| 11 | 118300240 | rs548877 | G | A | 0.985 | -7977.388 | -7978.907 | -1.519 | 8.47E-07 |
| 11 | 118412358 | rs61290697 | T | C | 0.987 | -7977.388 | -7978.907 | -1.519 | 8.47E-07 |
| 11 | 119989590 | rs7127949 | C | A | 1.026 | -7977.388 | -7978.907 | -1.519 | 8.47E-07 |
| 11 | 120325548 | rs11217874 | A | G | 0.98 | -7977.388 | -7978.907 | -1.519 | 8.47E-07 |
| 11 | 120898196 | rs12576574 | T | G | 1.019 | -7977.388 | -7978.907 | -1.519 | 8.47E-07 |
| 11 | 117769364 | rs11216609 | C | T | 1.005 | -7977.388 | -7978.909 | -1.52 | 8.46E-07 |
| 11 | 117769468 | rs11216610 | A | G | 1.003 | -7977.388 | -7978.908 | -1.52 | 8.46E-07 |
| 11 | 119750813 | rs73008690 | A | G | 1.014 | -7977.388 | -7978.908 | -1.52 | 8.46E-07 |
| 11 | 120053173 | rs11217755 | A | G | 0.991 | -7977.388 | -7978.908 | -1.52 | 8.46E-07 |
| 11 | 120076116 | rs562966 | C | T | 0.986 | -7977.388 | -7978.909 | -1.52 | 8.46E-07 |
| 11 | 120644424 | rs948027 | C | T | 0.987 | -7977.388 | -7978.908 | -1.52 | 8.46E-07 |
| 11 | 116448838 | rs34841529 | A | C | 0.977 | -7977.388 | -7978.909 | -1.521 | 8.45E-07 |
| 11 | 116697538 | rs2071522 | T | C | 1.026 | -7977.388 | -7978.909 | -1.521 | 8.45E-07 |
| 11 | 117131719 | rs11216338 | A | G | 1.027 | -7977.388 | -7978.909 | -1.521 | 8.45E-07 |
| 11 | 117319720 | rs11216396 | G | A | 0.974 | -7977.388 | -7978.91 | -1.521 | 8.45E-07 |
| 11 | 117348907 | rs552879 | T | C | 1.026 | -7977.388 | -7978.909 | -1.521 | 8.45E-07 |
| 11 | 118095148 | rs57495164 | T | C | 1.027 | -7977.388 | -7978.91 | -1.521 | 8.45E-07 |
| 11 | 118301066 | rs34815971 | G | A | 0.985 | -7977.388 | -7978.909 | -1.521 | 8.45E-07 |
| 11 | 118408471 | rs73022038 | T | C | 0.987 | -7977.388 | -7978.909 | -1.521 | 8.45E-07 |
| 11 | 118408653 | rs74404016 | T | C | 0.987 | -7977.388 | -7978.909 | -1.521 | 8.45E-07 |
| 11 | 118408789 | rs73022041 | C | T | 0.987 | -7977.388 | -7978.909 | -1.521 | 8.45E-07 |
| 11 | 118409302 | rs7128587 | A | G | 0.987 | -7977.388 | -7978.909 | -1.521 | 8.45E-07 |
| 11 | 118409374 | rs7131631 | A | G | 0.987 | -7977.388 | -7978.909 | -1.521 | 8.45E-07 |
| 11 | 118410468 | rs11216883 | G | A | 0.987 | -7977.388 | -7978.909 | -1.521 | 8.45E-07 |
| 11 | 118410488 | rs11216884 | C | T | 0.987 | -7977.388 | -7978.909 | -1.521 | 8.45E-07 |
| 11 | 118410545 | rs11216885 | G | A | 0.987 | -7977.388 | -7978.909 | -1.521 | 8.45E-07 |
| 11 | 118411424 | rs11216886 | C | A | 0.987 | -7977.388 | -7978.909 | -1.521 | 8.45E-07 |
| 11 | 119709638 | rs495981 | T | C | 0.976 | -7977.388 | -7978.909 | -1.521 | 8.45E-07 |
| 11 | 120034170 | rs626151 | T | G | 0.991 | -7977.388 | -7978.909 | -1.521 | 8.45E-07 |
| 11 | 120954911 | rs6589856 | A | T | 1.019 | -7977.388 | -7978.909 | -1.521 | 8.45E-07 |
| 11 | 121116720 | rs7129582 | G | A | 1.026 | -7977.388 | -7978.909 | -1.521 | 8.45E-07 |
| 11 | 116543111 | rs113264811 | T | C | 1.026 | -7977.388 | -7978.911 | -1.522 | 8.44E-07 |
| 11 | 116564408 | rs1787700 | T | C | 1.024 | -7977.388 | -7978.91 | -1.522 | 8.44E-07 |
| 11 | 117783176 | rs11216617 | T | G | 1.022 | -7977.388 | -7978.91 | -1.522 | 8.44E-07 |
| 11 | 117848645 | rs4505099 | G | A | 0.993 | -7977.388 | -7978.91 | -1.522 | 8.44E-07 |
| 11 | 118172520 | rs894678 | A | G | 0.974 | -7977.388 | -7978.911 | -1.522 | 8.44E-07 |
| 11 | 119362888 | rs549303 | C | T | 0.974 | -7977.388 | -7978.91 | -1.522 | 8.44E-07 |
| 11 | 120835068 | rs6589851 | T | C | 0.974 | -7977.388 | -7978.91 | -1.522 | 8.44E-07 |
| 11 | 117848129 | rs11216662 | C | A | 1 | -7977.388 | -7978.912 | -1.523 | 8.43E-07 |
| 11 | 117947433 | rs619585 | T | C | 1.019 | -7977.388 | -7978.911 | -1.523 | 8.43E-07 |
| 11 | 118408893 | rs73022042 | A | G | 0.987 | -7977.388 | -7978.911 | -1.523 | 8.43E-07 |
| 11 | 120006705 | rs620602 | G | A | 1.002 | -7977.388 | -7978.911 | -1.523 | 8.43E-07 |
| 11 | 120018743 | rs675647 | A | G | 1.001 | -7977.388 | -7978.911 | -1.523 | 8.43E-07 |
| 11 | 116458051 | rs7110616 | T | C | 0.974 | -7977.388 | -7978.912 | -1.524 | 8.43E-07 |
| 11 | 116540792 | rs3016354 | C | T | 1.026 | -7977.388 | -7978.913 | -1.524 | 8.43E-07 |
| 11 | 117100594 | rs500389 | G | A | 0.974 | -7977.388 | -7978.912 | -1.524 | 8.43E-07 |

| | | | | | | | |
|---|---|---|---|---|---|---|---|
| 11 | 117108318 | rs664011 | T | A | NULL | 0.681353734 | NULL&M_Disease |
| 11 | 117937131 | rs1786188 | A | G | NULL | 0.681353734 | NULL&M_Disease |
| 11 | 118415847 | rs3741320 | C | G | NULL | 0.681353734 | NULL&M_Disease |
| 11 | 118415848 | rs3741319 | C | A | NULL | 0.681353734 | NULL&M_Disease |
| 11 | 118747813 | rs6589706 | A | G | NULL | 0.681353734 | NULL&M_Disease |
| 11 | 118889378 | rs7113753 | A | G | NULL | 0.681353734 | NULL&M_Disease |
| 11 | 120010550 | rs56391092 | T | C | NULL | 0.681353734 | NULL&M_Disease |
| 11 | 120046336 | rs12419025 | T | A | NULL | 0.681353734 | NULL&M_Disease |
| 11 | 117084537 | rs12418744 | A | G | NULL | 0.681353734 | NULL&M_Disease |
| 11 | 117466151 | rs12803154 | T | C | NULL | 0.681353734 | NULL&M_Disease |
| 11 | 118424008 | rs11216892 | T | C | NULL | 0.681353734 | NULL&M_Disease |
| 11 | 118428791 | rs7929887 | G | A | NULL | 0.681353734 | NULL&M_Disease |
| 11 | 119856226 | rs4938746 | G | A | NULL | 0.681353734 | NULL&M_Disease |
| 11 | 116456659 | rs55751037 | T | C | NULL | 0.681353734 | NULL&M_Disease |
| 11 | 116547409 | rs1960036 | A | C | NULL | 0.681353734 | NULL&M_Disease |
| 11 | 118411713 | rs7932700 | T | G | NULL | 0.681353734 | NULL&M_Disease |
| 11 | 118419295 | rs59176377 | A | G | NULL | 0.681353734 | NULL&M_Disease |
| 11 | 120270249 | rs10892569 | G | A | NULL | 0.681353734 | NULL&M_Disease |
| 11 | 120750636 | rs74743296 | C | A | NULL | 0.681353734 | NULL&M_Disease |
| 11 | 120922228 | rs12576572 | T | G | NULL | 0.681353734 | NULL&M_Disease |
| 11 | 120927469 | rs7947565 | T | G | NULL | 0.681353734 | NULL&M_Disease |
| 11 | 120929197 | rs3758891 | A | G | NULL | 0.681353734 | NULL&M_Disease |
| 11 | 120933051 | rs57826163 | T | G | NULL | 0.681353734 | NULL&M_Disease |
| 11 | 116239244 | rs79099791 | C | T | NULL | 0.681353734 | NULL&M_Disease |
| 11 | 116584421 | rs1144036 | C | T | NULL | 0.681353734 | NULL&M_Disease |
| 11 | 116696921 | rs2849176 | T | C | NULL | 0.681353734 | NULL&M_Disease |
| 11 | 117133672 | rs552176 | A | G | NULL | 0.682438302 | NULL&M_Disease |
| 11 | 117327648 | rs9666820 | C | T | NULL | 0.682438302 | NULL&M_Disease |
| 11 | 117440404 | rs4936388 | G | T | NULL | 0.681353734 | NULL&M_Disease |
| 11 | 117768409 | rs755537 | G | A | NULL | 0.681353734 | NULL&M_Disease |
| 11 | 119505025 | rs1009364 | A | G | NULL | 0.681353734 | NULL&M_Disease |
| 11 | 119989380 | rs7128000 | A | G | NULL | 0.681353734 | NULL&M_Disease |
| 11 | 119989497 | rs7128133 | A | G | NULL | 0.681353734 | NULL&M_Disease |
| 11 | 116316813 | rs505390 | A | G | NULL | 0.682438302 | NULL&M_Disease |
| 11 | 116505492 | rs10892002 | C | G | NULL | 0.682438302 | NULL&M_Disease |
| 11 | 116540439 | rs2846015 | T | C | NULL | 0.682438302 | NULL&M_Disease |
| 11 | 116540807 | rs3017337 | T | C | NULL | 0.682438302 | NULL&M_Disease |
| 11 | 117459662 | rs498889 | C | T | NULL | 0.682438302 | NULL&M_Disease |
| 11 | 117758658 | rs580287 | G | C | NULL | 0.682438302 | NULL&M_Disease |
| 11 | 118099617 | rs11216825 | G | A | NULL | 0.682438302 | NULL&M_Disease |
| 11 | 118099713 | rs11216826 | A | T | NULL | 0.682438302 | NULL&M_Disease |
| 11 | 120110775 | rs10892553 | C | T | NULL | 0.682438302 | NULL&M_Disease |
| 11 | 120531435 | rs7939968 | A | T | NULL | 0.682438302 | NULL&M_Disease |
| 11 | 120955028 | rs2186601 | T | C | NULL | 0.682438302 | NULL&M_Disease |
| 11 | 117131664 | rs11216337 | C | T | NULL | 0.682438302 | NULL&M_Disease |
| 11 | 118407523 | rs7108845 | G | T | NULL | 0.682438302 | NULL&M_Disease |
| 11 | 119222706 | rs7122785 | G | A | NULL | 0.682438302 | NULL&M_Disease |
| 11 | 120000873 | rs477361 | T | G | NULL | 0.682438302 | NULL&M_Disease |
| 11 | 120000885 | rs477337 | T | C | NULL | 0.682438302 | NULL&M_Disease |
| 11 | 120898253 | rs12576759 | T | C | NULL | 0.682438302 | NULL&M_Disease |
| 11 | 120899492 | rs11218123 | T | G | NULL | 0.682438302 | NULL&M_Disease |
| 11 | 116429253 | rs11216005 | C | A | NULL | 0.682438302 | NULL&M_Disease |
| 11 | 116449283 | rs7122363 | T | C | NULL | 0.682438302 | NULL&M_Disease |
| 11 | 116504377 | rs11216067 | G | A | NULL | 0.682438302 | NULL&M_Disease |
| 11 | 117322782 | rs4938379 | T | C | NULL | 0.682438302 | NULL&M_Disease |
| 11 | 117359244 | rs1003261346 | A | G | NULL | 0.682438302 | NULL&M_Disease |
| 11 | 118028167 | rs615144 | A | C | NULL | 0.682438302 | NULL&M_Disease |
| 11 | 118422868 | rs2277293 | T | C | NULL | 0.682438302 | NULL&M_Disease |
| 11 | 118539151 | rs72999491 | A | G | NULL | 0.682438302 | NULL&M_Disease |
| 11 | 119692663 | rs542903 | T | C | NULL | 0.682438302 | NULL&M_Disease |
| 11 | 120000233 | rs504976 | C | A | NULL | 0.682438302 | NULL&M_Disease |
| 11 | 120268022 | rs4938803 | G | A | NULL | 0.682438302 | NULL&M_Disease |
| 11 | 120816284 | rs11824351 | C | T | NULL | 0.682438302 | NULL&M_Disease |
| 11 | 116697117 | rs2542050 | C | A | NULL | 0.682438302 | NULL&M_Disease |
| 11 | 116705100 | rs11216153 | T | G | NULL | 0.682438302 | NULL&M_Disease |
| 11 | 118303822 | rs3132893 | A | C | NULL | 0.682438302 | NULL&M_Disease |
| 11 | 118420101 | rs113739713 | G | A | NULL | 0.682438302 | NULL&M_Disease |
| 11 | 119686891 | rs79845538 | C | T | NULL | 0.682438302 | NULL&M_Disease |
| 11 | 119687133 | rs10892474 | C | T | NULL | 0.682438302 | NULL&M_Disease |
| 11 | 120148845 | rs12284449 | G | A | NULL | 0.682438302 | NULL&M_Disease |
| 11 | 120518929 | rs4935749 | A | G | NULL | 0.682438302 | NULL&M_Disease |
| 11 | 120836754 | rs12800734 | G | A | NULL | 0.682438302 | NULL&M_Disease |
| 11 | 116821918 | rs1473324 | G | T | NULL | 0.682438302 | NULL&M_Disease |
| 11 | 118096618 | rs10892226 | G | A | NULL | 0.682438302 | NULL&M_Disease |
| 11 | 118140372 | rs12793790 | A | G | NULL | 0.682438302 | NULL&M_Disease |
| 11 | 118409536 | rs11216880 | A | G | NULL | 0.682438302 | NULL&M_Disease |
| 11 | 118409583 | rs7101585 | A | G | NULL | 0.682438302 | NULL&M_Disease |
| 11 | 118971251 | rs649870 | G | A | NULL | 0.682438302 | NULL&M_Disease |
| 11 | 119680375 | rs73006796 | G | T | NULL | 0.682438302 | NULL&M_Disease |
| 11 | 116839535 | rs11823013 | C | T | NULL | 0.682438302 | NULL&M_Disease |
| 11 | 117344243 | rs488931 | A | G | NULL | 0.682438302 | NULL&M_Disease |
| 11 | 118130199 | rs61900882 | T | C | NULL | 0.682438302 | NULL&M_Disease |
| 11 | 116823996 | rs1473327 | T | A | NULL | 0.682438302 | NULL&M_Disease |
| 11 | 116948442 | rs4938336 | G | A | NULL | 0.682438302 | NULL&M_Disease |
| 11 | 117291975 | rs1902627 | T | C | NULL | 0.682438302 | NULL&M_Disease |
| 11 | 117314122 | rs61905305 | T | C | NULL | 0.682438302 | NULL&M_Disease |
| 11 | 118185919 | rs3782042 | A | G | NULL | 0.682438302 | NULL&M_Disease |
| 11 | 118222062 | rs4544037 | C | T | NULL | 0.682438302 | NULL&M_Disease |
| 11 | 118228968 | rs45489394 | G | C | NULL | 0.682438302 | NULL&M_Disease |
| 11 | 118409539 | rs11216881 | G | A | NULL | 0.682438302 | NULL&M_Disease |
| 11 | 118420736 | rs11216887 | C | T | NULL | 0.682438302 | NULL&M_Disease |
| 11 | 118421569 | rs11216890 | G | C | NULL | 0.682438302 | NULL&M_Disease |
| 11 | 118430519 | rs11552421 | T | C | NULL | 0.682438302 | NULL&M_Disease |
| 11 | 120000263 | rs504236 | T | C | NULL | 0.682438302 | NULL&M_Disease |
| 11 | 116205687 | rs10790137 | G | T | NULL | 0.682438302 | NULL&M_Disease |
| 11 | 116229759 | rs12808958 | A | G | NULL | 0.682438302 | NULL&M_Disease |

| | | | | | | | | | |
|---|---|---|---|---|---|---|---|---|---|
| 11 | 117108318 | rs664011 | T | A | 1.027 | -7977.388 | -7978.912 | -1.524 | 8.43E-07 |
| 11 | 117937131 | rs1786188 | A | G | 1.019 | -7977.388 | -7978.912 | -1.524 | 8.43E-07 |
| 11 | 118415847 | rs3741320 | C | G | 0.987 | -7977.388 | -7978.912 | -1.524 | 8.43E-07 |
| 11 | 118415848 | rs3741319 | C | A | 0.987 | -7977.388 | -7978.912 | -1.524 | 8.43E-07 |
| 11 | 118747813 | rs6589706 | A | G | 0.974 | -7977.388 | -7978.912 | -1.524 | 8.43E-07 |
| 11 | 118889378 | rs7113753 | A | G | 1.025 | -7977.388 | -7978.912 | -1.524 | 8.43E-07 |
| 11 | 120010550 | rs56391092 | T | C | 1.002 | -7977.388 | -7978.912 | -1.524 | 8.43E-07 |
| 11 | 120046336 | rs12419025 | T | A | 0.991 | -7977.388 | -7978.912 | -1.524 | 8.43E-07 |
| 11 | 117084537 | rs12418744 | A | G | 1.023 | -7977.388 | -7978.913 | -1.525 | 8.42E-07 |
| 11 | 117466151 | rs12803154 | T | C | 1.024 | -7977.388 | -7978.914 | -1.525 | 8.42E-07 |
| 11 | 118424008 | rs11216892 | T | C | 0.987 | -7977.388 | -7978.913 | -1.525 | 8.42E-07 |
| 11 | 118428791 | rs7929887 | G | A | 0.987 | -7977.388 | -7978.913 | -1.525 | 8.42E-07 |
| 11 | 119856226 | rs4938746 | G | A | 0.986 | -7977.388 | -7978.913 | -1.525 | 8.42E-07 |
| 11 | 116456659 | rs55751037 | T | C | 0.982 | -7977.388 | -7978.914 | -1.526 | 8.41E-07 |
| 11 | 116547409 | rs1960036 | A | C | 1.021 | -7977.388 | -7978.914 | -1.526 | 8.41E-07 |
| 11 | 118411713 | rs7932700 | T | G | 0.988 | -7977.388 | -7978.915 | -1.526 | 8.41E-07 |
| 11 | 118419295 | rs59176377 | A | G | 0.988 | -7977.388 | -7978.914 | -1.526 | 8.41E-07 |
| 11 | 120270249 | rs10892569 | G | A | 0.981 | -7977.388 | -7978.915 | -1.526 | 8.41E-07 |
| 11 | 120750636 | rs74743296 | C | A | 1.014 | -7977.388 | -7978.914 | -1.526 | 8.41E-07 |
| 11 | 120922228 | rs12576572 | T | G | 1.018 | -7977.388 | -7978.914 | -1.526 | 8.41E-07 |
| 11 | 120927469 | rs7947565 | T | G | 1.018 | -7977.388 | -7978.914 | -1.526 | 8.41E-07 |
| 11 | 120929197 | rs3758891 | A | G | 1.018 | -7977.388 | -7978.914 | -1.526 | 8.41E-07 |
| 11 | 120933051 | rs57826163 | T | G | 1.018 | -7977.388 | -7978.914 | -1.526 | 8.41E-07 |
| 11 | 116239244 | rs79099791 | C | T | 1.002 | -7977.388 | -7978.915 | -1.527 | 8.40E-07 |
| 11 | 116584421 | rs1144036 | C | T | 1.017 | -7977.388 | -7978.916 | -1.527 | 8.40E-07 |
| 11 | 116696921 | rs2849176 | T | C | 1.026 | -7977.388 | -7978.915 | -1.527 | 8.40E-07 |
| 11 | 117133672 | rs552176 | A | G | 1.026 | -7977.388 | -7978.916 | -1.527 | 8.40E-07 |
| 11 | 117327648 | rs9666820 | C | T | 0.974 | -7977.388 | -7978.916 | -1.527 | 8.40E-07 |
| 11 | 117440404 | rs4936388 | G | T | 1.025 | -7977.388 | -7978.916 | -1.527 | 8.40E-07 |
| 11 | 117768409 | rs755537 | G | A | 0.975 | -7977.388 | -7978.915 | -1.527 | 8.40E-07 |
| 11 | 119505025 | rs1009364 | A | G | 0.974 | -7977.388 | -7978.915 | -1.527 | 8.40E-07 |
| 11 | 119989380 | rs7128000 | A | G | 1.026 | -7977.388 | -7978.916 | -1.527 | 8.40E-07 |
| 11 | 119989497 | rs7128133 | A | G | 1.026 | -7977.388 | -7978.916 | -1.527 | 8.40E-07 |
| 11 | 116316813 | rs505390 | A | G | 1.026 | -7977.388 | -7978.916 | -1.528 | 8.39E-07 |
| 11 | 116505492 | rs10892002 | C | G | 1.026 | -7977.388 | -7978.916 | -1.528 | 8.39E-07 |
| 11 | 116540439 | rs2846015 | T | C | 1.025 | -7977.388 | -7978.916 | -1.528 | 8.39E-07 |
| 11 | 116540807 | rs3017337 | T | C | 1.025 | -7977.388 | -7978.916 | -1.528 | 8.39E-07 |
| 11 | 117459662 | rs498889 | C | T | 0.977 | -7977.388 | -7978.916 | -1.528 | 8.39E-07 |
| 11 | 117758658 | rs580287 | G | C | 1.026 | -7977.388 | -7978.916 | -1.528 | 8.39E-07 |
| 11 | 118099617 | rs11216825 | G | A | 1.026 | -7977.388 | -7978.916 | -1.528 | 8.39E-07 |
| 11 | 118099713 | rs11216826 | A | T | 1.026 | -7977.388 | -7978.916 | -1.528 | 8.39E-07 |
| 11 | 120110775 | rs10892553 | C | T | 1.022 | -7977.388 | -7978.916 | -1.528 | 8.39E-07 |
| 11 | 120531435 | rs7939968 | A | T | 1.026 | -7977.388 | -7978.916 | -1.528 | 8.39E-07 |
| 11 | 120955028 | rs2186601 | T | C | 1.018 | -7977.388 | -7978.917 | -1.528 | 8.39E-07 |
| 11 | 117131664 | rs11216337 | C | T | 1.026 | -7977.388 | -7978.917 | -1.529 | 8.38E-07 |
| 11 | 118407523 | rs7108845 | G | T | 0.988 | -7977.388 | -7978.917 | -1.529 | 8.38E-07 |
| 11 | 119222706 | rs7122785 | G | A | 0.975 | -7977.388 | -7978.918 | -1.529 | 8.38E-07 |
| 11 | 120000873 | rs477361 | T | G | 0.995 | -7977.388 | -7978.917 | -1.529 | 8.38E-07 |
| 11 | 120000885 | rs477337 | T | C | 0.995 | -7977.388 | -7978.917 | -1.529 | 8.38E-07 |
| 11 | 120898253 | rs12576759 | T | C | 1.018 | -7977.388 | -7978.917 | -1.529 | 8.38E-07 |
| 11 | 120899492 | rs11218123 | T | G | 1.018 | -7977.388 | -7978.917 | -1.529 | 8.38E-07 |
| 11 | 116429253 | rs11216005 | C | A | 1.025 | -7977.388 | -7978.918 | -1.53 | 8.38E-07 |
| 11 | 116449283 | rs7122363 | T | C | 0.978 | -7977.388 | -7978.918 | -1.53 | 8.38E-07 |
| 11 | 116504377 | rs11216067 | G | A | 0.986 | -7977.388 | -7978.919 | -1.53 | 8.38E-07 |
| 11 | 117322782 | rs4938379 | T | C | 0.974 | -7977.388 | -7978.918 | -1.53 | 8.38E-07 |
| 11 | 117359244 | rs1003261346 | A | G | 1.011 | -7977.388 | -7978.918 | -1.53 | 8.38E-07 |
| 11 | 118028167 | rs615144 | A | C | 1.026 | -7977.388 | -7978.918 | -1.53 | 8.38E-07 |
| 11 | 118422868 | rs2277293 | T | C | 0.988 | -7977.388 | -7978.918 | -1.53 | 8.38E-07 |
| 11 | 118539151 | rs72999491 | A | G | 1.012 | -7977.388 | -7978.918 | -1.53 | 8.38E-07 |
| 11 | 119692663 | rs542903 | T | C | 0.985 | -7977.388 | -7978.918 | -1.53 | 8.38E-07 |
| 11 | 120000233 | rs504976 | C | A | 0.993 | -7977.388 | -7978.918 | -1.53 | 8.38E-07 |
| 11 | 120268022 | rs4938803 | G | A | 0.981 | -7977.388 | -7978.918 | -1.53 | 8.38E-07 |
| 11 | 120816284 | rs11824351 | C | T | 1.026 | -7977.388 | -7978.919 | -1.53 | 8.38E-07 |
| 11 | 116697117 | rs2542050 | C | A | 1.026 | -7977.388 | -7978.919 | -1.531 | 8.37E-07 |
| 11 | 116705100 | rs11216153 | T | G | 0.982 | -7977.388 | -7978.919 | -1.531 | 8.37E-07 |
| 11 | 118303822 | rs3132893 | A | C | 0.987 | -7977.388 | -7978.919 | -1.531 | 8.37E-07 |
| 11 | 118420101 | rs113739713 | G | A | 0.988 | -7977.388 | -7978.919 | -1.531 | 8.37E-07 |
| 11 | 119686891 | rs79845538 | C | T | 0.976 | -7977.388 | -7978.919 | -1.531 | 8.37E-07 |
| 11 | 119687133 | rs10892474 | C | T | 0.976 | -7977.388 | -7978.919 | -1.531 | 8.37E-07 |
| 11 | 120148845 | rs12284449 | G | A | 1.025 | -7977.388 | -7978.92 | -1.531 | 8.37E-07 |
| 11 | 120518929 | rs4935749 | A | G | 1.026 | -7977.388 | -7978.919 | -1.531 | 8.37E-07 |
| 11 | 120836754 | rs12800734 | G | A | 0.974 | -7977.388 | -7978.919 | -1.531 | 8.37E-07 |
| 11 | 116821918 | rs1473324 | G | T | 0.988 | -7977.388 | -7978.92 | -1.532 | 8.36E-07 |
| 11 | 118096618 | rs10892226 | G | A | 1.026 | -7977.388 | -7978.92 | -1.532 | 8.36E-07 |
| 11 | 118140372 | rs12793790 | A | G | 0.975 | -7977.388 | -7978.92 | -1.532 | 8.36E-07 |
| 11 | 118409536 | rs11216880 | A | G | 0.989 | -7977.388 | -7978.921 | -1.532 | 8.36E-07 |
| 11 | 118409583 | rs7101585 | A | G | 0.989 | -7977.388 | -7978.921 | -1.532 | 8.36E-07 |
| 11 | 118971251 | rs649870 | G | A | 1.026 | -7977.388 | -7978.92 | -1.532 | 8.36E-07 |
| 11 | 119680375 | rs73006796 | G | T | 0.974 | -7977.388 | -7978.92 | -1.532 | 8.36E-07 |
| 11 | 116839535 | rs11823013 | C | T | 0.988 | -7977.388 | -7978.921 | -1.533 | 8.35E-07 |
| 11 | 117344243 | rs488931 | A | G | 1.026 | -7977.388 | -7978.922 | -1.533 | 8.35E-07 |
| 11 | 118130199 | rs61900882 | T | C | 1.026 | -7977.388 | -7978.921 | -1.533 | 8.35E-07 |
| 11 | 116823996 | rs1473327 | T | A | 0.988 | -7977.388 | -7978.922 | -1.534 | 8.34E-07 |
| 11 | 116948442 | rs4938336 | G | A | 0.988 | -7977.388 | -7978.922 | -1.534 | 8.34E-07 |
| 11 | 117291975 | rs1902627 | T | C | 0.975 | -7977.388 | -7978.923 | -1.534 | 8.34E-07 |
| 11 | 117314122 | rs61905305 | T | C | 1.008 | -7977.388 | -7978.922 | -1.534 | 8.34E-07 |
| 11 | 118185919 | rs3782042 | A | G | 0.975 | -7977.388 | -7978.922 | -1.534 | 8.34E-07 |
| 11 | 118222062 | rs4544037 | C | T | 1.021 | -7977.388 | -7978.923 | -1.534 | 8.34E-07 |
| 11 | 118228968 | rs45489394 | G | C | 1.021 | -7977.388 | -7978.923 | -1.534 | 8.34E-07 |
| 11 | 118409539 | rs11216881 | G | A | 0.989 | -7977.388 | -7978.922 | -1.534 | 8.34E-07 |
| 11 | 118420736 | rs11216887 | C | T | 0.989 | -7977.388 | -7978.922 | -1.534 | 8.34E-07 |
| 11 | 118421569 | rs11216890 | G | C | 0.989 | -7977.388 | -7978.923 | -1.534 | 8.34E-07 |
| 11 | 118430519 | rs11552421 | T | C | 0.989 | -7977.388 | -7978.923 | -1.534 | 8.34E-07 |
| 11 | 120000263 | rs504236 | T | C | 0.997 | -7977.388 | -7978.922 | -1.534 | 8.34E-07 |
| 11 | 116205687 | rs10790137 | G | T | 1.026 | -7977.388 | -7978.923 | -1.535 | 8.33E-07 |
| 11 | 116229759 | rs12808958 | A | G | 0.985 | -7977.388 | -7978.924 | -1.535 | 8.33E-07 |

| | | | | | | | |
|---|---|---|---|---|---|---|---|
| 11 | 116434082 | rs61904806 | C | T | NULL | 0.682438302 | NULL&M_Disease |
| 11 | 116457650 | rs10790152 | G | A | NULL | 0.682438302 | NULL&M_Disease |
| 11 | 116836209 | rs61905681 | C | A | NULL | 0.682438302 | NULL&M_Disease |
| 11 | 116911077 | rs59266239 | T | C | NULL | 0.682438302 | NULL&M_Disease |
| 11 | 117291936 | rs3891896 | G | T | NULL | 0.682438302 | NULL&M_Disease |
| 11 | 117377390 | rs2510852 | C | G | NULL | 0.682438302 | NULL&M_Disease |
| 11 | 118416253 | rs2277295 | T | C | NULL | 0.682438302 | NULL&M_Disease |
| 11 | 118429764 | rs111942423 | G | T | NULL | 0.682438302 | NULL&M_Disease |
| 11 | 119693573 | rs7395971 | C | T | NULL | 0.682438302 | NULL&M_Disease |
| 11 | 119989123 | rs10466596 | T | G | NULL | 0.682438302 | NULL&M_Disease |
| 11 | 120000872 | rs501021 | T | C | NULL | 0.682438302 | NULL&M_Disease |
| 11 | 120807747 | rs7926317 | C | T | NULL | 0.682438302 | NULL&M_Disease |
| 11 | 120825271 | rs11218080 | T | C | NULL | 0.682438302 | NULL&M_Disease |
| 11 | 120825575 | rs4936568 | A | G | NULL | 0.682438302 | NULL&M_Disease |
| 11 | 116697321 | rs2849173 | G | C | NULL | 0.682438302 | NULL&M_Disease |
| 11 | 116711141 | rs2727784 | C | T | NULL | 0.682438302 | NULL&M_Disease |
| 11 | 117133025 | rs637065 | A | G | NULL | 0.682438302 | NULL&M_Disease |
| 11 | 117368158 | rs497744 | T | G | NULL | 0.682438302 | NULL&M_Disease |
| 11 | 117802743 | rs3863316 | G | C | NULL | 0.682438302 | NULL&M_Disease |
| 11 | 118140597 | rs12795905 | T | C | NULL | 0.682438302 | NULL&M_Disease |
| 11 | 119656950 | rs7105596 | C | T | NULL | 0.682438302 | NULL&M_Disease |
| 11 | 116502267 | rs11216060 | A | G | NULL | 0.682438302 | NULL&M_Disease |
| 11 | 116833055 | rs56134734 | A | G | NULL | 0.682438302 | NULL&M_Disease |
| 11 | 116833655 | rs61905677 | G | T | NULL | 0.682438302 | NULL&M_Disease |
| 11 | 116834785 | rs61905678 | T | C | NULL | 0.682438302 | NULL&M_Disease |
| 11 | 116836031 | rs61905680 | A | G | NULL | 0.682438302 | NULL&M_Disease |
| 11 | 119224773 | rs12222464 | C | T | NULL | 0.682438302 | NULL&M_Disease |
| 11 | 119456168 | rs906828 | T | C | NULL | 0.683520894 | NULL&M_Disease |
| 11 | 119504257 | rs4394850 | C | G | NULL | 0.682438302 | NULL&M_Disease |
| 11 | 119858605 | rs2155993 | T | C | NULL | 0.682438302 | NULL&M_Disease |
| 11 | 119985050 | rs1317723 | A | G | NULL | 0.682438302 | NULL&M_Disease |
| 11 | 120621358 | rs3132781 | A | G | NULL | 0.682438302 | NULL&M_Disease |
| 11 | 120805056 | rs17124570 | A | G | NULL | 0.682438302 | NULL&M_Disease |
| 11 | 120930947 | rs12225989 | A | C | NULL | 0.682438302 | NULL&M_Disease |
| 11 | 116801811 | rs61907594 | T | C | NULL | 0.682438302 | NULL&M_Disease |
| 11 | 116802581 | rs79974099 | A | T | NULL | 0.682438302 | NULL&M_Disease |
| 11 | 116841446 | rs59294170 | T | A | NULL | 0.682438302 | NULL&M_Disease |
| 11 | 117137255 | rs507880 | G | T | NULL | 0.683520894 | NULL&M_Disease |
| 11 | 117137633 | rs2509211 | C | T | NULL | 0.683520894 | NULL&M_Disease |
| 11 | 117137647 | rs524085 | G | A | NULL | 0.683520894 | NULL&M_Disease |
| 11 | 117138291 | rs477269 | G | T | NULL | 0.683520894 | NULL&M_Disease |
| 11 | 117425003 | rs665218 | A | G | NULL | 0.682438302 | NULL&M_Disease |
| 11 | 117716453 | rs1145223 | T | C | NULL | 0.682438302 | NULL&M_Disease |
| 11 | 118425419 | rs2277292 | T | G | NULL | 0.682438302 | NULL&M_Disease |
| 11 | 120055170 | rs4938791 | A | G | NULL | 0.682438302 | NULL&M_Disease |
| 11 | 120526184 | rs4514426 | C | T | NULL | 0.683520894 | NULL&M_Disease |
| 11 | 120755218 | rs12577602 | T | G | NULL | 0.682438302 | NULL&M_Disease |
| 11 | 116959848 | rs111901638 | A | G | NULL | 0.683520894 | NULL&M_Disease |
| 11 | 118210106 | rs2276423 | G | C | NULL | 0.683520894 | NULL&M_Disease |
| 11 | 118235490 | rs12576486 | C | T | NULL | 0.683520894 | NULL&M_Disease |
| 11 | 118985151 | rs595782 | C | T | NULL | 0.683520894 | NULL&M_Disease |
| 11 | 119166651 | rs6589729 | A | G | NULL | 0.683520894 | NULL&M_Disease |
| 11 | 119360054 | rs607491 | G | C | NULL | 0.683520894 | NULL&M_Disease |
| 11 | 120517844 | rs6589827 | C | T | NULL | 0.683520894 | NULL&M_Disease |
| 11 | 120656572 | rs879602 | C | T | NULL | 0.683520894 | NULL&M_Disease |
| 11 | 116231844 | rs71478238 | G | T | NULL | 0.683520894 | NULL&M_Disease |
| 11 | 116809945 | rs61624438 | A | C | NULL | 0.683520894 | NULL&M_Disease |
| 11 | 117084091 | rs4938358 | T | C | NULL | 0.683520894 | NULL&M_Disease |
| 11 | 118422234 | rs11216891 | T | A | NULL | 0.683520894 | NULL&M_Disease |
| 11 | 118543589 | rs72999499 | G | C | NULL | 0.683520894 | NULL&M_Disease |
| 11 | 120749741 | rs4077910 | T | C | NULL | 0.683520894 | NULL&M_Disease |
| 11 | 121116786 | rs7129508 | C | A | NULL | 0.683520894 | NULL&M_Disease |
| 11 | 116327447 | rs59293325 | A | G | NULL | 0.683520894 | NULL&M_Disease |
| 11 | 116398667 | rs78972478 | T | C | NULL | 0.683520894 | NULL&M_Disease |
| 11 | 116457744 | rs7110192 | G | C | NULL | 0.683520894 | NULL&M_Disease |
| 11 | 116708413 | rs670 | T | C | NULL | 0.683520894 | NULL&M_Disease |
| 11 | 116796855 | rs61907591 | A | T | NULL | 0.683520894 | NULL&M_Disease |
| 11 | 116930888 | rs4938333 | G | A | NULL | 0.683520894 | NULL&M_Disease |
| 11 | 116931898 | rs61903394 | A | G | NULL | 0.683520894 | NULL&M_Disease |
| 11 | 117107775 | rs535602 | A | G | NULL | 0.683520894 | NULL&M_Disease |
| 11 | 117790794 | rs11826841 | G | A | NULL | 0.683520894 | NULL&M_Disease |
| 11 | 117960809 | rs7924314 | A | G | NULL | 0.683520894 | NULL&M_Disease |
| 11 | 118140203 | rs12789276 | G | A | NULL | 0.683520894 | NULL&M_Disease |
| 11 | 118423672 | rs73022058 | T | C | NULL | 0.683520894 | NULL&M_Disease |
| 11 | 118425051 | rs56403966 | A | G | NULL | 0.683520894 | NULL&M_Disease |
| 11 | 118425063 | rs55857590 | G | A | NULL | 0.683520894 | NULL&M_Disease |
| 11 | 120149741 | rs12798944 | C | T | NULL | 0.683520894 | NULL&M_Disease |
| 11 | 120622656 | rs3132782 | A | G | NULL | 0.683520894 | NULL&M_Disease |
| 11 | 121134395 | rs12419764 | G | A | NULL | 0.683520894 | NULL&M_Disease |
| 11 | 116224775 | rs17119495 | C | T | NULL | 0.683520894 | NULL&M_Disease |
| 11 | 116331957 | rs568466 | C | T | NULL | 0.683520894 | NULL&M_Disease |
| 11 | 116503644 | rs2096961 | A | G | NULL | 0.683520894 | NULL&M_Disease |
| 11 | 116822034 | rs1473325 | T | C | NULL | 0.683520894 | NULL&M_Disease |
| 11 | 117302038 | rs11606588 | C | G | NULL | 0.683520894 | NULL&M_Disease |
| 11 | 117322357 | rs4938378 | C | T | NULL | 0.683520894 | NULL&M_Disease |
| 11 | 117322454 | rs4936379 | A | C | NULL | 0.683520894 | NULL&M_Disease |
| 11 | 117426543 | rs490111 | G | A | NULL | 0.683520894 | NULL&M_Disease |
| 11 | 117552480 | rs76638771 | C | T | NULL | 0.683520894 | NULL&M_Disease |
| 11 | 117552758 | rs12295127 | A | G | NULL | 0.683520894 | NULL&M_Disease |
| 11 | 118405989 | rs3741325 | G | A | NULL | 0.683520894 | NULL&M_Disease |
| 11 | 120498337 | rs7928123 | T | C | NULL | 0.683520894 | NULL&M_Disease |
| 11 | 120623605 | rs3133865 | A | C | NULL | 0.683520894 | NULL&M_Disease |
| 11 | 116816630 | rs7924318 | G | C | NULL | 0.683520894 | NULL&M_Disease |
| 11 | 116972935 | rs4936363 | T | C | NULL | 0.683520894 | NULL&M_Disease |
| 11 | 117423538 | rs580714 | C | A | NULL | 0.683520894 | NULL&M_Disease |
| 11 | 117932630 | rs625267 | T | C | NULL | 0.683520894 | NULL&M_Disease |

| | | | | | | | | | |
|---|---|---|---|---|---|---|---|---|---|
| 11 | 116434082 | rs61904806 | C | T | 1.015 | -7977.388 | -7978.923 | -1.535 | 8.33E-07 |
| 11 | 116457650 | rs10790152 | G | A | 0.975 | -7977.388 | -7978.923 | -1.535 | 8.33E-07 |
| 11 | 116836209 | rs61905681 | C | A | 0.988 | -7977.388 | -7978.923 | -1.535 | 8.33E-07 |
| 11 | 116911077 | rs59266239 | T | C | 0.983 | -7977.388 | -7978.924 | -1.535 | 8.33E-07 |
| 11 | 117291936 | rs3891896 | G | T | 1.022 | -7977.388 | -7978.923 | -1.535 | 8.33E-07 |
| 11 | 117377390 | rs2510852 | C | G | 1.005 | -7977.388 | -7978.924 | -1.535 | 8.33E-07 |
| 11 | 118416253 | rs2277295 | T | C | 0.989 | -7977.388 | -7978.924 | -1.535 | 8.33E-07 |
| 11 | 118429764 | rs111942423 | G | T | 0.989 | -7977.388 | -7978.923 | -1.535 | 8.33E-07 |
| 11 | 119693573 | rs7395971 | C | T | 0.986 | -7977.388 | -7978.923 | -1.535 | 8.33E-07 |
| 11 | 119989123 | rs10466596 | T | G | 1.026 | -7977.388 | -7978.923 | -1.535 | 8.33E-07 |
| 11 | 120000872 | rs501021 | T | C | 0.997 | -7977.388 | -7978.923 | -1.535 | 8.33E-07 |
| 11 | 120807747 | rs7926317 | C | T | 0.975 | -7977.388 | -7978.923 | -1.535 | 8.33E-07 |
| 11 | 120825271 | rs11218080 | T | C | 0.976 | -7977.388 | -7978.923 | -1.535 | 8.33E-07 |
| 11 | 120825575 | rs4936568 | A | G | 0.976 | -7977.388 | -7978.923 | -1.535 | 8.33E-07 |
| 11 | 116697321 | rs2849173 | G | C | 1.026 | -7977.388 | -7978.925 | -1.536 | 8.33E-07 |
| 11 | 116711141 | rs2727784 | C | T | 0.975 | -7977.388 | -7978.924 | -1.536 | 8.33E-07 |
| 11 | 117133025 | rs637065 | A | G | 1.026 | -7977.388 | -7978.924 | -1.536 | 8.33E-07 |
| 11 | 117368158 | rs497744 | T | G | 0.997 | -7977.388 | -7978.925 | -1.536 | 8.33E-07 |
| 11 | 117802743 | rs3863316 | G | C | 1.026 | -7977.388 | -7978.924 | -1.536 | 8.33E-07 |
| 11 | 118140597 | rs12795905 | T | C | 0.975 | -7977.388 | -7978.924 | -1.536 | 8.33E-07 |
| 11 | 119656950 | rs7105596 | C | T | 0.985 | -7977.388 | -7978.924 | -1.536 | 8.33E-07 |
| 11 | 116502267 | rs11216060 | A | G | 0.986 | -7977.388 | -7978.926 | -1.537 | 8.32E-07 |
| 11 | 116833055 | rs56134734 | A | G | 0.988 | -7977.388 | -7978.925 | -1.537 | 8.32E-07 |
| 11 | 116833655 | rs61905677 | G | T | 0.988 | -7977.388 | -7978.925 | -1.537 | 8.32E-07 |
| 11 | 116834785 | rs61905678 | T | C | 0.988 | -7977.388 | -7978.925 | -1.537 | 8.32E-07 |
| 11 | 116836031 | rs61905680 | A | G | 0.988 | -7977.388 | -7978.925 | -1.537 | 8.32E-07 |
| 11 | 119224773 | rs12222464 | C | T | 0.976 | -7977.388 | -7978.926 | -1.537 | 8.32E-07 |
| 11 | 119456168 | rs906828 | T | C | 0.975 | -7977.388 | -7978.926 | -1.537 | 8.32E-07 |
| 11 | 119504257 | rs4394850 | C | G | 0.975 | -7977.388 | -7978.926 | -1.537 | 8.32E-07 |
| 11 | 119858605 | rs2155993 | T | C | 1.025 | -7977.388 | -7978.926 | -1.537 | 8.32E-07 |
| 11 | 119985050 | rs1317723 | A | G | 0.981 | -7977.388 | -7978.925 | -1.537 | 8.32E-07 |
| 11 | 120621358 | rs3132781 | A | G | 1.018 | -7977.388 | -7978.926 | -1.537 | 8.32E-07 |
| 11 | 120805056 | rs17124570 | A | G | 0.976 | -7977.388 | -7978.925 | -1.537 | 8.32E-07 |
| 11 | 120930947 | rs12225989 | A | C | 1.017 | -7977.388 | -7978.925 | -1.537 | 8.32E-07 |
| 11 | 116801811 | rs61907594 | T | C | 0.989 | -7977.388 | -7978.926 | -1.538 | 8.31E-07 |
| 11 | 116802581 | rs79974099 | A | T | 0.989 | -7977.388 | -7978.926 | -1.538 | 8.31E-07 |
| 11 | 116841446 | rs59294170 | T | A | 0.989 | -7977.388 | -7978.927 | -1.538 | 8.31E-07 |
| 11 | 117137255 | rs507880 | G | T | 1.026 | -7977.388 | -7978.926 | -1.538 | 8.31E-07 |
| 11 | 117137633 | rs2509211 | C | T | 1.026 | -7977.388 | -7978.926 | -1.538 | 8.31E-07 |
| 11 | 117137647 | rs524085 | G | A | 1.026 | -7977.388 | -7978.926 | -1.538 | 8.31E-07 |
| 11 | 117138291 | rs477269 | G | T | 1.026 | -7977.388 | -7978.926 | -1.538 | 8.31E-07 |
| 11 | 117425003 | rs665218 | A | G | 1.007 | -7977.388 | -7978.926 | -1.538 | 8.31E-07 |
| 11 | 117716453 | rs1145223 | T | C | 1.005 | -7977.388 | -7978.926 | -1.538 | 8.31E-07 |
| 11 | 118425419 | rs2277292 | T | G | 0.99 | -7977.388 | -7978.926 | -1.538 | 8.31E-07 |
| 11 | 120055170 | rs4938791 | A | G | 0.999 | -7977.388 | -7978.926 | -1.538 | 8.31E-07 |
| 11 | 120526184 | rs4514426 | C | T | 1.026 | -7977.388 | -7978.926 | -1.538 | 8.31E-07 |
| 11 | 120755218 | rs12577602 | T | G | 1.012 | -7977.388 | -7978.927 | -1.538 | 8.31E-07 |
| 11 | 116959848 | rs111901638 | A | G | 0.988 | -7977.388 | -7978.927 | -1.539 | 8.30E-07 |
| 11 | 118210106 | rs2276423 | G | C | 1.024 | -7977.388 | -7978.927 | -1.539 | 8.30E-07 |
| 11 | 118235490 | rs12576486 | C | T | 1.02 | -7977.388 | -7978.927 | -1.539 | 8.30E-07 |
| 11 | 118985151 | rs595782 | C | T | 1.026 | -7977.388 | -7978.927 | -1.539 | 8.30E-07 |
| 11 | 119166651 | rs6589729 | A | G | 1.026 | -7977.388 | -7978.927 | -1.539 | 8.30E-07 |
| 11 | 119360054 | rs607491 | G | C | 0.975 | -7977.388 | -7978.927 | -1.539 | 8.30E-07 |
| 11 | 120517844 | rs6589827 | C | T | 1.026 | -7977.388 | -7978.927 | -1.539 | 8.30E-07 |
| 11 | 120656572 | rs879602 | C | T | 0.994 | -7977.388 | -7978.927 | -1.539 | 8.30E-07 |
| 11 | 116231844 | rs71478238 | G | T | 0.985 | -7977.388 | -7978.928 | -1.54 | 8.29E-07 |
| 11 | 116809945 | rs61624438 | A | C | 0.989 | -7977.388 | -7978.928 | -1.54 | 8.29E-07 |
| 11 | 117084091 | rs4938358 | T | C | 1.022 | -7977.388 | -7978.928 | -1.54 | 8.29E-07 |
| 11 | 118422234 | rs11216891 | T | A | 0.99 | -7977.388 | -7978.928 | -1.54 | 8.29E-07 |
| 11 | 118543589 | rs72999499 | G | C | 1.011 | -7977.388 | -7978.929 | -1.54 | 8.29E-07 |
| 11 | 120749741 | rs4077910 | T | C | 1.011 | -7977.388 | -7978.929 | -1.54 | 8.29E-07 |
| 11 | 121116786 | rs7129508 | C | A | 1.026 | -7977.388 | -7978.928 | -1.54 | 8.29E-07 |
| 11 | 116327447 | rs59293325 | A | G | 1.013 | -7977.388 | -7978.929 | -1.541 | 8.28E-07 |
| 11 | 116398667 | rs78972478 | T | C | 1.022 | -7977.388 | -7978.929 | -1.541 | 8.28E-07 |
| 11 | 116457744 | rs7110192 | G | C | 0.975 | -7977.388 | -7978.929 | -1.541 | 8.28E-07 |
| 11 | 116708413 | rs670 | T | C | 0.983 | -7977.388 | -7978.929 | -1.541 | 8.28E-07 |
| 11 | 116796855 | rs61907591 | A | T | 0.989 | -7977.388 | -7978.93 | -1.541 | 8.28E-07 |
| 11 | 116930888 | rs4938333 | G | A | 0.989 | -7977.388 | -7978.929 | -1.541 | 8.28E-07 |
| 11 | 116931898 | rs61903394 | A | G | 0.989 | -7977.388 | -7978.929 | -1.541 | 8.28E-07 |
| 11 | 117107775 | rs535602 | A | G | 1.026 | -7977.388 | -7978.929 | -1.541 | 8.28E-07 |
| 11 | 117790794 | rs11826841 | G | A | 1.024 | -7977.388 | -7978.929 | -1.541 | 8.28E-07 |
| 11 | 117960809 | rs7924314 | A | G | 1.025 | -7977.388 | -7978.929 | -1.541 | 8.28E-07 |
| 11 | 118140203 | rs12789276 | G | A | 0.975 | -7977.388 | -7978.929 | -1.541 | 8.28E-07 |
| 11 | 118423672 | rs73022058 | T | C | 0.991 | -7977.388 | -7978.93 | -1.541 | 8.28E-07 |
| 11 | 118425051 | rs56403966 | A | G | 0.99 | -7977.388 | -7978.929 | -1.541 | 8.28E-07 |
| 11 | 118425063 | rs55857590 | G | A | 0.99 | -7977.388 | -7978.929 | -1.541 | 8.28E-07 |
| 11 | 120149741 | rs12798944 | C | T | 1.016 | -7977.388 | -7978.929 | -1.541 | 8.28E-07 |
| 11 | 120622656 | rs3132782 | A | G | 1.018 | -7977.388 | -7978.93 | -1.541 | 8.28E-07 |
| 11 | 121134395 | rs12419764 | G | A | 1.02 | -7977.388 | -7978.929 | -1.541 | 8.28E-07 |
| 11 | 116224775 | rs17119495 | C | T | 0.985 | -7977.388 | -7978.931 | -1.542 | 8.28E-07 |
| 11 | 116331957 | rs568466 | C | T | 0.991 | -7977.388 | -7978.931 | -1.542 | 8.28E-07 |
| 11 | 116503644 | rs2096961 | A | G | 0.987 | -7977.388 | -7978.93 | -1.542 | 8.28E-07 |
| 11 | 116822034 | rs1473325 | T | C | 0.989 | -7977.388 | -7978.93 | -1.542 | 8.28E-07 |
| 11 | 117302038 | rs11606588 | C | G | 0.975 | -7977.388 | -7978.93 | -1.542 | 8.28E-07 |
| 11 | 117322357 | rs4938378 | C | T | 0.975 | -7977.388 | -7978.93 | -1.542 | 8.28E-07 |
| 11 | 117322454 | rs4936379 | A | C | 0.975 | -7977.388 | -7978.93 | -1.542 | 8.28E-07 |
| 11 | 117426543 | rs490111 | G | A | 0.986 | -7977.388 | -7978.93 | -1.542 | 8.28E-07 |
| 11 | 117552480 | rs76638771 | C | T | 0.994 | -7977.388 | -7978.93 | -1.542 | 8.28E-07 |
| 11 | 117552758 | rs12295127 | A | G | 0.995 | -7977.388 | -7978.93 | -1.542 | 8.28E-07 |
| 11 | 118405989 | rs3741325 | G | A | 0.99 | -7977.388 | -7978.93 | -1.542 | 8.28E-07 |
| 11 | 120498337 | rs7928123 | T | C | 1.026 | -7977.388 | -7978.931 | -1.542 | 8.28E-07 |
| 11 | 120623605 | rs3133865 | A | C | 1.018 | -7977.388 | -7978.93 | -1.542 | 8.28E-07 |
| 11 | 116816630 | rs7924318 | G | C | 0.989 | -7977.388 | -7978.931 | -1.543 | 8.27E-07 |
| 11 | 116972935 | rs4936363 | T | C | 0.988 | -7977.388 | -7978.932 | -1.543 | 8.27E-07 |
| 11 | 117423538 | rs580714 | C | A | 1.004 | -7977.388 | -7978.931 | -1.543 | 8.27E-07 |
| 11 | 117932630 | rs625267 | T | C | 1.017 | -7977.388 | -7978.932 | -1.543 | 8.27E-07 |

| | | | | | | | |
|---|---|---|---|---|---|---|---|
| 11 | 118140136 | rs34786601 | C | T | NULL | 0.683520894 | NULL&M_Disease |
| 11 | 119505032 | rs1009363 | G | A | NULL | 0.683520894 | NULL&M_Disease |
| 11 | 119638088 | rs10750165 | G | A | NULL | 0.683520894 | NULL&M_Disease |
| 11 | 120180479 | rs717052 | A | G | NULL | 0.683520894 | NULL&M_Disease |
| 11 | 116920693 | rs11826651 | A | T | NULL | 0.683520894 | NULL&M_Disease |
| 11 | 117371824 | rs596707 | C | T | NULL | 0.683520894 | NULL&M_Disease |
| 11 | 117378713 | rs603858 | C | G | NULL | 0.683520894 | NULL&M_Disease |
| 11 | 117460123 | rs634406 | G | C | NULL | 0.683520894 | NULL&M_Disease |
| 11 | 118159684 | rs61900902 | C | T | NULL | 0.683520894 | NULL&M_Disease |
| 11 | 120959463 | rs17124764 | T | A | NULL | 0.683520894 | NULL&M_Disease |
| 11 | 121116638 | rs7129292 | A | G | NULL | 0.683520894 | NULL&M_Disease |
| 11 | 116834787 | rs61905679 | T | C | NULL | 0.683520894 | NULL&M_Disease |
| 11 | 117956943 | rs4938478 | G | A | NULL | 0.683520894 | NULL&M_Disease |
| 11 | 118430048 | rs11216896 | T | C | NULL | 0.683520894 | NULL&M_Disease |
| 11 | 118872741 | rs4573680 | C | T | NULL | 0.683520894 | NULL&M_Disease |
| 11 | 120221987 | rs12785387 | T | C | NULL | 0.683520894 | NULL&M_Disease |
| 11 | 120493360 | rs4936532 | A | G | NULL | 0.683520894 | NULL&M_Disease |
| 11 | 120527341 | rs4525243 | G | A | NULL | 0.683520894 | NULL&M_Disease |
| 11 | 116501072 | rs76055915 | T | C | NULL | 0.683520894 | NULL&M_Disease |
| 11 | 116501089 | rs74424292 | T | A | NULL | 0.683520894 | NULL&M_Disease |
| 11 | 116647120 | rs11216136 | G | T | NULL | 0.683520894 | NULL&M_Disease |
| 11 | 116862144 | rs61905687 | C | T | NULL | 0.683520894 | NULL&M_Disease |
| 11 | 116868654 | rs7950213 | A | C | NULL | 0.683520894 | NULL&M_Disease |
| 11 | 118951642 | rs2508948 | A | T | NULL | 0.683520894 | NULL&M_Disease |
| 11 | 120270170 | rs11217857 | G | A | NULL | 0.683520894 | NULL&M_Disease |
| 11 | 120959800 | rs2298491 | A | G | NULL | 0.683520894 | NULL&M_Disease |
| 11 | 116501053 | rs77649044 | C | T | NULL | 0.683520894 | NULL&M_Disease |
| 11 | 116863833 | rs7130531 | T | C | NULL | 0.683520894 | NULL&M_Disease |
| 11 | 116941613 | rs61903398 | T | C | NULL | 0.683520894 | NULL&M_Disease |
| 11 | 116942475 | rs61903399 | T | A | NULL | 0.683520894 | NULL&M_Disease |
| 11 | 118108003 | rs1793141 | G | A | NULL | 0.683520894 | NULL&M_Disease |
| 11 | 119938625 | rs11217669 | A | G | NULL | 0.683520894 | NULL&M_Disease |
| 11 | 119999755 | rs470941 | C | T | NULL | 0.683520894 | NULL&M_Disease |
| 11 | 120758691 | rs7112199 | A | G | NULL | 0.683520894 | NULL&M_Disease |
| 11 | 120835246 | rs7128320 | G | A | NULL | 0.683520894 | NULL&M_Disease |
| 11 | 120861967 | rs582800 | C | T | NULL | 0.683520894 | NULL&M_Disease |
| 11 | 116501987 | rs11216059 | A | C | NULL | 0.683520894 | NULL&M_Disease |
| 11 | 116542812 | rs1367711737 | T | G | NULL | 0.6846015 | NULL&M_Disease |
| 11 | 120009145 | rs2511023 | A | G | NULL | 0.683520894 | NULL&M_Disease |
| 11 | 120180657 | rs717051 | T | C | NULL | 0.683520894 | NULL&M_Disease |
| 11 | 120224789 | rs11824032 | A | G | NULL | 0.683520894 | NULL&M_Disease |
| 11 | 116228699 | rs531852 | C | A | NULL | 0.6846015 | NULL&M_Disease |
| 11 | 116454503 | rs7116825 | A | C | NULL | 0.6846015 | NULL&M_Disease |
| 11 | 118421512 | rs11216889 | A | G | NULL | 0.6846015 | NULL&M_Disease |
| 11 | 119782588 | rs536613 | G | A | NULL | 0.6846015 | NULL&M_Disease |
| 11 | 119977373 | rs77959607 | G | T | NULL | 0.6846015 | NULL&M_Disease |
| 11 | 121144673 | rs11218235 | T | C | NULL | 0.6846015 | NULL&M_Disease |
| 11 | 116574336 | rs1145198 | A | G | NULL | 0.6846015 | NULL&M_Disease |
| 11 | 117173560 | rs476469 | T | C | NULL | 0.6846015 | NULL&M_Disease |
| 11 | 117411041 | rs634867 | A | G | NULL | 0.6846015 | NULL&M_Disease |
| 11 | 117411392 | rs525110 | G | A | NULL | 0.6846015 | NULL&M_Disease |
| 11 | 117412453 | rs506836 | T | C | NULL | 0.6846015 | NULL&M_Disease |
| 11 | 117412621 | rs679223 | A | G | NULL | 0.6846015 | NULL&M_Disease |
| 11 | 117500940 | rs12273996 | A | G | NULL | 0.6846015 | NULL&M_Disease |
| 11 | 118100411 | rs869638 | G | C | NULL | 0.6846015 | NULL&M_Disease |
| 11 | 118135745 | rs3759001 | T | C | NULL | 0.6846015 | NULL&M_Disease |
| 11 | 120758479 | rs7111646 | T | A | NULL | 0.6846015 | NULL&M_Disease |
| 11 | 120907482 | rs2000525 | A | G | NULL | 0.6846015 | NULL&M_Disease |
| 11 | 116502752 | rs11216061 | C | T | NULL | 0.6846015 | NULL&M_Disease |
| 11 | 117135806 | rs7111678 | T | C | NULL | 0.6846015 | NULL&M_Disease |
| 11 | 117136070 | rs2509210 | T | C | NULL | 0.6846015 | NULL&M_Disease |
| 11 | 117136378 | rs672177 | G | C | NULL | 0.6846015 | NULL&M_Disease |
| 11 | 117313135 | rs11216390 | C | G | NULL | 0.6846015 | NULL&M_Disease |
| 11 | 117956579 | rs4938477 | G | A | NULL | 0.6846015 | NULL&M_Disease |
| 11 | 118113045 | rs1793146 | T | A | NULL | 0.6846015 | NULL&M_Disease |
| 11 | 118412296 | rs60067274 | T | A | NULL | 0.6846015 | NULL&M_Disease |
| 11 | 116502823 | rs11216064 | C | G | NULL | 0.6846015 | NULL&M_Disease |
| 11 | 116963965 | rs7950501 | T | C | NULL | 0.6846015 | NULL&M_Disease |
| 11 | 117431708 | rs475422 | C | T | NULL | 0.6846015 | NULL&M_Disease |
| 11 | 117499026 | rs7941794 | A | C | NULL | 0.6846015 | NULL&M_Disease |
| 11 | 119822879 | rs659471 | G | A | NULL | 0.6846015 | NULL&M_Disease |
| 11 | 120560994 | rs10790402 | A | G | NULL | 0.6846015 | NULL&M_Disease |
| 11 | 121010539 | rs4554895 | A | C | NULL | 0.6846015 | NULL&M_Disease |
| 11 | 116327971 | rs55837479 | A | G | NULL | 0.6846015 | NULL&M_Disease |
| 11 | 116699984 | rs2542052 | A | C | NULL | 0.6846015 | NULL&M_Disease |
| 11 | 117455669 | rs474749 | A | G | NULL | 0.6846015 | NULL&M_Disease |
| 11 | 118977911 | rs2509050 | G | A | NULL | 0.6846015 | NULL&M_Disease |
| 11 | 119885204 | rs949050 | T | C | NULL | 0.6846015 | NULL&M_Disease |
| 11 | 120326277 | rs476636 | G | A | NULL | 0.6846015 | NULL&M_Disease |
| 11 | 120727836 | rs10750177 | T | A | NULL | 0.6846015 | NULL&M_Disease |
| 11 | 117002596 | rs4938341 | A | G | NULL | 0.6846015 | NULL&M_Disease |
| 11 | 117091821 | rs7941150 | A | G | NULL | 0.6846015 | NULL&M_Disease |
| 11 | 117134761 | rs540922 | G | A | NULL | 0.6846015 | NULL&M_Disease |
| 11 | 117134766 | rs472785 | C | T | NULL | 0.6846015 | NULL&M_Disease |
| 11 | 117313136 | rs11216391 | T | C | NULL | 0.6846015 | NULL&M_Disease |
| 11 | 117726474 | rs553709 | C | T | NULL | 0.6846015 | NULL&M_Disease |
| 11 | 117764726 | rs11216608 | G | A | NULL | 0.6846015 | NULL&M_Disease |
| 11 | 118110977 | rs2853006 | T | C | NULL | 0.6846015 | NULL&M_Disease |
| 11 | 118896959 | rs57156480 | A | G | NULL | 0.6846015 | NULL&M_Disease |
| 11 | 119060963 | rs45535039 | C | T | NULL | 0.6846015 | NULL&M_Disease |
| 11 | 119504481 | rs11607829 | C | G | NULL | 0.6846015 | NULL&M_Disease |
| 11 | 119632975 | rs12290653 | A | G | NULL | 0.6846015 | NULL&M_Disease |
| 11 | 120156378 | rs10892559 | T | C | NULL | 0.6846015 | NULL&M_Disease |
| 11 | 120491438 | rs7128466 | A | G | NULL | 0.6846015 | NULL&M_Disease |
| 11 | 121099539 | rs4936595 | A | C | NULL | 0.6846015 | NULL&M_Disease |
| 11 | 116817784 | rs111327557 | T | C | NULL | 0.6846015 | NULL&M_Disease |

| | | | | | | | | | |
|---|---|---|---|---|---|---|---|---|---|
| 11 | 118140136 | rs34786601 | C | T | 0.975 | -7977.388 | -7978.931 | -1.543 | 8.27E-07 |
| 11 | 119505032 | rs1009363 | G | A | 0.975 | -7977.388 | -7978.931 | -1.543 | 8.27E-07 |
| 11 | 119638088 | rs10750165 | G | A | 1.026 | -7977.388 | -7978.932 | -1.543 | 8.27E-07 |
| 11 | 120180479 | rs717052 | A | G | 1.014 | -7977.388 | -7978.932 | -1.543 | 8.27E-07 |
| 11 | 116920693 | rs11826651 | A | T | 0.989 | -7977.388 | -7978.932 | -1.544 | 8.26E-07 |
| 11 | 117371824 | rs596707 | C | T | 1.002 | -7977.388 | -7978.932 | -1.544 | 8.26E-07 |
| 11 | 117378713 | rs603858 | C | G | 1.002 | -7977.388 | -7978.932 | -1.544 | 8.26E-07 |
| 11 | 117460123 | rs634406 | G | C | 0.977 | -7977.388 | -7978.932 | -1.544 | 8.26E-07 |
| 11 | 118159684 | rs61900902 | C | T | 0.975 | -7977.388 | -7978.933 | -1.544 | 8.26E-07 |
| 11 | 120959463 | rs17124764 | T | A | 1.017 | -7977.388 | -7978.932 | -1.544 | 8.26E-07 |
| 11 | 121116638 | rs7129292 | A | G | 1.026 | -7977.388 | -7978.932 | -1.544 | 8.26E-07 |
| 11 | 116834787 | rs61905679 | T | C | 0.989 | -7977.388 | -7978.933 | -1.545 | 8.25E-07 |
| 11 | 117956943 | rs4938478 | G | A | 0.975 | -7977.388 | -7978.934 | -1.545 | 8.25E-07 |
| 11 | 118430048 | rs11216896 | T | C | 0.991 | -7977.388 | -7978.933 | -1.545 | 8.25E-07 |
| 11 | 118872741 | rs4573680 | C | T | 1.024 | -7977.388 | -7978.933 | -1.545 | 8.25E-07 |
| 11 | 120221987 | rs12785387 | T | C | 0.982 | -7977.388 | -7978.934 | -1.545 | 8.25E-07 |
| 11 | 120493360 | rs4936532 | A | G | 0.988 | -7977.388 | -7978.933 | -1.545 | 8.25E-07 |
| 11 | 120527341 | rs4525243 | G | A | 1.025 | -7977.388 | -7978.934 | -1.545 | 8.25E-07 |
| 11 | 116501072 | rs76055915 | T | C | 0.988 | -7977.388 | -7978.934 | -1.546 | 8.24E-07 |
| 11 | 116501089 | rs74424292 | T | A | 0.988 | -7977.388 | -7978.934 | -1.546 | 8.24E-07 |
| 11 | 116647120 | rs11216136 | G | T | 1.017 | -7977.388 | -7978.934 | -1.546 | 8.24E-07 |
| 11 | 116862144 | rs61905687 | C | T | 0.99 | -7977.388 | -7978.934 | -1.546 | 8.24E-07 |
| 11 | 116868654 | rs7950213 | A | C | 0.99 | -7977.388 | -7978.934 | -1.546 | 8.24E-07 |
| 11 | 118951642 | rs2508948 | A | T | 0.975 | -7977.388 | -7978.934 | -1.546 | 8.24E-07 |
| 11 | 120270170 | rs11217857 | G | A | 0.982 | -7977.388 | -7978.935 | -1.546 | 8.24E-07 |
| 11 | 120959800 | rs2298491 | A | G | 1.016 | -7977.388 | -7978.935 | -1.546 | 8.24E-07 |
| 11 | 116501053 | rs77649044 | C | T | 0.988 | -7977.388 | -7978.935 | -1.547 | 8.23E-07 |
| 11 | 116863833 | rs7130531 | T | C | 0.989 | -7977.388 | -7978.935 | -1.547 | 8.23E-07 |
| 11 | 116941613 | rs61903398 | T | C | 0.99 | -7977.388 | -7978.935 | -1.547 | 8.23E-07 |
| 11 | 116942475 | rs61903399 | T | A | 0.99 | -7977.388 | -7978.935 | -1.547 | 8.23E-07 |
| 11 | 118108003 | rs1793141 | G | A | 1.024 | -7977.388 | -7978.935 | -1.547 | 8.23E-07 |
| 11 | 119938625 | rs11217669 | A | G | 0.977 | -7977.388 | -7978.935 | -1.547 | 8.23E-07 |
| 11 | 119999755 | rs470941 | C | T | 0.977 | -7977.388 | -7978.935 | -1.547 | 8.23E-07 |
| 11 | 120758691 | rs7112199 | A | G | 1.01 | -7977.388 | -7978.935 | -1.547 | 8.23E-07 |
| 11 | 120835246 | rs7128320 | G | A | 0.975 | -7977.388 | -7978.935 | -1.547 | 8.23E-07 |
| 11 | 120861967 | rs582800 | C | T | 1.021 | -7977.388 | -7978.936 | -1.547 | 8.23E-07 |
| 11 | 116501987 | rs11216059 | A | C | 0.988 | -7977.388 | -7978.936 | -1.548 | 8.23E-07 |
| 11 | 116542812 | rs1367711737 | T | G | 1.025 | -7977.388 | -7978.936 | -1.548 | 8.23E-07 |
| 11 | 120009145 | rs2511023 | A | G | 0.988 | -7977.388 | -7978.936 | -1.548 | 8.23E-07 |
| 11 | 120180657 | rs717051 | T | C | 1.013 | -7977.388 | -7978.936 | -1.548 | 8.23E-07 |
| 11 | 120224789 | rs11824032 | A | G | 0.983 | -7977.388 | -7978.936 | -1.548 | 8.23E-07 |
| 11 | 116228699 | rs531852 | C | A | 1.026 | -7977.388 | -7978.937 | -1.549 | 8.22E-07 |
| 11 | 116454503 | rs7116825 | A | C | 0.975 | -7977.388 | -7978.937 | -1.549 | 8.22E-07 |
| 11 | 118421512 | rs11216889 | A | G | 0.992 | -7977.388 | -7978.937 | -1.549 | 8.22E-07 |
| 11 | 119782588 | rs536613 | G | A | 0.975 | -7977.388 | -7978.937 | -1.549 | 8.22E-07 |
| 11 | 119977373 | rs77959607 | G | T | 0.985 | -7977.388 | -7978.937 | -1.549 | 8.22E-07 |
| 11 | 121144673 | rs11218235 | T | C | 1.019 | -7977.388 | -7978.937 | -1.549 | 8.22E-07 |
| 11 | 116574336 | rs1145198 | A | G | 1.023 | -7977.388 | -7978.938 | -1.55 | 8.21E-07 |
| 11 | 117173560 | rs476469 | T | C | 0.975 | -7977.388 | -7978.938 | -1.55 | 8.21E-07 |
| 11 | 117411041 | rs634867 | A | G | 0.999 | -7977.388 | -7978.938 | -1.55 | 8.21E-07 |
| 11 | 117411392 | rs525110 | G | A | 0.998 | -7977.388 | -7978.938 | -1.55 | 8.21E-07 |
| 11 | 117412453 | rs506836 | T | C | 0.998 | -7977.388 | -7978.939 | -1.55 | 8.21E-07 |
| 11 | 117412621 | rs679223 | A | G | 0.998 | -7977.388 | -7978.938 | -1.55 | 8.21E-07 |
| 11 | 117500940 | rs12273996 | A | G | 1.022 | -7977.388 | -7978.939 | -1.55 | 8.21E-07 |
| 11 | 118100411 | rs869638 | G | C | 1.026 | -7977.388 | -7978.939 | -1.55 | 8.21E-07 |
| 11 | 118135745 | rs3759001 | T | C | 1.025 | -7977.388 | -7978.939 | -1.55 | 8.21E-07 |
| 11 | 120758479 | rs7111646 | T | A | 1.01 | -7977.388 | -7978.938 | -1.55 | 8.21E-07 |
| 11 | 120907482 | rs2000525 | A | G | 1.016 | -7977.388 | -7978.938 | -1.55 | 8.21E-07 |
| 11 | 116502752 | rs11216061 | C | T | 0.988 | -7977.388 | -7978.939 | -1.551 | 8.20E-07 |
| 11 | 117135806 | rs7111678 | T | C | 1.026 | -7977.388 | -7978.939 | -1.551 | 8.20E-07 |
| 11 | 117136070 | rs2509210 | T | C | 1.026 | -7977.388 | -7978.939 | -1.551 | 8.20E-07 |
| 11 | 117136378 | rs672177 | G | C | 1.026 | -7977.388 | -7978.939 | -1.551 | 8.20E-07 |
| 11 | 117313135 | rs11216390 | C | G | 1.004 | -7977.388 | -7978.939 | -1.551 | 8.20E-07 |
| 11 | 117956579 | rs4938477 | G | A | 0.975 | -7977.388 | -7978.94 | -1.551 | 8.20E-07 |
| 11 | 118113045 | rs1793146 | T | A | 1.024 | -7977.388 | -7978.94 | -1.551 | 8.20E-07 |
| 11 | 118412296 | rs60067274 | T | A | 0.992 | -7977.388 | -7978.94 | -1.551 | 8.20E-07 |
| 11 | 116502823 | rs11216064 | C | G | 0.988 | -7977.388 | -7978.941 | -1.552 | 8.19E-07 |
| 11 | 116963965 | rs7950501 | T | C | 0.991 | -7977.388 | -7978.94 | -1.552 | 8.19E-07 |
| 11 | 117431708 | rs475422 | C | T | 0.978 | -7977.388 | -7978.941 | -1.552 | 8.19E-07 |
| 11 | 117499026 | rs7941794 | A | C | 0.975 | -7977.388 | -7978.94 | -1.552 | 8.19E-07 |
| 11 | 119822879 | rs659471 | G | A | 0.975 | -7977.388 | -7978.94 | -1.552 | 8.19E-07 |
| 11 | 120560994 | rs10790402 | A | G | 1.026 | -7977.388 | -7978.941 | -1.552 | 8.19E-07 |
| 11 | 121010539 | rs4554895 | A | C | 0.981 | -7977.388 | -7978.94 | -1.552 | 8.19E-07 |
| 11 | 116327971 | rs55837479 | A | G | 1.012 | -7977.388 | -7978.941 | -1.553 | 8.19E-07 |
| 11 | 116699984 | rs2542052 | A | C | 1.025 | -7977.388 | -7978.941 | -1.553 | 8.19E-07 |
| 11 | 117455669 | rs474749 | A | G | 1.025 | -7977.388 | -7978.941 | -1.553 | 8.19E-07 |
| 11 | 118977911 | rs2509050 | G | A | 1.025 | -7977.388 | -7978.941 | -1.553 | 8.19E-07 |
| 11 | 119885204 | rs949050 | T | C | 0.975 | -7977.388 | -7978.941 | -1.553 | 8.19E-07 |
| 11 | 120326277 | rs476636 | G | A | 0.99 | -7977.388 | -7978.941 | -1.553 | 8.19E-07 |
| 11 | 120727836 | rs10750177 | T | A | 1.026 | -7977.388 | -7978.941 | -1.553 | 8.19E-07 |
| 11 | 117002596 | rs4938341 | A | G | 0.985 | -7977.388 | -7978.942 | -1.554 | 8.18E-07 |
| 11 | 117091821 | rs7941150 | A | G | 0.986 | -7977.388 | -7978.943 | -1.554 | 8.18E-07 |
| 11 | 117134761 | rs540922 | G | A | 1.025 | -7977.388 | -7978.942 | -1.554 | 8.18E-07 |
| 11 | 117134766 | rs472785 | C | T | 1.025 | -7977.388 | -7978.942 | -1.554 | 8.18E-07 |
| 11 | 117313136 | rs11216391 | T | C | 1.002 | -7977.388 | -7978.942 | -1.554 | 8.18E-07 |
| 11 | 117726474 | rs553709 | C | T | 1.002 | -7977.388 | -7978.942 | -1.554 | 8.18E-07 |
| 11 | 117764726 | rs11216608 | G | A | 0.976 | -7977.388 | -7978.942 | -1.554 | 8.18E-07 |
| 11 | 118110977 | rs2853006 | T | C | 1.024 | -7977.388 | -7978.943 | -1.554 | 8.18E-07 |
| 11 | 118896959 | rs57156480 | A | G | 1.023 | -7977.388 | -7978.942 | -1.554 | 8.18E-07 |
| 11 | 119060963 | rs45535039 | C | T | 0.976 | -7977.388 | -7978.942 | -1.554 | 8.18E-07 |
| 11 | 119504481 | rs11607829 | C | G | 0.975 | -7977.388 | -7978.942 | -1.554 | 8.18E-07 |
| 11 | 119632975 | rs12290653 | A | G | 1.025 | -7977.388 | -7978.942 | -1.554 | 8.18E-07 |
| 11 | 120156378 | rs10892559 | T | C | 1.025 | -7977.388 | -7978.943 | -1.554 | 8.18E-07 |
| 11 | 120491438 | rs7128466 | A | G | 0.989 | -7977.388 | -7978.942 | -1.554 | 8.18E-07 |
| 11 | 121099539 | rs4936595 | A | C | 1.002 | -7977.388 | -7978.942 | -1.554 | 8.18E-07 |
| 11 | 116817784 | rs111327557 | T | C | 0.991 | -7977.388 | -7978.943 | -1.555 | 8.17E-07 |

| | | | | | | | |
|---|---|---|---|---|---|---|---|
| 11 | 116871223 | rs17120233 | G | A | NULL | 0.6846015 | NULL&M_Disease |
| 11 | 116872263 | rs1248452864 | G | A | NULL | 0.6846015 | NULL&M_Disease |
| 11 | 116877353 | rs77250716 | C | T | NULL | 0.6846015 | NULL&M_Disease |
| 11 | 116937159 | rs112895963 | T | A | NULL | 0.6846015 | NULL&M_Disease |
| 11 | 117785260 | rs4936410 | C | T | NULL | 0.6846015 | NULL&M_Disease |
| 11 | 118205702 | rs4938506 | A | G | NULL | 0.6846015 | NULL&M_Disease |
| 11 | 118216234 | rs3212264 | A | C | NULL | 0.6846015 | NULL&M_Disease |
| 11 | 118536950 | rs201359394 | T | A | NULL | 0.6846015 | NULL&M_Disease |
| 11 | 120812025 | rs10892648 | A | C | NULL | 0.6846015 | NULL&M_Disease |
| 11 | 116496744 | rs77984767 | G | A | NULL | 0.6846015 | NULL&M_Disease |
| 11 | 117473770 | rs12796722 | T | C | NULL | 0.6846015 | NULL&M_Disease |
| 11 | 118892940 | rs12280288 | C | T | NULL | 0.6846015 | NULL&M_Disease |
| 11 | 119383037 | rs7951668 | T | C | NULL | 0.6846015 | NULL&M_Disease |
| 11 | 121131331 | rs9666004 | T | C | NULL | 0.6846015 | NULL&M_Disease |
| 11 | 116500588 | rs11216055 | T | C | NULL | 0.6846015 | NULL&M_Disease |
| 11 | 116500629 | rs11216056 | A | C | NULL | 0.6846015 | NULL&M_Disease |
| 11 | 116500768 | rs11216057 | T | C | NULL | 0.6846015 | NULL&M_Disease |
| 11 | 116502776 | rs11216063 | A | T | NULL | 0.6846015 | NULL&M_Disease |
| 11 | 116952200 | rs4936362 | T | C | NULL | 0.6846015 | NULL&M_Disease |
| 11 | 117083004 | rs67201490 | G | T | NULL | 0.6846015 | NULL&M_Disease |
| 11 | 117084040 | rs4938357 | A | G | NULL | 0.6846015 | NULL&M_Disease |
| 11 | 117085305 | rs56081989 | A | G | NULL | 0.6846015 | NULL&M_Disease |
| 11 | 117764641 | rs11216607 | A | G | NULL | 0.6846015 | NULL&M_Disease |
| 11 | 120745874 | rs2230297 | T | C | NULL | 0.6846015 | NULL&M_Disease |
| 11 | 116493715 | rs11216047 | G | A | NULL | 0.685680114 | NULL&M_Disease |
| 11 | 116496309 | rs4340085 | T | C | NULL | 0.6846015 | NULL&M_Disease |
| 11 | 116498951 | rs11216054 | C | T | NULL | 0.685680114 | NULL&M_Disease |
| 11 | 116537962 | rs1784125 | C | G | NULL | 0.685680114 | NULL&M_Disease |
| 11 | 117003420 | rs4938342 | G | C | NULL | 0.6846015 | NULL&M_Disease |
| 11 | 117454157 | rs880812 | C | G | NULL | 0.685680114 | NULL&M_Disease |
| 11 | 117757345 | rs601122 | T | G | NULL | 0.685680114 | NULL&M_Disease |
| 11 | 119496472 | rs1355191 | C | T | NULL | 0.685680114 | NULL&M_Disease |
| 11 | 119951200 | rs6589790 | T | C | NULL | 0.6846015 | NULL&M_Disease |
| 11 | 121135004 | rs1278081233 | G | T | NULL | 0.685680114 | NULL&M_Disease |
| 11 | 116536418 | rs12803249 | G | A | NULL | 0.685680114 | NULL&M_Disease |
| 11 | 116961802 | rs61903421 | T | A | NULL | 0.685680114 | NULL&M_Disease |
| 11 | 116963897 | rs7950381 | A | C | NULL | 0.685680114 | NULL&M_Disease |
| 11 | 117053675 | rs4938355 | G | A | NULL | 0.685680114 | NULL&M_Disease |
| 11 | 117661100 | rs625447 | C | T | NULL | 0.685680114 | NULL&M_Disease |
| 11 | 120179563 | rs12787658 | G | A | NULL | 0.685680114 | NULL&M_Disease |
| 11 | 120821862 | rs12361041 | G | A | NULL | 0.685680114 | NULL&M_Disease |
| 11 | 116956665 | rs61903418 | A | G | NULL | 0.685680114 | NULL&M_Disease |
| 11 | 116960497 | rs11823918 | C | G | NULL | 0.685680114 | NULL&M_Disease |
| 11 | 117156552 | rs7083 | T | C | NULL | 0.685680114 | NULL&M_Disease |
| 11 | 117357085 | rs1315236635 | G | T | NULL | 0.685680114 | NULL&M_Disease |
| 11 | 118139755 | rs55852239 | A | G | NULL | 0.685680114 | NULL&M_Disease |
| 11 | 119568994 | rs4938713 | T | C | NULL | 0.685680114 | NULL&M_Disease |
| 11 | 119792847 | rs11217575 | G | A | NULL | 0.685680114 | NULL&M_Disease |
| 11 | 119951214 | rs6589792 | G | C | NULL | 0.685680114 | NULL&M_Disease |
| 11 | 119981785 | rs10736502 | G | C | NULL | 0.685680114 | NULL&M_Disease |
| 11 | 120198728 | rs11827818 | G | A | NULL | 0.685680114 | NULL&M_Disease |
| 11 | 121131224 | rs9665995 | T | C | NULL | 0.685680114 | NULL&M_Disease |
| 11 | 121140086 | rs10437696 | C | T | NULL | 0.685680114 | NULL&M_Disease |
| 11 | 116560610 | rs1787714 | A | G | NULL | 0.685680114 | NULL&M_Disease |
| 11 | 116880703 | rs61905691 | G | A | NULL | 0.685680114 | NULL&M_Disease |
| 11 | 116915020 | rs17120241 | C | T | NULL | 0.685680114 | NULL&M_Disease |
| 11 | 116919067 | rs61905716 | T | C | NULL | 0.685680114 | NULL&M_Disease |
| 11 | 116919368 | rs112151056 | A | G | NULL | 0.685680114 | NULL&M_Disease |
| 11 | 116922714 | rs61905719 | T | C | NULL | 0.685680114 | NULL&M_Disease |
| 11 | 117473944 | rs12797135 | T | C | NULL | 0.685680114 | NULL&M_Disease |
| 11 | 117692394 | rs501171 | T | C | NULL | 0.685680114 | NULL&M_Disease |
| 11 | 117791132 | rs7927300 | A | C | NULL | 0.685680114 | NULL&M_Disease |
| 11 | 119065044 | rs11217176 | A | G | NULL | 0.685680114 | NULL&M_Disease |
| 11 | 119620339 | rs34952981 | T | A | NULL | 0.685680114 | NULL&M_Disease |
| 11 | 119951213 | rs6589791 | A | C | NULL | 0.685680114 | NULL&M_Disease |
| 11 | 116219447 | rs569304 | A | G | NULL | 0.685680114 | NULL&M_Disease |
| 11 | 117319998 | rs11216397 | C | T | NULL | 0.685680114 | NULL&M_Disease |
| 11 | 117433965 | rs522742 | A | G | NULL | 0.685680114 | NULL&M_Disease |
| 11 | 118875727 | rs4936457 | G | A | NULL | 0.685680114 | NULL&M_Disease |
| 11 | 118875761 | rs4936458 | G | T | NULL | 0.685680114 | NULL&M_Disease |
| 11 | 119552057 | rs10790331 | C | A | NULL | 0.685680114 | NULL&M_Disease |
| 11 | 120248342 | rs12270397 | T | C | NULL | 0.685680114 | NULL&M_Disease |
| 11 | 121116824 | rs7115463 | T | C | NULL | 0.685680114 | NULL&M_Disease |
| 11 | 116495784 | rs112305884 | T | C | NULL | 0.685680114 | NULL&M_Disease |
| 11 | 116502768 | rs11216062 | A | C | NULL | 0.685680114 | NULL&M_Disease |
| 11 | 117126710 | rs512188 | A | G | NULL | 0.685680114 | NULL&M_Disease |
| 11 | 118880740 | rs11217122 | C | T | NULL | 0.685680114 | NULL&M_Disease |
| 11 | 119705210 | rs558034 | A | G | NULL | 0.685680114 | NULL&M_Disease |
| 11 | 120130512 | rs1941411 | G | A | NULL | 0.685680114 | NULL&M_Disease |
| 11 | 120542320 | rs10790392 | A | G | NULL | 0.685680114 | NULL&M_Disease |
| 11 | 120769478 | rs61409027 | T | C | NULL | 0.685680114 | NULL&M_Disease |
| 11 | 116464111 | rs11216025 | T | C | NULL | 0.685680114 | NULL&M_Disease |
| 11 | 117201097 | rs484553 | T | G | NULL | 0.685680114 | NULL&M_Disease |
| 11 | 120745967 | rs3824977 | G | A | NULL | 0.685680114 | NULL&M_Disease |
| 11 | 121136604 | rs10790422 | T | C | NULL | 0.685680114 | NULL&M_Disease |
| 11 | 116883396 | rs7125788 | C | T | NULL | 0.685680114 | NULL&M_Disease |
| 11 | 116883776 | rs61905705 | T | A | NULL | 0.685680114 | NULL&M_Disease |
| 11 | 116883777 | rs61905706 | T | G | NULL | 0.685680114 | NULL&M_Disease |
| 11 | 116893724 | rs4936359 | A | T | NULL | 0.685680114 | NULL&M_Disease |
| 11 | 116893769 | rs4936360 | C | G | NULL | 0.685680114 | NULL&M_Disease |
| 11 | 116897834 | rs61905709 | T | C | NULL | 0.685680114 | NULL&M_Disease |
| 11 | 116898479 | rs4938325 | A | G | NULL | 0.685680114 | NULL&M_Disease |
| 11 | 116902056 | rs7112191 | A | G | NULL | 0.685680114 | NULL&M_Disease |
| 11 | 119695491 | rs482090 | T | C | NULL | 0.685680114 | NULL&M_Disease |
| 11 | 116494264 | rs79801428 | A | G | NULL | 0.685680114 | NULL&M_Disease |
| 11 | 116909293 | rs61905712 | A | G | NULL | 0.685680114 | NULL&M_Disease |

| | | | | | | | | | |
|---|---|---|---|---|---|---|---|---|---|
| 11 | 116871223 | rs17120233 | G | A | 0.992 | -7977.388 | -7978.943 | -1.555 | 8.17E-07 |
| 11 | 116872263 | rs1248452864 | G | A | 0.992 | -7977.388 | -7978.943 | -1.555 | 8.17E-07 |
| 11 | 116877353 | rs77250716 | C | T | 0.992 | -7977.388 | -7978.943 | -1.555 | 8.17E-07 |
| 11 | 116937159 | rs112895963 | T | A | 0.991 | -7977.388 | -7978.943 | -1.555 | 8.17E-07 |
| 11 | 117785260 | rs4936410 | C | T | 0.976 | -7977.388 | -7978.944 | -1.555 | 8.17E-07 |
| 11 | 118205702 | rs4938506 | A | G | 1.023 | -7977.388 | -7978.944 | -1.555 | 8.17E-07 |
| 11 | 118216234 | rs3212264 | A | C | 1.02 | -7977.388 | -7978.943 | -1.555 | 8.17E-07 |
| 11 | 118536950 | rs201359394 | T | A | 0.977 | -7977.388 | -7978.943 | -1.555 | 8.17E-07 |
| 11 | 120812025 | rs10892648 | A | C | 0.977 | -7977.388 | -7978.944 | -1.555 | 8.17E-07 |
| 11 | 116496744 | rs77984767 | G | A | 0.989 | -7977.388 | -7978.945 | -1.556 | 8.16E-07 |
| 11 | 117473770 | rs12796722 | T | C | 1.023 | -7977.388 | -7978.944 | -1.556 | 8.16E-07 |
| 11 | 118892940 | rs12280288 | C | T | 1.022 | -7977.388 | -7978.944 | -1.556 | 8.16E-07 |
| 11 | 119383037 | rs7951668 | T | C | 1.025 | -7977.388 | -7978.944 | -1.556 | 8.16E-07 |
| 11 | 121131331 | rs9666004 | T | C | 0.975 | -7977.388 | -7978.945 | -1.556 | 8.16E-07 |
| 11 | 116500588 | rs11216055 | T | C | 0.989 | -7977.388 | -7978.945 | -1.557 | 8.15E-07 |
| 11 | 116500629 | rs11216056 | A | C | 0.989 | -7977.388 | -7978.945 | -1.557 | 8.15E-07 |
| 11 | 116500768 | rs11216057 | T | C | 0.989 | -7977.388 | -7978.945 | -1.557 | 8.15E-07 |
| 11 | 116502776 | rs11216063 | A | T | 0.989 | -7977.388 | -7978.945 | -1.557 | 8.15E-07 |
| 11 | 116952200 | rs4936362 | T | C | 0.992 | -7977.388 | -7978.945 | -1.557 | 8.15E-07 |
| 11 | 117083004 | rs67201490 | G | T | 0.986 | -7977.388 | -7978.946 | -1.557 | 8.15E-07 |
| 11 | 117084040 | rs4938357 | A | G | 0.986 | -7977.388 | -7978.946 | -1.557 | 8.15E-07 |
| 11 | 117085305 | rs56081989 | A | G | 0.986 | -7977.388 | -7978.946 | -1.557 | 8.15E-07 |
| 11 | 117764641 | rs11216607 | A | G | 0.976 | -7977.388 | -7978.945 | -1.557 | 8.15E-07 |
| 11 | 120745874 | rs2230297 | T | C | 1.007 | -7977.388 | -7978.945 | -1.557 | 8.15E-07 |
| 11 | 116493715 | rs11216047 | G | A | 1.022 | -7977.388 | -7978.947 | -1.558 | 8.14E-07 |
| 11 | 116496309 | rs4340085 | T | C | 0.989 | -7977.388 | -7978.946 | -1.558 | 8.14E-07 |
| 11 | 116498951 | rs11216054 | C | T | 0.978 | -7977.388 | -7978.946 | -1.558 | 8.14E-07 |
| 11 | 116537962 | rs1784125 | C | G | 1.024 | -7977.388 | -7978.946 | -1.558 | 8.14E-07 |
| 11 | 117003420 | rs4938342 | G | C | 0.985 | -7977.388 | -7978.946 | -1.558 | 8.14E-07 |
| 11 | 117454157 | rs880812 | C | G | 0.975 | -7977.388 | -7978.946 | -1.558 | 8.14E-07 |
| 11 | 117757345 | rs601122 | T | G | 1.025 | -7977.388 | -7978.946 | -1.558 | 8.14E-07 |
| 11 | 119496472 | rs1355191 | C | T | 1.025 | -7977.388 | -7978.946 | -1.558 | 8.14E-07 |
| 11 | 119951200 | rs6589790 | T | C | 0.994 | -7977.388 | -7978.946 | -1.558 | 8.14E-07 |
| 11 | 121135004 | rs1278081233 | G | T | 1.019 | -7977.388 | -7978.947 | -1.558 | 8.14E-07 |
| 11 | 116536418 | rs12803249 | G | A | 1.022 | -7977.388 | -7978.947 | -1.559 | 8.14E-07 |
| 11 | 116961802 | rs61903421 | T | A | 0.992 | -7977.388 | -7978.947 | -1.559 | 8.14E-07 |
| 11 | 116963897 | rs7950381 | A | C | 0.992 | -7977.388 | -7978.947 | -1.559 | 8.14E-07 |
| 11 | 117053675 | rs4938355 | G | A | 0.985 | -7977.388 | -7978.947 | -1.559 | 8.14E-07 |
| 11 | 117661100 | rs625447 | C | T | 0.975 | -7977.388 | -7978.947 | -1.559 | 8.14E-07 |
| 11 | 120179563 | rs12787658 | G | A | 1.025 | -7977.388 | -7978.947 | -1.559 | 8.14E-07 |
| 11 | 120821862 | rs12361041 | G | A | 0.977 | -7977.388 | -7978.947 | -1.559 | 8.14E-07 |
| 11 | 116956665 | rs61903418 | A | G | 0.992 | -7977.388 | -7978.948 | -1.56 | 8.13E-07 |
| 11 | 116960497 | rs11823918 | C | G | 0.992 | -7977.388 | -7978.948 | -1.56 | 8.13E-07 |
| 11 | 117156552 | rs7083 | T | C | 1.025 | -7977.388 | -7978.948 | -1.56 | 8.13E-07 |
| 11 | 117357085 | rs1315236635 | G | T | 1.025 | -7977.388 | -7978.948 | -1.56 | 8.13E-07 |
| 11 | 118139755 | rs55852239 | A | G | 0.976 | -7977.388 | -7978.949 | -1.56 | 8.13E-07 |
| 11 | 119568994 | rs4938713 | T | C | 1.009 | -7977.388 | -7978.948 | -1.56 | 8.13E-07 |
| 11 | 119792847 | rs11217575 | G | A | 0.99 | -7977.388 | -7978.948 | -1.56 | 8.13E-07 |
| 11 | 119951214 | rs6589792 | G | C | 0.995 | -7977.388 | -7978.948 | -1.56 | 8.13E-07 |
| 11 | 119981785 | rs10736502 | G | C | 0.978 | -7977.388 | -7978.949 | -1.56 | 8.13E-07 |
| 11 | 120198728 | rs11827818 | G | A | 0.983 | -7977.388 | -7978.949 | -1.56 | 8.13E-07 |
| 11 | 121131224 | rs9665995 | T | C | 0.975 | -7977.388 | -7978.948 | -1.56 | 8.13E-07 |
| 11 | 121140086 | rs10437696 | C | T | 1.018 | -7977.388 | -7978.948 | -1.56 | 8.13E-07 |
| 11 | 116560610 | rs1787714 | A | G | 0.975 | -7977.388 | -7978.949 | -1.561 | 8.12E-07 |
| 11 | 116880703 | rs61905691 | G | A | 0.993 | -7977.388 | -7978.95 | -1.561 | 8.12E-07 |
| 11 | 116915020 | rs17120241 | C | T | 0.993 | -7977.388 | -7978.949 | -1.561 | 8.12E-07 |
| 11 | 116919067 | rs61905716 | T | C | 0.993 | -7977.388 | -7978.949 | -1.561 | 8.12E-07 |
| 11 | 116919368 | rs112151056 | A | G | 0.993 | -7977.388 | -7978.949 | -1.561 | 8.12E-07 |
| 11 | 116922714 | rs61905719 | T | C | 0.993 | -7977.388 | -7978.949 | -1.561 | 8.12E-07 |
| 11 | 117473944 | rs12797135 | T | C | 1.023 | -7977.388 | -7978.949 | -1.561 | 8.12E-07 |
| 11 | 117692394 | rs501171 | T | C | 1.019 | -7977.388 | -7978.95 | -1.561 | 8.12E-07 |
| 11 | 117791132 | rs7927300 | A | C | 1.022 | -7977.388 | -7978.95 | -1.561 | 8.12E-07 |
| 11 | 119065044 | rs11217176 | A | G | 0.976 | -7977.388 | -7978.949 | -1.561 | 8.12E-07 |
| 11 | 119620339 | rs34952981 | T | A | 1.025 | -7977.388 | -7978.95 | -1.561 | 8.12E-07 |
| 11 | 119951213 | rs6589791 | A | C | 0.995 | -7977.388 | -7978.949 | -1.561 | 8.12E-07 |
| 11 | 116219447 | rs569304 | A | G | 1.025 | -7977.388 | -7978.95 | -1.562 | 8.11E-07 |
| 11 | 117319998 | rs11216397 | C | T | 0.976 | -7977.388 | -7978.951 | -1.562 | 8.11E-07 |
| 11 | 117433965 | rs522742 | A | G | 1.018 | -7977.388 | -7978.95 | -1.562 | 8.11E-07 |
| 11 | 118875727 | rs4936457 | G | A | 1.023 | -7977.388 | -7978.95 | -1.562 | 8.11E-07 |
| 11 | 118875761 | rs4936458 | G | T | 1.023 | -7977.388 | -7978.95 | -1.562 | 8.11E-07 |
| 11 | 119552057 | rs10790331 | C | A | 0.976 | -7977.388 | -7978.951 | -1.562 | 8.11E-07 |
| 11 | 120248342 | rs12270397 | T | C | 0.984 | -7977.388 | -7978.95 | -1.562 | 8.11E-07 |
| 11 | 121116824 | rs7115463 | T | C | 1.025 | -7977.388 | -7978.95 | -1.562 | 8.11E-07 |
| 11 | 116495784 | rs112305884 | T | C | 0.99 | -7977.388 | -7978.951 | -1.563 | 8.10E-07 |
| 11 | 116502768 | rs11216062 | A | C | 0.99 | -7977.388 | -7978.951 | -1.563 | 8.10E-07 |
| 11 | 117126710 | rs512188 | A | G | 1.025 | -7977.388 | -7978.952 | -1.563 | 8.10E-07 |
| 11 | 118880740 | rs11217122 | C | T | 1.022 | -7977.388 | -7978.951 | -1.563 | 8.10E-07 |
| 11 | 119705210 | rs558034 | A | G | 0.978 | -7977.388 | -7978.952 | -1.563 | 8.10E-07 |
| 11 | 120130512 | rs1941411 | G | A | 1.024 | -7977.388 | -7978.951 | -1.563 | 8.10E-07 |
| 11 | 120542320 | rs10790392 | A | G | 0.988 | -7977.388 | -7978.951 | -1.563 | 8.10E-07 |
| 11 | 120769478 | rs61409027 | T | C | 0.988 | -7977.388 | -7978.951 | -1.563 | 8.10E-07 |
| 11 | 116464111 | rs11216025 | T | C | 1.009 | -7977.388 | -7978.952 | -1.564 | 8.10E-07 |
| 11 | 117201097 | rs484553 | T | G | 0.976 | -7977.388 | -7978.952 | -1.564 | 8.10E-07 |
| 11 | 120745967 | rs3824977 | G | A | 1.006 | -7977.388 | -7978.953 | -1.564 | 8.10E-07 |
| 11 | 121136604 | rs10790422 | T | C | 1.018 | -7977.388 | -7978.952 | -1.564 | 8.10E-07 |
| 11 | 116883396 | rs7125788 | C | T | 0.994 | -7977.388 | -7978.953 | -1.565 | 8.09E-07 |
| 11 | 116883776 | rs61905705 | T | A | 0.994 | -7977.388 | -7978.954 | -1.565 | 8.09E-07 |
| 11 | 116883777 | rs61905706 | T | G | 0.994 | -7977.388 | -7978.954 | -1.565 | 8.09E-07 |
| 11 | 116893724 | rs4936359 | A | T | 0.994 | -7977.388 | -7978.953 | -1.565 | 8.09E-07 |
| 11 | 116893769 | rs4936360 | C | G | 0.994 | -7977.388 | -7978.953 | -1.565 | 8.09E-07 |
| 11 | 116897834 | rs61905709 | T | C | 0.994 | -7977.388 | -7978.953 | -1.565 | 8.09E-07 |
| 11 | 116898479 | rs4938325 | A | G | 0.994 | -7977.388 | -7978.953 | -1.565 | 8.09E-07 |
| 11 | 116902056 | rs7112191 | A | G | 0.994 | -7977.388 | -7978.953 | -1.565 | 8.09E-07 |
| 11 | 119695491 | rs482090 | T | C | 1.022 | -7977.388 | -7978.953 | -1.565 | 8.09E-07 |
| 11 | 116494264 | rs79801428 | A | G | 1.003 | -7977.388 | -7978.955 | -1.566 | 8.08E-07 |
| 11 | 116909293 | rs61905712 | A | G | 0.994 | -7977.388 | -7978.954 | -1.566 | 8.08E-07 |

| | | | | | | | |
|---|---|---|---|---|---|---|---|
| 11 | 116909429 | rs61905713 | T | C | NULL | 0.685680114 | NULL&M_Disease |
| 11 | 117699173 | rs76813465 | T | C | NULL | 0.685680114 | NULL&M_Disease |
| 11 | 117956409 | rs10892211 | C | T | NULL | 0.685680114 | NULL&M_Disease |
| 11 | 120212423 | rs10502235 | G | A | NULL | 0.685680114 | NULL&M_Disease |
| 11 | 120256337 | rs17123861 | C | T | NULL | 0.685680114 | NULL&M_Disease |
| 11 | 120518871 | rs4935748 | A | G | NULL | 0.685680114 | NULL&M_Disease |
| 11 | 121108865 | rs1941298 | T | C | NULL | 0.685680114 | NULL&M_Disease |
| 11 | 121128543 | rs4423192 | T | C | NULL | 0.685680114 | NULL&M_Disease |
| 11 | 116210133 | rs10736484 | T | C | NULL | 0.685680114 | NULL&M_Disease |
| 11 | 117793239 | rs10892194 | G | A | NULL | 0.685680114 | NULL&M_Disease |
| 11 | 118135973 | rs12798851 | G | T | NULL | 0.685680114 | NULL&M_Disease |
| 11 | 120101780 | rs2845706 | A | G | NULL | 0.685680114 | NULL&M_Disease |
| 11 | 120905771 | rs2000526 | C | T | NULL | 0.685680114 | NULL&M_Disease |
| 11 | 117266754 | rs693147 | A | C | NULL | 0.686756727 | NULL&M_Disease |
| 11 | 117902914 | rs590917 | T | G | NULL | 0.685680114 | NULL&M_Disease |
| 11 | 118117614 | rs17092740 | G | A | NULL | 0.686756727 | NULL&M_Disease |
| 11 | 118121421 | rs12577053 | G | A | NULL | 0.686756727 | NULL&M_Disease |
| 11 | 118121553 | rs12577061 | G | C | NULL | 0.686756727 | NULL&M_Disease |
| 11 | 119218530 | rs10892351 | T | G | NULL | 0.686756727 | NULL&M_Disease |
| 11 | 119507925 | rs4412764 | A | G | NULL | 0.686756727 | NULL&M_Disease |
| 11 | 119613788 | rs4505073 | G | C | NULL | 0.686756727 | NULL&M_Disease |
| 11 | 120149004 | rs12271393 | A | G | NULL | 0.685680114 | NULL&M_Disease |
| 11 | 120259430 | rs7932037 | A | T | NULL | 0.686756727 | NULL&M_Disease |
| 11 | 120261791 | rs7478880 | T | C | NULL | 0.686756727 | NULL&M_Disease |
| 11 | 120527541 | rs4622278 | C | T | NULL | 0.686756727 | NULL&M_Disease |
| 11 | 116303839 | rs12785979 | C | A | NULL | 0.686756727 | NULL&M_Disease |
| 11 | 116311642 | rs492517 | T | C | NULL | 0.686756727 | NULL&M_Disease |
| 11 | 116493328 | rs7104866 | T | C | NULL | 0.686756727 | NULL&M_Disease |
| 11 | 116567619 | rs4938300 | T | C | NULL | 0.686756727 | NULL&M_Disease |
| 11 | 116987342 | rs61903430 | C | G | NULL | 0.686756727 | NULL&M_Disease |
| 11 | 117042967 | rs56371319 | A | C | NULL | 0.686756727 | NULL&M_Disease |
| 11 | 117152540 | rs505045 | G | T | NULL | 0.686756727 | NULL&M_Disease |
| 11 | 117423936 | rs686598 | C | G | NULL | 0.686756727 | NULL&M_Disease |
| 11 | 117926158 | rs12272258 | A | G | NULL | 0.686756727 | NULL&M_Disease |
| 11 | 117956263 | rs10892209 | T | C | NULL | 0.686756727 | NULL&M_Disease |
| 11 | 117956338 | rs10892210 | T | G | NULL | 0.686756727 | NULL&M_Disease |
| 11 | 117127620 | rs655023 | C | T | NULL | 0.686756727 | NULL&M_Disease |
| 11 | 117751994 | rs584994 | G | T | NULL | 0.686756727 | NULL&M_Disease |
| 11 | 118835258 | rs11607835 | C | T | NULL | 0.686756727 | NULL&M_Disease |
| 11 | 118836638 | rs4310630 | G | A | NULL | 0.686756727 | NULL&M_Disease |
| 11 | 119949999 | rs11217675 | C | T | NULL | 0.686756727 | NULL&M_Disease |
| 11 | 120248249 | rs9633950 | G | T | NULL | 0.686756727 | NULL&M_Disease |
| 11 | 120406723 | rs11824600 | A | G | NULL | 0.686756727 | NULL&M_Disease |
| 11 | 121005824 | rs680511 | G | A | NULL | 0.686756727 | NULL&M_Disease |
| 11 | 116495059 | rs11216048 | G | A | NULL | 0.686756727 | NULL&M_Disease |
| 11 | 116495175 | rs11216049 | C | T | NULL | 0.686756727 | NULL&M_Disease |
| 11 | 116495314 | rs11216050 | T | C | NULL | 0.686756727 | NULL&M_Disease |
| 11 | 117311507 | rs79136761 | A | G | NULL | 0.686756727 | NULL&M_Disease |
| 11 | 118976719 | rs639373 | A | G | NULL | 0.686756727 | NULL&M_Disease |
| 11 | 119942571 | rs1790474 | A | G | NULL | 0.686756727 | NULL&M_Disease |
| 11 | 120768404 | rs10892636 | G | A | NULL | 0.686756727 | NULL&M_Disease |
| 11 | 120878492 | rs591671 | A | G | NULL | 0.686756727 | NULL&M_Disease |
| 11 | 116219158 | rs552104 | G | T | NULL | 0.686756727 | NULL&M_Disease |
| 11 | 116968798 | rs7128071 | A | G | NULL | 0.686756727 | NULL&M_Disease |
| 11 | 116980594 | rs4938339 | T | C | NULL | 0.686756727 | NULL&M_Disease |
| 11 | 119632722 | rs12277255 | G | A | NULL | 0.686756727 | NULL&M_Disease |
| 11 | 119951884 | rs12791310 | G | A | NULL | 0.686756727 | NULL&M_Disease |
| 11 | 116496360 | rs1240658 | C | T | NULL | 0.686756727 | NULL&M_Disease |
| 11 | 116507582 | rs74948313 | G | A | NULL | 0.686756727 | NULL&M_Disease |
| 11 | 117760253 | rs4938452 | A | T | NULL | 0.686756727 | NULL&M_Disease |
| 11 | 117908079 | rs685540 | C | G | NULL | 0.686756727 | NULL&M_Disease |
| 11 | 120074732 | rs1287392 | T | C | NULL | 0.686756727 | NULL&M_Disease |
| 11 | 121108679 | rs1941297 | T | C | NULL | 0.686756727 | NULL&M_Disease |
| 11 | 116311777 | rs529411 | C | T | NULL | 0.686756727 | NULL&M_Disease |
| 11 | 116986684 | rs61903429 | A | C | NULL | 0.686756727 | NULL&M_Disease |
| 11 | 117750740 | rs497768 | C | G | NULL | 0.686756727 | NULL&M_Disease |
| 11 | 117790769 | rs11826839 | G | A | NULL | 0.686756727 | NULL&M_Disease |
| 11 | 119772239 | rs515278 | C | G | NULL | 0.686756727 | NULL&M_Disease |
| 11 | 116968268 | rs61903426 | T | C | NULL | 0.686756727 | NULL&M_Disease |
| 11 | 116981335 | rs6589595 | A | G | NULL | 0.686756727 | NULL&M_Disease |
| 11 | 118848725 | rs10892318 | A | G | NULL | 0.686756727 | NULL&M_Disease |
| 11 | 116982928 | rs59700451 | A | G | NULL | 0.686756727 | NULL&M_Disease |
| 11 | 117555644 | rs10892161 | A | G | NULL | 0.686756727 | NULL&M_Disease |
| 11 | 119070949 | rs55771168 | C | T | NULL | 0.686756727 | NULL&M_Disease |
| 11 | 120400419 | rs2252222 | G | A | NULL | 0.686756727 | NULL&M_Disease |
| 11 | 120406694 | rs11827023 | C | T | NULL | 0.686756727 | NULL&M_Disease |
| 11 | 120605232 | rs3133223 | A | G | NULL | 0.686756727 | NULL&M_Disease |
| 11 | 116980622 | rs4938340 | A | G | NULL | 0.686756727 | NULL&M_Disease |
| 11 | 117326269 | rs61905307 | C | T | NULL | 0.686756727 | NULL&M_Disease |
| 11 | 118137667 | rs3759007 | T | C | NULL | 0.687831331 | NULL&M_Disease |
| 11 | 119864323 | rs10892509 | C | A | NULL | 0.686756727 | NULL&M_Disease |
| 11 | 119878605 | rs6589785 | T | G | NULL | 0.686756727 | NULL&M_Disease |
| 11 | 116232173 | rs12799683 | A | G | NULL | 0.686756727 | NULL&M_Disease |
| 11 | 117406111 | rs11216440 | A | G | NULL | 0.687831331 | NULL&M_Disease |
| 11 | 118138629 | rs34180117 | T | C | NULL | 0.687831331 | NULL&M_Disease |
| 11 | 118138922 | rs35850218 | C | T | NULL | 0.687831331 | NULL&M_Disease |
| 11 | 118146599 | rs11603267 | T | G | NULL | 0.687831331 | NULL&M_Disease |
| 11 | 118147496 | rs10502227 | C | T | NULL | 0.687831331 | NULL&M_Disease |
| 11 | 119919351 | rs145141827 | T | G | NULL | 0.686756727 | NULL&M_Disease |
| 11 | 119997599 | rs470725 | T | C | NULL | 0.687831331 | NULL&M_Disease |
| 11 | 120832264 | rs608891 | A | G | NULL | 0.686756727 | NULL&M_Disease |
| 11 | 116651115 | rs11604424 | C | T | NULL | 0.687831331 | NULL&M_Disease |
| 11 | 116988313 | rs75904569 | G | A | NULL | 0.687831331 | NULL&M_Disease |
| 11 | 117757547 | rs4938448 | C | A | NULL | 0.687831331 | NULL&M_Disease |
| 11 | 117760117 | rs4938450 | T | C | NULL | 0.687831331 | NULL&M_Disease |
| 11 | 118995353 | rs624588 | C | G | NULL | 0.687831331 | NULL&M_Disease |

| | | | | | | | | | |
|---|---|---|---|---|---|---|---|---|---|
| 11 | 116909429 | rs61905713 | T | C | 0.994 | -7977.388 | -7978.954 | -1.566 | 8.08E-07 |
| 11 | 117699173 | rs76813465 | T | C | 0.993 | -7977.388 | -7978.954 | -1.566 | 8.08E-07 |
| 11 | 117956409 | rs10892211 | C | T | 0.976 | -7977.388 | -7978.954 | -1.566 | 8.08E-07 |
| 11 | 120212423 | rs10502235 | G | A | 0.984 | -7977.388 | -7978.954 | -1.566 | 8.08E-07 |
| 11 | 120256337 | rs17123861 | C | T | 0.984 | -7977.388 | -7978.954 | -1.566 | 8.08E-07 |
| 11 | 120518871 | rs4935748 | A | G | 1.024 | -7977.388 | -7978.954 | -1.566 | 8.08E-07 |
| 11 | 121108865 | rs1941298 | T | C | 0.976 | -7977.388 | -7978.955 | -1.566 | 8.08E-07 |
| 11 | 121128543 | rs4423192 | T | C | 0.975 | -7977.388 | -7978.954 | -1.566 | 8.08E-07 |
| 11 | 116210133 | rs10736484 | T | C | 1.025 | -7977.388 | -7978.955 | -1.567 | 8.07E-07 |
| 11 | 117793239 | rs10892194 | G | A | 1.005 | -7977.388 | -7978.956 | -1.567 | 8.07E-07 |
| 11 | 118135973 | rs12798851 | G | T | 1.007 | -7977.388 | -7978.955 | -1.567 | 8.07E-07 |
| 11 | 120101780 | rs2845706 | A | G | 0.987 | -7977.388 | -7978.955 | -1.567 | 8.07E-07 |
| 11 | 120905771 | rs2000526 | C | T | 1.014 | -7977.388 | -7978.955 | -1.567 | 8.07E-07 |
| 11 | 117266754 | rs693147 | A | C | 0.976 | -7977.388 | -7978.956 | -1.568 | 8.06E-07 |
| 11 | 117902914 | rs590917 | T | G | 1.015 | -7977.388 | -7978.956 | -1.568 | 8.06E-07 |
| 11 | 118117614 | rs17092740 | G | A | 1.025 | -7977.388 | -7978.957 | -1.568 | 8.06E-07 |
| 11 | 118121421 | rs12577053 | G | A | 1.025 | -7977.388 | -7978.957 | -1.568 | 8.06E-07 |
| 11 | 118121553 | rs12577061 | G | C | 1.025 | -7977.388 | -7978.957 | -1.568 | 8.06E-07 |
| 11 | 119218530 | rs10892351 | T | G | 0.976 | -7977.388 | -7978.957 | -1.568 | 8.06E-07 |
| 11 | 119507925 | rs4412764 | A | G | 0.975 | -7977.388 | -7978.957 | -1.568 | 8.06E-07 |
| 11 | 119613788 | rs4505073 | G | C | 0.976 | -7977.388 | -7978.956 | -1.568 | 8.06E-07 |
| 11 | 120149004 | rs12271393 | A | G | 1.02 | -7977.388 | -7978.956 | -1.568 | 8.06E-07 |
| 11 | 120259430 | rs7932037 | A | T | 1.025 | -7977.388 | -7978.957 | -1.568 | 8.06E-07 |
| 11 | 120261791 | rs7478880 | T | C | 1.025 | -7977.388 | -7978.956 | -1.568 | 8.06E-07 |
| 11 | 120527541 | rs4622278 | C | T | 1.025 | -7977.388 | -7978.956 | -1.568 | 8.06E-07 |
| 11 | 116303839 | rs12785979 | C | A | 1.025 | -7977.388 | -7978.957 | -1.569 | 8.06E-07 |
| 11 | 116311642 | rs492517 | T | C | 0.976 | -7977.388 | -7978.957 | -1.569 | 8.06E-07 |
| 11 | 116493328 | rs7104866 | T | C | 0.976 | -7977.388 | -7978.957 | -1.569 | 8.06E-07 |
| 11 | 116567619 | rs4938300 | T | C | 1.018 | -7977.388 | -7978.957 | -1.569 | 8.06E-07 |
| 11 | 116987342 | rs61903430 | C | G | 1.006 | -7977.388 | -7978.957 | -1.569 | 8.06E-07 |
| 11 | 117042967 | rs56371319 | A | C | 0.986 | -7977.388 | -7978.957 | -1.569 | 8.06E-07 |
| 11 | 117152540 | rs505045 | G | T | 1.025 | -7977.388 | -7978.957 | -1.569 | 8.06E-07 |
| 11 | 117423936 | rs686598 | C | G | 1.025 | -7977.388 | -7978.957 | -1.569 | 8.06E-07 |
| 11 | 117926158 | rs12272258 | A | G | 1.024 | -7977.388 | -7978.957 | -1.569 | 8.06E-07 |
| 11 | 117956263 | rs10892209 | T | C | 0.976 | -7977.388 | -7978.957 | -1.569 | 8.06E-07 |
| 11 | 117956338 | rs10892210 | T | G | 0.976 | -7977.388 | -7978.957 | -1.569 | 8.06E-07 |
| 11 | 117127620 | rs655023 | C | T | 1.025 | -7977.388 | -7978.959 | -1.57 | 8.05E-07 |
| 11 | 117751994 | rs584994 | G | T | 0.976 | -7977.388 | -7978.958 | -1.57 | 8.05E-07 |
| 11 | 118835258 | rs11607835 | C | T | 1.025 | -7977.388 | -7978.958 | -1.57 | 8.05E-07 |
| 11 | 118836638 | rs4310630 | G | A | 1.025 | -7977.388 | -7978.958 | -1.57 | 8.05E-07 |
| 11 | 119949999 | rs11217675 | C | T | 0.998 | -7977.388 | -7978.958 | -1.57 | 8.05E-07 |
| 11 | 120248249 | rs9633950 | G | T | 1.025 | -7977.388 | -7978.959 | -1.57 | 8.05E-07 |
| 11 | 120406723 | rs11824600 | A | G | 1.025 | -7977.388 | -7978.959 | -1.57 | 8.05E-07 |
| 11 | 121005824 | rs680511 | G | A | 1.025 | -7977.388 | -7978.958 | -1.57 | 8.05E-07 |
| 11 | 116495059 | rs11216048 | G | A | 0.991 | -7977.388 | -7978.959 | -1.571 | 8.04E-07 |
| 11 | 116495175 | rs11216049 | C | T | 0.991 | -7977.388 | -7978.959 | -1.571 | 8.04E-07 |
| 11 | 116495314 | rs11216050 | T | C | 0.991 | -7977.388 | -7978.959 | -1.571 | 8.04E-07 |
| 11 | 117311507 | rs79136761 | A | G | 0.996 | -7977.388 | -7978.959 | -1.571 | 8.04E-07 |
| 11 | 118976719 | rs639373 | A | G | 1.025 | -7977.388 | -7978.959 | -1.571 | 8.04E-07 |
| 11 | 119942571 | rs1790474 | A | G | 1.019 | -7977.388 | -7978.959 | -1.571 | 8.04E-07 |
| 11 | 120768404 | rs10892636 | G | A | 0.989 | -7977.388 | -7978.96 | -1.571 | 8.04E-07 |
| 11 | 120878492 | rs591671 | A | G | 1.019 | -7977.388 | -7978.959 | -1.571 | 8.04E-07 |
| 11 | 116219158 | rs552104 | G | T | 1.024 | -7977.388 | -7978.96 | -1.572 | 8.03E-07 |
| 11 | 116968798 | rs7128071 | A | G | 0.993 | -7977.388 | -7978.96 | -1.572 | 8.03E-07 |
| 11 | 116980594 | rs4938339 | T | C | 1.005 | -7977.388 | -7978.96 | -1.572 | 8.03E-07 |
| 11 | 119632722 | rs12277255 | G | A | 1.025 | -7977.388 | -7978.96 | -1.572 | 8.03E-07 |
| 11 | 119951884 | rs12791310 | G | A | 1.024 | -7977.388 | -7978.96 | -1.572 | 8.03E-07 |
| 11 | 116496360 | rs1240658 | C | T | 1.016 | -7977.388 | -7978.962 | -1.573 | 8.02E-07 |
| 11 | 116507582 | rs74948313 | G | A | 1.003 | -7977.388 | -7978.961 | -1.573 | 8.02E-07 |
| 11 | 117760253 | rs4938452 | A | T | 1.025 | -7977.388 | -7978.962 | -1.573 | 8.02E-07 |
| 11 | 117908079 | rs685540 | C | G | 1.015 | -7977.388 | -7978.962 | -1.573 | 8.02E-07 |
| 11 | 120074732 | rs1287392 | T | C | 1.014 | -7977.388 | -7978.961 | -1.573 | 8.02E-07 |
| 11 | 121108679 | rs1941297 | T | C | 0.976 | -7977.388 | -7978.961 | -1.573 | 8.02E-07 |
| 11 | 116311777 | rs529411 | C | T | 0.976 | -7977.388 | -7978.962 | -1.574 | 8.02E-07 |
| 11 | 116986684 | rs61903429 | A | C | 1.004 | -7977.388 | -7978.962 | -1.574 | 8.02E-07 |
| 11 | 117750740 | rs497768 | C | G | 0.976 | -7977.388 | -7978.963 | -1.574 | 8.02E-07 |
| 11 | 117790769 | rs11826839 | G | A | 1.022 | -7977.388 | -7978.962 | -1.574 | 8.02E-07 |
| 11 | 119772239 | rs515278 | C | G | 0.977 | -7977.388 | -7978.962 | -1.574 | 8.02E-07 |
| 11 | 116968268 | rs61903426 | T | C | 0.994 | -7977.388 | -7978.963 | -1.575 | 8.01E-07 |
| 11 | 116981335 | rs6589595 | A | G | 1.004 | -7977.388 | -7978.964 | -1.575 | 8.01E-07 |
| 11 | 118848725 | rs10892318 | A | G | 1.025 | -7977.388 | -7978.963 | -1.575 | 8.01E-07 |
| 11 | 116982928 | rs59700451 | A | G | 1.003 | -7977.388 | -7978.964 | -1.576 | 8.00E-07 |
| 11 | 117555644 | rs10892161 | A | G | 0.98 | -7977.388 | -7978.964 | -1.576 | 8.00E-07 |
| 11 | 119070949 | rs55771168 | C | T | 0.977 | -7977.388 | -7978.964 | -1.576 | 8.00E-07 |
| 11 | 120400419 | rs2252222 | G | A | 0.984 | -7977.388 | -7978.964 | -1.576 | 8.00E-07 |
| 11 | 120406694 | rs11827023 | C | T | 1.025 | -7977.388 | -7978.965 | -1.576 | 8.00E-07 |
| 11 | 120605232 | rs3133223 | A | G | 0.979 | -7977.388 | -7978.965 | -1.576 | 8.00E-07 |
| 11 | 116980622 | rs4938340 | A | G | 0.999 | -7977.388 | -7978.966 | -1.577 | 7.99E-07 |
| 11 | 117326269 | rs61905307 | C | T | 1.002 | -7977.388 | -7978.965 | -1.577 | 7.99E-07 |
| 11 | 118137667 | rs3759007 | T | C | 0.976 | -7977.388 | -7978.966 | -1.577 | 7.99E-07 |
| 11 | 119864323 | rs10892509 | C | A | 0.995 | -7977.388 | -7978.965 | -1.577 | 7.99E-07 |
| 11 | 119878605 | rs6589785 | T | G | 1.023 | -7977.388 | -7978.966 | -1.577 | 7.99E-07 |
| 11 | 116232173 | rs12799683 | A | G | 0.99 | -7977.388 | -7978.966 | -1.578 | 7.98E-07 |
| 11 | 117406111 | rs11216440 | A | G | 0.978 | -7977.388 | -7978.967 | -1.578 | 7.98E-07 |
| 11 | 118138629 | rs34180117 | T | C | 0.976 | -7977.388 | -7978.966 | -1.578 | 7.98E-07 |
| 11 | 118138922 | rs35850218 | C | T | 0.976 | -7977.388 | -7978.966 | -1.578 | 7.98E-07 |
| 11 | 118146599 | rs11603267 | T | G | 1.02 | -7977.388 | -7978.966 | -1.578 | 7.98E-07 |
| 11 | 118147496 | rs10502227 | C | T | 1.02 | -7977.388 | -7978.966 | -1.578 | 7.98E-07 |
| 11 | 119919351 | rs145141827 | T | G | 1.014 | -7977.388 | -7978.966 | -1.578 | 7.98E-07 |
| 11 | 119997599 | rs470725 | T | C | 0.978 | -7977.388 | -7978.967 | -1.578 | 7.98E-07 |
| 11 | 120832264 | rs608891 | A | G | 1.019 | -7977.388 | -7978.966 | -1.578 | 7.98E-07 |
| 11 | 116651115 | rs11604424 | C | T | 0.98 | -7977.388 | -7978.967 | -1.579 | 7.98E-07 |
| 11 | 116988313 | rs75904569 | G | A | 1 | -7977.388 | -7978.967 | -1.579 | 7.98E-07 |
| 11 | 117757547 | rs4938448 | C | A | 1.025 | -7977.388 | -7978.968 | -1.579 | 7.98E-07 |
| 11 | 117760117 | rs4938450 | T | C | 0.977 | -7977.388 | -7978.968 | -1.579 | 7.98E-07 |
| 11 | 118995353 | rs624588 | C | G | 0.976 | -7977.388 | -7978.967 | -1.579 | 7.98E-07 |

| | | | | | | | |
|---|---|---|---|---|---|---|---|
| 11 | 120257495 | rs10790381 | G | A | NULL | 0.687831331 | NULL&M_Disease |
| 11 | 120545346 | rs11217962 | T | G | NULL | 0.687831331 | NULL&M_Disease |
| 11 | 121126285 | rs4144287 | G | A | NULL | 0.687831331 | NULL&M_Disease |
| 11 | 116312643 | rs500943 | T | C | NULL | 0.687831331 | NULL&M_Disease |
| 11 | 116454261 | rs7113362 | A | G | NULL | 0.687831331 | NULL&M_Disease |
| 11 | 116455707 | rs7121512 | G | T | NULL | 0.687831331 | NULL&M_Disease |
| 11 | 116498862 | rs10892000 | G | T | NULL | 0.687831331 | NULL&M_Disease |
| 11 | 117760135 | rs4938451 | T | C | NULL | 0.687831331 | NULL&M_Disease |
| 11 | 118139478 | rs56128937 | A | T | NULL | 0.687831331 | NULL&M_Disease |
| 11 | 121142453 | rs6589869 | G | C | NULL | 0.687831331 | NULL&M_Disease |
| 11 | 116223503 | rs565578 | C | T | NULL | 0.687831331 | NULL&M_Disease |
| 11 | 118137560 | rs3759006 | A | G | NULL | 0.687831331 | NULL&M_Disease |
| 11 | 118137911 | rs7948683 | A | G | NULL | 0.687831331 | NULL&M_Disease |
| 11 | 118143368 | rs4338560 | T | C | NULL | 0.687831331 | NULL&M_Disease |
| 11 | 119781157 | rs79163278 | C | T | NULL | 0.687831331 | NULL&M_Disease |
| 11 | 119814422 | rs12222499 | T | C | NULL | 0.687831331 | NULL&M_Disease |
| 11 | 120242348 | rs12807097 | A | G | NULL | 0.687831331 | NULL&M_Disease |
| 11 | 120540060 | rs11217955 | T | C | NULL | 0.687831331 | NULL&M_Disease |
| 11 | 120541873 | rs10892596 | T | C | NULL | 0.687831331 | NULL&M_Disease |
| 11 | 116493950 | rs1240659 | G | T | NULL | 0.687831331 | NULL&M_Disease |
| 11 | 117021426 | rs45625638 | C | G | NULL | 0.687831331 | NULL&M_Disease |
| 11 | 117628944 | rs34458464 | G | A | NULL | 0.687831331 | NULL&M_Disease |
| 11 | 117691386 | rs869789 | A | G | NULL | 0.687831331 | NULL&M_Disease |
| 11 | 118028094 | rs876479 | T | C | NULL | 0.687831331 | NULL&M_Disease |
| 11 | 118055217 | rs6589650 | C | T | NULL | 0.687831331 | NULL&M_Disease |
| 11 | 118055273 | rs7940757 | T | C | NULL | 0.687831331 | NULL&M_Disease |
| 11 | 118055733 | rs7944087 | T | C | NULL | 0.687831331 | NULL&M_Disease |
| 11 | 118056092 | rs7931206 | C | T | NULL | 0.687831331 | NULL&M_Disease |
| 11 | 118063479 | rs655499 | G | A | NULL | 0.687831331 | NULL&M_Disease |
| 11 | 118123517 | rs1715456 | A | G | NULL | 0.687831331 | NULL&M_Disease |
| 11 | 120541508 | rs12362348 | C | G | NULL | 0.687831331 | NULL&M_Disease |
| 11 | 120804291 | rs10892647 | G | A | NULL | 0.687831331 | NULL&M_Disease |
| 11 | 117322917 | rs10892110 | C | A | NULL | 0.687831331 | NULL&M_Disease |
| 11 | 117761400 | rs1939367 | A | C | NULL | 0.687831331 | NULL&M_Disease |
| 11 | 118119575 | rs12364013 | C | T | NULL | 0.687831331 | NULL&M_Disease |
| 11 | 119006300 | rs641382 | A | C | NULL | 0.687831331 | NULL&M_Disease |
| 11 | 120150059 | rs4936511 | C | A | NULL | 0.687831331 | NULL&M_Disease |
| 11 | 116269032 | rs9645661 | A | G | NULL | 0.687831331 | NULL&M_Disease |
| 11 | 117757443 | rs4938447 | C | A | NULL | 0.687831331 | NULL&M_Disease |
| 11 | 118055801 | rs7944321 | A | G | NULL | 0.687831331 | NULL&M_Disease |
| 11 | 118056047 | rs7944366 | G | A | NULL | 0.687831331 | NULL&M_Disease |
| 11 | 118138998 | rs12806041 | G | A | NULL | 0.687831331 | NULL&M_Disease |
| 11 | 119989636 | rs7128147 | T | C | NULL | 0.687831331 | NULL&M_Disease |
| 11 | 120249287 | rs11600703 | T | C | NULL | 0.687831331 | NULL&M_Disease |
| 11 | 120249493 | rs9633919 | A | G | NULL | 0.687831331 | NULL&M_Disease |
| 11 | 120250699 | rs12420378 | G | A | NULL | 0.687831331 | NULL&M_Disease |
| 11 | 120539410 | rs10892593 | T | C | NULL | 0.687831331 | NULL&M_Disease |
| 11 | 120769943 | rs7106413 | T | C | NULL | 0.687831331 | NULL&M_Disease |
| 11 | 121113812 | rs59868549 | T | A | NULL | 0.687831331 | NULL&M_Disease |
| 11 | 116314419 | rs480053 | A | G | NULL | 0.687831331 | NULL&M_Disease |
| 11 | 117010263 | rs145672593 | T | C | NULL | 0.687831331 | NULL&M_Disease |
| 11 | 117128971 | rs621858 | A | G | NULL | 0.687831331 | NULL&M_Disease |
| 11 | 117826191 | rs2512159 | G | C | NULL | 0.687831331 | NULL&M_Disease |
| 11 | 118053156 | rs4537793 | A | C | NULL | 0.687831331 | NULL&M_Disease |
| 11 | 118205636 | rs4938505 | A | G | NULL | 0.687831331 | NULL&M_Disease |
| 11 | 119972626 | rs1073635 | T | C | NULL | 0.687831331 | NULL&M_Disease |
| 11 | 120540410 | rs10892594 | C | A | NULL | 0.687831331 | NULL&M_Disease |
| 11 | 120541373 | rs12362311 | T | C | NULL | 0.687831331 | NULL&M_Disease |
| 11 | 120541660 | rs10082646 | C | T | NULL | 0.687831331 | NULL&M_Disease |
| 11 | 120551572 | rs10892610 | G | A | NULL | 0.687831331 | NULL&M_Disease |
| 11 | 120612223 | rs3018284 | G | T | NULL | 0.687831331 | NULL&M_Disease |
| 11 | 116311210 | rs534845 | A | G | NULL | 0.687831331 | NULL&M_Disease |
| 11 | 117143581 | rs673738 | T | C | NULL | 0.687831331 | NULL&M_Disease |
| 11 | 120540687 | rs10892595 | T | C | NULL | 0.687831331 | NULL&M_Disease |
| 11 | 120593695 | rs17311407 | T | C | NULL | 0.687831331 | NULL&M_Disease |
| 11 | 117025357 | rs68137714 | G | A | NULL | 0.687831331 | NULL&M_Disease |
| 11 | 117068534 | rs2075547 | A | C | NULL | 0.687831331 | NULL&M_Disease |
| 11 | 117328559 | rs491812 | T | C | NULL | 0.687831331 | NULL&M_Disease |
| 11 | 118055110 | rs7940717 | T | G | NULL | 0.687831331 | NULL&M_Disease |
| 11 | 118120351 | rs73024604 | C | T | NULL | 0.687831331 | NULL&M_Disease |
| 11 | 118120366 | rs73024606 | G | C | NULL | 0.687831331 | NULL&M_Disease |
| 11 | 118120802 | rs61900655 | G | C | NULL | 0.687831331 | NULL&M_Disease |
| 11 | 118184092 | rs3782041 | T | C | NULL | 0.687831331 | NULL&M_Disease |
| 11 | 118191274 | rs7937334 | T | C | NULL | 0.687831331 | NULL&M_Disease |
| 11 | 120063441 | rs12800456 | G | A | NULL | 0.687831331 | NULL&M_Disease |
| 11 | 120542279 | rs10790391 | A | G | NULL | 0.688903918 | NULL&M_Disease |
| 11 | 116505239 | rs11216068 | T | C | NULL | 0.688903918 | NULL&M_Disease |
| 11 | 116538689 | rs1784128 | A | G | NULL | 0.688903918 | NULL&M_Disease |
| 11 | 117087118 | rs56061158 | C | G | NULL | 0.688903918 | NULL&M_Disease |
| 11 | 117404769 | rs12806865 | G | A | NULL | 0.688903918 | NULL&M_Disease |
| 11 | 117759825 | rs6589632 | T | C | NULL | 0.688903918 | NULL&M_Disease |
| 11 | 118056722 | rs7935561 | A | G | NULL | 0.688903918 | NULL&M_Disease |
| 11 | 118362869 | rs537975 | T | C | NULL | 0.688903918 | NULL&M_Disease |
| 11 | 118869460 | rs3737504 | G | A | NULL | 0.688903918 | NULL&M_Disease |
| 11 | 118896116 | rs2276071 | C | A | NULL | 0.688903918 | NULL&M_Disease |
| 11 | 118980039 | rs510435 | T | C | NULL | 0.688903918 | NULL&M_Disease |
| 11 | 119978951 | rs4938782 | T | C | NULL | 0.688903918 | NULL&M_Disease |
| 11 | 117083373 | rs111765459 | C | T | NULL | 0.688903918 | NULL&M_Disease |
| 11 | 117138836 | rs619772 | T | A | NULL | 0.688903918 | NULL&M_Disease |
| 11 | 118140184 | rs12788933 | T | C | NULL | 0.688903918 | NULL&M_Disease |
| 11 | 118359161 | rs572126 | G | A | NULL | 0.688903918 | NULL&M_Disease |
| 11 | 119763417 | rs523223 | A | G | NULL | 0.688903918 | NULL&M_Disease |
| 11 | 120542603 | rs7934908 | A | G | NULL | 0.688903918 | NULL&M_Disease |
| 11 | 120551586 | rs10892611 | T | C | NULL | 0.688903918 | NULL&M_Disease |
| 11 | 121081632 | rs499443 | A | G | NULL | 0.688903918 | NULL&M_Disease |
| 11 | 121093676 | rs654329 | T | C | NULL | 0.688903918 | NULL&M_Disease |

| | | | | | | | | | |
|---|---|---|---|---|---|---|---|---|---|
| 11 | 120257495 | rs10790381 | G | A | 0.985 | -7977.388 | -7978.967 | -1.579 | 7.98E-07 |
| 11 | 120545346 | rs11217962 | T | G | 0.991 | -7977.388 | -7978.968 | -1.579 | 7.98E-07 |
| 11 | 121126285 | rs4144287 | G | A | 1.017 | -7977.388 | -7978.967 | -1.579 | 7.98E-07 |
| 11 | 116312643 | rs500943 | T | C | 0.977 | -7977.388 | -7978.969 | -1.58 | 7.97E-07 |
| 11 | 116454261 | rs7113362 | A | G | 0.976 | -7977.388 | -7978.968 | -1.58 | 7.97E-07 |
| 11 | 116455707 | rs7121512 | G | T | 0.976 | -7977.388 | -7978.968 | -1.58 | 7.97E-07 |
| 11 | 116498862 | rs10892000 | G | T | 0.979 | -7977.388 | -7978.968 | -1.58 | 7.97E-07 |
| 11 | 117760135 | rs4938451 | T | C | 1.025 | -7977.388 | -7978.968 | -1.58 | 7.97E-07 |
| 11 | 118139478 | rs56128937 | A | T | 0.976 | -7977.388 | -7978.969 | -1.58 | 7.97E-07 |
| 11 | 121142453 | rs6589869 | G | C | 1.017 | -7977.388 | -7978.969 | -1.58 | 7.97E-07 |
| 11 | 116223503 | rs565578 | C | T | 1.024 | -7977.388 | -7978.969 | -1.581 | 7.96E-07 |
| 11 | 118137560 | rs3759006 | A | G | 0.976 | -7977.388 | -7978.969 | -1.581 | 7.96E-07 |
| 11 | 118137911 | rs7948683 | A | G | 0.976 | -7977.388 | -7978.969 | -1.581 | 7.96E-07 |
| 11 | 118143368 | rs4338560 | T | C | 1.02 | -7977.388 | -7978.969 | -1.581 | 7.96E-07 |
| 11 | 119781157 | rs79163278 | C | T | 0.998 | -7977.388 | -7978.969 | -1.581 | 7.96E-07 |
| 11 | 119814422 | rs12222499 | T | C | 0.997 | -7977.388 | -7978.97 | -1.581 | 7.96E-07 |
| 11 | 120242348 | rs12807097 | A | G | 0.985 | -7977.388 | -7978.969 | -1.581 | 7.96E-07 |
| 11 | 120540060 | rs11217955 | T | C | 0.991 | -7977.388 | -7978.969 | -1.581 | 7.96E-07 |
| 11 | 120541873 | rs10892596 | T | C | 0.991 | -7977.388 | -7978.97 | -1.581 | 7.96E-07 |
| 11 | 116493950 | rs1240659 | G | T | 1.016 | -7977.388 | -7978.971 | -1.582 | 7.95E-07 |
| 11 | 117021426 | rs45625638 | C | G | 0.988 | -7977.388 | -7978.97 | -1.582 | 7.95E-07 |
| 11 | 117628944 | rs34458464 | G | A | 0.977 | -7977.388 | -7978.97 | -1.582 | 7.95E-07 |
| 11 | 117691386 | rs869789 | A | G | 1.001 | -7977.388 | -7978.97 | -1.582 | 7.95E-07 |
| 11 | 118028094 | rs876479 | T | C | 0.976 | -7977.388 | -7978.97 | -1.582 | 7.95E-07 |
| 11 | 118055217 | rs6589650 | C | T | 0.981 | -7977.388 | -7978.97 | -1.582 | 7.95E-07 |
| 11 | 118055273 | rs7940757 | T | C | 0.981 | -7977.388 | -7978.97 | -1.582 | 7.95E-07 |
| 11 | 118055733 | rs7944087 | T | C | 0.981 | -7977.388 | -7978.97 | -1.582 | 7.95E-07 |
| 11 | 118056092 | rs7931206 | C | T | 0.981 | -7977.388 | -7978.97 | -1.582 | 7.95E-07 |
| 11 | 118063479 | rs655499 | G | A | 0.996 | -7977.388 | -7978.97 | -1.582 | 7.95E-07 |
| 11 | 118123517 | rs1715456 | A | G | 1.025 | -7977.388 | -7978.971 | -1.582 | 7.95E-07 |
| 11 | 120541508 | rs12362348 | C | G | 0.99 | -7977.388 | -7978.971 | -1.582 | 7.95E-07 |
| 11 | 120804291 | rs10892647 | G | A | 0.977 | -7977.388 | -7978.97 | -1.582 | 7.95E-07 |
| 11 | 117322917 | rs10892110 | C | A | 0.976 | -7977.388 | -7978.971 | -1.583 | 7.94E-07 |
| 11 | 117761400 | rs1939367 | A | C | 1.025 | -7977.388 | -7978.971 | -1.583 | 7.94E-07 |
| 11 | 118119575 | rs12364013 | C | T | 1.025 | -7977.388 | -7978.972 | -1.583 | 7.94E-07 |
| 11 | 119006300 | rs641382 | A | C | 0.976 | -7977.388 | -7978.972 | -1.583 | 7.94E-07 |
| 11 | 120150059 | rs4936511 | C | A | 1.019 | -7977.388 | -7978.971 | -1.583 | 7.94E-07 |
| 11 | 116269032 | rs9645661 | A | G | 1.022 | -7977.388 | -7978.972 | -1.584 | 7.94E-07 |
| 11 | 117757443 | rs4938447 | C | A | 1.025 | -7977.388 | -7978.973 | -1.584 | 7.94E-07 |
| 11 | 118055801 | rs7944321 | A | G | 0.981 | -7977.388 | -7978.973 | -1.584 | 7.94E-07 |
| 11 | 118056047 | rs7944366 | G | A | 0.981 | -7977.388 | -7978.973 | -1.584 | 7.94E-07 |
| 11 | 118138998 | rs12806041 | G | A | 0.977 | -7977.388 | -7978.972 | -1.584 | 7.94E-07 |
| 11 | 119989636 | rs7128147 | T | C | 1.024 | -7977.388 | -7978.972 | -1.584 | 7.94E-07 |
| 11 | 120249287 | rs11600703 | T | C | 1.024 | -7977.388 | -7978.972 | -1.584 | 7.94E-07 |
| 11 | 120249493 | rs9633919 | A | G | 1.024 | -7977.388 | -7978.972 | -1.584 | 7.94E-07 |
| 11 | 120250699 | rs12420378 | G | A | 1.024 | -7977.388 | -7978.972 | -1.584 | 7.94E-07 |
| 11 | 120539410 | rs10892593 | T | C | 0.992 | -7977.388 | -7978.972 | -1.584 | 7.94E-07 |
| 11 | 120769943 | rs7106413 | T | C | 0.99 | -7977.388 | -7978.972 | -1.584 | 7.94E-07 |
| 11 | 121113812 | rs59868549 | T | A | 0.976 | -7977.388 | -7978.972 | -1.584 | 7.94E-07 |
| 11 | 116314419 | rs480053 | A | G | 0.978 | -7977.388 | -7978.973 | -1.585 | 7.93E-07 |
| 11 | 117010263 | rs145672593 | T | C | 0.99 | -7977.388 | -7978.974 | -1.585 | 7.93E-07 |
| 11 | 117128971 | rs621858 | A | G | 1.024 | -7977.388 | -7978.973 | -1.585 | 7.93E-07 |
| 11 | 117826191 | rs2512159 | G | C | 0.977 | -7977.388 | -7978.974 | -1.585 | 7.93E-07 |
| 11 | 118053156 | rs4537793 | A | C | 0.989 | -7977.388 | -7978.974 | -1.585 | 7.93E-07 |
| 11 | 118205636 | rs4938505 | A | G | 0.978 | -7977.388 | -7978.973 | -1.585 | 7.93E-07 |
| 11 | 119972626 | rs1073635 | T | C | 1.018 | -7977.388 | -7978.974 | -1.585 | 7.93E-07 |
| 11 | 120540410 | rs10892594 | C | A | 0.992 | -7977.388 | -7978.974 | -1.585 | 7.93E-07 |
| 11 | 120541373 | rs12362311 | T | C | 0.99 | -7977.388 | -7978.973 | -1.585 | 7.93E-07 |
| 11 | 120541660 | rs10082646 | C | T | 0.992 | -7977.388 | -7978.974 | -1.585 | 7.93E-07 |
| 11 | 120551572 | rs10892610 | G | A | 1.004 | -7977.388 | -7978.973 | -1.585 | 7.93E-07 |
| 11 | 120612223 | rs3018284 | G | T | 0.977 | -7977.388 | -7978.974 | -1.585 | 7.93E-07 |
| 11 | 116311210 | rs534845 | A | G | 1.024 | -7977.388 | -7978.974 | -1.586 | 7.92E-07 |
| 11 | 117143581 | rs673738 | T | C | 1.024 | -7977.388 | -7978.975 | -1.586 | 7.92E-07 |
| 11 | 120540687 | rs10892595 | T | C | 0.992 | -7977.388 | -7978.974 | -1.586 | 7.92E-07 |
| 11 | 120593695 | rs17311407 | T | C | 0.99 | -7977.388 | -7978.975 | -1.586 | 7.92E-07 |
| 11 | 117025357 | rs68137714 | G | A | 0.989 | -7977.388 | -7978.975 | -1.587 | 7.91E-07 |
| 11 | 117068534 | rs2075547 | A | C | 0.991 | -7977.388 | -7978.975 | -1.587 | 7.91E-07 |
| 11 | 117328559 | rs491812 | T | C | 0.976 | -7977.388 | -7978.975 | -1.587 | 7.91E-07 |
| 11 | 118055110 | rs7940717 | T | G | 0.981 | -7977.388 | -7978.975 | -1.587 | 7.91E-07 |
| 11 | 118120351 | rs73024604 | C | T | 1.024 | -7977.388 | -7978.975 | -1.587 | 7.91E-07 |
| 11 | 118120366 | rs73024606 | G | C | 1.024 | -7977.388 | -7978.975 | -1.587 | 7.91E-07 |
| 11 | 118120802 | rs61900655 | G | C | 1.024 | -7977.388 | -7978.975 | -1.587 | 7.91E-07 |
| 11 | 118184092 | rs3782041 | T | C | 0.977 | -7977.388 | -7978.975 | -1.587 | 7.91E-07 |
| 11 | 118191274 | rs7937334 | T | C | 1.021 | -7977.388 | -7978.975 | -1.587 | 7.91E-07 |
| 11 | 120063441 | rs12800456 | G | A | 1.023 | -7977.388 | -7978.975 | -1.587 | 7.91E-07 |
| 11 | 120542279 | rs10790391 | A | G | 0.993 | -7977.388 | -7978.977 | -1.588 | 7.90E-07 |
| 11 | 116505239 | rs11216068 | T | C | 1.024 | -7977.388 | -7978.977 | -1.589 | 7.90E-07 |
| 11 | 116538689 | rs1784128 | A | G | 0.976 | -7977.388 | -7978.977 | -1.589 | 7.90E-07 |
| 11 | 117087118 | rs56061158 | C | G | 0.991 | -7977.388 | -7978.977 | -1.589 | 7.90E-07 |
| 11 | 117404769 | rs12806865 | G | A | 0.979 | -7977.388 | -7978.978 | -1.589 | 7.90E-07 |
| 11 | 117759825 | rs6589632 | T | C | 0.977 | -7977.388 | -7978.978 | -1.589 | 7.90E-07 |
| 11 | 118056722 | rs7935561 | A | G | 0.981 | -7977.388 | -7978.977 | -1.589 | 7.90E-07 |
| 11 | 118362869 | rs537975 | T | C | 0.982 | -7977.388 | -7978.977 | -1.589 | 7.90E-07 |
| 11 | 118869460 | rs3737504 | G | A | 1.022 | -7977.388 | -7978.978 | -1.589 | 7.90E-07 |
| 11 | 118896116 | rs2276071 | C | A | 1.022 | -7977.388 | -7978.977 | -1.589 | 7.90E-07 |
| 11 | 118980039 | rs510435 | T | C | 1.024 | -7977.388 | -7978.977 | -1.589 | 7.90E-07 |
| 11 | 119978951 | rs4938782 | T | C | 0.979 | -7977.388 | -7978.978 | -1.589 | 7.90E-07 |
| 11 | 117083373 | rs111765459 | C | T | 1.019 | -7977.388 | -7978.978 | -1.59 | 7.89E-07 |
| 11 | 117138836 | rs619772 | T | A | 1.024 | -7977.388 | -7978.978 | -1.59 | 7.89E-07 |
| 11 | 118140184 | rs12788933 | T | C | 1.019 | -7977.388 | -7978.978 | -1.59 | 7.89E-07 |
| 11 | 118359161 | rs572126 | G | A | 0.982 | -7977.388 | -7978.979 | -1.59 | 7.89E-07 |
| 11 | 119763417 | rs523223 | A | G | 0.978 | -7977.388 | -7978.979 | -1.59 | 7.89E-07 |
| 11 | 120542603 | rs7934908 | A | G | 0.993 | -7977.388 | -7978.979 | -1.59 | 7.89E-07 |
| 11 | 120551586 | rs10892611 | T | C | 1.002 | -7977.388 | -7978.978 | -1.59 | 7.89E-07 |
| 11 | 121081632 | rs499443 | A | G | 0.996 | -7977.388 | -7978.978 | -1.59 | 7.89E-07 |
| 11 | 121093676 | rs654329 | T | C | 0.996 | -7977.388 | -7978.979 | -1.59 | 7.89E-07 |

| | | | | | | | |
|---|---|---|---|---|---|---|---|
| 11 | 121094021 | rs652916 | T | C | NULL | 0.688903918 | NULL&M_Disease |
| 11 | 121094059 | rs681231 | A | G | NULL | 0.688903918 | NULL&M_Disease |
| 11 | 116924366 | rs11826742 | G | C | NULL | 0.688903918 | NULL&M_Disease |
| 11 | 117552437 | rs11216498 | G | A | NULL | 0.688903918 | NULL&M_Disease |
| 11 | 117867397 | rs2508454 | T | C | NULL | 0.688903918 | NULL&M_Disease |
| 11 | 118114576 | rs56315433 | C | A | NULL | 0.688903918 | NULL&M_Disease |
| 11 | 118114893 | rs56356019 | A | G | NULL | 0.688903918 | NULL&M_Disease |
| 11 | 119974582 | rs61900463 | G | C | NULL | 0.688903918 | NULL&M_Disease |
| 11 | 120228989 | rs4938802 | G | A | NULL | 0.688903918 | NULL&M_Disease |
| 11 | 120542028 | rs10790390 | G | A | NULL | 0.688903918 | NULL&M_Disease |
| 11 | 120542339 | rs10892597 | T | C | NULL | 0.688903918 | NULL&M_Disease |
| 11 | 120542761 | rs7934830 | G | A | NULL | 0.688903918 | NULL&M_Disease |
| 11 | 120546394 | rs12225752 | G | A | NULL | 0.688903918 | NULL&M_Disease |
| 11 | 117145918 | rs644215 | T | A | NULL | 0.688903918 | NULL&M_Disease |
| 11 | 117145949 | rs493139 | T | C | NULL | 0.688903918 | NULL&M_Disease |
| 11 | 117720645 | rs61903804 | G | A | NULL | 0.688903918 | NULL&M_Disease |
| 11 | 118996781 | rs652872 | A | G | NULL | 0.688903918 | NULL&M_Disease |
| 11 | 119000044 | rs616458 | C | G | NULL | 0.688903918 | NULL&M_Disease |
| 11 | 119822450 | rs1046727798 | G | A | NULL | 0.688903918 | NULL&M_Disease |
| 11 | 120148717 | rs12284383 | G | A | NULL | 0.688903918 | NULL&M_Disease |
| 11 | 120542709 | rs7934825 | G | C | NULL | 0.688903918 | NULL&M_Disease |
| 11 | 120770075 | rs11218049 | G | A | NULL | 0.688903918 | NULL&M_Disease |
| 11 | 121074575 | rs571429 | A | C | NULL | 0.688903918 | NULL&M_Disease |
| 11 | 121092625 | rs590021 | T | C | NULL | 0.688903918 | NULL&M_Disease |
| 11 | 121139928 | rs59005463 | A | G | NULL | 0.688903918 | NULL&M_Disease |
| 11 | 117143605 | rs573956 | G | A | NULL | 0.688903918 | NULL&M_Disease |
| 11 | 117144727 | rs495438 | A | C | NULL | 0.688903918 | NULL&M_Disease |
| 11 | 117147987 | rs7925044 | G | A | NULL | 0.688903918 | NULL&M_Disease |
| 11 | 117954250 | rs10160375 | C | T | NULL | 0.688903918 | NULL&M_Disease |
| 11 | 118057227 | rs11216802 | T | C | NULL | 0.688903918 | NULL&M_Disease |
| 11 | 118057320 | rs11216804 | G | A | NULL | 0.688903918 | NULL&M_Disease |
| 11 | 118229061 | rs35865949 | A | T | NULL | 0.688903918 | NULL&M_Disease |
| 11 | 118230739 | rs3782046 | A | G | NULL | 0.688903918 | NULL&M_Disease |
| 11 | 119594525 | rs10790339 | C | T | NULL | 0.688903918 | NULL&M_Disease |
| 11 | 119884981 | rs652824 | C | T | NULL | 0.688903918 | NULL&M_Disease |
| 11 | 120544575 | rs11217958 | T | C | NULL | 0.688903918 | NULL&M_Disease |
| 11 | 120561883 | rs6589833 | G | C | NULL | 0.688903918 | NULL&M_Disease |
| 11 | 120831878 | rs607142 | T | C | NULL | 0.688903918 | NULL&M_Disease |
| 11 | 121090430 | rs514557 | T | C | NULL | 0.688903918 | NULL&M_Disease |
| 11 | 121090812 | rs489436 | A | C | NULL | 0.688903918 | NULL&M_Disease |
| 11 | 121090822 | rs618739 | A | G | NULL | 0.688903918 | NULL&M_Disease |
| 11 | 121090892 | rs543954 | A | G | NULL | 0.688903918 | NULL&M_Disease |
| 11 | 121091188 | rs588550 | A | G | NULL | 0.688903918 | NULL&M_Disease |
| 11 | 121093766 | rs475189 | T | C | NULL | 0.688903918 | NULL&M_Disease |
| 11 | 121098347 | rs11218211 | C | T | NULL | 0.688903918 | NULL&M_Disease |
| 11 | 116919208 | rs61905717 | A | G | NULL | 0.688903918 | NULL&M_Disease |
| 11 | 116919511 | rs17120244 | T | C | NULL | 0.688903918 | NULL&M_Disease |
| 11 | 117093486 | rs4936368 | T | C | NULL | 0.688903918 | NULL&M_Disease |
| 11 | 117093560 | rs4938359 | G | A | NULL | 0.688903918 | NULL&M_Disease |
| 11 | 119968219 | rs1790471 | T | C | NULL | 0.688903918 | NULL&M_Disease |
| 11 | 120562898 | rs9651695 | A | G | NULL | 0.688903918 | NULL&M_Disease |
| 11 | 121093832 | rs557827 | T | A | NULL | 0.688903918 | NULL&M_Disease |
| 11 | 117064506 | rs115787357 | G | T | NULL | 0.688903918 | NULL&M_Disease |
| 11 | 117791979 | rs10892193 | T | C | NULL | 0.688903918 | NULL&M_Disease |
| 11 | 118057244 | rs11216803 | A | C | NULL | 0.688903918 | NULL&M_Disease |
| 11 | 119003322 | rs662069 | A | C | NULL | 0.688903918 | NULL&M_Disease |
| 11 | 119005132 | rs100803 | T | C | NULL | 0.688903918 | NULL&M_Disease |
| 11 | 119005485 | rs7234 | T | C | NULL | 0.688903918 | NULL&M_Disease |
| 11 | 119751585 | rs895649 | G | A | NULL | 0.688903918 | NULL&M_Disease |
| 11 | 119822294 | rs11217593 | A | G | NULL | 0.688903918 | NULL&M_Disease |
| 11 | 120545811 | rs10892599 | T | G | NULL | 0.688903918 | NULL&M_Disease |
| 11 | 120548054 | rs10790393 | T | C | NULL | 0.688903918 | NULL&M_Disease |
| 11 | 121091284 | rs685710 | C | A | NULL | 0.688903918 | NULL&M_Disease |
| 11 | 116562805 | rs1784119 | T | C | NULL | 0.688903918 | NULL&M_Disease |
| 11 | 117729830 | rs7121573 | A | G | NULL | 0.688903918 | NULL&M_Disease |
| 11 | 121100193 | rs10790412 | C | G | NULL | 0.688903918 | NULL&M_Disease |
| 11 | 121143882 | rs7125177 | T | C | NULL | 0.688903918 | NULL&M_Disease |
| 11 | 118115526 | rs2368242 | T | C | NULL | 0.688903918 | NULL&M_Disease |
| 11 | 119004009 | rs672189 | T | G | NULL | 0.688903918 | NULL&M_Disease |
| 11 | 119964758 | rs1787028 | G | C | NULL | 0.688903918 | NULL&M_Disease |
| 11 | 120594279 | rs56129109 | C | A | NULL | 0.688903918 | NULL&M_Disease |
| 11 | 121100429 | rs9787799 | G | A | NULL | 0.688903918 | NULL&M_Disease |
| 11 | 121100648 | rs9787800 | G | A | NULL | 0.688903918 | NULL&M_Disease |
| 11 | 116599504 | rs180357 | C | T | NULL | 0.689974481 | NULL&M_Disease |
| 11 | 116961588 | rs56016514 | A | G | NULL | 0.689974481 | NULL&M_Disease |
| 11 | 116961892 | rs57582454 | A | G | NULL | 0.689974481 | NULL&M_Disease |
| 11 | 116962078 | rs61903423 | A | G | NULL | 0.689974481 | NULL&M_Disease |
| 11 | 116962563 | rs7932655 | A | T | NULL | 0.688903918 | NULL&M_Disease |
| 11 | 117011299 | rs7934690 | A | T | NULL | 0.689974481 | NULL&M_Disease |
| 11 | 118131049 | rs17121997 | T | C | NULL | 0.689974481 | NULL&M_Disease |
| 11 | 118214562 | rs12795615 | T | C | NULL | 0.688903918 | NULL&M_Disease |
| 11 | 120150533 | rs948404 | A | G | NULL | 0.689974481 | NULL&M_Disease |
| 11 | 120542648 | rs7934805 | T | C | NULL | 0.688903918 | NULL&M_Disease |
| 11 | 120542998 | rs7935087 | G | A | NULL | 0.689974481 | NULL&M_Disease |
| 11 | 120544237 | rs12360673 | G | A | NULL | 0.688903918 | NULL&M_Disease |
| 11 | 121100910 | rs9787912 | A | G | NULL | 0.689974481 | NULL&M_Disease |
| 11 | 116314641 | rs550646 | A | G | NULL | 0.689974481 | NULL&M_Disease |
| 11 | 116496310 | rs7114993 | T | G | NULL | 0.689974481 | NULL&M_Disease |
| 11 | 117145815 | rs492151 | G | A | NULL | 0.689974481 | NULL&M_Disease |
| 11 | 117327216 | rs10790189 | C | A | NULL | 0.689974481 | NULL&M_Disease |
| 11 | 118009674 | rs4936424 | T | C | NULL | 0.689974481 | NULL&M_Disease |
| 11 | 118226011 | rs12787202 | T | C | NULL | 0.689974481 | NULL&M_Disease |
| 11 | 118230264 | rs2276418 | T | A | NULL | 0.689974481 | NULL&M_Disease |
| 11 | 118233424 | rs34405565 | T | C | NULL | 0.689974481 | NULL&M_Disease |
| 11 | 120133494 | rs2715883 | G | A | NULL | 0.689974481 | NULL&M_Disease |
| 11 | 120544070 | rs12363187 | T | C | NULL | 0.689974481 | NULL&M_Disease |

| | | | | | | | | | |
|---|---|---|---|---|---|---|---|---|---|
| 11 | 121094021 | rs652916 | T | C | 0.997 | -7977.388 | -7978.979 | -1.59 | 7.89E-07 |
| 11 | 121094059 | rs681231 | A | G | 0.996 | -7977.388 | -7978.978 | -1.59 | 7.89E-07 |
| 11 | 116924366 | rs11826742 | G | C | 0.99 | -7977.388 | -7978.979 | -1.591 | 7.88E-07 |
| 11 | 117552437 | rs11216498 | G | A | 0.981 | -7977.388 | -7978.98 | -1.591 | 7.88E-07 |
| 11 | 117867397 | rs2508454 | T | C | 0.977 | -7977.388 | -7978.979 | -1.591 | 7.88E-07 |
| 11 | 118114576 | rs56315433 | C | A | 1.024 | -7977.388 | -7978.979 | -1.591 | 7.88E-07 |
| 11 | 118114893 | rs56356019 | A | G | 1.024 | -7977.388 | -7978.979 | -1.591 | 7.88E-07 |
| 11 | 119974582 | rs61900463 | G | C | 0.992 | -7977.388 | -7978.98 | -1.591 | 7.88E-07 |
| 11 | 120228989 | rs4938802 | G | A | 0.986 | -7977.388 | -7978.979 | -1.591 | 7.88E-07 |
| 11 | 120542028 | rs10790390 | G | A | 0.993 | -7977.388 | -7978.98 | -1.591 | 7.88E-07 |
| 11 | 120542339 | rs10892597 | T | C | 0.993 | -7977.388 | -7978.98 | -1.591 | 7.88E-07 |
| 11 | 120542761 | rs7934830 | G | A | 0.993 | -7977.388 | -7978.98 | -1.591 | 7.88E-07 |
| 11 | 120546394 | rs12225752 | G | A | 0.993 | -7977.388 | -7978.979 | -1.591 | 7.88E-07 |
| 11 | 117145918 | rs644215 | T | A | 1.024 | -7977.388 | -7978.981 | -1.592 | 7.87E-07 |
| 11 | 117145949 | rs493139 | T | C | 1.024 | -7977.388 | -7978.981 | -1.592 | 7.87E-07 |
| 11 | 117720645 | rs61903804 | G | A | 1.01 | -7977.388 | -7978.98 | -1.592 | 7.87E-07 |
| 11 | 118996781 | rs652872 | A | G | 0.976 | -7977.388 | -7978.981 | -1.592 | 7.87E-07 |
| 11 | 119000044 | rs616458 | C | G | 0.976 | -7977.388 | -7978.98 | -1.592 | 7.87E-07 |
| 11 | 119822450 | rs1046727798 | G | A | 0.998 | -7977.388 | -7978.981 | -1.592 | 7.87E-07 |
| 11 | 120148717 | rs12284383 | G | A | 1.018 | -7977.388 | -7978.981 | -1.592 | 7.87E-07 |
| 11 | 120542709 | rs7934825 | G | C | 0.994 | -7977.388 | -7978.981 | -1.592 | 7.87E-07 |
| 11 | 120770075 | rs11218049 | G | A | 0.992 | -7977.388 | -7978.98 | -1.592 | 7.87E-07 |
| 11 | 121074575 | rs571429 | A | C | 0.998 | -7977.388 | -7978.981 | -1.592 | 7.87E-07 |
| 11 | 121092625 | rs590021 | T | C | 0.997 | -7977.388 | -7978.981 | -1.592 | 7.87E-07 |
| 11 | 121139928 | rs59005463 | A | G | 1.022 | -7977.388 | -7978.98 | -1.592 | 7.87E-07 |
| 11 | 117143605 | rs573956 | G | A | 1.024 | -7977.388 | -7978.982 | -1.593 | 7.86E-07 |
| 11 | 117144727 | rs495438 | A | C | 1.024 | -7977.388 | -7978.982 | -1.593 | 7.86E-07 |
| 11 | 117147987 | rs7925044 | G | A | 1.024 | -7977.388 | -7978.981 | -1.593 | 7.86E-07 |
| 11 | 117954250 | rs10160375 | C | T | 0.977 | -7977.388 | -7978.981 | -1.593 | 7.86E-07 |
| 11 | 118057227 | rs11216802 | T | C | 0.981 | -7977.388 | -7978.981 | -1.593 | 7.86E-07 |
| 11 | 118057320 | rs11216804 | G | A | 0.981 | -7977.388 | -7978.982 | -1.593 | 7.86E-07 |
| 11 | 118229061 | rs35865949 | A | T | 1.009 | -7977.388 | -7978.981 | -1.593 | 7.86E-07 |
| 11 | 118230739 | rs3782046 | A | G | 1.009 | -7977.388 | -7978.981 | -1.593 | 7.86E-07 |
| 11 | 119594525 | rs10790339 | C | T | 0.976 | -7977.388 | -7978.981 | -1.593 | 7.86E-07 |
| 11 | 119884981 | rs652824 | C | T | 0.976 | -7977.388 | -7978.981 | -1.593 | 7.86E-07 |
| 11 | 120544575 | rs11217958 | T | C | 0.994 | -7977.388 | -7978.982 | -1.593 | 7.86E-07 |
| 11 | 120561883 | rs6589833 | G | C | 1.024 | -7977.388 | -7978.982 | -1.593 | 7.86E-07 |
| 11 | 120831878 | rs607142 | T | C | 1.018 | -7977.388 | -7978.982 | -1.593 | 7.86E-07 |
| 11 | 121090430 | rs514557 | T | C | 0.998 | -7977.388 | -7978.981 | -1.593 | 7.86E-07 |
| 11 | 121090812 | rs489436 | A | C | 0.997 | -7977.388 | -7978.981 | -1.593 | 7.86E-07 |
| 11 | 121090822 | rs618739 | A | G | 0.997 | -7977.388 | -7978.981 | -1.593 | 7.86E-07 |
| 11 | 121090892 | rs543954 | A | G | 0.997 | -7977.388 | -7978.981 | -1.593 | 7.86E-07 |
| 11 | 121091188 | rs588550 | A | G | 0.997 | -7977.388 | -7978.981 | -1.593 | 7.86E-07 |
| 11 | 121093766 | rs475189 | T | C | 0.998 | -7977.388 | -7978.982 | -1.593 | 7.86E-07 |
| 11 | 121098347 | rs11218211 | C | T | 0.999 | -7977.388 | -7978.981 | -1.593 | 7.86E-07 |
| 11 | 116919208 | rs61905717 | A | G | 1.016 | -7977.388 | -7978.982 | -1.594 | 7.86E-07 |
| 11 | 116919511 | rs17120244 | T | C | 1.016 | -7977.388 | -7978.982 | -1.594 | 7.86E-07 |
| 11 | 117093486 | rs4936368 | T | C | 1.019 | -7977.388 | -7978.982 | -1.594 | 7.86E-07 |
| 11 | 117093560 | rs4938359 | G | A | 1.019 | -7977.388 | -7978.982 | -1.594 | 7.86E-07 |
| 11 | 119968219 | rs1790471 | T | C | 0.978 | -7977.388 | -7978.982 | -1.594 | 7.86E-07 |
| 11 | 120562898 | rs9651695 | A | G | 1.024 | -7977.388 | -7978.982 | -1.594 | 7.86E-07 |
| 11 | 121093832 | rs557827 | T | A | 0.998 | -7977.388 | -7978.982 | -1.594 | 7.86E-07 |
| 11 | 117064506 | rs115787357 | G | T | 0.993 | -7977.388 | -7978.983 | -1.595 | 7.85E-07 |
| 11 | 117791979 | rs10892193 | T | C | 1.021 | -7977.388 | -7978.983 | -1.595 | 7.85E-07 |
| 11 | 118057244 | rs11216803 | A | C | 0.982 | -7977.388 | -7978.983 | -1.595 | 7.85E-07 |
| 11 | 119003322 | rs662069 | A | C | 0.976 | -7977.388 | -7978.983 | -1.595 | 7.85E-07 |
| 11 | 119005132 | rs100803 | T | C | 0.976 | -7977.388 | -7978.983 | -1.595 | 7.85E-07 |
| 11 | 119005485 | rs7234 | T | C | 0.976 | -7977.388 | -7978.983 | -1.595 | 7.85E-07 |
| 11 | 119751585 | rs895649 | G | A | 0.977 | -7977.388 | -7978.983 | -1.595 | 7.85E-07 |
| 11 | 119822294 | rs11217593 | A | G | 1.001 | -7977.388 | -7978.983 | -1.595 | 7.85E-07 |
| 11 | 120545811 | rs10892599 | T | G | 0.994 | -7977.388 | -7978.984 | -1.595 | 7.85E-07 |
| 11 | 120548054 | rs10790393 | T | C | 0.994 | -7977.388 | -7978.984 | -1.595 | 7.85E-07 |
| 11 | 121091284 | rs685710 | C | A | 0.998 | -7977.388 | -7978.983 | -1.595 | 7.85E-07 |
| 11 | 116562805 | rs1784119 | T | C | 0.977 | -7977.388 | -7978.984 | -1.596 | 7.84E-07 |
| 11 | 117729830 | rs7121573 | A | G | 0.988 | -7977.388 | -7978.985 | -1.596 | 7.84E-07 |
| 11 | 121100193 | rs10790412 | C | G | 0.996 | -7977.388 | -7978.984 | -1.596 | 7.84E-07 |
| 11 | 121143882 | rs7125177 | T | C | 0.976 | -7977.388 | -7978.985 | -1.596 | 7.84E-07 |
| 11 | 118115526 | rs2368242 | T | C | 1.024 | -7977.388 | -7978.985 | -1.597 | 7.83E-07 |
| 11 | 119004009 | rs672189 | T | G | 0.976 | -7977.388 | -7978.985 | -1.597 | 7.83E-07 |
| 11 | 119964758 | rs1787028 | G | C | 0.979 | -7977.388 | -7978.985 | -1.597 | 7.83E-07 |
| 11 | 120594279 | rs56129109 | C | A | 0.991 | -7977.388 | -7978.985 | -1.597 | 7.83E-07 |
| 11 | 121100429 | rs9787799 | G | A | 0.997 | -7977.388 | -7978.986 | -1.597 | 7.83E-07 |
| 11 | 121100648 | rs9787800 | G | A | 0.997 | -7977.388 | -7978.986 | -1.597 | 7.83E-07 |
| 11 | 116599504 | rs180357 | C | T | 1.022 | -7977.388 | -7978.987 | -1.598 | 7.83E-07 |
| 11 | 116961588 | rs56016514 | A | G | 1.016 | -7977.388 | -7978.987 | -1.598 | 7.83E-07 |
| 11 | 116961892 | rs57582454 | A | G | 1.016 | -7977.388 | -7978.987 | -1.598 | 7.83E-07 |
| 11 | 116962078 | rs61903423 | A | G | 1.016 | -7977.388 | -7978.987 | -1.598 | 7.83E-07 |
| 11 | 116962563 | rs7932655 | A | T | 1.016 | -7977.388 | -7978.986 | -1.598 | 7.83E-07 |
| 11 | 117011299 | rs7934690 | A | T | 0.978 | -7977.388 | -7978.987 | -1.598 | 7.83E-07 |
| 11 | 118131049 | rs17121997 | T | C | 1.024 | -7977.388 | -7978.987 | -1.598 | 7.83E-07 |
| 11 | 118214562 | rs12795615 | T | C | 1.007 | -7977.388 | -7978.986 | -1.598 | 7.83E-07 |
| 11 | 120150533 | rs948404 | A | G | 1.017 | -7977.388 | -7978.987 | -1.598 | 7.83E-07 |
| 11 | 120542648 | rs7934805 | T | C | 0.995 | -7977.388 | -7978.986 | -1.598 | 7.83E-07 |
| 11 | 120542998 | rs7935087 | G | A | 0.995 | -7977.388 | -7978.987 | -1.598 | 7.83E-07 |
| 11 | 120544237 | rs12360673 | G | A | 0.995 | -7977.388 | -7978.986 | -1.598 | 7.83E-07 |
| 11 | 121100910 | rs9787912 | A | G | 0.997 | -7977.388 | -7978.987 | -1.598 | 7.83E-07 |
| 11 | 116314641 | rs550646 | A | G | 0.979 | -7977.388 | -7978.987 | -1.599 | 7.82E-07 |
| 11 | 116496310 | rs7114993 | T | G | 0.98 | -7977.388 | -7978.987 | -1.599 | 7.82E-07 |
| 11 | 117145815 | rs492151 | G | A | 1.024 | -7977.388 | -7978.988 | -1.599 | 7.82E-07 |
| 11 | 117327216 | rs10790189 | C | A | 0.977 | -7977.388 | -7978.987 | -1.599 | 7.82E-07 |
| 11 | 118009674 | rs4936424 | T | C | 1.024 | -7977.388 | -7978.987 | -1.599 | 7.82E-07 |
| 11 | 118226011 | rs12787202 | T | C | 1.007 | -7977.388 | -7978.988 | -1.599 | 7.82E-07 |
| 11 | 118230264 | rs2276418 | T | A | 1.007 | -7977.388 | -7978.988 | -1.599 | 7.82E-07 |
| 11 | 118233424 | rs34405565 | T | C | 1.007 | -7977.388 | -7978.988 | -1.599 | 7.82E-07 |
| 11 | 120133494 | rs2715883 | G | A | 1.022 | -7977.388 | -7978.987 | -1.599 | 7.82E-07 |
| 11 | 120544070 | rs12363187 | T | C | 0.995 | -7977.388 | -7978.988 | -1.599 | 7.82E-07 |

| | | | | | | | |
|---|---|---|---|---|---|---|---|
| 11 | 116558525 | rs12799449 | G | A | NULL | 0.689974481 | NULL&M_Disease |
| 11 | 117065007 | rs73008568 | G | C | NULL | 0.689974481 | NULL&M_Disease |
| 11 | 117388932 | rs11216435 | T | C | NULL | 0.689974481 | NULL&M_Disease |
| 11 | 117909799 | rs652107 | G | A | NULL | 0.689974481 | NULL&M_Disease |
| 11 | 118221137 | rs3753060 | T | C | NULL | 0.689974481 | NULL&M_Disease |
| 11 | 118221349 | rs3753059 | C | T | NULL | 0.689974481 | NULL&M_Disease |
| 11 | 118221350 | rs3753058 | T | G | NULL | 0.689974481 | NULL&M_Disease |
| 11 | 118229679 | rs45474998 | A | G | NULL | 0.689974481 | NULL&M_Disease |
| 11 | 118233078 | rs3782045 | G | A | NULL | 0.689974481 | NULL&M_Disease |
| 11 | 120730638 | rs4414232 | C | T | NULL | 0.689974481 | NULL&M_Disease |
| 11 | 121103497 | rs11218215 | T | C | NULL | 0.689974481 | NULL&M_Disease |
| 11 | 121103527 | rs11218216 | C | T | NULL | 0.689974481 | NULL&M_Disease |
| 11 | 121126386 | rs7104501 | T | A | NULL | 0.689974481 | NULL&M_Disease |
| 11 | 117088362 | rs2239012 | G | A | NULL | 0.689974481 | NULL&M_Disease |
| 11 | 118111346 | rs10892228 | G | A | NULL | 0.689974481 | NULL&M_Disease |
| 11 | 118222634 | rs35902434 | C | T | NULL | 0.689974481 | NULL&M_Disease |
| 11 | 118239266 | rs2276421 | A | G | NULL | 0.689974481 | NULL&M_Disease |
| 11 | 118585146 | rs112799932 | G | T | NULL | 0.689974481 | NULL&M_Disease |
| 11 | 119004026 | rs537153 | T | C | NULL | 0.689974481 | NULL&M_Disease |
| 11 | 119382758 | rs7937927 | C | T | NULL | 0.689974481 | NULL&M_Disease |
| 11 | 120238534 | rs7131046 | A | G | NULL | 0.689974481 | NULL&M_Disease |
| 11 | 120830625 | rs688893 | C | T | NULL | 0.689974481 | NULL&M_Disease |
| 11 | 121101217 | rs11218212 | T | A | NULL | 0.689974481 | NULL&M_Disease |
| 11 | 116558427 | rs12799766 | A | G | NULL | 0.689974481 | NULL&M_Disease |
| 11 | 116570968 | rs1145194 | A | C | NULL | 0.689974481 | NULL&M_Disease |
| 11 | 118225452 | rs36061634 | A | T | NULL | 0.689974481 | NULL&M_Disease |
| 11 | 118228114 | rs35305304 | T | C | NULL | 0.689974481 | NULL&M_Disease |
| 11 | 118228624 | rs7110197 | C | G | NULL | 0.689974481 | NULL&M_Disease |
| 11 | 118234191 | rs3825054 | A | G | NULL | 0.689974481 | NULL&M_Disease |
| 11 | 118836821 | rs7112758 | A | G | NULL | 0.689974481 | NULL&M_Disease |
| 11 | 118848315 | rs58745438 | C | T | NULL | 0.689974481 | NULL&M_Disease |
| 11 | 119175075 | rs2509671 | A | C | NULL | 0.689974481 | NULL&M_Disease |
| 11 | 119865509 | rs657107 | T | C | NULL | 0.689974481 | NULL&M_Disease |
| 11 | 120239937 | rs11217843 | G | A | NULL | 0.689974481 | NULL&M_Disease |
| 11 | 120543750 | rs10892598 | T | G | NULL | 0.689974481 | NULL&M_Disease |
| 11 | 120769383 | rs57943482 | C | T | NULL | 0.689974481 | NULL&M_Disease |
| 11 | 121144114 | rs12363999 | T | C | NULL | 0.689974481 | NULL&M_Disease |
| 11 | 121145524 | rs2115926 | A | G | NULL | 0.689974481 | NULL&M_Disease |
| 11 | 116558245 | rs61906115 | A | T | NULL | 0.689974481 | NULL&M_Disease |
| 11 | 117721360 | rs79717347 | G | T | NULL | 0.689974481 | NULL&M_Disease |
| 11 | 118054431 | rs12295634 | T | C | NULL | 0.689974481 | NULL&M_Disease |
| 11 | 118867803 | rs4938618 | C | G | NULL | 0.689974481 | NULL&M_Disease |
| 11 | 118869539 | rs11822103 | A | G | NULL | 0.689974481 | NULL&M_Disease |
| 11 | 119368006 | rs10892372 | A | G | NULL | 0.689974481 | NULL&M_Disease |
| 11 | 119702432 | rs477756 | C | T | NULL | 0.689974481 | NULL&M_Disease |
| 11 | 120543440 | rs10082698 | G | A | NULL | 0.689974481 | NULL&M_Disease |
| 11 | 120543669 | rs11217957 | G | A | NULL | 0.689974481 | NULL&M_Disease |
| 11 | 120544735 | rs11217959 | C | T | NULL | 0.689974481 | NULL&M_Disease |
| 11 | 120545098 | rs61902672 | T | C | NULL | 0.689974481 | NULL&M_Disease |
| 11 | 120545103 | rs11217960 | T | C | NULL | 0.689974481 | NULL&M_Disease |
| 11 | 120545219 | rs11217961 | G | A | NULL | 0.689974481 | NULL&M_Disease |
| 11 | 120545502 | rs11217963 | T | C | NULL | 0.689974481 | NULL&M_Disease |
| 11 | 120545503 | rs11217964 | G | A | NULL | 0.689974481 | NULL&M_Disease |
| 11 | 120545929 | rs10892600 | G | A | NULL | 0.689974481 | NULL&M_Disease |
| 11 | 120546027 | rs10892601 | G | A | NULL | 0.689974481 | NULL&M_Disease |
| 11 | 120546082 | rs10892602 | A | G | NULL | 0.689974481 | NULL&M_Disease |
| 11 | 120546119 | rs10892603 | C | T | NULL | 0.689974481 | NULL&M_Disease |
| 11 | 120546223 | rs10892604 | T | C | NULL | 0.689974481 | NULL&M_Disease |
| 11 | 120546287 | rs12225737 | C | A | NULL | 0.689974481 | NULL&M_Disease |
| 11 | 120546914 | rs11217965 | T | C | NULL | 0.689974481 | NULL&M_Disease |
| 11 | 120546953 | rs10892605 | T | C | NULL | 0.689974481 | NULL&M_Disease |
| 11 | 120547052 | rs4628679 | C | T | NULL | 0.689974481 | NULL&M_Disease |
| 11 | 120547109 | rs4367954 | G | A | NULL | 0.689974481 | NULL&M_Disease |
| 11 | 120547112 | rs4268524 | T | C | NULL | 0.689974481 | NULL&M_Disease |
| 11 | 120547174 | rs4256992 | A | G | NULL | 0.689974481 | NULL&M_Disease |
| 11 | 120547912 | rs10502240 | T | C | NULL | 0.689974481 | NULL&M_Disease |
| 11 | 120548076 | rs10790394 | C | G | NULL | 0.689974481 | NULL&M_Disease |
| 11 | 120548318 | rs11607732 | G | A | NULL | 0.689974481 | NULL&M_Disease |
| 11 | 120548353 | rs11603586 | T | C | NULL | 0.689974481 | NULL&M_Disease |
| 11 | 121132370 | rs1941302 | T | A | NULL | 0.689974481 | NULL&M_Disease |
| 11 | 121132553 | rs7130538 | T | G | NULL | 0.689974481 | NULL&M_Disease |
| 11 | 116458862 | rs11216022 | T | C | NULL | 0.689974481 | NULL&M_Disease |
| 11 | 117014863 | rs76904522 | A | G | NULL | 0.689974481 | NULL&M_Disease |
| 11 | 117088994 | rs2239011 | G | A | NULL | 0.689974481 | NULL&M_Disease |
| 11 | 117089006 | rs2239010 | C | G | NULL | 0.689974481 | NULL&M_Disease |
| 11 | 117090545 | rs3882895 | C | T | NULL | 0.689974481 | NULL&M_Disease |
| 11 | 117139121 | rs558590 | G | A | NULL | 0.689974481 | NULL&M_Disease |
| 11 | 117554061 | rs1942923 | C | T | NULL | 0.689974481 | NULL&M_Disease |
| 11 | 117700323 | rs4472969 | A | G | NULL | 0.689974481 | NULL&M_Disease |
| 11 | 117775847 | rs4340086 | T | A | NULL | 0.689974481 | NULL&M_Disease |
| 11 | 117965530 | rs2276122 | G | A | NULL | 0.689974481 | NULL&M_Disease |
| 11 | 119194035 | rs10892348 | T | C | NULL | 0.689974481 | NULL&M_Disease |
| 11 | 120548902 | rs61902674 | A | C | NULL | 0.689974481 | NULL&M_Disease |
| 11 | 120769382 | rs57151589 | T | A | NULL | 0.689974481 | NULL&M_Disease |
| 11 | 120833767 | rs642280 | C | T | NULL | 0.689974481 | NULL&M_Disease |
| 11 | 117686194 | rs954319 | A | G | NULL | 0.689974481 | NULL&M_Disease |
| 11 | 118234678 | rs17122113 | G | A | NULL | 0.689974481 | NULL&M_Disease |
| 11 | 119866151 | rs633621 | A | G | NULL | 0.689974481 | NULL&M_Disease |
| 11 | 120221068 | rs1812974 | T | C | NULL | 0.689974481 | NULL&M_Disease |
| 11 | 120544352 | rs61902671 | A | C | NULL | 0.689974481 | NULL&M_Disease |
| 11 | 120548768 | rs61601696 | C | T | NULL | 0.689974481 | NULL&M_Disease |
| 11 | 120551626 | rs11217970 | A | G | NULL | 0.689974481 | NULL&M_Disease |
| 11 | 120551964 | rs12224951 | T | C | NULL | 0.689974481 | NULL&M_Disease |
| 11 | 117338555 | rs7113263 | A | G | NULL | 0.689974481 | NULL&M_Disease |
| 11 | 117716151 | rs12098917 | C | T | NULL | 0.689974481 | NULL&M_Disease |
| 11 | 117876035 | rs11216666 | C | T | NULL | 0.689974481 | NULL&M_Disease |

| | | | | | | | | | |
|---|---|---|---|---|---|---|---|---|---|
| 11 | 116558525 | rs12799449 | G | A | 1.02 | -7977.388 | -7978.989 | -1.6 | 7.81E-07 |
| 11 | 117065007 | rs73008568 | G | C | 0.993 | -7977.388 | -7978.988 | -1.6 | 7.81E-07 |
| 11 | 117388932 | rs11216435 | T | C | 1.018 | -7977.388 | -7978.988 | -1.6 | 7.81E-07 |
| 11 | 117909799 | rs652107 | G | A | 1.011 | -7977.388 | -7978.988 | -1.6 | 7.81E-07 |
| 11 | 118221137 | rs3753060 | T | C | 1.007 | -7977.388 | -7978.989 | -1.6 | 7.81E-07 |
| 11 | 118221349 | rs3753059 | C | T | 1.007 | -7977.388 | -7978.989 | -1.6 | 7.81E-07 |
| 11 | 118221350 | rs3753058 | T | G | 1.007 | -7977.388 | -7978.989 | -1.6 | 7.81E-07 |
| 11 | 118229679 | rs45474998 | A | G | 1.007 | -7977.388 | -7978.989 | -1.6 | 7.81E-07 |
| 11 | 118233078 | rs3782045 | G | A | 1.007 | -7977.388 | -7978.989 | -1.6 | 7.81E-07 |
| 11 | 120730638 | rs4414232 | C | T | 1.024 | -7977.388 | -7978.988 | -1.6 | 7.81E-07 |
| 11 | 121103497 | rs11218215 | T | C | 1.001 | -7977.388 | -7978.989 | -1.6 | 7.81E-07 |
| 11 | 121103527 | rs11218216 | C | T | 1.002 | -7977.388 | -7978.988 | -1.6 | 7.81E-07 |
| 11 | 121126386 | rs7104501 | T | A | 0.977 | -7977.388 | -7978.988 | -1.6 | 7.81E-07 |
| 11 | 117088362 | rs2239012 | G | A | 1.018 | -7977.388 | -7978.989 | -1.601 | 7.80E-07 |
| 11 | 118111346 | rs10892228 | G | A | 1.022 | -7977.388 | -7978.989 | -1.601 | 7.80E-07 |
| 11 | 118222634 | rs35902434 | C | T | 1.007 | -7977.388 | -7978.989 | -1.601 | 7.80E-07 |
| 11 | 118239266 | rs2276421 | A | G | 1.007 | -7977.388 | -7978.99 | -1.601 | 7.80E-07 |
| 11 | 118585146 | rs112799932 | G | T | 1.02 | -7977.388 | -7978.989 | -1.601 | 7.80E-07 |
| 11 | 119004026 | rs537153 | T | C | 0.977 | -7977.388 | -7978.989 | -1.601 | 7.80E-07 |
| 11 | 119382758 | rs7937927 | C | T | 1.024 | -7977.388 | -7978.989 | -1.601 | 7.80E-07 |
| 11 | 120238534 | rs7131046 | A | G | 1.024 | -7977.388 | -7978.989 | -1.601 | 7.80E-07 |
| 11 | 120830625 | rs688893 | C | T | 1.017 | -7977.388 | -7978.99 | -1.601 | 7.80E-07 |
| 11 | 121101217 | rs11218212 | T | A | 1 | -7977.388 | -7978.989 | -1.601 | 7.80E-07 |
| 11 | 116558427 | rs12799766 | A | G | 1.02 | -7977.388 | -7978.99 | -1.602 | 7.79E-07 |
| 11 | 116570968 | rs1145194 | A | C | 0.977 | -7977.388 | -7978.99 | -1.602 | 7.79E-07 |
| 11 | 118225452 | rs36061634 | A | T | 1.007 | -7977.388 | -7978.99 | -1.602 | 7.79E-07 |
| 11 | 118228114 | rs35305304 | T | C | 1.007 | -7977.388 | -7978.99 | -1.602 | 7.79E-07 |
| 11 | 118228624 | rs7110197 | C | G | 1.007 | -7977.388 | -7978.99 | -1.602 | 7.79E-07 |
| 11 | 118234191 | rs3825054 | A | G | 1.007 | -7977.388 | -7978.99 | -1.602 | 7.79E-07 |
| 11 | 118836821 | rs7112758 | A | G | 1.024 | -7977.388 | -7978.99 | -1.602 | 7.79E-07 |
| 11 | 118848315 | rs58745438 | C | T | 1.024 | -7977.388 | -7978.99 | -1.602 | 7.79E-07 |
| 11 | 119175075 | rs2509671 | A | C | 1.022 | -7977.388 | -7978.991 | -1.602 | 7.79E-07 |
| 11 | 119865509 | rs657107 | T | C | 0.995 | -7977.388 | -7978.99 | -1.602 | 7.79E-07 |
| 11 | 120239937 | rs11217843 | G | A | 0.987 | -7977.388 | -7978.99 | -1.602 | 7.79E-07 |
| 11 | 120543750 | rs10892598 | T | G | 0.996 | -7977.388 | -7978.99 | -1.602 | 7.79E-07 |
| 11 | 120769383 | rs57943482 | C | T | 0.995 | -7977.388 | -7978.99 | -1.602 | 7.79E-07 |
| 11 | 121144114 | rs12363999 | T | C | 0.977 | -7977.388 | -7978.991 | -1.602 | 7.79E-07 |
| 11 | 121145524 | rs2115926 | A | G | 0.977 | -7977.388 | -7978.99 | -1.602 | 7.79E-07 |
| 11 | 116558245 | rs61906115 | A | T | 1.02 | -7977.388 | -7978.991 | -1.603 | 7.79E-07 |
| 11 | 117721360 | rs79717347 | G | T | 1.009 | -7977.388 | -7978.991 | -1.603 | 7.79E-07 |
| 11 | 118054431 | rs12295634 | T | C | 0.982 | -7977.388 | -7978.992 | -1.603 | 7.79E-07 |
| 11 | 118867803 | rs4938618 | C | G | 1.021 | -7977.388 | -7978.991 | -1.603 | 7.79E-07 |
| 11 | 118869539 | rs11822103 | A | G | 1.021 | -7977.388 | -7978.991 | -1.603 | 7.79E-07 |
| 11 | 119368006 | rs10892372 | A | G | 0.977 | -7977.388 | -7978.991 | -1.603 | 7.79E-07 |
| 11 | 119702432 | rs477756 | C | T | 0.978 | -7977.388 | -7978.991 | -1.603 | 7.79E-07 |
| 11 | 120543440 | rs10082698 | G | A | 0.997 | -7977.388 | -7978.991 | -1.603 | 7.79E-07 |
| 11 | 120543669 | rs11217957 | G | A | 0.997 | -7977.388 | -7978.991 | -1.603 | 7.79E-07 |
| 11 | 120544735 | rs11217959 | C | T | 0.997 | -7977.388 | -7978.991 | -1.603 | 7.79E-07 |
| 11 | 120545098 | rs61902672 | T | C | 0.997 | -7977.388 | -7978.991 | -1.603 | 7.79E-07 |
| 11 | 120545103 | rs11217960 | T | C | 0.997 | -7977.388 | -7978.991 | -1.603 | 7.79E-07 |
| 11 | 120545219 | rs11217961 | G | A | 0.997 | -7977.388 | -7978.991 | -1.603 | 7.79E-07 |
| 11 | 120545502 | rs11217963 | T | C | 0.997 | -7977.388 | -7978.991 | -1.603 | 7.79E-07 |
| 11 | 120545503 | rs11217964 | G | A | 0.997 | -7977.388 | -7978.991 | -1.603 | 7.79E-07 |
| 11 | 120545929 | rs10892600 | G | A | 0.997 | -7977.388 | -7978.991 | -1.603 | 7.79E-07 |
| 11 | 120546027 | rs10892601 | G | A | 0.997 | -7977.388 | -7978.991 | -1.603 | 7.79E-07 |
| 11 | 120546082 | rs10892602 | A | G | 0.997 | -7977.388 | -7978.991 | -1.603 | 7.79E-07 |
| 11 | 120546119 | rs10892603 | C | T | 0.997 | -7977.388 | -7978.991 | -1.603 | 7.79E-07 |
| 11 | 120546223 | rs10892604 | T | C | 0.997 | -7977.388 | -7978.991 | -1.603 | 7.79E-07 |
| 11 | 120546287 | rs12225737 | C | A | 0.997 | -7977.388 | -7978.991 | -1.603 | 7.79E-07 |
| 11 | 120546914 | rs11217965 | T | C | 0.997 | -7977.388 | -7978.991 | -1.603 | 7.79E-07 |
| 11 | 120546953 | rs10892605 | T | C | 0.997 | -7977.388 | -7978.991 | -1.603 | 7.79E-07 |
| 11 | 120547052 | rs4628679 | C | T | 0.997 | -7977.388 | -7978.991 | -1.603 | 7.79E-07 |
| 11 | 120547109 | rs4367954 | G | A | 0.997 | -7977.388 | -7978.991 | -1.603 | 7.79E-07 |
| 11 | 120547112 | rs4268524 | T | C | 0.997 | -7977.388 | -7978.991 | -1.603 | 7.79E-07 |
| 11 | 120547174 | rs4256992 | A | G | 0.997 | -7977.388 | -7978.991 | -1.603 | 7.79E-07 |
| 11 | 120547912 | rs10502240 | T | C | 0.997 | -7977.388 | -7978.991 | -1.603 | 7.79E-07 |
| 11 | 120548076 | rs10790394 | C | G | 0.997 | -7977.388 | -7978.991 | -1.603 | 7.79E-07 |
| 11 | 120548318 | rs11607732 | G | A | 0.997 | -7977.388 | -7978.991 | -1.603 | 7.79E-07 |
| 11 | 120548353 | rs11603586 | T | C | 0.997 | -7977.388 | -7978.991 | -1.603 | 7.79E-07 |
| 11 | 121132370 | rs1941302 | T | A | 0.977 | -7977.388 | -7978.991 | -1.603 | 7.79E-07 |
| 11 | 121132553 | rs7130538 | T | G | 0.977 | -7977.388 | -7978.991 | -1.603 | 7.79E-07 |
| 11 | 116458862 | rs11216022 | T | C | 1.024 | -7977.388 | -7978.992 | -1.604 | 7.78E-07 |
| 11 | 117014863 | rs76904522 | A | G | 0.991 | -7977.388 | -7978.992 | -1.604 | 7.78E-07 |
| 11 | 117088994 | rs2239011 | G | A | 1.018 | -7977.388 | -7978.992 | -1.604 | 7.78E-07 |
| 11 | 117089006 | rs2239010 | C | G | 1.018 | -7977.388 | -7978.992 | -1.604 | 7.78E-07 |
| 11 | 117090545 | rs3882895 | C | T | 1.018 | -7977.388 | -7978.992 | -1.604 | 7.78E-07 |
| 11 | 117139121 | rs558590 | G | A | 1.024 | -7977.388 | -7978.992 | -1.604 | 7.78E-07 |
| 11 | 117554061 | rs1942923 | C | T | 0.982 | -7977.388 | -7978.992 | -1.604 | 7.78E-07 |
| 11 | 117700323 | rs4472969 | A | G | 0.986 | -7977.388 | -7978.993 | -1.604 | 7.78E-07 |
| 11 | 117775847 | rs4340086 | T | A | 1.023 | -7977.388 | -7978.992 | -1.604 | 7.78E-07 |
| 11 | 117965530 | rs2276122 | G | A | 0.977 | -7977.388 | -7978.992 | -1.604 | 7.78E-07 |
| 11 | 119194035 | rs10892348 | T | C | 1.02 | -7977.388 | -7978.992 | -1.604 | 7.78E-07 |
| 11 | 120548902 | rs61902674 | A | C | 0.997 | -7977.388 | -7978.992 | -1.604 | 7.78E-07 |
| 11 | 120769382 | rs57151589 | T | A | 0.996 | -7977.388 | -7978.992 | -1.604 | 7.78E-07 |
| 11 | 120833767 | rs642280 | C | T | 1.017 | -7977.388 | -7978.992 | -1.604 | 7.78E-07 |
| 11 | 117686194 | rs954319 | A | G | 1.024 | -7977.388 | -7978.993 | -1.605 | 7.77E-07 |
| 11 | 118234678 | rs17122113 | G | A | 1.006 | -7977.388 | -7978.993 | -1.605 | 7.77E-07 |
| 11 | 119866151 | rs633621 | A | G | 0.997 | -7977.388 | -7978.993 | -1.605 | 7.77E-07 |
| 11 | 120221068 | rs1812974 | T | C | 0.988 | -7977.388 | -7978.993 | -1.605 | 7.77E-07 |
| 11 | 120544352 | rs61902671 | A | C | 0.998 | -7977.388 | -7978.993 | -1.605 | 7.77E-07 |
| 11 | 120548768 | rs61601696 | C | T | 0.998 | -7977.388 | -7978.994 | -1.605 | 7.77E-07 |
| 11 | 120551626 | rs11217970 | A | G | 0.999 | -7977.388 | -7978.993 | -1.605 | 7.77E-07 |
| 11 | 120551964 | rs12224951 | T | C | 1.001 | -7977.388 | -7978.993 | -1.605 | 7.77E-07 |
| 11 | 117338555 | rs7113263 | A | G | 0.991 | -7977.388 | -7978.994 | -1.606 | 7.76E-07 |
| 11 | 117716151 | rs12098917 | C | T | 1.008 | -7977.388 | -7978.994 | -1.606 | 7.76E-07 |
| 11 | 117876035 | rs11216666 | C | T | 0.989 | -7977.388 | -7978.994 | -1.606 | 7.76E-07 |

| | | | | | | | |
|---|---|---|---|---|---|---|---|
| 11 | 117960427 | rs1893317 | A | G | NULL | 0.689974481 | NULL&M_Disease |
| 11 | 118214067 | rs3181259 | A | G | NULL | 0.689974481 | NULL&M_Disease |
| 11 | 119767614 | rs504485 | A | C | NULL | 0.689974481 | NULL&M_Disease |
| 11 | 119967615 | rs1632452 | C | T | NULL | 0.689974481 | NULL&M_Disease |
| 11 | 120137818 | rs2015914 | A | G | NULL | 0.689974481 | NULL&M_Disease |
| 11 | 120552193 | rs12224987 | T | C | NULL | 0.689974481 | NULL&M_Disease |
| 11 | 120552212 | rs11217972 | T | C | NULL | 0.689974481 | NULL&M_Disease |
| 11 | 117700725 | rs2004507 | G | T | NULL | 0.689974481 | NULL&M_Disease |
| 11 | 118213632 | rs2239695 | T | C | NULL | 0.689974481 | NULL&M_Disease |
| 11 | 118365956 | rs3132786 | C | T | NULL | 0.689974481 | NULL&M_Disease |
| 11 | 118896115 | rs2276072 | G | A | NULL | 0.689974481 | NULL&M_Disease |
| 11 | 119975789 | rs12802344 | G | T | NULL | 0.689974481 | NULL&M_Disease |
| 11 | 120145180 | rs12800691 | C | G | NULL | 0.689974481 | NULL&M_Disease |
| 11 | 121133458 | rs4936600 | C | T | NULL | 0.689974481 | NULL&M_Disease |
| 11 | 116464209 | rs76376430 | T | C | NULL | 0.689974481 | NULL&M_Disease |
| 11 | 116496187 | rs7114844 | A | G | NULL | 0.691043012 | NULL&M_Disease |
| 11 | 117212908 | rs495473 | G | A | NULL | 0.689974481 | NULL&M_Disease |
| 11 | 117954309 | rs656199 | A | G | NULL | 0.689974481 | NULL&M_Disease |
| 11 | 117990553 | rs12366111 | C | T | NULL | 0.689974481 | NULL&M_Disease |
| 11 | 118054128 | rs12274740 | C | T | NULL | 0.689974481 | NULL&M_Disease |
| 11 | 118109097 | rs12275546 | G | A | NULL | 0.691043012 | NULL&M_Disease |
| 11 | 118125672 | rs10790248 | G | A | NULL | 0.691043012 | NULL&M_Disease |
| 11 | 120702378 | rs58359402 | G | A | NULL | 0.691043012 | NULL&M_Disease |
| 11 | 116705516 | rs525028 | G | A | NULL | 0.691043012 | NULL&M_Disease |
| 11 | 116958943 | rs7930409 | G | A | NULL | 0.691043012 | NULL&M_Disease |
| 11 | 116959039 | rs7930783 | A | G | NULL | 0.691043012 | NULL&M_Disease |
| 11 | 117649675 | rs7924993 | A | G | NULL | 0.691043012 | NULL&M_Disease |
| 11 | 119384079 | rs606630 | T | C | NULL | 0.691043012 | NULL&M_Disease |
| 11 | 119633448 | rs35633147 | T | C | NULL | 0.691043012 | NULL&M_Disease |
| 11 | 120836290 | rs684613 | T | C | NULL | 0.691043012 | NULL&M_Disease |
| 11 | 116957219 | rs61903419 | A | G | NULL | 0.691043012 | NULL&M_Disease |
| 11 | 117775760 | rs4259862 | C | T | NULL | 0.691043012 | NULL&M_Disease |
| 11 | 117963766 | rs11216740 | A | G | NULL | 0.691043012 | NULL&M_Disease |
| 11 | 119384351 | rs4938661 | C | G | NULL | 0.691043012 | NULL&M_Disease |
| 11 | 119971934 | rs12798518 | A | G | NULL | 0.691043012 | NULL&M_Disease |
| 11 | 120184678 | rs4938796 | A | G | NULL | 0.691043012 | NULL&M_Disease |
| 11 | 120767546 | rs12574454 | G | A | NULL | 0.691043012 | NULL&M_Disease |
| 11 | 120770004 | rs7106442 | T | C | NULL | 0.691043012 | NULL&M_Disease |
| 11 | 116962759 | rs7946219 | T | C | NULL | 0.691043012 | NULL&M_Disease |
| 11 | 116963180 | rs7946729 | T | C | NULL | 0.691043012 | NULL&M_Disease |
| 11 | 117318324 | rs12419440 | C | T | NULL | 0.691043012 | NULL&M_Disease |
| 11 | 119238381 | rs10892354 | T | C | NULL | 0.691043012 | NULL&M_Disease |
| 11 | 119367674 | rs10892371 | G | A | NULL | 0.691043012 | NULL&M_Disease |
| 11 | 120150430 | rs1941408 | A | G | NULL | 0.691043012 | NULL&M_Disease |
| 11 | 120598054 | rs61901402 | A | G | NULL | 0.691043012 | NULL&M_Disease |
| 11 | 120731495 | rs7937438 | G | A | NULL | 0.691043012 | NULL&M_Disease |
| 11 | 121006345 | rs11218155 | A | T | NULL | 0.691043012 | NULL&M_Disease |
| 11 | 116375253 | rs7107783 | G | A | NULL | 0.691043012 | NULL&M_Disease |
| 11 | 116728277 | rs11216162 | A | G | NULL | 0.691043012 | NULL&M_Disease |
| 11 | 116919677 | rs61905718 | T | C | NULL | 0.691043012 | NULL&M_Disease |
| 11 | 118126675 | rs4938492 | T | C | NULL | 0.691043012 | NULL&M_Disease |
| 11 | 118129340 | rs1104542 | C | T | NULL | 0.691043012 | NULL&M_Disease |
| 11 | 119703507 | rs7395055 | C | T | NULL | 0.691043012 | NULL&M_Disease |
| 11 | 120273224 | rs7948928 | G | C | NULL | 0.691043012 | NULL&M_Disease |
| 11 | 116214668 | rs71478236 | A | G | NULL | 0.691043012 | NULL&M_Disease |
| 11 | 116251739 | rs1790047 | C | T | NULL | 0.691043012 | NULL&M_Disease |
| 11 | 117801262 | rs10892198 | G | C | NULL | 0.691043012 | NULL&M_Disease |
| 11 | 118846466 | rs4938602 | A | G | NULL | 0.691043012 | NULL&M_Disease |
| 11 | 119180316 | rs7914 | A | G | NULL | 0.691043012 | NULL&M_Disease |
| 11 | 119384277 | rs34980489 | A | T | NULL | 0.691043012 | NULL&M_Disease |
| 11 | 120000608 | rs12790585 | A | G | NULL | 0.691043012 | NULL&M_Disease |
| 11 | 120133107 | rs2155808 | T | A | NULL | 0.691043012 | NULL&M_Disease |
| 11 | 120530316 | rs4936538 | T | C | NULL | 0.691043012 | NULL&M_Disease |
| 11 | 120764218 | rs7103569 | G | A | NULL | 0.691043012 | NULL&M_Disease |
| 11 | 116314402 | rs480101 | T | C | NULL | 0.691043012 | NULL&M_Disease |
| 11 | 116518438 | rs1240770 | T | G | NULL | 0.691043012 | NULL&M_Disease |
| 11 | 116960403 | rs11823889 | A | G | NULL | 0.691043012 | NULL&M_Disease |
| 11 | 117361861 | rs597686 | T | C | NULL | 0.691043012 | NULL&M_Disease |
| 11 | 117706942 | rs73018483 | G | C | NULL | 0.691043012 | NULL&M_Disease |
| 11 | 117758431 | rs10892189 | T | C | NULL | 0.691043012 | NULL&M_Disease |
| 11 | 119784330 | rs611478 | A | G | NULL | 0.691043012 | NULL&M_Disease |
| 11 | 120146360 | rs4938793 | T | G | NULL | 0.691043012 | NULL&M_Disease |
| 11 | 120224544 | rs10892563 | C | T | NULL | 0.691043012 | NULL&M_Disease |
| 11 | 120594813 | rs4612814 | G | A | NULL | 0.691043012 | NULL&M_Disease |
| 11 | 117399918 | rs4938402 | A | G | NULL | 0.691043012 | NULL&M_Disease |
| 11 | 118126904 | rs4936425 | C | T | NULL | 0.691043012 | NULL&M_Disease |
| 11 | 118240715 | rs12792929 | C | G | NULL | 0.691043012 | NULL&M_Disease |
| 11 | 119366595 | rs617711 | T | C | NULL | 0.691043012 | NULL&M_Disease |
| 11 | 120726044 | rs7342199 | A | G | NULL | 0.691043012 | NULL&M_Disease |
| 11 | 120726066 | rs7342200 | A | G | NULL | 0.691043012 | NULL&M_Disease |
| 11 | 117087792 | rs2238006 | C | A | NULL | 0.691043012 | NULL&M_Disease |
| 11 | 117291793 | rs1902625 | C | T | NULL | 0.691043012 | NULL&M_Disease |
| 11 | 117305461 | rs7944427 | T | C | NULL | 0.691043012 | NULL&M_Disease |
| 11 | 117339276 | rs7114223 | A | G | NULL | 0.691043012 | NULL&M_Disease |
| 11 | 118110202 | rs11216827 | A | T | NULL | 0.691043012 | NULL&M_Disease |
| 11 | 118993549 | rs2508950 | T | C | NULL | 0.691043012 | NULL&M_Disease |
| 11 | 119364461 | rs656924 | T | G | NULL | 0.691043012 | NULL&M_Disease |
| 11 | 119860539 | rs10892508 | G | C | NULL | 0.691043012 | NULL&M_Disease |
| 11 | 119927888 | rs1545509 | A | G | NULL | 0.691043012 | NULL&M_Disease |
| 11 | 120133012 | rs2847494 | T | C | NULL | 0.691043012 | NULL&M_Disease |
| 11 | 116586423 | rs7926828 | C | T | NULL | 0.691043012 | NULL&M_Disease |
| 11 | 117328700 | rs648384 | G | A | NULL | 0.692109504 | NULL&M_Disease |
| 11 | 117550767 | rs10892158 | C | T | NULL | 0.691043012 | NULL&M_Disease |
| 11 | 118136745 | rs3759003 | G | T | NULL | 0.691043012 | NULL&M_Disease |
| 11 | 118149122 | rs1970750 | T | C | NULL | 0.691043012 | NULL&M_Disease |
| 11 | 116578223 | rs10892019 | T | C | NULL | 0.692109504 | NULL&M_Disease |

| | | | | | | | | | |
|---|---|---|---|---|---|---|---|---|---|
| 11 | 117960427 | rs1893317 | A | G | 1.022 | -7977.388 | -7978.995 | -1.606 | 7.76E-07 |
| 11 | 118214067 | rs3181259 | A | G | 1.006 | -7977.388 | -7978.994 | -1.606 | 7.76E-07 |
| 11 | 119767614 | rs504485 | A | C | 0.978 | -7977.388 | -7978.995 | -1.606 | 7.76E-07 |
| 11 | 119967615 | rs1632452 | C | T | 0.979 | -7977.388 | -7978.994 | -1.606 | 7.76E-07 |
| 11 | 120137818 | rs2015914 | A | G | 1.022 | -7977.388 | -7978.994 | -1.606 | 7.76E-07 |
| 11 | 120552193 | rs12224987 | T | C | 1 | -7977.388 | -7978.994 | -1.606 | 7.76E-07 |
| 11 | 120552212 | rs11217972 | T | C | 1 | -7977.388 | -7978.994 | -1.606 | 7.76E-07 |
| 11 | 117700725 | rs2004507 | G | T | 0.977 | -7977.388 | -7978.995 | -1.607 | 7.76E-07 |
| 11 | 118213632 | rs2239695 | T | C | 1.005 | -7977.388 | -7978.996 | -1.607 | 7.76E-07 |
| 11 | 118365956 | rs3132786 | C | T | 0.983 | -7977.388 | -7978.995 | -1.607 | 7.76E-07 |
| 11 | 118896115 | rs2276072 | G | A | 1.021 | -7977.388 | -7978.996 | -1.607 | 7.76E-07 |
| 11 | 119975789 | rs12802344 | G | T | 0.995 | -7977.388 | -7978.995 | -1.607 | 7.76E-07 |
| 11 | 120145180 | rs12800691 | C | G | 1.006 | -7977.388 | -7978.995 | -1.607 | 7.76E-07 |
| 11 | 121133458 | rs4936600 | C | T | 0.977 | -7977.388 | -7978.995 | -1.607 | 7.76E-07 |
| 11 | 116464209 | rs76376430 | T | C | 1.006 | -7977.388 | -7978.996 | -1.608 | 7.75E-07 |
| 11 | 116496187 | rs7114844 | A | G | 0.981 | -7977.388 | -7978.997 | -1.608 | 7.75E-07 |
| 11 | 117212908 | rs495473 | G | A | 0.984 | -7977.388 | -7978.996 | -1.608 | 7.75E-07 |
| 11 | 117954309 | rs656199 | A | G | 1.016 | -7977.388 | -7978.996 | -1.608 | 7.75E-07 |
| 11 | 117990553 | rs12366111 | C | T | 1.017 | -7977.388 | -7978.996 | -1.608 | 7.75E-07 |
| 11 | 118054128 | rs12274740 | C | T | 0.982 | -7977.388 | -7978.996 | -1.608 | 7.75E-07 |
| 11 | 118109097 | rs12275546 | G | A | 1.022 | -7977.388 | -7978.996 | -1.608 | 7.75E-07 |
| 11 | 118125672 | rs10790248 | G | A | 1.024 | -7977.388 | -7978.996 | -1.608 | 7.75E-07 |
| 11 | 120702378 | rs58359402 | G | A | 0.977 | -7977.388 | -7978.997 | -1.608 | 7.75E-07 |
| 11 | 116705516 | rs525028 | G | A | 0.978 | -7977.388 | -7978.997 | -1.609 | 7.74E-07 |
| 11 | 116958943 | rs7930409 | G | A | 1.015 | -7977.388 | -7978.997 | -1.609 | 7.74E-07 |
| 11 | 116959039 | rs7930783 | A | G | 1.015 | -7977.388 | -7978.997 | -1.609 | 7.74E-07 |
| 11 | 117649675 | rs7924993 | A | G | 1.008 | -7977.388 | -7978.998 | -1.609 | 7.74E-07 |
| 11 | 119384079 | rs606630 | T | C | 0.978 | -7977.388 | -7978.997 | -1.609 | 7.74E-07 |
| 11 | 119633448 | rs35633147 | T | C | 1.023 | -7977.388 | -7978.997 | -1.609 | 7.74E-07 |
| 11 | 120836290 | rs684613 | T | C | 1.017 | -7977.388 | -7978.997 | -1.609 | 7.74E-07 |
| 11 | 116957219 | rs61903419 | A | G | 1.015 | -7977.388 | -7978.999 | -1.61 | 7.73E-07 |
| 11 | 117775760 | rs4259862 | C | T | 1.023 | -7977.388 | -7978.999 | -1.61 | 7.73E-07 |
| 11 | 117963766 | rs11216740 | A | G | 0.977 | -7977.388 | -7978.998 | -1.61 | 7.73E-07 |
| 11 | 119384351 | rs4938661 | C | G | 1.016 | -7977.388 | -7978.998 | -1.61 | 7.73E-07 |
| 11 | 119971934 | rs12798518 | A | G | 1.017 | -7977.388 | -7978.998 | -1.61 | 7.73E-07 |
| 11 | 120184678 | rs4938796 | A | G | 0.977 | -7977.388 | -7978.998 | -1.61 | 7.73E-07 |
| 11 | 120767546 | rs12574454 | G | A | 0.996 | -7977.388 | -7978.998 | -1.61 | 7.73E-07 |
| 11 | 120770004 | rs7106442 | T | C | 0.996 | -7977.388 | -7978.998 | -1.61 | 7.73E-07 |
| 11 | 116962759 | rs7946219 | T | C | 1.015 | -7977.388 | -7979 | -1.611 | 7.72E-07 |
| 11 | 116963180 | rs7946729 | T | C | 1.015 | -7977.388 | -7979 | -1.611 | 7.72E-07 |
| 11 | 117318324 | rs12419440 | C | T | 0.978 | -7977.388 | -7978.999 | -1.611 | 7.72E-07 |
| 11 | 119238381 | rs10892354 | T | C | 0.977 | -7977.388 | -7978.999 | -1.611 | 7.72E-07 |
| 11 | 119367674 | rs10892371 | G | A | 1.023 | -7977.388 | -7978.999 | -1.611 | 7.72E-07 |
| 11 | 120150430 | rs1941408 | A | G | 1.016 | -7977.388 | -7978.999 | -1.611 | 7.72E-07 |
| 11 | 120598054 | rs61901402 | A | G | 0.997 | -7977.388 | -7978.999 | -1.611 | 7.72E-07 |
| 11 | 120731495 | rs7937438 | G | A | 1.023 | -7977.388 | -7978.999 | -1.611 | 7.72E-07 |
| 11 | 121006345 | rs11218155 | A | T | 0.986 | -7977.388 | -7978.999 | -1.611 | 7.72E-07 |
| 11 | 116375253 | rs7107783 | G | A | 0.977 | -7977.388 | -7979 | -1.612 | 7.72E-07 |
| 11 | 116728277 | rs11216162 | A | G | 1.012 | -7977.388 | -7979.001 | -1.612 | 7.72E-07 |
| 11 | 116919677 | rs61905718 | T | C | 1.003 | -7977.388 | -7979 | -1.612 | 7.72E-07 |
| 11 | 118126675 | rs4938492 | T | C | 1.024 | -7977.388 | -7979 | -1.612 | 7.72E-07 |
| 11 | 118129340 | rs1104542 | C | T | 1.024 | -7977.388 | -7979 | -1.612 | 7.72E-07 |
| 11 | 119703507 | rs7395055 | C | T | 0.977 | -7977.388 | -7979 | -1.612 | 7.72E-07 |
| 11 | 120273224 | rs7948928 | G | C | 1.023 | -7977.388 | -7979 | -1.612 | 7.72E-07 |
| 11 | 116214668 | rs71478236 | A | G | 1.002 | -7977.388 | -7979.001 | -1.613 | 7.71E-07 |
| 11 | 116251739 | rs1790047 | C | T | 1.02 | -7977.388 | -7979.001 | -1.613 | 7.71E-07 |
| 11 | 117801262 | rs10892198 | G | C | 0.977 | -7977.388 | -7979.002 | -1.613 | 7.71E-07 |
| 11 | 118846466 | rs4938602 | A | G | 1.023 | -7977.388 | -7979.001 | -1.613 | 7.71E-07 |
| 11 | 119180316 | rs7914 | A | G | 1.02 | -7977.388 | -7979.001 | -1.613 | 7.71E-07 |
| 11 | 119384277 | rs34980489 | A | T | 0.994 | -7977.388 | -7979.001 | -1.613 | 7.71E-07 |
| 11 | 120000608 | rs12790585 | A | G | 1.021 | -7977.388 | -7979.002 | -1.613 | 7.71E-07 |
| 11 | 120133107 | rs2155808 | T | A | 1.021 | -7977.388 | -7979.001 | -1.613 | 7.71E-07 |
| 11 | 120530316 | rs4936538 | T | C | 1.023 | -7977.388 | -7979.001 | -1.613 | 7.71E-07 |
| 11 | 120764218 | rs7103569 | G | A | 1.001 | -7977.388 | -7979.001 | -1.613 | 7.71E-07 |
| 11 | 116314402 | rs480101 | T | C | 0.989 | -7977.388 | -7979.002 | -1.614 | 7.70E-07 |
| 11 | 116518438 | rs1240770 | T | G | 1.016 | -7977.388 | -7979.002 | -1.614 | 7.70E-07 |
| 11 | 116960403 | rs11823889 | A | G | 1.015 | -7977.388 | -7979.002 | -1.614 | 7.70E-07 |
| 11 | 117361861 | rs597686 | T | C | 0.998 | -7977.388 | -7979.002 | -1.614 | 7.70E-07 |
| 11 | 117706942 | rs73018483 | G | C | 0.994 | -7977.388 | -7979.003 | -1.614 | 7.70E-07 |
| 11 | 117758431 | rs10892189 | T | C | 1.023 | -7977.388 | -7979.002 | -1.614 | 7.70E-07 |
| 11 | 119784330 | rs611478 | A | G | 0.978 | -7977.388 | -7979.002 | -1.614 | 7.70E-07 |
| 11 | 120146360 | rs4938793 | T | G | 1.015 | -7977.388 | -7979.003 | -1.614 | 7.70E-07 |
| 11 | 120224544 | rs10892563 | C | T | 1.023 | -7977.388 | -7979.002 | -1.614 | 7.70E-07 |
| 11 | 120594813 | rs4612814 | G | A | 0.995 | -7977.388 | -7979.002 | -1.614 | 7.70E-07 |
| 11 | 117399918 | rs4938402 | A | G | 1.016 | -7977.388 | -7979.004 | -1.615 | 7.69E-07 |
| 11 | 118126904 | rs4936425 | C | T | 1.023 | -7977.388 | -7979.004 | -1.615 | 7.69E-07 |
| 11 | 118240715 | rs12792929 | C | G | 1.001 | -7977.388 | -7979.003 | -1.615 | 7.69E-07 |
| 11 | 119366595 | rs617711 | T | C | 0.978 | -7977.388 | -7979.004 | -1.615 | 7.69E-07 |
| 11 | 120726044 | rs7342199 | A | G | 0.977 | -7977.388 | -7979.004 | -1.615 | 7.69E-07 |
| 11 | 120726066 | rs7342200 | A | G | 0.977 | -7977.388 | -7979.003 | -1.615 | 7.69E-07 |
| 11 | 117087792 | rs2238006 | C | A | 1.017 | -7977.388 | -7979.005 | -1.616 | 7.69E-07 |
| 11 | 117291793 | rs1902625 | C | T | 0.977 | -7977.388 | -7979.005 | -1.616 | 7.69E-07 |
| 11 | 117305461 | rs7944427 | T | C | 0.987 | -7977.388 | -7979.005 | -1.616 | 7.69E-07 |
| 11 | 117339276 | rs7114223 | A | G | 0.993 | -7977.388 | -7979.004 | -1.616 | 7.69E-07 |
| 11 | 118110202 | rs11216827 | A | T | 1.023 | -7977.388 | -7979.004 | -1.616 | 7.69E-07 |
| 11 | 118993549 | rs2508950 | T | C | 0.977 | -7977.388 | -7979.005 | -1.616 | 7.69E-07 |
| 11 | 119364461 | rs656924 | T | G | 0.978 | -7977.388 | -7979.004 | -1.616 | 7.69E-07 |
| 11 | 119860539 | rs10892508 | G | C | 0.978 | -7977.388 | -7979.005 | -1.616 | 7.69E-07 |
| 11 | 119927888 | rs1545509 | A | G | 1.022 | -7977.388 | -7979.004 | -1.616 | 7.69E-07 |
| 11 | 120133012 | rs2847494 | T | C | 1.021 | -7977.388 | -7979.005 | -1.616 | 7.69E-07 |
| 11 | 116586423 | rs7926828 | C | T | 1.015 | -7977.388 | -7979.006 | -1.617 | 7.68E-07 |
| 11 | 117328700 | rs648384 | G | A | 0.977 | -7977.388 | -7979.006 | -1.617 | 7.68E-07 |
| 11 | 117550767 | rs10892158 | C | T | 0.983 | -7977.388 | -7979.005 | -1.617 | 7.68E-07 |
| 11 | 118136745 | rs3759003 | G | T | 0.978 | -7977.388 | -7979.006 | -1.617 | 7.68E-07 |
| 11 | 118149122 | rs1970750 | T | C | 0.978 | -7977.388 | -7979.005 | -1.617 | 7.68E-07 |
| 11 | 116578223 | rs10892019 | T | C | 1.014 | -7977.388 | -7979.006 | -1.618 | 7.67E-07 |

| | | | | | | | |
|---|---|---|---|---|---|---|---|
| 11 | 117303242 | rs60580523 | T | C | NULL | 0.691043012 | NULL&M_Disease |
| 11 | 118363424 | rs477106 | A | C | NULL | 0.691043012 | NULL&M_Disease |
| 11 | 119362151 | rs521053 | T | C | NULL | 0.692109504 | NULL&M_Disease |
| 11 | 120235560 | rs10892565 | C | T | NULL | 0.691043012 | NULL&M_Disease |
| 11 | 116218868 | rs2581477 | A | G | NULL | 0.692109504 | NULL&M_Disease |
| 11 | 116218926 | rs693448 | C | A | NULL | 0.692109504 | NULL&M_Disease |
| 11 | 116700142 | rs2854117 | T | C | NULL | 0.692109504 | NULL&M_Disease |
| 11 | 116721201 | rs11216161 | T | C | NULL | 0.692109504 | NULL&M_Disease |
| 11 | 117091227 | rs3882897 | A | G | NULL | 0.692109504 | NULL&M_Disease |
| 11 | 117261488 | rs897837 | G | C | NULL | 0.692109504 | NULL&M_Disease |
| 11 | 117791353 | rs7930658 | G | A | NULL | 0.692109504 | NULL&M_Disease |
| 11 | 119358442 | rs534279 | G | A | NULL | 0.692109504 | NULL&M_Disease |
| 11 | 119846524 | rs633223 | A | G | NULL | 0.692109504 | NULL&M_Disease |
| 11 | 120269308 | rs7933086 | A | G | NULL | 0.692109504 | NULL&M_Disease |
| 11 | 116223405 | rs579706 | G | T | NULL | 0.692109504 | NULL&M_Disease |
| 11 | 116496623 | rs2156122 | A | C | NULL | 0.692109504 | NULL&M_Disease |
| 11 | 118068100 | rs1039377879 | T | A | NULL | 0.692109504 | NULL&M_Disease |
| 11 | 118136871 | rs3759005 | G | A | NULL | 0.692109504 | NULL&M_Disease |
| 11 | 119905908 | rs1530919 | A | G | NULL | 0.692109504 | NULL&M_Disease |
| 11 | 120193956 | rs12799299 | T | C | NULL | 0.692109504 | NULL&M_Disease |
| 11 | 120520364 | rs2000872 | G | A | NULL | 0.692109504 | NULL&M_Disease |
| 11 | 117876198 | rs11216667 | A | C | NULL | 0.692109504 | NULL&M_Disease |
| 11 | 117904029 | rs1786177 | A | G | NULL | 0.692109504 | NULL&M_Disease |
| 11 | 117913024 | rs638415 | C | A | NULL | 0.692109504 | NULL&M_Disease |
| 11 | 119366779 | rs618627 | C | T | NULL | 0.692109504 | NULL&M_Disease |
| 11 | 119857163 | rs2155169 | G | A | NULL | 0.692109504 | NULL&M_Disease |
| 11 | 120050515 | rs531260 | C | T | NULL | 0.692109504 | NULL&M_Disease |
| 11 | 116721826 | rs10047459 | C | T | NULL | 0.692109504 | NULL&M_Disease |
| 11 | 117904561 | rs1783850 | G | C | NULL | 0.692109504 | NULL&M_Disease |
| 11 | 118113629 | rs10790247 | T | G | NULL | 0.692109504 | NULL&M_Disease |
| 11 | 119364555 | rs668678 | A | G | NULL | 0.692109504 | NULL&M_Disease |
| 11 | 116566538 | rs111218525 | T | C | NULL | 0.692109504 | NULL&M_Disease |
| 11 | 117794456 | rs146556537 | A | G | NULL | 0.692109504 | NULL&M_Disease |
| 11 | 118072181 | rs1715441 | T | C | NULL | 0.692109504 | NULL&M_Disease |
| 11 | 119863827 | rs664511 | C | G | NULL | 0.692109504 | NULL&M_Disease |
| 11 | 120537660 | rs61900956 | G | A | NULL | 0.692109504 | NULL&M_Disease |
| 11 | 120827021 | rs1789661 | T | C | NULL | 0.692109504 | NULL&M_Disease |
| 11 | 120853317 | rs645545 | A | T | NULL | 0.692109504 | NULL&M_Disease |
| 11 | 117336280 | rs544153 | A | G | NULL | 0.692109504 | NULL&M_Disease |
| 11 | 117699586 | rs10892181 | G | T | NULL | 0.692109504 | NULL&M_Disease |
| 11 | 118112433 | rs1793145 | C | T | NULL | 0.692109504 | NULL&M_Disease |
| 11 | 116560967 | rs3017229 | A | G | NULL | 0.692109504 | NULL&M_Disease |
| 11 | 116749878 | rs11216172 | G | A | NULL | 0.692109504 | NULL&M_Disease |
| 11 | 117701783 | rs7130098 | C | G | NULL | 0.692109504 | NULL&M_Disease |
| 11 | 117710118 | rs562089 | C | A | NULL | 0.692109504 | NULL&M_Disease |
| 11 | 118096891 | rs1715437 | C | T | NULL | 0.692109504 | NULL&M_Disease |
| 11 | 119786013 | rs531411 | A | G | NULL | 0.692109504 | NULL&M_Disease |
| 11 | 120896545 | rs10892662 | T | G | NULL | 0.692109504 | NULL&M_Disease |
| 11 | 121112938 | rs12276890 | G | C | NULL | 0.692109504 | NULL&M_Disease |
| 11 | 116728630 | rs12225230 | C | G | NULL | 0.692109504 | NULL&M_Disease |
| 11 | 117906882 | rs653330 | A | G | NULL | 0.692109504 | NULL&M_Disease |
| 11 | 118120104 | rs2853005 | C | T | NULL | 0.692109504 | NULL&M_Disease |
| 11 | 119365287 | rs689580 | A | G | NULL | 0.692109504 | NULL&M_Disease |
| 11 | 119365632 | rs603075 | C | G | NULL | 0.692109504 | NULL&M_Disease |
| 11 | 119549825 | rs906830 | T | C | NULL | 0.692109504 | NULL&M_Disease |
| 11 | 119784002 | rs481400 | G | A | NULL | 0.692109504 | NULL&M_Disease |
| 11 | 119907605 | rs12420530 | T | G | NULL | 0.692109504 | NULL&M_Disease |
| 11 | 120538183 | rs11217951 | T | C | NULL | 0.692109504 | NULL&M_Disease |
| 11 | 120538217 | rs11217953 | A | G | NULL | 0.692109504 | NULL&M_Disease |
| 11 | 116314960 | rs547761 | C | T | NULL | 0.692109504 | NULL&M_Disease |
| 11 | 116398417 | rs4936354 | T | G | NULL | 0.692109504 | NULL&M_Disease |
| 11 | 116502834 | rs4502019 | G | C | NULL | 0.692109504 | NULL&M_Disease |
| 11 | 117694089 | rs635706 | G | A | NULL | 0.692109504 | NULL&M_Disease |
| 11 | 117791929 | rs11216622 | A | C | NULL | 0.692109504 | NULL&M_Disease |
| 11 | 120063980 | rs880487 | C | T | NULL | 0.692109504 | NULL&M_Disease |
| 11 | 120538215 | rs11217952 | T | G | NULL | 0.692109504 | NULL&M_Disease |
| 11 | 120628922 | rs1079071 | C | T | NULL | 0.692109504 | NULL&M_Disease |
| 11 | 120793387 | rs2850805 | G | A | NULL | 0.692109504 | NULL&M_Disease |
| 11 | 117304196 | rs2242133 | T | C | NULL | 0.692109504 | NULL&M_Disease |
| 11 | 119368120 | rs10892373 | T | G | NULL | 0.693173949 | NULL&M_Disease |
| 11 | 120070770 | rs2368940 | A | G | NULL | 0.693173949 | NULL&M_Disease |
| 11 | 120538307 | rs11217954 | C | G | NULL | 0.692109504 | NULL&M_Disease |
| 11 | 121125452 | rs7106859 | A | C | NULL | 0.693173949 | NULL&M_Disease |
| 11 | 117729616 | rs7121195 | A | G | NULL | 0.693173949 | NULL&M_Disease |
| 11 | 118119073 | rs11216832 | A | C | NULL | 0.693173949 | NULL&M_Disease |
| 11 | 118119957 | rs11216833 | T | G | NULL | 0.693173949 | NULL&M_Disease |
| 11 | 119364296 | rs67744990 | T | G | NULL | 0.693173949 | NULL&M_Disease |
| 11 | 120047745 | rs488672 | C | T | NULL | 0.693173949 | NULL&M_Disease |
| 11 | 120274501 | rs10892570 | T | G | NULL | 0.693173949 | NULL&M_Disease |
| 11 | 117554747 | rs12279393 | C | T | NULL | 0.693173949 | NULL&M_Disease |
| 11 | 117722292 | rs4525269 | C | T | NULL | 0.693173949 | NULL&M_Disease |
| 11 | 117722501 | rs4525270 | C | T | NULL | 0.693173949 | NULL&M_Disease |
| 11 | 119177938 | rs11217234 | G | A | NULL | 0.693173949 | NULL&M_Disease |
| 11 | 116223171 | rs556194 | C | A | NULL | 0.693173949 | NULL&M_Disease |
| 11 | 116329917 | rs518684 | A | C | NULL | 0.693173949 | NULL&M_Disease |
| 11 | 120150500 | rs1941409 | G | A | NULL | 0.693173949 | NULL&M_Disease |
| 11 | 120224650 | rs10892564 | G | A | NULL | 0.693173949 | NULL&M_Disease |
| 11 | 118106875 | rs1793139 | G | A | NULL | 0.693173949 | NULL&M_Disease |
| 11 | 119367238 | rs620566 | A | C | NULL | 0.693173949 | NULL&M_Disease |
| 11 | 120274211 | rs4938805 | G | A | NULL | 0.693173949 | NULL&M_Disease |
| 11 | 121164440 | rs751417 | G | A | NULL | 0.693173949 | NULL&M_Disease |
| 11 | 116705278 | rs12721030 | T | C | NULL | 0.693173949 | NULL&M_Disease |
| 11 | 117239173 | rs7115177 | C | A | NULL | 0.693173949 | NULL&M_Disease |
| 11 | 117468794 | rs622860 | G | A | NULL | 0.693173949 | NULL&M_Disease |
| 11 | 118066609 | rs12289120 | T | C | NULL | 0.693173949 | NULL&M_Disease |
| 11 | 118831824 | rs7113731 | G | A | NULL | 0.693173949 | NULL&M_Disease |

| | | | | | | | | | |
|---|---|---|---|---|---|---|---|---|---|
| 11 | 117303242 | rs60580523 | T | C | 0.987 | -7977.388 | -7979.006 | -1.618 | 7.67E-07 |
| 11 | 118363424 | rs477106 | A | C | 0.985 | -7977.388 | -7979.006 | -1.618 | 7.67E-07 |
| 11 | 119362151 | rs521053 | T | C | 0.978 | -7977.388 | -7979.006 | -1.618 | 7.67E-07 |
| 11 | 120235560 | rs10892565 | C | T | 0.989 | -7977.388 | -7979.006 | -1.618 | 7.67E-07 |
| 11 | 116218868 | rs2581477 | A | G | 1.022 | -7977.388 | -7979.008 | -1.619 | 7.66E-07 |
| 11 | 116218926 | rs693448 | C | A | 1.022 | -7977.388 | -7979.008 | -1.619 | 7.66E-07 |
| 11 | 116700142 | rs2854117 | T | C | 0.98 | -7977.388 | -7979.007 | -1.619 | 7.66E-07 |
| 11 | 116721201 | rs11216161 | T | C | 0.994 | -7977.388 | -7979.007 | -1.619 | 7.66E-07 |
| 11 | 117091227 | rs3882897 | A | G | 1.017 | -7977.388 | -7979.007 | -1.619 | 7.66E-07 |
| 11 | 117261488 | rs897837 | G | C | 0.989 | -7977.388 | -7979.007 | -1.619 | 7.66E-07 |
| 11 | 117791353 | rs7930658 | G | A | 1.019 | -7977.388 | -7979.007 | -1.619 | 7.66E-07 |
| 11 | 119358442 | rs534279 | G | A | 1.008 | -7977.388 | -7979.007 | -1.619 | 7.66E-07 |
| 11 | 119846524 | rs633223 | A | G | 0.977 | -7977.388 | -7979.007 | -1.619 | 7.66E-07 |
| 11 | 120269308 | rs7933086 | A | G | 1.023 | -7977.388 | -7979.007 | -1.619 | 7.66E-07 |
| 11 | 116223405 | rs579706 | G | T | 1.022 | -7977.388 | -7979.008 | -1.62 | 7.65E-07 |
| 11 | 116496623 | rs2156122 | A | C | 0.981 | -7977.388 | -7979.008 | -1.62 | 7.65E-07 |
| 11 | 118068100 | rs1039377879 | T | A | 0.98 | -7977.388 | -7979.008 | -1.62 | 7.65E-07 |
| 11 | 118136871 | rs3759005 | G | A | 0.978 | -7977.388 | -7979.009 | -1.62 | 7.65E-07 |
| 11 | 119905908 | rs1530919 | A | G | 1.022 | -7977.388 | -7979.009 | -1.62 | 7.65E-07 |
| 11 | 120193956 | rs12799299 | T | C | 0.977 | -7977.388 | -7979.009 | -1.62 | 7.65E-07 |
| 11 | 120520364 | rs2000872 | G | A | 1.023 | -7977.388 | -7979.008 | -1.62 | 7.65E-07 |
| 11 | 117876198 | rs11216667 | A | C | 0.991 | -7977.388 | -7979.01 | -1.621 | 7.65E-07 |
| 11 | 117904029 | rs1786177 | A | G | 1.009 | -7977.388 | -7979.009 | -1.621 | 7.65E-07 |
| 11 | 117913024 | rs638415 | C | A | 1.009 | -7977.388 | -7979.009 | -1.621 | 7.65E-07 |
| 11 | 119366779 | rs618627 | C | T | 0.978 | -7977.388 | -7979.009 | -1.621 | 7.65E-07 |
| 11 | 119857163 | rs2155169 | G | A | 1.022 | -7977.388 | -7979.009 | -1.621 | 7.65E-07 |
| 11 | 120050515 | rs531260 | C | T | 0.98 | -7977.388 | -7979.01 | -1.621 | 7.65E-07 |
| 11 | 116721826 | rs10047459 | C | T | 0.996 | -7977.388 | -7979.011 | -1.622 | 7.64E-07 |
| 11 | 117904561 | rs1783850 | G | C | 1.009 | -7977.388 | -7979.011 | -1.622 | 7.64E-07 |
| 11 | 118113629 | rs10790247 | T | G | 1.021 | -7977.388 | -7979.011 | -1.622 | 7.64E-07 |
| 11 | 119364555 | rs668678 | A | G | 0.978 | -7977.388 | -7979.01 | -1.622 | 7.64E-07 |
| 11 | 116566538 | rs111218525 | T | C | 1.013 | -7977.388 | -7979.011 | -1.623 | 7.63E-07 |
| 11 | 117794456 | rs146556537 | A | G | 1.019 | -7977.388 | -7979.011 | -1.623 | 7.63E-07 |
| 11 | 118072181 | rs1715441 | T | C | 0.98 | -7977.388 | -7979.012 | -1.623 | 7.63E-07 |
| 11 | 119863827 | rs664511 | C | G | 0.999 | -7977.388 | -7979.011 | -1.623 | 7.63E-07 |
| 11 | 120537660 | rs61900956 | G | A | 0.995 | -7977.388 | -7979.011 | -1.623 | 7.63E-07 |
| 11 | 120827021 | rs1789661 | T | C | 1.013 | -7977.388 | -7979.011 | -1.623 | 7.63E-07 |
| 11 | 120853317 | rs645545 | A | T | 1.016 | -7977.388 | -7979.011 | -1.623 | 7.63E-07 |
| 11 | 117336280 | rs544153 | A | G | 0.997 | -7977.388 | -7979.012 | -1.624 | 7.62E-07 |
| 11 | 117699586 | rs10892181 | G | T | 0.978 | -7977.388 | -7979.012 | -1.624 | 7.62E-07 |
| 11 | 118112433 | rs1793145 | C | T | 1.021 | -7977.388 | -7979.012 | -1.624 | 7.62E-07 |
| 11 | 116560967 | rs3017229 | A | G | 0.978 | -7977.388 | -7979.013 | -1.625 | 7.62E-07 |
| 11 | 116749878 | rs11216172 | G | A | 1.015 | -7977.388 | -7979.013 | -1.625 | 7.62E-07 |
| 11 | 117701783 | rs7130098 | C | G | 0.99 | -7977.388 | -7979.013 | -1.625 | 7.62E-07 |
| 11 | 117710118 | rs562089 | C | A | 1.012 | -7977.388 | -7979.013 | -1.625 | 7.62E-07 |
| 11 | 118096891 | rs1715437 | C | T | 1.021 | -7977.388 | -7979.014 | -1.625 | 7.62E-07 |
| 11 | 119786013 | rs531411 | A | G | 0.979 | -7977.388 | -7979.013 | -1.625 | 7.62E-07 |
| 11 | 120896545 | rs10892662 | T | G | 1.009 | -7977.388 | -7979.014 | -1.625 | 7.62E-07 |
| 11 | 121112938 | rs12276890 | G | C | 1.016 | -7977.388 | -7979.013 | -1.625 | 7.62E-07 |
| 11 | 116728630 | rs12225230 | C | G | 1.011 | -7977.388 | -7979.015 | -1.626 | 7.61E-07 |
| 11 | 117906882 | rs653330 | A | G | 1.008 | -7977.388 | -7979.014 | -1.626 | 7.61E-07 |
| 11 | 118120104 | rs2853005 | C | T | 1.021 | -7977.388 | -7979.014 | -1.626 | 7.61E-07 |
| 11 | 119365287 | rs689580 | A | G | 0.978 | -7977.388 | -7979.014 | -1.626 | 7.61E-07 |
| 11 | 119365632 | rs603075 | C | G | 0.978 | -7977.388 | -7979.014 | -1.626 | 7.61E-07 |
| 11 | 119549825 | rs906830 | T | C | 0.978 | -7977.388 | -7979.014 | -1.626 | 7.61E-07 |
| 11 | 119784002 | rs481400 | G | A | 0.979 | -7977.388 | -7979.014 | -1.626 | 7.61E-07 |
| 11 | 119907605 | rs12420530 | T | G | 1.021 | -7977.388 | -7979.015 | -1.626 | 7.61E-07 |
| 11 | 120538183 | rs11217951 | T | C | 0.998 | -7977.388 | -7979.015 | -1.626 | 7.61E-07 |
| 11 | 120538217 | rs11217953 | A | G | 0.997 | -7977.388 | -7979.015 | -1.626 | 7.61E-07 |
| 11 | 116314960 | rs547761 | C | T | 0.98 | -7977.388 | -7979.015 | -1.627 | 7.60E-07 |
| 11 | 116398417 | rs4936354 | T | G | 1.021 | -7977.388 | -7979.015 | -1.627 | 7.60E-07 |
| 11 | 116502834 | rs4502019 | G | C | 0.982 | -7977.388 | -7979.015 | -1.627 | 7.60E-07 |
| 11 | 117694089 | rs635706 | G | A | 1.015 | -7977.388 | -7979.015 | -1.627 | 7.60E-07 |
| 11 | 117791929 | rs11216622 | A | C | 1.019 | -7977.388 | -7979.016 | -1.627 | 7.60E-07 |
| 11 | 120063980 | rs880487 | C | T | 1.021 | -7977.388 | -7979.016 | -1.627 | 7.60E-07 |
| 11 | 120538215 | rs11217952 | T | G | 0.997 | -7977.388 | -7979.015 | -1.627 | 7.60E-07 |
| 11 | 120628922 | rs1079071 | C | T | 0.979 | -7977.388 | -7979.015 | -1.627 | 7.60E-07 |
| 11 | 120793387 | rs2850805 | G | A | 0.978 | -7977.388 | -7979.015 | -1.627 | 7.60E-07 |
| 11 | 117304196 | rs2242133 | T | C | 0.988 | -7977.388 | -7979.016 | -1.628 | 7.59E-07 |
| 11 | 119368120 | rs10892373 | T | G | 1.023 | -7977.388 | -7979.016 | -1.628 | 7.59E-07 |
| 11 | 120070770 | rs2368940 | A | G | 1.019 | -7977.388 | -7979.017 | -1.628 | 7.59E-07 |
| 11 | 120538307 | rs11217954 | C | G | 0.998 | -7977.388 | -7979.016 | -1.628 | 7.59E-07 |
| 11 | 121125452 | rs7106859 | A | C | 0.978 | -7977.388 | -7979.017 | -1.628 | 7.59E-07 |
| 11 | 117729616 | rs7121195 | A | G | 1.003 | -7977.388 | -7979.017 | -1.629 | 7.59E-07 |
| 11 | 118119073 | rs11216832 | A | C | 1.021 | -7977.388 | -7979.018 | -1.629 | 7.59E-07 |
| 11 | 118119957 | rs11216833 | T | G | 1.021 | -7977.388 | -7979.018 | -1.629 | 7.59E-07 |
| 11 | 119364296 | rs67744990 | T | G | 0.988 | -7977.388 | -7979.017 | -1.629 | 7.59E-07 |
| 11 | 120047745 | rs488672 | C | T | 0.98 | -7977.388 | -7979.018 | -1.629 | 7.59E-07 |
| 11 | 120274501 | rs10892570 | T | G | 1.022 | -7977.388 | -7979.018 | -1.629 | 7.59E-07 |
| 11 | 117554747 | rs12279393 | C | T | 0.984 | -7977.388 | -7979.018 | -1.63 | 7.58E-07 |
| 11 | 117722292 | rs4525269 | C | T | 1.001 | -7977.388 | -7979.018 | -1.63 | 7.58E-07 |
| 11 | 117722501 | rs4525270 | C | T | 1.001 | -7977.388 | -7979.018 | -1.63 | 7.58E-07 |
| 11 | 119177938 | rs11217234 | G | A | 0.981 | -7977.388 | -7979.018 | -1.63 | 7.58E-07 |
| 11 | 116223171 | rs556194 | C | A | 1.022 | -7977.388 | -7979.019 | -1.631 | 7.57E-07 |
| 11 | 116329917 | rs518684 | A | C | 1.023 | -7977.388 | -7979.019 | -1.631 | 7.57E-07 |
| 11 | 120150500 | rs1941409 | G | A | 1.02 | -7977.388 | -7979.019 | -1.631 | 7.57E-07 |
| 11 | 120224650 | rs10892564 | G | A | 1.022 | -7977.388 | -7979.019 | -1.631 | 7.57E-07 |
| 11 | 118106875 | rs1793139 | G | A | 1.02 | -7977.388 | -7979.02 | -1.632 | 7.56E-07 |
| 11 | 119367238 | rs620566 | A | C | 0.978 | -7977.388 | -7979.02 | -1.632 | 7.56E-07 |
| 11 | 120274211 | rs4938805 | G | A | 1.022 | -7977.388 | -7979.021 | -1.632 | 7.56E-07 |
| 11 | 121164440 | rs751417 | G | A | 1.021 | -7977.388 | -7979.02 | -1.632 | 7.56E-07 |
| 11 | 116705278 | rs12721030 | T | C | 1.014 | -7977.388 | -7979.021 | -1.633 | 7.56E-07 |
| 11 | 117239173 | rs7115177 | C | A | 1.02 | -7977.388 | -7979.022 | -1.633 | 7.56E-07 |
| 11 | 117468794 | rs622860 | G | A | 0.984 | -7977.388 | -7979.022 | -1.633 | 7.56E-07 |
| 11 | 118066609 | rs12289120 | T | C | 0.997 | -7977.388 | -7979.021 | -1.633 | 7.56E-07 |
| 11 | 118831824 | rs7113731 | G | A | 1.023 | -7977.388 | -7979.021 | -1.633 | 7.56E-07 |

| | | | | | | | |
|---|---|---|---|---|---|---|---|
| 11 | 119551498 | rs1533051 | T | C | NULL | 0.693173949 | NULL&M_Disease |
| 11 | 120110219 | rs2715890 | C | G | NULL | 0.693173949 | NULL&M_Disease |
| 11 | 116628401 | rs2075295 | C | T | NULL | 0.693173949 | NULL&M_Disease |
| 11 | 117403235 | rs3741280 | T | G | NULL | 0.693173949 | NULL&M_Disease |
| 11 | 117642382 | rs11825298 | T | C | NULL | 0.693173949 | NULL&M_Disease |
| 11 | 117646354 | rs7941246 | G | T | NULL | 0.693173949 | NULL&M_Disease |
| 11 | 117729680 | rs7121313 | A | G | NULL | 0.693173949 | NULL&M_Disease |
| 11 | 117784714 | rs11216618 | C | T | NULL | 0.693173949 | NULL&M_Disease |
| 11 | 118068752 | rs2298831 | C | T | NULL | 0.693173949 | NULL&M_Disease |
| 11 | 118098041 | rs1715438 | A | G | NULL | 0.693173949 | NULL&M_Disease |
| 11 | 119065310 | rs12575067 | A | G | NULL | 0.693173949 | NULL&M_Disease |
| 11 | 119366884 | rs598494 | C | G | NULL | 0.693173949 | NULL&M_Disease |
| 11 | 119366947 | rs619180 | T | C | NULL | 0.693173949 | NULL&M_Disease |
| 11 | 119367127 | rs620109 | T | C | NULL | 0.693173949 | NULL&M_Disease |
| 11 | 119969933 | rs7118773 | A | G | NULL | 0.693173949 | NULL&M_Disease |
| 11 | 120621602 | rs1945011 | A | G | NULL | 0.693173949 | NULL&M_Disease |
| 11 | 117093764 | rs3830008 | C | A | NULL | 0.693173949 | NULL&M_Disease |
| 11 | 117213189 | rs11216360 | A | G | NULL | 0.693173949 | NULL&M_Disease |
| 11 | 117256436 | rs3781981 | C | T | NULL | 0.693173949 | NULL&M_Disease |
| 11 | 117383716 | rs55975161 | A | G | NULL | 0.693173949 | NULL&M_Disease |
| 11 | 117557057 | rs11216502 | C | G | NULL | 0.693173949 | NULL&M_Disease |
| 11 | 118868750 | rs4445645 | C | G | NULL | 0.693173949 | NULL&M_Disease |
| 11 | 118993632 | rs1179920 | G | A | NULL | 0.693173949 | NULL&M_Disease |
| 11 | 119363369 | rs575021 | A | G | NULL | 0.693173949 | NULL&M_Disease |
| 11 | 119984202 | rs506127 | A | G | NULL | 0.693173949 | NULL&M_Disease |
| 11 | 120553706 | rs11217973 | T | C | NULL | 0.693173949 | NULL&M_Disease |
| 11 | 120946580 | rs7478744 | A | G | NULL | 0.693173949 | NULL&M_Disease |
| 11 | 116555765 | rs61906112 | A | T | NULL | 0.693173949 | NULL&M_Disease |
| 11 | 116964469 | rs56227539 | C | T | NULL | 0.693173949 | NULL&M_Disease |
| 11 | 117213893 | rs10892095 | G | A | NULL | 0.693173949 | NULL&M_Disease |
| 11 | 117729274 | rs10892186 | A | G | NULL | 0.693173949 | NULL&M_Disease |
| 11 | 118037813 | rs8192613 | T | G | NULL | 0.693173949 | NULL&M_Disease |
| 11 | 118091747 | rs1793187 | C | G | NULL | 0.693173949 | NULL&M_Disease |
| 11 | 118832924 | rs4635088 | C | G | NULL | 0.693173949 | NULL&M_Disease |
| 11 | 118843448 | rs11607245 | T | C | NULL | 0.693173949 | NULL&M_Disease |
| 11 | 119358692 | rs12785238 | C | T | NULL | 0.693173949 | NULL&M_Disease |
| 11 | 119367321 | rs596647 | A | C | NULL | 0.693173949 | NULL&M_Disease |
| 11 | 119827163 | rs4938741 | C | T | NULL | 0.693173949 | NULL&M_Disease |
| 11 | 119884781 | rs616889 | G | A | NULL | 0.693173949 | NULL&M_Disease |
| 11 | 120647226 | rs2852217 | C | T | NULL | 0.693173949 | NULL&M_Disease |
| 11 | 120936907 | rs10892667 | C | T | NULL | 0.693173949 | NULL&M_Disease |
| 11 | 116560738 | rs1787712 | G | C | NULL | 0.693173949 | NULL&M_Disease |
| 11 | 116686288 | rs7107757 | T | C | NULL | 0.693173949 | NULL&M_Disease |
| 11 | 117177603 | rs687740 | C | A | NULL | 0.693173949 | NULL&M_Disease |
| 11 | 118077689 | rs1715425 | C | T | NULL | 0.693173949 | NULL&M_Disease |
| 11 | 118125077 | rs1715453 | T | C | NULL | 0.693173949 | NULL&M_Disease |
| 11 | 116251791 | rs502483 | T | A | NULL | 0.69423634 | NULL&M_Disease |
| 11 | 116438159 | rs7350537 | T | C | NULL | 0.69423634 | NULL&M_Disease |
| 11 | 117638172 | rs3851576 | C | A | NULL | 0.69423634 | NULL&M_Disease |
| 11 | 119553678 | rs10790332 | T | C | NULL | 0.69423634 | NULL&M_Disease |
| 11 | 120598007 | rs17311414 | A | C | NULL | 0.69423634 | NULL&M_Disease |
| 11 | 121107345 | rs7129197 | T | G | NULL | 0.69423634 | NULL&M_Disease |
| 11 | 116964437 | rs59097294 | C | T | NULL | 0.69423634 | NULL&M_Disease |
| 11 | 118091067 | rs2097183 | T | C | NULL | 0.69423634 | NULL&M_Disease |
| 11 | 118091375 | rs11216822 | A | G | NULL | 0.69423634 | NULL&M_Disease |
| 11 | 118103972 | rs1793162 | A | C | NULL | 0.69423634 | NULL&M_Disease |
| 11 | 119363885 | rs579427 | T | C | NULL | 0.69423634 | NULL&M_Disease |
| 11 | 119365917 | rs604361 | T | C | NULL | 0.69423634 | NULL&M_Disease |
| 11 | 119628589 | rs10892451 | T | G | NULL | 0.69423634 | NULL&M_Disease |
| 11 | 119983799 | rs530792 | T | C | NULL | 0.69423634 | NULL&M_Disease |
| 11 | 120911771 | rs10892664 | A | T | NULL | 0.69423634 | NULL&M_Disease |
| 11 | 121111458 | rs948381 | C | A | NULL | 0.69423634 | NULL&M_Disease |
| 11 | 116319221 | rs505064 | C | T | NULL | 0.69423634 | NULL&M_Disease |
| 11 | 116432791 | rs4938283 | T | C | NULL | 0.69423634 | NULL&M_Disease |
| 11 | 117154091 | rs1158810726 | G | T | NULL | 0.69423634 | NULL&M_Disease |
| 11 | 117177741 | rs574603 | C | T | NULL | 0.69423634 | NULL&M_Disease |
| 11 | 117464063 | rs649233 | T | C | NULL | 0.69423634 | NULL&M_Disease |
| 11 | 117538453 | rs581580 | G | A | NULL | 0.69423634 | NULL&M_Disease |
| 11 | 117760895 | rs7942136 | C | T | NULL | 0.69423634 | NULL&M_Disease |
| 11 | 118125625 | rs1056562 | T | C | NULL | 0.69423634 | NULL&M_Disease |
| 11 | 119073886 | rs7104210 | G | T | NULL | 0.69423634 | NULL&M_Disease |
| 11 | 120606290 | rs2852232 | G | A | NULL | 0.69423634 | NULL&M_Disease |
| 11 | 120690057 | rs7940051 | A | G | NULL | 0.69423634 | NULL&M_Disease |
| 11 | 120690372 | rs56326825 | C | T | NULL | 0.69423634 | NULL&M_Disease |
| 11 | 120898232 | rs12576758 | T | C | NULL | 0.69423634 | NULL&M_Disease |
| 11 | 120898482 | rs68097814 | G | A | NULL | 0.69423634 | NULL&M_Disease |
| 11 | 116432066 | rs11216006 | T | C | NULL | 0.69423634 | NULL&M_Disease |
| 11 | 116499500 | rs61270867 | G | A | NULL | 0.69423634 | NULL&M_Disease |
| 11 | 116694938 | rs10892035 | A | G | NULL | 0.69423634 | NULL&M_Disease |
| 11 | 119365993 | rs629272 | A | G | NULL | 0.69423634 | NULL&M_Disease |
| 11 | 119983831 | rs530702 | A | G | NULL | 0.69423634 | NULL&M_Disease |
| 11 | 120899458 | rs11218122 | T | C | NULL | 0.69423634 | NULL&M_Disease |
| 11 | 120926467 | rs11218128 | A | T | NULL | 0.69423634 | NULL&M_Disease |
| 11 | 120932457 | rs67932968 | A | G | NULL | 0.69423634 | NULL&M_Disease |
| 11 | 116693464 | rs5092 | C | T | NULL | 0.69423634 | NULL&M_Disease |
| 11 | 117211240 | rs1258164206 | T | G | NULL | 0.69423634 | NULL&M_Disease |
| 11 | 117315191 | rs7125516 | C | T | NULL | 0.69423634 | NULL&M_Disease |
| 11 | 117677039 | rs12281156 | A | T | NULL | 0.69423634 | NULL&M_Disease |
| 11 | 117846083 | rs12420994 | G | C | NULL | 0.69423634 | NULL&M_Disease |
| 11 | 118080771 | rs1793176 | T | A | NULL | 0.69423634 | NULL&M_Disease |
| 11 | 118089434 | rs1793185 | A | C | NULL | 0.69423634 | NULL&M_Disease |
| 11 | 118091089 | rs1793186 | T | C | NULL | 0.69423634 | NULL&M_Disease |
| 11 | 120624339 | rs2156762 | C | T | NULL | 0.69423634 | NULL&M_Disease |
| 11 | 120898761 | rs11218120 | G | A | NULL | 0.69423634 | NULL&M_Disease |
| 11 | 120898784 | rs11218121 | C | T | NULL | 0.69423634 | NULL&M_Disease |
| 11 | 116322236 | rs7106296 | T | C | NULL | 0.69423634 | NULL&M_Disease |

| | | | | | | | | | |
|---|---|---|---|---|---|---|---|---|---|
| 11 | 119551498 | rs1533051 | T | C | 0.978 | -7977.388 | -7979.021 | -1.633 | 7.56E-07 |
| 11 | 120110219 | rs2715890 | C | G | 1.013 | -7977.388 | -7979.021 | -1.633 | 7.56E-07 |
| 11 | 116628401 | rs2075295 | C | T | 1.019 | -7977.388 | -7979.022 | -1.634 | 7.55E-07 |
| 11 | 117403235 | rs3741280 | T | G | 0.98 | -7977.388 | -7979.022 | -1.634 | 7.55E-07 |
| 11 | 117642382 | rs11825298 | T | C | 0.997 | -7977.388 | -7979.022 | -1.634 | 7.55E-07 |
| 11 | 117646354 | rs7941246 | G | T | 0.98 | -7977.388 | -7979.022 | -1.634 | 7.55E-07 |
| 11 | 117729680 | rs7121313 | A | G | 0.998 | -7977.388 | -7979.023 | -1.634 | 7.55E-07 |
| 11 | 117784714 | rs11216618 | C | T | 1.02 | -7977.388 | -7979.022 | -1.634 | 7.55E-07 |
| 11 | 118068752 | rs2298831 | C | T | 1.001 | -7977.388 | -7979.022 | -1.634 | 7.55E-07 |
| 11 | 118098041 | rs1715438 | A | G | 1.02 | -7977.388 | -7979.023 | -1.634 | 7.55E-07 |
| 11 | 119065310 | rs12575067 | A | G | 1.006 | -7977.388 | -7979.022 | -1.634 | 7.55E-07 |
| 11 | 119366884 | rs598494 | C | G | 0.978 | -7977.388 | -7979.023 | -1.634 | 7.55E-07 |
| 11 | 119366947 | rs619180 | T | C | 0.978 | -7977.388 | -7979.023 | -1.634 | 7.55E-07 |
| 11 | 119367127 | rs620109 | T | C | 0.978 | -7977.388 | -7979.023 | -1.634 | 7.55E-07 |
| 11 | 119969933 | rs7118773 | A | G | 0.979 | -7977.388 | -7979.023 | -1.634 | 7.55E-07 |
| 11 | 120621602 | rs1945011 | A | G | 0.98 | -7977.388 | -7979.022 | -1.634 | 7.55E-07 |
| 11 | 117093764 | rs3830008 | C | A | 1.016 | -7977.388 | -7979.023 | -1.635 | 7.54E-07 |
| 11 | 117213189 | rs11216360 | A | G | 0.991 | -7977.388 | -7979.023 | -1.635 | 7.54E-07 |
| 11 | 117256436 | rs3781981 | C | T | 1.022 | -7977.388 | -7979.023 | -1.635 | 7.54E-07 |
| 11 | 117383716 | rs55975161 | A | G | 0.99 | -7977.388 | -7979.024 | -1.635 | 7.54E-07 |
| 11 | 117557057 | rs11216502 | C | G | 0.984 | -7977.388 | -7979.023 | -1.635 | 7.54E-07 |
| 11 | 118868750 | rs4445645 | C | G | 1.017 | -7977.388 | -7979.023 | -1.635 | 7.54E-07 |
| 11 | 118993632 | rs1179920 | G | A | 0.978 | -7977.388 | -7979.024 | -1.635 | 7.54E-07 |
| 11 | 119363369 | rs575021 | A | G | 0.978 | -7977.388 | -7979.023 | -1.635 | 7.54E-07 |
| 11 | 119984202 | rs506127 | A | G | 0.995 | -7977.388 | -7979.024 | -1.635 | 7.54E-07 |
| 11 | 120553706 | rs11217973 | T | C | 0.98 | -7977.388 | -7979.023 | -1.635 | 7.54E-07 |
| 11 | 120946580 | rs7478744 | A | G | 1.007 | -7977.388 | -7979.023 | -1.635 | 7.54E-07 |
| 11 | 116555765 | rs61906112 | A | T | 1.018 | -7977.388 | -7979.024 | -1.636 | 7.53E-07 |
| 11 | 116964469 | rs56227539 | C | T | 1.012 | -7977.388 | -7979.024 | -1.636 | 7.53E-07 |
| 11 | 117213893 | rs10892095 | G | A | 0.991 | -7977.388 | -7979.025 | -1.636 | 7.53E-07 |
| 11 | 117729274 | rs10892186 | A | G | 1.016 | -7977.388 | -7979.024 | -1.636 | 7.53E-07 |
| 11 | 118037813 | rs8192613 | T | G | 0.978 | -7977.388 | -7979.024 | -1.636 | 7.53E-07 |
| 11 | 118091747 | rs1793187 | C | G | 1.02 | -7977.388 | -7979.024 | -1.636 | 7.53E-07 |
| 11 | 118832924 | rs4635088 | C | G | 1.023 | -7977.388 | -7979.024 | -1.636 | 7.53E-07 |
| 11 | 118843448 | rs11607245 | T | C | 1.023 | -7977.388 | -7979.024 | -1.636 | 7.53E-07 |
| 11 | 119358692 | rs12785238 | C | T | 0.992 | -7977.388 | -7979.024 | -1.636 | 7.53E-07 |
| 11 | 119367321 | rs596647 | A | C | 0.989 | -7977.388 | -7979.024 | -1.636 | 7.53E-07 |
| 11 | 119827163 | rs4938741 | C | T | 0.982 | -7977.388 | -7979.024 | -1.636 | 7.53E-07 |
| 11 | 119884781 | rs616889 | G | A | 0.978 | -7977.388 | -7979.024 | -1.636 | 7.53E-07 |
| 11 | 120647226 | rs2852217 | C | T | 0.981 | -7977.388 | -7979.024 | -1.636 | 7.53E-07 |
| 11 | 120936907 | rs10892667 | C | T | 1.007 | -7977.388 | -7979.024 | -1.636 | 7.53E-07 |
| 11 | 116560738 | rs1787712 | G | C | 1.02 | -7977.388 | -7979.025 | -1.637 | 7.53E-07 |
| 11 | 116686288 | rs7107757 | T | C | 1.005 | -7977.388 | -7979.026 | -1.637 | 7.53E-07 |
| 11 | 117177603 | rs687740 | C | A | 1.022 | -7977.388 | -7979.025 | -1.637 | 7.53E-07 |
| 11 | 118077689 | rs1715425 | C | T | 0.981 | -7977.388 | -7979.025 | -1.637 | 7.53E-07 |
| 11 | 118125077 | rs1715453 | T | C | 1.023 | -7977.388 | -7979.025 | -1.637 | 7.53E-07 |
| 11 | 116251791 | rs502483 | T | A | 1.018 | -7977.388 | -7979.026 | -1.638 | 7.52E-07 |
| 11 | 116438159 | rs7350537 | T | C | 0.985 | -7977.388 | -7979.026 | -1.638 | 7.52E-07 |
| 11 | 117638172 | rs3851576 | C | A | 0.98 | -7977.388 | -7979.026 | -1.638 | 7.52E-07 |
| 11 | 119553678 | rs10790332 | T | C | 0.978 | -7977.388 | -7979.027 | -1.638 | 7.52E-07 |
| 11 | 120598007 | rs17311414 | A | C | 0.999 | -7977.388 | -7979.027 | -1.638 | 7.52E-07 |
| 11 | 121107345 | rs7129197 | T | G | 0.978 | -7977.388 | -7979.026 | -1.638 | 7.52E-07 |
| 11 | 116964437 | rs59097294 | C | T | 1.012 | -7977.388 | -7979.028 | -1.639 | 7.51E-07 |
| 11 | 118091067 | rs2097183 | T | C | 1.016 | -7977.388 | -7979.028 | -1.639 | 7.51E-07 |
| 11 | 118091375 | rs11216822 | A | G | 1.016 | -7977.388 | -7979.028 | -1.639 | 7.51E-07 |
| 11 | 118103972 | rs1793162 | A | C | 1.02 | -7977.388 | -7979.028 | -1.639 | 7.51E-07 |
| 11 | 119363885 | rs579427 | T | C | 0.979 | -7977.388 | -7979.027 | -1.639 | 7.51E-07 |
| 11 | 119365917 | rs604361 | T | C | 0.978 | -7977.388 | -7979.027 | -1.639 | 7.51E-07 |
| 11 | 119628589 | rs10892451 | T | G | 1.022 | -7977.388 | -7979.027 | -1.639 | 7.51E-07 |
| 11 | 119983799 | rs530792 | T | C | 0.996 | -7977.388 | -7979.027 | -1.639 | 7.51E-07 |
| 11 | 120911771 | rs10892664 | A | T | 1.006 | -7977.388 | -7979.027 | -1.639 | 7.51E-07 |
| 11 | 121111458 | rs948381 | C | A | 0.978 | -7977.388 | -7979.027 | -1.639 | 7.51E-07 |
| 11 | 116319221 | rs505064 | C | T | 1.013 | -7977.388 | -7979.028 | -1.64 | 7.50E-07 |
| 11 | 116432791 | rs4938283 | T | C | 0.994 | -7977.388 | -7979.028 | -1.64 | 7.50E-07 |
| 11 | 117154091 | rs1158810726 | G | T | 0.993 | -7977.388 | -7979.029 | -1.64 | 7.50E-07 |
| 11 | 117177741 | rs574603 | C | T | 1.022 | -7977.388 | -7979.028 | -1.64 | 7.50E-07 |
| 11 | 117464063 | rs649233 | T | C | 0.986 | -7977.388 | -7979.029 | -1.64 | 7.50E-07 |
| 11 | 117538453 | rs581580 | G | A | 1.022 | -7977.388 | -7979.028 | -1.64 | 7.50E-07 |
| 11 | 117760895 | rs7942136 | C | T | 1.022 | -7977.388 | -7979.029 | -1.64 | 7.50E-07 |
| 11 | 118125625 | rs1056562 | T | C | 1.023 | -7977.388 | -7979.028 | -1.64 | 7.50E-07 |
| 11 | 119073886 | rs7104210 | G | T | 0.98 | -7977.388 | -7979.029 | -1.64 | 7.50E-07 |
| 11 | 120606290 | rs2852232 | G | A | 1.022 | -7977.388 | -7979.029 | -1.64 | 7.50E-07 |
| 11 | 120690057 | rs7940051 | A | G | 0.978 | -7977.388 | -7979.028 | -1.64 | 7.50E-07 |
| 11 | 120690372 | rs56326825 | C | T | 0.978 | -7977.388 | -7979.028 | -1.64 | 7.50E-07 |
| 11 | 120898232 | rs12576758 | T | C | 1.006 | -7977.388 | -7979.028 | -1.64 | 7.50E-07 |
| 11 | 120898482 | rs68097814 | G | A | 1.006 | -7977.388 | -7979.028 | -1.64 | 7.50E-07 |
| 11 | 116432066 | rs11216006 | T | C | 0.994 | -7977.388 | -7979.029 | -1.641 | 7.50E-07 |
| 11 | 116499500 | rs61270867 | G | A | 1.002 | -7977.388 | -7979.029 | -1.641 | 7.50E-07 |
| 11 | 116694938 | rs10892035 | A | G | 0.997 | -7977.388 | -7979.03 | -1.641 | 7.50E-07 |
| 11 | 119365993 | rs629272 | A | G | 0.979 | -7977.388 | -7979.029 | -1.641 | 7.50E-07 |
| 11 | 119983831 | rs530702 | A | G | 0.997 | -7977.388 | -7979.03 | -1.641 | 7.50E-07 |
| 11 | 120899458 | rs11218122 | T | C | 1.005 | -7977.388 | -7979.029 | -1.641 | 7.50E-07 |
| 11 | 120926467 | rs11218128 | A | T | 1.005 | -7977.388 | -7979.029 | -1.641 | 7.50E-07 |
| 11 | 120932457 | rs67932968 | A | G | 1.005 | -7977.388 | -7979.029 | -1.641 | 7.50E-07 |
| 11 | 116693464 | rs5092 | C | T | 1 | -7977.388 | -7979.03 | -1.642 | 7.49E-07 |
| 11 | 117211240 | rs1258164206 | T | G | 0.987 | -7977.388 | -7979.03 | -1.642 | 7.49E-07 |
| 11 | 117315191 | rs7125516 | C | T | 0.985 | -7977.388 | -7979.031 | -1.642 | 7.49E-07 |
| 11 | 117677039 | rs12281156 | A | T | 0.988 | -7977.388 | -7979.03 | -1.642 | 7.49E-07 |
| 11 | 117846083 | rs12420994 | G | C | 1.022 | -7977.388 | -7979.03 | -1.642 | 7.49E-07 |
| 11 | 118080771 | rs1793176 | T | A | 0.982 | -7977.388 | -7979.03 | -1.642 | 7.49E-07 |
| 11 | 118089434 | rs1793185 | A | C | 1.016 | -7977.388 | -7979.031 | -1.642 | 7.49E-07 |
| 11 | 118091089 | rs1793186 | T | C | 1.02 | -7977.388 | -7979.03 | -1.642 | 7.49E-07 |
| 11 | 120624339 | rs2156762 | C | T | 0.98 | -7977.388 | -7979.03 | -1.642 | 7.49E-07 |
| 11 | 120898761 | rs11218120 | G | A | 1.005 | -7977.388 | -7979.031 | -1.642 | 7.49E-07 |
| 11 | 120898784 | rs11218121 | C | T | 1.005 | -7977.388 | -7979.031 | -1.642 | 7.49E-07 |
| 11 | 116322236 | rs7106296 | T | C | 0.992 | -7977.388 | -7979.031 | -1.643 | 7.48E-07 |

| | | | | | | | |
|---|---|---|---|---|---|---|---|
| 11 | 116329043 | rs574223 | G | T | NULL | 0.69423634 | NULL&M_Disease |
| 11 | 117747110 | rs7119090 | C | G | NULL | 0.69423634 | NULL&M_Disease |
| 11 | 118884726 | rs58601794 | C | T | NULL | 0.69423634 | NULL&M_Disease |
| 11 | 120137465 | rs2096882 | T | G | NULL | 0.69423634 | NULL&M_Disease |
| 11 | 120690291 | rs7940347 | T | G | NULL | 0.69423634 | NULL&M_Disease |
| 11 | 117873048 | rs2512148 | A | C | NULL | 0.69423634 | NULL&M_Disease |
| 11 | 120789293 | rs2852213 | T | C | NULL | 0.69423634 | NULL&M_Disease |
| 11 | 118012310 | rs11216781 | A | G | NULL | 0.69423634 | NULL&M_Disease |
| 11 | 118884708 | rs61381062 | A | G | NULL | 0.69423634 | NULL&M_Disease |
| 11 | 119364258 | rs667319 | G | A | NULL | 0.69423634 | NULL&M_Disease |
| 11 | 119989630 | rs7128378 | A | G | NULL | 0.69423634 | NULL&M_Disease |
| 11 | 120857051 | rs11218092 | A | G | NULL | 0.69423634 | NULL&M_Disease |
| 11 | 116564553 | rs1787699 | G | A | NULL | 0.69423634 | NULL&M_Disease |
| 11 | 116564562 | rs1787698 | T | C | NULL | 0.69423634 | NULL&M_Disease |
| 11 | 117867008 | rs2512146 | A | G | NULL | 0.69423634 | NULL&M_Disease |
| 11 | 118074117 | rs1793173 | A | G | NULL | 0.69423634 | NULL&M_Disease |
| 11 | 118074337 | rs1793174 | A | G | NULL | 0.69423634 | NULL&M_Disease |
| 11 | 118114344 | rs2853033 | A | G | NULL | 0.69423634 | NULL&M_Disease |
| 11 | 118114813 | rs1793163 | T | C | NULL | 0.69423634 | NULL&M_Disease |
| 11 | 118114853 | rs1793164 | T | A | NULL | 0.69423634 | NULL&M_Disease |
| 11 | 118203642 | rs73612281 | T | C | NULL | 0.69423634 | NULL&M_Disease |
| 11 | 118841761 | rs11217108 | T | C | NULL | 0.69423634 | NULL&M_Disease |
| 11 | 118885163 | rs59608621 | G | A | NULL | 0.69423634 | NULL&M_Disease |
| 11 | 119560449 | rs4936492 | A | G | NULL | 0.69423634 | NULL&M_Disease |
| 11 | 116503403 | rs7946423 | T | A | NULL | 0.69423634 | NULL&M_Disease |
| 11 | 117232391 | rs17500548 | G | A | NULL | 0.69423634 | NULL&M_Disease |
| 11 | 117407301 | rs521609 | A | G | NULL | 0.69423634 | NULL&M_Disease |
| 11 | 117701057 | rs10790211 | A | G | NULL | 0.69423634 | NULL&M_Disease |
| 11 | 117776526 | rs3802872 | G | A | NULL | 0.69423634 | NULL&M_Disease |
| 11 | 117864063 | rs3135932 | G | A | NULL | 0.69423634 | NULL&M_Disease |
| 11 | 117955810 | rs10750123 | C | T | NULL | 0.695296669 | NULL&M_Disease |
| 11 | 118087040 | rs12280559 | T | C | NULL | 0.69423634 | NULL&M_Disease |
| 11 | 118885161 | rs59095933 | A | G | NULL | 0.69423634 | NULL&M_Disease |
| 11 | 118994516 | rs7103544 | G | A | NULL | 0.69423634 | NULL&M_Disease |
| 11 | 120136108 | rs10750169 | C | G | NULL | 0.69423634 | NULL&M_Disease |
| 11 | 121110846 | rs7112341 | A | T | NULL | 0.69423634 | NULL&M_Disease |
| 11 | 121115967 | rs10892697 | C | G | NULL | 0.69423634 | NULL&M_Disease |
| 11 | 117236060 | rs10892097 | T | C | NULL | 0.695296669 | NULL&M_Disease |
| 11 | 117236319 | rs10790179 | T | G | NULL | 0.695296669 | NULL&M_Disease |
| 11 | 117642030 | rs10892171 | A | T | NULL | 0.69423634 | NULL&M_Disease |
| 11 | 119490921 | rs4367956 | A | C | NULL | 0.695296669 | NULL&M_Disease |
| 11 | 116323001 | rs542671 | A | G | NULL | 0.695296669 | NULL&M_Disease |
| 11 | 116557248 | rs17092642 | C | T | NULL | 0.695296669 | NULL&M_Disease |
| 11 | 117225455 | rs542869 | A | T | NULL | 0.695296669 | NULL&M_Disease |
| 11 | 117232260 | rs12418678 | G | A | NULL | 0.695296669 | NULL&M_Disease |
| 11 | 117691774 | rs7483165 | C | T | NULL | 0.695296669 | NULL&M_Disease |
| 11 | 118078761 | rs7101554 | C | T | NULL | 0.695296669 | NULL&M_Disease |
| 11 | 118104627 | rs1573559 | G | A | NULL | 0.695296669 | NULL&M_Disease |
| 11 | 118107805 | rs1793140 | A | G | NULL | 0.695296669 | NULL&M_Disease |
| 11 | 118113907 | rs3018368 | G | C | NULL | 0.695296669 | NULL&M_Disease |
| 11 | 119369816 | rs35544562 | C | G | NULL | 0.695296669 | NULL&M_Disease |
| 11 | 119542000 | rs4576834 | T | C | NULL | 0.695296669 | NULL&M_Disease |
| 11 | 120074907 | rs12793410 | A | T | NULL | 0.695296669 | NULL&M_Disease |
| 11 | 120136078 | rs10790372 | T | G | NULL | 0.695296669 | NULL&M_Disease |
| 11 | 121108515 | rs12295531 | T | C | NULL | 0.695296669 | NULL&M_Disease |
| 11 | 117283808 | rs9163 | T | C | NULL | 0.695296669 | NULL&M_Disease |
| 11 | 118994357 | rs7119384 | T | A | NULL | 0.695296669 | NULL&M_Disease |
| 11 | 121110312 | rs10437694 | G | T | NULL | 0.695296669 | NULL&M_Disease |
| 11 | 116352339 | rs1354975 | T | C | NULL | 0.695296669 | NULL&M_Disease |
| 11 | 116626570 | rs180323 | G | A | NULL | 0.695296669 | NULL&M_Disease |
| 11 | 118393396 | rs573971 | A | G | NULL | 0.695296669 | NULL&M_Disease |
| 11 | 119628785 | rs11217467 | A | G | NULL | 0.695296669 | NULL&M_Disease |
| 11 | 121126349 | rs4144286 | A | G | NULL | 0.695296669 | NULL&M_Disease |
| 11 | 117177920 | rs573442 | T | C | NULL | 0.695296669 | NULL&M_Disease |
| 11 | 117192528 | rs56967930 | G | A | NULL | 0.695296669 | NULL&M_Disease |
| 11 | 117192701 | rs55986402 | C | A | NULL | 0.695296669 | NULL&M_Disease |
| 11 | 117213160 | rs493516 | A | G | NULL | 0.695296669 | NULL&M_Disease |
| 11 | 117398202 | rs7940527 | C | G | NULL | 0.695296669 | NULL&M_Disease |
| 11 | 118091636 | rs7103936 | C | A | NULL | 0.695296669 | NULL&M_Disease |
| 11 | 118127347 | rs1793149 | A | G | NULL | 0.695296669 | NULL&M_Disease |
| 11 | 119238455 | rs11217257 | G | A | NULL | 0.695296669 | NULL&M_Disease |
| 11 | 119365552 | rs683630 | T | C | NULL | 0.695296669 | NULL&M_Disease |
| 11 | 120513914 | rs1317514 | C | T | NULL | 0.695296669 | NULL&M_Disease |
| 11 | 120724121 | rs7104543 | G | A | NULL | 0.695296669 | NULL&M_Disease |
| 11 | 116219458 | rs554865 | T | C | NULL | 0.695296669 | NULL&M_Disease |
| 11 | 116438555 | rs12225221 | T | C | NULL | 0.695296669 | NULL&M_Disease |
| 11 | 116440163 | rs4936357 | T | C | NULL | 0.695296669 | NULL&M_Disease |
| 11 | 116440575 | rs10431039 | T | C | NULL | 0.695296669 | NULL&M_Disease |
| 11 | 117208747 | rs645305 | C | T | NULL | 0.695296669 | NULL&M_Disease |
| 11 | 117538059 | rs608811 | A | G | NULL | 0.695296669 | NULL&M_Disease |
| 11 | 118088362 | rs11216821 | C | T | NULL | 0.695296669 | NULL&M_Disease |
| 11 | 118112841 | rs1715418 | T | A | NULL | 0.695296669 | NULL&M_Disease |
| 11 | 119438208 | rs2663634 | T | C | NULL | 0.695296669 | NULL&M_Disease |
| 11 | 116440678 | rs10431079 | G | A | NULL | 0.695296669 | NULL&M_Disease |
| 11 | 117210901 | rs536021 | C | T | NULL | 0.695296669 | NULL&M_Disease |
| 11 | 117212892 | rs475385 | T | C | NULL | 0.695296669 | NULL&M_Disease |
| 11 | 116439792 | rs7943469 | G | T | NULL | 0.695296669 | NULL&M_Disease |
| 11 | 116440036 | rs1261967419 | A | T | NULL | 0.695296669 | NULL&M_Disease |
| 11 | 117189201 | rs61903708 | G | A | NULL | 0.695296669 | NULL&M_Disease |
| 11 | 117208270 | rs875209 | A | G | NULL | 0.695296669 | NULL&M_Disease |
| 11 | 117209268 | rs573801 | T | C | NULL | 0.695296669 | NULL&M_Disease |
| 11 | 117229891 | rs2508664 | T | C | NULL | 0.695296669 | NULL&M_Disease |
| 11 | 117231611 | rs11216366 | T | G | NULL | 0.695296669 | NULL&M_Disease |
| 11 | 117239190 | rs7115526 | T | G | NULL | 0.695296669 | NULL&M_Disease |
| 11 | 117248268 | rs7952470 | A | G | NULL | 0.695296669 | NULL&M_Disease |
| 11 | 118075152 | rs11216816 | A | C | NULL | 0.695296669 | NULL&M_Disease |

| | | | | | | | | | |
|---|---|---|---|---|---|---|---|---|---|
| 11 | 116329043 | rs574223 | G | T | 1.022 | -7977.388 | -7979.032 | -1.643 | 7.48E-07 |
| 11 | 117747110 | rs7119090 | C | G | 1.014 | -7977.388 | -7979.031 | -1.643 | 7.48E-07 |
| 11 | 118884726 | rs58601794 | C | T | 1.019 | -7977.388 | -7979.031 | -1.643 | 7.48E-07 |
| 11 | 120137465 | rs2096882 | T | G | 1.02 | -7977.388 | -7979.031 | -1.643 | 7.48E-07 |
| 11 | 120690291 | rs7940347 | T | G | 0.978 | -7977.388 | -7979.031 | -1.643 | 7.48E-07 |
| 11 | 117873048 | rs2512148 | A | C | 0.979 | -7977.388 | -7979.032 | -1.644 | 7.47E-07 |
| 11 | 120789293 | rs2852213 | T | C | 0.978 | -7977.388 | -7979.032 | -1.644 | 7.47E-07 |
| 11 | 118012310 | rs11216781 | A | G | 1.019 | -7977.388 | -7979.033 | -1.645 | 7.47E-07 |
| 11 | 118884708 | rs61381062 | A | G | 1.018 | -7977.388 | -7979.034 | -1.645 | 7.47E-07 |
| 11 | 119364258 | rs667319 | G | A | 0.979 | -7977.388 | -7979.033 | -1.645 | 7.47E-07 |
| 11 | 119989630 | rs7128378 | A | G | 1.022 | -7977.388 | -7979.033 | -1.645 | 7.47E-07 |
| 11 | 120857051 | rs11218092 | A | G | 1.013 | -7977.388 | -7979.034 | -1.645 | 7.47E-07 |
| 11 | 116564553 | rs1787699 | G | A | 0.978 | -7977.388 | -7979.034 | -1.646 | 7.46E-07 |
| 11 | 116564562 | rs1787698 | T | C | 0.978 | -7977.388 | -7979.034 | -1.646 | 7.46E-07 |
| 11 | 117867008 | rs2512146 | A | G | 0.979 | -7977.388 | -7979.034 | -1.646 | 7.46E-07 |
| 11 | 118074117 | rs1793173 | A | G | 0.982 | -7977.388 | -7979.034 | -1.646 | 7.46E-07 |
| 11 | 118074337 | rs1793174 | A | G | 0.982 | -7977.388 | -7979.034 | -1.646 | 7.46E-07 |
| 11 | 118114344 | rs2853033 | A | G | 1.019 | -7977.388 | -7979.034 | -1.646 | 7.46E-07 |
| 11 | 118114813 | rs1793163 | T | C | 1.019 | -7977.388 | -7979.034 | -1.646 | 7.46E-07 |
| 11 | 118114853 | rs1793164 | T | A | 1.019 | -7977.388 | -7979.034 | -1.646 | 7.46E-07 |
| 11 | 118203642 | rs73612281 | T | C | 0.981 | -7977.388 | -7979.035 | -1.646 | 7.46E-07 |
| 11 | 118841761 | rs11217108 | T | C | 1.022 | -7977.388 | -7979.035 | -1.646 | 7.46E-07 |
| 11 | 118885163 | rs59608621 | G | A | 1.018 | -7977.388 | -7979.034 | -1.646 | 7.46E-07 |
| 11 | 119560449 | rs4936492 | A | G | 0.979 | -7977.388 | -7979.034 | -1.646 | 7.46E-07 |
| 11 | 116503403 | rs7946423 | T | A | 0.983 | -7977.388 | -7979.035 | -1.647 | 7.45E-07 |
| 11 | 117232391 | rs17500548 | G | A | 0.988 | -7977.388 | -7979.036 | -1.647 | 7.45E-07 |
| 11 | 117407301 | rs521609 | A | G | 1.015 | -7977.388 | -7979.035 | -1.647 | 7.45E-07 |
| 11 | 117701057 | rs10790211 | A | G | 0.981 | -7977.388 | -7979.035 | -1.647 | 7.45E-07 |
| 11 | 117776526 | rs3802872 | G | A | 1.022 | -7977.388 | -7979.035 | -1.647 | 7.45E-07 |
| 11 | 117864063 | rs3135932 | G | A | 0.993 | -7977.388 | -7979.036 | -1.647 | 7.45E-07 |
| 11 | 117955810 | rs10750123 | C | T | 0.979 | -7977.388 | -7979.036 | -1.647 | 7.45E-07 |
| 11 | 118087040 | rs12280559 | T | C | 1.016 | -7977.388 | -7979.035 | -1.647 | 7.45E-07 |
| 11 | 118885161 | rs59095933 | A | G | 1.018 | -7977.388 | -7979.035 | -1.647 | 7.45E-07 |
| 11 | 118994516 | rs7103544 | G | A | 0.978 | -7977.388 | -7979.035 | -1.647 | 7.45E-07 |
| 11 | 120136108 | rs10750169 | C | G | 1.02 | -7977.388 | -7979.035 | -1.647 | 7.45E-07 |
| 11 | 121110846 | rs7112341 | A | T | 1.006 | -7977.388 | -7979.036 | -1.647 | 7.45E-07 |
| 11 | 121115967 | rs10892697 | C | G | 0.99 | -7977.388 | -7979.035 | -1.647 | 7.45E-07 |
| 11 | 117236060 | rs10892097 | T | C | 0.988 | -7977.388 | -7979.036 | -1.648 | 7.44E-07 |
| 11 | 117236319 | rs10790179 | T | G | 0.988 | -7977.388 | -7979.036 | -1.648 | 7.44E-07 |
| 11 | 117642030 | rs10892171 | A | T | 0.981 | -7977.388 | -7979.036 | -1.648 | 7.44E-07 |
| 11 | 119490921 | rs4367956 | A | C | 0.98 | -7977.388 | -7979.036 | -1.648 | 7.44E-07 |
| 11 | 116323001 | rs542671 | A | G | 0.992 | -7977.388 | -7979.037 | -1.649 | 7.44E-07 |
| 11 | 116557248 | rs17092642 | C | T | 1.017 | -7977.388 | -7979.037 | -1.649 | 7.44E-07 |
| 11 | 117225455 | rs542869 | A | T | 0.988 | -7977.388 | -7979.037 | -1.649 | 7.44E-07 |
| 11 | 117232260 | rs12418678 | G | A | 0.988 | -7977.388 | -7979.038 | -1.649 | 7.44E-07 |
| 11 | 117691774 | rs7483165 | C | T | 0.988 | -7977.388 | -7979.038 | -1.649 | 7.44E-07 |
| 11 | 118078761 | rs7101554 | C | T | 0.982 | -7977.388 | -7979.038 | -1.649 | 7.44E-07 |
| 11 | 118104627 | rs1573559 | G | A | 1.022 | -7977.388 | -7979.037 | -1.649 | 7.44E-07 |
| 11 | 118107805 | rs1793140 | A | G | 1.019 | -7977.388 | -7979.037 | -1.649 | 7.44E-07 |
| 11 | 118113907 | rs3018368 | G | C | 1.019 | -7977.388 | -7979.037 | -1.649 | 7.44E-07 |
| 11 | 119369816 | rs35544562 | C | G | 0.991 | -7977.388 | -7979.037 | -1.649 | 7.44E-07 |
| 11 | 119542000 | rs4576834 | T | C | 0.988 | -7977.388 | -7979.037 | -1.649 | 7.44E-07 |
| 11 | 120074907 | rs12793410 | A | T | 1.017 | -7977.388 | -7979.037 | -1.649 | 7.44E-07 |
| 11 | 120136078 | rs10790372 | T | G | 1.019 | -7977.388 | -7979.038 | -1.649 | 7.44E-07 |
| 11 | 121108515 | rs12295531 | T | C | 1.005 | -7977.388 | -7979.037 | -1.649 | 7.44E-07 |
| 11 | 117283808 | rs9163 | T | C | 0.986 | -7977.388 | -7979.038 | -1.65 | 7.43E-07 |
| 11 | 118994357 | rs7119384 | T | A | 0.978 | -7977.388 | -7979.039 | -1.65 | 7.43E-07 |
| 11 | 121110312 | rs10437694 | G | T | 1.005 | -7977.388 | -7979.038 | -1.65 | 7.43E-07 |
| 11 | 116352339 | rs1354975 | T | C | 1.007 | -7977.388 | -7979.039 | -1.651 | 7.42E-07 |
| 11 | 116626570 | rs180323 | G | A | 1.019 | -7977.388 | -7979.039 | -1.651 | 7.42E-07 |
| 11 | 118393396 | rs573971 | A | G | 0.987 | -7977.388 | -7979.04 | -1.651 | 7.42E-07 |
| 11 | 119628785 | rs11217467 | A | G | 1.022 | -7977.388 | -7979.039 | -1.651 | 7.42E-07 |
| 11 | 121126349 | rs4144286 | A | G | 0.978 | -7977.388 | -7979.039 | -1.651 | 7.42E-07 |
| 11 | 117177920 | rs573442 | T | C | 1.022 | -7977.388 | -7979.04 | -1.652 | 7.41E-07 |
| 11 | 117192528 | rs56967930 | G | A | 1.002 | -7977.388 | -7979.04 | -1.652 | 7.41E-07 |
| 11 | 117192701 | rs55986402 | C | A | 1.002 | -7977.388 | -7979.041 | -1.652 | 7.41E-07 |
| 11 | 117213160 | rs493516 | A | G | 0.985 | -7977.388 | -7979.041 | -1.652 | 7.41E-07 |
| 11 | 117398202 | rs7940527 | C | G | 1.015 | -7977.388 | -7979.04 | -1.652 | 7.41E-07 |
| 11 | 118091636 | rs7103936 | C | A | 1.015 | -7977.388 | -7979.04 | -1.652 | 7.41E-07 |
| 11 | 118127347 | rs1793149 | A | G | 1.022 | -7977.388 | -7979.04 | -1.652 | 7.41E-07 |
| 11 | 119238455 | rs11217257 | G | A | 0.979 | -7977.388 | -7979.041 | -1.652 | 7.41E-07 |
| 11 | 119365552 | rs683630 | T | C | 0.979 | -7977.388 | -7979.04 | -1.652 | 7.41E-07 |
| 11 | 120513914 | rs1317514 | C | T | 1.02 | -7977.388 | -7979.041 | -1.652 | 7.41E-07 |
| 11 | 120724121 | rs7104543 | G | A | 1.022 | -7977.388 | -7979.04 | -1.652 | 7.41E-07 |
| 11 | 116219458 | rs554865 | T | C | 1.02 | -7977.388 | -7979.042 | -1.653 | 7.41E-07 |
| 11 | 116438555 | rs12225221 | T | C | 0.998 | -7977.388 | -7979.041 | -1.653 | 7.41E-07 |
| 11 | 116440163 | rs4936357 | T | C | 1.001 | -7977.388 | -7979.042 | -1.653 | 7.41E-07 |
| 11 | 116440575 | rs10431039 | T | C | 1.001 | -7977.388 | -7979.041 | -1.653 | 7.41E-07 |
| 11 | 117208747 | rs645305 | C | T | 0.985 | -7977.388 | -7979.041 | -1.653 | 7.41E-07 |
| 11 | 117538059 | rs608811 | A | G | 1.022 | -7977.388 | -7979.042 | -1.653 | 7.41E-07 |
| 11 | 118088362 | rs11216821 | C | T | 1.014 | -7977.388 | -7979.041 | -1.653 | 7.41E-07 |
| 11 | 118112841 | rs1715418 | T | A | 1.019 | -7977.388 | -7979.041 | -1.653 | 7.41E-07 |
| 11 | 119438208 | rs2663634 | T | C | 1.007 | -7977.388 | -7979.042 | -1.653 | 7.41E-07 |
| 11 | 116440678 | rs10431079 | G | A | 1.009 | -7977.388 | -7979.042 | -1.654 | 7.40E-07 |
| 11 | 117210901 | rs536021 | C | T | 0.985 | -7977.388 | -7979.043 | -1.654 | 7.40E-07 |
| 11 | 117212892 | rs475385 | T | C | 0.986 | -7977.388 | -7979.042 | -1.654 | 7.40E-07 |
| 11 | 116439792 | rs7943469 | G | T | 1.008 | -7977.388 | -7979.043 | -1.655 | 7.39E-07 |
| 11 | 116440036 | rs1261967419 | A | T | 1.008 | -7977.388 | -7979.043 | -1.655 | 7.39E-07 |
| 11 | 117189201 | rs61903708 | G | A | 1.001 | -7977.388 | -7979.043 | -1.655 | 7.39E-07 |
| 11 | 117208270 | rs875209 | A | G | 0.985 | -7977.388 | -7979.043 | -1.655 | 7.39E-07 |
| 11 | 117209268 | rs573801 | T | C | 0.985 | -7977.388 | -7979.043 | -1.655 | 7.39E-07 |
| 11 | 117229891 | rs2508664 | T | C | 0.988 | -7977.388 | -7979.043 | -1.655 | 7.39E-07 |
| 11 | 117231611 | rs11216366 | T | G | 0.988 | -7977.388 | -7979.043 | -1.655 | 7.39E-07 |
| 11 | 117239190 | rs7115526 | T | G | 1.019 | -7977.388 | -7979.043 | -1.655 | 7.39E-07 |
| 11 | 117248268 | rs7952470 | A | G | 0.989 | -7977.388 | -7979.043 | -1.655 | 7.39E-07 |
| 11 | 118075152 | rs11216816 | A | C | 1.007 | -7977.388 | -7979.044 | -1.655 | 7.39E-07 |

| | | | | | | | |
|---|---|---|---|---|---|---|---|
| 11 | 118075474 | rs11216817 | T | C | NULL | 0.695296669 | NULL&M_Disease |
| 11 | 118083664 | rs1715429 | A | G | NULL | 0.695296669 | NULL&M_Disease |
| 11 | 118116522 | rs1614945 | C | A | NULL | 0.695296669 | NULL&M_Disease |
| 11 | 118837113 | rs3890562 | G | A | NULL | 0.695296669 | NULL&M_Disease |
| 11 | 120294861 | rs7115714 | A | G | NULL | 0.695296669 | NULL&M_Disease |
| 11 | 117212337 | rs474891 | T | G | NULL | 0.695296669 | NULL&M_Disease |
| 11 | 117212640 | rs561473 | C | T | NULL | 0.695296669 | NULL&M_Disease |
| 11 | 117237207 | rs11606575 | G | C | NULL | 0.695296669 | NULL&M_Disease |
| 11 | 118069979 | rs7116375 | A | C | NULL | 0.695296669 | NULL&M_Disease |
| 11 | 118070788 | rs11600134 | A | G | NULL | 0.695296669 | NULL&M_Disease |
| 11 | 118114258 | rs2853032 | G | C | NULL | 0.695296669 | NULL&M_Disease |
| 11 | 118126920 | rs1793151 | C | T | NULL | 0.695296669 | NULL&M_Disease |
| 11 | 118127314 | rs1793150 | G | A | NULL | 0.695296669 | NULL&M_Disease |
| 11 | 119361783 | rs689291 | C | G | NULL | 0.695296669 | NULL&M_Disease |
| 11 | 119362414 | rs686593 | G | C | NULL | 0.695296669 | NULL&M_Disease |
| 11 | 119383853 | rs592525 | A | G | NULL | 0.695296669 | NULL&M_Disease |
| 11 | 119480880 | rs1567378 | G | A | NULL | 0.695296669 | NULL&M_Disease |
| 11 | 120194024 | rs12799480 | T | C | NULL | 0.695296669 | NULL&M_Disease |
| 11 | 116368043 | rs1371711774 | T | C | NULL | 0.695296669 | NULL&M_Disease |
| 11 | 117178225 | rs558229 | G | A | NULL | 0.695296669 | NULL&M_Disease |
| 11 | 117187551 | rs11601511 | C | G | NULL | 0.695296669 | NULL&M_Disease |
| 11 | 117208949 | rs948181 | T | C | NULL | 0.695296669 | NULL&M_Disease |
| 11 | 117240357 | rs9651682 | A | G | NULL | 0.695296669 | NULL&M_Disease |
| 11 | 118291210 | rs58577143 | G | A | NULL | 0.695296669 | NULL&M_Disease |
| 11 | 118959331 | rs17075 | G | A | NULL | 0.695296669 | NULL&M_Disease |
| 11 | 116454342 | rs61904848 | C | A | NULL | 0.69635493 | NULL&M_Disease |
| 11 | 117210707 | rs578781 | A | G | NULL | 0.69635493 | NULL&M_Disease |
| 11 | 117307219 | rs11216385 | T | C | NULL | 0.69635493 | NULL&M_Disease |
| 11 | 118119020 | rs3018367 | T | C | NULL | 0.69635493 | NULL&M_Disease |
| 11 | 120190262 | rs11217815 | A | G | NULL | 0.69635493 | NULL&M_Disease |
| 11 | 120624775 | rs2852244 | C | G | NULL | 0.69635493 | NULL&M_Disease |
| 11 | 120846239 | rs570373 | G | A | NULL | 0.69635493 | NULL&M_Disease |
| 11 | 116568516 | rs1787690 | T | G | NULL | 0.69635493 | NULL&M_Disease |
| 11 | 118122737 | rs1715458 | A | G | NULL | 0.69635493 | NULL&M_Disease |
| 11 | 118123488 | rs1715457 | A | G | NULL | 0.69635493 | NULL&M_Disease |
| 11 | 118128445 | rs1715450 | C | G | NULL | 0.69635493 | NULL&M_Disease |
| 11 | 118128455 | rs1715449 | G | A | NULL | 0.69635493 | NULL&M_Disease |
| 11 | 117227694 | rs595367 | C | T | NULL | 0.69635493 | NULL&M_Disease |
| 11 | 117421799 | rs597583 | G | C | NULL | 0.69635493 | NULL&M_Disease |
| 11 | 118069240 | rs7112584 | A | G | NULL | 0.69635493 | NULL&M_Disease |
| 11 | 118105473 | rs12362596 | G | C | NULL | 0.69635493 | NULL&M_Disease |
| 11 | 116306907 | rs35724134 | A | C | NULL | 0.69635493 | NULL&M_Disease |
| 11 | 116569204 | rs947986 | A | G | NULL | 0.69635493 | NULL&M_Disease |
| 11 | 117214091 | rs573827 | A | G | NULL | 0.69635493 | NULL&M_Disease |
| 11 | 117238491 | rs10892098 | T | C | NULL | 0.69635493 | NULL&M_Disease |
| 11 | 117873372 | rs2508446 | A | G | NULL | 0.69635493 | NULL&M_Disease |
| 11 | 118071836 | rs11216813 | A | G | NULL | 0.69635493 | NULL&M_Disease |
| 11 | 118887581 | rs7105187 | T | C | NULL | 0.69635493 | NULL&M_Disease |
| 11 | 119365380 | rs1981988 | C | G | NULL | 0.69635493 | NULL&M_Disease |
| 11 | 119941819 | rs11608135 | A | G | NULL | 0.69635493 | NULL&M_Disease |
| 11 | 116565309 | rs12293222 | A | G | NULL | 0.69635493 | NULL&M_Disease |
| 11 | 117141469 | rs4936369 | C | T | NULL | 0.69635493 | NULL&M_Disease |
| 11 | 117955759 | rs10892208 | G | A | NULL | 0.69635493 | NULL&M_Disease |
| 11 | 118992177 | rs583713 | T | C | NULL | 0.69635493 | NULL&M_Disease |
| 11 | 119368153 | rs10892374 | G | A | NULL | 0.69635493 | NULL&M_Disease |
| 11 | 119368188 | rs10892375 | T | C | NULL | 0.69635493 | NULL&M_Disease |
| 11 | 116322769 | rs11602880 | A | G | NULL | 0.69635493 | NULL&M_Disease |
| 11 | 117447031 | rs505824 | T | C | NULL | 0.69635493 | NULL&M_Disease |
| 11 | 118080685 | rs12286021 | C | T | NULL | 0.69635493 | NULL&M_Disease |
| 11 | 119711982 | rs549406 | C | T | NULL | 0.69635493 | NULL&M_Disease |
| 11 | 120261187 | rs11217852 | G | A | NULL | 0.69635493 | NULL&M_Disease |
| 11 | 116536281 | rs1240776 | T | C | NULL | 0.69635493 | NULL&M_Disease |
| 11 | 117451094 | rs536031 | A | T | NULL | 0.69635493 | NULL&M_Disease |
| 11 | 117669418 | rs4528371 | T | A | NULL | 0.69635493 | NULL&M_Disease |
| 11 | 117841718 | rs34306783 | T | C | NULL | 0.69635493 | NULL&M_Disease |
| 11 | 120852324 | rs526397 | G | T | NULL | 0.69635493 | NULL&M_Disease |
| 11 | 118072232 | rs11216815 | A | G | NULL | 0.69635493 | NULL&M_Disease |
| 11 | 118129705 | rs1793135 | T | C | NULL | 0.69635493 | NULL&M_Disease |
| 11 | 118959732 | rs549893 | C | T | NULL | 0.69635493 | NULL&M_Disease |
| 11 | 120233564 | rs7924975 | C | G | NULL | 0.69635493 | NULL&M_Disease |
| 11 | 116439615 | rs7928307 | T | C | NULL | 0.69635493 | NULL&M_Disease |
| 11 | 116565055 | rs1784118 | T | C | NULL | 0.69635493 | NULL&M_Disease |
| 11 | 116571284 | rs11601584 | C | T | NULL | 0.69635493 | NULL&M_Disease |
| 11 | 117456121 | rs11216462 | C | G | NULL | 0.69635493 | NULL&M_Disease |
| 11 | 118224059 | rs45508093 | G | A | NULL | 0.69635493 | NULL&M_Disease |
| 11 | 120324080 | rs1373270 | A | G | NULL | 0.69635493 | NULL&M_Disease |
| 11 | 120741935 | rs4936561 | C | G | NULL | 0.69635493 | NULL&M_Disease |
| 11 | 120851125 | rs598804 | T | C | NULL | 0.69635493 | NULL&M_Disease |
| 11 | 120854479 | rs540787 | T | C | NULL | 0.69635493 | NULL&M_Disease |
| 11 | 116562288 | rs1787702 | A | G | NULL | 0.69635493 | NULL&M_Disease |
| 11 | 117228519 | rs12221818 | G | A | NULL | 0.69635493 | NULL&M_Disease |
| 11 | 118078718 | rs7101652 | A | G | NULL | 0.69635493 | NULL&M_Disease |
| 11 | 118080268 | rs11216818 | T | C | NULL | 0.69635493 | NULL&M_Disease |
| 11 | 118080520 | rs4306318 | T | C | NULL | 0.69635493 | NULL&M_Disease |
| 11 | 120148567 | rs2155809 | A | T | NULL | 0.69635493 | NULL&M_Disease |
| 11 | 120604533 | rs2852229 | A | G | NULL | 0.69635493 | NULL&M_Disease |
| 11 | 116218830 | rs2581476 | A | G | NULL | 0.697411114 | NULL&M_Disease |
| 11 | 117216599 | rs652346 | T | C | NULL | 0.697411114 | NULL&M_Disease |
| 11 | 117216603 | rs494722 | T | A | NULL | 0.697411114 | NULL&M_Disease |
| 11 | 117225467 | rs542900 | A | T | NULL | 0.697411114 | NULL&M_Disease |
| 11 | 117538766 | rs593380 | C | T | NULL | 0.697411114 | NULL&M_Disease |
| 11 | 117782638 | rs2277288 | G | A | NULL | 0.697411114 | NULL&M_Disease |
| 11 | 117824960 | rs2243935 | A | G | NULL | 0.697411114 | NULL&M_Disease |
| 11 | 119859475 | rs3740724 | T | C | NULL | 0.697411114 | NULL&M_Disease |
| 11 | 120852961 | rs499804 | A | C | NULL | 0.69635493 | NULL&M_Disease |
| 11 | 120853259 | rs645156 | T | C | NULL | 0.69635493 | NULL&M_Disease |

| | | | | | | | | | |
|---|---|---|---|---|---|---|---|---|---|
| 11 | 118075474 | rs11216817 | T | C | 1.007 | -7977.388 | -7979.044 | -1.655 | 7.39E-07 |
| 11 | 118083664 | rs1715429 | A | G | 0.982 | -7977.388 | -7979.044 | -1.655 | 7.39E-07 |
| 11 | 118116522 | rs1614945 | C | A | 1.019 | -7977.388 | -7979.044 | -1.655 | 7.39E-07 |
| 11 | 118837113 | rs3890562 | G | A | 1.022 | -7977.388 | -7979.043 | -1.655 | 7.39E-07 |
| 11 | 120294861 | rs7115714 | A | G | 1.021 | -7977.388 | -7979.043 | -1.655 | 7.39E-07 |
| 11 | 117212337 | rs474891 | T | G | 0.986 | -7977.388 | -7979.044 | -1.656 | 7.38E-07 |
| 11 | 117212640 | rs561473 | C | T | 0.986 | -7977.388 | -7979.044 | -1.656 | 7.38E-07 |
| 11 | 117237207 | rs11606575 | G | C | 1.006 | -7977.388 | -7979.044 | -1.656 | 7.38E-07 |
| 11 | 118069979 | rs7116375 | A | C | 1.008 | -7977.388 | -7979.044 | -1.656 | 7.38E-07 |
| 11 | 118070788 | rs11600134 | A | G | 1.008 | -7977.388 | -7979.044 | -1.656 | 7.38E-07 |
| 11 | 118114258 | rs2853032 | G | C | 1.019 | -7977.388 | -7979.044 | -1.656 | 7.38E-07 |
| 11 | 118126920 | rs1793151 | C | T | 1.022 | -7977.388 | -7979.044 | -1.656 | 7.38E-07 |
| 11 | 118127314 | rs1793150 | G | A | 1.022 | -7977.388 | -7979.044 | -1.656 | 7.38E-07 |
| 11 | 119361783 | rs689291 | C | G | 0.979 | -7977.388 | -7979.045 | -1.656 | 7.38E-07 |
| 11 | 119362414 | rs686593 | G | C | 0.979 | -7977.388 | -7979.045 | -1.656 | 7.38E-07 |
| 11 | 119383853 | rs592525 | A | G | 0.979 | -7977.388 | -7979.045 | -1.656 | 7.38E-07 |
| 11 | 119480880 | rs1567378 | G | A | 0.98 | -7977.388 | -7979.044 | -1.656 | 7.38E-07 |
| 11 | 120194024 | rs12799480 | T | C | 0.979 | -7977.388 | -7979.045 | -1.656 | 7.38E-07 |
| 11 | 116368043 | rs1371711774 | T | C | 0.979 | -7977.388 | -7979.045 | -1.657 | 7.38E-07 |
| 11 | 117178225 | rs558229 | G | A | 0.991 | -7977.388 | -7979.046 | -1.657 | 7.38E-07 |
| 11 | 117187551 | rs11601511 | C | G | 0.999 | -7977.388 | -7979.046 | -1.657 | 7.38E-07 |
| 11 | 117208949 | rs948181 | T | C | 0.986 | -7977.388 | -7979.046 | -1.657 | 7.38E-07 |
| 11 | 117240357 | rs9651682 | A | G | 1.019 | -7977.388 | -7979.046 | -1.657 | 7.38E-07 |
| 11 | 118291210 | rs58577143 | G | A | 0.989 | -7977.388 | -7979.045 | -1.657 | 7.38E-07 |
| 11 | 118959331 | rs17075 | G | A | 0.979 | -7977.388 | -7979.045 | -1.657 | 7.38E-07 |
| 11 | 116454342 | rs61904848 | C | A | 0.98 | -7977.388 | -7979.046 | -1.658 | 7.37E-07 |
| 11 | 117210707 | rs578781 | A | G | 0.986 | -7977.388 | -7979.046 | -1.658 | 7.37E-07 |
| 11 | 117307219 | rs11216385 | T | C | 0.99 | -7977.388 | -7979.047 | -1.658 | 7.37E-07 |
| 11 | 118119020 | rs3018367 | T | C | 1.019 | -7977.388 | -7979.046 | -1.658 | 7.37E-07 |
| 11 | 120190262 | rs11217815 | A | G | 0.979 | -7977.388 | -7979.046 | -1.658 | 7.37E-07 |
| 11 | 120624775 | rs2852244 | C | G | 0.981 | -7977.388 | -7979.047 | -1.658 | 7.37E-07 |
| 11 | 120846239 | rs570373 | G | A | 1.012 | -7977.388 | -7979.047 | -1.658 | 7.37E-07 |
| 11 | 116568516 | rs1787690 | T | G | 0.979 | -7977.388 | -7979.047 | -1.659 | 7.36E-07 |
| 11 | 118122737 | rs1715458 | A | G | 1.019 | -7977.388 | -7979.047 | -1.659 | 7.36E-07 |
| 11 | 118123488 | rs1715457 | A | G | 1.019 | -7977.388 | -7979.047 | -1.659 | 7.36E-07 |
| 11 | 118128445 | rs1715450 | C | G | 1.022 | -7977.388 | -7979.047 | -1.659 | 7.36E-07 |
| 11 | 118128455 | rs1715449 | G | A | 1.022 | -7977.388 | -7979.047 | -1.659 | 7.36E-07 |
| 11 | 117227694 | rs595367 | C | T | 0.989 | -7977.388 | -7979.048 | -1.66 | 7.35E-07 |
| 11 | 117421799 | rs597583 | G | C | 1.013 | -7977.388 | -7979.048 | -1.66 | 7.35E-07 |
| 11 | 118069240 | rs7112584 | A | G | 1.007 | -7977.388 | -7979.049 | -1.66 | 7.35E-07 |
| 11 | 118105473 | rs12362596 | G | C | 1.022 | -7977.388 | -7979.048 | -1.66 | 7.35E-07 |
| 11 | 116306907 | rs35724134 | A | C | 1.02 | -7977.388 | -7979.049 | -1.661 | 7.35E-07 |
| 11 | 116569204 | rs947986 | A | G | 0.979 | -7977.388 | -7979.05 | -1.661 | 7.35E-07 |
| 11 | 117214091 | rs573827 | A | G | 0.986 | -7977.388 | -7979.049 | -1.661 | 7.35E-07 |
| 11 | 117238491 | rs10892098 | T | C | 1.018 | -7977.388 | -7979.05 | -1.661 | 7.35E-07 |
| 11 | 117873372 | rs2508446 | A | G | 0.98 | -7977.388 | -7979.049 | -1.661 | 7.35E-07 |
| 11 | 118071836 | rs11216813 | A | G | 1.006 | -7977.388 | -7979.049 | -1.661 | 7.35E-07 |
| 11 | 118887581 | rs7105187 | T | C | 1.017 | -7977.388 | -7979.049 | -1.661 | 7.35E-07 |
| 11 | 119365380 | rs1981988 | C | G | 1.022 | -7977.388 | -7979.049 | -1.661 | 7.35E-07 |
| 11 | 119941819 | rs11608135 | A | G | 1.021 | -7977.388 | -7979.049 | -1.661 | 7.35E-07 |
| 11 | 116565309 | rs12293222 | A | G | 1.02 | -7977.388 | -7979.05 | -1.662 | 7.34E-07 |
| 11 | 117141469 | rs4936369 | C | T | 0.992 | -7977.388 | -7979.05 | -1.662 | 7.34E-07 |
| 11 | 117955759 | rs10892208 | G | A | 0.979 | -7977.388 | -7979.051 | -1.662 | 7.34E-07 |
| 11 | 118992177 | rs583713 | T | C | 0.979 | -7977.388 | -7979.05 | -1.662 | 7.34E-07 |
| 11 | 119368153 | rs10892374 | G | A | 1.021 | -7977.388 | -7979.051 | -1.662 | 7.34E-07 |
| 11 | 119368188 | rs10892375 | T | C | 1.022 | -7977.388 | -7979.05 | -1.662 | 7.34E-07 |
| 11 | 116322769 | rs11602880 | A | G | 0.993 | -7977.388 | -7979.051 | -1.663 | 7.33E-07 |
| 11 | 117447031 | rs505824 | T | C | 0.985 | -7977.388 | -7979.052 | -1.663 | 7.33E-07 |
| 11 | 118080685 | rs12286021 | C | T | 1.007 | -7977.388 | -7979.051 | -1.663 | 7.33E-07 |
| 11 | 119711982 | rs549406 | C | T | 0.985 | -7977.388 | -7979.051 | -1.663 | 7.33E-07 |
| 11 | 120261187 | rs11217852 | G | A | 1.021 | -7977.388 | -7979.052 | -1.663 | 7.33E-07 |
| 11 | 116536281 | rs1240776 | T | C | 0.979 | -7977.388 | -7979.052 | -1.664 | 7.33E-07 |
| 11 | 117451094 | rs536031 | A | T | 0.985 | -7977.388 | -7979.052 | -1.664 | 7.33E-07 |
| 11 | 117669418 | rs4528371 | T | A | 0.99 | -7977.388 | -7979.052 | -1.664 | 7.33E-07 |
| 11 | 117841718 | rs34306783 | T | C | 0.981 | -7977.388 | -7979.053 | -1.664 | 7.33E-07 |
| 11 | 120852324 | rs526397 | G | T | 1.012 | -7977.388 | -7979.052 | -1.664 | 7.33E-07 |
| 11 | 118072232 | rs11216815 | A | G | 1.006 | -7977.388 | -7979.053 | -1.665 | 7.32E-07 |
| 11 | 118129705 | rs1793135 | T | C | 1.019 | -7977.388 | -7979.053 | -1.665 | 7.32E-07 |
| 11 | 118959732 | rs549893 | C | T | 0.979 | -7977.388 | -7979.053 | -1.665 | 7.32E-07 |
| 11 | 120233564 | rs7924975 | C | G | 1.021 | -7977.388 | -7979.053 | -1.665 | 7.32E-07 |
| 11 | 116439615 | rs7928307 | T | C | 1.006 | -7977.388 | -7979.055 | -1.666 | 7.31E-07 |
| 11 | 116565055 | rs1784118 | T | C | 0.979 | -7977.388 | -7979.054 | -1.666 | 7.31E-07 |
| 11 | 116571284 | rs11601584 | C | T | 0.985 | -7977.388 | -7979.054 | -1.666 | 7.31E-07 |
| 11 | 117456121 | rs11216462 | C | G | 0.979 | -7977.388 | -7979.055 | -1.666 | 7.31E-07 |
| 11 | 118224059 | rs45508093 | G | A | 0.992 | -7977.388 | -7979.054 | -1.666 | 7.31E-07 |
| 11 | 120324080 | rs1373270 | A | G | 1.021 | -7977.388 | -7979.055 | -1.666 | 7.31E-07 |
| 11 | 120741935 | rs4936561 | C | G | 0.979 | -7977.388 | -7979.054 | -1.666 | 7.31E-07 |
| 11 | 120851125 | rs598804 | T | C | 1.012 | -7977.388 | -7979.054 | -1.666 | 7.31E-07 |
| 11 | 120854479 | rs540787 | T | C | 1.012 | -7977.388 | -7979.055 | -1.666 | 7.31E-07 |
| 11 | 116562288 | rs1787702 | A | G | 1.019 | -7977.388 | -7979.056 | -1.667 | 7.30E-07 |
| 11 | 117228519 | rs12221818 | G | A | 0.999 | -7977.388 | -7979.055 | -1.667 | 7.30E-07 |
| 11 | 118078718 | rs7101652 | A | G | 1.006 | -7977.388 | -7979.055 | -1.667 | 7.30E-07 |
| 11 | 118080268 | rs11216818 | T | C | 1.006 | -7977.388 | -7979.055 | -1.667 | 7.30E-07 |
| 11 | 118080520 | rs4306318 | T | C | 1.006 | -7977.388 | -7979.055 | -1.667 | 7.30E-07 |
| 11 | 120148567 | rs2155809 | A | T | 1.018 | -7977.388 | -7979.055 | -1.667 | 7.30E-07 |
| 11 | 120604533 | rs2852229 | A | G | 0.983 | -7977.388 | -7979.055 | -1.667 | 7.30E-07 |
| 11 | 116218830 | rs2581476 | A | G | 1.02 | -7977.388 | -7979.057 | -1.668 | 7.30E-07 |
| 11 | 117216599 | rs652346 | T | C | 0.99 | -7977.388 | -7979.056 | -1.668 | 7.30E-07 |
| 11 | 117216603 | rs494722 | T | A | 0.99 | -7977.388 | -7979.056 | -1.668 | 7.30E-07 |
| 11 | 117225467 | rs542900 | A | T | 0.991 | -7977.388 | -7979.057 | -1.668 | 7.30E-07 |
| 11 | 117538766 | rs593380 | C | T | 1.021 | -7977.388 | -7979.057 | -1.668 | 7.30E-07 |
| 11 | 117782638 | rs2277288 | G | A | 0.98 | -7977.388 | -7979.056 | -1.668 | 7.30E-07 |
| 11 | 117824960 | rs2243935 | A | G | 0.981 | -7977.388 | -7979.056 | -1.668 | 7.30E-07 |
| 11 | 119859475 | rs3740724 | T | C | 1.021 | -7977.388 | -7979.056 | -1.668 | 7.30E-07 |
| 11 | 120852961 | rs499804 | A | C | 1.012 | -7977.388 | -7979.056 | -1.668 | 7.30E-07 |
| 11 | 120853259 | rs645156 | T | C | 1.012 | -7977.388 | -7979.056 | -1.668 | 7.30E-07 |

| | | | | | | | |
|---|---|---|---|---|---|---|---|
| 11 | 120853941 | rs658696 | A | G | NULL | 0.69635493 | NULL&M_Disease |
| 11 | 116296641 | rs10750091 | A | G | NULL | 0.697411114 | NULL&M_Disease |
| 11 | 116299004 | rs1202587279 | T | C | NULL | 0.697411114 | NULL&M_Disease |
| 11 | 117497778 | rs11216474 | G | A | NULL | 0.697411114 | NULL&M_Disease |
| 11 | 118087841 | rs7939622 | A | G | NULL | 0.697411114 | NULL&M_Disease |
| 11 | 119482594 | rs7935369 | T | C | NULL | 0.697411114 | NULL&M_Disease |
| 11 | 120605140 | rs2846122 | A | G | NULL | 0.697411114 | NULL&M_Disease |
| 11 | 116293739 | rs501433 | T | C | NULL | 0.697411114 | NULL&M_Disease |
| 11 | 116326138 | rs12419822 | A | C | NULL | 0.697411114 | NULL&M_Disease |
| 11 | 116552766 | rs1787717 | T | C | NULL | 0.697411114 | NULL&M_Disease |
| 11 | 117233427 | rs11828506 | T | C | NULL | 0.697411114 | NULL&M_Disease |
| 11 | 117831809 | rs3020913 | G | A | NULL | 0.697411114 | NULL&M_Disease |
| 11 | 117927961 | rs585533 | A | G | NULL | 0.697411114 | NULL&M_Disease |
| 11 | 118072716 | rs12801003 | T | C | NULL | 0.697411114 | NULL&M_Disease |
| 11 | 120308290 | rs10892578 | G | A | NULL | 0.697411114 | NULL&M_Disease |
| 11 | 120496044 | rs12281635 | G | A | NULL | 0.697411114 | NULL&M_Disease |
| 11 | 116354930 | rs11826857 | C | T | NULL | 0.697411114 | NULL&M_Disease |
| 11 | 116354937 | rs11824358 | T | C | NULL | 0.697411114 | NULL&M_Disease |
| 11 | 116393980 | rs4938276 | T | C | NULL | 0.697411114 | NULL&M_Disease |
| 11 | 116426097 | rs1962593 | T | C | NULL | 0.697411114 | NULL&M_Disease |
| 11 | 116441277 | rs7110823 | G | C | NULL | 0.697411114 | NULL&M_Disease |
| 11 | 117323082 | rs11216398 | T | C | NULL | 0.697411114 | NULL&M_Disease |
| 11 | 117492654 | rs7116918 | G | A | NULL | 0.697411114 | NULL&M_Disease |
| 11 | 117709028 | rs516655 | G | A | NULL | 0.697411114 | NULL&M_Disease |
| 11 | 118840857 | rs12420418 | T | C | NULL | 0.697411114 | NULL&M_Disease |
| 11 | 119363485 | rs671182 | C | T | NULL | 0.697411114 | NULL&M_Disease |
| 11 | 119877773 | rs1944497 | G | A | NULL | 0.697411114 | NULL&M_Disease |
| 11 | 116538215 | rs1784126 | T | G | NULL | 0.697411114 | NULL&M_Disease |
| 11 | 117225441 | rs1041395195 | T | G | NULL | 0.697411114 | NULL&M_Disease |
| 11 | 117532543 | rs10892152 | G | A | NULL | 0.697411114 | NULL&M_Disease |
| 11 | 118115879 | rs1793165 | A | C | NULL | 0.697411114 | NULL&M_Disease |
| 11 | 119359257 | rs623989 | G | T | NULL | 0.697411114 | NULL&M_Disease |
| 11 | 119613090 | rs35351462 | A | C | NULL | 0.697411114 | NULL&M_Disease |
| 11 | 120855775 | rs61903170 | C | T | NULL | 0.697411114 | NULL&M_Disease |
| 11 | 116226662 | rs562236 | C | T | NULL | 0.697411114 | NULL&M_Disease |
| 11 | 116441138 | rs7110879 | A | G | NULL | 0.697411114 | NULL&M_Disease |
| 11 | 117192783 | rs60837248 | G | A | NULL | 0.697411114 | NULL&M_Disease |
| 11 | 120309036 | rs7951250 | G | A | NULL | 0.697411114 | NULL&M_Disease |
| 11 | 120685533 | rs3020103 | G | A | NULL | 0.697411114 | NULL&M_Disease |
| 11 | 117237487 | rs7109923 | G | A | NULL | 0.697411114 | NULL&M_Disease |
| 11 | 117872734 | rs2512147 | T | C | NULL | 0.697411114 | NULL&M_Disease |
| 11 | 118071857 | rs11216814 | G | A | NULL | 0.697411114 | NULL&M_Disease |
| 11 | 118877563 | rs58709307 | T | C | NULL | 0.697411114 | NULL&M_Disease |
| 11 | 119367405 | rs1366878 | T | C | NULL | 0.697411114 | NULL&M_Disease |
| 11 | 119877246 | rs4938754 | T | C | NULL | 0.697411114 | NULL&M_Disease |
| 11 | 120145046 | rs34172923 | G | C | NULL | 0.697411114 | NULL&M_Disease |
| 11 | 120732508 | rs3824978 | T | C | NULL | 0.697411114 | NULL&M_Disease |
| 11 | 116562266 | rs1787703 | T | C | NULL | 0.697411114 | NULL&M_Disease |
| 11 | 116575473 | rs4938301 | A | G | NULL | 0.697411114 | NULL&M_Disease |
| 11 | 116699445 | rs595049 | C | A | NULL | 0.697411114 | NULL&M_Disease |
| 11 | 117118712 | rs7928525 | T | A | NULL | 0.697411114 | NULL&M_Disease |
| 11 | 117120998 | rs1056136 | G | T | NULL | 0.697411114 | NULL&M_Disease |
| 11 | 118071762 | rs11216812 | A | G | NULL | 0.697411114 | NULL&M_Disease |
| 11 | 118121399 | rs7926717 | T | G | NULL | 0.697411114 | NULL&M_Disease |
| 11 | 118889185 | rs11217125 | C | T | NULL | 0.697411114 | NULL&M_Disease |
| 11 | 118992919 | rs673454 | A | C | NULL | 0.697411114 | NULL&M_Disease |
| 11 | 120608816 | rs2004901 | C | T | NULL | 0.697411114 | NULL&M_Disease |
| 11 | 116292614 | rs571025 | T | C | NULL | 0.697411114 | NULL&M_Disease |
| 11 | 116293365 | rs505009 | A | G | NULL | 0.697411114 | NULL&M_Disease |
| 11 | 116312926 | rs486476 | G | C | NULL | 0.697411114 | NULL&M_Disease |
| 11 | 116538576 | rs1784127 | A | G | NULL | 0.697411114 | NULL&M_Disease |
| 11 | 118246071 | rs61059752 | A | C | NULL | 0.697411114 | NULL&M_Disease |
| 11 | 118258056 | rs1079118 | A | G | NULL | 0.697411114 | NULL&M_Disease |
| 11 | 118263054 | rs73613959 | A | G | NULL | 0.697411114 | NULL&M_Disease |
| 11 | 118281418 | rs60545357 | C | A | NULL | 0.697411114 | NULL&M_Disease |
| 11 | 118590864 | rs11216982 | A | G | NULL | 0.697411114 | NULL&M_Disease |
| 11 | 119841515 | rs656382 | T | C | NULL | 0.697411114 | NULL&M_Disease |
| 11 | 120710315 | rs7949440 | A | G | NULL | 0.697411114 | NULL&M_Disease |
| 11 | 120850070 | rs478978 | G | C | NULL | 0.697411114 | NULL&M_Disease |
| 11 | 120850185 | rs477389 | G | C | NULL | 0.697411114 | NULL&M_Disease |
| 11 | 116296520 | rs10790141 | A | G | NULL | 0.697411114 | NULL&M_Disease |
| 11 | 116572530 | rs11216100 | G | C | NULL | 0.697411114 | NULL&M_Disease |
| 11 | 116645149 | rs2041967 | G | A | NULL | 0.697411114 | NULL&M_Disease |
| 11 | 117222139 | rs688207 | T | C | NULL | 0.697411114 | NULL&M_Disease |
| 11 | 117222592 | rs1369553921 | A | G | NULL | 0.697411114 | NULL&M_Disease |
| 11 | 117222942 | rs514806 | T | G | NULL | 0.697411114 | NULL&M_Disease |
| 11 | 117285057 | rs10736488 | G | A | NULL | 0.697411114 | NULL&M_Disease |
| 11 | 118270715 | rs45480496 | A | G | NULL | 0.697411114 | NULL&M_Disease |
| 11 | 119481661 | rs4581455 | A | G | NULL | 0.697411114 | NULL&M_Disease |
| 11 | 119585681 | rs7931316 | C | T | NULL | 0.698465216 | NULL&M_Disease |
| 11 | 120741937 | rs4936562 | C | T | NULL | 0.697411114 | NULL&M_Disease |
| 11 | 117235412 | rs7124084 | T | C | NULL | 0.698465216 | NULL&M_Disease |
| 11 | 117454633 | rs638979 | T | C | NULL | 0.698465216 | NULL&M_Disease |
| 11 | 118129583 | rs11216839 | T | A | NULL | 0.697411114 | NULL&M_Disease |
| 11 | 118616253 | rs7124498 | A | G | NULL | 0.697411114 | NULL&M_Disease |
| 11 | 118626286 | rs11217014 | C | T | NULL | 0.698465216 | NULL&M_Disease |
| 11 | 118627149 | rs11217016 | C | A | NULL | 0.698465216 | NULL&M_Disease |
| 11 | 119196915 | rs2509390 | C | A | NULL | 0.697411114 | NULL&M_Disease |
| 11 | 116509339 | rs58990403 | G | A | NULL | 0.698465216 | NULL&M_Disease |
| 11 | 116510680 | rs58089984 | G | T | NULL | 0.698465216 | NULL&M_Disease |
| 11 | 117196583 | rs73014327 | C | T | NULL | 0.698465216 | NULL&M_Disease |
| 11 | 117232954 | rs7934067 | G | A | NULL | 0.698465216 | NULL&M_Disease |
| 11 | 117692170 | rs10750114 | G | T | NULL | 0.698465216 | NULL&M_Disease |
| 11 | 118246906 | rs7933963 | T | C | NULL | 0.698465216 | NULL&M_Disease |
| 11 | 118886117 | rs10111 | T | C | NULL | 0.698465216 | NULL&M_Disease |
| 11 | 118889132 | rs11217124 | G | A | NULL | 0.698465216 | NULL&M_Disease |

| | | | | | | | | | |
|---|---|---|---|---|---|---|---|---|---|
| 11 | 120853941 | rs658696 | A | G | 1.012 | -7977.388 | -7979.056 | -1.668 | 7.30E-07 |
| 11 | 116296641 | rs10750091 | A | G | 1.016 | -7977.388 | -7979.057 | -1.669 | 7.29E-07 |
| 11 | 116299004 | rs1202587279 | T | C | 0.98 | -7977.388 | -7979.057 | -1.669 | 7.29E-07 |
| 11 | 117497778 | rs11216474 | G | A | 1.021 | -7977.388 | -7979.058 | -1.669 | 7.29E-07 |
| 11 | 118087841 | rs7939622 | A | G | 1.012 | -7977.388 | -7979.057 | -1.669 | 7.29E-07 |
| 11 | 119482594 | rs7935369 | T | C | 0.981 | -7977.388 | -7979.057 | -1.669 | 7.29E-07 |
| 11 | 120605140 | rs2846122 | A | G | 0.983 | -7977.388 | -7979.057 | -1.669 | 7.29E-07 |
| 11 | 116293739 | rs501433 | T | C | 1.016 | -7977.388 | -7979.058 | -1.67 | 7.28E-07 |
| 11 | 116326138 | rs12419822 | A | C | 1.003 | -7977.388 | -7979.059 | -1.67 | 7.28E-07 |
| 11 | 116552766 | rs1787717 | T | C | 0.979 | -7977.388 | -7979.058 | -1.67 | 7.28E-07 |
| 11 | 117233427 | rs11828506 | T | C | 0.99 | -7977.388 | -7979.058 | -1.67 | 7.28E-07 |
| 11 | 117831809 | rs3020913 | G | A | 0.981 | -7977.388 | -7979.058 | -1.67 | 7.28E-07 |
| 11 | 117927961 | rs585533 | A | G | 1.016 | -7977.388 | -7979.058 | -1.67 | 7.28E-07 |
| 11 | 118072716 | rs12801003 | T | C | 1.005 | -7977.388 | -7979.059 | -1.67 | 7.28E-07 |
| 11 | 120308290 | rs10892578 | G | A | 1.021 | -7977.388 | -7979.058 | -1.67 | 7.28E-07 |
| 11 | 120496044 | rs12281635 | G | A | 0.98 | -7977.388 | -7979.058 | -1.67 | 7.28E-07 |
| 11 | 116354930 | rs11826857 | C | T | 1.002 | -7977.388 | -7979.059 | -1.671 | 7.27E-07 |
| 11 | 116354937 | rs11824358 | T | C | 1.002 | -7977.388 | -7979.059 | -1.671 | 7.27E-07 |
| 11 | 116393980 | rs4938276 | T | C | 1.019 | -7977.388 | -7979.059 | -1.671 | 7.27E-07 |
| 11 | 116426097 | rs1962593 | T | C | 1.004 | -7977.388 | -7979.059 | -1.671 | 7.27E-07 |
| 11 | 116441277 | rs7110823 | G | C | 1.005 | -7977.388 | -7979.059 | -1.671 | 7.27E-07 |
| 11 | 117323082 | rs11216398 | T | C | 1.001 | -7977.388 | -7979.059 | -1.671 | 7.27E-07 |
| 11 | 117492654 | rs7116918 | G | A | 1.021 | -7977.388 | -7979.06 | -1.671 | 7.27E-07 |
| 11 | 117709028 | rs516655 | G | A | 1.009 | -7977.388 | -7979.06 | -1.671 | 7.27E-07 |
| 11 | 118840857 | rs12420418 | T | C | 1.021 | -7977.388 | -7979.06 | -1.671 | 7.27E-07 |
| 11 | 119363485 | rs671182 | C | T | 0.98 | -7977.388 | -7979.059 | -1.671 | 7.27E-07 |
| 11 | 119877773 | rs1944497 | G | A | 1.021 | -7977.388 | -7979.06 | -1.671 | 7.27E-07 |
| 11 | 116538215 | rs1784126 | T | G | 0.979 | -7977.388 | -7979.061 | -1.672 | 7.27E-07 |
| 11 | 117225441 | rs1041395195 | T | G | 0.991 | -7977.388 | -7979.06 | -1.672 | 7.27E-07 |
| 11 | 117532543 | rs10892152 | G | A | 0.981 | -7977.388 | -7979.061 | -1.672 | 7.27E-07 |
| 11 | 118115879 | rs1793165 | A | C | 1.018 | -7977.388 | -7979.06 | -1.672 | 7.27E-07 |
| 11 | 119359257 | rs623989 | G | T | 0.98 | -7977.388 | -7979.061 | -1.672 | 7.27E-07 |
| 11 | 119613090 | rs35351462 | A | C | 1.011 | -7977.388 | -7979.06 | -1.672 | 7.27E-07 |
| 11 | 120855775 | rs61903170 | C | T | 1.011 | -7977.388 | -7979.06 | -1.672 | 7.27E-07 |
| 11 | 116226662 | rs562236 | C | T | 1.019 | -7977.388 | -7979.061 | -1.673 | 7.26E-07 |
| 11 | 116441138 | rs7110879 | A | G | 1.004 | -7977.388 | -7979.061 | -1.673 | 7.26E-07 |
| 11 | 117192783 | rs60837248 | G | A | 1 | -7977.388 | -7979.062 | -1.673 | 7.26E-07 |
| 11 | 120309036 | rs7951250 | G | A | 1.021 | -7977.388 | -7979.061 | -1.673 | 7.26E-07 |
| 11 | 120685533 | rs3020103 | G | A | 0.979 | -7977.388 | -7979.062 | -1.673 | 7.26E-07 |
| 11 | 117237487 | rs7109923 | G | A | 0.991 | -7977.388 | -7979.062 | -1.674 | 7.25E-07 |
| 11 | 117872734 | rs2512147 | T | C | 0.981 | -7977.388 | -7979.063 | -1.674 | 7.25E-07 |
| 11 | 118071857 | rs11216814 | G | A | 1.004 | -7977.388 | -7979.063 | -1.674 | 7.25E-07 |
| 11 | 118877563 | rs58709307 | T | C | 1.013 | -7977.388 | -7979.062 | -1.674 | 7.25E-07 |
| 11 | 119367405 | rs1366878 | T | C | 1.021 | -7977.388 | -7979.062 | -1.674 | 7.25E-07 |
| 11 | 119877246 | rs4938754 | T | C | 1.021 | -7977.388 | -7979.062 | -1.674 | 7.25E-07 |
| 11 | 120145046 | rs34172923 | G | C | 1.017 | -7977.388 | -7979.062 | -1.674 | 7.25E-07 |
| 11 | 120732508 | rs3824978 | T | C | 1.021 | -7977.388 | -7979.062 | -1.674 | 7.25E-07 |
| 11 | 116562266 | rs1787703 | T | C | 1.019 | -7977.388 | -7979.064 | -1.675 | 7.25E-07 |
| 11 | 116575473 | rs4938301 | A | G | 1.011 | -7977.388 | -7979.063 | -1.675 | 7.25E-07 |
| 11 | 116699445 | rs595049 | C | A | 1.02 | -7977.388 | -7979.064 | -1.675 | 7.25E-07 |
| 11 | 117118712 | rs7928525 | T | A | 1.002 | -7977.388 | -7979.063 | -1.675 | 7.25E-07 |
| 11 | 117120998 | rs1056136 | G | T | 1.001 | -7977.388 | -7979.063 | -1.675 | 7.25E-07 |
| 11 | 118071762 | rs11216812 | A | G | 1.004 | -7977.388 | -7979.063 | -1.675 | 7.25E-07 |
| 11 | 118121399 | rs7926717 | T | G | 1.011 | -7977.388 | -7979.063 | -1.675 | 7.25E-07 |
| 11 | 118889185 | rs11217125 | C | T | 1.017 | -7977.388 | -7979.063 | -1.675 | 7.25E-07 |
| 11 | 118992919 | rs673454 | A | C | 0.979 | -7977.388 | -7979.064 | -1.675 | 7.25E-07 |
| 11 | 120608816 | rs2004901 | C | T | 0.983 | -7977.388 | -7979.063 | -1.675 | 7.25E-07 |
| 11 | 116292614 | rs571025 | T | C | 1.016 | -7977.388 | -7979.064 | -1.676 | 7.24E-07 |
| 11 | 116293365 | rs505009 | A | G | 1.016 | -7977.388 | -7979.065 | -1.676 | 7.24E-07 |
| 11 | 116312926 | rs486476 | G | C | 1.012 | -7977.388 | -7979.064 | -1.676 | 7.24E-07 |
| 11 | 116538576 | rs1784127 | A | G | 0.98 | -7977.388 | -7979.064 | -1.676 | 7.24E-07 |
| 11 | 118246071 | rs61059752 | A | C | 0.997 | -7977.388 | -7979.064 | -1.676 | 7.24E-07 |
| 11 | 118258056 | rs1079118 | A | G | 0.997 | -7977.388 | -7979.065 | -1.676 | 7.24E-07 |
| 11 | 118263054 | rs73613959 | A | G | 0.997 | -7977.388 | -7979.064 | -1.676 | 7.24E-07 |
| 11 | 118281418 | rs60545357 | C | A | 0.997 | -7977.388 | -7979.064 | -1.676 | 7.24E-07 |
| 11 | 118590864 | rs11216982 | A | G | 1.016 | -7977.388 | -7979.064 | -1.676 | 7.24E-07 |
| 11 | 119841515 | rs656382 | T | C | 0.985 | -7977.388 | -7979.064 | -1.676 | 7.24E-07 |
| 11 | 120710315 | rs7949440 | A | G | 0.991 | -7977.388 | -7979.065 | -1.676 | 7.24E-07 |
| 11 | 120850070 | rs478978 | G | C | 1.011 | -7977.388 | -7979.064 | -1.676 | 7.24E-07 |
| 11 | 120850185 | rs477389 | G | C | 1.011 | -7977.388 | -7979.064 | -1.676 | 7.24E-07 |
| 11 | 116296520 | rs10790141 | A | G | 1.016 | -7977.388 | -7979.065 | -1.677 | 7.23E-07 |
| 11 | 116572530 | rs11216100 | G | C | 0.986 | -7977.388 | -7979.065 | -1.677 | 7.23E-07 |
| 11 | 116645149 | rs2041967 | G | A | 1.004 | -7977.388 | -7979.065 | -1.677 | 7.23E-07 |
| 11 | 117222139 | rs688207 | T | C | 0.991 | -7977.388 | -7979.066 | -1.677 | 7.23E-07 |
| 11 | 117222592 | rs1369553921 | A | G | 0.991 | -7977.388 | -7979.066 | -1.677 | 7.23E-07 |
| 11 | 117222942 | rs514806 | T | G | 0.991 | -7977.388 | -7979.066 | -1.677 | 7.23E-07 |
| 11 | 117285057 | rs10736488 | G | A | 0.989 | -7977.388 | -7979.066 | -1.677 | 7.23E-07 |
| 11 | 118270715 | rs45480496 | A | G | 0.997 | -7977.388 | -7979.065 | -1.677 | 7.23E-07 |
| 11 | 119481661 | rs4581455 | A | G | 0.981 | -7977.388 | -7979.065 | -1.677 | 7.23E-07 |
| 11 | 119585681 | rs7931316 | C | T | 1.021 | -7977.388 | -7979.066 | -1.677 | 7.23E-07 |
| 11 | 120741937 | rs4936562 | C | T | 0.98 | -7977.388 | -7979.066 | -1.677 | 7.23E-07 |
| 11 | 117235412 | rs7124084 | T | C | 0.991 | -7977.388 | -7979.066 | -1.678 | 7.22E-07 |
| 11 | 117454633 | rs638979 | T | C | 0.989 | -7977.388 | -7979.066 | -1.678 | 7.22E-07 |
| 11 | 118129583 | rs11216839 | T | A | 1.011 | -7977.388 | -7979.066 | -1.678 | 7.22E-07 |
| 11 | 118616253 | rs7124498 | A | G | 1.016 | -7977.388 | -7979.066 | -1.678 | 7.22E-07 |
| 11 | 118626286 | rs11217014 | C | T | 1.016 | -7977.388 | -7979.066 | -1.678 | 7.22E-07 |
| 11 | 118627149 | rs11217016 | C | A | 1.016 | -7977.388 | -7979.066 | -1.678 | 7.22E-07 |
| 11 | 119196915 | rs2509390 | C | A | 1.017 | -7977.388 | -7979.066 | -1.678 | 7.22E-07 |
| 11 | 116509339 | rs58990403 | G | A | 1.007 | -7977.388 | -7979.067 | -1.679 | 7.22E-07 |
| 11 | 116510680 | rs58089984 | G | T | 1.001 | -7977.388 | -7979.067 | -1.679 | 7.22E-07 |
| 11 | 117196583 | rs73014327 | C | T | 0.999 | -7977.388 | -7979.067 | -1.679 | 7.22E-07 |
| 11 | 117232954 | rs7934067 | G | A | 0.992 | -7977.388 | -7979.068 | -1.679 | 7.22E-07 |
| 11 | 117692170 | rs10750114 | G | T | 0.987 | -7977.388 | -7979.067 | -1.679 | 7.22E-07 |
| 11 | 118246906 | rs7933963 | T | C | 0.999 | -7977.388 | -7979.067 | -1.679 | 7.22E-07 |
| 11 | 118886117 | rs10111 | T | C | 1.013 | -7977.388 | -7979.067 | -1.679 | 7.22E-07 |
| 11 | 118889132 | rs11217124 | G | A | 1.016 | -7977.388 | -7979.067 | -1.679 | 7.22E-07 |

| | | | | | | | |
|---|---|---|---|---|---|---|---|
| 11 | 119695450 | rs525914 | T | C | NULL | 0.698465216 | NULL&M_Disease |
| 11 | 119847991 | rs621150 | T | C | NULL | 0.698465216 | NULL&M_Disease |
| 11 | 120101092 | rs2845705 | A | T | NULL | 0.698465216 | NULL&M_Disease |
| 11 | 120238824 | rs7117063 | C | T | NULL | 0.698465216 | NULL&M_Disease |
| 11 | 120303939 | rs7131597 | T | G | NULL | 0.698465216 | NULL&M_Disease |
| 11 | 116288558 | rs578780 | T | C | NULL | 0.698465216 | NULL&M_Disease |
| 11 | 117218460 | rs1261744 | C | T | NULL | 0.698465216 | NULL&M_Disease |
| 11 | 117219770 | rs508871 | A | G | NULL | 0.698465216 | NULL&M_Disease |
| 11 | 117441495 | rs657137 | T | C | NULL | 0.698465216 | NULL&M_Disease |
| 11 | 117451316 | rs537997 | T | C | NULL | 0.698465216 | NULL&M_Disease |
| 11 | 118603030 | rs7943039 | A | G | NULL | 0.698465216 | NULL&M_Disease |
| 11 | 118610463 | rs4499035 | T | C | NULL | 0.698465216 | NULL&M_Disease |
| 11 | 118610826 | rs1893029 | T | G | NULL | 0.698465216 | NULL&M_Disease |
| 11 | 118895686 | rs8192696 | T | C | NULL | 0.698465216 | NULL&M_Disease |
| 11 | 119697594 | rs476937 | A | G | NULL | 0.698465216 | NULL&M_Disease |
| 11 | 119829572 | rs11217601 | C | A | NULL | 0.698465216 | NULL&M_Disease |
| 11 | 120968162 | rs11218140 | A | G | NULL | 0.698465216 | NULL&M_Disease |
| 11 | 116223363 | rs564591 | G | A | NULL | 0.698465216 | NULL&M_Disease |
| 11 | 117232492 | rs2305825 | T | C | NULL | 0.698465216 | NULL&M_Disease |
| 11 | 117811548 | rs7118414 | C | G | NULL | 0.698465216 | NULL&M_Disease |
| 11 | 119070910 | rs7105861 | G | C | NULL | 0.698465216 | NULL&M_Disease |
| 11 | 119742279 | rs11217547 | A | G | NULL | 0.698465216 | NULL&M_Disease |
| 11 | 120099679 | rs2508490 | G | A | NULL | 0.698465216 | NULL&M_Disease |
| 11 | 120604315 | rs2846125 | C | T | NULL | 0.698465216 | NULL&M_Disease |
| 11 | 120605882 | rs2852231 | C | T | NULL | 0.698465216 | NULL&M_Disease |
| 11 | 120609495 | rs948031 | G | A | NULL | 0.698465216 | NULL&M_Disease |
| 11 | 117200753 | rs11216357 | G | T | NULL | 0.698465216 | NULL&M_Disease |
| 11 | 117217391 | rs1242696 | T | G | NULL | 0.698465216 | NULL&M_Disease |
| 11 | 117232156 | rs12421205 | G | C | NULL | 0.698465216 | NULL&M_Disease |
| 11 | 117232233 | rs12421494 | A | G | NULL | 0.698465216 | NULL&M_Disease |
| 11 | 117692334 | rs4579962 | A | G | NULL | 0.698465216 | NULL&M_Disease |
| 11 | 118609598 | rs112996178 | T | C | NULL | 0.698465216 | NULL&M_Disease |
| 11 | 119206375 | rs2511841 | G | A | NULL | 0.698465216 | NULL&M_Disease |
| 11 | 119935789 | rs2034643 | T | G | NULL | 0.698465216 | NULL&M_Disease |
| 11 | 120002727 | rs550164 | G | A | NULL | 0.698465216 | NULL&M_Disease |
| 11 | 117974937 | rs11216755 | C | T | NULL | 0.698465216 | NULL&M_Disease |
| 11 | 118635364 | rs11217018 | C | T | NULL | 0.698465216 | NULL&M_Disease |
| 11 | 119019877 | rs71482170 | A | G | NULL | 0.698465216 | NULL&M_Disease |
| 11 | 119361345 | rs2509668 | G | A | NULL | 0.698465216 | NULL&M_Disease |
| 11 | 119879217 | rs647437 | C | T | NULL | 0.698465216 | NULL&M_Disease |
| 11 | 119973299 | rs7131629 | A | G | NULL | 0.698465216 | NULL&M_Disease |
| 11 | 120244577 | rs7126328 | T | C | NULL | 0.698465216 | NULL&M_Disease |
| 11 | 117224360 | rs657666 | A | G | NULL | 0.698465216 | NULL&M_Disease |
| 11 | 117224818 | rs553303 | T | C | NULL | 0.698465216 | NULL&M_Disease |
| 11 | 117224878 | rs655389 | A | G | NULL | 0.698465216 | NULL&M_Disease |
| 11 | 117224887 | rs553506 | T | C | NULL | 0.698465216 | NULL&M_Disease |
| 11 | 117226616 | rs535366 | C | T | NULL | 0.698465216 | NULL&M_Disease |
| 11 | 117228295 | rs534296 | G | A | NULL | 0.698465216 | NULL&M_Disease |
| 11 | 117228664 | rs558854 | A | G | NULL | 0.698465216 | NULL&M_Disease |
| 11 | 117228709 | rs486366 | G | A | NULL | 0.698465216 | NULL&M_Disease |
| 11 | 117229902 | rs2510450 | T | C | NULL | 0.698465216 | NULL&M_Disease |
| 11 | 117231057 | rs512730 | A | T | NULL | 0.698465216 | NULL&M_Disease |
| 11 | 117692121 | rs1053087454 | G | A | NULL | 0.698465216 | NULL&M_Disease |
| 11 | 117766606 | rs73020459 | T | C | NULL | 0.698465216 | NULL&M_Disease |
| 11 | 119361057 | rs35574604 | A | G | NULL | 0.698465216 | NULL&M_Disease |
| 11 | 119480810 | rs1567377 | C | T | NULL | 0.698465216 | NULL&M_Disease |
| 11 | 120211212 | rs7932658 | A | G | NULL | 0.698465216 | NULL&M_Disease |
| 11 | 120704659 | rs7114615 | T | G | NULL | 0.698465216 | NULL&M_Disease |
| 11 | 116297397 | rs483367 | T | C | NULL | 0.698465216 | NULL&M_Disease |
| 11 | 116325990 | rs552043 | T | C | NULL | 0.698465216 | NULL&M_Disease |
| 11 | 116553778 | rs11826999 | A | G | NULL | 0.698465216 | NULL&M_Disease |
| 11 | 116563822 | rs12420888 | G | C | NULL | 0.698465216 | NULL&M_Disease |
| 11 | 117225134 | rs654479 | A | G | NULL | 0.698465216 | NULL&M_Disease |
| 11 | 117845195 | rs36119355 | T | G | NULL | 0.698465216 | NULL&M_Disease |
| 11 | 118200657 | rs67859744 | A | T | NULL | 0.698465216 | NULL&M_Disease |
| 11 | 118624919 | rs11217013 | C | G | NULL | 0.698465216 | NULL&M_Disease |
| 11 | 118627778 | rs10892282 | C | G | NULL | 0.698465216 | NULL&M_Disease |
| 11 | 118832035 | rs10892317 | C | T | NULL | 0.698465216 | NULL&M_Disease |
| 11 | 119837112 | rs626466 | T | C | NULL | 0.698465216 | NULL&M_Disease |
| 11 | 120605867 | rs2846119 | C | G | NULL | 0.698465216 | NULL&M_Disease |
| 11 | 120724018 | rs6589848 | T | C | NULL | 0.698465216 | NULL&M_Disease |
| 11 | 116297866 | rs493602 | C | G | NULL | 0.698465216 | NULL&M_Disease |
| 11 | 116567881 | rs10892017 | C | G | NULL | 0.698465216 | NULL&M_Disease |
| 11 | 116580815 | rs11216109 | G | C | NULL | 0.698465216 | NULL&M_Disease |
| 11 | 117745418 | rs540780 | T | G | NULL | 0.698465216 | NULL&M_Disease |
| 11 | 118622985 | rs3825057 | C | T | NULL | 0.698465216 | NULL&M_Disease |
| 11 | 119701127 | rs555255 | G | A | NULL | 0.698465216 | NULL&M_Disease |
| 11 | 120208257 | rs4938799 | G | T | NULL | 0.698465216 | NULL&M_Disease |
| 11 | 120238721 | rs11217841 | A | C | NULL | 0.698465216 | NULL&M_Disease |
| 11 | 120530973 | rs6589829 | C | T | NULL | 0.698465216 | NULL&M_Disease |
| 11 | 120596053 | rs3133845 | A | G | NULL | 0.698465216 | NULL&M_Disease |
| 11 | 120597136 | rs3133846 | C | T | NULL | 0.698465216 | NULL&M_Disease |
| 11 | 116552815 | rs12806224 | C | T | NULL | 0.698465216 | NULL&M_Disease |
| 11 | 116553025 | rs12805061 | G | A | NULL | 0.698465216 | NULL&M_Disease |
| 11 | 117227686 | rs528720 | A | G | NULL | 0.698465216 | NULL&M_Disease |
| 11 | 119480660 | rs1459783673 | A | T | NULL | 0.698465216 | NULL&M_Disease |
| 11 | 119970538 | rs6589796 | A | C | NULL | 0.698465216 | NULL&M_Disease |
| 11 | 120254684 | rs1892894 | T | C | NULL | 0.698465216 | NULL&M_Disease |
| 11 | 116563608 | rs12785772 | A | G | NULL | 0.699517228 | NULL&M_Disease |
| 11 | 116568168 | rs11605568 | A | G | NULL | 0.698465216 | NULL&M_Disease |
| 11 | 117226849 | rs537323 | T | C | NULL | 0.698465216 | NULL&M_Disease |
| 11 | 117509143 | rs587099 | G | A | NULL | 0.699517228 | NULL&M_Disease |
| 11 | 118177115 | rs55942271 | T | C | NULL | 0.699517228 | NULL&M_Disease |
| 11 | 118613674 | rs7130875 | C | T | NULL | 0.699517228 | NULL&M_Disease |
| 11 | 118878222 | rs11606357 | C | T | NULL | 0.699517228 | NULL&M_Disease |
| 11 | 116317663 | rs497960 | C | T | NULL | 0.699517228 | NULL&M_Disease |

| | | | | | | | | | |
|---|---|---|---|---|---|---|---|---|---|
| 11 | 119695450 | rs525914 | T | C | 1.001 | -7977.388 | -7979.067 | -1.679 | 7.22E-07 |
| 11 | 119847991 | rs621150 | T | C | 1.021 | -7977.388 | -7979.067 | -1.679 | 7.22E-07 |
| 11 | 120101092 | rs2845705 | A | T | 0.989 | -7977.388 | -7979.067 | -1.679 | 7.22E-07 |
| 11 | 120238824 | rs7117063 | C | T | 1.02 | -7977.388 | -7979.067 | -1.679 | 7.22E-07 |
| 11 | 120303939 | rs7131597 | T | G | 1.02 | -7977.388 | -7979.067 | -1.679 | 7.22E-07 |
| 11 | 116288558 | rs578780 | T | C | 1.016 | -7977.388 | -7979.069 | -1.68 | 7.21E-07 |
| 11 | 117218460 | rs1261744 | C | T | 0.992 | -7977.388 | -7979.068 | -1.68 | 7.21E-07 |
| 11 | 117219770 | rs508871 | A | G | 0.992 | -7977.388 | -7979.068 | -1.68 | 7.21E-07 |
| 11 | 117441495 | rs657137 | T | C | 1.019 | -7977.388 | -7979.068 | -1.68 | 7.21E-07 |
| 11 | 117451316 | rs537997 | T | C | 0.989 | -7977.388 | -7979.069 | -1.68 | 7.21E-07 |
| 11 | 118603030 | rs7943039 | A | G | 1.015 | -7977.388 | -7979.069 | -1.68 | 7.21E-07 |
| 11 | 118610463 | rs4499035 | T | C | 1.015 | -7977.388 | -7979.068 | -1.68 | 7.21E-07 |
| 11 | 118610826 | rs1893029 | T | G | 1.015 | -7977.388 | -7979.068 | -1.68 | 7.21E-07 |
| 11 | 118895686 | rs8192696 | T | C | 1.012 | -7977.388 | -7979.068 | -1.68 | 7.21E-07 |
| 11 | 119697594 | rs476937 | A | G | 1 | -7977.388 | -7979.069 | -1.68 | 7.21E-07 |
| 11 | 119829572 | rs11217601 | C | A | 0.985 | -7977.388 | -7979.069 | -1.68 | 7.21E-07 |
| 11 | 120968162 | rs11218140 | A | G | 1.017 | -7977.388 | -7979.068 | -1.68 | 7.21E-07 |
| 11 | 116223363 | rs564591 | G | A | 1.02 | -7977.388 | -7979.069 | -1.681 | 7.20E-07 |
| 11 | 117232492 | rs2305825 | T | C | 0.992 | -7977.388 | -7979.069 | -1.681 | 7.20E-07 |
| 11 | 117811548 | rs7118414 | C | G | 1.021 | -7977.388 | -7979.069 | -1.681 | 7.20E-07 |
| 11 | 119070910 | rs7105861 | G | C | 1.021 | -7977.388 | -7979.069 | -1.681 | 7.20E-07 |
| 11 | 119742279 | rs11217547 | A | G | 0.98 | -7977.388 | -7979.069 | -1.681 | 7.20E-07 |
| 11 | 120099679 | rs2508490 | G | A | 0.989 | -7977.388 | -7979.069 | -1.681 | 7.20E-07 |
| 11 | 120604315 | rs2846125 | C | T | 0.984 | -7977.388 | -7979.069 | -1.681 | 7.20E-07 |
| 11 | 120605882 | rs2852231 | C | T | 0.984 | -7977.388 | -7979.069 | -1.681 | 7.20E-07 |
| 11 | 120609495 | rs948031 | G | A | 0.984 | -7977.388 | -7979.07 | -1.681 | 7.20E-07 |
| 11 | 117200753 | rs11216357 | G | T | 1 | -7977.388 | -7979.07 | -1.682 | 7.19E-07 |
| 11 | 117217391 | rs1242696 | T | G | 0.98 | -7977.388 | -7979.07 | -1.682 | 7.19E-07 |
| 11 | 117232156 | rs12421205 | G | C | 0.992 | -7977.388 | -7979.07 | -1.682 | 7.19E-07 |
| 11 | 117232233 | rs12421494 | A | G | 0.992 | -7977.388 | -7979.07 | -1.682 | 7.19E-07 |
| 11 | 117692334 | rs4579962 | A | G | 0.983 | -7977.388 | -7979.071 | -1.682 | 7.19E-07 |
| 11 | 118609598 | rs112996178 | T | C | 1.015 | -7977.388 | -7979.07 | -1.682 | 7.19E-07 |
| 11 | 119206375 | rs2511841 | G | A | 1.019 | -7977.388 | -7979.071 | -1.682 | 7.19E-07 |
| 11 | 119935789 | rs2034643 | T | G | 1.02 | -7977.388 | -7979.07 | -1.682 | 7.19E-07 |
| 11 | 120002727 | rs550164 | G | A | 0.984 | -7977.388 | -7979.07 | -1.682 | 7.19E-07 |
| 11 | 117974937 | rs11216755 | C | T | 1.021 | -7977.388 | -7979.071 | -1.683 | 7.19E-07 |
| 11 | 118635364 | rs11217018 | C | T | 1.015 | -7977.388 | -7979.072 | -1.683 | 7.19E-07 |
| 11 | 119019877 | rs71482170 | A | G | 0.98 | -7977.388 | -7979.072 | -1.683 | 7.19E-07 |
| 11 | 119361345 | rs2509668 | G | A | 0.98 | -7977.388 | -7979.071 | -1.683 | 7.19E-07 |
| 11 | 119879217 | rs647437 | C | T | 0.983 | -7977.388 | -7979.072 | -1.683 | 7.19E-07 |
| 11 | 119973299 | rs7131629 | A | G | 0.981 | -7977.388 | -7979.071 | -1.683 | 7.19E-07 |
| 11 | 120244577 | rs7126328 | T | C | 1.02 | -7977.388 | -7979.071 | -1.683 | 7.19E-07 |
| 11 | 117224360 | rs657666 | A | G | 0.992 | -7977.388 | -7979.072 | -1.684 | 7.18E-07 |
| 11 | 117224818 | rs553303 | T | C | 0.992 | -7977.388 | -7979.072 | -1.684 | 7.18E-07 |
| 11 | 117224878 | rs655389 | A | G | 0.992 | -7977.388 | -7979.072 | -1.684 | 7.18E-07 |
| 11 | 117224887 | rs553506 | T | C | 0.992 | -7977.388 | -7979.072 | -1.684 | 7.18E-07 |
| 11 | 117226616 | rs535366 | C | T | 0.992 | -7977.388 | -7979.072 | -1.684 | 7.18E-07 |
| 11 | 117228295 | rs534296 | G | A | 0.992 | -7977.388 | -7979.072 | -1.684 | 7.18E-07 |
| 11 | 117228664 | rs558854 | A | G | 0.992 | -7977.388 | -7979.072 | -1.684 | 7.18E-07 |
| 11 | 117228709 | rs486366 | G | A | 0.992 | -7977.388 | -7979.072 | -1.684 | 7.18E-07 |
| 11 | 117229902 | rs2510450 | T | C | 0.992 | -7977.388 | -7979.072 | -1.684 | 7.18E-07 |
| 11 | 117231057 | rs512730 | A | T | 0.992 | -7977.388 | -7979.072 | -1.684 | 7.18E-07 |
| 11 | 117692121 | rs1053087454 | G | A | 0.987 | -7977.388 | -7979.072 | -1.684 | 7.18E-07 |
| 11 | 117766606 | rs73020459 | T | C | 1.002 | -7977.388 | -7979.072 | -1.684 | 7.18E-07 |
| 11 | 119361057 | rs35574604 | A | G | 0.98 | -7977.388 | -7979.073 | -1.684 | 7.18E-07 |
| 11 | 119480810 | rs1567377 | C | T | 0.981 | -7977.388 | -7979.073 | -1.684 | 7.18E-07 |
| 11 | 120211212 | rs7932658 | A | G | 1.02 | -7977.388 | -7979.073 | -1.684 | 7.18E-07 |
| 11 | 120704659 | rs7114615 | T | G | 0.992 | -7977.388 | -7979.072 | -1.684 | 7.18E-07 |
| 11 | 116297397 | rs483367 | T | C | 1.015 | -7977.388 | -7979.073 | -1.685 | 7.17E-07 |
| 11 | 116325990 | rs552043 | T | C | 1 | -7977.388 | -7979.073 | -1.685 | 7.17E-07 |
| 11 | 116553778 | rs11826999 | A | G | 1.014 | -7977.388 | -7979.073 | -1.685 | 7.17E-07 |
| 11 | 116563822 | rs12420888 | G | C | 0.987 | -7977.388 | -7979.074 | -1.685 | 7.17E-07 |
| 11 | 117225134 | rs654479 | A | G | 0.993 | -7977.388 | -7979.074 | -1.685 | 7.17E-07 |
| 11 | 117845195 | rs36119355 | T | G | 0.982 | -7977.388 | -7979.074 | -1.685 | 7.17E-07 |
| 11 | 118200657 | rs67859744 | A | T | 0.983 | -7977.388 | -7979.074 | -1.685 | 7.17E-07 |
| 11 | 118624919 | rs11217013 | C | G | 1.015 | -7977.388 | -7979.073 | -1.685 | 7.17E-07 |
| 11 | 118627778 | rs10892282 | C | G | 1.015 | -7977.388 | -7979.073 | -1.685 | 7.17E-07 |
| 11 | 118832035 | rs10892317 | C | T | 0.982 | -7977.388 | -7979.073 | -1.685 | 7.17E-07 |
| 11 | 119837112 | rs626466 | T | C | 0.986 | -7977.388 | -7979.074 | -1.685 | 7.17E-07 |
| 11 | 120605867 | rs2846119 | C | G | 0.984 | -7977.388 | -7979.074 | -1.685 | 7.17E-07 |
| 11 | 120724018 | rs6589848 | T | C | 0.992 | -7977.388 | -7979.073 | -1.685 | 7.17E-07 |
| 11 | 116297866 | rs493602 | C | G | 1.015 | -7977.388 | -7979.074 | -1.686 | 7.17E-07 |
| 11 | 116567881 | rs10892017 | C | G | 0.987 | -7977.388 | -7979.074 | -1.686 | 7.17E-07 |
| 11 | 116580815 | rs11216109 | G | C | 1.01 | -7977.388 | -7979.074 | -1.686 | 7.17E-07 |
| 11 | 117745418 | rs540780 | T | G | 0.98 | -7977.388 | -7979.074 | -1.686 | 7.17E-07 |
| 11 | 118622985 | rs3825057 | C | T | 1.015 | -7977.388 | -7979.075 | -1.686 | 7.17E-07 |
| 11 | 119701127 | rs555255 | G | A | 0.99 | -7977.388 | -7979.075 | -1.686 | 7.17E-07 |
| 11 | 120208257 | rs4938799 | G | T | 1.02 | -7977.388 | -7979.074 | -1.686 | 7.17E-07 |
| 11 | 120238721 | rs11217841 | A | C | 1.02 | -7977.388 | -7979.074 | -1.686 | 7.17E-07 |
| 11 | 120530973 | rs6589829 | C | T | 1.02 | -7977.388 | -7979.074 | -1.686 | 7.17E-07 |
| 11 | 120596053 | rs3133845 | A | G | 0.982 | -7977.388 | -7979.074 | -1.686 | 7.17E-07 |
| 11 | 120597136 | rs3133846 | C | T | 0.982 | -7977.388 | -7979.075 | -1.686 | 7.17E-07 |
| 11 | 116552815 | rs12806224 | C | T | 1.014 | -7977.388 | -7979.075 | -1.687 | 7.16E-07 |
| 11 | 116553025 | rs12805061 | G | A | 1.014 | -7977.388 | -7979.075 | -1.687 | 7.16E-07 |
| 11 | 117227686 | rs528720 | A | G | 0.993 | -7977.388 | -7979.075 | -1.687 | 7.16E-07 |
| 11 | 119480660 | rs1459783673 | A | T | 0.982 | -7977.388 | -7979.076 | -1.687 | 7.16E-07 |
| 11 | 119970538 | rs6589796 | A | C | 0.98 | -7977.388 | -7979.075 | -1.687 | 7.16E-07 |
| 11 | 120254684 | rs1892894 | T | C | 1.02 | -7977.388 | -7979.075 | -1.687 | 7.16E-07 |
| 11 | 116563608 | rs12785772 | A | G | 0.987 | -7977.388 | -7979.077 | -1.688 | 7.15E-07 |
| 11 | 116568168 | rs11605568 | A | G | 0.987 | -7977.388 | -7979.076 | -1.688 | 7.15E-07 |
| 11 | 117226849 | rs537323 | T | C | 0.993 | -7977.388 | -7979.076 | -1.688 | 7.15E-07 |
| 11 | 117509143 | rs587099 | G | A | 1.02 | -7977.388 | -7979.076 | -1.688 | 7.15E-07 |
| 11 | 118177115 | rs55942271 | T | C | 0.982 | -7977.388 | -7979.077 | -1.688 | 7.15E-07 |
| 11 | 118613674 | rs7130875 | C | T | 1.015 | -7977.388 | -7979.076 | -1.688 | 7.15E-07 |
| 11 | 118878222 | rs11606357 | C | T | 1.012 | -7977.388 | -7979.077 | -1.688 | 7.15E-07 |
| 11 | 116317663 | rs497960 | C | T | 1.008 | -7977.388 | -7979.078 | -1.689 | 7.14E-07 |

| | | | | | | | |
|---|---|---|---|---|---|---|---|
| 11 | 116319242 | rs505123 | T | G | NULL | 0.699517228 | NULL&M_Disease |
| 11 | 118094756 | rs12275955 | C | T | NULL | 0.699517228 | NULL&M_Disease |
| 11 | 120854938 | rs673351 | C | G | NULL | 0.699517228 | NULL&M_Disease |
| 11 | 117905405 | rs1783839 | C | T | NULL | 0.699517228 | NULL&M_Disease |
| 11 | 117947208 | rs1786191 | C | T | NULL | 0.699517228 | NULL&M_Disease |
| 11 | 118013034 | rs10790241 | A | G | NULL | 0.699517228 | NULL&M_Disease |
| 11 | 118084728 | rs12286604 | A | G | NULL | 0.699517228 | NULL&M_Disease |
| 11 | 118629364 | rs73001486 | A | G | NULL | 0.699517228 | NULL&M_Disease |
| 11 | 118893878 | rs111322300 | C | T | NULL | 0.699517228 | NULL&M_Disease |
| 11 | 119482705 | rs7935488 | T | C | NULL | 0.699517228 | NULL&M_Disease |
| 11 | 119548698 | rs2292293 | G | T | NULL | 0.699517228 | NULL&M_Disease |
| 11 | 119695011 | rs566880 | T | C | NULL | 0.699517228 | NULL&M_Disease |
| 11 | 120312746 | rs723937 | A | G | NULL | 0.699517228 | NULL&M_Disease |
| 11 | 120524884 | rs1893907 | G | C | NULL | 0.699517228 | NULL&M_Disease |
| 11 | 120610584 | rs751368 | T | G | NULL | 0.699517228 | NULL&M_Disease |
| 11 | 120611896 | rs2511064 | A | G | NULL | 0.699517228 | NULL&M_Disease |
| 11 | 120787231 | rs11218059 | T | C | NULL | 0.699517228 | NULL&M_Disease |
| 11 | 120854936 | rs673346 | G | A | NULL | 0.699517228 | NULL&M_Disease |
| 11 | 116573838 | rs882398 | C | T | NULL | 0.699517228 | NULL&M_Disease |
| 11 | 117784434 | rs11605749 | G | C | NULL | 0.699517228 | NULL&M_Disease |
| 11 | 117947698 | rs620924 | A | G | NULL | 0.699517228 | NULL&M_Disease |
| 11 | 118136160 | rs3759002 | A | G | NULL | 0.699517228 | NULL&M_Disease |
| 11 | 119043500 | rs644252 | T | G | NULL | 0.699517228 | NULL&M_Disease |
| 11 | 120124953 | rs2715888 | A | G | NULL | 0.699517228 | NULL&M_Disease |
| 11 | 120608085 | rs949298 | A | G | NULL | 0.699517228 | NULL&M_Disease |
| 11 | 120608142 | rs949299 | A | T | NULL | 0.699517228 | NULL&M_Disease |
| 11 | 120608222 | rs948033 | G | A | NULL | 0.699517228 | NULL&M_Disease |
| 11 | 120608661 | rs948032 | A | G | NULL | 0.699517228 | NULL&M_Disease |
| 11 | 120608912 | rs1320648 | C | T | NULL | 0.699517228 | NULL&M_Disease |
| 11 | 120608964 | rs752979 | A | G | NULL | 0.699517228 | NULL&M_Disease |
| 11 | 116295758 | rs574233 | T | G | NULL | 0.699517228 | NULL&M_Disease |
| 11 | 116561328 | rs1787709 | G | C | NULL | 0.699517228 | NULL&M_Disease |
| 11 | 117558989 | rs11216503 | G | A | NULL | 0.699517228 | NULL&M_Disease |
| 11 | 118011860 | rs12804550 | A | G | NULL | 0.699517228 | NULL&M_Disease |
| 11 | 118883491 | rs60065241 | A | C | NULL | 0.699517228 | NULL&M_Disease |
| 11 | 118887887 | rs369816532 | T | G | NULL | 0.699517228 | NULL&M_Disease |
| 11 | 118887888 | rs3916582 | T | C | NULL | 0.699517228 | NULL&M_Disease |
| 11 | 118895202 | rs11006 | G | A | NULL | 0.699517228 | NULL&M_Disease |
| 11 | 120238655 | rs11217840 | G | A | NULL | 0.699517228 | NULL&M_Disease |
| 11 | 120827609 | rs644057 | A | G | NULL | 0.699517228 | NULL&M_Disease |
| 11 | 120855192 | rs674352 | T | C | NULL | 0.699517228 | NULL&M_Disease |
| 11 | 116562580 | rs11216096 | A | C | NULL | 0.699517228 | NULL&M_Disease |
| 11 | 116596752 | rs180364 | G | A | NULL | 0.699517228 | NULL&M_Disease |
| 11 | 118196399 | rs67435655 | C | T | NULL | 0.699517228 | NULL&M_Disease |
| 11 | 119878901 | rs10892512 | A | G | NULL | 0.699517228 | NULL&M_Disease |
| 11 | 120611914 | rs2508806 | C | T | NULL | 0.699517228 | NULL&M_Disease |
| 11 | 120738221 | rs3802913 | G | C | NULL | 0.699517228 | NULL&M_Disease |
| 11 | 120855386 | rs73585281 | G | A | NULL | 0.699517228 | NULL&M_Disease |
| 11 | 120855433 | rs73585283 | A | G | NULL | 0.699517228 | NULL&M_Disease |
| 11 | 121120755 | rs1893248 | T | C | NULL | 0.699517228 | NULL&M_Disease |
| 11 | 117692842 | rs12274595 | C | T | NULL | 0.699517228 | NULL&M_Disease |
| 11 | 117745003 | rs4938445 | A | G | NULL | 0.699517228 | NULL&M_Disease |
| 11 | 117757875 | rs7121851 | T | C | NULL | 0.699517228 | NULL&M_Disease |
| 11 | 117988654 | rs7123209 | C | G | NULL | 0.699517228 | NULL&M_Disease |
| 11 | 118640828 | rs11217021 | C | T | NULL | 0.699517228 | NULL&M_Disease |
| 11 | 119482832 | rs7949154 | C | T | NULL | 0.699517228 | NULL&M_Disease |
| 11 | 119603847 | rs12421432 | T | C | NULL | 0.699517228 | NULL&M_Disease |
| 11 | 119969030 | rs592347 | T | C | NULL | 0.699517228 | NULL&M_Disease |
| 11 | 120244192 | rs10892567 | G | A | NULL | 0.699517228 | NULL&M_Disease |
| 11 | 120600950 | rs3133851 | A | G | NULL | 0.699517228 | NULL&M_Disease |
| 11 | 120605524 | rs2846121 | A | G | NULL | 0.699517228 | NULL&M_Disease |
| 11 | 116329185 | rs517599 | C | T | NULL | 0.699517228 | NULL&M_Disease |
| 11 | 118197486 | rs66962317 | C | T | NULL | 0.699517228 | NULL&M_Disease |
| 11 | 118197645 | rs12575015 | T | A | NULL | 0.699517228 | NULL&M_Disease |
| 11 | 118655356 | rs7934890 | T | C | NULL | 0.699517228 | NULL&M_Disease |
| 11 | 119878269 | rs4938756 | C | T | NULL | 0.699517228 | NULL&M_Disease |
| 11 | 120183794 | rs7108217 | T | C | NULL | 0.699517228 | NULL&M_Disease |
| 11 | 120443983 | rs4298914 | T | C | NULL | 0.699517228 | NULL&M_Disease |
| 11 | 120606683 | rs2155263 | A | G | NULL | 0.699517228 | NULL&M_Disease |
| 11 | 120855705 | rs61903169 | T | A | NULL | 0.699517228 | NULL&M_Disease |
| 11 | 117863032 | rs2512145 | A | G | NULL | 0.699517228 | NULL&M_Disease |
| 11 | 117872544 | rs2508445 | T | G | NULL | 0.699517228 | NULL&M_Disease |
| 11 | 118633533 | rs7110526 | C | T | NULL | 0.699517228 | NULL&M_Disease |
| 11 | 118854372 | rs75067211 | T | C | NULL | 0.699517228 | NULL&M_Disease |
| 11 | 118892835 | rs61179449 | C | G | NULL | 0.699517228 | NULL&M_Disease |
| 11 | 120084056 | rs12793354 | C | A | NULL | 0.699517228 | NULL&M_Disease |
| 11 | 120605736 | rs2852230 | C | G | NULL | 0.699517228 | NULL&M_Disease |
| 11 | 120605759 | rs2846120 | T | C | NULL | 0.699517228 | NULL&M_Disease |
| 11 | 116298367 | rs490592 | A | C | NULL | 0.699517228 | NULL&M_Disease |
| 11 | 116329750 | rs541563 | A | G | NULL | 0.699517228 | NULL&M_Disease |
| 11 | 116561381 | rs1784123 | C | T | NULL | 0.699517228 | NULL&M_Disease |
| 11 | 116576400 | rs12802944 | C | T | NULL | 0.699517228 | NULL&M_Disease |
| 11 | 117173019 | rs681120 | C | A | NULL | 0.699517228 | NULL&M_Disease |
| 11 | 117241429 | rs12417440 | T | C | NULL | 0.699517228 | NULL&M_Disease |
| 11 | 117446945 | rs504935 | C | T | NULL | 0.699517228 | NULL&M_Disease |
| 11 | 117662248 | rs534761 | T | C | NULL | 0.699517228 | NULL&M_Disease |
| 11 | 117947454 | rs651013 | C | T | NULL | 0.699517228 | NULL&M_Disease |
| 11 | 120610692 | rs2155262 | A | G | NULL | 0.699517228 | NULL&M_Disease |
| 11 | 120738397 | rs3802912 | A | G | NULL | 0.699517228 | NULL&M_Disease |
| 11 | 117330736 | rs747061 | G | C | NULL | 0.700567142 | NULL&M_Disease |
| 11 | 117486829 | rs527005 | G | C | NULL | 0.699517228 | NULL&M_Disease |
| 11 | 118637475 | rs112910608 | T | C | NULL | 0.700567142 | NULL&M_Disease |
| 11 | 118892872 | rs58955701 | T | C | NULL | 0.699517228 | NULL&M_Disease |
| 11 | 119742360 | rs10892481 | T | C | NULL | 0.700567142 | NULL&M_Disease |
| 11 | 120213360 | rs7121779 | T | A | NULL | 0.700567142 | NULL&M_Disease |
| 11 | 120216260 | rs7952529 | C | A | NULL | 0.700567142 | NULL&M_Disease |

| | | | | | | | | | |
|---|---|---|---|---|---|---|---|---|---|
| 11 | 116319242 | rs505123 | T | G | 1.007 | -7977.388 | -7979.077 | -1.689 | 7.14E-07 |
| 11 | 118094756 | rs12275955 | C | T | 1.013 | -7977.388 | -7979.078 | -1.689 | 7.14E-07 |
| 11 | 120854938 | rs673351 | C | G | 1.01 | -7977.388 | -7979.077 | -1.689 | 7.14E-07 |
| 11 | 117905405 | rs1783839 | C | T | 0.996 | -7977.388 | -7979.078 | -1.69 | 7.14E-07 |
| 11 | 117947208 | rs1786191 | C | T | 1.014 | -7977.388 | -7979.078 | -1.69 | 7.14E-07 |
| 11 | 118013034 | rs10790241 | A | G | 1.006 | -7977.388 | -7979.078 | -1.69 | 7.14E-07 |
| 11 | 118084728 | rs12286604 | A | G | 1.01 | -7977.388 | -7979.078 | -1.69 | 7.14E-07 |
| 11 | 118629364 | rs73001486 | A | G | 1.015 | -7977.388 | -7979.078 | -1.69 | 7.14E-07 |
| 11 | 118893878 | rs111322300 | C | T | 1.012 | -7977.388 | -7979.078 | -1.69 | 7.14E-07 |
| 11 | 119482705 | rs7935488 | T | C | 0.982 | -7977.388 | -7979.079 | -1.69 | 7.14E-07 |
| 11 | 119548698 | rs2292293 | G | T | 0.98 | -7977.388 | -7979.079 | -1.69 | 7.14E-07 |
| 11 | 119695011 | rs566880 | T | C | 1.001 | -7977.388 | -7979.078 | -1.69 | 7.14E-07 |
| 11 | 120312746 | rs723937 | A | G | 1.02 | -7977.388 | -7979.078 | -1.69 | 7.14E-07 |
| 11 | 120524884 | rs1893907 | G | C | 1.019 | -7977.388 | -7979.078 | -1.69 | 7.14E-07 |
| 11 | 120610584 | rs751368 | T | G | 0.98 | -7977.388 | -7979.078 | -1.69 | 7.14E-07 |
| 11 | 120611896 | rs2511064 | A | G | 0.984 | -7977.388 | -7979.079 | -1.69 | 7.14E-07 |
| 11 | 120787231 | rs11218059 | T | C | 0.988 | -7977.388 | -7979.079 | -1.69 | 7.14E-07 |
| 11 | 120854936 | rs673346 | G | A | 1.009 | -7977.388 | -7979.078 | -1.69 | 7.14E-07 |
| 11 | 116573838 | rs882398 | C | T | 1.018 | -7977.388 | -7979.079 | -1.691 | 7.13E-07 |
| 11 | 117784434 | rs11605749 | G | C | 0.982 | -7977.388 | -7979.079 | -1.691 | 7.13E-07 |
| 11 | 117947698 | rs620924 | A | G | 1.014 | -7977.388 | -7979.079 | -1.691 | 7.13E-07 |
| 11 | 118136160 | rs3759002 | A | G | 1.008 | -7977.388 | -7979.079 | -1.691 | 7.13E-07 |
| 11 | 119043500 | rs644252 | T | G | 1.021 | -7977.388 | -7979.079 | -1.691 | 7.13E-07 |
| 11 | 120124953 | rs2715888 | A | G | 1.017 | -7977.388 | -7979.079 | -1.691 | 7.13E-07 |
| 11 | 120608085 | rs949298 | A | G | 0.985 | -7977.388 | -7979.079 | -1.691 | 7.13E-07 |
| 11 | 120608142 | rs949299 | A | T | 0.985 | -7977.388 | -7979.079 | -1.691 | 7.13E-07 |
| 11 | 120608222 | rs948033 | G | A | 0.985 | -7977.388 | -7979.079 | -1.691 | 7.13E-07 |
| 11 | 120608661 | rs948032 | A | G | 0.985 | -7977.388 | -7979.079 | -1.691 | 7.13E-07 |
| 11 | 120608912 | rs1320648 | C | T | 0.985 | -7977.388 | -7979.079 | -1.691 | 7.13E-07 |
| 11 | 120608964 | rs752979 | A | G | 0.985 | -7977.388 | -7979.079 | -1.691 | 7.13E-07 |
| 11 | 116295758 | rs574233 | T | G | 1.015 | -7977.388 | -7979.08 | -1.692 | 7.12E-07 |
| 11 | 116561328 | rs1787709 | G | C | 1.018 | -7977.388 | -7979.08 | -1.692 | 7.12E-07 |
| 11 | 117558989 | rs11216503 | G | A | 1.003 | -7977.388 | -7979.08 | -1.692 | 7.12E-07 |
| 11 | 118011860 | rs12804550 | A | G | 1.006 | -7977.388 | -7979.08 | -1.692 | 7.12E-07 |
| 11 | 118883491 | rs60065241 | A | C | 1.012 | -7977.388 | -7979.081 | -1.692 | 7.12E-07 |
| 11 | 118887887 | rs369816532 | T | G | 1.015 | -7977.388 | -7979.08 | -1.692 | 7.12E-07 |
| 11 | 118887888 | rs3916582 | T | C | 1.015 | -7977.388 | -7979.08 | -1.692 | 7.12E-07 |
| 11 | 118895202 | rs11006 | G | A | 1.012 | -7977.388 | -7979.08 | -1.692 | 7.12E-07 |
| 11 | 120238655 | rs11217840 | G | A | 1.02 | -7977.388 | -7979.081 | -1.692 | 7.12E-07 |
| 11 | 120827609 | rs644057 | A | G | 0.982 | -7977.388 | -7979.081 | -1.692 | 7.12E-07 |
| 11 | 120855192 | rs674352 | T | C | 1.009 | -7977.388 | -7979.08 | -1.692 | 7.12E-07 |
| 11 | 116562580 | rs11216096 | A | C | 0.988 | -7977.388 | -7979.081 | -1.693 | 7.12E-07 |
| 11 | 116596752 | rs180364 | G | A | 1.019 | -7977.388 | -7979.081 | -1.693 | 7.12E-07 |
| 11 | 118196399 | rs67435655 | C | T | 0.983 | -7977.388 | -7979.081 | -1.693 | 7.12E-07 |
| 11 | 119878901 | rs10892512 | A | G | 1.016 | -7977.388 | -7979.081 | -1.693 | 7.12E-07 |
| 11 | 120611914 | rs2508806 | C | T | 0.985 | -7977.388 | -7979.082 | -1.693 | 7.12E-07 |
| 11 | 120738221 | rs3802913 | G | C | 0.98 | -7977.388 | -7979.082 | -1.693 | 7.12E-07 |
| 11 | 120855386 | rs73585281 | G | A | 1.009 | -7977.388 | -7979.082 | -1.693 | 7.12E-07 |
| 11 | 120855433 | rs73585283 | A | G | 1.009 | -7977.388 | -7979.082 | -1.693 | 7.12E-07 |
| 11 | 121120755 | rs1893248 | T | C | 0.98 | -7977.388 | -7979.082 | -1.693 | 7.12E-07 |
| 11 | 117692842 | rs12274595 | C | T | 0.988 | -7977.388 | -7979.082 | -1.694 | 7.11E-07 |
| 11 | 117745003 | rs4938445 | A | G | 0.982 | -7977.388 | -7979.082 | -1.694 | 7.11E-07 |
| 11 | 117757875 | rs7121851 | T | C | 1.007 | -7977.388 | -7979.082 | -1.694 | 7.11E-07 |
| 11 | 117988654 | rs7123209 | C | G | 1.009 | -7977.388 | -7979.082 | -1.694 | 7.11E-07 |
| 11 | 118640828 | rs11217021 | C | T | 1.014 | -7977.388 | -7979.082 | -1.694 | 7.11E-07 |
| 11 | 119482832 | rs7949154 | C | T | 0.982 | -7977.388 | -7979.083 | -1.694 | 7.11E-07 |
| 11 | 119603847 | rs12421432 | T | C | 1.019 | -7977.388 | -7979.083 | -1.694 | 7.11E-07 |
| 11 | 119969030 | rs592347 | T | C | 1.017 | -7977.388 | -7979.082 | -1.694 | 7.11E-07 |
| 11 | 120244192 | rs10892567 | G | A | 1.02 | -7977.388 | -7979.082 | -1.694 | 7.11E-07 |
| 11 | 120600950 | rs3133851 | A | G | 0.98 | -7977.388 | -7979.083 | -1.694 | 7.11E-07 |
| 11 | 120605524 | rs2846121 | A | G | 0.985 | -7977.388 | -7979.082 | -1.694 | 7.11E-07 |
| 11 | 116329185 | rs517599 | C | T | 1.02 | -7977.388 | -7979.083 | -1.695 | 7.10E-07 |
| 11 | 118197486 | rs66962317 | C | T | 0.984 | -7977.388 | -7979.083 | -1.695 | 7.10E-07 |
| 11 | 118197645 | rs12575015 | T | A | 0.984 | -7977.388 | -7979.083 | -1.695 | 7.10E-07 |
| 11 | 118655356 | rs7934890 | T | C | 1.014 | -7977.388 | -7979.083 | -1.695 | 7.10E-07 |
| 11 | 119878269 | rs4938756 | C | T | 1.02 | -7977.388 | -7979.084 | -1.695 | 7.10E-07 |
| 11 | 120183794 | rs7108217 | T | C | 1.02 | -7977.388 | -7979.083 | -1.695 | 7.10E-07 |
| 11 | 120443983 | rs4298914 | T | C | 1.02 | -7977.388 | -7979.083 | -1.695 | 7.10E-07 |
| 11 | 120606683 | rs2155263 | A | G | 0.985 | -7977.388 | -7979.084 | -1.695 | 7.10E-07 |
| 11 | 120855705 | rs61903169 | T | A | 1.009 | -7977.388 | -7979.083 | -1.695 | 7.10E-07 |
| 11 | 117863032 | rs2512145 | A | G | 0.982 | -7977.388 | -7979.084 | -1.696 | 7.09E-07 |
| 11 | 117872544 | rs2508445 | T | G | 0.982 | -7977.388 | -7979.084 | -1.696 | 7.09E-07 |
| 11 | 118633533 | rs7110526 | C | T | 1.014 | -7977.388 | -7979.085 | -1.696 | 7.09E-07 |
| 11 | 118854372 | rs75067211 | T | C | 1.015 | -7977.388 | -7979.084 | -1.696 | 7.09E-07 |
| 11 | 118892835 | rs61179449 | C | G | 1.011 | -7977.388 | -7979.084 | -1.696 | 7.09E-07 |
| 11 | 120084056 | rs12793354 | C | A | 1.017 | -7977.388 | -7979.085 | -1.696 | 7.09E-07 |
| 11 | 120605736 | rs2852230 | C | G | 0.985 | -7977.388 | -7979.084 | -1.696 | 7.09E-07 |
| 11 | 120605759 | rs2846120 | T | C | 0.985 | -7977.388 | -7979.084 | -1.696 | 7.09E-07 |
| 11 | 116298367 | rs490592 | A | C | 0.999 | -7977.388 | -7979.085 | -1.697 | 7.09E-07 |
| 11 | 116329750 | rs541563 | A | G | 1.02 | -7977.388 | -7979.085 | -1.697 | 7.09E-07 |
| 11 | 116561381 | rs1784123 | C | T | 1.018 | -7977.388 | -7979.085 | -1.697 | 7.09E-07 |
| 11 | 116576400 | rs12802944 | C | T | 1.005 | -7977.388 | -7979.085 | -1.697 | 7.09E-07 |
| 11 | 117173019 | rs681120 | C | A | 1.019 | -7977.388 | -7979.086 | -1.697 | 7.09E-07 |
| 11 | 117241429 | rs12417440 | T | C | 1.012 | -7977.388 | -7979.085 | -1.697 | 7.09E-07 |
| 11 | 117446945 | rs504935 | C | T | 0.988 | -7977.388 | -7979.085 | -1.697 | 7.09E-07 |
| 11 | 117662248 | rs534761 | T | C | 1.018 | -7977.388 | -7979.086 | -1.697 | 7.09E-07 |
| 11 | 117947454 | rs651013 | C | T | 1.014 | -7977.388 | -7979.085 | -1.697 | 7.09E-07 |
| 11 | 120610692 | rs2155262 | A | G | 0.985 | -7977.388 | -7979.086 | -1.697 | 7.09E-07 |
| 11 | 120738397 | rs3802912 | A | G | 0.981 | -7977.388 | -7979.085 | -1.697 | 7.09E-07 |
| 11 | 117330736 | rs747061 | G | C | 0.98 | -7977.388 | -7979.086 | -1.698 | 7.08E-07 |
| 11 | 117486829 | rs527005 | G | C | 1.001 | -7977.388 | -7979.086 | -1.698 | 7.08E-07 |
| 11 | 118637475 | rs112910608 | T | C | 1.014 | -7977.388 | -7979.086 | -1.698 | 7.08E-07 |
| 11 | 118892872 | rs58955701 | T | C | 1.011 | -7977.388 | -7979.086 | -1.698 | 7.08E-07 |
| 11 | 119742360 | rs10892481 | T | C | 0.981 | -7977.388 | -7979.087 | -1.698 | 7.08E-07 |
| 11 | 120213360 | rs7121779 | T | A | 1.019 | -7977.388 | -7979.086 | -1.698 | 7.08E-07 |
| 11 | 120216260 | rs7952529 | C | A | 1.019 | -7977.388 | -7979.087 | -1.698 | 7.08E-07 |

| | | | | | | | |
|---|---|---|---|---|---|---|---|
| 11 | 120235703 | rs6589811 | T | C | NULL | 0.700567142 | NULL&M_Disease |
| 11 | 116213060 | rs12574090 | A | G | NULL | 0.700567142 | NULL&M_Disease |
| 11 | 116292263 | rs574778 | C | A | NULL | 0.700567142 | NULL&M_Disease |
| 11 | 117535285 | rs11216489 | A | G | NULL | 0.700567142 | NULL&M_Disease |
| 11 | 117745049 | rs4938446 | A | T | NULL | 0.700567142 | NULL&M_Disease |
| 11 | 120241923 | rs10892566 | C | T | NULL | 0.700567142 | NULL&M_Disease |
| 11 | 120516717 | rs2000874 | G | A | NULL | 0.700567142 | NULL&M_Disease |
| 11 | 121002972 | rs10892674 | A | G | NULL | 0.700567142 | NULL&M_Disease |
| 11 | 121003224 | rs11218153 | A | G | NULL | 0.700567142 | NULL&M_Disease |
| 11 | 121003358 | rs7105712 | C | T | NULL | 0.700567142 | NULL&M_Disease |
| 11 | 116321311 | rs503361 | C | T | NULL | 0.700567142 | NULL&M_Disease |
| 11 | 117324393 | rs7936274 | T | C | NULL | 0.700567142 | NULL&M_Disease |
| 11 | 117775056 | rs477408 | G | A | NULL | 0.700567142 | NULL&M_Disease |
| 11 | 118892277 | rs4938621 | T | C | NULL | 0.700567142 | NULL&M_Disease |
| 11 | 120604347 | rs2846124 | C | A | NULL | 0.700567142 | NULL&M_Disease |
| 11 | 120987937 | rs628721 | G | A | NULL | 0.700567142 | NULL&M_Disease |
| 11 | 116321980 | rs573052 | A | G | NULL | 0.700567142 | NULL&M_Disease |
| 11 | 117552640 | rs10892160 | G | A | NULL | 0.700567142 | NULL&M_Disease |
| 11 | 119236794 | rs4589313 | A | G | NULL | 0.700567142 | NULL&M_Disease |
| 11 | 119492095 | rs4500497 | T | C | NULL | 0.700567142 | NULL&M_Disease |
| 11 | 119950310 | rs3887268 | G | C | NULL | 0.700567142 | NULL&M_Disease |
| 11 | 120233536 | rs7924963 | A | G | NULL | 0.700567142 | NULL&M_Disease |
| 11 | 120516014 | rs1343789 | C | T | NULL | 0.700567142 | NULL&M_Disease |
| 11 | 120855998 | rs542378 | T | C | NULL | 0.700567142 | NULL&M_Disease |
| 11 | 116490970 | rs12365322 | G | T | NULL | 0.700567142 | NULL&M_Disease |
| 11 | 117997722 | rs11216777 | G | T | NULL | 0.700567142 | NULL&M_Disease |
| 11 | 119598653 | rs2048246 | C | T | NULL | 0.700567142 | NULL&M_Disease |
| 11 | 119949822 | rs722058 | A | G | NULL | 0.700567142 | NULL&M_Disease |
| 11 | 120562930 | rs9651696 | C | T | NULL | 0.700567142 | NULL&M_Disease |
| 11 | 116401884 | rs11600715 | G | A | NULL | 0.700567142 | NULL&M_Disease |
| 11 | 116727936 | rs625145 | T | A | NULL | 0.700567142 | NULL&M_Disease |
| 11 | 117138403 | rs551826 | G | C | NULL | 0.700567142 | NULL&M_Disease |
| 11 | 117646824 | rs10160554 | G | A | NULL | 0.700567142 | NULL&M_Disease |
| 11 | 118892757 | rs117289748 | G | C | NULL | 0.700567142 | NULL&M_Disease |
| 11 | 118892759 | rs143660533 | C | A | NULL | 0.700567142 | NULL&M_Disease |
| 11 | 120207405 | rs7126413 | G | A | NULL | 0.700567142 | NULL&M_Disease |
| 11 | 120233158 | rs10790380 | A | G | NULL | 0.700567142 | NULL&M_Disease |
| 11 | 120789852 | rs949295 | C | T | NULL | 0.700567142 | NULL&M_Disease |
| 11 | 120855930 | rs543181 | A | G | NULL | 0.700567142 | NULL&M_Disease |
| 11 | 121192287 | rs2370113 | T | G | NULL | 0.700567142 | NULL&M_Disease |
| 11 | 116319329 | rs12574404 | G | C | NULL | 0.700567142 | NULL&M_Disease |
| 11 | 118893023 | rs7127188 | C | A | NULL | 0.700567142 | NULL&M_Disease |
| 11 | 120202099 | rs11217821 | C | T | NULL | 0.700567142 | NULL&M_Disease |
| 11 | 120520629 | rs4245040 | T | C | NULL | 0.700567142 | NULL&M_Disease |
| 11 | 120521766 | rs10892591 | T | C | NULL | 0.700567142 | NULL&M_Disease |
| 11 | 120606577 | rs2155264 | G | C | NULL | 0.700567142 | NULL&M_Disease |
| 11 | 120731363 | rs7937284 | G | A | NULL | 0.700567142 | NULL&M_Disease |
| 11 | 120829249 | rs672176 | A | G | NULL | 0.700567142 | NULL&M_Disease |
| 11 | 116664040 | rs10750097 | G | A | NULL | 0.700567142 | NULL&M_Disease |
| 11 | 118148271 | rs2187557 | T | C | NULL | 0.700567142 | NULL&M_Disease |
| 11 | 118657297 | rs487177 | G | C | NULL | 0.700567142 | NULL&M_Disease |
| 11 | 120262922 | rs7107211 | A | G | NULL | 0.700567142 | NULL&M_Disease |
| 11 | 120396485 | rs7129707 | T | C | NULL | 0.700567142 | NULL&M_Disease |
| 11 | 120602425 | rs3133852 | G | T | NULL | 0.700567142 | NULL&M_Disease |
| 11 | 116561393 | rs1787708 | A | G | NULL | 0.700567142 | NULL&M_Disease |
| 11 | 117775034 | rs686992 | C | T | NULL | 0.700567142 | NULL&M_Disease |
| 11 | 118174629 | rs3825051 | G | A | NULL | 0.700567142 | NULL&M_Disease |
| 11 | 118195962 | rs55843215 | C | T | NULL | 0.700567142 | NULL&M_Disease |
| 11 | 118856582 | rs3887652 | C | G | NULL | 0.700567142 | NULL&M_Disease |
| 11 | 119172536 | rs1047417 | G | A | NULL | 0.700567142 | NULL&M_Disease |
| 11 | 120090735 | rs11217770 | C | A | NULL | 0.700567142 | NULL&M_Disease |
| 11 | 120615801 | rs948030 | C | G | NULL | 0.700567142 | NULL&M_Disease |
| 11 | 121095669 | rs694490 | C | A | NULL | 0.700567142 | NULL&M_Disease |
| 11 | 116397532 | rs4938277 | T | C | NULL | 0.700567142 | NULL&M_Disease |
| 11 | 116583439 | rs6589562 | A | G | NULL | 0.700567142 | NULL&M_Disease |
| 11 | 117523275 | rs4938425 | A | G | NULL | 0.700567142 | NULL&M_Disease |
| 11 | 118001445 | rs10790239 | T | C | NULL | 0.700567142 | NULL&M_Disease |
| 11 | 118879824 | rs73559119 | C | T | NULL | 0.700567142 | NULL&M_Disease |
| 11 | 118880012 | rs11825284 | A | G | NULL | 0.700567142 | NULL&M_Disease |
| 11 | 118894287 | rs569 | G | A | NULL | 0.700567142 | NULL&M_Disease |
| 11 | 119370899 | rs10892376 | A | C | NULL | 0.700567142 | NULL&M_Disease |
| 11 | 119575724 | rs715849 | A | G | NULL | 0.700567142 | NULL&M_Disease |
| 11 | 119948566 | rs4938769 | T | C | NULL | 0.700567142 | NULL&M_Disease |
| 11 | 120344526 | rs10790383 | T | C | NULL | 0.701614954 | NULL&M_Disease |
| 11 | 120523139 | rs4570590 | C | G | NULL | 0.700567142 | NULL&M_Disease |
| 11 | 120562489 | rs11217974 | G | A | NULL | 0.700567142 | NULL&M_Disease |
| 11 | 116308090 | rs11215931 | G | A | NULL | 0.701614954 | NULL&M_Disease |
| 11 | 117662138 | rs12795949 | A | G | NULL | 0.700567142 | NULL&M_Disease |
| 11 | 118102058 | rs1793167 | T | C | NULL | 0.701614954 | NULL&M_Disease |
| 11 | 118879285 | rs7120445 | C | A | NULL | 0.700567142 | NULL&M_Disease |
| 11 | 119062889 | rs7945884 | C | T | NULL | 0.701614954 | NULL&M_Disease |
| 11 | 119696100 | rs1789792 | G | T | NULL | 0.701614954 | NULL&M_Disease |
| 11 | 120615697 | rs948029 | T | A | NULL | 0.701614954 | NULL&M_Disease |
| 11 | 116226263 | rs577853 | T | C | NULL | 0.701614954 | NULL&M_Disease |
| 11 | 117440171 | rs479601 | G | A | NULL | 0.701614954 | NULL&M_Disease |
| 11 | 117732007 | rs11216591 | T | A | NULL | 0.701614954 | NULL&M_Disease |
| 11 | 118886656 | rs7131534 | A | G | NULL | 0.701614954 | NULL&M_Disease |
| 11 | 119697182 | rs473219 | C | T | NULL | 0.701614954 | NULL&M_Disease |
| 11 | 119742989 | rs10892482 | C | T | NULL | 0.701614954 | NULL&M_Disease |
| 11 | 119744600 | rs2044544 | T | C | NULL | 0.701614954 | NULL&M_Disease |
| 11 | 119874301 | rs7929916 | A | G | NULL | 0.701614954 | NULL&M_Disease |
| 11 | 119902167 | rs949189 | C | T | NULL | 0.701614954 | NULL&M_Disease |
| 11 | 120614390 | rs1892974 | T | A | NULL | 0.701614954 | NULL&M_Disease |
| 11 | 120730669 | rs4331110 | A | G | NULL | 0.701614954 | NULL&M_Disease |
| 11 | 116227259 | rs497898 | A | G | NULL | 0.701614954 | NULL&M_Disease |
| 11 | 117307002 | rs11216384 | C | T | NULL | 0.701614954 | NULL&M_Disease |

| | | | | | | | | | |
|---|---|---|---|---|---|---|---|---|---|
| 11 | 120235703 | rs6589811 | T | C | 1.02 | -7977.388 | -7979.086 | -1.698 | 7.08E-07 |
| 11 | 116213060 | rs12574090 | A | G | 1.006 | -7977.388 | -7979.088 | -1.699 | 7.07E-07 |
| 11 | 116292263 | rs574778 | C | A | 0.998 | -7977.388 | -7979.087 | -1.699 | 7.07E-07 |
| 11 | 117535285 | rs11216489 | A | G | 0.981 | -7977.388 | -7979.088 | -1.699 | 7.07E-07 |
| 11 | 117745049 | rs4938446 | A | T | 0.982 | -7977.388 | -7979.087 | -1.699 | 7.07E-07 |
| 11 | 120241923 | rs10892566 | C | T | 1.019 | -7977.388 | -7979.087 | -1.699 | 7.07E-07 |
| 11 | 120516717 | rs2000874 | G | A | 1.018 | -7977.388 | -7979.087 | -1.699 | 7.07E-07 |
| 11 | 121002972 | rs10892674 | A | G | 0.983 | -7977.388 | -7979.087 | -1.699 | 7.07E-07 |
| 11 | 121003224 | rs11218153 | A | G | 0.983 | -7977.388 | -7979.087 | -1.699 | 7.07E-07 |
| 11 | 121003358 | rs7105712 | C | T | 0.983 | -7977.388 | -7979.087 | -1.699 | 7.07E-07 |
| 11 | 116321311 | rs503361 | C | T | 1.004 | -7977.388 | -7979.088 | -1.7 | 7.07E-07 |
| 11 | 117324393 | rs7936274 | T | C | 0.98 | -7977.388 | -7979.088 | -1.7 | 7.07E-07 |
| 11 | 117775056 | rs477408 | G | A | 1.019 | -7977.388 | -7979.088 | -1.7 | 7.07E-07 |
| 11 | 118892277 | rs4938621 | T | C | 1.011 | -7977.388 | -7979.088 | -1.7 | 7.07E-07 |
| 11 | 120604347 | rs2846124 | C | A | 0.985 | -7977.388 | -7979.088 | -1.7 | 7.07E-07 |
| 11 | 120987937 | rs628721 | G | A | 1.008 | -7977.388 | -7979.088 | -1.7 | 7.07E-07 |
| 11 | 116321980 | rs573052 | A | G | 1.002 | -7977.388 | -7979.089 | -1.701 | 7.06E-07 |
| 11 | 117552640 | rs10892160 | G | A | 1.001 | -7977.388 | -7979.089 | -1.701 | 7.06E-07 |
| 11 | 119236794 | rs4589313 | A | G | 0.982 | -7977.388 | -7979.09 | -1.701 | 7.06E-07 |
| 11 | 119492095 | rs4500497 | T | C | 0.982 | -7977.388 | -7979.09 | -1.701 | 7.06E-07 |
| 11 | 119950310 | rs3887268 | G | C | 1.001 | -7977.388 | -7979.089 | -1.701 | 7.06E-07 |
| 11 | 120233536 | rs7924963 | A | G | 1.019 | -7977.388 | -7979.089 | -1.701 | 7.06E-07 |
| 11 | 120516014 | rs1343789 | C | T | 1.018 | -7977.388 | -7979.089 | -1.701 | 7.06E-07 |
| 11 | 120855998 | rs542378 | T | C | 1.008 | -7977.388 | -7979.09 | -1.701 | 7.06E-07 |
| 11 | 116490970 | rs12365322 | G | T | 1.01 | -7977.388 | -7979.09 | -1.702 | 7.05E-07 |
| 11 | 117997722 | rs11216777 | G | T | 1.008 | -7977.388 | -7979.09 | -1.702 | 7.05E-07 |
| 11 | 119598653 | rs2048246 | C | T | 1.007 | -7977.388 | -7979.09 | -1.702 | 7.05E-07 |
| 11 | 119949822 | rs722058 | A | G | 1.003 | -7977.388 | -7979.091 | -1.702 | 7.05E-07 |
| 11 | 120562930 | rs9651696 | C | T | 1.02 | -7977.388 | -7979.09 | -1.702 | 7.05E-07 |
| 11 | 116401884 | rs11600715 | G | A | 1.009 | -7977.388 | -7979.092 | -1.703 | 7.05E-07 |
| 11 | 116727936 | rs625145 | T | A | 1.003 | -7977.388 | -7979.091 | -1.703 | 7.05E-07 |
| 11 | 117138403 | rs551826 | G | C | 0.983 | -7977.388 | -7979.092 | -1.703 | 7.05E-07 |
| 11 | 117646824 | rs10160554 | G | A | 0.984 | -7977.388 | -7979.091 | -1.703 | 7.05E-07 |
| 11 | 118892757 | rs117289748 | G | C | 1.011 | -7977.388 | -7979.091 | -1.703 | 7.05E-07 |
| 11 | 118892759 | rs143660533 | C | A | 1.011 | -7977.388 | -7979.091 | -1.703 | 7.05E-07 |
| 11 | 120207405 | rs7126413 | G | A | 1.019 | -7977.388 | -7979.091 | -1.703 | 7.05E-07 |
| 11 | 120233158 | rs10790380 | A | G | 1.019 | -7977.388 | -7979.092 | -1.703 | 7.05E-07 |
| 11 | 120789852 | rs949295 | C | T | 0.98 | -7977.388 | -7979.091 | -1.703 | 7.05E-07 |
| 11 | 120855930 | rs543181 | A | G | 1.007 | -7977.388 | -7979.092 | -1.703 | 7.05E-07 |
| 11 | 121192287 | rs2370113 | T | G | 1.019 | -7977.388 | -7979.092 | -1.703 | 7.05E-07 |
| 11 | 116319329 | rs12574404 | G | C | 1.003 | -7977.388 | -7979.093 | -1.704 | 7.04E-07 |
| 11 | 118893023 | rs7127188 | C | A | 1.011 | -7977.388 | -7979.092 | -1.704 | 7.04E-07 |
| 11 | 120202099 | rs11217821 | C | T | 1.019 | -7977.388 | -7979.092 | -1.704 | 7.04E-07 |
| 11 | 120520629 | rs4245040 | T | C | 1.017 | -7977.388 | -7979.093 | -1.704 | 7.04E-07 |
| 11 | 120521766 | rs10892591 | T | C | 1.019 | -7977.388 | -7979.092 | -1.704 | 7.04E-07 |
| 11 | 120606577 | rs2155264 | G | C | 0.985 | -7977.388 | -7979.092 | -1.704 | 7.04E-07 |
| 11 | 120731363 | rs7937284 | G | A | 1.019 | -7977.388 | -7979.093 | -1.704 | 7.04E-07 |
| 11 | 120829249 | rs672176 | A | G | 0.983 | -7977.388 | -7979.093 | -1.704 | 7.04E-07 |
| 11 | 116664040 | rs10750097 | G | A | 0.991 | -7977.388 | -7979.093 | -1.705 | 7.03E-07 |
| 11 | 118148271 | rs2187557 | T | C | 0.984 | -7977.388 | -7979.093 | -1.705 | 7.03E-07 |
| 11 | 118657297 | rs487177 | G | C | 1.013 | -7977.388 | -7979.094 | -1.705 | 7.03E-07 |
| 11 | 120262922 | rs7107211 | A | G | 1.019 | -7977.388 | -7979.093 | -1.705 | 7.03E-07 |
| 11 | 120396485 | rs7129707 | T | C | 0.983 | -7977.388 | -7979.093 | -1.705 | 7.03E-07 |
| 11 | 120602425 | rs3133852 | G | T | 1.012 | -7977.388 | -7979.093 | -1.705 | 7.03E-07 |
| 11 | 116561393 | rs1787708 | A | G | 1.017 | -7977.388 | -7979.095 | -1.706 | 7.02E-07 |
| 11 | 117775034 | rs686992 | C | T | 1.019 | -7977.388 | -7979.094 | -1.706 | 7.02E-07 |
| 11 | 118174629 | rs3825051 | G | A | 0.982 | -7977.388 | -7979.094 | -1.706 | 7.02E-07 |
| 11 | 118195962 | rs55843215 | C | T | 0.984 | -7977.388 | -7979.095 | -1.706 | 7.02E-07 |
| 11 | 118856582 | rs3887652 | C | G | 1.014 | -7977.388 | -7979.094 | -1.706 | 7.02E-07 |
| 11 | 119172536 | rs1047417 | G | A | 0.986 | -7977.388 | -7979.094 | -1.706 | 7.02E-07 |
| 11 | 120090735 | rs11217770 | C | A | 1.017 | -7977.388 | -7979.095 | -1.706 | 7.02E-07 |
| 11 | 120615801 | rs948030 | C | G | 0.986 | -7977.388 | -7979.094 | -1.706 | 7.02E-07 |
| 11 | 121095669 | rs694490 | C | A | 1.008 | -7977.388 | -7979.094 | -1.706 | 7.02E-07 |
| 11 | 116397532 | rs4938277 | T | C | 1.017 | -7977.388 | -7979.095 | -1.707 | 7.02E-07 |
| 11 | 116583439 | rs6589562 | A | G | 1.007 | -7977.388 | -7979.095 | -1.707 | 7.02E-07 |
| 11 | 117523275 | rs4938425 | A | G | 0.983 | -7977.388 | -7979.096 | -1.707 | 7.02E-07 |
| 11 | 118001445 | rs10790239 | T | C | 1.007 | -7977.388 | -7979.095 | -1.707 | 7.02E-07 |
| 11 | 118879824 | rs73559119 | C | T | 1.01 | -7977.388 | -7979.096 | -1.707 | 7.02E-07 |
| 11 | 118880012 | rs11825284 | A | G | 1.01 | -7977.388 | -7979.096 | -1.707 | 7.02E-07 |
| 11 | 118894287 | rs569 | G | A | 1.01 | -7977.388 | -7979.095 | -1.707 | 7.02E-07 |
| 11 | 119370899 | rs10892376 | A | C | 1.02 | -7977.388 | -7979.095 | -1.707 | 7.02E-07 |
| 11 | 119575724 | rs715849 | A | G | 1.019 | -7977.388 | -7979.095 | -1.707 | 7.02E-07 |
| 11 | 119948566 | rs4938769 | T | C | 0.999 | -7977.388 | -7979.096 | -1.707 | 7.02E-07 |
| 11 | 120344526 | rs10790383 | T | C | 1.019 | -7977.388 | -7979.096 | -1.707 | 7.02E-07 |
| 11 | 120523139 | rs4570590 | C | G | 1.019 | -7977.388 | -7979.095 | -1.707 | 7.02E-07 |
| 11 | 120562489 | rs11217974 | G | A | 1.018 | -7977.388 | -7979.096 | -1.707 | 7.02E-07 |
| 11 | 116308090 | rs11215931 | G | A | 1.018 | -7977.388 | -7979.096 | -1.708 | 7.01E-07 |
| 11 | 117662138 | rs12795949 | A | G | 0.998 | -7977.388 | -7979.096 | -1.708 | 7.01E-07 |
| 11 | 118102058 | rs1793167 | T | C | 1.011 | -7977.388 | -7979.096 | -1.708 | 7.01E-07 |
| 11 | 118879285 | rs7120445 | C | A | 1.01 | -7977.388 | -7979.096 | -1.708 | 7.01E-07 |
| 11 | 119062889 | rs7945884 | C | T | 1.02 | -7977.388 | -7979.096 | -1.708 | 7.01E-07 |
| 11 | 119696100 | rs1789792 | G | T | 0.995 | -7977.388 | -7979.096 | -1.708 | 7.01E-07 |
| 11 | 120615697 | rs948029 | T | A | 0.986 | -7977.388 | -7979.097 | -1.708 | 7.01E-07 |
| 11 | 116226263 | rs577853 | T | C | 1.008 | -7977.388 | -7979.097 | -1.709 | 7.00E-07 |
| 11 | 117440171 | rs479601 | G | A | 0.993 | -7977.388 | -7979.097 | -1.709 | 7.00E-07 |
| 11 | 117732007 | rs11216591 | T | A | 1.011 | -7977.388 | -7979.097 | -1.709 | 7.00E-07 |
| 11 | 118886656 | rs7131534 | A | G | 1.01 | -7977.388 | -7979.097 | -1.709 | 7.00E-07 |
| 11 | 119697182 | rs473219 | C | T | 0.996 | -7977.388 | -7979.098 | -1.709 | 7.00E-07 |
| 11 | 119742989 | rs10892482 | C | T | 0.981 | -7977.388 | -7979.098 | -1.709 | 7.00E-07 |
| 11 | 119744600 | rs2044544 | T | C | 0.981 | -7977.388 | -7979.097 | -1.709 | 7.00E-07 |
| 11 | 119874301 | rs7929916 | A | G | 1.019 | -7977.388 | -7979.097 | -1.709 | 7.00E-07 |
| 11 | 119902167 | rs949189 | C | T | 0.982 | -7977.388 | -7979.098 | -1.709 | 7.00E-07 |
| 11 | 120614390 | rs1892974 | T | A | 0.986 | -7977.388 | -7979.097 | -1.709 | 7.00E-07 |
| 11 | 120730669 | rs4331110 | A | G | 1.019 | -7977.388 | -7979.098 | -1.709 | 7.00E-07 |
| 11 | 116227259 | rs497898 | A | G | 1.008 | -7977.388 | -7979.098 | -1.71 | 7.00E-07 |
| 11 | 117307002 | rs11216384 | C | T | 0.982 | -7977.388 | -7979.098 | -1.71 | 7.00E-07 |

| | | | | | | | |
|---|---|---|---|---|---|---|---|
| 11 | 117732806 | rs10892187 | A | G | NULL | 0.701614954 | NULL&M_Disease |
| 11 | 119708875 | rs11217530 | A | G | NULL | 0.701614954 | NULL&M_Disease |
| 11 | 120233626 | rs7924772 | G | A | NULL | 0.701614954 | NULL&M_Disease |
| 11 | 120233908 | rs6589810 | A | G | NULL | 0.701614954 | NULL&M_Disease |
| 11 | 120611308 | rs2187495 | T | C | NULL | 0.701614954 | NULL&M_Disease |
| 11 | 120611792 | rs1073388 | G | C | NULL | 0.701614954 | NULL&M_Disease |
| 11 | 120731903 | rs10790404 | T | C | NULL | 0.701614954 | NULL&M_Disease |
| 11 | 120988792 | rs642548 | C | T | NULL | 0.701614954 | NULL&M_Disease |
| 11 | 121096292 | rs2155368 | C | A | NULL | 0.701614954 | NULL&M_Disease |
| 11 | 116302248 | rs11601357 | A | G | NULL | 0.701614954 | NULL&M_Disease |
| 11 | 116563706 | rs12785990 | A | G | NULL | 0.701614954 | NULL&M_Disease |
| 11 | 118122077 | rs11216834 | C | T | NULL | 0.701614954 | NULL&M_Disease |
| 11 | 118126682 | rs10892230 | A | C | NULL | 0.701614954 | NULL&M_Disease |
| 11 | 118880668 | rs11607569 | C | T | NULL | 0.701614954 | NULL&M_Disease |
| 11 | 119483293 | rs6589766 | A | G | NULL | 0.701614954 | NULL&M_Disease |
| 11 | 120199924 | rs61898351 | C | T | NULL | 0.701614954 | NULL&M_Disease |
| 11 | 120270709 | rs7118391 | T | C | NULL | 0.701614954 | NULL&M_Disease |
| 11 | 120735982 | rs4935755 | T | C | NULL | 0.701614954 | NULL&M_Disease |
| 11 | 121095824 | rs524990 | G | T | NULL | 0.701614954 | NULL&M_Disease |
| 11 | 116464362 | rs75214151 | A | G | NULL | 0.701614954 | NULL&M_Disease |
| 11 | 116471044 | rs11216037 | A | G | NULL | 0.701614954 | NULL&M_Disease |
| 11 | 116585533 | rs2000571 | A | G | NULL | 0.701614954 | NULL&M_Disease |
| 11 | 117442684 | rs491300 | G | T | NULL | 0.701614954 | NULL&M_Disease |
| 11 | 118717708 | rs7107756 | A | G | NULL | 0.701614954 | NULL&M_Disease |
| 11 | 119737648 | rs10892480 | A | G | NULL | 0.701614954 | NULL&M_Disease |
| 11 | 119939519 | rs2100358 | A | C | NULL | 0.701614954 | NULL&M_Disease |
| 11 | 120239051 | rs7117321 | C | T | NULL | 0.701614954 | NULL&M_Disease |
| 11 | 120273865 | rs4938804 | A | G | NULL | 0.701614954 | NULL&M_Disease |
| 11 | 120347715 | rs10892582 | T | C | NULL | 0.701614954 | NULL&M_Disease |
| 11 | 120522003 | rs10892592 | T | C | NULL | 0.701614954 | NULL&M_Disease |
| 11 | 121095863 | rs623952 | A | T | NULL | 0.701614954 | NULL&M_Disease |
| 11 | 116560778 | rs1787711 | G | A | NULL | 0.701614954 | NULL&M_Disease |
| 11 | 116561202 | rs1787710 | A | G | NULL | 0.701614954 | NULL&M_Disease |
| 11 | 117862298 | rs2512144 | A | G | NULL | 0.701614954 | NULL&M_Disease |
| 11 | 118831335 | rs61506726 | G | A | NULL | 0.701614954 | NULL&M_Disease |
| 11 | 120218243 | rs4539328 | C | T | NULL | 0.701614954 | NULL&M_Disease |
| 11 | 120607495 | rs2077623 | T | C | NULL | 0.701614954 | NULL&M_Disease |
| 11 | 120687529 | rs7124930 | C | T | NULL | 0.701614954 | NULL&M_Disease |
| 11 | 121095137 | rs637525 | G | A | NULL | 0.701614954 | NULL&M_Disease |
| 11 | 121178302 | rs1061332 | A | C | NULL | 0.701614954 | NULL&M_Disease |
| 11 | 116722551 | rs688456 | T | G | NULL | 0.701614954 | NULL&M_Disease |
| 11 | 117734224 | rs4938442 | T | C | NULL | 0.701614954 | NULL&M_Disease |
| 11 | 118064582 | rs619250 | A | T | NULL | 0.701614954 | NULL&M_Disease |
| 11 | 118605914 | rs7926282 | C | A | NULL | 0.701614954 | NULL&M_Disease |
| 11 | 120221618 | rs10892562 | G | A | NULL | 0.701614954 | NULL&M_Disease |
| 11 | 120558738 | rs6589831 | T | C | NULL | 0.701614954 | NULL&M_Disease |
| 11 | 116224496 | rs2510872 | A | G | NULL | 0.701614954 | NULL&M_Disease |
| 11 | 116560883 | rs3017232 | T | C | NULL | 0.701614954 | NULL&M_Disease |
| 11 | 116561701 | rs1784122 | A | T | NULL | 0.701614954 | NULL&M_Disease |
| 11 | 116561859 | rs1784120 | G | C | NULL | 0.701614954 | NULL&M_Disease |
| 11 | 116561938 | rs1787705 | A | G | NULL | 0.701614954 | NULL&M_Disease |
| 11 | 116579663 | rs11216107 | C | T | NULL | 0.701614954 | NULL&M_Disease |
| 11 | 117440267 | rs631120 | A | G | NULL | 0.701614954 | NULL&M_Disease |
| 11 | 117739616 | rs1006160 | A | T | NULL | 0.701614954 | NULL&M_Disease |
| 11 | 118115147 | rs11216830 | A | G | NULL | 0.701614954 | NULL&M_Disease |
| 11 | 118119211 | rs12271064 | G | A | NULL | 0.701614954 | NULL&M_Disease |
| 11 | 118605451 | rs2186780 | T | C | NULL | 0.701614954 | NULL&M_Disease |
| 11 | 118765600 | rs3922 | G | A | NULL | 0.701614954 | NULL&M_Disease |
| 11 | 119331254 | rs34708086 | C | G | NULL | 0.701614954 | NULL&M_Disease |
| 11 | 119702490 | rs477908 | T | G | NULL | 0.701614954 | NULL&M_Disease |
| 11 | 120191872 | rs11217817 | G | C | NULL | 0.701614954 | NULL&M_Disease |
| 11 | 120203559 | rs34298380 | G | A | NULL | 0.701614954 | NULL&M_Disease |
| 11 | 120239456 | rs7104639 | G | C | NULL | 0.701614954 | NULL&M_Disease |
| 11 | 120733632 | rs4569018 | C | T | NULL | 0.701614954 | NULL&M_Disease |
| 11 | 120855348 | rs11607159 | A | G | NULL | 0.701614954 | NULL&M_Disease |
| 11 | 116622761 | rs180330 | A | G | NULL | 0.701614954 | NULL&M_Disease |
| 11 | 118099186 | rs3802863 | T | C | NULL | 0.701614954 | NULL&M_Disease |
| 11 | 118116496 | rs11605435 | C | T | NULL | 0.701614954 | NULL&M_Disease |
| 11 | 118117118 | rs12280557 | T | C | NULL | 0.701614954 | NULL&M_Disease |
| 11 | 118117938 | rs7117544 | C | A | NULL | 0.701614954 | NULL&M_Disease |
| 11 | 118836606 | rs113975105 | G | C | NULL | 0.701614954 | NULL&M_Disease |
| 11 | 119332549 | rs34198109 | G | C | NULL | 0.701614954 | NULL&M_Disease |
| 11 | 120220268 | rs10790378 | C | T | NULL | 0.701614954 | NULL&M_Disease |
| 11 | 120223399 | rs4938801 | T | C | NULL | 0.701614954 | NULL&M_Disease |
| 11 | 120855313 | rs12796457 | T | C | NULL | 0.701614954 | NULL&M_Disease |
| 11 | 121098423 | rs7948840 | G | A | NULL | 0.701614954 | NULL&M_Disease |
| 11 | 117922143 | rs1786185 | G | A | NULL | 0.701614954 | NULL&M_Disease |
| 11 | 117969163 | rs948462 | C | T | NULL | 0.701614954 | NULL&M_Disease |
| 11 | 119200202 | rs55987240 | C | A | NULL | 0.701614954 | NULL&M_Disease |
| 11 | 119750091 | rs11217552 | T | C | NULL | 0.701614954 | NULL&M_Disease |
| 11 | 119875060 | rs7127194 | A | T | NULL | 0.701614954 | NULL&M_Disease |
| 11 | 120278359 | rs6589812 | C | T | NULL | 0.701614954 | NULL&M_Disease |
| 11 | 120278688 | rs7937612 | C | T | NULL | 0.701614954 | NULL&M_Disease |
| 11 | 120676208 | rs3016415 | G | T | NULL | 0.701614954 | NULL&M_Disease |
| 11 | 120707644 | rs3934628 | T | C | NULL | 0.701614954 | NULL&M_Disease |
| 11 | 120855341 | rs12796298 | G | A | NULL | 0.701614954 | NULL&M_Disease |
| 11 | 120890823 | rs12283211 | A | G | NULL | 0.701614954 | NULL&M_Disease |
| 11 | 116299000 | rs12362811 | A | G | NULL | 0.702660654 | NULL&M_Disease |
| 11 | 116348066 | rs11215954 | T | G | NULL | 0.701614954 | NULL&M_Disease |
| 11 | 116560667 | rs12291024 | G | T | NULL | 0.701614954 | NULL&M_Disease |
| 11 | 116561928 | rs1787706 | A | G | NULL | 0.702660654 | NULL&M_Disease |
| 11 | 118095996 | rs6589656 | C | G | NULL | 0.702660654 | NULL&M_Disease |
| 11 | 118132705 | rs2120268 | A | G | NULL | 0.702660654 | NULL&M_Disease |
| 11 | 119569802 | rs7945424 | G | T | NULL | 0.702660654 | NULL&M_Disease |
| 11 | 120203628 | rs12806740 | A | G | NULL | 0.702660654 | NULL&M_Disease |
| 11 | 120687069 | rs7105363 | T | G | NULL | 0.702660654 | NULL&M_Disease |

| | | | | | | | | | |
|---|---|---|---|---|---|---|---|---|---|
| 11 | 117732806 | rs10892187 | A | G | 1.011 | -7977.388 | -7979.098 | -1.71 | 7.00E-07 |
| 11 | 119708875 | rs11217530 | A | G | 0.983 | -7977.388 | -7979.098 | -1.71 | 7.00E-07 |
| 11 | 120233626 | rs7924772 | G | A | 1.019 | -7977.388 | -7979.099 | -1.71 | 7.00E-07 |
| 11 | 120233908 | rs6589810 | A | G | 1.019 | -7977.388 | -7979.099 | -1.71 | 7.00E-07 |
| 11 | 120611308 | rs2187495 | T | C | 0.986 | -7977.388 | -7979.098 | -1.71 | 7.00E-07 |
| 11 | 120611792 | rs1073388 | G | C | 0.986 | -7977.388 | -7979.098 | -1.71 | 7.00E-07 |
| 11 | 120731903 | rs10790404 | T | C | 1.019 | -7977.388 | -7979.098 | -1.71 | 7.00E-07 |
| 11 | 120988792 | rs642548 | C | T | 1.006 | -7977.388 | -7979.098 | -1.71 | 7.00E-07 |
| 11 | 121096292 | rs2155368 | C | A | 1.007 | -7977.388 | -7979.098 | -1.71 | 7.00E-07 |
| 11 | 116302248 | rs11601357 | A | G | 0.981 | -7977.388 | -7979.099 | -1.711 | 6.99E-07 |
| 11 | 116563706 | rs12785990 | A | G | 0.99 | -7977.388 | -7979.1 | -1.711 | 6.99E-07 |
| 11 | 118122077 | rs11216834 | C | T | 1.008 | -7977.388 | -7979.099 | -1.711 | 6.99E-07 |
| 11 | 118126682 | rs10892230 | A | C | 1.008 | -7977.388 | -7979.099 | -1.711 | 6.99E-07 |
| 11 | 118880668 | rs11607569 | C | T | 1.01 | -7977.388 | -7979.099 | -1.711 | 6.99E-07 |
| 11 | 119483293 | rs6589766 | A | G | 0.983 | -7977.388 | -7979.099 | -1.711 | 6.99E-07 |
| 11 | 120199924 | rs61898351 | C | T | 1.019 | -7977.388 | -7979.099 | -1.711 | 6.99E-07 |
| 11 | 120270709 | rs7118391 | T | C | 1.019 | -7977.388 | -7979.1 | -1.711 | 6.99E-07 |
| 11 | 120735982 | rs4935755 | T | C | 0.981 | -7977.388 | -7979.099 | -1.711 | 6.99E-07 |
| 11 | 121095824 | rs524990 | G | T | 1.007 | -7977.388 | -7979.099 | -1.711 | 6.99E-07 |
| 11 | 116464362 | rs75214151 | A | G | 1.014 | -7977.388 | -7979.101 | -1.712 | 6.98E-07 |
| 11 | 116471044 | rs11216037 | A | G | 1.017 | -7977.388 | -7979.1 | -1.712 | 6.98E-07 |
| 11 | 116585533 | rs2000571 | A | G | 1.006 | -7977.388 | -7979.1 | -1.712 | 6.98E-07 |
| 11 | 117442684 | rs491300 | G | T | 0.994 | -7977.388 | -7979.101 | -1.712 | 6.98E-07 |
| 11 | 118717708 | rs7107756 | A | G | 0.981 | -7977.388 | -7979.101 | -1.712 | 6.98E-07 |
| 11 | 119737648 | rs10892480 | A | G | 0.986 | -7977.388 | -7979.1 | -1.712 | 6.98E-07 |
| 11 | 119939519 | rs2100358 | A | C | 1.019 | -7977.388 | -7979.1 | -1.712 | 6.98E-07 |
| 11 | 120239051 | rs7117321 | C | T | 1.019 | -7977.388 | -7979.101 | -1.712 | 6.98E-07 |
| 11 | 120273865 | rs4938804 | A | G | 1.019 | -7977.388 | -7979.101 | -1.712 | 6.98E-07 |
| 11 | 120347715 | rs10892582 | T | C | 1.019 | -7977.388 | -7979.1 | -1.712 | 6.98E-07 |
| 11 | 120522003 | rs10892592 | T | C | 1.018 | -7977.388 | -7979.101 | -1.712 | 6.98E-07 |
| 11 | 121095863 | rs623952 | A | T | 1.007 | -7977.388 | -7979.1 | -1.712 | 6.98E-07 |
| 11 | 116560778 | rs1787711 | G | A | 1.017 | -7977.388 | -7979.101 | -1.713 | 6.98E-07 |
| 11 | 116561202 | rs1787710 | A | G | 1.017 | -7977.388 | -7979.102 | -1.713 | 6.98E-07 |
| 11 | 117862298 | rs2512144 | A | G | 0.983 | -7977.388 | -7979.101 | -1.713 | 6.98E-07 |
| 11 | 118831335 | rs61506726 | G | A | 0.984 | -7977.388 | -7979.101 | -1.713 | 6.98E-07 |
| 11 | 120218243 | rs4539328 | C | T | 1.019 | -7977.388 | -7979.101 | -1.713 | 6.98E-07 |
| 11 | 120607495 | rs2077623 | T | C | 0.986 | -7977.388 | -7979.101 | -1.713 | 6.98E-07 |
| 11 | 120687529 | rs7124930 | C | T | 0.981 | -7977.388 | -7979.101 | -1.713 | 6.98E-07 |
| 11 | 121095137 | rs637525 | G | A | 1.007 | -7977.388 | -7979.101 | -1.713 | 6.98E-07 |
| 11 | 121178302 | rs1061332 | A | C | 1.017 | -7977.388 | -7979.102 | -1.713 | 6.98E-07 |
| 11 | 116722551 | rs688456 | T | G | 0.998 | -7977.388 | -7979.102 | -1.714 | 6.97E-07 |
| 11 | 117734224 | rs4938442 | T | C | 1.011 | -7977.388 | -7979.103 | -1.714 | 6.97E-07 |
| 11 | 118064582 | rs619250 | A | T | 0.981 | -7977.388 | -7979.103 | -1.714 | 6.97E-07 |
| 11 | 118605914 | rs7926282 | C | A | 0.998 | -7977.388 | -7979.102 | -1.714 | 6.97E-07 |
| 11 | 120221618 | rs10892562 | G | A | 1.019 | -7977.388 | -7979.102 | -1.714 | 6.97E-07 |
| 11 | 120558738 | rs6589831 | T | C | 1.019 | -7977.388 | -7979.103 | -1.714 | 6.97E-07 |
| 11 | 116224496 | rs2510872 | A | G | 1.008 | -7977.388 | -7979.103 | -1.715 | 6.96E-07 |
| 11 | 116560883 | rs3017232 | T | C | 1.017 | -7977.388 | -7979.103 | -1.715 | 6.96E-07 |
| 11 | 116561701 | rs1784122 | A | T | 1.017 | -7977.388 | -7979.103 | -1.715 | 6.96E-07 |
| 11 | 116561859 | rs1784120 | G | C | 1.017 | -7977.388 | -7979.103 | -1.715 | 6.96E-07 |
| 11 | 116561938 | rs1787705 | A | G | 1.017 | -7977.388 | -7979.103 | -1.715 | 6.96E-07 |
| 11 | 116579663 | rs11216107 | C | T | 1.017 | -7977.388 | -7979.103 | -1.715 | 6.96E-07 |
| 11 | 117440267 | rs631120 | A | G | 0.998 | -7977.388 | -7979.103 | -1.715 | 6.96E-07 |
| 11 | 117739616 | rs1006160 | A | T | 0.982 | -7977.388 | -7979.103 | -1.715 | 6.96E-07 |
| 11 | 118115147 | rs11216830 | A | G | 1.008 | -7977.388 | -7979.103 | -1.715 | 6.96E-07 |
| 11 | 118119211 | rs12271064 | G | A | 1.007 | -7977.388 | -7979.103 | -1.715 | 6.96E-07 |
| 11 | 118605451 | rs2186780 | T | C | 0.999 | -7977.388 | -7979.103 | -1.715 | 6.96E-07 |
| 11 | 118765600 | rs3922 | G | A | 1.019 | -7977.388 | -7979.104 | -1.715 | 6.96E-07 |
| 11 | 119331254 | rs34708086 | C | G | 1.016 | -7977.388 | -7979.103 | -1.715 | 6.96E-07 |
| 11 | 119702490 | rs477908 | T | G | 0.983 | -7977.388 | -7979.104 | -1.715 | 6.96E-07 |
| 11 | 120191872 | rs11217817 | G | C | 0.982 | -7977.388 | -7979.103 | -1.715 | 6.96E-07 |
| 11 | 120203559 | rs34298380 | G | A | 1.019 | -7977.388 | -7979.103 | -1.715 | 6.96E-07 |
| 11 | 120239456 | rs7104639 | G | C | 1.019 | -7977.388 | -7979.103 | -1.715 | 6.96E-07 |
| 11 | 120733632 | rs4569018 | C | T | 1.019 | -7977.388 | -7979.103 | -1.715 | 6.96E-07 |
| 11 | 120855348 | rs11607159 | A | G | 1.005 | -7977.388 | -7979.103 | -1.715 | 6.96E-07 |
| 11 | 116622761 | rs180330 | A | G | 1.015 | -7977.388 | -7979.104 | -1.716 | 6.95E-07 |
| 11 | 118099186 | rs3802863 | T | C | 1.01 | -7977.388 | -7979.105 | -1.716 | 6.95E-07 |
| 11 | 118116496 | rs11605435 | C | T | 1.008 | -7977.388 | -7979.105 | -1.716 | 6.95E-07 |
| 11 | 118117118 | rs12280557 | T | C | 1.008 | -7977.388 | -7979.105 | -1.716 | 6.95E-07 |
| 11 | 118117938 | rs7117544 | C | A | 1.008 | -7977.388 | -7979.105 | -1.716 | 6.95E-07 |
| 11 | 118836606 | rs113975105 | G | C | 0.984 | -7977.388 | -7979.105 | -1.716 | 6.95E-07 |
| 11 | 119332549 | rs34198109 | G | C | 1.016 | -7977.388 | -7979.104 | -1.716 | 6.95E-07 |
| 11 | 120220268 | rs10790378 | C | T | 1.019 | -7977.388 | -7979.105 | -1.716 | 6.95E-07 |
| 11 | 120223399 | rs4938801 | T | C | 1.019 | -7977.388 | -7979.105 | -1.716 | 6.95E-07 |
| 11 | 120855313 | rs12796457 | T | C | 1.004 | -7977.388 | -7979.104 | -1.716 | 6.95E-07 |
| 11 | 121098423 | rs7948840 | G | A | 1.006 | -7977.388 | -7979.104 | -1.716 | 6.95E-07 |
| 11 | 117922143 | rs1786185 | G | A | 0.984 | -7977.388 | -7979.105 | -1.717 | 6.95E-07 |
| 11 | 117969163 | rs948462 | C | T | 1.009 | -7977.388 | -7979.105 | -1.717 | 6.95E-07 |
| 11 | 119200202 | rs55987240 | C | A | 1.014 | -7977.388 | -7979.105 | -1.717 | 6.95E-07 |
| 11 | 119750091 | rs11217552 | T | C | 0.981 | -7977.388 | -7979.105 | -1.717 | 6.95E-07 |
| 11 | 119875060 | rs7127194 | A | T | 1.019 | -7977.388 | -7979.105 | -1.717 | 6.95E-07 |
| 11 | 120278359 | rs6589812 | C | T | 1.019 | -7977.388 | -7979.105 | -1.717 | 6.95E-07 |
| 11 | 120278688 | rs7937612 | C | T | 1.019 | -7977.388 | -7979.105 | -1.717 | 6.95E-07 |
| 11 | 120676208 | rs3016415 | G | T | 1.001 | -7977.388 | -7979.105 | -1.717 | 6.95E-07 |
| 11 | 120707644 | rs3934628 | T | C | 0.981 | -7977.388 | -7979.105 | -1.717 | 6.95E-07 |
| 11 | 120855341 | rs12796298 | G | A | 1.004 | -7977.388 | -7979.105 | -1.717 | 6.95E-07 |
| 11 | 120890823 | rs12283211 | A | G | 1.01 | -7977.388 | -7979.105 | -1.717 | 6.95E-07 |
| 11 | 116299000 | rs12362811 | A | G | 0.983 | -7977.388 | -7979.106 | -1.718 | 6.94E-07 |
| 11 | 116348066 | rs11215954 | T | G | 0.988 | -7977.388 | -7979.106 | -1.718 | 6.94E-07 |
| 11 | 116560667 | rs12291024 | G | T | 0.99 | -7977.388 | -7979.106 | -1.718 | 6.94E-07 |
| 11 | 116561928 | rs1787706 | A | G | 1.017 | -7977.388 | -7979.106 | -1.718 | 6.94E-07 |
| 11 | 118095996 | rs6589656 | C | G | 1.01 | -7977.388 | -7979.106 | -1.718 | 6.94E-07 |
| 11 | 118132705 | rs2120268 | A | G | 1.006 | -7977.388 | -7979.107 | -1.718 | 6.94E-07 |
| 11 | 119569802 | rs7945424 | G | T | 1.018 | -7977.388 | -7979.106 | -1.718 | 6.94E-07 |
| 11 | 120203628 | rs12806740 | A | G | 1.018 | -7977.388 | -7979.106 | -1.718 | 6.94E-07 |
| 11 | 120687069 | rs7105363 | T | G | 0.981 | -7977.388 | -7979.106 | -1.718 | 6.94E-07 |

| | | | | | | | |
|---|---|---|---|---|---|---|---|
| 11 | 116568225 | rs11606013 | T | G | NULL | 0.702660654 | NULL&M_Disease |
| 11 | 117817358 | rs2515796 | G | A | NULL | 0.702660654 | NULL&M_Disease |
| 11 | 118098890 | rs1793156 | C | T | NULL | 0.702660654 | NULL&M_Disease |
| 11 | 118098962 | rs1793155 | A | G | NULL | 0.702660654 | NULL&M_Disease |
| 11 | 118135449 | rs3759000 | C | T | NULL | 0.702660654 | NULL&M_Disease |
| 11 | 118708564 | rs484232 | T | G | NULL | 0.702660654 | NULL&M_Disease |
| 11 | 120239088 | rs7101415 | A | G | NULL | 0.702660654 | NULL&M_Disease |
| 11 | 120435854 | rs4297478 | C | T | NULL | 0.702660654 | NULL&M_Disease |
| 11 | 120494034 | rs4631889 | A | G | NULL | 0.702660654 | NULL&M_Disease |
| 11 | 120507687 | rs35801991 | C | T | NULL | 0.702660654 | NULL&M_Disease |
| 11 | 121094411 | rs531410 | A | T | NULL | 0.702660654 | NULL&M_Disease |
| 11 | 121095030 | rs649880 | C | G | NULL | 0.702660654 | NULL&M_Disease |
| 11 | 116568364 | rs11608173 | A | T | NULL | 0.702660654 | NULL&M_Disease |
| 11 | 117476142 | rs666290 | T | C | NULL | 0.702660654 | NULL&M_Disease |
| 11 | 117737747 | rs876797 | A | C | NULL | 0.702660654 | NULL&M_Disease |
| 11 | 117741051 | rs7937921 | A | G | NULL | 0.702660654 | NULL&M_Disease |
| 11 | 118093359 | rs3018383 | A | C | NULL | 0.702660654 | NULL&M_Disease |
| 11 | 118112911 | rs11216829 | A | G | NULL | 0.702660654 | NULL&M_Disease |
| 11 | 118113583 | rs10502228 | T | C | NULL | 0.702660654 | NULL&M_Disease |
| 11 | 119936695 | rs1893392 | T | C | NULL | 0.702660654 | NULL&M_Disease |
| 11 | 120521066 | rs4935750 | C | T | NULL | 0.702660654 | NULL&M_Disease |
| 11 | 120703432 | rs1245791313 | G | A | NULL | 0.702660654 | NULL&M_Disease |
| 11 | 116245350 | rs487962 | A | G | NULL | 0.702660654 | NULL&M_Disease |
| 11 | 116575754 | rs1263057 | T | C | NULL | 0.702660654 | NULL&M_Disease |
| 11 | 116581641 | rs2367970 | A | G | NULL | 0.702660654 | NULL&M_Disease |
| 11 | 117287794 | rs474490 | T | G | NULL | 0.702660654 | NULL&M_Disease |
| 11 | 117522237 | rs1893517 | A | G | NULL | 0.702660654 | NULL&M_Disease |
| 11 | 117776648 | rs3802871 | C | A | NULL | 0.702660654 | NULL&M_Disease |
| 11 | 118114473 | rs12286303 | T | C | NULL | 0.702660654 | NULL&M_Disease |
| 11 | 118114788 | rs12286567 | T | G | NULL | 0.702660654 | NULL&M_Disease |
| 11 | 120322079 | rs496034 | A | G | NULL | 0.702660654 | NULL&M_Disease |
| 11 | 120619548 | rs3133862 | G | T | NULL | 0.702660654 | NULL&M_Disease |
| 11 | 117869670 | rs2229113 | A | G | NULL | 0.702660654 | NULL&M_Disease |
| 11 | 119488812 | rs4354701 | A | G | NULL | 0.702660654 | NULL&M_Disease |
| 11 | 119566228 | rs7129848 | T | C | NULL | 0.702660654 | NULL&M_Disease |
| 11 | 119573389 | rs10892434 | T | C | NULL | 0.702660654 | NULL&M_Disease |
| 11 | 119974755 | rs1110703 | G | A | NULL | 0.702660654 | NULL&M_Disease |
| 11 | 121094778 | rs527813 | G | T | NULL | 0.702660654 | NULL&M_Disease |
| 11 | 117333214 | rs11216403 | C | T | NULL | 0.702660654 | NULL&M_Disease |
| 11 | 118103542 | rs10790246 | T | A | NULL | 0.702660654 | NULL&M_Disease |
| 11 | 120191828 | rs2212641 | C | T | NULL | 0.702660654 | NULL&M_Disease |
| 11 | 120343271 | rs7939137 | T | C | NULL | 0.702660654 | NULL&M_Disease |
| 11 | 120686999 | rs7104999 | G | C | NULL | 0.702660654 | NULL&M_Disease |
| 11 | 116623659 | rs180327 | C | T | NULL | 0.702660654 | NULL&M_Disease |
| 11 | 117662864 | rs12294117 | A | G | NULL | 0.702660654 | NULL&M_Disease |
| 11 | 117763868 | rs559545 | A | C | NULL | 0.702660654 | NULL&M_Disease |
| 11 | 118096165 | rs7943511 | T | C | NULL | 0.702660654 | NULL&M_Disease |
| 11 | 119043656 | rs643423 | C | T | NULL | 0.702660654 | NULL&M_Disease |
| 11 | 120480291 | rs11604139 | A | T | NULL | 0.702660654 | NULL&M_Disease |
| 11 | 120687234 | rs7105286 | G | A | NULL | 0.702660654 | NULL&M_Disease |
| 11 | 120737184 | rs7115168 | G | A | NULL | 0.702660654 | NULL&M_Disease |
| 11 | 120987966 | rs628756 | C | T | NULL | 0.702660654 | NULL&M_Disease |
| 11 | 116488539 | rs11216040 | A | G | NULL | 0.702660654 | NULL&M_Disease |
| 11 | 116560948 | rs3017231 | A | G | NULL | 0.702660654 | NULL&M_Disease |
| 11 | 116758101 | rs7106641 | G | A | NULL | 0.702660654 | NULL&M_Disease |
| 11 | 117331864 | rs529796 | A | G | NULL | 0.702660654 | NULL&M_Disease |
| 11 | 118094921 | rs12295438 | T | C | NULL | 0.702660654 | NULL&M_Disease |
| 11 | 118095803 | rs3802866 | T | C | NULL | 0.702660654 | NULL&M_Disease |
| 11 | 118096028 | rs7943272 | C | A | NULL | 0.702660654 | NULL&M_Disease |
| 11 | 118111676 | rs11216828 | A | T | NULL | 0.702660654 | NULL&M_Disease |
| 11 | 119331500 | rs1941453 | T | C | NULL | 0.702660654 | NULL&M_Disease |
| 11 | 119490180 | rs7928238 | G | A | NULL | 0.702660654 | NULL&M_Disease |
| 11 | 119746467 | rs562993 | A | T | NULL | 0.702660654 | NULL&M_Disease |
| 11 | 119854514 | rs7107144 | C | T | NULL | 0.702660654 | NULL&M_Disease |
| 11 | 119877101 | rs4938752 | G | C | NULL | 0.702660654 | NULL&M_Disease |
| 11 | 120480136 | rs1025130228 | T | A | NULL | 0.702660654 | NULL&M_Disease |
| 11 | 120480191 | rs11601552 | A | C | NULL | 0.702660654 | NULL&M_Disease |
| 11 | 120731470 | rs7937697 | A | G | NULL | 0.702660654 | NULL&M_Disease |
| 11 | 120981195 | rs592529 | T | C | NULL | 0.702660654 | NULL&M_Disease |
| 11 | 117638627 | rs4936407 | T | C | NULL | 0.702660654 | NULL&M_Disease |
| 11 | 117662605 | rs34336740 | G | C | NULL | 0.702660654 | NULL&M_Disease |
| 11 | 117790674 | rs12418028 | A | G | NULL | 0.702660654 | NULL&M_Disease |
| 11 | 118831727 | rs10892315 | C | T | NULL | 0.702660654 | NULL&M_Disease |
| 11 | 120216906 | rs12417581 | A | T | NULL | 0.702660654 | NULL&M_Disease |
| 11 | 116560961 | rs3017230 | C | G | NULL | 0.702660654 | NULL&M_Disease |
| 11 | 117664373 | rs12271220 | A | G | NULL | 0.702660654 | NULL&M_Disease |
| 11 | 119875343 | rs4482041 | T | C | NULL | 0.702660654 | NULL&M_Disease |
| 11 | 120228286 | rs4936517 | G | C | NULL | 0.702660654 | NULL&M_Disease |
| 11 | 120718009 | rs4936560 | C | T | NULL | 0.702660654 | NULL&M_Disease |
| 11 | 117741995 | rs10790218 | A | G | NULL | 0.703704238 | NULL&M_Disease |
| 11 | 117959216 | rs2276434 | T | C | NULL | 0.703704238 | NULL&M_Disease |
| 11 | 117959372 | rs2276435 | A | G | NULL | 0.703704238 | NULL&M_Disease |
| 11 | 118096818 | rs11216824 | A | G | NULL | 0.703704238 | NULL&M_Disease |
| 11 | 119842786 | rs7944850 | A | G | NULL | 0.703704238 | NULL&M_Disease |
| 11 | 119843219 | rs1944504 | T | C | NULL | 0.703704238 | NULL&M_Disease |
| 11 | 120137544 | rs2096883 | T | C | NULL | 0.702660654 | NULL&M_Disease |
| 11 | 120215843 | rs12361632 | G | C | NULL | 0.703704238 | NULL&M_Disease |
| 11 | 117666413 | rs11216547 | T | C | NULL | 0.703704238 | NULL&M_Disease |
| 11 | 117739916 | rs873713 | T | A | NULL | 0.703704238 | NULL&M_Disease |
| 11 | 118011805 | rs644287 | T | C | NULL | 0.703704238 | NULL&M_Disease |
| 11 | 118105222 | rs1547885 | T | C | NULL | 0.703704238 | NULL&M_Disease |
| 11 | 118596444 | rs68097885 | A | G | NULL | 0.703704238 | NULL&M_Disease |
| 11 | 116464824 | rs73011181 | T | A | NULL | 0.703704238 | NULL&M_Disease |
| 11 | 116489036 | rs12365864 | G | A | NULL | 0.703704238 | NULL&M_Disease |
| 11 | 117816152 | rs2515794 | C | A | NULL | 0.703704238 | NULL&M_Disease |
| 11 | 118104569 | rs1793159 | A | C | NULL | 0.703704238 | NULL&M_Disease |

| | | | | | | | | | |
|---|---|---|---|---|---|---|---|---|---|
| 11 | 116568225 | rs11606013 | T | G | 0.991 | -7977.388 | -7979.107 | -1.719 | 6.93E-07 |
| 11 | 117817358 | rs2515796 | G | A | 1.019 | -7977.388 | -7979.108 | -1.719 | 6.93E-07 |
| 11 | 118098890 | rs1793156 | C | T | 1.01 | -7977.388 | -7979.107 | -1.719 | 6.93E-07 |
| 11 | 118098962 | rs1793155 | A | G | 1.01 | -7977.388 | -7979.107 | -1.719 | 6.93E-07 |
| 11 | 118135449 | rs3759000 | C | T | 1.014 | -7977.388 | -7979.108 | -1.719 | 6.93E-07 |
| 11 | 118708564 | rs484232 | T | G | 0.981 | -7977.388 | -7979.108 | -1.719 | 6.93E-07 |
| 11 | 120239088 | rs7101415 | A | G | 1.019 | -7977.388 | -7979.107 | -1.719 | 6.93E-07 |
| 11 | 120435854 | rs4297478 | C | T | 1.019 | -7977.388 | -7979.107 | -1.719 | 6.93E-07 |
| 11 | 120494034 | rs4631889 | A | G | 0.982 | -7977.388 | -7979.107 | -1.719 | 6.93E-07 |
| 11 | 120507687 | rs35801991 | C | T | 0.981 | -7977.388 | -7979.107 | -1.719 | 6.93E-07 |
| 11 | 121094411 | rs531410 | A | T | 1.006 | -7977.388 | -7979.108 | -1.719 | 6.93E-07 |
| 11 | 121095030 | rs649880 | C | G | 1.006 | -7977.388 | -7979.107 | -1.719 | 6.93E-07 |
| 11 | 116568364 | rs11608173 | A | T | 0.991 | -7977.388 | -7979.108 | -1.72 | 6.93E-07 |
| 11 | 117476142 | rs666290 | T | C | 0.999 | -7977.388 | -7979.108 | -1.72 | 6.93E-07 |
| 11 | 117737747 | rs876797 | A | C | 1.01 | -7977.388 | -7979.108 | -1.72 | 6.93E-07 |
| 11 | 117741051 | rs7937921 | A | G | 1.011 | -7977.388 | -7979.108 | -1.72 | 6.93E-07 |
| 11 | 118093359 | rs3018383 | A | C | 1.009 | -7977.388 | -7979.108 | -1.72 | 6.93E-07 |
| 11 | 118112911 | rs11216829 | A | G | 1.007 | -7977.388 | -7979.109 | -1.72 | 6.93E-07 |
| 11 | 118113583 | rs10502228 | T | C | 1.007 | -7977.388 | -7979.109 | -1.72 | 6.93E-07 |
| 11 | 119936695 | rs1893392 | T | C | 1.018 | -7977.388 | -7979.108 | -1.72 | 6.93E-07 |
| 11 | 120521066 | rs4935750 | C | T | 1.016 | -7977.388 | -7979.108 | -1.72 | 6.93E-07 |
| 11 | 120703432 | rs1245791313 | G | A | 0.982 | -7977.388 | -7979.108 | -1.72 | 6.93E-07 |
| 11 | 116245350 | rs487962 | A | G | 1.012 | -7977.388 | -7979.109 | -1.721 | 6.92E-07 |
| 11 | 116575754 | rs1263057 | T | C | 1.016 | -7977.388 | -7979.109 | -1.721 | 6.92E-07 |
| 11 | 116581641 | rs2367970 | A | G | 1.005 | -7977.388 | -7979.109 | -1.721 | 6.92E-07 |
| 11 | 117287794 | rs474490 | T | G | 0.99 | -7977.388 | -7979.11 | -1.721 | 6.92E-07 |
| 11 | 117522237 | rs1893517 | A | G | 0.984 | -7977.388 | -7979.109 | -1.721 | 6.92E-07 |
| 11 | 117776648 | rs3802871 | C | A | 1.019 | -7977.388 | -7979.11 | -1.721 | 6.92E-07 |
| 11 | 118114473 | rs12286303 | T | C | 1.007 | -7977.388 | -7979.109 | -1.721 | 6.92E-07 |
| 11 | 118114788 | rs12286567 | T | G | 1.007 | -7977.388 | -7979.109 | -1.721 | 6.92E-07 |
| 11 | 120322079 | rs496034 | A | G | 0.981 | -7977.388 | -7979.11 | -1.721 | 6.92E-07 |
| 11 | 120619548 | rs3133862 | G | T | 1.008 | -7977.388 | -7979.109 | -1.721 | 6.92E-07 |
| 11 | 117869670 | rs2229113 | A | G | 0.983 | -7977.388 | -7979.11 | -1.722 | 6.91E-07 |
| 11 | 119488812 | rs4354701 | A | G | 1.019 | -7977.388 | -7979.111 | -1.722 | 6.91E-07 |
| 11 | 119566228 | rs7129848 | T | C | 0.988 | -7977.388 | -7979.11 | -1.722 | 6.91E-07 |
| 11 | 119573389 | rs10892434 | T | C | 1.018 | -7977.388 | -7979.11 | -1.722 | 6.91E-07 |
| 11 | 119974755 | rs1110703 | G | A | 0.984 | -7977.388 | -7979.11 | -1.722 | 6.91E-07 |
| 11 | 121094778 | rs527813 | G | T | 1.005 | -7977.388 | -7979.11 | -1.722 | 6.91E-07 |
| 11 | 117333214 | rs11216403 | C | T | 1.016 | -7977.388 | -7979.111 | -1.723 | 6.91E-07 |
| 11 | 118103542 | rs10790246 | T | A | 1.007 | -7977.388 | -7979.111 | -1.723 | 6.91E-07 |
| 11 | 120191828 | rs2212641 | C | T | 0.982 | -7977.388 | -7979.111 | -1.723 | 6.91E-07 |
| 11 | 120343271 | rs7939137 | T | C | 1.018 | -7977.388 | -7979.111 | -1.723 | 6.91E-07 |
| 11 | 120686999 | rs7104999 | G | C | 0.981 | -7977.388 | -7979.111 | -1.723 | 6.91E-07 |
| 11 | 116623659 | rs180327 | C | T | 1.018 | -7977.388 | -7979.112 | -1.724 | 6.90E-07 |
| 11 | 117662864 | rs12294117 | A | G | 0.994 | -7977.388 | -7979.113 | -1.724 | 6.90E-07 |
| 11 | 117763868 | rs559545 | A | C | 1.019 | -7977.388 | -7979.112 | -1.724 | 6.90E-07 |
| 11 | 118096165 | rs7943511 | T | C | 1.006 | -7977.388 | -7979.113 | -1.724 | 6.90E-07 |
| 11 | 119043656 | rs643423 | C | T | 1.019 | -7977.388 | -7979.113 | -1.724 | 6.90E-07 |
| 11 | 120480291 | rs11604139 | A | T | 0.982 | -7977.388 | -7979.112 | -1.724 | 6.90E-07 |
| 11 | 120687234 | rs7105286 | G | A | 0.982 | -7977.388 | -7979.112 | -1.724 | 6.90E-07 |
| 11 | 120737184 | rs7115168 | G | A | 0.982 | -7977.388 | -7979.113 | -1.724 | 6.90E-07 |
| 11 | 120987966 | rs628756 | C | T | 0.985 | -7977.388 | -7979.113 | -1.724 | 6.90E-07 |
| 11 | 116488539 | rs11216040 | A | G | 1.006 | -7977.388 | -7979.113 | -1.725 | 6.89E-07 |
| 11 | 116560948 | rs3017231 | A | G | 1.016 | -7977.388 | -7979.113 | -1.725 | 6.89E-07 |
| 11 | 116758101 | rs7106641 | G | A | 1.007 | -7977.388 | -7979.114 | -1.725 | 6.89E-07 |
| 11 | 117331864 | rs529796 | A | G | 1.019 | -7977.388 | -7979.113 | -1.725 | 6.89E-07 |
| 11 | 118094921 | rs12295438 | T | C | 1.006 | -7977.388 | -7979.114 | -1.725 | 6.89E-07 |
| 11 | 118095803 | rs3802866 | T | C | 1.006 | -7977.388 | -7979.114 | -1.725 | 6.89E-07 |
| 11 | 118096028 | rs7943272 | C | A | 1.006 | -7977.388 | -7979.114 | -1.725 | 6.89E-07 |
| 11 | 118111676 | rs11216828 | A | T | 1.006 | -7977.388 | -7979.113 | -1.725 | 6.89E-07 |
| 11 | 119331500 | rs1941453 | T | C | 1.016 | -7977.388 | -7979.113 | -1.725 | 6.89E-07 |
| 11 | 119490180 | rs7928238 | G | A | 1.019 | -7977.388 | -7979.113 | -1.725 | 6.89E-07 |
| 11 | 119746467 | rs562993 | A | T | 0.986 | -7977.388 | -7979.114 | -1.725 | 6.89E-07 |
| 11 | 119854514 | rs7107144 | C | T | 1.018 | -7977.388 | -7979.114 | -1.725 | 6.89E-07 |
| 11 | 119877101 | rs4938752 | G | C | 1.019 | -7977.388 | -7979.114 | -1.725 | 6.89E-07 |
| 11 | 120480136 | rs1025130228 | T | A | 0.982 | -7977.388 | -7979.113 | -1.725 | 6.89E-07 |
| 11 | 120480191 | rs11601552 | A | C | 0.982 | -7977.388 | -7979.113 | -1.725 | 6.89E-07 |
| 11 | 120731470 | rs7937697 | A | G | 1.019 | -7977.388 | -7979.113 | -1.725 | 6.89E-07 |
| 11 | 120981195 | rs592529 | T | C | 0.981 | -7977.388 | -7979.113 | -1.725 | 6.89E-07 |
| 11 | 117638627 | rs4936407 | T | C | 0.986 | -7977.388 | -7979.114 | -1.726 | 6.89E-07 |
| 11 | 117662605 | rs34336740 | G | C | 0.995 | -7977.388 | -7979.114 | -1.726 | 6.89E-07 |
| 11 | 117790674 | rs12418028 | A | G | 1.012 | -7977.388 | -7979.114 | -1.726 | 6.89E-07 |
| 11 | 118831727 | rs10892315 | C | T | 0.985 | -7977.388 | -7979.114 | -1.726 | 6.89E-07 |
| 11 | 120216906 | rs12417581 | A | T | 1.018 | -7977.388 | -7979.114 | -1.726 | 6.89E-07 |
| 11 | 116560961 | rs3017230 | C | G | 1.016 | -7977.388 | -7979.115 | -1.727 | 6.88E-07 |
| 11 | 117664373 | rs12271220 | A | G | 0.995 | -7977.388 | -7979.116 | -1.727 | 6.88E-07 |
| 11 | 119875343 | rs4482041 | T | C | 1.018 | -7977.388 | -7979.116 | -1.727 | 6.88E-07 |
| 11 | 120228286 | rs4936517 | G | C | 1.018 | -7977.388 | -7979.115 | -1.727 | 6.88E-07 |
| 11 | 120718009 | rs4936560 | C | T | 1.003 | -7977.388 | -7979.115 | -1.727 | 6.88E-07 |
| 11 | 117741995 | rs10790218 | A | G | 0.983 | -7977.388 | -7979.116 | -1.728 | 6.87E-07 |
| 11 | 117959216 | rs2276434 | T | C | 0.982 | -7977.388 | -7979.116 | -1.728 | 6.87E-07 |
| 11 | 117959372 | rs2276435 | A | G | 0.982 | -7977.388 | -7979.116 | -1.728 | 6.87E-07 |
| 11 | 118096818 | rs11216824 | A | G | 1.004 | -7977.388 | -7979.117 | -1.728 | 6.87E-07 |
| 11 | 119842786 | rs7944850 | A | G | 1.019 | -7977.388 | -7979.117 | -1.728 | 6.87E-07 |
| 11 | 119843219 | rs1944504 | T | C | 1.019 | -7977.388 | -7979.116 | -1.728 | 6.87E-07 |
| 11 | 120137544 | rs2096883 | T | C | 1.005 | -7977.388 | -7979.116 | -1.728 | 6.87E-07 |
| 11 | 120215843 | rs12361632 | G | C | 1.018 | -7977.388 | -7979.117 | -1.728 | 6.87E-07 |
| 11 | 117666413 | rs11216547 | T | C | 0.995 | -7977.388 | -7979.117 | -1.729 | 6.86E-07 |
| 11 | 117739916 | rs873713 | T | A | 1.009 | -7977.388 | -7979.117 | -1.729 | 6.86E-07 |
| 11 | 118011805 | rs644287 | T | C | 0.983 | -7977.388 | -7979.117 | -1.729 | 6.86E-07 |
| 11 | 118105222 | rs1547885 | T | C | 1.005 | -7977.388 | -7979.118 | -1.729 | 6.86E-07 |
| 11 | 118596444 | rs68097885 | A | G | 1.011 | -7977.388 | -7979.117 | -1.729 | 6.86E-07 |
| 11 | 116464824 | rs73011181 | T | A | 1.016 | -7977.388 | -7979.118 | -1.73 | 6.86E-07 |
| 11 | 116489036 | rs12365864 | G | A | 1.005 | -7977.388 | -7979.118 | -1.73 | 6.86E-07 |
| 11 | 117816152 | rs2515794 | C | A | 1.019 | -7977.388 | -7979.119 | -1.73 | 6.86E-07 |
| 11 | 118104569 | rs1793159 | A | C | 1.005 | -7977.388 | -7979.118 | -1.73 | 6.86E-07 |

| | | | | | | | |
|---|---|---|---|---|---|---|---|
| 11 | 118104577 | rs1715422 | C | T | NULL | 0.703704238 | NULL&M_Disease |
| 11 | 118107495 | rs9651688 | T | C | NULL | 0.703704238 | NULL&M_Disease |
| 11 | 118668296 | rs58868870 | T | C | NULL | 0.703704238 | NULL&M_Disease |
| 11 | 119488646 | rs4499013 | C | T | NULL | 0.703704238 | NULL&M_Disease |
| 11 | 119573259 | rs10892433 | T | C | NULL | 0.703704238 | NULL&M_Disease |
| 11 | 119607933 | rs10892445 | G | A | NULL | 0.703704238 | NULL&M_Disease |
| 11 | 119875093 | rs10892511 | T | C | NULL | 0.703704238 | NULL&M_Disease |
| 11 | 119878219 | rs4938755 | G | A | NULL | 0.703704238 | NULL&M_Disease |
| 11 | 116559002 | rs1145210 | T | C | NULL | 0.703704238 | NULL&M_Disease |
| 11 | 116560529 | rs79280888 | G | A | NULL | 0.703704238 | NULL&M_Disease |
| 11 | 117308936 | rs10892102 | T | G | NULL | 0.703704238 | NULL&M_Disease |
| 11 | 117763538 | rs10892192 | T | C | NULL | 0.703704238 | NULL&M_Disease |
| 11 | 118105233 | rs1715421 | C | T | NULL | 0.703704238 | NULL&M_Disease |
| 11 | 118108679 | rs9651667 | G | C | NULL | 0.703704238 | NULL&M_Disease |
| 11 | 118109196 | rs35409990 | A | G | NULL | 0.703704238 | NULL&M_Disease |
| 11 | 119163540 | rs12222094 | C | G | NULL | 0.703704238 | NULL&M_Disease |
| 11 | 119568715 | rs4936494 | A | G | NULL | 0.703704238 | NULL&M_Disease |
| 11 | 119568719 | rs4936495 | C | T | NULL | 0.703704238 | NULL&M_Disease |
| 11 | 120479125 | rs34884722 | G | A | NULL | 0.703704238 | NULL&M_Disease |
| 11 | 120688599 | rs4936553 | A | G | NULL | 0.703704238 | NULL&M_Disease |
| 11 | 120793442 | rs4936563 | T | C | NULL | 0.703704238 | NULL&M_Disease |
| 11 | 116560524 | rs34937367 | C | T | NULL | 0.703704238 | NULL&M_Disease |
| 11 | 116560992 | rs3017228 | G | A | NULL | 0.703704238 | NULL&M_Disease |
| 11 | 116763034 | rs499910 | G | A | NULL | 0.703704238 | NULL&M_Disease |
| 11 | 117762288 | rs10750117 | G | C | NULL | 0.703704238 | NULL&M_Disease |
| 11 | 117860582 | rs2512143 | A | G | NULL | 0.703704238 | NULL&M_Disease |
| 11 | 118187511 | rs4936428 | T | C | NULL | 0.703704238 | NULL&M_Disease |
| 11 | 119162720 | rs11217231 | C | G | NULL | 0.703704238 | NULL&M_Disease |
| 11 | 119572441 | rs885430 | G | A | NULL | 0.703704238 | NULL&M_Disease |
| 11 | 120712099 | rs10790403 | C | T | NULL | 0.703704238 | NULL&M_Disease |
| 11 | 120963898 | rs10892670 | T | C | NULL | 0.703704238 | NULL&M_Disease |
| 11 | 116526322 | rs486394 | C | A | NULL | 0.703704238 | NULL&M_Disease |
| 11 | 117308927 | rs10892101 | A | C | NULL | 0.703704238 | NULL&M_Disease |
| 11 | 117665021 | rs11216546 | T | C | NULL | 0.703704238 | NULL&M_Disease |
| 11 | 117731269 | rs11601324 | C | G | NULL | 0.703704238 | NULL&M_Disease |
| 11 | 117952097 | rs3825050 | T | C | NULL | 0.703704238 | NULL&M_Disease |
| 11 | 119197843 | rs9888142 | G | A | NULL | 0.703704238 | NULL&M_Disease |
| 11 | 119377274 | rs7928100 | A | G | NULL | 0.703704238 | NULL&M_Disease |
| 11 | 116246867 | rs512588 | T | G | NULL | 0.703704238 | NULL&M_Disease |
| 11 | 116339688 | rs11215948 | G | C | NULL | 0.703704238 | NULL&M_Disease |
| 11 | 118020057 | rs598746 | A | G | NULL | 0.703704238 | NULL&M_Disease |
| 11 | 118666289 | rs56104144 | C | G | NULL | 0.703704238 | NULL&M_Disease |
| 11 | 119870618 | rs61897870 | G | C | NULL | 0.703704238 | NULL&M_Disease |
| 11 | 120712591 | rs7110179 | G | A | NULL | 0.703704238 | NULL&M_Disease |
| 11 | 120715884 | rs6589847 | A | G | NULL | 0.703704238 | NULL&M_Disease |
| 11 | 121149050 | rs34805238 | T | C | NULL | 0.703704238 | NULL&M_Disease |
| 11 | 116574817 | rs1240781 | T | C | NULL | 0.703704238 | NULL&M_Disease |
| 11 | 118105599 | rs1715420 | A | T | NULL | 0.703704238 | NULL&M_Disease |
| 11 | 118108331 | rs55768116 | A | C | NULL | 0.703704238 | NULL&M_Disease |
| 11 | 119739796 | rs541612 | T | C | NULL | 0.703704238 | NULL&M_Disease |
| 11 | 116492107 | rs61904858 | G | A | NULL | 0.703704238 | NULL&M_Disease |
| 11 | 117198789 | rs2508665 | T | A | NULL | 0.703704238 | NULL&M_Disease |
| 11 | 118573743 | rs519294 | A | C | NULL | 0.703704238 | NULL&M_Disease |
| 11 | 119605480 | rs4489755 | T | G | NULL | 0.703704238 | NULL&M_Disease |
| 11 | 120094400 | rs12808181 | T | C | NULL | 0.703704238 | NULL&M_Disease |
| 11 | 120414851 | rs2252771 | G | C | NULL | 0.703704238 | NULL&M_Disease |
| 11 | 117308984 | rs58290590 | A | G | NULL | 0.703704238 | NULL&M_Disease |
| 11 | 117956971 | rs4938479 | A | G | NULL | 0.703704238 | NULL&M_Disease |
| 11 | 118173783 | rs59138930 | T | C | NULL | 0.703704238 | NULL&M_Disease |
| 11 | 120461856 | rs7950360 | C | G | NULL | 0.703704238 | NULL&M_Disease |
| 11 | 120462743 | rs1320720 | A | G | NULL | 0.703704238 | NULL&M_Disease |
| 11 | 116214441 | rs35777932 | C | T | NULL | 0.704745698 | NULL&M_Disease |
| 11 | 116214490 | rs71478235 | T | C | NULL | 0.704745698 | NULL&M_Disease |
| 11 | 116467589 | rs12361543 | G | A | NULL | 0.704745698 | NULL&M_Disease |
| 11 | 116507479 | rs10892003 | G | A | NULL | 0.704745698 | NULL&M_Disease |
| 11 | 117193369 | rs4936370 | T | G | NULL | 0.704745698 | NULL&M_Disease |
| 11 | 117317941 | rs12364375 | T | A | NULL | 0.704745698 | NULL&M_Disease |
| 11 | 117696958 | rs479991 | C | T | NULL | 0.704745698 | NULL&M_Disease |
| 11 | 119192956 | rs7944487 | G | A | NULL | 0.704745698 | NULL&M_Disease |
| 11 | 119822015 | rs17245340 | C | T | NULL | 0.704745698 | NULL&M_Disease |
| 11 | 120198093 | rs1893261 | A | G | NULL | 0.704745698 | NULL&M_Disease |
| 11 | 120270253 | rs10750172 | G | A | NULL | 0.704745698 | NULL&M_Disease |
| 11 | 121127685 | rs10219304 | A | G | NULL | 0.704745698 | NULL&M_Disease |
| 11 | 116492701 | rs11216046 | T | G | NULL | 0.704745698 | NULL&M_Disease |
| 11 | 117098436 | rs4938360 | T | C | NULL | 0.704745698 | NULL&M_Disease |
| 11 | 118901166 | rs2846281 | C | G | NULL | 0.704745698 | NULL&M_Disease |
| 11 | 119256303 | rs12420005 | T | C | NULL | 0.704745698 | NULL&M_Disease |
| 11 | 120353298 | rs12794946 | C | A | NULL | 0.704745698 | NULL&M_Disease |
| 11 | 116571250 | rs1145195 | A | G | NULL | 0.704745698 | NULL&M_Disease |
| 11 | 118176097 | rs3782040 | G | A | NULL | 0.704745698 | NULL&M_Disease |
| 11 | 121112467 | rs11218222 | G | C | NULL | 0.704745698 | NULL&M_Disease |
| 11 | 116315743 | rs575549 | A | G | NULL | 0.704745698 | NULL&M_Disease |
| 11 | 116574584 | rs1240782 | A | G | NULL | 0.704745698 | NULL&M_Disease |
| 11 | 117096652 | rs1232770834 | A | G | NULL | 0.704745698 | NULL&M_Disease |
| 11 | 117915659 | rs619744 | G | C | NULL | 0.704745698 | NULL&M_Disease |
| 11 | 118707249 | rs4938551 | C | G | NULL | 0.704745698 | NULL&M_Disease |
| 11 | 119597943 | rs7952366 | T | G | NULL | 0.704745698 | NULL&M_Disease |
| 11 | 119745598 | rs534427 | T | C | NULL | 0.704745698 | NULL&M_Disease |
| 11 | 119858093 | rs11217613 | G | A | NULL | 0.704745698 | NULL&M_Disease |
| 11 | 119876697 | rs4938750 | A | G | NULL | 0.704745698 | NULL&M_Disease |
| 11 | 119877108 | rs4938753 | T | C | NULL | 0.704745698 | NULL&M_Disease |
| 11 | 120142118 | rs7484249 | C | T | NULL | 0.704745698 | NULL&M_Disease |
| 11 | 121129412 | rs9943622 | T | G | NULL | 0.704745698 | NULL&M_Disease |
| 11 | 116468132 | rs12364542 | A | G | NULL | 0.704745698 | NULL&M_Disease |
| 11 | 117697982 | rs2845710 | A | G | NULL | 0.704745698 | NULL&M_Disease |
| 11 | 118475098 | rs604096 | G | A | NULL | 0.704745698 | NULL&M_Disease |

| | | | | | | | | | |
|---|---|---|---|---|---|---|---|---|---|
| 11 | 118104577 | rs1715422 | C | T | 1.005 | -7977.388 | -7979.118 | -1.73 | 6.86E-07 |
| 11 | 118107495 | rs9651688 | T | C | 1.005 | -7977.388 | -7979.119 | -1.73 | 6.86E-07 |
| 11 | 118668296 | rs58868870 | T | C | 1.011 | -7977.388 | -7979.118 | -1.73 | 6.86E-07 |
| 11 | 119488646 | rs4499013 | C | T | 1.018 | -7977.388 | -7979.118 | -1.73 | 6.86E-07 |
| 11 | 119573259 | rs10892433 | T | C | 1.018 | -7977.388 | -7979.118 | -1.73 | 6.86E-07 |
| 11 | 119607933 | rs10892445 | G | A | 0.982 | -7977.388 | -7979.118 | -1.73 | 6.86E-07 |
| 11 | 119875093 | rs10892511 | T | C | 1.018 | -7977.388 | -7979.118 | -1.73 | 6.86E-07 |
| 11 | 119878219 | rs4938755 | G | A | 1.018 | -7977.388 | -7979.118 | -1.73 | 6.86E-07 |
| 11 | 116559002 | rs1145210 | T | C | 1.016 | -7977.388 | -7979.119 | -1.731 | 6.85E-07 |
| 11 | 116560529 | rs79280888 | G | A | 1.016 | -7977.388 | -7979.119 | -1.731 | 6.85E-07 |
| 11 | 117308936 | rs10892102 | T | G | 1.014 | -7977.388 | -7979.119 | -1.731 | 6.85E-07 |
| 11 | 117763538 | rs10892192 | T | C | 0.982 | -7977.388 | -7979.119 | -1.731 | 6.85E-07 |
| 11 | 118105233 | rs1715421 | C | T | 1.005 | -7977.388 | -7979.12 | -1.731 | 6.85E-07 |
| 11 | 118108679 | rs9651667 | G | C | 1.005 | -7977.388 | -7979.119 | -1.731 | 6.85E-07 |
| 11 | 118109196 | rs35409990 | A | G | 1.005 | -7977.388 | -7979.119 | -1.731 | 6.85E-07 |
| 11 | 119163540 | rs12222094 | C | G | 0.988 | -7977.388 | -7979.12 | -1.731 | 6.85E-07 |
| 11 | 119568715 | rs4936494 | A | G | 1.018 | -7977.388 | -7979.12 | -1.731 | 6.85E-07 |
| 11 | 119568719 | rs4936495 | C | T | 1.018 | -7977.388 | -7979.12 | -1.731 | 6.85E-07 |
| 11 | 120479125 | rs34884722 | G | A | 0.982 | -7977.388 | -7979.12 | -1.731 | 6.85E-07 |
| 11 | 120688599 | rs4936553 | A | G | 0.983 | -7977.388 | -7979.12 | -1.731 | 6.85E-07 |
| 11 | 120793442 | rs4936563 | T | C | 0.993 | -7977.388 | -7979.12 | -1.731 | 6.85E-07 |
| 11 | 116560524 | rs34937367 | C | T | 1.016 | -7977.388 | -7979.121 | -1.732 | 6.84E-07 |
| 11 | 116560992 | rs3017228 | G | A | 1.016 | -7977.388 | -7979.12 | -1.732 | 6.84E-07 |
| 11 | 116763034 | rs499910 | G | A | 1.006 | -7977.388 | -7979.121 | -1.732 | 6.84E-07 |
| 11 | 117762288 | rs10750117 | G | C | 0.982 | -7977.388 | -7979.121 | -1.732 | 6.84E-07 |
| 11 | 117860582 | rs2512143 | A | G | 0.984 | -7977.388 | -7979.12 | -1.732 | 6.84E-07 |
| 11 | 118187511 | rs4936428 | T | C | 0.994 | -7977.388 | -7979.121 | -1.732 | 6.84E-07 |
| 11 | 119162720 | rs11217231 | C | G | 0.988 | -7977.388 | -7979.12 | -1.732 | 6.84E-07 |
| 11 | 119572441 | rs885430 | G | A | 1.018 | -7977.388 | -7979.12 | -1.732 | 6.84E-07 |
| 11 | 120712099 | rs10790403 | C | T | 0.996 | -7977.388 | -7979.12 | -1.732 | 6.84E-07 |
| 11 | 120963898 | rs10892670 | T | C | 1.013 | -7977.388 | -7979.12 | -1.732 | 6.84E-07 |
| 11 | 116526322 | rs486394 | C | A | 0.986 | -7977.388 | -7979.121 | -1.733 | 6.84E-07 |
| 11 | 117308927 | rs10892101 | A | C | 1.014 | -7977.388 | -7979.121 | -1.733 | 6.84E-07 |
| 11 | 117665021 | rs11216546 | T | C | 0.997 | -7977.388 | -7979.122 | -1.733 | 6.84E-07 |
| 11 | 117731269 | rs11601324 | C | G | 1.008 | -7977.388 | -7979.121 | -1.733 | 6.84E-07 |
| 11 | 117952097 | rs3825050 | T | C | 1.017 | -7977.388 | -7979.121 | -1.733 | 6.84E-07 |
| 11 | 119197843 | rs9888142 | G | A | 1.018 | -7977.388 | -7979.121 | -1.733 | 6.84E-07 |
| 11 | 119377274 | rs7928100 | A | G | 1.018 | -7977.388 | -7979.122 | -1.733 | 6.84E-07 |
| 11 | 116246867 | rs512588 | T | G | 1.01 | -7977.388 | -7979.122 | -1.734 | 6.83E-07 |
| 11 | 116339688 | rs11215948 | G | C | 1.01 | -7977.388 | -7979.122 | -1.734 | 6.83E-07 |
| 11 | 118020057 | rs598746 | A | G | 1.018 | -7977.388 | -7979.123 | -1.734 | 6.83E-07 |
| 11 | 118666289 | rs56104144 | C | G | 1.011 | -7977.388 | -7979.123 | -1.734 | 6.83E-07 |
| 11 | 119870618 | rs61897870 | G | C | 0.993 | -7977.388 | -7979.122 | -1.734 | 6.83E-07 |
| 11 | 120712591 | rs7110179 | G | A | 0.998 | -7977.388 | -7979.122 | -1.734 | 6.83E-07 |
| 11 | 120715884 | rs6589847 | A | G | 0.998 | -7977.388 | -7979.122 | -1.734 | 6.83E-07 |
| 11 | 121149050 | rs34805238 | T | C | 1.012 | -7977.388 | -7979.123 | -1.734 | 6.83E-07 |
| 11 | 116574817 | rs1240781 | T | C | 0.982 | -7977.388 | -7979.124 | -1.735 | 6.82E-07 |
| 11 | 118105599 | rs1715420 | A | T | 1.004 | -7977.388 | -7979.124 | -1.735 | 6.82E-07 |
| 11 | 118108331 | rs55768116 | A | C | 0.982 | -7977.388 | -7979.123 | -1.735 | 6.82E-07 |
| 11 | 119739796 | rs541612 | T | C | 0.986 | -7977.388 | -7979.123 | -1.735 | 6.82E-07 |
| 11 | 116492107 | rs61904858 | G | A | 1.004 | -7977.388 | -7979.124 | -1.736 | 6.82E-07 |
| 11 | 117198789 | rs2508665 | T | A | 0.988 | -7977.388 | -7979.124 | -1.736 | 6.82E-07 |
| 11 | 118573743 | rs519294 | A | C | 0.984 | -7977.388 | -7979.125 | -1.736 | 6.82E-07 |
| 11 | 119605480 | rs4489755 | T | G | 0.982 | -7977.388 | -7979.125 | -1.736 | 6.82E-07 |
| 11 | 120094400 | rs12808181 | T | C | 1.014 | -7977.388 | -7979.125 | -1.736 | 6.82E-07 |
| 11 | 120414851 | rs2252771 | G | C | 1.009 | -7977.388 | -7979.125 | -1.736 | 6.82E-07 |
| 11 | 117308984 | rs58290590 | A | G | 1.013 | -7977.388 | -7979.125 | -1.737 | 6.81E-07 |
| 11 | 117956971 | rs4938479 | A | G | 0.983 | -7977.388 | -7979.125 | -1.737 | 6.81E-07 |
| 11 | 118173783 | rs59138930 | T | C | 0.984 | -7977.388 | -7979.125 | -1.737 | 6.81E-07 |
| 11 | 120461856 | rs7950360 | C | G | 1.01 | -7977.388 | -7979.125 | -1.737 | 6.81E-07 |
| 11 | 120462743 | rs1320720 | A | G | 1.01 | -7977.388 | -7979.126 | -1.737 | 6.81E-07 |
| 11 | 116214441 | rs35777932 | C | T | 1.009 | -7977.388 | -7979.126 | -1.738 | 6.80E-07 |
| 11 | 116214490 | rs71478235 | T | C | 1.009 | -7977.388 | -7979.127 | -1.738 | 6.80E-07 |
| 11 | 116467589 | rs12361543 | G | A | 1.015 | -7977.388 | -7979.126 | -1.738 | 6.80E-07 |
| 11 | 116507479 | rs10892003 | G | A | 1.017 | -7977.388 | -7979.127 | -1.738 | 6.80E-07 |
| 11 | 117193369 | rs4936370 | T | G | 0.988 | -7977.388 | -7979.127 | -1.738 | 6.80E-07 |
| 11 | 117317941 | rs12364375 | T | A | 0.984 | -7977.388 | -7979.126 | -1.738 | 6.80E-07 |
| 11 | 117696958 | rs479991 | C | T | 0.991 | -7977.388 | -7979.126 | -1.738 | 6.80E-07 |
| 11 | 119192956 | rs7944487 | G | A | 0.985 | -7977.388 | -7979.126 | -1.738 | 6.80E-07 |
| 11 | 119822015 | rs17245340 | C | T | 0.984 | -7977.388 | -7979.127 | -1.738 | 6.80E-07 |
| 11 | 120198093 | rs1893261 | A | G | 1.017 | -7977.388 | -7979.126 | -1.738 | 6.80E-07 |
| 11 | 120270253 | rs10750172 | G | A | 1.018 | -7977.388 | -7979.126 | -1.738 | 6.80E-07 |
| 11 | 121127685 | rs10219304 | A | G | 1.014 | -7977.388 | -7979.126 | -1.738 | 6.80E-07 |
| 11 | 116492701 | rs11216046 | T | G | 1.003 | -7977.388 | -7979.128 | -1.739 | 6.80E-07 |
| 11 | 117098436 | rs4938360 | T | C | 1.004 | -7977.388 | -7979.127 | -1.739 | 6.80E-07 |
| 11 | 118901166 | rs2846281 | C | G | 1.004 | -7977.388 | -7979.127 | -1.739 | 6.80E-07 |
| 11 | 119256303 | rs12420005 | T | C | 1.015 | -7977.388 | -7979.127 | -1.739 | 6.80E-07 |
| 11 | 120353298 | rs12794946 | C | A | 1.018 | -7977.388 | -7979.128 | -1.739 | 6.80E-07 |
| 11 | 116571250 | rs1145195 | A | G | 0.982 | -7977.388 | -7979.128 | -1.74 | 6.79E-07 |
| 11 | 118176097 | rs3782040 | G | A | 0.984 | -7977.388 | -7979.128 | -1.74 | 6.79E-07 |
| 11 | 121112467 | rs11218222 | G | C | 0.982 | -7977.388 | -7979.128 | -1.74 | 6.79E-07 |
| 11 | 116315743 | rs575549 | A | G | 0.997 | -7977.388 | -7979.129 | -1.741 | 6.78E-07 |
| 11 | 116574584 | rs1240782 | A | G | 0.982 | -7977.388 | -7979.129 | -1.741 | 6.78E-07 |
| 11 | 117096652 | rs1232770834 | A | G | 1.004 | -7977.388 | -7979.129 | -1.741 | 6.78E-07 |
| 11 | 117915659 | rs619744 | G | C | 1.01 | -7977.388 | -7979.13 | -1.741 | 6.78E-07 |
| 11 | 118707249 | rs4938551 | C | G | 0.982 | -7977.388 | -7979.13 | -1.741 | 6.78E-07 |
| 11 | 119597943 | rs7952366 | T | G | 0.982 | -7977.388 | -7979.13 | -1.741 | 6.78E-07 |
| 11 | 119745598 | rs534427 | T | C | 0.987 | -7977.388 | -7979.129 | -1.741 | 6.78E-07 |
| 11 | 119858093 | rs11217613 | G | A | 1.018 | -7977.388 | -7979.13 | -1.741 | 6.78E-07 |
| 11 | 119876697 | rs4938750 | A | G | 1.018 | -7977.388 | -7979.129 | -1.741 | 6.78E-07 |
| 11 | 119877108 | rs4938753 | T | C | 1.018 | -7977.388 | -7979.129 | -1.741 | 6.78E-07 |
| 11 | 120142118 | rs7484249 | C | T | 1.014 | -7977.388 | -7979.129 | -1.741 | 6.78E-07 |
| 11 | 121129412 | rs9943622 | T | G | 1.012 | -7977.388 | -7979.129 | -1.741 | 6.78E-07 |
| 11 | 116468132 | rs12364542 | A | G | 1.015 | -7977.388 | -7979.131 | -1.742 | 6.78E-07 |
| 11 | 117697982 | rs2845710 | A | G | 0.992 | -7977.388 | -7979.13 | -1.742 | 6.78E-07 |
| 11 | 118475098 | rs604096 | G | A | 1.008 | -7977.388 | -7979.13 | -1.742 | 6.78E-07 |

| | | | | | | | |
|---|---|---|---|---|---|---|---|
| 11 | 118913993 | rs1003081 | T | C | NULL | 0.704745698 | NULL&M_Disease |
| 11 | 119972011 | rs6589802 | G | A | NULL | 0.704745698 | NULL&M_Disease |
| 11 | 120196433 | rs36011588 | G | C | NULL | 0.704745698 | NULL&M_Disease |
| 11 | 120330149 | rs2305010 | C | G | NULL | 0.704745698 | NULL&M_Disease |
| 11 | 116559743 | rs3016360 | C | T | NULL | 0.704745698 | NULL&M_Disease |
| 11 | 117172820 | rs680210 | T | C | NULL | 0.704745698 | NULL&M_Disease |
| 11 | 118124092 | rs1715455 | T | C | NULL | 0.704745698 | NULL&M_Disease |
| 11 | 118131223 | rs7120665 | C | T | NULL | 0.704745698 | NULL&M_Disease |
| 11 | 119876028 | rs2155168 | T | C | NULL | 0.704745698 | NULL&M_Disease |
| 11 | 120141165 | rs1941406 | A | G | NULL | 0.704745698 | NULL&M_Disease |
| 11 | 117320727 | rs10892108 | T | C | NULL | 0.704745698 | NULL&M_Disease |
| 11 | 117685220 | rs10892177 | C | G | NULL | 0.704745698 | NULL&M_Disease |
| 11 | 117685575 | rs11216561 | C | T | NULL | 0.704745698 | NULL&M_Disease |
| 11 | 118072373 | rs7949751 | G | A | NULL | 0.704745698 | NULL&M_Disease |
| 11 | 120610144 | rs2846114 | C | T | NULL | 0.704745698 | NULL&M_Disease |
| 11 | 121140993 | rs10790428 | C | T | NULL | 0.704745698 | NULL&M_Disease |
| 11 | 121149254 | rs11218238 | G | A | NULL | 0.704745698 | NULL&M_Disease |
| 11 | 121151411 | rs4462362 | C | G | NULL | 0.704745698 | NULL&M_Disease |
| 11 | 116680918 | rs1263172 | G | A | NULL | 0.704745698 | NULL&M_Disease |
| 11 | 117097009 | rs7939747 | G | A | NULL | 0.704745698 | NULL&M_Disease |
| 11 | 117914297 | rs1786180 | G | C | NULL | 0.704745698 | NULL&M_Disease |
| 11 | 117914304 | rs1786179 | A | T | NULL | 0.704745698 | NULL&M_Disease |
| 11 | 117914550 | rs1783853 | G | A | NULL | 0.704745698 | NULL&M_Disease |
| 11 | 119072976 | rs11217180 | T | C | NULL | 0.704745698 | NULL&M_Disease |
| 11 | 119166047 | rs1309012690 | A | G | NULL | 0.704745698 | NULL&M_Disease |
| 11 | 119330741 | rs12574794 | T | C | NULL | 0.704745698 | NULL&M_Disease |
| 11 | 119375180 | rs10750147 | A | G | NULL | 0.704745698 | NULL&M_Disease |
| 11 | 119488133 | rs7934906 | T | C | NULL | 0.704745698 | NULL&M_Disease |
| 11 | 119572103 | rs3935405 | T | C | NULL | 0.704745698 | NULL&M_Disease |
| 11 | 119596772 | rs12418662 | C | T | NULL | 0.704745698 | NULL&M_Disease |
| 11 | 120448992 | rs11602075 | C | T | NULL | 0.704745698 | NULL&M_Disease |
| 11 | 120494089 | rs6589821 | A | G | NULL | 0.704745698 | NULL&M_Disease |
| 11 | 121134420 | rs11218230 | G | T | NULL | 0.704745698 | NULL&M_Disease |
| 11 | 121136455 | rs10790419 | T | C | NULL | 0.704745698 | NULL&M_Disease |
| 11 | 116560325 | rs1787715 | T | C | NULL | 0.704745698 | NULL&M_Disease |
| 11 | 117193350 | rs4938370 | C | A | NULL | 0.704745698 | NULL&M_Disease |
| 11 | 117548726 | rs596440 | C | T | NULL | 0.704745698 | NULL&M_Disease |
| 11 | 117548858 | rs637454 | G | A | NULL | 0.704745698 | NULL&M_Disease |
| 11 | 117817276 | rs2515795 | A | G | NULL | 0.704745698 | NULL&M_Disease |
| 11 | 119874855 | rs7126918 | A | T | NULL | 0.704745698 | NULL&M_Disease |
| 11 | 120133153 | rs2715882 | T | C | NULL | 0.704745698 | NULL&M_Disease |
| 11 | 120722457 | rs11218031 | A | C | NULL | 0.704745698 | NULL&M_Disease |
| 11 | 121133278 | rs7937589 | C | G | NULL | 0.704745698 | NULL&M_Disease |
| 11 | 116572205 | rs1145196 | T | C | NULL | 0.704745698 | NULL&M_Disease |
| 11 | 117425009 | rs579239 | C | T | NULL | 0.704745698 | NULL&M_Disease |
| 11 | 118166212 | rs4938500 | A | C | NULL | 0.704745698 | NULL&M_Disease |
| 11 | 118262706 | rs61900932 | T | C | NULL | 0.704745698 | NULL&M_Disease |
| 11 | 119702015 | rs544201 | A | T | NULL | 0.704745698 | NULL&M_Disease |
| 11 | 120273316 | rs4381380 | A | G | NULL | 0.704745698 | NULL&M_Disease |
| 11 | 116560174 | rs1787716 | A | G | NULL | 0.705785028 | NULL&M_Disease |
| 11 | 116570942 | rs1145193 | G | A | NULL | 0.705785028 | NULL&M_Disease |
| 11 | 117072963 | rs634960 | C | T | NULL | 0.705785028 | NULL&M_Disease |
| 11 | 117697985 | rs11216566 | G | A | NULL | 0.705785028 | NULL&M_Disease |
| 11 | 117801589 | rs61900351 | A | T | NULL | 0.705785028 | NULL&M_Disease |
| 11 | 117972936 | rs881122 | A | C | NULL | 0.705785028 | NULL&M_Disease |
| 11 | 119012846 | rs659375 | A | T | NULL | 0.705785028 | NULL&M_Disease |
| 11 | 119969871 | rs623353 | C | A | NULL | 0.705785028 | NULL&M_Disease |
| 11 | 120479913 | rs34759019 | A | C | NULL | 0.705785028 | NULL&M_Disease |
| 11 | 121127518 | rs4533044 | C | T | NULL | 0.704745698 | NULL&M_Disease |
| 11 | 121134392 | rs11218229 | G | T | NULL | 0.705785028 | NULL&M_Disease |
| 11 | 121147950 | rs9971567 | G | A | NULL | 0.705785028 | NULL&M_Disease |
| 11 | 116310858 | rs580872 | A | G | NULL | 0.705785028 | NULL&M_Disease |
| 11 | 119868171 | rs7111799 | A | G | NULL | 0.705785028 | NULL&M_Disease |
| 11 | 121128562 | rs4517527 | G | T | NULL | 0.705785028 | NULL&M_Disease |
| 11 | 121128752 | rs4376915 | G | A | NULL | 0.705785028 | NULL&M_Disease |
| 11 | 116388314 | rs867899 | A | G | NULL | 0.705785028 | NULL&M_Disease |
| 11 | 116975704 | rs11216275 | G | T | NULL | 0.705785028 | NULL&M_Disease |
| 11 | 117627333 | rs10790204 | A | G | NULL | 0.705785028 | NULL&M_Disease |
| 11 | 118574185 | rs687664 | A | C | NULL | 0.705785028 | NULL&M_Disease |
| 11 | 118574188 | rs523715 | A | G | NULL | 0.705785028 | NULL&M_Disease |
| 11 | 119071897 | rs7932398 | G | C | NULL | 0.705785028 | NULL&M_Disease |
| 11 | 121127491 | rs4307734 | A | C | NULL | 0.705785028 | NULL&M_Disease |
| 11 | 121127870 | rs4370953 | G | A | NULL | 0.705785028 | NULL&M_Disease |
| 11 | 121127974 | rs12419859 | G | C | NULL | 0.705785028 | NULL&M_Disease |
| 11 | 121128034 | rs12421355 | A | T | NULL | 0.705785028 | NULL&M_Disease |
| 11 | 121129552 | rs9943676 | C | T | NULL | 0.705785028 | NULL&M_Disease |
| 11 | 121129931 | rs4936599 | T | C | NULL | 0.705785028 | NULL&M_Disease |
| 11 | 116570718 | rs1145191 | G | C | NULL | 0.705785028 | NULL&M_Disease |
| 11 | 117072525 | rs488962 | T | C | NULL | 0.705785028 | NULL&M_Disease |
| 11 | 117989951 | rs45615131 | T | A | NULL | 0.705785028 | NULL&M_Disease |
| 11 | 117998077 | rs17601549 | T | C | NULL | 0.705785028 | NULL&M_Disease |
| 11 | 119203939 | rs2511846 | C | G | NULL | 0.705785028 | NULL&M_Disease |
| 11 | 121132812 | rs10892701 | C | T | NULL | 0.705785028 | NULL&M_Disease |
| 11 | 121151102 | rs4415772 | G | T | NULL | 0.705785028 | NULL&M_Disease |
| 11 | 116312958 | rs485778 | A | G | NULL | 0.705785028 | NULL&M_Disease |
| 11 | 116470173 | rs11216034 | C | A | NULL | 0.705785028 | NULL&M_Disease |
| 11 | 116529468 | rs948690 | C | T | NULL | 0.705785028 | NULL&M_Disease |
| 11 | 118162796 | rs11216844 | G | A | NULL | 0.705785028 | NULL&M_Disease |
| 11 | 119236192 | rs45539534 | T | C | NULL | 0.705785028 | NULL&M_Disease |
| 11 | 120333801 | rs11601535 | G | A | NULL | 0.705785028 | NULL&M_Disease |
| 11 | 117309283 | rs10892103 | A | G | NULL | 0.705785028 | NULL&M_Disease |
| 11 | 119229964 | rs2241646 | G | A | NULL | 0.705785028 | NULL&M_Disease |
| 11 | 119612677 | rs12798188 | T | C | NULL | 0.705785028 | NULL&M_Disease |
| 11 | 119687318 | rs4938729 | C | T | NULL | 0.705785028 | NULL&M_Disease |
| 11 | 120256845 | rs622373 | G | A | NULL | 0.705785028 | NULL&M_Disease |
| 11 | 120993066 | rs577733 | A | G | NULL | 0.705785028 | NULL&M_Disease |

| | | | | | | | | | |
|---|---|---|---|---|---|---|---|---|---|
| 11 | 118913993 | rs1003081 | T | C | 0.982 | -7977.388 | -7979.131 | -1.742 | 6.78E-07 |
| 11 | 119972011 | rs6589802 | G | A | 0.982 | -7977.388 | -7979.13 | -1.742 | 6.78E-07 |
| 11 | 120196433 | rs36011588 | G | C | 1.017 | -7977.388 | -7979.13 | -1.742 | 6.78E-07 |
| 11 | 120330149 | rs2305010 | C | G | 1.017 | -7977.388 | -7979.13 | -1.742 | 6.78E-07 |
| 11 | 116559743 | rs3016360 | C | T | 1.015 | -7977.388 | -7979.132 | -1.743 | 6.77E-07 |
| 11 | 117172820 | rs680210 | T | C | 1.017 | -7977.388 | -7979.131 | -1.743 | 6.77E-07 |
| 11 | 118124092 | rs1715455 | T | C | 1.014 | -7977.388 | -7979.131 | -1.743 | 6.77E-07 |
| 11 | 118131223 | rs7120665 | C | T | 1.014 | -7977.388 | -7979.132 | -1.743 | 6.77E-07 |
| 11 | 119876028 | rs2155168 | T | C | 1.018 | -7977.388 | -7979.132 | -1.743 | 6.77E-07 |
| 11 | 120141165 | rs1941406 | A | G | 1.013 | -7977.388 | -7979.132 | -1.743 | 6.77E-07 |
| 11 | 117320727 | rs10892108 | T | C | 1.018 | -7977.388 | -7979.133 | -1.744 | 6.76E-07 |
| 11 | 117685220 | rs10892177 | C | G | 0.983 | -7977.388 | -7979.132 | -1.744 | 6.76E-07 |
| 11 | 117685575 | rs11216561 | C | T | 0.983 | -7977.388 | -7979.132 | -1.744 | 6.76E-07 |
| 11 | 118072373 | rs7949751 | G | A | 0.988 | -7977.388 | -7979.132 | -1.744 | 6.76E-07 |
| 11 | 120610144 | rs2846114 | C | T | 0.982 | -7977.388 | -7979.132 | -1.744 | 6.76E-07 |
| 11 | 121140993 | rs10790428 | C | T | 1.014 | -7977.388 | -7979.132 | -1.744 | 6.76E-07 |
| 11 | 121149254 | rs11218238 | G | A | 1.011 | -7977.388 | -7979.133 | -1.744 | 6.76E-07 |
| 11 | 121151411 | rs4462362 | C | G | 1.011 | -7977.388 | -7979.132 | -1.744 | 6.76E-07 |
| 11 | 116680918 | rs1263172 | G | A | 1.008 | -7977.388 | -7979.133 | -1.745 | 6.76E-07 |
| 11 | 117097009 | rs7939747 | G | A | 1.003 | -7977.388 | -7979.133 | -1.745 | 6.76E-07 |
| 11 | 117914297 | rs1786180 | G | C | 1.009 | -7977.388 | -7979.133 | -1.745 | 6.76E-07 |
| 11 | 117914304 | rs1786179 | A | T | 1.009 | -7977.388 | -7979.133 | -1.745 | 6.76E-07 |
| 11 | 117914550 | rs1783853 | G | A | 1.009 | -7977.388 | -7979.133 | -1.745 | 6.76E-07 |
| 11 | 119072976 | rs11217180 | T | C | 0.991 | -7977.388 | -7979.133 | -1.745 | 6.76E-07 |
| 11 | 119166047 | rs1309012690 | A | G | 0.99 | -7977.388 | -7979.133 | -1.745 | 6.76E-07 |
| 11 | 119330741 | rs12574794 | T | C | 1.014 | -7977.388 | -7979.133 | -1.745 | 6.76E-07 |
| 11 | 119375180 | rs10750147 | A | G | 1.018 | -7977.388 | -7979.133 | -1.745 | 6.76E-07 |
| 11 | 119488133 | rs7934906 | T | C | 1.018 | -7977.388 | -7979.134 | -1.745 | 6.76E-07 |
| 11 | 119572103 | rs3935405 | T | C | 1.017 | -7977.388 | -7979.133 | -1.745 | 6.76E-07 |
| 11 | 119596772 | rs12418662 | C | T | 0.982 | -7977.388 | -7979.134 | -1.745 | 6.76E-07 |
| 11 | 120448992 | rs11602075 | C | T | 1.018 | -7977.388 | -7979.133 | -1.745 | 6.76E-07 |
| 11 | 120494089 | rs6589821 | A | G | 0.983 | -7977.388 | -7979.133 | -1.745 | 6.76E-07 |
| 11 | 121134420 | rs11218230 | G | T | 1.011 | -7977.388 | -7979.133 | -1.745 | 6.76E-07 |
| 11 | 121136455 | rs10790419 | T | C | 1.014 | -7977.388 | -7979.133 | -1.745 | 6.76E-07 |
| 11 | 116560325 | rs1787715 | T | C | 1.015 | -7977.388 | -7979.134 | -1.746 | 6.75E-07 |
| 11 | 117193350 | rs4938370 | C | A | 0.989 | -7977.388 | -7979.135 | -1.746 | 6.75E-07 |
| 11 | 117548726 | rs596440 | C | T | 1 | -7977.388 | -7979.135 | -1.746 | 6.75E-07 |
| 11 | 117548858 | rs637454 | G | A | 1 | -7977.388 | -7979.135 | -1.746 | 6.75E-07 |
| 11 | 117817276 | rs2515795 | A | G | 1.018 | -7977.388 | -7979.134 | -1.746 | 6.75E-07 |
| 11 | 119874855 | rs7126918 | A | T | 1.018 | -7977.388 | -7979.134 | -1.746 | 6.75E-07 |
| 11 | 120133153 | rs2715882 | T | C | 1.009 | -7977.388 | -7979.134 | -1.746 | 6.75E-07 |
| 11 | 120722457 | rs11218031 | A | C | 0.983 | -7977.388 | -7979.134 | -1.746 | 6.75E-07 |
| 11 | 121133278 | rs7937589 | C | G | 1.013 | -7977.388 | -7979.134 | -1.746 | 6.75E-07 |
| 11 | 116572205 | rs1145196 | T | C | 0.982 | -7977.388 | -7979.136 | -1.747 | 6.74E-07 |
| 11 | 117425009 | rs579239 | C | T | 1.006 | -7977.388 | -7979.135 | -1.747 | 6.74E-07 |
| 11 | 118166212 | rs4938500 | A | C | 0.985 | -7977.388 | -7979.135 | -1.747 | 6.74E-07 |
| 11 | 118262706 | rs61900932 | T | C | 1.005 | -7977.388 | -7979.135 | -1.747 | 6.74E-07 |
| 11 | 119702015 | rs544201 | A | T | 0.985 | -7977.388 | -7979.135 | -1.747 | 6.74E-07 |
| 11 | 120273316 | rs4381380 | A | G | 1.017 | -7977.388 | -7979.135 | -1.747 | 6.74E-07 |
| 11 | 116560174 | rs1787716 | A | G | 1.015 | -7977.388 | -7979.137 | -1.748 | 6.74E-07 |
| 11 | 116570942 | rs1145193 | G | A | 0.982 | -7977.388 | -7979.137 | -1.748 | 6.74E-07 |
| 11 | 117072963 | rs634960 | C | T | 0.992 | -7977.388 | -7979.137 | -1.748 | 6.74E-07 |
| 11 | 117697985 | rs11216566 | G | A | 0.987 | -7977.388 | -7979.137 | -1.748 | 6.74E-07 |
| 11 | 117801589 | rs61900351 | A | T | 0.982 | -7977.388 | -7979.136 | -1.748 | 6.74E-07 |
| 11 | 117972936 | rs881122 | A | C | 0.998 | -7977.388 | -7979.136 | -1.748 | 6.74E-07 |
| 11 | 119012846 | rs659375 | A | T | 1.012 | -7977.388 | -7979.136 | -1.748 | 6.74E-07 |
| 11 | 119969871 | rs623353 | C | A | 0.983 | -7977.388 | -7979.136 | -1.748 | 6.74E-07 |
| 11 | 120479913 | rs34759019 | A | C | 0.983 | -7977.388 | -7979.136 | -1.748 | 6.74E-07 |
| 11 | 121127518 | rs4533044 | C | T | 1.011 | -7977.388 | -7979.136 | -1.748 | 6.74E-07 |
| 11 | 121134392 | rs11218229 | G | T | 1.011 | -7977.388 | -7979.136 | -1.748 | 6.74E-07 |
| 11 | 121147950 | rs9971567 | G | A | 1.011 | -7977.388 | -7979.136 | -1.748 | 6.74E-07 |
| 11 | 116310858 | rs580872 | A | G | 0.999 | -7977.388 | -7979.137 | -1.749 | 6.73E-07 |
| 11 | 119868171 | rs7111799 | A | G | 0.997 | -7977.388 | -7979.138 | -1.749 | 6.73E-07 |
| 11 | 121128562 | rs4517527 | G | T | 1.011 | -7977.388 | -7979.138 | -1.749 | 6.73E-07 |
| 11 | 121128752 | rs4376915 | G | A | 1.011 | -7977.388 | -7979.138 | -1.749 | 6.73E-07 |
| 11 | 116388314 | rs867899 | A | G | 1.018 | -7977.388 | -7979.139 | -1.75 | 6.72E-07 |
| 11 | 116975704 | rs11216275 | G | T | 1.003 | -7977.388 | -7979.138 | -1.75 | 6.72E-07 |
| 11 | 117627333 | rs10790204 | A | G | 0.987 | -7977.388 | -7979.139 | -1.75 | 6.72E-07 |
| 11 | 118574185 | rs687664 | A | C | 0.985 | -7977.388 | -7979.138 | -1.75 | 6.72E-07 |
| 11 | 118574188 | rs523715 | A | G | 0.985 | -7977.388 | -7979.138 | -1.75 | 6.72E-07 |
| 11 | 119071897 | rs7932398 | G | C | 0.992 | -7977.388 | -7979.138 | -1.75 | 6.72E-07 |
| 11 | 121127491 | rs4307734 | A | C | 1.011 | -7977.388 | -7979.139 | -1.75 | 6.72E-07 |
| 11 | 121127870 | rs4370953 | G | A | 1.011 | -7977.388 | -7979.139 | -1.75 | 6.72E-07 |
| 11 | 121127974 | rs12419859 | G | C | 1.011 | -7977.388 | -7979.139 | -1.75 | 6.72E-07 |
| 11 | 121128034 | rs12421355 | A | T | 1.011 | -7977.388 | -7979.139 | -1.75 | 6.72E-07 |
| 11 | 121129552 | rs9943676 | C | T | 1.011 | -7977.388 | -7979.139 | -1.75 | 6.72E-07 |
| 11 | 121129931 | rs4936599 | T | C | 1.011 | -7977.388 | -7979.139 | -1.75 | 6.72E-07 |
| 11 | 116570718 | rs1145191 | G | C | 0.983 | -7977.388 | -7979.139 | -1.751 | 6.72E-07 |
| 11 | 117072525 | rs488962 | T | C | 0.993 | -7977.388 | -7979.139 | -1.751 | 6.72E-07 |
| 11 | 117989951 | rs45615131 | T | A | 0.984 | -7977.388 | -7979.139 | -1.751 | 6.72E-07 |
| 11 | 117998077 | rs17601549 | T | C | 0.984 | -7977.388 | -7979.139 | -1.751 | 6.72E-07 |
| 11 | 119203939 | rs2511846 | C | G | 1.015 | -7977.388 | -7979.14 | -1.751 | 6.72E-07 |
| 11 | 121132812 | rs10892701 | C | T | 1.011 | -7977.388 | -7979.14 | -1.751 | 6.72E-07 |
| 11 | 121151102 | rs4415772 | G | T | 1.01 | -7977.388 | -7979.139 | -1.751 | 6.72E-07 |
| 11 | 116312958 | rs485778 | A | G | 1 | -7977.388 | -7979.14 | -1.752 | 6.71E-07 |
| 11 | 116470173 | rs11216034 | C | A | 1.014 | -7977.388 | -7979.141 | -1.752 | 6.71E-07 |
| 11 | 116529468 | rs948690 | C | T | 1.015 | -7977.388 | -7979.14 | -1.752 | 6.71E-07 |
| 11 | 118162796 | rs11216844 | G | A | 0.985 | -7977.388 | -7979.14 | -1.752 | 6.71E-07 |
| 11 | 119236192 | rs45539534 | T | C | 0.985 | -7977.388 | -7979.14 | -1.752 | 6.71E-07 |
| 11 | 120333801 | rs11601535 | G | A | 1.017 | -7977.388 | -7979.14 | -1.752 | 6.71E-07 |
| 11 | 117309283 | rs10892103 | A | G | 1.012 | -7977.388 | -7979.141 | -1.753 | 6.70E-07 |
| 11 | 119229964 | rs2241646 | G | A | 0.987 | -7977.388 | -7979.141 | -1.753 | 6.70E-07 |
| 11 | 119612677 | rs12798188 | T | C | 0.983 | -7977.388 | -7979.141 | -1.753 | 6.70E-07 |
| 11 | 119687318 | rs4938729 | C | T | 1.005 | -7977.388 | -7979.141 | -1.753 | 6.70E-07 |
| 11 | 120256845 | rs622373 | G | A | 1.016 | -7977.388 | -7979.142 | -1.753 | 6.70E-07 |
| 11 | 120993066 | rs577733 | A | G | 1.014 | -7977.388 | -7979.142 | -1.753 | 6.70E-07 |

| | | | | | | | |
|---|---|---|---|---|---|---|---|
| 11 | 121134099 | rs11218228 | G | A | NULL | 0.705785028 | NULL&M_Disease |
| 11 | 121140729 | rs10892715 | A | G | NULL | 0.705785028 | NULL&M_Disease |
| 11 | 116331960 | rs560801 | C | A | NULL | 0.705785028 | NULL&M_Disease |
| 11 | 116566763 | rs59854309 | T | C | NULL | 0.705785028 | NULL&M_Disease |
| 11 | 118163216 | rs11216845 | T | C | NULL | 0.705785028 | NULL&M_Disease |
| 11 | 118171058 | rs1318166 | G | A | NULL | 0.705785028 | NULL&M_Disease |
| 11 | 120494449 | rs10790388 | A | C | NULL | 0.705785028 | NULL&M_Disease |
| 11 | 120996292 | rs536069 | A | G | NULL | 0.705785028 | NULL&M_Disease |
| 11 | 121134434 | rs11218231 | G | A | NULL | 0.705785028 | NULL&M_Disease |
| 11 | 121134532 | rs4936602 | A | G | NULL | 0.705785028 | NULL&M_Disease |
| 11 | 121134781 | rs4935765 | C | T | NULL | 0.705785028 | NULL&M_Disease |
| 11 | 121148154 | rs9971517 | C | T | NULL | 0.705785028 | NULL&M_Disease |
| 11 | 117917622 | rs657671 | C | T | NULL | 0.705785028 | NULL&M_Disease |
| 11 | 120083648 | rs11600568 | G | C | NULL | 0.705785028 | NULL&M_Disease |
| 11 | 120445354 | rs4272790 | T | C | NULL | 0.705785028 | NULL&M_Disease |
| 11 | 120456829 | rs1320718 | C | T | NULL | 0.705785028 | NULL&M_Disease |
| 11 | 120792285 | rs2850806 | T | C | NULL | 0.705785028 | NULL&M_Disease |
| 11 | 121126530 | rs7119061 | G | A | NULL | 0.705785028 | NULL&M_Disease |
| 11 | 121134903 | rs4936604 | T | C | NULL | 0.705785028 | NULL&M_Disease |
| 11 | 121137520 | rs10790424 | T | C | NULL | 0.705785028 | NULL&M_Disease |
| 11 | 116427932 | rs7930491 | A | G | NULL | 0.705785028 | NULL&M_Disease |
| 11 | 117926593 | rs676812 | A | G | NULL | 0.705785028 | NULL&M_Disease |
| 11 | 120440360 | rs4310631 | G | C | NULL | 0.705785028 | NULL&M_Disease |
| 11 | 120450280 | rs10892587 | A | C | NULL | 0.705785028 | NULL&M_Disease |
| 11 | 121134854 | rs4936603 | A | C | NULL | 0.705785028 | NULL&M_Disease |
| 11 | 121135196 | rs10892702 | A | T | NULL | 0.705785028 | NULL&M_Disease |
| 11 | 121135199 | rs10892703 | A | T | NULL | 0.705785028 | NULL&M_Disease |
| 11 | 121135240 | rs10892704 | T | C | NULL | 0.705785028 | NULL&M_Disease |
| 11 | 121135286 | rs10892705 | T | A | NULL | 0.705785028 | NULL&M_Disease |
| 11 | 121135479 | rs1583125 | A | C | NULL | 0.705785028 | NULL&M_Disease |
| 11 | 121135578 | rs1583126 | T | G | NULL | 0.705785028 | NULL&M_Disease |
| 11 | 121135897 | rs4936607 | G | A | NULL | 0.705785028 | NULL&M_Disease |
| 11 | 121136069 | rs4936608 | T | C | NULL | 0.705785028 | NULL&M_Disease |
| 11 | 121136349 | rs10750180 | C | T | NULL | 0.705785028 | NULL&M_Disease |
| 11 | 121136428 | rs10790418 | T | C | NULL | 0.705785028 | NULL&M_Disease |
| 11 | 121136458 | rs10790420 | T | C | NULL | 0.705785028 | NULL&M_Disease |
| 11 | 121136529 | rs10790421 | C | T | NULL | 0.705785028 | NULL&M_Disease |
| 11 | 121136688 | rs10790423 | C | A | NULL | 0.705785028 | NULL&M_Disease |
| 11 | 121136915 | rs10892709 | C | T | NULL | 0.705785028 | NULL&M_Disease |
| 11 | 121137123 | rs10892710 | G | A | NULL | 0.705785028 | NULL&M_Disease |
| 11 | 121137332 | rs12293490 | T | C | NULL | 0.705785028 | NULL&M_Disease |
| 11 | 121137543 | rs10892711 | T | G | NULL | 0.705785028 | NULL&M_Disease |
| 11 | 121137603 | rs10892712 | C | T | NULL | 0.705785028 | NULL&M_Disease |
| 11 | 121137629 | rs10892713 | C | G | NULL | 0.705785028 | NULL&M_Disease |
| 11 | 121137690 | rs10892714 | A | G | NULL | 0.705785028 | NULL&M_Disease |
| 11 | 121137754 | rs7935472 | G | A | NULL | 0.705785028 | NULL&M_Disease |
| 11 | 121137859 | rs7950819 | A | T | NULL | 0.705785028 | NULL&M_Disease |
| 11 | 121138191 | rs7935970 | G | A | NULL | 0.705785028 | NULL&M_Disease |
| 11 | 121138260 | rs7951282 | C | T | NULL | 0.705785028 | NULL&M_Disease |
| 11 | 121138262 | rs7951286 | A | T | NULL | 0.705785028 | NULL&M_Disease |
| 11 | 121138302 | rs7936408 | T | G | NULL | 0.705785028 | NULL&M_Disease |
| 11 | 120078286 | rs10892551 | A | G | NULL | 0.705785028 | NULL&M_Disease |
| 11 | 121131225 | rs9667661 | A | G | NULL | 0.705785028 | NULL&M_Disease |
| 11 | 121132347 | rs1941303 | C | G | NULL | 0.705785028 | NULL&M_Disease |
| 11 | 121135037 | rs4936605 | G | A | NULL | 0.705785028 | NULL&M_Disease |
| 11 | 121135791 | rs1583127 | G | A | NULL | 0.705785028 | NULL&M_Disease |
| 11 | 121137458 | rs12280328 | G | A | NULL | 0.705785028 | NULL&M_Disease |
| 11 | 121137477 | rs12293637 | A | G | NULL | 0.705785028 | NULL&M_Disease |
| 11 | 121138891 | rs1560403 | A | G | NULL | 0.705785028 | NULL&M_Disease |
| 11 | 121139062 | rs1560404 | C | T | NULL | 0.705785028 | NULL&M_Disease |
| 11 | 116597889 | rs180363 | C | T | NULL | 0.706822221 | NULL&M_Disease |
| 11 | 117024264 | rs10790174 | G | A | NULL | 0.706822221 | NULL&M_Disease |
| 11 | 117296892 | rs12281346 | C | G | NULL | 0.706822221 | NULL&M_Disease |
| 11 | 117719605 | rs561394 | G | A | NULL | 0.706822221 | NULL&M_Disease |
| 11 | 117954657 | rs2187008 | G | C | NULL | 0.706822221 | NULL&M_Disease |
| 11 | 119083906 | rs11217186 | T | C | NULL | 0.705785028 | NULL&M_Disease |
| 11 | 119876516 | rs4938749 | G | A | NULL | 0.706822221 | NULL&M_Disease |
| 11 | 119973745 | rs1107149 | C | T | NULL | 0.706822221 | NULL&M_Disease |
| 11 | 120273597 | rs4245213 | A | G | NULL | 0.706822221 | NULL&M_Disease |
| 11 | 120319064 | rs2305008 | C | A | NULL | 0.706822221 | NULL&M_Disease |
| 11 | 120447136 | rs4938817 | C | A | NULL | 0.706822221 | NULL&M_Disease |
| 11 | 120456605 | rs4363614 | A | G | NULL | 0.706822221 | NULL&M_Disease |
| 11 | 120599866 | rs1939668 | A | G | NULL | 0.706822221 | NULL&M_Disease |
| 11 | 121150839 | rs12420651 | G | T | NULL | 0.706822221 | NULL&M_Disease |
| 11 | 116209581 | rs10891965 | A | G | NULL | 0.706822221 | NULL&M_Disease |
| 11 | 117180905 | rs4245169 | T | C | NULL | 0.706822221 | NULL&M_Disease |
| 11 | 117193741 | rs4938371 | T | C | NULL | 0.706822221 | NULL&M_Disease |
| 11 | 117805536 | rs10790224 | C | T | NULL | 0.706822221 | NULL&M_Disease |
| 11 | 119078886 | rs11217183 | G | A | NULL | 0.706822221 | NULL&M_Disease |
| 11 | 119078993 | rs11217184 | G | A | NULL | 0.706822221 | NULL&M_Disease |
| 11 | 119178050 | rs1052121 | G | C | NULL | 0.706822221 | NULL&M_Disease |
| 11 | 119216279 | rs36015759 | A | G | NULL | 0.706822221 | NULL&M_Disease |
| 11 | 119831773 | rs4245210 | A | G | NULL | 0.706822221 | NULL&M_Disease |
| 11 | 119941114 | rs7113913 | T | C | NULL | 0.706822221 | NULL&M_Disease |
| 11 | 119969402 | rs594081 | A | C | NULL | 0.706822221 | NULL&M_Disease |
| 11 | 120478676 | rs11600894 | T | C | NULL | 0.706822221 | NULL&M_Disease |
| 11 | 121131965 | rs12417414 | C | T | NULL | 0.706822221 | NULL&M_Disease |
| 11 | 121132211 | rs1941305 | A | G | NULL | 0.706822221 | NULL&M_Disease |
| 11 | 121132279 | rs1941304 | A | G | NULL | 0.706822221 | NULL&M_Disease |
| 11 | 117012210 | rs2735182 | A | T | NULL | 0.706822221 | NULL&M_Disease |
| 11 | 117019010 | rs2008908 | C | T | NULL | 0.706822221 | NULL&M_Disease |
| 11 | 119332065 | rs10750143 | G | A | NULL | 0.706822221 | NULL&M_Disease |
| 11 | 119656902 | rs7119690 | G | A | NULL | 0.706822221 | NULL&M_Disease |
| 11 | 119971523 | rs6589799 | C | T | NULL | 0.706822221 | NULL&M_Disease |
| 11 | 120715674 | rs9783385 | G | T | NULL | 0.706822221 | NULL&M_Disease |
| 11 | 121132675 | rs10892700 | T | C | NULL | 0.706822221 | NULL&M_Disease |

| | | | | | | | | | |
|---|---|---|---|---|---|---|---|---|---|
| 11 | 121134099 | rs11218228 | G | A | 1.01 | -7977.388 | -7979.141 | -1.753 | 6.70E-07 |
| 11 | 121140729 | rs10892715 | A | G | 1.013 | -7977.388 | -7979.141 | -1.753 | 6.70E-07 |
| 11 | 116331960 | rs560801 | C | A | 0.993 | -7977.388 | -7979.142 | -1.754 | 6.69E-07 |
| 11 | 116566763 | rs59854309 | T | C | 0.983 | -7977.388 | -7979.143 | -1.754 | 6.69E-07 |
| 11 | 118163216 | rs11216845 | T | C | 0.985 | -7977.388 | -7979.142 | -1.754 | 6.69E-07 |
| 11 | 118171058 | rs1318166 | G | A | 0.985 | -7977.388 | -7979.142 | -1.754 | 6.69E-07 |
| 11 | 120494449 | rs10790388 | A | C | 0.983 | -7977.388 | -7979.143 | -1.754 | 6.69E-07 |
| 11 | 120996292 | rs536069 | A | G | 1.014 | -7977.388 | -7979.142 | -1.754 | 6.69E-07 |
| 11 | 121134434 | rs11218231 | G | A | 1.01 | -7977.388 | -7979.142 | -1.754 | 6.69E-07 |
| 11 | 121134532 | rs4936602 | A | G | 1.01 | -7977.388 | -7979.143 | -1.754 | 6.69E-07 |
| 11 | 121134781 | rs4935765 | C | T | 1.01 | -7977.388 | -7979.143 | -1.754 | 6.69E-07 |
| 11 | 121148154 | rs9971517 | C | T | 1.01 | -7977.388 | -7979.143 | -1.754 | 6.69E-07 |
| 11 | 117917622 | rs657671 | C | T | 1.008 | -7977.388 | -7979.143 | -1.755 | 6.69E-07 |
| 11 | 120083648 | rs11600568 | G | C | 0.983 | -7977.388 | -7979.143 | -1.755 | 6.69E-07 |
| 11 | 120445354 | rs4272790 | T | C | 0.996 | -7977.388 | -7979.144 | -1.755 | 6.69E-07 |
| 11 | 120456829 | rs1320718 | C | T | 0.995 | -7977.388 | -7979.143 | -1.755 | 6.69E-07 |
| 11 | 120792285 | rs2850806 | T | C | 0.984 | -7977.388 | -7979.143 | -1.755 | 6.69E-07 |
| 11 | 121126530 | rs7119061 | G | A | 1.013 | -7977.388 | -7979.144 | -1.755 | 6.69E-07 |
| 11 | 121134903 | rs4936604 | T | C | 1.01 | -7977.388 | -7979.143 | -1.755 | 6.69E-07 |
| 11 | 121137520 | rs10790424 | T | C | 1.013 | -7977.388 | -7979.144 | -1.755 | 6.69E-07 |
| 11 | 116427932 | rs7930491 | A | G | 0.988 | -7977.388 | -7979.144 | -1.756 | 6.68E-07 |
| 11 | 117926593 | rs676812 | A | G | 1.007 | -7977.388 | -7979.144 | -1.756 | 6.68E-07 |
| 11 | 120440360 | rs4310631 | G | C | 0.996 | -7977.388 | -7979.144 | -1.756 | 6.68E-07 |
| 11 | 120450280 | rs10892587 | A | C | 0.999 | -7977.388 | -7979.144 | -1.756 | 6.68E-07 |
| 11 | 121134854 | rs4936603 | A | C | 1.01 | -7977.388 | -7979.144 | -1.756 | 6.68E-07 |
| 11 | 121135196 | rs10892702 | A | T | 1.01 | -7977.388 | -7979.144 | -1.756 | 6.68E-07 |
| 11 | 121135199 | rs10892703 | A | T | 1.01 | -7977.388 | -7979.144 | -1.756 | 6.68E-07 |
| 11 | 121135240 | rs10892704 | T | C | 1.01 | -7977.388 | -7979.144 | -1.756 | 6.68E-07 |
| 11 | 121135286 | rs10892705 | T | A | 1.01 | -7977.388 | -7979.144 | -1.756 | 6.68E-07 |
| 11 | 121135479 | rs1583125 | A | C | 1.01 | -7977.388 | -7979.144 | -1.756 | 6.68E-07 |
| 11 | 121135578 | rs1583126 | T | G | 1.01 | -7977.388 | -7979.144 | -1.756 | 6.68E-07 |
| 11 | 121135897 | rs4936607 | G | A | 1.01 | -7977.388 | -7979.144 | -1.756 | 6.68E-07 |
| 11 | 121136069 | rs4936608 | T | C | 1.01 | -7977.388 | -7979.144 | -1.756 | 6.68E-07 |
| 11 | 121136349 | rs10750180 | C | T | 1.01 | -7977.388 | -7979.144 | -1.756 | 6.68E-07 |
| 11 | 121136428 | rs10790418 | T | C | 1.01 | -7977.388 | -7979.144 | -1.756 | 6.68E-07 |
| 11 | 121136458 | rs10790420 | T | C | 1.01 | -7977.388 | -7979.144 | -1.756 | 6.68E-07 |
| 11 | 121136529 | rs10790421 | C | T | 1.01 | -7977.388 | -7979.144 | -1.756 | 6.68E-07 |
| 11 | 121136688 | rs10790423 | C | A | 1.01 | -7977.388 | -7979.144 | -1.756 | 6.68E-07 |
| 11 | 121136915 | rs10892709 | C | T | 1.01 | -7977.388 | -7979.144 | -1.756 | 6.68E-07 |
| 11 | 121137123 | rs10892710 | G | A | 1.01 | -7977.388 | -7979.144 | -1.756 | 6.68E-07 |
| 11 | 121137332 | rs12293490 | T | C | 1.01 | -7977.388 | -7979.144 | -1.756 | 6.68E-07 |
| 11 | 121137543 | rs10892711 | T | G | 1.01 | -7977.388 | -7979.144 | -1.756 | 6.68E-07 |
| 11 | 121137603 | rs10892712 | C | T | 1.01 | -7977.388 | -7979.144 | -1.756 | 6.68E-07 |
| 11 | 121137629 | rs10892713 | C | G | 1.01 | -7977.388 | -7979.144 | -1.756 | 6.68E-07 |
| 11 | 121137690 | rs10892714 | A | G | 1.01 | -7977.388 | -7979.144 | -1.756 | 6.68E-07 |
| 11 | 121137754 | rs7935472 | G | A | 1.01 | -7977.388 | -7979.144 | -1.756 | 6.68E-07 |
| 11 | 121137859 | rs7950819 | A | T | 1.01 | -7977.388 | -7979.144 | -1.756 | 6.68E-07 |
| 11 | 121138191 | rs7935970 | G | A | 1.01 | -7977.388 | -7979.144 | -1.756 | 6.68E-07 |
| 11 | 121138260 | rs7951282 | C | T | 1.01 | -7977.388 | -7979.144 | -1.756 | 6.68E-07 |
| 11 | 121138262 | rs7951286 | A | T | 1.01 | -7977.388 | -7979.144 | -1.756 | 6.68E-07 |
| 11 | 121138302 | rs7936408 | T | G | 1.01 | -7977.388 | -7979.144 | -1.756 | 6.68E-07 |
| 11 | 120078286 | rs10892551 | A | G | 0.995 | -7977.388 | -7979.146 | -1.757 | 6.67E-07 |
| 11 | 121131225 | rs9667661 | A | G | 1.01 | -7977.388 | -7979.145 | -1.757 | 6.67E-07 |
| 11 | 121132347 | rs1941303 | C | G | 1.01 | -7977.388 | -7979.145 | -1.757 | 6.67E-07 |
| 11 | 121135037 | rs4936605 | G | A | 1.01 | -7977.388 | -7979.146 | -1.757 | 6.67E-07 |
| 11 | 121135791 | rs1583127 | G | A | 1.01 | -7977.388 | -7979.146 | -1.757 | 6.67E-07 |
| 11 | 121137458 | rs12280328 | G | A | 1.01 | -7977.388 | -7979.145 | -1.757 | 6.67E-07 |
| 11 | 121137477 | rs12293637 | A | G | 1.01 | -7977.388 | -7979.145 | -1.757 | 6.67E-07 |
| 11 | 121138891 | rs1560403 | A | G | 1.01 | -7977.388 | -7979.146 | -1.757 | 6.67E-07 |
| 11 | 121139062 | rs1560404 | C | T | 1.01 | -7977.388 | -7979.146 | -1.757 | 6.67E-07 |
| 11 | 116597889 | rs180363 | C | T | 1.015 | -7977.388 | -7979.146 | -1.758 | 6.67E-07 |
| 11 | 117024264 | rs10790174 | G | A | 0.995 | -7977.388 | -7979.147 | -1.758 | 6.67E-07 |
| 11 | 117296892 | rs12281346 | C | G | 0.999 | -7977.388 | -7979.146 | -1.758 | 6.67E-07 |
| 11 | 117719605 | rs561394 | G | A | 1.013 | -7977.388 | -7979.146 | -1.758 | 6.67E-07 |
| 11 | 117954657 | rs2187008 | G | C | 0.984 | -7977.388 | -7979.147 | -1.758 | 6.67E-07 |
| 11 | 119083906 | rs11217186 | T | C | 0.991 | -7977.388 | -7979.146 | -1.758 | 6.67E-07 |
| 11 | 119876516 | rs4938749 | G | A | 1.017 | -7977.388 | -7979.146 | -1.758 | 6.67E-07 |
| 11 | 119973745 | rs1107149 | C | T | 0.983 | -7977.388 | -7979.146 | -1.758 | 6.67E-07 |
| 11 | 120273597 | rs4245213 | A | G | 1.017 | -7977.388 | -7979.146 | -1.758 | 6.67E-07 |
| 11 | 120319064 | rs2305008 | C | A | 1.017 | -7977.388 | -7979.146 | -1.758 | 6.67E-07 |
| 11 | 120447136 | rs4938817 | C | A | 0.996 | -7977.388 | -7979.146 | -1.758 | 6.67E-07 |
| 11 | 120456605 | rs4363614 | A | G | 1 | -7977.388 | -7979.146 | -1.758 | 6.67E-07 |
| 11 | 120599866 | rs1939668 | A | G | 1.005 | -7977.388 | -7979.146 | -1.758 | 6.67E-07 |
| 11 | 121150839 | rs12420651 | G | T | 1.01 | -7977.388 | -7979.146 | -1.758 | 6.67E-07 |
| 11 | 116209581 | rs10891965 | A | G | 0.988 | -7977.388 | -7979.147 | -1.759 | 6.66E-07 |
| 11 | 117180905 | rs4245169 | T | C | 1.004 | -7977.388 | -7979.147 | -1.759 | 6.66E-07 |
| 11 | 117193741 | rs4938371 | T | C | 0.99 | -7977.388 | -7979.148 | -1.759 | 6.66E-07 |
| 11 | 117805536 | rs10790224 | C | T | 1.017 | -7977.388 | -7979.147 | -1.759 | 6.66E-07 |
| 11 | 119078886 | rs11217183 | G | A | 0.992 | -7977.388 | -7979.147 | -1.759 | 6.66E-07 |
| 11 | 119078993 | rs11217184 | G | A | 0.992 | -7977.388 | -7979.147 | -1.759 | 6.66E-07 |
| 11 | 119178050 | rs1052121 | G | C | 1.014 | -7977.388 | -7979.148 | -1.759 | 6.66E-07 |
| 11 | 119216279 | rs36015759 | A | G | 0.997 | -7977.388 | -7979.148 | -1.759 | 6.66E-07 |
| 11 | 119831773 | rs4245210 | A | G | 1.017 | -7977.388 | -7979.147 | -1.759 | 6.66E-07 |
| 11 | 119941114 | rs7113913 | T | C | 1.016 | -7977.388 | -7979.148 | -1.759 | 6.66E-07 |
| 11 | 119969402 | rs594081 | A | C | 0.984 | -7977.388 | -7979.148 | -1.759 | 6.66E-07 |
| 11 | 120478676 | rs11600894 | T | C | 0.984 | -7977.388 | -7979.147 | -1.759 | 6.66E-07 |
| 11 | 121131965 | rs12417414 | C | T | 1.01 | -7977.388 | -7979.147 | -1.759 | 6.66E-07 |
| 11 | 121132211 | rs1941305 | A | G | 1.01 | -7977.388 | -7979.147 | -1.759 | 6.66E-07 |
| 11 | 121132279 | rs1941304 | A | G | 1.01 | -7977.388 | -7979.147 | -1.759 | 6.66E-07 |
| 11 | 117012210 | rs2735182 | A | T | 0.996 | -7977.388 | -7979.148 | -1.76 | 6.65E-07 |
| 11 | 117019010 | rs2008908 | C | T | 0.996 | -7977.388 | -7979.148 | -1.76 | 6.65E-07 |
| 11 | 119332065 | rs10750143 | G | A | 1.017 | -7977.388 | -7979.149 | -1.76 | 6.65E-07 |
| 11 | 119656902 | rs7119690 | G | A | 1.015 | -7977.388 | -7979.148 | -1.76 | 6.65E-07 |
| 11 | 119971523 | rs6589799 | C | T | 0.983 | -7977.388 | -7979.149 | -1.76 | 6.65E-07 |
| 11 | 120715674 | rs9783385 | G | T | 0.984 | -7977.388 | -7979.149 | -1.76 | 6.65E-07 |
| 11 | 121132675 | rs10892700 | T | C | 1.01 | -7977.388 | -7979.148 | -1.76 | 6.65E-07 |

| | | | | | | | |
|---|---|---|---|---|---|---|---|
| 11 | 117010979 | rs4277146 | G | A | NULL | 0.706822221 | NULL&M_Disease |
| 11 | 117016824 | rs10892079 | T | G | NULL | 0.706822221 | NULL&M_Disease |
| 11 | 117408156 | rs473343 | C | G | NULL | 0.706822221 | NULL&M_Disease |
| 11 | 117946691 | rs1786189 | T | C | NULL | 0.706822221 | NULL&M_Disease |
| 11 | 118573668 | rs574383 | C | T | NULL | 0.706822221 | NULL&M_Disease |
| 11 | 119065709 | rs10750139 | G | C | NULL | 0.706822221 | NULL&M_Disease |
| 11 | 119376478 | rs891291 | C | G | NULL | 0.706822221 | NULL&M_Disease |
| 11 | 120138885 | rs753309 | A | G | NULL | 0.706822221 | NULL&M_Disease |
| 11 | 120292097 | rs10750174 | T | C | NULL | 0.706822221 | NULL&M_Disease |
| 11 | 120294423 | rs10892576 | T | C | NULL | 0.706822221 | NULL&M_Disease |
| 11 | 120444179 | rs4561221 | T | C | NULL | 0.706822221 | NULL&M_Disease |
| 11 | 120525032 | rs1893906 | A | G | NULL | 0.706822221 | NULL&M_Disease |
| 11 | 121129528 | rs9943623 | A | C | NULL | 0.706822221 | NULL&M_Disease |
| 11 | 117021923 | rs4938349 | C | T | NULL | 0.706822221 | NULL&M_Disease |
| 11 | 117317664 | rs12361458 | T | C | NULL | 0.706822221 | NULL&M_Disease |
| 11 | 117823425 | rs2512158 | A | G | NULL | 0.706822221 | NULL&M_Disease |
| 11 | 117858983 | rs4252248 | G | A | NULL | 0.706822221 | NULL&M_Disease |
| 11 | 119075429 | rs4938638 | G | A | NULL | 0.706822221 | NULL&M_Disease |
| 11 | 119477261 | rs4938691 | C | T | NULL | 0.706822221 | NULL&M_Disease |
| 11 | 119940945 | rs7113973 | A | G | NULL | 0.706822221 | NULL&M_Disease |
| 11 | 120078028 | rs10892550 | T | G | NULL | 0.706822221 | NULL&M_Disease |
| 11 | 120274353 | rs4938806 | A | C | NULL | 0.706822221 | NULL&M_Disease |
| 11 | 120424000 | rs2248887 | A | G | NULL | 0.706822221 | NULL&M_Disease |
| 11 | 121062860 | rs11218195 | C | A | NULL | 0.706822221 | NULL&M_Disease |
| 11 | 121133909 | rs4936601 | T | G | NULL | 0.706822221 | NULL&M_Disease |
| 11 | 117003749 | rs4938344 | G | A | NULL | 0.706822221 | NULL&M_Disease |
| 11 | 117011440 | rs7951095 | C | A | NULL | 0.706822221 | NULL&M_Disease |
| 11 | 117016997 | rs4936365 | C | T | NULL | 0.706822221 | NULL&M_Disease |
| 11 | 117313086 | rs3863314 | A | G | NULL | 0.706822221 | NULL&M_Disease |
| 11 | 117927759 | rs584670 | A | G | NULL | 0.706822221 | NULL&M_Disease |
| 11 | 118027074 | rs630403 | A | G | NULL | 0.706822221 | NULL&M_Disease |
| 11 | 119376672 | rs891292 | G | A | NULL | 0.706822221 | NULL&M_Disease |
| 11 | 119453980 | rs11217330 | C | A | NULL | 0.706822221 | NULL&M_Disease |
| 11 | 120091272 | rs12786658 | A | G | NULL | 0.706822221 | NULL&M_Disease |
| 11 | 120135501 | rs2847495 | A | C | NULL | 0.706822221 | NULL&M_Disease |
| 11 | 120834573 | rs4936570 | T | C | NULL | 0.706822221 | NULL&M_Disease |
| 11 | 116417813 | rs4938278 | T | G | NULL | 0.706822221 | NULL&M_Disease |
| 11 | 116417966 | rs57903286 | T | C | NULL | 0.706822221 | NULL&M_Disease |
| 11 | 116428507 | rs2063176 | G | C | NULL | 0.706822221 | NULL&M_Disease |
| 11 | 116999225 | rs9326248 | T | G | NULL | 0.706822221 | NULL&M_Disease |
| 11 | 117323952 | rs7946391 | C | T | NULL | 0.706822221 | NULL&M_Disease |
| 11 | 117421858 | rs636530 | G | A | NULL | 0.706822221 | NULL&M_Disease |
| 11 | 117422262 | rs599411 | G | T | NULL | 0.706822221 | NULL&M_Disease |
| 11 | 117853613 | rs4936414 | C | T | NULL | 0.706822221 | NULL&M_Disease |
| 11 | 119595348 | rs11607610 | T | C | NULL | 0.706822221 | NULL&M_Disease |
| 11 | 119639182 | rs56303423 | A | G | NULL | 0.706822221 | NULL&M_Disease |
| 11 | 119974597 | rs1110705 | C | T | NULL | 0.706822221 | NULL&M_Disease |
| 11 | 121136531 | rs10750181 | T | C | NULL | 0.706822221 | NULL&M_Disease |
| 11 | 121136762 | rs10892707 | T | C | NULL | 0.706822221 | NULL&M_Disease |
| 11 | 121138583 | rs12276124 | C | T | NULL | 0.706822221 | NULL&M_Disease |
| 11 | 117032125 | rs3736120 | G | T | NULL | 0.706822221 | NULL&M_Disease |
| 11 | 117048417 | rs10790177 | A | G | NULL | 0.706822221 | NULL&M_Disease |
| 11 | 117081763 | rs171052 | G | C | NULL | 0.706822221 | NULL&M_Disease |
| 11 | 117324662 | rs10892114 | G | C | NULL | 0.706822221 | NULL&M_Disease |
| 11 | 117668104 | rs11216549 | G | C | NULL | 0.706822221 | NULL&M_Disease |
| 11 | 118174885 | rs2231439 | G | A | NULL | 0.706822221 | NULL&M_Disease |
| 11 | 118716563 | rs2851460 | A | G | NULL | 0.706822221 | NULL&M_Disease |
| 11 | 120199080 | rs11823300 | A | G | NULL | 0.706822221 | NULL&M_Disease |
| 11 | 121137448 | rs12280323 | G | A | NULL | 0.706822221 | NULL&M_Disease |
| 11 | 116700169 | rs2854116 | C | T | NULL | 0.706822221 | NULL&M_Disease |
| 11 | 116720089 | rs2289893 | T | C | NULL | 0.706822221 | NULL&M_Disease |
| 11 | 117051165 | rs4938354 | A | G | NULL | 0.706822221 | NULL&M_Disease |
| 11 | 117064693 | rs521171 | C | G | NULL | 0.706822221 | NULL&M_Disease |
| 11 | 117070857 | rs664971 | G | A | NULL | 0.706822221 | NULL&M_Disease |
| 11 | 117767498 | rs558247 | G | A | NULL | 0.706822221 | NULL&M_Disease |
| 11 | 119065102 | rs11217177 | C | T | NULL | 0.706822221 | NULL&M_Disease |
| 11 | 119683867 | rs7950804 | G | A | NULL | 0.706822221 | NULL&M_Disease |
| 11 | 119683955 | rs7950719 | C | T | NULL | 0.706822221 | NULL&M_Disease |
| 11 | 120709387 | rs11218024 | T | C | NULL | 0.706822221 | NULL&M_Disease |
| 11 | 121130758 | rs11218226 | C | T | NULL | 0.706822221 | NULL&M_Disease |
| 11 | 116388264 | rs867897 | G | C | NULL | 0.706822221 | NULL&M_Disease |
| 11 | 117003060 | rs11216284 | C | T | NULL | 0.706822221 | NULL&M_Disease |
| 11 | 117004087 | rs11216286 | G | A | NULL | 0.706822221 | NULL&M_Disease |
| 11 | 117034729 | rs10790175 | T | C | NULL | 0.706822221 | NULL&M_Disease |
| 11 | 117053165 | rs7946257 | A | G | NULL | 0.706822221 | NULL&M_Disease |
| 11 | 117056080 | rs7107152 | A | G | NULL | 0.706822221 | NULL&M_Disease |
| 11 | 117071070 | rs664082 | C | G | NULL | 0.706822221 | NULL&M_Disease |
| 11 | 117324868 | rs10790186 | G | A | NULL | 0.706822221 | NULL&M_Disease |
| 11 | 117922486 | rs624931 | C | T | NULL | 0.706822221 | NULL&M_Disease |
| 11 | 119063908 | rs2301574 | T | C | NULL | 0.706822221 | NULL&M_Disease |
| 11 | 119479391 | rs2271249 | C | T | NULL | 0.706822221 | NULL&M_Disease |
| 11 | 119590878 | rs11217426 | T | C | NULL | 0.706822221 | NULL&M_Disease |
| 11 | 119683473 | rs7950042 | A | C | NULL | 0.706822221 | NULL&M_Disease |
| 11 | 119689932 | rs4938731 | A | G | NULL | 0.706822221 | NULL&M_Disease |
| 11 | 121127540 | rs4604899 | C | T | NULL | 0.706822221 | NULL&M_Disease |
| 11 | 117069061 | rs585849 | A | G | NULL | 0.707857271 | NULL&M_Disease |
| 11 | 117193254 | rs61903732 | G | A | NULL | 0.707857271 | NULL&M_Disease |
| 11 | 117776819 | rs12224796 | G | A | NULL | 0.707857271 | NULL&M_Disease |
| 11 | 119479520 | rs2271250 | C | T | NULL | 0.707857271 | NULL&M_Disease |
| 11 | 119683777 | rs7950403 | A | G | NULL | 0.707857271 | NULL&M_Disease |
| 11 | 119957453 | rs20494 | A | G | NULL | 0.707857271 | NULL&M_Disease |
| 11 | 120288372 | rs10790382 | C | T | NULL | 0.707857271 | NULL&M_Disease |
| 11 | 120592483 | rs1939669 | A | T | NULL | 0.707857271 | NULL&M_Disease |
| 11 | 121112943 | rs10892695 | G | T | NULL | 0.707857271 | NULL&M_Disease |
| 11 | 117081974 | rs236910 | G | A | NULL | 0.707857271 | NULL&M_Disease |
| 11 | 117202437 | rs507133 | A | G | NULL | 0.707857271 | NULL&M_Disease |

| | | | | | | | | | |
|---|---|---|---|---|---|---|---|---|---|
| 11 | 117010979 | rs4277146 | G | A | 0.996 | -7977.388 | -7979.149 | -1.761 | 6.65E-07 |
| 11 | 117016824 | rs10892079 | T | G | 0.996 | -7977.388 | -7979.149 | -1.761 | 6.65E-07 |
| 11 | 117408156 | rs473343 | C | G | 1.002 | -7977.388 | -7979.15 | -1.761 | 6.65E-07 |
| 11 | 117946691 | rs1786189 | T | C | 1.006 | -7977.388 | -7979.149 | -1.761 | 6.65E-07 |
| 11 | 118573668 | rs574383 | C | T | 0.985 | -7977.388 | -7979.149 | -1.761 | 6.65E-07 |
| 11 | 119065709 | rs10750139 | G | C | 1.017 | -7977.388 | -7979.15 | -1.761 | 6.65E-07 |
| 11 | 119376478 | rs891291 | C | G | 1.017 | -7977.388 | -7979.149 | -1.761 | 6.65E-07 |
| 11 | 120138885 | rs753309 | A | G | 1.012 | -7977.388 | -7979.15 | -1.761 | 6.65E-07 |
| 11 | 120292097 | rs10750174 | T | C | 1.016 | -7977.388 | -7979.149 | -1.761 | 6.65E-07 |
| 11 | 120294423 | rs10892576 | T | C | 1.016 | -7977.388 | -7979.149 | -1.761 | 6.65E-07 |
| 11 | 120444179 | rs4561221 | T | C | 1.017 | -7977.388 | -7979.149 | -1.761 | 6.65E-07 |
| 11 | 120525032 | rs1893906 | A | G | 1.014 | -7977.388 | -7979.149 | -1.761 | 6.65E-07 |
| 11 | 121129528 | rs9943623 | A | C | 1.01 | -7977.388 | -7979.149 | -1.761 | 6.65E-07 |
| 11 | 117021923 | rs4938349 | C | T | 0.997 | -7977.388 | -7979.151 | -1.762 | 6.64E-07 |
| 11 | 117317664 | rs12361458 | T | C | 0.986 | -7977.388 | -7979.15 | -1.762 | 6.64E-07 |
| 11 | 117823425 | rs2512158 | A | G | 0.983 | -7977.388 | -7979.15 | -1.762 | 6.64E-07 |
| 11 | 117858983 | rs4252248 | G | A | 0.985 | -7977.388 | -7979.15 | -1.762 | 6.64E-07 |
| 11 | 119075429 | rs4938638 | G | A | 1.017 | -7977.388 | -7979.15 | -1.762 | 6.64E-07 |
| 11 | 119477261 | rs4938691 | C | T | 0.986 | -7977.388 | -7979.15 | -1.762 | 6.64E-07 |
| 11 | 119940945 | rs7113973 | A | G | 1.016 | -7977.388 | -7979.15 | -1.762 | 6.64E-07 |
| 11 | 120078028 | rs10892550 | T | G | 0.996 | -7977.388 | -7979.15 | -1.762 | 6.64E-07 |
| 11 | 120274353 | rs4938806 | A | C | 1.016 | -7977.388 | -7979.15 | -1.762 | 6.64E-07 |
| 11 | 120424000 | rs2248887 | A | G | 1.004 | -7977.388 | -7979.15 | -1.762 | 6.64E-07 |
| 11 | 121062860 | rs11218195 | C | A | 1.012 | -7977.388 | -7979.151 | -1.762 | 6.64E-07 |
| 11 | 121133909 | rs4936601 | T | G | 1.01 | -7977.388 | -7979.15 | -1.762 | 6.64E-07 |
| 11 | 117003749 | rs4938344 | G | A | 0.997 | -7977.388 | -7979.151 | -1.763 | 6.63E-07 |
| 11 | 117011440 | rs7951095 | C | A | 0.997 | -7977.388 | -7979.151 | -1.763 | 6.63E-07 |
| 11 | 117016997 | rs4936365 | C | T | 0.997 | -7977.388 | -7979.151 | -1.763 | 6.63E-07 |
| 11 | 117313086 | rs3863314 | A | G | 0.987 | -7977.388 | -7979.152 | -1.763 | 6.63E-07 |
| 11 | 117927759 | rs584670 | A | G | 1.006 | -7977.388 | -7979.152 | -1.763 | 6.63E-07 |
| 11 | 118027074 | rs630403 | A | G | 1.017 | -7977.388 | -7979.151 | -1.763 | 6.63E-07 |
| 11 | 119376672 | rs891292 | G | A | 1.017 | -7977.388 | -7979.151 | -1.763 | 6.63E-07 |
| 11 | 119453980 | rs11217330 | C | A | 1.012 | -7977.388 | -7979.151 | -1.763 | 6.63E-07 |
| 11 | 120091272 | rs12786658 | A | G | 1.013 | -7977.388 | -7979.151 | -1.763 | 6.63E-07 |
| 11 | 120135501 | rs2847495 | A | C | 1.006 | -7977.388 | -7979.152 | -1.763 | 6.63E-07 |
| 11 | 120834573 | rs4936570 | T | C | 1.007 | -7977.388 | -7979.152 | -1.763 | 6.63E-07 |
| 11 | 116417813 | rs4938278 | T | G | 0.99 | -7977.388 | -7979.152 | -1.764 | 6.63E-07 |
| 11 | 116417966 | rs57903286 | T | C | 0.99 | -7977.388 | -7979.152 | -1.764 | 6.63E-07 |
| 11 | 116428507 | rs2063176 | G | C | 0.989 | -7977.388 | -7979.153 | -1.764 | 6.63E-07 |
| 11 | 116999225 | rs9326248 | T | G | 0.998 | -7977.388 | -7979.153 | -1.764 | 6.63E-07 |
| 11 | 117323952 | rs7946391 | C | T | 0.983 | -7977.388 | -7979.152 | -1.764 | 6.63E-07 |
| 11 | 117421858 | rs636530 | G | A | 1 | -7977.388 | -7979.152 | -1.764 | 6.63E-07 |
| 11 | 117422262 | rs599411 | G | T | 1 | -7977.388 | -7979.152 | -1.764 | 6.63E-07 |
| 11 | 117853613 | rs4936414 | C | T | 0.985 | -7977.388 | -7979.152 | -1.764 | 6.63E-07 |
| 11 | 119595348 | rs11607610 | T | C | 0.983 | -7977.388 | -7979.152 | -1.764 | 6.63E-07 |
| 11 | 119639182 | rs56303423 | A | G | 0.997 | -7977.388 | -7979.152 | -1.764 | 6.63E-07 |
| 11 | 119974597 | rs1110705 | C | T | 0.983 | -7977.388 | -7979.153 | -1.764 | 6.63E-07 |
| 11 | 121136531 | rs10750181 | T | C | 1.009 | -7977.388 | -7979.153 | -1.764 | 6.63E-07 |
| 11 | 121136762 | rs10892707 | T | C | 1.009 | -7977.388 | -7979.152 | -1.764 | 6.63E-07 |
| 11 | 121138583 | rs12276124 | C | T | 1.009 | -7977.388 | -7979.153 | -1.764 | 6.63E-07 |
| 11 | 117032125 | rs3736120 | G | T | 0.998 | -7977.388 | -7979.154 | -1.765 | 6.62E-07 |
| 11 | 117048417 | rs10790177 | A | G | 0.998 | -7977.388 | -7979.154 | -1.765 | 6.62E-07 |
| 11 | 117081763 | rs171052 | G | C | 0.996 | -7977.388 | -7979.154 | -1.765 | 6.62E-07 |
| 11 | 117324662 | rs10892114 | G | C | 0.983 | -7977.388 | -7979.153 | -1.765 | 6.62E-07 |
| 11 | 117668104 | rs11216549 | G | C | 1.015 | -7977.388 | -7979.154 | -1.765 | 6.62E-07 |
| 11 | 118174885 | rs2231439 | G | A | 0.986 | -7977.388 | -7979.153 | -1.765 | 6.62E-07 |
| 11 | 118716563 | rs2851460 | A | G | 0.984 | -7977.388 | -7979.154 | -1.765 | 6.62E-07 |
| 11 | 120199080 | rs11823300 | A | G | 1.016 | -7977.388 | -7979.153 | -1.765 | 6.62E-07 |
| 11 | 121137448 | rs12280323 | G | A | 1.009 | -7977.388 | -7979.154 | -1.765 | 6.62E-07 |
| 11 | 116700169 | rs2854116 | C | T | 1.015 | -7977.388 | -7979.154 | -1.766 | 6.62E-07 |
| 11 | 116720089 | rs2289893 | T | C | 0.987 | -7977.388 | -7979.155 | -1.766 | 6.62E-07 |
| 11 | 117051165 | rs4938354 | A | G | 0.998 | -7977.388 | -7979.155 | -1.766 | 6.62E-07 |
| 11 | 117064693 | rs521171 | C | G | 1 | -7977.388 | -7979.154 | -1.766 | 6.62E-07 |
| 11 | 117070857 | rs664971 | G | A | 0.996 | -7977.388 | -7979.154 | -1.766 | 6.62E-07 |
| 11 | 117767498 | rs558247 | G | A | 0.984 | -7977.388 | -7979.154 | -1.766 | 6.62E-07 |
| 11 | 119065102 | rs11217177 | C | T | 0.996 | -7977.388 | -7979.155 | -1.766 | 6.62E-07 |
| 11 | 119683867 | rs7950804 | G | A | 0.983 | -7977.388 | -7979.154 | -1.766 | 6.62E-07 |
| 11 | 119683955 | rs7950719 | C | T | 0.983 | -7977.388 | -7979.154 | -1.766 | 6.62E-07 |
| 11 | 120709387 | rs11218024 | T | C | 0.984 | -7977.388 | -7979.154 | -1.766 | 6.62E-07 |
| 11 | 121130758 | rs11218226 | C | T | 1.009 | -7977.388 | -7979.155 | -1.766 | 6.62E-07 |
| 11 | 116388264 | rs867897 | G | C | 0.984 | -7977.388 | -7979.155 | -1.767 | 6.61E-07 |
| 11 | 117003060 | rs11216284 | C | T | 0.998 | -7977.388 | -7979.155 | -1.767 | 6.61E-07 |
| 11 | 117004087 | rs11216286 | G | A | 0.998 | -7977.388 | -7979.155 | -1.767 | 6.61E-07 |
| 11 | 117034729 | rs10790175 | T | C | 0.998 | -7977.388 | -7979.155 | -1.767 | 6.61E-07 |
| 11 | 117053165 | rs7946257 | A | G | 0.999 | -7977.388 | -7979.155 | -1.767 | 6.61E-07 |
| 11 | 117056080 | rs7107152 | A | G | 1 | -7977.388 | -7979.155 | -1.767 | 6.61E-07 |
| 11 | 117071070 | rs664082 | C | G | 0.996 | -7977.388 | -7979.156 | -1.767 | 6.61E-07 |
| 11 | 117324868 | rs10790186 | G | A | 0.984 | -7977.388 | -7979.155 | -1.767 | 6.61E-07 |
| 11 | 117922486 | rs624931 | C | T | 1.005 | -7977.388 | -7979.156 | -1.767 | 6.61E-07 |
| 11 | 119063908 | rs2301574 | T | C | 0.996 | -7977.388 | -7979.156 | -1.767 | 6.61E-07 |
| 11 | 119479391 | rs2271249 | C | T | 0.986 | -7977.388 | -7979.155 | -1.767 | 6.61E-07 |
| 11 | 119590878 | rs11217426 | T | C | 0.984 | -7977.388 | -7979.155 | -1.767 | 6.61E-07 |
| 11 | 119683473 | rs7950042 | A | C | 0.983 | -7977.388 | -7979.155 | -1.767 | 6.61E-07 |
| 11 | 119689932 | rs4938731 | A | G | 1.006 | -7977.388 | -7979.155 | -1.767 | 6.61E-07 |
| 11 | 121127540 | rs4604899 | C | T | 1.009 | -7977.388 | -7979.156 | -1.767 | 6.61E-07 |
| 11 | 117069061 | rs585849 | A | G | 0.997 | -7977.388 | -7979.156 | -1.768 | 6.60E-07 |
| 11 | 117193254 | rs61903732 | G | A | 0.992 | -7977.388 | -7979.156 | -1.768 | 6.60E-07 |
| 11 | 117776819 | rs12224796 | G | A | 1.016 | -7977.388 | -7979.156 | -1.768 | 6.60E-07 |
| 11 | 119479520 | rs2271250 | C | T | 0.986 | -7977.388 | -7979.157 | -1.768 | 6.60E-07 |
| 11 | 119683777 | rs7950403 | A | G | 0.983 | -7977.388 | -7979.157 | -1.768 | 6.60E-07 |
| 11 | 119957453 | rs20494 | A | G | 0.984 | -7977.388 | -7979.157 | -1.768 | 6.60E-07 |
| 11 | 120288372 | rs10790382 | C | T | 1.016 | -7977.388 | -7979.156 | -1.768 | 6.60E-07 |
| 11 | 120592483 | rs1939669 | A | T | 0.984 | -7977.388 | -7979.157 | -1.768 | 6.60E-07 |
| 11 | 121112943 | rs10892695 | G | T | 1.011 | -7977.388 | -7979.156 | -1.768 | 6.60E-07 |
| 11 | 117081974 | rs236910 | G | A | 0.997 | -7977.388 | -7979.157 | -1.769 | 6.60E-07 |
| 11 | 117202437 | rs507133 | A | G | 0.984 | -7977.388 | -7979.157 | -1.769 | 6.60E-07 |

| | | | | | | | |
|---|---|---|---|---|---|---|---|
| 11 | 117235368 | rs7945704 | A | G | NULL | 0.707857271 | NULL&M_Disease |
| 11 | 118594669 | rs140026975 | C | A | NULL | 0.707857271 | NULL&M_Disease |
| 11 | 119499592 | rs4278525 | A | G | NULL | 0.707857271 | NULL&M_Disease |
| 11 | 119589703 | rs9666263 | C | G | NULL | 0.707857271 | NULL&M_Disease |
| 11 | 119880792 | rs581491 | C | T | NULL | 0.707857271 | NULL&M_Disease |
| 11 | 120711869 | rs57053294 | C | T | NULL | 0.707857271 | NULL&M_Disease |
| 11 | 121127696 | rs4604900 | A | T | NULL | 0.707857271 | NULL&M_Disease |
| 11 | 121129841 | rs4936598 | A | G | NULL | 0.707857271 | NULL&M_Disease |
| 11 | 121130464 | rs12416965 | C | T | NULL | 0.707857271 | NULL&M_Disease |
| 11 | 121130611 | rs35255208 | G | A | NULL | 0.707857271 | NULL&M_Disease |
| 11 | 117325898 | rs7103239 | A | G | NULL | 0.707857271 | NULL&M_Disease |
| 11 | 117977949 | rs6589646 | T | C | NULL | 0.707857271 | NULL&M_Disease |
| 11 | 119198092 | rs4581461 | C | T | NULL | 0.707857271 | NULL&M_Disease |
| 11 | 120030227 | rs11217712 | G | T | NULL | 0.707857271 | NULL&M_Disease |
| 11 | 120336902 | rs4938808 | T | G | NULL | 0.707857271 | NULL&M_Disease |
| 11 | 120440043 | rs11601826 | G | A | NULL | 0.707857271 | NULL&M_Disease |
| 11 | 117067699 | rs8521 | C | T | NULL | 0.707857271 | NULL&M_Disease |
| 11 | 117508837 | rs695083 | G | A | NULL | 0.707857271 | NULL&M_Disease |
| 11 | 117695853 | rs11216564 | A | G | NULL | 0.707857271 | NULL&M_Disease |
| 11 | 117696744 | rs678776 | A | G | NULL | 0.707857271 | NULL&M_Disease |
| 11 | 119166280 | rs2155824 | A | T | NULL | 0.707857271 | NULL&M_Disease |
| 11 | 119634962 | rs59814528 | C | T | NULL | 0.707857271 | NULL&M_Disease |
| 11 | 119666454 | rs7946574 | T | C | NULL | 0.707857271 | NULL&M_Disease |
| 11 | 119976523 | rs7946597 | A | G | NULL | 0.707857271 | NULL&M_Disease |
| 11 | 119976805 | rs4938780 | T | C | NULL | 0.707857271 | NULL&M_Disease |
| 11 | 121136763 | rs10892708 | G | C | NULL | 0.707857271 | NULL&M_Disease |
| 11 | 117070895 | rs664922 | C | A | NULL | 0.707857271 | NULL&M_Disease |
| 11 | 117076564 | rs1242127 | G | A | NULL | 0.707857271 | NULL&M_Disease |
| 11 | 117309495 | rs10892104 | A | G | NULL | 0.707857271 | NULL&M_Disease |
| 11 | 117936146 | rs671567 | A | G | NULL | 0.707857271 | NULL&M_Disease |
| 11 | 118194599 | rs12577841 | T | C | NULL | 0.707857271 | NULL&M_Disease |
| 11 | 119561362 | rs7933848 | T | C | NULL | 0.707857271 | NULL&M_Disease |
| 11 | 121136756 | rs10892706 | A | T | NULL | 0.707857271 | NULL&M_Disease |
| 11 | 117940332 | rs654572 | C | T | NULL | 0.707857271 | NULL&M_Disease |
| 11 | 117940612 | rs624463 | G | A | NULL | 0.707857271 | NULL&M_Disease |
| 11 | 117978264 | rs1894177 | A | G | NULL | 0.707857271 | NULL&M_Disease |
| 11 | 120288368 | rs10750173 | C | T | NULL | 0.707857271 | NULL&M_Disease |
| 11 | 120494400 | rs7111020 | T | C | NULL | 0.707857271 | NULL&M_Disease |
| 11 | 121145976 | rs7939478 | C | A | NULL | 0.707857271 | NULL&M_Disease |
| 11 | 116338514 | rs1508099 | C | A | NULL | 0.707857271 | NULL&M_Disease |
| 11 | 116417048 | rs4635116 | C | A | NULL | 0.707857271 | NULL&M_Disease |
| 11 | 117941898 | rs640421 | T | C | NULL | 0.707857271 | NULL&M_Disease |
| 11 | 117942125 | rs652030 | C | T | NULL | 0.707857271 | NULL&M_Disease |
| 11 | 117942848 | rs655157 | T | G | NULL | 0.707857271 | NULL&M_Disease |
| 11 | 119455748 | rs7925188 | G | A | NULL | 0.707857271 | NULL&M_Disease |
| 11 | 119457828 | rs10892411 | G | C | NULL | 0.707857271 | NULL&M_Disease |
| 11 | 119684145 | rs895647 | T | G | NULL | 0.707857271 | NULL&M_Disease |
| 11 | 119684321 | rs895645 | G | A | NULL | 0.707857271 | NULL&M_Disease |
| 11 | 120142525 | rs7119027 | G | A | NULL | 0.707857271 | NULL&M_Disease |
| 11 | 120998942 | rs10502247 | T | C | NULL | 0.707857271 | NULL&M_Disease |
| 11 | 116311508 | rs532107 | G | A | NULL | 0.707857271 | NULL&M_Disease |
| 11 | 116568713 | rs1787691 | A | G | NULL | 0.707857271 | NULL&M_Disease |
| 11 | 117941554 | rs638997 | A | G | NULL | 0.707857271 | NULL&M_Disease |
| 11 | 119622391 | rs11217461 | A | G | NULL | 0.707857271 | NULL&M_Disease |
| 11 | 119987219 | rs576642 | C | T | NULL | 0.707857271 | NULL&M_Disease |
| 11 | 121145962 | rs7939570 | A | C | NULL | 0.707857271 | NULL&M_Disease |
| 11 | 121160519 | rs10790432 | C | T | NULL | 0.707857271 | NULL&M_Disease |
| 11 | 121161041 | rs7115581 | A | G | NULL | 0.707857271 | NULL&M_Disease |
| 11 | 121161106 | rs7115602 | A | G | NULL | 0.707857271 | NULL&M_Disease |
| 11 | 116301251 | rs12419991 | T | C | NULL | 0.707857271 | NULL&M_Disease |
| 11 | 116470227 | rs11216035 | C | T | NULL | 0.707857271 | NULL&M_Disease |
| 11 | 116567449 | rs1784106 | A | C | NULL | 0.707857271 | NULL&M_Disease |
| 11 | 116591553 | rs180373 | A | C | NULL | 0.707857271 | NULL&M_Disease |
| 11 | 117059552 | rs474339 | G | A | NULL | 0.707857271 | NULL&M_Disease |
| 11 | 118487673 | rs4938517 | C | T | NULL | 0.707857271 | NULL&M_Disease |
| 11 | 118840506 | rs55736521 | T | C | NULL | 0.707857271 | NULL&M_Disease |
| 11 | 118840705 | rs45534632 | G | A | NULL | 0.707857271 | NULL&M_Disease |
| 11 | 119181170 | rs2249466 | T | C | NULL | 0.707857271 | NULL&M_Disease |
| 11 | 119684301 | rs895646 | T | G | NULL | 0.707857271 | NULL&M_Disease |
| 11 | 120002927 | rs561498 | C | T | NULL | 0.707857271 | NULL&M_Disease |
| 11 | 121153455 | rs10750182 | A | T | NULL | 0.707857271 | NULL&M_Disease |
| 11 | 121156532 | rs7128047 | A | G | NULL | 0.707857271 | NULL&M_Disease |
| 11 | 121156809 | rs7128342 | A | T | NULL | 0.707857271 | NULL&M_Disease |
| 11 | 121158682 | rs2163281 | A | G | NULL | 0.707857271 | NULL&M_Disease |
| 11 | 116396188 | rs7107833 | C | T | NULL | 0.707857271 | NULL&M_Disease |
| 11 | 117941847 | rs671351 | A | C | NULL | 0.707857271 | NULL&M_Disease |
| 11 | 119479052 | rs3751020 | C | T | NULL | 0.707857271 | NULL&M_Disease |
| 11 | 119560847 | rs4938708 | T | C | NULL | 0.707857271 | NULL&M_Disease |
| 11 | 119561042 | rs4938709 | A | T | NULL | 0.707857271 | NULL&M_Disease |
| 11 | 119603642 | rs12361192 | C | T | NULL | 0.707857271 | NULL&M_Disease |
| 11 | 120481773 | rs12278532 | T | C | NULL | 0.707857271 | NULL&M_Disease |
| 11 | 120693133 | rs7952068 | G | C | NULL | 0.707857271 | NULL&M_Disease |
| 11 | 121141005 | rs7948174 | T | G | NULL | 0.707857271 | NULL&M_Disease |
| 11 | 117699878 | rs2019655 | T | C | NULL | 0.708890173 | NULL&M_Disease |
| 11 | 117960661 | rs7924498 | A | G | NULL | 0.708890173 | NULL&M_Disease |
| 11 | 119040463 | rs7483350 | G | C | NULL | 0.708890173 | NULL&M_Disease |
| 11 | 119575104 | rs2006949 | C | T | NULL | 0.708890173 | NULL&M_Disease |
| 11 | 119682999 | rs2121577 | G | A | NULL | 0.708890173 | NULL&M_Disease |
| 11 | 119683031 | rs2121576 | G | A | NULL | 0.708890173 | NULL&M_Disease |
| 11 | 119683964 | rs7937319 | T | C | NULL | 0.708890173 | NULL&M_Disease |
| 11 | 121142126 | rs7952199 | G | C | NULL | 0.708890173 | NULL&M_Disease |
| 11 | 121147301 | rs10790429 | A | T | NULL | 0.708890173 | NULL&M_Disease |
| 11 | 116572525 | rs1240783 | C | T | NULL | 0.708890173 | NULL&M_Disease |
| 11 | 118004863 | rs45584835 | G | A | NULL | 0.708890173 | NULL&M_Disease |
| 11 | 119684565 | rs4938727 | C | T | NULL | 0.708890173 | NULL&M_Disease |
| 11 | 120288944 | rs10892574 | C | G | NULL | 0.708890173 | NULL&M_Disease |

| | | | | | | | | | |
|---|---|---|---|---|---|---|---|---|---|
| 11 | 117235368 | rs7945704 | A | G | 0.997 | -7977.388 | -7979.157 | -1.769 | 6.60E-07 |
| 11 | 118594669 | rs140026975 | C | A | 1.006 | -7977.388 | -7979.157 | -1.769 | 6.60E-07 |
| 11 | 119499592 | rs4278525 | A | G | 0.991 | -7977.388 | -7979.157 | -1.769 | 6.60E-07 |
| 11 | 119589703 | rs9666263 | C | G | 0.984 | -7977.388 | -7979.158 | -1.769 | 6.60E-07 |
| 11 | 119880792 | rs581491 | C | T | 1.016 | -7977.388 | -7979.157 | -1.769 | 6.60E-07 |
| 11 | 120711869 | rs57053294 | C | T | 0.984 | -7977.388 | -7979.158 | -1.769 | 6.60E-07 |
| 11 | 121127696 | rs4604900 | A | T | 1.009 | -7977.388 | -7979.157 | -1.769 | 6.60E-07 |
| 11 | 121129841 | rs4936598 | A | G | 1.009 | -7977.388 | -7979.157 | -1.769 | 6.60E-07 |
| 11 | 121130464 | rs12416965 | C | T | 1.009 | -7977.388 | -7979.157 | -1.769 | 6.60E-07 |
| 11 | 121130611 | rs35255208 | G | A | 1.009 | -7977.388 | -7979.157 | -1.769 | 6.60E-07 |
| 11 | 117325898 | rs7103239 | A | G | 1.012 | -7977.388 | -7979.159 | -1.77 | 6.59E-07 |
| 11 | 117977949 | rs6589646 | T | C | 1.016 | -7977.388 | -7979.159 | -1.77 | 6.59E-07 |
| 11 | 119198092 | rs4581461 | C | T | 1.016 | -7977.388 | -7979.159 | -1.77 | 6.59E-07 |
| 11 | 120030227 | rs11217712 | G | T | 0.988 | -7977.388 | -7979.158 | -1.77 | 6.59E-07 |
| 11 | 120336902 | rs4938808 | T | G | 1.016 | -7977.388 | -7979.159 | -1.77 | 6.59E-07 |
| 11 | 120440043 | rs11601826 | G | A | 1.017 | -7977.388 | -7979.158 | -1.77 | 6.59E-07 |
| 11 | 117067699 | rs8521 | C | T | 0.998 | -7977.388 | -7979.159 | -1.771 | 6.58E-07 |
| 11 | 117508837 | rs695083 | G | A | 1.017 | -7977.388 | -7979.16 | -1.771 | 6.58E-07 |
| 11 | 117695853 | rs11216564 | A | G | 0.998 | -7977.388 | -7979.159 | -1.771 | 6.58E-07 |
| 11 | 117696744 | rs678776 | A | G | 0.987 | -7977.388 | -7979.159 | -1.771 | 6.58E-07 |
| 11 | 119166280 | rs2155824 | A | T | 0.995 | -7977.388 | -7979.159 | -1.771 | 6.58E-07 |
| 11 | 119634962 | rs59814528 | C | T | 1.016 | -7977.388 | -7979.16 | -1.771 | 6.58E-07 |
| 11 | 119666454 | rs7946574 | T | C | 1.002 | -7977.388 | -7979.159 | -1.771 | 6.58E-07 |
| 11 | 119976523 | rs7946597 | A | G | 0.987 | -7977.388 | -7979.159 | -1.771 | 6.58E-07 |
| 11 | 119976805 | rs4938780 | T | C | 0.987 | -7977.388 | -7979.159 | -1.771 | 6.58E-07 |
| 11 | 121136763 | rs10892708 | G | C | 1.008 | -7977.388 | -7979.16 | -1.771 | 6.58E-07 |
| 11 | 117070895 | rs664922 | C | A | 0.999 | -7977.388 | -7979.16 | -1.772 | 6.58E-07 |
| 11 | 117076564 | rs1242127 | G | A | 0.998 | -7977.388 | -7979.161 | -1.772 | 6.58E-07 |
| 11 | 117309495 | rs10892104 | A | G | 1.009 | -7977.388 | -7979.16 | -1.772 | 6.58E-07 |
| 11 | 117936146 | rs671567 | A | G | 1.003 | -7977.388 | -7979.16 | -1.772 | 6.58E-07 |
| 11 | 118194599 | rs12577841 | T | C | 0.989 | -7977.388 | -7979.16 | -1.772 | 6.58E-07 |
| 11 | 119561362 | rs7933848 | T | C | 0.985 | -7977.388 | -7979.161 | -1.772 | 6.58E-07 |
| 11 | 121136756 | rs10892706 | A | T | 1.008 | -7977.388 | -7979.161 | -1.772 | 6.58E-07 |
| 11 | 117940332 | rs654572 | C | T | 1.003 | -7977.388 | -7979.161 | -1.773 | 6.57E-07 |
| 11 | 117940612 | rs624463 | G | A | 1.003 | -7977.388 | -7979.161 | -1.773 | 6.57E-07 |
| 11 | 117978264 | rs1894177 | A | G | 1.016 | -7977.388 | -7979.161 | -1.773 | 6.57E-07 |
| 11 | 120288368 | rs10750173 | C | T | 1.016 | -7977.388 | -7979.161 | -1.773 | 6.57E-07 |
| 11 | 120494400 | rs7111020 | T | C | 0.984 | -7977.388 | -7979.161 | -1.773 | 6.57E-07 |
| 11 | 121145976 | rs7939478 | C | A | 1.008 | -7977.388 | -7979.161 | -1.773 | 6.57E-07 |
| 11 | 116338514 | rs1508099 | C | A | 0.988 | -7977.388 | -7979.162 | -1.774 | 6.56E-07 |
| 11 | 116417048 | rs4635116 | C | A | 0.991 | -7977.388 | -7979.162 | -1.774 | 6.56E-07 |
| 11 | 117941898 | rs640421 | T | C | 1.003 | -7977.388 | -7979.162 | -1.774 | 6.56E-07 |
| 11 | 117942125 | rs652030 | C | T | 1.003 | -7977.388 | -7979.162 | -1.774 | 6.56E-07 |
| 11 | 117942848 | rs655157 | T | G | 1.003 | -7977.388 | -7979.162 | -1.774 | 6.56E-07 |
| 11 | 119455748 | rs7925188 | G | A | 0.987 | -7977.388 | -7979.162 | -1.774 | 6.56E-07 |
| 11 | 119457828 | rs10892411 | G | C | 0.984 | -7977.388 | -7979.162 | -1.774 | 6.56E-07 |
| 11 | 119684145 | rs895647 | T | G | 0.984 | -7977.388 | -7979.162 | -1.774 | 6.56E-07 |
| 11 | 119684321 | rs895645 | G | A | 0.984 | -7977.388 | -7979.162 | -1.774 | 6.56E-07 |
| 11 | 120142525 | rs7119027 | G | A | 1.011 | -7977.388 | -7979.162 | -1.774 | 6.56E-07 |
| 11 | 120998942 | rs10502247 | T | C | 0.988 | -7977.388 | -7979.163 | -1.774 | 6.56E-07 |
| 11 | 116311508 | rs532107 | G | A | 1.003 | -7977.388 | -7979.164 | -1.775 | 6.56E-07 |
| 11 | 116568713 | rs1787691 | A | G | 0.984 | -7977.388 | -7979.163 | -1.775 | 6.56E-07 |
| 11 | 117941554 | rs638997 | A | G | 1.002 | -7977.388 | -7979.163 | -1.775 | 6.56E-07 |
| 11 | 119622391 | rs11217461 | A | G | 1.015 | -7977.388 | -7979.164 | -1.775 | 6.56E-07 |
| 11 | 119987219 | rs576642 | C | T | 1.017 | -7977.388 | -7979.163 | -1.775 | 6.56E-07 |
| 11 | 121145962 | rs7939570 | A | C | 1.007 | -7977.388 | -7979.163 | -1.775 | 6.56E-07 |
| 11 | 121160519 | rs10790432 | C | T | 1.012 | -7977.388 | -7979.163 | -1.775 | 6.56E-07 |
| 11 | 121161041 | rs7115581 | A | G | 1.012 | -7977.388 | -7979.163 | -1.775 | 6.56E-07 |
| 11 | 121161106 | rs7115602 | A | G | 1.012 | -7977.388 | -7979.163 | -1.775 | 6.56E-07 |
| 11 | 116301251 | rs12419991 | T | C | 1.014 | -7977.388 | -7979.165 | -1.776 | 6.55E-07 |
| 11 | 116470227 | rs11216035 | C | T | 1.012 | -7977.388 | -7979.164 | -1.776 | 6.55E-07 |
| 11 | 116567449 | rs1784106 | A | C | 0.984 | -7977.388 | -7979.164 | -1.776 | 6.55E-07 |
| 11 | 116591553 | rs180373 | A | C | 1.011 | -7977.388 | -7979.164 | -1.776 | 6.55E-07 |
| 11 | 117059552 | rs474339 | G | A | 0.998 | -7977.388 | -7979.164 | -1.776 | 6.55E-07 |
| 11 | 118487673 | rs4938517 | C | T | 1.014 | -7977.388 | -7979.164 | -1.776 | 6.55E-07 |
| 11 | 118840506 | rs55736521 | T | C | 0.987 | -7977.388 | -7979.164 | -1.776 | 6.55E-07 |
| 11 | 118840705 | rs45534632 | G | A | 0.987 | -7977.388 | -7979.164 | -1.776 | 6.55E-07 |
| 11 | 119181170 | rs2249466 | T | C | 1.011 | -7977.388 | -7979.164 | -1.776 | 6.55E-07 |
| 11 | 119684301 | rs895646 | T | G | 0.984 | -7977.388 | -7979.165 | -1.776 | 6.55E-07 |
| 11 | 120002927 | rs561498 | C | T | 0.994 | -7977.388 | -7979.164 | -1.776 | 6.55E-07 |
| 11 | 121153455 | rs10750182 | A | T | 1.012 | -7977.388 | -7979.165 | -1.776 | 6.55E-07 |
| 11 | 121156532 | rs7128047 | A | G | 1.012 | -7977.388 | -7979.164 | -1.776 | 6.55E-07 |
| 11 | 121156809 | rs7128342 | A | T | 1.012 | -7977.388 | -7979.164 | -1.776 | 6.55E-07 |
| 11 | 121158682 | rs2163281 | A | G | 1.012 | -7977.388 | -7979.164 | -1.776 | 6.55E-07 |
| 11 | 116396188 | rs7107833 | C | T | 1.013 | -7977.388 | -7979.165 | -1.777 | 6.54E-07 |
| 11 | 117941847 | rs671351 | A | C | 1 | -7977.388 | -7979.165 | -1.777 | 6.54E-07 |
| 11 | 119479052 | rs3751020 | C | T | 0.987 | -7977.388 | -7979.165 | -1.777 | 6.54E-07 |
| 11 | 119560847 | rs4938708 | T | C | 0.985 | -7977.388 | -7979.166 | -1.777 | 6.54E-07 |
| 11 | 119561042 | rs4938709 | A | T | 0.985 | -7977.388 | -7979.166 | -1.777 | 6.54E-07 |
| 11 | 119603642 | rs12361192 | C | T | 0.984 | -7977.388 | -7979.165 | -1.777 | 6.54E-07 |
| 11 | 120481773 | rs12278532 | T | C | 0.985 | -7977.388 | -7979.165 | -1.777 | 6.54E-07 |
| 11 | 120693133 | rs7952068 | G | C | 0.985 | -7977.388 | -7979.165 | -1.777 | 6.54E-07 |
| 11 | 121141005 | rs7948174 | T | G | 1.007 | -7977.388 | -7979.166 | -1.777 | 6.54E-07 |
| 11 | 117699878 | rs2019655 | T | C | 1.005 | -7977.388 | -7979.166 | -1.778 | 6.54E-07 |
| 11 | 117960661 | rs7924498 | A | G | 0.986 | -7977.388 | -7979.166 | -1.778 | 6.54E-07 |
| 11 | 119040463 | rs7483350 | G | C | 1.017 | -7977.388 | -7979.166 | -1.778 | 6.54E-07 |
| 11 | 119575104 | rs2006949 | C | T | 1.015 | -7977.388 | -7979.166 | -1.778 | 6.54E-07 |
| 11 | 119682999 | rs2121577 | G | A | 0.984 | -7977.388 | -7979.167 | -1.778 | 6.54E-07 |
| 11 | 119683031 | rs2121576 | G | A | 0.984 | -7977.388 | -7979.167 | -1.778 | 6.54E-07 |
| 11 | 119683964 | rs7937319 | T | C | 0.984 | -7977.388 | -7979.167 | -1.778 | 6.54E-07 |
| 11 | 121142126 | rs7952199 | G | C | 1.007 | -7977.388 | -7979.167 | -1.778 | 6.54E-07 |
| 11 | 121147301 | rs10790429 | A | T | 1.007 | -7977.388 | -7979.167 | -1.778 | 6.54E-07 |
| 11 | 116572525 | rs1240783 | C | T | 0.984 | -7977.388 | -7979.168 | -1.779 | 6.53E-07 |
| 11 | 118004863 | rs45584835 | G | A | 0.985 | -7977.388 | -7979.167 | -1.779 | 6.53E-07 |
| 11 | 119684565 | rs4938727 | C | T | 0.984 | -7977.388 | -7979.167 | -1.779 | 6.53E-07 |
| 11 | 120288944 | rs10892574 | C | G | 1.015 | -7977.388 | -7979.168 | -1.779 | 6.53E-07 |

| | | | | | | | |
|---|---|---|---|---|---|---|---|
| 11 | 120641809 | rs2846092 | A | G | NULL | 0.708890173 | NULL&M_Disease |
| 11 | 120710628 | rs11218025 | C | T | NULL | 0.708890173 | NULL&M_Disease |
| 11 | 116310757 | rs560320 | T | A | NULL | 0.708890173 | NULL&M_Disease |
| 11 | 117986152 | rs7925897 | C | A | NULL | 0.708890173 | NULL&M_Disease |
| 11 | 119358835 | rs625790 | C | T | NULL | 0.708890173 | NULL&M_Disease |
| 11 | 119479922 | rs4936488 | A | G | NULL | 0.708890173 | NULL&M_Disease |
| 11 | 119607659 | rs12786505 | A | G | NULL | 0.708890173 | NULL&M_Disease |
| 11 | 119683218 | rs2121575 | C | G | NULL | 0.708890173 | NULL&M_Disease |
| 11 | 119684646 | rs4938728 | T | C | NULL | 0.708890173 | NULL&M_Disease |
| 11 | 119903704 | rs1787023 | T | C | NULL | 0.708890173 | NULL&M_Disease |
| 11 | 120709874 | rs12223620 | G | T | NULL | 0.708890173 | NULL&M_Disease |
| 11 | 120831520 | rs493678 | T | C | NULL | 0.708890173 | NULL&M_Disease |
| 11 | 121139395 | rs10790425 | T | C | NULL | 0.708890173 | NULL&M_Disease |
| 11 | 121153921 | rs6589870 | G | A | NULL | 0.708890173 | NULL&M_Disease |
| 11 | 121154721 | rs10750183 | A | G | NULL | 0.708890173 | NULL&M_Disease |
| 11 | 121154941 | rs1583124 | G | A | NULL | 0.708890173 | NULL&M_Disease |
| 11 | 116272650 | rs4938269 | C | G | NULL | 0.708890173 | NULL&M_Disease |
| 11 | 116419981 | rs7947659 | C | A | NULL | 0.708890173 | NULL&M_Disease |
| 11 | 116575229 | rs1268832 | T | G | NULL | 0.708890173 | NULL&M_Disease |
| 11 | 118020568 | rs586053 | A | G | NULL | 0.708890173 | NULL&M_Disease |
| 11 | 118572021 | rs657539 | T | C | NULL | 0.708890173 | NULL&M_Disease |
| 11 | 119381784 | rs10892377 | A | G | NULL | 0.708890173 | NULL&M_Disease |
| 11 | 119700001 | rs579200 | G | T | NULL | 0.708890173 | NULL&M_Disease |
| 11 | 119879425 | rs648374 | C | T | NULL | 0.708890173 | NULL&M_Disease |
| 11 | 120190997 | rs11217816 | G | A | NULL | 0.708890173 | NULL&M_Disease |
| 11 | 120709008 | rs11218022 | A | G | NULL | 0.708890173 | NULL&M_Disease |
| 11 | 116227687 | rs482795 | G | A | NULL | 0.708890173 | NULL&M_Disease |
| 11 | 116575555 | rs1263058 | A | G | NULL | 0.708890173 | NULL&M_Disease |
| 11 | 116579099 | rs1262953 | A | G | NULL | 0.708890173 | NULL&M_Disease |
| 11 | 116668867 | rs7101592 | A | T | NULL | 0.708890173 | NULL&M_Disease |
| 11 | 117737359 | rs7101835 | G | A | NULL | 0.708890173 | NULL&M_Disease |
| 11 | 118486325 | rs12290522 | T | C | NULL | 0.708890173 | NULL&M_Disease |
| 11 | 119052826 | rs4245191 | C | A | NULL | 0.708890173 | NULL&M_Disease |
| 11 | 119713257 | rs471909 | T | A | NULL | 0.708890173 | NULL&M_Disease |
| 11 | 119947402 | rs664320 | A | G | NULL | 0.708890173 | NULL&M_Disease |
| 11 | 120273353 | rs4245212 | G | A | NULL | 0.708890173 | NULL&M_Disease |
| 11 | 120293490 | rs6589813 | G | A | NULL | 0.708890173 | NULL&M_Disease |
| 11 | 116344404 | rs17119743 | G | T | NULL | 0.708890173 | NULL&M_Disease |
| 11 | 117201749 | rs609663 | G | C | NULL | 0.708890173 | NULL&M_Disease |
| 11 | 118005622 | rs45460396 | G | T | NULL | 0.708890173 | NULL&M_Disease |
| 11 | 118176623 | rs56785719 | G | C | NULL | 0.708890173 | NULL&M_Disease |
| 11 | 119688712 | rs4245208 | C | T | NULL | 0.708890173 | NULL&M_Disease |
| 11 | 119693206 | rs557773 | C | T | NULL | 0.708890173 | NULL&M_Disease |
| 11 | 120689035 | rs6589845 | T | A | NULL | 0.708890173 | NULL&M_Disease |
| 11 | 121140526 | rs7126048 | G | A | NULL | 0.708890173 | NULL&M_Disease |
| 11 | 118012164 | rs656445 | A | G | NULL | 0.708890173 | NULL&M_Disease |
| 11 | 119478241 | rs4328225 | T | A | NULL | 0.708890173 | NULL&M_Disease |
| 11 | 119571342 | rs7924573 | C | G | NULL | 0.708890173 | NULL&M_Disease |
| 11 | 119579346 | rs10892438 | T | A | NULL | 0.708890173 | NULL&M_Disease |
| 11 | 119684574 | rs2368938 | C | T | NULL | 0.708890173 | NULL&M_Disease |
| 11 | 120481866 | rs12278678 | A | G | NULL | 0.708890173 | NULL&M_Disease |
| 11 | 117484392 | rs10892143 | C | G | NULL | 0.708890173 | NULL&M_Disease |
| 11 | 117839187 | rs11216657 | C | A | NULL | 0.708890173 | NULL&M_Disease |
| 11 | 118131813 | rs56083337 | T | A | NULL | 0.708890173 | NULL&M_Disease |
| 11 | 119749018 | rs35430963 | T | A | NULL | 0.708890173 | NULL&M_Disease |
| 11 | 121058189 | rs4280006 | C | T | NULL | 0.708890173 | NULL&M_Disease |
| 11 | 116426886 | rs906542 | G | A | NULL | 0.708890173 | NULL&M_Disease |
| 11 | 117201997 | rs563838 | T | C | NULL | 0.708890173 | NULL&M_Disease |
| 11 | 118005119 | rs868344 | C | T | NULL | 0.708890173 | NULL&M_Disease |
| 11 | 118492591 | rs2077173 | A | G | NULL | 0.708890173 | NULL&M_Disease |
| 11 | 118496066 | rs11603616 | A | G | NULL | 0.708890173 | NULL&M_Disease |
| 11 | 119684789 | rs2368939 | T | C | NULL | 0.708890173 | NULL&M_Disease |
| 11 | 119712224 | rs552040 | T | G | NULL | 0.708890173 | NULL&M_Disease |
| 11 | 119825798 | rs1350280779 | A | C | NULL | 0.708890173 | NULL&M_Disease |
| 11 | 119883753 | rs602168 | T | C | NULL | 0.708890173 | NULL&M_Disease |
| 11 | 120624732 | rs3133225 | A | G | NULL | 0.708890173 | NULL&M_Disease |
| 11 | 121037529 | rs2186747 | G | C | NULL | 0.708890173 | NULL&M_Disease |
| 11 | 121165092 | rs891441 | G | T | NULL | 0.708890173 | NULL&M_Disease |
| 11 | 116466894 | rs11216032 | A | G | NULL | 0.708890173 | NULL&M_Disease |
| 11 | 116568382 | rs1787689 | G | A | NULL | 0.708890173 | NULL&M_Disease |
| 11 | 116624703 | rs180326 | G | T | NULL | 0.708890173 | NULL&M_Disease |
| 11 | 117317593 | rs58777850 | C | T | NULL | 0.708890173 | NULL&M_Disease |
| 11 | 117533949 | rs12279593 | A | G | NULL | 0.708890173 | NULL&M_Disease |
| 11 | 117849268 | rs12364374 | A | G | NULL | 0.708890173 | NULL&M_Disease |
| 11 | 118522179 | rs1171461 | A | G | NULL | 0.708890173 | NULL&M_Disease |
| 11 | 119357305 | rs653106 | G | A | NULL | 0.708890173 | NULL&M_Disease |
| 11 | 116469461 | rs73011187 | T | C | NULL | 0.708890173 | NULL&M_Disease |
| 11 | 117357696 | rs4938389 | A | G | NULL | 0.709920918 | NULL&M_Disease |
| 11 | 117839307 | rs4372482 | C | T | NULL | 0.709920918 | NULL&M_Disease |
| 11 | 118497402 | rs12419204 | A | T | NULL | 0.709920918 | NULL&M_Disease |
| 11 | 118500173 | rs571001 | T | C | NULL | 0.709920918 | NULL&M_Disease |
| 11 | 119478022 | rs10892420 | G | A | NULL | 0.709920918 | NULL&M_Disease |
| 11 | 119883723 | rs621843 | A | G | NULL | 0.709920918 | NULL&M_Disease |
| 11 | 119883781 | rs668361 | A | G | NULL | 0.709920918 | NULL&M_Disease |
| 11 | 120592036 | rs3133226 | C | T | NULL | 0.709920918 | NULL&M_Disease |
| 11 | 120989780 | rs657405 | T | C | NULL | 0.709920918 | NULL&M_Disease |
| 11 | 120990070 | rs626517 | C | G | NULL | 0.709920918 | NULL&M_Disease |
| 11 | 117748844 | rs11605223 | G | C | NULL | 0.709920918 | NULL&M_Disease |
| 11 | 119989107 | rs10466595 | G | C | NULL | 0.709920918 | NULL&M_Disease |
| 11 | 120696839 | rs4317991 | A | G | NULL | 0.709920918 | NULL&M_Disease |
| 11 | 121040910 | rs1939935 | C | T | NULL | 0.709920918 | NULL&M_Disease |
| 11 | 118500193 | rs570952 | C | T | NULL | 0.709920918 | NULL&M_Disease |
| 11 | 118571790 | rs656287 | G | A | NULL | 0.709920918 | NULL&M_Disease |
| 11 | 119951702 | rs12362262 | T | C | NULL | 0.709920918 | NULL&M_Disease |
| 11 | 120559480 | rs6589832 | G | A | NULL | 0.709920918 | NULL&M_Disease |
| 11 | 121053640 | rs11218190 | A | G | NULL | 0.709920918 | NULL&M_Disease |

| | | | | | | | | | |
|---|---|---|---|---|---|---|---|---|---|
| 11 | 120641809 | rs2846092 | A | G | 0.99 | -7977.388 | -7979.168 | -1.779 | 6.53E-07 |
| 11 | 120710628 | rs11218025 | C | T | 0.985 | -7977.388 | -7979.167 | -1.779 | 6.53E-07 |
| 11 | 116310757 | rs560320 | T | A | 1.016 | -7977.388 | -7979.169 | -1.78 | 6.52E-07 |
| 11 | 117986152 | rs7925897 | C | A | 0.986 | -7977.388 | -7979.168 | -1.78 | 6.52E-07 |
| 11 | 119358835 | rs625790 | C | T | 0.985 | -7977.388 | -7979.168 | -1.78 | 6.52E-07 |
| 11 | 119479922 | rs4936488 | A | G | 0.987 | -7977.388 | -7979.168 | -1.78 | 6.52E-07 |
| 11 | 119607659 | rs12786505 | A | G | 0.984 | -7977.388 | -7979.169 | -1.78 | 6.52E-07 |
| 11 | 119683218 | rs2121575 | C | G | 0.984 | -7977.388 | -7979.169 | -1.78 | 6.52E-07 |
| 11 | 119684646 | rs4938728 | T | C | 0.984 | -7977.388 | -7979.168 | -1.78 | 6.52E-07 |
| 11 | 119903704 | rs1787023 | T | C | 1.015 | -7977.388 | -7979.168 | -1.78 | 6.52E-07 |
| 11 | 120709874 | rs12223620 | G | T | 0.985 | -7977.388 | -7979.169 | -1.78 | 6.52E-07 |
| 11 | 120831520 | rs493678 | T | C | 0.989 | -7977.388 | -7979.169 | -1.78 | 6.52E-07 |
| 11 | 121139395 | rs10790425 | T | C | 1.007 | -7977.388 | -7979.168 | -1.78 | 6.52E-07 |
| 11 | 121153921 | rs6589870 | G | A | 1.011 | -7977.388 | -7979.168 | -1.78 | 6.52E-07 |
| 11 | 121154721 | rs10750183 | A | G | 1.011 | -7977.388 | -7979.168 | -1.78 | 6.52E-07 |
| 11 | 121154941 | rs1583124 | G | A | 1.011 | -7977.388 | -7979.168 | -1.78 | 6.52E-07 |
| 11 | 116272650 | rs4938269 | C | G | 1.01 | -7977.388 | -7979.169 | -1.781 | 6.52E-07 |
| 11 | 116419981 | rs7947659 | C | A | 0.992 | -7977.388 | -7979.169 | -1.781 | 6.52E-07 |
| 11 | 116575229 | rs1268832 | T | G | 0.984 | -7977.388 | -7979.169 | -1.781 | 6.52E-07 |
| 11 | 118020568 | rs586053 | A | G | 1.016 | -7977.388 | -7979.169 | -1.781 | 6.52E-07 |
| 11 | 118572021 | rs657539 | T | C | 0.987 | -7977.388 | -7979.169 | -1.781 | 6.52E-07 |
| 11 | 119381784 | rs10892377 | A | G | 1.016 | -7977.388 | -7979.169 | -1.781 | 6.52E-07 |
| 11 | 119700001 | rs579200 | G | T | 0.996 | -7977.388 | -7979.169 | -1.781 | 6.52E-07 |
| 11 | 119879425 | rs648374 | C | T | 1.016 | -7977.388 | -7979.169 | -1.781 | 6.52E-07 |
| 11 | 120190997 | rs11217816 | G | A | 1.014 | -7977.388 | -7979.17 | -1.781 | 6.52E-07 |
| 11 | 120709008 | rs11218022 | A | G | 0.985 | -7977.388 | -7979.169 | -1.781 | 6.52E-07 |
| 11 | 116227687 | rs482795 | G | A | 1.009 | -7977.388 | -7979.17 | -1.782 | 6.51E-07 |
| 11 | 116575555 | rs1263058 | A | G | 0.985 | -7977.388 | -7979.17 | -1.782 | 6.51E-07 |
| 11 | 116579099 | rs1262953 | A | G | 0.99 | -7977.388 | -7979.17 | -1.782 | 6.51E-07 |
| 11 | 116668867 | rs7101592 | A | T | 1.004 | -7977.388 | -7979.17 | -1.782 | 6.51E-07 |
| 11 | 117737359 | rs7101835 | G | A | 0.985 | -7977.388 | -7979.171 | -1.782 | 6.51E-07 |
| 11 | 118486325 | rs12290522 | T | C | 1.014 | -7977.388 | -7979.17 | -1.782 | 6.51E-07 |
| 11 | 119052826 | rs4245191 | C | A | 1.017 | -7977.388 | -7979.171 | -1.782 | 6.51E-07 |
| 11 | 119713257 | rs471909 | T | A | 1.011 | -7977.388 | -7979.171 | -1.782 | 6.51E-07 |
| 11 | 119947402 | rs664320 | A | G | 1.005 | -7977.388 | -7979.17 | -1.782 | 6.51E-07 |
| 11 | 120273353 | rs4245212 | G | A | 0.984 | -7977.388 | -7979.171 | -1.782 | 6.51E-07 |
| 11 | 120293490 | rs6589813 | G | A | 1.015 | -7977.388 | -7979.17 | -1.782 | 6.51E-07 |
| 11 | 116344404 | rs17119743 | G | T | 1 | -7977.388 | -7979.171 | -1.783 | 6.50E-07 |
| 11 | 117201749 | rs609663 | G | C | 0.997 | -7977.388 | -7979.171 | -1.783 | 6.50E-07 |
| 11 | 118005622 | rs45460396 | G | T | 0.986 | -7977.388 | -7979.172 | -1.783 | 6.50E-07 |
| 11 | 118176623 | rs56785719 | G | C | 0.987 | -7977.388 | -7979.171 | -1.783 | 6.50E-07 |
| 11 | 119688712 | rs4245208 | C | T | 0.984 | -7977.388 | -7979.171 | -1.783 | 6.50E-07 |
| 11 | 119693206 | rs557773 | C | T | 0.984 | -7977.388 | -7979.172 | -1.783 | 6.50E-07 |
| 11 | 120689035 | rs6589845 | T | A | 0.984 | -7977.388 | -7979.171 | -1.783 | 6.50E-07 |
| 11 | 121140526 | rs7126048 | G | A | 1.007 | -7977.388 | -7979.171 | -1.783 | 6.50E-07 |
| 11 | 118012164 | rs656445 | A | G | 0.986 | -7977.388 | -7979.172 | -1.784 | 6.50E-07 |
| 11 | 119478241 | rs4328225 | T | A | 0.987 | -7977.388 | -7979.173 | -1.784 | 6.50E-07 |
| 11 | 119571342 | rs7924573 | C | G | 1.015 | -7977.388 | -7979.173 | -1.784 | 6.50E-07 |
| 11 | 119579346 | rs10892438 | T | A | 1.016 | -7977.388 | -7979.172 | -1.784 | 6.50E-07 |
| 11 | 119684574 | rs2368938 | C | T | 0.984 | -7977.388 | -7979.172 | -1.784 | 6.50E-07 |
| 11 | 120481866 | rs12278678 | A | G | 0.985 | -7977.388 | -7979.173 | -1.784 | 6.50E-07 |
| 11 | 117484392 | rs10892143 | C | G | 0.984 | -7977.388 | -7979.173 | -1.785 | 6.49E-07 |
| 11 | 117839187 | rs11216657 | C | A | 0.985 | -7977.388 | -7979.174 | -1.785 | 6.49E-07 |
| 11 | 118131813 | rs56083337 | T | A | 0.984 | -7977.388 | -7979.174 | -1.785 | 6.49E-07 |
| 11 | 119749018 | rs35430963 | T | A | 0.985 | -7977.388 | -7979.173 | -1.785 | 6.49E-07 |
| 11 | 121058189 | rs4280006 | C | T | 1.01 | -7977.388 | -7979.173 | -1.785 | 6.49E-07 |
| 11 | 116426886 | rs906542 | G | A | 0.984 | -7977.388 | -7979.174 | -1.786 | 6.48E-07 |
| 11 | 117201997 | rs563838 | T | C | 0.985 | -7977.388 | -7979.174 | -1.786 | 6.48E-07 |
| 11 | 118005119 | rs868344 | C | T | 0.986 | -7977.388 | -7979.175 | -1.786 | 6.48E-07 |
| 11 | 118492591 | rs2077173 | A | G | 1.013 | -7977.388 | -7979.174 | -1.786 | 6.48E-07 |
| 11 | 118496066 | rs11603616 | A | G | 1.013 | -7977.388 | -7979.175 | -1.786 | 6.48E-07 |
| 11 | 119684789 | rs2368939 | T | C | 0.984 | -7977.388 | -7979.175 | -1.786 | 6.48E-07 |
| 11 | 119712224 | rs552040 | T | G | 1.01 | -7977.388 | -7979.174 | -1.786 | 6.48E-07 |
| 11 | 119825798 | rs1350280779 | A | C | 0.985 | -7977.388 | -7979.174 | -1.786 | 6.48E-07 |
| 11 | 119883753 | rs602168 | T | C | 0.985 | -7977.388 | -7979.174 | -1.786 | 6.48E-07 |
| 11 | 120624732 | rs3133225 | A | G | 0.991 | -7977.388 | -7979.175 | -1.786 | 6.48E-07 |
| 11 | 121037529 | rs2186747 | G | C | 1.009 | -7977.388 | -7979.174 | -1.786 | 6.48E-07 |
| 11 | 121165092 | rs891441 | G | T | 1.016 | -7977.388 | -7979.175 | -1.786 | 6.48E-07 |
| 11 | 116466894 | rs11216032 | A | G | 1.012 | -7977.388 | -7979.175 | -1.787 | 6.48E-07 |
| 11 | 116568382 | rs1787689 | G | A | 0.984 | -7977.388 | -7979.175 | -1.787 | 6.48E-07 |
| 11 | 116624703 | rs180326 | G | T | 1.014 | -7977.388 | -7979.175 | -1.787 | 6.48E-07 |
| 11 | 117317593 | rs58777850 | C | T | 0.987 | -7977.388 | -7979.175 | -1.787 | 6.48E-07 |
| 11 | 117533949 | rs12279593 | A | G | 0.988 | -7977.388 | -7979.175 | -1.787 | 6.48E-07 |
| 11 | 117849268 | rs12364374 | A | G | 0.987 | -7977.388 | -7979.175 | -1.787 | 6.48E-07 |
| 11 | 118522179 | rs1171461 | A | G | 1.013 | -7977.388 | -7979.176 | -1.787 | 6.48E-07 |
| 11 | 119357305 | rs653106 | G | A | 1.015 | -7977.388 | -7979.175 | -1.787 | 6.48E-07 |
| 11 | 116469461 | rs73011187 | T | C | 1.011 | -7977.388 | -7979.176 | -1.788 | 6.47E-07 |
| 11 | 117357696 | rs4938389 | A | G | 1.014 | -7977.388 | -7979.176 | -1.788 | 6.47E-07 |
| 11 | 117839307 | rs4372482 | C | T | 0.985 | -7977.388 | -7979.177 | -1.788 | 6.47E-07 |
| 11 | 118497402 | rs12419204 | A | T | 1.013 | -7977.388 | -7979.176 | -1.788 | 6.47E-07 |
| 11 | 118500173 | rs571001 | T | C | 1.013 | -7977.388 | -7979.176 | -1.788 | 6.47E-07 |
| 11 | 119478022 | rs10892420 | G | A | 0.988 | -7977.388 | -7979.177 | -1.788 | 6.47E-07 |
| 11 | 119883723 | rs621843 | A | G | 0.985 | -7977.388 | -7979.176 | -1.788 | 6.47E-07 |
| 11 | 119883781 | rs668361 | A | G | 0.985 | -7977.388 | -7979.176 | -1.788 | 6.47E-07 |
| 11 | 120592036 | rs3133226 | C | T | 0.984 | -7977.388 | -7979.176 | -1.788 | 6.47E-07 |
| 11 | 120989780 | rs657405 | T | C | 1.012 | -7977.388 | -7979.176 | -1.788 | 6.47E-07 |
| 11 | 120990070 | rs626517 | C | G | 1.012 | -7977.388 | -7979.176 | -1.788 | 6.47E-07 |
| 11 | 117748844 | rs11605223 | G | C | 0.984 | -7977.388 | -7979.177 | -1.789 | 6.46E-07 |
| 11 | 119989107 | rs10466595 | G | C | 1.011 | -7977.388 | -7979.177 | -1.789 | 6.46E-07 |
| 11 | 120696839 | rs4317991 | A | G | 0.986 | -7977.388 | -7979.178 | -1.789 | 6.46E-07 |
| 11 | 121040910 | rs1939935 | C | T | 1.009 | -7977.388 | -7979.177 | -1.789 | 6.46E-07 |
| 11 | 118500193 | rs570952 | C | T | 1.013 | -7977.388 | -7979.178 | -1.79 | 6.46E-07 |
| 11 | 118571790 | rs656287 | G | A | 0.988 | -7977.388 | -7979.178 | -1.79 | 6.46E-07 |
| 11 | 119951702 | rs12362262 | T | C | 0.987 | -7977.388 | -7979.179 | -1.79 | 6.46E-07 |
| 11 | 120559480 | rs6589832 | G | A | 1.016 | -7977.388 | -7979.178 | -1.79 | 6.46E-07 |
| 11 | 121053640 | rs11218190 | A | G | 1.009 | -7977.388 | -7979.179 | -1.79 | 6.46E-07 |

| | | | | | | | |
|---|---|---|---|---|---|---|---|
| 11 | 116338899 | rs1508098 | G | A | NULL | 0.709920918 | NULL&M_Disease |
| 11 | 117970689 | rs11216748 | A | G | NULL | 0.709920918 | NULL&M_Disease |
| 11 | 119453770 | rs2368427 | C | T | NULL | 0.709920918 | NULL&M_Disease |
| 11 | 120602594 | rs2852228 | A | T | NULL | 0.709920918 | NULL&M_Disease |
| 11 | 121139585 | rs10790427 | G | A | NULL | 0.709920918 | NULL&M_Disease |
| 11 | 117845068 | rs4938466 | G | A | NULL | 0.709920918 | NULL&M_Disease |
| 11 | 120097498 | rs873059 | T | C | NULL | 0.709920918 | NULL&M_Disease |
| 11 | 117736735 | rs2155196 | T | C | NULL | 0.709920918 | NULL&M_Disease |
| 11 | 117736770 | rs1892937 | C | T | NULL | 0.709920918 | NULL&M_Disease |
| 11 | 118571357 | rs544452 | T | G | NULL | 0.709920918 | NULL&M_Disease |
| 11 | 118571911 | rs570949 | T | G | NULL | 0.709920918 | NULL&M_Disease |
| 11 | 118572171 | rs658040 | G | A | NULL | 0.709920918 | NULL&M_Disease |
| 11 | 118572611 | rs611217 | T | C | NULL | 0.709920918 | NULL&M_Disease |
| 11 | 119499014 | rs4582984 | C | A | NULL | 0.709920918 | NULL&M_Disease |
| 11 | 121049690 | rs7129267 | G | T | NULL | 0.709920918 | NULL&M_Disease |
| 11 | 116413688 | rs925152 | T | G | NULL | 0.709920918 | NULL&M_Disease |
| 11 | 117696720 | rs514527 | G | A | NULL | 0.709920918 | NULL&M_Disease |
| 11 | 117842292 | rs12790242 | A | G | NULL | 0.709920918 | NULL&M_Disease |
| 11 | 117970696 | rs11216749 | C | T | NULL | 0.709920918 | NULL&M_Disease |
| 11 | 119879255 | rs609066 | G | A | NULL | 0.709920918 | NULL&M_Disease |
| 11 | 120713654 | rs4482033 | G | A | NULL | 0.709920918 | NULL&M_Disease |
| 11 | 117316321 | rs4936378 | A | G | NULL | 0.709920918 | NULL&M_Disease |
| 11 | 117324521 | rs11605475 | A | G | NULL | 0.709920918 | NULL&M_Disease |
| 11 | 117736790 | rs2155197 | G | A | NULL | 0.709920918 | NULL&M_Disease |
| 11 | 118570914 | rs659564 | A | C | NULL | 0.709920918 | NULL&M_Disease |
| 11 | 118570919 | rs642224 | A | G | NULL | 0.709920918 | NULL&M_Disease |
| 11 | 118571023 | rs541631 | G | A | NULL | 0.709920918 | NULL&M_Disease |
| 11 | 118571123 | rs658676 | T | C | NULL | 0.709920918 | NULL&M_Disease |
| 11 | 118571168 | rs488141 | G | A | NULL | 0.709920918 | NULL&M_Disease |
| 11 | 118571316 | rs657769 | T | C | NULL | 0.709920918 | NULL&M_Disease |
| 11 | 118572747 | rs489126 | A | G | NULL | 0.709920918 | NULL&M_Disease |
| 11 | 118572760 | rs611733 | T | C | NULL | 0.709920918 | NULL&M_Disease |
| 11 | 118573022 | rs552079 | C | T | NULL | 0.709920918 | NULL&M_Disease |
| 11 | 118573339 | rs673547 | A | G | NULL | 0.709920918 | NULL&M_Disease |
| 11 | 118573535 | rs517445 | G | A | NULL | 0.709920918 | NULL&M_Disease |
| 11 | 119497488 | rs7116091 | C | T | NULL | 0.709920918 | NULL&M_Disease |
| 11 | 119498477 | rs10790324 | A | G | NULL | 0.709920918 | NULL&M_Disease |
| 11 | 119498618 | rs10750161 | A | G | NULL | 0.709920918 | NULL&M_Disease |
| 11 | 119945567 | rs10790365 | T | C | NULL | 0.709920918 | NULL&M_Disease |
| 11 | 120206163 | rs3809059 | A | T | NULL | 0.709920918 | NULL&M_Disease |
| 11 | 121007374 | rs663319 | G | A | NULL | 0.709920918 | NULL&M_Disease |
| 11 | 121139566 | rs10790426 | C | T | NULL | 0.709920918 | NULL&M_Disease |
| 11 | 116462788 | rs12420596 | A | G | NULL | 0.709920918 | NULL&M_Disease |
| 11 | 117690918 | rs11999 | C | A | NULL | 0.709920918 | NULL&M_Disease |
| 11 | 118569469 | rs7123811 | G | T | NULL | 0.709920918 | NULL&M_Disease |
| 11 | 119263703 | rs11217267 | C | T | NULL | 0.709920918 | NULL&M_Disease |
| 11 | 119498128 | rs10790323 | A | G | NULL | 0.709920918 | NULL&M_Disease |
| 11 | 119962278 | rs683743 | C | G | NULL | 0.709920918 | NULL&M_Disease |
| 11 | 120717153 | rs7121233 | C | T | NULL | 0.709920918 | NULL&M_Disease |
| 11 | 116422860 | rs12362410 | A | G | NULL | 0.709920918 | NULL&M_Disease |
| 11 | 117702690 | rs10790212 | T | C | NULL | 0.709920918 | NULL&M_Disease |
| 11 | 118767077 | rs581063 | C | T | NULL | 0.709920918 | NULL&M_Disease |
| 11 | 119468889 | rs11217344 | A | G | NULL | 0.709920918 | NULL&M_Disease |
| 11 | 119497757 | rs11217357 | T | G | NULL | 0.709920918 | NULL&M_Disease |
| 11 | 119497806 | rs11217358 | A | G | NULL | 0.709920918 | NULL&M_Disease |
| 11 | 117662909 | rs528431 | A | C | NULL | 0.710949503 | NULL&M_Disease |
| 11 | 117772848 | rs491822 | T | C | NULL | 0.710949503 | NULL&M_Disease |
| 11 | 118569694 | rs7108144 | A | G | NULL | 0.710949503 | NULL&M_Disease |
| 11 | 118569900 | rs12793594 | C | T | NULL | 0.710949503 | NULL&M_Disease |
| 11 | 118570062 | rs554456 | C | T | NULL | 0.710949503 | NULL&M_Disease |
| 11 | 118570098 | rs482889 | T | G | NULL | 0.710949503 | NULL&M_Disease |
| 11 | 118570697 | rs523793 | T | C | NULL | 0.710949503 | NULL&M_Disease |
| 11 | 119961694 | rs617320 | G | T | NULL | 0.710949503 | NULL&M_Disease |
| 11 | 120398613 | rs58418986 | A | G | NULL | 0.710949503 | NULL&M_Disease |
| 11 | 117973218 | rs4938482 | G | T | NULL | 0.710949503 | NULL&M_Disease |
| 11 | 119682612 | rs4936499 | C | T | NULL | 0.710949503 | NULL&M_Disease |
| 11 | 119716304 | rs514649 | G | A | NULL | 0.710949503 | NULL&M_Disease |
| 11 | 119934058 | rs671334 | A | G | NULL | 0.710949503 | NULL&M_Disease |
| 11 | 120488347 | rs10892588 | A | G | NULL | 0.710949503 | NULL&M_Disease |
| 11 | 120891614 | rs34929374 | G | A | NULL | 0.710949503 | NULL&M_Disease |
| 11 | 116421659 | rs884603 | T | C | NULL | 0.710949503 | NULL&M_Disease |
| 11 | 118066104 | rs590580 | A | G | NULL | 0.710949503 | NULL&M_Disease |
| 11 | 119291445 | rs3138092 | C | T | NULL | 0.710949503 | NULL&M_Disease |
| 11 | 120012390 | rs470411 | G | C | NULL | 0.710949503 | NULL&M_Disease |
| 11 | 118812512 | rs647769 | T | C | NULL | 0.710949503 | NULL&M_Disease |
| 11 | 119682874 | rs2166658 | T | C | NULL | 0.710949503 | NULL&M_Disease |
| 11 | 120348444 | rs7940963 | A | G | NULL | 0.710949503 | NULL&M_Disease |
| 11 | 120618079 | rs1939673 | C | T | NULL | 0.710949503 | NULL&M_Disease |
| 11 | 120708690 | rs4935754 | G | A | NULL | 0.710949503 | NULL&M_Disease |
| 11 | 121152173 | rs12419633 | A | G | NULL | 0.710949503 | NULL&M_Disease |
| 11 | 116272249 | rs1295453503 | T | C | NULL | 0.710949503 | NULL&M_Disease |
| 11 | 117736022 | rs11216594 | A | G | NULL | 0.710949503 | NULL&M_Disease |
| 11 | 117773893 | rs11216612 | T | C | NULL | 0.710949503 | NULL&M_Disease |
| 11 | 118182829 | rs1945765 | A | T | NULL | 0.710949503 | NULL&M_Disease |
| 11 | 118522338 | rs4938520 | G | A | NULL | 0.710949503 | NULL&M_Disease |
| 11 | 118568639 | rs511451 | T | C | NULL | 0.710949503 | NULL&M_Disease |
| 11 | 119668265 | rs4315086 | A | G | NULL | 0.710949503 | NULL&M_Disease |
| 11 | 120398707 | rs59912111 | A | C | NULL | 0.710949503 | NULL&M_Disease |
| 11 | 116400897 | rs17491207 | A | G | NULL | 0.710949503 | NULL&M_Disease |
| 11 | 116412160 | rs12802920 | C | A | NULL | 0.710949503 | NULL&M_Disease |
| 11 | 119685493 | rs4936501 | A | G | NULL | 0.710949503 | NULL&M_Disease |
| 11 | 119685596 | rs4936502 | T | C | NULL | 0.710949503 | NULL&M_Disease |
| 11 | 119686250 | rs11217520 | A | G | NULL | 0.710949503 | NULL&M_Disease |
| 11 | 119934081 | rs680630 | T | G | NULL | 0.710949503 | NULL&M_Disease |
| 11 | 120008957 | rs1790476 | G | C | NULL | 0.710949503 | NULL&M_Disease |
| 11 | 116215109 | rs11601506 | T | G | NULL | 0.710949503 | NULL&M_Disease |

| | | | | | | | | | |
|---|---|---|---|---|---|---|---|---|---|
| 11 | 116338899 | rs1508098 | G | A | 0.989 | -7977.388 | -7979.179 | -1.791 | 6.45E-07 |
| 11 | 117970689 | rs11216748 | A | G | 0.987 | -7977.388 | -7979.179 | -1.791 | 6.45E-07 |
| 11 | 119453770 | rs2368427 | C | T | 1.011 | -7977.388 | -7979.179 | -1.791 | 6.45E-07 |
| 11 | 120602594 | rs2852228 | A | T | 0.998 | -7977.388 | -7979.18 | -1.791 | 6.45E-07 |
| 11 | 121139585 | rs10790427 | G | A | 1.005 | -7977.388 | -7979.179 | -1.791 | 6.45E-07 |
| 11 | 117845068 | rs4938466 | G | A | 0.985 | -7977.388 | -7979.181 | -1.792 | 6.45E-07 |
| 11 | 120097498 | rs873059 | T | C | 1.009 | -7977.388 | -7979.18 | -1.792 | 6.45E-07 |
| 11 | 117736735 | rs2155196 | T | C | 0.986 | -7977.388 | -7979.181 | -1.793 | 6.44E-07 |
| 11 | 117736770 | rs1892937 | C | T | 0.986 | -7977.388 | -7979.182 | -1.793 | 6.44E-07 |
| 11 | 118571357 | rs544452 | T | G | 0.988 | -7977.388 | -7979.181 | -1.793 | 6.44E-07 |
| 11 | 118571911 | rs570949 | T | G | 0.988 | -7977.388 | -7979.181 | -1.793 | 6.44E-07 |
| 11 | 118572171 | rs658040 | G | A | 0.988 | -7977.388 | -7979.181 | -1.793 | 6.44E-07 |
| 11 | 118572611 | rs611217 | T | C | 0.988 | -7977.388 | -7979.182 | -1.793 | 6.44E-07 |
| 11 | 119499014 | rs4582984 | C | A | 0.996 | -7977.388 | -7979.182 | -1.793 | 6.44E-07 |
| 11 | 121049690 | rs7129267 | G | T | 1.009 | -7977.388 | -7979.181 | -1.793 | 6.44E-07 |
| 11 | 116413688 | rs925152 | T | G | 0.994 | -7977.388 | -7979.182 | -1.794 | 6.43E-07 |
| 11 | 117696720 | rs514527 | G | A | 0.999 | -7977.388 | -7979.182 | -1.794 | 6.43E-07 |
| 11 | 117842292 | rs12790242 | A | G | 0.985 | -7977.388 | -7979.182 | -1.794 | 6.43E-07 |
| 11 | 117970696 | rs11216749 | C | T | 0.987 | -7977.388 | -7979.182 | -1.794 | 6.43E-07 |
| 11 | 119879255 | rs609066 | G | A | 1.015 | -7977.388 | -7979.182 | -1.794 | 6.43E-07 |
| 11 | 120713654 | rs4482033 | G | A | 0.986 | -7977.388 | -7979.182 | -1.794 | 6.43E-07 |
| 11 | 117316321 | rs4936378 | A | G | 0.988 | -7977.388 | -7979.184 | -1.795 | 6.43E-07 |
| 11 | 117324521 | rs11605475 | A | G | 1.01 | -7977.388 | -7979.183 | -1.795 | 6.43E-07 |
| 11 | 117736790 | rs2155197 | G | A | 0.986 | -7977.388 | -7979.184 | -1.795 | 6.43E-07 |
| 11 | 118570914 | rs659564 | A | C | 0.988 | -7977.388 | -7979.183 | -1.795 | 6.43E-07 |
| 11 | 118570919 | rs642224 | A | G | 0.988 | -7977.388 | -7979.183 | -1.795 | 6.43E-07 |
| 11 | 118571023 | rs541631 | G | A | 0.988 | -7977.388 | -7979.183 | -1.795 | 6.43E-07 |
| 11 | 118571123 | rs658676 | T | C | 0.988 | -7977.388 | -7979.183 | -1.795 | 6.43E-07 |
| 11 | 118571168 | rs488141 | G | A | 0.988 | -7977.388 | -7979.183 | -1.795 | 6.43E-07 |
| 11 | 118571316 | rs657769 | T | C | 0.988 | -7977.388 | -7979.183 | -1.795 | 6.43E-07 |
| 11 | 118572747 | rs489126 | A | G | 0.988 | -7977.388 | -7979.184 | -1.795 | 6.43E-07 |
| 11 | 118572760 | rs611733 | T | C | 0.988 | -7977.388 | -7979.183 | -1.795 | 6.43E-07 |
| 11 | 118573022 | rs552079 | C | T | 0.988 | -7977.388 | -7979.183 | -1.795 | 6.43E-07 |
| 11 | 118573339 | rs673547 | A | G | 0.988 | -7977.388 | -7979.183 | -1.795 | 6.43E-07 |
| 11 | 118573535 | rs517445 | G | A | 0.988 | -7977.388 | -7979.183 | -1.795 | 6.43E-07 |
| 11 | 119497488 | rs7116091 | C | T | 0.997 | -7977.388 | -7979.184 | -1.795 | 6.43E-07 |
| 11 | 119498477 | rs10790324 | A | G | 0.997 | -7977.388 | -7979.183 | -1.795 | 6.43E-07 |
| 11 | 119498618 | rs10750161 | A | G | 0.997 | -7977.388 | -7979.183 | -1.795 | 6.43E-07 |
| 11 | 119945567 | rs10790365 | T | C | 1.015 | -7977.388 | -7979.183 | -1.795 | 6.43E-07 |
| 11 | 120206163 | rs3809059 | A | T | 0.985 | -7977.388 | -7979.183 | -1.795 | 6.43E-07 |
| 11 | 121007374 | rs663319 | G | A | 1.015 | -7977.388 | -7979.184 | -1.795 | 6.43E-07 |
| 11 | 121139566 | rs10790426 | C | T | 1.004 | -7977.388 | -7979.183 | -1.795 | 6.43E-07 |
| 11 | 116462788 | rs12420596 | A | G | 1.009 | -7977.388 | -7979.184 | -1.796 | 6.42E-07 |
| 11 | 117690918 | rs11999 | C | A | 0.987 | -7977.388 | -7979.185 | -1.796 | 6.42E-07 |
| 11 | 118569469 | rs7123811 | G | T | 0.988 | -7977.388 | -7979.185 | -1.796 | 6.42E-07 |
| 11 | 119263703 | rs11217267 | C | T | 1.011 | -7977.388 | -7979.184 | -1.796 | 6.42E-07 |
| 11 | 119498128 | rs10790323 | A | G | 0.998 | -7977.388 | -7979.185 | -1.796 | 6.42E-07 |
| 11 | 119962278 | rs683743 | C | G | 0.986 | -7977.388 | -7979.185 | -1.796 | 6.42E-07 |
| 11 | 120717153 | rs7121233 | C | T | 0.986 | -7977.388 | -7979.184 | -1.796 | 6.42E-07 |
| 11 | 116422860 | rs12362410 | A | G | 0.998 | -7977.388 | -7979.185 | -1.797 | 6.41E-07 |
| 11 | 117702690 | rs10790212 | T | C | 0.996 | -7977.388 | -7979.185 | -1.797 | 6.41E-07 |
| 11 | 118767077 | rs581063 | C | T | 1.015 | -7977.388 | -7979.185 | -1.797 | 6.41E-07 |
| 11 | 119468889 | rs11217344 | A | G | 0.992 | -7977.388 | -7979.185 | -1.797 | 6.41E-07 |
| 11 | 119497757 | rs11217357 | T | G | 0.998 | -7977.388 | -7979.185 | -1.797 | 6.41E-07 |
| 11 | 119497806 | rs11217358 | A | G | 0.998 | -7977.388 | -7979.185 | -1.797 | 6.41E-07 |
| 11 | 117662909 | rs528431 | A | C | 1.013 | -7977.388 | -7979.186 | -1.798 | 6.41E-07 |
| 11 | 117772848 | rs491822 | T | C | 1.014 | -7977.388 | -7979.186 | -1.798 | 6.41E-07 |
| 11 | 118569694 | rs7108144 | A | G | 0.988 | -7977.388 | -7979.187 | -1.798 | 6.41E-07 |
| 11 | 118569900 | rs12793594 | C | T | 0.988 | -7977.388 | -7979.187 | -1.798 | 6.41E-07 |
| 11 | 118570062 | rs554456 | C | T | 0.988 | -7977.388 | -7979.187 | -1.798 | 6.41E-07 |
| 11 | 118570098 | rs482889 | T | G | 0.988 | -7977.388 | -7979.187 | -1.798 | 6.41E-07 |
| 11 | 118570697 | rs523793 | T | C | 0.988 | -7977.388 | -7979.187 | -1.798 | 6.41E-07 |
| 11 | 119961694 | rs617320 | G | T | 0.986 | -7977.388 | -7979.187 | -1.798 | 6.41E-07 |
| 11 | 120398613 | rs58418986 | A | G | 1.005 | -7977.388 | -7979.187 | -1.798 | 6.41E-07 |
| 11 | 117973218 | rs4938482 | G | T | 1.007 | -7977.388 | -7979.188 | -1.799 | 6.40E-07 |
| 11 | 119682612 | rs4936499 | C | T | 0.985 | -7977.388 | -7979.187 | -1.799 | 6.40E-07 |
| 11 | 119716304 | rs514649 | G | A | 1.014 | -7977.388 | -7979.187 | -1.799 | 6.40E-07 |
| 11 | 119934058 | rs671334 | A | G | 0.993 | -7977.388 | -7979.188 | -1.799 | 6.40E-07 |
| 11 | 120488347 | rs10892588 | A | G | 0.986 | -7977.388 | -7979.187 | -1.799 | 6.40E-07 |
| 11 | 120891614 | rs34929374 | G | A | 0.995 | -7977.388 | -7979.187 | -1.799 | 6.40E-07 |
| 11 | 116421659 | rs884603 | T | C | 0.998 | -7977.388 | -7979.188 | -1.8 | 6.39E-07 |
| 11 | 118066104 | rs590580 | A | G | 0.985 | -7977.388 | -7979.188 | -1.8 | 6.39E-07 |
| 11 | 119291445 | rs3138092 | C | T | 1.007 | -7977.388 | -7979.188 | -1.8 | 6.39E-07 |
| 11 | 120012390 | rs470411 | G | C | 1.003 | -7977.388 | -7979.188 | -1.8 | 6.39E-07 |
| 11 | 118812512 | rs647769 | T | C | 0.986 | -7977.388 | -7979.189 | -1.801 | 6.39E-07 |
| 11 | 119682874 | rs2166658 | T | C | 0.985 | -7977.388 | -7979.189 | -1.801 | 6.39E-07 |
| 11 | 120348444 | rs7940963 | A | G | 0.985 | -7977.388 | -7979.189 | -1.801 | 6.39E-07 |
| 11 | 120618079 | rs1939673 | C | T | 0.986 | -7977.388 | -7979.189 | -1.801 | 6.39E-07 |
| 11 | 120708690 | rs4935754 | G | A | 0.986 | -7977.388 | -7979.19 | -1.801 | 6.39E-07 |
| 11 | 121152173 | rs12419633 | A | G | 0.993 | -7977.388 | -7979.189 | -1.801 | 6.39E-07 |
| 11 | 116272249 | rs1295453503 | T | C | 1 | -7977.388 | -7979.19 | -1.802 | 6.38E-07 |
| 11 | 117736022 | rs11216594 | A | G | 0.987 | -7977.388 | -7979.19 | -1.802 | 6.38E-07 |
| 11 | 117773893 | rs11216612 | T | C | 1.014 | -7977.388 | -7979.19 | -1.802 | 6.38E-07 |
| 11 | 118182829 | rs1945765 | A | T | 0.988 | -7977.388 | -7979.191 | -1.802 | 6.38E-07 |
| 11 | 118522338 | rs4938520 | G | A | 1.012 | -7977.388 | -7979.19 | -1.802 | 6.38E-07 |
| 11 | 118568639 | rs511451 | T | C | 0.988 | -7977.388 | -7979.19 | -1.802 | 6.38E-07 |
| 11 | 119668265 | rs4315086 | A | G | 1.002 | -7977.388 | -7979.191 | -1.802 | 6.38E-07 |
| 11 | 120398707 | rs59912111 | A | C | 1.004 | -7977.388 | -7979.19 | -1.802 | 6.38E-07 |
| 11 | 116400897 | rs17491207 | A | G | 1.011 | -7977.388 | -7979.191 | -1.803 | 6.37E-07 |
| 11 | 116412160 | rs12802920 | C | A | 0.996 | -7977.388 | -7979.191 | -1.803 | 6.37E-07 |
| 11 | 119685493 | rs4936501 | A | G | 0.985 | -7977.388 | -7979.191 | -1.803 | 6.37E-07 |
| 11 | 119685596 | rs4936502 | T | C | 0.985 | -7977.388 | -7979.191 | -1.803 | 6.37E-07 |
| 11 | 119686250 | rs11217520 | A | G | 0.985 | -7977.388 | -7979.191 | -1.803 | 6.37E-07 |
| 11 | 119934081 | rs680630 | T | G | 0.993 | -7977.388 | -7979.191 | -1.803 | 6.37E-07 |
| 11 | 120008957 | rs1790476 | G | C | 1.003 | -7977.388 | -7979.191 | -1.803 | 6.37E-07 |
| 11 | 116215109 | rs11601506 | T | G | 0.998 | -7977.388 | -7979.192 | -1.804 | 6.37E-07 |

| | | | | | | | |
|---|---|---|---|---|---|---|---|
| 11 | 118506402 | rs483283 | C | T | NULL | 0.710949503 | NULL&M_Disease |
| 11 | 118512162 | rs570836 | A | G | NULL | 0.710949503 | NULL&M_Disease |
| 11 | 118769442 | rs7104819 | G | A | NULL | 0.710949503 | NULL&M_Disease |
| 11 | 119263794 | rs10892359 | A | G | NULL | 0.710949503 | NULL&M_Disease |
| 11 | 119682703 | rs4936500 | G | C | NULL | 0.710949503 | NULL&M_Disease |
| 11 | 119686328 | rs11217521 | C | T | NULL | 0.710949503 | NULL&M_Disease |
| 11 | 119686548 | rs35676325 | A | G | NULL | 0.710949503 | NULL&M_Disease |
| 11 | 119963787 | rs590270 | C | T | NULL | 0.710949503 | NULL&M_Disease |
| 11 | 119964344 | rs664579 | G | A | NULL | 0.710949503 | NULL&M_Disease |
| 11 | 120089688 | rs11217769 | G | A | NULL | 0.710949503 | NULL&M_Disease |
| 11 | 120193477 | rs7934404 | A | G | NULL | 0.710949503 | NULL&M_Disease |
| 11 | 121179847 | rs2060008 | A | T | NULL | 0.710949503 | NULL&M_Disease |
| 11 | 116273569 | rs508023 | G | T | NULL | 0.710949503 | NULL&M_Disease |
| 11 | 116325442 | rs491575 | T | G | NULL | 0.710949503 | NULL&M_Disease |
| 11 | 116708020 | rs5070 | A | G | NULL | 0.710949503 | NULL&M_Disease |
| 11 | 118480223 | rs67307131 | C | T | NULL | 0.710949503 | NULL&M_Disease |
| 11 | 118568668 | rs11216949 | T | C | NULL | 0.710949503 | NULL&M_Disease |
| 11 | 118568813 | rs7479114 | T | C | NULL | 0.710949503 | NULL&M_Disease |
| 11 | 118569067 | rs4486664 | C | A | NULL | 0.710949503 | NULL&M_Disease |
| 11 | 119609314 | rs73006720 | C | A | NULL | 0.710949503 | NULL&M_Disease |
| 11 | 119614499 | rs6589770 | G | C | NULL | 0.710949503 | NULL&M_Disease |
| 11 | 119876939 | rs4938751 | C | T | NULL | 0.710949503 | NULL&M_Disease |
| 11 | 120126277 | rs2715881 | T | C | NULL | 0.710949503 | NULL&M_Disease |
| 11 | 120699625 | rs2004676 | T | C | NULL | 0.710949503 | NULL&M_Disease |
| 11 | 120704184 | rs4935753 | C | G | NULL | 0.710949503 | NULL&M_Disease |
| 11 | 120921866 | rs11218126 | T | C | NULL | 0.710949503 | NULL&M_Disease |
| 11 | 116273570 | rs508025 | A | T | NULL | 0.710949503 | NULL&M_Disease |
| 11 | 117313882 | rs57680824 | G | A | NULL | 0.710949503 | NULL&M_Disease |
| 11 | 117851174 | rs11216665 | G | T | NULL | 0.710949503 | NULL&M_Disease |
| 11 | 119686554 | rs35059164 | G | A | NULL | 0.710949503 | NULL&M_Disease |
| 11 | 119749812 | rs11217551 | T | G | NULL | 0.710949503 | NULL&M_Disease |
| 11 | 119964429 | rs665008 | A | G | NULL | 0.710949503 | NULL&M_Disease |
| 11 | 120005633 | rs605374 | T | G | NULL | 0.710949503 | NULL&M_Disease |
| 11 | 116412613 | rs10502219 | A | G | NULL | 0.710949503 | NULL&M_Disease |
| 11 | 119500417 | rs10790327 | T | C | NULL | 0.710949503 | NULL&M_Disease |
| 11 | 119681110 | rs4938726 | T | C | NULL | 0.710949503 | NULL&M_Disease |
| 11 | 119792443 | rs71484153 | C | T | NULL | 0.710949503 | NULL&M_Disease |
| 11 | 120252828 | rs10892568 | G | A | NULL | 0.710949503 | NULL&M_Disease |
| 11 | 117225033 | rs499744 | T | C | NULL | 0.711975919 | NULL&M_Disease |
| 11 | 117313577 | rs3894251 | C | G | NULL | 0.711975919 | NULL&M_Disease |
| 11 | 118480285 | rs12225399 | C | G | NULL | 0.711975919 | NULL&M_Disease |
| 11 | 118811684 | rs550999 | G | C | NULL | 0.711975919 | NULL&M_Disease |
| 11 | 119962750 | rs605230 | C | T | NULL | 0.711975919 | NULL&M_Disease |
| 11 | 119963001 | rs584653 | T | C | NULL | 0.711975919 | NULL&M_Disease |
| 11 | 119963324 | rs1628536 | G | A | NULL | 0.711975919 | NULL&M_Disease |
| 11 | 120337705 | rs894839 | G | A | NULL | 0.711975919 | NULL&M_Disease |
| 11 | 120706789 | rs7937037 | C | G | NULL | 0.711975919 | NULL&M_Disease |
| 11 | 116301080 | rs74508982 | T | C | NULL | 0.711975919 | NULL&M_Disease |
| 11 | 120398643 | rs17092876 | A | G | NULL | 0.711975919 | NULL&M_Disease |
| 11 | 120398747 | rs61226783 | G | A | NULL | 0.711975919 | NULL&M_Disease |
| 11 | 116276086 | rs573126 | C | A | NULL | 0.711975919 | NULL&M_Disease |
| 11 | 118528466 | rs7389 | G | T | NULL | 0.711975919 | NULL&M_Disease |
| 11 | 119961635 | rs617715 | A | T | NULL | 0.711975919 | NULL&M_Disease |
| 11 | 120309851 | rs538661 | T | C | NULL | 0.711975919 | NULL&M_Disease |
| 11 | 120704307 | rs4936558 | C | T | NULL | 0.711975919 | NULL&M_Disease |
| 11 | 120954769 | rs6589855 | C | T | NULL | 0.711975919 | NULL&M_Disease |
| 11 | 118568472 | rs570371 | G | A | NULL | 0.711975919 | NULL&M_Disease |
| 11 | 116343848 | rs7110240 | C | T | NULL | 0.711975919 | NULL&M_Disease |
| 11 | 117306738 | rs2305823 | C | A | NULL | 0.711975919 | NULL&M_Disease |
| 11 | 117463000 | rs498689 | G | A | NULL | 0.711975919 | NULL&M_Disease |
| 11 | 119610490 | rs58136657 | T | C | NULL | 0.711975919 | NULL&M_Disease |
| 11 | 119967026 | rs600484 | A | T | NULL | 0.711975919 | NULL&M_Disease |
| 11 | 119967247 | rs611656 | T | C | NULL | 0.711975919 | NULL&M_Disease |
| 11 | 119967383 | rs610075 | T | G | NULL | 0.711975919 | NULL&M_Disease |
| 11 | 119967980 | rs1629201 | C | T | NULL | 0.711975919 | NULL&M_Disease |
| 11 | 120238413 | rs7116481 | T | C | NULL | 0.711975919 | NULL&M_Disease |
| 11 | 120337519 | rs12417256 | C | A | NULL | 0.711975919 | NULL&M_Disease |
| 11 | 120531547 | rs7940234 | C | G | NULL | 0.711975919 | NULL&M_Disease |
| 11 | 116272339 | rs4938268 | G | A | NULL | 0.711975919 | NULL&M_Disease |
| 11 | 120521485 | rs4936536 | A | G | NULL | 0.711975919 | NULL&M_Disease |
| 11 | 120703415 | rs11218018 | G | A | NULL | 0.711975919 | NULL&M_Disease |
| 11 | 121015711 | rs11218161 | T | C | NULL | 0.711975919 | NULL&M_Disease |
| 11 | 116703671 | rs4225 | G | T | NULL | 0.711975919 | NULL&M_Disease |
| 11 | 117732028 | rs10790217 | G | A | NULL | 0.711975919 | NULL&M_Disease |
| 11 | 117800301 | rs11604677 | A | G | NULL | 0.711975919 | NULL&M_Disease |
| 11 | 117800326 | rs10790221 | G | A | NULL | 0.711975919 | NULL&M_Disease |
| 11 | 119452530 | rs12277021 | C | T | NULL | 0.711975919 | NULL&M_Disease |
| 11 | 119967484 | rs609650 | C | T | NULL | 0.711975919 | NULL&M_Disease |
| 11 | 119967918 | rs1629347 | A | C | NULL | 0.711975919 | NULL&M_Disease |
| 11 | 119967930 | rs1790472 | G | A | NULL | 0.711975919 | NULL&M_Disease |
| 11 | 119968311 | rs1670198 | C | T | NULL | 0.711975919 | NULL&M_Disease |
| 11 | 119968367 | rs1670197 | C | G | NULL | 0.711975919 | NULL&M_Disease |
| 11 | 119968392 | rs1787026 | C | G | NULL | 0.711975919 | NULL&M_Disease |
| 11 | 119968495 | rs1787025 | G | T | NULL | 0.711975919 | NULL&M_Disease |
| 11 | 119968506 | rs639775 | G | C | NULL | 0.711975919 | NULL&M_Disease |
| 11 | 119968598 | rs1670196 | C | T | NULL | 0.711975919 | NULL&M_Disease |
| 11 | 120221652 | rs10790379 | G | A | NULL | 0.711975919 | NULL&M_Disease |
| 11 | 120706759 | rs12226536 | G | C | NULL | 0.711975919 | NULL&M_Disease |
| 11 | 120991298 | rs519116 | T | C | NULL | 0.711975919 | NULL&M_Disease |
| 11 | 121101852 | rs11218213 | G | A | NULL | 0.711975919 | NULL&M_Disease |
| 11 | 121169193 | rs10892720 | T | G | NULL | 0.711975919 | NULL&M_Disease |
| 11 | 117319260 | rs10790184 | A | G | NULL | 0.711975919 | NULL&M_Disease |
| 11 | 117739122 | rs1320656 | C | A | NULL | 0.711975919 | NULL&M_Disease |
| 11 | 118179200 | rs1599809 | A | G | NULL | 0.711975919 | NULL&M_Disease |
| 11 | 118927907 | rs17552226 | G | C | NULL | 0.711975919 | NULL&M_Disease |
| 11 | 119439498 | rs755002 | T | C | NULL | 0.711975919 | NULL&M_Disease |

| | | | | | | | | | |
|---|---|---|---|---|---|---|---|---|---|
| 11 | 118506402 | rs483283 | C | T | 1.012 | -7977.388 | -7979.192 | -1.804 | 6.37E-07 |
| 11 | 118512162 | rs570836 | A | G | 1.012 | -7977.388 | -7979.192 | -1.804 | 6.37E-07 |
| 11 | 118769442 | rs7104819 | G | A | 1.014 | -7977.388 | -7979.192 | -1.804 | 6.37E-07 |
| 11 | 119263794 | rs10892359 | A | G | 1.01 | -7977.388 | -7979.192 | -1.804 | 6.37E-07 |
| 11 | 119682703 | rs4936500 | G | C | 0.985 | -7977.388 | -7979.192 | -1.804 | 6.37E-07 |
| 11 | 119686328 | rs11217521 | C | T | 0.985 | -7977.388 | -7979.193 | -1.804 | 6.37E-07 |
| 11 | 119686548 | rs35676325 | A | G | 0.985 | -7977.388 | -7979.193 | -1.804 | 6.37E-07 |
| 11 | 119963787 | rs590270 | C | T | 0.987 | -7977.388 | -7979.192 | -1.804 | 6.37E-07 |
| 11 | 119964344 | rs664579 | G | A | 0.987 | -7977.388 | -7979.192 | -1.804 | 6.37E-07 |
| 11 | 120089688 | rs11217769 | G | A | 0.986 | -7977.388 | -7979.192 | -1.804 | 6.37E-07 |
| 11 | 120193477 | rs7934404 | A | G | 1.003 | -7977.388 | -7979.192 | -1.804 | 6.37E-07 |
| 11 | 121179847 | rs2060008 | A | T | 1.015 | -7977.388 | -7979.192 | -1.804 | 6.37E-07 |
| 11 | 116273569 | rs508023 | G | T | 1.009 | -7977.388 | -7979.193 | -1.805 | 6.36E-07 |
| 11 | 116325442 | rs491575 | T | G | 0.986 | -7977.388 | -7979.193 | -1.805 | 6.36E-07 |
| 11 | 116708020 | rs5070 | A | G | 0.99 | -7977.388 | -7979.194 | -1.805 | 6.36E-07 |
| 11 | 118480223 | rs67307131 | C | T | 0.988 | -7977.388 | -7979.193 | -1.805 | 6.36E-07 |
| 11 | 118568668 | rs11216949 | T | C | 0.989 | -7977.388 | -7979.193 | -1.805 | 6.36E-07 |
| 11 | 118568813 | rs7479114 | T | C | 0.989 | -7977.388 | -7979.193 | -1.805 | 6.36E-07 |
| 11 | 118569067 | rs4486664 | C | A | 0.989 | -7977.388 | -7979.193 | -1.805 | 6.36E-07 |
| 11 | 119609314 | rs73006720 | C | A | 0.986 | -7977.388 | -7979.193 | -1.805 | 6.36E-07 |
| 11 | 119614499 | rs6589770 | G | C | 0.986 | -7977.388 | -7979.194 | -1.805 | 6.36E-07 |
| 11 | 119876939 | rs4938751 | C | T | 1.014 | -7977.388 | -7979.193 | -1.805 | 6.36E-07 |
| 11 | 120126277 | rs2715881 | T | C | 1.007 | -7977.388 | -7979.193 | -1.805 | 6.36E-07 |
| 11 | 120699625 | rs2004676 | T | C | 0.987 | -7977.388 | -7979.193 | -1.805 | 6.36E-07 |
| 11 | 120704184 | rs4935753 | C | G | 0.986 | -7977.388 | -7979.193 | -1.805 | 6.36E-07 |
| 11 | 120921866 | rs11218126 | T | C | 0.997 | -7977.388 | -7979.193 | -1.805 | 6.36E-07 |
| 11 | 116273570 | rs508025 | A | T | 1.009 | -7977.388 | -7979.194 | -1.806 | 6.36E-07 |
| 11 | 117313882 | rs57680824 | G | A | 0.991 | -7977.388 | -7979.195 | -1.806 | 6.36E-07 |
| 11 | 117851174 | rs11216665 | G | T | 0.989 | -7977.388 | -7979.194 | -1.806 | 6.36E-07 |
| 11 | 119686554 | rs35059164 | G | A | 0.985 | -7977.388 | -7979.194 | -1.806 | 6.36E-07 |
| 11 | 119749812 | rs11217551 | T | G | 0.986 | -7977.388 | -7979.194 | -1.806 | 6.36E-07 |
| 11 | 119964429 | rs665008 | A | G | 0.987 | -7977.388 | -7979.194 | -1.806 | 6.36E-07 |
| 11 | 120005633 | rs605374 | T | G | 0.997 | -7977.388 | -7979.194 | -1.806 | 6.36E-07 |
| 11 | 116412613 | rs10502219 | A | G | 0.997 | -7977.388 | -7979.195 | -1.807 | 6.35E-07 |
| 11 | 119500417 | rs10790327 | T | C | 1.011 | -7977.388 | -7979.195 | -1.807 | 6.35E-07 |
| 11 | 119681110 | rs4938726 | T | C | 0.985 | -7977.388 | -7979.195 | -1.807 | 6.35E-07 |
| 11 | 119792443 | rs71484153 | C | T | 0.992 | -7977.388 | -7979.196 | -1.807 | 6.35E-07 |
| 11 | 120252828 | rs10892568 | G | A | 0.985 | -7977.388 | -7979.195 | -1.807 | 6.35E-07 |
| 11 | 117225033 | rs499744 | T | C | 1.014 | -7977.388 | -7979.197 | -1.808 | 6.34E-07 |
| 11 | 117313577 | rs3894251 | C | G | 0.991 | -7977.388 | -7979.196 | -1.808 | 6.34E-07 |
| 11 | 118480285 | rs12225399 | C | G | 0.988 | -7977.388 | -7979.197 | -1.808 | 6.34E-07 |
| 11 | 118811684 | rs550999 | G | C | 0.987 | -7977.388 | -7979.196 | -1.808 | 6.34E-07 |
| 11 | 119962750 | rs605230 | C | T | 0.987 | -7977.388 | -7979.196 | -1.808 | 6.34E-07 |
| 11 | 119963001 | rs584653 | T | C | 0.987 | -7977.388 | -7979.196 | -1.808 | 6.34E-07 |
| 11 | 119963324 | rs1628536 | G | A | 0.987 | -7977.388 | -7979.196 | -1.808 | 6.34E-07 |
| 11 | 120337705 | rs894839 | G | A | 1.014 | -7977.388 | -7979.197 | -1.808 | 6.34E-07 |
| 11 | 120706789 | rs7937037 | C | G | 0.986 | -7977.388 | -7979.197 | -1.808 | 6.34E-07 |
| 11 | 116301080 | rs74508982 | T | C | 1.008 | -7977.388 | -7979.197 | -1.809 | 6.34E-07 |
| 11 | 120398643 | rs17092876 | A | G | 1.001 | -7977.388 | -7979.197 | -1.809 | 6.34E-07 |
| 11 | 120398747 | rs61226783 | G | A | 1.002 | -7977.388 | -7979.197 | -1.809 | 6.34E-07 |
| 11 | 116276086 | rs573126 | C | A | 1.009 | -7977.388 | -7979.198 | -1.81 | 6.33E-07 |
| 11 | 118528466 | rs7389 | G | T | 1.012 | -7977.388 | -7979.199 | -1.81 | 6.33E-07 |
| 11 | 119961635 | rs617715 | A | T | 0.987 | -7977.388 | -7979.198 | -1.81 | 6.33E-07 |
| 11 | 120309851 | rs538661 | T | C | 0.986 | -7977.388 | -7979.198 | -1.81 | 6.33E-07 |
| 11 | 120704307 | rs4936558 | C | T | 0.986 | -7977.388 | -7979.198 | -1.81 | 6.33E-07 |
| 11 | 120954769 | rs6589855 | C | T | 1.005 | -7977.388 | -7979.198 | -1.81 | 6.33E-07 |
| 11 | 118568472 | rs570371 | G | A | 0.989 | -7977.388 | -7979.199 | -1.811 | 6.32E-07 |
| 11 | 116343848 | rs7110240 | C | T | 1.007 | -7977.388 | -7979.201 | -1.812 | 6.32E-07 |
| 11 | 117306738 | rs2305823 | C | A | 0.989 | -7977.388 | -7979.2 | -1.812 | 6.32E-07 |
| 11 | 117463000 | rs498689 | G | A | 1.01 | -7977.388 | -7979.2 | -1.812 | 6.32E-07 |
| 11 | 119610490 | rs58136657 | T | C | 0.986 | -7977.388 | -7979.2 | -1.812 | 6.32E-07 |
| 11 | 119967026 | rs600484 | A | T | 0.987 | -7977.388 | -7979.201 | -1.812 | 6.32E-07 |
| 11 | 119967247 | rs611656 | T | C | 0.987 | -7977.388 | -7979.201 | -1.812 | 6.32E-07 |
| 11 | 119967383 | rs610075 | T | G | 0.987 | -7977.388 | -7979.201 | -1.812 | 6.32E-07 |
| 11 | 119967980 | rs1629201 | C | T | 0.987 | -7977.388 | -7979.2 | -1.812 | 6.32E-07 |
| 11 | 120238413 | rs7116481 | T | C | 0.986 | -7977.388 | -7979.2 | -1.812 | 6.32E-07 |
| 11 | 120337519 | rs12417256 | C | A | 1.013 | -7977.388 | -7979.201 | -1.812 | 6.32E-07 |
| 11 | 120531547 | rs7940234 | C | G | 1.012 | -7977.388 | -7979.201 | -1.812 | 6.32E-07 |
| 11 | 116272339 | rs4938268 | G | A | 0.986 | -7977.388 | -7979.201 | -1.813 | 6.31E-07 |
| 11 | 120521485 | rs4936536 | A | G | 1.009 | -7977.388 | -7979.201 | -1.813 | 6.31E-07 |
| 11 | 120703415 | rs11218018 | G | A | 0.986 | -7977.388 | -7979.201 | -1.813 | 6.31E-07 |
| 11 | 121015711 | rs11218161 | T | C | 1.006 | -7977.388 | -7979.201 | -1.813 | 6.31E-07 |
| 11 | 116703671 | rs4225 | G | T | 1.012 | -7977.388 | -7979.202 | -1.814 | 6.31E-07 |
| 11 | 117732028 | rs10790217 | G | A | 0.987 | -7977.388 | -7979.202 | -1.814 | 6.31E-07 |
| 11 | 117800301 | rs11604677 | A | G | 1.014 | -7977.388 | -7979.202 | -1.814 | 6.31E-07 |
| 11 | 117800326 | rs10790221 | G | A | 1.014 | -7977.388 | -7979.202 | -1.814 | 6.31E-07 |
| 11 | 119452530 | rs12277021 | C | T | 1.009 | -7977.388 | -7979.202 | -1.814 | 6.31E-07 |
| 11 | 119967484 | rs609650 | C | T | 0.987 | -7977.388 | -7979.202 | -1.814 | 6.31E-07 |
| 11 | 119967918 | rs1629347 | A | C | 0.987 | -7977.388 | -7979.202 | -1.814 | 6.31E-07 |
| 11 | 119967930 | rs1790472 | G | A | 0.987 | -7977.388 | -7979.202 | -1.814 | 6.31E-07 |
| 11 | 119968311 | rs1670198 | C | T | 0.987 | -7977.388 | -7979.202 | -1.814 | 6.31E-07 |
| 11 | 119968367 | rs1670197 | C | G | 0.987 | -7977.388 | -7979.202 | -1.814 | 6.31E-07 |
| 11 | 119968392 | rs1787026 | C | G | 0.987 | -7977.388 | -7979.202 | -1.814 | 6.31E-07 |
| 11 | 119968495 | rs1787025 | G | T | 0.987 | -7977.388 | -7979.202 | -1.814 | 6.31E-07 |
| 11 | 119968506 | rs639775 | G | C | 0.987 | -7977.388 | -7979.202 | -1.814 | 6.31E-07 |
| 11 | 119968598 | rs1670196 | C | T | 0.987 | -7977.388 | -7979.202 | -1.814 | 6.31E-07 |
| 11 | 120221652 | rs10790379 | G | A | 0.986 | -7977.388 | -7979.202 | -1.814 | 6.31E-07 |
| 11 | 120706759 | rs12226536 | G | C | 0.987 | -7977.388 | -7979.203 | -1.814 | 6.31E-07 |
| 11 | 120991298 | rs519116 | T | C | 1.009 | -7977.388 | -7979.202 | -1.814 | 6.31E-07 |
| 11 | 121101852 | rs11218213 | G | A | 1.005 | -7977.388 | -7979.202 | -1.814 | 6.31E-07 |
| 11 | 121169193 | rs10892720 | T | G | 1.014 | -7977.388 | -7979.202 | -1.814 | 6.31E-07 |
| 11 | 117319260 | rs10790184 | A | G | 1.013 | -7977.388 | -7979.203 | -1.815 | 6.30E-07 |
| 11 | 117739122 | rs1320656 | C | A | 0.987 | -7977.388 | -7979.203 | -1.815 | 6.30E-07 |
| 11 | 118179200 | rs1599809 | A | G | 0.99 | -7977.388 | -7979.203 | -1.815 | 6.30E-07 |
| 11 | 118927907 | rs17552226 | G | C | 0.995 | -7977.388 | -7979.203 | -1.815 | 6.30E-07 |
| 11 | 119439498 | rs755002 | T | C | 0.993 | -7977.388 | -7979.203 | -1.815 | 6.30E-07 |

| | | | | | | | |
|---|---|---|---|---|---|---|---|
| 11 | 119850718 | rs602030 | T | C | NULL | 0.711975919 | NULL&M_Disease |
| 11 | 120706839 | rs12223757 | G | A | NULL | 0.711975919 | NULL&M_Disease |
| 11 | 120995842 | rs560919 | C | T | NULL | 0.711975919 | NULL&M_Disease |
| 11 | 117738911 | rs948108 | A | G | NULL | 0.711975919 | NULL&M_Disease |
| 11 | 119451111 | rs10892409 | C | A | NULL | 0.711975919 | NULL&M_Disease |
| 11 | 119459317 | rs10892412 | C | T | NULL | 0.711975919 | NULL&M_Disease |
| 11 | 119647700 | rs34228131 | T | C | NULL | 0.711975919 | NULL&M_Disease |
| 11 | 119682166 | rs55728834 | C | T | NULL | 0.711975919 | NULL&M_Disease |
| 11 | 119967159 | rs654884 | T | C | NULL | 0.711975919 | NULL&M_Disease |
| 11 | 119968997 | rs637464 | G | T | NULL | 0.711975919 | NULL&M_Disease |
| 11 | 120994638 | rs584329 | T | C | NULL | 0.711975919 | NULL&M_Disease |
| 11 | 120994961 | rs659956 | A | G | NULL | 0.711975919 | NULL&M_Disease |
| 11 | 117312609 | rs3894250 | C | A | NULL | 0.711975919 | NULL&M_Disease |
| 11 | 117341163 | rs3888083 | C | T | NULL | 0.711975919 | NULL&M_Disease |
| 11 | 117797898 | rs34850032 | G | A | NULL | 0.711975919 | NULL&M_Disease |
| 11 | 118569097 | rs4393358 | T | C | NULL | 0.711975919 | NULL&M_Disease |
| 11 | 119451380 | rs56220308 | C | G | NULL | 0.711975919 | NULL&M_Disease |
| 11 | 119412474 | rs10750148 | A | T | NULL | 0.713000163 | NULL&M_Disease |
| 11 | 119428141 | rs11217316 | A | G | NULL | 0.713000163 | NULL&M_Disease |
| 11 | 119450113 | rs12277899 | T | A | NULL | 0.713000163 | NULL&M_Disease |
| 11 | 119451188 | rs12279777 | T | A | NULL | 0.713000163 | NULL&M_Disease |
| 11 | 119646732 | rs6589774 | A | G | NULL | 0.713000163 | NULL&M_Disease |
| 11 | 120996969 | rs661189 | G | C | NULL | 0.713000163 | NULL&M_Disease |
| 11 | 116416716 | rs4528369 | T | G | NULL | 0.713000163 | NULL&M_Disease |
| 11 | 116471690 | rs520176 | T | C | NULL | 0.713000163 | NULL&M_Disease |
| 11 | 116583017 | rs7480302 | G | A | NULL | 0.713000163 | NULL&M_Disease |
| 11 | 116584504 | rs4938302 | C | T | NULL | 0.713000163 | NULL&M_Disease |
| 11 | 116584987 | rs4938303 | C | T | NULL | 0.713000163 | NULL&M_Disease |
| 11 | 118191691 | rs6589659 | A | C | NULL | 0.713000163 | NULL&M_Disease |
| 11 | 118515616 | rs527290 | T | C | NULL | 0.713000163 | NULL&M_Disease |
| 11 | 118520480 | rs653008 | T | G | NULL | 0.713000163 | NULL&M_Disease |
| 11 | 119985475 | rs12807982 | A | C | NULL | 0.713000163 | NULL&M_Disease |
| 11 | 120704063 | rs4936557 | C | T | NULL | 0.713000163 | NULL&M_Disease |
| 11 | 117332343 | rs2298766 | T | C | NULL | 0.713000163 | NULL&M_Disease |
| 11 | 117685343 | rs10892178 | T | C | NULL | 0.713000163 | NULL&M_Disease |
| 11 | 118529595 | rs745663 | G | A | NULL | 0.713000163 | NULL&M_Disease |
| 11 | 118587347 | rs603486 | T | C | NULL | 0.713000163 | NULL&M_Disease |
| 11 | 119985570 | rs12808008 | G | A | NULL | 0.713000163 | NULL&M_Disease |
| 11 | 120034115 | rs626225 | G | A | NULL | 0.713000163 | NULL&M_Disease |
| 11 | 121151034 | rs73005128 | T | C | NULL | 0.713000163 | NULL&M_Disease |
| 11 | 117436095 | rs1940037 | T | C | NULL | 0.713000163 | NULL&M_Disease |
| 11 | 117838400 | rs11216656 | A | G | NULL | 0.713000163 | NULL&M_Disease |
| 11 | 118567733 | rs481903 | C | T | NULL | 0.713000163 | NULL&M_Disease |
| 11 | 119440533 | rs10750157 | T | C | NULL | 0.713000163 | NULL&M_Disease |
| 11 | 119850725 | rs615539 | G | A | NULL | 0.713000163 | NULL&M_Disease |
| 11 | 120107411 | rs882856 | G | A | NULL | 0.713000163 | NULL&M_Disease |
| 11 | 120497375 | rs4565903 | G | C | NULL | 0.713000163 | NULL&M_Disease |
| 11 | 121003046 | rs616120 | A | G | NULL | 0.713000163 | NULL&M_Disease |
| 11 | 121003514 | rs504148 | T | C | NULL | 0.713000163 | NULL&M_Disease |
| 11 | 121150625 | rs4457755 | G | A | NULL | 0.713000163 | NULL&M_Disease |
| 11 | 116303270 | rs11215929 | G | A | NULL | 0.713000163 | NULL&M_Disease |
| 11 | 117314320 | rs1304536246 | T | C | NULL | 0.713000163 | NULL&M_Disease |
| 11 | 119269557 | rs2096834 | T | C | NULL | 0.713000163 | NULL&M_Disease |
| 11 | 119320879 | rs11217290 | A | G | NULL | 0.713000163 | NULL&M_Disease |
| 11 | 119428144 | rs11217317 | T | C | NULL | 0.713000163 | NULL&M_Disease |
| 11 | 119966457 | rs624877 | T | C | NULL | 0.713000163 | NULL&M_Disease |
| 11 | 120483858 | rs11603831 | T | C | NULL | 0.713000163 | NULL&M_Disease |
| 11 | 116195089 | rs1866682 | T | C | NULL | 0.713000163 | NULL&M_Disease |
| 11 | 117772229 | rs575535 | C | G | NULL | 0.713000163 | NULL&M_Disease |
| 11 | 118042974 | rs679327 | G | A | NULL | 0.713000163 | NULL&M_Disease |
| 11 | 119181674 | rs2510146 | G | C | NULL | 0.713000163 | NULL&M_Disease |
| 11 | 119648152 | rs4388901 | C | T | NULL | 0.713000163 | NULL&M_Disease |
| 11 | 119648831 | rs34033074 | A | G | NULL | 0.713000163 | NULL&M_Disease |
| 11 | 119879327 | rs17245368 | T | C | NULL | 0.713000163 | NULL&M_Disease |
| 11 | 120557343 | rs10750176 | G | A | NULL | 0.713000163 | NULL&M_Disease |
| 11 | 121090790 | rs34271655 | C | T | NULL | 0.713000163 | NULL&M_Disease |
| 11 | 121183145 | rs35288685 | T | C | NULL | 0.713000163 | NULL&M_Disease |
| 11 | 117801635 | rs17453205 | G | A | NULL | 0.713000163 | NULL&M_Disease |
| 11 | 119643807 | rs12803325 | T | A | NULL | 0.713000163 | NULL&M_Disease |
| 11 | 119644249 | rs4420283 | C | T | NULL | 0.713000163 | NULL&M_Disease |
| 11 | 120023505 | rs12802402 | T | C | NULL | 0.713000163 | NULL&M_Disease |
| 11 | 121019288 | rs10892677 | T | C | NULL | 0.713000163 | NULL&M_Disease |
| 11 | 117845302 | rs7939408 | T | A | NULL | 0.713000163 | NULL&M_Disease |
| 11 | 119269112 | rs35720648 | C | G | NULL | 0.713000163 | NULL&M_Disease |
| 11 | 119299592 | rs12420302 | G | A | NULL | 0.713000163 | NULL&M_Disease |
| 11 | 119419568 | rs11217310 | C | G | NULL | 0.713000163 | NULL&M_Disease |
| 11 | 119444863 | rs11217326 | C | T | NULL | 0.713000163 | NULL&M_Disease |
| 11 | 119453252 | rs11217329 | A | G | NULL | 0.713000163 | NULL&M_Disease |
| 11 | 119453579 | rs1396776 | C | A | NULL | 0.713000163 | NULL&M_Disease |
| 11 | 120025443 | rs12791621 | G | A | NULL | 0.713000163 | NULL&M_Disease |
| 11 | 120355230 | rs503473 | G | A | NULL | 0.713000163 | NULL&M_Disease |
| 11 | 120477883 | rs11600354 | A | G | NULL | 0.713000163 | NULL&M_Disease |
| 11 | 116419268 | rs4938279 | A | G | NULL | 0.713000163 | NULL&M_Disease |
| 11 | 117331499 | rs10892119 | A | G | NULL | 0.713000163 | NULL&M_Disease |
| 11 | 117464173 | rs694005 | T | C | NULL | 0.713000163 | NULL&M_Disease |
| 11 | 117732718 | rs7103567 | T | C | NULL | 0.713000163 | NULL&M_Disease |
| 11 | 117799923 | rs7928668 | T | C | NULL | 0.713000163 | NULL&M_Disease |
| 11 | 117961908 | rs651474 | C | T | NULL | 0.713000163 | NULL&M_Disease |
| 11 | 118519550 | rs601318 | G | C | NULL | 0.713000163 | NULL&M_Disease |
| 11 | 119299408 | rs12417523 | T | C | NULL | 0.713000163 | NULL&M_Disease |
| 11 | 119299832 | rs12417611 | A | G | NULL | 0.713000163 | NULL&M_Disease |
| 11 | 119300681 | rs12807216 | A | C | NULL | 0.713000163 | NULL&M_Disease |
| 11 | 119425163 | rs11217315 | A | G | NULL | 0.713000163 | NULL&M_Disease |
| 11 | 119703617 | rs7395060 | G | A | NULL | 0.713000163 | NULL&M_Disease |
| 11 | 119799765 | rs12283803 | T | C | NULL | 0.713000163 | NULL&M_Disease |
| 11 | 119965319 | rs11217689 | G | C | NULL | 0.713000163 | NULL&M_Disease |

| | | | | | | | | | |
|---|---|---|---|---|---|---|---|---|---|
| 11 | 119850718 | rs602030 | T | C | 1.014 | -7977.388 | -7979.203 | -1.815 | 6.30E-07 |
| 11 | 120706839 | rs12223757 | G | A | 0.987 | -7977.388 | -7979.204 | -1.815 | 6.30E-07 |
| 11 | 120995842 | rs560919 | C | T | 1.009 | -7977.388 | -7979.203 | -1.815 | 6.30E-07 |
| 11 | 117738911 | rs948108 | A | G | 0.988 | -7977.388 | -7979.204 | -1.816 | 6.29E-07 |
| 11 | 119451111 | rs10892409 | C | A | 1.009 | -7977.388 | -7979.204 | -1.816 | 6.29E-07 |
| 11 | 119459317 | rs10892412 | C | T | 0.986 | -7977.388 | -7979.204 | -1.816 | 6.29E-07 |
| 11 | 119647700 | rs34228131 | T | C | 1.012 | -7977.388 | -7979.204 | -1.816 | 6.29E-07 |
| 11 | 119682166 | rs55728834 | C | T | 0.986 | -7977.388 | -7979.204 | -1.816 | 6.29E-07 |
| 11 | 119967159 | rs654884 | T | C | 0.988 | -7977.388 | -7979.204 | -1.816 | 6.29E-07 |
| 11 | 119968997 | rs637464 | G | T | 0.988 | -7977.388 | -7979.204 | -1.816 | 6.29E-07 |
| 11 | 120994638 | rs584329 | T | C | 1.009 | -7977.388 | -7979.204 | -1.816 | 6.29E-07 |
| 11 | 120994961 | rs659956 | A | G | 1.009 | -7977.388 | -7979.204 | -1.816 | 6.29E-07 |
| 11 | 117312609 | rs3894250 | C | A | 1.007 | -7977.388 | -7979.205 | -1.817 | 6.29E-07 |
| 11 | 117341163 | rs3888083 | C | T | 1.014 | -7977.388 | -7979.205 | -1.817 | 6.29E-07 |
| 11 | 117797898 | rs34850032 | G | A | 0.994 | -7977.388 | -7979.205 | -1.817 | 6.29E-07 |
| 11 | 118569097 | rs4393358 | T | C | 0.99 | -7977.388 | -7979.205 | -1.817 | 6.29E-07 |
| 11 | 119451380 | rs56220308 | C | G | 1.009 | -7977.388 | -7979.205 | -1.817 | 6.29E-07 |
| 11 | 119412474 | rs10750148 | A | T | 0.994 | -7977.388 | -7979.206 | -1.818 | 6.28E-07 |
| 11 | 119428141 | rs11217316 | A | G | 1.004 | -7977.388 | -7979.206 | -1.818 | 6.28E-07 |
| 11 | 119450113 | rs12277899 | T | A | 1.009 | -7977.388 | -7979.207 | -1.818 | 6.28E-07 |
| 11 | 119451188 | rs12279777 | T | A | 1.009 | -7977.388 | -7979.207 | -1.818 | 6.28E-07 |
| 11 | 119646732 | rs6589774 | A | G | 1.012 | -7977.388 | -7979.207 | -1.818 | 6.28E-07 |
| 11 | 120996969 | rs661189 | G | C | 1.009 | -7977.388 | -7979.206 | -1.818 | 6.28E-07 |
| 11 | 116416716 | rs4528369 | T | G | 0.992 | -7977.388 | -7979.207 | -1.819 | 6.27E-07 |
| 11 | 116471690 | rs520176 | T | C | 1.01 | -7977.388 | -7979.207 | -1.819 | 6.27E-07 |
| 11 | 116583017 | rs7480302 | G | A | 1.001 | -7977.388 | -7979.207 | -1.819 | 6.27E-07 |
| 11 | 116584504 | rs4938302 | C | T | 0.999 | -7977.388 | -7979.207 | -1.819 | 6.27E-07 |
| 11 | 116584987 | rs4938303 | C | T | 0.999 | -7977.388 | -7979.207 | -1.819 | 6.27E-07 |
| 11 | 118191691 | rs6589659 | A | C | 1.006 | -7977.388 | -7979.208 | -1.819 | 6.27E-07 |
| 11 | 118515616 | rs527290 | T | C | 1.011 | -7977.388 | -7979.208 | -1.819 | 6.27E-07 |
| 11 | 118520480 | rs653008 | T | G | 1.011 | -7977.388 | -7979.208 | -1.819 | 6.27E-07 |
| 11 | 119985475 | rs12807982 | A | C | 1.007 | -7977.388 | -7979.207 | -1.819 | 6.27E-07 |
| 11 | 120704063 | rs4936557 | C | T | 0.987 | -7977.388 | -7979.207 | -1.819 | 6.27E-07 |
| 11 | 117332343 | rs2298766 | T | C | 1.006 | -7977.388 | -7979.208 | -1.82 | 6.27E-07 |
| 11 | 117685343 | rs10892178 | T | C | 0.987 | -7977.388 | -7979.208 | -1.82 | 6.27E-07 |
| 11 | 118529595 | rs745663 | G | A | 1.011 | -7977.388 | -7979.208 | -1.82 | 6.27E-07 |
| 11 | 118587347 | rs603486 | T | C | 0.988 | -7977.388 | -7979.208 | -1.82 | 6.27E-07 |
| 11 | 119985570 | rs12808008 | G | A | 1.007 | -7977.388 | -7979.208 | -1.82 | 6.27E-07 |
| 11 | 120034115 | rs626225 | G | A | 1.014 | -7977.388 | -7979.208 | -1.82 | 6.27E-07 |
| 11 | 121151034 | rs73005128 | T | C | 0.997 | -7977.388 | -7979.208 | -1.82 | 6.27E-07 |
| 11 | 117436095 | rs1940037 | T | C | 0.994 | -7977.388 | -7979.209 | -1.821 | 6.26E-07 |
| 11 | 117838400 | rs11216656 | A | G | 0.987 | -7977.388 | -7979.209 | -1.821 | 6.26E-07 |
| 11 | 118567733 | rs481903 | C | T | 0.989 | -7977.388 | -7979.21 | -1.821 | 6.26E-07 |
| 11 | 119440533 | rs10750157 | T | C | 1.008 | -7977.388 | -7979.209 | -1.821 | 6.26E-07 |
| 11 | 119850725 | rs615539 | G | A | 1.013 | -7977.388 | -7979.209 | -1.821 | 6.26E-07 |
| 11 | 120107411 | rs882856 | G | A | 1.005 | -7977.388 | -7979.209 | -1.821 | 6.26E-07 |
| 11 | 120497375 | rs4565903 | G | C | 0.988 | -7977.388 | -7979.209 | -1.821 | 6.26E-07 |
| 11 | 121003046 | rs616120 | A | G | 1.009 | -7977.388 | -7979.209 | -1.821 | 6.26E-07 |
| 11 | 121003514 | rs504148 | T | C | 1.009 | -7977.388 | -7979.209 | -1.821 | 6.26E-07 |
| 11 | 121150625 | rs4457755 | G | A | 0.986 | -7977.388 | -7979.21 | -1.821 | 6.26E-07 |
| 11 | 116303270 | rs11215929 | G | A | 1.013 | -7977.388 | -7979.21 | -1.822 | 6.25E-07 |
| 11 | 117314320 | rs1304536246 | T | C | 0.992 | -7977.388 | -7979.21 | -1.822 | 6.25E-07 |
| 11 | 119269557 | rs2096834 | T | C | 1.007 | -7977.388 | -7979.21 | -1.822 | 6.25E-07 |
| 11 | 119320879 | rs11217290 | A | G | 0.996 | -7977.388 | -7979.21 | -1.822 | 6.25E-07 |
| 11 | 119428144 | rs11217317 | T | C | 1.003 | -7977.388 | -7979.21 | -1.822 | 6.25E-07 |
| 11 | 119966457 | rs624877 | T | C | 0.988 | -7977.388 | -7979.211 | -1.822 | 6.25E-07 |
| 11 | 120483858 | rs11603831 | T | C | 0.987 | -7977.388 | -7979.211 | -1.822 | 6.25E-07 |
| 11 | 116195089 | rs1866682 | T | C | 0.996 | -7977.388 | -7979.211 | -1.823 | 6.25E-07 |
| 11 | 117772229 | rs575535 | C | G | 1.013 | -7977.388 | -7979.211 | -1.823 | 6.25E-07 |
| 11 | 118042974 | rs679327 | G | A | 0.993 | -7977.388 | -7979.211 | -1.823 | 6.25E-07 |
| 11 | 119181674 | rs2510146 | G | C | 0.992 | -7977.388 | -7979.211 | -1.823 | 6.25E-07 |
| 11 | 119648152 | rs4388901 | C | T | 1.011 | -7977.388 | -7979.211 | -1.823 | 6.25E-07 |
| 11 | 119648831 | rs34033074 | A | G | 1.011 | -7977.388 | -7979.211 | -1.823 | 6.25E-07 |
| 11 | 119879327 | rs17245368 | T | C | 1.002 | -7977.388 | -7979.211 | -1.823 | 6.25E-07 |
| 11 | 120557343 | rs10750176 | G | A | 1.014 | -7977.388 | -7979.211 | -1.823 | 6.25E-07 |
| 11 | 121090790 | rs34271655 | C | T | 1.004 | -7977.388 | -7979.212 | -1.823 | 6.25E-07 |
| 11 | 121183145 | rs35288685 | T | C | 1.014 | -7977.388 | -7979.212 | -1.823 | 6.25E-07 |
| 11 | 117801635 | rs17453205 | G | A | 0.986 | -7977.388 | -7979.212 | -1.824 | 6.24E-07 |
| 11 | 119643807 | rs12803325 | T | A | 1.011 | -7977.388 | -7979.213 | -1.824 | 6.24E-07 |
| 11 | 119644249 | rs4420283 | C | T | 1.011 | -7977.388 | -7979.213 | -1.824 | 6.24E-07 |
| 11 | 120023505 | rs12802402 | T | C | 0.995 | -7977.388 | -7979.213 | -1.824 | 6.24E-07 |
| 11 | 121019288 | rs10892677 | T | C | 1.004 | -7977.388 | -7979.212 | -1.824 | 6.24E-07 |
| 11 | 117845302 | rs7939408 | T | A | 0.993 | -7977.388 | -7979.214 | -1.825 | 6.24E-07 |
| 11 | 119269112 | rs35720648 | C | G | 1.007 | -7977.388 | -7979.214 | -1.825 | 6.24E-07 |
| 11 | 119299592 | rs12420302 | G | A | 1.007 | -7977.388 | -7979.214 | -1.825 | 6.24E-07 |
| 11 | 119419568 | rs11217310 | C | G | 0.994 | -7977.388 | -7979.213 | -1.825 | 6.24E-07 |
| 11 | 119444863 | rs11217326 | C | T | 1.008 | -7977.388 | -7979.213 | -1.825 | 6.24E-07 |
| 11 | 119453252 | rs11217329 | A | G | 1.008 | -7977.388 | -7979.213 | -1.825 | 6.24E-07 |
| 11 | 119453579 | rs1396776 | C | A | 1.008 | -7977.388 | -7979.213 | -1.825 | 6.24E-07 |
| 11 | 120025443 | rs12791621 | G | A | 0.995 | -7977.388 | -7979.213 | -1.825 | 6.24E-07 |
| 11 | 120355230 | rs503473 | G | A | 0.986 | -7977.388 | -7979.213 | -1.825 | 6.24E-07 |
| 11 | 120477883 | rs11600354 | A | G | 0.988 | -7977.388 | -7979.213 | -1.825 | 6.24E-07 |
| 11 | 116419268 | rs4938279 | A | G | 0.996 | -7977.388 | -7979.214 | -1.826 | 6.23E-07 |
| 11 | 117331499 | rs10892119 | A | G | 1.006 | -7977.388 | -7979.215 | -1.826 | 6.23E-07 |
| 11 | 117464173 | rs694005 | T | C | 1.008 | -7977.388 | -7979.214 | -1.826 | 6.23E-07 |
| 11 | 117732718 | rs7103567 | T | C | 0.988 | -7977.388 | -7979.215 | -1.826 | 6.23E-07 |
| 11 | 117799923 | rs7928668 | T | C | 1.014 | -7977.388 | -7979.214 | -1.826 | 6.23E-07 |
| 11 | 117961908 | rs651474 | C | T | 1.013 | -7977.388 | -7979.214 | -1.826 | 6.23E-07 |
| 11 | 118519550 | rs601318 | G | C | 1.01 | -7977.388 | -7979.214 | -1.826 | 6.23E-07 |
| 11 | 119299408 | rs12417523 | T | C | 1.006 | -7977.388 | -7979.214 | -1.826 | 6.23E-07 |
| 11 | 119299832 | rs12417611 | A | G | 1.007 | -7977.388 | -7979.215 | -1.826 | 6.23E-07 |
| 11 | 119300681 | rs12807216 | A | C | 1.007 | -7977.388 | -7979.214 | -1.826 | 6.23E-07 |
| 11 | 119425163 | rs11217315 | A | G | 0.995 | -7977.388 | -7979.214 | -1.826 | 6.23E-07 |
| 11 | 119703617 | rs7395060 | G | A | 1.014 | -7977.388 | -7979.214 | -1.826 | 6.23E-07 |
| 11 | 119799765 | rs12283803 | T | C | 0.987 | -7977.388 | -7979.214 | -1.826 | 6.23E-07 |
| 11 | 119965319 | rs11217689 | G | C | 0.988 | -7977.388 | -7979.214 | -1.826 | 6.23E-07 |

| | | | | | | | |
|---|---|---|---|---|---|---|---|
| 11 | 119965383 | rs626302 | A | G | NULL | 0.713000163 | NULL&M_Disease |
| 11 | 119965482 | rs582951 | G | A | NULL | 0.713000163 | NULL&M_Disease |
| 11 | 119965639 | rs637685 | A | G | NULL | 0.713000163 | NULL&M_Disease |
| 11 | 121000784 | rs586473 | T | C | NULL | 0.713000163 | NULL&M_Disease |
| 11 | 121003848 | rs479299 | A | G | NULL | 0.713000163 | NULL&M_Disease |
| 11 | 119326630 | rs12421915 | T | C | NULL | 0.713000163 | NULL&M_Disease |
| 11 | 119412507 | rs10736498 | A | G | NULL | 0.713000163 | NULL&M_Disease |
| 11 | 119460505 | rs2887245 | T | C | NULL | 0.713000163 | NULL&M_Disease |
| 11 | 119570128 | rs11825206 | C | T | NULL | 0.713000163 | NULL&M_Disease |
| 11 | 116419442 | rs4938280 | C | T | NULL | 0.714022227 | NULL&M_Disease |
| 11 | 119417101 | rs10892388 | A | G | NULL | 0.714022227 | NULL&M_Disease |
| 11 | 119972861 | rs7938843 | T | G | NULL | 0.714022227 | NULL&M_Disease |
| 11 | 120038717 | rs12808627 | T | C | NULL | 0.714022227 | NULL&M_Disease |
| 11 | 120053941 | rs2076765 | G | A | NULL | 0.714022227 | NULL&M_Disease |
| 11 | 121001689 | rs670973 | A | C | NULL | 0.714022227 | NULL&M_Disease |
| 11 | 117444044 | rs746910 | G | A | NULL | 0.714022227 | NULL&M_Disease |
| 11 | 117963939 | rs7114565 | A | G | NULL | 0.714022227 | NULL&M_Disease |
| 11 | 118199295 | rs7110837 | A | G | NULL | 0.714022227 | NULL&M_Disease |
| 11 | 119297614 | rs7119828 | G | A | NULL | 0.714022227 | NULL&M_Disease |
| 11 | 119297711 | rs11217279 | A | G | NULL | 0.714022227 | NULL&M_Disease |
| 11 | 119299371 | rs57088249 | A | G | NULL | 0.714022227 | NULL&M_Disease |
| 11 | 119299377 | rs12417550 | A | G | NULL | 0.714022227 | NULL&M_Disease |
| 11 | 119416614 | rs12795183 | T | A | NULL | 0.714022227 | NULL&M_Disease |
| 11 | 121036315 | rs2276057 | A | G | NULL | 0.714022227 | NULL&M_Disease |
| 11 | 117961734 | rs11822370 | G | A | NULL | 0.714022227 | NULL&M_Disease |
| 11 | 117963635 | rs1783861 | T | A | NULL | 0.714022227 | NULL&M_Disease |
| 11 | 118200201 | rs11216853 | T | A | NULL | 0.714022227 | NULL&M_Disease |
| 11 | 118209960 | rs2276424 | A | C | NULL | 0.714022227 | NULL&M_Disease |
| 11 | 118505090 | rs645637 | A | G | NULL | 0.714022227 | NULL&M_Disease |
| 11 | 119422756 | rs10437690 | G | C | NULL | 0.714022227 | NULL&M_Disease |
| 11 | 119423690 | rs10790306 | C | T | NULL | 0.714022227 | NULL&M_Disease |
| 11 | 119968483 | rs10160467 | C | G | NULL | 0.714022227 | NULL&M_Disease |
| 11 | 120032185 | rs12799647 | G | T | NULL | 0.714022227 | NULL&M_Disease |
| 11 | 120037862 | rs12802652 | T | G | NULL | 0.714022227 | NULL&M_Disease |
| 11 | 120039696 | rs35652094 | A | G | NULL | 0.714022227 | NULL&M_Disease |
| 11 | 120490713 | rs4331108 | T | C | NULL | 0.714022227 | NULL&M_Disease |
| 11 | 120553943 | rs34612914 | G | A | NULL | 0.714022227 | NULL&M_Disease |
| 11 | 120717000 | rs11218028 | T | G | NULL | 0.714022227 | NULL&M_Disease |
| 11 | 120999732 | rs551199 | A | G | NULL | 0.714022227 | NULL&M_Disease |
| 11 | 121000212 | rs643744 | T | G | NULL | 0.714022227 | NULL&M_Disease |
| 11 | 117304947 | rs681160 | C | T | NULL | 0.714022227 | NULL&M_Disease |
| 11 | 119269952 | rs1583058 | C | T | NULL | 0.714022227 | NULL&M_Disease |
| 11 | 119424157 | rs10790307 | C | T | NULL | 0.714022227 | NULL&M_Disease |
| 11 | 119984638 | rs1901707 | T | C | NULL | 0.714022227 | NULL&M_Disease |
| 11 | 120036812 | rs12795599 | A | G | NULL | 0.714022227 | NULL&M_Disease |
| 11 | 120497350 | rs4445643 | T | G | NULL | 0.714022227 | NULL&M_Disease |
| 11 | 116329090 | rs570745 | A | G | NULL | 0.714022227 | NULL&M_Disease |
| 11 | 117267884 | rs573455 | A | G | NULL | 0.714022227 | NULL&M_Disease |
| 11 | 118067009 | rs647099 | A | G | NULL | 0.714022227 | NULL&M_Disease |
| 11 | 119262493 | rs35327319 | G | T | NULL | 0.714022227 | NULL&M_Disease |
| 11 | 119304053 | rs11217281 | T | C | NULL | 0.714022227 | NULL&M_Disease |
| 11 | 119308804 | rs12802321 | A | G | NULL | 0.714022227 | NULL&M_Disease |
| 11 | 119419133 | rs10790302 | A | G | NULL | 0.714022227 | NULL&M_Disease |
| 11 | 119419672 | rs11217311 | T | C | NULL | 0.714022227 | NULL&M_Disease |
| 11 | 119419976 | rs11217312 | C | A | NULL | 0.714022227 | NULL&M_Disease |
| 11 | 119437268 | rs10736501 | G | A | NULL | 0.714022227 | NULL&M_Disease |
| 11 | 119573524 | rs7932832 | G | A | NULL | 0.714022227 | NULL&M_Disease |
| 11 | 119703771 | rs1665777 | C | A | NULL | 0.714022227 | NULL&M_Disease |
| 11 | 120984087 | rs2298478 | G | C | NULL | 0.714022227 | NULL&M_Disease |
| 11 | 121082690 | rs12787035 | A | G | NULL | 0.714022227 | NULL&M_Disease |
| 11 | 117537799 | rs1145219 | T | G | NULL | 0.714022227 | NULL&M_Disease |
| 11 | 119233110 | rs59969944 | A | G | NULL | 0.714022227 | NULL&M_Disease |
| 11 | 119296520 | rs3862602 | G | A | NULL | 0.714022227 | NULL&M_Disease |
| 11 | 119444474 | rs11217325 | G | A | NULL | 0.714022227 | NULL&M_Disease |
| 11 | 119576555 | rs4938716 | T | G | NULL | 0.714022227 | NULL&M_Disease |
| 11 | 116537761 | rs6589557 | T | C | NULL | 0.714022227 | NULL&M_Disease |
| 11 | 116537942 | rs6589558 | A | G | NULL | 0.714022227 | NULL&M_Disease |
| 11 | 116563992 | rs1787701 | C | G | NULL | 0.714022227 | NULL&M_Disease |
| 11 | 117743822 | rs10750116 | C | G | NULL | 0.714022227 | NULL&M_Disease |
| 11 | 117800083 | rs10892196 | T | C | NULL | 0.714022227 | NULL&M_Disease |
| 11 | 117973673 | rs2276129 | A | G | NULL | 0.714022227 | NULL&M_Disease |
| 11 | 118066582 | rs648682 | T | C | NULL | 0.714022227 | NULL&M_Disease |
| 11 | 119290369 | rs1894006 | T | C | NULL | 0.714022227 | NULL&M_Disease |
| 11 | 119302637 | rs891293 | A | G | NULL | 0.714022227 | NULL&M_Disease |
| 11 | 119319811 | rs66461646 | C | T | NULL | 0.714022227 | NULL&M_Disease |
| 11 | 119413090 | rs10892386 | T | C | NULL | 0.714022227 | NULL&M_Disease |
| 11 | 119445417 | rs61898748 | T | C | NULL | 0.714022227 | NULL&M_Disease |
| 11 | 119445971 | rs12293402 | T | C | NULL | 0.714022227 | NULL&M_Disease |
| 11 | 119574060 | rs6589769 | G | C | NULL | 0.714022227 | NULL&M_Disease |
| 11 | 119660492 | rs34492753 | T | C | NULL | 0.714022227 | NULL&M_Disease |
| 11 | 119946041 | rs11217673 | T | G | NULL | 0.714022227 | NULL&M_Disease |
| 11 | 120055403 | rs12792820 | A | T | NULL | 0.714022227 | NULL&M_Disease |
| 11 | 117275561 | rs518368 | C | G | NULL | 0.714022227 | NULL&M_Disease |
| 11 | 117341808 | rs609341 | T | G | NULL | 0.714022227 | NULL&M_Disease |
| 11 | 117806937 | rs10790226 | T | C | NULL | 0.714022227 | NULL&M_Disease |
| 11 | 119274417 | rs12269936 | G | A | NULL | 0.714022227 | NULL&M_Disease |
| 11 | 119277583 | rs12421359 | G | A | NULL | 0.714022227 | NULL&M_Disease |
| 11 | 119304002 | rs11217280 | G | A | NULL | 0.714022227 | NULL&M_Disease |
| 11 | 119304238 | rs12283157 | C | G | NULL | 0.714022227 | NULL&M_Disease |
| 11 | 119306752 | rs61898859 | C | A | NULL | 0.714022227 | NULL&M_Disease |
| 11 | 119319701 | rs35514679 | A | C | NULL | 0.714022227 | NULL&M_Disease |
| 11 | 119428506 | rs58831762 | G | T | NULL | 0.714022227 | NULL&M_Disease |
| 11 | 119436709 | rs10750156 | G | A | NULL | 0.714022227 | NULL&M_Disease |
| 11 | 119436816 | rs10892408 | G | A | NULL | 0.714022227 | NULL&M_Disease |
| 11 | 119838883 | rs1944505 | A | G | NULL | 0.714022227 | NULL&M_Disease |
| 11 | 116197720 | rs930430 | A | G | NULL | 0.714022227 | NULL&M_Disease |

| | | | | | | | | | |
|---|---|---|---|---|---|---|---|---|---|
| 11 | 119965383 | rs626302 | A | G | 0.988 | -7977.388 | -7979.214 | -1.826 | 6.23E-07 |
| 11 | 119965482 | rs582951 | G | A | 0.988 | -7977.388 | -7979.214 | -1.826 | 6.23E-07 |
| 11 | 119965639 | rs637685 | A | G | 0.988 | -7977.388 | -7979.214 | -1.826 | 6.23E-07 |
| 11 | 121000784 | rs586473 | T | C | 1.008 | -7977.388 | -7979.215 | -1.826 | 6.23E-07 |
| 11 | 121003848 | rs479299 | A | G | 1.008 | -7977.388 | -7979.215 | -1.826 | 6.23E-07 |
| 11 | 119326630 | rs12421915 | T | C | 1.006 | -7977.388 | -7979.216 | -1.827 | 6.22E-07 |
| 11 | 119412507 | rs10736498 | A | G | 0.995 | -7977.388 | -7979.216 | -1.827 | 6.22E-07 |
| 11 | 119460505 | rs2887245 | T | C | 0.987 | -7977.388 | -7979.216 | -1.827 | 6.22E-07 |
| 11 | 119570128 | rs11825206 | C | T | 1.012 | -7977.388 | -7979.215 | -1.827 | 6.22E-07 |
| 11 | 116419442 | rs4938280 | C | T | 0.996 | -7977.388 | -7979.216 | -1.828 | 6.22E-07 |
| 11 | 119417101 | rs10892388 | A | G | 0.995 | -7977.388 | -7979.217 | -1.828 | 6.22E-07 |
| 11 | 119972861 | rs7938843 | T | G | 0.992 | -7977.388 | -7979.217 | -1.828 | 6.22E-07 |
| 11 | 120038717 | rs12808627 | T | C | 0.996 | -7977.388 | -7979.216 | -1.828 | 6.22E-07 |
| 11 | 120053941 | rs2076765 | G | A | 1.004 | -7977.388 | -7979.217 | -1.828 | 6.22E-07 |
| 11 | 121001689 | rs670973 | A | C | 1.008 | -7977.388 | -7979.216 | -1.828 | 6.22E-07 |
| 11 | 117444044 | rs746910 | G | A | 0.987 | -7977.388 | -7979.217 | -1.829 | 6.21E-07 |
| 11 | 117963939 | rs7114565 | A | G | 1.012 | -7977.388 | -7979.218 | -1.829 | 6.21E-07 |
| 11 | 118199295 | rs7110837 | A | G | 1 | -7977.388 | -7979.218 | -1.829 | 6.21E-07 |
| 11 | 119297614 | rs7119828 | G | A | 1.006 | -7977.388 | -7979.217 | -1.829 | 6.21E-07 |
| 11 | 119297711 | rs11217279 | A | G | 1.006 | -7977.388 | -7979.217 | -1.829 | 6.21E-07 |
| 11 | 119299371 | rs57088249 | A | G | 1.006 | -7977.388 | -7979.217 | -1.829 | 6.21E-07 |
| 11 | 119299377 | rs12417550 | A | G | 1.006 | -7977.388 | -7979.218 | -1.829 | 6.21E-07 |
| 11 | 119416614 | rs12795183 | T | A | 0.989 | -7977.388 | -7979.217 | -1.829 | 6.21E-07 |
| 11 | 121036315 | rs2276057 | A | G | 1.002 | -7977.388 | -7979.217 | -1.829 | 6.21E-07 |
| 11 | 117961734 | rs11822370 | G | A | 0.99 | -7977.388 | -7979.218 | -1.83 | 6.21E-07 |
| 11 | 117963635 | rs1783861 | T | A | 1.012 | -7977.388 | -7979.219 | -1.83 | 6.21E-07 |
| 11 | 118200201 | rs11216853 | T | A | 1 | -7977.388 | -7979.218 | -1.83 | 6.21E-07 |
| 11 | 118209960 | rs2276424 | A | C | 1.001 | -7977.388 | -7979.218 | -1.83 | 6.21E-07 |
| 11 | 118505090 | rs645637 | A | G | 1.01 | -7977.388 | -7979.218 | -1.83 | 6.21E-07 |
| 11 | 119422756 | rs10437690 | G | C | 0.995 | -7977.388 | -7979.218 | -1.83 | 6.21E-07 |
| 11 | 119423690 | rs10790306 | C | T | 0.995 | -7977.388 | -7979.218 | -1.83 | 6.21E-07 |
| 11 | 119968483 | rs10160467 | C | G | 0.99 | -7977.388 | -7979.218 | -1.83 | 6.21E-07 |
| 11 | 120032185 | rs12799647 | G | T | 0.996 | -7977.388 | -7979.219 | -1.83 | 6.21E-07 |
| 11 | 120037862 | rs12802652 | T | G | 0.996 | -7977.388 | -7979.218 | -1.83 | 6.21E-07 |
| 11 | 120039696 | rs35652094 | A | G | 0.996 | -7977.388 | -7979.219 | -1.83 | 6.21E-07 |
| 11 | 120490713 | rs4331108 | T | C | 0.988 | -7977.388 | -7979.218 | -1.83 | 6.21E-07 |
| 11 | 120553943 | rs34612914 | G | A | 0.988 | -7977.388 | -7979.218 | -1.83 | 6.21E-07 |
| 11 | 120717000 | rs11218028 | T | G | 0.988 | -7977.388 | -7979.219 | -1.83 | 6.21E-07 |
| 11 | 120999732 | rs551199 | A | G | 1.008 | -7977.388 | -7979.218 | -1.83 | 6.21E-07 |
| 11 | 121000212 | rs643744 | T | G | 1.008 | -7977.388 | -7979.218 | -1.83 | 6.21E-07 |
| 11 | 117304947 | rs681160 | C | T | 0.987 | -7977.388 | -7979.219 | -1.831 | 6.20E-07 |
| 11 | 119269952 | rs1583058 | C | T | 1.006 | -7977.388 | -7979.219 | -1.831 | 6.20E-07 |
| 11 | 119424157 | rs10790307 | C | T | 0.996 | -7977.388 | -7979.22 | -1.831 | 6.20E-07 |
| 11 | 119984638 | rs1901707 | T | C | 1.005 | -7977.388 | -7979.219 | -1.831 | 6.20E-07 |
| 11 | 120036812 | rs12795599 | A | G | 0.996 | -7977.388 | -7979.219 | -1.831 | 6.20E-07 |
| 11 | 120497350 | rs4445643 | T | G | 0.988 | -7977.388 | -7979.22 | -1.831 | 6.20E-07 |
| 11 | 116329090 | rs570745 | A | G | 1.013 | -7977.388 | -7979.22 | -1.832 | 6.19E-07 |
| 11 | 117267884 | rs573455 | A | G | 0.987 | -7977.388 | -7979.221 | -1.832 | 6.19E-07 |
| 11 | 118067009 | rs647099 | A | G | 0.987 | -7977.388 | -7979.22 | -1.832 | 6.19E-07 |
| 11 | 119262493 | rs35327319 | G | T | 1.007 | -7977.388 | -7979.221 | -1.832 | 6.19E-07 |
| 11 | 119304053 | rs11217281 | T | C | 1.005 | -7977.388 | -7979.221 | -1.832 | 6.19E-07 |
| 11 | 119308804 | rs12802321 | A | G | 1.006 | -7977.388 | -7979.221 | -1.832 | 6.19E-07 |
| 11 | 119419133 | rs10790302 | A | G | 0.996 | -7977.388 | -7979.22 | -1.832 | 6.19E-07 |
| 11 | 119419672 | rs11217311 | T | C | 0.996 | -7977.388 | -7979.22 | -1.832 | 6.19E-07 |
| 11 | 119419976 | rs11217312 | C | A | 0.996 | -7977.388 | -7979.22 | -1.832 | 6.19E-07 |
| 11 | 119437268 | rs10736501 | G | A | 0.996 | -7977.388 | -7979.22 | -1.832 | 6.19E-07 |
| 11 | 119573524 | rs7932832 | G | A | 1.012 | -7977.388 | -7979.22 | -1.832 | 6.19E-07 |
| 11 | 119703771 | rs1665777 | C | A | 1.005 | -7977.388 | -7979.22 | -1.832 | 6.19E-07 |
| 11 | 120984087 | rs2298478 | G | C | 1.003 | -7977.388 | -7979.22 | -1.832 | 6.19E-07 |
| 11 | 121082690 | rs12787035 | A | G | 1.001 | -7977.388 | -7979.22 | -1.832 | 6.19E-07 |
| 11 | 117537799 | rs1145219 | T | G | 1.013 | -7977.388 | -7979.221 | -1.833 | 6.19E-07 |
| 11 | 119233110 | rs59969944 | A | G | 0.991 | -7977.388 | -7979.221 | -1.833 | 6.19E-07 |
| 11 | 119296520 | rs3862602 | G | A | 1.005 | -7977.388 | -7979.221 | -1.833 | 6.19E-07 |
| 11 | 119444474 | rs11217325 | G | A | 1.007 | -7977.388 | -7979.221 | -1.833 | 6.19E-07 |
| 11 | 119576555 | rs4938716 | T | G | 1.009 | -7977.388 | -7979.222 | -1.833 | 6.19E-07 |
| 11 | 116537761 | rs6589557 | T | C | 0.995 | -7977.388 | -7979.223 | -1.834 | 6.18E-07 |
| 11 | 116537942 | rs6589558 | A | G | 0.995 | -7977.388 | -7979.223 | -1.834 | 6.18E-07 |
| 11 | 116563992 | rs1787701 | C | G | 0.987 | -7977.388 | -7979.222 | -1.834 | 6.18E-07 |
| 11 | 117743822 | rs10750116 | C | G | 0.988 | -7977.388 | -7979.222 | -1.834 | 6.18E-07 |
| 11 | 117800083 | rs10892196 | T | C | 1.013 | -7977.388 | -7979.223 | -1.834 | 6.18E-07 |
| 11 | 117973673 | rs2276129 | A | G | 0.99 | -7977.388 | -7979.222 | -1.834 | 6.18E-07 |
| 11 | 118066582 | rs648682 | T | C | 0.987 | -7977.388 | -7979.222 | -1.834 | 6.18E-07 |
| 11 | 119290369 | rs1894006 | T | C | 1.005 | -7977.388 | -7979.222 | -1.834 | 6.18E-07 |
| 11 | 119302637 | rs891293 | A | G | 1.005 | -7977.388 | -7979.222 | -1.834 | 6.18E-07 |
| 11 | 119319811 | rs66461646 | C | T | 1.005 | -7977.388 | -7979.223 | -1.834 | 6.18E-07 |
| 11 | 119413090 | rs10892386 | T | C | 0.998 | -7977.388 | -7979.223 | -1.834 | 6.18E-07 |
| 11 | 119445417 | rs61898748 | T | C | 1.006 | -7977.388 | -7979.222 | -1.834 | 6.18E-07 |
| 11 | 119445971 | rs12293402 | T | C | 1.006 | -7977.388 | -7979.222 | -1.834 | 6.18E-07 |
| 11 | 119574060 | rs6589769 | G | C | 1.012 | -7977.388 | -7979.223 | -1.834 | 6.18E-07 |
| 11 | 119660492 | rs34492753 | T | C | 1.009 | -7977.388 | -7979.222 | -1.834 | 6.18E-07 |
| 11 | 119946041 | rs11217673 | T | G | 1.01 | -7977.388 | -7979.222 | -1.834 | 6.18E-07 |
| 11 | 120055403 | rs12792820 | A | T | 1.002 | -7977.388 | -7979.223 | -1.834 | 6.18E-07 |
| 11 | 117275561 | rs518368 | C | G | 1.011 | -7977.388 | -7979.223 | -1.835 | 6.17E-07 |
| 11 | 117341808 | rs609341 | T | G | 1.012 | -7977.388 | -7979.223 | -1.835 | 6.17E-07 |
| 11 | 117806937 | rs10790226 | T | C | 1.011 | -7977.388 | -7979.223 | -1.835 | 6.17E-07 |
| 11 | 119274417 | rs12269936 | G | A | 1.005 | -7977.388 | -7979.223 | -1.835 | 6.17E-07 |
| 11 | 119277583 | rs12421359 | G | A | 1.005 | -7977.388 | -7979.223 | -1.835 | 6.17E-07 |
| 11 | 119304002 | rs11217280 | G | A | 1.005 | -7977.388 | -7979.224 | -1.835 | 6.17E-07 |
| 11 | 119304238 | rs12283157 | C | G | 1.005 | -7977.388 | -7979.224 | -1.835 | 6.17E-07 |
| 11 | 119306752 | rs61898859 | C | A | 1.005 | -7977.388 | -7979.224 | -1.835 | 6.17E-07 |
| 11 | 119319701 | rs35514679 | A | C | 1.005 | -7977.388 | -7979.223 | -1.835 | 6.17E-07 |
| 11 | 119428506 | rs58831762 | G | T | 0.996 | -7977.388 | -7979.223 | -1.835 | 6.17E-07 |
| 11 | 119436709 | rs10750156 | G | A | 0.996 | -7977.388 | -7979.223 | -1.835 | 6.17E-07 |
| 11 | 119436816 | rs10892408 | G | A | 0.996 | -7977.388 | -7979.223 | -1.835 | 6.17E-07 |
| 11 | 119838883 | rs1944505 | A | G | 0.998 | -7977.388 | -7979.223 | -1.835 | 6.17E-07 |
| 11 | 116197720 | rs930430 | A | G | 0.997 | -7977.388 | -7979.224 | -1.836 | 6.17E-07 |

| | | | | | | | |
|---|---|---|---|---|---|---|---|
| 11 | 117800574 | rs10790222 | A | G | NULL | 0.714022227 | NULL&M_Disease |
| 11 | 118067192 | rs605908 | C | A | NULL | 0.714022227 | NULL&M_Disease |
| 11 | 119257248 | rs56133216 | C | T | NULL | 0.714022227 | NULL&M_Disease |
| 11 | 119281397 | rs56027537 | C | T | NULL | 0.714022227 | NULL&M_Disease |
| 11 | 119317018 | rs735007 | G | A | NULL | 0.714022227 | NULL&M_Disease |
| 11 | 119412922 | rs10892385 | A | C | NULL | 0.714022227 | NULL&M_Disease |
| 11 | 119420468 | rs10892390 | T | C | NULL | 0.714022227 | NULL&M_Disease |
| 11 | 119420599 | rs10790304 | T | G | NULL | 0.714022227 | NULL&M_Disease |
| 11 | 119420615 | rs10790305 | A | G | NULL | 0.714022227 | NULL&M_Disease |
| 11 | 119425342 | rs10790308 | A | G | NULL | 0.714022227 | NULL&M_Disease |
| 11 | 119426055 | rs10790309 | A | G | NULL | 0.714022227 | NULL&M_Disease |
| 11 | 119426104 | rs10892391 | G | T | NULL | 0.714022227 | NULL&M_Disease |
| 11 | 119427370 | rs59209096 | A | G | NULL | 0.714022227 | NULL&M_Disease |
| 11 | 119973082 | rs7131339 | A | G | NULL | 0.714022227 | NULL&M_Disease |
| 11 | 120689119 | rs4935752 | C | A | NULL | 0.714022227 | NULL&M_Disease |
| 11 | 117966597 | rs1941631 | C | A | NULL | 0.714022227 | NULL&M_Disease |
| 11 | 119272485 | rs7111286 | C | G | NULL | 0.714022227 | NULL&M_Disease |
| 11 | 119276600 | rs12575027 | A | G | NULL | 0.714022227 | NULL&M_Disease |
| 11 | 119276834 | rs12577180 | C | A | NULL | 0.714022227 | NULL&M_Disease |
| 11 | 119284523 | rs12417301 | A | T | NULL | 0.714022227 | NULL&M_Disease |
| 11 | 119286275 | rs3138096 | G | C | NULL | 0.714022227 | NULL&M_Disease |
| 11 | 119288619 | rs3138094 | G | A | NULL | 0.714022227 | NULL&M_Disease |
| 11 | 119291403 | rs3138093 | C | T | NULL | 0.714022227 | NULL&M_Disease |
| 11 | 119292314 | rs1429512 | C | T | NULL | 0.714022227 | NULL&M_Disease |
| 11 | 119301580 | rs12418314 | T | C | NULL | 0.714022227 | NULL&M_Disease |
| 11 | 119301754 | rs12421120 | C | A | NULL | 0.714022227 | NULL&M_Disease |
| 11 | 119301942 | rs12418370 | A | G | NULL | 0.714022227 | NULL&M_Disease |
| 11 | 119418040 | rs1605853 | C | T | NULL | 0.714022227 | NULL&M_Disease |
| 11 | 119436098 | rs10892407 | A | C | NULL | 0.714022227 | NULL&M_Disease |
| 11 | 119492741 | rs7117705 | C | T | NULL | 0.714022227 | NULL&M_Disease |
| 11 | 119494843 | rs12797305 | G | A | NULL | 0.714022227 | NULL&M_Disease |
| 11 | 119569737 | rs7945395 | G | A | NULL | 0.714022227 | NULL&M_Disease |
| 11 | 119658670 | rs11217505 | C | G | NULL | 0.714022227 | NULL&M_Disease |
| 11 | 120323214 | rs496248 | A | G | NULL | 0.714022227 | NULL&M_Disease |
| 11 | 120879655 | rs668021 | C | G | NULL | 0.715042106 | NULL&M_Disease |
| 11 | 116561790 | rs1784121 | T | G | NULL | 0.715042106 | NULL&M_Disease |
| 11 | 116561996 | rs1787704 | T | C | NULL | 0.715042106 | NULL&M_Disease |
| 11 | 117442388 | rs2017098 | G | C | NULL | 0.715042106 | NULL&M_Disease |
| 11 | 117454506 | rs7948446 | T | C | NULL | 0.715042106 | NULL&M_Disease |
| 11 | 117969374 | rs4938481 | T | C | NULL | 0.715042106 | NULL&M_Disease |
| 11 | 119427371 | rs59427601 | A | G | NULL | 0.715042106 | NULL&M_Disease |
| 11 | 119704194 | rs527833 | A | G | NULL | 0.715042106 | NULL&M_Disease |
| 11 | 119883569 | rs622350 | A | C | NULL | 0.715042106 | NULL&M_Disease |
| 11 | 120058902 | rs12791393 | T | C | NULL | 0.715042106 | NULL&M_Disease |
| 11 | 120477398 | rs11600274 | A | G | NULL | 0.715042106 | NULL&M_Disease |
| 11 | 120486195 | rs12286173 | T | C | NULL | 0.715042106 | NULL&M_Disease |
| 11 | 120688502 | rs4936552 | T | A | NULL | 0.715042106 | NULL&M_Disease |
| 11 | 119306878 | rs12788547 | A | C | NULL | 0.715042106 | NULL&M_Disease |
| 11 | 119429736 | rs10750151 | T | C | NULL | 0.715042106 | NULL&M_Disease |
| 11 | 119431293 | rs10750152 | T | C | NULL | 0.715042106 | NULL&M_Disease |
| 11 | 119431487 | rs10892398 | C | T | NULL | 0.715042106 | NULL&M_Disease |
| 11 | 119431602 | rs55808044 | G | C | NULL | 0.715042106 | NULL&M_Disease |
| 11 | 119431655 | rs11217320 | G | A | NULL | 0.715042106 | NULL&M_Disease |
| 11 | 119432084 | rs11217321 | G | A | NULL | 0.715042106 | NULL&M_Disease |
| 11 | 119493539 | rs4519102 | C | A | NULL | 0.715042106 | NULL&M_Disease |
| 11 | 119647159 | rs7929505 | C | T | NULL | 0.715042106 | NULL&M_Disease |
| 11 | 119850450 | rs601052 | C | T | NULL | 0.715042106 | NULL&M_Disease |
| 11 | 120057972 | rs12808972 | A | G | NULL | 0.715042106 | NULL&M_Disease |
| 11 | 120544871 | rs2000869 | G | A | NULL | 0.715042106 | NULL&M_Disease |
| 11 | 121004172 | rs646371 | A | G | NULL | 0.715042106 | NULL&M_Disease |
| 11 | 117962275 | rs17528237 | T | C | NULL | 0.715042106 | NULL&M_Disease |
| 11 | 117973511 | rs4344527 | G | A | NULL | 0.715042106 | NULL&M_Disease |
| 11 | 118067132 | rs647573 | C | T | NULL | 0.715042106 | NULL&M_Disease |
| 11 | 119240809 | rs10892355 | C | T | NULL | 0.715042106 | NULL&M_Disease |
| 11 | 119308201 | rs56103633 | T | C | NULL | 0.715042106 | NULL&M_Disease |
| 11 | 119309000 | rs7130634 | A | T | NULL | 0.715042106 | NULL&M_Disease |
| 11 | 119426508 | rs10892392 | T | C | NULL | 0.715042106 | NULL&M_Disease |
| 11 | 119426651 | rs10892393 | T | C | NULL | 0.715042106 | NULL&M_Disease |
| 11 | 119429080 | rs10750150 | T | C | NULL | 0.715042106 | NULL&M_Disease |
| 11 | 119429098 | rs10736499 | C | T | NULL | 0.715042106 | NULL&M_Disease |
| 11 | 119429614 | rs10892394 | A | G | NULL | 0.715042106 | NULL&M_Disease |
| 11 | 119429656 | rs10892395 | G | A | NULL | 0.715042106 | NULL&M_Disease |
| 11 | 119429662 | rs10892396 | G | T | NULL | 0.715042106 | NULL&M_Disease |
| 11 | 119429848 | rs11217318 | A | G | NULL | 0.715042106 | NULL&M_Disease |
| 11 | 119429923 | rs11217319 | A | G | NULL | 0.715042106 | NULL&M_Disease |
| 11 | 119430203 | rs10892397 | G | A | NULL | 0.715042106 | NULL&M_Disease |
| 11 | 119430266 | rs12274984 | A | G | NULL | 0.715042106 | NULL&M_Disease |
| 11 | 119432144 | rs10892399 | C | T | NULL | 0.715042106 | NULL&M_Disease |
| 11 | 119432909 | rs10750154 | G | A | NULL | 0.715042106 | NULL&M_Disease |
| 11 | 119432935 | rs10790310 | T | C | NULL | 0.715042106 | NULL&M_Disease |
| 11 | 119433005 | rs10790311 | T | C | NULL | 0.715042106 | NULL&M_Disease |
| 11 | 119433521 | rs10892402 | G | A | NULL | 0.715042106 | NULL&M_Disease |
| 11 | 119434032 | rs10750155 | T | A | NULL | 0.715042106 | NULL&M_Disease |
| 11 | 119434275 | rs10790312 | T | C | NULL | 0.715042106 | NULL&M_Disease |
| 11 | 119434279 | rs2887244 | G | T | NULL | 0.715042106 | NULL&M_Disease |
| 11 | 119434468 | rs2368425 | T | G | NULL | 0.715042106 | NULL&M_Disease |
| 11 | 119434711 | rs4018655 | A | G | NULL | 0.715042106 | NULL&M_Disease |
| 11 | 119434843 | rs10892403 | A | C | NULL | 0.715042106 | NULL&M_Disease |
| 11 | 119630641 | rs11217471 | A | G | NULL | 0.715042106 | NULL&M_Disease |
| 11 | 119899963 | rs653278 | T | A | NULL | 0.715042106 | NULL&M_Disease |
| 11 | 117335722 | rs538847 | T | C | NULL | 0.715042106 | NULL&M_Disease |
| 11 | 117745659 | rs543312 | C | T | NULL | 0.715042106 | NULL&M_Disease |
| 11 | 117771069 | rs543480 | T | C | NULL | 0.715042106 | NULL&M_Disease |
| 11 | 117842533 | rs7122174 | A | T | NULL | 0.715042106 | NULL&M_Disease |
| 11 | 118067426 | rs648947 | A | G | NULL | 0.715042106 | NULL&M_Disease |
| 11 | 119260456 | rs56183153 | A | C | NULL | 0.715042106 | NULL&M_Disease |

| | | | | | | | | | |
|---|---|---|---|---|---|---|---|---|---|
| 11 | 117800574 | rs10790222 | A | G | 1.013 | -7977.388 | -7979.224 | -1.836 | 6.17E-07 |
| 11 | 118067192 | rs605908 | C | A | 0.987 | -7977.388 | -7979.224 | -1.836 | 6.17E-07 |
| 11 | 119257248 | rs56133216 | C | T | 1.006 | -7977.388 | -7979.225 | -1.836 | 6.17E-07 |
| 11 | 119281397 | rs56027537 | C | T | 1.005 | -7977.388 | -7979.225 | -1.836 | 6.17E-07 |
| 11 | 119317018 | rs735007 | G | A | 1.005 | -7977.388 | -7979.224 | -1.836 | 6.17E-07 |
| 11 | 119412922 | rs10892385 | A | C | 0.999 | -7977.388 | -7979.225 | -1.836 | 6.17E-07 |
| 11 | 119420468 | rs10892390 | T | C | 0.997 | -7977.388 | -7979.224 | -1.836 | 6.17E-07 |
| 11 | 119420599 | rs10790304 | T | G | 0.997 | -7977.388 | -7979.224 | -1.836 | 6.17E-07 |
| 11 | 119420615 | rs10790305 | A | G | 0.997 | -7977.388 | -7979.224 | -1.836 | 6.17E-07 |
| 11 | 119425342 | rs10790308 | A | G | 0.997 | -7977.388 | -7979.224 | -1.836 | 6.17E-07 |
| 11 | 119426055 | rs10790309 | A | G | 0.997 | -7977.388 | -7979.224 | -1.836 | 6.17E-07 |
| 11 | 119426104 | rs10892391 | G | T | 0.997 | -7977.388 | -7979.224 | -1.836 | 6.17E-07 |
| 11 | 119427370 | rs59209096 | A | G | 0.997 | -7977.388 | -7979.224 | -1.836 | 6.17E-07 |
| 11 | 119973082 | rs7131339 | A | G | 0.993 | -7977.388 | -7979.224 | -1.836 | 6.17E-07 |
| 11 | 120689119 | rs4935752 | C | A | 0.989 | -7977.388 | -7979.225 | -1.836 | 6.17E-07 |
| 11 | 117966597 | rs1941631 | C | A | 1.012 | -7977.388 | -7979.225 | -1.837 | 6.16E-07 |
| 11 | 119272485 | rs7111286 | C | G | 1.005 | -7977.388 | -7979.225 | -1.837 | 6.16E-07 |
| 11 | 119276600 | rs12575027 | A | G | 1.005 | -7977.388 | -7979.225 | -1.837 | 6.16E-07 |
| 11 | 119276834 | rs12577180 | C | A | 1.005 | -7977.388 | -7979.225 | -1.837 | 6.16E-07 |
| 11 | 119284523 | rs12417301 | A | T | 1.005 | -7977.388 | -7979.225 | -1.837 | 6.16E-07 |
| 11 | 119286275 | rs3138096 | G | C | 1.005 | -7977.388 | -7979.225 | -1.837 | 6.16E-07 |
| 11 | 119288619 | rs3138094 | G | A | 1.005 | -7977.388 | -7979.225 | -1.837 | 6.16E-07 |
| 11 | 119291403 | rs3138093 | C | T | 1.005 | -7977.388 | -7979.225 | -1.837 | 6.16E-07 |
| 11 | 119292314 | rs1429512 | C | T | 1.005 | -7977.388 | -7979.226 | -1.837 | 6.16E-07 |
| 11 | 119301580 | rs12418314 | T | C | 1.005 | -7977.388 | -7979.225 | -1.837 | 6.16E-07 |
| 11 | 119301754 | rs12421120 | C | A | 1.005 | -7977.388 | -7979.225 | -1.837 | 6.16E-07 |
| 11 | 119301942 | rs12418370 | A | G | 1.005 | -7977.388 | -7979.225 | -1.837 | 6.16E-07 |
| 11 | 119418040 | rs1605853 | C | T | 0.989 | -7977.388 | -7979.225 | -1.837 | 6.16E-07 |
| 11 | 119436098 | rs10892407 | A | C | 0.997 | -7977.388 | -7979.225 | -1.837 | 6.16E-07 |
| 11 | 119492741 | rs7117705 | C | T | 1.005 | -7977.388 | -7979.225 | -1.837 | 6.16E-07 |
| 11 | 119494843 | rs12797305 | G | A | 1.004 | -7977.388 | -7979.225 | -1.837 | 6.16E-07 |
| 11 | 119569737 | rs7945395 | G | A | 1.011 | -7977.388 | -7979.226 | -1.837 | 6.16E-07 |
| 11 | 119658670 | rs11217505 | C | G | 1.008 | -7977.388 | -7979.225 | -1.837 | 6.16E-07 |
| 11 | 120323214 | rs496248 | A | G | 0.987 | -7977.388 | -7979.225 | -1.837 | 6.16E-07 |
| 11 | 120879655 | rs668021 | C | G | 0.988 | -7977.388 | -7979.226 | -1.837 | 6.16E-07 |
| 11 | 116561790 | rs1784121 | T | G | 0.987 | -7977.388 | -7979.226 | -1.838 | 6.16E-07 |
| 11 | 116561996 | rs1787704 | T | C | 0.987 | -7977.388 | -7979.226 | -1.838 | 6.16E-07 |
| 11 | 117442388 | rs2017098 | G | C | 0.987 | -7977.388 | -7979.227 | -1.838 | 6.16E-07 |
| 11 | 117454506 | rs7948446 | T | C | 0.987 | -7977.388 | -7979.227 | -1.838 | 6.16E-07 |
| 11 | 117969374 | rs4938481 | T | C | 0.991 | -7977.388 | -7979.226 | -1.838 | 6.16E-07 |
| 11 | 119427371 | rs59427601 | A | G | 0.997 | -7977.388 | -7979.226 | -1.838 | 6.16E-07 |
| 11 | 119704194 | rs527833 | A | G | 0.998 | -7977.388 | -7979.227 | -1.838 | 6.16E-07 |
| 11 | 119883569 | rs622350 | A | C | 0.988 | -7977.388 | -7979.226 | -1.838 | 6.16E-07 |
| 11 | 120058902 | rs12791393 | T | C | 0.997 | -7977.388 | -7979.226 | -1.838 | 6.16E-07 |
| 11 | 120477398 | rs11600274 | A | G | 0.989 | -7977.388 | -7979.226 | -1.838 | 6.16E-07 |
| 11 | 120486195 | rs12286173 | T | C | 0.988 | -7977.388 | -7979.226 | -1.838 | 6.16E-07 |
| 11 | 120688502 | rs4936552 | T | A | 0.99 | -7977.388 | -7979.226 | -1.838 | 6.16E-07 |
| 11 | 119306878 | rs12788547 | A | C | 1.004 | -7977.388 | -7979.227 | -1.839 | 6.15E-07 |
| 11 | 119429736 | rs10750151 | T | C | 1.002 | -7977.388 | -7979.227 | -1.839 | 6.15E-07 |
| 11 | 119431293 | rs10750152 | T | C | 1.002 | -7977.388 | -7979.227 | -1.839 | 6.15E-07 |
| 11 | 119431487 | rs10892398 | C | T | 1.002 | -7977.388 | -7979.227 | -1.839 | 6.15E-07 |
| 11 | 119431602 | rs55808044 | G | C | 1.002 | -7977.388 | -7979.227 | -1.839 | 6.15E-07 |
| 11 | 119431655 | rs11217320 | G | A | 1.002 | -7977.388 | -7979.227 | -1.839 | 6.15E-07 |
| 11 | 119432084 | rs11217321 | G | A | 1.002 | -7977.388 | -7979.227 | -1.839 | 6.15E-07 |
| 11 | 119493539 | rs4519102 | C | A | 0.997 | -7977.388 | -7979.228 | -1.839 | 6.15E-07 |
| 11 | 119647159 | rs7929505 | C | T | 1.01 | -7977.388 | -7979.227 | -1.839 | 6.15E-07 |
| 11 | 119850450 | rs601052 | C | T | 1.012 | -7977.388 | -7979.227 | -1.839 | 6.15E-07 |
| 11 | 120057972 | rs12808972 | A | G | 1 | -7977.388 | -7979.227 | -1.839 | 6.15E-07 |
| 11 | 120544871 | rs2000869 | G | A | 0.99 | -7977.388 | -7979.228 | -1.839 | 6.15E-07 |
| 11 | 121004172 | rs646371 | A | G | 1.006 | -7977.388 | -7979.227 | -1.839 | 6.15E-07 |
| 11 | 117962275 | rs17528237 | T | C | 0.991 | -7977.388 | -7979.228 | -1.84 | 6.14E-07 |
| 11 | 117973511 | rs4344527 | G | A | 0.99 | -7977.388 | -7979.229 | -1.84 | 6.14E-07 |
| 11 | 118067132 | rs647573 | C | T | 0.988 | -7977.388 | -7979.228 | -1.84 | 6.14E-07 |
| 11 | 119240809 | rs10892355 | C | T | 0.989 | -7977.388 | -7979.229 | -1.84 | 6.14E-07 |
| 11 | 119308201 | rs56103633 | T | C | 1.004 | -7977.388 | -7979.228 | -1.84 | 6.14E-07 |
| 11 | 119309000 | rs7130634 | A | T | 1.004 | -7977.388 | -7979.228 | -1.84 | 6.14E-07 |
| 11 | 119426508 | rs10892392 | T | C | 0.998 | -7977.388 | -7979.229 | -1.84 | 6.14E-07 |
| 11 | 119426651 | rs10892393 | T | C | 0.998 | -7977.388 | -7979.229 | -1.84 | 6.14E-07 |
| 11 | 119429080 | rs10750150 | T | C | 0.999 | -7977.388 | -7979.228 | -1.84 | 6.14E-07 |
| 11 | 119429098 | rs10736499 | C | T | 0.999 | -7977.388 | -7979.228 | -1.84 | 6.14E-07 |
| 11 | 119429614 | rs10892394 | A | G | 1.001 | -7977.388 | -7979.229 | -1.84 | 6.14E-07 |
| 11 | 119429656 | rs10892395 | G | A | 1.001 | -7977.388 | -7979.229 | -1.84 | 6.14E-07 |
| 11 | 119429662 | rs10892396 | G | T | 1.001 | -7977.388 | -7979.229 | -1.84 | 6.14E-07 |
| 11 | 119429848 | rs11217318 | A | G | 0.999 | -7977.388 | -7979.228 | -1.84 | 6.14E-07 |
| 11 | 119429923 | rs11217319 | A | G | 1.001 | -7977.388 | -7979.229 | -1.84 | 6.14E-07 |
| 11 | 119430203 | rs10892397 | G | A | 1.001 | -7977.388 | -7979.229 | -1.84 | 6.14E-07 |
| 11 | 119430266 | rs12274984 | A | G | 1.001 | -7977.388 | -7979.229 | -1.84 | 6.14E-07 |
| 11 | 119432144 | rs10892399 | C | T | 1.001 | -7977.388 | -7979.229 | -1.84 | 6.14E-07 |
| 11 | 119432909 | rs10750154 | G | A | 1.002 | -7977.388 | -7979.228 | -1.84 | 6.14E-07 |
| 11 | 119432935 | rs10790310 | T | C | 1 | -7977.388 | -7979.229 | -1.84 | 6.14E-07 |
| 11 | 119433005 | rs10790311 | T | C | 1.001 | -7977.388 | -7979.229 | -1.84 | 6.14E-07 |
| 11 | 119433521 | rs10892402 | G | A | 1.001 | -7977.388 | -7979.229 | -1.84 | 6.14E-07 |
| 11 | 119434032 | rs10750155 | T | A | 1.001 | -7977.388 | -7979.229 | -1.84 | 6.14E-07 |
| 11 | 119434275 | rs10790312 | T | C | 1.001 | -7977.388 | -7979.229 | -1.84 | 6.14E-07 |
| 11 | 119434279 | rs2887244 | G | T | 1.001 | -7977.388 | -7979.229 | -1.84 | 6.14E-07 |
| 11 | 119434468 | rs2368425 | T | G | 1.001 | -7977.388 | -7979.229 | -1.84 | 6.14E-07 |
| 11 | 119434711 | rs4018655 | A | G | 1.001 | -7977.388 | -7979.229 | -1.84 | 6.14E-07 |
| 11 | 119434843 | rs10892403 | A | C | 1.001 | -7977.388 | -7979.229 | -1.84 | 6.14E-07 |
| 11 | 119630641 | rs11217471 | A | G | 0.987 | -7977.388 | -7979.228 | -1.84 | 6.14E-07 |
| 11 | 119899963 | rs653278 | T | A | 0.99 | -7977.388 | -7979.228 | -1.84 | 6.14E-07 |
| 11 | 117335722 | rs538847 | T | C | 1.013 | -7977.388 | -7979.23 | -1.841 | 6.14E-07 |
| 11 | 117745659 | rs543312 | C | T | 0.988 | -7977.388 | -7979.229 | -1.841 | 6.14E-07 |
| 11 | 117771069 | rs543480 | T | C | 1.011 | -7977.388 | -7979.23 | -1.841 | 6.14E-07 |
| 11 | 117842533 | rs7122174 | A | T | 0.988 | -7977.388 | -7979.229 | -1.841 | 6.14E-07 |
| 11 | 118067426 | rs648947 | A | G | 0.988 | -7977.388 | -7979.23 | -1.841 | 6.14E-07 |
| 11 | 119260456 | rs56183153 | A | C | 1.006 | -7977.388 | -7979.23 | -1.841 | 6.14E-07 |

| | | | | | | | |
|---|---|---|---|---|---|---|---|
| 11 | 119295063 | rs7130266 | G | A | NULL | 0.715042106 | NULL&M_Disease |
| 11 | 119422334 | rs2663625 | C | T | NULL | 0.715042106 | NULL&M_Disease |
| 11 | 119427346 | rs7103816 | C | T | NULL | 0.715042106 | NULL&M_Disease |
| 11 | 119427857 | rs7118777 | C | A | NULL | 0.715042106 | NULL&M_Disease |
| 11 | 119432823 | rs10750153 | T | C | NULL | 0.715042106 | NULL&M_Disease |
| 11 | 119435011 | rs10892404 | C | G | NULL | 0.715042106 | NULL&M_Disease |
| 11 | 119435047 | rs10892405 | G | A | NULL | 0.715042106 | NULL&M_Disease |
| 11 | 119435544 | rs10892406 | G | A | NULL | 0.715042106 | NULL&M_Disease |
| 11 | 119494489 | rs4938692 | T | C | NULL | 0.715042106 | NULL&M_Disease |
| 11 | 119572051 | rs3902806 | G | T | NULL | 0.715042106 | NULL&M_Disease |
| 11 | 119574609 | rs11217415 | C | T | NULL | 0.715042106 | NULL&M_Disease |
| 11 | 119899910 | rs583971 | A | G | NULL | 0.715042106 | NULL&M_Disease |
| 11 | 117095361 | rs236919 | T | C | NULL | 0.715042106 | NULL&M_Disease |
| 11 | 117476349 | rs568221 | T | C | NULL | 0.715042106 | NULL&M_Disease |
| 11 | 117733559 | rs7107910 | C | G | NULL | 0.715042106 | NULL&M_Disease |
| 11 | 117807414 | rs7116903 | T | C | NULL | 0.715042106 | NULL&M_Disease |
| 11 | 117808853 | rs7105537 | A | G | NULL | 0.715042106 | NULL&M_Disease |
| 11 | 118952371 | rs4614 | G | A | NULL | 0.715042106 | NULL&M_Disease |
| 11 | 119054989 | rs5012703 | C | T | NULL | 0.715042106 | NULL&M_Disease |
| 11 | 119312750 | rs7131641 | C | G | NULL | 0.715042106 | NULL&M_Disease |
| 11 | 119314499 | rs67851622 | G | C | NULL | 0.715042106 | NULL&M_Disease |
| 11 | 119317435 | rs735006 | G | A | NULL | 0.715042106 | NULL&M_Disease |
| 11 | 119494587 | rs4938693 | T | C | NULL | 0.715042106 | NULL&M_Disease |
| 11 | 119494885 | rs12798473 | T | C | NULL | 0.715042106 | NULL&M_Disease |
| 11 | 119495352 | rs7951065 | C | T | NULL | 0.715042106 | NULL&M_Disease |
| 11 | 119495527 | rs7951484 | G | A | NULL | 0.715042106 | NULL&M_Disease |
| 11 | 119495871 | rs4938694 | A | G | NULL | 0.715042106 | NULL&M_Disease |
| 11 | 119496085 | rs4938695 | T | C | NULL | 0.715042106 | NULL&M_Disease |
| 11 | 119895096 | rs595483 | G | A | NULL | 0.715042106 | NULL&M_Disease |
| 11 | 119974286 | rs2368961 | G | A | NULL | 0.715042106 | NULL&M_Disease |
| 11 | 117973558 | rs4388923 | A | C | NULL | 0.715042106 | NULL&M_Disease |
| 11 | 117973576 | rs3819108 | A | G | NULL | 0.715042106 | NULL&M_Disease |
| 11 | 118067621 | rs951657 | A | G | NULL | 0.715042106 | NULL&M_Disease |
| 11 | 119259720 | rs12792689 | G | A | NULL | 0.715042106 | NULL&M_Disease |
| 11 | 119438415 | rs2663635 | C | T | NULL | 0.715042106 | NULL&M_Disease |
| 11 | 119572905 | rs10790335 | C | T | NULL | 0.715042106 | NULL&M_Disease |
| 11 | 119574558 | rs11217414 | C | T | NULL | 0.715042106 | NULL&M_Disease |
| 11 | 119899904 | rs642475 | C | T | NULL | 0.715042106 | NULL&M_Disease |
| 11 | 119901095 | rs668400 | A | C | NULL | 0.715042106 | NULL&M_Disease |
| 11 | 119983919 | rs1075790 | A | C | NULL | 0.715042106 | NULL&M_Disease |
| 11 | 116300551 | rs11215927 | C | T | NULL | 0.715042106 | NULL&M_Disease |
| 11 | 119312046 | rs56152188 | C | T | NULL | 0.715042106 | NULL&M_Disease |
| 11 | 119312851 | rs61901262 | T | C | NULL | 0.715042106 | NULL&M_Disease |
| 11 | 119313221 | rs61901263 | T | C | NULL | 0.715042106 | NULL&M_Disease |
| 11 | 119314637 | rs55845693 | C | T | NULL | 0.715042106 | NULL&M_Disease |
| 11 | 119315316 | rs61901265 | A | G | NULL | 0.715042106 | NULL&M_Disease |
| 11 | 119315417 | rs17547501 | A | G | NULL | 0.715042106 | NULL&M_Disease |
| 11 | 119493873 | rs1190080879 | G | T | NULL | 0.715042106 | NULL&M_Disease |
| 11 | 119572571 | rs885429 | T | G | NULL | 0.715042106 | NULL&M_Disease |
| 11 | 120393776 | rs7931923 | G | A | NULL | 0.715042106 | NULL&M_Disease |
| 11 | 117835174 | rs11512467 | T | C | NULL | 0.715042106 | NULL&M_Disease |
| 11 | 119293216 | rs3138090 | A | C | NULL | 0.715042106 | NULL&M_Disease |
| 11 | 119293397 | rs3138089 | A | G | NULL | 0.715042106 | NULL&M_Disease |
| 11 | 119293441 | rs3138088 | G | A | NULL | 0.715042106 | NULL&M_Disease |
| 11 | 119294290 | rs11574653 | G | A | NULL | 0.715042106 | NULL&M_Disease |
| 11 | 119631325 | rs7131676 | G | A | NULL | 0.715042106 | NULL&M_Disease |
| 11 | 119636367 | rs10892453 | T | C | NULL | 0.715042106 | NULL&M_Disease |
| 11 | 119974641 | rs1110704 | A | G | NULL | 0.715042106 | NULL&M_Disease |
| 11 | 120184592 | rs4938795 | A | G | NULL | 0.715042106 | NULL&M_Disease |
| 11 | 117963955 | rs73010786 | T | C | NULL | 0.715042106 | NULL&M_Disease |
| 11 | 119412428 | rs7129877 | G | C | NULL | 0.715042106 | NULL&M_Disease |
| 11 | 119418419 | rs2663639 | G | A | NULL | 0.715042106 | NULL&M_Disease |
| 11 | 119446118 | rs4622279 | A | G | NULL | 0.715042106 | NULL&M_Disease |
| 11 | 119961097 | rs7102643 | T | C | NULL | 0.715042106 | NULL&M_Disease |
| 11 | 120498051 | rs7924856 | A | G | NULL | 0.715042106 | NULL&M_Disease |
| 11 | 119439459 | rs755003 | A | G | NULL | 0.715042106 | NULL&M_Disease |
| 11 | 117965646 | rs2276125 | A | G | NULL | 0.716059794 | NULL&M_Disease |
| 11 | 118068521 | rs1613643 | T | G | NULL | 0.716059794 | NULL&M_Disease |
| 11 | 119574140 | rs7111290 | G | A | NULL | 0.716059794 | NULL&M_Disease |
| 11 | 120397055 | rs11217904 | C | T | NULL | 0.716059794 | NULL&M_Disease |
| 11 | 120556389 | rs7121857 | G | T | NULL | 0.716059794 | NULL&M_Disease |
| 11 | 117966677 | rs1941633 | A | G | NULL | 0.716059794 | NULL&M_Disease |
| 11 | 119437438 | rs2845654 | C | A | NULL | 0.716059794 | NULL&M_Disease |
| 11 | 119816075 | rs10892500 | T | C | NULL | 0.716059794 | NULL&M_Disease |
| 11 | 119833857 | rs11217602 | T | C | NULL | 0.716059794 | NULL&M_Disease |
| 11 | 117535760 | rs73014416 | A | G | NULL | 0.716059794 | NULL&M_Disease |
| 11 | 117972543 | rs11216750 | T | A | NULL | 0.716059794 | NULL&M_Disease |
| 11 | 118567881 | rs497678 | C | T | NULL | 0.716059794 | NULL&M_Disease |
| 11 | 119492274 | rs4514434 | C | T | NULL | 0.716059794 | NULL&M_Disease |
| 11 | 119492297 | rs7130031 | T | C | NULL | 0.716059794 | NULL&M_Disease |
| 11 | 119575185 | rs2015078 | C | T | NULL | 0.716059794 | NULL&M_Disease |
| 11 | 119881024 | rs7936363 | T | G | NULL | 0.716059794 | NULL&M_Disease |
| 11 | 117536136 | rs11216491 | T | C | NULL | 0.716059794 | NULL&M_Disease |
| 11 | 117844785 | rs4938465 | T | A | NULL | 0.716059794 | NULL&M_Disease |
| 11 | 118188276 | rs56754533 | A | G | NULL | 0.716059794 | NULL&M_Disease |
| 11 | 119218648 | rs12577147 | A | G | NULL | 0.716059794 | NULL&M_Disease |
| 11 | 119438765 | rs1078762 | T | C | NULL | 0.716059794 | NULL&M_Disease |
| 11 | 119891597 | rs652449 | G | T | NULL | 0.716059794 | NULL&M_Disease |
| 11 | 119898559 | rs626440 | T | C | NULL | 0.716059794 | NULL&M_Disease |
| 11 | 117138456 | rs201365537 | A | G | NULL | 0.716059794 | NULL&M_Disease |
| 11 | 119572185 | rs3935406 | G | A | NULL | 0.716059794 | NULL&M_Disease |
| 11 | 119977712 | rs10892538 | A | G | NULL | 0.716059794 | NULL&M_Disease |
| 11 | 120983264 | rs638903 | A | C | NULL | 0.716059794 | NULL&M_Disease |
| 11 | 118480115 | rs12803321 | C | G | NULL | 0.716059794 | NULL&M_Disease |
| 11 | 119454670 | rs10466594 | C | T | NULL | 0.716059794 | NULL&M_Disease |
| 11 | 119640427 | rs34809912 | A | G | NULL | 0.716059794 | NULL&M_Disease |

| | | | | | | | | | |
|---|---|---|---|---|---|---|---|---|---|
| 11 | 119295063 | rs7130266 | G | A | 1.004 | -7977.388 | -7979.229 | -1.841 | 6.14E-07 |
| 11 | 119422334 | rs2663625 | C | T | 0.99 | -7977.388 | -7979.229 | -1.841 | 6.14E-07 |
| 11 | 119427346 | rs7103816 | C | T | 0.999 | -7977.388 | -7979.229 | -1.841 | 6.14E-07 |
| 11 | 119427857 | rs7118777 | C | A | 1 | -7977.388 | -7979.229 | -1.841 | 6.14E-07 |
| 11 | 119432823 | rs10750153 | T | C | 1 | -7977.388 | -7979.229 | -1.841 | 6.14E-07 |
| 11 | 119435011 | rs10892404 | C | G | 1 | -7977.388 | -7979.229 | -1.841 | 6.14E-07 |
| 11 | 119435047 | rs10892405 | G | A | 1 | -7977.388 | -7979.229 | -1.841 | 6.14E-07 |
| 11 | 119435544 | rs10892406 | G | A | 0.999 | -7977.388 | -7979.229 | -1.841 | 6.14E-07 |
| 11 | 119494489 | rs4938692 | T | C | 1 | -7977.388 | -7979.23 | -1.841 | 6.14E-07 |
| 11 | 119572051 | rs3902806 | G | T | 1.011 | -7977.388 | -7979.229 | -1.841 | 6.14E-07 |
| 11 | 119574609 | rs11217415 | C | T | 1.011 | -7977.388 | -7979.23 | -1.841 | 6.14E-07 |
| 11 | 119899910 | rs583971 | A | G | 0.99 | -7977.388 | -7979.23 | -1.841 | 6.14E-07 |
| 11 | 117095361 | rs236919 | T | C | 0.989 | -7977.388 | -7979.23 | -1.842 | 6.13E-07 |
| 11 | 117476349 | rs568221 | T | C | 1.002 | -7977.388 | -7979.23 | -1.842 | 6.13E-07 |
| 11 | 117733559 | rs7107910 | C | G | 0.99 | -7977.388 | -7979.23 | -1.842 | 6.13E-07 |
| 11 | 117807414 | rs7116903 | T | C | 1.01 | -7977.388 | -7979.23 | -1.842 | 6.13E-07 |
| 11 | 117808853 | rs7105537 | A | G | 1.01 | -7977.388 | -7979.23 | -1.842 | 6.13E-07 |
| 11 | 118952371 | rs4614 | G | A | 1.011 | -7977.388 | -7979.23 | -1.842 | 6.13E-07 |
| 11 | 119054989 | rs5012703 | C | T | 1.013 | -7977.388 | -7979.231 | -1.842 | 6.13E-07 |
| 11 | 119312750 | rs7131641 | C | G | 1.003 | -7977.388 | -7979.231 | -1.842 | 6.13E-07 |
| 11 | 119314499 | rs67851622 | G | C | 1.004 | -7977.388 | -7979.23 | -1.842 | 6.13E-07 |
| 11 | 119317435 | rs735006 | G | A | 1.004 | -7977.388 | -7979.23 | -1.842 | 6.13E-07 |
| 11 | 119494587 | rs4938693 | T | C | 1 | -7977.388 | -7979.23 | -1.842 | 6.13E-07 |
| 11 | 119494885 | rs12798473 | T | C | 1 | -7977.388 | -7979.23 | -1.842 | 6.13E-07 |
| 11 | 119495352 | rs7951065 | C | T | 1.001 | -7977.388 | -7979.23 | -1.842 | 6.13E-07 |
| 11 | 119495527 | rs7951484 | G | A | 1.001 | -7977.388 | -7979.23 | -1.842 | 6.13E-07 |
| 11 | 119495871 | rs4938694 | A | G | 1.001 | -7977.388 | -7979.23 | -1.842 | 6.13E-07 |
| 11 | 119496085 | rs4938695 | T | C | 1 | -7977.388 | -7979.23 | -1.842 | 6.13E-07 |
| 11 | 119895096 | rs595483 | G | A | 0.99 | -7977.388 | -7979.231 | -1.842 | 6.13E-07 |
| 11 | 119974286 | rs2368961 | G | A | 0.994 | -7977.388 | -7979.23 | -1.842 | 6.13E-07 |
| 11 | 117973558 | rs4388923 | A | C | 0.991 | -7977.388 | -7979.232 | -1.843 | 6.12E-07 |
| 11 | 117973576 | rs3819108 | A | G | 0.991 | -7977.388 | -7979.231 | -1.843 | 6.12E-07 |
| 11 | 118067621 | rs951657 | A | G | 0.988 | -7977.388 | -7979.231 | -1.843 | 6.12E-07 |
| 11 | 119259720 | rs12792689 | G | A | 1.005 | -7977.388 | -7979.231 | -1.843 | 6.12E-07 |
| 11 | 119438415 | rs2663635 | C | T | 0.99 | -7977.388 | -7979.232 | -1.843 | 6.12E-07 |
| 11 | 119572905 | rs10790335 | C | T | 1.011 | -7977.388 | -7979.231 | -1.843 | 6.12E-07 |
| 11 | 119574558 | rs11217414 | C | T | 1.011 | -7977.388 | -7979.231 | -1.843 | 6.12E-07 |
| 11 | 119899904 | rs642475 | C | T | 0.99 | -7977.388 | -7979.231 | -1.843 | 6.12E-07 |
| 11 | 119901095 | rs668400 | A | C | 0.988 | -7977.388 | -7979.231 | -1.843 | 6.12E-07 |
| 11 | 119983919 | rs1075790 | A | C | 1.003 | -7977.388 | -7979.231 | -1.843 | 6.12E-07 |
| 11 | 116300551 | rs11215927 | C | T | 1.008 | -7977.388 | -7979.233 | -1.844 | 6.12E-07 |
| 11 | 119312046 | rs56152188 | C | T | 1.003 | -7977.388 | -7979.232 | -1.844 | 6.12E-07 |
| 11 | 119312851 | rs61901262 | T | C | 1.003 | -7977.388 | -7979.233 | -1.844 | 6.12E-07 |
| 11 | 119313221 | rs61901263 | T | C | 1.003 | -7977.388 | -7979.233 | -1.844 | 6.12E-07 |
| 11 | 119314637 | rs55845693 | C | T | 1.003 | -7977.388 | -7979.233 | -1.844 | 6.12E-07 |
| 11 | 119315316 | rs61901265 | A | G | 1.003 | -7977.388 | -7979.233 | -1.844 | 6.12E-07 |
| 11 | 119315417 | rs17547501 | A | G | 1.003 | -7977.388 | -7979.233 | -1.844 | 6.12E-07 |
| 11 | 119493873 | rs1190080879 | G | T | 1 | -7977.388 | -7979.232 | -1.844 | 6.12E-07 |
| 11 | 119572571 | rs885429 | T | G | 1.011 | -7977.388 | -7979.232 | -1.844 | 6.12E-07 |
| 11 | 120393776 | rs7931923 | G | A | 0.993 | -7977.388 | -7979.232 | -1.844 | 6.12E-07 |
| 11 | 117835174 | rs11512467 | T | C | 0.988 | -7977.388 | -7979.233 | -1.845 | 6.11E-07 |
| 11 | 119293216 | rs3138090 | A | C | 1.002 | -7977.388 | -7979.233 | -1.845 | 6.11E-07 |
| 11 | 119293397 | rs3138089 | A | G | 1.002 | -7977.388 | -7979.233 | -1.845 | 6.11E-07 |
| 11 | 119293441 | rs3138088 | G | A | 1.002 | -7977.388 | -7979.233 | -1.845 | 6.11E-07 |
| 11 | 119294290 | rs11574653 | G | A | 1.002 | -7977.388 | -7979.233 | -1.845 | 6.11E-07 |
| 11 | 119631325 | rs7131676 | G | A | 0.988 | -7977.388 | -7979.234 | -1.845 | 6.11E-07 |
| 11 | 119636367 | rs10892453 | T | C | 1.011 | -7977.388 | -7979.233 | -1.845 | 6.11E-07 |
| 11 | 119974641 | rs1110704 | A | G | 0.995 | -7977.388 | -7979.234 | -1.845 | 6.11E-07 |
| 11 | 120184592 | rs4938795 | A | G | 1.008 | -7977.388 | -7979.233 | -1.845 | 6.11E-07 |
| 11 | 117963955 | rs73010786 | T | C | 0.992 | -7977.388 | -7979.234 | -1.846 | 6.11E-07 |
| 11 | 119412428 | rs7129877 | G | C | 0.99 | -7977.388 | -7979.234 | -1.846 | 6.11E-07 |
| 11 | 119418419 | rs2663639 | G | A | 0.99 | -7977.388 | -7979.234 | -1.846 | 6.11E-07 |
| 11 | 119446118 | rs4622279 | A | G | 1.004 | -7977.388 | -7979.234 | -1.846 | 6.11E-07 |
| 11 | 119961097 | rs7102643 | T | C | 0.991 | -7977.388 | -7979.234 | -1.846 | 6.11E-07 |
| 11 | 120498051 | rs7924856 | A | G | 0.99 | -7977.388 | -7979.235 | -1.846 | 6.11E-07 |
| 11 | 119439459 | rs755003 | A | G | 0.99 | -7977.388 | -7979.235 | -1.847 | 6.10E-07 |
| 11 | 117965646 | rs2276125 | A | G | 0.992 | -7977.388 | -7979.236 | -1.848 | 6.09E-07 |
| 11 | 118068521 | rs1613643 | T | G | 0.988 | -7977.388 | -7979.237 | -1.848 | 6.09E-07 |
| 11 | 119574140 | rs7111290 | G | A | 1.011 | -7977.388 | -7979.236 | -1.848 | 6.09E-07 |
| 11 | 120397055 | rs11217904 | C | T | 0.993 | -7977.388 | -7979.236 | -1.848 | 6.09E-07 |
| 11 | 120556389 | rs7121857 | G | T | 1.012 | -7977.388 | -7979.236 | -1.848 | 6.09E-07 |
| 11 | 117966677 | rs1941633 | A | G | 0.992 | -7977.388 | -7979.237 | -1.849 | 6.09E-07 |
| 11 | 119437438 | rs2845654 | C | A | 0.99 | -7977.388 | -7979.238 | -1.849 | 6.09E-07 |
| 11 | 119816075 | rs10892500 | T | C | 1.011 | -7977.388 | -7979.237 | -1.849 | 6.09E-07 |
| 11 | 119833857 | rs11217602 | T | C | 0.989 | -7977.388 | -7979.237 | -1.849 | 6.09E-07 |
| 11 | 117535760 | rs73014416 | A | G | 0.998 | -7977.388 | -7979.238 | -1.85 | 6.08E-07 |
| 11 | 117972543 | rs11216750 | T | A | 0.991 | -7977.388 | -7979.238 | -1.85 | 6.08E-07 |
| 11 | 118567881 | rs497678 | C | T | 0.992 | -7977.388 | -7979.238 | -1.85 | 6.08E-07 |
| 11 | 119492274 | rs4514434 | C | T | 1 | -7977.388 | -7979.239 | -1.85 | 6.08E-07 |
| 11 | 119492297 | rs7130031 | T | C | 0.999 | -7977.388 | -7979.238 | -1.85 | 6.08E-07 |
| 11 | 119575185 | rs2015078 | C | T | 1.01 | -7977.388 | -7979.239 | -1.85 | 6.08E-07 |
| 11 | 119881024 | rs7936363 | T | G | 0.99 | -7977.388 | -7979.238 | -1.85 | 6.08E-07 |
| 11 | 117536136 | rs11216491 | T | C | 1.002 | -7977.388 | -7979.239 | -1.851 | 6.08E-07 |
| 11 | 117844785 | rs4938465 | T | A | 0.988 | -7977.388 | -7979.239 | -1.851 | 6.08E-07 |
| 11 | 118188276 | rs56754533 | A | G | 0.996 | -7977.388 | -7979.239 | -1.851 | 6.08E-07 |
| 11 | 119218648 | rs12577147 | A | G | 0.992 | -7977.388 | -7979.239 | -1.851 | 6.08E-07 |
| 11 | 119438765 | rs1078762 | T | C | 0.99 | -7977.388 | -7979.239 | -1.851 | 6.08E-07 |
| 11 | 119891597 | rs652449 | G | T | 0.991 | -7977.388 | -7979.239 | -1.851 | 6.08E-07 |
| 11 | 119898559 | rs626440 | T | C | 0.991 | -7977.388 | -7979.24 | -1.851 | 6.08E-07 |
| 11 | 117138456 | rs201365537 | A | G | 1.012 | -7977.388 | -7979.24 | -1.852 | 6.07E-07 |
| 11 | 119572185 | rs3935406 | G | A | 1.01 | -7977.388 | -7979.24 | -1.852 | 6.07E-07 |
| 11 | 119977712 | rs10892538 | A | G | 0.997 | -7977.388 | -7979.24 | -1.852 | 6.07E-07 |
| 11 | 120983264 | rs638903 | A | C | 1.012 | -7977.388 | -7979.24 | -1.852 | 6.07E-07 |
| 11 | 118480115 | rs12803321 | C | G | 0.992 | -7977.388 | -7979.241 | -1.853 | 6.06E-07 |
| 11 | 119454670 | rs10466594 | C | T | 1.003 | -7977.388 | -7979.241 | -1.853 | 6.06E-07 |
| 11 | 119640427 | rs34809912 | A | G | 1.01 | -7977.388 | -7979.241 | -1.853 | 6.06E-07 |

| | | | | | | | |
|---|---|---|---|---|---|---|---|
| 11 | 119898995 | rs638399 | T | C | NULL | 0.716059794 | NULL&M_Disease |
| 11 | 119899143 | rs638948 | C | T | NULL | 0.716059794 | NULL&M_Disease |
| 11 | 120546088 | rs11820810 | G | A | NULL | 0.716059794 | NULL&M_Disease |
| 11 | 116991926 | rs7120515 | A | G | NULL | 0.716059794 | NULL&M_Disease |
| 11 | 116992076 | rs7106437 | T | C | NULL | 0.716059794 | NULL&M_Disease |
| 11 | 117805786 | rs10790225 | T | C | NULL | 0.716059794 | NULL&M_Disease |
| 11 | 117818041 | rs2187639 | A | C | NULL | 0.716059794 | NULL&M_Disease |
| 11 | 119406091 | rs10892384 | C | T | NULL | 0.716059794 | NULL&M_Disease |
| 11 | 119630418 | rs10790342 | A | G | NULL | 0.716059794 | NULL&M_Disease |
| 11 | 119792728 | rs1287836 | C | A | NULL | 0.716059794 | NULL&M_Disease |
| 11 | 119959585 | rs7949366 | T | C | NULL | 0.716059794 | NULL&M_Disease |
| 11 | 120040442 | rs2444240 | G | T | NULL | 0.716059794 | NULL&M_Disease |
| 11 | 120553937 | rs35407634 | T | G | NULL | 0.716059794 | NULL&M_Disease |
| 11 | 117954384 | rs10790230 | A | C | NULL | 0.716059794 | NULL&M_Disease |
| 11 | 119454588 | rs10466593 | T | C | NULL | 0.716059794 | NULL&M_Disease |
| 11 | 119750487 | rs543876 | C | T | NULL | 0.716059794 | NULL&M_Disease |
| 11 | 119896830 | rs10892516 | C | T | NULL | 0.716059794 | NULL&M_Disease |
| 11 | 116470373 | rs11216036 | T | A | NULL | 0.716059794 | NULL&M_Disease |
| 11 | 117530473 | rs10790199 | A | G | NULL | 0.716059794 | NULL&M_Disease |
| 11 | 117694392 | rs4936409 | G | A | NULL | 0.716059794 | NULL&M_Disease |
| 11 | 117973047 | rs881123 | T | C | NULL | 0.716059794 | NULL&M_Disease |
| 11 | 118189477 | rs67514601 | G | A | NULL | 0.716059794 | NULL&M_Disease |
| 11 | 119420469 | rs4322404 | C | T | NULL | 0.716059794 | NULL&M_Disease |
| 11 | 119437815 | rs2845653 | G | A | NULL | 0.716059794 | NULL&M_Disease |
| 11 | 119895416 | rs694662 | T | A | NULL | 0.716059794 | NULL&M_Disease |
| 11 | 119897949 | rs593598 | G | A | NULL | 0.716059794 | NULL&M_Disease |
| 11 | 119897964 | rs648009 | A | G | NULL | 0.716059794 | NULL&M_Disease |
| 11 | 120060793 | rs10892547 | T | G | NULL | 0.716059794 | NULL&M_Disease |
| 11 | 121161826 | rs12786590 | G | A | NULL | 0.716059794 | NULL&M_Disease |
| 11 | 121192985 | rs4537768 | T | G | NULL | 0.716059794 | NULL&M_Disease |
| 11 | 116560447 | rs1623797 | T | C | NULL | 0.716059794 | NULL&M_Disease |
| 11 | 117800700 | rs10790223 | C | T | NULL | 0.716059794 | NULL&M_Disease |
| 11 | 119055075 | rs10750138 | G | A | NULL | 0.716059794 | NULL&M_Disease |
| 11 | 119961707 | rs11217682 | G | A | NULL | 0.716059794 | NULL&M_Disease |
| 11 | 120183803 | rs7122401 | A | G | NULL | 0.716059794 | NULL&M_Disease |
| 11 | 120539693 | rs2000870 | G | C | NULL | 0.716059794 | NULL&M_Disease |
| 11 | 116389017 | rs883863 | A | G | NULL | 0.716059794 | NULL&M_Disease |
| 11 | 117312602 | rs3863313 | G | T | NULL | 0.717075285 | NULL&M_Disease |
| 11 | 117328639 | rs558582 | T | C | NULL | 0.717075285 | NULL&M_Disease |
| 11 | 119951870 | rs12365185 | G | A | NULL | 0.717075285 | NULL&M_Disease |
| 11 | 120645211 | rs2846095 | T | C | NULL | 0.717075285 | NULL&M_Disease |
| 11 | 121158100 | rs6589872 | C | T | NULL | 0.717075285 | NULL&M_Disease |
| 11 | 116430875 | rs4938281 | G | A | NULL | 0.717075285 | NULL&M_Disease |
| 11 | 117810129 | rs7110261 | A | G | NULL | 0.717075285 | NULL&M_Disease |
| 11 | 117986853 | rs10790236 | C | T | NULL | 0.717075285 | NULL&M_Disease |
| 11 | 119440066 | rs10790313 | A | G | NULL | 0.717075285 | NULL&M_Disease |
| 11 | 119469767 | rs12421460 | T | C | NULL | 0.717075285 | NULL&M_Disease |
| 11 | 119881715 | rs590282 | A | G | NULL | 0.717075285 | NULL&M_Disease |
| 11 | 119897453 | rs610831 | C | A | NULL | 0.717075285 | NULL&M_Disease |
| 11 | 119970508 | rs6589795 | T | C | NULL | 0.717075285 | NULL&M_Disease |
| 11 | 120705063 | rs4936559 | C | T | NULL | 0.717075285 | NULL&M_Disease |
| 11 | 117455699 | rs6589615 | C | G | NULL | 0.717075285 | NULL&M_Disease |
| 11 | 117529248 | rs73014407 | T | C | NULL | 0.717075285 | NULL&M_Disease |
| 11 | 117984995 | rs749688 | C | T | NULL | 0.717075285 | NULL&M_Disease |
| 11 | 118769032 | rs11217081 | G | A | NULL | 0.717075285 | NULL&M_Disease |
| 11 | 118938371 | rs1784460 | A | T | NULL | 0.717075285 | NULL&M_Disease |
| 11 | 118938388 | rs1784459 | A | C | NULL | 0.717075285 | NULL&M_Disease |
| 11 | 118938435 | rs1786684 | A | G | NULL | 0.717075285 | NULL&M_Disease |
| 11 | 119384214 | rs606151 | A | G | NULL | 0.717075285 | NULL&M_Disease |
| 11 | 119426636 | rs2663643 | A | G | NULL | 0.717075285 | NULL&M_Disease |
| 11 | 119427242 | rs2845662 | C | T | NULL | 0.717075285 | NULL&M_Disease |
| 11 | 119475996 | rs11217350 | C | T | NULL | 0.717075285 | NULL&M_Disease |
| 11 | 119630247 | rs7124166 | C | G | NULL | 0.717075285 | NULL&M_Disease |
| 11 | 119896745 | rs10892515 | A | T | NULL | 0.717075285 | NULL&M_Disease |
| 11 | 121158551 | rs7106203 | A | T | NULL | 0.717075285 | NULL&M_Disease |
| 11 | 116577441 | rs11216105 | G | A | NULL | 0.717075285 | NULL&M_Disease |
| 11 | 117686456 | rs4597100 | A | C | NULL | 0.717075285 | NULL&M_Disease |
| 11 | 119884319 | rs665758 | A | G | NULL | 0.717075285 | NULL&M_Disease |
| 11 | 119891625 | rs1243255182 | G | A | NULL | 0.717075285 | NULL&M_Disease |
| 11 | 120520218 | rs2000873 | C | A | NULL | 0.717075285 | NULL&M_Disease |
| 11 | 121192989 | rs1376996225 | T | G | NULL | 0.717075285 | NULL&M_Disease |
| 11 | 117341617 | rs549748 | A | T | NULL | 0.717075285 | NULL&M_Disease |
| 11 | 117811563 | rs7103799 | T | C | NULL | 0.717075285 | NULL&M_Disease |
| 11 | 117959224 | rs6589644 | T | C | NULL | 0.717075285 | NULL&M_Disease |
| 11 | 117960018 | rs948460 | T | A | NULL | 0.717075285 | NULL&M_Disease |
| 11 | 119419117 | rs1828279 | C | T | NULL | 0.717075285 | NULL&M_Disease |
| 11 | 119802591 | rs504542 | G | T | NULL | 0.717075285 | NULL&M_Disease |
| 11 | 119893208 | rs624736 | G | A | NULL | 0.717075285 | NULL&M_Disease |
| 11 | 119894303 | rs649451 | A | G | NULL | 0.717075285 | NULL&M_Disease |
| 11 | 116430949 | rs4938282 | T | C | NULL | 0.717075285 | NULL&M_Disease |
| 11 | 117324965 | rs10790187 | G | A | NULL | 0.717075285 | NULL&M_Disease |
| 11 | 117500025 | rs11216475 | T | C | NULL | 0.717075285 | NULL&M_Disease |
| 11 | 117960138 | rs948461 | T | G | NULL | 0.717075285 | NULL&M_Disease |
| 11 | 119455165 | rs7114739 | C | T | NULL | 0.717075285 | NULL&M_Disease |
| 11 | 120182578 | rs35238955 | T | G | NULL | 0.717075285 | NULL&M_Disease |
| 11 | 120557543 | rs10790401 | G | A | NULL | 0.717075285 | NULL&M_Disease |
| 11 | 120650611 | rs2846099 | T | A | NULL | 0.717075285 | NULL&M_Disease |
| 11 | 116313753 | rs541407 | C | T | NULL | 0.717075285 | NULL&M_Disease |
| 11 | 117312089 | rs7947950 | G | A | NULL | 0.717075285 | NULL&M_Disease |
| 11 | 118066695 | rs649138 | C | T | NULL | 0.717075285 | NULL&M_Disease |
| 11 | 119006722 | rs629345 | C | T | NULL | 0.717075285 | NULL&M_Disease |
| 11 | 119467192 | rs10892415 | T | A | NULL | 0.717075285 | NULL&M_Disease |
| 11 | 119468951 | rs10790315 | G | C | NULL | 0.717075285 | NULL&M_Disease |
| 11 | 119468994 | rs10790316 | T | C | NULL | 0.717075285 | NULL&M_Disease |
| 11 | 119469211 | rs7121550 | A | G | NULL | 0.717075285 | NULL&M_Disease |
| 11 | 119680383 | rs11607839 | C | A | NULL | 0.717075285 | NULL&M_Disease |

| | | | | | | | | | |
|---|---|---|---|---|---|---|---|---|---|
| 11 | 119898995 | rs638399 | T | C | 0.991 | -7977.388 | -7979.242 | -1.853 | 6.06E-07 |
| 11 | 119899143 | rs638948 | C | T | 0.991 | -7977.388 | -7979.241 | -1.853 | 6.06E-07 |
| 11 | 120546088 | rs11820810 | G | A | 0.992 | -7977.388 | -7979.241 | -1.853 | 6.06E-07 |
| 11 | 116991926 | rs7120515 | A | G | 1.006 | -7977.388 | -7979.243 | -1.854 | 6.06E-07 |
| 11 | 116992076 | rs7106437 | T | C | 1.006 | -7977.388 | -7979.243 | -1.854 | 6.06E-07 |
| 11 | 117805786 | rs10790225 | T | C | 0.988 | -7977.388 | -7979.242 | -1.854 | 6.06E-07 |
| 11 | 117818041 | rs2187639 | A | C | 1.007 | -7977.388 | -7979.242 | -1.854 | 6.06E-07 |
| 11 | 119406091 | rs10892384 | C | T | 1.011 | -7977.388 | -7979.243 | -1.854 | 6.06E-07 |
| 11 | 119630418 | rs10790342 | A | G | 0.988 | -7977.388 | -7979.242 | -1.854 | 6.06E-07 |
| 11 | 119792728 | rs1287836 | C | A | 1.004 | -7977.388 | -7979.243 | -1.854 | 6.06E-07 |
| 11 | 119959585 | rs7949366 | T | C | 0.992 | -7977.388 | -7979.242 | -1.854 | 6.06E-07 |
| 11 | 120040442 | rs2444240 | G | T | 1.011 | -7977.388 | -7979.243 | -1.854 | 6.06E-07 |
| 11 | 120553937 | rs35407634 | T | G | 0.991 | -7977.388 | -7979.243 | -1.854 | 6.06E-07 |
| 11 | 117954384 | rs10790230 | A | C | 0.991 | -7977.388 | -7979.244 | -1.855 | 6.05E-07 |
| 11 | 119454588 | rs10466593 | T | C | 1.002 | -7977.388 | -7979.244 | -1.855 | 6.05E-07 |
| 11 | 119750487 | rs543876 | C | T | 0.998 | -7977.388 | -7979.243 | -1.855 | 6.05E-07 |
| 11 | 119896830 | rs10892516 | C | T | 0.992 | -7977.388 | -7979.243 | -1.855 | 6.05E-07 |
| 11 | 116470373 | rs11216036 | T | A | 0.989 | -7977.388 | -7979.244 | -1.856 | 6.05E-07 |
| 11 | 117530473 | rs10790199 | A | G | 0.993 | -7977.388 | -7979.244 | -1.856 | 6.05E-07 |
| 11 | 117694392 | rs4936409 | G | A | 1.012 | -7977.388 | -7979.245 | -1.856 | 6.05E-07 |
| 11 | 117973047 | rs881123 | T | C | 0.992 | -7977.388 | -7979.244 | -1.856 | 6.05E-07 |
| 11 | 118189477 | rs67514601 | G | A | 1.002 | -7977.388 | -7979.244 | -1.856 | 6.05E-07 |
| 11 | 119420469 | rs4322404 | C | T | 0.991 | -7977.388 | -7979.244 | -1.856 | 6.05E-07 |
| 11 | 119437815 | rs2845653 | G | A | 0.991 | -7977.388 | -7979.245 | -1.856 | 6.05E-07 |
| 11 | 119895416 | rs694662 | T | A | 0.991 | -7977.388 | -7979.244 | -1.856 | 6.05E-07 |
| 11 | 119897949 | rs593598 | G | A | 0.992 | -7977.388 | -7979.244 | -1.856 | 6.05E-07 |
| 11 | 119897964 | rs648009 | A | G | 0.992 | -7977.388 | -7979.244 | -1.856 | 6.05E-07 |
| 11 | 120060793 | rs10892547 | T | G | 0.989 | -7977.388 | -7979.244 | -1.856 | 6.05E-07 |
| 11 | 121161826 | rs12786590 | G | A | 1.011 | -7977.388 | -7979.244 | -1.856 | 6.05E-07 |
| 11 | 121192985 | rs4537768 | T | G | 1.009 | -7977.388 | -7979.245 | -1.856 | 6.05E-07 |
| 11 | 116560447 | rs1623797 | T | C | 0.989 | -7977.388 | -7979.245 | -1.857 | 6.04E-07 |
| 11 | 117800700 | rs10790223 | C | T | 1.012 | -7977.388 | -7979.245 | -1.857 | 6.04E-07 |
| 11 | 119055075 | rs10750138 | G | A | 1.012 | -7977.388 | -7979.245 | -1.857 | 6.04E-07 |
| 11 | 119961707 | rs11217682 | G | A | 0.993 | -7977.388 | -7979.245 | -1.857 | 6.04E-07 |
| 11 | 120183803 | rs7122401 | A | G | 1.007 | -7977.388 | -7979.245 | -1.857 | 6.04E-07 |
| 11 | 120539693 | rs2000870 | G | C | 0.992 | -7977.388 | -7979.245 | -1.857 | 6.04E-07 |
| 11 | 116389017 | rs883863 | A | G | 1.004 | -7977.388 | -7979.246 | -1.858 | 6.03E-07 |
| 11 | 117312602 | rs3863313 | G | T | 1.011 | -7977.388 | -7979.247 | -1.858 | 6.03E-07 |
| 11 | 117328639 | rs558582 | T | C | 1.012 | -7977.388 | -7979.246 | -1.858 | 6.03E-07 |
| 11 | 119951870 | rs12365185 | G | A | 0.993 | -7977.388 | -7979.247 | -1.858 | 6.03E-07 |
| 11 | 120645211 | rs2846095 | T | C | 0.992 | -7977.388 | -7979.246 | -1.858 | 6.03E-07 |
| 11 | 121158100 | rs6589872 | C | T | 1.011 | -7977.388 | -7979.247 | -1.858 | 6.03E-07 |
| 11 | 116430875 | rs4938281 | G | A | 0.989 | -7977.388 | -7979.248 | -1.859 | 6.03E-07 |
| 11 | 117810129 | rs7110261 | A | G | 1.009 | -7977.388 | -7979.247 | -1.859 | 6.03E-07 |
| 11 | 117986853 | rs10790236 | C | T | 1.01 | -7977.388 | -7979.247 | -1.859 | 6.03E-07 |
| 11 | 119440066 | rs10790313 | A | G | 0.991 | -7977.388 | -7979.247 | -1.859 | 6.03E-07 |
| 11 | 119469767 | rs12421460 | T | C | 1.008 | -7977.388 | -7979.247 | -1.859 | 6.03E-07 |
| 11 | 119881715 | rs590282 | A | G | 0.991 | -7977.388 | -7979.247 | -1.859 | 6.03E-07 |
| 11 | 119897453 | rs610831 | C | A | 0.992 | -7977.388 | -7979.247 | -1.859 | 6.03E-07 |
| 11 | 119970508 | rs6589795 | T | C | 0.989 | -7977.388 | -7979.247 | -1.859 | 6.03E-07 |
| 11 | 120705063 | rs4936559 | C | T | 0.99 | -7977.388 | -7979.247 | -1.859 | 6.03E-07 |
| 11 | 117455699 | rs6589615 | C | G | 0.989 | -7977.388 | -7979.248 | -1.86 | 6.02E-07 |
| 11 | 117529248 | rs73014407 | T | C | 0.999 | -7977.388 | -7979.249 | -1.86 | 6.02E-07 |
| 11 | 117984995 | rs749688 | C | T | 1.01 | -7977.388 | -7979.248 | -1.86 | 6.02E-07 |
| 11 | 118769032 | rs11217081 | G | A | 1.011 | -7977.388 | -7979.248 | -1.86 | 6.02E-07 |
| 11 | 118938371 | rs1784460 | A | T | 0.99 | -7977.388 | -7979.248 | -1.86 | 6.02E-07 |
| 11 | 118938388 | rs1784459 | A | C | 0.99 | -7977.388 | -7979.248 | -1.86 | 6.02E-07 |
| 11 | 118938435 | rs1786684 | A | G | 0.99 | -7977.388 | -7979.248 | -1.86 | 6.02E-07 |
| 11 | 119384214 | rs606151 | A | G | 0.999 | -7977.388 | -7979.248 | -1.86 | 6.02E-07 |
| 11 | 119426636 | rs2663643 | A | G | 0.991 | -7977.388 | -7979.249 | -1.86 | 6.02E-07 |
| 11 | 119427242 | rs2845662 | C | T | 0.991 | -7977.388 | -7979.249 | -1.86 | 6.02E-07 |
| 11 | 119475996 | rs11217350 | C | T | 1.008 | -7977.388 | -7979.248 | -1.86 | 6.02E-07 |
| 11 | 119630247 | rs7124166 | C | G | 0.989 | -7977.388 | -7979.249 | -1.86 | 6.02E-07 |
| 11 | 119896745 | rs10892515 | A | T | 0.992 | -7977.388 | -7979.248 | -1.86 | 6.02E-07 |
| 11 | 121158551 | rs7106203 | A | T | 1.011 | -7977.388 | -7979.248 | -1.86 | 6.02E-07 |
| 11 | 116577441 | rs11216105 | G | A | 0.993 | -7977.388 | -7979.249 | -1.861 | 6.02E-07 |
| 11 | 117686456 | rs4597100 | A | C | 0.989 | -7977.388 | -7979.249 | -1.861 | 6.02E-07 |
| 11 | 119884319 | rs665758 | A | G | 0.989 | -7977.388 | -7979.249 | -1.861 | 6.02E-07 |
| 11 | 119891625 | rs1243255182 | G | A | 0.99 | -7977.388 | -7979.25 | -1.861 | 6.02E-07 |
| 11 | 120520218 | rs2000873 | C | A | 1.005 | -7977.388 | -7979.249 | -1.861 | 6.02E-07 |
| 11 | 121192989 | rs1376996225 | T | G | 1.008 | -7977.388 | -7979.249 | -1.861 | 6.02E-07 |
| 11 | 117341617 | rs549748 | A | T | 1.007 | -7977.388 | -7979.251 | -1.862 | 6.01E-07 |
| 11 | 117811563 | rs7103799 | T | C | 1.008 | -7977.388 | -7979.25 | -1.862 | 6.01E-07 |
| 11 | 117959224 | rs6589644 | T | C | 0.996 | -7977.388 | -7979.251 | -1.862 | 6.01E-07 |
| 11 | 117960018 | rs948460 | T | A | 0.996 | -7977.388 | -7979.251 | -1.862 | 6.01E-07 |
| 11 | 119419117 | rs1828279 | C | T | 0.991 | -7977.388 | -7979.25 | -1.862 | 6.01E-07 |
| 11 | 119802591 | rs504542 | G | T | 0.989 | -7977.388 | -7979.25 | -1.862 | 6.01E-07 |
| 11 | 119893208 | rs624736 | G | A | 0.992 | -7977.388 | -7979.25 | -1.862 | 6.01E-07 |
| 11 | 119894303 | rs649451 | A | G | 0.992 | -7977.388 | -7979.25 | -1.862 | 6.01E-07 |
| 11 | 116430949 | rs4938282 | T | C | 0.989 | -7977.388 | -7979.252 | -1.863 | 6.00E-07 |
| 11 | 117324965 | rs10790187 | G | A | 0.989 | -7977.388 | -7979.251 | -1.863 | 6.00E-07 |
| 11 | 117500025 | rs11216475 | T | C | 1.011 | -7977.388 | -7979.251 | -1.863 | 6.00E-07 |
| 11 | 117960138 | rs948461 | T | G | 0.997 | -7977.388 | -7979.252 | -1.863 | 6.00E-07 |
| 11 | 119455165 | rs7114739 | C | T | 0.99 | -7977.388 | -7979.251 | -1.863 | 6.00E-07 |
| 11 | 120182578 | rs35238955 | T | G | 1.006 | -7977.388 | -7979.251 | -1.863 | 6.00E-07 |
| 11 | 120557543 | rs10790401 | G | A | 1.011 | -7977.388 | -7979.251 | -1.863 | 6.00E-07 |
| 11 | 120650611 | rs2846099 | T | A | 0.993 | -7977.388 | -7979.252 | -1.863 | 6.00E-07 |
| 11 | 116313753 | rs541407 | C | T | 0.995 | -7977.388 | -7979.252 | -1.864 | 6.00E-07 |
| 11 | 117312089 | rs7947950 | G | A | 1.01 | -7977.388 | -7979.252 | -1.864 | 6.00E-07 |
| 11 | 118066695 | rs649138 | C | T | 0.989 | -7977.388 | -7979.252 | -1.864 | 6.00E-07 |
| 11 | 119006722 | rs629345 | C | T | 0.988 | -7977.388 | -7979.253 | -1.864 | 6.00E-07 |
| 11 | 119467192 | rs10892415 | T | A | 0.997 | -7977.388 | -7979.252 | -1.864 | 6.00E-07 |
| 11 | 119468951 | rs10790315 | G | C | 1.007 | -7977.388 | -7979.252 | -1.864 | 6.00E-07 |
| 11 | 119468994 | rs10790316 | T | C | 1.007 | -7977.388 | -7979.252 | -1.864 | 6.00E-07 |
| 11 | 119469211 | rs7121550 | A | G | 1.007 | -7977.388 | -7979.252 | -1.864 | 6.00E-07 |
| 11 | 119680383 | rs11607839 | C | A | 1.004 | -7977.388 | -7979.252 | -1.864 | 6.00E-07 |

| | | | | | | | |
|---|---|---|---|---|---|---|---|
| 11 | 119726873 | rs11602779 | A | G | NULL | 0.717075285 | NULL&M_Disease |
| 11 | 119732418 | rs4938733 | C | G | NULL | 0.717075285 | NULL&M_Disease |
| 11 | 119898379 | rs595377 | T | C | NULL | 0.717075285 | NULL&M_Disease |
| 11 | 116312872 | rs548890 | A | G | NULL | 0.717075285 | NULL&M_Disease |
| 11 | 117685037 | rs10892176 | C | T | NULL | 0.717075285 | NULL&M_Disease |
| 11 | 118477367 | rs498872 | A | G | NULL | 0.717075285 | NULL&M_Disease |
| 11 | 118938315 | rs1786141 | T | C | NULL | 0.717075285 | NULL&M_Disease |
| 11 | 119421928 | rs2845650 | C | T | NULL | 0.717075285 | NULL&M_Disease |
| 11 | 119459523 | rs35626144 | G | A | NULL | 0.717075285 | NULL&M_Disease |
| 11 | 119470258 | rs10790321 | A | G | NULL | 0.717075285 | NULL&M_Disease |
| 11 | 119470333 | rs10790322 | A | G | NULL | 0.717075285 | NULL&M_Disease |
| 11 | 119470389 | rs7950231 | G | A | NULL | 0.717075285 | NULL&M_Disease |
| 11 | 119472385 | rs906820 | T | C | NULL | 0.717075285 | NULL&M_Disease |
| 11 | 119895515 | rs587348 | T | C | NULL | 0.717075285 | NULL&M_Disease |
| 11 | 120060102 | rs12273399 | T | C | NULL | 0.717075285 | NULL&M_Disease |
| 11 | 120497794 | rs12807172 | T | G | NULL | 0.717075285 | NULL&M_Disease |
| 11 | 118826192 | rs12797009 | A | C | NULL | 0.717075285 | NULL&M_Disease |
| 11 | 118938957 | rs1786685 | A | G | NULL | 0.717075285 | NULL&M_Disease |
| 11 | 119008538 | rs11217164 | A | G | NULL | 0.717075285 | NULL&M_Disease |
| 11 | 119476676 | rs6589764 | C | T | NULL | 0.717075285 | NULL&M_Disease |
| 11 | 119701941 | rs11217529 | A | G | NULL | 0.717075285 | NULL&M_Disease |
| 11 | 119882223 | rs588052 | C | T | NULL | 0.717075285 | NULL&M_Disease |
| 11 | 119884311 | rs614677 | T | C | NULL | 0.717075285 | NULL&M_Disease |
| 11 | 119895939 | rs11217640 | C | T | NULL | 0.717075285 | NULL&M_Disease |
| 11 | 117521152 | rs12365289 | G | C | NULL | 0.717075285 | NULL&M_Disease |
| 11 | 118901262 | rs3759012 | T | C | NULL | 0.717075285 | NULL&M_Disease |
| 11 | 118939488 | rs10892327 | G | C | NULL | 0.717075285 | NULL&M_Disease |
| 11 | 118954265 | rs7924387 | A | G | NULL | 0.717075285 | NULL&M_Disease |
| 11 | 119414321 | rs4938680 | C | G | NULL | 0.717075285 | NULL&M_Disease |
| 11 | 119469009 | rs10790317 | A | G | NULL | 0.717075285 | NULL&M_Disease |
| 11 | 119884384 | rs665683 | C | G | NULL | 0.717075285 | NULL&M_Disease |
| 11 | 119898183 | rs624717 | G | T | NULL | 0.717075285 | NULL&M_Disease |
| 11 | 120603801 | rs3133854 | G | A | NULL | 0.717075285 | NULL&M_Disease |
| 11 | 116770517 | rs609177 | A | G | NULL | 0.718088575 | NULL&M_Disease |
| 11 | 117303393 | rs880615 | T | C | NULL | 0.718088575 | NULL&M_Disease |
| 11 | 117976568 | rs10790233 | A | G | NULL | 0.718088575 | NULL&M_Disease |
| 11 | 118942048 | rs1784305 | A | G | NULL | 0.718088575 | NULL&M_Disease |
| 11 | 119006462 | rs617290 | G | A | NULL | 0.718088575 | NULL&M_Disease |
| 11 | 120553182 | rs10892612 | A | G | NULL | 0.718088575 | NULL&M_Disease |
| 11 | 118567545 | rs644867 | G | A | NULL | 0.718088575 | NULL&M_Disease |
| 11 | 118770321 | rs11217083 | G | A | NULL | 0.718088575 | NULL&M_Disease |
| 11 | 118825762 | rs7110743 | A | C | NULL | 0.718088575 | NULL&M_Disease |
| 11 | 119012321 | rs620168 | G | C | NULL | 0.718088575 | NULL&M_Disease |
| 11 | 119427353 | rs1271978583 | G | C | NULL | 0.718088575 | NULL&M_Disease |
| 11 | 119429083 | rs2845657 | T | C | NULL | 0.718088575 | NULL&M_Disease |
| 11 | 119886924 | rs608298 | C | T | NULL | 0.718088575 | NULL&M_Disease |
| 11 | 120182869 | rs10892561 | T | C | NULL | 0.718088575 | NULL&M_Disease |
| 11 | 120182932 | rs10790376 | A | G | NULL | 0.718088575 | NULL&M_Disease |
| 11 | 116890993 | rs12808524 | T | C | NULL | 0.718088575 | NULL&M_Disease |
| 11 | 117524543 | rs10892150 | C | T | NULL | 0.718088575 | NULL&M_Disease |
| 11 | 117691148 | rs2014537 | C | T | NULL | 0.718088575 | NULL&M_Disease |
| 11 | 118942067 | rs1786690 | C | T | NULL | 0.718088575 | NULL&M_Disease |
| 11 | 118942638 | rs1614264 | C | A | NULL | 0.718088575 | NULL&M_Disease |
| 11 | 118995481 | rs1786142 | C | T | NULL | 0.718088575 | NULL&M_Disease |
| 11 | 118996948 | rs10790284 | C | A | NULL | 0.718088575 | NULL&M_Disease |
| 11 | 119216504 | rs3814762 | T | C | NULL | 0.718088575 | NULL&M_Disease |
| 11 | 119427309 | rs2663621 | C | T | NULL | 0.718088575 | NULL&M_Disease |
| 11 | 119469425 | rs12417207 | A | C | NULL | 0.718088575 | NULL&M_Disease |
| 11 | 119886078 | rs637463 | A | T | NULL | 0.718088575 | NULL&M_Disease |
| 11 | 119887725 | rs609467 | G | A | NULL | 0.718088575 | NULL&M_Disease |
| 11 | 120551569 | rs10790397 | A | C | NULL | 0.718088575 | NULL&M_Disease |
| 11 | 116314866 | rs565622 | C | A | NULL | 0.718088575 | NULL&M_Disease |
| 11 | 117691145 | rs2014536 | C | T | NULL | 0.718088575 | NULL&M_Disease |
| 11 | 118016622 | rs3018386 | G | C | NULL | 0.718088575 | NULL&M_Disease |
| 11 | 118478330 | rs7125115 | A | G | NULL | 0.718088575 | NULL&M_Disease |
| 11 | 118937985 | rs1784461 | A | G | NULL | 0.718088575 | NULL&M_Disease |
| 11 | 118989739 | rs533993 | G | A | NULL | 0.718088575 | NULL&M_Disease |
| 11 | 119000070 | rs526573 | A | G | NULL | 0.718088575 | NULL&M_Disease |
| 11 | 119595209 | rs11217430 | A | G | NULL | 0.718088575 | NULL&M_Disease |
| 11 | 119948132 | rs1551238 | C | A | NULL | 0.718088575 | NULL&M_Disease |
| 11 | 120645487 | rs2846096 | G | A | NULL | 0.718088575 | NULL&M_Disease |
| 11 | 121023906 | rs687329 | G | A | NULL | 0.718088575 | NULL&M_Disease |
| 11 | 116793231 | rs12225187 | G | A | NULL | 0.718088575 | NULL&M_Disease |
| 11 | 117037065 | rs6589602 | T | C | NULL | 0.718088575 | NULL&M_Disease |
| 11 | 118937106 | rs550974 | T | A | NULL | 0.718088575 | NULL&M_Disease |
| 11 | 118937444 | rs4454730 | A | G | NULL | 0.718088575 | NULL&M_Disease |
| 11 | 118937518 | rs4545564 | C | T | NULL | 0.718088575 | NULL&M_Disease |
| 11 | 119468661 | rs4938690 | G | A | NULL | 0.718088575 | NULL&M_Disease |
| 11 | 119472574 | rs10892419 | A | G | NULL | 0.718088575 | NULL&M_Disease |
| 11 | 119890949 | rs587304 | G | C | NULL | 0.718088575 | NULL&M_Disease |
| 11 | 119893574 | rs657706 | C | T | NULL | 0.718088575 | NULL&M_Disease |
| 11 | 119947953 | rs1551239 | A | G | NULL | 0.718088575 | NULL&M_Disease |
| 11 | 117331279 | rs680671 | A | T | NULL | 0.718088575 | NULL&M_Disease |
| 11 | 117488717 | rs4936389 | T | C | NULL | 0.718088575 | NULL&M_Disease |
| 11 | 118567486 | rs644498 | C | T | NULL | 0.718088575 | NULL&M_Disease |
| 11 | 118940957 | rs1784302 | G | C | NULL | 0.718088575 | NULL&M_Disease |
| 11 | 118950217 | rs1307145 | G | C | NULL | 0.718088575 | NULL&M_Disease |
| 11 | 118957091 | rs1799991 | A | G | NULL | 0.718088575 | NULL&M_Disease |
| 11 | 119004686 | rs658922 | A | T | NULL | 0.718088575 | NULL&M_Disease |
| 11 | 119882063 | rs639484 | T | C | NULL | 0.718088575 | NULL&M_Disease |
| 11 | 120537349 | rs11217950 | A | T | NULL | 0.718088575 | NULL&M_Disease |
| 11 | 120540795 | rs11217956 | T | C | NULL | 0.718088575 | NULL&M_Disease |
| 11 | 118928253 | rs2509121 | T | C | NULL | 0.718088575 | NULL&M_Disease |
| 11 | 120498338 | rs7928043 | A | G | NULL | 0.718088575 | NULL&M_Disease |
| 11 | 121194387 | rs10892722 | A | G | NULL | 0.718088575 | NULL&M_Disease |
| 11 | 117686567 | rs17121259 | G | C | NULL | 0.718088575 | NULL&M_Disease |

| | | | | | | | | | |
|---|---|---|---|---|---|---|---|---|---|
| 11 | 119726873 | rs11602779 | A | G | 0.999 | -7977.388 | -7979.252 | -1.864 | 6.00E-07 |
| 11 | 119732418 | rs4938733 | C | G | 1 | -7977.388 | -7979.252 | -1.864 | 6.00E-07 |
| 11 | 119898379 | rs595377 | T | C | 0.993 | -7977.388 | -7979.253 | -1.864 | 6.00E-07 |
| 11 | 116312872 | rs548890 | A | G | 0.996 | -7977.388 | -7979.253 | -1.865 | 5.99E-07 |
| 11 | 117685037 | rs10892176 | C | T | 0.99 | -7977.388 | -7979.253 | -1.865 | 5.99E-07 |
| 11 | 118477367 | rs498872 | A | G | 0.999 | -7977.388 | -7979.253 | -1.865 | 5.99E-07 |
| 11 | 118938315 | rs1786141 | T | C | 0.991 | -7977.388 | -7979.253 | -1.865 | 5.99E-07 |
| 11 | 119421928 | rs2845650 | C | T | 0.992 | -7977.388 | -7979.254 | -1.865 | 5.99E-07 |
| 11 | 119459523 | rs35626144 | G | A | 0.994 | -7977.388 | -7979.253 | -1.865 | 5.99E-07 |
| 11 | 119470258 | rs10790321 | A | G | 1.007 | -7977.388 | -7979.253 | -1.865 | 5.99E-07 |
| 11 | 119470333 | rs10790322 | A | G | 1.007 | -7977.388 | -7979.253 | -1.865 | 5.99E-07 |
| 11 | 119470389 | rs7950231 | G | A | 1.007 | -7977.388 | -7979.253 | -1.865 | 5.99E-07 |
| 11 | 119472385 | rs906820 | T | C | 1.007 | -7977.388 | -7979.253 | -1.865 | 5.99E-07 |
| 11 | 119895515 | rs587348 | T | C | 0.993 | -7977.388 | -7979.253 | -1.865 | 5.99E-07 |
| 11 | 120060102 | rs12273399 | T | C | 0.99 | -7977.388 | -7979.254 | -1.865 | 5.99E-07 |
| 11 | 120497794 | rs12807172 | T | G | 0.992 | -7977.388 | -7979.253 | -1.865 | 5.99E-07 |
| 11 | 118826192 | rs12797009 | A | C | 0.999 | -7977.388 | -7979.254 | -1.866 | 5.99E-07 |
| 11 | 118938957 | rs1786685 | A | G | 0.991 | -7977.388 | -7979.254 | -1.866 | 5.99E-07 |
| 11 | 119008538 | rs11217164 | A | G | 0.989 | -7977.388 | -7979.254 | -1.866 | 5.99E-07 |
| 11 | 119476676 | rs6589764 | C | T | 1.007 | -7977.388 | -7979.254 | -1.866 | 5.99E-07 |
| 11 | 119701941 | rs11217529 | A | G | 1.011 | -7977.388 | -7979.255 | -1.866 | 5.99E-07 |
| 11 | 119882223 | rs588052 | C | T | 0.992 | -7977.388 | -7979.254 | -1.866 | 5.99E-07 |
| 11 | 119884311 | rs614677 | T | C | 0.989 | -7977.388 | -7979.254 | -1.866 | 5.99E-07 |
| 11 | 119895939 | rs11217640 | C | T | 0.993 | -7977.388 | -7979.254 | -1.866 | 5.99E-07 |
| 11 | 117521152 | rs12365289 | G | C | 1.001 | -7977.388 | -7979.256 | -1.867 | 5.98E-07 |
| 11 | 118901262 | rs3759012 | T | C | 0.99 | -7977.388 | -7979.255 | -1.867 | 5.98E-07 |
| 11 | 118939488 | rs10892327 | G | C | 0.991 | -7977.388 | -7979.256 | -1.867 | 5.98E-07 |
| 11 | 118954265 | rs7924387 | A | G | 1.009 | -7977.388 | -7979.255 | -1.867 | 5.98E-07 |
| 11 | 119414321 | rs4938680 | C | G | 0.992 | -7977.388 | -7979.255 | -1.867 | 5.98E-07 |
| 11 | 119469009 | rs10790317 | A | G | 1.007 | -7977.388 | -7979.256 | -1.867 | 5.98E-07 |
| 11 | 119884384 | rs665683 | C | G | 0.989 | -7977.388 | -7979.255 | -1.867 | 5.98E-07 |
| 11 | 119898183 | rs624717 | G | T | 0.993 | -7977.388 | -7979.255 | -1.867 | 5.98E-07 |
| 11 | 120603801 | rs3133854 | G | A | 1.006 | -7977.388 | -7979.255 | -1.867 | 5.98E-07 |
| 11 | 116770517 | rs609177 | A | G | 0.993 | -7977.388 | -7979.256 | -1.868 | 5.97E-07 |
| 11 | 117303393 | rs880615 | T | C | 1.008 | -7977.388 | -7979.257 | -1.868 | 5.97E-07 |
| 11 | 117976568 | rs10790233 | A | G | 1.009 | -7977.388 | -7979.256 | -1.868 | 5.97E-07 |
| 11 | 118942048 | rs1784305 | A | G | 0.991 | -7977.388 | -7979.256 | -1.868 | 5.97E-07 |
| 11 | 119006462 | rs617290 | G | A | 0.989 | -7977.388 | -7979.256 | -1.868 | 5.97E-07 |
| 11 | 120553182 | rs10892612 | A | G | 1.011 | -7977.388 | -7979.256 | -1.868 | 5.97E-07 |
| 11 | 118567545 | rs644867 | G | A | 0.995 | -7977.388 | -7979.257 | -1.869 | 5.97E-07 |
| 11 | 118770321 | rs11217083 | G | A | 1.01 | -7977.388 | -7979.257 | -1.869 | 5.97E-07 |
| 11 | 118825762 | rs7110743 | A | C | 1.002 | -7977.388 | -7979.257 | -1.869 | 5.97E-07 |
| 11 | 119012321 | rs620168 | G | C | 0.989 | -7977.388 | -7979.258 | -1.869 | 5.97E-07 |
| 11 | 119427353 | rs1271978583 | G | C | 0.992 | -7977.388 | -7979.257 | -1.869 | 5.97E-07 |
| 11 | 119429083 | rs2845657 | T | C | 0.992 | -7977.388 | -7979.258 | -1.869 | 5.97E-07 |
| 11 | 119886924 | rs608298 | C | T | 0.993 | -7977.388 | -7979.257 | -1.869 | 5.97E-07 |
| 11 | 120182869 | rs10892561 | T | C | 1.005 | -7977.388 | -7979.257 | -1.869 | 5.97E-07 |
| 11 | 120182932 | rs10790376 | A | G | 1.005 | -7977.388 | -7979.257 | -1.869 | 5.97E-07 |
| 11 | 116890993 | rs12808524 | T | C | 0.995 | -7977.388 | -7979.258 | -1.87 | 5.96E-07 |
| 11 | 117524543 | rs10892150 | C | T | 1 | -7977.388 | -7979.258 | -1.87 | 5.96E-07 |
| 11 | 117691148 | rs2014537 | C | T | 1.001 | -7977.388 | -7979.259 | -1.87 | 5.96E-07 |
| 11 | 118942067 | rs1786690 | C | T | 0.991 | -7977.388 | -7979.259 | -1.87 | 5.96E-07 |
| 11 | 118942638 | rs1614264 | C | A | 0.991 | -7977.388 | -7979.258 | -1.87 | 5.96E-07 |
| 11 | 118995481 | rs1786142 | C | T | 0.989 | -7977.388 | -7979.258 | -1.87 | 5.96E-07 |
| 11 | 118996948 | rs10790284 | C | A | 0.989 | -7977.388 | -7979.258 | -1.87 | 5.96E-07 |
| 11 | 119216504 | rs3814762 | T | C | 1.005 | -7977.388 | -7979.258 | -1.87 | 5.96E-07 |
| 11 | 119427309 | rs2663621 | C | T | 0.992 | -7977.388 | -7979.258 | -1.87 | 5.96E-07 |
| 11 | 119469425 | rs12417207 | A | C | 1.006 | -7977.388 | -7979.258 | -1.87 | 5.96E-07 |
| 11 | 119886078 | rs637463 | A | T | 0.99 | -7977.388 | -7979.258 | -1.87 | 5.96E-07 |
| 11 | 119887725 | rs609467 | G | A | 0.993 | -7977.388 | -7979.258 | -1.87 | 5.96E-07 |
| 11 | 120551569 | rs10790397 | A | C | 1.011 | -7977.388 | -7979.258 | -1.87 | 5.96E-07 |
| 11 | 116314866 | rs565622 | C | A | 0.996 | -7977.388 | -7979.26 | -1.871 | 5.96E-07 |
| 11 | 117691145 | rs2014536 | C | T | 1 | -7977.388 | -7979.259 | -1.871 | 5.96E-07 |
| 11 | 118016622 | rs3018386 | G | C | 0.991 | -7977.388 | -7979.259 | -1.871 | 5.96E-07 |
| 11 | 118478330 | rs7125115 | A | G | 0.993 | -7977.388 | -7979.259 | -1.871 | 5.96E-07 |
| 11 | 118937985 | rs1784461 | A | G | 0.991 | -7977.388 | -7979.259 | -1.871 | 5.96E-07 |
| 11 | 118989739 | rs533993 | G | A | 0.989 | -7977.388 | -7979.259 | -1.871 | 5.96E-07 |
| 11 | 119000070 | rs526573 | A | G | 0.989 | -7977.388 | -7979.26 | -1.871 | 5.96E-07 |
| 11 | 119595209 | rs11217430 | A | G | 0.99 | -7977.388 | -7979.259 | -1.871 | 5.96E-07 |
| 11 | 119948132 | rs1551238 | C | A | 1.006 | -7977.388 | -7979.259 | -1.871 | 5.96E-07 |
| 11 | 120645487 | rs2846096 | G | A | 0.994 | -7977.388 | -7979.259 | -1.871 | 5.96E-07 |
| 11 | 121023906 | rs687329 | G | A | 1.008 | -7977.388 | -7979.259 | -1.871 | 5.96E-07 |
| 11 | 116793231 | rs12225187 | G | A | 0.994 | -7977.388 | -7979.261 | -1.872 | 5.95E-07 |
| 11 | 117037065 | rs6589602 | T | C | 0.992 | -7977.388 | -7979.26 | -1.872 | 5.95E-07 |
| 11 | 118937106 | rs550974 | T | A | 0.992 | -7977.388 | -7979.26 | -1.872 | 5.95E-07 |
| 11 | 118937444 | rs4454730 | A | G | 0.992 | -7977.388 | -7979.26 | -1.872 | 5.95E-07 |
| 11 | 118937518 | rs4545564 | C | T | 0.992 | -7977.388 | -7979.26 | -1.872 | 5.95E-07 |
| 11 | 119468661 | rs4938690 | G | A | 1.006 | -7977.388 | -7979.26 | -1.872 | 5.95E-07 |
| 11 | 119472574 | rs10892419 | A | G | 1.006 | -7977.388 | -7979.261 | -1.872 | 5.95E-07 |
| 11 | 119890949 | rs587304 | G | C | 0.994 | -7977.388 | -7979.26 | -1.872 | 5.95E-07 |
| 11 | 119893574 | rs657706 | C | T | 0.994 | -7977.388 | -7979.26 | -1.872 | 5.95E-07 |
| 11 | 119947953 | rs1551239 | A | G | 1.006 | -7977.388 | -7979.26 | -1.872 | 5.95E-07 |
| 11 | 117331279 | rs680671 | A | T | 1.01 | -7977.388 | -7979.261 | -1.873 | 5.94E-07 |
| 11 | 117488717 | rs4936389 | T | C | 0.99 | -7977.388 | -7979.261 | -1.873 | 5.94E-07 |
| 11 | 118567486 | rs644498 | C | T | 0.996 | -7977.388 | -7979.261 | -1.873 | 5.94E-07 |
| 11 | 118940957 | rs1784302 | G | C | 0.992 | -7977.388 | -7979.261 | -1.873 | 5.94E-07 |
| 11 | 118950217 | rs1307145 | G | C | 0.992 | -7977.388 | -7979.261 | -1.873 | 5.94E-07 |
| 11 | 118957091 | rs1799991 | A | G | 1.008 | -7977.388 | -7979.262 | -1.873 | 5.94E-07 |
| 11 | 119004686 | rs658922 | A | T | 0.989 | -7977.388 | -7979.261 | -1.873 | 5.94E-07 |
| 11 | 119882063 | rs639484 | T | C | 0.993 | -7977.388 | -7979.262 | -1.873 | 5.94E-07 |
| 11 | 120537349 | rs11217950 | A | T | 0.995 | -7977.388 | -7979.261 | -1.873 | 5.94E-07 |
| 11 | 120540795 | rs11217956 | T | C | 0.996 | -7977.388 | -7979.261 | -1.873 | 5.94E-07 |
| 11 | 118928253 | rs2509121 | T | C | 0.992 | -7977.388 | -7979.262 | -1.874 | 5.94E-07 |
| 11 | 120498338 | rs7928043 | A | G | 0.992 | -7977.388 | -7979.262 | -1.874 | 5.94E-07 |
| 11 | 121194387 | rs10892722 | A | G | 1.007 | -7977.388 | -7979.262 | -1.874 | 5.94E-07 |
| 11 | 117686567 | rs17121259 | G | C | 0.99 | -7977.388 | -7979.263 | -1.875 | 5.93E-07 |

| | | | | | | | |
|---|---|---|---|---|---|---|---|
| 11 | 117800169 | rs11600129 | C | T | NULL | 0.718088575 | NULL&M_Disease |
| 11 | 118566039 | rs6589674 | G | A | NULL | 0.718088575 | NULL&M_Disease |
| 11 | 118567257 | rs7126172 | G | A | NULL | 0.718088575 | NULL&M_Disease |
| 11 | 118997869 | rs547345 | C | T | NULL | 0.718088575 | NULL&M_Disease |
| 11 | 120187433 | rs7102217 | T | C | NULL | 0.718088575 | NULL&M_Disease |
| 11 | 120542687 | rs7934734 | G | A | NULL | 0.718088575 | NULL&M_Disease |
| 11 | 120556697 | rs7122193 | C | T | NULL | 0.718088575 | NULL&M_Disease |
| 11 | 116534288 | rs486915 | A | T | NULL | 0.718088575 | NULL&M_Disease |
| 11 | 116750874 | rs7396061 | A | C | NULL | 0.718088575 | NULL&M_Disease |
| 11 | 116987046 | rs11216277 | G | A | NULL | 0.718088575 | NULL&M_Disease |
| 11 | 116988936 | rs10892076 | G | A | NULL | 0.718088575 | NULL&M_Disease |
| 11 | 117035502 | rs10892080 | A | G | NULL | 0.718088575 | NULL&M_Disease |
| 11 | 117800682 | rs11216635 | G | A | NULL | 0.718088575 | NULL&M_Disease |
| 11 | 117979744 | rs10892216 | C | A | NULL | 0.718088575 | NULL&M_Disease |
| 11 | 117979814 | rs10892217 | C | T | NULL | 0.718088575 | NULL&M_Disease |
| 11 | 118949083 | rs1177563 | A | G | NULL | 0.718088575 | NULL&M_Disease |
| 11 | 118949799 | rs1168568 | A | G | NULL | 0.718088575 | NULL&M_Disease |
| 11 | 119412000 | rs11217309 | T | C | NULL | 0.718088575 | NULL&M_Disease |
| 11 | 119463270 | rs10892413 | A | G | NULL | 0.718088575 | NULL&M_Disease |
| 11 | 119476351 | rs2048247 | A | G | NULL | 0.718088575 | NULL&M_Disease |
| 11 | 119476354 | rs2048248 | A | G | NULL | 0.718088575 | NULL&M_Disease |
| 11 | 120055477 | rs566935 | C | G | NULL | 0.718088575 | NULL&M_Disease |
| 11 | 116520469 | rs506305 | T | C | NULL | 0.718088575 | NULL&M_Disease |
| 11 | 117000955 | rs10400366 | G | A | NULL | 0.718088575 | NULL&M_Disease |
| 11 | 118566313 | rs12794726 | T | C | NULL | 0.718088575 | NULL&M_Disease |
| 11 | 118567084 | rs542613 | A | G | NULL | 0.718088575 | NULL&M_Disease |
| 11 | 118567218 | rs11216947 | A | G | NULL | 0.718088575 | NULL&M_Disease |
| 11 | 119463904 | rs2368426 | A | G | NULL | 0.718088575 | NULL&M_Disease |
| 11 | 119472474 | rs4245196 | G | A | NULL | 0.718088575 | NULL&M_Disease |
| 11 | 119472670 | rs58254953 | A | G | NULL | 0.718088575 | NULL&M_Disease |
| 11 | 119727706 | rs10892478 | G | A | NULL | 0.718088575 | NULL&M_Disease |
| 11 | 120075574 | rs10892549 | T | C | NULL | 0.719099657 | NULL&M_Disease |
| 11 | 120980822 | rs538936 | A | G | NULL | 0.718088575 | NULL&M_Disease |
| 11 | 116330732 | rs513083 | T | C | NULL | 0.719099657 | NULL&M_Disease |
| 11 | 116473707 | rs9919543 | G | C | NULL | 0.719099657 | NULL&M_Disease |
| 11 | 116718858 | rs532815 | G | A | NULL | 0.719099657 | NULL&M_Disease |
| 11 | 118554259 | rs7106851 | A | G | NULL | 0.719099657 | NULL&M_Disease |
| 11 | 118566090 | rs7130076 | C | T | NULL | 0.719099657 | NULL&M_Disease |
| 11 | 118566175 | rs7114458 | A | G | NULL | 0.719099657 | NULL&M_Disease |
| 11 | 118566234 | rs7114216 | G | A | NULL | 0.719099657 | NULL&M_Disease |
| 11 | 118567375 | rs478296 | C | T | NULL | 0.719099657 | NULL&M_Disease |
| 11 | 118567444 | rs493834 | G | A | NULL | 0.719099657 | NULL&M_Disease |
| 11 | 119268984 | rs10892360 | A | G | NULL | 0.719099657 | NULL&M_Disease |
| 11 | 119429139 | rs2845656 | C | T | NULL | 0.719099657 | NULL&M_Disease |
| 11 | 119462854 | rs11608077 | G | A | NULL | 0.719099657 | NULL&M_Disease |
| 11 | 119571975 | rs1110166 | A | G | NULL | 0.719099657 | NULL&M_Disease |
| 11 | 119662942 | rs11217507 | G | T | NULL | 0.719099657 | NULL&M_Disease |
| 11 | 119887019 | rs607888 | A | G | NULL | 0.719099657 | NULL&M_Disease |
| 11 | 119969700 | rs605833 | C | T | NULL | 0.719099657 | NULL&M_Disease |
| 11 | 118943433 | rs1185460 | G | A | NULL | 0.719099657 | NULL&M_Disease |
| 11 | 119566184 | rs4938711 | T | C | NULL | 0.719099657 | NULL&M_Disease |
| 11 | 120539020 | rs7947520 | A | G | NULL | 0.719099657 | NULL&M_Disease |
| 11 | 120548795 | rs11217967 | A | C | NULL | 0.719099657 | NULL&M_Disease |
| 11 | 117965783 | rs2276126 | G | A | NULL | 0.719099657 | NULL&M_Disease |
| 11 | 117966994 | rs9651684 | C | G | NULL | 0.719099657 | NULL&M_Disease |
| 11 | 118539522 | rs2508991 | A | G | NULL | 0.719099657 | NULL&M_Disease |
| 11 | 118539566 | rs1783935 | T | A | NULL | 0.719099657 | NULL&M_Disease |
| 11 | 118546602 | rs526597 | G | A | NULL | 0.719099657 | NULL&M_Disease |
| 11 | 118548665 | rs1784298 | C | T | NULL | 0.719099657 | NULL&M_Disease |
| 11 | 118548671 | rs4938522 | T | A | NULL | 0.719099657 | NULL&M_Disease |
| 11 | 118548838 | rs4938523 | T | G | NULL | 0.719099657 | NULL&M_Disease |
| 11 | 118549582 | rs507080 | A | G | NULL | 0.719099657 | NULL&M_Disease |
| 11 | 118549803 | rs659969 | T | C | NULL | 0.719099657 | NULL&M_Disease |
| 11 | 118549908 | rs656575 | G | T | NULL | 0.719099657 | NULL&M_Disease |
| 11 | 118549985 | rs543327 | T | G | NULL | 0.719099657 | NULL&M_Disease |
| 11 | 118550017 | rs543413 | C | T | NULL | 0.719099657 | NULL&M_Disease |
| 11 | 118945185 | rs1169326 | A | G | NULL | 0.719099657 | NULL&M_Disease |
| 11 | 118945337 | rs1184902 | G | A | NULL | 0.719099657 | NULL&M_Disease |
| 11 | 118958127 | rs1799993 | A | C | NULL | 0.719099657 | NULL&M_Disease |
| 11 | 119702527 | rs570637 | T | C | NULL | 0.719099657 | NULL&M_Disease |
| 11 | 119848156 | rs651055 | C | G | NULL | 0.719099657 | NULL&M_Disease |
| 11 | 119950798 | rs12276260 | C | G | NULL | 0.719099657 | NULL&M_Disease |
| 11 | 120041444 | rs4936509 | C | T | NULL | 0.719099657 | NULL&M_Disease |
| 11 | 117009789 | rs2735181 | C | T | NULL | 0.719099657 | NULL&M_Disease |
| 11 | 117509797 | rs4938423 | T | C | NULL | 0.719099657 | NULL&M_Disease |
| 11 | 117965595 | rs2276123 | T | C | NULL | 0.719099657 | NULL&M_Disease |
| 11 | 118538157 | rs520543 | A | G | NULL | 0.719099657 | NULL&M_Disease |
| 11 | 118538187 | rs607527 | T | G | NULL | 0.719099657 | NULL&M_Disease |
| 11 | 118558449 | rs514143 | A | G | NULL | 0.719099657 | NULL&M_Disease |
| 11 | 118561410 | rs12798453 | G | T | NULL | 0.719099657 | NULL&M_Disease |
| 11 | 118564397 | rs558781 | T | G | NULL | 0.719099657 | NULL&M_Disease |
| 11 | 118564762 | rs516153 | A | C | NULL | 0.719099657 | NULL&M_Disease |
| 11 | 118564982 | rs518028 | C | T | NULL | 0.719099657 | NULL&M_Disease |
| 11 | 118566746 | rs533646 | G | C | NULL | 0.719099657 | NULL&M_Disease |
| 11 | 118567020 | rs642662 | G | A | NULL | 0.719099657 | NULL&M_Disease |
| 11 | 119565495 | rs7950059 | T | C | NULL | 0.719099657 | NULL&M_Disease |
| 11 | 119880367 | rs617124 | A | G | NULL | 0.719099657 | NULL&M_Disease |
| 11 | 119885822 | rs667670 | C | T | NULL | 0.719099657 | NULL&M_Disease |
| 11 | 119987517 | rs489059 | A | G | NULL | 0.719099657 | NULL&M_Disease |
| 11 | 116730738 | rs567312 | G | A | NULL | 0.719099657 | NULL&M_Disease |
| 11 | 116834313 | rs10892052 | G | A | NULL | 0.719099657 | NULL&M_Disease |
| 11 | 117042949 | rs7120565 | T | C | NULL | 0.719099657 | NULL&M_Disease |
| 11 | 117965082 | rs1080620 | A | G | NULL | 0.719099657 | NULL&M_Disease |
| 11 | 117965105 | rs1080619 | G | A | NULL | 0.719099657 | NULL&M_Disease |
| 11 | 118538308 | rs692750 | T | G | NULL | 0.719099657 | NULL&M_Disease |
| 11 | 118539140 | rs4589335 | T | C | NULL | 0.719099657 | NULL&M_Disease |

| | | | | | | | | | |
|---|---|---|---|---|---|---|---|---|---|
| 11 | 117800169 | rs11600129 | C | T | 1.01 | -7977.388 | -7979.264 | -1.875 | 5.93E-07 |
| 11 | 118566039 | rs6589674 | G | A | 0.996 | -7977.388 | -7979.264 | -1.875 | 5.93E-07 |
| 11 | 118567257 | rs7126172 | G | A | 0.996 | -7977.388 | -7979.263 | -1.875 | 5.93E-07 |
| 11 | 118997869 | rs547345 | C | T | 0.989 | -7977.388 | -7979.263 | -1.875 | 5.93E-07 |
| 11 | 120187433 | rs7102217 | T | C | 1.004 | -7977.388 | -7979.264 | -1.875 | 5.93E-07 |
| 11 | 120542687 | rs7934734 | G | A | 0.99 | -7977.388 | -7979.263 | -1.875 | 5.93E-07 |
| 11 | 120556697 | rs7122193 | C | T | 1.01 | -7977.388 | -7979.263 | -1.875 | 5.93E-07 |
| 11 | 116534288 | rs486915 | A | T | 1.004 | -7977.388 | -7979.264 | -1.876 | 5.93E-07 |
| 11 | 116750874 | rs7396061 | A | C | 0.998 | -7977.388 | -7979.264 | -1.876 | 5.93E-07 |
| 11 | 116987046 | rs11216277 | G | A | 1 | -7977.388 | -7979.264 | -1.876 | 5.93E-07 |
| 11 | 116988936 | rs10892076 | G | A | 1 | -7977.388 | -7979.264 | -1.876 | 5.93E-07 |
| 11 | 117035502 | rs10892080 | A | G | 0.993 | -7977.388 | -7979.264 | -1.876 | 5.93E-07 |
| 11 | 117800682 | rs11216635 | G | A | 1.01 | -7977.388 | -7979.265 | -1.876 | 5.93E-07 |
| 11 | 117979744 | rs10892216 | C | A | 1.009 | -7977.388 | -7979.264 | -1.876 | 5.93E-07 |
| 11 | 117979814 | rs10892217 | C | T | 1.009 | -7977.388 | -7979.264 | -1.876 | 5.93E-07 |
| 11 | 118949083 | rs1177563 | A | G | 0.992 | -7977.388 | -7979.265 | -1.876 | 5.93E-07 |
| 11 | 118949799 | rs1168568 | A | G | 0.992 | -7977.388 | -7979.265 | -1.876 | 5.93E-07 |
| 11 | 119412000 | rs11217309 | T | C | 0.993 | -7977.388 | -7979.264 | -1.876 | 5.93E-07 |
| 11 | 119463270 | rs10892413 | A | G | 1.004 | -7977.388 | -7979.264 | -1.876 | 5.93E-07 |
| 11 | 119476351 | rs2048247 | A | G | 1.005 | -7977.388 | -7979.265 | -1.876 | 5.93E-07 |
| 11 | 119476354 | rs2048248 | A | G | 1.005 | -7977.388 | -7979.265 | -1.876 | 5.93E-07 |
| 11 | 120055477 | rs566935 | C | G | 0.991 | -7977.388 | -7979.264 | -1.876 | 5.93E-07 |
| 11 | 116520469 | rs506305 | T | C | 1.007 | -7977.388 | -7979.265 | -1.877 | 5.92E-07 |
| 11 | 117000955 | rs10400366 | G | A | 0.993 | -7977.388 | -7979.266 | -1.877 | 5.92E-07 |
| 11 | 118566313 | rs12794726 | T | C | 0.996 | -7977.388 | -7979.265 | -1.877 | 5.92E-07 |
| 11 | 118567084 | rs542613 | A | G | 0.997 | -7977.388 | -7979.265 | -1.877 | 5.92E-07 |
| 11 | 118567218 | rs11216947 | A | G | 0.997 | -7977.388 | -7979.265 | -1.877 | 5.92E-07 |
| 11 | 119463904 | rs2368426 | A | G | 1.004 | -7977.388 | -7979.266 | -1.877 | 5.92E-07 |
| 11 | 119472474 | rs4245196 | G | A | 1.005 | -7977.388 | -7979.266 | -1.877 | 5.92E-07 |
| 11 | 119472670 | rs58254953 | A | G | 1.005 | -7977.388 | -7979.266 | -1.877 | 5.92E-07 |
| 11 | 119727706 | rs10892478 | G | A | 0.999 | -7977.388 | -7979.265 | -1.877 | 5.92E-07 |
| 11 | 120075574 | rs10892549 | T | C | 1.01 | -7977.388 | -7979.266 | -1.877 | 5.92E-07 |
| 11 | 120980822 | rs538936 | A | G | 1.01 | -7977.388 | -7979.265 | -1.877 | 5.92E-07 |
| 11 | 116330732 | rs513083 | T | C | 0.992 | -7977.388 | -7979.266 | -1.878 | 5.91E-07 |
| 11 | 116473707 | rs9919543 | G | C | 1.004 | -7977.388 | -7979.266 | -1.878 | 5.91E-07 |
| 11 | 116718858 | rs532815 | G | A | 1.005 | -7977.388 | -7979.266 | -1.878 | 5.91E-07 |
| 11 | 118554259 | rs7106851 | A | G | 0.997 | -7977.388 | -7979.266 | -1.878 | 5.91E-07 |
| 11 | 118566090 | rs7130076 | C | T | 0.997 | -7977.388 | -7979.266 | -1.878 | 5.91E-07 |
| 11 | 118566175 | rs7114458 | A | G | 0.997 | -7977.388 | -7979.266 | -1.878 | 5.91E-07 |
| 11 | 118566234 | rs7114216 | G | A | 0.997 | -7977.388 | -7979.266 | -1.878 | 5.91E-07 |
| 11 | 118567375 | rs478296 | C | T | 0.997 | -7977.388 | -7979.266 | -1.878 | 5.91E-07 |
| 11 | 118567444 | rs493834 | G | A | 0.997 | -7977.388 | -7979.267 | -1.878 | 5.91E-07 |
| 11 | 119268984 | rs10892360 | A | G | 1.009 | -7977.388 | -7979.266 | -1.878 | 5.91E-07 |
| 11 | 119429139 | rs2845656 | C | T | 0.993 | -7977.388 | -7979.267 | -1.878 | 5.91E-07 |
| 11 | 119462854 | rs11608077 | G | A | 1.004 | -7977.388 | -7979.267 | -1.878 | 5.91E-07 |
| 11 | 119571975 | rs1110166 | A | G | 1.007 | -7977.388 | -7979.266 | -1.878 | 5.91E-07 |
| 11 | 119662942 | rs11217507 | G | T | 1.003 | -7977.388 | -7979.266 | -1.878 | 5.91E-07 |
| 11 | 119887019 | rs607888 | A | G | 0.991 | -7977.388 | -7979.267 | -1.878 | 5.91E-07 |
| 11 | 119969700 | rs605833 | C | T | 0.993 | -7977.388 | -7979.266 | -1.878 | 5.91E-07 |
| 11 | 118943433 | rs1185460 | G | A | 0.992 | -7977.388 | -7979.267 | -1.879 | 5.91E-07 |
| 11 | 119566184 | rs4938711 | T | C | 0.992 | -7977.388 | -7979.267 | -1.879 | 5.91E-07 |
| 11 | 120539020 | rs7947520 | A | G | 1.01 | -7977.388 | -7979.267 | -1.879 | 5.91E-07 |
| 11 | 120548795 | rs11217967 | A | C | 1.002 | -7977.388 | -7979.267 | -1.879 | 5.91E-07 |
| 11 | 117965783 | rs2276126 | G | A | 0.997 | -7977.388 | -7979.268 | -1.88 | 5.90E-07 |
| 11 | 117966994 | rs9651684 | C | G | 0.997 | -7977.388 | -7979.268 | -1.88 | 5.90E-07 |
| 11 | 118539522 | rs2508991 | A | G | 1.003 | -7977.388 | -7979.268 | -1.88 | 5.90E-07 |
| 11 | 118539566 | rs1783935 | T | A | 1.003 | -7977.388 | -7979.269 | -1.88 | 5.90E-07 |
| 11 | 118546602 | rs526597 | G | A | 1.003 | -7977.388 | -7979.268 | -1.88 | 5.90E-07 |
| 11 | 118548665 | rs1784298 | C | T | 1.003 | -7977.388 | -7979.268 | -1.88 | 5.90E-07 |
| 11 | 118548671 | rs4938522 | T | A | 1.003 | -7977.388 | -7979.268 | -1.88 | 5.90E-07 |
| 11 | 118548838 | rs4938523 | T | G | 1.003 | -7977.388 | -7979.268 | -1.88 | 5.90E-07 |
| 11 | 118549582 | rs507080 | A | G | 1.003 | -7977.388 | -7979.268 | -1.88 | 5.90E-07 |
| 11 | 118549803 | rs659969 | T | C | 1.003 | -7977.388 | -7979.268 | -1.88 | 5.90E-07 |
| 11 | 118549908 | rs656575 | G | T | 1.003 | -7977.388 | -7979.268 | -1.88 | 5.90E-07 |
| 11 | 118549985 | rs543327 | T | G | 1.003 | -7977.388 | -7979.268 | -1.88 | 5.90E-07 |
| 11 | 118550017 | rs543413 | C | T | 1.003 | -7977.388 | -7979.268 | -1.88 | 5.90E-07 |
| 11 | 118945185 | rs1169326 | A | G | 0.992 | -7977.388 | -7979.268 | -1.88 | 5.90E-07 |
| 11 | 118945337 | rs1184902 | G | A | 0.992 | -7977.388 | -7979.268 | -1.88 | 5.90E-07 |
| 11 | 118958127 | rs1799993 | A | C | 1.008 | -7977.388 | -7979.268 | -1.88 | 5.90E-07 |
| 11 | 119702527 | rs570637 | T | C | 1.009 | -7977.388 | -7979.268 | -1.88 | 5.90E-07 |
| 11 | 119848156 | rs651055 | C | G | 1.008 | -7977.388 | -7979.269 | -1.88 | 5.90E-07 |
| 11 | 119950798 | rs12276260 | C | G | 0.996 | -7977.388 | -7979.268 | -1.88 | 5.90E-07 |
| 11 | 120041444 | rs4936509 | C | T | 1.009 | -7977.388 | -7979.268 | -1.88 | 5.90E-07 |
| 11 | 117009789 | rs2735181 | C | T | 0.993 | -7977.388 | -7979.269 | -1.881 | 5.90E-07 |
| 11 | 117509797 | rs4938423 | T | C | 1.009 | -7977.388 | -7979.269 | -1.881 | 5.90E-07 |
| 11 | 117965595 | rs2276123 | T | C | 0.997 | -7977.388 | -7979.269 | -1.881 | 5.90E-07 |
| 11 | 118538157 | rs520543 | A | G | 1.002 | -7977.388 | -7979.27 | -1.881 | 5.90E-07 |
| 11 | 118538187 | rs607527 | T | G | 1.003 | -7977.388 | -7979.27 | -1.881 | 5.90E-07 |
| 11 | 118558449 | rs514143 | A | G | 0.999 | -7977.388 | -7979.27 | -1.881 | 5.90E-07 |
| 11 | 118561410 | rs12798453 | G | T | 0.998 | -7977.388 | -7979.27 | -1.881 | 5.90E-07 |
| 11 | 118564397 | rs558781 | T | G | 0.998 | -7977.388 | -7979.269 | -1.881 | 5.90E-07 |
| 11 | 118564762 | rs516153 | A | C | 0.998 | -7977.388 | -7979.27 | -1.881 | 5.90E-07 |
| 11 | 118564982 | rs518028 | C | T | 0.998 | -7977.388 | -7979.269 | -1.881 | 5.90E-07 |
| 11 | 118566746 | rs533646 | G | C | 0.998 | -7977.388 | -7979.269 | -1.881 | 5.90E-07 |
| 11 | 118567020 | rs642662 | G | A | 0.998 | -7977.388 | -7979.27 | -1.881 | 5.90E-07 |
| 11 | 119565495 | rs7950059 | T | C | 0.992 | -7977.388 | -7979.27 | -1.881 | 5.90E-07 |
| 11 | 119880367 | rs617124 | A | G | 1.01 | -7977.388 | -7979.269 | -1.881 | 5.90E-07 |
| 11 | 119885822 | rs667670 | C | T | 0.991 | -7977.388 | -7979.269 | -1.881 | 5.90E-07 |
| 11 | 119987517 | rs489059 | A | G | 0.991 | -7977.388 | -7979.27 | -1.881 | 5.90E-07 |
| 11 | 116730738 | rs567312 | G | A | 0.994 | -7977.388 | -7979.27 | -1.882 | 5.89E-07 |
| 11 | 116834313 | rs10892052 | G | A | 0.997 | -7977.388 | -7979.27 | -1.882 | 5.89E-07 |
| 11 | 117042949 | rs7120565 | T | C | 0.993 | -7977.388 | -7979.27 | -1.882 | 5.89E-07 |
| 11 | 117965082 | rs1080620 | A | G | 0.997 | -7977.388 | -7979.271 | -1.882 | 5.89E-07 |
| 11 | 117965105 | rs1080619 | G | A | 0.997 | -7977.388 | -7979.271 | -1.882 | 5.89E-07 |
| 11 | 118538308 | rs692750 | T | G | 0.999 | -7977.388 | -7979.27 | -1.882 | 5.89E-07 |
| 11 | 118539140 | rs4589335 | T | C | 0.999 | -7977.388 | -7979.27 | -1.882 | 5.89E-07 |

| | | | | | | | |
|---|---|---|---|---|---|---|---|
| 11 | 118553326 | rs693037 | T | C | NULL | 0.719099657 | NULL&M_Disease |
| 11 | 118555719 | rs625513 | C | A | NULL | 0.719099657 | NULL&M_Disease |
| 11 | 118555754 | rs525485 | A | G | NULL | 0.719099657 | NULL&M_Disease |
| 11 | 118556084 | rs613377 | T | C | NULL | 0.719099657 | NULL&M_Disease |
| 11 | 118556691 | rs610664 | G | A | NULL | 0.719099657 | NULL&M_Disease |
| 11 | 118556797 | rs571751 | C | A | NULL | 0.719099657 | NULL&M_Disease |
| 11 | 118557198 | rs546890 | C | T | NULL | 0.719099657 | NULL&M_Disease |
| 11 | 118558078 | rs555356 | T | G | NULL | 0.719099657 | NULL&M_Disease |
| 11 | 118558333 | rs693306 | G | A | NULL | 0.719099657 | NULL&M_Disease |
| 11 | 118558455 | rs558907 | G | A | NULL | 0.719099657 | NULL&M_Disease |
| 11 | 118558827 | rs561845 | G | A | NULL | 0.719099657 | NULL&M_Disease |
| 11 | 118558889 | rs638805 | T | G | NULL | 0.719099657 | NULL&M_Disease |
| 11 | 118558901 | rs510408 | C | G | NULL | 0.719099657 | NULL&M_Disease |
| 11 | 118558952 | rs509641 | C | T | NULL | 0.719099657 | NULL&M_Disease |
| 11 | 118558953 | rs563549 | A | G | NULL | 0.719099657 | NULL&M_Disease |
| 11 | 118559003 | rs638301 | G | A | NULL | 0.719099657 | NULL&M_Disease |
| 11 | 118559084 | rs508579 | C | G | NULL | 0.719099657 | NULL&M_Disease |
| 11 | 118559178 | rs486462 | C | T | NULL | 0.719099657 | NULL&M_Disease |
| 11 | 118559247 | rs533144 | A | C | NULL | 0.719099657 | NULL&M_Disease |
| 11 | 118559542 | rs625918 | T | C | NULL | 0.719099657 | NULL&M_Disease |
| 11 | 118559709 | rs574217358 | A | G | NULL | 0.719099657 | NULL&M_Disease |
| 11 | 118559866 | rs7926959 | C | T | NULL | 0.719099657 | NULL&M_Disease |
| 11 | 118559881 | rs7926970 | G | T | NULL | 0.719099657 | NULL&M_Disease |
| 11 | 118560569 | rs4938529 | T | C | NULL | 0.719099657 | NULL&M_Disease |
| 11 | 118562520 | rs558593 | T | C | NULL | 0.719099657 | NULL&M_Disease |
| 11 | 118562767 | rs667982 | C | T | NULL | 0.719099657 | NULL&M_Disease |
| 11 | 118562799 | rs582630 | A | C | NULL | 0.719099657 | NULL&M_Disease |
| 11 | 118563381 | rs585039 | A | C | NULL | 0.719099657 | NULL&M_Disease |
| 11 | 118564413 | rs508640 | G | A | NULL | 0.719099657 | NULL&M_Disease |
| 11 | 118564601 | rs471227 | C | T | NULL | 0.719099657 | NULL&M_Disease |
| 11 | 118565231 | rs519982 | T | C | NULL | 0.719099657 | NULL&M_Disease |
| 11 | 118565336 | rs478371 | A | G | NULL | 0.719099657 | NULL&M_Disease |
| 11 | 118565679 | rs502601 | A | G | NULL | 0.719099657 | NULL&M_Disease |
| 11 | 119429855 | rs916493 | T | G | NULL | 0.719099657 | NULL&M_Disease |
| 11 | 119434182 | rs2663628 | C | A | NULL | 0.719099657 | NULL&M_Disease |
| 11 | 119648938 | rs11217495 | G | A | NULL | 0.719099657 | NULL&M_Disease |
| 11 | 119885840 | rs632039 | C | T | NULL | 0.719099657 | NULL&M_Disease |
| 11 | 116833730 | rs10502222 | C | T | NULL | 0.719099657 | NULL&M_Disease |
| 11 | 117698762 | rs7117314 | A | G | NULL | 0.719099657 | NULL&M_Disease |
| 11 | 118539170 | rs4457789 | T | G | NULL | 0.719099657 | NULL&M_Disease |
| 11 | 118545797 | rs600969 | T | C | NULL | 0.719099657 | NULL&M_Disease |
| 11 | 118548421 | rs562313 | A | G | NULL | 0.719099657 | NULL&M_Disease |
| 11 | 118550601 | rs673770 | G | T | NULL | 0.719099657 | NULL&M_Disease |
| 11 | 118550756 | rs642530 | C | T | NULL | 0.719099657 | NULL&M_Disease |
| 11 | 118550784 | rs642497 | A | C | NULL | 0.719099657 | NULL&M_Disease |
| 11 | 118551597 | rs12799314 | A | G | NULL | 0.719099657 | NULL&M_Disease |
| 11 | 118552074 | rs4938524 | T | G | NULL | 0.719099657 | NULL&M_Disease |
| 11 | 118552420 | rs4936433 | G | A | NULL | 0.719099657 | NULL&M_Disease |
| 11 | 118552715 | rs519848 | A | G | NULL | 0.719099657 | NULL&M_Disease |
| 11 | 118552842 | rs549370 | A | C | NULL | 0.719099657 | NULL&M_Disease |
| 11 | 118552996 | rs654792 | T | G | NULL | 0.719099657 | NULL&M_Disease |
| 11 | 118553218 | rs592280 | C | T | NULL | 0.719099657 | NULL&M_Disease |
| 11 | 118554100 | rs4331144 | G | A | NULL | 0.719099657 | NULL&M_Disease |
| 11 | 118554248 | rs7106652 | T | C | NULL | 0.719099657 | NULL&M_Disease |
| 11 | 118554436 | rs11605069 | G | T | NULL | 0.719099657 | NULL&M_Disease |
| 11 | 118554448 | rs11602869 | A | G | NULL | 0.719099657 | NULL&M_Disease |
| 11 | 118554939 | rs12796373 | A | G | NULL | 0.719099657 | NULL&M_Disease |
| 11 | 118584498 | rs569770 | T | C | NULL | 0.719099657 | NULL&M_Disease |
| 11 | 119633930 | rs11217476 | A | C | NULL | 0.719099657 | NULL&M_Disease |
| 11 | 119959144 | rs12285338 | G | A | NULL | 0.719099657 | NULL&M_Disease |
| 11 | 119959214 | rs12273529 | T | C | NULL | 0.719099657 | NULL&M_Disease |
| 11 | 119959362 | rs7949126 | T | C | NULL | 0.719099657 | NULL&M_Disease |
| 11 | 120044762 | rs573211 | A | G | NULL | 0.719099657 | NULL&M_Disease |
| 11 | 120563298 | rs7925270 | T | C | NULL | 0.719099657 | NULL&M_Disease |
| 11 | 116814596 | rs10892048 | A | G | NULL | 0.719099657 | NULL&M_Disease |
| 11 | 118949331 | rs1177562 | T | C | NULL | 0.719099657 | NULL&M_Disease |
| 11 | 118952173 | rs15818 | G | A | NULL | 0.719099657 | NULL&M_Disease |
| 11 | 119428421 | rs996889 | G | A | NULL | 0.719099657 | NULL&M_Disease |
| 11 | 119883621 | rs601366 | C | T | NULL | 0.719099657 | NULL&M_Disease |
| 11 | 119977230 | rs10892537 | T | C | NULL | 0.719099657 | NULL&M_Disease |
| 11 | 120980753 | rs538741 | T | C | NULL | 0.719099657 | NULL&M_Disease |
| 11 | 116798544 | rs1446105 | C | T | NULL | 0.719099657 | NULL&M_Disease |
| 11 | 116829200 | rs6589582 | A | C | NULL | 0.719099657 | NULL&M_Disease |
| 11 | 116830087 | rs7120706 | T | C | NULL | 0.719099657 | NULL&M_Disease |
| 11 | 116830261 | rs7120963 | T | C | NULL | 0.719099657 | NULL&M_Disease |
| 11 | 116930682 | rs4938331 | C | A | NULL | 0.719099657 | NULL&M_Disease |
| 11 | 116930798 | rs4938332 | A | G | NULL | 0.719099657 | NULL&M_Disease |
| 11 | 116931042 | rs4938334 | G | C | NULL | 0.719099657 | NULL&M_Disease |
| 11 | 116936081 | rs6589590 | A | C | NULL | 0.719099657 | NULL&M_Disease |
| 11 | 116952392 | rs11216267 | C | T | NULL | 0.719099657 | NULL&M_Disease |
| 11 | 116960313 | rs10892072 | G | A | NULL | 0.719099657 | NULL&M_Disease |
| 11 | 117227519 | rs564674 | T | C | NULL | 0.719099657 | NULL&M_Disease |
| 11 | 117266648 | rs484988 | C | G | NULL | 0.719099657 | NULL&M_Disease |
| 11 | 117436906 | rs1893900 | T | C | NULL | 0.719099657 | NULL&M_Disease |
| 11 | 118718061 | rs4938559 | T | C | NULL | 0.719099657 | NULL&M_Disease |
| 11 | 118718062 | rs4938560 | G | A | NULL | 0.719099657 | NULL&M_Disease |
| 11 | 119018023 | rs1168815 | C | T | NULL | 0.719099657 | NULL&M_Disease |
| 11 | 119971337 | rs6589797 | C | T | NULL | 0.719099657 | NULL&M_Disease |
| 11 | 120556557 | rs7106140 | T | G | NULL | 0.719099657 | NULL&M_Disease |
| 11 | 120556641 | rs7106261 | A | G | NULL | 0.719099657 | NULL&M_Disease |
| 11 | 120603644 | rs3133853 | C | T | NULL | 0.719099657 | NULL&M_Disease |
| 11 | 120603857 | rs3132777 | A | G | NULL | 0.719099657 | NULL&M_Disease |
| 11 | 116752163 | rs482371 | T | C | NULL | 0.719099657 | NULL&M_Disease |
| 11 | 116796846 | rs7121898 | A | T | NULL | 0.719099657 | NULL&M_Disease |
| 11 | 116853213 | rs3863306 | C | T | NULL | 0.719099657 | NULL&M_Disease |
| 11 | 116857003 | rs10892053 | C | T | NULL | 0.719099657 | NULL&M_Disease |

| | | | | | | | | | |
|---|---|---|---|---|---|---|---|---|---|
| 11 | 118553326 | rs693037 | T | C | 0.999 | -7977.388 | -7979.271 | -1.882 | 5.89E-07 |
| 11 | 118555719 | rs625513 | C | A | 0.999 | -7977.388 | -7979.27 | -1.882 | 5.89E-07 |
| 11 | 118555754 | rs525485 | A | G | 0.999 | -7977.388 | -7979.27 | -1.882 | 5.89E-07 |
| 11 | 118556084 | rs613377 | T | C | 0.999 | -7977.388 | -7979.271 | -1.882 | 5.89E-07 |
| 11 | 118556691 | rs610664 | G | A | 0.999 | -7977.388 | -7979.271 | -1.882 | 5.89E-07 |
| 11 | 118556797 | rs571751 | C | A | 0.999 | -7977.388 | -7979.271 | -1.882 | 5.89E-07 |
| 11 | 118557198 | rs546890 | C | T | 0.999 | -7977.388 | -7979.271 | -1.882 | 5.89E-07 |
| 11 | 118558078 | rs555356 | T | G | 0.999 | -7977.388 | -7979.271 | -1.882 | 5.89E-07 |
| 11 | 118558333 | rs693306 | G | A | 0.999 | -7977.388 | -7979.27 | -1.882 | 5.89E-07 |
| 11 | 118558455 | rs558907 | G | A | 0.999 | -7977.388 | -7979.27 | -1.882 | 5.89E-07 |
| 11 | 118558827 | rs561845 | G | A | 0.999 | -7977.388 | -7979.27 | -1.882 | 5.89E-07 |
| 11 | 118558889 | rs638805 | T | G | 0.999 | -7977.388 | -7979.27 | -1.882 | 5.89E-07 |
| 11 | 118558901 | rs510408 | C | G | 0.999 | -7977.388 | -7979.27 | -1.882 | 5.89E-07 |
| 11 | 118558952 | rs509641 | C | T | 0.999 | -7977.388 | -7979.27 | -1.882 | 5.89E-07 |
| 11 | 118558953 | rs563549 | A | G | 0.999 | -7977.388 | -7979.27 | -1.882 | 5.89E-07 |
| 11 | 118559003 | rs638301 | G | A | 0.999 | -7977.388 | -7979.27 | -1.882 | 5.89E-07 |
| 11 | 118559084 | rs508579 | C | G | 0.999 | -7977.388 | -7979.271 | -1.882 | 5.89E-07 |
| 11 | 118559178 | rs486462 | C | T | 0.999 | -7977.388 | -7979.271 | -1.882 | 5.89E-07 |
| 11 | 118559247 | rs533144 | A | C | 0.999 | -7977.388 | -7979.271 | -1.882 | 5.89E-07 |
| 11 | 118559542 | rs625918 | T | C | 0.999 | -7977.388 | -7979.271 | -1.882 | 5.89E-07 |
| 11 | 118559709 | rs574217358 | A | G | 0.999 | -7977.388 | -7979.271 | -1.882 | 5.89E-07 |
| 11 | 118559866 | rs7926959 | C | T | 0.999 | -7977.388 | -7979.271 | -1.882 | 5.89E-07 |
| 11 | 118559881 | rs7926970 | G | T | 0.999 | -7977.388 | -7979.271 | -1.882 | 5.89E-07 |
| 11 | 118560569 | rs4938529 | T | C | 0.999 | -7977.388 | -7979.27 | -1.882 | 5.89E-07 |
| 11 | 118562520 | rs558593 | T | C | 0.999 | -7977.388 | -7979.27 | -1.882 | 5.89E-07 |
| 11 | 118562767 | rs667982 | C | T | 0.999 | -7977.388 | -7979.271 | -1.882 | 5.89E-07 |
| 11 | 118562799 | rs582630 | A | C | 0.999 | -7977.388 | -7979.27 | -1.882 | 5.89E-07 |
| 11 | 118563381 | rs585039 | A | C | 0.999 | -7977.388 | -7979.27 | -1.882 | 5.89E-07 |
| 11 | 118564413 | rs508640 | G | A | 0.999 | -7977.388 | -7979.27 | -1.882 | 5.89E-07 |
| 11 | 118564601 | rs471227 | C | T | 0.999 | -7977.388 | -7979.27 | -1.882 | 5.89E-07 |
| 11 | 118565231 | rs519982 | T | C | 0.998 | -7977.388 | -7979.27 | -1.882 | 5.89E-07 |
| 11 | 118565336 | rs478371 | A | G | 0.999 | -7977.388 | -7979.271 | -1.882 | 5.89E-07 |
| 11 | 118565679 | rs502601 | A | G | 0.999 | -7977.388 | -7979.271 | -1.882 | 5.89E-07 |
| 11 | 119429855 | rs916493 | T | G | 0.994 | -7977.388 | -7979.271 | -1.882 | 5.89E-07 |
| 11 | 119434182 | rs2663628 | C | A | 0.994 | -7977.388 | -7979.271 | -1.882 | 5.89E-07 |
| 11 | 119648938 | rs11217495 | G | A | 0.991 | -7977.388 | -7979.271 | -1.882 | 5.89E-07 |
| 11 | 119885840 | rs632039 | C | T | 0.992 | -7977.388 | -7979.27 | -1.882 | 5.89E-07 |
| 11 | 116833730 | rs10502222 | C | T | 0.997 | -7977.388 | -7979.271 | -1.883 | 5.88E-07 |
| 11 | 117698762 | rs7117314 | A | G | 0.997 | -7977.388 | -7979.271 | -1.883 | 5.88E-07 |
| 11 | 118539170 | rs4457789 | T | G | 1.002 | -7977.388 | -7979.271 | -1.883 | 5.88E-07 |
| 11 | 118545797 | rs600969 | T | C | 1 | -7977.388 | -7979.271 | -1.883 | 5.88E-07 |
| 11 | 118548421 | rs562313 | A | G | 1 | -7977.388 | -7979.271 | -1.883 | 5.88E-07 |
| 11 | 118550601 | rs673770 | G | T | 1 | -7977.388 | -7979.271 | -1.883 | 5.88E-07 |
| 11 | 118550756 | rs642530 | C | T | 1 | -7977.388 | -7979.271 | -1.883 | 5.88E-07 |
| 11 | 118550784 | rs642497 | A | C | 1 | -7977.388 | -7979.271 | -1.883 | 5.88E-07 |
| 11 | 118551597 | rs12799314 | A | G | 1 | -7977.388 | -7979.271 | -1.883 | 5.88E-07 |
| 11 | 118552074 | rs4938524 | T | G | 0.999 | -7977.388 | -7979.271 | -1.883 | 5.88E-07 |
| 11 | 118552420 | rs4936433 | G | A | 0.999 | -7977.388 | -7979.271 | -1.883 | 5.88E-07 |
| 11 | 118552715 | rs519848 | A | G | 0.999 | -7977.388 | -7979.271 | -1.883 | 5.88E-07 |
| 11 | 118552842 | rs549370 | A | C | 0.999 | -7977.388 | -7979.271 | -1.883 | 5.88E-07 |
| 11 | 118552996 | rs654792 | T | G | 0.999 | -7977.388 | -7979.271 | -1.883 | 5.88E-07 |
| 11 | 118553218 | rs592280 | C | T | 0.999 | -7977.388 | -7979.271 | -1.883 | 5.88E-07 |
| 11 | 118554100 | rs4331144 | G | A | 1 | -7977.388 | -7979.271 | -1.883 | 5.88E-07 |
| 11 | 118554248 | rs7106652 | T | C | 0.999 | -7977.388 | -7979.271 | -1.883 | 5.88E-07 |
| 11 | 118554436 | rs11605069 | G | T | 0.999 | -7977.388 | -7979.271 | -1.883 | 5.88E-07 |
| 11 | 118554448 | rs11602869 | A | G | 0.999 | -7977.388 | -7979.271 | -1.883 | 5.88E-07 |
| 11 | 118554939 | rs12796373 | A | G | 0.999 | -7977.388 | -7979.271 | -1.883 | 5.88E-07 |
| 11 | 118584498 | rs569770 | T | C | 0.996 | -7977.388 | -7979.271 | -1.883 | 5.88E-07 |
| 11 | 119633930 | rs11217476 | A | C | 1.005 | -7977.388 | -7979.271 | -1.883 | 5.88E-07 |
| 11 | 119959144 | rs12285338 | G | A | 1.01 | -7977.388 | -7979.272 | -1.883 | 5.88E-07 |
| 11 | 119959214 | rs12273529 | T | C | 1.01 | -7977.388 | -7979.272 | -1.883 | 5.88E-07 |
| 11 | 119959362 | rs7949126 | T | C | 1.01 | -7977.388 | -7979.272 | -1.883 | 5.88E-07 |
| 11 | 120044762 | rs573211 | A | G | 1.008 | -7977.388 | -7979.271 | -1.883 | 5.88E-07 |
| 11 | 120563298 | rs7925270 | T | C | 1.009 | -7977.388 | -7979.271 | -1.883 | 5.88E-07 |
| 11 | 116814596 | rs10892048 | A | G | 0.998 | -7977.388 | -7979.272 | -1.884 | 5.88E-07 |
| 11 | 118949331 | rs1177562 | T | C | 0.993 | -7977.388 | -7979.273 | -1.884 | 5.88E-07 |
| 11 | 118952173 | rs15818 | G | A | 1.007 | -7977.388 | -7979.272 | -1.884 | 5.88E-07 |
| 11 | 119428421 | rs996889 | G | A | 0.994 | -7977.388 | -7979.272 | -1.884 | 5.88E-07 |
| 11 | 119883621 | rs601366 | C | T | 0.994 | -7977.388 | -7979.272 | -1.884 | 5.88E-07 |
| 11 | 119977230 | rs10892537 | T | C | 0.998 | -7977.388 | -7979.272 | -1.884 | 5.88E-07 |
| 11 | 120980753 | rs538741 | T | C | 1.01 | -7977.388 | -7979.272 | -1.884 | 5.88E-07 |
| 11 | 116798544 | rs1446105 | C | T | 0.998 | -7977.388 | -7979.273 | -1.885 | 5.87E-07 |
| 11 | 116829200 | rs6589582 | A | C | 0.998 | -7977.388 | -7979.274 | -1.885 | 5.87E-07 |
| 11 | 116830087 | rs7120706 | T | C | 0.998 | -7977.388 | -7979.273 | -1.885 | 5.87E-07 |
| 11 | 116830261 | rs7120963 | T | C | 0.998 | -7977.388 | -7979.273 | -1.885 | 5.87E-07 |
| 11 | 116930682 | rs4938331 | C | A | 0.998 | -7977.388 | -7979.273 | -1.885 | 5.87E-07 |
| 11 | 116930798 | rs4938332 | A | G | 0.998 | -7977.388 | -7979.273 | -1.885 | 5.87E-07 |
| 11 | 116931042 | rs4938334 | G | C | 0.998 | -7977.388 | -7979.273 | -1.885 | 5.87E-07 |
| 11 | 116936081 | rs6589590 | A | C | 0.998 | -7977.388 | -7979.274 | -1.885 | 5.87E-07 |
| 11 | 116952392 | rs11216267 | C | T | 0.998 | -7977.388 | -7979.273 | -1.885 | 5.87E-07 |
| 11 | 116960313 | rs10892072 | G | A | 0.998 | -7977.388 | -7979.273 | -1.885 | 5.87E-07 |
| 11 | 117227519 | rs564674 | T | C | 0.992 | -7977.388 | -7979.273 | -1.885 | 5.87E-07 |
| 11 | 117266648 | rs484988 | C | G | 1.006 | -7977.388 | -7979.274 | -1.885 | 5.87E-07 |
| 11 | 117436906 | rs1893900 | T | C | 1.009 | -7977.388 | -7979.273 | -1.885 | 5.87E-07 |
| 11 | 118718061 | rs4938559 | T | C | 0.991 | -7977.388 | -7979.273 | -1.885 | 5.87E-07 |
| 11 | 118718062 | rs4938560 | G | A | 0.991 | -7977.388 | -7979.273 | -1.885 | 5.87E-07 |
| 11 | 119018023 | rs1168815 | C | T | 0.99 | -7977.388 | -7979.273 | -1.885 | 5.87E-07 |
| 11 | 119971337 | rs6589797 | C | T | 1 | -7977.388 | -7979.273 | -1.885 | 5.87E-07 |
| 11 | 120556557 | rs7106140 | T | G | 1.009 | -7977.388 | -7979.274 | -1.885 | 5.87E-07 |
| 11 | 120556641 | rs7106261 | A | G | 1.009 | -7977.388 | -7979.274 | -1.885 | 5.87E-07 |
| 11 | 120603644 | rs3133853 | C | T | 0.997 | -7977.388 | -7979.273 | -1.885 | 5.87E-07 |
| 11 | 120603857 | rs3132777 | A | G | 1.001 | -7977.388 | -7979.273 | -1.885 | 5.87E-07 |
| 11 | 116752163 | rs482371 | T | C | 0.997 | -7977.388 | -7979.274 | -1.886 | 5.87E-07 |
| 11 | 116796846 | rs7121898 | A | T | 0.998 | -7977.388 | -7979.274 | -1.886 | 5.87E-07 |
| 11 | 116853213 | rs3863306 | C | T | 0.999 | -7977.388 | -7979.275 | -1.886 | 5.87E-07 |
| 11 | 116857003 | rs10892053 | C | T | 0.999 | -7977.388 | -7979.274 | -1.886 | 5.87E-07 |

| | | | | | | | |
|---|---|---|---|---|---|---|---|
| 11 | 117022352 | rs4938351 | A | C | NULL | 0.719099657 | NULL&M_Disease |
| 11 | 117042238 | rs10750103 | C | T | NULL | 0.719099657 | NULL&M_Disease |
| 11 | 117171337 | rs548381 | T | C | NULL | 0.719099657 | NULL&M_Disease |
| 11 | 118017365 | rs1715443 | G | T | NULL | 0.719099657 | NULL&M_Disease |
| 11 | 119637061 | rs6589772 | A | C | NULL | 0.719099657 | NULL&M_Disease |
| 11 | 119648830 | rs11217494 | C | T | NULL | 0.719099657 | NULL&M_Disease |
| 11 | 119656448 | rs6589775 | C | T | NULL | 0.719099657 | NULL&M_Disease |
| 11 | 120058623 | rs12790943 | T | C | NULL | 0.719099657 | NULL&M_Disease |
| 11 | 120058640 | rs12791292 | C | G | NULL | 0.719099657 | NULL&M_Disease |
| 11 | 120551969 | rs11217971 | A | G | NULL | 0.719099657 | NULL&M_Disease |
| 11 | 120556610 | rs7105943 | A | C | NULL | 0.719099657 | NULL&M_Disease |
| 11 | 116304152 | rs12787144 | A | G | NULL | 0.719099657 | NULL&M_Disease |
| 11 | 116317895 | rs567610 | T | C | NULL | 0.719099657 | NULL&M_Disease |
| 11 | 116800943 | rs10790167 | T | C | NULL | 0.719099657 | NULL&M_Disease |
| 11 | 116853123 | rs3863304 | G | A | NULL | 0.719099657 | NULL&M_Disease |
| 11 | 116853124 | rs3863305 | C | T | NULL | 0.719099657 | NULL&M_Disease |
| 11 | 116853606 | rs7131351 | G | A | NULL | 0.719099657 | NULL&M_Disease |
| 11 | 116886114 | rs1022783694 | C | T | NULL | 0.719099657 | NULL&M_Disease |
| 11 | 116896155 | rs10892063 | A | C | NULL | 0.719099657 | NULL&M_Disease |
| 11 | 116896553 | rs10892064 | G | C | NULL | 0.719099657 | NULL&M_Disease |
| 11 | 116897024 | rs10892065 | T | C | NULL | 0.719099657 | NULL&M_Disease |
| 11 | 116899782 | rs7926104 | G | A | NULL | 0.720108527 | NULL&M_Disease |
| 11 | 116901549 | rs11216242 | G | A | NULL | 0.720108527 | NULL&M_Disease |
| 11 | 116902616 | rs12804122 | C | G | NULL | 0.719099657 | NULL&M_Disease |
| 11 | 116921219 | rs4938328 | G | T | NULL | 0.719099657 | NULL&M_Disease |
| 11 | 117088450 | rs236915 | G | A | NULL | 0.719099657 | NULL&M_Disease |
| 11 | 118017093 | rs2946742 | G | A | NULL | 0.719099657 | NULL&M_Disease |
| 11 | 118018767 | rs658624 | G | A | NULL | 0.719099657 | NULL&M_Disease |
| 11 | 118917814 | rs501351 | G | T | NULL | 0.719099657 | NULL&M_Disease |
| 11 | 119016176 | rs671872 | G | A | NULL | 0.720108527 | NULL&M_Disease |
| 11 | 119326726 | rs6589750 | G | A | NULL | 0.719099657 | NULL&M_Disease |
| 11 | 119434753 | rs2845658 | A | G | NULL | 0.719099657 | NULL&M_Disease |
| 11 | 119885914 | rs638000 | T | C | NULL | 0.719099657 | NULL&M_Disease |
| 11 | 120652208 | rs2846103 | C | T | NULL | 0.719099657 | NULL&M_Disease |
| 11 | 116876621 | rs10892058 | C | T | NULL | 0.720108527 | NULL&M_Disease |
| 11 | 116881662 | rs10892060 | C | A | NULL | 0.720108527 | NULL&M_Disease |
| 11 | 116904357 | rs4575311 | G | A | NULL | 0.720108527 | NULL&M_Disease |
| 11 | 117688898 | rs540442 | G | A | NULL | 0.720108527 | NULL&M_Disease |
| 11 | 117689549 | rs527534 | T | G | NULL | 0.720108527 | NULL&M_Disease |
| 11 | 119231667 | rs11217255 | G | A | NULL | 0.720108527 | NULL&M_Disease |
| 11 | 116755041 | rs1269329 | T | C | NULL | 0.720108527 | NULL&M_Disease |
| 11 | 116772787 | rs518181 | C | A | NULL | 0.720108527 | NULL&M_Disease |
| 11 | 116878597 | rs11216227 | C | G | NULL | 0.720108527 | NULL&M_Disease |
| 11 | 116880798 | rs7127881 | G | A | NULL | 0.720108527 | NULL&M_Disease |
| 11 | 117699786 | rs10892182 | A | G | NULL | 0.720108527 | NULL&M_Disease |
| 11 | 119430659 | rs2845651 | G | A | NULL | 0.720108527 | NULL&M_Disease |
| 11 | 119630480 | rs10790343 | A | G | NULL | 0.720108527 | NULL&M_Disease |
| 11 | 119706311 | rs7118350 | G | C | NULL | 0.720108527 | NULL&M_Disease |
| 11 | 120649396 | rs949296 | A | T | NULL | 0.720108527 | NULL&M_Disease |
| 11 | 116881645 | rs10892059 | G | A | NULL | 0.720108527 | NULL&M_Disease |
| 11 | 117336108 | rs482287 | T | C | NULL | 0.720108527 | NULL&M_Disease |
| 11 | 117973454 | rs4996910 | A | G | NULL | 0.720108527 | NULL&M_Disease |
| 11 | 119649488 | rs10892458 | T | C | NULL | 0.720108527 | NULL&M_Disease |
| 11 | 119994421 | rs470392 | T | C | NULL | 0.720108527 | NULL&M_Disease |
| 11 | 116207032 | rs11215900 | T | C | NULL | 0.720108527 | NULL&M_Disease |
| 11 | 116713566 | rs591592 | C | T | NULL | 0.720108527 | NULL&M_Disease |
| 11 | 116713630 | rs579163 | C | G | NULL | 0.720108527 | NULL&M_Disease |
| 11 | 116720829 | rs660083 | C | G | NULL | 0.720108527 | NULL&M_Disease |
| 11 | 116732856 | rs484646 | A | G | NULL | 0.720108527 | NULL&M_Disease |
| 11 | 116760693 | rs660443 | T | C | NULL | 0.720108527 | NULL&M_Disease |
| 11 | 117015195 | rs2735183 | C | T | NULL | 0.720108527 | NULL&M_Disease |
| 11 | 117090558 | rs74830 | T | C | NULL | 0.720108527 | NULL&M_Disease |
| 11 | 117729618 | rs7121196 | A | G | NULL | 0.720108527 | NULL&M_Disease |
| 11 | 117973338 | rs35392431 | G | C | NULL | 0.720108527 | NULL&M_Disease |
| 11 | 117973439 | rs4996911 | A | G | NULL | 0.720108527 | NULL&M_Disease |
| 11 | 118008212 | rs2004235 | C | T | NULL | 0.720108527 | NULL&M_Disease |
| 11 | 118801640 | rs572608 | A | T | NULL | 0.720108527 | NULL&M_Disease |
| 11 | 118953202 | rs7127212 | T | C | NULL | 0.720108527 | NULL&M_Disease |
| 11 | 118954066 | rs7938944 | C | T | NULL | 0.720108527 | NULL&M_Disease |
| 11 | 118954194 | rs7952700 | C | G | NULL | 0.720108527 | NULL&M_Disease |
| 11 | 118965338 | rs1064193 | A | T | NULL | 0.720108527 | NULL&M_Disease |
| 11 | 119019520 | rs4301800 | A | C | NULL | 0.720108527 | NULL&M_Disease |
| 11 | 119564842 | rs7110874 | C | T | NULL | 0.720108527 | NULL&M_Disease |
| 11 | 119834038 | rs10790357 | C | G | NULL | 0.720108527 | NULL&M_Disease |
| 11 | 120552067 | rs10790399 | A | C | NULL | 0.720108527 | NULL&M_Disease |
| 11 | 116330989 | rs487611 | C | G | NULL | 0.720108527 | NULL&M_Disease |
| 11 | 117311608 | rs7947434 | G | C | NULL | 0.720108527 | NULL&M_Disease |
| 11 | 118577990 | rs480958 | G | A | NULL | 0.720108527 | NULL&M_Disease |
| 11 | 118802492 | rs688035 | G | C | NULL | 0.720108527 | NULL&M_Disease |
| 11 | 118928249 | rs11217133 | A | G | NULL | 0.720108527 | NULL&M_Disease |
| 11 | 119014474 | rs612132 | T | A | NULL | 0.720108527 | NULL&M_Disease |
| 11 | 119141812 | rs2511857 | A | T | NULL | 0.720108527 | NULL&M_Disease |
| 11 | 119630385 | rs10892452 | T | C | NULL | 0.720108527 | NULL&M_Disease |
| 11 | 120541263 | rs12365194 | A | G | NULL | 0.720108527 | NULL&M_Disease |
| 11 | 116556865 | rs1145212 | A | G | NULL | 0.720108527 | NULL&M_Disease |
| 11 | 118931030 | rs11217137 | G | A | NULL | 0.720108527 | NULL&M_Disease |
| 11 | 118993899 | rs28489214 | G | A | NULL | 0.720108527 | NULL&M_Disease |
| 11 | 119649342 | rs10892456 | G | A | NULL | 0.720108527 | NULL&M_Disease |
| 11 | 120001695 | rs670855 | T | C | NULL | 0.720108527 | NULL&M_Disease |
| 11 | 116315660 | rs520326 | G | T | NULL | 0.720108527 | NULL&M_Disease |
| 11 | 116722693 | rs689243 | G | C | NULL | 0.720108527 | NULL&M_Disease |
| 11 | 116733728 | rs476399 | T | C | NULL | 0.720108527 | NULL&M_Disease |
| 11 | 116741572 | rs533556 | A | C | NULL | 0.720108527 | NULL&M_Disease |
| 11 | 117989554 | rs3741311 | G | A | NULL | 0.720108527 | NULL&M_Disease |
| 11 | 118717732 | rs11217052 | G | A | NULL | 0.720108527 | NULL&M_Disease |
| 11 | 118947154 | rs1168570 | C | T | NULL | 0.720108527 | NULL&M_Disease |

| | | | | | | | | | |
|---|---|---|---|---|---|---|---|---|---|
| 11 | 117022352 | rs4938351 | A | C | 0.994 | -7977.388 | -7979.274 | -1.886 | 5.87E-07 |
| 11 | 117042238 | rs10750103 | C | T | 0.994 | -7977.388 | -7979.274 | -1.886 | 5.87E-07 |
| 11 | 117171337 | rs548381 | T | C | 1.007 | -7977.388 | -7979.274 | -1.886 | 5.87E-07 |
| 11 | 118017365 | rs1715443 | G | T | 0.992 | -7977.388 | -7979.274 | -1.886 | 5.87E-07 |
| 11 | 119637061 | rs6589772 | A | C | 0.991 | -7977.388 | -7979.275 | -1.886 | 5.87E-07 |
| 11 | 119648830 | rs11217494 | C | T | 0.991 | -7977.388 | -7979.274 | -1.886 | 5.87E-07 |
| 11 | 119656448 | rs6589775 | C | T | 0.991 | -7977.388 | -7979.274 | -1.886 | 5.87E-07 |
| 11 | 120058623 | rs12790943 | T | C | 1.008 | -7977.388 | -7979.275 | -1.886 | 5.87E-07 |
| 11 | 120058640 | rs12791292 | C | G | 1.008 | -7977.388 | -7979.275 | -1.886 | 5.87E-07 |
| 11 | 120551969 | rs11217971 | A | G | 1.01 | -7977.388 | -7979.274 | -1.886 | 5.87E-07 |
| 11 | 120556610 | rs7105943 | A | C | 1.009 | -7977.388 | -7979.275 | -1.886 | 5.87E-07 |
| 11 | 116304152 | rs12787144 | A | G | 0.992 | -7977.388 | -7979.275 | -1.887 | 5.86E-07 |
| 11 | 116317895 | rs567610 | T | C | 1.009 | -7977.388 | -7979.276 | -1.887 | 5.86E-07 |
| 11 | 116800943 | rs10790167 | T | C | 1 | -7977.388 | -7979.276 | -1.887 | 5.86E-07 |
| 11 | 116853123 | rs3863304 | G | A | 1.001 | -7977.388 | -7979.275 | -1.887 | 5.86E-07 |
| 11 | 116853124 | rs3863305 | C | T | 1.001 | -7977.388 | -7979.275 | -1.887 | 5.86E-07 |
| 11 | 116853606 | rs7131351 | G | A | 1 | -7977.388 | -7979.275 | -1.887 | 5.86E-07 |
| 11 | 116886114 | rs1022783694 | C | T | 0.999 | -7977.388 | -7979.275 | -1.887 | 5.86E-07 |
| 11 | 116896155 | rs10892063 | A | C | 0.999 | -7977.388 | -7979.275 | -1.887 | 5.86E-07 |
| 11 | 116896553 | rs10892064 | G | C | 0.999 | -7977.388 | -7979.275 | -1.887 | 5.86E-07 |
| 11 | 116897024 | rs10892065 | T | C | 0.999 | -7977.388 | -7979.275 | -1.887 | 5.86E-07 |
| 11 | 116899782 | rs7926104 | G | A | 1 | -7977.388 | -7979.276 | -1.887 | 5.86E-07 |
| 11 | 116901549 | rs11216242 | G | A | 1 | -7977.388 | -7979.276 | -1.887 | 5.86E-07 |
| 11 | 116902616 | rs12804122 | C | G | 1 | -7977.388 | -7979.276 | -1.887 | 5.86E-07 |
| 11 | 116921219 | rs4938328 | G | T | 0.999 | -7977.388 | -7979.275 | -1.887 | 5.86E-07 |
| 11 | 117088450 | rs236915 | G | A | 0.994 | -7977.388 | -7979.276 | -1.887 | 5.86E-07 |
| 11 | 118017093 | rs2946742 | G | A | 0.992 | -7977.388 | -7979.275 | -1.887 | 5.86E-07 |
| 11 | 118018767 | rs658624 | G | A | 0.992 | -7977.388 | -7979.275 | -1.887 | 5.86E-07 |
| 11 | 118917814 | rs501351 | G | T | 0.994 | -7977.388 | -7979.276 | -1.887 | 5.86E-07 |
| 11 | 119016176 | rs671872 | G | A | 0.99 | -7977.388 | -7979.276 | -1.887 | 5.86E-07 |
| 11 | 119326726 | rs6589750 | G | A | 1.008 | -7977.388 | -7979.275 | -1.887 | 5.86E-07 |
| 11 | 119434753 | rs2845658 | A | G | 0.995 | -7977.388 | -7979.276 | -1.887 | 5.86E-07 |
| 11 | 119885914 | rs638000 | T | C | 0.992 | -7977.388 | -7979.275 | -1.887 | 5.86E-07 |
| 11 | 120652208 | rs2846103 | C | T | 0.998 | -7977.388 | -7979.275 | -1.887 | 5.86E-07 |
| 11 | 116876621 | rs10892058 | C | T | 0.996 | -7977.388 | -7979.277 | -1.888 | 5.86E-07 |
| 11 | 116881662 | rs10892060 | C | A | 0.995 | -7977.388 | -7979.276 | -1.888 | 5.86E-07 |
| 11 | 116904357 | rs4575311 | G | A | 1 | -7977.388 | -7979.276 | -1.888 | 5.86E-07 |
| 11 | 117688898 | rs540442 | G | A | 0.999 | -7977.388 | -7979.276 | -1.888 | 5.86E-07 |
| 11 | 117689549 | rs527534 | T | G | 1.003 | -7977.388 | -7979.276 | -1.888 | 5.86E-07 |
| 11 | 119231667 | rs11217255 | G | A | 1.001 | -7977.388 | -7979.276 | -1.888 | 5.86E-07 |
| 11 | 116755041 | rs1269329 | T | C | 0.997 | -7977.388 | -7979.277 | -1.889 | 5.85E-07 |
| 11 | 116772787 | rs518181 | C | A | 0.997 | -7977.388 | -7979.277 | -1.889 | 5.85E-07 |
| 11 | 116878597 | rs11216227 | C | G | 0.996 | -7977.388 | -7979.277 | -1.889 | 5.85E-07 |
| 11 | 116880798 | rs7127881 | G | A | 0.996 | -7977.388 | -7979.277 | -1.889 | 5.85E-07 |
| 11 | 117699786 | rs10892182 | A | G | 0.999 | -7977.388 | -7979.278 | -1.889 | 5.85E-07 |
| 11 | 119430659 | rs2845651 | G | A | 0.995 | -7977.388 | -7979.277 | -1.889 | 5.85E-07 |
| 11 | 119630480 | rs10790343 | A | G | 0.991 | -7977.388 | -7979.277 | -1.889 | 5.85E-07 |
| 11 | 119706311 | rs7118350 | G | C | 1.009 | -7977.388 | -7979.277 | -1.889 | 5.85E-07 |
| 11 | 120649396 | rs949296 | A | T | 0.999 | -7977.388 | -7979.278 | -1.889 | 5.85E-07 |
| 11 | 116881645 | rs10892059 | G | A | 0.996 | -7977.388 | -7979.279 | -1.89 | 5.84E-07 |
| 11 | 117336108 | rs482287 | T | C | 0.992 | -7977.388 | -7979.278 | -1.89 | 5.84E-07 |
| 11 | 117973454 | rs4996910 | A | G | 0.997 | -7977.388 | -7979.279 | -1.89 | 5.84E-07 |
| 11 | 119649488 | rs10892458 | T | C | 0.991 | -7977.388 | -7979.279 | -1.89 | 5.84E-07 |
| 11 | 119994421 | rs470392 | T | C | 1.006 | -7977.388 | -7979.279 | -1.89 | 5.84E-07 |
| 11 | 116207032 | rs11215900 | T | C | 1.007 | -7977.388 | -7979.28 | -1.891 | 5.84E-07 |
| 11 | 116713566 | rs591592 | C | T | 1.002 | -7977.388 | -7979.279 | -1.891 | 5.84E-07 |
| 11 | 116713630 | rs579163 | C | G | 1.002 | -7977.388 | -7979.28 | -1.891 | 5.84E-07 |
| 11 | 116720829 | rs660083 | C | G | 1.001 | -7977.388 | -7979.28 | -1.891 | 5.84E-07 |
| 11 | 116732856 | rs484646 | A | G | 0.997 | -7977.388 | -7979.279 | -1.891 | 5.84E-07 |
| 11 | 116760693 | rs660443 | T | C | 0.998 | -7977.388 | -7979.279 | -1.891 | 5.84E-07 |
| 11 | 117015195 | rs2735183 | C | T | 0.995 | -7977.388 | -7979.28 | -1.891 | 5.84E-07 |
| 11 | 117090558 | rs74830 | T | C | 0.995 | -7977.388 | -7979.28 | -1.891 | 5.84E-07 |
| 11 | 117729618 | rs7121196 | A | G | 1.005 | -7977.388 | -7979.279 | -1.891 | 5.84E-07 |
| 11 | 117973338 | rs35392431 | G | C | 0.998 | -7977.388 | -7979.279 | -1.891 | 5.84E-07 |
| 11 | 117973439 | rs4996911 | A | G | 0.997 | -7977.388 | -7979.279 | -1.891 | 5.84E-07 |
| 11 | 118008212 | rs2004235 | C | T | 1.007 | -7977.388 | -7979.279 | -1.891 | 5.84E-07 |
| 11 | 118801640 | rs572608 | A | T | 0.992 | -7977.388 | -7979.28 | -1.891 | 5.84E-07 |
| 11 | 118953202 | rs7127212 | T | C | 1.006 | -7977.388 | -7979.279 | -1.891 | 5.84E-07 |
| 11 | 118954066 | rs7938944 | C | T | 1.006 | -7977.388 | -7979.279 | -1.891 | 5.84E-07 |
| 11 | 118954194 | rs7952700 | C | G | 1.006 | -7977.388 | -7979.279 | -1.891 | 5.84E-07 |
| 11 | 118965338 | rs1064193 | A | T | 0.999 | -7977.388 | -7979.28 | -1.891 | 5.84E-07 |
| 11 | 119019520 | rs4301800 | A | C | 0.99 | -7977.388 | -7979.279 | -1.891 | 5.84E-07 |
| 11 | 119564842 | rs7110874 | C | T | 0.994 | -7977.388 | -7979.28 | -1.891 | 5.84E-07 |
| 11 | 119834038 | rs10790357 | C | G | 1.008 | -7977.388 | -7979.279 | -1.891 | 5.84E-07 |
| 11 | 120552067 | rs10790399 | A | C | 1.009 | -7977.388 | -7979.279 | -1.891 | 5.84E-07 |
| 11 | 116330989 | rs487611 | C | G | 0.993 | -7977.388 | -7979.281 | -1.892 | 5.83E-07 |
| 11 | 117311608 | rs7947434 | G | C | 1.007 | -7977.388 | -7979.281 | -1.892 | 5.83E-07 |
| 11 | 118577990 | rs480958 | G | A | 1 | -7977.388 | -7979.28 | -1.892 | 5.83E-07 |
| 11 | 118802492 | rs688035 | G | C | 1.005 | -7977.388 | -7979.28 | -1.892 | 5.83E-07 |
| 11 | 118928249 | rs11217133 | A | G | 0.995 | -7977.388 | -7979.281 | -1.892 | 5.83E-07 |
| 11 | 119014474 | rs612132 | T | A | 0.991 | -7977.388 | -7979.28 | -1.892 | 5.83E-07 |
| 11 | 119141812 | rs2511857 | A | T | 1.008 | -7977.388 | -7979.281 | -1.892 | 5.83E-07 |
| 11 | 119630385 | rs10892452 | T | C | 0.991 | -7977.388 | -7979.28 | -1.892 | 5.83E-07 |
| 11 | 120541263 | rs12365194 | A | G | 1.009 | -7977.388 | -7979.28 | -1.892 | 5.83E-07 |
| 11 | 116556865 | rs1145212 | A | G | 1.002 | -7977.388 | -7979.282 | -1.893 | 5.83E-07 |
| 11 | 118931030 | rs11217137 | G | A | 0.995 | -7977.388 | -7979.282 | -1.893 | 5.83E-07 |
| 11 | 118993899 | rs28489214 | G | A | 0.991 | -7977.388 | -7979.281 | -1.893 | 5.83E-07 |
| 11 | 119649342 | rs10892456 | G | A | 0.992 | -7977.388 | -7979.282 | -1.893 | 5.83E-07 |
| 11 | 120001695 | rs670855 | T | C | 0.992 | -7977.388 | -7979.281 | -1.893 | 5.83E-07 |
| 11 | 116315660 | rs520326 | G | T | 0.994 | -7977.388 | -7979.282 | -1.894 | 5.82E-07 |
| 11 | 116722693 | rs689243 | G | C | 1.001 | -7977.388 | -7979.282 | -1.894 | 5.82E-07 |
| 11 | 116733728 | rs476399 | T | C | 0.998 | -7977.388 | -7979.282 | -1.894 | 5.82E-07 |
| 11 | 116741572 | rs533556 | A | C | 0.998 | -7977.388 | -7979.282 | -1.894 | 5.82E-07 |
| 11 | 117989554 | rs3741311 | G | A | 1.007 | -7977.388 | -7979.283 | -1.894 | 5.82E-07 |
| 11 | 118717732 | rs11217052 | G | A | 0.992 | -7977.388 | -7979.282 | -1.894 | 5.82E-07 |
| 11 | 118947154 | rs1168570 | C | T | 0.994 | -7977.388 | -7979.282 | -1.894 | 5.82E-07 |

| | | | | | | | |
|---|---|---|---|---|---|---|---|
| 11 | 118947168 | rs1168569 | G | A | NULL | 0.720108527 | NULL&M_Disease |
| 11 | 118958869 | rs1006195 | T | G | NULL | 0.720108527 | NULL&M_Disease |
| 11 | 119015932 | rs595412 | T | C | NULL | 0.720108527 | NULL&M_Disease |
| 11 | 116736679 | rs573549 | A | G | NULL | 0.720108527 | NULL&M_Disease |
| 11 | 117776747 | rs11601073 | T | C | NULL | 0.720108527 | NULL&M_Disease |
| 11 | 117990403 | rs7938794 | T | G | NULL | 0.720108527 | NULL&M_Disease |
| 11 | 117991403 | rs7928092 | G | C | NULL | 0.720108527 | NULL&M_Disease |
| 11 | 118012441 | rs11604096 | C | A | NULL | 0.720108527 | NULL&M_Disease |
| 11 | 118017088 | rs2946743 | G | A | NULL | 0.720108527 | NULL&M_Disease |
| 11 | 118800302 | rs12793214 | A | C | NULL | 0.720108527 | NULL&M_Disease |
| 11 | 119291480 | rs3138091 | G | A | NULL | 0.720108527 | NULL&M_Disease |
| 11 | 119448579 | rs4609614 | A | C | NULL | 0.720108527 | NULL&M_Disease |
| 11 | 119649969 | rs4608095 | A | G | NULL | 0.720108527 | NULL&M_Disease |
| 11 | 119650060 | rs12222723 | T | G | NULL | 0.720108527 | NULL&M_Disease |
| 11 | 119951111 | rs12276427 | A | G | NULL | 0.720108527 | NULL&M_Disease |
| 11 | 119971490 | rs6589798 | T | G | NULL | 0.720108527 | NULL&M_Disease |
| 11 | 116726769 | rs640621 | C | G | NULL | 0.720108527 | NULL&M_Disease |
| 11 | 117777599 | rs11216613 | G | T | NULL | 0.720108527 | NULL&M_Disease |
| 11 | 117992620 | rs4938485 | G | A | NULL | 0.720108527 | NULL&M_Disease |
| 11 | 118006957 | rs1814964 | C | T | NULL | 0.720108527 | NULL&M_Disease |
| 11 | 118989098 | rs1129013 | C | G | NULL | 0.720108527 | NULL&M_Disease |
| 11 | 119292712 | rs1001205 | T | C | NULL | 0.720108527 | NULL&M_Disease |
| 11 | 119649359 | rs10892457 | T | C | NULL | 0.720108527 | NULL&M_Disease |
| 11 | 119821850 | rs7931873 | G | A | NULL | 0.720108527 | NULL&M_Disease |
| 11 | 120013984 | rs470370 | C | T | NULL | 0.720108527 | NULL&M_Disease |
| 11 | 116521187 | rs534421 | G | A | NULL | 0.720108527 | NULL&M_Disease |
| 11 | 117014862 | rs2622934 | G | T | NULL | 0.720108527 | NULL&M_Disease |
| 11 | 118009279 | rs10790240 | G | T | NULL | 0.720108527 | NULL&M_Disease |
| 11 | 118017445 | rs1786183 | A | G | NULL | 0.720108527 | NULL&M_Disease |
| 11 | 119285008 | rs3138097 | C | T | NULL | 0.720108527 | NULL&M_Disease |
| 11 | 119290034 | rs1054735 | G | C | NULL | 0.721115178 | NULL&M_Disease |
| 11 | 119834547 | rs10750167 | G | A | NULL | 0.720108527 | NULL&M_Disease |
| 11 | 117776850 | rs12798929 | T | C | NULL | 0.721115178 | NULL&M_Disease |
| 11 | 118011776 | rs668468 | G | A | NULL | 0.721115178 | NULL&M_Disease |
| 11 | 118991019 | rs675866 | C | A | NULL | 0.721115178 | NULL&M_Disease |
| 11 | 119305126 | rs10790294 | T | C | NULL | 0.721115178 | NULL&M_Disease |
| 11 | 119309945 | rs4450179 | C | T | NULL | 0.721115178 | NULL&M_Disease |
| 11 | 119971706 | rs6589800 | C | A | NULL | 0.721115178 | NULL&M_Disease |
| 11 | 117702115 | rs7130303 | A | C | NULL | 0.721115178 | NULL&M_Disease |
| 11 | 117731136 | rs5014624 | A | G | NULL | 0.721115178 | NULL&M_Disease |
| 11 | 118076004 | rs1806 | C | T | NULL | 0.721115178 | NULL&M_Disease |
| 11 | 118990253 | rs1305565 | C | T | NULL | 0.721115178 | NULL&M_Disease |
| 11 | 119023107 | rs668033 | C | A | NULL | 0.721115178 | NULL&M_Disease |
| 11 | 119448580 | rs202125027 | A | T | NULL | 0.721115178 | NULL&M_Disease |
| 11 | 119972448 | rs1073636 | A | T | NULL | 0.721115178 | NULL&M_Disease |
| 11 | 119972642 | rs1073634 | T | C | NULL | 0.721115178 | NULL&M_Disease |
| 11 | 120551561 | rs10790396 | A | G | NULL | 0.721115178 | NULL&M_Disease |
| 11 | 121153421 | rs7949987 | G | A | NULL | 0.721115178 | NULL&M_Disease |
| 11 | 116317405 | rs556550 | G | A | NULL | 0.721115178 | NULL&M_Disease |
| 11 | 116465680 | rs11216029 | T | G | NULL | 0.721115178 | NULL&M_Disease |
| 11 | 117270818 | rs1243337 | C | A | NULL | 0.721115178 | NULL&M_Disease |
| 11 | 117992645 | rs4938486 | G | A | NULL | 0.721115178 | NULL&M_Disease |
| 11 | 118016981 | rs3018387 | T | A | NULL | 0.721115178 | NULL&M_Disease |
| 11 | 118018022 | rs628299 | G | A | NULL | 0.721115178 | NULL&M_Disease |
| 11 | 119414914 | rs7945998 | T | C | NULL | 0.721115178 | NULL&M_Disease |
| 11 | 119649844 | rs4258392 | C | T | NULL | 0.721115178 | NULL&M_Disease |
| 11 | 119649874 | rs5020633 | A | G | NULL | 0.721115178 | NULL&M_Disease |
| 11 | 119754895 | rs1358104428 | A | G | NULL | 0.721115178 | NULL&M_Disease |
| 11 | 119971987 | rs6589801 | T | C | NULL | 0.721115178 | NULL&M_Disease |
| 11 | 120976428 | rs504626 | A | G | NULL | 0.721115178 | NULL&M_Disease |
| 11 | 116465734 | rs11216030 | T | C | NULL | 0.721115178 | NULL&M_Disease |
| 11 | 118807735 | rs10892311 | T | C | NULL | 0.721115178 | NULL&M_Disease |
| 11 | 118942654 | rs10892328 | A | T | NULL | 0.721115178 | NULL&M_Disease |
| 11 | 118944675 | rs3825061 | T | C | NULL | 0.721115178 | NULL&M_Disease |
| 11 | 119649863 | rs5020632 | T | C | NULL | 0.721115178 | NULL&M_Disease |
| 11 | 119727494 | rs7113123 | T | C | NULL | 0.721115178 | NULL&M_Disease |
| 11 | 119846820 | rs636518 | G | A | NULL | 0.721115178 | NULL&M_Disease |
| 11 | 119964351 | rs664590 | T | G | NULL | 0.721115178 | NULL&M_Disease |
| 11 | 119967556 | rs613004 | T | C | NULL | 0.721115178 | NULL&M_Disease |
| 11 | 119967661 | rs1787027 | C | A | NULL | 0.721115178 | NULL&M_Disease |
| 11 | 119967716 | rs10892534 | A | C | NULL | 0.721115178 | NULL&M_Disease |
| 11 | 119967718 | rs10892535 | T | C | NULL | 0.721115178 | NULL&M_Disease |
| 11 | 121154437 | rs1982011 | G | A | NULL | 0.721115178 | NULL&M_Disease |
| 11 | 117315692 | rs11216394 | G | A | NULL | 0.721115178 | NULL&M_Disease |
| 11 | 118017434 | rs1793157 | C | A | NULL | 0.721115178 | NULL&M_Disease |
| 11 | 118735476 | rs10892301 | A | G | NULL | 0.721115178 | NULL&M_Disease |
| 11 | 118796570 | rs11217096 | T | C | NULL | 0.721115178 | NULL&M_Disease |
| 11 | 118941596 | rs1260808616 | A | G | NULL | 0.721115178 | NULL&M_Disease |
| 11 | 117730979 | rs6589627 | C | T | NULL | 0.721115178 | NULL&M_Disease |
| 11 | 117730983 | rs6589628 | A | G | NULL | 0.721115178 | NULL&M_Disease |
| 11 | 117731070 | rs6589629 | C | G | NULL | 0.721115178 | NULL&M_Disease |
| 11 | 117731085 | rs6589630 | C | T | NULL | 0.721115178 | NULL&M_Disease |
| 11 | 117731191 | rs5014623 | C | T | NULL | 0.721115178 | NULL&M_Disease |
| 11 | 117991333 | rs7927810 | A | G | NULL | 0.721115178 | NULL&M_Disease |
| 11 | 119306276 | rs4938655 | A | C | NULL | 0.721115178 | NULL&M_Disease |
| 11 | 119788216 | rs643153 | C | T | NULL | 0.721115178 | NULL&M_Disease |
| 11 | 120007889 | rs470645 | T | G | NULL | 0.721115178 | NULL&M_Disease |
| 11 | 116584317 | rs1144035 | C | T | NULL | 0.721115178 | NULL&M_Disease |
| 11 | 117731263 | rs5014622 | G | A | NULL | 0.721115178 | NULL&M_Disease |
| 11 | 119306183 | rs4938654 | T | C | NULL | 0.721115178 | NULL&M_Disease |
| 11 | 119748404 | rs10892483 | C | G | NULL | 0.721115178 | NULL&M_Disease |
| 11 | 119843707 | rs10790359 | G | A | NULL | 0.721115178 | NULL&M_Disease |
| 11 | 119843736 | rs10790360 | C | A | NULL | 0.721115178 | NULL&M_Disease |
| 11 | 119967769 | rs10790366 | T | G | NULL | 0.721115178 | NULL&M_Disease |
| 11 | 121158779 | rs2115923 | A | G | NULL | 0.721115178 | NULL&M_Disease |
| 11 | 117260075 | rs552498 | G | A | NULL | 0.721115178 | NULL&M_Disease |

| | | | | | | | | | |
|---|---|---|---|---|---|---|---|---|---|
| 11 | 118947168 | rs1168569 | G | A | 0.994 | -7977.388 | -7979.282 | -1.894 | 5.82E-07 |
| 11 | 118958869 | rs1006195 | T | G | 1.006 | -7977.388 | -7979.282 | -1.894 | 5.82E-07 |
| 11 | 119015932 | rs595412 | T | C | 0.991 | -7977.388 | -7979.282 | -1.894 | 5.82E-07 |
| 11 | 116736679 | rs573549 | A | G | 1.002 | -7977.388 | -7979.283 | -1.895 | 5.81E-07 |
| 11 | 117776747 | rs11601073 | T | C | 0.992 | -7977.388 | -7979.283 | -1.895 | 5.81E-07 |
| 11 | 117990403 | rs7938794 | T | G | 1.007 | -7977.388 | -7979.283 | -1.895 | 5.81E-07 |
| 11 | 117991403 | rs7928092 | G | C | 1.007 | -7977.388 | -7979.283 | -1.895 | 5.81E-07 |
| 11 | 118012441 | rs11604096 | C | A | 0.995 | -7977.388 | -7979.283 | -1.895 | 5.81E-07 |
| 11 | 118017088 | rs2946743 | G | A | 0.993 | -7977.388 | -7979.283 | -1.895 | 5.81E-07 |
| 11 | 118800302 | rs12793214 | A | C | 0.992 | -7977.388 | -7979.284 | -1.895 | 5.81E-07 |
| 11 | 119291480 | rs3138091 | G | A | 1.007 | -7977.388 | -7979.283 | -1.895 | 5.81E-07 |
| 11 | 119448579 | rs4609614 | A | C | 1.006 | -7977.388 | -7979.283 | -1.895 | 5.81E-07 |
| 11 | 119649969 | rs4608095 | A | G | 0.992 | -7977.388 | -7979.283 | -1.895 | 5.81E-07 |
| 11 | 119650060 | rs12222723 | T | G | 0.992 | -7977.388 | -7979.283 | -1.895 | 5.81E-07 |
| 11 | 119951111 | rs12276427 | A | G | 0.999 | -7977.388 | -7979.283 | -1.895 | 5.81E-07 |
| 11 | 119971490 | rs6589798 | T | G | 0.993 | -7977.388 | -7979.283 | -1.895 | 5.81E-07 |
| 11 | 116726769 | rs640621 | C | G | 0.999 | -7977.388 | -7979.284 | -1.896 | 5.81E-07 |
| 11 | 117777599 | rs11216613 | G | T | 1.008 | -7977.388 | -7979.284 | -1.896 | 5.81E-07 |
| 11 | 117992620 | rs4938485 | G | A | 1.006 | -7977.388 | -7979.285 | -1.896 | 5.81E-07 |
| 11 | 118006957 | rs1814964 | C | T | 1.007 | -7977.388 | -7979.284 | -1.896 | 5.81E-07 |
| 11 | 118989098 | rs1129013 | C | G | 0.991 | -7977.388 | -7979.284 | -1.896 | 5.81E-07 |
| 11 | 119292712 | rs1001205 | T | C | 1.007 | -7977.388 | -7979.284 | -1.896 | 5.81E-07 |
| 11 | 119649359 | rs10892457 | T | C | 0.992 | -7977.388 | -7979.284 | -1.896 | 5.81E-07 |
| 11 | 119821850 | rs7931873 | G | A | 0.992 | -7977.388 | -7979.284 | -1.896 | 5.81E-07 |
| 11 | 120013984 | rs470370 | C | T | 1.007 | -7977.388 | -7979.284 | -1.896 | 5.81E-07 |
| 11 | 116521187 | rs534421 | G | A | 1.001 | -7977.388 | -7979.286 | -1.897 | 5.80E-07 |
| 11 | 117014862 | rs2622934 | G | T | 0.996 | -7977.388 | -7979.286 | -1.897 | 5.80E-07 |
| 11 | 118009279 | rs10790240 | G | T | 1.007 | -7977.388 | -7979.286 | -1.897 | 5.80E-07 |
| 11 | 118017445 | rs1786183 | A | G | 0.993 | -7977.388 | -7979.285 | -1.897 | 5.80E-07 |
| 11 | 119285008 | rs3138097 | C | T | 1.007 | -7977.388 | -7979.285 | -1.897 | 5.80E-07 |
| 11 | 119290034 | rs1054735 | G | C | 1.006 | -7977.388 | -7979.286 | -1.897 | 5.80E-07 |
| 11 | 119834547 | rs10750167 | G | A | 1.008 | -7977.388 | -7979.285 | -1.897 | 5.80E-07 |
| 11 | 117776850 | rs12798929 | T | C | 0.992 | -7977.388 | -7979.286 | -1.898 | 5.80E-07 |
| 11 | 118011776 | rs668468 | G | A | 0.996 | -7977.388 | -7979.286 | -1.898 | 5.80E-07 |
| 11 | 118991019 | rs675866 | C | A | 0.991 | -7977.388 | -7979.286 | -1.898 | 5.80E-07 |
| 11 | 119305126 | rs10790294 | T | C | 1.006 | -7977.388 | -7979.286 | -1.898 | 5.80E-07 |
| 11 | 119309945 | rs4450179 | C | T | 1.006 | -7977.388 | -7979.286 | -1.898 | 5.80E-07 |
| 11 | 119971706 | rs6589800 | C | A | 0.993 | -7977.388 | -7979.286 | -1.898 | 5.80E-07 |
| 11 | 117702115 | rs7130303 | A | C | 0.993 | -7977.388 | -7979.287 | -1.899 | 5.79E-07 |
| 11 | 117731136 | rs5014624 | A | G | 0.997 | -7977.388 | -7979.287 | -1.899 | 5.79E-07 |
| 11 | 118076004 | rs1806 | C | T | 0.993 | -7977.388 | -7979.287 | -1.899 | 5.79E-07 |
| 11 | 118990253 | rs1305565 | C | T | 0.991 | -7977.388 | -7979.288 | -1.899 | 5.79E-07 |
| 11 | 119023107 | rs668033 | C | A | 0.991 | -7977.388 | -7979.287 | -1.899 | 5.79E-07 |
| 11 | 119448580 | rs202125027 | A | T | 1.005 | -7977.388 | -7979.288 | -1.899 | 5.79E-07 |
| 11 | 119972448 | rs1073636 | A | T | 0.993 | -7977.388 | -7979.287 | -1.899 | 5.79E-07 |
| 11 | 119972642 | rs1073634 | T | C | 0.993 | -7977.388 | -7979.287 | -1.899 | 5.79E-07 |
| 11 | 120551561 | rs10790396 | A | G | 1.008 | -7977.388 | -7979.287 | -1.899 | 5.79E-07 |
| 11 | 121153421 | rs7949987 | G | A | 1.003 | -7977.388 | -7979.287 | -1.899 | 5.79E-07 |
| 11 | 116317405 | rs556550 | G | A | 1.007 | -7977.388 | -7979.288 | -1.9 | 5.79E-07 |
| 11 | 116465680 | rs11216029 | T | G | 0.993 | -7977.388 | -7979.288 | -1.9 | 5.79E-07 |
| 11 | 117270818 | rs1243337 | C | A | 1.004 | -7977.388 | -7979.288 | -1.9 | 5.79E-07 |
| 11 | 117992645 | rs4938486 | G | A | 1.007 | -7977.388 | -7979.288 | -1.9 | 5.79E-07 |
| 11 | 118016981 | rs3018387 | T | A | 0.994 | -7977.388 | -7979.289 | -1.9 | 5.79E-07 |
| 11 | 118018022 | rs628299 | G | A | 0.994 | -7977.388 | -7979.288 | -1.9 | 5.79E-07 |
| 11 | 119414914 | rs7945998 | T | C | 0.998 | -7977.388 | -7979.289 | -1.9 | 5.79E-07 |
| 11 | 119649844 | rs4258392 | C | T | 0.992 | -7977.388 | -7979.288 | -1.9 | 5.79E-07 |
| 11 | 119649874 | rs5020633 | A | G | 0.992 | -7977.388 | -7979.288 | -1.9 | 5.79E-07 |
| 11 | 119754895 | rs1358104428 | A | G | 1.008 | -7977.388 | -7979.288 | -1.9 | 5.79E-07 |
| 11 | 119971987 | rs6589801 | T | C | 0.993 | -7977.388 | -7979.288 | -1.9 | 5.79E-07 |
| 11 | 120976428 | rs504626 | A | G | 1.008 | -7977.388 | -7979.288 | -1.9 | 5.79E-07 |
| 11 | 116465734 | rs11216030 | T | C | 0.993 | -7977.388 | -7979.29 | -1.901 | 5.78E-07 |
| 11 | 118807735 | rs10892311 | T | C | 0.993 | -7977.388 | -7979.289 | -1.901 | 5.78E-07 |
| 11 | 118942654 | rs10892328 | A | T | 0.996 | -7977.388 | -7979.29 | -1.901 | 5.78E-07 |
| 11 | 118944675 | rs3825061 | T | C | 0.996 | -7977.388 | -7979.29 | -1.901 | 5.78E-07 |
| 11 | 119649863 | rs5020632 | T | C | 0.993 | -7977.388 | -7979.289 | -1.901 | 5.78E-07 |
| 11 | 119727494 | rs7113123 | T | C | 0.998 | -7977.388 | -7979.289 | -1.901 | 5.78E-07 |
| 11 | 119846820 | rs636518 | G | A | 1.006 | -7977.388 | -7979.289 | -1.901 | 5.78E-07 |
| 11 | 119964351 | rs664590 | T | G | 0.997 | -7977.388 | -7979.289 | -1.901 | 5.78E-07 |
| 11 | 119967556 | rs613004 | T | C | 0.998 | -7977.388 | -7979.289 | -1.901 | 5.78E-07 |
| 11 | 119967661 | rs1787027 | C | A | 0.998 | -7977.388 | -7979.289 | -1.901 | 5.78E-07 |
| 11 | 119967716 | rs10892534 | A | C | 0.998 | -7977.388 | -7979.289 | -1.901 | 5.78E-07 |
| 11 | 119967718 | rs10892535 | T | C | 0.998 | -7977.388 | -7979.289 | -1.901 | 5.78E-07 |
| 11 | 121154437 | rs1982011 | G | A | 1.003 | -7977.388 | -7979.29 | -1.901 | 5.78E-07 |
| 11 | 117315692 | rs11216394 | G | A | 0.997 | -7977.388 | -7979.29 | -1.902 | 5.77E-07 |
| 11 | 118017434 | rs1793157 | C | A | 0.994 | -7977.388 | -7979.291 | -1.902 | 5.77E-07 |
| 11 | 118735476 | rs10892301 | A | G | 0.993 | -7977.388 | -7979.29 | -1.902 | 5.77E-07 |
| 11 | 118796570 | rs11217096 | T | C | 0.993 | -7977.388 | -7979.29 | -1.902 | 5.77E-07 |
| 11 | 118941596 | rs1260808616 | A | G | 0.996 | -7977.388 | -7979.29 | -1.902 | 5.77E-07 |
| 11 | 117730979 | rs6589627 | C | T | 0.998 | -7977.388 | -7979.291 | -1.903 | 5.77E-07 |
| 11 | 117730983 | rs6589628 | A | G | 0.998 | -7977.388 | -7979.291 | -1.903 | 5.77E-07 |
| 11 | 117731070 | rs6589629 | C | G | 0.998 | -7977.388 | -7979.291 | -1.903 | 5.77E-07 |
| 11 | 117731085 | rs6589630 | C | T | 0.998 | -7977.388 | -7979.291 | -1.903 | 5.77E-07 |
| 11 | 117731191 | rs5014623 | C | T | 0.998 | -7977.388 | -7979.291 | -1.903 | 5.77E-07 |
| 11 | 117991333 | rs7927810 | A | G | 1.006 | -7977.388 | -7979.291 | -1.903 | 5.77E-07 |
| 11 | 119306276 | rs4938655 | A | C | 1.005 | -7977.388 | -7979.291 | -1.903 | 5.77E-07 |
| 11 | 119788216 | rs643153 | C | T | 1.007 | -7977.388 | -7979.292 | -1.903 | 5.77E-07 |
| 11 | 120007889 | rs470645 | T | G | 1.006 | -7977.388 | -7979.292 | -1.903 | 5.77E-07 |
| 11 | 116584317 | rs1144035 | C | T | 1.007 | -7977.388 | -7979.292 | -1.904 | 5.76E-07 |
| 11 | 117731263 | rs5014622 | G | A | 0.998 | -7977.388 | -7979.292 | -1.904 | 5.76E-07 |
| 11 | 119306183 | rs4938654 | T | C | 1.005 | -7977.388 | -7979.292 | -1.904 | 5.76E-07 |
| 11 | 119748404 | rs10892483 | C | G | 0.995 | -7977.388 | -7979.292 | -1.904 | 5.76E-07 |
| 11 | 119843707 | rs10790359 | G | A | 1.008 | -7977.388 | -7979.292 | -1.904 | 5.76E-07 |
| 11 | 119843736 | rs10790360 | C | A | 1.008 | -7977.388 | -7979.292 | -1.904 | 5.76E-07 |
| 11 | 119967769 | rs10790366 | T | G | 1.007 | -7977.388 | -7979.292 | -1.904 | 5.76E-07 |
| 11 | 121158779 | rs2115923 | A | G | 1.001 | -7977.388 | -7979.293 | -1.904 | 5.76E-07 |
| 11 | 117260075 | rs552498 | G | A | 1.002 | -7977.388 | -7979.293 | -1.905 | 5.76E-07 |

| | | | | | | | |
|---|---|---|---|---|---|---|---|
| 11 | 117333521 | rs55871712 | T | C | NULL | 0.721115178 | NULL&M_Disease |
| 11 | 117539478 | rs947926 | C | A | NULL | 0.721115178 | NULL&M_Disease |
| 11 | 117699377 | rs11216569 | T | C | NULL | 0.721115178 | NULL&M_Disease |
| 11 | 117731284 | rs5014621 | A | G | NULL | 0.721115178 | NULL&M_Disease |
| 11 | 117731319 | rs5014620 | C | A | NULL | 0.721115178 | NULL&M_Disease |
| 11 | 117731439 | rs1815774 | C | G | NULL | 0.721115178 | NULL&M_Disease |
| 11 | 117731643 | rs948107 | A | C | NULL | 0.721115178 | NULL&M_Disease |
| 11 | 118006408 | rs7124127 | C | T | NULL | 0.721115178 | NULL&M_Disease |
| 11 | 118208750 | rs11216855 | C | T | NULL | 0.721115178 | NULL&M_Disease |
| 11 | 118992502 | rs595359 | T | G | NULL | 0.721115178 | NULL&M_Disease |
| 11 | 119443669 | rs10750158 | A | C | NULL | 0.721115178 | NULL&M_Disease |
| 11 | 119902255 | rs10892517 | T | C | NULL | 0.721115178 | NULL&M_Disease |
| 11 | 120060673 | rs7116997 | C | T | NULL | 0.721115178 | NULL&M_Disease |
| 11 | 120551684 | rs10790398 | T | C | NULL | 0.721115178 | NULL&M_Disease |
| 11 | 120617504 | rs1945010 | T | C | NULL | 0.721115178 | NULL&M_Disease |
| 11 | 121185726 | rs7102059 | T | A | NULL | 0.721115178 | NULL&M_Disease |
| 11 | 116316173 | rs490043 | G | A | NULL | 0.721115178 | NULL&M_Disease |
| 11 | 117334518 | rs4938386 | T | C | NULL | 0.721115178 | NULL&M_Disease |
| 11 | 117731927 | rs10790216 | C | T | NULL | 0.721115178 | NULL&M_Disease |
| 11 | 118016068 | rs1793137 | C | T | NULL | 0.721115178 | NULL&M_Disease |
| 11 | 119316659 | rs7111567 | T | G | NULL | 0.721115178 | NULL&M_Disease |
| 11 | 119566174 | rs4938710 | C | T | NULL | 0.721115178 | NULL&M_Disease |
| 11 | 119637414 | rs4341549 | G | A | NULL | 0.721115178 | NULL&M_Disease |
| 11 | 120540086 | rs10790389 | C | G | NULL | 0.721115178 | NULL&M_Disease |
| 11 | 121158509 | rs4936611 | T | C | NULL | 0.721115178 | NULL&M_Disease |
| 11 | 121160060 | rs1560409 | C | G | NULL | 0.721115178 | NULL&M_Disease |
| 11 | 117258198 | rs501776 | C | T | NULL | 0.721115178 | NULL&M_Disease |
| 11 | 117958396 | rs603407 | C | T | NULL | 0.721115178 | NULL&M_Disease |
| 11 | 117993422 | rs1941636 | C | T | NULL | 0.722119606 | NULL&M_Disease |
| 11 | 118801235 | rs602716 | A | G | NULL | 0.721115178 | NULL&M_Disease |
| 11 | 119444330 | rs1876793 | T | C | NULL | 0.721115178 | NULL&M_Disease |
| 11 | 119839344 | rs949096 | A | G | NULL | 0.721115178 | NULL&M_Disease |
| 11 | 119845535 | rs1787024 | C | T | NULL | 0.721115178 | NULL&M_Disease |
| 11 | 119903119 | rs11217646 | G | A | NULL | 0.721115178 | NULL&M_Disease |
| 11 | 119989892 | rs606970 | G | A | NULL | 0.721115178 | NULL&M_Disease |
| 11 | 116576415 | rs1263056 | G | A | NULL | 0.722119606 | NULL&M_Disease |
| 11 | 117323076 | rs4938380 | A | G | NULL | 0.722119606 | NULL&M_Disease |
| 11 | 117954081 | rs657118 | G | A | NULL | 0.722119606 | NULL&M_Disease |
| 11 | 117984623 | rs596134 | C | T | NULL | 0.722119606 | NULL&M_Disease |
| 11 | 118018725 | rs614825 | G | A | NULL | 0.722119606 | NULL&M_Disease |
| 11 | 118481431 | rs10892246 | A | G | NULL | 0.722119606 | NULL&M_Disease |
| 11 | 118803536 | rs615166 | A | G | NULL | 0.722119606 | NULL&M_Disease |
| 11 | 119311481 | rs6589741 | A | T | NULL | 0.722119606 | NULL&M_Disease |
| 11 | 119311933 | rs6589743 | T | A | NULL | 0.722119606 | NULL&M_Disease |
| 11 | 119312014 | rs6589744 | G | A | NULL | 0.722119606 | NULL&M_Disease |
| 11 | 119314765 | rs7932332 | G | A | NULL | 0.722119606 | NULL&M_Disease |
| 11 | 119410158 | rs11217308 | T | C | NULL | 0.722119606 | NULL&M_Disease |
| 11 | 119441311 | rs906823 | A | G | NULL | 0.722119606 | NULL&M_Disease |
| 11 | 119441696 | rs906824 | T | C | NULL | 0.722119606 | NULL&M_Disease |
| 11 | 119441804 | rs906825 | G | A | NULL | 0.722119606 | NULL&M_Disease |
| 11 | 119442188 | rs906826 | T | C | NULL | 0.722119606 | NULL&M_Disease |
| 11 | 119444978 | rs7108677 | G | C | NULL | 0.722119606 | NULL&M_Disease |
| 11 | 119444980 | rs7108679 | G | A | NULL | 0.722119606 | NULL&M_Disease |
| 11 | 119843952 | rs7342270 | A | T | NULL | 0.722119606 | NULL&M_Disease |
| 11 | 117469590 | rs748316 | C | T | NULL | 0.722119606 | NULL&M_Disease |
| 11 | 119315789 | rs4938659 | T | C | NULL | 0.722119606 | NULL&M_Disease |
| 11 | 119444016 | rs1876794 | C | G | NULL | 0.722119606 | NULL&M_Disease |
| 11 | 119444435 | rs1876792 | G | A | NULL | 0.722119606 | NULL&M_Disease |
| 11 | 119620456 | rs12365133 | G | A | NULL | 0.722119606 | NULL&M_Disease |
| 11 | 119840052 | rs200041620 | A | G | NULL | 0.722119606 | NULL&M_Disease |
| 11 | 119840067 | rs731392 | G | C | NULL | 0.722119606 | NULL&M_Disease |
| 11 | 119991166 | rs590843 | T | A | NULL | 0.722119606 | NULL&M_Disease |
| 11 | 120041110 | rs7949789 | A | T | NULL | 0.722119606 | NULL&M_Disease |
| 11 | 117811167 | rs3863318 | A | C | NULL | 0.722119606 | NULL&M_Disease |
| 11 | 117985080 | rs598061 | C | A | NULL | 0.722119606 | NULL&M_Disease |
| 11 | 117989246 | rs1941627 | G | A | NULL | 0.722119606 | NULL&M_Disease |
| 11 | 117989835 | rs10892220 | A | T | NULL | 0.722119606 | NULL&M_Disease |
| 11 | 118067187 | rs605560 | C | G | NULL | 0.722119606 | NULL&M_Disease |
| 11 | 119316614 | rs7111455 | T | C | NULL | 0.722119606 | NULL&M_Disease |
| 11 | 119317208 | rs2082049 | G | A | NULL | 0.722119606 | NULL&M_Disease |
| 11 | 119405750 | rs10892383 | T | C | NULL | 0.722119606 | NULL&M_Disease |
| 11 | 119568070 | rs4936493 | T | A | NULL | 0.722119606 | NULL&M_Disease |
| 11 | 119844270 | rs6589784 | C | A | NULL | 0.722119606 | NULL&M_Disease |
| 11 | 120067082 | rs896693 | A | G | NULL | 0.722119606 | NULL&M_Disease |
| 11 | 116583513 | rs886022 | A | C | NULL | 0.722119606 | NULL&M_Disease |
| 11 | 117958909 | rs1317876 | G | A | NULL | 0.722119606 | NULL&M_Disease |
| 11 | 117989989 | rs10750125 | C | T | NULL | 0.722119606 | NULL&M_Disease |
| 11 | 119406292 | rs4375449 | A | G | NULL | 0.722119606 | NULL&M_Disease |
| 11 | 119408686 | rs4938675 | A | G | NULL | 0.722119606 | NULL&M_Disease |
| 11 | 119446574 | rs1995822 | A | G | NULL | 0.722119606 | NULL&M_Disease |
| 11 | 119840401 | rs7951479 | T | C | NULL | 0.722119606 | NULL&M_Disease |
| 11 | 119840442 | rs7951593 | T | C | NULL | 0.722119606 | NULL&M_Disease |
| 11 | 118144445 | rs4245179 | G | A | NULL | 0.722119606 | NULL&M_Disease |
| 11 | 118144512 | rs4245180 | A | G | NULL | 0.722119606 | NULL&M_Disease |
| 11 | 118144804 | rs4938493 | G | T | NULL | 0.722119606 | NULL&M_Disease |
| 11 | 118145686 | rs17122021 | T | C | NULL | 0.722119606 | NULL&M_Disease |
| 11 | 118146197 | rs57461652 | A | C | NULL | 0.722119606 | NULL&M_Disease |
| 11 | 119451940 | rs4938687 | A | G | NULL | 0.722119606 | NULL&M_Disease |
| 11 | 119568301 | rs1467051 | T | C | NULL | 0.722119606 | NULL&M_Disease |
| 11 | 119836087 | rs10892504 | A | G | NULL | 0.722119606 | NULL&M_Disease |
| 11 | 116316963 | rs504339 | A | G | NULL | 0.722119606 | NULL&M_Disease |
| 11 | 117992718 | rs4936423 | C | G | NULL | 0.722119606 | NULL&M_Disease |
| 11 | 118068949 | rs1617182 | C | T | NULL | 0.722119606 | NULL&M_Disease |
| 11 | 118141993 | rs57506367 | C | T | NULL | 0.722119606 | NULL&M_Disease |
| 11 | 118142619 | rs7940047 | T | C | NULL | 0.722119606 | NULL&M_Disease |
| 11 | 118142930 | rs7940554 | T | C | NULL | 0.722119606 | NULL&M_Disease |

| | | | | | | | | | |
|---|---|---|---|---|---|---|---|---|---|
| 11 | 117333521 | rs55871712 | T | C | 1.007 | -7977.388 | -7979.293 | -1.905 | 5.76E-07 |
| 11 | 117539478 | rs947926 | C | A | 0.996 | -7977.388 | -7979.293 | -1.905 | 5.76E-07 |
| 11 | 117699377 | rs11216569 | T | C | 0.994 | -7977.388 | -7979.293 | -1.905 | 5.76E-07 |
| 11 | 117731284 | rs5014621 | A | G | 0.999 | -7977.388 | -7979.294 | -1.905 | 5.76E-07 |
| 11 | 117731319 | rs5014620 | C | A | 0.999 | -7977.388 | -7979.294 | -1.905 | 5.76E-07 |
| 11 | 117731439 | rs1815774 | C | G | 0.999 | -7977.388 | -7979.293 | -1.905 | 5.76E-07 |
| 11 | 117731643 | rs948107 | A | C | 0.998 | -7977.388 | -7979.293 | -1.905 | 5.76E-07 |
| 11 | 118006408 | rs7124127 | C | T | 1.007 | -7977.388 | -7979.293 | -1.905 | 5.76E-07 |
| 11 | 118208750 | rs11216855 | C | T | 1.007 | -7977.388 | -7979.293 | -1.905 | 5.76E-07 |
| 11 | 118992502 | rs595359 | T | G | 0.992 | -7977.388 | -7979.293 | -1.905 | 5.76E-07 |
| 11 | 119443669 | rs10750158 | A | C | 1.003 | -7977.388 | -7979.293 | -1.905 | 5.76E-07 |
| 11 | 119902255 | rs10892517 | T | C | 0.993 | -7977.388 | -7979.293 | -1.905 | 5.76E-07 |
| 11 | 120060673 | rs7116997 | C | T | 1.007 | -7977.388 | -7979.293 | -1.905 | 5.76E-07 |
| 11 | 120551684 | rs10790398 | T | C | 1.008 | -7977.388 | -7979.294 | -1.905 | 5.76E-07 |
| 11 | 120617504 | rs1945010 | T | C | 0.998 | -7977.388 | -7979.294 | -1.905 | 5.76E-07 |
| 11 | 121185726 | rs7102059 | T | A | 1.007 | -7977.388 | -7979.293 | -1.905 | 5.76E-07 |
| 11 | 116316173 | rs490043 | G | A | 1.004 | -7977.388 | -7979.295 | -1.906 | 5.75E-07 |
| 11 | 117334518 | rs4938386 | T | C | 1.007 | -7977.388 | -7979.295 | -1.906 | 5.75E-07 |
| 11 | 117731927 | rs10790216 | C | T | 0.999 | -7977.388 | -7979.294 | -1.906 | 5.75E-07 |
| 11 | 118016068 | rs1793137 | C | T | 0.997 | -7977.388 | -7979.294 | -1.906 | 5.75E-07 |
| 11 | 119316659 | rs7111567 | T | G | 1.005 | -7977.388 | -7979.294 | -1.906 | 5.75E-07 |
| 11 | 119566174 | rs4938710 | C | T | 0.997 | -7977.388 | -7979.295 | -1.906 | 5.75E-07 |
| 11 | 119637414 | rs4341549 | G | A | 0.993 | -7977.388 | -7979.295 | -1.906 | 5.75E-07 |
| 11 | 120540086 | rs10790389 | C | G | 0.992 | -7977.388 | -7979.294 | -1.906 | 5.75E-07 |
| 11 | 121158509 | rs4936611 | T | C | 1 | -7977.388 | -7979.294 | -1.906 | 5.75E-07 |
| 11 | 121160060 | rs1560409 | C | G | 1 | -7977.388 | -7979.294 | -1.906 | 5.75E-07 |
| 11 | 117258198 | rs501776 | C | T | 1.001 | -7977.388 | -7979.295 | -1.907 | 5.75E-07 |
| 11 | 117958396 | rs603407 | C | T | 1.006 | -7977.388 | -7979.295 | -1.907 | 5.75E-07 |
| 11 | 117993422 | rs1941636 | C | T | 1.005 | -7977.388 | -7979.296 | -1.907 | 5.75E-07 |
| 11 | 118801235 | rs602716 | A | G | 1.001 | -7977.388 | -7979.295 | -1.907 | 5.75E-07 |
| 11 | 119444330 | rs1876793 | T | C | 1.003 | -7977.388 | -7979.295 | -1.907 | 5.75E-07 |
| 11 | 119839344 | rs949096 | A | G | 1.005 | -7977.388 | -7979.296 | -1.907 | 5.75E-07 |
| 11 | 119845535 | rs1787024 | C | T | 1.005 | -7977.388 | -7979.296 | -1.907 | 5.75E-07 |
| 11 | 119903119 | rs11217646 | G | A | 0.994 | -7977.388 | -7979.296 | -1.907 | 5.75E-07 |
| 11 | 119989892 | rs606970 | G | A | 1.008 | -7977.388 | -7979.295 | -1.907 | 5.75E-07 |
| 11 | 116576415 | rs1263056 | G | A | 0.993 | -7977.388 | -7979.297 | -1.908 | 5.74E-07 |
| 11 | 117323076 | rs4938380 | A | G | 0.993 | -7977.388 | -7979.296 | -1.908 | 5.74E-07 |
| 11 | 117954081 | rs657118 | G | A | 0.994 | -7977.388 | -7979.296 | -1.908 | 5.74E-07 |
| 11 | 117984623 | rs596134 | C | T | 1.004 | -7977.388 | -7979.296 | -1.908 | 5.74E-07 |
| 11 | 118018725 | rs614825 | G | A | 0.995 | -7977.388 | -7979.296 | -1.908 | 5.74E-07 |
| 11 | 118481431 | rs10892246 | A | G | 0.996 | -7977.388 | -7979.296 | -1.908 | 5.74E-07 |
| 11 | 118803536 | rs615166 | A | G | 1 | -7977.388 | -7979.296 | -1.908 | 5.74E-07 |
| 11 | 119311481 | rs6589741 | A | T | 1.005 | -7977.388 | -7979.296 | -1.908 | 5.74E-07 |
| 11 | 119311933 | rs6589743 | T | A | 1.005 | -7977.388 | -7979.296 | -1.908 | 5.74E-07 |
| 11 | 119312014 | rs6589744 | G | A | 1.005 | -7977.388 | -7979.296 | -1.908 | 5.74E-07 |
| 11 | 119314765 | rs7932332 | G | A | 1.005 | -7977.388 | -7979.297 | -1.908 | 5.74E-07 |
| 11 | 119410158 | rs11217308 | T | C | 0.996 | -7977.388 | -7979.296 | -1.908 | 5.74E-07 |
| 11 | 119441311 | rs906823 | A | G | 1.002 | -7977.388 | -7979.297 | -1.908 | 5.74E-07 |
| 11 | 119441696 | rs906824 | T | C | 1.002 | -7977.388 | -7979.297 | -1.908 | 5.74E-07 |
| 11 | 119441804 | rs906825 | G | A | 1.002 | -7977.388 | -7979.297 | -1.908 | 5.74E-07 |
| 11 | 119442188 | rs906826 | T | C | 1.002 | -7977.388 | -7979.297 | -1.908 | 5.74E-07 |
| 11 | 119444978 | rs7108677 | G | C | 1.002 | -7977.388 | -7979.297 | -1.908 | 5.74E-07 |
| 11 | 119444980 | rs7108679 | G | A | 1.002 | -7977.388 | -7979.297 | -1.908 | 5.74E-07 |
| 11 | 119843952 | rs7342270 | A | T | 1.007 | -7977.388 | -7979.297 | -1.908 | 5.74E-07 |
| 11 | 117469590 | rs748316 | C | T | 0.993 | -7977.388 | -7979.297 | -1.909 | 5.73E-07 |
| 11 | 119315789 | rs4938659 | T | C | 1.005 | -7977.388 | -7979.297 | -1.909 | 5.73E-07 |
| 11 | 119444016 | rs1876794 | C | G | 1.002 | -7977.388 | -7979.297 | -1.909 | 5.73E-07 |
| 11 | 119444435 | rs1876792 | G | A | 1.002 | -7977.388 | -7979.297 | -1.909 | 5.73E-07 |
| 11 | 119620456 | rs12365133 | G | A | 1.007 | -7977.388 | -7979.297 | -1.909 | 5.73E-07 |
| 11 | 119840052 | rs200041620 | A | G | 1.005 | -7977.388 | -7979.297 | -1.909 | 5.73E-07 |
| 11 | 119840067 | rs731392 | G | C | 1.005 | -7977.388 | -7979.297 | -1.909 | 5.73E-07 |
| 11 | 119991166 | rs590843 | T | A | 1 | -7977.388 | -7979.297 | -1.909 | 5.73E-07 |
| 11 | 120041110 | rs7949789 | A | T | 0.996 | -7977.388 | -7979.297 | -1.909 | 5.73E-07 |
| 11 | 117811167 | rs3863318 | A | C | 0.993 | -7977.388 | -7979.298 | -1.91 | 5.73E-07 |
| 11 | 117985080 | rs598061 | C | A | 1.004 | -7977.388 | -7979.298 | -1.91 | 5.73E-07 |
| 11 | 117989246 | rs1941627 | G | A | 1.004 | -7977.388 | -7979.298 | -1.91 | 5.73E-07 |
| 11 | 117989835 | rs10892220 | A | T | 1.004 | -7977.388 | -7979.299 | -1.91 | 5.73E-07 |
| 11 | 118067187 | rs605560 | C | G | 0.994 | -7977.388 | -7979.298 | -1.91 | 5.73E-07 |
| 11 | 119316614 | rs7111455 | T | C | 1.004 | -7977.388 | -7979.298 | -1.91 | 5.73E-07 |
| 11 | 119317208 | rs2082049 | G | A | 1.004 | -7977.388 | -7979.298 | -1.91 | 5.73E-07 |
| 11 | 119405750 | rs10892383 | T | C | 1.003 | -7977.388 | -7979.298 | -1.91 | 5.73E-07 |
| 11 | 119568070 | rs4936493 | T | A | 1.002 | -7977.388 | -7979.299 | -1.91 | 5.73E-07 |
| 11 | 119844270 | rs6589784 | C | A | 1.007 | -7977.388 | -7979.299 | -1.91 | 5.73E-07 |
| 11 | 120067082 | rs896693 | A | G | 0.995 | -7977.388 | -7979.298 | -1.91 | 5.73E-07 |
| 11 | 116583513 | rs886022 | A | C | 1.006 | -7977.388 | -7979.299 | -1.911 | 5.72E-07 |
| 11 | 117958909 | rs1317876 | G | A | 1.006 | -7977.388 | -7979.3 | -1.911 | 5.72E-07 |
| 11 | 117989989 | rs10750125 | C | T | 1.004 | -7977.388 | -7979.299 | -1.911 | 5.72E-07 |
| 11 | 119406292 | rs4375449 | A | G | 1.003 | -7977.388 | -7979.3 | -1.911 | 5.72E-07 |
| 11 | 119408686 | rs4938675 | A | G | 0.997 | -7977.388 | -7979.3 | -1.911 | 5.72E-07 |
| 11 | 119446574 | rs1995822 | A | G | 1.001 | -7977.388 | -7979.299 | -1.911 | 5.72E-07 |
| 11 | 119840401 | rs7951479 | T | C | 1.004 | -7977.388 | -7979.299 | -1.911 | 5.72E-07 |
| 11 | 119840442 | rs7951593 | T | C | 1.007 | -7977.388 | -7979.299 | -1.911 | 5.72E-07 |
| 11 | 118144445 | rs4245179 | G | A | 1.005 | -7977.388 | -7979.301 | -1.912 | 5.72E-07 |
| 11 | 118144512 | rs4245180 | A | G | 1.005 | -7977.388 | -7979.301 | -1.912 | 5.72E-07 |
| 11 | 118144804 | rs4938493 | G | T | 1.005 | -7977.388 | -7979.301 | -1.912 | 5.72E-07 |
| 11 | 118145686 | rs17122021 | T | C | 1.005 | -7977.388 | -7979.301 | -1.912 | 5.72E-07 |
| 11 | 118146197 | rs57461652 | A | C | 1.005 | -7977.388 | -7979.3 | -1.912 | 5.72E-07 |
| 11 | 119451940 | rs4938687 | A | G | 1.001 | -7977.388 | -7979.3 | -1.912 | 5.72E-07 |
| 11 | 119568301 | rs1467051 | T | C | 1.001 | -7977.388 | -7979.3 | -1.912 | 5.72E-07 |
| 11 | 119836087 | rs10892504 | A | G | 1.004 | -7977.388 | -7979.3 | -1.912 | 5.72E-07 |
| 11 | 116316963 | rs504339 | A | G | 0.996 | -7977.388 | -7979.301 | -1.913 | 5.71E-07 |
| 11 | 117992718 | rs4936423 | C | G | 1.006 | -7977.388 | -7979.301 | -1.913 | 5.71E-07 |
| 11 | 118068949 | rs1617182 | C | T | 0.994 | -7977.388 | -7979.301 | -1.913 | 5.71E-07 |
| 11 | 118141993 | rs57506367 | C | T | 1.005 | -7977.388 | -7979.301 | -1.913 | 5.71E-07 |
| 11 | 118142619 | rs7940047 | T | C | 1.005 | -7977.388 | -7979.301 | -1.913 | 5.71E-07 |
| 11 | 118142930 | rs7940554 | T | C | 1.005 | -7977.388 | -7979.302 | -1.913 | 5.71E-07 |

| | | | | | | | |
|---|---|---|---|---|---|---|---|
| 11 | 118143586 | rs112590920 | A | G | NULL | 0.722119606 | NULL&M_Disease |
| 11 | 118148146 | rs2853009 | G | A | NULL | 0.722119606 | NULL&M_Disease |
| 11 | 119566825 | rs4623916 | T | A | NULL | 0.722119606 | NULL&M_Disease |
| 11 | 120028701 | rs618265 | A | G | NULL | 0.722119606 | NULL&M_Disease |
| 11 | 120556018 | rs7116106 | T | G | NULL | 0.722119606 | NULL&M_Disease |
| 11 | 121076285 | rs687560 | T | C | NULL | 0.722119606 | NULL&M_Disease |
| 11 | 121185128 | rs12226297 | A | G | NULL | 0.722119606 | NULL&M_Disease |
| 11 | 117996119 | rs7944738 | C | T | NULL | 0.722119606 | NULL&M_Disease |
| 11 | 118146199 | rs58162302 | T | A | NULL | 0.722119606 | NULL&M_Disease |
| 11 | 119411425 | rs4938679 | T | C | NULL | 0.722119606 | NULL&M_Disease |
| 11 | 119690671 | rs11217522 | T | G | NULL | 0.722119606 | NULL&M_Disease |
| 11 | 117989944 | rs10790237 | G | T | NULL | 0.722119606 | NULL&M_Disease |
| 11 | 117991034 | rs7924895 | G | A | NULL | 0.722119606 | NULL&M_Disease |
| 11 | 118022608 | rs583490 | A | G | NULL | 0.722119606 | NULL&M_Disease |
| 11 | 119485081 | rs871800 | C | G | NULL | 0.722119606 | NULL&M_Disease |
| 11 | 119485214 | rs884395 | G | A | NULL | 0.722119606 | NULL&M_Disease |
| 11 | 119789044 | rs538518 | G | T | NULL | 0.722119606 | NULL&M_Disease |
| 11 | 120074851 | rs10892548 | A | G | NULL | 0.722119606 | NULL&M_Disease |
| 11 | 117318148 | rs9326249 | A | G | NULL | 0.722119606 | NULL&M_Disease |
| 11 | 117989990 | rs10750126 | A | G | NULL | 0.722119606 | NULL&M_Disease |
| 11 | 119903434 | rs61897923 | C | T | NULL | 0.722119606 | NULL&M_Disease |
| 11 | 119979732 | rs1893391 | C | G | NULL | 0.722119606 | NULL&M_Disease |
| 11 | 120025243 | rs674637 | G | A | NULL | 0.722119606 | NULL&M_Disease |
| 11 | 120405843 | rs4084221 | A | G | NULL | 0.722119606 | NULL&M_Disease |
| 11 | 117991805 | rs7928275 | A | G | NULL | 0.722119606 | NULL&M_Disease |
| 11 | 117991817 | rs7943311 | T | G | NULL | 0.722119606 | NULL&M_Disease |
| 11 | 118070264 | rs1793169 | G | A | NULL | 0.722119606 | NULL&M_Disease |
| 11 | 118987971 | rs571445 | G | A | NULL | 0.722119606 | NULL&M_Disease |
| 11 | 119622781 | rs4317995 | A | G | NULL | 0.722119606 | NULL&M_Disease |
| 11 | 119654053 | rs10790345 | G | A | NULL | 0.722119606 | NULL&M_Disease |
| 11 | 116317399 | rs556514 | T | C | NULL | 0.723121805 | NULL&M_Disease |
| 11 | 117245133 | rs7110647 | T | C | NULL | 0.723121805 | NULL&M_Disease |
| 11 | 118017645 | rs1613495 | T | C | NULL | 0.723121805 | NULL&M_Disease |
| 11 | 118019906 | rs642936 | C | T | NULL | 0.723121805 | NULL&M_Disease |
| 11 | 119411610 | rs4936477 | G | A | NULL | 0.723121805 | NULL&M_Disease |
| 11 | 119622162 | rs11217460 | G | C | NULL | 0.723121805 | NULL&M_Disease |
| 11 | 119653947 | rs7942259 | T | C | NULL | 0.723121805 | NULL&M_Disease |
| 11 | 119658393 | rs10790346 | G | C | NULL | 0.723121805 | NULL&M_Disease |
| 11 | 119680923 | rs4936498 | G | T | NULL | 0.723121805 | NULL&M_Disease |
| 11 | 119719420 | rs1530604 | G | A | NULL | 0.723121805 | NULL&M_Disease |
| 11 | 119902691 | rs11605919 | C | T | NULL | 0.723121805 | NULL&M_Disease |
| 11 | 120543781 | rs4417279 | T | G | NULL | 0.723121805 | NULL&M_Disease |
| 11 | 116489629 | rs548207 | G | T | NULL | 0.723121805 | NULL&M_Disease |
| 11 | 117814506 | rs10160827 | T | C | NULL | 0.723121805 | NULL&M_Disease |
| 11 | 118146510 | rs58507982 | T | C | NULL | 0.723121805 | NULL&M_Disease |
| 11 | 119547005 | rs7122134 | G | A | NULL | 0.723121805 | NULL&M_Disease |
| 11 | 119639398 | rs10790344 | G | A | NULL | 0.723121805 | NULL&M_Disease |
| 11 | 116487699 | rs550767 | A | C | NULL | 0.723121805 | NULL&M_Disease |
| 11 | 118010890 | rs640323 | C | G | NULL | 0.723121805 | NULL&M_Disease |
| 11 | 119488176 | rs2136419 | T | C | NULL | 0.723121805 | NULL&M_Disease |
| 11 | 119564781 | rs7124934 | G | A | NULL | 0.723121805 | NULL&M_Disease |
| 11 | 120057343 | rs508205 | G | A | NULL | 0.723121805 | NULL&M_Disease |
| 11 | 120585300 | rs1939664 | A | T | NULL | 0.723121805 | NULL&M_Disease |
| 11 | 116464955 | rs11216027 | T | C | NULL | 0.723121805 | NULL&M_Disease |
| 11 | 117988784 | rs8924 | A | G | NULL | 0.723121805 | NULL&M_Disease |
| 11 | 119627450 | rs4351828 | A | G | NULL | 0.723121805 | NULL&M_Disease |
| 11 | 119978334 | rs1349689 | T | A | NULL | 0.723121805 | NULL&M_Disease |
| 11 | 120058685 | rs12278345 | G | T | NULL | 0.723121805 | NULL&M_Disease |
| 11 | 116577043 | rs1263055 | A | G | NULL | 0.723121805 | NULL&M_Disease |
| 11 | 119620940 | rs11217458 | A | G | NULL | 0.723121805 | NULL&M_Disease |
| 11 | 121085567 | rs508316 | C | A | NULL | 0.723121805 | NULL&M_Disease |
| 11 | 116316777 | rs489164 | C | T | NULL | 0.723121805 | NULL&M_Disease |
| 11 | 118004688 | rs3741315 | A | G | NULL | 0.723121805 | NULL&M_Disease |
| 11 | 119621691 | rs7949076 | T | C | NULL | 0.723121805 | NULL&M_Disease |
| 11 | 119655589 | rs7936400 | C | T | NULL | 0.723121805 | NULL&M_Disease |
| 11 | 119943275 | rs628944 | A | C | NULL | 0.723121805 | NULL&M_Disease |
| 11 | 120550286 | rs12574186 | G | A | NULL | 0.723121805 | NULL&M_Disease |
| 11 | 120584018 | rs10736504 | A | G | NULL | 0.723121805 | NULL&M_Disease |
| 11 | 120878916 | rs11218108 | T | C | NULL | 0.723121805 | NULL&M_Disease |
| 11 | 118016273 | rs1715444 | G | T | NULL | 0.723121805 | NULL&M_Disease |
| 11 | 118021740 | rs678262 | G | C | NULL | 0.723121805 | NULL&M_Disease |
| 11 | 119644451 | rs9326253 | C | T | NULL | 0.723121805 | NULL&M_Disease |
| 11 | 119727452 | rs6589781 | T | A | NULL | 0.723121805 | NULL&M_Disease |
| 11 | 120989335 | rs612969 | G | A | NULL | 0.723121805 | NULL&M_Disease |
| 11 | 116207302 | rs12420446 | A | G | NULL | 0.723121805 | NULL&M_Disease |
| 11 | 117473833 | rs1263889 | A | C | NULL | 0.723121805 | NULL&M_Disease |
| 11 | 118139403 | rs59049611 | C | T | NULL | 0.723121805 | NULL&M_Disease |
| 11 | 119777611 | rs509809 | G | A | NULL | 0.723121805 | NULL&M_Disease |
| 11 | 120026271 | rs470373 | T | C | NULL | 0.723121805 | NULL&M_Disease |
| 11 | 120028202 | rs470606 | G | T | NULL | 0.723121805 | NULL&M_Disease |
| 11 | 117317124 | rs7125517 | C | A | NULL | 0.723121805 | NULL&M_Disease |
| 11 | 118071097 | rs1626649 | T | C | NULL | 0.723121805 | NULL&M_Disease |
| 11 | 118961231 | rs494048 | T | C | NULL | 0.723121805 | NULL&M_Disease |
| 11 | 119696165 | rs477759 | T | C | NULL | 0.723121805 | NULL&M_Disease |
| 11 | 119773032 | rs486876 | A | G | NULL | 0.723121805 | NULL&M_Disease |
| 11 | 119786235 | rs529482 | G | A | NULL | 0.723121805 | NULL&M_Disease |
| 11 | 120542632 | rs7935000 | A | G | NULL | 0.723121805 | NULL&M_Disease |
| 11 | 120551419 | rs10790395 | A | C | NULL | 0.723121805 | NULL&M_Disease |
| 11 | 120569381 | rs12574240 | C | T | NULL | 0.723121805 | NULL&M_Disease |
| 11 | 121087285 | rs653478 | A | T | NULL | 0.723121805 | NULL&M_Disease |
| 11 | 117335074 | rs477890 | T | A | NULL | 0.723121805 | NULL&M_Disease |
| 11 | 117693255 | rs529623 | T | C | NULL | 0.723121805 | NULL&M_Disease |
| 11 | 117954724 | rs3949025 | A | G | NULL | 0.724121771 | NULL&M_Disease |
| 11 | 119785065 | rs472446 | G | A | NULL | 0.723121805 | NULL&M_Disease |
| 11 | 120056416 | rs2444239 | A | T | NULL | 0.724121771 | NULL&M_Disease |
| 11 | 120549404 | rs7119539 | G | A | NULL | 0.723121805 | NULL&M_Disease |

| | | | | | | | | | |
|---|---|---|---|---|---|---|---|---|---|
| 11 | 118143586 | rs112590920 | A | G | 1.005 | -7977.388 | -7979.302 | -1.913 | 5.71E-07 |
| 11 | 118148146 | rs2853009 | G | A | 1.006 | -7977.388 | -7979.301 | -1.913 | 5.71E-07 |
| 11 | 119566825 | rs4623916 | T | A | 0.998 | -7977.388 | -7979.302 | -1.913 | 5.71E-07 |
| 11 | 120028701 | rs618265 | A | G | 1.003 | -7977.388 | -7979.301 | -1.913 | 5.71E-07 |
| 11 | 120556018 | rs7116106 | T | G | 1.005 | -7977.388 | -7979.302 | -1.913 | 5.71E-07 |
| 11 | 121076285 | rs687560 | T | C | 1.005 | -7977.388 | -7979.301 | -1.913 | 5.71E-07 |
| 11 | 121185128 | rs12226297 | A | G | 1.006 | -7977.388 | -7979.302 | -1.913 | 5.71E-07 |
| 11 | 117996119 | rs7944738 | C | T | 1.003 | -7977.388 | -7979.303 | -1.914 | 5.71E-07 |
| 11 | 118146199 | rs58162302 | T | A | 1.005 | -7977.388 | -7979.302 | -1.914 | 5.71E-07 |
| 11 | 119411425 | rs4938679 | T | C | 0.997 | -7977.388 | -7979.302 | -1.914 | 5.71E-07 |
| 11 | 119690671 | rs11217522 | T | G | 1.006 | -7977.388 | -7979.303 | -1.914 | 5.71E-07 |
| 11 | 117989944 | rs10790237 | G | T | 1.003 | -7977.388 | -7979.303 | -1.915 | 5.70E-07 |
| 11 | 117991034 | rs7924895 | G | A | 1.005 | -7977.388 | -7979.303 | -1.915 | 5.70E-07 |
| 11 | 118022608 | rs583490 | A | G | 0.996 | -7977.388 | -7979.304 | -1.915 | 5.70E-07 |
| 11 | 119485081 | rs871800 | C | G | 1.004 | -7977.388 | -7979.303 | -1.915 | 5.70E-07 |
| 11 | 119485214 | rs884395 | G | A | 1.004 | -7977.388 | -7979.303 | -1.915 | 5.70E-07 |
| 11 | 119789044 | rs538518 | G | T | 1 | -7977.388 | -7979.304 | -1.915 | 5.70E-07 |
| 11 | 120074851 | rs10892548 | A | G | 0.996 | -7977.388 | -7979.303 | -1.915 | 5.70E-07 |
| 11 | 117318148 | rs9326249 | A | G | 1.005 | -7977.388 | -7979.304 | -1.916 | 5.69E-07 |
| 11 | 117989990 | rs10750126 | A | G | 1.003 | -7977.388 | -7979.305 | -1.916 | 5.69E-07 |
| 11 | 119903434 | rs61897923 | C | T | 0.995 | -7977.388 | -7979.304 | -1.916 | 5.69E-07 |
| 11 | 119979732 | rs1893391 | C | G | 1.005 | -7977.388 | -7979.304 | -1.916 | 5.69E-07 |
| 11 | 120025243 | rs674637 | G | A | 0.998 | -7977.388 | -7979.304 | -1.916 | 5.69E-07 |
| 11 | 120405843 | rs4084221 | A | G | 1.001 | -7977.388 | -7979.304 | -1.916 | 5.69E-07 |
| 11 | 117991805 | rs7928275 | A | G | 1.005 | -7977.388 | -7979.305 | -1.917 | 5.69E-07 |
| 11 | 117991817 | rs7943311 | T | G | 1.005 | -7977.388 | -7979.305 | -1.917 | 5.69E-07 |
| 11 | 118070264 | rs1793169 | G | A | 0.995 | -7977.388 | -7979.305 | -1.917 | 5.69E-07 |
| 11 | 118987971 | rs571445 | G | A | 0.994 | -7977.388 | -7979.305 | -1.917 | 5.69E-07 |
| 11 | 119622781 | rs4317995 | A | G | 0.994 | -7977.388 | -7979.305 | -1.917 | 5.69E-07 |
| 11 | 119654053 | rs10790345 | G | A | 0.994 | -7977.388 | -7979.305 | -1.917 | 5.69E-07 |
| 11 | 116317399 | rs556514 | T | C | 1.005 | -7977.388 | -7979.307 | -1.918 | 5.68E-07 |
| 11 | 117245133 | rs7110647 | T | C | 1.006 | -7977.388 | -7979.307 | -1.918 | 5.68E-07 |
| 11 | 118017645 | rs1613495 | T | C | 1 | -7977.388 | -7979.306 | -1.918 | 5.68E-07 |
| 11 | 118019906 | rs642936 | C | T | 0.996 | -7977.388 | -7979.306 | -1.918 | 5.68E-07 |
| 11 | 119411610 | rs4936477 | G | A | 0.998 | -7977.388 | -7979.307 | -1.918 | 5.68E-07 |
| 11 | 119622162 | rs11217460 | G | C | 0.994 | -7977.388 | -7979.307 | -1.918 | 5.68E-07 |
| 11 | 119653947 | rs7942259 | T | C | 0.994 | -7977.388 | -7979.307 | -1.918 | 5.68E-07 |
| 11 | 119658393 | rs10790346 | G | C | 0.994 | -7977.388 | -7979.307 | -1.918 | 5.68E-07 |
| 11 | 119680923 | rs4936498 | G | T | 1.005 | -7977.388 | -7979.306 | -1.918 | 5.68E-07 |
| 11 | 119719420 | rs1530604 | G | A | 1.006 | -7977.388 | -7979.306 | -1.918 | 5.68E-07 |
| 11 | 119902691 | rs11605919 | C | T | 0.995 | -7977.388 | -7979.306 | -1.918 | 5.68E-07 |
| 11 | 120543781 | rs4417279 | T | G | 1.006 | -7977.388 | -7979.306 | -1.918 | 5.68E-07 |
| 11 | 116489629 | rs548207 | G | T | 0.999 | -7977.388 | -7979.308 | -1.919 | 5.68E-07 |
| 11 | 117814506 | rs10160827 | T | C | 0.995 | -7977.388 | -7979.307 | -1.919 | 5.68E-07 |
| 11 | 118146510 | rs58507982 | T | C | 1.004 | -7977.388 | -7979.307 | -1.919 | 5.68E-07 |
| 11 | 119547005 | rs7122134 | G | A | 0.996 | -7977.388 | -7979.307 | -1.919 | 5.68E-07 |
| 11 | 119639398 | rs10790344 | G | A | 0.994 | -7977.388 | -7979.307 | -1.919 | 5.68E-07 |
| 11 | 116487699 | rs550767 | A | C | 1 | -7977.388 | -7979.308 | -1.92 | 5.67E-07 |
| 11 | 118010890 | rs640323 | C | G | 1 | -7977.388 | -7979.308 | -1.92 | 5.67E-07 |
| 11 | 119488176 | rs2136419 | T | C | 0.999 | -7977.388 | -7979.308 | -1.92 | 5.67E-07 |
| 11 | 119564781 | rs7124934 | G | A | 1.003 | -7977.388 | -7979.308 | -1.92 | 5.67E-07 |
| 11 | 120057343 | rs508205 | G | A | 0.997 | -7977.388 | -7979.308 | -1.92 | 5.67E-07 |
| 11 | 120585300 | rs1939664 | A | T | 0.994 | -7977.388 | -7979.308 | -1.92 | 5.67E-07 |
| 11 | 116464955 | rs11216027 | T | C | 0.997 | -7977.388 | -7979.309 | -1.921 | 5.67E-07 |
| 11 | 117988784 | rs8924 | A | G | 1.001 | -7977.388 | -7979.309 | -1.921 | 5.67E-07 |
| 11 | 119627450 | rs4351828 | A | G | 1.005 | -7977.388 | -7979.31 | -1.921 | 5.67E-07 |
| 11 | 119978334 | rs1349689 | T | A | 1.004 | -7977.388 | -7979.309 | -1.921 | 5.67E-07 |
| 11 | 120058685 | rs12278345 | G | T | 0.998 | -7977.388 | -7979.309 | -1.921 | 5.67E-07 |
| 11 | 116577043 | rs1263055 | A | G | 0.996 | -7977.388 | -7979.31 | -1.922 | 5.66E-07 |
| 11 | 119620940 | rs11217458 | A | G | 0.995 | -7977.388 | -7979.311 | -1.922 | 5.66E-07 |
| 11 | 121085567 | rs508316 | C | A | 1.004 | -7977.388 | -7979.31 | -1.922 | 5.66E-07 |
| 11 | 116316777 | rs489164 | C | T | 1.004 | -7977.388 | -7979.312 | -1.923 | 5.65E-07 |
| 11 | 118004688 | rs3741315 | A | G | 1.001 | -7977.388 | -7979.312 | -1.923 | 5.65E-07 |
| 11 | 119621691 | rs7949076 | T | C | 0.995 | -7977.388 | -7979.312 | -1.923 | 5.65E-07 |
| 11 | 119655589 | rs7936400 | C | T | 0.995 | -7977.388 | -7979.311 | -1.923 | 5.65E-07 |
| 11 | 119943275 | rs628944 | A | C | 0.997 | -7977.388 | -7979.312 | -1.923 | 5.65E-07 |
| 11 | 120550286 | rs12574186 | G | A | 0.995 | -7977.388 | -7979.311 | -1.923 | 5.65E-07 |
| 11 | 120584018 | rs10736504 | A | G | 0.995 | -7977.388 | -7979.311 | -1.923 | 5.65E-07 |
| 11 | 120878916 | rs11218108 | T | C | 0.995 | -7977.388 | -7979.312 | -1.923 | 5.65E-07 |
| 11 | 118016273 | rs1715444 | G | T | 0.998 | -7977.388 | -7979.312 | -1.924 | 5.65E-07 |
| 11 | 118021740 | rs678262 | G | C | 1.004 | -7977.388 | -7979.312 | -1.924 | 5.65E-07 |
| 11 | 119644451 | rs9326253 | C | T | 0.996 | -7977.388 | -7979.312 | -1.924 | 5.65E-07 |
| 11 | 119727452 | rs6589781 | T | A | 0.999 | -7977.388 | -7979.312 | -1.924 | 5.65E-07 |
| 11 | 120989335 | rs612969 | G | A | 1.002 | -7977.388 | -7979.313 | -1.924 | 5.65E-07 |
| 11 | 116207302 | rs12420446 | A | G | 1.002 | -7977.388 | -7979.314 | -1.925 | 5.64E-07 |
| 11 | 117473833 | rs1263889 | A | C | 0.997 | -7977.388 | -7979.313 | -1.925 | 5.64E-07 |
| 11 | 118139403 | rs59049611 | C | T | 1.003 | -7977.388 | -7979.313 | -1.925 | 5.64E-07 |
| 11 | 119777611 | rs509809 | G | A | 0.997 | -7977.388 | -7979.314 | -1.925 | 5.64E-07 |
| 11 | 120026271 | rs470373 | T | C | 1.001 | -7977.388 | -7979.313 | -1.925 | 5.64E-07 |
| 11 | 120028202 | rs470606 | G | T | 1.002 | -7977.388 | -7979.314 | -1.925 | 5.64E-07 |
| 11 | 117317124 | rs7125517 | C | A | 0.996 | -7977.388 | -7979.314 | -1.926 | 5.64E-07 |
| 11 | 118071097 | rs1626649 | T | C | 0.997 | -7977.388 | -7979.315 | -1.926 | 5.64E-07 |
| 11 | 118961231 | rs494048 | T | C | 0.996 | -7977.388 | -7979.315 | -1.926 | 5.64E-07 |
| 11 | 119696165 | rs477759 | T | C | 1.004 | -7977.388 | -7979.315 | -1.926 | 5.64E-07 |
| 11 | 119773032 | rs486876 | A | G | 0.997 | -7977.388 | -7979.315 | -1.926 | 5.64E-07 |
| 11 | 119786235 | rs529482 | G | A | 1.004 | -7977.388 | -7979.315 | -1.926 | 5.64E-07 |
| 11 | 120542632 | rs7935000 | A | G | 0.996 | -7977.388 | -7979.315 | -1.926 | 5.64E-07 |
| 11 | 120551419 | rs10790395 | A | C | 1.005 | -7977.388 | -7979.314 | -1.926 | 5.64E-07 |
| 11 | 120569381 | rs12574240 | C | T | 1.002 | -7977.388 | -7979.314 | -1.926 | 5.64E-07 |
| 11 | 121087285 | rs653478 | A | T | 1.003 | -7977.388 | -7979.314 | -1.926 | 5.64E-07 |
| 11 | 117335074 | rs477890 | T | A | 1.003 | -7977.388 | -7979.315 | -1.927 | 5.63E-07 |
| 11 | 117693255 | rs529623 | T | C | 1.004 | -7977.388 | -7979.315 | -1.927 | 5.63E-07 |
| 11 | 117954724 | rs3949025 | A | G | 0.997 | -7977.388 | -7979.316 | -1.927 | 5.63E-07 |
| 11 | 119785065 | rs472446 | G | A | 0.998 | -7977.388 | -7979.315 | -1.927 | 5.63E-07 |
| 11 | 120056416 | rs2444239 | A | T | 1 | -7977.388 | -7979.316 | -1.927 | 5.63E-07 |
| 11 | 120549404 | rs7119539 | G | A | 1.005 | -7977.388 | -7979.315 | -1.927 | 5.63E-07 |

| | | | | | | | |
|---|---|---|---|---|---|---|---|
| 11 | 120549745 | rs10892608 | A | G | NULL | 0.723121805 | NULL&M_Disease |
| 11 | 120551100 | rs10892609 | C | T | NULL | 0.723121805 | NULL&M_Disease |
| 11 | 120877650 | rs12419194 | A | G | NULL | 0.723121805 | NULL&M_Disease |
| 11 | 121082861 | rs679847 | G | A | NULL | 0.723121805 | NULL&M_Disease |
| 11 | 121083812 | rs665035 | T | C | NULL | 0.723121805 | NULL&M_Disease |
| 11 | 121084190 | rs681637 | C | T | NULL | 0.723121805 | NULL&M_Disease |
| 11 | 117779612 | rs947969 | A | G | NULL | 0.724121771 | NULL&M_Disease |
| 11 | 117996296 | rs7944957 | C | T | NULL | 0.724121771 | NULL&M_Disease |
| 11 | 118073344 | rs17121871 | T | C | NULL | 0.724121771 | NULL&M_Disease |
| 11 | 118194571 | rs10892234 | G | A | NULL | 0.724121771 | NULL&M_Disease |
| 11 | 118986630 | rs608444 | G | A | NULL | 0.724121771 | NULL&M_Disease |
| 11 | 119900629 | rs680472 | G | C | NULL | 0.724121771 | NULL&M_Disease |
| 11 | 119939826 | rs591744 | G | A | NULL | 0.724121771 | NULL&M_Disease |
| 11 | 120053650 | rs494965 | T | C | NULL | 0.724121771 | NULL&M_Disease |
| 11 | 121155632 | rs10892717 | G | C | NULL | 0.724121771 | NULL&M_Disease |
| 11 | 117502518 | rs36108816 | C | G | NULL | 0.724121771 | NULL&M_Disease |
| 11 | 118076069 | rs1805 | T | G | NULL | 0.724121771 | NULL&M_Disease |
| 11 | 118076954 | rs11600544 | C | G | NULL | 0.724121771 | NULL&M_Disease |
| 11 | 118079867 | rs4938489 | G | T | NULL | 0.724121771 | NULL&M_Disease |
| 11 | 118080133 | rs17121879 | G | T | NULL | 0.724121771 | NULL&M_Disease |
| 11 | 118080977 | rs4938490 | G | A | NULL | 0.724121771 | NULL&M_Disease |
| 11 | 118081345 | rs17121881 | A | T | NULL | 0.724121771 | NULL&M_Disease |
| 11 | 118081468 | rs17121882 | C | T | NULL | 0.724121771 | NULL&M_Disease |
| 11 | 119648005 | rs4418818 | A | G | NULL | 0.724121771 | NULL&M_Disease |
| 11 | 119648306 | rs4301801 | G | C | NULL | 0.724121771 | NULL&M_Disease |
| 11 | 119648368 | rs4269933 | G | A | NULL | 0.724121771 | NULL&M_Disease |
| 11 | 119648458 | rs4408323 | C | A | NULL | 0.724121771 | NULL&M_Disease |
| 11 | 119784660 | rs475724 | A | G | NULL | 0.724121771 | NULL&M_Disease |
| 11 | 119938682 | rs592979 | G | A | NULL | 0.724121771 | NULL&M_Disease |
| 11 | 119255795 | rs10892358 | G | A | NULL | 0.724121771 | NULL&M_Disease |
| 11 | 119650105 | rs7103360 | G | A | NULL | 0.724121771 | NULL&M_Disease |
| 11 | 119937400 | rs625573 | C | G | NULL | 0.724121771 | NULL&M_Disease |
| 11 | 120872976 | rs11218105 | G | A | NULL | 0.724121771 | NULL&M_Disease |
| 11 | 120877786 | rs1944516 | T | C | NULL | 0.724121771 | NULL&M_Disease |
| 11 | 119659720 | rs10892464 | C | T | NULL | 0.724121771 | NULL&M_Disease |
| 11 | 117511951 | rs657882 | C | G | NULL | 0.724121771 | NULL&M_Disease |
| 11 | 119633863 | rs7110679 | A | G | NULL | 0.724121771 | NULL&M_Disease |
| 11 | 119984535 | rs519672 | G | C | NULL | 0.724121771 | NULL&M_Disease |
| 11 | 119984863 | rs543584 | C | T | NULL | 0.724121771 | NULL&M_Disease |
| 11 | 116391537 | rs11215978 | C | T | NULL | 0.724121771 | NULL&M_Disease |
| 11 | 119185347 | rs6589732 | A | G | NULL | 0.724121771 | NULL&M_Disease |
| 11 | 119616190 | rs7109251 | A | G | NULL | 0.724121771 | NULL&M_Disease |
| 11 | 119642827 | rs10892454 | G | T | NULL | 0.724121771 | NULL&M_Disease |
| 11 | 119656449 | rs6589776 | A | G | NULL | 0.724121771 | NULL&M_Disease |
| 11 | 120557091 | rs10790400 | T | A | NULL | 0.724121771 | NULL&M_Disease |
| 11 | 120828855 | rs732481 | G | T | NULL | 0.724121771 | NULL&M_Disease |
| 11 | 119653781 | rs1466052 | C | T | NULL | 0.724121771 | NULL&M_Disease |
| 11 | 119653883 | rs10892459 | G | C | NULL | 0.724121771 | NULL&M_Disease |
| 11 | 119986167 | rs503559 | T | C | NULL | 0.724121771 | NULL&M_Disease |
| 11 | 120544816 | rs2000868 | T | C | NULL | 0.724121771 | NULL&M_Disease |
| 11 | 120550222 | rs35101469 | G | T | NULL | 0.724121771 | NULL&M_Disease |
| 11 | 118983681 | rs682795 | C | T | NULL | 0.724121771 | NULL&M_Disease |
| 11 | 119624823 | rs10892448 | A | C | NULL | 0.724121771 | NULL&M_Disease |
| 11 | 119624993 | rs11217466 | A | T | NULL | 0.724121771 | NULL&M_Disease |
| 11 | 119654058 | rs10892460 | C | T | NULL | 0.724121771 | NULL&M_Disease |
| 11 | 120869570 | rs11820008 | T | C | NULL | 0.724121771 | NULL&M_Disease |
| 11 | 119650848 | rs4331115 | T | C | NULL | 0.724121771 | NULL&M_Disease |
| 11 | 119652214 | rs5026369 | C | G | NULL | 0.724121771 | NULL&M_Disease |
| 11 | 119652686 | rs11217501 | C | T | NULL | 0.724121771 | NULL&M_Disease |
| 11 | 119653461 | rs1466051 | C | T | NULL | 0.724121771 | NULL&M_Disease |
| 11 | 119654208 | rs11606569 | C | T | NULL | 0.724121771 | NULL&M_Disease |
| 11 | 119654326 | rs35662046 | C | T | NULL | 0.724121771 | NULL&M_Disease |
| 11 | 119659753 | rs10892465 | C | T | NULL | 0.724121771 | NULL&M_Disease |
| 11 | 120865621 | rs11218096 | T | G | NULL | 0.724121771 | NULL&M_Disease |
| 11 | 120867855 | rs11218101 | A | G | NULL | 0.724121771 | NULL&M_Disease |
| 11 | 120868460 | rs11218102 | C | G | NULL | 0.724121771 | NULL&M_Disease |
| 11 | 120871452 | rs12799836 | G | A | NULL | 0.724121771 | NULL&M_Disease |
| 11 | 120872094 | rs4936571 | G | A | NULL | 0.724121771 | NULL&M_Disease |
| 11 | 120872214 | rs4936572 | A | G | NULL | 0.724121771 | NULL&M_Disease |
| 11 | 117324522 | rs10892113 | T | C | NULL | 0.725119498 | NULL&M_Disease |
| 11 | 118084980 | rs11216819 | G | A | NULL | 0.725119498 | NULL&M_Disease |
| 11 | 119652763 | rs11217502 | A | G | NULL | 0.725119498 | NULL&M_Disease |
| 11 | 119654807 | rs10892461 | C | T | NULL | 0.725119498 | NULL&M_Disease |
| 11 | 120549488 | rs7119655 | G | A | NULL | 0.725119498 | NULL&M_Disease |
| 11 | 120864922 | rs1944522 | T | C | NULL | 0.725119498 | NULL&M_Disease |
| 11 | 120865697 | rs11218097 | T | G | NULL | 0.725119498 | NULL&M_Disease |
| 11 | 120867747 | rs11218100 | T | C | NULL | 0.725119498 | NULL&M_Disease |
| 11 | 117323834 | rs11216400 | G | A | NULL | 0.725119498 | NULL&M_Disease |
| 11 | 119641049 | rs7926574 | C | T | NULL | 0.725119498 | NULL&M_Disease |
| 11 | 119794035 | rs4936506 | T | C | NULL | 0.725119498 | NULL&M_Disease |
| 11 | 119956652 | rs3915646 | C | T | NULL | 0.725119498 | NULL&M_Disease |
| 11 | 119958632 | rs7109733 | C | T | NULL | 0.725119498 | NULL&M_Disease |
| 11 | 120566535 | rs7126415 | C | G | NULL | 0.725119498 | NULL&M_Disease |
| 11 | 120585993 | rs3132775 | G | A | NULL | 0.725119498 | NULL&M_Disease |
| 11 | 119059404 | rs1815811 | G | A | NULL | 0.725119498 | NULL&M_Disease |
| 11 | 120550229 | rs11217968 | G | A | NULL | 0.725119498 | NULL&M_Disease |
| 11 | 120588584 | rs3132784 | C | G | NULL | 0.725119498 | NULL&M_Disease |
| 11 | 120550430 | rs7932158 | C | T | NULL | 0.725119498 | NULL&M_Disease |
| 11 | 118986659 | rs608793 | C | T | NULL | 0.725119498 | NULL&M_Disease |
| 11 | 118973133 | rs10790282 | G | A | NULL | 0.725119498 | NULL&M_Disease |
| 11 | 118981025 | rs617948 | A | G | NULL | 0.725119498 | NULL&M_Disease |

| | | | | | | | | | |
|---|---|---|---|---|---|---|---|---|---|
| 11 | 120549745 | rs10892608 | A | G | 1.005 | -7977.388 | -7979.315 | -1.927 | 5.63E-07 |
| 11 | 120551100 | rs10892609 | C | T | 1.005 | -7977.388 | -7979.316 | -1.927 | 5.63E-07 |
| 11 | 120877650 | rs12419194 | A | G | 0.996 | -7977.388 | -7979.315 | -1.927 | 5.63E-07 |
| 11 | 121082861 | rs679847 | G | A | 1.002 | -7977.388 | -7979.315 | -1.927 | 5.63E-07 |
| 11 | 121083812 | rs665035 | T | C | 1.002 | -7977.388 | -7979.315 | -1.927 | 5.63E-07 |
| 11 | 121084190 | rs681637 | C | T | 1.002 | -7977.388 | -7979.315 | -1.927 | 5.63E-07 |
| 11 | 117779612 | rs947969 | A | G | 0.999 | -7977.388 | -7979.317 | -1.928 | 5.63E-07 |
| 11 | 117996296 | rs7944957 | C | T | 1.003 | -7977.388 | -7979.317 | -1.928 | 5.63E-07 |
| 11 | 118073344 | rs17121871 | T | C | 0.998 | -7977.388 | -7979.317 | -1.928 | 5.63E-07 |
| 11 | 118194571 | rs10892234 | G | A | 1.003 | -7977.388 | -7979.316 | -1.928 | 5.63E-07 |
| 11 | 118986630 | rs608444 | G | A | 0.996 | -7977.388 | -7979.316 | -1.928 | 5.63E-07 |
| 11 | 119900629 | rs680472 | G | C | 1.003 | -7977.388 | -7979.316 | -1.928 | 5.63E-07 |
| 11 | 119939826 | rs591744 | G | A | 1.002 | -7977.388 | -7979.317 | -1.928 | 5.63E-07 |
| 11 | 120053650 | rs494965 | T | C | 1.001 | -7977.388 | -7979.316 | -1.928 | 5.63E-07 |
| 11 | 121155632 | rs10892717 | G | C | 1.001 | -7977.388 | -7979.317 | -1.928 | 5.63E-07 |
| 11 | 117502518 | rs36108816 | C | G | 0.997 | -7977.388 | -7979.317 | -1.929 | 5.62E-07 |
| 11 | 118076069 | rs1805 | T | G | 0.998 | -7977.388 | -7979.317 | -1.929 | 5.62E-07 |
| 11 | 118076954 | rs11600544 | C | G | 0.998 | -7977.388 | -7979.317 | -1.929 | 5.62E-07 |
| 11 | 118079867 | rs4938489 | G | T | 0.998 | -7977.388 | -7979.317 | -1.929 | 5.62E-07 |
| 11 | 118080133 | rs17121879 | G | T | 0.998 | -7977.388 | -7979.317 | -1.929 | 5.62E-07 |
| 11 | 118080977 | rs4938490 | G | A | 0.998 | -7977.388 | -7979.317 | -1.929 | 5.62E-07 |
| 11 | 118081345 | rs17121881 | A | T | 0.998 | -7977.388 | -7979.317 | -1.929 | 5.62E-07 |
| 11 | 118081468 | rs17121882 | C | T | 0.998 | -7977.388 | -7979.317 | -1.929 | 5.62E-07 |
| 11 | 119648005 | rs4418818 | A | G | 0.997 | -7977.388 | -7979.317 | -1.929 | 5.62E-07 |
| 11 | 119648306 | rs4301801 | G | C | 0.997 | -7977.388 | -7979.317 | -1.929 | 5.62E-07 |
| 11 | 119648368 | rs4269933 | G | A | 0.997 | -7977.388 | -7979.317 | -1.929 | 5.62E-07 |
| 11 | 119648458 | rs4408323 | C | A | 0.997 | -7977.388 | -7979.317 | -1.929 | 5.62E-07 |
| 11 | 119784660 | rs475724 | A | G | 0.998 | -7977.388 | -7979.317 | -1.929 | 5.62E-07 |
| 11 | 119938682 | rs592979 | G | A | 1.001 | -7977.388 | -7979.318 | -1.929 | 5.62E-07 |
| 11 | 119255795 | rs10892358 | G | A | 1 | -7977.388 | -7979.319 | -1.93 | 5.61E-07 |
| 11 | 119650105 | rs7103360 | G | A | 0.998 | -7977.388 | -7979.318 | -1.93 | 5.61E-07 |
| 11 | 119937400 | rs625573 | C | G | 1 | -7977.388 | -7979.318 | -1.93 | 5.61E-07 |
| 11 | 120872976 | rs11218105 | G | A | 1.004 | -7977.388 | -7979.318 | -1.93 | 5.61E-07 |
| 11 | 120877786 | rs1944516 | T | C | 1.003 | -7977.388 | -7979.319 | -1.93 | 5.61E-07 |
| 11 | 119659720 | rs10892464 | C | T | 0.997 | -7977.388 | -7979.319 | -1.931 | 5.61E-07 |
| 11 | 117511951 | rs657882 | C | G | 1 | -7977.388 | -7979.32 | -1.932 | 5.60E-07 |
| 11 | 119633863 | rs7110679 | A | G | 0.997 | -7977.388 | -7979.321 | -1.932 | 5.60E-07 |
| 11 | 119984535 | rs519672 | G | C | 0.997 | -7977.388 | -7979.321 | -1.932 | 5.60E-07 |
| 11 | 119984863 | rs543584 | C | T | 0.997 | -7977.388 | -7979.321 | -1.932 | 5.60E-07 |
| 11 | 116391537 | rs11215978 | C | T | 1.003 | -7977.388 | -7979.321 | -1.933 | 5.60E-07 |
| 11 | 119185347 | rs6589732 | A | G | 1.002 | -7977.388 | -7979.321 | -1.933 | 5.60E-07 |
| 11 | 119616190 | rs7109251 | A | G | 0.998 | -7977.388 | -7979.322 | -1.934 | 5.59E-07 |
| 11 | 119642827 | rs10892454 | G | T | 1.001 | -7977.388 | -7979.322 | -1.934 | 5.59E-07 |
| 11 | 119656449 | rs6589776 | A | G | 0.997 | -7977.388 | -7979.323 | -1.934 | 5.59E-07 |
| 11 | 120557091 | rs10790400 | T | A | 1 | -7977.388 | -7979.323 | -1.934 | 5.59E-07 |
| 11 | 120828855 | rs732481 | G | T | 0.997 | -7977.388 | -7979.322 | -1.934 | 5.59E-07 |
| 11 | 119653781 | rs1466052 | C | T | 0.998 | -7977.388 | -7979.324 | -1.935 | 5.59E-07 |
| 11 | 119653883 | rs10892459 | G | C | 0.998 | -7977.388 | -7979.324 | -1.935 | 5.59E-07 |
| 11 | 119986167 | rs503559 | T | C | 0.998 | -7977.388 | -7979.324 | -1.935 | 5.59E-07 |
| 11 | 120544816 | rs2000868 | T | C | 0.997 | -7977.388 | -7979.324 | -1.935 | 5.59E-07 |
| 11 | 120550222 | rs35101469 | G | T | 1.003 | -7977.388 | -7979.323 | -1.935 | 5.59E-07 |
| 11 | 118983681 | rs682795 | C | T | 1.002 | -7977.388 | -7979.324 | -1.936 | 5.58E-07 |
| 11 | 119624823 | rs10892448 | A | C | 0.998 | -7977.388 | -7979.325 | -1.936 | 5.58E-07 |
| 11 | 119624993 | rs11217466 | A | T | 1.002 | -7977.388 | -7979.324 | -1.936 | 5.58E-07 |
| 11 | 119654058 | rs10892460 | C | T | 0.998 | -7977.388 | -7979.325 | -1.936 | 5.58E-07 |
| 11 | 120869570 | rs11820008 | T | C | 0.998 | -7977.388 | -7979.324 | -1.936 | 5.58E-07 |
| 11 | 119650848 | rs4331115 | T | C | 0.998 | -7977.388 | -7979.325 | -1.937 | 5.58E-07 |
| 11 | 119652214 | rs5026369 | C | G | 0.998 | -7977.388 | -7979.325 | -1.937 | 5.58E-07 |
| 11 | 119652686 | rs11217501 | C | T | 0.998 | -7977.388 | -7979.325 | -1.937 | 5.58E-07 |
| 11 | 119653461 | rs1466051 | C | T | 0.998 | -7977.388 | -7979.325 | -1.937 | 5.58E-07 |
| 11 | 119654208 | rs11606569 | C | T | 0.998 | -7977.388 | -7979.325 | -1.937 | 5.58E-07 |
| 11 | 119654326 | rs35662046 | C | T | 0.998 | -7977.388 | -7979.325 | -1.937 | 5.58E-07 |
| 11 | 119659753 | rs10892465 | C | T | 1.001 | -7977.388 | -7979.325 | -1.937 | 5.58E-07 |
| 11 | 120865621 | rs11218096 | T | G | 1.001 | -7977.388 | -7979.326 | -1.937 | 5.58E-07 |
| 11 | 120867855 | rs11218101 | A | G | 0.999 | -7977.388 | -7979.325 | -1.937 | 5.58E-07 |
| 11 | 120868460 | rs11218102 | C | G | 1 | -7977.388 | -7979.326 | -1.937 | 5.58E-07 |
| 11 | 120871452 | rs12799836 | G | A | 1.001 | -7977.388 | -7979.325 | -1.937 | 5.58E-07 |
| 11 | 120872094 | rs4936571 | G | A | 0.999 | -7977.388 | -7979.325 | -1.937 | 5.58E-07 |
| 11 | 120872214 | rs4936572 | A | G | 0.999 | -7977.388 | -7979.325 | -1.937 | 5.58E-07 |
| 11 | 117324522 | rs10892113 | T | C | 0.999 | -7977.388 | -7979.327 | -1.938 | 5.57E-07 |
| 11 | 118084980 | rs11216819 | G | A | 0.999 | -7977.388 | -7979.326 | -1.938 | 5.57E-07 |
| 11 | 119652763 | rs11217502 | A | G | 0.999 | -7977.388 | -7979.326 | -1.938 | 5.57E-07 |
| 11 | 119654807 | rs10892461 | C | T | 0.999 | -7977.388 | -7979.326 | -1.938 | 5.57E-07 |
| 11 | 120549488 | rs7119655 | G | A | 1.002 | -7977.388 | -7979.326 | -1.938 | 5.57E-07 |
| 11 | 120864922 | rs1944522 | T | C | 1 | -7977.388 | -7979.326 | -1.938 | 5.57E-07 |
| 11 | 120865697 | rs11218097 | T | G | 1 | -7977.388 | -7979.326 | -1.938 | 5.57E-07 |
| 11 | 120867747 | rs11218100 | T | C | 1 | -7977.388 | -7979.326 | -1.938 | 5.57E-07 |
| 11 | 117323834 | rs11216400 | G | A | 0.999 | -7977.388 | -7979.327 | -1.939 | 5.56E-07 |
| 11 | 119641049 | rs7926574 | C | T | 1 | -7977.388 | -7979.327 | -1.939 | 5.56E-07 |
| 11 | 119794035 | rs4936506 | T | C | 1.002 | -7977.388 | -7979.327 | -1.939 | 5.56E-07 |
| 11 | 119956652 | rs3915646 | C | T | 0.999 | -7977.388 | -7979.327 | -1.939 | 5.56E-07 |
| 11 | 119958632 | rs7109733 | C | T | 0.999 | -7977.388 | -7979.327 | -1.939 | 5.56E-07 |
| 11 | 120566535 | rs7126415 | C | G | 1 | -7977.388 | -7979.327 | -1.939 | 5.56E-07 |
| 11 | 120585993 | rs3132775 | G | A | 1 | -7977.388 | -7979.327 | -1.939 | 5.56E-07 |
| 11 | 119059404 | rs1815811 | G | A | 1.002 | -7977.388 | -7979.328 | -1.94 | 5.56E-07 |
| 11 | 120550229 | rs11217968 | G | A | 1.001 | -7977.388 | -7979.329 | -1.94 | 5.56E-07 |
| 11 | 120588584 | rs3132784 | C | G | 0.999 | -7977.388 | -7979.329 | -1.94 | 5.56E-07 |
| 11 | 120550430 | rs7932158 | C | T | 1.001 | -7977.388 | -7979.329 | -1.941 | 5.55E-07 |
| 11 | 118986659 | rs608793 | C | T | 0.997 | -7977.388 | -7979.33 | -1.942 | 5.55E-07 |
| 11 | 118973133 | rs10790282 | G | A | 1.002 | -7977.388 | -7979.332 | -1.944 | 5.54E-07 |
| 11 | 118981025 | rs617948 | A | G | 0.998 | -7977.388 | -7979.333 | -1.945 | 5.53E-07 |

**Supplemental Data 5: Bayesian analysis of SjD Immunochip data in DDX6-CXCR5 locus (Dataset 3).**

Note: CHROM=chromosome; POS=basepair position; A1=major allele; A2=minor allele; OR=odd ratio; L+P0=The log null marginal likelihood; L+P1=The log alternative marginal likelihood; BF=Bayes factor; P_CAUSAL=The probability that this variant is the causal variant at this locus;

### CREDIBLE SET

| CHROM | POS | RSID | A1 | A2 | BestModel | BestModelProb | CredibleSet |
|---|---|---|---|---|---|---|---|
| 11 | 118724894 | rs9736016 | A | T | M_Disease | 0.99645971 | M_Disease |
| 11 | 118729391 | rs10790268 | A | G | M_Disease | 0.996370412 | M_Disease |
| 11 | 118729456 | rs10790269 | C | T | M_Disease | 0.996370412 | M_Disease |
| 11 | 118739056 | rs7122669 | A | G | M_Disease | 0.996370412 | M_Disease |
| 11 | 118721699 | rs7125066 | G | A | M_Disease | 0.996352285 | M_Disease |
| 11 | 118738298 | rs7119044 | G | C | M_Disease | 0.996352285 | M_Disease |
| 11 | 118734000 | rs11217074 | C | T | M_Disease | 0.995929862 | M_Disease |
| 11 | 118741842 | rs4938573 | C | T | M_Disease | 0.995806428 | M_Disease |
| 11 | 118740418 | rs7951740 | G | C | M_Disease | 0.995785496 | M_Disease |
| 11 | 118743286 | rs12365699 | A | G | M_Disease | 0.995570371 | M_Disease |
| 11 | 118741157 | rs7117261 | T | C | M_Disease | 0.995458744 | M_Disease |
| 11 | 118743338 | rs4936444 | T | C | M_Disease | 0.995390428 | M_Disease |
| 11 | 118743772 | rs6421571 | T | C | M_Disease | 0.99520321 | M_Disease |
| 11 | 118680648 | rs11217040 | A | C | M_Disease | 0.995106783 | M_Disease |
| 11 | 118683564 | rs17122453 | A | G | M_Disease | 0.995082376 | M_Disease |
| 11 | 118609790 | rs11217006 | A | G | M_Disease | 0.994933371 | M_Disease |
| 11 | 118681464 | rs7942535 | C | T | M_Disease | 0.994908103 | M_Disease |
| 11 | 118682350 | rs11217041 | G | A | M_Disease | 0.994908103 | M_Disease |
| 11 | 118682528 | rs11217042 | A | G | M_Disease | 0.994908103 | M_Disease |
| 11 | 118681083 | rs73005426 | G | A | M_Disease | 0.994831547 | M_Disease |
| 11 | 118740864 | rs4936443 | C | T | M_Disease | 0.994648433 | M_Disease |
| 11 | 118742800 | rs74541740 | A | G | M_Disease | 0.994648433 | M_Disease |
| 11 | 118696022 | rs11217044 | C | T | M_Disease | 0.994513701 | M_Disease |
| 11 | 118721565 | rs75249660 | A | G | M_Disease | 0.994403491 | M_Disease |
| 11 | 118745243 | rs7481797 | A | G | M_Disease | 0.994319391 | M_Disease |
| 11 | 118740931 | rs4938572 | C | T | M_Disease | 0.9942053 | M_Disease |
| 11 | 118745278 | rs7481819 | C | T | M_Disease | 0.994000133 | M_Disease |
| 11 | 118611781 | rs10892279 | A | G | M_Disease | 0.993910009 | M_Disease |
| 11 | 118610957 | rs6589684 | A | G | M_Disease | 0.993879669 | M_Disease |
| 11 | 118722149 | rs11217058 | A | G | M_Disease | 0.99384918 | M_Disease |
| 11 | 118725660 | rs4936441 | C | G | M_Disease | 0.993818539 | M_Disease |
| 11 | 118726843 | rs1037679725 | T | C | M_Disease | 0.993471317 | M_Disease |
| 11 | 118694547 | rs7123726 | C | T | M_Disease | 0.993438807 | M_Disease |
| 11 | 118701018 | rs79504890 | A | G | M_Disease | 0.993438807 | M_Disease |
| 11 | 118738281 | rs7119038 | G | A | M_Disease | 0.993340308 | M_Disease |
| 11 | 118667357 | rs10892294 | C | G | M_Disease | 0.993273827 | M_Disease |
| 11 | 118649512 | rs73003215 | A | G | M_Disease | 0.992931406 | M_Disease |
| 11 | 118665575 | rs10892293 | T | A | M_Disease | 0.992931406 | M_Disease |
| 11 | 118726753 | rs11217066 | G | T | M_Disease | 0.992860873 | M_Disease |
| 11 | 118612197 | rs6589685 | C | T | M_Disease | 0.992825344 | M_Disease |
| 11 | 118684610 | rs715412 | A | G | M_Disease | 0.992534727 | M_Disease |
| 11 | 118679304 | rs28409215 | A | G | M_Disease | 0.992460265 | M_Disease |
| 11 | 118698907 | rs11217046 | A | G | M_Disease | 0.992422759 | M_Disease |
| 11 | 118681079 | rs73005423 | G | A | M_Disease | 0.992193799 | M_Disease |
| 11 | 118697865 | rs11217045 | A | C | M_Disease | 0.992154977 | M_Disease |
| 11 | 118697906 | rs73005502 | G | A | M_Disease | 0.992154977 | M_Disease |
| 11 | 118669605 | rs11217032 | A | T | M_Disease | 0.991877809 | M_Disease |
| 11 | 118673706 | rs11217033 | C | T | M_Disease | 0.991877809 | M_Disease |
| 11 | 118675772 | rs11217036 | T | C | M_Disease | 0.991877809 | M_Disease |
| 11 | 118677086 | rs11217037 | A | G | M_Disease | 0.991877809 | M_Disease |
| 11 | 118677355 | rs11217038 | A | G | M_Disease | 0.991877809 | M_Disease |
| 11 | 118611817 | rs10892280 | T | C | M_Disease | 0.99146493 | M_Disease |
| 11 | 118642085 | rs10892286 | C | A | M_Disease | 0.99146493 | M_Disease |
| 11 | 118642999 | rs10892287 | C | T | M_Disease | 0.99146493 | M_Disease |
| 11 | 118734521 | rs1174717189 | T | A | M_Disease | 0.991337056 | M_Disease |
| 11 | 118653503 | rs10892290 | T | G | M_Disease | 0.991294011 | M_Disease |
| 11 | 118653517 | rs10892291 | T | C | M_Disease | 0.991294011 | M_Disease |
| 11 | 118744396 | rs7117313 | C | G | M_Disease | 0.991250754 | M_Disease |
| 11 | 118644694 | rs10892288 | G | A | M_Disease | 0.991207284 | M_Disease |
| 11 | 118618776 | rs1048024 | T | A | M_Disease | 0.991075585 | M_Disease |
| 11 | 118619047 | rs2077579 | G | T | M_Disease | 0.991075585 | M_Disease |
| 11 | 118639353 | rs11217020 | A | G | M_Disease | 0.991075585 | M_Disease |
| 11 | 118636980 | rs10892283 | G | A | M_Disease | 0.991031252 | M_Disease |
| 11 | 118655645 | rs874621 | A | G | M_Disease | 0.990986701 | M_Disease |
| 11 | 118610549 | rs11826521 | T | G | M_Disease | 0.990851729 | M_Disease |
| 11 | 118615785 | rs11217009 | T | C | M_Disease | 0.990851729 | M_Disease |
| 11 | 118697012 | rs571470 | A | G | M_Disease | 0.990851729 | M_Disease |
| 11 | 118655340 | rs10892292 | C | G | M_Disease | 0.990806295 | M_Disease |
| 11 | 118646003 | rs10892289 | T | C | M_Disease | 0.990760637 | M_Disease |
| 11 | 118619253 | rs2004781 | T | C | M_Disease | 0.990622311 | M_Disease |
| 11 | 118619960 | rs3889239 | T | C | M_Disease | 0.990622311 | M_Disease |
| 11 | 118628373 | rs74676029 | T | C | M_Disease | 0.990622311 | M_Disease |
| 11 | 118636343 | rs11217019 | A | C | M_Disease | 0.990528956 | M_Disease |
| 11 | 118612921 | rs2155433 | A | G | M_Disease | 0.990387196 | M_Disease |
| 11 | 118648373 | rs56043232 | T | C | M_Disease | 0.990387196 | M_Disease |
| 11 | 118615340 | rs56758835 | A | G | M_Disease | 0.990339477 | M_Disease |
| 11 | 118729210 | rs10892300 | T | C | M_Disease | 0.990048198 | M_Disease |
| 11 | 118729860 | rs11217070 | G | T | M_Disease | 0.989998813 | M_Disease |
| 11 | 118745884 | rs10790275 | G | C | M_Disease | 0.989998813 | M_Disease |
| 11 | 118729583 | rs12790489 | C | T | M_Disease | 0.989386938 | M_Disease |
| 11 | 118730141 | rs7929520 | G | A | M_Disease | 0.989386938 | M_Disease |
| 11 | 118731599 | rs34561290 | T | C | M_Disease | 0.989386938 | M_Disease |
| 11 | 118731762 | rs11217072 | G | A | M_Disease | 0.989386938 | M_Disease |
| 11 | 118661398 | rs57494551 | T | C | M_Disease | 0.989334307 | M_Disease |
| 11 | 118608716 | rs7945144 | G | A | M_Disease | 0.988569797 | M_Disease |
| 11 | 118724723 | rs57719838 | C | T | M_Disease | 0.987444978 | M_Disease |
| 11 | 118725942 | rs11217063 | C | G | M_Disease | 0.986874682 | M_Disease |

### FINEMAP

| CHROM | POS | RSID | A1 | A2 | OR_Disease | L+P0 | L+P1 | logBF | P_CAUSAL |
|---|---|---|---|---|---|---|---|---|---|
| 11 | 118724894 | rs9736016 | A | T | 1.219 | -4194.66 | -4183.377 | 11.282 | 0.035815633 |
| 11 | 118739056 | rs7122669 | A | G | 1.27 | -4194.66 | -4183.433 | 11.227 | 0.033898965 |
| 11 | 118729391 | rs10790268 | A | G | 1.271 | -4194.66 | -4183.433 | 11.226 | 0.033865083 |
| 11 | 118729456 | rs10790269 | C | T | 1.271 | -4194.66 | -4183.433 | 11.226 | 0.033865083 |
| 11 | 118721699 | rs7125066 | G | A | 1.219 | -4194.66 | -4183.439 | 11.221 | 0.03369618 |
| 11 | 118738298 | rs7119044 | G | C | 1.27 | -4194.66 | -4183.442 | 11.217 | 0.033561664 |
| 11 | 118734000 | rs11217074 | C | T | 1.268 | -4194.66 | -4183.667 | 10.993 | 0.026826346 |
| 11 | 118741842 | rs4938573 | C | T | 1.267 | -4194.66 | -4183.72 | 10.939 | 0.025416142 |
| 11 | 118740418 | rs7951740 | G | C | 1.267 | -4194.66 | -4183.733 | 10.926 | 0.02508787 |
| 11 | 118743286 | rs12365699 | A | G | 1.286 | -4194.66 | -4183.83 | 10.829 | 0.022768647 |
| 11 | 118741157 | rs7117261 | T | C | 1.265 | -4194.66 | -4183.884 | 10.776 | 0.02159333 |
| 11 | 118743338 | rs4936444 | T | C | 1.266 | -4194.66 | -4183.913 | 10.747 | 0.020976116 |
| 11 | 118743772 | rs6421571 | T | C | 1.264 | -4194.66 | -4183.991 | 10.669 | 0.019402161 |
| 11 | 118680648 | rs11217040 | A | C | 1.258 | -4194.66 | -4184.031 | 10.629 | 0.018641392 |
| 11 | 118683564 | rs17122453 | A | G | 1.256 | -4194.66 | -4184.044 | 10.616 | 0.018400622 |
| 11 | 118609790 | rs11217006 | A | G | 1.254 | -4194.66 | -4184.105 | 10.554 | 0.01729443 |
| 11 | 118681464 | rs7942535 | C | T | 1.255 | -4194.66 | -4184.111 | 10.549 | 0.017208174 |
| 11 | 118682350 | rs11217041 | G | A | 1.255 | -4194.66 | -4184.111 | 10.549 | 0.017208174 |
| 11 | 118682528 | rs11217042 | A | G | 1.255 | -4194.66 | -4184.111 | 10.549 | 0.017208174 |
| 11 | 118681083 | rs73005426 | G | A | 1.255 | -4194.66 | -4184.143 | 10.517 | 0.016666229 |
| 11 | 118740864 | rs4936443 | C | T | 1.26 | -4194.66 | -4184.21 | 10.449 | 0.01557059 |
| 11 | 118742800 | rs74541740 | A | G | 1.261 | -4194.66 | -4184.216 | 10.443 | 0.015477455 |
| 11 | 118696022 | rs11217044 | C | T | 1.254 | -4194.66 | -4184.263 | 10.396 | 0.014766845 |
| 11 | 118721565 | rs75249660 | A | G | 1.259 | -4194.66 | -4184.306 | 10.354 | 0.014159482 |
| 11 | 118745243 | rs7481797 | A | G | 1.259 | -4194.66 | -4184.335 | 10.324 | 0.013741006 |
| 11 | 118740931 | rs4938572 | C | T | 1.258 | -4194.66 | -4184.376 | 10.283 | 0.013189017 |
| 11 | 118745278 | rs7481819 | C | T | 1.258 | -4194.66 | -4184.439 | 10.221 | 0.012396132 |
| 11 | 118611781 | rs10892279 | A | G | 1.25 | -4194.66 | -4184.467 | 10.192 | 0.012041807 |
| 11 | 118610957 | rs6589684 | A | G | 1.25 | -4194.66 | -4184.485 | 10.175 | 0.011838826 |
| 11 | 118722149 | rs11217058 | A | G | 1.256 | -4194.66 | -4184.495 | 10.164 | 0.011709313 |
| 11 | 118725660 | rs4936441 | C | G | 1.256 | -4194.66 | -4184.506 | 10.154 | 0.011592803 |
| 11 | 118726843 | rs1037679725 | T | C | 1.254 | -4194.66 | -4184.617 | 10.043 | 0.010374849 |
| 11 | 118694547 | rs7123726 | C | T | 1.25 | -4194.66 | -4184.618 | 10.042 | 0.010364479 |
| 11 | 118701018 | rs79504890 | A | G | 1.252 | -4194.66 | -4184.626 | 10.034 | 0.010281894 |
| 11 | 118738281 | rs7119038 | G | A | 1.239 | -4194.66 | -4184.651 | 10.009 | 0.010028033 |
| 11 | 118667357 | rs10892294 | C | G | 1.25 | -4194.66 | -4184.671 | 9.989 | 0.009829465 |
| 11 | 118649512 | rs73003215 | A | G | 1.247 | -4194.66 | -4184.772 | 9.888 | 0.008885178 |
| 11 | 118665575 | rs10892293 | T | A | 1.249 | -4194.66 | -4184.775 | 9.885 | 0.008858562 |
| 11 | 118726753 | rs11217066 | G | T | 1.252 | -4194.66 | -4184.792 | 9.868 | 0.00870924 |
| 11 | 118612197 | rs6589685 | C | T | 1.246 | -4194.66 | -4184.8 | 9.86 | 0.008639844 |
| 11 | 118684610 | rs715412 | A | G | 1.247 | -4194.66 | -4184.884 | 9.776 | 0.007943742 |
| 11 | 118679304 | rs28409215 | A | G | 1.247 | -4194.66 | -4184.899 | 9.761 | 0.007825475 |
| 11 | 118698907 | rs11217046 | A | G | 1.249 | -4194.66 | -4184.911 | 9.749 | 0.007732131 |
| 11 | 118681079 | rs73005423 | G | A | 1.246 | -4194.66 | -4184.975 | 9.684 | 0.007245528 |
| 11 | 118697865 | rs11217045 | A | C | 1.248 | -4194.66 | -4184.979 | 9.681 | 0.007223824 |
| 11 | 118697906 | rs73005502 | G | A | 1.248 | -4194.66 | -4184.979 | 9.681 | 0.007223824 |
| 11 | 118669605 | rs11217032 | A | T | 1.245 | -4194.66 | -4185.049 | 9.611 | 0.006735449 |
| 11 | 118673706 | rs11217033 | C | T | 1.245 | -4194.66 | -4185.049 | 9.611 | 0.006735449 |
| 11 | 118675772 | rs11217036 | T | C | 1.245 | -4194.66 | -4185.049 | 9.611 | 0.006735449 |
| 11 | 118677086 | rs11217037 | A | G | 1.245 | -4194.66 | -4185.049 | 9.611 | 0.006735449 |
| 11 | 118677355 | rs11217038 | A | G | 1.245 | -4194.66 | -4185.049 | 9.611 | 0.006735449 |
| 11 | 118611817 | rs10892280 | T | C | 1.243 | -4194.66 | -4185.148 | 9.511 | 0.006094486 |
| 11 | 118642085 | rs10892286 | C | A | 1.244 | -4194.66 | -4185.151 | 9.509 | 0.006082309 |
| 11 | 118642999 | rs10892287 | C | T | 1.244 | -4194.66 | -4185.151 | 9.509 | 0.006082309 |
| 11 | 118734521 | rs1174717189 | T | A | 0.833 | -4194.66 | -4185.184 | 9.476 | 0.005884869 |
| 11 | 118653503 | rs10892290 | T | G | 1.244 | -4194.66 | -4185.188 | 9.472 | 0.005861377 |
| 11 | 118653517 | rs10892291 | T | C | 1.244 | -4194.66 | -4185.188 | 9.472 | 0.005861377 |
| 11 | 118744396 | rs7117313 | C | G | 1.248 | -4194.66 | -4185.2 | 9.459 | 0.005785672 |
| 11 | 118644694 | rs10892288 | G | A | 1.244 | -4194.66 | -4185.213 | 9.447 | 0.005716659 |
| 11 | 118618776 | rs1048024 | T | A | 1.243 | -4194.66 | -4185.239 | 9.42 | 0.005564374 |
| 11 | 118619047 | rs2077579 | G | T | 1.243 | -4194.66 | -4185.239 | 9.42 | 0.005564374 |
| 11 | 118639353 | rs11217020 | A | G | 1.243 | -4194.66 | -4185.241 | 9.419 | 0.005558812 |
| 11 | 118636980 | rs10892283 | G | A | 1.243 | -4194.66 | -4185.25 | 9.41 | 0.005509007 |
| 11 | 118655645 | rs874621 | A | G | 1.243 | -4194.66 | -4185.259 | 9.4 | 0.005454192 |
| 11 | 118610549 | rs11826521 | T | G | 1.242 | -4194.66 | -4185.292 | 9.367 | 0.005277141 |
| 11 | 118697012 | rs571470 | A | G | 1.198 | -4194.66 | -4185.293 | 9.366 | 0.005271867 |
| 11 | 118615785 | rs11217009 | T | C | 1.243 | -4194.66 | -4185.295 | 9.365 | 0.005266597 |
| 11 | 118655340 | rs10892292 | C | G | 1.242 | -4194.66 | -4185.302 | 9.358 | 0.00522986 |
| 11 | 118646003 | rs10892289 | T | C | 1.242 | -4194.66 | -4185.316 | 9.343 | 0.005151997 |
| 11 | 118628373 | rs74676029 | T | C | 1.242 | -4194.66 | -4185.341 | 9.318 | 0.005024794 |
| 11 | 118619253 | rs2004781 | T | C | 1.242 | -4194.66 | -4185.343 | 9.317 | 0.005019772 |
| 11 | 118619960 | rs3889239 | T | C | 1.242 | -4194.66 | -4185.343 | 9.317 | 0.005019772 |
| 11 | 118636343 | rs11217019 | A | C | 1.242 | -4194.66 | -4185.366 | 9.293 | 0.004900731 |
| 11 | 118612921 | rs2155433 | A | G | 1.239 | -4194.66 | -4185.387 | 9.272 | 0.004798889 |
| 11 | 118648373 | rs56043232 | T | C | 1.241 | -4194.66 | -4185.387 | 9.272 | 0.004798889 |
| 11 | 118615340 | rs56758835 | A | G | 1.241 | -4194.66 | -4185.398 | 9.262 | 0.004751139 |
| 11 | 118729210 | rs10892300 | T | C | 0.835 | -4194.66 | -4185.465 | 9.194 | 0.004438802 |
| 11 | 118729860 | rs11217070 | G | T | 0.836 | -4194.66 | -4185.469 | 9.19 | 0.004421082 |
| 11 | 118745884 | rs10790275 | G | C | 1.245 | -4194.66 | -4185.474 | 9.185 | 0.004399032 |
| 11 | 118729583 | rs12790489 | C | T | 0.836 | -4194.66 | -4185.595 | 9.064 | 0.003897692 |
| 11 | 118730141 | rs7929520 | G | A | 0.836 | -4194.66 | -4185.595 | 9.064 | 0.003897692 |
| 11 | 118731599 | rs34561290 | T | C | 0.836 | -4194.66 | -4185.595 | 9.064 | 0.003897692 |
| 11 | 118731762 | rs11217072 | G | A | 0.836 | -4194.66 | -4185.595 | 9.064 | 0.003897692 |
| 11 | 118661398 | rs57494551 | T | C | 1.239 | -4194.66 | -4185.6 | 9.059 | 0.003878252 |
| 11 | 118608716 | rs7945144 | G | A | 1.233 | -4194.66 | -4185.745 | 8.915 | 0.003358131 |
| 11 | 118724723 | rs57719838 | C | T | 0.839 | -4194.66 | -4185.931 | 8.728 | 0.00278538 |
| 11 | 118725942 | rs11217063 | C | G | 0.839 | -4194.66 | -4186.02 | 8.64 | 0.002550742 |

| | | | | | | | |
|---|---|---|---|---|---|---|---|
| 11 | 118726647 | rs11217065 | C | T | M_Disease | 0.986874682 | M_Disease |
| 11 | 118723379 | rs7103067 | A | G | M_Disease | 0.986211011 | M_Disease |
| 11 | 118740180 | rs7483509 | A | G | M_Disease | 0.985585482 | M_Disease |
| 11 | 118727169 | rs10790267 | T | C | M_Disease | 0.983373644 | M_Disease |
| 11 | 118728320 | rs4936442 | T | C | M_Disease | 0.982791609 | M_Disease |
| 11 | 118727332 | rs11217067 | T | G | M_Disease | 0.98236366 | M_Disease |
| 11 | 118607311 | rs56267528 | A | C | M_Disease | 0.981109038 | M_Disease |
| 11 | 118607871 | rs76409436 | A | G | M_Disease | 0.980922803 | M_Disease |
| 11 | 118607417 | rs55894437 | T | C | M_Disease | 0.980829011 | M_Disease |
| 11 | 118607794 | rs12364971 | G | T | M_Disease | 0.980734767 | M_Disease |
| 11 | 118711790 | rs622722 | A | G | M_Disease | 0.980449303 | M_Disease |
| 11 | 118750321 | rs11217078 | C | T | M_Disease | 0.977686561 | M_Disease |
| 11 | 118606521 | rs6589681 | T | C | M_Disease | 0.976454651 | M_Disease |
| 11 | 118607150 | rs11217002 | T | C | M_Disease | 0.975872979 | M_Disease |
| 11 | 118606722 | rs6589682 | T | C | M_Disease | 0.968323772 | M_Disease |
| 11 | 118607046 | rs11217001 | A | G | M_Disease | 0.966104836 | M_Disease |
| 11 | 118607569 | rs77209083 | G | A | M_Disease | 0.963910622 | M_Disease |
| 11 | 118612565 | rs6589686 | A | G | M_Disease | 0.962852366 | M_Disease |
| 11 | 118604837 | rs11216998 | T | C | M_Disease | 0.93991335 | M_Disease&NULL |
| 11 | 118604941 | rs11216999 | C | T | M_Disease | 0.915289434 | M_Disease&NULL |
| 11 | 118705753 | rs556437 | A | G | M_Disease | 0.903784458 | M_Disease&NULL |
| 11 | 118605463 | rs10892274 | T | C | M_Disease | 0.897981931 | M_Disease&NULL |
| 11 | 118605617 | rs10892275 | A | T | M_Disease | 0.897981931 | M_Disease&NULL |
| 11 | 118605841 | rs10892277 | A | G | M_Disease | 0.893785008 | M_Disease&NULL |
| 11 | 118605628 | rs10892276 | A | G | M_Disease | 0.891388203 | M_Disease&NULL |
| 11 | 118608532 | rs567680 | G | C | M_Disease | 0.878681162 | M_Disease&NULL |
| 11 | 118608625 | rs566733 | G | C | M_Disease | 0.868755531 | M_Disease&NULL |
| 11 | 118603520 | rs11216994 | T | C | M_Disease | 0.860566127 | M_Disease&NULL |
| 11 | 118603526 | rs11216995 | A | G | M_Disease | 0.860566127 | M_Disease&NULL |
| 11 | 118719984 | rs145680506 | T | C | M_Disease | 0.860566127 | M_Disease&NULL |
| 11 | 118657311 | rs1309473877 | A | C | M_Disease | 0.852582339 | M_Disease&NULL |
| 11 | 118603165 | rs73001429 | G | T | M_Disease | 0.851952802 | M_Disease&NULL |
| 11 | 118604638 | rs11216997 | T | C | M_Disease | 0.850050858 | M_Disease&NULL |
| 11 | 118598544 | rs10892271 | A | G | M_Disease | 0.848128836 | M_Disease&NULL |
| 11 | 118602707 | rs10892272 | G | C | M_Disease | 0.848128836 | M_Disease&NULL |
| 11 | 118602809 | rs10892273 | A | G | M_Disease | 0.848128836 | M_Disease&NULL |
| 11 | 118607622 | rs11217004 | A | G | M_Disease | 0.842904531 | M_Disease&NULL |
| 11 | 118617299 | rs607125 | C | T | M_Disease | 0.841575821 | M_Disease&NULL |
| 11 | 118603736 | rs112561833 | C | A | M_Disease | 0.83889105 | M_Disease&NULL |
| 11 | 118613235 | rs632124 | A | T | M_Disease | 0.83479513 | M_Disease&NULL |
| 11 | 118620697 | rs488219 | C | T | M_Disease | 0.829911475 | M_Disease&NULL |
| 11 | 118659896 | rs2105782 | C | T | M_Disease | 0.829911475 | M_Disease&NULL |
| 11 | 118624250 | rs546393 | C | T | M_Disease | 0.829204581 | M_Disease&NULL |
| 11 | 118699276 | rs625274 | T | C | M_Disease | 0.82849523 | M_Disease&NULL |
| 11 | 118644582 | rs2510897 | C | A | M_Disease | 0.827783608 | M_Disease&NULL |
| 11 | 118558425 | rs149509653 | A | G | M_Disease | 0.82706965 | M_Disease&NULL |
| 11 | 118651809 | rs555649 | C | T | M_Disease | 0.82706965 | M_Disease&NULL |
| 11 | 118680072 | rs607472 | G | C | M_Disease | 0.825634714 | M_Disease&NULL |
| 11 | 118658075 | rs515523 | G | A | M_Disease | 0.824913732 | M_Disease&NULL |
| 11 | 118700557 | rs4291686 | G | A | M_Disease | 0.822736696 | M_Disease&NULL |
| 11 | 118590668 | rs11216980 | A | C | M_Disease | 0.820538481 | M_Disease&NULL |
| 11 | 118585826 | rs11216972 | T | C | M_Disease | 0.819061207 | M_Disease&NULL |
| 11 | 118631288 | rs507890 | C | T | M_Disease | 0.819061207 | M_Disease&NULL |
| 11 | 118631653 | rs1784543 | T | A | M_Disease | 0.819061207 | M_Disease&NULL |
| 11 | 118587319 | rs10892263 | C | T | M_Disease | 0.815326614 | M_Disease&NULL |
| 11 | 118589012 | rs12421793 | A | C | M_Disease | 0.815326614 | M_Disease&NULL |
| 11 | 118632522 | rs581045 | T | C | M_Disease | 0.815326614 | M_Disease&NULL |
| 11 | 118629031 | rs555639 | A | G | M_Disease | 0.814572581 | M_Disease&NULL |
| 11 | 118588952 | rs12421790 | T | C | M_Disease | 0.813057386 | M_Disease&NULL |
| 11 | 118679629 | rs1784300 | T | A | M_Disease | 0.812296221 | M_Disease&NULL |
| 11 | 118662028 | rs12796102 | T | C | M_Disease | 0.802976114 | M_Disease&NULL |
| 11 | 118603495 | rs11216993 | A | T | M_Disease | 0.799792822 | M_Disease&NULL |
| 11 | 118587097 | rs10892262 | C | G | M_Disease | 0.790012373 | M_Disease&NULL |
| 11 | 118617920 | rs476246 | T | C | M_Disease | 0.787513156 | M_Disease&NULL |
| 11 | 118640518 | rs181603672 | G | A | M_Disease | 0.779026108 | M_Disease&NULL |
| 11 | 118617544 | rs472641 | G | A | M_Disease | 0.776433137 | M_Disease&NULL |
| 11 | 118619948 | rs112143902 | T | C | M_Disease | 0.776433137 | M_Disease&NULL |
| 11 | 118648297 | rs117422168 | G | A | M_Disease | 0.770298949 | M_Disease&NULL |
| 11 | 118640063 | rs524590 | T | A | M_Disease | 0.767634117 | M_Disease&NULL |
| 11 | 118621495 | rs487728 | G | C | M_Disease | 0.765845626 | M_Disease&NULL |
| 11 | 118653629 | rs540180 | T | G | M_Disease | 0.765845626 | M_Disease&NULL |
| 11 | 118654651 | rs2508916 | G | C | M_Disease | 0.765845626 | M_Disease&NULL |
| 11 | 118652326 | rs1790189 | G | T | M_Disease | 0.764947804 | M_Disease&NULL |
| 11 | 118652373 | rs1784293 | C | A | M_Disease | 0.764947804 | M_Disease&NULL |
| 11 | 118579747 | rs10790261 | G | A | M_Disease | 0.7640476 | M_Disease&NULL |
| 11 | 118769604 | rs61751532 | T | C | M_Disease | 0.761332715 | M_Disease&NULL |
| 11 | 118583525 | rs10892260 | C | A | M_Disease | 0.759510917 | M_Disease&NULL |
| 11 | 118624767 | rs581062 | A | G | M_Disease | 0.758596462 | M_Disease&NULL |
| 11 | 118664377 | rs598373 | T | C | M_Disease | 0.758596462 | M_Disease&NULL |
| 11 | 118579865 | rs10892258 | A | G | M_Disease | 0.757679639 | M_Disease&NULL |
| 11 | 118580338 | rs11216961 | G | C | M_Disease | 0.757679639 | M_Disease&NULL |
| 11 | 118580813 | rs11216962 | C | T | M_Disease | 0.757679639 | M_Disease&NULL |
| 11 | 118580814 | rs11216963 | A | G | M_Disease | 0.757679639 | M_Disease&NULL |
| 11 | 118581895 | rs11216964 | A | G | M_Disease | 0.757679639 | M_Disease&NULL |
| 11 | 118582613 | rs11216965 | A | C | M_Disease | 0.757679639 | M_Disease&NULL |
| 11 | 118583002 | rs10892259 | A | G | M_Disease | 0.757679639 | M_Disease&NULL |
| 11 | 118678881 | rs591756 | T | C | M_Disease | 0.757679639 | M_Disease&NULL |
| 11 | 118605520 | rs75598590 | C | T | M_Disease | 0.753988716 | M_Disease&NULL |
| 11 | 118677710 | rs688161 | G | A | M_Disease | 0.753988716 | M_Disease&NULL |
| 11 | 118678609 | rs2508570 | A | G | M_Disease | 0.753988716 | M_Disease&NULL |
| 11 | 118584340 | rs11216969 | C | T | M_Disease | 0.751195782 | M_Disease&NULL |
| 11 | 118679387 | rs473298 | G | A | M_Disease | 0.751195782 | M_Disease&NULL |
| 11 | 118679445 | rs516719 | C | T | M_Disease | 0.751195782 | M_Disease&NULL |

| | | | | | | | | | |
|---|---|---|---|---|---|---|---|---|---|
| 11 | 118726647 | rs11217065 | C | T | 0.839 | -4194.66 | -4186.02 | 8.64 | 0.002550742 |
| 11 | 118723379 | rs7103067 | A | G | 0.84 | -4194.66 | -4186.116 | 8.543 | 0.002314942 |
| 11 | 118740180 | rs7483509 | A | G | 0.84 | -4194.66 | -4186.208 | 8.452 | 0.002113583 |
| 11 | 118727169 | rs10790267 | T | C | 0.843 | -4194.66 | -4186.502 | 8.158 | 0.001575204 |
| 11 | 118728320 | rs4936442 | T | C | 0.843 | -4194.66 | -4186.569 | 8.09 | 0.00147165 |
| 11 | 118727332 | rs11217067 | T | G | 0.844 | -4194.66 | -4186.621 | 8.038 | 0.00139708 |
| 11 | 118607311 | rs56267528 | A | C | 1.219 | -4194.66 | -4186.764 | 7.896 | 0.001212136 |
| 11 | 118607871 | rs76409436 | A | G | 1.218 | -4194.66 | -4186.782 | 7.877 | 0.001189323 |
| 11 | 118607417 | rs55894437 | T | C | 1.219 | -4194.66 | -4186.789 | 7.87 | 0.001181027 |
| 11 | 118607794 | rs12364971 | G | T | 1.218 | -4194.66 | -4186.806 | 7.854 | 0.001162281 |
| 11 | 118711790 | rs622722 | A | G | 0.845 | -4194.66 | -4186.832 | 7.828 | 0.001132451 |
| 11 | 118750321 | rs11217078 | C | T | 1.187 | -4194.66 | -4187.097 | 7.563 | 0.000868823 |
| 11 | 118606521 | rs6589681 | T | C | 1.214 | -4194.66 | -4187.208 | 7.452 | 0.000777544 |
| 11 | 118607150 | rs11217002 | T | C | 1.212 | -4194.66 | -4187.265 | 7.394 | 0.000733729 |
| 11 | 118606722 | rs6589682 | T | C | 1.207 | -4194.66 | -4187.822 | 6.837 | 0.000420372 |
| 11 | 118607046 | rs11217001 | A | G | 1.204 | -4194.66 | -4187.966 | 6.693 | 0.000363995 |
| 11 | 118607569 | rs77209083 | G | A | 1.202 | -4194.66 | -4188.094 | 6.566 | 0.000320583 |
| 11 | 118612565 | rs6589686 | A | G | 1.209 | -4194.66 | -4188.154 | 6.506 | 0.000301913 |
| 11 | 118604837 | rs11216998 | T | C | 1.171 | -4194.66 | -4189.166 | 5.493 | 0.000109633 |
| 11 | 118604941 | rs11216999 | C | T | 1.162 | -4194.66 | -4189.901 | 4.758 | 5.26E-05 |
| 11 | 118705753 | rs556437 | A | G | 0.876 | -4194.66 | -4190.184 | 4.475 | 3.96E-05 |
| 11 | 118605463 | rs10892274 | T | C | 1.156 | -4194.66 | -4190.307 | 4.353 | 3.51E-05 |
| 11 | 118605617 | rs10892275 | A | T | 1.156 | -4194.66 | -4190.307 | 4.353 | 3.51E-05 |
| 11 | 118605841 | rs10892277 | A | G | 1.154 | -4194.66 | -4190.404 | 4.256 | 3.18E-05 |
| 11 | 118605628 | rs10892276 | A | G | 1.154 | -4194.66 | -4190.45 | 4.21 | 3.04E-05 |
| 11 | 118608532 | rs567680 | G | C | 0.881 | -4194.66 | -4190.705 | 3.954 | 2.35E-05 |
| 11 | 118608625 | rs566733 | G | C | 0.882 | -4194.66 | -4190.881 | 3.778 | 1.97E-05 |
| 11 | 118603520 | rs11216994 | T | C | 1.154 | -4194.66 | -4191.027 | 3.633 | 1.71E-05 |
| 11 | 118603526 | rs11216995 | A | G | 1.154 | -4194.66 | -4191.027 | 3.633 | 1.71E-05 |
| 11 | 118719984 | rs145680506 | T | C | 0.811 | -4194.66 | -4191.027 | 3.632 | 1.70E-05 |
| 11 | 118657311 | rs1309473877 | A | C | 0.885 | -4194.66 | -4191.153 | 3.507 | 1.50E-05 |
| 11 | 118603165 | rs73001429 | G | T | 1.152 | -4194.66 | -4191.164 | 3.496 | 1.49E-05 |
| 11 | 118604638 | rs11216997 | T | C | 1.152 | -4194.66 | -4191.19 | 3.47 | 1.45E-05 |
| 11 | 118598544 | rs10892271 | A | G | 1.151 | -4194.66 | -4191.219 | 3.441 | 1.41E-05 |
| 11 | 118602707 | rs10892272 | G | C | 1.151 | -4194.66 | -4191.219 | 3.44 | 1.41E-05 |
| 11 | 118602809 | rs10892273 | A | G | 1.151 | -4194.66 | -4191.219 | 3.44 | 1.41E-05 |
| 11 | 118607622 | rs11217004 | A | G | 0.887 | -4194.66 | -4191.304 | 3.355 | 1.29E-05 |
| 11 | 118617299 | rs607125 | C | T | 0.887 | -4194.66 | -4191.321 | 3.338 | 1.27E-05 |
| 11 | 118603736 | rs112561833 | C | A | 1.149 | -4194.66 | -4191.357 | 3.303 | 1.23E-05 |
| 11 | 118613235 | rs632124 | A | T | 0.888 | -4194.66 | -4191.416 | 3.243 | 1.16E-05 |
| 11 | 118659896 | rs2105782 | C | T | 0.889 | -4194.66 | -4191.489 | 3.171 | 1.08E-05 |
| 11 | 118620697 | rs488219 | C | T | 0.889 | -4194.66 | -4191.49 | 3.169 | 1.07E-05 |
| 11 | 118624250 | rs546393 | C | T | 0.889 | -4194.66 | -4191.506 | 3.154 | 1.06E-05 |
| 11 | 118699276 | rs625274 | T | C | 0.889 | -4194.66 | -4191.507 | 3.153 | 1.06E-05 |
| 11 | 118644582 | rs2510897 | C | A | 0.889 | -4194.66 | -4191.523 | 3.137 | 1.04E-05 |
| 11 | 118651809 | rs555649 | C | T | 0.889 | -4194.66 | -4191.527 | 3.133 | 1.04E-05 |
| 11 | 118558425 | rs149509653 | A | G | 0.869 | -4194.66 | -4191.531 | 3.129 | 1.03E-05 |
| 11 | 118680072 | rs607472 | G | C | 0.889 | -4194.66 | -4191.553 | 3.107 | 1.01E-05 |
| 11 | 118658075 | rs515523 | G | A | 0.889 | -4194.66 | -4191.558 | 3.102 | 1.00E-05 |
| 11 | 118700557 | rs4291686 | G | A | 0.89 | -4194.66 | -4191.595 | 3.065 | 9.67E-06 |
| 11 | 118590668 | rs11216980 | A | C | 1.146 | -4194.66 | -4191.621 | 3.039 | 9.42E-06 |
| 11 | 118585826 | rs11216972 | T | C | 1.145 | -4194.66 | -4191.64 | 3.019 | 9.24E-06 |
| 11 | 118631288 | rs507890 | C | T | 0.89 | -4194.66 | -4191.641 | 3.019 | 9.24E-06 |
| 11 | 118631653 | rs1784543 | T | A | 0.89 | -4194.66 | -4191.641 | 3.019 | 9.24E-06 |
| 11 | 118589012 | rs12421793 | A | C | 1.145 | -4194.66 | -4191.689 | 2.97 | 8.79E-06 |
| 11 | 118632522 | rs581045 | T | C | 0.891 | -4194.66 | -4191.689 | 2.97 | 8.79E-06 |
| 11 | 118587319 | rs10892263 | C | T | 1.145 | -4194.66 | -4191.694 | 2.966 | 8.76E-06 |
| 11 | 118629031 | rs555639 | A | G | 0.891 | -4194.66 | -4191.704 | 2.956 | 8.67E-06 |
| 11 | 118588952 | rs12421790 | T | C | 1.145 | -4194.66 | -4191.723 | 2.937 | 8.51E-06 |
| 11 | 118679629 | rs1784300 | T | A | 0.891 | -4194.66 | -4191.732 | 2.928 | 8.43E-06 |
| 11 | 118662028 | rs12796102 | T | C | 0.892 | -4194.66 | -4191.851 | 2.809 | 7.49E-06 |
| 11 | 118603495 | rs11216993 | A | T | 1.141 | -4194.66 | -4191.889 | 2.771 | 7.21E-06 |
| 11 | 118587097 | rs10892262 | C | G | 1.139 | -4194.66 | -4192.009 | 2.651 | 6.39E-06 |
| 11 | 118617920 | rs476246 | T | C | 1.123 | -4194.66 | -4192.045 | 2.614 | 6.16E-06 |
| 11 | 118640518 | rs181603672 | G | A | 1.249 | -4194.66 | -4192.146 | 2.514 | 5.57E-06 |
| 11 | 118617544 | rs472641 | G | A | 1.121 | -4194.66 | -4192.168 | 2.491 | 5.45E-06 |
| 11 | 118619948 | rs112143902 | T | C | 0.836 | -4194.66 | -4192.172 | 2.488 | 5.43E-06 |
| 11 | 118648297 | rs117422168 | G | A | 1.245 | -4194.66 | -4192.25 | 2.41 | 5.02E-06 |
| 11 | 118640063 | rs524590 | T | A | 1.119 | -4194.66 | -4192.269 | 2.391 | 4.93E-06 |
| 11 | 118653629 | rs540180 | T | G | 1.119 | -4194.66 | -4192.29 | 2.369 | 4.82E-06 |
| 11 | 118654651 | rs2508916 | G | C | 1.119 | -4194.66 | -4192.29 | 2.369 | 4.82E-06 |
| 11 | 118621495 | rs487728 | G | C | 1.119 | -4194.66 | -4192.294 | 2.366 | 4.81E-06 |
| 11 | 118652326 | rs1790189 | G | T | 1.119 | -4194.66 | -4192.299 | 2.361 | 4.78E-06 |
| 11 | 118652373 | rs1784293 | C | A | 1.119 | -4194.66 | -4192.299 | 2.361 | 4.78E-06 |
| 11 | 118579747 | rs10790261 | G | A | 1.131 | -4194.66 | -4192.314 | 2.346 | 4.71E-06 |
| 11 | 118769604 | rs61751532 | T | C | 0.83 | -4194.66 | -4192.349 | 2.311 | 4.55E-06 |
| 11 | 118583525 | rs10892260 | C | A | 1.13 | -4194.66 | -4192.36 | 2.3 | 4.50E-06 |
| 11 | 118624767 | rs581062 | A | G | 1.118 | -4194.66 | -4192.369 | 2.291 | 4.46E-06 |
| 11 | 118664377 | rs598373 | T | C | 1.118 | -4194.66 | -4192.37 | 2.289 | 4.45E-06 |
| 11 | 118678881 | rs591756 | T | C | 1.118 | -4194.66 | -4192.379 | 2.28 | 4.41E-06 |
| 11 | 118579865 | rs10892258 | A | G | 1.13 | -4194.66 | -4192.383 | 2.276 | 4.39E-06 |
| 11 | 118580338 | rs11216961 | G | C | 1.13 | -4194.66 | -4192.383 | 2.276 | 4.39E-06 |
| 11 | 118580813 | rs11216962 | C | T | 1.13 | -4194.66 | -4192.383 | 2.276 | 4.39E-06 |
| 11 | 118580814 | rs11216963 | A | G | 1.13 | -4194.66 | -4192.383 | 2.276 | 4.39E-06 |
| 11 | 118581895 | rs11216964 | A | G | 1.13 | -4194.66 | -4192.383 | 2.276 | 4.39E-06 |
| 11 | 118582613 | rs11216965 | A | C | 1.13 | -4194.66 | -4192.383 | 2.276 | 4.39E-06 |
| 11 | 118583002 | rs10892259 | A | G | 1.13 | -4194.66 | -4192.383 | 2.276 | 4.39E-06 |
| 11 | 118677710 | rs688161 | G | A | 1.117 | -4194.66 | -4192.425 | 2.235 | 4.22E-06 |
| 11 | 118678609 | rs2508570 | A | G | 1.117 | -4194.66 | -4192.425 | 2.235 | 4.22E-06 |
| 11 | 118605520 | rs75598590 | C | T | 1.239 | -4194.66 | -4192.428 | 2.232 | 4.20E-06 |
| 11 | 118679387 | rs473298 | G | A | 1.117 | -4194.66 | -4192.454 | 2.206 | 4.10E-06 |
| 11 | 118679445 | rs516719 | C | T | 1.117 | -4194.66 | -4192.454 | 2.206 | 4.10E-06 |
| 11 | 118584340 | rs11216969 | C | T | 1.129 | -4194.66 | -4192.456 | 2.203 | 4.08E-06 |

| | | | | | | | |
|---|---|---|---|---|---|---|---|
| 11 | 118578705 | rs10892256 | A | G | M_Disease | 0.750260106 | M_Disease&NULL |
| 11 | 118579327 | rs10892257 | A | G | M_Disease | 0.750260106 | M_Disease&NULL |
| 11 | 118678986 | rs477014 | C | T | M_Disease | 0.748381722 | M_Disease&NULL |
| 11 | 118680045 | rs2508569 | C | G | M_Disease | 0.748381722 | M_Disease&NULL |
| 11 | 118575326 | rs11216956 | A | G | M_Disease | 0.74743902 | M_Disease&NULL |
| 11 | 118626511 | rs499343 | T | C | M_Disease | 0.746493983 | M_Disease&NULL |
| 11 | 118577873 | rs11216960 | A | G | M_Disease | 0.743644892 | M_Disease&NULL |
| 11 | 118566069 | rs506255 | A | C | M_Disease | 0.737884101 | M_Disease&NULL |
| 11 | 118757219 | rs1623316 | G | C | M_Disease | 0.735945394 | M_Disease&NULL |
| 11 | 118563021 | rs117330495 | A | C | M_Disease | 0.733997517 | M_Disease&NULL |
| 11 | 118478681 | rs45578540 | T | C | M_Disease | 0.722119606 | M_Disease&NULL |
| 11 | 118590961 | rs111227443 | T | C | M_Disease | 0.720108527 | M_Disease&NULL |
| 11 | 118669178 | rs583213 | G | T | M_Disease | 0.720108527 | M_Disease&NULL |
| 11 | 118652569 | rs150792202 | A | G | M_Disease | 0.714022227 | M_Disease&NULL |
| 11 | 118695908 | rs583905 | G | A | M_Disease | 0.699517228 | M_Disease&NULL |
| 11 | 118685481 | rs515707 | T | C | M_Disease | 0.697411114 | M_Disease&NULL |
| 11 | 118689871 | rs587816 | A | C | M_Disease | 0.697411114 | M_Disease&NULL |
| 11 | 118690550 | rs577121 | C | T | M_Disease | 0.697411114 | M_Disease&NULL |
| 11 | 118707249 | rs4938551 | C | G | NULL | 0.69635493 | NULL&M_Disease |
| 11 | 118716563 | rs2851460 | A | G | NULL | 0.69635493 | NULL&M_Disease |
| 11 | 118708564 | rs484232 | T | G | NULL | 0.695296669 | NULL&M_Disease |
| 11 | 118717708 | rs7107756 | A | G | NULL | 0.695296669 | NULL&M_Disease |
| 11 | 118459069 | rs633308 | G | A | NULL | 0.69423634 | NULL&M_Disease |
| 11 | 118717732 | rs11217052 | G | A | NULL | 0.69423634 | NULL&M_Disease |
| 11 | 118718061 | rs4938559 | T | C | NULL | 0.69423634 | NULL&M_Disease |
| 11 | 118718062 | rs4938560 | G | A | NULL | 0.69423634 | NULL&M_Disease |
| 11 | 118735476 | rs10892301 | A | G | NULL | 0.69423634 | NULL&M_Disease |
| 11 | 118755738 | rs523604 | G | A | M_Disease | 0.69423634 | M_Disease&NULL |
| 11 | 118481431 | rs10892246 | A | G | NULL | 0.692109504 | NULL&M_Disease |
| 11 | 118696175 | rs584935 | A | G | M_Disease | 0.692109504 | M_Disease&NULL |
| 11 | 118480223 | rs67307131 | C | T | NULL | 0.691043012 | NULL&M_Disease |
| 11 | 118480285 | rs12225399 | C | G | NULL | 0.691043012 | NULL&M_Disease |
| 11 | 118486325 | rs12290522 | T | C | NULL | 0.691043012 | NULL&M_Disease |
| 11 | 118487673 | rs4938517 | C | T | NULL | 0.691043012 | NULL&M_Disease |
| 11 | 118492591 | rs2077173 | A | G | NULL | 0.691043012 | NULL&M_Disease |
| 11 | 118496066 | rs11603616 | A | G | NULL | 0.691043012 | NULL&M_Disease |
| 11 | 118497402 | rs12419204 | A | T | NULL | 0.691043012 | NULL&M_Disease |
| 11 | 118500173 | rs571001 | T | C | NULL | 0.691043012 | NULL&M_Disease |
| 11 | 118500193 | rs570952 | C | T | NULL | 0.691043012 | NULL&M_Disease |
| 11 | 118505090 | rs645637 | A | G | NULL | 0.691043012 | NULL&M_Disease |
| 11 | 118506402 | rs483283 | C | T | NULL | 0.691043012 | NULL&M_Disease |
| 11 | 118512162 | rs570836 | A | G | NULL | 0.691043012 | NULL&M_Disease |
| 11 | 118519550 | rs601318 | G | C | NULL | 0.691043012 | NULL&M_Disease |
| 11 | 118520480 | rs653008 | T | G | NULL | 0.691043012 | NULL&M_Disease |
| 11 | 118522179 | rs1171461 | A | G | NULL | 0.691043012 | NULL&M_Disease |
| 11 | 118522338 | rs4938520 | G | A | NULL | 0.691043012 | NULL&M_Disease |
| 11 | 118528466 | rs7389 | G | T | NULL | 0.691043012 | NULL&M_Disease |
| 11 | 118529595 | rs745663 | G | A | NULL | 0.691043012 | NULL&M_Disease |
| 11 | 118811684 | rs550999 | G | C | NULL | 0.691043012 | NULL&M_Disease |
| 11 | 118812512 | rs647769 | T | C | NULL | 0.691043012 | NULL&M_Disease |
| 11 | 118515616 | rs527290 | T | C | NULL | 0.689974481 | NULL&M_Disease |
| 11 | 118577990 | rs480958 | G | A | NULL | 0.689974481 | NULL&M_Disease |
| 11 | 118515779 | rs117679873 | A | G | M_Disease | 0.689974481 | M_Disease&NULL |
| 11 | 118480115 | rs12803321 | C | G | NULL | 0.687831331 | NULL&M_Disease |
| 11 | 118622985 | rs3825057 | C | T | NULL | 0.682438302 | NULL&M_Disease |
| 11 | 118603030 | rs7943039 | A | G | NULL | 0.681353734 | NULL&M_Disease |
| 11 | 118629364 | rs73001486 | A | G | NULL | 0.681353734 | NULL&M_Disease |
| 11 | 118640828 | rs11217021 | C | T | NULL | 0.681353734 | NULL&M_Disease |
| 11 | 118796570 | rs11217096 | T | C | NULL | 0.681353734 | NULL&M_Disease |
| 11 | 118477367 | rs498872 | A | G | NULL | 0.680267197 | NULL&M_Disease |
| 11 | 118594669 | rs140026975 | C | A | NULL | 0.680267197 | NULL&M_Disease |
| 11 | 118616253 | rs7124498 | A | G | NULL | 0.680267197 | NULL&M_Disease |
| 11 | 118624919 | rs11217013 | C | G | NULL | 0.680267197 | NULL&M_Disease |
| 11 | 118626286 | rs11217014 | C | T | NULL | 0.680267197 | NULL&M_Disease |
| 11 | 118627149 | rs11217016 | C | A | NULL | 0.680267197 | NULL&M_Disease |
| 11 | 118627778 | rs10892282 | C | G | NULL | 0.680267197 | NULL&M_Disease |
| 11 | 118633533 | rs7110526 | C | T | NULL | 0.680267197 | NULL&M_Disease |
| 11 | 118635364 | rs11217018 | C | T | NULL | 0.680267197 | NULL&M_Disease |
| 11 | 118637475 | rs112910608 | T | C | NULL | 0.680267197 | NULL&M_Disease |
| 11 | 118747576 | rs55756957 | G | A | NULL | 0.680267197 | NULL&M_Disease |
| 11 | 118770321 | rs11217083 | G | A | NULL | 0.680267197 | NULL&M_Disease |
| 11 | 118478330 | rs7125115 | A | G | NULL | 0.679178699 | NULL&M_Disease |
| 11 | 118596444 | rs68097885 | A | G | NULL | 0.679178699 | NULL&M_Disease |
| 11 | 118609598 | rs112996178 | T | C | NULL | 0.679178699 | NULL&M_Disease |
| 11 | 118610463 | rs4499035 | T | C | NULL | 0.67808825 | NULL&M_Disease |
| 11 | 118610826 | rs1893029 | T | G | NULL | 0.67808825 | NULL&M_Disease |
| 11 | 118613674 | rs7130875 | C | T | NULL | 0.67808825 | NULL&M_Disease |
| 11 | 118655356 | rs7934890 | T | C | NULL | 0.67808825 | NULL&M_Disease |
| 11 | 118657297 | rs487177 | G | C | NULL | 0.67808825 | NULL&M_Disease |
| 11 | 118666289 | rs56104144 | C | G | NULL | 0.676995856 | NULL&M_Disease |
| 11 | 118801235 | rs602716 | A | G | NULL | 0.676995856 | NULL&M_Disease |
| 11 | 118802492 | rs688035 | G | C | NULL | 0.676995856 | NULL&M_Disease |
| 11 | 118665324 | rs61901411 | T | C | M_Disease | 0.675901528 | M_Disease&NULL |
| 11 | 118668296 | rs58868870 | T | C | NULL | 0.675901528 | NULL&M_Disease |
| 11 | 118700264 | rs113641235 | T | C | M_Disease | 0.675901528 | M_Disease&NULL |
| 11 | 118769442 | rs7104819 | G | A | NULL | 0.675901528 | NULL&M_Disease |
| 11 | 118803536 | rs615166 | A | G | NULL | 0.675901528 | NULL&M_Disease |
| 11 | 118584498 | rs569770 | T | C | NULL | 0.674805273 | NULL&M_Disease |
| 11 | 118590864 | rs11216982 | A | G | NULL | 0.674805273 | NULL&M_Disease |
| 11 | 118605914 | rs7926282 | C | A | NULL | 0.674805273 | NULL&M_Disease |
| 11 | 118587347 | rs603486 | T | C | NULL | 0.673707099 | NULL&M_Disease |
| 11 | 118605451 | rs2186780 | T | C | NULL | 0.673707099 | NULL&M_Disease |
| 11 | 118539566 | rs1783935 | T | A | NULL | 0.672607017 | NULL&M_Disease |

| | | | | | | | | | |
|---|---|---|---|---|---|---|---|---|---|
| 11 | 118578705 | rs10892256 | A | G | 1.129 | -4194.66 | -4192.46 | 2.199 | 4.07E-06 |
| 11 | 118579327 | rs10892257 | A | G | 1.129 | -4194.66 | -4192.46 | 2.199 | 4.07E-06 |
| 11 | 118678986 | rs477014 | C | T | 1.116 | -4194.66 | -4192.483 | 2.177 | 3.98E-06 |
| 11 | 118680045 | rs2508569 | C | G | 1.116 | -4194.66 | -4192.483 | 2.177 | 3.98E-06 |
| 11 | 118575326 | rs11216956 | A | G | 1.128 | -4194.66 | -4192.493 | 2.166 | 3.94E-06 |
| 11 | 118626511 | rs499343 | T | C | 1.116 | -4194.66 | -4192.498 | 2.162 | 3.92E-06 |
| 11 | 118577873 | rs11216960 | A | G | 1.127 | -4194.66 | -4192.529 | 2.131 | 3.80E-06 |
| 11 | 118566069 | rs506255 | A | C | 0.862 | -4194.66 | -4192.589 | 2.071 | 3.58E-06 |
| 11 | 118757219 | rs1623316 | G | C | 0.901 | -4194.66 | -4192.615 | 2.044 | 3.48E-06 |
| 11 | 118563021 | rs117330495 | A | C | 1.241 | -4194.66 | -4192.642 | 2.018 | 3.39E-06 |
| 11 | 118478681 | rs45578540 | T | C | 1.237 | -4194.66 | -4192.759 | 1.9 | 3.02E-06 |
| 11 | 118669178 | rs583213 | G | T | 0.87 | -4194.66 | -4192.771 | 1.889 | 2.98E-06 |
| 11 | 118590961 | rs111227443 | T | C | 0.784 | -4194.66 | -4192.779 | 1.881 | 2.96E-06 |
| 11 | 118652569 | rs150792202 | A | G | 0.858 | -4194.66 | -4192.832 | 1.828 | 2.81E-06 |
| 11 | 118695908 | rs583905 | G | A | 0.885 | -4194.66 | -4192.97 | 1.689 | 2.44E-06 |
| 11 | 118685481 | rs515707 | T | C | 0.885 | -4194.66 | -4192.992 | 1.668 | 2.39E-06 |
| 11 | 118689871 | rs587816 | A | C | 0.885 | -4194.66 | -4192.992 | 1.668 | 2.39E-06 |
| 11 | 118690550 | rs577121 | C | T | 0.885 | -4194.66 | -4192.992 | 1.668 | 2.39E-06 |
| 11 | 118755738 | rs523604 | G | A | 1.101 | -4194.66 | -4193.024 | 1.635 | 2.31E-06 |
| 11 | 118696175 | rs584935 | A | G | 0.886 | -4194.66 | -4193.038 | 1.621 | 2.28E-06 |
| 11 | 118515779 | rs117679873 | A | G | 1.221 | -4194.66 | -4193.068 | 1.592 | 2.22E-06 |
| 11 | 118665324 | rs61901411 | T | C | 0.877 | -4194.66 | -4193.189 | 1.471 | 1.96E-06 |
| 11 | 118700264 | rs113641235 | T | C | 0.889 | -4194.66 | -4193.195 | 1.465 | 1.95E-06 |
| 11 | 118682462 | rs523148 | T | C | 0.89 | -4194.66 | -4193.247 | 1.413 | 1.85E-06 |
| 11 | 118668908 | rs582200 | G | A | 0.879 | -4194.66 | -4193.254 | 1.405 | 1.84E-06 |
| 11 | 118697375 | rs73005500 | C | T | 0.89 | -4194.66 | -4193.28 | 1.38 | 1.79E-06 |
| 11 | 118665009 | rs4938544 | A | G | 0.88 | -4194.66 | -4193.292 | 1.368 | 1.77E-06 |
| 11 | 118519213 | rs668628 | T | G | 0.89 | -4194.66 | -4193.292 | 1.367 | 1.77E-06 |
| 11 | 118462188 | rs4936431 | G | C | 1.19 | -4194.66 | -4193.331 | 1.328 | 1.70E-06 |
| 11 | 118469739 | rs143380378 | G | C | 1.19 | -4194.66 | -4193.331 | 1.328 | 1.70E-06 |
| 11 | 118677083 | rs675003 | T | C | 0.881 | -4194.66 | -4193.332 | 1.327 | 1.70E-06 |
| 11 | 118678780 | rs500254 | G | A | 0.881 | -4194.66 | -4193.332 | 1.327 | 1.70E-06 |
| 11 | 118662340 | rs5006102 | A | T | 0.881 | -4194.66 | -4193.333 | 1.326 | 1.70E-06 |
| 11 | 118700425 | rs10128600 | C | T | 0.892 | -4194.66 | -4193.34 | 1.32 | 1.69E-06 |
| 11 | 118662045 | rs7933859 | T | C | 0.881 | -4194.66 | -4193.365 | 1.294 | 1.65E-06 |
| 11 | 118662203 | rs4508234 | G | C | 0.881 | -4194.66 | -4193.365 | 1.294 | 1.65E-06 |
| 11 | 118820778 | rs139076649 | C | T | 0.819 | -4194.66 | -4193.417 | 1.242 | 1.56E-06 |
| 11 | 118615232 | rs139423236 | G | C | 1.237 | -4194.66 | -4193.43 | 1.23 | 1.54E-06 |
| 11 | 118500963 | rs680465 | A | G | 0.899 | -4194.66 | -4193.483 | 1.177 | 1.46E-06 |
| 11 | 118468037 | rs535774 | T | C | 0.869 | -4194.66 | -4193.522 | 1.138 | 1.41E-06 |
| 11 | 118762073 | rs77841686 | A | G | 1.217 | -4194.66 | -4193.56 | 1.1 | 1.36E-06 |
| 11 | 118763313 | rs78041268 | A | G | 1.217 | -4194.66 | -4193.56 | 1.1 | 1.36E-06 |
| 11 | 118767564 | rs75438046 | A | G | 1.217 | -4194.66 | -4193.56 | 1.1 | 1.36E-06 |
| 11 | 118805385 | rs55772432 | A | G | 0.873 | -4194.66 | -4193.63 | 1.029 | 1.26E-06 |
| 11 | 118783424 | rs149114341 | A | G | 1.212 | -4194.66 | -4193.634 | 1.025 | 1.26E-06 |
| 11 | 118573894 | rs576283 | G | A | 0.916 | -4194.66 | -4193.665 | 0.994 | 1.22E-06 |
| 11 | 118573916 | rs627391 | T | A | 0.916 | -4194.66 | -4193.665 | 0.994 | 1.22E-06 |
| 11 | 118574311 | rs524409 | T | C | 0.916 | -4194.66 | -4193.667 | 0.993 | 1.22E-06 |
| 11 | 118574675 | rs494459 | T | C | 0.916 | -4194.66 | -4193.667 | 0.993 | 1.22E-06 |
| 11 | 118575011 | rs530901 | C | T | 0.916 | -4194.66 | -4193.69 | 0.969 | 1.19E-06 |
| 11 | 118575180 | rs626645 | C | T | 0.916 | -4194.66 | -4193.69 | 0.969 | 1.19E-06 |
| 11 | 118575419 | rs625735 | A | G | 0.916 | -4194.66 | -4193.69 | 0.969 | 1.19E-06 |
| 11 | 118575479 | rs637563 | G | A | 0.916 | -4194.66 | -4193.69 | 0.969 | 1.19E-06 |
| 11 | 118575693 | rs638514 | C | T | 0.916 | -4194.66 | -4193.69 | 0.969 | 1.19E-06 |
| 11 | 118560857 | rs73001406 | A | G | 1.104 | -4194.66 | -4193.7 | 0.96 | 1.18E-06 |
| 11 | 118777978 | rs78755429 | A | C | 0.83 | -4194.66 | -4193.702 | 0.957 | 1.17E-06 |
| 11 | 118698160 | rs640373 | G | A | 0.899 | -4194.66 | -4193.72 | 0.94 | 1.16E-06 |
| 11 | 118698274 | rs639915 | A | G | 0.899 | -4194.66 | -4193.72 | 0.94 | 1.16E-06 |
| 11 | 118698294 | rs639885 | A | G | 0.899 | -4194.66 | -4193.72 | 0.94 | 1.16E-06 |
| 11 | 118698537 | rs638893 | G | A | 0.899 | -4194.66 | -4193.72 | 0.94 | 1.16E-06 |
| 11 | 118698777 | rs516332 | G | A | 0.899 | -4194.66 | -4193.72 | 0.94 | 1.16E-06 |
| 11 | 118699744 | rs2508564 | C | T | 0.899 | -4194.66 | -4193.72 | 0.94 | 1.16E-06 |
| 11 | 118699863 | rs2510901 | A | G | 0.899 | -4194.66 | -4193.72 | 0.94 | 1.16E-06 |
| 11 | 118599649 | rs182364290 | G | A | 1.217 | -4194.66 | -4193.724 | 0.935 | 1.15E-06 |
| 11 | 118895495 | rs8301 | G | C | 1.104 | -4194.66 | -4193.738 | 0.922 | 1.13E-06 |
| 11 | 118555692 | rs10892252 | G | A | 1.102 | -4194.66 | -4193.771 | 0.888 | 1.10E-06 |
| 11 | 118661197 | rs61901410 | G | C | 0.891 | -4194.66 | -4193.773 | 0.886 | 1.09E-06 |
| 11 | 118595207 | rs563537 | A | T | 0.918 | -4194.66 | -4193.784 | 0.876 | 1.08E-06 |
| 11 | 118580348 | rs516503 | T | C | 0.918 | -4194.66 | -4193.786 | 0.873 | 1.08E-06 |
| 11 | 118606197 | rs12420846 | T | C | 1.158 | -4194.66 | -4193.793 | 0.866 | 1.07E-06 |
| 11 | 118556398 | rs11216943 | A | G | 1.101 | -4194.66 | -4193.8 | 0.86 | 1.07E-06 |
| 11 | 118550524 | rs201134460 | T | C | 1.1 | -4194.66 | -4193.85 | 0.81 | 1.01E-06 |
| 11 | 118550522 | rs2277296 | C | T | 1.1 | -4194.66 | -4193.867 | 0.793 | 9.97E-07 |
| 11 | 118746769 | rs4938576 | T | G | 1.088 | -4194.66 | -4193.883 | 0.776 | 9.80E-07 |
| 11 | 118701898 | rs533254 | A | T | 0.902 | -4194.66 | -4193.886 | 0.773 | 9.77E-07 |
| 11 | 118580906 | rs496058 | G | A | 0.92 | -4194.66 | -4193.897 | 0.763 | 9.68E-07 |
| 11 | 118754353 | rs630923 | A | C | 1.111 | -4194.66 | -4193.901 | 0.759 | 9.64E-07 |
| 11 | 118588998 | rs570424 | C | T | 0.919 | -4194.66 | -4193.903 | 0.757 | 9.62E-07 |
| 11 | 118503442 | rs45621235 | T | C | 1.18 | -4194.66 | -4193.905 | 0.755 | 9.60E-07 |
| 11 | 118598612 | rs568688 | G | T | 0.92 | -4194.66 | -4193.96 | 0.699 | 9.08E-07 |
| 11 | 118855789 | rs4938609 | A | C | 1.211 | -4194.66 | -4193.968 | 0.692 | 9.01E-07 |
| 11 | 118441079 | rs7106316 | G | A | 0.882 | -4194.66 | -4193.972 | 0.687 | 8.97E-07 |
| 11 | 118543563 | rs10892251 | T | C | 1.097 | -4194.66 | -4193.989 | 0.671 | 8.83E-07 |
| 11 | 118598138 | rs543692 | C | T | 0.921 | -4194.66 | -4193.989 | 0.67 | 8.82E-07 |
| 11 | 118632893 | rs6589690 | A | G | 0.897 | -4194.66 | -4194.005 | 0.654 | 8.68E-07 |
| 11 | 118530869 | rs45447797 | G | C | 0.868 | -4194.66 | -4194.019 | 0.64 | 8.56E-07 |
| 11 | 118660932 | rs1784295 | A | G | 0.899 | -4194.66 | -4194.077 | 0.582 | 8.07E-07 |
| 11 | 118629143 | rs475946 | T | C | 0.899 | -4194.66 | -4194.101 | 0.559 | 7.89E-07 |
| 11 | 118758089 | rs497916 | T | C | 1.091 | -4194.66 | -4194.105 | 0.554 | 7.85E-07 |
| 11 | 118707171 | rs4938550 | G | A | 0.907 | -4194.66 | -4194.108 | 0.552 | 7.84E-07 |
| 11 | 118637666 | rs535996 | T | C | 0.9 | -4194.66 | -4194.123 | 0.537 | 7.72E-07 |
| 11 | 118661055 | rs1790188 | T | C | 0.9 | -4194.66 | -4194.126 | 0.533 | 7.69E-07 |
| 11 | 118702002 | rs534114 | A | G | 0.908 | -4194.66 | -4194.145 | 0.515 | 7.55E-07 |

| | | | | | | | |
|---|---|---|---|---|---|---|---|
| 11 | 118585146 | rs112799932 | G | T | NULL | 0.672607017 | NULL&M_Disease |
| 11 | 118767077 | rs581063 | C | T | NULL | 0.672607017 | NULL&M_Disease |
| 11 | 118807735 | rs10892311 | T | C | NULL | 0.672607017 | NULL&M_Disease |
| 11 | 118538187 | rs607527 | T | G | NULL | 0.671505034 | NULL&M_Disease |
| 11 | 118538157 | rs520543 | A | G | NULL | 0.67040116 | NULL&M_Disease |
| 11 | 118539170 | rs4457789 | T | G | NULL | 0.67040116 | NULL&M_Disease |
| 11 | 118539522 | rs2508991 | A | G | NULL | 0.67040116 | NULL&M_Disease |
| 11 | 118548665 | rs1784298 | C | T | NULL | 0.67040116 | NULL&M_Disease |
| 11 | 118548671 | rs4938522 | T | A | NULL | 0.67040116 | NULL&M_Disease |
| 11 | 118548838 | rs4938523 | T | G | NULL | 0.67040116 | NULL&M_Disease |
| 11 | 118682462 | rs523148 | T | C | M_Disease | 0.67040116 | M_Disease&NULL |
| 11 | 118769032 | rs11217081 | G | A | NULL | 0.67040116 | NULL&M_Disease |
| 11 | 118668908 | rs582200 | G | A | M_Disease | 0.669295403 | M_Disease&NULL |
| 11 | 118788081 | rs7929825 | G | C | NULL | 0.669295403 | NULL&M_Disease |
| 11 | 118831335 | rs61506726 | G | A | NULL | 0.669295403 | NULL&M_Disease |
| 11 | 118546602 | rs526597 | G | A | NULL | 0.668187772 | NULL&M_Disease |
| 11 | 118549582 | rs507080 | A | G | NULL | 0.668187772 | NULL&M_Disease |
| 11 | 118549803 | rs659969 | T | C | NULL | 0.668187772 | NULL&M_Disease |
| 11 | 118549908 | rs656575 | G | T | NULL | 0.668187772 | NULL&M_Disease |
| 11 | 118549985 | rs543327 | T | G | NULL | 0.668187772 | NULL&M_Disease |
| 11 | 118550017 | rs543413 | C | T | NULL | 0.668187772 | NULL&M_Disease |
| 11 | 118697375 | rs73005500 | C | T | M_Disease | 0.665966927 | M_Disease&NULL |
| 11 | 118765600 | rs3922 | G | A | NULL | 0.665966927 | NULL&M_Disease |
| 11 | 118519213 | rs668628 | T | G | M_Disease | 0.66485373 | M_Disease&NULL |
| 11 | 118562520 | rs558593 | T | C | NULL | 0.66485373 | NULL&M_Disease |
| 11 | 118567020 | rs642662 | G | A | NULL | 0.66485373 | NULL&M_Disease |
| 11 | 118665009 | rs4938544 | A | G | M_Disease | 0.66485373 | M_Disease&NULL |
| 11 | 118545797 | rs600969 | T | C | NULL | 0.663738697 | NULL&M_Disease |
| 11 | 118548421 | rs562313 | A | G | NULL | 0.663738697 | NULL&M_Disease |
| 11 | 118562767 | rs667982 | C | T | NULL | 0.663738697 | NULL&M_Disease |
| 11 | 118565679 | rs502601 | A | G | NULL | 0.663738697 | NULL&M_Disease |
| 11 | 118800302 | rs12793214 | A | C | NULL | 0.663738697 | NULL&M_Disease |
| 11 | 118801640 | rs572608 | A | T | NULL | 0.663738697 | NULL&M_Disease |
| 11 | 118538308 | rs692750 | T | G | NULL | 0.662621837 | NULL&M_Disease |
| 11 | 118539140 | rs4589335 | T | C | NULL | 0.662621837 | NULL&M_Disease |
| 11 | 118550601 | rs673770 | G | T | NULL | 0.662621837 | NULL&M_Disease |
| 11 | 118550756 | rs642530 | C | T | NULL | 0.662621837 | NULL&M_Disease |
| 11 | 118550784 | rs642497 | A | C | NULL | 0.662621837 | NULL&M_Disease |
| 11 | 118551597 | rs12799314 | A | G | NULL | 0.662621837 | NULL&M_Disease |
| 11 | 118552420 | rs4936433 | G | A | NULL | 0.662621837 | NULL&M_Disease |
| 11 | 118552715 | rs519848 | A | G | NULL | 0.662621837 | NULL&M_Disease |
| 11 | 118552842 | rs549370 | A | C | NULL | 0.662621837 | NULL&M_Disease |
| 11 | 118552996 | rs654792 | T | G | NULL | 0.662621837 | NULL&M_Disease |
| 11 | 118553218 | rs592280 | C | T | NULL | 0.662621837 | NULL&M_Disease |
| 11 | 118554248 | rs7106652 | T | C | NULL | 0.662621837 | NULL&M_Disease |
| 11 | 118554259 | rs7106851 | A | G | NULL | 0.662621837 | NULL&M_Disease |
| 11 | 118554436 | rs11605069 | G | T | NULL | 0.662621837 | NULL&M_Disease |
| 11 | 118554448 | rs11602869 | A | G | NULL | 0.662621837 | NULL&M_Disease |
| 11 | 118554939 | rs12796373 | A | G | NULL | 0.662621837 | NULL&M_Disease |
| 11 | 118561410 | rs12798453 | G | T | NULL | 0.662621837 | NULL&M_Disease |
| 11 | 118562799 | rs582630 | A | C | NULL | 0.662621837 | NULL&M_Disease |
| 11 | 118563381 | rs585039 | A | C | NULL | 0.662621837 | NULL&M_Disease |
| 11 | 118564397 | rs558781 | T | G | NULL | 0.662621837 | NULL&M_Disease |
| 11 | 118564413 | rs508640 | G | A | NULL | 0.662621837 | NULL&M_Disease |
| 11 | 118564601 | rs471227 | C | T | NULL | 0.662621837 | NULL&M_Disease |
| 11 | 118564762 | rs516153 | A | C | NULL | 0.662621837 | NULL&M_Disease |
| 11 | 118564982 | rs518028 | C | T | NULL | 0.662621837 | NULL&M_Disease |
| 11 | 118565231 | rs519982 | T | C | NULL | 0.662621837 | NULL&M_Disease |
| 11 | 118565336 | rs478371 | A | G | NULL | 0.662621837 | NULL&M_Disease |
| 11 | 118755630 | rs3176905 | C | T | NULL | 0.662621837 | NULL&M_Disease |
| 11 | 118462188 | rs4936431 | G | C | M_Disease | 0.661503159 | M_Disease&NULL |
| 11 | 118469739 | rs143380378 | G | C | M_Disease | 0.661503159 | M_Disease&NULL |
| 11 | 118554100 | rs4331144 | G | A | NULL | 0.661503159 | NULL&M_Disease |
| 11 | 118566090 | rs7130076 | C | T | NULL | 0.661503159 | NULL&M_Disease |
| 11 | 118566175 | rs7114458 | A | G | NULL | 0.661503159 | NULL&M_Disease |
| 11 | 118566234 | rs7114216 | G | A | NULL | 0.661503159 | NULL&M_Disease |
| 11 | 118566313 | rs12794726 | T | C | NULL | 0.661503159 | NULL&M_Disease |
| 11 | 118566746 | rs533646 | G | C | NULL | 0.661503159 | NULL&M_Disease |
| 11 | 118567444 | rs493834 | G | A | NULL | 0.661503159 | NULL&M_Disease |
| 11 | 118552074 | rs4938524 | T | G | NULL | 0.660382673 | NULL&M_Disease |
| 11 | 118553326 | rs693037 | T | C | NULL | 0.660382673 | NULL&M_Disease |
| 11 | 118555719 | rs625513 | C | A | NULL | 0.660382673 | NULL&M_Disease |
| 11 | 118556084 | rs613377 | T | C | NULL | 0.660382673 | NULL&M_Disease |
| 11 | 118556691 | rs610664 | G | A | NULL | 0.660382673 | NULL&M_Disease |
| 11 | 118556797 | rs571751 | C | A | NULL | 0.660382673 | NULL&M_Disease |
| 11 | 118557198 | rs546890 | C | T | NULL | 0.660382673 | NULL&M_Disease |
| 11 | 118558078 | rs555356 | T | G | NULL | 0.660382673 | NULL&M_Disease |
| 11 | 118558333 | rs693306 | G | A | NULL | 0.660382673 | NULL&M_Disease |
| 11 | 118558455 | rs558907 | G | A | NULL | 0.660382673 | NULL&M_Disease |
| 11 | 118558827 | rs561845 | G | A | NULL | 0.660382673 | NULL&M_Disease |
| 11 | 118558889 | rs638805 | T | G | NULL | 0.660382673 | NULL&M_Disease |
| 11 | 118558901 | rs510408 | C | G | NULL | 0.660382673 | NULL&M_Disease |
| 11 | 118558952 | rs509641 | C | T | NULL | 0.660382673 | NULL&M_Disease |
| 11 | 118558953 | rs563549 | A | G | NULL | 0.660382673 | NULL&M_Disease |
| 11 | 118559003 | rs638301 | G | A | NULL | 0.660382673 | NULL&M_Disease |
| 11 | 118559084 | rs508579 | C | G | NULL | 0.660382673 | NULL&M_Disease |
| 11 | 118559178 | rs486462 | C | T | NULL | 0.660382673 | NULL&M_Disease |
| 11 | 118559709 | rs574217358 | A | G | NULL | 0.660382673 | NULL&M_Disease |
| 11 | 118559866 | rs7926959 | C | T | NULL | 0.660382673 | NULL&M_Disease |
| 11 | 118559881 | rs7926970 | G | T | NULL | 0.660382673 | NULL&M_Disease |
| 11 | 118560569 | rs4938529 | T | C | NULL | 0.660382673 | NULL&M_Disease |
| 11 | 118560570 | rs4938530 | T | G | NULL | 0.660382673 | NULL&M_Disease |
| 11 | 118566039 | rs6589674 | G | A | NULL | 0.660382673 | NULL&M_Disease |

| | | | | | | | | | |
|---|---|---|---|---|---|---|---|---|---|
| 11 | 118702910 | rs7941321 | A | G | 0.908 | -4194.66 | -4194.145 | 0.515 | 7.55E-07 |
| 11 | 118703001 | rs7928228 | T | C | 0.908 | -4194.66 | -4194.145 | 0.515 | 7.55E-07 |
| 11 | 118703484 | rs2508921 | T | C | 0.908 | -4194.66 | -4194.145 | 0.515 | 7.55E-07 |
| 11 | 118703817 | rs1170396 | T | A | 0.908 | -4194.66 | -4194.145 | 0.515 | 7.55E-07 |
| 11 | 118704255 | rs480791 | G | A | 0.908 | -4194.66 | -4194.145 | 0.515 | 7.55E-07 |
| 11 | 118704265 | rs622690 | T | C | 0.908 | -4194.66 | -4194.145 | 0.515 | 7.55E-07 |
| 11 | 118640081 | rs529717 | A | G | 0.901 | -4194.66 | -4194.156 | 0.503 | 7.46E-07 |
| 11 | 118641451 | rs2508929 | G | T | 0.901 | -4194.66 | -4194.156 | 0.503 | 7.46E-07 |
| 11 | 118644213 | rs1790187 | A | G | 0.901 | -4194.66 | -4194.156 | 0.503 | 7.46E-07 |
| 11 | 118648396 | rs548167 | A | G | 0.901 | -4194.66 | -4194.156 | 0.503 | 7.46E-07 |
| 11 | 118652781 | rs2508915 | C | G | 0.901 | -4194.66 | -4194.156 | 0.503 | 7.46E-07 |
| 11 | 118655704 | rs6421568 | C | G | 0.901 | -4194.66 | -4194.156 | 0.503 | 7.46E-07 |
| 11 | 118655775 | rs6421569 | A | C | 0.901 | -4194.66 | -4194.156 | 0.503 | 7.46E-07 |
| 11 | 118701597 | rs4938549 | T | C | 0.908 | -4194.66 | -4194.166 | 0.493 | 7.39E-07 |
| 11 | 118701668 | rs530646 | A | G | 0.908 | -4194.66 | -4194.166 | 0.493 | 7.39E-07 |
| 11 | 118705654 | rs649338 | A | G | 0.908 | -4194.66 | -4194.176 | 0.483 | 7.31E-07 |
| 11 | 118748789 | rs145762171 | A | G | 0.837 | -4194.66 | -4194.177 | 0.482 | 7.31E-07 |
| 11 | 118609736 | rs1783937 | A | G | 0.901 | -4194.66 | -4194.191 | 0.469 | 7.21E-07 |
| 11 | 118812559 | rs74678032 | C | T | 0.855 | -4194.66 | -4194.195 | 0.465 | 7.18E-07 |
| 11 | 118725189 | rs112965917 | A | T | 0.875 | -4194.66 | -4194.224 | 0.436 | 6.98E-07 |
| 11 | 118899218 | rs4936459 | T | C | 1.108 | -4194.66 | -4194.226 | 0.433 | 6.96E-07 |
| 11 | 118702936 | rs7941696 | G | A | 0.91 | -4194.66 | -4194.228 | 0.431 | 6.94E-07 |
| 11 | 118614397 | rs10892281 | T | C | 0.903 | -4194.66 | -4194.242 | 0.417 | 6.85E-07 |
| 11 | 118615660 | rs6589687 | T | A | 0.903 | -4194.66 | -4194.242 | 0.417 | 6.85E-07 |
| 11 | 118622245 | rs485901 | G | A | 0.903 | -4194.66 | -4194.253 | 0.407 | 6.78E-07 |
| 11 | 118610629 | rs521910 | A | G | 0.903 | -4194.66 | -4194.256 | 0.404 | 6.76E-07 |
| 11 | 118798780 | rs77916839 | C | G | 1.153 | -4194.66 | -4194.256 | 0.403 | 6.75E-07 |
| 11 | 118618837 | rs640120 | A | C | 0.904 | -4194.66 | -4194.284 | 0.375 | 6.56E-07 |
| 11 | 118619423 | rs1961138 | C | T | 0.904 | -4194.66 | -4194.284 | 0.375 | 6.56E-07 |
| 11 | 118798824 | rs112119822 | G | A | 1.116 | -4194.66 | -4194.332 | 0.327 | 6.26E-07 |
| 11 | 118799206 | rs7101680 | A | G | 1.116 | -4194.66 | -4194.332 | 0.327 | 6.26E-07 |
| 11 | 118898319 | rs741811 | A | G | 1.104 | -4194.66 | -4194.363 | 0.296 | 6.07E-07 |
| 11 | 118545589 | rs113941645 | G | C | 0.867 | -4194.66 | -4194.374 | 0.285 | 6.00E-07 |
| 11 | 118705145 | rs636736 | G | A | 0.915 | -4194.66 | -4194.424 | 0.236 | 5.71E-07 |
| 11 | 118762701 | rs76457018 | A | G | 0.91 | -4194.66 | -4194.45 | 0.209 | 5.56E-07 |
| 11 | 118801440 | rs12280501 | T | C | 1.111 | -4194.66 | -4194.453 | 0.207 | 5.55E-07 |
| 11 | 118721562 | rs184045098 | T | C | 1.152 | -4194.66 | -4194.465 | 0.194 | 5.48E-07 |
| 11 | 118732181 | rs117079831 | A | G | 1.152 | -4194.66 | -4194.465 | 0.194 | 5.48E-07 |
| 11 | 118615738 | rs193090156 | T | G | 1.157 | -4194.66 | -4194.476 | 0.183 | 5.42E-07 |
| 11 | 118782222 | rs61900138 | C | T | 0.881 | -4194.66 | -4194.494 | 0.165 | 5.32E-07 |
| 11 | 118572267 | rs573905 | A | G | 0.93 | -4194.66 | -4194.495 | 0.164 | 5.32E-07 |
| 11 | 118518753 | rs117947426 | T | C | 1.152 | -4194.66 | -4194.509 | 0.151 | 5.25E-07 |
| 11 | 118503912 | rs45478298 | T | C | 0.885 | -4194.66 | -4194.516 | 0.144 | 5.21E-07 |
| 11 | 118607090 | rs10736495 | C | G | 0.91 | -4194.66 | -4194.518 | 0.142 | 5.20E-07 |
| 11 | 118789952 | rs11217090 | T | C | 1.107 | -4194.66 | -4194.524 | 0.136 | 5.17E-07 |
| 11 | 118602497 | rs478081 | G | A | 1.076 | -4194.66 | -4194.534 | 0.126 | 5.12E-07 |
| 11 | 118726340 | rs113786535 | T | C | 0.918 | -4194.66 | -4194.539 | 0.12 | 5.09E-07 |
| 11 | 118604678 | rs493720 | T | C | 0.912 | -4194.66 | -4194.558 | 0.102 | 5.00E-07 |
| 11 | 118803193 | rs117235696 | G | T | 1.13 | -4194.66 | -4194.566 | 0.094 | 4.96E-07 |
| 11 | 118605127 | rs526850 | G | T | 0.912 | -4194.66 | -4194.582 | 0.078 | 4.88E-07 |
| 11 | 118477746 | rs72999445 | G | T | 1.14 | -4194.66 | -4194.59 | 0.07 | 4.84E-07 |
| 11 | 118809882 | rs11217099 | A | C | 1.09 | -4194.66 | -4194.591 | 0.068 | 4.83E-07 |
| 11 | 118521706 | rs145106291 | G | C | 1.14 | -4194.66 | -4194.595 | 0.065 | 4.81E-07 |
| 11 | 118577614 | rs947931 | G | T | 1.088 | -4194.66 | -4194.594 | 0.065 | 4.81E-07 |
| 11 | 118803842 | rs11600378 | G | A | 1.127 | -4194.66 | -4194.596 | 0.064 | 4.81E-07 |
| 11 | 118804671 | rs143711539 | A | G | 1.127 | -4194.66 | -4194.596 | 0.064 | 4.81E-07 |
| 11 | 118602058 | rs562059 | G | T | 1.074 | -4194.66 | -4194.611 | 0.048 | 4.73E-07 |
| 11 | 118607694 | rs7119702 | G | A | 0.913 | -4194.66 | -4194.614 | 0.046 | 4.72E-07 |
| 11 | 118789657 | rs12294466 | T | C | 1.11 | -4194.66 | -4194.616 | 0.044 | 4.71E-07 |
| 11 | 118583971 | rs6589677 | A | G | 0.911 | -4194.66 | -4194.622 | 0.038 | 4.69E-07 |
| 11 | 118583712 | rs118008100 | A | G | 1.135 | -4194.66 | -4194.622 | 0.037 | 4.68E-07 |
| 11 | 118605447 | rs2186781 | T | C | 0.914 | -4194.66 | -4194.627 | 0.033 | 4.66E-07 |
| 11 | 118805144 | rs11605265 | A | G | 1.123 | -4194.66 | -4194.647 | 0.013 | 4.57E-07 |
| 11 | 118805220 | rs11605658 | T | C | 1.123 | -4194.66 | -4194.647 | 0.013 | 4.57E-07 |
| 11 | 118831755 | rs10892316 | T | C | 1.088 | -4194.66 | -4194.654 | 0.006 | 4.54E-07 |
| 11 | 118834718 | rs112939258 | T | C | 1.088 | -4194.66 | -4194.656 | 0.004 | 4.53E-07 |
| 11 | 118811641 | rs11606199 | A | G | 1.087 | -4194.66 | -4194.672 | -0.013 | 4.45E-07 |
| 11 | 118811315 | rs77638281 | A | G | 1.121 | -4194.66 | -4194.677 | -0.018 | 4.43E-07 |
| 11 | 118876552 | rs74884652 | C | T | 1.091 | -4194.66 | -4194.697 | -0.037 | 4.35E-07 |
| 11 | 118576463 | rs496547 | A | T | 1.072 | -4194.66 | -4194.704 | -0.044 | 4.32E-07 |
| 11 | 118876213 | rs76253444 | G | A | 1.086 | -4194.66 | -4194.71 | -0.05 | 4.29E-07 |
| 11 | 118877891 | rs75154936 | C | T | 1.09 | -4194.66 | -4194.727 | -0.068 | 4.22E-07 |
| 11 | 118890085 | rs4938619 | T | C | 1.09 | -4194.66 | -4194.727 | -0.068 | 4.22E-07 |
| 11 | 118893767 | rs111345266 | T | C | 1.09 | -4194.66 | -4194.727 | -0.068 | 4.22E-07 |
| 11 | 118559731 | rs113784020 | T | C | 0.897 | -4194.66 | -4194.75 | -0.091 | 4.12E-07 |
| 11 | 118799691 | rs191378986 | G | A | 1.117 | -4194.66 | -4194.752 | -0.092 | 4.12E-07 |
| 11 | 118433807 | rs146500266 | A | G | 1.114 | -4194.66 | -4194.753 | -0.093 | 4.11E-07 |
| 11 | 118834508 | rs111358668 | A | G | 1.085 | -4194.66 | -4194.755 | -0.095 | 4.10E-07 |
| 11 | 118896959 | rs57156480 | A | G | 1.079 | -4194.66 | -4194.756 | -0.097 | 4.09E-07 |
| 11 | 118797562 | rs142336193 | G | A | 0.896 | -4194.66 | -4194.771 | -0.111 | 4.04E-07 |
| 11 | 118712509 | rs75488969 | T | C | 0.914 | -4194.66 | -4194.778 | -0.118 | 4.01E-07 |
| 11 | 118856582 | rs3887652 | C | G | 1.077 | -4194.66 | -4194.784 | -0.125 | 3.98E-07 |
| 11 | 118873840 | rs9645663 | C | T | 1.084 | -4194.66 | -4194.784 | -0.125 | 3.98E-07 |
| 11 | 118789026 | rs11607239 | T | G | 0.899 | -4194.66 | -4194.798 | -0.138 | 3.93E-07 |
| 11 | 118601426 | rs186926311 | T | C | 0.899 | -4194.66 | -4194.8 | -0.141 | 3.92E-07 |
| 11 | 118875727 | rs4936457 | G | A | 1.076 | -4194.66 | -4194.807 | -0.148 | 3.89E-07 |
| 11 | 118875761 | rs4936458 | G | T | 1.076 | -4194.66 | -4194.807 | -0.148 | 3.89E-07 |
| 11 | 118539151 | rs72999491 | A | G | 1.086 | -4194.66 | -4194.812 | -0.152 | 3.88E-07 |
| 11 | 118853774 | rs11217109 | C | T | 1.083 | -4194.66 | -4194.813 | -0.154 | 3.87E-07 |
| 11 | 118572000 | rs182361280 | A | G | 0.906 | -4194.66 | -4194.819 | -0.16 | 3.84E-07 |
| 11 | 118873987 | rs9645664 | C | T | 1.076 | -4194.66 | -4194.819 | -0.16 | 3.84E-07 |
| 11 | 118621277 | rs117409833 | C | T | 1.108 | -4194.66 | -4194.823 | -0.163 | 3.83E-07 |
| 11 | 118870413 | rs10892319 | T | C | 1.082 | -4194.66 | -4194.838 | -0.179 | 3.77E-07 |

| | | | | | | | |
|---|---|---|---|---|---|---|---|
| 11 | 118754649 | rs10892307 | G | C | NULL | 0.660382673 | NULL&M_Disease |
| 11 | 118662340 | rs5006102 | A | T | M_Disease | 0.660382673 | M_Disease&NULL |
| 11 | 118677083 | rs675003 | T | C | M_Disease | 0.660382673 | M_Disease&NULL |
| 11 | 118678780 | rs500254 | G | A | M_Disease | 0.660382673 | M_Disease&NULL |
| 11 | 118559247 | rs533144 | A | C | NULL | 0.659260388 | NULL&M_Disease |
| 11 | 118559542 | rs625918 | T | C | NULL | 0.659260388 | NULL&M_Disease |
| 11 | 118567375 | rs478296 | C | T | NULL | 0.659260388 | NULL&M_Disease |
| 11 | 118567486 | rs644498 | C | T | NULL | 0.659260388 | NULL&M_Disease |
| 11 | 118700425 | rs10128600 | C | T | M_Disease | 0.659260388 | M_Disease&NULL |
| 11 | 118831727 | rs10892315 | C | T | NULL | 0.659260388 | NULL&M_Disease |
| 11 | 118555754 | rs525485 | A | G | NULL | 0.658136315 | NULL&M_Disease |
| 11 | 118558449 | rs514143 | A | G | NULL | 0.658136315 | NULL&M_Disease |
| 11 | 118567733 | rs481903 | C | T | NULL | 0.658136315 | NULL&M_Disease |
| 11 | 118567218 | rs11216947 | A | G | NULL | 0.657010463 | NULL&M_Disease |
| 11 | 118567257 | rs7126172 | G | A | NULL | 0.657010463 | NULL&M_Disease |
| 11 | 118662045 | rs7933859 | T | C | M_Disease | 0.657010463 | M_Disease&NULL |
| 11 | 118662203 | rs4508234 | G | C | M_Disease | 0.657010463 | M_Disease&NULL |
| 11 | 118832035 | rs10892317 | C | T | NULL | 0.654753461 | NULL&M_Disease |
| 11 | 118567084 | rs542613 | A | G | NULL | 0.653622331 | NULL&M_Disease |
| 11 | 118573668 | rs574383 | C | T | NULL | 0.653622331 | NULL&M_Disease |
| 11 | 118573743 | rs519294 | A | C | NULL | 0.653622331 | NULL&M_Disease |
| 11 | 118661376 | rs73003294 | T | C | NULL | 0.653622331 | NULL&M_Disease |
| 11 | 118429764 | rs111942423 | G | T | NULL | 0.652489462 | NULL&M_Disease |
| 11 | 118430048 | rs11216896 | T | C | NULL | 0.652489462 | NULL&M_Disease |
| 11 | 118430519 | rs11552421 | T | C | NULL | 0.652489462 | NULL&M_Disease |
| 11 | 118440863 | rs11216904 | G | A | NULL | 0.652489462 | NULL&M_Disease |
| 11 | 118441846 | rs10892243 | C | T | NULL | 0.652489462 | NULL&M_Disease |
| 11 | 118443944 | rs11216906 | A | G | NULL | 0.652489462 | NULL&M_Disease |
| 11 | 118445585 | rs73023341 | G | A | NULL | 0.652489462 | NULL&M_Disease |
| 11 | 118447580 | rs9736327 | G | T | NULL | 0.652489462 | NULL&M_Disease |
| 11 | 118447597 | rs9735421 | A | G | NULL | 0.652489462 | NULL&M_Disease |
| 11 | 118450046 | rs11216910 | G | A | NULL | 0.652489462 | NULL&M_Disease |
| 11 | 118453483 | rs60112336 | T | C | NULL | 0.652489462 | NULL&M_Disease |
| 11 | 118567545 | rs644867 | G | A | NULL | 0.652489462 | NULL&M_Disease |
| 11 | 118572760 | rs611733 | T | C | NULL | 0.652489462 | NULL&M_Disease |
| 11 | 118573022 | rs552079 | C | T | NULL | 0.652489462 | NULL&M_Disease |
| 11 | 118573339 | rs673547 | A | G | NULL | 0.652489462 | NULL&M_Disease |
| 11 | 118573535 | rs517445 | G | A | NULL | 0.652489462 | NULL&M_Disease |
| 11 | 118428791 | rs7929887 | G | A | NULL | 0.651354865 | NULL&M_Disease |
| 11 | 118432279 | rs11216897 | G | A | NULL | 0.651354865 | NULL&M_Disease |
| 11 | 118432404 | rs4326835 | T | G | NULL | 0.651354865 | NULL&M_Disease |
| 11 | 118432852 | rs11216898 | C | T | NULL | 0.651354865 | NULL&M_Disease |
| 11 | 118433729 | rs59108310 | G | A | NULL | 0.651354865 | NULL&M_Disease |
| 11 | 118434421 | rs11216899 | G | A | NULL | 0.651354865 | NULL&M_Disease |
| 11 | 118434835 | rs60753148 | T | C | NULL | 0.651354865 | NULL&M_Disease |
| 11 | 118436079 | rs11216900 | C | G | NULL | 0.651354865 | NULL&M_Disease |
| 11 | 118444525 | rs11216907 | A | G | NULL | 0.651354865 | NULL&M_Disease |
| 11 | 118445275 | rs17122269 | C | T | NULL | 0.651354865 | NULL&M_Disease |
| 11 | 118446298 | rs55844025 | T | C | NULL | 0.651354865 | NULL&M_Disease |
| 11 | 118448087 | rs9735449 | T | C | NULL | 0.651354865 | NULL&M_Disease |
| 11 | 118456414 | rs11216914 | C | T | NULL | 0.651354865 | NULL&M_Disease |
| 11 | 118463221 | rs201373950 | A | T | NULL | 0.651354865 | NULL&M_Disease |
| 11 | 118572747 | rs489126 | A | G | NULL | 0.651354865 | NULL&M_Disease |
| 11 | 118820778 | rs139076649 | C | T | M_Disease | 0.651354865 | M_Disease&NULL |
| 11 | 118435756 | rs2277291 | T | C | NULL | 0.650218549 | NULL&M_Disease |
| 11 | 118438448 | rs60837962 | T | A | NULL | 0.650218549 | NULL&M_Disease |
| 11 | 118439048 | rs12224844 | T | A | NULL | 0.650218549 | NULL&M_Disease |
| 11 | 118449370 | rs17122278 | G | A | NULL | 0.650218549 | NULL&M_Disease |
| 11 | 118449396 | rs73023350 | A | G | NULL | 0.650218549 | NULL&M_Disease |
| 11 | 118450875 | rs7126175 | T | C | NULL | 0.650218549 | NULL&M_Disease |
| 11 | 118615232 | rs139423236 | G | C | M_Disease | 0.650218549 | M_Disease&NULL |
| 11 | 118798757 | rs57879605 | T | C | NULL | 0.650218549 | NULL&M_Disease |
| 11 | 118437279 | rs11216902 | G | A | NULL | 0.649080524 | NULL&M_Disease |
| 11 | 118459039 | rs57438673 | A | G | NULL | 0.647940802 | NULL&M_Disease |
| 11 | 118459702 | rs113054880 | G | C | NULL | 0.647940802 | NULL&M_Disease |
| 11 | 118464815 | rs11216921 | C | G | NULL | 0.647940802 | NULL&M_Disease |
| 11 | 118466249 | rs7115634 | G | A | NULL | 0.647940802 | NULL&M_Disease |
| 11 | 118571357 | rs544452 | T | G | NULL | 0.647940802 | NULL&M_Disease |
| 11 | 118571790 | rs656287 | G | A | NULL | 0.647940802 | NULL&M_Disease |
| 11 | 118571911 | rs570949 | T | G | NULL | 0.647940802 | NULL&M_Disease |
| 11 | 118572171 | rs658040 | G | A | NULL | 0.647940802 | NULL&M_Disease |
| 11 | 118836606 | rs113975105 | G | C | NULL | 0.647940802 | NULL&M_Disease |
| 11 | 118451901 | rs11216911 | C | T | NULL | 0.646799393 | NULL&M_Disease |
| 11 | 118452334 | rs11216912 | C | G | NULL | 0.646799393 | NULL&M_Disease |
| 11 | 118453681 | rs11216913 | G | A | NULL | 0.646799393 | NULL&M_Disease |
| 11 | 118457630 | rs17092756 | C | G | NULL | 0.646799393 | NULL&M_Disease |
| 11 | 118462160 | rs11216917 | G | T | NULL | 0.646799393 | NULL&M_Disease |
| 11 | 118465295 | rs11216922 | G | A | NULL | 0.646799393 | NULL&M_Disease |
| 11 | 118465501 | rs11216923 | G | A | NULL | 0.646799393 | NULL&M_Disease |
| 11 | 118471996 | rs11216926 | T | C | NULL | 0.646799393 | NULL&M_Disease |
| 11 | 118572611 | rs611217 | T | C | NULL | 0.646799393 | NULL&M_Disease |
| 11 | 118718485 | rs4938561 | C | T | NULL | 0.646799393 | NULL&M_Disease |
| 11 | 118454177 | rs17842763 | G | A | NULL | 0.645656306 | NULL&M_Disease |
| 11 | 118461605 | rs11216916 | C | T | NULL | 0.645656306 | NULL&M_Disease |
| 11 | 118470092 | rs11216924 | G | C | NULL | 0.645656306 | NULL&M_Disease |
| 11 | 118473231 | rs1047079 | C | T | NULL | 0.645656306 | NULL&M_Disease |
| 11 | 118567881 | rs497678 | C | T | NULL | 0.645656306 | NULL&M_Disease |
| 11 | 118569900 | rs12793594 | C | T | NULL | 0.645656306 | NULL&M_Disease |
| 11 | 118570062 | rs554456 | C | T | NULL | 0.645656306 | NULL&M_Disease |
| 11 | 118570098 | rs482889 | T | G | NULL | 0.645656306 | NULL&M_Disease |
| 11 | 118570697 | rs523793 | T | C | NULL | 0.645656306 | NULL&M_Disease |
| 11 | 118570914 | rs659564 | A | C | NULL | 0.645656306 | NULL&M_Disease |
| 11 | 118570919 | rs642224 | A | G | NULL | 0.645656306 | NULL&M_Disease |

| | | | | | | | | | |
|---|---|---|---|---|---|---|---|---|---|
| 11 | 118556995 | rs72999502 | C | T | 1.101 | -4194.66 | -4194.844 | -0.185 | 3.75E-07 |
| 11 | 118619478 | rs189583461 | G | A | 1.104 | -4194.66 | -4194.844 | -0.185 | 3.75E-07 |
| 11 | 118867695 | rs4938617 | T | C | 1.085 | -4194.66 | -4194.854 | -0.194 | 3.72E-07 |
| 11 | 118853021 | rs113962716 | A | C | 1.082 | -4194.66 | -4194.86 | -0.2 | 3.69E-07 |
| 11 | 118543589 | rs72999499 | G | C | 1.084 | -4194.66 | -4194.863 | -0.203 | 3.68E-07 |
| 11 | 118749318 | rs78312641 | A | T | 1.085 | -4194.66 | -4194.874 | -0.215 | 3.64E-07 |
| 11 | 118872404 | rs6589713 | C | T | 1.073 | -4194.66 | -4194.874 | -0.215 | 3.64E-07 |
| 11 | 118598936 | rs184071181 | C | T | 0.908 | -4194.66 | -4194.876 | -0.216 | 3.64E-07 |
| 11 | 118652593 | rs139217641 | T | C | 1.088 | -4194.66 | -4194.877 | -0.217 | 3.63E-07 |
| 11 | 118809805 | rs142069912 | C | G | 0.907 | -4194.66 | -4194.876 | -0.217 | 3.63E-07 |
| 11 | 118793613 | rs11217091 | C | T | 1.091 | -4194.66 | -4194.883 | -0.223 | 3.61E-07 |
| 11 | 118501723 | rs61900957 | T | C | 0.918 | -4194.66 | -4194.885 | -0.225 | 3.60E-07 |
| 11 | 118892757 | rs117289748 | G | C | 1.076 | -4194.66 | -4194.888 | -0.228 | 3.59E-07 |
| 11 | 118892759 | rs143660533 | C | A | 1.076 | -4194.66 | -4194.888 | -0.228 | 3.59E-07 |
| 11 | 118893878 | rs111322300 | C | T | 1.076 | -4194.66 | -4194.888 | -0.228 | 3.59E-07 |
| 11 | 118894287 | rs569 | G | A | 1.076 | -4194.66 | -4194.888 | -0.228 | 3.59E-07 |
| 11 | 118895202 | rs11006 | G | A | 1.076 | -4194.66 | -4194.888 | -0.228 | 3.59E-07 |
| 11 | 118777378 | rs186296705 | G | T | 0.923 | -4194.66 | -4194.898 | -0.238 | 3.56E-07 |
| 11 | 118854372 | rs75067211 | T | C | 1.074 | -4194.66 | -4194.897 | -0.238 | 3.56E-07 |
| 11 | 118751326 | rs10892306 | A | T | 1.093 | -4194.66 | -4194.908 | -0.249 | 3.52E-07 |
| 11 | 118892940 | rs12280288 | C | T | 1.075 | -4194.66 | -4194.909 | -0.249 | 3.52E-07 |
| 11 | 118873385 | rs10892320 | T | C | 1.072 | -4194.66 | -4194.911 | -0.251 | 3.51E-07 |
| 11 | 118576762 | rs553188 | T | C | 0.922 | -4194.66 | -4194.915 | -0.255 | 3.50E-07 |
| 11 | 118852647 | rs80126726 | A | G | 1.079 | -4194.66 | -4194.925 | -0.265 | 3.46E-07 |
| 11 | 118872741 | rs4573680 | C | T | 1.073 | -4194.66 | -4194.927 | -0.268 | 3.45E-07 |
| 11 | 118895686 | rs8192696 | T | C | 1.076 | -4194.66 | -4194.927 | -0.268 | 3.45E-07 |
| 11 | 118576997 | rs498912 | C | T | 0.923 | -4194.66 | -4194.938 | -0.278 | 3.42E-07 |
| 11 | 118849368 | rs45491601 | A | G | 0.925 | -4194.66 | -4194.943 | -0.283 | 3.40E-07 |
| 11 | 118509741 | rs113313560 | T | C | 0.924 | -4194.66 | -4194.944 | -0.285 | 3.39E-07 |
| 11 | 118622744 | rs6589689 | G | A | 0.918 | -4194.66 | -4194.944 | -0.285 | 3.39E-07 |
| 11 | 118762521 | rs190509947 | T | A | 0.924 | -4194.66 | -4194.96 | -0.301 | 3.34E-07 |
| 11 | 118749563 | rs77643000 | C | T | 1.081 | -4194.66 | -4194.961 | -0.302 | 3.34E-07 |
| 11 | 118587349 | rs11216975 | C | A | 1.084 | -4194.66 | -4194.963 | -0.303 | 3.33E-07 |
| 11 | 118788517 | rs73575424 | A | C | 0.924 | -4194.66 | -4194.967 | -0.307 | 3.32E-07 |
| 11 | 118867803 | rs4938618 | C | G | 1.071 | -4194.66 | -4194.969 | -0.31 | 3.31E-07 |
| 11 | 118869539 | rs11822103 | A | G | 1.071 | -4194.66 | -4194.969 | -0.31 | 3.31E-07 |
| 11 | 118590743 | rs503542 | G | A | 0.94 | -4194.66 | -4194.972 | -0.312 | 3.30E-07 |
| 11 | 118601363 | rs557990 | T | C | 0.926 | -4194.66 | -4194.973 | -0.314 | 3.30E-07 |
| 11 | 118602392 | rs564892 | A | T | 0.926 | -4194.66 | -4194.973 | -0.314 | 3.30E-07 |
| 11 | 118828184 | rs45586434 | A | C | 1.077 | -4194.66 | -4194.977 | -0.317 | 3.29E-07 |
| 11 | 118424513 | rs144339774 | T | C | 1.079 | -4194.66 | -4194.984 | -0.325 | 3.26E-07 |
| 11 | 118896115 | rs2276072 | G | A | 1.071 | -4194.66 | -4194.986 | -0.326 | 3.26E-07 |
| 11 | 118896116 | rs2276071 | C | A | 1.071 | -4194.66 | -4194.986 | -0.326 | 3.26E-07 |
| 11 | 118722946 | rs148990923 | T | C | 1.083 | -4194.66 | -4194.986 | -0.327 | 3.25E-07 |
| 11 | 118709275 | rs187164533 | A | G | 1.08 | -4194.66 | -4194.99 | -0.331 | 3.24E-07 |
| 11 | 118765283 | rs665648 | A | G | 0.935 | -4194.66 | -4195.002 | -0.343 | 3.20E-07 |
| 11 | 118785982 | rs79157249 | T | A | 0.925 | -4194.66 | -4195.006 | -0.346 | 3.19E-07 |
| 11 | 118872581 | rs4558175 | A | G | 1.069 | -4194.66 | -4195.01 | -0.35 | 3.18E-07 |
| 11 | 118664181 | rs145462967 | A | G | 0.929 | -4194.66 | -4195.021 | -0.361 | 3.14E-07 |
| 11 | 118673968 | rs140269932 | A | G | 0.929 | -4194.66 | -4195.021 | -0.361 | 3.14E-07 |
| 11 | 118744615 | rs73009445 | A | G | 0.919 | -4194.66 | -4195.032 | -0.373 | 3.11E-07 |
| 11 | 118555383 | rs2508573 | T | C | 0.941 | -4194.66 | -4195.035 | -0.375 | 3.10E-07 |
| 11 | 118887568 | rs7121270 | G | T | 1.069 | -4194.66 | -4195.036 | -0.377 | 3.09E-07 |
| 11 | 118492713 | rs45457998 | G | A | 0.92 | -4194.66 | -4195.04 | -0.38 | 3.09E-07 |
| 11 | 118887581 | rs7105187 | T | C | 1.069 | -4194.66 | -4195.044 | -0.384 | 3.07E-07 |
| 11 | 118788769 | rs186418307 | T | G | 0.959 | -4194.66 | -4195.048 | -0.388 | 3.06E-07 |
| 11 | 118696055 | rs189417780 | C | A | 0.936 | -4194.66 | -4195.061 | -0.401 | 3.02E-07 |
| 11 | 118594326 | rs544096 | G | A | 0.929 | -4194.66 | -4195.064 | -0.404 | 3.01E-07 |
| 11 | 118791180 | rs146541591 | C | T | 0.927 | -4194.66 | -4195.064 | -0.404 | 3.01E-07 |
| 11 | 118725204 | rs55794678 | T | C | 1.064 | -4194.66 | -4195.069 | -0.409 | 3.00E-07 |
| 11 | 118781710 | rs6589707 | A | C | 1.03 | -4194.66 | -4195.069 | -0.41 | 2.99E-07 |
| 11 | 118787213 | rs7119222 | C | T | 1.03 | -4194.66 | -4195.069 | -0.41 | 2.99E-07 |
| 11 | 118759905 | rs118020807 | A | C | 1.052 | -4194.66 | -4195.072 | -0.412 | 2.99E-07 |
| 11 | 118880740 | rs11217122 | C | T | 1.07 | -4194.66 | -4195.071 | -0.412 | 2.99E-07 |
| 11 | 118613605 | rs139648410 | G | A | 0.949 | -4194.66 | -4195.079 | -0.419 | 2.97E-07 |
| 11 | 118765087 | rs2230321 | T | C | 1.017 | -4194.66 | -4195.082 | -0.422 | 2.96E-07 |
| 11 | 118694739 | rs188550348 | A | G | 1.031 | -4194.66 | -4195.085 | -0.426 | 2.95E-07 |
| 11 | 118797003 | rs73575428 | G | C | 0.927 | -4194.66 | -4195.087 | -0.428 | 2.94E-07 |
| 11 | 118797278 | rs74584888 | A | G | 0.927 | -4194.66 | -4195.087 | -0.428 | 2.94E-07 |
| 11 | 118797392 | rs73575429 | C | T | 0.927 | -4194.66 | -4195.087 | -0.428 | 2.94E-07 |
| 11 | 118797420 | rs111390501 | G | A | 0.927 | -4194.66 | -4195.087 | -0.428 | 2.94E-07 |
| 11 | 118797544 | rs56032119 | G | A | 0.927 | -4194.66 | -4195.087 | -0.428 | 2.94E-07 |
| 11 | 118798154 | rs60969913 | C | G | 0.927 | -4194.66 | -4195.087 | -0.428 | 2.94E-07 |
| 11 | 118786829 | rs7102433 | G | A | 1.017 | -4194.66 | -4195.089 | -0.43 | 2.94E-07 |
| 11 | 118413882 | rs146864992 | T | C | 0.976 | -4194.66 | -4195.091 | -0.431 | 2.93E-07 |
| 11 | 118891314 | rs117694374 | C | T | 0.991 | -4194.66 | -4195.091 | -0.431 | 2.93E-07 |
| 11 | 118761159 | rs57231586 | T | G | 0.985 | -4194.66 | -4195.091 | -0.432 | 2.93E-07 |
| 11 | 118761591 | rs58397111 | T | C | 0.986 | -4194.66 | -4195.093 | -0.433 | 2.93E-07 |
| 11 | 118887887 | rs369816532 | T | G | 1.068 | -4194.66 | -4195.093 | -0.433 | 2.93E-07 |
| 11 | 118762102 | rs58508419 | A | G | 0.995 | -4194.66 | -4195.095 | -0.435 | 2.92E-07 |
| 11 | 118792384 | rs182691948 | T | G | 0.953 | -4194.66 | -4195.098 | -0.438 | 2.91E-07 |
| 11 | 118797857 | rs150438978 | T | A | 0.953 | -4194.66 | -4195.098 | -0.438 | 2.91E-07 |
| 11 | 118419577 | rs191458787 | T | C | 0.977 | -4194.66 | -4195.098 | -0.439 | 2.91E-07 |
| 11 | 118421438 | rs145485933 | G | A | 0.977 | -4194.66 | -4195.098 | -0.439 | 2.91E-07 |
| 11 | 118510642 | rs141169729 | T | C | 1.051 | -4194.66 | -4195.1 | -0.44 | 2.91E-07 |
| 11 | 118666978 | rs1289369859 | A | G | 1.006 | -4194.66 | -4195.103 | -0.444 | 2.89E-07 |
| 11 | 118684537 | rs150583208 | G | A | 1.006 | -4194.66 | -4195.103 | -0.444 | 2.89E-07 |
| 11 | 118880668 | rs11607569 | C | T | 1.069 | -4194.66 | -4195.104 | -0.444 | 2.89E-07 |
| 11 | 118719332 | rs112073991 | G | A | 1.05 | -4194.66 | -4195.106 | -0.446 | 2.89E-07 |
| 11 | 118563286 | rs527619 | A | G | 0.943 | -4194.66 | -4195.106 | -0.447 | 2.89E-07 |
| 11 | 118565651 | rs540800 | A | G | 0.943 | -4194.66 | -4195.106 | -0.447 | 2.89E-07 |
| 11 | 118779681 | rs9971606 | C | T | 0.999 | -4194.66 | -4195.108 | -0.448 | 2.88E-07 |
| 11 | 118889378 | rs7113753 | A | G | 1.066 | -4194.66 | -4195.108 | -0.448 | 2.88E-07 |

| | | | | | | | |
|---|---|---|---|---|---|---|---|
| 11 | 118571023 | rs541631 | G | A | NULL | 0.645656306 | NULL&M_Disease |
| 11 | 118571123 | rs658676 | T | C | NULL | 0.645656306 | NULL&M_Disease |
| 11 | 118571168 | rs488141 | G | A | NULL | 0.645656306 | NULL&M_Disease |
| 11 | 118571316 | rs657769 | T | C | NULL | 0.645656306 | NULL&M_Disease |
| 11 | 118469571 | rs7931564 | G | A | NULL | 0.644511554 | NULL&M_Disease |
| 11 | 118720590 | rs4938564 | T | G | NULL | 0.644511554 | NULL&M_Disease |
| 11 | 118720684 | rs4936440 | G | C | NULL | 0.644511554 | NULL&M_Disease |
| 11 | 118720998 | rs4938565 | A | G | NULL | 0.644511554 | NULL&M_Disease |
| 11 | 118721189 | rs4938566 | T | A | NULL | 0.644511554 | NULL&M_Disease |
| 11 | 118721249 | rs4938567 | G | A | NULL | 0.644511554 | NULL&M_Disease |
| 11 | 118721850 | rs7946961 | G | A | NULL | 0.644511554 | NULL&M_Disease |
| 11 | 118722231 | rs4938570 | G | A | NULL | 0.644511554 | NULL&M_Disease |
| 11 | 118723087 | rs11217059 | T | C | NULL | 0.644511554 | NULL&M_Disease |
| 11 | 118724336 | rs10892298 | C | T | NULL | 0.644511554 | NULL&M_Disease |
| 11 | 118449746 | rs11216909 | T | C | NULL | 0.643365146 | NULL&M_Disease |
| 11 | 118471423 | rs11216925 | C | T | NULL | 0.643365146 | NULL&M_Disease |
| 11 | 118472354 | rs10892245 | A | C | NULL | 0.643365146 | NULL&M_Disease |
| 11 | 118500963 | rs680465 | A | G | M_Disease | 0.643365146 | M_Disease&NULL |
| 11 | 118504742 | rs633683 | T | C | NULL | 0.643365146 | NULL&M_Disease |
| 11 | 118572021 | rs657539 | T | C | NULL | 0.643365146 | NULL&M_Disease |
| 11 | 118681088 | rs2508922 | A | G | NULL | 0.643365146 | NULL&M_Disease |
| 11 | 118717929 | rs4938558 | T | C | NULL | 0.643365146 | NULL&M_Disease |
| 11 | 118718856 | rs2156751 | G | A | NULL | 0.643365146 | NULL&M_Disease |
| 11 | 118718972 | rs2187491 | C | T | NULL | 0.643365146 | NULL&M_Disease |
| 11 | 118719254 | rs2156750 | C | T | NULL | 0.643365146 | NULL&M_Disease |
| 11 | 118719405 | rs2156749 | G | C | NULL | 0.643365146 | NULL&M_Disease |
| 11 | 118719742 | rs7116715 | C | T | NULL | 0.643365146 | NULL&M_Disease |
| 11 | 118568668 | rs11216949 | T | C | NULL | 0.642217093 | NULL&M_Disease |
| 11 | 118568813 | rs7479114 | T | C | NULL | 0.642217093 | NULL&M_Disease |
| 11 | 118569067 | rs4486664 | C | A | NULL | 0.642217093 | NULL&M_Disease |
| 11 | 118569097 | rs4393358 | T | C | NULL | 0.642217093 | NULL&M_Disease |
| 11 | 118569469 | rs7123811 | G | T | NULL | 0.642217093 | NULL&M_Disease |
| 11 | 118569694 | rs7108144 | A | G | NULL | 0.642217093 | NULL&M_Disease |
| 11 | 118730669 | rs7933007 | C | T | NULL | 0.642217093 | NULL&M_Disease |
| 11 | 118473385 | rs17742 | C | T | NULL | 0.641067406 | NULL&M_Disease |
| 11 | 118514022 | rs613243 | C | G | NULL | 0.641067406 | NULL&M_Disease |
| 11 | 118685015 | rs540716 | T | C | NULL | 0.641067406 | NULL&M_Disease |
| 11 | 118723537 | rs7103314 | G | A | NULL | 0.641067406 | NULL&M_Disease |
| 11 | 118750967 | rs11602393 | A | T | NULL | 0.641067406 | NULL&M_Disease |
| 11 | 118816304 | rs586763 | T | C | NULL | 0.641067406 | NULL&M_Disease |
| 11 | 118692727 | rs503425 | C | T | NULL | 0.639916097 | NULL&M_Disease |
| 11 | 118725420 | rs11217062 | A | G | NULL | 0.639916097 | NULL&M_Disease |
| 11 | 118726230 | rs2156755 | A | G | NULL | 0.639916097 | NULL&M_Disease |
| 11 | 118700218 | rs2510902 | A | G | NULL | 0.638763175 | NULL&M_Disease |
| 11 | 118468037 | rs535774 | T | C | M_Disease | 0.638763175 | M_Disease&NULL |
| 11 | 118509668 | rs483598 | A | G | NULL | 0.63645254 | NULL&M_Disease |
| 11 | 118536950 | rs201359394 | T | A | NULL | 0.63645254 | NULL&M_Disease |
| 11 | 118568472 | rs570371 | G | A | NULL | 0.63645254 | NULL&M_Disease |
| 11 | 118568639 | rs511451 | T | C | NULL | 0.635294849 | NULL&M_Disease |
| 11 | 118765688 | rs676925 | G | C | NULL | 0.635294849 | NULL&M_Disease |
| 11 | 118762073 | rs77841686 | A | G | M_Disease | 0.635294849 | M_Disease&NULL |
| 11 | 118763313 | rs78041268 | A | G | M_Disease | 0.635294849 | M_Disease&NULL |
| 11 | 118767564 | rs75438046 | A | G | M_Disease | 0.635294849 | M_Disease&NULL |
| 11 | 118501459 | rs11216934 | A | G | NULL | 0.634135591 | NULL&M_Disease |
| 11 | 118816353 | rs586370 | A | G | NULL | 0.634135591 | NULL&M_Disease |
| 11 | 118462554 | rs11216920 | G | A | NULL | 0.630648526 | NULL&M_Disease |
| 11 | 118499469 | rs577396 | C | G | NULL | 0.630648526 | NULL&M_Disease |
| 11 | 118711830 | rs566848 | G | A | NULL | 0.630648526 | NULL&M_Disease |
| 11 | 118807637 | rs112974653 | T | G | NULL | 0.630648526 | NULL&M_Disease |
| 11 | 118807740 | rs10892312 | A | G | NULL | 0.628316188 | NULL&M_Disease |
| 11 | 118783424 | rs149114341 | A | G | M_Disease | 0.627147766 | M_Disease&NULL |
| 11 | 118806897 | rs10892310 | T | C | NULL | 0.627147766 | NULL&M_Disease |
| 11 | 118807876 | rs12282154 | A | G | NULL | 0.627147766 | NULL&M_Disease |
| 11 | 118809329 | rs10892313 | A | G | NULL | 0.627147766 | NULL&M_Disease |
| 11 | 118810405 | rs4938589 | T | C | NULL | 0.627147766 | NULL&M_Disease |
| 11 | 118805385 | rs55772432 | A | G | M_Disease | 0.625977858 | M_Disease&NULL |
| 11 | 118734640 | rs12286723 | T | C | NULL | 0.624806474 | NULL&M_Disease |
| 11 | 118808140 | rs7117408 | T | C | NULL | 0.624806474 | NULL&M_Disease |
| 11 | 118745664 | rs11217076 | G | T | NULL | 0.623633628 | NULL&M_Disease |
| 11 | 118573894 | rs576283 | G | A | M_Disease | 0.622459331 | M_Disease&NULL |
| 11 | 118573916 | rs627391 | T | A | M_Disease | 0.622459331 | M_Disease&NULL |
| 11 | 118574311 | rs524409 | T | C | M_Disease | 0.621283595 | M_Disease&NULL |
| 11 | 118574675 | rs494459 | T | C | M_Disease | 0.621283595 | M_Disease&NULL |
| 11 | 118535840 | rs12225548 | G | C | NULL | 0.618927855 | NULL&M_Disease |
| 11 | 118575011 | rs530901 | C | T | M_Disease | 0.618927855 | M_Disease&NULL |
| 11 | 118575180 | rs626645 | C | T | M_Disease | 0.618927855 | M_Disease&NULL |
| 11 | 118575419 | rs625735 | A | G | M_Disease | 0.618927855 | M_Disease&NULL |
| 11 | 118575479 | rs637563 | G | A | M_Disease | 0.618927855 | M_Disease&NULL |
| 11 | 118575693 | rs638514 | C | T | M_Disease | 0.618927855 | M_Disease&NULL |
| 11 | 118698170 | rs56296109 | A | C | NULL | 0.618927855 | NULL&M_Disease |
| 11 | 118762218 | rs543824 | G | A | NULL | 0.618927855 | NULL&M_Disease |
| 11 | 118777978 | rs78755429 | A | C | M_Disease | 0.618927855 | M_Disease&NULL |
| 11 | 118560857 | rs73001406 | A | G | M_Disease | 0.617747875 | M_Disease&NULL |
| 11 | 118763736 | rs12363277 | C | T | NULL | 0.617747875 | NULL&M_Disease |
| 11 | 118514625 | rs11216938 | T | C | NULL | 0.616566505 | NULL&M_Disease |
| 11 | 118529127 | rs7928371 | A | G | NULL | 0.616566505 | NULL&M_Disease |
| 11 | 118599649 | rs182364290 | G | A | M_Disease | 0.616566505 | M_Disease&NULL |
| 11 | 118490076 | rs10892247 | A | G | NULL | 0.615383756 | NULL&M_Disease |
| 11 | 118507478 | rs11216937 | T | C | NULL | 0.615383756 | NULL&M_Disease |
| 11 | 118516737 | rs12362152 | G | A | NULL | 0.615383756 | NULL&M_Disease |
| 11 | 118528424 | rs17748 | T | C | NULL | 0.615383756 | NULL&M_Disease |
| 11 | 118569414 | rs6589675 | C | G | NULL | 0.615383756 | NULL&M_Disease |
| 11 | 118698160 | rs640373 | G | A | M_Disease | 0.615383756 | M_Disease&NULL |

| | | | | | | | | | |
|---|---|---|---|---|---|---|---|---|---|
| 11 | 118759000 | rs77491736 | G | C | 0.974 | -4194.66 | -4195.109 | -0.449 | 2.88E-07 |
| 11 | 118887888 | rs3916582 | T | C | 1.067 | -4194.66 | -4195.108 | -0.449 | 2.88E-07 |
| 11 | 118518227 | rs10736494 | T | C | 1.059 | -4194.66 | -4195.115 | -0.456 | 2.86E-07 |
| 11 | 118884708 | rs61381062 | A | G | 1.067 | -4194.66 | -4195.116 | -0.456 | 2.86E-07 |
| 11 | 118801045 | rs75085081 | A | G | 0.935 | -4194.66 | -4195.118 | -0.458 | 2.85E-07 |
| 11 | 118755946 | rs76109832 | T | C | 0.926 | -4194.66 | -4195.12 | -0.46 | 2.85E-07 |
| 11 | 118540104 | rs506234 | G | C | 0.943 | -4194.66 | -4195.123 | -0.464 | 2.84E-07 |
| 11 | 118886656 | rs7131534 | A | G | 1.068 | -4194.66 | -4195.129 | -0.469 | 2.82E-07 |
| 11 | 118884726 | rs58601794 | C | T | 1.066 | -4194.66 | -4195.135 | -0.475 | 2.81E-07 |
| 11 | 118886117 | rs10111 | T | C | 1.068 | -4194.66 | -4195.136 | -0.476 | 2.80E-07 |
| 11 | 118650163 | rs73003218 | T | C | 0.97 | -4194.66 | -4195.14 | -0.48 | 2.79E-07 |
| 11 | 118530434 | rs117619140 | T | G | 1.066 | -4194.66 | -4195.144 | -0.484 | 2.78E-07 |
| 11 | 118888831 | rs11820805 | C | T | 1.065 | -4194.66 | -4195.144 | -0.484 | 2.78E-07 |
| 11 | 118533042 | rs142438429 | T | C | 1.031 | -4194.66 | -4195.146 | -0.487 | 2.77E-07 |
| 11 | 118752178 | rs61150131 | C | T | 0.928 | -4194.66 | -4195.149 | -0.489 | 2.77E-07 |
| 11 | 118591443 | rs111843292 | A | C | 0.96 | -4194.66 | -4195.151 | -0.491 | 2.76E-07 |
| 11 | 118825636 | rs1790191 | G | A | 1.059 | -4194.66 | -4195.152 | -0.492 | 2.76E-07 |
| 11 | 118885163 | rs59608621 | G | A | 1.066 | -4194.66 | -4195.157 | -0.497 | 2.74E-07 |
| 11 | 118877563 | rs58709307 | T | C | 1.067 | -4194.66 | -4195.161 | -0.501 | 2.73E-07 |
| 11 | 118832251 | rs79808604 | A | C | 0.998 | -4194.66 | -4195.165 | -0.506 | 2.72E-07 |
| 11 | 118751898 | rs60651507 | A | G | 0.93 | -4194.66 | -4195.168 | -0.509 | 2.71E-07 |
| 11 | 118819541 | rs150196102 | T | C | 1.011 | -4194.66 | -4195.169 | -0.51 | 2.71E-07 |
| 11 | 118557483 | rs6589673 | C | A | 1.043 | -4194.66 | -4195.17 | -0.511 | 2.71E-07 |
| 11 | 118869460 | rs3737504 | G | A | 1.065 | -4194.66 | -4195.173 | -0.514 | 2.70E-07 |
| 11 | 118885161 | rs59095933 | A | G | 1.065 | -4194.66 | -4195.175 | -0.515 | 2.70E-07 |
| 11 | 118879285 | rs7120445 | C | A | 1.066 | -4194.66 | -4195.177 | -0.518 | 2.69E-07 |
| 11 | 118578462 | rs516459 | C | T | 1.014 | -4194.66 | -4195.183 | -0.523 | 2.67E-07 |
| 11 | 118892277 | rs4938621 | T | C | 1.066 | -4194.66 | -4195.183 | -0.523 | 2.67E-07 |
| 11 | 118892835 | rs61179449 | C | G | 1.066 | -4194.66 | -4195.183 | -0.523 | 2.67E-07 |
| 11 | 118892872 | rs58955701 | T | C | 1.066 | -4194.66 | -4195.183 | -0.523 | 2.67E-07 |
| 11 | 118893023 | rs7127188 | C | A | 1.066 | -4194.66 | -4195.183 | -0.523 | 2.67E-07 |
| 11 | 118878222 | rs11606357 | C | T | 1.066 | -4194.66 | -4195.184 | -0.524 | 2.67E-07 |
| 11 | 118888619 | rs11602764 | G | A | 1.064 | -4194.66 | -4195.185 | -0.525 | 2.67E-07 |
| 11 | 118883491 | rs60065241 | A | C | 1.066 | -4194.66 | -4195.186 | -0.526 | 2.67E-07 |
| 11 | 118605352 | rs76478216 | C | T | 1.072 | -4194.66 | -4195.186 | -0.527 | 2.66E-07 |
| 11 | 118483912 | rs45621436 | G | T | 1.036 | -4194.66 | -4195.189 | -0.53 | 2.66E-07 |
| 11 | 118493319 | rs10790254 | A | G | 1.036 | -4194.66 | -4195.189 | -0.53 | 2.66E-07 |
| 11 | 118824439 | rs61900157 | C | A | 1.058 | -4194.66 | -4195.192 | -0.533 | 2.65E-07 |
| 11 | 118824996 | rs4426142 | A | G | 1.058 | -4194.66 | -4195.192 | -0.533 | 2.65E-07 |
| 11 | 118825069 | rs2097131 | A | G | 1.058 | -4194.66 | -4195.192 | -0.533 | 2.65E-07 |
| 11 | 118823148 | rs663003 | T | C | 1.058 | -4194.66 | -4195.196 | -0.536 | 2.64E-07 |
| 11 | 118436071 | rs73023329 | T | C | 0.934 | -4194.66 | -4195.198 | -0.539 | 2.63E-07 |
| 11 | 118559426 | rs10892253 | A | C | 1.034 | -4194.66 | -4195.2 | -0.54 | 2.63E-07 |
| 11 | 118561858 | rs10790257 | C | G | 1.034 | -4194.66 | -4195.2 | -0.54 | 2.63E-07 |
| 11 | 118561859 | rs10790258 | T | G | 1.034 | -4194.66 | -4195.2 | -0.54 | 2.63E-07 |
| 11 | 118748170 | rs77074463 | C | G | 1.073 | -4194.66 | -4195.201 | -0.541 | 2.63E-07 |
| 11 | 118691070 | rs140577174 | G | A | 1.029 | -4194.66 | -4195.203 | -0.544 | 2.62E-07 |
| 11 | 118782474 | rs11606719 | G | C | 1.019 | -4194.66 | -4195.203 | -0.544 | 2.62E-07 |
| 11 | 118477226 | rs45496391 | C | A | 1.028 | -4194.66 | -4195.204 | -0.545 | 2.62E-07 |
| 11 | 118455465 | rs45460493 | A | G | 1.027 | -4194.66 | -4195.206 | -0.546 | 2.61E-07 |
| 11 | 118507068 | rs10892249 | C | T | 1.027 | -4194.66 | -4195.207 | -0.547 | 2.61E-07 |
| 11 | 118868750 | rs4445645 | C | G | 1.065 | -4194.66 | -4195.207 | -0.547 | 2.61E-07 |
| 11 | 118879824 | rs73559119 | C | T | 1.065 | -4194.66 | -4195.207 | -0.547 | 2.61E-07 |
| 11 | 118880012 | rs11825284 | A | G | 1.065 | -4194.66 | -4195.207 | -0.547 | 2.61E-07 |
| 11 | 118518021 | rs10736493 | G | T | 1.024 | -4194.66 | -4195.211 | -0.551 | 2.60E-07 |
| 11 | 118421841 | rs145968026 | G | C | 1.023 | -4194.66 | -4195.213 | -0.554 | 2.59E-07 |
| 11 | 118487265 | rs76544852 | T | G | 1.023 | -4194.66 | -4195.214 | -0.554 | 2.59E-07 |
| 11 | 118580773 | rs675329 | T | C | 1.017 | -4194.66 | -4195.214 | -0.554 | 2.59E-07 |
| 11 | 118630532 | rs35888235 | T | G | 0.937 | -4194.66 | -4195.214 | -0.554 | 2.59E-07 |
| 11 | 118460957 | rs45600633 | A | G | 1.02 | -4194.66 | -4195.218 | -0.558 | 2.58E-07 |
| 11 | 118474283 | rs139202987 | G | A | 1.02 | -4194.66 | -4195.218 | -0.558 | 2.58E-07 |
| 11 | 118445559 | rs73555296 | T | G | 1.02 | -4194.66 | -4195.222 | -0.562 | 2.57E-07 |
| 11 | 118583261 | rs112246697 | A | G | 0.995 | -4194.66 | -4195.223 | -0.563 | 2.57E-07 |
| 11 | 118889132 | rs11217124 | G | A | 1.063 | -4194.66 | -4195.223 | -0.564 | 2.57E-07 |
| 11 | 118760443 | rs12786257 | A | G | 1.04 | -4194.66 | -4195.229 | -0.569 | 2.55E-07 |
| 11 | 118515579 | rs10790255 | G | T | 1.062 | -4194.66 | -4195.235 | -0.576 | 2.54E-07 |
| 11 | 118824683 | rs507763 | C | G | 1.065 | -4194.66 | -4195.236 | -0.576 | 2.54E-07 |
| 11 | 118824917 | rs11217102 | A | G | 1.065 | -4194.66 | -4195.236 | -0.576 | 2.54E-07 |
| 11 | 118475098 | rs604096 | G | A | 1.063 | -4194.66 | -4195.237 | -0.578 | 2.53E-07 |
| 11 | 118721767 | rs116884018 | C | T | 0.961 | -4194.66 | -4195.24 | -0.58 | 2.53E-07 |
| 11 | 118826045 | rs537080 | C | T | 1.065 | -4194.66 | -4195.24 | -0.58 | 2.53E-07 |
| 11 | 118889185 | rs11217125 | C | T | 1.063 | -4194.66 | -4195.24 | -0.581 | 2.52E-07 |
| 11 | 118793479 | rs138526953 | A | C | 0.992 | -4194.66 | -4195.244 | -0.585 | 2.51E-07 |
| 11 | 118633561 | rs12794288 | A | C | 0.942 | -4194.66 | -4195.248 | -0.588 | 2.51E-07 |
| 11 | 118466646 | rs184841209 | T | G | 0.939 | -4194.66 | -4195.253 | -0.593 | 2.49E-07 |
| 11 | 118603615 | rs74783030 | A | G | 0.961 | -4194.66 | -4195.271 | -0.612 | 2.45E-07 |
| 11 | 118782802 | rs76160025 | T | C | 0.965 | -4194.66 | -4195.276 | -0.617 | 2.43E-07 |
| 11 | 118539746 | rs72999492 | C | A | 1.063 | -4194.66 | -4195.282 | -0.622 | 2.42E-07 |
| 11 | 118614115 | rs73001451 | C | T | 0.94 | -4194.66 | -4195.281 | -0.622 | 2.42E-07 |
| 11 | 118826060 | rs625291 | T | C | 1.055 | -4194.66 | -4195.294 | -0.634 | 2.39E-07 |
| 11 | 118826063 | rs666191 | A | G | 1.055 | -4194.66 | -4195.31 | -0.65 | 2.36E-07 |
| 11 | 118729283 | rs11217068 | T | C | 1.061 | -4194.66 | -4195.319 | -0.659 | 2.33E-07 |
| 11 | 118712270 | rs7105658 | T | C | 1.061 | -4194.66 | -4195.324 | -0.664 | 2.32E-07 |
| 11 | 118700515 | rs112571283 | T | C | 0.985 | -4194.66 | -4195.328 | -0.669 | 2.31E-07 |
| 11 | 118874272 | rs192113495 | G | T | 1.003 | -4194.66 | -4195.334 | -0.675 | 2.30E-07 |
| 11 | 118757788 | rs3176912 | A | G | 0.986 | -4194.66 | -4195.353 | -0.693 | 2.26E-07 |
| 11 | 118568427 | rs117361624 | T | C | 0.946 | -4194.66 | -4195.354 | -0.694 | 2.25E-07 |
| 11 | 118729442 | rs12277355 | T | C | 1.06 | -4194.66 | -4195.356 | -0.697 | 2.25E-07 |
| 11 | 118543798 | rs76570573 | T | C | 0.946 | -4194.66 | -4195.357 | -0.698 | 2.25E-07 |
| 11 | 118711069 | rs538645 | C | T | 0.946 | -4194.66 | -4195.359 | -0.7 | 2.24E-07 |
| 11 | 118772160 | rs61730467 | A | G | 1.012 | -4194.66 | -4195.37 | -0.71 | 2.22E-07 |
| 11 | 118775524 | rs79108568 | G | T | 0.994 | -4194.66 | -4195.37 | -0.711 | 2.22E-07 |
| 11 | 118629959 | rs140110112 | A | G | 0.948 | -4194.66 | -4195.379 | -0.719 | 2.20E-07 |

| | | | | | | | |
|---|---|---|---|---|---|---|---|
| 11 | 118698274 | rs639915 | A | G | M_Disease | 0.615383756 | M_Disease&NULL |
| 11 | 118698294 | rs639885 | A | G | M_Disease | 0.615383756 | M_Disease&NULL |
| 11 | 118698537 | rs638893 | G | A | M_Disease | 0.615383756 | M_Disease&NULL |
| 11 | 118698777 | rs516332 | G | A | M_Disease | 0.615383756 | M_Disease&NULL |
| 11 | 118699744 | rs2508564 | C | T | M_Disease | 0.615383756 | M_Disease&NULL |
| 11 | 118699863 | rs2510901 | A | G | M_Disease | 0.615383756 | M_Disease&NULL |
| 11 | 118516774 | rs12365095 | T | C | NULL | 0.614199643 | NULL&M_Disease |
| 11 | 118516903 | rs80015904 | A | G | NULL | 0.614199643 | NULL&M_Disease |
| 11 | 118707559 | rs7118009 | T | C | NULL | 0.614199643 | NULL&M_Disease |
| 11 | 118727137 | rs117223320 | C | G | NULL | 0.614199643 | NULL&M_Disease |
| 11 | 118735876 | rs79843382 | T | C | NULL | 0.614199643 | NULL&M_Disease |
| 11 | 118757191 | rs1790192 | G | A | NULL | 0.614199643 | NULL&M_Disease |
| 11 | 118501022 | rs10892248 | A | G | NULL | 0.613014176 | NULL&M_Disease |
| 11 | 118689745 | rs492471 | T | C | NULL | 0.613014176 | NULL&M_Disease |
| 11 | 118715589 | rs116872113 | A | G | NULL | 0.613014176 | NULL&M_Disease |
| 11 | 118764051 | rs613791 | T | C | NULL | 0.613014176 | NULL&M_Disease |
| 11 | 118895495 | rs8301 | G | C | M_Disease | 0.613014176 | M_Disease&NULL |
| 11 | 118495287 | rs77683570 | C | T | NULL | 0.611827369 | NULL&M_Disease |
| 11 | 118496119 | rs45565037 | A | G | NULL | 0.611827369 | NULL&M_Disease |
| 11 | 118499394 | rs2236661 | G | C | NULL | 0.611827369 | NULL&M_Disease |
| 11 | 118521549 | rs494560 | A | G | NULL | 0.611827369 | NULL&M_Disease |
| 11 | 118728479 | rs7924647 | G | A | NULL | 0.611827369 | NULL&M_Disease |
| 11 | 118523799 | rs72999483 | C | T | NULL | 0.610639234 | NULL&M_Disease |
| 11 | 118723753 | rs7119568 | A | T | NULL | 0.610639234 | NULL&M_Disease |
| 11 | 118746223 | rs11217077 | C | T | NULL | 0.610639234 | NULL&M_Disease |
| 11 | 118759610 | rs566416 | G | T | NULL | 0.610639234 | NULL&M_Disease |
| 11 | 118811878 | rs74891547 | C | G | NULL | 0.610639234 | NULL&M_Disease |
| 11 | 118534082 | rs10790256 | T | C | NULL | 0.609449784 | NULL&M_Disease |
| 11 | 118583847 | rs12803363 | T | C | NULL | 0.609449784 | NULL&M_Disease |
| 11 | 118708785 | rs7122510 | A | C | NULL | 0.609449784 | NULL&M_Disease |
| 11 | 118708786 | rs7122511 | A | C | NULL | 0.609449784 | NULL&M_Disease |
| 11 | 118804266 | rs7123066 | T | C | NULL | 0.609449784 | NULL&M_Disease |
| 11 | 118804283 | rs7122959 | G | A | NULL | 0.609449784 | NULL&M_Disease |
| 11 | 118555692 | rs10892252 | G | A | M_Disease | 0.609449784 | M_Disease&NULL |
| 11 | 118661197 | rs61901410 | G | C | M_Disease | 0.609449784 | M_Disease&NULL |
| 11 | 118595207 | rs563537 | A | T | M_Disease | 0.608259031 | M_Disease&NULL |
| 11 | 118709600 | rs12272981 | G | A | NULL | 0.608259031 | NULL&M_Disease |
| 11 | 118510562 | rs4245184 | A | G | NULL | 0.607066988 | NULL&M_Disease |
| 11 | 118511352 | rs644273 | T | C | NULL | 0.607066988 | NULL&M_Disease |
| 11 | 118511757 | rs654423 | A | G | NULL | 0.607066988 | NULL&M_Disease |
| 11 | 118580348 | rs516503 | T | C | M_Disease | 0.607066988 | M_Disease&NULL |
| 11 | 118606197 | rs12420846 | T | C | M_Disease | 0.607066988 | M_Disease&NULL |
| 11 | 118791150 | rs12270280 | T | C | NULL | 0.607066988 | NULL&M_Disease |
| 11 | 118791532 | rs12271856 | T | G | NULL | 0.607066988 | NULL&M_Disease |
| 11 | 118792278 | rs77398170 | G | A | NULL | 0.607066988 | NULL&M_Disease |
| 11 | 118793918 | rs12282752 | C | T | NULL | 0.607066988 | NULL&M_Disease |
| 11 | 118486110 | rs45540840 | A | G | NULL | 0.605873668 | NULL&M_Disease |
| 11 | 118501299 | rs7936556 | A | G | NULL | 0.605873668 | NULL&M_Disease |
| 11 | 118615390 | rs78502545 | T | C | NULL | 0.605873668 | NULL&M_Disease |
| 11 | 118626825 | rs74337439 | A | G | NULL | 0.605873668 | NULL&M_Disease |
| 11 | 118556398 | rs11216943 | A | G | M_Disease | 0.605873668 | M_Disease&NULL |
| 11 | 118486067 | rs11603023 | T | C | NULL | 0.604679085 | NULL&M_Disease |
| 11 | 118806216 | rs1231742890 | G | T | NULL | 0.604679085 | NULL&M_Disease |
| 11 | 118480695 | rs34026809 | C | G | NULL | 0.60348325 | NULL&M_Disease |
| 11 | 118526691 | rs72999486 | T | C | NULL | 0.60348325 | NULL&M_Disease |
| 11 | 118724116 | rs12276533 | A | G | NULL | 0.60348325 | NULL&M_Disease |
| 11 | 118802128 | rs59069289 | T | G | NULL | 0.60348325 | NULL&M_Disease |
| 11 | 118802386 | rs59480472 | C | T | NULL | 0.60348325 | NULL&M_Disease |
| 11 | 118803160 | rs73575440 | A | G | NULL | 0.60348325 | NULL&M_Disease |
| 11 | 118803596 | rs111313044 | A | G | NULL | 0.60348325 | NULL&M_Disease |
| 11 | 118804643 | rs56933877 | A | G | NULL | 0.60348325 | NULL&M_Disease |
| 11 | 118804919 | rs73575441 | C | T | NULL | 0.60348325 | NULL&M_Disease |
| 11 | 118804989 | rs112045567 | G | A | NULL | 0.60348325 | NULL&M_Disease |
| 11 | 118502172 | rs35435609 | T | C | NULL | 0.602286177 | NULL&M_Disease |
| 11 | 118712427 | rs7108905 | G | C | NULL | 0.602286177 | NULL&M_Disease |
| 11 | 118805138 | rs7127427 | A | G | NULL | 0.602286177 | NULL&M_Disease |
| 11 | 118725383 | rs9735317 | T | C | NULL | 0.601087879 | NULL&M_Disease |
| 11 | 118488782 | rs11216930 | C | A | NULL | 0.599888369 | NULL&M_Disease |
| 11 | 118494790 | rs11825621 | C | T | NULL | 0.599888369 | NULL&M_Disease |
| 11 | 118644323 | rs6589695 | C | G | NULL | 0.599888369 | NULL&M_Disease |
| 11 | 118704751 | rs35262372 | G | A | NULL | 0.599888369 | NULL&M_Disease |
| 11 | 118765267 | rs598207 | C | G | NULL | 0.599888369 | NULL&M_Disease |
| 11 | 118801296 | rs75210328 | T | G | NULL | 0.599888369 | NULL&M_Disease |
| 11 | 118550524 | rs201134460 | T | C | M_Disease | 0.599888369 | M_Disease&NULL |
| 11 | 118550522 | rs2277296 | C | T | M_Disease | 0.59868766 | M_Disease&NULL |
| 11 | 118801577 | rs77875650 | A | G | NULL | 0.59868766 | NULL&M_Disease |
| 11 | 118691975 | rs77731159 | A | T | NULL | 0.597485766 | NULL&M_Disease |
| 11 | 118807639 | rs73575454 | A | G | NULL | 0.597485766 | NULL&M_Disease |
| 11 | 118809767 | rs75460330 | A | G | NULL | 0.597485766 | NULL&M_Disease |
| 11 | 118810054 | rs73557344 | C | G | NULL | 0.597485766 | NULL&M_Disease |
| 11 | 118483802 | rs72999455 | G | A | NULL | 0.596282699 | NULL&M_Disease |
| 11 | 118701898 | rs533254 | A | T | M_Disease | 0.596282699 | M_Disease&NULL |
| 11 | 118746769 | rs4938576 | T | G | M_Disease | 0.596282699 | M_Disease&NULL |
| 11 | 118530611 | rs2276065 | C | T | NULL | 0.595078474 | NULL&M_Disease |
| 11 | 118550841 | rs79699220 | C | G | NULL | 0.595078474 | NULL&M_Disease |
| 11 | 118711376 | rs118046730 | C | T | NULL | 0.595078474 | NULL&M_Disease |
| 11 | 118503442 | rs45621235 | T | C | M_Disease | 0.593873103 | M_Disease&NULL |
| 11 | 118542383 | rs687190 | C | T | NULL | 0.593873103 | NULL&M_Disease |
| 11 | 118579395 | rs11828220 | A | T | NULL | 0.593873103 | NULL&M_Disease |
| 11 | 118580906 | rs496058 | G | A | M_Disease | 0.593873103 | M_Disease&NULL |
| 11 | 118588998 | rs570424 | C | T | M_Disease | 0.593873103 | M_Disease&NULL |
| 11 | 118714492 | rs7117594 | T | C | NULL | 0.593873103 | NULL&M_Disease |
| 11 | 118754353 | rs630923 | A | C | M_Disease | 0.593873103 | M_Disease&NULL |

| | | | | | | | | | |
|---|---|---|---|---|---|---|---|---|---|
| 11 | 118812821 | rs74535623 | T | G | 1.02 | -4194.66 | -4195.38 | -0.721 | 2.19E-07 |
| 11 | 118813314 | rs73557345 | C | T | 1.02 | -4194.66 | -4195.38 | -0.721 | 2.19E-07 |
| 11 | 118509877 | rs112552940 | T | C | 1.016 | -4194.66 | -4195.39 | -0.73 | 2.17E-07 |
| 11 | 118713180 | rs2187490 | G | T | 0.943 | -4194.66 | -4195.391 | -0.731 | 2.17E-07 |
| 11 | 118765231 | rs12795395 | T | C | 0.983 | -4194.66 | -4195.391 | -0.732 | 2.17E-07 |
| 11 | 118773438 | rs35346926 | T | C | 0.983 | -4194.66 | -4195.391 | -0.732 | 2.17E-07 |
| 11 | 118707035 | rs143605660 | A | G | 0.95 | -4194.66 | -4195.402 | -0.743 | 2.15E-07 |
| 11 | 118464665 | rs113337722 | G | A | 0.999 | -4194.66 | -4195.409 | -0.749 | 2.13E-07 |
| 11 | 118496524 | rs35692099 | C | T | 0.987 | -4194.66 | -4195.41 | -0.75 | 2.13E-07 |
| 11 | 118530068 | rs112337443 | C | T | 1.003 | -4194.66 | -4195.41 | -0.751 | 2.13E-07 |
| 11 | 118502121 | rs17614795 | G | A | 0.99 | -4194.66 | -4195.413 | -0.754 | 2.12E-07 |
| 11 | 118714492 | rs7117594 | T | C | 1.057 | -4194.66 | -4195.418 | -0.758 | 2.11E-07 |
| 11 | 118542383 | rs687190 | C | T | 0.95 | -4194.66 | -4195.42 | -0.761 | 2.11E-07 |
| 11 | 118579395 | rs11828220 | A | T | 0.952 | -4194.66 | -4195.421 | -0.762 | 2.11E-07 |
| 11 | 118530611 | rs2276065 | C | T | 1.055 | -4194.66 | -4195.427 | -0.768 | 2.09E-07 |
| 11 | 118550841 | rs79699220 | C | G | 0.952 | -4194.66 | -4195.437 | -0.778 | 2.07E-07 |
| 11 | 118711376 | rs118046730 | C | T | 1.036 | -4194.66 | -4195.44 | -0.781 | 2.07E-07 |
| 11 | 118483802 | rs72999455 | G | A | 0.957 | -4194.66 | -4195.443 | -0.784 | 2.06E-07 |
| 11 | 118807639 | rs73575454 | A | G | 0.956 | -4194.66 | -4195.459 | -0.799 | 2.03E-07 |
| 11 | 118809767 | rs75460330 | A | G | 0.956 | -4194.66 | -4195.459 | -0.799 | 2.03E-07 |
| 11 | 118810054 | rs73557344 | C | G | 0.956 | -4194.66 | -4195.459 | -0.799 | 2.03E-07 |
| 11 | 118691975 | rs77731159 | A | T | 0.957 | -4194.66 | -4195.459 | -0.8 | 2.03E-07 |
| 11 | 118765267 | rs598207 | C | G | 1.053 | -4194.66 | -4195.472 | -0.812 | 2.00E-07 |
| 11 | 118801577 | rs77875650 | A | G | 1.03 | -4194.66 | -4195.472 | -0.812 | 2.00E-07 |
| 11 | 118494790 | rs11825621 | C | T | 0.96 | -4194.66 | -4195.472 | -0.813 | 2.00E-07 |
| 11 | 118488782 | rs11216930 | C | A | 1.054 | -4194.66 | -4195.474 | -0.815 | 2.00E-07 |
| 11 | 118644323 | rs6589695 | C | G | 1.007 | -4194.66 | -4195.475 | -0.815 | 2.00E-07 |
| 11 | 118801296 | rs75210328 | T | G | 1.029 | -4194.66 | -4195.477 | -0.817 | 1.99E-07 |
| 11 | 118704751 | rs35262372 | G | A | 1.026 | -4194.66 | -4195.48 | -0.82 | 1.99E-07 |
| 11 | 118725383 | rs9735317 | T | C | 1.054 | -4194.66 | -4195.479 | -0.82 | 1.99E-07 |
| 11 | 118712427 | rs7108905 | G | C | 1.054 | -4194.66 | -4195.496 | -0.836 | 1.96E-07 |
| 11 | 118502172 | rs35435609 | T | C | 0.965 | -4194.66 | -4195.5 | -0.841 | 1.95E-07 |
| 11 | 118805138 | rs7127427 | A | G | 1.019 | -4194.66 | -4195.501 | -0.841 | 1.95E-07 |
| 11 | 118526691 | rs72999486 | T | C | 0.966 | -4194.66 | -4195.501 | -0.842 | 1.94E-07 |
| 11 | 118724116 | rs12276533 | A | G | 1.053 | -4194.66 | -4195.502 | -0.842 | 1.94E-07 |
| 11 | 118486067 | rs11603023 | T | C | 0.953 | -4194.66 | -4195.508 | -0.848 | 1.93E-07 |
| 11 | 118802128 | rs59069289 | T | G | 0.963 | -4194.66 | -4195.508 | -0.848 | 1.93E-07 |
| 11 | 118802386 | rs59480472 | C | T | 0.963 | -4194.66 | -4195.508 | -0.848 | 1.93E-07 |
| 11 | 118803160 | rs73575440 | A | G | 0.963 | -4194.66 | -4195.508 | -0.848 | 1.93E-07 |
| 11 | 118803596 | rs111313044 | A | G | 0.963 | -4194.66 | -4195.508 | -0.848 | 1.93E-07 |
| 11 | 118804643 | rs56933877 | A | G | 0.963 | -4194.66 | -4195.508 | -0.848 | 1.93E-07 |
| 11 | 118804919 | rs73575441 | C | T | 0.963 | -4194.66 | -4195.508 | -0.848 | 1.93E-07 |
| 11 | 118804989 | rs112045567 | G | A | 0.963 | -4194.66 | -4195.508 | -0.848 | 1.93E-07 |
| 11 | 118480695 | rs34026809 | C | G | 0.992 | -4194.66 | -4195.508 | -0.849 | 1.93E-07 |
| 11 | 118486110 | rs45540840 | A | G | 1.052 | -4194.66 | -4195.518 | -0.858 | 1.91E-07 |
| 11 | 118501299 | rs7936556 | A | G | 0.953 | -4194.66 | -4195.52 | -0.86 | 1.91E-07 |
| 11 | 118806216 | rs1231742890 | G | T | 1.008 | -4194.66 | -4195.521 | -0.861 | 1.91E-07 |
| 11 | 118615390 | rs78502545 | T | C | 1.014 | -4194.66 | -4195.529 | -0.869 | 1.89E-07 |
| 11 | 118626825 | rs74337439 | A | G | 1.014 | -4194.66 | -4195.529 | -0.869 | 1.89E-07 |
| 11 | 118510562 | rs4245184 | A | G | 0.954 | -4194.66 | -4195.534 | -0.874 | 1.88E-07 |
| 11 | 118511352 | rs644273 | T | C | 0.954 | -4194.66 | -4195.534 | -0.874 | 1.88E-07 |
| 11 | 118511757 | rs654423 | A | G | 0.954 | -4194.66 | -4195.534 | -0.874 | 1.88E-07 |
| 11 | 118792278 | rs77398170 | G | A | 0.992 | -4194.66 | -4195.533 | -0.874 | 1.88E-07 |
| 11 | 118791150 | rs12270280 | T | C | 0.989 | -4194.66 | -4195.537 | -0.877 | 1.88E-07 |
| 11 | 118709600 | rs12272981 | G | A | 1.052 | -4194.66 | -4195.537 | -0.878 | 1.88E-07 |
| 11 | 118793918 | rs12282752 | C | T | 0.993 | -4194.66 | -4195.54 | -0.88 | 1.87E-07 |
| 11 | 118791532 | rs12271856 | T | G | 1.001 | -4194.66 | -4195.54 | -0.881 | 1.87E-07 |
| 11 | 118708785 | rs7122510 | A | C | 1.051 | -4194.66 | -4195.549 | -0.89 | 1.85E-07 |
| 11 | 118708786 | rs7122511 | A | C | 1.051 | -4194.66 | -4195.549 | -0.89 | 1.85E-07 |
| 11 | 118534082 | rs10790256 | T | C | 1.05 | -4194.66 | -4195.551 | -0.892 | 1.85E-07 |
| 11 | 118583847 | rs12803363 | T | C | 0.999 | -4194.66 | -4195.556 | -0.897 | 1.84E-07 |
| 11 | 118804266 | rs7123066 | T | C | 1.017 | -4194.66 | -4195.558 | -0.899 | 1.84E-07 |
| 11 | 118804283 | rs7122959 | G | A | 1.017 | -4194.66 | -4195.558 | -0.899 | 1.84E-07 |
| 11 | 118746223 | rs11217077 | C | T | 0.952 | -4194.66 | -4195.56 | -0.9 | 1.83E-07 |
| 11 | 118723753 | rs7119568 | A | T | 1.05 | -4194.66 | -4195.561 | -0.902 | 1.83E-07 |
| 11 | 118811878 | rs74891547 | C | G | 1.039 | -4194.66 | -4195.564 | -0.904 | 1.83E-07 |
| 11 | 118523799 | rs72999483 | C | T | 1.048 | -4194.66 | -4195.564 | -0.905 | 1.83E-07 |
| 11 | 118759610 | rs566416 | G | T | 1.05 | -4194.66 | -4195.566 | -0.907 | 1.82E-07 |
| 11 | 118495287 | rs77683570 | C | T | 1.05 | -4194.66 | -4195.567 | -0.908 | 1.82E-07 |
| 11 | 118496119 | rs45565037 | A | G | 1.05 | -4194.66 | -4195.568 | -0.909 | 1.82E-07 |
| 11 | 118499394 | rs2236661 | G | C | 1.05 | -4194.66 | -4195.568 | -0.909 | 1.82E-07 |
| 11 | 118521549 | rs494560 | A | G | 0.955 | -4194.66 | -4195.574 | -0.914 | 1.81E-07 |
| 11 | 118728479 | rs7924647 | G | A | 1.05 | -4194.66 | -4195.576 | -0.916 | 1.81E-07 |
| 11 | 118764051 | rs613791 | T | C | 1.049 | -4194.66 | -4195.585 | -0.925 | 1.79E-07 |
| 11 | 118501022 | rs10892248 | A | G | 1.049 | -4194.66 | -4195.587 | -0.927 | 1.79E-07 |
| 11 | 118715589 | rs116872113 | A | G | 1.018 | -4194.66 | -4195.587 | -0.927 | 1.79E-07 |
| 11 | 118516903 | rs80015904 | A | G | 1.049 | -4194.66 | -4195.588 | -0.928 | 1.78E-07 |
| 11 | 118689745 | rs492471 | T | C | 1.035 | -4194.66 | -4195.587 | -0.928 | 1.78E-07 |
| 11 | 118707559 | rs7118009 | T | C | 1.049 | -4194.66 | -4195.591 | -0.931 | 1.78E-07 |
| 11 | 118757191 | rs1790192 | G | A | 1.047 | -4194.66 | -4195.595 | -0.935 | 1.77E-07 |
| 11 | 118516774 | rs12365095 | T | C | 1.049 | -4194.66 | -4195.596 | -0.936 | 1.77E-07 |
| 11 | 118735876 | rs79843382 | T | C | 1.007 | -4194.66 | -4195.596 | -0.936 | 1.77E-07 |
| 11 | 118727137 | rs117223320 | C | G | 1.006 | -4194.66 | -4195.598 | -0.938 | 1.77E-07 |
| 11 | 118507478 | rs11216937 | T | C | 1.048 | -4194.66 | -4195.598 | -0.939 | 1.76E-07 |
| 11 | 118490076 | rs10892247 | A | G | 1.048 | -4194.66 | -4195.603 | -0.943 | 1.76E-07 |
| 11 | 118516737 | rs12362152 | G | A | 1.048 | -4194.66 | -4195.605 | -0.946 | 1.75E-07 |
| 11 | 118569414 | rs6589675 | C | G | 0.957 | -4194.66 | -4195.605 | -0.946 | 1.75E-07 |
| 11 | 118528424 | rs17748 | T | C | 1.048 | -4194.66 | -4195.607 | -0.947 | 1.75E-07 |
| 11 | 118514625 | rs11216938 | T | C | 1.048 | -4194.66 | -4195.607 | -0.948 | 1.75E-07 |
| 11 | 118529127 | rs7928371 | A | G | 1.048 | -4194.66 | -4195.616 | -0.956 | 1.73E-07 |
| 11 | 118763736 | rs12363277 | C | T | 1.013 | -4194.66 | -4195.624 | -0.965 | 1.72E-07 |
| 11 | 118762218 | rs543824 | G | A | 1.047 | -4194.66 | -4195.627 | -0.968 | 1.71E-07 |
| 11 | 118535840 | rs12225548 | G | C | 1.047 | -4194.66 | -4195.632 | -0.972 | 1.71E-07 |

| | | | | | | | |
|---|---|---|---|---|---|---|---|
| 11 | 118464665 | rs113337722 | G | A | NULL | 0.591458978 | NULL&M_Disease |
| 11 | 118496524 | rs35692099 | C | T | NULL | 0.591458978 | NULL&M_Disease |
| 11 | 118502121 | rs17614795 | G | A | NULL | 0.591458978 | NULL&M_Disease |
| 11 | 118530068 | rs112337443 | C | T | NULL | 0.591458978 | NULL&M_Disease |
| 11 | 118707035 | rs143605660 | A | G | NULL | 0.591458978 | NULL&M_Disease |
| 11 | 118713180 | rs2187490 | G | T | NULL | 0.590250252 | NULL&M_Disease |
| 11 | 118509877 | rs112552940 | T | C | NULL | 0.589040434 | NULL&M_Disease |
| 11 | 118765231 | rs12795395 | T | C | NULL | 0.589040434 | NULL&M_Disease |
| 11 | 118773438 | rs35346926 | T | C | NULL | 0.589040434 | NULL&M_Disease |
| 11 | 118855789 | rs4938609 | A | C | M_Disease | 0.587829538 | M_Disease&NULL |
| 11 | 118629959 | rs140110112 | A | G | NULL | 0.587829538 | NULL&M_Disease |
| 11 | 118812821 | rs74535623 | T | G | NULL | 0.587829538 | NULL&M_Disease |
| 11 | 118813314 | rs73557345 | C | T | NULL | 0.587829538 | NULL&M_Disease |
| 11 | 118711069 | rs538645 | C | T | NULL | 0.586617579 | NULL&M_Disease |
| 11 | 118772160 | rs61730467 | A | G | NULL | 0.586617579 | NULL&M_Disease |
| 11 | 118775524 | rs79108568 | G | T | NULL | 0.586617579 | NULL&M_Disease |
| 11 | 118598612 | rs568688 | G | T | M_Disease | 0.586617579 | M_Disease&NULL |
| 11 | 118543798 | rs76570573 | T | C | NULL | 0.585404569 | NULL&M_Disease |
| 11 | 118568427 | rs117361624 | T | C | NULL | 0.585404569 | NULL&M_Disease |
| 11 | 118729442 | rs12277355 | T | C | NULL | 0.585404569 | NULL&M_Disease |
| 11 | 118441079 | rs7106316 | G | A | M_Disease | 0.585404569 | M_Disease&NULL |
| 11 | 118757788 | rs3176912 | A | G | NULL | 0.584190523 | NULL&M_Disease |
| 11 | 118543563 | rs10892251 | T | C | M_Disease | 0.582975454 | M_Disease&NULL |
| 11 | 118598138 | rs543692 | C | T | M_Disease | 0.582975454 | M_Disease&NULL |
| 11 | 118632893 | rs6589690 | A | G | M_Disease | 0.581759377 | M_Disease&NULL |
| 11 | 118700515 | rs112571283 | T | C | NULL | 0.581759377 | NULL&M_Disease |
| 11 | 118712270 | rs7105658 | T | C | NULL | 0.581759377 | NULL&M_Disease |
| 11 | 118729283 | rs11217068 | T | C | NULL | 0.581759377 | NULL&M_Disease |
| 11 | 118874272 | rs192113495 | G | T | NULL | 0.581759377 | NULL&M_Disease |
| 11 | 118826063 | rs666191 | A | G | NULL | 0.580542305 | NULL&M_Disease |
| 11 | 118530869 | rs45447797 | G | C | M_Disease | 0.580542305 | M_Disease&NULL |
| 11 | 118826060 | rs625291 | T | C | NULL | 0.578105233 | NULL&M_Disease |
| 11 | 118539746 | rs72999492 | C | A | NULL | 0.576885261 | NULL&M_Disease |
| 11 | 118614115 | rs73001451 | C | T | NULL | 0.576885261 | NULL&M_Disease |
| 11 | 118782802 | rs76160025 | T | C | NULL | 0.575664351 | NULL&M_Disease |
| 11 | 118603615 | rs74783030 | A | G | NULL | 0.574442517 | NULL&M_Disease |
| 11 | 118466646 | rs184841209 | T | G | NULL | 0.573219773 | NULL&M_Disease |
| 11 | 118660932 | rs1784295 | A | G | M_Disease | 0.573219773 | M_Disease&NULL |
| 11 | 118475098 | rs604096 | G | A | NULL | 0.571996133 | NULL&M_Disease |
| 11 | 118633561 | rs12794288 | A | C | NULL | 0.571996133 | NULL&M_Disease |
| 11 | 118826045 | rs537080 | C | T | NULL | 0.571996133 | NULL&M_Disease |
| 11 | 118889185 | rs11217125 | C | T | NULL | 0.571996133 | NULL&M_Disease |
| 11 | 118515579 | rs10790255 | G | T | NULL | 0.570771612 | NULL&M_Disease |
| 11 | 118721767 | rs116884018 | C | T | NULL | 0.570771612 | NULL&M_Disease |
| 11 | 118793479 | rs138526953 | A | C | NULL | 0.570771612 | NULL&M_Disease |
| 11 | 118824683 | rs507763 | C | G | NULL | 0.570771612 | NULL&M_Disease |
| 11 | 118824917 | rs11217102 | A | G | NULL | 0.570771612 | NULL&M_Disease |
| 11 | 118760443 | rs12786257 | A | G | NULL | 0.569546224 | NULL&M_Disease |
| 11 | 118889132 | rs11217124 | G | A | NULL | 0.569546224 | NULL&M_Disease |
| 11 | 118629143 | rs475946 | T | C | M_Disease | 0.569546224 | M_Disease&NULL |
| 11 | 118758089 | rs497916 | T | C | M_Disease | 0.569546224 | M_Disease&NULL |
| 11 | 118445559 | rs73555296 | T | G | NULL | 0.568319983 | NULL&M_Disease |
| 11 | 118460957 | rs45600633 | A | G | NULL | 0.568319983 | NULL&M_Disease |
| 11 | 118474283 | rs139202987 | G | A | NULL | 0.568319983 | NULL&M_Disease |
| 11 | 118583261 | rs112246697 | A | G | NULL | 0.568319983 | NULL&M_Disease |
| 11 | 118630532 | rs35888235 | T | G | NULL | 0.568319983 | NULL&M_Disease |
| 11 | 118707171 | rs4938550 | G | A | M_Disease | 0.568319983 | M_Disease&NULL |
| 11 | 118421841 | rs145968026 | G | C | NULL | 0.567092905 | NULL&M_Disease |
| 11 | 118487265 | rs76544852 | T | G | NULL | 0.567092905 | NULL&M_Disease |
| 11 | 118518021 | rs10736493 | G | T | NULL | 0.567092905 | NULL&M_Disease |
| 11 | 118580773 | rs675329 | T | C | NULL | 0.567092905 | NULL&M_Disease |
| 11 | 118637666 | rs535996 | T | C | M_Disease | 0.567092905 | M_Disease&NULL |
| 11 | 118661055 | rs1790188 | T | C | M_Disease | 0.567092905 | M_Disease&NULL |
| 11 | 118748170 | rs77074463 | C | G | NULL | 0.567092905 | NULL&M_Disease |
| 11 | 118868750 | rs4445645 | C | G | NULL | 0.567092905 | NULL&M_Disease |
| 11 | 118879824 | rs73559119 | C | T | NULL | 0.567092905 | NULL&M_Disease |
| 11 | 118880012 | rs11825284 | A | G | NULL | 0.567092905 | NULL&M_Disease |
| 11 | 118436071 | rs73023329 | T | C | NULL | 0.565865003 | NULL&M_Disease |
| 11 | 118455465 | rs45460493 | A | G | NULL | 0.565865003 | NULL&M_Disease |
| 11 | 118477226 | rs45496391 | C | A | NULL | 0.565865003 | NULL&M_Disease |
| 11 | 118507068 | rs10892249 | C | T | NULL | 0.565865003 | NULL&M_Disease |
| 11 | 118559426 | rs10892253 | A | C | NULL | 0.565865003 | NULL&M_Disease |
| 11 | 118561858 | rs10790257 | C | G | NULL | 0.565865003 | NULL&M_Disease |
| 11 | 118561859 | rs10790258 | T | G | NULL | 0.565865003 | NULL&M_Disease |
| 11 | 118691070 | rs140577174 | G | A | NULL | 0.565865003 | NULL&M_Disease |
| 11 | 118782474 | rs11606719 | G | C | NULL | 0.565865003 | NULL&M_Disease |
| 11 | 118823148 | rs663003 | T | C | NULL | 0.565865003 | NULL&M_Disease |
| 11 | 118824439 | rs61900157 | C | A | NULL | 0.565865003 | NULL&M_Disease |
| 11 | 118824996 | rs4426142 | A | G | NULL | 0.565865003 | NULL&M_Disease |
| 11 | 118825069 | rs2097131 | A | G | NULL | 0.565865003 | NULL&M_Disease |
| 11 | 118483912 | rs45621436 | G | T | NULL | 0.564636292 | NULL&M_Disease |
| 11 | 118493319 | rs10790254 | A | G | NULL | 0.564636292 | NULL&M_Disease |
| 11 | 118605352 | rs76478216 | C | T | NULL | 0.564636292 | NULL&M_Disease |
| 11 | 118878222 | rs11606357 | C | T | NULL | 0.564636292 | NULL&M_Disease |
| 11 | 118879285 | rs7120445 | C | A | NULL | 0.564636292 | NULL&M_Disease |
| 11 | 118883491 | rs60065241 | A | C | NULL | 0.564636292 | NULL&M_Disease |
| 11 | 118888619 | rs11602764 | G | A | NULL | 0.564636292 | NULL&M_Disease |
| 11 | 118892277 | rs4938621 | T | C | NULL | 0.564636292 | NULL&M_Disease |
| 11 | 118892835 | rs61179449 | C | G | NULL | 0.564636292 | NULL&M_Disease |
| 11 | 118892872 | rs58955701 | T | C | NULL | 0.564636292 | NULL&M_Disease |
| 11 | 118893023 | rs7127188 | C | A | NULL | 0.564636292 | NULL&M_Disease |
| 11 | 118702002 | rs534114 | A | G | M_Disease | 0.564636292 | M_Disease&NULL |
| 11 | 118702910 | rs7941321 | A | G | M_Disease | 0.564636292 | M_Disease&NULL |

| | | | | | | | | | |
|---|---|---|---|---|---|---|---|---|---|
| 11 | 118698170 | rs56296109 | A | C | 1.031 | -4194.66 | -4195.637 | -0.977 | 1.70E-07 |
| 11 | 118745664 | rs11217076 | G | T | 0.957 | -4194.66 | -4195.673 | -1.014 | 1.64E-07 |
| 11 | 118734640 | rs12286723 | T | C | 1.044 | -4194.66 | -4195.682 | -1.023 | 1.62E-07 |
| 11 | 118808140 | rs7117408 | T | C | 1.038 | -4194.66 | -4195.682 | -1.023 | 1.62E-07 |
| 11 | 118807876 | rs12282154 | A | G | 1.037 | -4194.66 | -4195.7 | -1.041 | 1.59E-07 |
| 11 | 118810405 | rs4938589 | T | C | 1.036 | -4194.66 | -4195.703 | -1.043 | 1.59E-07 |
| 11 | 118806897 | rs10892310 | T | C | 0.958 | -4194.66 | -4195.704 | -1.044 | 1.59E-07 |
| 11 | 118809329 | rs10892313 | A | G | 1.036 | -4194.66 | -4195.707 | -1.047 | 1.58E-07 |
| 11 | 118807740 | rs10892312 | A | G | 1.035 | -4194.66 | -4195.712 | -1.053 | 1.57E-07 |
| 11 | 118499469 | rs577396 | C | G | 0.96 | -4194.66 | -4195.733 | -1.073 | 1.54E-07 |
| 11 | 118711830 | rs566848 | G | A | 0.959 | -4194.66 | -4195.733 | -1.074 | 1.54E-07 |
| 11 | 118807637 | rs112974653 | T | G | 1.033 | -4194.66 | -4195.737 | -1.077 | 1.54E-07 |
| 11 | 118462554 | rs11216920 | G | A | 1.011 | -4194.66 | -4195.739 | -1.08 | 1.53E-07 |
| 11 | 118501459 | rs11216934 | A | G | 0.96 | -4194.66 | -4195.756 | -1.097 | 1.51E-07 |
| 11 | 118816353 | rs586370 | A | G | 1.041 | -4194.66 | -4195.758 | -1.098 | 1.50E-07 |
| 11 | 118568639 | rs511451 | T | C | 0.961 | -4194.66 | -4195.77 | -1.111 | 1.49E-07 |
| 11 | 118536950 | rs201359394 | T | A | 0.961 | -4194.66 | -4195.777 | -1.117 | 1.48E-07 |
| 11 | 118765688 | rs676925 | G | C | 1.04 | -4194.66 | -4195.776 | -1.117 | 1.48E-07 |
| 11 | 118509668 | rs483598 | A | G | 0.961 | -4194.66 | -4195.781 | -1.122 | 1.47E-07 |
| 11 | 118568472 | rs570371 | G | A | 0.961 | -4194.66 | -4195.786 | -1.127 | 1.46E-07 |
| 11 | 118700218 | rs2510902 | A | G | 1.038 | -4194.66 | -4195.803 | -1.143 | 1.44E-07 |
| 11 | 118725420 | rs11217062 | A | G | 0.963 | -4194.66 | -4195.809 | -1.149 | 1.43E-07 |
| 11 | 118726230 | rs2156755 | A | G | 0.963 | -4194.66 | -4195.809 | -1.149 | 1.43E-07 |
| 11 | 118692727 | rs503425 | C | T | 1.037 | -4194.66 | -4195.814 | -1.154 | 1.42E-07 |
| 11 | 118816304 | rs586763 | T | C | 1.035 | -4194.66 | -4195.818 | -1.159 | 1.42E-07 |
| 11 | 118514022 | rs613243 | C | G | 0.963 | -4194.66 | -4195.821 | -1.162 | 1.41E-07 |
| 11 | 118685015 | rs540716 | T | C | 1.036 | -4194.66 | -4195.823 | -1.163 | 1.41E-07 |
| 11 | 118750967 | rs11602393 | A | T | 0.967 | -4194.66 | -4195.823 | -1.164 | 1.41E-07 |
| 11 | 118473385 | rs17742 | C | T | 1.026 | -4194.66 | -4195.824 | -1.165 | 1.41E-07 |
| 11 | 118723537 | rs7103314 | G | A | 0.964 | -4194.66 | -4195.824 | -1.165 | 1.41E-07 |
| 11 | 118730669 | rs7933007 | C | T | 0.964 | -4194.66 | -4195.828 | -1.168 | 1.40E-07 |
| 11 | 118568668 | rs11216949 | T | C | 0.963 | -4194.66 | -4195.828 | -1.169 | 1.40E-07 |
| 11 | 118568813 | rs7479114 | T | C | 0.963 | -4194.66 | -4195.828 | -1.169 | 1.40E-07 |
| 11 | 118569067 | rs4486664 | C | A | 0.963 | -4194.66 | -4195.828 | -1.169 | 1.40E-07 |
| 11 | 118569097 | rs4393358 | T | C | 0.963 | -4194.66 | -4195.834 | -1.174 | 1.39E-07 |
| 11 | 118569469 | rs7123811 | G | T | 0.963 | -4194.66 | -4195.834 | -1.175 | 1.39E-07 |
| 11 | 118569694 | rs7108144 | A | G | 0.963 | -4194.66 | -4195.834 | -1.175 | 1.39E-07 |
| 11 | 118717929 | rs4938558 | T | C | 0.964 | -4194.66 | -4195.839 | -1.179 | 1.39E-07 |
| 11 | 118719254 | rs2156750 | C | T | 0.964 | -4194.66 | -4195.84 | -1.18 | 1.39E-07 |
| 11 | 118719405 | rs2156749 | G | C | 0.964 | -4194.66 | -4195.84 | -1.18 | 1.39E-07 |
| 11 | 118719742 | rs7116715 | C | T | 0.964 | -4194.66 | -4195.84 | -1.18 | 1.39E-07 |
| 11 | 118681088 | rs2508922 | A | G | 1.035 | -4194.66 | -4195.842 | -1.182 | 1.38E-07 |
| 11 | 118471423 | rs11216925 | C | T | 1.024 | -4194.66 | -4195.843 | -1.183 | 1.38E-07 |
| 11 | 118472354 | rs10892245 | A | C | 1.024 | -4194.66 | -4195.843 | -1.183 | 1.38E-07 |
| 11 | 118449746 | rs11216909 | T | C | 1.024 | -4194.66 | -4195.843 | -1.184 | 1.38E-07 |
| 11 | 118572021 | rs657539 | T | C | 0.964 | -4194.66 | -4195.844 | -1.184 | 1.38E-07 |
| 11 | 118504742 | rs633683 | T | C | 0.964 | -4194.66 | -4195.844 | -1.185 | 1.38E-07 |
| 11 | 118718856 | rs2156751 | G | A | 0.965 | -4194.66 | -4195.845 | -1.186 | 1.38E-07 |
| 11 | 118718972 | rs2187491 | C | T | 0.965 | -4194.66 | -4195.845 | -1.186 | 1.38E-07 |
| 11 | 118720590 | rs4938564 | T | G | 0.965 | -4194.66 | -4195.851 | -1.192 | 1.37E-07 |
| 11 | 118720684 | rs4936440 | G | C | 0.965 | -4194.66 | -4195.851 | -1.192 | 1.37E-07 |
| 11 | 118720998 | rs4938565 | A | G | 0.965 | -4194.66 | -4195.851 | -1.192 | 1.37E-07 |
| 11 | 118721189 | rs4938566 | T | A | 0.965 | -4194.66 | -4195.851 | -1.192 | 1.37E-07 |
| 11 | 118721249 | rs4938567 | G | A | 0.965 | -4194.66 | -4195.851 | -1.192 | 1.37E-07 |
| 11 | 118721850 | rs7946961 | G | A | 0.965 | -4194.66 | -4195.851 | -1.192 | 1.37E-07 |
| 11 | 118722231 | rs4938570 | G | A | 0.965 | -4194.66 | -4195.851 | -1.192 | 1.37E-07 |
| 11 | 118723087 | rs11217059 | T | C | 0.965 | -4194.66 | -4195.851 | -1.192 | 1.37E-07 |
| 11 | 118724336 | rs10892298 | C | T | 0.965 | -4194.66 | -4195.851 | -1.192 | 1.37E-07 |
| 11 | 118469571 | rs7931564 | G | A | 1.022 | -4194.66 | -4195.855 | -1.195 | 1.37E-07 |
| 11 | 118569900 | rs12793594 | C | T | 0.964 | -4194.66 | -4195.861 | -1.201 | 1.36E-07 |
| 11 | 118570062 | rs554456 | C | T | 0.964 | -4194.66 | -4195.861 | -1.201 | 1.36E-07 |
| 11 | 118570098 | rs482889 | T | G | 0.964 | -4194.66 | -4195.861 | -1.201 | 1.36E-07 |
| 11 | 118570697 | rs523793 | T | C | 0.964 | -4194.66 | -4195.861 | -1.201 | 1.36E-07 |
| 11 | 118570914 | rs659564 | A | C | 0.964 | -4194.66 | -4195.861 | -1.201 | 1.36E-07 |
| 11 | 118570919 | rs642224 | A | G | 0.964 | -4194.66 | -4195.861 | -1.201 | 1.36E-07 |
| 11 | 118571023 | rs541631 | G | A | 0.964 | -4194.66 | -4195.861 | -1.201 | 1.36E-07 |
| 11 | 118571123 | rs658676 | T | C | 0.964 | -4194.66 | -4195.861 | -1.201 | 1.36E-07 |
| 11 | 118571168 | rs488141 | G | A | 0.964 | -4194.66 | -4195.861 | -1.201 | 1.36E-07 |
| 11 | 118571316 | rs657769 | T | C | 0.964 | -4194.66 | -4195.861 | -1.201 | 1.36E-07 |
| 11 | 118454177 | rs17842763 | G | A | 1.021 | -4194.66 | -4195.863 | -1.203 | 1.35E-07 |
| 11 | 118470092 | rs11216924 | G | C | 1.021 | -4194.66 | -4195.862 | -1.203 | 1.35E-07 |
| 11 | 118473231 | rs1047079 | C | T | 1.021 | -4194.66 | -4195.863 | -1.203 | 1.35E-07 |
| 11 | 118567881 | rs497678 | C | T | 0.964 | -4194.66 | -4195.863 | -1.204 | 1.35E-07 |
| 11 | 118461605 | rs11216916 | C | T | 1.021 | -4194.66 | -4195.866 | -1.207 | 1.35E-07 |
| 11 | 118465295 | rs11216922 | G | A | 1.02 | -4194.66 | -4195.868 | -1.209 | 1.35E-07 |
| 11 | 118465501 | rs11216923 | G | A | 1.02 | -4194.66 | -4195.868 | -1.209 | 1.35E-07 |
| 11 | 118718485 | rs4938561 | C | T | 0.966 | -4194.66 | -4195.869 | -1.209 | 1.35E-07 |
| 11 | 118451901 | rs11216911 | C | T | 1.019 | -4194.66 | -4195.873 | -1.213 | 1.34E-07 |
| 11 | 118452334 | rs11216912 | C | G | 1.019 | -4194.66 | -4195.873 | -1.213 | 1.34E-07 |
| 11 | 118453681 | rs11216913 | G | A | 1.019 | -4194.66 | -4195.873 | -1.213 | 1.34E-07 |
| 11 | 118462160 | rs11216917 | G | T | 1.019 | -4194.66 | -4195.874 | -1.214 | 1.34E-07 |
| 11 | 118572611 | rs611217 | T | C | 0.965 | -4194.66 | -4195.874 | -1.215 | 1.34E-07 |
| 11 | 118457630 | rs17092756 | C | G | 1.019 | -4194.66 | -4195.876 | -1.216 | 1.34E-07 |
| 11 | 118471996 | rs11216926 | T | C | 1.019 | -4194.66 | -4195.877 | -1.217 | 1.34E-07 |
| 11 | 118459702 | rs113054880 | G | C | 1.018 | -4194.66 | -4195.882 | -1.223 | 1.33E-07 |
| 11 | 118571357 | rs544452 | T | G | 0.965 | -4194.66 | -4195.883 | -1.224 | 1.33E-07 |
| 11 | 118571790 | rs656287 | G | A | 0.965 | -4194.66 | -4195.883 | -1.224 | 1.33E-07 |
| 11 | 118571911 | rs570949 | T | G | 0.965 | -4194.66 | -4195.883 | -1.224 | 1.33E-07 |
| 11 | 118572171 | rs658040 | G | A | 0.965 | -4194.66 | -4195.883 | -1.224 | 1.33E-07 |
| 11 | 118836606 | rs113975105 | G | C | 1.035 | -4194.66 | -4195.884 | -1.224 | 1.33E-07 |
| 11 | 118459039 | rs57438673 | A | G | 1.017 | -4194.66 | -4195.885 | -1.225 | 1.33E-07 |
| 11 | 118464815 | rs11216921 | C | G | 1.018 | -4194.66 | -4195.884 | -1.225 | 1.33E-07 |
| 11 | 118466249 | rs7115634 | G | A | 1.017 | -4194.66 | -4195.887 | -1.227 | 1.32E-07 |

| | | | | | | | |
|---|---|---|---|---|---|---|---|
| 11 | 118703001 | rs7928228 | T | C | M_Disease | 0.564636292 | M_Disease&NULL |
| 11 | 118703484 | rs2508921 | T | C | M_Disease | 0.564636292 | M_Disease&NULL |
| 11 | 118703817 | rs1170396 | T | A | M_Disease | 0.564636292 | M_Disease&NULL |
| 11 | 118704255 | rs480791 | G | A | M_Disease | 0.564636292 | M_Disease&NULL |
| 11 | 118704265 | rs622690 | T | C | M_Disease | 0.564636292 | M_Disease&NULL |
| 11 | 118578462 | rs516459 | C | T | NULL | 0.563406786 | NULL&M_Disease |
| 11 | 118640081 | rs529717 | A | G | M_Disease | 0.563406786 | M_Disease&NULL |
| 11 | 118641451 | rs2508929 | G | T | M_Disease | 0.563406786 | M_Disease&NULL |
| 11 | 118644213 | rs1790187 | A | G | M_Disease | 0.563406786 | M_Disease&NULL |
| 11 | 118648396 | rs548167 | A | G | M_Disease | 0.563406786 | M_Disease&NULL |
| 11 | 118652781 | rs2508915 | C | G | M_Disease | 0.563406786 | M_Disease&NULL |
| 11 | 118655704 | rs6421568 | C | G | M_Disease | 0.563406786 | M_Disease&NULL |
| 11 | 118655775 | rs6421569 | A | C | M_Disease | 0.563406786 | M_Disease&NULL |
| 11 | 118869460 | rs3737504 | G | A | NULL | 0.563406786 | NULL&M_Disease |
| 11 | 118885161 | rs59095933 | A | G | NULL | 0.563406786 | NULL&M_Disease |
| 11 | 118557483 | rs6589673 | C | A | NULL | 0.562176501 | NULL&M_Disease |
| 11 | 118701597 | rs4938549 | T | C | M_Disease | 0.562176501 | M_Disease&NULL |
| 11 | 118701668 | rs530646 | A | G | M_Disease | 0.562176501 | M_Disease&NULL |
| 11 | 118748789 | rs145762171 | A | G | M_Disease | 0.562176501 | M_Disease&NULL |
| 11 | 118751898 | rs60651507 | A | G | NULL | 0.562176501 | NULL&M_Disease |
| 11 | 118877563 | rs58709307 | T | C | NULL | 0.562176501 | NULL&M_Disease |
| 11 | 118885163 | rs59608621 | G | A | NULL | 0.562176501 | NULL&M_Disease |
| 11 | 118705654 | rs649338 | A | G | M_Disease | 0.56094545 | M_Disease&NULL |
| 11 | 118819541 | rs150196102 | T | C | NULL | 0.56094545 | NULL&M_Disease |
| 11 | 118825636 | rs1790191 | G | A | NULL | 0.56094545 | NULL&M_Disease |
| 11 | 118832251 | rs79808604 | A | C | NULL | 0.56094545 | NULL&M_Disease |
| 11 | 118591443 | rs111843292 | A | C | NULL | 0.559713649 | NULL&M_Disease |
| 11 | 118752178 | rs61150131 | C | T | NULL | 0.559713649 | NULL&M_Disease |
| 11 | 118888831 | rs11820805 | C | T | NULL | 0.559713649 | NULL&M_Disease |
| 11 | 118812559 | rs74678032 | C | T | M_Disease | 0.559713649 | M_Disease&NULL |
| 11 | 118530434 | rs117619140 | T | G | NULL | 0.558481112 | NULL&M_Disease |
| 11 | 118533042 | rs142438429 | T | C | NULL | 0.558481112 | NULL&M_Disease |
| 11 | 118609736 | rs1783937 | A | G | M_Disease | 0.558481112 | M_Disease&NULL |
| 11 | 118884726 | rs58601794 | C | T | NULL | 0.558481112 | NULL&M_Disease |
| 11 | 118886117 | rs10111 | T | C | NULL | 0.558481112 | NULL&M_Disease |
| 11 | 118886656 | rs7131534 | A | G | NULL | 0.558481112 | NULL&M_Disease |
| 11 | 118540104 | rs506234 | G | C | NULL | 0.557247855 | NULL&M_Disease |
| 11 | 118650163 | rs73003218 | T | C | NULL | 0.557247855 | NULL&M_Disease |
| 11 | 118563286 | rs527619 | A | G | NULL | 0.556013891 | NULL&M_Disease |
| 11 | 118565651 | rs540800 | A | G | NULL | 0.556013891 | NULL&M_Disease |
| 11 | 118755946 | rs76109832 | T | C | NULL | 0.556013891 | NULL&M_Disease |
| 11 | 118801045 | rs75085081 | A | G | NULL | 0.556013891 | NULL&M_Disease |
| 11 | 118884708 | rs61381062 | A | G | NULL | 0.556013891 | NULL&M_Disease |
| 11 | 118887888 | rs3916582 | T | C | NULL | 0.556013891 | NULL&M_Disease |
| 11 | 118889378 | rs7113753 | A | G | NULL | 0.556013891 | NULL&M_Disease |
| 11 | 118518227 | rs10736494 | T | C | NULL | 0.554779235 | NULL&M_Disease |
| 11 | 118880668 | rs11607569 | C | T | NULL | 0.554779235 | NULL&M_Disease |
| 11 | 118725189 | rs112965917 | A | T | M_Disease | 0.554779235 | M_Disease&NULL |
| 11 | 118899218 | rs4936459 | T | C | M_Disease | 0.554779235 | M_Disease&NULL |
| 11 | 118510642 | rs141169729 | T | C | NULL | 0.553543903 | NULL&M_Disease |
| 11 | 118666978 | rs1289369859 | A | G | NULL | 0.553543903 | NULL&M_Disease |
| 11 | 118684537 | rs150583208 | G | A | NULL | 0.553543903 | NULL&M_Disease |
| 11 | 118702936 | rs7941696 | G | A | M_Disease | 0.553543903 | M_Disease&NULL |
| 11 | 118719332 | rs112073991 | G | A | NULL | 0.553543903 | NULL&M_Disease |
| 11 | 118759000 | rs77491736 | G | C | NULL | 0.553543903 | NULL&M_Disease |
| 11 | 118779681 | rs9971606 | C | T | NULL | 0.553543903 | NULL&M_Disease |
| 11 | 118887887 | rs369816532 | T | G | NULL | 0.553543903 | NULL&M_Disease |
| 11 | 118419577 | rs191458787 | T | C | NULL | 0.55230791 | NULL&M_Disease |
| 11 | 118421438 | rs145485933 | G | A | NULL | 0.55230791 | NULL&M_Disease |
| 11 | 118614397 | rs10892281 | T | C | M_Disease | 0.55230791 | M_Disease&NULL |
| 11 | 118615660 | rs6589687 | T | A | M_Disease | 0.55230791 | M_Disease&NULL |
| 11 | 118761591 | rs58397111 | T | C | NULL | 0.55230791 | NULL&M_Disease |
| 11 | 118762102 | rs58508419 | A | G | NULL | 0.55230791 | NULL&M_Disease |
| 11 | 118792384 | rs182691948 | T | G | NULL | 0.55230791 | NULL&M_Disease |
| 11 | 118797003 | rs73575428 | G | C | NULL | 0.55230791 | NULL&M_Disease |
| 11 | 118797278 | rs74584888 | A | G | NULL | 0.55230791 | NULL&M_Disease |
| 11 | 118797392 | rs73575429 | C | T | NULL | 0.55230791 | NULL&M_Disease |
| 11 | 118797420 | rs111390501 | G | A | NULL | 0.55230791 | NULL&M_Disease |
| 11 | 118797544 | rs56032119 | G | A | NULL | 0.55230791 | NULL&M_Disease |
| 11 | 118797857 | rs150438978 | T | A | NULL | 0.55230791 | NULL&M_Disease |
| 11 | 118798154 | rs60969913 | C | G | NULL | 0.55230791 | NULL&M_Disease |
| 11 | 118413882 | rs146864992 | T | C | NULL | 0.551071269 | NULL&M_Disease |
| 11 | 118610629 | rs521910 | A | G | M_Disease | 0.551071269 | M_Disease&NULL |
| 11 | 118622245 | rs485901 | G | A | M_Disease | 0.551071269 | M_Disease&NULL |
| 11 | 118694739 | rs188550348 | A | G | NULL | 0.551071269 | NULL&M_Disease |
| 11 | 118761159 | rs57231586 | T | G | NULL | 0.551071269 | NULL&M_Disease |
| 11 | 118786829 | rs7102433 | G | A | NULL | 0.551071269 | NULL&M_Disease |
| 11 | 118798780 | rs77916839 | C | G | M_Disease | 0.551071269 | M_Disease&NULL |
| 11 | 118880740 | rs11217122 | C | T | NULL | 0.551071269 | NULL&M_Disease |
| 11 | 118891314 | rs117694374 | C | T | NULL | 0.551071269 | NULL&M_Disease |
| 11 | 118594326 | rs544096 | G | A | NULL | 0.549833997 | NULL&M_Disease |
| 11 | 118613605 | rs139648410 | G | A | NULL | 0.549833997 | NULL&M_Disease |
| 11 | 118725204 | rs55794678 | T | C | NULL | 0.549833997 | NULL&M_Disease |
| 11 | 118759905 | rs118020807 | A | C | NULL | 0.549833997 | NULL&M_Disease |
| 11 | 118765087 | rs2230321 | T | C | NULL | 0.549833997 | NULL&M_Disease |
| 11 | 118696055 | rs189417780 | C | A | NULL | 0.548596109 | NULL&M_Disease |
| 11 | 118781710 | rs6589707 | A | C | NULL | 0.548596109 | NULL&M_Disease |
| 11 | 118787213 | rs7119222 | C | T | NULL | 0.548596109 | NULL&M_Disease |
| 11 | 118791180 | rs146541591 | C | T | NULL | 0.548596109 | NULL&M_Disease |
| 11 | 118618837 | rs640120 | A | C | M_Disease | 0.547357618 | M_Disease&NULL |
| 11 | 118619423 | rs1961138 | C | T | M_Disease | 0.547357618 | M_Disease&NULL |
| 11 | 118887581 | rs7105187 | T | C | NULL | 0.547357618 | NULL&M_Disease |
| 11 | 118492713 | rs45457998 | G | A | NULL | 0.546118541 | NULL&M_Disease |

| | | | | | | | | | |
|---|---|---|---|---|---|---|---|---|---|
| 11 | 118437279 | rs11216902 | G | A | 1.013 | -4194.66 | -4195.891 | -1.231 | 1.32E-07 |
| 11 | 118435756 | rs2277291 | T | C | 1.009 | -4194.66 | -4195.902 | -1.242 | 1.30E-07 |
| 11 | 118798757 | rs57879605 | T | C | 1.03 | -4194.66 | -4195.903 | -1.243 | 1.30E-07 |
| 11 | 118438448 | rs60837962 | T | A | 1.009 | -4194.66 | -4195.905 | -1.245 | 1.30E-07 |
| 11 | 118439048 | rs12224844 | T | A | 1.01 | -4194.66 | -4195.906 | -1.246 | 1.30E-07 |
| 11 | 118449396 | rs73023350 | A | G | 1.012 | -4194.66 | -4195.905 | -1.246 | 1.30E-07 |
| 11 | 118450875 | rs7126175 | T | C | 1.012 | -4194.66 | -4195.905 | -1.246 | 1.30E-07 |
| 11 | 118449370 | rs17122278 | G | A | 1.009 | -4194.66 | -4195.908 | -1.248 | 1.30E-07 |
| 11 | 118463221 | rs201373950 | A | T | 1.023 | -4194.66 | -4195.908 | -1.249 | 1.29E-07 |
| 11 | 118432404 | rs4326835 | T | G | 1.006 | -4194.66 | -4195.91 | -1.25 | 1.29E-07 |
| 11 | 118432852 | rs11216898 | C | T | 1.007 | -4194.66 | -4195.91 | -1.25 | 1.29E-07 |
| 11 | 118436079 | rs11216900 | C | G | 1.007 | -4194.66 | -4195.91 | -1.25 | 1.29E-07 |
| 11 | 118446298 | rs55844025 | T | C | 1.008 | -4194.66 | -4195.91 | -1.25 | 1.29E-07 |
| 11 | 118456414 | rs11216914 | C | T | 1.011 | -4194.66 | -4195.909 | -1.25 | 1.29E-07 |
| 11 | 118433729 | rs59108310 | G | A | 1.006 | -4194.66 | -4195.911 | -1.251 | 1.29E-07 |
| 11 | 118432279 | rs11216897 | G | A | 1.006 | -4194.66 | -4195.912 | -1.252 | 1.29E-07 |
| 11 | 118434421 | rs11216899 | G | A | 1.005 | -4194.66 | -4195.912 | -1.252 | 1.29E-07 |
| 11 | 118434835 | rs60753148 | T | C | 1.005 | -4194.66 | -4195.912 | -1.252 | 1.29E-07 |
| 11 | 118444525 | rs11216907 | A | G | 1.007 | -4194.66 | -4195.914 | -1.254 | 1.29E-07 |
| 11 | 118445275 | rs17122269 | C | T | 1.006 | -4194.66 | -4195.913 | -1.254 | 1.29E-07 |
| 11 | 118572747 | rs489126 | A | G | 0.967 | -4194.66 | -4195.913 | -1.254 | 1.29E-07 |
| 11 | 118428791 | rs7929887 | G | A | 1.011 | -4194.66 | -4195.916 | -1.257 | 1.28E-07 |
| 11 | 118448087 | rs9735449 | T | C | 1.005 | -4194.66 | -4195.917 | -1.257 | 1.28E-07 |
| 11 | 118441846 | rs10892243 | C | T | 1.005 | -4194.66 | -4195.919 | -1.259 | 1.28E-07 |
| 11 | 118567545 | rs644867 | G | A | 0.967 | -4194.66 | -4195.918 | -1.259 | 1.28E-07 |
| 11 | 118440863 | rs11216904 | G | A | 1.004 | -4194.66 | -4195.92 | -1.26 | 1.28E-07 |
| 11 | 118447580 | rs9736327 | G | T | 1.006 | -4194.66 | -4195.92 | -1.26 | 1.28E-07 |
| 11 | 118447597 | rs9735421 | A | G | 1.006 | -4194.66 | -4195.92 | -1.26 | 1.28E-07 |
| 11 | 118572760 | rs611733 | T | C | 0.967 | -4194.66 | -4195.92 | -1.26 | 1.28E-07 |
| 11 | 118573022 | rs552079 | C | T | 0.967 | -4194.66 | -4195.92 | -1.26 | 1.28E-07 |
| 11 | 118573339 | rs673547 | A | G | 0.967 | -4194.66 | -4195.92 | -1.26 | 1.28E-07 |
| 11 | 118573535 | rs517445 | G | A | 0.967 | -4194.66 | -4195.92 | -1.26 | 1.28E-07 |
| 11 | 118445585 | rs73023341 | G | A | 1.005 | -4194.66 | -4195.922 | -1.262 | 1.28E-07 |
| 11 | 118429764 | rs111942423 | G | T | 1.009 | -4194.66 | -4195.923 | -1.263 | 1.28E-07 |
| 11 | 118443944 | rs11216906 | A | G | 1.004 | -4194.66 | -4195.923 | -1.264 | 1.27E-07 |
| 11 | 118450046 | rs11216910 | G | A | 1.004 | -4194.66 | -4195.923 | -1.264 | 1.27E-07 |
| 11 | 118430048 | rs11216896 | T | C | 1.008 | -4194.66 | -4195.925 | -1.265 | 1.27E-07 |
| 11 | 118430519 | rs11552421 | T | C | 1.009 | -4194.66 | -4195.924 | -1.265 | 1.27E-07 |
| 11 | 118453483 | rs60112336 | T | C | 1.002 | -4194.66 | -4195.924 | -1.265 | 1.27E-07 |
| 11 | 118567084 | rs542613 | A | G | 0.967 | -4194.66 | -4195.928 | -1.268 | 1.27E-07 |
| 11 | 118573668 | rs574383 | C | T | 0.967 | -4194.66 | -4195.929 | -1.269 | 1.27E-07 |
| 11 | 118573743 | rs519294 | A | C | 0.967 | -4194.66 | -4195.929 | -1.269 | 1.27E-07 |
| 11 | 118661376 | rs73003294 | T | C | 1.004 | -4194.66 | -4195.933 | -1.273 | 1.26E-07 |
| 11 | 118832035 | rs10892317 | C | T | 1.031 | -4194.66 | -4195.938 | -1.279 | 1.26E-07 |
| 11 | 118567218 | rs11216947 | A | G | 0.969 | -4194.66 | -4195.961 | -1.301 | 1.23E-07 |
| 11 | 118567257 | rs7126172 | G | A | 0.969 | -4194.66 | -4195.961 | -1.301 | 1.23E-07 |
| 11 | 118555754 | rs525485 | A | G | 0.97 | -4194.66 | -4195.973 | -1.313 | 1.21E-07 |
| 11 | 118558449 | rs514143 | A | G | 0.97 | -4194.66 | -4195.973 | -1.313 | 1.21E-07 |
| 11 | 118567733 | rs481903 | C | T | 0.969 | -4194.66 | -4195.972 | -1.313 | 1.21E-07 |
| 11 | 118567486 | rs644498 | C | T | 0.97 | -4194.66 | -4195.98 | -1.321 | 1.20E-07 |
| 11 | 118831727 | rs10892315 | C | T | 1.029 | -4194.66 | -4195.981 | -1.322 | 1.20E-07 |
| 11 | 118567375 | rs478296 | C | T | 0.97 | -4194.66 | -4195.982 | -1.323 | 1.20E-07 |
| 11 | 118559247 | rs533144 | A | C | 0.97 | -4194.66 | -4195.984 | -1.324 | 1.20E-07 |
| 11 | 118559542 | rs625918 | T | C | 0.97 | -4194.66 | -4195.984 | -1.324 | 1.20E-07 |
| 11 | 118754649 | rs10892307 | G | C | 0.985 | -4194.66 | -4195.987 | -1.328 | 1.20E-07 |
| 11 | 118552074 | rs4938524 | T | G | 0.97 | -4194.66 | -4195.988 | -1.329 | 1.19E-07 |
| 11 | 118555719 | rs625513 | C | A | 0.97 | -4194.66 | -4195.988 | -1.329 | 1.19E-07 |
| 11 | 118556691 | rs610664 | G | A | 0.97 | -4194.66 | -4195.988 | -1.329 | 1.19E-07 |
| 11 | 118556797 | rs571751 | C | A | 0.97 | -4194.66 | -4195.988 | -1.329 | 1.19E-07 |
| 11 | 118557198 | rs546890 | C | T | 0.97 | -4194.66 | -4195.988 | -1.329 | 1.19E-07 |
| 11 | 118558078 | rs555356 | T | G | 0.97 | -4194.66 | -4195.988 | -1.329 | 1.19E-07 |
| 11 | 118558333 | rs693306 | G | A | 0.97 | -4194.66 | -4195.988 | -1.329 | 1.19E-07 |
| 11 | 118558455 | rs558907 | G | A | 0.97 | -4194.66 | -4195.988 | -1.329 | 1.19E-07 |
| 11 | 118558827 | rs561845 | G | A | 0.97 | -4194.66 | -4195.988 | -1.329 | 1.19E-07 |
| 11 | 118558889 | rs638805 | T | G | 0.97 | -4194.66 | -4195.988 | -1.329 | 1.19E-07 |
| 11 | 118558901 | rs510408 | C | G | 0.97 | -4194.66 | -4195.988 | -1.329 | 1.19E-07 |
| 11 | 118558952 | rs509641 | C | T | 0.97 | -4194.66 | -4195.988 | -1.329 | 1.19E-07 |
| 11 | 118558953 | rs563549 | A | G | 0.97 | -4194.66 | -4195.988 | -1.329 | 1.19E-07 |
| 11 | 118559003 | rs638301 | G | A | 0.97 | -4194.66 | -4195.988 | -1.329 | 1.19E-07 |
| 11 | 118559084 | rs508579 | C | G | 0.97 | -4194.66 | -4195.988 | -1.329 | 1.19E-07 |
| 11 | 118559178 | rs486462 | C | T | 0.97 | -4194.66 | -4195.988 | -1.329 | 1.19E-07 |
| 11 | 118559709 | rs574217358 | A | G | 0.97 | -4194.66 | -4195.988 | -1.329 | 1.19E-07 |
| 11 | 118559866 | rs7926959 | C | T | 0.97 | -4194.66 | -4195.988 | -1.329 | 1.19E-07 |
| 11 | 118559881 | rs7926970 | G | T | 0.97 | -4194.66 | -4195.988 | -1.329 | 1.19E-07 |
| 11 | 118560569 | rs4938529 | T | C | 0.97 | -4194.66 | -4195.988 | -1.329 | 1.19E-07 |
| 11 | 118560570 | rs4938530 | T | G | 0.97 | -4194.66 | -4195.988 | -1.329 | 1.19E-07 |
| 11 | 118566039 | rs6589674 | G | A | 0.97 | -4194.66 | -4195.989 | -1.33 | 1.19E-07 |
| 11 | 118556084 | rs613377 | T | C | 0.971 | -4194.66 | -4195.996 | -1.336 | 1.19E-07 |
| 11 | 118553326 | rs693037 | T | C | 0.971 | -4194.66 | -4195.996 | -1.337 | 1.18E-07 |
| 11 | 118567444 | rs493834 | G | A | 0.971 | -4194.66 | -4195.999 | -1.339 | 1.18E-07 |
| 11 | 118554100 | rs4331144 | G | A | 0.971 | -4194.66 | -4196.006 | -1.346 | 1.17E-07 |
| 11 | 118566090 | rs7130076 | C | T | 0.971 | -4194.66 | -4196.006 | -1.346 | 1.17E-07 |
| 11 | 118566175 | rs7114458 | A | G | 0.971 | -4194.66 | -4196.006 | -1.346 | 1.17E-07 |
| 11 | 118566234 | rs7114216 | G | A | 0.971 | -4194.66 | -4196.006 | -1.346 | 1.17E-07 |
| 11 | 118566313 | rs12794726 | T | C | 0.971 | -4194.66 | -4196.006 | -1.346 | 1.17E-07 |
| 11 | 118566746 | rs533646 | G | C | 0.971 | -4194.66 | -4196.006 | -1.346 | 1.17E-07 |
| 11 | 118550601 | rs673770 | G | T | 0.971 | -4194.66 | -4196.008 | -1.348 | 1.17E-07 |
| 11 | 118550756 | rs642530 | C | T | 0.971 | -4194.66 | -4196.008 | -1.348 | 1.17E-07 |
| 11 | 118550784 | rs642497 | A | C | 0.971 | -4194.66 | -4196.008 | -1.348 | 1.17E-07 |
| 11 | 118551597 | rs12799314 | A | G | 0.971 | -4194.66 | -4196.008 | -1.348 | 1.17E-07 |
| 11 | 118561410 | rs12798453 | G | T | 0.971 | -4194.66 | -4196.007 | -1.348 | 1.17E-07 |
| 11 | 118562799 | rs582630 | A | C | 0.971 | -4194.66 | -4196.007 | -1.348 | 1.17E-07 |
| 11 | 118563381 | rs585039 | A | C | 0.971 | -4194.66 | -4196.007 | -1.348 | 1.17E-07 |

| | | | | | | | |
|---|---|---|---|---|---|---|---|
| 11 | 118555383 | rs2508573 | T | C | NULL | 0.546118541 | NULL&M_Disease |
| 11 | 118744615 | rs73009445 | A | G | NULL | 0.546118541 | NULL&M_Disease |
| 11 | 118788769 | rs186418307 | T | G | NULL | 0.546118541 | NULL&M_Disease |
| 11 | 118887568 | rs7121270 | G | T | NULL | 0.546118541 | NULL&M_Disease |
| 11 | 118664181 | rs145462967 | A | G | NULL | 0.543638687 | NULL&M_Disease |
| 11 | 118673968 | rs140269932 | A | G | NULL | 0.543638687 | NULL&M_Disease |
| 11 | 118872581 | rs4558175 | A | G | NULL | 0.543638687 | NULL&M_Disease |
| 11 | 118785982 | rs79157249 | T | A | NULL | 0.542397941 | NULL&M_Disease |
| 11 | 118765283 | rs665648 | A | G | NULL | 0.541156668 | NULL&M_Disease |
| 11 | 118798824 | rs112119822 | G | A | M_Disease | 0.541156668 | M_Disease&NULL |
| 11 | 118799206 | rs7101680 | A | G | M_Disease | 0.541156668 | M_Disease&NULL |
| 11 | 118709275 | rs187164533 | A | G | NULL | 0.539914885 | NULL&M_Disease |
| 11 | 118896115 | rs2276072 | G | A | NULL | 0.539914885 | NULL&M_Disease |
| 11 | 118896116 | rs2276071 | C | A | NULL | 0.539914885 | NULL&M_Disease |
| 11 | 118424513 | rs144339774 | T | C | NULL | 0.538672605 | NULL&M_Disease |
| 11 | 118590743 | rs503542 | G | A | NULL | 0.538672605 | NULL&M_Disease |
| 11 | 118601363 | rs557990 | T | C | NULL | 0.538672605 | NULL&M_Disease |
| 11 | 118602392 | rs564892 | A | T | NULL | 0.538672605 | NULL&M_Disease |
| 11 | 118722946 | rs148990923 | T | C | NULL | 0.538672605 | NULL&M_Disease |
| 11 | 118828184 | rs45586434 | A | C | NULL | 0.538672605 | NULL&M_Disease |
| 11 | 118867803 | rs4938618 | C | G | NULL | 0.538672605 | NULL&M_Disease |
| 11 | 118869539 | rs11822103 | A | G | NULL | 0.538672605 | NULL&M_Disease |
| 11 | 118545589 | rs113941645 | G | C | M_Disease | 0.537429845 | M_Disease&NULL |
| 11 | 118749563 | rs77643000 | C | T | NULL | 0.537429845 | NULL&M_Disease |
| 11 | 118788517 | rs73575424 | A | C | NULL | 0.537429845 | NULL&M_Disease |
| 11 | 118898319 | rs741811 | A | G | M_Disease | 0.537429845 | M_Disease&NULL |
| 11 | 118587349 | rs11216975 | C | A | NULL | 0.53618662 | NULL&M_Disease |
| 11 | 118762521 | rs190509947 | T | A | NULL | 0.53618662 | NULL&M_Disease |
| 11 | 118509741 | rs113313560 | T | C | NULL | 0.533698835 | NULL&M_Disease |
| 11 | 118576997 | rs498912 | C | T | NULL | 0.533698835 | NULL&M_Disease |
| 11 | 118622744 | rs6589689 | G | A | NULL | 0.533698835 | NULL&M_Disease |
| 11 | 118849368 | rs45491601 | A | G | NULL | 0.533698835 | NULL&M_Disease |
| 11 | 118872741 | rs4573680 | C | T | NULL | 0.533698835 | NULL&M_Disease |
| 11 | 118895686 | rs8192696 | T | C | NULL | 0.533698835 | NULL&M_Disease |
| 11 | 118852647 | rs80126726 | A | G | NULL | 0.532454306 | NULL&M_Disease |
| 11 | 118576762 | rs553188 | T | C | NULL | 0.531209373 | NULL&M_Disease |
| 11 | 118873385 | rs10892320 | T | C | NULL | 0.531209373 | NULL&M_Disease |
| 11 | 118892940 | rs12280288 | C | T | NULL | 0.531209373 | NULL&M_Disease |
| 11 | 118705145 | rs636736 | G | A | M_Disease | 0.529964052 | M_Disease&NULL |
| 11 | 118751326 | rs10892306 | A | T | NULL | 0.529964052 | NULL&M_Disease |
| 11 | 118854372 | rs75067211 | T | C | NULL | 0.529964052 | NULL&M_Disease |
| 11 | 118892757 | rs117289748 | G | C | NULL | 0.528718357 | NULL&M_Disease |
| 11 | 118892759 | rs143660533 | C | A | NULL | 0.528718357 | NULL&M_Disease |
| 11 | 118893878 | rs111322300 | C | T | NULL | 0.528718357 | NULL&M_Disease |
| 11 | 118894287 | rs569 | G | A | NULL | 0.528718357 | NULL&M_Disease |
| 11 | 118895202 | rs11006 | G | A | NULL | 0.528718357 | NULL&M_Disease |
| 11 | 118501723 | rs61900957 | T | C | NULL | 0.527472304 | NULL&M_Disease |
| 11 | 118777378 | rs186296705 | G | T | NULL | 0.527472304 | NULL&M_Disease |
| 11 | 118793613 | rs11217091 | C | T | NULL | 0.527472304 | NULL&M_Disease |
| 11 | 118598936 | rs184071181 | C | T | NULL | 0.526225909 | NULL&M_Disease |
| 11 | 118721562 | rs184045098 | T | C | M_Disease | 0.526225909 | M_Disease&NULL |
| 11 | 118732181 | rs117079831 | A | G | M_Disease | 0.526225909 | M_Disease&NULL |
| 11 | 118749318 | rs78312641 | A | T | NULL | 0.526225909 | NULL&M_Disease |
| 11 | 118762701 | rs76457018 | A | G | M_Disease | 0.526225909 | M_Disease&NULL |
| 11 | 118801440 | rs12280501 | T | C | M_Disease | 0.526225909 | M_Disease&NULL |
| 11 | 118872404 | rs6589713 | C | T | NULL | 0.526225909 | NULL&M_Disease |
| 11 | 118543589 | rs72999499 | G | C | NULL | 0.524979187 | NULL&M_Disease |
| 11 | 118615738 | rs193090156 | T | G | M_Disease | 0.524979187 | M_Disease&NULL |
| 11 | 118652593 | rs139217641 | T | C | NULL | 0.524979187 | NULL&M_Disease |
| 11 | 118809805 | rs142069912 | C | G | NULL | 0.524979187 | NULL&M_Disease |
| 11 | 118853021 | rs113962716 | A | C | NULL | 0.524979187 | NULL&M_Disease |
| 11 | 118867695 | rs4938617 | T | C | NULL | 0.523732154 | NULL&M_Disease |
| 11 | 118556995 | rs72999502 | C | T | NULL | 0.522484825 | NULL&M_Disease |
| 11 | 118870413 | rs10892319 | T | C | NULL | 0.522484825 | NULL&M_Disease |
| 11 | 118518753 | rs117947426 | T | C | M_Disease | 0.521237215 | M_Disease&NULL |
| 11 | 118572267 | rs573905 | A | G | M_Disease | 0.521237215 | M_Disease&NULL |
| 11 | 118619478 | rs189583461 | G | A | NULL | 0.521237215 | NULL&M_Disease |
| 11 | 118782222 | rs61900138 | C | T | M_Disease | 0.521237215 | M_Disease&NULL |
| 11 | 118873987 | rs9645664 | C | T | NULL | 0.51998934 | NULL&M_Disease |
| 11 | 118539151 | rs72999491 | A | G | NULL | 0.518741216 | NULL&M_Disease |
| 11 | 118621277 | rs117409833 | C | T | NULL | 0.518741216 | NULL&M_Disease |
| 11 | 118853774 | rs11217109 | C | T | NULL | 0.518741216 | NULL&M_Disease |
| 11 | 118875727 | rs4936457 | G | A | NULL | 0.518741216 | NULL&M_Disease |
| 11 | 118875761 | rs4936458 | G | T | NULL | 0.518741216 | NULL&M_Disease |
| 11 | 118503912 | rs45478298 | T | C | M_Disease | 0.518741216 | M_Disease&NULL |
| 11 | 118607090 | rs10736495 | C | G | M_Disease | 0.518741216 | M_Disease&NULL |
| 11 | 118572000 | rs182361280 | A | G | NULL | 0.517492858 | NULL&M_Disease |
| 11 | 118789952 | rs11217090 | T | C | M_Disease | 0.517492858 | M_Disease&NULL |
| 11 | 118601426 | rs186926311 | T | C | NULL | 0.516244281 | NULL&M_Disease |
| 11 | 118602497 | rs478081 | G | A | M_Disease | 0.516244281 | M_Disease&NULL |
| 11 | 118789026 | rs11607239 | T | G | NULL | 0.516244281 | NULL&M_Disease |
| 11 | 118726340 | rs113786535 | T | C | M_Disease | 0.514995502 | M_Disease&NULL |
| 11 | 118856582 | rs3887652 | C | G | NULL | 0.514995502 | NULL&M_Disease |
| 11 | 118873840 | rs9645663 | C | T | NULL | 0.514995502 | NULL&M_Disease |
| 11 | 118712509 | rs75488969 | T | C | NULL | 0.513746535 | NULL&M_Disease |
| 11 | 118604678 | rs493720 | T | C | M_Disease | 0.513746535 | M_Disease&NULL |
| 11 | 118797562 | rs142336193 | G | A | NULL | 0.512497396 | NULL&M_Disease |
| 11 | 118803193 | rs117235696 | G | T | M_Disease | 0.512497396 | M_Disease&NULL |
| 11 | 118477746 | rs72999445 | G | T | M_Disease | 0.511248102 | M_Disease&NULL |
| 11 | 118834508 | rs111358668 | A | G | NULL | 0.511248102 | NULL&M_Disease |
| 11 | 118896959 | rs57156480 | A | G | NULL | 0.511248102 | NULL&M_Disease |
| 11 | 118433807 | rs146500266 | A | G | NULL | 0.509998667 | NULL&M_Disease |
| 11 | 118521706 | rs145106291 | G | C | M_Disease | 0.509998667 | M_Disease&NULL |

| | | | | | | | | | |
|---|---|---|---|---|---|---|---|---|---|
| 11 | 118564397 | rs558781 | T | G | 0.971 | -4194.66 | -4196.007 | -1.348 | 1.17E-07 |
| 11 | 118564413 | rs508640 | G | A | 0.971 | -4194.66 | -4196.007 | -1.348 | 1.17E-07 |
| 11 | 118564601 | rs471227 | C | T | 0.971 | -4194.66 | -4196.007 | -1.348 | 1.17E-07 |
| 11 | 118564762 | rs516153 | A | C | 0.971 | -4194.66 | -4196.007 | -1.348 | 1.17E-07 |
| 11 | 118564982 | rs518028 | C | T | 0.971 | -4194.66 | -4196.007 | -1.348 | 1.17E-07 |
| 11 | 118565231 | rs519982 | T | C | 0.971 | -4194.66 | -4196.007 | -1.348 | 1.17E-07 |
| 11 | 118565336 | rs478371 | A | G | 0.971 | -4194.66 | -4196.007 | -1.348 | 1.17E-07 |
| 11 | 118538308 | rs692750 | T | G | 0.971 | -4194.66 | -4196.011 | -1.351 | 1.17E-07 |
| 11 | 118539140 | rs4589335 | T | C | 0.971 | -4194.66 | -4196.011 | -1.351 | 1.17E-07 |
| 11 | 118552420 | rs4936433 | G | A | 0.972 | -4194.66 | -4196.013 | -1.353 | 1.17E-07 |
| 11 | 118552715 | rs519848 | A | G | 0.972 | -4194.66 | -4196.013 | -1.353 | 1.17E-07 |
| 11 | 118552842 | rs549370 | A | C | 0.972 | -4194.66 | -4196.013 | -1.353 | 1.17E-07 |
| 11 | 118552996 | rs654792 | T | G | 0.972 | -4194.66 | -4196.013 | -1.353 | 1.17E-07 |
| 11 | 118553218 | rs592280 | C | T | 0.972 | -4194.66 | -4196.013 | -1.353 | 1.17E-07 |
| 11 | 118554248 | rs7106652 | T | C | 0.972 | -4194.66 | -4196.013 | -1.353 | 1.17E-07 |
| 11 | 118554259 | rs7106851 | A | G | 0.972 | -4194.66 | -4196.013 | -1.353 | 1.17E-07 |
| 11 | 118554436 | rs11605069 | G | T | 0.972 | -4194.66 | -4196.013 | -1.353 | 1.17E-07 |
| 11 | 118554448 | rs11602869 | A | G | 0.972 | -4194.66 | -4196.013 | -1.353 | 1.17E-07 |
| 11 | 118554939 | rs12796373 | A | G | 0.972 | -4194.66 | -4196.013 | -1.353 | 1.17E-07 |
| 11 | 118755630 | rs3176905 | C | T | 0.99 | -4194.66 | -4196.013 | -1.354 | 1.17E-07 |
| 11 | 118801640 | rs572608 | A | T | 0.971 | -4194.66 | -4196.018 | -1.358 | 1.16E-07 |
| 11 | 118800302 | rs12793214 | A | C | 0.971 | -4194.66 | -4196.019 | -1.36 | 1.16E-07 |
| 11 | 118565679 | rs502601 | A | G | 0.972 | -4194.66 | -4196.023 | -1.364 | 1.15E-07 |
| 11 | 118562767 | rs667982 | C | T | 0.972 | -4194.66 | -4196.024 | -1.365 | 1.15E-07 |
| 11 | 118545797 | rs600969 | T | C | 0.972 | -4194.66 | -4196.026 | -1.367 | 1.15E-07 |
| 11 | 118548421 | rs562313 | A | G | 0.972 | -4194.66 | -4196.026 | -1.367 | 1.15E-07 |
| 11 | 118562520 | rs558593 | T | C | 0.973 | -4194.66 | -4196.031 | -1.371 | 1.15E-07 |
| 11 | 118567020 | rs642662 | G | A | 0.973 | -4194.66 | -4196.033 | -1.373 | 1.14E-07 |
| 11 | 118765600 | rs3922 | G | A | 1.028 | -4194.66 | -4196.046 | -1.387 | 1.13E-07 |
| 11 | 118549582 | rs507080 | A | G | 0.974 | -4194.66 | -4196.06 | -1.401 | 1.11E-07 |
| 11 | 118549803 | rs659969 | T | C | 0.974 | -4194.66 | -4196.06 | -1.401 | 1.11E-07 |
| 11 | 118549908 | rs656575 | G | T | 0.974 | -4194.66 | -4196.06 | -1.401 | 1.11E-07 |
| 11 | 118549985 | rs543327 | T | G | 0.974 | -4194.66 | -4196.06 | -1.401 | 1.11E-07 |
| 11 | 118550017 | rs543413 | C | T | 0.974 | -4194.66 | -4196.06 | -1.401 | 1.11E-07 |
| 11 | 118546602 | rs526597 | G | A | 0.974 | -4194.66 | -4196.062 | -1.403 | 1.11E-07 |
| 11 | 118831335 | rs61506726 | G | A | 1.023 | -4194.66 | -4196.067 | -1.407 | 1.10E-07 |
| 11 | 118788081 | rs7929825 | G | C | 1.005 | -4194.66 | -4196.071 | -1.411 | 1.10E-07 |
| 11 | 118548665 | rs1784298 | C | T | 0.975 | -4194.66 | -4196.077 | -1.417 | 1.09E-07 |
| 11 | 118548671 | rs4938522 | T | A | 0.975 | -4194.66 | -4196.077 | -1.417 | 1.09E-07 |
| 11 | 118548838 | rs4938523 | T | G | 0.975 | -4194.66 | -4196.077 | -1.417 | 1.09E-07 |
| 11 | 118538157 | rs520543 | A | G | 0.975 | -4194.66 | -4196.082 | -1.423 | 1.09E-07 |
| 11 | 118539170 | rs4457789 | T | G | 0.975 | -4194.66 | -4196.082 | -1.423 | 1.09E-07 |
| 11 | 118539522 | rs2508991 | A | G | 0.975 | -4194.66 | -4196.082 | -1.423 | 1.09E-07 |
| 11 | 118769032 | rs11217081 | G | A | 1.027 | -4194.66 | -4196.083 | -1.423 | 1.09E-07 |
| 11 | 118538187 | rs607527 | T | G | 0.976 | -4194.66 | -4196.096 | -1.436 | 1.07E-07 |
| 11 | 118807735 | rs10892311 | T | C | 0.974 | -4194.66 | -4196.098 | -1.438 | 1.07E-07 |
| 11 | 118585146 | rs112799932 | G | T | 1.019 | -4194.66 | -4196.099 | -1.439 | 1.07E-07 |
| 11 | 118539566 | rs1783935 | T | A | 0.977 | -4194.66 | -4196.1 | -1.44 | 1.07E-07 |
| 11 | 118767077 | rs581063 | C | T | 1.026 | -4194.66 | -4196.1 | -1.44 | 1.07E-07 |
| 11 | 118587347 | rs603486 | T | C | 0.976 | -4194.66 | -4196.107 | -1.447 | 1.06E-07 |
| 11 | 118605451 | rs2186780 | T | C | 0.993 | -4194.66 | -4196.111 | -1.452 | 1.06E-07 |
| 11 | 118605914 | rs7926282 | C | A | 0.997 | -4194.66 | -4196.12 | -1.46 | 1.05E-07 |
| 11 | 118584498 | rs569770 | T | C | 0.978 | -4194.66 | -4196.124 | -1.465 | 1.04E-07 |
| 11 | 118590864 | rs11216982 | A | G | 1.016 | -4194.66 | -4196.125 | -1.466 | 1.04E-07 |
| 11 | 118668296 | rs58868870 | T | C | 1.015 | -4194.66 | -4196.13 | -1.47 | 1.04E-07 |
| 11 | 118803536 | rs615166 | A | G | 0.978 | -4194.66 | -4196.131 | -1.471 | 1.04E-07 |
| 11 | 118769442 | rs7104819 | G | A | 1.023 | -4194.66 | -4196.133 | -1.474 | 1.03E-07 |
| 11 | 118666289 | rs56104144 | C | G | 1.014 | -4194.66 | -4196.139 | -1.479 | 1.03E-07 |
| 11 | 118801235 | rs602716 | A | G | 0.979 | -4194.66 | -4196.143 | -1.483 | 1.02E-07 |
| 11 | 118802492 | rs688035 | G | C | 0.979 | -4194.66 | -4196.145 | -1.485 | 1.02E-07 |
| 11 | 118610463 | rs4499035 | T | C | 1.012 | -4194.66 | -4196.154 | -1.494 | 1.01E-07 |
| 11 | 118610826 | rs1893029 | T | G | 1.012 | -4194.66 | -4196.154 | -1.494 | 1.01E-07 |
| 11 | 118613674 | rs7130875 | C | T | 1.012 | -4194.66 | -4196.153 | -1.494 | 1.01E-07 |
| 11 | 118655356 | rs7934890 | T | C | 1.012 | -4194.66 | -4196.156 | -1.497 | 1.01E-07 |
| 11 | 118657297 | rs487177 | G | C | 1.012 | -4194.66 | -4196.156 | -1.497 | 1.01E-07 |
| 11 | 118609598 | rs112996178 | T | C | 1.011 | -4194.66 | -4196.159 | -1.5 | 1.01E-07 |
| 11 | 118478330 | rs7125115 | A | G | 1.02 | -4194.66 | -4196.161 | -1.501 | 1.01E-07 |
| 11 | 118596444 | rs68097885 | A | G | 1.011 | -4194.66 | -4196.161 | -1.502 | 1.00E-07 |
| 11 | 118624919 | rs11217013 | C | G | 1.009 | -4194.66 | -4196.17 | -1.51 | 9.97E-08 |
| 11 | 118770321 | rs11217083 | G | A | 1.021 | -4194.66 | -4196.171 | -1.511 | 9.96E-08 |
| 11 | 118626286 | rs11217014 | C | T | 1.009 | -4194.66 | -4196.171 | -1.512 | 9.95E-08 |
| 11 | 118627778 | rs10892282 | C | G | 1.009 | -4194.66 | -4196.171 | -1.512 | 9.95E-08 |
| 11 | 118627149 | rs11217016 | C | A | 1.008 | -4194.66 | -4196.172 | -1.513 | 9.94E-08 |
| 11 | 118747576 | rs55756957 | G | A | 0.992 | -4194.66 | -4196.172 | -1.513 | 9.94E-08 |
| 11 | 118477367 | rs498872 | A | G | 0.983 | -4194.66 | -4196.173 | -1.514 | 9.93E-08 |
| 11 | 118594669 | rs140026975 | C | A | 1.008 | -4194.66 | -4196.174 | -1.514 | 9.93E-08 |
| 11 | 118633533 | rs7110526 | C | T | 1.008 | -4194.66 | -4196.174 | -1.515 | 9.92E-08 |
| 11 | 118635364 | rs11217018 | C | T | 1.008 | -4194.66 | -4196.174 | -1.515 | 9.92E-08 |
| 11 | 118616253 | rs7124498 | A | G | 1.008 | -4194.66 | -4196.175 | -1.516 | 9.91E-08 |
| 11 | 118603030 | rs7943039 | A | G | 1.008 | -4194.66 | -4196.177 | -1.517 | 9.90E-08 |
| 11 | 118637475 | rs112910608 | T | C | 1.008 | -4194.66 | -4196.176 | -1.517 | 9.90E-08 |
| 11 | 118629364 | rs73001486 | A | G | 1.007 | -4194.66 | -4196.179 | -1.52 | 9.87E-08 |
| 11 | 118640828 | rs11217021 | C | T | 1.007 | -4194.66 | -4196.18 | -1.52 | 9.87E-08 |
| 11 | 118796570 | rs11217096 | T | C | 0.979 | -4194.66 | -4196.182 | -1.522 | 9.85E-08 |
| 11 | 118622985 | rs3825057 | C | T | 1.002 | -4194.66 | -4196.191 | -1.531 | 9.76E-08 |
| 11 | 118480115 | rs12803321 | C | G | 1.011 | -4194.66 | -4196.242 | -1.582 | 9.27E-08 |
| 11 | 118577990 | rs480958 | G | A | 0.99 | -4194.66 | -4196.259 | -1.599 | 9.12E-08 |
| 11 | 118515616 | rs527290 | T | C | 0.993 | -4194.66 | -4196.262 | -1.602 | 9.09E-08 |
| 11 | 118522179 | rs1171461 | A | G | 0.994 | -4194.66 | -4196.267 | -1.607 | 9.05E-08 |
| 11 | 118520480 | rs653008 | T | G | 0.995 | -4194.66 | -4196.267 | -1.608 | 9.04E-08 |
| 11 | 118519550 | rs601318 | G | C | 0.995 | -4194.66 | -4196.269 | -1.61 | 9.02E-08 |
| 11 | 118812512 | rs647769 | T | C | 0.995 | -4194.66 | -4196.27 | -1.61 | 9.02E-08 |
| 11 | 118480223 | rs67307131 | C | T | 1.006 | -4194.66 | -4196.271 | -1.611 | 9.01E-08 |

| | | | | | | | |
|---|---|---|---|---|---|---|---|
| 11 | 118559731 | rs113784020 | T | C | NULL | 0.509998667 | NULL&M_Disease |
| 11 | 118605127 | rs526850 | G | T | M_Disease | 0.509998667 | M_Disease&NULL |
| 11 | 118799691 | rs191378986 | G | A | NULL | 0.509998667 | NULL&M_Disease |
| 11 | 118577614 | rs947931 | G | T | M_Disease | 0.508749107 | M_Disease&NULL |
| 11 | 118803842 | rs11600378 | G | A | M_Disease | 0.508749107 | M_Disease&NULL |
| 11 | 118804671 | rs143711539 | A | G | M_Disease | 0.508749107 | M_Disease&NULL |
| 11 | 118809882 | rs11217099 | A | C | M_Disease | 0.508749107 | M_Disease&NULL |
| 11 | 118877891 | rs75154936 | C | T | NULL | 0.507499438 | NULL&M_Disease |
| 11 | 118890085 | rs4938619 | T | C | NULL | 0.507499438 | NULL&M_Disease |
| 11 | 118893767 | rs111345266 | T | C | NULL | 0.507499438 | NULL&M_Disease |
| 11 | 118583712 | rs118008100 | A | G | M_Disease | 0.506249674 | M_Disease&NULL |
| 11 | 118602058 | rs562059 | G | T | M_Disease | 0.506249674 | M_Disease&NULL |
| 11 | 118607694 | rs7119702 | G | A | M_Disease | 0.506249674 | M_Disease&NULL |
| 11 | 118789657 | rs12294466 | T | C | M_Disease | 0.506249674 | M_Disease&NULL |
| 11 | 118876213 | rs76253444 | G | A | NULL | 0.506249674 | NULL&M_Disease |
| 11 | 118576463 | rs496547 | A | T | NULL | 0.504999833 | NULL&M_Disease |
| 11 | 118583971 | rs6589677 | A | G | M_Disease | 0.504999833 | M_Disease&NULL |
| 11 | 118605447 | rs2186781 | T | C | M_Disease | 0.504999833 | M_Disease&NULL |
| 11 | 118876552 | rs74884652 | C | T | NULL | 0.50374993 | NULL&M_Disease |
| 11 | 118805144 | rs11605265 | A | G | M_Disease | 0.502499979 | M_Disease&NULL |
| 11 | 118805220 | rs11605658 | T | C | M_Disease | 0.502499979 | M_Disease&NULL |
| 11 | 118811315 | rs77638281 | A | G | NULL | 0.501249997 | NULL&M_Disease |
| 11 | 118811641 | rs11606199 | A | G | NULL | 0.501249997 | NULL&M_Disease |
| 11 | 118831755 | rs10892316 | T | C | M_Disease | 0.501249997 | M_Disease&NULL |
| 11 | 118834718 | rs112939258 | T | C | M_Disease | 0.501249997 | M_Disease&NULL |

| | | | | | | | | | |
|---|---|---|---|---|---|---|---|---|---|
| 11 | 118528466 | rs7389 | G | T | 0.996 | -4194.66 | -4196.271 | -1.611 | 9.01E-08 |
| 11 | 118529595 | rs745663 | G | A | 0.996 | -4194.66 | -4196.271 | -1.611 | 9.01E-08 |
| 11 | 118512162 | rs570836 | A | G | 0.996 | -4194.66 | -4196.271 | -1.612 | 9.00E-08 |
| 11 | 118505090 | rs645637 | A | G | 0.997 | -4194.66 | -4196.272 | -1.613 | 8.99E-08 |
| 11 | 118506402 | rs483283 | C | T | 0.996 | -4194.66 | -4196.272 | -1.613 | 8.99E-08 |
| 11 | 118522338 | rs4938520 | G | A | 0.996 | -4194.66 | -4196.273 | -1.613 | 8.99E-08 |
| 11 | 118480285 | rs12225399 | C | G | 1.005 | -4194.66 | -4196.273 | -1.614 | 8.98E-08 |
| 11 | 118492591 | rs2077173 | A | G | 0.997 | -4194.66 | -4196.273 | -1.614 | 8.98E-08 |
| 11 | 118497402 | rs12419204 | A | T | 0.997 | -4194.66 | -4196.273 | -1.614 | 8.98E-08 |
| 11 | 118500173 | rs571001 | T | C | 0.997 | -4194.66 | -4196.273 | -1.614 | 8.98E-08 |
| 11 | 118500193 | rs570952 | C | T | 0.997 | -4194.66 | -4196.273 | -1.614 | 8.98E-08 |
| 11 | 118496066 | rs11603616 | A | G | 0.998 | -4194.66 | -4196.274 | -1.615 | 8.97E-08 |
| 11 | 118486325 | rs12290522 | T | C | 0.998 | -4194.66 | -4196.275 | -1.616 | 8.96E-08 |
| 11 | 118487673 | rs4938517 | C | T | 0.998 | -4194.66 | -4196.275 | -1.616 | 8.96E-08 |
| 11 | 118811684 | rs550999 | G | C | 0.997 | -4194.66 | -4196.276 | -1.616 | 8.96E-08 |
| 11 | 118481431 | rs10892246 | A | G | 0.99 | -4194.66 | -4196.283 | -1.624 | 8.89E-08 |
| 11 | 118735476 | rs10892301 | A | G | 1.009 | -4194.66 | -4196.3 | -1.641 | 8.74E-08 |
| 11 | 118459069 | rs633308 | G | A | 0.998 | -4194.66 | -4196.302 | -1.642 | 8.73E-08 |
| 11 | 118717732 | rs11217052 | G | A | 1.006 | -4194.66 | -4196.306 | -1.647 | 8.69E-08 |
| 11 | 118718061 | rs4938559 | T | C | 1.006 | -4194.66 | -4196.306 | -1.647 | 8.69E-08 |
| 11 | 118718062 | rs4938560 | G | A | 1.006 | -4194.66 | -4196.306 | -1.647 | 8.69E-08 |
| 11 | 118708564 | rs484232 | T | G | 0.995 | -4194.66 | -4196.309 | -1.649 | 8.67E-08 |
| 11 | 118717708 | rs7107756 | A | G | 0.997 | -4194.66 | -4196.31 | -1.651 | 8.66E-08 |
| 11 | 118707249 | rs4938551 | C | G | 0.999 | -4194.66 | -4196.316 | -1.657 | 8.60E-08 |
| 11 | 118716563 | rs2851460 | A | G | 1 | -4194.66 | -4196.317 | -1.658 | 8.60E-08 |

**Supplemental Data 6: Bayesian analysis of SLE Immunochip data in DDX6-CXCR5 locus (Dataset 4).**

Note: CHROM=chromosome; POS=basepair position; A1=major allele; A2=minor allele; OR=odd ratio; L+P0=The log null marginal likelihood; L+P1=The log alternative marginal likelihood; BF=Bayes factor; P_CAUSAL=The probability that this variant is the causal variant at this locus;

### CREDIBLE SET

| CHROM | POS (GRCh37/hg19) | RSID | A1 | A2 | BestModel | BestModelProb | CredibleSet |
|---|---|---|---|---|---|---|---|
| 11 | 118607871 | rs76409436 | A | G | M_Disease | 0.966104836 | M_Disease |
| 11 | 118609790 | rs11217006 | A | G | M_Disease | 0.965108591 | M_Disease |
| 11 | 118603736 | rs112561833 | C | A | M_Disease | 0.964084154 | M_Disease |
| 11 | 118611817 | rs10892280 | T | C | M_Disease | 0.963561135 | M_Disease |
| 11 | 118610549 | rs11826521 | T | G | M_Disease | 0.962852366 | M_Disease |
| 11 | 118639353 | rs11217020 | A | G | M_Disease | 0.962673113 | M_Disease |
| 11 | 118603526 | rs11216995 | A | G | M_Disease | 0.961947753 | M_Disease |
| 11 | 118608716 | rs7945144 | G | A | M_Disease | 0.961022003 | M_Disease |
| 11 | 118611781 | rs10892279 | A | G | M_Disease | 0.960645684 | M_Disease |
| 11 | 118607794 | rs12364971 | G | T | M_Disease | 0.960265882 | M_Disease |
| 11 | 118618776 | rs1048024 | T | A | M_Disease | 0.960265882 | M_Disease |
| 11 | 118619047 | rs2077579 | G | T | M_Disease | 0.960265882 | M_Disease |
| 11 | 118619253 | rs2004781 | T | C | M_Disease | 0.960265882 | M_Disease |
| 11 | 118619960 | rs3889239 | T | C | M_Disease | 0.960265882 | M_Disease |
| 11 | 118598544 | rs10892271 | A | G | M_Disease | 0.959882568 | M_Disease |
| 11 | 118610957 | rs6589684 | A | G | M_Disease | 0.959882568 | M_Disease |
| 11 | 118636980 | rs10892283 | G | A | M_Disease | 0.959882568 | M_Disease |
| 11 | 118603520 | rs11216994 | T | C | M_Disease | 0.959689585 | M_Disease |
| 11 | 118636343 | rs11217019 | A | C | M_Disease | 0.959300946 | M_Disease |
| 11 | 118604638 | rs11216997 | T | C | M_Disease | 0.959105284 | M_Disease |
| 11 | 118628373 | rs74676029 | T | C | M_Disease | 0.959105284 | M_Disease |
| 11 | 118602707 | rs10892272 | G | C | M_Disease | 0.958908722 | M_Disease |
| 11 | 118602809 | rs10892273 | A | G | M_Disease | 0.958908722 | M_Disease |
| 11 | 118603495 | rs11216993 | A | T | M_Disease | 0.958908722 | M_Disease |
| 11 | 118607569 | rs77209083 | G | A | M_Disease | 0.958711255 | M_Disease |
| 11 | 118648373 | rs56043232 | T | C | M_Disease | 0.958313595 | M_Disease |
| 11 | 118655645 | rs874621 | A | G | M_Disease | 0.958113393 | M_Disease |
| 11 | 118607046 | rs11217001 | A | G | M_Disease | 0.957912272 | M_Disease |
| 11 | 118603165 | rs73001429 | G | T | M_Disease | 0.957507257 | M_Disease |
| 11 | 118646003 | rs10892289 | T | C | M_Disease | 0.957507257 | M_Disease |
| 11 | 118644694 | rs10892288 | G | A | M_Disease | 0.957303356 | M_Disease |
| 11 | 118661398 | rs57494551 | T | C | M_Disease | 0.957303356 | M_Disease |
| 11 | 118665575 | rs10892293 | T | A | M_Disease | 0.956892745 | M_Disease |
| 11 | 118615340 | rs56758835 | A | G | M_Disease | 0.956478365 | M_Disease |
| 11 | 118615785 | rs11217009 | T | C | M_Disease | 0.956478365 | M_Disease |
| 11 | 118612197 | rs6589685 | C | T | M_Disease | 0.956060185 | M_Disease |
| 11 | 118655340 | rs10892292 | C | G | M_Disease | 0.954997901 | M_Disease |
| 11 | 118642085 | rs10892286 | C | A | M_Disease | 0.954782527 | M_Disease |
| 11 | 118642999 | rs10892287 | C | T | M_Disease | 0.954782527 | M_Disease |
| 11 | 118587319 | rs10892263 | C | T | M_Disease | 0.954566171 | M_Disease |
| 11 | 118589012 | rs12421793 | A | C | M_Disease | 0.954348829 | M_Disease |
| 11 | 118653503 | rs10892290 | T | G | M_Disease | 0.954348829 | M_Disease |
| 11 | 118653517 | rs10892291 | T | C | M_Disease | 0.954348829 | M_Disease |
| 11 | 118649512 | rs73003215 | A | G | M_Disease | 0.953469525 | M_Disease |
| 11 | 118612921 | rs2155433 | A | G | M_Disease | 0.9527995 | M_Disease |
| 11 | 118550522 | rs2277296 | C | T | M_Disease | 0.952347732 | M_Disease |
| 11 | 118550524 | rs201134460 | T | C | M_Disease | 0.952347732 | M_Disease |
| 11 | 118543563 | rs10892251 | T | C | M_Disease | 0.951891858 | M_Disease |
| 11 | 118607417 | rs55894437 | T | C | M_Disease | 0.951662371 | M_Disease |
| 11 | 118588952 | rs12421790 | T | C | M_Disease | 0.951431846 | M_Disease |
| 11 | 118607150 | rs11217002 | T | C | M_Disease | 0.950967661 | M_Disease |
| 11 | 118607311 | rs56267528 | A | C | M_Disease | 0.950967661 | M_Disease |
| 11 | 118556398 | rs11216943 | A | G | M_Disease | 0.950733994 | M_Disease |
| 11 | 118675772 | rs11217036 | T | C | M_Disease | 0.950733994 | M_Disease |
| 11 | 118677086 | rs11217037 | A | G | M_Disease | 0.947846437 | M_Disease&NULL |
| 11 | 118677355 | rs11217038 | A | G | M_Disease | 0.947846437 | M_Disease&NULL |
| 11 | 118555692 | rs10892252 | G | A | M_Disease | 0.947598715 | M_Disease&NULL |
| 11 | 118669605 | rs11217032 | A | T | M_Disease | 0.947099933 | M_Disease&NULL |
| 11 | 118673706 | rs11217033 | C | T | M_Disease | 0.947099933 | M_Disease&NULL |
| 11 | 118684610 | rs715412 | A | G | M_Disease | 0.946848864 | M_Disease&NULL |
| 11 | 118606521 | rs6589681 | T | C | M_Disease | 0.946088893 | M_Disease&NULL |
| 11 | 118667357 | rs10892294 | C | G | M_Disease | 0.9458333 | M_Disease&NULL |
| 11 | 118683564 | rs17122453 | A | G | M_Disease | 0.9458333 | M_Disease&NULL |
| 11 | 118694547 | rs7123726 | C | T | M_Disease | 0.94505965 | M_Disease&NULL |
| 11 | 118606722 | rs6589682 | T | C | M_Disease | 0.942675824 | M_Disease&NULL |
| 11 | 118679304 | rs28409215 | A | G | M_Disease | 0.942675824 | M_Disease&NULL |
| 11 | 118680648 | rs11217040 | A | C | M_Disease | 0.942675824 | M_Disease&NULL |
| 11 | 118697865 | rs11217045 | A | C | M_Disease | 0.942675824 | M_Disease&NULL |
| 11 | 118697906 | rs73005502 | G | A | M_Disease | 0.942675824 | M_Disease&NULL |
| 11 | 118698907 | rs11217046 | A | G | M_Disease | 0.941859849 | M_Disease&NULL |
| 11 | 118696022 | rs11217044 | C | T | M_Disease | 0.941309825 | M_Disease&NULL |
| 11 | 118560857 | rs73001406 | A | G | M_Disease | 0.941032987 | M_Disease&NULL |
| 11 | 118682528 | rs11217042 | A | G | M_Disease | 0.941032987 | M_Disease&NULL |
| 11 | 118612565 | rs6589686 | A | G | M_Disease | 0.940475634 | M_Disease&NULL |
| 11 | 118701018 | rs79504890 | A | G | M_Disease | 0.939630347 | M_Disease&NULL |
| 11 | 118682350 | rs11217041 | G | A | M_Disease | 0.938773837 | M_Disease&NULL |
| 11 | 118587097 | rs10892262 | C | G | M_Disease | 0.937614149 | M_Disease&NULL |
| 11 | 118681464 | rs7942535 | C | T | M_Disease | 0.937026644 | M_Disease&NULL |
| 11 | 118590668 | rs11216980 | A | C | M_Disease | 0.935233031 | M_Disease&NULL |
| 11 | 118681079 | rs73005423 | G | A | M_Disease | 0.933702149 | M_Disease&NULL |
| 11 | 118681083 | rs73005426 | G | A | M_Disease | 0.933080433 | M_Disease&NULL |
| 11 | 118459069 | rs633308 | G | A | M_Disease | 0.928905696 | M_Disease&NULL |
| 11 | 118721699 | rs7125066 | G | A | M_Disease | 0.921651475 | M_Disease&NULL |
| 11 | 118724894 | rs9736016 | A | T | M_Disease | 0.919457587 | M_Disease&NULL |
| 11 | 118585826 | rs11216972 | T | C | M_Disease | 0.914900955 | M_Disease&NULL |
| 11 | 118721565 | rs75249660 | A | G | M_Disease | 0.914900955 | M_Disease&NULL |
| 11 | 118530611 | rs2276065 | C | T | M_Disease | 0.909701855 | M_Disease&NULL |
| 11 | 118722149 | rs11217058 | A | G | M_Disease | 0.908877039 | M_Disease&NULL |
| 11 | 118575326 | rs11216956 | A | G | M_Disease | 0.903348789 | M_Disease&NULL |
| 11 | 118725660 | rs4936441 | C | G | M_Disease | 0.898439072 | M_Disease&NULL |
| 11 | 118507478 | rs11216937 | T | C | M_Disease | 0.896135084 | M_Disease&NULL |
| 11 | 118516774 | rs12365095 | T | C | M_Disease | 0.896135084 | M_Disease&NULL |
| 11 | 118495287 | rs77683570 | C | T | M_Disease | 0.895200621 | M_Disease&NULL |
| 11 | 118486110 | rs45540840 | A | G | M_Disease | 0.894258741 | M_Disease&NULL |
| 11 | 118499394 | rs2236661 | G | C | M_Disease | 0.893785008 | M_Disease&NULL |
| 11 | 118726753 | rs11217066 | G | T | M_Disease | 0.893309406 | M_Disease&NULL |
| 11 | 118726843 | rs1037679725 | T | C | M_Disease | 0.893309406 | M_Disease&NULL |
| 11 | 118516903 | rs80015904 | A | G | M_Disease | 0.89283193 | M_Disease&NULL |

### FINEMAP

| CHROM | POS (GRCh37/hg19) | RSID | A1 | A2 | OR_Disease | L+P0 | L+P1 | logBF | P_CAUSAL |
|---|---|---|---|---|---|---|---|---|---|
| 11 | 118607871 | rs76409436 | A | G | 1.189 | -4813.425 | -4806.718 | 6.706 | 0.020245341 |
| 11 | 118609790 | rs11217006 | A | G | 1.189 | -4813.425 | -4806.778 | 6.647 | 0.01908542 |
| 11 | 118603736 | rs112561833 | C | A | 1.186 | -4813.425 | -4806.843 | 6.581 | 0.017866451 |
| 11 | 118611817 | rs10892280 | T | C | 1.19 | -4813.425 | -4806.87 | 6.555 | 0.01740791 |
| 11 | 118610549 | rs11826521 | T | G | 1.19 | -4813.425 | -4806.913 | 6.512 | 0.016675235 |
| 11 | 118639353 | rs11217020 | A | G | 1.19 | -4813.425 | -4806.926 | 6.499 | 0.01645986 |
| 11 | 118603526 | rs11216995 | A | G | 1.184 | -4813.425 | -4806.965 | 6.459 | 0.01581446 |
| 11 | 118608716 | rs7945144 | G | A | 1.186 | -4813.425 | -4807.011 | 6.413 | 0.015103473 |
| 11 | 118611781 | rs10892279 | A | G | 1.186 | -4813.425 | -4807.036 | 6.388 | 0.014730566 |
| 11 | 118618776 | rs1048024 | T | A | 1.188 | -4813.425 | -4807.053 | 6.372 | 0.014496753 |
| 11 | 118619047 | rs2077579 | G | T | 1.188 | -4813.425 | -4807.053 | 6.372 | 0.014496753 |
| 11 | 118619253 | rs2004781 | T | C | 1.188 | -4813.425 | -4807.053 | 6.372 | 0.014496753 |
| 11 | 118619960 | rs3889239 | T | C | 1.188 | -4813.425 | -4807.053 | 6.372 | 0.014496753 |
| 11 | 118607794 | rs12364971 | G | T | 1.185 | -4813.425 | -4807.057 | 6.368 | 0.014438882 |
| 11 | 118598544 | rs10892271 | A | G | 1.183 | -4813.425 | -4807.068 | 6.356 | 0.014266651 |
| 11 | 118636980 | rs10892283 | G | A | 1.188 | -4813.425 | -4807.072 | 6.352 | 0.014209698 |
| 11 | 118610957 | rs6589684 | A | G | 1.186 | -4813.425 | -4807.073 | 6.351 | 0.014195495 |
| 11 | 118603520 | rs11216994 | T | C | 1.182 | -4813.425 | -4807.079 | 6.346 | 0.014124695 |
| 11 | 118636343 | rs11217019 | A | C | 1.188 | -4813.425 | -4807.1 | 6.325 | 0.013831169 |
| 11 | 118604638 | rs11216997 | T | C | 1.182 | -4813.425 | -4807.107 | 6.317 | 0.013720961 |
| 11 | 118628373 | rs74676029 | T | C | 1.188 | -4813.425 | -4807.11 | 6.315 | 0.013693547 |
| 11 | 118602707 | rs10892272 | G | C | 1.182 | -4813.425 | -4807.119 | 6.306 | 0.013570858 |
| 11 | 118602809 | rs10892273 | A | G | 1.182 | -4813.425 | -4807.119 | 6.306 | 0.013570858 |
| 11 | 118603495 | rs11216993 | A | T | 1.183 | -4813.425 | -4807.122 | 6.302 | 0.013516683 |
| 11 | 118607569 | rs77209083 | G | A | 1.185 | -4813.425 | -4807.129 | 6.295 | 0.013422396 |
| 11 | 118648373 | rs56043232 | T | C | 1.187 | -4813.425 | -4807.153 | 6.271 | 0.013104094 |
| 11 | 118655645 | rs874621 | A | G | 1.187 | -4813.425 | -4807.163 | 6.262 | 0.012986686 |
| 11 | 118607046 | rs11217001 | A | G | 1.185 | -4813.425 | -4807.172 | 6.252 | 0.012857466 |
| 11 | 118603165 | rs73001429 | G | T | 1.181 | -4813.425 | -4807.189 | 6.235 | 0.012640737 |
| 11 | 118646003 | rs10892289 | T | C | 1.187 | -4813.425 | -4807.193 | 6.232 | 0.012602872 |
| 11 | 118644694 | rs10892288 | G | A | 1.186 | -4813.425 | -4807.2 | 6.224 | 0.012502451 |
| 11 | 118661398 | rs57494551 | T | C | 1.187 | -4813.425 | -4807.206 | 6.218 | 0.012427661 |
| 11 | 118665575 | rs10892293 | T | A | 1.185 | -4813.425 | -4807.22 | 6.204 | 0.012254886 |
| 11 | 118615340 | rs56758835 | A | G | 1.186 | -4813.425 | -4807.244 | 6.18 | 0.01196427 |
| 11 | 118615785 | rs11217009 | T | C | 1.186 | -4813.425 | -4807.244 | 6.18 | 0.01196427 |
| 11 | 118612197 | rs6589685 | C | T | 1.183 | -4813.425 | -4807.265 | 6.16 | 0.011727361 |
| 11 | 118655340 | rs10892292 | C | G | 1.185 | -4813.425 | -4807.311 | 6.113 | 0.011188928 |
| 11 | 118642085 | rs10892286 | C | A | 1.185 | -4813.425 | -4807.322 | 6.102 | 0.011066524 |
| 11 | 118642999 | rs10892287 | C | T | 1.185 | -4813.425 | -4807.322 | 6.102 | 0.011066524 |
| 11 | 118587319 | rs10892263 | C | T | 1.183 | -4813.425 | -4807.328 | 6.096 | 0.011000324 |
| 11 | 118589012 | rs12421793 | A | C | 1.182 | -4813.425 | -4807.338 | 6.086 | 0.010890869 |
| 11 | 118653503 | rs10892290 | T | G | 1.185 | -4813.425 | -4807.338 | 6.086 | 0.010890869 |
| 11 | 118653517 | rs10892291 | T | C | 1.185 | -4813.425 | -4807.338 | 6.086 | 0.010890869 |
| 11 | 118649512 | rs73003215 | A | G | 1.182 | -4813.425 | -4807.378 | 6.046 | 0.010463831 |
| 11 | 118612921 | rs2155433 | A | G | 1.182 | -4813.425 | -4807.414 | 6.01 | 0.010093833 |
| 11 | 118550522 | rs2277296 | C | T | 1.178 | -4813.425 | -4807.432 | 5.992 | 0.00991377 |
| 11 | 118550524 | rs201134460 | T | C | 1.178 | -4813.425 | -4807.432 | 5.992 | 0.00991377 |
| 11 | 118543563 | rs10892251 | T | C | 1.177 | -4813.425 | -4807.452 | 5.972 | 0.009717464 |
| 11 | 118607417 | rs55894437 | T | C | 1.18 | -4813.425 | -4807.458 | 5.967 | 0.009668998 |
| 11 | 118588952 | rs12421790 | T | C | 1.181 | -4813.425 | -4807.473 | 5.952 | 0.009525045 |
| 11 | 118607311 | rs56267528 | A | C | 1.179 | -4813.425 | -4807.487 | 5.937 | 0.009383236 |
| 11 | 118607150 | rs11217002 | T | C | 1.179 | -4813.425 | -4807.494 | 5.931 | 0.009327105 |
| 11 | 118675772 | rs11217036 | T | C | 1.181 | -4813.425 | -4807.502 | 5.923 | 0.009252786 |
| 11 | 118556398 | rs11216943 | A | G | 1.176 | -4813.425 | -4807.503 | 5.922 | 0.009243538 |
| 11 | 118677086 | rs11217037 | A | G | 1.18 | -4813.425 | -4807.617 | 5.808 | 0.00824762 |
| 11 | 118677355 | rs11217038 | A | G | 1.18 | -4813.425 | -4807.617 | 5.808 | 0.00824762 |
| 11 | 118555692 | rs10892252 | G | A | 1.175 | -4813.425 | -4807.631 | 5.794 | 0.008132958 |
| 11 | 118669605 | rs11217032 | A | T | 1.179 | -4813.425 | -4807.648 | 5.777 | 0.007995866 |
| 11 | 118673706 | rs11217033 | C | T | 1.179 | -4813.425 | -4807.648 | 5.777 | 0.007995866 |
| 11 | 118684610 | rs715412 | A | G | 1.179 | -4813.425 | -4807.657 | 5.767 | 0.007916306 |
| 11 | 118606521 | rs6589681 | T | C | 1.177 | -4813.425 | -4807.691 | 5.734 | 0.007659331 |
| 11 | 118683564 | rs17122453 | A | G | 1.177 | -4813.425 | -4807.699 | 5.725 | 0.007590707 |
| 11 | 118667357 | rs10892294 | C | G | 1.179 | -4813.425 | -4807.704 | 5.721 | 0.007560404 |
| 11 | 118694547 | rs7123726 | C | T | 1.177 | -4813.425 | -4807.73 | 5.694 | 0.007359005 |
| 11 | 118697865 | rs11217045 | A | C | 1.179 | -4813.425 | -4807.818 | 5.607 | 0.006745831 |
| 11 | 118697906 | rs73005502 | G | A | 1.179 | -4813.425 | -4807.818 | 5.607 | 0.006745831 |
| 11 | 118680648 | rs11217040 | A | C | 1.176 | -4813.425 | -4807.821 | 5.603 | 0.006718902 |
| 11 | 118606722 | rs6589682 | T | C | 1.179 | -4813.425 | -4807.823 | 5.601 | 0.006705477 |
| 11 | 118679304 | rs28409215 | A | G | 1.177 | -4813.425 | -4807.825 | 5.6 | 0.006698775 |
| 11 | 118698907 | rs11217046 | A | G | 1.178 | -4813.425 | -4807.851 | 5.573 | 0.006520328 |
| 11 | 118696022 | rs11217044 | C | T | 1.175 | -4813.425 | -4807.875 | 5.549 | 0.006365703 |
| 11 | 118682528 | rs11217042 | A | G | 1.175 | -4813.425 | -4807.877 | 5.547 | 0.006352984 |
| 11 | 118560857 | rs73001406 | A | G | 1.172 | -4813.425 | -4807.883 | 5.541 | 0.006314981 |
| 11 | 118612565 | rs6589686 | A | G | 1.181 | -4813.425 | -4807.902 | 5.523 | 0.006202328 |
| 11 | 118701018 | rs79504890 | A | G | 1.177 | -4813.425 | -4807.93 | 5.494 | 0.006025043 |
| 11 | 118682350 | rs11217041 | G | A | 1.173 | -4813.425 | -4807.966 | 5.458 | 0.005812 |
| 11 | 118587097 | rs10892262 | C | G | 1.173 | -4813.425 | -4808.002 | 5.423 | 0.005612098 |
| 11 | 118681464 | rs7942535 | C | T | 1.173 | -4813.425 | -4808.021 | 5.404 | 0.005506475 |
| 11 | 118590668 | rs11216980 | A | C | 1.173 | -4813.425 | -4808.081 | 5.344 | 0.005185803 |
| 11 | 118681079 | rs73005423 | G | A | 1.173 | -4813.425 | -4808.134 | 5.29 | 0.004913196 |
| 11 | 118681083 | rs73005426 | G | A | 1.171 | -4813.425 | -4808.155 | 5.27 | 0.004815908 |
| 11 | 118459069 | rs633308 | G | A | 1.143 | -4813.425 | -4808.284 | 5.141 | 0.004233058 |
| 11 | 118721699 | rs7125066 | G | A | 1.142 | -4813.425 | -4808.495 | 4.929 | 0.003424395 |
| 11 | 118724894 | rs9736016 | A | T | 1.141 | -4813.425 | -4808.548 | 4.876 | 0.003247627 |
| 11 | 118585826 | rs11216972 | T | C | 1.163 | -4813.425 | -4808.67 | 4.755 | 0.002877508 |
| 11 | 118721565 | rs75249660 | A | G | 1.169 | -4813.425 | -4808.676 | 4.748 | 0.002857436 |
| 11 | 118530611 | rs2276065 | C | T | 1.16 | -4813.425 | -4808.798 | 4.626 | 0.002529255 |
| 11 | 118722149 | rs11217058 | A | G | 1.167 | -4813.425 | -4808.825 | 4.599 | 0.002461879 |
| 11 | 118575326 | rs11216956 | A | G | 1.157 | -4813.425 | -4808.949 | 4.475 | 0.002174774 |
| 11 | 118725660 | rs4936441 | C | G | 1.163 | -4813.425 | -4809.065 | 4.36 | 0.00193852 |
| 11 | 118507478 | rs11216937 | T | C | 1.156 | -4813.425 | -4809.107 | 4.318 | 0.001858788 |
| 11 | 118516774 | rs12365095 | T | C | 1.156 | -4813.425 | -4809.114 | 4.311 | 0.001845822 |
| 11 | 118495287 | rs77683570 | C | T | 1.156 | -4813.425 | -4809.136 | 4.288 | 0.001803853 |
| 11 | 118486110 | rs45540840 | A | G | 1.156 | -4813.425 | -4809.149 | 4.276 | 0.001782336 |
| 11 | 118499394 | rs2236661 | G | C | 1.156 | -4813.425 | -4809.158 | 4.267 | 0.001766367 |
| 11 | 118726753 | rs11217066 | G | T | 1.162 | -4813.425 | -4809.168 | 4.256 | 0.001747043 |
| 11 | 118726843 | rs1037679725 | T | C | 1.162 | -4813.425 | -4809.168 | 4.256 | 0.001747043 |
| 11 | 118516903 | rs80015904 | A | G | 1.155 | -4813.425 | -4809.182 | 4.242 | 0.001722755 |

| | | | | | | | |
|---|---|---|---|---|---|---|---|
| 11 | 118514625 | rs11216938 | T | C | M_Disease | 0.889927426 | M_Disease&NULL |
| 11 | 118516737 | rs12362152 | G | A | M_Disease | 0.88844946 | M_Disease&NULL |
| 11 | 118534082 | rs10790256 | T | C | M_Disease | 0.88844946 | M_Disease&NULL |
| 11 | 118583525 | rs10892260 | C | A | M_Disease | 0.887952961 | M_Disease&NULL |
| 11 | 118741842 | rs4938573 | C | T | M_Disease | 0.887952961 | M_Disease&NULL |
| 11 | 118743772 | rs6421571 | T | C | M_Disease | 0.887454533 | M_Disease&NULL |
| 11 | 118496119 | rs45565037 | A | G | M_Disease | 0.88695417 | M_Disease&NULL |
| 11 | 118490076 | rs10892247 | A | G | M_Disease | 0.885441421 | M_Disease&NULL |
| 11 | 118501022 | rs10892248 | A | G | M_Disease | 0.885441421 | M_Disease&NULL |
| 11 | 118488782 | rs11216930 | C | A | M_Disease | 0.883911078 | M_Disease&NULL |
| 11 | 118535840 | rs12225548 | G | C | M_Disease | 0.883911078 | M_Disease&NULL |
| 11 | 118734000 | rs11217074 | C | T | M_Disease | 0.883911078 | M_Disease&NULL |
| 11 | 118529127 | rs7928371 | A | G | M_Disease | 0.881321047 | M_Disease&NULL |
| 11 | 118528424 | rs17748 | T | C | M_Disease | 0.880797078 | M_Disease&NULL |
| 11 | 118579747 | rs10790261 | G | A | M_Disease | 0.880797078 | M_Disease&NULL |
| 11 | 118579327 | rs10892257 | A | G | M_Disease | 0.88027111 | M_Disease&NULL |
| 11 | 118584340 | rs11216969 | C | T | M_Disease | 0.88027111 | M_Disease&NULL |
| 11 | 118743338 | rs4936444 | T | C | M_Disease | 0.88027111 | M_Disease&NULL |
| 11 | 118740418 | rs7951740 | G | C | M_Disease | 0.879743138 | M_Disease&NULL |
| 11 | 118581895 | rs11216964 | A | G | M_Disease | 0.879213157 | M_Disease&NULL |
| 11 | 118578705 | rs10892256 | A | G | M_Disease | 0.878147149 | M_Disease&NULL |
| 11 | 118579865 | rs10892258 | A | G | M_Disease | 0.877611113 | M_Disease&NULL |
| 11 | 118580338 | rs11216961 | G | C | M_Disease | 0.877611113 | M_Disease&NULL |
| 11 | 118580813 | rs11216962 | C | T | M_Disease | 0.877611113 | M_Disease&NULL |
| 11 | 118580814 | rs11216963 | A | G | M_Disease | 0.877611113 | M_Disease&NULL |
| 11 | 118582613 | rs11216965 | A | C | M_Disease | 0.877611113 | M_Disease&NULL |
| 11 | 118583002 | rs10892259 | A | G | M_Disease | 0.877611113 | M_Disease&NULL |
| 11 | 118577873 | rs11216960 | A | G | M_Disease | 0.875990818 | M_Disease&NULL |
| 11 | 118729391 | rs10790268 | A | G | M_Disease | 0.874352143 | M_Disease&NULL |
| 11 | 118729456 | rs10790269 | C | T | M_Disease | 0.874352143 | M_Disease&NULL |
| 11 | 118741157 | rs7117261 | T | C | M_Disease | 0.873801812 | M_Disease&NULL |
| 11 | 118742800 | rs74541740 | A | G | M_Disease | 0.872138434 | M_Disease&NULL |
| 11 | 118744396 | rs7117313 | C | G | M_Disease | 0.862948707 | M_Disease&NULL |
| 11 | 118745884 | rs10790275 | G | C | M_Disease | 0.861165006 | M_Disease&NULL |
| 11 | 118739056 | rs7122669 | A | G | M_Disease | 0.860566127 | M_Disease&NULL |
| 11 | 118738298 | rs7119044 | G | C | M_Disease | 0.859361874 | M_Disease&NULL |
| 11 | 118515579 | rs10790255 | G | T | M_Disease | 0.858148935 | M_Disease&NULL |
| 11 | 118740931 | rs4938572 | C | T | M_Disease | 0.856313167 | M_Disease&NULL |
| 11 | 118745243 | rs7481797 | A | G | M_Disease | 0.856313167 | M_Disease&NULL |
| 11 | 118745278 | rs7481819 | C | T | M_Disease | 0.85383477 | M_Disease&NULL |
| 11 | 118740864 | rs4936443 | C | T | M_Disease | 0.847483683 | M_Disease&NULL |
| 11 | 118640518 | rs181603672 | G | A | M_Disease | 0.825634714 | M_Disease&NULL |
| 11 | 118743286 | rs12365699 | A | G | M_Disease | 0.818319025 | M_Disease&NULL |
| 11 | 118469739 | rs143380378 | G | C | M_Disease | 0.817574476 | M_Disease&NULL |
| 11 | 118648297 | rs117422168 | G | A | M_Disease | 0.815326614 | M_Disease&NULL |
| 11 | 118605520 | rs75598590 | C | T | M_Disease | 0.802183889 | M_Disease&NULL |
| 11 | 118605463 | rs10892274 | T | C | M_Disease | 0.800592243 | M_Disease&NULL |
| 11 | 118605617 | rs10892275 | A | T | M_Disease | 0.800592243 | M_Disease&NULL |
| 11 | 118605841 | rs10892277 | A | G | M_Disease | 0.800592243 | M_Disease&NULL |
| 11 | 118604941 | rs11216999 | C | T | M_Disease | 0.796571127 | M_Disease&NULL |
| 11 | 118605628 | rs10892276 | A | G | M_Disease | 0.785834983 | M_Disease&NULL |
| 11 | 118462188 | rs4936431 | G | C | M_Disease | 0.781597464 | M_Disease&NULL |
| 11 | 118853774 | rs11217109 | C | T | M_Disease | 0.780742748 | M_Disease&NULL |
| 11 | 118763313 | rs78041268 | A | G | M_Disease | 0.777299861 | M_Disease&NULL |
| 11 | 118853021 | rs113962716 | A | C | M_Disease | 0.772942259 | M_Disease&NULL |
| 11 | 118852647 | rs80126726 | A | G | M_Disease | 0.766741064 | M_Disease&NULL |
| 11 | 118762073 | rs77841686 | A | G | M_Disease | 0.765845626 | M_Disease&NULL |
| 11 | 118809882 | rs11217099 | A | C | M_Disease | 0.765845626 | M_Disease&NULL |
| 11 | 118876213 | rs76253444 | G | A | M_Disease | 0.763145016 | M_Disease&NULL |
| 11 | 118604837 | rs11216998 | T | C | M_Disease | 0.760423002 | M_Disease&NULL |
| 11 | 118898319 | rs741811 | A | G | M_Disease | 0.756760451 | M_Disease&NULL |
| 11 | 118806897 | rs10892310 | T | C | M_Disease | 0.749322084 | M_Disease&NULL |
| 11 | 118811641 | rs11606199 | A | G | M_Disease | 0.748381722 | M_Disease&NULL |
| 11 | 118831755 | rs10892316 | T | C | M_Disease | 0.744596916 | M_Disease&NULL |
| 11 | 118873840 | rs9645663 | C | T | M_Disease | 0.744596916 | M_Disease&NULL |
| 11 | 118834508 | rs111358668 | A | G | M_Disease | 0.739813607 | M_Disease&NULL |
| 11 | 118834718 | rs112939258 | T | C | M_Disease | 0.739813607 | M_Disease&NULL |
| 11 | 118893767 | rs111345266 | T | C | M_Disease | 0.739813607 | M_Disease&NULL |
| 11 | 118767564 | rs75438046 | A | G | M_Disease | 0.733997517 | M_Disease&NULL |
| 11 | 118870413 | rs10892319 | T | C | M_Disease | 0.733997517 | M_Disease&NULL |
| 11 | 118617920 | rs476246 | T | C | M_Disease | 0.732040502 | M_Disease&NULL |
| 11 | 118890085 | rs4938619 | T | C | M_Disease | 0.732040502 | M_Disease&NULL |
| 11 | 118828184 | rs45586434 | A | C | M_Disease | 0.731058579 | M_Disease&NULL |
| 11 | 118606197 | rs12420846 | T | C | M_Disease | 0.729087922 | M_Disease&NULL |
| 11 | 118617544 | rs472641 | G | A | M_Disease | 0.728099198 | M_Disease&NULL |
| 11 | 118624767 | rs581062 | A | G | M_Disease | 0.723121805 | M_Disease&NULL |
| 11 | 118751326 | rs10892306 | A | T | M_Disease | 0.723121805 | M_Disease&NULL |
| 11 | 118876552 | rs74884652 | C | T | M_Disease | 0.721115178 | M_Disease&NULL |
| 11 | 118877891 | rs75154936 | C | T | M_Disease | 0.721115178 | M_Disease&NULL |
| 11 | 118478681 | rs45578540 | T | C | M_Disease | 0.719099657 | M_Disease&NULL |
| 11 | 118664377 | rs598373 | T | C | M_Disease | 0.717075285 | M_Disease&NULL |
| 11 | 118867695 | rs4938617 | T | C | M_Disease | 0.715042106 | M_Disease&NULL |
| 11 | 118810405 | rs4938589 | T | C | M_Disease | 0.710949503 | M_Disease&NULL |
| 11 | 118621495 | rs487728 | G | C | M_Disease | 0.709920918 | M_Disease&NULL |
| 11 | 118640063 | rs524590 | T | A | M_Disease | 0.709920918 | M_Disease&NULL |
| 11 | 118626511 | rs499343 | T | C | M_Disease | 0.708890173 | M_Disease&NULL |
| 11 | 118563021 | rs117330495 | A | C | M_Disease | 0.705785028 | M_Disease&NULL |
| 11 | 118816353 | rs586370 | A | G | NULL | 0.704745698 | NULL&M_Disease |
| 11 | 118770321 | rs11217083 | G | A | NULL | 0.703704238 | NULL&M_Disease |
| 11 | 118717708 | rs7107756 | A | G | NULL | 0.702660654 | NULL&M_Disease |
| 11 | 118765600 | rs3922 | G | A | NULL | 0.702660654 | NULL&M_Disease |
| 11 | 118767077 | rs581063 | C | T | NULL | 0.702660654 | NULL&M_Disease |
| 11 | 118769032 | rs11217081 | G | A | NULL | 0.702660654 | NULL&M_Disease |
| 11 | 118707249 | rs4938551 | C | G | NULL | 0.701614954 | NULL&M_Disease |
| 11 | 118716563 | rs2851460 | A | G | NULL | 0.701614954 | NULL&M_Disease |
| 11 | 118769442 | rs7104819 | G | A | NULL | 0.701614954 | NULL&M_Disease |
| 11 | 118515779 | rs117679873 | A | G | M_Disease | 0.700567142 | M_Disease&NULL |
| 11 | 118796570 | rs11217096 | T | C | NULL | 0.699517228 | NULL&M_Disease |
| 11 | 118807735 | rs10892311 | T | C | NULL | 0.699517228 | NULL&M_Disease |
| 11 | 118823148 | rs663003 | T | C | NULL | 0.699517228 | NULL&M_Disease |
| 11 | 118826060 | rs625291 | T | C | NULL | 0.699517228 | NULL&M_Disease |
| 11 | 118566069 | rs506255 | A | C | M_Disease | 0.698465216 | M_Disease&NULL |
| 11 | 118574185 | rs687664 | A | C | NULL | 0.698465216 | NULL&M_Disease |

| | | | | | | | | | |
|---|---|---|---|---|---|---|---|---|---|
| 11 | 118514625 | rs11216938 | T | C | 1.154 | -4813.425 | -4809.244 | 4.181 | 0.001620808 |
| 11 | 118516737 | rs12362152 | G | A | 1.154 | -4813.425 | -4809.27 | 4.155 | 0.00157921 |
| 11 | 118534082 | rs10790256 | T | C | 1.154 | -4813.425 | -4809.275 | 4.15 | 0.001571334 |
| 11 | 118741842 | rs4938573 | C | T | 1.16 | -4813.425 | -4809.282 | 4.142 | 0.001558813 |
| 11 | 118583525 | rs10892260 | C | A | 1.152 | -4813.425 | -4809.283 | 4.141 | 0.001557255 |
| 11 | 118743772 | rs6421571 | T | C | 1.161 | -4813.425 | -4809.292 | 4.132 | 0.001543303 |
| 11 | 118496119 | rs45565037 | A | G | 1.153 | -4813.425 | -4809.305 | 4.119 | 0.00152337 |
| 11 | 118490076 | rs10892247 | A | G | 1.153 | -4813.425 | -4809.333 | 4.092 | 0.001482789 |
| 11 | 118501022 | rs10892248 | A | G | 1.153 | -4813.425 | -4809.333 | 4.092 | 0.001482789 |
| 11 | 118488782 | rs11216930 | C | A | 1.153 | -4813.425 | -4809.36 | 4.064 | 0.001441847 |
| 11 | 118734000 | rs11217074 | C | T | 1.159 | -4813.425 | -4809.36 | 4.064 | 0.001441847 |
| 11 | 118535840 | rs12225548 | G | C | 1.153 | -4813.425 | -4809.362 | 4.063 | 0.001440406 |
| 11 | 118529127 | rs7928371 | A | G | 1.152 | -4813.425 | -4809.407 | 4.018 | 0.001377024 |
| 11 | 118528424 | rs17748 | T | C | 1.152 | -4813.425 | -4809.418 | 4.007 | 0.00136196 |
| 11 | 118579747 | rs10790261 | G | A | 1.15 | -4813.425 | -4809.422 | 4.003 | 0.001356523 |
| 11 | 118743338 | rs4936444 | T | C | 1.159 | -4813.425 | -4809.431 | 3.994 | 0.001344369 |
| 11 | 118579327 | rs10892257 | A | G | 1.15 | -4813.425 | -4809.432 | 3.992 | 0.001341683 |
| 11 | 118584340 | rs11216969 | C | T | 1.15 | -4813.425 | -4809.436 | 3.988 | 0.001336327 |
| 11 | 118740418 | rs7951740 | G | C | 1.158 | -4813.425 | -4809.44 | 3.985 | 0.001332324 |
| 11 | 118581895 | rs11216964 | A | G | 1.15 | -4813.425 | -4809.453 | 3.971 | 0.001313801 |
| 11 | 118578705 | rs10892256 | A | G | 1.149 | -4813.425 | -4809.472 | 3.952 | 0.001289075 |
| 11 | 118579865 | rs10892258 | A | G | 1.149 | -4813.425 | -4809.479 | 3.945 | 0.001280083 |
| 11 | 118580338 | rs11216961 | G | C | 1.149 | -4813.425 | -4809.479 | 3.945 | 0.001280083 |
| 11 | 118580813 | rs11216962 | C | T | 1.149 | -4813.425 | -4809.479 | 3.945 | 0.001280083 |
| 11 | 118580814 | rs11216963 | A | G | 1.149 | -4813.425 | -4809.479 | 3.945 | 0.001280083 |
| 11 | 118582613 | rs11216965 | A | C | 1.149 | -4813.425 | -4809.479 | 3.945 | 0.001280083 |
| 11 | 118583002 | rs10892259 | A | G | 1.149 | -4813.425 | -4809.479 | 3.945 | 0.001280083 |
| 11 | 118577873 | rs11216960 | A | G | 1.149 | -4813.425 | -4809.51 | 3.915 | 0.001242251 |
| 11 | 118729391 | rs10790268 | A | G | 1.156 | -4813.425 | -4809.546 | 3.878 | 0.001197127 |
| 11 | 118729456 | rs10790269 | C | T | 1.156 | -4813.425 | -4809.546 | 3.878 | 0.001197127 |
| 11 | 118741157 | rs7117261 | T | C | 1.156 | -4813.425 | -4809.556 | 3.869 | 0.001186401 |
| 11 | 118742800 | rs74541740 | A | G | 1.156 | -4813.425 | -4809.584 | 3.841 | 0.001153643 |
| 11 | 118744396 | rs7117313 | C | G | 1.153 | -4813.425 | -4809.745 | 3.68 | 0.000982087 |
| 11 | 118745884 | rs10790275 | G | C | 1.152 | -4813.425 | -4809.775 | 3.649 | 0.000952169 |
| 11 | 118739056 | rs7122669 | A | G | 1.153 | -4813.425 | -4809.78 | 3.645 | 0.000948309 |
| 11 | 118738298 | rs7119044 | G | C | 1.152 | -4813.425 | -4809.799 | 3.625 | 0.000929531 |
| 11 | 118515579 | rs10790255 | G | T | 1.142 | -4813.425 | -4809.817 | 3.608 | 0.000913862 |
| 11 | 118740931 | rs4938572 | C | T | 1.151 | -4813.425 | -4809.848 | 3.576 | 0.000885082 |
| 11 | 118745243 | rs7481797 | A | G | 1.151 | -4813.425 | -4809.849 | 3.576 | 0.000885082 |
| 11 | 118745278 | rs7481819 | C | T | 1.151 | -4813.425 | -4809.896 | 3.529 | 0.000844445 |
| 11 | 118740864 | rs4936443 | C | T | 1.149 | -4813.425 | -4809.994 | 3.431 | 0.000765615 |
| 11 | 118640518 | rs181603672 | G | A | 1.259 | -4813.425 | -4810.321 | 3.103 | 0.000551521 |
| 11 | 118743286 | rs12365699 | A | G | 1.152 | -4813.425 | -4810.414 | 3.011 | 0.000503045 |
| 11 | 118469739 | rs143380378 | G | C | 1.246 | -4813.425 | -4810.424 | 3.001 | 0.00049804 |
| 11 | 118648297 | rs117422168 | G | A | 1.255 | -4813.425 | -4810.454 | 2.971 | 0.00048332 |
| 11 | 118605520 | rs75598590 | C | T | 1.25 | -4813.425 | -4810.625 | 2.8 | 0.000407353 |
| 11 | 118605463 | rs10892274 | T | C | 1.124 | -4813.425 | -4810.645 | 2.78 | 0.000399287 |
| 11 | 118605617 | rs10892275 | A | T | 1.124 | -4813.425 | -4810.645 | 2.78 | 0.000399287 |
| 11 | 118605841 | rs10892277 | A | G | 1.124 | -4813.425 | -4810.645 | 2.78 | 0.000399287 |
| 11 | 118604941 | rs11216999 | C | T | 1.124 | -4813.425 | -4810.694 | 2.73 | 0.000379813 |
| 11 | 118605628 | rs10892276 | A | G | 1.122 | -4813.425 | -4810.818 | 2.607 | 0.000335855 |
| 11 | 118462188 | rs4936431 | G | C | 1.23 | -4813.425 | -4810.876 | 2.549 | 0.00031693 |
| 11 | 118853774 | rs11217109 | C | T | 1.145 | -4813.425 | -4810.887 | 2.538 | 0.000313463 |
| 11 | 118763313 | rs78041268 | A | G | 1.278 | -4813.425 | -4810.934 | 2.491 | 0.000299071 |
| 11 | 118853021 | rs113962716 | A | C | 1.143 | -4813.425 | -4810.973 | 2.451 | 0.000287344 |
| 11 | 118852647 | rs80126726 | A | G | 1.142 | -4813.425 | -4811.042 | 2.383 | 0.000268454 |
| 11 | 118809882 | rs11217099 | A | C | 1.14 | -4813.425 | -4811.056 | 2.369 | 0.000264722 |
| 11 | 118762073 | rs77841686 | A | G | 1.272 | -4813.425 | -4811.063 | 2.361 | 0.000262613 |
| 11 | 118876213 | rs76253444 | G | A | 1.14 | -4813.425 | -4811.079 | 2.346 | 0.000258703 |
| 11 | 118604837 | rs11216998 | T | C | 1.117 | -4813.425 | -4811.109 | 2.315 | 0.000250806 |
| 11 | 118898319 | rs741811 | A | G | 1.149 | -4813.425 | -4811.156 | 2.269 | 0.00023953 |
| 11 | 118806897 | rs10892310 | T | C | 0.897 | -4813.425 | -4811.232 | 2.192 | 0.000221779 |
| 11 | 118811641 | rs11606199 | A | G | 1.135 | -4813.425 | -4811.238 | 2.186 | 0.000220452 |
| 11 | 118873840 | rs9645663 | C | T | 1.136 | -4813.425 | -4811.284 | 2.141 | 0.000210752 |
| 11 | 118831755 | rs10892316 | T | C | 1.135 | -4813.425 | -4811.285 | 2.139 | 0.00021033 |
| 11 | 118893767 | rs111345266 | T | C | 1.143 | -4813.425 | -4811.328 | 2.096 | 0.000201478 |
| 11 | 118834508 | rs111358668 | A | G | 1.135 | -4813.425 | -4811.33 | 2.095 | 0.000201277 |
| 11 | 118834718 | rs112939258 | T | C | 1.135 | -4813.425 | -4811.33 | 2.095 | 0.000201277 |
| 11 | 118870413 | rs10892319 | T | C | 1.134 | -4813.425 | -4811.387 | 2.037 | 0.000189935 |
| 11 | 118767564 | rs75438046 | A | G | 1.254 | -4813.425 | -4811.402 | 2.022 | 0.000187107 |
| 11 | 118890085 | rs4938619 | T | C | 1.141 | -4813.425 | -4811.414 | 2.011 | 0.00018506 |
| 11 | 118617920 | rs476246 | T | C | 1.106 | -4813.425 | -4811.414 | 2.01 | 0.000184875 |
| 11 | 118828184 | rs45586434 | A | C | 1.132 | -4813.425 | -4811.422 | 2.003 | 0.000183585 |
| 11 | 118606197 | rs12420846 | T | C | 1.197 | -4813.425 | -4811.442 | 1.983 | 0.00017995 |
| 11 | 118617544 | rs472641 | G | A | 1.105 | -4813.425 | -4811.447 | 1.977 | 0.000178874 |
| 11 | 118624767 | rs581062 | A | G | 1.104 | -4813.425 | -4811.503 | 1.921 | 0.000169132 |
| 11 | 118751326 | rs10892306 | A | T | 1.186 | -4813.425 | -4811.506 | 1.918 | 0.000168625 |
| 11 | 118876552 | rs74884652 | C | T | 1.138 | -4813.425 | -4811.522 | 1.903 | 0.000166115 |
| 11 | 118877891 | rs75154936 | C | T | 1.138 | -4813.425 | -4811.522 | 1.903 | 0.000166115 |
| 11 | 118478681 | rs45578540 | T | C | 1.224 | -4813.425 | -4811.552 | 1.873 | 0.000161206 |
| 11 | 118664377 | rs598373 | T | C | 1.104 | -4813.425 | -4811.557 | 1.868 | 0.000160401 |
| 11 | 118867695 | rs4938617 | T | C | 1.137 | -4813.425 | -4811.582 | 1.843 | 0.000156441 |
| 11 | 118810405 | rs4938589 | T | C | 1.148 | -4813.425 | -4811.623 | 1.801 | 0.000150007 |
| 11 | 118640063 | rs524590 | T | A | 1.102 | -4813.425 | -4811.628 | 1.796 | 0.000149259 |
| 11 | 118621495 | rs487728 | G | C | 1.102 | -4813.425 | -4811.633 | 1.792 | 0.000148663 |
| 11 | 118626511 | rs499343 | T | C | 1.102 | -4813.425 | -4811.641 | 1.784 | 0.000147478 |
| 11 | 118563021 | rs117330495 | A | C | 1.214 | -4813.425 | -4811.674 | 1.751 | 0.000142691 |
| 11 | 118515779 | rs117679873 | A | G | 1.214 | -4813.425 | -4811.73 | 1.695 | 0.00013492 |
| 11 | 118566069 | rs506255 | A | C | 0.872 | -4813.425 | -4811.74 | 1.685 | 0.000133577 |
| 11 | 118725942 | rs11217063 | C | G | 0.911 | -4813.425 | -4811.793 | 1.631 | 0.000126555 |
| 11 | 118726647 | rs11217065 | C | T | 0.911 | -4813.425 | -4811.793 | 1.631 | 0.000126555 |
| 11 | 118723379 | rs7103067 | A | G | 0.911 | -4813.425 | -4811.798 | 1.626 | 0.000125924 |
| 11 | 118500963 | rs680465 | A | G | 0.897 | -4813.425 | -4811.822 | 1.603 | 0.000123061 |
| 11 | 118459269 | rs10736492 | G | A | 0.912 | -4813.425 | -4811.826 | 1.599 | 0.00012257 |
| 11 | 118677710 | rs688161 | G | A | 1.099 | -4813.425 | -4811.826 | 1.598 | 0.000122447 |
| 11 | 118783424 | rs149114341 | A | G | 1.232 | -4813.425 | -4811.829 | 1.595 | 0.00012208 |
| 11 | 118652326 | rs1790189 | G | T | 1.099 | -4813.425 | -4811.842 | 1.582 | 0.000120504 |
| 11 | 118652373 | rs1784293 | C | A | 1.099 | -4813.425 | -4811.842 | 1.582 | 0.000120504 |
| 11 | 118809329 | rs10892313 | A | G | 1.142 | -4813.425 | -4811.847 | 1.578 | 0.000120023 |
| 11 | 118678609 | rs2508570 | A | G | 1.099 | -4813.425 | -4811.85 | 1.574 | 0.000119543 |
| 11 | 118724723 | rs57719838 | C | T | 0.912 | -4813.425 | -4811.85 | 1.574 | 0.000119543 |
| 11 | 118542383 | rs687190 | C | T | 0.914 | -4813.425 | -4811.875 | 1.549 | 0.000116592 |
| 11 | 118653629 | rs540180 | T | G | 1.098 | -4813.425 | -4811.89 | 1.535 | 0.000114971 |

| | | | | | | | |
|---|---|---|---|---|---|---|---|
| 11 | 118574188 | rs523715 | A | G | NULL | 0.698465216 | NULL&M_Disease |
| 11 | 118708564 | rs484232 | T | G | NULL | 0.698465216 | NULL&M_Disease |
| 11 | 118717732 | rs11217052 | G | A | NULL | 0.698465216 | NULL&M_Disease |
| 11 | 118800302 | rs12793214 | A | C | NULL | 0.698465216 | NULL&M_Disease |
| 11 | 118801640 | rs572608 | A | T | NULL | 0.698465216 | NULL&M_Disease |
| 11 | 118826063 | rs666191 | A | G | NULL | 0.698465216 | NULL&M_Disease |
| 11 | 118718061 | rs4938559 | T | C | NULL | 0.697411114 | NULL&M_Disease |
| 11 | 118718062 | rs4938560 | G | A | NULL | 0.697411114 | NULL&M_Disease |
| 11 | 118735476 | rs10892301 | A | G | NULL | 0.69635493 | NULL&M_Disease |
| 11 | 118824439 | rs61900157 | C | A | NULL | 0.695296669 | NULL&M_Disease |
| 11 | 118824996 | rs4426142 | A | G | NULL | 0.695296669 | NULL&M_Disease |
| 11 | 118825069 | rs2097131 | A | G | NULL | 0.695296669 | NULL&M_Disease |
| 11 | 118725942 | rs11217063 | C | G | M_Disease | 0.693173949 | M_Disease&NULL |
| 11 | 118726647 | rs11217065 | C | T | M_Disease | 0.693173949 | M_Disease&NULL |
| 11 | 118750321 | rs11217078 | C | T | NULL | 0.693173949 | NULL&M_Disease |
| 11 | 118825636 | rs1790191 | G | A | NULL | 0.693173949 | NULL&M_Disease |
| 11 | 118723379 | rs7103067 | A | G | M_Disease | 0.692109504 | M_Disease&NULL |
| 11 | 118758089 | rs497916 | T | C | NULL | 0.691043012 | NULL&M_Disease |
| 11 | 118500963 | rs680465 | A | G | M_Disease | 0.689974481 | M_Disease&NULL |
| 11 | 118783424 | rs149114341 | A | G | M_Disease | 0.689974481 | M_Disease&NULL |
| 11 | 118459269 | rs10736492 | G | A | M_Disease | 0.688903918 | M_Disease&NULL |
| 11 | 118486325 | rs12290522 | T | C | NULL | 0.688903918 | NULL&M_Disease |
| 11 | 118487673 | rs4938517 | C | T | NULL | 0.688903918 | NULL&M_Disease |
| 11 | 118677710 | rs688161 | G | A | M_Disease | 0.688903918 | M_Disease&NULL |
| 11 | 118497402 | rs12419204 | A | T | NULL | 0.687831331 | NULL&M_Disease |
| 11 | 118500173 | rs571001 | T | C | NULL | 0.687831331 | NULL&M_Disease |
| 11 | 118500193 | rs570952 | C | T | NULL | 0.687831331 | NULL&M_Disease |
| 11 | 118652326 | rs1790189 | G | T | M_Disease | 0.687831331 | M_Disease&NULL |
| 11 | 118652373 | rs1784293 | C | A | M_Disease | 0.687831331 | M_Disease&NULL |
| 11 | 118809329 | rs10892313 | A | G | M_Disease | 0.687831331 | M_Disease&NULL |
| 11 | 118496066 | rs11603616 | A | G | NULL | 0.686756727 | NULL&M_Disease |
| 11 | 118505090 | rs645637 | A | G | NULL | 0.686756727 | NULL&M_Disease |
| 11 | 118529595 | rs745663 | G | A | NULL | 0.686756727 | NULL&M_Disease |
| 11 | 118678609 | rs2508570 | A | G | M_Disease | 0.686756727 | M_Disease&NULL |
| 11 | 118724723 | rs57719838 | C | T | M_Disease | 0.686756727 | M_Disease&NULL |
| 11 | 118895495 | rs8301 | G | C | NULL | 0.686756727 | NULL&M_Disease |
| 11 | 118492591 | rs2077173 | A | G | NULL | 0.685680114 | NULL&M_Disease |
| 11 | 118568472 | rs570371 | G | A | NULL | 0.685680114 | NULL&M_Disease |
| 11 | 118571790 | rs656287 | G | A | NULL | 0.685680114 | NULL&M_Disease |
| 11 | 118506402 | rs483283 | C | T | NULL | 0.6846015 | NULL&M_Disease |
| 11 | 118522338 | rs4938520 | G | A | NULL | 0.6846015 | NULL&M_Disease |
| 11 | 118542383 | rs687190 | C | T | M_Disease | 0.6846015 | M_Disease&NULL |
| 11 | 118568813 | rs7479114 | T | C | NULL | 0.6846015 | NULL&M_Disease |
| 11 | 118569067 | rs4486664 | C | A | NULL | 0.6846015 | NULL&M_Disease |
| 11 | 118569469 | rs7123811 | G | T | NULL | 0.6846015 | NULL&M_Disease |
| 11 | 118569694 | rs7108144 | A | G | NULL | 0.6846015 | NULL&M_Disease |
| 11 | 118569900 | rs12793594 | C | T | NULL | 0.6846015 | NULL&M_Disease |
| 11 | 118570062 | rs554456 | C | T | NULL | 0.6846015 | NULL&M_Disease |
| 11 | 118570098 | rs482889 | T | G | NULL | 0.6846015 | NULL&M_Disease |
| 11 | 118570697 | rs523793 | T | C | NULL | 0.6846015 | NULL&M_Disease |
| 11 | 118572747 | rs489126 | A | G | NULL | 0.6846015 | NULL&M_Disease |
| 11 | 118512162 | rs570836 | A | G | NULL | 0.683520894 | NULL&M_Disease |
| 11 | 118528466 | rs7389 | G | T | NULL | 0.683520894 | NULL&M_Disease |
| 11 | 118570914 | rs659564 | A | C | NULL | 0.683520894 | NULL&M_Disease |
| 11 | 118571357 | rs544452 | T | G | NULL | 0.683520894 | NULL&M_Disease |
| 11 | 118571911 | rs570949 | T | G | NULL | 0.683520894 | NULL&M_Disease |
| 11 | 118572171 | rs658040 | G | A | NULL | 0.683520894 | NULL&M_Disease |
| 11 | 118572760 | rs611733 | T | C | NULL | 0.683520894 | NULL&M_Disease |
| 11 | 118573022 | rs552079 | C | T | NULL | 0.683520894 | NULL&M_Disease |
| 11 | 118573339 | rs673547 | A | G | NULL | 0.683520894 | NULL&M_Disease |
| 11 | 118573535 | rs517445 | G | A | NULL | 0.683520894 | NULL&M_Disease |
| 11 | 118573668 | rs574383 | C | T | NULL | 0.683520894 | NULL&M_Disease |
| 11 | 118573743 | rs519294 | A | C | NULL | 0.683520894 | NULL&M_Disease |
| 11 | 118601426 | rs186926311 | T | C | M_Disease | 0.683520894 | M_Disease&NULL |
| 11 | 118605451 | rs2186780 | T | C | NULL | 0.683520894 | NULL&M_Disease |
| 11 | 118605914 | rs7926282 | C | A | NULL | 0.683520894 | NULL&M_Disease |
| 11 | 118757191 | rs1790192 | G | A | NULL | 0.683520894 | NULL&M_Disease |
| 11 | 118522179 | rs1171461 | A | G | NULL | 0.682438302 | NULL&M_Disease |
| 11 | 118539170 | rs4457789 | T | G | NULL | 0.682438302 | NULL&M_Disease |
| 11 | 118568668 | rs11216949 | T | C | NULL | 0.682438302 | NULL&M_Disease |
| 11 | 118572021 | rs657539 | T | C | NULL | 0.682438302 | NULL&M_Disease |
| 11 | 118653629 | rs540180 | T | G | M_Disease | 0.682438302 | M_Disease&NULL |
| 11 | 118654651 | rs2508916 | G | C | M_Disease | 0.682438302 | M_Disease&NULL |
| 11 | 118666289 | rs56104144 | C | G | NULL | 0.682438302 | NULL&M_Disease |
| 11 | 118569097 | rs4393358 | T | C | NULL | 0.681353734 | NULL&M_Disease |
| 11 | 118570919 | rs642224 | A | G | NULL | 0.681353734 | NULL&M_Disease |
| 11 | 118571023 | rs541631 | G | A | NULL | 0.681353734 | NULL&M_Disease |
| 11 | 118571123 | rs658676 | T | C | NULL | 0.681353734 | NULL&M_Disease |
| 11 | 118571168 | rs488141 | G | A | NULL | 0.681353734 | NULL&M_Disease |
| 11 | 118571316 | rs657769 | T | C | NULL | 0.681353734 | NULL&M_Disease |
| 11 | 118668296 | rs58868870 | T | C | NULL | 0.681353734 | NULL&M_Disease |
| 11 | 118539522 | rs2508991 | A | G | NULL | 0.680267197 | NULL&M_Disease |
| 11 | 118572611 | rs611217 | T | C | NULL | 0.680267197 | NULL&M_Disease |
| 11 | 118590864 | rs11216982 | A | G | NULL | 0.680267197 | NULL&M_Disease |
| 11 | 118678881 | rs591756 | T | C | M_Disease | 0.680267197 | M_Disease&NULL |
| 11 | 118538157 | rs520543 | A | G | NULL | 0.679178699 | NULL&M_Disease |
| 11 | 118539566 | rs1783935 | T | A | NULL | 0.679178699 | NULL&M_Disease |
| 11 | 118568639 | rs511451 | T | C | NULL | 0.679178699 | NULL&M_Disease |
| 11 | 118538187 | rs607527 | T | G | NULL | 0.67808825 | NULL&M_Disease |
| 11 | 118546602 | rs526597 | G | A | NULL | 0.67808825 | NULL&M_Disease |
| 11 | 118678986 | rs477014 | C | T | M_Disease | 0.676995856 | M_Disease&NULL |
| 11 | 118679387 | rs473298 | G | A | M_Disease | 0.676995856 | M_Disease&NULL |
| 11 | 118679445 | rs516719 | C | T | M_Disease | 0.676995856 | M_Disease&NULL |
| 11 | 118573519 | rs4639966 | C | T | NULL | 0.676995856 | NULL&M_Disease |
| 11 | 118655356 | rs7934890 | T | C | NULL | 0.675901528 | NULL&M_Disease |
| 11 | 118602058 | rs562059 | G | T | M_Disease | 0.674805273 | M_Disease&NULL |
| 11 | 118657297 | rs487177 | G | C | NULL | 0.674805273 | NULL&M_Disease |
| 11 | 118755738 | rs523604 | G | A | NULL | 0.674805273 | NULL&M_Disease |
| 11 | 118549985 | rs543327 | T | G | NULL | 0.673707099 | NULL&M_Disease |
| 11 | 118550017 | rs543413 | C | T | NULL | 0.673707099 | NULL&M_Disease |
| 11 | 118707171 | rs4938550 | G | A | NULL | 0.673707099 | NULL&M_Disease |
| 11 | 118808140 | rs7117408 | T | C | M_Disease | 0.673707099 | M_Disease&NULL |
| 11 | 118711790 | rs622722 | A | G | M_Disease | 0.672607017 | M_Disease&NULL |

| | | | | | | | | | |
|---|---|---|---|---|---|---|---|---|---|
| 11 | 118654651 | rs2508916 | G | C | 1.098 | -4813.425 | -4811.89 | 1.535 | 0.000114971 |
| 11 | 118601426 | rs186926311 | T | C | 1.28 | -4813.425 | -4811.894 | 1.53 | 0.000114398 |
| 11 | 118678881 | rs591756 | T | C | 1.098 | -4813.425 | -4811.91 | 1.514 | 0.000112582 |
| 11 | 118678986 | rs477014 | C | T | 1.097 | -4813.425 | -4811.942 | 1.483 | 0.000109145 |
| 11 | 118679387 | rs473298 | G | A | 1.097 | -4813.425 | -4811.942 | 1.483 | 0.000109145 |
| 11 | 118679445 | rs516719 | C | T | 1.097 | -4813.425 | -4811.942 | 1.483 | 0.000109145 |
| 11 | 118602058 | rs562059 | G | T | 1.096 | -4813.425 | -4811.963 | 1.461 | 0.00010677 |
| 11 | 118808140 | rs7117408 | T | C | 1.139 | -4813.425 | -4811.978 | 1.447 | 0.000105286 |
| 11 | 118711790 | rs622722 | A | G | 0.914 | -4813.425 | -4811.979 | 1.446 | 0.000105181 |
| 11 | 118807740 | rs10892312 | A | G | 1.138 | -4813.425 | -4812.007 | 1.417 | 0.000102174 |
| 11 | 118824683 | rs507763 | C | G | 1.114 | -4813.425 | -4812.016 | 1.409 | 0.00010136 |
| 11 | 118824917 | rs11217102 | A | G | 1.114 | -4813.425 | -4812.016 | 1.409 | 0.00010136 |
| 11 | 118807876 | rs12282154 | A | G | 1.137 | -4813.425 | -4812.021 | 1.403 | 0.000100754 |
| 11 | 118602497 | rs478081 | G | A | 1.095 | -4813.425 | -4812.023 | 1.402 | 0.000100653 |
| 11 | 118680045 | rs2508569 | C | G | 1.096 | -4813.425 | -4812.036 | 1.388 | 9.93E-05 |
| 11 | 118826045 | rs537080 | C | T | 1.115 | -4813.425 | -4812.037 | 1.388 | 9.93E-05 |
| 11 | 118728320 | rs4936442 | T | C | 0.914 | -4813.425 | -4812.041 | 1.383 | 9.88E-05 |
| 11 | 118727169 | rs10790267 | T | C | 0.915 | -4813.425 | -4812.067 | 1.358 | 9.63E-05 |
| 11 | 118727332 | rs11217067 | T | G | 0.915 | -4813.425 | -4812.067 | 1.358 | 9.63E-05 |
| 11 | 118477367 | rs498872 | A | G | 1.098 | -4813.425 | -4812.081 | 1.344 | 9.50E-05 |
| 11 | 118652569 | rs150792202 | A | G | 0.874 | -4813.425 | -4812.096 | 1.328 | 9.35E-05 |
| 11 | 118705753 | rs556437 | A | G | 0.917 | -4813.425 | -4812.113 | 1.312 | 9.20E-05 |
| 11 | 118499469 | rs577396 | C | G | 0.917 | -4813.425 | -4812.143 | 1.281 | 8.92E-05 |
| 11 | 118501459 | rs11216934 | A | G | 0.918 | -4813.425 | -4812.157 | 1.267 | 8.79E-05 |
| 11 | 118504742 | rs633683 | T | C | 0.916 | -4813.425 | -4812.157 | 1.267 | 8.79E-05 |
| 11 | 118697012 | rs571470 | A | G | 1.091 | -4813.425 | -4812.179 | 1.246 | 8.61E-05 |
| 11 | 118729210 | rs10892300 | T | C | 0.916 | -4813.425 | -4812.199 | 1.226 | 8.44E-05 |
| 11 | 118729583 | rs12790489 | C | T | 0.917 | -4813.425 | -4812.239 | 1.185 | 8.10E-05 |
| 11 | 118729860 | rs11217070 | G | T | 0.917 | -4813.425 | -4812.239 | 1.185 | 8.10E-05 |
| 11 | 118755630 | rs3176905 | C | T | 1.115 | -4813.425 | -4812.246 | 1.179 | 8.05E-05 |
| 11 | 118734521 | rs1174717189 | T | A | 0.917 | -4813.425 | -4812.249 | 1.176 | 8.03E-05 |
| 11 | 118509668 | rs483598 | A | G | 0.919 | -4813.425 | -4812.259 | 1.165 | 7.94E-05 |
| 11 | 118572267 | rs573905 | A | G | 0.919 | -4813.425 | -4812.267 | 1.157 | 7.88E-05 |
| 11 | 118731599 | rs34561290 | T | C | 0.918 | -4813.425 | -4812.277 | 1.148 | 7.81E-05 |
| 11 | 118731762 | rs11217072 | G | A | 0.918 | -4813.425 | -4812.277 | 1.148 | 7.81E-05 |
| 11 | 118530068 | rs112337443 | C | T | 0.836 | -4813.425 | -4812.282 | 1.142 | 7.76E-05 |
| 11 | 118540104 | rs506234 | G | C | 0.919 | -4813.425 | -4812.308 | 1.117 | 7.57E-05 |
| 11 | 118730141 | rs7929520 | G | A | 0.918 | -4813.425 | -4812.313 | 1.111 | 7.52E-05 |
| 11 | 118789952 | rs11217090 | T | C | 1.135 | -4813.425 | -4812.336 | 1.089 | 7.36E-05 |
| 11 | 118514022 | rs613243 | C | G | 0.92 | -4813.425 | -4812.352 | 1.072 | 7.24E-05 |
| 11 | 118555383 | rs2508573 | T | C | 0.92 | -4813.425 | -4812.36 | 1.065 | 7.19E-05 |
| 11 | 118807637 | rs112974653 | T | G | 1.128 | -4813.425 | -4812.376 | 1.048 | 7.06E-05 |
| 11 | 118899218 | rs4936459 | T | C | 1.116 | -4813.425 | -4812.435 | 0.99 | 6.67E-05 |
| 11 | 118719984 | rs145680506 | T | C | 0.873 | -4813.425 | -4812.451 | 0.974 | 6.56E-05 |
| 11 | 118740180 | rs7483509 | A | G | 0.92 | -4813.425 | -4812.491 | 0.933 | 6.30E-05 |
| 11 | 118801440 | rs12280501 | T | C | 1.13 | -4813.425 | -4812.507 | 0.918 | 6.20E-05 |
| 11 | 118699276 | rs625274 | T | C | 0.922 | -4813.425 | -4812.53 | 0.895 | 6.06E-05 |
| 11 | 118799206 | rs7101680 | A | G | 1.129 | -4813.425 | -4812.544 | 0.881 | 5.98E-05 |
| 11 | 118793613 | rs11217091 | C | T | 1.129 | -4813.425 | -4812.558 | 0.866 | 5.89E-05 |
| 11 | 118565651 | rs540800 | A | G | 0.923 | -4813.425 | -4812.563 | 0.862 | 5.87E-05 |
| 11 | 118563286 | rs527619 | A | G | 0.923 | -4813.425 | -4812.567 | 0.857 | 5.84E-05 |
| 11 | 118754649 | rs10892307 | G | C | 1.107 | -4813.425 | -4812.584 | 0.84 | 5.74E-05 |
| 11 | 118747576 | rs55756957 | G | A | 1.094 | -4813.425 | -4812.586 | 0.839 | 5.73E-05 |
| 11 | 118738281 | rs7119038 | G | A | 1.096 | -4813.425 | -4812.587 | 0.837 | 5.72E-05 |
| 11 | 118700557 | rs4291686 | G | A | 0.923 | -4813.425 | -4812.596 | 0.829 | 5.68E-05 |
| 11 | 118798824 | rs112119822 | G | A | 1.127 | -4813.425 | -4812.617 | 0.808 | 5.56E-05 |
| 11 | 118501299 | rs7936556 | A | G | 0.924 | -4813.425 | -4812.621 | 0.804 | 5.54E-05 |
| 11 | 118782222 | rs61900138 | C | T | 0.857 | -4813.425 | -4812.624 | 0.8 | 5.51E-05 |
| 11 | 118793918 | rs12282752 | C | T | 1.159 | -4813.425 | -4812.646 | 0.778 | 5.39E-05 |
| 11 | 118510562 | rs4245184 | A | G | 0.925 | -4813.425 | -4812.696 | 0.729 | 5.14E-05 |
| 11 | 118511352 | rs644273 | T | C | 0.925 | -4813.425 | -4812.696 | 0.729 | 5.14E-05 |
| 11 | 118511757 | rs654423 | A | G | 0.925 | -4813.425 | -4812.696 | 0.729 | 5.14E-05 |
| 11 | 118486067 | rs11603023 | T | C | 0.925 | -4813.425 | -4812.701 | 0.724 | 5.11E-05 |
| 11 | 118806216 | rs1231742890 | G | T | 1.156 | -4813.425 | -4812.729 | 0.695 | 4.96E-05 |
| 11 | 118725204 | rs55794678 | T | C | 1.196 | -4813.425 | -4812.731 | 0.694 | 4.96E-05 |
| 11 | 118509877 | rs112552940 | T | C | 0.855 | -4813.425 | -4812.733 | 0.691 | 4.94E-05 |
| 11 | 118679629 | rs1784300 | T | A | 0.925 | -4813.425 | -4812.738 | 0.687 | 4.92E-05 |
| 11 | 118521549 | rs494560 | A | G | 0.926 | -4813.425 | -4812.765 | 0.659 | 4.79E-05 |
| 11 | 118464665 | rs113337722 | G | A | 0.858 | -4813.425 | -4812.79 | 0.635 | 4.67E-05 |
| 11 | 118462367 | rs11216919 | G | A | 0.923 | -4813.425 | -4812.801 | 0.624 | 4.62E-05 |
| 11 | 118603615 | rs74783030 | A | G | 1.175 | -4813.425 | -4812.825 | 0.599 | 4.51E-05 |
| 11 | 118659896 | rs2105782 | C | T | 0.927 | -4813.425 | -4812.828 | 0.596 | 4.50E-05 |
| 11 | 118558425 | rs149509653 | A | G | 0.913 | -4813.425 | -4812.829 | 0.595 | 4.49E-05 |
| 11 | 118791150 | rs12270280 | T | C | 1.148 | -4813.425 | -4812.832 | 0.593 | 4.48E-05 |
| 11 | 118762521 | rs190509947 | T | A | 1.194 | -4813.425 | -4812.845 | 0.58 | 4.42E-05 |
| 11 | 118811878 | rs74891547 | C | G | 1.131 | -4813.425 | -4812.844 | 0.58 | 4.42E-05 |
| 11 | 118765283 | rs665648 | A | G | 0.84 | -4813.425 | -4812.881 | 0.544 | 4.27E-05 |
| 11 | 118680072 | rs607472 | G | C | 0.928 | -4813.425 | -4812.885 | 0.539 | 4.25E-05 |
| 11 | 118696055 | rs189417780 | C | A | 1.189 | -4813.425 | -4812.89 | 0.535 | 4.23E-05 |
| 11 | 118791532 | rs12271856 | T | G | 1.145 | -4813.425 | -4812.904 | 0.521 | 4.17E-05 |
| 11 | 118805138 | rs7127427 | A | G | 1.145 | -4813.425 | -4812.938 | 0.487 | 4.03E-05 |
| 11 | 118797562 | rs142336193 | G | A | 0.861 | -4813.425 | -4812.947 | 0.477 | 3.99E-05 |
| 11 | 118658075 | rs515523 | G | A | 0.929 | -4813.425 | -4812.955 | 0.469 | 3.96E-05 |
| 11 | 118651809 | rs555649 | C | T | 0.929 | -4813.425 | -4812.974 | 0.451 | 3.89E-05 |
| 11 | 118644582 | rs2510897 | C | A | 0.929 | -4813.425 | -4812.975 | 0.449 | 3.88E-05 |
| 11 | 118711376 | rs118046730 | C | T | 0.878 | -4813.425 | -4812.984 | 0.44 | 3.85E-05 |
| 11 | 118620697 | rs488219 | C | T | 0.929 | -4813.425 | -4812.986 | 0.439 | 3.84E-05 |
| 11 | 118750967 | rs11602393 | A | T | 1.095 | -4813.425 | -4812.985 | 0.439 | 3.84E-05 |
| 11 | 118801296 | rs75210328 | T | G | 1.141 | -4813.425 | -4812.985 | 0.439 | 3.84E-05 |
| 11 | 118801577 | rs77875650 | A | G | 1.141 | -4813.425 | -4812.985 | 0.439 | 3.84E-05 |
| 11 | 118629031 | rs555639 | A | G | 0.929 | -4813.425 | -4812.988 | 0.436 | 3.83E-05 |
| 11 | 118632522 | rs581045 | T | C | 0.929 | -4813.425 | -4812.988 | 0.436 | 3.83E-05 |
| 11 | 118624250 | rs546393 | C | T | 0.929 | -4813.425 | -4813.006 | 0.418 | 3.76E-05 |
| 11 | 118804266 | rs7123066 | T | C | 1.134 | -4813.425 | -4813.016 | 0.409 | 3.73E-05 |
| 11 | 118804283 | rs7122959 | G | A | 1.134 | -4813.425 | -4813.016 | 0.409 | 3.73E-05 |
| 11 | 118662028 | rs12796102 | T | C | 0.93 | -4813.425 | -4813.022 | 0.403 | 3.71E-05 |
| 11 | 118748170 | rs77074463 | C | G | 0.895 | -4813.425 | -4813.025 | 0.399 | 3.69E-05 |
| 11 | 118605352 | rs76478216 | C | T | 0.881 | -4813.425 | -4813.026 | 0.398 | 3.69E-05 |
| 11 | 118666978 | rs1289369859 | A | G | 0.853 | -4813.425 | -4813.078 | 0.347 | 3.50E-05 |
| 11 | 118477746 | rs72999445 | G | T | 1.173 | -4813.425 | -4813.085 | 0.339 | 3.48E-05 |
| 11 | 118746769 | rs4938576 | T | G | 1.075 | -4813.425 | -4813.086 | 0.339 | 3.48E-05 |
| 11 | 118617299 | rs607125 | C | T | 0.931 | -4813.425 | -4813.09 | 0.335 | 3.46E-05 |
| 11 | 118631288 | rs507890 | C | T | 0.931 | -4813.425 | -4813.094 | 0.33 | 3.45E-05 |

| | | | | | | | |
|---|---|---|---|---|---|---|---|
| 11 | 118519550 | rs601318 | G | C | NULL | 0.672607017 | NULL&M_Disease |
| 11 | 118520480 | rs653008 | T | G | NULL | 0.672607017 | NULL&M_Disease |
| 11 | 118548665 | rs1784298 | C | T | NULL | 0.672607017 | NULL&M_Disease |
| 11 | 118548671 | rs4938522 | T | A | NULL | 0.672607017 | NULL&M_Disease |
| 11 | 118548838 | rs4938523 | T | G | NULL | 0.672607017 | NULL&M_Disease |
| 11 | 118549582 | rs507080 | A | G | NULL | 0.672607017 | NULL&M_Disease |
| 11 | 118549803 | rs659969 | T | C | NULL | 0.672607017 | NULL&M_Disease |
| 11 | 118549908 | rs656575 | G | T | NULL | 0.672607017 | NULL&M_Disease |
| 11 | 118633533 | rs7110526 | C | T | NULL | 0.672607017 | NULL&M_Disease |
| 11 | 118635364 | rs11217018 | C | T | NULL | 0.672607017 | NULL&M_Disease |
| 11 | 118637475 | rs112910608 | T | C | NULL | 0.672607017 | NULL&M_Disease |
| 11 | 118640828 | rs11217021 | C | T | NULL | 0.672607017 | NULL&M_Disease |
| 11 | 118515616 | rs527290 | T | C | NULL | 0.671505034 | NULL&M_Disease |
| 11 | 118538308 | rs692750 | T | G | NULL | 0.67040116 | NULL&M_Disease |
| 11 | 118705145 | rs636736 | G | A | NULL | 0.67040116 | NULL&M_Disease |
| 11 | 118726340 | rs113786535 | T | C | NULL | 0.67040116 | NULL&M_Disease |
| 11 | 118807740 | rs10892312 | A | G | M_Disease | 0.67040116 | M_Disease&NULL |
| 11 | 118880740 | rs11217122 | C | T | NULL | 0.67040116 | NULL&M_Disease |
| 11 | 118609598 | rs112996178 | T | C | NULL | 0.669295403 | NULL&M_Disease |
| 11 | 118624919 | rs11217013 | C | G | NULL | 0.669295403 | NULL&M_Disease |
| 11 | 118705654 | rs649338 | A | G | NULL | 0.669295403 | NULL&M_Disease |
| 11 | 118824683 | rs507763 | C | G | M_Disease | 0.669295403 | M_Disease&NULL |
| 11 | 118824917 | rs11217102 | A | G | M_Disease | 0.669295403 | M_Disease&NULL |
| 11 | 118539140 | rs4589335 | T | C | NULL | 0.668187772 | NULL&M_Disease |
| 11 | 118602497 | rs478081 | G | A | M_Disease | 0.668187772 | M_Disease&NULL |
| 11 | 118613674 | rs7130875 | C | T | NULL | 0.668187772 | NULL&M_Disease |
| 11 | 118626286 | rs11217014 | C | T | NULL | 0.668187772 | NULL&M_Disease |
| 11 | 118627778 | rs10892282 | C | G | NULL | 0.668187772 | NULL&M_Disease |
| 11 | 118698160 | rs640373 | G | A | NULL | 0.668187772 | NULL&M_Disease |
| 11 | 118698274 | rs639915 | A | G | NULL | 0.668187772 | NULL&M_Disease |
| 11 | 118698294 | rs639885 | A | G | NULL | 0.668187772 | NULL&M_Disease |
| 11 | 118698537 | rs638893 | G | A | NULL | 0.668187772 | NULL&M_Disease |
| 11 | 118698777 | rs516332 | G | A | NULL | 0.668187772 | NULL&M_Disease |
| 11 | 118699863 | rs2510901 | A | G | NULL | 0.668187772 | NULL&M_Disease |
| 11 | 118701597 | rs4938549 | T | C | NULL | 0.668187772 | NULL&M_Disease |
| 11 | 118701668 | rs530646 | A | G | NULL | 0.668187772 | NULL&M_Disease |
| 11 | 118702002 | rs534114 | A | G | NULL | 0.668187772 | NULL&M_Disease |
| 11 | 118702910 | rs7941321 | A | G | NULL | 0.668187772 | NULL&M_Disease |
| 11 | 118703001 | rs7928228 | T | C | NULL | 0.668187772 | NULL&M_Disease |
| 11 | 118703484 | rs2508921 | T | C | NULL | 0.668187772 | NULL&M_Disease |
| 11 | 118703817 | rs1170298 | T | A | NULL | 0.668187772 | NULL&M_Disease |
| 11 | 118807876 | rs12282154 | A | G | M_Disease | 0.668187772 | M_Disease&NULL |
| 11 | 118831335 | rs61506726 | G | A | NULL | 0.668187772 | NULL&M_Disease |
| 11 | 118567375 | rs478296 | C | T | NULL | 0.667078277 | NULL&M_Disease |
| 11 | 118697375 | rs73005500 | C | T | NULL | 0.667078277 | NULL&M_Disease |
| 11 | 118699744 | rs2508564 | C | T | NULL | 0.667078277 | NULL&M_Disease |
| 11 | 118704255 | rs480791 | G | A | NULL | 0.667078277 | NULL&M_Disease |
| 11 | 118704265 | rs622690 | T | C | NULL | 0.667078277 | NULL&M_Disease |
| 11 | 118826045 | rs537080 | C | T | M_Disease | 0.667078277 | M_Disease&NULL |
| 11 | 118610826 | rs1893029 | T | G | NULL | 0.665966927 | NULL&M_Disease |
| 11 | 118616253 | rs7124498 | A | G | NULL | 0.665966927 | NULL&M_Disease |
| 11 | 118680045 | rs2508569 | C | G | M_Disease | 0.665966927 | M_Disease&NULL |
| 11 | 118728320 | rs4936442 | T | C | M_Disease | 0.665966927 | M_Disease&NULL |
| 11 | 118553326 | rs693037 | T | C | NULL | 0.66485373 | NULL&M_Disease |
| 11 | 118610463 | rs4499035 | T | C | NULL | 0.66485373 | NULL&M_Disease |
| 11 | 118700264 | rs113641235 | T | C | NULL | 0.66485373 | NULL&M_Disease |
| 11 | 118555719 | rs625513 | C | A | NULL | 0.663738697 | NULL&M_Disease |
| 11 | 118555754 | rs525485 | A | G | NULL | 0.663738697 | NULL&M_Disease |
| 11 | 118558449 | rs514143 | A | G | NULL | 0.663738697 | NULL&M_Disease |
| 11 | 118622985 | rs3825057 | C | T | NULL | 0.663738697 | NULL&M_Disease |
| 11 | 118701898 | rs533254 | A | T | NULL | 0.663738697 | NULL&M_Disease |
| 11 | 118702936 | rs7941696 | G | A | NULL | 0.663738697 | NULL&M_Disease |
| 11 | 118564397 | rs558781 | T | G | NULL | 0.662621837 | NULL&M_Disease |
| 11 | 118566039 | rs6589674 | G | A | NULL | 0.662621837 | NULL&M_Disease |
| 11 | 118567444 | rs493834 | G | A | NULL | 0.662621837 | NULL&M_Disease |
| 11 | 118727169 | rs10790267 | T | C | M_Disease | 0.662621837 | M_Disease&NULL |
| 11 | 118727332 | rs11217067 | T | G | M_Disease | 0.662621837 | M_Disease&NULL |
| 11 | 118895686 | rs8192696 | T | C | NULL | 0.662621837 | NULL&M_Disease |
| 11 | 118477367 | rs498872 | A | G | M_Disease | 0.661503159 | M_Disease&NULL |
| 11 | 118545797 | rs600969 | T | C | NULL | 0.661503159 | NULL&M_Disease |
| 11 | 118552074 | rs4938524 | T | G | NULL | 0.661503159 | NULL&M_Disease |
| 11 | 118567020 | rs642662 | G | A | NULL | 0.661503159 | NULL&M_Disease |
| 11 | 118585146 | rs112799932 | G | T | NULL | 0.661503159 | NULL&M_Disease |
| 11 | 118594669 | rs140026975 | C | A | NULL | 0.661503159 | NULL&M_Disease |
| 11 | 118627149 | rs11217016 | C | A | NULL | 0.661503159 | NULL&M_Disease |
| 11 | 118629364 | rs73001486 | A | G | NULL | 0.661503159 | NULL&M_Disease |
| 11 | 118700425 | rs10128600 | C | T | NULL | 0.661503159 | NULL&M_Disease |
| 11 | 118554100 | rs4331144 | G | A | NULL | 0.660382673 | NULL&M_Disease |
| 11 | 118556084 | rs613377 | T | C | NULL | 0.660382673 | NULL&M_Disease |
| 11 | 118556691 | rs610664 | G | A | NULL | 0.660382673 | NULL&M_Disease |
| 11 | 118556797 | rs571751 | C | A | NULL | 0.660382673 | NULL&M_Disease |
| 11 | 118557198 | rs546890 | C | T | NULL | 0.660382673 | NULL&M_Disease |
| 11 | 118558078 | rs555356 | T | G | NULL | 0.660382673 | NULL&M_Disease |
| 11 | 118558333 | rs693306 | G | A | NULL | 0.660382673 | NULL&M_Disease |
| 11 | 118558455 | rs558907 | G | A | NULL | 0.660382673 | NULL&M_Disease |
| 11 | 118558827 | rs561845 | G | A | NULL | 0.660382673 | NULL&M_Disease |
| 11 | 118558889 | rs638805 | T | G | NULL | 0.660382673 | NULL&M_Disease |
| 11 | 118558901 | rs510408 | C | G | NULL | 0.660382673 | NULL&M_Disease |
| 11 | 118558952 | rs509641 | C | T | NULL | 0.660382673 | NULL&M_Disease |
| 11 | 118558953 | rs563549 | A | G | NULL | 0.660382673 | NULL&M_Disease |
| 11 | 118559003 | rs638301 | G | A | NULL | 0.660382673 | NULL&M_Disease |
| 11 | 118559084 | rs508579 | C | G | NULL | 0.660382673 | NULL&M_Disease |
| 11 | 118559178 | rs486462 | C | T | NULL | 0.660382673 | NULL&M_Disease |
| 11 | 118559247 | rs533144 | A | C | NULL | 0.660382673 | NULL&M_Disease |
| 11 | 118559542 | rs625918 | T | C | NULL | 0.660382673 | NULL&M_Disease |
| 11 | 118559709 | rs574217358 | A | G | NULL | 0.660382673 | NULL&M_Disease |
| 11 | 118559866 | rs7926959 | C | T | NULL | 0.660382673 | NULL&M_Disease |
| 11 | 118559881 | rs7926970 | G | T | NULL | 0.660382673 | NULL&M_Disease |
| 11 | 118560569 | rs4938529 | T | C | NULL | 0.660382673 | NULL&M_Disease |
| 11 | 118560570 | rs4938530 | T | G | NULL | 0.660382673 | NULL&M_Disease |
| 11 | 118561410 | rs12798453 | G | T | NULL | 0.660382673 | NULL&M_Disease |
| 11 | 118562520 | rs558593 | T | C | NULL | 0.660382673 | NULL&M_Disease |
| 11 | 118562799 | rs582630 | A | C | NULL | 0.660382673 | NULL&M_Disease |

| | | | | | | | | | |
|---|---|---|---|---|---|---|---|---|---|
| 11 | 118631653 | rs1784543 | T | A | 0.931 | -4813.425 | -4813.094 | 0.33 | 3.45E-05 |
| 11 | 118684537 | rs150583208 | G | A | 0.855 | -4813.425 | -4813.102 | 0.323 | 3.42E-05 |
| 11 | 118781710 | rs6589707 | A | C | 0.856 | -4813.425 | -4813.119 | 0.306 | 3.36E-05 |
| 11 | 118523799 | rs72999483 | C | T | 1.103 | -4813.425 | -4813.13 | 0.295 | 3.33E-05 |
| 11 | 118747008 | rs75040111 | T | C | 1.165 | -4813.425 | -4813.162 | 0.263 | 3.22E-05 |
| 11 | 118787213 | rs7119222 | C | T | 0.861 | -4813.425 | -4813.181 | 0.244 | 3.16E-05 |
| 11 | 118689745 | rs492471 | T | C | 0.898 | -4813.425 | -4813.187 | 0.238 | 3.14E-05 |
| 11 | 118787897 | rs11602400 | C | A | 0.862 | -4813.425 | -4813.196 | 0.228 | 3.11E-05 |
| 11 | 118626825 | rs74337439 | A | G | 0.888 | -4813.425 | -4813.202 | 0.223 | 3.10E-05 |
| 11 | 118657311 | rs1309473877 | A | C | 0.933 | -4813.425 | -4813.201 | 0.223 | 3.10E-05 |
| 11 | 118761159 | rs57231586 | T | G | 0.864 | -4813.425 | -4813.209 | 0.215 | 3.07E-05 |
| 11 | 118576463 | rs496547 | A | T | 1.074 | -4813.425 | -4813.214 | 0.21 | 3.06E-05 |
| 11 | 118781530 | rs7926599 | C | G | 0.864 | -4813.425 | -4813.223 | 0.201 | 3.03E-05 |
| 11 | 118786829 | rs7102433 | G | A | 0.867 | -4813.425 | -4813.241 | 0.184 | 2.98E-05 |
| 11 | 118792278 | rs77398170 | G | A | 1.126 | -4813.425 | -4813.254 | 0.17 | 2.94E-05 |
| 11 | 118588998 | rs570424 | C | T | 0.934 | -4813.425 | -4813.286 | 0.138 | 2.84E-05 |
| 11 | 118615390 | rs78502545 | T | C | 0.892 | -4813.425 | -4813.302 | 0.122 | 2.80E-05 |
| 11 | 118819541 | rs150196102 | T | C | 1.146 | -4813.425 | -4813.306 | 0.118 | 2.79E-05 |
| 11 | 118721767 | rs116884018 | C | T | 1.138 | -4813.425 | -4813.321 | 0.103 | 2.75E-05 |
| 11 | 118789657 | rs12294466 | T | C | 1.107 | -4813.425 | -4813.322 | 0.103 | 2.75E-05 |
| 11 | 118849368 | rs45491601 | A | G | 1.146 | -4813.425 | -4813.323 | 0.102 | 2.74E-05 |
| 11 | 118761591 | rs58397111 | T | C | 0.874 | -4813.425 | -4813.323 | 0.101 | 2.74E-05 |
| 11 | 118762102 | rs58508419 | A | G | 0.874 | -4813.425 | -4813.323 | 0.101 | 2.74E-05 |
| 11 | 118622744 | rs6589689 | G | A | 0.877 | -4813.425 | -4813.331 | 0.093 | 2.72E-05 |
| 11 | 118782474 | rs11606719 | G | C | 0.88 | -4813.425 | -4813.337 | 0.088 | 2.70E-05 |
| 11 | 118608625 | rs566733 | G | C | 0.935 | -4813.425 | -4813.341 | 0.083 | 2.69E-05 |
| 11 | 118619948 | rs112143902 | T | C | 0.904 | -4813.425 | -4813.342 | 0.082 | 2.69E-05 |
| 11 | 118749318 | rs78312641 | A | T | 0.917 | -4813.425 | -4813.362 | 0.063 | 2.64E-05 |
| 11 | 118613235 | rs632124 | A | T | 0.935 | -4813.425 | -4813.363 | 0.062 | 2.64E-05 |
| 11 | 118749563 | rs77643000 | C | T | 0.917 | -4813.425 | -4813.362 | 0.062 | 2.64E-05 |
| 11 | 118816304 | rs586763 | T | C | 1.083 | -4813.425 | -4813.385 | 0.04 | 2.58E-05 |
| 11 | 118832251 | rs79808604 | A | C | 1.136 | -4813.425 | -4813.388 | 0.037 | 2.57E-05 |
| 11 | 118802492 | rs688035 | G | C | 0.934 | -4813.425 | -4813.394 | 0.03 | 2.55E-05 |
| 11 | 118798780 | rs77916839 | C | G | 1.12 | -4813.425 | -4813.407 | 0.018 | 2.52E-05 |
| 11 | 118413882 | rs146864992 | T | C | 0.883 | -4813.425 | -4813.414 | 0.01 | 2.50E-05 |
| 11 | 118672061 | rs183981091 | T | G | 0.885 | -4813.425 | -4813.429 | -0.004 | 2.47E-05 |
| 11 | 118468037 | rs535774 | T | C | 0.908 | -4813.425 | -4813.432 | -0.008 | 2.46E-05 |
| 11 | 118811684 | rs550999 | G | C | 0.934 | -4813.425 | -4813.44 | -0.015 | 2.44E-05 |
| 11 | 118764117 | rs73575405 | A | G | 0.886 | -4813.425 | -4813.443 | -0.019 | 2.43E-05 |
| 11 | 118608532 | rs567680 | G | C | 0.937 | -4813.425 | -4813.451 | -0.026 | 2.41E-05 |
| 11 | 118501723 | rs61900957 | T | C | 0.914 | -4813.425 | -4813.455 | -0.03 | 2.40E-05 |
| 11 | 118419577 | rs191458787 | T | C | 0.888 | -4813.425 | -4813.461 | -0.036 | 2.39E-05 |
| 11 | 118421438 | rs145485933 | G | A | 0.888 | -4813.425 | -4813.461 | -0.036 | 2.39E-05 |
| 11 | 118801235 | rs602716 | A | G | 0.935 | -4813.425 | -4813.462 | -0.037 | 2.39E-05 |
| 11 | 118812512 | rs647769 | T | C | 0.935 | -4813.425 | -4813.466 | -0.041 | 2.38E-05 |
| 11 | 118530434 | rs117619140 | T | G | 1.122 | -4813.425 | -4813.47 | -0.045 | 2.37E-05 |
| 11 | 118803193 | rs117235696 | G | T | 1.114 | -4813.425 | -4813.475 | -0.051 | 2.35E-05 |
| 11 | 118650163 | rs73003218 | T | C | 1.123 | -4813.425 | -4813.483 | -0.059 | 2.34E-05 |
| 11 | 118779681 | rs9971606 | C | T | 0.891 | -4813.425 | -4813.484 | -0.059 | 2.34E-05 |
| 11 | 118607622 | rs11217004 | A | G | 0.938 | -4813.425 | -4813.488 | -0.063 | 2.33E-05 |
| 11 | 118595207 | rs563537 | A | T | 0.938 | -4813.425 | -4813.494 | -0.069 | 2.31E-05 |
| 11 | 118577614 | rs947931 | G | T | 1.081 | -4813.425 | -4813.5 | -0.075 | 2.30E-05 |
| 11 | 118803842 | rs11600378 | G | A | 1.113 | -4813.425 | -4813.499 | -0.075 | 2.30E-05 |
| 11 | 118804671 | rs143711539 | A | G | 1.113 | -4813.425 | -4813.499 | -0.075 | 2.30E-05 |
| 11 | 118765087 | rs2230321 | T | C | 0.893 | -4813.425 | -4813.501 | -0.077 | 2.29E-05 |
| 11 | 118598138 | rs543692 | C | T | 0.938 | -4813.425 | -4813.531 | -0.107 | 2.23E-05 |
| 11 | 118759000 | rs77491736 | G | C | 0.898 | -4813.425 | -4813.547 | -0.122 | 2.19E-05 |
| 11 | 118721562 | rs184045098 | T | C | 1.114 | -4813.425 | -4813.551 | -0.126 | 2.18E-05 |
| 11 | 118803536 | rs615166 | A | G | 0.937 | -4813.425 | -4813.556 | -0.131 | 2.17E-05 |
| 11 | 118598612 | rs568688 | G | T | 0.939 | -4813.425 | -4813.562 | -0.137 | 2.16E-05 |
| 11 | 118805144 | rs11605265 | A | G | 1.108 | -4813.425 | -4813.562 | -0.137 | 2.16E-05 |
| 11 | 118805220 | rs11605658 | T | C | 1.108 | -4813.425 | -4813.562 | -0.137 | 2.16E-05 |
| 11 | 118735876 | rs79843382 | T | C | 1.1 | -4813.425 | -4813.579 | -0.154 | 2.12E-05 |
| 11 | 118732181 | rs117079831 | A | G | 1.107 | -4813.425 | -4813.601 | -0.177 | 2.08E-05 |
| 11 | 118487265 | rs76544852 | T | G | 0.903 | -4813.425 | -4813.606 | -0.181 | 2.07E-05 |
| 11 | 118460957 | rs45600633 | A | G | 0.904 | -4813.425 | -4813.615 | -0.19 | 2.05E-05 |
| 11 | 118474283 | rs139202987 | G | A | 0.904 | -4813.425 | -4813.615 | -0.19 | 2.05E-05 |
| 11 | 118619478 | rs189583461 | G | A | 1.104 | -4813.425 | -4813.623 | -0.199 | 2.03E-05 |
| 11 | 118621277 | rs117409833 | C | T | 1.104 | -4813.425 | -4813.623 | -0.199 | 2.03E-05 |
| 11 | 118421841 | rs145968026 | G | C | 0.905 | -4813.425 | -4813.628 | -0.204 | 2.02E-05 |
| 11 | 118507068 | rs10892249 | C | T | 0.905 | -4813.425 | -4813.632 | -0.208 | 2.01E-05 |
| 11 | 118518021 | rs10736493 | G | T | 0.905 | -4813.425 | -4813.632 | -0.208 | 2.01E-05 |
| 11 | 118764051 | rs613791 | T | C | 0.936 | -4813.425 | -4813.64 | -0.216 | 2.00E-05 |
| 11 | 118455465 | rs45460493 | A | G | 0.908 | -4813.425 | -4813.661 | -0.236 | 1.96E-05 |
| 11 | 118477226 | rs45496391 | C | A | 0.908 | -4813.425 | -4813.661 | -0.236 | 1.96E-05 |
| 11 | 118466646 | rs184841209 | T | G | 1.099 | -4813.425 | -4813.662 | -0.238 | 1.95E-05 |
| 11 | 118691070 | rs140577174 | G | A | 1.1 | -4813.425 | -4813.667 | -0.243 | 1.94E-05 |
| 11 | 118789026 | rs11607239 | T | G | 0.91 | -4813.425 | -4813.681 | -0.256 | 1.92E-05 |
| 11 | 118587349 | rs11216975 | C | A | 1.093 | -4813.425 | -4813.686 | -0.261 | 1.91E-05 |
| 11 | 118576762 | rs553188 | T | C | 0.926 | -4813.425 | -4813.696 | -0.272 | 1.89E-05 |
| 11 | 118727137 | rs117223320 | C | G | 1.092 | -4813.425 | -4813.703 | -0.279 | 1.87E-05 |
| 11 | 118759905 | rs118020807 | A | C | 1.083 | -4813.425 | -4813.724 | -0.3 | 1.84E-05 |
| 11 | 118811315 | rs77638281 | A | G | 1.094 | -4813.425 | -4813.726 | -0.301 | 1.83E-05 |
| 11 | 118576997 | rs498912 | C | T | 0.927 | -4813.425 | -4813.735 | -0.311 | 1.82E-05 |
| 11 | 118519213 | rs668628 | T | G | 0.931 | -4813.425 | -4813.737 | -0.312 | 1.81E-05 |
| 11 | 118782802 | rs76160025 | T | C | 1.092 | -4813.425 | -4813.74 | -0.315 | 1.81E-05 |
| 11 | 118715589 | rs116872113 | A | G | 1.09 | -4813.425 | -4813.74 | -0.316 | 1.81E-05 |
| 11 | 118777978 | rs78755429 | A | C | 0.915 | -4813.425 | -4813.743 | -0.318 | 1.80E-05 |
| 11 | 118673968 | rs140269932 | A | G | 1.082 | -4813.425 | -4813.749 | -0.325 | 1.79E-05 |
| 11 | 118805385 | rs55772432 | A | G | 0.923 | -4813.425 | -4813.759 | -0.334 | 1.77E-05 |
| 11 | 118664181 | rs145462967 | A | G | 1.077 | -4813.425 | -4813.775 | -0.35 | 1.75E-05 |
| 11 | 118483912 | rs45621436 | G | T | 0.922 | -4813.425 | -4813.779 | -0.355 | 1.74E-05 |
| 11 | 118493319 | rs10790254 | A | G | 0.922 | -4813.425 | -4813.779 | -0.355 | 1.74E-05 |
| 11 | 118518753 | rs117947426 | T | C | 1.058 | -4813.425 | -4813.782 | -0.357 | 1.73E-05 |
| 11 | 118601363 | rs557990 | T | C | 0.93 | -4813.425 | -4813.791 | -0.367 | 1.72E-05 |
| 11 | 118602392 | rs564892 | A | T | 0.93 | -4813.425 | -4813.791 | -0.367 | 1.72E-05 |
| 11 | 118445559 | rs73555296 | T | G | 0.924 | -4813.425 | -4813.797 | -0.372 | 1.71E-05 |
| 11 | 118503912 | rs45478298 | T | C | 1.086 | -4813.425 | -4813.807 | -0.382 | 1.69E-05 |
| 11 | 118607694 | rs7119702 | G | A | 0.931 | -4813.425 | -4813.813 | -0.389 | 1.68E-05 |
| 11 | 118748789 | rs145762171 | A | G | 0.944 | -4813.425 | -4813.817 | -0.392 | 1.67E-05 |
| 11 | 118798757 | rs57879605 | T | C | 1.069 | -4813.425 | -4813.827 | -0.402 | 1.66E-05 |
| 11 | 118583712 | rs118008100 | A | G | 1.064 | -4813.425 | -4813.834 | -0.409 | 1.65E-05 |
| 11 | 118610629 | rs521910 | A | G | 0.932 | -4813.425 | -4813.838 | -0.414 | 1.64E-05 |

| | | | | | | | |
|---|---|---|---|---|---|---|---|
| 11 | 118563381 | rs585039 | A | C | NULL | 0.660382673 | NULL&M_Disease |
| 11 | 118564413 | rs508640 | G | A | NULL | 0.660382673 | NULL&M_Disease |
| 11 | 118564601 | rs471227 | C | T | NULL | 0.660382673 | NULL&M_Disease |
| 11 | 118564762 | rs516153 | A | C | NULL | 0.660382673 | NULL&M_Disease |
| 11 | 118564982 | rs518028 | C | T | NULL | 0.660382673 | NULL&M_Disease |
| 11 | 118565231 | rs519982 | T | C | NULL | 0.660382673 | NULL&M_Disease |
| 11 | 118565336 | rs478371 | A | G | NULL | 0.660382673 | NULL&M_Disease |
| 11 | 118565679 | rs502601 | A | G | NULL | 0.660382673 | NULL&M_Disease |
| 11 | 118566090 | rs7130076 | C | T | NULL | 0.660382673 | NULL&M_Disease |
| 11 | 118566175 | rs7114458 | A | G | NULL | 0.660382673 | NULL&M_Disease |
| 11 | 118566234 | rs7114216 | G | A | NULL | 0.660382673 | NULL&M_Disease |
| 11 | 118566313 | rs12794726 | T | C | NULL | 0.660382673 | NULL&M_Disease |
| 11 | 118566746 | rs533646 | G | C | NULL | 0.660382673 | NULL&M_Disease |
| 11 | 118682462 | rs523148 | T | C | NULL | 0.660382673 | NULL&M_Disease |
| 11 | 118762701 | rs76457018 | A | G | NULL | 0.660382673 | NULL&M_Disease |
| 11 | 118879285 | rs7120445 | C | A | NULL | 0.660382673 | NULL&M_Disease |
| 11 | 118879824 | rs73559119 | C | T | NULL | 0.660382673 | NULL&M_Disease |
| 11 | 118880012 | rs11825284 | A | G | NULL | 0.660382673 | NULL&M_Disease |
| 11 | 118652569 | rs150792202 | A | G | M_Disease | 0.660382673 | M_Disease&NULL |
| 11 | 118552420 | rs4936433 | G | A | NULL | 0.659260388 | NULL&M_Disease |
| 11 | 118552715 | rs519848 | A | G | NULL | 0.659260388 | NULL&M_Disease |
| 11 | 118552842 | rs549370 | A | C | NULL | 0.659260388 | NULL&M_Disease |
| 11 | 118552996 | rs654792 | T | G | NULL | 0.659260388 | NULL&M_Disease |
| 11 | 118553218 | rs592280 | C | T | NULL | 0.659260388 | NULL&M_Disease |
| 11 | 118554248 | rs7106652 | T | C | NULL | 0.659260388 | NULL&M_Disease |
| 11 | 118554259 | rs7106851 | A | G | NULL | 0.659260388 | NULL&M_Disease |
| 11 | 118554436 | rs11605069 | G | T | NULL | 0.659260388 | NULL&M_Disease |
| 11 | 118554448 | rs11602869 | A | G | NULL | 0.659260388 | NULL&M_Disease |
| 11 | 118554939 | rs12796373 | A | G | NULL | 0.659260388 | NULL&M_Disease |
| 11 | 118562767 | rs667982 | C | T | NULL | 0.659260388 | NULL&M_Disease |
| 11 | 118567084 | rs542613 | A | G | NULL | 0.659260388 | NULL&M_Disease |
| 11 | 118567486 | rs644498 | C | T | NULL | 0.659260388 | NULL&M_Disease |
| 11 | 118700218 | rs2510902 | A | G | NULL | 0.659260388 | NULL&M_Disease |
| 11 | 118832035 | rs10892317 | C | T | NULL | 0.659260388 | NULL&M_Disease |
| 11 | 118877563 | rs58709307 | T | C | NULL | 0.659260388 | NULL&M_Disease |
| 11 | 118886656 | rs7131534 | A | G | NULL | 0.659260388 | NULL&M_Disease |
| 11 | 118705753 | rs556437 | A | G | M_Disease | 0.658136315 | M_Disease&NULL |
| 11 | 118548421 | rs562313 | A | G | NULL | 0.658136315 | NULL&M_Disease |
| 11 | 118550601 | rs673770 | G | T | NULL | 0.658136315 | NULL&M_Disease |
| 11 | 118550756 | rs642530 | C | T | NULL | 0.658136315 | NULL&M_Disease |
| 11 | 118550784 | rs642497 | A | C | NULL | 0.658136315 | NULL&M_Disease |
| 11 | 118551597 | rs12799314 | A | G | NULL | 0.658136315 | NULL&M_Disease |
| 11 | 118567545 | rs644867 | G | A | NULL | 0.658136315 | NULL&M_Disease |
| 11 | 118596444 | rs68097885 | A | G | NULL | 0.658136315 | NULL&M_Disease |
| 11 | 118689871 | rs587816 | A | C | NULL | 0.658136315 | NULL&M_Disease |
| 11 | 118695908 | rs583905 | G | A | NULL | 0.658136315 | NULL&M_Disease |
| 11 | 118696175 | rs584935 | A | G | NULL | 0.658136315 | NULL&M_Disease |
| 11 | 118894287 | rs569 | G | A | NULL | 0.658136315 | NULL&M_Disease |
| 11 | 118895202 | rs11006 | G | A | NULL | 0.658136315 | NULL&M_Disease |
| 11 | 118685481 | rs515707 | T | C | NULL | 0.657010463 | NULL&M_Disease |
| 11 | 118690550 | rs577121 | C | T | NULL | 0.657010463 | NULL&M_Disease |
| 11 | 118836606 | rs113975105 | G | C | NULL | 0.657010463 | NULL&M_Disease |
| 11 | 118868750 | rs4445645 | C | G | NULL | 0.657010463 | NULL&M_Disease |
| 11 | 118892277 | rs4938621 | T | C | NULL | 0.657010463 | NULL&M_Disease |
| 11 | 118892835 | rs61179449 | C | G | NULL | 0.657010463 | NULL&M_Disease |
| 11 | 118892872 | rs58955701 | T | C | NULL | 0.657010463 | NULL&M_Disease |
| 11 | 118892940 | rs12280288 | C | T | NULL | 0.657010463 | NULL&M_Disease |
| 11 | 118893023 | rs7127188 | C | A | NULL | 0.657010463 | NULL&M_Disease |
| 11 | 118878222 | rs11606357 | C | T | NULL | 0.655882841 | NULL&M_Disease |
| 11 | 118883491 | rs60065241 | A | C | NULL | 0.655882841 | NULL&M_Disease |
| 11 | 118886117 | rs10111 | T | C | NULL | 0.655882841 | NULL&M_Disease |
| 11 | 118892757 | rs117289748 | G | C | NULL | 0.655882841 | NULL&M_Disease |
| 11 | 118892759 | rs143660533 | C | A | NULL | 0.655882841 | NULL&M_Disease |
| 11 | 118893878 | rs111322300 | C | T | NULL | 0.655882841 | NULL&M_Disease |
| 11 | 118499469 | rs577396 | C | G | M_Disease | 0.654753461 | M_Disease&NULL |
| 11 | 118681088 | rs2508922 | A | G | NULL | 0.654753461 | NULL&M_Disease |
| 11 | 118785982 | rs79157249 | T | A | NULL | 0.654753461 | NULL&M_Disease |
| 11 | 118831727 | rs10892315 | C | T | NULL | 0.654753461 | NULL&M_Disease |
| 11 | 118567257 | rs7126172 | G | A | NULL | 0.653622331 | NULL&M_Disease |
| 11 | 118661376 | rs73003294 | T | C | NULL | 0.653622331 | NULL&M_Disease |
| 11 | 118685015 | rs540716 | T | C | NULL | 0.653622331 | NULL&M_Disease |
| 11 | 118501459 | rs11216934 | A | G | M_Disease | 0.652489462 | M_Disease&NULL |
| 11 | 118504742 | rs633683 | T | C | M_Disease | 0.652489462 | M_Disease&NULL |
| 11 | 118603030 | rs7943039 | A | G | NULL | 0.652489462 | NULL&M_Disease |
| 11 | 118788517 | rs73575424 | A | C | NULL | 0.652489462 | NULL&M_Disease |
| 11 | 118872581 | rs4558175 | A | G | NULL | 0.652489462 | NULL&M_Disease |
| 11 | 118880668 | rs11607569 | C | T | NULL | 0.652489462 | NULL&M_Disease |
| 11 | 118480115 | rs12803321 | C | G | NULL | 0.651354865 | NULL&M_Disease |
| 11 | 118567218 | rs11216947 | A | G | NULL | 0.651354865 | NULL&M_Disease |
| 11 | 118692727 | rs503425 | C | T | NULL | 0.651354865 | NULL&M_Disease |
| 11 | 118697012 | rs571470 | A | G | M_Disease | 0.650218549 | M_Disease&NULL |
| 11 | 118896115 | rs2276072 | G | A | NULL | 0.650218549 | NULL&M_Disease |
| 11 | 118896116 | rs2276071 | C | A | NULL | 0.650218549 | NULL&M_Disease |
| 11 | 118567733 | rs481903 | C | T | NULL | 0.649080524 | NULL&M_Disease |
| 11 | 118797003 | rs73575428 | G | C | NULL | 0.649080524 | NULL&M_Disease |
| 11 | 118797278 | rs74584888 | A | G | NULL | 0.649080524 | NULL&M_Disease |
| 11 | 118797392 | rs73575429 | C | T | NULL | 0.649080524 | NULL&M_Disease |
| 11 | 118797420 | rs111390501 | G | A | NULL | 0.649080524 | NULL&M_Disease |
| 11 | 118797544 | rs56032119 | G | A | NULL | 0.649080524 | NULL&M_Disease |
| 11 | 118798154 | rs60969913 | C | G | NULL | 0.649080524 | NULL&M_Disease |
| 11 | 118729210 | rs10892300 | T | C | M_Disease | 0.647940802 | M_Disease&NULL |
| 11 | 118481431 | rs10892246 | A | G | NULL | 0.646799393 | NULL&M_Disease |
| 11 | 118757219 | rs1623316 | G | C | NULL | 0.646799393 | NULL&M_Disease |
| 11 | 118896959 | rs57156480 | A | G | NULL | 0.646799393 | NULL&M_Disease |
| 11 | 118873385 | rs10892320 | T | C | NULL | 0.645656306 | NULL&M_Disease |
| 11 | 118754353 | rs630923 | A | C | NULL | 0.644511554 | NULL&M_Disease |
| 11 | 118729583 | rs12790489 | C | T | M_Disease | 0.643365146 | M_Disease&NULL |
| 11 | 118729860 | rs11217070 | G | T | M_Disease | 0.643365146 | M_Disease&NULL |
| 11 | 118755630 | rs3176905 | C | T | M_Disease | 0.643365146 | M_Disease&NULL |
| 11 | 118698170 | rs56296109 | A | C | NULL | 0.642217093 | NULL&M_Disease |
| 11 | 118734521 | rs1174717189 | T | A | M_Disease | 0.642217093 | M_Disease&NULL |
| 11 | 118462554 | rs11216920 | G | A | NULL | 0.641067406 | NULL&M_Disease |
| 11 | 118509668 | rs483598 | A | G | M_Disease | 0.641067406 | M_Disease&NULL |

| | | | | | | | | | |
|---|---|---|---|---|---|---|---|---|---|
| 11 | 118561858 | rs10790257 | C | G | 0.93 | -4813.425 | -4813.846 | -0.422 | 1.62E-05 |
| 11 | 118561859 | rs10790258 | T | G | 0.93 | -4813.425 | -4813.846 | -0.422 | 1.62E-05 |
| 11 | 118572000 | rs182361280 | A | G | 1.043 | -4813.425 | -4813.848 | -0.424 | 1.62E-05 |
| 11 | 118559426 | rs10892253 | A | C | 0.931 | -4813.425 | -4813.85 | -0.426 | 1.62E-05 |
| 11 | 118605127 | rs526850 | G | T | 0.932 | -4813.425 | -4813.85 | -0.426 | 1.62E-05 |
| 11 | 118757788 | rs3176912 | A | G | 1.08 | -4813.425 | -4813.85 | -0.426 | 1.62E-05 |
| 11 | 118539746 | rs72999492 | C | A | 1.082 | -4813.425 | -4813.853 | -0.429 | 1.61E-05 |
| 11 | 118615738 | rs193090156 | T | G | 1.02 | -4813.425 | -4813.858 | -0.433 | 1.61E-05 |
| 11 | 118590743 | rs503542 | G | A | 0.945 | -4813.425 | -4813.858 | -0.434 | 1.60E-05 |
| 11 | 118599649 | rs182364290 | G | A | 1.069 | -4813.425 | -4813.861 | -0.436 | 1.60E-05 |
| 11 | 118788769 | rs186418307 | T | G | 1.037 | -4813.425 | -4813.861 | -0.437 | 1.60E-05 |
| 11 | 118557483 | rs6589673 | C | A | 0.933 | -4813.425 | -4813.863 | -0.438 | 1.60E-05 |
| 11 | 118609736 | rs1783937 | A | G | 0.933 | -4813.425 | -4813.863 | -0.438 | 1.60E-05 |
| 11 | 118792384 | rs182691948 | T | G | 0.956 | -4813.425 | -4813.863 | -0.439 | 1.60E-05 |
| 11 | 118797857 | rs150438978 | T | A | 0.956 | -4813.425 | -4813.863 | -0.439 | 1.60E-05 |
| 11 | 118604678 | rs493720 | T | C | 0.933 | -4813.425 | -4813.869 | -0.445 | 1.59E-05 |
| 11 | 118605447 | rs2186781 | T | C | 0.933 | -4813.425 | -4813.877 | -0.453 | 1.57E-05 |
| 11 | 118594326 | rs544096 | G | A | 0.934 | -4813.425 | -4813.885 | -0.46 | 1.56E-05 |
| 11 | 118509741 | rs113313560 | T | C | 1.02 | -4813.425 | -4813.905 | -0.481 | 1.53E-05 |
| 11 | 118607090 | rs10736495 | C | G | 0.934 | -4813.425 | -4813.911 | -0.486 | 1.52E-05 |
| 11 | 118615232 | rs139423236 | G | C | 1.059 | -4813.425 | -4813.918 | -0.494 | 1.51E-05 |
| 11 | 118539151 | rs72999491 | A | G | 1.068 | -4813.425 | -4813.92 | -0.496 | 1.51E-05 |
| 11 | 118744615 | rs73009445 | A | G | 1.076 | -4813.425 | -4813.923 | -0.499 | 1.50E-05 |
| 11 | 118694739 | rs188550348 | A | G | 0.986 | -4813.425 | -4813.926 | -0.501 | 1.50E-05 |
| 11 | 118812559 | rs74678032 | C | T | 0.939 | -4813.425 | -4813.929 | -0.505 | 1.49E-05 |
| 11 | 118496524 | rs35692099 | C | T | 0.933 | -4813.425 | -4813.931 | -0.506 | 1.49E-05 |
| 11 | 118854372 | rs75067211 | T | C | 1.062 | -4813.425 | -4813.934 | -0.51 | 1.49E-05 |
| 11 | 118543589 | rs72999499 | G | C | 1.068 | -4813.425 | -4813.939 | -0.514 | 1.48E-05 |
| 11 | 118590961 | rs111227443 | T | C | 1.05 | -4813.425 | -4813.942 | -0.518 | 1.48E-05 |
| 11 | 118578462 | rs516459 | C | T | 0.975 | -4813.425 | -4813.955 | -0.531 | 1.46E-05 |
| 11 | 118502121 | rs17614795 | G | A | 0.936 | -4813.425 | -4813.958 | -0.534 | 1.45E-05 |
| 11 | 118556995 | rs72999502 | C | T | 1.073 | -4813.425 | -4813.968 | -0.543 | 1.44E-05 |
| 11 | 118799691 | rs191378986 | G | A | 0.976 | -4813.425 | -4813.97 | -0.546 | 1.43E-05 |
| 11 | 118772160 | rs61730467 | A | G | 1.064 | -4813.425 | -4813.972 | -0.547 | 1.43E-05 |
| 11 | 118855789 | rs4938609 | A | C | 1.007 | -4813.425 | -4813.978 | -0.553 | 1.42E-05 |
| 11 | 118533042 | rs142438429 | T | C | 0.976 | -4813.425 | -4813.978 | -0.554 | 1.42E-05 |
| 11 | 118791180 | rs146541591 | C | T | 0.951 | -4813.425 | -4813.997 | -0.573 | 1.40E-05 |
| 11 | 118583261 | rs112246697 | A | G | 0.97 | -4813.425 | -4814 | -0.575 | 1.39E-05 |
| 11 | 118510642 | rs141169729 | T | C | 0.989 | -4813.425 | -4814.003 | -0.579 | 1.39E-05 |
| 11 | 118793479 | rs138526953 | A | C | 1.031 | -4813.425 | -4814.01 | -0.586 | 1.38E-05 |
| 11 | 118580773 | rs675329 | T | C | 0.997 | -4813.425 | -4814.011 | -0.587 | 1.38E-05 |
| 11 | 118591443 | rs111843292 | A | C | 0.994 | -4813.425 | -4814.014 | -0.59 | 1.37E-05 |
| 11 | 118745664 | rs11217076 | G | T | 0.944 | -4813.425 | -4814.018 | -0.593 | 1.37E-05 |
| 11 | 118480285 | rs12225399 | C | G | 0.947 | -4813.425 | -4814.018 | -0.594 | 1.37E-05 |
| 11 | 118583971 | rs6589677 | A | G | 0.937 | -4813.425 | -4814.018 | -0.594 | 1.37E-05 |
| 11 | 118580906 | rs496058 | G | A | 0.948 | -4813.425 | -4814.022 | -0.598 | 1.36E-05 |
| 11 | 118574311 | rs524409 | T | C | 0.948 | -4813.425 | -4814.023 | -0.599 | 1.36E-05 |
| 11 | 118574675 | rs494459 | T | C | 0.948 | -4813.425 | -4814.023 | -0.599 | 1.36E-05 |
| 11 | 118709275 | rs187164533 | A | G | 0.984 | -4813.425 | -4814.032 | -0.607 | 1.35E-05 |
| 11 | 118575419 | rs625735 | A | G | 0.948 | -4813.425 | -4814.032 | -0.608 | 1.35E-05 |
| 11 | 118575479 | rs637563 | G | A | 0.948 | -4813.425 | -4814.032 | -0.608 | 1.35E-05 |
| 11 | 118575693 | rs638514 | C | T | 0.948 | -4813.425 | -4814.032 | -0.608 | 1.35E-05 |
| 11 | 118530869 | rs45447797 | G | C | 0.938 | -4813.425 | -4814.035 | -0.611 | 1.34E-05 |
| 11 | 118580348 | rs516503 | T | C | 0.949 | -4813.425 | -4814.039 | -0.615 | 1.34E-05 |
| 11 | 118820778 | rs139076649 | C | T | 1.02 | -4813.425 | -4814.043 | -0.619 | 1.33E-05 |
| 11 | 118722946 | rs148990923 | T | C | 0.978 | -4813.425 | -4814.046 | -0.622 | 1.33E-05 |
| 11 | 118573894 | rs576283 | G | A | 0.949 | -4813.425 | -4814.05 | -0.625 | 1.33E-05 |
| 11 | 118575011 | rs530901 | C | T | 0.949 | -4813.425 | -4814.05 | -0.625 | 1.33E-05 |
| 11 | 118575180 | rs626645 | C | T | 0.949 | -4813.425 | -4814.05 | -0.625 | 1.33E-05 |
| 11 | 118480695 | rs34026809 | C | G | 1.061 | -4813.425 | -4814.051 | -0.626 | 1.32E-05 |
| 11 | 118809805 | rs142069912 | C | G | 1 | -4813.425 | -4814.054 | -0.629 | 1.32E-05 |
| 11 | 118760443 | rs12786257 | A | G | 0.983 | -4813.425 | -4814.057 | -0.632 | 1.32E-05 |
| 11 | 118518227 | rs10736494 | T | C | 0.972 | -4813.425 | -4814.057 | -0.633 | 1.32E-05 |
| 11 | 118746223 | rs11217077 | C | T | 0.945 | -4813.425 | -4814.063 | -0.638 | 1.31E-05 |
| 11 | 118801045 | rs75085081 | A | G | 0.966 | -4813.425 | -4814.067 | -0.643 | 1.30E-05 |
| 11 | 118874272 | rs192113495 | G | T | 1.035 | -4813.425 | -4814.069 | -0.644 | 1.30E-05 |
| 11 | 118856582 | rs3887652 | C | G | 1.058 | -4813.425 | -4814.069 | -0.645 | 1.30E-05 |
| 11 | 118775524 | rs79108568 | G | T | 1.047 | -4813.425 | -4814.072 | -0.648 | 1.30E-05 |
| 11 | 118441079 | rs7106316 | G | A | 0.938 | -4813.425 | -4814.075 | -0.65 | 1.29E-05 |
| 11 | 118807639 | rs73575454 | A | G | 0.94 | -4813.425 | -4814.08 | -0.656 | 1.29E-05 |
| 11 | 118809767 | rs75460330 | A | G | 0.94 | -4813.425 | -4814.08 | -0.656 | 1.29E-05 |
| 11 | 118810054 | rs73557344 | C | G | 0.94 | -4813.425 | -4814.08 | -0.656 | 1.29E-05 |
| 11 | 118573916 | rs627391 | T | A | 0.95 | -4813.425 | -4814.083 | -0.659 | 1.28E-05 |
| 11 | 118788081 | rs7929825 | G | C | 1.06 | -4813.425 | -4814.09 | -0.665 | 1.27E-05 |
| 11 | 118700515 | rs112571283 | T | C | 1.029 | -4813.425 | -4814.092 | -0.667 | 1.27E-05 |
| 11 | 118545589 | rs113941645 | G | C | 1.014 | -4813.425 | -4814.094 | -0.669 | 1.27E-05 |
| 11 | 118730669 | rs7933007 | C | T | 0.946 | -4813.425 | -4814.094 | -0.669 | 1.27E-05 |
| 11 | 118577990 | rs480958 | G | A | 0.949 | -4813.425 | -4814.1 | -0.675 | 1.26E-05 |
| 11 | 118755946 | rs76109832 | T | C | 0.943 | -4813.425 | -4814.1 | -0.676 | 1.26E-05 |
| 11 | 118669178 | rs583213 | G | T | 0.942 | -4813.425 | -4814.101 | -0.677 | 1.26E-05 |
| 11 | 118614115 | rs73001451 | C | T | 1.058 | -4813.425 | -4814.106 | -0.682 | 1.25E-05 |
| 11 | 118717929 | rs4938558 | T | C | 0.947 | -4813.425 | -4814.131 | -0.706 | 1.22E-05 |
| 11 | 118583847 | rs12803363 | T | C | 1.054 | -4813.425 | -4814.132 | -0.708 | 1.22E-05 |
| 11 | 118559731 | rs113784020 | T | C | 0.959 | -4813.425 | -4814.133 | -0.709 | 1.22E-05 |
| 11 | 118802128 | rs59069289 | T | G | 0.945 | -4813.425 | -4814.144 | -0.719 | 1.21E-05 |
| 11 | 118802386 | rs59480472 | C | T | 0.945 | -4813.425 | -4814.144 | -0.719 | 1.21E-05 |
| 11 | 118803160 | rs73575440 | A | G | 0.945 | -4813.425 | -4814.144 | -0.719 | 1.21E-05 |
| 11 | 118803596 | rs111313044 | A | G | 0.945 | -4813.425 | -4814.144 | -0.719 | 1.21E-05 |
| 11 | 118804643 | rs56933877 | A | G | 0.945 | -4813.425 | -4814.144 | -0.719 | 1.21E-05 |
| 11 | 118804919 | rs73575441 | C | T | 0.945 | -4813.425 | -4814.144 | -0.719 | 1.21E-05 |
| 11 | 118804989 | rs112045567 | G | A | 0.945 | -4813.425 | -4814.144 | -0.719 | 1.21E-05 |
| 11 | 118584498 | rs569770 | T | C | 0.95 | -4813.425 | -4814.147 | -0.722 | 1.20E-05 |
| 11 | 118812821 | rs74535623 | T | G | 0.963 | -4813.425 | -4814.147 | -0.723 | 1.20E-05 |
| 11 | 118813314 | rs73557345 | C | T | 0.963 | -4813.425 | -4814.147 | -0.723 | 1.20E-05 |
| 11 | 118725420 | rs11217062 | A | G | 0.948 | -4813.425 | -4814.153 | -0.729 | 1.19E-05 |
| 11 | 118726230 | rs2156755 | A | G | 0.948 | -4813.425 | -4814.153 | -0.729 | 1.19E-05 |
| 11 | 118773438 | rs35346926 | T | C | 1.024 | -4813.425 | -4814.154 | -0.73 | 1.19E-05 |
| 11 | 118662045 | rs7933859 | T | C | 0.944 | -4813.425 | -4814.157 | -0.733 | 1.19E-05 |
| 11 | 118662203 | rs4508234 | G | C | 0.944 | -4813.425 | -4814.157 | -0.733 | 1.19E-05 |
| 11 | 118662340 | rs5006102 | A | T | 0.944 | -4813.425 | -4814.157 | -0.733 | 1.19E-05 |
| 11 | 118765231 | rs12795395 | T | C | 1.023 | -4813.425 | -4814.159 | -0.734 | 1.19E-05 |
| 11 | 118711830 | rs566848 | G | A | 0.948 | -4813.425 | -4814.165 | -0.741 | 1.18E-05 |
| 11 | 118598936 | rs184071181 | C | T | 1.043 | -4813.425 | -4814.169 | -0.745 | 1.18E-05 |

| | | | | | | | |
|---|---|---|---|---|---|---|---|
| 11 | 118712427 | rs7108905 | G | C | NULL | 0.641067406 | NULL&M_Disease |
| 11 | 118530068 | rs112337443 | C | T | M_Disease | 0.639916097 | M_Disease&NULL |
| 11 | 118572267 | rs573905 | A | G | M_Disease | 0.639916097 | M_Disease&NULL |
| 11 | 118438448 | rs60837962 | T | A | NULL | 0.638763175 | NULL&M_Disease |
| 11 | 118724116 | rs12276533 | A | G | NULL | 0.638763175 | NULL&M_Disease |
| 11 | 118731599 | rs34561290 | T | C | M_Disease | 0.638763175 | M_Disease&NULL |
| 11 | 118731762 | rs11217072 | G | A | M_Disease | 0.638763175 | M_Disease&NULL |
| 11 | 118889132 | rs11217124 | G | A | NULL | 0.638763175 | NULL&M_Disease |
| 11 | 118449746 | rs11216909 | T | C | NULL | 0.637608653 | NULL&M_Disease |
| 11 | 118872404 | rs6589713 | C | T | NULL | 0.637608653 | NULL&M_Disease |
| 11 | 118434421 | rs11216899 | G | A | NULL | 0.63645254 | NULL&M_Disease |
| 11 | 118434835 | rs60753148 | T | C | NULL | 0.63645254 | NULL&M_Disease |
| 11 | 118708785 | rs7122510 | A | C | NULL | 0.63645254 | NULL&M_Disease |
| 11 | 118708786 | rs7122511 | A | C | NULL | 0.63645254 | NULL&M_Disease |
| 11 | 118723753 | rs7119568 | A | T | NULL | 0.63645254 | NULL&M_Disease |
| 11 | 118540104 | rs506234 | G | C | M_Disease | 0.635294849 | M_Disease&NULL |
| 11 | 118713180 | rs2187490 | G | T | NULL | 0.635294849 | NULL&M_Disease |
| 11 | 118730141 | rs7929520 | G | A | M_Disease | 0.635294849 | M_Disease&NULL |
| 11 | 118885163 | rs59608621 | G | A | NULL | 0.635294849 | NULL&M_Disease |
| 11 | 118436079 | rs11216900 | C | G | NULL | 0.634135591 | NULL&M_Disease |
| 11 | 118445275 | rs17122269 | C | T | NULL | 0.634135591 | NULL&M_Disease |
| 11 | 118709600 | rs12272981 | G | A | NULL | 0.634135591 | NULL&M_Disease |
| 11 | 118728479 | rs7924647 | G | A | NULL | 0.634135591 | NULL&M_Disease |
| 11 | 118887888 | rs3916582 | T | C | NULL | 0.634135591 | NULL&M_Disease |
| 11 | 118789952 | rs11217090 | T | C | M_Disease | 0.632974777 | M_Disease&NULL |
| 11 | 118884708 | rs61381062 | A | G | NULL | 0.632974777 | NULL&M_Disease |
| 11 | 118884726 | rs58601794 | C | T | NULL | 0.632974777 | NULL&M_Disease |
| 11 | 118435756 | rs2277291 | T | C | NULL | 0.631812418 | NULL&M_Disease |
| 11 | 118456414 | rs11216914 | C | T | NULL | 0.631812418 | NULL&M_Disease |
| 11 | 118473231 | rs1047079 | C | T | NULL | 0.631812418 | NULL&M_Disease |
| 11 | 118712270 | rs7105658 | T | C | NULL | 0.631812418 | NULL&M_Disease |
| 11 | 118729283 | rs11217068 | T | C | NULL | 0.631812418 | NULL&M_Disease |
| 11 | 118729442 | rs12277355 | T | C | NULL | 0.631812418 | NULL&M_Disease |
| 11 | 118887887 | rs369816532 | T | G | NULL | 0.631812418 | NULL&M_Disease |
| 11 | 118888619 | rs11602764 | G | A | NULL | 0.631812418 | NULL&M_Disease |
| 11 | 118889185 | rs11217125 | C | T | NULL | 0.631812418 | NULL&M_Disease |
| 11 | 118459039 | rs57438673 | A | G | NULL | 0.630648526 | NULL&M_Disease |
| 11 | 118459702 | rs113054880 | G | C | NULL | 0.630648526 | NULL&M_Disease |
| 11 | 118514022 | rs613243 | C | G | M_Disease | 0.630648526 | M_Disease&NULL |
| 11 | 118707559 | rs7118009 | T | C | NULL | 0.630648526 | NULL&M_Disease |
| 11 | 118888831 | rs11820805 | C | T | NULL | 0.630648526 | NULL&M_Disease |
| 11 | 118555383 | rs2508573 | T | C | M_Disease | 0.629483112 | M_Disease&NULL |
| 11 | 118449396 | rs73023350 | A | G | NULL | 0.629483112 | NULL&M_Disease |
| 11 | 118450875 | rs7126175 | T | C | NULL | 0.629483112 | NULL&M_Disease |
| 11 | 118725383 | rs9735317 | T | C | NULL | 0.629483112 | NULL&M_Disease |
| 11 | 118432852 | rs11216898 | C | T | NULL | 0.628316188 | NULL&M_Disease |
| 11 | 118449370 | rs17122278 | G | A | NULL | 0.628316188 | NULL&M_Disease |
| 11 | 118734640 | rs12286723 | T | C | NULL | 0.628316188 | NULL&M_Disease |
| 11 | 118807637 | rs112974653 | T | G | M_Disease | 0.628316188 | M_Disease&NULL |
| 11 | 118885161 | rs59095933 | A | G | NULL | 0.628316188 | NULL&M_Disease |
| 11 | 118451901 | rs11216911 | C | T | NULL | 0.627147766 | NULL&M_Disease |
| 11 | 118452334 | rs11216912 | C | G | NULL | 0.627147766 | NULL&M_Disease |
| 11 | 118453681 | rs11216913 | G | A | NULL | 0.627147766 | NULL&M_Disease |
| 11 | 118454177 | rs17842763 | G | A | NULL | 0.627147766 | NULL&M_Disease |
| 11 | 118457630 | rs17092756 | C | G | NULL | 0.627147766 | NULL&M_Disease |
| 11 | 118470092 | rs11216924 | G | C | NULL | 0.627147766 | NULL&M_Disease |
| 11 | 118471996 | rs11216926 | T | C | NULL | 0.627147766 | NULL&M_Disease |
| 11 | 118550841 | rs79699220 | C | G | NULL | 0.627147766 | NULL&M_Disease |
| 11 | 118869460 | rs3737504 | G | A | NULL | 0.627147766 | NULL&M_Disease |
| 11 | 118453483 | rs60112336 | T | C | NULL | 0.625977858 | NULL&M_Disease |
| 11 | 118432404 | rs4326835 | T | G | NULL | 0.624806474 | NULL&M_Disease |
| 11 | 118478330 | rs7125115 | A | G | NULL | 0.624806474 | NULL&M_Disease |
| 11 | 118714492 | rs7117594 | T | C | NULL | 0.624806474 | NULL&M_Disease |
| 11 | 118444525 | rs11216907 | A | G | NULL | 0.623633628 | NULL&M_Disease |
| 11 | 118446298 | rs55844025 | T | C | NULL | 0.623633628 | NULL&M_Disease |
| 11 | 118466249 | rs7115634 | G | A | NULL | 0.623633628 | NULL&M_Disease |
| 11 | 118568427 | rs117361624 | T | C | NULL | 0.623633628 | NULL&M_Disease |
| 11 | 118579395 | rs11828220 | A | T | NULL | 0.623633628 | NULL&M_Disease |
| 11 | 118629959 | rs140110112 | A | G | NULL | 0.623633628 | NULL&M_Disease |
| 11 | 118691975 | rs77731159 | A | T | NULL | 0.623633628 | NULL&M_Disease |
| 11 | 118707035 | rs143605660 | A | G | NULL | 0.622459331 | NULL&M_Disease |
| 11 | 118872741 | rs4573680 | C | T | NULL | 0.622459331 | NULL&M_Disease |
| 11 | 118447580 | rs9736327 | G | T | NULL | 0.621283595 | NULL&M_Disease |
| 11 | 118447597 | rs9735421 | A | G | NULL | 0.621283595 | NULL&M_Disease |
| 11 | 118464815 | rs11216921 | C | G | NULL | 0.621283595 | NULL&M_Disease |
| 11 | 118469571 | rs7931564 | G | A | NULL | 0.621283595 | NULL&M_Disease |
| 11 | 118471423 | rs11216925 | C | T | NULL | 0.621283595 | NULL&M_Disease |
| 11 | 118472354 | rs10892245 | A | C | NULL | 0.621283595 | NULL&M_Disease |
| 11 | 118473385 | rs17742 | C | T | NULL | 0.621283595 | NULL&M_Disease |
| 11 | 118763736 | rs12363277 | C | T | NULL | 0.621283595 | NULL&M_Disease |
| 11 | 118769604 | rs61751532 | T | C | NULL | 0.621283595 | NULL&M_Disease |
| 11 | 118899218 | rs4936459 | T | C | M_Disease | 0.621283595 | M_Disease&NULL |
| 11 | 118432279 | rs11216897 | G | A | NULL | 0.620106432 | NULL&M_Disease |
| 11 | 118448087 | rs9735449 | T | C | NULL | 0.620106432 | NULL&M_Disease |
| 11 | 118483802 | rs72999455 | G | A | NULL | 0.620106432 | NULL&M_Disease |
| 11 | 118543798 | rs76570573 | T | C | NULL | 0.620106432 | NULL&M_Disease |
| 11 | 118765688 | rs676925 | G | C | NULL | 0.620106432 | NULL&M_Disease |
| 11 | 118450046 | rs11216910 | G | A | NULL | 0.618927855 | NULL&M_Disease |
| 11 | 118494790 | rs11825621 | C | T | NULL | 0.618927855 | NULL&M_Disease |
| 11 | 118502172 | rs35435609 | T | C | NULL | 0.618927855 | NULL&M_Disease |
| 11 | 118536950 | rs201359394 | T | A | NULL | 0.618927855 | NULL&M_Disease |
| 11 | 118719984 | rs145680506 | T | C | M_Disease | 0.618927855 | M_Disease&NULL |
| 11 | 118461605 | rs11216916 | C | T | NULL | 0.616566505 | NULL&M_Disease |
| 11 | 118462160 | rs11216917 | G | T | NULL | 0.616566505 | NULL&M_Disease |
| 11 | 118465295 | rs11216922 | G | A | NULL | 0.616566505 | NULL&M_Disease |
| 11 | 118465501 | rs11216923 | G | A | NULL | 0.616566505 | NULL&M_Disease |
| 11 | 118567881 | rs497678 | C | T | NULL | 0.616566505 | NULL&M_Disease |
| 11 | 118759610 | rs566416 | G | T | NULL | 0.616566505 | NULL&M_Disease |
| 11 | 118762218 | rs543824 | G | A | NULL | 0.616566505 | NULL&M_Disease |
| 11 | 118433729 | rs59108310 | G | A | NULL | 0.614199643 | NULL&M_Disease |
| 11 | 118437279 | rs11216902 | G | A | NULL | 0.614199643 | NULL&M_Disease |
| 11 | 118439048 | rs12224844 | T | A | NULL | 0.614199643 | NULL&M_Disease |
| 11 | 118443944 | rs11216906 | A | G | NULL | 0.614199643 | NULL&M_Disease |

| | | | | | | | | | |
|---|---|---|---|---|---|---|---|---|---|
| 11 | 118429764 | rs111942423 | G | T | 0.945 | -4813.425 | -4814.174 | -0.749 | 1.17E-05 |
| 11 | 118587347 | rs603486 | T | C | 0.951 | -4813.425 | -4814.174 | -0.75 | 1.17E-05 |
| 11 | 118678780 | rs500254 | G | A | 0.945 | -4813.425 | -4814.18 | -0.756 | 1.16E-05 |
| 11 | 118428791 | rs7929887 | G | A | 0.945 | -4813.425 | -4814.181 | -0.757 | 1.16E-05 |
| 11 | 118660932 | rs1784295 | A | G | 0.945 | -4813.425 | -4814.182 | -0.757 | 1.16E-05 |
| 11 | 118430519 | rs11552421 | T | C | 0.946 | -4813.425 | -4814.186 | -0.762 | 1.16E-05 |
| 11 | 118720590 | rs4938564 | T | G | 0.949 | -4813.425 | -4814.192 | -0.768 | 1.15E-05 |
| 11 | 118721850 | rs7946961 | G | A | 0.949 | -4813.425 | -4814.192 | -0.768 | 1.15E-05 |
| 11 | 118722231 | rs4938570 | G | A | 0.949 | -4813.425 | -4814.192 | -0.768 | 1.15E-05 |
| 11 | 118723087 | rs11217059 | T | C | 0.949 | -4813.425 | -4814.192 | -0.768 | 1.15E-05 |
| 11 | 118723537 | rs7103314 | G | A | 0.949 | -4813.425 | -4814.192 | -0.768 | 1.15E-05 |
| 11 | 118724336 | rs10892298 | C | T | 0.949 | -4813.425 | -4814.192 | -0.768 | 1.15E-05 |
| 11 | 118661055 | rs1790188 | T | C | 0.946 | -4813.425 | -4814.198 | -0.774 | 1.14E-05 |
| 11 | 118655704 | rs6421568 | C | G | 0.946 | -4813.425 | -4814.201 | -0.776 | 1.14E-05 |
| 11 | 118655775 | rs6421569 | A | C | 0.946 | -4813.425 | -4814.201 | -0.776 | 1.14E-05 |
| 11 | 118887581 | rs7105187 | T | C | 1.054 | -4813.425 | -4814.202 | -0.777 | 1.14E-05 |
| 11 | 118503442 | rs45621235 | T | C | 1.03 | -4813.425 | -4814.203 | -0.779 | 1.14E-05 |
| 11 | 118720684 | rs4936440 | G | C | 0.95 | -4813.425 | -4814.206 | -0.782 | 1.13E-05 |
| 11 | 118720998 | rs4938565 | A | G | 0.95 | -4813.425 | -4814.206 | -0.782 | 1.13E-05 |
| 11 | 118721189 | rs4938566 | T | A | 0.95 | -4813.425 | -4814.206 | -0.782 | 1.13E-05 |
| 11 | 118721249 | rs4938567 | G | A | 0.95 | -4813.425 | -4814.206 | -0.782 | 1.13E-05 |
| 11 | 118661197 | rs61901410 | G | C | 0.946 | -4813.425 | -4814.212 | -0.788 | 1.13E-05 |
| 11 | 118492713 | rs45457998 | G | A | 0.982 | -4813.425 | -4814.215 | -0.79 | 1.12E-05 |
| 11 | 118637666 | rs535996 | T | C | 0.947 | -4813.425 | -4814.219 | -0.794 | 1.12E-05 |
| 11 | 118648396 | rs548167 | A | G | 0.947 | -4813.425 | -4814.221 | -0.796 | 1.12E-05 |
| 11 | 118652781 | rs2508915 | C | G | 0.947 | -4813.425 | -4814.221 | -0.796 | 1.12E-05 |
| 11 | 118718485 | rs4938561 | C | T | 0.95 | -4813.425 | -4814.22 | -0.796 | 1.12E-05 |
| 11 | 118630532 | rs35888235 | T | G | 0.977 | -4813.425 | -4814.224 | -0.8 | 1.11E-05 |
| 11 | 118633561 | rs12794288 | A | C | 0.977 | -4813.425 | -4814.224 | -0.8 | 1.11E-05 |
| 11 | 118632893 | rs6589690 | A | G | 0.947 | -4813.425 | -4814.227 | -0.802 | 1.11E-05 |
| 11 | 118640081 | rs529717 | A | G | 0.947 | -4813.425 | -4814.228 | -0.803 | 1.11E-05 |
| 11 | 118641451 | rs2508929 | G | T | 0.947 | -4813.425 | -4814.228 | -0.803 | 1.11E-05 |
| 11 | 118644213 | rs1790187 | A | G | 0.947 | -4813.425 | -4814.228 | -0.803 | 1.11E-05 |
| 11 | 118665009 | rs4938544 | A | G | 0.947 | -4813.425 | -4814.228 | -0.804 | 1.11E-05 |
| 11 | 118751898 | rs60651507 | A | G | 0.956 | -4813.425 | -4814.229 | -0.805 | 1.11E-05 |
| 11 | 118719254 | rs2156750 | C | T | 0.95 | -4813.425 | -4814.231 | -0.806 | 1.11E-05 |
| 11 | 118719405 | rs2156749 | G | C | 0.95 | -4813.425 | -4814.231 | -0.806 | 1.11E-05 |
| 11 | 118719742 | rs7116715 | C | T | 0.95 | -4813.425 | -4814.231 | -0.806 | 1.11E-05 |
| 11 | 118665324 | rs61901411 | T | C | 0.947 | -4813.425 | -4814.236 | -0.811 | 1.10E-05 |
| 11 | 118677083 | rs675003 | T | C | 0.947 | -4813.425 | -4814.235 | -0.811 | 1.10E-05 |
| 11 | 118440863 | rs11216904 | G | A | 0.948 | -4813.425 | -4814.239 | -0.815 | 1.10E-05 |
| 11 | 118618837 | rs640120 | A | C | 0.948 | -4813.425 | -4814.242 | -0.818 | 1.09E-05 |
| 11 | 118619423 | rs1961138 | C | T | 0.948 | -4813.425 | -4814.242 | -0.818 | 1.09E-05 |
| 11 | 118622245 | rs485901 | G | A | 0.948 | -4813.425 | -4814.242 | -0.818 | 1.09E-05 |
| 11 | 118718856 | rs2156751 | G | A | 0.951 | -4813.425 | -4814.244 | -0.819 | 1.09E-05 |
| 11 | 118718972 | rs2187491 | C | T | 0.951 | -4813.425 | -4814.244 | -0.819 | 1.09E-05 |
| 11 | 118668908 | rs582200 | G | A | 0.948 | -4813.425 | -4814.254 | -0.829 | 1.08E-05 |
| 11 | 118441846 | rs10892243 | C | T | 0.948 | -4813.425 | -4814.255 | -0.831 | 1.08E-05 |
| 11 | 118712509 | rs75488969 | T | C | 0.95 | -4813.425 | -4814.275 | -0.85 | 1.06E-05 |
| 11 | 118887568 | rs7121270 | G | T | 1.051 | -4813.425 | -4814.275 | -0.85 | 1.06E-05 |
| 11 | 118869539 | rs11822103 | A | G | 1.051 | -4813.425 | -4814.279 | -0.854 | 1.05E-05 |
| 11 | 118480223 | rs67307131 | C | T | 0.954 | -4813.425 | -4814.283 | -0.858 | 1.05E-05 |
| 11 | 118569414 | rs6589675 | C | G | 0.95 | -4813.425 | -4814.284 | -0.86 | 1.05E-05 |
| 11 | 118752178 | rs61150131 | C | T | 0.961 | -4813.425 | -4814.285 | -0.86 | 1.05E-05 |
| 11 | 118614397 | rs10892281 | T | C | 0.95 | -4813.425 | -4814.285 | -0.861 | 1.05E-05 |
| 11 | 118615660 | rs6589687 | T | A | 0.95 | -4813.425 | -4814.285 | -0.861 | 1.05E-05 |
| 11 | 118765267 | rs598207 | C | G | 0.952 | -4813.425 | -4814.285 | -0.861 | 1.05E-05 |
| 11 | 118867803 | rs4938618 | C | G | 1.051 | -4813.425 | -4814.293 | -0.869 | 1.04E-05 |
| 11 | 118436071 | rs73023329 | T | C | 0.983 | -4813.425 | -4814.301 | -0.876 | 1.03E-05 |
| 11 | 118725189 | rs112965917 | A | T | 0.975 | -4813.425 | -4814.305 | -0.881 | 1.03E-05 |
| 11 | 118644323 | rs6589695 | C | G | 0.989 | -4813.425 | -4814.316 | -0.891 | 1.02E-05 |
| 11 | 118711069 | rs538645 | C | T | 0.953 | -4813.425 | -4814.32 | -0.895 | 1.01E-05 |
| 11 | 118875727 | rs4936457 | G | A | 1.049 | -4813.425 | -4814.323 | -0.898 | 1.01E-05 |
| 11 | 118875761 | rs4936458 | G | T | 1.049 | -4813.425 | -4814.323 | -0.898 | 1.01E-05 |
| 11 | 118704751 | rs35262372 | G | A | 1.005 | -4813.425 | -4814.337 | -0.912 | 9.95E-06 |
| 11 | 118430048 | rs11216896 | T | C | 0.952 | -4813.425 | -4814.338 | -0.913 | 9.94E-06 |
| 11 | 118889378 | rs7113753 | A | G | 1.049 | -4813.425 | -4814.337 | -0.913 | 9.94E-06 |
| 11 | 118873987 | rs9645664 | C | T | 1.049 | -4813.425 | -4814.344 | -0.92 | 9.87E-06 |
| 11 | 118629143 | rs475946 | T | C | 0.953 | -4813.425 | -4814.347 | -0.922 | 9.85E-06 |
| 11 | 118433729 | rs59108310 | G | A | 0.953 | -4813.425 | -4814.348 | -0.924 | 9.83E-06 |
| 11 | 118439048 | rs12224844 | T | A | 0.953 | -4813.425 | -4814.348 | -0.924 | 9.83E-06 |
| 11 | 118437279 | rs11216902 | G | A | 0.954 | -4813.425 | -4814.357 | -0.932 | 9.75E-06 |
| 11 | 118443944 | rs11216906 | A | G | 0.953 | -4813.425 | -4814.357 | -0.932 | 9.75E-06 |
| 11 | 118445585 | rs73023341 | G | A | 0.953 | -4813.425 | -4814.357 | -0.932 | 9.75E-06 |
| 11 | 118526691 | rs72999486 | T | C | 0.977 | -4813.425 | -4814.359 | -0.934 | 9.73E-06 |
| 11 | 118462160 | rs11216917 | G | T | 0.954 | -4813.425 | -4814.371 | -0.946 | 9.62E-06 |
| 11 | 118461605 | rs11216916 | C | T | 0.954 | -4813.425 | -4814.372 | -0.947 | 9.61E-06 |
| 11 | 118759610 | rs566416 | G | T | 0.955 | -4813.425 | -4814.373 | -0.948 | 9.60E-06 |
| 11 | 118567881 | rs497678 | C | T | 0.956 | -4813.425 | -4814.374 | -0.949 | 9.59E-06 |
| 11 | 118762218 | rs543824 | G | A | 0.955 | -4813.425 | -4814.374 | -0.95 | 9.58E-06 |
| 11 | 118465295 | rs11216922 | G | A | 0.954 | -4813.425 | -4814.376 | -0.952 | 9.56E-06 |
| 11 | 118465501 | rs11216923 | G | A | 0.954 | -4813.425 | -4814.376 | -0.952 | 9.56E-06 |
| 11 | 118450046 | rs11216910 | G | A | 0.955 | -4813.425 | -4814.393 | -0.969 | 9.40E-06 |
| 11 | 118494790 | rs11825621 | C | T | 0.985 | -4813.425 | -4814.394 | -0.969 | 9.40E-06 |
| 11 | 118502172 | rs35435609 | T | C | 0.985 | -4813.425 | -4814.394 | -0.969 | 9.40E-06 |
| 11 | 118536950 | rs201359394 | T | A | 0.956 | -4813.425 | -4814.395 | -0.97 | 9.39E-06 |
| 11 | 118765688 | rs676925 | G | C | 0.956 | -4813.425 | -4814.397 | -0.973 | 9.36E-06 |
| 11 | 118448087 | rs9735449 | T | C | 0.956 | -4813.425 | -4814.402 | -0.977 | 9.32E-06 |
| 11 | 118432279 | rs11216897 | G | A | 0.956 | -4813.425 | -4814.403 | -0.978 | 9.32E-06 |
| 11 | 118543798 | rs76570573 | T | C | 0.99 | -4813.425 | -4814.403 | -0.978 | 9.32E-06 |
| 11 | 118483802 | rs72999455 | G | A | 0.995 | -4813.425 | -4814.409 | -0.984 | 9.26E-06 |
| 11 | 118469571 | rs7931564 | G | A | 0.956 | -4813.425 | -4814.411 | -0.987 | 9.23E-06 |
| 11 | 118471423 | rs11216925 | C | T | 0.956 | -4813.425 | -4814.411 | -0.987 | 9.23E-06 |
| 11 | 118472354 | rs10892245 | A | C | 0.956 | -4813.425 | -4814.411 | -0.987 | 9.23E-06 |
| 11 | 118447580 | rs9736327 | G | T | 0.957 | -4813.425 | -4814.413 | -0.988 | 9.22E-06 |
| 11 | 118447597 | rs9735421 | A | G | 0.957 | -4813.425 | -4814.413 | -0.988 | 9.22E-06 |
| 11 | 118763736 | rs12363277 | C | T | 1.023 | -4813.425 | -4814.412 | -0.988 | 9.22E-06 |
| 11 | 118464815 | rs11216921 | C | G | 0.956 | -4813.425 | -4814.413 | -0.989 | 9.21E-06 |
| 11 | 118769604 | rs61751532 | T | C | 1.02 | -4813.425 | -4814.415 | -0.99 | 9.20E-06 |
| 11 | 118473385 | rs17742 | C | T | 0.957 | -4813.425 | -4814.417 | -0.993 | 9.18E-06 |
| 11 | 118872741 | rs4573680 | C | T | 1.046 | -4813.425 | -4814.418 | -0.994 | 9.17E-06 |
| 11 | 118707035 | rs143605660 | A | G | 1.013 | -4813.425 | -4814.428 | -1.004 | 9.08E-06 |
| 11 | 118466249 | rs7115634 | G | A | 0.958 | -4813.425 | -4814.432 | -1.007 | 9.05E-06 |

| | | | | | | | |
|---|---|---|---|---|---|---|---|
| 11 | 118445585 | rs73023341 | G | A | NULL | 0.614199643 | NULL&M_Disease |
| 11 | 118526691 | rs72999486 | T | C | NULL | 0.614199643 | NULL&M_Disease |
| 11 | 118740180 | rs7483509 | A | G | M_Disease | 0.614199643 | M_Disease&NULL |
| 11 | 118430048 | rs11216896 | T | C | NULL | 0.613014176 | NULL&M_Disease |
| 11 | 118629143 | rs475946 | T | C | NULL | 0.613014176 | NULL&M_Disease |
| 11 | 118801440 | rs12280501 | T | C | M_Disease | 0.613014176 | M_Disease&NULL |
| 11 | 118873987 | rs9645664 | C | T | NULL | 0.613014176 | NULL&M_Disease |
| 11 | 118889378 | rs7113753 | A | G | NULL | 0.613014176 | NULL&M_Disease |
| 11 | 118704751 | rs35262372 | G | A | NULL | 0.611827369 | NULL&M_Disease |
| 11 | 118711069 | rs538645 | C | T | NULL | 0.610639234 | NULL&M_Disease |
| 11 | 118875727 | rs4936457 | G | A | NULL | 0.610639234 | NULL&M_Disease |
| 11 | 118875761 | rs4936458 | G | T | NULL | 0.610639234 | NULL&M_Disease |
| 11 | 118644323 | rs6589695 | C | G | NULL | 0.609449784 | NULL&M_Disease |
| 11 | 118699276 | rs625274 | T | C | M_Disease | 0.609449784 | M_Disease&NULL |
| 11 | 118725189 | rs112965917 | A | T | NULL | 0.608259031 | NULL&M_Disease |
| 11 | 118799206 | rs7101680 | A | G | M_Disease | 0.608259031 | M_Disease&NULL |
| 11 | 118436071 | rs73023329 | T | C | NULL | 0.607066988 | NULL&M_Disease |
| 11 | 118793613 | rs11217091 | C | T | M_Disease | 0.607066988 | M_Disease&NULL |
| 11 | 118867803 | rs4938618 | C | G | NULL | 0.607066988 | NULL&M_Disease |
| 11 | 118480223 | rs67307131 | C | T | NULL | 0.605873668 | NULL&M_Disease |
| 11 | 118565651 | rs540800 | A | G | M_Disease | 0.605873668 | M_Disease&NULL |
| 11 | 118569414 | rs6589675 | C | G | NULL | 0.605873668 | NULL&M_Disease |
| 11 | 118614397 | rs10892281 | T | C | NULL | 0.605873668 | NULL&M_Disease |
| 11 | 118615660 | rs6589687 | T | A | NULL | 0.605873668 | NULL&M_Disease |
| 11 | 118752178 | rs61150131 | C | T | NULL | 0.605873668 | NULL&M_Disease |
| 11 | 118765267 | rs598207 | C | G | NULL | 0.605873668 | NULL&M_Disease |
| 11 | 118869539 | rs11822103 | A | G | NULL | 0.605873668 | NULL&M_Disease |
| 11 | 118563286 | rs527619 | A | G | M_Disease | 0.604679085 | M_Disease&NULL |
| 11 | 118712509 | rs75488969 | T | C | NULL | 0.604679085 | NULL&M_Disease |
| 11 | 118887568 | rs7121270 | G | T | NULL | 0.604679085 | NULL&M_Disease |
| 11 | 118747576 | rs55756957 | G | A | M_Disease | 0.60348325 | M_Disease&NULL |
| 11 | 118754649 | rs10892307 | G | C | M_Disease | 0.60348325 | M_Disease&NULL |
| 11 | 118441846 | rs10892243 | C | T | NULL | 0.602286177 | NULL&M_Disease |
| 11 | 118668908 | rs582200 | G | A | NULL | 0.602286177 | NULL&M_Disease |
| 11 | 118738281 | rs7119038 | G | A | M_Disease | 0.602286177 | M_Disease&NULL |
| 11 | 118440863 | rs11216904 | G | A | NULL | 0.601087879 | NULL&M_Disease |
| 11 | 118618837 | rs640120 | A | C | NULL | 0.601087879 | NULL&M_Disease |
| 11 | 118619423 | rs1961138 | C | T | NULL | 0.601087879 | NULL&M_Disease |
| 11 | 118622245 | rs485901 | G | A | NULL | 0.601087879 | NULL&M_Disease |
| 11 | 118700557 | rs4291686 | G | A | M_Disease | 0.601087879 | M_Disease&NULL |
| 11 | 118718856 | rs2156751 | G | A | NULL | 0.601087879 | NULL&M_Disease |
| 11 | 118718972 | rs2187491 | C | T | NULL | 0.601087879 | NULL&M_Disease |
| 11 | 118798824 | rs112119822 | G | A | M_Disease | 0.599888369 | M_Disease&NULL |
| 11 | 118640081 | rs529717 | A | G | NULL | 0.599888369 | NULL&M_Disease |
| 11 | 118641451 | rs2508929 | G | T | NULL | 0.599888369 | NULL&M_Disease |
| 11 | 118644213 | rs1790187 | A | G | NULL | 0.599888369 | NULL&M_Disease |
| 11 | 118665009 | rs4938544 | A | G | NULL | 0.599888369 | NULL&M_Disease |
| 11 | 118665324 | rs61901411 | T | C | NULL | 0.599888369 | NULL&M_Disease |
| 11 | 118677083 | rs675003 | T | C | NULL | 0.599888369 | NULL&M_Disease |
| 11 | 118719254 | rs2156750 | C | T | NULL | 0.599888369 | NULL&M_Disease |
| 11 | 118719405 | rs2156749 | G | C | NULL | 0.599888369 | NULL&M_Disease |
| 11 | 118719742 | rs7116715 | C | T | NULL | 0.599888369 | NULL&M_Disease |
| 11 | 118501299 | rs7936556 | A | G | M_Disease | 0.59868766 | M_Disease&NULL |
| 11 | 118630532 | rs35888235 | T | G | NULL | 0.59868766 | NULL&M_Disease |
| 11 | 118632893 | rs6589690 | A | G | NULL | 0.59868766 | NULL&M_Disease |
| 11 | 118633561 | rs12794288 | A | C | NULL | 0.59868766 | NULL&M_Disease |
| 11 | 118637666 | rs535996 | T | C | NULL | 0.59868766 | NULL&M_Disease |
| 11 | 118648396 | rs548167 | A | G | NULL | 0.59868766 | NULL&M_Disease |
| 11 | 118652781 | rs2508915 | C | G | NULL | 0.59868766 | NULL&M_Disease |
| 11 | 118718485 | rs4938561 | C | T | NULL | 0.59868766 | NULL&M_Disease |
| 11 | 118751898 | rs60651507 | A | G | NULL | 0.59868766 | NULL&M_Disease |
| 11 | 118782222 | rs61900138 | C | T | M_Disease | 0.59868766 | M_Disease&NULL |
| 11 | 118492713 | rs45457998 | G | A | NULL | 0.597485766 | NULL&M_Disease |
| 11 | 118661197 | rs61901410 | G | C | NULL | 0.597485766 | NULL&M_Disease |
| 11 | 118503442 | rs45621235 | T | C | NULL | 0.596282699 | NULL&M_Disease |
| 11 | 118655704 | rs6421568 | C | G | NULL | 0.596282699 | NULL&M_Disease |
| 11 | 118655775 | rs6421569 | A | C | NULL | 0.596282699 | NULL&M_Disease |
| 11 | 118661055 | rs1790188 | T | C | NULL | 0.596282699 | NULL&M_Disease |
| 11 | 118720684 | rs4936440 | G | C | NULL | 0.596282699 | NULL&M_Disease |
| 11 | 118720998 | rs4938565 | A | G | NULL | 0.596282699 | NULL&M_Disease |
| 11 | 118721189 | rs4938566 | T | A | NULL | 0.596282699 | NULL&M_Disease |
| 11 | 118721249 | rs4938567 | G | A | NULL | 0.596282699 | NULL&M_Disease |
| 11 | 118793918 | rs12282752 | C | T | M_Disease | 0.596282699 | M_Disease&NULL |
| 11 | 118887581 | rs7105187 | T | C | NULL | 0.596282699 | NULL&M_Disease |
| 11 | 118720590 | rs4938564 | T | G | NULL | 0.595078474 | NULL&M_Disease |
| 11 | 118721850 | rs7946961 | G | A | NULL | 0.595078474 | NULL&M_Disease |
| 11 | 118722231 | rs4938570 | G | A | NULL | 0.595078474 | NULL&M_Disease |
| 11 | 118723087 | rs11217059 | T | C | NULL | 0.595078474 | NULL&M_Disease |
| 11 | 118723537 | rs7103314 | G | A | NULL | 0.595078474 | NULL&M_Disease |
| 11 | 118724336 | rs10892298 | C | T | NULL | 0.595078474 | NULL&M_Disease |
| 11 | 118428791 | rs7929887 | G | A | NULL | 0.593873103 | NULL&M_Disease |
| 11 | 118430519 | rs11552421 | T | C | NULL | 0.593873103 | NULL&M_Disease |
| 11 | 118660932 | rs1784295 | A | G | NULL | 0.593873103 | NULL&M_Disease |
| 11 | 118678780 | rs500254 | G | A | NULL | 0.593873103 | NULL&M_Disease |
| 11 | 118429764 | rs111942423 | G | T | NULL | 0.5926666 | NULL&M_Disease |
| 11 | 118587347 | rs603486 | T | C | NULL | 0.5926666 | NULL&M_Disease |
| 11 | 118598936 | rs184071181 | C | T | NULL | 0.591458978 | NULL&M_Disease |
| 11 | 118711830 | rs566848 | G | A | NULL | 0.591458978 | NULL&M_Disease |
| 11 | 118584498 | rs569770 | T | C | NULL | 0.590250252 | NULL&M_Disease |
| 11 | 118662045 | rs7933859 | T | C | NULL | 0.590250252 | NULL&M_Disease |
| 11 | 118662203 | rs4508234 | G | C | NULL | 0.590250252 | NULL&M_Disease |
| 11 | 118662340 | rs5006102 | A | T | NULL | 0.590250252 | NULL&M_Disease |
| 11 | 118725420 | rs11217062 | A | G | NULL | 0.590250252 | NULL&M_Disease |
| 11 | 118726230 | rs2156755 | A | G | NULL | 0.590250252 | NULL&M_Disease |
| 11 | 118765231 | rs12795395 | T | C | NULL | 0.590250252 | NULL&M_Disease |
| 11 | 118773438 | rs35346926 | T | C | NULL | 0.590250252 | NULL&M_Disease |
| 11 | 118486067 | rs11603023 | T | C | M_Disease | 0.589040434 | M_Disease&NULL |
| 11 | 118510562 | rs4245184 | A | G | M_Disease | 0.589040434 | M_Disease&NULL |
| 11 | 118511352 | rs644273 | T | C | M_Disease | 0.589040434 | M_Disease&NULL |
| 11 | 118511757 | rs654423 | A | G | M_Disease | 0.589040434 | M_Disease&NULL |
| 11 | 118802128 | rs59069289 | T | G | NULL | 0.589040434 | NULL&M_Disease |
| 11 | 118802386 | rs59480472 | C | T | NULL | 0.589040434 | NULL&M_Disease |
| 11 | 118803160 | rs73575440 | A | G | NULL | 0.589040434 | NULL&M_Disease |

| | | | | | | | | | |
|---|---|---|---|---|---|---|---|---|---|
| 11 | 118446298 | rs55844025 | T | C | 0.958 | -4813.425 | -4814.432 | -1.008 | 9.04E-06 |
| 11 | 118579395 | rs11828220 | A | T | 0.987 | -4813.425 | -4814.432 | -1.008 | 9.04E-06 |
| 11 | 118629959 | rs140110112 | A | G | 0.988 | -4813.425 | -4814.432 | -1.008 | 9.04E-06 |
| 11 | 118568427 | rs117361624 | T | C | 0.998 | -4813.425 | -4814.434 | -1.009 | 9.03E-06 |
| 11 | 118691975 | rs77731159 | A | T | 1.01 | -4813.425 | -4814.435 | -1.01 | 9.02E-06 |
| 11 | 118444525 | rs11216907 | A | G | 0.958 | -4813.425 | -4814.436 | -1.012 | 9.00E-06 |
| 11 | 118478330 | rs7125115 | A | G | 0.958 | -4813.425 | -4814.442 | -1.017 | 8.96E-06 |
| 11 | 118714492 | rs7117594 | T | C | 1.045 | -4813.425 | -4814.442 | -1.017 | 8.96E-06 |
| 11 | 118432404 | rs4326835 | T | G | 0.959 | -4813.425 | -4814.443 | -1.018 | 8.95E-06 |
| 11 | 118453483 | rs60112336 | T | C | 0.959 | -4813.425 | -4814.448 | -1.024 | 8.90E-06 |
| 11 | 118470092 | rs11216924 | G | C | 0.959 | -4813.425 | -4814.458 | -1.034 | 8.81E-06 |
| 11 | 118471996 | rs11216926 | T | C | 0.959 | -4813.425 | -4814.458 | -1.034 | 8.81E-06 |
| 11 | 118451901 | rs11216911 | C | T | 0.959 | -4813.425 | -4814.459 | -1.035 | 8.80E-06 |
| 11 | 118452334 | rs11216912 | C | G | 0.959 | -4813.425 | -4814.459 | -1.035 | 8.80E-06 |
| 11 | 118453681 | rs11216913 | G | A | 0.959 | -4813.425 | -4814.459 | -1.035 | 8.80E-06 |
| 11 | 118550841 | rs79699220 | C | G | 0.991 | -4813.425 | -4814.461 | -1.037 | 8.78E-06 |
| 11 | 118454177 | rs17842763 | G | A | 0.959 | -4813.425 | -4814.463 | -1.038 | 8.77E-06 |
| 11 | 118457630 | rs17092756 | C | G | 0.959 | -4813.425 | -4814.463 | -1.038 | 8.77E-06 |
| 11 | 118734640 | rs12286723 | T | C | 1.044 | -4813.425 | -4814.467 | -1.042 | 8.74E-06 |
| 11 | 118869460 | rs3737504 | G | A | 1.044 | -4813.425 | -4814.466 | -1.042 | 8.74E-06 |
| 11 | 118449370 | rs17122278 | G | A | 0.96 | -4813.425 | -4814.473 | -1.048 | 8.69E-06 |
| 11 | 118885161 | rs59095933 | A | G | 1.043 | -4813.425 | -4814.473 | -1.048 | 8.69E-06 |
| 11 | 118432852 | rs11216898 | C | T | 0.961 | -4813.425 | -4814.475 | -1.051 | 8.66E-06 |
| 11 | 118725383 | rs9735317 | T | C | 1.043 | -4813.425 | -4814.478 | -1.053 | 8.64E-06 |
| 11 | 118449396 | rs73023350 | A | G | 0.961 | -4813.425 | -4814.481 | -1.056 | 8.62E-06 |
| 11 | 118450875 | rs7126175 | T | C | 0.961 | -4813.425 | -4814.484 | -1.059 | 8.59E-06 |
| 11 | 118888831 | rs11820805 | C | T | 1.043 | -4813.425 | -4814.492 | -1.067 | 8.52E-06 |
| 11 | 118459039 | rs57438673 | A | G | 0.961 | -4813.425 | -4814.495 | -1.07 | 8.50E-06 |
| 11 | 118459702 | rs113054880 | G | C | 0.961 | -4813.425 | -4814.495 | -1.07 | 8.50E-06 |
| 11 | 118707559 | rs7118009 | T | C | 1.042 | -4813.425 | -4814.495 | -1.071 | 8.49E-06 |
| 11 | 118887887 | rs369816532 | T | G | 1.042 | -4813.425 | -4814.497 | -1.073 | 8.47E-06 |
| 11 | 118712270 | rs7105658 | T | C | 1.042 | -4813.425 | -4814.498 | -1.074 | 8.46E-06 |
| 11 | 118435756 | rs2277291 | T | C | 0.962 | -4813.425 | -4814.5 | -1.075 | 8.45E-06 |
| 11 | 118729283 | rs11217068 | T | C | 1.042 | -4813.425 | -4814.5 | -1.075 | 8.45E-06 |
| 11 | 118729442 | rs12277355 | T | C | 1.042 | -4813.425 | -4814.5 | -1.075 | 8.45E-06 |
| 11 | 118456414 | rs11216914 | C | T | 0.962 | -4813.425 | -4814.5 | -1.076 | 8.45E-06 |
| 11 | 118473231 | rs1047079 | C | T | 0.962 | -4813.425 | -4814.5 | -1.076 | 8.45E-06 |
| 11 | 118888619 | rs11602764 | G | A | 1.042 | -4813.425 | -4814.502 | -1.077 | 8.44E-06 |
| 11 | 118889185 | rs11217125 | C | T | 1.042 | -4813.425 | -4814.506 | -1.081 | 8.40E-06 |
| 11 | 118884708 | rs61381062 | A | G | 1.042 | -4813.425 | -4814.513 | -1.089 | 8.34E-06 |
| 11 | 118884726 | rs58601794 | C | T | 1.042 | -4813.425 | -4814.513 | -1.089 | 8.34E-06 |
| 11 | 118728479 | rs7924647 | G | A | 1.041 | -4813.425 | -4814.518 | -1.094 | 8.30E-06 |
| 11 | 118709600 | rs12272981 | G | A | 1.041 | -4813.425 | -4814.521 | -1.096 | 8.28E-06 |
| 11 | 118887888 | rs3916582 | T | C | 1.041 | -4813.425 | -4814.522 | -1.097 | 8.27E-06 |
| 11 | 118436079 | rs11216900 | C | G | 0.964 | -4813.425 | -4814.522 | -1.098 | 8.26E-06 |
| 11 | 118445275 | rs17122269 | C | T | 0.964 | -4813.425 | -4814.524 | -1.099 | 8.25E-06 |
| 11 | 118713180 | rs2187490 | G | T | 0.973 | -4813.425 | -4814.529 | -1.104 | 8.21E-06 |
| 11 | 118885163 | rs59608621 | G | A | 1.041 | -4813.425 | -4814.532 | -1.108 | 8.18E-06 |
| 11 | 118434421 | rs11216899 | G | A | 0.965 | -4813.425 | -4814.542 | -1.117 | 8.11E-06 |
| 11 | 118723753 | rs7119568 | A | T | 1.04 | -4813.425 | -4814.542 | -1.117 | 8.11E-06 |
| 11 | 118708785 | rs7122510 | A | C | 1.04 | -4813.425 | -4814.545 | -1.12 | 8.08E-06 |
| 11 | 118708786 | rs7122511 | A | C | 1.04 | -4813.425 | -4814.545 | -1.12 | 8.08E-06 |
| 11 | 118434835 | rs60753148 | T | C | 0.965 | -4813.425 | -4814.547 | -1.122 | 8.07E-06 |
| 11 | 118872404 | rs6589713 | C | T | 1.04 | -4813.425 | -4814.554 | -1.129 | 8.01E-06 |
| 11 | 118449746 | rs11216909 | T | C | 0.965 | -4813.425 | -4814.556 | -1.132 | 7.99E-06 |
| 11 | 118724116 | rs12276533 | A | G | 1.039 | -4813.425 | -4814.559 | -1.135 | 7.96E-06 |
| 11 | 118438448 | rs60837962 | T | A | 0.967 | -4813.425 | -4814.561 | -1.136 | 7.95E-06 |
| 11 | 118889132 | rs11217124 | G | A | 1.04 | -4813.425 | -4814.561 | -1.137 | 7.95E-06 |
| 11 | 118712427 | rs7108905 | G | C | 1.038 | -4813.425 | -4814.58 | -1.155 | 7.80E-06 |
| 11 | 118462554 | rs11216920 | G | A | 0.988 | -4813.425 | -4814.588 | -1.164 | 7.73E-06 |
| 11 | 118698170 | rs56296109 | A | C | 1 | -4813.425 | -4814.591 | -1.167 | 7.71E-06 |
| 11 | 118754353 | rs630923 | A | C | 0.966 | -4813.425 | -4814.611 | -1.186 | 7.57E-06 |
| 11 | 118873385 | rs10892320 | T | C | 1.037 | -4813.425 | -4814.62 | -1.196 | 7.49E-06 |
| 11 | 118757219 | rs1623316 | G | C | 0.964 | -4813.425 | -4814.628 | -1.204 | 7.43E-06 |
| 11 | 118481431 | rs10892246 | A | G | 1.037 | -4813.425 | -4814.63 | -1.206 | 7.42E-06 |
| 11 | 118896959 | rs57156480 | A | G | 1.036 | -4813.425 | -4814.633 | -1.208 | 7.40E-06 |
| 11 | 118567733 | rs481903 | C | T | 0.965 | -4813.425 | -4814.651 | -1.227 | 7.26E-06 |
| 11 | 118797003 | rs73575428 | G | C | 0.998 | -4813.425 | -4814.652 | -1.227 | 7.26E-06 |
| 11 | 118797278 | rs74584888 | A | G | 0.998 | -4813.425 | -4814.652 | -1.227 | 7.26E-06 |
| 11 | 118797392 | rs73575429 | C | T | 0.998 | -4813.425 | -4814.652 | -1.227 | 7.26E-06 |
| 11 | 118797420 | rs111390501 | G | A | 0.998 | -4813.425 | -4814.652 | -1.227 | 7.26E-06 |
| 11 | 118797544 | rs56032119 | G | A | 0.998 | -4813.425 | -4814.652 | -1.227 | 7.26E-06 |
| 11 | 118798154 | rs60969913 | C | G | 0.998 | -4813.425 | -4814.652 | -1.227 | 7.26E-06 |
| 11 | 118896116 | rs2276071 | C | A | 1.035 | -4813.425 | -4814.66 | -1.235 | 7.20E-06 |
| 11 | 118896115 | rs2276072 | G | A | 1.034 | -4813.425 | -4814.663 | -1.238 | 7.18E-06 |
| 11 | 118480115 | rs12803321 | C | G | 0.966 | -4813.425 | -4814.667 | -1.242 | 7.15E-06 |
| 11 | 118692727 | rs503425 | C | T | 0.968 | -4813.425 | -4814.671 | -1.246 | 7.13E-06 |
| 11 | 118567218 | rs11216947 | A | G | 0.966 | -4813.425 | -4814.675 | -1.25 | 7.10E-06 |
| 11 | 118872581 | rs4558175 | A | G | 1.034 | -4813.425 | -4814.677 | -1.252 | 7.08E-06 |
| 11 | 118880668 | rs11607569 | C | T | 1.033 | -4813.425 | -4814.678 | -1.253 | 7.08E-06 |
| 11 | 118603030 | rs7943039 | A | G | 0.967 | -4813.425 | -4814.681 | -1.257 | 7.05E-06 |
| 11 | 118567257 | rs7126172 | G | A | 0.966 | -4813.425 | -4814.687 | -1.262 | 7.01E-06 |
| 11 | 118788517 | rs73575424 | A | C | 0.986 | -4813.425 | -4814.687 | -1.262 | 7.01E-06 |
| 11 | 118685015 | rs540716 | T | C | 0.969 | -4813.425 | -4814.688 | -1.263 | 7.01E-06 |
| 11 | 118661376 | rs73003294 | T | C | 1.021 | -4813.425 | -4814.689 | -1.264 | 7.00E-06 |
| 11 | 118785982 | rs79157249 | T | A | 0.991 | -4813.425 | -4814.701 | -1.276 | 6.91E-06 |
| 11 | 118681088 | rs2508922 | A | G | 0.969 | -4813.425 | -4814.704 | -1.279 | 6.89E-06 |
| 11 | 118831727 | rs10892315 | C | T | 1.033 | -4813.425 | -4814.704 | -1.28 | 6.89E-06 |
| 11 | 118886117 | rs10111 | T | C | 1.031 | -4813.425 | -4814.71 | -1.285 | 6.85E-06 |
| 11 | 118883491 | rs60065241 | A | C | 1.031 | -4813.425 | -4814.71 | -1.286 | 6.85E-06 |
| 11 | 118878222 | rs11606357 | C | T | 1.031 | -4813.425 | -4814.712 | -1.288 | 6.83E-06 |
| 11 | 118892757 | rs117289748 | G | C | 1.03 | -4813.425 | -4814.714 | -1.289 | 6.83E-06 |
| 11 | 118892759 | rs143660533 | C | A | 1.03 | -4813.425 | -4814.714 | -1.289 | 6.83E-06 |
| 11 | 118893878 | rs111322300 | C | T | 1.03 | -4813.425 | -4814.715 | -1.291 | 6.81E-06 |
| 11 | 118836606 | rs113975105 | G | C | 1.032 | -4813.425 | -4814.717 | -1.292 | 6.81E-06 |
| 11 | 118690550 | rs577121 | C | T | 0.973 | -4813.425 | -4814.718 | -1.293 | 6.80E-06 |
| 11 | 118868750 | rs4445645 | C | G | 1.03 | -4813.425 | -4814.718 | -1.294 | 6.79E-06 |
| 11 | 118892277 | rs4938621 | T | C | 1.03 | -4813.425 | -4814.72 | -1.296 | 6.78E-06 |
| 11 | 118892835 | rs61179449 | C | G | 1.03 | -4813.425 | -4814.72 | -1.296 | 6.78E-06 |
| 11 | 118892872 | rs58955701 | T | C | 1.03 | -4813.425 | -4814.72 | -1.296 | 6.78E-06 |
| 11 | 118893023 | rs7127188 | C | A | 1.03 | -4813.425 | -4814.72 | -1.296 | 6.78E-06 |
| 11 | 118892940 | rs12280288 | C | T | 1.03 | -4813.425 | -4814.725 | -1.3 | 6.75E-06 |
| 11 | 118596444 | rs68097885 | A | G | 0.969 | -4813.425 | -4814.727 | -1.302 | 6.74E-06 |

| | | | | | | | |
|---|---|---|---|---|---|---|---|
| 11 | 118803596 | rs111313044 | A | G | NULL | 0.589040434 | NULL&M_Disease |
| 11 | 118804643 | rs56933877 | A | G | NULL | 0.589040434 | NULL&M_Disease |
| 11 | 118804919 | rs73575441 | C | T | NULL | 0.589040434 | NULL&M_Disease |
| 11 | 118804989 | rs112045567 | G | A | NULL | 0.589040434 | NULL&M_Disease |
| 11 | 118812821 | rs74535623 | T | G | NULL | 0.589040434 | NULL&M_Disease |
| 11 | 118813314 | rs73557345 | C | T | NULL | 0.589040434 | NULL&M_Disease |
| 11 | 118559731 | rs113784020 | T | C | NULL | 0.587829538 | NULL&M_Disease |
| 11 | 118583847 | rs12803363 | T | C | NULL | 0.587829538 | NULL&M_Disease |
| 11 | 118717929 | rs4938558 | T | C | NULL | 0.587829538 | NULL&M_Disease |
| 11 | 118725204 | rs55794678 | T | C | M_Disease | 0.586617579 | M_Disease&NULL |
| 11 | 118806216 | rs1231742890 | G | T | M_Disease | 0.586617579 | M_Disease&NULL |
| 11 | 118509877 | rs112552940 | T | C | M_Disease | 0.585404569 | M_Disease&NULL |
| 11 | 118577990 | rs480958 | G | A | NULL | 0.584190523 | NULL&M_Disease |
| 11 | 118614115 | rs73001451 | C | T | NULL | 0.584190523 | NULL&M_Disease |
| 11 | 118669178 | rs583213 | G | T | NULL | 0.584190523 | NULL&M_Disease |
| 11 | 118679629 | rs1784300 | T | A | M_Disease | 0.584190523 | M_Disease&NULL |
| 11 | 118730669 | rs7933007 | C | T | NULL | 0.582975454 | NULL&M_Disease |
| 11 | 118755946 | rs76109832 | T | C | NULL | 0.582975454 | NULL&M_Disease |
| 11 | 118788081 | rs7929825 | G | C | NULL | 0.582975454 | NULL&M_Disease |
| 11 | 118521549 | rs494560 | A | G | M_Disease | 0.581759377 | M_Disease&NULL |
| 11 | 118545589 | rs113941645 | G | C | NULL | 0.581759377 | NULL&M_Disease |
| 11 | 118573916 | rs627391 | T | A | NULL | 0.581759377 | NULL&M_Disease |
| 11 | 118700515 | rs112571283 | T | C | NULL | 0.581759377 | NULL&M_Disease |
| 11 | 118807639 | rs73575454 | A | G | NULL | 0.581759377 | NULL&M_Disease |
| 11 | 118809767 | rs75460330 | A | G | NULL | 0.581759377 | NULL&M_Disease |
| 11 | 118810054 | rs73557344 | C | G | NULL | 0.581759377 | NULL&M_Disease |
| 11 | 118441079 | rs7106316 | G | A | NULL | 0.580542305 | NULL&M_Disease |
| 11 | 118856582 | rs3887652 | C | G | NULL | 0.580542305 | NULL&M_Disease |
| 11 | 118464665 | rs113337722 | G | A | M_Disease | 0.579324252 | M_Disease&NULL |
| 11 | 118746223 | rs11217077 | C | T | NULL | 0.579324252 | NULL&M_Disease |
| 11 | 118775524 | rs79108568 | G | T | NULL | 0.579324252 | NULL&M_Disease |
| 11 | 118801045 | rs75085081 | A | G | NULL | 0.579324252 | NULL&M_Disease |
| 11 | 118874272 | rs192113495 | G | T | NULL | 0.579324252 | NULL&M_Disease |
| 11 | 118518227 | rs10736494 | T | C | NULL | 0.578105233 | NULL&M_Disease |
| 11 | 118573894 | rs576283 | G | A | NULL | 0.578105233 | NULL&M_Disease |
| 11 | 118575011 | rs530901 | C | T | NULL | 0.578105233 | NULL&M_Disease |
| 11 | 118575180 | rs626645 | C | T | NULL | 0.578105233 | NULL&M_Disease |
| 11 | 118760443 | rs12786257 | A | G | NULL | 0.578105233 | NULL&M_Disease |
| 11 | 118462367 | rs11216919 | G | A | M_Disease | 0.576885261 | M_Disease&NULL |
| 11 | 118480695 | rs34026809 | C | G | NULL | 0.576885261 | NULL&M_Disease |
| 11 | 118580348 | rs516503 | T | C | NULL | 0.576885261 | NULL&M_Disease |
| 11 | 118722946 | rs148990923 | T | C | NULL | 0.576885261 | NULL&M_Disease |
| 11 | 118809805 | rs142069912 | C | G | NULL | 0.576885261 | NULL&M_Disease |
| 11 | 118530869 | rs45447797 | G | C | NULL | 0.575664351 | NULL&M_Disease |
| 11 | 118575419 | rs625735 | A | G | NULL | 0.575664351 | NULL&M_Disease |
| 11 | 118575479 | rs637563 | G | A | NULL | 0.575664351 | NULL&M_Disease |
| 11 | 118575693 | rs638514 | C | T | NULL | 0.575664351 | NULL&M_Disease |
| 11 | 118820778 | rs139076649 | C | T | NULL | 0.575664351 | NULL&M_Disease |
| 11 | 118480285 | rs12225399 | C | G | NULL | 0.574442517 | NULL&M_Disease |
| 11 | 118574311 | rs524409 | T | C | NULL | 0.574442517 | NULL&M_Disease |
| 11 | 118574675 | rs494459 | T | C | NULL | 0.574442517 | NULL&M_Disease |
| 11 | 118580906 | rs496058 | G | A | NULL | 0.574442517 | NULL&M_Disease |
| 11 | 118583971 | rs6589677 | A | G | NULL | 0.574442517 | NULL&M_Disease |
| 11 | 118603615 | rs74783030 | A | G | M_Disease | 0.574442517 | M_Disease&NULL |
| 11 | 118709275 | rs187164533 | A | G | NULL | 0.574442517 | NULL&M_Disease |
| 11 | 118745664 | rs11217076 | G | T | NULL | 0.574442517 | NULL&M_Disease |
| 11 | 118558425 | rs149509653 | A | G | M_Disease | 0.573219773 | M_Disease&NULL |
| 11 | 118659896 | rs2105782 | C | T | M_Disease | 0.573219773 | M_Disease&NULL |
| 11 | 118762521 | rs190509947 | T | A | M_Disease | 0.573219773 | M_Disease&NULL |
| 11 | 118791150 | rs12270280 | T | C | M_Disease | 0.573219773 | M_Disease&NULL |
| 11 | 118580773 | rs675329 | T | C | NULL | 0.571996133 | NULL&M_Disease |
| 11 | 118591443 | rs111843292 | A | C | NULL | 0.571996133 | NULL&M_Disease |
| 11 | 118793479 | rs138526953 | A | C | NULL | 0.571996133 | NULL&M_Disease |
| 11 | 118811878 | rs74891547 | C | G | M_Disease | 0.571996133 | M_Disease&NULL |
| 11 | 118510642 | rs141169729 | T | C | NULL | 0.570771612 | NULL&M_Disease |
| 11 | 118583261 | rs112246697 | A | G | NULL | 0.570771612 | NULL&M_Disease |
| 11 | 118791180 | rs146541591 | C | T | NULL | 0.570771612 | NULL&M_Disease |
| 11 | 118533042 | rs142438429 | T | C | NULL | 0.568319983 | NULL&M_Disease |
| 11 | 118765283 | rs665648 | A | G | M_Disease | 0.568319983 | M_Disease&NULL |
| 11 | 118556995 | rs72999502 | C | T | NULL | 0.567092905 | NULL&M_Disease |
| 11 | 118680072 | rs607472 | G | C | M_Disease | 0.567092905 | M_Disease&NULL |
| 11 | 118696055 | rs189417780 | C | A | M_Disease | 0.567092905 | M_Disease&NULL |
| 11 | 118772160 | rs61730467 | A | G | NULL | 0.567092905 | NULL&M_Disease |
| 11 | 118799691 | rs191378986 | G | A | NULL | 0.567092905 | NULL&M_Disease |
| 11 | 118855789 | rs4938609 | A | C | NULL | 0.567092905 | NULL&M_Disease |
| 11 | 118502121 | rs17614795 | G | A | NULL | 0.565865003 | NULL&M_Disease |
| 11 | 118791532 | rs12271856 | T | G | M_Disease | 0.564636292 | M_Disease&NULL |
| 11 | 118543589 | rs72999499 | G | C | NULL | 0.564636292 | NULL&M_Disease |
| 11 | 118578462 | rs516459 | C | T | NULL | 0.564636292 | NULL&M_Disease |
| 11 | 118590961 | rs111227443 | T | C | NULL | 0.563406786 | NULL&M_Disease |
| 11 | 118854372 | rs75067211 | T | C | NULL | 0.563406786 | NULL&M_Disease |
| 11 | 118496524 | rs35692099 | C | T | NULL | 0.562176501 | NULL&M_Disease |
| 11 | 118539151 | rs72999491 | A | G | NULL | 0.562176501 | NULL&M_Disease |
| 11 | 118744615 | rs73009445 | A | G | NULL | 0.562176501 | NULL&M_Disease |
| 11 | 118812559 | rs74678032 | C | T | NULL | 0.562176501 | NULL&M_Disease |
| 11 | 118607090 | rs10736495 | C | G | NULL | 0.56094545 | NULL&M_Disease |
| 11 | 118615232 | rs139423236 | G | C | NULL | 0.56094545 | NULL&M_Disease |
| 11 | 118694739 | rs188550348 | A | G | NULL | 0.56094545 | NULL&M_Disease |
| 11 | 118805138 | rs7127427 | A | G | M_Disease | 0.56094545 | M_Disease&NULL |
| 11 | 118797562 | rs142336193 | G | A | M_Disease | 0.559713649 | M_Disease&NULL |
| 11 | 118509741 | rs113313560 | T | C | NULL | 0.558481112 | NULL&M_Disease |
| 11 | 118658075 | rs515523 | G | A | M_Disease | 0.558481112 | M_Disease&NULL |
| 11 | 118594326 | rs544096 | G | A | NULL | 0.557247855 | NULL&M_Disease |
| 11 | 118605447 | rs2186781 | T | C | NULL | 0.557247855 | NULL&M_Disease |
| 11 | 118604678 | rs493720 | T | C | NULL | 0.556013891 | NULL&M_Disease |
| 11 | 118644582 | rs2510897 | C | A | M_Disease | 0.556013891 | M_Disease&NULL |
| 11 | 118651809 | rs555649 | C | T | M_Disease | 0.556013891 | M_Disease&NULL |
| 11 | 118711376 | rs118046730 | C | T | M_Disease | 0.554779235 | M_Disease&NULL |
| 11 | 118750967 | rs11602393 | A | T | M_Disease | 0.554779235 | M_Disease&NULL |
| 11 | 118801296 | rs75210328 | T | G | M_Disease | 0.554779235 | M_Disease&NULL |
| 11 | 118801577 | rs77875650 | A | G | M_Disease | 0.554779235 | M_Disease&NULL |
| 11 | 118590743 | rs503542 | G | A | NULL | 0.554779235 | NULL&M_Disease |
| 11 | 118609736 | rs1783937 | A | G | NULL | 0.554779235 | NULL&M_Disease |

| | | | | | | | | | |
|---|---|---|---|---|---|---|---|---|---|
| 11 | 118685481 | rs515707 | T | C | 0.974 | -4813.425 | -4814.727 | -1.302 | 6.74E-06 |
| 11 | 118894287 | rs569 | G | A | 1.029 | -4813.425 | -4814.729 | -1.304 | 6.72E-06 |
| 11 | 118895202 | rs11006 | G | A | 1.029 | -4813.425 | -4814.729 | -1.304 | 6.72E-06 |
| 11 | 118695908 | rs583905 | G | A | 0.974 | -4813.425 | -4814.731 | -1.306 | 6.71E-06 |
| 11 | 118696175 | rs584935 | A | G | 0.974 | -4813.425 | -4814.731 | -1.306 | 6.71E-06 |
| 11 | 118567545 | rs644867 | G | A | 0.968 | -4813.425 | -4814.732 | -1.307 | 6.70E-06 |
| 11 | 118689871 | rs587816 | A | C | 0.974 | -4813.425 | -4814.733 | -1.308 | 6.70E-06 |
| 11 | 118548421 | rs562313 | A | G | 0.968 | -4813.425 | -4814.734 | -1.309 | 6.69E-06 |
| 11 | 118550601 | rs673770 | G | T | 0.968 | -4813.425 | -4814.734 | -1.309 | 6.69E-06 |
| 11 | 118550756 | rs642530 | C | T | 0.968 | -4813.425 | -4814.734 | -1.309 | 6.69E-06 |
| 11 | 118550784 | rs642497 | A | C | 0.968 | -4813.425 | -4814.734 | -1.309 | 6.69E-06 |
| 11 | 118551597 | rs12799314 | A | G | 0.968 | -4813.425 | -4814.734 | -1.309 | 6.69E-06 |
| 11 | 118832035 | rs10892317 | C | T | 1.031 | -4813.425 | -4814.738 | -1.314 | 6.66E-06 |
| 11 | 118567486 | rs644498 | C | T | 0.969 | -4813.425 | -4814.74 | -1.315 | 6.65E-06 |
| 11 | 118877563 | rs58709307 | T | C | 1.029 | -4813.425 | -4814.74 | -1.316 | 6.64E-06 |
| 11 | 118567084 | rs542613 | A | G | 0.969 | -4813.425 | -4814.743 | -1.318 | 6.63E-06 |
| 11 | 118700218 | rs2510902 | A | G | 0.972 | -4813.425 | -4814.743 | -1.318 | 6.63E-06 |
| 11 | 118552420 | rs4936433 | G | A | 0.969 | -4813.425 | -4814.744 | -1.32 | 6.62E-06 |
| 11 | 118552715 | rs519848 | A | G | 0.969 | -4813.425 | -4814.744 | -1.32 | 6.62E-06 |
| 11 | 118552842 | rs549370 | A | C | 0.969 | -4813.425 | -4814.744 | -1.32 | 6.62E-06 |
| 11 | 118552996 | rs654792 | T | G | 0.969 | -4813.425 | -4814.744 | -1.32 | 6.62E-06 |
| 11 | 118553218 | rs592280 | C | T | 0.969 | -4813.425 | -4814.744 | -1.32 | 6.62E-06 |
| 11 | 118554248 | rs7106652 | T | C | 0.969 | -4813.425 | -4814.744 | -1.32 | 6.62E-06 |
| 11 | 118554259 | rs7106851 | A | G | 0.969 | -4813.425 | -4814.744 | -1.32 | 6.62E-06 |
| 11 | 118554436 | rs11605069 | G | T | 0.969 | -4813.425 | -4814.744 | -1.32 | 6.62E-06 |
| 11 | 118554448 | rs11602869 | A | G | 0.969 | -4813.425 | -4814.744 | -1.32 | 6.62E-06 |
| 11 | 118554939 | rs12796373 | A | G | 0.969 | -4813.425 | -4814.744 | -1.32 | 6.62E-06 |
| 11 | 118562767 | rs667982 | C | T | 0.969 | -4813.425 | -4814.745 | -1.32 | 6.62E-06 |
| 11 | 118559709 | rs574217358 | A | G | 0.969 | -4813.425 | -4814.747 | -1.322 | 6.60E-06 |
| 11 | 118559866 | rs7926959 | C | T | 0.969 | -4813.425 | -4814.747 | -1.322 | 6.60E-06 |
| 11 | 118559881 | rs7926970 | G | T | 0.969 | -4813.425 | -4814.747 | -1.322 | 6.60E-06 |
| 11 | 118560569 | rs4938529 | T | C | 0.969 | -4813.425 | -4814.747 | -1.322 | 6.60E-06 |
| 11 | 118560570 | rs4938530 | T | G | 0.969 | -4813.425 | -4814.747 | -1.322 | 6.60E-06 |
| 11 | 118561410 | rs12798453 | G | T | 0.969 | -4813.425 | -4814.747 | -1.322 | 6.60E-06 |
| 11 | 118562799 | rs582630 | A | C | 0.969 | -4813.425 | -4814.747 | -1.322 | 6.60E-06 |
| 11 | 118563381 | rs585039 | A | C | 0.969 | -4813.425 | -4814.747 | -1.322 | 6.60E-06 |
| 11 | 118564413 | rs508640 | G | A | 0.969 | -4813.425 | -4814.747 | -1.322 | 6.60E-06 |
| 11 | 118564601 | rs471227 | C | T | 0.969 | -4813.425 | -4814.747 | -1.322 | 6.60E-06 |
| 11 | 118564762 | rs516153 | A | C | 0.969 | -4813.425 | -4814.747 | -1.322 | 6.60E-06 |
| 11 | 118564982 | rs518028 | C | T | 0.969 | -4813.425 | -4814.747 | -1.322 | 6.60E-06 |
| 11 | 118565231 | rs519982 | T | C | 0.969 | -4813.425 | -4814.747 | -1.322 | 6.60E-06 |
| 11 | 118565336 | rs478371 | A | G | 0.969 | -4813.425 | -4814.747 | -1.322 | 6.60E-06 |
| 11 | 118886656 | rs7131534 | A | G | 1.028 | -4813.425 | -4814.746 | -1.322 | 6.60E-06 |
| 11 | 118554100 | rs4331144 | G | A | 0.969 | -4813.425 | -4814.748 | -1.323 | 6.60E-06 |
| 11 | 118556084 | rs613377 | T | C | 0.969 | -4813.425 | -4814.75 | -1.325 | 6.58E-06 |
| 11 | 118682462 | rs523148 | T | C | 0.976 | -4813.425 | -4814.752 | -1.328 | 6.56E-06 |
| 11 | 118556691 | rs610664 | G | A | 0.969 | -4813.425 | -4814.753 | -1.329 | 6.56E-06 |
| 11 | 118556797 | rs571751 | C | A | 0.969 | -4813.425 | -4814.753 | -1.329 | 6.56E-06 |
| 11 | 118557198 | rs546890 | C | T | 0.969 | -4813.425 | -4814.753 | -1.329 | 6.56E-06 |
| 11 | 118558333 | rs693306 | G | A | 0.969 | -4813.425 | -4814.753 | -1.329 | 6.56E-06 |
| 11 | 118558455 | rs558907 | G | A | 0.969 | -4813.425 | -4814.753 | -1.329 | 6.56E-06 |
| 11 | 118558827 | rs561845 | G | A | 0.969 | -4813.425 | -4814.753 | -1.329 | 6.56E-06 |
| 11 | 118558889 | rs638805 | T | G | 0.969 | -4813.425 | -4814.753 | -1.329 | 6.56E-06 |
| 11 | 118558901 | rs510408 | C | G | 0.969 | -4813.425 | -4814.753 | -1.329 | 6.56E-06 |
| 11 | 118558952 | rs509641 | C | T | 0.969 | -4813.425 | -4814.753 | -1.329 | 6.56E-06 |
| 11 | 118558953 | rs563549 | A | G | 0.969 | -4813.425 | -4814.753 | -1.329 | 6.56E-06 |
| 11 | 118559003 | rs638301 | G | A | 0.969 | -4813.425 | -4814.753 | -1.329 | 6.56E-06 |
| 11 | 118559084 | rs508579 | C | G | 0.969 | -4813.425 | -4814.753 | -1.329 | 6.56E-06 |
| 11 | 118559178 | rs486462 | C | T | 0.969 | -4813.425 | -4814.753 | -1.329 | 6.56E-06 |
| 11 | 118559247 | rs533144 | A | C | 0.969 | -4813.425 | -4814.753 | -1.329 | 6.56E-06 |
| 11 | 118559542 | rs625918 | T | C | 0.969 | -4813.425 | -4814.753 | -1.329 | 6.56E-06 |
| 11 | 118562520 | rs558593 | T | C | 0.969 | -4813.425 | -4814.753 | -1.329 | 6.56E-06 |
| 11 | 118565679 | rs502601 | A | G | 0.969 | -4813.425 | -4814.753 | -1.329 | 6.56E-06 |
| 11 | 118566090 | rs7130076 | C | T | 0.969 | -4813.425 | -4814.753 | -1.329 | 6.56E-06 |
| 11 | 118566175 | rs7114458 | A | G | 0.969 | -4813.425 | -4814.753 | -1.329 | 6.56E-06 |
| 11 | 118566234 | rs7114216 | G | A | 0.969 | -4813.425 | -4814.753 | -1.329 | 6.56E-06 |
| 11 | 118566313 | rs12794726 | T | C | 0.969 | -4813.425 | -4814.753 | -1.329 | 6.56E-06 |
| 11 | 118566746 | rs533646 | G | C | 0.969 | -4813.425 | -4814.753 | -1.329 | 6.56E-06 |
| 11 | 118879285 | rs7120445 | C | A | 1.028 | -4813.425 | -4814.753 | -1.329 | 6.56E-06 |
| 11 | 118879824 | rs73559119 | C | T | 1.028 | -4813.425 | -4814.753 | -1.329 | 6.56E-06 |
| 11 | 118880012 | rs11825284 | A | G | 1.028 | -4813.425 | -4814.753 | -1.329 | 6.56E-06 |
| 11 | 118558078 | rs555356 | T | G | 0.969 | -4813.425 | -4814.755 | -1.331 | 6.54E-06 |
| 11 | 118762701 | rs76457018 | A | G | 0.989 | -4813.425 | -4814.756 | -1.331 | 6.54E-06 |
| 11 | 118545797 | rs600969 | T | C | 0.969 | -4813.425 | -4814.757 | -1.332 | 6.54E-06 |
| 11 | 118585146 | rs112799932 | G | T | 0.971 | -4813.425 | -4814.758 | -1.334 | 6.53E-06 |
| 11 | 118594669 | rs140026975 | C | A | 0.971 | -4813.425 | -4814.759 | -1.334 | 6.53E-06 |
| 11 | 118627149 | rs11217016 | C | A | 0.971 | -4813.425 | -4814.76 | -1.335 | 6.52E-06 |
| 11 | 118567020 | rs642662 | G | A | 0.969 | -4813.425 | -4814.762 | -1.337 | 6.51E-06 |
| 11 | 118552074 | rs4938524 | T | G | 0.97 | -4813.425 | -4814.764 | -1.339 | 6.49E-06 |
| 11 | 118629364 | rs73001486 | A | G | 0.971 | -4813.425 | -4814.764 | -1.34 | 6.49E-06 |
| 11 | 118700425 | rs10128600 | C | T | 0.977 | -4813.425 | -4814.767 | -1.342 | 6.47E-06 |
| 11 | 118895686 | rs8192696 | T | C | 1.026 | -4813.425 | -4814.768 | -1.343 | 6.47E-06 |
| 11 | 118566039 | rs6589674 | G | A | 0.97 | -4813.425 | -4814.773 | -1.348 | 6.43E-06 |
| 11 | 118564397 | rs558781 | T | G | 0.97 | -4813.425 | -4814.775 | -1.35 | 6.42E-06 |
| 11 | 118567444 | rs493834 | G | A | 0.97 | -4813.425 | -4814.776 | -1.351 | 6.42E-06 |
| 11 | 118701898 | rs533254 | A | T | 0.979 | -4813.425 | -4814.778 | -1.353 | 6.40E-06 |
| 11 | 118555719 | rs625513 | C | A | 0.97 | -4813.425 | -4814.781 | -1.356 | 6.38E-06 |
| 11 | 118555754 | rs525485 | A | G | 0.97 | -4813.425 | -4814.781 | -1.356 | 6.38E-06 |
| 11 | 118558449 | rs514143 | A | G | 0.97 | -4813.425 | -4814.781 | -1.356 | 6.38E-06 |
| 11 | 118702936 | rs7941696 | G | A | 0.978 | -4813.425 | -4814.781 | -1.356 | 6.38E-06 |
| 11 | 118622985 | rs3825057 | C | T | 0.972 | -4813.425 | -4814.786 | -1.361 | 6.35E-06 |
| 11 | 118553326 | rs693037 | T | C | 0.971 | -4813.425 | -4814.788 | -1.363 | 6.34E-06 |
| 11 | 118610463 | rs4499035 | T | C | 0.973 | -4813.425 | -4814.791 | -1.366 | 6.32E-06 |
| 11 | 118700264 | rs113641235 | T | C | 0.98 | -4813.425 | -4814.795 | -1.37 | 6.29E-06 |
| 11 | 118610826 | rs1893029 | T | G | 0.973 | -4813.425 | -4814.797 | -1.372 | 6.28E-06 |
| 11 | 118616253 | rs7124498 | A | G | 0.973 | -4813.425 | -4814.801 | -1.377 | 6.25E-06 |
| 11 | 118567375 | rs478296 | C | T | 0.971 | -4813.425 | -4814.806 | -1.382 | 6.22E-06 |
| 11 | 118704255 | rs480791 | G | A | 0.982 | -4813.425 | -4814.809 | -1.385 | 6.20E-06 |
| 11 | 118704265 | rs622690 | T | C | 0.982 | -4813.425 | -4814.809 | -1.385 | 6.20E-06 |
| 11 | 118699744 | rs2508564 | C | T | 0.981 | -4813.425 | -4814.811 | -1.386 | 6.19E-06 |
| 11 | 118697375 | rs73005500 | C | T | 0.982 | -4813.425 | -4814.812 | -1.388 | 6.18E-06 |
| 11 | 118626286 | rs11217014 | C | T | 0.974 | -4813.425 | -4814.817 | -1.393 | 6.15E-06 |
| 11 | 118627778 | rs10892282 | C | G | 0.974 | -4813.425 | -4814.817 | -1.393 | 6.15E-06 |

| | | | | | | | |
|---|---|---|---|---|---|---|---|
| 11 | 118539746 | rs72999492 | C | A | NULL | 0.553543903 | NULL&M_Disease |
| 11 | 118557483 | rs6589673 | C | A | NULL | 0.553543903 | NULL&M_Disease |
| 11 | 118599649 | rs182364290 | G | A | NULL | 0.553543903 | NULL&M_Disease |
| 11 | 118605127 | rs526850 | G | T | NULL | 0.553543903 | NULL&M_Disease |
| 11 | 118620697 | rs488219 | C | T | M_Disease | 0.553543903 | M_Disease&NULL |
| 11 | 118629031 | rs555639 | A | G | M_Disease | 0.553543903 | M_Disease&NULL |
| 11 | 118632522 | rs581045 | T | C | M_Disease | 0.553543903 | M_Disease&NULL |
| 11 | 118788769 | rs186418307 | T | G | NULL | 0.553543903 | NULL&M_Disease |
| 11 | 118792384 | rs182691948 | T | G | NULL | 0.553543903 | NULL&M_Disease |
| 11 | 118797857 | rs150438978 | T | A | NULL | 0.553543903 | NULL&M_Disease |
| 11 | 118559426 | rs10892253 | A | C | NULL | 0.55230791 | NULL&M_Disease |
| 11 | 118561858 | rs10790257 | C | G | NULL | 0.55230791 | NULL&M_Disease |
| 11 | 118561859 | rs10790258 | T | G | NULL | 0.55230791 | NULL&M_Disease |
| 11 | 118610629 | rs521910 | A | G | NULL | 0.55230791 | NULL&M_Disease |
| 11 | 118615738 | rs193090156 | T | G | NULL | 0.55230791 | NULL&M_Disease |
| 11 | 118757788 | rs3176912 | A | G | NULL | 0.55230791 | NULL&M_Disease |
| 11 | 118572000 | rs182361280 | A | G | NULL | 0.551071269 | NULL&M_Disease |
| 11 | 118624250 | rs546393 | C | T | M_Disease | 0.551071269 | M_Disease&NULL |
| 11 | 118804266 | rs7123066 | T | C | M_Disease | 0.551071269 | M_Disease&NULL |
| 11 | 118804283 | rs7122959 | G | A | M_Disease | 0.551071269 | M_Disease&NULL |
| 11 | 118583712 | rs118008100 | A | G | NULL | 0.549833997 | NULL&M_Disease |
| 11 | 118605352 | rs76478216 | C | T | M_Disease | 0.549833997 | M_Disease&NULL |
| 11 | 118662028 | rs12796102 | T | C | M_Disease | 0.549833997 | M_Disease&NULL |
| 11 | 118748170 | rs77074463 | C | G | M_Disease | 0.549833997 | M_Disease&NULL |
| 11 | 118798757 | rs57879605 | T | C | NULL | 0.549833997 | NULL&M_Disease |
| 11 | 118607694 | rs7119702 | G | A | NULL | 0.548596109 | NULL&M_Disease |
| 11 | 118503912 | rs45478298 | T | C | NULL | 0.547357618 | NULL&M_Disease |
| 11 | 118748789 | rs145762171 | A | G | NULL | 0.547357618 | NULL&M_Disease |
| 11 | 118445559 | rs73555296 | T | G | NULL | 0.546118541 | NULL&M_Disease |
| 11 | 118601363 | rs557990 | T | C | NULL | 0.546118541 | NULL&M_Disease |
| 11 | 118602392 | rs564892 | A | T | NULL | 0.546118541 | NULL&M_Disease |
| 11 | 118666978 | rs1289369859 | A | G | M_Disease | 0.544878892 | M_Disease&NULL |
| 11 | 118477746 | rs72999445 | G | T | M_Disease | 0.543638687 | M_Disease&NULL |
| 11 | 118483912 | rs45621436 | G | T | NULL | 0.543638687 | NULL&M_Disease |
| 11 | 118493319 | rs10790254 | A | G | NULL | 0.543638687 | NULL&M_Disease |
| 11 | 118518753 | rs117947426 | T | C | NULL | 0.543638687 | NULL&M_Disease |
| 11 | 118664181 | rs145462967 | A | G | NULL | 0.542397941 | NULL&M_Disease |
| 11 | 118617299 | rs607125 | C | T | M_Disease | 0.541156668 | M_Disease&NULL |
| 11 | 118631288 | rs507890 | C | T | M_Disease | 0.541156668 | M_Disease&NULL |
| 11 | 118631653 | rs1784543 | T | A | M_Disease | 0.541156668 | M_Disease&NULL |
| 11 | 118684537 | rs150583208 | G | A | M_Disease | 0.541156668 | M_Disease&NULL |
| 11 | 118746769 | rs4938576 | T | G | M_Disease | 0.541156668 | M_Disease&NULL |
| 11 | 118805385 | rs55772432 | A | G | NULL | 0.541156668 | NULL&M_Disease |
| 11 | 118673968 | rs140269932 | A | G | NULL | 0.539914885 | NULL&M_Disease |
| 11 | 118715589 | rs116872113 | A | G | NULL | 0.539914885 | NULL&M_Disease |
| 11 | 118781710 | rs6589707 | A | C | M_Disease | 0.539914885 | M_Disease&NULL |
| 11 | 118519213 | rs668628 | T | G | NULL | 0.538672605 | NULL&M_Disease |
| 11 | 118576997 | rs498912 | C | T | NULL | 0.538672605 | NULL&M_Disease |
| 11 | 118777978 | rs78755429 | A | C | NULL | 0.538672605 | NULL&M_Disease |
| 11 | 118782802 | rs76160025 | T | C | NULL | 0.538672605 | NULL&M_Disease |
| 11 | 118811315 | rs77638281 | A | G | NULL | 0.537429845 | NULL&M_Disease |
| 11 | 118523799 | rs72999483 | C | T | M_Disease | 0.53618662 | M_Disease&NULL |
| 11 | 118759905 | rs118020807 | A | C | NULL | 0.53618662 | NULL&M_Disease |
| 11 | 118727137 | rs117223320 | C | G | NULL | 0.534942945 | NULL&M_Disease |
| 11 | 118747008 | rs75040111 | T | C | M_Disease | 0.533698835 | M_Disease&NULL |
| 11 | 118576762 | rs553188 | T | C | NULL | 0.533698835 | NULL&M_Disease |
| 11 | 118587349 | rs11216975 | C | A | NULL | 0.531209373 | NULL&M_Disease |
| 11 | 118787213 | rs7119222 | C | T | M_Disease | 0.531209373 | M_Disease&NULL |
| 11 | 118789026 | rs11607239 | T | G | NULL | 0.531209373 | NULL&M_Disease |
| 11 | 118466646 | rs184841209 | T | G | NULL | 0.529964052 | NULL&M_Disease |
| 11 | 118689745 | rs492471 | T | C | M_Disease | 0.529964052 | M_Disease&NULL |
| 11 | 118691070 | rs140577174 | G | A | NULL | 0.529964052 | NULL&M_Disease |
| 11 | 118787897 | rs11602400 | C | A | M_Disease | 0.529964052 | M_Disease&NULL |
| 11 | 118455465 | rs45460493 | A | G | NULL | 0.528718357 | NULL&M_Disease |
| 11 | 118477226 | rs45496391 | C | A | NULL | 0.528718357 | NULL&M_Disease |
| 11 | 118626825 | rs74337439 | A | G | M_Disease | 0.527472304 | M_Disease&NULL |
| 11 | 118657311 | rs1309473877 | A | C | M_Disease | 0.527472304 | M_Disease&NULL |
| 11 | 118761159 | rs57231586 | T | G | M_Disease | 0.527472304 | M_Disease&NULL |
| 11 | 118764051 | rs613791 | T | C | NULL | 0.527472304 | NULL&M_Disease |
| 11 | 118576463 | rs496547 | A | T | M_Disease | 0.526225909 | M_Disease&NULL |
| 11 | 118781530 | rs7926599 | C | G | M_Disease | 0.526225909 | M_Disease&NULL |
| 11 | 118421841 | rs145968026 | G | C | NULL | 0.524979187 | NULL&M_Disease |
| 11 | 118507068 | rs10892249 | C | T | NULL | 0.524979187 | NULL&M_Disease |
| 11 | 118518021 | rs10736493 | G | T | NULL | 0.524979187 | NULL&M_Disease |
| 11 | 118786829 | rs7102433 | G | A | M_Disease | 0.523732154 | M_Disease&NULL |
| 11 | 118619478 | rs189583461 | G | A | NULL | 0.523732154 | NULL&M_Disease |
| 11 | 118621277 | rs117409833 | C | T | NULL | 0.523732154 | NULL&M_Disease |
| 11 | 118460957 | rs45600633 | A | G | NULL | 0.522484825 | NULL&M_Disease |
| 11 | 118474283 | rs139202987 | G | A | NULL | 0.522484825 | NULL&M_Disease |
| 11 | 118487265 | rs76544852 | T | G | NULL | 0.522484825 | NULL&M_Disease |
| 11 | 118732181 | rs117079831 | A | G | NULL | 0.521237215 | NULL&M_Disease |
| 11 | 118792278 | rs77398170 | G | A | M_Disease | 0.521237215 | M_Disease&NULL |
| 11 | 118735876 | rs79843382 | T | C | NULL | 0.518741216 | NULL&M_Disease |
| 11 | 118598612 | rs568688 | G | T | NULL | 0.517492858 | NULL&M_Disease |
| 11 | 118588998 | rs570424 | C | T | M_Disease | 0.516244281 | M_Disease&NULL |
| 11 | 118803536 | rs615166 | A | G | NULL | 0.516244281 | NULL&M_Disease |
| 11 | 118805144 | rs11605265 | A | G | NULL | 0.516244281 | NULL&M_Disease |
| 11 | 118805220 | rs11605658 | T | C | NULL | 0.516244281 | NULL&M_Disease |
| 11 | 118819541 | rs150196102 | T | C | M_Disease | 0.516244281 | M_Disease&NULL |
| 11 | 118615390 | rs78502545 | T | C | M_Disease | 0.514995502 | M_Disease&NULL |
| 11 | 118721562 | rs184045098 | T | C | NULL | 0.514995502 | NULL&M_Disease |
| 11 | 118598138 | rs543692 | C | T | NULL | 0.513746535 | NULL&M_Disease |
| 11 | 118721767 | rs116884018 | C | T | M_Disease | 0.513746535 | M_Disease&NULL |
| 11 | 118759000 | rs77491736 | G | C | NULL | 0.513746535 | NULL&M_Disease |
| 11 | 118761591 | rs58397111 | T | C | M_Disease | 0.513746535 | M_Disease&NULL |
| 11 | 118762102 | rs58508419 | A | G | M_Disease | 0.513746535 | M_Disease&NULL |
| 11 | 118849368 | rs45491601 | A | G | M_Disease | 0.513746535 | M_Disease&NULL |
| 11 | 118622744 | rs6589689 | G | A | M_Disease | 0.512497396 | M_Disease&NULL |
| 11 | 118789657 | rs12294466 | T | C | M_Disease | 0.512497396 | M_Disease&NULL |
| 11 | 118782474 | rs11606719 | G | C | M_Disease | 0.511248102 | M_Disease&NULL |
| 11 | 118577614 | rs947931 | G | T | NULL | 0.509998667 | NULL&M_Disease |
| 11 | 118608625 | rs566733 | G | C | M_Disease | 0.509998667 | M_Disease&NULL |
| 11 | 118619948 | rs112143902 | T | C | M_Disease | 0.509998667 | M_Disease&NULL |

| | | | | | | | | | |
|---|---|---|---|---|---|---|---|---|---|
| 11 | 118699863 | rs2510901 | A | G | 0.982 | -4813.425 | -4814.818 | -1.393 | 6.15E-06 |
| 11 | 118702910 | rs7941321 | A | G | 0.983 | -4813.425 | -4814.817 | -1.393 | 6.15E-06 |
| 11 | 118703001 | rs7928228 | T | C | 0.983 | -4813.425 | -4814.817 | -1.393 | 6.15E-06 |
| 11 | 118703484 | rs2508921 | T | C | 0.983 | -4813.425 | -4814.817 | -1.393 | 6.15E-06 |
| 11 | 118703817 | rs1170396 | T | A | 0.983 | -4813.425 | -4814.817 | -1.393 | 6.15E-06 |
| 11 | 118831335 | rs61506726 | G | A | 1.027 | -4813.425 | -4814.818 | -1.393 | 6.15E-06 |
| 11 | 118698160 | rs640373 | G | A | 0.982 | -4813.425 | -4814.82 | -1.395 | 6.14E-06 |
| 11 | 118698274 | rs639915 | A | G | 0.982 | -4813.425 | -4814.82 | -1.395 | 6.14E-06 |
| 11 | 118698294 | rs639885 | A | G | 0.982 | -4813.425 | -4814.82 | -1.395 | 6.14E-06 |
| 11 | 118698537 | rs638893 | G | A | 0.982 | -4813.425 | -4814.82 | -1.395 | 6.14E-06 |
| 11 | 118698777 | rs516332 | G | A | 0.982 | -4813.425 | -4814.82 | -1.395 | 6.14E-06 |
| 11 | 118539140 | rs4589335 | T | C | 0.972 | -4813.425 | -4814.822 | -1.397 | 6.13E-06 |
| 11 | 118613674 | rs7130875 | C | T | 0.975 | -4813.425 | -4814.825 | -1.401 | 6.10E-06 |
| 11 | 118701597 | rs4938549 | T | C | 0.984 | -4813.425 | -4814.826 | -1.401 | 6.10E-06 |
| 11 | 118701668 | rs530646 | A | G | 0.984 | -4813.425 | -4814.826 | -1.401 | 6.10E-06 |
| 11 | 118702002 | rs534114 | A | G | 0.984 | -4813.425 | -4814.826 | -1.401 | 6.10E-06 |
| 11 | 118609598 | rs112996178 | T | C | 0.975 | -4813.425 | -4814.827 | -1.402 | 6.10E-06 |
| 11 | 118624919 | rs11217013 | C | G | 0.975 | -4813.425 | -4814.835 | -1.41 | 6.05E-06 |
| 11 | 118705654 | rs649338 | A | G | 0.985 | -4813.425 | -4814.834 | -1.41 | 6.05E-06 |
| 11 | 118726340 | rs113786535 | T | C | 0.985 | -4813.425 | -4814.839 | -1.414 | 6.02E-06 |
| 11 | 118705145 | rs636736 | G | A | 0.985 | -4813.425 | -4814.84 | -1.415 | 6.02E-06 |
| 11 | 118538308 | rs692750 | T | G | 0.973 | -4813.425 | -4814.841 | -1.417 | 6.01E-06 |
| 11 | 118880740 | rs11217122 | C | T | 1.022 | -4813.425 | -4814.845 | -1.42 | 5.99E-06 |
| 11 | 118515616 | rs527290 | T | C | 0.974 | -4813.425 | -4814.852 | -1.427 | 5.95E-06 |
| 11 | 118520480 | rs653008 | T | G | 0.974 | -4813.425 | -4814.857 | -1.432 | 5.92E-06 |
| 11 | 118640828 | rs11217021 | C | T | 0.977 | -4813.425 | -4814.859 | -1.434 | 5.90E-06 |
| 11 | 118548665 | rs1784298 | C | T | 0.974 | -4813.425 | -4814.86 | -1.435 | 5.90E-06 |
| 11 | 118548671 | rs4938522 | T | A | 0.974 | -4813.425 | -4814.86 | -1.435 | 5.90E-06 |
| 11 | 118548838 | rs4938523 | T | G | 0.974 | -4813.425 | -4814.86 | -1.435 | 5.90E-06 |
| 11 | 118549582 | rs507080 | A | G | 0.974 | -4813.425 | -4814.86 | -1.435 | 5.90E-06 |
| 11 | 118549803 | rs659969 | T | C | 0.974 | -4813.425 | -4814.86 | -1.435 | 5.90E-06 |
| 11 | 118549908 | rs656575 | G | T | 0.974 | -4813.425 | -4814.86 | -1.435 | 5.90E-06 |
| 11 | 118519550 | rs601318 | G | C | 0.974 | -4813.425 | -4814.862 | -1.438 | 5.88E-06 |
| 11 | 118633533 | rs7110526 | C | T | 0.977 | -4813.425 | -4814.863 | -1.438 | 5.88E-06 |
| 11 | 118635364 | rs11217018 | C | T | 0.977 | -4813.425 | -4814.863 | -1.438 | 5.88E-06 |
| 11 | 118637475 | rs112910608 | T | C | 0.977 | -4813.425 | -4814.863 | -1.438 | 5.88E-06 |
| 11 | 118707171 | rs4938550 | G | A | 0.993 | -4813.425 | -4814.871 | -1.447 | 5.83E-06 |
| 11 | 118549985 | rs543327 | T | G | 0.975 | -4813.425 | -4814.873 | -1.448 | 5.82E-06 |
| 11 | 118550017 | rs543413 | C | T | 0.975 | -4813.425 | -4814.873 | -1.448 | 5.82E-06 |
| 11 | 118755738 | rs523604 | G | A | 1.027 | -4813.425 | -4814.881 | -1.456 | 5.78E-06 |
| 11 | 118657297 | rs487177 | G | C | 0.979 | -4813.425 | -4814.883 | -1.459 | 5.76E-06 |
| 11 | 118655356 | rs7934890 | T | C | 0.979 | -4813.425 | -4814.893 | -1.469 | 5.70E-06 |
| 11 | 118573519 | rs4639966 | C | T | 0.98 | -4813.425 | -4814.901 | -1.477 | 5.66E-06 |
| 11 | 118538187 | rs607527 | T | G | 0.977 | -4813.425 | -4814.91 | -1.486 | 5.61E-06 |
| 11 | 118546602 | rs526597 | G | A | 0.977 | -4813.425 | -4814.913 | -1.488 | 5.59E-06 |
| 11 | 118539566 | rs1783935 | T | A | 0.977 | -4813.425 | -4814.92 | -1.495 | 5.55E-06 |
| 11 | 118568639 | rs511451 | T | C | 0.978 | -4813.425 | -4814.923 | -1.499 | 5.53E-06 |
| 11 | 118538157 | rs520543 | A | G | 0.977 | -4813.425 | -4814.925 | -1.5 | 5.53E-06 |
| 11 | 118590864 | rs11216982 | A | G | 0.981 | -4813.425 | -4814.927 | -1.502 | 5.52E-06 |
| 11 | 118539522 | rs2508991 | A | G | 0.978 | -4813.425 | -4814.931 | -1.506 | 5.49E-06 |
| 11 | 118572611 | rs611217 | T | C | 0.978 | -4813.425 | -4814.932 | -1.507 | 5.49E-06 |
| 11 | 118569097 | rs4393358 | T | C | 0.979 | -4813.425 | -4814.942 | -1.517 | 5.43E-06 |
| 11 | 118668296 | rs58868870 | T | C | 0.984 | -4813.425 | -4814.943 | -1.519 | 5.42E-06 |
| 11 | 118570919 | rs642224 | A | G | 0.979 | -4813.425 | -4814.946 | -1.521 | 5.41E-06 |
| 11 | 118571023 | rs541631 | G | A | 0.979 | -4813.425 | -4814.946 | -1.521 | 5.41E-06 |
| 11 | 118571123 | rs658676 | T | C | 0.979 | -4813.425 | -4814.946 | -1.521 | 5.41E-06 |
| 11 | 118571168 | rs488141 | G | A | 0.979 | -4813.425 | -4814.946 | -1.521 | 5.41E-06 |
| 11 | 118571316 | rs657769 | T | C | 0.979 | -4813.425 | -4814.946 | -1.521 | 5.41E-06 |
| 11 | 118522179 | rs1171461 | A | G | 0.979 | -4813.425 | -4814.948 | -1.524 | 5.40E-06 |
| 11 | 118572021 | rs657539 | T | C | 0.979 | -4813.425 | -4814.948 | -1.524 | 5.40E-06 |
| 11 | 118539170 | rs4457789 | T | G | 0.979 | -4813.425 | -4814.951 | -1.526 | 5.39E-06 |
| 11 | 118568668 | rs11216949 | T | C | 0.979 | -4813.425 | -4814.951 | -1.526 | 5.39E-06 |
| 11 | 118666289 | rs56104144 | C | G | 0.985 | -4813.425 | -4814.955 | -1.531 | 5.36E-06 |
| 11 | 118573668 | rs574383 | C | T | 0.98 | -4813.425 | -4814.959 | -1.534 | 5.34E-06 |
| 11 | 118605451 | rs2186780 | T | C | 0.997 | -4813.425 | -4814.958 | -1.534 | 5.34E-06 |
| 11 | 118605914 | rs7926282 | C | A | 0.998 | -4813.425 | -4814.96 | -1.535 | 5.34E-06 |
| 11 | 118512162 | rs570836 | A | G | 0.98 | -4813.425 | -4814.961 | -1.536 | 5.33E-06 |
| 11 | 118573743 | rs519294 | A | C | 0.98 | -4813.425 | -4814.962 | -1.538 | 5.32E-06 |
| 11 | 118757191 | rs1790192 | G | A | 1.022 | -4813.425 | -4814.963 | -1.538 | 5.32E-06 |
| 11 | 118528466 | rs7389 | G | T | 0.98 | -4813.425 | -4814.965 | -1.54 | 5.31E-06 |
| 11 | 118570914 | rs659564 | A | C | 0.98 | -4813.425 | -4814.965 | -1.541 | 5.31E-06 |
| 11 | 118571357 | rs544452 | T | G | 0.98 | -4813.425 | -4814.965 | -1.541 | 5.31E-06 |
| 11 | 118571911 | rs570949 | T | G | 0.98 | -4813.425 | -4814.965 | -1.541 | 5.31E-06 |
| 11 | 118572171 | rs658040 | G | A | 0.98 | -4813.425 | -4814.965 | -1.541 | 5.31E-06 |
| 11 | 118572760 | rs611733 | T | C | 0.98 | -4813.425 | -4814.965 | -1.541 | 5.31E-06 |
| 11 | 118573022 | rs552079 | C | T | 0.98 | -4813.425 | -4814.965 | -1.541 | 5.31E-06 |
| 11 | 118573339 | rs673547 | A | G | 0.98 | -4813.425 | -4814.965 | -1.541 | 5.31E-06 |
| 11 | 118573535 | rs517445 | G | A | 0.98 | -4813.425 | -4814.965 | -1.541 | 5.31E-06 |
| 11 | 118568813 | rs7479114 | T | C | 0.981 | -4813.425 | -4814.968 | -1.543 | 5.29E-06 |
| 11 | 118569067 | rs4486664 | C | A | 0.981 | -4813.425 | -4814.968 | -1.543 | 5.29E-06 |
| 11 | 118569469 | rs7123811 | G | T | 0.981 | -4813.425 | -4814.968 | -1.543 | 5.29E-06 |
| 11 | 118569694 | rs7108144 | A | G | 0.981 | -4813.425 | -4814.968 | -1.543 | 5.29E-06 |
| 11 | 118569900 | rs12793594 | C | T | 0.981 | -4813.425 | -4814.968 | -1.543 | 5.29E-06 |
| 11 | 118570062 | rs554456 | C | T | 0.981 | -4813.425 | -4814.968 | -1.543 | 5.29E-06 |
| 11 | 118570098 | rs482889 | T | G | 0.981 | -4813.425 | -4814.968 | -1.543 | 5.29E-06 |
| 11 | 118570697 | rs523793 | T | C | 0.981 | -4813.425 | -4814.968 | -1.543 | 5.29E-06 |
| 11 | 118506402 | rs483283 | C | T | 0.98 | -4813.425 | -4814.969 | -1.544 | 5.29E-06 |
| 11 | 118572747 | rs489126 | A | G | 0.981 | -4813.425 | -4814.971 | -1.547 | 5.27E-06 |
| 11 | 118522338 | rs4938520 | G | A | 0.981 | -4813.425 | -4814.975 | -1.55 | 5.26E-06 |
| 11 | 118492591 | rs2077173 | A | G | 0.981 | -4813.425 | -4814.98 | -1.555 | 5.23E-06 |
| 11 | 118568472 | rs570371 | G | A | 0.982 | -4813.425 | -4814.98 | -1.555 | 5.23E-06 |
| 11 | 118571790 | rs656287 | G | A | 0.982 | -4813.425 | -4814.981 | -1.556 | 5.23E-06 |
| 11 | 118529595 | rs745663 | G | A | 0.982 | -4813.425 | -4814.989 | -1.564 | 5.18E-06 |
| 11 | 118505090 | rs645637 | A | G | 0.982 | -4813.425 | -4814.995 | -1.57 | 5.15E-06 |
| 11 | 118496066 | rs11603616 | A | G | 0.982 | -4813.425 | -4814.996 | -1.571 | 5.15E-06 |
| 11 | 118895495 | rs8301 | G | C | 1.007 | -4813.425 | -4814.995 | -1.571 | 5.15E-06 |
| 11 | 118497402 | rs12419204 | A | T | 0.983 | -4813.425 | -4815.002 | -1.578 | 5.11E-06 |
| 11 | 118500173 | rs571001 | T | C | 0.983 | -4813.425 | -4815.002 | -1.578 | 5.11E-06 |
| 11 | 118500193 | rs570952 | C | T | 0.983 | -4813.425 | -4815.002 | -1.578 | 5.11E-06 |
| 11 | 118486325 | rs12290522 | T | C | 0.983 | -4813.425 | -4815.009 | -1.584 | 5.08E-06 |
| 11 | 118487673 | rs4938517 | C | T | 0.983 | -4813.425 | -4815.009 | -1.584 | 5.08E-06 |
| 11 | 118758089 | rs497916 | T | C | 0.993 | -4813.425 | -4815.032 | -1.607 | 4.97E-06 |
| 11 | 118750321 | rs11217078 | C | T | 1.014 | -4813.425 | -4815.052 | -1.627 | 4.87E-06 |

| | | | | | | | |
|---|---|---|---|---|---|---|---|
| 11 | 118595207 | rs563537 | A | T | NULL | 0.508749107 | NULL&M_Disease |
| 11 | 118607622 | rs11217004 | A | G | NULL | 0.508749107 | NULL&M_Disease |
| 11 | 118765087 | rs2230321 | T | C | NULL | 0.508749107 | NULL&M_Disease |
| 11 | 118803842 | rs11600378 | G | A | NULL | 0.508749107 | NULL&M_Disease |
| 11 | 118804671 | rs143711539 | A | G | NULL | 0.508749107 | NULL&M_Disease |
| 11 | 118613235 | rs632124 | A | T | M_Disease | 0.507499438 | M_Disease&NULL |
| 11 | 118749318 | rs78312641 | A | T | M_Disease | 0.507499438 | M_Disease&NULL |
| 11 | 118749563 | rs77643000 | C | T | M_Disease | 0.507499438 | M_Disease&NULL |
| 11 | 118650163 | rs73003218 | T | C | NULL | 0.506249674 | NULL&M_Disease |
| 11 | 118779681 | rs9971606 | C | T | NULL | 0.506249674 | NULL&M_Disease |
| 11 | 118803193 | rs117235696 | G | T | NULL | 0.506249674 | NULL&M_Disease |
| 11 | 118832251 | rs79808604 | A | C | M_Disease | 0.506249674 | M_Disease&NULL |
| 11 | 118530434 | rs117619140 | T | G | NULL | 0.504999833 | NULL&M_Disease |
| 11 | 118801235 | rs602716 | A | G | NULL | 0.504999833 | NULL&M_Disease |
| 11 | 118812512 | rs647769 | T | C | NULL | 0.504999833 | NULL&M_Disease |
| 11 | 118816304 | rs586763 | T | C | M_Disease | 0.504999833 | M_Disease&NULL |
| 11 | 118419577 | rs191458787 | T | C | NULL | 0.50374993 | NULL&M_Disease |
| 11 | 118421438 | rs145485933 | G | A | NULL | 0.50374993 | NULL&M_Disease |
| 11 | 118501723 | rs61900957 | T | C | NULL | 0.50374993 | NULL&M_Disease |
| 11 | 118608532 | rs567680 | G | C | NULL | 0.50374993 | NULL&M_Disease |
| 11 | 118802492 | rs688035 | G | C | M_Disease | 0.50374993 | M_Disease&NULL |
| 11 | 118413882 | rs146864992 | T | C | M_Disease | 0.502499979 | M_Disease&NULL |
| 11 | 118798780 | rs77916839 | C | G | M_Disease | 0.502499979 | M_Disease&NULL |
| 11 | 118811684 | rs550999 | G | C | NULL | 0.502499979 | NULL&M_Disease |
| 11 | 118468037 | rs535774 | T | C | NULL | 0.501249997 | NULL&M_Disease |
| 11 | 118672061 | rs183981091 | T | G | M_Disease | 0.501249997 | M_Disease&NULL |
| 11 | 118764117 | rs73575405 | A | G | NULL | 0.501249997 | NULL&M_Disease |

| | | | | | | | | | |
|---|---|---|---|---|---|---|---|---|---|
| 11 | 118825636 | rs1790191 | G | A | 1.018 | -4813.425 | -4815.052 | -1.627 | 4.87E-06 |
| 11 | 118824439 | rs61900157 | C | A | 1.016 | -4813.425 | -4815.075 | -1.651 | 4.75E-06 |
| 11 | 118824996 | rs4426142 | A | G | 1.016 | -4813.425 | -4815.075 | -1.651 | 4.75E-06 |
| 11 | 118825069 | rs2097131 | A | G | 1.016 | -4813.425 | -4815.075 | -1.651 | 4.75E-06 |
| 11 | 118735476 | rs10892301 | A | G | 0.986 | -4813.425 | -4815.084 | -1.659 | 4.71E-06 |
| 11 | 118718061 | rs4938559 | T | C | 0.987 | -4813.425 | -4815.095 | -1.671 | 4.66E-06 |
| 11 | 118718062 | rs4938560 | G | A | 0.987 | -4813.425 | -4815.095 | -1.671 | 4.66E-06 |
| 11 | 118574188 | rs523715 | A | G | 0.994 | -4813.425 | -4815.097 | -1.672 | 4.65E-06 |
| 11 | 118801640 | rs572608 | A | T | 1.014 | -4813.425 | -4815.097 | -1.672 | 4.65E-06 |
| 11 | 118574185 | rs687664 | A | C | 0.994 | -4813.425 | -4815.098 | -1.673 | 4.65E-06 |
| 11 | 118708564 | rs484232 | T | G | 0.988 | -4813.425 | -4815.1 | -1.675 | 4.64E-06 |
| 11 | 118717732 | rs11217052 | G | A | 0.988 | -4813.425 | -4815.102 | -1.677 | 4.63E-06 |
| 11 | 118826063 | rs666191 | A | G | 1.014 | -4813.425 | -4815.102 | -1.678 | 4.63E-06 |
| 11 | 118800302 | rs12793214 | A | C | 1.013 | -4813.425 | -4815.103 | -1.679 | 4.62E-06 |
| 11 | 118807735 | rs10892311 | T | C | 1.013 | -4813.425 | -4815.107 | -1.682 | 4.61E-06 |
| 11 | 118826060 | rs625291 | T | C | 1.013 | -4813.425 | -4815.106 | -1.682 | 4.61E-06 |
| 11 | 118823148 | rs663003 | T | C | 1.013 | -4813.425 | -4815.109 | -1.685 | 4.59E-06 |
| 11 | 118796570 | rs11217096 | T | C | 1.012 | -4813.425 | -4815.114 | -1.689 | 4.58E-06 |
| 11 | 118707249 | rs4938551 | C | G | 0.991 | -4813.425 | -4815.126 | -1.702 | 4.52E-06 |
| 11 | 118716563 | rs2851460 | A | G | 0.992 | -4813.425 | -4815.128 | -1.703 | 4.51E-06 |
| 11 | 118769442 | rs7104819 | G | A | 0.996 | -4813.425 | -4815.135 | -1.711 | 4.48E-06 |
| 11 | 118765600 | rs3922 | G | A | 0.997 | -4813.425 | -4815.138 | -1.714 | 4.46E-06 |
| 11 | 118769032 | rs11217081 | G | A | 1.006 | -4813.425 | -4815.14 | -1.715 | 4.46E-06 |
| 11 | 118767077 | rs581063 | C | T | 1.005 | -4813.425 | -4815.143 | -1.718 | 4.44E-06 |
| 11 | 118717708 | rs7107756 | A | G | 1.005 | -4813.425 | -4815.144 | -1.719 | 4.44E-06 |
| 11 | 118770321 | rs11217083 | G | A | 1.003 | -4813.425 | -4815.15 | -1.725 | 4.41E-06 |
| 11 | 118816353 | rs586370 | A | G | 0.994 | -4813.425 | -4815.161 | -1.736 | 4.37E-06 |

**Supplemental Data 7: Bayesian analysis of merged SjD and SLE Immunochip data in DDX6-CXCR5 locus (Merged Datasets 3 and 4) with bioinformatic analysis for the top 100 SNPs (RegulomeDB Score, Haploreg epimarks, and publications)**

Note: CHROM=chromosome; POS=basepair position; A1=major allele; A2=minor allele; OR=odd ratio; L+P0=The log null marginal likelihood; L+P1=The log alternative marginal likelihood; BF=Bayes factor;
P_CAUSAL=The probability that this variant is the causal variant at this locus; Disease2=SjD and Disease1=SLE

**CREDIBLE SET**

| CHROM | POS (GRCh37/hg19) | RSID | A1 | A2 | BestModel | BestModelProb | CredibleSet | Regulome DB Sc | Haplogreg Epimarks | Publications |
|---|---|---|---|---|---|---|---|---|---|---|
| 11 | 118722149 | rs11217058 | A | G | M_Disease2_Disease1 | 0.963238649 | M_Disease2_Disease1 | 7 | > 10 | None |
| 11 | 118743772 | rs6421571 | T | C | M_Disease2_Disease1 | 0.959528177 | M_Disease2_Disease1 | 5 | > 100 | Cordell et al., Nat Commun. 2015 Sep 22;6:8019.; Mells et al., Nat Genet. 2011 Mar 13;43(4)329-32. |
| 11 | 118721565 | rs75249660 | A | G | M_Disease2_Disease1 | 0.958080511 | M_Disease2_Disease1 | 5 | < 5 | None |
| 11 | 118725660 | rs4936441 | C | G | M_Disease2_Disease1 | 0.956144888 | M_Disease2_Disease1 | 3a | <10 | Yin *et al.* Ann Rheum Dis., 2021 May: 80(5); 632-640. |
| 11 | 118726843 | rs1037679725 | T | C | M_Disease2_Disease1 | 0.955986471 | M_Disease2_Disease1 | 5 | Not found | None |
| 11 | 118665575 | rs10892293 | T | A | M_Disease2_Disease1 | 0.955801177 | M_Disease2_Disease1 | 6 | <50 | None |
| 11 | 118741842 | rs4938573 | C | T | M_Disease2_Disease1 | 0.954551172 | M_Disease2_Disease1 | 4 | >350 | Current Study; Skibola et al., Am J Hum Genet. 2014 Oct 2;95(4):462-71.; Eyre et al., Nat Genet. 2012 Dec;44(12):1336-40.; Aqwari et al., Clin Exp Immunol. 2018 Jun;192(3):259-270. |
| 11 | 118726753 | rs11217066 | G | T | M_Disease2_Disease1 | 0.954366323 | M_Disease2_Disease1 | 7 | <50 | None |
| 11 | 118743338 | rs4936444 | T | C | M_Disease2_Disease1 | 0.95403071 | M_Disease2_Disease1 | 4 | >100 | None |
| 11 | 118729391 | rs10790268 | A | G | M_Disease2_Disease1 | 0.952519943 | M_Disease2_Disease1 | 4 | >10 | Laufer et al. Hum Mol Genet. 2019 Mar 1;28(5):858-874. |
| 11 | 118729456 | rs10790269 | C | T | M_Disease2_Disease1 | 0.952519943 | M_Disease2_Disease1 | 5 | >10 | None |
| 11 | 118611817 | rs10892280 | T | C | M_Disease2_Disease1 | 0.951969981 | M_Disease2_Disease1 | 4 | <50 | None |
| 11 | 118734000 | rs11217074 | C | T | M_Disease2_Disease1 | 0.951721438 | M_Disease2_Disease1 | 6 | <5 | None |
| 11 | 118610549 | rs11826521 | T | G | M_Disease2_Disease1 | 0.950927565 | M_Disease2_Disease1 | 5 | >150 | None |
| 11 | 118684610 | rs715412 | A | G | M_Disease2_Disease1 | 0.950350126 | M_Disease2_Disease1 | 3a | <50 | None |
| 11 | 118683564 | rs17122453 | A | G | M_Disease2_Disease1 | 0.949451207 | M_Disease2_Disease1&M_Disease2 | 5 | <20 | None |
| 11 | 118740418 | rs7951740 | G | C | M_Disease2_Disease1 | 0.949302449 | M_Disease2_Disease1&M_Disease2 | 3a | >200 | None |
| 11 | 118639353 | rs11217020 | A | G | M_Disease2_Disease1 | 0.94872921 | M_Disease2_Disease1&M_Disease2 | 5 | >150 | Lopez-Isac *et al.*, Nat Commun. 2019 Oct 31;10(1):4955 |
| 11 | 118741157 | rs7117261 | T | C | M_Disease2_Disease1 | 0.948501838 | M_Disease2_Disease1&M_Disease2 | 2b | >500 | Current Study; Juran et al., Hum Mol Genet. 2012 Dec 1;21(23):5209-21. |
| 11 | 118675772 | rs11217036 | T | C | M_Disease2_Disease1 | 0.948017749 | M_Disease2_Disease1&M_Disease2 | 6 | <5 | Pickrell et al., Nat Genet. 2016 Jul;48(7);709-17. |
| 11 | 118738298 | rs7119044 | G | C | M_Disease2_Disease1 | 0.946727268 | M_Disease2_Disease1&M_Disease2 | 7 | <50 | Cordell et al., J Hepatol. 2021 Sep;75(3):572-581. |
| 11 | 118739056 | rs7122669 | A | G | M_Disease2_Disease1 | 0.946704554 | M_Disease2_Disease1&M_Disease2 | 5 | <100 | None |
| 11 | 118677086 | rs11217037 | A | G | M_Disease2_Disease1 | 0.946670873 | M_Disease2_Disease1&M_Disease2 | 3a | <50 | Kim *et al* ., Ann Rheum Dis. 2015 Mar;74(3):e13 |
| 11 | 118677355 | rs11217038 | A | G | M_Disease2_Disease1 | 0.946670873 | M_Disease2_Disease1&M_Disease2 | 5 | <20 | None |
| 11 | 118682528 | rs11217042 | A | G | M_Disease2_Disease1 | 0.946475336 | M_Disease2_Disease1&M_Disease2 | 7 | <10 | None |
| 11 | 118745243 | rs7481797 | A | G | M_Disease2_Disease1 | 0.946349212 | M_Disease2_Disease1&M_Disease2 | 5 | <100 | None |
| 11 | 118669605 | rs11217032 | A | T | M_Disease2_Disease1 | 0.946108604 | M_Disease2_Disease1&M_Disease2 | 5 | <10 | None |
| 11 | 118673706 | rs11217033 | C | T | M_Disease2_Disease1 | 0.946108604 | M_Disease2_Disease1&M_Disease2 | 5 | <5 | None |
| 11 | 118667357 | rs10892294 | C | G | M_Disease2_Disease1 | 0.945850546 | M_Disease2_Disease1&M_Disease2 | 3a | <50 | Lessard et al., Nat Genet. 2013 Nov;45(11):1284-92.; Joachims *et al* ., 2016 ACR/ARHP Annual Meeting, Abstract Number:670. |
| 11 | 118679304 | rs28409215 | A | G | M_Disease2_Disease1 | 0.945661939 | M_Disease2_Disease1&M_Disease2 | 5 | <50 | None |
| 11 | 118682350 | rs11217041 | G | A | M_Disease2_Disease1 | 0.945345808 | M_Disease2_Disease1&M_Disease2 | 5 | <10 | None |
| 11 | 118636980 | rs10892283 | G | A | M_Disease2_Disease1 | 0.944922711 | M_Disease2_Disease1&M_Disease2 | 5 | <50 | None |
| 11 | 118681464 | rs7942535 | C | T | M_Disease2_Disease1 | 0.944347007 | M_Disease2_Disease1&M_Disease2 | 5 | <10 | None |
| 11 | 118742800 | rs74541740 | A | G | M_Disease2_Disease1 | 0.944309616 | M_Disease2_Disease1&M_Disease2 | 6 | <100 | None |
| 11 | 118644694 | rs10892288 | G | A | M_Disease2_Disease1 | 0.943654881 | M_Disease2_Disease1&M_Disease2 | 5 | <200 | None |
| 11 | 118655645 | rs874621 | A | G | M_Disease2_Disease1 | 0.94361347 | M_Disease2_Disease1&M_Disease2 | 5 | >200 | None |
| 11 | 118618776 | rs1048024 | T | A | M_Disease2_Disease1 | 0.943558833 | M_Disease2_Disease1&M_Disease2 | 3a | <100 | None |
| 11 | 118619047 | rs2077579 | G | T | M_Disease2_Disease1 | 0.943558833 | M_Disease2_Disease1&M_Disease2 | 4 | >100 | None |
| 11 | 118743286 | rs12365699 | A | G | M_Disease2_Disease1 | 0.943477353 | M_Disease2_Disease1&M_Disease2 | 1f | <150 | Joachims et al., 2016 ACR/ARHP Annual Meeting, Abstract Number:670.; International Multiple Sclerosis Genetics Consortium, Science. 2019 27;365(6460).; and others. |
| 11 | 118661398 | rs57494551 | T | C | M_Disease2_Disease1 | 0.943396597 | M_Disease2_Disease1&M_Disease2 | 2b | >500 | Current Study |
| 11 | 118642085 | rs10892286 | C | A | M_Disease2_Disease1 | 0.943333587 | M_Disease2_Disease1&M_Disease2 | 5 | <200 | Langefeld *et al.* , Nat Commun., 2017 Jul17;816021. |
| 11 | 118642999 | rs10892287 | C | T | M_Disease2_Disease1 | 0.943333587 | M_Disease2_Disease1&M_Disease2 | 3a | <200 | None |
| 11 | 118681079 | rs73005423 | G | A | M_Disease2_Disease1 | 0.9431223 | M_Disease2_Disease1&M_Disease2 | 5 | <10 | None |
| 11 | 118745278 | rs7481819 | C | T | M_Disease2_Disease1 | 0.942914434 | M_Disease2_Disease1&M_Disease2 | 5 | <100 | Current Study |
| 11 | 118646003 | rs10892289 | T | C | M_Disease2_Disease1 | 0.942700706 | M_Disease2_Disease1&M_Disease2 | 7 | >100 | None |
| 11 | 118648373 | rs56043232 | T | C | M_Disease2_Disease1 | 0.942633494 | M_Disease2_Disease1&M_Disease2 | 6 | <150 | None |
| 11 | 118619253 | rs2004781 | T | C | M_Disease2_Disease1 | 0.942575419 | M_Disease2_Disease1&M_Disease2 | 5 | <100 | None |
| 11 | 118619960 | rs3889239 | T | C | M_Disease2_Disease1 | 0.942575419 | M_Disease2_Disease1&M_Disease2 | 5 | <100 | None |
| 11 | 118636343 | rs11217019 | A | C | M_Disease2_Disease1 | 0.942399136 | M_Disease2_Disease1&M_Disease2 | 5 | <50 | None |
| 11 | 118628373 | rs74676029 | T | C | M_Disease2_Disease1 | 0.942161903 | M_Disease2_Disease1&M_Disease2 | 5 | <50 | None |
| 11 | 118653503 | rs10892290 | T | G | M_Disease2_Disease1 | 0.942108775 | M_Disease2_Disease1&M_Disease2 | 5 | >100 | None |
| 11 | 118653517 | rs10892291 | T | C | M_Disease2_Disease1 | 0.942108775 | M_Disease2_Disease1&M_Disease2 | 5 | >100 | None |
| 11 | 118681083 | rs73005426 | G | A | M_Disease2_Disease1 | 0.941994768 | M_Disease2_Disease1&M_Disease2 | 5 | <10 | None |
| 11 | 118655340 | rs10892292 | C | G | M_Disease2_Disease1 | 0.941854281 | M_Disease2_Disease1&M_Disease2 | 6 | >100 | None |
| 11 | 118615785 | rs11217009 | T | C | M_Disease2_Disease1 | 0.939997369 | M_Disease2_Disease1&M_Disease2 | 5 | <5 | None |
| 11 | 118615340 | rs56758835 | A | G | M_Disease2_Disease1 | 0.938680777 | M_Disease2_Disease1&M_Disease2 | 4 | <50 | None |
| 11 | 118609790 | rs11217006 | A | G | M_Disease2_Disease1 | 0.937599563 | M_Disease2_Disease1&M_Disease2 | 6 | >10 | None |
| 11 | 118680648 | rs11217040 | A | C | M_Disease2_Disease1 | 0.937526607 | M_Disease2_Disease1&M_Disease2 | 5 | <10 | Gutierrez-Achury et al., Hum Mol Genet. 2016 Jan 1;25(1):180-90. |
| 11 | 118611781 | rs10892279 | A | G | M_Disease2_Disease1 | 0.930698529 | M_Disease2_Disease1&M_Disease2 | 4 | <50 | Trynka et al., Nat Genet. 2011 Nov 6;43(12):1193-201. |
| 11 | 118610957 | rs6589684 | A | G | M_Disease2_Disease1 | 0.930138555 | M_Disease2_Disease1&M_Disease2 | 4 | <100 | None |
| 11 | 118649512 | rs73003215 | A | G | M_Disease2_Disease1 | 0.92578598 | M_Disease2_Disease1&M_Disease2 | 5 | <150 | None |
| 11 | 118740931 | rs4938572 | C | T | M_Disease2_Disease1 | 0.924980565 | M_Disease2_Disease1&M_Disease2 | 1b | >500 | Current Study |
| 11 | 118744396 | rs7117313 | C | G | M_Disease2_Disease1 | 0.924743274 | M_Disease2_Disease1&M_Disease2 | 5 | <100 | None |
| 11 | 118698907 | rs11217046 | A | G | M_Disease2_Disease1 | 0.924527128 | M_Disease2_Disease1&M_Disease2 | 7 | <5 | None |
| 11 | 118697865 | rs11217045 | A | C | M_Disease2_Disease1 | 0.924418308 | M_Disease2_Disease1&M_Disease2 | 5 | <20 | Current Study |
| 11 | 118697906 | rs73005502 | G | A | M_Disease2_Disease1 | 0.924418308 | M_Disease2_Disease1&M_Disease2 | 5 | <20 | None |
| 11 | 118740864 | rs4936443 | C | T | M_Disease2_Disease1 | 0.920976375 | M_Disease2_Disease1&M_Disease2 | 4 | >500 | Current Study |
| 11 | 118612197 | rs6589685 | C | T | M_Disease2_Disease1 | 0.920166683 | M_Disease2_Disease1&M_Disease2 | 5 | <100 | None |
| 11 | 118701018 | rs79504890 | A | G | M_Disease2_Disease1 | 0.918726597 | M_Disease2_Disease1&M_Disease2 | 6 | <5 | None |
| 11 | 118745884 | rs10790275 | G | C | M_Disease2_Disease1 | 0.917391272 | M_Disease2_Disease1&M_Disease2 | 3a | <100 | Grosche et el., Nat Commun., 2021 Nov15;12(1):6618. |
| 11 | 118696022 | rs11217044 | C | T | M_Disease2_Disease1 | 0.910972354 | M_Disease2_Disease1&M_Disease2 | 7 | <10 | None |
| 11 | 118612921 | rs2155433 | A | G | M_Disease2_Disease1 | 0.909436735 | M_Disease2_Disease1&M_Disease2 | 1f | <100 | None |
| 11 | 118607569 | rs77209083 | G | A | M_Disease2_Disease1 | 0.904965281 | M_Disease2_Disease1&M_Disease1&M_Disease2 | 6 | <5 | None |
| 11 | 118608716 | rs7945144 | G | A | M_Disease2_Disease1 | 0.903600434 | M_Disease2_Disease1&M_Disease2&M_Disease1 | 5 | <5 | Sakaue et al., Nat Genet. 2021 Oct;53(10):1415-1424. |
| 11 | 118607046 | rs11217001 | A | G | M_Disease2_Disease1 | 0.903205796 | M_Disease2_Disease1&M_Disease1&M_Disease2 | 5 | <5 | None |
| 11 | 118694547 | rs7123726 | C | T | M_Disease2_Disease1 | 0.894814685 | M_Disease2_Disease1&M_Disease2&M_Disease1 | 5 | <5 | None |
| 11 | 118607871 | rs76409436 | A | G | M_Disease2_Disease1 | 0.889832484 | M_Disease2_Disease1&M_Disease2&M_Disease1 | 7 | 0 | None |
| 11 | 118607794 | rs12364971 | G | T | M_Disease2_Disease1 | 0.885612838 | M_Disease2_Disease1&M_Disease2&M_Disease1 | 7 | 0 | None |
| 11 | 118607417 | rs55894437 | T | C | M_Disease2_Disease1 | 0.881749768 | M_Disease2_Disease1&M_Disease2&M_Disease1 | 5 | <5 | None |
| 11 | 118607311 | rs56267528 | A | C | M_Disease2_Disease1 | 0.880901953 | M_Disease2_Disease1&M_Disease2&M_Disease1 | 5 | <5 | None |
| 11 | 118738281 | rs7119038 | G | A | M_Disease2_Disease1 | 0.876812274 | M_Disease2_Disease1&M_Disease2 | 5 | <25 | Lessard et al., Nat Genet. 2013 Nov;45(11):1284-92.; Joachims et al., 2016 ACR/ARHP Annual Meeting, Abstract Number:670. |
| 11 | 118724894 | rs9736016 | A | T | M_Disease2_Disease1 | 0.867796485 | M_Disease2_Disease1&M_Disease2&M_Disease1 | 5 | <5 | International Multiple Sclerosis Genetics Consortium (IMSGC) et al., Nat Genet. 2013 Nov;45(11):1353-60. |
| 11 | 118606521 | rs6589681 | T | C | M_Disease2_Disease1 | 0.863889918 | M_Disease2_Disease1&M_Disease2&M_Disease1 | 7 | <5 | None |
| 11 | 118607150 | rs11217002 | T | C | M_Disease2_Disease1 | 0.862070397 | M_Disease2_Disease1&M_Disease2&M_Disease1 | 7 | <5 | None |
| 11 | 118721699 | rs7125066 | G | A | M_Disease2_Disease1 | 0.852192725 | M_Disease2_Disease1&M_Disease2&M_Disease1 | 5 | <5 | None |
| 11 | 118606722 | rs6589682 | T | C | M_Disease2_Disease1 | 0.824448997 | M_Disease2_Disease1&M_Disease2&M_Disease1 | 5 | <5 | None |
| 11 | 118589012 | rs12421793 | A | C | M_Disease2_Disease1 | 0.783530866 | M_Disease2_Disease1&M_Disease1&M_Disease2 | 5 | <100 | None |
| 11 | 118782222 | rs61900138 | C | T | M_Disease2_Disease1 | 0.78026013 | M_Disease2_Disease1&M_Disease1&M_Disease2 | 4 | >750 | None |
| 11 | 118603736 | rs112561833 | C | A | M_Disease2_Disease1 | 0.779920076 | M_Disease2_Disease1&M_Disease1&M_Disease2&NULL | 6 | <10 | None |
| 11 | 118604638 | rs11216997 | T | C | M_Disease2_Disease1 | 0.778170535 | M_Disease2_Disease1&M_Disease1&M_Disease2&NULL | 5 | <50 | None |
| 11 | 118588952 | rs12421790 | T | C | M_Disease2_Disease1 | 0.773921217 | M_Disease2_Disease1&M_Disease1&M_Disease2&NULL | 5 | <100 | None |
| 11 | 118603526 | rs11216995 | A | G | M_Disease2_Disease1 | 0.772468421 | M_Disease2_Disease1&M_Disease1&M_Disease2&NULL | 5 | <10 | None |
| 11 | 118587319 | rs10892263 | C | T | M_Disease2_Disease1 | 0.771949421 | M_Disease2_Disease1&M_Disease1&M_Disease2&NULL | 3a | <100 | None |
| 11 | 118603520 | rs11216994 | T | C | M_Disease2_Disease1 | 0.767402883 | M_Disease2_Disease1&M_Disease1&M_Disease2&NULL | 5 | <10 | None |
| 11 | 118598544 | rs10892271 | A | G | M_Disease2_Disease1 | 0.761943726 | M_Disease2_Disease1&M_Disease1&M_Disease2&NULL | 3a | <100 | None |
| 11 | 118602707 | rs10892272 | G | C | M_Disease2_Disease1 | 0.754019372 | M_Disease2_Disease1&M_Disease1&M_Disease2&NULL | 5 | <20 | None |
| 11 | 118602809 | rs10892273 | A | G | M_Disease2_Disease1 | 0.754019372 | M_Disease2_Disease1&M_Disease1&M_Disease2&NULL | 5 | <20 | None |
| 11 | 118603165 | rs73001429 | G | T | M_Disease2_Disease1 | 0.75378869 | M_Disease2_Disease1&M_Disease1&M_Disease2&NULL | 5 | <10 | None |
| 11 | 118612565 | rs6589686 | A | G | M_Disease2_Disease1 | 0.747215358 | M_Disease2_Disease1&M_Disease2&M_Disease1&NULL | 2b | <100 | None |
| 11 | 118590668 | rs11216980 | A | C | M_Disease2_Disease1 | 0.742987336 | M_Disease2_Disease1&M_Disease1&M_Disease2&NULL | 5 | <100 | None |
| 11 | 118587097 | rs10892262 | C | G | M_Disease2_Disease1 | 0.732943092 | M_Disease2_Disease1&M_Disease1&M_Disease2&NULL | | | |
| 11 | 118585826 | rs11216972 | T | C | M_Disease2_Disease1 | 0.690314199 | M_Disease2_Disease1&M_Disease1&M_Disease2&NULL | | | |
| 11 | 118797562 | rs142336193 | G | A | M_Disease2_Disease1 | 0.687244653 | M_Disease2_Disease1&M_Disease1&M_Disease2&NULL | | | |
| 11 | 118575326 | rs11216956 | A | G | M_Disease2_Disease1 | 0.673011405 | M_Disease2_Disease1&M_Disease1&M_Disease2&NULL | | | |
| 11 | 118583525 | rs10892260 | C | A | M_Disease2_Disease1 | 0.665523623 | M_Disease2_Disease1&M_Disease1&M_Disease2&NULL | | | |
| 11 | 118579747 | rs10790261 | G | A | M_Disease2_Disease1 | 0.663161413 | M_Disease2_Disease1&M_Disease1&M_Disease2&NULL | | | |
| 11 | 118603495 | rs11216993 | A | T | M_Disease2_Disease1 | 0.659847136 | M_Disease2_Disease1&M_Disease1&M_Disease2&NULL | | | |
| 11 | 118577873 | rs11216960 | A | G | M_Disease2_Disease1 | 0.654437695 | M_Disease2_Disease1&M_Disease1&M_Disease2&NULL | | | |
| 11 | 118581895 | rs11216964 | A | G | M_Disease2_Disease1 | 0.654051939 | M_Disease2_Disease1&M_Disease1&M_Disease2&NULL | | | |
| 11 | 118579865 | rs10892258 | A | G | M_Disease2_Disease1 | 0.652541941 | M_Disease2_Disease1&M_Disease1&M_Disease2&NULL | | | |
| 11 | 118580338 | rs11216961 | G | C | M_Disease2_Disease1 | 0.652541941 | M_Disease2_Disease1&M_Disease1&M_Disease2&NULL | | | |
| 11 | 118580813 | rs11216962 | C | T | M_Disease2_Disease1 | 0.652541941 | M_Disease2_Disease1&M_Disease1&M_Disease2&NULL | | | |
| 11 | 118580814 | rs11216963 | A | G | M_Disease2_Disease1 | 0.652541941 | M_Disease2_Disease1&M_Disease1&M_Disease2&NULL | | | |
| 11 | 118582613 | rs11216965 | A | C | M_Disease2_Disease1 | 0.652541941 | M_Disease2_Disease1&M_Disease1&M_Disease2&NULL | | | |
| 11 | 118583002 | rs10892259 | A | G | M_Disease2_Disease1 | 0.652541941 | M_Disease2_Disease1&M_Disease1&M_Disease2&NULL | | | |
| 11 | 118578705 | rs10892256 | A | G | M_Disease2_Disease1 | 0.649103025 | M_Disease2_Disease1&M_Disease1&M_Disease2&NULL | | | |
| 11 | 118579327 | rs10892257 | A | G | M_Disease2_Disease1 | 0.646516457 | M_Disease2_Disease1&M_Disease1&M_Disease2&NULL | | | |
| 11 | 118584340 | rs11216969 | C | T | M_Disease2_Disease1 | 0.646157512 | M_Disease2_Disease1&M_Disease1&M_Disease2&NULL | | | |
| 11 | 118590961 | rs111227443 | T | C | M_Disease2_Disease1 | 0.632166428 | M_Disease2_Disease1&M_Disease2&NULL&M_Disease1 | | | |
| 11 | 118604837 | rs11216998 | T | C | M_Disease2_Disease1 | 0.629334801 | M_Disease2_Disease1&M_Disease2&M_Disease1&NULL | | | |
| 11 | 118443944 | rs11216906 | A | G | M_Disease2_Disease1 | 0.608753729 | M_Disease2_Disease1&M_Disease1&NULL&M_Disease2 | | | |
| 11 | 118445585 | rs73023341 | G | A | M_Disease2_Disease1 | 0.607361906 | M_Disease2_Disease1&M_Disease1&NULL&M_Disease2 | | | |
| 11 | 118450046 | rs11216910 | G | A | M_Disease2_Disease1 | 0.602090113 | M_Disease2_Disease1&M_Disease1&NULL&M_Disease2 | | | |
| 11 | 118734521 | rs1174717189 | T | A | M_Disease2_Disease1 | 0.59773277 | M_Disease2_Disease1&M_Disease2&NULL&M_Disease1 | | | |
| 11 | 118604941 | rs11216999 | C | T | M_Disease2_Disease1 | 0.59323083 | M_Disease2_Disease1&M_Disease2&M_Disease1&NULL | | | |
| 11 | 118447580 | rs9736327 | G | T | M_Disease2_Disease1 | 0.590889596 | M_Disease2_Disease1&M_Disease1&NULL&M_Disease2 | | | |
| 11 | 118447597 | rs9735421 | A | G | M_Disease2_Disease1 | 0.590889596 | M_Disease2_Disease1&M_Disease1&NULL&M_Disease2 | | | |
| 11 | 118440863 | rs11216904 | G | A | M_Disease2_Disease1 | 0.589695862 | M_Disease2_Disease1&M_Disease1&NULL&M_Disease2 | | | |
| 11 | 118441846 | rs10892243 | C | T | M_Disease2_Disease1 | 0.588073947 | M_Disease2_Disease1&M_Disease1&NULL&M_Disease2 | | | |
| 11 | 118465295 | rs11216922 | G | A | M_Disease2_Disease1 | 0.57885693 | M_Disease2_Disease1&M_Disease1&NULL&M_Disease2 | | | |
| 11 | 118465501 | rs11216923 | G | A | M_Disease2_Disease1 | 0.57885693 | M_Disease2_Disease1&M_Disease1&NULL&M_Disease2 | | | |
| 11 | 118453483 | rs60112336 | T | C | M_Disease2_Disease1 | 0.578283734 | M_Disease2_Disease1&M_Disease1&NULL&M_Disease2 | | | |
| 11 | 118462160 | rs11216917 | G | T | M_Disease2_Disease1 | 0.574237214 | M_Disease2_Disease1&M_Disease1&NULL&M_Disease2 | | | |
| 11 | 118543563 | rs10892251 | T | C | M_Disease2_Disease1 | 0.572405119 | M_Disease2_Disease1&M_Disease1&NULL&M_Disease2 | | | |
| 11 | 118439048 | rs12224844 | T | A | M_Disease2_Disease1 | 0.571539591 | M_Disease2_Disease1&M_Disease1&NULL&M_Disease2 | | | |
| 11 | 118555692 | rs10892252 | G | A | M_Disease2_Disease1 | 0.570708233 | M_Disease2_Disease1&M_Disease1&NULL&M_Disease2 | | | |
| 11 | 118469571 | rs7931564 | G | A | M_Disease2_Disease1 | 0.569670636 | M_Disease2_Disease1&M_Disease1&NULL&M_Disease2 | | | |
| 11 | 118471423 | rs11216925 | C | T | M_Disease2_Disease1 | 0.569670636 | M_Disease2_Disease1&M_Disease1&NULL&M_Disease2 | | | |
| 11 | 118472354 | rs10892245 | A | C | M_Disease2_Disease1 | 0.569670636 | M_Disease2_Disease1&M_Disease1&NULL&M_Disease2 | | | |
| 11 | 118448087 | rs9735449 | T | C | M_Disease2_Disease1 | 0.568872669 | M_Disease2_Disease1&M_Disease1&NULL&M_Disease2 | | | |
| 11 | 118461605 | rs11216916 | C | T | M_Disease2_Disease1 | 0.567680638 | M_Disease2_Disease1&M_Disease1&NULL&M_Disease2 | | | |
| 11 | 118560857 | rs73001406 | A | G | M_Disease2_Disease1 | 0.564036828 | M_Disease2_Disease1&M_Disease1&NULL&M_Disease2 | | | |
| 11 | 118473385 | rs17742 | C | T | M_Disease2_Disease1 | 0.562266462 | M_Disease2_Disease1&M_Disease1&NULL&M_Disease2 | | | |
| 11 | 118724723 | rs57719838 | C | T | M_Disease2_Disease1 | 0.560658269 | M_Disease2_Disease1&M_Disease2&NULL&M_Disease1 | | | |
| 11 | 118725942 | rs11217063 | C | G | M_Disease2_Disease1 | 0.557065553 | M_Disease2_Disease1&M_Disease2&NULL&M_Disease1 | | | |
| 11 | 118726647 | rs11217065 | C | T | M_Disease2_Disease1 | 0.557065553 | M_Disease2_Disease1&M_Disease2&NULL&M_Disease1 | | | |
| 11 | 118437279 | rs11216902 | G | A | M_Disease2_Disease1 | 0.55647769 | M_Disease2_Disease1&M_Disease1&NULL&M_Disease2 | | | |
| 11 | 118433729 | rs59108310 | G | A | M_Disease2_Disease1 | 0.555161712 | M_Disease2_Disease1&M_Disease1&NULL&M_Disease2 | | | |
| 11 | 118446298 | rs55844025 | T | C | M_Disease2_Disease1 | 0.553862153 | M_Disease2_Disease1&M_Disease1&NULL&M_Disease2 | | | |
| 11 | 118464815 | rs11216921 | C | G | M_Disease2_Disease1 | 0.552538554 | M_Disease2_Disease1&M_Disease1&NULL&M_Disease2 | | | |
| 11 | 118605463 | rs10892274 | T | C | M_Disease2_Disease1 | 0.550498426 | M_Disease2_Disease1&M_Disease2&M_Disease1&NULL | | | |
| 11 | 118605617 | rs10892275 | A | T | M_Disease2_Disease1 | 0.550498426 | M_Disease2_Disease1&M_Disease2&M_Disease1&NULL | | | |
| 11 | 118471996 | rs11216926 | T | C | M_Disease2_Disease1 | 0.550250338 | M_Disease2_Disease1&M_Disease1&NULL&M_Disease2 | | | |
| 11 | 118550524 | rs201134460 | T | C | M_Disease2_Disease1 | 0.550040336 | M_Disease2_Disease1&M_Disease1&NULL&M_Disease2 | | | |
| 11 | 118470092 | rs11216924 | G | C | M_Disease2_Disease1 | 0.549420922 | M_Disease2_Disease1&M_Disease1&NULL&M_Disease2 | | | |
| 11 | 118457630 | rs17092756 | C | G | M_Disease2_Disease1 | 0.549028892 | M_Disease2_Disease1&M_Disease1&NULL&M_Disease2 | | | |
| 11 | 118723379 | rs7103067 | A | G | M_Disease2_Disease1 | 0.549023723 | M_Disease2_Disease1&M_Disease2&NULL&M_Disease1 | | | |
| 11 | 118550522 | rs2277296 | C | T | M_Disease2_Disease1 | 0.548612694 | M_Disease2_Disease1&M_Disease1&NULL&M_Disease2 | | | |
| 11 | 118556398 | rs11216943 | A | G | M_Disease2_Disease1 | 0.548479437 | M_Disease2_Disease1&M_Disease1&NULL&M_Disease2 | | | |
| 11 | 118451901 | rs11216911 | C | T | M_Disease2_Disease1 | 0.54845171 | M_Disease2_Disease1&M_Disease1&NULL&M_Disease2 | | | |
| 11 | 118452334 | rs11216912 | C | G | M_Disease2_Disease1 | 0.54845171 | M_Disease2_Disease1&M_Disease1&NULL&M_Disease2 | | | |
| 11 | 118453681 | rs11216913 | G | A | M_Disease2_Disease1 | 0.54845171 | M_Disease2_Disease1&M_Disease1&NULL&M_Disease2 | | | |
| 11 | 118466249 | rs7115634 | G | A | M_Disease2_Disease1 | 0.548281415 | M_Disease2_Disease1&M_Disease1&NULL&M_Disease2 | | | |
| 11 | 118444525 | rs11216907 | A | G | M_Disease2_Disease1 | 0.547886497 | M_Disease2_Disease1&M_Disease1&NULL&M_Disease2 | | | |
| 11 | 118729210 | rs10892300 | T | C | M_Disease2_Disease1 | 0.546748913 | M_Disease2_Disease1&M_Disease2&NULL&M_Disease1 | | | |
| 11 | 118454177 | rs17842763 | G | A | M_Disease2_Disease1 | 0.546407247 | M_Disease2_Disease1&M_Disease1&NULL&M_Disease2 | | | |
| 11 | 118729860 | rs11217070 | G | T | M_Disease2_Disease1 | 0.545640768 | M_Disease2_Disease1&M_Disease2&NULL&M_Disease1 | | | |
| 11 | 118605841 | rs10892277 | A | G | M_Disease2_Disease1 | 0.543417372 | M_Disease2_Disease1&M_Disease2&M_Disease1&NULL | | | |
| 11 | 118456414 | rs11216914 | C | T | M_Disease2_Disease1 | 0.543355783 | M_Disease2_Disease1&M_Disease1&NULL&M_Disease2 | | | |
| 11 | 118449396 | rs73023350 | A | G | M_Disease2_Disease1 | 0.54303355 | M_Disease2_Disease1&M_Disease1&NULL&M_Disease2 | | | |
| 11 | 118450875 | rs7126175 | T | C | M_Disease2_Disease1 | 0.541629883 | M_Disease2_Disease1&M_Disease1&NULL&M_Disease2 | | | |
| 11 | 118432279 | rs11216897 | G | A | M_Disease2_Disease1 | 0.541411169 | M_Disease2_Disease1&M_Disease1&NULL&M_Disease2 | | | |
| 11 | 118729583 | rs12790489 | C | T | M_Disease2_Disease1 | 0.539302199 | M_Disease2_Disease1&M_Disease2&NULL&M_Disease1 | | | |
| 11 | 118731599 | rs34561290 | T | C | M_Disease2_Disease1 | 0.538558248 | M_Disease2_Disease1&M_Disease2&NULL&M_Disease1 | | | |
| 11 | 118731762 | rs11217072 | G | A | M_Disease2_Disease1 | 0.538558248 | M_Disease2_Disease1&M_Disease2&NULL&M_Disease1 | | | |
| 11 | 118730141 | rs7929520 | G | A | M_Disease2_Disease1 | 0.536620971 | M_Disease2_Disease1&M_Disease2&NULL&M_Disease1 | | | |
| 11 | 118449746 | rs11216909 | T | C | M_Disease2_Disease1 | 0.534435855 | M_Disease2_Disease1&M_Disease1&NULL&M_Disease2 | | | |
| 11 | 118459039 | rs57438673 | A | G | M_Disease2_Disease1 | 0.533855688 | M_Disease2_Disease1&M_Disease1&NULL&M_Disease2 | | | |
| 11 | 118605628 | rs10892276 | A | G | M_Disease2_Disease1 | 0.532715281 | M_Disease2_Disease1&M_Disease2&M_Disease1&NULL | | | |
| 11 | 118459702 | rs113054880 | G | C | M_Disease2_Disease1 | 0.53237106 | M_Disease2_Disease1&M_Disease1&NULL&M_Disease2 | | | |
| 11 | 118805385 | rs55772432 | A | G | M_Disease2_Disease1 | 0.523090535 | M_Disease2_Disease1&M_Disease2&NULL&M_Disease1 | | | |
| 11 | 118449370 | rs17122278 | G | A | M_Disease2_Disease1 | 0.522971398 | M_Disease2_Disease1&M_Disease1&NULL&M_Disease2 | | | |
| 11 | 118740180 | rs7483509 | A | G | M_Disease2_Disease1 | 0.518331672 | M_Disease2_Disease1&M_Disease2&NULL&M_Disease1 | | | |
| 11 | 118473231 | rs1047079 | C | T | M_Disease2_Disease1 | 0.508733945 | M_Disease2_Disease1&M_Disease1&NULL&M_Disease2 | | | |
| 11 | 118436079 | rs11216900 | C | G | M_Disease2_Disease1 | 0.507720906 | M_Disease2_Disease1&M_Disease1&NULL&M_Disease2 | | | |
| 11 | 118445275 | rs17122269 | C | T | M_Disease2_Disease1 | 0.505731958 | M_Disease2_Disease1&M_Disease1&NULL&M_Disease2 | | | |
| 11 | 118432404 | rs4326835 | T | G | M_Disease2_Disease1 | 0.50181406 | M_Disease2_Disease1&M_Disease1&NULL&M_Disease2 | | | |
| 11 | 118711790 | rs622722 | A | G | M_Disease2_Disease1 | 0.499816148 | M_Disease2_Disease1&M_Disease2&NULL&M_Disease1 | | | |
| 11 | 118432852 | rs11216898 | C | T | M_Disease2_Disease1 | 0.499687336 | M_Disease2_Disease1&M_Disease1&NULL&M_Disease2 | | | |
| 11 | 118727169 | rs10790267 | T | C | M_Disease2_Disease1 | 0.497734883 | M_Disease2_Disease1&M_Disease2&NULL&M_Disease1 | | | |
| 11 | 118789026 | rs11607239 | T | G | M_Disease2_Disease1 | 0.496659458 | M_Disease2_Disease1&M_Disease2&NULL&M_Disease1 | | | |
| 11 | 118434421 | rs11216899 | G | A | M_Disease2_Disease1 | 0.495935101 | M_Disease2_Disease1&M_Disease1&NULL&M_Disease2 | | | |
| 11 | 118434835 | rs60753148 | T | C | M_Disease2_Disease1 | 0.495935101 | M_Disease2_Disease1&M_Disease1&NULL&M_Disease2 | | | |
| 11 | 118459069 | rs633308 | G | A | M_Disease2_Disease1 | 0.495638536 | M_Disease2_Disease1&M_Disease1&NULL&M_Disease2 | | | |
| 11 | 118728320 | rs4936442 | T | C | M_Disease2_Disease1 | 0.494101529 | M_Disease2_Disease1&M_Disease2&NULL&M_Disease1 | | | |
| 11 | 118727332 | rs11217067 | T | G | M_Disease2_Disease1 | 0.491278066 | M_Disease2_Disease1&M_Disease2&NULL&M_Disease1 | | | |
| 11 | 118809805 | rs142069912 | C | G | M_Disease2_Disease1 | 0.49117041 | M_Disease2_Disease1&M_Disease2&M_Disease1&NULL | | | |
| 11 | 118438448 | rs60837962 | T | A | M_Disease2_Disease1 | 0.486817336 | M_Disease2_Disease1&M_Disease1&NULL&M_Disease2 | | | |
| 11 | 118769604 | rs61751532 | T | C | M_Disease2_Disease1 | 0.485819712 | M_Disease2_Disease1&M_Disease2&NULL&M_Disease1 | | | |
| 11 | 118435756 | rs2277291 | T | C | M_Disease2_Disease1 | 0.481100188 | M_Disease2_Disease1&M_Disease1&NULL&M_Disease2 | | | |
| 11 | 118669178 | rs583213 | G | T | M_Disease2_Disease1 | 0.436426767 | M_Disease2_Disease1&M_Disease2&NULL&M_Disease1 | | | |
| 11 | 118750321 | rs11217078 | C | T | M_Disease2_Disease1 | 0.428988107 | M_Disease2_Disease1&M_Disease2&NULL&M_Disease1 | | | |
| 11 | 118587347 | rs603486 | T | C | M_Disease2_Disease1 | 0.426687758 | M_Disease2_Disease1&M_Disease1&NULL&M_Disease2 | | | |
| 11 | 118577990 | rs480958 | G | A | M_Disease2_Disease1 | 0.413006775 | M_Disease2_Disease1&M_Disease1&NULL&M_Disease2 | | | |
| 11 | 118584498 | rs569770 | T | C | M_Disease2_Disease1 | 0.406029182 | M_Disease2_Disease1&M_Disease1&NULL&M_Disease2 | | | |
| 11 | 118504742 | rs633683 | T | C | M_Disease2_Disease1 | 0.40507142 | M_Disease2_Disease1&M_Disease1&NULL&M_Disease2 | | | |
| 11 | 118530611 | rs2276065 | C | T | M_Disease2_Disease1 | 0.400992048 | M_Disease2_Disease1&M_Disease1&NULL&M_Disease2 | | | |
| 11 | 118697012 | rs571470 | A | G | M_Disease2_Disease1 | 0.400382205 | M_Disease2_Disease1&M_Disease2&NULL&M_Disease1 | | | |
| 11 | 118807639 | rs73575454 | A | G | M_Disease2_Disease1 | 0.394572754 | M_Disease2_Disease1&M_Disease1&M_Disease2&NULL | | | |
| 11 | 118809767 | rs75460330 | A | G | M_Disease2_Disease1 | 0.394572754 | M_Disease2_Disease1&M_Disease1&M_Disease2&NULL | | | |
| 11 | 118810054 | rs73557344 | C | G | M_Disease2_Disease1 | 0.394572754 | M_Disease2_Disease1&M_Disease1&M_Disease2&NULL | | | |
| 11 | 118749318 | rs78312641 | A | T | M_Disease2_Disease1 | 0.393249088 | M_Disease2_Disease1&M_Disease1&M_Disease2&NULL | | | |
| 11 | 118749563 | rs77643000 | C | T | M_Disease2_Disease1 | 0.391800553 | M_Disease2_Disease1&M_Disease1&M_Disease2&NULL | | | |
| 11 | 118748170 | rs77074463 | C | G | M_Disease2_Disease1 | 0.387955013 | M_Disease2_Disease1&M_Disease1&NULL&M_Disease2 | | | |
| 11 | 118509741 | rs113313560 | T | C | M_Disease2_Disease1 | 0.385934233 | M_Disease2_Disease1&M_Disease2&NULL&M_Disease1 | | | |
| 11 | 118662340 | rs5006102 | A | T | M_Disease2_Disease1 | 0.384801961 | M_Disease2_Disease1&M_Disease2&NULL&M_Disease1 | | | |
| 11 | 118662045 | rs7933859 | T | C | M_Disease2_Disease1 | 0.382152913 | M_Disease2_Disease1&M_Disease2&NULL&M_Disease1 | | | |
| 11 | 118662203 | rs4508234 | G | C | M_Disease2_Disease1 | 0.382152913 | M_Disease2_Disease1&M_Disease2&NULL&M_Disease1 | | | |
| 11 | 118668908 | rs582200 | G | A | M_Disease2_Disease1 | 0.380074586 | M_Disease2_Disease1&M_Disease2&NULL&M_Disease1 | | | |
| 11 | 118678780 | rs500254 | G | A | M_Disease2_Disease1 | 0.378756161 | M_Disease2_Disease1&M_Disease2&NULL&M_Disease1 | | | |
| 11 | 118665009 | rs4938544 | A | G | M_Disease2_Disease1 | 0.377308267 | M_Disease2_Disease1&M_Disease2&NULL&M_Disease1 | | | |
| 11 | 118601426 | rs186926311 | T | C | M_Disease2_Disease1 | 0.375757708 | M_Disease2_Disease1&M_Disease1&M_Disease2&NULL | | | |
| 11 | 118677083 | rs675003 | T | C | M_Disease2_Disease1 | 0.374787163 | M_Disease2_Disease1&M_Disease2&NULL&M_Disease1 | | | |
| 11 | 118802128 | rs59069289 | T | G | M_Disease2_Disease1 | 0.374738527 | M_Disease2_Disease1&M_Disease1&NULL&M_Disease2 | | | |
| 11 | 118802386 | rs59480472 | C | T | M_Disease2_Disease1 | 0.374738527 | M_Disease2_Disease1&M_Disease1&NULL&M_Disease2 | | | |
| 11 | 118803160 | rs73575440 | A | G | M_Disease2_Disease1 | 0.374738527 | M_Disease2_Disease1&M_Disease1&NULL&M_Disease2 | | | |
| 11 | 118803596 | rs111313044 | A | G | M_Disease2_Disease1 | 0.374738527 | M_Disease2_Disease1&M_Disease1&NULL&M_Disease2 | | | |
| 11 | 118804643 | rs56933877 | A | G | M_Disease2_Disease1 | 0.374738527 | M_Disease2_Disease1&M_Disease1&NULL&M_Disease2 | | | |
| 11 | 118804919 | rs73575441 | C | T | M_Disease2_Disease1 | 0.374738527 | M_Disease2_Disease1&M_Disease1&NULL&M_Disease2 | | | |
| 11 | 118804989 | rs112045567 | G | A | M_Disease2_Disease1 | 0.374738527 | M_Disease2_Disease1&M_Disease1&NULL&M_Disease2 | | | |
| 11 | 118665324 | rs61901411 | T | C | M_Disease2_Disease1 | 0.370387079 | M_Disease2_Disease1&M_Disease2&NULL&M_Disease1 | | | |
| 11 | 118820778 | rs139076649 | C | T | M_Disease2_Disease1 | 0.366299508 | M_Disease2_Disease1&M_Disease2&M_Disease1&NULL | | | |
| 11 | 118507478 | rs11216937 | T | C | M_Disease2_Disease1 | 0.363288595 | M_Disease2_Disease1&M_Disease1&NULL&M_Disease2 | | | |
| 11 | 118516903 | rs80015904 | A | G | M_Disease2_Disease1 | 0.362325805 | M_Disease2_Disease1&M_Disease1&NULL&M_Disease2 | | | |

**FINEMAP**

| CHROM | POS (GRCh37/hg19) | RSID | A1 | A2 | OR_Disease1 | OR_Disease2 | L+P0 | L+P1 | logBF | P_CAUSAL |
|---|---|---|---|---|---|---|---|---|---|---|
| 11 | 118722149 | rs11217058 | A | G | 1.198 | 1.284 | -9407.082 | -9391.613 | 15.469 | 0.058809446 |
| 11 | 118743772 | rs6421571 | T | C | 1.187 | 1.292 | -9407.706 | -9392.367 | 15.338 | 0.051588691 |
| 11 | 118721565 | rs75249660 | A | G | 1.193 | 1.281 | -9403.846 | -9388.853 | 14.993 | 0.036536161 |
| 11 | 118741842 | rs4938573 | C | T | 1.18 | 1.291 | -9407.283 | -9392.295 | 14.989 | 0.036390308 |
| 11 | 118743338 | rs4936444 | T | C | 1.181 | 1.292 | -9404.41 | -9389.483 | 14.927 | 0.034202628 |
| 11 | 118729391 | rs10790268 | A | G | 1.176 | 1.293 | -9405.615 | -9390.694 | 14.921 | 0.033998027 |
| 11 | 118729456 | rs10790269 | C | T | 1.176 | 1.293 | -9405.615 | -9390.694 | 14.921 | 0.033998027 |
| 11 | 118725660 | rs4936441 | C | G | 1.189 | 1.281 | -9407.766 | -9392.892 | 14.874 | 0.032437089 |
| 11 | 118726843 | rs1037679725 | T | C | 1.19 | 1.28 | -9407.766 | -9392.924 | 14.842 | 0.031415534 |
| 11 | 118734000 | rs11217074 | C | T | 1.179 | 1.29 | -9407.635 | -9392.893 | 14.742 | 0.028425951 |
| 11 | 118726753 | rs11217066 | G | T | 1.19 | 1.278 | -9407.766 | -9393.08 | 14.686 | 0.026877849 |
| 11 | 118740418 | rs7951740 | G | C | 1.177 | 1.288 | -9408.077 | -9393.528 | 14.549 | 0.023436684 |
| 11 | 118741157 | rs7117261 | T | C | 1.176 | 1.289 | -9408.077 | -9393.554 | 14.523 | 0.022835184 |
| 11 | 118738298 | rs7119044 | G | C | 1.171 | 1.291 | -9406.727 | -9392.227 | 14.5 | 0.022315968 |
| 11 | 118739056 | rs7122669 | A | G | 1.171 | 1.29 | -9407.169 | -9392.669 | 14.5 | 0.022315968 |
| 11 | 118665575 | rs10892293 | T | A | 1.203 | 1.258 | -9405.948 | -9391.462 | 14.485 | 0.021983727 |
| 11 | 118745243 | rs7481797 | A | G | 1.175 | 1.285 | -9398.925 | -9384.585 | 14.34 | 0.019016414 |
| 11 | 118743286 | rs12365699 | A | G | 1.179 | 1.313 | -9406.882 | -9392.555 | 14.328 | 0.01878958 |
| 11 | 118742800 | rs74541740 | A | G | 1.176 | 1.283 | -9404.747 | -9390.598 | 14.148 | 0.015694377 |
| 11 | 118611817 | rs10892280 | T | C | 1.208 | 1.249 | -9408.077 | -9393.969 | 14.108 | 0.015078992 |
| 11 | 118684610 | rs715412 | A | G | 1.197 | 1.258 | -9408.077 | -9393.98 | 14.097 | 0.014914032 |
| 11 | 118745278 | rs7481819 | C | T | 1.175 | 1.283 | -9397.235 | -9383.165 | 14.07 | 0.01451674 |
| 11 | 118683564 | rs17122453 | A | G | 1.192 | 1.258 | -9408.077 | -9394.013 | 14.064 | 0.014429901 |
| 11 | 118610549 | rs11826521 | T | G | 1.207 | 1.248 | -9408.077 | -9394.05 | 14.028 | 0.013919664 |
| 11 | 118675772 | rs11217036 | T | C | 1.198 | 1.254 | -9407.547 | -9393.66 | 13.887 | 0.012089079 |
| 11 | 118639353 | rs11217020 | A | G | 1.206 | 1.248 | -9408.077 | -9394.213 | 13.865 | 0.011826024 |
| 11 | 118682528 | rs11217042 | A | G | 1.19 | 1.257 | -9408.077 | -9394.212 | 13.865 | 0.011826024 |
| 11 | 118677086 | rs11217037 | A | G | 1.196 | 1.254 | -9408.077 | -9394.271 | 13.806 | 0.011148472 |
| 11 | 118677355 | rs11217038 | A | G | 1.196 | 1.254 | -9408.077 | -9394.271 | 13.806 | 0.011148472 |
| 11 | 118682350 | rs11217041 | G | A | 1.188 | 1.257 | -9408.077 | -9394.287 | 13.791 | 0.010982493 |
| 11 | 118667357 | rs10892294 | C | G | 1.194 | 1.255 | -9405.225 | -9391.45 | 13.775 | 0.010808172 |
| 11 | 118669605 | rs11217032 | A | T | 1.196 | 1.254 | -9407.547 | -9393.781 | 13.766 | 0.010711335 |
| 11 | 118673706 | rs11217033 | C | T | 1.196 | 1.254 | -9407.547 | -9393.781 | 13.766 | 0.010711335 |
| 11 | 118679304 | rs28409215 | A | G | 1.193 | 1.256 | -9408.077 | -9394.318 | 13.759 | 0.010636617 |
| 11 | 118681464 | rs7942535 | C | T | 1.187 | 1.257 | -9408.077 | -9394.341 | 13.736 | 0.010394767 |
| 11 | 118681079 | rs73005423 | G | A | 1.19 | 1.257 | -9405.811 | -9392.183 | 13.628 | 0.00933063 |
| 11 | 118636980 | rs10892283 | G | A | 1.204 | 1.246 | -9405.711 | -9392.11 | 13.601 | 0.009082073 |
| 11 | 118681083 | rs73005426 | G | A | 1.185 | 1.257 | -9407.063 | -9393.467 | 13.596 | 0.009036776 |
| 11 | 118644694 | rs10892288 | G | A | 1.203 | 1.246 | -9403.942 | -9390.424 | 13.518 | 0.008358696 |
| 11 | 118655645 | rs874621 | A | G | 1.204 | 1.245 | -9404.11 | -9390.602 | 13.508 | 0.008275526 |
| 11 | 118642085 | rs10892286 | C | A | 1.202 | 1.247 | -9403.942 | -9390.437 | 13.505 | 0.008250737 |
| 11 | 118642999 | rs10892287 | C | T | 1.202 | 1.247 | -9403.942 | -9390.437 | 13.505 | 0.008250737 |
| 11 | 118618776 | rs1048024 | T | A | 1.204 | 1.244 | -9405.711 | -9392.217 | 13.494 | 0.008160476 |
| 11 | 118619047 | rs2077579 | G | T | 1.204 | 1.244 | -9405.711 | -9392.217 | 13.494 | 0.008160476 |
| 11 | 118661398 | rs57494551 | T | C | 1.204 | 1.246 | -9404.13 | -9390.637 | 13.494 | 0.008160476 |
| 11 | 118646003 | rs10892289 | T | C | 1.202 | 1.246 | -9406.31 | -9392.849 | 13.46 | 0.007887683 |
| 11 | 118648373 | rs56043232 | T | C | 1.203 | 1.245 | -9406.31 | -9392.864 | 13.445 | 0.007770251 |
| 11 | 118619253 | rs2004781 | T | C | 1.204 | 1.244 | -9408.077 | -9394.648 | 13.429 | 0.007646916 |
| 11 | 118619960 | rs3889239 | T | C | 1.204 | 1.244 | -9408.077 | -9394.648 | 13.429 | 0.007646916 |
| 11 | 118636343 | rs11217019 | A | C | 1.203 | 1.244 | -9404.817 | -9391.396 | 13.42 | 0.007578403 |
| 11 | 118653503 | rs10892290 | T | G | 1.202 | 1.245 | -9402.857 | -9389.445 | 13.412 | 0.007518018 |
| 11 | 118653517 | rs10892291 | T | C | 1.202 | 1.245 | -9402.857 | -9389.445 | 13.412 | 0.007518018 |
| 11 | 118628373 | rs74676029 | T | C | 1.204 | 1.244 | -9405.711 | -9392.302 | 13.409 | 0.007495497 |
| 11 | 118655340 | rs10892292 | C | G | 1.202 | 1.246 | -9406.478 | -9393.071 | 13.407 | 0.007480521 |
| 11 | 118680648 | rs11217040 | A | C | 1.187 | 1.254 | -9402.521 | -9389.243 | 13.278 | 0.006575183 |
| 11 | 118615785 | rs11217009 | T | C | 1.202 | 1.244 | -9403.917 | -9390.648 | 13.269 | 0.006516272 |
| 11 | 118615340 | rs56758835 | A | G | 1.201 | 1.243 | -9406.284 | -9393.079 | 13.205 | 0.006112296 |
| 11 | 118609790 | rs11217006 | A | G | 1.193 | 1.245 | -9401.922 | -9388.734 | 13.188 | 0.006009265 |
| 11 | 118740931 | rs4938572 | C | T | 1.164 | 1.275 | -9404.889 | -9391.83 | 13.059 | 0.005281988 |
| 11 | 118744396 | rs7117313 | C | G | 1.168 | 1.271 | -9402.381 | -9389.422 | 12.96 | 0.004784122 |
| 11 | 118740864 | rs4936443 | C | T | 1.16 | 1.275 | -9404.544 | -9391.62 | 12.924 | 0.004614957 |
| 11 | 118611781 | rs10892279 | A | G | 1.191 | 1.241 | -9408.077 | -9395.301 | 12.777 | 0.003984064 |
| 11 | 118610957 | rs6589684 | A | G | 1.191 | 1.241 | -9406.825 | -9394.088 | 12.737 | 0.003827847 |
| 11 | 118745884 | rs10790275 | G | C | 1.165 | 1.268 | -9397.139 | -9384.576 | 12.563 | 0.003216528 |
| 11 | 118698907 | rs11217046 | A | G | 1.186 | 1.249 | -9397.116 | -9384.589 | 12.527 | 0.003102792 |
| 11 | 118697865 | rs11217045 | A | C | 1.186 | 1.249 | -9397.938 | -9385.424 | 12.513 | 0.003059656 |
| 11 | 118697906 | rs73005502 | G | A | 1.186 | 1.249 | -9397.938 | -9385.424 | 12.513 | 0.003059656 |
| 11 | 118649512 | rs73003215 | A | G | 1.19 | 1.239 | -9407.3 | -9394.794 | 12.505 | 0.003035276 |
| 11 | 118701018 | rs79504890 | A | G | 1.179 | 1.252 | -9394.215 | -9381.896 | 12.319 | 0.00252011 |
| 11 | 118612197 | rs6589685 | C | T | 1.188 | 1.236 | -9408.077 | -9395.855 | 12.222 | 0.002287141 |
| 11 | 118696022 | rs11217044 | C | T | 1.18 | 1.24 | -9387.535 | -9375.671 | 11.863 | 0.00159728 |
| 11 | 118612921 | rs2155433 | A | G | 1.186 | 1.231 | -9406.72 | -9395 | 11.72 | 0.001384449 |
| 11 | 118738281 | rs7119038 | G | A | 1.118 | 1.261 | -9392.164 | -9380.502 | 11.662 | 0.001306436 |
| 11 | 118607569 | rs77209083 | G | A | 1.206 | 1.206 | -9381.664 | -9370.196 | 11.467 | 0.00107498 |
| 11 | 118608716 | rs7945144 | G | A | 1.188 | 1.223 | -9399.001 | -9387.558 | 11.443 | 0.001049488 |
| 11 | 118607046 | rs11217001 | A | G | 1.206 | 1.207 | -9358.695 | -9347.303 | 11.392 | 0.000997306 |
| 11 | 118694547 | rs7123726 | C | T | 1.173 | 1.236 | -9390.695 | -9379.472 | 11.224 | 0.000843077 |
| 11 | 118607871 | rs76409436 | A | G | 1.193 | 1.208 | -9386.594 | -9375.709 | 10.886 | 0.000601278 |
| 11 | 118607794 | rs12364971 | G | T | 1.191 | 1.207 | -9386.592 | -9375.866 | 10.726 | 0.000512376 |
| 11 | 118607417 | rs55894437 | T | C | 1.19 | 1.207 | -9363.351 | -9352.75 | 10.601 | 0.00045217 |
| 11 | 118607311 | rs56267528 | A | C | 1.19 | 1.207 | -9363.351 | -9352.783 | 10.568 | 0.000437492 |
| 11 | 118724894 | rs9736016 | A | T | 1.132 | 1.2 | -9405.966 | -9395.52 | 10.446 | 0.000387245 |
| 11 | 118606521 | rs6589681 | T | C | 1.188 | 1.202 | -9344.14 | -9334.117 | 10.023 | 0.000253676 |
| 11 | 118607150 | rs11217002 | T | C | 1.187 | 1.201 | -9357.888 | -9347.909 | 9.979 | 0.000242756 |
| 11 | 118721699 | rs7125066 | G | A | 1.13 | 1.196 | -9405.966 | -9395.99 | 9.977 | 0.000242271 |
| 11 | 118606722 | rs6589682 | T | C | 1.183 | 1.195 | -9318.2 | -9309.219 | 8.982 | 8.96E-05 |
| 11 | 118589012 | rs12421793 | A | C | 1.203 | 1.145 | -9380.994 | -9372.652 | 8.342 | 4.72E-05 |
| 11 | 118782222 | rs61900138 | C | T | 0.704 | 0.766 | -9307.813 | -9299.505 | 8.309 | 4.57E-05 |
| 11 | 118603736 | rs112561833 | C | A | 1.197 | 1.147 | -9401.989 | -9393.769 | 8.22 | 4.18E-05 |
| 11 | 118604638 | rs11216997 | T | C | 1.195 | 1.147 | -9403.408 | -9395.241 | 8.166 | 3.96E-05 |
| 11 | 118588952 | rs12421790 | T | C | 1.201 | 1.144 | -9384.784 | -9376.644 | 8.14 | 3.86E-05 |
| 11 | 118587319 | rs10892263 | C | T | 1.2 | 1.144 | -9383.106 | -9375.027 | 8.079 | 3.63E-05 |
| 11 | 118603526 | rs11216995 | A | G | 1.194 | 1.146 | -9406.738 | -9398.691 | 8.048 | 3.52E-05 |
| 11 | 118603520 | rs11216994 | T | C | 1.193 | 1.146 | -9406.738 | -9398.802 | 7.936 | 3.15E-05 |
| 11 | 118598544 | rs10892271 | A | G | 1.192 | 1.145 | -9406.738 | -9398.905 | 7.833 | 2.84E-05 |
| 11 | 118602707 | rs10892272 | G | C | 1.191 | 1.143 | -9406.738 | -9399.05 | 7.688 | 2.46E-05 |
| 11 | 118602809 | rs10892273 | A | G | 1.191 | 1.143 | -9406.738 | -9399.05 | 7.688 | 2.46E-05 |
| 11 | 118603165 | rs73001429 | G | T | 1.19 | 1.143 | -9406.738 | -9399.066 | 7.673 | 2.42E-05 |
| 11 | 118590668 | rs11216980 | A | C | 1.193 | 1.143 | -9387.759 | -9380.275 | 7.485 | 2.00E-05 |
| 11 | 118612565 | rs6589686 | A | G | 1.172 | 1.184 | -9402.874 | -9395.508 | 7.367 | 1.78E-05 |
| 11 | 118587097 | rs10892262 | C | G | 1.191 | 1.141 | -9382.451 | -9375.153 | 7.298 | 1.66E-05 |
| 11 | 118585826 | rs11216972 | T | C | 1.177 | 1.141 | -9393.182 | -9386.693 | 6.489 | 7.40E-06 |
| 11 | 118797562 | rs142336193 | G | A | 0.727 | 0.738 | -9314.554 | -9308.083 | 6.471 | 7.27E-06 |
| 11 | 118590961 | rs111227443 | T | C | 0.893 | 0.648 | -9354.982 | -9348.664 | 6.317 | 6.23E-06 |
| 11 | 118443944 | rs11216906 | A | G | 0.805 | 0.967 | -9385.815 | -9379.516 | 6.299 | 6.12E-06 |
| 11 | 118445585 | rs73023341 | G | A | 0.805 | 0.968 | -9386.211 | -9379.931 | 6.28 | 6.01E-06 |
| 11 | 118575326 | rs11216956 | A | G | 1.177 | 1.13 | -9407.752 | -9401.48 | 6.273 | 5.97E-06 |
| 11 | 118450046 | rs11216910 | G | A | 0.805 | 0.966 | -9385.023 | -9378.855 | 6.168 | 5.37E-06 |
| 11 | 118583525 | rs10892260 | C | A | 1.173 | 1.133 | -9406.682 | -9400.576 | 6.106 | 5.05E-06 |
| 11 | 118579747 | rs10790261 | G | A | 1.171 | 1.134 | -9408.077 | -9402.026 | 6.052 | 4.78E-06 |
| 11 | 118603495 | rs11216993 | A | T | 1.176 | 1.134 | -9339.45 | -9333.425 | 6.025 | 4.66E-06 |
| 11 | 118447580 | rs9736327 | G | T | 0.808 | 0.967 | -9386.041 | -9380.078 | 5.963 | 4.38E-06 |
| 11 | 118447597 | rs9735421 | A | G | 0.808 | 0.967 | -9386.041 | -9380.078 | 5.963 | 4.38E-06 |
| 11 | 118577873 | rs11216960 | A | G | 1.171 | 1.131 | -9408.077 | -9402.147 | 5.93 | 4.23E-06 |
| 11 | 118581895 | rs11216964 | A | G | 1.17 | 1.133 | -9407.542 | -9401.621 | 5.922 | 4.20E-06 |
| 11 | 118440863 | rs11216904 | G | A | 0.807 | 0.963 | -9387.388 | -9381.478 | 5.91 | 4.15E-06 |
| 11 | 118579865 | rs10892258 | A | G | 1.17 | 1.133 | -9408.077 | -9402.182 | 5.896 | 4.09E-06 |
| 11 | 118580338 | rs11216961 | G | C | 1.17 | 1.133 | -9408.077 | -9402.182 | 5.896 | 4.09E-06 |
| 11 | 118580813 | rs11216962 | C | T | 1.17 | 1.133 | -9408.077 | -9402.182 | 5.896 | 4.09E-06 |
| 11 | 118580814 | rs11216963 | A | G | 1.17 | 1.133 | -9408.077 | -9402.182 | 5.896 | 4.09E-06 |
| 11 | 118582613 | rs11216965 | A | C | 1.17 | 1.133 | -9408.077 | -9402.182 | 5.896 | 4.09E-06 |
| 11 | 118583002 | rs10892259 | A | G | 1.17 | 1.133 | -9408.077 | -9402.182 | 5.896 | 4.09E-06 |
| 11 | 118441846 | rs10892243 | C | T | 0.808 | 0.961 | -9387.388 | -9381.516 | 5.872 | 4.00E-06 |
| 11 | 118578705 | rs10892256 | A | G | 1.17 | 1.131 | -9408.077 | -9402.235 | 5.842 | 3.88E-06 |
| 11 | 118579327 | rs10892257 | A | G | 1.17 | 1.131 | -9407.491 | -9401.684 | 5.807 | 3.74E-06 |
| 11 | 118584340 | rs11216969 | C | T | 1.17 | 1.13 | -9406.682 | -9400.875 | 5.807 | 3.74E-06 |
| 11 | 118465295 | rs11216922 | G | A | 0.811 | 0.968 | -9373.316 | -9367.564 | 5.752 | 3.54E-06 |
| 11 | 118465501 | rs11216923 | G | A | 0.811 | 0.968 | -9373.316 | -9367.564 | 5.752 | 3.54E-06 |
| 11 | 118453483 | rs60112336 | T | C | 0.81 | 0.965 | -9388.387 | -9382.666 | 5.721 | 3.44E-06 |
| 11 | 118462160 | rs11216917 | G | T | 0.812 | 0.967 | -9374.504 | -9368.834 | 5.669 | 3.26E-06 |
| 11 | 118439048 | rs12224844 | T | A | 0.81 | 0.968 | -9386.463 | -9380.832 | 5.631 | 3.14E-06 |
| 11 | 118469571 | rs7931564 | G | A | 0.813 | 0.968 | -9373.316 | -9367.721 | 5.595 | 3.03E-06 |
| 11 | 118471423 | rs11216925 | C | T | 0.813 | 0.968 | -9373.316 | -9367.721 | 5.595 | 3.03E-06 |
| 11 | 118472354 | rs10892245 | A | C | 0.813 | 0.968 | -9373.316 | -9367.721 | 5.595 | 3.03E-06 |
| 11 | 118461605 | rs11216916 | C | T | 0.813 | 0.969 | -9375.342 | -9369.788 | 5.555 | 2.91E-06 |
| 11 | 118448087 | rs9735449 | T | C | 0.81 | 0.961 | -9385.295 | -9379.761 | 5.535 | 2.85E-06 |
| 11 | 118734521 | rs1174717189 | T | A | 0.934 | 0.854 | -9403.112 | -9397.584 | 5.528 | 2.83E-06 |
| 11 | 118604837 | rs11216998 | T | C | 1.117 | 1.168 | -9391.999 | -9386.472 | 5.526 | 2.83E-06 |
| 11 | 118473385 | rs17742 | C | T | 0.814 | 0.971 | -9375.313 | -9369.835 | 5.478 | 2.69E-06 |
| 11 | 118437279 | rs11216902 | G | A | 0.813 | 0.976 | -9387.986 | -9382.593 | 5.393 | 2.47E-06 |
| 11 | 118433729 | rs59108310 | G | A | 0.813 | 0.969 | -9387 | -9381.664 | 5.336 | 2.34E-06 |
| 11 | 118446298 | rs55844025 | T | C | 0.813 | 0.965 | -9387.404 | -9382.117 | 5.287 | 2.23E-06 |
| 11 | 118464815 | rs11216921 | C | G | 0.815 | 0.965 | -9376.856 | -9371.595 | 5.261 | 2.17E-06 |
| 11 | 118471996 | rs11216926 | T | C | 0.815 | 0.965 | -9375.843 | -9370.612 | 5.231 | 2.10E-06 |
| 11 | 118470092 | rs11216924 | G | C | 0.816 | 0.967 | -9375.843 | -9370.615 | 5.228 | 2.10E-06 |
| 11 | 118451901 | rs11216911 | C | T | 0.816 | 0.969 | -9377.513 | -9372.296 | 5.217 | 2.08E-06 |
| 11 | 118452334 | rs11216912 | C | G | 0.816 | 0.969 | -9377.513 | -9372.296 | 5.217 | 2.08E-06 |
| 11 | 118453681 | rs11216913 | G | A | 0.816 | 0.969 | -9377.513 | -9372.296 | 5.217 | 2.08E-06 |
| 11 | 118457630 | rs17092756 | C | G | 0.816 | 0.967 | -9376.108 | -9370.897 | 5.21 | 2.06E-06 |
| 11 | 118444525 | rs11216907 | A | G | 0.814 | 0.963 | -9387.404 | -9382.221 | 5.183 | 2.01E-06 |
| 11 | 118454177 | rs17842763 | G | A | 0.816 | 0.97 | -9376.759 | -9371.578 | 5.181 | 2.00E-06 |
| 11 | 118466249 | rs7115634 | G | A | 0.816 | 0.964 | -9376.856 | -9371.675 | 5.181 | 2.00E-06 |
| 11 | 118449396 | rs73023350 | A | G | 0.817 | 0.966 | -9377.842 | -9372.738 | 5.105 | 1.86E-06 |
| 11 | 118456414 | rs11216914 | C | T | 0.816 | 0.963 | -9378.33 | -9373.234 | 5.097 | 1.84E-06 |
| 11 | 118432279 | rs11216897 | G | A | 0.815 | 0.969 | -9387 | -9381.906 | 5.094 | 1.84E-06 |
| 11 | 118450875 | rs7126175 | T | C | 0.817 | 0.967 | -9378.191 | -9373.102 | 5.088 | 1.82E-06 |
| 11 | 118543563 | rs10892251 | T | C | 1.179 | 1.084 | -9408.077 | -9402.993 | 5.084 | 1.82E-06 |
| 11 | 118555692 | rs10892252 | G | A | 1.177 | 1.089 | -9408.077 | -9403.069 | 5.008 | 1.68E-06 |
| 11 | 118449746 | rs11216909 | T | C | 0.818 | 0.966 | -9377.513 | -9372.558 | 4.955 | 1.60E-06 |
| 11 | 118459039 | rs57438673 | A | G | 0.818 | 0.966 | -9377.839 | -9372.9 | 4.939 | 1.57E-06 |
| 11 | 118604941 | rs11216999 | C | T | 1.121 | 1.156 | -9399.26 | -9394.324 | 4.936 | 1.57E-06 |
| 11 | 118459702 | rs113054880 | G | C | 0.819 | 0.967 | -9378.2 | -9373.279 | 4.921 | 1.54E-06 |
| 11 | 118560857 | rs73001406 | A | G | 1.175 | 1.092 | -9408.077 | -9403.212 | 4.865 | 1.46E-06 |
| 11 | 118449370 | rs17122278 | G | A | 0.818 | 0.967 | -9386.499 | -9381.733 | 4.765 | 1.32E-06 |
| 11 | 118724723 | rs57719838 | C | T | 0.925 | 0.863 | -9405.966 | -9401.221 | 4.745 | 1.29E-06 |
| 11 | 118550524 | rs201134460 | T | C | 1.174 | 1.083 | -9408.077 | -9403.354 | 4.723 | 1.27E-06 |
| 11 | 118550522 | rs2277296 | C | T | 1.174 | 1.083 | -9407.677 | -9402.971 | 4.705 | 1.24E-06 |
| 11 | 118725942 | rs11217063 | C | G | 0.925 | 0.864 | -9407.595 | -9402.899 | 4.696 | 1.23E-06 |
| 11 | 118726647 | rs11217065 | C | T | 0.925 | 0.864 | -9407.595 | -9402.899 | 4.696 | 1.23E-06 |
| 11 | 118556398 | rs11216943 | A | G | 1.173 | 1.085 | -9408.077 | -9403.393 | 4.684 | 1.22E-06 |
| 11 | 118729210 | rs10892300 | T | C | 0.934 | 0.862 | -9407.595 | -9402.93 | 4.665 | 1.19E-06 |
| 11 | 118729860 | rs11217070 | G | T | 0.934 | 0.862 | -9407.595 | -9402.94 | 4.655 | 1.18E-06 |
| 11 | 118723379 | rs7103067 | A | G | 0.925 | 0.865 | -9405.966 | -9401.404 | 4.562 | 1.08E-06 |
| 11 | 118729583 | rs12790489 | C | T | 0.934 | 0.863 | -9407.595 | -9403.042 | 4.553 | 1.07E-06 |
| 11 | 118731599 | rs34561290 | T | C | 0.935 | 0.863 | -9407.595 | -9403.056 | 4.539 | 1.05E-06 |
| 11 | 118731762 | rs11217072 | G | A | 0.935 | 0.863 | -9407.595 | -9403.056 | 4.539 | 1.05E-06 |
| 11 | 118473231 | rs1047079 | C | T | 0.823 | 0.969 | -9376.856 | -9372.322 | 4.534 | 1.05E-06 |
| 11 | 118730141 | rs7929520 | G | A | 0.935 | 0.862 | -9407.595 | -9403.067 | 4.528 | 1.04E-06 |
| 11 | 118436079 | rs11216900 | C | G | 0.821 | 0.969 | -9386.246 | -9381.738 | 4.509 | 1.02E-06 |
| 11 | 118445275 | rs17122269 | C | T | 0.822 | 0.966 | -9386.088 | -9381.624 | 4.464 | 9.77E-07 |
| 11 | 118432404 | rs4326835 | T | G | 0.822 | 0.97 | -9384.907 | -9380.499 | 4.408 | 9.24E-07 |
| 11 | 118459069 | rs633308 | G | A | 1.154 | 1.027 | -9105.923 | -9101.53 | 4.393 | 9.10E-07 |
| 11 | 118432852 | rs11216898 | C | T | 0.823 | 0.969 | -9386.246 | -9381.88 | 4.367 | 8.87E-07 |
| 11 | 118605463 | rs10892274 | T | C | 1.118 | 1.147 | -9396.088 | -9391.77 | 4.318 | 8.45E-07 |
| 11 | 118605617 | rs10892275 | A | T | 1.118 | 1.147 | -9396.088 | -9391.77 | 4.318 | 8.45E-07 |
| 11 | 118434421 | rs11216899 | G | A | 0.823 | 0.967 | -9386.246 | -9381.943 | 4.304 | 8.33E-07 |
| 11 | 118434835 | rs60753148 | T | C | 0.823 | 0.967 | -9386.246 | -9381.943 | 4.304 | 8.33E-07 |
| 11 | 118740180 | rs7483509 | A | G | 0.937 | 0.864 | -9402.633 | -9398.373 | 4.259 | 7.96E-07 |
| 11 | 118605841 | rs10892277 | A | G | 1.118 | 1.146 | -9396.088 | -9391.868 | 4.22 | 7.66E-07 |
| 11 | 118438448 | rs60837962 | T | A | 0.825 | 0.97 | -9384.393 | -9380.225 | 4.168 | 7.27E-07 |
| 11 | 118435756 | rs2277291 | T | C | 0.826 | 0.974 | -9384 | -9379.917 | 4.084 | 6.68E-07 |
| 11 | 118605628 | rs10892276 | A | G | 1.116 | 1.144 | -9394 | -9389.937 | 4.064 | 6.55E-07 |
| 11 | 118805385 | rs55772432 | A | G | 0.864 | 0.808 | -9230.439 | -9226.415 | 4.024 | 6.29E-07 |
| 11 | 118769604 | rs61751532 | T | C | 0.937 | 0.777 | -9262.41 | -9258.443 | 3.967 | 5.95E-07 |
| 11 | 118711790 | rs622722 | A | G | 0.931 | 0.869 | -9397.96 | -9394.077 | 3.883 | 5.47E-07 |
| 11 | 118727169 | rs10790267 | T | C | 0.932 | 0.87 | -9405.316 | -9401.474 | 3.843 | 5.25E-07 |
| 11 | 118728320 | rs4936442 | T | C | 0.931 | 0.87 | -9405.316 | -9401.527 | 3.789 | 4.98E-07 |
| 11 | 118789026 | rs11607239 | T | G | 0.834 | 0.737 | -9366.164 | -9362.398 | 3.766 | 4.86E-07 |
| 11 | 118727332 | rs11217067 | T | G | 0.932 | 0.871 | -9405.316 | -9401.569 | 3.748 | 4.78E-07 |
| 11 | 118809805 | rs142069912 | C | G | 0.824 | 0.717 | -9386.646 | -9382.928 | 3.718 | 4.64E-07 |
| 11 | 118750321 | rs11217078 | C | T | 0.967 | 1.134 | -9405.113 | -9401.671 | 3.442 | 3.52E-07 |
| 11 | 118669178 | rs583213 | G | T | 0.912 | 0.838 | -9407.18 | -9404.262 | 2.917 | 2.08E-07 |
| 11 | 118749318 | rs78312641 | A | T | 0.869 | 1.063 | -9404.454 | -9401.728 | 2.727 | 1.72E-07 |
| 11 | 118530611 | rs2276065 | C | T | 1.153 | 1.035 | -9407.272 | -9404.553 | 2.719 | 1.71E-07 |
| 11 | 118749563 | rs77643000 | C | T | 0.867 | 1.058 | -9405.718 | -9403.018 | 2.7 | 1.67E-07 |
| 11 | 118587347 | rs603486 | T | C | 0.893 | 0.917 | -9381.051 | -9378.418 | 2.632 | 1.56E-07 |
| 11 | 118504742 | rs633683 | T | C | 0.884 | 0.951 | -9372.295 | -9369.665 | 2.63 | 1.56E-07 |
| 11 | 118697012 | rs571470 | A | G | 1.044 | 1.14 | -9396.993 | -9394.384 | 2.609 | 1.53E-07 |
| 11 | 118748170 | rs77074463 | C | G | 0.832 | 1.022 | -9375.784 | -9373.201 | 2.583 | 1.49E-07 |
| 11 | 118577990 | rs480958 | G | A | 0.889 | 0.928 | -9406.526 | -9404.003 | 2.524 | 1.40E-07 |
| 11 | 118584498 | rs569770 | T | C | 0.892 | 0.923 | -9399.893 | -9397.517 | 2.376 | 1.21E-07 |
| 11 | 118509741 | rs113313560 | T | C | 0.874 | 0.726 | -9348.941 | -9346.703 | 2.238 | 1.06E-07 |
| 11 | 118601426 | rs186926311 | T | C | 1.281 | 0.866 | -9385.049 | -9382.821 | 2.229 | 1.05E-07 |
| 11 | 118807639 | rs73575454 | A | G | 0.835 | 0.855 | -9348.139 | -9345.944 | 2.195 | 1.01E-07 |
| 11 | 118809767 | rs75460330 | A | G | 0.835 | 0.855 | -9348.139 | -9345.944 | 2.195 | 1.01E-07 |
| 11 | 118810054 | rs73557344 | C | G | 0.835 | 0.855 | -9348.139 | -9345.944 | 2.195 | 1.01E-07 |
| 11 | 118507478 | rs11216937 | T | C | 1.145 | 1.025 | -9408.077 | -9405.95 | 2.127 | 9.44E-08 |
| 11 | 118820778 | rs139076649 | C | T | 1.136 | 0.811 | -9373.157 | -9371.041 | 2.116 | 9.34E-08 |
| 11 | 118662340 | rs5006102 | A | T | 0.913 | 0.85 | -9407.45 | -9405.337 | 2.114 | 9.32E-08 |
| 11 | 118516903 | rs80015904 | A | G | 1.145 | 1.026 | -9406.626 | -9404.522 | 2.104 | 9.23E-08 |
| 11 | 118503912 | rs45478298 | T | C | 1.228 | 0.927 | -9393.296 | -9391.217 | 2.08 | 9.01E-08 |
| 11 | 118668908 | rs582200 | G | A | 0.919 | 0.849 | -9408.077 | -9406.005 | 2.073 | 8.95E-08 |
| 11 | 118662045 | rs7933859 | T | C | 0.914 | 0.851 | -9408.077 | -9406.01 | 2.067 | 8.89E-08 |
| 11 | 118662203 | rs4508234 | G | C | 0.914 | 0.851 | -9408.077 | -9406.01 | 2.067 | 8.89E-08 |
| 11 | 118495287 | rs77683570 | C | T | 1.144 | 1.026 | -9408.077 | -9406.011 | 2.066 | 8.88E-08 |
| 11 | 118678780 | rs500254 | G | A | 0.916 | 0.851 | -9408.077 | -9406.045 | 2.033 | 8.60E-08 |
| 11 | 118665009 | rs4938544 | A | G | 0.918 | 0.85 | -9408.077 | -9406.052 | 2.025 | 8.53E-08 |
| 11 | 118534082 | rs10790256 | T | C | 1.143 | 1.026 | -9408.077 | -9406.079 | 1.998 | 8.30E-08 |
| 11 | 118490076 | rs10892247 | A | G | 1.143 | 1.024 | -9407.579 | -9405.588 | 1.991 | 8.24E-08 |
| 11 | 118677083 | rs675003 | T | C | 0.918 | 0.851 | -9408.077 | -9406.094 | 1.983 | 8.18E-08 |
| 11 | 118665324 | rs61901411 | T | C | 0.921 | 0.85 | -9408.077 | -9406.143 | 1.934 | 7.79E-08 |
| 11 | 118528424 | rs17748 | T | C | 1.142 | 1.023 | -9407.677 | -9405.753 | 1.923 | 7.70E-08 |
| 11 | 118802128 | rs59069289 | T | G | 0.84 | 0.86 | -9351.244 | -9349.332 | 1.912 | 7.62E-08 |
| 11 | 118802386 | rs59480472 | C | T | 0.84 | 0.86 | -9351.244 | -9349.332 | 1.912 | 7.62E-08 |

| | | | | | | | |
|---|---|---|---|---|---|---|---|
| 11 | 118503912 | rs45478298 | T | C | M_Disease2_Disease1 | 0.361456994 | M_Disease2_Disease1&M_Disease1&M_Disease2&NULL |
| 11 | 118664377 | rs598373 | T | C | M_Disease2_Disease1 | 0.361372521 | M_Disease2_Disease1&M_Disease1&NULL&M_Disease2 |
| 11 | 118777978 | rs78755429 | A | C | M_Disease2_Disease1 | 0.360759079 | M_Disease2_Disease1&M_Disease2&NULL&M_Disease1 |
| 11 | 118495287 | rs77683570 | C | T | M_Disease2_Disease1 | 0.360016901 | M_Disease2_Disease1&M_Disease1&NULL&M_Disease2 |
| 11 | 118615232 | rs139423236 | G | C | M_Disease2_Disease1 | 0.358125055 | M_Disease2_Disease1&M_Disease2&NULL&M_Disease1 |
| 11 | 118782474 | rs11606719 | G | C | M_Disease2_Disease1 | 0.356814822 | M_Disease2_Disease1&M_Disease1&NULL&M_Disease2 |
| 11 | 118762521 | rs190509947 | T | A | M_Disease2_Disease1 | 0.355979015 | M_Disease2_Disease1&M_Disease2&NULL&M_Disease1 |
| 11 | 118534082 | rs10790256 | T | C | M_Disease2_Disease1 | 0.355796522 | M_Disease2_Disease1&M_Disease1&NULL&M_Disease2 |
| 11 | 118490076 | rs10892247 | A | G | M_Disease2_Disease1 | 0.354759433 | M_Disease2_Disease1&M_Disease1&NULL&M_Disease2 |
| 11 | 118590743 | rs503542 | G | A | NULL | 0.353607383 | NULL&M_Disease2&M_Disease1&M_Disease2_Disease1 |
| 11 | 118812821 | rs74535623 | T | G | M_Disease2_Disease1 | 0.35270219 | M_Disease2_Disease1&M_Disease1&NULL&M_Disease2 |
| 11 | 118813314 | rs73557345 | C | T | M_Disease2_Disease1 | 0.35270219 | M_Disease2_Disease1&M_Disease1&NULL&M_Disease2 |
| 11 | 118752178 | rs61150131 | C | T | M_Disease2_Disease1 | 0.35138273 | M_Disease2_Disease1&M_Disease2&M_Disease1&NULL |
| 11 | 118528424 | rs17748 | T | C | M_Disease2_Disease1 | 0.351321038 | M_Disease2_Disease1&M_Disease1&NULL&M_Disease2 |
| 11 | 118594326 | rs544096 | G | A | M_Disease2_Disease1 | 0.350712104 | M_Disease2_Disease1&M_Disease1&M_Disease2&NULL |
| 11 | 118580906 | rs496058 | G | A | NULL | 0.350201768 | NULL&M_Disease2&M_Disease1&M_Disease2_Disease1 |
| 11 | 118580348 | rs516503 | T | C | NULL | 0.350043577 | NULL&M_Disease2&M_Disease1&M_Disease2_Disease1 |
| 11 | 118595207 | rs563537 | A | T | NULL | 0.349356198 | NULL&M_Disease2&M_Disease1&M_Disease2_Disease1 |
| 11 | 118617920 | rs476246 | T | C | M_Disease2_Disease1 | 0.349335712 | M_Disease2_Disease1&M_Disease1&M_Disease2&NULL |
| 11 | 118598612 | rs568688 | G | T | NULL | 0.349064832 | NULL&M_Disease2&M_Disease1&M_Disease2_Disease1 |
| 11 | 118598138 | rs543692 | C | T | NULL | 0.348773234 | NULL&M_Disease2&M_Disease1&M_Disease2_Disease1 |
| 11 | 118486110 | rs45540840 | A | G | M_Disease2_Disease1 | 0.348577311 | M_Disease2_Disease1&M_Disease1&NULL&M_Disease2 |
| 11 | 118516774 | rs12365095 | T | C | M_Disease2_Disease1 | 0.348006864 | M_Disease2_Disease1&M_Disease1&NULL&M_Disease2 |
| 11 | 118573916 | rs627391 | T | A | NULL | 0.347520005 | NULL&M_Disease2&M_Disease1&M_Disease2_Disease1 |
| 11 | 118617544 | rs472641 | G | A | M_Disease2_Disease1 | 0.347319381 | M_Disease2_Disease1&M_Disease1&M_Disease2&NULL |
| 11 | 118575419 | rs625735 | A | G | NULL | 0.347072207 | NULL&M_Disease2&M_Disease1&M_Disease2_Disease1 |
| 11 | 118575479 | rs637563 | G | A | NULL | 0.347072207 | NULL&M_Disease2&M_Disease1&M_Disease2_Disease1 |
| 11 | 118575693 | rs638514 | C | T | NULL | 0.347072207 | NULL&M_Disease2&M_Disease1&M_Disease2_Disease1 |
| 11 | 118573894 | rs576283 | G | A | NULL | 0.346931987 | NULL&M_Disease2&M_Disease1&M_Disease2_Disease1 |
| 11 | 118575011 | rs530901 | C | T | NULL | 0.346931987 | NULL&M_Disease2&M_Disease1&M_Disease2_Disease1 |
| 11 | 118575180 | rs626645 | C | T | NULL | 0.346931987 | NULL&M_Disease2&M_Disease1&M_Disease2_Disease1 |
| 11 | 118609736 | rs1783937 | A | G | M_Disease2_Disease1 | 0.346891909 | M_Disease2_Disease1&M_Disease2&NULL&M_Disease1 |
| 11 | 118786829 | rs7102433 | G | A | M_Disease2_Disease1 | 0.346856394 | M_Disease2_Disease1&M_Disease1&NULL&M_Disease2 |
| 11 | 118574311 | rs524409 | T | C | NULL | 0.346773865 | NULL&M_Disease2&M_Disease1&M_Disease2_Disease1 |
| 11 | 118574675 | rs494459 | T | C | NULL | 0.346773865 | NULL&M_Disease2&M_Disease1&M_Disease2_Disease1 |
| 11 | 118765600 | rs3922 | G | A | NULL | 0.345682974 | NULL&M_Disease1&M_Disease2&M_Disease2_Disease1 |
| 11 | 118588998 | rs570424 | C | T | NULL | 0.345435355 | NULL&M_Disease2&M_Disease1&M_Disease2_Disease1 |
| 11 | 118836606 | rs113975105 | G | C | NULL | 0.344942384 | NULL&M_Disease2&M_Disease1&M_Disease2_Disease1 |
| 11 | 118884708 | rs61381062 | A | G | NULL | 0.344491747 | NULL&M_Disease2&M_Disease1&M_Disease2_Disease1 |
| 11 | 118884726 | rs58601794 | C | T | NULL | 0.344491747 | NULL&M_Disease2&M_Disease1&M_Disease2_Disease1 |
| 11 | 118885163 | rs59608621 | G | A | NULL | 0.344491747 | NULL&M_Disease2&M_Disease1&M_Disease2_Disease1 |
| 11 | 118514625 | rs11216938 | T | C | M_Disease2_Disease1 | 0.344405812 | M_Disease2_Disease1&M_Disease1&NULL&M_Disease2 |
| 11 | 118769442 | rs7104819 | G | A | NULL | 0.344366774 | NULL&M_Disease1&M_Disease2&M_Disease2_Disease1 |
| 11 | 118885161 | rs59095933 | A | G | NULL | 0.344195683 | NULL&M_Disease2&M_Disease1&M_Disease2_Disease1 |
| 11 | 118889132 | rs11217124 | G | A | NULL | 0.344195683 | NULL&M_Disease2&M_Disease1&M_Disease2_Disease1 |
| 11 | 118889185 | rs11217125 | C | T | NULL | 0.344195683 | NULL&M_Disease2&M_Disease1&M_Disease2_Disease1 |
| 11 | 118869460 | rs3737504 | G | A | NULL | 0.343597213 | NULL&M_Disease2&M_Disease1&M_Disease2_Disease1 |
| 11 | 118832035 | rs10892317 | C | T | NULL | 0.343175201 | NULL&M_Disease2&M_Disease1&M_Disease2_Disease1 |
| 11 | 118872741 | rs4573680 | C | T | NULL | 0.343152358 | NULL&M_Disease2&M_Disease1&M_Disease2_Disease1 |
| 11 | 118867803 | rs4938618 | C | G | NULL | 0.342861926 | NULL&M_Disease2&M_Disease1&M_Disease2_Disease1 |
| 11 | 118869539 | rs11822103 | A | G | NULL | 0.342861926 | NULL&M_Disease2&M_Disease1&M_Disease2_Disease1 |
| 11 | 118887888 | rs3916582 | T | C | NULL | 0.342712053 | NULL&M_Disease2&M_Disease1&M_Disease2_Disease1 |
| 11 | 118831727 | rs10892315 | C | T | NULL | 0.342425 | NULL&M_Disease2&M_Disease1&M_Disease2_Disease1 |
| 11 | 118854372 | rs75067211 | T | C | NULL | 0.342414668 | NULL&M_Disease2&M_Disease1&M_Disease2_Disease1 |
| 11 | 118887887 | rs369816532 | T | G | NULL | 0.342414668 | NULL&M_Disease2&M_Disease1&M_Disease2_Disease1 |
| 11 | 118875727 | rs4936457 | G | A | NULL | 0.34241037 | NULL&M_Disease2&M_Disease1&M_Disease2_Disease1 |
| 11 | 118875761 | rs4936458 | G | T | NULL | 0.34241037 | NULL&M_Disease2&M_Disease1&M_Disease2_Disease1 |
| 11 | 118887568 | rs7121270 | G | T | NULL | 0.342270712 | NULL&M_Disease2&M_Disease1&M_Disease2_Disease1 |
| 11 | 118873987 | rs9645664 | C | T | NULL | 0.342111294 | NULL&M_Disease2&M_Disease1&M_Disease2_Disease1 |
| 11 | 118761159 | rs57231586 | T | G | M_Disease2_Disease1 | 0.341877145 | M_Disease2_Disease1&M_Disease1&NULL&M_Disease2 |
| 11 | 118887581 | rs7105187 | T | C | NULL | 0.34168746 | NULL&M_Disease2&M_Disease1&M_Disease2_Disease1 |
| 11 | 118883491 | rs60065241 | A | C | NULL | 0.341685173 | NULL&M_Disease2&M_Disease1&M_Disease2_Disease1 |
| 11 | 118878222 | rs11606357 | C | T | NULL | 0.341390406 | NULL&M_Disease2&M_Disease1&M_Disease2_Disease1 |
| 11 | 118856582 | rs3887652 | C | G | NULL | 0.341371521 | NULL&M_Disease2&M_Disease1&M_Disease2_Disease1 |
| 11 | 118535840 | rs12225548 | G | C | M_Disease2_Disease1 | 0.341300854 | M_Disease2_Disease1&M_Disease1&NULL&M_Disease2 |
| 11 | 118892277 | rs4938621 | T | C | NULL | 0.34123831 | NULL&M_Disease2&M_Disease1&M_Disease2_Disease1 |
| 11 | 118892835 | rs61179449 | C | G | NULL | 0.34123831 | NULL&M_Disease2&M_Disease1&M_Disease2_Disease1 |
| 11 | 118892872 | rs58955701 | T | C | NULL | 0.34123831 | NULL&M_Disease2&M_Disease1&M_Disease2_Disease1 |
| 11 | 118893023 | rs7127188 | C | A | NULL | 0.34123831 | NULL&M_Disease2&M_Disease1&M_Disease2_Disease1 |
| 11 | 118877563 | rs58709307 | T | C | NULL | 0.341095416 | NULL&M_Disease2&M_Disease1&M_Disease2_Disease1 |
| 11 | 118880668 | rs11607569 | C | T | NULL | 0.34108937 | NULL&M_Disease2&M_Disease1&M_Disease2_Disease1 |
| 11 | 118723753 | rs7119568 | A | T | NULL | 0.340947126 | NULL&M_Disease2&M_Disease1&M_Disease2_Disease1 |
| 11 | 118879285 | rs7120445 | C | A | NULL | 0.340944842 | NULL&M_Disease2&M_Disease1&M_Disease2_Disease1 |
| 11 | 118879824 | rs73559119 | C | T | NULL | 0.340943582 | NULL&M_Disease2&M_Disease1&M_Disease2_Disease1 |
| 11 | 118880012 | rs11825284 | A | G | NULL | 0.340943582 | NULL&M_Disease2&M_Disease1&M_Disease2_Disease1 |
| 11 | 118734640 | rs12286723 | T | C | NULL | 0.340804015 | NULL&M_Disease2&M_Disease1&M_Disease2_Disease1 |
| 11 | 118886656 | rs7131534 | A | G | NULL | 0.340798945 | NULL&M_Disease2&M_Disease1&M_Disease2_Disease1 |
| 11 | 118728479 | rs7924647 | G | A | NULL | 0.340797184 | NULL&M_Disease2&M_Disease1&M_Disease2_Disease1 |
| 11 | 118729442 | rs12277355 | T | C | NULL | 0.340647159 | NULL&M_Disease2&M_Disease1&M_Disease2_Disease1 |
| 11 | 118886117 | rs10111 | T | C | NULL | 0.340647123 | NULL&M_Disease2&M_Disease1&M_Disease2_Disease1 |
| 11 | 118714492 | rs7117594 | T | C | NULL | 0.34064511 | NULL&M_Disease2&M_Disease1&M_Disease2_Disease1 |
| 11 | 118888619 | rs11602764 | G | A | NULL | 0.340627473 | NULL&M_Disease2&M_Disease1&M_Disease2_Disease1 |
| 11 | 118889378 | rs7113753 | A | G | NULL | 0.340618407 | NULL&M_Disease2&M_Disease1&M_Disease2_Disease1 |
| 11 | 118725383 | rs9735317 | T | C | NULL | 0.340352506 | NULL&M_Disease2&M_Disease1&M_Disease2_Disease1 |
| 11 | 118729283 | rs11217068 | T | C | NULL | 0.340349491 | NULL&M_Disease2&M_Disease1&M_Disease2_Disease1 |
| 11 | 118724116 | rs12276533 | A | G | NULL | 0.340203086 | NULL&M_Disease2&M_Disease1&M_Disease2_Disease1 |
| 11 | 118499394 | rs2236661 | G | C | M_Disease2_Disease1 | 0.340134845 | M_Disease2_Disease1&M_Disease1&NULL&M_Disease2 |
| 11 | 118888831 | rs11820805 | C | T | NULL | 0.340027616 | NULL&M_Disease2&M_Disease1&M_Disease2_Disease1 |
| 11 | 118496119 | rs45565037 | A | G | M_Disease2_Disease1 | 0.339568745 | M_Disease2_Disease1&M_Disease1&NULL&M_Disease2 |
| 11 | 118868750 | rs4445645 | C | G | NULL | 0.339470216 | NULL&M_Disease1&M_Disease2&M_Disease2_Disease1 |
| 11 | 118707559 | rs7118009 | T | C | NULL | 0.339313311 | NULL&M_Disease2&M_Disease1&M_Disease2_Disease1 |
| 11 | 118709600 | rs12272981 | G | A | NULL | 0.339310305 | NULL&M_Disease2&M_Disease1&M_Disease2_Disease1 |
| 11 | 118872404 | rs6589713 | C | T | NULL | 0.33930037 | NULL&M_Disease2&M_Disease1&M_Disease2_Disease1 |
| 11 | 118712270 | rs7105658 | T | C | NULL | 0.338863784 | NULL&M_Disease2&M_Disease1&M_Disease2_Disease1 |
| 11 | 118893878 | rs111322300 | C | T | NULL | 0.338856275 | NULL&M_Disease2&M_Disease1&M_Disease2_Disease1 |
| 11 | 118896115 | rs2276072 | G | A | NULL | 0.338546425 | NULL&M_Disease2&M_Disease1&M_Disease2_Disease1 |
| 11 | 118896116 | rs2276071 | C | A | NULL | 0.338546425 | NULL&M_Disease2&M_Disease1&M_Disease2_Disease1 |
| 11 | 118712427 | rs7108905 | G | C | NULL | 0.338419321 | NULL&M_Disease2&M_Disease1&M_Disease2_Disease1 |
| 11 | 118892757 | rs117289748 | G | C | NULL | 0.338409951 | NULL&M_Disease2&M_Disease1&M_Disease2_Disease1 |
| 11 | 118892759 | rs143660533 | C | A | NULL | 0.338409951 | NULL&M_Disease2&M_Disease1&M_Disease2_Disease1 |
| 11 | 118894287 | rs569 | G | A | NULL | 0.338409951 | NULL&M_Disease2&M_Disease1&M_Disease2_Disease1 |
| 11 | 118895202 | rs11006 | G | A | NULL | 0.338409951 | NULL&M_Disease2&M_Disease1&M_Disease2_Disease1 |
| 11 | 118708785 | rs7122510 | A | C | NULL | 0.338119515 | NULL&M_Disease2&M_Disease1&M_Disease2_Disease1 |
| 11 | 118708786 | rs7122511 | A | C | NULL | 0.338119515 | NULL&M_Disease2&M_Disease1&M_Disease2_Disease1 |
| 11 | 118679387 | rs473298 | G | A | M_Disease2_Disease1 | 0.338061113 | M_Disease2_Disease1&M_Disease1&NULL&M_Disease2 |
| 11 | 118679445 | rs516719 | C | T | M_Disease2_Disease1 | 0.338061113 | M_Disease2_Disease1&M_Disease1&NULL&M_Disease2 |
| 11 | 118516737 | rs12362152 | G | A | M_Disease2_Disease1 | 0.337991182 | M_Disease2_Disease1&M_Disease1&NULL&M_Disease2 |
| 11 | 118831335 | rs61506726 | G | A | NULL | 0.337950041 | NULL&M_Disease2&M_Disease1&M_Disease2_Disease1 |
| 11 | 118501022 | rs10892248 | A | G | M_Disease2_Disease1 | 0.337788002 | M_Disease2_Disease1&M_Disease1&NULL&M_Disease2 |
| 11 | 118678986 | rs477014 | C | T | M_Disease2_Disease1 | 0.337563944 | M_Disease2_Disease1&NULL&M_Disease1&M_Disease2 |
| 11 | 118895686 | rs8192696 | T | C | NULL | 0.337532719 | NULL&M_Disease2&M_Disease1&M_Disease2_Disease1 |
| 11 | 118872581 | rs4558175 | A | G | NULL | 0.337084466 | NULL&M_Disease2&M_Disease1&M_Disease2_Disease1 |
| 11 | 118873385 | rs10892320 | T | C | NULL | 0.33692945 | NULL&M_Disease2&M_Disease1&M_Disease2_Disease1 |
| 11 | 118677710 | rs688161 | G | A | M_Disease2_Disease1 | 0.336817313 | M_Disease2_Disease1&M_Disease1&NULL&M_Disease2 |
| 11 | 118678609 | rs2508570 | A | G | M_Disease2_Disease1 | 0.336817313 | M_Disease2_Disease1&M_Disease1&NULL&M_Disease2 |
| 11 | 118678881 | rs591756 | T | C | M_Disease2_Disease1 | 0.335452873 | M_Disease2_Disease1&M_Disease2&NULL&M_Disease1 |
| 11 | 118896959 | rs57156480 | A | G | NULL | 0.335249154 | NULL&M_Disease2&M_Disease1&M_Disease2_Disease1 |
| 11 | 118735476 | rs10892301 | A | G | NULL | 0.33516528 | NULL&M_Disease1&M_Disease2&M_Disease2_Disease1 |
| 11 | 118892940 | rs12280288 | C | T | NULL | 0.33500945 | NULL&M_Disease2&M_Disease1&M_Disease2_Disease1 |
| 11 | 118680045 | rs2508569 | C | G | M_Disease2_Disease1 | 0.333722614 | M_Disease2_Disease1&NULL&M_Disease2&M_Disease1 |
| 11 | 118661197 | rs61901410 | G | C | M_Disease2_Disease1 | 0.33333834 | M_Disease2_Disease1&M_Disease2&NULL&M_Disease1 |
| 11 | 118529127 | rs7928371 | A | G | M_Disease2_Disease1 | 0.333220456 | M_Disease2_Disease1&M_Disease1&NULL&M_Disease2 |
| 11 | 118781710 | rs6589707 | A | C | M_Disease2_Disease1 | 0.332226122 | M_Disease2_Disease1&M_Disease1&NULL&M_Disease2 |
| 11 | 118769032 | rs11217081 | G | A | NULL | 0.331764736 | NULL&M_Disease2&M_Disease1&M_Disease2_Disease1 |
| 11 | 118604678 | rs493720 | T | C | M_Disease2_Disease1 | 0.330663335 | M_Disease2_Disease1&M_Disease2&NULL&M_Disease1 |
| 11 | 118640063 | rs524590 | T | A | M_Disease2_Disease1 | 0.330395081 | M_Disease2_Disease1&NULL&M_Disease2&M_Disease1 |
| 11 | 118661376 | rs73003294 | T | C | NULL | 0.330374091 | NULL&M_Disease2&M_Disease1&M_Disease2_Disease1 |
| 11 | 118880740 | rs11217122 | C | T | NULL | 0.33024999 | NULL&M_Disease1&M_Disease2&M_Disease2_Disease1 |
| 11 | 118757219 | rs1623316 | G | C | M_Disease2 | 0.33008318 | M_Disease2&M_Disease2_Disease1&NULL&M_Disease1 |
| 11 | 118610629 | rs521910 | A | G | M_Disease2_Disease1 | 0.330053503 | M_Disease2_Disease1&M_Disease2&NULL&M_Disease1 |
| 11 | 118767077 | rs581063 | C | T | NULL | 0.32994037 | NULL&M_Disease2&M_Disease1&M_Disease2_Disease1 |
| 11 | 118825636 | rs1790191 | G | A | NULL | 0.329777465 | NULL&M_Disease1&M_Disease2&M_Disease2_Disease1 |
| 11 | 118760443 | rs12786257 | A | G | M_Disease2_Disease1 | 0.329650593 | M_Disease2_Disease1&M_Disease2&M_Disease1&NULL |
| 11 | 118824439 | rs61900157 | C | A | NULL | 0.329613261 | NULL&M_Disease1&M_Disease2&M_Disease2_Disease1 |
| 11 | 118824996 | rs4426142 | A | G | NULL | 0.329613261 | NULL&M_Disease1&M_Disease2&M_Disease2_Disease1 |
| 11 | 118825069 | rs2097131 | A | G | NULL | 0.329613261 | NULL&M_Disease1&M_Disease2&M_Disease2_Disease1 |
| 11 | 118605127 | rs526850 | G | T | M_Disease2_Disease1 | 0.329202824 | M_Disease2_Disease1&M_Disease2&NULL&M_Disease1 |
| 11 | 118826060 | rs625291 | T | C | NULL | 0.32914368 | NULL&M_Disease1&M_Disease2&M_Disease2_Disease1 |
| 11 | 118826063 | rs666191 | A | G | NULL | 0.32914368 | NULL&M_Disease1&M_Disease2&M_Disease2_Disease1 |
| 11 | 118601363 | rs557990 | T | C | M_Disease2_Disease1 | 0.329027089 | M_Disease2_Disease1&M_Disease2&NULL&M_Disease1 |
| 11 | 118602392 | rs564892 | A | T | M_Disease2_Disease1 | 0.329027089 | M_Disease2_Disease1&M_Disease2&NULL&M_Disease1 |
| 11 | 118607694 | rs7119702 | G | A | M_Disease2_Disease1 | 0.328541731 | M_Disease2_Disease1&M_Disease2&NULL&M_Disease1 |
| 11 | 118624767 | rs581062 | A | G | M_Disease2_Disease1 | 0.328290479 | M_Disease2_Disease1&M_Disease1&NULL&M_Disease2 |
| 11 | 118652326 | rs1790189 | G | T | M_Disease2_Disease1 | 0.327814053 | M_Disease2_Disease1&NULL&M_Disease2&M_Disease1 |
| 11 | 118652373 | rs1784293 | C | A | M_Disease2_Disease1 | 0.327814053 | M_Disease2_Disease1&NULL&M_Disease2&M_Disease1 |
| 11 | 118488782 | rs11216930 | C | A | M_Disease2_Disease1 | 0.327802562 | M_Disease2_Disease1&M_Disease1&NULL&M_Disease2 |
| 11 | 118660932 | rs1784295 | A | G | M_Disease2_Disease1 | 0.327527815 | M_Disease2_Disease1&M_Disease2&NULL&M_Disease1 |
| 11 | 118751898 | rs60651507 | A | G | M_Disease2_Disease1 | 0.327073175 | M_Disease2_Disease1&M_Disease2&NULL&M_Disease1 |
| 11 | 118770321 | rs11217083 | G | A | NULL | 0.326379583 | NULL&M_Disease2&M_Disease1&M_Disease2_Disease1 |
| 11 | 118823148 | rs663003 | T | C | NULL | 0.326223651 | NULL&M_Disease1&M_Disease2&M_Disease2_Disease1 |
| 11 | 118605447 | rs2186781 | T | C | M_Disease2_Disease1 | 0.326175949 | M_Disease2_Disease1&M_Disease2&NULL&M_Disease1 |
| 11 | 118787213 | rs7119222 | C | T | M_Disease2_Disease1 | 0.325757221 | M_Disease2_Disease1&M_Disease1&NULL&M_Disease2 |
| 11 | 118717708 | rs7107756 | A | G | NULL | 0.325470095 | NULL&M_Disease2&M_Disease1&M_Disease2_Disease1 |
| 11 | 118626511 | rs499343 | T | C | M_Disease2_Disease1 | 0.325251087 | M_Disease2_Disease1&NULL&M_Disease1&M_Disease2 |
| 11 | 118605352 | rs76478216 | C | T | M_Disease2_Disease1 | 0.324701624 | M_Disease2_Disease1&M_Disease1&M_Disease2&NULL |
| 11 | 118816304 | rs586763 | T | C | NULL | 0.324492706 | NULL&M_Disease1&M_Disease2&M_Disease2_Disease1 |
| 11 | 118695908 | rs583905 | G | A | M_Disease2_Disease1 | 0.324037306 | M_Disease2_Disease1&M_Disease2&NULL&M_Disease1 |
| 11 | 118621495 | rs487728 | G | C | M_Disease2_Disease1 | 0.323910347 | M_Disease2_Disease1&NULL&M_Disease2&M_Disease1 |
| 11 | 118717732 | rs11217052 | G | A | NULL | 0.32363022 | NULL&M_Disease1&M_Disease2&M_Disease2_Disease1 |
| 11 | 118718061 | rs4938559 | T | C | NULL | 0.322857119 | NULL&M_Disease1&M_Disease2&M_Disease2_Disease1 |
| 11 | 118718062 | rs4938560 | G | A | NULL | 0.322857119 | NULL&M_Disease1&M_Disease2&M_Disease2_Disease1 |
| 11 | 118653629 | rs540180 | T | G | M_Disease2_Disease1 | 0.322566117 | M_Disease2_Disease1&NULL&M_Disease2&M_Disease1 |
| 11 | 118654651 | rs2508916 | G | C | M_Disease2_Disease1 | 0.322566117 | M_Disease2_Disease1&NULL&M_Disease2&M_Disease1 |
| 11 | 118599649 | rs182364290 | G | A | M_Disease2_Disease1 | 0.32200112 | M_Disease2_Disease1&M_Disease2&NULL&M_Disease1 |
| 11 | 118661055 | rs1790188 | T | C | M_Disease2_Disease1 | 0.320693461 | M_Disease2_Disease1&M_Disease2&NULL&M_Disease1 |
| 11 | 118685481 | rs515707 | T | C | M_Disease2_Disease1 | 0.319768935 | M_Disease2_Disease1&M_Disease2&NULL&M_Disease1 |
| 11 | 118700557 | rs4291686 | G | A | NULL | 0.319006605 | NULL&M_Disease2&M_Disease1&M_Disease2_Disease1 |
| 11 | 118716563 | rs2851460 | A | G | NULL | 0.318885617 | NULL&M_Disease1&M_Disease2&M_Disease2_Disease1 |
| 11 | 118761591 | rs58397111 | T | C | M_Disease2_Disease1 | 0.318885355 | M_Disease2_Disease1&M_Disease1&NULL&M_Disease2 |
| 11 | 118755738 | rs523604 | G | A | NULL | 0.318836881 | NULL&M_Disease2&M_Disease1&M_Disease2_Disease1 |
| 11 | 118690550 | rs577121 | C | T | M_Disease2_Disease1 | 0.318810946 | M_Disease2_Disease1&M_Disease2&NULL&M_Disease1 |
| 11 | 118696175 | rs584935 | A | G | M_Disease2_Disease1 | 0.318810946 | M_Disease2_Disease1&M_Disease2&NULL&M_Disease1 |
| 11 | 118809882 | rs11217099 | A | C | NULL | 0.318726903 | NULL&M_Disease1&M_Disease2&M_Disease2_Disease1 |
| 11 | 118637666 | rs535996 | T | C | M_Disease2_Disease1 | 0.318622251 | M_Disease2_Disease1&M_Disease2&NULL&M_Disease1 |
| 11 | 118689871 | rs587816 | A | C | M_Disease2_Disease1 | 0.318570879 | M_Disease2_Disease1&M_Disease2&NULL&M_Disease1 |
| 11 | 118899218 | rs4936459 | T | C | NULL | 0.318482134 | NULL&M_Disease1&M_Disease2&M_Disease2_Disease1 |
| 11 | 118478330 | rs7125115 | A | G | M_Disease1 | 0.318350261 | M_Disease1&M_Disease2_Disease1&NULL&M_Disease2 |
| 11 | 118655704 | rs6421568 | C | G | M_Disease2_Disease1 | 0.318248904 | M_Disease2_Disease1&M_Disease2&NULL&M_Disease1 |
| 11 | 118655775 | rs6421569 | A | C | M_Disease2_Disease1 | 0.318248904 | M_Disease2_Disease1&M_Disease2&NULL&M_Disease1 |
| 11 | 118477367 | rs498872 | A | G | M_Disease1 | 0.31803855 | M_Disease1&M_Disease2_Disease1&NULL&M_Disease2 |
| 11 | 118500963 | rs680465 | A | G | NULL | 0.317870743 | NULL&M_Disease2&M_Disease1&M_Disease2_Disease1 |
| 11 | 118699276 | rs625274 | T | C | NULL | 0.317419943 | NULL&M_Disease2&M_Disease1&M_Disease2_Disease1 |
| 11 | 118826045 | rs537080 | C | T | NULL | 0.317115874 | NULL&M_Disease1&M_Disease2&M_Disease2_Disease1 |
| 11 | 118806897 | rs10892310 | T | C | M_Disease2_Disease1 | 0.316977981 | M_Disease2_Disease1&M_Disease1&NULL&M_Disease2 |
| 11 | 118789657 | rs12294466 | T | C | NULL | 0.316972829 | NULL&M_Disease2&M_Disease1&M_Disease2_Disease1 |
| 11 | 118798824 | rs112119822 | G | A | NULL | 0.316393849 | NULL&M_Disease2&M_Disease1&M_Disease2_Disease1 |
| 11 | 118707249 | rs4938551 | C | G | NULL | 0.316367842 | NULL&M_Disease1&M_Disease2&M_Disease2_Disease1 |
| 11 | 118648396 | rs548167 | A | G | M_Disease2_Disease1 | 0.316289744 | M_Disease2_Disease1&M_Disease2&NULL&M_Disease1 |
| 11 | 118652781 | rs2508915 | C | G | M_Disease2_Disease1 | 0.316289744 | M_Disease2_Disease1&M_Disease2&NULL&M_Disease1 |
| 11 | 118558425 | rs149509653 | A | G | M_Disease2 | 0.316145547 | M_Disease2&M_Disease2_Disease1&NULL&M_Disease1 |
| 11 | 118662028 | rs12796102 | T | C | NULL | 0.316056896 | NULL&M_Disease2&M_Disease1&M_Disease2_Disease1 |
| 11 | 118828184 | rs45586434 | A | C | NULL | 0.315896387 | NULL&M_Disease1&M_Disease2&M_Disease2_Disease1 |
| 11 | 118798757 | rs57879605 | T | C | NULL | 0.315749313 | NULL&M_Disease2&M_Disease1&M_Disease2_Disease1 |
| 11 | 118631288 | rs507890 | C | T | NULL | 0.315575705 | NULL&M_Disease2&M_Disease1&M_Disease2_Disease1 |
| 11 | 118631653 | rs1784543 | T | A | NULL | 0.315575705 | NULL&M_Disease2&M_Disease1&M_Disease2_Disease1 |
| 11 | 118640081 | rs529717 | A | G | M_Disease2_Disease1 | 0.315533797 | M_Disease2_Disease1&M_Disease2&NULL&M_Disease1 |
| 11 | 118641451 | rs2508929 | G | T | M_Disease2_Disease1 | 0.315533797 | M_Disease2_Disease1&M_Disease2&NULL&M_Disease1 |
| 11 | 118644213 | rs1790187 | A | G | M_Disease2_Disease1 | 0.315533797 | M_Disease2_Disease1&M_Disease2&NULL&M_Disease1 |
| 11 | 118799206 | rs7101680 | A | G | NULL | 0.315347051 | NULL&M_Disease2&M_Disease1&M_Disease2_Disease1 |
| 11 | 118607090 | rs10736495 | C | G | M_Disease2_Disease1 | 0.314890762 | M_Disease2_Disease1&M_Disease2&NULL&M_Disease1 |
| 11 | 118801440 | rs12280501 | T | C | NULL | 0.314730221 | NULL&M_Disease2&M_Disease1&M_Disease2_Disease1 |
| 11 | 118700264 | rs113641235 | T | C | M_Disease2 | 0.314404232 | M_Disease2&M_Disease2_Disease1&NULL&M_Disease1 |
| 11 | 118632893 | rs6589690 | A | G | M_Disease2_Disease1 | 0.314351804 | M_Disease2_Disease1&M_Disease2&NULL&M_Disease1 |
| 11 | 118697375 | rs73005500 | C | T | M_Disease2 | 0.314212393 | M_Disease2&M_Disease2_Disease1&NULL&M_Disease1 |
| 11 | 118824683 | rs507763 | C | G | NULL | 0.314177807 | NULL&M_Disease1&M_Disease2&M_Disease2_Disease1 |
| 11 | 118824917 | rs11217102 | A | G | NULL | 0.314177807 | NULL&M_Disease1&M_Disease2&M_Disease2_Disease1 |
| 11 | 118515579 | rs10790255 | G | T | M_Disease1 | 0.313918081 | M_Disease1&M_Disease2_Disease1&NULL&M_Disease2 |
| 11 | 118629031 | rs555639 | A | G | NULL | 0.313671055 | NULL&M_Disease2&M_Disease1&M_Disease2_Disease1 |
| 11 | 118632522 | rs581045 | T | C | NULL | 0.313520632 | NULL&M_Disease2&M_Disease1&M_Disease2_Disease1 |
| 11 | 118679629 | rs1784300 | T | A | NULL | 0.313263812 | NULL&M_Disease2&M_Disease1&M_Disease2_Disease1 |
| 11 | 118755946 | rs76109832 | T | C | M_Disease2_Disease1 | 0.313171924 | M_Disease2_Disease1&M_Disease2&M_Disease1&NULL |
| 11 | 118789952 | rs11217090 | T | C | NULL | 0.312998521 | NULL&M_Disease2&M_Disease1&M_Disease2_Disease1 |
| 11 | 118658075 | rs515523 | G | A | NULL | 0.312829593 | NULL&M_Disease2&M_Disease1&M_Disease2_Disease1 |
| 11 | 118762102 | rs58508419 | A | G | M_Disease2_Disease1 | 0.312468537 | M_Disease2_Disease1&M_Disease1&NULL&M_Disease2 |
| 11 | 118630532 | rs35888235 | T | G | M_Disease2_Disease1 | 0.312331371 | M_Disease2_Disease1&M_Disease2&NULL&M_Disease1 |
| 11 | 118644582 | rs2510897 | C | A | NULL | 0.312035531 | NULL&M_Disease2&M_Disease1&M_Disease2_Disease1 |
| 11 | 118763736 | rs12363277 | C | T | NULL | 0.311977431 | NULL&M_Disease2&M_Disease1&M_Disease2_Disease1 |
| 11 | 118811641 | rs11606199 | A | G | NULL | 0.31182189 | NULL&M_Disease1&M_Disease2&M_Disease2_Disease1 |
| 11 | 118707171 | rs4938550 | G | A | M_Disease2 | 0.311631461 | M_Disease2&NULL&M_Disease2_Disease1&M_Disease1 |
| 11 | 118700425 | rs10128600 | C | T | M_Disease2 | 0.311498966 | M_Disease2&M_Disease2_Disease1&NULL&M_Disease1 |
| 11 | 118651809 | rs555649 | C | T | NULL | 0.311391201 | NULL&M_Disease2&M_Disease1&M_Disease2_Disease1 |
| 11 | 118699863 | rs2510901 | A | G | M_Disease2 | 0.311180376 | M_Disease2&M_Disease2_Disease1&NULL&M_Disease1 |
| 11 | 118682462 | rs523148 | T | C | M_Disease2 | 0.31089498 | M_Disease2&M_Disease2_Disease1&NULL&M_Disease1 |
| 11 | 118698160 | rs640373 | G | A | M_Disease2 | 0.310844939 | M_Disease2&M_Disease2_Disease1&NULL&M_Disease1 |
| 11 | 118698274 | rs639915 | A | G | M_Disease2 | 0.310844939 | M_Disease2&M_Disease2_Disease1&NULL&M_Disease1 |
| 11 | 118698294 | rs639885 | A | G | M_Disease2 | 0.310844939 | M_Disease2&M_Disease2_Disease1&NULL&M_Disease1 |
| 11 | 118698537 | rs638893 | G | A | M_Disease2 | 0.310844939 | M_Disease2&M_Disease2_Disease1&NULL&M_Disease1 |
| 11 | 118698777 | rs516332 | G | A | M_Disease2 | 0.310844939 | M_Disease2&M_Disease2_Disease1&NULL&M_Disease1 |
| 11 | 118699744 | rs2508564 | C | T | M_Disease2 | 0.310823243 | M_Disease2&M_Disease2_Disease1&NULL&M_Disease1 |
| 11 | 118717929 | rs4938558 | T | C | NULL | 0.310325301 | NULL&M_Disease1&M_Disease2&M_Disease2_Disease1 |
| 11 | 118624250 | rs546393 | C | T | NULL | 0.310273484 | NULL&M_Disease2&M_Disease1&M_Disease2_Disease1 |
| 11 | 118718856 | rs2156751 | G | A | NULL | 0.310155786 | NULL&M_Disease1&M_Disease2&M_Disease2_Disease1 |
| 11 | 118718972 | rs2187491 | C | T | NULL | 0.310155786 | NULL&M_Disease1&M_Disease2&M_Disease2_Disease1 |
| 11 | 118719254 | rs2156750 | C | T | NULL | 0.310155786 | NULL&M_Disease1&M_Disease2&M_Disease2_Disease1 |
| 11 | 118719405 | rs2156749 | G | C | NULL | 0.310155786 | NULL&M_Disease1&M_Disease2&M_Disease2_Disease1 |
| 11 | 118719742 | rs7116715 | C | T | NULL | 0.310155786 | NULL&M_Disease1&M_Disease2&M_Disease2_Disease1 |
| 11 | 118659896 | rs2105782 | C | T | NULL | 0.310081949 | NULL&M_Disease2&M_Disease1&M_Disease2_Disease1 |
| 11 | 118718485 | rs4938561 | C | T | NULL | 0.309662009 | NULL&M_Disease1&M_Disease2&M_Disease2_Disease1 |
| 11 | 118807637 | rs112974653 | T | G | NULL | 0.309625984 | NULL&M_Disease1&M_Disease2&M_Disease2_Disease1 |

| | | | | | | | | | | |
|---|---|---|---|---|---|---|---|---|---|---|
| 11 | 118803160 | rs73575440 | A | G | 0.84 | 0.86 | -9351.244 | -9349.332 | 1.912 | 7.62E-08 |
| 11 | 118803596 | rs111313044 | A | G | 0.84 | 0.86 | -9351.244 | -9349.332 | 1.912 | 7.62E-08 |
| 11 | 118804643 | rs56933877 | A | G | 0.84 | 0.86 | -9351.244 | -9349.332 | 1.912 | 7.62E-08 |
| 11 | 118804919 | rs73575441 | C | T | 0.84 | 0.86 | -9351.244 | -9349.332 | 1.912 | 7.62E-08 |
| 11 | 118804989 | rs112045567 | G | A | 0.84 | 0.86 | -9351.244 | -9349.332 | 1.912 | 7.62E-08 |
| 11 | 118762521 | rs190509947 | T | A | 1.056 | 0.753 | -9389.18 | -9387.269 | 1.911 | 7.61E-08 |
| 11 | 118486110 | rs45540840 | A | G | 1.141 | 1.022 | -9407.579 | -9405.695 | 1.884 | 7.41E-08 |
| 11 | 118516774 | rs12365095 | T | C | 1.141 | 1.022 | -9406.98 | -9405.106 | 1.874 | 7.33E-08 |
| 11 | 118615232 | rs139423236 | G | C | 1.085 | 1.311 | -9402.874 | -9401.016 | 1.859 | 7.22E-08 |
| 11 | 118514625 | rs11216938 | T | C | 1.14 | 1.021 | -9407.579 | -9405.762 | 1.817 | 6.93E-08 |
| 11 | 118782474 | rs11606719 | G | C | 0.775 | 0.887 | -9393.279 | -9391.505 | 1.774 | 6.63E-08 |
| 11 | 118535840 | rs12225548 | G | C | 1.139 | 1.02 | -9395.813 | -9394.041 | 1.772 | 6.62E-08 |
| 11 | 118777978 | rs78755429 | A | C | 0.862 | 0.789 | -9380.1 | -9378.329 | 1.771 | 6.61E-08 |
| 11 | 118499394 | rs2236661 | G | C | 1.139 | 1.022 | -9408.077 | -9406.341 | 1.737 | 6.39E-08 |
| 11 | 118496119 | rs45565037 | A | G | 1.138 | 1.022 | -9407.579 | -9405.857 | 1.722 | 6.30E-08 |
| 11 | 118516737 | rs12362152 | G | A | 1.138 | 1.021 | -9406.98 | -9405.27 | 1.709 | 6.22E-08 |
| 11 | 118812821 | rs74535623 | T | G | 0.8 | 0.889 | -9367.694 | -9365.997 | 1.696 | 6.14E-08 |
| 11 | 118813314 | rs73557345 | C | T | 0.8 | 0.889 | -9367.694 | -9365.997 | 1.696 | 6.14E-08 |
| 11 | 118501022 | rs10892248 | A | G | 1.138 | 1.022 | -9407.579 | -9405.884 | 1.694 | 6.12E-08 |
| 11 | 118529127 | rs7928371 | A | G | 1.137 | 1.02 | -9408.077 | -9406.449 | 1.629 | 5.74E-08 |
| 11 | 118664377 | rs598373 | T | C | 1.1 | 1.101 | -9406.478 | -9404.85 | 1.628 | 5.73E-08 |
| 11 | 118786829 | rs7102433 | G | A | 0.765 | 0.882 | -9391.946 | -9390.334 | 1.612 | 5.64E-08 |
| 11 | 118752178 | rs61150131 | C | T | 0.865 | 0.842 | -9404.679 | -9403.119 | 1.56 | 5.36E-08 |
| 11 | 118761159 | rs57231586 | T | G | 0.775 | 0.862 | -9401.168 | -9399.654 | 1.514 | 5.12E-08 |
| 11 | 118488782 | rs11216930 | C | A | 1.135 | 1.024 | -9398.429 | -9396.928 | 1.501 | 5.05E-08 |
| 11 | 118594326 | rs544096 | G | A | 0.888 | 0.882 | -9398.516 | -9397.022 | 1.494 | 5.01E-08 |
| 11 | 118605352 | rs76478216 | C | T | 0.854 | 1.089 | -9403.137 | -9401.672 | 1.465 | 4.87E-08 |
| 11 | 118609736 | rs1783937 | A | G | 0.9 | 0.871 | -9405.964 | -9404.507 | 1.457 | 4.83E-08 |
| 11 | 118617920 | rs476246 | T | C | 1.096 | 1.101 | -9408.077 | -9406.636 | 1.442 | 4.76E-08 |
| 11 | 118617544 | rs472641 | G | A | 1.095 | 1.101 | -9408.077 | -9406.66 | 1.417 | 4.64E-08 |
| 11 | 118781710 | rs6589707 | A | C | 0.773 | 0.891 | -9391.946 | -9390.558 | 1.387 | 4.51E-08 |
| 11 | 118695908 | rs583905 | G | A | 0.957 | 0.868 | -9407.257 | -9405.922 | 1.335 | 4.28E-08 |
| 11 | 118760443 | rs12786257 | A | G | 1.168 | 1.226 | -9400.212 | -9398.906 | 1.306 | 4.15E-08 |
| 11 | 118661197 | rs61901410 | G | C | 0.918 | 0.865 | -9407.198 | -9405.898 | 1.3 | 4.13E-08 |
| 11 | 118679387 | rs473298 | G | A | 1.094 | 1.099 | -9408.077 | -9406.799 | 1.279 | 4.04E-08 |
| 11 | 118679445 | rs516719 | C | T | 1.094 | 1.099 | -9408.077 | -9406.799 | 1.279 | 4.04E-08 |
| 11 | 118787213 | rs7119222 | C | T | 0.779 | 0.891 | -9391.946 | -9390.676 | 1.27 | 4.01E-08 |
| 11 | 118685481 | rs515707 | T | C | 0.958 | 0.869 | -9407.537 | -9406.27 | 1.267 | 4.00E-08 |
| 11 | 118599649 | rs182364290 | G | A | 1.069 | 1.279 | -9394.688 | -9393.423 | 1.265 | 3.99E-08 |
| 11 | 118678986 | rs477014 | C | T | 1.094 | 1.099 | -9407.167 | -9405.907 | 1.26 | 3.97E-08 |
| 11 | 118678609 | rs2508570 | A | G | 1.094 | 1.099 | -9408.077 | -9406.821 | 1.257 | 3.96E-08 |
| 11 | 118677710 | rs688161 | G | A | 1.094 | 1.099 | -9408.077 | -9406.824 | 1.253 | 3.94E-08 |
| 11 | 118696175 | rs584935 | A | G | 0.958 | 0.869 | -9407.257 | -9406.011 | 1.246 | 3.91E-08 |
| 11 | 118690550 | rs577121 | C | T | 0.958 | 0.87 | -9408.077 | -9406.833 | 1.244 | 3.90E-08 |
| 11 | 118689871 | rs587816 | A | C | 0.959 | 0.87 | -9408.077 | -9406.837 | 1.241 | 3.89E-08 |
| 11 | 118678881 | rs591756 | T | C | 1.093 | 1.1 | -9407.167 | -9405.929 | 1.238 | 3.88E-08 |
| 11 | 118680045 | rs2508569 | C | G | 1.093 | 1.099 | -9406.232 | -9405.027 | 1.205 | 3.76E-08 |
| 11 | 118604678 | rs493720 | T | C | 0.898 | 0.878 | -9407.273 | -9406.078 | 1.196 | 3.72E-08 |
| 11 | 118610629 | rs521910 | A | G | 0.902 | 0.876 | -9408.077 | -9406.883 | 1.195 | 3.72E-08 |
| 11 | 118751898 | rs60651507 | A | G | 0.869 | 0.847 | -9407.224 | -9406.03 | 1.194 | 3.71E-08 |
| 11 | 118660932 | rs1784295 | A | G | 0.914 | 0.869 | -9408.077 | -9406.887 | 1.191 | 3.70E-08 |
| 11 | 118605127 | rs526850 | G | T | 0.898 | 0.879 | -9408.077 | -9406.9 | 1.177 | 3.65E-08 |
| 11 | 118607694 | rs7119702 | G | A | 0.899 | 0.878 | -9404.743 | -9403.571 | 1.172 | 3.63E-08 |
| 11 | 118601363 | rs557990 | T | C | 0.894 | 0.884 | -9405.061 | -9403.897 | 1.165 | 3.61E-08 |
| 11 | 118602392 | rs564892 | A | T | 0.894 | 0.884 | -9405.061 | -9403.897 | 1.165 | 3.61E-08 |
| 11 | 118640063 | rs524590 | T | A | 1.092 | 1.098 | -9408.077 | -9406.92 | 1.157 | 3.58E-08 |
| 11 | 118605447 | rs2186781 | T | C | 0.899 | 0.88 | -9407.273 | -9406.14 | 1.133 | 3.49E-08 |
| 11 | 118761591 | rs58397111 | T | C | 0.795 | 0.86 | -9401.168 | -9400.035 | 1.133 | 3.49E-08 |
| 11 | 118624767 | rs581062 | A | G | 1.094 | 1.096 | -9408.077 | -9406.948 | 1.129 | 3.48E-08 |
| 11 | 118477367 | rs498872 | A | G | 1.109 | 0.982 | -9288.845 | -9287.72 | 1.125 | 3.47E-08 |
| 11 | 118652326 | rs1790189 | G | T | 1.091 | 1.098 | -9406.478 | -9405.364 | 1.114 | 3.43E-08 |
| 11 | 118652373 | rs1784293 | C | A | 1.091 | 1.098 | -9406.478 | -9405.364 | 1.114 | 3.43E-08 |
| 11 | 118757219 | rs1623316 | G | C | 0.989 | 0.893 | -9193.855 | -9192.754 | 1.101 | 3.38E-08 |
| 11 | 118626511 | rs499343 | T | C | 1.092 | 1.097 | -9406.922 | -9405.841 | 1.082 | 3.32E-08 |
| 11 | 118661055 | rs1790188 | T | C | 0.915 | 0.87 | -9408.077 | -9406.999 | 1.078 | 3.31E-08 |
| 11 | 118630532 | rs35888235 | T | G | 0.916 | 0.812 | -9345.024 | -9343.962 | 1.062 | 3.26E-08 |
| 11 | 118621495 | rs487728 | G | C | 1.091 | 1.097 | -9407.282 | -9406.22 | 1.061 | 3.25E-08 |
| 11 | 118558425 | rs149509653 | A | G | 0.97 | 0.877 | -9346.064 | -9345.005 | 1.059 | 3.25E-08 |
| 11 | 118637666 | rs535996 | T | C | 0.916 | 0.87 | -9408.077 | -9407.026 | 1.051 | 3.22E-08 |
| 11 | 118762102 | rs58508419 | A | G | 0.797 | 0.87 | -9401.168 | -9400.128 | 1.04 | 3.18E-08 |
| 11 | 118653629 | rs540180 | T | G | 1.09 | 1.098 | -9406.478 | -9405.439 | 1.039 | 3.18E-08 |
| 11 | 118654651 | rs2508916 | G | C | 1.09 | 1.098 | -9406.478 | -9405.439 | 1.039 | 3.18E-08 |
| 11 | 118655704 | rs6421568 | C | G | 0.915 | 0.871 | -9408.077 | -9407.04 | 1.038 | 3.18E-08 |
| 11 | 118655775 | rs6421569 | A | C | 0.915 | 0.871 | -9408.077 | -9407.04 | 1.038 | 3.18E-08 |
| 11 | 118806897 | rs10892310 | T | C | 0.896 | 0.933 | -9193.536 | -9192.501 | 1.035 | 3.17E-08 |
| 11 | 118648396 | rs548167 | A | G | 0.916 | 0.871 | -9408.077 | -9407.065 | 1.013 | 3.10E-08 |
| 11 | 118652781 | rs2508915 | C | G | 0.916 | 0.871 | -9408.077 | -9407.065 | 1.013 | 3.10E-08 |
| 11 | 118640081 | rs529717 | A | G | 0.917 | 0.871 | -9408.077 | -9407.072 | 1.005 | 3.07E-08 |
| 11 | 118641451 | rs2508929 | G | T | 0.917 | 0.871 | -9408.077 | -9407.072 | 1.005 | 3.07E-08 |
| 11 | 118644213 | rs1790187 | A | G | 0.917 | 0.871 | -9408.077 | -9407.072 | 1.005 | 3.07E-08 |
| 11 | 118632893 | rs6589690 | A | G | 0.919 | 0.87 | -9407.198 | -9406.201 | 0.996 | 3.05E-08 |
| 11 | 118755946 | rs76109832 | T | C | 0.867 | 0.859 | -9403.637 | -9402.666 | 0.97 | 2.97E-08 |
| 11 | 118622744 | rs6589689 | G | A | 0.8 | 0.9 | -9401.626 | -9400.658 | 0.968 | 2.96E-08 |
| 11 | 118607090 | rs10736495 | C | G | 0.904 | 0.88 | -9407.198 | -9406.245 | 0.952 | 2.92E-08 |
| 11 | 118633561 | rs12794288 | A | C | 0.917 | 0.816 | -9345.024 | -9344.096 | 0.928 | 2.85E-08 |
| 11 | 118629143 | rs475946 | T | C | 0.922 | 0.871 | -9407.198 | -9406.317 | 0.881 | 2.72E-08 |
| 11 | 118622245 | rs485901 | G | A | 0.917 | 0.874 | -9407.198 | -9406.336 | 0.862 | 2.67E-08 |
| 11 | 118759000 | rs77491736 | G | C | 0.806 | 0.883 | -9391.276 | -9390.426 | 0.85 | 2.63E-08 |
| 11 | 118618837 | rs640120 | A | C | 0.918 | 0.875 | -9408.077 | -9407.261 | 0.817 | 2.55E-08 |
| 11 | 118619423 | rs1961138 | C | T | 0.918 | 0.875 | -9408.077 | -9407.261 | 0.817 | 2.55E-08 |
| 11 | 118614397 | rs10892281 | T | C | 0.92 | 0.874 | -9408.077 | -9407.273 | 0.804 | 2.51E-08 |
| 11 | 118615660 | rs6589687 | T | A | 0.92 | 0.874 | -9408.077 | -9407.273 | 0.804 | 2.51E-08 |
| 11 | 118682462 | rs523148 | T | C | 0.966 | 0.876 | -9407.537 | -9406.748 | 0.789 | 2.48E-08 |
| 11 | 118700425 | rs10128600 | C | T | 0.967 | 0.876 | -9406.73 | -9405.949 | 0.781 | 2.46E-08 |
| 11 | 118812559 | rs74678032 | C | T | 0.905 | 0.812 | -9379.324 | -9378.558 | 0.765 | 2.42E-08 |
| 11 | 118700264 | rs113641235 | T | C | 0.973 | 0.877 | -9405.806 | -9405.077 | 0.73 | 2.34E-08 |
| 11 | 118763313 | rs78041268 | A | G | 1.204 | 1.123 | -9405.051 | -9404.33 | 0.72 | 2.31E-08 |
| 11 | 118666978 | rs1289369859 | A | G | 0.806 | 0.985 | -9401.346 | -9400.698 | 0.649 | 2.15E-08 |
| 11 | 118711376 | rs118046730 | C | T | 0.873 | 1.084 | -9405.028 | -9404.388 | 0.641 | 2.14E-08 |
| 11 | 118697375 | rs73005500 | C | T | 0.974 | 0.879 | -9406.727 | -9406.086 | 0.64 | 2.13E-08 |
| 11 | 118779681 | rs9971606 | C | T | 0.821 | 0.877 | -9401.563 | -9400.929 | 0.634 | 2.12E-08 |
| 11 | 118762073 | rs77841686 | A | G | 1.198 | 1.124 | -9405.051 | -9404.418 | 0.633 | 2.12E-08 |
| 11 | 118699863 | rs2510901 | A | G | 0.968 | 0.879 | -9406.337 | -9405.734 | 0.603 | 2.06E-08 |
| 11 | 118699744 | rs2508564 | C | T | 0.967 | 0.879 | -9406.337 | -9405.735 | 0.601 | 2.05E-08 |
| 11 | 118576762 | rs553188 | T | C | 0.896 | 0.893 | -9408.077 | -9407.483 | 0.595 | 2.04E-08 |
| 11 | 118684537 | rs150583208 | G | A | 0.808 | 0.979 | -9399.841 | -9399.249 | 0.592 | 2.03E-08 |
| 11 | 118698160 | rs640373 | G | A | 0.968 | 0.88 | -9407.257 | -9406.702 | 0.555 | 1.96E-08 |
| 11 | 118698274 | rs639915 | A | G | 0.968 | 0.88 | -9407.257 | -9406.702 | 0.555 | 1.96E-08 |
| 11 | 118698294 | rs639885 | A | G | 0.968 | 0.88 | -9407.257 | -9406.702 | 0.555 | 1.96E-08 |
| 11 | 118698537 | rs638893 | G | A | 0.968 | 0.88 | -9407.257 | -9406.702 | 0.555 | 1.96E-08 |
| 11 | 118698777 | rs516332 | G | A | 0.968 | 0.88 | -9407.257 | -9406.702 | 0.555 | 1.96E-08 |
| 11 | 118719984 | rs145680506 | T | C | 0.901 | 0.868 | -9351.192 | -9350.647 | 0.545 | 1.94E-08 |
| 11 | 118591443 | rs111843292 | A | C | 0.882 | 0.822 | -9349.102 | -9348.569 | 0.532 | 1.92E-08 |
| 11 | 118576997 | rs498912 | C | T | 0.897 | 0.894 | -9408.077 | -9407.552 | 0.526 | 1.90E-08 |
| 11 | 118478330 | rs7125115 | A | G | 0.904 | 0.973 | -9328.99 | -9328.47 | 0.52 | 1.89E-08 |
| 11 | 118673968 | rs140269932 | A | G | 0.97 | 0.809 | -9344.982 | -9344.482 | 0.5 | 1.86E-08 |
| 11 | 118702936 | rs7941696 | G | A | 0.958 | 0.881 | -9408.077 | -9407.581 | 0.496 | 1.85E-08 |
| 11 | 118664181 | rs145462967 | A | G | 0.966 | 0.809 | -9344.982 | -9344.496 | 0.486 | 1.83E-08 |
| 11 | 118762701 | rs76457018 | A | G | 0.961 | 0.874 | -9408.077 | -9407.602 | 0.475 | 1.81E-08 |
| 11 | 118583971 | rs6589677 | A | G | 0.91 | 0.884 | -9407.683 | -9407.23 | 0.453 | 1.77E-08 |
| 11 | 118563021 | rs117330495 | A | C | 1.114 | 1.182 | -9350.291 | -9349.851 | 0.44 | 1.75E-08 |
| 11 | 118480285 | rs12225399 | C | G | 0.905 | 0.962 | -9330.2 | -9329.766 | 0.434 | 1.74E-08 |
| 11 | 118765087 | rs2230321 | T | C | 0.828 | 0.891 | -9401.168 | -9400.755 | 0.413 | 1.70E-08 |
| 11 | 118767564 | rs75438046 | A | G | 1.181 | 1.125 | -9405.085 | -9404.677 | 0.408 | 1.69E-08 |
| 11 | 118791180 | rs146541591 | C | T | 0.864 | 0.858 | -9385.059 | -9384.679 | 0.381 | 1.65E-08 |
| 11 | 118801045 | rs75085081 | A | G | 0.866 | 0.858 | -9381.047 | -9380.679 | 0.369 | 1.63E-08 |
| 11 | 118583261 | rs112246697 | A | G | 0.861 | 0.851 | -9342.428 | -9342.07 | 0.358 | 1.61E-08 |
| 11 | 118689745 | rs492471 | T | C | 0.866 | 0.997 | -9399.372 | -9399.014 | 0.358 | 1.61E-08 |
| 11 | 118757788 | rs3176912 | A | G | 0.963 | 0.824 | -9398.479 | -9398.127 | 0.352 | 1.60E-08 |
| 11 | 118545589 | rs113941645 | G | C | 0.991 | 0.828 | -9360.629 | -9360.29 | 0.339 | 1.58E-08 |
| 11 | 118559731 | rs113784020 | T | C | 0.962 | 0.834 | -9342.213 | -9341.876 | 0.337 | 1.58E-08 |
| 11 | 118503442 | rs45621235 | T | C | 1.001 | 1.198 | -9376.023 | -9375.709 | 0.314 | 1.54E-08 |
| 11 | 118515579 | rs10790255 | G | T | 1.113 | 1.027 | -9408.077 | -9407.791 | 0.286 | 1.50E-08 |
| 11 | 118487265 | rs76544852 | T | G | 0.834 | 0.989 | -9407.166 | -9406.893 | 0.273 | 1.48E-08 |
| 11 | 118421841 | rs145968026 | G | C | 0.835 | 0.992 | -9407.166 | -9406.911 | 0.256 | 1.45E-08 |
| 11 | 118460957 | rs45600633 | A | G | 0.835 | 0.992 | -9407.166 | -9406.911 | 0.256 | 1.45E-08 |
| 11 | 118474283 | rs139202987 | G | A | 0.835 | 0.992 | -9407.166 | -9406.911 | 0.256 | 1.45E-08 |
| 11 | 118652569 | rs150792202 | A | G | 0.891 | 0.898 | -9365.219 | -9364.966 | 0.254 | 1.45E-08 |
| 11 | 118518021 | rs10736493 | G | T | 0.835 | 0.989 | -9406.793 | -9406.543 | 0.249 | 1.44E-08 |
| 11 | 118696055 | rs189417780 | C | A | 1.182 | 0.932 | -9391.139 | -9390.898 | 0.241 | 1.43E-08 |
| 11 | 118507068 | rs10892249 | C | T | 0.836 | 0.992 | -9407.166 | -9406.936 | 0.231 | 1.42E-08 |
| 11 | 118626825 | rs74337439 | A | G | 0.878 | 1.065 | -9407.133 | -9406.919 | 0.214 | 1.39E-08 |
| 11 | 118455465 | rs45460493 | A | G | 0.839 | 0.995 | -9407.166 | -9406.984 | 0.183 | 1.35E-08 |
| 11 | 118477226 | rs45496391 | C | A | 0.839 | 0.995 | -9407.166 | -9406.984 | 0.183 | 1.35E-08 |
| 11 | 118701898 | rs533254 | A | T | 0.971 | 0.886 | -9408.077 | -9407.909 | 0.168 | 1.33E-08 |
| 11 | 118748789 | rs145762171 | A | G | 0.959 | 0.824 | -9403.123 | -9402.968 | 0.155 | 1.31E-08 |
| 11 | 118775524 | rs79108568 | G | T | 0.934 | 0.837 | -9389.623 | -9389.479 | 0.143 | 1.30E-08 |
| 11 | 118566069 | rs506255 | A | C | 0.935 | 0.88 | -9327.085 | -9326.959 | 0.126 | 1.28E-08 |
| 11 | 118483912 | rs45621436 | G | T | 0.842 | 0.995 | -9408.077 | -9407.958 | 0.119 | 1.27E-08 |
| 11 | 118493319 | rs10790254 | A | G | 0.842 | 0.995 | -9408.077 | -9407.958 | 0.119 | 1.27E-08 |
| 11 | 118466646 | rs184841209 | T | G | 1.148 | 0.952 | -9390.311 | -9390.193 | 0.118 | 1.27E-08 |
| 11 | 118701597 | rs4938549 | T | C | 0.973 | 0.887 | -9408.077 | -9407.979 | 0.099 | 1.24E-08 |
| 11 | 118701668 | rs530646 | A | G | 0.973 | 0.887 | -9408.077 | -9407.979 | 0.099 | 1.24E-08 |
| 11 | 118702002 | rs534114 | A | G | 0.973 | 0.887 | -9408.077 | -9407.979 | 0.099 | 1.24E-08 |
| 11 | 118702910 | rs7941321 | A | G | 0.972 | 0.887 | -9408.077 | -9407.98 | 0.097 | 1.24E-08 |
| 11 | 118703001 | rs7928228 | T | C | 0.972 | 0.887 | -9408.077 | -9407.98 | 0.097 | 1.24E-08 |
| 11 | 118703484 | rs2508921 | T | C | 0.972 | 0.887 | -9408.077 | -9407.98 | 0.097 | 1.24E-08 |
| 11 | 118703817 | rs1170396 | T | A | 0.972 | 0.887 | -9408.077 | -9407.98 | 0.097 | 1.24E-08 |
| 11 | 118855789 | rs4938609 | A | C | 0.866 | 1.087 | -9364.192 | -9364.094 | 0.097 | 1.24E-08 |
| 11 | 118704255 | rs480791 | G | A | 0.971 | 0.887 | -9408.077 | -9407.981 | 0.096 | 1.24E-08 |
| 11 | 118704265 | rs622690 | T | C | 0.971 | 0.887 | -9408.077 | -9407.981 | 0.096 | 1.24E-08 |
| 11 | 118615390 | rs78502545 | T | C | 0.883 | 1.064 | -9407.133 | -9407.063 | 0.07 | 1.21E-08 |
| 11 | 118750967 | rs11602393 | A | T | 1.05 | 0.914 | -9383.82 | -9383.787 | 0.033 | 1.16E-08 |
| 11 | 118705654 | rs649338 | A | G | 0.974 | 0.888 | -9408.077 | -9408.05 | 0.027 | 1.16E-08 |
| 11 | 118480223 | rs67307131 | C | T | 0.91 | 0.961 | -9329.055 | -9329.035 | 0.02 | 1.15E-08 |
| 11 | 118783424 | rs149114341 | A | G | 1.155 | 1.118 | -9400.09 | -9400.085 | 0.005 | 1.13E-08 |
| 11 | 118557483 | rs6589673 | C | A | 0.851 | 0.997 | -9408.077 | -9408.12 | -0.043 | 1.08E-08 |
| 11 | 118561858 | rs10790257 | C | G | 0.85 | 0.987 | -9408.077 | -9408.121 | -0.043 | 1.08E-08 |
| 11 | 118561859 | rs10790258 | T | G | 0.85 | 0.987 | -9408.077 | -9408.121 | -0.043 | 1.08E-08 |
| 11 | 118782802 | rs76160025 | T | C | 0.975 | 0.842 | -9391.759 | -9391.803 | -0.043 | 1.08E-08 |
| 11 | 118755630 | rs3176905 | C | T | 1.071 | 0.931 | -9401.554 | -9401.605 | -0.051 | 1.07E-08 |
| 11 | 118559426 | rs10892253 | A | C | 0.851 | 0.987 | -9408.077 | -9408.131 | -0.054 | 1.07E-08 |
| 11 | 118598936 | rs184071181 | C | T | 1.136 | 0.945 | -9396.012 | -9396.069 | -0.056 | 1.06E-08 |
| 11 | 118705145 | rs636736 | G | A | 0.972 | 0.89 | -9406.03 | -9406.095 | -0.065 | 1.05E-08 |
| 11 | 118754649 | rs10892307 | G | C | 1.065 | 0.926 | -9396.988 | -9397.058 | -0.07 | 1.05E-08 |
| 11 | 118445559 | rs73555296 | T | G | 0.852 | 0.987 | -9405.733 | -9405.817 | -0.084 | 1.03E-08 |
| 11 | 118707171 | rs4938550 | G | A | 0.986 | 0.891 | -9406.703 | -9406.792 | -0.089 | 1.03E-08 |
| 11 | 118692727 | rs503425 | C | T | 0.901 | 0.972 | -9404.757 | -9404.852 | -0.095 | 1.02E-08 |
| 11 | 118788517 | rs73575424 | A | C | 0.94 | 0.884 | -9387.017 | -9387.125 | -0.108 | 1.01E-08 |
| 11 | 118721767 | rs116884018 | C | T | 1.152 | 0.949 | -9362.943 | -9363.056 | -0.113 | 1.01E-08 |
| 11 | 118691070 | rs140577174 | G | A | 0.892 | 0.868 | -9338.22 | -9338.338 | -0.118 | 1.00E-08 |
| 11 | 118765283 | rs665648 | A | G | 0.881 | 0.88 | -9382.684 | -9382.808 | -0.124 | 9.94E-09 |
| 11 | 118772160 | rs61730467 | A | G | 0.954 | 0.848 | -9393.727 | -9393.868 | -0.141 | 9.77E-09 |
| 11 | 118685015 | rs540716 | T | C | 0.902 | 0.972 | -9404.112 | -9404.271 | -0.159 | 9.60E-09 |
| 11 | 118603615 | rs74783030 | A | G | 1.143 | 0.945 | -9387.218 | -9387.385 | -0.167 | 9.52E-09 |
| 11 | 118793918 | rs12282752 | C | T | 1.041 | 0.876 | -9362.392 | -9362.571 | -0.179 | 9.41E-09 |
| 11 | 118681088 | rs2508922 | A | G | 0.902 | 0.97 | -9404.112 | -9404.303 | -0.191 | 9.30E-09 |
| 11 | 118413882 | rs146864992 | T | C | 0.858 | 0.978 | -9399.512 | -9399.704 | -0.192 | 9.29E-09 |
| 11 | 118478681 | rs45578540 | T | C | 1.119 | 1.132 | -9338.393 | -9338.597 | -0.203 | 9.19E-09 |
| 11 | 118791150 | rs12270280 | T | C | 1.039 | 0.878 | -9364.432 | -9364.668 | -0.237 | 8.88E-09 |
| 11 | 118785982 | rs79157249 | T | A | 0.946 | 0.886 | -9388.043 | -9388.296 | -0.253 | 8.74E-09 |
| 11 | 118799691 | rs191378986 | G | A | 0.866 | 1.004 | -9367.27 | -9367.535 | -0.266 | 8.63E-09 |
| 11 | 118754353 | rs630923 | A | C | 0.953 | 1.089 | -9380.559 | -9380.849 | -0.291 | 8.41E-09 |
| 11 | 118419577 | rs191458787 | T | C | 0.868 | 0.979 | -9399.512 | -9399.828 | -0.316 | 8.21E-09 |
| 11 | 118421438 | rs145485933 | G | A | 0.868 | 0.979 | -9399.512 | -9399.828 | -0.316 | 8.21E-09 |
| 11 | 118700218 | rs2510902 | A | G | 0.905 | 0.973 | -9397.962 | -9398.281 | -0.319 | 8.18E-09 |
| 11 | 118700515 | rs112571283 | T | C | 0.932 | 0.864 | -9332.07 | -9332.389 | -0.319 | 8.18E-09 |
| 11 | 118711069 | rs538645 | C | T | 0.933 | 0.914 | -9399.972 | -9400.293 | -0.32 | 8.17E-09 |
| 11 | 118694739 | rs188550348 | A | G | 0.888 | 0.918 | -9363.958 | -9364.318 | -0.361 | 7.84E-09 |
| 11 | 118492713 | rs45457998 | G | A | 0.984 | 0.867 | -9338.801 | -9339.201 | -0.4 | 7.54E-09 |
| 11 | 118496524 | rs35692099 | C | T | 0.895 | 0.894 | -9346.699 | -9347.101 | -0.402 | 7.53E-09 |
| 11 | 118502121 | rs17614795 | G | A | 0.895 | 0.894 | -9346.699 | -9347.101 | -0.402 | 7.53E-09 |
| 11 | 118644323 | rs6589695 | C | G | 0.877 | 0.936 | -9383.415 | -9383.835 | -0.419 | 7.40E-09 |
| 11 | 118515779 | rs117679873 | A | G | 1.107 | 1.119 | -9339.362 | -9339.814 | -0.452 | 7.16E-09 |
| 11 | 118744615 | rs73009445 | A | G | 1.101 | 0.935 | -9400.033 | -9400.486 | -0.452 | 7.16E-09 |
| 11 | 118747576 | rs55756957 | G | A | 1.026 | 0.918 | -9294.415 | -9294.867 | -0.452 | 7.16E-09 |
| 11 | 118650163 | rs73003218 | T | C | 0.936 | 0.889 | -9356.325 | -9356.83 | -0.505 | 6.79E-09 |
| 11 | 118480115 | rs12803321 | C | G | 0.917 | 0.969 | -9328.464 | -9328.975 | -0.511 | 6.75E-09 |
| 11 | 118727137 | rs117223320 | C | G | 0.914 | 0.891 | -9269.862 | -9270.389 | -0.527 | 6.64E-09 |
| 11 | 118735876 | rs79843382 | T | C | 0.918 | 0.889 | -9267.306 | -9267.852 | -0.546 | 6.52E-09 |
| 11 | 118898319 | rs741811 | A | G | 1.11 | 1.049 | -9408.077 | -9408.624 | -0.547 | 6.51E-09 |
| 11 | 118568427 | rs117361624 | T | C | 0.912 | 0.896 | -9381.777 | -9382.338 | -0.56 | 6.43E-09 |
| 11 | 118791532 | rs12271856 | T | G | 1.032 | 0.889 | -9362.372 | -9362.941 | -0.569 | 6.37E-09 |
| 11 | 118832251 | rs79808604 | A | C | 1.068 | 0.919 | -9388.834 | -9389.402 | -0.569 | 6.37E-09 |
| 11 | 118792278 | rs77398170 | G | A | 1.012 | 0.882 | -9362.372 | -9362.954 | -0.582 | 6.29E-09 |
| 11 | 118819541 | rs150196102 | T | C | 1.088 | 0.938 | -9389.679 | -9390.266 | -0.587 | 6.26E-09 |
| 11 | 118518227 | rs10736494 | T | C | 0.891 | 1.011 | -9407.704 | -9408.302 | -0.599 | 6.18E-09 |
| 11 | 118746223 | rs11217077 | C | T | 0.927 | 0.927 | -9382.848 | -9383.454 | -0.606 | 6.14E-09 |
| 11 | 118807735 | rs10892311 | T | C | 1.082 | 1.055 | -9164.094 | -9164.724 | -0.629 | 6.00E-09 |
| 11 | 118501723 | rs61900957 | T | C | 0.934 | 0.898 | -9392.76 | -9393.391 | -0.631 | 5.99E-09 |
| 11 | 118530434 | rs117619140 | T | G | 0.941 | 0.891 | -9299.397 | -9300.063 | -0.666 | 5.78E-09 |
| 11 | 118477746 | rs72999445 | G | T | 1.082 | 1.062 | -9377.421 | -9378.11 | -0.69 | 5.65E-09 |
| 11 | 118585146 | rs112799932 | G | T | 0.915 | 0.966 | -9400.288 | -9400.985 | -0.697 | 5.61E-09 |
| 11 | 118806216 | rs1231742890 | G | T | 1.026 | 0.893 | -9356.854 | -9357.57 | -0.716 | 5.50E-09 |
| 11 | 118745664 | rs11217076 | G | T | 0.927 | 0.932 | -9382.546 | -9383.279 | -0.734 | 5.40E-09 |
| 11 | 118804266 | rs7123066 | T | C | 0.986 | 0.886 | -9349.066 | -9349.831 | -0.764 | 5.24E-09 |
| 11 | 118804283 | rs7122959 | G | A | 0.986 | 0.886 | -9349.066 | -9349.831 | -0.764 | 5.24E-09 |
| 11 | 118725189 | rs112965917 | A | T | 0.989 | 0.888 | -9373.722 | -9374.504 | -0.783 | 5.14E-09 |
| 11 | 118722946 | rs148990923 | T | C | 0.906 | 0.959 | -9397.992 | -9398.781 | -0.789 | 5.11E-09 |
| 11 | 118629959 | rs140110112 | A | G | 0.924 | 0.902 | -9405.943 | -9406.743 | -0.8 | 5.06E-09 |
| 11 | 118764051 | rs613791 | T | C | 0.933 | 1.034 | -9402.209 | -9403.013 | -0.804 | 5.04E-09 |
| 11 | 118759905 | rs118020807 | A | C | 1.084 | 1.026 | -9404.163 | -9404.977 | -0.814 | 4.99E-09 |
| 11 | 118579395 | rs11828220 | A | T | 0.928 | 0.9 | -9404.724 | -9405.539 | -0.815 | 4.98E-09 |
| 11 | 118602058 | rs562059 | G | T | 1.082 | 1.047 | -9403.722 | -9404.539 | -0.818 | 4.97E-09 |
| 11 | 118621277 | rs117409833 | C | T | 1.091 | 1.038 | -9399.472 | -9400.307 | -0.835 | 4.88E-09 |
| 11 | 118569414 | rs6589675 | C | G | 0.917 | 0.922 | -9374.805 | -9375.64 | -0.836 | 4.88E-09 |
| 11 | 118709275 | rs187164533 | A | G | 0.919 | 0.954 | -9392.613 | -9393.474 | -0.861 | 4.76E-09 |
| 11 | 118533042 | rs142438429 | T | C | 0.924 | 0.962 | -9392.102 | -9392.969 | -0.866 | 4.73E-09 |
| 11 | 118602497 | rs478081 | G | A | 1.081 | 1.047 | -9403.722 | -9404.589 | -0.867 | 4.73E-09 |

| | | | | | | | |
|---|---|---|---|---|---|---|---|
| 11 | 118705654 | rs649338 | A | G | M_Disease2 | 0.309534526 | M_Disease2&M_Disease2_Disease1&NULL&M_Disease1 |
| 11 | 118701898 | rs533254 | A | T | M_Disease2 | 0.309430367 | M_Disease2&M_Disease2_Disease1&NULL&M_Disease1 |
| 11 | 118720590 | rs4938564 | T | G | NULL | 0.309368883 | NULL&M_Disease1&M_Disease2&M_Disease2_Disease1 |
| 11 | 118720684 | rs4936440 | G | C | NULL | 0.309368883 | NULL&M_Disease1&M_Disease2&M_Disease2_Disease1 |
| 11 | 118720998 | rs4938565 | A | G | NULL | 0.309368883 | NULL&M_Disease1&M_Disease2&M_Disease2_Disease1 |
| 11 | 118721189 | rs4938566 | T | A | NULL | 0.309368883 | NULL&M_Disease1&M_Disease2&M_Disease2_Disease1 |
| 11 | 118721249 | rs4938567 | G | A | NULL | 0.309368883 | NULL&M_Disease1&M_Disease2&M_Disease2_Disease1 |
| 11 | 118721850 | rs7946961 | G | A | NULL | 0.309368883 | NULL&M_Disease1&M_Disease2&M_Disease2_Disease1 |
| 11 | 118722231 | rs4938570 | G | A | NULL | 0.309368883 | NULL&M_Disease1&M_Disease2&M_Disease2_Disease1 |
| 11 | 118723087 | rs11217059 | T | C | NULL | 0.309368883 | NULL&M_Disease1&M_Disease2&M_Disease2_Disease1 |
| 11 | 118724336 | rs10892298 | C | T | NULL | 0.309368883 | NULL&M_Disease1&M_Disease2&M_Disease2_Disease1 |
| 11 | 118701597 | rs4938549 | T | C | M_Disease2 | 0.309233902 | M_Disease2&M_Disease2_Disease1&NULL&M_Disease1 |
| 11 | 118701668 | rs530646 | A | G | M_Disease2 | 0.309233902 | M_Disease2&M_Disease2_Disease1&NULL&M_Disease1 |
| 11 | 118702002 | rs534114 | A | G | M_Disease2 | 0.309233902 | M_Disease2&M_Disease2_Disease1&NULL&M_Disease1 |
| 11 | 118702910 | rs7941321 | A | G | M_Disease2 | 0.308784059 | M_Disease2&M_Disease2_Disease1&NULL&M_Disease1 |
| 11 | 118703001 | rs7928228 | T | C | M_Disease2 | 0.308784059 | M_Disease2&M_Disease2_Disease1&NULL&M_Disease1 |
| 11 | 118703484 | rs2508921 | T | C | M_Disease2 | 0.308784059 | M_Disease2&M_Disease2_Disease1&NULL&M_Disease1 |
| 11 | 118703817 | rs1170396 | T | A | M_Disease2 | 0.308784059 | M_Disease2&M_Disease2_Disease1&NULL&M_Disease1 |
| 11 | 118617299 | rs607125 | C | T | NULL | 0.308768941 | NULL&M_Disease2&M_Disease1&M_Disease2_Disease1 |
| 11 | 118723537 | rs7103314 | G | A | NULL | 0.308713434 | NULL&M_Disease1&M_Disease2&M_Disease2_Disease1 |
| 11 | 118620697 | rs488219 | C | T | NULL | 0.308633604 | NULL&M_Disease2&M_Disease1&M_Disease2_Disease1 |
| 11 | 118480285 | rs12225399 | C | G | M_Disease1 | 0.308443222 | M_Disease1&M_Disease2_Disease1&NULL&M_Disease2 |
| 11 | 118704255 | rs480791 | G | A | M_Disease2 | 0.308428445 | M_Disease2&M_Disease2_Disease1&NULL&M_Disease1 |
| 11 | 118704265 | rs622690 | T | C | M_Disease2 | 0.308428445 | M_Disease2&M_Disease2_Disease1&NULL&M_Disease1 |
| 11 | 118713180 | rs2187490 | G | T | NULL | 0.308345822 | NULL&M_Disease2&M_Disease1&M_Disease2_Disease1 |
| 11 | 118622744 | rs6589689 | G | A | M_Disease2_Disease1 | 0.307831503 | M_Disease2_Disease1&M_Disease1&NULL&M_Disease2 |
| 11 | 118705145 | rs636736 | G | A | M_Disease2 | 0.307663685 | M_Disease2&NULL&M_Disease2_Disease1&M_Disease1 |
| 11 | 118556995 | rs72999502 | C | T | NULL | 0.307355032 | NULL&M_Disease2&M_Disease1&M_Disease2_Disease1 |
| 11 | 118622245 | rs485901 | G | A | M_Disease2_Disease1 | 0.307336585 | M_Disease2_Disease1&M_Disease2&NULL&M_Disease1 |
| 11 | 118725420 | rs11217062 | A | G | NULL | 0.307284094 | NULL&M_Disease1&M_Disease2&M_Disease2_Disease1 |
| 11 | 118726230 | rs2156755 | A | G | NULL | 0.307284094 | NULL&M_Disease1&M_Disease2&M_Disease2_Disease1 |
| 11 | 118692727 | rs503425 | C | T | M_Disease1 | 0.307252972 | M_Disease1&NULL&M_Disease2_Disease1&M_Disease2 |
| 11 | 118629143 | rs475946 | T | C | M_Disease2_Disease1 | 0.30724056 | M_Disease2_Disease1&M_Disease2&NULL&M_Disease1 |
| 11 | 118605914 | rs7926282 | C | A | NULL | 0.306415199 | NULL&M_Disease2&M_Disease1&M_Disease2_Disease1 |
| 11 | 118685015 | rs540716 | T | C | M_Disease1 | 0.306257039 | M_Disease1&NULL&M_Disease2_Disease1&M_Disease2 |
| 11 | 118657311 | rs1309473877 | A | C | NULL | 0.305841329 | NULL&M_Disease2&M_Disease1&M_Disease2_Disease1 |
| 11 | 118793613 | rs11217091 | C | T | NULL | 0.305653774 | NULL&M_Disease2&M_Disease1&M_Disease2_Disease1 |
| 11 | 118765688 | rs676925 | G | C | NULL | 0.305619267 | NULL&M_Disease2&M_Disease1&M_Disease2_Disease1 |
| 11 | 118480115 | rs12803321 | C | G | M_Disease1 | 0.305243658 | M_Disease1&NULL&M_Disease2_Disease1&M_Disease2 |
| 11 | 118808140 | rs7117408 | T | C | NULL | 0.305228279 | NULL&M_Disease1&M_Disease2&M_Disease2_Disease1 |
| 11 | 118681088 | rs2508922 | A | G | M_Disease1 | 0.305224112 | M_Disease1&NULL&M_Disease2_Disease1&M_Disease2 |
| 11 | 118668296 | rs58868870 | T | C | NULL | 0.305199373 | NULL&M_Disease1&M_Disease2&M_Disease2_Disease1 |
| 11 | 118700218 | rs2510902 | A | G | M_Disease1 | 0.305173979 | M_Disease1&NULL&M_Disease2_Disease1&M_Disease2 |
| 11 | 118633561 | rs12794288 | A | C | M_Disease2_Disease1 | 0.30492941 | M_Disease2_Disease1&M_Disease2&NULL&M_Disease1 |
| 11 | 118680072 | rs607472 | G | C | NULL | 0.304690164 | NULL&M_Disease2&M_Disease1&M_Disease2_Disease1 |
| 11 | 118539746 | rs72999492 | C | A | NULL | 0.304582181 | NULL&M_Disease2&M_Disease1&M_Disease2_Disease1 |
| 11 | 118730669 | rs7933007 | C | T | NULL | 0.304513353 | NULL&M_Disease1&M_Disease2&M_Disease2_Disease1 |
| 11 | 118666289 | rs56104144 | C | G | NULL | 0.304466365 | NULL&M_Disease1&M_Disease2&M_Disease2_Disease1 |
| 11 | 118810405 | rs4938589 | T | C | NULL | 0.30432192 | NULL&M_Disease1&M_Disease2&M_Disease2_Disease1 |
| 11 | 118480223 | rs67307131 | C | T | M_Disease1 | 0.304076682 | M_Disease1&M_Disease2_Disease1&NULL&M_Disease2 |
| 11 | 118618837 | rs640120 | A | C | M_Disease2_Disease1 | 0.30403547 | M_Disease2_Disease1&M_Disease2&NULL&M_Disease1 |
| 11 | 118619423 | rs1961138 | C | T | M_Disease2_Disease1 | 0.30403547 | M_Disease2_Disease1&M_Disease2&NULL&M_Disease1 |
| 11 | 118702936 | rs7941696 | G | A | M_Disease2 | 0.303853211 | M_Disease2&M_Disease2_Disease1&NULL&M_Disease1 |
| 11 | 118809329 | rs10892313 | A | G | NULL | 0.303757588 | NULL&M_Disease1&M_Disease2&M_Disease2_Disease1 |
| 11 | 118607622 | rs11217004 | A | G | NULL | 0.30371361 | NULL&M_Disease2&M_Disease1&M_Disease2_Disease1 |
| 11 | 118762701 | rs76457018 | A | G | M_Disease2 | 0.303477895 | M_Disease2&M_Disease2_Disease1&NULL&M_Disease1 |
| 11 | 118708564 | rs484232 | T | G | NULL | 0.303472482 | NULL&M_Disease1&M_Disease2&M_Disease2_Disease1 |
| 11 | 118614397 | rs10892281 | T | C | M_Disease2_Disease1 | 0.303166108 | M_Disease2_Disease1&M_Disease2&NULL&M_Disease1 |
| 11 | 118615660 | rs6589687 | T | A | M_Disease2_Disease1 | 0.303166108 | M_Disease2_Disease1&M_Disease2&NULL&M_Disease1 |
| 11 | 118613235 | rs632124 | A | T | NULL | 0.303041372 | NULL&M_Disease2&M_Disease1&M_Disease2_Disease1 |
| 11 | 118605451 | rs2186780 | T | C | NULL | 0.30302387 | NULL&M_Disease2&M_Disease1&M_Disease2_Disease1 |
| 11 | 118757191 | rs1790192 | G | A | NULL | 0.302984555 | NULL&M_Disease2&M_Disease1&M_Disease2_Disease1 |
| 11 | 118816353 | rs586370 | A | G | NULL | 0.301933523 | NULL&M_Disease1&M_Disease2&M_Disease2_Disease1 |
| 11 | 118526691 | rs72999486 | T | C | NULL | 0.301278642 | NULL&M_Disease2&M_Disease1&M_Disease2_Disease1 |
| 11 | 118609598 | rs112996178 | T | C | NULL | 0.301269671 | NULL&M_Disease1&M_Disease2&M_Disease2_Disease1 |
| 11 | 118805144 | rs11605265 | A | G | NULL | 0.3009814 | NULL&M_Disease2&M_Disease1&M_Disease2_Disease1 |
| 11 | 118805220 | rs11605658 | T | C | NULL | 0.3009814 | NULL&M_Disease2&M_Disease1&M_Disease2_Disease1 |
| 11 | 118759000 | rs77491736 | G | C | M_Disease2_Disease1 | 0.300931733 | M_Disease2_Disease1&M_Disease1&NULL&M_Disease2 |
| 11 | 118803842 | rs11600378 | G | A | NULL | 0.300675322 | NULL&M_Disease2&M_Disease1&M_Disease2_Disease1 |
| 11 | 118804671 | rs143711539 | A | G | NULL | 0.300675322 | NULL&M_Disease2&M_Disease1&M_Disease2_Disease1 |
| 11 | 118811315 | rs77638281 | A | G | NULL | 0.300673571 | NULL&M_Disease2&M_Disease1&M_Disease2_Disease1 |
| 11 | 118481431 | rs10892246 | A | G | NULL | 0.300624307 | NULL&M_Disease1&M_Disease2&M_Disease2_Disease1 |
| 11 | 118803193 | rs117235696 | G | T | NULL | 0.300499519 | NULL&M_Disease2&M_Disease1&M_Disease2_Disease1 |
| 11 | 118608532 | rs567680 | G | C | M_Disease2 | 0.300484723 | M_Disease2&NULL&M_Disease1&M_Disease2_Disease1 |
| 11 | 118834718 | rs112939258 | T | C | NULL | 0.30047114 | NULL&M_Disease1&M_Disease2&M_Disease2_Disease1 |
| 11 | 118895495 | rs8301 | G | C | NULL | 0.300298365 | NULL&M_Disease1&M_Disease2&M_Disease2_Disease1 |
| 11 | 118747576 | rs55756957 | G | A | M_Disease2 | 0.300129822 | M_Disease2&NULL&M_Disease1&M_Disease2_Disease1 |
| 11 | 118834508 | rs111358668 | A | G | NULL | 0.300025365 | NULL&M_Disease1&M_Disease2&M_Disease2_Disease1 |
| 11 | 118619948 | rs112143902 | T | C | NULL | 0.299865421 | NULL&M_Disease2&M_Disease1&M_Disease2_Disease1 |
| 11 | 118441079 | rs7106316 | G | A | NULL | 0.299731175 | NULL&M_Disease2&M_Disease1&M_Disease2_Disease1 |
| 11 | 118608625 | rs566733 | G | C | M_Disease2 | 0.299475368 | M_Disease2&NULL&M_Disease1&M_Disease2_Disease1 |
| 11 | 118502172 | rs35435609 | T | C | NULL | 0.299400151 | NULL&M_Disease2&M_Disease1&M_Disease2_Disease1 |
| 11 | 118519213 | rs668628 | T | G | M_Disease2 | 0.299204572 | M_Disease2&NULL&M_Disease1&M_Disease2_Disease1 |
| 11 | 118798780 | rs77916839 | C | G | NULL | 0.298954886 | NULL&M_Disease2&M_Disease1&M_Disease2_Disease1 |
| 11 | 118585146 | rs112799932 | G | T | M_Disease1 | 0.298625173 | M_Disease1&NULL&M_Disease2_Disease1&M_Disease2 |
| 11 | 118689745 | rs492471 | T | C | M_Disease1 | 0.298256737 | M_Disease1&M_Disease2_Disease1&NULL&M_Disease2 |
| 11 | 118796570 | rs11217096 | T | C | NULL | 0.298087506 | NULL&M_Disease1&M_Disease2&M_Disease2_Disease1 |
| 11 | 118483802 | rs72999455 | G | A | NULL | 0.297836219 | NULL&M_Disease2&M_Disease1&M_Disease2_Disease1 |
| 11 | 118494790 | rs11825621 | C | T | NULL | 0.297836219 | NULL&M_Disease2&M_Disease1&M_Disease2_Disease1 |
| 11 | 118831755 | rs10892316 | T | C | NULL | 0.297701599 | NULL&M_Disease1&M_Disease2&M_Disease2_Disease1 |
| 11 | 118590864 | rs11216982 | A | G | NULL | 0.297641249 | NULL&M_Disease1&M_Disease2&M_Disease2_Disease1 |
| 11 | 118610826 | rs1893029 | T | G | NULL | 0.297489644 | NULL&M_Disease1&M_Disease2&M_Disease2_Disease1 |
| 11 | 118596444 | rs68097885 | A | G | M_Disease1 | 0.29722546 | M_Disease1&NULL&M_Disease2&M_Disease2_Disease1 |
| 11 | 118610463 | rs4499035 | T | C | NULL | 0.297164025 | NULL&M_Disease1&M_Disease2&M_Disease2_Disease1 |
| 11 | 118711830 | rs566848 | G | A | NULL | 0.297091747 | NULL&M_Disease1&M_Disease2&M_Disease2_Disease1 |
| 11 | 118624919 | rs11217013 | C | G | NULL | 0.296674932 | NULL&M_Disease1&M_Disease2&M_Disease2_Disease1 |
| 11 | 118801640 | rs572608 | A | T | NULL | 0.296063102 | NULL&M_Disease1&M_Disease2&M_Disease2_Disease1 |
| 11 | 118637475 | rs112910608 | T | C | NULL | 0.295882082 | NULL&M_Disease1&M_Disease2&M_Disease2_Disease1 |
| 11 | 118853774 | rs11217109 | C | T | NULL | 0.295805055 | NULL&M_Disease1&M_Disease2&M_Disease2_Disease1 |
| 11 | 118812559 | rs74678032 | C | T | M_Disease2_Disease1 | 0.295748334 | M_Disease2_Disease1&M_Disease2&NULL&M_Disease1 |
| 11 | 118633533 | rs7110526 | C | T | NULL | 0.295624749 | NULL&M_Disease1&M_Disease2&M_Disease2_Disease1 |
| 11 | 118635364 | rs11217018 | C | T | NULL | 0.295624749 | NULL&M_Disease1&M_Disease2&M_Disease2_Disease1 |
| 11 | 118468037 | rs535774 | T | C | NULL | 0.295619537 | NULL&M_Disease2&M_Disease1&M_Disease2_Disease1 |
| 11 | 118655356 | rs7934890 | T | C | NULL | 0.295498799 | NULL&M_Disease1&M_Disease2&M_Disease2_Disease1 |
| 11 | 118853021 | rs113962716 | A | C | NULL | 0.295061058 | NULL&M_Disease1&M_Disease2&M_Disease2_Disease1 |
| 11 | 118603030 | rs7943039 | A | G | M_Disease1 | 0.294934103 | M_Disease1&NULL&M_Disease2&M_Disease2_Disease1 |
| 11 | 118803536 | rs615166 | A | G | NULL | 0.294771291 | NULL&M_Disease1&M_Disease2&M_Disease2_Disease1 |
| 11 | 118870413 | rs10892319 | T | C | NULL | 0.294738428 | NULL&M_Disease1&M_Disease2&M_Disease2_Disease1 |
| 11 | 118765267 | rs598207 | C | G | NULL | 0.294724202 | NULL&M_Disease2&M_Disease1&M_Disease2_Disease1 |
| 11 | 118436071 | rs73023329 | T | C | NULL | 0.294660532 | NULL&M_Disease2&M_Disease1&M_Disease2_Disease1 |
| 11 | 118640828 | rs11217021 | C | T | NULL | 0.294559357 | NULL&M_Disease1&M_Disease2&M_Disease2_Disease1 |
| 11 | 118657297 | rs487177 | G | C | NULL | 0.294518185 | NULL&M_Disease1&M_Disease2&M_Disease2_Disease1 |
| 11 | 118616253 | rs7124498 | A | G | NULL | 0.29438999 | NULL&M_Disease1&M_Disease2&M_Disease2_Disease1 |
| 11 | 118543798 | rs76570573 | T | C | NULL | 0.294247688 | NULL&M_Disease2&M_Disease1&M_Disease2_Disease1 |
| 11 | 118852647 | rs80126726 | A | G | NULL | 0.294092764 | NULL&M_Disease1&M_Disease2&M_Disease2_Disease1 |
| 11 | 118801235 | rs602716 | A | G | NULL | 0.294092454 | NULL&M_Disease1&M_Disease2&M_Disease2_Disease1 |
| 11 | 118763313 | rs78041268 | A | G | M_Disease2_Disease1 | 0.294070326 | M_Disease2_Disease1&M_Disease1&NULL&M_Disease2 |
| 11 | 118576463 | rs496547 | A | T | NULL | 0.293915057 | NULL&M_Disease1&M_Disease2&M_Disease2_Disease1 |
| 11 | 118758089 | rs497916 | T | C | NULL | 0.293909127 | NULL&M_Disease2&M_Disease1&M_Disease2_Disease1 |
| 11 | 118726340 | rs113786535 | T | C | M_Disease2 | 0.293869383 | M_Disease2&NULL&M_Disease1&M_Disease2_Disease1 |
| 11 | 118788081 | rs7929825 | G | C | M_Disease2 | 0.293789031 | M_Disease2&NULL&M_Disease1&M_Disease2_Disease1 |
| 11 | 118550841 | rs79699220 | C | G | NULL | 0.293771685 | NULL&M_Disease2&M_Disease1&M_Disease2_Disease1 |
| 11 | 118613674 | rs7130875 | C | T | NULL | 0.293747803 | NULL&M_Disease1&M_Disease2&M_Disease2_Disease1 |
| 11 | 118698170 | rs56296109 | A | C | NULL | 0.29361678 | NULL&M_Disease1&M_Disease2&M_Disease2_Disease1 |
| 11 | 118762218 | rs543824 | G | A | NULL | 0.29346109 | NULL&M_Disease2&M_Disease1&M_Disease2_Disease1 |
| 11 | 118802492 | rs688035 | G | C | NULL | 0.293401605 | NULL&M_Disease1&M_Disease2&M_Disease2_Disease1 |
| 11 | 118759610 | rs566416 | G | T | NULL | 0.293393078 | NULL&M_Disease2&M_Disease1&M_Disease2_Disease1 |
| 11 | 118800302 | rs12793214 | A | C | NULL | 0.292988493 | NULL&M_Disease1&M_Disease2&M_Disease2_Disease1 |
| 11 | 118606197 | rs12420846 | T | C | NULL | 0.292920453 | NULL&M_Disease1&M_Disease2&M_Disease2_Disease1 |
| 11 | 118576762 | rs553188 | T | C | M_Disease2_Disease1 | 0.292863712 | M_Disease2_Disease1&NULL&M_Disease1&M_Disease2 |
| 11 | 118874272 | rs192113495 | G | T | NULL | 0.292729232 | NULL&M_Disease2&M_Disease1&M_Disease2_Disease1 |
| 11 | 118626286 | rs11217014 | C | T | NULL | 0.292225645 | NULL&M_Disease1&M_Disease2&M_Disease2_Disease1 |
| 11 | 118627778 | rs10892282 | C | G | NULL | 0.292225645 | NULL&M_Disease1&M_Disease2&M_Disease2_Disease1 |
| 11 | 118530869 | rs45447797 | G | C | NULL | 0.29222315 | NULL&M_Disease2&M_Disease1&M_Disease2_Disease1 |
| 11 | 118594669 | rs140026975 | C | A | M_Disease1 | 0.292143089 | M_Disease1&NULL&M_Disease2&M_Disease2_Disease1 |
| 11 | 118873840 | rs9645663 | C | T | NULL | 0.291756939 | NULL&M_Disease1&M_Disease2&M_Disease2_Disease1 |
| 11 | 118627149 | rs11217016 | C | A | M_Disease1 | 0.291398687 | M_Disease1&NULL&M_Disease2&M_Disease2_Disease1 |
| 11 | 118811878 | rs74891547 | C | G | NULL | 0.290206882 | NULL&M_Disease1&M_Disease2&M_Disease2_Disease1 |
| 11 | 118705753 | rs556437 | A | G | M_Disease2 | 0.290151859 | M_Disease2&NULL&M_Disease1&M_Disease2_Disease1 |
| 11 | 118629364 | rs73001486 | A | G | NULL | 0.289878576 | NULL&M_Disease1&M_Disease2&M_Disease2_Disease1 |
| 11 | 118890085 | rs4938619 | T | C | M_Disease1 | 0.289814582 | M_Disease1&NULL&M_Disease2&M_Disease2_Disease1 |
| 11 | 118893767 | rs111345266 | T | C | M_Disease1 | 0.289328457 | M_Disease1&NULL&M_Disease2&M_Disease2_Disease1 |
| 11 | 118746769 | rs4938576 | T | G | NULL | 0.289149156 | NULL&M_Disease2&M_Disease1&M_Disease2_Disease1 |
| 11 | 118764051 | rs613791 | T | C | M_Disease1 | 0.289148966 | M_Disease1&NULL&M_Disease2&M_Disease2_Disease1 |
| 11 | 118543589 | rs72999499 | G | C | NULL | 0.289067047 | NULL&M_Disease1&M_Disease2&M_Disease2_Disease1 |
| 11 | 118797003 | rs73575428 | G | C | M_Disease2 | 0.288964844 | M_Disease2&NULL&M_Disease1&M_Disease2_Disease1 |
| 11 | 118797278 | rs74584888 | A | G | M_Disease2 | 0.288964844 | M_Disease2&NULL&M_Disease1&M_Disease2_Disease1 |
| 11 | 118797392 | rs73575429 | C | T | M_Disease2 | 0.288964844 | M_Disease2&NULL&M_Disease1&M_Disease2_Disease1 |
| 11 | 118797420 | rs111390501 | G | A | M_Disease2 | 0.288964844 | M_Disease2&NULL&M_Disease1&M_Disease2_Disease1 |
| 11 | 118797544 | rs56032119 | G | A | M_Disease2 | 0.288964844 | M_Disease2&NULL&M_Disease1&M_Disease2_Disease1 |
| 11 | 118798154 | rs60969913 | C | G | M_Disease2 | 0.288964844 | M_Disease2&NULL&M_Disease1&M_Disease2_Disease1 |
| 11 | 118762073 | rs77841686 | A | G | M_Disease2_Disease1 | 0.288908242 | M_Disease2_Disease1&M_Disease1&NULL&M_Disease2 |
| 11 | 118539151 | rs72999491 | A | G | NULL | 0.28890204 | NULL&M_Disease1&M_Disease2&M_Disease2_Disease1 |
| 11 | 118867695 | rs4938617 | T | C | NULL | 0.288659918 | NULL&M_Disease1&M_Disease2&M_Disease2_Disease1 |
| 11 | 118576997 | rs498912 | C | T | M_Disease2_Disease1 | 0.28862346 | M_Disease2_Disease1&NULL&M_Disease1&M_Disease2 |
| 11 | 118779681 | rs9971606 | C | T | M_Disease2_Disease1 | 0.28856147 | M_Disease2_Disease1&M_Disease1&NULL&M_Disease2 |
| 11 | 118583847 | rs12803363 | T | C | NULL | 0.288471199 | NULL&M_Disease1&M_Disease2&M_Disease2_Disease1 |
| 11 | 118503442 | rs45621235 | T | C | M_Disease2 | 0.288251608 | M_Disease2&M_Disease2_Disease1&NULL&M_Disease1 |
| 11 | 118876213 | rs76253444 | G | A | NULL | 0.287409572 | NULL&M_Disease1&M_Disease2&M_Disease2_Disease1 |
| 11 | 118719984 | rs145680506 | T | C | M_Disease2_Disease1 | 0.287300215 | M_Disease2_Disease1&M_Disease2&NULL&M_Disease1 |
| 11 | 118876552 | rs74884652 | C | T | M_Disease1 | 0.287020251 | M_Disease1&NULL&M_Disease2&M_Disease2_Disease1 |
| 11 | 118877891 | rs75154936 | C | T | M_Disease1 | 0.287020251 | M_Disease1&NULL&M_Disease2&M_Disease2_Disease1 |
| 11 | 118898319 | rs741811 | A | G | M_Disease1 | 0.286728288 | M_Disease1&NULL&M_Disease2_Disease1&M_Disease2 |
| 11 | 118712509 | rs75488969 | T | C | NULL | 0.286675671 | NULL&M_Disease2&M_Disease1&M_Disease2_Disease1 |
| 11 | 118577614 | rs947931 | G | T | NULL | 0.286531577 | NULL&M_Disease1&M_Disease2&M_Disease2_Disease1 |
| 11 | 118602058 | rs562059 | G | T | M_Disease1 | 0.286036514 | M_Disease1&NULL&M_Disease2&M_Disease2_Disease1 |
| 11 | 118666978 | rs1289369859 | A | G | M_Disease2_Disease1 | 0.285943519 | M_Disease2_Disease1&M_Disease1&NULL&M_Disease2 |
| 11 | 118622985 | rs3825057 | C | T | NULL | 0.285699271 | NULL&M_Disease1&M_Disease2&M_Disease2_Disease1 |
| 11 | 118559731 | rs113784020 | T | C | M_Disease2 | 0.285468884 | M_Disease2&M_Disease2_Disease1&NULL&M_Disease1 |
| 11 | 118602497 | rs478081 | G | A | M_Disease1 | 0.285347891 | M_Disease1&NULL&M_Disease2&M_Disease2_Disease1 |
| 11 | 118785982 | rs79157249 | T | A | M_Disease2 | 0.285328436 | M_Disease2&NULL&M_Disease2_Disease1&M_Disease1 |
| 11 | 118793479 | rs138526953 | A | C | NULL | 0.285127386 | NULL&M_Disease1&M_Disease2&M_Disease2_Disease1 |
| 11 | 118725204 | rs55794678 | T | C | NULL | 0.284508751 | NULL&M_Disease2&M_Disease1&M_Disease2_Disease1 |
| 11 | 118765231 | rs12795395 | T | C | NULL | 0.284463868 | NULL&M_Disease2&M_Disease1&M_Disease2_Disease1 |
| 11 | 118583971 | rs6589677 | A | G | M_Disease2_Disease1 | 0.283862213 | M_Disease2_Disease1&M_Disease2&NULL&M_Disease1 |
| 11 | 118591443 | rs111843292 | A | C | M_Disease2_Disease1 | 0.2834801 | M_Disease2_Disease1&M_Disease2&NULL&M_Disease1 |
| 11 | 118580773 | rs675329 | T | C | NULL | 0.28339059 | NULL&M_Disease1&M_Disease2&M_Disease2_Disease1 |
| 11 | 118757788 | rs3176912 | A | G | M_Disease2 | 0.283152062 | M_Disease2&M_Disease2_Disease1&NULL&M_Disease1 |
| 11 | 118462554 | rs11216920 | G | A | NULL | 0.283140926 | NULL&M_Disease1&M_Disease2&M_Disease2_Disease1 |
| 11 | 118773438 | rs35346926 | T | C | NULL | 0.283102155 | NULL&M_Disease2&M_Disease1&M_Disease2_Disease1 |
| 11 | 118691975 | rs77731159 | A | T | NULL | 0.283061136 | NULL&M_Disease2&M_Disease1&M_Disease2_Disease1 |
| 11 | 118684537 | rs150583208 | G | A | M_Disease2_Disease1 | 0.282753827 | M_Disease2_Disease1&M_Disease1&NULL&M_Disease2 |
| 11 | 118804266 | rs7123066 | T | C | M_Disease2 | 0.28222124 | M_Disease2&NULL&M_Disease1&M_Disease2_Disease1 |
| 11 | 118804283 | rs7122959 | G | A | M_Disease2 | 0.28222124 | M_Disease2&NULL&M_Disease1&M_Disease2_Disease1 |
| 11 | 118750967 | rs11602393 | A | T | M_Disease2 | 0.282034827 | M_Disease2&M_Disease2_Disease1&NULL&M_Disease1 |
| 11 | 118569414 | rs6589675 | C | G | NULL | 0.281905145 | NULL&M_Disease1&M_Disease2&M_Disease2_Disease1 |
| 11 | 118545589 | rs113941645 | G | C | M_Disease2 | 0.281873638 | M_Disease2&M_Disease2_Disease1&NULL&M_Disease1 |
| 11 | 118788517 | rs73575424 | A | C | M_Disease2 | 0.281835044 | M_Disease2&NULL&M_Disease2_Disease1&M_Disease1 |
| 11 | 118469739 | rs143380378 | G | C | NULL | 0.281438206 | NULL&M_Disease1&M_Disease2&M_Disease2_Disease1 |
| 11 | 118792278 | rs77398170 | G | A | M_Disease2 | 0.28120308 | M_Disease2&NULL&M_Disease1&M_Disease2_Disease1 |
| 11 | 118704751 | rs35262372 | G | A | NULL | 0.281180316 | NULL&M_Disease1&M_Disease2&M_Disease2_Disease1 |
| 11 | 118462188 | rs4936431 | G | C | NULL | 0.280684276 | NULL&M_Disease1&M_Disease2&M_Disease2_Disease1 |
| 11 | 118745664 | rs11217076 | G | T | NULL | 0.280658614 | NULL&M_Disease1&M_Disease2&M_Disease2_Disease1 |
| 11 | 118725189 | rs112965917 | A | T | M_Disease2 | 0.280606107 | M_Disease2&NULL&M_Disease1&M_Disease2_Disease1 |
| 11 | 118754353 | rs630923 | A | C | M_Disease2 | 0.280461156 | M_Disease2&NULL&M_Disease1&M_Disease2_Disease1 |
| 11 | 118480695 | rs34026809 | C | G | NULL | 0.280383388 | NULL&M_Disease1&M_Disease2&M_Disease2_Disease1 |
| 11 | 118587349 | rs11216975 | C | A | NULL | 0.280354405 | NULL&M_Disease1&M_Disease2&M_Disease2_Disease1 |
| 11 | 118792384 | rs182691948 | T | G | NULL | 0.280233027 | NULL&M_Disease2&M_Disease1&M_Disease2_Disease1 |
| 11 | 118797857 | rs150438978 | T | A | NULL | 0.280233027 | NULL&M_Disease2&M_Disease1&M_Disease2_Disease1 |
| 11 | 118492713 | rs45457998 | G | A | M_Disease2 | 0.280214644 | M_Disease2&NULL&M_Disease2_Disease1&M_Disease1 |
| 11 | 118801296 | rs75210328 | T | G | NULL | 0.280189202 | NULL&M_Disease2&M_Disease1&M_Disease2_Disease1 |
| 11 | 118801577 | rs77875650 | A | G | NULL | 0.280189202 | NULL&M_Disease2&M_Disease1&M_Disease2_Disease1 |
| 11 | 118566069 | rs506255 | A | C | M_Disease2 | 0.28017654 | M_Disease2&M_Disease2_Disease1&NULL&M_Disease1 |
| 11 | 118791150 | rs12270280 | T | C | M_Disease2 | 0.279775227 | M_Disease2&NULL&M_Disease2_Disease1&M_Disease1 |
| 11 | 118707035 | rs143605660 | A | G | NULL | 0.279664169 | NULL&M_Disease2&M_Disease1&M_Disease2_Disease1 |
| 11 | 118793918 | rs12282752 | C | T | M_Disease2 | 0.279471898 | M_Disease2&NULL&M_Disease2_Disease1&M_Disease1 |
| 11 | 118782802 | rs76160025 | T | C | M_Disease2 | 0.279413403 | M_Disease2&NULL&M_Disease2_Disease1&M_Disease1 |
| 11 | 118583712 | rs118008100 | A | G | NULL | 0.279302956 | NULL&M_Disease2&M_Disease1&M_Disease2_Disease1 |
| 11 | 118711376 | rs118046730 | C | T | M_Disease2_Disease1 | 0.279192733 | M_Disease2_Disease1&M_Disease1&M_Disease2&NULL |
| 11 | 118510642 | rs141169729 | T | C | NULL | 0.279035285 | NULL&M_Disease1&M_Disease2&M_Disease2_Disease1 |
| 11 | 118579395 | rs11828220 | A | T | NULL | 0.27892763 | NULL&M_Disease2&M_Disease1&M_Disease2_Disease1 |
| 11 | 118466646 | rs184841209 | T | G | M_Disease1 | 0.278617255 | M_Disease1&M_Disease2_Disease1&NULL&M_Disease2 |
| 11 | 118629959 | rs140110112 | A | G | NULL | 0.278467356 | NULL&M_Disease2&M_Disease1&M_Disease2_Disease1 |
| 11 | 118563021 | rs117330495 | A | C | M_Disease2_Disease1 | 0.278269823 | M_Disease2_Disease1&M_Disease2&NULL&M_Disease1 |
| 11 | 118578462 | rs516459 | C | T | NULL | 0.27825311 | NULL&M_Disease2&M_Disease1&M_Disease2_Disease1 |
| 11 | 118732181 | rs117079831 | A | G | NULL | 0.278157278 | NULL&M_Disease1&M_Disease2&M_Disease2_Disease1 |
| 11 | 118791532 | rs12271856 | T | G | M_Disease2 | 0.27789829 | M_Disease2&NULL&M_Disease1&M_Disease2_Disease1 |
| 11 | 118807735 | rs10892311 | T | C | M_Disease1 | 0.277615625 | M_Disease1&NULL&M_Disease2_Disease1&M_Disease2 |
| 11 | 118772160 | rs61730467 | A | G | M_Disease2 | 0.277530442 | M_Disease2&NULL&M_Disease2_Disease1&M_Disease1 |
| 11 | 118619478 | rs189583461 | G | A | NULL | 0.277406969 | NULL&M_Disease1&M_Disease2&M_Disease2_Disease1 |
| 11 | 118673968 | rs140269932 | A | G | M_Disease2_Disease1 | 0.277383251 | M_Disease2_Disease1&M_Disease2&NULL&M_Disease1 |
| 11 | 118721562 | rs184045098 | T | C | NULL | 0.277339513 | NULL&M_Disease1&M_Disease2&M_Disease2_Disease1 |
| 11 | 118709275 | rs187164533 | A | G | NULL | 0.277135326 | NULL&M_Disease1&M_Disease2&M_Disease2_Disease1 |
| 11 | 118664181 | rs145462967 | A | G | M_Disease2_Disease1 | 0.277098676 | M_Disease2_Disease1&M_Disease2&NULL&M_Disease1 |
| 11 | 118746223 | rs11217077 | C | T | NULL | 0.277081499 | NULL&M_Disease1&M_Disease2&M_Disease2_Disease1 |
| 11 | 118533042 | rs142438429 | T | C | NULL | 0.276901156 | NULL&M_Disease1&M_Disease2&M_Disease2_Disease1 |
| 11 | 118806216 | rs1231742890 | G | T | M_Disease2 | 0.276777484 | M_Disease2&NULL&M_Disease1&M_Disease2_Disease1 |
| 11 | 118487265 | rs76544852 | T | G | M_Disease1 | 0.276754427 | M_Disease1&M_Disease2_Disease1&NULL&M_Disease2 |
| 11 | 118767564 | rs75438046 | A | G | M_Disease2_Disease1 | 0.276531362 | M_Disease2_Disease1&M_Disease1&NULL&M_Disease2 |
| 11 | 118518021 | rs10736493 | G | T | M_Disease1 | 0.276484828 | M_Disease1&M_Disease2_Disease1&NULL&M_Disease2 |
| 11 | 118421841 | rs145968026 | G | C | M_Disease1 | 0.276474187 | M_Disease1&M_Disease2_Disease1&NULL&M_Disease2 |
| 11 | 118460957 | rs45600633 | A | G | M_Disease1 | 0.276474187 | M_Disease1&M_Disease2_Disease1&NULL&M_Disease2 |
| 11 | 118474283 | rs139202987 | G | A | M_Disease1 | 0.276474187 | M_Disease1&M_Disease2_Disease1&NULL&M_Disease2 |
| 11 | 118507068 | rs10892249 | C | T | M_Disease1 | 0.276292759 | M_Disease1&M_Disease2_Disease1&NULL&M_Disease2 |
| 11 | 118765087 | rs2230321 | T | C | M_Disease2_Disease1 | 0.276238444 | M_Disease2_Disease1&M_Disease1&NULL&M_Disease2 |
| 11 | 118805138 | rs7127427 | A | G | NULL | 0.275978321 | NULL&M_Disease2&M_Disease1&M_Disease2_Disease1 |
| 11 | 118621277 | rs117409833 | C | T | NULL | 0.275833564 | NULL&M_Disease1&M_Disease2&M_Disease2_Disease1 |
| 11 | 118455465 | rs45460493 | A | G | M_Disease1 | 0.275728727 | M_Disease1&M_Disease2_Disease1&NULL&M_Disease2 |
| 11 | 118477226 | rs45496391 | C | A | M_Disease1 | 0.275728727 | M_Disease1&M_Disease2_Disease1&NULL&M_Disease2 |

| | | | | | | | | | | |
|---|---|---|---|---|---|---|---|---|---|---|
| 11 | 118721562 | rs184045098 | T | C | 1.078 | 1.06 | -9396.222 | -9397.093 | -0.87 | 4.72E-09 |
| 11 | 118619478 | rs189583461 | G | A | 1.08 | 1.045 | -9397.104 | -9397.989 | -0.885 | 4.65E-09 |
| 11 | 118707035 | rs143605660 | A | G | 0.946 | 0.895 | -9404.007 | -9404.894 | -0.886 | 4.64E-09 |
| 11 | 118797003 | rs73575428 | G | C | 0.971 | 0.897 | -9370.343 | -9371.232 | -0.889 | 4.63E-09 |
| 11 | 118797278 | rs74584888 | A | G | 0.971 | 0.897 | -9370.343 | -9371.232 | -0.889 | 4.63E-09 |
| 11 | 118797392 | rs73575429 | C | T | 0.971 | 0.897 | -9370.343 | -9371.232 | -0.889 | 4.63E-09 |
| 11 | 118797420 | rs111390501 | G | A | 0.971 | 0.897 | -9370.343 | -9371.232 | -0.889 | 4.63E-09 |
| 11 | 118797544 | rs56032119 | G | A | 0.971 | 0.897 | -9370.343 | -9371.232 | -0.889 | 4.63E-09 |
| 11 | 118798154 | rs60969913 | C | G | 0.971 | 0.897 | -9370.343 | -9371.232 | -0.889 | 4.63E-09 |
| 11 | 118805138 | rs7127427 | A | G | 1.025 | 0.902 | -9358.773 | -9359.662 | -0.889 | 4.63E-09 |
| 11 | 118614115 | rs73001451 | C | T | 1.098 | 0.964 | -9408.077 | -9408.969 | -0.892 | 4.61E-09 |
| 11 | 118603030 | rs7943039 | A | G | 0.918 | 0.965 | -9398.825 | -9399.723 | -0.898 | 4.59E-09 |
| 11 | 118732181 | rs117079831 | A | G | 1.071 | 1.06 | -9396.222 | -9397.124 | -0.901 | 4.57E-09 |
| 11 | 118890085 | rs4938619 | T | C | 1.1 | 1.034 | -9408.077 | -9408.98 | -0.903 | 4.56E-09 |
| 11 | 118462188 | rs4936431 | G | C | 1.096 | 1.084 | -9287.534 | -9288.442 | -0.908 | 4.54E-09 |
| 11 | 118893767 | rs111345266 | T | C | 1.1 | 1.034 | -9408.077 | -9408.997 | -0.919 | 4.49E-09 |
| 11 | 118578462 | rs516459 | C | T | 0.959 | 1.049 | -9402.93 | -9403.865 | -0.935 | 4.42E-09 |
| 11 | 118469739 | rs143380378 | G | C | 1.092 | 1.085 | -9280.629 | -9281.565 | -0.936 | 4.41E-09 |
| 11 | 118704751 | rs35262372 | G | A | 1.096 | 1.078 | -9405.606 | -9406.545 | -0.939 | 4.40E-09 |
| 11 | 118510642 | rs141169729 | T | C | 0.934 | 0.979 | -9391.316 | -9392.267 | -0.951 | 4.35E-09 |
| 11 | 118788081 | rs7929825 | G | C | 0.967 | 0.908 | -9353.634 | -9354.587 | -0.954 | 4.34E-09 |
| 11 | 118583712 | rs118008100 | A | G | 0.971 | 1.03 | -9384.761 | -9385.725 | -0.964 | 4.29E-09 |
| 11 | 118691975 | rs77731159 | A | T | 0.942 | 0.902 | -9404.257 | -9405.24 | -0.983 | 4.21E-09 |
| 11 | 118746769 | rs4938576 | T | G | 1.061 | 1.072 | -9404.26 | -9405.248 | -0.988 | 4.19E-09 |
| 11 | 118587349 | rs11216975 | C | A | 1.052 | 0.983 | -9389.764 | -9390.755 | -0.991 | 4.18E-09 |
| 11 | 118792384 | rs182691948 | T | G | 1.009 | 0.96 | -9398.379 | -9399.375 | -0.996 | 4.16E-09 |
| 11 | 118797857 | rs150438978 | T | A | 1.009 | 0.96 | -9398.379 | -9399.375 | -0.996 | 4.16E-09 |
| 11 | 118480695 | rs34026809 | C | G | 1.108 | 1.01 | -9350.306 | -9351.303 | -0.997 | 4.15E-09 |
| 11 | 118519213 | rs668628 | T | G | 0.981 | 0.909 | -9385.631 | -9386.631 | -0.999 | 4.14E-09 |
| 11 | 118622985 | rs3825057 | C | T | 0.922 | 0.956 | -9406.151 | -9407.15 | -0.999 | 4.14E-09 |
| 11 | 118577614 | rs947931 | G | T | 1.088 | 1.059 | -9290.677 | -9291.695 | -1.017 | 4.07E-09 |
| 11 | 118596444 | rs68097885 | A | G | 0.92 | 0.971 | -9398.257 | -9399.278 | -1.021 | 4.05E-09 |
| 11 | 118462554 | rs11216920 | G | A | 0.905 | 0.966 | -9366.957 | -9367.984 | -1.027 | 4.03E-09 |
| 11 | 118801296 | rs75210328 | T | G | 1.025 | 0.91 | -9356.396 | -9357.425 | -1.029 | 4.02E-09 |
| 11 | 118801577 | rs77875650 | A | G | 1.025 | 0.91 | -9356.396 | -9357.425 | -1.029 | 4.02E-09 |
| 11 | 118594669 | rs140026975 | C | A | 0.921 | 0.964 | -9394.868 | -9395.9 | -1.033 | 4.01E-09 |
| 11 | 118712509 | rs75488969 | T | C | 0.94 | 0.909 | -9407.039 | -9408.083 | -1.044 | 3.96E-09 |
| 11 | 118876552 | rs74884652 | C | T | 1.095 | 1.034 | -9408.077 | -9409.147 | -1.07 | 3.86E-09 |
| 11 | 118877891 | rs75154936 | C | T | 1.095 | 1.034 | -9408.077 | -9409.147 | -1.07 | 3.86E-09 |
| 11 | 118627149 | rs11217016 | C | A | 0.922 | 0.964 | -9408.077 | -9409.16 | -1.083 | 3.81E-09 |
| 11 | 118705753 | rs556437 | A | G | 0.962 | 0.924 | -9388.633 | -9389.717 | -1.083 | 3.81E-09 |
| 11 | 118580773 | rs675329 | T | C | 0.954 | 0.983 | -9391.511 | -9392.597 | -1.086 | 3.80E-09 |
| 11 | 118773438 | rs35346926 | T | C | 1.026 | 0.924 | -9390.325 | -9391.417 | -1.092 | 3.78E-09 |
| 11 | 118725204 | rs55794678 | T | C | 1.01 | 0.982 | -9361.884 | -9363.004 | -1.12 | 3.67E-09 |
| 11 | 118876213 | rs76253444 | G | A | 1.091 | 1.036 | -9390.282 | -9391.411 | -1.129 | 3.64E-09 |
| 11 | 118765231 | rs12795395 | T | C | 1.024 | 0.927 | -9390.864 | -9391.997 | -1.133 | 3.62E-09 |
| 11 | 118793479 | rs138526953 | A | C | 0.957 | 0.997 | -9390.7 | -9391.844 | -1.144 | 3.59E-09 |
| 11 | 118629364 | rs73001486 | A | G | 0.924 | 0.964 | -9408.077 | -9409.265 | -1.188 | 3.43E-09 |
| 11 | 118802492 | rs688035 | G | C | 0.93 | 0.953 | -9193.36 | -9194.557 | -1.196 | 3.40E-09 |
| 11 | 118576463 | rs496547 | A | T | 1.072 | 1.052 | -9405.568 | -9406.772 | -1.203 | 3.38E-09 |
| 11 | 118800302 | rs12793214 | A | C | 1.075 | 1.044 | -9177.441 | -9178.645 | -1.204 | 3.38E-09 |
| 11 | 118867695 | rs4938617 | T | C | 1.091 | 1.027 | -9404.515 | -9405.733 | -1.218 | 3.33E-09 |
| 11 | 118801235 | rs602716 | A | G | 0.93 | 0.953 | -9196.458 | -9197.679 | -1.221 | 3.32E-09 |
| 11 | 118583847 | rs12803363 | T | C | 1.092 | 1.01 | -9350.485 | -9351.721 | -1.237 | 3.27E-09 |
| 11 | 118803536 | rs615166 | A | G | 0.931 | 0.953 | -9194.217 | -9195.458 | -1.241 | 3.25E-09 |
| 11 | 118626286 | rs11217014 | C | T | 0.925 | 0.965 | -9408.077 | -9409.338 | -1.261 | 3.19E-09 |
| 11 | 118627778 | rs10892282 | C | G | 0.925 | 0.965 | -9408.077 | -9409.338 | -1.261 | 3.19E-09 |
| 11 | 118873840 | rs9645663 | C | T | 1.087 | 1.03 | -9387.564 | -9388.854 | -1.29 | 3.10E-09 |
| 11 | 118801640 | rs572608 | A | T | 1.072 | 1.045 | -9175.387 | -9176.682 | -1.295 | 3.08E-09 |
| 11 | 118711830 | rs566848 | G | A | 0.936 | 0.941 | -9399.316 | -9400.62 | -1.304 | 3.06E-09 |
| 11 | 118657297 | rs487177 | G | C | 0.926 | 0.965 | -9404.073 | -9405.398 | -1.325 | 2.99E-09 |
| 11 | 118606197 | rs12420846 | T | C | 1.081 | 1.053 | -9299.818 | -9301.147 | -1.33 | 2.98E-09 |
| 11 | 118640828 | rs11217021 | C | T | 0.927 | 0.963 | -9406.825 | -9408.157 | -1.331 | 2.97E-09 |
| 11 | 118796570 | rs11217096 | T | C | 1.07 | 1.048 | -9182.462 | -9183.797 | -1.334 | 2.96E-09 |
| 11 | 118613674 | rs7130875 | C | T | 0.926 | 0.969 | -9408.077 | -9409.412 | -1.335 | 2.96E-09 |
| 11 | 118616253 | rs7124498 | A | G | 0.927 | 0.967 | -9406.926 | -9408.269 | -1.343 | 2.94E-09 |
| 11 | 118530869 | rs45447797 | G | C | 0.975 | 0.918 | -9350.298 | -9351.642 | -1.344 | 2.94E-09 |
| 11 | 118811878 | rs74891547 | C | G | 1.082 | 0.984 | -9327.668 | -9329.014 | -1.345 | 2.93E-09 |
| 11 | 118633533 | rs7110526 | C | T | 0.927 | 0.964 | -9407.183 | -9408.536 | -1.352 | 2.91E-09 |
| 11 | 118635364 | rs11217018 | C | T | 0.927 | 0.964 | -9407.183 | -9408.536 | -1.352 | 2.91E-09 |
| 11 | 118655356 | rs7934890 | T | C | 0.927 | 0.965 | -9404.073 | -9405.435 | -1.362 | 2.88E-09 |
| 11 | 118698170 | rs56296109 | A | C | 0.919 | 0.967 | -9391.949 | -9393.312 | -1.363 | 2.88E-09 |
| 11 | 118852647 | rs80126726 | A | G | 1.085 | 1.028 | -9361.491 | -9362.863 | -1.371 | 2.86E-09 |
| 11 | 118637475 | rs112910608 | T | C | 0.928 | 0.964 | -9408.077 | -9409.449 | -1.372 | 2.85E-09 |
| 11 | 118726340 | rs113786535 | T | C | 0.981 | 0.918 | -9393.913 | -9395.286 | -1.374 | 2.85E-09 |
| 11 | 118539151 | rs72999491 | A | G | 0.939 | 1.037 | -9332.623 | -9333.998 | -1.375 | 2.85E-09 |
| 11 | 118543589 | rs72999499 | G | C | 0.938 | 1.036 | -9332.402 | -9333.78 | -1.378 | 2.84E-09 |
| 11 | 118468037 | rs535774 | T | C | 0.952 | 0.919 | -9343.283 | -9344.662 | -1.379 | 2.83E-09 |
| 11 | 118550841 | rs79699220 | C | G | 0.972 | 0.916 | -9352.388 | -9353.769 | -1.381 | 2.83E-09 |
| 11 | 118874272 | rs192113495 | G | T | 0.993 | 1.013 | -9389.181 | -9390.568 | -1.387 | 2.81E-09 |
| 11 | 118853021 | rs113962716 | A | C | 1.084 | 1.029 | -9362.344 | -9363.739 | -1.395 | 2.79E-09 |
| 11 | 118870413 | rs10892319 | T | C | 1.084 | 1.029 | -9386.706 | -9388.107 | -1.4 | 2.78E-09 |
| 11 | 118543798 | rs76570573 | T | C | 0.978 | 0.92 | -9373.528 | -9374.936 | -1.408 | 2.75E-09 |
| 11 | 118853774 | rs11217109 | C | T | 1.084 | 1.031 | -9365.983 | -9367.395 | -1.412 | 2.74E-09 |
| 11 | 118624919 | rs11217013 | C | G | 0.928 | 0.968 | -9408.077 | -9409.492 | -1.415 | 2.73E-09 |
| 11 | 118436071 | rs73023329 | T | C | 0.976 | 0.929 | -9358.531 | -9359.956 | -1.424 | 2.71E-09 |
| 11 | 118608625 | rs566733 | G | C | 0.976 | 0.928 | -9393.557 | -9394.992 | -1.435 | 2.68E-09 |
| 11 | 118610463 | rs4499035 | T | C | 0.928 | 0.97 | -9406.622 | -9408.06 | -1.438 | 2.67E-09 |
| 11 | 118610826 | rs1893029 | T | G | 0.928 | 0.97 | -9406.622 | -9408.075 | -1.452 | 2.63E-09 |
| 11 | 118831755 | rs10892316 | T | C | 1.081 | 1.036 | -9363.991 | -9365.444 | -1.454 | 2.63E-09 |
| 11 | 118608532 | rs567680 | G | C | 0.978 | 0.928 | -9394.356 | -9395.824 | -1.468 | 2.59E-09 |
| 11 | 118590864 | rs11216982 | A | G | 0.929 | 0.971 | -9388.478 | -9389.947 | -1.469 | 2.59E-09 |
| 11 | 118708564 | rs484232 | T | G | 0.944 | 0.944 | -9406.511 | -9407.993 | -1.483 | 2.55E-09 |
| 11 | 118483802 | rs72999455 | G | A | 0.975 | 0.928 | -9370.78 | -9372.299 | -1.519 | 2.46E-09 |
| 11 | 118494790 | rs11825621 | C | T | 0.976 | 0.928 | -9375.279 | -9376.798 | -1.519 | 2.46E-09 |
| 11 | 118605451 | rs2186780 | T | C | 0.945 | 0.935 | -9402.466 | -9403.995 | -1.529 | 2.44E-09 |
| 11 | 118619948 | rs112143902 | T | C | 0.963 | 0.925 | -9403.993 | -9405.541 | -1.548 | 2.39E-09 |
| 11 | 118758089 | rs497916 | T | C | 0.983 | 1.069 | -9403.704 | -9405.253 | -1.549 | 2.39E-09 |
| 11 | 118834718 | rs112939258 | T | C | 1.078 | 1.033 | -9366.504 | -9368.058 | -1.554 | 2.38E-09 |
| 11 | 118757191 | rs1790192 | G | A | 1.039 | 1.072 | -9392.806 | -9394.363 | -1.557 | 2.37E-09 |
| 11 | 118441079 | rs7106316 | G | A | 0.97 | 0.924 | -9337.84 | -9339.399 | -1.558 | 2.37E-09 |
| 11 | 118759610 | rs566416 | G | T | 0.971 | 1.062 | -9407.355 | -9408.914 | -1.559 | 2.37E-09 |
| 11 | 118762218 | rs543824 | G | A | 0.966 | 1.058 | -9403.62 | -9405.182 | -1.561 | 2.36E-09 |
| 11 | 118834508 | rs111358668 | A | G | 1.079 | 1.029 | -9366.956 | -9368.517 | -1.561 | 2.36E-09 |
| 11 | 118609598 | rs112996178 | T | C | 0.931 | 0.97 | -9403.592 | -9405.158 | -1.566 | 2.35E-09 |
| 11 | 118502172 | rs35435609 | T | C | 0.976 | 0.932 | -9375.279 | -9376.85 | -1.571 | 2.34E-09 |
| 11 | 118730669 | rs7933007 | C | T | 0.938 | 0.949 | -9406.759 | -9408.336 | -1.577 | 2.33E-09 |
| 11 | 118798780 | rs77916839 | C | G | 1.015 | 1.034 | -9364.127 | -9365.708 | -1.581 | 2.32E-09 |
| 11 | 118680072 | rs607472 | G | C | 0.964 | 0.934 | -9401.469 | -9403.079 | -1.61 | 2.25E-09 |
| 11 | 118765267 | rs598207 | C | G | 0.96 | 1.048 | -9400.848 | -9402.465 | -1.617 | 2.23E-09 |
| 11 | 118803193 | rs117235696 | G | T | 1.021 | 1.017 | -9363.915 | -9365.535 | -1.62 | 2.23E-09 |
| 11 | 118803842 | rs11600378 | G | A | 1.019 | 1.015 | -9363.915 | -9365.541 | -1.626 | 2.21E-09 |
| 11 | 118804671 | rs143711539 | A | G | 1.019 | 1.015 | -9363.915 | -9365.541 | -1.626 | 2.21E-09 |
| 11 | 118526691 | rs72999486 | T | C | 0.974 | 0.938 | -9370.939 | -9372.571 | -1.632 | 2.20E-09 |
| 11 | 118811315 | rs77638281 | A | G | 0.997 | 0.991 | -9361.459 | -9363.095 | -1.636 | 2.19E-09 |
| 11 | 118805144 | rs11605265 | A | G | 1.016 | 1.009 | -9363.25 | -9364.892 | -1.642 | 2.18E-09 |
| 11 | 118805220 | rs11605658 | T | C | 1.016 | 1.009 | -9363.25 | -9364.892 | -1.642 | 2.18E-09 |
| 11 | 118605914 | rs7926282 | C | A | 0.946 | 0.939 | -9402.466 | -9404.109 | -1.643 | 2.18E-09 |
| 11 | 118666289 | rs56104144 | C | G | 0.933 | 0.969 | -9404.515 | -9406.173 | -1.659 | 2.14E-09 |
| 11 | 118607622 | rs11217004 | A | G | 0.976 | 0.932 | -9379.278 | -9380.947 | -1.669 | 2.12E-09 |
| 11 | 118613235 | rs632124 | A | T | 0.98 | 0.932 | -9406.031 | -9407.703 | -1.672 | 2.11E-09 |
| 11 | 118725420 | rs11217062 | A | G | 0.94 | 0.95 | -9408.077 | -9409.751 | -1.674 | 2.11E-09 |
| 11 | 118726230 | rs2156755 | A | G | 0.94 | 0.95 | -9408.077 | -9409.751 | -1.674 | 2.11E-09 |
| 11 | 118668296 | rs58868870 | T | C | 0.934 | 0.97 | -9406.926 | -9408.628 | -1.703 | 2.05E-09 |
| 11 | 118723537 | rs7103314 | G | A | 0.941 | 0.951 | -9408.077 | -9409.804 | -1.727 | 2.00E-09 |
| 11 | 118539746 | rs72999492 | C | A | 1.04 | 1.033 | -9376.144 | -9377.883 | -1.74 | 1.98E-09 |
| 11 | 118657311 | rs1309473877 | A | C | 0.977 | 0.934 | -9406.278 | -9408.023 | -1.745 | 1.97E-09 |
| 11 | 118720590 | rs4938564 | T | G | 0.941 | 0.952 | -9408.077 | -9409.829 | -1.752 | 1.95E-09 |
| 11 | 118720684 | rs4936440 | G | C | 0.941 | 0.952 | -9408.077 | -9409.829 | -1.752 | 1.95E-09 |
| 11 | 118720998 | rs4938565 | A | G | 0.941 | 0.952 | -9408.077 | -9409.829 | -1.752 | 1.95E-09 |
| 11 | 118721189 | rs4938566 | T | A | 0.941 | 0.952 | -9408.077 | -9409.829 | -1.752 | 1.95E-09 |
| 11 | 118721249 | rs4938567 | G | A | 0.941 | 0.952 | -9408.077 | -9409.829 | -1.752 | 1.95E-09 |
| 11 | 118721850 | rs7946961 | G | A | 0.941 | 0.952 | -9408.077 | -9409.829 | -1.752 | 1.95E-09 |
| 11 | 118722231 | rs4938570 | G | A | 0.941 | 0.952 | -9408.077 | -9409.829 | -1.752 | 1.95E-09 |
| 11 | 118723087 | rs11217059 | T | C | 0.941 | 0.952 | -9408.077 | -9409.829 | -1.752 | 1.95E-09 |
| 11 | 118724336 | rs10892298 | C | T | 0.941 | 0.952 | -9408.077 | -9409.829 | -1.752 | 1.95E-09 |
| 11 | 118718485 | rs4938561 | C | T | 0.942 | 0.951 | -9408.077 | -9409.836 | -1.759 | 1.94E-09 |
| 11 | 118816353 | rs586370 | A | G | 0.94 | 1.002 | -9081.913 | -9083.676 | -1.763 | 1.93E-09 |
| 11 | 118895495 | rs8301 | G | C | 0.958 | 1.043 | -9343.759 | -9345.526 | -1.767 | 1.92E-09 |
| 11 | 118718856 | rs2156751 | G | A | 0.942 | 0.951 | -9408.077 | -9409.85 | -1.772 | 1.91E-09 |
| 11 | 118718972 | rs2187491 | C | T | 0.942 | 0.951 | -9408.077 | -9409.85 | -1.772 | 1.91E-09 |
| 11 | 118719254 | rs2156750 | C | T | 0.942 | 0.951 | -9408.077 | -9409.85 | -1.772 | 1.91E-09 |
| 11 | 118719405 | rs2156749 | G | C | 0.942 | 0.951 | -9408.077 | -9409.85 | -1.772 | 1.91E-09 |
| 11 | 118719742 | rs7116715 | C | T | 0.942 | 0.951 | -9408.077 | -9409.85 | -1.772 | 1.91E-09 |
| 11 | 118717929 | rs4938558 | T | C | 0.942 | 0.952 | -9407.669 | -9409.448 | -1.78 | 1.90E-09 |
| 11 | 118481431 | rs10892246 | A | G | 1.046 | 0.967 | -9332.042 | -9333.845 | -1.803 | 1.85E-09 |
| 11 | 118620697 | rs488219 | C | T | 0.973 | 0.935 | -9406.926 | -9408.736 | -1.811 | 1.84E-09 |
| 11 | 118713180 | rs2187490 | G | T | 0.957 | 0.941 | -9406.618 | -9408.433 | -1.815 | 1.83E-09 |
| 11 | 118809329 | rs10892313 | A | G | 1.051 | 0.968 | -9307.906 | -9309.72 | -1.815 | 1.83E-09 |
| 11 | 118810405 | rs4938589 | T | C | 1.052 | 0.969 | -9305.085 | -9306.9 | -1.816 | 1.83E-09 |
| 11 | 118617299 | rs607125 | C | T | 0.976 | 0.935 | -9407.198 | -9409.019 | -1.822 | 1.82E-09 |
| 11 | 118659896 | rs2105782 | C | T | 0.972 | 0.936 | -9405.139 | -9406.975 | -1.836 | 1.79E-09 |
| 11 | 118793613 | rs11217091 | C | T | 1.017 | 0.943 | -9316.248 | -9318.083 | -1.836 | 1.79E-09 |
| 11 | 118808140 | rs7117408 | T | C | 1.051 | 0.971 | -9310.762 | -9312.607 | -1.845 | 1.78E-09 |
| 11 | 118556995 | rs72999502 | C | T | 1.012 | 1.041 | -9360.258 | -9362.106 | -1.848 | 1.77E-09 |
| 11 | 118624250 | rs546393 | C | T | 0.974 | 0.936 | -9406.926 | -9408.789 | -1.864 | 1.75E-09 |
| 11 | 118651809 | rs555649 | C | T | 0.973 | 0.937 | -9408.077 | -9409.968 | -1.891 | 1.70E-09 |
| 11 | 118679629 | rs1784300 | T | A | 0.968 | 0.939 | -9405.914 | -9407.818 | -1.905 | 1.67E-09 |
| 11 | 118644582 | rs2510897 | C | A | 0.973 | 0.937 | -9406.709 | -9408.618 | -1.909 | 1.67E-09 |
| 11 | 118765688 | rs676925 | G | C | 0.977 | 1.056 | -9398.257 | -9400.178 | -1.922 | 1.65E-09 |
| 11 | 118658075 | rs515523 | G | A | 0.973 | 0.938 | -9406.682 | -9408.613 | -1.931 | 1.63E-09 |
| 11 | 118811641 | rs11606199 | A | G | 1.065 | 1.022 | -9350.254 | -9352.205 | -1.952 | 1.60E-09 |
| 11 | 118707249 | rs4938551 | C | G | 0.952 | 0.951 | -9404.178 | -9406.137 | -1.959 | 1.59E-09 |
| 11 | 118632522 | rs581045 | T | C | 0.973 | 0.938 | -9406.926 | -9408.887 | -1.961 | 1.58E-09 |
| 11 | 118629031 | rs555639 | A | G | 0.973 | 0.939 | -9406.926 | -9408.894 | -1.968 | 1.57E-09 |
| 11 | 118807637 | rs112974653 | T | G | 1.041 | 0.968 | -9309.326 | -9311.314 | -1.988 | 1.54E-09 |
| 11 | 118763736 | rs12363277 | C | T | 1 | 0.974 | -9388.154 | -9390.147 | -1.993 | 1.53E-09 |
| 11 | 118662028 | rs12796102 | T | C | 0.974 | 0.94 | -9405.731 | -9407.768 | -2.037 | 1.47E-09 |
| 11 | 118716563 | rs2851460 | A | G | 0.953 | 0.952 | -9405.21 | -9407.252 | -2.042 | 1.46E-09 |
| 11 | 118631288 | rs507890 | C | T | 0.976 | 0.94 | -9406.046 | -9408.094 | -2.048 | 1.45E-09 |
| 11 | 118631653 | rs1784543 | T | A | 0.976 | 0.94 | -9406.046 | -9408.094 | -2.048 | 1.45E-09 |
| 11 | 118699276 | rs625274 | T | C | 0.97 | 0.942 | -9406.409 | -9408.471 | -2.062 | 1.43E-09 |
| 11 | 118824683 | rs507763 | C | G | 1.061 | 1.012 | -9380.958 | -9383.022 | -2.063 | 1.43E-09 |
| 11 | 118824917 | rs11217102 | A | G | 1.061 | 1.012 | -9380.958 | -9383.022 | -2.063 | 1.43E-09 |
| 11 | 118789952 | rs11217090 | T | C | 1.026 | 0.966 | -9319.057 | -9321.129 | -2.073 | 1.42E-09 |
| 11 | 118828184 | rs45586434 | A | C | 1.06 | 1.026 | -9357.873 | -9359.946 | -2.073 | 1.42E-09 |
| 11 | 118500963 | rs680465 | A | G | 0.964 | 0.942 | -9385.911 | -9387.991 | -2.081 | 1.40E-09 |
| 11 | 118801440 | rs12280501 | T | C | 1.009 | 0.959 | -9304.486 | -9306.6 | -2.113 | 1.36E-09 |
| 11 | 118700557 | rs4291686 | G | A | 0.971 | 0.943 | -9406.409 | -9408.527 | -2.119 | 1.35E-09 |
| 11 | 118799206 | rs7101680 | A | G | 1.012 | 0.962 | -9304.142 | -9306.271 | -2.129 | 1.34E-09 |
| 11 | 118826045 | rs537080 | C | T | 1.059 | 1.015 | -9367.044 | -9369.186 | -2.142 | 1.32E-09 |
| 11 | 118899218 | rs4936459 | T | C | 1.049 | 1.039 | -9378.807 | -9380.956 | -2.149 | 1.31E-09 |
| 11 | 118798824 | rs112119822 | G | A | 1.01 | 0.963 | -9304.525 | -9306.677 | -2.152 | 1.31E-09 |
| 11 | 118789657 | rs12294466 | T | C | 0.995 | 0.98 | -9351.232 | -9353.391 | -2.159 | 1.30E-09 |
| 11 | 118798757 | rs57879605 | T | C | 1.007 | 0.946 | -9275.067 | -9277.243 | -2.176 | 1.28E-09 |
| 11 | 118718061 | rs4938559 | T | C | 0.95 | 0.965 | -9402.844 | -9405.059 | -2.215 | 1.23E-09 |
| 11 | 118718062 | rs4938560 | G | A | 0.95 | 0.965 | -9402.844 | -9405.059 | -2.215 | 1.23E-09 |
| 11 | 118717732 | rs11217052 | G | A | 0.95 | 0.965 | -9402.844 | -9405.083 | -2.239 | 1.20E-09 |
| 11 | 118809882 | rs11217099 | A | C | 1.051 | 0.999 | -9313.936 | -9316.184 | -2.249 | 1.19E-09 |
| 11 | 118717708 | rs7107756 | A | G | 0.967 | 0.949 | -9395.688 | -9398.005 | -2.316 | 1.11E-09 |
| 11 | 118770321 | rs11217083 | G | A | 0.959 | 0.955 | -9392.892 | -9395.21 | -2.318 | 1.11E-09 |
| 11 | 118755738 | rs523604 | G | A | 0.984 | 1.048 | -9406.217 | -9408.558 | -2.341 | 1.08E-09 |
| 11 | 118816304 | rs586763 | T | C | 1.044 | 0.997 | -9408.077 | -9410.509 | -2.432 | 9.89E-10 |
| 11 | 118767077 | rs581063 | C | T | 0.961 | 0.959 | -9397.325 | -9399.787 | -2.462 | 9.60E-10 |
| 11 | 118769032 | rs11217081 | G | A | 0.962 | 0.96 | -9396.903 | -9399.417 | -2.514 | 9.11E-10 |
| 11 | 118823148 | rs663003 | T | C | 0.956 | 1.005 | -9080.025 | -9082.546 | -2.521 | 9.05E-10 |
| 11 | 118661376 | rs73003294 | T | C | 0.989 | 0.979 | -9400.286 | -9402.88 | -2.594 | 8.41E-10 |
| 11 | 118826060 | rs625291 | T | C | 0.959 | 1.007 | -9076.385 | -9079.006 | -2.621 | 8.19E-10 |
| 11 | 118826063 | rs666191 | A | G | 0.959 | 1.007 | -9075.723 | -9078.344 | -2.621 | 8.19E-10 |
| 11 | 118880740 | rs11217122 | C | T | 0.969 | 1.014 | -9381.429 | -9384.071 | -2.642 | 8.02E-10 |
| 11 | 118824439 | rs61900157 | C | A | 0.96 | 1.007 | -9080.786 | -9083.433 | -2.647 | 7.98E-10 |
| 11 | 118824996 | rs4426142 | A | G | 0.96 | 1.007 | -9080.786 | -9083.433 | -2.647 | 7.98E-10 |
| 11 | 118825069 | rs2097131 | A | G | 0.96 | 1.007 | -9080.786 | -9083.433 | -2.647 | 7.98E-10 |
| 11 | 118825636 | rs1790191 | G | A | 0.961 | 1.008 | -9080.786 | -9083.441 | -2.655 | 7.91E-10 |
| 11 | 118735476 | rs10892301 | A | G | 0.959 | 0.976 | -9406.438 | -9409.123 | -2.684 | 7.69E-10 |
| 11 | 118896959 | rs57156480 | A | G | 0.998 | 1.033 | -9380.468 | -9383.252 | -2.784 | 6.95E-10 |
| 11 | 118892940 | rs12280288 | C | T | 0.982 | 1.021 | -9383.168 | -9385.963 | -2.795 | 6.88E-10 |
| 11 | 118831335 | rs61506726 | G | A | 0.978 | 0.965 | -9280.55 | -9283.355 | -2.805 | 6.81E-10 |
| 11 | 118895686 | rs8192696 | T | C | 0.988 | 1.017 | -9395.524 | -9398.374 | -2.85 | 6.51E-10 |
| 11 | 118872581 | rs4558175 | A | G | 0.981 | 1.02 | -9350.498 | -9353.36 | -2.862 | 6.43E-10 |
| 11 | 118873385 | rs10892320 | T | C | 0.984 | 1.023 | -9352.066 | -9354.929 | -2.863 | 6.43E-10 |
| 11 | 118712427 | rs7108905 | G | C | 0.996 | 1.018 | -9401.123 | -9403.996 | -2.873 | 6.36E-10 |
| 11 | 118708785 | rs7122510 | A | C | 0.997 | 1.018 | -9399.815 | -9402.691 | -2.876 | 6.34E-10 |
| 11 | 118708786 | rs7122511 | A | C | 0.997 | 1.018 | -9399.815 | -9402.691 | -2.876 | 6.34E-10 |
| 11 | 118894287 | rs569 | G | A | 0.994 | 1.021 | -9399.915 | -9402.793 | -2.878 | 6.33E-10 |
| 11 | 118895202 | rs11006 | G | A | 0.994 | 1.021 | -9399.915 | -9402.793 | -2.878 | 6.33E-10 |
| 11 | 118893878 | rs111322300 | C | T | 0.995 | 1.021 | -9399.915 | -9402.8 | -2.885 | 6.29E-10 |
| 11 | 118896115 | rs2276072 | G | A | 0.994 | 1.026 | -9376.674 | -9379.559 | -2.886 | 6.28E-10 |
| 11 | 118896116 | rs2276071 | C | A | 0.994 | 1.026 | -9376.674 | -9379.559 | -2.886 | 6.28E-10 |
| 11 | 118892757 | rs117289748 | G | C | 0.995 | 1.021 | -9400.309 | -9403.197 | -2.888 | 6.27E-10 |
| 11 | 118892759 | rs143660533 | C | A | 0.995 | 1.021 | -9400.309 | -9403.197 | -2.888 | 6.27E-10 |
| 11 | 118712270 | rs7105658 | T | C | 0.996 | 1.017 | -9402.75 | -9405.643 | -2.893 | 6.24E-10 |
| 11 | 118709600 | rs12272981 | G | A | 0.998 | 1.016 | -9399.914 | -9402.81 | -2.896 | 6.22E-10 |
| 11 | 118868750 | rs4445645 | C | G | 0.98 | 1 | -9391.275 | -9394.178 | -2.903 | 6.17E-10 |
| 11 | 118707559 | rs7118009 | T | C | 0.999 | 1.015 | -9399.815 | -9402.728 | -2.912 | 6.12E-10 |
| 11 | 118872404 | rs6589713 | C | T | 0.987 | 1.021 | -9348.45 | -9351.375 | -2.924 | 6.05E-10 |
| 11 | 118729283 | rs11217068 | T | C | 1.002 | 1.017 | -9405.241 | -9408.169 | -2.928 | 6.02E-10 |
| 11 | 118724116 | rs12276533 | A | G | 1 | 1.013 | -9404.598 | -9407.532 | -2.934 | 5.99E-10 |
| 11 | 118725383 | rs9735317 | T | C | 1.004 | 1.014 | -9405.281 | -9408.215 | -2.934 | 5.99E-10 |
| 11 | 118734640 | rs12286723 | T | C | 1.011 | 1.014 | -9406.383 | -9409.32 | -2.937 | 5.97E-10 |
| 11 | 118729442 | rs12277355 | T | C | 1.002 | 1.015 | -9405.241 | -9408.181 | -2.94 | 5.95E-10 |
| 11 | 118888831 | rs11820805 | C | T | 0.993 | 1.024 | -9358.609 | -9361.557 | -2.949 | 5.90E-10 |
| 11 | 118886656 | rs7131534 | A | G | 0.989 | 1.009 | -9399.264 | -9402.216 | -2.952 | 5.88E-10 |
| 11 | 118728479 | rs7924647 | G | A | 1.001 | 1.01 | -9407.357 | -9410.31 | -2.953 | 5.87E-10 |
| 11 | 118714492 | rs7117594 | T | C | 0.999 | 1.01 | -9407.766 | -9410.722 | -2.956 | 5.86E-10 |
| 11 | 118723753 | rs7119568 | A | T | 1 | 1.009 | -9404.598 | -9407.556 | -2.959 | 5.84E-10 |
| 11 | 118886117 | rs10111 | T | C | 0.991 | 1.01 | -9400.347 | -9403.306 | -2.959 | 5.84E-10 |
| 11 | 118889378 | rs7113753 | A | G | 0.995 | 1.024 | -9356.408 | -9359.368 | -2.96 | 5.83E-10 |
| 11 | 118877563 | rs58709307 | T | C | 0.989 | 1.008 | -9398.593 | -9401.554 | -2.961 | 5.83E-10 |

| | | | | | | | |
|---|---|---|---|---|---|---|---|
| 11 | 118791180 | rs146541591 | C | T | M_Disease2_Disease1 | 0.275710379 | M_Disease2_Disease1&M_Disease2&NULL&M_Disease1 |
| 11 | 118483912 | rs45621436 | G | T | M_Disease1 | 0.275495357 | M_Disease1&M_Disease2_Disease1&NULL&M_Disease2 |
| 11 | 118493319 | rs10790254 | A | G | M_Disease1 | 0.275495357 | M_Disease1&M_Disease2_Disease1&NULL&M_Disease2 |
| 11 | 118614115 | rs73001451 | C | T | NULL | 0.275231206 | NULL&M_Disease1&M_Disease2&M_Disease2_Disease1 |
| 11 | 118801045 | rs75085081 | A | G | M_Disease2_Disease1 | 0.275154234 | M_Disease2_Disease1&M_Disease2&NULL&M_Disease1 |
| 11 | 118759905 | rs118020807 | A | C | NULL | 0.274982576 | NULL&M_Disease1&M_Disease2&M_Disease2_Disease1 |
| 11 | 118775524 | rs79108568 | G | T | M_Disease2 | 0.274850076 | M_Disease2&M_Disease2_Disease1&NULL&M_Disease1 |
| 11 | 118722946 | rs148990923 | T | C | NULL | 0.274801037 | NULL&M_Disease1&M_Disease2&M_Disease2_Disease1 |
| 11 | 118501723 | rs61900957 | T | C | NULL | 0.274791119 | NULL&M_Disease2&M_Disease1&M_Disease2_Disease1 |
| 11 | 118598936 | rs184071181 | C | T | M_Disease1 | 0.274540095 | M_Disease1&NULL&M_Disease2_Disease1&M_Disease2 |
| 11 | 118583261 | rs112246697 | A | G | M_Disease2_Disease1 | 0.27448936 | M_Disease2_Disease1&M_Disease2&NULL&M_Disease1 |
| 11 | 118561858 | rs10790257 | C | G | M_Disease1 | 0.274087026 | M_Disease1&NULL&M_Disease2_Disease1&M_Disease2 |
| 11 | 118561859 | rs10790258 | T | G | M_Disease1 | 0.274087026 | M_Disease1&NULL&M_Disease2_Disease1&M_Disease2 |
| 11 | 118626825 | rs74337439 | A | G | M_Disease1 | 0.273946447 | M_Disease1&M_Disease2_Disease1&NULL&M_Disease2 |
| 11 | 118557483 | rs6589673 | C | A | M_Disease1 | 0.273879065 | M_Disease1&NULL&M_Disease2_Disease1&M_Disease2 |
| 11 | 118559426 | rs10892253 | A | C | M_Disease1 | 0.273650743 | M_Disease1&NULL&M_Disease2_Disease1&M_Disease2 |
| 11 | 118445559 | rs73555296 | T | G | M_Disease1 | 0.273437643 | M_Disease1&NULL&M_Disease2_Disease1&M_Disease2 |
| 11 | 118615390 | rs78502545 | T | C | M_Disease1 | 0.272758203 | M_Disease1&M_Disease2_Disease1&NULL&M_Disease2 |
| 11 | 118652569 | rs150792202 | A | G | M_Disease2_Disease1 | 0.271646383 | M_Disease2_Disease1&NULL&M_Disease1&M_Disease2 |
| 11 | 118568427 | rs117361624 | T | C | NULL | 0.271462774 | NULL&M_Disease2&M_Disease1&M_Disease2_Disease1 |
| 11 | 118748789 | rs145762171 | A | G | M_Disease2 | 0.27126901 | M_Disease2&M_Disease2_Disease1&NULL&M_Disease1 |
| 11 | 118530434 | rs117619140 | T | G | NULL | 0.271266628 | NULL&M_Disease2&M_Disease1&M_Disease2_Disease1 |
| 11 | 118477746 | rs72999445 | G | T | NULL | 0.271060501 | NULL&M_Disease1&M_Disease2&M_Disease2_Disease1 |
| 11 | 118644323 | rs6589695 | C | G | M_Disease1 | 0.270585501 | M_Disease1&NULL&M_Disease2&M_Disease2_Disease1 |
| 11 | 118735876 | rs79843382 | T | C | NULL | 0.270506391 | NULL&M_Disease2&M_Disease1&M_Disease2_Disease1 |
| 11 | 118727137 | rs117223320 | C | G | NULL | 0.270462139 | NULL&M_Disease2&M_Disease1&M_Disease2_Disease1 |
| 11 | 118721767 | rs116884018 | C | T | M_Disease1 | 0.269155337 | M_Disease1&NULL&M_Disease2_Disease1&M_Disease2 |
| 11 | 118603615 | rs74783030 | A | G | M_Disease1 | 0.268932587 | M_Disease1&NULL&M_Disease2_Disease1&M_Disease2 |
| 11 | 118711069 | rs538645 | C | T | NULL | 0.268378838 | NULL&M_Disease2&M_Disease2_Disease1&M_Disease1 |
| 11 | 118518227 | rs10736494 | T | C | NULL | 0.267741119 | NULL&M_Disease1&M_Disease2&M_Disease2_Disease1 |
| 11 | 118700515 | rs112571283 | T | C | M_Disease2 | 0.267346376 | M_Disease2&NULL&M_Disease2_Disease1&M_Disease1 |
| 11 | 118799691 | rs191378986 | G | A | M_Disease1 | 0.267225481 | M_Disease1&NULL&M_Disease2&M_Disease2_Disease1 |
| 11 | 118413882 | rs146864992 | T | C | M_Disease1 | 0.266929099 | M_Disease1&NULL&M_Disease2_Disease1&M_Disease2 |
| 11 | 118819541 | rs150196102 | T | C | NULL | 0.266797163 | NULL&M_Disease1&M_Disease2&M_Disease2_Disease1 |
| 11 | 118515779 | rs117679873 | A | G | NULL | 0.266752615 | NULL&M_Disease2&M_Disease1&M_Disease2_Disease1 |
| 11 | 118696055 | rs189417780 | C | A | M_Disease1 | 0.266549035 | M_Disease1&M_Disease2_Disease1&NULL&M_Disease2 |
| 11 | 118832251 | rs79808604 | A | C | NULL | 0.266290137 | NULL&M_Disease2&M_Disease1&M_Disease2_Disease1 |
| 11 | 118419577 | rs191458787 | T | C | M_Disease1 | 0.265775101 | M_Disease1&NULL&M_Disease2&M_Disease2_Disease1 |
| 11 | 118421438 | rs145485933 | G | A | M_Disease1 | 0.265775101 | M_Disease1&NULL&M_Disease2&M_Disease2_Disease1 |
| 11 | 118744615 | rs73009445 | A | G | M_Disease1 | 0.265692612 | M_Disease1&NULL&M_Disease2&M_Disease2_Disease1 |
| 11 | 118650163 | rs73003218 | T | C | NULL | 0.26564103 | NULL&M_Disease2&M_Disease1&M_Disease2_Disease1 |
| 11 | 118754649 | rs10892307 | G | C | M_Disease2 | 0.265153885 | M_Disease2&M_Disease1&NULL&M_Disease2_Disease1 |
| 11 | 118496524 | rs35692099 | C | T | NULL | 0.26508173 | NULL&M_Disease1&M_Disease2&M_Disease2_Disease1 |
| 11 | 118502121 | rs17614795 | G | A | NULL | 0.26508173 | NULL&M_Disease1&M_Disease2&M_Disease2_Disease1 |
| 11 | 118694739 | rs188550348 | A | G | NULL | 0.261391564 | NULL&M_Disease1&M_Disease2&M_Disease2_Disease1 |
| 11 | 118855789 | rs4938609 | A | C | M_Disease1 | 0.259435204 | M_Disease1&M_Disease2_Disease1&NULL&M_Disease2 |
| 11 | 118478681 | rs45578540 | T | C | NULL | 0.259365613 | NULL&M_Disease2&M_Disease1&M_Disease2_Disease1 |
| 11 | 118755630 | rs3176905 | C | T | M_Disease2 | 0.258770077 | M_Disease2&M_Disease1&NULL&M_Disease2_Disease1 |
| 11 | 118783424 | rs149114341 | A | G | M_Disease1 | 0.256428678 | M_Disease1&M_Disease2_Disease1&NULL&M_Disease2 |
| 11 | 118691070 | rs140577174 | G | A | NULL | 0.255214404 | NULL&M_Disease2&M_Disease2_Disease1&M_Disease1 |
| 11 | 118765283 | rs665648 | A | G | NULL | 0.255137005 | NULL&M_Disease1&M_Disease2&M_Disease2_Disease1 |

| | | | | | | | | | | |
|---|---|---|---|---|---|---|---|---|---|---|
| 11 | 118879285 | rs7120445 | C | A | 0.988 | 1.006 | -9397.509 | -9400.474 | -2.964 | 5.81E-10 |
| 11 | 118856582 | rs3887652 | C | G | 1.005 | 1.025 | -9362.188 | -9365.153 | -2.965 | 5.80E-10 |
| 11 | 118880668 | rs11607569 | C | T | 0.996 | 1.013 | -9400.889 | -9403.858 | -2.968 | 5.79E-10 |
| 11 | 118887581 | rs7105187 | T | C | 1.012 | 1.023 | -9373.938 | -9376.907 | -2.968 | 5.79E-10 |
| 11 | 118888619 | rs11602764 | G | A | 0.993 | 1.022 | -9361.787 | -9364.756 | -2.969 | 5.78E-10 |
| 11 | 118879824 | rs73559119 | C | T | 0.989 | 1.007 | -9399.806 | -9402.776 | -2.97 | 5.77E-10 |
| 11 | 118880012 | rs11825284 | A | G | 0.989 | 1.007 | -9399.806 | -9402.776 | -2.97 | 5.77E-10 |
| 11 | 118892277 | rs4938621 | T | C | 0.991 | 1.008 | -9400.347 | -9403.322 | -2.974 | 5.75E-10 |
| 11 | 118892835 | rs61179449 | C | G | 0.991 | 1.008 | -9400.347 | -9403.322 | -2.974 | 5.75E-10 |
| 11 | 118892872 | rs58955701 | T | C | 0.991 | 1.008 | -9400.347 | -9403.322 | -2.974 | 5.75E-10 |
| 11 | 118893023 | rs7127188 | C | A | 0.991 | 1.008 | -9400.347 | -9403.322 | -2.974 | 5.75E-10 |
| 11 | 118878222 | rs11606357 | C | T | 0.991 | 1.008 | -9398.593 | -9401.568 | -2.975 | 5.75E-10 |
| 11 | 118831727 | rs10892315 | C | T | 0.985 | 0.974 | -9293.361 | -9296.338 | -2.977 | 5.73E-10 |
| 11 | 118883491 | rs60065241 | A | C | 0.993 | 1.007 | -9399.806 | -9402.791 | -2.985 | 5.69E-10 |
| 11 | 118887568 | rs7121270 | G | T | 1.01 | 1.022 | -9372.851 | -9375.841 | -2.99 | 5.66E-10 |
| 11 | 118875727 | rs4936457 | G | A | 1.002 | 1.022 | -9372.333 | -9375.334 | -3 | 5.60E-10 |
| 11 | 118875761 | rs4936458 | G | T | 1.002 | 1.022 | -9372.333 | -9375.334 | -3 | 5.60E-10 |
| 11 | 118854372 | rs75067211 | T | C | 1.007 | 1.021 | -9350.458 | -9353.46 | -3.002 | 5.59E-10 |
| 11 | 118887887 | rs369816532 | T | G | 1.005 | 1.02 | -9381.98 | -9384.982 | -3.002 | 5.59E-10 |
| 11 | 118873987 | rs9645664 | C | T | 0.999 | 1.02 | -9368.077 | -9371.081 | -3.004 | 5.58E-10 |
| 11 | 118832035 | rs10892317 | C | T | 0.985 | 0.976 | -9294.529 | -9297.536 | -3.007 | 5.56E-10 |
| 11 | 118887888 | rs3916582 | T | C | 1.005 | 1.019 | -9381.712 | -9384.719 | -3.007 | 5.56E-10 |
| 11 | 118769442 | rs7104819 | G | A | 0.97 | 0.981 | -9402.011 | -9405.027 | -3.016 | 5.51E-10 |
| 11 | 118867803 | rs4938618 | C | G | 1.005 | 1.02 | -9371.49 | -9374.506 | -3.016 | 5.51E-10 |
| 11 | 118869539 | rs11822103 | A | G | 1.006 | 1.02 | -9371.49 | -9374.507 | -3.017 | 5.51E-10 |
| 11 | 118872741 | rs4573680 | C | T | 0.999 | 1.018 | -9371.401 | -9374.435 | -3.034 | 5.42E-10 |
| 11 | 118588998 | rs570424 | C | T | 0.975 | 0.973 | -9393.803 | -9396.839 | -3.036 | 5.41E-10 |
| 11 | 118869460 | rs3737504 | G | A | 0.993 | 1.012 | -9366.531 | -9369.583 | -3.052 | 5.32E-10 |
| 11 | 118889132 | rs11217124 | G | A | 0.999 | 1.012 | -9380.893 | -9383.953 | -3.06 | 5.28E-10 |
| 11 | 118885161 | rs59095933 | A | G | 1.003 | 1.012 | -9380.801 | -9383.862 | -3.061 | 5.27E-10 |
| 11 | 118884708 | rs61381062 | A | G | 1.001 | 1.011 | -9379.713 | -9382.781 | -3.068 | 5.24E-10 |
| 11 | 118884726 | rs58601794 | C | T | 1.001 | 1.011 | -9379.713 | -9382.781 | -3.068 | 5.24E-10 |
| 11 | 118885163 | rs59608621 | G | A | 1.001 | 1.011 | -9379.713 | -9382.781 | -3.068 | 5.24E-10 |
| 11 | 118889185 | rs11217125 | C | T | 1 | 1.011 | -9381.19 | -9384.257 | -3.068 | 5.24E-10 |
| 11 | 118765600 | rs3922 | G | A | 0.971 | 0.985 | -9407.383 | -9410.457 | -3.074 | 5.20E-10 |
| 11 | 118836606 | rs113975105 | G | C | 0.989 | 0.98 | -9296.3 | -9299.376 | -3.076 | 5.19E-10 |
| 11 | 118574311 | rs524409 | T | C | 0.994 | 0.971 | -9408.077 | -9411.229 | -3.152 | 4.81E-10 |
| 11 | 118574675 | rs494459 | T | C | 0.994 | 0.971 | -9408.077 | -9411.229 | -3.152 | 4.81E-10 |
| 11 | 118573894 | rs576283 | G | A | 0.994 | 0.972 | -9406.827 | -9409.985 | -3.158 | 4.78E-10 |
| 11 | 118575011 | rs530901 | C | T | 0.995 | 0.972 | -9408.077 | -9411.236 | -3.159 | 4.78E-10 |
| 11 | 118575180 | rs626645 | C | T | 0.995 | 0.972 | -9408.077 | -9411.236 | -3.159 | 4.78E-10 |
| 11 | 118575419 | rs625735 | A | G | 0.994 | 0.972 | -9408.077 | -9411.237 | -3.16 | 4.78E-10 |
| 11 | 118575479 | rs637563 | G | A | 0.994 | 0.972 | -9408.077 | -9411.237 | -3.16 | 4.78E-10 |
| 11 | 118575693 | rs638514 | C | T | 0.994 | 0.972 | -9408.077 | -9411.237 | -3.16 | 4.78E-10 |
| 11 | 118573916 | rs627391 | T | A | 0.995 | 0.972 | -9405.575 | -9408.738 | -3.163 | 4.76E-10 |
| 11 | 118598138 | rs543692 | C | T | 0.986 | 0.975 | -9396.25 | -9399.441 | -3.191 | 4.63E-10 |
| 11 | 118598612 | rs568688 | G | T | 0.986 | 0.975 | -9395.454 | -9398.648 | -3.195 | 4.61E-10 |
| 11 | 118595207 | rs563537 | A | T | 0.988 | 0.975 | -9388.704 | -9391.914 | -3.21 | 4.54E-10 |
| 11 | 118580348 | rs516503 | T | C | 0.995 | 0.977 | -9404.948 | -9408.21 | -3.262 | 4.31E-10 |
| 11 | 118580906 | rs496058 | G | A | 0.994 | 0.977 | -9404.948 | -9408.214 | -3.265 | 4.30E-10 |
| 11 | 118590743 | rs503542 | G | A | 0.988 | 0.986 | -9352.886 | -9356.257 | -3.371 | 3.87E-10 |
